# Sorting receptor SORCS2 facilitates a protective stress response in pancreatic islets

**DOI:** 10.1101/2023.05.15.540791

**Authors:** Oleksandra Kalnytska, Per Qvist, Séverine Kunz, Thomas Conrad, Thomas E. Willnow, Vanessa Schmidt

**Author notes:** **Correspondence to:** Thomas E. Willnow, Max-Delbrück-Center for Molecular Medicine Robert-Rössle-Str. 10, D-13125 Berlin, Germany, Vanessa Schmidt, Max-Delbrück-Center for Molecular Medicine Robert-Roessle-Str. 10, D-13125 Berlin, Germany.

## Abstract

**Objective:** SORCS2 is an intracellular sorting receptor genetically associated with body mass index (BMI) in humans, yet its mode of action remains unknown. Elucidating the receptor function that defines its role in metabolic health is the objective of this work.

**Methods:** Combining *in vivo* metabolic studies in SORCS2-deficient mouse models with *ex vivo* structural and functional analyses as well as single-cell transcriptomics of murine pancreatic tissues, we studied the pathophysiological consequences of receptor dysfunction for metabolism.

**Results:** Our studies identified an important role for SORCS2 in islet stress response essential to sustain glucose-stimulated insulin release. In detail, we show that SORCS2 is predominantly expressed in islet alpha cells. Loss of receptor expression coincides with the inability of these cells to produce osteopontin, a secreted factor that facilitates insulin release from beta cells under stress. In line with diminished osteopontin levels, beta cells in SORCS2- deficient islets show changes in gene expression patterns related to aggravated ER stress, protein misfolding, as well as mitochondrial dysfunction; and they exhibit defects in insulin granule maturation and a blunted response to glucose stimulation *in vivo* and *ex vivo*. Impaired glucose tolerance in receptor mutant mice coincides with alterations in body weight and composition.

**Conclusion:** Our data identified a novel concept in protective islet stress response involving the alpha cell receptor SORCS2 and provide experimental support for association of *SORCS2* with metabolic control in humans.

## 1. Introduction

Vacuolar protein sorting 10 protein (VPS10P) domain receptors are a group of type 1 transmembrane proteins expressed in various mammalian cell types. The gene family comprises five members designated SORLA, sortilin, as well as sortilin-related receptors CNS expressed (SORCS) −1, −2, and −3. Mainly, VPS10P domain receptors act as intracellular sorting factors that direct cargo proteins between cell surface and intracellular compartments, defining secretory and endocytic capacities of target cells (reviewed in [1]).

Early studies have mainly focused on functions of VPS10P domain receptors in neuronal protein trafficking, providing molecular explanations for their causal roles in psychiatric and neurodegenerative diseases (reviewed in [1; 2]). Surprisingly, recent work now uncovered genetic association of these receptors with metabolic traits in humans and mouse models, including obesity [3], hypercholesterolemia [4–6], and type 2 diabetes (T2D) [7; 8]. Corroborating important functions for VPS10P domain receptors in metabolic health, SORCS1 has been implicated in biogenesis and trafficking of insulin secretory granules in pancreatic islet beta cells. In mice, targeted *Sorcs1* gene disruption [9] or expression of a mutant receptor variant [10] impairs the ability of beta cells to sustain release of insulin during a glucose challenge.

Interestingly, SORCS1 shares close structural similarity with SORCS2, a VPS10P domain receptor genetically associated with human body mass index (BMI) [11]. Still, the role of SORCS2 in metabolic control remains unknown. In this study, we identified a SORCS1-related function for SORCS2 in facilitating glucose-stimulated insulin release, possibly by providing a protective islet stress response that strengthens secretory granule maturation and insulin release from beta cells.

## 2. Materials and methods

### 2.1 Animal experimentation

The *Sorcs2^-/-^* mouse model was described previously [12]. Male mice and wildtype controls (*Sorcs2^+/+^*) inbred on C57BL/6N background and 20-30 weeks of age were used for the study. Animals were housed in a controlled environment (12 h light/dark cycle) and fed a normal mouse chow (4.5% crude fat, 39% carbohydrates; Sniff Deutschland, V1124-3). Animal experimentation was approved by the Berlin State Office for Health and Social Affairs (X9009/22, G0230/17).

### 2.2 Metabolic phenotyping of mice

Nuclear magnetic resonance imaging was applied to assess body composition. Gas exchange was recorded in metabolic cages (TSE Phenomaster System, TSE Systems). In short, mice were kept individually in a metabolic cage for 4 days, and parameters were recorded at an interval of 8 min. The first day was considered equilibration, and quantification would start on day 2. The respiratory exchange ratio was quantified as the ratio between consumed O_2_ and produced CO_2_ [13]. For glucose tolerance tests (GTT), mice were fasted overnight and intraperitoneally (i.p.) injected with a dose of D-glucose of 2 g/kg body weight. Blood glucose levels were measured with a glucometer from tail tip blood before and every 15 min for 120 min following glucose injection. For insulin tolerance tests (ITT), animals were fasted overnight before i.p. injection with a human recombinant insulin dose of 0.75 U/kg body weight. Blood glucose was measured with a glucometer from tail tip blood before and every 15 min for 120 min following insulin injection. To test glucose-stimulated insulin secretion (GSIS), mice were fasted overnight and i.p. injected with a D-glucose dose of 2 g/kg body weight the next morning. Blood was collected from the facial vein into EDTA-coated tubes with 600 KIU/ml aprotinin (Serva, USA) before and at 2 min or 30 min after glucose injection. Plasma levels of insulin and other hormones were measured using commercially available ELISA kits (Supplementary table S1).

### 2.3 Histological analysis of mouse pancreatic tissue

Immunohistological analyses were performed on 4 µm cryo- or paraffin sections of tissue fixed in 4% paraformaldehyde using standard protocols. Primary antibodies were applied overnight in 1% NDS, 1% BSA, and 0.5 % Triton X-100 in TBS at 4°C. Secondary antibodies were applied for 1 h at room temperature. Source and concentrations of antibodies used are given as supplementary information (Supplementary table S2). For co-staining of SORCS2 with pancreatic cell markers, tissue sections were rehydrated in PBS for 15 min, subjected to antigen retrieval in 10 mM sodium citrate, 0.05% Tween-20 in PBS (pH 8.5) at 80°C for 30 min, and blocked in 10% NDS, 1% BSA, 0.3% Triton X-100, 0.05 % Tween-20 in TBS for 1 h at room temperature. For islet morphological analyses, tissues were stained for pancreatic cell markers glucagon, insulin, somatostatin, and pancreatic polypeptide Y, and images were obtained on a Leica SP8 DLS confocal microscope with a 63X objective. CellProfiler software [14] was used to measure tissue areas covered by the various cell populations in the islets and given as percent of the total islet area (defined manually). The total beta cell mass was quantified as the percentage of insulin-positive tissue area multiplied by pancreas weight, as described before [15].

For electron microscopic analysis of beta cell vesicles, isolated islets were incubated for 2 hours in 1.6 mM D-glucose in KRBH, washed with PBS, and fixed by immersion in 4 % (w/v) paraformaldehyde and 2.5% (v/v) glutaraldehyde in 0.1 M phosphate buffer for 2h at room temperature. Samples were postfixed in 1% (v/v) osmium tetroxide for 2h at room temperature, followed by incubation in 0.5% uranyl acetate overnight at 4°C. After dehydration through a graded ethanol series, embedding was done in PolyBed® 812 resin (Polysciences, Germany), and 60-80 nm sections were cut and stained with uranyl acetate and lead citrate and examined at 80 kV with a Zeiss EM 910 electron microscope (Zeiss, Germany) using a Quemesa CCD camera and iTEM software (Emsis GmbH, Germany). Fifteen beta cell images per sample (n = 4) were analyzed by manual quantification of mature, immature, crystal-containing, and empty vesicles in the entire cell (total) or situated 0.2 μm from the plasma membrane (docked). Numbers are given as % of total number of vesicles per cell [16; 17].

### 2.4 Metabolic analysis of isolated pancreatic islets

Pancreatic islets were collected as previously described [18] and cultured overnight in RPMI 1640 (PAN-Biotech, Germany) supplemented with 2 mM L-Glutamine, 100 U/ml penicillin, 100 mg/ml streptomycin, and 10% FBS (Gibco 10270-106) before further experiments. For GSIS, 30 islets per sample were handpicked into low protein-binding tubes (ThermoFisher, USA) and washed with KRBH containing 11 mM D-glucose. After pre-incubation with KRBH buffer containing 1.6 mM glucose for 1h at 37°C, islets of Langerhans were incubated consequentially in 1.6 mM glucose KRBH, 16 mM glucose KRBH for 1h, and 30 mM KCl KRBH for 30 min at 37°C. After each incubation, supernatants were collected into new tubes coated with 600 KIU/ml aprotinin, and total protein concentration and hormone content were determined using the Pierce BCA protein assay kit (ThermoFisher, USA) and commercially available ELISA (Supplementary table S1), respectively. To analyze total islet hormone content, 30 handpicked islets after overnight recovery were collected into low-protein binding tubes, lysed, and hormone content, as well as total protein concentrations were determined (Supplementary table S1). Hormone levels were normalized to total islet protein content.

### 2.5 Single-cell RNA sequencing of pancreatic islets

After overnight recovery, islets from 3 animals per genotype (400-600 islets per sample) were washed once in PBS and digested in 200 µl Accutase (Innovative Cell Technologies, USA) with the addition of 50 units/ml DNase-1 (Roche, Switzerland) for 10 min at 37°C [19; 20]. The digestion reaction was stopped by adding 1 ml of 0.5 mM EDTA, 2% FBS in HBSS. The cell suspension was centrifuged for 3 min at 200 x g and pellets resuspended in 1% BSA in PBS by pipetting 4 times up and down and filtering through 40 μm Flowmi strainer (Merck, Germany). Apoptotic and duplet cells were removed by staining the cell suspension with propidium iodide (Invitrogen, USA) and performing flow cytometry with BD FACS Aria III (BD Biosciences, USA).

Libraries were generated using the Chromium Next GEM Single Cell 3ʹ Reagent Kits v3.1 (10X Genomics Inc., USA). Briefly, a droplet emulsion targeting 10,000 cells was generated in a microfluidic Next GEM Chip G, followed by barcoded cDNA generation inside the droplets. Purified and amplified cDNA was then subjected to library preparation and sequenced on a NovaSeq 6000 instrument (Illumina, USA) to a depth of 40,000 mean read pairs per cell. Sequence data were aligned with mouse genome GRCm39 using STAR version 2.7.8a [21] and gene-level counts for cells in each sample were aggregated into a single count matrix using the DropletUtils tool [22] on Galaxy [23]. All subsequent analysis was performed using Automated Single Cell Analysis Platform VI [24]. Following pre-treatment, 8,846 cells and 22,106 genes were included in the analysis. Data were normalized using Seurat [25] and dimensional reduction with UMAP [26]. Clusters of cells were identified using Seurat [27]. Differentially expressed genes (DEGs) and cell-type (cluster) markers were determined using Limma [28]. GO ontology was performed with cluster Profiler R package [29] and visualized with Genekitr (https://genekitr.top/genekitr/).

### 2.6 Expression analysis

Pancreatic islets were isolated and cultured overnight as described above. Total RNA was isolated using the RNeasy Micro kit (Qiagen, USA) and reversely transcribed to cDNA with high-capacity RNA to cDNA kit (Applied Biosystems, USA). cDNA was subjected to qRT-PCR with TaqMan Gene Expression Assays: *SPP1* (Mm00436767_m1) and *Gapdh* (Mm99999915_g1). Relative gene expression was quantified with the cycle threshold (CT) comparative method (2^-ddCT^) with normalization to CT values. Determination of protein concentrations in islet supernatants or lysates was performed using standard Western blotting or ELISA.

### 2.7 Statistical methods

Data were analyzed using GraphPad Prism 7.0 (GraphPad Software, USA). Normal distribution was tested using D’Adostino-Pearson omnibus normality test or Shapiro-Wilk normality test, depending on the sample size. Two-group analysis was performed using Student’s t-test or Mann-Whitney U test, depending on a normal distribution. ROUT outlier test was applied where indicated. Data with two variables was analyzed using Two-way ANOVA and Sidak’s (for repeated measures) or Turkey’s multiple comparison tests. Data are represented as mean ± standard deviation (SD).

## 3. Results

### 3.1 SORCS2 deficiency impairs glucose-stimulated insulin release from murine pancreatic islets *in vivo*

Initially, we interrogated expression of SORCS2 in the pancreas, in line with the reported expression of the related diabetes risk factor SORCS1 in this tissue [9]. Western blot analysis confirmed the presence of SORCS2 in islets purified from pancreata of wildtype mice (*Sorcs2^+/+^*), and its absence in this tissue from mice carrying a targeted *Sorcs2* gene disruption (*Sorcs2^-/-^*) [12] (Fig. 1A). Immunohistological analyses refined the cell-type specific expression of SORCS2 to alpha, delta, and pancreatic polypeptide (PP) cells, but not beta cells of the murine pancreas (Fig. 1B, C). This expression pattern contrasts with that of SORCS1, which is reported to localize to beta cells [9; 10].

**Figure 1.**
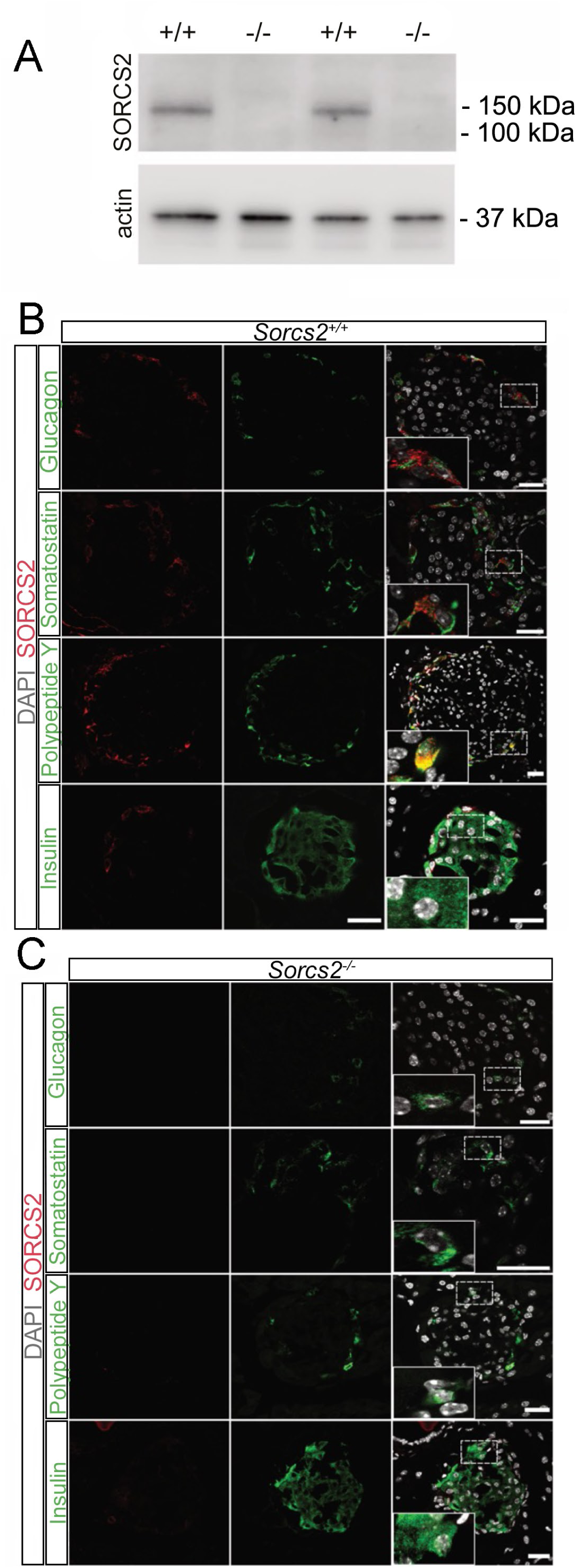
SORCS2 expression in mouse pancreatic islets. **(A)** Analysis of SORCS2 expression in total lysate from murine islets using Western blot analysis. The receptor is detected in samples from two *Sorcs2*^+/+^ but not two *Sorcs2^-/-^* mice. Detection of actin served as loading control. The migration of marker proteins of the indicated molecular weights in the gel is given to the right of the blot. **(B, C**) Immunofluorescence staining of pancreatic sections from *Sorcs2*^+/+^ (B) and *Sorcs2^-/-^* (C) animals for SORCS2 (red), as well as glucagon, somatostatin, pancreatic polypeptide Y, or insulin (green). Nuclei were counterstained with DAPI (grey). Individual as well as merged channel configurations are given. Stippled boxes indicate the area of sections shown in the higher magnification insets of the respective panels. SORCS2 is most prominently expressed in alpha (glucagon+), delta (somatostatin+), and PP (pancreatic polypeptide Y+) cells of wildtype pancreatic islets. The receptor is not expressed in SORCS2-deficient pancreatic tissue. Scale bars: 25 μm.

Given the expression of SORCS2 in the pancreas, we explored the consequences of receptor dysfunction for glucose homeostasis and insulin action in *Sorcs2^-/-^* mice. These studies were carried out in males at 20-26 weeks of age, fed a normal chow diet (see methods for details). When compared to wildtype controls, adult mutant animals exhibited a significant decrease in body weight before and after overnight fasting (Fig. 2A), albeit at normal body size (Fig. 2B). An impact of SORCS2 activity on body homeostasis was corroborated by analysis of body composition using magnetic resonance spectroscopy, documenting a decreased fat and an increased lean mass as compared to controls (Fig. 2C, D).

**Figure 2.**
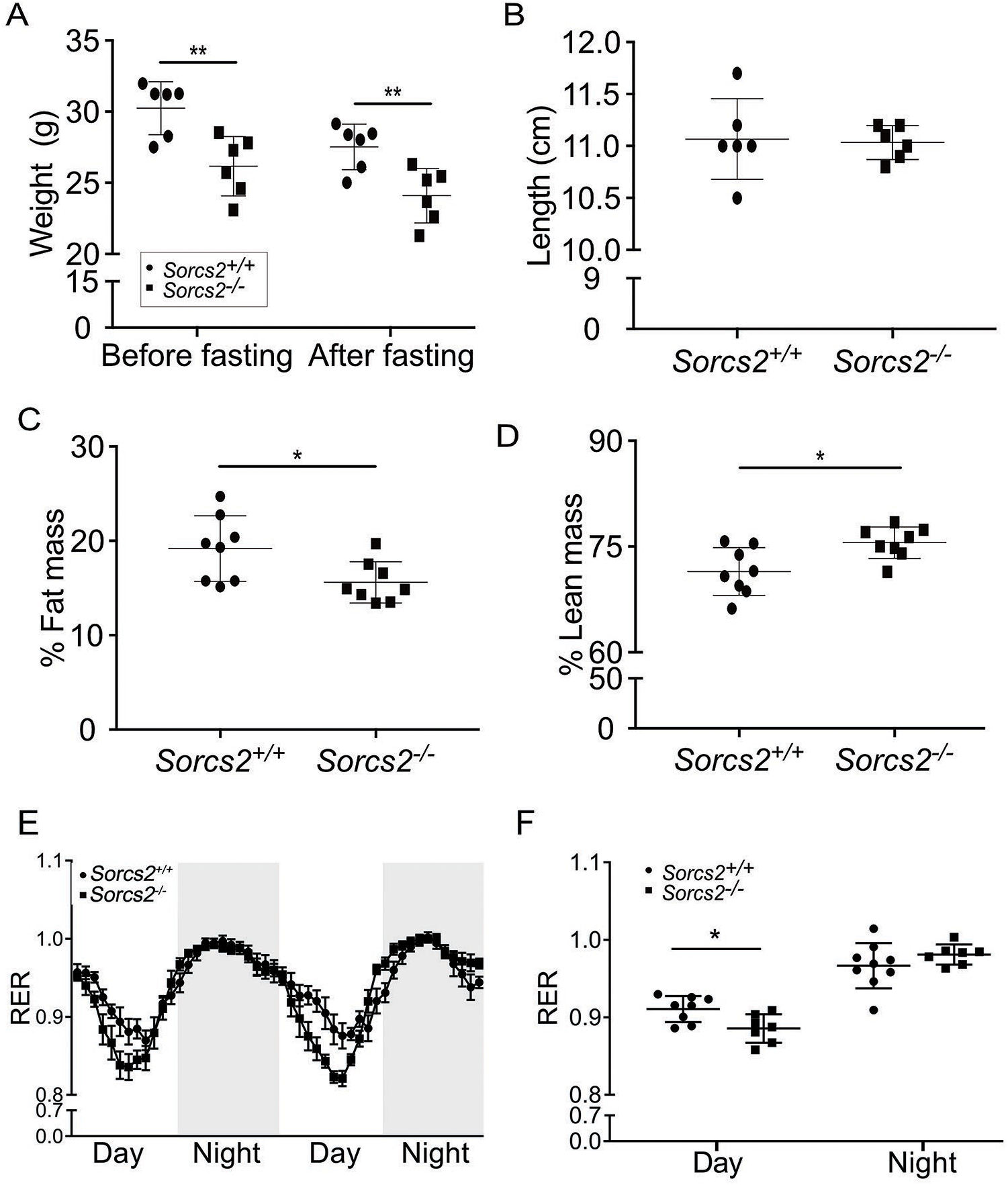
Effects of SORCS2 deficiency on whole body metabolism of mice. **(A)** Body weights of *Sorcs2*^+/+^ (circles) and *Sorcs2^-/-^* (squares) mice measured before and after 16 hours fasting (n=6 animals per genotype). **(B)** Length of *Sorcs2*^+/+^ and *Sorcs2^-/-^* animals measured from nose tip to beginning of the tail (n=6 animals per genotype). **(C, D)** Fat (C) and lean (D) mass of *Sorcs2*^+/+^ and *Sorcs2^-/-^* mice as analyzed by nuclear magnetic resonance spectroscopy (n = 8 animals per genotype). **(E-F)** Basic metabolic rates were determined in *Sorcs2*^+/+^ (circles) and *Sorcs2^-/-^* (squares) mice during day (6 am – 6 pm) or night (6 pm – 6 am) using indirect gas calorimetry and expressed as respiratory exchange ratio (RER). RER curves (E) as well as RER quantification for day and night (F) are given (n = 7 - 8 animals per genotype). Significance of data was determined using two-way ANOVA followed by Sidak’s or Turkey multiple comparisons tests (A) or using unpaired Student’s t-test (B-D and F). *, p < 0.05

Metabolic rates in *Sorcs2^-/-^* mice, expressed as respiratory exchange ratio (RER), were lower during day time when compared to wildtypes, indicating a relative shift in metabolic fuel consumption from carbohydrates to lipids (Fig. 2E, F). In line with this observation, *in vivo* metabolic stress tests showed a blunted response of *Sorcs2^-/-^* mice to a glucose bolus (glucose tolerance test, GTT; Fig. 3A, B). By contrast, the sensitivity to insulin (insulin tolerance test, ITT) was normal (Fig. 3C). Also, fasting plasma levels of glucagon, glucagon- like peptide 1 (GLP-1), somatostatin, somatostatin-28, neuropeptide Y (NPY), and peptide YY (PYY) were comparable to wildtypes (Supplementary table S3). A defect in glucose-stimulated insulin release in *Sorcs2^-/-^* mice was substantiated by decreased levels of plasma insulin both 2 min and 30 min after an intraperitoneal (i.p.) dose of glucose (Fig. 3D, E). A similar decrease was seen for plasma C-peptide levels (Fig. 3F).

**Figure 3.**
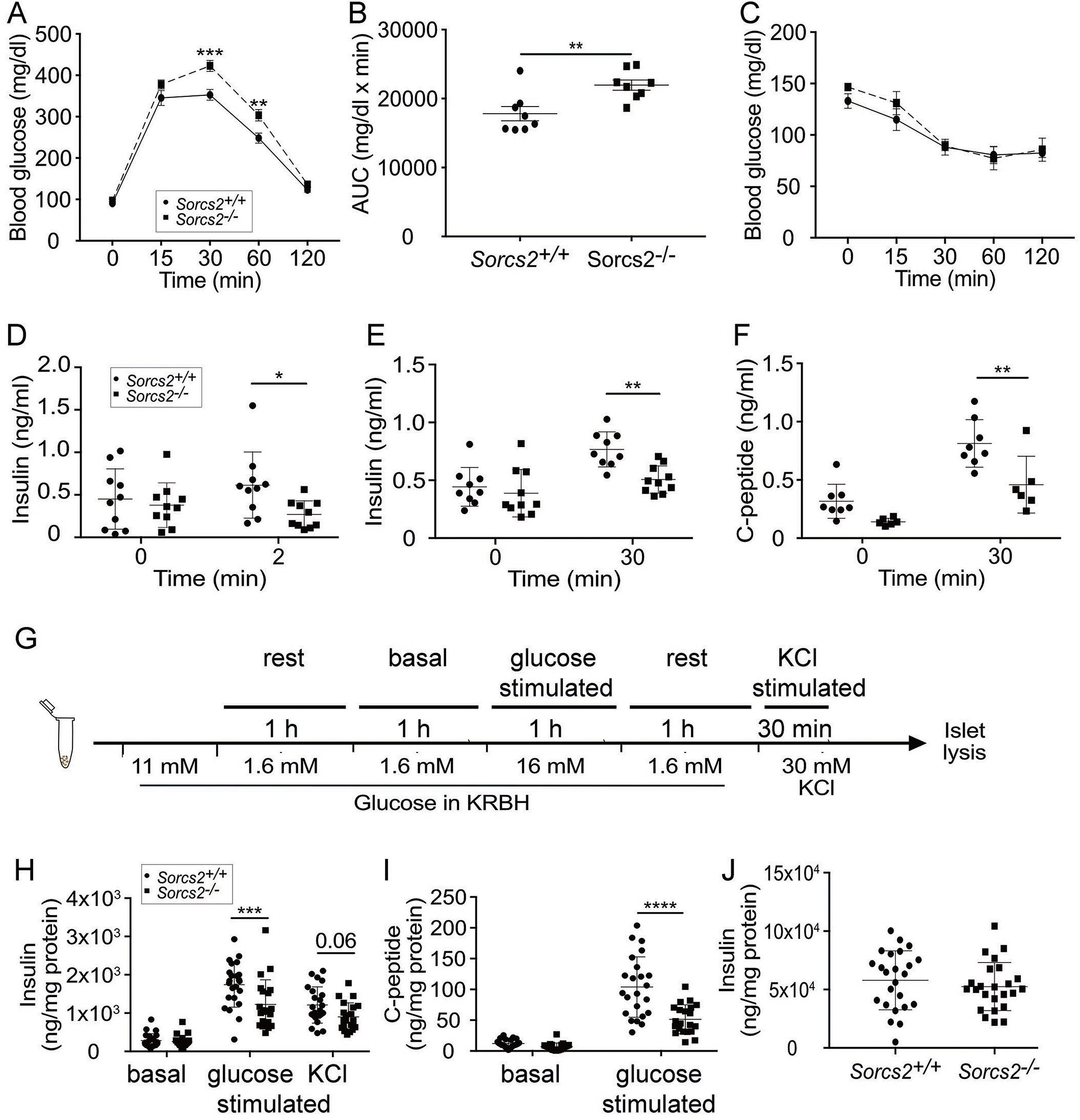
Effects of SORCS2 deficiency on glucose homeostasis *in vivo* and *in vitro*. **(A)** Glucose tolerance test (GTT) performed in *Sorcs2*^+/+^ and *Sorcs2^-/-^* mice (n=8 animals per genotype). Blood glucose levels were measured every 15 min for 120 min after i.p. injection of 2 g/kg body weight glucose following 16 hours fasting (n=8 animals per genotype). **(B)** Area under the curve (AUC) for the GTT shown in panel A. **(C)** Insulin tolerance test (ITT) performed in *Sorcs2*^+/+^ (filled circles) and *Sorcs2^-/-^* (filled squares) mice by determination of blood glucose levels every 15 min for 120 min following an intraperitoneal (i.p.) injection of 0.75 U/kg body weight of insulin (n = 8 animals per genotype). Animals had been fasted for 16 hours before insulin application. **(D-F)** Plasma levels of insulin (D, E) and C-peptide (F) levels were determined by ELISA under basal conditions, or 2 or 30 minutes after i.p. injection of 2 g/kg body weight of glucose (n = 6 - 12 animals per group). ROUT 10% outlier test was applied to D and E. Significance of data was determined using two-way ANOVA followed by Sidak’s or Turkey multiple comparisons tests (A, C, D-F), or Mann-Whitney U test (B). **(G)** Protocol for glucose and KCl stimulation in isolated pancreatic islets. **(H - I)** Insulin and C-peptide levels normalized to total islet protein as determined by ELISA in supernatants of isolated *Sorcs2^+/+^*(filled circles) and *Sorcs2^-/-^* (filled squares) islets treated consequentially for 1 hour with 1.6 mM (basal) and 16 mM glucose (glucose-stimulated), followed by 30 min with 30 mM KCl in Krebs-Ringer-Bicarbonate Hepes buffer (KRBH) (n = 21-23 animals per genotype). **(J)** Insulin level determined by ELISA in lysates of isolated islets from *Sorcs2^+/+^*and *Sorcs2^-/-^* animals cultured overnight (n = 24 animals per genotype). ROUT outlier test 10% was applied to J and 0.1% applied to H and I. Significance of data was determined using two-way ANOVA followed by Sidak’s or Turkey’s multiple comparisons tests (H, I), or unpaired Student’s t-test (J). *, p < 0.05; **, p < 0.01; ***, p < 0.001; ****, p < 0.0001.

### 3.2 SORCS2 deficiency impairs glucose-stimulated insulin release from pancreatic islets *in vitro*

To exclude a confounding effect of SORCS2 deficiency in non-pancreatic tissues on insulin secretion *in vivo*, we tested insulin release from isolated islets stimulated with glucose *ex vivo* (Fig. 3G). SORCS2-deficient islets showed a blunted response in release of insulin and C- peptide when challenged with 16 mM glucose (Fig. 3H, I), albeit at normal insulin islet content (Fig. 3J). A trend towards decreased release of insulin was also seen when treating mutant islets with KCl as a secretagogue (p=0.06, Fig. 3H). This observation argued for a global release defect, rather than insensitivity to glucose uptake and catabolism, to underlie the impaired insulin secretion in SORCS2-deficient islets. This release defect was specific to beta cells as islet content (Supplementary table S4) and secreted levels (Supplementary table S5) of non-beta cell hormones glucagon, GLP-1, somatostatin, somatostatin-28, NPY, and PYY were not affected by receptor deficiency under basal or glucose-stimulated conditions.

Taken together, our data documented a similar phenotype in SORCS2 mutant mice as reported in SORCS1-deficient animals before, namely an inability to sustain insulin release upon glucose stress [9]. Of note, while an impact of receptor deficiency on metabolic phenotypes was only reported in SORCS1 KO mice made diabetic by deletion of the leptin gene (ob/ob) [9], metabolic disturbances were apparent in SORCS2-deficient animals without additional genetic or dietary stressors.

### 3.3 SORCS2 deficiency affects secretory granule formation in islet beta cells

Given the unimpaired release of hormones from non-beta cells, we reasoned that SORCS2 deficiency may impact insulin secretion from beta cells by a non-endocrine mechanism, perhaps by affecting structural integrity of pancreatic islets or islet cell types. Morphometric analysis of immunostainings for insulin documented comparable levels of beta cell area and mass but a slight decrease in islet number in mutant versus control islets (Fig. 4A, B). Similarly, islet areas covered by alpha (glucagon+), delta (somatostatin+), and PP (PPY+) cells were similar comparing genotypes (Fig. 4C, D). However, transmission electron microscopy showed distinct alterations in the composition of secretory granules in mutant beta cells with a relative increase in immature and a decrease in crystallized vesicles (Fig. 4E, F). The levels of vesicles docked at the plasma membrane were not changed, arguing that SORCS2 deficiency affects vesicle maturation rather than membrane docking (Fig. 4E, F). Defects in secretory granule biogenesis are shared by beta cells in SORCS1-deficient mice [9; 10], supporting a paracrine effect of SORCS2 on functional integrity of the insulin release machinery as well.

**Figure 4.**
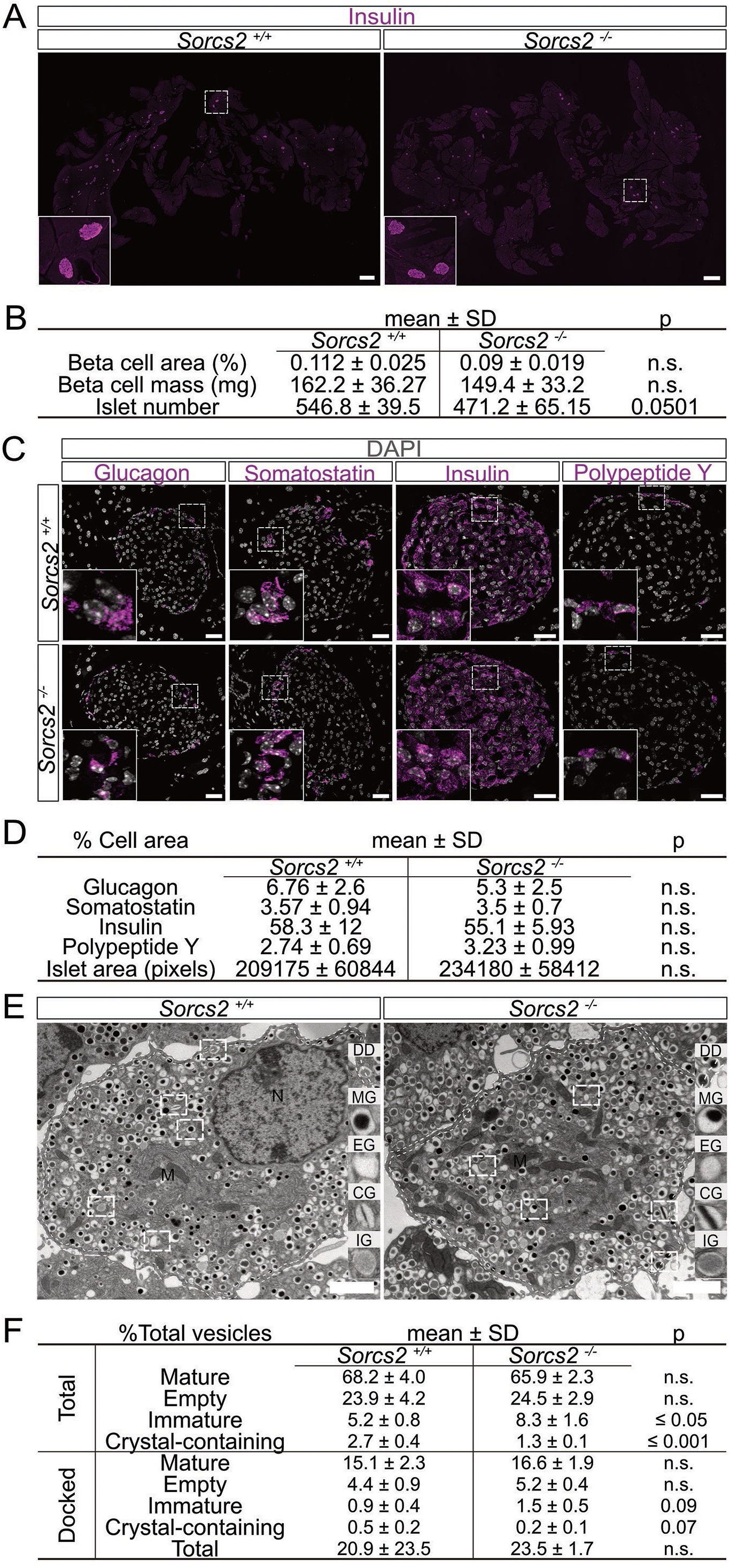
Composition and structure of cell types in wildtype and SORCS2-deficient murine pancreatic islets. **(A)** Representative images of murine pancreatic sections of the indicted *Sorcs2* genotypes immunostained for insulin (purple). (**B**) Quantitative analysis of beta cell area and mass in wildtype and SORCS2-deficient islets based on immunostainings for insulin (as exemplified in A). The beta cell area was quantified as insulin+ area per total pancreas area in each section. The beta cell mass was quantified as beta cell area multiplied by weight of the respective pancreatic tissue. Islet numbers were manually counted on six histological sections per pancreas, spaced 200 µm apart (n = 5-6 animals per genotype). (**C**) Representative images of murine islets of the indicted *Sorcs2* genotypes immunostained for insulin, glucagon, somatostatin, and pancreatic polypeptide Y (purple). Nuclei were counterstained with DAPI (grey). (**D**) Area covered by cells expressing insulin, glucagon, pancreatic polypeptide Y, or somatostatin in *Sorcs2*^+/+^ and *Sorcs2^-/-^* pancreata were quantified based on immunohistological stainings for the various hormones (as exemplified in C) and expressed as % of the total islet cell area. For each mouse, 30-35 pancreatic islets were analyzed (n = 6 animals per genotype). Stippled boxes in A and C indicate the area of sections shown in the higher magnification insets in the respective panels. Scale bars: 1000 µM (A) and 25 µm (C). Statistical significance was tested by unpaired Student’s t-test with p < 0.05 considered significant. n.s., not significant. (**E**) Representative transmission electron microscopic (EM) images of beta cells in isolated *Sorcs2*^+/+^ and *Sorcs2^-/-^* islets treated for 1 hour with 1.6 mM glucose. Stippled boxes highlight the different types of secretory granules shown in the higher magnification insets and identified as mature granules (MG), empty granules (EG), crystal-containing granules (CG), as well as immature granules (IG). Inset DD indicates exemplary vesicles with a docking distance of 0.2 µm from the plasma membrane. Scale bar: 2 µm. (**F**) Quantification of different types of granules per cell identified on EM sections from *Sorcs2*^+/+^ and *Sorcs2^-/-^* islets treated for one hour with 1.6 mM glucose (n = 4 animals, 14-27 quantified cells per animal). Statistical significance was determined using unpaired Student’s t-test or Mann-Whitney U test with p < 0.05 considered significant. n.s., not significant.

### 3.4 Single-cell RNA sequencing identifies increased cell stress in SORCS2-deficient islet cell types

To explore the cellular basis of SORCS2 deficiency phenotypes, we performed single-cell RNA sequencing (scRNAseq) in pools of islets isolated from *Sorcs2^+/+^* or *Sorcs2^-/-^*pancreata (400-600 islets per genotype). Unsupervised clustering of the scRNAseq data identified a total of 18 distinct cell clusters in wildtype and receptor mutant islets (Fig. 5A). Based on expression of their respective markers, 14 cell clusters were annotated as beta cells, expressing e.g., *Ins1*, *Ins2*, and *Chromogranin* (*Chg*) *a* and *Chgb* (Fig. 5B; Supplementary figure S1). Single cell clusters were identified as alpha cells (cluster #6; *Glucagon/GcG*, *Transthyretin/Ttr*), delta cells (cluster #11; *Somatostatin/Sst*), and PP cells (cluster #16; *Pancreatic polypeptide Y/Ppy*) (Fig. 5A, B; Supplementary figure S1), respectively. Cell cluster #14 showed characteristics of both beta (*Ins1, Ins2*) and non-beta cell types (*Gcg, Ttr, Ppy, Pyy, Chga, Chgb*), suggesting a progenitor or transdifferentiating cell type [30; 31]. In line with immunohistology, *Sorcs2* transcripts in wildtype islets were largely confined to alpha, delta, and PP cell types (Fig. 5C). Of note, *Sorcs2* transcripts were also found in cluster 14 with both beta and non-beta cell type characteristics (Fig. 5C).

**Figure 5.**
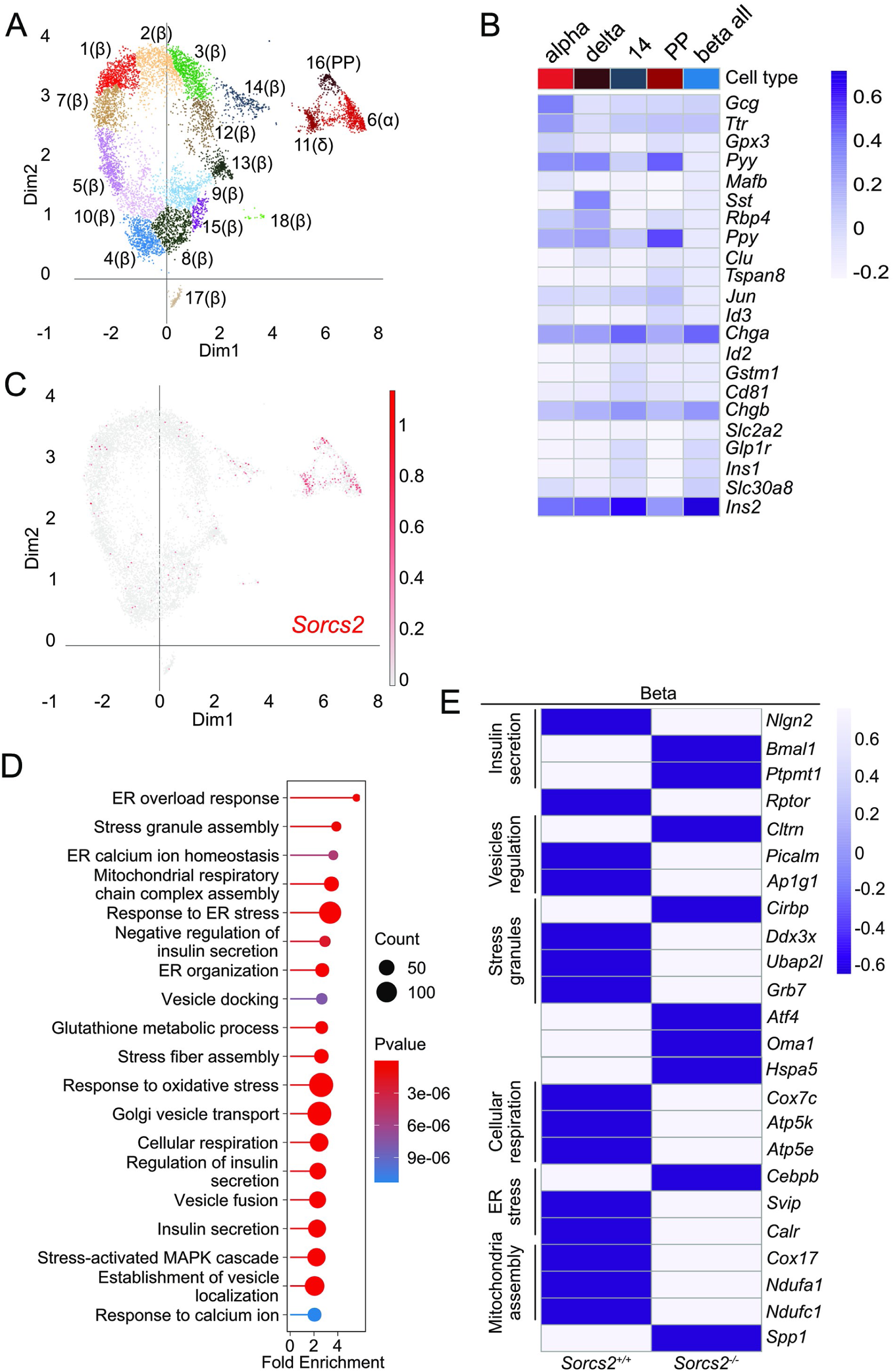
Single-cell RNA sequence analysis identifies impact of SORCS2 deficiency on islet beta cell function. (**A**) Cells from pancreatic islets of *Sorcs2^+/+^* and *Sorcs2^-/-^* mice were analyzed by unsupervised clustering and visualized using uniform manifold approximation and projection for dimension reduction (UMAP) plot. A total of 18 distinct cell clusters were identified in the pooled data set. (**B**) Heatmap showing the top five most unique identifiers per cluster according to the adjusted p-value calculated with linear models for microarray data (Limma) moderated t-statistics for alpha, delta, all beta, beta cluster 14, and PP clusters. (**C**) UMAP plot localizing *Sorcs2* transcripts (red) to the indicated clusters of alpha, delta, PP cells, as well as cluster 14. (**D)** Graph representing biological processes impacted in beta cells of *Sorcs2^+/+^*versus *Sorcs2^-/-^* islets. Fold enrichment shown in the graphs is calculated by dividing the percentage of DEGs in the respective GO ontology term by the corresponding percentage in the background gene list. **(E)** Heatmaps of normalized levels of differentially expressed genes (DEGs) associated with biological processes in GO analysis in beta cell populations of pancreatic islets from *Sorcs2^+/+^* and *Sorcs2^-/-^* animals. Gene ontology analysis was performed using clusterProfiler R package [29] and visualized with Genekitr (https://genekitr.top/genekitr/). Benjamini-Hochberg (BH) test was applied to calculate adjusted p value of GO terms. Significance of DEGs was determined using Limma moderated t-statistics.

Comparative analysis of differentially expressed genes (DEGs) in the combined dataset of beta cell clusters (Fig. 5D-E; Supplementary data file 1) identified enrichment in genes related to the gene ontology (GO) term “insulin secretion”, including *Nlgn2*, *Bmal1*, and *Ptpmt1*, regulators of insulin release [32 ; 33 ; 34] as well as mTOR-associated protein *Rptor*, implicated in maturation of beta cells [35]. Also, genes related to the GO terms “vesicle transport/fusion/docking“, including *Cltrn* [36], *Picalm* [37], and *Ap1g1* [38] were differentially expressed. These alterations constituted a molecular signature of insulin secretion defects, possibly due to faulty granule vesicle maturation seen in SORCS2-deficient islets. Interestingly, enrichment in DEGs related to “stress granule assembly“ was also obvious, including genes encoding stress factors CIRBP [39] and DDX3X [40], as well as stress granule associated proteins UBAP2L and GRB7 [41]. Enhanced stress of mutant islet beta cells was further supported by DEGs related to stress response mechanisms, such as stress-induced transcription factor ATF4 [42], mitochondrial stress relay molecule OMA1 [43], endoplasmic reticulum stress regulator C/EBPB [44], and heat shock protein HSPA5 [45]. Finally, enrichment of DEGs related to mitochondrial and respiratory activities argued for changes in cellular energy homeostasis, possibly as a consequence of enhanced beta cell stress [46–48]. Aggravated beta cell stress was supported by changes in size of some beta cell clusters in *Sorcs2^-/-^* islets. While cells from both *Sorcs2* genotypes were represented in all clusters (Fig. 6A), a shift in relative size of some beta cell clusters in mutants was noted (Fig. 6B). Particularly, more than 10% of cells derived from *Sorcs2*^-/-^ islets were found in beta cluster 13 that is characterized by high expression of stress-induced genes (e.g., *MT1*, *MT2*, and genes from the heat shock protein-encoding family; Supplementary data file 2), and an overall change in gene expression patterns in GO terms related to protein (mis)folding (Fig. 6C). Less than 1% of cells derived from *Sorcs2*^+/+^ islets were found in this cluster.

**Figure 6.**
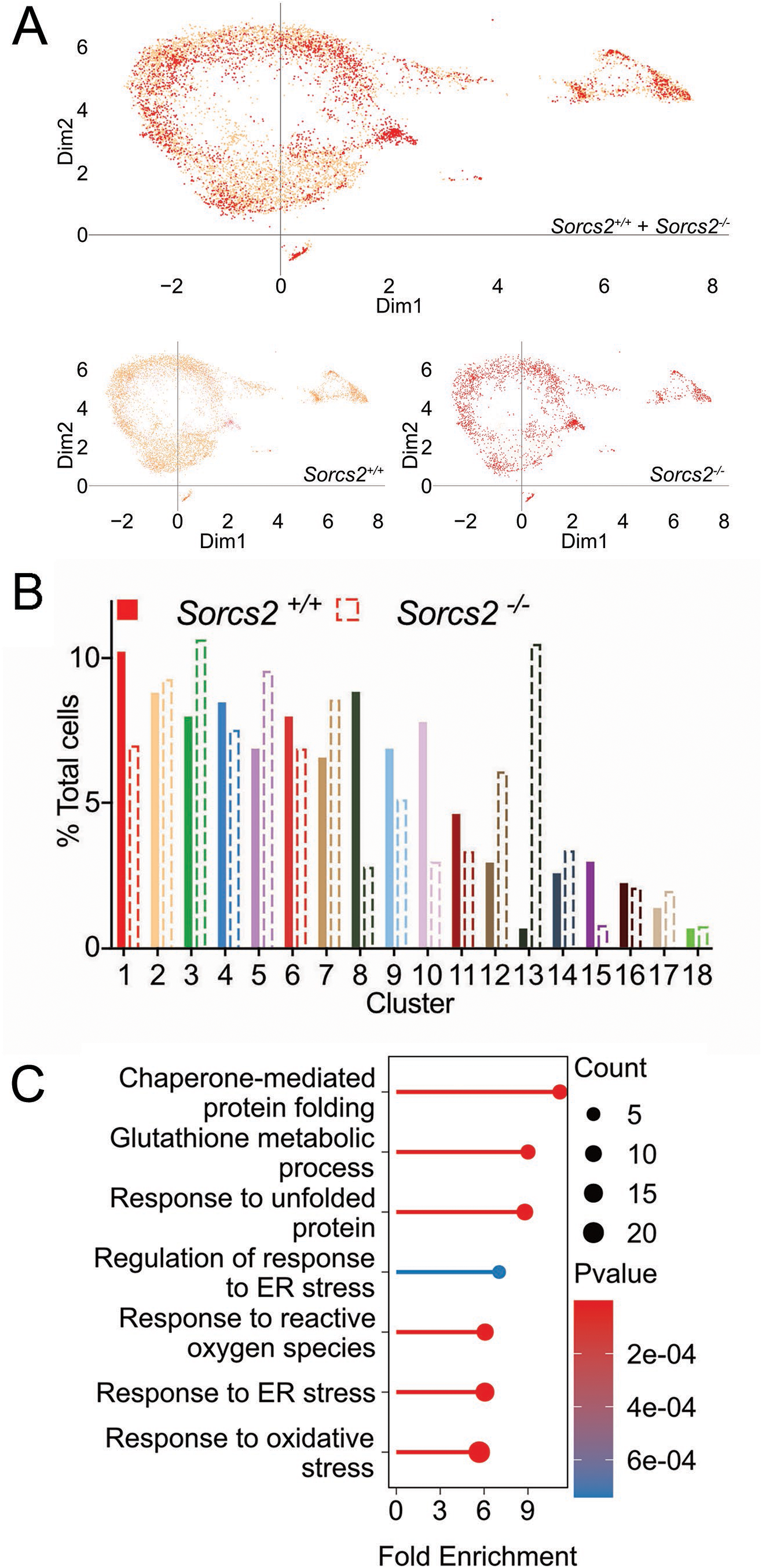
Effect of SORCS2 deficiency on size of pancreatic islets beta cell clusters. (**A**) Unsupervised clustering and visualized using uniform manifold approximation and projection for dimension reduction (UMAP) plots for single-cell RNA sequencing of cells from isolated *Sorcs2^+/+^* or *Sorcs2^-/-^* islets are given as individual genotypes (lower panels) or combined (upper panel). (**B**) Quantitative contribution of individual cell clusters to the total cell counts in *Sorcs2^+/+^* (solid bars) and *Sorcs2^-/-^* (stippled bars) islets. (**C**) Graph representing biological processes impacted in beta cell cluster 13 of *Sorcs2^+/+^*versus *Sorcs2^-/-^* islets. Gene ontology analysis was performed using clusterProfiler R package [29] and visualized with Genekitr (https://genekitr.top/genekitr/). Benjamini-Hochberg (BH) test was applied to calculate adjusted p value of GO terms.

To identify possible primary causes of SORCS2 deficiency underlying enhanced beta cell stress, we focused on DEGs in alpha cells as receptor-expressing islet cell type (Fig. 7; Supplementary data file 3). Similar to beta cells, alpha cells showed enrichment of DEGs related to the GO terms “mitochondria” and “respiratory function”. Furthermore, expression changes related to “Cellular response to calcium” and in genes encoding transcription factors implicated in islet cell proliferation and maturation were noteworthy. The latter included *Jund*, *Fosb*, *Zfp800*, *Klf6*, *Gabpb1*, and *Klf4* (Fig. 7B, Supplementary data file 3).

**Figure 7.**
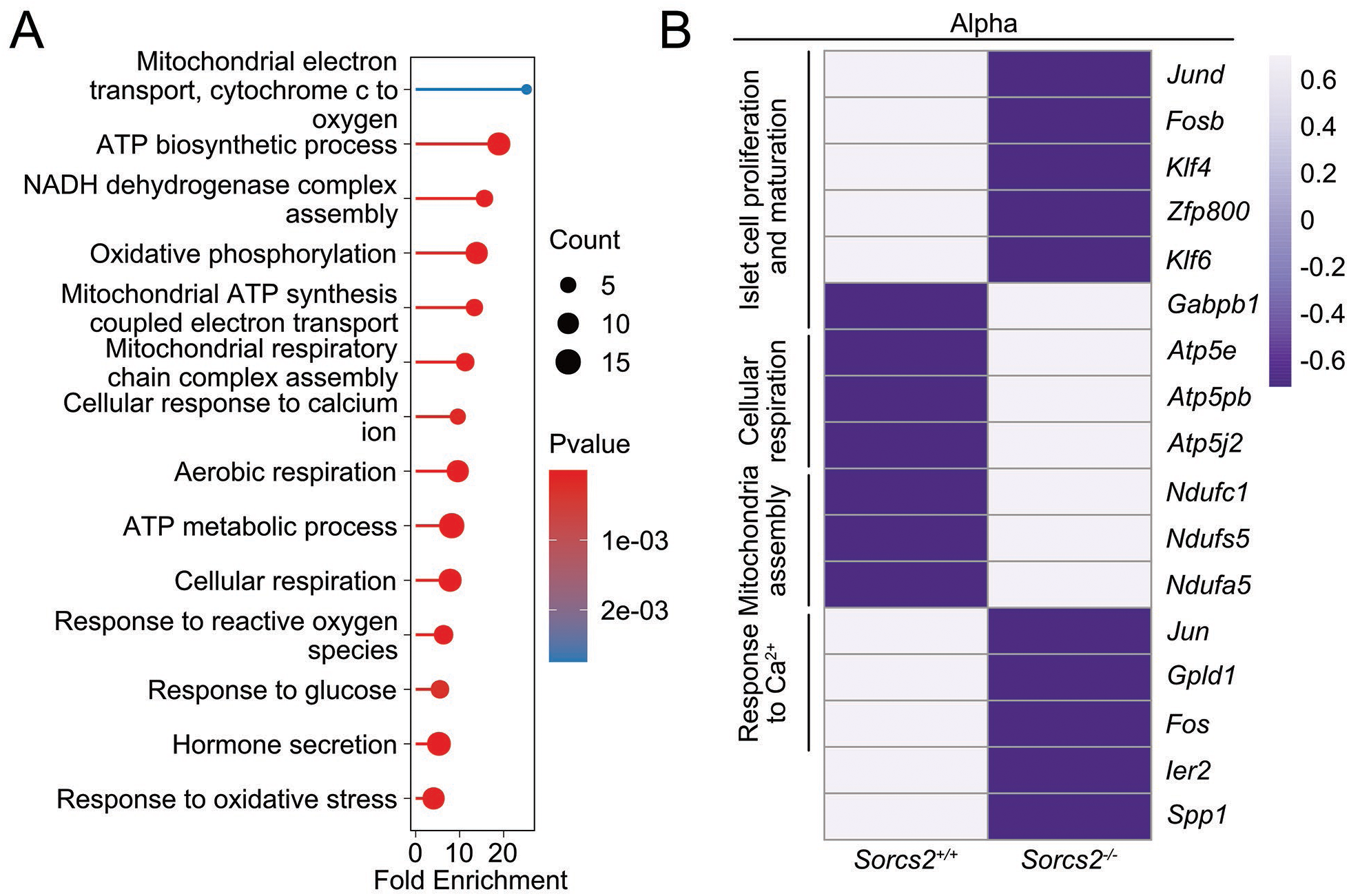
Effect of SORCS2 deficiency on gene expression in pancreatic islets alpha cells. (**A**) Graph representing biological processes impacted in alpha cells of *Sorcs2^+/+^* versus *Sorcs2^-/-^* islets. **(B)** Heatmaps of normalized levels of differentially expressed genes (DEGs) associated with biological processes in GO analysis in the alpha cell population of pancreatic islets from *Sorcs2^+/+^* and *Sorcs2^-/-^* animals. Gene ontology analysis was performed using clusterProfiler R package [29] and visualized with Genekitr (https://genekitr.top/genekitr/). Benjamini-Hochberg (BH) test was applied to calculate adjusted p value of GO terms. Significance of DEGs was determined using Limma moderated t-statistics.

### 3.5 Loss of SORCS2 activity attenuates expression of osteopontin in islets

One gene differentially expressed in receptor mutant islet alpha cells was *Spp1* (Fig. 7B). It encodes osteopontin, a secreted protein that acts in cell stress response in various metabolic tissues, including adipose tissue and pancreas [49]. With relevance to islet function, osteopontin is released from various islet cell types upon stress imposed by inflammatory cytokines or glucose. The secreted factor acts on beta cells to facilitate secretory granule docking and insulin release in a calcium-dependent manner [49; 50]. *Spp1* transcript levels were significantly reduced in SORCS2-deficient alpha cells (Fig. 7B).

Reduced osteopontin expression was confirmed in isolated islets, documenting decreased levels of *Spp1* transcripts (Fig. 8A), as well as total and released osteopontin protein in mutant compared to wildtype tissue (Fig. 8B, C). Cellular defects related to a diminished expression of osteopontin were supported by decreased expression of *Klf4, Jund,* and *Fosb*, transcription factors that induce *Spp1* [51–53], in mutant alpha cells (Fig. 7B). Also, transcripts encoding IER2, an immediate early response factor that stimulates osteopontin secretion [54] were among the significantly changed transcripts (Fig. 7B).

**Figure 8.**
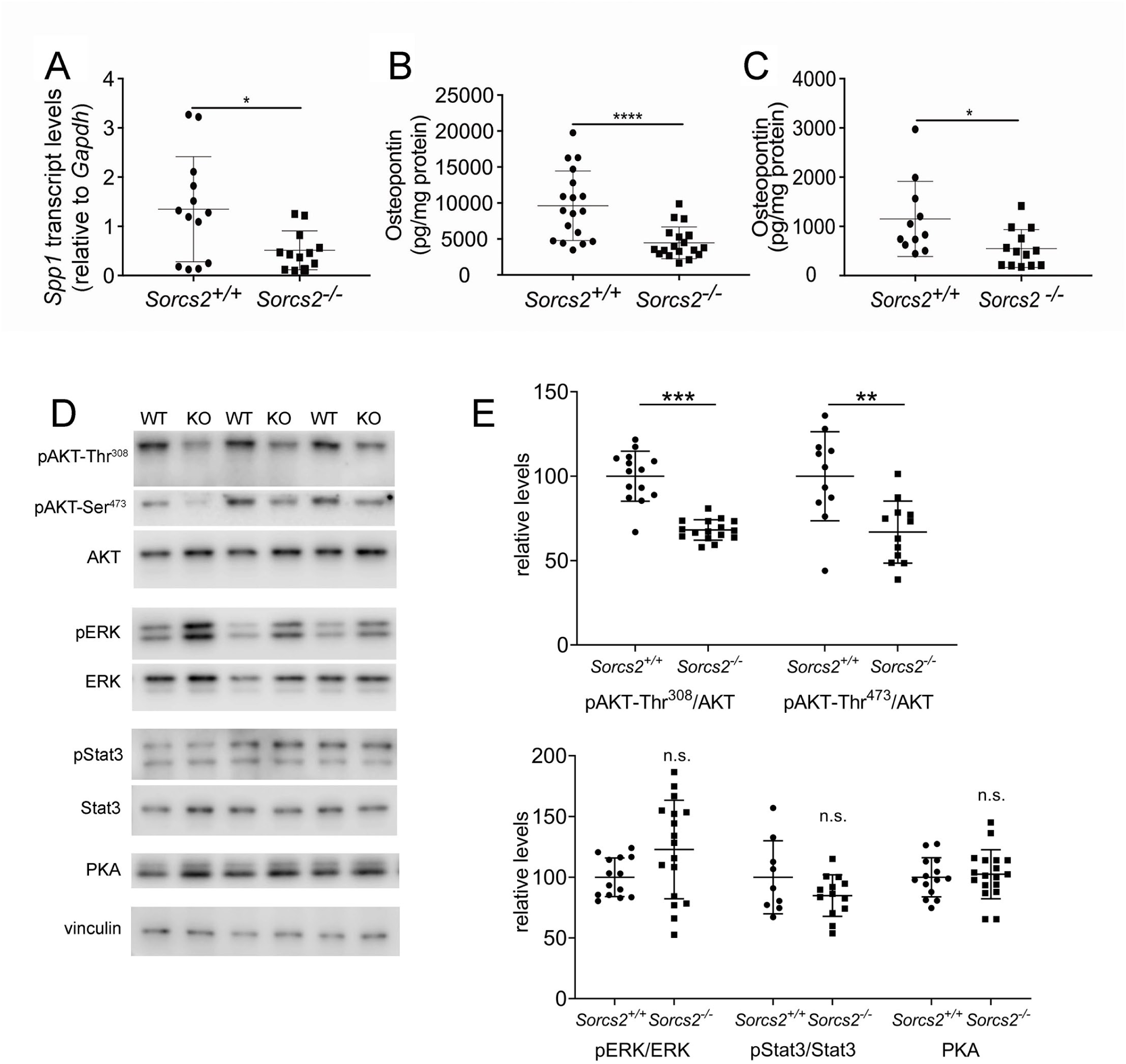
Impaired osteopontin expression in SORCS2-deficient islets. **(A)** Relative transcript levels for *Spp1* were determined by quantitative RT-PCR in *Sorcs2^+/+^*and *Sorcs2^-/-^* islets cultured overnight (n = 12-13 animals per genotype). **(B)** Osteopontin levels were determined by ELISA in lysates of isolated *Sorcs2^+/+^* and *Sorcs2^-/-^* islets cultured overnight (n = 18 animals per genotype). (**C)** Secreted osteopontin levels were determined by ELISA in the supernatant of isolated *Sorcs2^+/+^*and *Sorcs2^-/-^* islets cultured overnight (n = 11- 13 animals per genotype). (**D**) Levels of the indicated proteins in lysates of isolated *Sorcs2^+/+^* (WT) and *Sorcs2^-/-^* (KO) islets were detected using Western blot analysis. The blots show representative data of three individual pools of islet preparation for each genotype (approximately 250 islets per pool). Detection of vinculin served as loading control. p, phosphorylated. **(E**) Quantitative analysis of protein levels based on densitometric scanning of replicate blots (as exemplified in D) of a total of 13-18 pools of isolated islets per genotypes. Significance of data was determined using unpaired Student’s t-test (A, E) or Mann-Whitney U test (B, C). *, p <0.05; **, p < 0.01; ***, p < 0.001; ****, p < 0.0001; n.s., not significant.

Several signaling pathways are implicated in induction of *Spp1* transcription, including ERK, AKT, PKA, and Stat3 [55; 56]. While activation levels of ERK, Stat3, and PKA were not impacted, Western blot analysis documented a defect in AKT signaling in SORCS2-deficient islets, as suggested by a relative decrease in levels of phosphorylated (p) variants AKT-Ser^473^ and AKT-Thr^308^ compared to control tissue (Fig. 8D, E). These findings implicated impaired AKT signaling in loss of osteopontin expression in stressed islets lacking the receptor SORCS2.

## 4. Discussion

Our study identified an important function for the VPS10P domain SORCS2 in control of insulin secretion from murine islets. Loss of the receptor impairs the ability of islet beta cells to release the hormone during a glucose challenge, a defect coinciding with aggravated beta cell stress. These findings provide experimental support for a presumed role of SORCS2 in control of systemic metabolism suggested by association studies [11]. Interestingly, insulin secretion defects are also seen in mice lacking the diabetes risk factor *Sorcs1* [9], suggesting related functions for both receptors in glucose homeostasis.

Beta cells from SORCS2-deficient islets are characterized by a relative increase in immature and a corresponding decrease in crystal-containing vesicles, arguing for impaired maturation of secretory granules as underlying cause for the insulin release defect. This assumption is supported by scRNAseq data documenting changes in gene signatures related to vesicle formation and insulin secretion. While insulin granule defects are phenotypes shared by SORCS1- and SORCS2-deficient pancreata, the molecular mode of action of both receptors must be different. According to current hypotheses, SORCS1 acts as an intracellular sorting receptor that facilitates replenishment of secretory granules during glucose stress [9; 10]. By contrast, distinct expression of SORCS2 in non-beta cells implies a non-cell autonomous mechanism of receptor action in insulin secretion. Based on our scRNAseq data, this receptor action likely involves a protective islet stress response, essential to sustain insulin release under adverse conditions imposed by various stressors, such as hyperglycemia or inflammation [57]. Elevated cell stress in beta cells from SORCS2-deficient islets is obvious from DEGs related to GO terms such as “ER overload”, “Response to ER stress”, “Response to oxidate stress”, “Stress granule assembly” or “Stress response signaling”, just to name a few terms. Also, changes in mitochondrial gene expression argue for enhanced cell stress in beta cells as documented in numerous studies [58–60].

Taken together, the above findings suggest that insulin granule defects in SORCS2 mutant islets are a secondary consequence of aggravated cell stress caused by loss of protective receptor activities in this tissue. In support of this hypothesis, SORCS2 has been identified as a stress response factor in the brain before. In neurons, SORCS2 acts as sorting receptor that sustains cell surface expression of the neuronal amino acid transporter EAAT3 to facilitate import of cysteine, required for synthesis of the reactive oxygen species scavenger glutathione. Lack of SORCS2 impairs neuronal cysteine uptake, resulting in oxidative brain damage [61]. A second mechanism of neuroprotective receptor action concerns astrocyte- mediated stress response in stroke when SORCS2 controls astrocytic release of endostatin, a growth factor linked to post-stroke angiogenesis [62]. The lost ability of astrocytes to release endostatin in SORCS2 mutant mice results in a blunted endostatin response and in impaired vascularization of the ischemic brain [62]. Reduced expression of osteopontin in SORCS2- deficient islets supports our hypothesis about a role for SORCS2 in protective stress response in the pancreas as well. Secreted osteopontin strengthens the insulin release machinery in stressed islets in a calcium-dependent manner [50] and improves glucose-stimulated insulin release [49], while loss of the protein is associated with changes in granule structure and function [50]. Reduced *Spp1* transcript levels are seen in both alpha and beta cells of SORCS2-deficient islets (Figs. 5E and 7B). Thus, it is unclear whether SORCS2 directly or indirectly facilitates osteopontin expression in alpha cells. However, exogenous addition of osteopontin promotes glucose-stimulated insulin release from islets [49] while disruption of expression in beta cells [50; 63] does not decrease insulin secretion. These findings point to an essential role for osteopontin from non-beta cells to strengthen the insulin release capabilities of beta cells, a mechanism whereby SORCS2 may protect functional integrity of stressed islet *in trans*.

Currently, the stress signals that act through SORCS2 and the molecular mode of receptor action in islet stress response remain unknown. These actions include, but may not be restricted to, secretion of stress factors such as osteopontin from alpha cells. Increased cell stress and reduced *Spp1* expression are apparent in islets freshly isolated from mouse pancreata without prior glucose challenge, suggesting chronic insult *in vivo*. Multiple stressors are known to induce *Spp1* transcription and also to impact islet functions, including Il-1β [64–66] or glucose [67]. Because the release of various hormones from non-beta cells is not impacted by SORCS2 deficiency, the receptor is unlikely to have a generalized role in secretory cell function. More likely, SORCS2 is involved in relaying stress signals into stress factor production, as shown for endostatin in the ischemic brain [62]. Based on our data, AKT signaling, a pathway known to induce *Spp1* transcription, may be a promising target pathway to interrogate SORCS2 actions in the future.

Although much still needs to be learned about the mode of SORCS2 (and SORCS1) in glucose homeostasis, yet the identification of two related receptors that may act in concert *in cis* or *trans* to sustain insulin release capabilities in stressed islets points towards a new paradigm in control of metabolism, relevant to human health.

## Acknowledgement

We are indebted to K. Kampf, C. Schiel, and S.C. Carneiro Raimundo for expert technical assistance, as well as to A. Shih for sharing protocols. Also, we are grateful for assistance from the Bioinformatics Core Facility of Aarhus University Health.

## Data availability

All data are available on reasonable request from the authors. The scRNAsq data are available from the GEO database (accession number GSE231551).

## Authors’ relationships and activities

The authors declare no competing interest related to this manuscript.

## Contribution statement

OK and VS designed the study, performed experiments, and analyzed data. TC performed scRNAseq, and PQ and OK carried out the bioinformatical analyses. SK performed the EM studies. VS and TEW designed experiments and interpreted data. OK, VS, and TEW wrote the manuscript with editorial input, review and approval for publication from all authors. VS and TEW are guarantors of this work.

## Abbreviations

GLP-1: glucagon-like peptide 1
GSIS: glucose-stimulated insulin secretion
GTT: glucose tolerance test
ITT: insulin tolerance test
i.p.: intraperitoneal
NPY: neuropeptide Y
PPY: pancreatic polypeptide Y
SORCS: sorting-related receptor CNS expressed
VPS10P: vacuolar protein sorting 10 protein

## SUPPLEMENTARY TABLES

**Supplementary table S1:** ELISA kits used for determination of hormone levels.

| Hormone | Manufacturer | Catalog number |
| --- | --- | --- |
| Insulin | Crystal Chemicals, USA | 90080 |
| C-peptide | Crystal Chemicals, USA | 90050 |
| Glucagon | Crystal Chemicals, USA | 81518 |
| GLP-1 | Crystal Chemicals, USA | 81508 |
| Somatostatin | Phoenix Pharmaceuticals Inc, USA | EKE-060-03 |
| Somatostatin-28 | Phoenix Pharmaceuticals Inc, USA | FEK-060-14 |
| Neuropeptide Y (NPY) | Phoenix Pharmaceuticals Inc, USA | EK-049-03 |
| Peptide YY | Phoenix Pharmaceuticals. Inc, USA | EK-059-04 |
| Osteopontin | Abcam, USA | Ab100734-1001 |

**Supplementary table S2:** Primary and secondary antibodies used for immunodetections.

| Antibody | Manufacturer | Catalog number | Host | Dilution |
| --- | --- | --- | --- | --- |
| SORCS2 | R&D systems, USA | AF4237 | Sheep | 1:20 |
| SORCS2 | in-house |  | Rabbit | 1:1000 |
| Insulin | Agilent, USA | IR002 | Guinea pig | 1:3 |
| Insulin | ProteinTech, Germany | 15848 | Mouse | 1:500 |
| Glucagon | Abcam, USA | Ab10988 | Mouse | 1:500 |
| Somatostatin | Abcam, USA | Ab111912 | Rabbit | 1:250 |
| Pancreatic polypeptide | Sigma-Aldrich, USA | AB939-I | Rabbit | 1:500 |
| Anti-mouse Alexa 555 | Abcam, USA | Ab150106 | Donkey | 1:1000 |
| Anti-mouse Alexa 647 | Invitrogen, USA | A31571 | Donkey | 1:1000 |
| Anti-guinea pig Alexa 647 | Merck Millipore | AP193SA6 | Donkey | 1:1000 |
| Anti-rabbit Alexa 555 | Invitrogen, USA | A31572 | Donkey | 1:1000 |
| Anti-rabbit Alexa 647 | Abcam, USA | Ab150075 | Donkey | 1:1000 |
| DAPI | Roche | 10236276001 |  | 1:2000 |
| AKT | Cell Signaling Tech, USA | 4691 | Rabbit | 1:1000 |
| pAKT-T | Cell Signaling Tech, USA | 2965 | Rabbit | 1:1000 |
| pAKT-S | Cell Signaling Tech, USA | 9271 | Rabbit | 1:1000 |
| ERK | Cell Signaling Tech, USA | 9102 | Rabbit | 1:1000 |
| pERK | Cell Signaling Tech, USA | 4370 | Rabbit | 1:2000 |
| PKA (cAMP-Protein Kinase Catalytic subunit) | Abcam, USA | ab136960 | Rabbit | 1:1000 |
| Stat3 | Cell Signaling Tech, USA | 9139 | Mouse | 1:1000 |
| pStat3 | Cell Signaling Tech, USA | 9145 | Rabbit | 1:2000 |
| pPI3K | Cell Signaling Tech, USA | 4228 | Rabbit | 1:1000 |
| pJNK | Cell Signaling Tech, USA | 9251 | Rabbit | 1:1000 |
| Vinculin | Abcam, USA | 700062 | Rabbit | 1:1000 |
| Alpha-tubulin | Sigma-Aldrich, USA | 14-4502-82 | Mouse | 1:1000 |
| Anti-Rabbit IgG peroxidase | Sigma-Aldrich, USA | A0545 | Goat | 1:10000 |
| Anti-Mouse IgG peroxidase | Sigma-Aldrich, USA | A9917 | Goat | 1:10000 |

**Supplementary table S3.**
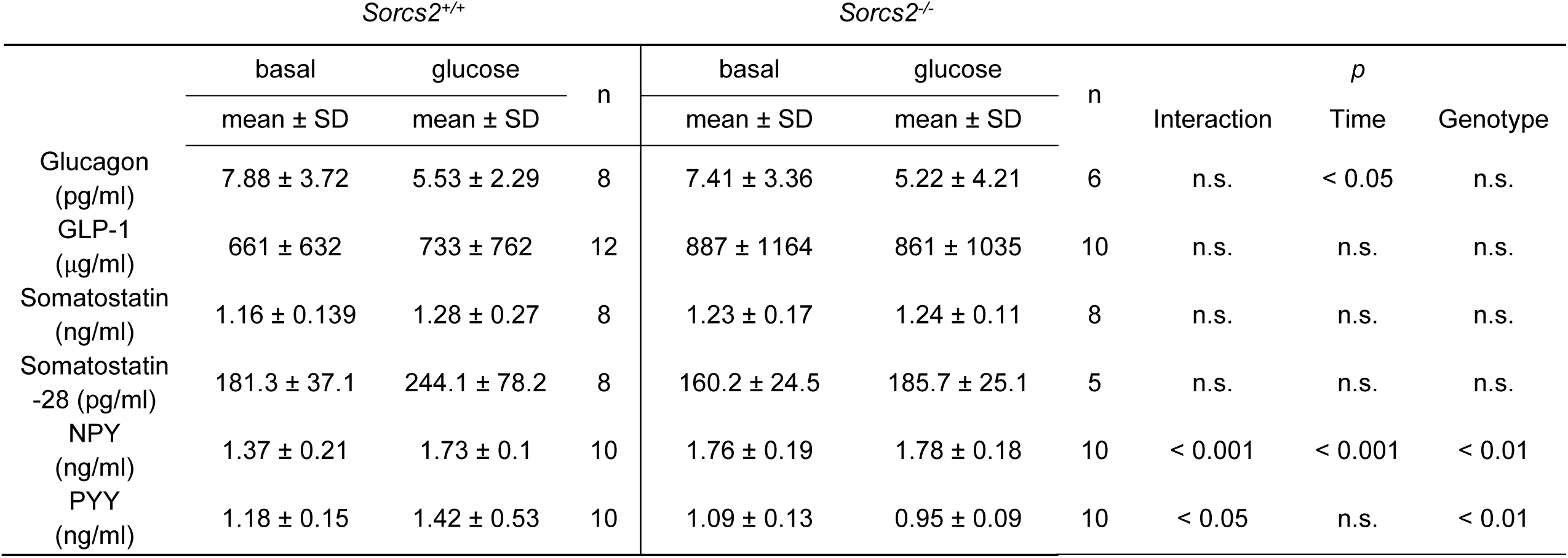
Plasma levels of pancreatic hormones in wildtype and SORCS2-deficient mice. Plasma hormone levels were determined by ELISA under basal conditions (basal) or 30 minutes after intraperitoneal injection of 2 g/kg body weight of glucose. Data are given as mean ± standard deviation (SD). Significance of data was evaluated using two-way ANOVA followed by Sidak’s or Turkey’s multiple comparisons tests. n, number of animals; n.s., not significant.

**Supplementary table S4.** Total hormone content in isolated wildtype and SORCS2-deficient islets. Hormone levels were determined by ELISA in lysates of isolated pancreatic islets from *Sorcs2^+/+^* and *Sorcs2^-/-^* mice kept overnight in culture medium. Data are given as mean ± standard deviation (SD). Significance of data was determined using unpaired Student’s t-test or Mann-Whitney U test. n, number of animals; n.s. not significant.

|  | *Sorcs2*<sup>+/+</sup> |  | *Sorcs2*<sup>-/-</sup> |  | p |
| --- | --- | --- | --- | --- | --- |
| | mean ± SD | n | mean ± SD | n | |
| Glucagon<br>(ng/mg protein) | 576.6 ± 439.3 | 12 | 653.2 ± 337.6 | 12 | n.s. |
| GLP-1<br>(mg/mg protein) | 11.74 ± 5.12 | 11 | 15.06 ± 5.26 | 12 | < 0.05 |
| Total somatostatin<br>(ng/mg protein) | 403.5 ± 315.5 | 13 | 462.8 ± 386 | 13 | n.s. |
| Somatostatin-28<br>(ng/mg protein) | 20.78 ± 9.63 | 11 | 26.74 ± 11.59 | 11 | n.s. |
| NPY<br>(ng/mg protein) | 1.20 ± 0.19 | 8 | 1.36 ± 0.37 | 8 | n.s. |
| PYY<br>(ng/mg protein) | 1.62 ± 0.55 | 8 | 0.99 ± 0.16 | 8 | < 0.01 |

**Supplementary table S5.** Levels of hormones released from wildtype and SORCS2-deficient pancreatic islets. Isolated pancreatic islets from *Sorcs2^+/+^* and *Sorcs2^-/-^* mice were treated for 1 hour with 1.6 mM or 16 mM glucose in Krebs-Ringer-Bicarbonate Hepes buffer. Levels of indicated hormones released into the supernatant were determined by ELISA thereafter. Data are given as mean ± standard deviation (SD). Significance of data was evaluated using two-way ANOVA followed by Sidak’s or Turkey’s multiple comparisons tests. n, number of animals; n.s., not significant.

|  | *Sorcs2*<sup>+/+</sup> |  |  | *Sorcs2*<sup>-/-</sup> |  |  |  |  |  |
| --- | --- | --- | --- | --- | --- | --- | --- | --- | --- |
|  | 1.6 mM | 16 mM | n | 1.6 mM | 16 mM | n | interaction | *p*<br>time | genotype |
|  | mean ± SD | mean ± SD |  | mean ± SD | mean ± SD |  |  |  |  |
| Glucagon<br>(ng/mg protein) | 1.59 ± 1.24 | 1.87 ± 1.71 | 23 | 1.47 ± 1.15 | 2.27 ± 3.69 | 23 | n.s. | n.s. | n.s. |
| GLP-1<br>(µg/mg protein) | 512 ± 317 | 590 ± 443 | 21 | 471 ± 196 | 531 ± 332 | 21 | n.s. | n.s. | n.s. |
| Somatostatin<br>(ng/mg protein) | 0.81 ± 0.49 | 1.18 ± 0.51 | 16 | 0.81 ± 0.51 | 1.316 ± 0.54 | 16 | n.s. | < 0.0001 | n.s. |
| Somatostatin-28<br>(pg/mg protein) | 580.9 ± 131.42 | 858.3 ± 353.56 | 7 | 475.4 ± 453.61 | 1306 ± 1218.63 | 7 | n.s. | < 0.01 | n.s. |
| NPY<br>(ng/mg protein) | 0.03 ± 0.03 | 0.07 ± 0.05 | 12 | 0.06 ± 0.05 | 0.111 ± 0.09 | 13 | n.s. | < 0.0001 | n.s. |
| PYY<br>(ng/mg protein) | 0.64 ± 0.26 | 0.46 ± 0.21 | 18 | 0.76 ± 0.41 | 0.574 ± 0.28 | 18 | n.s. | ≤ 0.001 | n.s. |

## SUPPLEMENTARY FIGURES

**Supplementary figure S1.**
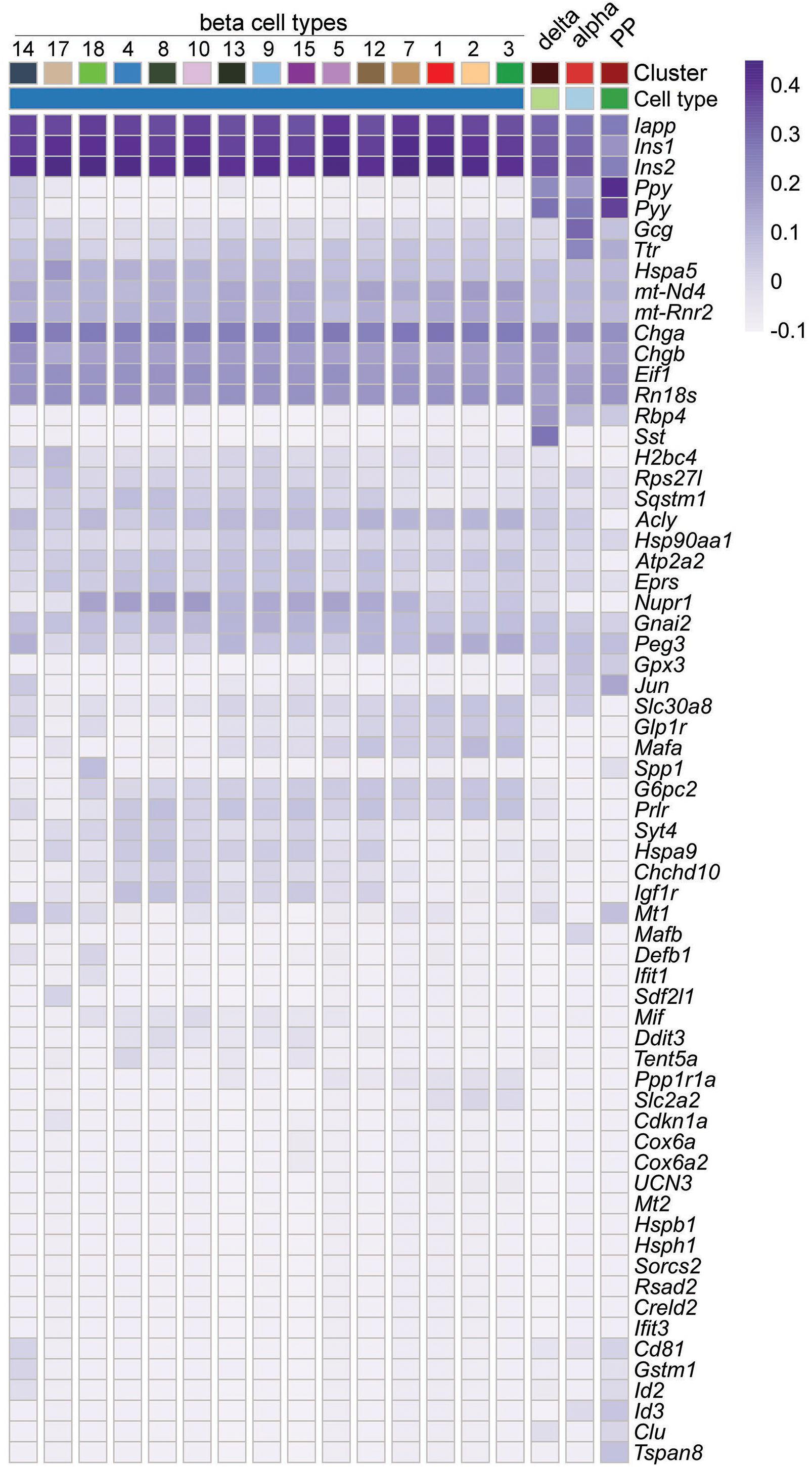
Annotation of islet cell types from single-cell RNA sequencing. Heatmap showing the top 20 most unique identifiers per cluster according to the adjusted p-value calculated with linear models for microarray data (Limma) moderated t-statistics for all 18 cell clusters identified by single-cell RNA sequencing of *Sorcs2^+/+^* and *Sorcs2^-/-^*islets.

## SUPPLEMENTARY DATA FILES

**File 1:** Differentially expressed genes in all beta cell clusters comparing *Sorcs2^+/+^* and *Sorcs2^-/-^* islets.

**File 2:** Differentially expressed genes in beta cell cluster #13 comparing *Sorcs2^+/+^* and *Sorcs2^-/-^* islets.

**File 3:** Differentially expressed genes in alpha cells comparing *Sorcs2^+/+^* and *Sorcs2^-/-^* islets.

