## Supplementary data file 1 for "Sorting receptor SORCS2 facilitates a protective stress response in pancreatic islets"

| GeneID | $Log_2FC$ | p-value | FDR | Average (*Sorcs2* $^{-/-}$ ) | Average (*Sorcs2* $^{+/+}$ ) |
|---|---|---|---|---|---|
| 0610009B22Rik | -0.002672007 | 0.773200955 | 0.840082665 | 0.402485709 | 0.385289692 |
| 0610009E02Rik | 0.006456701 | 0.235631855 | 0.349469721 | 0.08829633 | 0.081386867 |
| 0610009L18Rik | 0.005938828 | 0.379160621 | 0.500706269 | 0.133734409 | 0.130780324 |
| 0610010F05Rik | -0.014745313 | 0.021566663 | 0.049544211 | 0.11330532 | 0.123370286 |
| 0610010K14Rik | 0.021345576 | 0.000493344 | 0.001918518 | 0.139508623 | 0.104193517 |
| 0610012G03Rik | 0.04166493 | 2.60E-05 | 0.000136499 | 0.618557437 | 0.569741604 |
| 0610030E20Rik | 0.011513348 | 0.076075158 | 0.141740645 | 0.130977428 | 0.11910731 |
| 0610040B10Rik | 0.020618756 | 0.000105054 | 0.000488208 | 0.092729313 | 0.072685509 |
| 0610040J01Rik | 0.044938809 | 1.25E-06 | 8.60E-06 | 0.421584992 | 0.359231037 |
| 1010001B22Rik | 0.002963273 | 0.499767403 | 0.614175034 | 0.05483195 | 0.051515851 |
| 1110002L01Rik | -0.012880411 | 0.00728457 | 0.0199158 | 0.053911667 | 0.064951749 |
| 1110004F10Rik | 1.44E-05 | 0.998851497 | 0.998954249 | 0.608601597 | 0.598695272 |
| 1110008P14Rik | -0.044309478 | 6.01E-06 | 3.63E-05 | 0.662518553 | 0.705249904 |
| 1110018N20Rik | -0.017486821 | 0.007441161 | 0.020258462 | 0.120194756 | 0.132710379 |
| 1110020A21Rik | 0.003636865 | 0.571252846 | 0.678275546 | 0.124268847 | 0.115635514 |
| 1110032A03Rik | -0.030865384 | 0.000482092 | 0.00187927 | 0.300759583 | 0.313859479 |
| 1110035H17Rik | -0.002642572 | 0.502646086 | 0.616854993 | 0.039993579 | 0.041342265 |
| 1110038B12Rik | -0.025673844 | 0.000144675 | 0.000649069 | 0.108035933 | 0.134383247 |
| 1110038F14Rik | 0.007780016 | 0.375409295 | 0.497306059 | 0.317676255 | 0.302414038 |
| 1110046J04Rik | 0.001420386 | 0.7262063 | 0.802566516 | 0.047043151 | 0.042173799 |
| 1110051M20Rik | 4.61E-05 | 0.993710231 | 0.996138326 | 0.104371969 | 0.099062155 |
| 1110059E24Rik | 0.001823084 | 0.79145684 | 0.854000377 | 0.152686159 | 0.144808065 |
| 1110059G10Rik | 0.017151898 | 0.01214375 | 0.030562138 | 0.158373618 | 0.140675223 |
| 1110065P20Rik | -0.012721793 | 0.206083592 | 0.314380147 | 0.5866716 | 0.592522863 |
| 1190007I07Rik | -0.030121978 | 7.12E-05 | 0.000341605 | 0.171658328 | 0.197359959 |
| 1300002E11Rik | -0.004646373 | 0.591033709 | 0.695471874 | 0.294775563 | 0.289586174 |
| 1500002C15Rik | -0.012288259 | 0.004484818 | 0.013164673 | 0.043157889 | 0.0533594 |
| 1500009L16Rik | -0.021548938 | 0.018014389 | 0.042664041 | 0.304056187 | 0.319031999 |
| 1500011B03Rik | -0.06504942 | 6.00E-11 | 7.94E-10 | 0.718181494 | 0.791282352 |
| 1500015A07Rik | 0.011992218 | 0.045653416 | 0.09243735 | 0.10873101 | 0.099313906 |
| 1500026H17Rik | 0.015053957 | 0.026250367 | 0.058412925 | 0.145830073 | 0.136508909 |
| 1600012H06Rik | 0.011745421 | 0.036731559 | 0.077195032 | 0.096874931 | 0.086988565 |
| 1600014C10Rik | 0.012481959 | 0.001921387 | 0.006310719 | 0.050394212 | 0.039896418 |
| 1600020E01Rik | -0.014979511 | 0.025379943 | 0.056866514 | 0.12273842 | 0.135569652 |
| 1700007L15Rik | -0.009413935 | 0.07196666 | 0.135240333 | 0.076535298 | 0.078915268 |
| 1700008J07Rik | -0.002797504 | 0.505463094 | 0.619635912 | 0.04901089 | 0.047667496 |
| 1700016K19Rik | -0.020729121 | 4.96E-06 | 3.05E-05 | 0.03789598 | 0.05848957 |
| 1700019D03Rik | 0.042313602 | 5.57E-07 | 4.13E-06 | 0.28348618 | 0.231074028 |
| 1700029J07Rik | -0.019127554 | 9.53E-05 | 0.000447676 | 0.054785349 | 0.069792752 |
| 1700030C10Rik | 0.013762324 | 0.001108219 | 0.003913587 | 0.055957611 | 0.041363971 |
| 1700030K09Rik | -0.004391655 | 0.265692187 | 0.382165918 | 0.040799035 | 0.041926826 |
| 1700037C18Rik | -0.010001 | 0.050464218 | 0.100762606 | 0.071728494 | 0.075938111 |
| 1700037H04Rik | 0.004229213 | 0.345210973 | 0.467559358 | 0.058304823 | 0.05334548 |
| 1700052K11Rik | -0.015592303 | 0.000281283 | 0.001161696 | 0.045210832 | 0.053892728 |
| 1700056E22Rik | 0.001118747 | 0.84261389 | 0.892362989 | 0.090347753 | 0.087724161 |
| 1700063J08Rik | -0.004604678 | 0.320708558 | 0.442264273 | 0.054375828 | 0.058218455 |
| 1700084E18Rik | 0.009031212 | 0.03113588 | 0.067462229 | 0.049004575 | 0.044674552 |
| 1700086L19Rik | -0.019861657 | 0.074224571 | 0.13887823 | 0.461222784 | 0.502221623 |
| 1700086O06Rik | -0.001115597 | 0.793912795 | 0.856080323 | 0.049207983 | 0.049302371 |
| 1700086P04Rik | -0.000480205 | 0.905367107 | 0.93560051 | 0.046841968 | 0.043176816 |
| 1700088E04Rik | 0.002322492 | 0.706344351 | 0.788141832 | 0.112434016 | 0.110184803 |
| 1700096K18Rik | -0.000392378 | 0.951938268 | 0.967765747 | 0.132398922 | 0.125776554 |
| 1700110K17Rik | -0.067098028 | 1.16E-22 | 4.07E-21 | 0.0911017 | 0.153802561 |
| 1700113A16Rik | -0.011989309 | 0.024764698 | 0.055835435 | 0.076171442 | 0.082246429 |
| 1700123O20Rik | 0.009427977 | 0.220494323 | 0.331423285 | 0.20990997 | 0.195531775 |
| 1810008I18Rik | -0.144856289 | 6.48E-38 | 4.85E-36 | 0.409677886 | 0.568919841 |
| 1810010H24Rik | -0.000283264 | 0.950716298 | 0.967130255 | 0.059694555 | 0.057032562 |
| 1810013L24Rik | 0.012325915 | 0.082707274 | 0.151427518 | 0.166678356 | 0.15431791 |
| 1810014B01Rik | -0.00426134 | 0.351060496 | 0.47343739 | 0.057438569 | 0.058655685 |
| 1810021B22Rik | 0.004698353 | 0.235870972 | 0.349670264 | 0.046100582 | 0.040265449 |
| 1810024B03Rik | -0.008341481 | 0.07910619 | 0.146127757 | 0.056622104 | 0.065290357 |
| 1810030O07Rik | -0.019515954 | 0.004743533 | 0.013790859 | 0.141962622 | 0.153595515 |
| 1810037I17Rik | 0.001932046 | 0.821834907 | 0.877541103 | 1.197567688 | 1.224764435 |
| 1810044D09Rik | 0.011024687 | 0.10263466 | 0.180599848 | 0.147373283 | 0.134901066 |
| 1810058I24Rik | 0.047181821 | 2.76E-06 | 1.78E-05 | 0.5659162 | 0.5118669 |
| 1810059H22Rik | 0.008671197 | 0.136828481 | 0.227859968 | 0.106649546 | 0.094255775 |
| 1810062G17Rik | 0.000588021 | 0.912256064 | 0.939654252 | 0.084701985 | 0.078986882 |
| 2010001A14Rik | -0.002001638 | 0.646939091 | 0.740381618 | 0.055871275 | 0.051601185 |
| 2010204K13Rik | -0.027658266 | 6.80E-05 | 0.000328046 | 0.121097325 | 0.146429207 |
| 2010315B03Rik | 0.000192925 | 0.975636688 | 0.983731579 | 0.118263292 | 0.115490727 |

| | | | | | |
|---|---|---|---|---|---|
| 2010320M18Rik | -0.001760836 | 0.823070943 | 0.878074806 | 0.218572526 | 0.213272183 |
| 2200002J24Rik | 0.011049859 | 0.018495875 | 0.043549744 | 0.061405834 | 0.050935988 |
| 2210016F16Rik | -0.02651334 | 0.002567502 | 0.008112204 | 0.312417569 | 0.324103179 |
| 2210016L21Rik | 0.020281678 | 0.040208754 | 0.083359144 | 0.536114848 | 0.501232616 |
| 2210408F21Rik | -0.028704112 | 0.000277874 | 0.001150058 | 0.182165524 | 0.212838601 |
| 2210408I21Rik | -0.004022335 | 0.309966956 | 0.431362546 | 0.04083561 | 0.042083545 |
| 2300009A05Rik | 0.0351629 | 0.000381277 | 0.001523542 | 0.751630765 | 0.701071734 |
| 2310009B15Rik | -0.012697297 | 0.075626896 | 0.141181117 | 0.149007989 | 0.161265901 |
| 2310010J17Rik | 0.052692064 | 6.39E-11 | 8.40E-10 | 0.260347273 | 0.207209862 |
| 2310011J03Rik | 0.002835424 | 0.720041654 | 0.798658866 | 0.226442719 | 0.214548717 |
| 2310015A10Rik | -0.01032136 | 0.062667755 | 0.120956108 | 0.085517257 | 0.089358351 |
| 2310022A10Rik | 0.01322604 | 0.005485225 | 0.015606484 | 0.068643647 | 0.058813266 |
| 2310022B05Rik | 0.046448225 | 4.79E-08 | 4.27E-07 | 0.314629258 | 0.24579858 |
| 2310030G06Rik | -0.014844637 | 0.076173851 | 0.141897333 | 0.256259385 | 0.26490475 |
| 2310033P09Rik | -0.012498683 | 0.113353063 | 0.195419442 | 0.197757507 | 0.207399856 |
| 2310034G01Rik | -0.009726634 | 0.110457917 | 0.191625959 | 0.11286845 | 0.111479108 |
| 2310039H08Rik | 0.020931397 | 0.043009795 | 0.088011203 | 0.677582999 | 0.67225308 |
| 2310040G24Rik | -0.016130801 | 0.011427723 | 0.029030657 | 0.105975987 | 0.119211854 |
| 2310057M21Rik | 0.004596983 | 0.38760898 | 0.508547166 | 0.085993097 | 0.079084116 |
| 2310061I04Rik | 0.02445966 | 0.001802197 | 0.005961538 | 0.220039158 | 0.194131239 |
| 2310068J16Rik | 0.012286248 | 0.001011342 | 0.003616133 | 0.043678309 | 0.032772418 |
| 2410002F23Rik | -0.005047021 | 0.438276539 | 0.556547089 | 0.124724478 | 0.127192894 |
| 2410004B18Rik | -0.015901694 | 0.010807476 | 0.027657353 | 0.10937263 | 0.119303735 |
| 2410006H16Rik | -0.042591158 | 0.000914585 | 0.003312863 | 0.943245937 | 1.068364663 |
| 2410021H03Rik | -0.034846689 | 7.07E-07 | 5.12E-06 | 0.108423483 | 0.139274172 |
| 2410022M11Rik | 0.002855975 | 0.626696609 | 0.724895233 | 0.107391004 | 0.098793068 |
| 2510002D24Rik | -0.020099629 | 0.019965692 | 0.046400058 | 0.270797225 | 0.285419575 |
| 2510009E07Rik | 0.004614747 | 0.531475256 | 0.6428219 | 0.181513097 | 0.169023601 |
| 2510039O18Rik | 0.017404453 | 0.021972885 | 0.050287285 | 0.210421666 | 0.188491173 |
| 2510046G10Rik | -0.004142358 | 0.535758974 | 0.646796069 | 0.144287956 | 0.135694176 |
| 2610001J05Rik | -0.013578184 | 0.153580456 | 0.248810063 | 0.372427366 | 0.386961461 |
| 2610002M06Rik | 8.20E-05 | 0.989753993 | 0.993241689 | 0.125532953 | 0.121107395 |
| 2610005L07Rik | -0.006432271 | 0.187669868 | 0.291411349 | 0.059806424 | 0.065919857 |
| 2610008E11Rik | -0.00802959 | 0.188713574 | 0.292798175 | 0.103572269 | 0.109937658 |
| 2610021A01Rik | 0.0060317 | 0.101374627 | 0.178699534 | 0.039454706 | 0.033533267 |
| 2610037D02Rik | -0.017242674 | 7.42E-05 | 0.000355819 | 0.044652253 | 0.053831001 |
| 2610301B20Rik | 0.006232979 | 0.292886589 | 0.412374137 | 0.101434812 | 0.097602232 |
| 2610306M01Rik | -0.008956118 | 0.136960615 | 0.228001901 | 0.107204044 | 0.106997321 |
| 2610507B11Rik | -0.011450293 | 0.194818418 | 0.300448074 | 0.292997246 | 0.292368045 |
| 2700033N17Rik | 0.01300407 | 0.001358504 | 0.004671871 | 0.049691982 | 0.039514201 |
| 2700046G09Rik | -0.022747481 | 0.000352498 | 0.001423169 | 0.115027839 | 0.126439909 |
| 2700049A03Rik | 0.000981597 | 0.838013528 | 0.889040541 | 0.065375517 | 0.061234065 |
| 2700062C07Rik | -0.002316777 | 0.713450301 | 0.794065693 | 0.12443514 | 0.121511286 |
| 2700081O15Rik | 0.004645918 | 0.212570205 | 0.32225285 | 0.038736425 | 0.034491845 |
| 2700097O09Rik | -0.005295542 | 0.207811608 | 0.316172841 | 0.046352218 | 0.048352086 |
| 2810001G20Rik | -0.008929608 | 0.099972735 | 0.176747577 | 0.081935993 | 0.086910549 |
| 2810004N23Rik | 0.027861323 | 0.001157924 | 0.004064021 | 0.277675308 | 0.261002727 |
| 2810006K23Rik | -0.002593352 | 0.67773566 | 0.765973853 | 0.112032841 | 0.115589153 |
| 2810013P06Rik | -0.011811481 | 0.14635777 | 0.240271908 | 0.220292992 | 0.225628813 |
| 2810021J22Rik | 0.00858782 | 0.03870666 | 0.080613999 | 0.051677633 | 0.042785585 |
| 2810025M15Rik | -0.055051589 | 2.49E-12 | 3.86E-11 | 0.182090637 | 0.221629389 |
| 2810402E24Rik | -0.009881373 | 0.108536475 | 0.189238093 | 0.10247609 | 0.111945782 |
| 2810403D21Rik | -0.006453855 | 0.146320836 | 0.240271908 | 0.050708955 | 0.054719469 |
| 2900005J15Rik | 0.001157843 | 0.852297563 | 0.898680132 | 0.119322724 | 0.110665855 |
| 2900026A02Rik | 0.059055174 | 7.26E-13 | 1.18E-11 | 0.257393578 | 0.210335637 |
| 2900027M19Rik | -0.014378002 | 0.031295683 | 0.067717923 | 0.120720095 | 0.129786985 |
| 2900052L18Rik | -0.016560285 | 0.004119233 | 0.01221323 | 0.087553106 | 0.095136636 |
| 2900076A07Rik | -0.019694622 | 0.000724942 | 0.00270087 | 0.085149205 | 0.09966245 |
| 2900092N22Rik | -0.008696413 | 0.041120912 | 0.084858151 | 0.039992217 | 0.048516065 |
| 2900093K20Rik | 0.041369064 | 3.67E-07 | 2.81E-06 | 0.241603462 | 0.205920856 |
| 3110009E18Rik | 0.013737688 | 0.006481979 | 0.018041167 | 0.078792421 | 0.067732741 |
| 3110040N11Rik | 0.004815484 | 0.512664776 | 0.626540158 | 0.183712152 | 0.173338061 |
| 3110056K07Rik | 0.00527934 | 0.31232182 | 0.433708432 | 0.08120306 | 0.075184812 |
| 3110082I17Rik | 0.002836716 | 0.582430065 | 0.688519588 | 0.072218209 | 0.071935378 |
| 3222401L13Rik | -0.042026009 | 8.63E-10 | 9.95E-09 | 0.109494185 | 0.146366523 |
| 3830406C13Rik | -0.02154714 | 0.008756183 | 0.023220843 | 0.244666915 | 0.24736551 |
| 4632415L05Rik | -0.006746857 | 0.264612421 | 0.380950978 | 0.104031319 | 0.108369761 |
| 4632427E13Rik | -0.00149796 | 0.852505296 | 0.898680132 | 0.212049425 | 0.201550964 |
| 4632428C04Rik | 0.014785505 | 0.00039055 | 0.001555479 | 0.059096972 | 0.042278994 |
| 4732414G09Rik | -0.004534164 | 0.26994521 | 0.386567585 | 0.042786892 | 0.046759244 |
| 4833420G17Rik | -0.005660031 | 0.454854465 | 0.573553191 | 0.186079547 | 0.184656921 |
| 4833438C02Rik | 0.005811663 | 0.186764217 | 0.290329664 | 0.055622498 | 0.049491104 |

| | | | | | |
|---|---|---|---|---|---|
| 4833439L19Rik | 0.09287384 | 2.06E-19 | 5.70E-18 | 0.723304984 | 0.608949284 |
| 4921524J17Rik | 0.033990694 | 5.26E-06 | 3.22E-05 | 0.195876854 | 0.169652474 |
| 4930402H24Rik | 0.049976273 | 1.43E-09 | 1.60E-08 | 0.275866483 | 0.212994502 |
| 4930430F08Rik | -0.003546486 | 0.509547847 | 0.623471538 | 0.082847334 | 0.082203399 |
| 4930431P19Rik | -0.008812783 | 0.155871764 | 0.252018175 | 0.116897095 | 0.117324988 |
| 4930453N24Rik | 0.007425827 | 0.337431099 | 0.459454503 | 0.22459003 | 0.20950227 |
| 4930481A15Rik | -0.013075784 | 0.004761499 | 0.013834817 | 0.056341767 | 0.061683437 |
| 4930503L19Rik | -0.005655282 | 0.172421428 | 0.272256151 | 0.043225249 | 0.047440993 |
| 4930522L14Rik | 0.008812431 | 0.017398403 | 0.041471517 | 0.04059127 | 0.033987512 |
| 4930550C14Rik | -0.0066184 | 0.168441614 | 0.267143454 | 0.056086906 | 0.061224813 |
| 4930570G19Rik | -0.000817347 | 0.841551933 | 0.89162684 | 0.046954613 | 0.044580862 |
| 4930578M01Rik | 0.009624425 | 0.077882185 | 0.144305433 | 0.091172632 | 0.076805992 |
| 4930579K19Rik | -0.017917212 | 0.001715922 | 0.005701363 | 0.085493045 | 0.098136161 |
| 4931406C07Rik | -0.025739529 | 0.001767474 | 0.00585664 | 0.206799933 | 0.236359846 |
| 4932438A13Rik | 0.017567822 | 0.078628349 | 0.145466186 | 0.43918998 | 0.420268298 |
| 4933404O12Rik | -0.008210157 | 0.038847564 | 0.080855494 | 0.036224987 | 0.043315487 |
| 4933421O10Rik | 0.015767638 | 0.047561351 | 0.095911938 | 0.224177779 | 0.210128095 |
| 4933434E20Rik | -0.018797198 | 0.031019073 | 0.067271749 | 0.284761368 | 0.291996048 |
| 5031425E22Rik | -0.000212866 | 0.977866703 | 0.985162703 | 0.197173764 | 0.192555968 |
| 5031439G07Rik | 0.028550671 | 4.72E-07 | 3.55E-06 | 0.107799678 | 0.084747387 |
| 5033430I15Rik | 0.007071904 | 0.153093443 | 0.248145124 | 0.075773502 | 0.063727591 |
| 5330406M23Rik | -0.008939272 | 0.057172393 | 0.111882047 | 0.054558654 | 0.05988722 |
| 5330431K02Rik | -0.00635635 | 0.13614379 | 0.227069811 | 0.042013737 | 0.048064878 |
| 5330434G04Rik | -0.05136552 | 1.13E-12 | 1.82E-11 | 0.11132024 | 0.15579928 |
| 5330438D12Rik | -0.026893772 | 4.65E-06 | 2.87E-05 | 0.091458861 | 0.104486634 |
| 5330439M10Rik | -0.006671794 | 0.323947657 | 0.446030183 | 0.086464063 | 0.109748432 |
| 5430403G16Rik | -0.003520451 | 0.400939361 | 0.521019522 | 0.044873777 | 0.047029946 |
| 5430405H02Rik | -0.001243635 | 0.824135198 | 0.878474013 | 0.088832092 | 0.08890099 |
| 5430416N02Rik | -0.040721813 | 2.97E-07 | 2.31E-06 | 0.145604063 | 0.194305634 |
| 5530601H04Rik | -0.006618532 | 0.151990849 | 0.246893072 | 0.056156526 | 0.05859415 |
| 5730409E04Rik | 0.007935581 | 0.100908813 | 0.178078686 | 0.067089648 | 0.061364729 |
| 5730455P16Rik | 0.016352621 | 0.040325447 | 0.083529591 | 0.220991659 | 0.202817422 |
| 5730480H06Rik | -0.00971236 | 0.044337215 | 0.090271498 | 0.060856537 | 0.066111912 |
| 5830444B04Rik | 0.00217411 | 0.630154045 | 0.727336772 | 0.058029471 | 0.051801868 |
| 5830448L01Rik | -0.000404464 | 0.916280472 | 0.942055706 | 0.040204008 | 0.038849931 |
| 5830454E08Rik | -0.000915095 | 0.862031442 | 0.905724595 | 0.080870485 | 0.076963087 |
| 5930430L01Rik | 0.000664107 | 0.892694906 | 0.927319145 | 0.066799554 | 0.064031137 |
| 6030458C11Rik | -0.002633291 | 0.604673956 | 0.706992207 | 0.070516368 | 0.072722926 |
| 6330403K07Rik | 0.004195085 | 0.365151939 | 0.487973492 | 0.053763624 | 0.050947613 |
| 6330403L08Rik | 0.00211354 | 0.603545136 | 0.706303637 | 0.043363575 | 0.042892458 |
| 6330418K02Rik | -0.010639717 | 0.024238764 | 0.054751224 | 0.058086026 | 0.063424348 |
| 6330419E04Rik | -0.014521336 | 0.005163418 | 0.014834145 | 0.061391187 | 0.073579195 |
| 6430503K07Rik | -0.006517333 | 0.141704723 | 0.234175304 | 0.050230338 | 0.051804941 |
| 6430548M08Rik | 0.018829625 | 0.013846992 | 0.034115677 | 0.190142051 | 0.17976896 |
| 6430590A07Rik | -0.013477121 | 0.010048951 | 0.025990383 | 0.072786965 | 0.077383629 |
| 6820431F20Rik | -0.047425255 | 2.10E-06 | 1.38E-05 | 0.399634779 | 0.448401672 |
| 7SK | 0.036340528 | 1.34E-10 | 1.70E-09 | 0.117657997 | 0.074876038 |
| 8030462N17Rik | 0.02061527 | 0.001313541 | 0.004534889 | 0.134912114 | 0.114140737 |
| 8430429K09Rik | 0.014043209 | 0.00890551 | 0.023559011 | 0.089545952 | 0.076712295 |
| 9130023H24Rik | 0.020051007 | 8.80E-05 | 0.000416084 | 0.086909488 | 0.065920193 |
| 9130401M01Rik | -0.000284299 | 0.969748189 | 0.979724815 | 0.197032877 | 0.188652647 |
| 9230114K14Rik | 0.001895392 | 0.755850186 | 0.826310076 | 0.106224682 | 0.103643889 |
| 9330020H09Rik | 0.022896346 | 0.000125602 | 0.000572211 | 0.111788374 | 0.097882332 |
| 9330151L19Rik | -0.008967597 | 0.015276881 | 0.037111904 | 0.034441749 | 0.038762554 |
| 9330159F19Rik | 0.00376402 | 0.320712109 | 0.442264273 | 0.039692161 | 0.036253191 |
| 9330160F10Rik | -0.005565656 | 0.235940893 | 0.349720592 | 0.055785268 | 0.060486213 |
| 9330162G02Rik | 0.01801164 | 1.19E-05 | 6.70E-05 | 0.052518837 | 0.039084261 |
| 9430038I01Rik | -0.0031331 | 0.422786843 | 0.541534567 | 0.038872608 | 0.040444767 |
| 9430065F17Rik | -0.004607206 | 0.30271148 | 0.422899987 | 0.052108997 | 0.055449204 |
| 9430081H08Rik | 0.010292099 | 0.021008607 | 0.048468361 | 0.056808489 | 0.050137054 |
| 9530068E07Rik | 0.035038946 | 0.000198084 | 0.000850983 | 0.440357023 | 0.400674602 |
| 9530077C05Rik | 0.004473719 | 0.318671117 | 0.440449332 | 0.055289198 | 0.051978672 |
| 9930017N22Rik | -0.01190549 | 0.005427452 | 0.015478832 | 0.039212928 | 0.049483009 |
| 9930021J03Rik | 0.018779308 | 0.035567463 | 0.075252856 | 0.317921393 | 0.293582221 |
| A030001D20Rik | -0.015904952 | 0.005850263 | 0.016483753 | 0.088027364 | 0.098836489 |
| A130010J15Rik | 0.004330996 | 0.365348544 | 0.488169123 | 0.06279708 | 0.060325028 |
| A130014A01Rik | -0.009921879 | 0.017071147 | 0.040777812 | 0.041409521 | 0.047840319 |
| A1cf | -0.005839294 | 0.119672295 | 0.203935855 | 0.037552699 | 0.036980688 |
| A230051N06Rik | -0.038349956 | 2.30E-07 | 1.82E-06 | 0.150089098 | 0.182590258 |
| A230057D06Rik | -0.101935618 | 1.98E-21 | 6.19E-20 | 0.380886114 | 0.460928903 |
| A2ml1 | 0.029492839 | 3.55E-10 | 4.27E-09 | 0.078455672 | 0.04844716 |
| A330023F24Rik | -0.01226476 | 0.043326343 | 0.088547131 | 0.096483703 | 0.104238076 |

| | | | | | |
|---|---|---|---|---|---|
| A330076H08Rik | -0.142273613 | 2.96E-31 | 1.61E-29 | 0.812962421 | 0.970581297 |
| A430005L14Rik | 0.01102641 | 0.228402987 | 0.34104344 | 0.363560176 | 0.337923696 |
| A430018G15Rik | 0.005455521 | 0.319895957 | 0.44145188 | 0.091288067 | 0.083080421 |
| A430033K04Rik | 0.006364781 | 0.132276517 | 0.221761045 | 0.052403637 | 0.045696301 |
| A530058N18Rik | -0.024778498 | 0.010306754 | 0.026536617 | 0.394334042 | 0.39962609 |
| A730020E08Rik | 0.003774618 | 0.393329555 | 0.513902692 | 0.057339013 | 0.051923354 |
| A730063M14Rik | -0.003200944 | 0.5579254 | 0.666521349 | 0.074561177 | 0.084122169 |
| A930004J17Rik | -0.012365474 | 0.002901946 | 0.00903096 | 0.036563714 | 0.047690621 |
| A930005H10Rik | -0.014158347 | 0.019293106 | 0.045034231 | 0.099720078 | 0.108102241 |
| AA414768 | 0.005780496 | 0.150573226 | 0.245616259 | 0.045477588 | 0.040955151 |
| Aacs | 0.031201631 | 1.62E-07 | 1.32E-06 | 0.126726881 | 0.091321955 |
| Aagab | 0.009322859 | 0.169974514 | 0.269223236 | 0.156924065 | 0.140638947 |
| Aak1 | 0.054057138 | 1.59E-09 | 1.76E-08 | 0.336969705 | 0.289180761 |
| Aamdc | -0.008141006 | 0.178061229 | 0.279707751 | 0.105439952 | 0.111060922 |
| Aamp | -0.047538472 | 8.55E-07 | 6.06E-06 | 0.495428121 | 0.533146272 |
| Aar2 | 0.013489973 | 0.014352139 | 0.035190793 | 0.095731925 | 0.08219394 |
| Aars | 0.017488682 | 0.093076303 | 0.166431455 | 0.672751684 | 0.645519874 |
| Aasdh | 0.009770677 | 0.021713763 | 0.049835034 | 0.05309985 | 0.045410635 |
| Aasdhppt | -0.013880936 | 0.075867649 | 0.14156523 | 0.202749094 | 0.20835124 |
| Aatf | -0.005109881 | 0.449533655 | 0.567961121 | 0.137115685 | 0.139474017 |
| Aatk | -0.006879624 | 0.186523435 | 0.290048119 | 0.073320322 | 0.074200527 |
| Abat | 0.021553024 | 0.009079919 | 0.023922756 | 0.265002193 | 0.224639548 |
| Abca1 | 0.017794847 | 0.076843161 | 0.142948117 | 0.407523412 | 0.3844471 |
| Abca2 | 0.007141173 | 0.292943136 | 0.41239403 | 0.154377619 | 0.13838658 |
| Abca3 | 0.038411943 | 3.95E-06 | 2.48E-05 | 0.269745362 | 0.229361973 |
| Abca5 | 0.013045062 | 0.147345006 | 0.241444152 | 0.310615661 | 0.30176274 |
| Abca7 | -0.017338195 | 0.005921822 | 0.016668197 | 0.11394602 | 0.120453769 |
| Abcb10 | 0.042762731 | 1.50E-07 | 1.23E-06 | 0.254283087 | 0.209424079 |
| Abcb4 | 0.03655686 | 1.24E-08 | 1.21E-07 | 0.136006548 | 0.106217917 |
| Abcb7 | 0.009582518 | 0.104840697 | 0.183650677 | 0.110102353 | 0.098408426 |
| Abcb8 | -0.011061226 | 0.026881312 | 0.059585071 | 0.066810706 | 0.07165962 |
| Abcb9 | -0.012865368 | 0.041648712 | 0.085767586 | 0.117060444 | 0.121365574 |
| Abcc1 | 0.009423341 | 0.069447367 | 0.131509018 | 0.079872364 | 0.071481014 |
| Abcc4 | -0.016259235 | 0.00308134 | 0.009513112 | 0.071210615 | 0.087814487 |
| Abcc5 | 0.00550163 | 0.410424854 | 0.529824018 | 0.137725779 | 0.13086245 |
| Abcc8 | -0.105136156 | 2.62E-26 | 1.15E-24 | 1.261598096 | 1.373788023 |
| Abcd1 | 0.019274253 | 0.000192906 | 0.000831679 | 0.085880752 | 0.069110215 |
| Abcd3 | 0.052376775 | 3.46E-07 | 2.66E-06 | 0.775174592 | 0.690827758 |
| Abcd4 | -0.002805632 | 0.466108699 | 0.583656462 | 0.040504482 | 0.039938791 |
| Abce1 | 0.043749283 | 8.04E-07 | 5.74E-06 | 0.306038472 | 0.271520634 |
| Abcf1 | 0.02652658 | 0.006249617 | 0.017474482 | 0.45135935 | 0.42650362 |
| Abcf2 | -0.012642282 | 0.06719268 | 0.128112814 | 0.144998752 | 0.151005811 |
| Abcf3 | 0.008030963 | 0.292019696 | 0.411495378 | 0.1976883 | 0.191488661 |
| Abcg1 | 0.031435655 | 0.000144162 | 0.000647663 | 0.261543221 | 0.2229315 |
| Abhd10 | -0.004551846 | 0.417748833 | 0.537003062 | 0.08809331 | 0.090874115 |
| Abhd11 | 0.006611834 | 0.337430483 | 0.459454503 | 0.151303594 | 0.14434835 |
| Abhd12 | -0.010651698 | 0.147098547 | 0.241202913 | 0.177891879 | 0.178032257 |
| Abhd13 | 0.017349388 | 0.009126028 | 0.024011702 | 0.142800307 | 0.126806255 |
| Abhd14a | -0.049824484 | 7.43E-13 | 1.20E-11 | 0.131265353 | 0.165275115 |
| Abhd14b | -0.018646873 | 0.008025851 | 0.021626199 | 0.155725011 | 0.160138946 |
| Abhd15 | 0.010979569 | 0.012344581 | 0.030979354 | 0.059240094 | 0.047909812 |
| Abhd16a | 0.005665817 | 0.523900217 | 0.63595429 | 0.327266944 | 0.314200312 |
| Abhd17a | -0.002901895 | 0.745415786 | 0.818215228 | 0.359021448 | 0.344001719 |
| Abhd17b | -0.003138478 | 0.631605222 | 0.728493867 | 0.127243294 | 0.128629561 |
| Abhd17c | 0.017249638 | 0.084236189 | 0.153763468 | 0.384487181 | 0.390788611 |
| Abhd18 | -0.008330599 | 0.109193998 | 0.189950319 | 0.069166212 | 0.073191273 |
| Abhd2 | 0.122290247 | 5.36E-49 | 6.52E-47 | 0.311924109 | 0.194968562 |
| Abhd3 | -0.011799827 | 0.004895371 | 0.014163093 | 0.044564101 | 0.050218768 |
| Abhd4 | 0.009028923 | 0.040803744 | 0.084295367 | 0.060365911 | 0.049273845 |
| Abhd5 | 0.008697231 | 0.196752504 | 0.302901295 | 0.145455269 | 0.137510898 |
| Abhd6 | -0.012077038 | 0.092051274 | 0.164841127 | 0.156319347 | 0.165853269 |
| Abhd8 | 0.011864943 | 0.136552392 | 0.227595124 | 0.246094181 | 0.219684752 |
| Abi1 | 0.028539707 | 0.000865899 | 0.003157641 | 0.275668617 | 0.260801079 |
| Abi2 | 0.014697729 | 0.068747662 | 0.130463551 | 0.228042167 | 0.216396296 |
| Abitram | 0.008268765 | 0.15791921 | 0.254597305 | 0.105492938 | 0.095156737 |
| Abl1 | 0.01053403 | 0.034357666 | 0.073170915 | 0.074178568 | 0.064212777 |
| Abl2 | 0.016702087 | 0.009647135 | 0.025131149 | 0.129621913 | 0.116052764 |
| Ablim1 | 0.025297403 | 0.000293416 | 0.0012077 | 0.156042891 | 0.132229665 |
| Abr | 0.038388646 | 3.06E-07 | 2.37E-06 | 0.212392544 | 0.169758643 |
| Abracl | 0.030357279 | 0.001908961 | 0.006276265 | 0.959741218 | 0.92823297 |
| Abraxas2 | -0.027492597 | 0.003419754 | 0.01042221 | 0.345318786 | 0.372358915 |
| Abt1 | -0.008873667 | 0.151009538 | 0.246006139 | 0.113539715 | 0.11650327 |

| | | | | | |
|---|---|---|---|---|---|
| Abtb1 | -0.012480513 | 0.021506113 | 0.04944016 | 0.084969494 | 0.086381836 |
| Acaa1a | -0.060286416 | 6.87E-11 | 9.00E-10 | 0.351261754 | 0.380827326 |
| Acaa2 | -0.019055192 | 0.002603912 | 0.008211232 | 0.117582964 | 0.123484692 |
| Acaca | 0.006422502 | 0.284415655 | 0.403603707 | 0.10824031 | 0.099973652 |
| Acad11 | -0.021416294 | 8.56E-05 | 0.000405287 | 0.075682351 | 0.089431133 |
| Acad12 | 0.008521127 | 0.035355661 | 0.074843158 | 0.051231753 | 0.040067304 |
| Acad8 | -0.00561824 | 0.285640649 | 0.404928315 | 0.07740865 | 0.078294233 |
| Acad9 | -0.003030059 | 0.537390127 | 0.648041033 | 0.06507703 | 0.066713011 |
| Acadl | -0.01223905 | 0.163170076 | 0.2610399 | 0.301948143 | 0.301434875 |
| Acadm | 0.008554135 | 0.188765085 | 0.292831363 | 0.135425353 | 0.125261856 |
| Acads | -0.003517768 | 0.587399689 | 0.692206034 | 0.131272475 | 0.12855644 |
| Acadsb | 0.019802321 | 0.026170641 | 0.058262188 | 0.312227643 | 0.291671202 |
| Acadvl | -0.038829501 | 2.68E-06 | 1.73E-05 | 0.247614942 | 0.265745661 |
| Acap2 | 0.022036893 | 0.005567075 | 0.01577933 | 0.225319999 | 0.201624437 |
| Acap3 | 0.001006185 | 0.797281185 | 0.858450165 | 0.041687538 | 0.040146951 |
| Acat1 | -0.080270407 | 1.05E-15 | 2.13E-14 | 0.515862667 | 0.584830519 |
| Acbd3 | 0.044444847 | 2.00E-05 | 0.000108438 | 0.738914034 | 0.691578478 |
| Acbd4 | -0.007292359 | 0.09873523 | 0.174814041 | 0.05094797 | 0.05492446 |
| Acbd5 | 0.011484487 | 0.24809398 | 0.362484171 | 0.495394703 | 0.491875845 |
| Acbd6 | 0.021711912 | 0.023175385 | 0.05261819 | 0.478559194 | 0.440536605 |
| Acd | -0.005570854 | 0.441310318 | 0.55988763 | 0.168961309 | 0.167867997 |
| Acer2 | -0.002987773 | 0.463555121 | 0.581207492 | 0.044749397 | 0.044301119 |
| Acer3 | -0.001868205 | 0.643800559 | 0.737745053 | 0.042046729 | 0.044444652 |
| Acin1 | 0.034720827 | 0.000639248 | 0.002414439 | 0.620317098 | 0.584543119 |
| Acly | -0.060209961 | 1.63E-14 | 3.07E-13 | 2.144269776 | 2.286502893 |
| Aco1 | 0.002490148 | 0.667278593 | 0.757064125 | 0.102146237 | 0.095656548 |
| Aco2 | 0.09585732 | 4.83E-22 | 1.60E-20 | 0.644986232 | 0.525397695 |
| Acot11 | 0.002543996 | 0.515932093 | 0.628715444 | 0.041836139 | 0.039105444 |
| Acot13 | -0.013805257 | 0.167084479 | 0.265554243 | 0.567040362 | 0.553002957 |
| Acot2 | -0.000717913 | 0.864011925 | 0.906923335 | 0.050175607 | 0.046824503 |
| Acot8 | -0.016337415 | 0.01371869 | 0.033851042 | 0.126080121 | 0.136284518 |
| Acot9 | 0.000898479 | 0.824471808 | 0.878605165 | 0.042433004 | 0.043242733 |
| Acox1 | -0.000169683 | 0.984792943 | 0.990703331 | 0.302914106 | 0.302839473 |
| Acox3 | 0.005902458 | 0.155019079 | 0.250806371 | 0.049950168 | 0.043869838 |
| Acp1 | 0.00164209 | 0.870303942 | 0.910970599 | 0.553507155 | 0.552356955 |
| Acp2 | 0.005689532 | 0.420033556 | 0.539155827 | 0.15723533 | 0.151991288 |
| Acp6 | -0.007634367 | 0.083648698 | 0.152853985 | 0.051249653 | 0.05601389 |
| Acsf3 | 0.006310701 | 0.289012162 | 0.408457078 | 0.105625386 | 0.098554772 |
| Acsl1 | -0.006984635 | 0.217268659 | 0.327740249 | 0.086749739 | 0.091956655 |
| Acsl3 | -0.000609313 | 0.904944594 | 0.935446182 | 0.069302858 | 0.069672808 |
| Acsl4 | -0.00547612 | 0.507430231 | 0.621549289 | 0.21335238 | 0.227795961 |
| Acsl5 | -0.00910799 | 0.268009946 | 0.384759701 | 0.229634843 | 0.240076696 |
| Acsl6 | -0.003958162 | 0.392751705 | 0.513354676 | 0.056376399 | 0.058709412 |
| Acss2 | 0.006599613 | 0.250407286 | 0.365205466 | 0.103650821 | 0.093080117 |
| Acta1 | -0.007215247 | 0.151839089 | 0.246735782 | 0.066701292 | 0.071564109 |
| Actb | 0.009265901 | 0.334120202 | 0.456994458 | 1.657389005 | 1.684449156 |
| Actg1 | 0.038246929 | 0.000181719 | 0.000789044 | 1.724410244 | 1.688132805 |
| Actl6a | -0.013825054 | 0.077615898 | 0.144004152 | 0.208839709 | 0.208375007 |
| Actl6b | -0.009659442 | 0.139930692 | 0.231935147 | 0.136728681 | 0.133959602 |
| Actn1 | 0.01713012 | 0.036542243 | 0.076896902 | 0.230773252 | 0.222787397 |
| Actn4 | 0.086046863 | 4.28E-22 | 1.43E-20 | 0.345743898 | 0.266932268 |
| Actr10 | -0.059659426 | 7.29E-10 | 8.47E-09 | 0.478717495 | 0.522380351 |
| Actr1a | 0.018158676 | 0.007547724 | 0.020491196 | 0.152950597 | 0.13570918 |
| Actr1b | 0.028935591 | 0.001438427 | 0.004906803 | 0.319061202 | 0.301805771 |
| Actr2 | 0.01175484 | 0.217409556 | 0.327901908 | 0.448391714 | 0.436244349 |
| Actr3 | -0.006951355 | 0.485355473 | 0.601328648 | 0.559567259 | 0.547469811 |
| Actr3b | 0.006012098 | 0.132167743 | 0.221616902 | 0.046092676 | 0.040038286 |
| Actr5 | -0.007555402 | 0.054940421 | 0.108014312 | 0.038005005 | 0.042838613 |
| Actr6 | -0.016661516 | 0.000973343 | 0.003494401 | 0.062476195 | 0.073997588 |
| Actr8 | -0.005215065 | 0.435779317 | 0.554317222 | 0.13904902 | 0.136908686 |
| Acvr1 | 0.005347406 | 0.239214245 | 0.353027067 | 0.060764484 | 0.054886482 |
| Acvr1b | 0.007988495 | 0.084557933 | 0.154234939 | 0.063626815 | 0.055909508 |
| Acvr1c | 0.012588344 | 0.022704097 | 0.051632569 | 0.093821135 | 0.072208405 |
| Acvr2a | 0.017151288 | 0.004931537 | 0.014252201 | 0.11649349 | 0.099800444 |
| Acvr2b | 0.003568902 | 0.49340498 | 0.608486807 | 0.084543683 | 0.075227896 |
| Acyp1 | -0.001458008 | 0.873050832 | 0.912863002 | 0.36268896 | 0.358643183 |
| Acyp2 | -0.044685452 | 1.68E-08 | 1.60E-07 | 0.185485072 | 0.219116635 |
| Adal | 0.002028133 | 0.630416782 | 0.727553651 | 0.046842292 | 0.046150168 |
| Adam10 | 0.038954032 | 1.15E-07 | 9.65E-07 | 0.192026531 | 0.159222241 |
| Adam15 | 0.001028052 | 0.853042899 | 0.898806011 | 0.082609645 | 0.086262752 |
| Adam17 | 0.004183778 | 0.391172367 | 0.512075844 | 0.067450404 | 0.063491884 |
| Adam22 | 0.068947741 | 7.63E-19 | 2.01E-17 | 0.232034973 | 0.172105065 |

| | | | | | |
|---|---|---|---|---|---|
| Adam32 | 0.029965271 | 6.32E-09 | 6.46E-08 | 0.098453094 | 0.063571675 |
| Adam9 | 0.004653633 | 0.534614102 | 0.645574251 | 0.171072856 | 0.176798823 |
| Adar | 0.007106126 | 0.2728901 | 0.390267365 | 0.128266864 | 0.120646542 |
| Adarb1 | 0.016429478 | 0.026680676 | 0.059234879 | 0.18822366 | 0.161561181 |
| Adat1 | -0.002134978 | 0.583027132 | 0.688846454 | 0.038479468 | 0.040850743 |
| Adck1 | 0.006452653 | 0.134281301 | 0.224734517 | 0.055050765 | 0.049848185 |
| Adck2 | 0.007837661 | 0.038558891 | 0.080357887 | 0.047259751 | 0.035781798 |
| Adck5 | -0.01486829 | 0.00999349 | 0.025873956 | 0.092822955 | 0.102181292 |
| Adcy6 | 0.00523214 | 0.36549604 | 0.488299093 | 0.099832379 | 0.092205999 |
| Adcy9 | 0.040770335 | 4.98E-07 | 3.73E-06 | 0.258340424 | 0.210149159 |
| Add1 | 0.022879612 | 0.001855931 | 0.006118467 | 0.190455691 | 0.165117096 |
| Add3 | 0.029262841 | 1.84E-05 | 0.000100397 | 0.162839145 | 0.129186649 |
| Adgb | -0.004675909 | 0.248067791 | 0.362484171 | 0.040968367 | 0.043482752 |
| Adgrb3 | -0.01542838 | 0.001907637 | 0.006276265 | 0.058718437 | 0.067854726 |
| Adgrg1 | 0.025273316 | 0.006228007 | 0.01741907 | 0.399554241 | 0.354726364 |
| Adgrl1 | 0.023946827 | 0.004037328 | 0.012007006 | 0.265754858 | 0.229164449 |
| Adh1 | 0.044543029 | 2.03E-08 | 1.90E-07 | 0.24689752 | 0.175938447 |
| Adh5 | -0.033533744 | 0.000117008 | 0.000538105 | 0.297155463 | 0.312673886 |
| Adi1 | 0.014032373 | 0.147249017 | 0.241343563 | 0.412369921 | 0.414509707 |
| Adipor1 | 0.031121871 | 0.000891276 | 0.003241671 | 0.396459903 | 0.365347284 |
| Adipor2 | 0.018488873 | 0.024863712 | 0.05598078 | 0.256854372 | 0.235581348 |
| Adk | 0.013193088 | 0.143902102 | 0.237201803 | 0.362498144 | 0.333108692 |
| Adm2 | -0.097179086 | 6.59E-28 | 3.14E-26 | 0.148086314 | 0.242717861 |
| Adnp | 0.002999437 | 0.549551787 | 0.658865763 | 0.068153054 | 0.067275491 |
| Adnp2 | 0.004115985 | 0.409138255 | 0.529012118 | 0.071681324 | 0.06609821 |
| Ado | 0.003511585 | 0.548583185 | 0.658029084 | 0.10249111 | 0.097286451 |
| Adora1 | 0.047501968 | 8.73E-18 | 2.08E-16 | 0.11123381 | 0.06738021 |
| Adora2b | 0.047842453 | 9.69E-15 | 1.86E-13 | 0.135595622 | 0.087916686 |
| Adpgk | -0.004086997 | 0.38220188 | 0.503355009 | 0.055415082 | 0.061070324 |
| Adprh | -0.013610725 | 0.116981114 | 0.200227181 | 0.264124521 | 0.280757721 |
| Adprm | 0.001153257 | 0.842390942 | 0.892270806 | 0.104300825 | 0.097122092 |
| Adprs | -0.003491228 | 0.605043779 | 0.707254493 | 0.149241688 | 0.143038694 |
| Adra2a | 0.014270493 | 0.156513296 | 0.252761173 | 0.442452702 | 0.423617986 |
| Adrm1 | -0.00956751 | 0.301530662 | 0.42173516 | 0.385021113 | 0.371695131 |
| Adsl | 0.009227364 | 0.098005659 | 0.173617167 | 0.095140167 | 0.08611796 |
| Adss | 0.023323063 | 0.000215691 | 0.000917947 | 0.134873114 | 0.108330763 |
| Adssl1 | -0.000195186 | 0.962377629 | 0.974028092 | 0.045947327 | 0.045466126 |
| Aebp2 | -0.000625377 | 0.933660986 | 0.95498886 | 0.170846618 | 0.176549468 |
| Aen | -0.031688118 | 0.00024833 | 0.001039633 | 0.199707928 | 0.233083245 |
| Afdn | 0.023331016 | 0.016053174 | 0.038726788 | 0.444932413 | 0.428100506 |
| Aff1 | 0.006381483 | 0.462263154 | 0.580036446 | 0.279031465 | 0.265078986 |
| Aff2 | -0.0278266 | 7.55E-06 | 4.46E-05 | 0.094391637 | 0.114016529 |
| Aff4 | 0.036375982 | 0.000404861 | 0.00160721 | 0.718889923 | 0.695506711 |
| Afg1l | 0.008332951 | 0.029066894 | 0.063746524 | 0.044304716 | 0.036144976 |
| Afg3l1 | 0.009630184 | 0.217587626 | 0.328068688 | 0.21368443 | 0.204782416 |
| Afg3l2 | 0.035573873 | 2.33E-05 | 0.000124074 | 0.266900956 | 0.232042172 |
| Aftph | 0.02054047 | 0.026382828 | 0.058667396 | 0.375609792 | 0.352855608 |
| Aga | -0.156098428 | 5.68E-52 | 7.26E-50 | 0.548862206 | 0.703729895 |
| Agap1 | -0.063642369 | 1.42E-09 | 1.59E-08 | 0.590192496 | 0.671302115 |
| Agap3 | 0.006868348 | 0.330998195 | 0.453617769 | 0.169569301 | 0.153875074 |
| Agbl3 | 0.002105446 | 0.731094959 | 0.806347775 | 0.117098189 | 0.108592911 |
| Agbl4 | -0.012387809 | 0.003620174 | 0.010947227 | 0.04250446 | 0.050595492 |
| Agfg1 | 0.035780218 | 0.000119454 | 0.000548055 | 0.391040377 | 0.350326497 |
| Agfg2 | 0.003370089 | 0.430340776 | 0.549051577 | 0.052086284 | 0.047766014 |
| Aggf1 | 0.039649708 | 2.02E-06 | 1.33E-05 | 0.273435795 | 0.228912617 |
| Agk | -0.012237212 | 0.007454547 | 0.020283544 | 0.049930856 | 0.059077482 |
| Agl | 0.01828676 | 0.02552102 | 0.05710365 | 0.234557843 | 0.218433468 |
| Ago1 | 0.027507616 | 1.56E-05 | 8.64E-05 | 0.133876574 | 0.110099512 |
| Ago2 | 0.061461054 | 1.35E-10 | 1.71E-09 | 0.412172352 | 0.365838248 |
| Ago3 | 0.010314495 | 0.1492397 | 0.243963729 | 0.159499349 | 0.147948651 |
| Ago4 | -0.000694953 | 0.883519327 | 0.920245864 | 0.062013656 | 0.060341912 |
| Agpat1 | 0.012648713 | 0.070525518 | 0.133084062 | 0.166059885 | 0.150321788 |
| Agpat2 | -0.007065582 | 0.248517296 | 0.362939034 | 0.113043262 | 0.11311059 |
| Agpat3 | -0.008556104 | 0.382929031 | 0.503992294 | 0.576950751 | 0.553566362 |
| Agpat4 | -0.038509335 | 1.43E-13 | 2.53E-12 | 0.047004339 | 0.078849968 |
| Agpat5 | -0.013458682 | 0.088623794 | 0.16017857 | 0.197790244 | 0.209167963 |
| Agps | 0.002276726 | 0.702477578 | 0.784998507 | 0.10386963 | 0.101535314 |
| Agt | -0.008700694 | 0.067470155 | 0.1284655 | 0.061754911 | 0.061699136 |
| Agtpbp1 | -0.015174264 | 0.064202149 | 0.123354405 | 0.218818897 | 0.230179127 |
| Agtrap | 0.017304752 | 0.01227881 | 0.03085412 | 0.163656932 | 0.143130871 |
| Ahctf1 | 0.010557875 | 0.137977517 | 0.229341326 | 0.161001651 | 0.151348605 |
| Ahcyl1 | 0.060799324 | 1.24E-09 | 1.39E-08 | 0.643211428 | 0.565812947 |

| | | | | | |
|---|---|---|---|---|---|
| Ahcyl2 | 0.014922959 | 0.097803255 | 0.173290185 | 0.328993982 | 0.311732138 |
| Ahdc1 | 0.012591898 | 0.034772162 | 0.073827246 | 0.113173095 | 0.096212624 |
| Ahi1 | -0.024029351 | 0.030446597 | 0.066175233 | 0.727089254 | 0.817528836 |
| Ahsa1 | -0.01300698 | 0.153785819 | 0.249059758 | 0.35297024 | 0.344525272 |
| Ahsa2 | -0.005742368 | 0.371156464 | 0.493622866 | 0.12167307 | 0.123925318 |
| AI413582 | 0.02517933 | 0.006835756 | 0.01887453 | 0.350721838 | 0.334581256 |
| AI480526 | 0.002155805 | 0.694248224 | 0.77857668 | 0.091067237 | 0.081139184 |
| AI506816 | -0.008056568 | 0.072247407 | 0.135674964 | 0.04715747 | 0.054937587 |
| AI597479 | 0.013374461 | 0.00236853 | 0.007572131 | 0.061346486 | 0.048068173 |
| AI661453 | -0.004764293 | 0.372369206 | 0.494559211 | 0.076129646 | 0.078959731 |
| AI837181 | 0.005594298 | 0.382944978 | 0.503992294 | 0.133465198 | 0.122967121 |
| AI987944 | 0.015293989 | 0.00748247 | 0.020336756 | 0.104715424 | 0.087818569 |
| Aida | 0.013631481 | 0.122466305 | 0.207677903 | 0.299643483 | 0.281957657 |
| Aifm1 | 0.013769658 | 0.019659942 | 0.045743386 | 0.107894954 | 0.096706619 |
| Aifm2 | -0.005178195 | 0.251869657 | 0.366842383 | 0.053244278 | 0.056123293 |
| Aig1 | -0.061123698 | 1.58E-09 | 1.75E-08 | 0.538034207 | 0.589687208 |
| Aimp1 | -0.002216261 | 0.825471372 | 0.879207675 | 0.575473714 | 0.574544829 |
| Aimp2 | -0.02618183 | 8.11E-06 | 4.76E-05 | 0.087759783 | 0.10460882 |
| Aip | -0.014648226 | 0.068273439 | 0.129817011 | 0.232640806 | 0.232621433 |
| Airn | -0.009515129 | 0.125403326 | 0.211802012 | 0.106317919 | 0.112437621 |
| Ak1 | 0.002187432 | 0.721803092 | 0.799609122 | 0.121410919 | 0.107232207 |
| Ak2 | 0.008019326 | 0.360021379 | 0.482842853 | 0.307654777 | 0.289898322 |
| Ak3 | -0.004039697 | 0.500970163 | 0.615264265 | 0.103948607 | 0.10606015 |
| Ak5 | -0.0024914 | 0.537356019 | 0.648041033 | 0.043270481 | 0.042382412 |
| Ak6 | -0.015759377 | 0.038384694 | 0.080012006 | 0.164049181 | 0.181765553 |
| Akain1 | -0.015387512 | 0.085113375 | 0.155102574 | 0.287239796 | 0.306080102 |
| Akap1 | 0.002095038 | 0.644299502 | 0.738229789 | 0.054632239 | 0.054288951 |
| Akap10 | 0.003789008 | 0.391375659 | 0.512075844 | 0.05325058 | 0.051580077 |
| Akap11 | 0.028671472 | 0.004600451 | 0.013439178 | 0.515299347 | 0.486634804 |
| Akap13 | -0.001155616 | 0.917056914 | 0.942575366 | 0.650769739 | 0.639724301 |
| Akap17b | -0.007048082 | 0.122751362 | 0.20808361 | 0.051705301 | 0.054805779 |
| Akap6 | 0.014296539 | 0.00656028 | 0.018233191 | 0.07810527 | 0.068954838 |
| Akap7 | 0.028011665 | 6.71E-07 | 4.88E-06 | 0.106754281 | 0.080585761 |
| Akap8 | -0.004020966 | 0.567561267 | 0.675046567 | 0.14867843 | 0.15369755 |
| Akap8l | -0.010564388 | 0.258635238 | 0.374810009 | 0.331048494 | 0.338049285 |
| Akap9 | 0.034549526 | 0.000458589 | 0.001798086 | 1.06413289 | 1.02642607 |
| Akip1 | -0.013205195 | 0.011197096 | 0.028541732 | 0.070578378 | 0.07931688 |
| Akirin1 | -0.001895884 | 0.852497806 | 0.898680132 | 0.556860755 | 0.575927971 |
| Akirin2 | 0.030706521 | 0.000795784 | 0.002930536 | 0.374346207 | 0.33248852 |
| Akr1a1 | 0.007444513 | 0.43158105 | 0.550200757 | 0.919080829 | 0.864417727 |
| Akr1b10 | -0.008699602 | 0.109035765 | 0.189814646 | 0.079364062 | 0.083976735 |
| Akr1b3 | -0.01219219 | 0.174472053 | 0.275181261 | 0.316737854 | 0.319520234 |
| Akr1c12 | -0.001202596 | 0.882850492 | 0.919773195 | 0.262981455 | 0.234022018 |
| Akr1c13 | 0.0269421 | 0.001307576 | 0.004517503 | 0.280334429 | 0.231902685 |
| Akr1c14 | -0.017894904 | 0.01508763 | 0.036698007 | 0.154497403 | 0.16272304 |
| Akr1c19 | -0.025298073 | 0.00346772 | 0.010532074 | 0.272148594 | 0.258396249 |
| Akr1e1 | -0.028005769 | 8.13E-05 | 0.000386941 | 0.153975061 | 0.167123336 |
| Akr7a5 | -0.036548354 | 5.07E-06 | 3.11E-05 | 0.214926756 | 0.238004388 |
| Akt1 | 0.033361164 | 0.000148396 | 0.000662095 | 0.348064669 | 0.296261025 |
| Akt1s1 | 0.005124763 | 0.536819532 | 0.647593931 | 0.25155749 | 0.242206097 |
| Akt2 | 0.021526625 | 0.016123053 | 0.038867743 | 0.304268419 | 0.287749103 |
| Akt3 | 0.027315039 | 0.00027695 | 0.001147211 | 0.196419652 | 0.170904249 |
| Aktip | -0.004946691 | 0.565067376 | 0.672656426 | 0.293910264 | 0.281796408 |
| Alad | -0.000209712 | 0.975267733 | 0.98346156 | 0.149628547 | 0.142658933 |
| Alas1 | 0.006296952 | 0.268695759 | 0.385345946 | 0.096991423 | 0.090588062 |
| Alcam | -0.064325509 | 5.07E-10 | 6.00E-09 | 0.374316458 | 0.468503406 |
| Aldh18a1 | 0.008670502 | 0.34263176 | 0.464778285 | 0.341225371 | 0.324532603 |
| Aldh1l2 | -0.0551601 | 9.38E-08 | 7.99E-07 | 0.379947074 | 0.430509842 |
| Aldh2 | -0.012111817 | 0.07099833 | 0.133820427 | 0.138047625 | 0.140244922 |
| Aldh3a2 | 0.002508434 | 0.725557323 | 0.802122845 | 0.159977577 | 0.157228883 |
| Aldh5a1 | -0.054876747 | 2.87E-10 | 3.50E-09 | 0.220359659 | 0.24776352 |
| Aldh6a1 | -0.013512878 | 0.008867798 | 0.023478413 | 0.068530168 | 0.076066855 |
| Aldh7a1 | -0.015017268 | 0.018320178 | 0.043219793 | 0.123294362 | 0.126847521 |
| Aldh9a1 | -0.029472411 | 6.51E-05 | 0.000314668 | 0.170846474 | 0.179316659 |
| Aldoa | -0.063706614 | 3.41E-16 | 7.20E-15 | 1.648763777 | 1.712213469 |
| Aldoc | 0.007257907 | 0.082576912 | 0.151217317 | 0.054788956 | 0.043834348 |
| Alg1 | -0.00994952 | 0.140396555 | 0.232487703 | 0.142829314 | 0.144330845 |
| Alg10b | 0.009684823 | 0.162955068 | 0.260824691 | 0.146415961 | 0.143806811 |
| Alg11 | -0.005242618 | 0.380026611 | 0.501305117 | 0.099231627 | 0.103333416 |
| Alg12 | -0.002523189 | 0.572346746 | 0.679242562 | 0.052517355 | 0.054455988 |
| Alg13 | 0.000642076 | 0.862391898 | 0.905809641 | 0.036116835 | 0.036389535 |
| Alg14 | -0.005188182 | 0.368166982 | 0.49085565 | 0.095963988 | 0.095638766 |

| | | | | | |
|---|---|---|---|---|---|
| Alg2 | -0.03590131 | 1.54E-05 | 8.52E-05 | 0.236416352 | 0.257458251 |
| Alg3 | -0.019530409 | 0.001556649 | 0.005238402 | 0.100723058 | 0.118719152 |
| Alg5 | -0.003092839 | 0.70995037 | 0.791075931 | 0.235122933 | 0.242486966 |
| Alg6 | -0.007320759 | 0.12166122 | 0.206637034 | 0.060092658 | 0.061904191 |
| Alg9 | 0.006368935 | 0.149499867 | 0.244234197 | 0.055228034 | 0.050535616 |
| Alkbh1 | 0.00692685 | 0.431350014 | 0.550122634 | 0.260470534 | 0.275016399 |
| Alkbh2 | -0.025431806 | 5.09E-06 | 3.12E-05 | 0.075264309 | 0.093255826 |
| Alkbh3 | 0.00099418 | 0.868939179 | 0.910400898 | 0.108629124 | 0.105118768 |
| Alkbh5 | 0.059799051 | 3.76E-09 | 3.97E-08 | 0.669608192 | 0.595568218 |
| Alkbh6 | -0.051982891 | 7.39E-10 | 8.58E-09 | 0.2545979 | 0.289131912 |
| Alkbh7 | -0.047027342 | 1.01E-10 | 1.29E-09 | 0.151904262 | 0.183255252 |
| Alkbh8 | 0.010527887 | 0.077999617 | 0.14449548 | 0.10411256 | 0.096658922 |
| Alms1 | -0.003964817 | 0.335142498 | 0.458006096 | 0.042196213 | 0.044413564 |
| Alox5ap | -0.014501244 | 0.016693733 | 0.040045141 | 0.093625678 | 0.103653034 |
| Alyref | 0.013013828 | 0.160329219 | 0.25741849 | 0.380083824 | 0.353739521 |
| Alyref2 | -0.011154506 | 0.042036582 | 0.086383355 | 0.084495487 | 0.08885249 |
| Amacr | -0.013238932 | 0.008820482 | 0.023365865 | 0.065951626 | 0.07467661 |
| Ambp | -0.019674929 | 0.00020045 | 0.000860005 | 0.05422915 | 0.070772251 |
| Ambra1 | 0.007198465 | 0.163395406 | 0.261185487 | 0.079043692 | 0.070999883 |
| Amd1 | -0.007608654 | 0.224903227 | 0.336748679 | 0.119229603 | 0.116786512 |
| Amd2 | 0.007495305 | 0.116881081 | 0.200091191 | 0.071298679 | 0.06077027 |
| Amdhd2 | 0.001553385 | 0.781198514 | 0.846218603 | 0.095761912 | 0.089539288 |
| Amer1 | 0.010426206 | 0.005264951 | 0.015076834 | 0.04199706 | 0.033394575 |
| Amer2 | 0.000485492 | 0.947234076 | 0.964799339 | 0.17349634 | 0.16774884 |
| Amfr | 0.041201207 | 1.10E-05 | 6.24E-05 | 0.414002142 | 0.369009031 |
| Amigo1 | 0.009892095 | 0.16676087 | 0.265169968 | 0.175619566 | 0.159183334 |
| Amigo2 | 0.059720572 | 3.25E-19 | 8.81E-18 | 0.162080561 | 0.107519548 |
| Ammecr1l | -0.006374776 | 0.215494849 | 0.325607862 | 0.069983512 | 0.074909865 |
| Amn1 | -0.012670447 | 0.086033397 | 0.156398034 | 0.163671156 | 0.175495087 |
| Amotl1 | -0.011614536 | 0.088666981 | 0.160226837 | 0.129153387 | 0.136691952 |
| Amotl2 | 0.006652931 | 0.10398204 | 0.18244241 | 0.049570519 | 0.040859037 |
| Ampd2 | 0.025360048 | 0.002551493 | 0.00806949 | 0.280219471 | 0.246517902 |
| Amph | 0.01735034 | 0.000383221 | 0.001528168 | 0.073556327 | 0.060433955 |
| Amz2 | -0.004780814 | 0.56859833 | 0.675894928 | 0.26441718 | 0.26066569 |
| Anapc1 | 0.005711942 | 0.436559971 | 0.555019752 | 0.170374722 | 0.163069923 |
| Anapc10 | -0.006662926 | 0.218035138 | 0.328590546 | 0.084225739 | 0.085854502 |
| Anapc11 | 0.02732464 | 0.003883036 | 0.011622807 | 0.468047352 | 0.424449428 |
| Anapc13 | 0.125189656 | 2.28E-37 | 1.64E-35 | 0.899689169 | 0.773590858 |
| Anapc15 | 0.002871038 | 0.601106249 | 0.70404388 | 0.087392885 | 0.084246264 |
| Anapc16 | -0.045299995 | 7.32E-06 | 4.34E-05 | 0.566388661 | 0.603838405 |
| Anapc2 | 0.00390417 | 0.554971024 | 0.66443127 | 0.140859614 | 0.133155311 |
| Anapc4 | -0.007172454 | 0.380502482 | 0.501660581 | 0.232503014 | 0.233026163 |
| Anapc5 | 0.017247324 | 0.079895887 | 0.147306621 | 0.597344355 | 0.536410599 |
| Anapc7 | -0.008980773 | 0.179212146 | 0.281279261 | 0.131990923 | 0.13782479 |
| Ang | -0.020519075 | 0.058735458 | 0.114548871 | 0.566995873 | 0.557895684 |
| Angel1 | 0.011760594 | 0.006881708 | 0.018969085 | 0.054207038 | 0.04853719 |
| Angel2 | 0.014991393 | 0.08146753 | 0.149495531 | 0.279259029 | 0.260817397 |
| Angptl6 | -0.189413913 | 2.91E-68 | 7.65E-66 | 0.285188963 | 0.477963171 |
| Ank | 0.056494122 | 1.97E-07 | 1.58E-06 | 0.688665894 | 0.619610073 |
| Ank2 | 0.032725444 | 0.001744813 | 0.005787469 | 0.536756069 | 0.503545552 |
| Ank3 | 0.040456688 | 7.37E-07 | 5.30E-06 | 0.244236339 | 0.205489651 |
| Ankfy1 | 0.026940895 | 1.32E-05 | 7.38E-05 | 0.127302934 | 0.103557852 |
| Ankhd1 | 0.013485798 | 0.142757231 | 0.235674275 | 0.339952692 | 0.323747316 |
| Ankib1 | 0.003941422 | 0.584703963 | 0.689948444 | 0.161109766 | 0.15798806 |
| Ankle2 | 0.003836967 | 0.593728091 | 0.697718421 | 0.16510148 | 0.160441122 |
| Ankmy2 | 0.007737954 | 0.242598862 | 0.356495965 | 0.14540704 | 0.131853083 |
| Ankra2 | 0.028290767 | 0.000915464 | 0.003313138 | 0.269204544 | 0.264729929 |
| Ankrd10 | -0.107528726 | 3.33E-22 | 1.13E-20 | 0.601341281 | 0.732205112 |
| Ankrd11 | 0.01177895 | 0.273792357 | 0.391500116 | 0.767678712 | 0.766307742 |
| Ankrd12 | 0.009377242 | 0.386050406 | 0.507186764 | 0.735085973 | 0.718717559 |
| Ankrd13a | 0.007882547 | 0.176709937 | 0.278124334 | 0.101627705 | 0.093148759 |
| Ankrd13c | 0.007194843 | 0.380395423 | 0.501655495 | 0.250451483 | 0.230101597 |
| Ankrd16 | -0.008530937 | 0.041939739 | 0.086257276 | 0.043371462 | 0.0491171 |
| Ankrd17 | 0.026044575 | 0.008059381 | 0.021699191 | 0.488014251 | 0.451838371 |
| Ankrd24 | 0.002433583 | 0.760997689 | 0.830536544 | 0.224913722 | 0.217505809 |
| Ankrd26 | 0.014369967 | 0.013539874 | 0.033486302 | 0.098791074 | 0.086687809 |
| Ankrd27 | 0.009416772 | 0.048540522 | 0.097623284 | 0.06989675 | 0.059437969 |
| Ankrd28 | 0.0047675 | 0.342313516 | 0.464670763 | 0.070776558 | 0.067691553 |
| Ankrd33b | 0.017278237 | 0.022084013 | 0.050494068 | 0.186050364 | 0.16437239 |
| Ankrd37 | 0.014054139 | 0.009467438 | 0.024769224 | 0.093587782 | 0.074984084 |
| Ankrd39 | -0.010243473 | 0.111100045 | 0.192568129 | 0.120726263 | 0.126297214 |
| Ankrd40 | 0.013001205 | 0.149642899 | 0.244426792 | 0.345996204 | 0.32268899 |

| | | | | | |
|---|---|---|---|---|---|
| Ankrd44 | 0.041920116 | 1.73E-07 | 1.40E-06 | 0.239202665 | 0.187706721 |
| Ankrd46 | 0.007486415 | 0.158902397 | 0.255939215 | 0.085738029 | 0.077527475 |
| Ankrd49 | 0.010823959 | 0.051309803 | 0.102064183 | 0.088599271 | 0.082742278 |
| Ankrd50 | 0.008193368 | 0.042081203 | 0.086416926 | 0.046838472 | 0.041054958 |
| Ankrd54 | -0.034016591 | 0.000129518 | 0.000588398 | 0.310391548 | 0.321765517 |
| Anks1 | 0.027143597 | 2.04E-05 | 0.00011038 | 0.13346808 | 0.109268123 |
| Anks3 | -0.010227693 | 0.069808678 | 0.132038904 | 0.08756675 | 0.091750334 |
| Anks4b | 0.016132872 | 0.035585756 | 0.075258804 | 0.18557291 | 0.168540366 |
| Ankzf1 | -0.010754468 | 0.024111485 | 0.054489041 | 0.057594527 | 0.06393516 |
| Ano10 | 0.009655577 | 0.050305151 | 0.100548248 | 0.071772028 | 0.063619749 |
| Ano6 | 0.044177428 | 6.85E-12 | 9.94E-11 | 0.145698835 | 0.108913185 |
| Ano8 | 0.010355798 | 0.022301124 | 0.050870842 | 0.0655505 | 0.053562496 |
| Anp32a | 0.001770429 | 0.857219067 | 0.902033096 | 0.709342708 | 0.667427272 |
| Anp32b | -0.011446015 | 0.245442542 | 0.359366325 | 0.939322493 | 0.937928709 |
| Anp32e | -0.049314252 | 4.00E-07 | 3.04E-06 | 0.475617239 | 0.509084744 |
| Anpep | 0.020425401 | 0.001677932 | 0.005590424 | 0.130385675 | 0.10329034 |
| Antkmt | -0.006471252 | 0.242102306 | 0.356030649 | 0.085320681 | 0.08964156 |
| Anxa11 | 0.028332211 | 0.00025866 | 0.001079267 | 0.223462982 | 0.18744788 |
| Anxa4 | 0.030409781 | 0.001741373 | 0.005780003 | 0.446135504 | 0.4256943 |
| Anxa5 | -0.005125008 | 0.562314678 | 0.670446812 | 1.33296035 | 1.285108931 |
| Anxa6 | 0.008586995 | 0.307759213 | 0.428842636 | 0.266286471 | 0.252521362 |
| Anxa7 | -0.00442247 | 0.622901052 | 0.72170707 | 0.344034739 | 0.333950412 |
| Aopep | 0.009050833 | 0.379142792 | 0.500706269 | 0.530107541 | 0.516187653 |
| Aox1 | -0.010759018 | 0.012378699 | 0.031031492 | 0.042723825 | 0.050403637 |
| Ap1ar | -0.011562988 | 0.099278366 | 0.175679701 | 0.14289869 | 0.154849043 |
| Ap1b1 | 0.05347346 | 2.09E-11 | 2.89E-10 | 0.256389302 | 0.205235784 |
| Ap1g1 | 0.032992908 | 2.36E-05 | 0.000125441 | 0.215230938 | 0.189586982 |
| Ap1m1 | 0.005588926 | 0.336443535 | 0.459074252 | 0.104047601 | 0.095994215 |
| Ap1m2 | -0.008721407 | 0.202012521 | 0.309480331 | 0.144080967 | 0.149489237 |
| Ap1s1 | 0.026529836 | 0.003983701 | 0.011876586 | 0.382727848 | 0.347311621 |
| Ap1s2 | 0.057124192 | 8.62E-08 | 7.37E-07 | 0.663031075 | 0.562525197 |
| Ap1s3 | 0.004768863 | 0.205680854 | 0.313913541 | 0.042345632 | 0.035897535 |
| Ap2a1 | 0.022408025 | 0.008647933 | 0.023015387 | 0.290867289 | 0.267235194 |
| Ap2a2 | 0.054284257 | 1.65E-09 | 1.82E-08 | 0.355053619 | 0.302776993 |
| Ap2b1 | 0.026780116 | 0.002504899 | 0.007947987 | 0.321715761 | 0.294970365 |
| Ap2m1 | -0.021159973 | 0.032398467 | 0.069608375 | 0.55069215 | 0.555170825 |
| Ap2s1 | 0.009343097 | 0.335430504 | 0.458270848 | 0.536543118 | 0.526536188 |
| Ap3b1 | 0.004369066 | 0.595807466 | 0.699570071 | 0.25000755 | 0.23743199 |
| Ap3b2 | 0.025843713 | 0.008986687 | 0.023722122 | 0.531811824 | 0.487716281 |
| Ap3d1 | 0.037084049 | 0.000122754 | 0.000561079 | 0.433653323 | 0.392377479 |
| Ap3m1 | 0.007309466 | 0.343334735 | 0.465472081 | 0.204372116 | 0.192981505 |
| Ap3m2 | -0.008784995 | 0.107484842 | 0.187606398 | 0.079455831 | 0.084173727 |
| Ap3s1 | 0.014088828 | 0.140570092 | 0.232695801 | 0.45718901 | 0.434945442 |
| Ap3s2 | -0.00098199 | 0.891155586 | 0.926115939 | 0.170414257 | 0.164603471 |
| Ap4b1 | 0.003501101 | 0.367416468 | 0.490101161 | 0.047023067 | 0.038999872 |
| Ap4m1 | -0.014579837 | 0.001036564 | 0.00369409 | 0.047796208 | 0.056827439 |
| Ap4s1 | -0.02489927 | 0.001091489 | 0.003861518 | 0.177581443 | 0.193191131 |
| Ap5m1 | 0.004685789 | 0.451098651 | 0.569408087 | 0.12015561 | 0.112246297 |
| Ap5s1 | -0.000633901 | 0.893892731 | 0.927936305 | 0.066271296 | 0.062545905 |
| Apba1 | -0.002066578 | 0.671545846 | 0.760397009 | 0.062134772 | 0.066225836 |
| Apba3 | 0.003795031 | 0.393205062 | 0.513809088 | 0.055557831 | 0.052553552 |
| Apbb1 | -0.023348543 | 0.002642229 | 0.008318573 | 0.186232475 | 0.201899007 |
| Apbb2 | 0.008006136 | 0.258453561 | 0.374748028 | 0.162008991 | 0.151648518 |
| Apbb3 | 0.010966405 | 0.009527246 | 0.024872149 | 0.055447201 | 0.044411823 |
| Apc | 0.040008991 | 6.51E-06 | 3.90E-05 | 0.308486986 | 0.27423537 |
| Apeh | 0.009774971 | 0.12022263 | 0.20465845 | 0.130232027 | 0.116172953 |
| Apex1 | 0.002208475 | 0.706622088 | 0.788351314 | 0.100561001 | 0.094482985 |
| Aph1a | -0.008933197 | 0.279933544 | 0.398815052 | 0.250317876 | 0.248515684 |
| Aph1b | 0.005346549 | 0.249056088 | 0.363452911 | 0.062605508 | 0.056409563 |
| Api5 | 0.011656364 | 0.197882144 | 0.304494422 | 0.330047289 | 0.318987431 |
| Apip | -0.075534313 | 2.17E-22 | 7.45E-21 | 0.163345857 | 0.223183181 |
| Aplp1 | -0.078383055 | 1.11E-16 | 2.43E-15 | 1.001137641 | 1.058436225 |
| Aplp2 | 0.00883892 | 0.314106172 | 0.435564143 | 0.287943768 | 0.279154207 |
| Apmap | -0.041362945 | 9.87E-10 | 1.13E-08 | 0.115128461 | 0.150068702 |
| Apoa2 | -0.004217804 | 0.314691564 | 0.436127069 | 0.044864716 | 0.047728214 |
| Apobec1 | 0.030286343 | 9.70E-05 | 0.000453994 | 0.213965974 | 0.180077449 |
| Apobec3 | 0.011023509 | 0.024848801 | 0.055961609 | 0.068255009 | 0.060581351 |
| Apoo | -0.011250736 | 0.131181176 | 0.220228526 | 0.188929104 | 0.186535633 |
| Apool | 0.007160272 | 0.201974138 | 0.309480331 | 0.100217805 | 0.0877806 |
| App | -0.099186573 | 3.08E-20 | 8.99E-19 | 0.485523693 | 0.604345901 |
| Appbp2 | 0.027578818 | 0.002479861 | 0.007883979 | 0.364992999 | 0.333845033 |
| Appl1 | -0.000411889 | 0.963379639 | 0.974810247 | 0.313788254 | 0.31662625 |

| | | | | | |
|---|---|---|---|---|---|
| Appl2 | 0.011531438 | 0.075932357 | 0.141637448 | 0.13588306 | 0.122132867 |
| Aprt | -0.001972607 | 0.841956475 | 0.891958249 | 0.470170959 | 0.473695871 |
| Aptx | 0.000283631 | 0.964295157 | 0.975376522 | 0.123640171 | 0.119277333 |
| Aqp11 | 0.035441799 | 8.71E-08 | 7.44E-07 | 0.170097893 | 0.120412281 |
| Aqp4 | 0.052094537 | 6.79E-22 | 2.23E-20 | 0.105546181 | 0.051917207 |
| Aqr | 0.026880352 | 0.000893265 | 0.003247688 | 0.238152244 | 0.210626272 |
| Araf | -0.040789577 | 3.30E-05 | 0.000169722 | 0.502911196 | 0.545027077 |
| Arap1 | 0.012176737 | 0.022214816 | 0.050745404 | 0.08714312 | 0.0753104 |
| Arc | 0.002245951 | 0.616933635 | 0.7170148 | 0.048949166 | 0.047991775 |
| Arcn1 | -0.007248436 | 0.479462796 | 0.596002723 | 0.680289447 | 0.673881173 |
| Arel1 | 0.003509323 | 0.568622158 | 0.675894928 | 0.10868203 | 0.106777143 |
| Arf1 | -0.009909481 | 0.256538578 | 0.372360115 | 1.248330705 | 1.252926764 |
| Arf2 | -0.002005518 | 0.735663879 | 0.810530852 | 0.101468597 | 0.10247835 |
| Arf3 | 0.035517249 | 0.000272967 | 0.001132161 | 0.565706859 | 0.518344167 |
| Arf4 | -0.03557895 | 0.000544434 | 0.002092918 | 0.794924767 | 0.839969232 |
| Arf5 | 0.003107347 | 0.703895308 | 0.786130981 | 1.359610995 | 1.356285647 |
| Arf6 | 0.006634819 | 0.359139447 | 0.481859467 | 0.165252456 | 0.156953326 |
| Arfgap1 | 0.005533984 | 0.509576717 | 0.623471538 | 0.262925027 | 0.250818244 |
| Arfgap2 | 0.011598521 | 0.167212505 | 0.265704662 | 0.264164799 | 0.249867467 |
| Arfgap3 | -0.010144248 | 0.280389085 | 0.399230036 | 0.36378603 | 0.363303393 |
| Arfgef1 | 0.032577198 | 0.001479279 | 0.005023597 | 0.737514359 | 0.693014362 |
| Arfgef2 | 0.033185199 | 2.52E-05 | 0.000132882 | 0.224637326 | 0.19135303 |
| Arfgef3 | 0.003880447 | 0.692808143 | 0.777410061 | 0.921011679 | 0.933050015 |
| Arfip1 | 0.008210952 | 0.341133758 | 0.463498867 | 0.289369897 | 0.276821082 |
| Arfip2 | -0.020947099 | 0.019370555 | 0.045160801 | 0.314018982 | 0.320695537 |
| Arfrp1 | -0.020709126 | 0.018004567 | 0.04265117 | 0.271107624 | 0.29144878 |
| Arglu1 | -0.051739983 | 1.18E-06 | 8.17E-06 | 0.683341221 | 0.715944531 |
| Arhgap1 | -0.009264007 | 0.220455585 | 0.331416298 | 0.172183181 | 0.185959531 |
| Arhgap12 | 0.012168976 | 0.091934716 | 0.164784164 | 0.171488102 | 0.161329492 |
| Arhgap17 | -0.003220252 | 0.60542239 | 0.707323309 | 0.108113718 | 0.111820359 |
| Arhgap18 | -0.005291779 | 0.248514495 | 0.362939034 | 0.055739616 | 0.057877491 |
| Arhgap21 | 0.011117332 | 0.130468739 | 0.219070307 | 0.172761238 | 0.165806023 |
| Arhgap23 | -0.001747361 | 0.659668187 | 0.750967433 | 0.041320154 | 0.041841093 |
| Arhgap24 | 0.022337044 | 0.005841825 | 0.016466867 | 0.233373851 | 0.210378036 |
| Arhgap26 | 0.020235243 | 4.57E-05 | 0.000226421 | 0.078248208 | 0.061988986 |
| Arhgap32 | 0.002620427 | 0.710317273 | 0.791303372 | 0.155079501 | 0.150141332 |
| Arhgap35 | 0.031077171 | 3.93E-05 | 0.000197637 | 0.206937141 | 0.173711573 |
| Arhgap39 | -0.012300757 | 0.002282642 | 0.007336146 | 0.037824438 | 0.046023012 |
| Arhgap44 | -0.000969582 | 0.870411146 | 0.91098473 | 0.100566656 | 0.102670559 |
| Arhgap5 | 0.036180811 | 0.000221711 | 0.000940026 | 1.126006395 | 1.055845034 |
| Arhgap6 | 0.026438555 | 1.21E-08 | 1.19E-07 | 0.066660423 | 0.048268465 |
| Arhgdia | -0.006283522 | 0.529108456 | 0.640517048 | 0.634395621 | 0.619049525 |
| Arhgdig | -0.083037276 | 2.95E-18 | 7.40E-17 | 0.329825209 | 0.409972209 |
| Arhgef1 | 0.032759184 | 3.02E-06 | 1.94E-05 | 0.171776404 | 0.14211095 |
| Arhgef11 | -0.002253951 | 0.713080911 | 0.793745434 | 0.107923083 | 0.108989365 |
| Arhgef12 | 0.050220975 | 1.32E-06 | 9.00E-06 | 0.747452989 | 0.701242414 |
| Arhgef16 | 0.03958973 | 6.21E-07 | 4.55E-06 | 0.242688963 | 0.200379061 |
| Arhgef17 | 0.054766463 | 3.38E-13 | 5.77E-12 | 0.209597194 | 0.161738088 |
| Arhgef18 | 0.019620997 | 0.000310052 | 0.001267721 | 0.093900517 | 0.077264659 |
| Arhgef2 | -0.021918189 | 0.024795108 | 0.055878081 | 0.298287889 | 0.326988883 |
| Arhgef25 | -0.012518062 | 0.001356793 | 0.004667638 | 0.033874453 | 0.042884433 |
| Arhgef28 | 0.015373944 | 0.010165045 | 0.026248225 | 0.114389026 | 0.097689864 |
| Arhgef3 | 0.006368781 | 0.08019611 | 0.147617671 | 0.039661084 | 0.032438877 |
| Arhgef5 | 0.0301067 | 0.000101856 | 0.000475165 | 0.203973306 | 0.18482718 |
| Arhgef7 | 0.002067762 | 0.729535441 | 0.805147413 | 0.102365903 | 0.102749232 |
| Arhgef9 | -0.004071456 | 0.38072555 | 0.501886616 | 0.061003507 | 0.058395476 |
| Arid1a | 0.057730183 | 2.79E-09 | 3.00E-08 | 0.500301132 | 0.432807645 |
| Arid1b | 0.035172146 | 9.71E-06 | 5.59E-05 | 0.237484777 | 0.20166674 |
| Arid2 | 0.024147024 | 0.002984665 | 0.009258746 | 0.247868938 | 0.217887491 |
| Arid3a | -0.017126713 | 0.003408292 | 0.01040032 | 0.083564408 | 0.098153022 |
| Arid3b | 0.008658878 | 0.046496681 | 0.094018454 | 0.054274711 | 0.047439974 |
| Arid4a | 0.019889663 | 0.010239325 | 0.026404964 | 0.204959401 | 0.181819375 |
| Arid4b | 0.00117381 | 0.907080397 | 0.935957559 | 0.472309619 | 0.468358808 |
| Arid5a | 0.006702668 | 0.132507586 | 0.22211013 | 0.046127499 | 0.048467877 |
| Arid5b | 0.055097108 | 1.80E-13 | 3.16E-12 | 0.186941026 | 0.14144157 |
| Arih1 | 0.037159469 | 1.61E-05 | 8.87E-05 | 0.296052178 | 0.257946951 |
| Arih2 | 0.002679546 | 0.720739213 | 0.798794646 | 0.178968342 | 0.177142959 |
| Arl1 | -0.051083735 | 9.81E-08 | 8.30E-07 | 0.948420558 | 1.000620531 |
| Arl13b | 0.011888073 | 0.061959349 | 0.119922116 | 0.126396735 | 0.116702884 |
| Arl14ep | -0.012901559 | 0.085499496 | 0.155631174 | 0.169726113 | 0.178601886 |
| Arl15 | 0.004499355 | 0.213683602 | 0.323508882 | 0.037478151 | 0.033127894 |
| Arl16 | -0.003476794 | 0.542212562 | 0.652491762 | 0.097038848 | 0.093965059 |

| | | | | | |
|---|---|---|---|---|---|
| Arl2 | -0.002261906 | 0.810178728 | 0.86908944 | 0.399958617 | 0.398603843 |
| Arl2bp | -0.006881158 | 0.442419158 | 0.561001573 | 0.335219133 | 0.331551235 |
| Arl3 | -0.010224918 | 0.292990158 | 0.412400509 | 0.833228571 | 0.834387482 |
| Arl4a | -0.025253372 | 0.013910879 | 0.034255717 | 0.435059042 | 0.489608022 |
| Arl5a | 0.03427133 | 6.34E-05 | 0.000306851 | 0.277829717 | 0.250742162 |
| Arl5b | 0.003405523 | 0.475482656 | 0.592342694 | 0.060253444 | 0.060604039 |
| Arl6 | -0.050041665 | 1.20E-10 | 1.53E-09 | 0.195451712 | 0.22466054 |
| Arl6ip1 | -0.05990241 | 2.54E-09 | 2.74E-08 | 0.628025262 | 0.686766267 |
| Arl6ip4 | -0.012558061 | 0.168215492 | 0.266915459 | 0.323820147 | 0.334813758 |
| Arl6ip5 | -0.005888201 | 0.530013059 | 0.641532241 | 0.383852653 | 0.37242285 |
| Arl6ip6 | 0.006508297 | 0.307976042 | 0.429083273 | 0.126990294 | 0.118508528 |
| Arl8a | 0.008247837 | 0.334753673 | 0.45757468 | 0.284000529 | 0.271668622 |
| Arl8b | 0.049508167 | 9.17E-07 | 6.46E-06 | 0.58080553 | 0.536254519 |
| Armc1 | 0.028014943 | 0.001020955 | 0.003645144 | 0.305044943 | 0.266505969 |
| Armc10 | 0.015783577 | 0.089177973 | 0.160929011 | 0.341639578 | 0.347461365 |
| Armc5 | 0.03673228 | 2.06E-07 | 1.65E-06 | 0.186724553 | 0.145306463 |
| Armc8 | 0.024479108 | 0.001349489 | 0.004649091 | 0.207321763 | 0.183593446 |
| Armc9 | -0.004326167 | 0.40153648 | 0.521611125 | 0.075953126 | 0.075386886 |
| Armcx1 | 0.007735067 | 0.272133238 | 0.389299491 | 0.158319478 | 0.151082937 |
| Armcx2 | 0.007401072 | 0.333987994 | 0.456877905 | 0.201336521 | 0.191886361 |
| Armcx3 | -0.045483079 | 1.68E-06 | 1.12E-05 | 0.363105364 | 0.40731187 |
| Armcx4 | 0.000252545 | 0.967201139 | 0.977761201 | 0.104759971 | 0.105869179 |
| Armcx5 | -0.010767393 | 0.133787638 | 0.224062604 | 0.148888244 | 0.15838445 |
| Armcx6 | -0.001449958 | 0.758063789 | 0.827985188 | 0.058675327 | 0.059794873 |
| Armh3 | 0.009741453 | 0.017182476 | 0.041023582 | 0.050972866 | 0.041852791 |
| Armt1 | -0.013147656 | 0.008910916 | 0.0235669 | 0.063023755 | 0.072137487 |
| Arnt | 0.024274276 | 0.0017192 | 0.005710304 | 0.198858212 | 0.184367525 |
| Arnt2 | 0.001686497 | 0.696473955 | 0.780172807 | 0.047966736 | 0.049182837 |
| Arntl | -0.021279527 | 9.21E-06 | 5.33E-05 | 0.048025137 | 0.067272106 |
| Arpc1a | 0.03509725 | 0.000273607 | 0.001134332 | 0.946388723 | 0.852989797 |
| Arpc1b | 0.002442921 | 0.763877032 | 0.832930967 | 0.231875713 | 0.224786495 |
| Arpc2 | 0.007477938 | 0.392598217 | 0.513292074 | 1.208851656 | 1.173147905 |
| Arpc3 | -0.050101208 | 2.27E-07 | 1.80E-06 | 0.833443667 | 0.872584399 |
| Arpc4 | -0.004315982 | 0.565216442 | 0.672751499 | 0.185861431 | 0.184459683 |
| Arpc5 | -0.008176431 | 0.409782509 | 0.529699889 | 0.764457106 | 0.768422191 |
| Arpc5l | -0.115731047 | 4.30E-25 | 1.74E-23 | 1.198409847 | 1.416330182 |
| Arpin | 0.016246568 | 0.004202288 | 0.012421599 | 0.104274916 | 0.088717823 |
| Arpp19 | 0.084021875 | 6.80E-18 | 1.65E-16 | 0.88236654 | 0.835197593 |
| Arrb2 | 0.007953063 | 0.165096487 | 0.263242628 | 0.103746149 | 0.090049295 |
| Arrdc1 | 0.014022388 | 0.008059948 | 0.021699191 | 0.088609523 | 0.076165322 |
| Arrdc3 | -0.006460849 | 0.246166585 | 0.360317907 | 0.080735762 | 0.08600923 |
| Arrdc4 | 0.00268612 | 0.722009504 | 0.799740259 | 0.166483001 | 0.157659228 |
| Arsa | 1.99E-05 | 0.996893909 | 0.997612206 | 0.073507879 | 0.072497498 |
| Arsb | -0.002623627 | 0.741161355 | 0.814741146 | 0.223942814 | 0.211539945 |
| Arxes1 | -0.004864296 | 0.361548602 | 0.484423306 | 0.083438432 | 0.081495859 |
| Arxes2 | -0.025694145 | 0.000933128 | 0.003364938 | 0.194558709 | 0.210625667 |
| As3mt | -0.036783319 | 2.61E-05 | 0.000137094 | 0.29008872 | 0.306455215 |
| Asah1 | -0.008441586 | 0.407551874 | 0.527732994 | 0.674155242 | 0.670105329 |
| Asah2 | 0.026126674 | 0.000654657 | 0.00246307 | 0.201038672 | 0.179227975 |
| Asap1 | 0.024514227 | 0.0007751 | 0.002863039 | 0.174407499 | 0.152368486 |
| Asap2 | 0.005260033 | 0.165030617 | 0.263193514 | 0.040854815 | 0.035067876 |
| Asb1 | 0.008059578 | 0.116299383 | 0.199376229 | 0.081440141 | 0.071974587 |
| Asb11 | -0.028430723 | 1.57E-06 | 1.06E-05 | 0.069941947 | 0.096270965 |
| Asb13 | -0.003807171 | 0.554240971 | 0.663832786 | 0.123536481 | 0.124245377 |
| Asb3 | -0.01114907 | 0.025202618 | 0.056573505 | 0.068103878 | 0.072156195 |
| Asb6 | -0.003785344 | 0.393410056 | 0.513938803 | 0.054312614 | 0.054077964 |
| Asb7 | 0.025654928 | 8.91E-06 | 5.17E-05 | 0.107505022 | 0.084605617 |
| Asb8 | -0.002674777 | 0.687465801 | 0.773021341 | 0.138960117 | 0.134396826 |
| Ascc1 | -0.01412253 | 0.079774935 | 0.147139426 | 0.221918026 | 0.226868605 |
| Ascc2 | -0.028811919 | 9.94E-05 | 0.000464319 | 0.141129424 | 0.171859133 |
| Ascc3 | 0.091168731 | 6.45E-19 | 1.71E-17 | 0.594744815 | 0.521978726 |
| Asf1a | -0.022490284 | 0.001324696 | 0.004568533 | 0.149487454 | 0.158917272 |
| Ash1l | 0.029128861 | 0.004928738 | 0.014251264 | 0.786173571 | 0.752769279 |
| Ash2l | 0.006591106 | 0.389198109 | 0.510287797 | 0.196955147 | 0.187885679 |
| Asic1 | -0.000687729 | 0.899159094 | 0.931148776 | 0.079806611 | 0.079576041 |
| Asl | -0.009706159 | 0.087966238 | 0.159286229 | 0.091619252 | 0.095315616 |
| Asns | -0.06521892 | 3.32E-11 | 4.52E-10 | 1.214810793 | 1.295994001 |
| Asnsd1 | 0.009033074 | 0.176754964 | 0.278150172 | 0.146138817 | 0.134006677 |
| Asph | -0.014175849 | 0.031030806 | 0.067279548 | 0.120299524 | 0.13190239 |
| Asphd1 | -0.012258187 | 0.019003131 | 0.044506404 | 0.070445536 | 0.079121475 |
| Aspscr1 | -0.019362797 | 0.005313282 | 0.015182805 | 0.146492751 | 0.160181691 |
| Asrgl1 | -0.035672393 | 3.47E-06 | 2.19E-05 | 0.180000921 | 0.205175931 |

| | | | | | |
|---|---|---|---|---|---|
| Asxl1 | 0.009479568 | 0.076926021 | 0.143051794 | 0.085792492 | 0.074933388 |
| Asxl2 | 0.016890412 | 0.070157003 | 0.132594553 | 0.361159919 | 0.336788342 |
| Atad1 | 0.01431602 | 0.113404776 | 0.195419442 | 0.342639548 | 0.325272172 |
| Atad2 | 0.002377366 | 0.627485218 | 0.725634744 | 0.064575145 | 0.061843541 |
| Atad2b | 0.000361277 | 0.959246356 | 0.971967751 | 0.157713189 | 0.151334812 |
| Atad3a | -0.002777759 | 0.639686688 | 0.735284226 | 0.094095891 | 0.100898784 |
| Atat1 | -7.31E-05 | 0.988730825 | 0.993079296 | 0.074567768 | 0.073448469 |
| Ate1 | -0.003125924 | 0.746786374 | 0.819349637 | 0.414361046 | 0.423155691 |
| Atf1 | -0.019310777 | 0.024927502 | 0.056072461 | 0.25555606 | 0.274826641 |
| Atf2 | 0.054223565 | 1.13E-09 | 1.28E-08 | 0.350722087 | 0.298081554 |
| Atf3 | -0.032588055 | 6.43E-06 | 3.86E-05 | 0.081628509 | 0.138495473 |
| Atf4 | -0.049452211 | 1.32E-07 | 1.10E-06 | 1.197928193 | 1.263891695 |
| Atf5 | -0.172511089 | 6.90E-67 | 1.64E-64 | 1.491103177 | 1.681534409 |
| Atf6 | -0.044983394 | 7.49E-06 | 4.43E-05 | 0.420304542 | 0.474758185 |
| Atf6b | -0.017376737 | 0.005208137 | 0.014946138 | 0.105710698 | 0.118103643 |
| Atf7 | 0.005980783 | 0.339105258 | 0.461475548 | 0.118126655 | 0.110495976 |
| Atf7ip | -0.002140217 | 0.785378647 | 0.84989439 | 0.198817495 | 0.19907061 |
| Atg10 | -0.017775443 | 0.01380028 | 0.034026457 | 0.170140623 | 0.172662022 |
| Atg101 | -0.014443911 | 0.040507698 | 0.083857546 | 0.156640856 | 0.159066711 |
| Atg12 | -0.006563648 | 0.380455261 | 0.501660581 | 0.175829398 | 0.180302148 |
| Atg13 | -0.028008301 | 0.000337109 | 0.00136671 | 0.20000124 | 0.212828085 |
| Atg14 | 0.014591152 | 0.003937898 | 0.011765287 | 0.079682084 | 0.067634507 |
| Atg16l1 | 0.018503268 | 0.002502593 | 0.00794326 | 0.118237637 | 0.102229136 |
| Atg16l2 | 0.018011199 | 0.001046704 | 0.003726128 | 0.09924718 | 0.079991582 |
| Atg2a | -0.005557147 | 0.24007218 | 0.354062763 | 0.056732152 | 0.061961772 |
| Atg2b | 0.008078683 | 0.189159876 | 0.293350186 | 0.114695273 | 0.106948514 |
| Atg3 | -0.009747921 | 0.290640548 | 0.410102671 | 0.342207137 | 0.346632683 |
| Atg4a | 0.00173282 | 0.64972314 | 0.742955583 | 0.043981695 | 0.038703223 |
| Atg4b | 0.002396559 | 0.698869036 | 0.782224818 | 0.117741849 | 0.112558264 |
| Atg4c | 0.000888413 | 0.847906773 | 0.896211094 | 0.061896587 | 0.058082697 |
| Atg4d | -0.012186043 | 0.114317479 | 0.196637391 | 0.187247153 | 0.195595036 |
| Atg5 | -0.006522555 | 0.452505195 | 0.57096113 | 0.301367611 | 0.300890376 |
| Atg7 | -4.97E-05 | 0.991228116 | 0.994193721 | 0.05631957 | 0.055658494 |
| Atic | -0.030990983 | 7.46E-05 | 0.000357109 | 0.183154017 | 0.211088004 |
| Atl1 | -0.011220126 | 0.00987233 | 0.025594345 | 0.045111124 | 0.052056739 |
| Atl2 | 0.017628923 | 0.012331229 | 0.03096983 | 0.166282814 | 0.148009545 |
| Atl3 | 0.023458603 | 0.001354802 | 0.004662438 | 0.182513288 | 0.16503846 |
| Atm | 0.007974497 | 0.288238511 | 0.407659994 | 0.182650447 | 0.169176068 |
| Atmin | 0.001303078 | 0.823972953 | 0.878458718 | 0.100748118 | 0.097643016 |
| Atn1 | 0.017617418 | 0.013460781 | 0.033367086 | 0.170601238 | 0.152359673 |
| Atox1 | 0.037648021 | 0.000152985 | 0.000679762 | 0.954746992 | 0.946648634 |
| Atp11a | -0.005646282 | 0.362661714 | 0.485446397 | 0.102365436 | 0.113885337 |
| Atp11b | -0.001822266 | 0.835961661 | 0.887832561 | 0.273547475 | 0.285212516 |
| Atp13a1 | 0.008853171 | 0.215520119 | 0.325607862 | 0.166281854 | 0.157202734 |
| Atp13a2 | 0.005584754 | 0.389930706 | 0.511179385 | 0.13363615 | 0.12329666 |
| Atp13a3 | 0.048579575 | 2.27E-08 | 2.11E-07 | 0.288203407 | 0.256659333 |
| Atp1a1 | 0.130730842 | 9.96E-49 | 1.19E-46 | 1.575936906 | 1.428522721 |
| Atp1b1 | 0.104375782 | 3.52E-28 | 1.69E-26 | 1.378670932 | 1.243108599 |
| Atp1b2 | 0.006770349 | 0.177045065 | 0.278516524 | 0.075894381 | 0.065739776 |
| Atp1b3 | 0.025165646 | 0.004399182 | 0.012944565 | 0.313650564 | 0.295129157 |
| Atp23 | 0.001258537 | 0.74741909 | 0.81976629 | 0.043946878 | 0.041035637 |
| Atp2a2 | 0.084215891 | 9.45E-24 | 3.52E-22 | 2.174279887 | 1.999260542 |
| Atp2a3 | 0.048063297 | 3.51E-06 | 2.21E-05 | 0.791466609 | 0.714350017 |
| Atp2b1 | 0.04125379 | 4.06E-05 | 0.000203699 | 0.925023581 | 0.878996567 |
| Atp2b2 | 0.009214162 | 0.201470784 | 0.308991792 | 0.169361178 | 0.1558975 |
| Atp2b4 | 0.051318069 | 4.36E-18 | 1.08E-16 | 0.130367218 | 0.082231678 |
| Atp2c1 | 0.006836764 | 0.479886844 | 0.596301111 | 0.458872279 | 0.440976578 |
| Atp5a1 | 0.015206401 | 0.040289023 | 0.083480368 | 1.549645497 | 1.505086564 |
| Atp5b | 0.010237708 | 0.213410264 | 0.323173611 | 1.491459706 | 1.465234427 |
| Atp5c1 | -0.049527475 | 1.50E-08 | 1.44E-07 | 1.288035655 | 1.362442026 |
| Atp5d | 0.017871386 | 0.022418829 | 0.051055483 | 1.509467164 | 1.462110338 |
| Atp5e | 0.11611226 | 2.21E-103 | 1.53E-100 | 2.41289628 | 2.233275477 |
| Atp5g1 | -0.027713158 | 0.001134689 | 0.003991117 | 1.812000416 | 1.880421923 |
| Atp5g2 | 0.04075184 | 7.39E-08 | 6.39E-07 | 0.218100974 | 0.181641541 |
| Atp5g3 | 0.029433385 | 0.001583661 | 0.005310141 | 1.042204709 | 1.002850318 |
| Atp5h | -0.026252167 | 0.000240414 | 0.001010757 | 1.599088427 | 1.61199162 |
| Atp5j | 0.006568655 | 0.363899994 | 0.486705917 | 1.601124297 | 1.566580627 |
| Atp5j2 | 0.114515873 | 1.27E-60 | 2.37E-58 | 1.723815407 | 1.58581498 |
| Atp5k | 0.124696385 | 2.98E-38 | 2.28E-36 | 0.97864293 | 0.858431703 |
| Atp5l | 0.172684878 | 1.77E-72 | 5.51E-70 | 1.260143706 | 1.077923588 |
| Atp5md | 0.127590623 | 3.39E-55 | 5.15E-53 | 1.494470787 | 1.345241971 |
| Atp5mpl | 0.096964979 | 1.42E-23 | 5.26E-22 | 0.905822336 | 0.791294952 |

| | | | | | |
|---|---|---|---|---|---|
| Atp5o | -0.021630802 | 0.029164852 | 0.0639181 | 0.711424033 | 0.697618829 |
| Atp5pb | -0.056860796 | 2.00E-10 | 2.47E-09 | 1.258119955 | 1.33689852 |
| Atp6ap1 | 0.030080045 | 0.002101567 | 0.00680821 | 0.640800929 | 0.610955007 |
| Atp6ap2 | -0.040333966 | 6.06E-06 | 3.65E-05 | 1.262402463 | 1.317830476 |
| Atp6v0a1 | 0.047185262 | 2.28E-07 | 1.80E-06 | 0.364217691 | 0.323037347 |
| Atp6v0a2 | -0.002541297 | 0.726530485 | 0.802742286 | 0.165372335 | 0.168910966 |
| Atp6v0b | -0.006651712 | 0.395769292 | 0.516327034 | 1.486870499 | 1.52942319 |
| Atp6v0c | -0.020547054 | 0.001611714 | 0.005391978 | 1.853099916 | 1.868946511 |
| Atp6v0d1 | 0.00141516 | 0.885525189 | 0.921841299 | 0.557302142 | 0.55810524 |
| Atp6v0e | 0.049951932 | 3.94E-08 | 3.55E-07 | 1.390994813 | 1.327857367 |
| Atp6v0e2 | 0.028342808 | 0.001934244 | 0.006344373 | 0.377128047 | 0.342131738 |
| Atp6v1a | 0.075089184 | 2.78E-13 | 4.77E-12 | 0.652811652 | 0.589867035 |
| Atp6v1b2 | 0.038611535 | 0.000174307 | 0.000762995 | 0.764308308 | 0.712272738 |
| Atp6v1c1 | 0.032767418 | 0.000859169 | 0.003137807 | 0.581868881 | 0.537060569 |
| Atp6v1d | 0.027025027 | 0.006410039 | 0.017871638 | 0.808537542 | 0.771999165 |
| Atp6v1e1 | 0.105891993 | 8.81E-31 | 4.60E-29 | 1.272149704 | 1.103214624 |
| Atp6v1f | -0.022940675 | 0.003338769 | 0.010217032 | 1.398723698 | 1.419482924 |
| Atp6v1g1 | -0.049188237 | 4.24E-09 | 4.46E-08 | 1.369407888 | 1.436160979 |
| Atp6v1h | -0.004115487 | 0.65274465 | 0.745235587 | 0.340092823 | 0.34298689 |
| Atp7a | 0.038762238 | 9.22E-07 | 6.48E-06 | 0.238544095 | 0.190249619 |
| Atp8a1 | 0.022612764 | 0.002329961 | 0.007466013 | 0.191825268 | 0.168110409 |
| Atp8a2 | 0.003311094 | 0.414401785 | 0.533901015 | 0.045616872 | 0.041269638 |
| Atp8b1 | 0.112238258 | 2.74E-27 | 1.24E-25 | 0.769660126 | 0.640801622 |
| Atp8b2 | -0.00082989 | 0.865338384 | 0.908021562 | 0.067460389 | 0.065254186 |
| Atp9a | 0.020125046 | 0.021305675 | 0.049048964 | 0.292329843 | 0.273671208 |
| Atp9b | 0.00839717 | 0.114340417 | 0.196642054 | 0.08273079 | 0.075545921 |
| Atpaf1 | 0.001190675 | 0.839307091 | 0.889840537 | 0.105451067 | 0.099372538 |
| Atpaf2 | -0.000643965 | 0.897101431 | 0.929909384 | 0.071185292 | 0.069053635 |
| Atpif1 | 0.002524413 | 0.714927344 | 0.795030596 | 1.762334623 | 1.775896217 |
| Atpsckmt | -0.012085876 | 0.081267228 | 0.149268844 | 0.148217588 | 0.153496948 |
| Atr | 1.49E-05 | 0.997545222 | 0.997853138 | 0.06423151 | 0.062226344 |
| Atraid | -0.23520807 | 1.62E-122 | 1.43E-119 | 0.696162209 | 0.971381379 |
| Atrip | 0.009638226 | 0.062302005 | 0.120417513 | 0.076443512 | 0.070115793 |
| Atrn | 0.013872379 | 0.024551238 | 0.055392697 | 0.118223441 | 0.104947757 |
| Atrnl1 | 0.01322452 | 0.0028225 | 0.008806273 | 0.062441236 | 0.047911433 |
| Atrx | -0.002035988 | 0.844878396 | 0.894079434 | 0.989951649 | 0.977516771 |
| Atxn1 | 0.015817859 | 0.023914187 | 0.054131251 | 0.155440645 | 0.135490652 |
| Atxn10 | -0.037790063 | 0.000121285 | 0.00055508 | 0.60992205 | 0.614291488 |
| Atxn1l | 0.010833306 | 0.023698383 | 0.05369277 | 0.068745165 | 0.05950766 |
| Atxn2 | 0.03356404 | 0.000114148 | 0.000526697 | 0.295364772 | 0.266790178 |
| Atxn2l | -0.038541397 | 5.97E-05 | 0.000290129 | 0.378035191 | 0.41815298 |
| Atxn3 | -0.002929304 | 0.569567241 | 0.676852795 | 0.07193486 | 0.073774353 |
| Atxn7 | 0.019103501 | 0.007086374 | 0.019428574 | 0.16507349 | 0.147244652 |
| Atxn7l3 | 0.01531224 | 0.021201658 | 0.048855776 | 0.138605823 | 0.127712759 |
| Atxn7l3b | 0.002286213 | 0.804427164 | 0.864254712 | 0.954119496 | 0.954714306 |
| AU022252 | -0.00225307 | 0.591363263 | 0.695527902 | 0.048091409 | 0.047632019 |
| AU040320 | 0.013510596 | 0.074983368 | 0.140109227 | 0.188089537 | 0.181418199 |
| AU041133 | 0.003022931 | 0.592358913 | 0.6965304 | 0.095875746 | 0.088363065 |
| Auh | -0.008383612 | 0.410207986 | 0.529794153 | 0.537939414 | 0.529435345 |
| Aup1 | 0.018073325 | 0.028842298 | 0.063339694 | 0.244303923 | 0.240641674 |
| Aurkaip1 | -0.05598589 | 1.84E-08 | 1.74E-07 | 0.725650048 | 0.779179489 |
| Auts2 | 0.064630663 | 1.03E-09 | 1.17E-08 | 0.543451953 | 0.465755977 |
| AV356131 | -0.014295377 | 0.043478851 | 0.088821473 | 0.155789616 | 0.158685552 |
| Aven | -0.025166887 | 8.21E-07 | 5.85E-06 | 0.064606831 | 0.079481364 |
| Avl9 | 0.020830103 | 0.022354964 | 0.050957786 | 0.330402308 | 0.318360544 |
| Avpi1 | -0.114588342 | 6.43E-25 | 2.56E-23 | 0.288773386 | 0.435919388 |
| AW146154 | -0.004423771 | 0.248590795 | 0.362991846 | 0.037750207 | 0.039242073 |
| AW209491 | 0.008759261 | 0.160718174 | 0.257881183 | 0.124754461 | 0.113187983 |
| AW549877 | 0.009488993 | 0.219131408 | 0.329963906 | 0.196402882 | 0.186866893 |
| AW554918 | -0.001645905 | 0.678544063 | 0.766446615 | 0.040535613 | 0.042122981 |
| Axin1 | 0.018085633 | 0.015072232 | 0.03666973 | 0.198806616 | 0.171966736 |
| Azi2 | 0.008452112 | 0.362223169 | 0.484926143 | 0.376791725 | 0.36171719 |
| Azin1 | -0.023019135 | 0.018040725 | 0.042705606 | 0.344454059 | 0.382862716 |
| B230112I24Rik | -0.018747274 | 4.20E-05 | 0.000209554 | 0.04334794 | 0.059629833 |
| B230216N24Rik | -0.002277932 | 0.634612873 | 0.730520142 | 0.044803463 | 0.054828838 |
| B230217C12Rik | -0.008484045 | 0.180771773 | 0.283005342 | 0.121587016 | 0.120400113 |
| B230219D22Rik | 0.073642747 | 1.96E-13 | 3.42E-12 | 0.608271807 | 0.539914845 |
| B230307C23Rik | -0.000314016 | 0.947778442 | 0.965075865 | 0.060557341 | 0.06350652 |
| B230322F03Rik | 0.004166489 | 0.352191102 | 0.474501371 | 0.061109863 | 0.052954956 |
| B230354K17Rik | 0.008824222 | 0.101502464 | 0.178866587 | 0.085141813 | 0.076578408 |
| B2m | 0.009666166 | 0.381853243 | 0.50309616 | 0.36998996 | 0.372180295 |
| B330016D10Rik | 0.003583204 | 0.548217321 | 0.657681466 | 0.11211568 | 0.101397111 |

| | | | | | |
|---|---|---|---|---|---|
| B3galnt1 | 0.011389745 | 0.093583587 | 0.167123371 | 0.152981246 | 0.136674654 |
| B3galnt2 | 0.000632548 | 0.897064261 | 0.929909384 | 0.063832147 | 0.06544352 |
| B3galt2 | 0.026622604 | 7.88E-07 | 5.63E-06 | 0.0903257 | 0.069029794 |
| B3galt4 | -0.003953099 | 0.335913763 | 0.4586326 | 0.044388905 | 0.046468131 |
| B3galt5 | -0.006436294 | 0.201386337 | 0.308959755 | 0.069708975 | 0.070491911 |
| B3galt6 | -0.001408039 | 0.798511849 | 0.859382621 | 0.086228337 | 0.086671654 |
| B3gat3 | -0.040993236 | 2.53E-06 | 1.64E-05 | 0.292307464 | 0.319086717 |
| B3gnt3 | -0.004340042 | 0.370478777 | 0.49291447 | 0.059619135 | 0.064410611 |
| B4galt1 | 0.007775443 | 0.078983608 | 0.145929046 | 0.05687164 | 0.050435169 |
| B4galt3 | -0.006485651 | 0.450003221 | 0.56839435 | 0.270674461 | 0.277970784 |
| B4galt5 | 0.022838993 | 3.40E-05 | 0.000174429 | 0.095027934 | 0.076903716 |
| B4galt6 | 0.038767249 | 2.22E-05 | 0.000118917 | 0.359272424 | 0.318771004 |
| B4galt7 | 0.003980124 | 0.384909918 | 0.5058935 | 0.060926697 | 0.056230422 |
| B630019K06Rik | -0.016097196 | 0.004225394 | 0.012478517 | 0.088239436 | 0.092965157 |
| B830012L14Rik | -0.000350519 | 0.958833992 | 0.971740148 | 0.126350153 | 0.123121933 |
| B830017H08Rik | -0.043387791 | 1.88E-05 | 0.000102495 | 1.438122841 | 1.469771176 |
| B930095G15Rik | 0.009740523 | 0.07725732 | 0.143475771 | 0.087503424 | 0.08031288 |
| B9d1 | -0.04061842 | 9.01E-07 | 6.36E-06 | 0.234074314 | 0.264994558 |
| B9d2 | -0.034325853 | 1.13E-06 | 7.80E-06 | 0.142302091 | 0.167261977 |
| Babam1 | -0.052142289 | 9.82E-10 | 1.12E-08 | 0.263570858 | 0.295870407 |
| Babam2 | -0.040912655 | 8.13E-06 | 4.76E-05 | 0.310711021 | 0.344267356 |
| Bace1 | 0.028620622 | 2.91E-05 | 0.000151493 | 0.162107692 | 0.135006132 |
| Bace2 | -0.131821876 | 6.57E-63 | 1.33E-60 | 1.694374373 | 1.818528489 |
| Bach1 | 0.015782449 | 0.054964348 | 0.108039504 | 0.233425632 | 0.213040484 |
| Bad | -0.036989266 | 1.74E-05 | 9.57E-05 | 0.26933329 | 0.295103215 |
| Bag1 | -0.026934368 | 0.004540783 | 0.013308862 | 0.84031386 | 0.85451895 |
| Bag2 | -0.023494943 | 0.002135114 | 0.006905381 | 0.174646374 | 0.18932908 |
| Bag3 | 0.002944385 | 0.638260923 | 0.733732138 | 0.104513714 | 0.109791256 |
| Bag4 | 0.026692107 | 0.000853272 | 0.00312096 | 0.242336023 | 0.209929316 |
| Bag5 | -0.006980464 | 0.404923831 | 0.524679436 | 0.257159123 | 0.257621857 |
| Bag6 | 0.030151249 | 0.000248885 | 0.00104116 | 0.267769017 | 0.230097294 |
| Bahd1 | 0.00234866 | 0.641554642 | 0.736088758 | 0.072340963 | 0.068303413 |
| Baiap2 | 0.064557235 | 6.65E-38 | 4.93E-36 | 0.106807654 | 0.047090788 |
| Baiap2l1 | 0.033932868 | 6.60E-07 | 4.81E-06 | 0.155512295 | 0.129873805 |
| Baiap3 | -2.40E-05 | 0.997084121 | 0.997699859 | 0.133216968 | 0.118071851 |
| Bak1 | 0.0008851 | 0.817611517 | 0.874553765 | 0.041217487 | 0.038805632 |
| Bambi | 0.014495708 | 0.095980016 | 0.170713084 | 0.238399529 | 0.216926268 |
| Banf1 | -0.004555865 | 0.624220581 | 0.722815316 | 0.348414616 | 0.352487427 |
| Banp | 0.03121239 | 0.000974847 | 0.003498509 | 0.386148857 | 0.321302567 |
| Bap1 | 0.015928078 | 0.004807953 | 0.013940627 | 0.103931351 | 0.086852331 |
| Barhl1 | -0.015060661 | 0.003416998 | 0.010417075 | 0.05594095 | 0.067481019 |
| Bax | -0.046686206 | 7.94E-06 | 4.67E-05 | 0.790327588 | 0.845169678 |
| Baz1a | 0.003923122 | 0.46845946 | 0.585801925 | 0.076781618 | 0.078095035 |
| Baz1b | 0.024340415 | 0.017322513 | 0.041324947 | 0.492343993 | 0.457374679 |
| Baz2a | 0.031379082 | 0.000747223 | 0.002771653 | 0.379789422 | 0.336326589 |
| Baz2b | 0.049955135 | 1.47E-06 | 9.94E-06 | 0.721191828 | 0.646491434 |
| BB218582 | 0.000536931 | 0.910466881 | 0.938619225 | 0.073386314 | 0.063540429 |
| Bbc3 | -0.018874844 | 0.002357195 | 0.007543334 | 0.104935712 | 0.116183938 |
| Bbip1 | -0.035534501 | 0.000357036 | 0.001439694 | 0.509028796 | 0.538927752 |
| Bbs1 | 0.007536739 | 0.074527402 | 0.139337578 | 0.050026668 | 0.044868811 |
| Bbs2 | 0.002566298 | 0.620614356 | 0.719915615 | 0.083266278 | 0.07472092 |
| Bbs4 | -0.006466254 | 0.291038818 | 0.410485912 | 0.109310484 | 0.110652449 |
| Bbs5 | 0.001917588 | 0.631315826 | 0.728331845 | 0.045059673 | 0.042643777 |
| Bbs7 | 0.007130318 | 0.180133362 | 0.282369647 | 0.083579637 | 0.077053705 |
| Bbx | 0.026713641 | 0.008557925 | 0.022807211 | 0.650286729 | 0.60359932 |
| BC003965 | -0.0100626 | 0.157119828 | 0.25365642 | 0.16092178 | 0.165170623 |
| BC004004 | 0.00222645 | 0.81881652 | 0.875168669 | 0.492824592 | 0.4663038 |
| BC005537 | 0.001910612 | 0.851755491 | 0.898618218 | 0.616094959 | 0.600564499 |
| BC005624 | -0.005454487 | 0.562940988 | 0.670879032 | 0.406366842 | 0.405750466 |
| BC029722 | 0.00991856 | 0.186228753 | 0.289728906 | 0.195546206 | 0.181853709 |
| BC031181 | 0.011654695 | 0.240480837 | 0.354396649 | 0.733151473 | 0.720733329 |
| BC037704 | -0.002762904 | 0.504409595 | 0.618706798 | 0.045402021 | 0.045647328 |
| BC051142 | -0.018038739 | 4.06E-05 | 0.000203696 | 0.039486408 | 0.054752273 |
| BC051226 | -0.013498534 | 0.031020289 | 0.067271749 | 0.108594587 | 0.118752375 |
| Bcap29 | -0.029488591 | 1.63E-05 | 8.97E-05 | 0.122892384 | 0.149968658 |
| Bcap31 | -0.043112622 | 5.25E-06 | 3.21E-05 | 0.378345673 | 0.417278232 |
| Bcar1 | 0.03899918 | 4.76E-07 | 3.57E-06 | 0.227311859 | 0.182963648 |
| Bcas2 | 0.002277297 | 0.827337348 | 0.880886398 | 0.792684086 | 0.784456955 |
| Bcas3 | -0.006515049 | 0.111617381 | 0.1933614 | 0.043518103 | 0.046445005 |
| Bcat2 | -0.023509659 | 1.66E-06 | 1.11E-05 | 0.055128693 | 0.073444552 |
| Bccip | -0.053089951 | 8.89E-08 | 7.59E-07 | 0.589455712 | 0.625617332 |
| Bcdin3d | 0.001353774 | 0.811702878 | 0.870340286 | 0.097338166 | 0.092748733 |

| | | | | | |
|---|---|---|---|---|---|
| Bckdha | -0.016757091 | 0.014814959 | 0.036170526 | 0.149223547 | 0.150664378 |
| Bckdhb | 0.012193881 | 0.012042985 | 0.030363565 | 0.073604181 | 0.062345202 |
| Bckdk | 0.024690943 | 0.005086744 | 0.014648497 | 0.338408732 | 0.303194637 |
| Bcl10 | -0.010333186 | 0.121303191 | 0.206245125 | 0.133566157 | 0.13983331 |
| Bcl11a | -0.013993452 | 0.000893666 | 0.003247931 | 0.039882711 | 0.04792619 |
| Bcl2l1 | -0.001403552 | 0.845637221 | 0.894439423 | 0.157505834 | 0.160569364 |
| Bcl2l11 | 0.012111899 | 0.025946023 | 0.05785487 | 0.093663171 | 0.076826681 |
| Bcl2l13 | 0.016557708 | 0.009628331 | 0.025102341 | 0.128101707 | 0.116047629 |
| Bcl2l2 | 0.006917959 | 0.325350824 | 0.447518493 | 0.157816139 | 0.153142003 |
| Bcl3 | -0.01077696 | 0.109218503 | 0.189950319 | 0.104464283 | 0.127991951 |
| Bcl6b | 0.016156052 | 3.28E-05 | 0.000168864 | 0.048885697 | 0.034181946 |
| Bcl7a | 0.0085606 | 0.23824978 | 0.352138515 | 0.169779084 | 0.160766804 |
| Bcl7b | 0.008436907 | 0.382160221 | 0.503355009 | 0.441094467 | 0.430389206 |
| Bcl7c | 0.005942901 | 0.460909895 | 0.579010983 | 0.250230922 | 0.230349259 |
| Bcl9 | 0.066209526 | 3.24E-15 | 6.39E-14 | 0.287784467 | 0.224392308 |
| Bcl9l | 0.005853695 | 0.242906688 | 0.356665305 | 0.070658688 | 0.066791368 |
| Bclaf1 | -0.003657739 | 0.728484421 | 0.804444064 | 0.608450458 | 0.619071558 |
| Bclaf3 | -0.002803481 | 0.598110191 | 0.701595955 | 0.075758694 | 0.078571717 |
| Bcor | 0.034659718 | 0.00021237 | 0.000905155 | 0.385872253 | 0.351677336 |
| Bcorl1 | 0.006610605 | 0.181652181 | 0.283839673 | 0.073479271 | 0.064217246 |
| Bcr | 0.045447065 | 5.07E-08 | 4.50E-07 | 0.268783155 | 0.222814966 |
| Bcs1l | 0.003935239 | 0.433739486 | 0.552138484 | 0.072433557 | 0.069033544 |
| Bdh1 | 0.020453091 | 2.17E-05 | 0.000116774 | 0.072996104 | 0.05828551 |
| Bdp1 | 0.01310919 | 0.119868203 | 0.204126585 | 0.248903213 | 0.232843836 |
| Becn1 | -0.002358096 | 0.799598942 | 0.860208134 | 0.395260322 | 0.397151212 |
| Bend5 | -0.000639161 | 0.870261425 | 0.910970599 | 0.039952523 | 0.040532862 |
| Bend7 | 0.002801496 | 0.733517926 | 0.808441365 | 0.219460487 | 0.220061454 |
| Bet1 | -0.063122532 | 8.51E-11 | 1.10E-09 | 0.406107247 | 0.476440971 |
| Bet1l | 0.023731197 | 0.006365297 | 0.017762175 | 0.300738593 | 0.272281861 |
| Bex1 | -0.113837277 | 4.16E-26 | 1.81E-24 | 0.742624471 | 0.860142475 |
| Bex2 | -0.074686657 | 1.82E-16 | 3.91E-15 | 1.494155012 | 1.63182 |
| Bex3 | -0.056255667 | 3.94E-15 | 7.70E-14 | 2.065571142 | 2.223170021 |
| Bex4 | -0.139428946 | 1.85E-41 | 1.66E-39 | 0.725332386 | 0.868303426 |
| Bfar | -0.008187539 | 0.278247187 | 0.396703204 | 0.17711587 | 0.180966122 |
| Bhlha15 | -0.011994127 | 0.022113991 | 0.050550722 | 0.067732391 | 0.077607417 |
| Bhlhb9 | -0.011263555 | 0.149110991 | 0.243885777 | 0.200809156 | 0.207335945 |
| Bhlhe40 | 0.01921946 | 0.001932671 | 0.006341352 | 0.103585598 | 0.097141751 |
| Bhlhe41 | 0.10547542 | 5.81E-30 | 2.99E-28 | 0.413162112 | 0.279898418 |
| Bicc1 | 0.014944463 | 0.015936224 | 0.038473297 | 0.118811072 | 0.102846445 |
| Bicd1 | -0.003176057 | 0.74098038 | 0.814634315 | 0.391198404 | 0.384448048 |
| Bicd2 | 0.020466435 | 1.77E-05 | 9.67E-05 | 0.070542701 | 0.056245818 |
| Bicdl1 | -0.006775067 | 0.171154315 | 0.270606968 | 0.0651938 | 0.069079038 |
| Bicral | 0.016035659 | 0.007852033 | 0.021204852 | 0.113547703 | 0.098599009 |
| Bik | -0.023302058 | 7.30E-08 | 6.32E-07 | 0.036283651 | 0.052737726 |
| Bin1 | -0.006734448 | 0.316096457 | 0.437824441 | 0.137750446 | 0.139819479 |
| Bin3 | 0.002983513 | 0.566809134 | 0.674307618 | 0.084727313 | 0.076762215 |
| Birc2 | -0.007551565 | 0.339200487 | 0.46147595 | 0.215240101 | 0.209178989 |
| Birc6 | 0.028990932 | 0.00419087 | 0.012391618 | 0.590202814 | 0.5531747 |
| Bivm | -0.003234979 | 0.432911868 | 0.551318992 | 0.045493112 | 0.046083506 |
| Blcap | 0.032176366 | 4.00E-05 | 0.000200941 | 0.233610707 | 0.198014678 |
| Blmh | 0.003024401 | 0.730407108 | 0.806017924 | 0.300223432 | 0.288997818 |
| Bloc1s1 | 0.029110883 | 2.29E-13 | 3.96E-12 | 0.058917517 | 0.034394919 |
| Bloc1s2 | -0.001642621 | 0.838190721 | 0.889131499 | 0.224964491 | 0.221495344 |
| Bloc1s3 | -0.00142847 | 0.713079095 | 0.793745434 | 0.041689348 | 0.040764374 |
| Bloc1s4 | 0.005359204 | 0.313958917 | 0.435422053 | 0.090354341 | 0.079968165 |
| Bloc1s5 | -0.003311516 | 0.526720337 | 0.638500638 | 0.079319182 | 0.077895499 |
| Bloc1s6 | 0.009155421 | 0.166240946 | 0.264602894 | 0.136945507 | 0.129916776 |
| Blvra | 0.027399973 | 6.74E-07 | 4.90E-06 | 0.10863384 | 0.077822765 |
| Blvrb | -0.038436283 | 9.19E-09 | 9.22E-08 | 0.116018256 | 0.144856384 |
| Blzf1 | -0.016501358 | 0.0106468 | 0.027296464 | 0.115114587 | 0.127236202 |
| Bmi1 | -0.003259672 | 0.712541991 | 0.793327215 | 0.291876602 | 0.288803094 |
| Bmp1 | 0.005962215 | 0.468423638 | 0.585801925 | 0.251400931 | 0.229124083 |
| Bmp2k | 0.001172526 | 0.763060713 | 0.832320907 | 0.039404715 | 0.038483052 |
| Bmpr1a | 0.022371673 | 0.010048079 | 0.025990383 | 0.301375263 | 0.277591362 |
| Bmpr2 | 0.006539936 | 0.32552363 | 0.447692846 | 0.134841284 | 0.129067427 |
| Bms1 | 0.030691565 | 9.80E-07 | 6.86E-06 | 0.125245505 | 0.103995656 |
| Bmt2 | 0.023975673 | 0.000237645 | 0.001001033 | 0.138053745 | 0.117196558 |
| Bmyc | -0.012906258 | 0.167322794 | 0.265802648 | 0.399375384 | 0.387166589 |
| Bnip1 | 9.44E-05 | 0.986880442 | 0.992187348 | 0.093976518 | 0.093593869 |
| Bnip2 | -0.019048079 | 0.000661159 | 0.002484648 | 0.073852938 | 0.091636972 |
| Bnip3 | -0.05486959 | 4.97E-08 | 4.42E-07 | 0.486958747 | 0.543081383 |
| Bnip3l | -0.000122189 | 0.988892754 | 0.993079296 | 0.333556335 | 0.303241839 |

| | | | | | |
|---|---|---|---|---|---|
| Bod1 | 0.030407103 | 0.001294764 | 0.004478012 | 0.457346841 | 0.411346335 |
| Bod1l | -0.014097264 | 0.165817069 | 0.264101171 | 0.536155831 | 0.545495452 |
| Bola1 | -0.004035868 | 0.686370653 | 0.772594129 | 0.488874949 | 0.503579541 |
| Bola2 | 0.077032417 | 1.47E-14 | 2.78E-13 | 0.655501996 | 0.623155235 |
| Bola3 | 0.04060864 | 4.37E-05 | 0.000217665 | 0.640536317 | 0.581947396 |
| Bop1 | 0.003609641 | 0.511236352 | 0.625030157 | 0.089478723 | 0.084609265 |
| Borcs5 | 0.005456017 | 0.234701853 | 0.348521676 | 0.064381327 | 0.057054001 |
| Borcs6 | 0.001180053 | 0.849376712 | 0.896952957 | 0.11918257 | 0.112019228 |
| Borcs7 | 0.00225376 | 0.796981717 | 0.858342334 | 0.298401214 | 0.293292649 |
| Borcs8 | -0.032676467 | 2.01E-05 | 0.000108984 | 0.181646241 | 0.203502138 |
| Bpgm | -0.005993674 | 0.363045799 | 0.485826738 | 0.131870687 | 0.135780675 |
| Bphl | -0.04769645 | 5.47E-09 | 5.63E-08 | 0.238190487 | 0.254157149 |
| Bpnt1 | -0.005786678 | 0.519862768 | 0.632158328 | 0.329757156 | 0.315429609 |
| Bpnt2 | 0.046609384 | 1.28E-06 | 8.77E-06 | 0.461162273 | 0.405484137 |
| Bptf | 0.030452875 | 0.003722943 | 0.011209184 | 0.701472283 | 0.663793529 |
| Braf | 0.048441058 | 1.68E-06 | 1.12E-05 | 0.59157227 | 0.533079324 |
| Brap | 0.012120072 | 0.084495227 | 0.154149483 | 0.164300533 | 0.148082694 |
| Brat1 | 0.008719965 | 0.013793036 | 0.034017225 | 0.037782836 | 0.030187539 |
| Brcc3 | -0.022277858 | 0.002547134 | 0.00806095 | 0.165690807 | 0.178474708 |
| Brd1 | 0.003512572 | 0.71998831 | 0.798658866 | 0.472643809 | 0.458142944 |
| Brd2 | -0.049241494 | 1.33E-06 | 9.10E-06 | 0.689599028 | 0.768861234 |
| Brd3 | 0.038938455 | 3.08E-05 | 0.000159272 | 0.386145125 | 0.346863698 |
| Brd3os | -0.002767836 | 0.583066687 | 0.688846454 | 0.073050543 | 0.072619359 |
| Brd4 | 0.048993515 | 4.04E-07 | 3.07E-06 | 0.450918098 | 0.40760325 |
| Brd7 | 0.036074392 | 8.06E-06 | 4.73E-05 | 0.242309899 | 0.212076823 |
| Brd8 | 0.024824416 | 0.009097352 | 0.023962194 | 0.391142717 | 0.369471169 |
| Brd9 | 0.008818108 | 0.18034648 | 0.282544264 | 0.134083859 | 0.128227631 |
| Brdt | -0.002815573 | 0.488419116 | 0.60396981 | 0.043785536 | 0.044652417 |
| Brf1 | 0.004311376 | 0.438088635 | 0.556381151 | 0.092573589 | 0.085874087 |
| Brf2 | 0.003184723 | 0.456614412 | 0.575175604 | 0.055913014 | 0.048895206 |
| Bri3 | 0.001816332 | 0.852797743 | 0.89874251 | 0.907081794 | 0.862186615 |
| Bri3bp | 0.009391612 | 0.100563693 | 0.177534087 | 0.099575847 | 0.090584099 |
| Brip1os | -0.001360659 | 0.761471204 | 0.830680324 | 0.05868981 | 0.055151703 |
| Brix1 | -0.014839536 | 0.085829893 | 0.156144876 | 0.256496792 | 0.265979145 |
| Brk1 | -0.003094784 | 0.748668423 | 0.820766172 | 0.817766034 | 0.816692316 |
| Brms1l | -0.004891505 | 0.335365285 | 0.458246142 | 0.067111333 | 0.071000137 |
| Brox | 0.010514469 | 0.149284121 | 0.243963729 | 0.178811535 | 0.164934737 |
| Brpf1 | 0.011130526 | 0.040522809 | 0.083857546 | 0.085735094 | 0.077815796 |
| Brpf3 | -0.000377798 | 0.938156381 | 0.957861409 | 0.064318845 | 0.065108644 |
| Brsk1 | 0.00701761 | 0.123168215 | 0.208649833 | 0.057199027 | 0.054069095 |
| Brsk2 | 0.007031063 | 0.189542499 | 0.293662499 | 0.081871416 | 0.077415149 |
| Brwd1 | -0.004335909 | 0.629798846 | 0.72709944 | 0.305442976 | 0.305231723 |
| Brwd3 | 0.007908268 | 0.127218923 | 0.214353964 | 0.074792848 | 0.069413638 |
| Bscl2 | -0.040874227 | 8.35E-07 | 5.94E-06 | 0.224143557 | 0.255418129 |
| Bsdc1 | 0.011031888 | 0.150996497 | 0.246006139 | 0.204484934 | 0.190547307 |
| Bsg | -0.076694876 | 7.22E-31 | 3.80E-29 | 1.799401148 | 1.890759411 |
| Bsn | 0.016755401 | 0.001423914 | 0.004864121 | 0.086602254 | 0.071929771 |
| Bspry | -0.014826884 | 0.003132094 | 0.009646031 | 0.063478812 | 0.073939826 |
| Bst2 | 0.100810443 | 1.80E-37 | 1.30E-35 | 0.180284442 | 0.106715736 |
| Btaf1 | 0.007005996 | 0.282686958 | 0.401619553 | 0.125214889 | 0.12241244 |
| Btbd1 | 0.005235516 | 0.532899992 | 0.643984303 | 0.293885341 | 0.267482721 |
| Btbd10 | -0.008141712 | 0.163893638 | 0.261637759 | 0.099849472 | 0.100871289 |
| Btbd17 | 0.013984695 | 0.073740144 | 0.138132097 | 0.234295406 | 0.196187589 |
| Btbd2 | 0.017336659 | 0.004371582 | 0.012878946 | 0.123760077 | 0.103149103 |
| Btbd3 | 0.020824145 | 3.39E-05 | 0.000174207 | 0.079473509 | 0.063762379 |
| Btbd6 | 0.004466159 | 0.386545703 | 0.507494575 | 0.082349502 | 0.073758072 |
| Btbd7 | 0.02872378 | 0.000910534 | 0.00330223 | 0.28693649 | 0.267479763 |
| Btbd8 | 0.03920228 | 3.35E-09 | 3.55E-08 | 0.13302616 | 0.110253784 |
| Btbd9 | 0.032773202 | 2.23E-05 | 0.000119231 | 0.218058599 | 0.188618864 |
| Btf3 | -0.09148147 | 2.26E-28 | 1.10E-26 | 1.646632792 | 1.800860892 |
| Btf3l4 | -0.006149606 | 0.413187376 | 0.532900991 | 0.195724817 | 0.187094369 |
| Btf3-ps18 | -0.015418788 | 0.000888346 | 0.003233434 | 0.053481222 | 0.063408757 |
| Btg1 | 0.041522656 | 0.000123332 | 0.000563457 | 0.675722592 | 0.604924051 |
| Btg2 | -0.115652547 | 1.46E-22 | 5.07E-21 | 1.256893672 | 1.486141745 |
| Btg3 | 0.084138971 | 7.21E-16 | 1.49E-14 | 0.676971257 | 0.583890042 |
| Btrc | 0.015802787 | 0.045455346 | 0.092162018 | 0.227997976 | 0.202944804 |
| Bub3 | -0.023505308 | 0.007353165 | 0.020075111 | 0.309497437 | 0.31809207 |
| Bud13 | 0.003655487 | 0.413245873 | 0.532905741 | 0.057044132 | 0.052287323 |
| Bud23 | -0.029656498 | 1.21E-05 | 6.83E-05 | 0.132244891 | 0.152884475 |
| Bud31 | -0.009528299 | 0.327070234 | 0.449509103 | 0.427384648 | 0.429347139 |
| Bysl | -9.52E-05 | 0.985282546 | 0.990990784 | 0.074272147 | 0.073927499 |
| Bzw1 | -0.006935111 | 0.480612097 | 0.596973401 | 0.511523607 | 0.508243899 |

| | | | | | |
|---|---|---|---|---|---|
| Bzw2 | -0.025185321 | 0.001161535 | 0.004075224 | 0.16975509 | 0.193715817 |
| C030006K11Rik | -0.007989364 | 0.198073917 | 0.304647147 | 0.114113703 | 0.114645643 |
| C030014I23Rik | 0.003497082 | 0.553678751 | 0.663241106 | 0.108779187 | 0.100918059 |
| C030034I22Rik | -0.001464662 | 0.774592843 | 0.841155029 | 0.072531779 | 0.071287931 |
| C130036L24Rik | 0.005490193 | 0.19153917 | 0.296095375 | 0.052291652 | 0.045975962 |
| C130089K02Rik | -0.010037503 | 0.131882058 | 0.22125235 | 0.117163751 | 0.123693424 |
| C1d | 0.036845004 | 4.47E-05 | 0.000221872 | 0.396919767 | 0.340884832 |
| C1galt1 | 0.04116415 | 3.67E-05 | 0.000186667 | 0.463168581 | 0.431484826 |
| C1galt1c1 | -0.020351474 | 0.00500397 | 0.014444357 | 0.167090966 | 0.17697523 |
| C1qa | 0.120470842 | 5.41E-74 | 1.75E-71 | 0.197739227 | 0.079347415 |
| C1qbp | -0.021773335 | 0.017780045 | 0.042201561 | 0.311416256 | 0.326721882 |
| C1qtnf12 | 0.001907986 | 0.713734639 | 0.794152828 | 0.076930103 | 0.074847729 |
| C1qtnf4 | -0.03604656 | 0.000431795 | 0.001702316 | 0.893019506 | 0.911031769 |
| C1qtnf6 | 0.001847269 | 0.617849222 | 0.717821479 | 0.036970682 | 0.035818726 |
| C2cd2l | 0.006144803 | 0.338575122 | 0.460947674 | 0.128856806 | 0.118371812 |
| C2cd3 | 0.009169777 | 0.12170029 | 0.206667287 | 0.108425179 | 0.097581449 |
| C2cd4a | 0.000475378 | 0.938760257 | 0.958276692 | 0.080584036 | 0.092376126 |
| C2cd4b | -0.003421699 | 0.800166866 | 0.860628639 | 1.171208342 | 1.200732097 |
| C2cd4c | -0.001397936 | 0.731120719 | 0.806347775 | 0.044833489 | 0.044050708 |
| C2cd5 | -0.027624707 | 9.94E-06 | 5.71E-05 | 0.097011196 | 0.119297836 |
| C330007P06Rik | -0.015494729 | 0.031890194 | 0.068713755 | 0.161309835 | 0.168484988 |
| C330013E15Rik | 0.004606391 | 0.315586294 | 0.437180102 | 0.060655759 | 0.055557294 |
| C330018D20Rik | -0.004449937 | 0.27626806 | 0.394286271 | 0.046779137 | 0.045973248 |
| C77080 | 0.034025714 | 4.63E-08 | 4.14E-07 | 0.13148044 | 0.100633391 |
| C87436 | -0.007548498 | 0.257040636 | 0.372921812 | 0.131701931 | 0.135536787 |
| C8b | -0.006296524 | 0.168346956 | 0.267040676 | 0.050435515 | 0.055622826 |
| C920006O11Rik | -0.00082302 | 0.881050952 | 0.918560575 | 0.085464736 | 0.08478924 |
| C920021L13Rik | 0.011920128 | 0.007258299 | 0.019849559 | 0.056692284 | 0.050208138 |
| C9orf72 | -0.019991398 | 0.001272733 | 0.004414381 | 0.089552644 | 0.114792212 |
| Caap1 | -0.009910097 | 0.151238515 | 0.246247002 | 0.143647134 | 0.148612873 |
| Cab39 | 0.045087132 | 5.60E-07 | 4.14E-06 | 0.358821927 | 0.309094192 |
| Cab39l | 0.011220231 | 0.118638355 | 0.2024578 | 0.17533465 | 0.159881605 |
| Cabcoco1 | -0.009056243 | 0.025569278 | 0.057198463 | 0.041324884 | 0.045794097 |
| Cabin1 | -0.004896597 | 0.491624191 | 0.60677547 | 0.155247565 | 0.158440983 |
| Cables2 | 0.003276398 | 0.464489911 | 0.581929242 | 0.054286184 | 0.05294703 |
| Cabp1 | -0.038367392 | 3.85E-05 | 0.000194218 | 0.310447174 | 0.345642057 |
| Cabp7 | 0.063024367 | 4.88E-18 | 1.20E-16 | 0.216426678 | 0.139238141 |
| Cacfd1 | -0.001674172 | 0.837210821 | 0.888382842 | 0.224824695 | 0.231024846 |
| Cacna1a | 0.01423403 | 0.110874695 | 0.192226307 | 0.277732151 | 0.263159096 |
| Cacna1c | 0.024243676 | 0.00046936 | 0.001833314 | 0.155107364 | 0.13685284 |
| Cacna1d | -0.001707325 | 0.865792446 | 0.908203945 | 0.475071617 | 0.473597839 |
| Cacna2d1 | 0.071495016 | 3.14E-12 | 4.75E-11 | 0.744253903 | 0.641487478 |
| Cacnb2 | 0.012046764 | 0.077277451 | 0.143485748 | 0.1442614 | 0.133354763 |
| Cacnb3 | 0.015783414 | 0.030963026 | 0.067177535 | 0.180939456 | 0.15645254 |
| Cactin | 0.002514612 | 0.601138475 | 0.70404388 | 0.068476293 | 0.063406785 |
| Cacul1 | 0.024282415 | 0.000695699 | 0.002602379 | 0.174671527 | 0.152611189 |
| Cacybp | 0.002721887 | 0.794083874 | 0.856169838 | 0.728435563 | 0.6963169 |
| Cad | 0.017921467 | 0.006983051 | 0.01919944 | 0.144328225 | 0.126456526 |
| Cadm1 | 0.019006549 | 0.001080351 | 0.003833275 | 0.089342432 | 0.079719291 |
| Cadps | 0.050103145 | 7.25E-07 | 5.23E-06 | 0.492827411 | 0.411525669 |
| Cadps2 | -0.008802094 | 0.029714487 | 0.064932399 | 0.037709854 | 0.044968783 |
| Calb1 | 0.025820103 | 9.28E-07 | 6.52E-06 | 0.085589437 | 0.057164568 |
| Calcoco1 | 0.003186199 | 0.57541351 | 0.681800139 | 0.100773486 | 0.089292585 |
| Cald1 | -0.057610229 | 9.76E-19 | 2.53E-17 | 0.066581752 | 0.11647521 |
| Calm1 | 0.007756404 | 0.111262543 | 0.192781045 | 2.817890536 | 2.817025312 |
| Calm2 | -0.096948181 | 1.58E-32 | 9.22E-31 | 1.306805451 | 1.42805838 |
| Calm3 | 0.007938273 | 0.400233585 | 0.520405365 | 1.016578898 | 0.983951773 |
| Calr | 0.087976249 | 3.57E-22 | 1.20E-20 | 1.57368512 | 1.459286265 |
| Calu | 0.006445004 | 0.246407896 | 0.360574837 | 0.088181247 | 0.084790844 |
| Camk1 | -0.017539598 | 0.015139065 | 0.036813905 | 0.157067368 | 0.170208215 |
| Camk2b | 0.050600983 | 6.48E-07 | 4.73E-06 | 0.669789033 | 0.595597068 |
| Camk2d | -0.017667255 | 0.020885829 | 0.048230886 | 0.169328585 | 0.187112059 |
| Camk2g | 0.006318906 | 0.22501473 | 0.336863751 | 0.080794932 | 0.07318386 |
| Camk2n1 | 0.061750009 | 1.13E-20 | 3.38E-19 | 2.515934661 | 2.294867234 |
| Camk2n2 | -0.046213277 | 9.90E-06 | 5.69E-05 | 0.696143044 | 0.744595684 |
| Camkk2 | 0.029446984 | 5.78E-07 | 4.26E-06 | 0.12198213 | 0.089174274 |
| Camkmt | -0.00105164 | 0.857557202 | 0.902240254 | 0.097022758 | 0.099564903 |
| Caml | -0.018261413 | 0.022453346 | 0.051122116 | 0.216587783 | 0.228378151 |
| Camsap1 | 0.000814619 | 0.907338861 | 0.935957559 | 0.147202489 | 0.148963348 |
| Camsap2 | 0.026326878 | 4.86E-05 | 0.000239836 | 0.135199704 | 0.11352685 |
| Camsap3 | 0.013574463 | 0.000735712 | 0.002734774 | 0.050126004 | 0.038399552 |
| Camta1 | -0.010503934 | 0.22793812 | 0.340453895 | 0.283174276 | 0.295264354 |

| | | | | | |
|---|---|---|---|---|---|
| Camta2 | 0.019143547 | 0.003410928 | 0.010405097 | 0.138667101 | 0.124988522 |
| Cand1 | 0.055840839 | 1.57E-10 | 1.97E-09 | 0.320394334 | 0.269056779 |
| Cand2 | 0.003672256 | 0.387129387 | 0.508123653 | 0.05179256 | 0.046586307 |
| Cant1 | 0.00213232 | 0.76431544 | 0.833222102 | 0.15901016 | 0.153310693 |
| Canx | -0.008297338 | 0.346427941 | 0.468554875 | 1.266601114 | 1.275980745 |
| Cap1 | 0.010022508 | 0.210335119 | 0.319412375 | 0.231618796 | 0.220063623 |
| Capn1 | 0.006349702 | 0.252413549 | 0.367469979 | 0.092778616 | 0.087537197 |
| Capn10 | -0.004433334 | 0.417208297 | 0.536461488 | 0.087476795 | 0.085081403 |
| Capn15 | 0.00710767 | 0.235609124 | 0.349469721 | 0.105524173 | 0.100067454 |
| Capn2 | 0.012837187 | 0.068711337 | 0.130434151 | 0.164605298 | 0.149343313 |
| Capn5 | 0.019932493 | 0.000139238 | 0.000628152 | 0.086891202 | 0.069814766 |
| Capn7 | 0.024625286 | 0.001059438 | 0.003767322 | 0.197284054 | 0.175924833 |
| Capns1 | -0.032298374 | 0.000963758 | 0.003466393 | 0.734898295 | 0.765775494 |
| Caprin1 | 0.034501941 | 0.000725549 | 0.00270087 | 0.613132094 | 0.566170502 |
| Capsl | -0.019219477 | 0.009040022 | 0.023830557 | 0.170199887 | 0.179356609 |
| Capza1 | -0.000726984 | 0.852653241 | 0.898687642 | 0.041293975 | 0.040974877 |
| Capza2 | -0.000478345 | 0.961461141 | 0.973476902 | 0.599830078 | 0.585200648 |
| Capzb | 0.067724479 | 2.62E-12 | 4.02E-11 | 0.57578752 | 0.494497265 |
| Car10 | 0.036878297 | 2.41E-05 | 0.000127517 | 0.312653009 | 0.265887461 |
| Car8 | -0.038032212 | 3.21E-06 | 2.04E-05 | 0.149268509 | 0.19303812 |
| Card19 | 0.008971278 | 0.28625343 | 0.405619567 | 0.25708332 | 0.244219048 |
| Card6 | 0.00577115 | 0.134406709 | 0.224866981 | 0.042693538 | 0.03719126 |
| Carf | -0.00868789 | 0.12545769 | 0.211826965 | 0.082141431 | 0.088716757 |
| Carhsp1 | -0.00755699 | 0.230282782 | 0.343322989 | 0.112749906 | 0.117892051 |
| Carm1 | 0.012440709 | 0.030341042 | 0.066004835 | 0.101537066 | 0.090134945 |
| Carmil1 | 0.014855079 | 0.004438209 | 0.013037801 | 0.083767708 | 0.071422159 |
| Carmil2 | -2.69E-05 | 0.995149352 | 0.996584467 | 0.057540912 | 0.051850607 |
| Carmil3 | 0.020676966 | 0.000196244 | 0.000844198 | 0.105834498 | 0.078431279 |
| Carnmt1 | -0.006200337 | 0.423594357 | 0.542008994 | 0.202332197 | 0.200457738 |
| Cars | 0.018718562 | 0.051132046 | 0.101845063 | 0.416040698 | 0.394741626 |
| Cars2 | -0.015566564 | 0.006283112 | 0.017558039 | 0.089504271 | 0.09660511 |
| Cartpt | -0.016908553 | 0.003818544 | 0.011450921 | 0.043552291 | 0.071859501 |
| Casc3 | 0.039496262 | 3.43E-07 | 2.64E-06 | 0.208658949 | 0.182206446 |
| Casd1 | 0.003383953 | 0.603383684 | 0.706248034 | 0.127763227 | 0.122974842 |
| Cask | -0.003543379 | 0.635456486 | 0.7311999 | 0.169230529 | 0.176262894 |
| Caskin1 | -0.017408481 | 1.67E-05 | 9.19E-05 | 0.033048776 | 0.047421448 |
| Casp2 | 0.001101124 | 0.780192543 | 0.845411491 | 0.042669852 | 0.040387119 |
| Casp3 | 0.006028377 | 0.275535313 | 0.393587174 | 0.089046295 | 0.081377205 |
| Casp6 | -0.010215739 | 0.010829445 | 0.027691706 | 0.039089829 | 0.045425426 |
| Casp8 | 0.006633654 | 0.134774737 | 0.225288857 | 0.059570436 | 0.051714231 |
| Casp8ap2 | 0.015055303 | 0.037704469 | 0.07883072 | 0.166934027 | 0.15367881 |
| Casp9 | -0.003974449 | 0.287487543 | 0.406775417 | 0.035991892 | 0.038085548 |
| Casr | 0.010363216 | 0.013538542 | 0.033486302 | 0.053596253 | 0.044009345 |
| Cast | 0.026587476 | 0.000635147 | 0.002399883 | 0.198754679 | 0.182041396 |
| Castor2 | 0.092228037 | 6.22E-18 | 1.52E-16 | 0.665918214 | 0.553190444 |
| Casz1 | 0.027687024 | 0.001474055 | 0.005010757 | 0.283816762 | 0.254011463 |
| Cbarp | 0.105100569 | 5.26E-32 | 3.01E-30 | 0.338296927 | 0.255178702 |
| Cbfa2t2 | 0.014671965 | 0.011288282 | 0.028738698 | 0.10829285 | 0.091488111 |
| Cbfa2t3 | 0.016049526 | 0.097042193 | 0.172224206 | 0.37637495 | 0.355567122 |
| Cbfb | 0.008446428 | 0.158048517 | 0.254690483 | 0.108607008 | 0.100087869 |
| Cbl | 0.011821364 | 0.13448696 | 0.224885143 | 0.208325199 | 0.194922751 |
| Cblb | 0.017969068 | 0.000153522 | 0.000681212 | 0.070750756 | 0.055346322 |
| Cblc | -0.007374682 | 0.171701793 | 0.271340188 | 0.069455052 | 0.08041587 |
| Cbll1 | 0.037657391 | 3.83E-07 | 2.92E-06 | 0.192432275 | 0.164368031 |
| Cbr1 | -0.037106552 | 3.82E-05 | 0.000192973 | 0.332370488 | 0.343241633 |
| Cbr3 | -0.006045615 | 0.291680431 | 0.411271524 | 0.09065418 | 0.092983742 |
| Cbr4 | -0.002368322 | 0.661663679 | 0.752625985 | 0.090277626 | 0.085983818 |
| Cbs | -0.0082845 | 0.300155989 | 0.420114674 | 0.191879487 | 0.190261163 |
| Cbwd1 | 0.00037836 | 0.951315357 | 0.967407817 | 0.114966215 | 0.113691763 |
| Cbx1 | 0.014260853 | 0.141865922 | 0.234401851 | 0.512757678 | 0.491160996 |
| Cbx3 | 0.012487432 | 0.195307326 | 0.301053488 | 0.48853289 | 0.446017132 |
| Cbx4 | -0.029445435 | 0.000530977 | 0.00204685 | 0.189814921 | 0.230863851 |
| Cbx5 | -0.027340474 | 0.004626994 | 0.013496439 | 0.404979296 | 0.426021214 |
| Cbx6 | 0.037402273 | 7.12E-05 | 0.000341605 | 0.399717016 | 0.347927871 |
| Cbx7 | -0.00252301 | 0.705118488 | 0.787225763 | 0.131494452 | 0.132063029 |
| Cbx8 | -0.003000106 | 0.437066291 | 0.555445553 | 0.037747925 | 0.039019753 |
| Cby1 | 0.001408296 | 0.836508459 | 0.888187078 | 0.15071738 | 0.146012074 |
| Cc2d1a | -0.003563561 | 0.493448878 | 0.608486807 | 0.075558073 | 0.076306804 |
| Cc2d1b | 0.017212194 | 0.005479999 | 0.015596178 | 0.122947477 | 0.107064108 |
| Cc2d2a | 0.037907975 | 8.36E-09 | 8.43E-08 | 0.138066047 | 0.113253524 |
| Ccar1 | 0.008444278 | 0.384216043 | 0.505391472 | 0.396888486 | 0.384439231 |
| Ccar2 | -0.002044497 | 0.683713631 | 0.770495412 | 0.071976061 | 0.069338776 |

| | | | | | |
|---|---|---|---|---|---|
| Ccdc107 | -0.017414198 | 0.045554879 | 0.092306072 | 0.250427397 | 0.276074063 |
| Ccdc112 | -0.004848237 | 0.486158685 | 0.602016907 | 0.135046835 | 0.143965272 |
| Ccdc113 | -0.009869836 | 0.016123597 | 0.038867743 | 0.04241217 | 0.046371831 |
| Ccdc115 | -0.011611553 | 0.122433316 | 0.207677903 | 0.181441852 | 0.183349715 |
| Ccdc117 | 0.000892299 | 0.868353314 | 0.91010467 | 0.086447498 | 0.080238295 |
| Ccdc12 | -0.01211108 | 0.213059619 | 0.322894095 | 0.553744661 | 0.551353706 |
| Ccdc122 | -0.010443802 | 0.04698262 | 0.094862934 | 0.073789133 | 0.077709156 |
| Ccdc124 | -0.004528056 | 0.633019482 | 0.729345272 | 0.436566712 | 0.434835247 |
| Ccdc126 | 0.001202984 | 0.750017222 | 0.821874147 | 0.040074756 | 0.037728701 |
| Ccdc127 | -0.009905855 | 0.267004681 | 0.38359975 | 0.301447352 | 0.314377845 |
| Ccdc130 | 0.006762604 | 0.227622924 | 0.34019217 | 0.097695415 | 0.086093843 |
| Ccdc134 | -0.004588894 | 0.231926178 | 0.345032334 | 0.036877313 | 0.039952036 |
| Ccdc136 | -0.013936116 | 0.002417363 | 0.007705443 | 0.049197371 | 0.060688629 |
| Ccdc137 | 0.003030703 | 0.524724485 | 0.636795836 | 0.062950586 | 0.06060048 |
| Ccdc148 | 0.010570485 | 0.020258605 | 0.04694974 | 0.061461793 | 0.049248679 |
| Ccdc149 | 0.006327002 | 0.127277574 | 0.214378478 | 0.051652872 | 0.044485455 |
| Ccdc157 | -0.005471893 | 0.202599045 | 0.310176511 | 0.043314609 | 0.049827061 |
| Ccdc160 | -0.00350228 | 0.396635942 | 0.517180073 | 0.043465267 | 0.045936153 |
| Ccdc162 | 0.007530749 | 0.235026517 | 0.348843938 | 0.120838504 | 0.111471391 |
| Ccdc163 | -0.001173705 | 0.78898723 | 0.852281539 | 0.053635683 | 0.051512596 |
| Ccdc166 | 0.008339616 | 0.113202805 | 0.195238188 | 0.090286164 | 0.076882261 |
| Ccdc167 | -0.011357684 | 0.111233107 | 0.192764397 | 0.146165578 | 0.159804501 |
| Ccdc173 | 0.005153861 | 0.291942557 | 0.411462096 | 0.067912124 | 0.063000555 |
| Ccdc174 | 0.011051921 | 0.185935362 | 0.289411397 | 0.259579961 | 0.246908752 |
| Ccdc181 | 0.012012197 | 0.053646622 | 0.105813037 | 0.122029696 | 0.110219457 |
| Ccdc184 | 0.037734983 | 6.78E-12 | 9.85E-11 | 0.108973338 | 0.074561319 |
| Ccdc186 | 0.002508949 | 0.79524397 | 0.856945453 | 0.391420334 | 0.387954647 |
| Ccdc189 | -0.01952284 | 3.92E-05 | 0.000197342 | 0.054104573 | 0.066823873 |
| Ccdc191 | 0.00990933 | 0.136722008 | 0.227721666 | 0.136816598 | 0.126673613 |
| Ccdc22 | -0.013279162 | 0.000914598 | 0.003312863 | 0.040480596 | 0.046591641 |
| Ccdc25 | 0.014278259 | 0.110573253 | 0.191791823 | 0.323292636 | 0.317812936 |
| Ccdc28a | -0.016458978 | 0.003678941 | 0.011093879 | 0.088050494 | 0.094415539 |
| Ccdc28b | 0.006619589 | 0.318331926 | 0.440108514 | 0.141641301 | 0.130773669 |
| Ccdc30 | -0.00190936 | 0.757740411 | 0.827724975 | 0.112792928 | 0.113141493 |
| Ccdc32 | -0.001223563 | 0.849409256 | 0.896952957 | 0.125574546 | 0.126565472 |
| Ccdc34 | -0.019810515 | 0.012446431 | 0.031162555 | 0.196868461 | 0.208652232 |
| Ccdc43 | -0.011474605 | 0.061480214 | 0.119142093 | 0.106820093 | 0.115308003 |
| Ccdc47 | -0.102952254 | 8.20E-19 | 2.14E-17 | 0.641044782 | 0.809638393 |
| Ccdc50 | 0.013890894 | 0.096919965 | 0.172082247 | 0.24602612 | 0.236571435 |
| Ccdc51 | -0.004898139 | 0.192287397 | 0.29711031 | 0.036591367 | 0.038766102 |
| Ccdc58 | -0.011623025 | 0.073564583 | 0.137882182 | 0.124336446 | 0.129305299 |
| Ccdc59 | 0.002256046 | 0.827822775 | 0.881306726 | 0.601700751 | 0.628939736 |
| Ccdc6 | 0.044319492 | 6.89E-08 | 5.99E-07 | 0.255242366 | 0.213746676 |
| Ccdc66 | 0.003895114 | 0.538866886 | 0.649580145 | 0.122862777 | 0.115144487 |
| Ccdc71 | 0.013434885 | 0.013586732 | 0.033593645 | 0.097702024 | 0.080789476 |
| Ccdc71l | 0.000704405 | 0.857601542 | 0.902240254 | 0.042233799 | 0.040152622 |
| Ccdc73 | 0.009362894 | 0.015436947 | 0.037416605 | 0.046358033 | 0.038158386 |
| Ccdc77 | -0.008817685 | 0.034513751 | 0.073438978 | 0.043053328 | 0.048634198 |
| Ccdc82 | -0.003720246 | 0.560299379 | 0.668744137 | 0.114390876 | 0.118013553 |
| Ccdc84 | -0.008712894 | 0.060405136 | 0.117360496 | 0.053524298 | 0.061130083 |
| Ccdc85b | 0.077197086 | 5.26E-16 | 1.10E-14 | 0.478290387 | 0.38826022 |
| Ccdc86 | 0.003615094 | 0.591152833 | 0.695471874 | 0.138721674 | 0.13333629 |
| Ccdc88a | 0.002695405 | 0.797329262 | 0.858450165 | 0.552031064 | 0.550698475 |
| Ccdc9 | 0.006539049 | 0.251760316 | 0.366842383 | 0.089990035 | 0.085824321 |
| Ccdc90b | -0.00889046 | 0.15381337 | 0.249062888 | 0.115680233 | 0.116094009 |
| Ccdc91 | -0.024924604 | 0.001083851 | 0.00384429 | 0.187219354 | 0.195587566 |
| Ccdc92 | 0.019963241 | 0.00847361 | 0.022607145 | 0.201201987 | 0.18279593 |
| Ccdc92b | 0.016582588 | 0.000391918 | 0.001560291 | 0.071210888 | 0.054449551 |
| Ccdc93 | 0.006067892 | 0.151718356 | 0.246730735 | 0.051956688 | 0.047174752 |
| Ccdc97 | 0.008564054 | 0.124205359 | 0.210119963 | 0.091856193 | 0.085049511 |
| Ccdc9b | 0.010874647 | 0.012128341 | 0.030539168 | 0.057354909 | 0.045532617 |
| Ccl25 | -0.006999876 | 0.129902299 | 0.218269988 | 0.05555388 | 0.059877758 |
| Ccl27a | -0.001120131 | 0.878478553 | 0.917255772 | 0.170299948 | 0.169995098 |
| Ccl28 | -0.024537544 | 2.46E-07 | 1.93E-06 | 0.040029952 | 0.062906075 |
| Ccm2 | 0.008234242 | 0.260628222 | 0.376665316 | 0.192023175 | 0.173022002 |
| Ccnc | -0.007720206 | 0.255957499 | 0.371572167 | 0.146065452 | 0.145352779 |
| Ccnd1 | -0.018449721 | 0.108697356 | 0.189436287 | 0.718090138 | 0.755511031 |
| Ccnd2 | 0.086062033 | 3.13E-31 | 1.69E-29 | 2.521886029 | 2.319157775 |
| Ccnd3 | 0.00957685 | 0.202916002 | 0.310473618 | 0.192464525 | 0.180690749 |
| Ccndbp1 | -0.018924047 | 0.010233582 | 0.026397157 | 0.169476024 | 0.1789215 |
| Ccng1 | 0.039315643 | 0.000114415 | 0.000527467 | 0.378950185 | 0.341842398 |
| Ccng2 | 0.009270307 | 0.031833644 | 0.068622325 | 0.057578193 | 0.046915236 |

| | | | | | |
|---|---|---|---|---|---|
| Ccnh | -0.011990424 | 0.1517598 | 0.246730735 | 0.26100146 | 0.260106337 |
| Ccni | -0.01772698 | 0.074543299 | 0.139340502 | 0.573088602 | 0.566132222 |
| Ccnk | 0.014480271 | 0.021891364 | 0.050136122 | 0.119240683 | 0.109851804 |
| Ccnl1 | 0.008197752 | 0.382589972 | 0.503686843 | 0.342816448 | 0.345386269 |
| Ccnl2 | -0.00542476 | 0.568768622 | 0.675986374 | 0.389667447 | 0.389583707 |
| Ccnq | -0.008015106 | 0.216865859 | 0.327207137 | 0.130422843 | 0.129599386 |
| Ccnt1 | 0.028606591 | 0.00140553 | 0.004815705 | 0.326388523 | 0.297215305 |
| Ccnt2 | 0.004194505 | 0.628845317 | 0.726715297 | 0.274699053 | 0.264943238 |
| Ccny | -0.018387414 | 0.044061256 | 0.089841344 | 0.335518994 | 0.34905898 |
| Ccp110 | -0.004121313 | 0.366836485 | 0.489551724 | 0.052440946 | 0.056492961 |
| Ccpg1 | -0.026270335 | 0.001742091 | 0.005780413 | 0.243996903 | 0.259379539 |
| Ccpg1os | -0.035368509 | 2.11E-06 | 1.39E-05 | 0.17635832 | 0.190467044 |
| Ccs | -0.034069321 | 4.40E-06 | 2.73E-05 | 0.168850534 | 0.191317345 |
| Ccser1 | 0.004601588 | 0.431165196 | 0.54995907 | 0.099651559 | 0.094940142 |
| Ccser2 | 0.022594433 | 0.000588297 | 0.002242031 | 0.141209336 | 0.118943263 |
| Cct2 | -0.009545256 | 0.339153333 | 0.46147595 | 0.635208142 | 0.630869936 |
| Cct3 | -0.034781235 | 0.0002989 | 0.001227676 | 0.41437641 | 0.435649698 |
| Cct4 | -0.049195467 | 8.96E-07 | 6.34E-06 | 0.509624441 | 0.538751649 |
| Cct5 | -0.039130942 | 9.21E-05 | 0.000433306 | 0.612693641 | 0.629426792 |
| Cct6a | 0.023960992 | 0.018669417 | 0.043873357 | 0.616524002 | 0.578814983 |
| Cct7 | -0.057801016 | 3.91E-09 | 4.11E-08 | 0.780910436 | 0.838126534 |
| Cct8 | -0.019775104 | 0.045825678 | 0.09273881 | 0.534874023 | 0.546568489 |
| Ccz1 | 0.00208732 | 0.790371439 | 0.853302736 | 0.221387774 | 0.21062543 |
| Cd151 | -0.011306645 | 0.057880352 | 0.113153585 | 0.09972168 | 0.105529739 |
| Cd164 | -0.041476351 | 7.91E-11 | 1.03E-09 | 2.048797972 | 2.167809071 |
| Cd164l2 | 0.023740356 | 4.26E-06 | 2.66E-05 | 0.093687922 | 0.066412143 |
| Cd200 | 0.046427563 | 1.25E-06 | 8.60E-06 | 0.375440683 | 0.342213077 |
| Cd24a | 0.003813995 | 0.542606171 | 0.652792624 | 0.077073343 | 0.085263806 |
| Cd274 | -0.006097693 | 0.254349137 | 0.369457987 | 0.06773538 | 0.077000808 |
| Cd276 | 0.002702085 | 0.561355208 | 0.669713502 | 0.059464438 | 0.057930873 |
| Cd2ap | 0.072792931 | 2.29E-12 | 3.55E-11 | 0.640417395 | 0.556867439 |
| Cd2bp2 | 0.00574632 | 0.404540858 | 0.52439283 | 0.153504165 | 0.145329864 |
| Cd320 | -0.016035214 | 0.00851115 | 0.022694843 | 0.102321049 | 0.113738426 |
| Cd3eap | 0.009673848 | 0.143134449 | 0.236256896 | 0.135324324 | 0.124371974 |
| Cd44 | 0.050250153 | 2.72E-13 | 4.67E-12 | 0.146818319 | 0.110028487 |
| Cd47 | 0.03502684 | 0.00030841 | 0.001261943 | 1.020116557 | 1.004473635 |
| Cd59a | 0.008419873 | 0.290378954 | 0.409793031 | 0.215907218 | 0.202021469 |
| Cd63 | -0.03848088 | 1.83E-08 | 1.73E-07 | 2.364044195 | 2.456845256 |
| Cd79a | 0.050297496 | 8.29E-16 | 1.70E-14 | 0.119924315 | 0.060655209 |
| Cd81 | -0.128503354 | 1.27E-19 | 3.57E-18 | 0.460979473 | 0.608025089 |
| Cd82 | -0.003525769 | 0.686700826 | 0.772786831 | 0.252800167 | 0.268859783 |
| Cd9 | 0.05028184 | 3.84E-75 | 1.38E-72 | 0.045661504 | 0.00535703 |
| Cd99l2 | -0.008252325 | 0.14575935 | 0.23961319 | 0.086399398 | 0.092952243 |
| Cda | 0.022546368 | 0.01926444 | 0.044999733 | 0.466755672 | 0.410495697 |
| Cdadc1 | 0.007010497 | 0.351979388 | 0.474393093 | 0.192185834 | 0.183777686 |
| Cdc123 | -0.02354058 | 0.00871034 | 0.023137138 | 0.330280658 | 0.341230287 |
| Cdc14b | 0.007196258 | 0.319393947 | 0.441055367 | 0.166207295 | 0.156922368 |
| Cdc16 | 0.020391864 | 0.011730662 | 0.029699349 | 0.2445044 | 0.219689425 |
| Cdc23 | 0.008084762 | 0.229382737 | 0.34224355 | 0.148058065 | 0.135186484 |
| Cdc25a | 0.011258883 | 0.090154341 | 0.162324934 | 0.134638651 | 0.126955333 |
| Cdc26 | -0.008911715 | 0.328458806 | 0.451027756 | 0.3808515 | 0.368661995 |
| Cdc27 | 0.00463737 | 0.479009235 | 0.595591225 | 0.136069327 | 0.126616702 |
| Cdc34 | 0.025649562 | 0.007121253 | 0.019502204 | 0.392183132 | 0.366286779 |
| Cdc37 | -0.008591328 | 0.374457227 | 0.49648874 | 0.535913832 | 0.537197491 |
| Cdc37l1 | 0.012978783 | 0.185529261 | 0.288825536 | 0.482883295 | 0.47646874 |
| Cdc40 | 0.006015564 | 0.466371651 | 0.58391052 | 0.244627117 | 0.235205596 |
| Cdc42 | 0.016403397 | 0.097174739 | 0.172333603 | 0.839243474 | 0.792319129 |
| Cdc42bpa | 0.004263669 | 0.640403077 | 0.73568096 | 0.323909107 | 0.313287336 |
| Cdc42bpb | 0.015436797 | 0.024788847 | 0.055876924 | 0.161474605 | 0.139395174 |
| Cdc42bpg | 0.000662966 | 0.893865727 | 0.927936305 | 0.068488642 | 0.067826529 |
| Cdc42ep3 | 0.007580064 | 0.039378492 | 0.08182041 | 0.039026665 | 0.03274002 |
| Cdc42ep5 | 0.001689726 | 0.677666004 | 0.765973853 | 0.045212651 | 0.042525145 |
| Cdc42se1 | 0.043736408 | 3.10E-11 | 4.22E-10 | 0.152182825 | 0.117712039 |
| Cdc42se2 | -0.017761343 | 0.06774795 | 0.128969173 | 0.338572343 | 0.37778662 |
| Cdc5l | 0.031690664 | 0.001163235 | 0.004079717 | 0.523235588 | 0.451860045 |
| Cdc73 | -0.01226686 | 0.186048556 | 0.289494888 | 0.352786607 | 0.354109541 |
| Cdcp1 | 0.010676319 | 0.090847474 | 0.163319685 | 0.121388955 | 0.110784306 |
| Cdh1 | 0.102446956 | 3.69E-24 | 1.41E-22 | 0.600267711 | 0.484746038 |
| Cdh2 | 0.0825442 | 9.15E-17 | 2.01E-15 | 0.533248164 | 0.434258311 |
| Cdh22 | 0.004924733 | 0.34286317 | 0.464962441 | 0.083847143 | 0.071094506 |
| Cdh4 | -0.022025823 | 1.88E-05 | 0.000102304 | 0.053738518 | 0.072548317 |
| Cdh7 | 0.027059455 | 3.54E-08 | 3.21E-07 | 0.084721031 | 0.056866522 |

| | | | | | |
|---|---|---|---|---|---|
| Cdh8 | 0.030365517 | 1.87E-05 | 0.000101903 | 0.174826815 | 0.143359434 |
| Cdhr1 | 0.020192406 | 0.001851752 | 0.006106761 | 0.1408376 | 0.116668483 |
| Cdip1 | 0.016505543 | 0.026006015 | 0.057962053 | 0.185104944 | 0.169911758 |
| Cdipt | -0.015389399 | 0.093491133 | 0.16698894 | 0.340527614 | 0.354840119 |
| Cdk10 | -0.012901749 | 0.015445414 | 0.037427796 | 0.077404179 | 0.082798872 |
| Cdk11b | 0.021499065 | 0.025287256 | 0.05671112 | 0.396935124 | 0.378506746 |
| Cdk12 | -0.002263267 | 0.812959383 | 0.871111112 | 0.365743173 | 0.369333644 |
| Cdk13 | 0.030615531 | 0.000683444 | 0.002560479 | 0.337105237 | 0.299627594 |
| Cdk14 | -0.0602012 | 5.83E-12 | 8.56E-11 | 0.242624756 | 0.280158211 |
| Cdk16 | 0.003061306 | 0.715269115 | 0.795270623 | 0.260668926 | 0.258293992 |
| Cdk17 | 0.006343961 | 0.192883442 | 0.297889249 | 0.070978115 | 0.064009277 |
| Cdk2ap1 | 0.001749355 | 0.8293326 | 0.882430944 | 0.276504223 | 0.24174458 |
| Cdk2ap2 | -0.078199915 | 2.17E-15 | 4.34E-14 | 0.401505109 | 0.463794772 |
| Cdk4 | -0.015275034 | 0.117264277 | 0.200605895 | 0.486445915 | 0.476803381 |
| Cdk5 | -0.001293684 | 0.852387826 | 0.898680132 | 0.155563751 | 0.151399735 |
| Cdk5r2 | 0.00780751 | 0.230546569 | 0.343610876 | 0.134315288 | 0.123788702 |
| Cdk5rap2 | 0.008804953 | 0.076703286 | 0.142746811 | 0.070870436 | 0.062133648 |
| Cdk5rap3 | -0.084067114 | 6.75E-19 | 1.79E-17 | 0.385316933 | 0.44857051 |
| Cdk7 | -0.005258639 | 0.415958466 | 0.535267797 | 0.125022023 | 0.129863862 |
| Cdk8 | 0.031950152 | 0.001018126 | 0.003636377 | 0.444386104 | 0.392061227 |
| Cdk9 | -0.018567332 | 0.019662786 | 0.045743386 | 0.226769962 | 0.229586938 |
| Cdkal1 | -0.003494113 | 0.480845572 | 0.597187105 | 0.06967371 | 0.068608795 |
| Cdkl1 | 0.018447415 | 0.001248774 | 0.00433601 | 0.111262351 | 0.087275671 |
| Cdkl2 | -0.009272681 | 0.206263776 | 0.314506969 | 0.161826025 | 0.169835409 |
| Cdkl3 | -0.001805514 | 0.660970967 | 0.75210203 | 0.043205437 | 0.044137122 |
| Cdkl5 | 0.020491645 | 0.000174267 | 0.000762995 | 0.093125027 | 0.074593872 |
| Cdkn1a | -0.060781714 | 8.38E-06 | 4.90E-05 | 0.635365238 | 0.709466556 |
| Cdkn1b | -0.006778103 | 0.487932966 | 0.603598969 | 0.409989692 | 0.396640047 |
| Cdkn2aip | -0.024197835 | 0.000596871 | 0.002272036 | 0.142170991 | 0.158155372 |
| Cdkn2aipnl | 0.00803127 | 0.181927131 | 0.284082166 | 0.109870954 | 0.101570194 |
| Cdkn2b | -0.00334504 | 0.634495638 | 0.730520142 | 0.153978109 | 0.148168286 |
| Cdkn2c | -0.032368839 | 1.95E-09 | 2.13E-08 | 0.071861181 | 0.088618629 |
| Cdkn2d | -0.007806844 | 0.258646826 | 0.374810009 | 0.154878318 | 0.151309185 |
| Cdon | 0.007794134 | 0.077668926 | 0.144021581 | 0.059375576 | 0.048939614 |
| Cdpf1 | 0.000677736 | 0.925168401 | 0.949286247 | 0.173393821 | 0.169598306 |
| Cdr2 | 0.006955981 | 0.292495274 | 0.411882829 | 0.133626127 | 0.127488365 |
| Cds1 | 0.0102683 | 0.118418694 | 0.202118424 | 0.136824057 | 0.122258686 |
| Cds2 | 0.036625725 | 4.60E-05 | 0.000228062 | 0.351055103 | 0.314704085 |
| Cdv3 | 0.06649275 | 1.73E-11 | 2.43E-10 | 0.488653258 | 0.431633212 |
| Cdyl | 0.019535985 | 0.000683432 | 0.002560479 | 0.096545093 | 0.087745383 |
| Cebpb | -0.096087029 | 2.15E-17 | 4.99E-16 | 0.976142955 | 1.113658286 |
| Cebpd | 0.049229092 | 5.00E-05 | 0.000245704 | 0.409610874 | 0.408229125 |
| Cebpg | -0.033543121 | 0.001549318 | 0.005215536 | 0.710594749 | 0.753353107 |
| Cebpz | 0.060964366 | 3.82E-10 | 4.59E-09 | 0.468822115 | 0.428036154 |
| Cebpzos | -0.020395735 | 0.031937592 | 0.06878913 | 0.439710033 | 0.444912028 |
| Celf1 | 0.060367041 | 1.01E-09 | 1.14E-08 | 0.542732584 | 0.479341684 |
| Celf2 | 0.01887195 | 0.007222926 | 0.01975838 | 0.168858359 | 0.145385415 |
| Celf3 | 0.03631564 | 0.000247659 | 0.001037371 | 0.509982292 | 0.496186872 |
| Celf4 | -0.037596261 | 0.000209367 | 0.000894712 | 0.450340404 | 0.485259714 |
| Celf6 | -0.028173014 | 8.56E-05 | 0.000405287 | 0.14659358 | 0.170802109 |
| Celsr2 | 0.008722597 | 0.091395088 | 0.164149833 | 0.0812357 | 0.069815034 |
| Celsr3 | 0.00044595 | 0.936653592 | 0.957014679 | 0.088565821 | 0.089542848 |
| Cemip2 | 0.010210654 | 0.031105987 | 0.067412485 | 0.062805941 | 0.057096074 |
| Cenpb | 0.003925806 | 0.664310701 | 0.754313085 | 0.356344725 | 0.333132728 |
| Cenpc1 | 0.003712158 | 0.517109649 | 0.629840273 | 0.095453805 | 0.091241483 |
| Cenpj | -0.001220826 | 0.803093969 | 0.863108508 | 0.063066802 | 0.06479547 |
| Cenpq | -0.001070319 | 0.849553257 | 0.896952957 | 0.088441787 | 0.089074868 |
| Cenps | 0.007150689 | 0.2094423 | 0.318260593 | 0.094588942 | 0.086936793 |
| Cenpv | -0.024278046 | 0.011693383 | 0.029620393 | 0.485766027 | 0.485755439 |
| Cenpx | -0.022300804 | 0.016849597 | 0.04035767 | 0.373647715 | 0.387062282 |
| Cep104 | 0.00498444 | 0.484150424 | 0.600447815 | 0.161627751 | 0.156266886 |
| Cep120 | 0.018327979 | 0.003548519 | 0.010748916 | 0.12850524 | 0.111448921 |
| Cep126 | -0.05574301 | 9.85E-14 | 1.77E-12 | 0.128226933 | 0.179245246 |
| Cep131 | -0.001088258 | 0.764865943 | 0.833541833 | 0.035708854 | 0.034939891 |
| Cep135 | 0.006826962 | 0.096926803 | 0.172082247 | 0.046207098 | 0.041034835 |
| Cep162 | 0.00602519 | 0.379776706 | 0.501220354 | 0.143588408 | 0.135343019 |
| Cep164 | 0.011975775 | 0.112960145 | 0.194923418 | 0.189756095 | 0.179307057 |
| Cep170 | 0.032934776 | 8.25E-05 | 0.000391903 | 0.258723228 | 0.223042858 |
| Cep170b | 0.046484 | 5.40E-08 | 4.77E-07 | 0.285974836 | 0.248240702 |
| Cep19 | -0.008231412 | 0.247630152 | 0.36196968 | 0.158420641 | 0.162883478 |
| Cep192 | 0.002549425 | 0.632523832 | 0.729098641 | 0.080094962 | 0.077466574 |
| Cep20 | -0.01972495 | 0.015228106 | 0.03700266 | 0.234286282 | 0.243599003 |

| | | | | | |
|---|---|---|---|---|---|
| Cep250 | 0.003539137 | 0.36893693 | 0.491343128 | 0.042065048 | 0.039703887 |
| Cep290 | 0.006416114 | 0.506099248 | 0.61999917 | 0.360104786 | 0.353779091 |
| Cep295 | 0.004071435 | 0.399797187 | 0.519974184 | 0.063362487 | 0.061325819 |
| Cep350 | 0.003639356 | 0.687018979 | 0.772787055 | 0.288683431 | 0.283044588 |
| Cep43 | 0.011762364 | 0.005531087 | 0.015695886 | 0.053104014 | 0.044281915 |
| Cep44 | -0.000520894 | 0.931483083 | 0.953551493 | 0.106048404 | 0.106898433 |
| Cep57 | -0.005070091 | 0.539941045 | 0.650632975 | 0.241167273 | 0.236487467 |
| Cep63 | -0.01738386 | 0.007691924 | 0.020836133 | 0.123118231 | 0.13036173 |
| Cep68 | -0.010619947 | 0.029866235 | 0.065205374 | 0.060536421 | 0.068581641 |
| Cep70 | -0.014132809 | 0.057959236 | 0.113262249 | 0.171196582 | 0.177885407 |
| Cep76 | 0.006718234 | 0.064541147 | 0.12385887 | 0.037981945 | 0.032996172 |
| Cep78 | 0.002468786 | 0.641324479 | 0.736074138 | 0.080371534 | 0.077189391 |
| Cep83 | 0.021181512 | 0.008089453 | 0.021755369 | 0.217284638 | 0.199527146 |
| Cep83os | -0.01069469 | 0.033626886 | 0.071819109 | 0.065219594 | 0.071692579 |
| Cep85 | -0.013860977 | 0.032336092 | 0.06951647 | 0.117300438 | 0.128191124 |
| Cep89 | -0.008680546 | 0.095871622 | 0.170613932 | 0.074099874 | 0.078586923 |
| Cep97 | -0.001796133 | 0.71476402 | 0.794981787 | 0.065751387 | 0.066496664 |
| Cept1 | -0.002559073 | 0.650432332 | 0.74341678 | 0.092289096 | 0.09273153 |
| Cerk | 0.044122536 | 5.24E-07 | 3.90E-06 | 0.320706131 | 0.277341408 |
| Cerkl | 0.003094903 | 0.684740256 | 0.771473493 | 0.18661695 | 0.183361322 |
| Cers2 | -0.024587694 | 0.002462433 | 0.007833191 | 0.217958419 | 0.238330622 |
| Cers4 | -0.013810727 | 0.018786854 | 0.044074758 | 0.091112881 | 0.101342612 |
| Cers5 | -0.022412419 | 0.001318231 | 0.00454785 | 0.143386474 | 0.15907026 |
| Cers6 | 0.02127538 | 0.000895568 | 0.00325363 | 0.130283593 | 0.111642466 |
| Cert1 | 0.014910423 | 0.099502716 | 0.176044661 | 0.326264044 | 0.302616228 |
| Cetn2 | 0.011866984 | 0.170437881 | 0.269737438 | 0.322232558 | 0.292998276 |
| Cetn3 | -0.02034062 | 0.027256953 | 0.060280278 | 0.365166104 | 0.371910317 |
| Cetn4 | -0.016003329 | 0.015643325 | 0.037860196 | 0.1190874 | 0.133498097 |
| Cfap126 | -0.003581589 | 0.557323931 | 0.665995796 | 0.119147715 | 0.110074944 |
| Cfap20 | 0.004189058 | 0.662280135 | 0.752974795 | 0.487588263 | 0.47417799 |
| Cfap298 | 0.0123403 | 0.153853601 | 0.249086546 | 0.293336965 | 0.280820667 |
| Cfap36 | -0.04588067 | 3.25E-06 | 2.06E-05 | 0.879155206 | 0.924270632 |
| Cfap410 | 0.00091739 | 0.8975683 | 0.930194969 | 0.176683183 | 0.16133278 |
| Cfap97 | 0.019130847 | 0.023288652 | 0.05286301 | 0.277258093 | 0.246517215 |
| Cfdp1 | -0.021025424 | 0.028309121 | 0.062323659 | 0.493013509 | 0.495838727 |
| Cfl1 | 0.032556545 | 0.000474726 | 0.001852784 | 1.144424015 | 1.072293698 |
| Cfl2 | 0.008351023 | 0.381901837 | 0.50309616 | 0.391749642 | 0.376918177 |
| Cflar | 0.005483825 | 0.510404966 | 0.624366775 | 0.243279572 | 0.23841508 |
| Cggbp1 | 0.043512969 | 5.42E-06 | 3.30E-05 | 0.414362706 | 0.384591116 |
| Cgn | 0.019533044 | 0.00228093 | 0.007333069 | 0.12940904 | 0.112645333 |
| Cgref1 | -0.031287976 | 2.03E-10 | 2.50E-09 | 0.051608026 | 0.073660152 |
| Cgrrf1 | 0.047581622 | 1.56E-09 | 1.74E-08 | 0.230927538 | 0.191662482 |
| Chac1 | -0.002399556 | 0.830791372 | 0.883789662 | 0.58314218 | 0.568435731 |
| Chac2 | 0.02118868 | 1.19E-05 | 6.69E-05 | 0.07540816 | 0.058861374 |
| Chadl | -0.005238322 | 0.238223284 | 0.352138515 | 0.057421078 | 0.055869794 |
| Champ1 | 0.004875729 | 0.350979518 | 0.473433388 | 0.081536787 | 0.073764326 |
| Chaserr | -0.023814393 | 0.008931647 | 0.02359638 | 0.323574517 | 0.337929488 |
| Chchd1 | 0.017887688 | 0.072926396 | 0.136791515 | 0.808610542 | 0.804295341 |
| Chchd10 | -0.118883741 | 2.25E-25 | 9.27E-24 | 1.176215363 | 1.423514036 |
| Chchd2 | 0.015535137 | 0.010692285 | 0.027398628 | 2.328241453 | 2.19359858 |
| Chchd3 | 0.011679341 | 0.150702536 | 0.245744725 | 0.25529494 | 0.235317709 |
| Chchd4 | -0.023197948 | 0.004839159 | 0.014018564 | 0.23439485 | 0.250607276 |
| Chchd5 | -0.006947705 | 0.336978249 | 0.459288173 | 0.167175161 | 0.166561311 |
| Chchd6 | -0.032772015 | 0.000308412 | 0.001261943 | 0.318477547 | 0.347118966 |
| Chchd7 | 0.009575971 | 0.337396005 | 0.459454503 | 0.603031865 | 0.580225525 |
| Chd1 | 0.012181969 | 0.122316337 | 0.207554223 | 0.212849853 | 0.200667588 |
| Chd1l | -0.000398604 | 0.924330258 | 0.948726644 | 0.046845461 | 0.047231211 |
| Chd2 | 0.121329179 | 7.06E-39 | 5.81E-37 | 0.344241805 | 0.233621694 |
| Chd3 | -0.031281477 | 0.002320351 | 0.007438443 | 0.491581163 | 0.507391244 |
| Chd4 | 0.063651799 | 5.05E-10 | 5.98E-09 | 1.022550956 | 0.935221989 |
| Chd5 | -0.03015342 | 8.57E-05 | 0.000405754 | 0.151646182 | 0.177530899 |
| Chd6 | 0.019891129 | 0.017424938 | 0.041520894 | 0.253033346 | 0.230736061 |
| Chd7 | 0.013947277 | 0.166185692 | 0.264601588 | 0.431866067 | 0.42096773 |
| Chd8 | 0.043466666 | 1.23E-06 | 8.42E-06 | 0.320878717 | 0.274770605 |
| Chd9 | 0.021636893 | 0.031570959 | 0.068222463 | 0.512246171 | 0.480965947 |
| Cherp | 0.002631766 | 0.681616447 | 0.768755812 | 0.124634117 | 0.121962446 |
| Chfr | -0.005972194 | 0.318426041 | 0.440173179 | 0.102829574 | 0.105289085 |
| Chga | -0.010248351 | 0.019029109 | 0.044524909 | 5.328309275 | 5.287266421 |
| Chgb | -0.098590419 | 2.23E-60 | 4.01E-58 | 2.993444953 | 3.356842913 |
| Chic1 | 0.050076021 | 5.25E-07 | 3.91E-06 | 1.726062633 | 1.589602661 |
| Chic2 | -0.011319572 | 0.203722511 | 0.311560523 | 0.313734099 | 0.309768446 |
| Chid1 | -0.024186155 | 0.003424014 | 0.010431676 | 0.241406137 | 0.254994227 |

| | | | | | |
|---|---|---|---|---|---|
| Chka | -0.072570758 | 1.89E-11 | 2.65E-10 | 0.51794266 | 0.5976725 |
| Chkb | -0.001073712 | 0.823405072 | 0.87823852 | 0.065513555 | 0.063581481 |
| Chm | 0.01905777 | 0.006810733 | 0.018816126 | 0.164790196 | 0.146570848 |
| Chmp1a | -0.0026199 | 0.752541385 | 0.823759774 | 0.269919745 | 0.258845794 |
| Chmp1b | 0.005565243 | 0.378745032 | 0.500519364 | 0.120689842 | 0.115584373 |
| Chmp2a | -0.021724081 | 0.023418239 | 0.053119952 | 1.011354024 | 1.048552357 |
| Chmp2b | 0.009259176 | 0.240825132 | 0.354796475 | 0.228508967 | 0.209240192 |
| Chmp3 | -0.008046252 | 0.404539874 | 0.52439283 | 0.480133127 | 0.475242555 |
| Chmp4b | 0.020641 | 0.038723038 | 0.080630837 | 0.92212829 | 0.895890613 |
| Chmp4c | 0.081809483 | 5.73E-23 | 2.05E-21 | 0.274576073 | 0.196662243 |
| Chmp5 | -0.024992275 | 0.008242234 | 0.022062498 | 0.788712933 | 0.797457769 |
| Chmp6 | 0.001180359 | 0.844504487 | 0.893878348 | 0.112631658 | 0.106085593 |
| Chmp7 | 0.015492885 | 0.022549388 | 0.051304739 | 0.163817804 | 0.139556287 |
| Chn1 | 0.012580655 | 0.020039504 | 0.046518961 | 0.094920977 | 0.078426595 |
| Chordc1 | -0.025578969 | 0.005823477 | 0.016423646 | 0.337042342 | 0.365088438 |
| Chp1 | 0.027643903 | 0.000800821 | 0.002946852 | 0.258744302 | 0.229079867 |
| Chpf | 0.01035863 | 0.220433976 | 0.331416298 | 0.256482639 | 0.247898835 |
| Chpf2 | -0.005857135 | 0.345326115 | 0.467650159 | 0.110649292 | 0.113632423 |
| Chpt1 | -0.015274222 | 0.018541854 | 0.043628434 | 0.126074278 | 0.131279636 |
| Chrac1 | 0.024842864 | 0.009984834 | 0.025858433 | 0.488504459 | 0.458708327 |
| Chrm3 | 0.033340028 | 1.29E-07 | 1.07E-06 | 0.133506335 | 0.103316733 |
| Chrna1os | -0.002706113 | 0.596481306 | 0.700023087 | 0.068145256 | 0.071323585 |
| Chrnb2 | 0.014871543 | 0.007287528 | 0.019918288 | 0.095781858 | 0.081947996 |
| Chst11 | -0.017251651 | 0.035603653 | 0.075272591 | 0.220235496 | 0.240634956 |
| Chst12 | 0.013229761 | 0.020046189 | 0.046518961 | 0.110993269 | 0.086619743 |
| Chtop | 0.019287517 | 0.034425736 | 0.073283777 | 0.354863247 | 0.332358376 |
| Chuk | 0.012818467 | 0.130264265 | 0.21877971 | 0.265171939 | 0.250641738 |
| Churc1 | -0.036016043 | 0.000232212 | 0.000980273 | 0.660585664 | 0.668265813 |
| Ciao1 | 0.001376267 | 0.828544108 | 0.881869732 | 0.130547615 | 0.119181748 |
| Ciao2a | -0.019039883 | 0.016430865 | 0.039490945 | 0.221346036 | 0.231960451 |
| Ciao2b | -0.005742559 | 0.480171662 | 0.596502543 | 0.247610662 | 0.245742495 |
| Ciao3 | 0.008121138 | 0.156311105 | 0.252560505 | 0.098794787 | 0.091746125 |
| Ciapin1 | -0.048590143 | 8.23E-10 | 9.51E-09 | 0.158852717 | 0.204096221 |
| Ciart | -0.002164696 | 0.629059249 | 0.726715297 | 0.052258073 | 0.0521805 |
| Cib1 | -0.040227379 | 8.59E-06 | 5.01E-05 | 0.311786769 | 0.33998865 |
| Cib2 | 0.002303243 | 0.813587735 | 0.871618127 | 0.505105333 | 0.463178655 |
| Cib3 | 0.009917784 | 0.259770894 | 0.375817355 | 0.271996991 | 0.221510558 |
| Cibar1 | -0.036623586 | 0.000180104 | 0.000782659 | 0.490669024 | 0.509346953 |
| Cibar2 | 0.005083283 | 0.305898792 | 0.426678344 | 0.076481214 | 0.064267411 |
| Cic | 0.025948375 | 0.000106361 | 0.000493813 | 0.144199514 | 0.126658525 |
| Cinp | -0.00368212 | 0.528203961 | 0.63966105 | 0.096594718 | 0.098979449 |
| Cipc | 0.005977778 | 0.260241714 | 0.376301228 | 0.079699294 | 0.076300648 |
| Cir1 | 0.011647686 | 0.070857656 | 0.133607085 | 0.133102998 | 0.119824199 |
| Cirbp | -0.055997389 | 4.43E-09 | 4.63E-08 | 1.031096003 | 1.074551256 |
| Cisd1 | 0.012609143 | 0.180892039 | 0.283102447 | 0.992597308 | 0.965678019 |
| Cisd2 | -0.007210894 | 0.459241086 | 0.577586266 | 0.4933576 | 0.500612274 |
| Cisd3 | -0.037745767 | 5.43E-07 | 4.04E-06 | 0.166764977 | 0.196518084 |
| Cish | 0.018999291 | 0.003945438 | 0.011777481 | 0.140449379 | 0.118003745 |
| Cited2 | 0.111640855 | 1.21E-24 | 4.80E-23 | 0.543006827 | 0.396955522 |
| Cited4 | -0.008292032 | 0.147600907 | 0.241741242 | 0.09182947 | 0.092946205 |
| Ciz1 | 0.001711274 | 0.772452555 | 0.839551004 | 0.10467885 | 0.10075519 |
| Ckap4 | 0.000631569 | 0.945009028 | 0.96313846 | 0.333269691 | 0.326876591 |
| Ckap5 | 0.014314437 | 0.083350557 | 0.152433054 | 0.259372904 | 0.234839403 |
| Ckb | -0.036201297 | 0.012464438 | 0.031199606 | 0.790483828 | 0.851175409 |
| Ckmt1 | -0.00582418 | 0.353561906 | 0.475952485 | 0.122014797 | 0.117222315 |
| Cks1b | -0.036255013 | 1.42E-11 | 2.01E-10 | 0.056792871 | 0.088548983 |
| Clasp1 | -0.055497278 | 8.33E-14 | 1.50E-12 | 0.13535499 | 0.185765977 |
| Clasp2 | 0.003834363 | 0.652866848 | 0.745235587 | 0.243923394 | 0.24405268 |
| Clasrp | 0.003328136 | 0.448010702 | 0.566905739 | 0.05677559 | 0.051443977 |
| Clba1 | 0.002573533 | 0.641542024 | 0.736088758 | 0.091967258 | 0.085338458 |
| Clcc1 | 0.026652445 | 3.48E-05 | 0.000178232 | 0.140372733 | 0.116574752 |
| Clcn3 | -0.008063269 | 0.42938178 | 0.548115764 | 0.705126797 | 0.71018627 |
| Clcn4 | 0.020838367 | 0.017018065 | 0.040681001 | 0.30101486 | 0.277212659 |
| Clcn5 | -0.004794689 | 0.482116568 | 0.598459816 | 0.1266867 | 0.13361411 |
| Clcn7 | 0.015749519 | 0.001879648 | 0.00619036 | 0.078752505 | 0.065812528 |
| Cldn12 | 0.003369749 | 0.580903121 | 0.687048679 | 0.106670666 | 0.107516102 |
| Cldn3 | 0.033090275 | 0.000853973 | 0.003122349 | 0.348473449 | 0.320376139 |
| Cldn4 | 0.088769089 | 1.67E-23 | 6.18E-22 | 0.322183562 | 0.221012729 |
| Cldn7 | 0.053963569 | 1.07E-06 | 7.42E-06 | 1.087958537 | 1.088590645 |
| Cldnd1 | 0.004753934 | 0.587988357 | 0.692815756 | 0.302684897 | 0.291987914 |
| Clec11a | -0.016511587 | 0.000915697 | 0.003313138 | 0.065185759 | 0.073574909 |
| Clec16a | -0.001165845 | 0.779712891 | 0.845157198 | 0.045603474 | 0.046674187 |

| | | | | | |
|---|---|---|---|---|---|
| Clic1 | 0.001245754 | 0.901981409 | 0.933176892 | 0.800149411 | 0.796594038 |
| Clic4 | 0.019570514 | 0.061450349 | 0.119126679 | 0.648312179 | 0.642860248 |
| Clic5 | 0.051303709 | 1.39E-37 | 1.02E-35 | 0.069734363 | 0.027323143 |
| Clint1 | 0.040855529 | 4.65E-05 | 0.000230001 | 0.528449645 | 0.492057762 |
| Clip1 | 0.018455441 | 0.071740576 | 0.134931685 | 0.54090666 | 0.54187991 |
| Clip2 | 0.008916335 | 0.030253668 | 0.06585897 | 0.048292545 | 0.042148534 |
| Clip3 | 0.017237953 | 0.001857898 | 0.006122874 | 0.09783433 | 0.081355741 |
| Clip4 | 0.026062881 | 2.78E-11 | 3.80E-10 | 0.052952992 | 0.030670481 |
| Clk1 | -0.005983141 | 0.543051532 | 0.65297562 | 0.366639826 | 0.373921436 |
| Clk2 | 0.007398798 | 0.177600565 | 0.2790743 | 0.088777651 | 0.082256705 |
| Clk3 | 0.005668191 | 0.41426864 | 0.533870589 | 0.153687154 | 0.146089826 |
| Clk4 | 0.003865521 | 0.650222771 | 0.743264642 | 0.239536569 | 0.242148599 |
| Cln3 | -0.001919243 | 0.752585303 | 0.823759774 | 0.11015913 | 0.107336205 |
| Cln5 | -0.012146036 | 0.066841055 | 0.127542441 | 0.122941879 | 0.13511548 |
| Cln6 | 0.056829695 | 1.31E-09 | 1.47E-08 | 0.409436799 | 0.346595667 |
| Cln8 | -0.002776314 | 0.550729247 | 0.660033251 | 0.056732639 | 0.059400559 |
| Clns1a | -0.009582609 | 0.288989398 | 0.408457078 | 0.345372206 | 0.347545896 |
| Clock | 0.003551843 | 0.710891855 | 0.791762013 | 0.396598271 | 0.392232524 |
| Clp1 | 0.00632189 | 0.145405007 | 0.239192466 | 0.056362634 | 0.048804312 |
| Clpb | -0.032897449 | 8.66E-06 | 5.04E-05 | 0.15610115 | 0.183212677 |
| Clpp | -0.052214803 | 1.48E-07 | 1.22E-06 | 0.535217215 | 0.582013579 |
| Clptm1 | -0.004580868 | 0.58584433 | 0.69095943 | 0.257618013 | 0.258170703 |
| Clptm1l | 0.003145831 | 0.747230469 | 0.819651881 | 0.437039188 | 0.442826927 |
| Clpx | 0.024577844 | 0.009338455 | 0.024477881 | 0.378403312 | 0.354734727 |
| Clstn1 | 0.019350405 | 0.024575115 | 0.055433705 | 0.296616933 | 0.266310031 |
| Clstn3 | -0.008717538 | 0.07998804 | 0.147380307 | 0.061482504 | 0.068486873 |
| Clta | 0.028805669 | 0.001103286 | 0.003898998 | 1.233337625 | 1.163714881 |
| Cltb | 0.024026677 | 0.010231695 | 0.026397157 | 1.014591878 | 0.990638946 |
| Cltc | 0.04629962 | 9.31E-07 | 6.53E-06 | 1.154788403 | 1.093400396 |
| Cltrn | -0.125124565 | 1.42E-38 | 1.12E-36 | 1.575474057 | 1.760692181 |
| Clu | 0.079679232 | 6.94E-14 | 1.25E-12 | 0.499585301 | 0.406194561 |
| Cluap1 | -0.019823967 | 0.001364087 | 0.004689412 | 0.106631536 | 0.117549456 |
| Cluh | 0.013462914 | 0.063505329 | 0.122236957 | 0.177567198 | 0.160213401 |
| Clybl | -0.012803732 | 0.00507697 | 0.014626537 | 0.051121807 | 0.059748655 |
| Cmas | -0.054758245 | 8.38E-08 | 7.18E-07 | 0.604574673 | 0.677499178 |
| Cmbl | -0.042083771 | 5.63E-07 | 4.16E-06 | 0.215778644 | 0.245932223 |
| Cmc1 | 0.007958366 | 0.350825962 | 0.473318068 | 0.288530939 | 0.27902758 |
| Cmc2 | -0.007223653 | 0.283642397 | 0.402682738 | 0.135878993 | 0.140612927 |
| Cmc4 | -0.020794883 | 0.000794772 | 0.002927919 | 0.10829611 | 0.121627569 |
| Cmip | 0.022100394 | 0.027681909 | 0.061108882 | 0.537020934 | 0.49862888 |
| Cmpk1 | -0.005397392 | 0.579694134 | 0.686036077 | 0.719136081 | 0.710625503 |
| Cmpk2 | 0.016524206 | 0.015770438 | 0.038139353 | 0.142280591 | 0.121892397 |
| Cmss1 | -0.031376555 | 5.72E-08 | 5.03E-07 | 0.080453607 | 0.105743879 |
| Cmtm4 | 0.039946607 | 4.05E-05 | 0.000203344 | 0.485760445 | 0.442953013 |
| Cmtm6 | 0.005304148 | 0.394795593 | 0.515402545 | 0.121677131 | 0.111352351 |
| Cmtr2 | 0.008544569 | 0.035213718 | 0.074618084 | 0.046245523 | 0.041493467 |
| Cnbp | -0.017070754 | 0.084330974 | 0.15390759 | 0.894658636 | 0.889547503 |
| Cndp2 | -0.006313219 | 0.369021097 | 0.491377775 | 0.155176368 | 0.156098896 |
| Cnep1r1 | -0.002062287 | 0.765722642 | 0.833914587 | 0.154046626 | 0.151015564 |
| Cnih1 | 0.008609285 | 0.31523277 | 0.436752599 | 0.273986131 | 0.262404605 |
| Cnih4 | 0.007005231 | 0.340108411 | 0.462452304 | 0.176413582 | 0.166231732 |
| Cnksr1 | 0.011706475 | 0.017616801 | 0.041895923 | 0.074430559 | 0.064244814 |
| Cnksr3 | 0.011534764 | 0.001041965 | 0.003711975 | 0.040345166 | 0.028582955 |
| Cnn3 | 0.074610749 | 2.45E-14 | 4.57E-13 | 0.443580111 | 0.378814073 |
| Cnnm1 | -0.000523052 | 0.900159248 | 0.93179678 | 0.044259636 | 0.044884124 |
| Cnnm2 | -0.010094659 | 0.087305192 | 0.158354679 | 0.098124418 | 0.103079219 |
| Cnnm3 | -0.001866077 | 0.72391496 | 0.80090399 | 0.075657577 | 0.077254666 |
| Cnot1 | 0.037352581 | 2.43E-05 | 0.000128717 | 0.323381045 | 0.282936922 |
| Cnot10 | 0.008085649 | 0.283918792 | 0.403016279 | 0.19297949 | 0.18513834 |
| Cnot11 | -0.001576719 | 0.8106439 | 0.869396579 | 0.136150901 | 0.131910773 |
| Cnot2 | 0.004231 | 0.572332068 | 0.679242562 | 0.192153065 | 0.181453857 |
| Cnot3 | 0.017098302 | 0.032966291 | 0.070649891 | 0.226988468 | 0.210563893 |
| Cnot4 | -0.00093434 | 0.914991745 | 0.941328016 | 0.297904871 | 0.292687193 |
| Cnot6 | -0.007515916 | 0.3279457 | 0.450541855 | 0.189641828 | 0.193706885 |
| Cnot6l | 0.07855024 | 4.54E-14 | 8.29E-13 | 0.797167111 | 0.665402131 |
| Cnot7 | 0.024706385 | 0.007365948 | 0.020093082 | 0.384391389 | 0.341734753 |
| Cnot8 | 0.002770514 | 0.617429088 | 0.717419098 | 0.095985226 | 0.088219079 |
| Cnot9 | -0.000426489 | 0.92914308 | 0.951828279 | 0.063874085 | 0.06377465 |
| Cnppd1 | -0.008365731 | 0.189422719 | 0.293617294 | 0.115091095 | 0.119391062 |
| Cnpy2 | -0.105098109 | 1.99E-31 | 1.09E-29 | 1.134961749 | 1.292827077 |
| Cnpy3 | -0.020067272 | 0.003670149 | 0.01107207 | 0.150331705 | 0.156144729 |
| Cnpy4 | -0.001150377 | 0.764825916 | 0.833541833 | 0.039339478 | 0.039095589 |

| | | | | | |
|---|---|---|---|---|---|
| Cnrip1 | -0.022452384 | 0.000324499 | 0.001321099 | 0.089389771 | 0.112083601 |
| Cnst | 0.013858374 | 0.017525517 | 0.041729875 | 0.104985172 | 0.092393798 |
| Cntfr | 0.036344119 | 1.43E-07 | 1.18E-06 | 0.16955508 | 0.121941641 |
| Cntln | -0.030875308 | 0.000124836 | 0.000569524 | 0.190508636 | 0.211282797 |
| Cntn1 | -0.097903966 | 1.13E-21 | 3.64E-20 | 0.340866104 | 0.444497074 |
| Cntnap2 | -0.04470793 | 2.65E-08 | 2.44E-07 | 0.180029437 | 0.215660975 |
| Cntrl | 0.003869317 | 0.522866638 | 0.635096134 | 0.104521486 | 0.098954024 |
| Cntrob | 0.012257245 | 0.004125779 | 0.012222023 | 0.055262002 | 0.044865485 |
| Coa3 | 0.034931559 | 4.37E-05 | 0.000217665 | 1.345442893 | 1.26331475 |
| Coa4 | 0.015875599 | 0.015174304 | 0.036890369 | 0.144013812 | 0.125373744 |
| Coa5 | 0.014342935 | 0.078226917 | 0.144888948 | 0.246589254 | 0.230665317 |
| Coa6 | 0.007739592 | 0.286708953 | 0.405991296 | 0.183060712 | 0.16791223 |
| Coa7 | 0.021015766 | 0.000306241 | 0.001254119 | 0.113515223 | 0.091261603 |
| Coa8 | -0.026470314 | 0.005391035 | 0.015392553 | 0.405685941 | 0.422531463 |
| Coasy | -0.006455355 | 0.448947477 | 0.567724685 | 0.260019303 | 0.26643017 |
| Cobll1 | 0.03464907 | 1.96E-05 | 0.000106687 | 0.240919788 | 0.208155414 |
| Cog1 | 0.015024036 | 0.040329963 | 0.083529591 | 0.194746286 | 0.169630838 |
| Cog2 | 0.025049003 | 0.000141918 | 0.000639058 | 0.141844499 | 0.119821726 |
| Cog3 | -0.001373314 | 0.791081505 | 0.853690131 | 0.074143615 | 0.073701051 |
| Cog4 | 0.00819647 | 0.212422911 | 0.322079778 | 0.137467241 | 0.126170271 |
| Cog5 | 0.018676444 | 0.005477169 | 0.01559269 | 0.149945934 | 0.132924182 |
| Cog6 | 0.021026774 | 0.01251882 | 0.03130349 | 0.265269922 | 0.250699249 |
| Cog7 | -0.002153692 | 0.789839561 | 0.852850586 | 0.246383001 | 0.230393812 |
| Cog8 | -0.005167331 | 0.449543389 | 0.567961121 | 0.150171861 | 0.142866175 |
| Coil | 0.012649689 | 0.006850948 | 0.018900374 | 0.066516857 | 0.055415352 |
| Col27a1 | -0.059586134 | 1.29E-27 | 5.99E-26 | 0.036079682 | 0.092633376 |
| Col4a1 | 0.02126374 | 6.80E-06 | 4.05E-05 | 0.076636724 | 0.052293558 |
| Col6a6 | 0.035815296 | 5.23E-08 | 4.63E-07 | 0.147213746 | 0.106220162 |
| Colgalt1 | -0.003550983 | 0.629092362 | 0.726715297 | 0.167220524 | 0.172922109 |
| Commd1 | -0.044295175 | 2.13E-06 | 1.40E-05 | 0.349044058 | 0.37192939 |
| Commd10 | 0.00134382 | 0.765372961 | 0.833720552 | 0.060209218 | 0.054381511 |
| Commd2 | -0.021546079 | 0.003966067 | 0.011831269 | 0.186211088 | 0.193330508 |
| Commd3 | 0.007577523 | 0.455947294 | 0.574409692 | 0.529973242 | 0.512419819 |
| Commd4 | -0.012905172 | 0.12696332 | 0.214071695 | 0.266968357 | 0.271141245 |
| Commd6 | 0.013409701 | 0.169237645 | 0.268274643 | 0.573834144 | 0.561460317 |
| Commd7 | -0.013792257 | 0.164183606 | 0.261971609 | 0.460174549 | 0.46994529 |
| Commd8 | -0.003464272 | 0.62890324 | 0.726715297 | 0.169650255 | 0.164656692 |
| Commd9 | -0.021046961 | 0.014186341 | 0.034828184 | 0.290707803 | 0.291092731 |
| Comt | -0.007580318 | 0.460832668 | 0.578988782 | 0.643073788 | 0.616453108 |
| Comtd1 | -0.023266428 | 0.000228801 | 0.000968395 | 0.102842039 | 0.12113271 |
| Cop1 | 0.008512208 | 0.265742204 | 0.382181317 | 0.202315535 | 0.186091215 |
| Copa | 0.021444649 | 0.029212099 | 0.063955957 | 0.866068719 | 0.83473864 |
| Copb1 | 0.005274869 | 0.601076785 | 0.70404388 | 0.909585767 | 0.898184287 |
| Copb2 | 0.026666717 | 0.006652106 | 0.018414516 | 1.02281659 | 0.966506764 |
| Cope | -0.095375047 | 2.32E-23 | 8.55E-22 | 1.054321793 | 1.185941955 |
| Copg1 | 0.033409596 | 0.000720583 | 0.00268719 | 0.590513252 | 0.541318929 |
| Copg2 | 0.004393302 | 0.525450192 | 0.63751738 | 0.15844337 | 0.147198431 |
| Coprs | -0.018771568 | 0.016552293 | 0.039753309 | 0.206319508 | 0.217501336 |
| Cops2 | 0.04238074 | 1.34E-06 | 9.14E-06 | 0.328042212 | 0.277781864 |
| Cops3 | -0.009785478 | 0.306472344 | 0.427355728 | 0.442380112 | 0.441580253 |
| Cops4 | -0.033226232 | 0.000409308 | 0.001622215 | 0.39156925 | 0.40741361 |
| Cops5 | -0.017169807 | 0.067880519 | 0.129163801 | 0.370269585 | 0.37883481 |
| Cops6 | -0.013913987 | 0.115056658 | 0.197738808 | 0.319255652 | 0.322571031 |
| Cops7a | -0.002142716 | 0.813909812 | 0.871649173 | 0.364677081 | 0.351155141 |
| Cops7b | 0.005175258 | 0.219990273 | 0.330870271 | 0.050447627 | 0.046328552 |
| Cops8 | -0.008512642 | 0.316519286 | 0.4382228 | 0.264841498 | 0.268993408 |
| Cops9 | 0.089858882 | 1.92E-21 | 6.01E-20 | 0.990760524 | 0.930199388 |
| Copz1 | 0.003484637 | 0.70943467 | 0.790591914 | 0.411830285 | 0.394195663 |
| Copz2 | -0.024974912 | 0.010287251 | 0.026500438 | 0.444797446 | 0.454801404 |
| Coq10a | -0.003794189 | 0.659275965 | 0.750614746 | 0.274852211 | 0.274811358 |
| Coq10b | 0.031213691 | 0.000350429 | 0.001417169 | 0.276277975 | 0.246197908 |
| Coq2 | -0.072389832 | 1.14E-15 | 2.33E-14 | 0.283959864 | 0.34497427 |
| Coq4 | -0.008072222 | 0.108598225 | 0.189311807 | 0.071819542 | 0.072118309 |
| Coq5 | -0.010203457 | 0.166891136 | 0.265290324 | 0.168277136 | 0.176419018 |
| Coq6 | -0.010866379 | 0.026401374 | 0.058695211 | 0.06219043 | 0.069391903 |
| Coq7 | 0.000936742 | 0.901427631 | 0.932901792 | 0.194947266 | 0.189825112 |
| Coq8a | -0.006150527 | 0.310579543 | 0.43202952 | 0.110518116 | 0.103726403 |
| Coq8b | -0.017918289 | 0.00609416 | 0.017098824 | 0.117310815 | 0.132657679 |
| Coq9 | -0.005442219 | 0.494084419 | 0.609193243 | 0.2262303 | 0.220970745 |
| Coro1c | 0.0313864 | 1.42E-05 | 7.92E-05 | 0.186604006 | 0.156209971 |
| Coro2b | 0.045409582 | 4.64E-09 | 4.83E-08 | 0.23052353 | 0.184009389 |
| Coro7 | 0.008032952 | 0.121450041 | 0.206386523 | 0.079872914 | 0.07235285 |

| | | | | | |
|---|---|---|---|---|---|
| Cotl1 | 0.039255735 | 3.49E-07 | 2.67E-06 | 0.232751425 | 0.182189926 |
| Cox10 | 0.002553282 | 0.600051517 | 0.70329764 | 0.065402595 | 0.064057346 |
| Cox11 | -0.006511294 | 0.150565528 | 0.245616259 | 0.052667038 | 0.056975916 |
| Cox14 | 0.025016494 | 0.007679716 | 0.020808862 | 0.943319038 | 0.916633202 |
| Cox15 | 0.001591741 | 0.719803132 | 0.79857652 | 0.056777382 | 0.053017367 |
| Cox16 | 0.016231987 | 0.028853268 | 0.06334212 | 0.177400253 | 0.175630622 |
| Cox17 | 0.120847099 | 4.33E-38 | 3.26E-36 | 1.096959659 | 0.980058261 |
| Cox18 | -0.027445336 | 0.000528352 | 0.00203854 | 0.202773668 | 0.219289215 |
| Cox19 | -0.005633812 | 0.472933768 | 0.589998985 | 0.21002776 | 0.212302096 |
| Cox20 | 0.040344348 | 2.27E-05 | 0.000121444 | 0.418798243 | 0.381632654 |
| Cox4i1 | -0.011898382 | 0.046678853 | 0.094308355 | 1.997150632 | 2.002946455 |
| Cox5a | 0.013856192 | 0.159567917 | 0.256543623 | 1.005640871 | 0.983844571 |
| Cox5b | 0.052258329 | 5.40E-10 | 6.35E-09 | 1.444822561 | 1.376068106 |
| Cox6a1 | 0.032412449 | 0.000179746 | 0.000782577 | 1.253702813 | 1.230220589 |
| Cox6a2 | -0.080649166 | 7.13E-07 | 5.15E-06 | 0.691536269 | 0.781738439 |
| Cox6b1 | 0.043864432 | 1.02E-09 | 1.16E-08 | 1.636778671 | 1.588988492 |
| Cox6c | 0.142892987 | 5.08E-88 | 2.90E-85 | 1.807458678 | 1.611922819 |
| Cox7a2 | 0.10872559 | 6.06E-58 | 9.99E-56 | 1.823657266 | 1.667394467 |
| Cox7a2l | 0.052117444 | 1.46E-10 | 1.85E-09 | 1.843070111 | 1.675286646 |
| Cox7b | 0.043221225 | 1.31E-06 | 8.94E-06 | 1.186699564 | 1.172625401 |
| Cox7c | 0.137335161 | 3.68E-87 | 1.99E-84 | 1.892997505 | 1.723105174 |
| Cox8a | 0.067512401 | 1.41E-48 | 1.67E-46 | 2.660524938 | 2.530557563 |
| Cpd | 0.087012097 | 1.27E-19 | 3.56E-18 | 0.447668566 | 0.365075456 |
| Cpe | -0.017004684 | 3.26E-05 | 0.000168046 | 3.036790634 | 3.079230306 |
| Cpeb1 | 0.057306596 | 5.67E-10 | 6.64E-09 | 0.360458818 | 0.303008884 |
| Cpeb2 | -0.000176295 | 0.970933466 | 0.980449395 | 0.063350076 | 0.061947053 |
| Cpeb3 | 0.007705111 | 0.268134609 | 0.384875711 | 0.148910742 | 0.141540945 |
| Cpeb4 | 0.013329532 | 0.163017205 | 0.260881196 | 0.360667896 | 0.35877558 |
| Cplane1 | 0.000578929 | 0.934447523 | 0.955579975 | 0.154788365 | 0.148854925 |
| Cplx2 | 0.086975452 | 1.55E-15 | 3.12E-14 | 1.243448769 | 1.062729194 |
| Cpm | 0.015848038 | 0.005470946 | 0.015584102 | 0.097191825 | 0.080493636 |
| Cpn1 | -0.062551151 | 5.04E-13 | 8.34E-12 | 0.251379964 | 0.294099858 |
| Cpne3 | -0.005004838 | 0.566273454 | 0.67376215 | 0.293451044 | 0.289488188 |
| Cpne4 | -0.047230964 | 2.34E-16 | 5.01E-15 | 0.056919822 | 0.098509165 |
| Cpne8 | 0.008415647 | 0.033518311 | 0.071632731 | 0.042032237 | 0.038290506 |
| Cpox | -0.028806234 | 0.000160639 | 0.000708913 | 0.168053183 | 0.193427105 |
| Cpped1 | -0.004424202 | 0.340859635 | 0.463344152 | 0.059035873 | 0.059139686 |
| Cpq | -0.068149488 | 3.83E-12 | 5.77E-11 | 0.468775562 | 0.511420587 |
| Cpsf1 | 0.013454049 | 0.011586917 | 0.029388991 | 0.090331874 | 0.075655129 |
| Cpsf2 | -0.005386668 | 0.485520955 | 0.601457024 | 0.191232008 | 0.195540155 |
| Cpsf3 | -0.003615298 | 0.60455414 | 0.706992207 | 0.15248647 | 0.149688987 |
| Cpsf6 | -0.018622355 | 0.031449517 | 0.067990261 | 0.257547835 | 0.270033559 |
| Cpsf7 | 0.004946954 | 0.517203413 | 0.629869921 | 0.191199079 | 0.18323752 |
| Cpt1a | 0.003903503 | 0.496863581 | 0.611921426 | 0.094128531 | 0.089206094 |
| Cpt2 | 0.020463708 | 0.004267541 | 0.012599161 | 0.184167323 | 0.153949579 |
| Cptp | 0.001887126 | 0.627782373 | 0.725892035 | 0.040976417 | 0.039660896 |
| Cr1l | -0.08744405 | 1.73E-22 | 5.98E-21 | 0.250808522 | 0.328016492 |
| Cracdl | 0.012189962 | 0.006562223 | 0.018233191 | 0.060525086 | 0.049653473 |
| Cradd | -0.007990746 | 0.104831004 | 0.183650677 | 0.067756335 | 0.06920382 |
| Cramp1l | 0.013145902 | 0.009829766 | 0.025491457 | 0.079181656 | 0.066237621 |
| Crat | 0.001265077 | 0.753974433 | 0.824723159 | 0.045506683 | 0.042751756 |
| Crb3 | 0.0160765 | 0.087859624 | 0.159123374 | 0.398682574 | 0.387369861 |
| Crbn | -0.020144135 | 0.004119034 | 0.01221323 | 0.146541546 | 0.156296257 |
| Crcp | -0.008262846 | 0.219288916 | 0.33012184 | 0.138081029 | 0.140429087 |
| Creb1 | 0.013269415 | 0.112458038 | 0.194570177 | 0.253681665 | 0.235697125 |
| Creb3 | -0.0432884 | 5.87E-06 | 3.55E-05 | 0.333527659 | 0.376161906 |
| Creb3l1 | 0.043189089 | 1.12E-06 | 7.75E-06 | 0.327655797 | 0.286061687 |
| Creb3l2 | -0.003255658 | 0.739779823 | 0.813958741 | 0.437285712 | 0.431614488 |
| Crebbp | 0.036089507 | 0.000136531 | 0.000617086 | 0.400724446 | 0.360555439 |
| Crebl2 | 0.008757198 | 0.193831687 | 0.299211124 | 0.145594331 | 0.131708989 |
| Crebrf | 0.006723785 | 0.463959852 | 0.581564941 | 0.328628338 | 0.303676399 |
| Crebzf | -0.039816097 | 1.85E-07 | 1.49E-06 | 0.165765446 | 0.201044075 |
| Creg1 | -0.150056005 | 1.81E-72 | 5.51E-70 | 1.396747244 | 1.636255536 |
| Creld1 | -0.062284902 | 7.35E-18 | 1.78E-16 | 0.136417643 | 0.184335837 |
| Creld2 | 0.040366571 | 0.000146598 | 0.000655578 | 0.471705592 | 0.437518463 |
| Crem | -0.011761262 | 0.268901673 | 0.385470667 | 0.41276094 | 0.464535585 |
| Crip1 | 0.032089225 | 0.001573517 | 0.005282366 | 0.475861447 | 0.457294365 |
| Cript | -0.008477002 | 0.361447771 | 0.48435496 | 0.407667744 | 0.39917446 |
| Crk | 0.025633961 | 0.00124395 | 0.004323804 | 0.22381363 | 0.207515534 |
| Crkl | 0.011499928 | 0.063635817 | 0.122392515 | 0.116458624 | 0.105252003 |
| Crlf1 | -0.030587992 | 2.32E-05 | 0.000123787 | 0.127818337 | 0.159072854 |
| Crlf2 | -0.044714849 | 2.45E-09 | 2.65E-08 | 0.164564176 | 0.197063629 |

| | | | | | |
|---|---|---|---|---|---|
| Crlf3 | 0.003288983 | 0.40975608 | 0.529699889 | 0.043706195 | 0.040957056 |
| Crls1 | -0.016730338 | 0.053029587 | 0.104702203 | 0.253848764 | 0.266332913 |
| Crmp1 | 0.004382922 | 0.3976697 | 0.517972243 | 0.075363219 | 0.071856676 |
| Crnkl1 | 0.007274584 | 0.283367798 | 0.402469208 | 0.149048485 | 0.137709941 |
| Crocc | 0.011380328 | 0.013017215 | 0.03236659 | 0.066447935 | 0.052129367 |
| Crot | 0.063177292 | 9.12E-17 | 2.01E-15 | 0.221573528 | 0.162786705 |
| Crtap | -0.000672553 | 0.887807675 | 0.923228817 | 0.063081244 | 0.060540499 |
| Crtc1 | 0.013693958 | 0.012381316 | 0.031031492 | 0.093803539 | 0.079839016 |
| Crtc2 | 0.008501857 | 0.123708807 | 0.209419645 | 0.091442499 | 0.082412362 |
| Crtc3 | 0.007450497 | 0.354520237 | 0.476978376 | 0.219176679 | 0.219450717 |
| Cry1 | 0.009578988 | 0.098842676 | 0.17494056 | 0.089920741 | 0.09132123 |
| Cry2 | 0.031539127 | 0.000148286 | 0.00066193 | 0.257645033 | 0.225347144 |
| Cryba2 | -0.003030882 | 0.72072918 | 0.798794646 | 0.21421675 | 0.195754254 |
| Crybb3 | -0.05542635 | 1.56E-06 | 1.05E-05 | 0.166830811 | 0.275436178 |
| Crybg1 | 0.008812014 | 0.044111241 | 0.089886709 | 0.054582999 | 0.048242468 |
| Crybg3 | 0.028074463 | 4.20E-05 | 0.000209554 | 0.153303052 | 0.124203644 |
| Cryl1 | 0.028965436 | 3.73E-05 | 0.000189372 | 0.179646929 | 0.135419914 |
| Cryz | -0.014607423 | 0.004640984 | 0.01352913 | 0.069388019 | 0.077569212 |
| Cryzl1 | -0.012597364 | 0.129388641 | 0.217632589 | 0.232456607 | 0.240878774 |
| Cryzl2 | -0.065200047 | 2.72E-21 | 8.42E-20 | 0.11701581 | 0.161561305 |
| Cs | 0.007891142 | 0.263166474 | 0.379362525 | 0.160266837 | 0.154655332 |
| Csad | -0.014548756 | 0.042080445 | 0.086416926 | 0.155696166 | 0.160139729 |
| Csde1 | 0.059406015 | 4.44E-10 | 5.29E-09 | 1.125915366 | 1.036323951 |
| Cse1l | 0.026477152 | 5.09E-05 | 0.000249699 | 0.14070077 | 0.11965556 |
| Csf1 | 0.028068476 | 2.16E-10 | 2.66E-09 | 0.063655742 | 0.041162414 |
| Csf2ra | -0.029587432 | 0.000133336 | 0.00060405 | 0.195626684 | 0.20776367 |
| Csgalnact2 | 0.005436354 | 0.319085931 | 0.440789078 | 0.084816565 | 0.081447284 |
| Csk | 0.010327684 | 0.085424613 | 0.155582257 | 0.111840404 | 0.101483506 |
| Csnk1a1 | 0.040730017 | 1.14E-06 | 7.85E-06 | 1.417516113 | 1.341659671 |
| Csnk1d | 0.027286124 | 0.001509904 | 0.005107356 | 0.290967517 | 0.265988128 |
| Csnk1e | 0.019599488 | 0.007058018 | 0.019367218 | 0.17854 | 0.157622216 |
| Csnk1g1 | -0.010619518 | 0.133155156 | 0.223080204 | 0.14529492 | 0.156478935 |
| Csnk1g2 | 0.011471914 | 0.196730565 | 0.302901295 | 0.326918598 | 0.312096971 |
| Csnk1g3 | 0.00623956 | 0.446980037 | 0.56582551 | 0.244568293 | 0.234172151 |
| Csnk2a1 | 0.05215968 | 3.36E-08 | 3.06E-07 | 0.416642989 | 0.37671997 |
| Csnk2a2 | 0.050070215 | 3.40E-07 | 2.62E-06 | 0.480138023 | 0.437123298 |
| Csnk2b | -0.044225042 | 6.08E-06 | 3.66E-05 | 0.431841375 | 0.462449127 |
| Cspp1 | 0.006080759 | 0.55833805 | 0.666850433 | 0.463993148 | 0.459716778 |
| Csrnp1 | 0.016075988 | 0.000843274 | 0.003090203 | 0.06306963 | 0.056679084 |
| Csrp1 | 0.0023057 | 0.589803486 | 0.694449496 | 0.053941282 | 0.04719283 |
| Cst3 | -0.036532223 | 1.16E-11 | 1.65E-10 | 2.253817701 | 2.304665968 |
| Cst6 | -0.016262111 | 0.008789311 | 0.023295988 | 0.10701465 | 0.118244243 |
| Cstb | 0.008428004 | 0.403501892 | 0.523394983 | 0.697398022 | 0.693900808 |
| Cstf1 | 0.00695514 | 0.135438096 | 0.226242126 | 0.065829923 | 0.05742903 |
| Cstf2 | 0.006530179 | 0.203407488 | 0.311176648 | 0.075387685 | 0.07241177 |
| Cstf2t | 0.011645908 | 0.037236262 | 0.078036417 | 0.094274568 | 0.084273251 |
| Cstf3 | -0.028342609 | 0.0002396 | 0.001007959 | 0.183693681 | 0.205146981 |
| Ctbp1 | 0.008537677 | 0.374732484 | 0.496680192 | 0.473735625 | 0.456561106 |
| Ctbp2 | 0.008594711 | 0.136913304 | 0.227962175 | 0.102461345 | 0.091248353 |
| Ctbs | -0.012078157 | 0.022889686 | 0.052030284 | 0.075324517 | 0.083224681 |
| Ctc1 | 0.00952067 | 0.012169688 | 0.030611568 | 0.0445489 | 0.035137468 |
| Ctcf | 0.008338137 | 0.394317455 | 0.5149858 | 0.494418902 | 0.477417369 |
| Ctdnep1 | 0.018990553 | 0.030367961 | 0.066048617 | 0.317933728 | 0.294426628 |
| Ctdp1 | 0.00772841 | 0.063001716 | 0.121430184 | 0.052143651 | 0.042593243 |
| Ctdsp1 | 0.009652303 | 0.15152535 | 0.246631417 | 0.146008212 | 0.132011113 |
| Ctdsp2 | 0.048952979 | 1.68E-09 | 1.85E-08 | 0.254384183 | 0.204120187 |
| Ctdspl | 0.080765737 | 2.52E-18 | 6.35E-17 | 0.457790934 | 0.341672457 |
| Ctdspl2 | 0.020991038 | 0.005462192 | 0.015563725 | 0.187199147 | 0.167218694 |
| Ctf1 | -0.027194172 | 3.06E-06 | 1.96E-05 | 0.081038347 | 0.099565321 |
| Ctif | 0.016177239 | 0.000218783 | 0.000929231 | 0.058019748 | 0.046425539 |
| Ctnna1 | 0.051433453 | 1.60E-07 | 1.31E-06 | 0.486374665 | 0.437879574 |
| Ctnnal1 | 0.009142857 | 0.314478718 | 0.435956381 | 0.218666171 | 0.227795352 |
| Ctnnb1 | -0.019826631 | 0.047182058 | 0.095226068 | 0.788110391 | 0.789711575 |
| Ctnnbip1 | -0.011232547 | 0.221153649 | 0.332022932 | 0.357016442 | 0.336819015 |
| Ctnnbl1 | 0.016645979 | 0.014948638 | 0.03645991 | 0.15974611 | 0.138534855 |
| Ctnnd1 | 0.024909316 | 0.002095388 | 0.006797253 | 0.239583093 | 0.21517339 |
| Ctnnd2 | 0.000134312 | 0.986394228 | 0.99190367 | 0.211946371 | 0.205021402 |
| Ctns | 0.00480633 | 0.2515264 | 0.366562683 | 0.048683097 | 0.04591447 |
| Ctps | -0.009139626 | 0.096211222 | 0.171073934 | 0.080198848 | 0.08710271 |
| Ctps2 | -0.010902603 | 0.105622139 | 0.184765976 | 0.141249393 | 0.144222342 |
| Ctr9 | 0.030700869 | 0.00018826 | 0.000814535 | 0.256032176 | 0.221274822 |
| Ctsa | -0.033604265 | 3.65E-05 | 0.000185964 | 0.220963799 | 0.242835556 |

| | | | | | |
|---|---|---|---|---|---|
| Ctsb | -0.030919253 | 0.003355674 | 0.010262304 | 0.907392382 | 0.977353397 |
| Ctsd | -0.02290894 | 0.021727182 | 0.049848644 | 0.689582709 | 0.692208295 |
| Ctsf | -0.062287774 | 2.10E-12 | 3.27E-11 | 0.284015085 | 0.322552085 |
| Ctsl | -0.10065613 | 2.54E-25 | 1.04E-23 | 1.087157894 | 1.246665495 |
| Ctso | -0.004826599 | 0.586000806 | 0.691060145 | 0.306864229 | 0.302392679 |
| Ctsz | -0.06840183 | 1.98E-12 | 3.10E-11 | 0.298301814 | 0.376959849 |
| Cttn | 0.016054566 | 0.052324248 | 0.103667483 | 0.252637643 | 0.235653561 |
| Cttnbp2 | -0.076032664 | 9.60E-16 | 1.96E-14 | 0.252233448 | 0.331636703 |
| Ctu1 | 0.007779006 | 0.026697538 | 0.059258782 | 0.037283907 | 0.029738994 |
| Ctu2 | -0.005819481 | 0.269702836 | 0.386277397 | 0.076524816 | 0.079838697 |
| Ctxn1 | -0.012961699 | 0.001977974 | 0.006472521 | 0.040655358 | 0.049994243 |
| Cuedc1 | -0.006161744 | 0.437196848 | 0.555538851 | 0.209572235 | 0.208743596 |
| Cuedc2 | -0.038452829 | 3.03E-06 | 1.94E-05 | 0.247806375 | 0.262433377 |
| Cul1 | 0.018341299 | 0.060633768 | 0.117731475 | 0.488895908 | 0.467255186 |
| Cul2 | -0.001440935 | 0.851137057 | 0.898160693 | 0.183323026 | 0.189076313 |
| Cul3 | 0.027145889 | 0.007135535 | 0.019530313 | 0.672473644 | 0.619537551 |
| Cul4a | 0.01228851 | 0.125421126 | 0.211802012 | 0.236744664 | 0.222139425 |
| Cul4b | -0.001972535 | 0.770930104 | 0.838364929 | 0.144325986 | 0.138003766 |
| Cul5 | 0.002917839 | 0.760685329 | 0.830288847 | 0.419486049 | 0.410605429 |
| Cul7 | 0.012874353 | 0.021257237 | 0.048972242 | 0.099538517 | 0.084889501 |
| Cul9 | -0.006081658 | 0.27762617 | 0.396050129 | 0.087998307 | 0.089905193 |
| Cuta | -0.006230517 | 0.536719948 | 0.647554149 | 0.692459966 | 0.68807869 |
| Cutc | 0.00564281 | 0.334423318 | 0.457344703 | 0.106016948 | 0.096200792 |
| Cux1 | -0.026991871 | 0.001277982 | 0.004431005 | 0.231483075 | 0.256342439 |
| Cux2 | 0.008227009 | 0.186909786 | 0.290476142 | 0.126312188 | 0.109788515 |
| Cwc15 | -0.037968367 | 0.00012916 | 0.000587319 | 0.506335405 | 0.541828062 |
| Cwc22 | 0.009940041 | 0.059003725 | 0.11500285 | 0.082074543 | 0.073263814 |
| Cwc25 | 0.027428161 | 0.00010418 | 0.000484723 | 0.171238956 | 0.148340596 |
| Cwc27 | -0.005878494 | 0.40330384 | 0.52320789 | 0.149308967 | 0.149896006 |
| Cwf19l1 | 0.00833355 | 0.016033062 | 0.038687871 | 0.036531132 | 0.028951788 |
| Cwf19l2 | -0.013864571 | 0.091093782 | 0.163699398 | 0.214704431 | 0.227454722 |
| Cxadr | 0.006919103 | 0.312590672 | 0.433895847 | 0.143541515 | 0.140331046 |
| Cxcl16 | 0.035084026 | 1.05E-20 | 3.16E-19 | 0.056617317 | 0.024626951 |
| Cxxc1 | 0.001674144 | 0.784170849 | 0.848870838 | 0.112390672 | 0.110118622 |
| Cxxc4 | 0.028472201 | 0.00688887 | 0.018978065 | 0.587423152 | 0.525100519 |
| Cxxc5 | 0.017316386 | 0.082011437 | 0.15040845 | 0.552718255 | 0.514356006 |
| Cyb561 | -0.024340864 | 0.010159803 | 0.026241659 | 0.353074891 | 0.376658438 |
| Cyb561a3 | -0.002086982 | 0.583131693 | 0.688846454 | 0.036543437 | 0.038589232 |
| Cyb561d1 | 0.009254756 | 0.021016921 | 0.048476038 | 0.046503354 | 0.040346423 |
| Cyb561d2 | -0.014817387 | 0.012851599 | 0.032028518 | 0.101592392 | 0.108504067 |
| Cyb5a | -0.080187742 | 4.07E-17 | 9.29E-16 | 0.364492021 | 0.434874669 |
| Cyb5b | 0.051453503 | 1.79E-07 | 1.44E-06 | 0.547324313 | 0.482180779 |
| Cyb5d2 | -0.005616212 | 0.260224416 | 0.376301228 | 0.069569988 | 0.070419891 |
| Cyb5r1 | -0.196388022 | 1.23E-66 | 2.78E-64 | 0.290852105 | 0.509258381 |
| Cyb5r3 | 0.011400629 | 0.254637283 | 0.369821282 | 1.119767011 | 1.050008225 |
| Cyb5r4 | 0.014234216 | 0.053623673 | 0.105789234 | 0.184368876 | 0.169311564 |
| Cyb5rl | 0.00162771 | 0.673525256 | 0.762283182 | 0.042641495 | 0.039053052 |
| Cybc1 | 0.001530787 | 0.72404291 | 0.80090399 | 0.05195356 | 0.048491634 |
| Cyc1 | -0.013740541 | 0.186261812 | 0.289733974 | 0.570093716 | 0.586419953 |
| Cycs | 0.004852047 | 0.648075658 | 0.741507773 | 0.701194809 | 0.688946475 |
| Cyfip1 | 0.02195915 | 0.003415134 | 0.01041466 | 0.198881731 | 0.174196971 |
| Cyfip2 | 0.085589553 | 3.46E-17 | 7.93E-16 | 0.991109154 | 0.888105661 |
| Cyhr1 | 0.005608897 | 0.555024349 | 0.66443127 | 0.491809904 | 0.466084912 |
| Cyld | 0.023384108 | 0.005852902 | 0.016483753 | 0.278506155 | 0.250728309 |
| Cyp20a1 | -0.015476167 | 0.03348586 | 0.071596555 | 0.157212537 | 0.171129542 |
| Cyp2j6 | -0.014009891 | 0.002418777 | 0.007707424 | 0.054250068 | 0.062147881 |
| Cyp2u1 | 0.008168747 | 0.147863977 | 0.242049769 | 0.099984337 | 0.087241275 |
| Cyp4f13 | -0.020321941 | 0.00016933 | 0.000744561 | 0.074944744 | 0.087203733 |
| Cyp4f16 | -0.008481343 | 0.043157475 | 0.088276241 | 0.042698131 | 0.049512395 |
| Cyp4f17 | -0.006895145 | 0.114675611 | 0.197183638 | 0.047034471 | 0.053221108 |
| Cyp4f39 | -0.014459701 | 0.001317839 | 0.00454785 | 0.046245638 | 0.056240528 |
| Cyp4v3 | -0.00356963 | 0.584396067 | 0.689837125 | 0.132057597 | 0.129499621 |
| Cyp51 | 0.03548832 | 2.12E-06 | 1.39E-05 | 0.190617722 | 0.163327846 |
| Cyren | 0.008655009 | 0.108227911 | 0.188801678 | 0.084644071 | 0.079553119 |
| Cyrib | -3.85E-05 | 0.996247187 | 0.997175452 | 0.252274907 | 0.242728855 |
| Cystm1 | 0.06255279 | 9.22E-10 | 1.06E-08 | 0.804528233 | 0.714342959 |
| Cyth1 | 0.01756529 | 0.006973632 | 0.019178969 | 0.136701421 | 0.121721862 |
| Cyth2 | -0.002054323 | 0.806422467 | 0.865824321 | 0.274280996 | 0.261922809 |
| Cyth3 | -0.006085086 | 0.238117966 | 0.352138515 | 0.064207058 | 0.072583451 |
| Czib | -0.038499717 | 5.28E-06 | 3.23E-05 | 0.244066155 | 0.274069726 |
| D030056L22Rik | -0.007914552 | 0.10887745 | 0.18962855 | 0.066953185 | 0.068791071 |
| D10Wsu102e | 0.006007997 | 0.372195525 | 0.494396077 | 0.127240289 | 0.126773704 |

| | | | | | |
|---|---|---|---|---|---|
| D11Wsu47e | -0.001193055 | 0.747212936 | 0.819651881 | 0.037204663 | 0.036372817 |
| D130020L05Rik | -0.0164379 | 0.008147194 | 0.021864736 | 0.10321888 | 0.116382652 |
| D17H6S53E | 0.004810322 | 0.242955691 | 0.356665305 | 0.052875185 | 0.044416377 |
| D230025D16Rik | 0.006524638 | 0.171049872 | 0.270485825 | 0.064543212 | 0.0583869 |
| D330023K18Rik | 0.002805029 | 0.679343521 | 0.767082197 | 0.146149175 | 0.140254274 |
| D330050I16Rik | 0.006680895 | 0.085221246 | 0.155240951 | 0.044303706 | 0.038259262 |
| D3Ertd751e | -0.003994012 | 0.576764192 | 0.683067545 | 0.1574144 | 0.154872612 |
| D5Ertd579e | 0.020391318 | 0.036550657 | 0.076897962 | 0.43534255 | 0.410530932 |
| D630024D03Rik | -0.001182063 | 0.880364083 | 0.918041362 | 0.224941666 | 0.200874415 |
| D630039A03Rik | 0.005065405 | 0.272238277 | 0.389392457 | 0.06558832 | 0.054321425 |
| D630045J12Rik | 0.001978029 | 0.641088025 | 0.736074138 | 0.050219325 | 0.046417411 |
| D6Wsu163e | -0.00570377 | 0.214254528 | 0.323997904 | 0.055805959 | 0.05798394 |
| D730003I15Rik | -0.047230559 | 7.86E-06 | 4.63E-05 | 0.514728836 | 0.57185481 |
| D830050J10Rik | 0.000707709 | 0.875346709 | 0.914476758 | 0.057273653 | 0.05444715 |
| D8Ertd738e | 0.046617341 | 7.45E-07 | 5.35E-06 | 1.121997217 | 1.067350247 |
| Daam1 | 0.052601846 | 9.97E-07 | 6.96E-06 | 0.570883624 | 0.531832056 |
| Dab2ip | 0.018857697 | 0.002677918 | 0.008425476 | 0.127971155 | 0.108255298 |
| Dach1 | 0.053532604 | 2.33E-09 | 2.52E-08 | 0.347614248 | 0.273192585 |
| Dact1 | 0.01612003 | 5.16E-05 | 0.000252618 | 0.050155995 | 0.03636223 |
| Dact2 | 0.015903127 | 0.000759436 | 0.002809405 | 0.079404952 | 0.054771553 |
| Dad1 | -0.018406654 | 0.011263742 | 0.028696568 | 1.806330255 | 1.883962331 |
| Dag1 | 0.041419376 | 7.55E-07 | 5.42E-06 | 0.267161466 | 0.233136192 |
| Dagla | 0.005372211 | 0.271616282 | 0.388781937 | 0.068630878 | 0.063754695 |
| Daglb | 0.001810189 | 0.697333046 | 0.780865223 | 0.063400578 | 0.057797662 |
| Dalrd3 | -0.02163514 | 0.004722529 | 0.013733901 | 0.191949513 | 0.20064775 |
| Dancr | -0.002227083 | 0.633612249 | 0.729941733 | 0.058489053 | 0.059192003 |
| Dand5 | -0.002268788 | 0.634306486 | 0.730481836 | 0.066476925 | 0.063986541 |
| Dap | -0.021236842 | 0.028325887 | 0.06234645 | 1.080152937 | 1.065886146 |
| Dap3 | -0.001137658 | 0.898366686 | 0.930724736 | 0.3164429 | 0.308681336 |
| Dapk1 | 0.024746649 | 0.000113243 | 0.000523015 | 0.13749318 | 0.112592585 |
| Dapk3 | -0.013572926 | 0.108425999 | 0.189079384 | 0.262331873 | 0.267081948 |
| Dapl1 | 0.222801879 | 2.85E-70 | 7.70E-68 | 0.575187436 | 0.347622269 |
| Dapp1 | -0.006380246 | 0.310573896 | 0.43202952 | 0.099258426 | 0.110378427 |
| Dars | -0.000565664 | 0.94146033 | 0.96028615 | 0.205420981 | 0.199594946 |
| Daxx | 0.007044409 | 0.226372569 | 0.338635808 | 0.102933904 | 0.095549016 |
| Dazap1 | -0.008970889 | 0.321980089 | 0.443624869 | 0.356587091 | 0.354429487 |
| Dazap2 | 0.023983674 | 0.009465365 | 0.024769224 | 0.380210711 | 0.34454398 |
| Dbi | 0.052038142 | 2.08E-08 | 1.95E-07 | 0.392041996 | 0.343747112 |
| Dbnl | 0.002571333 | 0.729265322 | 0.804940674 | 0.189278197 | 0.179681126 |
| Dbp | 0.107878725 | 5.89E-36 | 4.01E-34 | 0.30978763 | 0.153307434 |
| Dbpht2 | -0.017325991 | 0.017325703 | 0.041324947 | 0.139188761 | 0.155402275 |
| Dbr1 | 0.010299187 | 0.092214285 | 0.16510263 | 0.116080818 | 0.103326715 |
| Dbt | 0.010766597 | 0.035143679 | 0.074485905 | 0.075634771 | 0.067497044 |
| Dcaf1 | 0.02359326 | 0.001087306 | 0.003852328 | 0.171470282 | 0.154191101 |
| Dcaf10 | 0.000886919 | 0.891847641 | 0.926538017 | 0.128693094 | 0.123737071 |
| Dcaf11 | -0.021363286 | 0.015064473 | 0.036660026 | 0.31119567 | 0.316035621 |
| Dcaf12 | 0.001937486 | 0.74491969 | 0.817769739 | 0.106991796 | 0.104443071 |
| Dcaf12l1 | 0.025635148 | 0.00157697 | 0.005292132 | 0.246831386 | 0.210133463 |
| Dcaf13 | -0.000351898 | 0.960676233 | 0.97298618 | 0.163570416 | 0.156934051 |
| Dcaf15 | 0.010985548 | 0.009714977 | 0.025273119 | 0.054767335 | 0.045527101 |
| Dcaf17 | 0.019599365 | 0.002028241 | 0.006614096 | 0.126946977 | 0.109712733 |
| Dcaf4 | 0.006242248 | 0.083615122 | 0.152830649 | 0.038750733 | 0.032472036 |
| Dcaf5 | 0.018053392 | 0.006112075 | 0.017139198 | 0.144524276 | 0.123692489 |
| Dcaf6 | 0.028840964 | 0.000217712 | 0.00092549 | 0.219971973 | 0.189763092 |
| Dcaf7 | 0.043963139 | 9.36E-07 | 6.56E-06 | 0.341654289 | 0.30563896 |
| Dcaf8 | -0.003984199 | 0.688478417 | 0.773801985 | 0.566677137 | 0.536401176 |
| Dcakd | -0.007034811 | 0.117489731 | 0.200827171 | 0.056681716 | 0.056685798 |
| Dcbld1 | 0.004384946 | 0.277866007 | 0.396272664 | 0.042638293 | 0.041573026 |
| Dcbld2 | 0.004191415 | 0.258713418 | 0.374845283 | 0.038543674 | 0.035443486 |
| Dcdc2a | 0.020652938 | 0.008158382 | 0.021880218 | 0.204962826 | 0.172440774 |
| Dclre1a | 0.005897122 | 0.157200409 | 0.253703017 | 0.048692312 | 0.043413959 |
| Dcp1a | 0.000783494 | 0.863927859 | 0.906923335 | 0.054796659 | 0.055431049 |
| Dcp1b | 0.001681464 | 0.720384792 | 0.798776488 | 0.0636406 | 0.059739151 |
| Dcp2 | 0.018036452 | 0.001353577 | 0.004659871 | 0.09935731 | 0.085407584 |
| Dcps | -0.022681393 | 0.000244785 | 0.001026661 | 0.106489335 | 0.119480136 |
| Dct | 0.01834038 | 1.19E-05 | 6.74E-05 | 0.058139078 | 0.036998033 |
| Dctn1 | 0.017988334 | 0.032624346 | 0.070047237 | 0.272040844 | 0.250387551 |
| Dctn2 | -0.026305476 | 0.006575854 | 0.018260627 | 0.513335207 | 0.525839843 |
| Dctn3 | -0.018636983 | 0.052579757 | 0.10406767 | 0.92103637 | 0.929937036 |
| Dctn4 | 0.019455876 | 0.018450556 | 0.043464092 | 0.255621612 | 0.232579676 |
| Dctn5 | 0.011812476 | 0.03744806 | 0.07837891 | 0.101361401 | 0.087514247 |
| Dctn6 | -0.017920068 | 0.039449684 | 0.081950817 | 0.303656848 | 0.299395845 |

| | | | | | |
|---|---|---|---|---|---|
| Dctpp1 | -0.006911143 | 0.349348566 | 0.47178313 | 0.17483547 | 0.178254612 |
| Dcun1d1 | 0.011317957 | 0.06862341 | 0.130324718 | 0.126380211 | 0.111156453 |
| Dcun1d2 | -0.002808198 | 0.582726246 | 0.688702232 | 0.070867908 | 0.072521469 |
| Dcun1d3 | 0.023299016 | 0.000375827 | 0.001503618 | 0.144856748 | 0.120039293 |
| Dcun1d4 | -0.00087913 | 0.869308773 | 0.910400898 | 0.083238494 | 0.079839022 |
| Dcun1d5 | -0.015901295 | 0.091717446 | 0.164513629 | 0.399068632 | 0.407834426 |
| Dcx | 0.001733812 | 0.765498602 | 0.833763994 | 0.089842675 | 0.088226503 |
| Dcxr | -0.020213337 | 0.000929636 | 0.003356079 | 0.094803561 | 0.11037301 |
| Dda1 | 0.021577576 | 0.012936854 | 0.032199717 | 0.30399075 | 0.279939535 |
| Ddb1 | 0.008005538 | 0.427928863 | 0.546578805 | 0.608710314 | 0.578835436 |
| Ddc | 0.00424136 | 0.722464983 | 0.79997774 | 0.91067554 | 0.811402171 |
| Ddhd1 | -0.005689907 | 0.329244974 | 0.451915803 | 0.090716913 | 0.095152513 |
| Ddhd2 | 0.003090875 | 0.724189554 | 0.800975068 | 0.299348412 | 0.28414978 |
| Ddi2 | -0.017740907 | 0.042088474 | 0.086416926 | 0.272418524 | 0.282185007 |
| Ddit3 | -0.068852177 | 4.87E-07 | 3.65E-06 | 0.845702934 | 0.975399143 |
| Ddit4 | -0.052739492 | 2.14E-09 | 2.33E-08 | 0.207168084 | 0.267862645 |
| Ddo | 0.002625181 | 0.513786287 | 0.627145942 | 0.045492711 | 0.041384477 |
| Ddost | -0.094053155 | 4.06E-23 | 1.47E-21 | 1.018055715 | 1.158812043 |
| Ddr1 | 0.000386163 | 0.966405143 | 0.977159729 | 0.346327161 | 0.32802585 |
| Ddrgk1 | -0.105662915 | 4.39E-26 | 1.90E-24 | 0.663760222 | 0.786049785 |
| Ddt | -0.056075542 | 1.79E-08 | 1.70E-07 | 0.524928517 | 0.564161327 |
| Ddx1 | 0.048732111 | 2.48E-07 | 1.94E-06 | 0.422581246 | 0.367767404 |
| Ddx10 | -0.004824851 | 0.402133736 | 0.52210793 | 0.092940804 | 0.095582471 |
| Ddx17 | -0.079234904 | 2.05E-14 | 3.86E-13 | 0.516837433 | 0.595922789 |
| Ddx18 | 0.033587779 | 0.000154137 | 0.000683321 | 0.302875988 | 0.284860575 |
| Ddx19a | 0.011468329 | 0.053494305 | 0.105555437 | 0.11305907 | 0.098287364 |
| Ddx19b | -0.002249885 | 0.689379276 | 0.774456358 | 0.083389693 | 0.089614844 |
| Ddx20 | 0.01143114 | 0.006416213 | 0.017883722 | 0.054010723 | 0.043952478 |
| Ddx21 | 0.042388866 | 3.22E-06 | 2.05E-05 | 0.302158563 | 0.277295014 |
| Ddx23 | 0.022864804 | 0.001993886 | 0.006520201 | 0.177146711 | 0.158844444 |
| Ddx24 | 0.022298503 | 0.021267651 | 0.048984625 | 0.923241513 | 0.907174797 |
| Ddx25 | -0.023168549 | 0.000135772 | 0.000613941 | 0.103987868 | 0.114939056 |
| Ddx27 | 0.03383447 | 5.13E-06 | 3.15E-05 | 0.188270841 | 0.160367835 |
| Ddx28 | 0.004080531 | 0.37279165 | 0.494985034 | 0.060083028 | 0.055539737 |
| Ddx31 | 0.020088002 | 0.00309643 | 0.009556664 | 0.154794799 | 0.126421887 |
| Ddx39a | -0.013617828 | 0.008221929 | 0.02202634 | 0.065661403 | 0.077911234 |
| Ddx39b | -0.049797911 | 3.33E-07 | 2.57E-06 | 0.431358357 | 0.465503701 |
| Ddx3x | 0.069921018 | 5.79E-12 | 8.51E-11 | 0.76393301 | 0.696283081 |
| Ddx3y | 0.048164105 | 8.41E-07 | 5.97E-06 | 0.422546469 | 0.37323796 |
| Ddx41 | -0.013672835 | 0.143164926 | 0.236267087 | 0.35985642 | 0.375324661 |
| Ddx42 | 0.021493181 | 0.028966031 | 0.063553996 | 0.467381648 | 0.445970212 |
| Ddx46 | 0.03344414 | 0.000542811 | 0.002087503 | 0.412721253 | 0.374737824 |
| Ddx47 | 0.036376881 | 9.92E-07 | 6.93E-06 | 0.197830746 | 0.168179503 |
| Ddx49 | -0.012994466 | 0.05423447 | 0.106805134 | 0.13103959 | 0.141163751 |
| Ddx5 | 0.007349456 | 0.231824687 | 0.344993556 | 2.229582019 | 2.222580841 |
| Ddx50 | -0.026790467 | 0.00636138 | 0.01775634 | 0.852215208 | 0.886439723 |
| Ddx52 | 0.028256763 | 0.000186099 | 0.000805902 | 0.19789833 | 0.171601663 |
| Ddx54 | 0.011628683 | 0.116167958 | 0.199221183 | 0.181410735 | 0.169863974 |
| Ddx55 | -0.01136583 | 0.090913677 | 0.163405947 | 0.127938991 | 0.135912165 |
| Ddx56 | -0.018692939 | 0.004762963 | 0.013834935 | 0.123931191 | 0.136330501 |
| Ddx58 | 0.031682336 | 1.04E-06 | 7.28E-06 | 0.13238426 | 0.109142495 |
| Ddx6 | 0.008007332 | 0.44391043 | 0.562525704 | 0.839022004 | 0.794294693 |
| Ddx60 | 0.059257656 | 1.44E-26 | 6.36E-25 | 0.104394386 | 0.056592211 |
| Deaf1 | 0.018570784 | 0.000467443 | 0.001827295 | 0.095143856 | 0.074737587 |
| Decr1 | -0.003149539 | 0.612359977 | 0.712783653 | 0.12194895 | 0.117040045 |
| Decr2 | -0.000494233 | 0.927316135 | 0.950628786 | 0.086717846 | 0.083571335 |
| Dedd | 0.002586593 | 0.669718004 | 0.758924183 | 0.114665622 | 0.107080401 |
| Dedd2 | 0.000798538 | 0.879986799 | 0.917943312 | 0.08273937 | 0.077043503 |
| Def8 | 0.002745649 | 0.630961779 | 0.728009781 | 0.095985023 | 0.090419133 |
| Defb1 | 0.080340952 | 9.85E-09 | 9.80E-08 | 1.04003182 | 0.990306976 |
| Degs1 | 0.117285764 | 2.55E-28 | 1.23E-26 | 0.807779077 | 0.668513703 |
| Degs2 | -0.024947563 | 1.02E-05 | 5.84E-05 | 0.073798714 | 0.093076651 |
| Dek | 0.025548537 | 0.006914839 | 0.019038819 | 0.390320897 | 0.362990733 |
| Dele1 | 0.015338308 | 0.029793126 | 0.065074987 | 0.167144382 | 0.147212058 |
| Denn2b | 0.032705123 | 5.53E-06 | 3.36E-05 | 0.179316023 | 0.1465385 |
| Dennd10 | -0.02882196 | 4.04E-05 | 0.000202813 | 0.136290574 | 0.15923561 |
| Dennd11 | 0.012014951 | 0.160150392 | 0.257289775 | 0.257967049 | 0.257460408 |
| Dennd1a | 0.011782765 | 0.06733075 | 0.128300579 | 0.132370545 | 0.117588816 |
| Dennd1b | 0.008975755 | 0.199110501 | 0.305999413 | 0.153258825 | 0.144390686 |
| Dennd2d | -0.003250263 | 0.534443359 | 0.645448241 | 0.075277543 | 0.076412718 |
| Dennd4a | -0.009120565 | 0.260298885 | 0.376301228 | 0.195184794 | 0.212896386 |
| Dennd4c | 0.07024362 | 1.19E-12 | 1.92E-11 | 0.473308222 | 0.379747404 |

| | | | | | |
|---|---|---|---|---|---|
| Dennd5a | 0.016933334 | 0.019375743 | 0.045162065 | 0.174348765 | 0.16081489 |
| Dennd5b | 0.01115733 | 0.135899954 | 0.22677984 | 0.176919338 | 0.169025212 |
| Dennd6a | 0.029060222 | 9.60E-05 | 0.00044998 | 0.19678506 | 0.172143481 |
| Denr | 0.058887105 | 9.01E-09 | 9.04E-08 | 0.932035689 | 0.886359669 |
| Depdc5 | -0.003135695 | 0.475033187 | 0.592010337 | 0.052959349 | 0.05223193 |
| Depp1 | 0.015814983 | 0.015942328 | 0.038478479 | 0.103538549 | 0.093584928 |
| Deptor | 0.036762731 | 0.000324066 | 0.001319888 | 0.587581905 | 0.530760425 |
| Dera | 0.008849752 | 0.136706794 | 0.227721666 | 0.110852462 | 0.101061777 |
| Derl1 | -0.10966273 | 5.24E-25 | 2.09E-23 | 0.746506809 | 0.879677597 |
| Derl2 | -0.044110974 | 6.69E-06 | 3.99E-05 | 0.46558316 | 0.487227251 |
| Desi1 | 0.044131242 | 8.80E-06 | 5.12E-05 | 0.578055035 | 0.502763454 |
| Desi2 | 0.02267466 | 0.01580627 | 0.038207 | 0.374066532 | 0.352447591 |
| Dexi | -0.002852038 | 0.654527329 | 0.746517444 | 0.127646211 | 0.122524817 |
| Dffa | 0.009716731 | 0.070303947 | 0.132769032 | 0.088901915 | 0.077515764 |
| Dffb | -0.003409105 | 0.522486272 | 0.634754399 | 0.078237772 | 0.078531615 |
| Dgat1 | -0.000393622 | 0.931101148 | 0.953375563 | 0.057543097 | 0.057049399 |
| Dgcr2 | 0.007530819 | 0.302220003 | 0.422577717 | 0.182708044 | 0.173472768 |
| Dgcr6 | -0.032764457 | 0.000272744 | 0.001131717 | 0.342520489 | 0.362145532 |
| Dgcr8 | 0.002661808 | 0.573589153 | 0.680467816 | 0.06397955 | 0.060251147 |
| Dgkd | 0.019807119 | 0.016176022 | 0.038984454 | 0.258326121 | 0.230235842 |
| Dgke | 0.024951353 | 0.00024315 | 0.001020683 | 0.147888573 | 0.12702816 |
| Dgkg | -0.032650832 | 3.83E-05 | 0.000193293 | 0.146377703 | 0.18857464 |
| Dgkh | 0.022157625 | 0.003341994 | 0.010223684 | 0.178690707 | 0.166923 |
| Dgkq | -0.006956803 | 0.087170028 | 0.158198061 | 0.039810856 | 0.045152936 |
| Dgkz | 0.005745174 | 0.43410422 | 0.552475616 | 0.169406619 | 0.164865614 |
| Dguok | -0.000473425 | 0.959370761 | 0.971967751 | 0.38251516 | 0.372741074 |
| Dhcr7 | 0.007789665 | 0.024871532 | 0.05598542 | 0.03707493 | 0.029672316 |
| Dhdds | -0.031385866 | 5.35E-05 | 0.000261434 | 0.184001701 | 0.212644205 |
| Dhdh | -0.001422402 | 0.801087576 | 0.861333047 | 0.094494909 | 0.089152377 |
| Dhps | -0.031670693 | 4.06E-05 | 0.000203699 | 0.183055473 | 0.209835354 |
| Dhrs1 | -0.021474403 | 0.000382426 | 0.001526251 | 0.099962348 | 0.113707609 |
| Dhrs4 | 0.000428666 | 0.956404047 | 0.970443551 | 0.226060465 | 0.208583557 |
| Dhrs7 | -0.024000956 | 0.022388268 | 0.05099783 | 0.587508465 | 0.628214971 |
| Dhrs7b | -0.00203073 | 0.732780053 | 0.807811279 | 0.104786541 | 0.104192203 |
| Dhx15 | 0.055930118 | 4.35E-10 | 5.19E-09 | 0.337046296 | 0.299467426 |
| Dhx16 | 0.017472172 | 0.001012935 | 0.003620499 | 0.086591894 | 0.073754301 |
| Dhx29 | -0.001088434 | 0.877400777 | 0.916228824 | 0.150482014 | 0.151584339 |
| Dhx30 | 0.007411245 | 0.359462276 | 0.482226058 | 0.243126195 | 0.228827482 |
| Dhx32 | 0.039573506 | 2.15E-05 | 0.000115864 | 0.406435031 | 0.364782995 |
| Dhx34 | 0.012195366 | 0.011480401 | 0.029149244 | 0.074221203 | 0.059626219 |
| Dhx36 | 0.030144475 | 0.002778636 | 0.008705735 | 0.552846433 | 0.52728442 |
| Dhx38 | 0.012815835 | 0.014927079 | 0.036416829 | 0.084014728 | 0.073658104 |
| Dhx40 | -0.020418874 | 0.020954838 | 0.048367268 | 0.286051367 | 0.292770001 |
| Dhx57 | -0.001734884 | 0.746456598 | 0.819080253 | 0.083981841 | 0.080983778 |
| Dhx8 | 0.021946118 | 0.000682596 | 0.002559276 | 0.133251694 | 0.116109158 |
| Dhx9 | -0.009170045 | 0.306813927 | 0.427648029 | 0.294773833 | 0.296978961 |
| Diablo | -0.043206456 | 4.40E-08 | 3.94E-07 | 0.20578328 | 0.228887966 |
| Diaph1 | 0.044487805 | 4.04E-07 | 3.07E-06 | 0.30985438 | 0.261036766 |
| Diaph2 | -0.000279215 | 0.970969633 | 0.980449395 | 0.185125337 | 0.184736861 |
| Dicer1 | -0.011004134 | 0.260892655 | 0.376935412 | 0.445813986 | 0.443018807 |
| Dido1 | -0.001255496 | 0.871459496 | 0.911787475 | 0.197123201 | 0.194711194 |
| Dimt1 | -0.01612592 | 0.006188201 | 0.017327675 | 0.092837209 | 0.103657128 |
| Dio1 | 0.088971675 | 1.17E-22 | 4.09E-21 | 0.356166174 | 0.261293488 |
| Dip2b | 0.042332169 | 2.12E-08 | 1.98E-07 | 0.194374768 | 0.163313072 |
| Dip2c | 0.070536776 | 1.02E-17 | 2.41E-16 | 0.264864702 | 0.203259522 |
| Dipk1b | -0.015067014 | 0.0217741 | 0.049926368 | 0.129519904 | 0.136174664 |
| Dipk2a | 0.002091196 | 0.717591954 | 0.796941852 | 0.093948373 | 0.09212996 |
| Diras1 | 0.01859988 | 5.06E-06 | 3.11E-05 | 0.052833839 | 0.038962583 |
| Dis3 | -0.002805902 | 0.522520002 | 0.634754399 | 0.049644868 | 0.052094514 |
| Dis3l | -0.006409409 | 0.182318675 | 0.284602145 | 0.061865419 | 0.064346244 |
| Dis3l2 | 0.016203547 | 0.000406755 | 0.001613413 | 0.065406475 | 0.051822777 |
| Disp2 | 0.012575268 | 0.213731465 | 0.323508882 | 0.565587275 | 0.541231568 |
| Dixdc1 | 0.003892553 | 0.452143568 | 0.570652963 | 0.077002294 | 0.069646897 |
| Dkc1 | -0.010525221 | 0.12955567 | 0.217838158 | 0.146889637 | 0.153156818 |
| Dlat | 0.029368312 | 0.000422767 | 0.001670111 | 0.262002401 | 0.237395584 |
| Dld | 0.012100691 | 0.184094919 | 0.2870983 | 0.344198937 | 0.323376507 |
| Dleu2 | -0.024169768 | 0.000600473 | 0.00228396 | 0.131096873 | 0.153094518 |
| Dlg1 | 0.010627386 | 0.152747359 | 0.247790726 | 0.185185496 | 0.172371745 |
| Dlg3 | 0.007893132 | 0.23144077 | 0.344626614 | 0.137409032 | 0.125172875 |
| Dlg4 | 0.021117311 | 0.001667141 | 0.005560188 | 0.151123389 | 0.129502447 |
| Dlg5 | 0.012795781 | 0.012892959 | 0.032115128 | 0.079276575 | 0.068828311 |
| Dlgap1 | -0.022644642 | 0.001535026 | 0.005177051 | 0.133444618 | 0.14853128 |

| | | | | | |
|---|---|---|---|---|---|
| Dlgap4 | 0.045743105 | 8.14E-09 | 8.22E-08 | 0.243061023 | 0.199180017 |
| Dlk1 | 0.046669599 | 1.58E-32 | 9.22E-31 | 0.060254364 | 0.014694939 |
| Dlst | 0.008378223 | 0.293146687 | 0.412561102 | 0.233570977 | 0.212488998 |
| Dmac1 | -0.014707623 | 0.072198249 | 0.135608844 | 0.225789893 | 0.242610812 |
| Dmac2 | -0.006026782 | 0.220870175 | 0.331825091 | 0.062279538 | 0.066738756 |
| Dmac2l | -0.020436195 | 0.000261105 | 0.001088068 | 0.084710589 | 0.094984901 |
| Dmap1 | -0.013715114 | 0.008773957 | 0.023261633 | 0.073958835 | 0.081246443 |
| Dmpk | 0.019687886 | 0.0166944 | 0.040045141 | 0.226823991 | 0.210451368 |
| Dmrtc1a | -0.015391688 | 0.000119322 | 0.000547707 | 0.038523046 | 0.046786084 |
| Dmtf1 | 0.04470171 | 2.18E-10 | 2.67E-09 | 0.177041215 | 0.135132651 |
| Dmwd | 0.027454211 | 9.87E-06 | 5.67E-05 | 0.12848792 | 0.105157404 |
| Dmxl1 | 0.040142177 | 1.57E-05 | 8.70E-05 | 0.351876752 | 0.319758172 |
| Dmxl2 | 0.002969816 | 0.731726852 | 0.806924734 | 0.278734806 | 0.275437951 |
| Dnaaf5 | 0.006311587 | 0.071037887 | 0.133869033 | 0.035753901 | 0.030520304 |
| Dnaja1 | -8.01E-05 | 0.993781693 | 0.996138326 | 0.698114755 | 0.668965999 |
| Dnaja2 | -0.0003263 | 0.973552511 | 0.982139412 | 0.556508017 | 0.541614735 |
| Dnaja3 | -0.018560275 | 0.023991922 | 0.054269304 | 0.217961594 | 0.23296507 |
| Dnaja4 | -0.0069697 | 0.469932661 | 0.587159148 | 0.409096776 | 0.414323324 |
| Dnajb1 | -0.027648143 | 0.000756853 | 0.002800962 | 0.170297465 | 0.207837133 |
| Dnajb11 | -0.040383435 | 6.33E-05 | 0.000306749 | 0.444850663 | 0.49052571 |
| Dnajb12 | 0.005482685 | 0.461414038 | 0.579344863 | 0.188077365 | 0.179581575 |
| Dnajb14 | 0.032028999 | 0.001280651 | 0.004438676 | 0.547236166 | 0.51599593 |
| Dnajb2 | 0.057704348 | 7.03E-11 | 9.18E-10 | 0.326794103 | 0.271677307 |
| Dnajb4 | -0.006373755 | 0.415096814 | 0.534466274 | 0.184465841 | 0.191457236 |
| Dnajb6 | 0.013117304 | 0.184955784 | 0.288107583 | 0.723905844 | 0.69884591 |
| Dnajb9 | -0.172796409 | 3.84E-53 | 5.18E-51 | 0.781700061 | 0.990501306 |
| Dnajc1 | -0.006998158 | 0.494588155 | 0.609737008 | 0.570758994 | 0.591987418 |
| Dnajc10 | -0.013992544 | 0.132598056 | 0.222223462 | 0.325384578 | 0.342100255 |
| Dnajc11 | 0.015528359 | 0.004883235 | 0.014137823 | 0.094642779 | 0.08256144 |
| Dnajc12 | 0.027625541 | 0.004663235 | 0.013585846 | 0.502631132 | 0.459156038 |
| Dnajc13 | 0.034323236 | 3.08E-06 | 1.97E-05 | 0.193001796 | 0.158128271 |
| Dnajc14 | 0.018934311 | 0.011323777 | 0.028796693 | 0.19893141 | 0.175344763 |
| Dnajc15 | -0.032776545 | 0.000675198 | 0.002532515 | 0.419133034 | 0.454320883 |
| Dnajc16 | 0.005147172 | 0.246975267 | 0.361175323 | 0.056158659 | 0.052319783 |
| Dnajc17 | -0.009250406 | 0.122791161 | 0.20808361 | 0.101112082 | 0.106588393 |
| Dnajc18 | -0.008057071 | 0.224322343 | 0.33598241 | 0.13269233 | 0.135778886 |
| Dnajc19 | 0.010914525 | 0.26743604 | 0.384105951 | 0.546350042 | 0.537554215 |
| Dnajc2 | -0.024830331 | 0.007721743 | 0.020893622 | 0.325916521 | 0.347046174 |
| Dnajc21 | 0.063909147 | 9.89E-10 | 1.13E-08 | 0.804522131 | 0.742320226 |
| Dnajc24 | 0.113758899 | 2.06E-30 | 1.07E-28 | 0.526623821 | 0.381450325 |
| Dnajc25 | -0.008943154 | 0.143329755 | 0.23641871 | 0.106312385 | 0.109403132 |
| Dnajc27 | -0.002135035 | 0.588175949 | 0.692934477 | 0.041766134 | 0.042101935 |
| Dnajc3 | -0.100372112 | 1.17E-33 | 7.16E-32 | 1.38380427 | 1.572874499 |
| Dnajc30 | -0.008382167 | 0.236585105 | 0.35024827 | 0.159566458 | 0.16169391 |
| Dnajc4 | -0.038928031 | 3.91E-06 | 2.45E-05 | 0.250428682 | 0.276068062 |
| Dnajc5 | 0.019248126 | 0.043391392 | 0.088661436 | 0.408790296 | 0.391914549 |
| Dnajc6 | -0.004763036 | 0.42588994 | 0.544444707 | 0.100552858 | 0.105568462 |
| Dnajc7 | -0.013085428 | 0.165115567 | 0.263242628 | 0.400933648 | 0.401865904 |
| Dnajc8 | 0.022555144 | 0.020904403 | 0.048262315 | 0.495042595 | 0.473051117 |
| Dnajc9 | -0.011626128 | 0.16262048 | 0.260417774 | 0.247859346 | 0.250119348 |
| Dnal1 | -0.001985403 | 0.699303249 | 0.782503788 | 0.074114371 | 0.073750508 |
| Dnal4 | 0.006222124 | 0.402395824 | 0.522308705 | 0.192727092 | 0.175909408 |
| Dnase2a | 0.004394925 | 0.501859703 | 0.616123252 | 0.119896665 | 0.120115749 |
| Dner | -0.012718827 | 0.020450608 | 0.04732702 | 0.076662878 | 0.087535634 |
| Dnlz | -0.03565845 | 0.00011787 | 0.00054181 | 0.354261425 | 0.385716245 |
| Dnm1l | 0.048316777 | 4.78E-09 | 4.97E-08 | 0.261176642 | 0.223724831 |
| Dnm2 | 0.014169514 | 0.129808588 | 0.218150231 | 0.408061639 | 0.379825739 |
| Dnm3 | -0.015599694 | 0.062425395 | 0.120574473 | 0.19775838 | 0.214890722 |
| Dnmt1 | 0.008103292 | 0.191401276 | 0.295976333 | 0.112227744 | 0.106934265 |
| Dnmt3a | 0.048804116 | 9.76E-07 | 6.83E-06 | 0.465797302 | 0.393909272 |
| Dnmt3bos | 0.007796345 | 0.037817926 | 0.079033937 | 0.041657202 | 0.033993987 |
| Dnpep | -0.010537787 | 0.146885929 | 0.241016878 | 0.1696689 | 0.171912078 |
| Dnph1 | -0.029763264 | 1.45E-11 | 2.05E-10 | 0.035298319 | 0.060249468 |
| Dnttip1 | 0.002148709 | 0.78726446 | 0.851256775 | 0.228169312 | 0.221699249 |
| Dnttip2 | 0.022914477 | 0.010172982 | 0.026261744 | 0.307906114 | 0.283868073 |
| Dock10 | 0.022621654 | 7.05E-09 | 7.14E-08 | 0.051832748 | 0.032352951 |
| Dock11 | 0.003161653 | 0.560336324 | 0.668744137 | 0.086558106 | 0.080081918 |
| Dock5 | -0.029153127 | 4.75E-07 | 3.56E-06 | 0.073601659 | 0.099335857 |
| Dock7 | 0.003904027 | 0.565400458 | 0.672818551 | 0.148528637 | 0.140596253 |
| Dock8 | 0.001511755 | 0.711102074 | 0.791905425 | 0.047860796 | 0.043355543 |
| Dock9 | -0.001457386 | 0.796019109 | 0.857495598 | 0.087497054 | 0.089656881 |
| Dohh | -0.004080602 | 0.643224313 | 0.737432403 | 0.329039206 | 0.315667924 |

| | | | | | |
|---|---|---|---|---|---|
| Dolk | -0.010742187 | 0.020157855 | 0.046742455 | 0.054440509 | 0.061902915 |
| Dop1a | -0.014415389 | 0.051315545 | 0.102064183 | 0.156698624 | 0.168566437 |
| Dop1b | 0.006346657 | 0.373426328 | 0.495557024 | 0.162707405 | 0.157742831 |
| Dot1l | 0.023674278 | 3.87E-05 | 0.000195149 | 0.104854654 | 0.085160208 |
| Dpagt1 | -0.027807031 | 3.90E-06 | 2.45E-05 | 0.09012489 | 0.111787126 |
| Dpf2 | -0.008095428 | 0.354757077 | 0.477164956 | 0.299202802 | 0.29932079 |
| Dph3 | 0.017187394 | 0.082909863 | 0.151769852 | 0.55132715 | 0.532771231 |
| Dph5 | -0.007160972 | 0.150410057 | 0.245432456 | 0.066119401 | 0.070018554 |
| Dph6 | -0.014309302 | 0.052760016 | 0.10431558 | 0.152874026 | 0.174413462 |
| Dph7 | -0.002742608 | 0.561437959 | 0.669730041 | 0.060881942 | 0.062228205 |
| Dpm1 | -0.010241194 | 0.242974112 | 0.356665305 | 0.317374483 | 0.300670718 |
| Dpm2 | -0.009932612 | 0.259003893 | 0.374930889 | 0.313767483 | 0.312291256 |
| Dpm3 | 0.115516466 | 2.52E-46 | 2.60E-44 | 1.497170578 | 1.421045871 |
| Dpp3 | 0.000556358 | 0.937492389 | 0.95738456 | 0.161350484 | 0.157802122 |
| Dpp4 | 0.014493424 | 0.08599867 | 0.156393391 | 0.263279449 | 0.235040795 |
| Dpp7 | -0.042493888 | 6.90E-09 | 7.01E-08 | 0.164598873 | 0.186444636 |
| Dpp8 | -0.008310827 | 0.307269893 | 0.428222176 | 0.219874473 | 0.22348385 |
| Dpp9 | -0.005636273 | 0.511483429 | 0.625253602 | 0.279415488 | 0.273562064 |
| Dpy19l4 | 0.019341959 | 0.00206647 | 0.006714645 | 0.128211745 | 0.109530667 |
| Dpy30 | -0.027654432 | 0.003828462 | 0.011473584 | 0.457200429 | 0.471018635 |
| Dpysl2 | 0.027384753 | 0.002543165 | 0.008051008 | 0.384739668 | 0.329865484 |
| Dr1 | -0.003300789 | 0.626473023 | 0.724722838 | 0.137785798 | 0.136837323 |
| Dram2 | -0.008497287 | 0.232362936 | 0.345629202 | 0.158583478 | 0.161932549 |
| Drap1 | -0.069919739 | 4.11E-13 | 6.87E-12 | 0.884294808 | 0.954376816 |
| Drg1 | -0.004349853 | 0.612740896 | 0.712908927 | 0.283253312 | 0.284877929 |
| Drg2 | -0.005738048 | 0.367188458 | 0.489954184 | 0.12673782 | 0.122120755 |
| Drosha | 0.01471688 | 0.014977596 | 0.036509365 | 0.119556559 | 0.102599767 |
| Dsc2 | 0.007937195 | 0.050941957 | 0.101528845 | 0.046244923 | 0.041085898 |
| Dsg2 | 0.038086823 | 2.28E-07 | 1.81E-06 | 0.180218236 | 0.158006909 |
| Dsp | 0.071157772 | 2.72E-12 | 4.16E-11 | 0.458365549 | 0.384368914 |
| Dst | -0.056216931 | 1.56E-08 | 1.50E-07 | 1.221078844 | 1.346716234 |
| Dstn | 0.09263953 | 1.51E-24 | 5.90E-23 | 1.312762153 | 1.158760384 |
| Dstyk | 0.005365525 | 0.369114574 | 0.491377775 | 0.108186131 | 0.10150277 |
| Dtd1 | -0.058155514 | 7.86E-12 | 1.14E-10 | 0.236849069 | 0.284715357 |
| Dtd2 | -0.031253239 | 0.00010224 | 0.000476727 | 0.214879983 | 0.232918755 |
| Dtnb | 0.011297309 | 0.105712444 | 0.184877924 | 0.157324153 | 0.144282447 |
| Dtnbos | 0.019039015 | 0.000525888 | 0.00203126 | 0.098280067 | 0.068819756 |
| Dtnbp1 | -0.010217598 | 0.195930641 | 0.301828187 | 0.217551692 | 0.21584971 |
| Dtwd1 | -0.00922371 | 0.042189526 | 0.086551292 | 0.053441161 | 0.057984853 |
| Dtx2 | 0.00386325 | 0.38447084 | 0.505589816 | 0.061006975 | 0.052828666 |
| Dtx3 | -0.028985647 | 0.000350946 | 0.001418667 | 0.219460312 | 0.236546418 |
| Dtx3l | 0.033862852 | 4.25E-09 | 4.46E-08 | 0.1084033 | 0.08263674 |
| Dtymk | -0.0311389 | 0.000154821 | 0.000686038 | 0.223048743 | 0.250605506 |
| Dus1l | 0.008042817 | 0.198923636 | 0.305760568 | 0.124437203 | 0.113482164 |
| Dus2 | -0.009583787 | 0.078786642 | 0.145703583 | 0.080298216 | 0.085730866 |
| Dus3l | -0.015444127 | 0.000616515 | 0.002336749 | 0.04943892 | 0.059062848 |
| Dusp1 | -0.014263848 | 0.251516685 | 0.366562683 | 0.483960322 | 0.570662118 |
| Dusp10 | 0.026888219 | 7.40E-06 | 4.38E-05 | 0.116188909 | 0.092909388 |
| Dusp11 | 0.011105961 | 0.244939573 | 0.358792003 | 0.425670936 | 0.417887791 |
| Dusp12 | -0.006266225 | 0.286588257 | 0.405975672 | 0.099779905 | 0.102187421 |
| Dusp14 | -0.003252032 | 0.543094102 | 0.65297562 | 0.081471528 | 0.080023775 |
| Dusp16 | 0.022644659 | 1.04E-05 | 5.92E-05 | 0.083404402 | 0.065313328 |
| Dusp18 | -0.004912778 | 0.49910972 | 0.613832346 | 0.158053941 | 0.162612309 |
| Dusp19 | 0.013448528 | 0.054086272 | 0.106593702 | 0.172145576 | 0.149491364 |
| Dusp22 | -0.01586428 | 0.025460933 | 0.057008565 | 0.149909589 | 0.159322392 |
| Dusp23 | -0.007834332 | 0.289354395 | 0.408762486 | 0.172417038 | 0.168167034 |
| Dusp26 | 0.045710086 | 2.01E-13 | 3.49E-12 | 0.131521137 | 0.084589277 |
| Dusp28 | -0.025296101 | 0.000399505 | 0.001587602 | 0.156364378 | 0.16780471 |
| Dusp3 | 0.062472711 | 5.96E-15 | 1.15E-13 | 0.245890435 | 0.190366165 |
| Dusp4 | 0.010154702 | 0.057066947 | 0.11169818 | 0.070035928 | 0.070478948 |
| Dusp5 | -0.026797436 | 0.004046071 | 0.012029328 | 0.253470229 | 0.293569154 |
| Dusp7 | 0.024854166 | 1.09E-06 | 7.59E-06 | 0.083861517 | 0.063107614 |
| Dusp8 | 0.026114831 | 0.000421864 | 0.001667219 | 0.171410074 | 0.157128227 |
| Dut | -0.051917904 | 2.24E-07 | 1.78E-06 | 0.438415332 | 0.481463669 |
| Dvl1 | -0.000971502 | 0.870047071 | 0.910909159 | 0.102044248 | 0.102043422 |
| Dvl2 | 0.000247817 | 0.950530399 | 0.967042334 | 0.042767297 | 0.041244969 |
| Dvl3 | 0.012360883 | 0.062948576 | 0.12137764 | 0.135082739 | 0.127539952 |
| Dxo | -0.024612131 | 3.02E-05 | 0.000156835 | 0.090861732 | 0.107148128 |
| Dym | -8.40E-05 | 0.989768102 | 0.993241689 | 0.129209583 | 0.128685764 |
| Dync1h1 | 0.010147037 | 0.320860899 | 0.442406702 | 0.630694055 | 0.615088187 |
| Dync1i2 | 0.033540945 | 0.000718359 | 0.002679926 | 0.568846834 | 0.53450948 |
| Dync1li1 | 0.011385109 | 0.152511483 | 0.247565295 | 0.217748799 | 0.20767248 |

| | | | | | |
|---|---|---|---|---|---|
| Dync1li2 | -0.004691427 | 0.548225735 | 0.657681466 | 0.192898209 | 0.196168355 |
| Dync2h1 | -0.023699193 | 0.008144911 | 0.021864736 | 0.282389963 | 0.288806556 |
| Dync2i1 | 0.007490019 | 0.132748326 | 0.222436958 | 0.071255768 | 0.065145655 |
| Dync2i2 | -0.005227372 | 0.364207086 | 0.486978585 | 0.096777529 | 0.097545724 |
| Dync2li1 | -0.025467374 | 0.001515134 | 0.005119963 | 0.214238266 | 0.232663985 |
| Dynll1 | -0.050416362 | 7.21E-07 | 5.21E-06 | 0.538285753 | 0.593991578 |
| Dynll2 | 0.04320826 | 2.30E-06 | 1.50E-05 | 1.225369849 | 1.140870159 |
| Dynlrb1 | -0.011476768 | 0.230051866 | 0.343032972 | 0.818217389 | 0.808313177 |
| Dynlt1a | 0.00270248 | 0.583341612 | 0.689010709 | 0.075691774 | 0.065124045 |
| Dynlt1b | -0.005000404 | 0.29745599 | 0.417236637 | 0.055906873 | 0.064139794 |
| Dynlt1f | -0.009881142 | 0.145060376 | 0.238746737 | 0.15486505 | 0.145518093 |
| Dynlt2a2 | 0.001274053 | 0.772778996 | 0.839718051 | 0.052539247 | 0.050976113 |
| Dynlt2b | -0.02862536 | 0.000270397 | 0.001123899 | 0.20123118 | 0.221671728 |
| Dynlt3 | -0.007471357 | 0.447822071 | 0.566743839 | 0.509007061 | 0.510508097 |
| Dyrk1a | 0.031919183 | 5.86E-06 | 3.55E-05 | 0.176116318 | 0.142842499 |
| Dyrk2 | 0.07106727 | 8.28E-15 | 1.60E-13 | 0.390209881 | 0.308174329 |
| Dyrk3 | -0.018803319 | 0.001309293 | 0.004521829 | 0.078852999 | 0.101385818 |
| Dzank1 | -0.000665323 | 0.860338717 | 0.90433701 | 0.037645159 | 0.037447657 |
| Dzip1l | 0.022592731 | 2.21E-08 | 2.05E-07 | 0.055215777 | 0.035849688 |
| Dzip3 | -0.00417474 | 0.49942032 | 0.613935435 | 0.10491826 | 0.106511721 |
| E030030I06Rik | -0.010914132 | 0.009737971 | 0.025306751 | 0.041064076 | 0.049245539 |
| E130102H24Rik | 0.013027692 | 0.007371748 | 0.020102696 | 0.066895827 | 0.060547355 |
| E130308A19Rik | 0.014631087 | 0.000362583 | 0.001456024 | 0.05277198 | 0.039462298 |
| E130309D02Rik | 0.005538752 | 0.263466123 | 0.37963801 | 0.073518806 | 0.066682243 |
| E130311K13Rik | -0.003259277 | 0.547629823 | 0.657372162 | 0.084060187 | 0.083104507 |
| E130317F20Rik | 0.008073427 | 0.096912664 | 0.172082247 | 0.067165538 | 0.062548602 |
| E230020A03Rik | -0.000934698 | 0.879458172 | 0.917821182 | 0.100089865 | 0.099879586 |
| E2f1 | 0.000874988 | 0.820241391 | 0.876306242 | 0.038849793 | 0.036843703 |
| E2f4 | -0.001362882 | 0.850326706 | 0.897597854 | 0.17162084 | 0.16305817 |
| E2f5 | 0.005873692 | 0.348481539 | 0.4710048 | 0.115824414 | 0.114867468 |
| E2f6 | -0.006453226 | 0.296191795 | 0.416003558 | 0.10808503 | 0.113694886 |
| E4f1 | -0.004717383 | 0.311479844 | 0.432910228 | 0.059792299 | 0.060075211 |
| Eaf1 | 0.01080534 | 0.191351173 | 0.295945928 | 0.22648554 | 0.230597697 |
| Eapp | 0.013571863 | 0.110396323 | 0.191587478 | 0.276520653 | 0.264105577 |
| Ears2 | -0.00848474 | 0.094757715 | 0.168786095 | 0.069292579 | 0.073450185 |
| Ebag9 | 0.008447592 | 0.238735248 | 0.352625962 | 0.172600512 | 0.161948527 |
| Ebna1bp2 | -0.062116252 | 2.61E-11 | 3.58E-10 | 0.345406355 | 0.400464357 |
| Ebp | -0.015095335 | 0.060683624 | 0.11780475 | 0.229695958 | 0.235768532 |
| Ebpl | -0.049720656 | 1.25E-10 | 1.59E-09 | 0.183399862 | 0.218366684 |
| Ecd | 0.043656356 | 2.13E-07 | 1.70E-06 | 0.278473468 | 0.240493444 |
| Ece1 | 0.054625044 | 3.75E-07 | 2.87E-06 | 1.034221873 | 0.919227969 |
| Ech1 | -0.039224499 | 8.99E-06 | 5.22E-05 | 0.306035931 | 0.323342727 |
| Echdc1 | -0.007036802 | 0.240002067 | 0.354035993 | 0.105634116 | 0.107476057 |
| Echdc2 | 0.012201112 | 0.097004802 | 0.17218928 | 0.187558549 | 0.166928265 |
| Echdc3 | -0.008513703 | 0.022307552 | 0.050873567 | 0.035582786 | 0.038870792 |
| Echs1 | -0.082972446 | 1.80E-19 | 5.00E-18 | 0.307888546 | 0.37362326 |
| Eci1 | -0.046422138 | 2.03E-08 | 1.90E-07 | 0.239497668 | 0.262369592 |
| Eci2 | -0.038091728 | 9.35E-05 | 0.000439702 | 0.464482464 | 0.476879024 |
| Ecpas | 0.016572423 | 0.091733016 | 0.164513629 | 0.485209019 | 0.452627904 |
| Ecsit | 0.02240673 | 0.000989135 | 0.003543247 | 0.158256109 | 0.136032657 |
| Edc3 | 0.007199887 | 0.137833293 | 0.229140779 | 0.072197867 | 0.062321878 |
| Edc4 | 0.015414868 | 0.017478442 | 0.04162798 | 0.134195807 | 0.118930242 |
| Edem1 | -0.016213342 | 0.075138657 | 0.140372411 | 0.283197937 | 0.310732649 |
| Edem2 | -0.082253386 | 9.38E-19 | 2.44E-17 | 0.304096061 | 0.371159341 |
| Edem3 | 0.003302475 | 0.63283253 | 0.729216292 | 0.146160288 | 0.144582636 |
| Edf1 | -0.01251691 | 0.126772321 | 0.21382382 | 1.357120367 | 1.374108219 |
| Edrf1 | 0.023928731 | 3.16E-05 | 0.000163441 | 0.104113393 | 0.084849698 |
| Eea1 | 0.037213465 | 0.000261529 | 0.001089369 | 0.564901916 | 0.527171442 |
| Eed | -0.005604983 | 0.399848488 | 0.519974184 | 0.131338247 | 0.132700985 |
| Eef1a1 | -0.04451281 | 1.18E-32 | 6.99E-31 | 3.597770755 | 3.665558821 |
| Eef1akmt1 | -0.025211271 | 0.001442087 | 0.004912393 | 0.22197667 | 0.231509139 |
| Eef1akmt2 | -0.021002846 | 0.002495917 | 0.007927246 | 0.14572534 | 0.158413228 |
| Eef1akmt3 | -0.01224795 | 0.005284698 | 0.015115573 | 0.041721737 | 0.054242303 |
| Eef1akmt4 | -0.01467136 | 0.000177917 | 0.000777047 | 0.033590218 | 0.044201929 |
| Eef1aknmt | 0.005590488 | 0.166720304 | 0.26514883 | 0.047022435 | 0.041793946 |
| Eef1b2 | -0.044543125 | 9.80E-09 | 9.77E-08 | 1.620909828 | 1.704136248 |
| Eef1d | -0.067883699 | 6.87E-13 | 1.12E-11 | 0.866060189 | 0.915928064 |
| Eef1e1 | -0.0169411 | 0.012751367 | 0.031786868 | 0.132086416 | 0.148622702 |
| Eef1g | -0.034671003 | 0.000349752 | 0.001415017 | 0.973350395 | 0.995997572 |
| Eef2 | 0.000471004 | 0.942056979 | 0.960652643 | 2.088880451 | 2.026450285 |
| Eef2k | -0.007847059 | 0.048639089 | 0.097760847 | 0.038097806 | 0.043549385 |
| Eef2kmt | -0.024231493 | 0.005514367 | 0.015661898 | 0.28211721 | 0.296854899 |

| | | | | | |
|---|---|---|---|---|---|
| Eefsec | 0.010567529 | 0.03659871 | 0.0769824 | 0.072334435 | 0.066800111 |
| Eepd1 | 0.016856966 | 2.91E-05 | 0.000151493 | 0.056103652 | 0.037672338 |
| Efcab1 | -0.034211283 | 0.000746339 | 0.002769431 | 0.607255805 | 0.627021617 |
| Efcab10 | -0.008983943 | 0.054188048 | 0.106765856 | 0.053330364 | 0.061356101 |
| Efcab14 | 0.032305459 | 0.000867903 | 0.003163763 | 0.507068832 | 0.45863467 |
| Efcab15 | 0.078338161 | 4.20E-16 | 8.81E-15 | 0.489216428 | 0.402972117 |
| Efcab2 | -0.007175233 | 0.106543199 | 0.186129915 | 0.052014763 | 0.055492416 |
| Efemp2 | 0.003333918 | 0.563594641 | 0.671478811 | 0.094659609 | 0.087497853 |
| Efhc2 | 0.001151677 | 0.771023068 | 0.838372248 | 0.042795558 | 0.041101393 |
| Efhd2 | 0.014383075 | 0.021791698 | 0.049934096 | 0.123517169 | 0.113069165 |
| Efl1 | 0.0061072 | 0.244645517 | 0.358469287 | 0.079305 | 0.075834254 |
| Efna2 | -0.005902634 | 0.109198486 | 0.189950319 | 0.034860017 | 0.037315408 |
| Efna5 | -0.001268817 | 0.854618112 | 0.899880714 | 0.131568526 | 0.139665698 |
| Efnb1 | 0.012715962 | 0.010939721 | 0.027951633 | 0.079045403 | 0.06416543 |
| Efr3a | 0.047724587 | 4.82E-07 | 3.61E-06 | 0.441801601 | 0.385145083 |
| Eftud2 | 0.006400652 | 0.281321835 | 0.40026502 | 0.108598963 | 0.101044502 |
| Egf | -0.00520498 | 0.302311264 | 0.422644537 | 0.065294558 | 0.069954486 |
| Egfr | 0.030801571 | 6.27E-10 | 7.34E-09 | 0.084476633 | 0.057933947 |
| Egln1 | -0.005681211 | 0.518048477 | 0.630702927 | 0.308348529 | 0.299919904 |
| Egln2 | -0.017296252 | 0.054195587 | 0.106765856 | 0.346628558 | 0.348111163 |
| Egr1 | 0.038817601 | 0.006471966 | 0.018020213 | 1.110163266 | 1.077259784 |
| Ehbp1 | 0.005630415 | 0.297396378 | 0.417213216 | 0.085024271 | 0.078744483 |
| Ehd1 | -0.002102724 | 0.731078043 | 0.806347775 | 0.117455802 | 0.109717606 |
| Ehd3 | -0.010379707 | 0.029358573 | 0.064226833 | 0.056632732 | 0.064149294 |
| Ehd4 | 0.010612045 | 0.056348924 | 0.110537579 | 0.087178799 | 0.080203647 |
| Ehhadh | -0.002590597 | 0.498086905 | 0.612806997 | 0.037860433 | 0.039365579 |
| Ehmt1 | 0.010166255 | 0.155724677 | 0.251906042 | 0.16489266 | 0.152812331 |
| Ehmt2 | 0.028256633 | 0.001499238 | 0.005080372 | 0.338666838 | 0.2996727 |
| Ei24 | 0.002518689 | 0.779807482 | 0.845157198 | 0.380815497 | 0.352892084 |
| Eid1 | 0.019257033 | 0.040515038 | 0.083857546 | 1.240487453 | 1.183744048 |
| Eid2 | -0.03994795 | 7.35E-07 | 5.29E-06 | 0.225144225 | 0.245708099 |
| Eid2b | -0.000984758 | 0.885680035 | 0.92190378 | 0.149065902 | 0.145276464 |
| Eif1 | -0.017795355 | 6.86E-06 | 4.08E-05 | 3.693307811 | 3.724069296 |
| Eif1a | -0.006940452 | 0.485318724 | 0.601328648 | 0.410852992 | 0.444692694 |
| Eif1ad | 0.002838262 | 0.744551853 | 0.817636181 | 0.25714657 | 0.266703493 |
| Eif1ax | -0.024508559 | 0.008128336 | 0.021829748 | 0.353566009 | 0.379199259 |
| Eif1b | 0.031938148 | 0.001406273 | 0.004815705 | 0.776056097 | 0.730028989 |
| Eif2a | -0.01475404 | 0.080207181 | 0.147617671 | 0.266727046 | 0.265089618 |
| Eif2ak1 | 0.002617183 | 0.674447954 | 0.763149791 | 0.118868721 | 0.112143343 |
| Eif2ak2 | 0.009521705 | 0.168349342 | 0.267040676 | 0.146861456 | 0.139691365 |
| Eif2ak3 | 0.013173563 | 0.038322163 | 0.079915931 | 0.124086199 | 0.114783358 |
| Eif2ak4 | 0.013630937 | 0.002789789 | 0.008721006 | 0.062103511 | 0.052619509 |
| Eif2b1 | -0.004283789 | 0.468484876 | 0.585801925 | 0.104212584 | 0.103856585 |
| Eif2b2 | -0.000642361 | 0.935187547 | 0.956003506 | 0.213634135 | 0.209872882 |
| Eif2b3 | -0.003320369 | 0.648257088 | 0.741628078 | 0.163154514 | 0.167970936 |
| Eif2b4 | 0.003595694 | 0.562953214 | 0.670879032 | 0.119287393 | 0.112414892 |
| Eif2b5 | 0.031888568 | 5.33E-05 | 0.000260533 | 0.23758323 | 0.202551962 |
| Eif2d | -0.011006237 | 0.089364785 | 0.16112857 | 0.124710827 | 0.128900506 |
| Eif2s1 | -0.006096533 | 0.514807804 | 0.627896308 | 0.349647135 | 0.351879935 |
| Eif2s2 | 0.053209973 | 1.88E-10 | 2.34E-09 | 1.641688839 | 1.570720036 |
| Eif2s3x | 0.038142297 | 3.39E-06 | 2.15E-05 | 0.265038348 | 0.225737368 |
| Eif2s3y | -0.030554771 | 0.000561973 | 0.00215353 | 0.266042187 | 0.290265722 |
| Eif3a | 0.034222835 | 0.00032299 | 0.001316055 | 1.250229691 | 1.199471672 |
| Eif3b | 0.002162721 | 0.822507802 | 0.877955737 | 0.409883009 | 0.408045437 |
| Eif3c | 0.011307374 | 0.235371944 | 0.349303319 | 1.713906626 | 1.768499355 |
| Eif3d | -0.000429991 | 0.964440908 | 0.975376522 | 0.440106505 | 0.430683554 |
| Eif3e | -0.057229505 | 1.07E-08 | 1.06E-07 | 0.660192112 | 0.70217711 |
| Eif3f | -0.021667631 | 0.026181935 | 0.058273985 | 0.998199047 | 0.960158532 |
| Eif3g | -0.040172528 | 8.03E-05 | 0.00038265 | 0.52124834 | 0.560801545 |
| Eif3h | -0.061797658 | 7.54E-11 | 9.82E-10 | 0.90107093 | 0.95106492 |
| Eif3i | -0.08071818 | 7.20E-16 | 1.49E-14 | 0.621471087 | 0.693086164 |
| Eif3j1 | 0.057097572 | 6.66E-09 | 6.78E-08 | 0.455070817 | 0.412702176 |
| Eif3k | -0.080915356 | 2.96E-17 | 6.81E-16 | 0.871609146 | 0.953609787 |
| Eif3l | -0.028543418 | 0.001147281 | 0.004031032 | 0.318408918 | 0.325852245 |
| Eif3m | -0.015969842 | 0.114240709 | 0.196540112 | 0.682565036 | 0.666531738 |
| Eif4a1 | -0.031811658 | 0.001582359 | 0.005308383 | 0.918946847 | 0.942744429 |
| Eif4a2 | 0.083565837 | 1.91E-20 | 5.65E-19 | 1.281684429 | 1.123718269 |
| Eif4a3 | -0.004429548 | 0.627218021 | 0.725412039 | 0.355962335 | 0.345211266 |
| Eif4b | 0.034303681 | 0.000618924 | 0.002344963 | 0.614561372 | 0.548170399 |
| Eif4e | 0.023675907 | 0.019324946 | 0.045076087 | 0.661439851 | 0.628736154 |
| Eif4e2 | -0.007110391 | 0.369506731 | 0.491765153 | 0.214633947 | 0.213108222 |
| Eif4e3 | 0.006425232 | 0.289271402 | 0.408723488 | 0.114655943 | 0.103093419 |

| | | | | | |
|---|---|---|---|---|---|
| Eif4ebp1 | 0.00070585 | 0.952875378 | 0.968313414 | 1.034012779 | 1.040796071 |
| Eif4ebp2 | 0.046070354 | 1.10E-07 | 9.24E-07 | 0.316936797 | 0.264615384 |
| Eif4enif1 | 0.007594456 | 0.269987221 | 0.386570804 | 0.148082529 | 0.142332526 |
| Eif4g1 | 0.003941145 | 0.695372175 | 0.779477492 | 0.753322682 | 0.747200171 |
| Eif4g2 | -0.005957885 | 0.489986744 | 0.605369314 | 1.423096608 | 1.440471035 |
| Eif4g3 | -0.062920709 | 2.36E-09 | 2.55E-08 | 0.674036194 | 0.744623688 |
| Eif4h | 0.033065052 | 0.000858602 | 0.003136912 | 0.504694126 | 0.458374956 |
| Eif5 | 0.021133091 | 0.018459817 | 0.043475374 | 1.417198068 | 1.398759185 |
| Eif5a | -0.03361043 | 0.000149435 | 0.00066612 | 1.461024071 | 1.52450226 |
| Eif5a2 | 0.002034535 | 0.641358159 | 0.736074138 | 0.0495094 | 0.050517379 |
| Eif5b | 0.094776304 | 5.22E-21 | 1.60E-19 | 0.886024127 | 0.794780043 |
| Eif6 | -0.022806385 | 0.027145304 | 0.060129107 | 0.521546801 | 0.546132134 |
| Eipr1 | 0.008318153 | 0.178055674 | 0.279707751 | 0.120968794 | 0.113993086 |
| Elac1 | -0.00085085 | 0.848217703 | 0.896442278 | 0.05242184 | 0.051968691 |
| Elac2 | 0.001452552 | 0.756183147 | 0.826566212 | 0.061627432 | 0.059648216 |
| Elapor1 | -0.091776067 | 4.77E-19 | 1.29E-17 | 0.584294501 | 0.685750894 |
| Elapor2 | 0.141906645 | 5.39E-54 | 7.48E-52 | 0.428195509 | 0.288046237 |
| Elavl1 | -0.005971743 | 0.490888375 | 0.606252132 | 0.293427464 | 0.286242577 |
| Elavl4 | 0.023971271 | 0.012313562 | 0.03093345 | 0.330947726 | 0.304009289 |
| Elf1 | 0.008410671 | 0.339877915 | 0.4622682 | 0.295703213 | 0.286149731 |
| Elf2 | 0.011099952 | 0.210046271 | 0.319023567 | 0.303144253 | 0.295827445 |
| Elf3 | -0.004943892 | 0.419051821 | 0.538114347 | 0.096815818 | 0.104240925 |
| Elk3 | -0.005719632 | 0.377028292 | 0.498703273 | 0.122594141 | 0.12432332 |
| Elk4 | -0.0031321 | 0.575968844 | 0.68235835 | 0.087320252 | 0.089391654 |
| Ell | 0.010452365 | 0.048608125 | 0.097739027 | 0.084918572 | 0.07576728 |
| Ell2 | -0.004016386 | 0.668769563 | 0.758047999 | 0.341706426 | 0.365284211 |
| Elmo1 | 0.00277784 | 0.675462351 | 0.764119732 | 0.123286198 | 0.126078141 |
| Elmo2 | 0.001568381 | 0.720684008 | 0.798794646 | 0.054062905 | 0.052400588 |
| Elmo3 | 0.004946363 | 0.400461732 | 0.52056277 | 0.107045234 | 0.098216218 |
| Elmod1 | -0.003993614 | 0.469359145 | 0.586593342 | 0.081121287 | 0.083134082 |
| Elmod2 | 0.00698839 | 0.187386216 | 0.291110385 | 0.081437186 | 0.075444167 |
| Elmod3 | 0.006878422 | 0.110275551 | 0.191480426 | 0.05552718 | 0.048725993 |
| Eloa | 0.021969793 | 0.006871921 | 0.018947481 | 0.245927064 | 0.220962887 |
| Elob | 0.083606444 | 1.41E-27 | 6.51E-26 | 1.572098802 | 1.473212722 |
| Eloc | -0.004824649 | 0.616262956 | 0.716406608 | 0.789967398 | 0.806197973 |
| Elof1 | -0.007352305 | 0.455872765 | 0.57439023 | 0.63343464 | 0.6172278 |
| Elovl1 | 0.005206717 | 0.334473342 | 0.457348781 | 0.085135573 | 0.07962267 |
| Elovl2 | 0.004949798 | 0.360306847 | 0.482959212 | 0.083484161 | 0.078504603 |
| Elovl4 | 0.004332245 | 0.336875733 | 0.459277223 | 0.058130562 | 0.053338751 |
| Elovl5 | 0.043360052 | 1.33E-07 | 1.10E-06 | 0.269208003 | 0.217497076 |
| Elovl6 | 0.015623667 | 0.020048801 | 0.046518961 | 0.133461545 | 0.126583116 |
| Elovl7 | 0.007749697 | 0.122756208 | 0.20808361 | 0.070384883 | 0.067028093 |
| Elp1 | 0.014532919 | 0.034853462 | 0.073943268 | 0.155213748 | 0.138882249 |
| Elp2 | 0.00658865 | 0.302615542 | 0.422826717 | 0.134590107 | 0.12024597 |
| Elp3 | 0.00949853 | 0.279654671 | 0.39859298 | 0.309941714 | 0.290478611 |
| Elp4 | 0.007017401 | 0.323601925 | 0.445680395 | 0.167305491 | 0.154753014 |
| Elp5 | -0.002182787 | 0.749502502 | 0.821587927 | 0.146364424 | 0.14585271 |
| Elp6 | -0.00260482 | 0.563577892 | 0.671478811 | 0.055914092 | 0.056261795 |
| Emb | 0.021716765 | 0.021618274 | 0.049627595 | 1.195172276 | 1.12793079 |
| Emc1 | -0.001636172 | 0.810101108 | 0.86908944 | 0.140514785 | 0.141381923 |
| Emc10 | -0.054419028 | 3.19E-08 | 2.91E-07 | 0.559413314 | 0.614620741 |
| Emc2 | -0.055459486 | 1.51E-10 | 1.90E-09 | 0.264610555 | 0.306008179 |
| Emc3 | -0.008126918 | 0.289284748 | 0.408723488 | 0.184639155 | 0.192416686 |
| Emc4 | -0.04194828 | 6.35E-06 | 3.81E-05 | 0.344676081 | 0.37552811 |
| Emc6 | -0.006209433 | 0.510436858 | 0.624366775 | 0.386130385 | 0.395402747 |
| Emc7 | -0.055738409 | 5.98E-08 | 5.25E-07 | 0.796354012 | 0.878070774 |
| Emc8 | -0.003378205 | 0.641175551 | 0.736074138 | 0.179150269 | 0.175224184 |
| Emc9 | -0.018197858 | 0.019383704 | 0.045169792 | 0.204523525 | 0.213490858 |
| Emd | 0.001598776 | 0.869385099 | 0.910400898 | 0.429214578 | 0.421542561 |
| Emg1 | -0.019510393 | 0.043724875 | 0.089249052 | 0.488084167 | 0.500529073 |
| Emid1 | 0.079514521 | 1.65E-52 | 2.14E-50 | 0.122101119 | 0.045502235 |
| Emilin1 | 0.030399421 | 3.37E-05 | 0.00017302 | 0.176423445 | 0.144575922 |
| Eml1 | -4.21E-05 | 0.992536837 | 0.995198343 | 0.058397934 | 0.054692719 |
| Eml2 | -0.005272228 | 0.432062747 | 0.550529472 | 0.144020486 | 0.142089028 |
| Eml3 | 0.004738354 | 0.33674756 | 0.459193399 | 0.067609599 | 0.064341662 |
| Eml4 | 0.025497318 | 0.000525466 | 0.002030439 | 0.185465672 | 0.162523687 |
| Eml5 | -0.032890913 | 0.000879073 | 0.003200879 | 0.381063642 | 0.413464174 |
| Eml6 | -0.004601926 | 0.363116622 | 0.485854638 | 0.068190121 | 0.070750072 |
| Emsy | 0.006982959 | 0.321777479 | 0.443418944 | 0.158368813 | 0.149268544 |
| Enah | 0.076033921 | 1.89E-14 | 3.56E-13 | 0.525116493 | 0.435442799 |
| Enc1 | 0.032076657 | 3.69E-10 | 4.44E-09 | 0.08901397 | 0.059933445 |
| Endod1 | -0.013089801 | 0.01931013 | 0.04506315 | 0.083345244 | 0.091314708 |

| | | | | | |
|---|---|---|---|---|---|
| Endog | -0.039062351 | 2.01E-06 | 1.33E-05 | 0.218193602 | 0.247195532 |
| Endov | 0.001691131 | 0.717492525 | 0.796922465 | 0.059463608 | 0.058481082 |
| Enho | -0.050562013 | 2.09E-08 | 1.95E-07 | 1.561990945 | 1.586333178 |
| Eno1 | 0.033369172 | 0.000581039 | 0.002219591 | 0.421689276 | 0.37235557 |
| Eno2 | 0.005773475 | 0.414532352 | 0.53399393 | 0.152601928 | 0.148606571 |
| Eno3 | -0.012515788 | 0.003278487 | 0.010048378 | 0.045747783 | 0.051168523 |
| Enoph1 | -0.00428059 | 0.36028015 | 0.482959212 | 0.060625453 | 0.060421325 |
| Enox2 | 0.004099478 | 0.39863227 | 0.518947902 | 0.067455048 | 0.061472373 |
| Enpep | -0.023318064 | 0.000284414 | 0.001173133 | 0.097303159 | 0.121326621 |
| Enpp2 | -0.041901425 | 0.000362405 | 0.001456024 | 0.645385461 | 0.695164549 |
| Enpp4 | 0.014588111 | 0.045149532 | 0.091675803 | 0.182021321 | 0.162649586 |
| Enpp5 | -0.008879993 | 0.349488336 | 0.471906334 | 0.412195024 | 0.402193171 |
| Ensa | 0.00197864 | 0.813905449 | 0.871649173 | 0.258363512 | 0.258879807 |
| ENSMUSG00000095 | -0.091400906 | 2.81E-18 | 7.05E-17 | 0.490200493 | 0.575958875 |
| ENSMUSG00000115 | 0.02232303 | 0.006037256 | 0.016953842 | 0.177957123 | 0.181036753 |
| ENSMUSG00000117 | 0.0002438 | 0.967016683 | 0.977676392 | 0.101511507 | 0.096063946 |
| Entpd2 | 0.011891021 | 0.009903197 | 0.025667524 | 0.066180065 | 0.053568412 |
| Entpd3 | -0.114463832 | 3.57E-26 | 1.56E-24 | 0.634040908 | 0.800573533 |
| Entpd5 | -0.007118973 | 0.15796399 | 0.254597305 | 0.064407022 | 0.070833172 |
| Entpd7 | 0.021714373 | 0.00696325 | 0.019155835 | 0.215983409 | 0.204254444 |
| Entr1 | -0.004466831 | 0.584471517 | 0.689842429 | 0.242780459 | 0.237545822 |
| Eny2 | -0.009745151 | 0.329777559 | 0.452455183 | 0.697534607 | 0.719023331 |
| Eola1 | 0.008300184 | 0.173098266 | 0.273280504 | 0.120841275 | 0.107820927 |
| Ep300 | 0.02175094 | 0.012872721 | 0.032072936 | 0.303724394 | 0.274749063 |
| Ep400 | 0.027595633 | 0.00106763 | 0.00379229 | 0.271022102 | 0.246810155 |
| Epb41 | 0.027825869 | 0.001104537 | 0.003902003 | 0.273884103 | 0.244721647 |
| Epb41l1 | 0.011003201 | 0.037461074 | 0.078389274 | 0.08762421 | 0.074465321 |
| Epb41l3 | 0.084897537 | 9.27E-16 | 1.90E-14 | 0.825499237 | 0.684610775 |
| Epb41l4a | 0.019349122 | 2.23E-06 | 1.46E-05 | 0.048316595 | 0.036039186 |
| Epb41l4aos | 0.015497272 | 0.011762365 | 0.029756366 | 0.113203783 | 0.103696411 |
| Epb41l4b | 0.048930449 | 8.47E-09 | 8.53E-08 | 0.29606859 | 0.242313876 |
| Epb41l5 | 0.018546586 | 0.00808863 | 0.021755369 | 0.161759151 | 0.143325869 |
| Epc1 | 0.039731241 | 2.16E-07 | 1.72E-06 | 0.213201194 | 0.177917307 |
| Epc2 | 0.020882665 | 0.008231214 | 0.022039069 | 0.211334042 | 0.196031794 |
| Epcam | 0.021568918 | 0.011961645 | 0.030205485 | 1.477158802 | 1.476959134 |
| Epg5 | 0.008723355 | 0.063803369 | 0.122660937 | 0.063806343 | 0.058116109 |
| Ephx1 | -0.011284216 | 0.057006409 | 0.111624634 | 0.096499079 | 0.101224502 |
| Epm2a | 0.0065312 | 0.061750767 | 0.119589832 | 0.036991551 | 0.029772105 |
| Epm2aip1 | -0.023073874 | 0.010827348 | 0.027691706 | 0.298734343 | 0.319447877 |
| Epn1 | 0.012557067 | 0.188147664 | 0.29205997 | 0.468410244 | 0.437837927 |
| Epn2 | 0.013288663 | 0.016980464 | 0.040611088 | 0.096464503 | 0.081901983 |
| Eprs | -0.077357833 | 2.95E-16 | 6.26E-15 | 1.692321728 | 1.847819996 |
| Eps15 | -0.010341887 | 0.087367527 | 0.15840362 | 0.098651013 | 0.107899754 |
| Eps15l1 | 0.002950964 | 0.600296953 | 0.703482037 | 0.093885615 | 0.088420158 |
| Eps8 | 0.024818094 | 0.005237963 | 0.015017245 | 0.309019033 | 0.279563097 |
| Eps8l2 | 0.053691636 | 2.81E-12 | 4.29E-11 | 0.231542158 | 0.170047947 |
| Erap1 | 0.000142668 | 0.985257363 | 0.990990784 | 0.196740688 | 0.196487429 |
| Erbin | 0.023281573 | 0.009610814 | 0.025070119 | 0.318907215 | 0.29075069 |
| Erc1 | 0.006524734 | 0.288907385 | 0.408427744 | 0.108970656 | 0.103583288 |
| Erc2 | 0.03137109 | 7.48E-09 | 7.57E-08 | 0.100827152 | 0.071040278 |
| Ercc1 | -0.008959726 | 0.070978281 | 0.133808581 | 0.065859799 | 0.071284487 |
| Ercc2 | 0.001273686 | 0.74991258 | 0.821852114 | 0.044315505 | 0.041802109 |
| Ercc3 | 0.009545563 | 0.054873662 | 0.10793517 | 0.072360414 | 0.064426795 |
| Ercc4 | 0.001321597 | 0.743397621 | 0.816737673 | 0.046544095 | 0.042590626 |
| Ercc5 | 0.021313386 | 0.00065411 | 0.002461967 | 0.125640884 | 0.108485361 |
| Ercc6 | 0.005823521 | 0.24419675 | 0.357973583 | 0.071223572 | 0.065838695 |
| Ercc6l2 | 0.018351705 | 0.012097591 | 0.030485427 | 0.172813792 | 0.157580097 |
| Erf | 0.005769673 | 0.177405343 | 0.278857678 | 0.051520176 | 0.046611617 |
| Erg28 | 0.001725465 | 0.810385744 | 0.8692156 | 0.174830668 | 0.166794154 |
| Ergic1 | -0.001942631 | 0.8437706 | 0.893296066 | 0.455991796 | 0.458467553 |
| Ergic2 | 0.011556596 | 0.24178041 | 0.355826392 | 0.508291347 | 0.503220415 |
| Ergic3 | -0.07609171 | 3.19E-15 | 6.31E-14 | 0.457716864 | 0.512141658 |
| Erh | -0.003700485 | 0.696086835 | 0.77991889 | 0.364506606 | 0.350866898 |
| Eri1 | 0.002807163 | 0.618546363 | 0.718422269 | 0.09355161 | 0.089329998 |
| Eri3 | -0.015294846 | 0.044529249 | 0.090567648 | 0.192499687 | 0.197267318 |
| Erich1 | 0.020325227 | 0.001953006 | 0.006399435 | 0.139708191 | 0.124121755 |
| Erich3 | -0.004614711 | 0.495561406 | 0.610627122 | 0.133751609 | 0.134240281 |
| Erlec1 | -0.087726216 | 8.59E-18 | 2.05E-16 | 0.690260456 | 0.793295403 |
| Erlin1 | -0.006175714 | 0.337261411 | 0.45944503 | 0.105390118 | 0.118123277 |
| Erlin2 | 0.021985791 | 0.001610517 | 0.005389826 | 0.161532399 | 0.144983307 |
| Ermard | -0.002587626 | 0.561527056 | 0.669754145 | 0.052204876 | 0.053387838 |
| Ermp1 | 0.006752536 | 0.178751326 | 0.280655749 | 0.0728685 | 0.066629751 |

| | | | | | |
|---|---|---|---|---|---|
| Ern1 | -0.031908321 | 0.001568588 | 0.005271279 | 0.43094387 | 0.476428883 |
| Ero1a | -0.04075745 | 9.81E-05 | 0.000458207 | 0.422898474 | 0.485644835 |
| Ero1b | 0.047957155 | 5.34E-08 | 4.72E-07 | 1.841906508 | 1.705340022 |
| Erp29 | -0.037914534 | 0.000143869 | 0.000646643 | 0.685791035 | 0.727230885 |
| Erp44 | -0.032729034 | 0.001178248 | 0.004129388 | 0.495019808 | 0.538896948 |
| Errfi1 | 0.034682239 | 0.000116437 | 0.000535986 | 0.24272463 | 0.236912985 |
| Esco1 | 0.019970149 | 0.010420435 | 0.026803664 | 0.211476801 | 0.189316629 |
| Esd | -0.073362903 | 6.32E-13 | 1.04E-11 | 0.876178231 | 0.960560315 |
| Esf1 | 0.011936464 | 0.178381581 | 0.280120454 | 0.296875025 | 0.296143213 |
| Esrp1 | -0.01226825 | 0.216057423 | 0.326216845 | 0.417575683 | 0.443591668 |
| Esrra | -0.000327662 | 0.949598114 | 0.966498416 | 0.07500091 | 0.074407461 |
| Ess2 | -0.002483849 | 0.596163365 | 0.699818913 | 0.060911903 | 0.060354027 |
| Esyt1 | 0.038090259 | 3.49E-05 | 0.000178551 | 0.351615632 | 0.289608419 |
| Esyt2 | 0.029965005 | 0.0001464 | 0.000654993 | 0.215850964 | 0.194916645 |
| Etaa1 | 0.007201683 | 0.15439673 | 0.249882638 | 0.075815336 | 0.067587513 |
| Etf1 | 0.029112114 | 0.003906739 | 0.011690157 | 0.533951897 | 0.507205465 |
| Etfa | -0.019865917 | 0.02274889 | 0.051722337 | 0.313802432 | 0.310112952 |
| Etfb | -0.080896916 | 2.78E-17 | 6.41E-16 | 0.477558622 | 0.508966051 |
| Etfdh | -0.013586116 | 0.026449464 | 0.05878868 | 0.104818667 | 0.112209167 |
| Etfrf1 | -0.013210327 | 0.076629562 | 0.142636915 | 0.179507279 | 0.186856638 |
| Ethe1 | -0.016793801 | 0.004483729 | 0.013164673 | 0.096053594 | 0.106752916 |
| Etl4 | 0.014025496 | 0.060893425 | 0.118141265 | 0.189716397 | 0.170278629 |
| Etnk1 | -0.001928232 | 0.847476188 | 0.895886971 | 0.523542878 | 0.527594285 |
| Etohd2 | -0.009611424 | 0.019322903 | 0.045076087 | 0.043829763 | 0.047772554 |
| Ets2 | 0.023473323 | 0.011139679 | 0.028410273 | 0.332146702 | 0.324946207 |
| Etv1 | -0.087718832 | 4.66E-17 | 1.05E-15 | 0.256275582 | 0.363131844 |
| Etv3 | 0.016642453 | 0.004412453 | 0.012975763 | 0.105845899 | 0.093529969 |
| Etv5 | -0.017147952 | 0.004163528 | 0.012318266 | 0.080051023 | 0.10242867 |
| Etv6 | 0.035910732 | 4.66E-12 | 6.93E-11 | 0.089240603 | 0.063942814 |
| Eva1b | -0.018855214 | 0.060906697 | 0.118143438 | 0.490217343 | 0.494690014 |
| Eva1c | -0.006865757 | 0.110439457 | 0.191625959 | 0.048100253 | 0.051410465 |
| Evi5 | 0.024834006 | 0.001526262 | 0.005153985 | 0.22494264 | 0.199013254 |
| Evi5l | 0.00886794 | 0.029989989 | 0.065416798 | 0.049398885 | 0.04233351 |
| Evl | -0.027829805 | 1.79E-05 | 9.77E-05 | 0.11569175 | 0.132886742 |
| Ewsr1 | 0.018953401 | 0.05015758 | 0.100294527 | 0.521463997 | 0.482641731 |
| Exd2 | -0.011048905 | 0.0371854 | 0.077963438 | 0.075538969 | 0.082858581 |
| Exoc1 | 0.001858544 | 0.749603014 | 0.821605468 | 0.098677765 | 0.096068389 |
| Exoc2 | 0.006714656 | 0.296786505 | 0.416598095 | 0.122168917 | 0.118269756 |
| Exoc3 | 0.007223651 | 0.415590289 | 0.534864812 | 0.327840236 | 0.310961639 |
| Exoc4 | 0.027885905 | 0.000707899 | 0.002643946 | 0.241609968 | 0.218177888 |
| Exoc5 | 0.060715646 | 1.24E-12 | 1.98E-11 | 0.306486217 | 0.24423682 |
| Exoc6 | 0.012333197 | 0.003791862 | 0.011381438 | 0.05528948 | 0.045073478 |
| Exoc6b | 0.03201962 | 0.000114921 | 0.000529509 | 0.245086046 | 0.224928258 |
| Exoc7 | -0.059730476 | 4.18E-12 | 6.22E-11 | 0.245403862 | 0.279126793 |
| Exoc8 | -0.003595959 | 0.464050316 | 0.581603348 | 0.067598909 | 0.066825031 |
| Exosc1 | -0.022560578 | 0.001960471 | 0.006417407 | 0.156445507 | 0.173768771 |
| Exosc10 | 0.006808693 | 0.265297103 | 0.381654104 | 0.113343886 | 0.106283932 |
| Exosc2 | -0.002565505 | 0.58414274 | 0.689705577 | 0.05736756 | 0.059003601 |
| Exosc3 | -0.015422913 | 0.05825456 | 0.113725067 | 0.229310303 | 0.237463139 |
| Exosc4 | -0.022815763 | 0.010456847 | 0.026866138 | 0.308607723 | 0.329184094 |
| Exosc5 | -0.076775026 | 8.36E-21 | 2.53E-19 | 0.194091273 | 0.256218788 |
| Exosc7 | -0.010232102 | 0.23828693 | 0.352138515 | 0.305265843 | 0.305002292 |
| Exosc8 | 0.005020555 | 0.341545239 | 0.463947667 | 0.08242256 | 0.077678676 |
| Exosc9 | -0.008455198 | 0.183619713 | 0.286495081 | 0.118758982 | 0.122866881 |
| Exph5 | 0.017289322 | 0.038989449 | 0.081116076 | 0.246198077 | 0.220510735 |
| Ext1 | 0.012970972 | 0.027933371 | 0.061580098 | 0.100029831 | 0.093713036 |
| Ext2 | 0.014629497 | 0.029640465 | 0.064785207 | 0.14832545 | 0.135511147 |
| Extl2 | 0.023109983 | 0.000191025 | 0.000824663 | 0.12679246 | 0.105546313 |
| Extl3 | 0.017683572 | 0.001192314 | 0.004169667 | 0.095455082 | 0.07983723 |
| Eya1 | 0.006623875 | 0.108783743 | 0.189499292 | 0.04960369 | 0.042449229 |
| Eya3 | 0.011919568 | 0.03906495 | 0.081238384 | 0.100220161 | 0.091696717 |
| Ezh1 | 0.02024808 | 0.004222554 | 0.01247771 | 0.165762018 | 0.14779351 |
| Ezh2 | 1.57E-05 | 0.996757721 | 0.997578605 | 0.038914284 | 0.03836361 |
| Ezr | 0.006709638 | 0.325660654 | 0.447817946 | 0.14829627 | 0.137037034 |
| F11r | 0.024220246 | 0.001193927 | 0.004173807 | 0.188979955 | 0.167407492 |
| F2rl1 | 0.008041085 | 0.026578462 | 0.059034911 | 0.039853078 | 0.031506199 |
| F3 | 0.005647805 | 0.28352637 | 0.402635608 | 0.08424156 | 0.074276643 |
| F8a | 0.030391954 | 4.65E-05 | 0.000229924 | 0.197211721 | 0.17181405 |
| Faah | 0.004313625 | 0.49555097 | 0.610627122 | 0.122447927 | 0.116813552 |
| Faap100 | 0.000181465 | 0.969518871 | 0.979594935 | 0.061774854 | 0.06136219 |
| Faap20 | -0.02366943 | 0.004316805 | 0.012736867 | 0.236916603 | 0.252933108 |
| Faap24 | 0.001856259 | 0.633957409 | 0.730166323 | 0.042657369 | 0.03953213 |

| | | | | | |
|---|---|---|---|---|---|
| Fabp3 | -0.071343409 | 2.81E-20 | 8.23E-19 | 0.147996273 | 0.200845536 |
| Fabp5 | -0.026891204 | 1.11E-06 | 7.70E-06 | 0.051539316 | 0.075935431 |
| Fadd | -0.003301254 | 0.4151157 | 0.534466274 | 0.043770445 | 0.044936112 |
| Fads1 | 0.001049675 | 0.846268437 | 0.894771261 | 0.083285623 | 0.08198526 |
| Fads6 | -0.011409006 | 0.076855565 | 0.142948117 | 0.116482328 | 0.1252057 |
| Faf1 | 0.000913404 | 0.917173726 | 0.942575366 | 0.307494651 | 0.293619552 |
| Faf2 | 0.024045424 | 0.008597756 | 0.022900654 | 0.363908237 | 0.336939968 |
| Fah | -0.005102738 | 0.192465442 | 0.297290917 | 0.036810767 | 0.041501071 |
| Fahd1 | -0.010794569 | 0.116207759 | 0.199254292 | 0.150139056 | 0.150058483 |
| Fahd2a | -0.050988769 | 3.69E-13 | 6.27E-12 | 0.137145651 | 0.171158515 |
| Faim | -0.027476498 | 0.000232077 | 0.000980273 | 0.173031712 | 0.191417013 |
| Fam102a | -0.001284739 | 0.883671565 | 0.920305834 | 0.275671878 | 0.276263053 |
| Fam104a | 0.066029698 | 2.18E-11 | 3.02E-10 | 0.830489307 | 0.732800492 |
| Fam110a | 0.01212798 | 0.041963086 | 0.086268793 | 0.10955547 | 0.100177487 |
| Fam110b | 0.033609787 | 6.63E-07 | 4.83E-06 | 0.167740424 | 0.128864334 |
| Fam114a1 | 0.043983341 | 2.21E-07 | 1.76E-06 | 0.292794187 | 0.24623785 |
| Fam114a2 | 0.00254382 | 0.775542204 | 0.841614164 | 0.328618427 | 0.316426128 |
| Fam117a | -0.002304983 | 0.772673801 | 0.839697596 | 0.233175711 | 0.219248204 |
| Fam117b | 0.007955321 | 0.239037422 | 0.352911438 | 0.147078175 | 0.13462071 |
| Fam118a | 0.003043933 | 0.576220642 | 0.682506955 | 0.086181273 | 0.082959011 |
| Fam118b | -0.007558374 | 0.135765412 | 0.226672048 | 0.069732847 | 0.073599196 |
| Fam120a | 0.021358732 | 0.031190201 | 0.067534773 | 0.554875174 | 0.529082112 |
| Fam120b | -0.003692247 | 0.633751927 | 0.730016142 | 0.195377853 | 0.19752196 |
| Fam120c | -0.002641565 | 0.565411187 | 0.672818551 | 0.054447363 | 0.056578544 |
| Fam122a | -0.011965216 | 0.243580222 | 0.357285293 | 0.568377368 | 0.580528903 |
| Fam126b | 0.032021056 | 0.000170878 | 0.000749689 | 0.272445208 | 0.239788505 |
| Fam131a | 0.006716471 | 0.246780232 | 0.361048316 | 0.10249044 | 0.093688655 |
| Fam133b | 0.020815868 | 0.007653043 | 0.020748156 | 0.207329267 | 0.188799136 |
| Fam135a | -0.056150166 | 1.35E-07 | 1.11E-06 | 0.673207864 | 0.765650758 |
| Fam135b | -0.02994794 | 1.82E-05 | 9.93E-05 | 0.121658668 | 0.144826947 |
| Fam136a | 0.014969234 | 0.025333263 | 0.056775008 | 0.142885181 | 0.130132798 |
| Fam13b | -0.016078558 | 0.025960185 | 0.057873175 | 0.144916388 | 0.157407223 |
| Fam149a | -0.002998611 | 0.568129078 | 0.675556616 | 0.072753878 | 0.076580379 |
| Fam149b | -0.000597467 | 0.926127992 | 0.950170572 | 0.132224916 | 0.125131003 |
| Fam151a | 0.067047583 | 7.78E-10 | 9.01E-09 | 0.726630879 | 0.593273969 |
| Fam155a | 0.020174315 | 0.002236475 | 0.007192527 | 0.146288246 | 0.121363846 |
| Fam160a1 | 0.009821407 | 0.009799127 | 0.025431691 | 0.044698877 | 0.035211052 |
| Fam160a2 | -0.00136428 | 0.811571013 | 0.870294881 | 0.099142048 | 0.09283838 |
| Fam160b1 | 0.025389921 | 0.001587286 | 0.005319404 | 0.238439793 | 0.214500481 |
| Fam160b2 | 0.004453404 | 0.39026036 | 0.511391726 | 0.074289882 | 0.07219206 |
| Fam161a | -0.005809533 | 0.129761275 | 0.218138194 | 0.037425264 | 0.039955834 |
| Fam162a | -0.070757048 | 4.52E-16 | 9.47E-15 | 0.240263281 | 0.29650929 |
| Fam163a | -0.015925643 | 0.000158112 | 0.000699029 | 0.036495915 | 0.049759924 |
| Fam168a | 0.010268468 | 0.17215544 | 0.271973451 | 0.184737825 | 0.17544052 |
| Fam168b | 0.006870398 | 0.268783711 | 0.385379095 | 0.119091808 | 0.112241321 |
| Fam169a | 0.018122089 | 0.00203767 | 0.006638816 | 0.109183361 | 0.091008003 |
| Fam171a1 | 0.0205008 | 0.000148095 | 0.000661662 | 0.097396692 | 0.076989551 |
| Fam171b | -0.01373666 | 0.118414153 | 0.202118424 | 0.260604748 | 0.271235838 |
| Fam172a | -0.003084337 | 0.681813164 | 0.768888479 | 0.186469343 | 0.184078518 |
| Fam174a | -0.052353225 | 1.57E-07 | 1.28E-06 | 0.601802249 | 0.632813715 |
| Fam174b | 0.026817314 | 0.007724389 | 0.020894966 | 0.580577993 | 0.531592717 |
| Fam174c | -0.018251361 | 0.022946482 | 0.052135007 | 0.188656175 | 0.214903758 |
| Fam183b | 0.025281804 | 0.01692631 | 0.040511469 | 0.866382763 | 0.786500833 |
| Fam184a | 0.026360153 | 0.000171887 | 0.000753418 | 0.156442774 | 0.138491235 |
| Fam185a | 0.001172751 | 0.764522428 | 0.833354304 | 0.041581695 | 0.040360737 |
| Fam187b | -0.014599426 | 0.021068721 | 0.04858399 | 0.11227389 | 0.123906713 |
| Fam189b | 0.000837221 | 0.828623741 | 0.881869732 | 0.041034094 | 0.038885342 |
| Fam193a | 0.016948213 | 0.029737696 | 0.064968512 | 0.204098397 | 0.192134756 |
| Fam193b | -0.015282226 | 0.014666707 | 0.035862607 | 0.101987252 | 0.114403196 |
| Fam199x | 0.011713563 | 0.055095237 | 0.10825301 | 0.113640268 | 0.103185397 |
| Fam204a | 0.009568878 | 0.308790991 | 0.430095417 | 0.39150658 | 0.382553115 |
| Fam207a | 0.021456197 | 0.003144727 | 0.009675012 | 0.182238956 | 0.157012654 |
| Fam20a | 0.018297549 | 0.001791699 | 0.005932866 | 0.11772055 | 0.092499103 |
| Fam20b | 0.027996088 | 0.001287762 | 0.004460142 | 0.318236123 | 0.280579392 |
| Fam20c | 0.040749695 | 8.24E-06 | 4.83E-05 | 0.382613443 | 0.325615316 |
| Fam210a | -0.005772385 | 0.29831543 | 0.417959737 | 0.085107358 | 0.08757699 |
| Fam210b | -0.186854833 | 2.37E-57 | 3.78E-55 | 0.435467256 | 0.641550416 |
| Fam214a | 0.028122343 | 0.00014508 | 0.000650284 | 0.181661582 | 0.15887388 |
| Fam214b | 0.017718625 | 0.002796609 | 0.008739146 | 0.110119252 | 0.095342964 |
| Fam216a | -0.02694606 | 0.002012263 | 0.006569249 | 0.28942082 | 0.300333906 |
| Fam217b | 0.006857959 | 0.088374083 | 0.159816375 | 0.050302636 | 0.040695909 |
| Fam219a | 0.016727171 | 0.017996684 | 0.042642886 | 0.160181836 | 0.149068376 |

| | | | | | |
|---|---|---|---|---|---|
| Fam219b | -0.003377284 | 0.544432039 | 0.65416464 | 0.090916204 | 0.088657482 |
| Fam220a | 8.40E-05 | 0.984632552 | 0.990703331 | 0.048148921 | 0.048831081 |
| Fam222a | 0.016515846 | 0.00965783 | 0.025145533 | 0.128120769 | 0.109561867 |
| Fam222b | 0.019807091 | 0.00408628 | 0.012130324 | 0.162809386 | 0.137546332 |
| Fam229b | -0.025125371 | 1.77E-05 | 9.67E-05 | 0.091041936 | 0.106045509 |
| Fam234a | 0.043015249 | 4.62E-06 | 2.85E-05 | 0.39869289 | 0.330671721 |
| Fam234b | 0.016244348 | 0.003548422 | 0.010748916 | 0.101341466 | 0.083175142 |
| Fam241a | 0.004686521 | 0.250191685 | 0.364945771 | 0.046126413 | 0.042671463 |
| Fam241b | -0.01314726 | 0.077494744 | 0.143806814 | 0.184311129 | 0.181762532 |
| Fam32a | -0.011397 | 0.204141503 | 0.312152201 | 0.339317405 | 0.333409548 |
| Fam3a | -0.011016101 | 0.147162249 | 0.241266675 | 0.196900412 | 0.19896627 |
| Fam3c | -0.028960498 | 4.18E-05 | 0.000208646 | 0.143314882 | 0.166204 |
| Fam50a | 0.014633254 | 0.109042872 | 0.189814646 | 0.303891223 | 0.298231858 |
| Fam53a | -0.004141169 | 0.543580375 | 0.653398666 | 0.147017303 | 0.146532064 |
| Fam53c | 0.010407615 | 0.071169425 | 0.134087725 | 0.100277184 | 0.091660974 |
| Fam71e1 | -0.017445496 | 0.006388268 | 0.017816046 | 0.116863078 | 0.126224306 |
| Fam76a | 0.019259333 | 0.037423977 | 0.078362246 | 0.38604565 | 0.359862079 |
| Fam76b | -0.007866707 | 0.252970227 | 0.367949813 | 0.136587412 | 0.143855869 |
| Fam83g | 0.022916304 | 0.001347275 | 0.004643107 | 0.169668929 | 0.145821527 |
| Fam83h | 0.001640939 | 0.723006326 | 0.800420062 | 0.059470011 | 0.055878168 |
| Fam8a1 | 0.01885035 | 0.007110028 | 0.01947695 | 0.169413551 | 0.148918638 |
| Fam91a1 | 0.037904289 | 2.32E-05 | 0.000123629 | 0.329025752 | 0.293843353 |
| Fam98a | 0.009139474 | 0.262881187 | 0.379076075 | 0.233736347 | 0.223647723 |
| Fam98b | -0.007222247 | 0.358121501 | 0.480759077 | 0.192849897 | 0.209501048 |
| Fam98c | -0.020398573 | 0.003008266 | 0.009314128 | 0.137380372 | 0.150974025 |
| Fance | 0.006137409 | 0.236940393 | 0.35061408 | 0.083834788 | 0.074239514 |
| Fancf | 0.008333127 | 0.051784984 | 0.102724672 | 0.057768708 | 0.045536286 |
| Fancl | -0.007773549 | 0.048734037 | 0.097931441 | 0.041230216 | 0.044314522 |
| Fancm | 0.004420576 | 0.413685394 | 0.533331043 | 0.080086903 | 0.077267897 |
| Far1 | -0.009027913 | 0.269078177 | 0.385609971 | 0.229714139 | 0.236733003 |
| Farp1 | 0.018572097 | 0.006049457 | 0.016983199 | 0.147863796 | 0.133971193 |
| Farp2 | 0.002913387 | 0.430654501 | 0.549379748 | 0.038016102 | 0.034791942 |
| Fars2 | 0.004484753 | 0.531142633 | 0.642579477 | 0.17853388 | 0.163723505 |
| Farsa | -0.017226693 | 0.009717231 | 0.025273119 | 0.127143097 | 0.137657987 |
| Farsb | -0.00106588 | 0.894630535 | 0.928392142 | 0.228291604 | 0.224032531 |
| Fasn | 0.019345189 | 0.001197082 | 0.004183333 | 0.118692716 | 0.096925468 |
| Fastk | 0.007840353 | 0.319450723 | 0.441055367 | 0.21476895 | 0.204492653 |
| Fastkd1 | -0.003864446 | 0.412563435 | 0.53228743 | 0.05804125 | 0.061301869 |
| Fastkd2 | -0.004715943 | 0.346168305 | 0.468399202 | 0.066557845 | 0.070091031 |
| Fat1 | 0.00410395 | 0.397988555 | 0.518318115 | 0.065067002 | 0.061916966 |
| Fau | 0.013230464 | 0.001423642 | 0.004864121 | 3.110106619 | 3.108819297 |
| Fbf1 | 0.008570929 | 0.031851482 | 0.068645558 | 0.04689085 | 0.03910717 |
| Fbh1 | 0.009092836 | 0.115464334 | 0.198259317 | 0.108968304 | 0.093198562 |
| Fbl | 0.00231253 | 0.769904157 | 0.837717763 | 0.211293265 | 0.2038051 |
| Fbrs | 0.010871613 | 0.065347603 | 0.125208789 | 0.106462066 | 0.096774092 |
| Fbrsl1 | 0.008888565 | 0.240294278 | 0.354175405 | 0.197303513 | 0.182872267 |
| Fbxl12 | -0.00418793 | 0.462865726 | 0.580567745 | 0.088772607 | 0.091747641 |
| Fbxl14 | 0.02524494 | 0.000134474 | 0.000608924 | 0.150835421 | 0.125284823 |
| Fbxl15 | -0.028422551 | 0.000197891 | 0.00085053 | 0.179775303 | 0.197763179 |
| Fbxl16 | 0.023630846 | 0.011493734 | 0.029167862 | 0.398635455 | 0.371835813 |
| Fbxl17 | 0.006466051 | 0.364537355 | 0.48728615 | 0.169520587 | 0.156369457 |
| Fbxl18 | 0.012964496 | 0.003837771 | 0.011497938 | 0.061700943 | 0.050854992 |
| Fbxl2 | -0.009187064 | 0.068112907 | 0.129537106 | 0.069223081 | 0.072400042 |
| Fbxl20 | 0.004694223 | 0.371094565 | 0.493608066 | 0.080146802 | 0.075458971 |
| Fbxl3 | 0.001444059 | 0.823589777 | 0.878251308 | 0.12439993 | 0.121466406 |
| Fbxl4 | 0.007320622 | 0.160279444 | 0.257389619 | 0.084594403 | 0.07297733 |
| Fbxl5 | 0.021486516 | 0.020836366 | 0.048128094 | 0.363575607 | 0.349042625 |
| Fbxl6 | -0.018790999 | 0.001248799 | 0.00433601 | 0.092969204 | 0.104660884 |
| Fbxo11 | 0.033477061 | 1.34E-05 | 7.52E-05 | 0.214792057 | 0.179313982 |
| Fbxo16 | -0.008481763 | 0.084193666 | 0.153714709 | 0.065374871 | 0.069485394 |
| Fbxo17 | -0.005319828 | 0.209300548 | 0.318139451 | 0.04827379 | 0.049998617 |
| Fbxo2 | -0.047058913 | 8.40E-07 | 5.97E-06 | 0.359450479 | 0.393087596 |
| Fbxo21 | 0.03673696 | 2.12E-06 | 1.39E-05 | 0.226454172 | 0.182721067 |
| Fbxo22 | -0.009788981 | 0.180361006 | 0.282544264 | 0.171289026 | 0.173991735 |
| Fbxo25 | 0.001043985 | 0.84852757 | 0.896531337 | 0.091125707 | 0.085826303 |
| Fbxo28 | 0.008407026 | 0.210657705 | 0.319802343 | 0.145635595 | 0.133499767 |
| Fbxo3 | 0.03815922 | 1.72E-05 | 9.46E-05 | 0.346624708 | 0.303423127 |
| Fbxo30 | 0.001234306 | 0.751293362 | 0.822808839 | 0.040124264 | 0.039303642 |
| Fbxo31 | -0.00646741 | 0.304932503 | 0.425452611 | 0.128009044 | 0.121457202 |
| Fbxo32 | 0.004510433 | 0.526344891 | 0.638284275 | 0.158288222 | 0.143319823 |
| Fbxo33 | 0.026090378 | 0.000294284 | 0.001210762 | 0.171143814 | 0.15086656 |
| Fbxo34 | 0.025742647 | 4.08E-05 | 0.000204028 | 0.131957423 | 0.109134074 |

| | | | | | |
|---|---|---|---|---|---|
| Fbxo36 | -0.014899978 | 0.024316142 | 0.054900496 | 0.127914784 | 0.13763761 |
| Fbxo38 | 0.025684414 | 5.53E-05 | 0.000269699 | 0.136178205 | 0.112509795 |
| Fbxo4 | 0.015268367 | 0.012101395 | 0.030487111 | 0.117075055 | 0.101508433 |
| Fbxo42 | -0.000243973 | 0.966066541 | 0.976918963 | 0.092227817 | 0.093871921 |
| Fbxo44 | 0.000461782 | 0.943540549 | 0.962045224 | 0.138343916 | 0.130023387 |
| Fbxo45 | 0.001646397 | 0.750531598 | 0.822252445 | 0.076655505 | 0.074046209 |
| Fbxo46 | -0.002946236 | 0.446622378 | 0.565571095 | 0.040506372 | 0.04039405 |
| Fbxo6 | 0.005858698 | 0.39027931 | 0.511391726 | 0.146438322 | 0.137776601 |
| Fbxo7 | -0.003271383 | 0.605466424 | 0.707323309 | 0.123871225 | 0.121119466 |
| Fbxo8 | -0.004971337 | 0.49950388 | 0.613935435 | 0.17269743 | 0.175895625 |
| Fbxo9 | -0.032227067 | 0.000270853 | 0.001125312 | 0.29644396 | 0.309740628 |
| Fbxw11 | 0.052761679 | 3.09E-12 | 4.70E-11 | 0.21599429 | 0.171360713 |
| Fbxw2 | -0.016273999 | 0.060308984 | 0.117261067 | 0.288882359 | 0.292347574 |
| Fbxw4 | 0.00866131 | 0.12133229 | 0.206258528 | 0.097544933 | 0.086399992 |
| Fbxw5 | 0.01333929 | 0.073627151 | 0.137972852 | 0.200166073 | 0.177451329 |
| Fbxw7 | 0.050124483 | 5.22E-08 | 4.62E-07 | 0.341600153 | 0.297060183 |
| Fbxw8 | 0.00412383 | 0.499356337 | 0.613935435 | 0.113452245 | 0.107425421 |
| Fbxw9 | -0.012691035 | 0.014386921 | 0.035267184 | 0.069540185 | 0.077497745 |
| Fcer1g | 0.014472396 | 0.001702606 | 0.005662928 | 0.068876544 | 0.050477143 |
| Fcf1 | -0.031588521 | 4.98E-05 | 0.000244655 | 0.184238479 | 0.207417457 |
| Fcho2 | 0.019162804 | 0.024083967 | 0.054439508 | 0.259359398 | 0.241278544 |
| Fchsd2 | -0.005257025 | 0.274963698 | 0.393059413 | 0.061596809 | 0.063467579 |
| Fdft1 | 0.007287973 | 0.36218162 | 0.484926143 | 0.201326062 | 0.204792791 |
| Fdps | 0.006957993 | 0.386418361 | 0.507472647 | 0.227627989 | 0.213844101 |
| Fdx1 | -0.020085253 | 0.028524562 | 0.062712752 | 0.365512546 | 0.375103256 |
| Fdxacb1 | -0.002911025 | 0.42232847 | 0.541101395 | 0.036599948 | 0.035386231 |
| Fdxr | -0.00951857 | 0.042126739 | 0.086440724 | 0.057350387 | 0.062602059 |
| Fech | 0.001414896 | 0.845678583 | 0.894439423 | 0.175212834 | 0.167481115 |
| Fem1a | 0.005941726 | 0.37211555 | 0.49435739 | 0.141492927 | 0.13241843 |
| Fem1b | 0.062959582 | 1.82E-13 | 3.18E-12 | 0.301058612 | 0.240530224 |
| Fem1c | 0.000320832 | 0.956667938 | 0.970443551 | 0.101147748 | 0.099285619 |
| Fer | 8.38E-05 | 0.989024047 | 0.993079296 | 0.107144518 | 0.107226238 |
| Fer1l6 | 0.030999184 | 1.42E-08 | 1.38E-07 | 0.100893659 | 0.066565615 |
| Fermt2 | -0.00927571 | 0.34158994 | 0.463947667 | 0.426685731 | 0.451277116 |
| Fez1 | -0.032789597 | 1.60E-06 | 1.07E-05 | 0.122920911 | 0.147139978 |
| Fez2 | 0.000250954 | 0.957942721 | 0.971432058 | 0.062824262 | 0.062225274 |
| Ffar1 | 0.056513884 | 5.38E-09 | 5.55E-08 | 0.42499332 | 0.347794864 |
| Ffar2 | -0.041660876 | 2.08E-08 | 1.95E-07 | 0.138059448 | 0.171311961 |
| Ffar3 | 0.012176609 | 0.003450009 | 0.010488114 | 0.056535483 | 0.040866548 |
| Fgd1 | 0.003408561 | 0.402720602 | 0.522608746 | 0.04754377 | 0.042870172 |
| Fgd2 | 0.007134903 | 0.337216117 | 0.45944503 | 0.204026154 | 0.176447699 |
| Fgd4 | 0.00977903 | 0.073773256 | 0.138166749 | 0.089145578 | 0.077556447 |
| Fgd6 | 0.033090953 | 3.33E-07 | 2.57E-06 | 0.141746532 | 0.109613501 |
| Fgf12 | 0.002411138 | 0.774516974 | 0.841155029 | 0.253914019 | 0.245133569 |
| Fgf9 | 0.004336995 | 0.491197074 | 0.606325286 | 0.115209076 | 0.111856478 |
| Fgfbp3 | 0.005052264 | 0.207759819 | 0.316143522 | 0.048079892 | 0.04177755 |
| Fgfr1 | 0.000729731 | 0.898573207 | 0.930839501 | 0.089140564 | 0.089925901 |
| Fgfr1op2 | 0.016680839 | 0.094579613 | 0.168530608 | 0.556014754 | 0.54048288 |
| Fggy | -0.001561227 | 0.736212401 | 0.810859517 | 0.063459803 | 0.059793944 |
| Fgl2 | 0.016183526 | 0.051218301 | 0.101967416 | 0.215666761 | 0.209225554 |
| Fh1 | 0.059519943 | 3.96E-06 | 2.48E-05 | 0.818125827 | 0.778758086 |
| Fhad1 | 0.004366284 | 0.398841875 | 0.519012275 | 0.077373015 | 0.07162568 |
| Fhdc1 | 0.017395743 | 0.024001503 | 0.054278346 | 0.208601733 | 0.182863642 |
| Fhit | -0.0117822 | 0.003876453 | 0.011606675 | 0.041594316 | 0.048034172 |
| Fhl1 | -0.000839139 | 0.905048766 | 0.935454402 | 0.14144169 | 0.150902391 |
| Fhod3 | 0.081044486 | 1.63E-21 | 5.15E-20 | 0.284031515 | 0.202714692 |
| Fibp | -0.052312902 | 1.37E-09 | 1.54E-08 | 0.285577823 | 0.32047184 |
| Ficd | -0.011179762 | 0.104383153 | 0.18298107 | 0.134192848 | 0.144704962 |
| Fig4 | 0.003930283 | 0.417647096 | 0.536943278 | 0.067745641 | 0.062964833 |
| Filip1 | 0.030453674 | 0.000374057 | 0.001497769 | 0.251641895 | 0.224338938 |
| Fip1l1 | 0.051837805 | 3.84E-07 | 2.93E-06 | 0.663722033 | 0.611577389 |
| Firre | -0.010012229 | 0.094740025 | 0.16878551 | 0.088924787 | 0.103780593 |
| Fis1 | -0.025931262 | 0.006641912 | 0.018398134 | 0.777426009 | 0.770933422 |
| Fitm2 | 0.020847487 | 0.016199812 | 0.039012775 | 0.283426295 | 0.265845146 |
| Fiz1 | -0.005748721 | 0.289817826 | 0.409244019 | 0.081477827 | 0.083555658 |
| Fkbp11 | -0.031497834 | 0.004170617 | 0.012335486 | 0.400361011 | 0.476564543 |
| Fkbp15 | 0.009725064 | 0.150647779 | 0.245696646 | 0.142609501 | 0.134281305 |
| Fkbp1a | 0.025311178 | 0.004940792 | 0.014270465 | 1.17098696 | 1.089609388 |
| Fkbp1b | 0.03977881 | 0.000508186 | 0.001970718 | 0.830969854 | 0.727695549 |
| Fkbp2 | 0.037906542 | 9.21E-06 | 5.33E-05 | 1.486968876 | 1.480234437 |
| Fkbp3 | -0.002108287 | 0.832602993 | 0.885415589 | 0.539274443 | 0.533167717 |
| Fkbp4 | 0.007010242 | 0.483937016 | 0.60033631 | 0.695989213 | 0.654242538 |

| | | | | | |
|---|---|---|---|---|---|
| Fkbp5 | 0.003127292 | 0.693287141 | 0.777782279 | 0.210515151 | 0.21017123 |
| Fkbp8 | -0.02182367 | 0.021954202 | 0.050256357 | 0.474480384 | 0.468731699 |
| Fkbp9 | 0.01058759 | 0.289491731 | 0.408897081 | 0.684336243 | 0.625532491 |
| Fkbpl | 0.009849733 | 0.031966798 | 0.068817806 | 0.066236155 | 0.055042472 |
| Fkrp | 0.005799162 | 0.282416175 | 0.401485512 | 0.085357145 | 0.077746193 |
| Fktn | 0.011844474 | 0.094549932 | 0.168508605 | 0.164426834 | 0.148797283 |
| Flad1 | -0.018835978 | 0.002126121 | 0.006880876 | 0.101409248 | 0.115132315 |
| Flcn | 0.023237195 | 0.000345284 | 0.001397523 | 0.132989622 | 0.115764806 |
| Flii | 0.022689809 | 0.003425006 | 0.010431676 | 0.217002206 | 0.189908364 |
| Flnb | 0.057132956 | 2.52E-08 | 2.32E-07 | 0.528166022 | 0.479921344 |
| Flot1 | -0.044406148 | 6.65E-07 | 4.84E-06 | 0.253521305 | 0.29789413 |
| Flot2 | 3.41E-06 | 0.999626811 | 0.999626811 | 0.166143083 | 0.169375494 |
| Flrt1 | -0.000795652 | 0.93159631 | 0.953566996 | 0.296690284 | 0.303703675 |
| Flt3l | -0.022174296 | 2.86E-06 | 1.84E-05 | 0.049175051 | 0.066834157 |
| Flvcr1 | 0.003484924 | 0.416688565 | 0.535994473 | 0.050946503 | 0.047528567 |
| Flywch1 | -0.045018988 | 1.81E-06 | 1.20E-05 | 0.339805856 | 0.373287689 |
| Flywch2 | -0.047900066 | 9.52E-10 | 1.09E-08 | 0.182129051 | 0.215355028 |
| Fmc1 | -0.022521189 | 0.016977369 | 0.040611088 | 0.34354923 | 0.383232228 |
| Fmn1 | -0.034530738 | 6.77E-13 | 1.11E-11 | 0.033068538 | 0.068098626 |
| Fmn2 | 0.018326343 | 0.042220429 | 0.086596416 | 0.328856239 | 0.289407876 |
| Fmnl2 | 0.06080419 | 2.89E-10 | 3.52E-09 | 0.380042535 | 0.321251755 |
| Fmo1 | 0.013285614 | 0.005531177 | 0.015695886 | 0.07312739 | 0.05058531 |
| Fmo5 | -0.012319631 | 0.001572856 | 0.00528197 | 0.037737995 | 0.041504364 |
| Fmr1 | 0.034223737 | 1.93E-05 | 0.000104863 | 0.230899921 | 0.20079537 |
| Fn3k | -0.001379602 | 0.775184306 | 0.841391295 | 0.071097463 | 0.064658936 |
| Fnbp1 | 0.001374912 | 0.761167016 | 0.83057396 | 0.055559439 | 0.054677291 |
| Fnbp1l | 0.028139167 | 3.70E-06 | 2.33E-05 | 0.118367155 | 0.098631525 |
| Fnbp4 | 0.008894927 | 0.25793195 | 0.374047496 | 0.202362898 | 0.19278165 |
| Fndc10 | 0.000569809 | 0.916139095 | 0.942009972 | 0.090084542 | 0.081255382 |
| Fndc3a | 0.004781282 | 0.598345178 | 0.70170227 | 1.166005199 | 1.159304363 |
| Fndc3b | 0.002246685 | 0.820542122 | 0.876531207 | 0.445590218 | 0.465535535 |
| Fndc5 | -0.008217669 | 0.048964515 | 0.098313304 | 0.041568471 | 0.04720082 |
| Fnip1 | 0.041945423 | 9.21E-06 | 5.33E-05 | 0.381991844 | 0.338513773 |
| Fnip2 | 0.051902287 | 2.04E-11 | 2.83E-10 | 0.223471925 | 0.170408349 |
| Fnta | -0.017250563 | 0.041735516 | 0.085909949 | 0.272762068 | 0.271259607 |
| Fntb | 0.000826547 | 0.865561204 | 0.908059353 | 0.064383773 | 0.065168365 |
| Focad | 0.008370771 | 0.103362239 | 0.18161715 | 0.077886764 | 0.069859243 |
| Folr1 | 0.004298588 | 0.609114372 | 0.710049151 | 0.189116502 | 0.182800946 |
| Fos | -0.069612583 | 1.56E-06 | 1.05E-05 | 1.395773358 | 1.604653485 |
| Fosb | -0.110834392 | 5.15E-26 | 2.19E-24 | 0.202416966 | 0.361805499 |
| Fosl2 | 0.020907559 | 0.000333987 | 0.001355184 | 0.087103352 | 0.083539772 |
| Foxa2 | 0.057880034 | 7.48E-09 | 7.57E-08 | 0.524191043 | 0.452850454 |
| Foxa3 | 0.01897177 | 0.001539602 | 0.00518822 | 0.120069294 | 0.099981387 |
| Foxj2 | 0.02610513 | 1.76E-05 | 9.65E-05 | 0.116740897 | 0.098610117 |
| Foxj3 | 0.016296179 | 0.025519043 | 0.05710365 | 0.173880085 | 0.159690114 |
| Foxk1 | 0.003613544 | 0.457640356 | 0.576318593 | 0.066352148 | 0.063420613 |
| Foxk2 | 0.023511465 | 0.000644365 | 0.002431879 | 0.160433959 | 0.140239681 |
| Foxn2 | 0.008681403 | 0.105382042 | 0.184466009 | 0.081109574 | 0.076638489 |
| Foxn3 | 0.018921032 | 0.019340617 | 0.045101818 | 0.230942384 | 0.207895698 |
| Foxo1 | 0.055584091 | 1.04E-09 | 1.18E-08 | 0.364655101 | 0.29497001 |
| Foxo3 | 0.020580907 | 0.00210039 | 0.006806989 | 0.155938092 | 0.126946305 |
| Foxo6 | 0.016165189 | 0.001066775 | 0.003790639 | 0.078751516 | 0.061135665 |
| Foxp1 | 0.058113034 | 1.02E-08 | 1.02E-07 | 0.546265162 | 0.484625244 |
| Foxp2 | -0.00891616 | 0.068385369 | 0.129953589 | 0.05187374 | 0.059538531 |
| Foxp4 | -0.015849279 | 0.031633003 | 0.068295816 | 0.167412313 | 0.173955502 |
| Foxred1 | -0.010503288 | 0.021089006 | 0.048619236 | 0.051870492 | 0.058533934 |
| Foxred2 | 0.008233429 | 0.057029088 | 0.111646556 | 0.056253622 | 0.04665637 |
| Fpgs | 5.53E-05 | 0.989934259 | 0.9933059 | 0.049943337 | 0.05123728 |
| Fpgt | 0.013098361 | 0.042705549 | 0.087467833 | 0.132779255 | 0.117020567 |
| Fra10ac1 | 0.031840109 | 0.000182752 | 0.000792822 | 0.27330999 | 0.233029067 |
| Fras1 | 0.012280581 | 0.012422649 | 0.031111023 | 0.076464501 | 0.060321905 |
| Frg1 | -0.013271207 | 0.076787462 | 0.142876115 | 0.19270868 | 0.19110477 |
| Frk | 0.001289532 | 0.840865469 | 0.891093753 | 0.119926859 | 0.119148587 |
| Frmd4a | 0.003131251 | 0.605337575 | 0.707323309 | 0.104321372 | 0.102421922 |
| Frmd4b | 0.00856663 | 0.156165956 | 0.252409879 | 0.112572298 | 0.096915104 |
| Frmd5 | 0.007461389 | 0.205153831 | 0.313404391 | 0.104678777 | 0.095613316 |
| Frrs1 | -0.005923737 | 0.231870761 | 0.345002684 | 0.060199901 | 0.068188192 |
| Frrs1l | -0.005327427 | 0.185297703 | 0.288511253 | 0.039117433 | 0.042409534 |
| Frs2 | 0.001310129 | 0.832479901 | 0.885415589 | 0.114941837 | 0.113918323 |
| Fry | 0.006047672 | 0.417905405 | 0.537133309 | 0.169489197 | 0.167212501 |
| Fryl | -0.004330777 | 0.614564024 | 0.714602493 | 0.269799438 | 0.267415418 |
| Frzb | 0.007992774 | 0.422501251 | 0.541251438 | 0.535238644 | 0.487707444 |

| | | | | | |
|---|---|---|---|---|---|
| Fsd1l | 0.002709503 | 0.592864382 | 0.696871904 | 0.072655906 | 0.069741748 |
| Fth1 | -0.092188515 | 2.67E-90 | 1.62E-87 | 3.001885921 | 3.200411293 |
| Ftl1 | -0.049810365 | 1.13E-18 | 2.90E-17 | 3.279318916 | 3.329985448 |
| Ftl1-ps1 | -0.005854301 | 0.217715589 | 0.328210723 | 0.058368863 | 0.06415337 |
| Fto | -0.001617108 | 0.816152105 | 0.873280392 | 0.150258577 | 0.150393624 |
| Ftsj1 | -0.00274784 | 0.519040708 | 0.63147463 | 0.050004231 | 0.049726331 |
| Ftsj3 | -0.01508433 | 0.033422325 | 0.071523628 | 0.140746701 | 0.15430047 |
| Ftx | -0.019860951 | 0.001133656 | 0.003988927 | 0.096080354 | 0.109124255 |
| Fubp1 | -0.038533434 | 0.000121324 | 0.00055508 | 0.489490681 | 0.509171426 |
| Fubp3 | 0.013096116 | 0.065020915 | 0.124705728 | 0.167080453 | 0.151416209 |
| Fuca1 | -0.047366834 | 6.08E-07 | 4.46E-06 | 0.449491667 | 0.475914534 |
| Fuca2 | -0.012446432 | 0.053161966 | 0.104942261 | 0.11469983 | 0.125655028 |
| Fundc1 | -0.01265342 | 0.102711522 | 0.180669698 | 0.206486056 | 0.206603871 |
| Fundc2 | 0.011272949 | 0.240199707 | 0.354089697 | 0.517987886 | 0.493603068 |
| Fuom | -0.084444273 | 1.52E-16 | 3.28E-15 | 0.626909811 | 0.722094227 |
| Furin | 0.017146122 | 0.009818646 | 0.025475548 | 0.148334317 | 0.12917322 |
| Fus | -0.048968232 | 3.30E-06 | 2.10E-05 | 0.635701498 | 0.690788361 |
| Fut1 | -0.025017891 | 1.40E-09 | 1.56E-08 | 0.031044631 | 0.049360579 |
| Fut11 | 0.008337456 | 0.102540127 | 0.180466169 | 0.07744408 | 0.070142546 |
| Fut8 | 0.001540578 | 0.794446039 | 0.856370372 | 0.102259872 | 0.099539379 |
| Fuz | -0.014034262 | 0.010138852 | 0.026201467 | 0.081155163 | 0.088038526 |
| Fxn | -0.03037515 | 5.02E-05 | 0.000246802 | 0.172590888 | 0.191066178 |
| Fxr1 | 0.047603324 | 1.12E-06 | 7.73E-06 | 0.511467957 | 0.449256947 |
| Fxr2 | 0.033003314 | 0.000153134 | 0.000680113 | 0.323127286 | 0.281649707 |
| Fxyd3 | 0.058668283 | 1.25E-11 | 1.78E-10 | 0.200602172 | 0.159005612 |
| Fxyd6 | 0.109397208 | 3.23E-22 | 1.10E-20 | 0.73550633 | 0.572275628 |
| Fyco1 | 0.033943467 | 1.55E-05 | 8.60E-05 | 0.229784696 | 0.193638128 |
| Fyn | -0.000996064 | 0.824290322 | 0.878508059 | 0.050517963 | 0.053180894 |
| Fyttd1 | 0.020937808 | 0.030416573 | 0.066139549 | 0.47077451 | 0.438315798 |
| Fzd3 | 0.005965178 | 0.488713647 | 0.604129615 | 0.269138576 | 0.262785268 |
| Fzd6 | 0.008854648 | 0.03650189 | 0.076845251 | 0.051038149 | 0.045411564 |
| Fzd7 | -0.001474874 | 0.687457619 | 0.773021341 | 0.036202441 | 0.034079756 |
| Fzd8 | 0.00280884 | 0.479413129 | 0.596002723 | 0.043498248 | 0.03986702 |
| Fzr1 | 0.020034773 | 0.000917132 | 0.003317098 | 0.121878147 | 0.100952657 |
| G2e3 | 0.011253593 | 0.052976732 | 0.104640347 | 0.101250106 | 0.089622135 |
| G3bp1 | 0.015819485 | 0.109914127 | 0.19088695 | 0.595290668 | 0.561982509 |
| G3bp2 | 0.030539502 | 0.001908812 | 0.006276265 | 0.791005507 | 0.751851961 |
| G6pc2 | 0.178381689 | 2.10E-57 | 3.41E-55 | 2.126017715 | 1.738015491 |
| G6pc3 | -0.02550449 | 0.001101454 | 0.003893942 | 0.205996357 | 0.215732872 |
| G6pdx | -0.010512785 | 0.070643877 | 0.133281539 | 0.09178668 | 0.09846518 |
| G730003C15Rik | -0.001427637 | 0.748667996 | 0.820766172 | 0.056386975 | 0.054288679 |
| Gaa | -0.056689562 | 1.91E-08 | 1.80E-07 | 0.461046152 | 0.512494495 |
| Gab2 | 0.002713274 | 0.589985523 | 0.694579711 | 0.065911587 | 0.068837936 |
| Gabarap | 0.0140352 | 0.024849441 | 0.055961609 | 2.026456062 | 1.978038402 |
| Gabarapl1 | 0.019100905 | 0.058767995 | 0.114589339 | 0.631787116 | 0.605002113 |
| Gabarapl2 | 0.046194271 | 3.33E-08 | 3.03E-07 | 1.29959127 | 1.173793571 |
| Gabbr1 | -0.013998243 | 0.039768266 | 0.082542075 | 0.134880527 | 0.14533826 |
| Gabbr2 | 0.073163718 | 1.75E-13 | 3.07E-12 | 0.348775536 | 0.31798999 |
| Gabpa | 0.003624196 | 0.518602351 | 0.631167118 | 0.090475732 | 0.086900955 |
| Gabpb1 | 0.007622653 | 0.112727695 | 0.194833539 | 0.067167177 | 0.060339911 |
| Gabpb2 | 0.004260978 | 0.549233166 | 0.658564977 | 0.154875166 | 0.150969348 |
| Gabrb3 | -0.015470847 | 0.048319039 | 0.097278463 | 0.193420005 | 0.205428278 |
| Gad1 | 0.089535421 | 4.54E-19 | 1.23E-17 | 0.385291758 | 0.277836861 |
| Gadd45a | -0.022236631 | 0.018683671 | 0.043896243 | 0.294580598 | 0.307485979 |
| Gadd45b | 0.066238985 | 6.55E-20 | 1.90E-18 | 0.154977937 | 0.102671917 |
| Gadd45g | -0.042013328 | 0.001125691 | 0.003965206 | 0.89955929 | 0.946333212 |
| Gadd45gip1 | 0.022835727 | 0.022074203 | 0.05048351 | 0.715886056 | 0.662618828 |
| Gak | 0.041970667 | 4.13E-07 | 3.13E-06 | 0.261624853 | 0.227010187 |
| Gal3st1 | 0.01362054 | 0.003986793 | 0.011882157 | 0.075704459 | 0.05683453 |
| Galc | -0.012092927 | 0.056633448 | 0.111006125 | 0.111880681 | 0.122051213 |
| Gale | 0.022245382 | 0.000207378 | 0.000886989 | 0.114473071 | 0.095949039 |
| Galk2 | 0.000659342 | 0.926663903 | 0.950401057 | 0.177352025 | 0.163480187 |
| Galns | -0.013535513 | 0.029273286 | 0.064069088 | 0.101976682 | 0.114499826 |
| Galnt1 | 0.013978532 | 0.115346682 | 0.198127286 | 0.317899446 | 0.29990721 |
| Galnt10 | 0.023054338 | 8.52E-06 | 4.98E-05 | 0.090643514 | 0.067344536 |
| Galnt11 | 0.002073369 | 0.771656653 | 0.838779738 | 0.162508876 | 0.159980811 |
| Galnt12 | -0.00152052 | 0.7885847 | 0.852144587 | 0.094912431 | 0.09204099 |
| Galnt14 | 0.001532521 | 0.853981972 | 0.899697956 | 0.261708189 | 0.234794628 |
| Galnt18 | 0.067736888 | 1.50E-24 | 5.90E-23 | 0.177398881 | 0.106477397 |
| Galnt7 | 0.002597947 | 0.625778727 | 0.724172101 | 0.078658001 | 0.077060616 |
| Galr1 | 0.014157533 | 0.018440189 | 0.043460732 | 0.11017057 | 0.093719767 |
| Galr3 | -0.006644934 | 0.140840977 | 0.233025183 | 0.051611135 | 0.056620919 |

| | | | | | |
|---|---|---|---|---|---|
| Galt | -0.011577892 | 0.008323816 | 0.022250244 | 0.048684474 | 0.05504991 |
| Gan | 0.007729793 | 0.084610692 | 0.154302222 | 0.058924005 | 0.051392168 |
| Ganab | -0.024792991 | 0.008002505 | 0.021569269 | 0.376210734 | 0.397002494 |
| Ganc | -0.008850519 | 0.186918201 | 0.290476142 | 0.131386093 | 0.135616651 |
| Gapdh | -0.073349606 | 8.07E-19 | 2.11E-17 | 1.715499655 | 1.757046855 |
| Gapvd1 | -0.000891703 | 0.918918318 | 0.944268458 | 0.291851974 | 0.290199723 |
| Gar1 | -0.030224585 | 6.84E-05 | 0.000329438 | 0.161587082 | 0.187547936 |
| Garre1 | 0.011376705 | 0.050429309 | 0.100713587 | 0.101598898 | 0.091831199 |
| Gars | -0.008288918 | 0.39495232 | 0.515537924 | 1.166724927 | 1.160718693 |
| Gart | -0.000736224 | 0.90553008 | 0.93560051 | 0.117152258 | 0.112320139 |
| Gas2 | -0.017324723 | 0.00819216 | 0.021964749 | 0.129164043 | 0.131481791 |
| Gas2l1 | 0.002978579 | 0.515804327 | 0.628715444 | 0.058172486 | 0.056613903 |
| Gas5 | -0.025143815 | 0.008755149 | 0.023220843 | 1.461159223 | 1.560079421 |
| Gas8 | -0.003098643 | 0.58458071 | 0.689887553 | 0.094907705 | 0.093068153 |
| Gata6 | 0.01349989 | 0.007902842 | 0.021318377 | 0.072335761 | 0.059961268 |
| Gatad1 | -0.032332038 | 0.000587095 | 0.002238584 | 0.392758737 | 0.408621104 |
| Gatad2a | 0.048499509 | 1.01E-08 | 1.00E-07 | 0.273150349 | 0.238793977 |
| Gatad2b | 0.048183631 | 3.04E-07 | 2.35E-06 | 0.413974607 | 0.353324723 |
| Gatb | 0.00435904 | 0.241918968 | 0.355843696 | 0.040526726 | 0.035203376 |
| Gatc | 0.006878144 | 0.218083641 | 0.328612702 | 0.094382728 | 0.086085301 |
| Gatd1 | -0.020807383 | 0.015629479 | 0.037836105 | 0.286482256 | 0.291125284 |
| Gatd3a | -0.00763907 | 0.21327727 | 0.323086014 | 0.118009912 | 0.116029269 |
| Gatm | 0.038446824 | 7.18E-07 | 5.18E-06 | 0.190739307 | 0.148721159 |
| Gba | -0.004202029 | 0.464473247 | 0.581929242 | 0.087918324 | 0.095197152 |
| Gba2 | -0.008686189 | 0.017707726 | 0.042081172 | 0.03262177 | 0.037897166 |
| Gbe1 | 0.006028269 | 0.288032268 | 0.407427573 | 0.095346306 | 0.08794152 |
| Gbf1 | 0.039457834 | 8.38E-05 | 0.000397787 | 0.570825713 | 0.524956985 |
| Gbp7 | 0.015138155 | 0.01364833 | 0.033711653 | 0.107248404 | 0.096796179 |
| Gc | -0.158142579 | 9.38E-46 | 9.50E-44 | 0.269097 | 0.459375252 |
| Gca | -0.021675829 | 3.72E-05 | 0.000188972 | 0.063172633 | 0.082776523 |
| Gcat | -0.000451734 | 0.935255025 | 0.956003506 | 0.089501332 | 0.084234289 |
| Gcc1 | 0.011719158 | 0.036013286 | 0.076030656 | 0.091882238 | 0.084600455 |
| Gcc2 | 0.047605233 | 3.23E-06 | 2.05E-05 | 0.54100127 | 0.497409621 |
| Gcdh | -0.016516693 | 0.006844921 | 0.018889105 | 0.105273 | 0.113358222 |
| Gcfc2 | -0.007831868 | 0.044587899 | 0.090667967 | 0.036554953 | 0.041211869 |
| Gcg | 0.019210914 | 0.061080396 | 0.118433109 | 1.67453534 | 1.727444691 |
| Gcgr | 0.001010505 | 0.891760997 | 0.926538017 | 0.183995846 | 0.16170766 |
| Gch1 | -0.027221395 | 0.000979012 | 0.00351216 | 1.565507114 | 1.609155154 |
| Gck | 0.035252167 | 2.42E-06 | 1.58E-05 | 0.212026369 | 0.171247172 |
| Gclc | -0.044811849 | 2.70E-08 | 2.48E-07 | 0.179294771 | 0.225365855 |
| Gclm | -0.130856368 | 1.70E-39 | 1.44E-37 | 0.996392323 | 1.158304233 |
| Gcn1 | 0.012673703 | 0.112024251 | 0.193893496 | 0.221420043 | 0.208211815 |
| Gcnt2 | 0.022491684 | 0.006962896 | 0.019155835 | 0.236275414 | 0.229367235 |
| Gcsh | -0.02919511 | 0.000513863 | 0.001989558 | 0.266469623 | 0.274723404 |
| Gdap1 | -0.031378453 | 0.001669442 | 0.005564043 | 0.462458641 | 0.487453684 |
| Gdap1l1 | 0.034222096 | 0.000139179 | 0.000628152 | 0.333162311 | 0.288986917 |
| Gdap2 | -0.015059174 | 0.053324956 | 0.105242636 | 0.179432017 | 0.199148397 |
| Gde1 | -0.045276159 | 2.04E-05 | 0.000110074 | 0.701766007 | 0.767159917 |
| Gdf15 | -0.09181582 | 1.12E-25 | 4.69E-24 | 0.094451825 | 0.190242162 |
| Gdi1 | 0.000198025 | 0.983700474 | 0.989915381 | 0.507934669 | 0.499074006 |
| Gdi2 | 0.032836001 | 0.000796583 | 0.002932366 | 0.876747113 | 0.81130708 |
| Gdpd1 | -0.027095682 | 0.000281224 | 0.001161696 | 0.163759738 | 0.18371386 |
| Gdpgp1 | 0.004997119 | 0.253128686 | 0.368125218 | 0.058301271 | 0.0497831 |
| Gem | 0.00558724 | 0.312683733 | 0.433911297 | 0.063297158 | 0.069728427 |
| Gemin2 | 0.004787304 | 0.359938214 | 0.482797919 | 0.081580585 | 0.0753178 |
| Gemin5 | 0.001190903 | 0.788826443 | 0.852202543 | 0.054434881 | 0.05285064 |
| Gemin6 | -0.011249753 | 0.116703821 | 0.199893331 | 0.151462668 | 0.15792348 |
| Gemin7 | 0.044518464 | 2.05E-07 | 1.64E-06 | 0.336506817 | 0.277849322 |
| Gemin8 | -0.000596271 | 0.886533061 | 0.922199275 | 0.049554718 | 0.047039799 |
| Get1 | 0.000489506 | 0.902129216 | 0.9332305 | 0.045177439 | 0.041804766 |
| Get3 | -0.01677321 | 0.052702933 | 0.104226591 | 0.300353035 | 0.298570953 |
| Get4 | 0.009756814 | 0.245501124 | 0.359397971 | 0.264378531 | 0.250931024 |
| Gfer | -0.022431923 | 0.006436825 | 0.017936031 | 0.209689331 | 0.234980345 |
| Gfm1 | 0.006050812 | 0.26867332 | 0.385345946 | 0.090394156 | 0.083304474 |
| Gfm2 | 0.004160247 | 0.470882837 | 0.58817945 | 0.095408537 | 0.093206849 |
| Gfod2 | 0.010266576 | 0.070252397 | 0.132723242 | 0.103948386 | 0.09024386 |
| Gfpt1 | -0.031115994 | 0.001291465 | 0.004471376 | 1.103927581 | 1.18509748 |
| Gfus | -0.025157884 | 0.001189176 | 0.004160192 | 0.184066333 | 0.20651622 |
| Gga1 | 0.002837386 | 0.682223145 | 0.769199066 | 0.155865666 | 0.147761215 |
| Gga2 | 0.024038983 | 0.007526132 | 0.020443994 | 0.333682612 | 0.307906536 |
| Gga3 | 0.007698286 | 0.091914498 | 0.164778305 | 0.060370914 | 0.053875703 |
| Ggact | 0.060830487 | 3.52E-11 | 4.77E-10 | 0.389984955 | 0.310149696 |

| | | | | | |
|---|---|---|---|---|---|
| Ggct | -0.000608476 | 0.929212322 | 0.951828279 | 0.144005624 | 0.143601571 |
| Ggcx | -0.008473461 | 0.169679077 | 0.268799085 | 0.109575734 | 0.11154087 |
| Ggh | -0.066928273 | 1.55E-17 | 3.64E-16 | 0.178152394 | 0.230518948 |
| Ggnbp1 | 0.003280029 | 0.436067682 | 0.55461146 | 0.047798651 | 0.046476768 |
| Ggnbp2 | 0.035786066 | 0.000232163 | 0.000980273 | 0.515133514 | 0.4721642 |
| Ggps1 | -0.029319703 | 0.000279281 | 0.001154902 | 0.21498827 | 0.236122535 |
| Ggt7 | 0.005449091 | 0.312691066 | 0.433911297 | 0.089706277 | 0.079964601 |
| Ghdc | -0.007760845 | 0.114057529 | 0.196321875 | 0.065230921 | 0.067337695 |
| Ghitm | -0.123851235 | 6.92E-30 | 3.52E-28 | 1.044941999 | 1.268412321 |
| Ghr | -0.041685455 | 5.16E-05 | 0.000252928 | 0.703394704 | 0.719041509 |
| Gid4 | 0.004327932 | 0.515223321 | 0.628166683 | 0.141493348 | 0.132187378 |
| Gid8 | -0.004136813 | 0.583581072 | 0.6891261 | 0.195613611 | 0.189264026 |
| Gigyf1 | 0.022391976 | 0.004067815 | 0.01208289 | 0.229005851 | 0.190306087 |
| Gigyf2 | 0.019631408 | 0.025327565 | 0.056775008 | 0.302510946 | 0.279752151 |
| Gin1 | 0.005179421 | 0.28774771 | 0.40708429 | 0.068802887 | 0.062988678 |
| Ginm1 | -0.011999896 | 0.211666351 | 0.321183122 | 0.462975126 | 0.464508245 |
| Gins4 | -0.00677151 | 0.195537706 | 0.301270615 | 0.076412109 | 0.079588771 |
| Gipc1 | -0.011560944 | 0.06148281 | 0.119142093 | 0.113511237 | 0.116837759 |
| Gipr | -0.000333653 | 0.963888649 | 0.975139143 | 0.178100214 | 0.162148445 |
| Git1 | 0.013397504 | 0.063005898 | 0.121430184 | 0.168146032 | 0.158900051 |
| Git2 | -0.016688757 | 0.020439507 | 0.047312593 | 0.145462299 | 0.15741917 |
| Gjd2 | 0.108944589 | 4.87E-24 | 1.84E-22 | 0.664024493 | 0.519522346 |
| Gkap1 | 0.011703132 | 0.200918892 | 0.308388611 | 0.33827201 | 0.323280169 |
| Gla | -0.012528437 | 0.037022013 | 0.07765437 | 0.086026952 | 0.101864849 |
| Glb1 | -0.027649225 | 9.66E-06 | 5.57E-05 | 0.10944046 | 0.124772477 |
| Glb1l2 | 0.003257352 | 0.677266098 | 0.765625698 | 0.217131754 | 0.201285872 |
| Glce | 0.006060636 | 0.436180708 | 0.554682648 | 0.19145437 | 0.192116085 |
| Gle1 | -0.010363518 | 0.125986113 | 0.212608399 | 0.132999042 | 0.138727307 |
| Glg1 | 0.026307974 | 0.008814496 | 0.023356372 | 0.767956031 | 0.720899315 |
| Glis2 | 0.0081507 | 0.026727456 | 0.059298112 | 0.040792204 | 0.03387335 |
| Glis3 | 0.031547443 | 5.95E-06 | 3.60E-05 | 0.167218171 | 0.136746286 |
| Glmn | -0.006963101 | 0.115059498 | 0.197738808 | 0.048672502 | 0.054427583 |
| Glmp | -0.016387774 | 0.072161276 | 0.13556559 | 0.367843125 | 0.36337077 |
| Glo1 | -0.000573942 | 0.952188356 | 0.967851339 | 0.468694132 | 0.444824062 |
| Glod4 | -0.037100687 | 2.42E-05 | 0.000128185 | 0.298183448 | 0.319665661 |
| Glp1r | 0.021822745 | 0.07165802 | 0.13480249 | 1.482983498 | 1.367568868 |
| Glrb | -0.030008699 | 3.03E-05 | 0.00015715 | 0.144890089 | 0.168754633 |
| Glrx | -0.001960308 | 0.823597734 | 0.878251308 | 0.318899582 | 0.289215372 |
| Glrx2 | -0.039532112 | 5.54E-05 | 0.000269994 | 0.500078912 | 0.544320025 |
| Glrx3 | -0.040858558 | 1.16E-05 | 6.54E-05 | 0.371523482 | 0.388239245 |
| Glrx5 | -0.039208122 | 7.11E-05 | 0.000341544 | 0.817984418 | 0.831290791 |
| Gls | 0.007397832 | 0.422236109 | 0.541089139 | 0.327210261 | 0.330053943 |
| Glt1d1 | -0.010271295 | 0.033750354 | 0.072019522 | 0.056270305 | 0.065548865 |
| Glt8d1 | -0.040181114 | 4.50E-07 | 3.39E-06 | 0.207583005 | 0.235447981 |
| Gltp | -0.002858906 | 0.64446447 | 0.738331791 | 0.119902348 | 0.11513584 |
| Glud1 | 0.018517194 | 0.044233162 | 0.090078508 | 0.363672477 | 0.337892175 |
| Glul | 0.130923151 | 1.71E-30 | 8.89E-29 | 0.773291546 | 0.58244186 |
| Glyr1 | 0.036296144 | 0.000169637 | 0.000745333 | 0.455153547 | 0.422371586 |
| Gm10033 | 0.000506835 | 0.931734675 | 0.953608223 | 0.104883051 | 0.099524156 |
| Gm10039 | -0.014178123 | 0.001836004 | 0.006062095 | 0.045436824 | 0.060945315 |
| Gm10076 | 0.143804214 | 3.43E-125 | 3.33E-122 | 2.526961248 | 2.374428245 |
| Gm10138 | 0.000980935 | 0.872671086 | 0.912662252 | 0.104788865 | 0.10875582 |
| Gm10177 | 0.018291057 | 0.001818763 | 0.006008159 | 0.112893726 | 0.094333167 |
| Gm10275 | -0.007517157 | 0.081010836 | 0.14882603 | 0.037069681 | 0.052989858 |
| Gm10399 | -0.003927173 | 0.360757104 | 0.483496081 | 0.049993722 | 0.050472604 |
| Gm10524 | 0.004248975 | 0.376719581 | 0.498430561 | 0.067320839 | 0.062728168 |
| Gm10762 | -0.009627763 | 0.052045766 | 0.103136759 | 0.067049993 | 0.07048178 |
| Gm10941 | -0.008952423 | 0.331479293 | 0.454213063 | 0.294435661 | 0.284145196 |
| Gm11423 | -0.000386611 | 0.940136856 | 0.959566866 | 0.076070015 | 0.073836306 |
| Gm11478 | 0.001552275 | 0.753795009 | 0.824619678 | 0.07719219 | 0.06858067 |
| Gm11789 | 0.026682766 | 5.09E-05 | 0.000249699 | 0.151839255 | 0.107783567 |
| Gm11837 | -0.014844636 | 0.007509919 | 0.020405655 | 0.081086923 | 0.089908481 |
| Gm12254 | 0.00046246 | 0.8894972 | 0.924590162 | 0.033419934 | 0.029280188 |
| Gm12258 | -0.029701942 | 4.60E-08 | 4.12E-07 | 0.069207224 | 0.089251065 |
| Gm12350 | 0.005706486 | 0.194039613 | 0.29948454 | 0.053591377 | 0.051588994 |
| Gm12976 | -0.037293967 | 2.96E-09 | 3.16E-08 | 0.09558158 | 0.126555319 |
| Gm13436 | -0.00531944 | 0.147372799 | 0.241448998 | 0.034656583 | 0.037058024 |
| Gm13594 | 0.019446323 | 4.84E-09 | 5.03E-08 | 0.040421359 | 0.023657106 |
| Gm13611 | -0.000596627 | 0.907155939 | 0.935957559 | 0.073703986 | 0.073994208 |
| Gm14167 | -0.006921819 | 0.158797724 | 0.255813003 | 0.064853906 | 0.068620494 |
| Gm14295 | 0.00354915 | 0.373241048 | 0.495446405 | 0.045108923 | 0.041364253 |
| Gm14296 | -0.005561243 | 0.151304721 | 0.246313546 | 0.037627414 | 0.041748056 |

| | | | | | |
|---|---|---|---|---|---|
| Gm14303 | -0.000264347 | 0.949299861 | 0.966296016 | 0.043082591 | 0.046340282 |
| Gm14305 | -0.000201908 | 0.958614315 | 0.971740148 | 0.04581141 | 0.040673224 |
| Gm14966 | -0.005877245 | 0.111944167 | 0.193823898 | 0.035448647 | 0.03809992 |
| Gm15327 | -0.000584417 | 0.874322644 | 0.91389794 | 0.038047215 | 0.035940369 |
| Gm15417 | -0.02144977 | 0.013674743 | 0.033761903 | 0.24878303 | 0.279441671 |
| Gm15427 | -0.01506205 | 0.002707671 | 0.008499832 | 0.058540536 | 0.074470611 |
| Gm15441 | -0.000783376 | 0.835114042 | 0.887291259 | 0.039395209 | 0.035633514 |
| Gm15446 | 0.003795518 | 0.301733577 | 0.421958261 | 0.037349299 | 0.034351493 |
| Gm15500 | -0.038318214 | 1.76E-08 | 1.67E-07 | 0.121138195 | 0.150633524 |
| Gm15706 | -0.064359848 | 4.31E-11 | 5.77E-10 | 0.363461076 | 0.41227431 |
| Gm15860 | 0.002799795 | 0.521255145 | 0.633705339 | 0.056276211 | 0.050348274 |
| Gm15972 | 0.010951278 | 0.013310188 | 0.033035907 | 0.057806594 | 0.050565452 |
| Gm16062 | 0.07558493 | 3.89E-37 | 2.76E-35 | 0.1103467 | 0.062770447 |
| Gm16124 | -0.012123749 | 0.040119223 | 0.083217215 | 0.100174378 | 0.103334964 |
| Gm16174 | 0.07607819 | 2.19E-16 | 4.71E-15 | 0.319466702 | 0.287491386 |
| Gm16253 | 0.019053893 | 2.06E-06 | 1.36E-05 | 0.054760394 | 0.038349428 |
| Gm16286 | -0.022728352 | 0.018579544 | 0.043693838 | 0.466470955 | 0.471413855 |
| Gm16315 | -0.018932577 | 0.001026466 | 0.003660785 | 0.079738049 | 0.095992204 |
| Gm16536 | 0.00648844 | 0.102893609 | 0.180957248 | 0.044726669 | 0.040065114 |
| Gm1673 | 0.060802371 | 2.24E-12 | 3.48E-11 | 0.282143276 | 0.23249691 |
| Gm16740 | 0.005004734 | 0.241935027 | 0.355843696 | 0.052880961 | 0.047540739 |
| Gm17018 | -0.034262511 | 0.00014055 | 0.000633484 | 0.315352787 | 0.344825513 |
| Gm17066 | -0.001223829 | 0.745575501 | 0.81829815 | 0.038168045 | 0.037079189 |
| Gm17251 | -0.01651443 | 0.004438914 | 0.013037801 | 0.091914463 | 0.101503014 |
| Gm17491 | -0.004379685 | 0.308551678 | 0.429823673 | 0.048908602 | 0.04981885 |
| Gm17767 | 0.03061244 | 2.26E-10 | 2.77E-09 | 0.076386109 | 0.047376678 |
| Gm18811 | -0.008737773 | 0.039175855 | 0.081434181 | 0.037462931 | 0.047411279 |
| Gm19196 | 0.006147553 | 0.112823964 | 0.194880198 | 0.043151892 | 0.037982005 |
| Gm19412 | 0.000381231 | 0.933660259 | 0.95498886 | 0.05614274 | 0.056300365 |
| Gm19531 | -0.002920497 | 0.441816987 | 0.560450332 | 0.036801942 | 0.038780784 |
| Gm2011 | -0.004359904 | 0.478267607 | 0.594973471 | 0.116734828 | 0.113706903 |
| Gm20208 | 0.044882958 | 3.21E-08 | 2.93E-07 | 0.249501994 | 0.201302344 |
| Gm20257 | -0.017605288 | 0.000715867 | 0.00267268 | 0.072008802 | 0.081146707 |
| Gm20300 | -0.001943213 | 0.751203181 | 0.822802764 | 0.110617306 | 0.108095159 |
| Gm20342 | -0.006086856 | 0.14825138 | 0.242602242 | 0.04527838 | 0.045098062 |
| Gm20400 | -0.007240766 | 0.124528253 | 0.210440409 | 0.059570316 | 0.060433726 |
| Gm20604 | 0.005524929 | 0.336638196 | 0.459193399 | 0.10155529 | 0.095132841 |
| Gm20634 | -0.00430115 | 0.264137083 | 0.380435663 | 0.037693474 | 0.039832209 |
| Gm20712 | 0.019720277 | 1.62E-06 | 1.09E-05 | 0.057298455 | 0.040205396 |
| Gm21926 | -0.012094593 | 0.015016445 | 0.036585681 | 0.064538361 | 0.072526795 |
| Gm26559 | -0.00066078 | 0.881254912 | 0.9186747 | 0.05386724 | 0.053118316 |
| Gm26670 | 0.005815553 | 0.151661527 | 0.246729144 | 0.047583795 | 0.042474284 |
| Gm26782 | -0.024298569 | 0.000490808 | 0.001909418 | 0.150712104 | 0.156235846 |
| Gm26812 | 0.005186664 | 0.209195014 | 0.318028761 | 0.051575474 | 0.042799203 |
| Gm26881 | 0.003786257 | 0.499252575 | 0.613930374 | 0.093171678 | 0.09025573 |
| Gm27033 | 0.053562271 | 5.00E-10 | 5.94E-09 | 0.29611255 | 0.220452156 |
| Gm28438 | -0.426252374 | 0 | 0 | 0.00399016 | 0.448948105 |
| Gm29054 | 0.003628167 | 0.34716411 | 0.469419955 | 0.040487701 | 0.036546693 |
| Gm2a | -0.027489203 | 3.79E-05 | 0.000192167 | 0.130905225 | 0.142639868 |
| Gm30025 | 0.025496187 | 7.91E-05 | 0.000377525 | 0.143339354 | 0.118310423 |
| Gm30173 | 0.003913296 | 0.464968403 | 0.582453654 | 0.090653136 | 0.075838811 |
| Gm30648 | 0.002234807 | 0.795831821 | 0.857388848 | 0.265998979 | 0.252683428 |
| Gm32031 | 0.031161865 | 0.000172431 | 0.000755462 | 0.249406252 | 0.235231243 |
| Gm32856 | -0.020201438 | 1.86E-05 | 0.000101662 | 0.044695168 | 0.063385427 |
| Gm3294 | 0.000254611 | 0.961941618 | 0.973760559 | 0.076915946 | 0.077114359 |
| Gm33447 | -0.002402611 | 0.571535306 | 0.678528055 | 0.049812836 | 0.04714483 |
| Gm33989 | -0.013373277 | 0.038038634 | 0.07942689 | 0.113473985 | 0.119418714 |
| Gm34220 | -0.007588359 | 0.124505731 | 0.210438928 | 0.057970798 | 0.068005072 |
| Gm34552 | 0.001787646 | 0.632469501 | 0.729098641 | 0.037071748 | 0.035942371 |
| Gm34759 | -0.015054409 | 0.000815972 | 0.00299354 | 0.047555532 | 0.055372758 |
| Gm3625 | 0.013106721 | 0.010049151 | 0.025990383 | 0.083203435 | 0.067490333 |
| Gm36371 | -0.00189123 | 0.621564458 | 0.720845719 | 0.041500862 | 0.038427428 |
| Gm36738 | 0.06836803 | 3.91E-14 | 7.19E-13 | 0.342954738 | 0.28547077 |
| Gm36839 | -0.0091781 | 0.060387605 | 0.117360496 | 0.062303499 | 0.066616344 |
| Gm37033 | 0.003584666 | 0.482476984 | 0.598830746 | 0.072598037 | 0.064145232 |
| Gm37140 | 0.001120617 | 0.797353501 | 0.858450165 | 0.043340628 | 0.04806306 |
| Gm37254 | -0.001015152 | 0.833699691 | 0.886301628 | 0.063998494 | 0.061164696 |
| Gm37297 | -0.014535797 | 0.000619649 | 0.002346796 | 0.038527487 | 0.05012541 |
| Gm37376 | 0.017430198 | 0.007477999 | 0.020335975 | 0.102077239 | 0.096543485 |
| Gm37452 | -0.003647405 | 0.548112766 | 0.657681466 | 0.10155013 | 0.099520301 |
| Gm37629 | 0.012158903 | 0.006584208 | 0.018276464 | 0.058844275 | 0.048432903 |
| Gm37728 | -0.028857285 | 5.58E-10 | 6.56E-09 | 0.038783091 | 0.063058622 |

| | | | | | |
|---|---|---|---|---|---|
| Gm37834 | 0.005115418 | 0.24499892 | 0.358824872 | 0.052731196 | 0.047206395 |
| Gm37859 | -0.015868482 | 0.000447805 | 0.001759014 | 0.040177433 | 0.055291422 |
| Gm37899 | -0.002039337 | 0.767825443 | 0.835923736 | 0.127938718 | 0.12557796 |
| Gm379 | -0.080919931 | 2.37E-23 | 8.68E-22 | 0.137323531 | 0.221758234 |
| Gm38020 | -0.013602512 | 0.000580953 | 0.002219591 | 0.034791608 | 0.04366321 |
| Gm38070 | -0.166801304 | 5.87E-30 | 3.00E-28 | 0.466163829 | 0.733463083 |
| Gm38157 | -0.01277696 | 0.001665545 | 0.00555677 | 0.03405426 | 0.045922401 |
| Gm38285 | -0.002978705 | 0.474719904 | 0.591695758 | 0.043478121 | 0.044383038 |
| Gm38340 | -0.013110677 | 0.011262285 | 0.028696568 | 0.062125481 | 0.070492526 |
| Gm38394 | -0.007966507 | 0.238561272 | 0.352476092 | 0.129937598 | 0.1346947 |
| Gm3924 | -0.004546138 | 0.516709113 | 0.62950451 | 0.142875814 | 0.142219086 |
| Gm4013 | -0.005982436 | 0.492113244 | 0.607147837 | 0.262435767 | 0.275403949 |
| Gm40309 | -0.007466515 | 0.069722066 | 0.131952098 | 0.041734184 | 0.045898379 |
| Gm40477 | -2.37E-05 | 0.996252331 | 0.997175452 | 0.051297366 | 0.062886449 |
| Gm4204 | 0.009377395 | 0.052802686 | 0.104360177 | 0.069644327 | 0.06195929 |
| Gm42067 | -0.018121238 | 0.00292929 | 0.00910731 | 0.089932511 | 0.111717703 |
| Gm42444 | -0.006047115 | 0.23619714 | 0.349833729 | 0.068308701 | 0.070178146 |
| Gm42446 | -0.002554671 | 0.644918522 | 0.738677883 | 0.08298681 | 0.081502805 |
| Gm42447 | 0.002506916 | 0.689943239 | 0.77482132 | 0.11308899 | 0.105365136 |
| Gm42448 | 0.003522095 | 0.51334161 | 0.626894502 | 0.08290704 | 0.075084936 |
| Gm42685 | 0.021464136 | 7.37E-07 | 5.30E-06 | 0.05783904 | 0.041862394 |
| Gm42769 | -0.00611983 | 0.150098382 | 0.245006123 | 0.043139954 | 0.047997581 |
| Gm42770 | -0.001927407 | 0.662599429 | 0.753249725 | 0.047941147 | 0.050354764 |
| Gm42826 | 0.005401768 | 0.22768999 | 0.340224972 | 0.049811963 | 0.052929489 |
| Gm4285 | -0.002070873 | 0.57444437 | 0.680734784 | 0.035778108 | 0.035479394 |
| Gm42970 | 0.00732499 | 0.123399515 | 0.209005241 | 0.059495084 | 0.0566177 |
| Gm42992 | -0.002942572 | 0.484125195 | 0.600447815 | 0.045307079 | 0.044287011 |
| Gm43305 | -0.012767792 | 0.012971873 | 0.032262099 | 0.070970208 | 0.073994805 |
| Gm4332 | -0.00978941 | 0.0456577 | 0.09243735 | 0.058632139 | 0.069458895 |
| Gm43359 | 0.005075644 | 0.418073884 | 0.537207811 | 0.115549956 | 0.101351694 |
| Gm43360 | 0.001429724 | 0.73325823 | 0.80824677 | 0.043785019 | 0.042722243 |
| Gm43362 | 0.005594253 | 0.4626559 | 0.580454337 | 0.182606511 | 0.180796632 |
| Gm43843 | -0.019083363 | 1.01E-05 | 5.78E-05 | 0.037456786 | 0.054143961 |
| Gm43872 | -0.016423564 | 0.000185093 | 0.000801903 | 0.038505634 | 0.053240927 |
| Gm43874 | -0.011974251 | 0.008286955 | 0.022163901 | 0.046184229 | 0.055258647 |
| Gm43879 | -0.023961409 | 0.001075613 | 0.00381925 | 0.130586842 | 0.15254119 |
| Gm43880 | -0.010606524 | 0.036938953 | 0.077513598 | 0.062672552 | 0.069366406 |
| Gm43882 | -0.013854493 | 0.000645311 | 0.002434504 | 0.034146859 | 0.045071536 |
| Gm43885 | -0.015400311 | 0.00277335 | 0.008691975 | 0.061859635 | 0.072985654 |
| Gm43889 | -0.00136383 | 0.817881862 | 0.874657334 | 0.090402056 | 0.093380166 |
| Gm44109 | 0.029348346 | 7.54E-11 | 9.82E-10 | 0.072211676 | 0.045475627 |
| Gm44164 | -0.014388463 | 0.000450137 | 0.001767461 | 0.034684103 | 0.047059849 |
| Gm4419 | 0.029179308 | 4.49E-06 | 2.78E-05 | 0.143546369 | 0.109715193 |
| Gm44250 | -0.025828482 | 3.69E-06 | 2.32E-05 | 0.066072453 | 0.090825929 |
| Gm44509 | 0.004990737 | 0.160051312 | 0.257235717 | 0.036514352 | 0.03107362 |
| Gm44797 | -0.005053202 | 0.241449986 | 0.355497738 | 0.0471814 | 0.049514828 |
| Gm44799 | -0.003957813 | 0.327965923 | 0.450541855 | 0.040025391 | 0.042191418 |
| Gm44800 | -0.015013737 | 0.001407932 | 0.004819689 | 0.049107642 | 0.060260778 |
| Gm44829 | -0.003418904 | 0.398705903 | 0.518974265 | 0.040459215 | 0.041080666 |
| Gm44950 | 0.006456963 | 0.13482061 | 0.225326796 | 0.052511078 | 0.047119233 |
| Gm45051 | 0.008924531 | 0.023435384 | 0.053146444 | 0.044550345 | 0.038588189 |
| Gm45716 | 0.042463742 | 3.40E-11 | 4.61E-10 | 0.135631365 | 0.096399942 |
| Gm45847 | 0.010577154 | 0.007769539 | 0.021011255 | 0.047655716 | 0.036564306 |
| Gm45871 | 0.005413776 | 0.194320775 | 0.299823294 | 0.051568025 | 0.044430596 |
| Gm46123 | 0.008999944 | 0.017962 | 0.042581458 | 0.045462826 | 0.032993322 |
| Gm46404 | 0.011992708 | 0.001525569 | 0.005153435 | 0.044815304 | 0.033620579 |
| Gm46430 | -0.003676697 | 0.337282368 | 0.45944503 | 0.038043803 | 0.039122964 |
| Gm46565 | 0.008186013 | 0.078376603 | 0.145134265 | 0.061436223 | 0.055403371 |
| Gm47163 | -0.003169256 | 0.630616697 | 0.727697985 | 0.131484305 | 0.127513876 |
| Gm47283 | 0.005820527 | 0.253506367 | 0.368509106 | 0.078709873 | 0.06722276 |
| Gm47586 | -0.004682618 | 0.408796735 | 0.528781514 | 0.084723864 | 0.086626732 |
| Gm47889 | 0.014060941 | 0.000517567 | 0.002002304 | 0.051178732 | 0.040177861 |
| Gm4791 | -0.000497004 | 0.937698933 | 0.957494909 | 0.127213221 | 0.108983093 |
| Gm48226 | 0.010811221 | 0.113796246 | 0.195948831 | 0.15554273 | 0.136592351 |
| Gm48266 | 0.00176299 | 0.715509308 | 0.795446711 | 0.062816605 | 0.059453852 |
| Gm48267 | -0.004406798 | 0.354974131 | 0.477390857 | 0.057435574 | 0.05880116 |
| Gm4890 | -0.002737828 | 0.461050434 | 0.579085718 | 0.036475275 | 0.036598956 |
| Gm49041 | -0.010487367 | 0.021444671 | 0.04931057 | 0.051321399 | 0.058007011 |
| Gm49066 | 0.002723499 | 0.590799867 | 0.695454208 | 0.070715862 | 0.067827536 |
| Gm4924 | -0.013682999 | 0.012126732 | 0.030539168 | 0.076453071 | 0.086748295 |
| Gm49336 | 0.009920666 | 0.097096194 | 0.172288592 | 0.109262579 | 0.098223449 |
| Gm49338 | 0.00081725 | 0.853010789 | 0.898806011 | 0.055914433 | 0.051723761 |

| | | | | | |
|---|---|---|---|---|---|
| Gm49463 | -0.001936055 | 0.606631262 | 0.708088501 | 0.037526628 | 0.037455968 |
| Gm49492 | -0.00013862 | 0.971366335 | 0.980646262 | 0.038541015 | 0.038179881 |
| Gm49494 | 0.000920903 | 0.827332206 | 0.880886398 | 0.045711055 | 0.046488412 |
| Gm49542 | -0.017874933 | 9.81E-05 | 0.000458207 | 0.050891975 | 0.057195407 |
| Gm49599 | -0.04647185 | 1.08E-06 | 7.51E-06 | 0.322928885 | 0.360784181 |
| Gm49797 | 0.006919467 | 0.137830167 | 0.229140779 | 0.061798756 | 0.055106039 |
| Gm49980 | 0.003182512 | 0.700904305 | 0.783674188 | 0.229070922 | 0.217166913 |
| Gm50240 | 0.005963125 | 0.392836875 | 0.513396975 | 0.161351 | 0.150586902 |
| Gm50340 | -0.007523053 | 0.130041169 | 0.218465568 | 0.064842668 | 0.069650432 |
| Gm50353 | 0.021676051 | 1.81E-10 | 2.26E-09 | 0.043957358 | 0.022386175 |
| Gm50394 | -0.000368361 | 0.937302113 | 0.957290802 | 0.055426244 | 0.05965183 |
| Gm5113 | -0.008488226 | 0.034527464 | 0.07345208 | 0.039051645 | 0.044684729 |
| Gm52969 | -0.035950765 | 2.15E-18 | 5.46E-17 | 0.019027127 | 0.051061207 |
| Gm5617 | 0.004686241 | 0.645203968 | 0.738863141 | 0.658544033 | 0.645441687 |
| Gm5860 | -0.053584585 | 1.67E-10 | 2.09E-09 | 0.166948427 | 0.229869304 |
| Gm5870 | -0.003632791 | 0.393837427 | 0.514427982 | 0.043094467 | 0.049901568 |
| Gm5914 | 0.024155272 | 0.000151664 | 0.000674509 | 0.129550921 | 0.113060076 |
| Gm6563 | -0.012815628 | 0.001145871 | 0.004027535 | 0.032792341 | 0.044924177 |
| Gm6658 | 0.000866038 | 0.825210336 | 0.879103099 | 0.042570006 | 0.040684352 |
| Gm6710 | -0.009200002 | 0.011893145 | 0.03004031 | 0.034386594 | 0.038077592 |
| Gm7072 | 0.009421218 | 0.151755861 | 0.246730735 | 0.137128497 | 0.126117084 |
| Gm7160 | 0.000436223 | 0.926434668 | 0.950353357 | 0.065831164 | 0.05987074 |
| Gm7240 | 0.017830513 | 0.001594408 | 0.005341432 | 0.097698608 | 0.086553862 |
| Gm7467 | -0.000507851 | 0.914807695 | 0.941238269 | 0.066531588 | 0.061955823 |
| Gm8186 | 0.015085745 | 3.81E-05 | 0.000192645 | 0.042878527 | 0.031801011 |
| Gm8624 | -0.007310107 | 0.113823579 | 0.19596119 | 0.053299978 | 0.060835225 |
| Gm8730 | -0.016585955 | 0.000369058 | 0.001480801 | 0.051384818 | 0.064806508 |
| Gm8797 | 0.000424641 | 0.912402026 | 0.939654252 | 0.040363936 | 0.040610361 |
| Gm9385 | 0.01517261 | 0.000200774 | 0.000861018 | 0.054020047 | 0.040966672 |
| Gm9616 | -0.016714633 | 0.001956246 | 0.006407892 | 0.071077326 | 0.087120492 |
| Gm9794 | -0.027022906 | 0.000296527 | 0.001219473 | 0.155297245 | 0.191183021 |
| Gm9843 | -0.007017515 | 0.253323506 | 0.368353444 | 0.104181981 | 0.113251661 |
| Gm9856 | -0.009466125 | 0.115891896 | 0.198923202 | 0.103213691 | 0.104993974 |
| Gmcl1 | 0.012737924 | 0.029439632 | 0.064389674 | 0.1034427 | 0.091662094 |
| Gmds | -0.049782631 | 1.28E-07 | 1.07E-06 | 0.32843819 | 0.379989092 |
| Gmeb1 | 0.006958071 | 0.063013297 | 0.121430184 | 0.04301356 | 0.034149824 |
| Gmeb2 | 0.015910475 | 0.000759713 | 0.002809405 | 0.067443001 | 0.055067131 |
| Gmfb | 0.022486316 | 0.006642404 | 0.018398134 | 0.249803708 | 0.232492223 |
| Gmnn | -0.007984215 | 0.042121236 | 0.086440724 | 0.038237681 | 0.041978574 |
| Gmppa | 0.007485363 | 0.37825751 | 0.499989057 | 0.280567549 | 0.262402761 |
| Gmppb | -0.061646995 | 3.36E-12 | 5.08E-11 | 0.240503788 | 0.294881649 |
| Gmpr | -0.022833651 | 0.026293051 | 0.058494519 | 0.794867114 | 0.762142799 |
| Gmpr2 | -0.01186868 | 0.030026879 | 0.065482574 | 0.08346002 | 0.089806603 |
| Gmps | -0.016900777 | 0.068365876 | 0.129941945 | 0.359919319 | 0.3718629 |
| Gna11 | 0.020683453 | 0.035075308 | 0.074373423 | 0.800323135 | 0.747630555 |
| Gna12 | -0.007725599 | 0.164068864 | 0.2618315 | 0.086483845 | 0.089040486 |
| Gna13 | -0.002986761 | 0.73748219 | 0.812021349 | 0.312832527 | 0.312124094 |
| Gnai1 | 0.015463561 | 0.008218267 | 0.022022599 | 0.096619941 | 0.079288308 |
| Gnai2 | -0.095629451 | 8.70E-39 | 7.05E-37 | 2.052253099 | 2.233919913 |
| Gnai3 | 0.066678985 | 1.97E-11 | 2.74E-10 | 0.531360736 | 0.452295848 |
| Gnao1 | 0.090749722 | 1.03E-18 | 2.65E-17 | 0.808249683 | 0.696123023 |
| Gnaq | 0.071877177 | 2.27E-13 | 3.94E-12 | 0.484660321 | 0.415240429 |
| Gnas | -0.034455341 | 4.97E-14 | 9.05E-13 | 2.665643612 | 2.706489994 |
| Gnat2 | -0.019163005 | 1.74E-05 | 9.56E-05 | 0.047533742 | 0.059193798 |
| Gnaz | 0.036156455 | 0.000178057 | 0.000777313 | 0.445727501 | 0.396824112 |
| Gnb1 | 0.051316944 | 1.89E-07 | 1.52E-06 | 0.839249623 | 0.746710348 |
| Gnb2 | -0.033049783 | 0.000155985 | 0.00069025 | 1.229932713 | 1.234805147 |
| Gnb5 | -0.011271677 | 0.103450094 | 0.181738673 | 0.151299454 | 0.15524615 |
| Gne | 0.012281607 | 0.061576166 | 0.119299221 | 0.137362665 | 0.125956623 |
| Gng12 | 0.111054175 | 6.98E-29 | 3.48E-27 | 1.860977006 | 1.65850293 |
| Gng2 | -0.021112846 | 4.66E-08 | 4.16E-07 | 0.025302729 | 0.0439104 |
| Gng4 | 0.082704822 | 7.89E-16 | 1.63E-14 | 0.621770719 | 0.536245171 |
| Gng5 | -0.09258984 | 6.03E-35 | 3.88E-33 | 2.160379685 | 2.436156863 |
| Gnl1 | -0.004350346 | 0.599382836 | 0.70274963 | 0.253402742 | 0.250134524 |
| Gnl2 | -0.004278625 | 0.592689834 | 0.696750976 | 0.223816518 | 0.214650462 |
| Gnl3 | -0.009643993 | 0.335649911 | 0.458377361 | 0.375916061 | 0.420568714 |
| Gnl3l | 0.017253245 | 0.063624735 | 0.122392515 | 0.387388277 | 0.371632032 |
| Gnpat | 0.008265238 | 0.180116538 | 0.282369647 | 0.114207615 | 0.107813344 |
| Gnpda1 | 0.00934578 | 0.183565264 | 0.286456099 | 0.161112953 | 0.146182688 |
| Gnpda2 | -0.044649178 | 1.41E-07 | 1.16E-06 | 0.25243287 | 0.277984193 |
| Gnpnat1 | -0.054510679 | 6.67E-08 | 5.81E-07 | 0.300481437 | 0.366065351 |
| Gnptab | 0.002161829 | 0.606314755 | 0.707874182 | 0.047660356 | 0.046359584 |

| | | | | | |
|---|---|---|---|---|---|
| Gnptg | -0.124296062 | 4.61E-36 | 3.18E-34 | 0.578165548 | 0.707600268 |
| Gns | -0.006992613 | 0.483340504 | 0.599749378 | 0.775005344 | 0.789890367 |
| Golga1 | -0.014751682 | 0.049395763 | 0.098974775 | 0.175713793 | 0.183245992 |
| Golga2 | 0.030337044 | 0.000164044 | 0.000722861 | 0.246633084 | 0.213850441 |
| Golga3 | 0.040981133 | 2.30E-05 | 0.000122922 | 0.42233057 | 0.378516104 |
| Golga4 | -0.009267449 | 0.330121588 | 0.452671661 | 1.218734752 | 1.235211139 |
| Golga5 | 0.003907265 | 0.649188022 | 0.742518347 | 0.272962431 | 0.260490813 |
| Golga7 | 0.005698794 | 0.489505599 | 0.604928617 | 0.251814189 | 0.236387836 |
| Golga7b | -0.003504511 | 0.356344369 | 0.478703876 | 0.040163226 | 0.037641278 |
| Golgb1 | -0.026265783 | 0.003006353 | 0.009312213 | 1.403816325 | 1.458606045 |
| Golim4 | -0.027114581 | 0.009918091 | 0.025699275 | 0.653698285 | 0.687761749 |
| Golm1 | -0.028107557 | 1.18E-07 | 9.90E-07 | 0.063956198 | 0.084456377 |
| Golm2 | -0.00588402 | 0.564634951 | 0.672223977 | 0.69859523 | 0.699920601 |
| Golph3 | 0.031512664 | 0.001535223 | 0.005177051 | 0.613859984 | 0.586151708 |
| Golph3l | 0.001699313 | 0.75729668 | 0.8275192 | 0.089658604 | 0.084300317 |
| Golt1b | -0.003979363 | 0.680577437 | 0.767851206 | 0.418715098 | 0.410954352 |
| Gon4l | 0.022277733 | 0.000124017 | 0.00056632 | 0.111456017 | 0.088905753 |
| Gon7 | 0.001986336 | 0.711582176 | 0.79234932 | 0.083160016 | 0.079069586 |
| Gopc | 0.019505384 | 0.004666187 | 0.013590375 | 0.159988384 | 0.13956479 |
| Gorab | 0.000233741 | 0.970185834 | 0.979963291 | 0.116469768 | 0.115999548 |
| Gorasp1 | -0.00097796 | 0.832679838 | 0.885415589 | 0.060606044 | 0.059061942 |
| Gorasp2 | 0.027021554 | 0.006609677 | 0.018338836 | 0.630158632 | 0.614514309 |
| Gosr1 | 0.015942412 | 0.018547245 | 0.043628434 | 0.146343238 | 0.13335631 |
| Gosr2 | -0.014275957 | 0.128538331 | 0.21638955 | 0.403736777 | 0.411938982 |
| Got1 | -0.072531545 | 1.27E-14 | 2.42E-13 | 0.251210822 | 0.325915869 |
| Got2 | 0.033015453 | 0.000359414 | 0.001447484 | 0.385134699 | 0.349461973 |
| Gpaa1 | -0.012852316 | 0.034812885 | 0.073897569 | 0.106126227 | 0.112012616 |
| Gpalpp1 | 0.010173142 | 0.112576323 | 0.194675741 | 0.129791718 | 0.121465539 |
| Gpam | 0.002351419 | 0.557222683 | 0.665995796 | 0.045502475 | 0.041253261 |
| Gpank1 | 0.002794194 | 0.628736002 | 0.726715297 | 0.102033981 | 0.094790966 |
| Gpat4 | 0.03687969 | 2.31E-07 | 1.83E-06 | 0.177976551 | 0.145574632 |
| Gpatch1 | 0.012681371 | 0.042688247 | 0.087463675 | 0.123256925 | 0.110020783 |
| Gpatch11 | 0.011194939 | 0.140700904 | 0.232833054 | 0.20103854 | 0.190341809 |
| Gpatch2 | 0.0220863 | 0.009403248 | 0.024621163 | 0.262454702 | 0.241240547 |
| Gpatch2l | 0.001191121 | 0.789388669 | 0.852620446 | 0.0547277 | 0.053865972 |
| Gpatch4 | -0.00652053 | 0.266350399 | 0.382886083 | 0.086150485 | 0.096320535 |
| Gpatch8 | 0.024071809 | 0.007414106 | 0.020201776 | 0.321379837 | 0.298614263 |
| Gpbp1 | 0.013109996 | 0.152873638 | 0.247912846 | 0.364396388 | 0.344022424 |
| Gpbp1l1 | 0.013062339 | 0.103101712 | 0.181253369 | 0.233415763 | 0.217507154 |
| Gpc4 | -0.006389489 | 0.468379461 | 0.585801925 | 0.299990268 | 0.296304466 |
| Gpcpd1 | -0.012667784 | 0.058688948 | 0.114504104 | 0.117907673 | 0.137737646 |
| Gpd1 | -0.010691355 | 0.033950795 | 0.072399568 | 0.071870659 | 0.072028095 |
| Gpd1l | 0.001103212 | 0.788342837 | 0.852058817 | 0.045915424 | 0.044584903 |
| Gpd2 | 0.159038876 | 1.56E-61 | 2.98E-59 | 0.481179244 | 0.30294399 |
| Gphn | 0.043981168 | 3.05E-09 | 3.25E-08 | 0.209603554 | 0.163632066 |
| Gpi1 | -0.031410158 | 0.001454407 | 0.004949157 | 0.542564325 | 0.543708392 |
| Gpkow | 0.008181892 | 0.100149445 | 0.176899146 | 0.072521503 | 0.065047631 |
| Gpld1 | -0.022376926 | 1.23E-07 | 1.03E-06 | 0.030833331 | 0.051171624 |
| Gpn1 | -0.007252124 | 0.219455672 | 0.330270594 | 0.099570152 | 0.103720751 |
| Gpn2 | 0.002024553 | 0.679934515 | 0.767393006 | 0.07077781 | 0.067134014 |
| Gpn3 | -0.003374452 | 0.545866435 | 0.655579183 | 0.094936394 | 0.090326409 |
| Gpr107 | 0.005442067 | 0.514611601 | 0.627814529 | 0.235342562 | 0.236358776 |
| Gpr108 | -0.012038985 | 0.138161456 | 0.22949012 | 0.225258237 | 0.234317028 |
| Gpr119 | 0.00865157 | 0.041984412 | 0.086294389 | 0.057319949 | 0.040148595 |
| Gpr137 | 0.005999803 | 0.316989867 | 0.438686902 | 0.111747314 | 0.102649412 |
| Gpr137b | -0.006010515 | 0.261152025 | 0.377198038 | 0.075625586 | 0.080977391 |
| Gpr137b-ps | -0.008937316 | 0.124504525 | 0.210438928 | 0.087538406 | 0.096912572 |
| Gpr155 | 0.007247971 | 0.054421066 | 0.107123224 | 0.040274647 | 0.034342005 |
| Gpr158 | 0.014809658 | 0.091253085 | 0.163955367 | 0.297308391 | 0.279835975 |
| Gpr180 | -0.015932685 | 0.037978223 | 0.079334827 | 0.192812085 | 0.199775823 |
| Gpr19 | -0.000172989 | 0.968950117 | 0.979122029 | 0.054275657 | 0.053249167 |
| Gpr27 | 0.004483939 | 0.593953257 | 0.697898667 | 0.274773727 | 0.25640985 |
| Gpr45 | -0.038667517 | 9.56E-05 | 0.000448765 | 0.459692374 | 0.47921954 |
| Gpr85 | -0.010613488 | 0.093341384 | 0.166844077 | 0.107122296 | 0.117558304 |
| Gpr89 | 0.00188527 | 0.777703361 | 0.843842866 | 0.133994557 | 0.133571367 |
| Gprasp1 | -0.062303414 | 1.64E-09 | 1.81E-08 | 0.395311201 | 0.463521597 |
| Gprasp2 | -0.040048985 | 1.50E-06 | 1.02E-05 | 0.204627157 | 0.235100935 |
| Gprc5c | 0.03622328 | 5.21E-13 | 8.60E-12 | 0.08936488 | 0.055845975 |
| Gprin1 | 0.016361672 | 5.93E-05 | 0.000288372 | 0.054431159 | 0.039746454 |
| Gprin3 | 0.028239922 | 9.21E-06 | 5.33E-05 | 0.132612472 | 0.106850407 |
| Gps1 | -0.029970752 | 0.001528276 | 0.005158992 | 0.398368465 | 0.41860226 |
| Gps2 | 0.021395303 | 0.005672699 | 0.016050634 | 0.218786228 | 0.194035556 |

| | | | | | |
|---|---|---|---|---|---|
| Gpsm1 | 0.027749375 | 0.000764882 | 0.002827447 | 0.254463973 | 0.204997335 |
| Gpt2 | -0.012496036 | 0.064331845 | 0.123554761 | 0.121893186 | 0.136832641 |
| Gpx1 | 0.147435495 | 4.66E-44 | 4.44E-42 | 0.706780251 | 0.56099726 |
| Gpx2 | -0.118106767 | 4.32E-23 | 1.56E-21 | 0.604155564 | 0.772629554 |
| Gpx3 | 0.078770255 | 4.16E-17 | 9.41E-16 | 0.230881417 | 0.169560494 |
| Gpx4 | -0.135819497 | 7.51E-48 | 8.40E-46 | 1.247063904 | 1.416224425 |
| Gpx4-ps2 | -0.044226095 | 6.57E-09 | 6.69E-08 | 0.153669594 | 0.202193318 |
| Gramd1a | 0.024003258 | 0.001223437 | 0.004263173 | 0.198644183 | 0.170764822 |
| Gramd1b | -0.008465571 | 0.163697197 | 0.26151241 | 0.098429662 | 0.106426844 |
| Gramd1c | -0.000629525 | 0.898351446 | 0.930724736 | 0.068141242 | 0.067139106 |
| Gramd3 | -0.027042032 | 0.007713704 | 0.020882291 | 0.521115514 | 0.520295781 |
| Grb10 | -0.004951483 | 0.518658106 | 0.631167118 | 0.179086267 | 0.184010017 |
| Grb14 | 0.004971109 | 0.492112122 | 0.607147837 | 0.178033967 | 0.159532516 |
| Grb2 | 0.036048373 | 0.000130325 | 0.000591791 | 0.47941078 | 0.421032027 |
| Grb7 | 0.015815349 | 0.010951826 | 0.027975211 | 0.131276803 | 0.108816686 |
| Grcc10 | 0.025124217 | 0.004097012 | 0.012154761 | 0.29961012 | 0.281767277 |
| Grhpr | -0.025347572 | 0.001111426 | 0.003922061 | 0.188149072 | 0.209615534 |
| Gria2 | -0.071744876 | 4.19E-18 | 1.04E-16 | 0.137281557 | 0.198844569 |
| Grik5 | -0.004384883 | 0.451036875 | 0.569404039 | 0.100826373 | 0.097182626 |
| Grin1 | 0.028316338 | 8.24E-06 | 4.82E-05 | 0.130929558 | 0.10855389 |
| Grina | 0.032142259 | 0.001327088 | 0.00457516 | 0.545082339 | 0.507913638 |
| Grip1 | 0.008986611 | 0.149252945 | 0.243963729 | 0.116137165 | 0.107373489 |
| Gripap1 | -0.003384791 | 0.718789499 | 0.797872331 | 0.36381236 | 0.37474745 |
| Grk2 | 0.039713126 | 8.80E-07 | 6.23E-06 | 0.250074453 | 0.215555404 |
| Grk3 | 0.049092381 | 5.95E-08 | 5.23E-07 | 0.329406698 | 0.286148986 |
| Grk5 | 0.063805323 | 3.19E-25 | 1.29E-23 | 0.150882451 | 0.08734097 |
| Grk6 | 0.003298367 | 0.51083159 | 0.624613849 | 0.07101207 | 0.068103621 |
| Grn | 0.028560291 | 0.000229884 | 0.000972556 | 0.211865679 | 0.18731613 |
| Grpel1 | -0.017465495 | 0.091341445 | 0.164083801 | 0.659677756 | 0.670837552 |
| Grpel2 | 0.016390008 | 0.003302281 | 0.010114926 | 0.096642688 | 0.083082998 |
| Grsf1 | 0.031087643 | 0.000734358 | 0.002731228 | 0.37948748 | 0.345892304 |
| Grtp1 | -0.015691714 | 0.027154757 | 0.060136343 | 0.158157421 | 0.162759915 |
| Grwd1 | -0.003536908 | 0.373412218 | 0.495557024 | 0.040436921 | 0.042394272 |
| Gsap | 0.008392701 | 0.04504997 | 0.09151187 | 0.051709637 | 0.043421783 |
| Gsdma | -0.123159685 | 7.76E-26 | 3.27E-24 | 0.431811325 | 0.60931428 |
| Gse1 | 0.046327074 | 2.74E-06 | 1.77E-05 | 0.466623396 | 0.415254677 |
| Gsk3a | -0.004644274 | 0.556022575 | 0.664983574 | 0.224293104 | 0.219571314 |
| Gsk3b | 0.037363945 | 0.000110599 | 0.000512266 | 1.003317686 | 0.963233241 |
| Gskip | -0.005931665 | 0.380005925 | 0.501305117 | 0.136046736 | 0.141731199 |
| Gsn | 0.042819595 | 5.32E-06 | 3.25E-05 | 0.420841271 | 0.340124278 |
| Gspt1 | 0.074865182 | 1.39E-12 | 2.20E-11 | 0.74823775 | 0.689202647 |
| Gspt2 | -0.002054731 | 0.695464216 | 0.779490789 | 0.075052809 | 0.076435502 |
| Gsr | -0.024297595 | 0.009482333 | 0.024801274 | 0.365425679 | 0.384850413 |
| Gss | 0.010272648 | 0.068973853 | 0.130790677 | 0.098069212 | 0.088261237 |
| Gstm1 | -0.068764157 | 1.22E-08 | 1.19E-07 | 0.658046902 | 0.687423187 |
| Gstm2 | -0.054052448 | 4.10E-13 | 6.87E-12 | 0.116996215 | 0.154766837 |
| Gstm4 | -0.012978038 | 0.010820434 | 0.027683226 | 0.067871342 | 0.075472395 |
| Gstm5 | -0.045102095 | 7.26E-07 | 5.23E-06 | 0.306113372 | 0.344924128 |
| Gsto1 | -0.06381259 | 5.36E-19 | 1.43E-17 | 0.093316475 | 0.161211231 |
| Gsto2 | -0.009532504 | 0.01871675 | 0.043963336 | 0.040974937 | 0.045646564 |
| Gstp1 | -0.104781177 | 5.76E-19 | 1.53E-17 | 0.48516873 | 0.605201803 |
| Gstp-ps | -0.023222536 | 9.44E-07 | 6.61E-06 | 0.049394349 | 0.066156333 |
| Gstt1 | -0.059999496 | 1.59E-10 | 2.00E-09 | 0.347841986 | 0.379171816 |
| Gstt2 | -0.020849778 | 0.00054116 | 0.00208198 | 0.093838524 | 0.109872847 |
| Gstz1 | -0.029220429 | 0.001646495 | 0.005498875 | 0.358500734 | 0.364873543 |
| Gt(ROSA)26Sor | -0.014554261 | 0.041612425 | 0.085735526 | 0.154846608 | 0.162518797 |
| Gtf2a1 | 0.005106242 | 0.51368576 | 0.627145942 | 0.203356109 | 0.197351457 |
| Gtf2a2 | -0.029248842 | 0.003805312 | 0.011414762 | 0.597580174 | 0.620696581 |
| Gtf2b | -0.047776438 | 5.76E-07 | 4.24E-06 | 0.371815152 | 0.422939401 |
| Gtf2e1 | 0.01017121 | 0.010960901 | 0.027991038 | 0.048941239 | 0.039255416 |
| Gtf2e2 | 0.000909711 | 0.908910262 | 0.937253454 | 0.232358386 | 0.221444819 |
| Gtf2f1 | 0.009444559 | 0.20097064 | 0.308419347 | 0.179457016 | 0.163955661 |
| Gtf2f2 | 0.005179388 | 0.484888081 | 0.601132609 | 0.187440321 | 0.171574775 |
| Gtf2h1 | 0.003775406 | 0.591142452 | 0.695471874 | 0.152809273 | 0.147613278 |
| Gtf2h2 | 0.000664791 | 0.882126808 | 0.919287901 | 0.057082607 | 0.054148945 |
| Gtf2h3 | -0.007919428 | 0.12397854 | 0.209839723 | 0.075703148 | 0.077938833 |
| Gtf2h4 | -0.002998856 | 0.55508937 | 0.66443127 | 0.072573709 | 0.072813672 |
| Gtf2h5 | 0.009859541 | 0.316147639 | 0.437832956 | 0.712644873 | 0.711158431 |
| Gtf2i | 0.023696726 | 0.016209768 | 0.039027084 | 0.560632181 | 0.515742676 |
| Gtf2ird1 | 0.018468277 | 0.000499975 | 0.001941974 | 0.087895629 | 0.072493503 |
| Gtf3a | 0.000300753 | 0.971557465 | 0.980737377 | 0.272271152 | 0.2667258 |
| Gtf3c1 | 0.048580109 | 1.06E-09 | 1.20E-08 | 0.249272742 | 0.199358842 |

| | | | | | |
|---|---|---|---|---|---|
| Gtf3c2 | 0.008871427 | 0.202626108 | 0.310176511 | 0.15629264 | 0.147127552 |
| Gtf3c3 | 0.012844009 | 0.042521313 | 0.08714001 | 0.126574927 | 0.115823551 |
| Gtf3c4 | 0.022805485 | 0.000549446 | 0.00210968 | 0.14247949 | 0.121250905 |
| Gtf3c5 | 0.00199627 | 0.60939981 | 0.710296721 | 0.043200889 | 0.040037461 |
| Gtf3c6 | -0.007758023 | 0.295751272 | 0.415625017 | 0.176820727 | 0.179288426 |
| Gtpbp1 | 0.018750007 | 0.000445599 | 0.001752473 | 0.089446979 | 0.073901885 |
| Gtpbp10 | 0.004601199 | 0.290706302 | 0.410135927 | 0.054793084 | 0.050403495 |
| Gtpbp2 | -0.071021443 | 9.10E-11 | 1.17E-09 | 0.529589758 | 0.624781149 |
| Gtpbp3 | 0.00259742 | 0.497083391 | 0.612114595 | 0.042324999 | 0.038333413 |
| Gtpbp4 | 0.018180439 | 0.047301328 | 0.095446972 | 0.328726881 | 0.308667625 |
| Gtpbp6 | -0.009906587 | 0.138833816 | 0.2304101 | 0.133982524 | 0.14092932 |
| Gtpbp8 | 0.001175817 | 0.854436854 | 0.899880714 | 0.127729382 | 0.122501376 |
| Gucd1 | 0.024089578 | 4.41E-05 | 0.000218984 | 0.118577793 | 0.094086589 |
| Gucy1b1 | 0.000302412 | 0.946562921 | 0.964317795 | 0.048955273 | 0.052359894 |
| Gucy2c | 0.01345847 | 0.032264827 | 0.069413289 | 0.104404305 | 0.103240249 |
| Guf1 | -0.004892591 | 0.468668459 | 0.585896833 | 0.140760874 | 0.13764532 |
| Guk1 | -0.004382383 | 0.622778677 | 0.721706572 | 1.191607176 | 1.166165933 |
| Gusb | -0.02424565 | 0.003729093 | 0.011220749 | 0.2510219 | 0.264844998 |
| Gxylt1 | 0.029084103 | 9.32E-06 | 5.38E-05 | 0.143439784 | 0.116764578 |
| Gyg | -0.000538569 | 0.936736442 | 0.957014679 | 0.146049655 | 0.141269978 |
| Gys1 | -0.009673876 | 0.059991602 | 0.116671006 | 0.062171512 | 0.073729601 |
| Gzf1 | 0.014620821 | 0.031681244 | 0.068354429 | 0.151999819 | 0.13653723 |
| H13 | -0.075582556 | 1.35E-13 | 2.39E-12 | 0.567620245 | 0.662531033 |
| H1f0 | 0.006824474 | 0.524541065 | 0.636652713 | 0.580795301 | 0.552297132 |
| H1f10 | -0.018427368 | 0.077153509 | 0.143337744 | 0.374025595 | 0.348165016 |
| H1f2 | -0.011288258 | 0.388153806 | 0.509087441 | 1.143479646 | 1.178571074 |
| H1f3 | 0.004521037 | 0.368855738 | 0.491324858 | 0.065929889 | 0.063184965 |
| H1f4 | 0.020331698 | 0.044108264 | 0.089886709 | 0.372230319 | 0.350220488 |
| H2ac18 | 0.09941776 | 1.20E-21 | 3.85E-20 | 0.476873519 | 0.430550836 |
| H2ac24 | 0.122396258 | 4.42E-47 | 4.83E-45 | 0.278621721 | 0.188514064 |
| H2ac8 | 0.02618048 | 1.22E-05 | 6.89E-05 | 0.104391513 | 0.092708539 |
| H2aj | 0.047198106 | 2.88E-09 | 3.08E-08 | 1.544533456 | 1.54457953 |
| H2aw | 0.008917969 | 0.150033007 | 0.244940536 | 0.122131512 | 0.106214152 |
| H2ax | -0.013359444 | 0.080861888 | 0.148608559 | 0.197656748 | 0.192792389 |
| H2az1 | -0.087717781 | 1.20E-34 | 7.61E-33 | 2.078589516 | 2.223258251 |
| H2az2 | 0.007742169 | 0.441869142 | 0.560450332 | 0.634151373 | 0.60011351 |
| H2bc4 | 0.067225241 | 9.41E-11 | 1.21E-09 | 1.520658593 | 1.388752894 |
| H2bc6 | 0.001432914 | 0.803212716 | 0.863140712 | 0.093037081 | 0.088166235 |
| H2bu2 | 0.008685907 | 0.216881587 | 0.327207137 | 0.127953576 | 0.115385802 |
| H2-D1 | -0.076977553 | 9.61E-16 | 1.96E-14 | 1.365443484 | 1.490522872 |
| H2-K1 | -0.006838025 | 0.369090344 | 0.491377775 | 1.933972127 | 1.878443953 |
| H2-K2 | 0.003960665 | 0.327075437 | 0.449509103 | 0.050107381 | 0.042491122 |
| H2-Ke6 | 0.003098121 | 0.553317689 | 0.662890273 | 0.084159407 | 0.076335109 |
| H2-M3 | -0.023094222 | 0.000986331 | 0.00353711 | 0.139883707 | 0.156064204 |
| H2-Q2 | -0.033025323 | 0.000785373 | 0.002896417 | 0.471037907 | 0.480198145 |
| H2-Q4 | 0.034776608 | 5.86E-09 | 6.01E-08 | 0.117140488 | 0.088329507 |
| H2-T22 | -0.006488566 | 0.33008621 | 0.452671661 | 0.139049255 | 0.137303747 |
| H2-T23 | -0.06754508 | 6.14E-12 | 8.95E-11 | 0.344811873 | 0.374673875 |
| H3f3a | -0.039408808 | 1.77E-09 | 1.94E-08 | 1.900181109 | 1.900966122 |
| H3f3b | -0.064415329 | 1.40E-20 | 4.18E-19 | 2.178033689 | 2.314585015 |
| H4c14 | -0.019172378 | 4.45E-05 | 0.000220998 | 0.049988561 | 0.061852511 |
| H4c4 | 0.022519916 | 0.000669633 | 0.002515521 | 0.132104091 | 0.120594771 |
| H4c8 | 0.092068841 | 1.05E-27 | 4.89E-26 | 0.250880428 | 0.186380689 |
| H4c9 | 0.038761331 | 5.53E-07 | 4.10E-06 | 0.201283722 | 0.163431159 |
| H6pd | 0.012665114 | 0.005550774 | 0.015742307 | 0.067799149 | 0.05244597 |
| Habp4 | -0.003024181 | 0.67885884 | 0.766713016 | 0.174696025 | 0.172930335 |
| Hacd1 | 0.000230806 | 0.958701737 | 0.971740148 | 0.055396629 | 0.052830434 |
| Hacd2 | 0.012724793 | 0.028142038 | 0.06198389 | 0.109951422 | 0.094333751 |
| Hacd3 | 0.02291664 | 0.000694542 | 0.002599051 | 0.153637229 | 0.133016867 |
| Hace1 | 0.000363958 | 0.943997964 | 0.962309763 | 0.075453834 | 0.073237903 |
| Hadh | 0.057367783 | 1.08E-08 | 1.07E-07 | 1.924463924 | 1.689073811 |
| Hadha | 0.00283526 | 0.760463507 | 0.830139929 | 0.370409026 | 0.361748104 |
| Hadhb | -0.034863496 | 0.000231093 | 0.000977245 | 0.431228069 | 0.437116681 |
| Hagh | -0.127382715 | 1.57E-38 | 1.22E-36 | 0.671729211 | 0.785176408 |
| Haghl | -0.042435223 | 7.72E-08 | 6.64E-07 | 0.20314095 | 0.226161247 |
| Hap1 | -0.048188114 | 1.60E-05 | 8.81E-05 | 0.794137042 | 0.893050053 |
| Hapln1 | 0.004480047 | 0.35655394 | 0.478919233 | 0.061501715 | 0.059289061 |
| Hapln4 | -0.046235638 | 6.22E-09 | 6.36E-08 | 0.181070903 | 0.223798846 |
| Harbi1 | -0.000343845 | 0.95404698 | 0.968946518 | 0.105418113 | 0.103005865 |
| Hars | -0.01643462 | 0.0338713 | 0.072245893 | 0.186312626 | 0.20199853 |
| Hars2 | -0.009410677 | 0.052769391 | 0.10431558 | 0.062568656 | 0.066257563 |
| Hat1 | -0.015589364 | 0.007000331 | 0.01923607 | 0.09048617 | 0.100919976 |

| | | | | | |
|---|---|---|---|---|---|
| Haus1 | 0.003173175 | 0.457908674 | 0.576432491 | 0.052720036 | 0.048428999 |
| Haus2 | 0.013913259 | 0.021572114 | 0.049545025 | 0.115643825 | 0.100637242 |
| Haus7 | -0.009500316 | 0.032203793 | 0.069312658 | 0.050842513 | 0.056036032 |
| Hax1 | -0.069886793 | 1.22E-10 | 1.55E-09 | 0.57295374 | 0.643917827 |
| Hbegf | -0.018360982 | 0.002431968 | 0.00774438 | 0.082910701 | 0.103450047 |
| Hbp1 | -0.016124958 | 0.051928886 | 0.10294711 | 0.246269997 | 0.245908653 |
| Hbs1l | 0.002201773 | 0.8051486 | 0.864676911 | 0.284371735 | 0.283522134 |
| Hccs | 0.006530903 | 0.335978335 | 0.4586326 | 0.144325613 | 0.138182466 |
| Hcfc1 | -0.016807704 | 0.01780137 | 0.042241864 | 0.14390437 | 0.158923195 |
| Hcfc1r1 | 0.038057472 | 0.000150365 | 0.000669342 | 0.930657835 | 0.869870211 |
| Hcfc2 | -0.00292662 | 0.622826387 | 0.721706572 | 0.100393696 | 0.103787554 |
| Hcn2 | 0.01514008 | 0.01441539 | 0.035319158 | 0.127531375 | 0.107106226 |
| Hcn4 | 0.022185626 | 0.000106523 | 0.000494326 | 0.105385836 | 0.084439149 |
| Hdac1 | -0.037970037 | 1.44E-06 | 9.76E-06 | 0.198663837 | 0.221448417 |
| Hdac11 | 0.014087654 | 0.033712331 | 0.071969979 | 0.14920795 | 0.130929511 |
| Hdac2 | -0.011211258 | 0.191434916 | 0.295981274 | 0.290244858 | 0.28605455 |
| Hdac3 | -0.003762278 | 0.569954004 | 0.677229629 | 0.141144742 | 0.13521449 |
| Hdac4 | 0.020799485 | 0.00144504 | 0.004920729 | 0.138038345 | 0.118235822 |
| Hdac5 | 0.022652921 | 0.008952753 | 0.023645385 | 0.285684874 | 0.26337673 |
| Hdac6 | 0.065255125 | 2.37E-11 | 3.27E-10 | 0.435477225 | 0.345861779 |
| Hdac8 | -0.011522485 | 0.006619949 | 0.01836074 | 0.042556895 | 0.05152392 |
| Hddc2 | -0.023085936 | 0.000769861 | 0.002844769 | 0.129947753 | 0.149064446 |
| Hddc3 | 0.006483436 | 0.137240406 | 0.228311298 | 0.060454297 | 0.049742663 |
| Hdgf | 0.05763411 | 9.59E-09 | 9.60E-08 | 0.640043342 | 0.581514351 |
| Hdgfl2 | -0.011161117 | 0.104664436 | 0.183408011 | 0.147613117 | 0.149564831 |
| Hdgfl3 | 0.004690899 | 0.595972627 | 0.699679492 | 0.285234161 | 0.282250729 |
| Hdhd3 | -0.001110972 | 0.756254521 | 0.826566212 | 0.036239646 | 0.034328676 |
| Hdhd5 | -0.017166998 | 0.000748842 | 0.002776598 | 0.069150953 | 0.075755316 |
| Hdlbp | 0.031672232 | 0.000968889 | 0.003483558 | 1.182118577 | 1.140698449 |
| Heatr1 | -0.01269506 | 0.008258001 | 0.02209254 | 0.055822056 | 0.066579891 |
| Heatr3 | 0.002147304 | 0.687413014 | 0.773021341 | 0.078610132 | 0.076736074 |
| Heatr5a | 0.001229359 | 0.818536057 | 0.874965096 | 0.077502954 | 0.078245465 |
| Heatr5b | 0.006085373 | 0.212380903 | 0.322066314 | 0.069280305 | 0.063460885 |
| Heatr6 | -0.009666615 | 0.165524945 | 0.263765533 | 0.144399644 | 0.148201867 |
| Heca | 0.022612831 | 0.005185615 | 0.014889117 | 0.234014888 | 0.214330676 |
| Hectd1 | 0.00653426 | 0.51588209 | 0.628715444 | 0.609512635 | 0.591646016 |
| Hectd3 | 0.006414941 | 0.181684251 | 0.283839673 | 0.067599821 | 0.061543665 |
| Hectd4 | 0.020461977 | 0.001401996 | 0.004807834 | 0.138795787 | 0.113175914 |
| Heg1 | 0.028260962 | 2.54E-05 | 0.000133794 | 0.150140066 | 0.123886571 |
| Helb | 0.004986 | 0.215897208 | 0.326076225 | 0.04594733 | 0.041046144 |
| Helq | -0.007340387 | 0.2145172 | 0.324344669 | 0.096389246 | 0.101118109 |
| Helz | 0.038457578 | 1.78E-05 | 9.74E-05 | 0.334407807 | 0.291172507 |
| Hemk1 | -0.007571723 | 0.055935247 | 0.109748229 | 0.040926579 | 0.043270082 |
| Hepacam2 | 0.098099511 | 4.20E-21 | 1.29E-19 | 0.695324114 | 0.547596195 |
| Herc1 | -0.010774294 | 0.28576659 | 0.40504779 | 0.508371537 | 0.518177906 |
| Herc2 | 0.00700012 | 0.474198881 | 0.591349606 | 0.444160734 | 0.424391811 |
| Herc3 | -0.010608749 | 0.07136679 | 0.13433261 | 0.096446324 | 0.101386822 |
| Herc4 | 0.00233584 | 0.717925634 | 0.797221361 | 0.113169633 | 0.118883669 |
| Herc6 | 0.019121921 | 0.000148291 | 0.00066193 | 0.072846326 | 0.060301566 |
| Herpud1 | -0.02117696 | 0.038340631 | 0.079937297 | 1.162131327 | 1.191182154 |
| Herpud2 | 0.000121007 | 0.987863817 | 0.992728006 | 0.228879911 | 0.21721353 |
| Hes1 | -0.003750496 | 0.514104807 | 0.627341191 | 0.086228313 | 0.085719887 |
| Hes6 | -0.012032084 | 0.19010125 | 0.294246832 | 0.34544223 | 0.332560487 |
| Hexa | -0.107808435 | 2.08E-28 | 1.01E-26 | 0.833747818 | 0.924288283 |
| Hexb | -0.018729766 | 0.003369264 | 0.010300626 | 0.119891453 | 0.127143947 |
| Hexim1 | -0.012181299 | 0.101080523 | 0.178284624 | 0.156744904 | 0.167032588 |
| Hfe | -0.099015696 | 7.10E-32 | 3.99E-30 | 0.168800268 | 0.249086198 |
| Hgfac | 0.000759026 | 0.913414939 | 0.940302842 | 0.163278369 | 0.148839548 |
| Hgs | 0.005984496 | 0.394646085 | 0.515276556 | 0.159593557 | 0.152875958 |
| Hgsnat | 0.005091857 | 0.398301338 | 0.518586528 | 0.108507574 | 0.104848551 |
| Hhat | -0.021583906 | 3.27E-06 | 2.08E-05 | 0.042668029 | 0.060882741 |
| Hibadh | 0.004260948 | 0.631712017 | 0.728529565 | 0.348690036 | 0.330517227 |
| Hibch | -0.012593863 | 0.041873468 | 0.086139199 | 0.106106125 | 0.114877658 |
| Hic2 | 0.006152886 | 0.112814075 | 0.194880198 | 0.041374263 | 0.037595437 |
| Hid1 | -0.022884806 | 0.00712614 | 0.01951009 | 0.252396591 | 0.268505842 |
| Hif1a | 0.015913099 | 0.092001695 | 0.164841127 | 0.34154556 | 0.342445519 |
| Hif1an | 0.006087954 | 0.309774477 | 0.43119981 | 0.109750315 | 0.103765843 |
| Higd2a | -0.017147217 | 0.090149765 | 0.162324934 | 0.582553157 | 0.603014423 |
| Hikeshi | -0.023351994 | 0.002987137 | 0.009262731 | 0.214640961 | 0.221314368 |
| Hilpda | -0.005608882 | 0.241829205 | 0.355843696 | 0.060202694 | 0.064313616 |
| Hinfp | -0.00540537 | 0.2394266 | 0.353271423 | 0.054188659 | 0.05746237 |
| Hint1 | -0.032968277 | 2.90E-05 | 0.000151252 | 1.48426001 | 1.551304441 |

| | | | | | |
|---|---|---|---|---|---|
| Hint2 | -0.082074351 | 8.21E-17 | 1.81E-15 | 0.580776645 | 0.661569522 |
| Hint3 | 0.040643139 | 4.67E-05 | 0.000230507 | 0.64813295 | 0.610113284 |
| Hip1 | 0.016979224 | 0.000271165 | 0.001126128 | 0.065983398 | 0.050214106 |
| Hip1r | -0.049981865 | 1.44E-08 | 1.39E-07 | 0.250511303 | 0.290136174 |
| Hipk1 | 0.000638276 | 0.935227133 | 0.956003506 | 0.208830474 | 0.204692755 |
| Hipk2 | 0.053201946 | 1.09E-08 | 1.08E-07 | 0.386573916 | 0.332186968 |
| Hipk3 | 0.039909589 | 1.12E-05 | 6.34E-05 | 0.351494965 | 0.313504085 |
| Hira | 0.011140971 | 0.034857231 | 0.073943268 | 0.085881071 | 0.075737629 |
| Hirip3 | -0.007543478 | 0.216840733 | 0.327207137 | 0.095798877 | 0.107973929 |
| Hivep1 | 0.051170497 | 2.52E-12 | 3.89E-11 | 0.190108764 | 0.147428189 |
| Hivep2 | 0.068158937 | 5.97E-18 | 1.47E-16 | 0.222013752 | 0.175566147 |
| Hivep3 | 0.053304693 | 3.05E-10 | 3.70E-09 | 0.290614573 | 0.231252313 |
| Hjurp | -0.006778111 | 0.252679149 | 0.367612273 | 0.09779823 | 0.101034826 |
| Hlcs | -0.006880053 | 0.282330847 | 0.401465627 | 0.119840824 | 0.120747999 |
| Hlf | 0.019228034 | 3.00E-07 | 2.33E-06 | 0.04601083 | 0.028046031 |
| Hltf | -0.009619499 | 0.250554085 | 0.365364754 | 0.225928261 | 0.233057726 |
| Hmbox1 | 8.02E-05 | 0.989323519 | 0.993079296 | 0.105565154 | 0.103586868 |
| Hmbs | -0.007925035 | 0.294436202 | 0.414076198 | 0.198782822 | 0.192666304 |
| Hmces | -0.006635696 | 0.409112569 | 0.529012118 | 0.218590844 | 0.221277463 |
| Hmg20a | 0.006318057 | 0.334779747 | 0.45757468 | 0.131729242 | 0.125436632 |
| Hmg20b | -0.011024876 | 0.213290334 | 0.323086014 | 0.299666402 | 0.303978618 |
| Hmga1 | 0.01648798 | 3.49E-05 | 0.000178772 | 0.050556157 | 0.037817388 |
| Hmgb1 | -0.053579801 | 5.36E-10 | 6.33E-09 | 1.241737885 | 1.236750029 |
| Hmgb2 | 0.0125232 | 0.142454315 | 0.23521414 | 0.268698194 | 0.248637467 |
| Hmgcl | 0.012662916 | 0.086280143 | 0.156817265 | 0.194382092 | 0.176854203 |
| Hmgcll1 | 0.002257764 | 0.673166202 | 0.761965516 | 0.083198081 | 0.078870995 |
| Hmgcr | 0.047033546 | 1.69E-06 | 1.13E-05 | 0.450770722 | 0.403175784 |
| Hmgcs1 | 0.03140144 | 1.18E-05 | 6.64E-05 | 0.179675173 | 0.14801097 |
| Hmgn1 | 0.05433983 | 1.28E-07 | 1.07E-06 | 0.814480715 | 0.724333036 |
| Hmgn2 | -0.031500759 | 0.000971625 | 0.003490813 | 0.394182674 | 0.398684938 |
| Hmgn3 | 0.059365901 | 9.72E-11 | 1.25E-09 | 1.293921897 | 1.145663587 |
| Hmgn5 | -0.012403273 | 0.102460281 | 0.180358294 | 0.176612053 | 0.186310901 |
| Hmgxb3 | -0.005616484 | 0.356249415 | 0.478642456 | 0.102408213 | 0.107219773 |
| Hmgxb4 | -0.01053464 | 0.179406596 | 0.281502731 | 0.194091164 | 0.201477661 |
| Hmox1 | -0.128823365 | 1.73E-43 | 1.62E-41 | 0.144576721 | 0.275744589 |
| Hmox2 | -0.055531258 | 1.78E-08 | 1.69E-07 | 0.467563514 | 0.505112176 |
| Hnf1a | 0.01169553 | 0.015871904 | 0.038337056 | 0.06760662 | 0.060796891 |
| Hnf1aos1 | -0.039145552 | 4.40E-07 | 3.32E-06 | 0.186279533 | 0.203981811 |
| Hnf1b | 0.010404731 | 0.025852184 | 0.057672082 | 0.065113225 | 0.056059309 |
| Hnf4a | 0.006854353 | 0.095944793 | 0.170681661 | 0.050927622 | 0.043725843 |
| Hnf4g | 0.018102533 | 0.000177044 | 0.000773583 | 0.071835725 | 0.058286778 |
| Hnrnpa0 | -0.051626678 | 1.28E-07 | 1.07E-06 | 0.450642304 | 0.471667445 |
| Hnrnpa1 | 0.064027912 | 2.49E-10 | 3.04E-09 | 0.480990534 | 0.398177921 |
| Hnrnpa2b1 | -0.013283347 | 0.144834041 | 0.238414586 | 1.286545859 | 1.28231771 |
| Hnrnpa3 | -0.020066569 | 0.046569996 | 0.094127546 | 1.102476006 | 1.130962924 |
| Hnrnpab | 0.029488477 | 0.004011148 | 0.011940105 | 0.71574775 | 0.712120252 |
| Hnrnpc | 0.042348257 | 2.12E-05 | 0.000114343 | 0.719278745 | 0.66021755 |
| Hnrnpd | -0.013694282 | 0.180170224 | 0.282381898 | 0.733417285 | 0.744317396 |
| Hnrnpdl | 0.007787613 | 0.450503095 | 0.568951817 | 0.741944764 | 0.744114557 |
| Hnrnpf | -0.018426032 | 0.072838973 | 0.136680273 | 0.735619989 | 0.737647911 |
| Hnrnph1 | -0.008043642 | 0.421999917 | 0.540965611 | 0.556435497 | 0.571514874 |
| Hnrnph2 | 0.014732621 | 0.104592181 | 0.183314437 | 0.345188916 | 0.334803612 |
| Hnrnph3 | -0.016082399 | 0.093642338 | 0.167197578 | 0.44406701 | 0.43966585 |
| Hnrnpk | -0.013806957 | 0.118895923 | 0.202861735 | 1.177857688 | 1.169704701 |
| Hnrnpl | 0.035378553 | 0.000165654 | 0.000729386 | 0.427119786 | 0.382322972 |
| Hnrnpll | 0.002317968 | 0.753351353 | 0.824227082 | 0.17727782 | 0.171435382 |
| Hnrnpm | 0.014027239 | 0.136189535 | 0.227107145 | 0.394208841 | 0.375022314 |
| Hnrnpr | 0.003456186 | 0.721674733 | 0.799558034 | 0.479415075 | 0.450818748 |
| Hnrnpu | 0.021545498 | 0.032442525 | 0.069687632 | 1.055034795 | 1.008147199 |
| Hnrnpul1 | 0.02500817 | 0.004549559 | 0.01331742 | 0.318122438 | 0.28984775 |
| Hnrnpul2 | 0.034196144 | 0.000777586 | 0.002871131 | 0.572959446 | 0.534394447 |
| Homer1 | 0.001670777 | 0.72019628 | 0.798739247 | 0.059482938 | 0.058132239 |
| Homez | 0.006538093 | 0.119211724 | 0.203257871 | 0.054847313 | 0.045296171 |
| Hook1 | 0.007414406 | 0.470295306 | 0.587536746 | 0.584038704 | 0.581372185 |
| Hook2 | -0.017030568 | 0.016340439 | 0.039302758 | 0.146264762 | 0.156180191 |
| Hook3 | -0.042987144 | 1.49E-05 | 8.31E-05 | 0.430042912 | 0.455824722 |
| Hopx | -0.019665976 | 0.027108169 | 0.060060534 | 0.305037131 | 0.30584976 |
| Hp1bp3 | 0.030233031 | 0.002740972 | 0.00859604 | 0.593382438 | 0.56183051 |
| Hpcal1 | -0.003019313 | 0.708067928 | 0.789340259 | 0.225023375 | 0.216534369 |
| Hpf1 | 0.008784726 | 0.33475558 | 0.45757468 | 0.335124233 | 0.330366583 |
| Hpgds | 0.015952497 | 0.004491587 | 0.013176587 | 0.078420651 | 0.074402625 |
| Hprt | -0.055130363 | 2.44E-08 | 2.26E-07 | 0.605683334 | 0.635285202 |

| | | | | | |
|---|---|---|---|---|---|
| Hps1 | -0.000704959 | 0.898983742 | 0.931066362 | 0.092893714 | 0.088016623 |
| Hps3 | -7.00E-05 | 0.989242578 | 0.993079296 | 0.072129852 | 0.072508008 |
| Hps4 | 0.03296644 | 1.30E-07 | 1.08E-06 | 0.129106818 | 0.107140266 |
| Hps5 | 0.002139413 | 0.611791335 | 0.712486267 | 0.048158388 | 0.046827912 |
| Hras | 0.00420779 | 0.652167768 | 0.744874887 | 0.394782681 | 0.383071529 |
| Hs1bp3 | 0.007112273 | 0.105629428 | 0.184765976 | 0.058062503 | 0.050417158 |
| Hs2st1 | -0.007356509 | 0.268798153 | 0.385379095 | 0.128717591 | 0.134595433 |
| Hs3st6 | 0.006876396 | 0.371517197 | 0.493832402 | 0.224085137 | 0.191225112 |
| Hs6st1 | -0.001374135 | 0.857832333 | 0.902287779 | 0.187806876 | 0.18896249 |
| Hsbp1 | 0.051465779 | 1.10E-07 | 9.25E-07 | 0.940250311 | 0.884356593 |
| Hsbp1l1 | -0.01771314 | 0.015039022 | 0.036625594 | 0.166967951 | 0.17399715 |
| Hscb | -0.01569696 | 0.0153882 | 0.037317056 | 0.119239422 | 0.127507756 |
| Hsd17b10 | -0.022593436 | 0.005959957 | 0.016756133 | 0.229766999 | 0.247445729 |
| Hsd17b11 | 0.018784918 | 0.005415297 | 0.015457285 | 0.147989834 | 0.12969151 |
| Hsd17b12 | 0.004514625 | 0.634505083 | 0.730520142 | 0.351030496 | 0.359944929 |
| Hsd17b4 | -0.004394308 | 0.502326897 | 0.616541105 | 0.135983072 | 0.1327746 |
| Hsd17b7 | -0.005515386 | 0.189530069 | 0.293662499 | 0.041152296 | 0.047494676 |
| Hsdl1 | 0.010834617 | 0.079466541 | 0.146681928 | 0.122432654 | 0.107078629 |
| Hsdl2 | -0.00979576 | 0.189496738 | 0.293662499 | 0.189306917 | 0.185366032 |
| Hsf1 | 0.007046755 | 0.234787748 | 0.348595981 | 0.108112573 | 0.100316041 |
| Hsf2 | 0.039084126 | 1.59E-08 | 1.52E-07 | 0.168966539 | 0.132499525 |
| Hsp90aa1 | 0.039063434 | 2.18E-05 | 0.000116988 | 1.671565489 | 1.551056113 |
| Hsp90ab1 | 0.005129941 | 0.298880774 | 0.418570856 | 3.052068194 | 3.040755674 |
| Hsp90b1 | -0.01411327 | 0.030845625 | 0.066952705 | 2.531039559 | 2.619912684 |
| Hspa12a | 0.041437166 | 5.03E-14 | 9.12E-13 | 0.111988458 | 0.068479934 |
| Hspa13 | -0.076489973 | 3.49E-16 | 7.37E-15 | 0.304754044 | 0.37828454 |
| Hspa14 | -0.003400955 | 0.623876389 | 0.722578777 | 0.15234698 | 0.147744683 |
| Hspa1a | -0.036070548 | 7.63E-06 | 4.50E-05 | 0.102130049 | 0.154647282 |
| Hspa1b | -0.012008615 | 0.067100441 | 0.127987147 | 0.075274741 | 0.096724546 |
| Hspa2 | -0.067483697 | 1.38E-17 | 3.26E-16 | 0.148178952 | 0.204252279 |
| Hspa4 | 0.019069584 | 0.070103788 | 0.13251974 | 0.709214137 | 0.68355694 |
| Hspa4l | 0.089292816 | 1.09E-21 | 3.50E-20 | 0.365395915 | 0.278021887 |
| Hspa5 | -0.068644303 | 1.56E-15 | 3.15E-14 | 2.197656837 | 2.41311181 |
| Hspa8 | -0.049530885 | 1.91E-07 | 1.53E-06 | 1.483231362 | 1.464191214 |
| Hspa9 | -0.047473602 | 2.39E-05 | 0.000126741 | 1.368682138 | 1.50165455 |
| Hspb1 | 0.083254693 | 7.42E-22 | 2.42E-20 | 0.20139977 | 0.09878825 |
| Hspb11 | -0.004622926 | 0.53155041 | 0.642832825 | 0.179762387 | 0.176649607 |
| Hspb6 | 0.021525043 | 0.002205715 | 0.007105354 | 0.155441769 | 0.137543533 |
| Hspb8 | 0.00871026 | 0.054021286 | 0.106487214 | 0.055811252 | 0.048284931 |
| Hspbp1 | 0.001742766 | 0.79878164 | 0.859519104 | 0.148199306 | 0.143364099 |
| Hspd1 | 0.060438889 | 1.96E-08 | 1.84E-07 | 1.015099412 | 0.911667039 |
| Hspe1 | 0.135149072 | 5.78E-36 | 3.95E-34 | 1.214153506 | 1.00098876 |
| Hsph1 | 0.109020384 | 7.40E-24 | 2.77E-22 | 0.496592173 | 0.360382049 |
| Htatip2 | -0.162364966 | 7.75E-59 | 1.34E-56 | 0.307747845 | 0.464091856 |
| Htatsf1 | 0.007313711 | 0.400310062 | 0.520435199 | 0.29534752 | 0.279735992 |
| Htra2 | -0.007179523 | 0.146845587 | 0.240991357 | 0.064670735 | 0.070173042 |
| Htt | 0.032410388 | 3.07E-06 | 1.97E-05 | 0.168448037 | 0.137492905 |
| Hus1 | 0.013597927 | 0.003285166 | 0.010065675 | 0.065413596 | 0.053914434 |
| Huwe1 | -0.021214013 | 0.039713869 | 0.082464382 | 0.56676315 | 0.580780579 |
| Hyal2 | -0.013937671 | 0.005664425 | 0.016031888 | 0.066146561 | 0.075095509 |
| Hyi | -0.026579373 | 0.00455042 | 0.01331742 | 0.325008057 | 0.350074839 |
| Hyou1 | -0.022588483 | 0.031739102 | 0.068448879 | 0.619920075 | 0.666781202 |
| Hypk | 0.002731552 | 0.73650666 | 0.811091726 | 0.250628073 | 0.227030736 |
| Iah1 | -0.029122687 | 0.000918382 | 0.003319149 | 0.269812687 | 0.286521505 |
| Iapp | -0.024300765 | 3.93E-48 | 4.55E-46 | 9.220726894 | 9.351862273 |
| Iars | 0.012595854 | 0.175710977 | 0.276776105 | 0.358494274 | 0.345339455 |
| Iars2 | 0.0299281 | 2.30E-05 | 0.000122791 | 0.171375242 | 0.146931877 |
| Ibtk | -0.015013199 | 0.068066415 | 0.129474014 | 0.218012949 | 0.227154866 |
| Ica1 | 0.026884748 | 0.004672247 | 0.01360395 | 0.40649399 | 0.386960985 |
| Ica1l | -0.042433654 | 1.73E-08 | 1.64E-07 | 0.160731912 | 0.186330632 |
| Ice1 | 0.007300476 | 0.311848237 | 0.433245901 | 0.162151588 | 0.157425403 |
| Ice2 | -0.004807179 | 0.246867923 | 0.361072656 | 0.044966224 | 0.046735616 |
| Icmt | 0.004681387 | 0.382429693 | 0.503586818 | 0.086384059 | 0.080170772 |
| Id2 | 0.072752599 | 1.51E-08 | 1.45E-07 | 0.545890249 | 0.492109146 |
| Id3 | -0.016944176 | 0.06239469 | 0.120572685 | 0.140514599 | 0.17008826 |
| Ide | 0.005358299 | 0.448813952 | 0.567629666 | 0.159265351 | 0.15343376 |
| Idh1 | 0.024821589 | 0.000446912 | 0.001756216 | 0.168118694 | 0.143135539 |
| Idh2 | 0.045547115 | 4.70E-06 | 2.90E-05 | 0.552910716 | 0.464810888 |
| Idh3a | -0.000162162 | 0.984723997 | 0.990703331 | 0.268319709 | 0.261942119 |
| Idh3b | -0.003017903 | 0.720827105 | 0.798800993 | 0.274906982 | 0.266748309 |
| Idh3g | -0.003798511 | 0.669854805 | 0.758924183 | 0.336836101 | 0.32736143 |
| Idi1 | 0.011187575 | 0.009830022 | 0.025491457 | 0.056387334 | 0.046567884 |

| | | | | | |
|---|---|---|---|---|---|
| Idnk | 0.039381157 | 2.59E-05 | 0.000136439 | 0.370391886 | 0.338386224 |
| Ids | -0.011645759 | 0.260116583 | 0.376261482 | 0.687663987 | 0.707110564 |
| Idua | 0.00233867 | 0.618672632 | 0.718434702 | 0.060884308 | 0.058447601 |
| Ier2 | -0.11254254 | 9.75E-20 | 2.78E-18 | 0.585587057 | 0.734603369 |
| Ier3 | -0.026383814 | 0.000102598 | 0.000478169 | 0.10357635 | 0.129429705 |
| Ier3ip1 | 0.101156718 | 1.00E-24 | 3.98E-23 | 0.846055544 | 0.759040593 |
| Ier5 | -0.033598534 | 0.000215372 | 0.000917149 | 0.277225423 | 0.288092361 |
| Iffo1 | 0.002368882 | 0.688618257 | 0.773833088 | 0.106630317 | 0.099353809 |
| Ifi27 | 0.032592665 | 0.008656716 | 0.023026153 | 0.486843702 | 0.494557983 |
| Ifi30 | -0.011818663 | 0.153631592 | 0.248851438 | 0.224166219 | 0.226325458 |
| Ifi35 | 0.000250725 | 0.972330866 | 0.981212444 | 0.171851449 | 0.162371804 |
| Ifih1 | 0.016829579 | 0.001187566 | 0.004157292 | 0.078286513 | 0.066979425 |
| Ifit1 | 0.040717689 | 7.12E-13 | 1.16E-11 | 0.084440807 | 0.055934772 |
| Ifitm2 | -0.058930354 | 1.96E-11 | 2.74E-10 | 0.205030199 | 0.248500804 |
| Ifnar1 | 0.005689386 | 0.267714597 | 0.384449233 | 0.073287449 | 0.069548881 |
| Ifnar2 | -0.021222975 | 0.00112808 | 0.003972181 | 0.115444234 | 0.132441511 |
| Ifngr1 | -0.003300406 | 0.525953793 | 0.637889568 | 0.076133029 | 0.076104974 |
| Ifngr2 | -0.014146322 | 0.003942316 | 0.011774868 | 0.057930469 | 0.068593805 |
| Ifrd1 | -0.019205418 | 0.059395135 | 0.115656023 | 0.374728089 | 0.409007719 |
| Ifrd2 | -0.004712392 | 0.287383277 | 0.406687077 | 0.052280195 | 0.05449809 |
| Ift122 | -0.005478252 | 0.342536789 | 0.464735061 | 0.095100661 | 0.095668522 |
| Ift140 | -0.002415013 | 0.508049432 | 0.622150974 | 0.035675267 | 0.035079463 |
| Ift172 | -0.018295682 | 0.000988157 | 0.003542355 | 0.078799005 | 0.090221507 |
| Ift20 | -0.065988485 | 5.27E-12 | 7.78E-11 | 1.020687432 | 1.100449452 |
| Ift22 | -0.006634346 | 0.397532172 | 0.517931892 | 0.20788108 | 0.209434431 |
| Ift27 | -0.022644123 | 0.001414737 | 0.004839575 | 0.159546252 | 0.169968474 |
| Ift43 | -0.016405493 | 0.001420566 | 0.004856097 | 0.070710805 | 0.080466659 |
| Ift46 | -0.007685815 | 0.226189761 | 0.338414413 | 0.121888374 | 0.122310613 |
| Ift52 | -0.008347676 | 0.266647945 | 0.383226787 | 0.193483712 | 0.190123018 |
| Ift57 | 0.045993795 | 3.40E-05 | 0.000174609 | 0.674928889 | 0.636952971 |
| Ift74 | -0.029592154 | 0.001234064 | 0.004290977 | 0.314640676 | 0.343545088 |
| Ift80 | -0.011662425 | 0.041115487 | 0.084858151 | 0.090335913 | 0.094476188 |
| Ift81 | 0.035000255 | 0.000160948 | 0.000709955 | 0.365740437 | 0.324297601 |
| Ift88 | 0.012358188 | 0.089069904 | 0.160775642 | 0.180120079 | 0.158022687 |
| Iftap | -0.044060806 | 8.17E-08 | 7.02E-07 | 0.236838114 | 0.26533314 |
| Igbp1 | 0.005775297 | 0.412602147 | 0.53228743 | 0.170028798 | 0.155141913 |
| Igf1r | -0.01845096 | 0.150925734 | 0.246006139 | 1.404442697 | 1.550428573 |
| Igf2r | 0.056552246 | 3.38E-09 | 3.59E-08 | 0.408401327 | 0.368115981 |
| Igfals | 0.01512889 | 0.028271177 | 0.062254221 | 0.163421575 | 0.136465212 |
| Igfbp7 | 0.050350685 | 5.93E-06 | 3.59E-05 | 0.611399627 | 0.518474452 |
| Ighmbp2 | 0.009864931 | 0.01011239 | 0.026140032 | 0.043227859 | 0.036196375 |
| Igip | 0.021131193 | 0.001485473 | 0.005040758 | 0.142468607 | 0.123085852 |
| Igsf1 | 0.006879374 | 0.161170577 | 0.258479108 | 0.070655132 | 0.062308814 |
| Igsf11 | -0.003011701 | 0.463050993 | 0.5807252 | 0.041628334 | 0.04473189 |
| Igsf8 | 0.013919781 | 0.03160962 | 0.068275655 | 0.136032096 | 0.119672449 |
| Igsf9b | 0.020699392 | 0.001089436 | 0.003857065 | 0.128973039 | 0.10869464 |
| Igtp | -0.006655441 | 0.088533187 | 0.160044561 | 0.037057709 | 0.041242205 |
| Ik | 0.041793277 | 1.52E-05 | 8.45E-05 | 0.448292763 | 0.403758391 |
| Ikbip | -0.027865328 | 0.000372614 | 0.001492607 | 0.199160398 | 0.215477272 |
| Ikbkb | -0.007156606 | 0.260736763 | 0.376766173 | 0.114201528 | 0.120988181 |
| Ikbkg | -0.020723792 | 0.000672445 | 0.002525112 | 0.089478779 | 0.110315294 |
| Ikzf2 | 0.011273861 | 0.098825744 | 0.17494056 | 0.151465681 | 0.131523163 |
| Ikzf3 | 0.005856432 | 0.238777443 | 0.352634711 | 0.072359687 | 0.066415472 |
| Ikzf4 | 0.011522568 | 0.049989401 | 0.100019954 | 0.096433249 | 0.091782092 |
| Ikzf5 | 0.010590532 | 0.067265992 | 0.128203425 | 0.100330524 | 0.090126433 |
| Il10rb | -0.00226111 | 0.639998366 | 0.735555516 | 0.062850501 | 0.065266784 |
| Il11ra1 | -0.014429026 | 0.015286575 | 0.037126176 | 0.100177346 | 0.105212288 |
| Il13ra1 | 0.009446128 | 0.087269132 | 0.158318809 | 0.086734347 | 0.082918919 |
| Il15ra | -0.002226512 | 0.55515071 | 0.66443127 | 0.038053645 | 0.038104566 |
| Il17ra | 0.012544743 | 0.015326437 | 0.03718583 | 0.081851986 | 0.070350275 |
| Il17rc | 0.000688414 | 0.883071567 | 0.919878043 | 0.057626883 | 0.058094803 |
| Il17re | 0.012287923 | 0.004933493 | 0.014253616 | 0.056851992 | 0.043092839 |
| Il18 | 0.00218069 | 0.543389649 | 0.653250175 | 0.036553611 | 0.032103477 |
| Il18bp | -0.017325818 | 0.00412597 | 0.012222023 | 0.077938019 | 0.102212931 |
| Il1r1 | 0.024257834 | 0.018356645 | 0.043295319 | 1.694152317 | 1.625768981 |
| Il3ra | -0.025751644 | 1.22E-06 | 8.39E-06 | 0.065023576 | 0.084502827 |
| Il6ra | 0.018539952 | 0.048163376 | 0.096985158 | 0.355025791 | 0.307497318 |
| Il6st | 0.001834097 | 0.844860114 | 0.894079434 | 0.318094548 | 0.327159629 |
| Ildr1 | 0.021921123 | 0.005441185 | 0.015508414 | 0.22894446 | 0.205407497 |
| Ilf2 | -0.003434695 | 0.695025397 | 0.77921346 | 0.296473001 | 0.29003393 |
| Ilf3 | 0.03387641 | 3.93E-05 | 0.000197993 | 0.26661631 | 0.224520683 |
| Ilk | 0.014565696 | 0.06098355 | 0.118268915 | 0.220936117 | 0.198036101 |

| | | | | | |
|---|---|---|---|---|---|
| Ilkap | -0.031283388 | 0.000865884 | 0.003157641 | 0.387976265 | 0.403782475 |
| Ilrun | 0.04585888 | 9.16E-07 | 6.46E-06 | 0.39530223 | 0.353866279 |
| Ilvbl | 0.00758614 | 0.162031096 | 0.259645017 | 0.087294396 | 0.082067277 |
| Immp1l | -0.061841426 | 9.67E-10 | 1.11E-08 | 0.60866511 | 0.631287438 |
| Immp2l | -0.033275086 | 5.60E-10 | 6.58E-09 | 0.067220342 | 0.090268283 |
| Immt | -0.016297795 | 0.091640197 | 0.164468524 | 0.441277786 | 0.451522309 |
| Imp3 | -0.019371721 | 0.056929336 | 0.111541114 | 0.650376559 | 0.655982085 |
| Imp4 | -0.009712335 | 0.137168079 | 0.228230031 | 0.129170095 | 0.134044895 |
| Impa1 | -0.007782838 | 0.367798538 | 0.490431681 | 0.300736337 | 0.292999171 |
| Impact | -0.000545837 | 0.954900071 | 0.969560156 | 0.373562914 | 0.385936496 |
| Impdh1 | 0.000397658 | 0.96240628 | 0.974028092 | 0.283838296 | 0.266190835 |
| Impdh2 | 0.002482391 | 0.707685235 | 0.789133848 | 0.137448585 | 0.127293457 |
| Ina | 0.007765999 | 0.384324674 | 0.505465975 | 0.294259387 | 0.278769659 |
| Inafm1 | 0.005281581 | 0.50512696 | 0.61935229 | 0.220944106 | 0.212810716 |
| Inafm2 | 0.011767237 | 0.089295249 | 0.161062784 | 0.154161459 | 0.143371504 |
| Ing1 | -0.004707973 | 0.527819631 | 0.639275253 | 0.176593347 | 0.173252942 |
| Ing2 | 0.066964445 | 8.33E-11 | 1.08E-09 | 0.626713393 | 0.55611987 |
| Ing3 | -0.010474909 | 0.065132326 | 0.124894769 | 0.088582891 | 0.093323806 |
| Ing5 | 0.006571741 | 0.316593718 | 0.438263438 | 0.131145121 | 0.123768967 |
| Inha | 0.009468798 | 0.084089761 | 0.15358269 | 0.089898094 | 0.080608596 |
| Inip | 0.009504297 | 0.03560776 | 0.075272591 | 0.061046118 | 0.051565648 |
| Ino80 | -0.001825983 | 0.78588316 | 0.850251066 | 0.139598945 | 0.134230593 |
| Ino80b | 0.009371196 | 0.16013897 | 0.257289775 | 0.142746956 | 0.131600771 |
| Ino80d | 0.022831962 | 0.00367057 | 0.01107207 | 0.211628781 | 0.187149605 |
| Ino80e | -0.004531305 | 0.525805716 | 0.637869125 | 0.163476259 | 0.16202841 |
| Inpp1 | 0.017558056 | 0.000500383 | 0.001942779 | 0.080821236 | 0.064104638 |
| Inpp4a | 0.031928423 | 0.000283607 | 0.001170299 | 0.332232379 | 0.284820142 |
| Inpp5a | 0.025300728 | 0.000549329 | 0.00210968 | 0.179104306 | 0.160225044 |
| Inpp5b | -0.015603895 | 0.00553384 | 0.015698861 | 0.0764441 | 0.092879455 |
| Inpp5e | 0.01049998 | 0.005947126 | 0.016724895 | 0.045069796 | 0.035181599 |
| Inpp5f | -0.011219607 | 0.304576508 | 0.425138953 | 0.522553381 | 0.608982224 |
| Inpp5j | 0.01017652 | 0.019571385 | 0.04556346 | 0.05715896 | 0.047389511 |
| Inpp5k | -0.013567388 | 0.01167565 | 0.029583183 | 0.082009014 | 0.085117692 |
| Inppl1 | -0.011627841 | 0.085666738 | 0.155877227 | 0.136103228 | 0.142507307 |
| Ins1 | 0.008039468 | 9.78E-10 | 1.12E-08 | 10.69431767 | 10.62831239 |
| Ins2 | -0.003267042 | 1.00E-04 | 0.000466715 | 11.99720871 | 12.02750287 |
| Insig1 | 0.02435066 | 0.000862133 | 0.00314745 | 0.175356657 | 0.148822061 |
| Insig2 | 0.025543537 | 0.004714587 | 0.013719011 | 0.34406653 | 0.320457724 |
| Insm1 | 0.027376837 | 0.008061868 | 0.021699191 | 1.142224089 | 1.06277842 |
| Insr | -0.001315969 | 0.850805791 | 0.897908587 | 0.14730981 | 0.146151704 |
| Insrr | -0.092872353 | 4.96E-19 | 1.33E-17 | 0.465603388 | 0.498018382 |
| Ints1 | -0.016399297 | 0.005489738 | 0.015610187 | 0.090136835 | 0.102883622 |
| Ints10 | 0.000810315 | 0.878799545 | 0.917492394 | 0.082469689 | 0.080395557 |
| Ints11 | -0.006598514 | 0.235663803 | 0.349469721 | 0.087703862 | 0.09032134 |
| Ints12 | -0.002769057 | 0.591230676 | 0.695471874 | 0.078839094 | 0.075653485 |
| Ints13 | -0.006114899 | 0.332890424 | 0.45563293 | 0.113452921 | 0.120473971 |
| Ints14 | -0.003433404 | 0.438716546 | 0.557033075 | 0.05312245 | 0.053607272 |
| Ints2 | 0.014840187 | 0.004929747 | 0.014251264 | 0.083715533 | 0.071343521 |
| Ints3 | 0.007255042 | 0.351770261 | 0.474262998 | 0.202076726 | 0.193492649 |
| Ints4 | 0.009100744 | 0.079286741 | 0.146433453 | 0.079733023 | 0.072762665 |
| Ints5 | 0.015195761 | 0.001206536 | 0.004210319 | 0.06987716 | 0.055432515 |
| Ints6 | 0.014041547 | 0.07250105 | 0.136072434 | 0.205376337 | 0.188674576 |
| Ints6l | -0.031520328 | 6.56E-06 | 3.92E-05 | 0.125010903 | 0.154815801 |
| Ints7 | 0.004184072 | 0.366266529 | 0.488992474 | 0.060697199 | 0.057208839 |
| Ints8 | 0.005339053 | 0.324393873 | 0.446459888 | 0.087983068 | 0.080237684 |
| Ints9 | 0.007287223 | 0.156397671 | 0.252658384 | 0.079085655 | 0.071099403 |
| Intu | -0.000622868 | 0.904250816 | 0.935126735 | 0.074369375 | 0.073636103 |
| Invs | -0.01099447 | 0.033513519 | 0.071632731 | 0.071905195 | 0.075939348 |
| Ip6k1 | 0.040416943 | 2.99E-06 | 1.92E-05 | 0.313305899 | 0.268851654 |
| Ip6k2 | 0.035211824 | 0.000257279 | 0.001073966 | 0.481491398 | 0.434401931 |
| Ipmk | 0.057786762 | 1.24E-12 | 1.99E-11 | 0.256483065 | 0.200426363 |
| Ipo13 | 0.005388353 | 0.235997492 | 0.349751161 | 0.060518312 | 0.054420476 |
| Ipo4 | 0.006686295 | 0.329728021 | 0.452455183 | 0.141129812 | 0.142731555 |
| Ipo5 | 0.033597189 | 4.83E-05 | 0.000238396 | 0.247388473 | 0.220302919 |
| Ipo7 | 0.030623378 | 0.000465974 | 0.001823018 | 0.300102144 | 0.269926758 |
| Ipo8 | -0.011703345 | 0.076059994 | 0.141739555 | 0.121028692 | 0.132522736 |
| Ipo9 | -0.003560909 | 0.540422598 | 0.651132544 | 0.095648758 | 0.097162782 |
| Ipp | 0.008789168 | 0.072358411 | 0.135830946 | 0.074153666 | 0.063337822 |
| Ippk | 0.01910666 | 1.59E-05 | 8.76E-05 | 0.062975029 | 0.0475947 |
| Iqcb1 | 0.00065177 | 0.924584313 | 0.948887226 | 0.152576842 | 0.143382345 |
| Iqcc | -0.004223969 | 0.492006941 | 0.607147837 | 0.116951912 | 0.115288334 |
| Iqce | 2.99E-05 | 0.994972421 | 0.996584467 | 0.060077649 | 0.060143772 |

| | | | | | |
|---|---|---|---|---|---|
| Iqgap1 | 0.044845344 | 1.25E-05 | 7.01E-05 | 0.869598212 | 0.809553019 |
| Iqgap2 | 0.02144251 | 1.18E-08 | 1.16E-07 | 0.04637223 | 0.029016062 |
| Irak1 | -0.038261466 | 0.000144839 | 0.000649503 | 0.562369585 | 0.596229952 |
| Irak1bp1 | -0.00840269 | 0.332014171 | 0.454817777 | 0.276162084 | 0.274754654 |
| Ireb2 | -0.01958642 | 0.049238918 | 0.098762277 | 0.397472264 | 0.419753214 |
| Irf1 | 0.019702421 | 0.00593341 | 0.016695979 | 0.158653966 | 0.139481402 |
| Irf2 | 0.037874015 | 1.41E-05 | 7.88E-05 | 0.307504045 | 0.266502766 |
| Irf2bp1 | 0.02324084 | 0.002051317 | 0.006674331 | 0.211367778 | 0.178141218 |
| Irf2bp2 | 0.048864672 | 3.50E-06 | 2.21E-05 | 0.883119146 | 0.781174453 |
| Irf2bpl | 0.059108713 | 1.38E-10 | 1.75E-09 | 0.356743996 | 0.302051027 |
| Irf3 | -0.019701788 | 0.001583975 | 0.005310141 | 0.10521734 | 0.120693107 |
| Irf6 | 0.048484219 | 3.67E-09 | 3.88E-08 | 0.264545262 | 0.216178161 |
| Irf7 | 0.015159339 | 0.004084499 | 0.012128741 | 0.059368055 | 0.054419701 |
| Irf9 | 0.009452812 | 0.059457282 | 0.115747636 | 0.067636686 | 0.064487785 |
| Irgm1 | 0.006476882 | 0.178098302 | 0.27972079 | 0.062787523 | 0.059044249 |
| Irgq | 0.014809988 | 0.025721007 | 0.05743216 | 0.145923609 | 0.128557726 |
| Irs2 | 0.024331584 | 0.013100258 | 0.032556418 | 0.327553374 | 0.34801115 |
| Isca1 | 0.00198451 | 0.770626511 | 0.838150543 | 0.146399773 | 0.142181578 |
| Isca2 | -0.016873242 | 0.04443732 | 0.09039975 | 0.26882708 | 0.275288842 |
| Iscu | -0.043819626 | 6.82E-06 | 4.06E-05 | 0.611552057 | 0.620178224 |
| Isg20 | -0.001823556 | 0.86297283 | 0.906028278 | 0.744557903 | 0.744701652 |
| Isg20l2 | 0.004083756 | 0.330781851 | 0.45338519 | 0.048279189 | 0.045435524 |
| Isl1 | 0.024355276 | 0.03071563 | 0.066685429 | 0.869171361 | 0.814207193 |
| Isoc1 | 0.091964476 | 2.16E-21 | 6.72E-20 | 0.506847717 | 0.394995884 |
| Isoc2b | 0.001198489 | 0.879178855 | 0.917691307 | 0.237639392 | 0.222613201 |
| Ist1 | 0.008534435 | 0.242896654 | 0.356665305 | 0.17905235 | 0.167691425 |
| Isy1 | 0.010620098 | 0.094370703 | 0.168312599 | 0.122096135 | 0.115889 |
| Isyna1 | -0.003295633 | 0.508856824 | 0.62282589 | 0.068605377 | 0.068102217 |
| Itch | 0.019363353 | 0.018050777 | 0.042708605 | 0.246807085 | 0.22882465 |
| Itfg1 | -0.027518866 | 0.003011072 | 0.009319847 | 0.368317243 | 0.385977988 |
| Itfg2 | -0.005982829 | 0.340725515 | 0.463226605 | 0.117391203 | 0.118132096 |
| Itga2 | 0.012951995 | 0.000578021 | 0.002209798 | 0.045122998 | 0.030713555 |
| Itga3 | 0.003619458 | 0.446871088 | 0.56576126 | 0.064717244 | 0.061097032 |
| Itga6 | -0.040026472 | 3.15E-06 | 2.01E-05 | 0.199700689 | 0.24521861 |
| Itgav | 0.014758254 | 0.144808923 | 0.238413608 | 0.544267413 | 0.535807663 |
| Itgb1 | 0.056536217 | 1.44E-08 | 1.39E-07 | 0.552172554 | 0.51211835 |
| Itgb1bp1 | -0.007331408 | 0.342451007 | 0.464735061 | 0.200450796 | 0.202379725 |
| Itgb3bp | -0.023276291 | 0.006627544 | 0.018367441 | 0.271798037 | 0.282165585 |
| Itgb5 | -0.002644043 | 0.72326558 | 0.800420062 | 0.176106645 | 0.1811519 |
| Itgb8 | 0.04514358 | 5.94E-13 | 9.79E-12 | 0.13034075 | 0.093690227 |
| Itm2b | -0.072978136 | 3.16E-32 | 1.82E-30 | 2.372892463 | 2.504181096 |
| Itm2c | -0.05475081 | 9.44E-08 | 8.03E-07 | 0.840264003 | 0.89459201 |
| Itpa | -0.029391363 | 0.000242834 | 0.001019796 | 0.207403338 | 0.229712731 |
| Itpk1 | 0.013412574 | 0.005639717 | 0.015976065 | 0.071356782 | 0.056248906 |
| Itpkb | 0.065240426 | 7.01E-13 | 1.14E-11 | 0.362108228 | 0.275786881 |
| Itpr1 | 0.048168341 | 3.57E-13 | 6.08E-12 | 0.157862167 | 0.100821099 |
| Itpr2 | 0.005228313 | 0.422263248 | 0.541089139 | 0.123715333 | 0.113208163 |
| Itpr3 | 0.021109884 | 0.001204403 | 0.004205397 | 0.138295768 | 0.116749865 |
| Itprid2 | 0.04250827 | 2.15E-07 | 1.71E-06 | 0.255367556 | 0.218518279 |
| Itsn1 | 0.007050975 | 0.097150877 | 0.17232272 | 0.051127572 | 0.045627632 |
| Itsn2 | 0.036742619 | 1.32E-06 | 9.00E-06 | 0.204796118 | 0.169405847 |
| Ivd | 0.013058594 | 0.128998809 | 0.217089566 | 0.278403605 | 0.259899148 |
| Ivns1abp | -0.026952011 | 0.005721459 | 0.016169776 | 0.371248579 | 0.416007977 |
| Iws1 | 0.025365404 | 0.00205612 | 0.006685483 | 0.249195248 | 0.219783308 |
| Izumo4 | -0.009739575 | 0.161699829 | 0.259156897 | 0.146603443 | 0.152013546 |
| Jade1 | 0.024534309 | 0.021781055 | 0.049930538 | 0.664185612 | 0.6568248 |
| Jade2 | 0.013204885 | 0.036833988 | 0.077360128 | 0.13110274 | 0.110062735 |
| Jade3 | 0.034136494 | 2.63E-07 | 2.05E-06 | 0.15029028 | 0.116626262 |
| Jag2 | 0.010491248 | 0.062422118 | 0.120574473 | 0.098646252 | 0.086252526 |
| Jagn1 | -0.048995325 | 4.25E-07 | 3.22E-06 | 0.404231263 | 0.446899387 |
| Jak1 | 0.055524237 | 3.43E-08 | 3.12E-07 | 0.688212745 | 0.605864953 |
| Jak2 | 0.015192067 | 0.020637687 | 0.047725879 | 0.128816501 | 0.119282648 |
| Jakmip1 | -0.099412418 | 1.57E-40 | 1.36E-38 | 0.135169406 | 0.203756323 |
| Jam2 | -0.059334498 | 5.11E-18 | 1.26E-16 | 0.094958063 | 0.147301143 |
| Jam3 | -0.002966562 | 0.539271367 | 0.649906561 | 0.06679739 | 0.065356949 |
| Jarid2 | 0.011681045 | 0.07562963 | 0.141181117 | 0.134128261 | 0.121583815 |
| Jazf1 | 0.005745578 | 0.150801761 | 0.245865289 | 0.04613513 | 0.040356952 |
| Jkamp | 0.014708675 | 0.080942059 | 0.148727783 | 0.276128184 | 0.252262129 |
| Jmjd1c | 0.009479341 | 0.32162789 | 0.44332188 | 0.367052575 | 0.363155915 |
| Jmjd6 | -0.016116764 | 0.033462608 | 0.071578321 | 0.17972666 | 0.190267858 |
| Jmjd8 | 0.002645361 | 0.692801313 | 0.777410061 | 0.138945297 | 0.133652467 |
| Jmy | 0.002031829 | 0.782923864 | 0.847898608 | 0.179782415 | 0.171029517 |

| | | | | | |
|---|---|---|---|---|---|
| Josd1 | 0.026661161 | 0.000105795 | 0.00049142 | 0.153061703 | 0.13545184 |
| Josd2 | -0.01474056 | 0.063776292 | 0.122633131 | 0.218187032 | 0.220661701 |
| Jph3 | 0.003769116 | 0.455242191 | 0.573888166 | 0.072763836 | 0.068261308 |
| Jpt1 | -0.041066831 | 5.64E-08 | 4.96E-07 | 0.159456375 | 0.195035066 |
| Jpt2 | -0.008373198 | 0.205609331 | 0.31390294 | 0.12803967 | 0.134678493 |
| Jpx | 0.003693378 | 0.295590363 | 0.41549132 | 0.036839005 | 0.031237715 |
| Jrkl | -0.00492459 | 0.262228892 | 0.378303796 | 0.051806049 | 0.052890717 |
| Jtb | -0.064479419 | 2.06E-10 | 2.53E-09 | 0.726347381 | 0.796879172 |
| Jun | -0.134897808 | 6.19E-21 | 1.88E-19 | 0.71083471 | 1.048304278 |
| Junb | 0.004873618 | 0.724025681 | 0.80090399 | 1.015871161 | 1.018807164 |
| Jund | -0.007657057 | 0.342235792 | 0.464630131 | 1.795539344 | 1.815691883 |
| Jup | 0.035884994 | 4.64E-08 | 4.15E-07 | 0.143269309 | 0.109995422 |
| Kalrn | 0.01114406 | 0.017609043 | 0.041887722 | 0.069062283 | 0.054217934 |
| Kank1 | 0.015774742 | 0.000164097 | 0.000722861 | 0.053366162 | 0.04233841 |
| Kansl1 | 0.028339811 | 0.000658985 | 0.002478395 | 0.261796365 | 0.237860648 |
| Kansl1l | 0.006239603 | 0.222250275 | 0.333444008 | 0.0773937 | 0.068386239 |
| Kansl2 | 0.001043325 | 0.880213131 | 0.917982413 | 0.147200161 | 0.144769958 |
| Kansl3 | 0.007555293 | 0.252700352 | 0.367612273 | 0.13834016 | 0.1277004 |
| Kantr | 0.014297272 | 0.019131219 | 0.044742293 | 0.117507108 | 0.101633636 |
| Kars | 0.025746111 | 0.001231085 | 0.004285216 | 0.234527524 | 0.211188966 |
| Kat14 | 0.0339035 | 1.86E-07 | 1.50E-06 | 0.147472595 | 0.114085925 |
| Kat2a | -0.006141005 | 0.198623018 | 0.305395062 | 0.061960648 | 0.065153667 |
| Kat2b | 0.016400636 | 0.024843089 | 0.055961609 | 0.176837083 | 0.161196627 |
| Kat5 | 0.004036364 | 0.521928145 | 0.634193904 | 0.122464449 | 0.117043868 |
| Kat6a | 0.046736786 | 1.09E-08 | 1.08E-07 | 0.248360801 | 0.199899134 |
| Kat6b | -0.004291967 | 0.583511546 | 0.6891261 | 0.194330104 | 0.194870165 |
| Kat7 | 0.027953643 | 2.71E-05 | 0.00014187 | 0.148696745 | 0.12462729 |
| Kat8 | -0.027043838 | 1.39E-06 | 9.46E-06 | 0.078157952 | 0.09597067 |
| Katna1 | 0.004578556 | 0.564622442 | 0.672223977 | 0.224696289 | 0.214625338 |
| Katnal1 | 0.00526654 | 0.281111122 | 0.400023759 | 0.06968549 | 0.064298462 |
| Katnb1 | -0.007144719 | 0.087381055 | 0.15840362 | 0.044923555 | 0.048663951 |
| Katnbl1 | 0.002092225 | 0.699742606 | 0.782751998 | 0.083304305 | 0.082995706 |
| Katnip | 0.001443049 | 0.822971887 | 0.878074806 | 0.119357577 | 0.11857278 |
| Kazald1 | -0.025629938 | 2.73E-10 | 3.32E-09 | 0.034897404 | 0.050468496 |
| Kazn | -0.000387451 | 0.926501658 | 0.950353357 | 0.048789168 | 0.047475442 |
| Kbtbd2 | 0.007143996 | 0.310872115 | 0.432374636 | 0.158008843 | 0.151296528 |
| Kbtbd3 | 0.007978914 | 0.209090286 | 0.317919263 | 0.13175999 | 0.117949328 |
| Kbtbd4 | 0.008614891 | 0.091787789 | 0.164581499 | 0.078206674 | 0.070888103 |
| Kbtbd7 | -0.007362678 | 0.221494098 | 0.332412105 | 0.097645744 | 0.105284951 |
| Kcmf1 | 0.023295611 | 0.018215747 | 0.04301518 | 0.489255802 | 0.457661649 |
| Kcna2 | -0.000703114 | 0.867620641 | 0.909630958 | 0.042948711 | 0.044852033 |
| Kcnab3 | 0.011454141 | 0.018127211 | 0.042879013 | 0.073558356 | 0.058777661 |
| Kcnb1 | 0.037182015 | 3.94E-05 | 0.000198375 | 0.351693921 | 0.307261978 |
| Kcnb2 | 0.011077021 | 0.08920453 | 0.160929011 | 0.11961305 | 0.115424468 |
| Kcnc3 | -0.014745106 | 0.018530709 | 0.0436212 | 0.103371544 | 0.115624673 |
| Kcnh1 | -0.02880329 | 4.35E-05 | 0.000216638 | 0.109637097 | 0.144874957 |
| Kcnh2 | -0.005931165 | 0.46171208 | 0.579507788 | 0.225511494 | 0.222643073 |
| Kcnh6 | -0.0016081 | 0.682261888 | 0.769199066 | 0.040647721 | 0.040861 |
| Kcnh8 | 0.015213119 | 2.63E-05 | 0.000138235 | 0.041492788 | 0.029738455 |
| Kcnip1 | -0.03766283 | 0.000202447 | 0.000867427 | 0.507808768 | 0.544123846 |
| Kcnip4 | -0.003461559 | 0.383675026 | 0.504816431 | 0.038579711 | 0.042209458 |
| Kcnj11 | 0.012088545 | 0.15306989 | 0.248145124 | 0.260368726 | 0.236975869 |
| Kcnj12 | 0.028999434 | 4.74E-11 | 6.31E-10 | 0.068930459 | 0.041967841 |
| Kcnk1 | 0.00862597 | 0.28246872 | 0.401485512 | 0.216000452 | 0.213565697 |
| Kcnk12 | 0.00088554 | 0.894873767 | 0.928392142 | 0.142834442 | 0.136136245 |
| Kcnk16 | 0.113886706 | 2.15E-34 | 1.34E-32 | 0.396782937 | 0.271156328 |
| Kcnma1 | 0.001442747 | 0.880111515 | 0.917974912 | 0.36340194 | 0.361776004 |
| Kcnmb2 | 0.066340138 | 4.08E-11 | 5.49E-10 | 0.517739496 | 0.421434479 |
| Kcnn3 | 0.013476641 | 0.035358427 | 0.074843158 | 0.121171617 | 0.110729421 |
| Kcnq1ot1 | -0.058266087 | 9.55E-06 | 5.51E-05 | 0.747664541 | 0.823298174 |
| Kctd10 | 0.007100973 | 0.11343872 | 0.195437043 | 0.059173424 | 0.052799612 |
| Kctd11 | 0.018256725 | 0.00018009 | 0.000782659 | 0.079565348 | 0.059741074 |
| Kctd12 | 0.016579373 | 0.00764012 | 0.020718897 | 0.116127338 | 0.101550959 |
| Kctd12b | 0.029381681 | 7.77E-07 | 5.56E-06 | 0.118943063 | 0.080233367 |
| Kctd13 | -0.021448149 | 0.009343212 | 0.024483749 | 0.25039186 | 0.255723048 |
| Kctd15 | 0.000182225 | 0.971953968 | 0.980933923 | 0.072503723 | 0.073097195 |
| Kctd18 | 2.59E-05 | 0.995443044 | 0.996700732 | 0.055664568 | 0.055350774 |
| Kctd2 | -0.007869977 | 0.289847172 | 0.409244019 | 0.185182877 | 0.18284886 |
| Kctd20 | -0.011232276 | 0.205938041 | 0.314207413 | 0.289360691 | 0.290732483 |
| Kctd3 | 0.00885238 | 0.07587908 | 0.14156523 | 0.076577736 | 0.06664988 |
| Kctd5 | -0.001718573 | 0.70772043 | 0.789133848 | 0.056854601 | 0.0583933 |
| Kctd6 | 0.014168721 | 0.014246308 | 0.03495775 | 0.112594795 | 0.093531608 |

| | | | | | |
|---|---|---|---|---|---|
| Kctd8 | 0.02626305 | 3.44E-09 | 3.65E-08 | 0.068137785 | 0.042695108 |
| Kctd9 | -0.003845309 | 0.391276688 | 0.512075844 | 0.053522688 | 0.055006313 |
| Kdelr1 | 0.040266593 | 3.30E-05 | 0.00016996 | 0.773973905 | 0.704871439 |
| Kdelr2 | -0.014907503 | 0.14579098 | 0.239624667 | 0.87605173 | 0.86532258 |
| Kdelr3 | -0.01212327 | 0.012134098 | 0.030545755 | 0.054582134 | 0.063816448 |
| Kdf1 | -0.006644442 | 0.229641899 | 0.34257765 | 0.090067372 | 0.090132585 |
| Kdm1a | 0.052496071 | 3.66E-07 | 2.81E-06 | 0.684965016 | 0.621795014 |
| Kdm1b | -0.002750557 | 0.487496021 | 0.603211953 | 0.040176253 | 0.041944107 |
| Kdm2a | 0.030306245 | 0.001379626 | 0.004737803 | 0.391696941 | 0.356856422 |
| Kdm2b | -0.039176786 | 0.00019006 | 0.000820863 | 0.422976537 | 0.440589481 |
| Kdm3a | 0.007442834 | 0.160617404 | 0.257762035 | 0.081210823 | 0.072749125 |
| Kdm3b | 0.007425212 | 0.205093752 | 0.313381557 | 0.101982367 | 0.091517328 |
| Kdm4a | -0.002452261 | 0.748409853 | 0.820667786 | 0.188281044 | 0.190394476 |
| Kdm4b | 0.011467255 | 0.013612166 | 0.033639421 | 0.068653271 | 0.056320756 |
| Kdm4c | 0.017252916 | 0.011831278 | 0.029907354 | 0.149126799 | 0.135937287 |
| Kdm5a | 0.017330242 | 0.05203616 | 0.103136759 | 0.312476488 | 0.28761657 |
| Kdm5b | -0.009242198 | 0.328957402 | 0.451584844 | 0.398338336 | 0.395130047 |
| Kdm5c | 0.029192891 | 0.000178168 | 0.000777445 | 0.220212073 | 0.192198563 |
| Kdm5d | -0.008361959 | 0.135246596 | 0.225999898 | 0.084501359 | 0.08855756 |
| Kdm6a | 0.010219466 | 0.146226953 | 0.240219405 | 0.155133796 | 0.142635243 |
| Kdm6b | 0.033891456 | 0.000957791 | 0.00344748 | 0.514261566 | 0.479171691 |
| Kdm7a | -0.015005897 | 0.092990675 | 0.166339529 | 0.270567834 | 0.288882676 |
| Kdsr | -0.011108829 | 0.044671397 | 0.090799775 | 0.08127803 | 0.088892568 |
| Keap1 | 0.012472095 | 0.195045841 | 0.300734677 | 0.466901091 | 0.449069369 |
| Khdc4 | -0.002984991 | 0.68537162 | 0.77195284 | 0.165799298 | 0.167799281 |
| Khdrbs1 | 0.021384638 | 0.021128589 | 0.048698942 | 0.349083262 | 0.326719824 |
| Khdrbs3 | 0.023316473 | 0.017056067 | 0.040751803 | 0.565755071 | 0.512616312 |
| Khk | -0.007205857 | 0.296079427 | 0.415905822 | 0.16025089 | 0.147308969 |
| Khnyn | 0.008240087 | 0.166300845 | 0.2646425 | 0.111169165 | 0.099526169 |
| Khsrp | 0.010836961 | 0.117244476 | 0.200605895 | 0.154447925 | 0.139714739 |
| Kidins220 | 0.034183645 | 0.00051339 | 0.001988518 | 0.496059167 | 0.466446461 |
| Kif12 | -0.031397769 | 0.000735874 | 0.002734774 | 0.357087864 | 0.357426563 |
| Kif13a | -0.000387764 | 0.927100428 | 0.950566374 | 0.046762746 | 0.046549517 |
| Kif13b | 0.031890155 | 0.000107738 | 0.000499726 | 0.256428349 | 0.223134289 |
| Kif16b | -0.007475743 | 0.214070171 | 0.323870247 | 0.100328916 | 0.103898201 |
| Kif19a | 0.007306414 | 0.118268493 | 0.201970043 | 0.063281572 | 0.056608666 |
| Kif1a | 0.022522335 | 0.025943022 | 0.05785487 | 0.620466012 | 0.587018887 |
| Kif1b | 0.073141774 | 1.93E-12 | 3.03E-11 | 0.885473646 | 0.76717992 |
| Kif1c | 0.020561523 | 6.23E-05 | 0.000302437 | 0.087654473 | 0.067108795 |
| Kif21a | 0.111466927 | 8.04E-28 | 3.78E-26 | 0.942194701 | 0.818824467 |
| Kif2a | -0.008827088 | 0.299396105 | 0.419232167 | 0.268612508 | 0.267902888 |
| Kif3a | -0.023114205 | 0.001057219 | 0.003760805 | 0.14158192 | 0.159442226 |
| Kif3b | 0.022987485 | 0.001176117 | 0.004123409 | 0.171218392 | 0.148472673 |
| Kif5b | -0.033391478 | 5.79E-06 | 3.51E-05 | 1.910952083 | 2.00675496 |
| Kif5c | 0.011064942 | 0.248966812 | 0.363377173 | 0.352082194 | 0.345644182 |
| Kifap3 | 0.023068012 | 0.004035594 | 0.012005523 | 0.234845897 | 0.218339727 |
| Kifbp | -0.000587042 | 0.94147065 | 0.96028615 | 0.226844128 | 0.217068552 |
| Kifc3 | 0.012528784 | 0.018868387 | 0.044255359 | 0.089119289 | 0.075396047 |
| Kin | 0.001452445 | 0.872287267 | 0.91235899 | 0.300717925 | 0.307889428 |
| Kiz | -0.020918874 | 0.001558176 | 0.005241726 | 0.135110276 | 0.14128215 |
| Kl | -0.018246741 | 0.001110525 | 0.003920307 | 0.074804734 | 0.089493244 |
| Klc1 | 0.001590981 | 0.83662786 | 0.888187078 | 0.185524184 | 0.189625238 |
| Klc2 | -0.001721352 | 0.791002081 | 0.853690131 | 0.130959675 | 0.12542684 |
| Klc3 | 0.000466579 | 0.935962994 | 0.956626601 | 0.100504738 | 0.098213066 |
| Klc4 | 0.000749277 | 0.871889094 | 0.912080018 | 0.061147843 | 0.058635335 |
| Klf10 | 0.009698297 | 0.17710268 | 0.278517027 | 0.153398862 | 0.142949061 |
| Klf11 | 0.00385411 | 0.455287709 | 0.573888166 | 0.079179853 | 0.07212729 |
| Klf13 | 0.052502186 | 9.78E-09 | 9.76E-08 | 0.366231019 | 0.314919211 |
| Klf16 | 0.00209981 | 0.563931329 | 0.671715313 | 0.03833961 | 0.033842733 |
| Klf3 | 0.005286386 | 0.472136228 | 0.589382179 | 0.170417346 | 0.163814983 |
| Klf4 | -0.018319763 | 0.003606208 | 0.010908386 | 0.065875847 | 0.10337325 |
| Klf5 | 0.049499755 | 1.39E-11 | 1.97E-10 | 0.18375564 | 0.140242622 |
| Klf6 | -0.005642244 | 0.374454959 | 0.49648874 | 0.098171666 | 0.107268388 |
| Klf7 | 0.004606111 | 0.589678205 | 0.694386083 | 0.246778887 | 0.239256097 |
| Klf9 | 0.048262884 | 4.24E-06 | 2.65E-05 | 0.76063799 | 0.68784664 |
| Klhdc1 | 0.007727432 | 0.261636778 | 0.377561638 | 0.148812262 | 0.143044192 |
| Klhdc10 | 0.000874893 | 0.914432148 | 0.941138874 | 0.239220282 | 0.236459547 |
| Klhdc2 | -0.045518375 | 4.09E-06 | 2.56E-05 | 0.504491356 | 0.537188643 |
| Klhdc3 | 0.012627872 | 0.032970434 | 0.070649891 | 0.107848566 | 0.096592134 |
| Klhdc4 | -0.002587939 | 0.620325877 | 0.719745187 | 0.074256615 | 0.076956455 |
| Klhdc8b | -0.000232028 | 0.959738461 | 0.972238962 | 0.057689726 | 0.055488643 |
| Klhl11 | 0.003057286 | 0.489844975 | 0.605271079 | 0.05172668 | 0.051262162 |

| | | | | | |
|---|---|---|---|---|---|
| Klhl12 | -0.007947568 | 0.144587387 | 0.238169871 | 0.08017068 | 0.084715884 |
| Klhl15 | 0.004569003 | 0.370522418 | 0.49291447 | 0.078341455 | 0.071402021 |
| Klhl2 | -0.00172395 | 0.765289263 | 0.833720552 | 0.095754626 | 0.093659395 |
| Klhl20 | 0.010816902 | 0.007072082 | 0.01939486 | 0.0477553 | 0.03940059 |
| Klhl21 | 0.002024657 | 0.724307559 | 0.800998625 | 0.09636906 | 0.093477467 |
| Klhl22 | 0.001705686 | 0.819194163 | 0.875476053 | 0.176295557 | 0.171396013 |
| Klhl24 | 0.00082194 | 0.921451838 | 0.946271762 | 0.241815621 | 0.231771067 |
| Klhl26 | 0.001542324 | 0.73869224 | 0.812946112 | 0.062089007 | 0.05791789 |
| Klhl28 | 0.014399607 | 0.010454321 | 0.026866138 | 0.096284006 | 0.083720276 |
| Klhl32 | 0.047804875 | 3.24E-07 | 2.51E-06 | 0.417850749 | 0.346865405 |
| Klhl36 | 0.008457423 | 0.068524046 | 0.130166232 | 0.063067893 | 0.056340508 |
| Klhl42 | 0.011373011 | 0.031316537 | 0.067717926 | 0.087897875 | 0.07423135 |
| Klhl7 | 0.007760681 | 0.376239123 | 0.497862632 | 0.318146094 | 0.281204069 |
| Klhl8 | -0.010341807 | 0.054877962 | 0.10793517 | 0.08163206 | 0.084687231 |
| Klhl9 | 0.024750629 | 0.004356007 | 0.012840843 | 0.3084614 | 0.27002731 |
| Kmt2a | 0.039260764 | 0.000248413 | 0.001039633 | 0.671764258 | 0.612878483 |
| Kmt2b | 0.017491822 | 0.01501887 | 0.036585681 | 0.170915786 | 0.152941152 |
| Kmt2c | 0.02661849 | 0.008148111 | 0.021864736 | 0.504261147 | 0.471058459 |
| Kmt2d | 0.017430539 | 0.025707446 | 0.057415067 | 0.217637247 | 0.197733552 |
| Kmt2e | 0.005513769 | 0.594696065 | 0.6985182 | 0.692628441 | 0.667816243 |
| Kmt5a | 0.06565826 | 3.19E-14 | 5.92E-13 | 0.294506328 | 0.235316667 |
| Kmt5b | -0.017137392 | 0.043973766 | 0.089700577 | 0.246273198 | 0.25787054 |
| Kmt5c | 0.003872539 | 0.366502445 | 0.489240254 | 0.054207034 | 0.047151337 |
| Knop1 | 0.018438767 | 0.033537506 | 0.071643954 | 0.274987774 | 0.266553377 |
| Kpna1 | 0.040489457 | 6.77E-07 | 4.91E-06 | 0.241865853 | 0.21264544 |
| Kpna2 | 0.019273644 | 0.000259484 | 0.00108224 | 0.0920933 | 0.072372366 |
| Kpna3 | 0.023365298 | 0.00405768 | 0.012059879 | 0.25784359 | 0.220697337 |
| Kpna4 | 0.028705356 | 0.000622927 | 0.002355539 | 0.266274064 | 0.241304523 |
| Kpna6 | 0.023853353 | 0.000310084 | 0.001267721 | 0.149850879 | 0.127813689 |
| Kpnb1 | 0.023583577 | 0.00779528 | 0.021063288 | 0.318584655 | 0.295202259 |
| Kptn | -0.013843642 | 0.001613662 | 0.005396637 | 0.047586331 | 0.056068654 |
| Kras | -0.013757211 | 0.172169537 | 0.271973451 | 0.876952501 | 0.902879207 |
| Krba1 | 0.004605628 | 0.268209519 | 0.384875711 | 0.051248218 | 0.044543622 |
| Krcc1 | -0.015979581 | 0.119744506 | 0.203993043 | 0.585528944 | 0.592125226 |
| Kremen1 | 0.010839254 | 0.003748234 | 0.011271368 | 0.043708379 | 0.033674589 |
| Kri1 | 0.006364255 | 0.193207718 | 0.298342668 | 0.06962783 | 0.06179613 |
| Krit1 | -0.015669766 | 0.082570642 | 0.151217317 | 0.303267899 | 0.314832773 |
| Krr1 | 0.012243851 | 0.091443973 | 0.164176973 | 0.177827578 | 0.160685366 |
| Krt10 | 0.00531515 | 0.523256937 | 0.635332077 | 0.251911714 | 0.242612046 |
| Krt18 | -0.075118781 | 1.21E-13 | 2.17E-12 | 1.115909042 | 1.221330636 |
| Krt222 | -0.005293408 | 0.241421123 | 0.355497738 | 0.050730821 | 0.056772065 |
| Krt7 | -0.139925345 | 9.36E-56 | 1.44E-53 | 1.525792266 | 1.687757986 |
| Krt8 | -0.018906037 | 0.08415175 | 0.153667039 | 0.716249201 | 0.731327343 |
| Krtcap2 | -0.11220752 | 1.76E-38 | 1.36E-36 | 1.788708538 | 2.114141124 |
| Krtcap3 | -0.040187322 | 4.56E-07 | 3.43E-06 | 0.186219205 | 0.225210399 |
| Ksr2 | 0.003806978 | 0.64856746 | 0.741895852 | 0.235499905 | 0.232777551 |
| Kti12 | -0.010465591 | 0.071188338 | 0.134087725 | 0.093518323 | 0.09674899 |
| Ktn1 | 0.087092865 | 7.57E-17 | 1.67E-15 | 0.82575431 | 0.744477701 |
| Kxd1 | -0.004071728 | 0.615046086 | 0.715077499 | 0.249593708 | 0.238255503 |
| Ky | 0.021576935 | 0.002309709 | 0.007408443 | 0.165728015 | 0.138142032 |
| Kyat1 | -0.00951726 | 0.043154198 | 0.088276241 | 0.056220213 | 0.062970001 |
| Kyat3 | -0.004941834 | 0.209086132 | 0.317919263 | 0.03643227 | 0.04175665 |
| L2hgdh | -0.002418965 | 0.597160168 | 0.700566086 | 0.054004524 | 0.057374479 |
| L3mbtl2 | 0.013110587 | 0.013503772 | 0.033433387 | 0.086508533 | 0.074188387 |
| Lacc1 | 0.004159612 | 0.327675163 | 0.450269673 | 0.049067649 | 0.045598734 |
| Lactb | -0.006617339 | 0.31309934 | 0.434415839 | 0.12562052 | 0.130377021 |
| Lactb2 | 0.005687511 | 0.495950528 | 0.611029151 | 0.24211817 | 0.239861847 |
| Lage3 | -0.003678381 | 0.660889764 | 0.752097657 | 0.268277113 | 0.265472988 |
| Lamc1 | 0.006223164 | 0.138878675 | 0.230442841 | 0.050090824 | 0.045427771 |
| Lamc2 | 0.006093198 | 0.181030883 | 0.283182983 | 0.055475671 | 0.054418037 |
| Lamp1 | 0.003600438 | 0.665631984 | 0.755636869 | 1.306334108 | 1.279688397 |
| Lamp2 | -0.030116079 | 0.003049264 | 0.009420065 | 0.780684688 | 0.837262119 |
| Lamtor1 | -0.0237084 | 0.016565415 | 0.039774997 | 0.543712649 | 0.561234293 |
| Lamtor2 | -0.014203783 | 0.119013936 | 0.202991839 | 0.982837809 | 1.022425928 |
| Lamtor3 | -0.004926369 | 0.602064346 | 0.705043312 | 0.383606974 | 0.379773247 |
| Lamtor4 | 0.004586996 | 0.641638779 | 0.736088758 | 0.75205398 | 0.739760822 |
| Lamtor5 | -0.006899865 | 0.467593672 | 0.585063794 | 0.45015761 | 0.449909188 |
| Lancl1 | -0.032527453 | 1.16E-05 | 6.59E-05 | 0.173265332 | 0.189373376 |
| Lancl2 | 0.005957239 | 0.169659072 | 0.268799085 | 0.054317415 | 0.049212657 |
| Lap3 | -0.015806837 | 0.036040192 | 0.076037923 | 0.180559353 | 0.188731925 |
| Laptm4a | 0.03752325 | 1.02E-05 | 5.81E-05 | 1.262263858 | 1.208480234 |
| Laptm4b | 0.027156144 | 0.000807263 | 0.002964946 | 0.250064072 | 0.221166184 |

| | | | | | |
|---|---|---|---|---|---|
| Large2 | 0.00454081 | 0.296493668 | 0.416307256 | 0.052357387 | 0.049472496 |
| Larp1 | 0.069156129 | 3.98E-12 | 5.95E-11 | 0.549036338 | 0.465814122 |
| Larp1b | 0.039305949 | 2.40E-05 | 0.000127292 | 0.38917152 | 0.360688635 |
| Larp4 | 0.020784743 | 0.031593634 | 0.06825629 | 0.429558667 | 0.412582056 |
| Larp4b | 0.050010038 | 5.50E-07 | 4.09E-06 | 0.54410254 | 0.492236155 |
| Larp7 | 0.007554708 | 0.202171201 | 0.309626405 | 0.107228713 | 0.095925171 |
| Lars | -0.038517268 | 0.00034465 | 0.001395537 | 0.844350319 | 0.922105335 |
| Lars2 | 0.013238874 | 0.170544569 | 0.269862353 | 0.334921969 | 0.310402187 |
| Las1l | -0.027266958 | 0.000583849 | 0.002228576 | 0.192357251 | 0.217886786 |
| Lasp1 | 0.024837746 | 0.002977687 | 0.009240047 | 0.273596835 | 0.247724123 |
| Lats1 | -0.001537769 | 0.85136716 | 0.898306005 | 0.237390874 | 0.238548056 |
| Lbh | 0.039110409 | 0.000210112 | 0.000897499 | 0.52249093 | 0.502614281 |
| Lbr | 0.033541095 | 3.96E-08 | 3.56E-07 | 0.128417591 | 0.097770865 |
| Lclat1 | 0.00593433 | 0.205106649 | 0.313381557 | 0.062183663 | 0.057828987 |
| Lcmt1 | -0.011955028 | 0.086025611 | 0.156398034 | 0.157984963 | 0.157396296 |
| Lcmt2 | 0.014505274 | 0.009100657 | 0.023964405 | 0.100560437 | 0.08502108 |
| Lcor | -0.005463108 | 0.408587779 | 0.528581555 | 0.126180258 | 0.130444278 |
| Lcorl | -0.000565721 | 0.953555422 | 0.968700712 | 0.384918584 | 0.389759893 |
| Ldah | 0.003087979 | 0.645232263 | 0.738863141 | 0.144535346 | 0.137401534 |
| Ldb1 | 0.03008763 | 9.60E-05 | 0.00044998 | 0.221407264 | 0.182726185 |
| Ldhb | 0.011961474 | 0.025080456 | 0.056377386 | 0.095449283 | 0.076732057 |
| Ldlr | 0.038987975 | 9.81E-06 | 5.64E-05 | 0.27970899 | 0.245815557 |
| Ldlrad4 | -0.008690085 | 0.080141554 | 0.147591625 | 0.064890866 | 0.068854894 |
| Ldlrap1 | 0.008881199 | 0.04186417 | 0.086138299 | 0.054514109 | 0.047615018 |
| Lefty1 | -0.001293672 | 0.830357418 | 0.88342469 | 0.099293661 | 0.0930407 |
| Lekr1 | 0.00354276 | 0.395641126 | 0.516235921 | 0.049544582 | 0.044925754 |
| Lemd2 | -0.001141186 | 0.886141154 | 0.921988902 | 0.215617814 | 0.214108566 |
| Lemd3 | 0.006538745 | 0.216872471 | 0.327207137 | 0.081389715 | 0.075490289 |
| Leng1 | 0.001689327 | 0.752940466 | 0.823959179 | 0.082093551 | 0.079849825 |
| Leng8 | -0.016139675 | 0.020755563 | 0.047952847 | 0.137889666 | 0.152667216 |
| Leng9 | 0.003088831 | 0.556333831 | 0.665192167 | 0.078852263 | 0.075241054 |
| Leo1 | -0.034530386 | 2.86E-05 | 0.000149474 | 0.204208695 | 0.23809285 |
| Leprot | 0.001979654 | 0.787338091 | 0.851256775 | 0.174606444 | 0.172956903 |
| Leprotl1 | -0.06598316 | 2.85E-09 | 3.06E-08 | 0.99571769 | 1.127506085 |
| Letm1 | 0.025781721 | 0.000960829 | 0.003457135 | 0.226253222 | 0.201263716 |
| Letm2 | 0.014008853 | 0.014584978 | 0.035689695 | 0.106700476 | 0.088493446 |
| Letmd1 | 0.007454109 | 0.101656218 | 0.179105066 | 0.061159255 | 0.053791207 |
| Lgals3bp | -0.014743227 | 0.159302694 | 0.256319544 | 0.859517719 | 0.900546869 |
| Lgals8 | -0.00504643 | 0.55454836 | 0.664037339 | 0.252559419 | 0.258385316 |
| Lgalsl | -0.045003784 | 3.96E-07 | 3.01E-06 | 0.249012242 | 0.279304369 |
| Lgi1 | 0.035104481 | 2.37E-07 | 1.87E-06 | 0.152192368 | 0.113086748 |
| Lgi2 | 0.006980174 | 0.337227156 | 0.45944503 | 0.146070684 | 0.139094802 |
| Lgmn | -0.025790769 | 4.33E-07 | 3.27E-06 | 0.061515002 | 0.080822048 |
| Lgr4 | 0.013937627 | 0.000648507 | 0.002443714 | 0.053604801 | 0.040029049 |
| Lhfpl4 | 0.017757129 | 0.014012742 | 0.03446291 | 0.183548666 | 0.159490099 |
| Lhfpl5 | -0.013972727 | 0.003262652 | 0.010006151 | 0.056716763 | 0.063803076 |
| Lhx1 | 0.004175612 | 0.518928492 | 0.631417121 | 0.115400078 | 0.111716033 |
| Lhx1os | -0.010617186 | 0.006764098 | 0.018692598 | 0.03557346 | 0.043393367 |
| Lias | 0.006432475 | 0.458913447 | 0.57740075 | 0.303111509 | 0.282321571 |
| Lifr | 0.074725309 | 4.80E-20 | 1.40E-18 | 0.271816065 | 0.192347901 |
| Lig1 | 0.002603734 | 0.536255537 | 0.647148672 | 0.046911045 | 0.04517055 |
| Lig3 | 0.002566296 | 0.662272299 | 0.752974795 | 0.096719124 | 0.095386251 |
| Lig4 | 0.012333984 | 0.005233544 | 0.015009001 | 0.058647114 | 0.04811108 |
| Lima1 | 0.006714174 | 0.308991044 | 0.430253596 | 0.125354507 | 0.125011326 |
| Limch1 | 0.007589308 | 0.287205073 | 0.406538818 | 0.148665988 | 0.141664986 |
| Limd1 | 0.026138561 | 0.00315252 | 0.00969592 | 0.308414858 | 0.276789763 |
| Limd2 | 0.048101346 | 1.48E-08 | 1.43E-07 | 0.301259793 | 0.234011465 |
| Limk2 | 0.008518932 | 0.115212642 | 0.197967003 | 0.087944994 | 0.079471212 |
| Lims1 | 0.000917283 | 0.888665417 | 0.924021945 | 0.138106772 | 0.130232353 |
| Lin37 | -0.003434432 | 0.573226985 | 0.680124538 | 0.108746024 | 0.106826925 |
| Lin52 | 0.001582107 | 0.687609596 | 0.773093616 | 0.040713242 | 0.039592064 |
| LIN54 | 0.018789636 | 0.000180168 | 0.000782659 | 0.077395421 | 0.062727726 |
| Lin7c | 0.017621381 | 0.054257597 | 0.106823077 | 0.38025759 | 0.35473341 |
| Lins1 | 0.000155991 | 0.968456462 | 0.978928439 | 0.041067914 | 0.040518141 |
| Lipa | 0.015526536 | 0.000472525 | 0.001844936 | 0.061277012 | 0.047872588 |
| Lipo3 | 0.001147296 | 0.83586958 | 0.887831756 | 0.090629507 | 0.084731711 |
| Litaf | -0.003922548 | 0.464464723 | 0.581929242 | 0.05864186 | 0.07156133 |
| Lix1l | 0.003959821 | 0.446422639 | 0.565488064 | 0.081164546 | 0.072453124 |
| Llgl1 | 0.000860053 | 0.821901204 | 0.877541103 | 0.038115044 | 0.037914854 |
| Llgl2 | 0.023885333 | 0.000398814 | 0.001585796 | 0.162033379 | 0.12945413 |
| Llph | 0.009302977 | 0.33695471 | 0.459288173 | 0.416111206 | 0.405933046 |
| Lman1 | -0.096814236 | 6.23E-29 | 3.12E-27 | 1.37328299 | 1.57366527 |

| | | | | | |
|---|---|---|---|---|---|
| Lman2 | -0.062631069 | 2.97E-10 | 3.61E-09 | 0.611123661 | 0.663746705 |
| Lman2l | -0.040316526 | 6.00E-08 | 5.26E-07 | 0.161571628 | 0.190700148 |
| Lmbr1 | -0.001731825 | 0.728685312 | 0.804574515 | 0.066652261 | 0.069532662 |
| Lmbr1l | -0.001528435 | 0.685425967 | 0.77195284 | 0.037155018 | 0.038276166 |
| Lmbrd1 | 0.011972819 | 0.167233782 | 0.265704662 | 0.281625978 | 0.274356296 |
| Lmbrd2 | -0.013465115 | 0.089364869 | 0.16112857 | 0.203450268 | 0.21544676 |
| Lmf1 | -0.018436038 | 0.002221641 | 0.007154289 | 0.095228561 | 0.110578205 |
| Lmf2 | -2.79E-05 | 0.995470491 | 0.996700732 | 0.068226875 | 0.06701399 |
| Lmna | -0.003900553 | 0.691533063 | 0.776248058 | 0.4217914 | 0.430376793 |
| Lmnb1 | 0.005307159 | 0.201883704 | 0.309429824 | 0.050084008 | 0.04466102 |
| Lmnb2 | 0.006969263 | 0.085634099 | 0.155847007 | 0.048087574 | 0.042329246 |
| Lmo1 | -0.04571861 | 1.41E-06 | 9.58E-06 | 0.256463592 | 0.321562755 |
| Lmo4 | -0.018895434 | 0.000465689 | 0.001822635 | 0.059861889 | 0.077056361 |
| Lmo7 | 0.054220198 | 1.29E-08 | 1.26E-07 | 0.3893938 | 0.332892772 |
| Lmtk2 | 0.0189793 | 0.002051061 | 0.006674331 | 0.120689622 | 0.104675558 |
| Lncpint | -0.031360295 | 5.45E-07 | 4.05E-06 | 0.084994933 | 0.114526732 |
| Lnpep | 0.011414932 | 0.214037323 | 0.323870247 | 0.352612536 | 0.334136821 |
| Lnpk | 0.001339782 | 0.835632557 | 0.88767698 | 0.119833441 | 0.12229334 |
| Lnx1 | 0.010322533 | 0.019279145 | 0.045023263 | 0.061187574 | 0.049042506 |
| Lonp1 | -0.084413445 | 2.25E-14 | 4.21E-13 | 0.426512376 | 0.533583421 |
| Lonp2 | 0.011019216 | 0.211279867 | 0.320646716 | 0.315055738 | 0.290311079 |
| Lonrf1 | 0.018709168 | 2.58E-05 | 0.000135968 | 0.060422523 | 0.046189418 |
| Lonrf2 | 0.010181104 | 0.01996975 | 0.046400058 | 0.056816875 | 0.047361814 |
| Loxl2 | 0.03256279 | 7.02E-08 | 6.09E-07 | 0.125718697 | 0.092032586 |
| Lpar6 | 0.006003218 | 0.138491515 | 0.229881254 | 0.045088667 | 0.040483978 |
| Lpcat1 | 0.006589046 | 0.226753727 | 0.339101635 | 0.090069381 | 0.080530846 |
| Lpcat3 | -0.003147709 | 0.697449098 | 0.780905233 | 0.23644858 | 0.230050332 |
| Lpgat1 | 0.090191831 | 1.22E-18 | 3.13E-17 | 0.504878727 | 0.41797075 |
| Lpin1 | 0.030708763 | 3.58E-06 | 2.26E-05 | 0.157567887 | 0.121659877 |
| Lpin2 | 0.026007592 | 1.43E-05 | 7.95E-05 | 0.115588725 | 0.091923765 |
| Lpl | -0.050181096 | 8.78E-06 | 5.11E-05 | 0.408328934 | 0.446682611 |
| Lpp | 0.02661957 | 0.011625771 | 0.02947985 | 0.633109856 | 0.586443449 |
| Lratd2 | 5.01E-05 | 0.994127772 | 0.996209693 | 0.133731066 | 0.138888027 |
| Lrba | -0.013432272 | 0.120087514 | 0.20446424 | 0.256393843 | 0.262418984 |
| Lrch1 | 0.015531792 | 0.003030605 | 0.009374337 | 0.088823059 | 0.070471415 |
| Lrch2 | 0.003766933 | 0.391839283 | 0.512506593 | 0.052294253 | 0.04914188 |
| Lrch3 | 0.00222285 | 0.702708506 | 0.785076085 | 0.098686422 | 0.095066177 |
| Lrfn3 | 0.030245344 | 0.00014775 | 0.000660424 | 0.242837174 | 0.204266564 |
| Lrfn4 | 0.023371478 | 3.63E-06 | 2.29E-05 | 0.085079981 | 0.061597749 |
| Lrif1 | 0.00900341 | 0.29696807 | 0.416732618 | 0.260131124 | 0.253342776 |
| Lrig2 | 0.013449282 | 0.005161666 | 0.014834145 | 0.072576045 | 0.059706459 |
| Lrp10 | 0.031455693 | 0.000325532 | 0.001324747 | 0.306175092 | 0.282328645 |
| Lrp11 | 0.036852037 | 0.000142587 | 0.000641772 | 0.55911023 | 0.504320444 |
| Lrp12 | -0.000706091 | 0.904246678 | 0.935126735 | 0.100745369 | 0.098204046 |
| Lrp3 | -0.001224161 | 0.84354027 | 0.893149477 | 0.116251419 | 0.112177569 |
| Lrp5 | 0.005822033 | 0.129778987 | 0.218138194 | 0.04273625 | 0.036599856 |
| Lrp6 | 0.041651615 | 1.23E-07 | 1.03E-06 | 0.232435657 | 0.191277349 |
| Lrp8 | 0.005798038 | 0.379096126 | 0.500706269 | 0.128417363 | 0.120558097 |
| Lrpap1 | -0.095407755 | 9.85E-23 | 3.48E-21 | 0.51468811 | 0.589696713 |
| Lrpprc | 0.029779123 | 0.00056163 | 0.002153062 | 0.266023969 | 0.240271645 |
| Lrrc1 | 0.032600965 | 1.61E-08 | 1.53E-07 | 0.116897003 | 0.084642752 |
| Lrrc10b | 0.100968638 | 2.05E-21 | 6.38E-20 | 0.627389088 | 0.522979625 |
| Lrrc14 | 0.0147307 | 0.01127871 | 0.028727173 | 0.102429034 | 0.093222082 |
| Lrrc20 | 0.006187511 | 0.238604152 | 0.352485878 | 0.078688015 | 0.075747765 |
| Lrrc24 | -0.000162178 | 0.975744707 | 0.983738468 | 0.079932841 | 0.076356792 |
| Lrrc27 | -0.000873308 | 0.807621634 | 0.86692034 | 0.035819778 | 0.03376216 |
| Lrrc40 | 0.012065757 | 0.019155506 | 0.04478832 | 0.083585718 | 0.07150111 |
| Lrrc41 | 0.042520553 | 5.38E-07 | 4.00E-06 | 0.288761013 | 0.249268766 |
| Lrrc42 | 0.003175446 | 0.631606131 | 0.728493867 | 0.144808327 | 0.133900194 |
| Lrrc45 | -0.009530393 | 0.034723395 | 0.07375592 | 0.052349004 | 0.058389504 |
| Lrrc47 | 0.028067539 | 8.50E-05 | 0.00040344 | 0.174987329 | 0.150366811 |
| Lrrc49 | 0.004742724 | 0.258960128 | 0.374930889 | 0.053530293 | 0.045258891 |
| Lrrc51 | -0.022952953 | 1.67E-06 | 1.12E-05 | 0.053420415 | 0.068972328 |
| Lrrc57 | -0.001169788 | 0.821938828 | 0.877541103 | 0.076695414 | 0.075620237 |
| Lrrc58 | 0.01007657 | 0.315218117 | 0.436752599 | 0.539894555 | 0.537679312 |
| Lrrc59 | 0.016571381 | 0.06248677 | 0.120630734 | 0.299604517 | 0.288363499 |
| Lrrc61 | 0.010169671 | 0.01671293 | 0.0400797 | 0.057187684 | 0.045436077 |
| Lrrc73 | -0.009585504 | 0.263785518 | 0.38004191 | 0.269915045 | 0.273882513 |
| Lrrc8a | -0.002213802 | 0.632530223 | 0.729098641 | 0.060696821 | 0.058261765 |
| Lrrc8d | 0.019663676 | 0.027458714 | 0.060671276 | 0.332686495 | 0.311254396 |
| Lrrcc1 | -0.01713168 | 0.007027354 | 0.019292423 | 0.110318268 | 0.121026264 |
| Lrrfip1 | 0.11162412 | 1.42E-132 | 1.73E-129 | 0.123655223 | 0.024231137 |

| | | | | | |
|---|---|---|---|---|---|
| Lrrfip2 | 0.064993057 | 1.64E-08 | 1.56E-07 | 0.781005124 | 0.723328693 |
| Lrrn1 | 0.011213947 | 0.22586929 | 0.337986953 | 0.347568365 | 0.3290859 |
| Lrrn2 | 0.009097894 | 0.058710968 | 0.114524083 | 0.06896472 | 0.058010166 |
| Lrwd1 | -0.002139542 | 0.589426951 | 0.694174296 | 0.042859777 | 0.042577608 |
| Lsg1 | 0.000629516 | 0.930019636 | 0.952465454 | 0.162998273 | 0.157580503 |
| Lsm1 | -0.000219499 | 0.979473817 | 0.986270787 | 0.283746542 | 0.27976436 |
| Lsm10 | 0.010730406 | 0.157555662 | 0.254064712 | 0.208487526 | 0.192036217 |
| Lsm12 | -0.02491322 | 0.009523336 | 0.024872149 | 0.419832071 | 0.43763767 |
| Lsm14a | -0.00228405 | 0.78635096 | 0.850567872 | 0.252584685 | 0.253663522 |
| Lsm14b | -0.009989018 | 0.217561392 | 0.328068688 | 0.222598269 | 0.224637372 |
| Lsm2 | -0.024332135 | 0.002657672 | 0.008364481 | 0.215170799 | 0.233261781 |
| Lsm3 | 0.008748188 | 0.24070009 | 0.354666001 | 0.196995728 | 0.177082149 |
| Lsm4 | -0.032317245 | 0.001430976 | 0.0048831 | 0.662595331 | 0.706843796 |
| Lsm5 | 0.015067856 | 0.069962527 | 0.132278429 | 0.254219391 | 0.245676357 |
| Lsm6 | 0.031084618 | 0.001383337 | 0.00474887 | 0.864450651 | 0.829781062 |
| Lsm7 | 0.020652964 | 0.014755762 | 0.036035046 | 0.281472509 | 0.266303343 |
| Lsm8 | -0.009319335 | 0.264944738 | 0.381260027 | 0.263161731 | 0.258898388 |
| Lsr | -0.004097054 | 0.611085834 | 0.711896017 | 0.235251845 | 0.232865316 |
| Lss | 0.008606558 | 0.062810015 | 0.12113449 | 0.061743911 | 0.055107433 |
| Lst1 | 0.00459978 | 0.230966883 | 0.344079073 | 0.038714395 | 0.036805243 |
| Lta4h | 0.008499077 | 0.180599212 | 0.282780727 | 0.126253685 | 0.116198405 |
| Ltbp3 | -0.02531122 | 0.000233601 | 0.000985277 | 0.135501813 | 0.151629489 |
| Ltbr | -0.012828688 | 0.023403822 | 0.05309964 | 0.083342703 | 0.094696922 |
| Ltn1 | 0.009112948 | 0.215773342 | 0.325939781 | 0.18089176 | 0.170503033 |
| LTO1 | 0.003654768 | 0.514159083 | 0.627341191 | 0.090568411 | 0.088751428 |
| Ltv1 | -0.020250276 | 0.004089486 | 0.012136136 | 0.142360366 | 0.157295106 |
| Luc7l | -0.03384094 | 0.000268464 | 0.001116341 | 0.322777958 | 0.353570811 |
| Luc7l2 | -0.05036081 | 1.24E-06 | 8.54E-06 | 0.970480026 | 1.030530155 |
| Luc7l3 | -0.003049985 | 0.765166525 | 0.833720552 | 0.53267249 | 0.538621298 |
| Lurap1l | 0.051012912 | 1.25E-12 | 2.00E-11 | 0.199894829 | 0.142951722 |
| Luzp1 | 0.038937184 | 2.45E-07 | 1.93E-06 | 0.206959555 | 0.169357286 |
| Lxn | -0.011853115 | 0.015880174 | 0.038347505 | 0.060864433 | 0.068936812 |
| Ly6e | 0.010647731 | 0.311671328 | 0.433114444 | 0.522932685 | 0.529478883 |
| Ly96 | -0.029862633 | 6.30E-05 | 0.000305383 | 0.150909871 | 0.181010456 |
| Lyar | -0.009645885 | 0.24148396 | 0.355497738 | 0.222998517 | 0.235861167 |
| Lypla1 | 0.004766383 | 0.487034907 | 0.602871577 | 0.152135923 | 0.144452003 |
| Lypla2 | 0.017885537 | 0.019545302 | 0.045513636 | 0.202897836 | 0.191311844 |
| Lyplal1 | 0.008117184 | 0.274747684 | 0.39280838 | 0.206680615 | 0.18170522 |
| Lyrm1 | -0.003170916 | 0.480060509 | 0.596440673 | 0.056862261 | 0.055687725 |
| Lyrm2 | -0.016041677 | 0.063979495 | 0.122950909 | 0.271928928 | 0.291607452 |
| Lyrm4 | -0.009415236 | 0.148359053 | 0.242737582 | 0.133294322 | 0.132137868 |
| Lyrm9 | -0.001252366 | 0.886053639 | 0.921988902 | 0.277393903 | 0.29048726 |
| Lysmd1 | -0.000145746 | 0.970038698 | 0.979916482 | 0.041816984 | 0.040056784 |
| Lysmd2 | 0.008923073 | 0.355064486 | 0.477446325 | 0.489709779 | 0.456669694 |
| Lysmd3 | 0.001243826 | 0.877109449 | 0.916022995 | 0.211654717 | 0.209973824 |
| Lysmd4 | 0.002817195 | 0.528953245 | 0.640408898 | 0.055261234 | 0.053191648 |
| Lyst | -0.000472303 | 0.95100337 | 0.967219873 | 0.179584943 | 0.182864664 |
| Lzic | 0.006211203 | 0.16580804 | 0.264101171 | 0.056143735 | 0.051448724 |
| Lztfl1 | -0.031025161 | 0.000528648 | 0.00203868 | 0.313033673 | 0.32780744 |
| Lztr1 | -0.056842586 | 4.08E-13 | 6.86E-12 | 0.167702161 | 0.219040028 |
| Lzts2 | 0.037060505 | 3.18E-07 | 2.46E-06 | 0.200267522 | 0.152808181 |
| M6pr | 0.001372141 | 0.882328842 | 0.919399893 | 0.364665736 | 0.36436882 |
| Macf1 | 0.021272259 | 0.049712533 | 0.09954794 | 0.852619346 | 0.82330778 |
| Macir | -0.004898678 | 0.510564322 | 0.624444123 | 0.168042421 | 0.172580621 |
| Maco1 | 0.017223479 | 0.082060913 | 0.150470802 | 0.510868194 | 0.49118058 |
| Macrod1 | -0.000238019 | 0.961202309 | 0.973316201 | 0.076327129 | 0.066943256 |
| Macrod2 | 0.01066129 | 0.146112493 | 0.240112519 | 0.184616344 | 0.165003412 |
| Macroh2a1 | 0.075784395 | 3.41E-15 | 6.72E-14 | 0.980437864 | 0.877119491 |
| Macroh2a2 | 0.013722095 | 0.073205037 | 0.137287687 | 0.193708098 | 0.182790965 |
| Mad2l1bp | -0.001671769 | 0.6470292 | 0.740397585 | 0.036786194 | 0.035401673 |
| Mad2l2 | 0.008447883 | 0.141581268 | 0.234011065 | 0.10425101 | 0.092626953 |
| Madd | -0.097257417 | 4.51E-21 | 1.38E-19 | 0.759376958 | 0.888766728 |
| Maea | 0.012912892 | 0.067058329 | 0.127931921 | 0.173364091 | 0.154040306 |
| Maf1 | 0.004764385 | 0.505504606 | 0.619635912 | 0.167931027 | 0.155439786 |
| Mafa | 0.258472594 | 1.25E-61 | 2.43E-59 | 2.148718574 | 1.593754017 |
| Mafb | 0.030844982 | 2.25E-08 | 2.09E-07 | 0.081350159 | 0.054637968 |
| Maff | -0.009140923 | 0.071359355 | 0.13433261 | 0.057571667 | 0.071215898 |
| Mafg | -0.03382228 | 0.000613116 | 0.002324772 | 0.432956273 | 0.469889908 |
| Mafk | 0.010400046 | 0.075454832 | 0.140935999 | 0.095728388 | 0.093255613 |
| Maged1 | 0.023515986 | 0.001864431 | 0.006142324 | 1.764019715 | 1.686764645 |
| Maged2 | -0.012099696 | 0.181241583 | 0.283375791 | 0.331881362 | 0.334134981 |
| Magee1 | -0.003564155 | 0.513051036 | 0.626840185 | 0.085776668 | 0.083526302 |

| | | | | | |
|---|---|---|---|---|---|
| Magef1 | -0.008923736 | 0.066276467 | 0.126838546 | 0.062810241 | 0.066623141 |
| Mageh1 | 0.001829915 | 0.854618251 | 0.899880714 | 0.515930876 | 0.501245634 |
| Magi1 | 0.019557219 | 1.25E-05 | 7.01E-05 | 0.065754807 | 0.047252399 |
| Magi2 | -0.014696707 | 0.059384694 | 0.115656023 | 0.187519785 | 0.201943636 |
| Magoh | -0.015320546 | 0.09666637 | 0.171747383 | 0.36037359 | 0.364486379 |
| Magohb | -0.003238334 | 0.618905081 | 0.718618798 | 0.135224439 | 0.131187379 |
| Magt1 | -0.069602738 | 5.05E-11 | 6.72E-10 | 0.666152512 | 0.758634112 |
| Maip1 | 0.003560029 | 0.603912037 | 0.706611244 | 0.150923066 | 0.14715623 |
| Mak16 | 0.022820474 | 0.00664477 | 0.018399446 | 0.252769035 | 0.235462332 |
| Mal2 | -0.010117386 | 0.024082821 | 0.054439508 | 0.048633736 | 0.056808404 |
| Malat1 | -0.06361302 | 2.60E-12 | 4.00E-11 | 5.277592936 | 5.697134751 |
| Malsu1 | 0.018005613 | 0.038162666 | 0.079617477 | 0.295410928 | 0.279393441 |
| Malt1 | 0.004011607 | 0.353553891 | 0.475952485 | 0.052786337 | 0.0475929 |
| Maml1 | 0.010380749 | 0.008736545 | 0.023187741 | 0.048304244 | 0.038751524 |
| Maml3 | 0.048202875 | 7.12E-08 | 6.17E-07 | 0.313528351 | 0.269126409 |
| Man1a | -0.003523266 | 0.7267589 | 0.802903412 | 0.506302624 | 0.530031606 |
| Man1a2 | 0.005942373 | 0.526570167 | 0.638398199 | 0.386509773 | 0.375017387 |
| Man1b1 | -0.003651316 | 0.536072374 | 0.647013733 | 0.102991135 | 0.10345913 |
| Man1c1 | 0.007636617 | 0.087488824 | 0.158527947 | 0.059227311 | 0.051254682 |
| Man2a1 | 0.002147642 | 0.72208599 | 0.799740259 | 0.096927448 | 0.099591519 |
| Man2a2 | -0.010352764 | 0.15899176 | 0.255955935 | 0.162254953 | 0.168858768 |
| Man2b1 | -0.013031275 | 0.062250483 | 0.120341857 | 0.152622264 | 0.153519098 |
| Man2b2 | -0.005938416 | 0.365889734 | 0.488665782 | 0.136847024 | 0.132084154 |
| Man2c1 | -0.012362069 | 0.048637016 | 0.097760847 | 0.113650142 | 0.119696616 |
| Man2c1os | 0.00097625 | 0.806727613 | 0.8660563 | 0.042902445 | 0.042072317 |
| Manba | -0.004127491 | 0.408282304 | 0.528256662 | 0.071182045 | 0.069512022 |
| Manbal | -0.001969995 | 0.801196737 | 0.861355157 | 0.223690661 | 0.212383442 |
| Manea | 0.004903288 | 0.326726119 | 0.449156013 | 0.069192671 | 0.066318385 |
| Maneal | 0.002942405 | 0.532587679 | 0.643734017 | 0.064621492 | 0.060711666 |
| Manf | -0.103006411 | 3.99E-22 | 1.33E-20 | 1.337108134 | 1.534603205 |
| Maoa | -0.004886851 | 0.350453124 | 0.472880677 | 0.0763997 | 0.077572551 |
| Maob | 0.117889189 | 2.47E-31 | 1.35E-29 | 0.595725756 | 0.444445191 |
| Map10 | -0.004011822 | 0.343597663 | 0.465763591 | 0.045798726 | 0.048597888 |
| Map11 | 0.011800221 | 0.019190844 | 0.044860155 | 0.079142257 | 0.068069798 |
| Map1b | -0.03316856 | 0.000597145 | 0.002272191 | 1.188472196 | 1.254635348 |
| Map1lc3a | -0.013993387 | 0.170078256 | 0.269299805 | 0.754631289 | 0.763282116 |
| Map1lc3b | -0.004454668 | 0.654134034 | 0.74639906 | 0.900641042 | 0.877959533 |
| Map1s | 0.010841992 | 0.04891071 | 0.098225558 | 0.093824123 | 0.082090618 |
| Map2 | 0.096041435 | 5.54E-21 | 1.69E-19 | 0.535013178 | 0.418298472 |
| Map2k1 | 0.028724693 | 7.75E-05 | 0.000370685 | 0.18769241 | 0.158979658 |
| Map2k2 | -0.000123583 | 0.990155638 | 0.993322992 | 0.67106123 | 0.646331645 |
| Map2k3 | -0.014405985 | 0.017661454 | 0.041991846 | 0.094724414 | 0.108726973 |
| Map2k4 | 0.023840614 | 0.000218278 | 0.000927489 | 0.139552696 | 0.118709626 |
| Map2k5 | 0.015685611 | 0.004494553 | 0.013181311 | 0.099907147 | 0.083029418 |
| Map2k7 | 0.006825861 | 0.303287326 | 0.42364359 | 0.135702052 | 0.130018176 |
| Map3k1 | 0.036888137 | 9.72E-08 | 8.23E-07 | 0.155967401 | 0.12825925 |
| Map3k10 | 0.010357577 | 0.104100591 | 0.182584512 | 0.13267667 | 0.118740541 |
| Map3k11 | 0.013320297 | 0.005123582 | 0.014732761 | 0.067760426 | 0.058054588 |
| Map3k12 | 0.004198364 | 0.368745009 | 0.491289431 | 0.062052667 | 0.056625843 |
| Map3k15 | 0.006330826 | 0.463144041 | 0.580766976 | 0.26893904 | 0.255619392 |
| Map3k2 | 0.028906198 | 0.0028765 | 0.008966122 | 0.412263339 | 0.391043701 |
| Map3k20 | 0.018548669 | 2.97E-06 | 1.91E-05 | 0.04868575 | 0.034775372 |
| Map3k3 | 0.014076399 | 0.004549111 | 0.01331742 | 0.075773286 | 0.062691119 |
| Map3k4 | 0.064616148 | 2.87E-14 | 5.34E-13 | 0.286896529 | 0.231323236 |
| Map3k5 | 0.01456607 | 0.008101486 | 0.021775683 | 0.096159632 | 0.079685896 |
| Map3k7 | 0.015865972 | 0.03585058 | 0.075752952 | 0.197370003 | 0.18171186 |
| Map4 | -0.004928855 | 0.523122941 | 0.635328074 | 0.184164418 | 0.188667248 |
| Map4k2 | -0.005065193 | 0.277049305 | 0.395285198 | 0.056724835 | 0.060001684 |
| Map4k3 | 0.002162276 | 0.697868662 | 0.781285022 | 0.090378582 | 0.088422892 |
| Map4k4 | 0.014241333 | 0.096667525 | 0.171747383 | 0.260422635 | 0.249126023 |
| Map4k5 | 0.018801237 | 0.002828212 | 0.008821262 | 0.126807851 | 0.107954493 |
| Map6 | 0.012309443 | 0.0039902 | 0.011888668 | 0.055196949 | 0.044850663 |
| Map7 | 0.026730738 | 0.002884469 | 0.0089852 | 0.328776556 | 0.300062846 |
| Map7d1 | 0.051884015 | 4.05E-10 | 4.86E-09 | 0.263315514 | 0.211083277 |
| Map7d2 | -0.025910251 | 0.003826886 | 0.011472398 | 0.246051402 | 0.272295294 |
| Map9 | -0.006466785 | 0.427952877 | 0.546578805 | 0.220373068 | 0.222956621 |
| Mapk1 | 0.026393602 | 0.005793785 | 0.016350415 | 0.452773305 | 0.421968249 |
| Mapk10 | -0.046574059 | 5.32E-09 | 5.49E-08 | 0.185572242 | 0.219906807 |
| Mapk13 | -0.014525965 | 0.001086966 | 0.003852328 | 0.048736853 | 0.056396264 |
| Mapk14 | 0.01648967 | 0.077870017 | 0.144305433 | 0.360105229 | 0.347696469 |
| Mapk15 | -0.023587358 | 0.005106643 | 0.014692744 | 0.220262863 | 0.21319973 |
| Mapk1ip1 | 0.006010169 | 0.371888233 | 0.49412292 | 0.153123757 | 0.140301857 |

| | | | | | |
|---|---|---|---|---|---|
| Mapk1ip1l | 0.00707574 | 0.374106956 | 0.496189335 | 0.21715115 | 0.209720867 |
| Mapk3 | 0.047662321 | 9.27E-07 | 6.52E-06 | 0.570178433 | 0.486165534 |
| Mapk6 | 0.006105026 | 0.409096451 | 0.529012118 | 0.158587493 | 0.164769109 |
| Mapk8 | 0.002328832 | 0.688687831 | 0.773833088 | 0.09442354 | 0.094057969 |
| Mapk8ip1 | 0.029682534 | 0.000528402 | 0.00203854 | 0.319481347 | 0.270829936 |
| Mapk8ip2 | 0.02626119 | 0.003742476 | 0.011257534 | 0.35273166 | 0.317786293 |
| Mapk8ip3 | 0.023751389 | 0.003479728 | 0.010565244 | 0.233198039 | 0.213125451 |
| Mapk9 | -0.002917262 | 0.686914262 | 0.772787055 | 0.165910955 | 0.167057869 |
| Mapkap1 | 0.011078825 | 0.100201572 | 0.176959071 | 0.144623809 | 0.13282917 |
| Mapkapk2 | 0.017494818 | 0.051563619 | 0.102390014 | 0.282042164 | 0.278991531 |
| Mapkapk3 | 0.017241135 | 0.000109538 | 0.000507835 | 0.059634742 | 0.046063151 |
| Mapkbp1 | 0.004755463 | 0.223152731 | 0.334591433 | 0.042748483 | 0.038844209 |
| Mapre1 | 0.024669004 | 0.008197124 | 0.021971999 | 0.383964411 | 0.356706525 |
| Mapre2 | 0.042776756 | 1.18E-05 | 6.64E-05 | 0.494792548 | 0.447622909 |
| Mapre3 | 0.030640599 | 0.000318593 | 0.001301094 | 0.305424891 | 0.257630018 |
| Mapt | 0.014301128 | 0.081650201 | 0.149802464 | 0.238010976 | 0.213410063 |
| Marchf2 | -0.005814918 | 0.392704724 | 0.513354676 | 0.146518688 | 0.144441245 |
| Marchf4 | 0.011064319 | 0.034612919 | 0.073585567 | 0.088514872 | 0.072732095 |
| Marchf5 | -0.016384893 | 0.126484904 | 0.21337606 | 0.72463332 | 0.756805626 |
| Marchf6 | -0.044041498 | 1.56E-05 | 8.64E-05 | 0.561579935 | 0.604440573 |
| Marchf7 | 0.064076093 | 2.50E-11 | 3.43E-10 | 0.450501276 | 0.381486594 |
| Marchf8 | 0.010286987 | 0.291814356 | 0.411341043 | 0.482654286 | 0.466630588 |
| Marchf9 | -0.020387238 | 0.004956244 | 0.014310843 | 0.175241989 | 0.177080364 |
| Marcks | 0.052517351 | 5.61E-07 | 4.15E-06 | 0.594236708 | 0.532991802 |
| Marf1 | 0.010619934 | 0.190083407 | 0.294246832 | 0.255621547 | 0.224372155 |
| Mark1 | 0.005902595 | 0.302610926 | 0.422826717 | 0.092432798 | 0.090182553 |
| Mark2 | 0.054976957 | 4.19E-09 | 4.40E-08 | 0.399893893 | 0.344034184 |
| Mark3 | 0.010830964 | 0.249426783 | 0.363939244 | 0.37922098 | 0.365343863 |
| Mark4 | 0.009208351 | 0.08212979 | 0.1505687 | 0.084033165 | 0.075986806 |
| Mars1 | -0.023020696 | 0.014895721 | 0.036358575 | 0.408932057 | 0.414566211 |
| Marveld1 | -0.016974997 | 0.004135791 | 0.012247383 | 0.095512561 | 0.106374499 |
| Marveld2 | 0.011211981 | 0.005297811 | 0.015144169 | 0.048272021 | 0.039990388 |
| Marveld3 | -0.000840648 | 0.911698121 | 0.939231656 | 0.190457883 | 0.186242098 |
| Mast1 | 0.008391627 | 0.032378276 | 0.069580371 | 0.047970633 | 0.037796066 |
| Mast2 | 0.007560008 | 0.183831202 | 0.286733025 | 0.094712796 | 0.089776872 |
| Mast3 | 0.019881938 | 0.0021606 | 0.006980843 | 0.132625456 | 0.115759855 |
| Mast4 | 0.040925285 | 0.000145661 | 0.000652285 | 0.769582856 | 0.730006489 |
| Mat1a | 0.002113432 | 0.753021735 | 0.823959179 | 0.133504218 | 0.132435804 |
| Mat2a | -0.126444222 | 7.43E-29 | 3.68E-27 | 0.365988908 | 0.518335022 |
| Mat2b | -0.059533016 | 4.37E-11 | 5.84E-10 | 0.307070998 | 0.349009602 |
| Matr3 | 0.021751315 | 0.027720235 | 0.061179597 | 0.520327418 | 0.483912598 |
| Mau2 | -0.00739166 | 0.35144315 | 0.473887699 | 0.205625552 | 0.201704834 |
| Max | -0.014712397 | 0.056781394 | 0.111273677 | 0.200869298 | 0.205077859 |
| Maz | 0.002224277 | 0.805161828 | 0.864676911 | 0.345905517 | 0.325337866 |
| Mbd1 | 0.015688745 | 0.010755892 | 0.027539841 | 0.116508509 | 0.105992855 |
| Mbd2 | 0.007261357 | 0.455297205 | 0.573888166 | 0.826925588 | 0.792131908 |
| Mbd3 | 0.020064914 | 0.028622907 | 0.06290052 | 0.388854082 | 0.355358321 |
| Mbd5 | -0.002443925 | 0.67064304 | 0.759634816 | 0.092213476 | 0.092783683 |
| Mbd6 | -0.001082242 | 0.876798309 | 0.915894827 | 0.147014046 | 0.150035807 |
| Mbip | -0.002579955 | 0.607886578 | 0.708872893 | 0.071255754 | 0.070630182 |
| Mblac2 | -0.00440846 | 0.404977832 | 0.524679436 | 0.079094809 | 0.079515772 |
| Mbnl1 | 0.065976294 | 3.35E-11 | 4.55E-10 | 0.503481222 | 0.438368688 |
| Mbnl2 | 0.009337352 | 0.364143173 | 0.486960101 | 0.697878095 | 0.693874592 |
| Mboat2 | 0.046610332 | 8.37E-07 | 5.96E-06 | 0.447407585 | 0.373842488 |
| Mboat7 | 0.006179697 | 0.252025289 | 0.366959691 | 0.084164928 | 0.080462617 |
| Mbp | 0.041299071 | 4.20E-06 | 2.63E-05 | 0.331742216 | 0.296221256 |
| Mbtd1 | -0.000327436 | 0.958679679 | 0.971740148 | 0.1177344 | 0.115171556 |
| Mbtps1 | -0.002594056 | 0.743308832 | 0.816732422 | 0.219371229 | 0.216619153 |
| Mbtps2 | -0.003992469 | 0.45302008 | 0.571536623 | 0.076654488 | 0.079650124 |
| Mcat | -0.001073255 | 0.828059034 | 0.881461725 | 0.075103925 | 0.068791939 |
| Mcc | 0.012542283 | 0.034636247 | 0.073612336 | 0.108211405 | 0.096069643 |
| Mccc1 | -0.007476288 | 0.145543769 | 0.239380227 | 0.07110517 | 0.074869467 |
| Mccc2 | -0.013541862 | 0.001754385 | 0.005817235 | 0.046444415 | 0.054220196 |
| Mcee | 0.002302979 | 0.787698466 | 0.851551705 | 0.300683593 | 0.286030298 |
| Mcf2l | 0.054585085 | 5.29E-09 | 5.47E-08 | 0.383639433 | 0.319626714 |
| Mcfd2 | -0.029523777 | 0.000176211 | 0.000770983 | 0.194541307 | 0.217549124 |
| Mcl1 | -0.011317342 | 0.231409141 | 0.344626614 | 0.354892599 | 0.373852164 |
| Mcm3ap | 0.012151283 | 0.025298445 | 0.056723128 | 0.08928141 | 0.077792316 |
| Mcm4 | 0.007372045 | 0.054688926 | 0.10760691 | 0.041531017 | 0.035706732 |
| Mcm7 | -0.009057528 | 0.019292741 | 0.045034231 | 0.036578232 | 0.041834804 |
| Mcm9 | -0.002094648 | 0.610222825 | 0.711085497 | 0.04406539 | 0.045409791 |
| Mcmbp | 0.005736055 | 0.367278003 | 0.490006415 | 0.129922676 | 0.118175141 |

| | | | | | |
|---|---|---|---|---|---|
| Mcph1 | -3.39E-05 | 0.993121186 | 0.995681569 | 0.038356193 | 0.040505088 |
| Mcrip1 | 0.014003203 | 0.170708004 | 0.269989135 | 0.728075585 | 0.69484323 |
| Mcrip2 | -0.009332941 | 0.093478842 | 0.16698894 | 0.081061324 | 0.089319802 |
| Mcrs1 | -0.005702727 | 0.402734491 | 0.522608746 | 0.153872609 | 0.147661364 |
| Mcts1 | -0.030218368 | 0.000612409 | 0.002323903 | 0.296200551 | 0.313428711 |
| Mcts2 | 0.007628505 | 0.120474301 | 0.20501508 | 0.070392785 | 0.063155365 |
| Mcu | -0.002086176 | 0.654370297 | 0.74642591 | 0.05684948 | 0.060114314 |
| Mcub | -0.009258665 | 0.08243874 | 0.151021185 | 0.067902313 | 0.078349863 |
| Mcur1 | 0.00158649 | 0.841301059 | 0.89145819 | 0.210451101 | 0.209013076 |
| Mdc1 | 0.012300791 | 0.017875402 | 0.042407189 | 0.08144504 | 0.070983418 |
| Mdh1 | -0.022549489 | 0.012600643 | 0.031467622 | 1.332385859 | 1.369643568 |
| Mdh2 | -0.028586035 | 0.003754715 | 0.01128707 | 0.81914766 | 0.818028789 |
| Mdm1 | -2.49E-05 | 0.997443836 | 0.997853138 | 0.188329616 | 0.189030752 |
| Mdm2 | 0.007332091 | 0.379807689 | 0.501220354 | 0.251064807 | 0.236759531 |
| Mdm4 | -0.036571307 | 8.57E-06 | 5.00E-05 | 0.207875784 | 0.240884991 |
| Mdn1 | 0.009338335 | 0.101389285 | 0.178699534 | 0.094009548 | 0.087541153 |
| Mdp1 | -0.019394674 | 0.020159547 | 0.046742455 | 0.246889605 | 0.257523016 |
| Me1 | -0.040016897 | 1.74E-07 | 1.42E-06 | 0.162884324 | 0.19641955 |
| Me2 | 0.010544411 | 0.068398906 | 0.129953912 | 0.099576507 | 0.093233871 |
| Mea1 | -0.043160814 | 4.49E-06 | 2.78E-05 | 0.346666089 | 0.381096 |
| Meaf6 | -0.023607173 | 0.002179547 | 0.007030378 | 0.193740714 | 0.206930482 |
| Mecp2 | 0.022070993 | 0.011160752 | 0.028456552 | 0.290614762 | 0.270868007 |
| Mecr | -0.010294244 | 0.046629045 | 0.094227306 | 0.071518867 | 0.077330271 |
| Med1 | 0.015247279 | 0.034853309 | 0.073943268 | 0.172496991 | 0.157395099 |
| Med10 | -0.029023151 | 0.00080544 | 0.002959369 | 0.279865965 | 0.302725241 |
| Med11 | 0.011307695 | 0.04021337 | 0.083359144 | 0.095019286 | 0.083199225 |
| Med12 | 0.00874924 | 0.028139642 | 0.06198389 | 0.04439713 | 0.039670557 |
| Med12l | 0.001872131 | 0.74001347 | 0.814123679 | 0.089871265 | 0.087918775 |
| Med13 | 0.021748843 | 0.019845512 | 0.046146392 | 0.363849434 | 0.327918135 |
| Med13l | 0.038507313 | 1.22E-06 | 8.39E-06 | 0.236076257 | 0.194911162 |
| Med14 | 0.005516614 | 0.351008831 | 0.473433388 | 0.103793493 | 0.098671757 |
| Med15 | 0.024830846 | 0.000605472 | 0.002301172 | 0.171377096 | 0.155303013 |
| Med16 | 0.001648137 | 0.764271232 | 0.833222102 | 0.087905358 | 0.083056888 |
| Med17 | -0.002909661 | 0.525396115 | 0.63751738 | 0.057155488 | 0.058115351 |
| Med19 | -0.010994242 | 0.127032236 | 0.214150754 | 0.157263223 | 0.168112522 |
| Med20 | -0.000998004 | 0.848361089 | 0.89649636 | 0.077264715 | 0.075844358 |
| Med21 | -0.040433311 | 3.46E-06 | 2.19E-05 | 0.283136738 | 0.308492001 |
| Med22 | 0.002938071 | 0.625059936 | 0.723504481 | 0.107154914 | 0.104214496 |
| Med23 | 0.015514766 | 0.002953423 | 0.009173539 | 0.084716331 | 0.070192654 |
| Med24 | 0.002733763 | 0.542901316 | 0.65297562 | 0.058595269 | 0.054559057 |
| Med25 | 0.020726919 | 0.012553473 | 0.031382069 | 0.27399265 | 0.240066705 |
| Med26 | 0.004962638 | 0.226482181 | 0.338747655 | 0.048695099 | 0.043976362 |
| Med27 | -3.62E-05 | 0.995144852 | 0.996584467 | 0.109774784 | 0.103609801 |
| Med28 | 0.002607129 | 0.789707154 | 0.852850586 | 0.800612606 | 0.769612884 |
| Med29 | -0.004182148 | 0.652672813 | 0.745235587 | 0.376856982 | 0.380807043 |
| Med30 | -0.050050916 | 9.78E-09 | 9.76E-08 | 0.280552573 | 0.316720904 |
| Med31 | -0.003607848 | 0.658285546 | 0.749846076 | 0.24929102 | 0.243353225 |
| Med4 | -0.005895089 | 0.353758391 | 0.476085144 | 0.118794736 | 0.121401953 |
| Med6 | -0.004957281 | 0.41019553 | 0.529794153 | 0.111646445 | 0.110472097 |
| Med7 | -0.005518393 | 0.452305242 | 0.570782913 | 0.17851665 | 0.175993201 |
| Med8 | -0.001398439 | 0.865456917 | 0.908047933 | 0.246997246 | 0.242283135 |
| Med9 | -0.023544251 | 0.002285342 | 0.007342396 | 0.194091977 | 0.205652384 |
| Med9os | 0.001363524 | 0.774815955 | 0.841266309 | 0.059328437 | 0.060531378 |
| Mef2a | 0.019054709 | 0.025148825 | 0.056504941 | 0.265475439 | 0.248401954 |
| Mef2d | 0.029386374 | 0.000878411 | 0.003199669 | 0.306831954 | 0.290916929 |
| Meg3 | -0.036488981 | 0.000132755 | 0.000601979 | 2.530699584 | 2.596213199 |
| Megf8 | -0.004954234 | 0.44084785 | 0.559373897 | 0.126174861 | 0.122895366 |
| Megf9 | 0.075741836 | 7.42E-16 | 1.54E-14 | 0.383531981 | 0.326015612 |
| Meig1 | 0.042044853 | 2.64E-09 | 2.84E-08 | 0.1661405 | 0.136016403 |
| Meis1 | 0.0026648 | 0.628952112 | 0.726715297 | 0.086103172 | 0.082112597 |
| Meis2 | 0.062869408 | 2.02E-09 | 2.19E-08 | 0.94190422 | 0.870593528 |
| Meis3 | -0.023200843 | 0.001795399 | 0.005943095 | 0.178549612 | 0.188050564 |
| Memo1 | -0.006963501 | 0.280165756 | 0.399028931 | 0.1231538 | 0.125429317 |
| Men1 | 0.009801795 | 0.111972015 | 0.193837594 | 0.11883596 | 0.107831525 |
| Meox1 | -0.001345443 | 0.727310931 | 0.803421983 | 0.03908256 | 0.039098441 |
| Mepce | 0.006962732 | 0.319573363 | 0.441069312 | 0.159879869 | 0.147825936 |
| Mesd | 0.005913316 | 0.405504638 | 0.525291951 | 0.1703361 | 0.156794637 |
| Met | 0.043309264 | 2.59E-16 | 5.53E-15 | 0.098842846 | 0.063349407 |
| Metap1 | 0.013523614 | 0.05078726 | 0.101262047 | 0.156664313 | 0.144316327 |
| Metap1d | -0.013715142 | 0.023009091 | 0.052252834 | 0.106893497 | 0.109882978 |
| Metap2 | 0.134715552 | 1.43E-44 | 1.37E-42 | 0.992834702 | 0.854710636 |
| Metrn | -0.037301275 | 4.22E-24 | 1.61E-22 | 0.013957482 | 0.044378508 |

| | | | | | |
|---|---|---|---|---|---|
| Mettl1 | -0.085106821 | 6.05E-17 | 1.35E-15 | 0.41530643 | 0.500508763 |
| Mettl14 | 0.008646627 | 0.066227427 | 0.126819588 | 0.064362265 | 0.057345869 |
| Mettl16 | -0.003051873 | 0.673942031 | 0.762666095 | 0.165333007 | 0.164413885 |
| Mettl18 | 0.004454395 | 0.330118317 | 0.452671661 | 0.067151786 | 0.055376656 |
| Mettl21a | -0.002318927 | 0.610093659 | 0.711020205 | 0.056837772 | 0.056337445 |
| Mettl22 | -0.009793491 | 0.049387289 | 0.098974775 | 0.063536011 | 0.071633272 |
| Mettl23 | -0.042958014 | 5.63E-07 | 4.16E-06 | 0.271165956 | 0.297781638 |
| Mettl25 | -0.008197745 | 0.048818538 | 0.098080973 | 0.041453736 | 0.047861114 |
| Mettl26 | -0.079859883 | 1.65E-15 | 3.31E-14 | 0.81531577 | 0.877375585 |
| Mettl27 | -0.014440979 | 0.03011711 | 0.06563518 | 0.124921171 | 0.132203839 |
| Mettl3 | -0.007388093 | 0.153250762 | 0.248358711 | 0.073638935 | 0.075932769 |
| Mettl4 | -0.017162069 | 0.001930401 | 0.006336042 | 0.077929476 | 0.089330583 |
| Mettl5 | -0.023440574 | 0.000570385 | 0.002182324 | 0.137327529 | 0.149362014 |
| Mettl6 | -0.005725117 | 0.29209315 | 0.411495378 | 0.078230554 | 0.084116919 |
| Mettl7a1 | -0.014382234 | 0.117301464 | 0.200634207 | 0.268573535 | 0.294216815 |
| Mettl8 | 0.002856787 | 0.463713897 | 0.581331594 | 0.042419058 | 0.038787328 |
| Mettl9 | -0.049672112 | 5.07E-07 | 3.79E-06 | 0.583471618 | 0.620533079 |
| Mex3c | 0.009306494 | 0.100039516 | 0.176833485 | 0.093431042 | 0.087291629 |
| Mex3d | 0.010224334 | 0.009009859 | 0.02377038 | 0.047525779 | 0.038039454 |
| Mfap1a | 0.010038686 | 0.170009037 | 0.269234055 | 0.178367234 | 0.1643998 |
| Mfap1b | 0.029897679 | 0.000113549 | 0.000524179 | 0.215261506 | 0.184255307 |
| Mfap3 | 0.003207447 | 0.57190327 | 0.678882002 | 0.094119147 | 0.089777961 |
| Mff | -0.021053163 | 0.031184814 | 0.067534773 | 0.562173042 | 0.565889596 |
| Mfhas1 | 0.014135041 | 0.001032924 | 0.003682466 | 0.055826475 | 0.04524956 |
| Mfn1 | 0.000451341 | 0.940221556 | 0.959566866 | 0.105998398 | 0.104302047 |
| Mfn2 | 0.019187409 | 0.009156439 | 0.024085201 | 0.189343202 | 0.1688489 |
| Mfsd1 | 0.008361283 | 0.23447315 | 0.348291025 | 0.161082334 | 0.153928637 |
| Mfsd10 | -0.00766159 | 0.174262888 | 0.274985196 | 0.086733456 | 0.092703378 |
| Mfsd11 | 0.028246762 | 0.000102792 | 0.000478841 | 0.186404493 | 0.160186985 |
| Mfsd13a | 0.014830991 | 0.001640222 | 0.00548169 | 0.070988003 | 0.056917736 |
| Mfsd14a | 0.007251216 | 0.332368393 | 0.455033632 | 0.188867197 | 0.178037414 |
| Mfsd14b | 0.011783187 | 0.075987045 | 0.141661966 | 0.139125472 | 0.129399915 |
| Mfsd5 | 0.018134412 | 0.001207611 | 0.004212555 | 0.09630269 | 0.08357378 |
| Mfsd6 | 0.007322184 | 0.336056612 | 0.45867505 | 0.193987876 | 0.184828988 |
| Mfsd6l | 0.001862394 | 0.667545378 | 0.757190079 | 0.052126104 | 0.049792491 |
| Mfsd8 | -0.005034061 | 0.306713494 | 0.427569341 | 0.063825175 | 0.06718971 |
| Mga | 0.045806111 | 1.87E-06 | 1.24E-05 | 0.41929992 | 0.365742336 |
| Mgat1 | 0.003392442 | 0.618588028 | 0.718422269 | 0.1501954 | 0.143593854 |
| Mgat2 | -0.039497515 | 2.07E-06 | 1.36E-05 | 0.221497681 | 0.253106115 |
| Mgat3 | 0.023790783 | 4.96E-05 | 0.000244065 | 0.117394321 | 0.092477615 |
| Mgat4b | 0.00722631 | 0.280938192 | 0.399836203 | 0.141229402 | 0.131280168 |
| Mgat5 | 0.004591883 | 0.333496298 | 0.45633371 | 0.062758956 | 0.059934146 |
| Mgme1 | 0.010809697 | 0.010483996 | 0.026921661 | 0.054613945 | 0.041925253 |
| Mgmt | -0.020586405 | 0.000144252 | 0.00064777 | 0.073889467 | 0.085511906 |
| Mgrn1 | 0.030945759 | 5.00E-06 | 3.08E-05 | 0.161993952 | 0.132321348 |
| Mgst3 | -0.001154787 | 0.879492094 | 0.917821182 | 0.192346713 | 0.187864482 |
| Mia2 | -0.046147992 | 7.26E-06 | 4.31E-05 | 0.767437165 | 0.833698186 |
| Mia3 | 0.003270763 | 0.713537083 | 0.794071373 | 1.285317812 | 1.307402024 |
| Mib1 | 0.018732046 | 0.041624324 | 0.085735526 | 0.35379418 | 0.332537264 |
| Mib2 | -0.007499893 | 0.300370597 | 0.420354533 | 0.161807417 | 0.164439569 |
| Mical2 | 0.029986185 | 2.56E-06 | 1.66E-05 | 0.134547286 | 0.106257677 |
| Mical3 | 0.032184932 | 4.42E-05 | 0.00021939 | 0.218216615 | 0.185266226 |
| Micall1 | 0.006398642 | 0.138066546 | 0.229397379 | 0.052595859 | 0.047049952 |
| Micos10 | 0.086710277 | 3.19E-31 | 1.72E-29 | 1.62383899 | 1.517668736 |
| Micos13 | -0.018099304 | 0.021792879 | 0.049934096 | 1.348319874 | 1.407782371 |
| Micu1 | 0.012000375 | 0.076367171 | 0.142230198 | 0.153071401 | 0.137270272 |
| Micu2 | 0.018300533 | 0.03234139 | 0.06951647 | 0.28991902 | 0.258340585 |
| Micu3 | -0.003269146 | 0.725692395 | 0.802166645 | 0.379832021 | 0.372104371 |
| Mid1 | 0.018523699 | 1.60E-06 | 1.07E-05 | 0.047967123 | 0.033276252 |
| Mid1ip1 | -0.010998757 | 0.275338101 | 0.393455962 | 0.476277698 | 0.472228737 |
| Mid2 | -0.002665597 | 0.65419321 | 0.74639906 | 0.10198521 | 0.103374137 |
| Mideas | -0.015783771 | 0.011784434 | 0.029796689 | 0.095279325 | 0.114970663 |
| Midn | 0.035032778 | 9.21E-06 | 5.33E-05 | 0.198446938 | 0.178201719 |
| Mief1 | -0.001013001 | 0.85478245 | 0.899956138 | 0.087822828 | 0.086847351 |
| Mien1 | -0.016823652 | 0.07696429 | 0.143095587 | 0.985098112 | 0.992366096 |
| Mier1 | 0.02558198 | 0.009106054 | 0.023966363 | 0.481309337 | 0.446598199 |
| Mier2 | 0.012601542 | 0.00239548 | 0.007643208 | 0.054341828 | 0.043182505 |
| Mier3 | 0.006504598 | 0.284386835 | 0.403603707 | 0.110499388 | 0.104491118 |
| Mif | 0.01029119 | 0.372603359 | 0.494802603 | 1.04384739 | 1.08401562 |
| Mif4gd | -0.008328166 | 0.161685709 | 0.259156897 | 0.107883294 | 0.105475626 |
| Miga1 | -0.010787151 | 0.134131345 | 0.224560864 | 0.155380323 | 0.163150993 |
| Miip | -0.004671469 | 0.252511953 | 0.367556839 | 0.043096649 | 0.045312966 |

| | | | | | |
|---|---|---|---|---|---|
| Mindy1 | 0.004417416 | 0.437828178 | 0.556195682 | 0.091553882 | 0.089465886 |
| Mindy2 | 0.057675818 | 1.49E-09 | 1.66E-08 | 0.438444213 | 0.379056195 |
| Mindy3 | -0.008953067 | 0.201599483 | 0.309140406 | 0.15039569 | 0.154925808 |
| Mink1 | 0.009609492 | 0.152669826 | 0.247747629 | 0.143149398 | 0.131805104 |
| Minpp1 | 0.000938542 | 0.862810006 | 0.905955166 | 0.088455518 | 0.082177264 |
| Mios | 0.013162668 | 0.01533811 | 0.037204866 | 0.090094733 | 0.076493138 |
| Mipep | -0.011832533 | 0.010724484 | 0.027473895 | 0.055887042 | 0.060978302 |
| Mipol1 | 0.005155693 | 0.383380102 | 0.504496664 | 0.103261121 | 0.097319373 |
| Mir22hg | -0.065144739 | 1.94E-15 | 3.89E-14 | 0.170151492 | 0.242213561 |
| Mir6236 | -0.248049102 | 4.37E-85 | 2.12E-82 | 0.576609843 | 0.765902927 |
| Mir670hg | 0.110661892 | 4.32E-22 | 1.43E-20 | 0.613598767 | 0.529194666 |
| Mir703 | -0.022225103 | 2.95E-05 | 0.00015351 | 0.071728949 | 0.085695267 |
| Mirg | 0.011737588 | 0.050148814 | 0.100294527 | 0.103354404 | 0.094118164 |
| Mis12 | 0.020992327 | 0.000121327 | 0.00055508 | 0.09536216 | 0.077884967 |
| Mitd1 | 0.021138155 | 0.000608341 | 0.002310271 | 0.11757247 | 0.101792137 |
| Mitf | -0.008738563 | 0.03607074 | 0.076085862 | 0.040729745 | 0.047170502 |
| Mkks | 0.005062358 | 0.179235242 | 0.281279261 | 0.043157406 | 0.035883316 |
| Mkln1 | 0.020335493 | 0.030134289 | 0.065657901 | 0.383073683 | 0.354725497 |
| Mknk1 | -0.013495117 | 0.067448845 | 0.128450082 | 0.167988771 | 0.176135683 |
| Mknk2 | 0.006700186 | 0.164973486 | 0.263145568 | 0.065867161 | 0.059587971 |
| Mkrn1 | -0.013505773 | 0.162694338 | 0.260493141 | 0.516596781 | 0.50818753 |
| Mkrn2 | 0.025715379 | 0.000103316 | 0.000481052 | 0.150709978 | 0.123036662 |
| Mkrn2os | 0.017922081 | 0.013533571 | 0.033486302 | 0.174925596 | 0.157632361 |
| Mks1 | -0.002664767 | 0.508293551 | 0.622371524 | 0.0446702 | 0.043343216 |
| Mlec | -0.021871631 | 0.030202833 | 0.065777764 | 0.794836404 | 0.835266684 |
| Mlf2 | 0.031298244 | 0.001373085 | 0.004717007 | 0.642815884 | 0.595005658 |
| Mlh1 | -0.003478344 | 0.403860002 | 0.523719746 | 0.044651921 | 0.047165711 |
| Mlh3 | -0.012011782 | 0.091723509 | 0.164513629 | 0.14719243 | 0.153777062 |
| Mllt10 | -0.00480346 | 0.588294266 | 0.692934477 | 0.307064365 | 0.300908676 |
| Mllt11 | -0.009108009 | 0.213820909 | 0.323593848 | 0.166308575 | 0.169767777 |
| Mllt3 | 0.024209587 | 0.001090565 | 0.003859654 | 0.189273587 | 0.162402701 |
| Mllt6 | -0.015888081 | 0.017747546 | 0.0421553 | 0.129827253 | 0.13996904 |
| Mlx | -0.001571314 | 0.846173967 | 0.894768687 | 0.226187762 | 0.228930037 |
| Mlxip | 0.016877621 | 0.009335757 | 0.024477409 | 0.136898565 | 0.119653337 |
| Mlxipl | -0.035544162 | 0.001714406 | 0.005698276 | 0.497848078 | 0.477901247 |
| Mlycd | 0.001781413 | 0.821545461 | 0.877506589 | 0.221571515 | 0.211479639 |
| Mmaa | 0.00376294 | 0.377388995 | 0.498976718 | 0.055542025 | 0.046868374 |
| Mmab | -0.001430923 | 0.700525585 | 0.783537706 | 0.035966698 | 0.036805191 |
| Mmachc | -5.22E-05 | 0.989401982 | 0.993079296 | 0.04194925 | 0.041515235 |
| Mmadhc | -0.007833102 | 0.341057853 | 0.463483987 | 0.243825503 | 0.242476583 |
| Mmgt1 | -0.004669429 | 0.417217168 | 0.536461488 | 0.091331183 | 0.094400739 |
| Mmgt2 | -0.007697119 | 0.341647344 | 0.463960815 | 0.237219094 | 0.235460954 |
| Mmp24 | 0.001581465 | 0.683156357 | 0.769956655 | 0.041230706 | 0.038090275 |
| Mms19 | -0.005829725 | 0.285604501 | 0.404928315 | 0.08092403 | 0.085626309 |
| Mmut | -0.001514464 | 0.836281425 | 0.888075152 | 0.171433544 | 0.170074454 |
| Mnat1 | 0.028793945 | 0.000278696 | 0.001152971 | 0.229829825 | 0.204969463 |
| Mns1 | -0.020068809 | 0.017340882 | 0.041351007 | 0.247008789 | 0.242494469 |
| Mnt | 0.000694261 | 0.914697901 | 0.941224914 | 0.124998932 | 0.121233555 |
| Mnx1 | 0.009937373 | 0.147051097 | 0.241202913 | 0.150601136 | 0.127544852 |
| Mob1a | 0.023451438 | 0.000150925 | 0.000671531 | 0.12274522 | 0.104727973 |
| Mob1b | 0.005430365 | 0.50956125 | 0.623471538 | 0.235572498 | 0.231073282 |
| Mob2 | -0.023340005 | 0.001510353 | 0.005107356 | 0.171211269 | 0.18160793 |
| Mob3c | 0.00365815 | 0.555899628 | 0.66491833 | 0.109558719 | 0.109749633 |
| Mob4 | -0.024996972 | 0.00836765 | 0.022348982 | 0.40571025 | 0.413046843 |
| Mocs2 | 0.01008801 | 0.313289844 | 0.434618131 | 0.587713907 | 0.571433321 |
| Mocs3 | -0.00143408 | 0.763736026 | 0.832930967 | 0.066718856 | 0.063183185 |
| Mogs | -0.005147697 | 0.512838668 | 0.626673897 | 0.198979107 | 0.204900081 |
| Mon1a | -0.000502381 | 0.919404352 | 0.944568224 | 0.067906777 | 0.068415021 |
| Mon1b | 0.006482116 | 0.113528277 | 0.195532054 | 0.048759145 | 0.042919741 |
| Mon2 | 0.01815405 | 0.045397059 | 0.092063038 | 0.329152618 | 0.312692462 |
| Morc2a | -0.004899288 | 0.420461266 | 0.539531226 | 0.102442291 | 0.106963269 |
| Morc3 | -0.020910472 | 0.001689633 | 0.005625553 | 0.122491294 | 0.137870742 |
| Morc4 | 0.010248893 | 0.040845583 | 0.084363874 | 0.073070426 | 0.063579603 |
| Morf4l1 | 0.051910927 | 3.97E-12 | 5.94E-11 | 1.650862734 | 1.576749393 |
| Morf4l2 | 0.036148838 | 6.11E-05 | 0.000296592 | 1.255890977 | 1.2011438 |
| Morn2 | -0.014504866 | 0.109007066 | 0.189814646 | 0.360171983 | 0.351880902 |
| Morn4 | -0.003867345 | 0.397638164 | 0.517972243 | 0.053444515 | 0.058569243 |
| Mosmo | -0.01078193 | 0.12724728 | 0.214364591 | 0.149327347 | 0.156971052 |
| Mospd2 | 0.006785354 | 0.311012831 | 0.432480362 | 0.139394085 | 0.129739807 |
| Mospd3 | 0.001408583 | 0.794209791 | 0.856210644 | 0.09158087 | 0.082710921 |
| Mov10 | 0.008411527 | 0.032802731 | 0.070352558 | 0.046572387 | 0.038580846 |
| Mpc1 | 0.029321336 | 0.004151066 | 0.012288875 | 0.672324102 | 0.632675372 |

| | | | | | |
|---|---|---|---|---|---|
| Mpc2 | -0.062320499 | 4.06E-10 | 4.86E-09 | 0.858830085 | 0.937923052 |
| Mpdu1 | -0.006356594 | 0.338644669 | 0.460977803 | 0.132826664 | 0.135160656 |
| Mpdz | 0.018333013 | 0.007218111 | 0.019750767 | 0.151313259 | 0.136783868 |
| Mpg | -0.019083777 | 0.000778614 | 0.002873835 | 0.083294618 | 0.097203414 |
| Mphosph10 | -0.002085564 | 0.822969914 | 0.878074806 | 0.318829083 | 0.333153371 |
| Mphosph6 | 0.028583906 | 4.07E-05 | 0.000203877 | 0.163257351 | 0.1423324 |
| Mphosph8 | 0.011748753 | 0.241938575 | 0.355843696 | 0.488232142 | 0.482860645 |
| Mphosph9 | -0.007325969 | 0.261617482 | 0.377561638 | 0.122180689 | 0.123105303 |
| Mpi | -0.013113681 | 0.002028722 | 0.006614096 | 0.043485682 | 0.052431077 |
| Mplkip | 0.014504537 | 0.078624012 | 0.145466186 | 0.262188558 | 0.240306393 |
| Mpnd | -0.009429918 | 0.176565471 | 0.277986965 | 0.15059549 | 0.153441154 |
| Mpp2 | 0.014660929 | 0.005337595 | 0.015244446 | 0.077559386 | 0.070921235 |
| Mpp3 | 0.008514334 | 0.057540497 | 0.112557086 | 0.061627632 | 0.051144376 |
| Mpp5 | 0.030197154 | 1.05E-05 | 6.01E-05 | 0.165595515 | 0.13609584 |
| Mpp6 | -0.059980039 | 1.13E-11 | 1.62E-10 | 0.234334536 | 0.283954923 |
| Mpp7 | 0.023170156 | 0.004122175 | 0.012218226 | 0.221094456 | 0.198011282 |
| Mppe1 | -0.006702597 | 0.163707522 | 0.26151241 | 0.064162909 | 0.063582835 |
| Mpped2 | -0.005744545 | 0.382609271 | 0.503686843 | 0.138128558 | 0.133374536 |
| Mprip | 0.02274582 | 0.010010325 | 0.025910644 | 0.305103455 | 0.282500742 |
| Mpst | -3.58E-05 | 0.99436524 | 0.996209693 | 0.075814206 | 0.071977925 |
| Mpv17 | -0.022254643 | 0.007305536 | 0.019961893 | 0.226025562 | 0.24759055 |
| Mpv17l | -0.022490463 | 5.94E-06 | 3.59E-05 | 0.0543217 | 0.073458506 |
| Mpv17l2 | -0.022787896 | 0.001668798 | 0.005563803 | 0.151858081 | 0.168998368 |
| Mpz | -0.013514152 | 0.00281725 | 0.008795539 | 0.05177792 | 0.059228992 |
| Mpzl1 | 0.036734632 | 8.28E-08 | 7.11E-07 | 0.16233544 | 0.128979431 |
| Mras | 0.015508077 | 0.060526996 | 0.117571121 | 0.250491584 | 0.226479953 |
| Mre11a | -0.002139748 | 0.637486988 | 0.733110899 | 0.056174589 | 0.055696364 |
| Mreg | 0.010425268 | 0.04414221 | 0.089930966 | 0.083911149 | 0.069437594 |
| Mrfap1 | -0.020951156 | 0.000458676 | 0.001798086 | 2.037593481 | 2.031465325 |
| Mrgbp | 0.004669311 | 0.374555889 | 0.496513819 | 0.081634203 | 0.076724771 |
| Mrgpre | 0.009942212 | 0.038138511 | 0.079590614 | 0.067909982 | 0.059457829 |
| Mri1 | -0.006914322 | 0.244529966 | 0.358353984 | 0.098147364 | 0.103159492 |
| Mrln | 0.146918119 | 2.51E-39 | 2.08E-37 | 0.930818048 | 0.774834202 |
| Mrm2 | -0.003650624 | 0.409959183 | 0.529721316 | 0.056128187 | 0.053817266 |
| Mrm3 | -0.009642062 | 0.020990907 | 0.048439021 | 0.041539143 | 0.048680734 |
| Mroh1 | 0.00586295 | 0.376178564 | 0.497862632 | 0.135325216 | 0.127063171 |
| Mrpl1 | 0.039511331 | 7.71E-07 | 5.52E-06 | 0.245862763 | 0.201530492 |
| Mrpl10 | 0.008561327 | 0.261318857 | 0.377270854 | 0.208029907 | 0.190727422 |
| Mrpl11 | -0.02881545 | 0.001408859 | 0.004821164 | 0.343762845 | 0.361586308 |
| Mrpl12 | 0.000889956 | 0.922684058 | 0.947242062 | 0.355566159 | 0.346252019 |
| Mrpl13 | -0.062121572 | 1.92E-11 | 2.69E-10 | 0.341702619 | 0.395664376 |
| Mrpl14 | 0.004659577 | 0.628225607 | 0.726318154 | 0.507344059 | 0.49065782 |
| Mrpl15 | -0.01317851 | 0.13604621 | 0.226962048 | 0.279322859 | 0.293318635 |
| Mrpl16 | 0.004251624 | 0.541394327 | 0.651737946 | 0.164402611 | 0.15283623 |
| Mrpl17 | -0.014546428 | 0.150976565 | 0.246006139 | 0.621858 | 0.625006878 |
| Mrpl18 | 0.002252432 | 0.783851758 | 0.848714422 | 0.244927157 | 0.234247915 |
| Mrpl19 | -9.56E-05 | 0.989089068 | 0.993079296 | 0.156379738 | 0.158025283 |
| Mrpl2 | -0.028188429 | 0.002518307 | 0.007985317 | 0.355084832 | 0.377854259 |
| Mrpl20 | -0.00728732 | 0.468304327 | 0.585801925 | 0.785526587 | 0.795549044 |
| Mrpl21 | -0.049329707 | 4.35E-07 | 3.28E-06 | 0.408334537 | 0.456508497 |
| Mrpl22 | -0.004075789 | 0.515509448 | 0.628436722 | 0.124941168 | 0.119346001 |
| Mrpl23 | -0.001493339 | 0.774982623 | 0.841266309 | 0.078807544 | 0.077624007 |
| Mrpl23-ps1 | 0.005983343 | 0.078963116 | 0.145929046 | 0.036808083 | 0.029159734 |
| Mrpl24 | -0.100138022 | 4.86E-23 | 1.74E-21 | 0.480146603 | 0.580168825 |
| Mrpl27 | -0.025867172 | 0.005487912 | 0.01560956 | 0.372913259 | 0.384302611 |
| Mrpl28 | -0.013159224 | 0.166695916 | 0.26514883 | 0.469080669 | 0.46199201 |
| Mrpl3 | -0.025226945 | 0.000557696 | 0.002138825 | 0.166140651 | 0.180207083 |
| Mrpl30 | 0.002615096 | 0.793832453 | 0.856080323 | 0.710934893 | 0.687072836 |
| Mrpl32 | -0.022620389 | 0.007867782 | 0.021241482 | 0.268705457 | 0.278301437 |
| Mrpl33 | 0.114100917 | 1.72E-28 | 8.49E-27 | 0.869858522 | 0.754083184 |
| Mrpl34 | 0.003250661 | 0.740641361 | 0.814515637 | 0.528703715 | 0.520766458 |
| Mrpl35 | -0.008573585 | 0.275045751 | 0.393118904 | 0.207095958 | 0.2114329 |
| Mrpl36 | -0.016423371 | 0.101058078 | 0.17827738 | 0.553396576 | 0.56645577 |
| Mrpl37 | 0.000645225 | 0.904725841 | 0.935418975 | 0.082781021 | 0.082039966 |
| Mrpl38 | -0.029181249 | 0.00049935 | 0.001940319 | 0.25113421 | 0.268222941 |
| Mrpl39 | -0.017063877 | 0.009164509 | 0.024099906 | 0.123548278 | 0.134527386 |
| Mrpl4 | -0.049225326 | 1.13E-07 | 9.48E-07 | 0.340538315 | 0.375094166 |
| Mrpl40 | -0.002243766 | 0.781175171 | 0.846218603 | 0.240612788 | 0.228599169 |
| Mrpl41 | -0.000960345 | 0.915510323 | 0.941562611 | 0.362062759 | 0.352567781 |
| Mrpl42 | -0.049681906 | 5.82E-07 | 4.28E-06 | 0.601939719 | 0.632711948 |
| Mrpl43 | -0.016948633 | 0.070195915 | 0.132642311 | 0.405902587 | 0.410091046 |
| Mrpl44 | -0.020332177 | 0.002599471 | 0.00820255 | 0.136049229 | 0.148463365 |

| | | | | | |
|---|---|---|---|---|---|
| Mrpl45 | 0.019234308 | 0.001076134 | 0.003819707 | 0.116773157 | 0.095517106 |
| Mrpl46 | -0.00137989 | 0.858028017 | 0.902395974 | 0.206826019 | 0.20280862 |
| Mrpl47 | -0.018688922 | 0.000785626 | 0.002896417 | 0.075292812 | 0.091902583 |
| Mrpl48 | -0.008955308 | 0.332345556 | 0.455033632 | 0.377048181 | 0.370577878 |
| Mrpl49 | -0.000904643 | 0.894752007 | 0.928392142 | 0.153231661 | 0.146650685 |
| Mrpl50 | 0.02178374 | 0.007882203 | 0.021268604 | 0.263666985 | 0.23203949 |
| Mrpl51 | -0.00118913 | 0.902943209 | 0.933973176 | 0.480009066 | 0.477111551 |
| Mrpl52 | 0.038267226 | 0.000139578 | 0.000629395 | 0.774195066 | 0.788152329 |
| Mrpl54 | 0.004402219 | 0.661360133 | 0.752456784 | 0.779967129 | 0.784299907 |
| Mrpl55 | -0.024586117 | 0.005500688 | 0.015636752 | 0.297827668 | 0.318971856 |
| Mrpl57 | 0.020374395 | 0.045617721 | 0.092394892 | 0.830066752 | 0.838820121 |
| Mrpl58 | -0.055108449 | 1.15E-08 | 1.13E-07 | 0.454936425 | 0.502337806 |
| Mrpl9 | -0.015918913 | 0.041128534 | 0.084858151 | 0.203591605 | 0.208993358 |
| Mrps10 | -0.018512238 | 0.014702214 | 0.035922324 | 0.180140989 | 0.195449573 |
| Mrps11 | -0.024871803 | 0.003440777 | 0.010469869 | 0.279263246 | 0.293354625 |
| Mrps12 | 0.041504446 | 2.20E-05 | 0.000118086 | 0.841783443 | 0.779577084 |
| Mrps14 | -0.036955673 | 0.000193444 | 0.000833627 | 0.719945319 | 0.754190522 |
| Mrps15 | -0.013646802 | 0.11656111 | 0.199719266 | 0.307341261 | 0.305725749 |
| Mrps16 | 0.016003266 | 0.114854759 | 0.197456758 | 0.728444823 | 0.714895024 |
| Mrps17 | -0.047824885 | 3.27E-07 | 2.52E-06 | 0.370441179 | 0.412838468 |
| Mrps18a | -0.00387429 | 0.679532164 | 0.767191835 | 0.412150932 | 0.404689398 |
| Mrps18b | -0.02565258 | 0.002191296 | 0.007063588 | 0.220627334 | 0.237093191 |
| Mrps18c | 0.019014622 | 0.036026936 | 0.076037923 | 0.348899587 | 0.332499133 |
| Mrps2 | 0.00086853 | 0.852453282 | 0.898680132 | 0.061820846 | 0.059537301 |
| Mrps21 | 0.095851885 | 4.70E-23 | 1.69E-21 | 0.862338891 | 0.764047782 |
| Mrps22 | -0.006396994 | 0.241677248 | 0.355728418 | 0.082923734 | 0.084657336 |
| Mrps23 | -0.010912441 | 0.2209323 | 0.331825091 | 0.322587189 | 0.323230513 |
| Mrps24 | -0.019805464 | 0.045390907 | 0.092063038 | 0.654308788 | 0.664252913 |
| Mrps25 | -0.005667073 | 0.440331285 | 0.558791378 | 0.173401949 | 0.176072728 |
| Mrps26 | -0.039910094 | 4.41E-05 | 0.000218984 | 0.449278285 | 0.486051107 |
| Mrps27 | -0.003327925 | 0.379932926 | 0.501305117 | 0.036294756 | 0.038310147 |
| Mrps28 | -0.03283708 | 0.000642542 | 0.002425938 | 0.39865632 | 0.417056703 |
| Mrps30 | -0.002036923 | 0.739708476 | 0.813958741 | 0.112694916 | 0.109357306 |
| Mrps31 | -0.006992373 | 0.348778043 | 0.471340024 | 0.183547626 | 0.182199964 |
| Mrps33 | 0.008440539 | 0.37508832 | 0.496948576 | 1.034766324 | 0.97706166 |
| Mrps34 | 0.000450818 | 0.963905283 | 0.975139143 | 0.539567709 | 0.53934518 |
| Mrps35 | -0.026979614 | 0.000193872 | 0.000835101 | 0.158943122 | 0.175158838 |
| Mrps36 | 0.030102657 | 0.000442289 | 0.001740161 | 0.311527729 | 0.270635418 |
| Mrps5 | 0.040535072 | 1.34E-06 | 9.14E-06 | 0.261356649 | 0.236550298 |
| Mrps6 | -0.010953061 | 0.059785264 | 0.116316257 | 0.098687067 | 0.101661291 |
| Mrps7 | -0.002648063 | 0.763825189 | 0.832930967 | 0.3293226 | 0.320050706 |
| Mrps9 | -0.008380458 | 0.261466609 | 0.377428118 | 0.182065021 | 0.184093678 |
| Mrrf | -0.018778587 | 0.00201397 | 0.006572612 | 0.100504896 | 0.114299984 |
| Mrs2 | 0.012420048 | 0.105025712 | 0.183908497 | 0.193703673 | 0.18498093 |
| Mrtfb | 0.015762561 | 0.001227082 | 0.004272811 | 0.073925438 | 0.060942705 |
| Mrto4 | -0.02253813 | 0.001186161 | 0.004154129 | 0.138482988 | 0.154514484 |
| Msantd2 | -0.005207225 | 0.205825173 | 0.314084497 | 0.041264375 | 0.045551172 |
| Msantd3 | -0.005837329 | 0.231666411 | 0.344856966 | 0.063824222 | 0.067096543 |
| Msantd4 | 0.028437782 | 0.000302563 | 0.001241148 | 0.2284527 | 0.197859661 |
| Msh2 | 0.000808211 | 0.872930286 | 0.912835134 | 0.074526718 | 0.070614297 |
| Msh3 | -0.006862261 | 0.105533929 | 0.184698624 | 0.043268732 | 0.048803359 |
| Msh6 | 0.011884797 | 0.004775538 | 0.013859038 | 0.055071399 | 0.043920548 |
| Msi1 | -0.005604131 | 0.196722858 | 0.302901295 | 0.051472755 | 0.051337144 |
| Msi2 | 0.039940355 | 0.000155678 | 0.000689207 | 0.913639452 | 0.873806528 |
| Msl1 | 0.020402518 | 0.02568901 | 0.057393548 | 0.37441234 | 0.340376773 |
| Msl2 | 0.016823541 | 0.027423124 | 0.060606414 | 0.196745101 | 0.178745444 |
| Msl3 | 0.023737245 | 0.000563012 | 0.002156639 | 0.156969301 | 0.139423648 |
| Msl3l2 | 0.002178461 | 0.705269742 | 0.787304217 | 0.098795484 | 0.094253332 |
| Msmo1 | 0.006575928 | 0.485336489 | 0.601328648 | 0.335323089 | 0.346495855 |
| Msra | 0.005393749 | 0.147077002 | 0.241202913 | 0.040911573 | 0.034896581 |
| Msrb1 | -0.017398129 | 0.010851615 | 0.027733807 | 0.145401226 | 0.151988917 |
| Msrb2 | -0.019359942 | 0.000255303 | 0.001066174 | 0.071781496 | 0.082864491 |
| Msto1 | 0.001527406 | 0.69272872 | 0.777410061 | 0.040716183 | 0.039673086 |
| Mt1 | 0.331487939 | 2.89E-78 | 1.22E-75 | 1.882350436 | 1.501125299 |
| Mt2 | 0.282985633 | 3.89E-82 | 1.80E-79 | 0.721159085 | 0.410813098 |
| Mt3 | -0.00415941 | 0.499834822 | 0.61418025 | 0.079371617 | 0.093381957 |
| Mta1 | 0.026773715 | 0.000215749 | 0.000917947 | 0.192670887 | 0.162600762 |
| Mta2 | 0.017136411 | 0.004569783 | 0.013356647 | 0.11401609 | 0.101562619 |
| Mta3 | -0.031039643 | 8.52E-05 | 0.000403984 | 0.206965275 | 0.223135941 |
| Mtap | -0.003856771 | 0.54394178 | 0.653671444 | 0.121573871 | 0.123084096 |
| Mtarc2 | -0.036917898 | 0.000143449 | 0.000645057 | 0.380010323 | 0.423501854 |
| mt-Atp6 | -0.009193083 | 0.163152043 | 0.2610399 | 0.109269775 | 0.117913472 |

| | | | | | |
|---|---|---|---|---|---|
| mt-Atp8 | 0.026233563 | 0.003373705 | 0.010310959 | 0.227734322 | 0.238010218 |
| Mtch1 | 0.001308594 | 0.886079582 | 0.921988902 | 1.162445605 | 1.104074478 |
| Mtch2 | 0.004025113 | 0.677810436 | 0.765973853 | 0.499206387 | 0.491844213 |
| Mtcl1 | 0.021923578 | 0.0062179 | 0.017395805 | 0.223451056 | 0.203451318 |
| mt-Co1 | -0.019415817 | 0.19889534 | 0.305760568 | 1.391615027 | 1.555440146 |
| mt-Cytb | -0.171938164 | 2.37E-28 | 1.15E-26 | 3.371601488 | 4.014595436 |
| Mtdh | 0.012748951 | 0.180252971 | 0.282466051 | 1.019313722 | 1.018922514 |
| Mterf1a | 0.002900786 | 0.455583034 | 0.574174003 | 0.041213322 | 0.038813685 |
| Mterf2 | 0.001372042 | 0.724831149 | 0.801411171 | 0.044055858 | 0.03997738 |
| Mterf3 | -0.004770542 | 0.516310265 | 0.629097431 | 0.173766964 | 0.172741002 |
| Mterf4 | -0.002154387 | 0.643187001 | 0.737432403 | 0.060893727 | 0.060239336 |
| Mtf1 | 0.016141036 | 0.03677838 | 0.077276726 | 0.217940014 | 0.194892428 |
| Mtf2 | -0.002614534 | 0.66413503 | 0.754201701 | 0.102977319 | 0.103457053 |
| Mtfmt | -0.001800947 | 0.676518895 | 0.764958909 | 0.048863525 | 0.050733433 |
| Mtfp1 | -0.013919696 | 0.00505941 | 0.014582742 | 0.064266274 | 0.071473611 |
| Mtfr1 | -0.017363666 | 0.021431993 | 0.049293077 | 0.182260441 | 0.190100354 |
| Mtfr1l | -0.031202338 | 0.000260424 | 0.001085694 | 0.268654216 | 0.288243188 |
| Mtg1 | -0.01363454 | 0.028348839 | 0.062364735 | 0.11076565 | 0.119175953 |
| Mtg2 | -0.018526769 | 0.003177685 | 0.009770226 | 0.113391762 | 0.123221316 |
| Mthfd1 | 0.031754254 | 3.77E-05 | 0.000191264 | 0.234029486 | 0.184463411 |
| Mthfd1l | 0.007856521 | 0.236076092 | 0.34979625 | 0.134198744 | 0.126874599 |
| Mthfd2 | 0.008076387 | 0.448066622 | 0.566905739 | 1.183678837 | 1.160404997 |
| Mthfd2l | -0.02085008 | 0.003844721 | 0.011515211 | 0.157908474 | 0.165765115 |
| Mthfr | -0.002535891 | 0.659742115 | 0.750967433 | 0.094894532 | 0.094427053 |
| Mthfs | -0.008399643 | 0.076386516 | 0.142238979 | 0.066715956 | 0.064807988 |
| Mthfsd | 0.002563865 | 0.654344535 | 0.74642591 | 0.097493712 | 0.094271589 |
| Mthfsl | -0.008245078 | 0.153033037 | 0.248129951 | 0.103913876 | 0.099624705 |
| Mtif2 | 0.01610107 | 0.031152278 | 0.06748272 | 0.188894719 | 0.171983157 |
| Mtif3 | 0.004030392 | 0.541710222 | 0.652037487 | 0.138854834 | 0.131072487 |
| Mtln | 0.029196966 | 0.003143243 | 0.009673506 | 0.495847365 | 0.486554005 |
| Mtm1 | 0.000946485 | 0.832684091 | 0.885415589 | 0.053667751 | 0.053583942 |
| Mtmr1 | 0.006396038 | 0.278945178 | 0.397640032 | 0.096717875 | 0.095201703 |
| Mtmr10 | -0.004467269 | 0.36206402 | 0.484846612 | 0.060891993 | 0.066202854 |
| Mtmr11 | -0.018981289 | 0.000457745 | 0.001795884 | 0.071412135 | 0.086648151 |
| Mtmr12 | -2.64E-05 | 0.99464955 | 0.996391852 | 0.041045506 | 0.040654824 |
| Mtmr14 | -0.002168963 | 0.55552225 | 0.664630484 | 0.036221369 | 0.036106085 |
| Mtmr2 | 0.020422679 | 0.002533908 | 0.008028923 | 0.152723051 | 0.135588201 |
| Mtmr3 | 0.040634799 | 6.33E-07 | 4.63E-06 | 0.258204636 | 0.216607529 |
| Mtmr4 | 0.005901006 | 0.253942475 | 0.369032696 | 0.074098171 | 0.070815332 |
| Mtmr6 | 0.044059907 | 2.32E-07 | 1.83E-06 | 0.289765735 | 0.247040704 |
| Mtmr7 | 0.028825563 | 0.000889709 | 0.003237183 | 0.310696056 | 0.273583282 |
| Mtmr9 | 0.016754618 | 0.000358475 | 0.001444298 | 0.071415787 | 0.055259589 |
| mt-Nd1 | -0.096462912 | 3.14E-09 | 3.34E-08 | 2.184308774 | 2.57785588 |
| mt-Nd2 | -0.117503506 | 6.65E-13 | 1.09E-11 | 1.855034056 | 2.232598369 |
| mt-Nd3 | 0.623596308 | 0 | 0 | 0.612167385 | 0.002268718 |
| mt-Nd4 | -0.156266871 | 3.41E-22 | 1.15E-20 | 2.377285554 | 2.854176536 |
| mt-Nd5 | -0.028011504 | 0.061792268 | 0.119646371 | 1.044332374 | 1.181281079 |
| mt-Nd6 | 0.008792911 | 0.156426527 | 0.252663016 | 0.087028245 | 0.090140267 |
| Mto1 | -0.012025559 | 0.012642539 | 0.031556037 | 0.059793299 | 0.067229268 |
| Mtor | 0.005601877 | 0.399575399 | 0.519758099 | 0.140739113 | 0.131858999 |
| Mtpap | 0.002583663 | 0.646812517 | 0.74032391 | 0.091926214 | 0.088346988 |
| Mtpn | 0.06029578 | 3.14E-10 | 3.79E-09 | 0.46580963 | 0.407507868 |
| Mtres1 | -0.00388825 | 0.59124409 | 0.695471874 | 0.168482786 | 0.166041771 |
| Mtrex | 0.007918871 | 0.206513878 | 0.314740229 | 0.122790565 | 0.111950822 |
| Mtrf1 | -0.004518696 | 0.236102785 | 0.34979625 | 0.036837724 | 0.039880748 |
| Mtrf1l | 0.002035809 | 0.662887019 | 0.753400468 | 0.0627817 | 0.058295173 |
| mt-Rnr1 | 0.067990304 | 5.68E-07 | 4.19E-06 | 0.810800552 | 0.727664725 |
| mt-Rnr2 | 0.00078981 | 0.96103465 | 0.973247798 | 2.706380388 | 2.836245495 |
| Mtrr | -0.001667507 | 0.668077224 | 0.757439857 | 0.038116618 | 0.040200023 |
| Mtss1 | 0.028889624 | 0.001426288 | 0.004870521 | 0.332917231 | 0.295515923 |
| Mtss2 | 0.007681084 | 0.366622281 | 0.489333034 | 0.240977578 | 0.245527853 |
| Mttp | -0.012657291 | 0.02753764 | 0.060818022 | 0.086509803 | 0.095540618 |
| Mturn | -0.006778871 | 0.136619591 | 0.227668094 | 0.053962356 | 0.057368348 |
| Mtus1 | 0.044781346 | 1.89E-10 | 2.34E-09 | 0.176670628 | 0.13743146 |
| Mtus2 | -0.009807186 | 0.023386995 | 0.053073848 | 0.037690853 | 0.048705839 |
| Mtx1 | -0.016776424 | 0.025159828 | 0.05650715 | 0.171480715 | 0.183331273 |
| Mtx2 | -0.021626508 | 0.013983068 | 0.034407337 | 0.307039443 | 0.315556431 |
| Mtx3 | -0.005654214 | 0.181388818 | 0.283514805 | 0.043597138 | 0.047782235 |
| Muc4 | 0.009310554 | 0.189793175 | 0.293944989 | 0.148355515 | 0.139304198 |
| Mul1 | -0.001247318 | 0.854148576 | 0.899775973 | 0.143544615 | 0.142984764 |
| Mus81 | -0.004208994 | 0.425533694 | 0.544178889 | 0.076038165 | 0.078992616 |
| Mvb12a | -0.01451762 | 0.13836168 | 0.229704961 | 0.49377224 | 0.504774657 |

| | | | | | |
|---|---|---|---|---|---|
| Mvk | 0.00075426 | 0.850617832 | 0.897807682 | 0.045937063 | 0.042677607 |
| Mvp | -0.009726545 | 0.287036709 | 0.406374091 | 0.315389802 | 0.322522732 |
| Mxd1 | 0.017525826 | 0.002354609 | 0.007537539 | 0.102446868 | 0.086965018 |
| Mxd4 | -0.048823077 | 1.70E-06 | 1.14E-05 | 0.541352603 | 0.556396898 |
| Mxi1 | 0.022272276 | 0.000612687 | 0.002324049 | 0.142235999 | 0.119489263 |
| Mxra7 | -0.011125524 | 0.016541574 | 0.039737381 | 0.057650052 | 0.060781876 |
| Mxra8 | -0.001055865 | 0.803507106 | 0.863361636 | 0.049276628 | 0.048608346 |
| Myadm | 0.009925874 | 0.067890046 | 0.129163801 | 0.087179051 | 0.080958542 |
| Mybbp1a | 0.026287828 | 0.002442475 | 0.007775292 | 0.277280702 | 0.257147781 |
| Mycbp | 0.041695194 | 1.39E-08 | 1.35E-07 | 0.202442987 | 0.161274219 |
| Mycbp2 | 0.046608394 | 5.64E-06 | 3.43E-05 | 0.84720092 | 0.792403515 |
| Myd88 | -0.003516844 | 0.377378293 | 0.498976718 | 0.040646197 | 0.043471171 |
| Mydgf | -0.045057828 | 6.97E-06 | 4.14E-05 | 0.650662284 | 0.711462844 |
| Myef2 | 0.016776418 | 0.05292206 | 0.104553601 | 0.286002155 | 0.264057499 |
| Myg1 | -0.054476977 | 3.99E-12 | 5.96E-11 | 0.191654162 | 0.228211957 |
| Myh10 | 0.026024066 | 2.03E-05 | 0.000109732 | 0.120311627 | 0.098739171 |
| Myh14 | 0.012116235 | 0.069200362 | 0.131169023 | 0.130058459 | 0.12081663 |
| Myh9 | 0.024843667 | 0.007772949 | 0.021014629 | 0.325187795 | 0.315841009 |
| Myl12a | -0.007714716 | 0.353703764 | 0.476077529 | 0.252794472 | 0.250103137 |
| Myl12b | 0.023288537 | 0.002033592 | 0.006627751 | 0.248173306 | 0.193790069 |
| Myl6 | 0.0250768 | 0.00036208 | 0.001455806 | 1.807426674 | 1.733968266 |
| Mylip | 0.00433345 | 0.291651884 | 0.411271524 | 0.046082599 | 0.041929677 |
| Mynn | 0.018331933 | 0.002717178 | 0.00852417 | 0.118901566 | 0.103251862 |
| Myo10 | 0.064751833 | 2.62E-13 | 4.51E-12 | 0.31423719 | 0.243877303 |
| Myo15b | 0.013786974 | 0.001390822 | 0.004772882 | 0.053289132 | 0.042137306 |
| Myo18a | 0.03716123 | 2.41E-06 | 1.57E-05 | 0.227085428 | 0.190085277 |
| Myo1b | 0.042571996 | 1.35E-15 | 2.74E-14 | 0.100474252 | 0.063525696 |
| Myo1c | 0.043728847 | 5.66E-10 | 6.64E-09 | 0.166537107 | 0.138389151 |
| Myo1d | 0.003124897 | 0.543760751 | 0.65353468 | 0.076712802 | 0.072536198 |
| Myo5a | 0.001355772 | 0.870958502 | 0.911361378 | 0.238654319 | 0.231558219 |
| Myo5b | 0.006920377 | 0.124448082 | 0.210414654 | 0.05678636 | 0.052055972 |
| Myo6 | 0.005205618 | 0.607162723 | 0.708504807 | 0.542164231 | 0.527992859 |
| Myo9a | 0.002249401 | 0.716818949 | 0.796462125 | 0.111576417 | 0.108047758 |
| Myo9b | -0.001914109 | 0.676774023 | 0.765158397 | 0.055518759 | 0.056534458 |
| Mypop | 0.008723636 | 0.128659261 | 0.216555633 | 0.104611002 | 0.094038299 |
| Mysm1 | 0.040590049 | 1.07E-05 | 6.11E-05 | 0.356612503 | 0.315948164 |
| Myt1 | 0.013693076 | 0.042060382 | 0.086413997 | 0.143750514 | 0.127676155 |
| Myt1l | -0.017954365 | 0.035940005 | 0.075892426 | 0.23352027 | 0.242767237 |
| Mzt1 | -0.002040431 | 0.784666906 | 0.849313255 | 0.184453049 | 0.179241127 |
| Mzt2 | -0.009132974 | 0.18110153 | 0.283202361 | 0.145800873 | 0.148254582 |
| N4bp1 | 0.017243625 | 0.057627638 | 0.112704868 | 0.340224501 | 0.314178441 |
| N4bp2 | -0.006469381 | 0.253680004 | 0.368706384 | 0.087956855 | 0.089892465 |
| N4bp2l1 | -0.046299832 | 2.98E-08 | 2.74E-07 | 0.205359577 | 0.244187487 |
| N4bp2l2 | -0.009024237 | 0.35542827 | 0.477803324 | 0.438869032 | 0.461023579 |
| N4bp2os | 0.000587918 | 0.866271793 | 0.908608736 | 0.034544024 | 0.031350169 |
| N6amt1 | -0.014669409 | 0.075612502 | 0.141181117 | 0.226480673 | 0.2362601 |
| Naa10 | -0.020936282 | 0.002173586 | 0.007015804 | 0.137387101 | 0.151928612 |
| Naa15 | 0.034137302 | 0.000240861 | 0.00101195 | 0.378575617 | 0.338468292 |
| Naa16 | -0.007467677 | 0.245413196 | 0.359366325 | 0.117684431 | 0.121255287 |
| Naa20 | 0.020520565 | 0.047081775 | 0.0950434 | 0.623602531 | 0.589769058 |
| Naa25 | 0.000749817 | 0.907711624 | 0.936216042 | 0.123466833 | 0.121899004 |
| Naa30 | 0.00646811 | 0.34299692 | 0.465078947 | 0.148644301 | 0.137474706 |
| Naa35 | -0.010818757 | 0.227128378 | 0.339557449 | 0.322592376 | 0.323618365 |
| Naa38 | 0.071968996 | 2.48E-13 | 4.28E-12 | 0.538818381 | 0.475153709 |
| Naa40 | 0.005014562 | 0.355707228 | 0.478046125 | 0.083246903 | 0.080102221 |
| Naa50 | 0.023790348 | 0.010747239 | 0.027524936 | 0.370536951 | 0.341462817 |
| Naa60 | 0.025462592 | 0.000155486 | 0.00068867 | 0.152139235 | 0.129666251 |
| Naa80 | -0.007373925 | 0.160277812 | 0.257389619 | 0.074846716 | 0.079507458 |
| Naaa | -0.066163859 | 2.27E-24 | 8.78E-23 | 0.068207328 | 0.127982226 |
| Nab1 | 0.024781827 | 0.000918005 | 0.00331902 | 0.175909959 | 0.162601413 |
| Nabp1 | -0.01140937 | 0.090178664 | 0.162324934 | 0.115369502 | 0.128245528 |
| Nabp2 | -0.033480328 | 0.000334613 | 0.001357158 | 0.35191591 | 0.377292919 |
| Naca | -0.046269328 | 3.85E-12 | 5.79E-11 | 1.990013013 | 2.030833852 |
| Nacc1 | 0.022349158 | 0.000297328 | 0.001221736 | 0.126210941 | 0.104225287 |
| Nacc2 | -0.000247295 | 0.956502812 | 0.970443551 | 0.055828439 | 0.0557133 |
| Nadk | -0.00597929 | 0.363903035 | 0.486705917 | 0.129321404 | 0.132899857 |
| Nadk2 | 0.00384961 | 0.420492502 | 0.539531226 | 0.066185326 | 0.061067336 |
| Nae1 | -0.020303386 | 0.020312566 | 0.047052362 | 0.294685163 | 0.305698393 |
| Naf1 | 0.001641422 | 0.754165519 | 0.824746589 | 0.075433312 | 0.07648928 |
| Naga | -0.018921393 | 0.049337206 | 0.09889821 | 0.444750255 | 0.450128493 |
| Nagk | -0.017030826 | 0.060881514 | 0.118141265 | 0.332324458 | 0.340507464 |
| Naglu | 0.001916494 | 0.74275462 | 0.816215713 | 0.100826738 | 0.096885299 |

| | | | | | |
|---|---|---|---|---|---|
| Nagpa | -0.00498109 | 0.222455058 | 0.33369975 | 0.043670627 | 0.045994811 |
| Naip2 | 0.004324054 | 0.49105268 | 0.606325286 | 0.117261241 | 0.112839489 |
| Nalcn | -0.002684115 | 0.657022606 | 0.748984152 | 0.106579563 | 0.103448036 |
| Nampt | 0.040791476 | 4.76E-08 | 4.25E-07 | 0.200315706 | 0.16539679 |
| Nanp | 0.015117431 | 0.000315297 | 0.001288489 | 0.05563246 | 0.042944388 |
| Nans | -0.037466433 | 3.71E-05 | 0.000188368 | 0.309974379 | 0.34149 |
| Nap1l1 | 0.021990651 | 0.025410705 | 0.056909209 | 0.791025959 | 0.737056314 |
| Nap1l2 | -0.007800513 | 0.189670663 | 0.293814242 | 0.098908223 | 0.103364605 |
| Nap1l3 | 0.006103024 | 0.219420062 | 0.330268128 | 0.071640851 | 0.065125992 |
| Nap1l4 | 0.010396888 | 0.251105436 | 0.366113835 | 0.351174173 | 0.330720091 |
| Nap1l5 | -0.130483749 | 1.73E-34 | 1.08E-32 | 0.251157814 | 0.392916951 |
| Napa | -0.003943636 | 0.690636823 | 0.775510648 | 0.866931056 | 0.847206914 |
| Napb | -0.001104654 | 0.869172068 | 0.910400898 | 0.116171584 | 0.126154977 |
| Napepld | 0.023703873 | 1.42E-05 | 7.94E-05 | 0.092754316 | 0.075325341 |
| Napg | -0.022455124 | 0.011671194 | 0.029579599 | 0.298309084 | 0.319797951 |
| Naprt | -0.022505635 | 1.96E-07 | 1.57E-06 | 0.037642663 | 0.055489902 |
| Narf | 0.008268731 | 0.019507479 | 0.045436443 | 0.037749693 | 0.030260703 |
| Nars | 0.016730868 | 0.094525557 | 0.168496052 | 1.048796728 | 1.064462215 |
| Nars2 | 0.008420855 | 0.090404224 | 0.162700827 | 0.073255527 | 0.065032533 |
| Nasp | 0.000918014 | 0.895545849 | 0.928990263 | 0.143944634 | 0.145743375 |
| Nat10 | 0.00268411 | 0.592456817 | 0.696561274 | 0.071349162 | 0.067940504 |
| Nat14 | 0.002600242 | 0.637494295 | 0.733110899 | 0.088865554 | 0.083706036 |
| Nat8f1 | 0.001831507 | 0.744370127 | 0.817528963 | 0.091923475 | 0.091364009 |
| Nat9 | -0.014710186 | 0.075989216 | 0.141661966 | 0.251570783 | 0.252231472 |
| Natd1 | 0.007459539 | 0.040268873 | 0.083456402 | 0.039787595 | 0.032677155 |
| Nav1 | 0.024948158 | 0.00952544 | 0.024872149 | 0.385141646 | 0.351751601 |
| Nav2 | -0.035651244 | 0.000380005 | 0.001519081 | 0.419299027 | 0.435818879 |
| Nav3 | 0.003189413 | 0.504887241 | 0.619136447 | 0.060272032 | 0.056925151 |
| Naxd | -0.05375464 | 1.35E-08 | 1.31E-07 | 0.395609648 | 0.436135376 |
| Naxe | -0.080534018 | 2.85E-16 | 6.04E-15 | 0.466272855 | 0.527135844 |
| Nbas | -0.044519914 | 1.79E-06 | 1.19E-05 | 0.350534473 | 0.380568191 |
| Nbdy | 0.006552838 | 0.46705972 | 0.584470923 | 0.367853198 | 0.338592812 |
| Nbea | 0.00138647 | 0.891328791 | 0.926196933 | 0.477715299 | 0.472431579 |
| Nbeal1 | 0.016465075 | 0.087498556 | 0.158527947 | 0.408109473 | 0.385605825 |
| Nbeal2 | 0.001473002 | 0.761288269 | 0.83057396 | 0.067926183 | 0.062385539 |
| Nbl1 | -0.016456459 | 0.018319537 | 0.043219793 | 0.130963698 | 0.142437705 |
| Nbn | 0.016334361 | 0.00673941 | 0.018634967 | 0.113043303 | 0.099357518 |
| Nbr1 | 0.017050186 | 0.068634289 | 0.130324718 | 0.397264538 | 0.370144607 |
| Ncald | 0.024436696 | 0.014244038 | 0.03495775 | 0.490766638 | 0.481699999 |
| Ncam1 | 0.015768116 | 0.121564963 | 0.206545713 | 0.514372588 | 0.475971679 |
| Ncapd3 | -0.015393037 | 0.005176646 | 0.014867756 | 0.078073224 | 0.087969176 |
| Ncaph2 | 0.005694638 | 0.375039362 | 0.496948576 | 0.132446035 | 0.12173645 |
| Ncbp1 | 0.016550727 | 0.022157612 | 0.05063853 | 0.176026504 | 0.159829272 |
| Ncbp2 | -0.012083274 | 0.160889825 | 0.258114006 | 0.288788857 | 0.289250718 |
| Ncbp3 | 0.005335383 | 0.460634333 | 0.578889218 | 0.170231796 | 0.163698468 |
| Ncdn | 0.013643861 | 0.009274232 | 0.024355506 | 0.086342996 | 0.072088178 |
| Nceh1 | -0.046772786 | 1.38E-08 | 1.34E-07 | 0.212443396 | 0.25108095 |
| Nck1 | 0.05169969 | 6.91E-09 | 7.01E-08 | 0.338756102 | 0.297980159 |
| Nck2 | 0.002969415 | 0.540813772 | 0.651374368 | 0.073078999 | 0.062587691 |
| Nckap1 | 0.012677952 | 0.200842693 | 0.308320332 | 0.828199863 | 0.801272081 |
| Nckap5 | -0.003440241 | 0.475189226 | 0.592128897 | 0.058100308 | 0.060090494 |
| Nckipsd | 0.006270374 | 0.121108725 | 0.206022577 | 0.046042949 | 0.041604949 |
| Ncl | 0.029930829 | 0.002376106 | 0.007591357 | 1.282125908 | 1.240405374 |
| Ncln | 0.0023152 | 0.725761963 | 0.802166645 | 0.138038677 | 0.13190403 |
| Ncoa1 | 0.052273912 | 1.83E-08 | 1.73E-07 | 0.392782693 | 0.326428727 |
| Ncoa2 | 0.031534661 | 4.94E-05 | 0.000243573 | 0.221275313 | 0.188404349 |
| Ncoa3 | 0.03202998 | 2.01E-06 | 1.33E-05 | 0.159790195 | 0.129096846 |
| Ncoa4 | 0.027905906 | 4.97E-05 | 0.000244341 | 0.167409183 | 0.136227711 |
| Ncoa5 | 0.011314514 | 0.03275633 | 0.070284052 | 0.087516327 | 0.075475104 |
| Ncoa6 | -0.000203476 | 0.976333885 | 0.98412836 | 0.143656219 | 0.143001057 |
| Ncoa7 | 0.097468579 | 9.06E-23 | 3.21E-21 | 0.442255274 | 0.357116116 |
| Ncor1 | 0.025631821 | 0.005650696 | 0.015997691 | 1.244990352 | 1.180974358 |
| Ncor2 | 0.052069922 | 1.84E-10 | 2.30E-09 | 0.254481157 | 0.210729693 |
| Ncstn | -0.027299034 | 0.00122588 | 0.004270155 | 0.241575248 | 0.270955488 |
| Ndel1 | 0.012799692 | 0.137158591 | 0.228230031 | 0.272982406 | 0.265611407 |
| Ndfip1 | -0.085147098 | 4.90E-25 | 1.97E-23 | 1.366126604 | 1.47968994 |
| Ndfip2 | -0.003931736 | 0.686146374 | 0.772486348 | 0.520474526 | 0.521432479 |
| Ndn | -0.027389652 | 0.001493173 | 0.00506335 | 0.256486155 | 0.276594872 |
| Ndrg1 | -0.029292546 | 0.004560749 | 0.013335217 | 0.485764937 | 0.50524048 |
| Ndrg3 | 0.036316591 | 8.85E-05 | 0.000418095 | 0.426199152 | 0.367447636 |
| Ndrg4 | -0.013572466 | 0.012379548 | 0.031031492 | 0.073143172 | 0.08531387 |
| Ndst1 | 0.034957707 | 1.31E-05 | 7.37E-05 | 0.237682536 | 0.205071584 |

| | | | | | |
|---|---|---|---|---|---|
| Ndufa1 | 0.120431362 | 1.98E-43 | 1.84E-41 | 1.227751507 | 1.128150432 |
| Ndufa10 | 0.007914544 | 0.409925069 | 0.529721316 | 0.460783914 | 0.447933937 |
| Ndufa11 | -0.00825096 | 0.306186905 | 0.427018949 | 1.396065471 | 1.431962997 |
| Ndufa12 | 0.020873521 | 0.033661705 | 0.071877684 | 0.485171307 | 0.463703912 |
| Ndufa13 | 0.017445575 | 0.028906955 | 0.063438694 | 1.466143789 | 1.484444888 |
| Ndufa2 | 0.090852257 | 3.95E-40 | 3.37E-38 | 1.760853925 | 1.681340642 |
| Ndufa3 | 0.064191474 | 1.18E-10 | 1.50E-09 | 0.633316253 | 0.583111363 |
| Ndufa4 | 0.099986235 | 1.61E-31 | 8.88E-30 | 1.539598064 | 1.378992793 |
| Ndufa5 | 0.067504331 | 2.44E-12 | 3.79E-11 | 0.966813901 | 0.921179854 |
| Ndufa6 | 0.047351538 | 6.49E-09 | 6.62E-08 | 1.486576617 | 1.41114542 |
| Ndufa7 | 0.023258743 | 0.001402894 | 0.004809215 | 1.598308861 | 1.595933861 |
| Ndufa8 | 0.003479764 | 0.720527904 | 0.798794646 | 0.696207646 | 0.676697191 |
| Ndufa9 | 0.023367821 | 0.006148612 | 0.017231711 | 0.300045096 | 0.269331576 |
| Ndufab1 | -0.041837941 | 2.37E-05 | 0.000126018 | 0.55600417 | 0.549348354 |
| Ndufaf1 | -0.00060325 | 0.892992166 | 0.927528829 | 0.057439498 | 0.054697108 |
| Ndufaf2 | -0.006579819 | 0.390261999 | 0.511391726 | 0.189389347 | 0.197663561 |
| Ndufaf3 | -0.036756058 | 0.000112153 | 0.000518228 | 0.417081785 | 0.4426684 |
| Ndufaf4 | 0.005244788 | 0.542576763 | 0.652792624 | 0.25614129 | 0.253673466 |
| Ndufaf5 | 0.000528807 | 0.901689461 | 0.933073436 | 0.051154803 | 0.049142216 |
| Ndufaf6 | 0.008272417 | 0.155844806 | 0.252016501 | 0.105570474 | 0.095662703 |
| Ndufaf7 | -0.004694324 | 0.490806515 | 0.606228044 | 0.14788025 | 0.14532938 |
| Ndufaf8 | 0.057288391 | 4.28E-09 | 4.49E-08 | 0.555341418 | 0.510615764 |
| Ndufb1 | 0.095286367 | 5.32E-32 | 3.03E-30 | 1.374423594 | 1.293218479 |
| Ndufb10 | -0.028681717 | 0.00119862 | 0.004187202 | 1.146121606 | 1.169216415 |
| Ndufb11 | -0.03172927 | 1.25E-05 | 7.01E-05 | 1.575263546 | 1.603048069 |
| Ndufb2 | 0.070178359 | 5.00E-14 | 9.08E-13 | 1.052282544 | 0.997740384 |
| Ndufb3 | 0.040054764 | 3.32E-05 | 0.000170819 | 0.865295444 | 0.826623847 |
| Ndufb4 | 0.027984075 | 0.002712228 | 0.008511388 | 0.363705528 | 0.356204737 |
| Ndufb5 | -0.058159307 | 6.39E-10 | 7.47E-09 | 0.961611384 | 1.020621621 |
| Ndufb6 | -0.018378019 | 0.069806215 | 0.132038904 | 0.74045091 | 0.7669222 |
| Ndufb7 | 0.009222312 | 0.286712716 | 0.405991296 | 1.1366591 | 1.145658716 |
| Ndufb8 | -0.015908045 | 0.071209106 | 0.134087725 | 1.22007667 | 1.194964225 |
| Ndufb9 | -0.006170958 | 0.432040447 | 0.550529472 | 1.552238607 | 1.546562267 |
| Ndufc1 | 0.091143173 | 5.42E-27 | 2.43E-25 | 1.306527495 | 1.208411409 |
| Ndufc2 | -0.021414119 | 0.027209645 | 0.060199869 | 1.585690214 | 1.668407384 |
| Ndufs1 | 0.025074767 | 0.002482334 | 0.007889263 | 0.26476014 | 0.236135582 |
| Ndufs2 | -0.043774891 | 0.000111176 | 0.000514449 | 1.055917641 | 1.084255777 |
| Ndufs3 | -0.029878994 | 0.001818154 | 0.006008159 | 0.501103688 | 0.494255445 |
| Ndufs4 | 0.020664416 | 0.039027221 | 0.081177289 | 0.65352609 | 0.629411552 |
| Ndufs5 | 0.071319294 | 4.80E-14 | 8.76E-13 | 0.996810184 | 0.916337712 |
| Ndufs6 | -0.001260573 | 0.89381322 | 0.927936305 | 0.902808081 | 0.923064113 |
| Ndufs7 | -0.019011317 | 0.041752712 | 0.085927151 | 0.897643583 | 0.913785498 |
| Ndufs8 | -0.068411765 | 3.55E-12 | 5.35E-11 | 0.491686315 | 0.546046753 |
| Ndufv1 | -0.087757313 | 5.07E-19 | 1.36E-17 | 0.482903895 | 0.566393196 |
| Ndufv2 | -0.022111591 | 0.025845011 | 0.057669313 | 0.692560595 | 0.678075793 |
| Ndufv3 | 0.091221676 | 7.78E-28 | 3.69E-26 | 1.435778508 | 1.317435351 |
| Neat1 | -0.100866251 | 5.98E-16 | 1.24E-14 | 0.615472339 | 0.797546315 |
| Nebl | 0.00985771 | 0.026144802 | 0.058217995 | 0.053079493 | 0.047159119 |
| Necab2 | -0.067904949 | 1.13E-19 | 3.20E-18 | 0.120210134 | 0.178152088 |
| Necap1 | 0.006854417 | 0.371453037 | 0.493814634 | 0.195970928 | 0.189709355 |
| Necap2 | -0.025762251 | 1.47E-05 | 8.18E-05 | 0.089051717 | 0.108750626 |
| Nectin1 | 0.029321654 | 4.61E-07 | 3.47E-06 | 0.113898378 | 0.085621859 |
| Nectin2 | 0.015470075 | 0.006036728 | 0.016953842 | 0.098419233 | 0.086870446 |
| Nectin3 | 0.056469842 | 1.49E-08 | 1.43E-07 | 0.528871382 | 0.451104018 |
| Nedd4 | 0.047072239 | 1.36E-06 | 9.27E-06 | 1.047851743 | 0.959831201 |
| Nedd4l | 0.033838574 | 1.75E-07 | 1.42E-06 | 0.140421057 | 0.107798648 |
| Nedd8 | 0.033605553 | 0.00024619 | 0.001032109 | 0.982754196 | 0.93934956 |
| Nedd9 | -0.012127488 | 0.047408741 | 0.095643864 | 0.094928134 | 0.106447538 |
| Nefm | 0.009734177 | 0.093319991 | 0.166836513 | 0.083197297 | 0.078812804 |
| Negr1 | 0.009645756 | 0.184979175 | 0.288107583 | 0.172509558 | 0.158730663 |
| Neil1 | 0.005238025 | 0.147046402 | 0.241202913 | 0.037281126 | 0.032910851 |
| Neil2 | -0.005964461 | 0.219511504 | 0.330303489 | 0.062851479 | 0.066951125 |
| Nek1 | -0.005965694 | 0.303480061 | 0.42379103 | 0.094069597 | 0.097018234 |
| Nek4 | -0.000594175 | 0.910984908 | 0.938894867 | 0.077347482 | 0.078173689 |
| Nek7 | 0.024186217 | 0.005742955 | 0.016221095 | 0.30205102 | 0.276053516 |
| Nek9 | 0.022207662 | 0.002694619 | 0.008467062 | 0.188871928 | 0.164300228 |
| Nelfa | 0.016889431 | 0.010555 | 0.027082531 | 0.145570392 | 0.128598559 |
| Nelfb | -0.0110577 | 0.138018757 | 0.229370659 | 0.187424019 | 0.189951376 |
| Nelfcd | -0.023585833 | 9.56E-05 | 0.000448765 | 0.094679477 | 0.11209168 |
| Nelfe | -0.033499616 | 0.000124563 | 0.000568547 | 0.277980076 | 0.302910695 |
| Nell1 | -0.01296499 | 0.184589978 | 0.287731885 | 0.46118256 | 0.45170081 |
| Nemf | -0.013146192 | 0.160350378 | 0.25741849 | 0.347399221 | 0.356882385 |

| | | | | | |
|---|---|---|---|---|---|
| Nemp1 | 0.002748814 | 0.567826242 | 0.67527911 | 0.063102951 | 0.062580908 |
| Nenf | -0.097548445 | 6.16E-36 | 4.16E-34 | 1.363904867 | 1.49368974 |
| Neo1 | 0.024418926 | 0.001470212 | 0.004999441 | 0.211390391 | 0.187305844 |
| Net1 | 0.023266496 | 2.43E-07 | 1.91E-06 | 0.067774532 | 0.046534613 |
| Neu1 | -0.013299997 | 0.16742331 | 0.265918873 | 0.397815425 | 0.420416535 |
| Neurl1a | 0.046449113 | 3.30E-17 | 7.59E-16 | 0.109647989 | 0.07131039 |
| Neurl4 | 0.007544793 | 0.152699222 | 0.247753977 | 0.081003188 | 0.074632301 |
| Neurod1 | 0.061983251 | 5.29E-10 | 6.25E-09 | 1.278792178 | 1.178187889 |
| Nf1 | -0.001994869 | 0.798253968 | 0.859236611 | 0.196302467 | 0.197309869 |
| Nf2 | 0.015097721 | 0.041956679 | 0.086268793 | 0.188477962 | 0.171657318 |
| Nfasc | -0.027285751 | 0.001915056 | 0.006292049 | 0.254013309 | 0.28754178 |
| Nfat5 | 0.0149456 | 0.116518616 | 0.199681647 | 0.33542103 | 0.32386786 |
| Nfatc2 | 0.014912878 | 0.008081138 | 0.021745039 | 0.101060769 | 0.081993183 |
| Nfatc3 | 0.006593014 | 0.30926637 | 0.430572483 | 0.138900999 | 0.124740608 |
| Nfe2l1 | 0.023442083 | 0.018211405 | 0.04301518 | 0.891908514 | 0.885243068 |
| Nfe2l2 | 0.019634706 | 0.012478765 | 0.031227427 | 0.220233667 | 0.19384675 |
| Nfia | 0.027009874 | 8.13E-06 | 4.76E-05 | 0.1206832 | 0.093892877 |
| Nfic | 0.058281911 | 1.65E-08 | 1.57E-07 | 0.647865557 | 0.589653855 |
| Nfil3 | -0.025778386 | 0.002705373 | 0.008495362 | 0.200201359 | 0.228525108 |
| Nfix | 0.031090759 | 8.53E-05 | 0.000404374 | 0.203808292 | 0.168712582 |
| Nfkb1 | -0.002779034 | 0.728822491 | 0.804628497 | 0.216419396 | 0.215212588 |
| Nfkb2 | 0.004661149 | 0.228989797 | 0.341762175 | 0.039949 | 0.036428696 |
| Nfkbia | 0.046263169 | 7.85E-05 | 0.000374999 | 0.403740939 | 0.363582367 |
| Nfkbib | 0.003802641 | 0.582852599 | 0.688767833 | 0.149021909 | 0.145254152 |
| Nfkbil1 | 0.023673992 | 0.001904616 | 0.006270463 | 0.211444636 | 0.181785955 |
| Nfkbiz | 0.024231643 | 0.000898499 | 0.003263058 | 0.156198017 | 0.146257056 |
| Nfrkb | -0.003579267 | 0.549868556 | 0.659164254 | 0.101762977 | 0.103668656 |
| Nfs1 | -0.029569493 | 2.79E-05 | 0.000145872 | 0.150907854 | 0.165825638 |
| Nfu1 | -0.061648418 | 9.58E-11 | 1.23E-09 | 0.307302739 | 0.367712473 |
| Nfx1 | 0.04212152 | 6.34E-08 | 5.54E-07 | 0.221583217 | 0.185209053 |
| Nfxl1 | -0.03014733 | 9.65E-05 | 0.000452293 | 0.173301198 | 0.20259793 |
| Nfyb | -0.015906879 | 0.062057497 | 0.120088174 | 0.240025975 | 0.251658305 |
| Nfyc | 0.018968744 | 0.000353546 | 0.001426213 | 0.094856695 | 0.074745326 |
| Ngdn | -0.008285788 | 0.31214115 | 0.433581405 | 0.236377344 | 0.235542645 |
| Ngly1 | -0.017357707 | 0.014099571 | 0.03465016 | 0.151758962 | 0.161743081 |
| Ngrn | -0.027700041 | 0.004433328 | 0.013033206 | 0.445169705 | 0.454674622 |
| Nhlrc2 | 0.013012754 | 0.061729942 | 0.11957332 | 0.155921936 | 0.143492812 |
| Nhlrc3 | 0.002861199 | 0.574209376 | 0.680705225 | 0.070419708 | 0.068743513 |
| Nhp2 | -0.019288004 | 0.012224437 | 0.030733379 | 0.176577486 | 0.196253537 |
| Nhsl2 | -0.008626378 | 0.025161433 | 0.05650715 | 0.035327783 | 0.03995133 |
| Niban1 | -0.050447248 | 2.50E-19 | 6.88E-18 | 0.042370332 | 0.092306476 |
| Niban2 | 0.019215492 | 0.0008556 | 0.003127121 | 0.114081859 | 0.087175388 |
| Nicn1 | -0.006045616 | 0.349162817 | 0.471597793 | 0.125402664 | 0.123999941 |
| Nif3l1 | 0.015645933 | 0.00067272 | 0.002525166 | 0.067167867 | 0.053447037 |
| Nifk | 0.006339809 | 0.374830174 | 0.49674195 | 0.164777153 | 0.15546202 |
| Nim1k | 0.011572451 | 0.006585317 | 0.018276464 | 0.054191325 | 0.04462148 |
| Nin | 0.001136205 | 0.794716936 | 0.856551155 | 0.049167733 | 0.04836332 |
| Ninj1 | -0.000920954 | 0.927356862 | 0.950628786 | 0.798416167 | 0.808096449 |
| Nip7 | -0.013454989 | 0.092697276 | 0.165886958 | 0.204176083 | 0.215727895 |
| Nipa1 | 0.015019204 | 0.002781794 | 0.008710017 | 0.080171174 | 0.065099859 |
| Nipa2 | 0.001779047 | 0.827291794 | 0.880886398 | 0.228654117 | 0.230700542 |
| Nipal1 | 0.02576719 | 0.011700859 | 0.02963161 | 0.485385783 | 0.429986104 |
| Nipal2 | 0.001222834 | 0.886639066 | 0.922210869 | 0.250248581 | 0.255241506 |
| Nipal3 | 0.013914799 | 0.139943589 | 0.231935147 | 0.385618685 | 0.354657764 |
| Nipbl | 0.047341731 | 3.57E-06 | 2.25E-05 | 0.669169758 | 0.604248384 |
| Nipsnap1 | -0.015411252 | 0.031693191 | 0.068365033 | 0.172176004 | 0.172681553 |
| Nipsnap2 | -0.019216506 | 0.014395506 | 0.035279332 | 0.202603869 | 0.214193577 |
| Nipsnap3b | -0.013719094 | 0.070325087 | 0.13278316 | 0.18331066 | 0.188445258 |
| Nisch | 0.021696054 | 0.026765432 | 0.059368818 | 0.934019533 | 0.871979544 |
| Nit1 | -0.002165802 | 0.769268534 | 0.837307287 | 0.184670165 | 0.1762478 |
| Nit2 | -0.000960458 | 0.81361186 | 0.871618127 | 0.046892009 | 0.044553088 |
| Nkap | 7.69E-05 | 0.991786198 | 0.994650858 | 0.185960199 | 0.183754342 |
| Nkapd1 | -0.012126767 | 0.09083805 | 0.163319685 | 0.155697203 | 0.162586157 |
| Nkiras1 | -0.02938454 | 0.000730247 | 0.002716976 | 0.281488576 | 0.301832948 |
| Nkiras2 | 0.010769644 | 0.073901323 | 0.138353295 | 0.118532995 | 0.101830647 |
| Nktr | -0.081830559 | 9.11E-15 | 1.75E-13 | 1.042431951 | 1.158638155 |
| Nkx2-2 | 0.050425715 | 9.13E-07 | 6.44E-06 | 0.60459806 | 0.53640341 |
| Nkx6-1 | 0.105807382 | 7.77E-22 | 2.53E-20 | 1.411441054 | 1.181503808 |
| Nlgn2 | 0.027113576 | 3.88E-06 | 2.44E-05 | 0.113106609 | 0.087184434 |
| Nlk | 0.045872559 | 1.10E-07 | 9.25E-07 | 0.297141257 | 0.254541182 |
| Nln | 0.001824668 | 0.632655434 | 0.729098641 | 0.038741282 | 0.037194619 |
| Nmd3 | -0.013808865 | 0.051698496 | 0.102576477 | 0.149708117 | 0.158480921 |

| | | | | | |
|---|---|---|---|---|---|
| Nme1 | -0.041071211 | 5.42E-05 | 0.000264755 | 1.093655731 | 1.165025985 |
| Nme2 | 0.009634687 | 0.137063383 | 0.228133918 | 0.118165973 | 0.122019767 |
| Nme4 | -0.01487488 | 0.002171557 | 0.007011584 | 0.066294742 | 0.068659405 |
| Nme5 | -0.023832179 | 1.39E-07 | 1.15E-06 | 0.04702946 | 0.061756995 |
| Nme6 | -0.007672573 | 0.114093663 | 0.196321875 | 0.062029311 | 0.067091623 |
| Nme7 | 0.020034407 | 2.88E-06 | 1.85E-05 | 0.057765419 | 0.04293926 |
| Nmi | -0.014966445 | 0.02611469 | 0.058164265 | 0.129085001 | 0.140215242 |
| Nmnat1 | -0.002521103 | 0.519760092 | 0.632112536 | 0.039749201 | 0.040449761 |
| Nmnat2 | -0.027588952 | 3.26E-07 | 2.52E-06 | 0.063793449 | 0.087112913 |
| Nmnat3 | -0.009141932 | 0.083560379 | 0.152788039 | 0.075123448 | 0.079106673 |
| Nmral1 | -0.005364215 | 0.223561935 | 0.334998324 | 0.05125175 | 0.053138701 |
| Nmt1 | 0.021595543 | 0.012008676 | 0.030301787 | 0.306080964 | 0.273694767 |
| Nmt2 | 0.043411243 | 2.13E-07 | 1.70E-06 | 0.276057302 | 0.232294139 |
| Nnat | 0.102014356 | 3.12E-65 | 6.74E-63 | 0.14348278 | 0.035871548 |
| Noa1 | -0.002943365 | 0.556911604 | 0.665719244 | 0.070625781 | 0.07037453 |
| Nob1 | 0.006252117 | 0.344834315 | 0.467216936 | 0.139623985 | 0.128755715 |
| Noc2l | 0.004515518 | 0.521406784 | 0.633718809 | 0.153364797 | 0.149477522 |
| Noc3l | 0.013476231 | 0.00977294 | 0.025377276 | 0.081291802 | 0.068894025 |
| Noc4l | -0.005989621 | 0.172278604 | 0.272074819 | 0.049203841 | 0.054022524 |
| Noct | 0.029222932 | 2.48E-09 | 2.67E-08 | 0.079959095 | 0.056444039 |
| Nod1 | 0.019164268 | 2.20E-05 | 0.000118086 | 0.07071738 | 0.049975541 |
| Nol10 | 0.005590492 | 0.127166815 | 0.214337042 | 0.037291797 | 0.033741102 |
| Nol11 | -0.002962203 | 0.696042453 | 0.77991889 | 0.187301576 | 0.188469099 |
| Nol12 | -0.019772683 | 0.020033988 | 0.046518961 | 0.25406359 | 0.266238018 |
| Nol3 | -0.038982149 | 1.41E-08 | 1.37E-07 | 0.117189007 | 0.14939945 |
| Nol4 | 0.004082541 | 0.638210707 | 0.733732138 | 0.286821068 | 0.272201636 |
| Nol4l | 0.008199025 | 0.031939327 | 0.06878913 | 0.042843431 | 0.03584884 |
| Nol6 | 0.012978135 | 0.007876922 | 0.021260255 | 0.072177055 | 0.062395553 |
| Nol7 | -0.023345279 | 0.019879126 | 0.0462135 | 0.723190992 | 0.740404196 |
| Nol8 | -0.008548088 | 0.259278811 | 0.375272978 | 0.17429784 | 0.17903175 |
| Nol9 | -0.001400578 | 0.740788651 | 0.814515637 | 0.048284624 | 0.047094416 |
| Nolc1 | 0.01833721 | 0.048330054 | 0.097280493 | 0.336382123 | 0.327422421 |
| Nom1 | 0.01504812 | 0.03043345 | 0.066161449 | 0.154964249 | 0.14203742 |
| Nomo1 | -0.020971931 | 0.002346151 | 0.007512938 | 0.132718055 | 0.151267813 |
| Nono | 0.029037552 | 0.0026007 | 0.008203765 | 0.485164638 | 0.443600512 |
| Nop10 | 0.011058868 | 0.243793414 | 0.35754406 | 0.989849359 | 1.008837674 |
| Nop14 | -0.009935366 | 0.243979518 | 0.357763027 | 0.233960234 | 0.24874026 |
| Nop16 | -0.02727353 | 2.10E-05 | 0.000113496 | 0.108436879 | 0.130170106 |
| Nop2 | -0.001331343 | 0.774620549 | 0.841155029 | 0.059904284 | 0.059089977 |
| Nop53 | 0.062303975 | 3.08E-11 | 4.20E-10 | 0.442925922 | 0.361290513 |
| Nop56 | -0.014740534 | 0.05806782 | 0.113451638 | 0.172392589 | 0.191662823 |
| Nop58 | -0.025002795 | 0.012341306 | 0.030979354 | 0.368389539 | 0.407453894 |
| Nop9 | 0.00029084 | 0.949089415 | 0.966182963 | 0.057410176 | 0.05673092 |
| Norad | 0.013760641 | 0.177090679 | 0.278517027 | 0.758020531 | 0.727376112 |
| Nosip | -0.016342379 | 0.008461745 | 0.022581688 | 0.106434734 | 0.117891298 |
| Nostrin | -0.006023769 | 0.247092715 | 0.361292733 | 0.048992926 | 0.06018407 |
| Notch1 | 0.017525338 | 0.00200938 | 0.006562041 | 0.098691631 | 0.082224786 |
| Nova2 | -0.006160431 | 0.18807917 | 0.29200027 | 0.059341031 | 0.060153125 |
| Npas3 | -0.004331977 | 0.293792185 | 0.413290063 | 0.040614436 | 0.044237767 |
| Npas4 | 0.004786833 | 0.360252598 | 0.482959212 | 0.056983234 | 0.061937386 |
| Npat | 0.012416207 | 0.055845089 | 0.109615578 | 0.134329972 | 0.11983985 |
| Npc1 | 0.008083756 | 0.152451509 | 0.247517296 | 0.094985219 | 0.087011367 |
| Npc2 | -0.04609951 | 7.01E-07 | 5.08E-06 | 1.028714649 | 1.052196021 |
| Npdc1 | -0.035233847 | 0.000389974 | 0.001553822 | 0.69624415 | 0.721224768 |
| Npepl1 | -0.012645796 | 0.228645967 | 0.341353822 | 0.454789206 | 0.462562951 |
| Npepps | 0.06820234 | 1.81E-11 | 2.54E-10 | 0.579522351 | 0.499750507 |
| Nphp1 | -0.019440416 | 0.002893307 | 0.009007773 | 0.123058044 | 0.13442563 |
| Nploc4 | 0.018708586 | 0.009500793 | 0.024836438 | 0.170188098 | 0.156296655 |
| Npm1 | -0.028451936 | 0.002883867 | 0.0089852 | 1.354472137 | 1.388993629 |
| Npm3 | -0.01322235 | 0.084866779 | 0.154682194 | 0.18272163 | 0.19208378 |
| Npnt | 0.026841685 | 0.000225515 | 0.000954905 | 0.17854154 | 0.144672795 |
| Npr1 | -0.014841151 | 0.00252798 | 0.008013377 | 0.061611134 | 0.068895593 |
| Nprl2 | -0.019424032 | 0.000739475 | 0.002745009 | 0.086919337 | 0.100130125 |
| Nprl3 | 0.009534058 | 0.081697805 | 0.14986152 | 0.089455651 | 0.082327073 |
| Nptn | -0.030134844 | 0.000813947 | 0.002987237 | 0.326701893 | 0.34178478 |
| Nptxr | 0.080426703 | 2.24E-22 | 7.67E-21 | 0.280911567 | 0.199420169 |
| Nqo1 | -0.012728433 | 0.008998281 | 0.023746277 | 0.051994099 | 0.059696767 |
| Nqo2 | -0.001931071 | 0.680076293 | 0.767402409 | 0.061478449 | 0.059318778 |
| Nr1d1 | 0.067975233 | 5.05E-26 | 2.15E-24 | 0.173490212 | 0.087351455 |
| Nr1d2 | 0.080363992 | 2.51E-24 | 9.66E-23 | 0.26171397 | 0.167562925 |
| Nr1h2 | 0.003063595 | 0.659071083 | 0.750555121 | 0.159704918 | 0.146247896 |
| Nr1h3 | -0.037378204 | 2.96E-09 | 3.16E-08 | 0.09566981 | 0.120109188 |

| | | | | | |
|---|---|---|---|---|---|
| Nr1h4 | 0.008753461 | 0.043179917 | 0.088303567 | 0.056931401 | 0.046947662 |
| Nr2c1 | -0.000566169 | 0.894850311 | 0.928392142 | 0.048793094 | 0.048648003 |
| Nr2c2 | 0.012842203 | 0.079978648 | 0.147380307 | 0.182268872 | 0.164570544 |
| Nr2c2ap | -0.019540649 | 0.006328137 | 0.017668623 | 0.152775121 | 0.168793808 |
| Nr2f6 | -0.007538284 | 0.358457479 | 0.481077252 | 0.25810025 | 0.245967905 |
| Nr3c1 | 0.045679549 | 5.70E-06 | 3.46E-05 | 0.515030058 | 0.454061082 |
| Nr3c2 | 0.002694082 | 0.486009086 | 0.601957987 | 0.041541839 | 0.038340283 |
| Nr4a1 | 0.08405683 | 6.73E-20 | 1.94E-18 | 0.259896753 | 0.218086941 |
| Nr4a2 | -0.014342265 | 0.133931227 | 0.224264449 | 0.242011554 | 0.287398951 |
| Nr6a1 | 0.005680642 | 0.233823266 | 0.34748239 | 0.063507937 | 0.060356711 |
| Nras | 0.001074642 | 0.871926732 | 0.912080018 | 0.134115487 | 0.13141501 |
| Nrbf2 | 0.007505049 | 0.17975848 | 0.28200935 | 0.095010693 | 0.087193264 |
| Nrbp1 | 0.005468572 | 0.465390936 | 0.582907844 | 0.183119382 | 0.178670001 |
| Nrbp2 | -0.010199011 | 0.020255177 | 0.04694974 | 0.042876563 | 0.053241115 |
| Nrcam | -0.046959267 | 4.04E-14 | 7.41E-13 | 0.056176902 | 0.105459661 |
| Nrd1 | 0.031831902 | 0.002083281 | 0.006763385 | 0.599041575 | 0.585951694 |
| Nrde2 | 0.000981239 | 0.816102439 | 0.873280392 | 0.049689903 | 0.047584757 |
| Nrep | 0.001641576 | 0.77064675 | 0.838150543 | 0.09055233 | 0.086930947 |
| Nrf1 | -0.079112733 | 1.22E-16 | 2.67E-15 | 0.237484233 | 0.317746729 |
| Nrip1 | -0.00780413 | 0.364345174 | 0.48709623 | 0.235274022 | 0.247182658 |
| Nrip3 | 0.010649465 | 0.011362473 | 0.02887999 | 0.049964029 | 0.042853927 |
| Nrsn1 | -0.022839512 | 4.15E-08 | 3.73E-07 | 0.032431981 | 0.049683252 |
| Nrtn | -0.021367411 | 0.035894716 | 0.075829733 | 0.382475333 | 0.402720777 |
| Nrxn1 | -0.043422615 | 7.95E-06 | 4.67E-05 | 0.380530342 | 0.400166346 |
| Nsa2 | 0.016204175 | 0.100907933 | 0.178078686 | 0.68192437 | 0.651311872 |
| Nsd1 | -0.020085887 | 0.050612983 | 0.100997417 | 0.544802836 | 0.555826138 |
| Nsd2 | 0.010660182 | 0.206350174 | 0.314589366 | 0.249779809 | 0.23983977 |
| Nsd3 | 0.01042405 | 0.30153027 | 0.42173516 | 0.589573851 | 0.578322838 |
| Nsf | 0.004468772 | 0.604634526 | 0.706992207 | 0.27754788 | 0.265359179 |
| Nsfl1c | -0.028217755 | 0.001044698 | 0.003720351 | 0.284363533 | 0.292545243 |
| Nsmaf | 0.019932911 | 0.002426976 | 0.007731016 | 0.141595361 | 0.122869534 |
| Nsmce1 | -0.015300018 | 0.051423366 | 0.1022159 | 0.19584592 | 0.210171428 |
| Nsmce2 | -0.013176015 | 0.069101453 | 0.131007083 | 0.168955741 | 0.172523011 |
| Nsmce3 | 0.003199149 | 0.723101864 | 0.800420062 | 0.345000424 | 0.328981933 |
| Nsmce4a | -0.020977902 | 0.024307529 | 0.054893798 | 0.38902013 | 0.388337735 |
| Nsmf | 0.006940787 | 0.145640855 | 0.239499389 | 0.067863727 | 0.059587643 |
| Nsrp1 | 0.061238336 | 2.08E-12 | 3.25E-11 | 0.314419442 | 0.259076554 |
| Nsun2 | 0.010101891 | 0.18133869 | 0.28348203 | 0.199393091 | 0.183904238 |
| Nsun3 | 0.008828773 | 0.083237864 | 0.152255601 | 0.076981502 | 0.067830113 |
| Nsun4 | -0.010248675 | 0.01739995 | 0.041471517 | 0.044836018 | 0.052740004 |
| Nsun6 | -0.006632967 | 0.111738918 | 0.193502986 | 0.041233176 | 0.046601437 |
| Nt5c | -0.019734118 | 0.012927027 | 0.032183498 | 0.21075698 | 0.220856914 |
| Nt5c2 | 0.013083635 | 0.118269436 | 0.201970043 | 0.268087709 | 0.244327718 |
| Nt5c3 | 0.021135269 | 0.037555362 | 0.07855276 | 0.588890796 | 0.537811314 |
| Nt5c3b | -0.010144475 | 0.062703482 | 0.120975038 | 0.08212553 | 0.087700246 |
| Nt5dc1 | -0.009222979 | 0.036990654 | 0.077605338 | 0.050111022 | 0.054378737 |
| Nt5dc2 | 0.02570536 | 0.000408756 | 0.001620689 | 0.196948665 | 0.155720896 |
| Nt5dc3 | 0.027561466 | 2.74E-06 | 1.77E-05 | 0.114908954 | 0.092183142 |
| Nt5m | -0.020385659 | 0.013606897 | 0.033634948 | 0.257520239 | 0.252889945 |
| Ntan1 | -0.031663992 | 0.000168684 | 0.000742057 | 0.261694647 | 0.274667592 |
| Ntmt1 | -0.004557122 | 0.486049188 | 0.601957987 | 0.132191984 | 0.12983497 |
| Ntpcr | 0.000798907 | 0.900168212 | 0.93179678 | 0.120615317 | 0.119282865 |
| Ntrk2 | -0.094181865 | 6.60E-20 | 1.91E-18 | 0.590792702 | 0.675563213 |
| Nuak1 | 0.019790759 | 0.001505385 | 0.005097649 | 0.129482397 | 0.100475504 |
| Nub1 | 0.010871166 | 0.213319186 | 0.323086014 | 0.2995279 | 0.280300096 |
| Nubp1 | 0.004479573 | 0.497920387 | 0.612757215 | 0.138399745 | 0.130319733 |
| Nubp2 | -0.002827833 | 0.594174496 | 0.697989905 | 0.080620974 | 0.078312491 |
| Nubpl | -0.015294683 | 0.002395175 | 0.007643208 | 0.068669082 | 0.074792041 |
| Nucb1 | -0.045235317 | 4.81E-06 | 2.96E-05 | 0.522576165 | 0.570251056 |
| Nucb2 | -0.125392558 | 1.69E-33 | 1.03E-31 | 0.962932786 | 1.188439192 |
| Nucks1 | 0.0022711 | 0.814892838 | 0.87250971 | 0.841841965 | 0.828979 |
| Nudc | 0.008077663 | 0.423169838 | 0.541822358 | 0.705279099 | 0.679461049 |
| Nudcd1 | -0.014531729 | 0.02154905 | 0.049515449 | 0.103936668 | 0.11605449 |
| Nudcd2 | -0.007178062 | 0.397469706 | 0.517919914 | 0.262262825 | 0.262067012 |
| Nudcd3 | 0.052533384 | 2.37E-08 | 2.20E-07 | 0.424050839 | 0.389401621 |
| Nudt13 | -0.003573088 | 0.556462219 | 0.665263857 | 0.102710784 | 0.105530951 |
| Nudt14 | -0.021020846 | 0.006271806 | 0.017531484 | 0.202651072 | 0.207727918 |
| Nudt16 | 0.011896482 | 0.038141604 | 0.079590614 | 0.104087475 | 0.090154621 |
| Nudt16l1 | 0.002937493 | 0.667101465 | 0.757039856 | 0.151992398 | 0.144647206 |
| Nudt18 | 0.023770958 | 0.000865545 | 0.003157641 | 0.178953125 | 0.155168557 |
| Nudt19 | 0.019335803 | 0.011861372 | 0.029967843 | 0.216935911 | 0.189692858 |
| Nudt2 | -0.015304458 | 0.025503244 | 0.057090154 | 0.146308994 | 0.153157961 |

| | | | | | |
|---|---|---|---|---|---|
| Nudt21 | 0.013496606 | 0.059480768 | 0.115770176 | 0.18338405 | 0.158987211 |
| Nudt22 | -0.018321317 | 0.002146335 | 0.006939363 | 0.096157997 | 0.108829514 |
| Nudt3 | 0.021015701 | 0.012573992 | 0.031425283 | 0.267262736 | 0.249558733 |
| Nudt4 | 0.071055899 | 6.61E-14 | 1.20E-12 | 0.404282731 | 0.339042464 |
| Nudt5 | -0.022183639 | 0.012485991 | 0.031237469 | 0.289805819 | 0.30580315 |
| Nudt7 | 0.004522243 | 0.574391909 | 0.680734784 | 0.214970322 | 0.20713547 |
| Nudt9 | -0.009958608 | 0.195066126 | 0.300734677 | 0.205202487 | 0.205100164 |
| Nufip1 | 0.009935226 | 0.087144919 | 0.158182021 | 0.104059537 | 0.092888575 |
| Nufip2 | 0.005075468 | 0.584961385 | 0.690015817 | 0.344219641 | 0.339833386 |
| Numa1 | 0.029067316 | 0.001982032 | 0.006483618 | 0.379278612 | 0.340696116 |
| Numb | 0.002162123 | 0.670716443 | 0.759634816 | 0.073539966 | 0.071101963 |
| Numbl | 0.004496766 | 0.292493517 | 0.411882829 | 0.051986387 | 0.047839551 |
| Nup133 | -0.007037101 | 0.115412523 | 0.198205361 | 0.051093517 | 0.055353529 |
| Nup153 | -0.001532471 | 0.821690132 | 0.877541103 | 0.141650956 | 0.140700957 |
| Nup155 | -0.005669438 | 0.158961085 | 0.255955935 | 0.040993727 | 0.044491516 |
| Nup160 | -0.001941173 | 0.724375634 | 0.800998625 | 0.081191326 | 0.08544004 |
| Nup188 | 0.003000319 | 0.484400694 | 0.600681574 | 0.051305592 | 0.049347078 |
| Nup205 | 0.006648802 | 0.153362019 | 0.248497591 | 0.062343528 | 0.057720312 |
| Nup210 | -0.018151506 | 0.008931771 | 0.02359638 | 0.136781853 | 0.150353015 |
| Nup214 | 0.003687086 | 0.608651654 | 0.709594841 | 0.168494064 | 0.162643541 |
| Nup35 | 0.004222085 | 0.530569526 | 0.642045921 | 0.143437035 | 0.13249922 |
| Nup37 | -0.013670924 | 0.005253005 | 0.015047058 | 0.060539561 | 0.07039352 |
| Nup43 | -0.006331968 | 0.165691825 | 0.263988188 | 0.054712898 | 0.057781287 |
| Nup50 | 0.02740699 | 0.001088258 | 0.003854296 | 0.26223794 | 0.235759941 |
| Nup54 | -0.002062216 | 0.714971457 | 0.795030596 | 0.090035858 | 0.08969121 |
| Nup62 | 0.0068884 | 0.254926694 | 0.370186306 | 0.111005263 | 0.100569163 |
| Nup85 | 0.001035938 | 0.795708188 | 0.85735066 | 0.046016954 | 0.042806435 |
| Nup88 | 0.021529816 | 0.004618602 | 0.013480051 | 0.205413606 | 0.183255808 |
| Nup93 | -0.036136391 | 5.47E-06 | 3.33E-05 | 0.167614038 | 0.202339555 |
| Nup98 | 0.003959564 | 0.62512216 | 0.723504481 | 0.226834749 | 0.220945164 |
| Nupr1 | -0.556882525 | 1.26E-176 | 4.08E-173 | 1.483060125 | 2.743180604 |
| Nupr1l | -0.022147816 | 0.00030414 | 0.001246709 | 0.103598342 | 0.115949949 |
| Nus1 | 0.026245064 | 0.004486764 | 0.013166412 | 0.36334331 | 0.337909742 |
| Nvl | -0.001308806 | 0.848578622 | 0.896531337 | 0.140182726 | 0.140419506 |
| Nxf1 | 0.004024597 | 0.47790404 | 0.594673374 | 0.093053561 | 0.089289555 |
| Nxn | -0.004959892 | 0.330368391 | 0.452946199 | 0.073758535 | 0.073618043 |
| Nxt1 | 0.009891146 | 0.131369625 | 0.220468755 | 0.145842096 | 0.131831785 |
| Nxt2 | 0.002105885 | 0.683796081 | 0.770499014 | 0.079445216 | 0.074046308 |
| Oaf | -0.003507497 | 0.462864572 | 0.580567745 | 0.068841862 | 0.062150625 |
| Oard1 | -0.035803562 | 8.20E-05 | 0.000389844 | 0.333860594 | 0.356878768 |
| Oasl2 | 0.024921885 | 1.76E-09 | 1.93E-08 | 0.050041453 | 0.032252861 |
| Oaz1 | -0.004019828 | 0.506076157 | 0.61999917 | 2.097011437 | 2.042727428 |
| Oaz2 | 0.006903533 | 0.423451279 | 0.54189724 | 0.300828867 | 0.283039403 |
| Obi1 | 0.003712809 | 0.385228446 | 0.506243708 | 0.051991099 | 0.048379661 |
| Obsl1 | -0.008329428 | 0.079996491 | 0.147380307 | 0.057853055 | 0.063909491 |
| Ocel1 | -0.004127219 | 0.498266009 | 0.612949784 | 0.11086527 | 0.109320333 |
| Ociad1 | 0.001254778 | 0.899637433 | 0.931544906 | 0.688549958 | 0.667129788 |
| Ociad2 | 0.017631973 | 0.082175284 | 0.150623045 | 0.944785525 | 0.869743164 |
| Ocln | 0.017568019 | 7.50E-05 | 0.000359081 | 0.060550594 | 0.047279819 |
| Ocrl | 0.004114786 | 0.558114508 | 0.666665345 | 0.138193093 | 0.13786383 |
| Odc1 | -0.005917385 | 0.605663644 | 0.707383704 | 0.424029824 | 0.493100959 |
| Odf2 | 0.002862064 | 0.681555157 | 0.768755812 | 0.146809825 | 0.146662318 |
| Odf2l | 0.009061014 | 0.139995926 | 0.231982341 | 0.111339051 | 0.104373737 |
| Odr4 | -0.002094615 | 0.822712814 | 0.878074806 | 0.355118135 | 0.361660356 |
| Ofd1 | -0.004833148 | 0.521496005 | 0.63374802 | 0.176700098 | 0.18061275 |
| Oga | 0.030651917 | 0.001148868 | 0.004035151 | 0.38600168 | 0.34029431 |
| Ogdh | 0.067256268 | 2.39E-11 | 3.29E-10 | 0.610528335 | 0.544453315 |
| Ogdhl | 0.046396157 | 1.29E-10 | 1.64E-09 | 0.18160897 | 0.141489816 |
| Ogfod1 | 0.005112793 | 0.433782529 | 0.552138484 | 0.125211148 | 0.124749538 |
| Ogfod2 | 0.002923187 | 0.704304031 | 0.786406775 | 0.203794722 | 0.195118606 |
| Ogfod3 | -0.032886845 | 2.95E-07 | 2.29E-06 | 0.109189095 | 0.131963854 |
| Ogfr | -0.005159265 | 0.497849758 | 0.612757215 | 0.195560331 | 0.189458193 |
| Ogfrl1 | 0.009321256 | 0.009331306 | 0.024472339 | 0.039058125 | 0.029911745 |
| Ogg1 | 0.001568544 | 0.649500789 | 0.742788691 | 0.034766052 | 0.030735169 |
| Ogt | -0.066694414 | 2.71E-12 | 4.15E-11 | 0.318805391 | 0.373515728 |
| Oip5os1 | -0.004220451 | 0.607777674 | 0.70883092 | 0.234253862 | 0.232013893 |
| Ola1 | -0.040701129 | 1.83E-05 | 0.000100085 | 0.380515918 | 0.412242741 |
| Olfm1 | 0.010938875 | 0.020436824 | 0.047312593 | 0.06516605 | 0.057397919 |
| Olfm4 | -0.007107087 | 0.206984218 | 0.315259371 | 0.093280898 | 0.088260029 |
| Oma1 | -0.013057162 | 0.009220706 | 0.024238822 | 0.063973746 | 0.072603012 |
| Opa1 | 0.01953265 | 0.013966635 | 0.034375602 | 0.220218248 | 0.19990129 |
| Opa3 | 0.021219266 | 0.001219883 | 0.00425384 | 0.141433883 | 0.119091044 |

| | | | | | |
|---|---|---|---|---|---|
| Ophn1 | 0.001025191 | 0.812120315 | 0.870539744 | 0.04958431 | 0.048686871 |
| Optn | 0.003020216 | 0.570827632 | 0.677936262 | 0.079243357 | 0.077188316 |
| Orai1 | 0.001789948 | 0.716703954 | 0.796462125 | 0.068611852 | 0.066920198 |
| Orai2 | 0.009542758 | 0.026985829 | 0.059803108 | 0.054466867 | 0.047233998 |
| Orai3 | 0.003038193 | 0.666967072 | 0.756975703 | 0.166594417 | 0.156735734 |
| Orc2 | 0.000603479 | 0.93656426 | 0.957014679 | 0.182364272 | 0.18033303 |
| Orc3 | -0.002331196 | 0.707185037 | 0.788717785 | 0.113562814 | 0.112529353 |
| Orc4 | 0.009292747 | 0.221166479 | 0.332022932 | 0.193208935 | 0.18292415 |
| Orc5 | 0.017224349 | 0.000810871 | 0.002977071 | 0.08341893 | 0.067396701 |
| Orc6 | 0.026327435 | 2.14E-05 | 0.000115197 | 0.12922028 | 0.098488639 |
| Ormdl1 | -0.007589268 | 0.157285594 | 0.253764597 | 0.080722318 | 0.082979652 |
| Ormdl2 | -0.016198334 | 0.044165754 | 0.08996008 | 0.224062956 | 0.232877271 |
| Ormdl3 | -0.057286809 | 8.77E-09 | 8.82E-08 | 0.472577398 | 0.536023891 |
| Os9 | -0.064907575 | 3.14E-10 | 3.79E-09 | 0.592902997 | 0.675040337 |
| Osbp | 0.000406929 | 0.96443082 | 0.975376522 | 0.330421188 | 0.325582752 |
| Osbpl11 | 0.016460119 | 0.0001306 | 0.000592759 | 0.058588243 | 0.043593927 |
| Osbpl1a | 0.059453373 | 4.50E-15 | 8.72E-14 | 0.216676211 | 0.163819827 |
| Osbpl2 | 0.013201658 | 0.095907554 | 0.17064664 | 0.221741201 | 0.205592932 |
| Osbpl3 | 0.01015601 | 0.015472719 | 0.037475281 | 0.052659312 | 0.041780539 |
| Osbpl5 | 0.004901137 | 0.280701555 | 0.399557909 | 0.061573268 | 0.051978255 |
| Osbpl6 | 0.017198236 | 0.004624382 | 0.01349287 | 0.116805888 | 0.098653866 |
| Osbpl7 | 0.003531126 | 0.390955353 | 0.511909478 | 0.048525989 | 0.043924539 |
| Osbpl8 | 0.029327563 | 5.46E-05 | 0.000266152 | 0.179889231 | 0.152174247 |
| Osbpl9 | 0.017718305 | 0.0133901 | 0.033208814 | 0.174026311 | 0.156872182 |
| Oscp1 | -0.016795131 | 0.000361802 | 0.00145529 | 0.0559668 | 0.066149238 |
| Oser1 | -0.008716926 | 0.246416295 | 0.360574837 | 0.180593672 | 0.179997273 |
| Osgep | -0.035891419 | 8.65E-06 | 5.04E-05 | 0.225762139 | 0.243517148 |
| Osgepl1 | -0.004673487 | 0.296537271 | 0.416308353 | 0.053350032 | 0.05430987 |
| Osgin1 | -0.022495335 | 0.000699205 | 0.002613482 | 0.100457029 | 0.128994175 |
| Osgin2 | 0.004722948 | 0.289905735 | 0.409244019 | 0.055475436 | 0.052924446 |
| Ost4 | 0.078498652 | 2.18E-19 | 6.03E-18 | 1.304744537 | 1.208113259 |
| Ostc | -0.096980261 | 1.24E-25 | 5.15E-24 | 1.265203117 | 1.418179639 |
| Ostf1 | -0.011295676 | 0.138082026 | 0.229397379 | 0.1988273 | 0.203157806 |
| Ostm1 | -0.004349604 | 0.530103879 | 0.641562295 | 0.137896755 | 0.144058128 |
| Otub1 | 0.010509343 | 0.244783718 | 0.358617737 | 0.350260405 | 0.337173944 |
| Otub2 | -0.001031002 | 0.862711557 | 0.905949638 | 0.106952423 | 0.102829634 |
| Otud3 | 0.015057496 | 0.005023286 | 0.014495811 | 0.087775441 | 0.074591999 |
| Otud4 | 0.020778303 | 0.013165012 | 0.032700626 | 0.246759919 | 0.233092516 |
| Otud5 | 0.035509012 | 2.23E-06 | 1.46E-05 | 0.206453591 | 0.172082545 |
| Otud6b | 0.020085992 | 0.002618991 | 0.008256106 | 0.143443487 | 0.127145693 |
| Otud7b | 0.016303054 | 0.029529082 | 0.064570792 | 0.186376709 | 0.169703921 |
| Otulin | 0.034650701 | 3.87E-05 | 0.000195432 | 0.275804462 | 0.243005578 |
| Otulinl | 0.010604287 | 0.007612751 | 0.020656199 | 0.048461173 | 0.038956215 |
| Ovol2 | 0.027569856 | 0.000111394 | 0.000515209 | 0.157042568 | 0.1288442 |
| Oxa1l | 0.005276216 | 0.42040253 | 0.539531226 | 0.139459958 | 0.127467532 |
| Oxct1 | 0.04576643 | 4.50E-06 | 2.78E-05 | 0.581955906 | 0.513036881 |
| Oxld1 | 0.003729402 | 0.545331222 | 0.655017314 | 0.123111278 | 0.112000606 |
| Oxnad1 | -0.012673002 | 0.031269786 | 0.06767695 | 0.09712303 | 0.103392612 |
| Oxr1 | 0.010277391 | 0.210003723 | 0.319008781 | 0.252897812 | 0.231143309 |
| Oxsm | 0.015678287 | 0.000149577 | 0.000666445 | 0.05167755 | 0.0402684 |
| Oxsr1 | 0.022959109 | 0.00054861 | 0.002108137 | 0.148093489 | 0.126302884 |
| P2rx4 | 0.006896543 | 0.364777465 | 0.487540076 | 0.191600092 | 0.178277043 |
| P2ry1 | 0.032002989 | 3.73E-07 | 2.85E-06 | 0.131810151 | 0.101055648 |
| P3h1 | -0.003031446 | 0.459314054 | 0.577603316 | 0.040340467 | 0.04536732 |
| P3h4 | -0.027659846 | 0.000329861 | 0.00134068 | 0.199263251 | 0.207263083 |
| P4ha1 | -0.007304658 | 0.377175969 | 0.498830741 | 0.212146678 | 0.224786038 |
| P4ha2 | -0.00581816 | 0.402005449 | 0.522011082 | 0.136104648 | 0.144479778 |
| P4hb | 0.06512495 | 1.93E-11 | 2.70E-10 | 1.1239784 | 1.080379868 |
| P4htm | -0.005013801 | 0.487159444 | 0.602948964 | 0.160254553 | 0.163696231 |
| Pa2g4 | -0.059113747 | 1.61E-09 | 1.78E-08 | 0.394138021 | 0.449854834 |
| Pabpc1 | 0.031773654 | 0.000850859 | 0.00311565 | 1.098979993 | 1.054629221 |
| Pabpc1l | -0.008719015 | 0.018728798 | 0.043974199 | 0.036793415 | 0.037818177 |
| Pabpc4 | 0.005721376 | 0.358849042 | 0.481536285 | 0.129501473 | 0.110777853 |
| Pabpn1 | -0.008414818 | 0.24730155 | 0.361543709 | 0.171790588 | 0.169151979 |
| Pacc1 | 0.017800631 | 0.013375481 | 0.033181022 | 0.184290416 | 0.156069512 |
| Pacrg | -0.004013801 | 0.449392058 | 0.567961121 | 0.084890613 | 0.081238792 |
| Pacs1 | 0.01004473 | 0.064439022 | 0.123711725 | 0.08967647 | 0.080020022 |
| Pacs2 | -0.000855499 | 0.85874132 | 0.902950802 | 0.06341632 | 0.063261109 |
| Pacsin1 | 0.01914477 | 0.006020389 | 0.01691625 | 0.162781517 | 0.143500346 |
| Pacsin2 | -0.017527337 | 0.010074652 | 0.026049406 | 0.122526385 | 0.141752723 |
| Paf1 | 0.007132975 | 0.328555235 | 0.451096455 | 0.173390407 | 0.167516889 |
| Pafah1b1 | 0.047756243 | 5.73E-07 | 4.22E-06 | 1.083676974 | 1.038619796 |

| | | | | | |
|---|---|---|---|---|---|
| Pafah1b2 | 0.010848433 | 0.257105936 | 0.372960894 | 0.490964867 | 0.471311034 |
| Pafah1b3 | -0.00139068 | 0.862125171 | 0.90572519 | 0.213952319 | 0.216145188 |
| Pag1 | 0.009813496 | 0.058945014 | 0.114911456 | 0.076410068 | 0.070884764 |
| Paics | -0.001207589 | 0.88743411 | 0.922939069 | 0.268281067 | 0.266709858 |
| Paip1 | -0.004089971 | 0.630108641 | 0.727336772 | 0.274335561 | 0.267686873 |
| Paip2 | -0.042184767 | 4.54E-06 | 2.81E-05 | 0.975381578 | 1.005236181 |
| Paip2b | 0.00993508 | 0.223737496 | 0.33520973 | 0.233370695 | 0.227405184 |
| Pak1 | 0.015529128 | 2.95E-05 | 0.000153194 | 0.043579947 | 0.030949478 |
| Pak1ip1 | -0.027377776 | 0.003326937 | 0.010184031 | 0.349654884 | 0.372098678 |
| Pak2 | 0.041725196 | 2.68E-05 | 0.00014053 | 0.596390428 | 0.560359159 |
| Pak3 | -0.001464624 | 0.893574085 | 0.927936305 | 0.458277368 | 0.464794431 |
| Pak4 | 0.026093249 | 1.49E-05 | 8.28E-05 | 0.121747767 | 0.098718168 |
| Pam | -0.12035346 | 4.92E-48 | 5.63E-46 | 1.790203211 | 2.108256428 |
| Pam16 | -0.01181215 | 0.243301946 | 0.35703872 | 0.634812478 | 0.649074904 |
| Pamr1 | -0.001830915 | 0.812869638 | 0.871110959 | 0.183170596 | 0.185602626 |
| Pan2 | 0.000686963 | 0.869193144 | 0.910400898 | 0.046874119 | 0.044770875 |
| Pan3 | 0.020273947 | 0.00522812 | 0.01499787 | 0.173071978 | 0.154723898 |
| Pank1 | 0.003469912 | 0.533075895 | 0.644111274 | 0.093457565 | 0.084634826 |
| Pank2 | 0.016577719 | 0.011735358 | 0.029703502 | 0.140166731 | 0.124091847 |
| Pank3 | 0.023861207 | 0.00528024 | 0.015111713 | 0.281443445 | 0.253185309 |
| Pank4 | 0.004055676 | 0.26862466 | 0.385345946 | 0.039167334 | 0.034291091 |
| Papola | 0.038139932 | 4.23E-05 | 0.000210774 | 1.120718694 | 1.055682885 |
| Papolg | -0.000187991 | 0.961871584 | 0.973760559 | 0.039923333 | 0.039803149 |
| Pappa2 | -0.113336662 | 7.12E-22 | 2.33E-20 | 0.245592957 | 0.394386197 |
| Papss1 | 0.006657315 | 0.348404404 | 0.470966019 | 0.173124338 | 0.159379637 |
| Papss2 | 0.020863757 | 0.048859346 | 0.098142678 | 0.960994662 | 0.867383534 |
| Paqr3 | -0.017984632 | 2.87E-05 | 0.000149906 | 0.03523723 | 0.052234042 |
| Paqr9 | 0.006327949 | 0.105998355 | 0.185344606 | 0.044222642 | 0.038053655 |
| Pard3 | 0.023297242 | 0.002989285 | 0.009264211 | 0.218086338 | 0.189203605 |
| Pard3b | -0.026863003 | 3.14E-06 | 2.00E-05 | 0.075307695 | 0.096942882 |
| Pard6a | 0.00772166 | 0.325796678 | 0.447941636 | 0.210728784 | 0.200405759 |
| Pard6g | 0.01178663 | 0.020037492 | 0.046518961 | 0.073968723 | 0.063655935 |
| Parg | 0.006276977 | 0.400663762 | 0.520755762 | 0.180328411 | 0.172723831 |
| Park7 | 0.009662357 | 0.265861611 | 0.382296491 | 1.621294445 | 1.583910613 |
| Parl | -0.059754479 | 2.34E-13 | 4.04E-12 | 0.206844343 | 0.254611806 |
| Parm1 | 0.053548123 | 1.32E-21 | 4.19E-20 | 0.116793311 | 0.070173187 |
| Parn | 0.012727322 | 0.078389223 | 0.145134265 | 0.170069299 | 0.160322441 |
| Parp1 | 0.021291885 | 0.008745771 | 0.023205892 | 0.231746148 | 0.209454291 |
| Parp10 | 0.002546217 | 0.6045107 | 0.706992207 | 0.068835265 | 0.065331602 |
| Parp11 | -0.001683321 | 0.659277856 | 0.750614746 | 0.040168343 | 0.038512868 |
| Parp12 | 0.015794585 | 0.011375793 | 0.028906288 | 0.125069551 | 0.107783433 |
| Parp14 | 0.027555234 | 2.46E-06 | 1.60E-05 | 0.104769847 | 0.083147304 |
| Parp16 | 0.001677106 | 0.714428862 | 0.794790868 | 0.058780348 | 0.055844235 |
| Parp2 | -0.002102267 | 0.7793809 | 0.845157198 | 0.179009849 | 0.181066915 |
| Parp4 | 0.001911645 | 0.742222869 | 0.815723573 | 0.099791504 | 0.095332914 |
| Parp6 | 0.005879512 | 0.403126627 | 0.523047787 | 0.17026999 | 0.150301383 |
| Parp8 | 0.007567882 | 0.362751946 | 0.485500334 | 0.253229705 | 0.24016717 |
| Parp9 | 0.005290532 | 0.289892966 | 0.409244019 | 0.072638109 | 0.064592056 |
| Particl | 0.000425465 | 0.91961464 | 0.944684439 | 0.04946831 | 0.047278925 |
| Parva | 0.023482027 | 0.00455697 | 0.013328178 | 0.268713839 | 0.238960491 |
| Patj | 0.015053583 | 0.010545806 | 0.027073232 | 0.10757328 | 0.092922103 |
| Patl1 | 0.014869899 | 0.003498294 | 0.010614986 | 0.078574196 | 0.066919961 |
| Patz1 | 0.009731304 | 0.065477062 | 0.125432118 | 0.084877415 | 0.07424252 |
| Paupar | -0.00533347 | 0.199262787 | 0.30618505 | 0.041865414 | 0.04376107 |
| Pawr | 0.009391195 | 0.174428543 | 0.275157277 | 0.150790662 | 0.139834381 |
| Pax6 | 0.043647328 | 7.14E-06 | 4.24E-05 | 1.270663244 | 1.199300221 |
| Pax6os1 | -0.004528109 | 0.419994138 | 0.539155827 | 0.09165854 | 0.08750767 |
| Paxbp1 | 0.016700036 | 0.030186173 | 0.065756212 | 0.188877282 | 0.181267321 |
| Paxip1 | 0.011907447 | 0.118299535 | 0.201985964 | 0.197443693 | 0.178171321 |
| Paxx | -0.045839865 | 4.85E-09 | 5.04E-08 | 0.197380427 | 0.222274996 |
| Pbdc1 | -0.015827927 | 0.066552441 | 0.127150558 | 0.265148849 | 0.28092114 |
| Pbrm1 | 0.039826245 | 0.000111569 | 0.000515776 | 0.766868679 | 0.720159391 |
| Pbx1 | 0.012686697 | 0.171190548 | 0.270620245 | 0.360148965 | 0.33807891 |
| Pbx2 | 0.016899187 | 0.031518843 | 0.068124986 | 0.233157985 | 0.2034132 |
| Pbx3 | 0.003523779 | 0.671931781 | 0.76065682 | 0.251254507 | 0.237807849 |
| Pbxip1 | -0.052400699 | 2.06E-07 | 1.65E-06 | 0.339884137 | 0.411687313 |
| Pcbd1 | -0.032082764 | 0.000785108 | 0.002896417 | 1.054051478 | 1.03234753 |
| Pcbd2 | -0.080494167 | 1.00E-17 | 2.38E-16 | 0.358195963 | 0.421503906 |
| Pcbp1 | 0.001103671 | 0.911574243 | 0.93920356 | 0.887729121 | 0.886119155 |
| Pcbp2 | 0.066461679 | 3.00E-15 | 5.93E-14 | 1.428733569 | 1.342518978 |
| Pcbp3 | 0.004991688 | 0.493423082 | 0.608486807 | 0.167743194 | 0.162910943 |
| Pcbp4 | 0.005495295 | 0.340078924 | 0.462452304 | 0.104870819 | 0.094827339 |

| | | | | | |
|---|---|---|---|---|---|
| Pcca | -0.014504939 | 0.044078574 | 0.08985781 | 0.163738358 | 0.165878384 |
| Pccb | 0.012837735 | 0.081308568 | 0.149316566 | 0.189363922 | 0.168083181 |
| Pcdh1 | 0.016877904 | 0.015323652 | 0.03718583 | 0.155539892 | 0.143240294 |
| Pcdh15 | 0.029773611 | 2.43E-19 | 6.71E-18 | 0.045622603 | 0.018877592 |
| Pcdhga12 | -0.010144409 | 0.014563186 | 0.035645341 | 0.041301151 | 0.046713062 |
| Pcdhga5 | -0.003061717 | 0.474540103 | 0.591644962 | 0.040214817 | 0.047007808 |
| Pced1a | 0.004553364 | 0.317695426 | 0.439413136 | 0.060252521 | 0.05424249 |
| Pcf11 | 0.02359573 | 0.005938769 | 0.016706225 | 0.271588461 | 0.248017963 |
| Pcgf1 | -0.005565672 | 0.266705662 | 0.383226787 | 0.068318465 | 0.071064082 |
| Pcgf2 | 0.010938885 | 0.038066352 | 0.079467699 | 0.08193439 | 0.074583413 |
| Pcgf3 | 0.009329334 | 0.135894989 | 0.22677984 | 0.123300886 | 0.111942754 |
| Pcgf5 | 0.011653918 | 0.189338873 | 0.293573066 | 0.309752695 | 0.295957581 |
| Pcgf6 | -0.006408423 | 0.319387619 | 0.441055367 | 0.117859393 | 0.123980394 |
| Pcid2 | -0.002333885 | 0.663486523 | 0.753729373 | 0.080980738 | 0.080320519 |
| Pcif1 | 0.043527889 | 6.29E-06 | 3.79E-05 | 0.468019513 | 0.427562173 |
| Pck2 | -0.032550378 | 0.001941918 | 0.006365247 | 0.847622554 | 0.859339269 |
| Pclo | -0.003189871 | 0.771426415 | 0.838623237 | 1.061669774 | 1.045952241 |
| Pcm1 | 0.048655936 | 2.89E-06 | 1.86E-05 | 0.701808676 | 0.637000624 |
| Pcmt1 | -0.044259756 | 6.44E-06 | 3.86E-05 | 0.481160483 | 0.519320498 |
| Pcmtd1 | 0.009055406 | 0.374485901 | 0.49648874 | 0.74151558 | 0.726741225 |
| Pcmtd2 | 0.00123971 | 0.884273645 | 0.92083423 | 0.271207991 | 0.259190413 |
| Pcna | -0.011533894 | 0.160015572 | 0.257220798 | 0.227844637 | 0.241430079 |
| Pcnp | 0.004117273 | 0.675907241 | 0.764534055 | 0.599599287 | 0.588057958 |
| Pcnt | 0.012219004 | 0.016765351 | 0.040185588 | 0.081354112 | 0.06640466 |
| Pcnx | 0.015531908 | 0.044746233 | 0.090913872 | 0.201257523 | 0.184543544 |
| Pcnx3 | -0.000528169 | 0.934690571 | 0.95572799 | 0.127486158 | 0.125710563 |
| Pcnx4 | 0.00361477 | 0.474619402 | 0.591646343 | 0.071859294 | 0.069296327 |
| Pcp4 | 0.170592833 | 2.84E-75 | 1.06E-72 | 0.333483899 | 0.178366379 |
| Pcsk1 | -0.038528077 | 0.000194036 | 0.000835438 | 1.335098586 | 1.416829575 |
| Pcsk1n | -0.07625913 | 3.15E-71 | 8.76E-69 | 3.494805121 | 3.617143451 |
| Pcsk2 | -0.018400235 | 2.35E-06 | 1.53E-05 | 3.557790488 | 3.574240633 |
| Pcsk2os1 | -0.04177438 | 4.29E-06 | 2.68E-05 | 0.322114969 | 0.34486146 |
| Pcsk4 | -0.012818386 | 0.00531445 | 0.015182805 | 0.053909711 | 0.060507249 |
| Pcsk6 | -0.026969722 | 4.38E-06 | 2.73E-05 | 0.079169556 | 0.101443269 |
| Pcsk7 | -0.003999517 | 0.397165255 | 0.517699696 | 0.056461451 | 0.06155768 |
| Pcsk9 | 0.024526052 | 7.09E-08 | 6.15E-07 | 0.069854261 | 0.048463286 |
| Pctp | 0.009986478 | 0.00963404 | 0.025110492 | 0.046066028 | 0.036052983 |
| Pcx | -0.013672185 | 0.106110394 | 0.185473795 | 0.286571875 | 0.270195016 |
| Pcyox1 | 0.014227398 | 0.065202232 | 0.124954878 | 0.208987881 | 0.18919866 |
| Pcyox1l | -0.001098066 | 0.760300956 | 0.830055687 | 0.036734305 | 0.034126303 |
| Pcyt1a | 0.034465331 | 5.18E-06 | 3.17E-05 | 0.200844524 | 0.17176154 |
| Pcyt1b | 0.010263397 | 0.065155605 | 0.124905352 | 0.092442482 | 0.081521719 |
| Pcyt2 | 0.010208397 | 0.090848885 | 0.163319685 | 0.114922159 | 0.103627269 |
| Pdap1 | 0.102641157 | 4.69E-26 | 2.02E-24 | 1.049921576 | 0.933516411 |
| Pdcd10 | 0.022180517 | 0.020594397 | 0.0476371 | 0.482477096 | 0.449956908 |
| Pdcd11 | 0.020057798 | 0.00529257 | 0.015133637 | 0.169050189 | 0.15520218 |
| Pdcd2 | -0.017350521 | 0.01330392 | 0.033028781 | 0.151832141 | 0.15657389 |
| Pdcd2l | -0.012187241 | 0.041623239 | 0.085735526 | 0.102279492 | 0.106500881 |
| Pdcd4 | 0.147405078 | 2.30E-45 | 2.28E-43 | 1.017468155 | 0.822143776 |
| Pdcd5 | 0.025453481 | 0.009484791 | 0.024801274 | 0.570264849 | 0.54065683 |
| Pdcd6 | -0.021404536 | 0.030614977 | 0.066496604 | 0.616784715 | 0.623810043 |
| Pdcd6ip | 0.043219725 | 1.51E-05 | 8.39E-05 | 0.556605137 | 0.518704861 |
| Pdcd7 | 0.023656727 | 0.000461526 | 0.00180822 | 0.157653098 | 0.1285595 |
| Pdcl | -0.032503741 | 1.53E-06 | 1.03E-05 | 0.126903549 | 0.146177573 |
| Pdcl3 | -0.00852044 | 0.258626655 | 0.374810009 | 0.193336596 | 0.191017274 |
| Pde10a | 0.025215147 | 0.032330797 | 0.06951647 | 0.617642853 | 0.661889025 |
| Pde12 | 0.023683404 | 9.61E-06 | 5.54E-05 | 0.091685985 | 0.073847564 |
| Pde1c | 0.073624645 | 2.07E-24 | 8.04E-23 | 0.180079218 | 0.129677636 |
| Pde3b | 0.016366499 | 0.094001639 | 0.167715899 | 0.447945673 | 0.421615038 |
| Pde4b | 0.016542703 | 0.025647132 | 0.057320476 | 0.188078188 | 0.165484891 |
| Pde4d | -0.028043182 | 4.51E-09 | 4.70E-08 | 0.040143653 | 0.06663858 |
| Pde4dip | 0.018881509 | 0.051201952 | 0.101963412 | 0.422108799 | 0.404377649 |
| Pde5a | -0.039137678 | 0.000290764 | 0.001197292 | 0.601150373 | 0.637774259 |
| Pde6d | -0.021564339 | 0.006193144 | 0.017331532 | 0.210021508 | 0.219559629 |
| Pde7a | -0.00814993 | 0.04292885 | 0.087864059 | 0.037919049 | 0.044787899 |
| Pde8a | 0.004533959 | 0.613264733 | 0.713347659 | 0.329774598 | 0.313459782 |
| Pdgfa | 0.015997361 | 0.110328389 | 0.191537964 | 0.46849529 | 0.448722786 |
| Pdgfrl | -0.002350154 | 0.671634748 | 0.760409109 | 0.093486744 | 0.08773899 |
| Pdha1 | 0.004891916 | 0.574088964 | 0.680705225 | 0.310799612 | 0.296117179 |
| Pdhb | 0.006631989 | 0.461411811 | 0.579344863 | 0.33978157 | 0.319433429 |
| Pdhx | 0.00835848 | 0.066617837 | 0.1271914 | 0.062007717 | 0.054930131 |
| Pdia3 | 0.011484831 | 0.149309908 | 0.243964861 | 1.716417643 | 1.719954263 |

| | | | | | |
|---|---|---|---|---|---|
| Pdia4 | -0.028596735 | 0.004896334 | 0.014163093 | 0.460243719 | 0.493801412 |
| Pdia5 | -0.005637372 | 0.349881155 | 0.472305552 | 0.100576048 | 0.106570504 |
| Pdia6 | 0.021217538 | 0.038660303 | 0.080534705 | 1.301851769 | 1.303954809 |
| Pdik1l | 0.026551534 | 4.68E-06 | 2.89E-05 | 0.114592516 | 0.088517337 |
| Pdk1 | 0.023353541 | 3.61E-05 | 0.000183992 | 0.105644342 | 0.082128706 |
| Pdk2 | 0.006996312 | 0.064486518 | 0.123778466 | 0.045702214 | 0.035041419 |
| Pdk3 | -0.002369961 | 0.661613787 | 0.752625985 | 0.081323773 | 0.081720207 |
| Pdlim5 | 0.004502991 | 0.450982959 | 0.569404039 | 0.10181357 | 0.098460436 |
| Pdlim7 | 0.029788077 | 1.51E-09 | 1.68E-08 | 0.083736204 | 0.057342396 |
| Pdp2 | 0.003849429 | 0.385445701 | 0.506460752 | 0.055007751 | 0.050578564 |
| Pdpk1 | 0.050022838 | 1.76E-07 | 1.43E-06 | 0.456666247 | 0.408121188 |
| Pdrg1 | -0.065571586 | 4.60E-11 | 6.14E-10 | 0.364722785 | 0.42697328 |
| Pds5a | 0.012856919 | 0.136415133 | 0.227444336 | 0.274592673 | 0.263044299 |
| Pds5b | -0.001275283 | 0.874495802 | 0.913954363 | 0.219466983 | 0.21470715 |
| Pdss2 | 0.000814664 | 0.861672642 | 0.905445463 | 0.061932369 | 0.059507598 |
| Pdx1 | 0.059385545 | 6.04E-08 | 5.28E-07 | 0.96031385 | 0.822892952 |
| Pdxdc1 | 0.033875718 | 0.000463702 | 0.00181559 | 0.465329937 | 0.433178444 |
| Pdxk | 0.025014426 | 5.91E-06 | 3.58E-05 | 0.099808301 | 0.077195167 |
| Pdxp | -0.002174494 | 0.582644587 | 0.688689443 | 0.045922474 | 0.041606986 |
| Pdyn | -0.017634833 | 0.190333409 | 0.294559281 | 0.360464584 | 0.468749566 |
| Pdzd11 | -0.023528567 | 0.005760825 | 0.016266844 | 0.265018038 | 0.273864209 |
| Pdzd4 | -0.008632019 | 0.120442199 | 0.204996334 | 0.085266345 | 0.089674294 |
| Pdzd8 | -0.000998775 | 0.905415553 | 0.93560051 | 0.24510838 | 0.245621668 |
| Pdzk1ip1 | 0.020054824 | 0.035472883 | 0.075069084 | 0.439966676 | 0.403776158 |
| Pea15a | 0.082902097 | 4.23E-18 | 1.05E-16 | 0.437646714 | 0.342719856 |
| Pebp1 | 0.00820117 | 0.162074571 | 0.259671881 | 2.276557393 | 2.159438437 |
| Pecr | -0.022740042 | 0.001142498 | 0.004017132 | 0.148525159 | 0.163502227 |
| Pef1 | -0.012411514 | 0.06718092 | 0.128112814 | 0.147435524 | 0.146552241 |
| Peg13 | 0.022149866 | 0.002566127 | 0.008110497 | 0.192487884 | 0.161257437 |
| Peg3 | 0.008936376 | 0.368390032 | 0.491085684 | 2.216632698 | 2.169328679 |
| Peli1 | 0.02042414 | 0.002083557 | 0.006763385 | 0.144590803 | 0.125144538 |
| Peli3 | 0.003125694 | 0.48263453 | 0.598949821 | 0.055753839 | 0.051521326 |
| Pelp1 | 0.012376294 | 0.03688193 | 0.077410648 | 0.109148713 | 0.096482302 |
| Pemt | -0.006492069 | 0.165242845 | 0.26340235 | 0.057041778 | 0.059951862 |
| Pepd | 0.012238477 | 0.070427198 | 0.132950139 | 0.153668726 | 0.139183643 |
| Per1 | 0.02041194 | 0.00021037 | 0.000897966 | 0.092588748 | 0.077150398 |
| Per2 | 0.010080788 | 0.033473039 | 0.071584885 | 0.063417168 | 0.057491061 |
| Per3 | 0.044860161 | 3.62E-14 | 6.69E-13 | 0.126467542 | 0.073554092 |
| Perp | 0.016434837 | 0.000121283 | 0.00055508 | 0.058454638 | 0.044676295 |
| Pes1 | 0.012657888 | 0.041658577 | 0.085769734 | 0.119388622 | 0.107962683 |
| Pet100 | 0.06065231 | 4.95E-12 | 7.33E-11 | 0.327569223 | 0.283685047 |
| Pex1 | 0.002887616 | 0.584774242 | 0.689948444 | 0.079670834 | 0.07615683 |
| Pex10 | 0.001156833 | 0.758815125 | 0.828619639 | 0.039394748 | 0.037706993 |
| Pex11a | -0.008674188 | 0.220221241 | 0.331166419 | 0.154284688 | 0.160192082 |
| Pex11b | -0.007189287 | 0.319517893 | 0.441055367 | 0.17140026 | 0.169745152 |
| Pex11g | -0.035563521 | 3.73E-08 | 3.38E-07 | 0.102393272 | 0.135204468 |
| Pex12 | -0.002838323 | 0.591196611 | 0.695471874 | 0.079286072 | 0.078541406 |
| Pex13 | -0.005824026 | 0.396512863 | 0.517088941 | 0.148290263 | 0.148846602 |
| Pex14 | 0.009070061 | 0.149804968 | 0.244650412 | 0.127579513 | 0.117111422 |
| Pex16 | -0.021585681 | 0.003799254 | 0.011400108 | 0.169899656 | 0.185741365 |
| Pex19 | 0.010060472 | 0.118971996 | 0.202955913 | 0.135684377 | 0.121540513 |
| Pex2 | -0.027391911 | 0.000620343 | 0.00234851 | 0.220666162 | 0.236161666 |
| Pex26 | -0.00018823 | 0.972474433 | 0.981255468 | 0.08807103 | 0.08233969 |
| Pex3 | 0.00746539 | 0.325118438 | 0.447262127 | 0.191555173 | 0.186869829 |
| Pex5 | 0.007661106 | 0.151810392 | 0.246735782 | 0.090061797 | 0.078954369 |
| Pex5l | 0.000767134 | 0.92765817 | 0.950687696 | 0.24916667 | 0.249580096 |
| Pex6 | 0.004347469 | 0.555400995 | 0.664630484 | 0.193663579 | 0.178645787 |
| Pex7 | -0.010429971 | 0.251845004 | 0.366842383 | 0.354239182 | 0.34661084 |
| Pfas | 0.003464588 | 0.391357642 | 0.512075844 | 0.045066284 | 0.041732405 |
| Pfdn1 | 0.007245547 | 0.462153377 | 0.579973554 | 0.652198846 | 0.640443421 |
| Pfdn2 | 0.029284128 | 0.001838539 | 0.006067304 | 0.902240509 | 0.86341791 |
| Pfdn4 | 0.006983976 | 0.414402084 | 0.533901015 | 0.296424456 | 0.277441908 |
| Pfdn5 | -0.02603412 | 3.54E-05 | 0.000180852 | 1.844132233 | 1.866821576 |
| Pfdn6 | -0.039482828 | 6.40E-05 | 0.000309655 | 0.537868317 | 0.578070109 |
| Pfkfb2 | 0.012770039 | 0.052416844 | 0.103808629 | 0.143117716 | 0.121023916 |
| Pfkl | 0.00349661 | 0.629209617 | 0.726742 | 0.177056232 | 0.16250936 |
| Pfkm | 0.004157254 | 0.492582968 | 0.607650249 | 0.114184114 | 0.105825187 |
| Pfkp | -0.006887217 | 0.345590662 | 0.467812923 | 0.152043773 | 0.161602445 |
| Pfn1 | 0.026457663 | 0.005088417 | 0.014648975 | 1.057119975 | 1.016993573 |
| Pfn2 | -0.017854396 | 0.011995946 | 0.030284235 | 0.156386805 | 0.163269662 |
| Pgam1 | -0.04001024 | 7.04E-05 | 0.000338729 | 0.746933589 | 0.768603238 |
| Pgam5 | -0.004681545 | 0.395464637 | 0.516137361 | 0.08402298 | 0.086644038 |

| | | | | | |
|---|---|---|---|---|---|
| Pgap1 | 0.011550856 | 0.164841638 | 0.262978406 | 0.237396103 | 0.220865051 |
| Pgap2 | -0.017843825 | 0.004791653 | 0.013901656 | 0.112387911 | 0.122391427 |
| Pgap3 | 0.004690851 | 0.18054482 | 0.282748601 | 0.035892472 | 0.03050266 |
| Pgap4 | -0.00106315 | 0.842435068 | 0.892270806 | 0.085473094 | 0.081522104 |
| Pgap6 | 0.00933852 | 0.159323082 | 0.256319544 | 0.143469682 | 0.12817288 |
| Pgbd5 | 0.007217791 | 0.183712692 | 0.286594159 | 0.089300893 | 0.079579547 |
| Pgd | -0.044099611 | 7.85E-11 | 1.02E-09 | 0.107398889 | 0.150530178 |
| Pgf | 0.011313574 | 0.002177723 | 0.007026824 | 0.042801542 | 0.030113601 |
| Pgghg | -0.003426148 | 0.483556726 | 0.599941104 | 0.066696719 | 0.065916228 |
| Pggt1b | 0.013280947 | 0.084777395 | 0.154548254 | 0.195907627 | 0.18228482 |
| Pgk1 | 0.005474763 | 0.605658534 | 0.707383704 | 0.548270164 | 0.559636828 |
| Pgls | -0.09091128 | 1.44E-20 | 4.28E-19 | 0.871792478 | 0.996900144 |
| Pgm1 | -0.002592955 | 0.690991064 | 0.775764703 | 0.123091762 | 0.129085161 |
| Pgm2 | -0.00504869 | 0.349141043 | 0.471597793 | 0.083334245 | 0.083617069 |
| Pgm3 | -0.057634851 | 2.49E-11 | 3.43E-10 | 0.23124922 | 0.279568051 |
| Pgp | 0.005852551 | 0.548714573 | 0.658105487 | 0.498861065 | 0.47531956 |
| Pgpep1 | 0.002949499 | 0.634640313 | 0.730520142 | 0.124930026 | 0.113541495 |
| Pgrmc1 | -0.045096855 | 1.10E-05 | 6.25E-05 | 0.566611521 | 0.602509423 |
| Pgrmc2 | -0.021949704 | 0.035929015 | 0.075885702 | 0.613117875 | 0.664725637 |
| Pgs1 | 0.00389468 | 0.490073302 | 0.605399319 | 0.091824547 | 0.088894901 |
| Phactr1 | 0.091796984 | 5.47E-17 | 1.22E-15 | 0.900652792 | 0.772391022 |
| Phactr2 | -0.004502433 | 0.266676388 | 0.383226787 | 0.04262995 | 0.044119479 |
| Phactr4 | 0.015630614 | 0.019973693 | 0.046400058 | 0.141135242 | 0.130460325 |
| Phax | 0.006150059 | 0.486971945 | 0.602870399 | 0.327225437 | 0.312836951 |
| Phb | -0.007660774 | 0.173902318 | 0.274460769 | 0.08818085 | 0.093861619 |
| Phb2 | 0.004913733 | 0.619054704 | 0.718706691 | 0.484776806 | 0.462566058 |
| Phc1 | 0.00361673 | 0.500890636 | 0.615244316 | 0.085561197 | 0.0796045 |
| Phc2 | 0.010700268 | 0.199989753 | 0.30705944 | 0.26279245 | 0.242678401 |
| Phc3 | 0.01536282 | 0.028632739 | 0.062907906 | 0.155415647 | 0.141688676 |
| Pheta1 | 0.005652025 | 0.103086751 | 0.181253369 | 0.038658621 | 0.030294398 |
| Phf1 | -0.008603116 | 0.269420724 | 0.385987073 | 0.203404265 | 0.204885411 |
| Phf10 | -0.05363977 | 1.05E-06 | 7.32E-06 | 0.498883363 | 0.561788086 |
| Phf12 | 0.013176469 | 0.088878773 | 0.160532533 | 0.203681479 | 0.188817906 |
| Phf13 | 0.014275423 | 0.006528107 | 0.018159158 | 0.083591628 | 0.070555064 |
| Phf14 | 0.018704972 | 0.045508204 | 0.09223072 | 0.38636993 | 0.36465335 |
| Phf2 | 0.014996717 | 0.053816993 | 0.106127548 | 0.21235725 | 0.193318803 |
| Phf20 | 0.011719949 | 0.141038185 | 0.233272072 | 0.220498349 | 0.203849917 |
| Phf20l1 | 0.049220777 | 2.48E-06 | 1.61E-05 | 0.622754802 | 0.576526803 |
| Phf21a | 0.030960993 | 0.000142944 | 0.000643081 | 0.24255839 | 0.207452427 |
| Phf21b | 0.007575703 | 0.189815536 | 0.293944989 | 0.102231578 | 0.094452518 |
| Phf23 | 0.013400557 | 0.045378788 | 0.092063038 | 0.147301735 | 0.13219795 |
| Phf3 | 0.025992281 | 0.007715407 | 0.020882291 | 0.445346983 | 0.418577423 |
| Phf5a | -0.028978111 | 0.002581007 | 0.008152224 | 0.398469424 | 0.423828225 |
| Phf6 | 0.008097472 | 0.143517085 | 0.236647405 | 0.089357262 | 0.084107309 |
| Phf7 | 0.010071848 | 0.040172597 | 0.083310151 | 0.071160207 | 0.06452059 |
| Phf8 | 0.000864881 | 0.89642143 | 0.929501828 | 0.132228814 | 0.130636378 |
| Phgdh | -0.107242426 | 1.02E-19 | 2.90E-18 | 0.565817181 | 0.676904914 |
| Phip | 0.002258767 | 0.835078735 | 0.887291259 | 0.663058181 | 0.642535226 |
| Phka1 | -0.000157705 | 0.974824371 | 0.983204937 | 0.070847084 | 0.069995209 |
| Phka2 | 0.006277332 | 0.132122501 | 0.221579258 | 0.051862306 | 0.044158321 |
| Phkb | -0.025001084 | 0.000125154 | 0.000570709 | 0.121606046 | 0.134819461 |
| Phkg2 | -0.006241799 | 0.320099697 | 0.441669081 | 0.114215245 | 0.116024807 |
| Phlda1 | 0.000771002 | 0.895992665 | 0.929191473 | 0.093197679 | 0.090992496 |
| Phlda3 | 0.004231495 | 0.397247453 | 0.517699696 | 0.057894672 | 0.053168529 |
| Phldb2 | 0.043595637 | 5.88E-07 | 4.32E-06 | 0.300845033 | 0.250248582 |
| Phlpp1 | 0.005623478 | 0.209956837 | 0.3189874 | 0.058227697 | 0.052748615 |
| Phlpp2 | 0.014332285 | 0.00395246 | 0.011794295 | 0.075177297 | 0.063036446 |
| Phospho2 | 0.019552078 | 0.008557983 | 0.022807211 | 0.183397888 | 0.165581299 |
| Phpt1 | 0.008644561 | 0.123574829 | 0.209229274 | 0.103899227 | 0.08989545 |
| Phrf1 | 0.005999728 | 0.36197569 | 0.484795103 | 0.132745434 | 0.125966557 |
| Phtf1 | 0.008669459 | 0.147993729 | 0.242221385 | 0.109045019 | 0.10001872 |
| Phyh | -0.021248551 | 0.035332594 | 0.07483736 | 0.728733226 | 0.747002188 |
| Phyhip | 0.010052992 | 0.006546244 | 0.0182044 | 0.044869093 | 0.033446263 |
| Phyhipl | 0.013762366 | 0.024339014 | 0.054939377 | 0.120656835 | 0.104449705 |
| Phykpl | -0.010286218 | 0.038835254 | 0.080847183 | 0.063166886 | 0.070512279 |
| Pi4k2a | 0.013592607 | 0.140504336 | 0.23262656 | 0.32214097 | 0.315762154 |
| Pi4k2b | -0.009885909 | 0.14414531 | 0.237522153 | 0.132306859 | 0.139146252 |
| Pi4ka | 0.054779853 | 1.72E-09 | 1.88E-08 | 0.370458382 | 0.311249612 |
| Pi4kb | 0.004813677 | 0.418438071 | 0.537604721 | 0.108267049 | 0.100461709 |
| Pias1 | 0.02794581 | 0.000127854 | 0.000581926 | 0.197099203 | 0.165092126 |
| Pias2 | 0.009947737 | 0.147817483 | 0.242014411 | 0.152578079 | 0.146507396 |
| Pias3 | 0.017551733 | 6.96E-05 | 0.000334989 | 0.063341313 | 0.047853713 |

| | | | | | |
|---|---|---|---|---|---|
| Pias4 | 0.012974945 | 0.052523847 | 0.10397818 | 0.141815205 | 0.128724154 |
| Pibf1 | 0.016768275 | 0.005058773 | 0.014582742 | 0.111123832 | 0.09745828 |
| Picalm | 0.035012555 | 0.000238051 | 0.001002308 | 0.413523942 | 0.379101252 |
| Pick1 | -0.007986383 | 0.260406067 | 0.376400206 | 0.157990158 | 0.159644621 |
| Pid1 | 0.012469453 | 0.013504722 | 0.033433387 | 0.079696531 | 0.063920704 |
| Pigbos1 | 0.027601641 | 0.002820066 | 0.008801502 | 0.385858689 | 0.36072909 |
| Pigc | 0.006546988 | 0.159218473 | 0.256236053 | 0.066351784 | 0.057289812 |
| Pigf | -0.025270618 | 3.21E-05 | 0.000165711 | 0.100423616 | 0.114709469 |
| Pigg | 0.01017045 | 0.033831897 | 0.072177682 | 0.070108082 | 0.060171635 |
| Pigh | 0.008547953 | 0.11925943 | 0.203303556 | 0.093092698 | 0.082232785 |
| Pigk | -0.049831144 | 1.45E-08 | 1.40E-07 | 0.293253988 | 0.326762087 |
| Pigm | -0.001958825 | 0.652378173 | 0.745027675 | 0.049194723 | 0.050832802 |
| Pigo | 0.009779575 | 0.015192276 | 0.036924826 | 0.04750974 | 0.040616838 |
| Pigp | -0.049194543 | 8.50E-07 | 6.03E-06 | 0.692528082 | 0.709410612 |
| Pigq | 0.001441941 | 0.839897863 | 0.890262432 | 0.16864334 | 0.162323971 |
| Pigs | -0.000446246 | 0.945117636 | 0.963148182 | 0.129930456 | 0.126592454 |
| Pigt | -0.005402708 | 0.47192754 | 0.589197321 | 0.195722273 | 0.190494938 |
| Pigu | 0.004183273 | 0.478534998 | 0.595153799 | 0.103380984 | 0.100201297 |
| Pigv | 0.001449393 | 0.786562993 | 0.850702571 | 0.083645945 | 0.078270069 |
| Pigx | -0.059793941 | 7.93E-13 | 1.28E-11 | 0.213608229 | 0.262645444 |
| Pigyl | 0.020180574 | 0.027163832 | 0.060142741 | 0.410156051 | 0.376453318 |
| Pih1d1 | -0.007930816 | 0.146329371 | 0.240271908 | 0.086281924 | 0.087885789 |
| Pih1d2 | -0.012852768 | 0.001959241 | 0.006415541 | 0.042991952 | 0.049734036 |
| Pik3c2a | 0.029053761 | 0.002634225 | 0.008298749 | 0.387743707 | 0.356674322 |
| Pik3c3 | 0.021500108 | 0.0001535 | 0.000681212 | 0.107060994 | 0.084462118 |
| Pik3ca | 0.027135565 | 0.000520655 | 0.002013448 | 0.220314831 | 0.188552167 |
| Pik3ip1 | 0.008307162 | 0.023444434 | 0.053154569 | 0.042904154 | 0.031510275 |
| Pik3r1 | 0.042767749 | 2.41E-05 | 0.000127517 | 0.471947299 | 0.441204183 |
| Pik3r2 | 0.004630061 | 0.42323774 | 0.541833626 | 0.10535665 | 0.095507745 |
| Pik3r3 | -0.000758898 | 0.905579968 | 0.93560051 | 0.119424082 | 0.118539324 |
| Pik3r4 | 0.012060907 | 0.050382088 | 0.100681327 | 0.120511678 | 0.106236287 |
| Pikfyve | 0.0337049 | 1.27E-05 | 7.10E-05 | 0.212350441 | 0.181886175 |
| Pim1 | 0.008818309 | 0.10413081 | 0.18260457 | 0.088824292 | 0.07806021 |
| Pim2 | 0.031284637 | 1.01E-06 | 7.05E-06 | 0.124950515 | 0.103306839 |
| Pim3 | -0.026346307 | 0.00018389 | 0.000797049 | 0.133337748 | 0.154583614 |
| Pin1 | -0.017337689 | 0.081371631 | 0.149404155 | 0.548563696 | 0.556781427 |
| Pin4 | 0.046440504 | 4.26E-07 | 3.23E-06 | 0.352743119 | 0.335921843 |
| Pink1 | -0.008951186 | 0.288656006 | 0.408131717 | 0.269638174 | 0.256351606 |
| Pinx1 | 0.009948884 | 0.021591705 | 0.049578309 | 0.054521574 | 0.047288172 |
| Pip4k2a | 0.010815154 | 0.085496297 | 0.155631174 | 0.120305891 | 0.110888396 |
| Pip4k2b | 0.011583527 | 0.005267308 | 0.015079142 | 0.052832552 | 0.042226395 |
| Pip4k2c | -0.003310383 | 0.704161491 | 0.78633793 | 0.283974704 | 0.279553829 |
| Pip4p1 | 0.000724351 | 0.929077261 | 0.951828279 | 0.258275144 | 0.240528632 |
| Pip4p2 | -0.000237569 | 0.970815829 | 0.980449395 | 0.141306737 | 0.131301402 |
| Pip5k1a | 0.025818818 | 0.001003803 | 0.003591821 | 0.218259444 | 0.197720215 |
| Pip5k1b | -0.006424578 | 0.513140279 | 0.626840185 | 0.437248832 | 0.441686657 |
| Pip5k1c | 0.019923653 | 0.001537004 | 0.00518126 | 0.132392411 | 0.110216859 |
| Pir | -0.045604668 | 2.87E-21 | 8.87E-20 | 0.037814033 | 0.07450081 |
| Pisd | -0.013915268 | 0.018049402 | 0.042708605 | 0.092184933 | 0.101697052 |
| Pithd1 | -0.036868993 | 0.000297151 | 0.001221524 | 0.458644359 | 0.487199323 |
| Pitpna | -0.013985656 | 0.125601688 | 0.212033271 | 0.340932938 | 0.354694687 |
| Pitpnb | -0.027531977 | 0.004031079 | 0.011995759 | 0.460887547 | 0.466683767 |
| Pitpnc1 | -0.023540676 | 0.002397899 | 0.007646724 | 0.166869294 | 0.192733592 |
| Pitpnm1 | 0.009181341 | 0.13665626 | 0.227690173 | 0.110839218 | 0.106503861 |
| Pitpnm2 | 0.064303603 | 6.78E-17 | 1.50E-15 | 0.220644182 | 0.161449627 |
| Pitpnm3 | 0.012738855 | 0.062698891 | 0.120975038 | 0.143464602 | 0.132032822 |
| Pitrm1 | 0.003171379 | 0.640290807 | 0.73568096 | 0.144315454 | 0.138929765 |
| Pja1 | -0.019373373 | 0.023672782 | 0.053647269 | 0.278873637 | 0.278733039 |
| Pja2 | 0.007463612 | 0.454786929 | 0.573542421 | 0.644625298 | 0.620216109 |
| Pkd1 | 0.034703899 | 2.66E-05 | 0.000139382 | 0.248676299 | 0.220077624 |
| Pkd2 | -0.004475894 | 0.574296062 | 0.680724984 | 0.202600928 | 0.207623484 |
| Pkdcc | -0.01034685 | 0.324397516 | 0.446459888 | 0.609919068 | 0.582773635 |
| Pkia | 0.007166223 | 0.229272043 | 0.342130898 | 0.105083351 | 0.098961913 |
| Pkib | 0.004105966 | 0.580777957 | 0.686984219 | 0.178299538 | 0.170284414 |
| Pkig | -0.014104208 | 0.10731389 | 0.187341648 | 0.324639124 | 0.31747562 |
| Pkm | 0.042827963 | 1.52E-06 | 1.02E-05 | 1.533779218 | 1.371491219 |
| Pkn1 | 0.012688709 | 0.109080112 | 0.18984548 | 0.21358074 | 0.203948075 |
| Pkn2 | 0.075482441 | 1.24E-13 | 2.22E-12 | 0.628450868 | 0.551770434 |
| Pknox1 | -0.011431147 | 0.098071408 | 0.173701991 | 0.133869581 | 0.141971508 |
| Pkp3 | -0.00024772 | 0.950840838 | 0.967155747 | 0.042475989 | 0.042858619 |
| Pkp4 | 0.124389692 | 1.30E-34 | 8.18E-33 | 0.58449742 | 0.434711731 |
| Pla2g12a | -0.141321532 | 8.38E-39 | 6.85E-37 | 0.48092064 | 0.656636576 |

| | | | | | |
|---|---|---|---|---|---|
| Pla2g15 | 0.00246552 | 0.646316281 | 0.739843033 | 0.084307199 | 0.08032357 |
| Pla2g2f | 0.07312946 | 1.40E-14 | 2.67E-13 | 0.359864482 | 0.273961011 |
| Pla2g6 | -0.007631269 | 0.233256826 | 0.346746615 | 0.119582082 | 0.119255103 |
| Plaa | 0.016291725 | 0.04348859 | 0.088822704 | 0.228842525 | 0.214348547 |
| Plaat3 | 0.082345161 | 1.81E-24 | 7.05E-23 | 0.256759147 | 0.171054002 |
| Plag1 | 0.002189636 | 0.620393404 | 0.719745187 | 0.053094877 | 0.050565572 |
| Plagl2 | 0.004436552 | 0.356061012 | 0.478455447 | 0.06592513 | 0.062142078 |
| Plbd2 | -0.00222326 | 0.624889644 | 0.723407612 | 0.053133107 | 0.056975371 |
| Plcb1 | 0.017866466 | 0.008675864 | 0.023070773 | 0.146297026 | 0.130205265 |
| Plcb4 | 0.027579431 | 0.000208172 | 0.000889994 | 0.184571222 | 0.158573733 |
| Plcg1 | 0.010405698 | 0.063495369 | 0.122236957 | 0.091209019 | 0.084277054 |
| Plch1 | 0.011228761 | 0.018165866 | 0.042944863 | 0.070342264 | 0.058005052 |
| Plcl2 | 0.010803221 | 0.101020098 | 0.178242721 | 0.131989782 | 0.124808456 |
| Plcxd2 | -0.000427133 | 0.947625372 | 0.965075865 | 0.124003236 | 0.118270725 |
| Plcxd3 | 0.006426968 | 0.356854778 | 0.479190905 | 0.162686262 | 0.144758728 |
| Pld3 | -0.034912714 | 8.61E-05 | 0.000407225 | 0.29553211 | 0.32446965 |
| Plec | -0.00312398 | 0.64585506 | 0.739489211 | 0.131105257 | 0.136844936 |
| Plekha1 | 0.049220834 | 1.62E-07 | 1.32E-06 | 0.379391957 | 0.333750267 |
| Plekha2 | 0.01509126 | 0.003710835 | 0.011179651 | 0.083391465 | 0.06848746 |
| Plekha3 | -0.022396253 | 0.001658174 | 0.005535977 | 0.148405472 | 0.166967448 |
| Plekha5 | -0.029216857 | 0.000167876 | 0.000738838 | 0.185958956 | 0.201380887 |
| Plekha6 | 0.002398928 | 0.785163259 | 0.849755895 | 0.285679583 | 0.28489994 |
| Plekha8 | 0.006792703 | 0.190100058 | 0.294246832 | 0.079322057 | 0.073933677 |
| Plekhb2 | 0.002086969 | 0.798566406 | 0.859382621 | 0.245467865 | 0.239200373 |
| Plekhf2 | 0.012144797 | 0.011450488 | 0.029080888 | 0.069681875 | 0.059516596 |
| Plekhg3 | 0.031583539 | 1.56E-07 | 1.27E-06 | 0.127929786 | 0.094948176 |
| Plekhh1 | 0.013315443 | 0.002788618 | 0.008721006 | 0.060480128 | 0.048026362 |
| Plekhh2 | 0.008496465 | 0.049475307 | 0.09911373 | 0.051632309 | 0.046247851 |
| Plekhh3 | -0.003162758 | 0.562860599 | 0.670879032 | 0.084878359 | 0.084656565 |
| Plekhj1 | -0.028111503 | 0.002336072 | 0.007483128 | 0.376325143 | 0.392005904 |
| Plekhm1 | 0.005146942 | 0.240135773 | 0.354089697 | 0.052737286 | 0.050383365 |
| Plekhm2 | 0.000987687 | 0.849107749 | 0.89689543 | 0.077403117 | 0.072873944 |
| Plekhm3 | 0.011879845 | 0.013517165 | 0.033455671 | 0.069809942 | 0.059614706 |
| Plgrkt | -0.022447369 | 0.007099079 | 0.019457921 | 0.229080862 | 0.251454503 |
| Plin2 | -0.000806909 | 0.879718064 | 0.917845858 | 0.08064303 | 0.078495876 |
| Plk3 | -0.021588058 | 0.013635174 | 0.033687716 | 0.219104658 | 0.245458639 |
| Plod1 | -0.009808509 | 0.044652709 | 0.09078077 | 0.060879368 | 0.06816235 |
| Plod3 | -0.020456457 | 0.001513267 | 0.00511543 | 0.108607342 | 0.129999136 |
| Plp2 | 0.005058442 | 0.375807207 | 0.497697543 | 0.103021929 | 0.090176589 |
| Plpbp | 0.014305471 | 0.103611184 | 0.181988787 | 0.310497913 | 0.288432922 |
| Plpp2 | 0.010638173 | 0.036433822 | 0.076735185 | 0.073182494 | 0.067867157 |
| Plpp5 | -0.097116866 | 2.95E-25 | 1.20E-23 | 0.291923093 | 0.391510826 |
| Plpp6 | -0.000734594 | 0.905863773 | 0.93560051 | 0.113531813 | 0.112831212 |
| Plpp7 | -0.008638147 | 0.037777458 | 0.078966339 | 0.043874727 | 0.049301688 |
| Plppr1 | -0.006264895 | 0.202834829 | 0.310447136 | 0.069596543 | 0.067260234 |
| Plppr2 | 0.000718528 | 0.862658382 | 0.905949638 | 0.046916113 | 0.045987154 |
| Plppr4 | 0.007585175 | 0.21873597 | 0.329544569 | 0.107267482 | 0.103103838 |
| Plrg1 | -0.006521102 | 0.227714863 | 0.340224972 | 0.084839981 | 0.084138715 |
| Pls1 | 0.00311831 | 0.469482515 | 0.58667211 | 0.051297273 | 0.048268027 |
| Pls3 | -0.0212548 | 2.73E-05 | 0.000142901 | 0.055865123 | 0.076062085 |
| Plscr1 | -0.000852773 | 0.898677175 | 0.930848018 | 0.139200101 | 0.132226359 |
| Plscr3 | -0.00399544 | 0.435642798 | 0.55421608 | 0.077093815 | 0.075037757 |
| Plut | -0.001890413 | 0.822294538 | 0.877824476 | 0.196627108 | 0.163816618 |
| Plxna2 | 0.022052315 | 0.003189356 | 0.009796306 | 0.184862971 | 0.163213118 |
| Plxnb1 | -0.011066997 | 0.037275486 | 0.078084954 | 0.072108436 | 0.081437139 |
| Plxnb2 | 0.041945989 | 2.50E-05 | 0.000132063 | 0.488404688 | 0.442412136 |
| Pmepa1 | 0.018127405 | 0.014912677 | 0.036390824 | 0.159090685 | 0.150206915 |
| Pml | 0.009841375 | 0.03973453 | 0.082489665 | 0.06885125 | 0.058927548 |
| Pmm1 | -0.06211644 | 2.20E-15 | 4.38E-14 | 0.162302357 | 0.213815769 |
| Pmm2 | -0.004119474 | 0.51317797 | 0.626840185 | 0.117949388 | 0.117690205 |
| Pmpca | 0.00666238 | 0.331973125 | 0.454817777 | 0.153122464 | 0.143251272 |
| Pmpcb | -0.017232879 | 0.049925122 | 0.099911905 | 0.310159132 | 0.309385838 |
| Pms2 | 0.000350184 | 0.92819686 | 0.95108873 | 0.040573541 | 0.038969426 |
| Pmvk | -0.00058958 | 0.957433215 | 0.971016661 | 0.467634327 | 0.506016531 |
| Pnck | -0.005587702 | 0.206432639 | 0.314665745 | 0.053933963 | 0.053476222 |
| Pnisr | -0.007337738 | 0.485098422 | 0.601240038 | 0.537618771 | 0.534673385 |
| Pnkd | -0.01975444 | 0.040789452 | 0.084291513 | 0.461772084 | 0.479388015 |
| Pnkp | -0.014378269 | 0.015866558 | 0.038333667 | 0.100462674 | 0.108576536 |
| Pnmal1 | -0.045957865 | 1.41E-06 | 9.58E-06 | 0.337499988 | 0.375283186 |
| Pnmal2 | -0.00827131 | 0.437580703 | 0.555953945 | 0.741837233 | 0.733312209 |
| Pnn | -0.02267931 | 0.034180725 | 0.072841957 | 0.580162469 | 0.603965608 |
| Pno1 | -0.012767641 | 0.099846828 | 0.176557086 | 0.180090416 | 0.19393275 |

| | | | | | |
|---|---|---|---|---|---|
| Pnp | -0.029892903 | 6.84E-05 | 0.000329438 | 0.17865388 | 0.192446331 |
| Pnpla2 | 0.001244157 | 0.867805046 | 0.909647852 | 0.183075387 | 0.180290554 |
| Pnpla6 | -0.007715147 | 0.089415312 | 0.161189629 | 0.052587741 | 0.057981381 |
| Pnpla8 | -0.017447993 | 0.077395413 | 0.143677335 | 0.476833522 | 0.496653266 |
| Pnpo | 0.023607026 | 0.003714168 | 0.011186228 | 0.226568765 | 0.214174925 |
| Pnpt1 | -0.005025209 | 0.279845376 | 0.398758408 | 0.055790039 | 0.060431854 |
| Pnrc1 | -0.062319154 | 1.61E-09 | 1.78E-08 | 0.739830505 | 0.797629245 |
| Pnrc2 | -0.037621875 | 6.90E-05 | 0.00033231 | 0.329895473 | 0.375105946 |
| Poc1b | -9.20E-05 | 0.987177483 | 0.992383361 | 0.093165432 | 0.090694582 |
| Poc5 | 0.007245619 | 0.127186833 | 0.214337042 | 0.06523431 | 0.058813803 |
| Podxl2 | -0.029360527 | 0.000341138 | 0.00138247 | 0.225700086 | 0.235663659 |
| Pofut1 | -0.013249181 | 0.008388565 | 0.02239869 | 0.063234159 | 0.072527024 |
| Pofut2 | 0.136002585 | 3.37E-43 | 3.09E-41 | 0.984615538 | 0.850950857 |
| Pogk | 0.020925726 | 0.000842349 | 0.003087978 | 0.121072743 | 0.107253346 |
| Poglut1 | -0.014592463 | 0.011743049 | 0.029715232 | 0.094114161 | 0.101865674 |
| Poglut3 | 0.012896397 | 0.003532872 | 0.01070988 | 0.058890458 | 0.048387935 |
| Pogz | 0.013401543 | 0.026051357 | 0.058049803 | 0.117591663 | 0.100054015 |
| Polb | -0.015453424 | 0.061812378 | 0.119661478 | 0.238316517 | 0.243633436 |
| Pold2 | -0.007807446 | 0.07771261 | 0.144046139 | 0.049809344 | 0.055928 |
| Pold3 | -0.009212486 | 0.168932307 | 0.267877979 | 0.136556913 | 0.139458391 |
| Pold4 | -0.019067285 | 0.033733616 | 0.071999609 | 0.314926669 | 0.324440566 |
| Poldip2 | -0.008746087 | 0.261227748 | 0.377243871 | 0.206178874 | 0.205116258 |
| Poldip3 | 0.017802876 | 0.023789059 | 0.053860558 | 0.23884037 | 0.209452542 |
| Pole3 | 0.007030694 | 0.264670172 | 0.380977705 | 0.118669278 | 0.11115918 |
| Pole4 | -0.010056371 | 0.059910975 | 0.116537514 | 0.074073026 | 0.081580965 |
| Polg | -0.002786366 | 0.577673782 | 0.68381158 | 0.069125556 | 0.069238133 |
| Polk | 0.012183668 | 0.009626882 | 0.025102341 | 0.062895728 | 0.054068236 |
| Poll | -0.000696494 | 0.867823887 | 0.909647852 | 0.048369735 | 0.04718768 |
| Polr1a | -0.005547205 | 0.275517136 | 0.393587174 | 0.068481773 | 0.072054484 |
| Polr1b | -0.004823552 | 0.19632116 | 0.302381863 | 0.035447533 | 0.037427521 |
| Polr1c | 0.006977554 | 0.310039821 | 0.431402195 | 0.151861756 | 0.140748949 |
| Polr1d | 0.03060874 | 0.002008786 | 0.006562041 | 0.817909063 | 0.756482515 |
| Polr1f | -0.000516472 | 0.942075941 | 0.960652643 | 0.156809172 | 0.154462216 |
| Polr1h | -0.015100026 | 0.102409082 | 0.180300814 | 0.373364538 | 0.382352795 |
| Polr1has | 0.014275339 | 0.016298157 | 0.03922236 | 0.115949299 | 0.098132593 |
| Polr2a | 0.009402859 | 0.29777776 | 0.417567486 | 0.316487115 | 0.31485082 |
| Polr2b | -0.00232861 | 0.771271826 | 0.838548948 | 0.226875599 | 0.220657886 |
| Polr2c | -0.013594434 | 0.159515941 | 0.256513166 | 0.481653491 | 0.483635422 |
| Polr2d | -0.00251821 | 0.66304412 | 0.753402868 | 0.102765457 | 0.097502084 |
| Polr2e | -0.050837042 | 1.42E-07 | 1.17E-06 | 0.493189238 | 0.533828405 |
| Polr2f | -0.030252853 | 0.00206606 | 0.006714645 | 0.601197075 | 0.658417211 |
| Polr2g | -0.034595431 | 0.000527344 | 0.002036076 | 0.60969873 | 0.64157328 |
| Polr2h | -0.038212094 | 7.77E-07 | 5.56E-06 | 0.176537908 | 0.208217915 |
| Polr2i | 0.025002428 | 0.009412441 | 0.024638598 | 0.473213736 | 0.464486931 |
| Polr2j | 0.032369169 | 0.00072564 | 0.00270087 | 0.609580349 | 0.577869462 |
| Polr2k | 0.023220294 | 0.019025246 | 0.044524909 | 0.54186199 | 0.537887249 |
| Polr2l | 0.028741746 | 3.47E-05 | 0.000177626 | 0.162116406 | 0.142745015 |
| Polr2m | -0.010124032 | 0.295840567 | 0.415690416 | 0.474024048 | 0.467962045 |
| Polr3a | -0.007907491 | 0.112475251 | 0.194570177 | 0.065106997 | 0.069158937 |
| Polr3b | 0.000690182 | 0.860874482 | 0.904802347 | 0.041805674 | 0.040866525 |
| Polr3c | -0.010866307 | 0.049235343 | 0.098762277 | 0.089239352 | 0.090742463 |
| Polr3d | 0.022382981 | 0.007027867 | 0.019292423 | 0.240383438 | 0.230365155 |
| Polr3e | 0.016820518 | 0.000134682 | 0.000609578 | 0.060477933 | 0.047372847 |
| Polr3f | 0.008979988 | 0.117429023 | 0.200817056 | 0.096998642 | 0.090276251 |
| Polr3g | -0.00807591 | 0.055521536 | 0.109024515 | 0.043014594 | 0.048647724 |
| Polr3gl | -0.022637042 | 0.003464492 | 0.01052885 | 0.195067552 | 0.205551801 |
| Polr3h | -0.027724744 | 7.37E-06 | 4.37E-05 | 0.102540636 | 0.121165766 |
| Polr3k | -0.018575108 | 0.029105984 | 0.063803468 | 0.261590814 | 0.27289569 |
| Pom121 | 0.042396018 | 1.73E-06 | 1.15E-05 | 0.313249702 | 0.281723653 |
| Pomc | -0.026934156 | 0.001095123 | 0.003872965 | 0.207459918 | 0.234532066 |
| Pomgnt1 | -0.00863137 | 0.201463635 | 0.308991792 | 0.135525433 | 0.139848558 |
| Pomgnt2 | 0.014368359 | 0.014980031 | 0.036509365 | 0.118078732 | 0.096018308 |
| Pomp | -0.004277096 | 0.649875255 | 0.74304213 | 1.01991616 | 1.021115418 |
| Pomt2 | -0.005638194 | 0.368559543 | 0.491109633 | 0.111934194 | 0.115174033 |
| Pon2 | -0.031440658 | 7.19E-06 | 4.27E-05 | 0.119032524 | 0.147189202 |
| Pon3 | -0.033905438 | 4.85E-09 | 5.03E-08 | 0.069703252 | 0.101823849 |
| Pop4 | 0.006569861 | 0.106448973 | 0.185998726 | 0.049912738 | 0.043359339 |
| Pop5 | 0.024632466 | 0.014037133 | 0.034514165 | 0.733320359 | 0.709363288 |
| Pop7 | 0.006422688 | 0.423290104 | 0.541833626 | 0.242908632 | 0.229895708 |
| Por | 0.049871099 | 5.94E-07 | 4.36E-06 | 0.549698555 | 0.506638009 |
| Pot1a | 0.001968967 | 0.650808508 | 0.743701361 | 0.052651815 | 0.049992134 |
| Pou2f1 | 0.005246778 | 0.442238667 | 0.560845854 | 0.141579082 | 0.131672189 |

| | | | | | |
|---|---|---|---|---|---|
| Pou6f1 | 0.007811996 | 0.074212579 | 0.13887823 | 0.053013717 | 0.048886872 |
| Ppa1 | -0.028083584 | 0.003382355 | 0.010334146 | 0.360037229 | 0.392152684 |
| Ppa2 | -0.035428175 | 4.42E-06 | 2.74E-05 | 0.193389678 | 0.211829406 |
| Ppan | -0.001467453 | 0.816378473 | 0.873330932 | 0.116421652 | 0.116087871 |
| Ppard | 0.015105024 | 0.00638646 | 0.017816046 | 0.093412476 | 0.083431031 |
| Pparg | -0.036325598 | 4.99E-09 | 5.17E-08 | 0.09355758 | 0.120269398 |
| Ppat | 0.075164173 | 5.04E-13 | 8.34E-12 | 0.476265141 | 0.416530193 |
| Ppcdc | -0.013378017 | 0.003971469 | 0.011843749 | 0.056905732 | 0.061472908 |
| Ppcs | 0.006262867 | 0.365645011 | 0.488430997 | 0.157586565 | 0.146725651 |
| Ppdpf | -0.021355047 | 0.014952238 | 0.03645991 | 0.315762996 | 0.317640783 |
| Ppfia1 | 0.023028996 | 0.015052128 | 0.036639155 | 0.369453647 | 0.35660814 |
| Ppfibp1 | -0.013741368 | 0.106213904 | 0.185621351 | 0.218438597 | 0.242134125 |
| Pphln1 | 0.030730572 | 4.26E-07 | 3.23E-06 | 0.129355838 | 0.097984596 |
| Ppia | -0.026895187 | 7.90E-05 | 0.000377038 | 1.985559074 | 1.942489774 |
| Ppib | -0.068396531 | 1.96E-11 | 2.74E-10 | 0.732513447 | 0.81290654 |
| Ppid | -0.039732673 | 2.72E-06 | 1.76E-05 | 0.243961415 | 0.267023742 |
| Ppie | -0.035315851 | 1.41E-07 | 1.16E-06 | 0.121361289 | 0.147611264 |
| Ppif | -0.004507362 | 0.318333992 | 0.440108514 | 0.057705436 | 0.057043175 |
| Ppig | 0.049999302 | 7.26E-07 | 5.23E-06 | 0.852655172 | 0.791133379 |
| Ppih | -0.010786664 | 0.011635178 | 0.02949601 | 0.046661032 | 0.051516399 |
| Ppil1 | -0.007158011 | 0.176690408 | 0.278124334 | 0.078979546 | 0.080638941 |
| Ppil2 | -0.019349091 | 0.008823833 | 0.023368375 | 0.180660659 | 0.186042299 |
| Ppil3 | -0.007998811 | 0.145737602 | 0.23961319 | 0.086170152 | 0.089355999 |
| Ppil4 | 0.022896237 | 0.005428012 | 0.015478832 | 0.250176756 | 0.227195168 |
| Ppil6 | 0.003016319 | 0.388177864 | 0.509087441 | 0.036371982 | 0.030951537 |
| Ppip5k1 | 0.00906634 | 0.167533088 | 0.266049768 | 0.138763717 | 0.119614252 |
| Ppip5k2 | -0.012208927 | 0.095133497 | 0.169362362 | 0.147417001 | 0.165414163 |
| Ppl | 0.014514702 | 0.001129143 | 0.003974485 | 0.059761617 | 0.047084519 |
| Ppm1a | 0.004576717 | 0.625847973 | 0.724172101 | 0.400422023 | 0.387802135 |
| Ppm1b | -0.00560795 | 0.586940504 | 0.691916525 | 0.614980331 | 0.606516717 |
| Ppm1d | 0.006890695 | 0.21413427 | 0.323916815 | 0.09231992 | 0.084388962 |
| Ppm1e | -0.00463398 | 0.269169908 | 0.385684575 | 0.043065511 | 0.047367302 |
| Ppm1g | -0.015321476 | 0.097579286 | 0.172956394 | 0.339901464 | 0.349535115 |
| Ppm1h | 0.016355337 | 0.008251398 | 0.022080949 | 0.121892365 | 0.105979468 |
| Ppm1k | -0.002583933 | 0.686195443 | 0.772486348 | 0.11701392 | 0.119310595 |
| Ppm1l | 0.054862381 | 5.31E-08 | 4.70E-07 | 0.555489527 | 0.479342751 |
| Ppme1 | 0.030451079 | 0.000541095 | 0.00208198 | 0.312137777 | 0.282145075 |
| Ppox | -0.010613382 | 0.058318744 | 0.113827511 | 0.088187948 | 0.093173766 |
| Ppp1ca | -0.019293609 | 0.051234774 | 0.101967416 | 0.77895407 | 0.798670652 |
| Ppp1cb | 0.028978745 | 0.003665794 | 0.01106453 | 0.827409185 | 0.766830364 |
| Ppp1cc | -0.009362055 | 0.238194145 | 0.352138515 | 0.209867977 | 0.213495659 |
| Ppp1r10 | 0.050665695 | 2.62E-07 | 2.05E-06 | 0.431764714 | 0.375625725 |
| Ppp1r11 | -0.034082358 | 0.000649514 | 0.002446561 | 0.575528555 | 0.607973014 |
| Ppp1r12a | 0.037420214 | 8.09E-05 | 0.000385127 | 0.407484613 | 0.359110396 |
| Ppp1r12b | -0.001870614 | 0.708819792 | 0.790087826 | 0.066043865 | 0.068620727 |
| Ppp1r12c | 0.015865649 | 0.032764358 | 0.070285765 | 0.186219677 | 0.170680878 |
| Ppp1r13b | 0.036909267 | 3.47E-07 | 2.66E-06 | 0.183003227 | 0.152288106 |
| Ppp1r14b | -0.097989238 | 1.24E-21 | 3.96E-20 | 0.641200128 | 0.744452211 |
| Ppp1r15a | -0.121364591 | 1.80E-21 | 5.67E-20 | 0.597535913 | 0.786012944 |
| Ppp1r15b | 0.023293124 | 0.005040224 | 0.014540372 | 0.250812282 | 0.230693118 |
| Ppp1r16a | 0.009250993 | 0.113065103 | 0.195069908 | 0.111668466 | 0.097121993 |
| Ppp1r1a | 0.192050401 | 1.65E-60 | 3.03E-58 | 1.469867322 | 1.053634746 |
| Ppp1r2 | 0.023557757 | 0.018590375 | 0.043698168 | 0.789095156 | 0.741261474 |
| Ppp1r21 | 0.013948395 | 0.019837169 | 0.046138028 | 0.10434892 | 0.098022891 |
| Ppp1r35 | -0.014872137 | 0.054237497 | 0.106805134 | 0.199572043 | 0.205273077 |
| Ppp1r37 | -0.002564685 | 0.77980263 | 0.845157198 | 0.399042032 | 0.363716384 |
| Ppp1r3e | -0.005847493 | 0.16244344 | 0.260177121 | 0.044688221 | 0.047968664 |
| Ppp1r42 | -0.017967423 | 4.39E-05 | 0.000218493 | 0.041215893 | 0.057073246 |
| Ppp1r7 | -0.019891941 | 0.010144834 | 0.026209958 | 0.193614755 | 0.207292592 |
| Ppp1r8 | 0.011666204 | 0.052411613 | 0.103808629 | 0.109627286 | 0.100799627 |
| Ppp1r9a | 0.009737275 | 0.194163627 | 0.299628378 | 0.183537197 | 0.169620611 |
| Ppp1r9b | 0.017013219 | 0.009939415 | 0.025747667 | 0.140254737 | 0.124710627 |
| Ppp2ca | 0.008832133 | 0.367449842 | 0.490101161 | 0.69355059 | 0.667812864 |
| Ppp2cb | 0.00079806 | 0.912494938 | 0.939654252 | 0.166442633 | 0.162460167 |
| Ppp2r1a | 0.019634787 | 0.03195164 | 0.068800409 | 0.383239671 | 0.347329574 |
| Ppp2r1b | 0.006508517 | 0.208882329 | 0.317702441 | 0.079226956 | 0.072206946 |
| Ppp2r2a | 0.013668533 | 0.088152263 | 0.159533936 | 0.226191048 | 0.209746485 |
| Ppp2r2d | 0.076175286 | 1.27E-14 | 2.42E-13 | 0.492596603 | 0.437190299 |
| Ppp2r3a | 0.058636388 | 3.72E-13 | 6.31E-12 | 0.22663234 | 0.184278027 |
| Ppp2r3c | 0.008652758 | 0.169497859 | 0.26864333 | 0.127995098 | 0.115753458 |
| Ppp2r3d | 0.005904068 | 0.303387768 | 0.423723012 | 0.100439137 | 0.091033003 |
| Ppp2r5a | -0.002465714 | 0.807834831 | 0.867053458 | 0.599619445 | 0.582026502 |

| | | | | | |
|---|---|---|---|---|---|
| Ppp2r5b | 0.005706253 | 0.408239469 | 0.528256662 | 0.158834663 | 0.145220312 |
| Ppp2r5c | 0.025325898 | 0.00902136 | 0.023794266 | 0.515091638 | 0.484702925 |
| Ppp2r5d | 0.005134454 | 0.242912092 | 0.356665305 | 0.055022948 | 0.05005722 |
| Ppp2r5e | 0.015895688 | 0.064826733 | 0.124382375 | 0.275692671 | 0.262183208 |
| Ppp3ca | 0.023336115 | 0.008692378 | 0.023095737 | 0.314565534 | 0.283646846 |
| Ppp3cb | -0.001708603 | 0.856746253 | 0.901633154 | 0.39941973 | 0.391492926 |
| Ppp3cc | -0.006851899 | 0.094500565 | 0.168482393 | 0.04287967 | 0.046322433 |
| Ppp3r1 | 0.033457701 | 0.000120608 | 0.000552829 | 0.307580742 | 0.273569134 |
| Ppp4c | -0.000274655 | 0.974912116 | 0.983204937 | 0.311770228 | 0.297709431 |
| Ppp4r1 | 0.009660234 | 0.033301457 | 0.071280661 | 0.05987821 | 0.052302585 |
| Ppp4r1l-ps | -0.032306724 | 0.000900139 | 0.003267793 | 0.392451899 | 0.411329779 |
| Ppp4r2 | 0.028574229 | 0.001301293 | 0.004498993 | 0.327481299 | 0.287878895 |
| Ppp4r3a | 0.040127152 | 3.06E-06 | 1.96E-05 | 0.288679644 | 0.252897985 |
| Ppp4r3b | 0.015479541 | 0.092051254 | 0.164841127 | 0.341999536 | 0.316728514 |
| Ppp4r4 | 0.009056339 | 0.022557077 | 0.051310224 | 0.045202285 | 0.0380685 |
| Ppp5c | -0.004485235 | 0.533137566 | 0.644111274 | 0.166445071 | 0.16763011 |
| Ppp6c | 0.000602984 | 0.945954277 | 0.96379873 | 0.330134958 | 0.315321502 |
| Ppp6r1 | 0.028272307 | 0.000104204 | 0.000484723 | 0.178387789 | 0.154851848 |
| Ppp6r2 | 0.011599396 | 0.069304808 | 0.131290206 | 0.131236411 | 0.117181528 |
| Ppp6r3 | 0.008841453 | 0.296384181 | 0.416213637 | 0.274498703 | 0.258066538 |
| Pprc1 | 0.005194047 | 0.227796379 | 0.340294468 | 0.053062648 | 0.048533902 |
| Ppt1 | 0.00600034 | 0.384734101 | 0.505730791 | 0.154588456 | 0.142862317 |
| Pptc7 | 0.015038542 | 0.021363903 | 0.049148098 | 0.137396098 | 0.122174448 |
| Ppwd1 | 0.023437999 | 0.00060617 | 0.002302924 | 0.145734044 | 0.132570416 |
| Ppy | 0.179615107 | 4.13E-46 | 4.23E-44 | 1.366759037 | 1.145152482 |
| Pqbp1 | -0.005044671 | 0.496775445 | 0.611890393 | 0.178694148 | 0.182113456 |
| Pradc1 | -0.01443774 | 0.00378953 | 0.011377954 | 0.063091133 | 0.07249606 |
| Praf2 | -0.001992187 | 0.619686061 | 0.719182032 | 0.043479137 | 0.043947041 |
| Pramel12 | 0.005292407 | 0.224510823 | 0.336212911 | 0.056019378 | 0.049458723 |
| Prcc | 0.015155061 | 0.006841727 | 0.018885654 | 0.104580509 | 0.086680322 |
| Prdm2 | 0.015421443 | 0.069294116 | 0.131290206 | 0.259318808 | 0.244786876 |
| Prdm4 | 0.006094255 | 0.112555351 | 0.194674101 | 0.043158998 | 0.03786666 |
| Prdx1 | -0.124313014 | 1.84E-45 | 1.84E-43 | 1.317194808 | 1.487800354 |
| Prdx2 | -0.015830435 | 0.078745237 | 0.145654718 | 1.150573953 | 1.140075117 |
| Prdx3 | -0.027195492 | 0.001495496 | 0.00506946 | 0.272538571 | 0.284171982 |
| Prdx4 | -0.061800609 | 1.52E-16 | 3.28E-15 | 0.142633204 | 0.192883407 |
| Prdx5 | -0.060266428 | 2.05E-10 | 2.53E-09 | 0.906047427 | 0.979454775 |
| Prdx6 | -0.020885919 | 0.03166543 | 0.068335475 | 0.9969694 | 0.984665822 |
| Preb | -0.028061716 | 0.00210049 | 0.006806989 | 0.326093215 | 0.346133817 |
| Prelid1 | 0.003456566 | 0.700671548 | 0.783610812 | 0.351743104 | 0.331541439 |
| Prelid3b | -0.009906358 | 0.243371958 | 0.357087561 | 0.267679206 | 0.266029614 |
| Prep | 0.012235002 | 0.00690652 | 0.0190213 | 0.062875152 | 0.052123856 |
| Prepl | 0.018949964 | 0.05000756 | 0.100035698 | 0.438694924 | 0.408620778 |
| Prickle1 | 0.015320288 | 0.003256208 | 0.00998954 | 0.087376432 | 0.071243952 |
| Prim1 | -0.009705489 | 0.038569172 | 0.080362085 | 0.058408723 | 0.063248043 |
| Prkaa1 | 0.043853473 | 5.35E-08 | 4.73E-07 | 0.257935402 | 0.208167602 |
| Prkaa2 | -0.000586438 | 0.895785506 | 0.92913973 | 0.052074887 | 0.054084402 |
| Prkab1 | 0.01115926 | 0.029094721 | 0.06379316 | 0.082751553 | 0.068311342 |
| Prkab2 | 0.011463097 | 0.041586852 | 0.085723668 | 0.095078395 | 0.082951665 |
| Prkaca | 0.021309093 | 0.013032279 | 0.03239576 | 0.311991715 | 0.272702112 |
| Prkacb | 0.070270647 | 1.71E-13 | 3.02E-12 | 1.151252665 | 1.047207956 |
| Prkag1 | -0.013106822 | 0.080029712 | 0.147413578 | 0.182054688 | 0.184764778 |
| Prkag2 | 0.03596974 | 1.30E-07 | 1.08E-06 | 0.159916715 | 0.128423668 |
| Prkar1a | -0.084397753 | 2.92E-19 | 7.99E-18 | 1.552607651 | 1.811452735 |
| Prkar1b | 0.042051917 | 2.20E-06 | 1.44E-05 | 0.355586935 | 0.295137673 |
| Prkar2a | 0.051076874 | 1.88E-09 | 2.05E-08 | 0.277023795 | 0.234929221 |
| Prkar2b | 0.038677325 | 1.60E-06 | 1.07E-05 | 0.250095787 | 0.205776027 |
| Prkca | 0.087415302 | 8.95E-22 | 2.89E-20 | 0.386874683 | 0.288134744 |
| Prkcb | 0.052351753 | 2.15E-09 | 2.33E-08 | 0.330825935 | 0.273957202 |
| Prkcd | -0.003403128 | 0.436893429 | 0.555339697 | 0.050727216 | 0.053541123 |
| Prkce | 0.027501987 | 0.001232101 | 0.004285682 | 0.278512891 | 0.248782138 |
| Prkci | 0.012652222 | 0.013475835 | 0.033387377 | 0.079759756 | 0.068434196 |
| Prkcsh | -0.074694814 | 6.78E-13 | 1.11E-11 | 0.623701658 | 0.726876908 |
| Prkcz | -0.003009362 | 0.624229315 | 0.722815316 | 0.113319594 | 0.111762771 |
| Prkd1 | -0.003162749 | 0.640169282 | 0.735664983 | 0.145368571 | 0.137985532 |
| Prkd2 | 0.007960866 | 0.040711971 | 0.084195231 | 0.046384904 | 0.038160068 |
| Prkdc | -0.008353649 | 0.127370735 | 0.21449823 | 0.079726361 | 0.084164815 |
| Prkn | -0.012651628 | 0.003466251 | 0.010530904 | 0.043909739 | 0.053496461 |
| Prkra | -0.010977292 | 0.063638569 | 0.122392515 | 0.097624725 | 0.105469536 |
| Prkrip1 | 0.019824268 | 0.004774595 | 0.013859038 | 0.163244084 | 0.147245439 |
| Prkx | 0.008343883 | 0.097697438 | 0.17313425 | 0.073911815 | 0.067628503 |
| Prlr | 0.081819356 | 1.99E-15 | 3.98E-14 | 2.081691248 | 1.861424464 |

| | | | | | |
|---|---|---|---|---|---|
| Prmt1 | -0.063406843 | 9.22E-12 | 1.32E-10 | 0.391404892 | 0.433302268 |
| Prmt2 | 0.003073867 | 0.544486759 | 0.65416464 | 0.081490763 | 0.07247234 |
| Prmt3 | -0.01124177 | 0.021756144 | 0.049896965 | 0.059205448 | 0.068008717 |
| Prmt5 | 0.00644157 | 0.240970213 | 0.354902653 | 0.090475309 | 0.083194343 |
| Prmt7 | -0.01246838 | 0.025647405 | 0.057320476 | 0.082959114 | 0.091793648 |
| Prmt9 | -0.005760386 | 0.219089803 | 0.329963906 | 0.058722116 | 0.060815435 |
| Prnp | -0.077567679 | 2.04E-29 | 1.04E-27 | 1.75666857 | 1.907664883 |
| Prob1 | 0.001652735 | 0.650834311 | 0.743701361 | 0.036904443 | 0.034127377 |
| Prom1 | -0.013422753 | 0.159505688 | 0.256513166 | 0.281401016 | 0.304864853 |
| Prorp | 0.001215983 | 0.783064018 | 0.847955935 | 0.053913992 | 0.052526011 |
| Prorsd1 | 0.013151701 | 0.169593398 | 0.26875094 | 0.412663573 | 0.400404673 |
| Proscos | 0.021953594 | 1.04E-10 | 1.34E-09 | 0.041675598 | 0.023225414 |
| Proser1 | 0.007932184 | 0.156574673 | 0.252818297 | 0.093088212 | 0.084933399 |
| Prox1 | 0.043484447 | 1.67E-06 | 1.12E-05 | 0.308697991 | 0.276703705 |
| Prpf18 | 0.002784477 | 0.680100716 | 0.767402409 | 0.140584507 | 0.135724127 |
| Prpf19 | 0.021768476 | 0.018445964 | 0.043463806 | 0.410884026 | 0.368910781 |
| Prpf3 | 0.0151141 | 0.000803468 | 0.00295352 | 0.063776947 | 0.049586809 |
| Prpf31 | -0.005163558 | 0.407939689 | 0.528164823 | 0.114277772 | 0.117341516 |
| Prpf38a | 0.00614358 | 0.398113979 | 0.518412015 | 0.171627844 | 0.163313289 |
| Prpf38b | -0.002565797 | 0.786780528 | 0.850843191 | 0.351582743 | 0.350888531 |
| Prpf39 | -0.018271438 | 0.019257284 | 0.044993827 | 0.180611533 | 0.195297542 |
| Prpf4 | 0.010277304 | 0.145256535 | 0.238988666 | 0.160410134 | 0.147633986 |
| Prpf40a | -0.011210219 | 0.269005557 | 0.385562734 | 0.656207208 | 0.659049361 |
| Prpf40b | -0.013036951 | 0.021546859 | 0.049515449 | 0.085807109 | 0.093922282 |
| Prpf4b | -0.019593129 | 0.067795585 | 0.129034588 | 0.890199847 | 0.938610451 |
| Prpf6 | 0.077358813 | 2.13E-20 | 6.29E-19 | 0.277475966 | 0.211771002 |
| Prpf8 | 0.052511827 | 4.19E-08 | 3.76E-07 | 0.496192413 | 0.428629772 |
| Prps1 | -0.007074288 | 0.461722621 | 0.579507788 | 0.464384611 | 0.450356449 |
| Prps1l3 | -0.001467409 | 0.716756278 | 0.796462125 | 0.044219174 | 0.043783357 |
| Prps2 | -0.003612695 | 0.540824305 | 0.651374368 | 0.105434058 | 0.104374307 |
| Prpsap1 | -0.006601436 | 0.378049199 | 0.499781658 | 0.184177325 | 0.182421339 |
| Prpsap2 | 0.008796749 | 0.195361162 | 0.301053488 | 0.150041271 | 0.138399559 |
| Prr12 | 0.00746692 | 0.257469418 | 0.373432446 | 0.137677166 | 0.127230164 |
| Prr13 | -0.02503588 | 0.012344108 | 0.030979354 | 0.525363026 | 0.535117102 |
| Prr14 | -0.002668563 | 0.6673608 | 0.757069043 | 0.111514623 | 0.112091807 |
| Prr14l | 0.02963142 | 1.08E-05 | 6.17E-05 | 0.158765701 | 0.128118739 |
| Prr15l | 0.039992604 | 1.11E-07 | 9.36E-07 | 0.1977683 | 0.153528585 |
| Prr3 | -0.007699895 | 0.142022245 | 0.234580406 | 0.071720512 | 0.077657988 |
| Prr7 | 0.009273297 | 0.041592115 | 0.085723668 | 0.063335824 | 0.053213457 |
| Prrc1 | 0.020379726 | 0.043771738 | 0.089325952 | 0.626365137 | 0.592978116 |
| Prrc2a | 0.001502947 | 0.873771613 | 0.913518402 | 0.399068623 | 0.394170614 |
| Prrc2b | 0.033703024 | 0.000922927 | 0.003334335 | 0.798557422 | 0.733066288 |
| Prrc2c | 0.030957597 | 0.002307438 | 0.007406045 | 1.039923726 | 1.012315433 |
| Prrg2 | -0.006743496 | 0.418713572 | 0.537887599 | 0.246597641 | 0.250225256 |
| Prrt1 | -0.009276989 | 0.040731352 | 0.084217398 | 0.046782518 | 0.054714553 |
| Prss36 | 0.004235714 | 0.336706639 | 0.459193399 | 0.059839229 | 0.051614931 |
| Prss53 | -0.004824848 | 0.632497171 | 0.729098641 | 1.393158223 | 1.297017852 |
| Prune1 | 0.009491242 | 0.062178289 | 0.120295945 | 0.077176783 | 0.068390028 |
| Prune2 | -0.040745135 | 3.92E-05 | 0.000197342 | 0.513675442 | 0.511809644 |
| Prxl2a | 0.025682717 | 0.002190478 | 0.007063293 | 0.280248 | 0.24926942 |
| Prxl2b | -0.030317688 | 0.000510562 | 0.00197835 | 0.295245816 | 0.312377232 |
| Prxl2c | -0.038397894 | 1.22E-05 | 6.88E-05 | 0.285156193 | 0.304917981 |
| Psap | -0.023553258 | 0.001462341 | 0.004974414 | 1.709052362 | 1.684138406 |
| Psat1 | -0.002528663 | 0.785518298 | 0.849950907 | 0.287409086 | 0.301556587 |
| Psd | 0.003707105 | 0.477704355 | 0.594500991 | 0.08246983 | 0.073109928 |
| Psd3 | 0.041574064 | 1.58E-11 | 2.24E-10 | 0.134079077 | 0.095690063 |
| Psd4 | 0.010308804 | 0.003130969 | 0.009646031 | 0.038252695 | 0.028338049 |
| Psen1 | 0.008178432 | 0.191604084 | 0.296148634 | 0.118554318 | 0.1128328 |
| Psen2 | -0.016740489 | 0.034111501 | 0.072710373 | 0.201871207 | 0.219055957 |
| Psenen | -0.020177314 | 0.037105938 | 0.077813618 | 0.819118199 | 0.832044643 |
| Psip1 | -0.030907282 | 0.002001049 | 0.006541426 | 0.749110199 | 0.762275004 |
| Pskh1 | 0.010405496 | 0.050401274 | 0.10069897 | 0.090330453 | 0.077502395 |
| Psma1 | -0.046312284 | 2.98E-06 | 1.91E-05 | 0.620524335 | 0.656534758 |
| Psma2 | -0.036917158 | 0.000176885 | 0.000773343 | 0.960315097 | 1.005391921 |
| Psma3 | -0.087996103 | 8.14E-20 | 2.33E-18 | 0.950747174 | 1.037213149 |
| Psma4 | -0.037677908 | 0.000202169 | 0.000866615 | 0.58281434 | 0.612893534 |
| Psma5 | -0.022959056 | 0.018902893 | 0.044314908 | 0.449288509 | 0.458449206 |
| Psma6 | -0.037334581 | 0.000222457 | 0.000942778 | 0.589815846 | 0.62379877 |
| Psma7 | -0.081834765 | 1.11E-22 | 3.88E-21 | 1.3539631 | 1.5016865 |
| Psmb1 | -0.057694357 | 2.98E-10 | 3.61E-09 | 1.064579316 | 1.131794491 |
| Psmb10 | 0.030194829 | 0.000707503 | 0.002643485 | 0.262424527 | 0.239440529 |
| Psmb2 | -0.025608458 | 0.009784903 | 0.025401557 | 0.927176601 | 0.93549369 |

| | | | | | |
|---|---|---|---|---|---|
| Psmb3 | -0.073606505 | 1.56E-12 | 2.46E-11 | 0.727310876 | 0.790042002 |
| Psmb4 | -0.069580442 | 3.78E-13 | 6.41E-12 | 1.007286274 | 1.10354606 |
| Psmb5 | -0.121531109 | 3.78E-31 | 2.02E-29 | 0.704583878 | 0.830252563 |
| Psmb6 | -0.081476116 | 7.24E-17 | 1.60E-15 | 0.851424756 | 0.947410793 |
| Psmb7 | 0.022614144 | 0.017608896 | 0.041887722 | 0.472953903 | 0.441572923 |
| Psmb8 | 0.065382003 | 4.12E-11 | 5.52E-10 | 0.383145126 | 0.340775175 |
| Psmb9 | 0.004059654 | 0.581211256 | 0.687329501 | 0.166664417 | 0.159028715 |
| Psmc1 | -0.001118582 | 0.906668624 | 0.935897369 | 0.418553331 | 0.409124585 |
| Psmc2 | -0.055296513 | 6.47E-08 | 5.64E-07 | 0.587822488 | 0.640451243 |
| Psmc3 | -0.024578827 | 0.007961277 | 0.02147005 | 0.359155659 | 0.377107231 |
| Psmc4 | -0.034146603 | 0.000365077 | 0.001465433 | 0.414928993 | 0.437101557 |
| Psmc5 | -0.100493848 | 6.26E-23 | 2.23E-21 | 0.674429132 | 0.779646043 |
| Psmc6 | -0.016980534 | 0.089980839 | 0.162118924 | 0.571833187 | 0.578664968 |
| Psmd1 | 0.022425971 | 0.022514243 | 0.051236768 | 0.505465552 | 0.478558815 |
| Psmd10 | -0.011623572 | 0.114076689 | 0.196321875 | 0.17774027 | 0.182764043 |
| Psmd11 | -0.038294191 | 0.000110111 | 0.000510248 | 0.653710705 | 0.68221129 |
| Psmd12 | 0.008209637 | 0.413659591 | 0.533331043 | 0.648752511 | 0.636598729 |
| Psmd13 | -0.007781823 | 0.423724554 | 0.542104239 | 0.481842252 | 0.47739895 |
| Psmd14 | -0.015040802 | 0.07653506 | 0.142488291 | 0.262537004 | 0.271534542 |
| Psmd2 | 0.009727395 | 0.32149485 | 0.443217943 | 0.537470729 | 0.520298811 |
| Psmd3 | -0.011441217 | 0.236897213 | 0.350603548 | 0.455701938 | 0.449378847 |
| Psmd4 | 0.010130559 | 0.316367615 | 0.438075196 | 0.780961731 | 0.782597653 |
| Psmd5 | -0.004906055 | 0.466707811 | 0.584105733 | 0.135267602 | 0.137498776 |
| Psmd6 | -0.054562934 | 7.59E-08 | 6.55E-07 | 0.583882616 | 0.629254274 |
| Psmd7 | -0.054315834 | 3.86E-08 | 3.48E-07 | 0.7940458 | 0.871402516 |
| Psmd8 | -0.045180718 | 7.55E-06 | 4.46E-05 | 0.643311845 | 0.673857628 |
| Psmd9 | -0.034057148 | 2.14E-05 | 0.000115276 | 0.207657296 | 0.234610368 |
| Psme1 | 0.065650033 | 1.82E-10 | 2.28E-09 | 0.585254072 | 0.526932017 |
| Psme2 | -0.017877795 | 0.084765279 | 0.154548254 | 0.818990654 | 0.839461863 |
| Psme3 | 0.027630279 | 0.000153841 | 0.000682318 | 0.184700527 | 0.162243824 |
| Psme3ip1 | 0.000413523 | 0.955475915 | 0.969840973 | 0.181867219 | 0.17581793 |
| Psme4 | 0.017106596 | 0.080482733 | 0.148023673 | 0.460628558 | 0.431128561 |
| Psmf1 | 0.011666134 | 0.122245095 | 0.207519961 | 0.202681092 | 0.183284921 |
| Psmg1 | -0.035377291 | 2.23E-06 | 1.46E-05 | 0.166748568 | 0.19397879 |
| Psmg2 | -0.055264454 | 3.77E-11 | 5.10E-10 | 0.20417163 | 0.259577336 |
| Psmg3 | -0.006387987 | 0.379489369 | 0.500936272 | 0.176614255 | 0.177691356 |
| Psmg4 | 0.041532896 | 1.14E-05 | 6.48E-05 | 0.404180703 | 0.388734194 |
| Pspc1 | -0.007877012 | 0.117743089 | 0.201177208 | 0.064296261 | 0.070556726 |
| Psph | -0.11267725 | 3.18E-19 | 8.65E-18 | 0.599997059 | 0.746990022 |
| Pstk | 0.002329449 | 0.621767375 | 0.72099504 | 0.064241108 | 0.059953141 |
| Ptar1 | -0.007738138 | 0.20440785 | 0.312510319 | 0.105147341 | 0.108743851 |
| Ptbp1 | 0.019238513 | 0.04569455 | 0.092492696 | 0.436375689 | 0.410771809 |
| Ptbp2 | 0.01584051 | 0.01532294 | 0.03718583 | 0.13340621 | 0.121114005 |
| Ptbp3 | 0.018803149 | 0.057238275 | 0.111988431 | 1.082594217 | 1.08290926 |
| Ptcd1 | 0.004911247 | 0.198218111 | 0.304820702 | 0.042951812 | 0.037022074 |
| Ptcd2 | 0.006540391 | 0.3455749 | 0.467812923 | 0.159381294 | 0.150320765 |
| Ptcd3 | -0.007722718 | 0.116851474 | 0.200075736 | 0.065066307 | 0.069215816 |
| Ptch1 | 0.002695882 | 0.641285037 | 0.736074138 | 0.098486008 | 0.091731867 |
| Ptdss1 | 0.001166304 | 0.843012933 | 0.89268835 | 0.10219686 | 0.099502654 |
| Ptdss2 | -0.001725009 | 0.714654207 | 0.794950595 | 0.062111386 | 0.060583207 |
| Pten | 0.030510275 | 0.002169826 | 0.00700832 | 0.658757387 | 0.599983814 |
| Ptges2 | 0.013485057 | 0.022296419 | 0.050870842 | 0.110564866 | 0.0972445 |
| Ptges3 | -0.024275048 | 0.014636691 | 0.035807225 | 0.817282175 | 0.835157382 |
| Ptgr2 | 0.006134748 | 0.41801416 | 0.537202071 | 0.201851532 | 0.188847612 |
| Ptk2 | 0.002051127 | 0.740189975 | 0.814133605 | 0.112488373 | 0.110850271 |
| Ptk2b | 0.01364313 | 0.00118792 | 0.004157292 | 0.056152999 | 0.042451884 |
| Ptma | 0.04750664 | 1.57E-17 | 3.67E-16 | 2.469242222 | 2.371180003 |
| Ptms | 0.088256052 | 1.61E-19 | 4.50E-18 | 1.240231652 | 1.090153938 |
| Ptov1 | 0.003910015 | 0.695056483 | 0.77921346 | 0.692525032 | 0.635472614 |
| Ptp4a2 | 0.035567525 | 0.000351653 | 0.001420937 | 0.577099048 | 0.530290356 |
| Ptp4a3 | -0.0391242 | 3.00E-07 | 2.32E-06 | 0.171696357 | 0.196879129 |
| Ptpa | 0.029496532 | 0.001180311 | 0.004135131 | 0.37254012 | 0.330940388 |
| Ptpdc1 | 0.00471332 | 0.345495202 | 0.467812923 | 0.074236211 | 0.06656801 |
| Ptpmt1 | -0.024788794 | 0.002694564 | 0.008467062 | 0.245288541 | 0.259999714 |
| Ptpn1 | 0.056381164 | 9.26E-09 | 9.28E-08 | 0.449908335 | 0.398770612 |
| Ptpn11 | 0.048950994 | 1.48E-06 | 9.99E-06 | 0.741554212 | 0.66468909 |
| Ptpn12 | 0.013080711 | 0.025771759 | 0.057532271 | 0.099287089 | 0.093936674 |
| Ptpn13 | 0.013541962 | 0.003643387 | 0.011010571 | 0.064771822 | 0.052796518 |
| Ptpn2 | 0.01534865 | 0.052603822 | 0.104093482 | 0.199329779 | 0.190148088 |
| Ptpn21 | 0.037136418 | 1.20E-08 | 1.18E-07 | 0.145122334 | 0.11414682 |
| Ptpn23 | 0.031633537 | 1.90E-07 | 1.53E-06 | 0.122034361 | 0.098525226 |
| Ptpn3 | 0.013950533 | 0.000332182 | 0.001348987 | 0.048136198 | 0.035908081 |

| | | | | | |
|---|---|---|---|---|---|
| Ptpn4 | 0.006797578 | 0.335964355 | 0.4586326 | 0.160939382 | 0.151623164 |
| Ptpn6 | -0.004310648 | 0.505549481 | 0.619635912 | 0.128236422 | 0.127239564 |
| Ptpn9 | 0.016134999 | 0.025691907 | 0.057393548 | 0.165007451 | 0.15960284 |
| Ptpra | -0.002407095 | 0.779870934 | 0.845157198 | 0.291273823 | 0.283963129 |
| Ptprd | -0.005907568 | 0.256858151 | 0.372768315 | 0.068960984 | 0.073252447 |
| Ptprf | 0.05233316 | 8.49E-08 | 7.27E-07 | 1.056979456 | 0.979463869 |
| Ptprj | -0.008598101 | 0.244518982 | 0.358353984 | 0.167135594 | 0.173722693 |
| Ptprm | -0.021531011 | 0.000853122 | 0.00312096 | 0.101457906 | 0.126317002 |
| Ptprn | -0.125498548 | 3.57E-43 | 3.25E-41 | 1.335626596 | 1.610103692 |
| Ptprn2 | 0.054025172 | 9.95E-09 | 9.89E-08 | 0.364541484 | 0.311037153 |
| Ptprr | 0.012454652 | 0.003932229 | 0.011751961 | 0.058499652 | 0.044879251 |
| Ptprs | 0.003003004 | 0.562066615 | 0.670307289 | 0.0711382 | 0.071051662 |
| Ptprt | 0.007015511 | 0.391283565 | 0.512075844 | 0.236445748 | 0.208034712 |
| Ptpru | 0.016844763 | 0.000249868 | 0.001044376 | 0.070999273 | 0.052679411 |
| Ptrh2 | -0.057465191 | 3.13E-13 | 5.36E-12 | 0.169047714 | 0.217579853 |
| Ptrhd1 | 0.020329813 | 0.00973357 | 0.025302078 | 0.220895168 | 0.20136992 |
| Pts | -0.062401672 | 2.03E-10 | 2.50E-09 | 0.462047347 | 0.488586286 |
| Pttg1 | -0.034095418 | 1.74E-06 | 1.15E-05 | 0.146228295 | 0.166697906 |
| Pttg1ip | 0.025184257 | 0.003022647 | 0.009352697 | 0.291810637 | 0.262890995 |
| Puf60 | -0.042954393 | 4.39E-06 | 2.73E-05 | 0.353710256 | 0.383371852 |
| Pum1 | 0.042602906 | 1.11E-05 | 6.30E-05 | 0.481654277 | 0.440655513 |
| Pum2 | 0.052501911 | 9.55E-08 | 8.10E-07 | 0.573283442 | 0.509090305 |
| Pum3 | 0.030929724 | 5.03E-05 | 0.000246802 | 0.207658139 | 0.182690439 |
| Pura | 0.000129764 | 0.989145315 | 0.993079296 | 1.049183334 | 1.014722011 |
| Purb | 0.024699682 | 0.016485708 | 0.039612963 | 0.739904388 | 0.692496476 |
| Pus1 | -0.001341435 | 0.815501331 | 0.872866347 | 0.095511365 | 0.095297746 |
| Pus10 | 0.000673259 | 0.876895059 | 0.915897482 | 0.051534039 | 0.05029881 |
| Pus3 | -0.017008202 | 0.002132546 | 0.006899372 | 0.079415728 | 0.092370846 |
| Pus7 | -0.002321306 | 0.678544128 | 0.766446615 | 0.085207939 | 0.087623052 |
| Pusl1 | -0.010281909 | 0.008493479 | 0.022653938 | 0.036221364 | 0.043360871 |
| Pwp1 | 0.006630065 | 0.285994852 | 0.405312238 | 0.119585812 | 0.110235603 |
| Pwwp2a | 0.003318209 | 0.595036044 | 0.698833102 | 0.119092469 | 0.111553724 |
| Pwwp3a | 0.017514345 | 0.003132331 | 0.009646031 | 0.114448722 | 0.094514908 |
| Pwwp3b | -0.024531576 | 0.006933638 | 0.019085173 | 0.178094188 | 0.233978113 |
| Pxk | 0.041829016 | 2.49E-06 | 1.62E-05 | 0.335294044 | 0.291592944 |
| Pxmp4 | 0.010625379 | 0.051534255 | 0.102353775 | 0.090943266 | 0.079002472 |
| Pxn | -0.001966938 | 0.606113397 | 0.707738943 | 0.039796935 | 0.038546057 |
| Pxylp1 | 0.000451913 | 0.920625387 | 0.945622822 | 0.052623142 | 0.053515292 |
| Pycard | 0.018627294 | 0.000425396 | 0.001679815 | 0.086308533 | 0.069228663 |
| Pycr1 | -0.072689012 | 7.63E-15 | 1.48E-13 | 0.312733433 | 0.36186054 |
| Pycr2 | -0.066879093 | 7.63E-12 | 1.11E-10 | 0.792065396 | 0.853872061 |
| Pycrl | -0.033966916 | 1.53E-07 | 1.26E-06 | 0.109266597 | 0.133795545 |
| Pygo1 | -0.002894552 | 0.49539496 | 0.6105768 | 0.045221554 | 0.04623525 |
| Pygo2 | 0.002458967 | 0.607380572 | 0.708537788 | 0.063437622 | 0.060814509 |
| Pym1 | -0.000542543 | 0.904940141 | 0.935446182 | 0.060245266 | 0.056459022 |
| Pyroxd1 | -0.012735622 | 0.059130449 | 0.115226743 | 0.125915714 | 0.1389837 |
| Pyurf | -0.004995649 | 0.458656923 | 0.577224933 | 0.146403063 | 0.142128106 |
| Pyy | 0.067266019 | 2.01E-05 | 0.000108932 | 1.023011818 | 0.998776416 |
| Qars | 0.00487199 | 0.494979664 | 0.610142296 | 0.169644629 | 0.15702114 |
| Qdpr | -0.00314593 | 0.754767875 | 0.825312482 | 0.628932445 | 0.624135927 |
| Qk | 0.002661852 | 0.650082614 | 0.743191813 | 0.099499048 | 0.094868605 |
| Qpctl | -0.017812405 | 0.00585236 | 0.016483753 | 0.115150178 | 0.129359713 |
| Qrich1 | 0.008816923 | 0.222793065 | 0.334103683 | 0.175452392 | 0.161653996 |
| Qrsl1 | -0.005146096 | 0.207450487 | 0.315821114 | 0.041432029 | 0.044629261 |
| Qser1 | 0.012504411 | 0.08072716 | 0.148417068 | 0.164004606 | 0.150079546 |
| Qsox1 | -0.027083718 | 0.003549066 | 0.010748916 | 0.347466108 | 0.35223709 |
| Qsox2 | 0.003604909 | 0.655883241 | 0.747888444 | 0.237208907 | 0.230979739 |
| Qtrt1 | -0.008154749 | 0.182250174 | 0.284540901 | 0.102817128 | 0.110256157 |
| Qtrt2 | -0.002674553 | 0.527217494 | 0.638780869 | 0.0461106 | 0.048117852 |
| R3hcc1 | -0.008309152 | 0.164003172 | 0.261769634 | 0.100135549 | 0.106578757 |
| R3hcc1l | 0.004880631 | 0.259739287 | 0.375817355 | 0.052511899 | 0.048901055 |
| R3hdm1 | 0.005700194 | 0.5707372 | 0.677911675 | 0.492434892 | 0.484376371 |
| R3hdm2 | 0.057634437 | 1.40E-08 | 1.36E-07 | 0.581819245 | 0.533086557 |
| R3hdm4 | -0.003644354 | 0.653252576 | 0.745500827 | 0.238261311 | 0.238091965 |
| Rab10 | 0.029677598 | 0.003138154 | 0.009660903 | 0.668184128 | 0.63738648 |
| Rab10os | 0.035333195 | 1.42E-08 | 1.37E-07 | 0.127182803 | 0.101610439 |
| Rab11a | -0.000230575 | 0.981201479 | 0.987908118 | 0.579239298 | 0.564481165 |
| Rab11b | 0.013687129 | 0.13184363 | 0.221226057 | 0.367951747 | 0.337788038 |
| Rab11fip1 | 0.002746468 | 0.539156081 | 0.64984818 | 0.055122955 | 0.05218784 |
| Rab11fip2 | 0.03421127 | 8.91E-05 | 0.000420017 | 0.306135435 | 0.270145608 |
| Rab11fip3 | 0.005544937 | 0.223558356 | 0.334998324 | 0.059835236 | 0.053528844 |
| Rab12 | 0.036601405 | 3.07E-05 | 0.000158855 | 0.314593929 | 0.268379733 |

| | | | | | |
|---|---|---|---|---|---|
| Rab13 | -0.00734217 | 0.093708516 | 0.167285016 | 0.049967394 | 0.053613414 |
| Rab14 | 0.046066731 | 2.60E-07 | 2.03E-06 | 1.209343908 | 1.145710667 |
| Rab18 | -0.01239048 | 0.207232223 | 0.315587667 | 0.564109317 | 0.577281081 |
| Rab19 | 0.007589136 | 0.089859733 | 0.161930736 | 0.061894088 | 0.052303095 |
| Rab1a | 0.034169653 | 0.000411938 | 0.001631971 | 0.856592917 | 0.812110712 |
| Rab1b | 0.022191683 | 0.00951377 | 0.024863676 | 0.292945173 | 0.271339302 |
| Rab20 | 0.012469417 | 0.000179881 | 0.000782659 | 0.037789903 | 0.025159248 |
| Rab21 | 0.044637357 | 5.66E-06 | 3.43E-05 | 0.530893057 | 0.47041611 |
| Rab22a | 0.007456387 | 0.449347311 | 0.567961121 | 0.495420315 | 0.480919334 |
| Rab23 | 0.003620802 | 0.466661133 | 0.584105733 | 0.070319461 | 0.067104121 |
| Rab24 | -0.03354731 | 0.000381661 | 0.001524451 | 0.399962331 | 0.42158346 |
| Rab25 | -0.071009486 | 1.84E-13 | 3.21E-12 | 0.451995296 | 0.510292711 |
| Rab27a | 0.07552571 | 1.82E-14 | 3.43E-13 | 0.510030845 | 0.425354282 |
| Rab28 | -0.015605435 | 0.092437214 | 0.165471294 | 0.373392579 | 0.374717527 |
| Rab2a | 0.026656359 | 0.000612399 | 0.002323903 | 1.469390438 | 1.426059496 |
| Rab2b | -0.006122389 | 0.165494996 | 0.263761042 | 0.047407373 | 0.053208832 |
| Rab33b | 0.003266827 | 0.657075276 | 0.748984152 | 0.180340158 | 0.172035735 |
| Rab34 | -0.028022485 | 0.00021739 | 0.000924524 | 0.178166116 | 0.195827494 |
| Rab35 | 0.008370025 | 0.166707161 | 0.26514883 | 0.117292088 | 0.105207584 |
| Rab36 | -0.011098645 | 0.048107133 | 0.096912049 | 0.082050595 | 0.091536833 |
| Rab37 | 0.037768963 | 1.39E-05 | 7.78E-05 | 0.306143077 | 0.244105608 |
| Rab39b | -0.041225713 | 6.74E-06 | 4.02E-05 | 0.283706202 | 0.331229379 |
| Rab3a | 0.006025222 | 0.54625509 | 0.655964919 | 0.594462715 | 0.558788601 |
| Rab3b | 0.081366637 | 4.84E-24 | 1.83E-22 | 0.265325647 | 0.182221064 |
| Rab3d | 0.025479488 | 0.007480478 | 0.020336756 | 0.395668883 | 0.376909687 |
| Rab3gap1 | 0.000576373 | 0.951927898 | 0.967765747 | 0.408379508 | 0.411752785 |
| Rab3gap2 | 0.035253525 | 2.21E-05 | 0.000118328 | 0.256169406 | 0.227355397 |
| Rab3ip | -0.005893712 | 0.425570057 | 0.544178889 | 0.186517013 | 0.18035942 |
| Rab40b | -0.020924331 | 7.53E-06 | 4.45E-05 | 0.049660537 | 0.064840771 |
| Rab40c | 0.010921817 | 0.0444374 | 0.09039975 | 0.09484642 | 0.080112832 |
| Rab4a | -0.020033883 | 0.007102713 | 0.019462395 | 0.171511288 | 0.180614564 |
| Rab5a | 0.000263693 | 0.978899974 | 0.98589884 | 0.548030641 | 0.553685722 |
| Rab5b | 0.021802341 | 0.003908093 | 0.011690609 | 0.212999351 | 0.184426148 |
| Rab5c | 0.029673814 | 0.000587162 | 0.002238584 | 0.298537912 | 0.265300645 |
| Rab5if | -0.014297686 | 0.117496909 | 0.200827171 | 0.338585983 | 0.348640942 |
| Rab6a | 0.069755179 | 1.65E-11 | 2.33E-10 | 0.670402724 | 0.581604119 |
| Rab6b | -0.006915758 | 0.312233063 | 0.43364712 | 0.147916272 | 0.142873968 |
| Rab7 | 0.035227526 | 0.000415115 | 0.00164389 | 0.865265602 | 0.800760737 |
| Rab8a | -0.000438873 | 0.959284012 | 0.971967751 | 0.288764625 | 0.280272156 |
| Rab8b | 0.011209629 | 0.010282681 | 0.026495687 | 0.055902613 | 0.048283641 |
| Rab9 | -0.002436652 | 0.717420124 | 0.796922465 | 0.151899064 | 0.14440922 |
| Rabac1 | -0.05458291 | 3.37E-12 | 5.09E-11 | 1.491332859 | 1.591907812 |
| Rabep1 | 0.01749283 | 0.066589905 | 0.127163044 | 0.432420009 | 0.404705305 |
| Rabep2 | 0.00491094 | 0.220753034 | 0.331724416 | 0.047594857 | 0.040413118 |
| Rabepk | -0.013106758 | 0.018304179 | 0.043203018 | 0.086641415 | 0.093260979 |
| Rabgap1 | 0.007586492 | 0.376934224 | 0.49864669 | 0.276111538 | 0.265128243 |
| Rabgap1l | 0.027546735 | 4.80E-05 | 0.000236745 | 0.154038712 | 0.125551698 |
| Rabgef1 | 0.015661384 | 0.007357275 | 0.020080692 | 0.106644553 | 0.091949538 |
| Rabggta | 0.002275229 | 0.623401782 | 0.722115111 | 0.059503391 | 0.0586044 |
| Rabggtb | 0.012619328 | 0.152532001 | 0.247565295 | 0.309943843 | 0.287911677 |
| Rabif | 0.002727165 | 0.603238698 | 0.706248034 | 0.082406235 | 0.075139524 |
| Rabl2 | -0.001700584 | 0.667778917 | 0.757278272 | 0.043364201 | 0.042351074 |
| Rabl3 | 0.000259336 | 0.955400782 | 0.969840973 | 0.059984979 | 0.056986196 |
| Rabl6 | 0.028059677 | 0.00378602 | 0.011370924 | 0.44062869 | 0.414506702 |
| Rac1 | 0.033247116 | 0.000723007 | 0.002695198 | 0.628422259 | 0.581688313 |
| Racgap1 | 0.012033678 | 0.002690169 | 0.008461282 | 0.053433474 | 0.038486583 |
| Rack1 | -0.085825804 | 1.70E-33 | 1.03E-31 | 1.897003292 | 2.063131136 |
| Rad1 | -0.000179801 | 0.972733832 | 0.98141535 | 0.084309306 | 0.079331528 |
| Rad17 | 0.011038593 | 0.180971784 | 0.283144625 | 0.256287212 | 0.244228617 |
| Rad21 | 0.019259451 | 0.05590974 | 0.109720325 | 0.575820477 | 0.551077093 |
| Rad23a | -0.030931719 | 0.001694846 | 0.005640976 | 0.482011535 | 0.495026734 |
| Rad23b | 0.014608225 | 0.141314271 | 0.233688952 | 0.506322362 | 0.490517798 |
| Rad50 | 0.02077774 | 0.001836337 | 0.006062095 | 0.137532967 | 0.121114134 |
| Rad51ap2 | -0.060675804 | 2.00E-12 | 3.13E-11 | 0.199272673 | 0.240698181 |
| Rad51d | 0.00658747 | 0.180977237 | 0.283144625 | 0.071865921 | 0.065462684 |
| Rad54l2 | 0.022852539 | 4.07E-05 | 0.000203877 | 0.098212341 | 0.079800999 |
| Rae1 | -0.000593987 | 0.933673021 | 0.95498886 | 0.167128218 | 0.162248861 |
| Raf1 | 0.002948444 | 0.66218342 | 0.752974795 | 0.145408378 | 0.138781238 |
| Rai1 | 0.014131808 | 0.028521246 | 0.062712752 | 0.132267827 | 0.117165571 |
| Rai2 | 0.022548689 | 1.36E-05 | 7.62E-05 | 0.09676319 | 0.070470645 |
| Rala | 0.016490203 | 0.112932269 | 0.194923418 | 0.640199513 | 0.633458496 |
| Ralb | 0.020865098 | 0.000396864 | 0.001579335 | 0.115364728 | 0.093607575 |

| | | | | | |
|---|---|---|---|---|---|
| Ralbp1 | 0.058475698 | 2.87E-09 | 3.08E-08 | 0.828431076 | 0.769297042 |
| Ralgapa1 | 0.004056432 | 0.637766498 | 0.733337184 | 0.265995066 | 0.26191664 |
| Ralgapa2 | -0.005693054 | 0.433547116 | 0.551983377 | 0.154442478 | 0.159402664 |
| Ralgapb | 0.012576854 | 0.091415354 | 0.164155905 | 0.183273162 | 0.171934215 |
| Ralgds | 0.01004504 | 0.116055577 | 0.199112272 | 0.125814829 | 0.115864623 |
| Ralgps1 | 0.007221107 | 0.109203409 | 0.189950319 | 0.059242382 | 0.052414239 |
| Ralgps2 | 0.02324597 | 0.000304652 | 0.001248137 | 0.133179219 | 0.117654162 |
| Raly | 0.011674212 | 0.162282064 | 0.259961481 | 0.283716182 | 0.255013932 |
| Ramac | -0.014205106 | 0.09621809 | 0.171073934 | 0.293506732 | 0.286358324 |
| Ramp1 | 0.085718452 | 1.03E-54 | 1.50E-52 | 0.129237192 | 0.052102217 |
| Ramp2 | -0.052929602 | 3.86E-13 | 6.52E-12 | 0.116497949 | 0.169499097 |
| Ran | -0.037374334 | 0.000204413 | 0.000875078 | 0.928591199 | 0.978651734 |
| Ranbp1 | -0.054220901 | 5.18E-08 | 4.59E-07 | 0.495928317 | 0.540199849 |
| Ranbp10 | -0.002867557 | 0.617363503 | 0.717419098 | 0.094070787 | 0.093290059 |
| Ranbp2 | -0.015632911 | 0.134215622 | 0.224663272 | 0.552865986 | 0.564787675 |
| Ranbp3 | -0.002612718 | 0.705603917 | 0.787542987 | 0.160063631 | 0.148555795 |
| Ranbp6 | 0.010313223 | 0.018382454 | 0.043345674 | 0.057517666 | 0.048148319 |
| Ranbp9 | 0.030071051 | 3.77E-05 | 0.000191123 | 0.187299981 | 0.157775684 |
| Rangap1 | 0.061363028 | 1.89E-10 | 2.35E-09 | 0.440849705 | 0.383117134 |
| Rangrf | -0.005596915 | 0.206118213 | 0.314383632 | 0.053476743 | 0.055224429 |
| Rap1a | 0.030185303 | 0.002483449 | 0.007890224 | 0.716009481 | 0.654549808 |
| Rap1b | 0.005943212 | 0.560786173 | 0.669116737 | 1.163022272 | 1.17741209 |
| Rap1gap | 0.003433459 | 0.619191465 | 0.718733395 | 0.149929947 | 0.138999364 |
| Rap1gap2 | 0.111752628 | 6.67E-26 | 2.82E-24 | 0.735519523 | 0.613543523 |
| Rap1gapos | 0.01350785 | 0.000754164 | 0.002792074 | 0.046936965 | 0.037367064 |
| Rap1gds1 | 0.010033663 | 0.051357478 | 0.102126693 | 0.078511403 | 0.070363585 |
| Rap2a | 0.009060452 | 0.13992991 | 0.231935147 | 0.108604264 | 0.103684661 |
| Rap2b | 0.001504558 | 0.752516391 | 0.823759774 | 0.061057959 | 0.060497981 |
| Rap2c | -0.001799069 | 0.700967052 | 0.783674188 | 0.057476967 | 0.059860837 |
| Rapgef1 | 0.025234637 | 8.87E-05 | 0.00041868 | 0.138298993 | 0.116194301 |
| Rapgef2 | 0.003188829 | 0.552539254 | 0.662039269 | 0.08176899 | 0.078982053 |
| Rapgef4 | -0.001463742 | 0.866533083 | 0.90878475 | 0.27481872 | 0.270790001 |
| Rapgef6 | -0.003275523 | 0.651757337 | 0.744493577 | 0.16158899 | 0.164183872 |
| Raph1 | -0.017082692 | 0.008108215 | 0.021781725 | 0.105199391 | 0.120255727 |
| Rara | 0.031120397 | 1.38E-14 | 2.62E-13 | 0.059983054 | 0.034527723 |
| Rarg | 0.011256748 | 0.002109508 | 0.006831658 | 0.04315573 | 0.031584417 |
| Rars | -0.027276963 | 0.002893569 | 0.009007773 | 0.26335711 | 0.291123117 |
| Rars2 | -0.012032046 | 0.060610404 | 0.117709618 | 0.122216308 | 0.127767027 |
| Rasa1 | 0.030674008 | 0.000498505 | 0.001937813 | 0.317570264 | 0.285674825 |
| Rasa2 | 0.023720514 | 0.000947858 | 0.00341299 | 0.166881083 | 0.147362339 |
| Rasal2 | -0.008408757 | 0.232969126 | 0.346424876 | 0.145668103 | 0.153768503 |
| Rasd1 | -0.062242815 | 9.22E-07 | 6.48E-06 | 0.442261176 | 0.511002559 |
| Rasd2 | -0.024074803 | 6.42E-05 | 0.000310325 | 0.06937529 | 0.099588233 |
| Rasgef1b | -0.009166978 | 0.108708604 | 0.189436287 | 0.075219534 | 0.09137756 |
| Rasgrf1 | 0.041157618 | 0.000188803 | 0.000816186 | 0.658714232 | 0.600192096 |
| Rasgrf2 | -0.085544295 | 7.11E-32 | 3.99E-30 | 0.082142404 | 0.170775597 |
| Rassf1 | -0.013764299 | 0.046719627 | 0.094371122 | 0.129433905 | 0.143940446 |
| Rassf10 | -0.017163461 | 0.006440132 | 0.017940104 | 0.101525263 | 0.115217388 |
| Rassf3 | 0.009368633 | 0.246814144 | 0.361048316 | 0.231431094 | 0.222895389 |
| Rassf4 | -0.033318019 | 3.89E-06 | 2.44E-05 | 0.141355648 | 0.164242259 |
| Rassf7 | -0.000921337 | 0.855065657 | 0.900156829 | 0.075026621 | 0.070260713 |
| Rassf8 | 0.01074233 | 0.027188687 | 0.060170365 | 0.072333482 | 0.061269185 |
| Rb1 | 0.024256399 | 0.000116501 | 0.000536027 | 0.126334243 | 0.107428397 |
| Rb1cc1 | -0.010422691 | 0.302397961 | 0.422704958 | 0.467989845 | 0.470848423 |
| Rbbp4 | 0.000549108 | 0.954096381 | 0.968946518 | 0.465891318 | 0.444030672 |
| Rbbp5 | 0.015096278 | 0.002089734 | 0.006781173 | 0.074185321 | 0.061151347 |
| Rbbp6 | 0.019351623 | 0.055773625 | 0.109497412 | 0.639251553 | 0.604860693 |
| Rbbp7 | 0.007563383 | 0.459095842 | 0.577478299 | 0.662887529 | 0.648357705 |
| Rbbp8 | 0.008713197 | 0.060319131 | 0.117261067 | 0.059987853 | 0.054148111 |
| Rbbp9 | -0.004678018 | 0.282199792 | 0.40139669 | 0.050622426 | 0.05183753 |
| Rbck1 | -0.015583506 | 0.040673616 | 0.084133808 | 0.196922969 | 0.197162952 |
| Rbfa | -0.00410432 | 0.568503036 | 0.675894928 | 0.168764795 | 0.167622766 |
| Rbfox2 | 0.03141252 | 0.000240473 | 0.001010757 | 0.284573121 | 0.248627705 |
| Rbis | 0.022428095 | 0.013946853 | 0.034335605 | 0.328227729 | 0.33597656 |
| Rbks | -0.022146049 | 3.88E-09 | 4.08E-08 | 0.026297903 | 0.043167682 |
| Rbl2 | 0.02264487 | 0.000266915 | 0.001110374 | 0.125970098 | 0.103857213 |
| Rbm10 | -0.001455433 | 0.836803035 | 0.888187078 | 0.167565558 | 0.15609766 |
| Rbm12 | 0.01295616 | 0.003755778 | 0.01128707 | 0.062447691 | 0.049905453 |
| Rbm12b2 | 0.009706462 | 0.014147584 | 0.034759366 | 0.047631463 | 0.038100157 |
| Rbm14 | 0.036367128 | 7.33E-11 | 9.56E-10 | 0.104873033 | 0.073600479 |
| Rbm15 | 0.002744581 | 0.622557298 | 0.721566768 | 0.089546044 | 0.085665243 |
| Rbm15b | -0.002679363 | 0.662992896 | 0.753402868 | 0.10847491 | 0.108475306 |

| | | | | | |
|---|---|---|---|---|---|
| Rbm17 | 0.0057287 | 0.541033775 | 0.651465242 | 0.370060255 | 0.36349657 |
| Rbm18 | -0.01732269 | 0.017436156 | 0.041537445 | 0.162495774 | 0.172202014 |
| Rbm19 | 0.008739148 | 0.036729841 | 0.077195032 | 0.049416204 | 0.044083645 |
| Rbm22 | -0.000832422 | 0.901287748 | 0.932856328 | 0.139854498 | 0.137276006 |
| Rbm25 | -0.004583191 | 0.663854607 | 0.754059409 | 0.674130345 | 0.685972498 |
| Rbm26 | -0.012085798 | 0.222069104 | 0.33322362 | 0.429236607 | 0.453780241 |
| Rbm27 | 0.04953291 | 2.79E-08 | 2.56E-07 | 0.336801305 | 0.288097939 |
| Rbm28 | 0.030814279 | 0.000358338 | 0.001444298 | 0.278526588 | 0.257118906 |
| Rbm3 | -0.035991535 | 0.000563929 | 0.002158471 | 0.663182112 | 0.730219599 |
| Rbm33 | 0.028362012 | 0.000446546 | 0.001755486 | 0.231088256 | 0.207384622 |
| Rbm34 | -0.006621079 | 0.314321834 | 0.435801037 | 0.121771139 | 0.12813061 |
| Rbm39 | -0.00998917 | 0.201711854 | 0.309215176 | 1.55938096 | 1.584199632 |
| Rbm43 | 0.016175436 | 0.023562272 | 0.053409282 | 0.17184775 | 0.155914814 |
| Rbm47 | 0.037381107 | 3.89E-05 | 0.000196205 | 0.35806873 | 0.314537821 |
| Rbm48 | 0.005384903 | 0.376091654 | 0.497862632 | 0.10943513 | 0.106668132 |
| Rbm4b | -0.012464518 | 0.088060927 | 0.159398312 | 0.172574161 | 0.171710239 |
| Rbm5 | 0.003275906 | 0.744768721 | 0.817769739 | 0.459986075 | 0.449871991 |
| Rbm6 | 0.006087951 | 0.472218825 | 0.589409605 | 0.262053488 | 0.250039496 |
| Rbm7 | -0.010335479 | 0.212968082 | 0.322805689 | 0.257911989 | 0.251379549 |
| Rbm8a | -0.058326029 | 3.82E-09 | 4.03E-08 | 0.50962988 | 0.564147696 |
| Rbms1 | 0.021296547 | 0.000110698 | 0.000512481 | 0.095901259 | 0.077210778 |
| Rbms2 | -0.007133822 | 0.157443509 | 0.253968109 | 0.06893745 | 0.072826974 |
| Rbmx | -0.008102746 | 0.235622076 | 0.349469721 | 0.143237668 | 0.14552676 |
| Rbmx2 | 0.021875214 | 0.000869718 | 0.003169188 | 0.14607898 | 0.121484613 |
| Rbmxl1 | -0.005802635 | 0.345658521 | 0.46783964 | 0.117498055 | 0.113144437 |
| Rbp4 | 0.175784713 | 2.82E-64 | 5.83E-62 | 0.315517459 | 0.132515748 |
| Rbpj | 0.008229435 | 0.197933514 | 0.304494422 | 0.127881689 | 0.121723309 |
| Rbpms | 0.029739096 | 0.003065961 | 0.009468638 | 0.565868077 | 0.519322511 |
| Rbsn | 0.008555586 | 0.211807165 | 0.321346639 | 0.151351827 | 0.14205619 |
| Rbx1 | 0.020438245 | 0.01794674 | 0.04255566 | 1.252632225 | 1.241768952 |
| Rc3h1 | 0.034844103 | 6.72E-06 | 4.01E-05 | 0.211677369 | 0.179522732 |
| Rc3h2 | 0.029497827 | 0.001508295 | 0.005103947 | 0.375817305 | 0.339663322 |
| Rcan1 | 0.006046451 | 0.215337221 | 0.325432684 | 0.068632244 | 0.063604538 |
| Rcan2 | 0.061297645 | 1.24E-13 | 2.22E-12 | 0.294282234 | 0.218299992 |
| Rcan3 | 0.029290672 | 0.003998983 | 0.011907538 | 0.609219205 | 0.562661099 |
| Rcbtb1 | 0.033631901 | 0.000136682 | 0.000617483 | 0.321349678 | 0.280693437 |
| Rcbtb2 | 0.004644685 | 0.458443764 | 0.577031366 | 0.122582691 | 0.113796361 |
| Rcc1l | -0.00828822 | 0.08723157 | 0.158280202 | 0.060083472 | 0.065993321 |
| Rcc2 | -0.080639306 | 3.65E-15 | 7.16E-14 | 0.404468647 | 0.510577784 |
| Rce1 | -0.004020755 | 0.587157427 | 0.692088325 | 0.181513943 | 0.177516323 |
| Rchy1 | 0.000516706 | 0.953338872 | 0.968581933 | 0.307510093 | 0.296978052 |
| Rcl1 | -0.011156948 | 0.034006066 | 0.072501529 | 0.072495561 | 0.081291861 |
| Rcn1 | -0.014928947 | 0.004824346 | 0.013983987 | 0.070009763 | 0.080726976 |
| Rcn2 | -0.011857452 | 0.206551886 | 0.314748814 | 0.403719559 | 0.412748887 |
| Rcor1 | 0.010922804 | 0.16332007 | 0.261185487 | 0.210886995 | 0.195363189 |
| Rcor2 | 0.007197729 | 0.026679495 | 0.059234879 | 0.033843663 | 0.025962023 |
| Rcor3 | 0.0265171 | 0.000479891 | 0.00187144 | 0.202543153 | 0.173844118 |
| Rdh11 | 0.00465269 | 0.288640282 | 0.408131717 | 0.054416995 | 0.050352624 |
| Rdh13 | -0.00087949 | 0.823290391 | 0.87821255 | 0.039005216 | 0.040023619 |
| Rdh14 | -0.004492656 | 0.577195584 | 0.683328722 | 0.240643472 | 0.237041263 |
| Rdm1 | -0.083894986 | 8.29E-41 | 7.26E-39 | 0.068067428 | 0.131784409 |
| Rdx | 0.061640439 | 7.99E-10 | 9.25E-09 | 0.744899855 | 0.672229226 |
| Rec8 | -0.016208136 | 0.008154681 | 0.021876328 | 0.098515788 | 0.107723091 |
| Recql | -0.007842465 | 0.109581503 | 0.190409867 | 0.061265289 | 0.067325397 |
| Reep2 | 0.014082999 | 0.001369932 | 0.004707839 | 0.060247962 | 0.048038571 |
| Reep3 | 0.025420081 | 0.004570825 | 0.013356647 | 0.3196552 | 0.293095471 |
| Reep4 | -0.0032185 | 0.368535956 | 0.491109633 | 0.034744912 | 0.034759924 |
| Reep5 | -0.048674583 | 1.78E-07 | 1.44E-06 | 1.237238886 | 1.339806107 |
| Reep6 | -0.08646662 | 2.72E-17 | 6.29E-16 | 0.494689225 | 0.569538801 |
| Rel | 0.007484423 | 0.139031759 | 0.230620502 | 0.06718571 | 0.064547912 |
| Rela | 0.013632723 | 0.025800365 | 0.057582908 | 0.115180221 | 0.105489668 |
| Relb | 0.003085428 | 0.477186808 | 0.593962623 | 0.047901824 | 0.046678355 |
| Relch | 0.007391223 | 0.324234593 | 0.446362038 | 0.18920086 | 0.178148557 |
| Rell1 | 0.03883419 | 1.59E-10 | 1.99E-09 | 0.12328595 | 0.092776431 |
| Rell2 | 0.002266333 | 0.537228095 | 0.648006395 | 0.036555061 | 0.034626659 |
| Relt | 0.004160069 | 0.437917471 | 0.556236433 | 0.086224024 | 0.078326065 |
| Rem2 | 0.009558305 | 0.012419753 | 0.031111023 | 0.042299286 | 0.034805767 |
| Repin1 | 0.014709719 | 0.000932005 | 0.003362134 | 0.062911544 | 0.0489421 |
| Reps1 | 0.031927024 | 4.08E-05 | 0.000204028 | 0.22056714 | 0.191417039 |
| Reps2 | -0.011216745 | 0.156229537 | 0.252470672 | 0.186000585 | 0.202555951 |
| Rer1 | -0.019678868 | 0.056955187 | 0.111569279 | 0.689201588 | 0.703289598 |
| Rere | 0.084156011 | 8.47E-18 | 2.03E-16 | 0.494009385 | 0.406851364 |

| | | | | | |
|---|---|---|---|---|---|
| Resf1 | 0.007307273 | 0.384604694 | 0.505697435 | 0.214973254 | 0.214019723 |
| Resp18 | -0.109102149 | 1.02E-17 | 2.41E-16 | 2.696032913 | 2.94569951 |
| Ret | 0.004225763 | 0.603359646 | 0.706248034 | 0.242663543 | 0.221186155 |
| Retreg1 | 0.057829271 | 2.00E-11 | 2.77E-10 | 0.24368924 | 0.200443459 |
| Retreg2 | 0.003417533 | 0.718667233 | 0.797862605 | 0.447290555 | 0.428688462 |
| Retreg3 | 0.012481319 | 0.090175883 | 0.162324934 | 0.194624415 | 0.171191259 |
| Retsat | -0.029093507 | 3.81E-11 | 5.13E-10 | 0.032250718 | 0.057835648 |
| Rev1 | 0.011351034 | 0.051890377 | 0.102891749 | 0.102536125 | 0.093234761 |
| Rev3l | 0.009903148 | 0.262458145 | 0.378522191 | 0.278753603 | 0.266917082 |
| Rex1bd | -0.005800218 | 0.558805579 | 0.667244883 | 0.746021626 | 0.758653005 |
| Rexo1 | 0.012178946 | 0.102249148 | 0.180094443 | 0.182173074 | 0.175324185 |
| Rexo2 | -0.087241839 | 1.23E-17 | 2.89E-16 | 0.78182463 | 0.90152316 |
| Rexo4 | 0.000153586 | 0.979003024 | 0.98589884 | 0.09572393 | 0.096804836 |
| Rfc1 | 0.021546635 | 0.01367562 | 0.033761903 | 0.30717098 | 0.286342207 |
| Rfc2 | -0.005906632 | 0.426649671 | 0.545344215 | 0.185330257 | 0.182762752 |
| Rfc3 | 0.018462311 | 0.000170635 | 0.000749283 | 0.074687632 | 0.059662057 |
| Rfc4 | -0.006898093 | 0.110883062 | 0.192226307 | 0.048662114 | 0.05188537 |
| Rfc5 | 0.004211753 | 0.268183922 | 0.384875711 | 0.040488116 | 0.036912787 |
| Rfesd | -0.007087201 | 0.401361096 | 0.52145297 | 0.270114686 | 0.264004948 |
| Rffl | -0.005694195 | 0.154969691 | 0.250768198 | 0.04083534 | 0.043991761 |
| Rfk | 0.005955213 | 0.54096353 | 0.651461345 | 0.535838447 | 0.50988963 |
| Rflna | 0.074711872 | 1.83E-13 | 3.19E-12 | 0.378677443 | 0.306417609 |
| Rflnb | 0.02333338 | 0.000249136 | 0.001041764 | 0.126901146 | 0.103291592 |
| Rfng | 0.013737455 | 0.001570989 | 0.005277526 | 0.061732081 | 0.047110506 |
| Rft1 | -0.009398576 | 0.02734175 | 0.060454058 | 0.045352493 | 0.051378339 |
| Rfwd3 | 0.006055221 | 0.167803983 | 0.266392933 | 0.05828058 | 0.050880869 |
| Rfx2 | -0.00054496 | 0.882876307 | 0.919773195 | 0.036980711 | 0.035725795 |
| Rfx3 | 0.008965576 | 0.177473024 | 0.278918968 | 0.134403356 | 0.125617984 |
| Rfx5 | 0.002350578 | 0.680911564 | 0.768139038 | 0.090371596 | 0.090924312 |
| Rfx6 | 0.017250516 | 0.030090438 | 0.065591757 | 0.219864026 | 0.194503041 |
| Rfx7 | -0.007253326 | 0.258979746 | 0.374930889 | 0.120707424 | 0.120748996 |
| Rfxank | -0.006598121 | 0.104046513 | 0.182522592 | 0.041132005 | 0.045031014 |
| Rfxap | -0.005932871 | 0.408184375 | 0.528256662 | 0.16839804 | 0.169367207 |
| Rgl1 | 0.001732956 | 0.761279174 | 0.83057396 | 0.090987686 | 0.089982365 |
| Rgl2 | -0.010893385 | 0.016722447 | 0.040092633 | 0.053770813 | 0.059248721 |
| Rgl3 | -0.005608607 | 0.228157999 | 0.340729965 | 0.057919015 | 0.059775695 |
| Rgmb | 0.012585465 | 0.0150495 | 0.036639155 | 0.084992396 | 0.071862827 |
| Rgp1 | 0.007773247 | 0.095726418 | 0.170386715 | 0.065465578 | 0.056659776 |
| Rgs11 | -0.060280951 | 1.35E-12 | 2.15E-11 | 0.212711752 | 0.259660877 |
| Rgs16 | 0.021856864 | 0.008640585 | 0.023002126 | 0.214312082 | 0.183051102 |
| Rgs17 | 0.037068999 | 8.24E-05 | 0.000391332 | 0.285390686 | 0.276905444 |
| Rgs2 | 0.102911644 | 2.33E-33 | 1.40E-31 | 0.236911634 | 0.158843427 |
| Rgs3 | 0.004762342 | 0.264429904 | 0.380764303 | 0.051346682 | 0.045340836 |
| Rgs4 | -0.066181135 | 9.37E-08 | 7.98E-07 | 0.264179159 | 0.426061047 |
| Rgs7 | 0.016819633 | 0.019018695 | 0.044521974 | 0.16121782 | 0.150565835 |
| Rgs7bp | 0.010847592 | 0.122449436 | 0.207677903 | 0.143417569 | 0.134957676 |
| Rgs9 | -0.086411169 | 4.71E-27 | 2.12E-25 | 0.14401164 | 0.228569528 |
| Rhbdd1 | 0.008690408 | 0.231829141 | 0.344993556 | 0.165438674 | 0.162667229 |
| Rhbdd2 | -0.004689216 | 0.562722163 | 0.670850383 | 0.221465231 | 0.22796175 |
| Rhbdd3 | 0.003842961 | 0.506100948 | 0.61999917 | 0.098990063 | 0.095448884 |
| Rhbdl2 | 0.005323189 | 0.34248334 | 0.464735061 | 0.092034393 | 0.085717496 |
| Rheb | 0.022574295 | 0.032850776 | 0.070440063 | 0.888154691 | 0.858497985 |
| Rhno1 | -0.001425348 | 0.73612753 | 0.810857903 | 0.049825447 | 0.04833496 |
| Rhoa | -0.042260135 | 7.69E-07 | 5.52E-06 | 1.201509888 | 1.221509514 |
| Rhob | -0.056150052 | 2.02E-12 | 3.15E-11 | 0.159855465 | 0.216882656 |
| Rhobtb1 | 0.026059274 | 0.006104544 | 0.017123016 | 0.37065917 | 0.340896353 |
| Rhobtb2 | 0.013009761 | 0.001604779 | 0.005372472 | 0.054452506 | 0.041371575 |
| Rhobtb3 | 0.009505533 | 0.236398256 | 0.35007827 | 0.23212351 | 0.216236425 |
| Rhoc | -0.002384568 | 0.657552855 | 0.749440663 | 0.076893135 | 0.079695761 |
| Rhog | 0.001809523 | 0.693417849 | 0.77782489 | 0.063680855 | 0.056472999 |
| Rhoq | -0.020155418 | 0.013497653 | 0.033432912 | 0.202877554 | 0.228626899 |
| Rhot1 | 0.007484619 | 0.26318701 | 0.379362525 | 0.143186262 | 0.134280217 |
| Rhot2 | 0.006883298 | 0.227340106 | 0.339821727 | 0.098756874 | 0.090218057 |
| Rhou | -0.028275807 | 0.000104846 | 0.000487478 | 0.122568135 | 0.16159372 |
| Rhpn2 | 0.016440076 | 0.019007467 | 0.044506404 | 0.16354655 | 0.14551413 |
| Rian | 0.036829159 | 0.000581475 | 0.002220385 | 1.01110921 | 0.965852321 |
| Ric1 | 0.009539945 | 0.082190419 | 0.150623045 | 0.087510134 | 0.080328883 |
| Ric3 | 0.010648528 | 0.122780088 | 0.20808361 | 0.153794702 | 0.143710817 |
| Ric8a | -0.002094209 | 0.773682131 | 0.84051153 | 0.171694754 | 0.168077397 |
| Ric8b | 0.019222682 | 0.007778622 | 0.021024121 | 0.167246312 | 0.155443336 |
| Rictor | -0.003785814 | 0.629264982 | 0.726742 | 0.199645484 | 0.194787712 |
| Rida | -0.041239284 | 7.73E-10 | 8.97E-09 | 0.12708135 | 0.150947309 |

| | | | | | |
|---|---|---|---|---|---|
| Rif1 | 0.000249432 | 0.9789813 | 0.98589884 | 0.350055542 | 0.354629383 |
| Riiad1 | -0.034613176 | 2.83E-07 | 2.20E-06 | 0.100519971 | 0.137125242 |
| Rilpl1 | -0.008934722 | 0.091678487 | 0.164506875 | 0.079403925 | 0.081666886 |
| Rilpl2 | 0.007357908 | 0.311432352 | 0.432906109 | 0.179763485 | 0.165998023 |
| Rimbp2 | 0.027997962 | 0.002963635 | 0.00919938 | 0.382357172 | 0.348656214 |
| Rims2 | 0.007718872 | 0.386911028 | 0.507905619 | 0.296212992 | 0.283440597 |
| Rims3 | 0.047700247 | 1.04E-08 | 1.03E-07 | 0.285465011 | 0.223953126 |
| Ring1 | -0.019174246 | 0.007712686 | 0.020882291 | 0.162409843 | 0.174145304 |
| Rint1 | 0.009869887 | 0.233224894 | 0.346746615 | 0.241729606 | 0.235600943 |
| Riok1 | -0.012711519 | 0.108745994 | 0.189467482 | 0.192000102 | 0.205138365 |
| Riok2 | 0.006205614 | 0.26114283 | 0.377198038 | 0.091749213 | 0.084144521 |
| Riok3 | -0.006124478 | 0.548815413 | 0.658145238 | 0.612726974 | 0.620227026 |
| Riox1 | 0.003922989 | 0.291758279 | 0.411321634 | 0.040299337 | 0.035985223 |
| Riox2 | -0.000418969 | 0.941518986 | 0.96028615 | 0.091827782 | 0.092550865 |
| Ripk1 | 0.004159772 | 0.513788121 | 0.627145942 | 0.119188332 | 0.116350257 |
| Ripk2 | 0.011491191 | 0.006308117 | 0.017617787 | 0.051876627 | 0.0435413 |
| Ripk4 | 0.011527345 | 0.052614212 | 0.104093482 | 0.111662786 | 0.098014125 |
| Ripor1 | 0.01099709 | 0.069325306 | 0.131303453 | 0.115379891 | 0.102668033 |
| Ripply3 | 0.024563209 | 0.000178735 | 0.00077887 | 0.152974623 | 0.117626096 |
| Rit1 | -0.041580669 | 7.43E-08 | 6.42E-07 | 0.168798537 | 0.204747433 |
| Rita1 | -0.013914548 | 0.001026246 | 0.003660785 | 0.042054097 | 0.05140771 |
| Rlf | -0.002486772 | 0.748269107 | 0.820606007 | 0.185339738 | 0.189818166 |
| Rlim | -0.005105098 | 0.525912175 | 0.637889568 | 0.203927639 | 0.21389906 |
| Rmc1 | -0.007093108 | 0.163835088 | 0.261630211 | 0.072600982 | 0.07580608 |
| Rmdn1 | 0.011431972 | 0.086938161 | 0.15783619 | 0.142618658 | 0.132436005 |
| Rmdn3 | 0.016931186 | 0.014292068 | 0.035052343 | 0.156955584 | 0.145413895 |
| Rmi1 | -0.000642167 | 0.905829621 | 0.93560051 | 0.084085265 | 0.081966146 |
| Rmnd1 | -0.006237832 | 0.151014672 | 0.246006139 | 0.051190762 | 0.051910248 |
| Rmnd5a | -0.033050723 | 0.000285671 | 0.001177819 | 0.315694775 | 0.340945745 |
| Rmnd5b | 0.010319943 | 0.069644468 | 0.131830903 | 0.102436909 | 0.08953418 |
| Rn18s | -0.042607512 | 4.00E-13 | 6.74E-12 | 3.807369158 | 3.875386283 |
| Rnase4 | -0.148031628 | 9.38E-38 | 6.91E-36 | 0.911055227 | 1.0721585 |
| Rnaseh1 | -0.002472847 | 0.685243672 | 0.771951214 | 0.110343329 | 0.11169039 |
| Rnaseh2a | -0.015515352 | 0.009287321 | 0.024383293 | 0.102760661 | 0.108073559 |
| Rnaseh2b | 0.009436234 | 0.086443172 | 0.157054854 | 0.09945398 | 0.084084331 |
| Rnaseh2c | 0.018918719 | 0.022350568 | 0.050957786 | 0.239761598 | 0.230243269 |
| Rnaset2a | -0.02614974 | 3.12E-06 | 1.99E-05 | 0.079645422 | 0.096346478 |
| Rnaset2b | -0.007994375 | 0.079840622 | 0.147232649 | 0.057580172 | 0.060017081 |
| Rnd2 | 0.002403458 | 0.504594668 | 0.618855729 | 0.037142175 | 0.033327771 |
| Rnf10 | 0.019988797 | 0.044411468 | 0.090384822 | 0.701872123 | 0.672095349 |
| Rnf103 | -0.000532506 | 0.948416999 | 0.965599546 | 0.226145532 | 0.232463599 |
| Rnf11 | -0.037994583 | 6.59E-05 | 0.000318153 | 0.935346419 | 0.959785899 |
| Rnf111 | 0.024318769 | 0.000915169 | 0.003313138 | 0.18042164 | 0.163032942 |
| Rnf113a1 | 0.003735754 | 0.354588781 | 0.477004584 | 0.047843982 | 0.042591131 |
| Rnf113a2 | -0.007357427 | 0.271686945 | 0.388781937 | 0.139225375 | 0.140479004 |
| Rnf114 | 0.027506094 | 0.000299627 | 0.001230143 | 0.209633879 | 0.181725034 |
| Rnf115 | 0.002417972 | 0.798926763 | 0.859580123 | 0.448942895 | 0.422764561 |
| Rnf121 | 0.001896765 | 0.770531992 | 0.838150543 | 0.129112874 | 0.127323697 |
| Rnf123 | 0.001269983 | 0.779666737 | 0.845157198 | 0.057597821 | 0.055478071 |
| Rnf126 | 0.002952505 | 0.686844526 | 0.772787055 | 0.189152252 | 0.175372182 |
| Rnf128 | 0.043972705 | 3.88E-07 | 2.95E-06 | 0.294868133 | 0.25200158 |
| Rnf13 | -0.01935673 | 0.005685331 | 0.016081694 | 0.140359695 | 0.155279138 |
| Rnf130 | 0.049297021 | 6.81E-07 | 4.94E-06 | 0.953319224 | 0.876150487 |
| Rnf135 | -0.003954808 | 0.296847741 | 0.416623897 | 0.038571851 | 0.038269624 |
| Rnf138 | 0.01224277 | 0.129228746 | 0.217438884 | 0.228969857 | 0.217258662 |
| Rnf138rt1 | -0.003284088 | 0.662042946 | 0.752969294 | 0.158358762 | 0.1665635 |
| Rnf139 | 0.010706211 | 0.093485387 | 0.16698894 | 0.132799027 | 0.118555672 |
| Rnf14 | 0.019623203 | 0.043272479 | 0.088469299 | 0.461678943 | 0.443206351 |
| Rnf141 | 0.020097326 | 0.010351956 | 0.026645941 | 0.218372267 | 0.195298191 |
| Rnf145 | 0.011573691 | 0.081390695 | 0.14941094 | 0.138974997 | 0.126753025 |
| Rnf146 | -0.007136005 | 0.352013143 | 0.474393093 | 0.19860355 | 0.197787014 |
| Rnf149 | 0.012233164 | 0.016298944 | 0.03922236 | 0.081531751 | 0.068553766 |
| Rnf150 | 0.018734892 | 0.010421503 | 0.026803664 | 0.16140043 | 0.146642374 |
| Rnf157 | 0.003978883 | 0.371318473 | 0.493770783 | 0.057820346 | 0.051855361 |
| Rnf166 | 0.004830343 | 0.484974872 | 0.601163548 | 0.149280136 | 0.145306757 |
| Rnf167 | 0.017030449 | 0.047457503 | 0.095722374 | 0.303597162 | 0.275310656 |
| Rnf168 | 0.00675676 | 0.332405766 | 0.455033632 | 0.150997253 | 0.145015281 |
| Rnf169 | 0.018330533 | 0.000277852 | 0.001150058 | 0.080776978 | 0.064068197 |
| Rnf170 | 0.001276779 | 0.854511057 | 0.899880714 | 0.150216841 | 0.145019122 |
| Rnf181 | -0.035255178 | 2.07E-05 | 0.000111963 | 0.238609032 | 0.255118614 |
| Rnf182 | 0.020302787 | 0.003130417 | 0.009646031 | 0.160038681 | 0.128073078 |
| Rnf185 | -0.00422471 | 0.527280032 | 0.638780869 | 0.134214373 | 0.135674604 |

| | | | | | |
|---|---|---|---|---|---|
| Rnf186 | 0.030954901 | 1.56E-08 | 1.50E-07 | 0.110787533 | 0.07327223 |
| Rnf187 | -0.023220237 | 0.01608388 | 0.038791239 | 0.464336303 | 0.462180113 |
| Rnf19a | 0.01377006 | 0.00358499 | 0.01084758 | 0.070084989 | 0.056514275 |
| Rnf19b | 0.026626947 | 8.93E-06 | 5.18E-05 | 0.118897336 | 0.095590133 |
| Rnf2 | 0.015140666 | 0.029886301 | 0.065219892 | 0.165820079 | 0.145741159 |
| Rnf20 | 0.021582907 | 0.018286214 | 0.043171096 | 0.347144024 | 0.327843189 |
| Rnf208 | -0.005811168 | 0.248457318 | 0.362939034 | 0.068894634 | 0.070088658 |
| Rnf213 | 0.011614987 | 0.014467853 | 0.035438767 | 0.064426282 | 0.056776532 |
| Rnf214 | -0.001899278 | 0.791988373 | 0.854384261 | 0.160895798 | 0.163186671 |
| Rnf215 | -0.01729982 | 0.002508995 | 0.007958385 | 0.090117708 | 0.101497879 |
| Rnf216 | 0.019797771 | 0.006003755 | 0.016874388 | 0.178020251 | 0.155686359 |
| Rnf217 | 0.033898306 | 7.84E-06 | 4.62E-05 | 0.208470245 | 0.170956791 |
| Rnf220 | 0.017465826 | 0.025256436 | 0.05665507 | 0.212607324 | 0.199765435 |
| Rnf227 | 0.060765053 | 1.76E-12 | 2.77E-11 | 0.298297458 | 0.239955111 |
| Rnf25 | 0.012258941 | 0.038879599 | 0.080904851 | 0.11163571 | 0.098435627 |
| Rnf26 | 0.004377746 | 0.258968983 | 0.374930889 | 0.044938809 | 0.03856165 |
| Rnf31 | 0.006814546 | 0.19092036 | 0.295373606 | 0.079794354 | 0.073568058 |
| Rnf34 | 0.020762914 | 0.00318062 | 0.009776159 | 0.175988557 | 0.149637287 |
| Rnf38 | 0.00439571 | 0.416375388 | 0.535662501 | 0.086551726 | 0.080315396 |
| Rnf4 | 0.011194469 | 0.151872919 | 0.246742733 | 0.208365268 | 0.197419621 |
| Rnf40 | 0.004212702 | 0.427073827 | 0.545671146 | 0.080530459 | 0.07630749 |
| Rnf41 | 0.009959965 | 0.108378546 | 0.18903054 | 0.116570714 | 0.109754899 |
| Rnf44 | 0.01472634 | 0.017770302 | 0.042191724 | 0.119950106 | 0.108907507 |
| Rnf5 | -0.05632053 | 3.39E-11 | 4.60E-10 | 0.263170814 | 0.297650065 |
| Rnf6 | 0.059866877 | 3.02E-10 | 3.66E-09 | 0.432302506 | 0.363368513 |
| Rnf7 | 0.034166982 | 0.000454972 | 0.001785722 | 0.847933233 | 0.777695241 |
| Rnf8 | -0.00392343 | 0.580104681 | 0.686438378 | 0.169853923 | 0.163028709 |
| Rnft1 | 0.001962044 | 0.723144227 | 0.800420062 | 0.089729944 | 0.086521327 |
| Rngtt | 0.001778207 | 0.752991726 | 0.823959179 | 0.093282701 | 0.089906571 |
| Rnh1 | -0.012257445 | 0.225736483 | 0.337840223 | 0.564354194 | 0.567712233 |
| Rnmt | -0.019286033 | 0.013266079 | 0.032943248 | 0.191298294 | 0.205700329 |
| Rnpc3 | -0.004853287 | 0.43516039 | 0.553674822 | 0.109042236 | 0.110511192 |
| Rnpep | 0.000999352 | 0.880647169 | 0.918238071 | 0.138121372 | 0.131470377 |
| Rnpepl1 | 0.025684708 | 0.002960057 | 0.009191209 | 0.305044766 | 0.269285534 |
| Rnps1 | 0.022956193 | 0.012958652 | 0.032237465 | 0.349245387 | 0.339628484 |
| Ro60 | 0.041440235 | 2.35E-05 | 0.000125065 | 0.4762832 | 0.416886758 |
| Robo1 | 0.03324006 | 2.27E-07 | 1.80E-06 | 0.140724395 | 0.108197034 |
| Robo2 | 0.06878179 | 7.78E-12 | 1.13E-10 | 0.485039725 | 0.39162926 |
| Rock1 | 0.04336024 | 1.48E-05 | 8.21E-05 | 0.948592075 | 0.890841846 |
| Rock2 | 0.025846628 | 0.010606198 | 0.027206718 | 0.470272966 | 0.451303532 |
| Rogdi | -0.019526896 | 0.034420149 | 0.073283777 | 0.357356508 | 0.354803196 |
| Rom1 | -0.006986635 | 0.214240232 | 0.323997904 | 0.09285567 | 0.093110728 |
| Romo1 | 0.161210196 | 2.11E-72 | 6.21E-70 | 1.255407676 | 1.095063367 |
| Ropn1l | 0.008284283 | 0.079695857 | 0.147021465 | 0.070033312 | 0.058367888 |
| Ror2 | -0.00064197 | 0.911501634 | 0.93920356 | 0.09483266 | 0.095624991 |
| Rora | 0.005486265 | 0.57366594 | 0.680475875 | 0.422087828 | 0.406989 |
| Rorc | -0.007822582 | 0.207326165 | 0.31568128 | 0.102934352 | 0.112301783 |
| Rp2 | 0.008809944 | 0.192407295 | 0.297248327 | 0.141713944 | 0.133489025 |
| Rp9 | 0.037422725 | 0.000189009 | 0.000816688 | 0.713735323 | 0.664004101 |
| Rpa1 | 0.010329435 | 0.167744545 | 0.266342065 | 0.197355986 | 0.172334588 |
| Rpa2 | -0.001759714 | 0.646189626 | 0.739785156 | 0.040362908 | 0.038824617 |
| Rpa3 | -0.012365919 | 0.077016851 | 0.143165932 | 0.146980691 | 0.153628156 |
| Rpain | 0.001194272 | 0.755164207 | 0.825652994 | 0.041286366 | 0.038100732 |
| Rpap1 | 0.013232037 | 0.007996718 | 0.021559647 | 0.077798976 | 0.06491068 |
| Rpap2 | -0.011014563 | 0.042794761 | 0.087608057 | 0.080309893 | 0.086332195 |
| Rpap3 | 0.001357296 | 0.804871135 | 0.864636152 | 0.087174598 | 0.08537154 |
| Rpe | -0.004249872 | 0.535985099 | 0.646988718 | 0.150633854 | 0.147491569 |
| Rpf1 | -0.014649533 | 0.036651394 | 0.077076542 | 0.149935653 | 0.156864064 |
| Rpf2 | -0.016347916 | 0.008094702 | 0.021763467 | 0.104420527 | 0.115904797 |
| Rpgr | -0.002259767 | 0.689736689 | 0.774678846 | 0.088658703 | 0.089854831 |
| Rpgrip1 | 0.020992836 | 2.00E-05 | 0.000108444 | 0.086331054 | 0.058911944 |
| Rpgrip1l | -0.002858104 | 0.52724257 | 0.638780869 | 0.054945276 | 0.054441336 |
| Rph3al | 0.016701615 | 0.06739186 | 0.128366704 | 0.361533601 | 0.326666737 |
| Rpia | 0.041921015 | 7.48E-08 | 6.45E-07 | 0.248757585 | 0.188023189 |
| Rpl10 | -0.038290525 | 5.98E-18 | 1.47E-16 | 2.947740267 | 2.956354249 |
| Rpl10a | -0.107889629 | 1.16E-31 | 6.42E-30 | 1.249669102 | 1.309078841 |
| Rpl10a-ps1 | -0.015100214 | 0.006726601 | 0.018604841 | 0.085199464 | 0.092922544 |
| Rpl10-ps3 | -0.013743697 | 0.002535723 | 0.008030066 | 0.05262965 | 0.060230679 |
| Rpl11 | -0.009821434 | 0.037565709 | 0.078557502 | 2.705763296 | 2.696513648 |
| Rpl12 | -0.10484622 | 2.09E-40 | 1.80E-38 | 1.913626795 | 2.197784948 |
| Rpl13 | -0.028663354 | 1.14E-08 | 1.13E-07 | 2.769131883 | 2.810807588 |
| Rpl13a | 0.019938486 | 0.055300087 | 0.10863355 | 0.616390977 | 0.589419773 |

| | | | | | |
|---|---|---|---|---|---|
| Rpl14 | -0.038833803 | 1.78E-11 | 2.51E-10 | 2.307563076 | 2.377718486 |
| Rpl15 | -0.001259194 | 0.890560988 | 0.925596955 | 1.275674223 | 1.241841982 |
| Rpl17 | -0.024428636 | 6.52E-06 | 3.90E-05 | 2.581214458 | 2.604463372 |
| Rpl18 | -0.013261533 | 0.029910874 | 0.065258869 | 2.248711642 | 2.26941789 |
| Rpl18a | -0.033877832 | 1.61E-08 | 1.54E-07 | 2.166753565 | 2.218196039 |
| Rpl19 | -0.035806425 | 1.02E-19 | 2.90E-18 | 3.268828535 | 3.344102649 |
| Rpl19-ps11 | -0.013891757 | 0.000428949 | 0.001691782 | 0.033465054 | 0.044846953 |
| Rpl21 | -0.053701391 | 1.16E-26 | 5.15E-25 | 2.670973814 | 2.758047535 |
| Rpl22 | 0.000562155 | 0.93660875 | 0.957014679 | 1.762216612 | 1.801264542 |
| Rpl22l1 | 0.043862173 | 7.64E-08 | 6.58E-07 | 1.801046115 | 1.788969134 |
| Rpl23 | -0.023945547 | 1.94E-06 | 1.28E-05 | 2.885104787 | 2.997057979 |
| Rpl23a | -0.001070012 | 0.917129824 | 0.942575366 | 0.910481095 | 0.937114911 |
| Rpl24 | 0.009540825 | 0.350351598 | 0.472809306 | 0.71799321 | 0.653630746 |
| Rpl26 | -0.020219733 | 2.48E-05 | 0.000131186 | 2.660493944 | 2.757351904 |
| Rpl27 | 0.096651989 | 5.00E-26 | 2.14E-24 | 1.301891718 | 1.234113443 |
| Rpl27a | 0.105384992 | 1.88E-48 | 2.20E-46 | 2.044962583 | 1.926554791 |
| Rpl28 | 0.026058262 | 1.06E-07 | 8.93E-07 | 2.669037709 | 2.647221484 |
| Rpl29 | -0.050035614 | 1.65E-12 | 2.60E-11 | 1.86697852 | 1.868215131 |
| Rpl3 | -0.0980073 | 7.88E-53 | 1.05E-50 | 2.118184685 | 2.283272022 |
| Rpl30 | 0.061436563 | 8.04E-28 | 3.78E-26 | 2.452514209 | 2.396243044 |
| Rpl31 | 0.057686139 | 5.00E-12 | 7.40E-11 | 1.411366168 | 1.412275238 |
| Rpl32 | -0.040545637 | 1.98E-18 | 5.03E-17 | 3.145236242 | 3.356225251 |
| Rpl34 | 0.045792524 | 4.97E-19 | 1.33E-17 | 2.607369796 | 2.605059772 |
| Rpl35 | 0.054380595 | 3.26E-14 | 6.04E-13 | 1.906114159 | 1.917518627 |
| Rpl35a | 0.083007653 | 1.42E-54 | 2.03E-52 | 2.638872577 | 2.533625019 |
| Rpl36 | 0.061751236 | 1.38E-36 | 9.64E-35 | 2.71384579 | 2.692887904 |
| Rpl36a | 0.006392177 | 0.428262639 | 0.546902585 | 1.553785504 | 1.639262832 |
| Rpl36al | 0.04596899 | 1.19E-06 | 8.18E-06 | 1.144038929 | 1.141052858 |
| Rpl37 | 0.078737785 | 3.57E-54 | 5.02E-52 | 3.022257341 | 2.966060479 |
| Rpl37a | 0.109030617 | 2.06E-162 | 5.01E-159 | 3.165338729 | 2.977714232 |
| Rpl37rt | -0.012403584 | 0.04843159 | 0.097444519 | 0.106151946 | 0.120307427 |
| Rpl38 | 0.096462109 | 1.96E-76 | 7.95E-74 | 2.622402175 | 2.521274897 |
| Rpl39 | 0.098641926 | 7.05E-76 | 2.74E-73 | 2.650332287 | 2.539390474 |
| Rpl3-ps1 | -0.023564859 | 2.66E-05 | 0.000139382 | 0.077282181 | 0.096187311 |
| Rpl4 | -0.02822509 | 0.000159649 | 0.000705502 | 1.733211926 | 1.788157522 |
| Rpl41 | 0.084853219 | 4.78E-128 | 5.17E-125 | 3.854484028 | 3.656497898 |
| Rpl5 | -0.088361761 | 3.92E-22 | 1.31E-20 | 1.322157705 | 1.401166576 |
| Rpl6 | -0.008954207 | 0.102254817 | 0.180094443 | 2.409115931 | 2.423268882 |
| Rpl6l | -0.018121925 | 5.35E-06 | 3.27E-05 | 0.0374658 | 0.04755496 |
| Rpl7 | -0.048049185 | 1.54E-09 | 1.71E-08 | 1.595310847 | 1.639703811 |
| Rpl7a | -0.047087732 | 6.03E-11 | 7.96E-10 | 1.900856889 | 1.9326757 |
| Rpl7l1 | -0.022528172 | 0.008316689 | 0.022237307 | 0.267172972 | 0.280874149 |
| Rpl8 | -0.074769697 | 4.33E-67 | 1.05E-64 | 2.977000367 | 3.183192921 |
| Rpl9 | -0.064475099 | 5.00E-49 | 6.15E-47 | 3.101999622 | 3.222911744 |
| Rpl9-ps6 | -0.018386607 | 0.000556932 | 0.002136739 | 0.070506634 | 0.085207695 |
| Rplp0 | -0.046538912 | 5.18E-16 | 1.08E-14 | 2.406491958 | 2.489172409 |
| Rplp1 | 0.042719457 | 1.74E-18 | 4.46E-17 | 3.047689218 | 3.010937874 |
| Rplp2 | 0.06209444 | 9.79E-34 | 6.02E-32 | 2.536101114 | 2.504873715 |
| Rpn1 | 0.068060186 | 7.94E-12 | 1.14E-10 | 0.966172229 | 0.910628661 |
| Rpn2 | -0.063030468 | 5.11E-11 | 6.78E-10 | 0.975613843 | 1.07179934 |
| Rpp14 | 0.00077998 | 0.900132444 | 0.93179678 | 0.124370899 | 0.117347092 |
| Rpp21 | 0.021985211 | 0.019053798 | 0.04457195 | 0.362203543 | 0.36219057 |
| Rpp25l | 0.009246455 | 0.236135856 | 0.34979625 | 0.229103495 | 0.214213673 |
| Rpp30 | -0.010398916 | 0.051584287 | 0.102410137 | 0.080866539 | 0.083333213 |
| Rpp38 | 0.014693114 | 0.005282712 | 0.01511434 | 0.08487427 | 0.07249074 |
| Rprd1a | 0.00927496 | 0.221210738 | 0.332038103 | 0.190159388 | 0.178864817 |
| Rprd1b | 0.005582516 | 0.554407181 | 0.663950063 | 0.409990941 | 0.394324672 |
| Rprd2 | 0.030040297 | 0.000196498 | 0.000844914 | 0.241501185 | 0.21368727 |
| Rprml | -0.005207921 | 0.39062428 | 0.511605719 | 0.096692652 | 0.095276203 |
| Rps10 | 0.009116254 | 0.341038127 | 0.463483987 | 0.972553786 | 0.936829469 |
| Rps10-ps1 | -0.003988989 | 0.419622857 | 0.538770922 | 0.078681162 | 0.070465995 |
| Rps10-ps2 | -0.013941233 | 0.026300461 | 0.058497616 | 0.123148987 | 0.124927411 |
| Rps11 | -0.043801817 | 4.33E-13 | 7.23E-12 | 2.392657377 | 2.540103986 |
| Rps11-ps1 | -0.009367672 | 0.07163484 | 0.134784965 | 0.068529332 | 0.077404991 |
| Rps11-ps2 | -0.01506094 | 0.002987855 | 0.009262731 | 0.059202121 | 0.075946614 |
| Rps12 | 0.014958513 | 0.006571391 | 0.018253446 | 2.553043261 | 2.58753912 |
| Rps13 | -0.038553619 | 1.29E-12 | 2.06E-11 | 2.554994306 | 2.680986523 |
| Rps14 | -0.020429687 | 3.89E-07 | 2.96E-06 | 3.139182657 | 3.275744964 |
| Rps15 | -0.013519759 | 0.031221143 | 0.067586717 | 2.186132015 | 2.347181875 |
| Rps15a | 0.002375891 | 0.718450488 | 0.79771307 | 2.233688235 | 2.278407988 |
| Rps16 | -0.031848294 | 1.40E-08 | 1.36E-07 | 2.449332006 | 2.532370282 |
| Rps17 | 0.02647808 | 3.96E-05 | 0.000199203 | 2.038334812 | 2.118997576 |

| | | | | | |
|---|---|---|---|---|---|
| Rps18 | -0.078953913 | 1.50E-38 | 1.18E-36 | 2.374254954 | 2.606235743 |
| Rps19 | 0.062544057 | 1.19E-15 | 2.42E-14 | 1.829438513 | 1.756208446 |
| Rps19bp1 | 0.010215466 | 0.223928146 | 0.335443672 | 0.249469654 | 0.243855267 |
| Rps2 | -0.046583706 | 5.02E-13 | 8.32E-12 | 2.569558398 | 2.594347446 |
| Rps20 | 0.000138519 | 0.978473112 | 0.985671494 | 2.799706518 | 2.863054526 |
| Rps21 | 0.1132895 | 1.43E-151 | 2.78E-148 | 3.152097802 | 2.966796463 |
| Rps23 | 0.019658218 | 0.000353305 | 0.001425832 | 2.516567902 | 2.536215597 |
| Rps23-ps1 | -0.009988838 | 0.074356911 | 0.139093614 | 0.08146028 | 0.092033245 |
| Rps24 | -0.047588784 | 8.15E-32 | 4.56E-30 | 3.411170859 | 3.56001786 |
| Rps25 | 0.034776467 | 1.85E-06 | 1.23E-05 | 1.800011159 | 1.835033546 |
| Rps26 | 0.025969662 | 0.000360736 | 0.001451602 | 1.899961285 | 1.872299356 |
| Rps26-ps1 | -0.002071177 | 0.612559907 | 0.712783653 | 0.043678172 | 0.044654611 |
| Rps27 | 0.070115925 | 1.73E-49 | 2.15E-47 | 2.81370176 | 2.757638944 |
| Rps27a | -0.009835259 | 0.03131658 | 0.067717926 | 3.109787969 | 3.146743966 |
| Rps27a-ps3 | -0.020125558 | 0.000304175 | 0.001246709 | 0.08111894 | 0.095643493 |
| Rps27l | 0.048275744 | 1.21E-07 | 1.01E-06 | 1.499179251 | 1.504338005 |
| Rps27rt | -0.010266943 | 0.170423836 | 0.269737438 | 0.165268892 | 0.182278815 |
| Rps28 | 0.108408501 | 5.84E-31 | 3.08E-29 | 1.198359471 | 1.126833178 |
| Rps29 | 0.074487159 | 7.91E-67 | 1.83E-64 | 2.963281246 | 2.909281353 |
| Rps2-ps10 | -0.006868784 | 0.101099674 | 0.178286058 | 0.04282907 | 0.049368443 |
| Rps3 | -0.093311877 | 3.19E-65 | 6.74E-63 | 2.500713908 | 2.72983013 |
| Rps3a1 | -0.091918347 | 1.63E-74 | 5.67E-72 | 3.04837606 | 3.20840373 |
| Rps4l | -0.037507762 | 0.001439862 | 0.004908253 | 0.668700424 | 0.694387336 |
| Rps4x | -0.082507487 | 8.04E-47 | 8.49E-45 | 2.410989829 | 2.539833668 |
| Rps5 | -0.057619051 | 4.06E-23 | 1.47E-21 | 2.473291242 | 2.610056024 |
| Rps6 | -0.067524307 | 1.33E-16 | 2.88E-15 | 1.604652899 | 1.691640979 |
| Rps6ka2 | 0.019262091 | 0.025216456 | 0.05657844 | 0.28543047 | 0.249468339 |
| Rps6ka3 | 0.023170788 | 0.002393011 | 0.007642856 | 0.197262905 | 0.179127413 |
| Rps6ka5 | 0.010610002 | 0.003181837 | 0.009776807 | 0.043382439 | 0.030839428 |
| Rps6kb1 | 0.006994618 | 0.410317044 | 0.529794153 | 0.265260108 | 0.258054925 |
| Rps6kb2 | -0.002816529 | 0.632635383 | 0.729098641 | 0.096241276 | 0.100442954 |
| Rps6kc1 | -0.001384459 | 0.718922199 | 0.797872331 | 0.039684174 | 0.039452864 |
| Rps6-ps4 | -0.02955964 | 2.81E-07 | 2.19E-06 | 0.087913235 | 0.10314209 |
| Rps7 | -0.032442371 | 4.34E-06 | 2.71E-05 | 1.946083001 | 1.953624735 |
| Rps8 | 0.003322234 | 0.461524735 | 0.579409023 | 3.336450458 | 3.411308811 |
| Rps9 | -0.044597741 | 3.11E-14 | 5.77E-13 | 2.255148838 | 2.336054557 |
| Rpsa | -0.017630634 | 0.001685412 | 0.005613422 | 2.423272209 | 2.400053378 |
| Rpsa-ps10 | -0.019632212 | 9.08E-05 | 0.000427506 | 0.0618157 | 0.075636066 |
| Rptor | 0.017133636 | 0.00246308 | 0.007833191 | 0.10135065 | 0.0851301 |
| Rpusd1 | 0.001631125 | 0.790927736 | 0.853690131 | 0.11460098 | 0.106984504 |
| Rpusd4 | -0.005123359 | 0.266145158 | 0.382647622 | 0.057003836 | 0.059593059 |
| Rraga | -0.015855519 | 0.109438121 | 0.190230183 | 0.531948974 | 0.532874781 |
| Rragb | 0.012993837 | 0.00279739 | 0.008739146 | 0.055122873 | 0.045713526 |
| Rragc | 0.017831515 | 0.04435734 | 0.090293563 | 0.324906174 | 0.299205871 |
| Rragd | 0.000793539 | 0.933857793 | 0.955077368 | 0.36448499 | 0.365808992 |
| Rras | -0.015932474 | 0.008684337 | 0.023086992 | 0.100896506 | 0.111234447 |
| Rras2 | 0.01144289 | 0.022059092 | 0.050460822 | 0.072618765 | 0.064629813 |
| Rrbp1 | 0.056785603 | 4.49E-10 | 5.34E-09 | 1.496624894 | 1.481857723 |
| Rreb1 | 0.015942896 | 0.011839491 | 0.029920335 | 0.127598137 | 0.110895999 |
| Rrm1 | 0.000928219 | 0.838437875 | 0.889163645 | 0.056299708 | 0.055593632 |
| Rrm2b | 0.003941317 | 0.679774369 | 0.767301337 | 0.408549351 | 0.394855866 |
| Rrn3 | 0.00819043 | 0.174292992 | 0.274988067 | 0.112262502 | 0.103363557 |
| Rrnad1 | -0.002367337 | 0.611138053 | 0.711896017 | 0.059361026 | 0.059022287 |
| Rrp1 | -0.014997135 | 0.131244333 | 0.220296513 | 0.71108808 | 0.716678853 |
| Rrp12 | -0.003143655 | 0.472747163 | 0.589954905 | 0.045634615 | 0.05173311 |
| Rrp15 | -0.002839181 | 0.604995837 | 0.707254493 | 0.085722508 | 0.087134463 |
| Rrp1b | 0.005934599 | 0.352129563 | 0.474484215 | 0.121608909 | 0.116892252 |
| Rrp36 | 0.002306221 | 0.699512848 | 0.78258503 | 0.10799846 | 0.103863442 |
| Rrp7a | 0.017315998 | 0.019255873 | 0.044993827 | 0.192790328 | 0.168314146 |
| Rrp8 | 0.024625577 | 0.000198266 | 0.000851389 | 0.138055224 | 0.123728774 |
| Rrp9 | -0.001463493 | 0.751096062 | 0.822778132 | 0.057549008 | 0.057830405 |
| Rrs1 | -0.013797088 | 0.028665168 | 0.062964927 | 0.104359022 | 0.119024188 |
| Rsad2 | 0.068928312 | 5.79E-22 | 1.91E-20 | 0.142727471 | 0.086754392 |
| Rsbn1 | -0.021296466 | 0.020336543 | 0.047096683 | 0.308754473 | 0.33896081 |
| Rsbn1l | 0.010898933 | 0.258650519 | 0.374810009 | 0.431170728 | 0.426073529 |
| Rsf1 | 0.051460658 | 2.79E-07 | 2.18E-06 | 0.521993784 | 0.455899909 |
| Rsl1d1 | -0.011564698 | 0.239162677 | 0.353027067 | 0.428902323 | 0.431179095 |
| Rsl24d1 | -0.00801684 | 0.240017613 | 0.354035993 | 0.145201884 | 0.143280941 |
| Rsph1 | -0.013598115 | 0.006708549 | 0.018565475 | 0.058213783 | 0.072373953 |
| Rsph3a | 0.007965453 | 0.043279159 | 0.088469299 | 0.048628873 | 0.03946372 |
| Rsph3b | 0.003157256 | 0.371710463 | 0.493954227 | 0.036145976 | 0.032067672 |
| Rsph9 | -0.024586257 | 0.000801517 | 0.002948298 | 0.158240073 | 0.178263819 |

| | | | | | |
|---|---|---|---|---|---|
| Rspo4 | 0.004936208 | 0.580583212 | 0.686908675 | 0.277422658 | 0.265278598 |
| Rspry1 | 0.003709088 | 0.612532869 | 0.712783653 | 0.173353608 | 0.165673378 |
| Rsrc1 | 0.004997695 | 0.564123909 | 0.671862384 | 0.282765899 | 0.280142861 |
| Rsrc2 | 0.052032095 | 2.72E-07 | 2.13E-06 | 0.818680894 | 0.763223166 |
| Rsrp1 | -0.138163741 | 4.29E-41 | 3.83E-39 | 0.872613013 | 1.004541913 |
| Rsu1 | -0.007465333 | 0.238296574 | 0.352138515 | 0.121716566 | 0.121987369 |
| Rtca | 0.003803405 | 0.52757604 | 0.639059838 | 0.104485903 | 0.104942295 |
| Rtcb | 0.02701796 | 0.005055409 | 0.014579853 | 0.49988105 | 0.456625132 |
| Rtf1 | 0.03869212 | 2.77E-05 | 0.000145019 | 0.375483545 | 0.337035452 |
| Rtf2 | 0.004748464 | 0.570514276 | 0.677729691 | 0.266671646 | 0.253675047 |
| Rtl5 | 0.01716507 | 0.015737177 | 0.038068383 | 0.156840939 | 0.146801663 |
| Rtl6 | 0.0150825 | 0.009053835 | 0.023860501 | 0.105708682 | 0.090905781 |
| Rtl8a | 0.011199263 | 0.157964508 | 0.254597305 | 0.253049489 | 0.21503593 |
| Rtl8b | 0.006692848 | 0.322612695 | 0.444380932 | 0.165079812 | 0.141949401 |
| Rtn1 | -0.020502433 | 0.000993992 | 0.003559332 | 0.094574005 | 0.114105657 |
| Rtn2 | 0.0036254 | 0.605781998 | 0.707436947 | 0.170387578 | 0.154423138 |
| Rtn3 | 0.048534251 | 1.11E-06 | 7.71E-06 | 0.679048782 | 0.621648249 |
| Rtn4 | 0.006481709 | 0.486416613 | 0.602259591 | 1.156328371 | 1.130043115 |
| Rtn4ip1 | 0.007570515 | 0.051391538 | 0.102173524 | 0.042762205 | 0.038068196 |
| Rtn4rl1 | 0.005744955 | 0.432148133 | 0.550562725 | 0.178901135 | 0.162929462 |
| Rtraf | -0.043145716 | 1.78E-05 | 9.75E-05 | 0.721074203 | 0.736940633 |
| Rubcn | 0.02884953 | 1.58E-05 | 8.76E-05 | 0.146900121 | 0.125490135 |
| Rufy1 | 0.002078109 | 0.765237542 | 0.833720552 | 0.157273058 | 0.149806708 |
| Rufy2 | 0.001612174 | 0.823049198 | 0.878074806 | 0.16357989 | 0.158264342 |
| Rufy3 | 0.022093508 | 0.023739035 | 0.053772343 | 0.457525332 | 0.414281775 |
| Rundc1 | 0.001718834 | 0.793316645 | 0.855627294 | 0.132871017 | 0.130210064 |
| Rundc3a | -0.027431507 | 0.005474691 | 0.015590201 | 0.763576778 | 0.768471427 |
| Rundc3b | 0.074742128 | 2.55E-25 | 1.04E-23 | 0.194327147 | 0.134431554 |
| Runx1t1 | 0.023663977 | 0.012508941 | 0.031286833 | 0.349908423 | 0.314387916 |
| Rusc1 | -0.019645417 | 0.00865276 | 0.023021931 | 0.169009502 | 0.184026318 |
| Rusf1 | 4.07E-05 | 0.992033056 | 0.994795809 | 0.045546936 | 0.043917096 |
| Ruvbl1 | -0.00813996 | 0.186364501 | 0.289847333 | 0.114633106 | 0.115054875 |
| Ruvbl2 | 0.004151884 | 0.488734769 | 0.604129615 | 0.109992723 | 0.103103151 |
| Rwdd1 | 0.061704369 | 5.46E-12 | 8.04E-11 | 1.114499954 | 1.030656417 |
| Rwdd3 | -0.003594019 | 0.373622614 | 0.495682186 | 0.045052517 | 0.044677047 |
| Rwdd4a | -0.012599357 | 0.215109134 | 0.325138525 | 0.522867604 | 0.547761666 |
| Rxra | 0.001442489 | 0.802169161 | 0.862305239 | 0.093652397 | 0.093203098 |
| Rxrb | -0.015278743 | 0.036728988 | 0.077195032 | 0.169421521 | 0.176533246 |
| Rxylt1 | 0.023368305 | 0.008341399 | 0.022284991 | 0.301729892 | 0.292739117 |
| Rybp | 0.061630099 | 3.90E-12 | 5.85E-11 | 0.327404437 | 0.267252591 |
| Ryk | 0.007120489 | 0.390961517 | 0.511909478 | 0.257139777 | 0.246286286 |
| S100a1 | -0.044919394 | 2.39E-05 | 0.00012703 | 0.676799369 | 0.709873778 |
| S100a10 | 0.279301901 | 4.38E-148 | 7.10E-145 | 0.615758178 | 0.328658365 |
| S100a11 | 0.025800585 | 0.009262609 | 0.024331555 | 0.897325486 | 0.819837348 |
| S100a13 | -0.022803131 | 0.016615948 | 0.039886481 | 0.388764933 | 0.41481864 |
| S100a4 | 0.013204854 | 0.002043988 | 0.00665717 | 0.056911289 | 0.042059774 |
| S100a6 | 0.006300248 | 0.434429615 | 0.552817371 | 0.125032017 | 0.123536514 |
| S100pbp | 0.0015268 | 0.793233602 | 0.855627294 | 0.099178642 | 0.094208539 |
| S100z | 0.013000976 | 0.012589919 | 0.031448918 | 0.074035785 | 0.062009127 |
| Saal1 | 0.005111675 | 0.152849183 | 0.247912846 | 0.038134857 | 0.032284581 |
| Sac3d1 | -0.003534831 | 0.600067263 | 0.70329764 | 0.142890293 | 0.139147133 |
| Sacm1l | 0.02875309 | 0.001548814 | 0.005215536 | 0.351110125 | 0.313909708 |
| Sacs | 0.012645668 | 0.006127242 | 0.017176773 | 0.059325083 | 0.051697271 |
| Sae1 | -0.005074855 | 0.504195928 | 0.618600809 | 0.199036743 | 0.194084191 |
| Safb | 0.031839365 | 0.000594971 | 0.002265692 | 0.372990081 | 0.33265235 |
| Safb2 | 0.020687752 | 0.027840371 | 0.061430797 | 0.350055896 | 0.330211329 |
| Sall2 | 0.008316504 | 0.051534841 | 0.102353775 | 0.053368126 | 0.046674021 |
| Sall4 | -0.003218082 | 0.47541386 | 0.59233289 | 0.054036238 | 0.054409018 |
| Samd1 | -0.004666102 | 0.426882164 | 0.545569659 | 0.106052176 | 0.099575409 |
| Samd10 | 0.011388946 | 0.086695575 | 0.157425173 | 0.147991955 | 0.131119089 |
| Samd12 | -0.005910522 | 0.259388149 | 0.375375348 | 0.07028003 | 0.076474746 |
| Samd4b | 0.066840368 | 2.46E-12 | 3.80E-11 | 0.445454617 | 0.388159473 |
| Samd8 | 0.010552705 | 0.202873294 | 0.310457132 | 0.238807665 | 0.230084008 |
| Samd9l | 0.0174279 | 0.002017523 | 0.006582 | 0.091575477 | 0.079816066 |
| Samhd1 | -0.006298196 | 0.236795651 | 0.350506596 | 0.075886536 | 0.079813407 |
| Samm50 | -0.021544168 | 0.016882179 | 0.040425749 | 0.332830052 | 0.342644595 |
| Sap130 | 0.005237647 | 0.297151487 | 0.416929825 | 0.069266437 | 0.066798813 |
| Sap18 | -0.003189484 | 0.735917595 | 0.810718511 | 0.454725719 | 0.432428345 |
| Sap30 | 0.0007835 | 0.889336691 | 0.924522167 | 0.09698398 | 0.09036001 |
| Sap30bp | 0.006539437 | 0.304728007 | 0.42528936 | 0.122895955 | 0.117647131 |
| Sap30l | 0.006396846 | 0.459036298 | 0.577478117 | 0.299907909 | 0.286312683 |
| Sar1a | 0.019414303 | 0.047666086 | 0.096083285 | 0.897777029 | 0.87213639 |

| | | | | | |
|---|---|---|---|---|---|
| Sar1b | -0.028982385 | 0.004058828 | 0.012059879 | 0.869601088 | 0.887682745 |
| Saraf | -0.079584542 | 6.26E-17 | 1.39E-15 | 1.295592028 | 1.455937949 |
| Sarnp | -0.028211 | 0.002787545 | 0.008721006 | 0.380582742 | 0.395321053 |
| Sars | -0.067652054 | 2.10E-14 | 3.95E-13 | 1.264390499 | 1.350067191 |
| Sart1 | -0.006433324 | 0.390421399 | 0.511408898 | 0.182337994 | 0.179636709 |
| Sart3 | 0.009393881 | 0.090814359 | 0.163319685 | 0.095115461 | 0.085918524 |
| Sat1 | -0.038327773 | 3.77E-05 | 0.000191325 | 0.229792478 | 0.294703171 |
| Sat2 | -0.010716398 | 0.031628184 | 0.068295816 | 0.070150361 | 0.073811261 |
| Satb1 | 0.00576144 | 0.470991259 | 0.58817945 | 0.212115457 | 0.202848177 |
| Sav1 | -0.005119507 | 0.352482674 | 0.474828399 | 0.085423404 | 0.087098804 |
| Saysd1 | 0.00444657 | 0.60304148 | 0.706102525 | 0.274775556 | 0.268893616 |
| Sbds | -0.040746084 | 5.27E-05 | 0.000258028 | 0.555113993 | 0.590578203 |
| Sbf1 | 0.00112319 | 0.833264254 | 0.885935595 | 0.078205298 | 0.079077105 |
| Sbf2 | -0.006620075 | 0.271691471 | 0.388781937 | 0.100951217 | 0.104691947 |
| Sbno1 | 0.008182716 | 0.413996093 | 0.533590086 | 0.598662897 | 0.578064545 |
| Sbno2 | -0.006549165 | 0.103819565 | 0.182288932 | 0.036908344 | 0.044312001 |
| Sc5d | 0.005268013 | 0.39145728 | 0.512075844 | 0.11207848 | 0.108647085 |
| Scaf1 | 0.031652273 | 2.02E-05 | 0.000109395 | 0.202769539 | 0.168730743 |
| Scaf11 | 0.006237164 | 0.542221862 | 0.652491762 | 0.894280676 | 0.876723185 |
| Scaf4 | 0.019190977 | 0.002953071 | 0.009173539 | 0.129736197 | 0.116260025 |
| Scaf8 | 0.005963813 | 0.396155904 | 0.516692741 | 0.157393364 | 0.149175264 |
| Scai | -0.008841092 | 0.115733327 | 0.198686103 | 0.080988695 | 0.090589457 |
| Scamp1 | 0.017025391 | 0.034593885 | 0.073561187 | 0.237566397 | 0.220877771 |
| Scamp2 | 0.006942715 | 0.454266628 | 0.57296058 | 0.395670976 | 0.380353646 |
| Scamp3 | 0.016748636 | 0.044047304 | 0.089831737 | 0.263015381 | 0.243296976 |
| Scamp5 | 0.010522146 | 0.152088641 | 0.247010653 | 0.165246512 | 0.163082763 |
| Scand1 | 0.03340377 | 5.39E-05 | 0.000263087 | 1.379566787 | 1.352482749 |
| Scap | 0.008066109 | 0.243553708 | 0.357285293 | 0.160196096 | 0.145066625 |
| Scaper | 0.011046888 | 0.166229465 | 0.264602894 | 0.221317505 | 0.202290884 |
| Scarb1 | 0.000880132 | 0.81807377 | 0.874663278 | 0.039610144 | 0.037576671 |
| Scarb2 | -0.042795015 | 2.08E-05 | 0.000112231 | 0.616998893 | 0.66399037 |
| Scarf2 | -0.002060843 | 0.658636738 | 0.750148355 | 0.055098623 | 0.053684743 |
| Sccpdh | -0.005143727 | 0.361733216 | 0.484537107 | 0.087192879 | 0.092444058 |
| Scd2 | 0.056002674 | 4.02E-15 | 7.82E-14 | 0.165354021 | 0.121129905 |
| Scfd1 | -0.012757475 | 0.210498375 | 0.319610371 | 0.566568416 | 0.589541318 |
| Scfd2 | -0.007589671 | 0.093420726 | 0.166955202 | 0.049714985 | 0.057199366 |
| Scg2 | -0.084524837 | 5.65E-106 | 4.22E-103 | 4.091761181 | 4.442067978 |
| Scg3 | 0.004632669 | 0.443974831 | 0.562533991 | 3.16259815 | 3.226123294 |
| Scg5 | -0.117110515 | 7.07E-142 | 9.82E-139 | 2.657338122 | 2.928609629 |
| Scgn | 0.156502959 | 4.56E-45 | 4.48E-43 | 0.756305917 | 0.543723091 |
| Sclt1 | 0.006091166 | 0.373616668 | 0.495682186 | 0.140540361 | 0.136454735 |
| Scly | 0.025598086 | 0.00845472 | 0.022569135 | 0.424583728 | 0.389368689 |
| Scmh1 | -0.006365829 | 0.355690972 | 0.478046125 | 0.142137324 | 0.14610491 |
| Scn1b | 0.00860061 | 0.080823197 | 0.148565536 | 0.075767113 | 0.061290249 |
| Scn3a | -0.024160659 | 6.87E-05 | 0.00033062 | 0.074744141 | 0.103673679 |
| Scn8a | -0.011390407 | 0.08803632 | 0.159383445 | 0.123176389 | 0.135406097 |
| Scn9a | 0.005443763 | 0.498069516 | 0.612806997 | 0.198542201 | 0.193070176 |
| Scnm1 | -0.013714076 | 0.034286771 | 0.073051938 | 0.123773763 | 0.126926573 |
| Scnn1b | 0.033618124 | 0.001061369 | 0.003772807 | 0.557471265 | 0.498500552 |
| Sco2 | -0.008515076 | 0.094831725 | 0.168886982 | 0.068348842 | 0.073885621 |
| Scoc | -0.01361636 | 0.150127745 | 0.245012916 | 0.414043766 | 0.413890507 |
| Scp2 | -0.002677765 | 0.780376546 | 0.845516637 | 0.571170483 | 0.530350763 |
| Scpep1 | -0.033439435 | 0.000320999 | 0.001310141 | 0.27868612 | 0.31992331 |
| Scrib | -0.010922883 | 0.040641906 | 0.084086106 | 0.073766693 | 0.080795168 |
| Scrn1 | 0.021389539 | 0.011563218 | 0.029336535 | 0.28604454 | 0.250670337 |
| Scrn2 | -0.027963836 | 6.99E-10 | 8.14E-09 | 0.043744175 | 0.063917614 |
| Scrn3 | 0.004574998 | 0.511626688 | 0.625350095 | 0.156919637 | 0.147577867 |
| Scrt1 | -0.01530764 | 0.008870775 | 0.023479899 | 0.086056073 | 0.099356764 |
| Scyl1 | 0.003388104 | 0.667888219 | 0.757313887 | 0.223024874 | 0.212141738 |
| Scyl2 | 0.017155516 | 0.005429214 | 0.015478832 | 0.119152272 | 0.104322146 |
| Scyl3 | -0.005485309 | 0.332388816 | 0.455033632 | 0.088576654 | 0.091193636 |
| Sdad1 | 0.001065299 | 0.870717491 | 0.911207261 | 0.125343838 | 0.123613531 |
| Sdc2 | -0.041108925 | 8.95E-08 | 7.63E-07 | 0.168028074 | 0.202447393 |
| Sdc3 | 0.018678409 | 0.00036061 | 0.001451602 | 0.087845364 | 0.06915069 |
| Sdc4 | 0.030907638 | 0.003383869 | 0.010335524 | 0.54950123 | 0.504804228 |
| Sdcbp | 0.001313944 | 0.893957152 | 0.927936305 | 0.483329844 | 0.488312189 |
| Sdcbp2 | 0.005993102 | 0.223337186 | 0.334764743 | 0.063630643 | 0.054743018 |
| Sdccag8 | 0.030586836 | 1.01E-05 | 5.78E-05 | 0.172538046 | 0.140860302 |
| Sde2 | 0.001717855 | 0.838495402 | 0.889163645 | 0.256423874 | 0.253953811 |
| Sdf2 | -0.036255636 | 0.000129397 | 0.000588124 | 0.433174192 | 0.458282551 |
| Sdf2l1 | 0.053523459 | 4.98E-06 | 3.06E-05 | 0.765778316 | 0.744988447 |
| Sdf4 | -0.06827624 | 3.87E-13 | 6.53E-12 | 0.89134308 | 0.961322435 |

| | | | | | |
|---|---|---|---|---|---|
| Sdha | 0.008638622 | 0.371674225 | 0.493954227 | 0.599480407 | 0.567139235 |
| Sdhaf1 | -0.019366177 | 0.038220462 | 0.079720946 | 0.349663365 | 0.365233525 |
| Sdhaf2 | 0.005283607 | 0.432722723 | 0.551222525 | 0.147709112 | 0.137732971 |
| Sdhaf3 | 0.009765509 | 0.070823121 | 0.133567873 | 0.090289878 | 0.08048138 |
| Sdhaf4 | 6.21E-05 | 0.994330482 | 0.996209693 | 0.327606677 | 0.302590845 |
| Sdhb | -0.051588237 | 2.46E-07 | 1.93E-06 | 0.654218713 | 0.680901919 |
| Sdhc | 0.030930118 | 0.001442043 | 0.004912393 | 0.53592442 | 0.489787234 |
| Sdhd | 0.006257314 | 0.514076339 | 0.627341191 | 0.465541356 | 0.426776362 |
| Sdk2 | 0.028616205 | 0.006718014 | 0.018586379 | 0.681274906 | 0.660031263 |
| Sdr39u1 | 0.018221576 | 0.012014643 | 0.030301787 | 0.182603822 | 0.161901861 |
| Sdsl | -0.006246216 | 0.140185455 | 0.232217241 | 0.046666615 | 0.049608057 |
| Sec11a | -0.063579505 | 9.15E-10 | 1.05E-08 | 0.549839672 | 0.625762582 |
| Sec11c | -0.114247886 | 4.26E-38 | 3.24E-36 | 1.377091119 | 1.61797315 |
| Sec13 | -0.007898003 | 0.422889368 | 0.541534567 | 0.48476776 | 0.4836638 |
| Sec14l1 | 0.0446574 | 1.17E-07 | 9.77E-07 | 0.287027961 | 0.239498442 |
| Sec16a | 0.014460235 | 0.112393066 | 0.194497221 | 0.327125997 | 0.314016862 |
| Sec16b | -0.007632017 | 0.202043989 | 0.309480331 | 0.091646583 | 0.102063897 |
| Sec22a | -0.002636749 | 0.679078793 | 0.766872345 | 0.121852408 | 0.120342505 |
| Sec22b | 0.008771069 | 0.381174855 | 0.502342678 | 0.634899669 | 0.614188206 |
| Sec22c | -0.003187498 | 0.521265336 | 0.633705339 | 0.063754599 | 0.066817313 |
| Sec23a | 0.010883058 | 0.068718754 | 0.130434151 | 0.107129268 | 0.098370414 |
| Sec23b | -0.038433441 | 3.63E-05 | 0.00018531 | 0.330255154 | 0.362286882 |
| Sec23ip | 0.006894111 | 0.396150642 | 0.516692741 | 0.234960375 | 0.220540037 |
| Sec24a | 0.006598528 | 0.40097388 | 0.521019522 | 0.200056389 | 0.1972929 |
| Sec24b | 0.015256821 | 0.020710537 | 0.047860196 | 0.139997692 | 0.126461511 |
| Sec24c | 0.026232789 | 0.001453874 | 0.004949078 | 0.252237195 | 0.224794891 |
| Sec24d | -0.005322926 | 0.519189353 | 0.631576438 | 0.236285809 | 0.236192523 |
| Sec31a | 0.038626691 | 0.000125224 | 0.000570757 | 0.789549097 | 0.721109656 |
| Sec61a1 | -0.036990309 | 0.000352499 | 0.001423169 | 0.632273329 | 0.65629507 |
| Sec61a2 | -0.005614378 | 0.290270997 | 0.40970015 | 0.076052092 | 0.079002012 |
| Sec61b | -0.021875743 | 0.000328782 | 0.001337412 | 2.243238624 | 2.331390317 |
| Sec61g | 0.160184305 | 4.88E-55 | 7.19E-53 | 0.934995939 | 0.816654503 |
| Sec62 | 0.085523246 | 1.64E-27 | 7.51E-26 | 1.501681315 | 1.414404044 |
| Sec63 | 0.005646676 | 0.576024992 | 0.68235835 | 0.6118362 | 0.616506237 |
| Secisbp2 | -0.008268437 | 0.174565702 | 0.275239662 | 0.108162635 | 0.110457613 |
| Secisbp2l | 0.011361263 | 0.06647009 | 0.127109012 | 0.120067055 | 0.107968379 |
| Seh1l | 0.018831279 | 0.018912625 | 0.044327035 | 0.235254327 | 0.217075605 |
| Sel1l | 0.044872349 | 1.65E-05 | 9.09E-05 | 0.728618758 | 0.697937344 |
| Sel1l3 | 0.009869269 | 0.098583949 | 0.174577988 | 0.10451954 | 0.098069932 |
| Selenof | -0.075349617 | 3.81E-18 | 9.50E-17 | 1.171506697 | 1.275265903 |
| Selenoh | -0.019066198 | 0.031650016 | 0.068317375 | 0.33467686 | 0.328775226 |
| Selenoi | 0.013415233 | 0.039939525 | 0.082862156 | 0.135688327 | 0.123003066 |
| Selenok | -0.006933447 | 0.349147108 | 0.471597793 | 1.745025701 | 1.754299631 |
| Selenom | 0.047583633 | 1.66E-09 | 1.83E-08 | 1.655434157 | 1.577691441 |
| Selenon | 0.011437748 | 0.037933095 | 0.079257586 | 0.095299109 | 0.081922975 |
| Selenoo | 0.008078856 | 0.163890912 | 0.261637759 | 0.104236252 | 0.094911576 |
| Selenop | 0.118401696 | 1.79E-25 | 7.42E-24 | 1.143308204 | 0.922655848 |
| Selenos | -0.038791462 | 8.16E-05 | 0.000388143 | 1.06263529 | 1.138616121 |
| Selenot | 0.063117494 | 1.07E-09 | 1.21E-08 | 0.8514701 | 0.776368994 |
| Selenow | 0.163882937 | 3.46E-55 | 5.17E-53 | 1.293997454 | 1.055952536 |
| Sem1 | 0.127823175 | 6.63E-59 | 1.17E-56 | 1.550267329 | 1.387615043 |
| Sema4b | 0.017644901 | 0.000839749 | 0.003079608 | 0.088732457 | 0.070931069 |
| Sema4d | -0.004660144 | 0.349559247 | 0.471936537 | 0.06614768 | 0.070348059 |
| Sema6d | -0.001078525 | 0.779179767 | 0.845157198 | 0.03861578 | 0.039135379 |
| Senp1 | 0.005189396 | 0.368872674 | 0.491324858 | 0.097748798 | 0.092898741 |
| Senp2 | 0.003075195 | 0.685602815 | 0.771997981 | 0.18669709 | 0.18495113 |
| Senp3 | -0.002796458 | 0.660070508 | 0.751253276 | 0.118922889 | 0.120990783 |
| Senp5 | 0.035520541 | 1.52E-06 | 1.02E-05 | 0.194257101 | 0.165649252 |
| Senp6 | 0.003873731 | 0.689668131 | 0.774678846 | 0.914238782 | 0.900278651 |
| Senp7 | 0.008180175 | 0.265199728 | 0.381570483 | 0.174507697 | 0.165385229 |
| Senp8 | 0.010401944 | 0.034562501 | 0.07351053 | 0.076169085 | 0.063466805 |
| Sephs1 | -0.010966452 | 0.030041948 | 0.065500745 | 0.069475148 | 0.073509187 |
| Sephs2 | -0.005105245 | 0.547938058 | 0.65757978 | 0.27826507 | 0.266951376 |
| Sepsecs | -0.006078441 | 0.421267202 | 0.540382602 | 0.191656893 | 0.182880184 |
| Septin10 | 0.02041771 | 3.81E-05 | 0.000192645 | 0.079550164 | 0.061108573 |
| Septin11 | -0.02506007 | 0.004550552 | 0.01331742 | 0.266228215 | 0.286237502 |
| Septin7 | -0.030870601 | 0.0018014 | 0.005960929 | 0.674980419 | 0.697041661 |
| Septin8 | -0.004632032 | 0.318014864 | 0.439792391 | 0.052795609 | 0.058505937 |
| Septin9 | 0.01301166 | 0.135485301 | 0.226282098 | 0.244901152 | 0.248232786 |
| Serbp1 | 0.019191749 | 0.030333475 | 0.066003144 | 1.292866614 | 1.270706723 |
| Serf1 | -0.026056947 | 0.000141064 | 0.000635509 | 0.132399772 | 0.150187945 |
| Serf2 | 0.190911668 | 5.54E-101 | 3.59E-98 | 1.417550665 | 1.171800896 |

| | | | | | |
|---|---|---|---|---|---|
| Sergef | -0.016406142 | 0.008691623 | 0.023095737 | 0.101416866 | 0.120475225 |
| Serhl | -0.002151462 | 0.651694188 | 0.744493577 | 0.065120108 | 0.062627766 |
| Serinc1 | -0.041663122 | 6.41E-06 | 3.85E-05 | 1.03982436 | 1.100729557 |
| Serinc3 | 0.010051068 | 0.281364436 | 0.400267054 | 0.388524589 | 0.369605051 |
| Serinc5 | 0.001985522 | 0.706744809 | 0.788351314 | 0.079059573 | 0.075215086 |
| Serp1 | -0.042755236 | 2.28E-05 | 0.000121789 | 1.056954034 | 1.134439816 |
| Serp2 | -0.008222317 | 0.129308574 | 0.217535552 | 0.078707007 | 0.083858177 |
| Serpina10 | -0.035881173 | 0.001809054 | 0.005982185 | 0.533847964 | 0.605097384 |
| Serpina7 | -0.182254775 | 2.80E-51 | 3.53E-49 | 0.125571149 | 0.394694669 |
| Serpinb6a | 0.024190754 | 0.013807569 | 0.034027169 | 0.531025612 | 0.504421019 |
| Serpinb9 | -0.002801951 | 0.570368503 | 0.677639324 | 0.067731266 | 0.065092408 |
| Serping1 | -0.050366396 | 2.56E-09 | 2.76E-08 | 0.156059188 | 0.224395979 |
| Serpini1 | -0.057119687 | 5.06E-11 | 6.72E-10 | 0.225472008 | 0.280808252 |
| Sertad1 | -0.020351544 | 0.002451485 | 0.007801421 | 0.11462921 | 0.134939606 |
| Sertad2 | -0.009305425 | 0.099588648 | 0.176132589 | 0.082799865 | 0.090472721 |
| Sertad3 | 0.00149525 | 0.757722511 | 0.827724975 | 0.068836587 | 0.06352255 |
| Sesn1 | 0.013319298 | 0.008893381 | 0.023533329 | 0.079332311 | 0.06659603 |
| Sesn2 | -0.024150125 | 0.01815891 | 0.042943546 | 0.455425055 | 0.481054747 |
| Sesn3 | 0.063334805 | 1.23E-09 | 1.39E-08 | 0.609746751 | 0.517090905 |
| Sestd1 | 0.002993212 | 0.472777069 | 0.589954905 | 0.047965546 | 0.044537867 |
| Set | -0.013515429 | 0.170649518 | 0.269940548 | 0.970515256 | 0.999294257 |
| Setbp1 | 0.014881493 | 0.024492887 | 0.055273874 | 0.138783482 | 0.119081217 |
| Setd1a | 0.021772423 | 1.94E-05 | 0.000105262 | 0.081918449 | 0.065478219 |
| Setd1b | -0.011795738 | 0.134737505 | 0.225265353 | 0.198922338 | 0.200508153 |
| Setd2 | 0.027005259 | 0.001603938 | 0.005371507 | 0.265345978 | 0.241715144 |
| Setd3 | -0.005998298 | 0.538111668 | 0.648830663 | 0.575185407 | 0.562317888 |
| Setd5 | 0.015932109 | 0.109677829 | 0.190510604 | 0.484413101 | 0.461507746 |
| Setd6 | -0.007479664 | 0.223303731 | 0.334764743 | 0.109343874 | 0.113357893 |
| Setd7 | 0.048796995 | 2.13E-08 | 1.98E-07 | 0.296800055 | 0.254220093 |
| Setdb1 | 0.004127605 | 0.461088515 | 0.579085718 | 0.092926319 | 0.086204155 |
| Setx | -0.015627465 | 0.051686857 | 0.102576477 | 0.191378454 | 0.20888329 |
| Sez6l | -0.04212582 | 0.00015266 | 0.00067863 | 0.493107958 | 0.536673451 |
| Sez6l2 | -0.078976796 | 8.12E-18 | 1.95E-16 | 1.020416982 | 1.136292497 |
| Sf1 | 0.035503045 | 9.78E-05 | 0.000457469 | 0.360803151 | 0.319747618 |
| Sf3a1 | 0.038741914 | 3.31E-08 | 3.02E-07 | 0.178202557 | 0.142546408 |
| Sf3a2 | 0.002109802 | 0.726300149 | 0.802579 | 0.113626232 | 0.105476899 |
| Sf3a3 | 0.002260903 | 0.751729816 | 0.823194106 | 0.171565739 | 0.16297489 |
| Sf3b1 | -0.052319086 | 2.38E-07 | 1.87E-06 | 0.642703359 | 0.692065399 |
| Sf3b2 | 0.012509751 | 0.184904286 | 0.288106507 | 1.02188319 | 0.97369372 |
| Sf3b3 | 0.005646187 | 0.523191168 | 0.635331569 | 0.302937669 | 0.303376101 |
| Sf3b4 | -0.005269433 | 0.403566123 | 0.523408464 | 0.122167878 | 0.117984385 |
| Sf3b5 | 0.008676361 | 0.396724116 | 0.517225675 | 0.615093674 | 0.608177797 |
| Sf3b6 | -0.052706625 | 1.66E-08 | 1.58E-07 | 1.037591167 | 1.088008966 |
| Sfi1 | 0.002714546 | 0.542962725 | 0.65297562 | 0.054197489 | 0.051485679 |
| Sfmbt1 | 0.012902085 | 0.11083279 | 0.192207703 | 0.235378225 | 0.216223342 |
| Sfpq | -0.015590851 | 0.124308254 | 0.210251365 | 0.485300093 | 0.49355077 |
| Sfr1 | -0.037972564 | 0.000146279 | 0.000654752 | 0.603419097 | 0.623566222 |
| Sfrp5 | -0.035351395 | 1.29E-16 | 2.82E-15 | 0.028308212 | 0.050406801 |
| Sfswap | 0.013008303 | 0.078567878 | 0.145409653 | 0.179723732 | 0.168557899 |
| Sft2d1 | 0.006680251 | 0.075776518 | 0.141428165 | 0.04272037 | 0.035250192 |
| Sft2d2 | 0.007681726 | 0.234268528 | 0.34809088 | 0.124353319 | 0.123850034 |
| Sft2d3 | 0.018983835 | 0.018730421 | 0.043974199 | 0.235097802 | 0.221466329 |
| Sfxn1 | 0.017539412 | 0.07769248 | 0.144036287 | 0.677086934 | 0.651431951 |
| Sfxn2 | 0.014707628 | 0.051038369 | 0.101700148 | 0.189390935 | 0.173405497 |
| Sfxn3 | -3.43E-05 | 0.994227554 | 0.996209693 | 0.062994857 | 0.06030009 |
| Sgcb | -0.009521373 | 0.08736961 | 0.15840362 | 0.079373285 | 0.089740126 |
| Sgcz | -0.008978567 | 0.133372444 | 0.223405738 | 0.091314091 | 0.101737048 |
| Sgf29 | -0.015673584 | 0.004362625 | 0.012856454 | 0.082279433 | 0.088941097 |
| Sgk2 | 0.00046972 | 0.915181211 | 0.941423313 | 0.052453783 | 0.050888103 |
| Sgk3 | 0.008666219 | 0.138302585 | 0.229685298 | 0.105348751 | 0.095602384 |
| Sgms1 | 0.001816497 | 0.770561134 | 0.838150543 | 0.107927338 | 0.110870007 |
| Sgms2 | 0.03517208 | 1.53E-05 | 8.50E-05 | 0.235041242 | 0.198693036 |
| Sgpl1 | 0.048994577 | 6.79E-09 | 6.91E-08 | 0.290919917 | 0.235166095 |
| Sgpp1 | -0.016214477 | 0.063363804 | 0.122033064 | 0.279494555 | 0.300048454 |
| Sgpp2 | 0.031607012 | 1.19E-06 | 8.21E-06 | 0.146625565 | 0.11672209 |
| Sgsm1 | 0.021769723 | 0.001723603 | 0.005722974 | 0.160955296 | 0.141410251 |
| Sgsm2 | 0.036780719 | 2.52E-08 | 2.32E-07 | 0.146692606 | 0.120101609 |
| Sgsm3 | -0.001346091 | 0.846046131 | 0.894730827 | 0.154846479 | 0.149221705 |
| Sgta | -0.00171551 | 0.84521301 | 0.894324935 | 0.323531331 | 0.311764443 |
| Sgtb | 0.009587603 | 0.068951176 | 0.130774907 | 0.079108093 | 0.073909086 |
| Sh2b1 | 0.015136911 | 0.003274411 | 0.01003905 | 0.082857087 | 0.069352289 |
| Sh2d3c | -0.012393952 | 0.007426656 | 0.020230303 | 0.056386387 | 0.062938782 |

| | | | | | |
|---|---|---|---|---|---|
| Sh2d5 | -0.000246724 | 0.94780337 | 0.965075865 | 0.03973655 | 0.036048013 |
| Sh3bgrl | -0.002744575 | 0.767760073 | 0.835923736 | 0.369697172 | 0.364214533 |
| Sh3bgrl3 | 0.047266925 | 1.75E-07 | 1.42E-06 | 0.330152511 | 0.282055453 |
| Sh3bp4 | 0.002780076 | 0.598225178 | 0.70164618 | 0.083533055 | 0.075353448 |
| Sh3bp5l | -0.005676758 | 0.263205208 | 0.379362525 | 0.072459095 | 0.073646158 |
| Sh3d19 | 0.048498052 | 9.39E-12 | 1.35E-10 | 0.181921635 | 0.138351249 |
| Sh3gl1 | -0.029321518 | 0.000128251 | 0.000583463 | 0.173042929 | 0.200416499 |
| Sh3gl2 | 0.022120637 | 0.000132888 | 0.000602303 | 0.110500206 | 0.088822881 |
| Sh3glb1 | 0.02516888 | 0.00763257 | 0.020704198 | 1.052983968 | 1.009599009 |
| Sh3glb2 | -0.011323562 | 0.096732882 | 0.171832099 | 0.143849269 | 0.147051491 |
| Sh3kbp1 | 0.000691021 | 0.931115097 | 0.953375563 | 0.208445226 | 0.20339805 |
| Sh3pxd2a | 0.066657061 | 3.19E-10 | 3.85E-09 | 0.575403698 | 0.491209392 |
| Sh3rf1 | 0.008400947 | 0.021353949 | 0.049136826 | 0.039514301 | 0.032773757 |
| Sh3yl1 | 0.001777741 | 0.799792858 | 0.86032155 | 0.158859786 | 0.152960047 |
| Sharpin | 0.01503505 | 0.014524286 | 0.03555908 | 0.124050946 | 0.107346287 |
| Shb | 0.006724801 | 0.328393075 | 0.451001197 | 0.150362339 | 0.141057495 |
| Shc1 | -0.115418129 | 4.28E-31 | 2.28E-29 | 0.317820523 | 0.43298975 |
| Shc2 | -0.00656478 | 0.199815557 | 0.306840443 | 0.074466009 | 0.072644417 |
| Shfl | -0.014886301 | 0.058182486 | 0.113607177 | 0.196420757 | 0.211109007 |
| Shisa2 | 0.026542473 | 4.77E-08 | 4.25E-07 | 0.075466153 | 0.055248558 |
| Shisa5 | 0.028177569 | 0.002193294 | 0.007067683 | 0.325627457 | 0.310446345 |
| Shisal1 | 0.016952866 | 0.000466211 | 0.001823211 | 0.065911006 | 0.054433894 |
| Shisal2b | 0.146895852 | 5.74E-35 | 3.72E-33 | 0.517594191 | 0.381821626 |
| Shkbp1 | 0.002952465 | 0.560251213 | 0.668744137 | 0.07789126 | 0.071095482 |
| Shld1 | -0.000504614 | 0.933534402 | 0.95498886 | 0.114643644 | 0.108661308 |
| Shld2 | 0.004275736 | 0.38470061 | 0.505730791 | 0.069055922 | 0.06412389 |
| Shld3 | 0.005369263 | 0.163264368 | 0.261147777 | 0.043104332 | 0.03712435 |
| Shmt2 | 0.03036528 | 0.00036252 | 0.001456024 | 0.26868227 | 0.234121376 |
| Shoc2 | 0.035761096 | 6.24E-05 | 0.000302673 | 0.327144029 | 0.286917713 |
| Shprh | 0.005443508 | 0.417018009 | 0.536347279 | 0.136657273 | 0.129237933 |
| Shq1 | -0.005212617 | 0.174843405 | 0.275543456 | 0.038038627 | 0.041061166 |
| Shroom2 | 0.00508973 | 0.214925104 | 0.324910878 | 0.048147929 | 0.0425597 |
| Shroom3 | 0.018712862 | 0.002050408 | 0.006674331 | 0.118478054 | 0.09961934 |
| Shtn1 | 0.018720844 | 0.01235956 | 0.031001511 | 0.179479295 | 0.166302252 |
| Siae | 0.013528716 | 0.01272349 | 0.031725512 | 0.093464972 | 0.07948909 |
| Siah1a | -0.003665982 | 0.59571076 | 0.699541008 | 0.154760983 | 0.149829244 |
| Siah2 | 0.016892578 | 0.079969972 | 0.147380307 | 0.397979713 | 0.387457739 |
| Sidt2 | 0.005976003 | 0.324492429 | 0.446527302 | 0.108135634 | 0.104586299 |
| Sigirr | 0.000490251 | 0.91451748 | 0.941138874 | 0.05659889 | 0.055107227 |
| Sigmar1 | -0.015238414 | 0.01220346 | 0.030688577 | 0.103583732 | 0.112327287 |
| Sik1 | -0.012195111 | 0.029280746 | 0.06407099 | 0.063232258 | 0.089179981 |
| Sik2 | 0.00634208 | 0.184812017 | 0.28803181 | 0.059440977 | 0.057607559 |
| Sik3 | 0.04318984 | 1.52E-07 | 1.24E-06 | 0.255672342 | 0.221805669 |
| Sike1 | 0.014767719 | 0.058099885 | 0.113491476 | 0.210592146 | 0.196960095 |
| Sil1 | -0.038929656 | 4.65E-05 | 0.000229924 | 0.422090835 | 0.445448036 |
| Sim1 | 0.004003564 | 0.345916298 | 0.468123364 | 0.045933375 | 0.043201514 |
| Simc1 | 0.004170212 | 0.30479154 | 0.425316973 | 0.045885592 | 0.042103839 |
| Sin3a | 0.013093911 | 0.072861828 | 0.136696775 | 0.177471068 | 0.161661446 |
| Sin3b | 0.004784227 | 0.62972956 | 0.72709944 | 0.554876482 | 0.542826946 |
| Sinhcaf | -0.001731027 | 0.779559208 | 0.845157198 | 0.105219298 | 0.109439459 |
| Sipa1l1 | -0.002098969 | 0.75728418 | 0.8275192 | 0.134470382 | 0.138163794 |
| Sipa1l3 | 0.028162648 | 0.000170999 | 0.000749867 | 0.200814078 | 0.167441054 |
| Sirpa | -0.001617265 | 0.859140621 | 0.903272966 | 0.349706769 | 0.326114089 |
| Sirt1 | 0.001671851 | 0.759804167 | 0.829606481 | 0.082021017 | 0.081294577 |
| Sirt2 | -0.014249991 | 0.081444708 | 0.149481868 | 0.24548803 | 0.245691513 |
| Sirt3 | -0.026998799 | 9.66E-05 | 0.00045242 | 0.146545209 | 0.159277837 |
| Sirt4 | -0.017903448 | 2.66E-05 | 0.000139382 | 0.042183074 | 0.053774711 |
| Sirt5 | -0.010404795 | 0.025409724 | 0.056909209 | 0.059282029 | 0.0615674 |
| Sirt6 | 0.005635663 | 0.234901124 | 0.348711059 | 0.065610906 | 0.0601921 |
| Sirt7 | 0.000661543 | 0.923155189 | 0.947620605 | 0.149985109 | 0.142335525 |
| Siva1 | 0.004097646 | 0.635972456 | 0.731620426 | 0.300937396 | 0.284059144 |
| Six4 | 0.004665462 | 0.460364364 | 0.578699553 | 0.11976608 | 0.108545318 |
| Six5 | -0.000143784 | 0.971908952 | 0.980933923 | 0.046098138 | 0.043856396 |
| Ska2 | 0.006771621 | 0.06831671 | 0.129873886 | 0.040899379 | 0.034182171 |
| Skap1 | -0.048134832 | 1.73E-06 | 1.15E-05 | 0.487871994 | 0.514560463 |
| Skap2 | -0.014931284 | 0.044703601 | 0.090846239 | 0.167514803 | 0.179503985 |
| Ski | 0.011738726 | 0.088330479 | 0.159772817 | 0.160260487 | 0.139329661 |
| Skil | -0.007502891 | 0.381551847 | 0.502771356 | 0.226999832 | 0.244449152 |
| Skiv2l | 0.003637546 | 0.573775673 | 0.680523008 | 0.132976357 | 0.124285201 |
| Skp1 | 0.037986948 | 0.000115107 | 0.000530116 | 0.57131243 | 0.528401629 |
| Slain1 | 0.007091449 | 0.184919214 | 0.288106507 | 0.086734064 | 0.077133117 |
| Slain2 | 0.023572285 | 0.007587644 | 0.020593824 | 0.319250871 | 0.290970198 |

| | | | | | |
|---|---|---|---|---|---|
| Slbp | -0.01152349 | 0.103717173 | 0.182142044 | 0.148789507 | 0.156299959 |
| Slc10a3 | 0.007291568 | 0.063529198 | 0.122254723 | 0.04460187 | 0.039353228 |
| Slc10a7 | 0.004403444 | 0.379443814 | 0.500936272 | 0.071952272 | 0.068319883 |
| Slc11a2 | 0.018630488 | 0.001405928 | 0.004815705 | 0.106097318 | 0.092218615 |
| Slc12a2 | -0.000401028 | 0.937075882 | 0.957160299 | 0.067974398 | 0.067913851 |
| Slc12a6 | -0.002151773 | 0.730502656 | 0.806031872 | 0.115043933 | 0.112740429 |
| Slc12a7 | 0.026751539 | 0.000220194 | 0.000933998 | 0.179399261 | 0.144439384 |
| Slc12a9 | 0.001440338 | 0.713773648 | 0.794152828 | 0.042108369 | 0.040144837 |
| Slc15a4 | 0.007421959 | 0.097221766 | 0.172385557 | 0.0581638 | 0.051094928 |
| Slc16a10 | 0.073453435 | 1.12E-13 | 2.02E-12 | 0.49800515 | 0.416571971 |
| Slc16a11 | -0.000245156 | 0.963014784 | 0.974542493 | 0.081850851 | 0.077575877 |
| Slc16a12 | 0.046185196 | 2.17E-07 | 1.72E-06 | 0.319839729 | 0.269171964 |
| Slc16a6 | -0.012052491 | 0.104873902 | 0.183675747 | 0.135521898 | 0.161029393 |
| Slc16a9 | 0.038112579 | 1.13E-10 | 1.45E-09 | 0.126529719 | 0.084033909 |
| Slc17a5 | 0.009245592 | 0.151764019 | 0.246730735 | 0.129979493 | 0.120813505 |
| Slc1a4 | 0.016850806 | 0.012604068 | 0.031468093 | 0.143453562 | 0.13113633 |
| Slc20a1 | -0.014955018 | 0.003925392 | 0.011735136 | 0.063234449 | 0.076117409 |
| Slc20a2 | 0.006694217 | 0.107233217 | 0.187234435 | 0.05001872 | 0.044019046 |
| Slc22a15 | -0.001153462 | 0.797435038 | 0.858450165 | 0.053358744 | 0.053562585 |
| Slc22a17 | 0.019692944 | 0.006077531 | 0.017057089 | 0.174538226 | 0.154581579 |
| Slc22a23 | 0.01865587 | 0.029812001 | 0.065101588 | 0.260827986 | 0.244113565 |
| Slc23a2 | 0.01535795 | 0.019000314 | 0.044506404 | 0.119309381 | 0.118469939 |
| Slc24a2 | 0.045963253 | 4.39E-06 | 2.73E-05 | 0.491774447 | 0.429375761 |
| Slc24a5 | 0.009221855 | 0.083190771 | 0.152198095 | 0.096582486 | 0.076506456 |
| Slc25a1 | 0.082426408 | 1.35E-16 | 2.93E-15 | 0.530341816 | 0.464384341 |
| Slc25a10 | 0.003445699 | 0.35299138 | 0.475316093 | 0.037290739 | 0.035674771 |
| Slc25a11 | -0.015849472 | 0.066447882 | 0.127102912 | 0.293227621 | 0.293308674 |
| Slc25a12 | 0.001085389 | 0.888971865 | 0.924241736 | 0.205708032 | 0.203729386 |
| Slc25a14 | -0.005988554 | 0.372909964 | 0.495074515 | 0.143501688 | 0.141999958 |
| Slc25a16 | 0.0023266 | 0.750262193 | 0.822049932 | 0.174334557 | 0.170221387 |
| Slc25a17 | -0.018555106 | 0.022933256 | 0.052117138 | 0.23591554 | 0.242572713 |
| Slc25a19 | 0.002201858 | 0.657752329 | 0.749580136 | 0.073962703 | 0.068419741 |
| Slc25a20 | -0.007478752 | 0.122985253 | 0.208376199 | 0.064945644 | 0.067112569 |
| Slc25a22 | -0.003303291 | 0.547401143 | 0.657178798 | 0.083324729 | 0.085638877 |
| Slc25a23 | 0.016381286 | 0.025071248 | 0.056369722 | 0.177058935 | 0.161772046 |
| Slc25a25 | 0.029770386 | 2.79E-06 | 1.80E-05 | 0.124944161 | 0.10879403 |
| Slc25a27 | -0.010700348 | 0.047647842 | 0.096066429 | 0.079481426 | 0.084219706 |
| Slc25a28 | -0.014956733 | 0.103897262 | 0.182352841 | 0.346355571 | 0.365404337 |
| Slc25a29 | 0.002299768 | 0.701937328 | 0.784484964 | 0.103304972 | 0.102022999 |
| Slc25a3 | -0.0510529 | 1.71E-10 | 2.14E-09 | 1.67905772 | 1.757029525 |
| Slc25a32 | 0.01384544 | 0.016185282 | 0.038997104 | 0.099663169 | 0.089369452 |
| Slc25a35 | -0.006994633 | 0.092796462 | 0.166022673 | 0.043055175 | 0.048519112 |
| Slc25a36 | 0.025324131 | 0.005526322 | 0.015691269 | 0.350824567 | 0.322336372 |
| Slc25a37 | -0.001630651 | 0.70677536 | 0.788351314 | 0.046568893 | 0.049530561 |
| Slc25a38 | -0.003151965 | 0.488083807 | 0.603631951 | 0.05560253 | 0.056580771 |
| Slc25a39 | -0.052435448 | 1.03E-07 | 8.68E-07 | 0.911079857 | 0.945604208 |
| Slc25a4 | 0.047400631 | 6.36E-08 | 5.55E-07 | 1.35791761 | 1.259263336 |
| Slc25a40 | 0.033602227 | 4.16E-10 | 4.98E-09 | 0.095224582 | 0.071394515 |
| Slc25a42 | 0.007251938 | 0.126915627 | 0.2140284 | 0.064675484 | 0.058950743 |
| Slc25a44 | 0.030712427 | 0.000138841 | 0.000626946 | 0.245861601 | 0.208843024 |
| Slc25a46 | 0.014406017 | 0.075981731 | 0.141661966 | 0.238081296 | 0.225257854 |
| Slc25a5 | -0.052540162 | 1.47E-07 | 1.21E-06 | 0.935825587 | 0.988034248 |
| Slc25a51 | 0.030410167 | 0.001546643 | 0.005210141 | 0.455518617 | 0.423355148 |
| Slc25a53 | 0.003243723 | 0.470940501 | 0.58817945 | 0.054596225 | 0.052512281 |
| Slc26a2 | 0.014974483 | 0.002078634 | 0.006751914 | 0.072924758 | 0.059051725 |
| Slc27a1 | -0.000783329 | 0.862244722 | 0.90575291 | 0.055699007 | 0.054389893 |
| Slc27a4 | 0.000167802 | 0.97633305 | 0.98412836 | 0.091675378 | 0.090286631 |
| Slc29a1 | 0.006734967 | 0.432820692 | 0.551275091 | 0.299807895 | 0.266650225 |
| Slc29a3 | 0.002199586 | 0.564474401 | 0.672197468 | 0.038222454 | 0.037790975 |
| Slc29a4 | 0.024193961 | 5.58E-08 | 4.91E-07 | 0.067662589 | 0.045058216 |
| Slc2a13 | 0.013872446 | 9.78E-05 | 0.000457469 | 0.039776757 | 0.028517187 |
| Slc2a2 | 0.272128857 | 2.91E-74 | 9.76E-72 | 1.34325911 | 0.826565295 |
| Slc2a5 | 0.024368666 | 0.014285731 | 0.035045642 | 0.983460346 | 0.900238353 |
| Slc2a8 | -0.000304263 | 0.947204728 | 0.964799339 | 0.063312386 | 0.057844 |
| Slc30a1 | 0.010903033 | 0.029179089 | 0.06393489 | 0.069150768 | 0.06187743 |
| Slc30a4 | 0.006364251 | 0.311081585 | 0.432480362 | 0.116022014 | 0.113556674 |
| Slc30a5 | -0.007247217 | 0.351899258 | 0.474371129 | 0.200334532 | 0.201985371 |
| Slc30a6 | -0.003568849 | 0.408010431 | 0.528186073 | 0.048560776 | 0.051696196 |
| Slc30a7 | 0.001328377 | 0.839507272 | 0.889945448 | 0.130125391 | 0.129185248 |
| Slc30a8 | 0.083240628 | 4.47E-15 | 8.69E-14 | 1.78845903 | 1.487983685 |
| Slc30a9 | -0.03222538 | 0.000576383 | 0.002204404 | 0.350685209 | 0.383291042 |
| Slc31a1 | 0.015671491 | 0.024678934 | 0.055654974 | 0.156113686 | 0.144548669 |

| | | | | | |
|---|---|---|---|---|---|
| Slc31a2 | 0.008658466 | 0.188697885 | 0.292798175 | 0.149944708 | 0.129110301 |
| Slc33a1 | -0.056914634 | 1.00E-09 | 1.14E-08 | 0.330801423 | 0.380868605 |
| Slc35a1 | -0.009433739 | 0.180116771 | 0.282369647 | 0.154654387 | 0.160593632 |
| Slc35a2 | 0.003190472 | 0.640910731 | 0.736074138 | 0.142035645 | 0.138794596 |
| Slc35a3 | 0.006141944 | 0.276052915 | 0.394094926 | 0.092543861 | 0.089250014 |
| Slc35a4 | -0.001115511 | 0.867544491 | 0.909630958 | 0.137484624 | 0.134173083 |
| Slc35a5 | 0.010079238 | 0.11814048 | 0.201820725 | 0.132749914 | 0.121945099 |
| Slc35b1 | -0.044294577 | 1.77E-05 | 9.71E-05 | 0.528687374 | 0.588877196 |
| Slc35b2 | 0.033686775 | 7.90E-06 | 4.65E-05 | 0.211592607 | 0.17812289 |
| Slc35b3 | -0.004739642 | 0.398974946 | 0.519115956 | 0.088406775 | 0.091728654 |
| Slc35b4 | 0.039609187 | 1.61E-07 | 1.31E-06 | 0.208319056 | 0.159274132 |
| Slc35c1 | 0.001734201 | 0.717158879 | 0.79673393 | 0.063405679 | 0.062594537 |
| Slc35c2 | -0.031248121 | 1.00E-05 | 5.75E-05 | 0.147179862 | 0.166580498 |
| Slc35d1 | 0.002895426 | 0.508685885 | 0.622695061 | 0.051818064 | 0.050650956 |
| Slc35d3 | 0.019826978 | 0.001247563 | 0.004334813 | 0.118501465 | 0.099581046 |
| Slc35e1 | 0.020487776 | 0.000910644 | 0.00330223 | 0.120767854 | 0.106601658 |
| Slc35e2 | 0.008307412 | 0.071946479 | 0.135240333 | 0.062121752 | 0.054910755 |
| Slc35e4 | 0.014085178 | 0.028587827 | 0.062837635 | 0.129388283 | 0.11785174 |
| Slc35f5 | -0.007070666 | 0.429786876 | 0.548416854 | 0.315205049 | 0.318509859 |
| Slc35f6 | 0.003104514 | 0.408204254 | 0.528256662 | 0.040299628 | 0.035986689 |
| Slc35g1 | 0.0226201 | 0.00017691 | 0.000773343 | 0.117906305 | 0.094113134 |
| Slc35g2 | 0.00341976 | 0.701510599 | 0.784188346 | 0.300743327 | 0.298808451 |
| Slc36a1 | 0.018155775 | 0.005094589 | 0.014662403 | 0.13963307 | 0.118111313 |
| Slc36a4 | 0.004329768 | 0.476203412 | 0.593012626 | 0.108601571 | 0.103145751 |
| Slc37a1 | 0.001863304 | 0.791663897 | 0.854128999 | 0.14191115 | 0.143294045 |
| Slc37a3 | 0.01613124 | 0.012359788 | 0.031001511 | 0.133430412 | 0.120630857 |
| Slc37a4 | 0.006248961 | 0.390367594 | 0.511407324 | 0.185611696 | 0.163614558 |
| Slc38a10 | 0.008624571 | 0.388954016 | 0.510036545 | 0.553183592 | 0.529620042 |
| Slc38a2 | -0.014643103 | 0.163833409 | 0.261630211 | 0.71134078 | 0.733827068 |
| Slc38a4 | 0.023334837 | 0.027527468 | 0.060809372 | 0.59268187 | 0.57143164 |
| Slc38a6 | -0.000254771 | 0.956668483 | 0.970443551 | 0.05672777 | 0.058406726 |
| Slc38a7 | -0.003436771 | 0.521396171 | 0.633718809 | 0.075848307 | 0.079751996 |
| Slc38a9 | -0.005537983 | 0.242752225 | 0.356608814 | 0.058382396 | 0.060880203 |
| Slc39a1 | 0.033267439 | 6.93E-07 | 5.02E-06 | 0.156792288 | 0.124856125 |
| Slc39a10 | -0.007237936 | 0.294890098 | 0.414594582 | 0.139915868 | 0.141491925 |
| Slc39a13 | 0.003666089 | 0.560686165 | 0.669079525 | 0.120496469 | 0.115377699 |
| Slc39a14 | 0.001659773 | 0.722417836 | 0.79997774 | 0.059419847 | 0.05893217 |
| Slc39a3 | 0.019634775 | 0.001924219 | 0.006317886 | 0.134922592 | 0.114041942 |
| Slc39a6 | -0.00615853 | 0.424311041 | 0.54271174 | 0.186647523 | 0.194256223 |
| Slc39a9 | 0.01630062 | 0.034761215 | 0.073820126 | 0.206752986 | 0.190285926 |
| Slc3a2 | -0.03699207 | 0.00032973 | 0.00134068 | 1.105147212 | 1.209968508 |
| Slc40a1 | -0.012930232 | 0.106019981 | 0.185349084 | 0.180585408 | 0.184367355 |
| Slc41a1 | 0.001773956 | 0.735642034 | 0.810530852 | 0.074685423 | 0.07617882 |
| Slc41a3 | -0.006560361 | 0.083659335 | 0.152853985 | 0.036679737 | 0.038786812 |
| Slc43a2 | 0.025353813 | 0.013704988 | 0.033825817 | 0.462983802 | 0.410625189 |
| Slc44a1 | 0.030178745 | 0.000188809 | 0.000816186 | 0.242369025 | 0.208834837 |
| Slc44a3 | 0.002735351 | 0.744925819 | 0.817769739 | 0.260666456 | 0.248873945 |
| Slc45a4 | 0.00390123 | 0.479844521 | 0.596301111 | 0.089441969 | 0.083720346 |
| Slc46a3 | 0.005496419 | 0.116063489 | 0.199112272 | 0.036231825 | 0.030504257 |
| Slc48a1 | -0.016847055 | 0.045489562 | 0.09221216 | 0.257066085 | 0.260415133 |
| Slc49a4 | 0.034567625 | 1.26E-09 | 1.41E-08 | 0.10906308 | 0.078417563 |
| Slc4a10 | 0.035638093 | 4.48E-07 | 3.37E-06 | 0.162581228 | 0.139146712 |
| Slc4a1ap | 0.0200936 | 0.001789929 | 0.005929026 | 0.13194151 | 0.115633675 |
| Slc4a2 | 0.001070116 | 0.867436743 | 0.909630958 | 0.128357687 | 0.121027803 |
| Slc4a3 | -0.020222972 | 0.001400622 | 0.004804816 | 0.106233846 | 0.120332723 |
| Slc4a4 | -0.000151505 | 0.976816648 | 0.984410849 | 0.068632127 | 0.070653645 |
| Slc4a7 | 0.052196068 | 1.61E-06 | 1.08E-05 | 0.791791399 | 0.738855565 |
| Slc4a8 | -0.000628889 | 0.913943342 | 0.940747186 | 0.097327832 | 0.096723622 |
| Slc50a1 | -0.070420128 | 8.41E-12 | 1.21E-10 | 0.599699569 | 0.680633623 |
| Slc52a2 | -0.009065556 | 0.039869043 | 0.082733583 | 0.048806418 | 0.055366245 |
| Slc5a6 | 0.011632546 | 0.012009429 | 0.030301787 | 0.065677982 | 0.055279235 |
| Slc66a2 | -0.008099719 | 0.247755223 | 0.362098058 | 0.147223922 | 0.154371113 |
| Slc6a17 | -0.017813242 | 0.033524884 | 0.071632731 | 0.213847683 | 0.239939254 |
| Slc6a19 | 0.01581309 | 0.001125293 | 0.003965206 | 0.072082385 | 0.056653723 |
| Slc6a6 | 0.076934847 | 2.96E-13 | 5.08E-12 | 0.828086466 | 0.723606442 |
| Slc6a8 | 0.004873623 | 0.421499139 | 0.54060879 | 0.101947141 | 0.099671727 |
| Slc6a9 | 0.003507098 | 0.448593724 | 0.567498788 | 0.059967078 | 0.056935861 |
| Slc7a1 | 0.029854966 | 0.001204694 | 0.004205397 | 0.318193659 | 0.298626059 |
| Slc7a11 | -0.019098672 | 0.000539307 | 0.002076491 | 0.065353448 | 0.084774249 |
| Slc7a14 | 0.132749427 | 6.56E-36 | 4.40E-34 | 0.835520391 | 0.697079868 |
| Slc7a2 | 0.047235017 | 6.51E-07 | 4.75E-06 | 0.393333436 | 0.312047022 |
| Slc7a4 | 0.00211861 | 0.705645645 | 0.787542987 | 0.095875788 | 0.088531224 |

| | | | | | |
|---|---|---|---|---|---|
| Slc7a5 | 0.117708957 | 3.52E-36 | 2.44E-34 | 0.42058996 | 0.318809028 |
| Slc7a6 | -0.002509643 | 0.5500325 | 0.659279492 | 0.04561005 | 0.048199861 |
| Slc7a6os | -0.008164565 | 0.138335447 | 0.229700635 | 0.076725984 | 0.087292857 |
| Slc7a8 | 0.011219739 | 0.18054961 | 0.282748601 | 0.252723296 | 0.24232389 |
| Slc8a1 | -0.029252085 | 0.002229107 | 0.007175953 | 0.335787312 | 0.363529144 |
| Slc9a1 | 0.003738315 | 0.412011868 | 0.531667027 | 0.059292252 | 0.055429141 |
| Slc9a3r1 | 0.01520921 | 0.005162106 | 0.014834145 | 0.095360829 | 0.076163989 |
| Slc9a3r2 | -0.009150427 | 0.04243774 | 0.086987077 | 0.052043156 | 0.057510881 |
| Slc9a6 | 0.010479084 | 0.058409109 | 0.113981003 | 0.090868696 | 0.084048052 |
| Slc9a7 | 0.010083142 | 0.031046425 | 0.067298404 | 0.067095319 | 0.055591008 |
| Slc9a8 | 0.008316222 | 0.181807424 | 0.283940847 | 0.122810779 | 0.113243164 |
| Slc9a9 | -0.011678645 | 0.026708806 | 0.05927026 | 0.0725953 | 0.080376079 |
| Slco1a6 | -0.018208036 | 0.002555295 | 0.008078886 | 0.085380256 | 0.098034519 |
| Slco2a1 | 0.027368217 | 3.89E-10 | 4.67E-09 | 0.067778133 | 0.042075302 |
| Slco3a1 | 0.030859213 | 3.46E-06 | 2.19E-05 | 0.150944698 | 0.122382667 |
| Slf1 | -0.007914295 | 0.177373971 | 0.278857678 | 0.097679128 | 0.101323555 |
| Slf2 | -0.002307247 | 0.737517847 | 0.812021349 | 0.139421147 | 0.142698349 |
| Slirp | 0.053795445 | 2.27E-08 | 2.11E-07 | 0.462150145 | 0.444945935 |
| Slitrk6 | 0.097896776 | 9.27E-45 | 9.01E-43 | 0.195393362 | 0.10558334 |
| Slk | 0.00781376 | 0.391446476 | 0.512075844 | 0.316785285 | 0.305968759 |
| Slmap | 0.005984625 | 0.402296627 | 0.522249674 | 0.165623562 | 0.156528925 |
| Sltm | 0.070056992 | 3.61E-11 | 4.88E-10 | 0.618671875 | 0.550285914 |
| Slu7 | -0.014322303 | 0.089060664 | 0.160775642 | 0.25192905 | 0.256298281 |
| Slx1b | 6.15E-05 | 0.990074779 | 0.993322992 | 0.069841176 | 0.066319879 |
| Slx4ip | -0.005029891 | 0.318853923 | 0.440576726 | 0.066593902 | 0.070326296 |
| Smad1 | 0.043369542 | 1.42E-06 | 9.66E-06 | 0.331469415 | 0.294406444 |
| Smad2 | 0.01458739 | 0.021300973 | 0.049048964 | 0.129086618 | 0.113320535 |
| Smad4 | 0.040577404 | 8.58E-06 | 5.00E-05 | 0.381030657 | 0.332658246 |
| Smad5 | 0.033827613 | 4.34E-06 | 2.71E-05 | 0.193222222 | 0.159349237 |
| Smad7 | 0.000116419 | 0.988288807 | 0.992977827 | 0.188260184 | 0.185160867 |
| Smap1 | 0.00027802 | 0.977329325 | 0.984825424 | 0.507539393 | 0.501737402 |
| Smap2 | 0.135533022 | 9.40E-39 | 7.55E-37 | 0.981230735 | 0.812993001 |
| Smarca1 | 0.010015988 | 0.03610345 | 0.076138338 | 0.067286542 | 0.055727436 |
| Smarca2 | 0.02990297 | 0.002231786 | 0.007182198 | 0.431620559 | 0.404082523 |
| Smarca4 | 0.059980331 | 3.41E-09 | 3.61E-08 | 0.668026789 | 0.593136171 |
| Smarca5 | 0.09005585 | 2.37E-18 | 5.99E-17 | 0.730969123 | 0.631353545 |
| Smarcad1 | 0.017194513 | 0.025173934 | 0.056522167 | 0.201931615 | 0.184778447 |
| Smarcal1 | 0.004006827 | 0.475876509 | 0.592681453 | 0.096728175 | 0.088631359 |
| Smarcb1 | 0.000148263 | 0.985881682 | 0.991490816 | 0.250011413 | 0.247692569 |
| Smarcc1 | -0.007484884 | 0.390303086 | 0.511391726 | 0.275739905 | 0.27350245 |
| Smarcc2 | 0.000577277 | 0.953332266 | 0.968581933 | 0.500507303 | 0.492822756 |
| Smarcd1 | 0.02282091 | 0.002320592 | 0.007438443 | 0.188965122 | 0.175537811 |
| Smarcd2 | -0.008373325 | 0.270836148 | 0.387729205 | 0.192858271 | 0.19069253 |
| Smarce1 | 0.045130522 | 5.52E-09 | 5.67E-08 | 0.229907801 | 0.18779683 |
| Smc1a | 0.004100866 | 0.663420028 | 0.753729373 | 0.358062849 | 0.350607547 |
| Smc3 | 0.025375918 | 0.00873617 | 0.023187741 | 0.473918589 | 0.432621039 |
| Smc4 | 0.002753991 | 0.657953043 | 0.749690707 | 0.110491315 | 0.104459024 |
| Smc5 | -0.001589121 | 0.81413773 | 0.871797226 | 0.135614925 | 0.134285785 |
| Smc6 | 0.013381241 | 0.187564114 | 0.29129366 | 0.553812659 | 0.534746424 |
| Smchd1 | -0.003299658 | 0.71074363 | 0.791687622 | 0.299587073 | 0.29495972 |
| Smcr8 | 0.010782914 | 0.018023473 | 0.042675159 | 0.061654593 | 0.053163048 |
| Smdt1 | 0.045943569 | 6.81E-10 | 7.93E-09 | 1.649879795 | 1.578211623 |
| Smg1 | 0.022888041 | 0.009563832 | 0.024954259 | 0.285553396 | 0.263820449 |
| Smg5 | 0.015578998 | 0.009770318 | 0.025377247 | 0.112490278 | 0.098853102 |
| Smg6 | 0.013875748 | 0.062432822 | 0.120574473 | 0.18091496 | 0.169159869 |
| Smg7 | 0.026792453 | 0.001153022 | 0.004048279 | 0.257180648 | 0.227506513 |
| Smg9 | 0.001227397 | 0.811894618 | 0.870449876 | 0.08000006 | 0.072467269 |
| Smim1 | 0.032328652 | 0.001629595 | 0.005448047 | 0.601177227 | 0.568237158 |
| Smim10l1 | -0.013132034 | 0.190494485 | 0.29476164 | 0.838909523 | 0.842634375 |
| Smim11 | 0.000469019 | 0.960193001 | 0.972598078 | 0.427708631 | 0.433164255 |
| Smim12 | 0.017606488 | 0.001220906 | 0.004255881 | 0.099087102 | 0.081364716 |
| Smim13 | -0.003665698 | 0.64291037 | 0.737246357 | 0.203784185 | 0.212471754 |
| Smim14 | -0.081033128 | 1.18E-19 | 3.34E-18 | 1.077589726 | 1.179502937 |
| Smim15 | 0.012678606 | 0.071209093 | 0.134087725 | 0.161952487 | 0.15170144 |
| Smim19 | -0.007557425 | 0.426985519 | 0.545630023 | 0.405822041 | 0.411635072 |
| Smim20 | 0.025679347 | 0.006519407 | 0.018140147 | 0.453665162 | 0.402772459 |
| Smim22 | 0.126977823 | 5.35E-35 | 3.49E-33 | 0.733705524 | 0.623195216 |
| Smim26 | 0.019003549 | 0.056409132 | 0.110611048 | 0.589635 | 0.600152309 |
| Smim27 | 0.061009002 | 1.20E-12 | 1.92E-11 | 0.308059178 | 0.257613317 |
| Smim4 | 0.082024352 | 4.16E-17 | 9.41E-16 | 0.525871396 | 0.470462163 |
| Smim7 | -0.033191466 | 0.000717638 | 0.002678263 | 0.553233519 | 0.580776895 |
| Smim8 | -0.019293237 | 0.020291403 | 0.047014542 | 0.244032895 | 0.249134147 |

| | | | | | |
|---|---|---|---|---|---|
| Smn1 | -0.012204883 | 0.088885684 | 0.160532533 | 0.157546354 | 0.163060802 |
| Smndc1 | -0.009755298 | 0.275958241 | 0.394071597 | 0.30180395 | 0.303193359 |
| Smoc1 | -0.017796755 | 0.000149711 | 0.000666738 | 0.044880279 | 0.061011467 |
| Smpd1 | 0.012460297 | 0.140089096 | 0.232097169 | 0.277342262 | 0.260057961 |
| Smpd2 | 0.001554218 | 0.812157527 | 0.870539744 | 0.139065233 | 0.128947597 |
| Smpd3 | 0.012401963 | 0.030299471 | 0.065943913 | 0.096504612 | 0.085858042 |
| Smpd4 | 0.007827699 | 0.041448547 | 0.085500269 | 0.04295266 | 0.036606471 |
| Smpdl3a | -0.013729116 | 0.037500897 | 0.078455718 | 0.131940903 | 0.135217902 |
| Sms | 0.011317631 | 0.174934477 | 0.2756423 | 0.238218333 | 0.240516221 |
| Smu1 | 0.007818237 | 0.329852693 | 0.45249441 | 0.237431985 | 0.221617598 |
| Smug1 | 0.00509127 | 0.322018175 | 0.443624869 | 0.078042954 | 0.070810977 |
| Smurf1 | 0.000888299 | 0.819890365 | 0.876123778 | 0.039663141 | 0.039946547 |
| Smurf2 | 0.014175066 | 0.028122153 | 0.061968172 | 0.13098451 | 0.117630313 |
| Smyd2 | -0.019426903 | 0.020660868 | 0.047768124 | 0.253808221 | 0.266579553 |
| Smyd3 | 0.010543654 | 0.088810281 | 0.160455966 | 0.129198305 | 0.111399244 |
| Smyd4 | 0.009869698 | 0.079458892 | 0.146681928 | 0.09770886 | 0.08475298 |
| Snap23 | -0.007507563 | 0.296025577 | 0.415890269 | 0.163293302 | 0.165007532 |
| Snap25 | -0.072598613 | 2.95E-12 | 4.50E-11 | 0.630947038 | 0.731947563 |
| Snap29 | -0.004610627 | 0.478249066 | 0.594973471 | 0.125821906 | 0.127901651 |
| Snap47 | -0.009075979 | 0.31463649 | 0.43611291 | 0.35512742 | 0.348015491 |
| Snap91 | -0.023950899 | 3.05E-05 | 0.000158155 | 0.075380814 | 0.099287606 |
| Snapc1 | 0.017406933 | 0.023971045 | 0.054234698 | 0.203598663 | 0.188245423 |
| Snapc2 | 0.006302707 | 0.219150066 | 0.329963906 | 0.078467571 | 0.071146786 |
| Snapc3 | -0.012969841 | 0.008916361 | 0.023574887 | 0.062155496 | 0.069846649 |
| Snapc4 | 0.003749489 | 0.335628062 | 0.458377361 | 0.041934569 | 0.038803613 |
| Snapc5 | -0.01296751 | 0.105106002 | 0.184015946 | 0.217872214 | 0.22294821 |
| Snapin | -0.007902319 | 0.21373149 | 0.323508882 | 0.119608443 | 0.122142206 |
| Sncb | 0.037524534 | 1.28E-05 | 7.19E-05 | 0.298092714 | 0.239986781 |
| Snd1 | -0.041105358 | 5.64E-05 | 0.000274655 | 0.528291446 | 0.565485598 |
| Snf8 | -0.021439309 | 0.027178818 | 0.06016222 | 0.526158701 | 0.535454651 |
| Snhg1 | 0.045779001 | 3.65E-05 | 0.000186035 | 0.519727352 | 0.479760147 |
| Snhg10 | -0.001296292 | 0.758720034 | 0.828608871 | 0.048493096 | 0.048581688 |
| Snhg11 | -0.208495117 | 4.45E-62 | 8.83E-60 | 0.227338424 | 0.459646258 |
| Snhg12 | -0.101562771 | 8.27E-18 | 1.98E-16 | 0.428307699 | 0.578167896 |
| Snhg14 | -0.011798607 | 0.002394843 | 0.007643208 | 0.032462704 | 0.042512277 |
| Snhg15 | -0.021936856 | 0.00102275 | 0.003650212 | 0.100373776 | 0.12826795 |
| Snhg16 | -0.013774382 | 0.000935375 | 0.00337179 | 0.03831318 | 0.049444014 |
| Snhg17 | -0.008325906 | 0.066543767 | 0.127150558 | 0.051466346 | 0.05747916 |
| Snhg18 | -0.011384166 | 0.151843261 | 0.246735782 | 0.171758303 | 0.193904679 |
| Snhg20 | 0.000914451 | 0.910378816 | 0.938619225 | 0.235259719 | 0.228784944 |
| Snhg3 | 0.029472924 | 0.001646302 | 0.005498875 | 0.366346812 | 0.356882069 |
| Snhg4 | -0.035018844 | 1.16E-05 | 6.54E-05 | 0.184925752 | 0.211435107 |
| Snhg5 | -0.010433154 | 0.171274195 | 0.270708459 | 0.146599405 | 0.166595848 |
| Snhg6 | 0.062264733 | 1.64E-12 | 2.59E-11 | 0.280188777 | 0.256165027 |
| Snhg8 | 0.019535607 | 0.066484933 | 0.127112393 | 0.607022732 | 0.645962625 |
| Snhg9 | -0.009173795 | 0.195365815 | 0.301053488 | 0.126089691 | 0.161240695 |
| Snip1 | 0.004283394 | 0.433124159 | 0.551517102 | 0.088137506 | 0.082949111 |
| Snph | 0.019946194 | 0.001567283 | 0.005268714 | 0.132906253 | 0.111926405 |
| Snrk | 0.027556777 | 3.54E-05 | 0.000180852 | 0.15358901 | 0.127041323 |
| Snrnp200 | 0.06244894 | 4.87E-12 | 7.23E-11 | 0.354118818 | 0.290058306 |
| Snrnp25 | -0.01099058 | 0.030471269 | 0.066199257 | 0.069523948 | 0.075948685 |
| Snrnp27 | 0.011406945 | 0.237554917 | 0.351469928 | 0.466934283 | 0.451863832 |
| Snrnp35 | -0.003592736 | 0.501271751 | 0.615556899 | 0.077120541 | 0.079062337 |
| Snrnp40 | -0.005842802 | 0.365921302 | 0.488665782 | 0.13111544 | 0.128169354 |
| Snrnp48 | 0.024503951 | 0.004885034 | 0.014138822 | 0.288189669 | 0.262404092 |
| Snrnp70 | -0.079737338 | 9.36E-15 | 1.80E-13 | 0.627572224 | 0.728348999 |
| Snrpa | -0.033787456 | 1.44E-05 | 8.02E-05 | 0.196051919 | 0.213809725 |
| Snrpa1 | 0.020027221 | 0.010437246 | 0.026829958 | 0.20097205 | 0.186592281 |
| Snrpb | -0.018386632 | 0.063244237 | 0.121851065 | 0.585098636 | 0.591646084 |
| Snrpb2 | 0.03243756 | 2.56E-05 | 0.000134631 | 0.216005874 | 0.187499325 |
| Snrpc | -0.006876998 | 0.414584055 | 0.53399393 | 0.259779869 | 0.252539199 |
| Snrpd1 | 0.035398866 | 0.00030246 | 0.001241148 | 0.499560471 | 0.471832461 |
| Snrpd2 | 0.060472113 | 9.30E-10 | 1.07E-08 | 0.690814586 | 0.639290249 |
| Snrpd3 | 0.017275197 | 0.066579503 | 0.127163044 | 0.428173877 | 0.410226167 |
| Snrpe | 0.036718647 | 0.000283598 | 0.001170299 | 0.663575658 | 0.642375265 |
| Snrpf | 0.122363688 | 7.76E-36 | 5.17E-34 | 0.562231691 | 0.469164708 |
| Snrpg | 0.13261168 | 6.12E-41 | 5.41E-39 | 0.697635766 | 0.594277372 |
| Snrpn | -0.049253807 | 2.12E-09 | 2.30E-08 | 0.245941907 | 0.261026829 |
| Sntb2 | 0.014694512 | 0.022196643 | 0.050715808 | 0.13306027 | 0.115336798 |
| Sntg1 | 0.00355045 | 0.546750192 | 0.656478371 | 0.098356917 | 0.094228657 |
| Snu13 | -0.009944499 | 0.308993068 | 0.430253596 | 0.506397892 | 0.497728998 |
| Snupn | -0.002041957 | 0.696395497 | 0.780172807 | 0.075367414 | 0.076844512 |

| | | | | | |
|---|---|---|---|---|---|
| Snw1 | 0.012838654 | 0.184191028 | 0.287202113 | 0.477290305 | 0.450253416 |
| Snx1 | 0.003307863 | 0.651489575 | 0.744362633 | 0.18746706 | 0.174794743 |
| Snx10 | -0.022438464 | 0.035624476 | 0.075291555 | 0.498343385 | 0.548152929 |
| Snx11 | -0.012399538 | 0.005513228 | 0.015661898 | 0.049849974 | 0.057768651 |
| Snx12 | -0.025494548 | 0.000169659 | 0.000745333 | 0.131430258 | 0.149767464 |
| Snx13 | 0.027523555 | 0.000187111 | 0.000809924 | 0.186101713 | 0.162161899 |
| Snx14 | -0.013158321 | 0.071974232 | 0.135240333 | 0.163152393 | 0.174286569 |
| Snx15 | -0.022099381 | 0.004224156 | 0.012478517 | 0.198781183 | 0.207424204 |
| Snx16 | -0.005759833 | 0.392089493 | 0.51276487 | 0.13520345 | 0.137149553 |
| Snx17 | -0.03009832 | 0.000725126 | 0.00270087 | 0.31617498 | 0.332715527 |
| Snx18 | 0.02223641 | 2.76E-05 | 0.000144338 | 0.093317471 | 0.07234627 |
| Snx19 | 0.004778218 | 0.427413442 | 0.54603331 | 0.1049013 | 0.10099289 |
| Snx2 | 0.004569709 | 0.612209443 | 0.712783653 | 0.360806589 | 0.342141037 |
| Snx21 | -0.010141532 | 0.116819741 | 0.200056635 | 0.131686292 | 0.130094872 |
| Snx24 | 0.011226095 | 0.100100949 | 0.176877757 | 0.157013575 | 0.144178485 |
| Snx25 | 0.006308025 | 0.199465003 | 0.306378873 | 0.075751376 | 0.064288825 |
| Snx27 | 0.020450845 | 0.009304042 | 0.024414006 | 0.227702068 | 0.201508649 |
| Snx3 | -0.003221529 | 0.740183788 | 0.814133605 | 0.809724965 | 0.789904771 |
| Snx30 | 0.029800486 | 0.000219258 | 0.000930436 | 0.237837521 | 0.203584977 |
| Snx32 | -0.022421071 | 0.000120199 | 0.000551217 | 0.084129486 | 0.101814291 |
| Snx4 | 0.008619383 | 0.293679383 | 0.413191167 | 0.253964451 | 0.234742996 |
| Snx5 | 0.015267868 | 0.109600248 | 0.190409867 | 0.4670224 | 0.425196301 |
| Snx6 | 0.011567874 | 0.23005297 | 0.343032972 | 0.453489236 | 0.435513672 |
| Snx7 | -0.011039076 | 0.00964008 | 0.0251195 | 0.046189246 | 0.052146377 |
| Snx8 | -0.000552647 | 0.917107199 | 0.942575366 | 0.083011158 | 0.079392296 |
| Snx9 | 0.005887399 | 0.418781805 | 0.537904176 | 0.176678511 | 0.16586015 |
| Soat1 | -0.001862628 | 0.808432654 | 0.867503561 | 0.178445521 | 0.18295916 |
| Sobp | 0.058720364 | 2.46E-11 | 3.39E-10 | 0.312454191 | 0.256323632 |
| Socs2 | 0.017978699 | 0.041547311 | 0.085667647 | 0.312899235 | 0.272591757 |
| Socs3 | -0.014277938 | 0.050655483 | 0.101061484 | 0.124232115 | 0.146504718 |
| Socs4 | 0.010815686 | 0.123488115 | 0.209118874 | 0.153805226 | 0.146778501 |
| Socs5 | 0.003179453 | 0.600894569 | 0.70404388 | 0.108565045 | 0.103447096 |
| Socs6 | 0.011313866 | 0.076019475 | 0.141691207 | 0.124710165 | 0.117015527 |
| Socs7 | 0.009910782 | 0.172186442 | 0.271973451 | 0.17391457 | 0.161969198 |
| Sod1 | -0.032286887 | 0.001491068 | 0.005057979 | 0.811800983 | 0.8947192 |
| Sod2 | -0.066384824 | 2.66E-11 | 3.64E-10 | 0.722617065 | 0.787201445 |
| Soga3 | 0.020743783 | 0.013465711 | 0.033370798 | 0.26495323 | 0.237524282 |
| Son | 0.028946779 | 0.000594445 | 0.002264576 | 1.41846592 | 1.416478079 |
| Sorbs2 | 0.048250945 | 6.20E-06 | 3.73E-05 | 0.667379739 | 0.612828772 |
| Sord | 0.001595276 | 0.716832297 | 0.796462125 | 0.058893064 | 0.051925887 |
| Sorl1 | 0.022529018 | 0.015657653 | 0.03788544 | 0.28997763 | 0.28783074 |
| Sort1 | 0.003616992 | 0.688433794 | 0.773801985 | 0.327734096 | 0.317283472 |
| Sos1 | 0.020050373 | 0.007847754 | 0.021199183 | 0.191856852 | 0.173797434 |
| Sos2 | 0.015408312 | 0.041076358 | 0.084804492 | 0.185077801 | 0.173937797 |
| Sowaha | 0.002198138 | 0.610589817 | 0.711427876 | 0.053239313 | 0.048313299 |
| Sowahb | 0.007086833 | 0.051699726 | 0.102576477 | 0.038601123 | 0.033044556 |
| Sowahc | 0.009102358 | 0.369966679 | 0.492309889 | 0.364465675 | 0.385407035 |
| Sox12 | -0.002525311 | 0.671397564 | 0.760317661 | 0.099247872 | 0.101461743 |
| Sox4 | 0.081651071 | 7.76E-18 | 1.88E-16 | 0.353657791 | 0.259552593 |
| Sp1 | 0.007902486 | 0.276200204 | 0.394247304 | 0.177183245 | 0.162418011 |
| Sp2 | 0.004767228 | 0.251906989 | 0.366842383 | 0.051540025 | 0.044338383 |
| Sp3 | 0.024109524 | 0.003727364 | 0.011219018 | 0.259398533 | 0.22662638 |
| Sp3os | -0.018944172 | 0.017050082 | 0.040747518 | 0.191638259 | 0.212272924 |
| Sp4 | 0.004098057 | 0.517114283 | 0.629840273 | 0.121217834 | 0.117235456 |
| Spa17 | -0.005128138 | 0.506053209 | 0.61999917 | 0.202773714 | 0.196912179 |
| Spaca6 | -0.005823167 | 0.285067367 | 0.404292478 | 0.076008128 | 0.083677017 |
| Spag7 | 0.017236812 | 0.04362613 | 0.089066198 | 0.290575286 | 0.264307545 |
| Spag9 | 0.047592149 | 1.92E-06 | 1.27E-05 | 0.481243852 | 0.432922553 |
| Spast | 0.009503893 | 0.213157971 | 0.322992798 | 0.195076544 | 0.183956393 |
| Spata13 | 0.012327131 | 0.000697986 | 0.002609931 | 0.040579545 | 0.031499384 |
| Spata2 | 0.009535367 | 0.048386023 | 0.097372991 | 0.06564389 | 0.060426083 |
| Spata24 | -0.018893437 | 0.000145539 | 0.000652043 | 0.062113682 | 0.072675022 |
| Spata5 | 0.014296263 | 0.009106331 | 0.023966363 | 0.091272824 | 0.078233781 |
| SPATA5L1 | 0.007568115 | 0.03096211 | 0.067177535 | 0.037142426 | 0.029749005 |
| Spata6 | -0.006530593 | 0.147747435 | 0.241940469 | 0.052330234 | 0.056609591 |
| Spata7 | -0.007868432 | 0.207745395 | 0.316143522 | 0.109819201 | 0.112404603 |
| Spats2 | -0.004020793 | 0.429280242 | 0.548058111 | 0.069042729 | 0.072077553 |
| Spc24 | -0.014088525 | 0.000421469 | 0.001666335 | 0.0325241 | 0.045376364 |
| Spc25 | 0.061061888 | 5.21E-07 | 3.89E-06 | 0.847411668 | 0.760739474 |
| Spcs1 | -0.000986343 | 0.915287235 | 0.941432766 | 1.356592613 | 1.346375937 |
| Spcs2 | -0.167552942 | 2.71E-85 | 1.39E-82 | 1.348430101 | 1.615925781 |
| Spcs3 | -0.012857448 | 0.199483467 | 0.306378873 | 0.701531036 | 0.728454307 |

| | | | | | |
|---|---|---|---|---|---|
| Specc1 | 0.022244247 | 2.88E-09 | 3.08E-08 | 0.049517231 | 0.030353309 |
| Specc1l | 0.030414959 | 0.000750848 | 0.002782976 | 0.338734677 | 0.299837437 |
| Spen | 0.026069029 | 0.004615831 | 0.013476009 | 0.33140131 | 0.301595599 |
| Spg11 | -0.000346857 | 0.93294564 | 0.954747107 | 0.04593778 | 0.045117189 |
| Spg20 | -0.003290154 | 0.604417997 | 0.706992207 | 0.120607073 | 0.121296967 |
| Spg21 | -0.013336756 | 0.116335087 | 0.199402277 | 0.266362588 | 0.268576817 |
| Spg7 | -0.01091113 | 0.146493164 | 0.240453578 | 0.183698842 | 0.188990399 |
| Sphk2 | -0.004568568 | 0.517923388 | 0.630667733 | 0.152034816 | 0.155082386 |
| Sphkap | 0.074995914 | 1.37E-12 | 2.17E-11 | 1.250018956 | 1.123795187 |
| Spice1 | 0.009991647 | 0.025592331 | 0.057236862 | 0.059801371 | 0.050207343 |
| Spin1 | 0.023468756 | 3.53E-05 | 0.000180234 | 0.107042968 | 0.084843482 |
| Spindoc | 0.002859934 | 0.488063121 | 0.603631951 | 0.049144893 | 0.044257795 |
| Spint1 | -0.000550713 | 0.955834807 | 0.970103976 | 0.597292928 | 0.583280721 |
| Spint2 | -0.069125411 | 1.42E-16 | 3.08E-15 | 1.285183421 | 1.347804147 |
| Spire1 | 0.013204107 | 0.069797883 | 0.132038904 | 0.169277567 | 0.160167417 |
| Spire2 | -0.002311779 | 0.557537903 | 0.666140284 | 0.042912067 | 0.041621199 |
| Spns1 | -0.001201093 | 0.845294366 | 0.894324935 | 0.112785558 | 0.109828316 |
| Spock1 | -0.093119438 | 2.99E-32 | 1.73E-30 | 0.139034 | 0.214554369 |
| Spock2 | 0.057972418 | 3.58E-08 | 3.25E-07 | 1.099803144 | 1.023697233 |
| Spon2 | -0.047601959 | 4.65E-17 | 1.05E-15 | 0.061499319 | 0.097340134 |
| Spop | -0.022607515 | 0.018753545 | 0.044017602 | 0.441084271 | 0.448502165 |
| Spopl | 0.0089638 | 0.178923588 | 0.280880852 | 0.133049561 | 0.124660934 |
| Spout1 | -0.003885939 | 0.4789251 | 0.595562781 | 0.084902534 | 0.086373233 |
| Spp1 | -0.380089249 | 8.46E-72 | 2.42E-69 | 0.456507273 | 1.222374831 |
| Sppl2a | -0.018138244 | 0.049533153 | 0.099209171 | 0.320379887 | 0.340945109 |
| Sppl2b | 0.002508871 | 0.478421554 | 0.595088837 | 0.037316163 | 0.031730883 |
| Sppl3 | 0.001749974 | 0.809421268 | 0.868468554 | 0.173818284 | 0.16761268 |
| Spr | -0.017269431 | 0.029553102 | 0.064608783 | 0.216591336 | 0.219839983 |
| Spred2 | 0.002469047 | 0.653192538 | 0.745500827 | 0.080010648 | 0.083313585 |
| Spred3 | -0.01174891 | 0.003447965 | 0.010485179 | 0.03295843 | 0.045152561 |
| Spring1 | 0.001017401 | 0.864348057 | 0.907178216 | 0.099567057 | 0.102544523 |
| Sprtn | 0.0041552 | 0.378761671 | 0.500519364 | 0.062779498 | 0.059283734 |
| Spry2 | 0.011245383 | 0.010846747 | 0.027728655 | 0.056624928 | 0.048804134 |
| Spryd3 | 0.029961744 | 8.04E-05 | 0.000383199 | 0.212269088 | 0.177816599 |
| Spryd4 | -0.003557718 | 0.415561445 | 0.534864812 | 0.051101777 | 0.052945821 |
| Spryd7 | 0.027052326 | 0.000231499 | 0.000978535 | 0.185647155 | 0.16676968 |
| Spsb1 | -0.012481675 | 0.005804468 | 0.016375809 | 0.048666814 | 0.054073013 |
| Spsb2 | -0.025029757 | 0.002586202 | 0.008165981 | 0.218790593 | 0.2418641 |
| Spsb3 | -0.001794455 | 0.788276962 | 0.852058817 | 0.149174336 | 0.138642433 |
| Sptan1 | 0.038445766 | 1.26E-05 | 7.07E-05 | 0.311256027 | 0.264988104 |
| Sptbn1 | 0.054001073 | 1.62E-07 | 1.32E-06 | 0.631375165 | 0.56211525 |
| Sptbn2 | 0.014347865 | 0.04803987 | 0.096796604 | 0.164927383 | 0.15485899 |
| Sptbn4 | 0.013939464 | 0.066557209 | 0.127150558 | 0.19872345 | 0.177391718 |
| Sptlc1 | -0.008340731 | 0.215965747 | 0.326129077 | 0.13977687 | 0.143600008 |
| Sptlc2 | 0.016860441 | 0.009322465 | 0.024455749 | 0.134867044 | 0.121203172 |
| Sptssa | -0.022325857 | 0.016985762 | 0.040613768 | 0.432853659 | 0.431197065 |
| Spty2d1 | 0.03137957 | 0.000419695 | 0.001660672 | 0.305384316 | 0.278585208 |
| Sqle | -0.000237375 | 0.981953479 | 0.988562879 | 0.547646975 | 0.57535506 |
| Sqor | 0.012605717 | 0.144577623 | 0.238169871 | 0.303220834 | 0.279913948 |
| Sqstm1 | -0.113452756 | 2.51E-24 | 9.66E-23 | 1.456263689 | 1.704597191 |
| Sra1 | 0.04330559 | 2.52E-05 | 0.000132683 | 0.920116802 | 0.857715423 |
| Srbd1 | 0.013523297 | 0.008226698 | 0.022033046 | 0.079057843 | 0.067196199 |
| Src | 0.014238696 | 0.003913631 | 0.011703575 | 0.071862371 | 0.062708941 |
| Srcap | 0.003005341 | 0.504316272 | 0.618670385 | 0.054959802 | 0.052287163 |
| Srd5a3 | 0.008425183 | 0.119862843 | 0.204126585 | 0.090368123 | 0.081016052 |
| Srebf1 | 0.031030413 | 2.02E-06 | 1.33E-05 | 0.145294057 | 0.114378576 |
| Srebf2 | 0.051729211 | 3.96E-11 | 5.33E-10 | 0.234332667 | 0.184349324 |
| Srek1 | -0.00056523 | 0.953690019 | 0.968736221 | 0.405190445 | 0.401027912 |
| Srek1ip1 | 0.074413389 | 5.29E-17 | 1.18E-15 | 0.343029538 | 0.277107214 |
| Srf | 0.004168405 | 0.275996041 | 0.394071597 | 0.041065209 | 0.036730892 |
| Srfbp1 | 0.005521662 | 0.298029758 | 0.417770934 | 0.085946065 | 0.078071935 |
| Srgap1 | 0.033320006 | 5.30E-09 | 5.48E-08 | 0.109854172 | 0.080036111 |
| Sri | 0.120095607 | 2.47E-35 | 1.64E-33 | 0.531073444 | 0.399953092 |
| Srm | -0.017041599 | 0.05086503 | 0.101396314 | 0.248473898 | 0.260102446 |
| Srp14 | -0.067791654 | 2.82E-16 | 5.99E-15 | 1.239591568 | 1.323950968 |
| Srp19 | -0.023443274 | 0.01796939 | 0.042588592 | 0.461777772 | 0.483487602 |
| Srp54a | 0.016491708 | 0.03679859 | 0.077302484 | 0.202300271 | 0.196327498 |
| Srp68 | 0.032178998 | 0.000442239 | 0.001740161 | 0.373975591 | 0.327497744 |
| Srp72 | -0.073329056 | 3.36E-13 | 5.73E-12 | 0.705982081 | 0.789135117 |
| Srp9 | -0.019755823 | 0.026512674 | 0.058902244 | 1.268584095 | 1.288100823 |
| Srpk1 | -0.03423506 | 0.000322882 | 0.001316055 | 0.375598965 | 0.40913347 |
| Srpk2 | 0.030656471 | 0.002808719 | 0.008771719 | 0.749342446 | 0.692172929 |

| | | | | | |
|---|---|---|---|---|---|
| Srpr | -0.048790442 | 1.16E-07 | 9.75E-07 | 1.195059561 | 1.292137891 |
| Srprb | -0.004483297 | 0.582063423 | 0.68816984 | 0.230627657 | 0.229124438 |
| Srr | -0.00654963 | 0.208713697 | 0.317495628 | 0.075131669 | 0.077392906 |
| Srrd | 0.003592153 | 0.497900408 | 0.612757215 | 0.084283596 | 0.077619311 |
| Srrm1 | 0.026945221 | 0.007373608 | 0.020102696 | 0.829779096 | 0.792139742 |
| Srrm2 | -0.047825306 | 5.24E-09 | 5.43E-08 | 1.471461347 | 1.539360351 |
| Srrm3 | -0.007202176 | 0.280585103 | 0.399450633 | 0.129110302 | 0.134001809 |
| Srrm4 | -0.01406896 | 0.000660619 | 0.002483582 | 0.032693412 | 0.046994188 |
| Srrt | 0.004569714 | 0.473785167 | 0.590985295 | 0.12641826 | 0.116197166 |
| Srsf1 | -0.019243943 | 0.030235925 | 0.065835086 | 0.2909657 | 0.301154429 |
| Srsf10 | -0.006845317 | 0.477139164 | 0.593962623 | 0.438484791 | 0.451320249 |
| Srsf11 | -0.038793251 | 0.000156055 | 0.00069025 | 0.706774782 | 0.745343346 |
| Srsf2 | -0.059960323 | 1.71E-09 | 1.88E-08 | 0.441671884 | 0.504258423 |
| Srsf3 | -0.135077509 | 1.05E-38 | 8.36E-37 | 0.62498845 | 0.736264229 |
| Srsf4 | 0.014072537 | 0.069614073 | 0.13179903 | 0.212176283 | 0.193930183 |
| Srsf5 | -0.103895422 | 5.99E-27 | 2.67E-25 | 0.874988489 | 1.002668298 |
| Srsf6 | -0.070097745 | 3.85E-13 | 6.51E-12 | 0.431810457 | 0.481420299 |
| Srsf7 | -0.089824935 | 2.46E-17 | 5.71E-16 | 0.585303305 | 0.695426209 |
| Srsf9 | -0.030907326 | 0.00166118 | 0.00554411 | 0.586298645 | 0.606107275 |
| Srxn1 | -0.066774011 | 7.77E-11 | 1.01E-09 | 0.329514228 | 0.416607536 |
| Ss18 | -0.003736388 | 0.497398795 | 0.612425416 | 0.085247851 | 0.086794659 |
| Ss18l2 | -0.036000194 | 0.001506321 | 0.005099042 | 1.03282213 | 1.154863527 |
| Ssb | -0.032698447 | 0.000930277 | 0.003357145 | 0.854277545 | 0.903038112 |
| Ssbp1 | -0.033833836 | 0.000674717 | 0.002531685 | 0.518562248 | 0.557816165 |
| Ssbp2 | 0.00072316 | 0.910664536 | 0.938664188 | 0.130684349 | 0.123558626 |
| Ssbp3 | 0.050989801 | 5.47E-08 | 4.82E-07 | 0.420426586 | 0.355580185 |
| Ssbp4 | -0.065712755 | 1.99E-11 | 2.77E-10 | 0.451984685 | 0.494601635 |
| Ssh1 | 0.024515243 | 0.000160325 | 0.000707849 | 0.137004863 | 0.1151448 |
| Ssh2 | 0.001532307 | 0.775562775 | 0.841614164 | 0.080095053 | 0.078608753 |
| Ssna1 | -0.028300436 | 0.001439386 | 0.004908253 | 0.292963392 | 0.30996203 |
| Sspn | 0.006629459 | 0.096641915 | 0.171747383 | 0.047887117 | 0.038978281 |
| Ssr1 | -0.029393137 | 0.003665604 | 0.01106453 | 0.640036138 | 0.665629374 |
| Ssr2 | -0.075179269 | 1.71E-13 | 3.02E-12 | 0.889359139 | 0.987314139 |
| Ssr3 | -0.024383376 | 0.016957704 | 0.040576617 | 0.614574725 | 0.655496422 |
| Ssr4 | -0.053841289 | 1.25E-13 | 2.23E-12 | 2.203867057 | 2.422780741 |
| Ssrp1 | -0.007391563 | 0.44666271 | 0.565571095 | 0.468499532 | 0.468296797 |
| Sst | 0.023193596 | 0.053000062 | 0.104665165 | 0.407916485 | 0.389769537 |
| Sstr3 | 0.056689297 | 9.16E-13 | 1.48E-11 | 0.244910386 | 0.185417956 |
| Ssu72 | 0.011737145 | 0.235624024 | 0.349469721 | 0.547554805 | 0.515589829 |
| Ssx2ip | 0.029647914 | 0.000114424 | 0.000527467 | 0.204796143 | 0.182351592 |
| St13 | 0.084908399 | 4.16E-17 | 9.41E-16 | 0.978895314 | 0.859367662 |
| St14 | -0.000729636 | 0.897401907 | 0.930121678 | 0.085326224 | 0.08598822 |
| St18 | 0.063841894 | 5.01E-10 | 5.95E-09 | 0.526465617 | 0.470349261 |
| St3gal1 | 0.015789695 | 0.000925215 | 0.00334136 | 0.068004765 | 0.055350941 |
| St3gal3 | 0.009152672 | 0.124208952 | 0.210119963 | 0.109573556 | 0.097336627 |
| St3gal5 | 0.018006477 | 0.050608179 | 0.100997417 | 0.329480557 | 0.308633902 |
| St3gal6 | 0.007149948 | 0.136536301 | 0.227595124 | 0.065642458 | 0.058117923 |
| St6gal1 | -0.00982022 | 0.07175495 | 0.134932615 | 0.07835392 | 0.081913407 |
| St6galnac4 | 0.000512615 | 0.912615254 | 0.939678617 | 0.059586096 | 0.059668946 |
| St6galnac6 | 0.00478023 | 0.394559293 | 0.515232431 | 0.10217339 | 0.086769545 |
| St7l | 0.001422364 | 0.834967646 | 0.887291259 | 0.141857646 | 0.137751083 |
| St8sia1 | -0.009195372 | 0.124939864 | 0.211099297 | 0.095406149 | 0.097292428 |
| St8sia3 | 0.005127144 | 0.407495461 | 0.527730235 | 0.107772224 | 0.102970566 |
| St8sia5 | -0.040224945 | 9.97E-07 | 6.96E-06 | 0.160305185 | 0.213341857 |
| Stag1 | 0.025781119 | 0.002098844 | 0.00680619 | 0.236889421 | 0.227132273 |
| Stag2 | 0.028284176 | 0.00278179 | 0.008710017 | 0.397569081 | 0.364374451 |
| Stam | 0.004003594 | 0.449499275 | 0.567961121 | 0.081989724 | 0.077539849 |
| Stam2 | 0.05239709 | 6.79E-10 | 7.93E-09 | 0.287761691 | 0.240704842 |
| Stambp | 0.021767458 | 0.005241908 | 0.015021418 | 0.204039685 | 0.19145847 |
| Star | -0.056285919 | 2.60E-16 | 5.54E-15 | 0.05109295 | 0.116135664 |
| Stard10 | 0.039799729 | 7.19E-05 | 0.000344954 | 0.82183046 | 0.713633433 |
| Stard13 | 0.010826537 | 0.041460889 | 0.085507586 | 0.084715475 | 0.075966713 |
| Stard3 | -0.003689789 | 0.460781828 | 0.578988782 | 0.070604896 | 0.070843634 |
| Stard3nl | -0.036126439 | 0.000118979 | 0.000546394 | 0.334685879 | 0.367368556 |
| Stard4 | 0.023701882 | 4.48E-09 | 4.69E-08 | 0.055734837 | 0.036201278 |
| Stard5 | -0.027965936 | 0.000403497 | 0.00160245 | 0.160523763 | 0.191911336 |
| Stard7 | 0.032282835 | 0.000243921 | 0.001023477 | 0.314301697 | 0.282739436 |
| Stat1 | 0.027189868 | 8.56E-07 | 6.07E-06 | 0.094548841 | 0.074086045 |
| Stat2 | 0.010766159 | 0.018757964 | 0.044017602 | 0.057671175 | 0.050768161 |
| Stat3 | 0.04875415 | 3.33E-06 | 2.11E-05 | 0.685714268 | 0.662929611 |
| Stat5a | 0.007566415 | 0.070460282 | 0.132986771 | 0.053447073 | 0.043774097 |
| Stat5b | 0.022004351 | 0.001485361 | 0.005040758 | 0.167706798 | 0.140254239 |

| | | | | | |
|---|---|---|---|---|---|
| Stat6 | -0.001628669 | 0.743753251 | 0.817036058 | 0.066056931 | 0.067224253 |
| Stau1 | 0.031263656 | 7.74E-06 | 4.56E-05 | 0.169782401 | 0.142272683 |
| Stau2 | -0.031094152 | 0.000417011 | 0.001650726 | 0.282906546 | 0.298520419 |
| Stbd1 | -0.003422224 | 0.536317306 | 0.647148672 | 0.074087311 | 0.079939245 |
| Steap1 | -0.101368629 | 1.19E-32 | 6.99E-31 | 0.102078521 | 0.209366983 |
| Steap2 | -0.003537931 | 0.663938745 | 0.754066879 | 0.21863278 | 0.219662466 |
| Stim1 | 0.033185205 | 0.00050436 | 0.001956666 | 0.422233579 | 0.38619551 |
| Stim2 | 0.036612618 | 1.40E-07 | 1.15E-06 | 0.172240394 | 0.137320052 |
| Stimate | 0.009557799 | 0.017144411 | 0.040942758 | 0.047293156 | 0.040654672 |
| Stip1 | 0.017225876 | 0.045194065 | 0.091727912 | 0.289313945 | 0.259709168 |
| Stk11 | 0.00240716 | 0.800759702 | 0.861075747 | 0.428023188 | 0.413125501 |
| Stk11ip | 0.003709279 | 0.352930917 | 0.475300509 | 0.043782689 | 0.040839004 |
| Stk16 | -0.006557234 | 0.458915408 | 0.57740075 | 0.329301961 | 0.321622131 |
| Stk19 | -0.043210246 | 8.15E-08 | 7.01E-07 | 0.198421511 | 0.233259091 |
| Stk24 | -0.000621397 | 0.924785553 | 0.948993576 | 0.133492471 | 0.131480307 |
| Stk25 | 0.047869621 | 2.14E-09 | 2.33E-08 | 0.269877272 | 0.211937141 |
| Stk26 | 0.030428029 | 0.000322795 | 0.001316055 | 0.252845403 | 0.225169529 |
| Stk3 | 0.003293532 | 0.497785174 | 0.612757215 | 0.062973867 | 0.062476797 |
| Stk32a | -0.074047582 | 5.31E-12 | 7.84E-11 | 0.50317753 | 0.608871447 |
| Stk35 | 0.025241386 | 0.000973271 | 0.003494401 | 0.163769906 | 0.163110419 |
| Stk38 | 0.004338771 | 0.605172614 | 0.707320048 | 0.239732087 | 0.238037702 |
| Stk38l | 0.021870046 | 0.000739232 | 0.002745009 | 0.128986642 | 0.115217768 |
| Stk39 | -0.009029503 | 0.166800601 | 0.26518977 | 0.125958493 | 0.129737737 |
| Stk4 | -0.000824893 | 0.906056291 | 0.935699943 | 0.150320542 | 0.152242997 |
| Stk40 | 0.017778845 | 0.04685441 | 0.094623716 | 0.298461479 | 0.29711867 |
| Stmn3 | -0.111014923 | 1.73E-28 | 8.51E-27 | 0.328281592 | 0.431527074 |
| Stoml1 | -0.007175746 | 0.083162396 | 0.152174818 | 0.044328831 | 0.048005561 |
| Stoml2 | -0.027989024 | 0.000483198 | 0.001882826 | 0.222758818 | 0.237787802 |
| Stox1 | 0.009479001 | 0.13562917 | 0.226483475 | 0.126701079 | 0.11716775 |
| Stox2 | 0.002426798 | 0.740761411 | 0.814515637 | 0.16550052 | 0.163448555 |
| Strada | 0.008397444 | 0.049314873 | 0.098873829 | 0.055113527 | 0.046060259 |
| Stradb | 0.014460215 | 0.000684531 | 0.002563563 | 0.060070816 | 0.045102804 |
| Strap | 0.054223613 | 2.70E-08 | 2.48E-07 | 0.508924571 | 0.439815445 |
| Strbp | -0.010530218 | 0.286525974 | 0.405946592 | 0.47096069 | 0.481816208 |
| Strip1 | 0.003017113 | 0.596882738 | 0.700325124 | 0.096463619 | 0.0919629 |
| Strn | 0.02591187 | 8.99E-07 | 6.35E-06 | 0.090853809 | 0.069751334 |
| Strn3 | 0.061148771 | 2.65E-10 | 3.23E-09 | 0.48073719 | 0.411089189 |
| Strn4 | 0.016569437 | 0.016377544 | 0.039382262 | 0.16160895 | 0.142480163 |
| Stt3a | -0.01376396 | 0.175186462 | 0.275994618 | 0.529135954 | 0.54704353 |
| Stt3b | -0.01005815 | 0.234474878 | 0.348291025 | 0.249015461 | 0.261411505 |
| Stub1 | -0.027570507 | 0.005510887 | 0.015661163 | 0.805759837 | 0.830053565 |
| Stx12 | 0.037516958 | 3.19E-05 | 0.000164485 | 0.330326148 | 0.299378543 |
| Stx16 | -0.016857394 | 0.051235427 | 0.101967416 | 0.2098673 | 0.227368414 |
| Stx17 | -0.012313248 | 0.05269395 | 0.104226591 | 0.115347047 | 0.122903253 |
| Stx18 | -0.043346521 | 2.41E-07 | 1.89E-06 | 0.220732403 | 0.259252403 |
| Stx3 | 0.025281196 | 0.002234736 | 0.007189312 | 0.258057103 | 0.231678017 |
| Stx4a | 0.009778658 | 0.228838803 | 0.341589259 | 0.231670958 | 0.220678299 |
| Stx5a | -0.025071377 | 0.004545742 | 0.01331742 | 0.297807858 | 0.307819578 |
| Stx6 | 0.002889009 | 0.700970041 | 0.783674188 | 0.201685796 | 0.188971143 |
| Stx7 | 0.031720578 | 0.001292618 | 0.004473774 | 0.618541106 | 0.57015043 |
| Stx8 | -0.006450202 | 0.459929359 | 0.578302279 | 0.313179534 | 0.311322266 |
| Stxbp1 | 0.057949041 | 8.58E-09 | 8.63E-08 | 0.600620389 | 0.549983678 |
| Stxbp2 | -0.002755631 | 0.703011662 | 0.785234303 | 0.168227559 | 0.168217475 |
| Stxbp3 | 0.00309716 | 0.686558467 | 0.77271607 | 0.204117996 | 0.190155554 |
| Stxbp4 | 0.007915043 | 0.074599196 | 0.139418182 | 0.058404298 | 0.050401874 |
| Stxbp5 | 0.014814217 | 0.037245248 | 0.078038427 | 0.162406231 | 0.150928239 |
| Stxbp5l | 0.010608181 | 0.006560971 | 0.018233191 | 0.044423435 | 0.032497835 |
| Stxbp6 | 0.009866997 | 0.174709037 | 0.275420992 | 0.181159389 | 0.163962678 |
| Sub1 | -0.022389006 | 0.010975461 | 0.028020859 | 1.269745442 | 1.257990883 |
| Sucla2 | 0.022231083 | 0.010304397 | 0.026536617 | 0.309323861 | 0.279932504 |
| Suclg1 | -0.023862922 | 0.015512018 | 0.037561105 | 0.641783358 | 0.646728074 |
| Suclg2 | 0.002242156 | 0.779214687 | 0.845157198 | 0.222141439 | 0.215443873 |
| Suco | 0.036790537 | 0.000183668 | 0.000796441 | 0.500620777 | 0.459111702 |
| Suds3 | 0.009779215 | 0.298051759 | 0.417770934 | 0.452481805 | 0.418972553 |
| Sugp1 | -0.011772217 | 0.10122164 | 0.178468768 | 0.158394504 | 0.162587252 |
| Sugp2 | 0.008912477 | 0.032242615 | 0.069380854 | 0.049931169 | 0.043361208 |
| Sugt1 | -0.039892589 | 2.92E-05 | 0.000151984 | 0.37753057 | 0.413909517 |
| Sult1c2 | -0.001169002 | 0.789864912 | 0.852850586 | 0.055464989 | 0.050228712 |
| Sult1d1 | 0.069041585 | 2.06E-17 | 4.81E-16 | 0.218853482 | 0.146414092 |
| Sult2b1 | -0.018478688 | 0.000420042 | 0.001661369 | 0.067906435 | 0.081814784 |
| Sult4a1 | 0.010418528 | 0.006291754 | 0.017577135 | 0.045366871 | 0.035101109 |
| Sumf1 | 0.013213718 | 0.036336691 | 0.076547196 | 0.131605094 | 0.11341963 |

| | | | | | |
|---|---|---|---|---|---|
| Sumf2 | -0.010401159 | 0.017214402 | 0.041089716 | 0.047042187 | 0.054645558 |
| Sumo1 | 0.021593865 | 0.028856434 | 0.06334212 | 0.884547537 | 0.829338947 |
| Sumo2 | 0.038445749 | 7.39E-06 | 4.37E-05 | 1.389643334 | 1.298692481 |
| Sumo3 | 0.016491061 | 0.100372163 | 0.177228145 | 0.570813546 | 0.545265073 |
| Sun1 | 0.027044613 | 0.000426278 | 0.001682613 | 0.216750007 | 0.18886638 |
| Suox | -0.026049632 | 0.001054461 | 0.00375237 | 0.191780289 | 0.213168958 |
| Supt16 | 0.019213721 | 0.051078202 | 0.101758663 | 0.478146153 | 0.448453059 |
| Supt20 | 0.00731073 | 0.232463937 | 0.345726541 | 0.112462123 | 0.106838665 |
| Supt3 | -0.000984932 | 0.839316779 | 0.889840537 | 0.069024852 | 0.065433112 |
| Supt4a | -0.018291587 | 0.045297728 | 0.091919121 | 0.339576643 | 0.342484314 |
| Supt5 | 0.037765285 | 5.28E-05 | 0.000258375 | 0.379210515 | 0.353955818 |
| Supt6 | 0.043717677 | 1.05E-05 | 5.97E-05 | 0.406862061 | 0.378412789 |
| Supt7l | -0.004295645 | 0.333184517 | 0.455971266 | 0.054670345 | 0.054648596 |
| Supv3l1 | 0.006445904 | 0.125352072 | 0.211758966 | 0.053107456 | 0.045746355 |
| Surf1 | -0.042940277 | 1.29E-06 | 8.84E-06 | 0.284812366 | 0.317192514 |
| Surf2 | -0.023409104 | 0.001761703 | 0.005839506 | 0.167296842 | 0.185453038 |
| Surf4 | 0.001313258 | 0.893829218 | 0.927936305 | 0.823846173 | 0.83588452 |
| Surf6 | 0.008968216 | 0.051495393 | 0.102326209 | 0.060955802 | 0.054878384 |
| Susd4 | 0.008225723 | 0.034640654 | 0.073612336 | 0.045675246 | 0.037367079 |
| Susd6 | 0.039840059 | 6.45E-07 | 4.72E-06 | 0.219574544 | 0.196084479 |
| Suz12 | 0.023552828 | 0.012071985 | 0.030428788 | 0.384504538 | 0.356035624 |
| Sv2a | 0.025378411 | 0.008330257 | 0.022261341 | 0.401913503 | 0.380070623 |
| Svbp | -0.003563627 | 0.500296411 | 0.614592078 | 0.081504698 | 0.080584829 |
| Svil | -0.000446521 | 0.951387177 | 0.967407817 | 0.15157484 | 0.153723377 |
| Svip | 0.089996662 | 8.18E-20 | 2.34E-18 | 0.567166218 | 0.464157917 |
| Svop | 0.008596073 | 0.0146918 | 0.03591493 | 0.038774823 | 0.029948862 |
| Swi5 | -0.031940598 | 0.000265274 | 0.001104023 | 1.244170811 | 1.259821701 |
| Swsap1 | -0.007440291 | 0.070703874 | 0.133368852 | 0.043657128 | 0.046392837 |
| Swt1 | -0.004830271 | 0.557348467 | 0.665995796 | 0.227692317 | 0.225740315 |
| Syap1 | 0.013162103 | 0.134376076 | 0.224854425 | 0.305766802 | 0.300499071 |
| Sybu | -0.000436148 | 0.922688987 | 0.947242062 | 0.052368838 | 0.053668271 |
| Syce2 | -0.000177681 | 0.976805301 | 0.984410849 | 0.11727603 | 0.107797697 |
| Syf2 | -0.031815443 | 0.001085382 | 0.003848316 | 0.513601205 | 0.534708104 |
| Sympk | 0.021708327 | 0.007452339 | 0.020283214 | 0.249571842 | 0.21561105 |
| Syn1 | 0.040771462 | 1.82E-07 | 1.46E-06 | 0.227823205 | 0.190783192 |
| Syn2 | 0.004264888 | 0.629564987 | 0.727002114 | 0.30819772 | 0.290368286 |
| Syncrip | 0.010952818 | 0.230957977 | 0.344079073 | 0.333099248 | 0.31616503 |
| Syndig1l | 0.057220331 | 1.09E-08 | 1.07E-07 | 0.497710183 | 0.417919586 |
| Syne1 | -0.071770249 | 1.86E-19 | 5.16E-18 | 0.148229437 | 0.208022729 |
| Syne2 | 0.055965486 | 3.05E-19 | 8.32E-18 | 0.135584666 | 0.083792726 |
| Syne4 | -0.036744165 | 7.82E-05 | 0.000373599 | 0.319178547 | 0.356289181 |
| Syngr1 | 0.026656203 | 6.23E-05 | 0.000302437 | 0.143837665 | 0.122154023 |
| Syngr2 | 0.014131662 | 0.078976813 | 0.145929046 | 0.239929277 | 0.221981934 |
| Synj1 | 0.043210247 | 2.19E-08 | 2.04E-07 | 0.221476921 | 0.183256653 |
| Synj2 | 0.01230635 | 0.003038816 | 0.009393757 | 0.054701861 | 0.042123724 |
| Synj2bp | 0.011917803 | 0.174532728 | 0.275232308 | 0.316626398 | 0.297994559 |
| Synpo | 0.016941025 | 0.000238537 | 0.001003922 | 0.068169899 | 0.050715754 |
| Synpr | -0.003001657 | 0.643765214 | 0.737745053 | 0.123756257 | 0.119472396 |
| Synrg | 0.095179733 | 1.90E-21 | 5.98E-20 | 0.615755784 | 0.528275567 |
| Syp | -0.008819009 | 0.375628605 | 0.497528787 | 0.716162196 | 0.691809235 |
| Sypl | 0.015566384 | 0.066707545 | 0.127312673 | 0.278370371 | 0.257549022 |
| Sys1 | -0.005186986 | 0.594169388 | 0.697989905 | 0.898001939 | 0.901594184 |
| Syt11 | 0.038977184 | 3.50E-05 | 0.000179166 | 0.356780082 | 0.326166758 |
| Syt13 | -0.05962755 | 6.27E-11 | 8.25E-10 | 1.397785016 | 1.445691338 |
| Syt14 | 0.041329909 | 4.84E-06 | 2.98E-05 | 0.339401092 | 0.296443023 |
| Syt4 | -0.054448252 | 4.61E-06 | 2.85E-05 | 1.212108991 | 1.392177117 |
| Syt5 | -0.002147289 | 0.796408051 | 0.857819529 | 0.203809979 | 0.219994256 |
| Syt7 | 0.092008958 | 1.08E-18 | 2.78E-17 | 0.98791806 | 0.862739053 |
| Syt9 | 0.004583815 | 0.276878743 | 0.395099829 | 0.048817937 | 0.045283358 |
| Sytl1 | -0.063243124 | 2.69E-22 | 9.18E-21 | 0.094279656 | 0.144050099 |
| Sytl4 | 0.144680776 | 2.11E-39 | 1.76E-37 | 0.825994186 | 0.627580025 |
| Sytl5 | -0.009843073 | 0.086407803 | 0.157019936 | 0.086122706 | 0.095036805 |
| Syvn1 | -0.003570458 | 0.5815406 | 0.687635334 | 0.120888202 | 0.124092458 |
| Szrd1 | 0.030675966 | 3.04E-05 | 0.000157721 | 0.19355047 | 0.166311754 |
| Szt2 | 0.001331114 | 0.719155483 | 0.798040134 | 0.037495742 | 0.034979087 |
| T2 | 0.02899475 | 8.97E-07 | 6.34E-06 | 0.124621564 | 0.077376455 |
| Tab1 | 0.013179981 | 0.004320916 | 0.01274513 | 0.067095257 | 0.052779948 |
| Tab2 | 0.040743723 | 1.00E-05 | 5.74E-05 | 0.361763959 | 0.32743339 |
| Tab3 | 0.002194819 | 0.584315338 | 0.689825588 | 0.04410514 | 0.042440191 |
| Tacc1 | 0.005237124 | 0.596404925 | 0.70001795 | 0.48578892 | 0.480084832 |
| Tacc2 | -0.009838506 | 0.290938723 | 0.410404276 | 0.28853291 | 0.29892824 |
| Taco1 | 0.001687854 | 0.662849218 | 0.753400468 | 0.043801739 | 0.039485958 |

| | | | | | |
|---|---|---|---|---|---|
| Tada1 | 0.003639822 | 0.515207688 | 0.628166683 | 0.091473162 | 0.086338221 |
| Tada2a | 0.007955263 | 0.124160252 | 0.2101107 | 0.081150981 | 0.072385862 |
| Tada2b | 0.004886564 | 0.429466576 | 0.548152036 | 0.112212993 | 0.108298368 |
| Tada3 | -0.009650431 | 0.160505008 | 0.257624184 | 0.151648788 | 0.151505745 |
| Taf1 | -0.005879744 | 0.532318354 | 0.643523822 | 0.352839262 | 0.355863093 |
| Taf10 | -0.013484611 | 0.169020627 | 0.26797432 | 0.878898272 | 0.876690441 |
| Taf11 | -0.009981867 | 0.238050267 | 0.352138515 | 0.257642957 | 0.261164639 |
| Taf12 | -0.023117139 | 0.000844347 | 0.003092969 | 0.141460346 | 0.15690425 |
| Taf13 | -0.011611057 | 0.04077333 | 0.084286268 | 0.087488094 | 0.097137192 |
| Taf15 | -0.075423783 | 6.07E-12 | 8.86E-11 | 0.491431334 | 0.592020482 |
| Taf1a | -0.006844171 | 0.097130819 | 0.172318581 | 0.04161787 | 0.04613782 |
| Taf1b | 0.008304015 | 0.086582995 | 0.157279499 | 0.069485232 | 0.061116791 |
| Taf1d | 0.020092862 | 0.008036236 | 0.021648182 | 0.179466826 | 0.174541843 |
| Taf2 | 0.010955477 | 0.035246548 | 0.074671376 | 0.081477183 | 0.071620818 |
| Taf3 | 0.033233796 | 2.19E-05 | 0.000117725 | 0.21245343 | 0.186916858 |
| Taf4 | 0.00753321 | 0.091530747 | 0.164302422 | 0.057719501 | 0.050817382 |
| Taf5 | -0.006556679 | 0.159137985 | 0.256148922 | 0.055508091 | 0.060242704 |
| Taf5l | 0.011233776 | 0.046487853 | 0.094018454 | 0.098603978 | 0.087562355 |
| Taf6 | -0.005970019 | 0.349094396 | 0.471597793 | 0.117903342 | 0.122219875 |
| Taf7 | -0.03075532 | 0.000222964 | 0.000944513 | 0.205199448 | 0.232071775 |
| Taf8 | -0.001863794 | 0.64331469 | 0.737449053 | 0.04719402 | 0.043939865 |
| Taf9b | -0.020120061 | 0.000970745 | 0.00348894 | 0.106450785 | 0.11516133 |
| Tafa1 | -0.014180558 | 0.000944145 | 0.003402143 | 0.042995848 | 0.051781795 |
| Tafa5 | 0.032032887 | 3.29E-05 | 0.000169296 | 0.225330444 | 0.18631807 |
| Tagap1 | -0.003448651 | 0.40397029 | 0.5237929 | 0.044766385 | 0.045684589 |
| Tagln2 | -0.094093695 | 3.81E-17 | 8.71E-16 | 0.944840809 | 1.132078722 |
| Taldo1 | -0.096248967 | 2.15E-22 | 7.41E-21 | 0.619472666 | 0.694060795 |
| Tanc1 | 0.012660908 | 0.01266117 | 0.031586322 | 0.074797212 | 0.06585312 |
| Tanc2 | 0.015098676 | 0.03046117 | 0.066192109 | 0.150997814 | 0.137535863 |
| Tango2 | -0.000277036 | 0.955346338 | 0.969840973 | 0.071825805 | 0.067778667 |
| Tank | 0.003400898 | 0.577117407 | 0.683319381 | 0.111755495 | 0.105656988 |
| Taok1 | 0.021818878 | 0.025105197 | 0.056419956 | 0.466196775 | 0.433037962 |
| Taok2 | 0.047040075 | 1.88E-07 | 1.51E-06 | 0.339411521 | 0.277834173 |
| Taok3 | 0.010272447 | 0.317653805 | 0.439413136 | 0.524237297 | 0.503654878 |
| Tap1 | 0.014205467 | 0.001285665 | 0.004454466 | 0.057728409 | 0.047720207 |
| Tap2 | -0.001711999 | 0.808160709 | 0.867307475 | 0.154743727 | 0.152826667 |
| Tapbp | -0.01731574 | 0.089197052 | 0.160929011 | 0.715821261 | 0.733883247 |
| Tapbpl | -0.01088509 | 0.207700853 | 0.316143522 | 0.228307464 | 0.251531423 |
| Tapt1 | 0.018199423 | 0.001531935 | 0.00516955 | 0.105730619 | 0.087197666 |
| Tarbp2 | -0.006818081 | 0.180098476 | 0.282369647 | 0.073004808 | 0.073397758 |
| Tardbp | -0.007853988 | 0.406070641 | 0.525955072 | 0.377069002 | 0.387509143 |
| Tars | -0.0184259 | 0.079587864 | 0.146877982 | 0.503047721 | 0.531804087 |
| Tars2 | -0.00033387 | 0.9441679 | 0.962382085 | 0.061484516 | 0.061140323 |
| Tarsl2 | 0.016588571 | 0.001707116 | 0.005675985 | 0.088105251 | 0.071621949 |
| Tasor | -0.014206881 | 0.14097001 | 0.233198985 | 0.367346921 | 0.375518185 |
| Tasor2 | -0.01806402 | 0.038032642 | 0.07942689 | 0.24990605 | 0.268168907 |
| Tasp1 | -0.012409922 | 0.016811308 | 0.040285812 | 0.068842508 | 0.077870134 |
| Tat | 0.000179577 | 0.971362609 | 0.980646262 | 0.058543661 | 0.065379665 |
| Tatdn1 | -0.01735423 | 0.009358171 | 0.02451634 | 0.126127742 | 0.139283047 |
| Tatdn2 | 0.007145583 | 0.143381337 | 0.236463675 | 0.07313486 | 0.062515016 |
| Tatdn3 | -0.007467296 | 0.083114867 | 0.15211648 | 0.049532987 | 0.052781366 |
| Tax1bp1 | -0.009903877 | 0.300936867 | 0.421025791 | 1.029399832 | 1.013840969 |
| Tax1bp3 | 0.005653436 | 0.188983633 | 0.293123626 | 0.053633839 | 0.048875319 |
| Taz | -0.047289414 | 9.81E-09 | 9.77E-08 | 0.216926295 | 0.254065703 |
| Tbc1d1 | 0.054219789 | 1.05E-09 | 1.19E-08 | 0.323593493 | 0.270287658 |
| Tbc1d10a | 0.002296099 | 0.667254305 | 0.757064125 | 0.086648167 | 0.080958058 |
| Tbc1d10b | 0.035002138 | 1.38E-08 | 1.35E-07 | 0.13443605 | 0.101285069 |
| Tbc1d12 | 0.007995622 | 0.138900951 | 0.230442841 | 0.086390769 | 0.07903336 |
| Tbc1d13 | -0.003739467 | 0.379278095 | 0.500793377 | 0.045997047 | 0.049177923 |
| Tbc1d14 | 0.01672366 | 0.004795412 | 0.013908412 | 0.110883537 | 0.095367846 |
| Tbc1d15 | -0.026110219 | 0.009561339 | 0.024954259 | 0.478168769 | 0.507458144 |
| Tbc1d16 | 0.033350282 | 0.000149357 | 0.000666076 | 0.306139137 | 0.260382963 |
| Tbc1d17 | 0.011040551 | 0.112665496 | 0.194795297 | 0.167798085 | 0.1484454 |
| Tbc1d19 | -0.006367226 | 0.336767016 | 0.459193399 | 0.130335705 | 0.134012548 |
| Tbc1d20 | 0.031450113 | 0.000122682 | 0.000561016 | 0.243172225 | 0.220419507 |
| Tbc1d22a | 0.009883172 | 0.023923684 | 0.054140144 | 0.059519229 | 0.049341156 |
| Tbc1d22b | 0.020885996 | 0.001231652 | 0.004285654 | 0.13792772 | 0.1160913 |
| Tbc1d23 | 0.023958443 | 0.001350778 | 0.004651882 | 0.187903469 | 0.167617537 |
| Tbc1d24 | 0.023899582 | 4.10E-05 | 0.000204821 | 0.108143741 | 0.086742491 |
| Tbc1d30 | 0.012327298 | 0.009387569 | 0.024586732 | 0.068661023 | 0.057392825 |
| Tbc1d31 | -0.007915965 | 0.122329325 | 0.207554223 | 0.068233524 | 0.072133951 |
| Tbc1d32 | 0.001797682 | 0.728899729 | 0.804628497 | 0.075614062 | 0.0735546 |

| | | | | | |
|---|---|---|---|---|---|
| Tbc1d4 | 0.009643477 | 0.018588914 | 0.043698168 | 0.050434359 | 0.040067425 |
| Tbc1d5 | 0.008493713 | 0.147258796 | 0.241343563 | 0.10499221 | 0.095683864 |
| Tbc1d7 | -0.003305655 | 0.510651688 | 0.624472416 | 0.074772578 | 0.07235153 |
| Tbc1d8 | -0.015619658 | 0.012240767 | 0.030766478 | 0.102317808 | 0.115217597 |
| Tbc1d8b | 0.009332558 | 0.151567439 | 0.246658627 | 0.132084018 | 0.123094331 |
| Tbc1d9 | 0.014920347 | 0.0223722 | 0.050973174 | 0.131410079 | 0.120959167 |
| Tbc1d9b | -0.003711173 | 0.634870794 | 0.730698929 | 0.200710905 | 0.202158119 |
| Tbca | 0.009789362 | 0.303606575 | 0.423906811 | 0.983742811 | 0.976738163 |
| Tbcb | -0.075021682 | 4.26E-14 | 7.80E-13 | 0.54717791 | 0.624185629 |
| Tbcc | 0.003021897 | 0.694495452 | 0.778764104 | 0.209684211 | 0.195550069 |
| Tbccd1 | 0.001761547 | 0.719318351 | 0.798129766 | 0.070257811 | 0.064750345 |
| Tbcd | 0.033408591 | 1.10E-05 | 6.27E-05 | 0.212736322 | 0.172980471 |
| Tbce | -0.017896223 | 0.012408075 | 0.031090543 | 0.151899098 | 0.165049678 |
| Tbcel | -0.003467042 | 0.48761691 | 0.603284755 | 0.066030329 | 0.068355576 |
| Tbck | 0.00605945 | 0.37900634 | 0.500706269 | 0.154777905 | 0.140811639 |
| Tbk1 | 0.017290549 | 0.004282237 | 0.01263871 | 0.114053487 | 0.100926844 |
| Tbl1x | 0.058199649 | 1.25E-08 | 1.22E-07 | 0.754287534 | 0.662878121 |
| Tbl1xr1 | -0.002306811 | 0.67046511 | 0.759527126 | 0.08090259 | 0.080060281 |
| Tbl2 | 0.003635929 | 0.562128711 | 0.670307289 | 0.115497593 | 0.112957189 |
| Tbl3 | -0.011466019 | 0.033049583 | 0.070788289 | 0.073418819 | 0.083980195 |
| Tbp | 0.005158669 | 0.239224678 | 0.353027067 | 0.052659803 | 0.048779229 |
| Tbpl1 | 0.000438398 | 0.95264329 | 0.968178765 | 0.172219313 | 0.1712163 |
| Tbrg1 | 0.046001149 | 5.53E-06 | 3.36E-05 | 0.96421396 | 0.922484964 |
| Tbrg4 | -0.006283297 | 0.116131345 | 0.199193531 | 0.03987612 | 0.04376903 |
| Tcaf1 | 0.037734858 | 4.62E-05 | 0.00022852 | 0.359704828 | 0.320831157 |
| Tcaim | 0.009360835 | 0.057001625 | 0.111624634 | 0.06990912 | 0.063429536 |
| Tcea1 | -0.060754602 | 2.34E-08 | 2.17E-07 | 0.555624404 | 0.637809223 |
| Tceal1 | -0.022812205 | 0.003554778 | 0.010762863 | 0.209546193 | 0.218030928 |
| Tceal3 | -0.056525769 | 3.16E-08 | 2.89E-07 | 0.497182531 | 0.547732529 |
| Tceal8 | -0.009413153 | 0.339727687 | 0.462128525 | 0.489273771 | 0.485489793 |
| Tceal9 | -0.000380606 | 0.950421099 | 0.967032331 | 2.118995792 | 2.142692747 |
| Tceanc | 0.004838327 | 0.299679263 | 0.419568231 | 0.060725459 | 0.057250938 |
| Tceanc2 | 0.012552958 | 0.0140036 | 0.03444914 | 0.079907728 | 0.068777444 |
| Tcerg1 | 0.035386153 | 8.56E-06 | 5.00E-05 | 0.226291518 | 0.196505549 |
| Tcf12 | 0.010618658 | 0.185966575 | 0.289413646 | 0.222903101 | 0.212029624 |
| Tcf20 | 0.03544655 | 1.50E-06 | 1.01E-05 | 0.184556883 | 0.156388606 |
| Tcf24 | -0.050005655 | 6.20E-13 | 1.02E-11 | 0.1066214 | 0.149416855 |
| Tcf25 | 0.028201436 | 0.001119792 | 0.003947288 | 1.271720766 | 1.278254091 |
| Tcf3 | -0.002806465 | 0.607311045 | 0.708537788 | 0.087534502 | 0.08416728 |
| Tcf4 | 0.015585571 | 0.113408014 | 0.195419442 | 0.411605302 | 0.389560538 |
| Tcf7l2 | 0.002191802 | 0.685483323 | 0.77195284 | 0.082678103 | 0.077389439 |
| Tchp | 0.010308585 | 0.059398326 | 0.115656023 | 0.089427005 | 0.080217638 |
| Tcim | -0.018460832 | 0.001815901 | 0.006002783 | 0.062658973 | 0.086159015 |
| Tcn2 | -0.0508876 | 2.46E-08 | 2.27E-07 | 0.335253091 | 0.367672511 |
| Tcof1 | 0.01152535 | 0.093801514 | 0.16742029 | 0.138713939 | 0.138162197 |
| Tcp1 | -0.00654352 | 0.519713384 | 0.632112536 | 0.620234216 | 0.614256026 |
| Tcp11l1 | -0.008942211 | 0.233297312 | 0.346753778 | 0.161155894 | 0.172834095 |
| Tcp11l2 | -0.015947188 | 0.004750185 | 0.013806069 | 0.088358097 | 0.09435435 |
| Tcta | -0.001834683 | 0.837847314 | 0.888961212 | 0.326381086 | 0.321381968 |
| Tctn2 | 0.003418347 | 0.415323442 | 0.534662937 | 0.05072598 | 0.046926532 |
| Tctn3 | -0.006521058 | 0.150002943 | 0.244932585 | 0.051044068 | 0.056578415 |
| Tdg | 0.028205235 | 1.89E-06 | 1.25E-05 | 0.116030785 | 0.092845828 |
| Tdp1 | -0.010484517 | 0.014741771 | 0.036009923 | 0.047070555 | 0.05281831 |
| Tdp2 | -0.01479894 | 0.017691455 | 0.042052892 | 0.110640581 | 0.117665103 |
| Tdrd3 | 0.006924632 | 0.146341259 | 0.240271908 | 0.068104942 | 0.060077835 |
| Tdrd7 | 0.022548804 | 3.57E-05 | 0.000182004 | 0.097644445 | 0.077380836 |
| Tdrkh | -0.035462156 | 4.80E-08 | 4.27E-07 | 0.110134066 | 0.135383599 |
| Tead1 | 0.04608322 | 1.78E-07 | 1.43E-06 | 0.302097987 | 0.252287535 |
| Tec | 0.020266693 | 4.55E-06 | 2.81E-05 | 0.065872937 | 0.045870511 |
| Tecpr1 | 0.002970704 | 0.65800358 | 0.749690707 | 0.141439323 | 0.132717615 |
| Tecpr2 | -0.00557824 | 0.277904446 | 0.396272664 | 0.066864942 | 0.07361674 |
| Tecr | -0.075708289 | 3.94E-14 | 7.24E-13 | 0.750392979 | 0.817656954 |
| Tef | 0.045847761 | 3.06E-12 | 4.65E-11 | 0.160346161 | 0.108345582 |
| Tefm | -0.003379698 | 0.454248702 | 0.57296058 | 0.057596895 | 0.05668321 |
| Ten1 | -0.029943516 | 0.001842377 | 0.006077907 | 0.402367516 | 0.417602904 |
| Tenm2 | 1.87E-05 | 0.998109217 | 0.998314589 | 0.195249342 | 0.193387413 |
| Tenm3 | 0.016559845 | 0.015810836 | 0.038208537 | 0.156743764 | 0.134154642 |
| Tenm4 | -0.063660093 | 7.75E-19 | 2.04E-17 | 0.088886695 | 0.150293763 |
| Tent2 | 0.004376284 | 0.376075284 | 0.497862632 | 0.072143641 | 0.064894944 |
| Tent4a | 0.007386633 | 0.113534092 | 0.195532054 | 0.061862315 | 0.056393193 |
| Tent4b | -0.002149495 | 0.723998133 | 0.80090399 | 0.104006493 | 0.106969019 |
| Tent5a | 0.024448684 | 0.078812098 | 0.145722939 | 0.924887859 | 1.021351044 |

| | | | | | |
|---|---|---|---|---|---|
| Tent5c | 0.038475757 | 1.68E-05 | 9.25E-05 | 0.330476235 | 0.277627675 |
| Tep1 | 0.000860228 | 0.864965387 | 0.907728141 | 0.072896127 | 0.070057963 |
| Terf1 | -0.011900025 | 0.039581343 | 0.082206754 | 0.091912524 | 0.097749604 |
| Terf2 | 0.013233736 | 0.080303914 | 0.147750691 | 0.189787842 | 0.178279864 |
| Terf2ip | 0.002454095 | 0.728170652 | 0.804188922 | 0.161847908 | 0.153155141 |
| Tes | -0.004430147 | 0.626337508 | 0.724652297 | 0.312126812 | 0.315596234 |
| Tesk1 | 0.008178497 | 0.271827823 | 0.388919807 | 0.187069949 | 0.177982168 |
| Tet1 | -0.00273554 | 0.501692581 | 0.615995867 | 0.045101657 | 0.041804312 |
| Tet2 | 0.026997588 | 0.000179972 | 0.000782659 | 0.17435798 | 0.14766894 |
| Tet3 | -0.000113006 | 0.987928766 | 0.992728006 | 0.175994688 | 0.174066703 |
| Tex10 | -0.00163156 | 0.769803333 | 0.837701814 | 0.087228529 | 0.088395219 |
| Tex2 | 0.002969333 | 0.532320364 | 0.643523822 | 0.065386049 | 0.061362318 |
| Tex261 | -0.004462315 | 0.635335855 | 0.731147631 | 0.413993644 | 0.417755176 |
| Tex264 | -0.021893686 | 0.009764062 | 0.025367773 | 0.262182634 | 0.268296259 |
| Tex30 | -0.00854031 | 0.185201737 | 0.288408023 | 0.117957422 | 0.124664634 |
| Tex9 | -0.014289782 | 0.008578079 | 0.022854504 | 0.076785355 | 0.085645753 |
| Tfam | -0.019585115 | 0.020681721 | 0.047804966 | 0.251775291 | 0.26380138 |
| Tfb2m | -0.001652101 | 0.788597495 | 0.852144587 | 0.112242826 | 0.109699884 |
| Tfcp2 | -0.007581606 | 0.135825005 | 0.226732607 | 0.065838003 | 0.072044005 |
| Tfdp1 | -0.023110925 | 0.001264694 | 0.004388062 | 0.148683881 | 0.167512672 |
| Tfdp2 | -0.008718702 | 0.26243241 | 0.378522191 | 0.200176638 | 0.199376726 |
| Tfe3 | 0.010767292 | 0.121175722 | 0.206100485 | 0.145674505 | 0.140790825 |
| Tfg | -0.027431096 | 0.002907508 | 0.009045373 | 0.367450162 | 0.377742345 |
| Tfip11 | 0.004294913 | 0.404791935 | 0.524648339 | 0.078881022 | 0.073285787 |
| Tfpt | -0.007858016 | 0.187350715 | 0.29110175 | 0.102619808 | 0.104953967 |
| Tfrc | 0.049477241 | 1.97E-06 | 1.30E-05 | 0.413583655 | 0.364840164 |
| Tgds | 0.005693463 | 0.251172941 | 0.366157345 | 0.078500922 | 0.066064849 |
| Tgfbr1 | 0.026772339 | 2.42E-05 | 0.000128021 | 0.140474964 | 0.11171539 |
| Tgfbr3 | 0.019314601 | 1.19E-06 | 8.21E-06 | 0.053371546 | 0.034592592 |
| Tgfbrap1 | 0.016012628 | 0.016845462 | 0.04035767 | 0.144225311 | 0.129468457 |
| Tgoln1 | -0.018298881 | 0.033434738 | 0.071534445 | 1.289982375 | 1.331823152 |
| Tgs1 | 0.003160995 | 0.636268995 | 0.731874961 | 0.133212906 | 0.130066593 |
| Th | 0.022333958 | 1.82E-09 | 1.99E-08 | 0.04872827 | 0.024115654 |
| Tha1 | -0.013551995 | 0.00731152 | 0.019972633 | 0.068248839 | 0.074220262 |
| Thada | 0.002056927 | 0.655225915 | 0.74722655 | 0.060030262 | 0.055905403 |
| Thap1 | 0.000701333 | 0.907871073 | 0.936281168 | 0.108518018 | 0.105997685 |
| Thap11 | 0.009951575 | 0.163506542 | 0.261320171 | 0.187525359 | 0.162707561 |
| Thap12 | 0.024006844 | 0.000199369 | 0.000855748 | 0.136611015 | 0.116754949 |
| Thap2 | 0.012564654 | 0.073545171 | 0.137872378 | 0.164960962 | 0.14709334 |
| Thap3 | -0.009932048 | 0.201229736 | 0.308768229 | 0.205149243 | 0.204715401 |
| Thap4 | -0.003281563 | 0.528330164 | 0.639734195 | 0.076481153 | 0.076046428 |
| Thap6 | 0.00543685 | 0.229997762 | 0.343032972 | 0.058024694 | 0.053186084 |
| Thap7 | -0.001655243 | 0.82548937 | 0.879207675 | 0.20097558 | 0.186869594 |
| Thbs3 | -0.007842902 | 0.043902076 | 0.089573133 | 0.034870646 | 0.042109542 |
| Them4 | -0.01423637 | 0.029002548 | 0.06361976 | 0.114871811 | 0.126149552 |
| Them6 | -0.004532241 | 0.428595564 | 0.547255854 | 0.097967421 | 0.095303265 |
| Thnsl1 | 0.011899974 | 0.029215018 | 0.063955957 | 0.089861927 | 0.079260132 |
| Thoc1 | 0.014189804 | 0.04270827 | 0.087467833 | 0.154271242 | 0.146162736 |
| Thoc2 | 0.018697464 | 0.049217429 | 0.098762277 | 0.39504116 | 0.370902218 |
| Thoc2l | -0.009092281 | 0.381086488 | 0.50229431 | 0.536519782 | 0.561098374 |
| Thoc3 | -0.011031689 | 0.159522588 | 0.256513166 | 0.209892385 | 0.210810368 |
| Thoc5 | -0.006818542 | 0.212169219 | 0.321845709 | 0.082219322 | 0.087260672 |
| Thoc6 | -0.003547789 | 0.419056133 | 0.538114347 | 0.050427127 | 0.052243552 |
| Thoc7 | -0.000155639 | 0.98748451 | 0.992589372 | 0.602157379 | 0.5900301 |
| Thop1 | -0.020141104 | 0.00023494 | 0.000990499 | 0.076660297 | 0.090402946 |
| Thra | 0.032459915 | 2.17E-05 | 0.000116774 | 0.23294153 | 0.178201449 |
| Thrap3 | 0.019765286 | 0.045917144 | 0.092904573 | 0.46459168 | 0.44352734 |
| Thrb | 0.023700853 | 9.25E-06 | 5.35E-05 | 0.090846899 | 0.070405692 |
| Thsd4 | 0.003611998 | 0.335544484 | 0.458362157 | 0.039079829 | 0.03365539 |
| Thumpd1 | -0.003361097 | 0.665948224 | 0.755907605 | 0.196305568 | 0.200850424 |
| Thumpd3 | -0.003640087 | 0.566870619 | 0.674307618 | 0.119936397 | 0.119137885 |
| Thyn1 | -0.056348897 | 1.49E-12 | 2.36E-11 | 0.171398209 | 0.217878367 |
| Tia1 | -0.029238939 | 0.002743574 | 0.008601429 | 0.379953602 | 0.404306618 |
| Tial1 | -0.016217839 | 0.07196517 | 0.135240333 | 0.308633241 | 0.324580614 |
| Ticam1 | 0.0051019 | 0.205642726 | 0.313904628 | 0.04750531 | 0.04181231 |
| Tigar | 0.008270555 | 0.135437172 | 0.226242126 | 0.092010779 | 0.084233769 |
| Tigd2 | 0.003240442 | 0.702842179 | 0.785135202 | 0.262116038 | 0.25730229 |
| Timm10 | -0.016008038 | 0.035028194 | 0.074289726 | 0.175879696 | 0.189221418 |
| Timm10b | 0.024141916 | 0.015434144 | 0.037416605 | 0.521696539 | 0.488661142 |
| Timm13 | -0.000929147 | 0.916051255 | 0.942009972 | 1.157001419 | 1.181654431 |
| Timm17a | -0.020895859 | 0.031315828 | 0.067717926 | 0.45690976 | 0.461905452 |
| Timm17b | -0.055716528 | 5.49E-09 | 5.65E-08 | 0.424010912 | 0.46720897 |

| | | | | | |
|---|---|---|---|---|---|
| Timm21 | -0.002559858 | 0.533949589 | 0.64493203 | 0.047031782 | 0.04663631 |
| Timm22 | 0.003547979 | 0.652814618 | 0.745235587 | 0.206496356 | 0.210435467 |
| Timm23 | -0.019905737 | 0.04926014 | 0.098784464 | 0.533015085 | 0.54451401 |
| Timm29 | 0.009519618 | 0.183155158 | 0.285862008 | 0.163751515 | 0.156503242 |
| Timm44 | 0.008684374 | 0.219912186 | 0.330804003 | 0.16020772 | 0.153711617 |
| Timm50 | -0.013846309 | 0.087859933 | 0.159123374 | 0.211223864 | 0.227298699 |
| Timm8a1 | -0.011623429 | 0.036875486 | 0.077410648 | 0.078732195 | 0.090251843 |
| Timm8b | 0.108366727 | 2.95E-27 | 1.34E-25 | 0.919796725 | 0.797096331 |
| Timm9 | -0.013428846 | 0.106570386 | 0.186143962 | 0.241653844 | 0.253932933 |
| Timmdc1 | 0.007930198 | 0.175747596 | 0.27678894 | 0.108546449 | 0.097755326 |
| Timp2 | -0.021497232 | 0.000382113 | 0.001525627 | 0.094347157 | 0.109553402 |
| Tinf2 | -1.99E-05 | 0.996010659 | 0.997138877 | 0.040247365 | 0.040829708 |
| Tiparp | 0.04830202 | 4.85E-13 | 8.08E-12 | 0.141540382 | 0.110099168 |
| Tipin | -0.000426928 | 0.952221565 | 0.967851339 | 0.166140616 | 0.161414957 |
| Tiprl | -0.007239458 | 0.425801147 | 0.544402781 | 0.352268142 | 0.344332941 |
| Tjap1 | 0.006928592 | 0.112944007 | 0.194923418 | 0.057322652 | 0.04899836 |
| Tjp1 | 0.065374749 | 3.26E-12 | 4.94E-11 | 0.397182609 | 0.332663674 |
| Tjp2 | 0.043660421 | 6.92E-11 | 9.05E-10 | 0.159285359 | 0.116731112 |
| Tjp3 | 0.001428654 | 0.738312991 | 0.812620729 | 0.051967152 | 0.048293335 |
| Tk2 | -0.001329233 | 0.818523022 | 0.874965096 | 0.09933328 | 0.094277607 |
| Tkfc | -0.009695171 | 0.036036173 | 0.076037923 | 0.057495049 | 0.060798724 |
| Tkt | -0.040880702 | 1.44E-05 | 8.01E-05 | 0.353194216 | 0.38415215 |
| Tlcd1 | -0.007769328 | 0.103117916 | 0.181253369 | 0.058358884 | 0.063789913 |
| Tlcd3b | -0.003773575 | 0.489339326 | 0.604800016 | 0.085455019 | 0.084363129 |
| Tlcd4 | -0.009206985 | 0.297579184 | 0.417349224 | 0.28038328 | 0.292414426 |
| Tle1 | -0.021517644 | 0.00229005 | 0.007352664 | 0.140020047 | 0.159164677 |
| Tle2 | 0.000509607 | 0.906668144 | 0.935897369 | 0.052786845 | 0.050232616 |
| Tle3 | 0.067521262 | 1.92E-15 | 3.86E-14 | 0.286176763 | 0.215849858 |
| Tle4 | 0.002525571 | 0.558737443 | 0.667244883 | 0.046560893 | 0.045984625 |
| Tle5 | -0.00487579 | 0.421129209 | 0.540276876 | 2.071867904 | 2.001091307 |
| Tle6 | 0.007814473 | 0.027932265 | 0.061580098 | 0.038596626 | 0.030451139 |
| Tlk1 | 0.022900078 | 0.025022123 | 0.056272282 | 0.579980153 | 0.553074043 |
| Tlk2 | 0.043971501 | 4.49E-08 | 4.02E-07 | 0.23975934 | 0.197733687 |
| Tln1 | 0.035497867 | 9.01E-05 | 0.00042472 | 0.334174835 | 0.301707266 |
| Tln2 | 0.005151022 | 0.220762906 | 0.331724416 | 0.053088886 | 0.044510888 |
| Tlnrd1 | 0.016945968 | 0.000427403 | 0.001686369 | 0.07610806 | 0.058222631 |
| Tlr2 | 0.009671513 | 0.050491185 | 0.100795749 | 0.073814329 | 0.063745226 |
| Tm2d1 | -0.063850064 | 1.45E-10 | 1.84E-09 | 0.556565206 | 0.604012313 |
| Tm2d2 | -0.022130439 | 0.023750053 | 0.053784771 | 0.538724283 | 0.5401004 |
| Tm2d3 | -0.024898284 | 0.00965012 | 0.02513219 | 0.445323796 | 0.460443568 |
| Tm4sf4 | 0.014274603 | 0.115257287 | 0.198008719 | 1.661027087 | 1.641762035 |
| Tm7sf3 | 0.003292096 | 0.669830483 | 0.758924183 | 0.208607916 | 0.194008394 |
| Tm9sf2 | 0.013242272 | 0.184353225 | 0.287408924 | 0.648353776 | 0.664742565 |
| Tm9sf3 | -0.022516901 | 0.021729948 | 0.049848644 | 0.835537235 | 0.864065265 |
| Tm9sf4 | 0.019526496 | 0.033244685 | 0.071174814 | 0.359270905 | 0.343438964 |
| Tma16 | -0.013595956 | 0.016430136 | 0.039490945 | 0.082820793 | 0.09377452 |
| Tma7 | 0.098427317 | 2.97E-34 | 1.84E-32 | 1.39105787 | 1.273055394 |
| Tmbim1 | 0.002949441 | 0.644581387 | 0.738378725 | 0.122349057 | 0.120167402 |
| Tmbim4 | -0.051224619 | 4.94E-07 | 3.70E-06 | 0.786926662 | 0.84598517 |
| Tmbim6 | -0.152529978 | 4.20E-66 | 9.28E-64 | 1.326451937 | 1.583985472 |
| Tmc4 | -0.030402555 | 0.000488207 | 0.001900819 | 0.274757427 | 0.295723517 |
| Tmc6 | -0.003745406 | 0.387523429 | 0.508547166 | 0.047155263 | 0.05195091 |
| Tmcc1 | -0.005844695 | 0.38025019 | 0.501531997 | 0.128304364 | 0.131785905 |
| Tmcc2 | 0.006825049 | 0.176416066 | 0.277796727 | 0.075094759 | 0.068541533 |
| Tmcc3 | 0.039007843 | 9.33E-07 | 6.54E-06 | 0.222867168 | 0.192248918 |
| Tmco1 | -0.027858647 | 0.004411056 | 0.012975579 | 0.638770648 | 0.666378044 |
| Tmco3 | -0.014934093 | 0.066419031 | 0.127086365 | 0.230806492 | 0.235419173 |
| Tmco6 | -0.001126968 | 0.815585055 | 0.872866347 | 0.068531132 | 0.06331508 |
| Tmed1 | -0.000476802 | 0.907270735 | 0.935957559 | 0.045757651 | 0.044030451 |
| Tmed10 | -0.105217365 | 3.79E-47 | 4.19E-45 | 1.577571221 | 1.765864284 |
| Tmed2 | -0.03933692 | 1.98E-05 | 0.000107609 | 1.093701072 | 1.133192975 |
| Tmed3 | -0.08640966 | 8.51E-22 | 2.76E-20 | 1.387216284 | 1.554380425 |
| Tmed4 | -0.042531289 | 1.15E-05 | 6.52E-05 | 0.634269486 | 0.655098024 |
| Tmed5 | -0.036991544 | 0.000160276 | 0.000707849 | 0.365075239 | 0.420029789 |
| Tmed6 | -0.030356318 | 6.50E-07 | 4.75E-06 | 0.079332328 | 0.105556358 |
| Tmed7 | -0.00925514 | 0.336236061 | 0.458855557 | 0.447753725 | 0.448698926 |
| Tmed8 | 0.040213134 | 5.25E-06 | 3.21E-05 | 0.312704067 | 0.271511362 |
| Tmed9 | -0.102906959 | 2.07E-29 | 1.04E-27 | 1.147414769 | 1.294600928 |
| Tmem101 | -0.004806638 | 0.472880315 | 0.589998985 | 0.136386332 | 0.138794565 |
| Tmem102 | 0.008850945 | 0.057847244 | 0.113111606 | 0.065131419 | 0.056980252 |
| Tmem104 | 0.016791315 | 7.88E-05 | 0.000376281 | 0.060130973 | 0.0445204 |
| Tmem106a | 0.013361392 | 5.58E-05 | 0.000271661 | 0.037230164 | 0.024890075 |

| | | | | | |
|---|---|---|---|---|---|
| Tmem106b | -0.000248901 | 0.977487467 | 0.984882698 | 0.296362516 | 0.296990634 |
| Tmem106c | -0.006534389 | 0.422187159 | 0.541089139 | 0.246806726 | 0.237521553 |
| Tmem107 | 0.009459129 | 0.321661442 | 0.44332188 | 0.450057549 | 0.424387237 |
| Tmem108 | -0.027958449 | 0.002055256 | 0.006684911 | 0.262265492 | 0.295480628 |
| Tmem109 | -0.002272542 | 0.766865803 | 0.83506601 | 0.206871338 | 0.196404229 |
| Tmem11 | -0.013213862 | 0.10796298 | 0.188407125 | 0.228402258 | 0.231772005 |
| Tmem115 | -0.004780331 | 0.499509287 | 0.613935435 | 0.154458318 | 0.15812239 |
| Tmem117 | 0.012890976 | 0.021907877 | 0.050162123 | 0.094956466 | 0.084010464 |
| Tmem120a | -0.003676573 | 0.383746486 | 0.50484213 | 0.046040212 | 0.048647114 |
| Tmem121 | -0.007323251 | 0.147595163 | 0.241741242 | 0.0699208 | 0.074297604 |
| Tmem123 | 0.004601826 | 0.471883676 | 0.589197321 | 0.126134608 | 0.120662611 |
| Tmem126a | -0.050221545 | 2.69E-06 | 1.74E-05 | 0.622525185 | 0.689874374 |
| Tmem126b | -0.004942778 | 0.449679288 | 0.568058997 | 0.131128116 | 0.128821152 |
| Tmem127 | -0.001015837 | 0.907364739 | 0.935957559 | 0.274193812 | 0.278176337 |
| Tmem128 | -0.030295814 | 0.000909721 | 0.003301345 | 0.321795174 | 0.352016053 |
| Tmem129 | 0.011870023 | 0.015303514 | 0.037158033 | 0.074962562 | 0.062805038 |
| Tmem130 | -0.012872009 | 0.068773423 | 0.130486966 | 0.139310858 | 0.156468152 |
| Tmem131 | 0.022016179 | 0.004838752 | 0.014018564 | 0.217489907 | 0.19927236 |
| Tmem131l | 0.014892997 | 0.013754033 | 0.03392964 | 0.113703057 | 0.098096887 |
| Tmem132a | 0.023799523 | 6.25E-09 | 6.39E-08 | 0.057068231 | 0.037172033 |
| Tmem132b | 0.050721388 | 6.10E-13 | 1.00E-11 | 0.179572695 | 0.130371334 |
| Tmem134 | -0.018785215 | 0.049898997 | 0.099880184 | 0.55874329 | 0.573016639 |
| Tmem135 | 0.012797158 | 0.060406493 | 0.117360496 | 0.152772084 | 0.139942736 |
| Tmem138 | -0.005487338 | 0.282656108 | 0.401619553 | 0.071177792 | 0.073065772 |
| Tmem140 | 0.000812378 | 0.855916983 | 0.900955491 | 0.053700428 | 0.052047268 |
| Tmem141 | -0.019313898 | 0.007004171 | 0.019241183 | 0.155449034 | 0.168260222 |
| Tmem143 | 0.008382196 | 0.082336265 | 0.150861886 | 0.070976359 | 0.061662107 |
| Tmem147 | -0.046198615 | 7.97E-06 | 4.68E-05 | 0.704514058 | 0.750961072 |
| Tmem14c | -0.000825561 | 0.933478795 | 0.95498886 | 0.597187281 | 0.594716446 |
| Tmem150a | -0.011527303 | 0.005692251 | 0.016092058 | 0.039641467 | 0.049984645 |
| Tmem150c | -0.006297659 | 0.336586709 | 0.459193399 | 0.129853597 | 0.131490447 |
| Tmem151a | 0.001428441 | 0.754151446 | 0.824746589 | 0.058192982 | 0.05538439 |
| Tmem159 | -0.008479651 | 0.191009386 | 0.29546432 | 0.130973727 | 0.1297518 |
| Tmem160 | -0.075002036 | 3.75E-14 | 6.93E-13 | 0.562759157 | 0.664057546 |
| Tmem161a | -0.015949211 | 0.018543916 | 0.043628434 | 0.134549136 | 0.146742998 |
| Tmem161b | -0.000380593 | 0.927513844 | 0.950687696 | 0.047423232 | 0.047180827 |
| Tmem163 | 0.022311278 | 0.027214403 | 0.060199869 | 0.54443609 | 0.501936339 |
| Tmem164 | 0.006617268 | 0.199366969 | 0.306296725 | 0.081564567 | 0.072774465 |
| Tmem165 | -0.021740722 | 0.005891397 | 0.016587362 | 0.203243125 | 0.22135156 |
| Tmem167 | 0.028988973 | 0.003430681 | 0.010445687 | 0.894962655 | 0.876533333 |
| Tmem167b | 0.015308666 | 0.064984172 | 0.124659849 | 0.254149152 | 0.241139526 |
| Tmem168 | 0.000487478 | 0.919052713 | 0.944306751 | 0.063854152 | 0.062495709 |
| Tmem17 | -0.004621402 | 0.225535804 | 0.337591854 | 0.038179558 | 0.039940359 |
| Tmem170 | -0.004348769 | 0.319098286 | 0.440789078 | 0.051411682 | 0.053210217 |
| Tmem170b | 0.008940852 | 0.067382392 | 0.128366704 | 0.069650829 | 0.062755986 |
| Tmem175 | 0.005371183 | 0.275362514 | 0.393455962 | 0.072946787 | 0.064427036 |
| Tmem176a | -0.067233871 | 6.33E-11 | 8.33E-10 | 0.781468182 | 0.882806822 |
| Tmem176b | -0.083319892 | 2.28E-18 | 5.76E-17 | 0.976186437 | 1.082593613 |
| Tmem178b | 0.039892494 | 5.17E-10 | 6.12E-09 | 0.143489189 | 0.107701285 |
| Tmem179 | -0.046660021 | 2.06E-17 | 4.81E-16 | 0.045040342 | 0.084861366 |
| Tmem179b | -0.014635289 | 0.107118565 | 0.187067844 | 0.332533915 | 0.33190805 |
| Tmem18 | -0.001550114 | 0.731956814 | 0.807086781 | 0.053984412 | 0.056508606 |
| Tmem181a | -0.001109245 | 0.852199069 | 0.898680132 | 0.105904539 | 0.101467195 |
| Tmem181b-ps | 0.00127393 | 0.834296873 | 0.886839515 | 0.11626518 | 0.108779048 |
| Tmem183a | 0.00327114 | 0.682325936 | 0.769199066 | 0.224630535 | 0.220758827 |
| Tmem184a | -0.015366072 | 0.084416181 | 0.15403418 | 0.310840947 | 0.314140091 |
| Tmem184b | 0.039789017 | 1.99E-08 | 1.87E-07 | 0.179613813 | 0.146232891 |
| Tmem184c | 0.002502477 | 0.743902798 | 0.817108011 | 0.20279347 | 0.193238549 |
| Tmem185a | 0.005172806 | 0.421563716 | 0.540620294 | 0.12772812 | 0.121727209 |
| Tmem185b | 0.000329766 | 0.9385464 | 0.958158994 | 0.048663905 | 0.04857979 |
| Tmem186 | 0.006307349 | 0.134484778 | 0.224885143 | 0.053416307 | 0.045915567 |
| Tmem189 | 0.005935826 | 0.263849583 | 0.380077885 | 0.080674191 | 0.075906596 |
| Tmem19 | -0.015046657 | 0.028353438 | 0.062364735 | 0.143908232 | 0.152426519 |
| Tmem191c | -0.014724328 | 0.009678062 | 0.025191463 | 0.087962484 | 0.094131454 |
| Tmem192 | -0.014172671 | 0.073435573 | 0.137693471 | 0.217250714 | 0.224334123 |
| Tmem198 | -0.000606186 | 0.910524369 | 0.938619225 | 0.082357132 | 0.080713364 |
| Tmem199 | -0.004928826 | 0.541213566 | 0.651601026 | 0.229259157 | 0.228597091 |
| Tmem203 | 0.001358008 | 0.859661736 | 0.903723118 | 0.190747503 | 0.190846363 |
| Tmem205 | -0.055225252 | 3.31E-09 | 3.52E-08 | 0.348906648 | 0.405349952 |
| Tmem208 | -0.030268033 | 0.002116085 | 0.006850676 | 0.474982388 | 0.507501043 |
| Tmem209 | -0.002471267 | 0.574179544 | 0.680705225 | 0.04977328 | 0.051348437 |
| Tmem212 | -0.008393718 | 0.264739944 | 0.381021722 | 0.150951386 | 0.165114274 |

| | | | | | |
|---|---|---|---|---|---|
| Tmem214 | -0.008064875 | 0.31106583 | 0.432480362 | 0.21162865 | 0.217089045 |
| Tmem215 | 0.099513825 | 8.77E-21 | 2.65E-19 | 0.756351107 | 0.593535508 |
| Tmem216 | -0.016367605 | 0.005769684 | 0.016287127 | 0.096288663 | 0.107552112 |
| Tmem218 | -0.019545764 | 0.032655177 | 0.070097954 | 0.336290828 | 0.349193897 |
| Tmem219 | -0.022053974 | 0.022296046 | 0.050870842 | 0.446223277 | 0.463532504 |
| Tmem220 | -0.00525851 | 0.163568041 | 0.261375492 | 0.03691178 | 0.038481472 |
| Tmem222 | -0.020227455 | 0.010212814 | 0.026357573 | 0.210438649 | 0.219050746 |
| Tmem223 | -0.01002372 | 0.317473903 | 0.439231719 | 0.456065981 | 0.450123685 |
| Tmem229b | 0.033054502 | 5.67E-07 | 4.18E-06 | 0.157493224 | 0.117534433 |
| Tmem230 | -0.027215625 | 0.000537819 | 0.002071582 | 0.199531002 | 0.22012023 |
| Tmem231 | -0.006603888 | 0.204940873 | 0.31322672 | 0.074625959 | 0.078459267 |
| Tmem234 | 0.017548423 | 0.073740546 | 0.138132097 | 0.81514549 | 0.801767741 |
| Tmem237 | -0.003676052 | 0.523575926 | 0.635640004 | 0.093784585 | 0.096340544 |
| Tmem238 | -0.030119958 | 0.002934861 | 0.009121714 | 0.549070169 | 0.593795757 |
| Tmem242 | 0.0151684 | 0.036161784 | 0.076244821 | 0.187988816 | 0.165432868 |
| Tmem243 | 0.109082007 | 2.58E-35 | 1.69E-33 | 0.371340819 | 0.258318992 |
| Tmem245 | 0.037685601 | 3.14E-05 | 0.000162457 | 0.317526132 | 0.278188562 |
| Tmem248 | -0.003318341 | 0.737858611 | 0.81221257 | 0.576958683 | 0.576045733 |
| Tmem250-ps | 0.032025454 | 1.69E-05 | 9.30E-05 | 0.21416377 | 0.17510973 |
| Tmem251 | 0.016697194 | 0.02791878 | 0.061575858 | 0.193238318 | 0.178537254 |
| Tmem254a | 0.123663426 | 1.57E-79 | 6.93E-77 | 0.210691054 | 0.095769093 |
| Tmem255a | 0.019465373 | 2.43E-05 | 0.000128469 | 0.069120114 | 0.048636176 |
| Tmem256 | 0.108078063 | 1.42E-27 | 6.52E-26 | 1.169499336 | 1.086931398 |
| Tmem258 | 0.019052053 | 0.064225209 | 0.123374329 | 0.857348469 | 0.902393347 |
| Tmem259 | -0.022674245 | 0.005422223 | 0.015472512 | 0.224054624 | 0.238240613 |
| Tmem260 | -0.005822376 | 0.32014421 | 0.441669081 | 0.095580229 | 0.097716976 |
| Tmem263 | 0.008389722 | 0.34632544 | 0.468481415 | 0.326037623 | 0.305263404 |
| Tmem267 | -0.023756778 | 5.68E-09 | 5.83E-08 | 0.031446414 | 0.048826466 |
| Tmem268 | 0.000951402 | 0.836843616 | 0.888187078 | 0.060166975 | 0.057427704 |
| Tmem28 | 0.01234181 | 0.003655577 | 0.011043978 | 0.054822755 | 0.045247582 |
| Tmem29 | 0.005083947 | 0.282284288 | 0.401458141 | 0.065030928 | 0.059317292 |
| Tmem30a | -0.006425607 | 0.515150103 | 0.628166683 | 0.853273047 | 0.844646824 |
| Tmem30b | 0.00417664 | 0.498627083 | 0.613316359 | 0.119495749 | 0.109544216 |
| Tmem33 | -0.015937015 | 0.090664421 | 0.163108716 | 0.38893281 | 0.399440666 |
| Tmem35b | -0.006195484 | 0.112834873 | 0.194880198 | 0.04009399 | 0.040870614 |
| Tmem37 | -0.018019837 | 2.48E-06 | 1.61E-05 | 0.028318562 | 0.042522017 |
| Tmem38a | -0.000764157 | 0.901834781 | 0.933124494 | 0.120003429 | 0.11359944 |
| Tmem38b | -0.0110777 | 0.035344074 | 0.074843158 | 0.074398163 | 0.080635109 |
| Tmem39a | -0.001782744 | 0.815172207 | 0.872712718 | 0.18916554 | 0.191176657 |
| Tmem39b | -0.003282666 | 0.436925873 | 0.555339697 | 0.047783005 | 0.049309738 |
| Tmem41a | 0.002524202 | 0.473973701 | 0.591144622 | 0.035997916 | 0.032765284 |
| Tmem41b | -0.007496125 | 0.332401913 | 0.455033632 | 0.190257224 | 0.19846163 |
| Tmem42 | -0.018193504 | 0.025625042 | 0.057296839 | 0.239753724 | 0.236479153 |
| Tmem43 | 0.006696886 | 0.211036847 | 0.320327904 | 0.085362144 | 0.078165353 |
| Tmem50a | -0.039534519 | 5.95E-05 | 0.000289447 | 0.67956651 | 0.711627739 |
| Tmem50b | 0.032070858 | 4.18E-06 | 2.62E-05 | 0.168893842 | 0.140384529 |
| Tmem51 | -2.06E-05 | 0.997399799 | 0.997853138 | 0.122429078 | 0.116536551 |
| Tmem53 | -0.001033903 | 0.806116463 | 0.865591369 | 0.048581151 | 0.047624413 |
| Tmem59 | -0.098946745 | 8.60E-33 | 5.13E-31 | 1.37751291 | 1.501070154 |
| Tmem59l | -0.072623126 | 5.88E-12 | 8.63E-11 | 0.270028395 | 0.349275035 |
| Tmem60 | 0.036927688 | 4.23E-06 | 2.65E-05 | 0.248293569 | 0.212138759 |
| Tmem62 | -0.014803778 | 0.009295877 | 0.024399168 | 0.083498601 | 0.097192827 |
| Tmem63a | 0.008260124 | 0.21785232 | 0.328365931 | 0.145932119 | 0.133585414 |
| Tmem63b | 0.028902836 | 0.000399595 | 0.001587602 | 0.241511933 | 0.222144813 |
| Tmem64 | 0.0045076 | 0.442961286 | 0.561615757 | 0.101191958 | 0.097700072 |
| Tmem65 | 0.081329926 | 3.46E-15 | 6.80E-14 | 1.161216484 | 1.040868917 |
| Tmem67 | 0.009584508 | 0.017575701 | 0.041828878 | 0.047523934 | 0.040155836 |
| Tmem68 | 0.002944807 | 0.586155722 | 0.691158997 | 0.084284721 | 0.081485304 |
| Tmem70 | -0.02029963 | 0.010282655 | 0.026495687 | 0.208913306 | 0.21940634 |
| Tmem71 | 0.049500417 | 7.47E-19 | 1.97E-17 | 0.10474452 | 0.0699824 |
| Tmem80 | -0.002998142 | 0.667642896 | 0.757212347 | 0.15733555 | 0.154075739 |
| Tmem86a | -0.011693202 | 0.013331652 | 0.033080736 | 0.060437014 | 0.062666859 |
| Tmem87a | -0.001095067 | 0.825207941 | 0.879103099 | 0.063925763 | 0.067048107 |
| Tmem87b | 0.00859155 | 0.248756942 | 0.363179905 | 0.1848766 | 0.177822667 |
| Tmem8b | 0.003366141 | 0.432065405 | 0.550529472 | 0.050532657 | 0.047769817 |
| Tmem9 | -0.008321582 | 0.251905901 | 0.366842383 | 0.174835632 | 0.17248785 |
| Tmem94 | 0.030531475 | 1.70E-09 | 1.87E-08 | 0.086319722 | 0.063010686 |
| Tmem97 | -0.05003442 | 2.23E-10 | 2.74E-09 | 0.152844388 | 0.19743561 |
| Tmem98 | 0.010294234 | 0.006815163 | 0.018823016 | 0.046529733 | 0.035214046 |
| Tmem9b | 0.023824115 | 0.003710804 | 0.011179651 | 0.268778515 | 0.235590519 |
| Tmf1 | 0.049598555 | 6.08E-07 | 4.46E-06 | 0.493504438 | 0.44162344 |
| Tmigd3 | -0.032647113 | 1.00E-05 | 5.74E-05 | 0.147733049 | 0.158478189 |

| | | | | | |
|---|---|---|---|---|---|
| Tmod1 | 0.055095087 | 6.84E-11 | 8.98E-10 | 0.287281614 | 0.216805082 |
| Tmod2 | -0.014766367 | 0.148515174 | 0.242952132 | 0.667919692 | 0.672539831 |
| Tmod3 | -0.008247279 | 0.206962442 | 0.315259371 | 0.120281338 | 0.127641262 |
| Tmpo | 0.035460021 | 0.000131928 | 0.000598508 | 0.377683726 | 0.340356748 |
| Tmprss2 | 0.005260788 | 0.242366293 | 0.356364957 | 0.056149737 | 0.052898548 |
| Tmprss4 | 0.027018514 | 1.99E-05 | 0.00010812 | 0.128841119 | 0.105226759 |
| Tmsb10 | 0.163519127 | 1.21E-52 | 1.58E-50 | 0.691765218 | 0.521898857 |
| Tmsb15b1 | 0.027641047 | 1.92E-06 | 1.27E-05 | 0.117174543 | 0.08487886 |
| Tmsb15b2 | 0.259745432 | 1.05E-114 | 8.51E-112 | 0.584247729 | 0.27253247 |
| Tmsb15l | 0.024227254 | 6.00E-12 | 8.78E-11 | 0.051851808 | 0.023019422 |
| Tmsb4x | 0.146104635 | 1.24E-36 | 8.72E-35 | 1.771635769 | 1.433835373 |
| Tmtc1 | -0.015415832 | 0.018178092 | 0.042957561 | 0.096464042 | 0.122428001 |
| Tmtc2 | 0.008078556 | 0.134438779 | 0.224881935 | 0.080476376 | 0.076922024 |
| Tmtc3 | 0.036716527 | 3.78E-07 | 2.89E-06 | 0.186209896 | 0.149850373 |
| Tmtc4 | 0.003455049 | 0.502225763 | 0.616494807 | 0.076359504 | 0.071364853 |
| Tmub1 | 0.005443296 | 0.431467168 | 0.550199869 | 0.159360162 | 0.149298284 |
| Tmub2 | -0.00163453 | 0.813887855 | 0.871649173 | 0.150062863 | 0.151446717 |
| Tmx1 | -0.030319156 | 0.001475273 | 0.005013142 | 0.394833488 | 0.423459561 |
| Tmx2 | -0.015606286 | 0.045094302 | 0.091582787 | 0.196949732 | 0.205145158 |
| Tmx3 | -0.019454661 | 0.02833559 | 0.062353692 | 0.275364634 | 0.294866433 |
| Tmx4 | -0.018368894 | 0.069269027 | 0.131273584 | 0.54440895 | 0.571831937 |
| Tnfaip1 | -0.011698968 | 0.100084995 | 0.176877757 | 0.154909468 | 0.161134553 |
| Tnfaip8 | 0.03693267 | 2.10E-07 | 1.68E-06 | 0.170080256 | 0.130475216 |
| Tnfrsf11a | 0.002029309 | 0.61828572 | 0.718242774 | 0.044402769 | 0.04336411 |
| Tnfrsf12a | 0.040102021 | 1.40E-06 | 9.48E-06 | 0.200462894 | 0.172965775 |
| Tnfrsf1a | -0.018074158 | 0.036457175 | 0.07676774 | 0.232854043 | 0.254503433 |
| Tnfrsf21 | -0.011860281 | 0.085180699 | 0.155196168 | 0.130507122 | 0.142347405 |
| Tnfrsf23 | -0.011293096 | 0.05654994 | 0.110864794 | 0.077937924 | 0.096851748 |
| Tnfrsf9 | 0.028677367 | 4.10E-11 | 5.51E-10 | 0.06768478 | 0.04024325 |
| Tnfsf12 | -0.014880648 | 0.026489576 | 0.05886438 | 0.138867961 | 0.142145469 |
| Tnik | 0.017025832 | 0.010472218 | 0.026898523 | 0.143887471 | 0.122991589 |
| Tnip1 | 0.005339769 | 0.240194279 | 0.354089697 | 0.057993869 | 0.052555152 |
| Tnip2 | 0.005420678 | 0.219621402 | 0.330417714 | 0.052239692 | 0.05005224 |
| Tnk1 | 0.004811248 | 0.317352911 | 0.439126814 | 0.06719694 | 0.061787419 |
| Tnk2 | 0.03251218 | 7.31E-08 | 6.32E-07 | 0.129874222 | 0.095405858 |
| Tnks | -0.009204993 | 0.311855052 | 0.433245901 | 0.31367326 | 0.319040687 |
| Tnks1bp1 | 0.01500003 | 0.001907918 | 0.006276265 | 0.072358281 | 0.059331804 |
| Tnks2 | 0.047683494 | 2.48E-06 | 1.61E-05 | 0.593577076 | 0.536564414 |
| Tnnc1 | 0.014854623 | 0.000406669 | 0.001613413 | 0.051754895 | 0.041079202 |
| Tnpo1 | 0.017198393 | 0.079434511 | 0.146678503 | 0.471170381 | 0.447646308 |
| Tnpo2 | -0.004144174 | 0.531876102 | 0.6431467 | 0.13684812 | 0.133176429 |
| Tnpo3 | 0.008846539 | 0.298196891 | 0.417853873 | 0.2743952 | 0.25884964 |
| Tnr | 0.019539435 | 0.004157258 | 0.012303459 | 0.153069914 | 0.127848986 |
| Tnrc18 | 0.022061751 | 0.005560422 | 0.015765068 | 0.224838523 | 0.201683917 |
| Tnrc6a | 0.048519458 | 1.69E-06 | 1.13E-05 | 0.533099288 | 0.487855509 |
| Tnrc6b | 0.036333289 | 0.000271491 | 0.001126999 | 0.482467002 | 0.433370829 |
| Tnrc6c | -0.001105015 | 0.912399639 | 0.939654252 | 0.483037855 | 0.483967889 |
| Tns2 | 0.027284109 | 7.00E-08 | 6.08E-07 | 0.091484705 | 0.06274278 |
| Tns3 | 0.031534976 | 2.98E-05 | 0.000154637 | 0.199588584 | 0.16719653 |
| Tob1 | -0.004924965 | 0.612098651 | 0.712758783 | 0.401190017 | 0.413289183 |
| Tob2 | 0.012354978 | 0.122276687 | 0.207537351 | 0.206689695 | 0.198141414 |
| Toe1 | -0.009166332 | 0.077040733 | 0.143182949 | 0.072407013 | 0.077300653 |
| Togaram1 | 0.027337277 | 0.000853234 | 0.00312096 | 0.240873792 | 0.216167205 |
| Tollip | 0.054771336 | 6.06E-12 | 8.86E-11 | 0.248014143 | 0.192484831 |
| Tom1 | 0.002719008 | 0.62249779 | 0.721566768 | 0.090860811 | 0.085979721 |
| Tom1l1 | 0.013813255 | 0.006193061 | 0.017331532 | 0.082734116 | 0.067112408 |
| Tom1l2 | 0.016485596 | 0.014173186 | 0.034804676 | 0.143580145 | 0.126508896 |
| Tomm20 | 0.016591095 | 0.077082791 | 0.143233734 | 1.073058812 | 1.04905281 |
| Tomm22 | 0.006255796 | 0.513805536 | 0.627145942 | 0.483552607 | 0.460602481 |
| Tomm34 | -0.020622972 | 0.006172968 | 0.017294984 | 0.201343926 | 0.198476441 |
| Tomm40 | -0.01163552 | 0.17989305 | 0.282174933 | 0.26235771 | 0.27429106 |
| Tomm40l | 0.004229545 | 0.443375759 | 0.562067952 | 0.08955934 | 0.08419507 |
| Tomm5 | 0.005155736 | 0.586558087 | 0.691549566 | 0.430867855 | 0.422724089 |
| Tomm6 | 0.019333439 | 0.000118358 | 0.000543799 | 0.089392333 | 0.066304995 |
| Tomm7 | 0.148559979 | 1.63E-67 | 4.07E-65 | 1.326436445 | 1.162647679 |
| Tomm70a | 0.010409248 | 0.256948097 | 0.372843193 | 0.349290767 | 0.33438588 |
| Top1 | 0.05042827 | 1.13E-07 | 9.50E-07 | 1.006320663 | 0.942334399 |
| Top2b | 0.022409149 | 0.01619615 | 0.039012775 | 0.368330272 | 0.348324591 |
| Top3b | 0.012188659 | 0.036517638 | 0.076861761 | 0.105380626 | 0.092607822 |
| Topbp1 | 0.015010818 | 0.000552387 | 0.002120136 | 0.057647956 | 0.045087396 |
| Topors | 0.027237476 | 0.000803545 | 0.00295352 | 0.232270714 | 0.205446489 |
| Tor1a | -0.024198634 | 0.003116538 | 0.009609571 | 0.224945391 | 0.241034533 |

| | | | | | |
|---|---|---|---|---|---|
| Tor1aip1 | 0.006652343 | 0.465565032 | 0.583050784 | 0.327588304 | 0.31789022 |
| Tor1aip2 | 0.026361101 | 0.003945614 | 0.011777481 | 0.364132031 | 0.336717089 |
| Tor1b | -0.043355797 | 2.11E-05 | 0.000113696 | 0.518283875 | 0.578270678 |
| Tor2a | -0.036202538 | 4.71E-05 | 0.000232756 | 0.288142734 | 0.317147395 |
| Tor3a | 0.015782791 | 0.094419257 | 0.168368308 | 0.369667446 | 0.353060407 |
| Tox | 0.025092385 | 0.000163076 | 0.000719012 | 0.139092786 | 0.121741205 |
| Tox3 | -0.002240531 | 0.836947686 | 0.888200568 | 0.657769467 | 0.638115376 |
| Tox4 | -0.00482711 | 0.593609992 | 0.697663968 | 0.334720691 | 0.336070376 |
| Tpcn1 | 0.001954765 | 0.730844797 | 0.806317876 | 0.096354386 | 0.089774312 |
| Tpd52 | -0.003243084 | 0.727758684 | 0.803825258 | 0.93000667 | 0.911846086 |
| Tpd52l1 | -0.034872747 | 6.38E-08 | 5.57E-07 | 0.100127129 | 0.130496207 |
| Tpd52l2 | 0.042279787 | 2.30E-06 | 1.50E-05 | 0.348736617 | 0.305679292 |
| Tpgs1 | -0.004000041 | 0.641673753 | 0.736088758 | 0.284562384 | 0.281527495 |
| Tpgs2 | -0.003366288 | 0.508590584 | 0.622656802 | 0.075091235 | 0.073194286 |
| Tpi1 | 0.010129553 | 0.330707048 | 0.453346577 | 0.644821307 | 0.634439066 |
| Tpk1 | 0.008427432 | 0.015453116 | 0.037437127 | 0.037359239 | 0.029075038 |
| Tpm1 | 0.086139526 | 9.45E-21 | 2.84E-19 | 0.353650393 | 0.285526353 |
| Tpm3 | 0.027849529 | 0.002398155 | 0.007646724 | 0.372187997 | 0.341023028 |
| Tpm4 | 0.067694806 | 1.01E-11 | 1.45E-10 | 0.450872562 | 0.404886452 |
| Tpp1 | 0.02333041 | 0.004554961 | 0.013326311 | 0.242409304 | 0.220805771 |
| Tpp2 | 0.028766644 | 0.000343692 | 0.001392237 | 0.241360116 | 0.208258276 |
| Tpr | 0.022888272 | 0.022269499 | 0.050858366 | 1.028049382 | 0.996245493 |
| Tpra1 | 0.00060803 | 0.896811437 | 0.929807059 | 0.062051417 | 0.059362938 |
| Tprgl | -0.048445976 | 1.29E-07 | 1.07E-06 | 0.399335706 | 0.41631669 |
| Tprkb | -0.005742609 | 0.43639755 | 0.55488582 | 0.169842674 | 0.172277073 |
| Tpst1 | 0.006420053 | 0.244045052 | 0.357805157 | 0.090760765 | 0.083455995 |
| Tpst2 | -0.006976496 | 0.43153235 | 0.550200757 | 0.356611998 | 0.334438475 |
| Tpt1 | -0.056514843 | 6.46E-47 | 6.91E-45 | 3.594749159 | 3.763378523 |
| Tpt1-ps3 | -0.026132564 | 0.001018043 | 0.003636377 | 0.18543322 | 0.22265743 |
| Tra2a | -0.045303614 | 6.59E-06 | 3.94E-05 | 0.412548009 | 0.47074505 |
| Tra2b | -0.002279229 | 0.824898578 | 0.878963609 | 0.620118261 | 0.646113018 |
| Trabd | -0.070377395 | 4.16E-12 | 6.20E-11 | 0.465975223 | 0.535594541 |
| Tradd | -0.008523284 | 0.077669733 | 0.144021581 | 0.065602349 | 0.066841503 |
| Traf2 | -0.003923009 | 0.344862038 | 0.467216936 | 0.045260555 | 0.047687823 |
| Traf3 | 0.007827395 | 0.242411411 | 0.356368787 | 0.133550569 | 0.130259715 |
| Traf3ip1 | 0.005357611 | 0.230364069 | 0.343391518 | 0.056345965 | 0.051823594 |
| Traf3ip2 | -0.007994832 | 0.143274005 | 0.236406971 | 0.071469234 | 0.083981953 |
| Traf4 | 0.001317069 | 0.710078229 | 0.791127727 | 0.036412269 | 0.032810759 |
| Traf6 | 0.016463024 | 0.002829828 | 0.008823474 | 0.093699482 | 0.080731882 |
| Traf7 | 0.006533405 | 0.26842918 | 0.385134075 | 0.111708431 | 0.099806709 |
| Trafd1 | 0.043637089 | 1.26E-08 | 1.23E-07 | 0.20581587 | 0.169377797 |
| Trak1 | 0.035031953 | 8.48E-06 | 4.96E-05 | 0.217276354 | 0.199863111 |
| Trak2 | 0.015845556 | 0.007360311 | 0.020083341 | 0.110820627 | 0.094456513 |
| Tram1 | -0.01333133 | 0.143635116 | 0.236801865 | 1.081906962 | 1.106838165 |
| Trap1 | -0.013142201 | 0.033004123 | 0.070706496 | 0.108600286 | 0.114880387 |
| Trappc1 | 7.14E-05 | 0.994065816 | 0.996209693 | 0.468198961 | 0.450455621 |
| Trappc10 | 0.036579067 | 3.27E-05 | 0.000168501 | 0.301215059 | 0.274133499 |
| Trappc11 | 0.022571712 | 0.005641981 | 0.015977671 | 0.241860963 | 0.22194015 |
| Trappc12 | 0.003888878 | 0.552319899 | 0.661858013 | 0.132849285 | 0.124627256 |
| Trappc13 | -0.015167462 | 0.059182139 | 0.115304359 | 0.228571012 | 0.232316027 |
| Trappc2 | -0.01533928 | 0.031822346 | 0.068613184 | 0.155867045 | 0.166268319 |
| Trappc2l | -0.029121295 | 0.002361696 | 0.007555253 | 0.863724493 | 0.895095543 |
| Trappc3 | -0.000968167 | 0.913326412 | 0.940302842 | 0.349899319 | 0.333315754 |
| Trappc4 | -0.034932394 | 0.000286838 | 0.001182129 | 0.439503265 | 0.461211777 |
| Trappc5 | -0.021926535 | 0.008923506 | 0.023587364 | 0.248896633 | 0.268972156 |
| Trappc6a | -0.060511655 | 1.18E-10 | 1.51E-09 | 0.389649006 | 0.439007744 |
| Trappc6b | -0.026466898 | 0.007064425 | 0.019379328 | 0.642575707 | 0.656079181 |
| Trappc8 | 0.012792595 | 0.048479827 | 0.097521389 | 0.130249293 | 0.11770678 |
| Trappc9 | 0.008819577 | 0.206819093 | 0.315106602 | 0.160156936 | 0.148476332 |
| Trerf1 | 0.020704253 | 0.000191227 | 0.000825172 | 0.099913397 | 0.078721459 |
| Triap1 | -0.016150949 | 0.062714893 | 0.120975038 | 0.296667484 | 0.305465527 |
| Trib1 | 0.120258178 | 3.37E-37 | 2.41E-35 | 0.405961265 | 0.276561926 |
| Trib3 | -0.105892814 | 8.12E-20 | 2.33E-18 | 0.42980995 | 0.558367019 |
| Trim11 | -0.002991943 | 0.538253467 | 0.648921156 | 0.063780534 | 0.065554632 |
| Trim2 | -0.001926966 | 0.757445468 | 0.827588766 | 0.107998584 | 0.107848716 |
| Trim23 | 0.000484648 | 0.931348032 | 0.953513644 | 0.091106461 | 0.08849158 |
| Trim24 | 0.029231779 | 0.000318649 | 0.001301094 | 0.228150879 | 0.205991035 |
| Trim25 | 0.018771461 | 0.013805565 | 0.034027169 | 0.203619523 | 0.184791176 |
| Trim26 | 0.014386336 | 0.049804885 | 0.099712334 | 0.17871088 | 0.167036925 |
| Trim27 | -0.016329355 | 0.042398638 | 0.086925254 | 0.219047143 | 0.228847512 |
| Trim28 | 0.028243294 | 0.000371239 | 0.001488325 | 0.242909301 | 0.207911143 |
| Trim3 | 0.01438984 | 0.003640121 | 0.011004121 | 0.0755611 | 0.062961498 |

| | | | | | |
|---|---|---|---|---|---|
| Trim32 | 0.004435906 | 0.280237115 | 0.3990721 | 0.04897806 | 0.044290553 |
| Trim33 | 0.009057859 | 0.262181865 | 0.378292089 | 0.235355749 | 0.220762853 |
| Trim35 | 0.035940217 | 0.000524505 | 0.00202753 | 0.522981927 | 0.481187729 |
| Trim37 | 0.011451571 | 0.181673309 | 0.283839673 | 0.27134407 | 0.258214139 |
| Trim39 | -0.022731517 | 0.000436098 | 0.001717887 | 0.111393489 | 0.130256927 |
| Trim41 | 0.00624859 | 0.404974116 | 0.524679436 | 0.187367564 | 0.181541106 |
| Trim44 | 0.010276823 | 0.312497621 | 0.433828627 | 0.488052675 | 0.495603564 |
| Trim46 | -0.005425261 | 0.252661673 | 0.367612273 | 0.055437603 | 0.062757814 |
| Trim56 | 0.003807281 | 0.669016495 | 0.758239492 | 0.283741408 | 0.281457631 |
| Trim8 | 0.036328902 | 1.65E-07 | 1.34E-06 | 0.179329905 | 0.139119292 |
| Trim9 | 0.006732328 | 0.163366551 | 0.261185487 | 0.06365511 | 0.060199239 |
| Trio | 0.027352449 | 0.001503917 | 0.005094455 | 0.285891467 | 0.252612783 |
| Triobp | 0.000638463 | 0.905694541 | 0.93560051 | 0.084098606 | 0.080088042 |
| Trip11 | 0.018747694 | 0.078947381 | 0.145929046 | 0.789249668 | 0.776047813 |
| Trip12 | 0.035566614 | 0.000250838 | 0.00104798 | 0.497208027 | 0.445918157 |
| Trip4 | 0.007262169 | 0.300445308 | 0.420398573 | 0.157567194 | 0.148925301 |
| Triqk | -0.036314349 | 3.86E-08 | 3.48E-07 | 0.118889923 | 0.142602663 |
| Trir | -0.033153961 | 0.000535989 | 0.002065354 | 0.765917218 | 0.777318055 |
| Trit1 | 0.003033775 | 0.518082038 | 0.630702927 | 0.062812019 | 0.058041693 |
| Trmt1 | -0.018980631 | 0.00262605 | 0.008275676 | 0.107679079 | 0.121710565 |
| Trmt10a | 0.004584856 | 0.37609497 | 0.497862632 | 0.074602107 | 0.071746686 |
| Trmt10b | 0.002210861 | 0.696668974 | 0.780301356 | 0.09698058 | 0.092543333 |
| Trmt10c | 0.006795144 | 0.313528016 | 0.434886484 | 0.141162223 | 0.134804566 |
| Trmt11 | 0.000853581 | 0.869223041 | 0.910400898 | 0.075461581 | 0.074232881 |
| Trmt112 | -0.007983937 | 0.429771396 | 0.548416854 | 0.646936002 | 0.635270926 |
| Trmt12 | -0.007714798 | 0.119172644 | 0.20322688 | 0.068551572 | 0.069198502 |
| Trmt13 | -0.013507433 | 0.005734211 | 0.016201103 | 0.05512883 | 0.068500084 |
| Trmt1l | 0.007566189 | 0.235777135 | 0.34958446 | 0.126335818 | 0.115681409 |
| Trmt2a | 0.009641482 | 0.105554064 | 0.184700614 | 0.108786234 | 0.1009028 |
| Trmt2b | 0.012399686 | 0.016685787 | 0.040044242 | 0.086311821 | 0.07026207 |
| Trmt5 | 0.006247272 | 0.083826877 | 0.153131323 | 0.03974601 | 0.032718981 |
| Trmt6 | -0.008631067 | 0.368490822 | 0.491109633 | 0.370450081 | 0.392242367 |
| Trmt61a | 0.007506818 | 0.092018646 | 0.164841127 | 0.055857168 | 0.050397724 |
| Trmt61b | -0.034862156 | 8.29E-08 | 7.11E-07 | 0.105352892 | 0.134336776 |
| Trmt9b | 0.008398697 | 0.050226645 | 0.100411976 | 0.054828084 | 0.045433137 |
| Trmu | 0.000176233 | 0.968745471 | 0.979118772 | 0.054607447 | 0.053955611 |
| Trnau1ap | -0.02847691 | 0.000170882 | 0.000749689 | 0.1819074 | 0.199805519 |
| Trnt1 | -0.00856036 | 0.261261364 | 0.377243871 | 0.18148189 | 0.188132699 |
| Trp53 | -0.017933522 | 0.011319903 | 0.028794373 | 0.143193301 | 0.156186382 |
| Trp53bp1 | -0.000884256 | 0.91136808 | 0.939190213 | 0.210650492 | 0.213375224 |
| Trp53bp2 | 0.018713156 | 0.00230921 | 0.007408443 | 0.120493848 | 0.103781219 |
| Trp53cor1 | 0.002351308 | 0.71890568 | 0.797872331 | 0.12358543 | 0.124313165 |
| Trp53i11 | 0.013515526 | 0.02420212 | 0.054681154 | 0.111284727 | 0.096666591 |
| Trp53i13 | -0.049254595 | 2.57E-08 | 2.37E-07 | 0.299632821 | 0.328852085 |
| Trp53inp1 | -0.017792369 | 0.052464885 | 0.10388261 | 0.264053076 | 0.28599424 |
| Trp53inp2 | 0.044525239 | 9.52E-06 | 5.50E-05 | 0.637015723 | 0.5918835 |
| Trp53rka | 0.005811048 | 0.292225406 | 0.411622051 | 0.092435232 | 0.082170674 |
| Trp53rkb | 0.005334543 | 0.318815079 | 0.440576726 | 0.082454578 | 0.078431578 |
| Trpc1 | 0.015399243 | 0.091998808 | 0.164841127 | 0.31887062 | 0.307883779 |
| Trpc4 | -0.01119491 | 0.006853957 | 0.01890331 | 0.039130937 | 0.047448325 |
| Trpc4ap | 0.025537845 | 0.003444894 | 0.010479116 | 0.310340929 | 0.27679198 |
| Trpm3 | -0.010876373 | 0.103850848 | 0.182310933 | 0.121206119 | 0.130049147 |
| Trpm4 | -0.006818812 | 0.36622572 | 0.488992474 | 0.191375215 | 0.189423395 |
| Trpm5 | 0.09262588 | 5.47E-24 | 2.06E-22 | 0.382262639 | 0.240897574 |
| Trpm7 | -0.006049413 | 0.378826563 | 0.500537081 | 0.140963376 | 0.140904227 |
| Trps1 | 0.008126208 | 0.254288178 | 0.369424636 | 0.157013935 | 0.148730738 |
| Trpt1 | -0.00487878 | 0.248928354 | 0.363375594 | 0.048954171 | 0.049786983 |
| Trrap | 0.011285861 | 0.13730673 | 0.228346839 | 0.190228468 | 0.178473697 |
| Trub1 | -0.003267635 | 0.622470795 | 0.721566768 | 0.139420189 | 0.134996688 |
| Trub2 | -0.001827041 | 0.735053614 | 0.810042081 | 0.07939327 | 0.080465829 |
| Tsc1 | 0.021181969 | 0.003102254 | 0.009571601 | 0.175689485 | 0.150642466 |
| Tsc2 | -0.005665402 | 0.386476803 | 0.507472647 | 0.130723075 | 0.130329231 |
| Tsc22d1 | 0.039108661 | 0.00023345 | 0.00098507 | 1.204993353 | 1.16149612 |
| Tsc22d2 | 0.005186855 | 0.500267928 | 0.614592078 | 0.183938364 | 0.18069957 |
| Tsc22d3 | -0.002104718 | 0.769406979 | 0.837364228 | 0.166468055 | 0.156899027 |
| Tsc22d4 | 0.05395891 | 1.98E-09 | 2.16E-08 | 0.338237845 | 0.307181743 |
| Tsen15 | -0.020640904 | 0.003106794 | 0.009582566 | 0.151830636 | 0.162447866 |
| Tsen2 | -0.021803781 | 2.28E-05 | 0.000121955 | 0.062903928 | 0.079229208 |
| Tsen34 | -0.023326143 | 0.004322323 | 0.012745411 | 0.256986318 | 0.255610247 |
| Tsen54 | 0.002852615 | 0.514694601 | 0.627837002 | 0.05838092 | 0.051510006 |
| Tsfm | -0.030874402 | 3.13E-05 | 0.00016184 | 0.168100147 | 0.187955919 |
| Tsg101 | -0.037528776 | 4.57E-05 | 0.000226421 | 0.365371309 | 0.382743005 |

| | | | | | |
|---|---|---|---|---|---|
| Tsga10 | 0.032629943 | 1.46E-13 | 2.59E-12 | 0.071890794 | 0.040708653 |
| Tshz1 | 0.10649755 | 4.78E-24 | 1.82E-22 | 0.723874636 | 0.600710954 |
| Tshz2 | -0.000353767 | 0.97366435 | 0.982150322 | 0.6206829 | 0.612885424 |
| Tshz3 | 0.029999246 | 3.65E-05 | 0.000185947 | 0.181180376 | 0.148171202 |
| Tsn | 0.029753356 | 0.002597665 | 0.008199514 | 0.708565732 | 0.660976216 |
| Tsnax | -0.032497007 | 0.000178271 | 0.000777545 | 0.285119735 | 0.302894617 |
| Tspan13 | 0.059817519 | 1.87E-10 | 2.33E-09 | 1.067453469 | 0.98525394 |
| Tspan15 | 0.013884032 | 0.139815711 | 0.231841777 | 0.383103646 | 0.369248893 |
| Tspan17 | 0.004394173 | 0.518321955 | 0.630915994 | 0.155109631 | 0.143073503 |
| Tspan2 | -0.020146889 | 0.022038742 | 0.050426137 | 0.276977767 | 0.283528398 |
| Tspan3 | 0.013384794 | 0.173409154 | 0.273726871 | 0.752253022 | 0.698374152 |
| Tspan31 | -0.016071519 | 0.080660237 | 0.148322077 | 0.371130599 | 0.374143287 |
| Tspan33 | -0.011313766 | 0.1394423 | 0.231262034 | 0.195325861 | 0.196012952 |
| Tspan5 | 0.018860727 | 0.063507392 | 0.122236957 | 0.633886744 | 0.586567403 |
| Tspan7 | 0.037542336 | 0.000194694 | 0.000837898 | 0.746112185 | 0.694345112 |
| Tspan8 | 0.032361118 | 2.07E-05 | 0.000111963 | 0.13801451 | 0.115458741 |
| Tspo | -0.078549186 | 7.93E-18 | 1.91E-16 | 0.288575495 | 0.357880187 |
| Tspyl1 | 0.01650426 | 0.06220511 | 0.120295945 | 0.320676571 | 0.298225394 |
| Tspyl2 | -0.033975471 | 6.64E-05 | 0.000320327 | 0.225793811 | 0.255205565 |
| Tspyl4 | 0.047835084 | 5.37E-06 | 3.27E-05 | 0.925389596 | 0.873021409 |
| Tspyl5 | 0.003665591 | 0.392403862 | 0.513106974 | 0.049436075 | 0.047212937 |
| Tsr1 | 0.003785659 | 0.599836297 | 0.703196489 | 0.160343165 | 0.158037723 |
| Tsr2 | 0.006879367 | 0.304075721 | 0.424500885 | 0.142415064 | 0.132207642 |
| Tsr3 | -0.015413248 | 0.046266437 | 0.093591823 | 0.198677388 | 0.205038286 |
| Tssc4 | -0.052116204 | 5.33E-07 | 3.97E-06 | 0.749493902 | 0.734599675 |
| Tstd2 | 0.012451707 | 0.011487805 | 0.029160428 | 0.07355256 | 0.06175001 |
| Tstd3 | 0.002663032 | 0.620391992 | 0.719745187 | 0.086414535 | 0.0810565 |
| Ttbk2 | 0.016801301 | 0.004602628 | 0.013441499 | 0.107822962 | 0.0917851 |
| Ttc1 | 0.00157149 | 0.852553678 | 0.898680132 | 0.251561617 | 0.25286414 |
| Ttc13 | -0.005545408 | 0.231397164 | 0.344626614 | 0.05502766 | 0.05922279 |
| Ttc14 | -0.044949477 | 1.53E-05 | 8.48E-05 | 0.5355992 | 0.583547639 |
| Ttc17 | -0.0001047 | 0.982536142 | 0.98894465 | 0.063276806 | 0.061858914 |
| Ttc19 | 0.015371535 | 0.074071354 | 0.138644918 | 0.280997983 | 0.270351904 |
| Ttc21b | -0.000888626 | 0.848739782 | 0.896604168 | 0.058149597 | 0.057348141 |
| Ttc26 | -0.000751375 | 0.868810392 | 0.910400898 | 0.055633558 | 0.055255708 |
| Ttc28 | 0.088359244 | 4.01E-16 | 8.43E-15 | 0.565603551 | 0.451959518 |
| Ttc3 | 0.050976649 | 1.30E-06 | 8.92E-06 | 0.746843205 | 0.672739219 |
| Ttc30b | 6.42E-05 | 0.987753943 | 0.992728006 | 0.05096431 | 0.0460336 |
| Ttc32 | -0.032534907 | 6.84E-06 | 4.07E-05 | 0.14822756 | 0.174188827 |
| Ttc33 | -0.016205247 | 0.062214361 | 0.120295945 | 0.304798977 | 0.29981649 |
| Ttc37 | 0.021613866 | 0.004526881 | 0.013272116 | 0.194983464 | 0.178422977 |
| Ttc38 | 0.008323797 | 0.033232782 | 0.071165002 | 0.047164776 | 0.037046114 |
| Ttc39a | -0.008624269 | 0.287236694 | 0.406538818 | 0.242337735 | 0.233168778 |
| Ttc39b | -0.002152232 | 0.824168018 | 0.878474013 | 0.416306318 | 0.419931654 |
| Ttc39c | -0.006779271 | 0.12941294 | 0.217635808 | 0.042060868 | 0.053885212 |
| Ttc4 | 0.018993853 | 0.006884075 | 0.018970233 | 0.168905156 | 0.146879901 |
| Ttc5 | -0.016471931 | 0.00972213 | 0.025279099 | 0.121571039 | 0.127846459 |
| Ttc7 | 0.014129998 | 0.003660604 | 0.011055728 | 0.073786792 | 0.058842776 |
| Ttc7b | 0.014151892 | 0.0081062 | 0.021781725 | 0.088808847 | 0.077544687 |
| Ttc8 | -0.030130169 | 6.45E-05 | 0.000311892 | 0.171165643 | 0.191477152 |
| Ttc9 | -0.014021883 | 0.020517923 | 0.047471501 | 0.088641377 | 0.10865122 |
| Ttc9c | -0.000142338 | 0.982454057 | 0.98894465 | 0.126738644 | 0.124067118 |
| Ttf1 | 0.01260035 | 0.06984152 | 0.132075328 | 0.151296098 | 0.139729705 |
| Tti1 | 0.01377906 | 0.009222321 | 0.024238822 | 0.087004033 | 0.075526738 |
| Tti2 | -0.001795653 | 0.720394183 | 0.798776488 | 0.072223999 | 0.069814656 |
| Ttl | 0.005777163 | 0.220903743 | 0.331825091 | 0.066601019 | 0.057983745 |
| Ttll1 | -0.001885905 | 0.668523832 | 0.757857823 | 0.053872855 | 0.052877768 |
| Ttll5 | -0.00810504 | 0.063343075 | 0.122017313 | 0.045704869 | 0.052572335 |
| Ttll7 | 0.052662108 | 2.33E-07 | 1.84E-06 | 0.664563568 | 0.617586592 |
| Ttpal | -0.006711019 | 0.143942099 | 0.237227511 | 0.056602878 | 0.058917775 |
| Ttr | 0.171975267 | 7.13E-48 | 8.06E-46 | 2.249044835 | 1.902689677 |
| Ttyh1 | -0.077144458 | 1.74E-13 | 3.06E-12 | 0.189306448 | 0.288422843 |
| Ttyh2 | 0.04119281 | 3.79E-08 | 3.43E-07 | 0.209808199 | 0.163618308 |
| Tuba1a | -0.013680952 | 0.163378698 | 0.261185487 | 0.385793711 | 0.373649351 |
| Tuba1b | 0.020000505 | 0.004844924 | 0.014031085 | 0.171582117 | 0.143860966 |
| Tuba4a | -0.030657014 | 0.00156206 | 0.005252974 | 0.308305708 | 0.35714707 |
| Tubb2a | 0.015666076 | 0.068952085 | 0.130774907 | 0.2498261 | 0.232362444 |
| Tubb3 | -0.037004277 | 2.51E-06 | 1.63E-05 | 0.142778057 | 0.175921973 |
| Tubb4b | -0.095091197 | 7.83E-19 | 2.05E-17 | 0.485056628 | 0.59398788 |
| Tubb5 | 0.023377937 | 0.00903012 | 0.023810909 | 0.280188926 | 0.266862545 |
| Tubg1 | 0.006363983 | 0.387909369 | 0.508872606 | 0.173423203 | 0.170046821 |
| Tubg2 | -0.005741949 | 0.170626654 | 0.269940548 | 0.045829858 | 0.048216102 |

| | | | | | |
|---|---|---|---|---|---|
| Tubgcp2 | 0.000104683 | 0.983704228 | 0.989915381 | 0.071867798 | 0.071970481 |
| Tubgcp3 | 0.00210562 | 0.699359742 | 0.782503788 | 0.082303029 | 0.081989334 |
| Tubgcp4 | -0.003796333 | 0.457760626 | 0.576320747 | 0.072329077 | 0.07291476 |
| Tubgcp5 | 0.005411713 | 0.355277953 | 0.477667302 | 0.100997563 | 0.094866758 |
| Tufm | 0.005320353 | 0.360222679 | 0.482959212 | 0.109714372 | 0.096111514 |
| Tuft1 | 0.022901267 | 9.58E-06 | 5.52E-05 | 0.0885893 | 0.067834511 |
| Tug1 | 0.011298537 | 0.158966065 | 0.255955935 | 0.234309477 | 0.213389411 |
| Tulp3 | 0.012221465 | 0.008629787 | 0.022979674 | 0.067379231 | 0.054946171 |
| Tulp4 | 0.068723218 | 1.24E-11 | 1.77E-10 | 0.708196545 | 0.6332313 |
| Tusc1 | -0.006287819 | 0.242442209 | 0.356368787 | 0.082552022 | 0.082646813 |
| Tusc2 | 0.011036754 | 0.168058432 | 0.266709774 | 0.240490359 | 0.222116039 |
| TUSC3 | -0.008059572 | 0.422887283 | 0.541534567 | 0.738326003 | 0.731477821 |
| Tut1 | 0.001448239 | 0.732563656 | 0.807664307 | 0.051680882 | 0.0469287 |
| Tut4 | 0.025599554 | 0.010996249 | 0.028059195 | 0.519540002 | 0.490270597 |
| Tut7 | 0.023550376 | 0.011289147 | 0.028738698 | 0.331377727 | 0.311045085 |
| Tvp23b | -0.036341439 | 0.000210406 | 0.000897966 | 0.522502705 | 0.534179271 |
| Twf1 | -0.012555647 | 0.205518865 | 0.313814105 | 0.494481399 | 0.505205293 |
| Twf2 | -0.004122834 | 0.592131161 | 0.696346818 | 0.195420635 | 0.185184699 |
| Twnk | -0.008225899 | 0.049236019 | 0.098762277 | 0.041238913 | 0.047687698 |
| Twsg1 | 0.168441256 | 2.49E-56 | 3.90E-54 | 0.83737372 | 0.648517747 |
| Txlna | 0.037074383 | 6.53E-06 | 3.90E-05 | 0.254541793 | 0.221431786 |
| Txlng | -0.000737378 | 0.907126067 | 0.935957559 | 0.117323892 | 0.114649521 |
| Txn1 | -0.056817673 | 1.98E-08 | 1.86E-07 | 0.799368389 | 0.887185068 |
| Txn2 | 0.009726937 | 0.312431667 | 0.43379901 | 0.417125379 | 0.400489392 |
| Txndc11 | -0.011489592 | 0.141439109 | 0.233855616 | 0.188159914 | 0.195754957 |
| Txndc12 | -0.038234062 | 2.46E-05 | 0.000129815 | 0.29340412 | 0.327208381 |
| Txndc15 | -0.064962134 | 3.03E-12 | 4.62E-11 | 0.369921534 | 0.410640606 |
| Txndc16 | -0.008658056 | 0.089493285 | 0.161300281 | 0.070631056 | 0.073626667 |
| Txndc17 | 0.004715076 | 0.612875309 | 0.712979987 | 0.948158358 | 0.912326553 |
| Txndc5 | -0.023882848 | 0.005077614 | 0.014626537 | 0.248136634 | 0.264633406 |
| Txndc9 | 0.01195643 | 0.188652639 | 0.292797087 | 0.371016726 | 0.343170322 |
| Txnip | 0.031725998 | 0.037196424 | 0.077969737 | 1.224311938 | 1.101814143 |
| Txnl1 | -0.038047226 | 0.000127204 | 0.00057924 | 0.995844202 | 1.031843018 |
| Txnl4a | 0.025177868 | 0.009688869 | 0.025212844 | 0.434071532 | 0.399092202 |
| Txnrd1 | -0.084794637 | 7.74E-16 | 1.60E-14 | 0.944261136 | 1.106397971 |
| Txnrd2 | -0.000528619 | 0.927707691 | 0.950687696 | 0.096901436 | 0.098252284 |
| Tyk2 | -0.004563913 | 0.252548826 | 0.367556839 | 0.041093228 | 0.043065293 |
| Tyms | -0.014327642 | 0.000486516 | 0.001894997 | 0.03943193 | 0.049290655 |
| Tysnd1 | -0.002379243 | 0.678419164 | 0.766446615 | 0.095938738 | 0.094574262 |
| Tyw1 | -0.007903709 | 0.161599021 | 0.259080753 | 0.086796086 | 0.092509876 |
| Tyw5 | -0.00883119 | 0.062485253 | 0.120630734 | 0.058994804 | 0.062982092 |
| U2af1 | -0.030100803 | 0.002785375 | 0.00871842 | 0.484716111 | 0.506496749 |
| U2af2 | 0.007625469 | 0.373980893 | 0.496089813 | 0.274104838 | 0.266457203 |
| U2surp | 0.074358641 | 4.98E-13 | 8.27E-12 | 0.719888512 | 0.632473494 |
| Uaca | 0.03088548 | 0.000383011 | 0.001527957 | 0.285773679 | 0.250842562 |
| Uap1 | 0.000925761 | 0.886367338 | 0.922125549 | 0.133534875 | 0.126458578 |
| Uap1l1 | 0.035647219 | 3.13E-05 | 0.00016184 | 0.276407329 | 0.233811438 |
| Uba1 | 0.023840643 | 0.012647327 | 0.031559885 | 0.459023165 | 0.425106073 |
| Uba2 | 0.008830008 | 0.258851784 | 0.374930889 | 0.222392244 | 0.208928273 |
| Uba3 | 0.016648145 | 0.019589269 | 0.045594176 | 0.170609671 | 0.154988415 |
| Uba5 | -0.041952289 | 1.14E-05 | 6.48E-05 | 0.344918479 | 0.382526962 |
| Uba52 | 0.158113609 | 3.69E-58 | 6.18E-56 | 1.265027756 | 1.164412465 |
| Uba6 | 0.026274342 | 0.000625082 | 0.002362772 | 0.208878893 | 0.178960097 |
| Ubac1 | -0.000440111 | 0.940415391 | 0.959663948 | 0.101270189 | 0.098508656 |
| Ubac2 | -0.002321305 | 0.641952591 | 0.736234882 | 0.067590496 | 0.070609453 |
| Ubald1 | 0.006160648 | 0.358365035 | 0.481019587 | 0.14771716 | 0.134920354 |
| Ubald2 | -0.009392171 | 0.219000648 | 0.329892206 | 0.203384997 | 0.198038937 |
| Ubap1 | 0.017442729 | 0.003510222 | 0.010644535 | 0.117306896 | 0.097712588 |
| Ubap2 | 0.037477129 | 5.12E-07 | 3.83E-06 | 0.192866691 | 0.167727363 |
| Ubap2l | 0.06223619 | 1.26E-09 | 1.41E-08 | 0.664309416 | 0.615867025 |
| Ubb | -0.00323857 | 0.555472478 | 0.664630484 | 2.72993992 | 2.651834123 |
| Ubb-ps | 0.002397232 | 0.686942312 | 0.772787055 | 0.109734956 | 0.10350787 |
| Ubc | -0.027406152 | 2.30E-05 | 0.000122633 | 2.348342231 | 2.364992759 |
| Ube2a | -0.027648536 | 0.002694421 | 0.008467062 | 0.331215938 | 0.35250088 |
| Ube2b | 0.007852027 | 0.415046991 | 0.534466274 | 0.999377924 | 0.978940098 |
| Ube2d1 | -0.007316033 | 0.25013552 | 0.364918596 | 0.120186149 | 0.12407773 |
| Ube2d2a | -0.008530759 | 0.38758465 | 0.508547166 | 0.631006847 | 0.621954673 |
| Ube2d3 | -0.024494609 | 0.007028776 | 0.019292423 | 1.155414096 | 1.166201329 |
| Ube2d-ps | -0.000331067 | 0.956491652 | 0.970443551 | 0.105722417 | 0.106368854 |
| Ube2e1 | -0.04441437 | 3.17E-06 | 2.02E-05 | 0.427953487 | 0.462524643 |
| Ube2e2 | -0.006250509 | 0.342854841 | 0.464962441 | 0.134153458 | 0.132879052 |
| Ube2e3 | -0.025218089 | 0.00896147 | 0.023661979 | 0.502927259 | 0.498429754 |

| | | | | | |
|---|---|---|---|---|---|
| Ube2f | 0.036909416 | 2.43E-06 | 1.58E-05 | 0.229690968 | 0.193912651 |
| Ube2g1 | -0.004254759 | 0.530728729 | 0.642158644 | 0.143430902 | 0.142506099 |
| Ube2g2 | -0.022997913 | 0.00617553 | 0.017297176 | 0.242240253 | 0.258187919 |
| Ube2h | 0.031507311 | 0.000620964 | 0.002349946 | 0.373398571 | 0.340305124 |
| Ube2i | 0.004155079 | 0.674887314 | 0.763558067 | 0.520383822 | 0.507555861 |
| Ube2j1 | 0.006564414 | 0.45694386 | 0.575516026 | 0.316758379 | 0.301647547 |
| Ube2j2 | 0.000650099 | 0.940849544 | 0.96000622 | 0.294630327 | 0.287940111 |
| Ube2k | 0.014263562 | 0.142403659 | 0.235170438 | 0.704995498 | 0.690384893 |
| Ube2l3 | -0.016356912 | 0.096518815 | 0.171577239 | 0.606653441 | 0.602126439 |
| Ube2l6 | -6.57E-05 | 0.988381653 | 0.992977827 | 0.048777988 | 0.050996064 |
| Ube2m | -0.028707502 | 0.003440248 | 0.010469869 | 0.516715048 | 0.535403944 |
| Ube2n | -0.014811174 | 0.116662426 | 0.19985764 | 0.442633808 | 0.43578502 |
| Ube2o | 0.034650634 | 8.83E-05 | 0.000417086 | 0.309449715 | 0.280119325 |
| Ube2q1 | -0.016857118 | 0.063924554 | 0.122869615 | 0.335246889 | 0.342725403 |
| Ube2q2 | -0.051328403 | 3.91E-08 | 3.52E-07 | 0.3424331 | 0.385096548 |
| Ube2r2 | 0.040691899 | 2.81E-05 | 0.000146678 | 0.556835783 | 0.496799725 |
| Ube2s | 0.006019186 | 0.143314699 | 0.23641871 | 0.048057052 | 0.044550645 |
| Ube2v2 | -0.006614858 | 0.449096759 | 0.567839601 | 0.31165112 | 0.305364858 |
| Ube2w | -0.031067503 | 0.001009889 | 0.003612266 | 0.376428417 | 0.398373044 |
| Ube2z | 0.015979528 | 0.056383693 | 0.110583471 | 0.264949937 | 0.245332064 |
| Ube3a | 0.05396 | 6.03E-08 | 5.28E-07 | 0.85591465 | 0.780647084 |
| Ube3b | 0.023223118 | 1.94E-06 | 1.28E-05 | 0.078354328 | 0.058659152 |
| Ube3c | 0.042739198 | 8.05E-07 | 5.74E-06 | 0.291108215 | 0.249159177 |
| Ube4a | -0.005243462 | 0.477210661 | 0.593962623 | 0.171165203 | 0.170261449 |
| Ube4b | 0.055910028 | 4.27E-10 | 5.10E-09 | 0.346569769 | 0.295656399 |
| Ubfd1 | -0.020405257 | 0.012015366 | 0.030301787 | 0.218978523 | 0.231778163 |
| Ubl3 | 0.0349156 | 9.56E-05 | 0.000448765 | 1.250267953 | 1.206373846 |
| Ubl4a | 0.011332288 | 0.203535316 | 0.311323213 | 0.317803477 | 0.306048879 |
| Ubl5 | 0.113561688 | 1.19E-46 | 1.24E-44 | 1.430462522 | 1.327161668 |
| Ubl7 | -0.00568799 | 0.344361383 | 0.466668716 | 0.112499003 | 0.109192162 |
| Ublcp1 | -0.012460035 | 0.040513907 | 0.083857546 | 0.110761028 | 0.113009676 |
| Ubn1 | 0.059964073 | 9.29E-10 | 1.07E-08 | 0.524773634 | 0.46013813 |
| Ubn2 | -0.034741507 | 0.000753348 | 0.002791179 | 0.656140365 | 0.681864414 |
| Ubp1 | -0.007420017 | 0.305345575 | 0.425967812 | 0.161262932 | 0.168629771 |
| Ubqln1 | 0.002545585 | 0.779836979 | 0.845157198 | 0.354254471 | 0.350889636 |
| Ubqln2 | -0.025042176 | 0.010783344 | 0.027602862 | 0.490919643 | 0.481640714 |
| Ubqln4 | -0.007080949 | 0.154031036 | 0.24933229 | 0.063499157 | 0.069222217 |
| Ubr1 | 0.003502088 | 0.656513264 | 0.748519051 | 0.208530309 | 0.201688175 |
| Ubr2 | 0.025764739 | 0.011773553 | 0.029776921 | 0.63668473 | 0.610383305 |
| Ubr3 | -0.002437026 | 0.800311697 | 0.860689194 | 0.434481405 | 0.43127421 |
| Ubr4 | -0.049671217 | 1.69E-05 | 9.28E-05 | 0.824997775 | 0.971499127 |
| Ubr5 | 0.002101275 | 0.835178184 | 0.887291259 | 0.523525556 | 0.51011344 |
| Ubr7 | 0.014416321 | 0.032948037 | 0.070633036 | 0.148285485 | 0.134329717 |
| Ubtd1 | -0.000592173 | 0.879033799 | 0.917638419 | 0.04201419 | 0.040006502 |
| Ubtd2 | 0.013354456 | 0.000437011 | 0.001720787 | 0.047469458 | 0.034233297 |
| Ubtf | 0.006256615 | 0.534727068 | 0.645630471 | 0.465296409 | 0.436789421 |
| Ubxn1 | -0.028307738 | 0.00300669 | 0.009312213 | 0.856540919 | 0.873493238 |
| Ubxn10 | 0.013681388 | 0.018211773 | 0.04301518 | 0.108255361 | 0.089158116 |
| Ubxn2a | -0.001532969 | 0.879798966 | 0.917845858 | 0.550463563 | 0.547672232 |
| Ubxn2b | -0.009450266 | 0.097395447 | 0.172662023 | 0.090541775 | 0.095774885 |
| Ubxn4 | -0.05453177 | 4.50E-09 | 4.70E-08 | 1.046399144 | 1.120285321 |
| Ubxn6 | -0.049410011 | 1.75E-07 | 1.42E-06 | 0.400868179 | 0.428140953 |
| Ubxn7 | 0.003523653 | 0.596685049 | 0.700177676 | 0.140322707 | 0.130997942 |
| Ubxn8 | 0.008018118 | 0.10934155 | 0.190130308 | 0.076319909 | 0.066927717 |
| Uchl3 | -0.040342526 | 4.48E-07 | 3.37E-06 | 0.19473783 | 0.224454856 |
| Uchl5 | 0.012913464 | 0.111712385 | 0.193491502 | 0.227395192 | 0.21203495 |
| Uck1 | -0.005769245 | 0.341164153 | 0.463498867 | 0.110394478 | 0.107691107 |
| Uck2 | 0.035768174 | 2.18E-06 | 1.43E-05 | 0.203683267 | 0.171164705 |
| Uckl1 | -0.022913343 | 0.003308508 | 0.010130807 | 0.203446698 | 0.21550616 |
| Ucn3 | 0.207628556 | 1.30E-58 | 2.22E-56 | 1.099677684 | 0.74819979 |
| Ucp2 | -0.012318918 | 0.254065582 | 0.369156418 | 0.700020627 | 0.686190371 |
| Ufc1 | -0.039739414 | 1.37E-05 | 7.68E-05 | 0.999392568 | 1.028905377 |
| Ufd1 | -0.023649916 | 0.006472597 | 0.018020213 | 0.2695833 | 0.288525384 |
| Ufl1 | -0.041620581 | 2.73E-05 | 0.000143027 | 0.445281255 | 0.479556044 |
| Ufm1 | 0.039379574 | 3.80E-05 | 0.000192543 | 0.989319683 | 0.959732321 |
| Ufsp1 | -0.010847432 | 0.029885835 | 0.065219892 | 0.063701974 | 0.071248091 |
| Ufsp2 | 0.005871473 | 0.556119499 | 0.665017683 | 0.545461566 | 0.557696218 |
| Ugcg | 0.00033182 | 0.95884625 | 0.971740148 | 0.122830708 | 0.123253598 |
| Ugdh | -0.046144415 | 5.38E-10 | 6.34E-09 | 0.150849697 | 0.18506027 |
| Uggt1 | -0.015546499 | 0.112690431 | 0.194803764 | 0.437581815 | 0.463649998 |
| Uggt2 | -0.012832835 | 0.016913065 | 0.040489737 | 0.076635423 | 0.083103963 |
| Ugp2 | 0.006631596 | 0.369286209 | 0.491538954 | 0.183709506 | 0.171983151 |

| | | | | | |
|---|---|---|---|---|---|
| Ugt2b34 | -0.079446756 | 4.62E-17 | 1.04E-15 | 0.244525751 | 0.335763254 |
| Uhmk1 | 0.00858634 | 0.269613049 | 0.386205697 | 0.21124309 | 0.199814409 |
| Uhrf1bp1 | 0.015730858 | 0.003504081 | 0.010629229 | 0.090474396 | 0.076326411 |
| Uhrf1bp1l | 0.026198053 | 0.000432025 | 0.001702531 | 0.183637836 | 0.16175803 |
| Uhrf2 | 0.019942579 | 0.018401062 | 0.043379031 | 0.271076276 | 0.24634574 |
| Uimc1 | 0.003551192 | 0.612395387 | 0.712783653 | 0.156313501 | 0.152934928 |
| Ulk1 | -0.000317047 | 0.958306147 | 0.971699245 | 0.107623683 | 0.104952982 |
| Ulk2 | 0.007142531 | 0.323745548 | 0.445815045 | 0.167582804 | 0.162772077 |
| Ulk3 | 0.007206775 | 0.03431194 | 0.073089545 | 0.036424318 | 0.028224494 |
| Ulk4 | 0.003241399 | 0.413893565 | 0.533528672 | 0.047629505 | 0.04014698 |
| Umad1 | 0.004018209 | 0.479856924 | 0.596301111 | 0.094479849 | 0.091764023 |
| Umps | -0.004412367 | 0.559797662 | 0.668347399 | 0.196499471 | 0.192788394 |
| Unc119 | -0.015547182 | 0.029197313 | 0.063945996 | 0.163060877 | 0.166766309 |
| Unc119b | 0.008489454 | 0.193529777 | 0.298792519 | 0.12971636 | 0.124347245 |
| Unc13a | 0.024819706 | 0.000290099 | 0.001195061 | 0.148910848 | 0.136280326 |
| Unc13b | -0.019041303 | 0.00048929 | 0.001904275 | 0.071388434 | 0.086712189 |
| Unc45a | 0.003216537 | 0.682818274 | 0.769664841 | 0.217239661 | 0.205970128 |
| Unc50 | -0.050964602 | 9.49E-08 | 8.06E-07 | 0.416478325 | 0.448094036 |
| Unc5a | 0.004967949 | 0.409833631 | 0.529699889 | 0.108782291 | 0.102825222 |
| Unc5b | -0.024964276 | 9.47E-08 | 8.05E-07 | 0.043701594 | 0.063931331 |
| Unc79 | -0.000415592 | 0.943649913 | 0.962055836 | 0.095911957 | 0.096994741 |
| Unc80 | -0.011287506 | 0.285213384 | 0.404440566 | 0.666428789 | 0.64977404 |
| Unc93b1 | -0.004111326 | 0.491141009 | 0.606325286 | 0.103191481 | 0.103078846 |
| Unk | 0.003951918 | 0.357641408 | 0.480180882 | 0.051853074 | 0.047437976 |
| Upb1 | -0.023402252 | 0.026816549 | 0.059465103 | 0.508387858 | 0.52991944 |
| Upf1 | 0.034670045 | 1.03E-05 | 5.88E-05 | 0.227732577 | 0.191101612 |
| Upf2 | 0.035564018 | 7.17E-06 | 4.26E-05 | 0.224375098 | 0.195642381 |
| Upf3a | 0.008999564 | 0.181084484 | 0.283202361 | 0.1457842 | 0.13473164 |
| Upf3b | 0.016853672 | 0.108003986 | 0.188444858 | 0.627257083 | 0.6003161 |
| Upp1 | 0.080472947 | 1.83E-27 | 8.37E-26 | 0.22649483 | 0.141748751 |
| Upp2 | -0.000406963 | 0.954599759 | 0.969356472 | 0.171559411 | 0.163269028 |
| Uqcc1 | 0.020211494 | 0.001119771 | 0.003947288 | 0.126179897 | 0.108208075 |
| Uqcc2 | 0.061160037 | 2.63E-20 | 7.74E-19 | 2.037199984 | 1.931187255 |
| Uqcc3 | 0.021967905 | 0.018922557 | 0.044339625 | 0.422325077 | 0.399046673 |
| Uqcr10 | 0.065400539 | 4.12E-17 | 9.38E-16 | 1.565357131 | 1.514285611 |
| Uqcr11 | 0.139059871 | 4.44E-68 | 1.14E-65 | 1.530101949 | 1.346244263 |
| Uqcrb | 0.011033447 | 0.177382606 | 0.278857678 | 1.465364594 | 1.465564497 |
| Uqcrc1 | -0.035772423 | 0.000178704 | 0.00077887 | 0.446610932 | 0.472832646 |
| Uqcrc2 | -0.029790895 | 0.003190198 | 0.009796306 | 0.710047064 | 0.728854452 |
| Uqcrfs1 | -0.032034725 | 0.001116059 | 0.003936982 | 0.700305065 | 0.696295573 |
| Uqcrh | 0.064934386 | 6.45E-18 | 1.57E-16 | 1.561906534 | 1.471000628 |
| Uqcrq | 0.00810616 | 0.319503383 | 0.441055367 | 2.311213422 | 2.460080074 |
| Urb1 | 0.009835345 | 0.007675649 | 0.020803642 | 0.041029686 | 0.032681841 |
| Urgcp | 0.006009292 | 0.367563433 | 0.490185417 | 0.14744424 | 0.134197102 |
| Uri1 | -0.027878635 | 0.007462668 | 0.02029996 | 0.550849594 | 0.587485157 |
| Urm1 | -0.011329159 | 0.103912234 | 0.182352841 | 0.155679915 | 0.15492194 |
| Urod | -0.037662226 | 7.08E-07 | 5.12E-06 | 0.187326636 | 0.208650376 |
| Uros | -0.00531835 | 0.299791785 | 0.419665332 | 0.076625653 | 0.074862009 |
| Use1 | -0.056927221 | 1.22E-08 | 1.20E-07 | 0.576211739 | 0.645080552 |
| Usf1 | -0.023294006 | 0.001419073 | 0.004852702 | 0.170101422 | 0.178825361 |
| Usf2 | -0.011572914 | 0.205297696 | 0.313525637 | 0.373114747 | 0.364125418 |
| Usf3 | 0.01936988 | 0.001430789 | 0.0048831 | 0.116644473 | 0.099939601 |
| Ush1c | 0.015217127 | 0.058152116 | 0.113570685 | 0.237693078 | 0.210308663 |
| Uso1 | 0.034956374 | 0.000739365 | 0.002745009 | 0.716492108 | 0.679548875 |
| Usp1 | -0.011032661 | 0.146143216 | 0.240122417 | 0.179838159 | 0.190234506 |
| Usp10 | 0.019250654 | 0.01038969 | 0.026735988 | 0.19225911 | 0.179346494 |
| Usp11 | 0.001681858 | 0.698706433 | 0.782132866 | 0.053117378 | 0.048579386 |
| Usp12 | 0.012036649 | 0.087466619 | 0.158527947 | 0.165626997 | 0.153694692 |
| Usp14 | 0.037994895 | 3.69E-05 | 0.000187463 | 0.363924156 | 0.327186045 |
| Usp15 | 0.01926807 | 0.035581804 | 0.075258804 | 0.352273433 | 0.32880059 |
| Usp16 | 0.025027197 | 0.012585342 | 0.031445565 | 0.574154014 | 0.559030503 |
| Usp18 | 8.73E-05 | 0.986665955 | 0.992074301 | 0.070521836 | 0.067243749 |
| Usp19 | 0.021517914 | 0.012696863 | 0.031667239 | 0.289370929 | 0.269291182 |
| Usp20 | -0.012228454 | 0.080382034 | 0.14786644 | 0.147052295 | 0.155470957 |
| Usp21 | 0.000361587 | 0.926743676 | 0.950401057 | 0.041660366 | 0.040456686 |
| Usp22 | 0.014529818 | 0.085466201 | 0.155628845 | 0.269752487 | 0.250047887 |
| Usp24 | 0.029461157 | 2.35E-05 | 0.000125127 | 0.161105981 | 0.136189961 |
| Usp25 | 0.011921493 | 0.014647799 | 0.035825385 | 0.070205134 | 0.060578477 |
| Usp29 | 0.046700673 | 6.02E-09 | 6.17E-08 | 0.234563337 | 0.191870474 |
| Usp3 | 0.042885675 | 9.52E-08 | 8.08E-07 | 0.238700302 | 0.206170355 |
| Usp30 | 0.007062328 | 0.137308745 | 0.228346839 | 0.067187321 | 0.059726525 |
| Usp31 | 0.006990495 | 0.283585142 | 0.402660253 | 0.124756943 | 0.120930954 |

| | | | | | |
|---|---|---|---|---|---|
| Usp32 | 0.017758152 | 0.00237357 | 0.00758575 | 0.106630289 | 0.092445619 |
| Usp33 | 0.027212178 | 0.004373483 | 0.012880643 | 0.409877393 | 0.378972036 |
| Usp34 | 0.025505105 | 0.013897101 | 0.034230458 | 0.671980163 | 0.640648316 |
| Usp35 | 0.020268722 | 0.007387747 | 0.020135598 | 0.186889633 | 0.172774579 |
| Usp36 | 0.006219763 | 0.431855403 | 0.550478331 | 0.198535197 | 0.19932612 |
| Usp37 | 0.016704186 | 0.010980414 | 0.028026144 | 0.14227498 | 0.124247403 |
| Usp38 | 0.007542443 | 0.201678197 | 0.309212337 | 0.104048215 | 0.097666763 |
| Usp39 | 0.007741344 | 0.170236438 | 0.269506375 | 0.095442642 | 0.087706802 |
| Usp4 | 0.035248848 | 3.10E-06 | 1.98E-05 | 0.215010962 | 0.176424344 |
| Usp40 | -0.010490283 | 0.072294699 | 0.135737556 | 0.090878247 | 0.09767435 |
| Usp42 | 0.012454472 | 0.032297171 | 0.069467499 | 0.101337814 | 0.09206753 |
| Usp45 | 0.0339862 | 1.01E-05 | 5.79E-05 | 0.211735076 | 0.178461278 |
| Usp46 | 0.024282686 | 0.002288108 | 0.007348857 | 0.225681995 | 0.205186462 |
| Usp47 | 0.0476399 | 3.54E-07 | 2.72E-06 | 0.401676611 | 0.353027398 |
| Usp48 | -0.006896741 | 0.439866005 | 0.55834669 | 0.312610973 | 0.310686163 |
| Usp5 | -0.005502485 | 0.410474646 | 0.529824018 | 0.13769927 | 0.137580367 |
| Usp50 | 0.064041606 | 2.20E-14 | 4.11E-13 | 0.258558145 | 0.234316835 |
| Usp51 | 0.00835806 | 0.055424772 | 0.108856491 | 0.056263495 | 0.046567414 |
| Usp53 | 0.013347374 | 0.024917896 | 0.056064793 | 0.109588416 | 0.097242266 |
| Usp6nl | 0.003039302 | 0.607072993 | 0.708504807 | 0.099342909 | 0.098594194 |
| Usp7 | -0.008991939 | 0.338897574 | 0.461257485 | 0.37502436 | 0.377300825 |
| Usp8 | 0.011565687 | 0.201986391 | 0.309480331 | 0.333514476 | 0.320448656 |
| Usp9x | 0.010172933 | 0.320197267 | 0.441679601 | 0.940126211 | 0.895662214 |
| Uspl1 | 0.018177687 | 0.009517463 | 0.024866642 | 0.150422912 | 0.139693623 |
| Ust | 0.011886417 | 0.002500781 | 0.007940101 | 0.046520042 | 0.03712818 |
| Utp11 | -0.00137163 | 0.857773811 | 0.902287779 | 0.198610118 | 0.193855666 |
| Utp14a | -0.002602038 | 0.688744878 | 0.773833088 | 0.116404334 | 0.121405535 |
| Utp15 | -0.000452249 | 0.930030298 | 0.952465454 | 0.072412007 | 0.073743789 |
| Utp18 | 0.026170465 | 0.000563228 | 0.002156639 | 0.197818573 | 0.173829832 |
| Utp20 | 0.002272267 | 0.619225562 | 0.718733395 | 0.05583396 | 0.054932276 |
| Utp23 | -0.005292798 | 0.342222965 | 0.464630131 | 0.084656827 | 0.088628995 |
| Utp25 | -0.00322479 | 0.447157685 | 0.565976697 | 0.045310624 | 0.047787601 |
| Utp3 | 0.019322485 | 0.036170092 | 0.076245801 | 0.346531087 | 0.331974452 |
| Utp4 | 0.001952738 | 0.7232761 | 0.800420062 | 0.088891929 | 0.084487036 |
| Utp6 | 0.015567253 | 0.006623238 | 0.01836074 | 0.10339527 | 0.087644462 |
| Utrn | 0.095610788 | 2.53E-23 | 9.25E-22 | 0.42954618 | 0.316600748 |
| Uty | -0.008545535 | 0.205270332 | 0.313525637 | 0.134617794 | 0.136719386 |
| Uvrag | 0.032877169 | 5.95E-06 | 3.60E-05 | 0.188114015 | 0.155742535 |
| Uvssa | -0.004636158 | 0.532626665 | 0.643734017 | 0.161359572 | 0.168184478 |
| Uxs1 | -0.00612454 | 0.298115012 | 0.417799358 | 0.094864984 | 0.101155996 |
| Uxt | -0.03201918 | 0.000178985 | 0.000779612 | 0.241023231 | 0.272385078 |
| Vac14 | -0.002408961 | 0.6404102 | 0.73568096 | 0.072073834 | 0.072711671 |
| Vamp2 | -0.001552304 | 0.874091845 | 0.913754937 | 0.647963982 | 0.634749503 |
| Vamp3 | 0.017488301 | 0.032567565 | 0.06994077 | 0.248738889 | 0.224997912 |
| Vamp4 | 0.015216884 | 0.104221714 | 0.18273102 | 0.421743837 | 0.394049126 |
| Vamp5 | -0.0025104 | 0.50288112 | 0.617065537 | 0.03841492 | 0.037563057 |
| Vamp8 | 0.015235355 | 0.109437691 | 0.190230183 | 0.940204482 | 0.908550141 |
| Vapa | -0.025483586 | 0.007546951 | 0.020491196 | 0.861388105 | 0.859894561 |
| Vapb | -0.011262327 | 0.258792847 | 0.374904494 | 0.967686076 | 0.961866491 |
| Vars | -0.019231243 | 0.027602668 | 0.060947794 | 0.25973604 | 0.273413818 |
| Vars2 | 0.006297673 | 0.102374816 | 0.180273132 | 0.043454604 | 0.037435794 |
| Vasp | -0.005099234 | 0.481951972 | 0.598377069 | 0.169285237 | 0.17202963 |
| Vat1 | 0.038941325 | 1.65E-06 | 1.10E-05 | 0.244858138 | 0.209615213 |
| Vat1l | 0.012355217 | 0.066259207 | 0.12683048 | 0.146694575 | 0.131721595 |
| Vbp1 | -0.001057673 | 0.904394125 | 0.935175461 | 0.306860916 | 0.297821787 |
| Vcl | 0.02125297 | 0.000205551 | 0.000879562 | 0.102359925 | 0.0834976 |
| Vcp | 0.026256757 | 0.005692305 | 0.016092058 | 1.07591583 | 1.017419531 |
| Vcpip1 | 0.01708914 | 0.034449961 | 0.073319291 | 0.229635642 | 0.21306515 |
| Vcpkmt | -0.004598009 | 0.202496836 | 0.31007627 | 0.034953859 | 0.035401903 |
| Vdac1 | -0.012232252 | 0.238968693 | 0.352863553 | 0.795835256 | 0.804985156 |
| Vdac2 | -0.055259015 | 3.40E-08 | 3.10E-07 | 0.738804961 | 0.780448032 |
| Vdac3 | -0.155436107 | 6.07E-47 | 6.56E-45 | 0.837009635 | 1.02280245 |
| Vdr | 0.052499153 | 2.74E-07 | 2.14E-06 | 0.477156991 | 0.403390005 |
| Vegfa | 0.008332487 | 0.39724463 | 0.517699696 | 1.157361565 | 1.128208269 |
| Vegfb | -0.020304799 | 0.021889586 | 0.050136122 | 0.291914394 | 0.299030757 |
| Vezf1 | 0.056294939 | 3.58E-09 | 3.79E-08 | 0.443801507 | 0.379688265 |
| Vezt | -0.005406988 | 0.352807337 | 0.475199907 | 0.093331633 | 0.096616697 |
| Vgf | -0.172891167 | 6.13E-54 | 8.39E-52 | 0.212213651 | 0.434687722 |
| Vgll4 | 0.021654171 | 6.74E-05 | 0.000325177 | 0.096419438 | 0.076075135 |
| Vhl | 0.002571695 | 0.640673194 | 0.735896123 | 0.08756042 | 0.083810334 |
| Vil1 | 0.071144621 | 1.87E-18 | 4.78E-17 | 0.261123494 | 0.185251838 |
| Vipas39 | 0.008766635 | 0.167924815 | 0.266541233 | 0.124705371 | 0.115495128 |

| | | | | | |
|---|---|---|---|---|---|
| Vipr1 | 0.013644729 | 0.042767339 | 0.08757036 | 0.134977228 | 0.128610176 |
| Virma | 0.039123539 | 1.43E-07 | 1.18E-06 | 0.203841895 | 0.164028346 |
| Vkorc1 | -0.010069503 | 0.298841322 | 0.418570856 | 0.527729851 | 0.474710938 |
| Vkorc1l1 | -0.000623055 | 0.937004375 | 0.957160299 | 0.21510559 | 0.209457719 |
| Vldlr | 0.065231652 | 1.98E-10 | 2.45E-09 | 0.54628525 | 0.480166225 |
| Vma21 | 0.019313432 | 0.026080327 | 0.058101041 | 0.300249203 | 0.279989367 |
| Vmac | -0.024063639 | 0.001695711 | 0.005641926 | 0.196230431 | 0.202013269 |
| Vmp1 | -0.068357281 | 7.88E-12 | 1.14E-10 | 0.328652193 | 0.413930657 |
| Vopp1 | -0.000609892 | 0.884681617 | 0.921160402 | 0.046869434 | 0.04762189 |
| Vps11 | 0.014243492 | 0.050677766 | 0.101085194 | 0.1865511 | 0.168301695 |
| Vps13a | 0.0171945 | 0.062197637 | 0.120295945 | 0.345190297 | 0.320436792 |
| Vps13b | -0.016058459 | 0.040793208 | 0.084291513 | 0.195198276 | 0.202270814 |
| Vps13c | 0.007458884 | 0.44028176 | 0.558791378 | 0.370382601 | 0.373535591 |
| Vps13d | 0.023662309 | 0.000192077 | 0.000828471 | 0.13591736 | 0.109915287 |
| Vps16 | -0.008561885 | 0.088461734 | 0.159945132 | 0.069634728 | 0.072874325 |
| Vps18 | 0.005441343 | 0.324545712 | 0.446537421 | 0.090201957 | 0.084392854 |
| Vps26a | -0.007589451 | 0.23821986 | 0.352138515 | 0.120297412 | 0.124544154 |
| Vps26b | 0.012739106 | 0.051888635 | 0.102891749 | 0.13773713 | 0.125879142 |
| Vps26c | 0.014062909 | 0.018898901 | 0.044314908 | 0.112134235 | 0.09895512 |
| Vps28 | -0.105103775 | 3.42E-26 | 1.50E-24 | 0.645796791 | 0.744311304 |
| Vps29 | -0.002896868 | 0.767956731 | 0.835973053 | 0.577314366 | 0.570177418 |
| Vps33a | 0.000979079 | 0.840629304 | 0.890940596 | 0.066586591 | 0.063882902 |
| Vps33b | 0.001953557 | 0.628742095 | 0.726715297 | 0.045915415 | 0.043353977 |
| Vps35 | 0.078164787 | 1.20E-14 | 2.29E-13 | 0.683788718 | 0.586444097 |
| Vps35l | -0.006628434 | 0.328020984 | 0.45055383 | 0.141760234 | 0.142685103 |
| Vps36 | 0.017746192 | 0.045578107 | 0.092333894 | 0.322330423 | 0.308439905 |
| Vps37a | 0.08877574 | 3.78E-18 | 9.45E-17 | 0.779503832 | 0.648721174 |
| Vps37b | -0.016583339 | 0.010258505 | 0.026447412 | 0.114571675 | 0.125440826 |
| Vps37c | 0.01599197 | 0.010639052 | 0.027283795 | 0.127465596 | 0.110174736 |
| Vps37d | 0.004915883 | 0.267258955 | 0.383908328 | 0.059455776 | 0.051360494 |
| Vps39 | 0.020382143 | 0.000333191 | 0.00135252 | 0.100917202 | 0.084942688 |
| Vps41 | 0.015174798 | 0.067266578 | 0.128203425 | 0.263764138 | 0.239584154 |
| Vps45 | 0.000665362 | 0.921146596 | 0.94605823 | 0.139126734 | 0.139460063 |
| Vps4a | 0.029188548 | 9.34E-05 | 0.000439373 | 0.194828204 | 0.173643761 |
| Vps4b | 0.032996231 | 0.000646459 | 0.002437889 | 0.4427599 | 0.427312566 |
| Vps50 | 0.004964058 | 0.46025942 | 0.578642452 | 0.142895898 | 0.134226302 |
| Vps51 | -0.004954891 | 0.480929436 | 0.597214967 | 0.153212123 | 0.155696947 |
| Vps52 | 0.017949565 | 0.00471652 | 0.013720529 | 0.129774041 | 0.115369512 |
| Vps53 | 0.024197902 | 0.004650287 | 0.013552185 | 0.294485401 | 0.269686773 |
| Vps54 | 0.012214929 | 0.047707496 | 0.096146823 | 0.115399265 | 0.105877897 |
| Vps72 | 0.010265307 | 0.14023295 | 0.232256344 | 0.168715364 | 0.15057748 |
| Vps8 | 0.103844322 | 1.36E-43 | 1.28E-41 | 0.249758321 | 0.131776946 |
| Vps9d1 | 0.003637032 | 0.457728448 | 0.576320747 | 0.070764546 | 0.06517247 |
| Vrk1 | 0.031400691 | 0.000279471 | 0.001155195 | 0.27723085 | 0.245042504 |
| Vrk3 | -0.016905565 | 0.017276521 | 0.041227869 | 0.155287287 | 0.165519151 |
| Vsig10 | 0.014557961 | 0.006622536 | 0.01836074 | 0.091636481 | 0.076400847 |
| Vsnl1 | 0.005909413 | 0.160164341 | 0.257289775 | 0.060675069 | 0.044391406 |
| Vstm2l | 0.049026867 | 5.86E-07 | 4.31E-06 | 0.611901038 | 0.464590418 |
| Vta1 | -0.013035414 | 0.121377929 | 0.206300039 | 0.263291026 | 0.26506353 |
| Vti1a | -0.013243856 | 0.022478312 | 0.051166976 | 0.093100911 | 0.101164271 |
| Vti1b | -0.040638707 | 4.10E-05 | 0.000205035 | 0.587972026 | 0.611754088 |
| Vwa5a | 0.001164262 | 0.852203379 | 0.898680132 | 0.115517681 | 0.112233011 |
| Vwa5b2 | 0.008677895 | 0.284805013 | 0.404038281 | 0.230163047 | 0.20550367 |
| Vwa8 | -0.005977892 | 0.401618119 | 0.521647476 | 0.162156984 | 0.159580082 |
| Wac | 0.065758801 | 1.12E-11 | 1.60E-10 | 0.485393499 | 0.414030013 |
| Wapl | 0.018062934 | 0.066240773 | 0.126820165 | 0.536147014 | 0.50762412 |
| Wars | -0.023738713 | 0.017940593 | 0.042551463 | 0.34454214 | 0.40008611 |
| Wasf2 | 0.024376994 | 0.005118189 | 0.014721609 | 0.301699504 | 0.275795299 |
| Washc1 | 0.002048888 | 0.737660301 | 0.812086224 | 0.122794958 | 0.111039082 |
| Washc2 | 0.012708378 | 0.11942678 | 0.203553148 | 0.23841037 | 0.218520872 |
| Washc3 | -0.032625177 | 0.000264883 | 0.001102866 | 0.313953919 | 0.337348717 |
| Washc4 | 0.027181128 | 6.06E-05 | 0.000294541 | 0.152799427 | 0.129067859 |
| Washc5 | 0.019767612 | 0.004393512 | 0.012935712 | 0.153647289 | 0.143906128 |
| Wasl | 0.027049681 | 0.005242509 | 0.015021418 | 0.558676931 | 0.522341781 |
| Wbp1 | 0.004880651 | 0.531442428 | 0.6428219 | 0.225630253 | 0.207557812 |
| Wbp11 | 0.068209347 | 2.58E-12 | 3.97E-11 | 0.490361444 | 0.425820105 |
| Wbp1l | 0.012413282 | 0.02810949 | 0.061954309 | 0.09852798 | 0.086858837 |
| Wbp2 | 0.020170234 | 0.036250773 | 0.076382752 | 0.44604845 | 0.413657749 |
| Wbp4 | -0.005800022 | 0.563805035 | 0.67164717 | 0.525696403 | 0.537943883 |
| Wdcp | -0.006756588 | 0.267853502 | 0.384591898 | 0.105669174 | 0.109409269 |
| Wdfy1 | -0.113492122 | 1.10E-34 | 7.05E-33 | 0.252817746 | 0.360097512 |
| Wdfy2 | 0.010802517 | 0.039134212 | 0.081365015 | 0.082161749 | 0.071468189 |

| | | | | | |
|---|---|---|---|---|---|
| Wdfy3 | 0.02083986 | 0.002789098 | 0.008721006 | 0.163941395 | 0.142664976 |
| Wdpcp | -0.010797444 | 0.02741558 | 0.060603516 | 0.067552357 | 0.069204036 |
| Wdr1 | 0.008790518 | 0.295613398 | 0.41549132 | 0.249797949 | 0.249262715 |
| Wdr11 | 0.016076955 | 0.010691055 | 0.027398628 | 0.130376348 | 0.112263018 |
| Wdr12 | -0.034218987 | 1.13E-05 | 6.40E-05 | 0.173376711 | 0.201336727 |
| Wdr13 | -0.014290855 | 0.034720926 | 0.07375592 | 0.133645306 | 0.142251818 |
| Wdr18 | -0.026304732 | 0.00037118 | 0.001488325 | 0.164213477 | 0.1824059 |
| Wdr20 | -0.004853928 | 0.371444639 | 0.493814634 | 0.080620022 | 0.083141021 |
| Wdr24 | 0.008971788 | 0.059775796 | 0.116316257 | 0.06705543 | 0.059886876 |
| Wdr26 | 0.03764064 | 9.02E-05 | 0.000424797 | 0.468659003 | 0.41807192 |
| Wdr27 | 0.002334595 | 0.641919637 | 0.736234882 | 0.06981328 | 0.068544043 |
| Wdr3 | 0.001355213 | 0.790827926 | 0.853690131 | 0.073904337 | 0.070154212 |
| Wdr33 | 0.021767273 | 0.022297592 | 0.050870842 | 0.436041388 | 0.407214672 |
| Wdr35 | 0.017542114 | 0.008717725 | 0.023150428 | 0.148557343 | 0.129224607 |
| Wdr36 | 0.003352608 | 0.607206547 | 0.708504807 | 0.124507635 | 0.123439757 |
| Wdr37 | 0.012805041 | 0.051499912 | 0.102326209 | 0.139448831 | 0.125791647 |
| Wdr4 | 0.00603953 | 0.141569684 | 0.234011065 | 0.049935111 | 0.043271014 |
| Wdr41 | 0.004919296 | 0.399233288 | 0.51938258 | 0.104553812 | 0.09596768 |
| Wdr43 | 0.025990983 | 0.002914119 | 0.00906304 | 0.27002557 | 0.259204822 |
| Wdr44 | -0.004108336 | 0.421643233 | 0.540650951 | 0.070417632 | 0.071817885 |
| Wdr45 | -0.011249424 | 0.042396457 | 0.086925254 | 0.087979433 | 0.089002489 |
| Wdr45b | 0.031039636 | 0.00020284 | 0.000868727 | 0.267341488 | 0.238608639 |
| Wdr46 | -0.016139218 | 0.003680248 | 0.011094379 | 0.082621906 | 0.09253688 |
| Wdr47 | 0.008450895 | 0.156099921 | 0.2523451 | 0.108555389 | 0.099662847 |
| Wdr48 | -0.004599794 | 0.450656064 | 0.569071081 | 0.109072657 | 0.108558142 |
| Wdr5 | 0.000702024 | 0.885462599 | 0.921841299 | 0.06675236 | 0.064652804 |
| Wdr53 | -0.000763189 | 0.879607132 | 0.917842711 | 0.077656853 | 0.072007538 |
| Wdr54 | 0.001573648 | 0.679598445 | 0.767191835 | 0.038862919 | 0.038606005 |
| Wdr55 | 0.001038026 | 0.812373315 | 0.870675049 | 0.056819462 | 0.05226101 |
| Wdr59 | -0.005381171 | 0.194408363 | 0.299910838 | 0.043235196 | 0.04685456 |
| Wdr6 | 0.050919155 | 1.92E-08 | 1.81E-07 | 0.332939229 | 0.276056228 |
| Wdr61 | 0.007297044 | 0.42342459 | 0.54189724 | 0.357488056 | 0.343076118 |
| Wdr7 | 0.023630003 | 0.000386082 | 0.001538946 | 0.144613811 | 0.124640888 |
| Wdr70 | 0.008920338 | 0.12570947 | 0.212178381 | 0.101588482 | 0.092770976 |
| Wdr73 | -0.012568389 | 0.014513953 | 0.035542734 | 0.07073189 | 0.077374503 |
| Wdr74 | -0.026529353 | 0.000374429 | 0.001498641 | 0.163390752 | 0.182108641 |
| Wdr75 | -0.007199598 | 0.356835735 | 0.479190905 | 0.185379914 | 0.19728752 |
| Wdr77 | 0.001176813 | 0.816241403 | 0.873280392 | 0.073577235 | 0.070829356 |
| Wdr81 | 0.002148064 | 0.579289124 | 0.685640232 | 0.042339454 | 0.037579562 |
| Wdr82 | 0.003818729 | 0.54792527 | 0.65757978 | 0.120040749 | 0.116501902 |
| Wdr83 | -0.010361956 | 0.060705262 | 0.117823229 | 0.087941082 | 0.091197811 |
| Wdr83os | -0.074161864 | 1.21E-13 | 2.17E-12 | 0.640039362 | 0.711423045 |
| Wdr89 | -0.001351862 | 0.843959892 | 0.89339918 | 0.127401923 | 0.147389231 |
| Wdr90 | 0.00391034 | 0.424247979 | 0.542702481 | 0.064819947 | 0.064680627 |
| Wdr92 | 0.001436089 | 0.778431111 | 0.844538251 | 0.076855286 | 0.071682865 |
| Wdsub1 | 0.020072858 | 0.003403768 | 0.010389775 | 0.149771747 | 0.132008172 |
| Wdtc1 | 0.004895073 | 0.421812939 | 0.54079723 | 0.114809058 | 0.108131967 |
| Wdyhv1 | -0.010230576 | 0.092703543 | 0.165886958 | 0.101554352 | 0.11140234 |
| Wee1 | -0.002198433 | 0.607708066 | 0.70883092 | 0.042121026 | 0.049292342 |
| Wfs1 | -0.031550374 | 0.00091183 | 0.003305297 | 0.376953513 | 0.403866392 |
| Whamm | -0.00363179 | 0.613930579 | 0.714035985 | 0.157034019 | 0.158911788 |
| Whrn | 0.00389349 | 0.488674063 | 0.604129615 | 0.094232737 | 0.087261598 |
| Wipf2 | 0.017897962 | 0.013123812 | 0.032606619 | 0.172355079 | 0.155226133 |
| Wipi1 | -0.124389186 | 1.94E-25 | 8.01E-24 | 0.386405768 | 0.571984108 |
| Wipi2 | -0.000552579 | 0.950250524 | 0.967032331 | 0.319496803 | 0.309941423 |
| Wiz | 0.010907849 | 0.074376189 | 0.139093614 | 0.115940047 | 0.105321329 |
| Wls | -0.053778973 | 4.04E-08 | 3.63E-07 | 1.003891211 | 1.069396414 |
| Wnk1 | 0.030280084 | 0.002534536 | 0.008028923 | 0.775125243 | 0.73545358 |
| Wnk2 | 0.01386248 | 0.000274847 | 0.001138985 | 0.045885785 | 0.034094103 |
| Wnk3 | 0.004649887 | 0.27557759 | 0.393589736 | 0.048856994 | 0.044792849 |
| Wnk4 | 0.023748107 | 0.00018182 | 0.000789132 | 0.127784727 | 0.106730639 |
| Wnt4 | -0.009097835 | 0.410757736 | 0.530119038 | 1.650197849 | 1.615012516 |
| Wrap53 | -0.003536693 | 0.333845286 | 0.456746956 | 0.035567401 | 0.035721813 |
| Wrap73 | -0.008762119 | 0.093073513 | 0.166431455 | 0.078187116 | 0.080938816 |
| Wrn | 0.006119773 | 0.292085669 | 0.411495378 | 0.10153854 | 0.092847142 |
| Wrnip1 | 0.03048723 | 4.32E-06 | 2.69E-05 | 0.153312633 | 0.124539762 |
| Wsb2 | 0.039587635 | 7.94E-05 | 0.000378495 | 0.629279383 | 0.572163083 |
| Wtap | 0.030624196 | 0.000144569 | 0.000648891 | 0.256627256 | 0.215536508 |
| Wtip | -0.003272376 | 0.484807593 | 0.601109478 | 0.062551114 | 0.06117238 |
| Wwox | -0.012252817 | 0.02009968 | 0.046625886 | 0.069063441 | 0.080556678 |
| Wwp1 | 0.007442394 | 0.196783026 | 0.302901295 | 0.099840572 | 0.091504097 |
| Wwp2 | 0.007677689 | 0.242602068 | 0.356495965 | 0.133517884 | 0.124766809 |

| | | | | | |
|---|---|---|---|---|---|
| Xab2 | 0.014620301 | 0.011312555 | 0.028783215 | 0.114128654 | 0.093060418 |
| Xaf1 | 0.034996837 | 3.94E-15 | 7.70E-14 | 0.061596396 | 0.037674915 |
| Xbp1 | -0.051821568 | 1.17E-07 | 9.77E-07 | 0.957588011 | 0.994518224 |
| Xiap | 0.018766842 | 0.014696435 | 0.035917229 | 0.203875193 | 0.184981281 |
| Xk | 0.004723878 | 0.398809648 | 0.519012275 | 0.095786494 | 0.086303006 |
| Xkr6 | -0.011680663 | 0.077248118 | 0.143475771 | 0.121407542 | 0.130838912 |
| Xpa | -0.027311095 | 0.000213437 | 0.000909304 | 0.164252556 | 0.18228014 |
| Xpc | 0.010229377 | 0.128449176 | 0.216276912 | 0.14402731 | 0.132376334 |
| Xpnpep1 | -0.002511485 | 0.678075335 | 0.766184148 | 0.106131888 | 0.106453906 |
| Xpnpep3 | 0.016578836 | 0.001294347 | 0.004478012 | 0.081865231 | 0.067166111 |
| Xpo1 | 0.03434314 | 3.20E-06 | 2.04E-05 | 0.195331591 | 0.15828109 |
| Xpo4 | -0.002031176 | 0.752023419 | 0.823422872 | 0.121918503 | 0.12089864 |
| Xpo5 | 0.007271668 | 0.231549258 | 0.344735358 | 0.10939642 | 0.103494018 |
| Xpo6 | 0.008485216 | 0.23466911 | 0.348521676 | 0.168385368 | 0.157673319 |
| Xpo7 | 0.022238379 | 0.007314855 | 0.019976129 | 0.257968425 | 0.23703938 |
| Xpot | -0.005074545 | 0.605383199 | 0.707323309 | 0.409729369 | 0.41545937 |
| Xpr1 | 0.01240389 | 0.113168016 | 0.19521282 | 0.201884759 | 0.192358263 |
| Xrcc1 | 0.011422829 | 0.057931276 | 0.113230372 | 0.119627612 | 0.102076241 |
| Xrcc3 | 0.003678677 | 0.286724567 | 0.405991296 | 0.037653681 | 0.029678447 |
| Xrcc4 | -0.008295504 | 0.144686542 | 0.238292828 | 0.092685545 | 0.093786379 |
| Xrcc5 | 0.000974847 | 0.897899326 | 0.930438846 | 0.190907584 | 0.186734229 |
| Xrcc6 | -0.014908378 | 0.050423333 | 0.100713587 | 0.184366478 | 0.192206393 |
| Xrn1 | 0.02766882 | 0.002005584 | 0.006554047 | 0.303456567 | 0.279146279 |
| Xrn2 | 0.010351105 | 0.23094019 | 0.344079073 | 0.27342186 | 0.266500154 |
| Xxylt1 | -0.0136757 | 0.00050919 | 0.001973823 | 0.034346464 | 0.04352461 |
| Yae1d1 | 0.005595368 | 0.475625499 | 0.592444728 | 0.210728937 | 0.20446956 |
| Yaf2 | 0.040312663 | 6.35E-06 | 3.81E-05 | 0.35298294 | 0.305500625 |
| Yars | -0.001375294 | 0.881571804 | 0.918906517 | 0.346814514 | 0.336483117 |
| Yars2 | -0.007863479 | 0.088333562 | 0.159772817 | 0.056664238 | 0.059485782 |
| Ybey | -0.001020848 | 0.823735905 | 0.87830231 | 0.055365525 | 0.05670757 |
| Ybx1 | -0.027300681 | 0.000647314 | 0.002440165 | 1.482876411 | 1.53190061 |
| Ybx3 | 0.023911915 | 0.021341382 | 0.049119536 | 0.5498676 | 0.531670944 |
| Ydjc | -0.029925664 | 3.79E-08 | 3.43E-07 | 0.069400362 | 0.091581612 |
| Yeats2 | 0.014172244 | 0.003995181 | 0.011899863 | 0.073194772 | 0.061826068 |
| Yeats4 | -0.007344076 | 0.410018182 | 0.529727145 | 0.322054541 | 0.318794074 |
| Yes1 | -0.020426647 | 0.000946724 | 0.003410169 | 0.097368656 | 0.114053119 |
| Yif1a | -0.035249679 | 0.000372275 | 0.001491861 | 0.456053196 | 0.489742353 |
| Yif1b | 0.009363791 | 0.264443589 | 0.380764303 | 0.259374042 | 0.248524041 |
| Yipf1 | -0.025875972 | 0.003777983 | 0.011350293 | 0.329638612 | 0.347437374 |
| Yipf2 | -0.027501945 | 0.000587016 | 0.002238584 | 0.214003115 | 0.228970478 |
| Yipf3 | -0.001123604 | 0.906728792 | 0.935897369 | 0.456893373 | 0.455909841 |
| Yipf4 | -0.067143206 | 6.58E-12 | 9.58E-11 | 0.47297191 | 0.523007633 |
| Yipf5 | 0.00918597 | 0.342552093 | 0.464735061 | 0.461619759 | 0.44860574 |
| Yipf6 | 0.017549715 | 0.050786161 | 0.101262047 | 0.333499133 | 0.313907549 |
| Yju2 | 0.002588804 | 0.624727348 | 0.723305856 | 0.079246125 | 0.076061751 |
| Ykt6 | 0.026807887 | 0.000219172 | 0.000930436 | 0.172954701 | 0.153489537 |
| Ylpm1 | 0.024587757 | 0.006742131 | 0.01863719 | 0.315232581 | 0.299576197 |
| Yme1l1 | 0.024180959 | 0.011294178 | 0.028743978 | 0.427068577 | 0.396967911 |
| Yod1 | 0.006348295 | 0.316684318 | 0.438326444 | 0.124460885 | 0.113773926 |
| Ypel2 | -0.020945876 | 0.022364562 | 0.050967714 | 0.282520721 | 0.303168908 |
| Ypel3 | -0.019838123 | 0.046538449 | 0.094083345 | 0.829961189 | 0.786284339 |
| Ypel4 | -0.00119249 | 0.849607807 | 0.896952957 | 0.117708414 | 0.114976235 |
| Ypel5 | 0.002058976 | 0.820013965 | 0.876159552 | 0.33732959 | 0.333657613 |
| Yrdc | -0.004155976 | 0.641238501 | 0.736074138 | 0.281916786 | 0.285500453 |
| Ythdc1 | 0.09283001 | 1.95E-20 | 5.77E-19 | 0.544720001 | 0.463795872 |
| Ythdc2 | -0.000313726 | 0.956847377 | 0.970523756 | 0.095938821 | 0.093702854 |
| Ythdf1 | 0.015640001 | 0.039223326 | 0.081515429 | 0.203645218 | 0.183121854 |
| Ythdf2 | 0.008597775 | 0.255589269 | 0.371093022 | 0.201542856 | 0.182974939 |
| Ythdf3 | 0.04594519 | 2.50E-08 | 2.31E-07 | 0.264304376 | 0.224664693 |
| Ywhab | -0.028615065 | 0.00443539 | 0.013035327 | 0.612065032 | 0.626755494 |
| Ywhae | 0.027491209 | 0.000517036 | 0.002001044 | 1.426507559 | 1.367083783 |
| Ywhag | 0.014328756 | 0.155800135 | 0.251986177 | 0.486117508 | 0.490789495 |
| Ywhah | -0.01855781 | 0.078560752 | 0.145409653 | 0.671052097 | 0.693087144 |
| Ywhaq | -0.021528752 | 0.003220449 | 0.009882958 | 1.702711248 | 1.686207607 |
| Ywhaz | 0.055330721 | 1.63E-08 | 1.56E-07 | 0.888670581 | 0.857343307 |
| Yy1 | 0.024246278 | 0.015793869 | 0.038186519 | 0.587036703 | 0.540791669 |
| Zadh2 | 0.016217966 | 0.004765589 | 0.013838427 | 0.10531776 | 0.088257488 |
| Zbed3 | -0.008734437 | 0.157291243 | 0.253764597 | 0.109934289 | 0.113638162 |
| Zbtb1 | -0.003558879 | 0.555893706 | 0.66491833 | 0.100732584 | 0.103905848 |
| Zbtb10 | 0.006282904 | 0.221047032 | 0.331946129 | 0.077118511 | 0.068445021 |
| Zbtb11 | -0.012590403 | 0.112915616 | 0.194923418 | 0.195605748 | 0.204021268 |
| Zbtb11os1 | -0.0027762 | 0.601096019 | 0.70404388 | 0.078569384 | 0.078365222 |

| | | | | | |
|---|---|---|---|---|---|
| Zbtb12 | 0.009217601 | 0.055049081 | 0.108184185 | 0.071418017 | 0.060247215 |
| Zbtb14 | 0.044972472 | 1.03E-10 | 1.32E-09 | 0.186061059 | 0.136039343 |
| Zbtb17 | 0.014086177 | 0.000212167 | 0.000904688 | 0.045616618 | 0.034766518 |
| Zbtb18 | -0.005399867 | 0.443609573 | 0.562291039 | 0.138798557 | 0.146131445 |
| Zbtb2 | -0.004714913 | 0.24094075 | 0.354902653 | 0.039420525 | 0.04284484 |
| Zbtb20 | 0.035585386 | 0.000650393 | 0.002448924 | 1.245082134 | 1.177297802 |
| Zbtb21 | 0.014622543 | 0.01757413 | 0.041828878 | 0.114422717 | 0.104584585 |
| Zbtb22 | -0.003811715 | 0.446222318 | 0.565307971 | 0.07054251 | 0.070306575 |
| Zbtb24 | -0.000399006 | 0.945857323 | 0.96379873 | 0.097874371 | 0.099066913 |
| Zbtb25 | 0.007924598 | 0.043601297 | 0.089034197 | 0.047833817 | 0.038615447 |
| Zbtb26 | -0.002346103 | 0.544650888 | 0.654280976 | 0.040105061 | 0.039799486 |
| Zbtb33 | 0.023309081 | 4.35E-05 | 0.000216638 | 0.104927387 | 0.085140148 |
| Zbtb37 | -0.007159171 | 0.115976502 | 0.199033283 | 0.050356471 | 0.055603477 |
| Zbtb38 | 0.013083593 | 0.140635994 | 0.23276526 | 0.310024935 | 0.289900256 |
| Zbtb4 | 0.031515456 | 0.000378318 | 0.00151296 | 0.327636183 | 0.289141749 |
| Zbtb40 | 0.012333291 | 0.006988051 | 0.019207756 | 0.064111069 | 0.052787391 |
| Zbtb41 | 0.016420792 | 0.012922928 | 0.032181533 | 0.137808912 | 0.127185579 |
| Zbtb43 | -0.002636781 | 0.623357036 | 0.722115111 | 0.078646202 | 0.080481622 |
| Zbtb44 | -0.002927675 | 0.701866461 | 0.784484964 | 0.190464676 | 0.184561536 |
| Zbtb45 | 0.013368323 | 0.001479383 | 0.005023597 | 0.059224732 | 0.044182178 |
| Zbtb48 | 0.005729348 | 0.112771676 | 0.194874909 | 0.037764853 | 0.032383705 |
| Zbtb5 | -0.002507028 | 0.588302939 | 0.692934477 | 0.058714719 | 0.057665565 |
| Zbtb6 | 0.005894092 | 0.306659672 | 0.42755562 | 0.096008609 | 0.09082609 |
| Zbtb7a | 0.055001839 | 6.74E-08 | 5.87E-07 | 0.643644318 | 0.575754469 |
| Zbtb7b | -0.010920356 | 0.11719644 | 0.200560427 | 0.136121498 | 0.150375502 |
| Zbtb7c | 0.018257226 | 1.09E-06 | 7.59E-06 | 0.047506141 | 0.03122962 |
| Zbtb8a | 0.001724583 | 0.635787759 | 0.731494508 | 0.036711615 | 0.034029474 |
| Zbtb8os | -0.047837619 | 3.63E-08 | 3.29E-07 | 0.25619405 | 0.295588232 |
| Zc2hc1a | 0.003624259 | 0.542951355 | 0.65297562 | 0.110831818 | 0.101947734 |
| Zc2hc1c | 0.023503586 | 9.97E-07 | 6.96E-06 | 0.080039387 | 0.056336229 |
| Zc3h10 | 0.025628254 | 0.000247056 | 0.001035292 | 0.166128725 | 0.140759315 |
| Zc3h12b | 0.005302704 | 0.162943999 | 0.260824691 | 0.040357171 | 0.036187334 |
| Zc3h12c | 0.016546689 | 0.005824803 | 0.016423646 | 0.108145277 | 0.095478185 |
| Zc3h13 | 0.035531416 | 8.91E-05 | 0.000420017 | 0.320823821 | 0.285647049 |
| Zc3h14 | 0.013654446 | 0.14927382 | 0.243963729 | 0.432020715 | 0.407101554 |
| Zc3h15 | 0.056767446 | 9.32E-09 | 9.33E-08 | 0.844847146 | 0.78788856 |
| Zc3h18 | 0.004469088 | 0.584973294 | 0.690015817 | 0.22337207 | 0.219350791 |
| Zc3h3 | 0.063347253 | 4.16E-11 | 5.57E-10 | 0.381954805 | 0.268122175 |
| Zc3h4 | 0.015767332 | 0.00348557 | 0.01057968 | 0.092780415 | 0.077244936 |
| Zc3h6 | 0.025559695 | 0.000461634 | 0.00180822 | 0.17660651 | 0.152397958 |
| Zc3h7a | -0.01465053 | 0.086664339 | 0.157397853 | 0.240202263 | 0.250041581 |
| Zc3h7b | 0.022219045 | 0.002550059 | 0.00806758 | 0.189163187 | 0.167488825 |
| Zc3h8 | 0.007841488 | 0.100132999 | 0.176899146 | 0.066844593 | 0.057522788 |
| Zc3hav1 | -0.003391259 | 0.699336071 | 0.782503788 | 0.293332174 | 0.289574253 |
| Zc3hav1l | -0.000726431 | 0.858525431 | 0.902821443 | 0.044356597 | 0.043127773 |
| Zcchc10 | 0.008549788 | 0.149204763 | 0.243963729 | 0.107812114 | 0.098603181 |
| Zcchc12 | -0.089484151 | 1.10E-22 | 3.87E-21 | 0.144234311 | 0.240235198 |
| Zcchc14 | 0.027859382 | 5.28E-05 | 0.000258375 | 0.157561583 | 0.133328674 |
| Zcchc17 | -0.002249376 | 0.805175897 | 0.864676911 | 0.349242316 | 0.346700464 |
| Zcchc18 | 0.013693381 | 0.177034863 | 0.278516524 | 0.66125022 | 0.622463301 |
| Zcchc2 | 0.003561215 | 0.621027836 | 0.720309308 | 0.168162113 | 0.157962349 |
| Zcchc3 | 0.008854983 | 0.12066704 | 0.205307134 | 0.103502307 | 0.089136685 |
| Zcchc4 | -0.00919756 | 0.022999716 | 0.052243747 | 0.037829012 | 0.043983126 |
| Zcchc7 | 0.009518263 | 0.309805664 | 0.43119981 | 0.352752939 | 0.356153093 |
| Zcchc8 | 0.02151813 | 0.000621289 | 0.002350263 | 0.129080503 | 0.109556485 |
| Zcchc9 | 0.04019601 | 7.99E-07 | 5.70E-06 | 0.247172003 | 0.220239875 |
| Zcrb1 | 0.053513437 | 7.86E-09 | 7.95E-08 | 1.170705242 | 1.089145344 |
| Zdbf2 | 0.013990628 | 0.151195218 | 0.24621774 | 0.361218039 | 0.347427216 |
| Zdhhc1 | -0.009830852 | 0.070301046 | 0.132769032 | 0.085509291 | 0.085190252 |
| Zdhhc12 | -0.009154861 | 0.121605387 | 0.206578294 | 0.099380456 | 0.10474008 |
| Zdhhc14 | 0.018133214 | 2.96E-05 | 0.000153601 | 0.059012882 | 0.044686075 |
| Zdhhc16 | -0.005497937 | 0.243214496 | 0.356964271 | 0.06220065 | 0.063388182 |
| Zdhhc17 | 0.005958985 | 0.345135845 | 0.467522738 | 0.122387155 | 0.114923865 |
| Zdhhc18 | 0.007590861 | 0.064404079 | 0.12366906 | 0.048366257 | 0.043123886 |
| Zdhhc2 | 0.068216558 | 5.79E-11 | 7.67E-10 | 1.001746129 | 0.883074368 |
| Zdhhc20 | 0.002078357 | 0.802528205 | 0.862595822 | 0.23568909 | 0.231093402 |
| Zdhhc21 | 0.007178713 | 0.191978802 | 0.296680642 | 0.08434397 | 0.079773903 |
| Zdhhc22 | -0.00830431 | 0.109584175 | 0.190409867 | 0.063556801 | 0.070718695 |
| Zdhhc23 | 0.00974635 | 0.066453826 | 0.127102912 | 0.081123553 | 0.073645188 |
| Zdhhc24 | 0.002111992 | 0.612460008 | 0.712783653 | 0.046608389 | 0.046129082 |
| Zdhhc3 | 0.027823765 | 0.002640041 | 0.008314375 | 0.380129843 | 0.337150509 |
| Zdhhc4 | -0.013108339 | 0.01771201 | 0.042081172 | 0.082692894 | 0.090037362 |

| | | | | | |
|---|---|---|---|---|---|
| Zdhhc5 | 0.003665976 | 0.622309909 | 0.721538096 | 0.183071732 | 0.174853318 |
| Zdhhc6 | -0.007610618 | 0.20444676 | 0.312520661 | 0.106000732 | 0.108741707 |
| Zdhhc7 | 0.006561789 | 0.180807098 | 0.283015072 | 0.07063575 | 0.062947475 |
| Zdhhc8 | 0.025344762 | 0.000330457 | 0.001342544 | 0.16714332 | 0.14627301 |
| Zeb1 | -0.004926831 | 0.439549803 | 0.558018175 | 0.112647405 | 0.116340544 |
| Zer1 | 0.000396682 | 0.950387857 | 0.967032331 | 0.117028454 | 0.114842605 |
| Zfand1 | -0.013796526 | 0.018168301 | 0.042944863 | 0.098669843 | 0.10198785 |
| Zfand2a | -0.000396376 | 0.927073838 | 0.950566374 | 0.046273837 | 0.050664742 |
| Zfand2b | -0.001277777 | 0.870057854 | 0.910909159 | 0.20920718 | 0.204451261 |
| Zfand3 | 0.0161901 | 0.052842327 | 0.104417297 | 0.269388269 | 0.253682902 |
| Zfand5 | -0.023845751 | 0.019443673 | 0.04529868 | 0.540768171 | 0.564799191 |
| Zfand6 | 0.011964061 | 0.216627042 | 0.327026104 | 0.551577336 | 0.519046771 |
| Zfas1 | 0.029840675 | 0.005208549 | 0.014946138 | 0.449235185 | 0.453956649 |
| Zfc3h1 | -0.011720652 | 0.094339684 | 0.168288149 | 0.137670695 | 0.147214557 |
| Zfhx2 | 0.050170466 | 1.74E-06 | 1.15E-05 | 0.712945602 | 0.676031593 |
| Zfhx3 | 0.040977578 | 4.43E-06 | 2.75E-05 | 0.294683542 | 0.255827332 |
| Zfp1 | -0.002918794 | 0.60637484 | 0.707874182 | 0.088602324 | 0.090671474 |
| Zfp106 | 0.008214509 | 0.412851689 | 0.532538692 | 0.544862156 | 0.545094174 |
| Zfp110 | -0.01708096 | 0.010550376 | 0.027077812 | 0.128888969 | 0.138908415 |
| Zfp113 | 0.00902654 | 0.117571009 | 0.2009185 | 0.103874846 | 0.090966408 |
| Zfp12 | 0.012628101 | 0.007434131 | 0.020244991 | 0.06914093 | 0.056276971 |
| Zfp120 | -0.000855037 | 0.896026543 | 0.929191473 | 0.132481688 | 0.126467797 |
| Zfp13 | -0.004111242 | 0.293490935 | 0.412985796 | 0.040961235 | 0.041378238 |
| Zfp131 | 0.008987883 | 0.279852769 | 0.398758408 | 0.235702199 | 0.232843931 |
| Zfp142 | 0.002506205 | 0.643642661 | 0.737738027 | 0.082785928 | 0.080185925 |
| Zfp143 | 0.006337541 | 0.161004487 | 0.258255342 | 0.05717647 | 0.051921497 |
| Zfp146 | -0.007121823 | 0.158530461 | 0.255424783 | 0.068953108 | 0.071119117 |
| Zfp148 | 0.036663516 | 0.000211859 | 0.000903772 | 0.510086396 | 0.464025764 |
| Zfp157 | 0.003165283 | 0.573229836 | 0.680124538 | 0.093768775 | 0.087781653 |
| Zfp160 | 0.006087653 | 0.23652719 | 0.350215861 | 0.07365266 | 0.07055227 |
| Zfp174 | -0.006261415 | 0.161398107 | 0.258801319 | 0.050052493 | 0.056041418 |
| Zfp180 | 0.010062752 | 0.048127193 | 0.096932374 | 0.075739355 | 0.06800861 |
| Zfp182 | -0.009401478 | 0.027878193 | 0.061500293 | 0.043186518 | 0.050967666 |
| Zfp2 | -0.001734818 | 0.708030674 | 0.789340259 | 0.057064668 | 0.05855878 |
| Zfp207 | 0.001044433 | 0.906443473 | 0.935897369 | 0.316265517 | 0.302791631 |
| Zfp212 | -0.001822835 | 0.696757369 | 0.780310464 | 0.062047324 | 0.060140327 |
| Zfp213 | -0.005925071 | 0.141932784 | 0.234472477 | 0.042959544 | 0.045007327 |
| Zfp219 | 0.005613349 | 0.29792926 | 0.417719681 | 0.086550379 | 0.079224021 |
| Zfp235 | 0.002968827 | 0.468681307 | 0.585896833 | 0.046884895 | 0.043390286 |
| Zfp236 | 0.012354393 | 0.024918326 | 0.056064793 | 0.091381434 | 0.081535586 |
| Zfp24 | 0.033137192 | 7.44E-05 | 0.000356405 | 0.28177585 | 0.245058055 |
| Zfp251 | 0.00835689 | 0.212238081 | 0.321899941 | 0.141011114 | 0.131284717 |
| Zfp26 | -0.001464229 | 0.774434453 | 0.841155029 | 0.072013624 | 0.070765951 |
| Zfp260 | 0.005099286 | 0.52691373 | 0.638575826 | 0.220820005 | 0.214458731 |
| Zfp263 | 0.017028565 | 0.002153722 | 0.006960933 | 0.096375137 | 0.081212709 |
| Zfp266 | 0.012858699 | 0.112891851 | 0.194923418 | 0.238389828 | 0.222263776 |
| Zfp267 | 0.006759311 | 0.187438963 | 0.291145806 | 0.076290448 | 0.068038613 |
| Zfp275 | 0.001033582 | 0.856020084 | 0.900966467 | 0.091253095 | 0.091776879 |
| Zfp277 | -0.00395918 | 0.481988359 | 0.598377069 | 0.09427852 | 0.09144757 |
| Zfp280b | 0.006567962 | 0.071543622 | 0.134639391 | 0.03845197 | 0.033328361 |
| Zfp280d | -0.004260516 | 0.616529064 | 0.716630268 | 0.245848425 | 0.248065746 |
| Zfp281 | 0.002132491 | 0.693299859 | 0.777782279 | 0.083041878 | 0.079790929 |
| Zfp282 | 0.005262798 | 0.343730021 | 0.465878051 | 0.093732744 | 0.085692872 |
| Zfp292 | 0.014465495 | 0.145233912 | 0.238988666 | 0.450901482 | 0.427273544 |
| Zfp30 | 0.004342876 | 0.476959009 | 0.593877496 | 0.108983665 | 0.104087806 |
| Zfp316 | 0.001573026 | 0.70594443 | 0.787786014 | 0.048219397 | 0.045857905 |
| Zfp317 | 0.006731425 | 0.124399932 | 0.210369827 | 0.053730949 | 0.048459839 |
| Zfp318 | 0.006071672 | 0.416347147 | 0.535662501 | 0.171660678 | 0.166278345 |
| Zfp319 | 0.007955447 | 0.054460808 | 0.107179752 | 0.050608822 | 0.043503762 |
| Zfp322a | -0.006077585 | 0.354016175 | 0.476366125 | 0.126278459 | 0.127414979 |
| Zfp324 | 0.005050235 | 0.157200882 | 0.253703017 | 0.037109738 | 0.032461468 |
| Zfp326 | 0.001280987 | 0.87473026 | 0.913977643 | 0.221172055 | 0.217848325 |
| Zfp329 | 0.002606428 | 0.745834271 | 0.81848976 | 0.218526451 | 0.215251474 |
| Zfp330 | -0.015186994 | 0.094995099 | 0.169146952 | 0.308643125 | 0.317709755 |
| Zfp335 | 0.007800803 | 0.102673214 | 0.180634996 | 0.065014755 | 0.059535767 |
| Zfp346 | 0.001755195 | 0.741549299 | 0.815075442 | 0.079982995 | 0.075990173 |
| Zfp35 | 0.003946609 | 0.302443218 | 0.422707442 | 0.04146931 | 0.036692329 |
| Zfp358 | 0.002952818 | 0.608482716 | 0.709482965 | 0.104192512 | 0.095650812 |
| Zfp36 | -0.003517607 | 0.599126674 | 0.702534016 | 0.111577189 | 0.124719904 |
| Zfp362 | 0.011941048 | 0.004634285 | 0.013513652 | 0.055270416 | 0.04327453 |
| Zfp367 | 0.000728974 | 0.861041627 | 0.904880197 | 0.047258532 | 0.044642044 |
| Zfp369 | -0.000624443 | 0.87477494 | 0.913977643 | 0.040918749 | 0.042162759 |

| | | | | | |
|---|---|---|---|---|---|
| Zfp36l2 | -0.036449025 | 7.26E-05 | 0.00034843 | 0.268976923 | 0.309432382 |
| Zfp37 | 0.001026208 | 0.847507968 | 0.895886971 | 0.078612012 | 0.079328076 |
| Zfp384 | 0.000837468 | 0.845511384 | 0.894439423 | 0.053007945 | 0.048674124 |
| Zfp386 | 0.001776376 | 0.784003421 | 0.848784105 | 0.130807906 | 0.124769925 |
| Zfp395 | 0.016506525 | 0.004685595 | 0.013638727 | 0.110701164 | 0.090366298 |
| Zfp397 | -0.005187114 | 0.496502388 | 0.611631553 | 0.181767075 | 0.188375848 |
| Zfp398 | 0.015499594 | 0.036863605 | 0.077405608 | 0.185714946 | 0.164516356 |
| Zfp40 | 0.004451321 | 0.282768811 | 0.401677145 | 0.048078285 | 0.043489924 |
| Zfp407 | -0.004203175 | 0.395646353 | 0.516235921 | 0.059929864 | 0.065043041 |
| Zfp410 | -0.002099269 | 0.709400429 | 0.790591914 | 0.090890858 | 0.090358095 |
| Zfp414 | 0.004626877 | 0.284603223 | 0.403810936 | 0.059012302 | 0.048902611 |
| Zfp420 | 0.0023225 | 0.526850714 | 0.638575826 | 0.037702606 | 0.034645499 |
| Zfp422 | 0.006445421 | 0.376217048 | 0.497862632 | 0.176350046 | 0.164149257 |
| Zfp426 | 0.015116528 | 0.011359988 | 0.02887999 | 0.10657252 | 0.097218039 |
| Zfp428 | -0.006812565 | 0.110381397 | 0.191587478 | 0.049165444 | 0.051081382 |
| Zfp438 | 0.007380511 | 0.074382607 | 0.139093614 | 0.051368595 | 0.042932257 |
| Zfp442 | -0.001563315 | 0.691022663 | 0.775764703 | 0.042381731 | 0.04135509 |
| Zfp444 | 0.006140369 | 0.282459998 | 0.401485512 | 0.103862786 | 0.092121135 |
| Zfp445 | 0.011284063 | 0.19462668 | 0.300199998 | 0.288708386 | 0.275164496 |
| Zfp451 | 0.016931986 | 0.035129307 | 0.074471681 | 0.224366538 | 0.206324868 |
| Zfp462 | 0.014781365 | 0.036185234 | 0.076261186 | 0.155619769 | 0.142951263 |
| Zfp467 | -0.012417804 | 0.066032081 | 0.126470428 | 0.128847228 | 0.134909971 |
| Zfp507 | 0.004601476 | 0.386426997 | 0.507472647 | 0.079659276 | 0.077783053 |
| Zfp51 | 0.008467112 | 0.029196794 | 0.063945996 | 0.045038433 | 0.037031566 |
| Zfp511 | -0.008868034 | 0.209445718 | 0.318260593 | 0.156940594 | 0.156046119 |
| Zfp512 | 0.00315878 | 0.603576488 | 0.706303637 | 0.11431547 | 0.1062818 |
| Zfp512b | -0.004942803 | 0.410276689 | 0.529794153 | 0.103402871 | 0.104101146 |
| Zfp513 | 0.007893553 | 0.085902959 | 0.156248563 | 0.064408909 | 0.054900675 |
| Zfp516 | 0.034768415 | 0.000685155 | 0.002564912 | 0.569734098 | 0.50945032 |
| Zfp518a | 0.008846826 | 0.122192203 | 0.207466399 | 0.097846245 | 0.086872037 |
| Zfp518b | -0.000916974 | 0.815420362 | 0.872866347 | 0.040088012 | 0.040475665 |
| Zfp52 | 0.006959546 | 0.080216433 | 0.147617671 | 0.044585968 | 0.040864776 |
| Zfp523 | 0.003888534 | 0.513232655 | 0.626840185 | 0.113860975 | 0.102494922 |
| Zfp524 | 0.010034217 | 0.073889386 | 0.138353295 | 0.101044323 | 0.086323487 |
| Zfp53 | 0.005184119 | 0.14471136 | 0.238293334 | 0.038432238 | 0.03160373 |
| Zfp553 | 0.022385503 | 7.64E-05 | 0.000365723 | 0.107537358 | 0.084007038 |
| Zfp560 | -0.024163824 | 0.009226915 | 0.024244344 | 0.367197278 | 0.384786671 |
| Zfp563 | 0.006524886 | 0.09956157 | 0.176116736 | 0.045698083 | 0.039346703 |
| Zfp568 | -0.004682012 | 0.540592642 | 0.651256712 | 0.177514798 | 0.175534392 |
| Zfp574 | 0.007095271 | 0.113408403 | 0.195419442 | 0.058169921 | 0.052196637 |
| Zfp579 | 0.011445997 | 0.025210558 | 0.056578266 | 0.08344151 | 0.069504506 |
| Zfp580 | -0.007862688 | 0.079651677 | 0.146967851 | 0.054361502 | 0.055914529 |
| Zfp592 | 0.024932598 | 1.65E-05 | 9.09E-05 | 0.110583712 | 0.087640404 |
| Zfp593 | -0.008999718 | 0.206208566 | 0.314472107 | 0.137737312 | 0.155632508 |
| Zfp597 | -0.002402846 | 0.664673201 | 0.75463656 | 0.088090423 | 0.085812358 |
| Zfp598 | -0.003489399 | 0.614003377 | 0.714035985 | 0.137230442 | 0.147350283 |
| Zfp60 | 0.018365249 | 0.001911705 | 0.006283162 | 0.108884489 | 0.092847866 |
| Zfp605 | -0.00543325 | 0.346322536 | 0.468481415 | 0.092975283 | 0.096774439 |
| Zfp606 | -0.010045742 | 0.037434556 | 0.078367518 | 0.061001283 | 0.066438016 |
| Zfp608 | 0.022413019 | 0.005639771 | 0.015976065 | 0.225913567 | 0.200296135 |
| Zfp609 | 0.017065118 | 0.01215202 | 0.030575035 | 0.148265512 | 0.132085715 |
| Zfp61 | 0.011057321 | 0.012944001 | 0.032209259 | 0.05865009 | 0.048988228 |
| Zfp612 | 0.017972647 | 0.051250293 | 0.101976125 | 0.332167666 | 0.317922233 |
| Zfp617 | 0.001874886 | 0.721400358 | 0.799345142 | 0.081827942 | 0.076509894 |
| Zfp618 | 0.003248171 | 0.46655142 | 0.58406038 | 0.055255245 | 0.051588422 |
| Zfp62 | 0.010160315 | 0.126365242 | 0.21321119 | 0.138362873 | 0.126022896 |
| Zfp622 | 0.037583663 | 4.95E-05 | 0.000243995 | 0.356647132 | 0.316669762 |
| Zfp626 | 0.00742244 | 0.049050848 | 0.098466311 | 0.04182849 | 0.034962783 |
| Zfp628 | 0.008004839 | 0.037315178 | 0.078151264 | 0.042640772 | 0.036225758 |
| Zfp629 | 0.002752116 | 0.507540312 | 0.62160581 | 0.04793614 | 0.045044309 |
| Zfp637 | 0.011176693 | 0.121227871 | 0.206153116 | 0.184742407 | 0.166452601 |
| Zfp638 | -0.024298935 | 0.014166467 | 0.034796967 | 0.4439689 | 0.450553888 |
| Zfp639 | 0.047003013 | 8.38E-10 | 9.68E-09 | 0.210322957 | 0.179441581 |
| Zfp64 | 0.010460682 | 0.032663814 | 0.070101015 | 0.073658366 | 0.063505059 |
| Zfp641 | 0.007694357 | 0.045179083 | 0.091716652 | 0.041431928 | 0.03668109 |
| Zfp644 | -0.00584606 | 0.533386051 | 0.64433141 | 0.366451994 | 0.370543601 |
| Zfp646 | 0.016836131 | 0.011125583 | 0.028381768 | 0.139480463 | 0.122323435 |
| Zfp65 | 0.004445313 | 0.348227354 | 0.470792149 | 0.063524038 | 0.058792238 |
| Zfp651 | 0.023400695 | 5.30E-06 | 3.24E-05 | 0.087710893 | 0.062997944 |
| Zfp652 | 0.026337965 | 0.004396759 | 0.012941354 | 0.353747051 | 0.329047539 |
| Zfp654 | 0.028281372 | 6.30E-06 | 3.79E-05 | 0.13106963 | 0.104342234 |
| Zfp655 | 0.0143035 | 0.01777156 | 0.042191724 | 0.10882294 | 0.100373304 |

| | | | | | |
|---|---|---|---|---|---|
| Zfp664 | 0.031614046 | 0.000753676 | 0.002791329 | 0.39200066 | 0.369365738 |
| Zfp667 | 0.008605798 | 0.294772518 | 0.414489214 | 0.236478174 | 0.22221815 |
| Zfp672 | 0.024520938 | 0.000603762 | 0.002295572 | 0.166931808 | 0.150777511 |
| Zfp68 | 0.001667283 | 0.817978244 | 0.874657334 | 0.168459309 | 0.162101356 |
| Zfp687 | 0.00423305 | 0.460484641 | 0.578775912 | 0.097403433 | 0.090538149 |
| Zfp688 | 0.001798375 | 0.794789958 | 0.856551155 | 0.158462792 | 0.14968411 |
| Zfp692 | -0.004288337 | 0.263235918 | 0.379362525 | 0.037776721 | 0.039353495 |
| Zfp697 | -0.014479165 | 0.000322086 | 0.001314024 | 0.03731165 | 0.046872809 |
| Zfp704 | 0.015786752 | 0.065163541 | 0.124905352 | 0.269409168 | 0.247518415 |
| Zfp706 | 0.028836726 | 0.003571267 | 0.010809421 | 0.743701852 | 0.69624696 |
| Zfp707 | 0.007308164 | 0.090593492 | 0.163011278 | 0.055173391 | 0.047889549 |
| Zfp710 | 0.017989577 | 0.00046874 | 0.001831627 | 0.084341097 | 0.069463926 |
| Zfp715 | 0.007061309 | 0.053990989 | 0.106449076 | 0.03870428 | 0.033348965 |
| Zfp719 | 0.007218623 | 0.166064638 | 0.264452156 | 0.080161518 | 0.073348634 |
| Zfp729a | 0.009383002 | 0.07747822 | 0.143803599 | 0.079624262 | 0.072605378 |
| Zfp729b | 0.012684506 | 0.003042761 | 0.009402962 | 0.057807537 | 0.04535503 |
| Zfp738 | 0.011686885 | 0.066550866 | 0.127150558 | 0.125872136 | 0.113516218 |
| Zfp74 | 0.006918212 | 0.197943298 | 0.304494422 | 0.087535319 | 0.078462136 |
| Zfp740 | 0.002784658 | 0.658294205 | 0.749846076 | 0.12386481 | 0.115281976 |
| Zfp746 | 0.004861937 | 0.44950248 | 0.567961121 | 0.122560098 | 0.119397048 |
| Zfp748 | 0.006467386 | 0.113334757 | 0.195419442 | 0.048519427 | 0.042662509 |
| Zfp758 | 0.007513463 | 0.130273168 | 0.21877971 | 0.070860399 | 0.063626964 |
| Zfp768 | 0.006440188 | 0.151643509 | 0.246729144 | 0.060477261 | 0.053454392 |
| Zfp770 | -0.005870899 | 0.181774691 | 0.283935338 | 0.050880702 | 0.052856271 |
| Zfp771 | 0.01770587 | 0.026821073 | 0.059465103 | 0.235767109 | 0.217965785 |
| Zfp777 | 0.007535314 | 0.093829152 | 0.16743888 | 0.059357952 | 0.052623247 |
| Zfp780b | 0.002984927 | 0.574066279 | 0.680705225 | 0.080995583 | 0.078018817 |
| Zfp787 | 0.025105648 | 0.000989111 | 0.003543247 | 0.203807359 | 0.18211743 |
| Zfp788 | -0.003110154 | 0.603998687 | 0.706627586 | 0.103985161 | 0.105017858 |
| Zfp790 | -0.000310833 | 0.94095145 | 0.960009444 | 0.047561067 | 0.046818061 |
| Zfp799 | 0.018114813 | 5.11E-05 | 0.000250687 | 0.064145155 | 0.049744158 |
| Zfp800 | -0.010073454 | 0.222673696 | 0.333976191 | 0.239397375 | 0.243603431 |
| Zfp809 | 0.019310997 | 8.82E-05 | 0.000416749 | 0.076193993 | 0.059768507 |
| Zfp81 | 0.004708536 | 0.277872981 | 0.396272664 | 0.051793292 | 0.048252983 |
| Zfp810 | 0.001214656 | 0.834681365 | 0.887151222 | 0.096558139 | 0.094885433 |
| Zfp821 | 0.008287125 | 0.151128033 | 0.246149562 | 0.10589927 | 0.09440377 |
| Zfp827 | 0.002437736 | 0.676457195 | 0.764958909 | 0.097131624 | 0.095411512 |
| Zfp830 | -0.003286081 | 0.611733936 | 0.712486267 | 0.125409955 | 0.127162294 |
| Zfp839 | -0.003060272 | 0.468834614 | 0.586013129 | 0.045559291 | 0.04751754 |
| Zfp84 | 0.002717713 | 0.587338822 | 0.692206034 | 0.070007123 | 0.068650543 |
| Zfp846 | 0.000653653 | 0.885844276 | 0.921976025 | 0.057415166 | 0.055086628 |
| Zfp862-ps | -0.001261689 | 0.788697496 | 0.85215793 | 0.064688202 | 0.059247904 |
| Zfp865 | 0.005845614 | 0.260267564 | 0.376301228 | 0.077825578 | 0.072237251 |
| Zfp866 | 0.003323799 | 0.448677432 | 0.567530835 | 0.055418649 | 0.050877677 |
| Zfp868 | 0.006431509 | 0.119747819 | 0.203993043 | 0.048595803 | 0.043560059 |
| Zfp869 | 0.009920175 | 0.253486423 | 0.368509106 | 0.276198254 | 0.268945478 |
| Zfp87 | 0.01605451 | 0.000479444 | 0.001870448 | 0.066224459 | 0.052857398 |
| Zfp871 | 0.045769695 | 1.27E-06 | 8.68E-06 | 0.415501736 | 0.361952576 |
| Zfp874a | 0.002688711 | 0.487434877 | 0.603211953 | 0.041072285 | 0.038177305 |
| Zfp9 | 0.009168731 | 0.226947488 | 0.339339199 | 0.186867308 | 0.178631579 |
| Zfp90 | -0.003989773 | 0.363424356 | 0.486199475 | 0.051291198 | 0.052265088 |
| Zfp91 | 0.056466229 | 6.22E-11 | 8.21E-10 | 0.311060902 | 0.257941843 |
| Zfp92 | 0.005438471 | 0.350260333 | 0.472751764 | 0.090370863 | 0.090865697 |
| Zfp930 | -0.004151371 | 0.298491714 | 0.418146462 | 0.041612975 | 0.043569897 |
| Zfp931 | 0.010215058 | 0.041033819 | 0.084734662 | 0.077597711 | 0.065708589 |
| Zfp932 | 0.006116255 | 0.189363989 | 0.293573066 | 0.065651597 | 0.057749739 |
| Zfp938 | 0.015074484 | 0.054937508 | 0.108014312 | 0.208065351 | 0.200376879 |
| Zfp941 | -0.001905949 | 0.70256515 | 0.785006136 | 0.066553902 | 0.067621614 |
| Zfp942 | 0.000988402 | 0.836782411 | 0.888187078 | 0.064013894 | 0.061248733 |
| Zfp943 | -0.005458482 | 0.293961747 | 0.413468765 | 0.075090611 | 0.076178606 |
| Zfp944 | -0.000643902 | 0.894697448 | 0.928392142 | 0.066315353 | 0.064996918 |
| Zfp945 | -0.062009033 | 2.30E-15 | 4.57E-14 | 0.168755044 | 0.219186725 |
| Zfp946 | -0.002775446 | 0.580643437 | 0.686908675 | 0.066711299 | 0.069713999 |
| Zfp948 | 0.013543607 | 0.001002992 | 0.00359024 | 0.051078739 | 0.040699276 |
| Zfp949 | 0.00452857 | 0.361724671 | 0.484537107 | 0.074140902 | 0.065871861 |
| Zfp950 | -0.026985145 | 0.003035086 | 0.009385213 | 0.295334803 | 0.318163394 |
| Zfp952 | 0.007572244 | 0.117452641 | 0.20082212 | 0.068699236 | 0.062253503 |
| Zfp954 | 0.008658731 | 0.042113099 | 0.086440724 | 0.053935531 | 0.047209005 |
| Zfp955b | 0.003569238 | 0.455737329 | 0.574294013 | 0.068004256 | 0.061339592 |
| Zfp958 | 0.006449234 | 0.174789625 | 0.275503361 | 0.065533096 | 0.058835942 |
| Zfp959 | 0.0149118 | 0.000135648 | 0.000613665 | 0.049612706 | 0.036322384 |
| Zfp961 | 0.00061336 | 0.881773082 | 0.919017786 | 0.046490997 | 0.044190102 |

| | | | | | |
|---|---|---|---|---|---|
| Zfp974 | -0.001166883 | 0.774908915 | 0.841266309 | 0.044060486 | 0.044055479 |
| Zfp976 | -0.006536461 | 0.152307551 | 0.247324872 | 0.050086365 | 0.056200781 |
| Zfp979 | -0.020478343 | 7.76E-07 | 5.55E-06 | 0.035777663 | 0.049481856 |
| Zfp983 | 0.006797796 | 0.132021191 | 0.221447552 | 0.05953362 | 0.054102308 |
| Zfp994 | 0.000999054 | 0.869151576 | 0.910400898 | 0.109083949 | 0.103211877 |
| Zfp995 | -0.003854649 | 0.346653201 | 0.468794327 | 0.044634439 | 0.046464341 |
| Zfpl1 | -0.011204385 | 0.166320271 | 0.2646425 | 0.225608547 | 0.224858073 |
| Zfr | 0.008329882 | 0.410342519 | 0.529794153 | 0.794804974 | 0.772036177 |
| Zfr2 | -0.003500962 | 0.491122695 | 0.606325286 | 0.070375255 | 0.072068423 |
| Zfx | 0.005821265 | 0.401802772 | 0.521817599 | 0.155799247 | 0.14587721 |
| Zfyve1 | 0.003825357 | 0.386230979 | 0.507355435 | 0.057793642 | 0.05150054 |
| Zfyve16 | 0.002729751 | 0.695972145 | 0.77991889 | 0.145111589 | 0.146193378 |
| Zfyve19 | -0.013305795 | 0.01344181 | 0.033328559 | 0.080225234 | 0.085434113 |
| Zfyve21 | -0.002338212 | 0.782515071 | 0.847550303 | 0.285231644 | 0.265669998 |
| Zfyve26 | -0.00256368 | 0.576939275 | 0.683191673 | 0.055593654 | 0.056220232 |
| Zfyve27 | 0.000228716 | 0.968900949 | 0.979122029 | 0.09859133 | 0.098207858 |
| Zfyve9 | -0.008088592 | 0.329760566 | 0.452455183 | 0.227424835 | 0.235079239 |
| Zgpat | -7.62E-05 | 0.988933084 | 0.993079296 | 0.089632378 | 0.085415947 |
| Zhx1 | 0.006475053 | 0.446648608 | 0.565571095 | 0.264947781 | 0.258715424 |
| Zhx2 | 0.018071272 | 0.000504362 | 0.001956666 | 0.08161898 | 0.067867251 |
| Zhx3 | 0.030128908 | 1.55E-07 | 1.27E-06 | 0.111596163 | 0.085088153 |
| Zim1 | 0.017290541 | 0.00125264 | 0.004347792 | 0.092698528 | 0.069450425 |
| Zkscan1 | 0.027688623 | 0.000787002 | 0.002900392 | 0.255212611 | 0.229849319 |
| Zkscan14 | -0.005128143 | 0.280067324 | 0.398947184 | 0.061956663 | 0.062242381 |
| Zkscan17 | 0.024848301 | 2.75E-07 | 2.15E-06 | 0.075687566 | 0.056859264 |
| Zkscan3 | 0.014790377 | 0.090113996 | 0.162324934 | 0.301189299 | 0.282735437 |
| Zkscan6 | 0.003180458 | 0.474557438 | 0.591644962 | 0.05678887 | 0.052253769 |
| Zkscan7 | -0.001720379 | 0.716329051 | 0.796266983 | 0.05943872 | 0.060328984 |
| Zkscan8 | 0.00194253 | 0.693733126 | 0.77808877 | 0.066743709 | 0.064366288 |
| Zmat1 | 0.003059369 | 0.526543029 | 0.638398199 | 0.067066957 | 0.060135619 |
| Zmat2 | 0.043266653 | 8.82E-06 | 5.12E-05 | 0.5271335 | 0.483747223 |
| Zmat3 | 0.00415107 | 0.571091395 | 0.678166672 | 0.16201735 | 0.162584217 |
| Zmat5 | -0.02501783 | 0.001306247 | 0.004514515 | 0.208519475 | 0.218813727 |
| Zmiz1 | 0.072186585 | 2.23E-15 | 4.44E-14 | 0.354698537 | 0.277804588 |
| Zmiz2 | 0.013267078 | 0.083587735 | 0.152809319 | 0.204000283 | 0.189564065 |
| Zmpste24 | 9.48E-05 | 0.991130957 | 0.994193721 | 0.269072549 | 0.266306733 |
| Zmym1 | 0.001748078 | 0.68827067 | 0.773747393 | 0.05135261 | 0.049983748 |
| Zmym2 | 0.010182463 | 0.186560975 | 0.290060098 | 0.201233354 | 0.188538036 |
| Zmym3 | 0.014184508 | 0.003192885 | 0.00980146 | 0.074111235 | 0.059505052 |
| Zmym4 | 0.014486975 | 0.089058049 | 0.160775642 | 0.270288915 | 0.253325287 |
| Zmym5 | 0.001700603 | 0.849623512 | 0.896952957 | 0.306185186 | 0.296018146 |
| Zmym6 | -0.011002931 | 0.016332015 | 0.039292216 | 0.050420884 | 0.058356018 |
| Zmynd11 | 0.040601367 | 3.41E-05 | 0.000175034 | 0.480388988 | 0.433410899 |
| Zmynd19 | -0.000526014 | 0.92177109 | 0.946499635 | 0.081557052 | 0.080757135 |
| Zmynd8 | 0.073803977 | 4.97E-13 | 8.27E-12 | 0.593145155 | 0.508690441 |
| Znfx1 | 0.028692454 | 0.00023515 | 0.000990955 | 0.213611415 | 0.182791884 |
| Znhit1 | 0.012144628 | 0.197530035 | 0.304003007 | 0.40974492 | 0.387437032 |
| Znhit2 | 0.001210764 | 0.874564641 | 0.913954363 | 0.202578733 | 0.192736621 |
| Znhit3 | -0.020552592 | 0.003391904 | 0.010356813 | 0.1478356 | 0.159169107 |
| Znhit6 | -0.007737932 | 0.242682616 | 0.356560434 | 0.125074703 | 0.131836696 |
| Znrf1 | -0.000681715 | 0.906513359 | 0.935897369 | 0.098523116 | 0.095400385 |
| Znrf2 | 0.00193405 | 0.838382219 | 0.889163645 | 0.376326794 | 0.374933993 |
| Znrf3 | 0.006220343 | 0.103293475 | 0.181529133 | 0.042906769 | 0.036256274 |
| Zpr1 | -0.012468861 | 0.077638645 | 0.144018872 | 0.152214678 | 0.158007973 |
| Zranb1 | 0.024925137 | 0.0140922 | 0.0346408 | 0.542766416 | 0.514906895 |
| Zranb2 | -0.032220788 | 0.001936508 | 0.006349656 | 0.696989191 | 0.746175603 |
| Zrsr1 | 0.031254132 | 5.97E-06 | 3.60E-05 | 0.153852937 | 0.130929941 |
| Zrsr2 | 0.006885208 | 0.443706368 | 0.562340413 | 0.315931094 | 0.32044296 |
| Zscan12 | 0.006821075 | 0.088211781 | 0.159611936 | 0.046613706 | 0.040688205 |
| Zscan2 | 0.006798997 | 0.08017133 | 0.147617671 | 0.046006972 | 0.037542999 |
| Zscan21 | 0.001069937 | 0.786104676 | 0.85039609 | 0.041109346 | 0.041373414 |
| Zscan22 | 0.007430547 | 0.066657811 | 0.127242732 | 0.046315884 | 0.041350198 |
| Zscan26 | 0.001711464 | 0.828763606 | 0.881922042 | 0.217451147 | 0.203884494 |
| Zscan29 | 0.013956869 | 0.00236464 | 0.00756218 | 0.065694694 | 0.054250316 |
| Zswim1 | 0.005629211 | 0.284882037 | 0.404088586 | 0.078451026 | 0.075488921 |
| Zswim5 | 0.013902371 | 0.000397536 | 0.001581361 | 0.048978007 | 0.037410659 |
| Zswim7 | -0.019465842 | 0.022282222 | 0.050870842 | 0.252484912 | 0.26114935 |
| Zswim8 | 0.017911945 | 0.030703648 | 0.066674306 | 0.25443846 | 0.237580343 |
| Zup1 | -0.001673946 | 0.817976451 | 0.874657334 | 0.157448052 | 0.162506553 |
| Zw10 | 0.002105089 | 0.676172864 | 0.764745531 | 0.073216799 | 0.069057486 |
| Zwint | -0.059791409 | 1.98E-08 | 1.86E-07 | 0.601890759 | 0.678832086 |
| Zxdb | 0.006717567 | 0.282587381 | 0.401595456 | 0.12977945 | 0.113683671 |

| | | | | | |
|---|---|---|---|---|---|
| Zxdc | 0.01019592 | 0.136055834 | 0.226962048 | 0.147092915 | 0.135688625 |
| Zyg11b | -0.013887231 | 0.157529373 | 0.25406446 | 0.363769017 | 0.389087292 |
| Zzef1 | 0.022808522 | 0.002698527 | 0.0084766 | 0.202460023 | 0.176471215 |
| Zzz3 | 0.022745535 | 0.010428207 | 0.026813813 | 0.315399288 | 0.291498995 |
