## Supplementary data file 2 for "Sorting receptor SORCS2 facilitates a protective stress response in pancreatic islets"

| GeneID | $Log_2FC$ | p-value | FDR | Average (*Sorcs2* $^{-/-}$ ) | Average (*Sorcs2* $^{+/+}$ ) |
|---|---|---|---|---|---|
| 0610009B22Rik | 0.126313347 | 0.048424836 | 0.344953921 | 0.431289068 | 0.321563272 |
| 0610009E02Rik | -0.052824462 | 0.128223977 | 0.508260084 | 0.071562265 | 0.111640131 |
| 0610009L18Rik | -0.042826107 | 0.410650017 | 0.780328327 | 0.190524769 | 0.229015899 |
| 0610010F05Rik | 0.001909528 | 0.964234092 | 0.991970325 | 0.120570864 | 0.120533619 |
| 0610010K14Rik | 0.047363278 | 0.310098954 | 0.712953768 | 0.156515432 | 0.130785565 |
| 0610012G03Rik | 0.186597438 | 0.005894083 | 0.113767137 | 0.848644618 | 0.646839077 |
| 0610030E20Rik | -0.033190396 | 0.513593362 | 0.836550101 | 0.182985944 | 0.209378771 |
| 0610040B10Rik | 0.041164967 | 0.318211731 | 0.717175454 | 0.114538834 | 0.076849465 |
| 0610040J01Rik | 0.134520789 | 0.041058862 | 0.31946341 | 0.571322906 | 0.44672394 |
| 1010001B22Rik | -0.066715117 | 0.027865422 | 0.26510449 | 0.04762392 | 0.110710849 |
| 1110002L01Rik | 0.027439647 | 0.37563648 | 0.760593776 | 0.066324697 | 0.043107726 |
| 1110004F10Rik | 0.093539489 | 0.169410444 | 0.57227323 | 0.743485483 | 0.637305561 |
| 1110008P14Rik | -0.103054311 | 0.116465708 | 0.488986751 | 0.640223681 | 0.757439061 |
| 1110018N20Rik | 0.026314529 | 0.552678059 | 0.852623515 | 0.132184629 | 0.109971177 |
| 1110019D14Rik | -0.022754407 | 0.317437374 | 0.716863138 | 0.027660639 | 0.045099641 |
| 1110020A21Rik | 0.022705511 | 0.635715481 | 0.890285515 | 0.148110277 | 0.12532713 |
| 1110032A03Rik | -0.050065825 | 0.405343113 | 0.777010853 | 0.302883799 | 0.36431863 |
| 1110035H17Rik | 0.04587134 | 0.091417581 | 0.4462832 | 0.051558915 | 0.010512739 |
| 1110038B12Rik | -0.086881037 | 0.078275787 | 0.419462934 | 0.15153031 | 0.229994368 |
| 1110038F14Rik | 0.098380139 | 0.119240973 | 0.492433827 | 0.406081645 | 0.31725061 |
| 1110046J04Rik | 0.031081435 | 0.330524799 | 0.725898152 | 0.066517769 | 0.049027679 |
| 1110051M20Rik | 0.004593974 | 0.900635338 | 0.974070444 | 0.089110201 | 0.08115221 |
| 1110059E24Rik | -0.047031559 | 0.37709931 | 0.761412185 | 0.196778182 | 0.237777106 |
| 1110059G10Rik | -0.027605098 | 0.57950622 | 0.865234762 | 0.175590623 | 0.198100748 |
| 1110065P20Rik | 0.030382711 | 0.660821876 | 0.89868852 | 0.634586134 | 0.589673654 |
| 1190007I07Rik | -0.01829252 | 0.699770301 | 0.911507617 | 0.14620423 | 0.169549392 |
| 1300002E11Rik | 0.065845167 | 0.27570607 | 0.682937271 | 0.353348271 | 0.284727665 |
| 1500002C15Rik | -0.023828008 | 0.376004544 | 0.760642399 | 0.040332874 | 0.063849009 |
| 1500009L16Rik | -0.030730499 | 0.632474637 | 0.889078599 | 0.351239394 | 0.386191548 |
| 1500011B03Rik | -0.115084022 | 0.093786285 | 0.45227176 | 0.693030815 | 0.827449509 |
| 1500015A07Rik | 0.006307464 | 0.897359675 | 0.972761535 | 0.169957401 | 0.163019344 |
| 1500026H17Rik | -0.021803743 | 0.663437747 | 0.899593114 | 0.170715411 | 0.178823726 |
| 1600012H06Rik | -0.023239804 | 0.607760481 | 0.878733629 | 0.143570833 | 0.167944467 |
| 1600014C10Rik | 1.74E-05 | 0.999521019 | 0.999919513 | 0.054408725 | 0.05188877 |
| 1600020E01Rik | 0.01351103 | 0.769527504 | 0.931397082 | 0.141585085 | 0.127692421 |
| 1700007K13Rik | 0.037196629 | 0.202537695 | 0.61304911 | 0.055649358 | 0.02517262 |
| 1700007L15Rik | -0.028863026 | 0.42532652 | 0.790344696 | 0.084736748 | 0.107807011 |
| 1700008J07Rik | 0.003568098 | 0.897599531 | 0.972811413 | 0.04837566 | 0.044921774 |
| 1700017B05Rik | 0.034710273 | 0.236735291 | 0.647618456 | 0.05805981 | 0.026912155 |
| 1700019D03Rik | 0.055199985 | 0.37372681 | 0.759023874 | 0.336456995 | 0.281703322 |
| 1700029J07Rik | -0.011981876 | 0.731860708 | 0.920770724 | 0.077074447 | 0.086337808 |
| 1700030C10Rik | 0.046773076 | 0.162976741 | 0.562993622 | 0.076127291 | 0.035891262 |
| 1700030K09Rik | -0.052745779 | 0.067592946 | 0.394989542 | 0.044642551 | 0.09363783 |
| 1700037C18Rik | 0.039310018 | 0.221025354 | 0.633858837 | 0.070213977 | 0.034408445 |
| 1700037H04Rik | -0.064474867 | 0.053622129 | 0.361325633 | 0.065280289 | 0.123042536 |
| 1700052K11Rik | -0.012436398 | 0.615225455 | 0.882439418 | 0.033970331 | 0.045858355 |
| 1700056E22Rik | 0.058920598 | 0.135650547 | 0.519071499 | 0.10655809 | 0.050861419 |
| 1700063J08Rik | 0.000288587 | 0.992306513 | 0.998695538 | 0.05602731 | 0.055098261 |
| 1700066M21Rik | -0.019830023 | 0.435656043 | 0.796975049 | 0.036234609 | 0.050999601 |
| 1700084E18Rik | 0.033049263 | 0.31784559 | 0.717175454 | 0.071426597 | 0.041544368 |
| 1700086L19Rik | -0.164161258 | 0.030706514 | 0.279219789 | 0.563588452 | 0.760525344 |
| 1700086O06Rik | -0.029054831 | 0.366171345 | 0.753107222 | 0.06237268 | 0.08917434 |
| 1700086P04Rik | 0.02013013 | 0.489828151 | 0.826459641 | 0.055010716 | 0.035192264 |
| 1700088E04Rik | -0.037408882 | 0.336762081 | 0.731477644 | 0.098004557 | 0.128823243 |
| 1700096K18Rik | 0.02822479 | 0.537733067 | 0.847820734 | 0.147485564 | 0.125762872 |
| 1700110K17Rik | 0.003680587 | 0.925241489 | 0.981467853 | 0.09810626 | 0.092643505 |
| 1700113A16Rik | -0.041665612 | 0.20048721 | 0.610932587 | 0.0663944 | 0.104739224 |
| 1700123O20Rik | 0.007036323 | 0.899827227 | 0.973983939 | 0.234595075 | 0.235039424 |
| 1810008I18Rik | -0.181483921 | 0.005618548 | 0.111670034 | 0.35190838 | 0.533159566 |
| 1810010H24Rik | 0.026543334 | 0.408929253 | 0.779567504 | 0.069905216 | 0.048512617 |
| 1810013L24Rik | -0.062503373 | 0.258140217 | 0.665948842 | 0.215842983 | 0.27803348 |
| 1810014B01Rik | -0.038548044 | 0.175762932 | 0.579734651 | 0.0489257 | 0.083830253 |
| 1810024B03Rik | -0.04807666 | 0.111520096 | 0.482676674 | 0.05149632 | 0.089451664 |
| 1810030O07Rik | 0.026181952 | 0.610281934 | 0.879490606 | 0.19180973 | 0.164292594 |

| | | | | | |
|---|---|---|---|---|---|
| 1810037I17Rik | -0.032448773 | 0.572679851 | 0.861775874 | 1.26111986 | 1.289373661 |
| 1810044D09Rik | -0.091399023 | 0.050481982 | 0.354499194 | 0.141516351 | 0.223874538 |
| 1810055G02Rik | -0.018225366 | 0.469446991 | 0.814050152 | 0.035358589 | 0.049595635 |
| 1810058I24Rik | 0.153985187 | 0.029991447 | 0.274935828 | 0.75958193 | 0.591280576 |
| 1810059H22Rik | -0.019770934 | 0.610190203 | 0.879490606 | 0.096379761 | 0.111370142 |
| 1810062G17Rik | 0.031159714 | 0.380668262 | 0.762006446 | 0.083898779 | 0.061697061 |
| 2010001A14Rik | -0.03129552 | 0.303834398 | 0.708427238 | 0.053724984 | 0.081460395 |
| 2010204K13Rik | -0.046598422 | 0.271673925 | 0.67876599 | 0.110440432 | 0.15345601 |
| 2010315B03Rik | -0.004018397 | 0.929813091 | 0.982502239 | 0.1410279 | 0.144064989 |
| 2010320M18Rik | 0.006566485 | 0.901423471 | 0.974269668 | 0.203094771 | 0.185362575 |
| 2210016F16Rik | -0.130548318 | 0.028190265 | 0.26718195 | 0.314335012 | 0.434738364 |
| 2210016L21Rik | -0.015798212 | 0.813633211 | 0.948397417 | 0.540798598 | 0.547913922 |
| 2210408F21Rik | -0.109840064 | 0.034416454 | 0.293212179 | 0.191153957 | 0.281910735 |
| 2210408I21Rik | 0.051758342 | 0.064539274 | 0.387197068 | 0.053915156 | 0.011026637 |
| 2300009A05Rik | 0.013119735 | 0.840175051 | 0.954911647 | 0.894312883 | 0.893998457 |
| 2310009B15Rik | -0.041521941 | 0.370863631 | 0.75760494 | 0.142057525 | 0.173462472 |
| 2310010J17Rik | -0.004755832 | 0.9307326 | 0.982803527 | 0.221800663 | 0.237900849 |
| 2310011J03Rik | 0.060204964 | 0.322348749 | 0.719780733 | 0.312475808 | 0.259805726 |
| 2310015A10Rik | -0.01670478 | 0.634438157 | 0.88965971 | 0.075422626 | 0.085112507 |
| 2310022A10Rik | -0.025177858 | 0.519762696 | 0.838588359 | 0.101374264 | 0.121130225 |
| 2310022B05Rik | -0.00856035 | 0.888885373 | 0.970176435 | 0.358843872 | 0.383599969 |
| 2310030G06Rik | 0.008689445 | 0.88342737 | 0.968793593 | 0.317949437 | 0.317264197 |
| 2310033P09Rik | 0.020384655 | 0.719723941 | 0.918833528 | 0.242571739 | 0.223414894 |
| 2310034G01Rik | 0.003029184 | 0.936014104 | 0.984394449 | 0.094597851 | 0.094554021 |
| 2310039H08Rik | 0.001271544 | 0.985668389 | 0.996791297 | 0.726517926 | 0.713171848 |
| 2310040G24Rik | -0.052695744 | 0.207188045 | 0.618605371 | 0.107237478 | 0.157025631 |
| 2310057M21Rik | 0.108462305 | 0.009706842 | 0.152947522 | 0.131033592 | 0.03822914 |
| 2310061I04Rik | 0.101098583 | 0.091324838 | 0.446264555 | 0.309539048 | 0.212965914 |
| 2410002F23Rik | -0.046846702 | 0.344648206 | 0.737445936 | 0.165921864 | 0.19500508 |
| 2410004B18Rik | -0.012384399 | 0.787732295 | 0.937804093 | 0.136817181 | 0.14156397 |
| 2410006H16Rik | -0.070335807 | 0.326364806 | 0.720866629 | 1.167405733 | 1.322348979 |
| 2410021H03Rik | -0.177572921 | 2.65E-06 | 0.000474809 | 0.076219299 | 0.238322725 |
| 2410022M11Rik | -0.059583767 | 0.112786938 | 0.483221516 | 0.0898582 | 0.141421464 |
| 2510002D24Rik | -0.0409253 | 0.50728519 | 0.833462342 | 0.33964764 | 0.352822149 |
| 2510009E07Rik | 0.014592517 | 0.793033841 | 0.939604453 | 0.226687198 | 0.211234597 |
| 2510039O18Rik | 0.048228962 | 0.412118873 | 0.781344376 | 0.276294686 | 0.235896695 |
| 2510046G10Rik | -0.101511217 | 0.012513998 | 0.178420124 | 0.103455309 | 0.200827665 |
| 2610001J05Rik | -0.041450801 | 0.531578212 | 0.845612734 | 0.460493754 | 0.494200136 |
| 2610002M06Rik | -0.007191438 | 0.874802949 | 0.96500153 | 0.135711645 | 0.136456857 |
| 2610005L07Rik | 0.06603184 | 0.055273041 | 0.365283656 | 0.083869752 | 0.031570842 |
| 2610008E11Rik | 0.012472524 | 0.775342888 | 0.933636662 | 0.131638222 | 0.121362932 |
| 2610021A01Rik | 0.050517719 | 0.125030337 | 0.503785425 | 0.074698115 | 0.0364859 |
| 2610037D02Rik | -0.00276261 | 0.931308457 | 0.98302061 | 0.063783822 | 0.068694897 |
| 2610044O15Rik8 | -0.06405169 | 0.0267588 | 0.260502501 | 0.043974105 | 0.095453289 |
| 2610301B20Rik | 0.000595543 | 0.987720282 | 0.997268589 | 0.098624162 | 0.096594976 |
| 2610306M01Rik | -0.103873665 | 0.00673889 | 0.125488383 | 0.084616971 | 0.176574594 |
| 2610507B11Rik | 0.00575522 | 0.925683119 | 0.981625089 | 0.32322514 | 0.317588623 |
| 2700033N17Rik | -0.010194298 | 0.745411381 | 0.924566547 | 0.060422041 | 0.064695892 |
| 2700046G09Rik | -0.099963651 | 0.005846581 | 0.113654038 | 0.074574268 | 0.163402022 |
| 2700049A03Rik | 0.032761133 | 0.405943702 | 0.777419755 | 0.103236319 | 0.075363317 |
| 2700062C07Rik | -0.052267334 | 0.261438609 | 0.668468258 | 0.139285334 | 0.183369737 |
| 2700097O09Rik | 0.01396489 | 0.643931875 | 0.893069536 | 0.058021277 | 0.048277592 |
| 2810001G20Rik | -0.045284814 | 0.250221335 | 0.662228053 | 0.096719575 | 0.13701318 |
| 2810004N23Rik | -0.114285924 | 0.069744362 | 0.400867525 | 0.346438239 | 0.446651156 |
| 2810006K23Rik | 0.01430451 | 0.73913231 | 0.922843014 | 0.124399824 | 0.111504159 |
| 2810013P06Rik | -0.024560143 | 0.661791378 | 0.898920864 | 0.235061903 | 0.267543452 |
| 2810021J22Rik | 0.025862309 | 0.357938678 | 0.746823177 | 0.053365579 | 0.032528324 |
| 2810025M15Rik | -0.092900234 | 0.083211132 | 0.431177145 | 0.200596329 | 0.29579266 |
| 2810402E24Rik | 0.007332287 | 0.875480986 | 0.965096393 | 0.146785757 | 0.141027296 |
| 2810403D21Rik | -0.021536507 | 0.418824691 | 0.7851594 | 0.041033054 | 0.058320514 |
| 2810454H06Rik | -0.036588606 | 0.125368498 | 0.503839619 | 0.027529558 | 0.060584124 |
| 2900005J15Rik | -0.051327219 | 0.221750433 | 0.633858837 | 0.11552213 | 0.163014256 |
| 2900026A02Rik | 0.086344145 | 0.170587283 | 0.572831081 | 0.391715128 | 0.292076908 |
| 2900027M19Rik | -0.049731815 | 0.297808347 | 0.702656882 | 0.150440233 | 0.19113387 |
| 2900052L18Rik | -0.024112353 | 0.572096783 | 0.861494041 | 0.11602073 | 0.13865438 |

| | | | | | |
|---|---|---|---|---|---|
| 2900076A07Rik | -0.085501221 | 0.038933958 | 0.311933003 | 0.099717727 | 0.178642261 |
| 2900092N22Rik | -0.011482966 | 0.705976779 | 0.912844672 | 0.055817754 | 0.061940597 |
| 2900093K20Rik | 0.12515637 | 0.065252274 | 0.389310714 | 0.451032902 | 0.333135421 |
| 3010003L21Rik | 0.018048215 | 0.529783731 | 0.8444401 | 0.053216694 | 0.037589078 |
| 3110009E18Rik | 0.026406257 | 0.515364093 | 0.837141169 | 0.111553967 | 0.084079172 |
| 3110040N11Rik | -0.015016847 | 0.778953551 | 0.934651141 | 0.219393877 | 0.243690225 |
| 3110056K07Rik | 0.025234344 | 0.501802339 | 0.832218079 | 0.097581107 | 0.076916248 |
| 3110082I17Rik | -0.057883697 | 0.183307135 | 0.590066231 | 0.118902556 | 0.176450364 |
| 3222401L13Rik | -0.040017802 | 0.384683079 | 0.764861331 | 0.135641167 | 0.168138418 |
| 3830406C13Rik | -0.163771201 | 0.003771303 | 0.086973441 | 0.22013194 | 0.359640654 |
| 4632415L05Rik | -0.07848998 | 0.068894472 | 0.399753362 | 0.118280542 | 0.181250097 |
| 4632427E13Rik | -0.004960367 | 0.932549992 | 0.983607006 | 0.289137805 | 0.300400186 |
| 4632428C04Rik | -0.026454872 | 0.395239044 | 0.771942846 | 0.058873004 | 0.082165231 |
| 4732414G09Rik | -0.000653658 | 0.980398183 | 0.995540893 | 0.042875814 | 0.041429761 |
| 4732440D04Rik | -0.00707014 | 0.823171125 | 0.950930859 | 0.058782284 | 0.068859688 |
| 4732491K20Rik | -0.041477341 | 0.09902849 | 0.463315751 | 0.033280164 | 0.069057124 |
| 4833420G17Rik | -0.05277439 | 0.339115706 | 0.732701236 | 0.211107312 | 0.264945543 |
| 4833439L19Rik | -0.004893474 | 0.943660304 | 0.986918206 | 0.726738613 | 0.712248011 |
| 4921524J17Rik | 0.025770266 | 0.656527874 | 0.8981287 | 0.266607317 | 0.236454428 |
| 4930402H24Rik | 0.046849934 | 0.44062604 | 0.80031914 | 0.319423354 | 0.279091393 |
| 4930430F08Rik | 0.005871382 | 0.885302629 | 0.969534369 | 0.110128023 | 0.109307167 |
| 4930431P19Rik | 0.017372527 | 0.688725195 | 0.909796728 | 0.13066348 | 0.11443243 |
| 4930453N24Rik | 0.040116975 | 0.486785835 | 0.825733812 | 0.285223845 | 0.247243187 |
| 4930481A15Rik | -0.050871828 | 0.042491375 | 0.325502204 | 0.03284082 | 0.074812819 |
| 4930488L21Rik | -0.043669707 | 0.041130551 | 0.319536229 | 0.0197503 | 0.056528642 |
| 4930503L19Rik | -0.015121862 | 0.63996262 | 0.891345382 | 0.066542455 | 0.081978388 |
| 4930522L14Rik | 0.042582237 | 0.136684122 | 0.520249727 | 0.056498914 | 0.018558178 |
| 4930539J05Rik | -0.061839877 | 0.008948383 | 0.146996602 | 0.026164637 | 0.078071461 |
| 4930550C14Rik | -0.053103131 | 0.144123317 | 0.533279057 | 0.074137878 | 0.123334397 |
| 4930563E22Rik | -0.040869584 | 0.083578037 | 0.431740995 | 0.027498552 | 0.062859897 |
| 4930570G19Rik | 0.007144151 | 0.807007483 | 0.945532285 | 0.053806749 | 0.045392132 |
| 4930578M01Rik | -0.002473884 | 0.947721774 | 0.987651726 | 0.095515696 | 0.095804514 |
| 4930579K19Rik | -0.008043848 | 0.813414907 | 0.948397417 | 0.073457624 | 0.083772461 |
| 4930581F22Rik | -0.043533193 | 0.124009757 | 0.501565251 | 0.042650621 | 0.077869833 |
| 4931406C07Rik | -0.046927255 | 0.389685536 | 0.768272192 | 0.216233825 | 0.261862548 |
| 4931414P19Rik | -0.054098365 | 0.015090502 | 0.197980673 | 0.02022622 | 0.069394214 |
| 4931428F04Rik | -0.015144058 | 0.589326153 | 0.870246667 | 0.046535934 | 0.060730159 |
| 4932438A13Rik | -0.139710958 | 0.039134288 | 0.312482778 | 0.518236341 | 0.665420295 |
| 4933404O12Rik | -0.013845759 | 0.581454216 | 0.866173066 | 0.036691988 | 0.049262355 |
| 4933421O10Rik | 0.030872623 | 0.589005032 | 0.86992264 | 0.251010264 | 0.233102198 |
| 4933434E20Rik | 0.114503252 | 0.058355675 | 0.372056206 | 0.370402848 | 0.273436727 |
| 4933440N22Rik | 0.044461159 | 0.105529323 | 0.472434351 | 0.05135835 | 0.014267956 |
| 5031425E22Rik | -0.012817188 | 0.807336962 | 0.945756429 | 0.199067519 | 0.219547585 |
| 5031439G07Rik | 0.070820898 | 0.160867306 | 0.559080731 | 0.187137396 | 0.121507329 |
| 5033430I15Rik | -0.024630353 | 0.549659931 | 0.852455047 | 0.104615896 | 0.122444975 |
| 5330406M23Rik | -0.019845161 | 0.526242371 | 0.842199436 | 0.060085759 | 0.073521792 |
| 5330431K02Rik | -0.024502963 | 0.422819489 | 0.788524566 | 0.056103848 | 0.07829362 |
| 5330434G04Rik | -0.075921204 | 0.118106097 | 0.491192747 | 0.140269379 | 0.206488512 |
| 5330438D12Rik | 0.011776198 | 0.760275902 | 0.927880499 | 0.098870206 | 0.100825002 |
| 5330439M10Rik | -0.034149082 | 0.467433513 | 0.813188482 | 0.127608561 | 0.144822099 |
| 5430402O13Rik | -0.051980069 | 0.030565494 | 0.278642931 | 0.027699065 | 0.074523969 |
| 5430403G16Rik | -0.049322 | 0.111974929 | 0.483186743 | 0.051360183 | 0.097864152 |
| 5430405H02Rik | -0.117423666 | 0.005537827 | 0.111389124 | 0.103606639 | 0.204357654 |
| 5430416N02Rik | -0.146580313 | 0.002536414 | 0.068435438 | 0.145284941 | 0.288353617 |
| 5530601H04Rik | -0.034060106 | 0.242639918 | 0.653091138 | 0.050328399 | 0.078689022 |
| 5730409E04Rik | 0.00653967 | 0.86691552 | 0.962125879 | 0.102468756 | 0.105989884 |
| 5730455P16Rik | 0.071606382 | 0.215366906 | 0.629611185 | 0.27922896 | 0.204365193 |
| 5730480H06Rik | -0.036162025 | 0.274422775 | 0.682000399 | 0.06592489 | 0.093133499 |
| 5830444B04Rik | -0.011508646 | 0.710396486 | 0.914486281 | 0.057761345 | 0.063220069 |
| 5830448L01Rik | -0.050633644 | 0.073048981 | 0.410160881 | 0.041822936 | 0.085546855 |
| 5830454E08Rik | 0.023893314 | 0.469126236 | 0.813940204 | 0.074247676 | 0.055180595 |
| 5930430L01Rik | 0.039103055 | 0.299367477 | 0.704513802 | 0.096529507 | 0.062333886 |
| 6030458C11Rik | 0.056175655 | 0.123735788 | 0.501387205 | 0.091328403 | 0.04259934 |
| 6330403K07Rik | 0.015044278 | 0.68793928 | 0.909796728 | 0.08536165 | 0.070083999 |
| 6330403L08Rik | 0.042484917 | 0.221387176 | 0.633858837 | 0.079881343 | 0.042238486 |

| | | | | | |
|---|---|---|---|---|---|
| 6330419E04Rik | -0.023543767 | 0.547329979 | 0.851449318 | 0.094166979 | 0.122178469 |
| 6430548M08Rik | 0.087250647 | 0.153833105 | 0.550257617 | 0.318571974 | 0.237673483 |
| 6430590A07Rik | -0.015890559 | 0.646329243 | 0.89474313 | 0.075572034 | 0.102195064 |
| 6720427I07Rik | 0.024151219 | 0.409405569 | 0.779735046 | 0.054809003 | 0.031822305 |
| 6820431F20Rik | -0.091125723 | 0.172577581 | 0.575294174 | 0.495314304 | 0.606675973 |
| 7SK | 0.015097546 | 0.719560332 | 0.918741515 | 0.10898514 | 0.114868086 |
| 8030462N17Rik | -0.012102909 | 0.807051418 | 0.945532285 | 0.1771633 | 0.19457511 |
| 8430429K09Rik | -0.007292675 | 0.865297948 | 0.961687023 | 0.119695239 | 0.124606814 |
| 9130023H24Rik | 0.014453666 | 0.722752697 | 0.919483582 | 0.109553833 | 0.092336903 |
| 9130230N09Rik | -0.044077987 | 0.034076733 | 0.291879728 | 0.019312796 | 0.060006072 |
| 9130401M01Rik | -0.017109106 | 0.740590569 | 0.922901128 | 0.206391112 | 0.222200442 |
| 9130604C24Rik | -0.065216521 | 0.001640824 | 0.053297575 | 0.01635385 | 0.073180655 |
| 9230114K14Rik | 0.042798684 | 0.357630715 | 0.746576432 | 0.150771352 | 0.108910861 |
| 9330020H09Rik | 0.013488723 | 0.781764524 | 0.936001977 | 0.168669514 | 0.161109846 |
| 9330151L19Rik | -0.033308038 | 0.210342282 | 0.623103727 | 0.039625518 | 0.070638246 |
| 9330159F19Rik | -0.003683344 | 0.916124056 | 0.980132949 | 0.075691068 | 0.075235097 |
| 9330160F10Rik | 0.002987911 | 0.918763477 | 0.980385833 | 0.05324228 | 0.054808601 |
| 9330162G02Rik | 0.005132019 | 0.891857153 | 0.971118429 | 0.088319903 | 0.080111989 |
| 9430038I01Rik | -0.029945441 | 0.31582363 | 0.714975653 | 0.051476702 | 0.082638683 |
| 9430065F17Rik | -0.02369446 | 0.409245193 | 0.779735046 | 0.048075102 | 0.064563943 |
| 9430081H08Rik | 0.043295413 | 0.192947035 | 0.60325934 | 0.074335755 | 0.040733033 |
| 9530068E07Rik | 0.057561241 | 0.389269895 | 0.768056209 | 0.496104607 | 0.443809256 |
| 9530077C05Rik | 0.004454096 | 0.886629701 | 0.969612367 | 0.062614996 | 0.061103198 |
| 9930017N22Rik | -0.029517502 | 0.351707048 | 0.74208191 | 0.059647604 | 0.082494399 |
| 9930021J03Rik | -0.100381445 | 0.110114751 | 0.479280902 | 0.412377985 | 0.524809576 |
| A030001D20Rik | -0.02320694 | 0.590794722 | 0.871005673 | 0.118312851 | 0.145565315 |
| A130010J15Rik | -0.005859781 | 0.878179651 | 0.966691068 | 0.09482204 | 0.096720783 |
| A130014A01Rik | -0.011280558 | 0.732456144 | 0.921260942 | 0.065432431 | 0.078923733 |
| A230051N06Rik | -0.162573073 | 0.000285598 | 0.016862021 | 0.116881637 | 0.269782102 |
| A230057D06Rik | -0.294038133 | 9.48E-06 | 0.001260277 | 0.359955738 | 0.677914968 |
| A2ml1 | 0.122625725 | 0.006846337 | 0.126783553 | 0.147651226 | 0.036479163 |
| A330023F24Rik | -0.062755056 | 0.125232525 | 0.503791124 | 0.110812067 | 0.154261558 |
| A330076H08Rik | -0.196435351 | 0.00708933 | 0.128637347 | 0.821382314 | 1.074641381 |
| A430005L14Rik | 0.014576496 | 0.826921317 | 0.952687014 | 0.493093818 | 0.473300717 |
| A430018G15Rik | -0.034708243 | 0.329569278 | 0.724619244 | 0.080262151 | 0.108437909 |
| A430019L02Rik | -0.039551017 | 0.043612365 | 0.329086044 | 0.015108651 | 0.049507874 |
| A430027C01Rik | -0.018985353 | 0.442098871 | 0.800580622 | 0.03358444 | 0.049334466 |
| A430033K04Rik | 0.011417835 | 0.737589872 | 0.922843014 | 0.073816731 | 0.068937204 |
| A530017D24Rik | -0.03074937 | 0.269617628 | 0.677215248 | 0.044533177 | 0.071505531 |
| A530058N18Rik | -0.033255978 | 0.613981395 | 0.881747211 | 0.483317573 | 0.526495305 |
| A730020E08Rik | 0.040635483 | 0.181572574 | 0.589059475 | 0.062605016 | 0.024954362 |
| A730063M14Rik | -0.05219306 | 0.122572102 | 0.498887344 | 0.071526066 | 0.116911691 |
| A930004J17Rik | -0.010237934 | 0.724995806 | 0.92014727 | 0.051904279 | 0.05852099 |
| A930005H10Rik | -0.044179864 | 0.202344537 | 0.612939912 | 0.077256164 | 0.112390429 |
| A930019D19Rik | -0.022355528 | 0.321607816 | 0.719227395 | 0.026314151 | 0.046545353 |
| A930029G22Rik | -0.065134916 | 0.007771283 | 0.135652807 | 0.02488405 | 0.07961081 |
| AA414768 | 0.001435787 | 0.965413521 | 0.992256509 | 0.068923435 | 0.068523628 |
| Aacs | 0.111722913 | 0.016697428 | 0.207483393 | 0.156414255 | 0.054090357 |
| Aagab | -0.035311763 | 0.503732252 | 0.832249514 | 0.188751186 | 0.220565759 |
| Aak1 | 0.084247054 | 0.192273572 | 0.60309438 | 0.438223618 | 0.360725641 |
| Aamdc | 0.036408826 | 0.337555057 | 0.732352032 | 0.100395081 | 0.068411313 |
| Aamp | -0.091952285 | 0.15866722 | 0.556789205 | 0.507280922 | 0.597503329 |
| Aar2 | -0.021935264 | 0.599872591 | 0.875897759 | 0.115806349 | 0.134155893 |
| Aars | -0.023907585 | 0.727968789 | 0.920519505 | 0.800253178 | 0.833532069 |
| Aars2 | 0.031803183 | 0.276985604 | 0.684249202 | 0.057666532 | 0.031076448 |
| Aarsd1 | -0.013154144 | 0.603052467 | 0.876723966 | 0.037012262 | 0.050110382 |
| Aasdh | 0.026447358 | 0.44624812 | 0.803695026 | 0.079549569 | 0.055413798 |
| Aasdhppt | -0.074645691 | 0.194060282 | 0.60325934 | 0.243754334 | 0.317819756 |
| Aatf | 0.062852019 | 0.226499028 | 0.638587289 | 0.208609665 | 0.168919639 |
| Aatk | -0.076129239 | 0.016385896 | 0.206355375 | 0.055315318 | 0.119338922 |
| Abat | -0.079414381 | 0.120088825 | 0.493583757 | 0.173843268 | 0.251168052 |
| Abca1 | -0.122574514 | 0.073561844 | 0.411102578 | 0.437121789 | 0.549091434 |
| Abca2 | -0.054585001 | 0.217813329 | 0.632272789 | 0.125287135 | 0.175247843 |
| Abca3 | 0.081299399 | 0.163067141 | 0.562993622 | 0.301680628 | 0.221780756 |
| Abca5 | -0.080733488 | 0.204127163 | 0.615817354 | 0.389222146 | 0.457383399 |

| | | | | | |
|---|---|---|---|---|---|
| Abca7 | -0.055519816 | 0.136658157 | 0.520249727 | 0.082849008 | 0.126449037 |
| Abca8b | -0.050657421 | 0.036565273 | 0.301071077 | 0.028162015 | 0.07026672 |
| Abcb10 | 0.030933116 | 0.595409623 | 0.873342705 | 0.276447123 | 0.246905122 |
| Abcb4 | 0.029640911 | 0.561915434 | 0.857134531 | 0.18566249 | 0.152465773 |
| Abcb7 | 0.012718521 | 0.763760911 | 0.928730554 | 0.118571427 | 0.104526144 |
| Abcb8 | -0.084289448 | 0.003836709 | 0.087719938 | 0.043951965 | 0.120513148 |
| Abcb9 | -0.035936373 | 0.36623656 | 0.753107222 | 0.09779603 | 0.137350408 |
| Abcc1 | -0.09165648 | 0.020628017 | 0.227971987 | 0.094070531 | 0.180570289 |
| Abcc4 | -0.01304229 | 0.710601718 | 0.914509508 | 0.079063599 | 0.088894625 |
| Abcc5 | -0.016403538 | 0.75429247 | 0.926726954 | 0.201949031 | 0.208466217 |
| Abcc8 | -0.312913497 | 8.94E-06 | 0.001234663 | 1.061212248 | 1.500749707 |
| Abcd1 | -0.020652772 | 0.617482773 | 0.883238152 | 0.113322979 | 0.131852255 |
| Abcd3 | -0.036626706 | 0.594174316 | 0.873038736 | 0.869523093 | 0.89666912 |
| Abcd4 | -0.020974796 | 0.392963551 | 0.769745347 | 0.033821881 | 0.049735901 |
| Abce1 | 0.136843103 | 0.032351099 | 0.285006239 | 0.443987509 | 0.321272119 |
| Abcf1 | 0.004466071 | 0.946370758 | 0.987084873 | 0.54987954 | 0.53424787 |
| Abcf2 | -0.024450415 | 0.609435809 | 0.878855197 | 0.155306821 | 0.188033575 |
| Abcf3 | 0.05029683 | 0.379202894 | 0.762006446 | 0.256781748 | 0.218905888 |
| Abcg1 | -0.064053598 | 0.313773168 | 0.714906079 | 0.339207306 | 0.40201887 |
| Abhd10 | -0.009159471 | 0.812016197 | 0.948397417 | 0.097575187 | 0.109166147 |
| Abhd11 | 0.001044154 | 0.983702217 | 0.99608406 | 0.183797706 | 0.177777193 |
| Abhd12 | -0.077462209 | 0.104430739 | 0.471724268 | 0.153243764 | 0.217731904 |
| Abhd13 | 1.99E-05 | 0.999690599 | 0.99998562 | 0.180313003 | 0.188359932 |
| Abhd14a | -0.217099379 | 1.63E-08 | 8.40E-06 | 0.077200596 | 0.267073779 |
| Abhd14b | -0.036325139 | 0.386833209 | 0.766593284 | 0.11850522 | 0.147923536 |
| Abhd15 | -0.005560446 | 0.862023743 | 0.960216905 | 0.063663671 | 0.071095692 |
| Abhd16a | -0.027327752 | 0.661393018 | 0.898920864 | 0.397956789 | 0.42963309 |
| Abhd17a | 0.101751204 | 0.107397526 | 0.47507667 | 0.397628034 | 0.30987625 |
| Abhd17b | 0.015780915 | 0.759293673 | 0.927719223 | 0.183380672 | 0.166409438 |
| Abhd17c | 0.127978496 | 0.050630641 | 0.354499194 | 0.498853169 | 0.365489908 |
| Abhd18 | -0.038844099 | 0.259639278 | 0.667102369 | 0.071369503 | 0.101878935 |
| Abhd2 | 0.20605693 | 0.001572227 | 0.052080673 | 0.466385932 | 0.27120205 |
| Abhd3 | -0.003064658 | 0.917538028 | 0.980132949 | 0.054187609 | 0.05967035 |
| Abhd4 | -0.060956243 | 0.093156249 | 0.45188518 | 0.075858553 | 0.132265239 |
| Abhd5 | 0.116777266 | 0.025014215 | 0.251569802 | 0.212556996 | 0.111640559 |
| Abhd6 | 0.052663213 | 0.293653017 | 0.699263423 | 0.192108976 | 0.140636689 |
| Abhd8 | 0.035119474 | 0.541926627 | 0.850208191 | 0.294096046 | 0.287462829 |
| Abi1 | 0.102704989 | 0.102946893 | 0.468979611 | 0.409358029 | 0.308208124 |
| Abi2 | -0.10177852 | 0.075582544 | 0.414321131 | 0.302091031 | 0.39207858 |
| Abitram | -0.020585967 | 0.659702685 | 0.898227584 | 0.147507441 | 0.170523991 |
| Abl1 | 0.000258046 | 0.995109753 | 0.99872501 | 0.114713159 | 0.110720515 |
| Abl2 | 0.031209466 | 0.530530001 | 0.84508505 | 0.177616926 | 0.149830735 |
| Ablim1 | 0.139932938 | 0.017655041 | 0.21172951 | 0.276724427 | 0.142219153 |
| Abr | 0.04409985 | 0.445977741 | 0.803559158 | 0.259619213 | 0.230258373 |
| Abracl | 0.115244844 | 0.071460702 | 0.404769223 | 1.22235624 | 1.045416009 |
| Abraxas2 | -0.027178295 | 0.677843537 | 0.907377379 | 0.453803483 | 0.478877683 |
| Abt1 | 0.031760252 | 0.516701416 | 0.837438925 | 0.168749705 | 0.149775512 |
| Abtb1 | -0.092980493 | 0.005586555 | 0.111475656 | 0.06019392 | 0.14139965 |
| Acaa1a | -0.09978488 | 0.084811508 | 0.433957325 | 0.258570522 | 0.358273207 |
| Acaa2 | -0.002025166 | 0.961880768 | 0.991441074 | 0.116134522 | 0.117335531 |
| Acaca | -0.079718082 | 0.073961694 | 0.412240835 | 0.127605371 | 0.197418602 |
| Acad11 | -0.02726002 | 0.380737504 | 0.762006446 | 0.058138134 | 0.083886873 |
| Acad8 | -0.02919778 | 0.415935789 | 0.78353657 | 0.079246854 | 0.108249621 |
| Acad9 | 0.003169972 | 0.92146895 | 0.98085171 | 0.068898237 | 0.065844987 |
| Acadl | 0.052440275 | 0.401239137 | 0.774740795 | 0.370780552 | 0.321504559 |
| Acadm | -0.02655355 | 0.602974305 | 0.876723966 | 0.179254203 | 0.190272868 |
| Acads | 0.028827106 | 0.51434095 | 0.83659041 | 0.134044104 | 0.110663908 |
| Acadsb | -0.025579259 | 0.685094416 | 0.909796728 | 0.370534922 | 0.38508736 |
| Acadvl | -0.094644867 | 0.08758542 | 0.43990675 | 0.21368135 | 0.304448532 |
| Acap2 | 0.040199761 | 0.514356821 | 0.83659041 | 0.328674878 | 0.297128764 |
| Acap3 | 0.031847794 | 0.286001655 | 0.693090656 | 0.058935712 | 0.033383113 |
| Acat1 | -0.124928918 | 0.062205229 | 0.380250374 | 0.563891468 | 0.719696936 |
| Acbd3 | 0.046617449 | 0.496420683 | 0.830056214 | 0.956871876 | 0.926302234 |
| Acbd4 | -0.014234779 | 0.621181269 | 0.884242858 | 0.050038554 | 0.06282512 |
| Acbd5 | -0.067570794 | 0.305234702 | 0.709411642 | 0.563502974 | 0.633900183 |

| | | | | | |
|---|---|---|---|---|---|
| Acbd6 | -0.021924376 | 0.740795855 | 0.922901128 | 0.547257381 | 0.567678756 |
| Accs | -0.05242861 | 0.021619578 | 0.233831581 | 0.024265714 | 0.068604936 |
| Acd | 0.029192843 | 0.574137636 | 0.862168892 | 0.19358293 | 0.173056726 |
| Acer2 | -0.045153918 | 0.099855312 | 0.46464894 | 0.040589013 | 0.083980081 |
| Acer3 | -0.01030435 | 0.747281059 | 0.924930563 | 0.064922182 | 0.076777369 |
| Acin1 | -0.116866236 | 0.090862479 | 0.445385778 | 0.700555453 | 0.809393709 |
| Acly | -0.182340445 | 0.000591825 | 0.028286403 | 2.155005531 | 2.49037248 |
| Aco1 | 0.023747938 | 0.558929383 | 0.856615394 | 0.117294839 | 0.098039518 |
| Aco2 | 0.064583405 | 0.339097049 | 0.732701236 | 0.733688059 | 0.666005457 |
| Acot11 | -0.030522645 | 0.259742558 | 0.667102369 | 0.039717094 | 0.064201457 |
| Acot13 | -0.164291016 | 0.013379526 | 0.187294708 | 0.517124939 | 0.712668559 |
| Acot2 | 0.005259821 | 0.849230376 | 0.957111129 | 0.047758124 | 0.043260316 |
| Acot6 | -0.010955917 | 0.641319273 | 0.891787413 | 0.031622047 | 0.044916085 |
| Acot8 | -0.068787096 | 0.130717253 | 0.510509367 | 0.131751308 | 0.187852999 |
| Acot9 | 0.040398449 | 0.200860714 | 0.611438197 | 0.066326425 | 0.030555943 |
| Acox1 | -0.008847497 | 0.886223091 | 0.969590273 | 0.362436359 | 0.376083897 |
| Acp1 | 0.064028495 | 0.334587584 | 0.730169996 | 0.798300668 | 0.712163205 |
| Acp2 | 0.072625463 | 0.153339698 | 0.549817979 | 0.195056093 | 0.117463845 |
| Acp6 | -0.046132407 | 0.118480491 | 0.491192747 | 0.049716927 | 0.086335991 |
| Acsf3 | 0.009559589 | 0.812251412 | 0.948397417 | 0.10654284 | 0.090241345 |
| Acsl1 | -0.002655943 | 0.95069334 | 0.988114084 | 0.12257908 | 0.129763496 |
| Acsl3 | 0.040136495 | 0.28194903 | 0.688868362 | 0.094058733 | 0.059153838 |
| Acsl4 | -0.039062451 | 0.47525058 | 0.817387411 | 0.211289417 | 0.249788094 |
| Acsl5 | -0.012102993 | 0.832515009 | 0.953797263 | 0.28392668 | 0.289982694 |
| Acsl6 | -0.018623162 | 0.533978935 | 0.846288737 | 0.053697933 | 0.067479857 |
| Acss2 | 0.019258549 | 0.637647742 | 0.891210491 | 0.113656168 | 0.096855145 |
| Acta1 | 0.027049349 | 0.424514164 | 0.789923742 | 0.076556397 | 0.050671802 |
| Actb | 0.077822728 | 0.164323872 | 0.563869641 | 2.218671843 | 2.060976407 |
| Actg1 | 0.107030081 | 0.072063564 | 0.406577847 | 2.530428956 | 2.298878125 |
| Actl6a | 0.074630656 | 0.183508373 | 0.590066231 | 0.260488131 | 0.197629796 |
| Actl6b | -0.107857438 | 0.015780554 | 0.202803361 | 0.124726863 | 0.211392613 |
| Actn1 | -0.051438093 | 0.365185506 | 0.752802229 | 0.259353271 | 0.297352322 |
| Actn4 | 0.220744409 | 0.00122111 | 0.043929327 | 0.622234489 | 0.397215948 |
| Actr10 | -0.079973756 | 0.228737646 | 0.640401604 | 0.496142398 | 0.60662014 |
| Actr1a | -0.025278178 | 0.634250613 | 0.88965971 | 0.211094766 | 0.244608324 |
| Actr1b | 0.133382305 | 0.040919779 | 0.319308264 | 0.452668126 | 0.309333787 |
| Actr2 | -0.146329989 | 0.026472859 | 0.258974747 | 0.529980227 | 0.658354237 |
| Actr3 | 0.025145359 | 0.70435747 | 0.912507936 | 0.714584138 | 0.696680288 |
| Actr3b | 0.039479762 | 0.231976454 | 0.643345304 | 0.073788788 | 0.040202987 |
| Actr6 | -0.004707705 | 0.887962975 | 0.970045467 | 0.068695356 | 0.078552722 |
| Actr8 | -0.002105858 | 0.966571862 | 0.992504747 | 0.175331658 | 0.18856983 |
| Acvr1 | 0.016513035 | 0.653250429 | 0.897191373 | 0.092856293 | 0.078464423 |
| Acvr1b | 0.031991724 | 0.321701787 | 0.719227395 | 0.072298817 | 0.039858907 |
| Acvr1c | -0.05843878 | 0.113781349 | 0.484874023 | 0.076041596 | 0.136042539 |
| Acvr2a | 0.054001221 | 0.304593373 | 0.709410087 | 0.193355557 | 0.148287845 |
| Acvr2b | -0.039185595 | 0.229276619 | 0.640979764 | 0.063167409 | 0.101300269 |
| Acyp1 | -0.048914261 | 0.426168878 | 0.790724712 | 0.377441565 | 0.428697701 |
| Acyp2 | -0.052144771 | 0.260576669 | 0.667877433 | 0.139763269 | 0.191383859 |
| Adal | -0.031314124 | 0.261091522 | 0.66834369 | 0.04450629 | 0.072165004 |
| Adam10 | 0.068461802 | 0.24073087 | 0.650744882 | 0.293866328 | 0.22512699 |
| Adam15 | 0.02136625 | 0.571725209 | 0.861404566 | 0.094425401 | 0.075211446 |
| Adam17 | 0.017490281 | 0.640743747 | 0.891357586 | 0.092240792 | 0.080313661 |
| Adam22 | 0.051604469 | 0.342259855 | 0.735068249 | 0.229737696 | 0.163229475 |
| Adam32 | 0.091751973 | 0.019063882 | 0.218929273 | 0.108656789 | 0.032863202 |
| Adam9 | -0.044104891 | 0.415817207 | 0.783472369 | 0.217732015 | 0.255578711 |
| Adar | 0.033611131 | 0.526559757 | 0.842243869 | 0.207723513 | 0.167683601 |
| Adarb1 | 0.052150065 | 0.306229606 | 0.709680069 | 0.176949576 | 0.131763528 |
| Adat2 | -0.022690443 | 0.340958417 | 0.734220045 | 0.029794319 | 0.048554695 |
| Adck1 | -0.016562468 | 0.578057213 | 0.864673658 | 0.055556142 | 0.071018141 |
| Adck5 | -0.007729424 | 0.827569283 | 0.952851099 | 0.083890428 | 0.09394734 |
| Adcy6 | -0.018451006 | 0.641450835 | 0.891846798 | 0.102154359 | 0.1146902 |
| Adcy9 | 0.036047655 | 0.554463765 | 0.853469818 | 0.331277633 | 0.303659021 |
| Add1 | -0.085405436 | 0.11950063 | 0.492780536 | 0.227197703 | 0.306572775 |
| Add3 | 0.034964421 | 0.542201879 | 0.850208191 | 0.261249787 | 0.228371456 |
| Adgb | -0.042759054 | 0.101593223 | 0.467719786 | 0.034816104 | 0.072441929 |

| | | | | | |
|---|---|---|---|---|---|
| Adgrb3 | 0.007967272 | 0.820408598 | 0.950668356 | 0.081642374 | 0.073898472 |
| Adgrg1 | -0.105781562 | 0.099174861 | 0.463315751 | 0.474819644 | 0.564337227 |
| Adgrl1 | 0.019679943 | 0.732821449 | 0.921306823 | 0.254615182 | 0.23900006 |
| Adh1 | 0.022847214 | 0.672205944 | 0.905281039 | 0.20522263 | 0.196196543 |
| Adh5 | -0.006215327 | 0.915355854 | 0.980132949 | 0.282326695 | 0.300524068 |
| Adi1 | -0.150260888 | 0.021785736 | 0.235078507 | 0.432640261 | 0.566067346 |
| Adipor1 | 0.010523941 | 0.869030861 | 0.963382235 | 0.47327949 | 0.464610326 |
| Adipor2 | 0.040552418 | 0.497076184 | 0.830056214 | 0.304355847 | 0.267489694 |
| Adk | -0.060399397 | 0.332370852 | 0.72806771 | 0.386711685 | 0.433242326 |
| Adm2 | -0.226982988 | 9.71E-08 | 3.36E-05 | 0.077443157 | 0.293577256 |
| Adnp | -0.057112579 | 0.15253639 | 0.549597327 | 0.101480141 | 0.148777925 |
| Adnp2 | 0.017853315 | 0.633560176 | 0.889501117 | 0.089356415 | 0.071999492 |
| Ado | -0.001515636 | 0.971435559 | 0.993892166 | 0.121345547 | 0.128390075 |
| Adora1 | 0.033119269 | 0.499833928 | 0.830876638 | 0.152173817 | 0.123838746 |
| Adora2b | 0.081232073 | 0.081601588 | 0.42777546 | 0.150783401 | 0.088739608 |
| Adpgk | -0.020599439 | 0.553603219 | 0.852880354 | 0.078025814 | 0.096655265 |
| Adprh | -0.137077487 | 0.021476889 | 0.232789997 | 0.303820574 | 0.434878293 |
| Adprm | -0.054590231 | 0.215990142 | 0.629652353 | 0.121298318 | 0.169933448 |
| Adprs | -0.107572396 | 0.042150466 | 0.324597482 | 0.186405589 | 0.288364907 |
| Adra2a | -0.172557424 | 0.010450997 | 0.160622874 | 0.402543603 | 0.565726137 |
| Adrm1 | 0.138290462 | 0.037785725 | 0.306840872 | 0.441769984 | 0.312468475 |
| Adsl | 0.022129803 | 0.573953163 | 0.862168892 | 0.102086206 | 0.088034872 |
| Adss | -0.035619587 | 0.469597527 | 0.814050152 | 0.169523756 | 0.199713973 |
| Adssl1 | 0.037649941 | 0.204387494 | 0.616256046 | 0.057554942 | 0.025677804 |
| Aebp2 | -0.047026072 | 0.406890748 | 0.777736088 | 0.249353566 | 0.295977647 |
| Aen | -0.018226221 | 0.770606515 | 0.931764557 | 0.277764324 | 0.286434446 |
| Afdn | 0.069721594 | 0.303390056 | 0.708181201 | 0.622924247 | 0.519733729 |
| Aff1 | -0.132714099 | 0.028968508 | 0.271130568 | 0.299442021 | 0.425163816 |
| Aff2 | -0.016303532 | 0.703521372 | 0.912277105 | 0.110998592 | 0.133550491 |
| Aff4 | -0.071709756 | 0.269230769 | 0.676978829 | 0.920443757 | 1.008996397 |
| Afg1l | 0.020263715 | 0.50752604 | 0.833674307 | 0.060020934 | 0.040558887 |
| Afg3l1 | -0.079414808 | 0.15100743 | 0.546523037 | 0.226718831 | 0.309107023 |
| Afg3l2 | -0.004802073 | 0.938341336 | 0.985197198 | 0.348225952 | 0.346668663 |
| Aftph | -0.025347111 | 0.68559725 | 0.909796728 | 0.435896932 | 0.451888247 |
| Aga | -0.211012934 | 0.001272378 | 0.044967808 | 0.465957115 | 0.674067616 |
| Agap1 | -0.247333805 | 0.000657709 | 0.029470643 | 0.646346726 | 0.921623608 |
| Agap3 | 0.026765308 | 0.605984238 | 0.878178429 | 0.194033404 | 0.177384122 |
| Agbl3 | -0.048875204 | 0.261522633 | 0.668468258 | 0.119832361 | 0.160239433 |
| Agbl4 | -0.041896481 | 0.100933646 | 0.466401306 | 0.032974349 | 0.073053556 |
| Agfg1 | 0.080336361 | 0.225041788 | 0.637343235 | 0.539939856 | 0.457033658 |
| Agfg2 | 0.045127259 | 0.222508538 | 0.634095002 | 0.092985227 | 0.052490172 |
| Aggf1 | 0.005440642 | 0.929838474 | 0.982502239 | 0.363691922 | 0.361548536 |
| Agk | -0.049146652 | 0.13756572 | 0.522790545 | 0.065744557 | 0.107656484 |
| Agl | 0.016296019 | 0.777879723 | 0.934339474 | 0.278523863 | 0.267676628 |
| Ago1 | 0.027120171 | 0.58062983 | 0.866173066 | 0.17442579 | 0.15188536 |
| Ago2 | -0.042465653 | 0.532808522 | 0.84612966 | 0.576674899 | 0.608953004 |
| Ago3 | 0.053292786 | 0.359107479 | 0.747455833 | 0.273206031 | 0.223127479 |
| Ago4 | -0.003599703 | 0.917509442 | 0.980132949 | 0.0803507 | 0.080687612 |
| Agpat1 | -0.030148907 | 0.547064328 | 0.851446616 | 0.173188462 | 0.195243137 |
| Agpat2 | 0.033524426 | 0.440417353 | 0.80031914 | 0.134344221 | 0.102452758 |
| Agpat3 | -0.109970432 | 0.104410023 | 0.471724268 | 0.570892086 | 0.698449009 |
| Agpat4 | -0.025704536 | 0.326497927 | 0.720866629 | 0.036375559 | 0.054678652 |
| Agpat5 | -0.034593828 | 0.544798493 | 0.851259637 | 0.242804161 | 0.272282855 |
| Agps | 0.018396976 | 0.686433623 | 0.909796728 | 0.145587156 | 0.128389645 |
| Agt | 0.017180967 | 0.520197347 | 0.838864153 | 0.047913697 | 0.037905555 |
| Agtpbp1 | -0.087536689 | 0.141226784 | 0.528664246 | 0.271670291 | 0.350323058 |
| Agtrap | -0.049665299 | 0.302057028 | 0.707525413 | 0.150337778 | 0.194851531 |
| Ahctf1 | 0.023938056 | 0.653816311 | 0.897454058 | 0.212398034 | 0.201540955 |
| Ahcyl1 | 0.073864501 | 0.257021648 | 0.665256942 | 0.754101066 | 0.664194892 |
| Ahcyl2 | -0.006528482 | 0.920004966 | 0.980436936 | 0.403158042 | 0.419396977 |
| Ahdc1 | 0.032856265 | 0.483979637 | 0.822768706 | 0.146827964 | 0.122799591 |
| Ahi1 | -0.090450589 | 0.197395323 | 0.607747501 | 0.715584995 | 0.809100919 |
| Ahnak | 0.063645414 | 0.030992098 | 0.280493857 | 0.058545508 | 0 |
| Ahsa1 | 0.031965453 | 0.635112812 | 0.889813971 | 0.483227797 | 0.437059369 |
| Ahsa2 | -0.054830738 | 0.296831402 | 0.702656882 | 0.184020807 | 0.21848066 |

| | | | | | |
|---|---|---|---|---|---|
| AI413582 | 0.10121512 | 0.119460897 | 0.492780536 | 0.487577925 | 0.394161965 |
| AI480526 | -0.035192357 | 0.375198649 | 0.760147021 | 0.100768461 | 0.132058579 |
| AI506816 | -0.023708318 | 0.432049088 | 0.794226501 | 0.053151148 | 0.072357849 |
| AI597479 | -0.003447889 | 0.921844102 | 0.980859278 | 0.080632063 | 0.082294208 |
| AI661453 | -0.040465024 | 0.241990119 | 0.652555608 | 0.071605282 | 0.110800236 |
| AI837181 | 0.042302299 | 0.360683667 | 0.749054824 | 0.152850515 | 0.121472663 |
| AI987944 | 0.04984639 | 0.29353738 | 0.69925611 | 0.160152249 | 0.118293219 |
| Aida | 0.003735466 | 0.951073506 | 0.988114084 | 0.344069125 | 0.330967957 |
| Aifm1 | 0.029536003 | 0.478650154 | 0.819081437 | 0.118194389 | 0.095999912 |
| Aifm2 | 0.050170057 | 0.109008086 | 0.477972625 | 0.066437487 | 0.023371654 |
| Aig1 | -0.041230066 | 0.525326496 | 0.842015656 | 0.533294471 | 0.583512543 |
| Aimp1 | 0.063167427 | 0.359675067 | 0.747719327 | 0.683537019 | 0.60776843 |
| Aimp2 | 0.053054801 | 0.199443624 | 0.610108047 | 0.113131251 | 0.068355247 |
| Aip | 0.046825278 | 0.404858355 | 0.776552851 | 0.247468097 | 0.212039069 |
| Airn | 0.014653159 | 0.747564944 | 0.924930563 | 0.141608515 | 0.135785515 |
| Ak1 | -0.01907806 | 0.639103445 | 0.891345382 | 0.105383283 | 0.125630201 |
| Ak2 | 0.059495676 | 0.365338373 | 0.752802229 | 0.426942181 | 0.369542407 |
| Ak3 | -0.005914082 | 0.884164033 | 0.968954079 | 0.108905863 | 0.107956338 |
| Ak5 | -0.027248145 | 0.315615016 | 0.714975653 | 0.039540873 | 0.066807607 |
| Ak6 | -0.007078576 | 0.900814236 | 0.974086672 | 0.223551068 | 0.236531466 |
| Akain1 | 0.019299164 | 0.759775209 | 0.927720653 | 0.38073696 | 0.363753471 |
| Akap1 | 0.055421831 | 0.135903583 | 0.519431685 | 0.09825554 | 0.053977852 |
| Akap10 | 0.012131026 | 0.729758244 | 0.920522999 | 0.083542628 | 0.073019168 |
| Akap11 | 0.079328129 | 0.253328465 | 0.663146569 | 0.675131545 | 0.576306772 |
| Akap13 | -0.08855173 | 0.215112373 | 0.629611185 | 0.790564896 | 0.925317294 |
| Akap17b | -0.036198876 | 0.224711425 | 0.637067124 | 0.050921307 | 0.077383046 |
| Akap6 | -0.00634759 | 0.882662857 | 0.96861827 | 0.119508027 | 0.117291859 |
| Akap7 | 0.016266973 | 0.691253426 | 0.909796728 | 0.111944835 | 0.100799052 |
| Akap8 | -0.05719088 | 0.281786658 | 0.688819456 | 0.1917286 | 0.243586576 |
| Akap8l | 0.064579529 | 0.334955101 | 0.73038987 | 0.455246247 | 0.406060217 |
| Akap9 | -0.094265426 | 0.132821582 | 0.513464375 | 1.25653207 | 1.386225751 |
| Akip1 | -0.106431325 | 0.003829772 | 0.087719938 | 0.080210204 | 0.172579161 |
| Akirin1 | 0.019281929 | 0.780765318 | 0.935482467 | 0.743679542 | 0.705313667 |
| Akirin2 | 0.104926972 | 0.105635506 | 0.472566726 | 0.499381463 | 0.407768556 |
| Akna | -0.0479454 | 0.010912186 | 0.165446543 | 0.01236195 | 0.054740124 |
| Akr1a1 | 0.035344303 | 0.587779343 | 0.869466748 | 0.871956325 | 0.843580941 |
| Akr1b10 | -0.001538962 | 0.968367365 | 0.993505636 | 0.096557816 | 0.092573695 |
| Akr1b3 | -0.10981038 | 0.061703443 | 0.380250374 | 0.290808872 | 0.379729741 |
| Akr1c12 | -0.007946297 | 0.888279485 | 0.970148116 | 0.232262986 | 0.252438619 |
| Akr1c13 | -0.060970434 | 0.296561782 | 0.702656882 | 0.248185641 | 0.313794422 |
| Akr1c14 | -0.145177509 | 0.002581007 | 0.069451932 | 0.13540505 | 0.282055851 |
| Akr1c19 | -0.186692073 | 0.001270571 | 0.044967808 | 0.236227053 | 0.438528785 |
| Akr1e1 | -0.09529103 | 0.033644514 | 0.290545455 | 0.118296408 | 0.200661369 |
| Akr7a5 | -0.099585036 | 0.0560196 | 0.367897032 | 0.189018132 | 0.287545613 |
| Akt1 | 0.035220678 | 0.562978782 | 0.857583554 | 0.345004208 | 0.291847328 |
| Akt1s1 | 0.029975452 | 0.6084444 | 0.878855197 | 0.300472506 | 0.280473591 |
| Akt2 | -0.036287084 | 0.553692768 | 0.852887401 | 0.326609366 | 0.357411805 |
| Akt3 | 0.021275382 | 0.702526623 | 0.912194012 | 0.257424079 | 0.234253978 |
| Aktip | 0.018959485 | 0.756555331 | 0.926833377 | 0.340754706 | 0.330695286 |
| Alad | 0.006966297 | 0.883426129 | 0.968793593 | 0.159625334 | 0.157498669 |
| Alas1 | 0.004260747 | 0.92751387 | 0.982010202 | 0.154625734 | 0.158502948 |
| Alcam | -0.147033966 | 0.033891982 | 0.291744277 | 0.483304775 | 0.629158373 |
| Aldh18a1 | -0.076909543 | 0.200631455 | 0.610964171 | 0.332569371 | 0.399627061 |
| Aldh1l2 | -0.269908971 | 8.98E-06 | 0.001234663 | 0.263507048 | 0.532434003 |
| Aldh2 | -0.045956993 | 0.253345292 | 0.663146569 | 0.103666417 | 0.148429337 |
| Aldh3a2 | -0.022547001 | 0.667421127 | 0.901808587 | 0.199894598 | 0.210620717 |
| Aldh4a1 | 0.011351189 | 0.679862215 | 0.907783945 | 0.048605038 | 0.037957179 |
| Aldh5a1 | -0.177933721 | 0.001104731 | 0.040616055 | 0.193097385 | 0.370261231 |
| Aldh6a1 | -0.055640552 | 0.102742135 | 0.468979611 | 0.066316726 | 0.120832443 |
| Aldh7a1 | -0.021960413 | 0.599987278 | 0.875927602 | 0.113846367 | 0.127126237 |
| Aldh9a1 | -0.104109783 | 0.036186079 | 0.299721418 | 0.161655746 | 0.250874819 |
| Aldoa | -0.116118264 | 0.043803693 | 0.329086044 | 1.570224817 | 1.770363259 |
| Aldoc | -0.043850017 | 0.086587252 | 0.437601331 | 0.031287045 | 0.065155776 |
| Alg1 | 0.011675846 | 0.800602861 | 0.941935402 | 0.150155641 | 0.143620359 |
| Alg10b | 0.052462174 | 0.307673788 | 0.710239606 | 0.199881389 | 0.148007358 |

| | | | | | |
|---|---|---|---|---|---|
| Alg11 | -0.039331809 | 0.371382582 | 0.75760494 | 0.126662441 | 0.165358698 |
| Alg12 | -0.005593562 | 0.850366694 | 0.957175116 | 0.053014921 | 0.060965847 |
| Alg13 | -0.050207534 | 0.104059026 | 0.471105296 | 0.054675766 | 0.096221631 |
| Alg14 | 0.005835709 | 0.879239829 | 0.967221631 | 0.09724179 | 0.090137008 |
| Alg2 | -0.101338398 | 0.069425413 | 0.400472912 | 0.254514299 | 0.35460321 |
| Alg3 | 0.03792565 | 0.391993281 | 0.769704269 | 0.139308258 | 0.112060091 |
| Alg5 | -0.000137781 | 0.99811526 | 0.99938181 | 0.299494988 | 0.297595137 |
| Alg6 | -0.000787923 | 0.981704368 | 0.995792496 | 0.074693595 | 0.080683413 |
| Alg9 | -0.018190779 | 0.556027951 | 0.854910009 | 0.056664955 | 0.072383073 |
| Alkbh1 | -0.03275534 | 0.606692193 | 0.878443384 | 0.362330013 | 0.391519066 |
| Alkbh2 | 0.027639264 | 0.420879688 | 0.787234019 | 0.07871191 | 0.060384064 |
| Alkbh3 | 0.053708215 | 0.200333688 | 0.610852219 | 0.126199781 | 0.08062269 |
| Alkbh5 | -0.018342352 | 0.775639838 | 0.933636662 | 0.89888571 | 0.929450082 |
| Alkbh6 | -0.131633293 | 0.010987135 | 0.165713947 | 0.179995554 | 0.301376676 |
| Alkbh7 | -0.045343836 | 0.344444065 | 0.737445936 | 0.150874728 | 0.19852197 |
| Alkbh8 | -0.03627851 | 0.422438344 | 0.788202829 | 0.129835729 | 0.160795289 |
| Alms1 | -0.035200897 | 0.236389639 | 0.647025581 | 0.050379427 | 0.079460551 |
| Alox5ap | 0.033136431 | 0.384383244 | 0.764576818 | 0.096672841 | 0.068320496 |
| Als2 | -0.010195779 | 0.72332444 | 0.919686776 | 0.050748314 | 0.063772722 |
| Alyref | 0.031548064 | 0.636046196 | 0.890500163 | 0.417151418 | 0.381485069 |
| Alyref2 | -0.001429301 | 0.970988808 | 0.993892166 | 0.102730188 | 0.107367255 |
| Amacr | -0.018463271 | 0.535017351 | 0.846460359 | 0.054575561 | 0.071327371 |
| Ambp | 0.044446151 | 0.194032432 | 0.60325934 | 0.076224425 | 0.032044535 |
| Ambra1 | -0.009722125 | 0.809538975 | 0.947672346 | 0.106397614 | 0.115492508 |
| Amd1 | -0.007537337 | 0.858215186 | 0.958698478 | 0.120077465 | 0.130777913 |
| Amd2 | -0.046469414 | 0.143606714 | 0.532857888 | 0.060890554 | 0.101787488 |
| Amdhd2 | 0.025647115 | 0.532837198 | 0.84612966 | 0.117761299 | 0.10159501 |
| Amer1 | 0.010744927 | 0.741588261 | 0.923030925 | 0.069301858 | 0.064223749 |
| Amer2 | 0.036848282 | 0.509787609 | 0.834904701 | 0.236379392 | 0.209330889 |
| Amfr | -0.009826208 | 0.883445049 | 0.968793593 | 0.526877052 | 0.552003782 |
| Amigo1 | -0.046286605 | 0.392202099 | 0.769704269 | 0.211681457 | 0.243996161 |
| Amigo2 | 0.05415736 | 0.296724708 | 0.702656882 | 0.193194188 | 0.139404829 |
| Ammecr1l | -0.031203103 | 0.412359835 | 0.781358105 | 0.094301318 | 0.128444199 |
| Amn1 | -0.00956003 | 0.856856061 | 0.958302663 | 0.208742095 | 0.21755041 |
| Amotl1 | -0.004741077 | 0.915906379 | 0.980132949 | 0.129057839 | 0.131498046 |
| Amotl2 | -0.009961539 | 0.729943464 | 0.920522999 | 0.048714612 | 0.056618891 |
| Ampd2 | 0.093251282 | 0.109145865 | 0.477972625 | 0.30173556 | 0.196464354 |
| Amph | -0.004335948 | 0.916940747 | 0.980132949 | 0.114185014 | 0.111133732 |
| Amt | -0.079528548 | 0.001728403 | 0.055248355 | 0.028565586 | 0.097290336 |
| Amz2 | -0.132591431 | 0.021767748 | 0.235078507 | 0.276685982 | 0.400786166 |
| Anapc1 | -0.083322698 | 0.11989308 | 0.493385339 | 0.196647316 | 0.272055085 |
| Anapc10 | -0.013885466 | 0.733594115 | 0.921306823 | 0.104965108 | 0.119025951 |
| Anapc11 | -0.015302993 | 0.816207715 | 0.949036482 | 0.505306132 | 0.538865472 |
| Anapc13 | 0.048331467 | 0.472105658 | 0.815861654 | 0.896782066 | 0.813749466 |
| Anapc15 | 0.046563639 | 0.265190076 | 0.672177989 | 0.124144281 | 0.084229252 |
| Anapc16 | -0.077301507 | 0.244809459 | 0.654809396 | 0.579365867 | 0.671510264 |
| Anapc2 | 0.023338627 | 0.636749333 | 0.890755879 | 0.176298944 | 0.155296656 |
| Anapc4 | 0.042017502 | 0.485946388 | 0.824728423 | 0.305076026 | 0.276433961 |
| Anapc5 | -0.044996152 | 0.501002385 | 0.831764926 | 0.531222452 | 0.573906736 |
| Anapc7 | 0.026748923 | 0.548020042 | 0.851862654 | 0.136451204 | 0.108421518 |
| Ang | -0.064909617 | 0.376756597 | 0.761269164 | 0.578218582 | 0.621620489 |
| Angel1 | -0.012746675 | 0.697033743 | 0.910970095 | 0.067891183 | 0.076185058 |
| Angel2 | 0.023396862 | 0.703033021 | 0.912277105 | 0.31631969 | 0.287245535 |
| Angptl6 | -0.402057841 | 6.30E-12 | 6.32E-09 | 0.224563996 | 0.631108669 |
| Ank | -0.074261663 | 0.291691271 | 0.697566301 | 0.679244605 | 0.767193792 |
| Ank2 | -0.037084458 | 0.590068363 | 0.87075199 | 0.577123495 | 0.612883523 |
| Ank3 | 0.000360483 | 0.995349569 | 0.99872501 | 0.332980378 | 0.312585359 |
| Ankfy1 | 0.046998187 | 0.336122635 | 0.73102121 | 0.176455897 | 0.129757845 |
| Ankhd1 | -0.014041779 | 0.83618507 | 0.954911647 | 0.461931787 | 0.480871901 |
| Ankib1 | -0.113675159 | 0.028130134 | 0.26686404 | 0.187366811 | 0.294726684 |
| Ankle2 | -0.04178975 | 0.418252217 | 0.784672431 | 0.195037354 | 0.243793629 |
| Ankmy2 | 0.053014459 | 0.274279434 | 0.681927836 | 0.171022698 | 0.140354213 |
| Ankra2 | -0.002657466 | 0.96497375 | 0.991970325 | 0.368711332 | 0.352376186 |
| Ankrd10 | -0.234789815 | 0.000819 | 0.033552272 | 0.62572771 | 0.890498679 |
| Ankrd11 | -0.069158393 | 0.301727199 | 0.706960943 | 1.007260084 | 1.089566813 |

| | | | | | |
|---|---|---|---|---|---|
| Ankrd12 | -0.037320872 | 0.591915058 | 0.871888969 | 1.009857719 | 1.06288976 |
| Ankrd13a | -0.128917247 | 0.002063584 | 0.062198787 | 0.107005922 | 0.223620852 |
| Ankrd13b | -0.042201756 | 0.113254031 | 0.483626277 | 0.037202475 | 0.070319151 |
| Ankrd13c | -0.115791542 | 0.04911018 | 0.347612176 | 0.283637603 | 0.40424724 |
| Ankrd13d | -0.024777414 | 0.278022642 | 0.684996416 | 0.02725584 | 0.048237218 |
| Ankrd16 | -0.026526829 | 0.343697024 | 0.73648314 | 0.044507843 | 0.064749933 |
| Ankrd17 | -0.083735132 | 0.226656383 | 0.638720711 | 0.657039865 | 0.737301331 |
| Ankrd24 | 0.096732914 | 0.084608646 | 0.433706763 | 0.253959839 | 0.173969332 |
| Ankrd26 | -0.029121585 | 0.505220314 | 0.832249514 | 0.120228197 | 0.145177244 |
| Ankrd27 | 0.009109148 | 0.785205082 | 0.936442895 | 0.073425173 | 0.065091497 |
| Ankrd28 | 0.012987091 | 0.698866465 | 0.911319029 | 0.07554001 | 0.066410202 |
| Ankrd33b | 0.090457389 | 0.168692078 | 0.57227323 | 0.368501304 | 0.288986218 |
| Ankrd37 | 0.005510368 | 0.890673554 | 0.970579007 | 0.107656002 | 0.103510448 |
| Ankrd39 | -0.021830813 | 0.613024112 | 0.881106572 | 0.124516361 | 0.145180605 |
| Ankrd40 | 0.129389865 | 0.044088007 | 0.329446347 | 0.448249347 | 0.318247766 |
| Ankrd44 | 0.023974569 | 0.691530652 | 0.909796728 | 0.288540494 | 0.262588613 |
| Ankrd46 | 0.041050776 | 0.315741757 | 0.714975653 | 0.115314496 | 0.077989938 |
| Ankrd49 | 0.066623988 | 0.150540968 | 0.545732529 | 0.152179865 | 0.091103904 |
| Ankrd50 | -0.014454069 | 0.655680297 | 0.897825804 | 0.063436417 | 0.080760243 |
| Ankrd54 | -0.080047119 | 0.19122373 | 0.601854055 | 0.337591185 | 0.410250296 |
| Anks1 | 0.032281119 | 0.532533929 | 0.84612966 | 0.187283775 | 0.163729079 |
| Anks3 | -0.08926341 | 0.036308843 | 0.300252265 | 0.105675537 | 0.190597432 |
| Anks4b | -0.010002914 | 0.86324853 | 0.960676137 | 0.257249677 | 0.26415009 |
| Ankzf1 | -0.073545397 | 0.009527972 | 0.151556661 | 0.042417128 | 0.101500071 |
| Ano10 | -0.004127546 | 0.90098132 | 0.974162395 | 0.069745604 | 0.074923436 |
| Ano6 | 0.040396178 | 0.441641914 | 0.800354711 | 0.204940784 | 0.167588141 |
| Ano8 | 0.012084711 | 0.716417527 | 0.916579082 | 0.074400734 | 0.064460108 |
| Anp32a | 0.003907502 | 0.955324442 | 0.989278596 | 0.783990323 | 0.782951874 |
| Anp32b | -0.152926964 | 0.015139303 | 0.198113669 | 1.069974263 | 1.288445373 |
| Anp32e | -0.12083811 | 0.071128451 | 0.404027315 | 0.540340713 | 0.676184142 |
| Anpep | 0.005939892 | 0.882319481 | 0.968472149 | 0.099999948 | 0.103068713 |
| Antkmt | 0.007421686 | 0.832220454 | 0.953756189 | 0.079167993 | 0.069091621 |
| Anxa11 | -0.013595115 | 0.814701542 | 0.94873644 | 0.264129816 | 0.270270836 |
| Anxa4 | 0.064422869 | 0.339794366 | 0.733286616 | 0.552086703 | 0.463294822 |
| Anxa5 | -0.03576114 | 0.546335627 | 0.851446616 | 1.293759377 | 1.361060392 |
| Anxa6 | 0.032947757 | 0.573218511 | 0.861931715 | 0.283447182 | 0.24752984 |
| Anxa7 | -0.0243525 | 0.70448936 | 0.912507936 | 0.384898269 | 0.410279777 |
| Aopep | 0.05322264 | 0.447526741 | 0.804698299 | 0.607834088 | 0.553231955 |
| Ap1ar | 0.033882968 | 0.50417185 | 0.832249514 | 0.191201193 | 0.17523797 |
| Ap1b1 | 0.057685634 | 0.320862974 | 0.718701556 | 0.309826797 | 0.254048898 |
| Ap1g1 | 0.024259266 | 0.683823034 | 0.909438425 | 0.299956367 | 0.274110211 |
| Ap1m1 | 0.052864183 | 0.240706603 | 0.650744882 | 0.143331904 | 0.100603375 |
| Ap1m2 | -0.033496633 | 0.430912036 | 0.793421321 | 0.11735282 | 0.14766909 |
| Ap1s1 | 0.05034763 | 0.444117328 | 0.801932586 | 0.493436151 | 0.44836183 |
| Ap1s2 | -0.034312113 | 0.615634139 | 0.882439418 | 0.613242644 | 0.630638446 |
| Ap1s3 | -0.003669682 | 0.902538848 | 0.974796343 | 0.057868607 | 0.063990192 |
| Ap2a1 | -0.002332592 | 0.969530474 | 0.993839856 | 0.332971277 | 0.338778125 |
| Ap2a2 | 0.061849679 | 0.32586891 | 0.720866629 | 0.430148649 | 0.380221194 |
| Ap2b1 | -0.120176188 | 0.058848601 | 0.373129125 | 0.360358046 | 0.470570192 |
| Ap2m1 | 0.045197273 | 0.507746442 | 0.833674307 | 0.608081202 | 0.550436085 |
| Ap2s1 | 0.012799121 | 0.851953509 | 0.957461155 | 0.59300001 | 0.565287724 |
| Ap3b1 | -0.059548231 | 0.303117456 | 0.708026508 | 0.248235239 | 0.298380527 |
| Ap3b2 | -0.012110711 | 0.854886467 | 0.957723375 | 0.534420823 | 0.528484567 |
| Ap3d1 | 0.007309733 | 0.913361314 | 0.979979308 | 0.491389814 | 0.493415402 |
| Ap3m1 | -0.002114298 | 0.968660571 | 0.993505636 | 0.213532791 | 0.214545521 |
| Ap3m2 | -0.009463614 | 0.81987999 | 0.950575887 | 0.109613028 | 0.114103146 |
| Ap3s1 | 0.096708502 | 0.140078358 | 0.528161714 | 0.585193119 | 0.471373039 |
| Ap3s2 | -0.011377467 | 0.824836882 | 0.951815105 | 0.189550593 | 0.216190781 |
| Ap4b1 | -0.014322482 | 0.637054912 | 0.890794114 | 0.057199104 | 0.073991232 |
| Ap4e1 | 0.028790212 | 0.29139842 | 0.697147764 | 0.049140941 | 0.022404858 |
| Ap4m1 | -0.025108353 | 0.372885877 | 0.75825687 | 0.046815817 | 0.064033647 |
| Ap4s1 | -0.050372586 | 0.334021862 | 0.729929513 | 0.184772718 | 0.22623344 |
| Ap5m1 | 0.003816316 | 0.933103612 | 0.983777411 | 0.145447462 | 0.146637853 |
| Ap5s1 | -0.026463885 | 0.456537491 | 0.808445095 | 0.077175217 | 0.101381223 |
| Apba1 | -0.080457857 | 0.019703007 | 0.223934391 | 0.068099437 | 0.140245895 |

| | | | | | |
|---|---|---|---|---|---|
| Apba3 | 0.014775323 | 0.643149204 | 0.892561795 | 0.069162588 | 0.054993979 |
| Apbb1 | -0.07802088 | 0.172162726 | 0.575294174 | 0.249999063 | 0.324486521 |
| Apbb2 | -0.016876664 | 0.741093899 | 0.922901128 | 0.185648366 | 0.203716901 |
| Apbb3 | 0.084647702 | 0.017475278 | 0.210563469 | 0.092897973 | 0.025474914 |
| Apc | 0.06890423 | 0.288742175 | 0.695246414 | 0.437250191 | 0.35805867 |
| Apeh | -0.005605645 | 0.899223149 | 0.973834997 | 0.133716707 | 0.136139346 |
| Apex1 | 0.019723033 | 0.629028632 | 0.887864718 | 0.111925627 | 0.106822831 |
| Aph1a | 0.030211262 | 0.598801227 | 0.875574105 | 0.276242385 | 0.24022301 |
| Aph1b | 0.018332914 | 0.582796884 | 0.866383905 | 0.070213202 | 0.054108285 |
| Api5 | -0.023005598 | 0.726282292 | 0.92014727 | 0.445764624 | 0.465880562 |
| Apip | -0.09148701 | 0.047201447 | 0.340345488 | 0.133756725 | 0.215529005 |
| Aplp1 | -0.194372991 | 0.004357852 | 0.096131339 | 0.891637239 | 1.134087786 |
| Aplp2 | 0.072892013 | 0.251916588 | 0.663127858 | 0.402346771 | 0.336997528 |
| Apmap | -0.004193184 | 0.918131446 | 0.980385833 | 0.10763608 | 0.108790784 |
| Apoa2 | -0.021181242 | 0.426231357 | 0.790724712 | 0.039383478 | 0.056411564 |
| Apobec1 | 0.050745246 | 0.311777524 | 0.71405691 | 0.183389423 | 0.140328586 |
| Apobec3 | 0.038817541 | 0.318882714 | 0.717175454 | 0.103585435 | 0.063010795 |
| Apoo | -0.04314831 | 0.361668345 | 0.750013467 | 0.154388632 | 0.196981926 |
| Apool | 0.053289103 | 0.219054873 | 0.632622127 | 0.125106366 | 0.078205546 |
| App | -0.200516151 | 0.00329183 | 0.080573298 | 0.441594144 | 0.681863153 |
| Appbp2 | -0.031786201 | 0.617944951 | 0.883238152 | 0.457963468 | 0.473995471 |
| Appl1 | -0.167259046 | 0.008924191 | 0.146996602 | 0.418904726 | 0.584046171 |
| Appl2 | 0.010965638 | 0.815355709 | 0.948993269 | 0.152512785 | 0.141381494 |
| Aprt | 0.118032617 | 0.082473258 | 0.429166311 | 0.671666529 | 0.549001984 |
| Aptx | -0.027365806 | 0.535520242 | 0.846859409 | 0.127640248 | 0.156989455 |
| Aqp11 | 0.124368397 | 0.014434357 | 0.193677739 | 0.194344275 | 0.086458139 |
| Aqp4 | -0.044516413 | 0.195125278 | 0.604840152 | 0.067310936 | 0.105001864 |
| Aqr | -0.0116115 | 0.851293051 | 0.957461155 | 0.319846989 | 0.330291733 |
| Araf | -0.049249915 | 0.468229451 | 0.813213564 | 0.574622015 | 0.599750874 |
| Arap1 | 0.003080938 | 0.936042704 | 0.984394449 | 0.097450339 | 0.090692442 |
| Arcn1 | -0.092568799 | 0.181526963 | 0.589059475 | 0.783133338 | 0.90677774 |
| Arel1 | -0.019834172 | 0.662787606 | 0.899270729 | 0.134902362 | 0.14896038 |
| Arf1 | 0.067884149 | 0.224986547 | 0.637343235 | 1.448326134 | 1.343416044 |
| Arf2 | -0.021484862 | 0.640028527 | 0.891345382 | 0.145725823 | 0.167344432 |
| Arf3 | -0.025536621 | 0.710886839 | 0.914531044 | 0.598162688 | 0.621711967 |
| Arf4 | 0.014864121 | 0.830705496 | 0.95325696 | 0.952823585 | 0.92720259 |
| Arf5 | -0.061038334 | 0.232511714 | 0.643543798 | 1.547472747 | 1.654874166 |
| Arf6 | 0.004253938 | 0.9418274 | 0.986168593 | 0.248926785 | 0.245523085 |
| Arfgap1 | -0.023468832 | 0.697813436 | 0.910970095 | 0.294793175 | 0.311680657 |
| Arfgap2 | -0.000819784 | 0.989103157 | 0.997411191 | 0.300643379 | 0.291580731 |
| Arfgap3 | -0.115420942 | 0.0850478 | 0.434272336 | 0.405789954 | 0.533870694 |
| Arfgef1 | -0.115907123 | 0.086330432 | 0.43696958 | 0.854421514 | 0.973872264 |
| Arfgef2 | 0.027210801 | 0.652530872 | 0.896875696 | 0.317382154 | 0.286614384 |
| Arfgef3 | -0.080259286 | 0.206549258 | 0.617923966 | 1.039995537 | 1.132611422 |
| Arfip1 | 0.012907292 | 0.829085887 | 0.952883894 | 0.338131564 | 0.338769039 |
| Arfip2 | -0.030667491 | 0.632058482 | 0.888980506 | 0.398185421 | 0.425662369 |
| Arfrp1 | -0.106819932 | 0.08548171 | 0.435299206 | 0.324591376 | 0.437919824 |
| Arglu1 | -0.157956949 | 0.026231488 | 0.257330576 | 0.728481572 | 0.890914753 |
| Arhgap1 | 0.054023433 | 0.307567618 | 0.710199288 | 0.213222588 | 0.161705907 |
| Arhgap12 | -0.085641072 | 0.09874128 | 0.463315751 | 0.19266873 | 0.279486468 |
| Arhgap17 | 0.031871621 | 0.490091762 | 0.82659234 | 0.148754235 | 0.117183736 |
| Arhgap18 | -0.002971421 | 0.923555518 | 0.980869592 | 0.059245243 | 0.059342808 |
| Arhgap21 | -0.073743971 | 0.18061477 | 0.588008578 | 0.2423814 | 0.314999502 |
| Arhgap23 | 0.013447412 | 0.651756637 | 0.896489156 | 0.055314987 | 0.044556513 |
| Arhgap24 | 0.079505513 | 0.180980696 | 0.588627104 | 0.291627881 | 0.212452619 |
| Arhgap26 | -0.026297533 | 0.544854378 | 0.851259637 | 0.120722856 | 0.143382484 |
| Arhgap32 | -0.010313924 | 0.85140509 | 0.957461155 | 0.22443042 | 0.22615819 |
| Arhgap35 | 0.057963207 | 0.29747799 | 0.702656882 | 0.233161582 | 0.176750692 |
| Arhgap44 | 0.00135755 | 0.974488419 | 0.994250968 | 0.119189444 | 0.11796526 |
| Arhgap5 | 0.107158403 | 0.07184361 | 0.40592424 | 1.402650564 | 1.265349715 |
| Arhgap6 | 0.058680673 | 0.158727562 | 0.556789205 | 0.11634128 | 0.063472511 |
| Arhgdia | -0.028129608 | 0.683241807 | 0.909173483 | 0.765778774 | 0.786971206 |
| Arhgdig | -0.081031874 | 0.192384996 | 0.60309438 | 0.381385652 | 0.447477302 |
| Arhgef1 | 0.069628718 | 0.166531038 | 0.568127031 | 0.19513357 | 0.127467589 |
| Arhgef10l | -0.025554332 | 0.326101511 | 0.720866629 | 0.038102263 | 0.060577156 |

| | | | | | |
|---|---|---|---|---|---|
| Arhgef11 | -0.046847956 | 0.231820678 | 0.643291165 | 0.097637209 | 0.133383136 |
| Arhgef12 | 0.07562918 | 0.259259679 | 0.666715193 | 0.927017302 | 0.815994808 |
| Arhgef16 | 0.037301194 | 0.552926816 | 0.852623515 | 0.348887065 | 0.311164704 |
| Arhgef17 | 0.024906781 | 0.659408476 | 0.898227584 | 0.238334884 | 0.199719797 |
| Arhgef18 | 0.002733242 | 0.952463203 | 0.98838165 | 0.138132618 | 0.13210538 |
| Arhgef2 | -0.086564377 | 0.199709148 | 0.610108047 | 0.450682211 | 0.535960878 |
| Arhgef25 | -0.039240643 | 0.173455293 | 0.575630234 | 0.048407015 | 0.080784888 |
| Arhgef28 | 0.064630808 | 0.153102341 | 0.549602359 | 0.147099971 | 0.096597422 |
| Arhgef5 | 0.049379684 | 0.430047334 | 0.793295051 | 0.350164478 | 0.298675784 |
| Arhgef7 | -0.025647099 | 0.559101831 | 0.856748867 | 0.128063042 | 0.152510702 |
| Arhgef9 | -0.015700447 | 0.54976004 | 0.852455047 | 0.039164293 | 0.053395069 |
| Arid1a | -0.013010368 | 0.845458457 | 0.955950886 | 0.580049045 | 0.583451932 |
| Arid1b | 0.083975187 | 0.142718025 | 0.531525349 | 0.275318803 | 0.197544238 |
| Arid2 | -0.040334118 | 0.483867549 | 0.822768706 | 0.273538012 | 0.317428227 |
| Arid3a | -0.051018513 | 0.214982044 | 0.629611185 | 0.104896437 | 0.149747216 |
| Arid3b | -0.036997436 | 0.232463487 | 0.643543798 | 0.058842808 | 0.089077771 |
| Arid4a | 0.046121888 | 0.431209626 | 0.793555374 | 0.274085275 | 0.22184388 |
| Arid4b | 0.004516101 | 0.948415603 | 0.987651726 | 0.579958192 | 0.565992856 |
| Arid5a | 0.059784649 | 0.15574062 | 0.552356396 | 0.117717142 | 0.056654087 |
| Arid5b | 0.059100482 | 0.338389806 | 0.732353334 | 0.26467157 | 0.194196214 |
| Arih1 | 0.00293307 | 0.963525649 | 0.991970325 | 0.424108338 | 0.427054397 |
| Arih2 | 0.013295855 | 0.813937965 | 0.948397417 | 0.244168458 | 0.242632371 |
| Arl1 | -0.019012792 | 0.760166274 | 0.927859527 | 1.14536357 | 1.178380423 |
| Arl13b | 0.031642587 | 0.518478153 | 0.838109801 | 0.171363653 | 0.14321837 |
| Arl14ep | -0.125419323 | 0.01423856 | 0.193060412 | 0.169632607 | 0.281954209 |
| Arl15 | -0.026731071 | 0.324067178 | 0.720406991 | 0.043072831 | 0.066124384 |
| Arl16 | -0.011973281 | 0.753620738 | 0.926726954 | 0.093488907 | 0.109405074 |
| Arl2 | -0.099234802 | 0.12616987 | 0.505736405 | 0.421453522 | 0.515664911 |
| Arl2bp | -0.059872584 | 0.344807589 | 0.737445936 | 0.374224466 | 0.438396438 |
| Arl3 | -0.074204073 | 0.254138503 | 0.663146569 | 0.860422192 | 0.948586794 |
| Arl4a | -0.18675343 | 0.008827796 | 0.146996602 | 0.571482427 | 0.767923709 |
| Arl5a | 0.093487161 | 0.13551677 | 0.518757369 | 0.380208768 | 0.288231216 |
| Arl5b | 0.040778011 | 0.307907425 | 0.710440053 | 0.109917055 | 0.075594774 |
| Arl6 | -0.015961596 | 0.747702996 | 0.924930563 | 0.16925455 | 0.187221651 |
| Arl6ip1 | 0.005091455 | 0.940605475 | 0.986168593 | 0.851474335 | 0.841314418 |
| Arl6ip4 | 0.093612073 | 0.136621869 | 0.520249727 | 0.407833439 | 0.321501768 |
| Arl6ip5 | -0.036675416 | 0.597632615 | 0.874521003 | 0.531021886 | 0.580592477 |
| Arl6ip6 | -0.000240548 | 0.995904872 | 0.998890496 | 0.154500426 | 0.158600528 |
| Arl8a | 0.082176974 | 0.198446324 | 0.609389286 | 0.416633734 | 0.339353569 |
| Arl8b | 0.044702188 | 0.49825186 | 0.830722138 | 0.837781465 | 0.756371117 |
| Armc1 | 0.068003104 | 0.290449517 | 0.696593024 | 0.395749727 | 0.340716153 |
| Armc10 | -0.013699329 | 0.830167641 | 0.95325696 | 0.404268385 | 0.404276478 |
| Armc5 | -0.012933837 | 0.793682353 | 0.939757086 | 0.167341795 | 0.179218317 |
| Armc6 | 0.001907315 | 0.951097954 | 0.988114084 | 0.056782389 | 0.052218023 |
| Armc8 | 0.003242736 | 0.956390844 | 0.989397512 | 0.287350328 | 0.279158139 |
| Armc9 | 0.006956933 | 0.849533153 | 0.957111129 | 0.086936665 | 0.08171049 |
| Armcx1 | -0.00547653 | 0.915249061 | 0.980132949 | 0.17632331 | 0.189852439 |
| Armcx2 | -0.011160945 | 0.838278717 | 0.954911647 | 0.2237742 | 0.228909781 |
| Armcx3 | -0.124125726 | 0.065642841 | 0.389687514 | 0.408733191 | 0.509477482 |
| Armcx4 | 0.035466911 | 0.376804344 | 0.761269164 | 0.111541991 | 0.076546407 |
| Armcx5 | 0.016244644 | 0.754272867 | 0.926726954 | 0.188155167 | 0.186470928 |
| Armcx6 | -0.0533827 | 0.101937173 | 0.467829632 | 0.059506978 | 0.110048532 |
| Armh3 | 0.02253211 | 0.412365791 | 0.781358105 | 0.049359411 | 0.030606608 |
| Armt1 | 0.031712012 | 0.341554694 | 0.734873412 | 0.076770976 | 0.052288459 |
| Arnt | 0.032829098 | 0.600151419 | 0.875927602 | 0.345252431 | 0.306054637 |
| Arnt2 | -0.025391292 | 0.417347018 | 0.784482356 | 0.061212637 | 0.085149144 |
| Arntl | -0.068184574 | 0.026163918 | 0.257205918 | 0.047975008 | 0.106323514 |
| Arpc1a | 0.047965246 | 0.458825672 | 0.809370658 | 0.986130143 | 0.942188653 |
| Arpc1b | -0.026077773 | 0.651628262 | 0.896468516 | 0.259075017 | 0.293879082 |
| Arpc2 | 0.066028208 | 0.255341067 | 0.664253092 | 1.43472698 | 1.337986843 |
| Arpc3 | -0.035268018 | 0.600381196 | 0.876003207 | 0.843404783 | 0.901142656 |
| Arpc4 | 0.014810709 | 0.794975731 | 0.939757086 | 0.25129753 | 0.245699444 |
| Arpc5 | -0.000450557 | 0.994803566 | 0.998704266 | 0.972658799 | 0.963675069 |
| Arpc5l | -0.072503778 | 0.207984582 | 0.620020309 | 1.363786354 | 1.489545344 |
| Arpin | -0.024948148 | 0.561259702 | 0.857134531 | 0.120517413 | 0.142736685 |

| | | | | | |
|---|---|---|---|---|---|
| Arpp19 | 0.004595952 | 0.941360793 | 0.986168593 | 1.092255407 | 1.081794931 |
| Arrb2 | 0.028330304 | 0.538594315 | 0.848357012 | 0.145124665 | 0.120181647 |
| Arrdc1 | -0.040852423 | 0.338413347 | 0.732353334 | 0.119033735 | 0.154156904 |
| Arrdc2 | -0.036357148 | 0.173543128 | 0.575630234 | 0.038575432 | 0.074503096 |
| Arrdc3 | 0.021811621 | 0.60273371 | 0.876632118 | 0.117036654 | 0.110323241 |
| Arrdc4 | 0.043553504 | 0.451048812 | 0.806495742 | 0.246834694 | 0.204621004 |
| Arsa | 0.033788061 | 0.364523835 | 0.752802229 | 0.095956252 | 0.063520583 |
| Arsb | -0.048633568 | 0.381329212 | 0.762430537 | 0.2204281 | 0.256218753 |
| Arxes1 | -0.008687816 | 0.774183292 | 0.933157531 | 0.056024865 | 0.061469293 |
| Arxes2 | -0.193423022 | 0.000151674 | 0.010687095 | 0.167926349 | 0.338801981 |
| As3mt | -0.069252881 | 0.230115682 | 0.641131395 | 0.254638873 | 0.317022889 |
| Asah1 | -0.05944707 | 0.394960841 | 0.771549623 | 0.726020265 | 0.785835099 |
| Asah2 | -0.103874019 | 0.052978963 | 0.359290441 | 0.196657837 | 0.279334092 |
| Asap1 | 0.063460767 | 0.285570739 | 0.692407818 | 0.278063622 | 0.219640842 |
| Asb1 | -0.026656088 | 0.465692131 | 0.812791331 | 0.082566392 | 0.106513997 |
| Asb11 | -0.060765991 | 0.069141361 | 0.399753362 | 0.063610067 | 0.116182299 |
| Asb13 | 0.030482207 | 0.561641497 | 0.857134531 | 0.196641586 | 0.174365171 |
| Asb3 | -0.022016031 | 0.443359499 | 0.801368502 | 0.050707549 | 0.071563764 |
| Asb6 | -0.044293192 | 0.215247816 | 0.629611185 | 0.077070215 | 0.119461162 |
| Asb7 | 0.055849896 | 0.192596092 | 0.60325934 | 0.133777636 | 0.08979437 |
| Asb8 | -0.027945706 | 0.546029329 | 0.851405371 | 0.141409665 | 0.162799273 |
| Ascc1 | 0.007159763 | 0.894382295 | 0.972016422 | 0.215558307 | 0.203170124 |
| Ascc2 | 0.033123439 | 0.476537757 | 0.818027958 | 0.144943683 | 0.11699465 |
| Ascc3 | 0.159565588 | 0.017861061 | 0.212658918 | 0.923518113 | 0.726433994 |
| Asf1a | 0.006660098 | 0.895918995 | 0.97233366 | 0.186090325 | 0.187536066 |
| Ash1l | -0.085024148 | 0.196267588 | 0.606881632 | 1.004197278 | 1.087979568 |
| Ash2l | -0.023646569 | 0.675660216 | 0.906025597 | 0.235651969 | 0.247291458 |
| Asic1 | -0.022714504 | 0.542083381 | 0.850208191 | 0.088726046 | 0.107776358 |
| Asl | -0.007783031 | 0.826583241 | 0.952625559 | 0.083203752 | 0.091204504 |
| Asns | -0.137071206 | 0.038501873 | 0.310147112 | 1.119418112 | 1.291113287 |
| Asnsd1 | -0.014627222 | 0.766094872 | 0.92882091 | 0.176388949 | 0.204391342 |
| Asph | 0.052390097 | 0.260669902 | 0.667945828 | 0.153403886 | 0.105707142 |
| Asphd1 | -0.036728546 | 0.259854228 | 0.66721844 | 0.065866558 | 0.096001061 |
| Aspscr1 | -0.056782996 | 0.23004766 | 0.641131395 | 0.147070598 | 0.200083205 |
| Asrgl1 | -0.09916136 | 0.053711053 | 0.361325633 | 0.176965912 | 0.268535426 |
| Asxl1 | -0.004173654 | 0.909059196 | 0.978154712 | 0.083017912 | 0.087444829 |
| Asxl2 | -0.093998845 | 0.147908789 | 0.539745621 | 0.473090696 | 0.544542944 |
| Atad1 | -0.044051553 | 0.499572107 | 0.830722138 | 0.412695159 | 0.445086542 |
| Atad2 | 0.031465511 | 0.479938805 | 0.819939708 | 0.134336274 | 0.115885386 |
| Atad2b | -0.109127409 | 0.043649783 | 0.329086044 | 0.213015501 | 0.313304651 |
| Atad3a | -0.02530923 | 0.573880261 | 0.862168892 | 0.138326912 | 0.15429439 |
| Atat1 | 0.004238284 | 0.900038572 | 0.973983939 | 0.077725629 | 0.073045816 |
| Ate1 | 0.02750198 | 0.684331981 | 0.909679945 | 0.568384505 | 0.518015332 |
| Atf1 | 0.086080099 | 0.173543598 | 0.575630234 | 0.380318528 | 0.315473092 |
| Atf2 | 0.012202338 | 0.841783459 | 0.955118764 | 0.392953152 | 0.385130944 |
| Atf3 | -0.102788702 | 0.030121104 | 0.275342007 | 0.110690869 | 0.213227364 |
| Atf4 | -0.057873054 | 0.36362028 | 0.751765374 | 1.316170906 | 1.398375139 |
| Atf5 | -0.403140152 | 2.47E-08 | 1.03E-05 | 1.207792877 | 1.820765711 |
| Atf6 | -0.081583144 | 0.228039676 | 0.639451668 | 0.492348528 | 0.572573608 |
| Atf6b | 0.021799592 | 0.581915923 | 0.866173066 | 0.111060151 | 0.088158517 |
| Atf7 | -0.02159863 | 0.629852972 | 0.888022795 | 0.140577866 | 0.163359351 |
| Atf7ip | -0.143079731 | 0.016027927 | 0.203940048 | 0.272943822 | 0.416410589 |
| Atg10 | -0.042179943 | 0.368387932 | 0.755518937 | 0.14694963 | 0.186242629 |
| Atg101 | 0.001704089 | 0.973974802 | 0.994149348 | 0.193560799 | 0.188993473 |
| Atg12 | -0.011665004 | 0.840438077 | 0.954911647 | 0.255728236 | 0.272889574 |
| Atg13 | -0.039759418 | 0.451045867 | 0.806495742 | 0.199310904 | 0.229860229 |
| Atg14 | -0.022160266 | 0.592193198 | 0.872005806 | 0.109561535 | 0.126808501 |
| Atg16l1 | -0.06917273 | 0.165429819 | 0.566504119 | 0.167075383 | 0.23508967 |
| Atg16l2 | 0.026101851 | 0.581153767 | 0.866173066 | 0.155480662 | 0.13677268 |
| Atg2a | -0.068424473 | 0.036510915 | 0.301071077 | 0.064329345 | 0.115546405 |
| Atg2b | 0.056890988 | 0.241051663 | 0.651287862 | 0.16469957 | 0.112898497 |
| Atg3 | -0.058650944 | 0.359062779 | 0.747455833 | 0.453865724 | 0.546866166 |
| Atg4a | -0.008889107 | 0.743155061 | 0.924253661 | 0.042635043 | 0.049097964 |
| Atg4b | -0.053647231 | 0.235017741 | 0.64504573 | 0.134406581 | 0.184458564 |
| Atg4c | -0.002132404 | 0.948179291 | 0.987651726 | 0.066498664 | 0.064760291 |

| | | | | | |
|---|---|---|---|---|---|
| Atg4d | -0.07677144 | 0.145189835 | 0.533628684 | 0.198260975 | 0.264761811 |
| Atg5 | -0.051088425 | 0.392332749 | 0.769704269 | 0.320183048 | 0.385810357 |
| Atg7 | -0.039406548 | 0.208114817 | 0.620020309 | 0.056770214 | 0.090020617 |
| Atic | -0.003631994 | 0.945201763 | 0.987084873 | 0.201768751 | 0.203146634 |
| Atl2 | 0.02179195 | 0.697139838 | 0.910970095 | 0.226791127 | 0.20999239 |
| Atl3 | -0.027580264 | 0.617690598 | 0.883238152 | 0.226696613 | 0.249681277 |
| Atm | -0.084478645 | 0.110409292 | 0.479938529 | 0.180121967 | 0.249784323 |
| Atmin | -0.028572573 | 0.506849819 | 0.833276466 | 0.124663245 | 0.15036651 |
| Atn1 | -0.013968579 | 0.798349535 | 0.940958114 | 0.215133406 | 0.217220916 |
| Atox1 | 0.090667477 | 0.162276997 | 0.561259206 | 1.038082688 | 0.886648141 |
| Atp11a | 0.022597149 | 0.61824753 | 0.883323909 | 0.133179895 | 0.105708833 |
| Atp11b | -0.023996574 | 0.701222591 | 0.912152193 | 0.366547851 | 0.401130958 |
| Atp11c | -0.009324125 | 0.733586986 | 0.921306823 | 0.045038258 | 0.049802766 |
| Atp13a1 | 0.020855308 | 0.691102712 | 0.909796728 | 0.204764347 | 0.184646646 |
| Atp13a2 | 0.027291541 | 0.500574148 | 0.831556227 | 0.11133608 | 0.089162419 |
| Atp13a3 | 0.08547619 | 0.196721825 | 0.60713211 | 0.502618535 | 0.412646768 |
| Atp1a1 | 0.039152994 | 0.455897205 | 0.808152996 | 1.910734116 | 1.808399382 |
| Atp1b1 | 0.18136197 | 0.00046543 | 0.023956493 | 1.842989208 | 1.486115638 |
| Atp1b2 | 0.018749756 | 0.531136576 | 0.845476197 | 0.061067633 | 0.047134075 |
| Atp1b3 | 0.055787915 | 0.377473344 | 0.76154773 | 0.413553083 | 0.364114641 |
| Atp23 | 0.010632124 | 0.717906802 | 0.917797805 | 0.056932892 | 0.051171958 |
| Atp2a2 | 0.02183088 | 0.660236921 | 0.898481506 | 2.4318199 | 2.356327703 |
| Atp2a3 | -0.064897437 | 0.347507255 | 0.739124045 | 0.701295754 | 0.761209559 |
| Atp2b1 | 0.014253057 | 0.820831266 | 0.950892784 | 1.216571946 | 1.1930071 |
| Atp2b2 | -0.030320448 | 0.567281972 | 0.859183514 | 0.195084429 | 0.220221735 |
| Atp2b4 | 0.01581652 | 0.746662338 | 0.924645267 | 0.167552953 | 0.147425166 |
| Atp2c1 | 0.06838762 | 0.310710422 | 0.71348779 | 0.554441637 | 0.464508377 |
| Atp5a1 | -0.05380431 | 0.313766743 | 0.714906079 | 1.585225434 | 1.690275308 |
| Atp5b | -0.034263305 | 0.516260946 | 0.837427561 | 1.563603936 | 1.61682193 |
| Atp5c1 | -0.148317245 | 0.015946111 | 0.20388677 | 1.306870486 | 1.50788912 |
| Atp5d | 0.022809695 | 0.657450879 | 0.898224427 | 1.629154479 | 1.590092678 |
| Atp5e | 0.140057389 | 9.25E-05 | 0.007469648 | 2.57567659 | 2.242595132 |
| Atp5g1 | -0.020927506 | 0.719503946 | 0.918741515 | 1.746876484 | 1.765919832 |
| Atp5g2 | 0.034151765 | 0.535072067 | 0.846460359 | 0.239212823 | 0.215972 |
| Atp5g3 | -0.011420682 | 0.851899832 | 0.957461155 | 1.216130667 | 1.234599194 |
| Atp5h | -0.091338177 | 0.076992115 | 0.418199494 | 1.533391119 | 1.682630099 |
| Atp5j | 0.001028345 | 0.985639416 | 0.996791297 | 1.497115043 | 1.525612342 |
| Atp5j2 | 0.103200695 | 0.033092944 | 0.287968471 | 1.873450852 | 1.669318537 |
| Atp5k | 0.099958489 | 0.102836604 | 0.468979611 | 1.051499976 | 0.87928968 |
| Atp5l | 0.059481201 | 0.315605689 | 0.714975653 | 1.362553056 | 1.22829022 |
| Atp5md | 0.091702285 | 0.056515226 | 0.368100794 | 1.686504776 | 1.521805037 |
| Atp5mpl | 0.022360584 | 0.739884921 | 0.922901128 | 0.871616983 | 0.844015333 |
| Atp5o | -0.040510712 | 0.554825621 | 0.853469818 | 0.676288642 | 0.738654339 |
| Atp5pb | 0.016369095 | 0.771432248 | 0.932251553 | 1.435993782 | 1.398495713 |
| Atp6ap1 | -0.029534131 | 0.654869233 | 0.897454058 | 0.616519397 | 0.664760641 |
| Atp6ap2 | -0.025690089 | 0.660255267 | 0.898481506 | 1.383778214 | 1.442274402 |
| Atp6v0a1 | -0.136845387 | 0.029902875 | 0.274935828 | 0.379184629 | 0.517627048 |
| Atp6v0a2 | 0.054423725 | 0.297550887 | 0.702656882 | 0.199308954 | 0.158203643 |
| Atp6v0b | 0.010162737 | 0.831988142 | 0.953695326 | 1.787111774 | 1.73542239 |
| Atp6v0c | -0.044209375 | 0.353095599 | 0.743880213 | 1.94941952 | 2.060099732 |
| Atp6v0d1 | -0.077284911 | 0.250466673 | 0.662258694 | 0.673748122 | 0.735440234 |
| Atp6v0e | 0.238360427 | 2.72E-06 | 0.000479487 | 1.813317327 | 1.411836072 |
| Atp6v0e2 | 0.028404479 | 0.657811887 | 0.898224427 | 0.420139394 | 0.400848644 |
| Atp6v1a | 0.202563494 | 0.003086196 | 0.077130473 | 0.939406351 | 0.681267491 |
| Atp6v1b2 | 0.168504673 | 0.010929227 | 0.165454977 | 1.092998382 | 0.885217663 |
| Atp6v1c1 | 0.058770674 | 0.372749398 | 0.75825687 | 0.708910328 | 0.650611697 |
| Atp6v1d | 0.056051692 | 0.38708228 | 0.766754459 | 1.024864173 | 0.926730166 |
| Atp6v1e1 | 0.1356369 | 0.014031249 | 0.191923581 | 1.583963375 | 1.350781747 |
| Atp6v1f | -0.006143797 | 0.90820441 | 0.977744305 | 1.442044747 | 1.437861485 |
| Atp6v1g1 | -0.110341654 | 0.070188511 | 0.402558548 | 1.331751753 | 1.512205063 |
| Atp6v1h | -0.09439607 | 0.138933728 | 0.525848983 | 0.381397621 | 0.485313421 |
| Atp7a | 0.081264384 | 0.157351694 | 0.554924229 | 0.266096569 | 0.198699814 |
| Atp8a1 | 0.017754121 | 0.730406618 | 0.920770724 | 0.194760535 | 0.174093721 |
| Atp8a2 | -0.035261893 | 0.198840623 | 0.609389286 | 0.041530247 | 0.075089749 |
| Atp8b1 | 0.037902424 | 0.562845609 | 0.857583554 | 1.003863206 | 0.915958019 |

| | | | | | |
|---|---|---|---|---|---|
| Atp8b2 | -0.080070214 | 0.022626639 | 0.238404929 | 0.075838161 | 0.148094781 |
| Atp9a | -0.002039232 | 0.973440204 | 0.994079882 | 0.332163034 | 0.320304113 |
| Atp9b | -0.051422761 | 0.199742457 | 0.610108047 | 0.100912307 | 0.139356471 |
| Atpaf1 | 0.043681895 | 0.319743198 | 0.717256849 | 0.135762511 | 0.098324663 |
| Atpaf2 | -0.012405488 | 0.746160715 | 0.924566547 | 0.094696568 | 0.106843221 |
| Atpif1 | -0.143475923 | 0.003622454 | 0.084949926 | 1.756691765 | 2.004479313 |
| Atpsckmt | 0.000293663 | 0.994938481 | 0.998719626 | 0.152532321 | 0.147835839 |
| Atr | -0.014811568 | 0.703953988 | 0.912277105 | 0.097137421 | 0.116358302 |
| Atraid | -0.372257135 | 2.76E-08 | 1.11E-05 | 0.642595084 | 1.085369026 |
| Atrip | 0.008126148 | 0.853096186 | 0.957461155 | 0.12482023 | 0.116044694 |
| Atrn | 0.042165071 | 0.353686791 | 0.744256656 | 0.146818674 | 0.107408375 |
| Atrnl1 | 0.039770275 | 0.243599463 | 0.653570652 | 0.077641257 | 0.042453561 |
| Atrx | -0.125243661 | 0.073457285 | 0.410998661 | 1.085345803 | 1.238958298 |
| Atxn1 | -0.001824477 | 0.974239443 | 0.994170647 | 0.21997368 | 0.222357524 |
| Atxn10 | -0.123162499 | 0.061895365 | 0.380250374 | 0.54071258 | 0.685306967 |
| Atxn1l | 0.005535107 | 0.889685228 | 0.970458414 | 0.108091406 | 0.10273087 |
| Atxn2 | 0.074510472 | 0.237303697 | 0.648466434 | 0.379632711 | 0.316623682 |
| Atxn2l | -0.042312471 | 0.504747567 | 0.832249514 | 0.414474551 | 0.441490481 |
| Atxn3 | 0.052883932 | 0.170230484 | 0.572762895 | 0.103571605 | 0.055675268 |
| Atxn7 | 0.050517488 | 0.350486776 | 0.740908966 | 0.20362677 | 0.165768702 |
| Atxn7l1 | -0.002302996 | 0.930927994 | 0.982825736 | 0.043221141 | 0.046126623 |
| Atxn7l3 | -0.013520783 | 0.788814879 | 0.938142932 | 0.179711195 | 0.204193893 |
| Atxn7l3b | 0.050482568 | 0.396028457 | 0.772781583 | 1.207929108 | 1.142151893 |
| AU022252 | -0.01218704 | 0.691264283 | 0.909796728 | 0.059542675 | 0.06499569 |
| AU040320 | -0.086253653 | 0.131622148 | 0.512050022 | 0.235461622 | 0.304004084 |
| AU041133 | 0.052287914 | 0.193837312 | 0.60325934 | 0.113999252 | 0.071333901 |
| Auh | 0.137945849 | 0.04549703 | 0.334195605 | 0.650915205 | 0.501718482 |
| Aup1 | -0.032492924 | 0.584172394 | 0.8671012 | 0.302343327 | 0.303291508 |
| Aurkaip1 | -0.034353103 | 0.619283665 | 0.883797831 | 0.793348212 | 0.830153409 |
| Auts2 | 0.120016048 | 0.086117069 | 0.436930819 | 0.715506955 | 0.54158761 |
| AV356131 | 0.02667723 | 0.588318702 | 0.86992264 | 0.182239339 | 0.163791189 |
| Aven | -0.00636385 | 0.848968018 | 0.957111129 | 0.071300476 | 0.078905822 |
| Avl9 | -0.045051474 | 0.487831768 | 0.826112275 | 0.455170787 | 0.504019922 |
| Avpi1 | -0.185905047 | 0.003703305 | 0.085962309 | 0.320242739 | 0.509768668 |
| AW146154 | -0.022753639 | 0.391967371 | 0.769704269 | 0.041591868 | 0.063171568 |
| AW209491 | 0.041163123 | 0.353998334 | 0.744256656 | 0.137282868 | 0.102668977 |
| AW549877 | -0.049362292 | 0.410045253 | 0.780101937 | 0.292504035 | 0.326176138 |
| AW554918 | -0.01634044 | 0.559983817 | 0.856904336 | 0.046496238 | 0.062650088 |
| Axin1 | 0.048776309 | 0.389599229 | 0.768258286 | 0.259122 | 0.222748609 |
| Axin2 | -0.078325786 | 0.000464067 | 0.023956493 | 0.017899551 | 0.086629321 |
| Azi2 | 0.047076536 | 0.470201857 | 0.814364464 | 0.538548687 | 0.49106044 |
| Azin1 | -0.040679481 | 0.539540086 | 0.84899357 | 0.432354635 | 0.466594162 |
| B230112I24Rik | -0.047345577 | 0.100305016 | 0.465319642 | 0.045699452 | 0.085378108 |
| B230216N24Rik | -0.04233174 | 0.218122444 | 0.632396893 | 0.06511964 | 0.101872106 |
| B230217C12Rik | 0.01301932 | 0.784772724 | 0.936423418 | 0.15111272 | 0.138142928 |
| B230219D22Rik | 0.223242846 | 0.00097641 | 0.037405445 | 0.832193471 | 0.583071505 |
| B230307C23Rik | -0.013120936 | 0.660853942 | 0.89868852 | 0.054177508 | 0.061542155 |
| B230322F03Rik | -0.040747914 | 0.213014691 | 0.627539904 | 0.061280341 | 0.093413283 |
| B230354K17Rik | 0.000881929 | 0.980827218 | 0.995540893 | 0.088640287 | 0.089933259 |
| B230362B09Rik | -0.079471384 | 0.000505187 | 0.025226667 | 0.018985689 | 0.089954437 |
| B230369F24Rik | -0.012617809 | 0.644851652 | 0.893466123 | 0.042260807 | 0.052685634 |
| B230377A18Rik | -0.037526169 | 0.168551735 | 0.57227323 | 0.041604101 | 0.072903262 |
| B2m | -0.16813829 | 0.011300636 | 0.168327452 | 0.332969748 | 0.486709774 |
| B330016D10Rik | 0.010936002 | 0.801307536 | 0.942433061 | 0.128053634 | 0.118333535 |
| B3galnt1 | 0.066689058 | 0.168869419 | 0.57227323 | 0.169766519 | 0.110564734 |
| B3galnt2 | -0.08921643 | 0.01375067 | 0.189915708 | 0.0770724 | 0.148771728 |
| B3galt2 | -0.031548645 | 0.40530882 | 0.777010853 | 0.089571657 | 0.11223589 |
| B3galt4 | 0.032457509 | 0.326180164 | 0.720866629 | 0.07214431 | 0.045588064 |
| B3galt5 | 0.028113876 | 0.442052958 | 0.800580622 | 0.089427676 | 0.058958661 |
| B3galt6 | -0.011464554 | 0.769082833 | 0.931268476 | 0.101591521 | 0.115508741 |
| B3gat3 | -0.219462339 | 0.000195548 | 0.012828237 | 0.266807662 | 0.47317159 |
| B3gnt3 | 0.015519806 | 0.649761359 | 0.895585658 | 0.078361752 | 0.064743162 |
| B4galt1 | 0.002763003 | 0.927043407 | 0.981818446 | 0.057953064 | 0.055975666 |
| B4galt3 | -0.101174919 | 0.092975943 | 0.451693871 | 0.310008733 | 0.410042638 |
| B4galt5 | -0.000968395 | 0.980813214 | 0.995540893 | 0.103560148 | 0.102430987 |

| | | | | | |
|---|---|---|---|---|---|
| B4galt6 | -0.055386467 | 0.397399569 | 0.774053847 | 0.451283412 | 0.498054731 |
| B4galt7 | -0.061793099 | 0.075345635 | 0.414153416 | 0.074743669 | 0.130646753 |
| B630019K06Rik | -0.046198992 | 0.172665456 | 0.575294174 | 0.06935424 | 0.110901264 |
| B830012L14Rik | -0.049622792 | 0.30560346 | 0.709411642 | 0.147090155 | 0.173230047 |
| B830017H08Rik | -0.143595347 | 0.021368564 | 0.232117183 | 1.326529077 | 1.577378854 |
| B930095G15Rik | -0.002496879 | 0.954741997 | 0.989066902 | 0.124602614 | 0.127515758 |
| B9d1 | -0.085305332 | 0.112834407 | 0.483221516 | 0.218564879 | 0.287563411 |
| B9d2 | -0.024350357 | 0.608378998 | 0.878855197 | 0.153234564 | 0.171863241 |
| Babam1 | -0.069173676 | 0.212577303 | 0.626803288 | 0.231218341 | 0.293882675 |
| Babam2 | -0.085208861 | 0.146428971 | 0.536193937 | 0.276586169 | 0.377720512 |
| Bace1 | 0.017665445 | 0.723832881 | 0.919830015 | 0.180537338 | 0.160837205 |
| Bace2 | -0.252689906 | 1.21E-05 | 0.001459181 | 1.433759511 | 1.905551717 |
| Bach1 | -0.034937179 | 0.582032068 | 0.866173066 | 0.33691923 | 0.361557317 |
| Bad | -0.033113893 | 0.585690857 | 0.868327789 | 0.303346514 | 0.329255893 |
| Bag1 | -0.089999271 | 0.187455042 | 0.594820883 | 0.835834143 | 0.979117725 |
| Bag2 | -0.076953555 | 0.165430265 | 0.566504119 | 0.205486403 | 0.286361051 |
| Bag3 | 0.028739868 | 0.609193739 | 0.878855197 | 0.221817602 | 0.185664676 |
| Bag4 | 0.051752847 | 0.372895876 | 0.75825687 | 0.28948526 | 0.251466693 |
| Bag5 | -0.002594366 | 0.966108527 | 0.992409302 | 0.34927545 | 0.351182897 |
| Bag6 | 0.106820187 | 0.073015648 | 0.410160881 | 0.331008226 | 0.237429325 |
| Bahd1 | 0.034518479 | 0.371551874 | 0.75760494 | 0.100295251 | 0.069712225 |
| Baiap2 | 0.148685307 | 0.003089215 | 0.077130473 | 0.182479469 | 0.057711061 |
| Baiap2l1 | 0.090310029 | 0.112909167 | 0.483221516 | 0.263061272 | 0.178903002 |
| Baiap3 | -0.007668318 | 0.834860124 | 0.954911647 | 0.086567125 | 0.092343133 |
| Bak1 | -0.042740555 | 0.154250164 | 0.550976877 | 0.049535491 | 0.084483148 |
| Bambi | 0.066543363 | 0.324517862 | 0.720456931 | 0.354366536 | 0.291925354 |
| Banf1 | 0.057420396 | 0.373105238 | 0.758519786 | 0.444940917 | 0.388281129 |
| Banp | -0.026264911 | 0.673731896 | 0.905654932 | 0.372237508 | 0.398123368 |
| Bap1 | 0.008645942 | 0.849493088 | 0.957111129 | 0.148194989 | 0.146984161 |
| Barhl1 | -0.06398394 | 0.044811603 | 0.331447358 | 0.050855764 | 0.118197225 |
| Bax | -0.101849414 | 0.144782815 | 0.533279057 | 0.799794404 | 0.905068882 |
| Baz1a | 0.059836161 | 0.195655752 | 0.605923103 | 0.145495719 | 0.099438151 |
| Baz1b | 0.073731411 | 0.306501854 | 0.709680069 | 0.67029606 | 0.579018295 |
| Baz2a | 0.103743562 | 0.106656157 | 0.474004524 | 0.472567911 | 0.376325888 |
| Baz2b | 0.066456313 | 0.33707423 | 0.731980538 | 0.860085003 | 0.763330711 |
| BB218582 | -0.036633662 | 0.212086926 | 0.625908991 | 0.050383835 | 0.079648516 |
| Bbc3 | -0.049996582 | 0.254186556 | 0.663146569 | 0.113276229 | 0.161725605 |
| Bbip1 | -0.049772707 | 0.454442253 | 0.807876964 | 0.541610469 | 0.586478648 |
| Bbof1 | -0.01916522 | 0.450542172 | 0.806495742 | 0.036547015 | 0.053260284 |
| Bbs1 | 0.018125977 | 0.517735383 | 0.837743618 | 0.052196864 | 0.035136237 |
| Bbs10 | -0.012879964 | 0.608299752 | 0.878855197 | 0.036148476 | 0.048093692 |
| Bbs2 | -0.076468176 | 0.020410846 | 0.227815517 | 0.062066389 | 0.128150759 |
| Bbs4 | -0.038224364 | 0.374922906 | 0.760042036 | 0.123842937 | 0.150651665 |
| Bbs5 | -0.006584645 | 0.839369653 | 0.954911647 | 0.068522474 | 0.068722871 |
| Bbs7 | 0.010790879 | 0.76882323 | 0.931268476 | 0.089850542 | 0.075958677 |
| Bbs9 | -0.03153174 | 0.243346144 | 0.653166476 | 0.040640737 | 0.069686294 |
| Bbx | 0.041688761 | 0.560284935 | 0.856904336 | 0.764514253 | 0.743820179 |
| BC002059 | -0.011095327 | 0.658313089 | 0.898224427 | 0.034881032 | 0.047874875 |
| BC003965 | -0.006692758 | 0.904768886 | 0.975652137 | 0.244556927 | 0.262259642 |
| BC004004 | 0.05252424 | 0.427019521 | 0.79105114 | 0.473073521 | 0.411669387 |
| BC005537 | -0.087853327 | 0.205289664 | 0.616549498 | 0.642562222 | 0.757629465 |
| BC005624 | -0.17313918 | 0.007100314 | 0.128637347 | 0.479681764 | 0.663224853 |
| BC024063 | -0.045493119 | 0.01448501 | 0.193677739 | 0.011916506 | 0.053303064 |
| BC029722 | 0.134128496 | 0.015063928 | 0.197901365 | 0.258757136 | 0.136711582 |
| BC031181 | -0.018461187 | 0.794580121 | 0.939757086 | 0.859667951 | 0.880990472 |
| BC037032 | -0.050232753 | 0.003508848 | 0.08334591 | 0.00783683 | 0.050413082 |
| BC037704 | -0.016397255 | 0.590357123 | 0.870796216 | 0.054052934 | 0.070201131 |
| BC051142 | -0.009541524 | 0.750394579 | 0.92612569 | 0.054743635 | 0.06068918 |
| BC051226 | 0.019738223 | 0.637650878 | 0.891210491 | 0.118295578 | 0.09599978 |
| Bcap29 | -0.055813767 | 0.234162597 | 0.644459522 | 0.143742368 | 0.198948467 |
| Bcap31 | -0.032853578 | 0.610717502 | 0.879829419 | 0.432699238 | 0.448763496 |
| Bcar1 | 0.016753447 | 0.765683258 | 0.92882091 | 0.235125304 | 0.218329862 |
| Bcas2 | 0.103474097 | 0.106309627 | 0.473814268 | 1.064589957 | 0.945178637 |
| Bcas3 | -0.024722457 | 0.325621132 | 0.720833548 | 0.035491925 | 0.055265563 |
| Bcat2 | -0.028774739 | 0.344592201 | 0.737445936 | 0.054947372 | 0.082552655 |

| | | | | | |
|---|---|---|---|---|---|
| Bccip | -0.06825867 | 0.319818683 | 0.717256849 | 0.60883978 | 0.678150879 |
| Bcdin3d | 0.094219685 | 0.024508284 | 0.248116801 | 0.126485694 | 0.044349607 |
| Bckdha | -0.077466296 | 0.04700234 | 0.339397474 | 0.091539972 | 0.165020109 |
| Bckdhb | -0.007699522 | 0.817073132 | 0.949185535 | 0.07215008 | 0.078944055 |
| Bckdk | -0.004328339 | 0.944572279 | 0.987070339 | 0.37661989 | 0.371078597 |
| Bcl10 | -0.044867986 | 0.389866073 | 0.768386831 | 0.195616205 | 0.236045767 |
| Bcl2l1 | 0.009356285 | 0.864872448 | 0.961442912 | 0.221765916 | 0.208349236 |
| Bcl2l11 | 0.048137452 | 0.22600109 | 0.638587289 | 0.105097701 | 0.065722432 |
| Bcl2l13 | 0.002998507 | 0.949122233 | 0.987767085 | 0.15402994 | 0.152128864 |
| Bcl2l2 | 0.011843386 | 0.817759311 | 0.949543059 | 0.195633624 | 0.175492661 |
| Bcl3 | -0.029713471 | 0.545541807 | 0.851259637 | 0.157490155 | 0.190037064 |
| Bcl6b | 0.002930406 | 0.92483154 | 0.981375217 | 0.060887886 | 0.057516786 |
| Bcl7a | 0.094031014 | 0.079748199 | 0.423285392 | 0.229860509 | 0.146451088 |
| Bcl7b | -0.089126938 | 0.177621825 | 0.583182944 | 0.479783215 | 0.571126608 |
| Bcl7c | 0.01834522 | 0.748873511 | 0.925186829 | 0.267726931 | 0.255171918 |
| Bcl9 | 0.10741666 | 0.087740011 | 0.440323248 | 0.365294488 | 0.251535948 |
| Bcl9l | 0.006288604 | 0.863093352 | 0.960676137 | 0.086334427 | 0.078677456 |
| Bclaf1 | -0.195332416 | 0.005110232 | 0.105973959 | 0.674827796 | 0.908635183 |
| Bclaf3 | -0.04615076 | 0.261510126 | 0.668468258 | 0.109237741 | 0.159518318 |
| Bcor | -0.005493214 | 0.9335959 | 0.983869133 | 0.472161706 | 0.494173504 |
| Bcorl1 | -0.015403841 | 0.644515986 | 0.893466123 | 0.070036975 | 0.083316871 |
| Bcr | 0.028542972 | 0.662539257 | 0.899122029 | 0.378468969 | 0.349081816 |
| Bcs1l | -0.004648092 | 0.90722602 | 0.977187883 | 0.106061093 | 0.108483574 |
| Bdh1 | 0.07591589 | 0.042836052 | 0.327353972 | 0.102210489 | 0.040171784 |
| Bdp1 | -0.019485658 | 0.759130213 | 0.927719223 | 0.366260829 | 0.392051031 |
| Becn1 | -0.027568711 | 0.675023506 | 0.906025597 | 0.470284515 | 0.493023693 |
| Bend7 | -0.060547884 | 0.315242125 | 0.714975653 | 0.288522984 | 0.357277731 |
| Bet1 | -0.128416893 | 0.04462812 | 0.330723271 | 0.402111255 | 0.522048085 |
| Bet1l | -0.084528662 | 0.183657159 | 0.590066231 | 0.368431825 | 0.467691204 |
| Bex1 | -0.21019682 | 0.00211571 | 0.063200525 | 0.714914822 | 0.955454334 |
| Bex2 | -0.004468686 | 0.931890776 | 0.98340088 | 1.625597246 | 1.638769887 |
| Bex3 | -0.136392527 | 0.003243898 | 0.079642162 | 2.168877838 | 2.454179509 |
| Bex4 | -0.32237566 | 2.95E-06 | 0.000502094 | 0.637652442 | 1.0321389 |
| Bfar | -0.007620244 | 0.897264731 | 0.972761535 | 0.253312895 | 0.250995464 |
| Bhlha15 | -0.093407716 | 0.001891739 | 0.059335578 | 0.046679342 | 0.130211356 |
| Bhlhb9 | -0.125049846 | 0.023863106 | 0.245655381 | 0.229487316 | 0.335168703 |
| Bhlhe40 | -0.054812675 | 0.270094583 | 0.677226912 | 0.163794124 | 0.204730726 |
| Bhlhe41 | 0.096010308 | 0.183933561 | 0.590326848 | 0.57799686 | 0.477503306 |
| Bicc1 | -0.11277738 | 0.022142285 | 0.236913658 | 0.151902721 | 0.262054663 |
| Bicd1 | -0.038395275 | 0.575206435 | 0.862723699 | 0.477975729 | 0.509514597 |
| Bicd2 | -0.012220244 | 0.756499889 | 0.926833377 | 0.10122133 | 0.106747665 |
| Bicdl1 | -0.037352735 | 0.297386554 | 0.702656882 | 0.080889617 | 0.111627322 |
| Bicra | -0.006034822 | 0.816510219 | 0.949036482 | 0.039642377 | 0.044058655 |
| Bicral | 0.013356149 | 0.779151276 | 0.934664917 | 0.152499799 | 0.140792544 |
| Bin1 | -0.025363746 | 0.604630027 | 0.877736706 | 0.165209508 | 0.188478683 |
| Bin3 | 0.039342219 | 0.255820413 | 0.664689244 | 0.086377072 | 0.05329259 |
| Birc2 | -0.020228786 | 0.725846464 | 0.92014727 | 0.266484253 | 0.284218468 |
| Birc3 | -0.020842537 | 0.542261128 | 0.850208191 | 0.069719136 | 0.094314922 |
| Birc6 | -0.110532315 | 0.097981314 | 0.461275068 | 0.676442826 | 0.788837163 |
| Bivm | -0.019019073 | 0.529037471 | 0.844323583 | 0.058321878 | 0.077707996 |
| Blcap | -0.004574426 | 0.935690887 | 0.984289378 | 0.268505538 | 0.263837055 |
| Blmh | 0.011480448 | 0.853390584 | 0.957461155 | 0.376800145 | 0.368991243 |
| Bloc1s1 | 0.005741792 | 0.873669884 | 0.964940674 | 0.08167979 | 0.076133223 |
| Bloc1s2 | -0.042775752 | 0.461591504 | 0.810421416 | 0.262099794 | 0.308477932 |
| Bloc1s3 | -0.044347124 | 0.051134607 | 0.355933715 | 0.025078184 | 0.061489148 |
| Bloc1s4 | 0.049674744 | 0.272310503 | 0.680042969 | 0.143060302 | 0.100450324 |
| Bloc1s5 | -0.042925774 | 0.170740016 | 0.572958054 | 0.059018315 | 0.09365367 |
| Bloc1s6 | 0.015209153 | 0.763453563 | 0.928730554 | 0.186407525 | 0.177671859 |
| Blvra | 0.058364558 | 0.140207102 | 0.528448624 | 0.109330068 | 0.059141111 |
| Blvrb | -0.155106025 | 0.000772518 | 0.032691088 | 0.125860627 | 0.269982529 |
| Blzf1 | -0.073500745 | 0.095652263 | 0.455869783 | 0.11798892 | 0.183335421 |
| Bmi1 | -0.035431571 | 0.557866936 | 0.855384435 | 0.342094334 | 0.38649036 |
| Bmp1 | -0.06501195 | 0.21875589 | 0.632622127 | 0.196447762 | 0.243075467 |
| Bmp2k | 0.013540907 | 0.665899085 | 0.901122977 | 0.061836799 | 0.05008741 |
| Bmpr1a | 0.029732831 | 0.633526304 | 0.889501117 | 0.406315758 | 0.379390995 |

| | | | | | |
|---|---|---|---|---|---|
| Bmpr2 | 0.005825713 | 0.908798876 | 0.978084314 | 0.1870717 | 0.176349174 |
| Bms1 | 0.045950176 | 0.380174866 | 0.762006446 | 0.198821001 | 0.154034799 |
| Bmt2 | -0.047719551 | 0.349365519 | 0.740480699 | 0.176986734 | 0.215497988 |
| Bmyc | 0.079099888 | 0.220107348 | 0.633234352 | 0.457096343 | 0.388190824 |
| Bnip1 | 0.013485582 | 0.741495484 | 0.923029911 | 0.112624395 | 0.103172757 |
| Bnip2 | -0.09128611 | 0.037958121 | 0.307458537 | 0.115325668 | 0.213574571 |
| Bnip3 | 0.011073886 | 0.869918335 | 0.964130426 | 0.573716838 | 0.541299362 |
| Bnip3l | 0.00083182 | 0.988819526 | 0.997411191 | 0.305983231 | 0.317834676 |
| Bod1 | 0.126359907 | 0.057719249 | 0.370772606 | 0.586526816 | 0.467175755 |
| Bod1l | -0.106859118 | 0.124503706 | 0.502565678 | 0.635042735 | 0.73244394 |
| Bola1 | 0.0565549 | 0.415600807 | 0.783472369 | 0.580891328 | 0.520263473 |
| Bola2 | 0.078585545 | 0.248303416 | 0.659783035 | 0.798373478 | 0.679673925 |
| Bola3 | 0.017586115 | 0.799398469 | 0.941260986 | 0.668356011 | 0.641586643 |
| Bop1 | -0.041926352 | 0.269321158 | 0.676978829 | 0.09074833 | 0.127081365 |
| Borcs5 | -0.016362875 | 0.612612266 | 0.880789186 | 0.067092059 | 0.081465462 |
| Borcs6 | 0.048341073 | 0.361901724 | 0.750032543 | 0.203771419 | 0.16291952 |
| Borcs7 | -0.049186771 | 0.391239833 | 0.769704269 | 0.263329013 | 0.303613524 |
| Borcs8 | -0.134629468 | 0.002698017 | 0.071134541 | 0.127764777 | 0.238858128 |
| Bpgm | 0.034705228 | 0.482644501 | 0.822183105 | 0.172663883 | 0.139951177 |
| Bphl | -0.098981134 | 0.065155217 | 0.389263637 | 0.198707889 | 0.294698785 |
| Bpnt1 | -0.139892298 | 0.015865728 | 0.203377154 | 0.239204151 | 0.385480226 |
| Bpnt2 | -0.025540629 | 0.701746319 | 0.912152193 | 0.60123184 | 0.630287264 |
| Bptf | -0.031337255 | 0.656385965 | 0.89805697 | 0.90644812 | 0.934326882 |
| Braf | 0.008392625 | 0.899958649 | 0.973983939 | 0.695813116 | 0.678034291 |
| Brap | 0.042577364 | 0.420881998 | 0.787234019 | 0.216486105 | 0.189891783 |
| Brcc3 | -0.019088771 | 0.724504696 | 0.919830015 | 0.201217024 | 0.212786948 |
| Brd1 | -0.096626646 | 0.166441542 | 0.568127031 | 0.582431582 | 0.70330649 |
| Brd2 | -0.119757922 | 0.082087195 | 0.427831307 | 0.828009732 | 0.9725641 |
| Brd3 | 0.030031931 | 0.643167676 | 0.892561795 | 0.493723743 | 0.450583029 |
| Brd3os | -0.00210305 | 0.953103639 | 0.988662797 | 0.084776844 | 0.084638469 |
| Brd4 | -0.049734776 | 0.454996694 | 0.807876964 | 0.553021725 | 0.604102913 |
| Brd7 | -0.00621839 | 0.916412317 | 0.980132949 | 0.319581669 | 0.315883619 |
| Brd8 | 0.013071092 | 0.845301579 | 0.955950886 | 0.539412035 | 0.532197222 |
| Brd9 | 0.04626856 | 0.342304374 | 0.735068249 | 0.173359305 | 0.122886026 |
| Brdt | -0.021553214 | 0.423003216 | 0.78859241 | 0.041368648 | 0.057408154 |
| Brf1 | 0.018574452 | 0.63349687 | 0.889501117 | 0.102587371 | 0.091143929 |
| Brf2 | 0.010162757 | 0.776190908 | 0.933751738 | 0.084712695 | 0.078839911 |
| Bri3 | -0.026557887 | 0.690837342 | 0.909796728 | 1.104281086 | 1.142794259 |
| Bri3bp | 0.037073021 | 0.398507112 | 0.774321615 | 0.129869915 | 0.090106882 |
| Brip1os | -0.000877733 | 0.976704103 | 0.994468977 | 0.056289285 | 0.053759971 |
| Brix1 | -0.110069426 | 0.090891676 | 0.445385778 | 0.325055152 | 0.433861253 |
| Brk1 | -0.057920146 | 0.362149224 | 0.750073837 | 0.962261973 | 1.034963916 |
| Brms1l | -0.029179497 | 0.38010612 | 0.762006446 | 0.066516391 | 0.092395427 |
| Brox | 0.037145345 | 0.479783359 | 0.819939708 | 0.210202911 | 0.174413402 |
| Brpf1 | 0.013021818 | 0.754234803 | 0.926726954 | 0.113997955 | 0.10054553 |
| Brpf3 | -0.023386837 | 0.511426515 | 0.835198424 | 0.079091805 | 0.093394308 |
| Brsk1 | -0.009339869 | 0.790858855 | 0.938824685 | 0.078059057 | 0.087581166 |
| Brsk2 | 0.0642504 | 0.156622986 | 0.554127149 | 0.143766831 | 0.089742267 |
| Brwd1 | -0.048486762 | 0.455965581 | 0.808152996 | 0.361107857 | 0.391543598 |
| Brwd3 | -0.044406068 | 0.294059496 | 0.69977963 | 0.110560964 | 0.153971644 |
| Bscl2 | -0.038007459 | 0.463284703 | 0.811254677 | 0.199282277 | 0.225376656 |
| Bsdc1 | 0.049919393 | 0.384848061 | 0.764895046 | 0.259826181 | 0.214242038 |
| Bsg | -0.072583437 | 0.144560316 | 0.533279057 | 1.734594085 | 1.885895665 |
| Bsn | 0.063237565 | 0.137171608 | 0.521907288 | 0.122278161 | 0.056542112 |
| Bspry | -0.010756101 | 0.762724108 | 0.92872278 | 0.08210181 | 0.099078821 |
| Bst2 | 0.104527727 | 0.061415438 | 0.380250374 | 0.215827033 | 0.097732251 |
| Btaf1 | 0.006313654 | 0.902318115 | 0.974696557 | 0.180061271 | 0.183139261 |
| Btbd1 | 0.123931675 | 0.041351434 | 0.320745244 | 0.329113765 | 0.218861053 |
| Btbd10 | -0.022897281 | 0.625001548 | 0.88668351 | 0.143261596 | 0.167164371 |
| Btbd17 | -0.005088686 | 0.929241178 | 0.982502239 | 0.24614468 | 0.257499142 |
| Btbd2 | -0.122843125 | 0.008295697 | 0.141359743 | 0.136244921 | 0.247429766 |
| Btbd3 | 0.009079278 | 0.837895989 | 0.954911647 | 0.131721437 | 0.120812037 |
| Btbd6 | -0.016880569 | 0.615137939 | 0.882439418 | 0.070727037 | 0.091146719 |
| Btbd7 | 0.024022928 | 0.700376169 | 0.911997354 | 0.420907943 | 0.402411869 |
| Btbd8 | 0.076343989 | 0.183387935 | 0.590066231 | 0.253190568 | 0.176646768 |

| | | | | | |
|---|---|---|---|---|---|
| Btbd9 | 0.033010719 | 0.557444941 | 0.855384435 | 0.254141764 | 0.227975961 |
| Btd | -0.022003989 | 0.37215772 | 0.757676884 | 0.033564909 | 0.050157052 |
| Btf3 | -0.134186695 | 0.00574972 | 0.11293531 | 1.726431486 | 1.994178836 |
| Btf3l4 | -0.022798554 | 0.649205762 | 0.895557756 | 0.181778929 | 0.202055726 |
| Btf3-ps18 | -0.046468691 | 0.130120667 | 0.509514066 | 0.056363006 | 0.0974696 |
| Btg1 | -0.000617386 | 0.993256363 | 0.998695538 | 0.846183654 | 0.854735467 |
| Btg2 | -0.470058186 | 9.88E-10 | 7.08E-07 | 1.09975278 | 1.813627024 |
| Btg3 | 0.140519921 | 0.048088798 | 0.343780101 | 0.994665456 | 0.830405204 |
| Btrc | 0.005623677 | 0.922425887 | 0.980859278 | 0.262559218 | 0.268512648 |
| Bub3 | -0.087907124 | 0.138781137 | 0.525640103 | 0.30201878 | 0.376695522 |
| Bud13 | 0.008812708 | 0.798667749 | 0.940982298 | 0.078014746 | 0.069458254 |
| Bud23 | -0.154806013 | 0.000716142 | 0.031383916 | 0.120964403 | 0.261018677 |
| Bud31 | 0.057093842 | 0.400955124 | 0.774740795 | 0.523246402 | 0.472324163 |
| Bysl | -0.039940282 | 0.308750233 | 0.711182121 | 0.092737367 | 0.133928705 |
| Bzw1 | -0.022176141 | 0.754294542 | 0.926726954 | 0.69193589 | 0.733500659 |
| Bzw2 | -0.054206379 | 0.31937728 | 0.717256849 | 0.205644277 | 0.264650914 |
| C030006K11Rik | 0.049280506 | 0.282292627 | 0.689384696 | 0.139262432 | 0.092386795 |
| C030014I23Rik | -0.001464829 | 0.970016951 | 0.993892166 | 0.102828516 | 0.099341414 |
| C030015E24Rik | -0.032632663 | 0.118363017 | 0.491192747 | 0.019734235 | 0.047428701 |
| C030034I22Rik | 0.00031716 | 0.993723864 | 0.998695538 | 0.10491176 | 0.104452036 |
| C130036L24Rik | 0.023611971 | 0.396827305 | 0.773576286 | 0.050726761 | 0.032330434 |
| C130051F05Rik | -0.035638455 | 0.149139222 | 0.542986195 | 0.029728242 | 0.061935722 |
| C130089K02Rik | -0.136723063 | 0.006907097 | 0.126971675 | 0.165564904 | 0.293144135 |
| C1d | 0.057565808 | 0.365633556 | 0.752886169 | 0.419076735 | 0.359633366 |
| C1galt1 | -0.016452829 | 0.809799362 | 0.947866612 | 0.626943154 | 0.634177104 |
| C1galt1c1 | 0.059983052 | 0.243167975 | 0.653166476 | 0.198557269 | 0.148570785 |
| C1qa | 0.226346151 | 0.000230365 | 0.014633967 | 0.31135482 | 0.103411367 |
| C1qbp | 0.06542474 | 0.315584709 | 0.714975653 | 0.406010019 | 0.346535926 |
| C1qtnf12 | -0.007337509 | 0.834296442 | 0.954794181 | 0.081688819 | 0.08394981 |
| C1qtnf4 | -0.111540834 | 0.102221489 | 0.468127803 | 0.960938757 | 1.123715832 |
| C230037L18Rik | -0.05453503 | 0.002175941 | 0.064192029 | 0.00790314 | 0.057110257 |
| C230096K16Rik | 0.032195413 | 0.311505205 | 0.71405691 | 0.063900117 | 0.03946798 |
| C2cd2l | -0.058392115 | 0.210515566 | 0.623103727 | 0.139784641 | 0.202516713 |
| C2cd3 | -0.064327564 | 0.116506695 | 0.488986751 | 0.104705519 | 0.164951627 |
| C2cd4a | 0.021289969 | 0.599496463 | 0.875897759 | 0.097746689 | 0.082953328 |
| C2cd4b | -0.166550808 | 0.031323645 | 0.281212368 | 1.086461572 | 1.305305368 |
| C2cd4c | -0.011008265 | 0.716508748 | 0.916579082 | 0.055569242 | 0.06096943 |
| C2cd5 | -0.074341327 | 0.064469505 | 0.387009825 | 0.100762407 | 0.170546648 |
| C330007P06Rik | -0.081163344 | 0.112945868 | 0.483221516 | 0.184183482 | 0.259562045 |
| C330013E15Rik | -0.015819344 | 0.620614141 | 0.883937014 | 0.064475482 | 0.081491948 |
| C330018D20Rik | 0.001915762 | 0.947731068 | 0.987651726 | 0.053657451 | 0.048444335 |
| C630043F03Rik | -0.013751797 | 0.601750753 | 0.876557788 | 0.038016533 | 0.051100478 |
| C77080 | -0.011194967 | 0.828709254 | 0.952874813 | 0.184417701 | 0.182821884 |
| C87436 | -0.050775951 | 0.296244429 | 0.702600505 | 0.163951422 | 0.214186979 |
| C8b | -0.041452679 | 0.263661496 | 0.670246461 | 0.079450595 | 0.111932912 |
| C920006O11Rik | -0.053225883 | 0.168959807 | 0.57227323 | 0.093756217 | 0.144509804 |
| C920021L13Rik | 0.012990211 | 0.672321602 | 0.905281039 | 0.061031596 | 0.049281632 |
| C9orf72 | 0.038624642 | 0.405173934 | 0.776983336 | 0.145942916 | 0.10552819 |
| Caap1 | 0.004060567 | 0.937550163 | 0.985021197 | 0.196883677 | 0.19236374 |
| Cab39 | 0.044319589 | 0.48839039 | 0.826256745 | 0.498958685 | 0.454250138 |
| Cab39l | 0.044489071 | 0.399564323 | 0.774544012 | 0.215740455 | 0.17745758 |
| Cabin1 | -0.014088897 | 0.791965117 | 0.939347252 | 0.208596861 | 0.224476746 |
| Cables2 | -0.085816275 | 0.016891528 | 0.208334711 | 0.072437985 | 0.145855896 |
| Cabp1 | -0.034848936 | 0.573092568 | 0.861889183 | 0.321916518 | 0.354779103 |
| Cabp7 | 0.120578832 | 0.025947168 | 0.256582976 | 0.225989155 | 0.117902393 |
| Cacfd1 | -0.076201739 | 0.185936707 | 0.592701341 | 0.249704095 | 0.317661695 |
| Cacna1a | -0.079865718 | 0.200014535 | 0.61058019 | 0.336103051 | 0.402784214 |
| Cacna1b | -0.025895802 | 0.22491026 | 0.6373304 | 0.022918521 | 0.044701488 |
| Cacna1c | 0.004451162 | 0.934921847 | 0.984081583 | 0.223278111 | 0.213063484 |
| Cacna1d | -0.159409663 | 0.014778203 | 0.195426645 | 0.489515263 | 0.656512483 |
| Cacna2d1 | -0.090431649 | 0.188680859 | 0.596845187 | 0.721594386 | 0.836960812 |
| Cacnb2 | -0.039333595 | 0.439321429 | 0.800048525 | 0.177430489 | 0.217303046 |
| Cacnb3 | 0.000235143 | 0.996521935 | 0.998992348 | 0.208714651 | 0.199257285 |
| Cactin | -0.01036094 | 0.772862182 | 0.932469974 | 0.087077158 | 0.105079941 |
| Cacul1 | 0.121306756 | 0.034766781 | 0.29382176 | 0.287876242 | 0.175399525 |

| | | | | | |
|---|---|---|---|---|---|
| Cacybp | 0.232481531 | 0.003446761 | 0.082961956 | 1.275067076 | 0.916843803 |
| Cad | -0.019330766 | 0.654880357 | 0.897454058 | 0.124988791 | 0.144220422 |
| Cadm1 | 0.074112783 | 0.139341396 | 0.526569876 | 0.166268781 | 0.105405972 |
| Cadm4 | -0.01879424 | 0.434834862 | 0.796720977 | 0.031619391 | 0.047348383 |
| Cadps | -0.071536694 | 0.296990731 | 0.702656882 | 0.511238415 | 0.584798822 |
| Cadps2 | -0.039393459 | 0.14054848 | 0.528664246 | 0.037644485 | 0.070446627 |
| Calb1 | -0.021855559 | 0.533534752 | 0.84612966 | 0.06609426 | 0.082258371 |
| Calcoco1 | -0.015059593 | 0.702379979 | 0.912152193 | 0.099488447 | 0.115570578 |
| Cald1 | -0.084006844 | 0.022694679 | 0.238404929 | 0.078853419 | 0.14520046 |
| Calm1 | 0.051812337 | 0.075809809 | 0.415113505 | 3.138710086 | 2.981006315 |
| Calm2 | -0.165296967 | 0.003144072 | 0.078305325 | 1.359900301 | 1.628362575 |
| Calm3 | 0.013920125 | 0.821968718 | 0.950930859 | 1.089360926 | 1.060334784 |
| Calr | 0.355319017 | 1.76E-12 | 1.96E-09 | 2.122916174 | 1.45922184 |
| Calu | 0.040093113 | 0.404492787 | 0.776552851 | 0.157558748 | 0.124003556 |
| Caly | -0.049856253 | 0.031099096 | 0.280770404 | 0.024167919 | 0.07042772 |
| Camk1 | -0.015868134 | 0.757994024 | 0.927237784 | 0.188529109 | 0.20168613 |
| Camk2b | 0.004061845 | 0.952975129 | 0.988658594 | 0.782263162 | 0.76450969 |
| Camk2d | 0.047515274 | 0.436205514 | 0.797485381 | 0.302577016 | 0.246012688 |
| Camk2g | -0.010289123 | 0.795755253 | 0.939757086 | 0.100623065 | 0.116825337 |
| Camk2n1 | 0.083526238 | 0.027742135 | 0.264936073 | 2.803038853 | 2.608004038 |
| Camk2n2 | -0.142987512 | 0.042961227 | 0.327660967 | 0.802785899 | 1.016561274 |
| Camkk2 | -0.00989792 | 0.837989187 | 0.954911647 | 0.162491809 | 0.171536436 |
| Camkmt | -0.018486398 | 0.638165755 | 0.89125563 | 0.096785776 | 0.110416106 |
| Caml | -0.128091716 | 0.027210458 | 0.262622702 | 0.256491123 | 0.375908496 |
| Camsap1 | 0.057216421 | 0.275225212 | 0.68241738 | 0.213776919 | 0.165605929 |
| Camsap2 | -0.019637032 | 0.697283756 | 0.910970095 | 0.184206071 | 0.203397555 |
| Camsap3 | 0.019937873 | 0.556509168 | 0.85521955 | 0.073339057 | 0.052986277 |
| Camta1 | -0.054287311 | 0.34287265 | 0.735501771 | 0.254091042 | 0.307677139 |
| Camta2 | -0.089901431 | 0.055013384 | 0.364775849 | 0.148456665 | 0.215425324 |
| Cand1 | -0.123032561 | 0.052779686 | 0.359036238 | 0.400257037 | 0.519937221 |
| Cand2 | -0.062567827 | 0.035552979 | 0.296383099 | 0.048488325 | 0.101735343 |
| Cant1 | 0.039303056 | 0.459607951 | 0.809611113 | 0.205560959 | 0.165743669 |
| Canx | 0.033235064 | 0.559975401 | 0.856904336 | 1.496907291 | 1.425458753 |
| Cap1 | -0.000471735 | 0.993409096 | 0.998695538 | 0.274834812 | 0.275033983 |
| Capn1 | -0.014854264 | 0.705554613 | 0.912844672 | 0.101273157 | 0.11125009 |
| Capn10 | 0.110929326 | 0.00830948 | 0.141359743 | 0.131337146 | 0.033820044 |
| Capn15 | 0.039831767 | 0.415761273 | 0.783472369 | 0.173145177 | 0.133801016 |
| Capn2 | 0.101038446 | 0.077324498 | 0.418285047 | 0.281961378 | 0.188828207 |
| Capn5 | -0.042720983 | 0.284574112 | 0.691925959 | 0.099710495 | 0.135711848 |
| Capn7 | 0.020840841 | 0.708725578 | 0.913767764 | 0.240841875 | 0.209524739 |
| Capns1 | -0.217769676 | 0.001656375 | 0.053629162 | 0.723909566 | 0.987152239 |
| Caprin1 | -0.122637551 | 0.069773395 | 0.400867525 | 0.728707278 | 0.842091104 |
| Capsl | 0.046736659 | 0.381088477 | 0.762430261 | 0.223666345 | 0.167372487 |
| Capza1 | 0.020977355 | 0.498174513 | 0.830722138 | 0.062948781 | 0.053001272 |
| Capza2 | 0.15122555 | 0.025979929 | 0.256654086 | 0.811478523 | 0.662012713 |
| Capzb | 0.053676256 | 0.429737909 | 0.79317385 | 0.665476937 | 0.610100362 |
| Car10 | -0.008518312 | 0.890303709 | 0.970579007 | 0.330511238 | 0.348389589 |
| Car8 | -0.065920018 | 0.196901036 | 0.607156897 | 0.176757366 | 0.216111627 |
| Card19 | 0.155514989 | 0.014274476 | 0.193090184 | 0.384881031 | 0.234232915 |
| Card6 | 0.037979097 | 0.234377494 | 0.644859348 | 0.069810589 | 0.03492937 |
| Carf | 0.030434039 | 0.453891465 | 0.807844867 | 0.11060654 | 0.081937485 |
| Carhsp1 | -0.008044056 | 0.85760745 | 0.958476825 | 0.136746283 | 0.147695475 |
| Carm1 | 0.016705395 | 0.703094195 | 0.912277105 | 0.130553298 | 0.112590285 |
| Carmil1 | -0.033263055 | 0.377254438 | 0.761412185 | 0.092162233 | 0.113888687 |
| Carmil2 | 0.033333292 | 0.250935863 | 0.663121659 | 0.056601894 | 0.035015006 |
| Carmil3 | -0.011770075 | 0.757777751 | 0.927237784 | 0.093284569 | 0.108921687 |
| Carnmt1 | -0.100224957 | 0.075945573 | 0.41563016 | 0.232405351 | 0.32602092 |
| Cars | -0.051283902 | 0.456479029 | 0.808445095 | 0.500536163 | 0.53510924 |
| Cars2 | 0.000141801 | 0.997179461 | 0.999265301 | 0.10802171 | 0.107455652 |
| Cartpt | 0.063707664 | 0.08170313 | 0.42777546 | 0.083222597 | 0.023339107 |
| Casc3 | 0.006900742 | 0.910833826 | 0.978717991 | 0.342148397 | 0.327912849 |
| Casd1 | -0.130017173 | 0.002109727 | 0.063200525 | 0.107638299 | 0.213444857 |
| Cask | 0.039565555 | 0.471706944 | 0.815453427 | 0.223620713 | 0.181110457 |
| Caskin1 | -0.01408499 | 0.559381717 | 0.856904336 | 0.033644691 | 0.046998929 |
| Casp2 | -0.035168771 | 0.239564551 | 0.650744882 | 0.049680682 | 0.075346198 |

| | | | | | |
|---|---|---|---|---|---|
| Casp3 | 0.023524337 | 0.588892954 | 0.86992264 | 0.130924406 | 0.111328795 |
| Casp6 | -0.022686921 | 0.436659583 | 0.797879526 | 0.05090888 | 0.072093088 |
| Casp8 | 0.02244589 | 0.481524041 | 0.821110567 | 0.069035972 | 0.050572803 |
| Casp8ap2 | -0.015605426 | 0.782566184 | 0.936026395 | 0.24335224 | 0.250038836 |
| Casp9 | 0.043255259 | 0.108695981 | 0.477972625 | 0.049679242 | 0.017074904 |
| Casr | -0.002504258 | 0.936602771 | 0.98453816 | 0.061784885 | 0.064911651 |
| Cast | 0.116132764 | 0.06591446 | 0.390314712 | 0.388007179 | 0.27921102 |
| Castor2 | 0.002841892 | 0.966792686 | 0.992504747 | 0.660327704 | 0.64700411 |
| Casz1 | -0.074096572 | 0.252725773 | 0.663146569 | 0.372334641 | 0.440674243 |
| Cbarp | 0.148019005 | 0.023281996 | 0.241886829 | 0.526891152 | 0.360448032 |
| Cbfa2t2 | 0.071804533 | 0.119284703 | 0.492433827 | 0.153055981 | 0.094683654 |
| Cbfa2t3 | -0.002019229 | 0.976093299 | 0.994468977 | 0.425531108 | 0.431541285 |
| Cbfb | -0.029496492 | 0.524199118 | 0.841062594 | 0.142488474 | 0.181177529 |
| Cbl | -0.012273471 | 0.830860893 | 0.95325696 | 0.278961041 | 0.286755171 |
| Cblb | 0.054801193 | 0.159015214 | 0.556789205 | 0.107954487 | 0.058455084 |
| Cblc | 0.062005815 | 0.14063495 | 0.528664246 | 0.122327952 | 0.068999363 |
| Cbll1 | 0.022182978 | 0.703375106 | 0.912277105 | 0.297778005 | 0.269151453 |
| Cbr1 | -0.118234222 | 0.058720774 | 0.372687182 | 0.324249155 | 0.475085841 |
| Cbr3 | 0.020205798 | 0.662226542 | 0.898950007 | 0.143360014 | 0.137400187 |
| Cbr4 | 0.046015403 | 0.179219202 | 0.585554405 | 0.0813789 | 0.040439583 |
| Cbs | 0.078723895 | 0.132624935 | 0.513250014 | 0.196152214 | 0.122055437 |
| Cbwd1 | 0.014538459 | 0.753072511 | 0.926506029 | 0.153287791 | 0.144123358 |
| Cbx1 | -0.160226546 | 0.017828653 | 0.212525165 | 0.529023494 | 0.705457162 |
| Cbx3 | 0.017639432 | 0.783956458 | 0.936288787 | 0.554547006 | 0.561994915 |
| Cbx4 | -0.009963705 | 0.870472203 | 0.964236463 | 0.271036236 | 0.278059682 |
| Cbx5 | -0.21477168 | 0.000895263 | 0.035657774 | 0.359060007 | 0.567018777 |
| Cbx6 | -0.050431103 | 0.440005411 | 0.800219853 | 0.517973106 | 0.58568314 |
| Cbx7 | -0.055704866 | 0.244022049 | 0.653830568 | 0.150542906 | 0.200657195 |
| Cby1 | 0.035976095 | 0.469446924 | 0.814050152 | 0.181825735 | 0.143974194 |
| Cc2d1a | -0.005301883 | 0.873301608 | 0.964940674 | 0.068835895 | 0.0721513 |
| Cc2d1b | 0.018327442 | 0.703920269 | 0.912277105 | 0.166278869 | 0.149857095 |
| Cc2d2a | 0.015850559 | 0.744568537 | 0.924566547 | 0.160590433 | 0.140628662 |
| Ccar1 | -0.070597982 | 0.302923914 | 0.708026508 | 0.579388766 | 0.668983259 |
| Ccar2 | -0.011678648 | 0.739068104 | 0.922843014 | 0.08071712 | 0.086298084 |
| Ccdc107 | -0.01683951 | 0.769069541 | 0.931268476 | 0.262034701 | 0.280806224 |
| Ccdc112 | 0.000953965 | 0.985254691 | 0.996674192 | 0.192279586 | 0.188251899 |
| Ccdc113 | -0.014178501 | 0.569744326 | 0.860834533 | 0.035125037 | 0.048560716 |
| Ccdc115 | 0.051550987 | 0.355917966 | 0.745326231 | 0.231222188 | 0.193038187 |
| Ccdc117 | 0.004208623 | 0.915018796 | 0.980132949 | 0.105091818 | 0.101732149 |
| Ccdc12 | -0.057552328 | 0.41342599 | 0.781607961 | 0.592103007 | 0.626386436 |
| Ccdc122 | -0.018672414 | 0.608954482 | 0.878855197 | 0.084141921 | 0.099714002 |
| Ccdc124 | -0.064565801 | 0.326274099 | 0.720866629 | 0.500844527 | 0.571806743 |
| Ccdc126 | -0.006743504 | 0.802947498 | 0.943517638 | 0.04499769 | 0.049357304 |
| Ccdc127 | -0.015009892 | 0.815283315 | 0.948993269 | 0.392655 | 0.399485395 |
| Ccdc130 | 0.033567049 | 0.44168225 | 0.800354711 | 0.131988293 | 0.110110211 |
| Ccdc134 | 0.048875179 | 0.110561271 | 0.480391115 | 0.061814981 | 0.025175239 |
| Ccdc136 | -0.037909065 | 0.172918634 | 0.575371867 | 0.04349757 | 0.077989027 |
| Ccdc137 | 0.007529688 | 0.83851467 | 0.954911647 | 0.087306238 | 0.081936093 |
| Ccdc138 | -0.06613298 | 0.004770261 | 0.101438784 | 0.023679247 | 0.078043044 |
| Ccdc148 | 0.009963789 | 0.792506978 | 0.939347252 | 0.084830047 | 0.071122802 |
| Ccdc149 | -0.058614501 | 0.050623808 | 0.354499194 | 0.046529661 | 0.095155507 |
| Ccdc159 | -0.035764313 | 0.088606084 | 0.443118715 | 0.019856265 | 0.045715712 |
| Ccdc160 | -0.01110198 | 0.692161204 | 0.909796728 | 0.046269196 | 0.057519215 |
| Ccdc162 | 0.013877465 | 0.762854758 | 0.928730554 | 0.142655818 | 0.129399874 |
| Ccdc166 | 0.064910094 | 0.143292456 | 0.532282155 | 0.140423453 | 0.081267428 |
| Ccdc167 | -0.005751976 | 0.903841158 | 0.975048764 | 0.15612338 | 0.162278016 |
| Ccdc171 | -0.029087393 | 0.153458933 | 0.549817979 | 0.019424973 | 0.042963162 |
| Ccdc173 | 0.013382358 | 0.726345205 | 0.92014727 | 0.1005114 | 0.084727414 |
| Ccdc174 | -0.022404583 | 0.706921757 | 0.913266781 | 0.306644678 | 0.321549877 |
| Ccdc181 | -0.027560659 | 0.541620747 | 0.850208191 | 0.130162194 | 0.148935019 |
| Ccdc184 | 0.119268857 | 0.010815352 | 0.164475284 | 0.162728953 | 0.057920206 |
| Ccdc186 | 0.00758453 | 0.910479564 | 0.978693812 | 0.500249378 | 0.473509216 |
| Ccdc191 | -0.057017439 | 0.210184158 | 0.622857512 | 0.13483552 | 0.1745938 |
| Ccdc22 | -0.023411048 | 0.322588387 | 0.720155614 | 0.029728072 | 0.050778301 |
| Ccdc25 | 0.031332696 | 0.629213214 | 0.887864718 | 0.447577294 | 0.415824986 |

| | | | | | |
|---|---|---|---|---|---|
| Ccdc28a | -0.054935315 | 0.061154692 | 0.380250374 | 0.048518454 | 0.095239866 |
| Ccdc28b | 0.039480587 | 0.425975299 | 0.790724712 | 0.18115049 | 0.150136855 |
| Ccdc30 | 0.018354029 | 0.698881142 | 0.911319029 | 0.157817352 | 0.140712324 |
| Ccdc32 | -0.030296219 | 0.491253468 | 0.827270742 | 0.125972613 | 0.157969904 |
| Ccdc34 | -0.018512981 | 0.72585757 | 0.92014727 | 0.19244198 | 0.204894782 |
| Ccdc43 | -0.051951404 | 0.181700521 | 0.589059475 | 0.096964054 | 0.140493158 |
| Ccdc47 | -0.101215827 | 0.144524758 | 0.533279057 | 0.804339063 | 0.965639618 |
| Ccdc50 | 0.050866031 | 0.415602135 | 0.783472369 | 0.374221976 | 0.328283309 |
| Ccdc58 | 0.060715193 | 0.14098383 | 0.528664246 | 0.119463573 | 0.068060048 |
| Ccdc59 | -0.021931148 | 0.75305845 | 0.926506029 | 0.800518554 | 0.831167973 |
| Ccdc6 | 0.067019954 | 0.291833608 | 0.697566301 | 0.378203866 | 0.30007933 |
| Ccdc62 | -0.044977156 | 0.036561415 | 0.301071077 | 0.019021341 | 0.05883574 |
| Ccdc66 | -0.027623638 | 0.524270589 | 0.841062594 | 0.125171242 | 0.152784826 |
| Ccdc71 | 0.047270823 | 0.266836018 | 0.674278226 | 0.128770097 | 0.090559834 |
| Ccdc71l | 0.03038838 | 0.317647636 | 0.717070945 | 0.058612876 | 0.03229682 |
| Ccdc77 | -0.028747589 | 0.324141006 | 0.720406991 | 0.049755189 | 0.071756356 |
| Ccdc82 | 0.021852705 | 0.648756913 | 0.895430849 | 0.156792194 | 0.131487327 |
| Ccdc84 | -0.026571354 | 0.383943916 | 0.764308823 | 0.057798632 | 0.074914391 |
| Ccdc85b | 0.168123551 | 0.016768305 | 0.207525869 | 0.755013379 | 0.532736454 |
| Ccdc85c | -0.045758635 | 0.039497247 | 0.313386458 | 0.022823613 | 0.060420137 |
| Ccdc86 | 0.04962134 | 0.386728439 | 0.766593284 | 0.226817074 | 0.177252712 |
| Ccdc88a | -0.0103468 | 0.880546623 | 0.967810606 | 0.673874262 | 0.67524089 |
| Ccdc9 | 0.07937069 | 0.127991206 | 0.508260084 | 0.190081823 | 0.121735998 |
| Ccdc90b | -0.006055039 | 0.894693853 | 0.972181682 | 0.136934483 | 0.147174639 |
| Ccdc91 | -0.05235103 | 0.292912771 | 0.698685876 | 0.165163269 | 0.20303627 |
| Ccdc92 | 0.067747122 | 0.242616654 | 0.653091138 | 0.274087036 | 0.212535937 |
| Ccdc92b | 0.063385926 | 0.079478922 | 0.422526451 | 0.091111645 | 0.037435124 |
| Ccdc93 | -0.002715768 | 0.93668164 | 0.98453816 | 0.077384705 | 0.082117241 |
| Ccdc97 | -0.065987445 | 0.153255278 | 0.549817979 | 0.140084354 | 0.192957512 |
| Ccdc9b | 0.065294405 | 0.049012939 | 0.347612176 | 0.077765588 | 0.022586297 |
| Ccl25 | -0.056133419 | 0.109933429 | 0.478907045 | 0.07385336 | 0.120663666 |
| Ccl27a | -0.015206509 | 0.735266745 | 0.921955891 | 0.133266405 | 0.140497461 |
| Ccl28 | 0.041132408 | 0.217693714 | 0.632272789 | 0.070828754 | 0.030195027 |
| Ccm2 | 0.020313746 | 0.699401263 | 0.911319029 | 0.217536941 | 0.200254393 |
| Ccnc | -0.059529246 | 0.184289883 | 0.590567014 | 0.131922856 | 0.184066117 |
| Ccnd1 | -0.048891582 | 0.506780316 | 0.833276466 | 0.773208985 | 0.837884186 |
| Ccnd2 | -0.029508709 | 0.508123632 | 0.834020751 | 2.617413793 | 2.633574168 |
| Ccnd3 | -0.076801004 | 0.183571114 | 0.590066231 | 0.24416272 | 0.325143956 |
| Ccndbp1 | -0.073032073 | 0.110805004 | 0.481241811 | 0.137335133 | 0.211515499 |
| Ccne2 | -0.017791444 | 0.519975301 | 0.838796544 | 0.042950824 | 0.056940676 |
| Ccng1 | 0.173372355 | 0.015295184 | 0.198599946 | 0.551335203 | 0.361710505 |
| Ccng2 | -0.043417502 | 0.207208454 | 0.618605371 | 0.068405233 | 0.111008548 |
| Ccnh | -0.057756834 | 0.329460281 | 0.724538308 | 0.279543837 | 0.334320796 |
| Ccni | -0.275697867 | 3.80E-05 | 0.003895425 | 0.503137564 | 0.79296557 |
| Ccnk | 0.018205279 | 0.729940321 | 0.920522999 | 0.194823984 | 0.181411703 |
| Ccnl1 | -0.104936815 | 0.117334302 | 0.490497456 | 0.480580764 | 0.570954612 |
| Ccnl2 | 0.08088928 | 0.222568167 | 0.634095002 | 0.567629089 | 0.473748373 |
| Ccnq | -0.05218189 | 0.230114122 | 0.641131395 | 0.120731231 | 0.17734834 |
| Ccnt1 | 0.002325612 | 0.97180549 | 0.993892166 | 0.492831621 | 0.491875583 |
| Ccnt2 | -0.119969285 | 0.061148177 | 0.380250374 | 0.360503367 | 0.469698977 |
| Ccny | -0.180999689 | 0.003472352 | 0.083243072 | 0.363746581 | 0.542236234 |
| Ccp110 | -0.05011754 | 0.14989133 | 0.544321624 | 0.071971723 | 0.116542298 |
| Ccpg1 | -0.089198564 | 0.124786458 | 0.503409034 | 0.246099454 | 0.328665852 |
| Ccpg1os | -0.027052909 | 0.576121318 | 0.863450749 | 0.156082864 | 0.186680169 |
| Ccrl2 | -0.028725814 | 0.305648631 | 0.709411642 | 0.045513062 | 0.068824156 |
| Ccs | -0.072934289 | 0.125896557 | 0.504843686 | 0.15320823 | 0.229280547 |
| Ccser1 | 0.027108829 | 0.524549537 | 0.841229722 | 0.121663703 | 0.100617372 |
| Ccser2 | -0.040003257 | 0.42792586 | 0.791658813 | 0.168090165 | 0.198507294 |
| Cct2 | -0.116152749 | 0.074848629 | 0.41344108 | 0.761520718 | 0.906769679 |
| Cct3 | -0.054684918 | 0.426092187 | 0.790724712 | 0.489806297 | 0.54397349 |
| Cct4 | -0.027275945 | 0.685167479 | 0.909796728 | 0.582754762 | 0.633045376 |
| Cct5 | -0.028869825 | 0.670842295 | 0.904224039 | 0.764459068 | 0.819035812 |
| Cct6a | 0.052982343 | 0.446729804 | 0.804157614 | 0.782035315 | 0.715661133 |
| Cct7 | -0.146714623 | 0.027827919 | 0.26510449 | 0.82589812 | 0.977840332 |
| Cct8 | -0.010938893 | 0.875722985 | 0.965151159 | 0.708541016 | 0.703775427 |

| | | | | | |
|---|---|---|---|---|---|
| Ccz1 | -0.040277057 | 0.48003811 | 0.819943608 | 0.255802906 | 0.306180884 |
| Cd151 | -0.00438333 | 0.919704971 | 0.980436936 | 0.124223101 | 0.120356986 |
| Cd164 | -0.183117964 | 5.41E-05 | 0.004934051 | 2.085661571 | 2.422747533 |
| Cd164l2 | 0.003494729 | 0.91111302 | 0.978717991 | 0.059541912 | 0.055110665 |
| Cd200 | 0.157683211 | 0.025039178 | 0.251569802 | 0.612278662 | 0.425284343 |
| Cd24a | 0.033214429 | 0.522079715 | 0.840435301 | 0.168057082 | 0.140174757 |
| Cd274 | -0.025125547 | 0.526294742 | 0.842199436 | 0.099651107 | 0.126508646 |
| Cd276 | 0.011465497 | 0.757697399 | 0.927237784 | 0.091325071 | 0.074376221 |
| Cd2ap | 0.190832318 | 0.004839916 | 0.10205513 | 0.971827556 | 0.72295309 |
| Cd2bp2 | -0.005417934 | 0.915485237 | 0.980132949 | 0.18149044 | 0.197836003 |
| Cd320 | 0.001015538 | 0.979613572 | 0.995540893 | 0.105065393 | 0.108967317 |
| Cd3eap | -0.014819932 | 0.764129447 | 0.928730554 | 0.165498102 | 0.168852052 |
| Cd44 | 0.083595503 | 0.133566445 | 0.515221524 | 0.230623682 | 0.133294902 |
| Cd47 | 0.027537497 | 0.674400258 | 0.906025597 | 1.229036916 | 1.182538138 |
| Cd59a | 0.029695195 | 0.614881062 | 0.882439418 | 0.284314649 | 0.238882313 |
| Cd59b | 0.024065214 | 0.499529269 | 0.830722138 | 0.080090483 | 0.058926776 |
| Cd63 | 0.05923766 | 0.139642814 | 0.527113549 | 2.476834511 | 2.332156404 |
| Cd63-ps | 0.02647365 | 0.361088719 | 0.749638535 | 0.055600682 | 0.028070353 |
| Cd79a | 0.000958366 | 0.981630444 | 0.995792496 | 0.101102207 | 0.106886737 |
| Cd81 | -0.300789909 | 0.000119601 | 0.008958504 | 0.442558301 | 0.757921182 |
| Cd82 | -0.020868019 | 0.724127945 | 0.919830015 | 0.308847258 | 0.318109015 |
| Cd9 | 0.067501087 | 0.036793111 | 0.302698738 | 0.072000654 | 0.011219573 |
| Cd99l2 | -0.008258075 | 0.828525996 | 0.952874813 | 0.095985797 | 0.097522583 |
| Cda | 0.080441816 | 0.254687323 | 0.66334523 | 0.502910537 | 0.423466939 |
| Cdadc1 | -0.011005059 | 0.840785545 | 0.954960614 | 0.234889819 | 0.236754352 |
| Cdan1 | -0.029563597 | 0.194057172 | 0.60325934 | 0.026295614 | 0.054754917 |
| Cdc123 | -0.04296812 | 0.48137726 | 0.820999756 | 0.350652341 | 0.384392973 |
| Cdc14b | -0.121109283 | 0.018440452 | 0.2157189 | 0.165038236 | 0.265718638 |
| Cdc16 | -0.028872524 | 0.623198132 | 0.885857477 | 0.314566555 | 0.336611183 |
| Cdc23 | 0.035935532 | 0.462350043 | 0.810535897 | 0.171233226 | 0.138894367 |
| Cdc25a | -0.018315532 | 0.736163441 | 0.922302218 | 0.217683568 | 0.234220421 |
| Cdc25b | -0.050989265 | 0.02916581 | 0.272146451 | 0.025923703 | 0.067654679 |
| Cdc26 | -0.030123732 | 0.642443825 | 0.892561795 | 0.419385149 | 0.457015007 |
| Cdc27 | 0.041111008 | 0.414925758 | 0.783085706 | 0.18207265 | 0.144407238 |
| Cdc34 | 0.153096405 | 0.022257631 | 0.236913658 | 0.564681054 | 0.40908723 |
| Cdc37 | -0.050013293 | 0.459316249 | 0.809393611 | 0.582427365 | 0.627980958 |
| Cdc37l1 | 0.064802833 | 0.338035025 | 0.732352032 | 0.609581656 | 0.548534057 |
| Cdc40 | -0.107709407 | 0.074942329 | 0.41344108 | 0.335153415 | 0.435830677 |
| Cdc42 | -0.000695713 | 0.991381274 | 0.998321067 | 1.10942201 | 1.09827827 |
| Cdc42bpa | -0.012894457 | 0.837813703 | 0.954911647 | 0.373001152 | 0.374736628 |
| Cdc42bpb | -0.031024571 | 0.514276521 | 0.83659041 | 0.151281831 | 0.173608372 |
| Cdc42bpg | -0.06806498 | 0.076093332 | 0.416211866 | 0.090858673 | 0.150544073 |
| Cdc42ep3 | -0.008521122 | 0.79908727 | 0.941255596 | 0.069960215 | 0.076486437 |
| Cdc42ep5 | -0.030314532 | 0.400597702 | 0.774740795 | 0.077448589 | 0.109632796 |
| Cdc42se1 | 0.044186939 | 0.439640346 | 0.800048525 | 0.243982194 | 0.193980416 |
| Cdc42se2 | 0.021698221 | 0.745254641 | 0.924566547 | 0.465501448 | 0.436457453 |
| Cdc5l | -0.053619307 | 0.429698902 | 0.79317385 | 0.563377753 | 0.626204102 |
| Cdc73 | -0.082293493 | 0.187196034 | 0.594494963 | 0.381014846 | 0.467510801 |
| Cdca4 | 0.013534412 | 0.657746305 | 0.898224427 | 0.058784911 | 0.050825431 |
| Cdcp1 | -0.032975228 | 0.488738732 | 0.826256745 | 0.143564309 | 0.175602694 |
| Cdh1 | 0.330106059 | 1.41E-06 | 0.000283039 | 0.869608081 | 0.507577688 |
| Cdh2 | 0.056772289 | 0.410205277 | 0.780101937 | 0.69527617 | 0.640562949 |
| Cdh22 | 0.071774542 | 0.036192324 | 0.299721418 | 0.084032505 | 0.025812935 |
| Cdh4 | -0.054201096 | 0.079272294 | 0.422322725 | 0.050569465 | 0.094245793 |
| Cdh7 | -0.004008184 | 0.911441063 | 0.978933542 | 0.084261519 | 0.083373827 |
| Cdh8 | 0.016411249 | 0.742443359 | 0.923637085 | 0.178382669 | 0.171037044 |
| Cdhr1 | -0.042426684 | 0.358947485 | 0.747455833 | 0.1471236 | 0.188437389 |
| Cdip1 | 0.015468128 | 0.786022158 | 0.937083312 | 0.244766631 | 0.227436678 |
| Cdipt | -0.010254044 | 0.872909653 | 0.964911254 | 0.438358019 | 0.448483991 |
| Cdk10 | -0.030271671 | 0.392849982 | 0.769704269 | 0.075218142 | 0.098974965 |
| Cdk11b | 0.109460361 | 0.112944189 | 0.483221516 | 0.555199689 | 0.455152213 |
| Cdk12 | -0.02544925 | 0.71187172 | 0.915109107 | 0.49279331 | 0.522414782 |
| Cdk13 | 0.037344175 | 0.565189251 | 0.858087314 | 0.454525287 | 0.424593144 |
| Cdk14 | -0.126754157 | 0.035083025 | 0.294914843 | 0.250806411 | 0.376221107 |
| Cdk16 | 0.038671136 | 0.510216512 | 0.834957869 | 0.283494432 | 0.23871124 |

| | | | | | |
|---|---|---|---|---|---|
| Cdk17 | -0.025802029 | 0.51471253 | 0.836743564 | 0.103238712 | 0.130001483 |
| Cdk18 | -0.031477479 | 0.205292174 | 0.616549498 | 0.032030717 | 0.057260512 |
| Cdk2ap1 | -0.084249807 | 0.128668391 | 0.508260084 | 0.233760971 | 0.328879727 |
| Cdk2ap2 | -0.133162937 | 0.040116912 | 0.315774775 | 0.39940637 | 0.546947467 |
| Cdk4 | 0.053547525 | 0.411798895 | 0.781032788 | 0.629405404 | 0.579680162 |
| Cdk5 | -0.063816998 | 0.151249016 | 0.546523037 | 0.126920549 | 0.178971737 |
| Cdk5r1 | -0.019587726 | 0.490263855 | 0.826674417 | 0.04801745 | 0.066043864 |
| Cdk5r2 | 0.075015464 | 0.157789719 | 0.556086872 | 0.213128829 | 0.151819947 |
| Cdk5rap2 | 0.045595534 | 0.205407811 | 0.616549498 | 0.088792034 | 0.052212634 |
| Cdk5rap3 | -0.136833941 | 0.027500705 | 0.264360019 | 0.329846311 | 0.455850194 |
| Cdk7 | 0.037761391 | 0.40487208 | 0.776552851 | 0.140659292 | 0.116475848 |
| Cdk8 | 0.113946023 | 0.093779019 | 0.45227176 | 0.635625651 | 0.513197919 |
| Cdk9 | 0.0178803 | 0.751359212 | 0.926186851 | 0.278021771 | 0.271202076 |
| Cdkal1 | 0.008065795 | 0.818125617 | 0.94985851 | 0.083232865 | 0.074617847 |
| Cdkl1 | 0.064697175 | 0.113250891 | 0.483626277 | 0.117756877 | 0.065477759 |
| Cdkl2 | -0.080358166 | 0.100384608 | 0.465385827 | 0.157099121 | 0.225210053 |
| Cdkl3 | 0.021308766 | 0.510948595 | 0.834957869 | 0.066602412 | 0.047839006 |
| Cdkl5 | 0.031478058 | 0.502127415 | 0.832218079 | 0.151010203 | 0.117749858 |
| Cdkn1a | -0.028955188 | 0.728569936 | 0.920519505 | 0.96601625 | 1.026291998 |
| Cdkn1b | -0.108947494 | 0.108308578 | 0.477972625 | 0.506401412 | 0.646582279 |
| Cdkn2aip | -0.023718627 | 0.638413973 | 0.89125563 | 0.17611189 | 0.215858686 |
| Cdkn2aipnl | 0.062942353 | 0.155459115 | 0.552356396 | 0.141608288 | 0.08874912 |
| Cdkn2b | -0.024338784 | 0.640171977 | 0.891345382 | 0.174519874 | 0.198842247 |
| Cdkn2c | -0.012455664 | 0.72059425 | 0.919412687 | 0.077416028 | 0.089249673 |
| Cdkn2d | 0.032862284 | 0.52165496 | 0.840205104 | 0.190435209 | 0.165210669 |
| Cdon | 0.044020866 | 0.157019625 | 0.554909286 | 0.065975387 | 0.030415618 |
| Cdpf1 | -0.037145981 | 0.480692046 | 0.820388721 | 0.197555235 | 0.223838972 |
| Cdr2 | -0.013366419 | 0.792333011 | 0.939347252 | 0.18222431 | 0.209816656 |
| Cds1 | 0.123726539 | 0.012644225 | 0.179505076 | 0.187027844 | 0.074805454 |
| Cds2 | 0.041099111 | 0.5229574 | 0.841062594 | 0.429112 | 0.379750152 |
| Cdv3 | 0.054031863 | 0.443033804 | 0.801368502 | 0.686418609 | 0.629345459 |
| Cdyl | -0.018198914 | 0.712212981 | 0.915109107 | 0.161639422 | 0.182398897 |
| Cebpb | -0.266070066 | 9.55E-05 | 0.007605136 | 0.949250097 | 1.345125649 |
| Cebpd | -0.072517187 | 0.34119759 | 0.734419946 | 0.493364169 | 0.577409387 |
| Cebpg | -0.149480317 | 0.030102072 | 0.275342007 | 0.84531155 | 1.020001451 |
| Cebpz | -0.015073073 | 0.825480934 | 0.952011047 | 0.602607039 | 0.623452087 |
| Cebpzos | -0.018532676 | 0.778199122 | 0.934523337 | 0.450496176 | 0.464033777 |
| Celf1 | 0.001451057 | 0.982625421 | 0.995921574 | 0.748894665 | 0.727243175 |
| Celf2 | 0.000116716 | 0.998157794 | 0.99938181 | 0.182030197 | 0.187313073 |
| Celf3 | -0.039813004 | 0.545652272 | 0.851259637 | 0.548935344 | 0.577050085 |
| Celf4 | -0.098120038 | 0.151497056 | 0.546576546 | 0.505888497 | 0.597234715 |
| Celf6 | -0.019207332 | 0.693576092 | 0.909871028 | 0.16558989 | 0.176669833 |
| Celsr2 | -0.021100704 | 0.537631579 | 0.847793898 | 0.074693953 | 0.096319648 |
| Celsr3 | -0.031220333 | 0.458908373 | 0.809370658 | 0.115011821 | 0.144755835 |
| Cemip2 | -0.027484249 | 0.479543007 | 0.819939708 | 0.094952529 | 0.118330526 |
| Cenpb | -0.037886936 | 0.564893165 | 0.858087314 | 0.425329016 | 0.474027986 |
| Cenpc1 | -0.050810157 | 0.206780964 | 0.618432816 | 0.100822715 | 0.146647567 |
| Cenpj | -0.021385063 | 0.549301527 | 0.852402509 | 0.078920801 | 0.092055196 |
| Cenpo | -0.057896417 | 0.018535302 | 0.216324216 | 0.028307558 | 0.075784455 |
| Cenpq | 0.044169002 | 0.320094632 | 0.717489703 | 0.138216216 | 0.107772095 |
| Cenps | 0.006173656 | 0.884381115 | 0.968954079 | 0.117191452 | 0.10679934 |
| Cenpv | -0.153421628 | 0.020979753 | 0.229383208 | 0.50815835 | 0.648513033 |
| Cenpw | -0.024246994 | 0.358913309 | 0.747455833 | 0.039197729 | 0.061683998 |
| Cenpx | -0.10842418 | 0.084871797 | 0.433957325 | 0.327926214 | 0.43331301 |
| Cep104 | -0.072562449 | 0.171295231 | 0.574245234 | 0.204181442 | 0.266616959 |
| Cep112 | -0.046957848 | 0.080353281 | 0.425115437 | 0.039378617 | 0.076100061 |
| Cep120 | 0.019994321 | 0.676394533 | 0.906611913 | 0.165689976 | 0.147763893 |
| Cep126 | -0.10017244 | 0.028748962 | 0.270181023 | 0.127459223 | 0.219400175 |
| Cep135 | 0.015282503 | 0.69067682 | 0.909796728 | 0.097999157 | 0.072304829 |
| Cep162 | 0.013902429 | 0.772483071 | 0.932469974 | 0.159563842 | 0.155592995 |
| Cep164 | 0.002710419 | 0.961313875 | 0.991441074 | 0.227361567 | 0.224815109 |
| Cep170 | 0.037620695 | 0.556655307 | 0.85521955 | 0.366320536 | 0.316401366 |
| Cep170b | 0.053850312 | 0.388586566 | 0.767160379 | 0.36754392 | 0.316641223 |
| Cep19 | -0.041050987 | 0.432757212 | 0.795079432 | 0.205069715 | 0.236579392 |
| Cep192 | 0.016097296 | 0.70863957 | 0.913767764 | 0.123392377 | 0.110389717 |

| | | | | | |
|---|---|---|---|---|---|
| Cep20 | 0.006871337 | 0.906432922 | 0.976716534 | 0.31284237 | 0.318288804 |
| Cep250 | 0.019522189 | 0.560573468 | 0.856904336 | 0.075919852 | 0.057275709 |
| Cep290 | 0.001709866 | 0.979401244 | 0.995540893 | 0.469478671 | 0.450681666 |
| Cep295 | 0.036473448 | 0.289706226 | 0.695966637 | 0.080309394 | 0.048572655 |
| Cep350 | 0.001993145 | 0.975306244 | 0.994455356 | 0.374325196 | 0.361941561 |
| Cep43 | 0.010992222 | 0.764058537 | 0.928730554 | 0.086780324 | 0.077224131 |
| Cep44 | 0.034065753 | 0.476135551 | 0.817948631 | 0.159323304 | 0.121348599 |
| Cep57 | 0.044075271 | 0.462098125 | 0.810535897 | 0.320652542 | 0.281445254 |
| Cep57l1 | -0.02573317 | 0.373403035 | 0.758519786 | 0.048155028 | 0.072523424 |
| Cep63 | -0.099858746 | 0.017083683 | 0.208804661 | 0.106062085 | 0.19285329 |
| Cep68 | -0.00616618 | 0.862241965 | 0.960216905 | 0.084027791 | 0.091557506 |
| Cep70 | -0.106428475 | 0.043257725 | 0.328509254 | 0.191145181 | 0.284492646 |
| Cep76 | -0.038225606 | 0.24211588 | 0.652555608 | 0.063601848 | 0.091995558 |
| Cep78 | -0.001720794 | 0.964258397 | 0.991970325 | 0.097800187 | 0.097810231 |
| Cep83 | -0.057475096 | 0.313470392 | 0.714906079 | 0.252754396 | 0.300727196 |
| Cep83os | 0.022373121 | 0.474829649 | 0.817387411 | 0.064921548 | 0.042742157 |
| Cep85 | 0.051376374 | 0.293121332 | 0.698992351 | 0.169278421 | 0.123166247 |
| Cep89 | -0.012550102 | 0.734691377 | 0.921732448 | 0.08929618 | 0.097842875 |
| Cep95 | 0.0411629 | 0.181665118 | 0.589059475 | 0.065130538 | 0.030038545 |
| Cep97 | -0.053640777 | 0.097756946 | 0.460898239 | 0.060832069 | 0.112859264 |
| Cept1 | -0.048621326 | 0.262270593 | 0.668764912 | 0.129190052 | 0.171188958 |
| Cerk | 0.007416696 | 0.90334964 | 0.974950771 | 0.351421085 | 0.334569191 |
| Cerkl | -0.009412629 | 0.866509991 | 0.962125879 | 0.236369783 | 0.242827822 |
| Cers2 | -0.039725347 | 0.479801454 | 0.819939708 | 0.251458387 | 0.2975826 |
| Cers4 | 0.003130184 | 0.937995337 | 0.985059229 | 0.105395734 | 0.103047346 |
| Cers5 | -0.065251252 | 0.16395451 | 0.563565555 | 0.145109875 | 0.202247165 |
| Cers6 | 0.028354932 | 0.588931373 | 0.86992264 | 0.196248804 | 0.169634152 |
| Cert1 | 0.050880331 | 0.428142098 | 0.791658813 | 0.411613892 | 0.362107904 |
| Cetn2 | -0.032077465 | 0.591462083 | 0.871477529 | 0.295413414 | 0.326918148 |
| Cetn3 | -0.119581622 | 0.065457136 | 0.389491953 | 0.39648514 | 0.536531479 |
| Cetn4 | 0.036080668 | 0.42221074 | 0.787974934 | 0.141968385 | 0.101114737 |
| Cfap126 | -0.090367776 | 0.025572346 | 0.254380218 | 0.09936576 | 0.182524227 |
| Cfap20 | -0.054637074 | 0.400285773 | 0.774740795 | 0.564484504 | 0.618527127 |
| Cfap298 | 0.0576634 | 0.369254833 | 0.756523935 | 0.418775868 | 0.363847435 |
| Cfap36 | -0.122477027 | 0.077110131 | 0.418199494 | 0.881860165 | 1.020581881 |
| Cfap410 | -0.021331649 | 0.692480703 | 0.909796728 | 0.208249562 | 0.230039569 |
| Cfap97 | 0.015107069 | 0.800904085 | 0.942068945 | 0.321833303 | 0.309025083 |
| Cfdp1 | -0.0743368 | 0.246114711 | 0.655935044 | 0.477697557 | 0.567226351 |
| Cfl1 | 0.084527911 | 0.187227117 | 0.594494963 | 1.304312922 | 1.189650657 |
| Cfl2 | 0.062754325 | 0.375202319 | 0.760147021 | 0.611436971 | 0.556799171 |
| Cflar | 0.027775576 | 0.6396186 | 0.891345382 | 0.28970627 | 0.266138232 |
| Cggbp1 | 0.041666511 | 0.550000342 | 0.852459771 | 0.600550566 | 0.560153337 |
| Cgn | -0.032497419 | 0.515446092 | 0.837141169 | 0.158069793 | 0.183677147 |
| Cgref1 | -0.016548085 | 0.630924497 | 0.888561606 | 0.074862793 | 0.093128874 |
| Cgrrf1 | 0.170406188 | 0.005909168 | 0.113839386 | 0.340665171 | 0.18556075 |
| Chac1 | -0.159343143 | 0.024506946 | 0.248116801 | 0.560619932 | 0.718927705 |
| Chac2 | -0.008598542 | 0.830716033 | 0.95325696 | 0.10891218 | 0.11475351 |
| Chadl | -0.019005513 | 0.520267944 | 0.838864153 | 0.054227006 | 0.071579995 |
| Champ1 | -0.035548152 | 0.381723841 | 0.762597742 | 0.103278847 | 0.141150481 |
| Chaserr | -0.108471457 | 0.105832427 | 0.473025898 | 0.422526948 | 0.55787217 |
| Chchd1 | 0.079182646 | 0.221266843 | 0.633858837 | 0.964657158 | 0.852433035 |
| Chchd10 | -0.129875759 | 0.0581476 | 0.371463807 | 1.207480223 | 1.395007981 |
| Chchd2 | 0.014647699 | 0.709858821 | 0.914392244 | 2.400044345 | 2.399990659 |
| Chchd3 | 0.027287455 | 0.636603254 | 0.890755879 | 0.272371426 | 0.2444238 |
| Chchd4 | -0.006889191 | 0.910075236 | 0.978641543 | 0.317737254 | 0.330373838 |
| Chchd5 | -0.097173475 | 0.039170794 | 0.312494944 | 0.149367536 | 0.241946543 |
| Chchd6 | -0.123884276 | 0.034641438 | 0.293577356 | 0.279442067 | 0.385294659 |
| Chchd7 | -0.001670921 | 0.980571029 | 0.995540893 | 0.65058233 | 0.669927192 |
| Chd1 | 0.036250789 | 0.551557715 | 0.852505562 | 0.319216289 | 0.296910465 |
| Chd1l | -0.016072927 | 0.552454597 | 0.852623515 | 0.044832616 | 0.059035269 |
| Chd2 | 0.237599165 | 0.001112587 | 0.04075561 | 0.598167627 | 0.330448218 |
| Chd3 | -0.116587776 | 0.082096552 | 0.427831307 | 0.449530057 | 0.564778828 |
| Chd4 | -0.011106149 | 0.872279301 | 0.964745713 | 1.292591115 | 1.254360756 |
| Chd5 | -0.129882732 | 0.005865577 | 0.113654038 | 0.134629161 | 0.249348344 |
| Chd6 | 0.045948713 | 0.461670491 | 0.810421416 | 0.366506168 | 0.309637762 |

| | | | | | |
|---|---|---|---|---|---|
| Chd7 | -0.050625723 | 0.448900245 | 0.805868674 | 0.51121497 | 0.559267781 |
| Chd8 | 0.035308281 | 0.596948101 | 0.874171008 | 0.430685403 | 0.387923571 |
| Chd9 | -0.074126491 | 0.285018888 | 0.692407818 | 0.65151685 | 0.69733932 |
| Cherp | 0.02145654 | 0.658628179 | 0.898224427 | 0.165741371 | 0.145539778 |
| Chfr | -0.008978451 | 0.840210303 | 0.954911647 | 0.136653894 | 0.13696142 |
| Chga | -0.126700578 | 0.00075026 | 0.032181025 | 5.087149773 | 5.480266354 |
| Chgb | -0.26247239 | 8.19E-10 | 6.33E-07 | 3.049124412 | 3.7604862 |
| Chic1 | -0.121058849 | 0.043671233 | 0.329086044 | 1.737109011 | 1.972823757 |
| Chic2 | -0.035940735 | 0.57864937 | 0.865142348 | 0.390976158 | 0.435233505 |
| Chid1 | -0.040733055 | 0.475132076 | 0.817387411 | 0.243018396 | 0.283731739 |
| Chka | -0.259874612 | 0.000250145 | 0.015403105 | 0.64487684 | 0.939395823 |
| Chkb | -0.018243714 | 0.575582223 | 0.863006716 | 0.065489217 | 0.080802082 |
| Chm | -0.002385927 | 0.964097248 | 0.991970325 | 0.21225056 | 0.213787034 |
| Chmp1a | 0.052416639 | 0.388154185 | 0.767160379 | 0.339421172 | 0.288856742 |
| Chmp1b | -0.014722254 | 0.745926813 | 0.924566547 | 0.134185101 | 0.147820984 |
| Chmp2a | 0.012179482 | 0.844171918 | 0.955775921 | 1.269252273 | 1.268444235 |
| Chmp2b | 0.057442828 | 0.313762796 | 0.714906079 | 0.28530637 | 0.250941108 |
| Chmp3 | -0.123891159 | 0.054340495 | 0.36336812 | 0.495355624 | 0.617672075 |
| Chmp4b | 0.037072181 | 0.559528146 | 0.856904336 | 1.15290985 | 1.081228361 |
| Chmp4c | 0.057203312 | 0.363416343 | 0.751765374 | 0.360560821 | 0.307521682 |
| Chmp5 | -0.082150321 | 0.23154537 | 0.643122701 | 0.807814895 | 0.917463496 |
| Chmp6 | -0.0073108 | 0.866345748 | 0.962125879 | 0.127156326 | 0.134936817 |
| Chmp7 | -0.017697975 | 0.727054699 | 0.920455328 | 0.179057107 | 0.201948049 |
| Chn1 | 0.036944861 | 0.347011344 | 0.738383053 | 0.105351356 | 0.07399599 |
| Chordc1 | -0.0963801 | 0.145497164 | 0.533755497 | 0.430527793 | 0.525456121 |
| Chp1 | -0.109874849 | 0.071314493 | 0.404242678 | 0.300620854 | 0.409609282 |
| Chpf | -0.125641758 | 0.031400077 | 0.281646624 | 0.244147759 | 0.355160038 |
| Chpf2 | -0.047442176 | 0.260498405 | 0.667847379 | 0.113287396 | 0.159018178 |
| Chpt1 | -0.042420439 | 0.325974254 | 0.720866629 | 0.118883468 | 0.165472172 |
| Chrac1 | -0.0429219 | 0.523995848 | 0.841062594 | 0.578384592 | 0.610355537 |
| Chrm3 | 0.03979392 | 0.405765313 | 0.777374774 | 0.15436763 | 0.112717224 |
| Chrna1os | -0.077609853 | 0.025173398 | 0.252412985 | 0.066761577 | 0.137513198 |
| Chrnb2 | -0.015639206 | 0.670081145 | 0.903736153 | 0.086131038 | 0.097256581 |
| Chst11 | -0.027469156 | 0.629697434 | 0.888005452 | 0.268063984 | 0.288453728 |
| Chst12 | -0.015627015 | 0.694208023 | 0.909986409 | 0.100372093 | 0.112688885 |
| Chtop | 0.028031467 | 0.659016557 | 0.898224427 | 0.457986472 | 0.432096069 |
| Chuk | -0.011528403 | 0.849723797 | 0.957111129 | 0.329070944 | 0.345517372 |
| Churc1 | -0.015170995 | 0.826745318 | 0.952687014 | 0.637799806 | 0.674783844 |
| Ciao1 | -0.071797174 | 0.146608397 | 0.536416643 | 0.157555187 | 0.233889095 |
| Ciao2a | 0.004082676 | 0.939549098 | 0.985603501 | 0.219763807 | 0.226434516 |
| Ciao2b | -0.054945233 | 0.34327286 | 0.735897614 | 0.264127013 | 0.330131131 |
| Ciao3 | 0.070225459 | 0.102106805 | 0.468127803 | 0.127308663 | 0.066632789 |
| Ciapin1 | -0.084860072 | 0.10363468 | 0.469469748 | 0.183069982 | 0.261310593 |
| Ciart | -0.030974986 | 0.245497053 | 0.655159244 | 0.036260838 | 0.064524966 |
| Cib1 | -0.088796437 | 0.151224355 | 0.546523037 | 0.347345975 | 0.428114698 |
| Cib2 | -0.00700175 | 0.91436103 | 0.980132949 | 0.44046119 | 0.475484233 |
| Cib3 | -0.083010196 | 0.103522801 | 0.469469748 | 0.163774308 | 0.234472071 |
| Cibar1 | 0.023136474 | 0.725478757 | 0.92014727 | 0.537415998 | 0.525908568 |
| Cibar2 | 0.001740108 | 0.95842899 | 0.990297691 | 0.069381128 | 0.069193983 |
| Cic | 0.122370229 | 0.024371761 | 0.247768538 | 0.218714916 | 0.102217713 |
| Cilk1 | -0.027251564 | 0.340187587 | 0.733546237 | 0.047712279 | 0.070168837 |
| Cinp | 0.002758256 | 0.951808359 | 0.98838165 | 0.137974795 | 0.143589618 |
| Cipc | -0.011195647 | 0.803278113 | 0.943517638 | 0.131035768 | 0.149181934 |
| Cir1 | 0.018349246 | 0.690561576 | 0.909796728 | 0.141342451 | 0.130475384 |
| Cirbp | -0.256536686 | 0.000296007 | 0.017273394 | 0.874989683 | 1.209752144 |
| Cisd1 | 0.07799781 | 0.205035518 | 0.616549498 | 1.237810662 | 1.113754365 |
| Cisd2 | 0.056419673 | 0.39808495 | 0.774311756 | 0.547077095 | 0.495271038 |
| Cisd3 | -0.019201254 | 0.696490042 | 0.910970095 | 0.168983516 | 0.182576282 |
| Cish | -0.09782268 | 0.027550123 | 0.264360019 | 0.120130055 | 0.205531742 |
| Cited2 | -0.088717768 | 0.224753879 | 0.637067124 | 0.515749137 | 0.618040293 |
| Cited4 | -0.053850895 | 0.229355297 | 0.640979764 | 0.125912657 | 0.178738498 |
| Ciz1 | -0.05888691 | 0.182637114 | 0.590066231 | 0.125831448 | 0.190607748 |
| Ckap4 | -0.138993902 | 0.033975179 | 0.291747273 | 0.438548332 | 0.58347149 |
| Ckap5 | 0.021611135 | 0.719486784 | 0.918741515 | 0.313200322 | 0.292366267 |
| Ckb | -0.226426749 | 0.005480215 | 0.110673874 | 0.768981518 | 1.05657811 |

| | | | | | |
|---|---|---|---|---|---|
| Cklf | -0.005998571 | 0.818976673 | 0.950128492 | 0.040247969 | 0.046229317 |
| Ckmt1 | -0.046953024 | 0.228079802 | 0.639451668 | 0.100424895 | 0.149323397 |
| Cks1b | -0.012532542 | 0.690768307 | 0.909796728 | 0.059675644 | 0.065092876 |
| Clasp1 | -0.184971635 | 0.000107375 | 0.008226858 | 0.126718596 | 0.30004684 |
| Clasp2 | -0.089424469 | 0.141398176 | 0.528664246 | 0.292925799 | 0.389140804 |
| Clasrp | 0.026279168 | 0.39734867 | 0.774053847 | 0.063354922 | 0.04083448 |
| Clba1 | -0.021071357 | 0.618745758 | 0.883658391 | 0.114646319 | 0.130490632 |
| Clcc1 | 0.077684517 | 0.139163124 | 0.526292492 | 0.211767293 | 0.142001971 |
| Clcn3 | -0.023859991 | 0.722786698 | 0.919483582 | 0.881269276 | 0.908680572 |
| Clcn4 | -0.108833999 | 0.075484236 | 0.414292853 | 0.304458262 | 0.407925478 |
| Clcn5 | 0.024618052 | 0.605282768 | 0.87802905 | 0.149905898 | 0.122747819 |
| Clcn7 | 0.034805351 | 0.370129588 | 0.757234136 | 0.103108637 | 0.073676064 |
| Cldn12 | 0.047628842 | 0.274648362 | 0.682000399 | 0.136440785 | 0.09023342 |
| Cldn3 | 0.221385394 | 0.003245357 | 0.079642162 | 0.627573389 | 0.412730206 |
| Cldn4 | 0.232939216 | 0.001714791 | 0.054988351 | 0.488029248 | 0.258818096 |
| Cldn7 | 0.058505557 | 0.362208147 | 0.750073837 | 1.427797112 | 1.284775542 |
| Cldnd1 | 0.145541883 | 0.027654081 | 0.264598674 | 0.449742716 | 0.310448638 |
| Clec11a | 0.03010713 | 0.33520422 | 0.730736446 | 0.069245064 | 0.045726776 |
| Clec16a | 0.019113013 | 0.518675141 | 0.838109801 | 0.054657848 | 0.041195781 |
| Clic1 | 0.143009344 | 0.038262019 | 0.308710515 | 1.088669579 | 0.929611203 |
| Clic4 | -0.109395849 | 0.109200661 | 0.477972625 | 0.749895864 | 0.8568284 |
| Clic5 | 0.071976801 | 0.074907107 | 0.41344108 | 0.11628029 | 0.045062759 |
| Clint1 | 0.081481801 | 0.246041499 | 0.655935044 | 0.656979536 | 0.554175674 |
| Clip1 | -0.085277598 | 0.208268134 | 0.620292955 | 0.734036126 | 0.82974545 |
| Clip2 | -0.009923132 | 0.765551055 | 0.928784713 | 0.06877266 | 0.077294548 |
| Clip3 | -0.025205709 | 0.55787053 | 0.855384435 | 0.119871185 | 0.132316768 |
| Clk1 | -0.157617908 | 0.018583925 | 0.216639787 | 0.456711373 | 0.642315231 |
| Clk2 | 0.029413845 | 0.506380537 | 0.833269979 | 0.139068083 | 0.109631691 |
| Clk3 | -0.032645317 | 0.550672191 | 0.852505562 | 0.223879527 | 0.26943439 |
| Clk4 | 0.066487932 | 0.285317879 | 0.692407818 | 0.378019279 | 0.32105968 |
| Cln3 | -0.070360396 | 0.112200894 | 0.483221516 | 0.127207946 | 0.185278361 |
| Cln5 | 0.039980006 | 0.41788751 | 0.784482356 | 0.178194143 | 0.141412492 |
| Cln6 | 0.077693306 | 0.227502701 | 0.639441223 | 0.460000138 | 0.384106668 |
| Cln8 | 0.049809591 | 0.153070201 | 0.549602359 | 0.083342549 | 0.040053624 |
| Clns1a | -0.108627099 | 0.077582539 | 0.418285047 | 0.383312586 | 0.508702154 |
| Clock | -0.148185103 | 0.022387837 | 0.237282704 | 0.477714386 | 0.638868405 |
| Clp1 | 0.035040523 | 0.329379496 | 0.724538308 | 0.08673059 | 0.063996536 |
| Clpb | -0.016857227 | 0.709018443 | 0.913931846 | 0.145081951 | 0.155126275 |
| Clpp | -0.056056935 | 0.413188981 | 0.781607961 | 0.625172576 | 0.686514419 |
| Clptm1 | 0.045651425 | 0.439718436 | 0.800048525 | 0.28907955 | 0.245356329 |
| Clptm1l | -0.034088834 | 0.60222649 | 0.876569856 | 0.565450812 | 0.601152562 |
| Clpx | 0.106231827 | 0.112771809 | 0.483221516 | 0.52051344 | 0.399574593 |
| Clstn1 | -0.062010925 | 0.294819342 | 0.701208942 | 0.301582567 | 0.362298364 |
| Clstn3 | -0.108001965 | 0.000595829 | 0.028287387 | 0.051108024 | 0.147202138 |
| Clta | -0.015182084 | 0.80514905 | 0.94472738 | 1.345276434 | 1.399429633 |
| Cltb | 0.002769287 | 0.961749249 | 0.991441074 | 1.257323958 | 1.247445148 |
| Cltc | -0.093652084 | 0.094326778 | 0.453645362 | 1.350912442 | 1.506978989 |
| Cltrn | -0.418648997 | 5.65E-10 | 4.73E-07 | 1.299493826 | 2.019151215 |
| Clu | 0.395406576 | 1.64E-06 | 0.000316477 | 0.933269299 | 0.463356948 |
| Cluap1 | -0.044999279 | 0.296186149 | 0.702600505 | 0.122794626 | 0.159024711 |
| Cluh | -0.001819742 | 0.971068485 | 0.993892166 | 0.17869235 | 0.176112354 |
| Cmas | -0.142715761 | 0.028298886 | 0.267705863 | 0.636366512 | 0.816065942 |
| Cmbl | -0.127814443 | 0.022754999 | 0.238404929 | 0.224692744 | 0.348274707 |
| Cmc1 | -0.084078488 | 0.169609008 | 0.57227323 | 0.289838522 | 0.373044146 |
| Cmc2 | -0.113700093 | 0.008132817 | 0.139775831 | 0.115375821 | 0.215170158 |
| Cmc4 | -0.006305893 | 0.885899326 | 0.969590273 | 0.128627433 | 0.140883675 |
| Cmip | -0.006109219 | 0.927072987 | 0.981818446 | 0.672556558 | 0.693270428 |
| Cmpk1 | -0.106204321 | 0.116490915 | 0.488986751 | 0.769809698 | 0.902776162 |
| Cmpk2 | 0.061033163 | 0.257433293 | 0.665256942 | 0.203688441 | 0.139915803 |
| Cmss1 | -0.041585444 | 0.234520986 | 0.644900587 | 0.076663517 | 0.109414815 |
| Cmtm4 | 0.030377777 | 0.638373218 | 0.89125563 | 0.663283421 | 0.627491133 |
| Cmtm6 | -0.078234748 | 0.074820457 | 0.41344108 | 0.119166677 | 0.197898855 |
| Cmtm8 | 0.036664845 | 0.251083428 | 0.663127858 | 0.068681678 | 0.042176157 |
| Cmtr1 | 0.006673885 | 0.823026991 | 0.950930859 | 0.054370169 | 0.048079488 |
| Cmtr2 | 0.013889837 | 0.680975988 | 0.907937831 | 0.073782677 | 0.063401069 |

| | | | | | |
|---|---|---|---|---|---|
| Cnbd2 | -0.060368385 | 0.007468976 | 0.132683384 | 0.022113951 | 0.074096327 |
| Cnbp | -0.032545556 | 0.624791843 | 0.88668351 | 1.129835688 | 1.166152523 |
| Cndp2 | -0.061112989 | 0.195961587 | 0.606495975 | 0.152072986 | 0.216597209 |
| Cnep1r1 | 0.029462987 | 0.566197342 | 0.858706969 | 0.202307368 | 0.184947703 |
| Cnih1 | 0.122710579 | 0.047655463 | 0.341775906 | 0.336978462 | 0.215350427 |
| Cnih4 | 0.139596816 | 0.017501372 | 0.210625024 | 0.298948372 | 0.178337132 |
| Cnksr1 | 0.011384087 | 0.772707046 | 0.932469974 | 0.103899063 | 0.094206205 |
| Cnksr3 | 0.020226486 | 0.516344063 | 0.837427561 | 0.060961416 | 0.044750082 |
| Cnn3 | 0.181314188 | 0.008149052 | 0.139815444 | 0.732918584 | 0.545542229 |
| Cnnm1 | 0.014878514 | 0.658910678 | 0.898224427 | 0.072444488 | 0.058182006 |
| Cnnm2 | -0.033852274 | 0.392715279 | 0.769704269 | 0.104212852 | 0.132525838 |
| Cnnm3 | -0.039891183 | 0.278107044 | 0.684996416 | 0.082826657 | 0.112219015 |
| Cnot1 | 0.011548339 | 0.859802004 | 0.959220425 | 0.418573177 | 0.40015765 |
| Cnot10 | -0.035775232 | 0.515070308 | 0.836801664 | 0.227619773 | 0.260602712 |
| Cnot11 | -0.024461106 | 0.640043861 | 0.891345382 | 0.194235785 | 0.226797232 |
| Cnot2 | 0.034241977 | 0.545588304 | 0.851259637 | 0.246385601 | 0.229485258 |
| Cnot3 | 0.068174422 | 0.226800898 | 0.638720711 | 0.2616664 | 0.205119042 |
| Cnot4 | 0.02371412 | 0.703914521 | 0.912277105 | 0.364163668 | 0.346569589 |
| Cnot6 | -0.045607803 | 0.412506005 | 0.781358105 | 0.236697969 | 0.265766783 |
| Cnot6l | -0.091308642 | 0.173398227 | 0.575630234 | 0.922943747 | 1.036962035 |
| Cnot7 | 0.047919766 | 0.463022668 | 0.811092454 | 0.47966635 | 0.421947957 |
| Cnot8 | -0.007136366 | 0.842097466 | 0.955188334 | 0.086502932 | 0.089134865 |
| Cnot9 | -0.026590635 | 0.494811357 | 0.828842054 | 0.099403158 | 0.126334351 |
| Cnp | -0.029693875 | 0.224555179 | 0.637067124 | 0.032964152 | 0.055306667 |
| Cnppd1 | -0.008205495 | 0.85726971 | 0.958302663 | 0.139203456 | 0.154036776 |
| Cnpy2 | -0.05954015 | 0.337546449 | 0.732352032 | 1.16214191 | 1.223431376 |
| Cnpy3 | -0.016917681 | 0.73301994 | 0.921306823 | 0.172562773 | 0.187942001 |
| Cnpy4 | -0.003151011 | 0.924819512 | 0.981375217 | 0.068552504 | 0.068353556 |
| Cnrip1 | -0.104075685 | 0.009468758 | 0.151151979 | 0.08913205 | 0.183907317 |
| Cnst | -0.0265693 | 0.56389746 | 0.857926031 | 0.139942345 | 0.166367763 |
| Cntfr | 0.031371532 | 0.52882223 | 0.844309234 | 0.19028407 | 0.184759779 |
| Cntln | -0.15781105 | 0.002880537 | 0.074240929 | 0.179349142 | 0.323875383 |
| Cntn1 | -0.091338708 | 0.141475756 | 0.528664246 | 0.319602316 | 0.40600663 |
| Cntnap2 | -0.080345639 | 0.109433354 | 0.477972625 | 0.165689854 | 0.239027113 |
| Cntrl | -0.013604186 | 0.755919033 | 0.926726954 | 0.126339655 | 0.132715008 |
| Cntrob | -0.016563352 | 0.617636948 | 0.883238152 | 0.068673122 | 0.083618193 |
| Coa3 | 0.143715601 | 0.009487471 | 0.151151979 | 1.527296914 | 1.327010801 |
| Coa4 | -0.056373408 | 0.258552337 | 0.665948842 | 0.179315088 | 0.240957041 |
| Coa5 | -0.010641432 | 0.848985699 | 0.957111129 | 0.24921013 | 0.257364468 |
| Coa6 | 0.07383218 | 0.174889259 | 0.577992588 | 0.227694874 | 0.164907622 |
| Coa7 | 0.066458181 | 0.17268107 | 0.575294174 | 0.174232712 | 0.119469166 |
| Coa8 | -0.099879838 | 0.116408906 | 0.488986751 | 0.406900009 | 0.504172877 |
| Coasy | -0.039252108 | 0.517317576 | 0.837694133 | 0.328736304 | 0.358597496 |
| Cobl | -0.030694435 | 0.186837082 | 0.594008169 | 0.026892015 | 0.05793177 |
| Cobll1 | -0.009414132 | 0.881828171 | 0.96826489 | 0.387397319 | 0.418361549 |
| Cog1 | 0.053656097 | 0.311050527 | 0.713603232 | 0.200237668 | 0.144185544 |
| Cog2 | 0.09061911 | 0.078038419 | 0.418637956 | 0.195583023 | 0.118912204 |
| Cog3 | -0.038271007 | 0.324423593 | 0.720406991 | 0.095520236 | 0.12971587 |
| Cog4 | 0.05441568 | 0.254235908 | 0.663146569 | 0.171595747 | 0.136574494 |
| Cog5 | -0.063373851 | 0.205395414 | 0.616549498 | 0.161966909 | 0.224187975 |
| Cog6 | -0.05595894 | 0.363492863 | 0.751765374 | 0.324401384 | 0.362023794 |
| Cog7 | -0.088956332 | 0.086331641 | 0.43696958 | 0.188056012 | 0.268937091 |
| Cog8 | 0.045670113 | 0.378517426 | 0.762006446 | 0.191090012 | 0.160075804 |
| Coil | 0.018704934 | 0.634763638 | 0.889697337 | 0.104156923 | 0.088256841 |
| Col27a1 | -0.047353117 | 0.102235344 | 0.468127803 | 0.041122887 | 0.08537345 |
| Col4a1 | 0.067273285 | 0.095218459 | 0.455812225 | 0.115946714 | 0.054760787 |
| Col6a6 | -0.100104184 | 0.014955884 | 0.197256514 | 0.10025008 | 0.186186449 |
| Col9a2 | -0.050582227 | 0.002767026 | 0.072513428 | 0.00672673 | 0.052045063 |
| Colgalt1 | -0.032079356 | 0.546415531 | 0.851446616 | 0.205598687 | 0.227744055 |
| Commd1 | -0.017897511 | 0.761604208 | 0.928598328 | 0.30686244 | 0.340192855 |
| Commd10 | 0.006205033 | 0.84982394 | 0.957111129 | 0.068034742 | 0.065007356 |
| Commd2 | 0.012694644 | 0.812635852 | 0.948397417 | 0.206495852 | 0.195754728 |
| Commd3 | 0.091132028 | 0.182646146 | 0.590066231 | 0.737296744 | 0.662277808 |
| Commd4 | -0.002560227 | 0.964736517 | 0.991970325 | 0.300633827 | 0.312726347 |
| Commd6 | -0.132216803 | 0.051614303 | 0.355933715 | 0.647540329 | 0.793768369 |

| | | | | | |
|---|---|---|---|---|---|
| Commd7 | -0.097527285 | 0.136573686 | 0.520249727 | 0.481802965 | 0.560063966 |
| Commd8 | -0.000680032 | 0.989620403 | 0.9975952 | 0.207588588 | 0.214360831 |
| Commd9 | -0.120990734 | 0.045308344 | 0.333646255 | 0.308787151 | 0.423787653 |
| Comt | -0.045160793 | 0.525874902 | 0.842051614 | 0.755991827 | 0.817263238 |
| Comtd1 | 0.066477893 | 0.118633405 | 0.491192747 | 0.128651914 | 0.073913476 |
| Cop1 | -0.017393329 | 0.746230308 | 0.924566547 | 0.221932438 | 0.242310894 |
| Copa | -0.147128626 | 0.030791344 | 0.279445638 | 0.846422183 | 0.980997802 |
| Copb1 | -0.069843902 | 0.297716171 | 0.702656882 | 0.990871947 | 1.081725846 |
| Copb2 | -0.049022853 | 0.460440077 | 0.810111566 | 1.150653547 | 1.223026711 |
| Cope | -0.232218729 | 0.000639651 | 0.028919709 | 0.919386571 | 1.251102469 |
| Copg1 | -0.052085327 | 0.436885537 | 0.797968132 | 0.605225012 | 0.660869824 |
| Copg2 | -0.006546769 | 0.893472568 | 0.971676636 | 0.161456246 | 0.169837116 |
| Coprs | -0.136225556 | 0.009289818 | 0.149665973 | 0.182172459 | 0.299514119 |
| Cops2 | -0.027571538 | 0.673145742 | 0.905559579 | 0.437354976 | 0.48498767 |
| Cops3 | -0.161525287 | 0.017068682 | 0.208804661 | 0.490491084 | 0.643731947 |
| Cops4 | -0.058781851 | 0.377249448 | 0.761412185 | 0.489264249 | 0.543363983 |
| Cops5 | 0.093569235 | 0.152966428 | 0.549602359 | 0.49626925 | 0.411487944 |
| Cops6 | 0.032866468 | 0.601862216 | 0.876557788 | 0.380871342 | 0.351899768 |
| Cops7a | -0.037464463 | 0.553942359 | 0.853010042 | 0.384951423 | 0.416784072 |
| Cops7b | 0.052442954 | 0.144749882 | 0.533279057 | 0.091773338 | 0.044273973 |
| Cops8 | 0.013937765 | 0.818763835 | 0.950128492 | 0.351779448 | 0.345048945 |
| Cops9 | 0.164705699 | 0.00612213 | 0.117043465 | 1.17896375 | 0.950182705 |
| Copz1 | -0.067111083 | 0.30685259 | 0.709680069 | 0.436303481 | 0.502026918 |
| Copz2 | -0.049211758 | 0.429936593 | 0.793263728 | 0.389652249 | 0.436187185 |
| Coq10a | 0.0732822 | 0.217321194 | 0.631880888 | 0.343073714 | 0.258822201 |
| Coq10b | 0.042603066 | 0.496814623 | 0.830056214 | 0.357164224 | 0.315390642 |
| Coq2 | -0.119507047 | 0.036923306 | 0.303216817 | 0.247583303 | 0.355904974 |
| Coq3 | 0.008886689 | 0.789735453 | 0.938440574 | 0.07562117 | 0.074003344 |
| Coq4 | -0.062590536 | 0.06217629 | 0.380250374 | 0.065850612 | 0.119207721 |
| Coq5 | 0.006794485 | 0.901662942 | 0.974269668 | 0.214286608 | 0.210160877 |
| Coq6 | -0.100265586 | 0.000778253 | 0.032691088 | 0.046797291 | 0.131295861 |
| Coq7 | -0.062791967 | 0.236471578 | 0.647073398 | 0.205564177 | 0.262012401 |
| Coq8a | -0.02306351 | 0.480774071 | 0.820389213 | 0.065510337 | 0.086817177 |
| Coq8b | -0.01689919 | 0.711263643 | 0.914546911 | 0.138279784 | 0.165170291 |
| Coq9 | -0.069548582 | 0.214637698 | 0.629611185 | 0.216667841 | 0.281143263 |
| Coro1c | 0.033061126 | 0.562209804 | 0.857192739 | 0.265387282 | 0.248555391 |
| Coro2b | 0.138637547 | 0.02093616 | 0.229383208 | 0.321419465 | 0.190664895 |
| Coro7 | 0.031218324 | 0.383848432 | 0.764270326 | 0.086621717 | 0.059596631 |
| Cotl1 | 0.132372684 | 0.026913482 | 0.261501085 | 0.313003019 | 0.192371813 |
| Cox10 | 0.004145379 | 0.919785643 | 0.980436936 | 0.112868615 | 0.109722859 |
| Cox11 | 0.049723695 | 0.152819535 | 0.549602359 | 0.084310234 | 0.040447386 |
| Cox14 | 0.027083167 | 0.641955933 | 0.892178303 | 1.223144932 | 1.179608368 |
| Cox15 | 0.007244786 | 0.841087886 | 0.954960614 | 0.086621564 | 0.077952571 |
| Cox16 | 0.03124247 | 0.545988203 | 0.851405371 | 0.203103306 | 0.166943945 |
| Cox17 | 0.079512311 | 0.198394928 | 0.609389286 | 1.223210347 | 1.071882656 |
| Cox18 | -0.017397529 | 0.758094322 | 0.927247467 | 0.252231613 | 0.269350601 |
| Cox19 | 0.060040303 | 0.290861474 | 0.69667177 | 0.2609915 | 0.219060175 |
| Cox20 | 0.158325738 | 0.015890277 | 0.203432022 | 0.520859396 | 0.370887425 |
| Cox4i1 | -0.030196161 | 0.49663172 | 0.830056214 | 1.972356216 | 2.006443008 |
| Cox5a | 0.105629115 | 0.106225809 | 0.473814268 | 1.275865542 | 1.122639694 |
| Cox5b | -0.09553559 | 0.103028798 | 0.468979611 | 1.480811156 | 1.630225272 |
| Cox6a1 | -0.001199596 | 0.983708003 | 0.99608406 | 1.379753087 | 1.336706411 |
| Cox6a2 | 0.03138144 | 0.717789164 | 0.917764311 | 0.623862882 | 0.594833468 |
| Cox6b1 | -0.018612599 | 0.727540191 | 0.920455328 | 1.672678041 | 1.667259006 |
| Cox6c | 0.097119775 | 0.028808709 | 0.270430142 | 1.958532952 | 1.72891276 |
| Cox7a2 | 0.113448991 | 0.007972022 | 0.138153252 | 1.997433613 | 1.753175227 |
| Cox7a2l | -0.032514128 | 0.47412569 | 0.817387411 | 1.980414551 | 2.055719622 |
| Cox7b | 0.024973146 | 0.674873657 | 0.906025597 | 1.295109465 | 1.219890938 |
| Cox7c | 0.054338933 | 0.203630305 | 0.614950615 | 2.075012159 | 1.918617667 |
| Cox8a | -0.016591142 | 0.597709882 | 0.874521003 | 2.694221863 | 2.725897235 |
| Cpd | 0.185098991 | 0.00718413 | 0.128762694 | 0.624544648 | 0.427390196 |
| Cpe | -0.054604691 | 0.131460016 | 0.511891761 | 3.08383986 | 3.177048583 |
| Cpeb1 | 0.099700578 | 0.127326697 | 0.507536958 | 0.500688055 | 0.385985971 |
| Cpeb1os1 | -0.061358996 | 0.000499706 | 0.025103854 | 0.007230441 | 0.05929766 |
| Cpeb2 | -0.038478565 | 0.315774114 | 0.714975653 | 0.090686307 | 0.127965511 |

| | | | | | |
|---|---|---|---|---|---|
| Cpeb3 | -0.001494291 | 0.975828515 | 0.994455356 | 0.164826095 | 0.172041494 |
| Cpeb4 | -0.095152869 | 0.158676061 | 0.556789205 | 0.486459429 | 0.611249978 |
| Cplane1 | -0.085784087 | 0.07034848 | 0.402558548 | 0.147146136 | 0.219335669 |
| Cplx2 | -0.105710967 | 0.121316501 | 0.495787345 | 1.149651513 | 1.272696152 |
| Cpm | -0.066381736 | 0.103412676 | 0.469315384 | 0.104519164 | 0.160741089 |
| Cpn1 | -0.098242392 | 0.078561047 | 0.420542523 | 0.229098939 | 0.321661763 |
| Cpne3 | 0.017231349 | 0.773198935 | 0.93265205 | 0.350092386 | 0.345746918 |
| Cpne4 | -0.064582688 | 0.074855981 | 0.41344108 | 0.073801127 | 0.130994673 |
| Cpne8 | -0.042820423 | 0.169597729 | 0.57227323 | 0.054417134 | 0.092790097 |
| Cpox | -0.06075465 | 0.261926474 | 0.668764912 | 0.209530071 | 0.280205381 |
| Cpped1 | -0.047984824 | 0.169680695 | 0.57227323 | 0.071089376 | 0.121678302 |
| Cpq | -0.108597231 | 0.103222904 | 0.469315384 | 0.417494899 | 0.539357903 |
| Cpsf1 | -0.038170415 | 0.290720325 | 0.69667177 | 0.084463452 | 0.113881636 |
| Cpsf2 | -0.025368226 | 0.646386613 | 0.89474313 | 0.232046153 | 0.250815529 |
| Cpsf3 | -0.077227078 | 0.109351112 | 0.477972625 | 0.152288396 | 0.220923298 |
| Cpsf4 | 0.022927612 | 0.412689195 | 0.781358105 | 0.053033467 | 0.039005925 |
| Cpsf6 | 0.091554051 | 0.139215622 | 0.526292728 | 0.352868752 | 0.255960949 |
| Cpsf7 | -0.029620059 | 0.594330219 | 0.87313999 | 0.222980337 | 0.238394376 |
| Cpt1a | -0.00572406 | 0.887830875 | 0.970045467 | 0.111752407 | 0.110933938 |
| Cpt2 | 0.098312455 | 0.065448685 | 0.389491953 | 0.228816027 | 0.159670223 |
| Cptp | 0.049543126 | 0.103384323 | 0.469315384 | 0.062939248 | 0.027731894 |
| Cr1l | -0.191244271 | 0.000712217 | 0.031383916 | 0.221300513 | 0.405777727 |
| Cracdl | -0.050889708 | 0.1937459 | 0.60325934 | 0.096867205 | 0.139003411 |
| Cradd | -0.011868272 | 0.707991859 | 0.913767764 | 0.063669737 | 0.072531938 |
| Cramp1l | 0.012977607 | 0.743703541 | 0.924566547 | 0.104841954 | 0.095444085 |
| Crat | 0.062390586 | 0.031901626 | 0.283462667 | 0.059481538 | 0.007562972 |
| Crb3 | -0.019194049 | 0.770032928 | 0.931407628 | 0.50225065 | 0.53153902 |
| Crbn | -0.016476171 | 0.748147355 | 0.924930563 | 0.197992626 | 0.213692689 |
| Crcp | -0.043688957 | 0.365342752 | 0.752802229 | 0.145716155 | 0.183976114 |
| Creb1 | -0.082945294 | 0.179080493 | 0.585380409 | 0.322252827 | 0.410140306 |
| Creb3 | -0.135631999 | 0.037776341 | 0.306840872 | 0.406675213 | 0.534448609 |
| Creb3l1 | 0.040567201 | 0.514142125 | 0.83659041 | 0.401401012 | 0.366437245 |
| Creb3l2 | -0.113928881 | 0.077805936 | 0.418285047 | 0.456421384 | 0.573729697 |
| Crebbp | 0.010435527 | 0.873342267 | 0.964940674 | 0.485694642 | 0.472789361 |
| Crebl2 | 0.020361052 | 0.68951906 | 0.909796728 | 0.176483064 | 0.157481407 |
| Crebrf | -0.149213429 | 0.018158694 | 0.21492785 | 0.338660339 | 0.485602893 |
| Crebzf | -0.075846191 | 0.161523858 | 0.559811795 | 0.218229324 | 0.294779676 |
| Creg1 | -0.13394091 | 0.02538222 | 0.2537682 | 1.382725955 | 1.634291187 |
| Creld1 | -0.233041808 | 1.09E-07 | 3.66E-05 | 0.110633354 | 0.320944246 |
| Creld2 | 0.183237708 | 0.008840252 | 0.146996602 | 0.676440953 | 0.486057882 |
| Crem | -0.128771151 | 0.062652068 | 0.381808623 | 0.432194541 | 0.554400402 |
| Crip1 | -0.131310843 | 0.059354238 | 0.375386569 | 0.512655685 | 0.620384291 |
| Crip2 | 0.029980471 | 0.305752576 | 0.709411642 | 0.05406139 | 0.031415433 |
| Cript | -0.018041566 | 0.78455996 | 0.936423418 | 0.543959026 | 0.55543081 |
| Crk | 0.136961253 | 0.021828551 | 0.235078507 | 0.321662451 | 0.191276335 |
| Crkl | 0.031098588 | 0.537618413 | 0.847793898 | 0.177878012 | 0.147666774 |
| Crlf1 | -0.151122839 | 0.001328933 | 0.046438985 | 0.130434386 | 0.269047954 |
| Crlf2 | -0.140208708 | 0.002365303 | 0.066061828 | 0.127751328 | 0.262384303 |
| Crlf3 | 0.016665405 | 0.582857374 | 0.866383905 | 0.059633575 | 0.045727085 |
| Crls1 | 0.017054115 | 0.779997423 | 0.935229775 | 0.354824003 | 0.367990744 |
| Crmp1 | 0.012866589 | 0.760759987 | 0.928245562 | 0.116128713 | 0.105693214 |
| Crnkl1 | 0.00906538 | 0.862078747 | 0.960216905 | 0.199691411 | 0.193900569 |
| Crocc | 0.008244519 | 0.796244374 | 0.939955539 | 0.068893526 | 0.063141829 |
| Crot | 0.030284171 | 0.583296478 | 0.86669826 | 0.243662048 | 0.213133728 |
| Crtap | -0.024760653 | 0.484360231 | 0.823010604 | 0.077532038 | 0.09775559 |
| Crtc1 | 0.010988477 | 0.788873802 | 0.938142932 | 0.11515366 | 0.106375198 |
| Crtc2 | 0.010911122 | 0.799433184 | 0.941260986 | 0.124211962 | 0.115163648 |
| Crtc3 | 0.016649167 | 0.775983303 | 0.933654329 | 0.294260294 | 0.275022222 |
| Cry1 | 0.080496634 | 0.11457142 | 0.486032687 | 0.189033647 | 0.119959647 |
| Cry2 | -0.049541797 | 0.426749974 | 0.79105114 | 0.334091824 | 0.36396432 |
| Cryba2 | 0.017887258 | 0.763128893 | 0.928730554 | 0.241195164 | 0.217976821 |
| Crybb3 | 0.005115607 | 0.923397362 | 0.980859278 | 0.16857912 | 0.145144665 |
| Crybg1 | 0.025211934 | 0.465866583 | 0.812791331 | 0.076000073 | 0.052895076 |
| Crybg3 | 0.133424194 | 0.016723529 | 0.207483393 | 0.235182393 | 0.1200292 |
| Cryl1 | -0.035943556 | 0.379810411 | 0.762006446 | 0.109595392 | 0.133745933 |

| | | | | | |
|---|---|---|---|---|---|
| Cryz | -0.099995822 | 0.003525823 | 0.08334591 | 0.061968766 | 0.152970488 |
| Cryzl1 | 0.013382108 | 0.822626406 | 0.950930859 | 0.288830248 | 0.275499724 |
| Cryzl2 | -0.152641806 | 5.64E-05 | 0.00505496 | 0.077518503 | 0.212651347 |
| Cs | 0.042303645 | 0.423013005 | 0.78859241 | 0.206882128 | 0.168521608 |
| Csad | -0.039831482 | 0.313195553 | 0.714906079 | 0.095274571 | 0.130446853 |
| Csde1 | -0.000745695 | 0.989643858 | 0.9975952 | 1.324190091 | 1.327396119 |
| Cse1l | -0.01446107 | 0.782554045 | 0.936026395 | 0.192840526 | 0.205050343 |
| Csf1 | 0.051109463 | 0.132650433 | 0.513250014 | 0.077854182 | 0.029667109 |
| Csf2ra | -0.055768649 | 0.263761156 | 0.670246461 | 0.168133401 | 0.213961734 |
| Csgalnact2 | 0.026979572 | 0.516426432 | 0.837427561 | 0.119385993 | 0.091579119 |
| Csk | -0.088427658 | 0.070794936 | 0.403272567 | 0.156867161 | 0.233193305 |
| Csmd2 | -0.023352912 | 0.286801204 | 0.693329451 | 0.025160318 | 0.048442466 |
| Csnk1a1 | -0.003025412 | 0.953001086 | 0.988658594 | 1.677999509 | 1.699792759 |
| Csnk1d | -0.034287431 | 0.581760651 | 0.866173066 | 0.390061935 | 0.428443464 |
| Csnk1e | -0.070399261 | 0.219038641 | 0.632622127 | 0.228560565 | 0.294056966 |
| Csnk1g1 | -0.025437509 | 0.625564094 | 0.886834296 | 0.192540703 | 0.216097088 |
| Csnk1g2 | 0.047679048 | 0.45179186 | 0.806495742 | 0.396794436 | 0.349911882 |
| Csnk1g3 | 0.017437068 | 0.772382243 | 0.932469974 | 0.323889759 | 0.307348349 |
| Csnk2a1 | 0.091998715 | 0.179087768 | 0.585380409 | 0.61133504 | 0.495228501 |
| Csnk2a2 | 0.101822748 | 0.125307515 | 0.503839619 | 0.691077935 | 0.583879159 |
| Csnk2b | 0.019586451 | 0.764389382 | 0.928757959 | 0.475341933 | 0.450539156 |
| Cspg5 | -0.056876724 | 0.002431884 | 0.067057187 | 0.011267276 | 0.058796946 |
| Cspp1 | -0.009957845 | 0.888003239 | 0.970045467 | 0.626246566 | 0.599750619 |
| Csrnp1 | -0.015829874 | 0.692753148 | 0.909796728 | 0.097410739 | 0.107517025 |
| Csrp1 | 0.025671633 | 0.403441538 | 0.775884789 | 0.062380952 | 0.046906407 |
| Csrp2 | 0.059503649 | 0.161230753 | 0.559461938 | 0.123808388 | 0.07043799 |
| Cst3 | -0.047471792 | 0.277990962 | 0.684996416 | 2.16606893 | 2.251911255 |
| Cst6 | 0.002908593 | 0.941306468 | 0.986168593 | 0.105857786 | 0.102408813 |
| Cstb | 0.054401516 | 0.419872743 | 0.786284042 | 0.899777586 | 0.839206651 |
| Cstf1 | 0.040055139 | 0.296347068 | 0.702656882 | 0.100679476 | 0.064354154 |
| Cstf2 | -0.018935469 | 0.619755821 | 0.883866495 | 0.095912949 | 0.10660052 |
| Cstf2t | 0.015743725 | 0.728939425 | 0.920522999 | 0.142544006 | 0.129566214 |
| Cstf3 | -0.105340612 | 0.035278947 | 0.295084312 | 0.168011852 | 0.263067514 |
| Ctbp1 | -0.04861621 | 0.460178573 | 0.810111566 | 0.527434283 | 0.550571423 |
| Ctbp2 | 0.066846992 | 0.141246361 | 0.528664246 | 0.149576145 | 0.093423106 |
| Ctbs | -0.001784514 | 0.960762351 | 0.991375098 | 0.086720102 | 0.090826511 |
| Ctc1 | -0.000218167 | 0.993779752 | 0.998695538 | 0.048329362 | 0.050411905 |
| Ctcf | -0.066307649 | 0.334876216 | 0.73038987 | 0.485877291 | 0.568767256 |
| Ctdnep1 | 0.090016439 | 0.152264257 | 0.548949837 | 0.41602335 | 0.320486206 |
| Ctdp1 | -0.014952368 | 0.648656229 | 0.895430849 | 0.066966508 | 0.081570535 |
| Ctdsp1 | -0.026080138 | 0.598888331 | 0.875574105 | 0.161402991 | 0.179072824 |
| Ctdsp2 | 0.190536131 | 0.002420562 | 0.066938901 | 0.40879687 | 0.235616357 |
| Ctdspl | 0.12539791 | 0.054193582 | 0.362869235 | 0.523297623 | 0.394484111 |
| Ctdspl2 | 0.039758013 | 0.489107345 | 0.826256745 | 0.254513207 | 0.222579621 |
| Ctf1 | -0.070405336 | 0.054621715 | 0.363639613 | 0.074188934 | 0.136906962 |
| Ctif | 0.030618706 | 0.430219024 | 0.793295051 | 0.096917494 | 0.062144581 |
| Ctnna1 | 0.017597008 | 0.794142512 | 0.939757086 | 0.617409116 | 0.592797489 |
| Ctnnal1 | 0.092031263 | 0.161184404 | 0.559461938 | 0.36225298 | 0.269000204 |
| Ctnnb1 | -0.18821388 | 0.005583318 | 0.111475656 | 0.812020259 | 1.037949069 |
| Ctnnbip1 | -0.048336673 | 0.451356098 | 0.806495742 | 0.404101528 | 0.449932702 |
| Ctnnbl1 | 0.000594827 | 0.991225723 | 0.998287436 | 0.21924778 | 0.222167643 |
| Ctnnd1 | -0.035472246 | 0.562196273 | 0.857192739 | 0.32168259 | 0.342203721 |
| Ctnnd2 | -0.171491681 | 0.000678737 | 0.030277683 | 0.159246638 | 0.325672462 |
| Ctns | 0.035316112 | 0.266283126 | 0.673899077 | 0.070116897 | 0.042553143 |
| Ctps | 0.011381621 | 0.764995265 | 0.928757959 | 0.098486238 | 0.09499073 |
| Ctps2 | -0.104085948 | 0.021982416 | 0.23623533 | 0.127249024 | 0.218862244 |
| Ctr9 | -0.084118577 | 0.15969275 | 0.557786338 | 0.276165493 | 0.362922241 |
| Ctrb1 | 0.122701771 | 0.002200886 | 0.064525187 | 0.119866935 | 0 |
| Ctsa | -0.191829383 | 0.000555722 | 0.027208692 | 0.214160756 | 0.387222099 |
| Ctsb | 0.04216802 | 0.52643768 | 0.842199436 | 1.059888019 | 1.024633565 |
| Ctsd | -0.127489689 | 0.062089533 | 0.380250374 | 0.673898189 | 0.822198145 |
| Ctsf | -0.128081114 | 0.016554022 | 0.206965324 | 0.20753172 | 0.33951436 |
| Ctsl | -0.156964754 | 0.013613301 | 0.189247507 | 1.18458501 | 1.400809243 |
| Ctso | -0.018284486 | 0.762544125 | 0.92872278 | 0.318762111 | 0.339321056 |
| Ctsz | 0.056850183 | 0.403070129 | 0.775319066 | 0.424932212 | 0.354207051 |

| | | | | | |
|---|---|---|---|---|---|
| Cttn | 0.042569073 | 0.497932325 | 0.830722138 | 0.360038538 | 0.316753813 |
| Cttnbp2 | -0.080448856 | 0.178542462 | 0.584861057 | 0.275264145 | 0.33236294 |
| Ctu1 | 0.002408267 | 0.936257477 | 0.984517161 | 0.056267837 | 0.058765237 |
| Ctu2 | -0.052341457 | 0.155599796 | 0.552356396 | 0.081499165 | 0.127077752 |
| Ctxn1 | -0.043619681 | 0.101911361 | 0.467829632 | 0.03887867 | 0.07179416 |
| Cuedc1 | -0.058363917 | 0.335950586 | 0.730805382 | 0.285709041 | 0.340062694 |
| Cuedc2 | 0.034675106 | 0.52194514 | 0.840353444 | 0.221103781 | 0.19005861 |
| Cul1 | -0.039306857 | 0.56414326 | 0.857926031 | 0.649570808 | 0.696236512 |
| Cul2 | 0.052945844 | 0.329360112 | 0.724538308 | 0.235014573 | 0.196237199 |
| Cul3 | -0.089130328 | 0.188934026 | 0.596946223 | 0.846526668 | 0.968504738 |
| Cul4a | -0.018329595 | 0.752211831 | 0.926186851 | 0.274816404 | 0.284592506 |
| Cul4b | -0.052455896 | 0.308906077 | 0.711182121 | 0.185694573 | 0.225400433 |
| Cul5 | 0.039804005 | 0.551082836 | 0.852505562 | 0.524230263 | 0.494657312 |
| Cul7 | -0.029621607 | 0.457389834 | 0.808898849 | 0.099533345 | 0.133768395 |
| Cul9 | 0.018175606 | 0.575141927 | 0.862723699 | 0.069977853 | 0.055760039 |
| Cuta | -0.026295535 | 0.696180031 | 0.910970095 | 0.771356385 | 0.79223159 |
| Cutc | -0.01628006 | 0.715623877 | 0.916513571 | 0.129458599 | 0.151310598 |
| Cux1 | -0.065635879 | 0.226775283 | 0.638720711 | 0.21547781 | 0.268636485 |
| Cux2 | 0.05441409 | 0.15566613 | 0.552356396 | 0.101776049 | 0.048022298 |
| Cwc15 | -0.130978315 | 0.049305642 | 0.348262299 | 0.54987286 | 0.718749444 |
| Cwc22 | -0.01686265 | 0.709867925 | 0.914392244 | 0.134369618 | 0.146352123 |
| Cwc25 | 0.008416986 | 0.883638255 | 0.968874499 | 0.260821364 | 0.241354799 |
| Cwc27 | -0.048510834 | 0.316464286 | 0.715693767 | 0.159930397 | 0.20449628 |
| Cwf19l1 | -0.015565858 | 0.560058773 | 0.856904336 | 0.040640106 | 0.060765282 |
| Cwf19l2 | 0.02972936 | 0.626755627 | 0.887020055 | 0.34316196 | 0.315383188 |
| Cxadr | 0.0679976 | 0.202379483 | 0.612939912 | 0.213637869 | 0.150273338 |
| Cxcl16 | 0.080243783 | 0.053433361 | 0.360908913 | 0.118167515 | 0.048686651 |
| Cxxc1 | -0.038678687 | 0.40643198 | 0.777612997 | 0.148257587 | 0.187151287 |
| Cxxc4 | 0.11268111 | 0.124527333 | 0.502565678 | 0.689877833 | 0.549891192 |
| Cxxc5 | -0.031902391 | 0.634648154 | 0.88965971 | 0.634826766 | 0.680249149 |
| Cyb561 | -0.209366268 | 0.001088871 | 0.040180149 | 0.376825078 | 0.594795411 |
| Cyb561a3 | 0.026842969 | 0.337977641 | 0.732352032 | 0.051442911 | 0.028220241 |
| Cyb561d1 | 0.014975714 | 0.649157699 | 0.895557756 | 0.072438036 | 0.059445219 |
| Cyb561d2 | -0.059711746 | 0.09959283 | 0.463642501 | 0.079389796 | 0.136371355 |
| Cyb5a | -0.171118279 | 0.006237388 | 0.118794421 | 0.360736519 | 0.533081626 |
| Cyb5b | 0.035852953 | 0.608056795 | 0.87877121 | 0.630696728 | 0.59671146 |
| Cyb5d2 | -0.024633221 | 0.463929071 | 0.811254677 | 0.071251178 | 0.09081017 |
| Cyb5r1 | -0.386937684 | 1.78E-10 | 1.62E-07 | 0.214048896 | 0.618382898 |
| Cyb5r3 | -0.038207574 | 0.551836223 | 0.852505562 | 1.088328913 | 1.140977621 |
| Cyb5r4 | 0.009702435 | 0.866611798 | 0.962125879 | 0.268991108 | 0.25891378 |
| Cyb5rl | -0.016116816 | 0.545727055 | 0.851259637 | 0.040333403 | 0.055071563 |
| Cybc1 | -0.021241344 | 0.550176454 | 0.852505562 | 0.072445752 | 0.085823108 |
| Cyc1 | 0.098749185 | 0.154646652 | 0.551399093 | 0.652931455 | 0.539737291 |
| Cycs | 0.01014386 | 0.885488515 | 0.969590273 | 0.761264419 | 0.760353369 |
| Cyfip1 | 0.045781524 | 0.406305671 | 0.777612997 | 0.236499745 | 0.202194763 |
| Cyfip2 | 0.012020636 | 0.854828591 | 0.957723375 | 1.159274726 | 1.118358803 |
| Cyhr1 | 0.052549636 | 0.435551139 | 0.796975049 | 0.594440945 | 0.548396118 |
| Cyld | 0.02027484 | 0.743215234 | 0.924253661 | 0.372987517 | 0.356789199 |
| Cyp20a1 | -0.019220111 | 0.693105475 | 0.909848241 | 0.162461206 | 0.178116417 |
| Cyp2j6 | -0.051329946 | 0.09776351 | 0.460898239 | 0.053298364 | 0.104164802 |
| Cyp2u1 | -0.017169249 | 0.678046026 | 0.907527399 | 0.111753089 | 0.13493831 |
| Cyp4f13 | -0.054266172 | 0.090910211 | 0.445385778 | 0.057632781 | 0.101169963 |
| Cyp4f16 | -0.025231393 | 0.401116155 | 0.774740795 | 0.05438734 | 0.073831942 |
| Cyp4f17 | 0.029104976 | 0.341129685 | 0.734419946 | 0.062948913 | 0.037411086 |
| Cyp4f39 | -0.039920149 | 0.123037926 | 0.499567826 | 0.036466337 | 0.069614853 |
| Cyp4v3 | -0.107296295 | 0.01164169 | 0.171133529 | 0.114490237 | 0.207601255 |
| Cyp51 | 0.016879853 | 0.778607004 | 0.934641125 | 0.294181545 | 0.269998972 |
| Cyp7b1 | -0.046633413 | 0.074891978 | 0.41344108 | 0.03625418 | 0.077236903 |
| Cyren | 0.029377071 | 0.489816509 | 0.826459641 | 0.124739312 | 0.092699444 |
| Cyrib | -0.009618986 | 0.871223982 | 0.964535088 | 0.299332769 | 0.315089073 |
| Cystm1 | 0.173193252 | 0.012514473 | 0.178420124 | 1.072898996 | 0.887859244 |
| Cyth1 | -0.005368983 | 0.916816766 | 0.980132949 | 0.196472135 | 0.193474391 |
| Cyth2 | -0.00491125 | 0.9354326 | 0.984268477 | 0.307263107 | 0.31747792 |
| Cyth3 | -0.026489282 | 0.474286022 | 0.817387411 | 0.085723611 | 0.106447096 |
| Czib | -0.049316727 | 0.373376101 | 0.758519786 | 0.238645736 | 0.289633065 |

| | | | | | |
|---|---|---|---|---|---|
| D030056L22Rik | 0.039959028 | 0.278654148 | 0.68567092 | 0.091327523 | 0.063430968 |
| D10Wsu102e | 0.053491802 | 0.315475948 | 0.714975653 | 0.200971072 | 0.14885697 |
| D11Wsu47e | -0.031562081 | 0.27141862 | 0.67876599 | 0.045635221 | 0.073326551 |
| D130020L05Rik | -0.053267381 | 0.176924155 | 0.581653371 | 0.090316262 | 0.140445118 |
| D17H6S53E | 0.016293812 | 0.626167997 | 0.886938778 | 0.072666212 | 0.064024429 |
| D230025D16Rik | -0.092585261 | 0.013793818 | 0.189915708 | 0.078391482 | 0.163011667 |
| D230049E03Rik | -0.023471374 | 0.266182287 | 0.673899077 | 0.022136624 | 0.040473396 |
| D330023K18Rik | 0.013330356 | 0.769621299 | 0.931397082 | 0.140579018 | 0.121743987 |
| D3Ertd751e | -0.072755561 | 0.099339415 | 0.463315751 | 0.124078355 | 0.194310853 |
| D530018E20Rik | -0.062235435 | 0.00032064 | 0.018285574 | 0.006869419 | 0.058240679 |
| D5Ertd579e | 0.072210577 | 0.280791812 | 0.687895392 | 0.573418258 | 0.495823458 |
| D630024D03Rik | -0.047713561 | 0.332959186 | 0.728881491 | 0.159526671 | 0.19771049 |
| D630039A03Rik | 0.010483864 | 0.740554152 | 0.922901128 | 0.063542114 | 0.050980129 |
| D630045J12Rik | 0.02313547 | 0.392585455 | 0.769704269 | 0.049555134 | 0.033223977 |
| D6Wsu163e | -0.015252471 | 0.665554737 | 0.901006558 | 0.075816786 | 0.094987955 |
| D730003I15Rik | -0.145591592 | 0.030288062 | 0.276616272 | 0.496281875 | 0.642948899 |
| D830050J10Rik | -0.048744823 | 0.125395175 | 0.503839619 | 0.059039519 | 0.097150902 |
| D8Ertd738e | 0.063031214 | 0.312711719 | 0.714800165 | 1.412378585 | 1.32081336 |
| Daam1 | -0.077177335 | 0.271479829 | 0.67876599 | 0.650409053 | 0.71719203 |
| Dab2ip | 0.055799391 | 0.238873456 | 0.650445287 | 0.162191798 | 0.117260675 |
| Dach1 | 0.024623938 | 0.687740475 | 0.909796728 | 0.314822064 | 0.287994203 |
| Dach2 | -0.038390805 | 0.063301864 | 0.383440442 | 0.017662469 | 0.051917718 |
| Dact1 | 0.054791001 | 0.102998269 | 0.468979611 | 0.077375884 | 0.033687396 |
| Dact2 | -0.032617608 | 0.347118411 | 0.738454323 | 0.075595248 | 0.110305046 |
| Dad1 | -0.078619401 | 0.128747748 | 0.508260084 | 1.856956651 | 1.984883155 |
| Dag1 | 0.039642174 | 0.524053474 | 0.841062594 | 0.365371471 | 0.31553659 |
| Dagla | -0.029311237 | 0.415772606 | 0.783472369 | 0.0818024 | 0.101117528 |
| Daglb | -0.020171413 | 0.578042086 | 0.864673658 | 0.083990932 | 0.103284779 |
| Dalrd3 | 0.03621511 | 0.504462717 | 0.832249514 | 0.22522544 | 0.192090131 |
| Dancr | 0.008622453 | 0.783131894 | 0.936124803 | 0.064315417 | 0.057454587 |
| Dand5 | -0.012575874 | 0.686189371 | 0.909796728 | 0.059457604 | 0.071757844 |
| Dap | -0.002432895 | 0.969877754 | 0.993892166 | 1.063627364 | 1.10054254 |
| Dap3 | 0.083278612 | 0.193451816 | 0.60325934 | 0.395168982 | 0.329795337 |
| Dapk1 | -0.04234071 | 0.394079659 | 0.770629121 | 0.172997436 | 0.214543709 |
| Dapk3 | 0.048052861 | 0.414852256 | 0.783085706 | 0.291504146 | 0.24921345 |
| Dapl1 | 0.180309398 | 0.043836609 | 0.329086044 | 0.820038919 | 0.571325494 |
| Dapp1 | 0.084060436 | 0.064178214 | 0.386185094 | 0.143449854 | 0.068359525 |
| Dars | -0.077473903 | 0.168853754 | 0.57227323 | 0.223099413 | 0.292386444 |
| Dars2 | -0.027864425 | 0.193567161 | 0.60325934 | 0.022844758 | 0.047558829 |
| Daxx | -0.01608971 | 0.735305062 | 0.921955891 | 0.153951974 | 0.16866298 |
| Dazap1 | -0.004810314 | 0.938652413 | 0.985324082 | 0.390015437 | 0.398096398 |
| Dazap2 | 0.051625384 | 0.430460568 | 0.793311823 | 0.493278621 | 0.441350096 |
| Dbf4 | 0.074321461 | 0.043685745 | 0.329086044 | 0.089598324 | 0.017700417 |
| Dbi | 0.028270894 | 0.656970446 | 0.898224427 | 0.40267342 | 0.365114324 |
| Dbnl | 0.014837501 | 0.783631202 | 0.936124803 | 0.211796022 | 0.200467042 |
| Dbp | 0.176829641 | 0.009698865 | 0.152947522 | 0.37790185 | 0.214116031 |
| Dbpht2 | -0.014856773 | 0.784406882 | 0.936423418 | 0.201087937 | 0.206748494 |
| Dbr1 | -0.001391092 | 0.975791964 | 0.994455356 | 0.13797059 | 0.136627239 |
| Dbt | 0.037515367 | 0.334093473 | 0.729929513 | 0.102487322 | 0.067709061 |
| Dcaf1 | -0.104367239 | 0.07122925 | 0.404142444 | 0.229376858 | 0.329002178 |
| Dcaf10 | 0.006748749 | 0.900138181 | 0.973983939 | 0.204409838 | 0.201700967 |
| Dcaf11 | -0.095369105 | 0.128556337 | 0.508260084 | 0.312482965 | 0.40255827 |
| Dcaf12 | -0.022339388 | 0.565105496 | 0.858087314 | 0.098922204 | 0.122990286 |
| Dcaf12l1 | -0.060896905 | 0.286735806 | 0.693329451 | 0.245721387 | 0.303852197 |
| Dcaf13 | 0.018127773 | 0.755206405 | 0.926726954 | 0.254331439 | 0.234926601 |
| Dcaf15 | -0.003235356 | 0.911012862 | 0.978717991 | 0.05317002 | 0.052508596 |
| Dcaf17 | 0.003501544 | 0.944684435 | 0.987070339 | 0.181578163 | 0.175808263 |
| Dcaf5 | 0.035048276 | 0.491784224 | 0.827270742 | 0.182789876 | 0.1443307 |
| Dcaf6 | 0.041516024 | 0.466063668 | 0.812978978 | 0.258746747 | 0.212883386 |
| Dcaf7 | -0.005617587 | 0.929499376 | 0.982502239 | 0.435493548 | 0.420024022 |
| Dcaf8 | 0.022685946 | 0.735147028 | 0.921955891 | 0.588481788 | 0.56758375 |
| Dcakd | 0.000560156 | 0.985563763 | 0.996791297 | 0.060184573 | 0.061604417 |
| Dcbld1 | -0.057537546 | 0.088169746 | 0.441426313 | 0.065440216 | 0.110600734 |
| Dcdc2a | -0.083178992 | 0.108354048 | 0.477972625 | 0.17972951 | 0.273546924 |
| Dck | -0.059630226 | 0.031314862 | 0.281212368 | 0.041711312 | 0.095724102 |

| | | | | | |
|---|---|---|---|---|---|
| Dclre1a | -0.045230722 | 0.148334339 | 0.540999912 | 0.058716478 | 0.097199465 |
| Dclre1c | -0.017998972 | 0.48912918 | 0.826256745 | 0.039195269 | 0.053318097 |
| Dcp1a | -0.001067372 | 0.970913364 | 0.993892166 | 0.054511806 | 0.055511491 |
| Dcp1b | -0.029342734 | 0.4140702 | 0.782518584 | 0.080785268 | 0.105518838 |
| Dcp2 | -0.055556906 | 0.23470816 | 0.64504573 | 0.144993672 | 0.201876024 |
| Dcps | -0.036387419 | 0.324042675 | 0.720406991 | 0.086049295 | 0.121530154 |
| Dctn1 | -0.0647432 | 0.270679766 | 0.678123091 | 0.30129483 | 0.353584135 |
| Dctn2 | -0.027897628 | 0.674578474 | 0.906025597 | 0.532604419 | 0.569977856 |
| Dctn3 | -0.003148387 | 0.96186236 | 0.991441074 | 1.059965798 | 1.025521751 |
| Dctn4 | -0.068480478 | 0.263771398 | 0.670246461 | 0.321873406 | 0.379348318 |
| Dctn5 | 0.013388758 | 0.759542087 | 0.927719223 | 0.126908751 | 0.118235237 |
| Dctn6 | -0.027156658 | 0.653054023 | 0.897167155 | 0.296060994 | 0.333959834 |
| Dctpp1 | 0.036903642 | 0.503596531 | 0.832249514 | 0.255549933 | 0.228190485 |
| Dcun1d1 | -0.028145503 | 0.54419044 | 0.85118271 | 0.143307539 | 0.174775725 |
| Dcun1d2 | 0.007453935 | 0.840278391 | 0.954911647 | 0.089073933 | 0.085072939 |
| Dcun1d3 | 0.022230484 | 0.663169862 | 0.899491338 | 0.193281627 | 0.178373063 |
| Dcun1d4 | 0.013676719 | 0.714368707 | 0.915840939 | 0.091249989 | 0.080201397 |
| Dcun1d5 | -0.016654139 | 0.80311658 | 0.943517638 | 0.455122465 | 0.476778944 |
| Dcx | -0.033368449 | 0.458913471 | 0.809370658 | 0.133409944 | 0.154530613 |
| Dcxr | -0.015746159 | 0.714776956 | 0.916116931 | 0.122999564 | 0.133977618 |
| Dda1 | 0.143128046 | 0.022213273 | 0.236913658 | 0.405632582 | 0.274739782 |
| Ddb1 | -0.128372558 | 0.056766497 | 0.368804883 | 0.528850726 | 0.657514403 |
| Ddc | -0.262627111 | 0.000323607 | 0.018350512 | 0.666803636 | 1.005353538 |
| Ddhd1 | -0.048957388 | 0.276324415 | 0.683667283 | 0.136548895 | 0.170860183 |
| Ddhd2 | 0.013844597 | 0.82690121 | 0.952687014 | 0.371901959 | 0.354164854 |
| Ddi2 | 0.019801444 | 0.740381964 | 0.922901128 | 0.293342944 | 0.281901379 |
| Ddit3 | -0.043359383 | 0.592963104 | 0.872442739 | 1.252310487 | 1.272967566 |
| Ddit4 | -0.113559743 | 0.037169605 | 0.304267772 | 0.196109286 | 0.297850246 |
| Ddo | 0.013095852 | 0.650646117 | 0.895820999 | 0.053774229 | 0.041465159 |
| Ddost | -0.119351468 | 0.076782093 | 0.418199494 | 1.086877825 | 1.212765661 |
| Ddr1 | 0.021757219 | 0.732729534 | 0.921306823 | 0.363817581 | 0.326229282 |
| Ddrgk1 | -0.192174985 | 0.004828054 | 0.10205513 | 0.590938596 | 0.809967478 |
| Ddt | 0.077507054 | 0.253042638 | 0.663146569 | 0.50374863 | 0.429985309 |
| Ddx1 | 0.054545194 | 0.437135933 | 0.798116245 | 0.561568288 | 0.482856645 |
| Ddx10 | -0.011280223 | 0.798459725 | 0.940958114 | 0.131654312 | 0.145482349 |
| Ddx17 | -0.19867992 | 0.003786728 | 0.086973441 | 0.61304839 | 0.817571208 |
| Ddx18 | 0.043578192 | 0.509808763 | 0.834904701 | 0.409532303 | 0.356956485 |
| Ddx19a | 0.001141046 | 0.979845714 | 0.995540893 | 0.139646062 | 0.136075917 |
| Ddx19b | -0.038099125 | 0.350046709 | 0.7406656 | 0.110015824 | 0.145210699 |
| Ddx20 | 0.02762522 | 0.421057961 | 0.787234019 | 0.081157282 | 0.061784862 |
| Ddx21 | 0.017588335 | 0.795584036 | 0.939757086 | 0.427816002 | 0.40508728 |
| Ddx23 | 0.025980772 | 0.652068242 | 0.896672002 | 0.255092114 | 0.232618805 |
| Ddx24 | 0.033526049 | 0.604976835 | 0.87798619 | 1.11551728 | 1.079119427 |
| Ddx25 | -0.066185986 | 0.087353432 | 0.439557333 | 0.091564551 | 0.14627995 |
| Ddx27 | 0.156221026 | 0.007889573 | 0.137002848 | 0.301226075 | 0.154637494 |
| Ddx28 | 0.03685181 | 0.315750662 | 0.714975653 | 0.091786656 | 0.061292707 |
| Ddx31 | 0.033492203 | 0.496855715 | 0.830056214 | 0.185387729 | 0.15862282 |
| Ddx39a | -0.037365371 | 0.313213451 | 0.714906079 | 0.083414631 | 0.11380298 |
| Ddx39b | -0.157275996 | 0.017264761 | 0.209327864 | 0.399336546 | 0.539713153 |
| Ddx3x | 0.020788473 | 0.751267772 | 0.926186851 | 1.002142585 | 0.977395271 |
| Ddx3y | 0.042681256 | 0.533674141 | 0.84612966 | 0.626190208 | 0.592754573 |
| Ddx41 | -0.113371807 | 0.071018713 | 0.40386109 | 0.372817389 | 0.471030586 |
| Ddx42 | -0.006289264 | 0.925211972 | 0.981467853 | 0.577503235 | 0.593148087 |
| Ddx46 | -0.054517298 | 0.426946263 | 0.79105114 | 0.474949728 | 0.516601357 |
| Ddx47 | 0.012459273 | 0.830509077 | 0.95325696 | 0.284347827 | 0.265655351 |
| Ddx49 | 0.03634109 | 0.412826744 | 0.781358105 | 0.133226538 | 0.096601267 |
| Ddx5 | -0.091274146 | 0.024240146 | 0.246752889 | 2.367485809 | 2.569938453 |
| Ddx50 | -0.076006361 | 0.227849907 | 0.639451668 | 0.969909123 | 1.0664168 |
| Ddx52 | 0.100166925 | 0.0975962 | 0.460898239 | 0.331496168 | 0.254311202 |
| Ddx54 | 0.016282135 | 0.769584349 | 0.931397082 | 0.238804294 | 0.221071446 |
| Ddx55 | 0.079438309 | 0.130327073 | 0.509823306 | 0.197341544 | 0.134825499 |
| Ddx56 | -0.064052137 | 0.159970875 | 0.558431893 | 0.136191916 | 0.192925866 |
| Ddx58 | 0.082482262 | 0.138413944 | 0.524447246 | 0.228964436 | 0.141825843 |
| Ddx59 | -0.04634537 | 0.071892302 | 0.40592424 | 0.034948882 | 0.07336857 |
| Ddx6 | -0.124206901 | 0.073186436 | 0.410379808 | 0.916101624 | 1.060724209 |

| | | | | | |
|---|---|---|---|---|---|
| Ddx60 | 0.153501764 | 0.005310875 | 0.108344012 | 0.219119394 | 0.064412753 |
| Deaf1 | -0.012905552 | 0.75449985 | 0.926726954 | 0.109625999 | 0.120750366 |
| Decr1 | -0.025125848 | 0.550548395 | 0.852505562 | 0.116020185 | 0.132380801 |
| Decr2 | -0.01071225 | 0.754599368 | 0.926726954 | 0.077723251 | 0.08485787 |
| Dedd | -0.072682259 | 0.101537807 | 0.467719786 | 0.126633532 | 0.19807127 |
| Dedd2 | 0.031616273 | 0.397330546 | 0.774053847 | 0.093505806 | 0.073451737 |
| Def8 | 0.03682269 | 0.391997025 | 0.769704269 | 0.125153875 | 0.09213378 |
| Defb1 | 0.047197085 | 0.552885699 | 0.852623515 | 1.236004561 | 1.234647288 |
| Degs1 | 0.23374319 | 0.000787237 | 0.032786291 | 1.1919441 | 0.87144279 |
| Degs2 | -0.00393556 | 0.920056985 | 0.980436936 | 0.092870579 | 0.099183606 |
| Dek | 0.003755561 | 0.956325206 | 0.989397512 | 0.523132141 | 0.502204067 |
| Dele1 | 0.082264461 | 0.149438114 | 0.543502554 | 0.279517046 | 0.20355414 |
| Denn2b | 0.041152237 | 0.465639961 | 0.812791331 | 0.23724617 | 0.188268719 |
| Dennd10 | -0.144031155 | 0.004695005 | 0.10090742 | 0.162382228 | 0.299538922 |
| Dennd11 | 0.003018043 | 0.961973892 | 0.991441074 | 0.371812135 | 0.360364578 |
| Dennd1a | -0.031376917 | 0.51773674 | 0.837743618 | 0.163210909 | 0.182946674 |
| Dennd1b | 0.065167813 | 0.208074469 | 0.620020309 | 0.19589071 | 0.136737718 |
| Dennd2d | -0.032671095 | 0.361645237 | 0.750013467 | 0.078853903 | 0.104772983 |
| Dennd4a | -0.092615814 | 0.112827805 | 0.483221516 | 0.269347151 | 0.33811408 |
| Dennd4c | -0.050521105 | 0.448004004 | 0.804951652 | 0.536221597 | 0.574834729 |
| Dennd5a | 0.071670061 | 0.203717325 | 0.614950615 | 0.25733434 | 0.186168157 |
| Dennd5b | 0.001874298 | 0.973379341 | 0.994079882 | 0.242789181 | 0.24781126 |
| Dennd6a | -0.03734038 | 0.50246745 | 0.832222078 | 0.24726974 | 0.277763105 |
| Denr | 0.045506567 | 0.455912338 | 0.808152996 | 1.24607019 | 1.165858444 |
| Depdc5 | -0.024967545 | 0.417994584 | 0.784482356 | 0.055033149 | 0.077115918 |
| Depp1 | -0.0378514 | 0.426283891 | 0.790724712 | 0.122056853 | 0.178114292 |
| Deptor | -0.054148776 | 0.427672848 | 0.791545709 | 0.592312315 | 0.659662779 |
| Dera | 0.05776672 | 0.25485111 | 0.663366335 | 0.182922793 | 0.121705143 |
| Derl1 | -0.127760281 | 0.054608729 | 0.363639613 | 0.825254862 | 0.971803535 |
| Derl2 | 0.073292936 | 0.27849322 | 0.685442974 | 0.495821679 | 0.432576917 |
| Derl3 | 0.125894241 | 0.005287131 | 0.108299861 | 0.141828373 | 0.016379579 |
| Desi1 | 0.0296645 | 0.659250274 | 0.898224427 | 0.584673769 | 0.557081866 |
| Desi2 | -0.016396013 | 0.807036042 | 0.945532285 | 0.485541264 | 0.499218276 |
| Dexi | 0.02229046 | 0.64560309 | 0.894028451 | 0.165846565 | 0.143697697 |
| Dffa | -0.004257264 | 0.908182498 | 0.977744305 | 0.088474769 | 0.085354446 |
| Dffb | -0.071125484 | 0.042429754 | 0.325425723 | 0.070411212 | 0.140916132 |
| Dgat1 | 0.031807408 | 0.371491048 | 0.75760494 | 0.085047487 | 0.058748201 |
| Dgcr2 | 0.005962189 | 0.913158307 | 0.979979308 | 0.233943571 | 0.219379707 |
| Dgcr6 | -0.141305351 | 0.017268454 | 0.209327864 | 0.309829926 | 0.446944453 |
| Dgcr8 | -0.042110764 | 0.23767634 | 0.648778194 | 0.076192632 | 0.113997269 |
| Dgkb | 0.014086439 | 0.630476129 | 0.888402205 | 0.052687872 | 0.042172537 |
| Dgkd | 0.043618648 | 0.479934552 | 0.819939708 | 0.329866728 | 0.284528374 |
| Dgke | 0.034537696 | 0.565536015 | 0.858087314 | 0.281645863 | 0.244301944 |
| Dgkg | -0.033482718 | 0.540416957 | 0.849380522 | 0.207821417 | 0.2476195 |
| Dgkh | 0.115052588 | 0.054919559 | 0.364775849 | 0.30453501 | 0.187388272 |
| Dgkq | 0.01269101 | 0.689202251 | 0.909796728 | 0.063283786 | 0.053187337 |
| Dgkz | 0.051125988 | 0.346938378 | 0.738383053 | 0.216049575 | 0.172631417 |
| Dguok | -0.100422425 | 0.135887772 | 0.519431685 | 0.479083768 | 0.58732163 |
| Dhcr24 | -0.041050795 | 0.149035525 | 0.542986195 | 0.042893336 | 0.077636434 |
| Dhcr7 | 0.032119244 | 0.2580709 | 0.665948842 | 0.055731164 | 0.030902879 |
| Dhdds | -0.085306135 | 0.100015524 | 0.465178782 | 0.180118951 | 0.258875294 |
| Dhdh | -0.011573812 | 0.727811147 | 0.920494013 | 0.074317825 | 0.079499826 |
| Dhfr | -0.024468069 | 0.284265193 | 0.691925959 | 0.026907493 | 0.047933467 |
| Dhodh | -0.015933047 | 0.52119173 | 0.840083731 | 0.035770678 | 0.050255128 |
| Dhps | -0.014359446 | 0.78798419 | 0.937804093 | 0.204516862 | 0.23207767 |
| Dhrs1 | -0.006230406 | 0.849153416 | 0.957111129 | 0.069087619 | 0.073246037 |
| Dhrs13 | 0.009011889 | 0.776672808 | 0.934040758 | 0.066670803 | 0.067351109 |
| Dhrs4 | -0.111996293 | 0.029952271 | 0.274935828 | 0.184684444 | 0.277878071 |
| Dhrs7 | 0.060406281 | 0.377970222 | 0.761937562 | 0.667883151 | 0.619038153 |
| Dhrs7b | -0.003608939 | 0.933743973 | 0.983869133 | 0.126947896 | 0.123325162 |
| Dhx15 | 0.062977198 | 0.324371948 | 0.720406991 | 0.435453513 | 0.368195801 |
| Dhx16 | 0.014358194 | 0.740999846 | 0.922901128 | 0.129841564 | 0.11941144 |
| Dhx29 | 0.018432384 | 0.716411265 | 0.916579082 | 0.189948284 | 0.170583701 |
| Dhx30 | -0.003215863 | 0.955197185 | 0.989278596 | 0.258905735 | 0.283333806 |
| Dhx32 | 0.112675953 | 0.090607374 | 0.445385778 | 0.577361931 | 0.455460835 |

| | | | | | |
|---|---|---|---|---|---|
| Dhx33 | 0.001334743 | 0.96599712 | 0.992396427 | 0.060088651 | 0.057382365 |
| Dhx34 | -0.028774598 | 0.44906165 | 0.806014267 | 0.09463559 | 0.12440316 |
| Dhx36 | 0.149893809 | 0.029994493 | 0.274935828 | 0.717481216 | 0.5613375 |
| Dhx38 | -0.046028039 | 0.273534827 | 0.681426919 | 0.113953595 | 0.153167002 |
| Dhx40 | -0.098713077 | 0.100540299 | 0.46546263 | 0.316740359 | 0.416230968 |
| Dhx57 | -0.008302905 | 0.829623815 | 0.953065609 | 0.095779264 | 0.109883524 |
| Dhx8 | 0.070457993 | 0.194445403 | 0.603853652 | 0.215952168 | 0.150666306 |
| Dhx9 | -0.154637842 | 0.01162259 | 0.171133529 | 0.276753401 | 0.424203659 |
| Diablo | -0.05532175 | 0.290011642 | 0.69604181 | 0.199147668 | 0.248696886 |
| Diaph1 | 0.05534371 | 0.396380459 | 0.773119057 | 0.441078716 | 0.381264276 |
| Diaph2 | -0.038115423 | 0.510279793 | 0.834957869 | 0.240025205 | 0.271663415 |
| Dicer1 | -0.067380703 | 0.318354868 | 0.717175454 | 0.524971476 | 0.592450499 |
| Dido1 | -0.027009615 | 0.636863091 | 0.890755879 | 0.250695549 | 0.289088244 |
| Dimt1 | 0.031257244 | 0.489799185 | 0.826459641 | 0.134795762 | 0.10263488 |
| Dio1 | 0.146535613 | 0.028012242 | 0.266248935 | 0.449081806 | 0.289473606 |
| Dip2a | -0.016511986 | 0.551024775 | 0.852505562 | 0.044626303 | 0.058482522 |
| Dip2b | -0.038778083 | 0.519278047 | 0.838496482 | 0.28848401 | 0.336502276 |
| Dip2c | 0.023476827 | 0.702762263 | 0.912263947 | 0.361087115 | 0.325072583 |
| Dipk1b | 0.05462973 | 0.260134097 | 0.667410092 | 0.165305129 | 0.116470993 |
| Dipk2a | -0.062903109 | 0.186075684 | 0.592902108 | 0.141698911 | 0.198580077 |
| Diras1 | 0.049914957 | 0.141695164 | 0.529287072 | 0.081889031 | 0.035986927 |
| Dis3 | -0.068554552 | 0.069039671 | 0.399753362 | 0.086005443 | 0.145582695 |
| Dis3l | 0.008703643 | 0.784119505 | 0.936347328 | 0.063808229 | 0.056048555 |
| Dis3l2 | -0.002595927 | 0.9417175 | 0.986168593 | 0.085443571 | 0.079794978 |
| Disp2 | -0.121585192 | 0.074811481 | 0.41344108 | 0.560740991 | 0.663578507 |
| Dixdc1 | 0.043764903 | 0.319451344 | 0.717256849 | 0.133299932 | 0.094559454 |
| Dkc1 | -0.021764767 | 0.659732365 | 0.898227584 | 0.180180465 | 0.193719035 |
| Dlat | 0.019700887 | 0.748161391 | 0.924930563 | 0.334688751 | 0.299368583 |
| Dld | -0.003523142 | 0.955847878 | 0.989308832 | 0.439384344 | 0.457337422 |
| Dleu2 | -0.096549156 | 0.058666838 | 0.372682944 | 0.171083429 | 0.264619104 |
| Dlg1 | -0.066665997 | 0.225563991 | 0.638019508 | 0.214795181 | 0.261577976 |
| Dlg3 | -0.01886731 | 0.68043344 | 0.907783945 | 0.141008136 | 0.162288439 |
| Dlg4 | -0.014015178 | 0.79218395 | 0.939347252 | 0.188073512 | 0.192161016 |
| Dlg5 | 0.06545047 | 0.115766852 | 0.48800104 | 0.126839336 | 0.071646694 |
| Dlgap1 | -0.037163465 | 0.412800816 | 0.781358105 | 0.127128385 | 0.159086015 |
| Dlgap4 | 0.122303703 | 0.051346394 | 0.355933715 | 0.381613487 | 0.266717705 |
| Dlst | 0.019927654 | 0.734033205 | 0.921626545 | 0.285324253 | 0.25786189 |
| Dmac1 | -0.003735689 | 0.943842538 | 0.986918206 | 0.216977038 | 0.218578783 |
| Dmac2 | -0.055919296 | 0.132107497 | 0.513143556 | 0.084777946 | 0.130108437 |
| Dmac2l | -0.048578361 | 0.184892511 | 0.591385002 | 0.082917745 | 0.12502578 |
| Dmap1 | -0.027677883 | 0.38684816 | 0.766593284 | 0.064132146 | 0.090064673 |
| Dmpk | -0.093789003 | 0.077710266 | 0.418285047 | 0.1931202 | 0.273069166 |
| Dmrtc1a | -0.021044958 | 0.439362925 | 0.800048525 | 0.043621451 | 0.061820994 |
| Dmtf1 | 0.165947964 | 0.005636788 | 0.111811149 | 0.295712184 | 0.143967222 |
| Dmwd | 0.094927244 | 0.069103658 | 0.399753362 | 0.216373381 | 0.122832352 |
| Dmxl1 | 0.023219498 | 0.725755373 | 0.92014727 | 0.486490745 | 0.4581394 |
| Dmxl2 | -0.053793047 | 0.37498537 | 0.760042036 | 0.323572138 | 0.369853415 |
| Dnaja1 | -0.077919288 | 0.341299397 | 0.734481571 | 0.859819589 | 0.939340879 |
| Dnaja2 | -0.070197126 | 0.310713738 | 0.71348779 | 0.647878326 | 0.713910213 |
| Dnaja3 | -0.082946035 | 0.140755412 | 0.528664246 | 0.248378731 | 0.326229431 |
| Dnaja4 | 0.129778171 | 0.071595279 | 0.40513269 | 0.561598478 | 0.404768658 |
| Dnajb1 | 0.000243732 | 0.997274138 | 0.999265301 | 0.342729302 | 0.307098995 |
| Dnajb11 | -0.024105166 | 0.728257229 | 0.920519505 | 0.664493902 | 0.660485348 |
| Dnajb12 | -0.001761064 | 0.972234864 | 0.994070262 | 0.18655253 | 0.182704341 |
| Dnajb14 | 0.098840458 | 0.126439926 | 0.505886017 | 0.705152252 | 0.587932398 |
| Dnajb2 | 0.271840499 | 4.81E-05 | 0.004588036 | 0.550374475 | 0.274844825 |
| Dnajb4 | 0.015806972 | 0.77578258 | 0.933636662 | 0.222593848 | 0.215599022 |
| Dnajb6 | -0.043064567 | 0.513831048 | 0.836550101 | 0.876352569 | 0.929496874 |
| Dnajb9 | -0.114631688 | 0.112502273 | 0.483221516 | 0.952335991 | 1.107128519 |
| Dnajc1 | -0.083653888 | 0.218854206 | 0.632622127 | 0.740267697 | 0.819967761 |
| Dnajc10 | 0.035406844 | 0.581873816 | 0.866173066 | 0.417722883 | 0.377047798 |
| Dnajc11 | -0.00735798 | 0.854913406 | 0.957723375 | 0.108480388 | 0.113173546 |
| Dnajc12 | -0.032577477 | 0.636823167 | 0.890755879 | 0.561107702 | 0.599619914 |
| Dnajc13 | 0.045294806 | 0.428842454 | 0.792251095 | 0.261225587 | 0.215702174 |
| Dnajc14 | 0.081542626 | 0.146695766 | 0.536416643 | 0.265006108 | 0.196138198 |

| | | | | | |
|---|---|---|---|---|---|
| Dnajc15 | 0.014915299 | 0.820798763 | 0.950892784 | 0.43961566 | 0.40716436 |
| Dnajc16 | -0.012785058 | 0.684935781 | 0.909796728 | 0.062693298 | 0.079576819 |
| Dnajc17 | -0.041122939 | 0.324338787 | 0.720406991 | 0.111844318 | 0.15559622 |
| Dnajc18 | -0.05485642 | 0.23255289 | 0.643543798 | 0.136073109 | 0.178222917 |
| Dnajc19 | -0.014514671 | 0.825399332 | 0.952011047 | 0.636327693 | 0.645856419 |
| Dnajc2 | -0.005976353 | 0.925113061 | 0.981467853 | 0.39346519 | 0.403523275 |
| Dnajc21 | 0.101818659 | 0.128324945 | 0.508260084 | 1.133423258 | 0.988854747 |
| Dnajc24 | 0.210861715 | 0.003440033 | 0.082961956 | 0.631486436 | 0.391857468 |
| Dnajc25 | -0.121614975 | 0.003668016 | 0.085618324 | 0.103445894 | 0.222661174 |
| Dnajc27 | -0.026121097 | 0.324865738 | 0.720820254 | 0.039945112 | 0.060541065 |
| Dnajc3 | -0.133504673 | 0.022681769 | 0.238404929 | 1.481255677 | 1.683753184 |
| Dnajc30 | 0.035215901 | 0.508942175 | 0.8346818 | 0.213221564 | 0.192609808 |
| Dnajc4 | 0.06705296 | 0.223921716 | 0.63650588 | 0.258943306 | 0.199494263 |
| Dnajc5 | 0.048762644 | 0.467182106 | 0.813188482 | 0.579596373 | 0.53864589 |
| Dnajc6 | 0.040657701 | 0.347798111 | 0.739124045 | 0.135098356 | 0.099769913 |
| Dnajc7 | -0.056205749 | 0.38946361 | 0.768258286 | 0.401472383 | 0.440074029 |
| Dnajc8 | -0.068942218 | 0.299983715 | 0.70506682 | 0.570889256 | 0.653942729 |
| Dnajc9 | -0.071638937 | 0.234885467 | 0.64504573 | 0.309228006 | 0.391237872 |
| Dnal1 | -0.03088495 | 0.379214246 | 0.762006446 | 0.075126265 | 0.099721967 |
| Dnal4 | 0.041905116 | 0.451223168 | 0.806495742 | 0.229337564 | 0.192491013 |
| Dnase2a | -0.040408395 | 0.398275374 | 0.774311756 | 0.157996658 | 0.186911656 |
| Dner | -0.054792811 | 0.116671504 | 0.489152833 | 0.074005251 | 0.120828917 |
| Dnlz | -0.110052406 | 0.080071409 | 0.424327735 | 0.420997604 | 0.530504551 |
| Dnm1l | 0.141069922 | 0.022727864 | 0.238404929 | 0.385428202 | 0.247989264 |
| Dnm2 | 0.016591825 | 0.79591537 | 0.939835596 | 0.449781509 | 0.418984101 |
| Dnm3 | -0.023373368 | 0.686218199 | 0.909796728 | 0.244698259 | 0.25565873 |
| Dnmbp | 0.027037442 | 0.337511809 | 0.732352032 | 0.051808257 | 0.030879331 |
| Dnmt1 | 0.004969346 | 0.913376176 | 0.979979308 | 0.143870092 | 0.139358746 |
| Dnmt3a | 0.031671469 | 0.645269802 | 0.893813552 | 0.582250613 | 0.561528826 |
| Dnmt3b | -0.028806852 | 0.284565036 | 0.691925959 | 0.042167075 | 0.064242303 |
| Dnpep | 0.030763237 | 0.548987005 | 0.852402509 | 0.194154988 | 0.171436241 |
| Dnph1 | -0.028430914 | 0.232359093 | 0.643543798 | 0.028528179 | 0.056494342 |
| Dnttip1 | 0.055551131 | 0.335493908 | 0.730736446 | 0.260292934 | 0.213744696 |
| Dnttip2 | -0.002820749 | 0.967017064 | 0.992631446 | 0.478596643 | 0.500591711 |
| Dock10 | 0.072890192 | 0.038581366 | 0.310538229 | 0.083780475 | 0.021681417 |
| Dock11 | 0.028046378 | 0.504695706 | 0.832249514 | 0.118223583 | 0.09167488 |
| Dock5 | -0.162656732 | 2.11E-07 | 6.04E-05 | 0.046942929 | 0.186115489 |
| Dock7 | -0.101738957 | 0.037374146 | 0.304731357 | 0.158214426 | 0.244850714 |
| Dock8 | -0.026520338 | 0.293499128 | 0.69925611 | 0.034644776 | 0.058846571 |
| Dock9 | -0.000986396 | 0.981106697 | 0.995588708 | 0.117412215 | 0.119370426 |
| Dohh | 0.0879468 | 0.169194201 | 0.57227323 | 0.407434954 | 0.322941464 |
| Dolk | 0.009250528 | 0.758411404 | 0.927409267 | 0.057809825 | 0.052601773 |
| Dop1a | -0.038289533 | 0.486387013 | 0.825197169 | 0.222383813 | 0.259298374 |
| Dop1b | -0.008846238 | 0.861003821 | 0.959667007 | 0.186457275 | 0.195639651 |
| Dot1l | 0.089993885 | 0.083753372 | 0.431794059 | 0.199083423 | 0.113008789 |
| Dpagt1 | -0.013684654 | 0.71616399 | 0.916579082 | 0.092367439 | 0.106086083 |
| Dpf2 | 0.019504728 | 0.755670424 | 0.926726954 | 0.365686382 | 0.351302209 |
| Dph3 | 0.052465998 | 0.440437417 | 0.80031914 | 0.797317985 | 0.739503233 |
| Dph5 | 0.002868369 | 0.93804883 | 0.985059229 | 0.089077922 | 0.084254493 |
| Dph6 | -0.139828468 | 0.004031098 | 0.09071778 | 0.143456809 | 0.264374442 |
| Dph7 | -0.017394904 | 0.601069568 | 0.876370606 | 0.07050262 | 0.088920128 |
| Dpm1 | -0.010343151 | 0.858012717 | 0.9586857 | 0.279581074 | 0.313272899 |
| Dpm2 | -0.035985915 | 0.573670585 | 0.862168892 | 0.405578915 | 0.455664009 |
| Dpm3 | 0.117193088 | 0.020585762 | 0.227971987 | 1.832716454 | 1.611998975 |
| Dpp3 | 0.005590074 | 0.912411312 | 0.97958006 | 0.189571995 | 0.190556282 |
| Dpp4 | -0.1056746 | 0.056942805 | 0.368959821 | 0.229447979 | 0.318817302 |
| Dpp7 | 0.03448172 | 0.442125972 | 0.800580622 | 0.144581087 | 0.119603849 |
| Dpp8 | -0.051462274 | 0.356573948 | 0.745571198 | 0.243579653 | 0.284843016 |
| Dpp9 | -0.048855884 | 0.394852159 | 0.771549623 | 0.271065024 | 0.310323548 |
| Dpy19l4 | 0.053026979 | 0.242824861 | 0.653166476 | 0.149555255 | 0.100678623 |
| Dpy30 | 0.025700176 | 0.690107015 | 0.909796728 | 0.492664564 | 0.488244268 |
| Dpysl2 | 0.07738596 | 0.232096577 | 0.643345304 | 0.437766792 | 0.372341261 |
| Dr1 | -0.025016818 | 0.632159633 | 0.888980506 | 0.19216625 | 0.214528333 |
| Dram2 | -0.035925691 | 0.461985247 | 0.810513184 | 0.171265311 | 0.195898263 |
| Drap1 | -0.124308852 | 0.069420761 | 0.400472912 | 0.863405448 | 1.005323936 |

| | | | | | |
|---|---|---|---|---|---|
| Drg1 | 0.079091817 | 0.202918071 | 0.614015279 | 0.37667489 | 0.307681947 |
| Drg2 | 0.027507039 | 0.553831628 | 0.852970394 | 0.14890886 | 0.12741741 |
| Drosha | 0.007913059 | 0.838470122 | 0.954911647 | 0.103937383 | 0.093590799 |
| Dsc2 | 0.030159697 | 0.332500026 | 0.728191744 | 0.062872587 | 0.040086706 |
| Dsg2 | 0.047932683 | 0.416560902 | 0.784195194 | 0.282316723 | 0.236109101 |
| Dsp | 0.065613001 | 0.371714252 | 0.75760494 | 0.74536143 | 0.646616262 |
| Dst | -0.274656605 | 9.27E-06 | 0.00125691 | 1.314131586 | 1.747009736 |
| Dstn | 0.141601266 | 0.012686469 | 0.179596745 | 1.677694251 | 1.434594787 |
| Dstyk | 0.018039525 | 0.699289094 | 0.911319029 | 0.159544122 | 0.148711796 |
| Dtd1 | -0.054508431 | 0.318287701 | 0.717175454 | 0.218746213 | 0.265901676 |
| Dtd2 | 0.012965163 | 0.808963358 | 0.947108973 | 0.229314779 | 0.227206965 |
| Dtnb | 0.075393374 | 0.172681686 | 0.575294174 | 0.236758581 | 0.16059311 |
| Dtnbos | -0.068604323 | 0.119083039 | 0.492069355 | 0.110833199 | 0.199133613 |
| Dtnbp1 | -0.002493728 | 0.964125926 | 0.991970325 | 0.241675955 | 0.245794419 |
| Dtwd1 | 0.003995999 | 0.895127917 | 0.972229731 | 0.058114339 | 0.057149735 |
| Dtx2 | -0.035044669 | 0.269977311 | 0.677226912 | 0.060577083 | 0.092838588 |
| Dtx3 | -0.13279198 | 0.014298172 | 0.193150408 | 0.194470186 | 0.31933077 |
| Dtx3l | 0.071455357 | 0.172283465 | 0.575294174 | 0.206549944 | 0.1484944 |
| Dtymk | 0.086055621 | 0.135020344 | 0.518167175 | 0.292477861 | 0.214144931 |
| Dus1l | -0.072178513 | 0.124916688 | 0.503529639 | 0.135486892 | 0.201079693 |
| Dus2 | -0.053264747 | 0.140585932 | 0.528664246 | 0.079026335 | 0.124657006 |
| Dus3l | 0.02232517 | 0.41689062 | 0.784482356 | 0.049206487 | 0.031982503 |
| Dus4l | -0.030492507 | 0.244875298 | 0.654809396 | 0.039533586 | 0.066988421 |
| Dusp1 | -0.351837344 | 4.31E-07 | 0.000110955 | 0.372200445 | 0.780134631 |
| Dusp10 | 0.033142117 | 0.512791933 | 0.836077425 | 0.182301504 | 0.152749623 |
| Dusp11 | 0.051368786 | 0.457125716 | 0.808898849 | 0.551304927 | 0.489504896 |
| Dusp12 | 0.087550488 | 0.024137305 | 0.246431256 | 0.111978288 | 0.034101401 |
| Dusp14 | 0.024073906 | 0.531182484 | 0.845476197 | 0.101136306 | 0.081996797 |
| Dusp16 | 0.026106736 | 0.563105888 | 0.857617435 | 0.133615626 | 0.11004493 |
| Dusp18 | 0.021736341 | 0.657814267 | 0.898224427 | 0.174052207 | 0.15395719 |
| Dusp19 | 0.006676418 | 0.897313652 | 0.972761535 | 0.192678561 | 0.190033007 |
| Dusp22 | -0.007825267 | 0.874480484 | 0.964947297 | 0.173395184 | 0.168549811 |
| Dusp23 | 0.007645585 | 0.876882935 | 0.966020504 | 0.170659289 | 0.154317288 |
| Dusp26 | -0.08661532 | 0.025473742 | 0.254305979 | 0.091742025 | 0.164596254 |
| Dusp28 | -0.072498328 | 0.121754719 | 0.496977281 | 0.138925328 | 0.212631574 |
| Dusp3 | 0.22368737 | 0.000382491 | 0.020978469 | 0.385778773 | 0.172555074 |
| Dusp4 | 0.093051842 | 0.066452105 | 0.39159102 | 0.165546596 | 0.076393995 |
| Dusp5 | -0.180689347 | 0.004116883 | 0.092234709 | 0.309446747 | 0.502866775 |
| Dusp7 | 0.073681597 | 0.106470708 | 0.473900883 | 0.146317881 | 0.083555831 |
| Dusp8 | -0.031136385 | 0.603441035 | 0.876723966 | 0.274542863 | 0.296674235 |
| Dut | 0.130080831 | 0.056977954 | 0.368959821 | 0.521923343 | 0.398760511 |
| Dvl1 | -0.105717684 | 0.015166538 | 0.198211637 | 0.114748329 | 0.207466939 |
| Dvl2 | 0.015592915 | 0.629705654 | 0.888005452 | 0.068096311 | 0.058034399 |
| Dvl3 | -0.09840944 | 0.078489443 | 0.420383428 | 0.212499475 | 0.295245993 |
| Dxo | -0.017413871 | 0.626004161 | 0.886938778 | 0.083124979 | 0.097619709 |
| Dym | -0.097509969 | 0.039198081 | 0.312494944 | 0.150762131 | 0.237789557 |
| Dync1h1 | -0.179177747 | 0.009141829 | 0.1482335 | 0.695085772 | 0.899620505 |
| Dync1i2 | -0.023151886 | 0.739311293 | 0.922843014 | 0.71515603 | 0.736423282 |
| Dync1li1 | -0.084750704 | 0.163079021 | 0.562993622 | 0.300600167 | 0.393530664 |
| Dync1li2 | -0.071049218 | 0.218071885 | 0.632396893 | 0.239171462 | 0.312838118 |
| Dync2h1 | -0.090989213 | 0.115607239 | 0.487541958 | 0.245693441 | 0.328619681 |
| Dync2i1 | 0.043863169 | 0.254133747 | 0.663146569 | 0.102576507 | 0.060364941 |
| Dync2i2 | 0.027902358 | 0.457999231 | 0.809095362 | 0.097663712 | 0.071012948 |
| Dync2li1 | -0.104171396 | 0.056439337 | 0.367897032 | 0.209102614 | 0.30752843 |
| Dynll1 | 0.039172728 | 0.569174263 | 0.860620982 | 0.735695803 | 0.680954951 |
| Dynll2 | -0.010723611 | 0.860918764 | 0.959667007 | 1.430515024 | 1.43145795 |
| Dynlrb1 | -0.107285733 | 0.109447361 | 0.477972625 | 0.806244319 | 0.929682622 |
| Dynlt1a | -0.048484829 | 0.208100913 | 0.620020309 | 0.090236786 | 0.130302107 |
| Dynlt1b | 0.058030567 | 0.115407384 | 0.487313382 | 0.090642626 | 0.036069076 |
| Dynlt1f | -0.057125673 | 0.192376592 | 0.60309438 | 0.118026325 | 0.174466899 |
| Dynlt2a2 | -0.014429976 | 0.673088428 | 0.905559579 | 0.071734156 | 0.085873265 |
| Dynlt2b | 0.01660187 | 0.746850449 | 0.92465005 | 0.193129513 | 0.186797489 |
| Dynlt3 | -0.054723565 | 0.411727089 | 0.781032788 | 0.544518745 | 0.614552748 |
| Dyrk1a | 0.065829602 | 0.22743129 | 0.639441223 | 0.232478133 | 0.184728865 |
| Dyrk2 | 0.090646214 | 0.160670863 | 0.558784978 | 0.453190064 | 0.363849216 |

| | | | | | |
|---|---|---|---|---|---|
| Dyrk3 | -0.039778459 | 0.320822529 | 0.718701556 | 0.096144508 | 0.128828035 |
| Dzip1l | 0.005810279 | 0.888088851 | 0.970045467 | 0.108018942 | 0.104447755 |
| Dzip3 | -0.003319229 | 0.934345833 | 0.983960188 | 0.105239298 | 0.104904573 |
| E130102H24Rik | -0.00820907 | 0.833318561 | 0.954303562 | 0.097453234 | 0.101614643 |
| E130308A19Rik | 0.036859296 | 0.31459714 | 0.714975653 | 0.089344947 | 0.052016444 |
| E130309D02Rik | 0.016240747 | 0.679693435 | 0.907783945 | 0.102861489 | 0.089421795 |
| E130311K13Rik | -0.064011359 | 0.105353586 | 0.472434351 | 0.097681231 | 0.159567952 |
| E130317F20Rik | 0.025407489 | 0.493424982 | 0.82875113 | 0.090126296 | 0.068228914 |
| E230001N04Rik | -0.029063733 | 0.300673479 | 0.706227193 | 0.041534855 | 0.063934477 |
| E230020A03Rik | -0.07357031 | 0.124117662 | 0.501719281 | 0.151037238 | 0.207704051 |
| E2f4 | -0.052386054 | 0.331252138 | 0.727045202 | 0.208980918 | 0.272484712 |
| E2f5 | -0.073508293 | 0.15145015 | 0.546576546 | 0.177142047 | 0.240570988 |
| E2f6 | -0.098267307 | 0.027137273 | 0.262405402 | 0.123410405 | 0.206546932 |
| E330037G11Rik | -0.044527899 | 0.016536717 | 0.206965324 | 0.01185356 | 0.048915427 |
| E4f1 | 0.009739793 | 0.778705571 | 0.934641125 | 0.078909021 | 0.074244474 |
| Eaf1 | 0.142515581 | 0.015443273 | 0.199744575 | 0.308873016 | 0.175423617 |
| Eapp | -0.004725512 | 0.937785244 | 0.985021197 | 0.342044245 | 0.342251218 |
| Ears2 | -0.01168983 | 0.734603027 | 0.921732448 | 0.073550267 | 0.090390147 |
| Ebag9 | 0.034998811 | 0.513094337 | 0.836298776 | 0.217612906 | 0.190868153 |
| Ebi3 | -0.013092564 | 0.579541679 | 0.865234762 | 0.031281859 | 0.041747376 |
| Ebna1bp2 | -0.055846878 | 0.379527225 | 0.762006446 | 0.373696878 | 0.419844884 |
| Ebp | 0.108948471 | 0.062840446 | 0.382492148 | 0.283929709 | 0.182478546 |
| Ebpl | -0.156228617 | 0.002223496 | 0.064615296 | 0.177041684 | 0.316643263 |
| Ecd | -0.017115185 | 0.78373463 | 0.936135247 | 0.40721626 | 0.41967819 |
| Ece1 | -0.03599086 | 0.590643997 | 0.871005673 | 1.034908725 | 1.063024412 |
| Ece2 | -0.042544093 | 0.08357287 | 0.431740995 | 0.030790165 | 0.066956386 |
| Ech1 | 0.003477915 | 0.954124385 | 0.989066902 | 0.316783905 | 0.323893769 |
| Echdc1 | -0.015545665 | 0.692602246 | 0.909796728 | 0.101466497 | 0.115126461 |
| Echdc2 | -0.018468403 | 0.716050547 | 0.916579082 | 0.184876563 | 0.204389446 |
| Echdc3 | -0.038942444 | 0.126543039 | 0.505998346 | 0.035378376 | 0.072183675 |
| Echs1 | -0.210193655 | 0.000339411 | 0.019138566 | 0.272979637 | 0.485381917 |
| Eci1 | -0.108545117 | 0.045839677 | 0.335834191 | 0.210272937 | 0.310146501 |
| Eci2 | -0.115190605 | 0.082858793 | 0.430310873 | 0.431477846 | 0.550817252 |
| Ecpas | -0.129171491 | 0.052203844 | 0.357414722 | 0.475911574 | 0.591168792 |
| Ecsit | 0.061192807 | 0.237575268 | 0.648678718 | 0.206764422 | 0.153340442 |
| Edaradd | 0.041287419 | 0.154253764 | 0.550976877 | 0.056107312 | 0.020932688 |
| Edc3 | -0.044377836 | 0.191522727 | 0.602043906 | 0.070449326 | 0.112698025 |
| Edc4 | 0.054658865 | 0.289229023 | 0.695472148 | 0.203015146 | 0.158193065 |
| Edem1 | 0.218942573 | 0.000998353 | 0.038100628 | 0.475918803 | 0.254299751 |
| Edem2 | -0.164819466 | 0.004992527 | 0.104178779 | 0.270375601 | 0.431057284 |
| Edem3 | -0.035232816 | 0.50929915 | 0.834904701 | 0.207693658 | 0.242629311 |
| Edf1 | -0.087985783 | 0.137793096 | 0.522884425 | 1.405470667 | 1.526732918 |
| Edrf1 | -0.018644874 | 0.716031727 | 0.916579082 | 0.179429664 | 0.199706237 |
| Eea1 | 0.097437101 | 0.14118349 | 0.528664246 | 0.824748207 | 0.701040777 |
| Eed | -0.053351908 | 0.328451117 | 0.723428541 | 0.200859018 | 0.253060573 |
| Eef1a1 | -0.131455216 | 1.56E-07 | 4.73E-05 | 3.511192764 | 3.957800992 |
| Eef1akmt1 | -0.058573274 | 0.258442208 | 0.665948842 | 0.190781682 | 0.245873533 |
| Eef1akmt2 | -0.078852083 | 0.061478264 | 0.380250374 | 0.113226213 | 0.182351081 |
| Eef1akmt4 | -0.00872791 | 0.761237438 | 0.928506001 | 0.049342798 | 0.060473347 |
| Eef1aknmt | 0.016089464 | 0.605618722 | 0.87802905 | 0.064629101 | 0.052333936 |
| Eef1b2 | -0.07720545 | 0.095742655 | 0.456084021 | 1.709634595 | 1.836449544 |
| Eef1d | -0.118420323 | 0.068888028 | 0.399753362 | 0.861496963 | 1.00349289 |
| Eef1e1 | -0.012813818 | 0.783591124 | 0.936124803 | 0.153238245 | 0.163878675 |
| Eef1g | -0.158261945 | 0.017388736 | 0.210531661 | 0.97792443 | 1.181395067 |
| Eef2 | -0.156831564 | 0.000509172 | 0.025299777 | 2.039989654 | 2.384689464 |
| Eef2kmt | -0.099846278 | 0.092470121 | 0.450107955 | 0.299622283 | 0.39994589 |
| Eefsec | -0.059034496 | 0.124029586 | 0.501565251 | 0.092697695 | 0.141463835 |
| Eepd1 | 0.025639957 | 0.471139498 | 0.814612772 | 0.081424351 | 0.068783438 |
| Efcab1 | -0.077349154 | 0.248454965 | 0.659783035 | 0.575847976 | 0.630214698 |
| Efcab10 | -0.034849119 | 0.238955397 | 0.650445287 | 0.05030353 | 0.07231577 |
| Efcab14 | -0.090471787 | 0.160188775 | 0.558431893 | 0.526810056 | 0.629662205 |
| Efcab15 | 0.042188557 | 0.518181651 | 0.837916389 | 0.512611314 | 0.46823908 |
| Efcab7 | 0.02623775 | 0.355486018 | 0.745326231 | 0.052404016 | 0.030334111 |
| Efemp2 | 0.016934878 | 0.640106321 | 0.891345382 | 0.085221659 | 0.068605943 |
| Efhc2 | 0.001009339 | 0.973028808 | 0.994079882 | 0.05615456 | 0.051911394 |

| | | | | | |
|---|---|---|---|---|---|
| Efhd2 | -0.093550129 | 0.057470557 | 0.370238754 | 0.161674568 | 0.25370938 |
| Efl1 | 0.085266646 | 0.035200009 | 0.295084312 | 0.119805268 | 0.043398925 |
| Efna5 | 0.037058117 | 0.480130878 | 0.819943608 | 0.193668643 | 0.158312235 |
| Efnb1 | 0.0810307 | 0.043362935 | 0.328974889 | 0.113223542 | 0.041857112 |
| Efr3a | 0.118270719 | 0.069768575 | 0.400867525 | 0.483223604 | 0.38486191 |
| Eftud2 | 0.053760562 | 0.224553282 | 0.637067124 | 0.134664603 | 0.089242742 |
| Egf | -0.039875111 | 0.209180893 | 0.622272857 | 0.061584712 | 0.099150499 |
| Egfr | 0.12160635 | 0.003032909 | 0.076294 | 0.121741191 | 0.020588336 |
| Egln1 | 0.075832645 | 0.224097837 | 0.636826159 | 0.418654004 | 0.338433923 |
| Egln2 | -0.032452547 | 0.606989276 | 0.878576426 | 0.43138869 | 0.465202662 |
| Egr1 | -0.388883768 | 4.65E-06 | 0.00074156 | 0.892966175 | 1.446914822 |
| Ehbp1 | -0.008830986 | 0.825974427 | 0.952251932 | 0.105282313 | 0.107779626 |
| Ehd1 | -0.009046556 | 0.838448876 | 0.954911647 | 0.135222682 | 0.147951125 |
| Ehd3 | 0.012046567 | 0.707228296 | 0.913336388 | 0.068452361 | 0.055735625 |
| Ehd4 | 0.10601348 | 0.035806894 | 0.298004804 | 0.180696177 | 0.086764527 |
| Ehf | -0.016067887 | 0.576743464 | 0.863980874 | 0.045255357 | 0.056307497 |
| Ehhadh | 0.031179529 | 0.241474525 | 0.651877302 | 0.048052471 | 0.022823791 |
| Ehmt1 | 0.093252088 | 0.11132619 | 0.482253844 | 0.28389218 | 0.190520414 |
| Ehmt2 | -0.075794461 | 0.221780789 | 0.633858837 | 0.353602801 | 0.423711725 |
| Ei24 | 0.028640753 | 0.639554741 | 0.891345382 | 0.394116535 | 0.383683621 |
| Eid1 | -0.027190862 | 0.647987559 | 0.895376633 | 1.340445193 | 1.389272383 |
| Eid2 | 0.050243602 | 0.395696041 | 0.772384513 | 0.280659001 | 0.230644357 |
| Eid2b | 0.038564509 | 0.464601855 | 0.811698663 | 0.21768274 | 0.188038908 |
| Eif1 | -0.078374837 | 0.001030692 | 0.038850011 | 3.75780848 | 4.049189615 |
| Eif1a | 0.000863447 | 0.990367744 | 0.997733826 | 0.616403854 | 0.605177405 |
| Eif1ad | 0.03612763 | 0.556923026 | 0.855368999 | 0.345860054 | 0.320248217 |
| Eif1ax | 0.007588473 | 0.905578491 | 0.976167673 | 0.427713654 | 0.425011237 |
| Eif1b | -0.065222472 | 0.345424496 | 0.73792645 | 0.902534725 | 0.982540554 |
| Eif2a | -0.028101623 | 0.638766956 | 0.891336616 | 0.318370787 | 0.337719668 |
| Eif2ak1 | -0.069576736 | 0.127554833 | 0.508244485 | 0.137375882 | 0.19577106 |
| Eif2ak2 | 0.015389825 | 0.773881349 | 0.932949618 | 0.204224123 | 0.179530267 |
| Eif2ak3 | 0.019430525 | 0.69932231 | 0.911319029 | 0.181271953 | 0.167777734 |
| Eif2ak4 | -0.016532495 | 0.63877496 | 0.891336616 | 0.080659254 | 0.089178362 |
| Eif2b1 | 0.073513486 | 0.065127129 | 0.389263637 | 0.115356047 | 0.063805259 |
| Eif2b2 | -0.042544413 | 0.452081896 | 0.806552593 | 0.240045134 | 0.280216557 |
| Eif2b3 | 0.014406695 | 0.776676713 | 0.934040758 | 0.18264297 | 0.171076577 |
| Eif2b4 | 0.090259591 | 0.062917861 | 0.382731252 | 0.174795213 | 0.098581604 |
| Eif2b5 | 0.143280349 | 0.013662977 | 0.189413402 | 0.303565004 | 0.169867145 |
| Eif2d | -0.013088219 | 0.777652664 | 0.934318864 | 0.147861563 | 0.164142712 |
| Eif2s1 | 0.02719327 | 0.685533451 | 0.909796728 | 0.502355988 | 0.469203252 |
| Eif2s2 | 0.054680898 | 0.237094344 | 0.648247326 | 1.913934777 | 1.794413601 |
| Eif2s3x | -0.019732261 | 0.73953682 | 0.922843014 | 0.312873741 | 0.331048447 |
| Eif2s3y | -0.026682513 | 0.657751791 | 0.898224427 | 0.33540855 | 0.352753391 |
| Eif3a | -0.027810239 | 0.625303279 | 0.88668351 | 1.564817647 | 1.587744561 |
| Eif3b | 0.060848991 | 0.361840257 | 0.750032543 | 0.504328979 | 0.441367541 |
| Eif3c | -0.211984891 | 7.19E-05 | 0.006165423 | 1.834588551 | 2.211867694 |
| Eif3d | -0.012295123 | 0.859297495 | 0.959220425 | 0.559495424 | 0.571960756 |
| Eif3e | -0.138526 | 0.039665541 | 0.313977159 | 0.641580955 | 0.809529393 |
| Eif3f | -0.09608349 | 0.138993459 | 0.525848983 | 0.968493808 | 1.115401212 |
| Eif3g | -0.014087065 | 0.831127915 | 0.953266013 | 0.582232223 | 0.605263978 |
| Eif3h | -0.142762013 | 0.032571661 | 0.286271246 | 0.920887886 | 1.113379596 |
| Eif3i | -0.273899333 | 4.47E-05 | 0.004405397 | 0.575060062 | 0.890899144 |
| Eif3j1 | 0.133213807 | 0.047545347 | 0.341541257 | 0.734436884 | 0.565627446 |
| Eif3k | -0.12396272 | 0.059983832 | 0.378409279 | 0.767036806 | 0.912622578 |
| Eif3l | -0.042706051 | 0.492904507 | 0.828403211 | 0.32881926 | 0.379816169 |
| Eif3m | -0.094749903 | 0.173219801 | 0.575630234 | 0.781396471 | 0.925809923 |
| Eif4a1 | -0.120588994 | 0.081809485 | 0.42777546 | 1.053049375 | 1.21799604 |
| Eif4a2 | 0.058609102 | 0.323171718 | 0.720244873 | 1.473202192 | 1.349828449 |
| Eif4a3 | -0.004036673 | 0.95071487 | 0.988114084 | 0.462271169 | 0.469361823 |
| Eif4b | 0.009834213 | 0.878109109 | 0.966691068 | 0.767685924 | 0.766995603 |
| Eif4e | 0.110978307 | 0.099186039 | 0.463315751 | 0.850570541 | 0.715577265 |
| Eif4e2 | -0.0344099 | 0.548334606 | 0.852076539 | 0.264454717 | 0.303821924 |
| Eif4e3 | -0.001890733 | 0.969580539 | 0.993839856 | 0.166912758 | 0.172174983 |
| Eif4ebp1 | -0.157522543 | 0.02551421 | 0.254305979 | 0.980442926 | 1.150846813 |
| Eif4ebp2 | 0.065406483 | 0.302363125 | 0.707681683 | 0.393821929 | 0.331088593 |

| | | | | | |
|---|---|---|---|---|---|
| Eif4enif1 | 0.055615561 | 0.298391306 | 0.703205809 | 0.206832796 | 0.162339971 |
| Eif4g1 | -0.076641425 | 0.234151877 | 0.644459522 | 0.952339787 | 1.042145438 |
| Eif4g2 | -0.064232022 | 0.221260006 | 0.633858837 | 1.571955329 | 1.684669064 |
| Eif4g3 | -0.295901808 | 2.37E-05 | 0.002557803 | 0.664128419 | 1.014453033 |
| Eif4h | -0.007077299 | 0.916846042 | 0.980132949 | 0.617946769 | 0.634804761 |
| Eif5 | 0.190116105 | 0.000144481 | 0.010542562 | 1.963073683 | 1.635317438 |
| Eif5a | -0.013039307 | 0.812855531 | 0.948397417 | 1.621545645 | 1.657668589 |
| Eif5a2 | -0.000581638 | 0.986744788 | 0.997268589 | 0.07922312 | 0.083407606 |
| Eif5b | 0.128784066 | 0.058079046 | 0.37129897 | 1.258850517 | 1.031154911 |
| Eif6 | -0.114566164 | 0.115502423 | 0.487479818 | 0.746870235 | 0.875245671 |
| Eipr1 | 0.0285094 | 0.548412319 | 0.852076539 | 0.161511377 | 0.133322523 |
| Elac1 | 0.033663662 | 0.317471022 | 0.716863138 | 0.077232023 | 0.051155027 |
| Elac2 | 0.05402541 | 0.121049577 | 0.495099678 | 0.086079991 | 0.042400023 |
| Elapor1 | -0.162320807 | 0.017434924 | 0.210563469 | 0.571871892 | 0.782362521 |
| Elapor2 | 0.081090803 | 0.222477979 | 0.634095002 | 0.496829577 | 0.409376614 |
| Elavl1 | -0.037065984 | 0.543058572 | 0.850472599 | 0.384993249 | 0.426576533 |
| Elavl4 | -0.121551726 | 0.061935748 | 0.380250374 | 0.389639055 | 0.507250151 |
| Elf1 | -0.101302114 | 0.120943387 | 0.495048299 | 0.39220601 | 0.493405788 |
| Elf2 | -0.005980412 | 0.924860833 | 0.981375217 | 0.381426678 | 0.38313444 |
| Elf3 | -0.057078192 | 0.195012272 | 0.604676607 | 0.116836331 | 0.161959565 |
| Elk1 | -0.02057577 | 0.371877804 | 0.75760494 | 0.028090926 | 0.047951892 |
| Elk3 | -0.077216088 | 0.114850706 | 0.486793251 | 0.154727534 | 0.2234662 |
| Elk4 | -0.019042976 | 0.631707033 | 0.88887544 | 0.100074378 | 0.116980518 |
| Ell | 0.075945528 | 0.085498313 | 0.435299206 | 0.134245876 | 0.071308689 |
| Ell2 | 0.004556549 | 0.947724898 | 0.987651726 | 0.501917753 | 0.49558883 |
| Elmo1 | -0.013767017 | 0.801502687 | 0.942552134 | 0.221990068 | 0.23111276 |
| Elmo2 | 0.024156255 | 0.433414494 | 0.795625782 | 0.059985558 | 0.043342924 |
| Elmo3 | -0.042831512 | 0.28709086 | 0.693509257 | 0.104378039 | 0.139546776 |
| Elmod1 | -0.000594433 | 0.98626814 | 0.997196869 | 0.076599607 | 0.076117986 |
| Elmod2 | -0.01498459 | 0.699914282 | 0.911514162 | 0.091950341 | 0.110549051 |
| Elmod3 | 0.030336936 | 0.301063565 | 0.706666217 | 0.056713974 | 0.028103389 |
| Eloa | 0.056064465 | 0.368353263 | 0.755518937 | 0.344141522 | 0.291589765 |
| Elob | -0.063742856 | 0.22826858 | 0.639623601 | 1.571152908 | 1.667816475 |
| Eloc | -0.021786773 | 0.73166832 | 0.920770724 | 0.883562767 | 0.928384432 |
| Elof1 | -0.1121508 | 0.108516578 | 0.477972625 | 0.666972636 | 0.748076823 |
| Elovl1 | -0.023335998 | 0.563257361 | 0.857617435 | 0.106341764 | 0.119111892 |
| Elovl2 | 0.005325892 | 0.901585929 | 0.974269668 | 0.118734068 | 0.113924019 |
| Elovl4 | -0.031461237 | 0.315648765 | 0.714975653 | 0.060106073 | 0.089914467 |
| Elovl5 | 0.04028938 | 0.50478477 | 0.832249514 | 0.317856626 | 0.288805632 |
| Elovl6 | 0.008840524 | 0.860938701 | 0.959667007 | 0.172415301 | 0.172353598 |
| Elovl7 | 0.023285851 | 0.607521534 | 0.878733629 | 0.140850192 | 0.115311146 |
| Elp1 | -0.081680169 | 0.126693632 | 0.50621974 | 0.196622966 | 0.283091136 |
| Elp2 | -0.060539369 | 0.20566684 | 0.616737719 | 0.150702585 | 0.202714809 |
| Elp3 | -0.008151091 | 0.896145911 | 0.97233366 | 0.373166068 | 0.375412446 |
| Elp4 | -0.003736583 | 0.94136887 | 0.986168593 | 0.181008374 | 0.179758049 |
| Elp5 | 0.033610885 | 0.514625602 | 0.836743564 | 0.194867449 | 0.167267085 |
| Elp6 | 0.026913103 | 0.328524697 | 0.723431853 | 0.050849051 | 0.032093392 |
| Emb | -0.015030153 | 0.814441642 | 0.94873644 | 1.240075311 | 1.232759511 |
| Emc1 | 0.032235765 | 0.502464567 | 0.832222078 | 0.163873948 | 0.135309709 |
| Emc10 | -0.074797552 | 0.26550574 | 0.672779882 | 0.538354181 | 0.609011046 |
| Emc2 | -0.027161039 | 0.626933159 | 0.887105554 | 0.237347799 | 0.264924688 |
| Emc3 | 0.116764995 | 0.043989533 | 0.329446347 | 0.283035151 | 0.168964655 |
| Emc4 | 0.036209641 | 0.557721369 | 0.855384435 | 0.371593823 | 0.333300304 |
| Emc6 | 0.012163946 | 0.852175561 | 0.957461155 | 0.486230628 | 0.480400917 |
| Emc7 | 0.054432092 | 0.402014628 | 0.775220138 | 0.961632784 | 0.896411712 |
| Emc8 | -0.007533883 | 0.887949836 | 0.970045467 | 0.215319896 | 0.223732241 |
| Emc9 | 0.063741306 | 0.231591496 | 0.643122701 | 0.228272991 | 0.168234893 |
| Emd | 0.051361034 | 0.464790105 | 0.811886231 | 0.567956283 | 0.525869437 |
| Emg1 | -0.030382557 | 0.640646168 | 0.891345382 | 0.658691136 | 0.689618461 |
| Emid1 | 0.115420755 | 0.016576788 | 0.206965324 | 0.163764789 | 0.058991134 |
| Emilin1 | 0.018560416 | 0.701729759 | 0.912152193 | 0.165253145 | 0.140247121 |
| Eml1 | -0.020343602 | 0.493350745 | 0.82875113 | 0.052842806 | 0.06892635 |
| Eml2 | -0.149245343 | 0.001406162 | 0.048334428 | 0.145336302 | 0.283147069 |
| Eml3 | 0.01038951 | 0.753414256 | 0.926604446 | 0.070795354 | 0.060292449 |
| Eml4 | 0.033538423 | 0.547672316 | 0.851675805 | 0.249251163 | 0.209164567 |

| | | | | | |
|---|---|---|---|---|---|
| Eml5 | -0.040896115 | 0.539550519 | 0.84899357 | 0.496671888 | 0.515857744 |
| Eml6 | 0.00705081 | 0.816393613 | 0.949036482 | 0.059463968 | 0.048305253 |
| Emsy | 0.037866416 | 0.488122483 | 0.826256745 | 0.227468651 | 0.196729894 |
| Enah | 0.083795346 | 0.218711779 | 0.632622127 | 0.708612669 | 0.612693842 |
| Enc1 | 0.135551049 | 0.012048891 | 0.174761154 | 0.217654361 | 0.110647762 |
| Endod1 | -0.034276492 | 0.333444424 | 0.729464185 | 0.078767637 | 0.11084168 |
| Endog | -0.091050261 | 0.120877825 | 0.495002338 | 0.259685625 | 0.370043451 |
| Endov | -0.030377105 | 0.431688538 | 0.793999974 | 0.094141576 | 0.12289132 |
| Enho | -0.097909016 | 0.128193267 | 0.508260084 | 1.554955008 | 1.709493888 |
| Eno1 | 0.070345002 | 0.29238659 | 0.698069506 | 0.487871964 | 0.419734881 |
| Eno2 | -0.010288501 | 0.846260852 | 0.956199501 | 0.202403011 | 0.213445128 |
| Eno3 | -0.059913742 | 0.021592727 | 0.2337931 | 0.035179184 | 0.086517429 |
| Enoph1 | -0.02150498 | 0.479636721 | 0.819939708 | 0.057486615 | 0.074641205 |
| Enox2 | -0.035429975 | 0.233274237 | 0.643935869 | 0.04699017 | 0.08332071 |
| Enpep | -0.098849664 | 0.019153673 | 0.219709041 | 0.102355853 | 0.19313683 |
| Enpp2 | -0.169925103 | 0.013936598 | 0.191618681 | 0.531519342 | 0.694248005 |
| Enpp4 | -0.019583134 | 0.7141486 | 0.915840939 | 0.210685477 | 0.217813714 |
| Enpp5 | -0.012300336 | 0.848354121 | 0.957111129 | 0.409699046 | 0.402776181 |
| Ensa | 0.017976549 | 0.765226521 | 0.928757959 | 0.338866347 | 0.338437248 |
| ENSMUSG00000095041 | -0.112188101 | 0.093211528 | 0.45188518 | 0.553496941 | 0.660100934 |
| ENSMUSG00000115801 | -0.012958509 | 0.835386854 | 0.954911647 | 0.316948622 | 0.303098128 |
| ENSMUSG00000117901 | 0.012509774 | 0.770003557 | 0.931407628 | 0.122163368 | 0.104166573 |
| Entpd2 | 0.03213919 | 0.258233351 | 0.665948842 | 0.054603084 | 0.026402209 |
| Entpd3 | -0.204200038 | 0.00347586 | 0.083243072 | 0.632202762 | 0.855322007 |
| Entpd5 | -0.005966405 | 0.850142401 | 0.957111129 | 0.06034267 | 0.065941147 |
| Entpd6 | -0.015532375 | 0.545095123 | 0.851259637 | 0.037868936 | 0.05059301 |
| Entpd7 | 0.012275909 | 0.839803871 | 0.954911647 | 0.299994229 | 0.276468578 |
| Entr1 | 0.007828918 | 0.89224369 | 0.971118429 | 0.281274581 | 0.270766766 |
| Eny2 | -0.099298782 | 0.142524467 | 0.531198691 | 0.797364949 | 0.918539865 |
| Eola1 | 0.048302521 | 0.300799168 | 0.706227193 | 0.154539868 | 0.115121679 |
| Ep300 | 0.060488043 | 0.353943877 | 0.744256656 | 0.421968792 | 0.367784082 |
| Ep400 | -0.021420053 | 0.723073021 | 0.919483582 | 0.353935164 | 0.363641647 |
| Epb41 | 0.087804781 | 0.160235514 | 0.558431893 | 0.36337826 | 0.289629193 |
| Epb41l1 | 0.045563407 | 0.232477373 | 0.643543798 | 0.103193357 | 0.060877377 |
| Epb41l3 | -0.136170151 | 0.046785869 | 0.339117542 | 0.929252887 | 1.101479599 |
| Epb41l4a | 0.03381114 | 0.242770469 | 0.653166476 | 0.053936284 | 0.020457016 |
| Epb41l4aos | 0.049354075 | 0.310858036 | 0.71348779 | 0.166912491 | 0.122482047 |
| Epb41l4b | 0.09340039 | 0.142942656 | 0.531967163 | 0.383557488 | 0.282941237 |
| Epb41l5 | 0.083959583 | 0.107365596 | 0.47507667 | 0.207418228 | 0.126517478 |
| Epc1 | -0.090381016 | 0.120987693 | 0.495048299 | 0.278834222 | 0.358932028 |
| Epc2 | -0.105042221 | 0.070629137 | 0.402952703 | 0.240385838 | 0.333373188 |
| Epcam | 0.125920294 | 0.010276759 | 0.159021773 | 1.940137789 | 1.677463719 |
| Epg5 | -0.070390261 | 0.053469821 | 0.360912301 | 0.080213916 | 0.143470707 |
| Ephx1 | -0.101948526 | 0.01403526 | 0.191923581 | 0.095911443 | 0.195458172 |
| Ephx4 | 0.014038477 | 0.619948204 | 0.883866495 | 0.050915191 | 0.038690374 |
| Epm2aip1 | -0.002805205 | 0.965762064 | 0.992256509 | 0.407993449 | 0.408471024 |
| Epn1 | -0.085499424 | 0.187484203 | 0.594820883 | 0.511053977 | 0.577043488 |
| Epn2 | -0.026750964 | 0.545277141 | 0.851259637 | 0.126231478 | 0.14280105 |
| EPPK1 | 0.083913087 | 0.019901724 | 0.225455531 | 0.092254911 | 0.017039727 |
| Eprs | -0.262119134 | 1.91E-07 | 5.63E-05 | 1.824763349 | 2.356983254 |
| Eps15 | 0.007866551 | 0.859681301 | 0.959220425 | 0.134480364 | 0.130211994 |
| Eps15l1 | 0.029149633 | 0.474613756 | 0.817387411 | 0.115116218 | 0.088199292 |
| Eps8 | -0.008784164 | 0.893648204 | 0.971676636 | 0.391551922 | 0.400852889 |
| Eps8l2 | 0.121800483 | 0.056236792 | 0.367897032 | 0.335759374 | 0.231582614 |
| Eral1 | -0.077118991 | 0.000377208 | 0.020917314 | 0.016701103 | 0.085653228 |
| Erap1 | -0.017362713 | 0.74814509 | 0.924930563 | 0.216433731 | 0.23522623 |
| Erbin | 0.020260706 | 0.756387253 | 0.926833377 | 0.41873187 | 0.40678926 |
| Erc1 | 0.023065379 | 0.611292092 | 0.87989943 | 0.140862623 | 0.118280747 |
| Erc2 | -0.036169805 | 0.358080723 | 0.746894475 | 0.094437635 | 0.129774828 |
| Ercc1 | -0.071285881 | 0.023758852 | 0.245330129 | 0.054824861 | 0.111803559 |
| Ercc2 | -0.005765143 | 0.856887463 | 0.958302663 | 0.064824535 | 0.072953196 |
| Ercc3 | -0.016061651 | 0.678546709 | 0.907783945 | 0.095012863 | 0.10807771 |
| Ercc4 | 0.018228084 | 0.537016351 | 0.847793898 | 0.057589303 | 0.041618986 |
| Ercc5 | 0.001649037 | 0.973776368 | 0.994079882 | 0.176811988 | 0.175384227 |
| Ercc6 | 0.084138686 | 0.030971536 | 0.280493857 | 0.110604376 | 0.040064812 |

| | | | | | |
|---|---|---|---|---|---|
| Ercc6l2 | 0.074219508 | 0.16045933 | 0.558508561 | 0.208231296 | 0.137202244 |
| Ercc8 | -0.064685328 | 0.007254871 | 0.129345839 | 0.027179157 | 0.081813337 |
| Erf | 0.03691409 | 0.284009778 | 0.691894695 | 0.076205223 | 0.041759853 |
| Erg28 | -0.012966247 | 0.813206939 | 0.948397417 | 0.227810221 | 0.240359599 |
| Ergic1 | -0.054063846 | 0.428207203 | 0.791658813 | 0.499248941 | 0.576467431 |
| Ergic2 | 0.056922243 | 0.400996915 | 0.774740795 | 0.711062426 | 0.650160631 |
| Ergic3 | -0.216327079 | 0.000616059 | 0.028602902 | 0.365861694 | 0.584249948 |
| Erh | -0.013647192 | 0.835678743 | 0.954911647 | 0.388836127 | 0.421945971 |
| Eri1 | 0.008163602 | 0.858334077 | 0.958724586 | 0.142548347 | 0.129204519 |
| Eri3 | -0.01920729 | 0.688951042 | 0.909796728 | 0.1567278 | 0.163482953 |
| Erich1 | 0.118813926 | 0.024137505 | 0.246431256 | 0.210596047 | 0.107747798 |
| Erich3 | -0.008398498 | 0.847613225 | 0.956938604 | 0.124419452 | 0.129575381 |
| Erlec1 | -0.086056387 | 0.213687578 | 0.628127689 | 0.698108634 | 0.775570132 |
| Erlin1 | -0.018866584 | 0.694873356 | 0.910242368 | 0.154499997 | 0.160524809 |
| Erlin2 | -0.01280751 | 0.811156663 | 0.948397417 | 0.201266797 | 0.218265169 |
| Ermard | 0.024550751 | 0.454784928 | 0.807876964 | 0.072160326 | 0.051582379 |
| Ermp1 | 0.062541379 | 0.084477538 | 0.433265738 | 0.093048282 | 0.035650949 |
| Ern1 | -0.145151825 | 0.029644428 | 0.273726881 | 0.506050455 | 0.649266603 |
| Ero1a | 0.022969776 | 0.739434269 | 0.922843014 | 0.506594217 | 0.48285426 |
| Ero1b | 0.167274979 | 0.00154579 | 0.05174005 | 2.04176639 | 1.711960332 |
| Erp29 | -0.093865272 | 0.13515006 | 0.518330263 | 0.842446579 | 0.96466017 |
| Erp44 | 0.036827942 | 0.57565308 | 0.863006716 | 0.638717483 | 0.59512737 |
| Errfi1 | 0.040353033 | 0.537455103 | 0.847793898 | 0.349022939 | 0.299827968 |
| Esco1 | 0.044442182 | 0.451898648 | 0.806495742 | 0.295594504 | 0.265820025 |
| Esd | -0.044800016 | 0.497509324 | 0.830450871 | 0.959980022 | 1.077645422 |
| Esf1 | 0.048062973 | 0.446745328 | 0.804157614 | 0.373673062 | 0.319371576 |
| Esrp1 | -0.121205952 | 0.064954647 | 0.388652983 | 0.468600468 | 0.58191812 |
| Esrra | 0.036716432 | 0.350976861 | 0.741476479 | 0.10835176 | 0.075803031 |
| Ess2 | 0.02644922 | 0.431596897 | 0.793976916 | 0.075034943 | 0.054814757 |
| Esyt1 | 0.09291738 | 0.150266124 | 0.545074479 | 0.400626444 | 0.308022424 |
| Esyt2 | 0.081793425 | 0.178403696 | 0.584793566 | 0.355178172 | 0.263732153 |
| Etaa1 | 0.023391084 | 0.500813094 | 0.831764926 | 0.079687605 | 0.062748364 |
| Etf1 | 0.012954168 | 0.850132977 | 0.957111129 | 0.71536433 | 0.712758621 |
| Etfa | -0.086884714 | 0.141283175 | 0.528664246 | 0.281375136 | 0.36186074 |
| Etfb | -0.117869824 | 0.060728904 | 0.380221737 | 0.331289892 | 0.451499705 |
| Etfdh | -0.080724451 | 0.054413995 | 0.36337543 | 0.108258894 | 0.175040959 |
| Etfrf1 | 0.068864203 | 0.176913853 | 0.581653371 | 0.198273122 | 0.135009979 |
| Ethe1 | -0.00122752 | 0.976204437 | 0.994468977 | 0.111949821 | 0.114847983 |
| Etl4 | 0.000433962 | 0.993735538 | 0.998695538 | 0.224995497 | 0.217975877 |
| Etnk1 | -0.097727588 | 0.157068574 | 0.554909286 | 0.568909164 | 0.672607811 |
| Ets2 | 0.00415961 | 0.948537154 | 0.987651726 | 0.444887134 | 0.442040718 |
| Etv1 | -0.214817577 | 0.00160822 | 0.052831084 | 0.305808442 | 0.526550737 |
| Etv3 | 0.024484271 | 0.589599454 | 0.870394135 | 0.145014073 | 0.121544492 |
| Etv5 | -0.036016737 | 0.369484601 | 0.756685767 | 0.102930712 | 0.130329414 |
| Etv6 | 0.012229567 | 0.775674413 | 0.933636662 | 0.123540446 | 0.106981785 |
| Eva1b | -0.047901735 | 0.479885487 | 0.819939708 | 0.61996359 | 0.668259555 |
| Eva1c | -0.01479773 | 0.63837742 | 0.89125563 | 0.060631576 | 0.076708842 |
| Evi5 | -0.030069575 | 0.587048394 | 0.869181609 | 0.23355288 | 0.265985701 |
| Evi5l | 0.03244186 | 0.317462745 | 0.716863138 | 0.070678794 | 0.041175583 |
| Evl | -0.072795227 | 0.09372468 | 0.45227176 | 0.11724874 | 0.179985556 |
| Ewsr1 | -0.012338307 | 0.858108972 | 0.958686526 | 0.615645238 | 0.602445838 |
| Exd2 | -0.078682985 | 0.023371312 | 0.242332495 | 0.067413615 | 0.138753292 |
| Exo5 | 0.039974569 | 0.166882569 | 0.568374737 | 0.054247074 | 0.019380246 |
| Exoc1 | 0.053279535 | 0.200560215 | 0.610932587 | 0.121099721 | 0.083993775 |
| Exoc2 | -0.036475742 | 0.416246959 | 0.783840662 | 0.127423621 | 0.175475137 |
| Exoc3 | -0.046706362 | 0.474713047 | 0.817387411 | 0.425099701 | 0.463903307 |
| Exoc4 | 0.030974319 | 0.602017495 | 0.876569856 | 0.304480117 | 0.263008844 |
| Exoc5 | 0.045399662 | 0.473321875 | 0.816838319 | 0.435439737 | 0.409567494 |
| Exoc6 | 0.020599107 | 0.540520598 | 0.849380522 | 0.071778165 | 0.055078971 |
| Exoc6b | 0.084756048 | 0.168889771 | 0.57227323 | 0.37954615 | 0.287811797 |
| Exoc7 | -0.129620584 | 0.01658578 | 0.206965324 | 0.200026544 | 0.323439774 |
| Exoc8 | -0.019027373 | 0.606534319 | 0.878341504 | 0.084424736 | 0.104120467 |
| Exosc1 | -0.162188247 | 0.001738827 | 0.055405105 | 0.164984019 | 0.318900049 |
| Exosc10 | -0.039160725 | 0.385500552 | 0.765434033 | 0.139643614 | 0.169366249 |
| Exosc2 | -0.060106936 | 0.090651001 | 0.445385778 | 0.071438827 | 0.122710431 |

| | | | | | |
|---|---|---|---|---|---|
| Exosc3 | -0.016514674 | 0.77234246 | 0.932469974 | 0.248998264 | 0.26803367 |
| Exosc4 | 0.00283412 | 0.96573754 | 0.992256509 | 0.419692562 | 0.408586489 |
| Exosc5 | -0.168061262 | 0.000247224 | 0.015317211 | 0.118097853 | 0.271095124 |
| Exosc7 | 0.137301384 | 0.026153297 | 0.257205918 | 0.379655529 | 0.245356795 |
| Exosc8 | 0.026359392 | 0.524771217 | 0.841259975 | 0.119260869 | 0.098444169 |
| Exosc9 | 0.016181051 | 0.721792487 | 0.919475778 | 0.145471361 | 0.1387123 |
| Exph5 | -0.042340328 | 0.487829595 | 0.826112275 | 0.298739688 | 0.35062317 |
| Ext1 | 0.013684488 | 0.75536194 | 0.926726954 | 0.133860504 | 0.11399484 |
| Ext2 | -0.068224802 | 0.139433352 | 0.526719064 | 0.148888311 | 0.209075114 |
| Extl2 | 0.025127471 | 0.597185738 | 0.874263893 | 0.163042062 | 0.137430295 |
| Extl3 | -0.009336021 | 0.816417078 | 0.949036482 | 0.10938044 | 0.120891061 |
| Eya1 | 0.003019032 | 0.917666998 | 0.980166399 | 0.052602211 | 0.04767021 |
| Eya3 | 0.030492475 | 0.499064573 | 0.830722138 | 0.140703887 | 0.118199803 |
| Ezh1 | -0.040743603 | 0.470376644 | 0.814364464 | 0.237546668 | 0.264449085 |
| Ezh2 | -0.052126364 | 0.081048449 | 0.426199169 | 0.045430717 | 0.096959178 |
| Ezr | 0.130508551 | 0.015241584 | 0.198397782 | 0.223489221 | 0.115923336 |
| F11r | 0.155931442 | 0.009245179 | 0.149508372 | 0.314851217 | 0.169500223 |
| F3 | 0.062156613 | 0.14202412 | 0.53012127 | 0.124683928 | 0.077871861 |
| F8 | -0.043640137 | 0.026243183 | 0.257330576 | 0.015317641 | 0.052734253 |
| F8a | 0.019154775 | 0.741208129 | 0.922901128 | 0.291014882 | 0.282327952 |
| Faah | -0.079638673 | 0.090770158 | 0.445385778 | 0.151806884 | 0.217935489 |
| Faap100 | -0.028642415 | 0.407506222 | 0.777888922 | 0.076096751 | 0.102938914 |
| Faap20 | -0.124346514 | 0.032696127 | 0.286612247 | 0.26438768 | 0.383414341 |
| Faap24 | 0.05431523 | 0.096044837 | 0.456656576 | 0.072950597 | 0.025546485 |
| Fabp3 | -0.089982188 | 0.073502301 | 0.410998661 | 0.153568505 | 0.22503174 |
| Fabp5 | -0.08401297 | 0.011413825 | 0.16901784 | 0.058249397 | 0.130284505 |
| Fads1 | -0.0610209 | 0.081818383 | 0.42777546 | 0.073210632 | 0.128632653 |
| Fads6 | -0.042772917 | 0.345901159 | 0.738085711 | 0.131941549 | 0.173945455 |
| Faf1 | 0.071977076 | 0.235289052 | 0.645421211 | 0.348148312 | 0.276671047 |
| Faf2 | 0.109725864 | 0.099261348 | 0.463315751 | 0.567927944 | 0.481816812 |
| Fah | -0.02790407 | 0.209465769 | 0.622488392 | 0.025097817 | 0.049687078 |
| Fahd1 | -0.014206816 | 0.780905354 | 0.935482467 | 0.184704372 | 0.205213755 |
| Fahd2a | -0.143776874 | 0.00023328 | 0.014633967 | 0.087270496 | 0.213799681 |
| Faim | -0.053035085 | 0.349164372 | 0.740452736 | 0.234940976 | 0.279039944 |
| Fam102a | -0.085623761 | 0.143254823 | 0.532282155 | 0.291656348 | 0.360145491 |
| Fam104a | 0.05981729 | 0.338763893 | 0.732638051 | 1.0856393 | 0.998597276 |
| Fam110a | 0.026295187 | 0.550017881 | 0.852459771 | 0.127486659 | 0.104469047 |
| Fam110b | 0.094444087 | 0.082954997 | 0.430353379 | 0.231182752 | 0.15647855 |
| Fam114a1 | 0.092269527 | 0.133332542 | 0.514912939 | 0.359078765 | 0.259556526 |
| Fam114a2 | 0.081300844 | 0.192323238 | 0.60309438 | 0.369977153 | 0.288154944 |
| Fam117a | -0.019191764 | 0.711008848 | 0.914546911 | 0.188848154 | 0.206676409 |
| Fam117b | -0.03620307 | 0.461100248 | 0.810249429 | 0.173134822 | 0.214973205 |
| Fam118a | 0.029460214 | 0.463943593 | 0.811254677 | 0.11283056 | 0.093063986 |
| Fam118b | -0.039728445 | 0.284202757 | 0.691925959 | 0.086719804 | 0.119322614 |
| Fam120a | -0.154916009 | 0.026963627 | 0.261686913 | 0.584462166 | 0.750860568 |
| Fam120b | 0.024912603 | 0.66934382 | 0.903227201 | 0.272092421 | 0.255223187 |
| Fam120c | 0.03777314 | 0.1852709 | 0.591560356 | 0.056047875 | 0.029208936 |
| Fam122a | 0.057106779 | 0.403935947 | 0.77632362 | 0.696010566 | 0.624692333 |
| Fam126b | 0.120480569 | 0.06386089 | 0.38543852 | 0.396267964 | 0.280231003 |
| Fam131a | 0.049692453 | 0.281072509 | 0.68791143 | 0.155600195 | 0.114219947 |
| Fam133b | -0.043607909 | 0.451266617 | 0.806495742 | 0.23959776 | 0.27473201 |
| Fam135a | -0.081578969 | 0.231631819 | 0.643122701 | 0.828164103 | 0.932124833 |
| Fam135b | -0.06692627 | 0.118954424 | 0.491942957 | 0.112529846 | 0.16819155 |
| Fam136a | -0.06117363 | 0.268455097 | 0.676166273 | 0.219695559 | 0.283687013 |
| Fam13b | -0.039441851 | 0.416591489 | 0.784195194 | 0.152920436 | 0.186856059 |
| Fam149a | -0.013210852 | 0.708205514 | 0.913767764 | 0.081726881 | 0.094425499 |
| Fam149b | -0.045772301 | 0.316672112 | 0.715864412 | 0.136444918 | 0.175616985 |
| Fam151a | 0.073308607 | 0.288240316 | 0.694614177 | 0.598577205 | 0.513895505 |
| Fam155a | 0.004669786 | 0.913787564 | 0.980132949 | 0.124347082 | 0.129890305 |
| Fam160a1 | -0.016105938 | 0.577972017 | 0.864673658 | 0.050666319 | 0.063898583 |
| Fam160a2 | 0.029320528 | 0.485523388 | 0.824318763 | 0.124041337 | 0.10013777 |
| Fam160b1 | 0.055729756 | 0.34613624 | 0.738085711 | 0.302992112 | 0.260237337 |
| Fam160b2 | -0.017159945 | 0.63187469 | 0.88887544 | 0.082086584 | 0.090754543 |
| Fam161a | 0.001697691 | 0.949839456 | 0.98784658 | 0.045141487 | 0.044680826 |
| Fam162a | -0.086761031 | 0.100063846 | 0.465187968 | 0.203050412 | 0.282494615 |

| | | | | | |
|---|---|---|---|---|---|
| Fam163a | -0.025067451 | 0.381516268 | 0.762568749 | 0.047290399 | 0.06907526 |
| Fam167a | -0.030406168 | 0.144356647 | 0.533279057 | 0.020277648 | 0.044027922 |
| Fam168a | -0.009437998 | 0.864491881 | 0.961442912 | 0.240378047 | 0.247256708 |
| Fam168b | -0.054215054 | 0.275012428 | 0.68241738 | 0.164393013 | 0.208972997 |
| Fam169a | 0.002183907 | 0.961183558 | 0.991408835 | 0.140347888 | 0.134061584 |
| Fam171a1 | 0.011938245 | 0.784836752 | 0.936423418 | 0.130151649 | 0.123556345 |
| Fam171b | -0.046424056 | 0.455453565 | 0.808152996 | 0.34309491 | 0.379417923 |
| Fam172a | -0.003214644 | 0.952191666 | 0.98838165 | 0.215485298 | 0.205152661 |
| Fam174a | 0.02527217 | 0.706985902 | 0.913266781 | 0.627529055 | 0.6337897 |
| Fam174b | 0.106054666 | 0.130668668 | 0.510509367 | 0.734685862 | 0.617347187 |
| Fam174c | -0.025993821 | 0.643373568 | 0.892591387 | 0.226453331 | 0.249767552 |
| Fam177a | -0.032328666 | 0.336769771 | 0.731477644 | 0.06888927 | 0.097796031 |
| Fam183b | -0.086188176 | 0.222112069 | 0.633924544 | 0.804493535 | 0.905920122 |
| Fam184a | 0.082641153 | 0.17370047 | 0.575960231 | 0.311559207 | 0.239097746 |
| Fam185a | -0.00606325 | 0.831016235 | 0.95325696 | 0.048981062 | 0.056090927 |
| Fam187b | -0.112220209 | 0.003017311 | 0.076092349 | 0.085436521 | 0.181438952 |
| Fam189b | -0.003773429 | 0.90122013 | 0.974269668 | 0.053279231 | 0.052494291 |
| Fam193a | -0.000246603 | 0.996603605 | 0.998992348 | 0.28121782 | 0.279058157 |
| Fam193b | -0.14204203 | 0.002848623 | 0.073880191 | 0.143548981 | 0.271336558 |
| Fam199x | -0.026131586 | 0.591093905 | 0.871281744 | 0.155418749 | 0.185320848 |
| Fam204a | 0.059532068 | 0.356783482 | 0.745738403 | 0.452945356 | 0.388776227 |
| Fam207a | 0.025203253 | 0.659582907 | 0.898227584 | 0.256965543 | 0.233295433 |
| Fam20a | 0.026390931 | 0.525614245 | 0.842051614 | 0.120570886 | 0.095938327 |
| Fam20b | -0.010937822 | 0.861090073 | 0.959667007 | 0.369783119 | 0.382639143 |
| Fam20c | 0.041650238 | 0.524292554 | 0.841062594 | 0.432299154 | 0.381300681 |
| Fam210a | -0.039304888 | 0.286681339 | 0.693329451 | 0.084544949 | 0.120384563 |
| Fam210b | -0.15251504 | 0.032952981 | 0.287906455 | 0.495804705 | 0.67096449 |
| Fam214a | 0.053326359 | 0.388522801 | 0.767160379 | 0.337318886 | 0.266567431 |
| Fam214b | 0.071236387 | 0.116919691 | 0.489579867 | 0.149961056 | 0.083380662 |
| Fam216a | 0.006894852 | 0.911011444 | 0.978717991 | 0.330891895 | 0.327029195 |
| Fam217b | -0.001734958 | 0.94950442 | 0.987787714 | 0.045848676 | 0.047005345 |
| Fam219a | -0.001071723 | 0.984486652 | 0.996098037 | 0.222454177 | 0.210342799 |
| Fam219b | -0.020460464 | 0.613256152 | 0.88121002 | 0.107111424 | 0.126987963 |
| Fam220a | -0.060840866 | 0.060914472 | 0.380250374 | 0.056431331 | 0.107113569 |
| Fam220a | -0.026465476 | 0.305141655 | 0.709411642 | 0.038166872 | 0.055855763 |
| Fam222a | 0.022296702 | 0.647685063 | 0.895376633 | 0.163773416 | 0.146284648 |
| Fam222b | 0.045680227 | 0.391551923 | 0.769704269 | 0.218403782 | 0.182494442 |
| Fam229b | -0.000519196 | 0.98703967 | 0.997268589 | 0.063439244 | 0.062329086 |
| Fam234a | 0.025294149 | 0.710581828 | 0.914509508 | 0.457335876 | 0.42610344 |
| Fam234b | 0.034805663 | 0.415044758 | 0.783085706 | 0.127241289 | 0.097077709 |
| Fam241a | -0.030919085 | 0.349398883 | 0.740480699 | 0.067516066 | 0.09172537 |
| Fam241b | 0.052975057 | 0.315851554 | 0.714975653 | 0.215582398 | 0.166955227 |
| Fam32a | -0.052436579 | 0.398068176 | 0.774311756 | 0.367342621 | 0.420635275 |
| Fam3a | 0.009457006 | 0.851470247 | 0.957461155 | 0.178909602 | 0.172640022 |
| Fam3c | -0.032485797 | 0.504909223 | 0.832249514 | 0.168309435 | 0.202120142 |
| Fam50a | -0.066357745 | 0.313669395 | 0.714906079 | 0.390429727 | 0.468204684 |
| Fam53a | 0.006006909 | 0.899512787 | 0.973938494 | 0.169576426 | 0.170488857 |
| Fam53b | -0.025964416 | 0.282424501 | 0.689538973 | 0.030282099 | 0.050977615 |
| Fam53c | 0.01141847 | 0.785084967 | 0.936442895 | 0.122101402 | 0.115917608 |
| Fam71e1 | -0.049834276 | 0.194829894 | 0.60429779 | 0.09449318 | 0.137459654 |
| Fam76a | 0.076224813 | 0.250307911 | 0.662228053 | 0.552900676 | 0.470753661 |
| Fam76b | -0.055549628 | 0.274047057 | 0.681791282 | 0.175666209 | 0.222199745 |
| Fam83b | -0.035290926 | 0.078857971 | 0.421233343 | 0.016964283 | 0.048573395 |
| Fam83g | 0.118091338 | 0.038150417 | 0.308057713 | 0.248610983 | 0.141920641 |
| Fam83h | -0.011830249 | 0.724265301 | 0.919830015 | 0.071036652 | 0.087261448 |
| Fam8a1 | -0.007366626 | 0.879708548 | 0.96752517 | 0.175109644 | 0.189490261 |
| Fam91a1 | 0.007524274 | 0.908666346 | 0.978046554 | 0.422696872 | 0.399490646 |
| Fam98a | -0.045893225 | 0.445624895 | 0.803149051 | 0.307981053 | 0.348574205 |
| Fam98b | -0.070566574 | 0.234763942 | 0.64504573 | 0.259782172 | 0.330459513 |
| Fam98c | -0.118881878 | 0.01238343 | 0.177814713 | 0.139990713 | 0.245362375 |
| Fance | 0.01638805 | 0.660824358 | 0.89868852 | 0.090933253 | 0.076555251 |
| Fancf | -0.024459851 | 0.483723236 | 0.822768706 | 0.074664241 | 0.094442541 |
| Fancl | -0.071357037 | 0.001911871 | 0.059662472 | 0.022514626 | 0.082361296 |
| Fancm | 0.001156345 | 0.974993587 | 0.994455356 | 0.087955334 | 0.092071831 |
| Far1 | -0.016358542 | 0.77754984 | 0.934307164 | 0.267468124 | 0.279920451 |

| | | | | | |
|---|---|---|---|---|---|
| Farp1 | 0.044648887 | 0.367273316 | 0.754003328 | 0.176133744 | 0.130094945 |
| Farp2 | -0.009699496 | 0.717773949 | 0.917764311 | 0.041438505 | 0.050024541 |
| Fars2 | 0.001094594 | 0.984096993 | 0.99608406 | 0.21370364 | 0.216070248 |
| Farsa | -0.005173425 | 0.917382162 | 0.980132949 | 0.172945541 | 0.180451176 |
| Farsb | -0.032510097 | 0.562133778 | 0.857192739 | 0.250921316 | 0.287172748 |
| Fasn | -0.00032156 | 0.994212312 | 0.998695538 | 0.132090042 | 0.131039269 |
| Fastk | 0.070422185 | 0.216964216 | 0.631434486 | 0.271061384 | 0.203420659 |
| Fastkd1 | 0.002808595 | 0.939796852 | 0.985679047 | 0.090667475 | 0.087621858 |
| Fastkd2 | -0.047487006 | 0.141308717 | 0.528664246 | 0.062120265 | 0.103422759 |
| Fat1 | -0.05798608 | 0.119819486 | 0.493284734 | 0.08208778 | 0.138193143 |
| Fau | -0.058208997 | 0.025384405 | 0.2537682 | 3.172365171 | 3.350131904 |
| Faxc | -0.053548541 | 0.077873113 | 0.418318999 | 0.051000287 | 0.097013068 |
| Fbf1 | 0.004985462 | 0.863335534 | 0.960676137 | 0.052045919 | 0.043768761 |
| Fbh1 | -0.004859801 | 0.903057813 | 0.974950771 | 0.107811984 | 0.111724201 |
| Fbl | 0.072045092 | 0.210842833 | 0.623520777 | 0.25715795 | 0.19428454 |
| Fbp1 | -0.052945778 | 0.075490527 | 0.414292853 | 0.036396541 | 0.102310501 |
| Fbrs | -0.00101416 | 0.982135443 | 0.995827199 | 0.137968898 | 0.13560256 |
| Fbrsl1 | 0.021535136 | 0.690189846 | 0.909796728 | 0.223713794 | 0.209182682 |
| Fbxl12 | 0.031063899 | 0.494328724 | 0.828842054 | 0.144556698 | 0.119636189 |
| Fbxl14 | 0.048901928 | 0.36677074 | 0.753633555 | 0.213247454 | 0.179783483 |
| Fbxl15 | 0.044012508 | 0.410528058 | 0.780244294 | 0.210803281 | 0.164866553 |
| Fbxl16 | 0.004051429 | 0.95107638 | 0.988114084 | 0.495371417 | 0.482617809 |
| Fbxl17 | -0.02723895 | 0.615011229 | 0.882439418 | 0.21175545 | 0.233239014 |
| Fbxl18 | -0.010755338 | 0.771651271 | 0.9322553 | 0.088712544 | 0.098651017 |
| Fbxl19 | -0.034715224 | 0.129762616 | 0.509514066 | 0.026323143 | 0.053733447 |
| Fbxl2 | -0.055257132 | 0.123037316 | 0.499567826 | 0.078009748 | 0.121423048 |
| Fbxl20 | -0.007243754 | 0.843481157 | 0.95544362 | 0.087311946 | 0.0936427 |
| Fbxl3 | 0.071448972 | 0.142786556 | 0.531583332 | 0.168745488 | 0.109472705 |
| Fbxl4 | 0.051644368 | 0.151489685 | 0.546576546 | 0.090954083 | 0.046892699 |
| Fbxl5 | -0.042699986 | 0.523043086 | 0.841062594 | 0.480757428 | 0.502580698 |
| Fbxl6 | -0.030976699 | 0.379745803 | 0.762006446 | 0.076746885 | 0.109982556 |
| Fbxo10 | 0.048955367 | 0.080985078 | 0.426199169 | 0.052832113 | 0.010728959 |
| Fbxo11 | -0.04023208 | 0.487188739 | 0.82599888 | 0.282560337 | 0.326111453 |
| Fbxo16 | -0.034471688 | 0.314958602 | 0.714975653 | 0.074040401 | 0.102519263 |
| Fbxo17 | -5.31E-05 | 0.998769102 | 0.999466149 | 0.076598917 | 0.078559726 |
| Fbxo2 | -0.138558395 | 0.022743231 | 0.238404929 | 0.287098389 | 0.439943055 |
| Fbxo21 | 0.053614135 | 0.345102127 | 0.737702595 | 0.253352092 | 0.205873479 |
| Fbxo22 | -0.044420938 | 0.424879596 | 0.790344696 | 0.218708207 | 0.253888525 |
| Fbxo25 | -0.020281074 | 0.564537586 | 0.858087314 | 0.079290003 | 0.098462534 |
| Fbxo28 | 0.111864013 | 0.032624804 | 0.286486051 | 0.200654969 | 0.104544387 |
| Fbxo3 | 0.040657454 | 0.533252768 | 0.84612966 | 0.472729843 | 0.440554007 |
| Fbxo30 | -0.039724276 | 0.227884456 | 0.639451668 | 0.062604492 | 0.101047903 |
| Fbxo31 | -0.011911032 | 0.778381498 | 0.934523337 | 0.121207094 | 0.13308946 |
| Fbxo32 | -0.045453787 | 0.370803326 | 0.75760494 | 0.172565676 | 0.228783303 |
| Fbxo33 | 0.022824614 | 0.683736211 | 0.909438425 | 0.246293689 | 0.225708573 |
| Fbxo34 | 0.043208055 | 0.38804713 | 0.767160379 | 0.173003831 | 0.149937594 |
| Fbxo36 | 0.060583493 | 0.160575576 | 0.558647159 | 0.133169108 | 0.090930765 |
| Fbxo38 | 0.014664472 | 0.779825988 | 0.935139002 | 0.20677343 | 0.197814324 |
| Fbxo4 | 0.061519415 | 0.225337945 | 0.63782204 | 0.183558899 | 0.125525945 |
| Fbxo42 | -0.03551632 | 0.41157248 | 0.781032788 | 0.120372974 | 0.15478454 |
| Fbxo44 | -0.036610719 | 0.454818104 | 0.807876964 | 0.17237358 | 0.203083978 |
| Fbxo45 | 0.049824779 | 0.241303586 | 0.651739388 | 0.130805238 | 0.091479419 |
| Fbxo46 | -0.010576931 | 0.690915126 | 0.909796728 | 0.042122053 | 0.053263144 |
| Fbxo6 | 0.050424244 | 0.318583624 | 0.717175454 | 0.191498842 | 0.149437287 |
| Fbxo7 | -0.020377732 | 0.675568188 | 0.906025597 | 0.172150575 | 0.184809907 |
| Fbxo8 | -0.018862423 | 0.720778001 | 0.919412687 | 0.194088497 | 0.219777166 |
| Fbxo9 | -0.09850764 | 0.098906547 | 0.463315751 | 0.273340299 | 0.366139739 |
| Fbxw11 | 0.080593415 | 0.177155911 | 0.582224582 | 0.322916469 | 0.24713737 |
| Fbxw17 | -0.015696644 | 0.582477862 | 0.866379712 | 0.046809347 | 0.061005592 |
| Fbxw2 | -0.157209808 | 0.009861748 | 0.154418667 | 0.350610165 | 0.530642458 |
| Fbxw4 | -0.020333351 | 0.601732822 | 0.876557788 | 0.099319456 | 0.116651704 |
| Fbxw5 | 0.03249669 | 0.541647532 | 0.850208191 | 0.211492545 | 0.183817016 |
| Fbxw7 | 0.222769121 | 0.001236081 | 0.044151419 | 0.55134041 | 0.317476398 |
| Fbxw8 | -0.018448273 | 0.656278509 | 0.89805697 | 0.113155943 | 0.134106216 |
| Fbxw9 | -0.030730891 | 0.364983982 | 0.752802229 | 0.069415296 | 0.096669065 |

| | | | | | |
|---|---|---|---|---|---|
| Fcer1g | 0.011768644 | 0.761343468 | 0.928506001 | 0.089670979 | 0.084614238 |
| Fcf1 | -0.035179356 | 0.502609785 | 0.832249514 | 0.192550954 | 0.218971929 |
| Fcho2 | 0.026439458 | 0.670057379 | 0.903736153 | 0.34477853 | 0.319889571 |
| Fchsd2 | -0.006950254 | 0.859753888 | 0.959220425 | 0.100400104 | 0.104167238 |
| Fcsk | -0.063252487 | 0.003537447 | 0.08334591 | 0.018202119 | 0.06974967 |
| Fdft1 | 0.017885283 | 0.752428772 | 0.926186851 | 0.265153516 | 0.241321031 |
| Fdps | 0.013269427 | 0.804186172 | 0.944269608 | 0.205860779 | 0.18746468 |
| Fdx1 | -0.002547139 | 0.968400091 | 0.993505636 | 0.427203415 | 0.431422693 |
| Fdxacb1 | 0.001950618 | 0.944305644 | 0.987070339 | 0.048604556 | 0.049839358 |
| Fdxr | -0.016395819 | 0.532751983 | 0.84612966 | 0.038525217 | 0.051726525 |
| Fech | 0.018400313 | 0.731667573 | 0.920770724 | 0.2073697 | 0.188588166 |
| Fem1a | 0.094374869 | 0.07306642 | 0.410160881 | 0.207378455 | 0.125134317 |
| Fem1b | 0.103384307 | 0.113632363 | 0.484712294 | 0.427613539 | 0.308152937 |
| Fem1c | 0.021346078 | 0.620729487 | 0.88397586 | 0.128465307 | 0.107815679 |
| Fer | -0.021307526 | 0.627605275 | 0.887846955 | 0.125679867 | 0.13784055 |
| Fer1l6 | 0.041511025 | 0.32314131 | 0.720244873 | 0.116813809 | 0.081235877 |
| Fermt2 | 0.073257557 | 0.275292216 | 0.68241738 | 0.588755618 | 0.514905934 |
| Fez1 | -0.027987261 | 0.556894208 | 0.855368999 | 0.153278294 | 0.189549923 |
| Fez2 | 0.010894083 | 0.731882916 | 0.920770724 | 0.064635508 | 0.057387971 |
| Ffar1 | -0.083314047 | 0.211890137 | 0.625696178 | 0.447693578 | 0.526404156 |
| Ffar2 | 0.008337192 | 0.875887172 | 0.965226125 | 0.193892949 | 0.180905198 |
| Fgd1 | 0.025946009 | 0.407501933 | 0.777888922 | 0.063654459 | 0.040717474 |
| Fgd2 | -0.031396466 | 0.526982908 | 0.842652134 | 0.176826781 | 0.203485238 |
| Fgd4 | -0.065815623 | 0.132855134 | 0.513464375 | 0.11645643 | 0.182114709 |
| Fgd6 | 0.098606205 | 0.072938948 | 0.410133458 | 0.227401347 | 0.152569548 |
| Fgf12 | -0.039905421 | 0.506404659 | 0.833269979 | 0.286557092 | 0.311099949 |
| Fgf9 | -0.001019825 | 0.983124293 | 0.99608406 | 0.164984937 | 0.171161282 |
| Fgfbp3 | -0.048457308 | 0.152930553 | 0.549602359 | 0.066264482 | 0.108677007 |
| Fgfr1 | -0.064842889 | 0.105183414 | 0.472399448 | 0.099211935 | 0.157699711 |
| Fgfr1op2 | -0.103676443 | 0.12972671 | 0.509514066 | 0.682920947 | 0.794156303 |
| Fggy | -0.005526342 | 0.853484351 | 0.957461155 | 0.057485187 | 0.058674742 |
| Fgl2 | 0.025978825 | 0.683422758 | 0.909267527 | 0.330582674 | 0.28036645 |
| Fh1 | 0.17966478 | 0.024501336 | 0.248116801 | 0.947254776 | 0.752722789 |
| Fhad1 | 0.006822937 | 0.854956804 | 0.957723375 | 0.093793038 | 0.083403905 |
| Fhdc1 | 0.00604791 | 0.912571199 | 0.979623222 | 0.216846951 | 0.205665152 |
| Fhit | -0.017022315 | 0.488912913 | 0.826256745 | 0.03433945 | 0.046648185 |
| Fhl1 | -0.000839501 | 0.987096568 | 0.997268589 | 0.203501769 | 0.190037555 |
| Fhod3 | 0.154575472 | 0.027059786 | 0.26190846 | 0.516989294 | 0.362693889 |
| Fibp | -0.08594699 | 0.12658781 | 0.505998346 | 0.240790539 | 0.315475604 |
| Ficd | 0.019665168 | 0.704177642 | 0.912449135 | 0.186236817 | 0.163921846 |
| Fig4 | -0.040056167 | 0.277065578 | 0.684278349 | 0.083856415 | 0.111532649 |
| Filip1 | -0.044723055 | 0.482257742 | 0.821803219 | 0.321183027 | 0.370862503 |
| Fip1l1 | -0.093135675 | 0.170137584 | 0.572762895 | 0.858482823 | 0.956822793 |
| Firre | -0.068645891 | 0.106549289 | 0.473998412 | 0.109857653 | 0.162685982 |
| Fis1 | -0.139391944 | 0.033392315 | 0.289428903 | 0.728388483 | 0.907342946 |
| Fitm2 | -0.09366548 | 0.133387004 | 0.51492514 | 0.346107585 | 0.429695152 |
| Fiz1 | -0.021449423 | 0.611117454 | 0.87989943 | 0.117305985 | 0.139322364 |
| Fkbp11 | 0.049998 | 0.451901171 | 0.806495742 | 0.456919616 | 0.368374646 |
| Fkbp14 | -0.024253843 | 0.359729013 | 0.747719327 | 0.040559904 | 0.060295335 |
| Fkbp15 | 0.066295168 | 0.183253211 | 0.590066231 | 0.17742131 | 0.10978183 |
| Fkbp1a | 0.005201119 | 0.927214905 | 0.981818446 | 1.246084521 | 1.247419654 |
| Fkbp1b | -0.183285789 | 0.007560492 | 0.133456507 | 0.606545994 | 0.791934407 |
| Fkbp2 | 0.188737744 | 0.000887923 | 0.035506311 | 1.788314892 | 1.376288854 |
| Fkbp3 | -0.030637325 | 0.641865442 | 0.892178303 | 0.546501194 | 0.578625419 |
| Fkbp4 | 0.134523043 | 0.057848949 | 0.370772606 | 0.969393941 | 0.785656769 |
| Fkbp5 | 0.009549825 | 0.867274593 | 0.96228555 | 0.25998288 | 0.246550767 |
| Fkbp7 | -0.016582384 | 0.523545673 | 0.841062594 | 0.036922914 | 0.053985798 |
| Fkbp8 | -0.118264654 | 0.060133785 | 0.378409279 | 0.410388076 | 0.517447881 |
| Fkbp9 | -0.123306584 | 0.062126768 | 0.380250374 | 0.611113915 | 0.743904022 |
| Fkbpl | 0.018671845 | 0.595514348 | 0.873342705 | 0.086608537 | 0.079998893 |
| Fkrp | 0.007035833 | 0.829600379 | 0.953065609 | 0.070196207 | 0.063491762 |
| Fktn | 0.055390196 | 0.294078956 | 0.69977963 | 0.206234926 | 0.140654302 |
| Flad1 | 0.103432861 | 0.018504987 | 0.216221828 | 0.14246404 | 0.053392235 |
| Flcn | 0.080949997 | 0.095419381 | 0.455842134 | 0.172533763 | 0.09920584 |
| Flii | -0.065897249 | 0.231799087 | 0.643291165 | 0.231560238 | 0.280852174 |

| | | | | | |
|---|---|---|---|---|---|
| Flnb | 0.023109532 | 0.73867575 | 0.922843014 | 0.677730944 | 0.677821423 |
| Flot1 | -0.075816673 | 0.18391859 | 0.590326848 | 0.24478854 | 0.309401212 |
| Flot2 | -0.005702145 | 0.917992519 | 0.980385833 | 0.222141674 | 0.215706509 |
| Flrt1 | -0.020454584 | 0.751538216 | 0.926186851 | 0.368609904 | 0.384845465 |
| Flt3l | -0.019318449 | 0.477062651 | 0.818444859 | 0.042668062 | 0.054946966 |
| Flvcr1 | 0.01803117 | 0.663697928 | 0.899599744 | 0.110370601 | 0.097254288 |
| Flywch1 | -0.115953898 | 0.063728217 | 0.385325372 | 0.33371871 | 0.428633224 |
| Flywch2 | -0.080152474 | 0.075342706 | 0.414153416 | 0.121619904 | 0.191312091 |
| Fmc1 | -0.035989471 | 0.569486663 | 0.860833982 | 0.370992059 | 0.397948578 |
| Fmn1 | -0.048200066 | 0.060380898 | 0.379013803 | 0.032373148 | 0.071703206 |
| Fmn2 | -0.213324212 | 0.000403457 | 0.021771487 | 0.291640724 | 0.493131618 |
| Fmnl2 | 0.024357937 | 0.715934378 | 0.916579082 | 0.451494249 | 0.415818557 |
| Fmo1 | 0.069467294 | 0.07349529 | 0.410998661 | 0.09801044 | 0.038200068 |
| Fmr1 | 0.109573401 | 0.069467909 | 0.400487881 | 0.32709834 | 0.240278148 |
| Fnbp1 | -0.006552138 | 0.831367492 | 0.953431846 | 0.06055074 | 0.064161538 |
| Fnbp1l | 0.010068732 | 0.84128901 | 0.954960614 | 0.175022377 | 0.154263734 |
| Fnbp4 | 0.042601664 | 0.495032943 | 0.828936545 | 0.337583165 | 0.295702559 |
| Fndc10 | -0.045484472 | 0.253204883 | 0.663146569 | 0.100519205 | 0.14006067 |
| Fndc3a | -0.00312766 | 0.956792465 | 0.989625513 | 1.322121243 | 1.32563349 |
| Fndc3b | 0.007447838 | 0.911614266 | 0.979014807 | 0.592260577 | 0.579411791 |
| Fnip1 | 0.176891883 | 0.011417166 | 0.16901784 | 0.604034566 | 0.41440537 |
| Fnip2 | 0.099841201 | 0.11794308 | 0.491192747 | 0.355668319 | 0.258539429 |
| Fnta | 0.018927578 | 0.75786508 | 0.927237784 | 0.341633497 | 0.322961889 |
| Fntb | 6.95E-05 | 0.998239819 | 0.99938181 | 0.064823538 | 0.061552868 |
| Focad | 0.007005656 | 0.833363627 | 0.954303562 | 0.071474598 | 0.064537301 |
| Folr1 | -0.101958246 | 0.057806597 | 0.370772606 | 0.166982109 | 0.274235727 |
| Fos | -0.681817357 | 1.92E-15 | 3.21E-12 | 1.062275113 | 2.073157277 |
| Fosb | -0.314221294 | 2.39E-07 | 6.67E-05 | 0.191703739 | 0.509494555 |
| Fosl2 | -0.046362706 | 0.323591015 | 0.720311159 | 0.13218062 | 0.177328032 |
| Foxa2 | -0.052257614 | 0.45502327 | 0.807876964 | 0.597351222 | 0.642444602 |
| Foxa3 | 0.019813824 | 0.677131812 | 0.9069403 | 0.157400749 | 0.137388545 |
| Foxj2 | 0.065063274 | 0.190217942 | 0.599440342 | 0.177325656 | 0.119932338 |
| Foxj3 | 0.049310687 | 0.401096556 | 0.774740795 | 0.263811218 | 0.216587215 |
| Foxk1 | -0.024754036 | 0.531386062 | 0.845517106 | 0.100714862 | 0.124730965 |
| Foxk2 | 0.107954221 | 0.050765618 | 0.354829042 | 0.254385455 | 0.150557097 |
| Foxn2 | -0.017073217 | 0.68234404 | 0.909157989 | 0.114221063 | 0.12866792 |
| Foxn3 | 0.011850462 | 0.842320608 | 0.955188334 | 0.298262443 | 0.28868861 |
| Foxo1 | 0.043248467 | 0.532927939 | 0.84612966 | 0.449778053 | 0.404737806 |
| Foxo3 | -0.045235601 | 0.390233729 | 0.76844731 | 0.188525195 | 0.222837794 |
| Foxo6 | -0.009772253 | 0.811024422 | 0.948397417 | 0.106183211 | 0.117209242 |
| Foxp1 | 0.136466766 | 0.049144334 | 0.347612176 | 0.664283687 | 0.502416262 |
| Foxp2 | -0.0425632 | 0.221125602 | 0.633858837 | 0.068912574 | 0.106947717 |
| Foxp4 | -0.023421646 | 0.66907026 | 0.903100887 | 0.219620161 | 0.242446666 |
| Foxred1 | -0.025020547 | 0.414420396 | 0.782795478 | 0.055616204 | 0.076243534 |
| Foxred2 | 0.030227628 | 0.297665747 | 0.702656882 | 0.056772486 | 0.027481846 |
| Fpgs | -0.021769298 | 0.522165162 | 0.840438058 | 0.073186022 | 0.089552597 |
| Fpgt | 0.081336298 | 0.132323083 | 0.513250014 | 0.218796463 | 0.150563766 |
| Fra10ac1 | -0.006698859 | 0.914039036 | 0.980132949 | 0.334530575 | 0.329561314 |
| Fras1 | 0.02010849 | 0.583941662 | 0.867093874 | 0.088779638 | 0.075778759 |
| Frg1 | -0.007623931 | 0.886159636 | 0.969590273 | 0.212795585 | 0.221352999 |
| Frk | 0.018356921 | 0.724660145 | 0.919870226 | 0.185249934 | 0.159489442 |
| Frmd4a | -0.039846869 | 0.365179225 | 0.752802229 | 0.124579913 | 0.159296714 |
| Frmd4b | -0.006121245 | 0.88093458 | 0.967919035 | 0.108389452 | 0.117535228 |
| Frmd5 | -0.036747703 | 0.444390659 | 0.801932586 | 0.150630877 | 0.184627589 |
| Frrs1 | -0.028616452 | 0.417066762 | 0.784482356 | 0.074699023 | 0.104233079 |
| Frrs1l | -0.028283426 | 0.295939153 | 0.702458178 | 0.037930767 | 0.06334658 |
| Frs2 | 0.04321703 | 0.356532706 | 0.745571198 | 0.160241451 | 0.126251121 |
| Fry | -0.032654427 | 0.547821214 | 0.851685491 | 0.205805488 | 0.223799602 |
| Fryl | -0.182914346 | 0.002269044 | 0.065256147 | 0.294023415 | 0.463450312 |
| Frzb | -0.14483563 | 0.033128543 | 0.287968471 | 0.470554387 | 0.606281932 |
| Fsd1l | 0.018151977 | 0.654108063 | 0.897454058 | 0.110245128 | 0.096399346 |
| Fth1 | -0.206823308 | 2.26E-09 | 1.45E-06 | 2.881097709 | 3.452308554 |
| Ftl1 | 0.006655998 | 0.841547248 | 0.954960614 | 3.480574333 | 3.534390842 |
| Ftl1-ps1 | 0.025999945 | 0.470083574 | 0.814330141 | 0.087157111 | 0.070005922 |
| Fto | -0.019276279 | 0.70663347 | 0.913155676 | 0.192239026 | 0.201846229 |

| | | | | | |
|---|---|---|---|---|---|
| Ftsj1 | 0.049435766 | 0.089530717 | 0.444861289 | 0.058486811 | 0.017795444 |
| Ftsj3 | -0.105299383 | 0.031490426 | 0.281953085 | 0.148594345 | 0.246303813 |
| Ftx | -0.125466307 | 0.004715663 | 0.100919217 | 0.116525503 | 0.227546597 |
| Fubp1 | -0.061099916 | 0.380480801 | 0.762006446 | 0.603672136 | 0.659346066 |
| Fubp3 | -0.04947369 | 0.353129902 | 0.743880213 | 0.204968505 | 0.254003658 |
| Fuca1 | -0.065569544 | 0.306967484 | 0.709680069 | 0.413394148 | 0.466345296 |
| Fuca2 | -0.01802754 | 0.651610648 | 0.896468516 | 0.102455576 | 0.119322917 |
| Fundc1 | 0.003830524 | 0.943850539 | 0.986918206 | 0.219199649 | 0.218900551 |
| Fundc2 | -0.025999787 | 0.690341518 | 0.909796728 | 0.552234914 | 0.576764644 |
| Fuom | -0.261848774 | 7.15E-05 | 0.006165423 | 0.547406538 | 0.823004687 |
| Furin | -0.011639837 | 0.811559317 | 0.948397417 | 0.163766731 | 0.17390172 |
| Fus | -0.227063564 | 0.000950302 | 0.037078044 | 0.745026294 | 0.988736061 |
| Fut11 | -0.01352603 | 0.717730939 | 0.917764311 | 0.090583956 | 0.107597893 |
| Fut8 | -0.020883501 | 0.637707855 | 0.891210491 | 0.132391405 | 0.153491691 |
| Fuz | -0.035018931 | 0.313697385 | 0.714906079 | 0.073668174 | 0.108692576 |
| Fxn | -0.120006874 | 0.017254392 | 0.209327864 | 0.175389581 | 0.278680033 |
| Fxr1 | -0.082366734 | 0.229984299 | 0.641131395 | 0.664209505 | 0.770608566 |
| Fxr2 | 0.04022546 | 0.514245717 | 0.83659041 | 0.378277659 | 0.352327094 |
| Fxyd3 | -0.00725552 | 0.907284436 | 0.977187883 | 0.259143819 | 0.22911898 |
| Fxyd6 | -0.054091336 | 0.460626353 | 0.810111566 | 0.645425404 | 0.745190622 |
| Fyco1 | 0.06692793 | 0.22185828 | 0.633871779 | 0.235728747 | 0.172793513 |
| Fyn | 0.004502611 | 0.886850832 | 0.969712176 | 0.063928736 | 0.060355939 |
| Fyttd1 | -0.005978267 | 0.933728347 | 0.983869133 | 0.628120033 | 0.626735927 |
| Fzd3 | -0.118645356 | 0.050620274 | 0.354499194 | 0.334225611 | 0.436719143 |
| Fzd6 | 0.02813252 | 0.40681346 | 0.777736088 | 0.073752196 | 0.046415629 |
| Fzd8 | 0.006141103 | 0.828736985 | 0.952874813 | 0.050046623 | 0.045366891 |
| Fzr1 | 0.059952652 | 0.227329599 | 0.639441223 | 0.175311662 | 0.125713993 |
| G2e3 | 0.048765962 | 0.289175891 | 0.695472148 | 0.14314048 | 0.096080505 |
| G3bp1 | -0.024394539 | 0.721164588 | 0.919475778 | 0.734779293 | 0.744449342 |
| G3bp2 | 0.072810443 | 0.256755206 | 0.665256942 | 0.981340269 | 0.877218954 |
| G6pc2 | 0.188702296 | 0.00526861 | 0.108299861 | 2.213586253 | 1.770574782 |
| G6pc3 | -0.079805363 | 0.120206935 | 0.493866968 | 0.186035372 | 0.26533793 |
| G6pdx | -0.023506203 | 0.529615739 | 0.844323583 | 0.088182033 | 0.10740383 |
| G730003C15Rik | 0.028423598 | 0.417360299 | 0.784482356 | 0.083433742 | 0.057212185 |
| Gaa | -0.156948473 | 0.015462941 | 0.199744575 | 0.388956076 | 0.546898376 |
| Gab2 | 0.030666935 | 0.432909146 | 0.795079432 | 0.104908534 | 0.074189692 |
| Gabarap | -0.040725874 | 0.314606251 | 0.714975653 | 2.156743465 | 2.242820321 |
| Gabarapl1 | 0.030658552 | 0.653896686 | 0.897454058 | 0.798126057 | 0.752887925 |
| Gabarapl2 | -0.03933548 | 0.49638199 | 0.830056214 | 1.27539399 | 1.356242277 |
| Gabbr1 | 0.011958464 | 0.807941921 | 0.946354658 | 0.172184177 | 0.160356213 |
| Gabbr2 | 0.135746698 | 0.052911426 | 0.359257749 | 0.531692417 | 0.360387925 |
| Gabpa | 0.121670132 | 0.006808588 | 0.126317563 | 0.149960766 | 0.04178445 |
| Gabpb1 | 0.061862389 | 0.174189777 | 0.577186219 | 0.144595455 | 0.097582666 |
| Gabpb2 | -0.040628606 | 0.467280996 | 0.813188482 | 0.242210786 | 0.283653809 |
| Gabrb3 | -0.061616265 | 0.278057999 | 0.684996416 | 0.243981124 | 0.299207941 |
| Gad1 | 0.043098086 | 0.538967527 | 0.848567384 | 0.439488335 | 0.404109234 |
| Gadd45a | 0.00935597 | 0.884048762 | 0.968954079 | 0.332205242 | 0.310894671 |
| Gadd45b | 0.132765938 | 0.058014739 | 0.37129897 | 0.368624057 | 0.231829028 |
| Gadd45g | -0.112447973 | 0.144714297 | 0.533279057 | 0.875672901 | 1.034123633 |
| Gadd45gip1 | 0.011910577 | 0.861610018 | 0.960033276 | 0.764671752 | 0.733012891 |
| Gak | 0.045002796 | 0.469128632 | 0.813940204 | 0.327889362 | 0.286992759 |
| Gal3st1 | 0.018994588 | 0.565887167 | 0.858366253 | 0.074596207 | 0.057330038 |
| Galc | -0.036354673 | 0.399291639 | 0.774544012 | 0.123114387 | 0.151073158 |
| Gale | 0.027655532 | 0.523504029 | 0.841062594 | 0.129111363 | 0.104623567 |
| Galk2 | -0.014999079 | 0.763551698 | 0.928730554 | 0.170188368 | 0.177125196 |
| Galns | -0.030237161 | 0.430356835 | 0.793295051 | 0.094442759 | 0.112671068 |
| Galnt1 | -0.040559094 | 0.503617952 | 0.832249514 | 0.321104575 | 0.357251016 |
| Galnt10 | 0.027459701 | 0.488490072 | 0.826256745 | 0.103306334 | 0.082310564 |
| Galnt11 | 0.043671418 | 0.392373875 | 0.769704269 | 0.198854894 | 0.154053975 |
| Galnt12 | -0.042433099 | 0.243014801 | 0.653166476 | 0.079884036 | 0.113781971 |
| Galnt14 | -0.082867998 | 0.128012358 | 0.508260084 | 0.212285412 | 0.290478333 |
| Galnt18 | 0.010461566 | 0.834362834 | 0.954794181 | 0.172844329 | 0.168165776 |
| Galnt4 | -0.012209912 | 0.690529367 | 0.909796728 | 0.057286672 | 0.067448937 |
| Galnt7 | 0.007935893 | 0.842945157 | 0.955306089 | 0.105942814 | 0.09871209 |
| Galr1 | 0.041943051 | 0.399412965 | 0.774544012 | 0.171776359 | 0.129261884 |

| | | | | | |
|---|---|---|---|---|---|
| Galr3 | -0.001945353 | 0.94586838 | 0.987084873 | 0.051345276 | 0.052615928 |
| Galt | -0.002810583 | 0.922049544 | 0.980859278 | 0.049980134 | 0.049444566 |
| Gan | -0.019253676 | 0.614175121 | 0.88189924 | 0.096848616 | 0.110823654 |
| Ganab | -0.118264153 | 0.053701792 | 0.361325633 | 0.335459381 | 0.446837722 |
| Ganc | -0.00815978 | 0.862329913 | 0.960216905 | 0.14935296 | 0.16406913 |
| Gapdh | -0.201247852 | 0.000719169 | 0.031383916 | 1.489904117 | 1.859738837 |
| Gapvd1 | -0.173004755 | 0.005456551 | 0.110446789 | 0.346177033 | 0.513743401 |
| Gar1 | -0.041869139 | 0.455015951 | 0.807876964 | 0.213241081 | 0.250751058 |
| Garre1 | -0.011309665 | 0.789175188 | 0.938279005 | 0.121374591 | 0.13171348 |
| Gars | -0.176817153 | 0.005658969 | 0.112029722 | 1.102782805 | 1.373108933 |
| Gart | 0.000415739 | 0.992657456 | 0.998695538 | 0.132625713 | 0.144925502 |
| Gas2 | -0.054392506 | 0.182061133 | 0.589470435 | 0.102661826 | 0.150150995 |
| Gas2l1 | 0.036207636 | 0.323310576 | 0.720244873 | 0.08780696 | 0.052470566 |
| Gas5 | -0.106001759 | 0.051721061 | 0.355933715 | 1.61507863 | 1.786259531 |
| Gas6 | -0.02206431 | 0.327819428 | 0.722581934 | 0.028361579 | 0.047344743 |
| Gas8 | 0.014547719 | 0.73186906 | 0.920770724 | 0.123222489 | 0.112558371 |
| Gata6 | -0.002350221 | 0.951970056 | 0.98838165 | 0.093599569 | 0.092049511 |
| Gatad1 | -0.106851699 | 0.107289241 | 0.47507667 | 0.457546696 | 0.547823152 |
| Gatad2a | 0.029573652 | 0.636749457 | 0.890755879 | 0.358777167 | 0.334603547 |
| Gatad2b | 0.017704929 | 0.789727713 | 0.938440574 | 0.472149836 | 0.450289989 |
| Gatc | 0.029217739 | 0.504373448 | 0.832249514 | 0.130360663 | 0.103779631 |
| Gatd1 | -0.079099044 | 0.188049469 | 0.595974902 | 0.30876444 | 0.385436032 |
| Gatd3a | -0.021278628 | 0.630009352 | 0.888118521 | 0.127959288 | 0.152892717 |
| Gatm | -0.022436493 | 0.69009232 | 0.909796728 | 0.213692354 | 0.226451685 |
| Gba | -0.044763823 | 0.246310569 | 0.655935044 | 0.09706396 | 0.124411627 |
| Gbe1 | -0.064845211 | 0.068746906 | 0.399753362 | 0.075682564 | 0.127468772 |
| Gbf1 | -0.013763604 | 0.842991921 | 0.955306089 | 0.639862627 | 0.655616277 |
| Gbp7 | 0.026359265 | 0.594693719 | 0.873162794 | 0.168421543 | 0.142534878 |
| Gc | -0.226314661 | 0.000320441 | 0.018285574 | 0.274557405 | 0.488596761 |
| Gca | -0.056183295 | 0.102449804 | 0.468895886 | 0.07193926 | 0.121921123 |
| Gcat | 0.040378277 | 0.24023213 | 0.650744882 | 0.081223939 | 0.049064544 |
| Gcc1 | -0.020271058 | 0.619279628 | 0.883797831 | 0.107612883 | 0.128705216 |
| Gcc2 | 0.079252949 | 0.251263175 | 0.663127858 | 0.7303839 | 0.624460675 |
| Gcdh | -0.020238943 | 0.638241162 | 0.89125563 | 0.12303648 | 0.147812274 |
| Gcfc2 | -0.061644256 | 0.020923101 | 0.229383208 | 0.034638018 | 0.089691139 |
| Gcg | 0.089428095 | 0.145258277 | 0.533628684 | 2.58937983 | 2.107365951 |
| Gcgr | -0.187091961 | 2.01E-05 | 0.002221072 | 0.112646189 | 0.287896064 |
| Gch1 | -0.059414986 | 0.271722801 | 0.67876599 | 1.810333111 | 1.936309915 |
| Gck | 0.055812151 | 0.345647151 | 0.73792645 | 0.2978603 | 0.246010624 |
| Gclc | -0.208493508 | 0.000276058 | 0.016492796 | 0.221973902 | 0.438877635 |
| Gclm | -0.309108354 | 2.42E-06 | 0.000449067 | 1.071519226 | 1.551882483 |
| Gcn1 | 0.064901969 | 0.26669388 | 0.674088763 | 0.292560436 | 0.236215988 |
| Gcnt2 | -0.032958152 | 0.593767393 | 0.872824154 | 0.325253106 | 0.335157342 |
| Gcsh | -0.163660871 | 0.003201507 | 0.079146623 | 0.230560954 | 0.401907944 |
| Gdap1 | -0.165963143 | 0.014462384 | 0.193677739 | 0.429153701 | 0.600013905 |
| Gdap1l1 | 0.108151494 | 0.114000887 | 0.485047438 | 0.502520435 | 0.409924226 |
| Gdap2 | -0.071800962 | 0.217156511 | 0.631640679 | 0.256298077 | 0.319645994 |
| Gde1 | 0.066468952 | 0.347745163 | 0.739124045 | 0.943512116 | 0.859821869 |
| Gdf15 | -0.052486379 | 0.267590206 | 0.675673686 | 0.122081227 | 0.183524547 |
| Gdi1 | -0.108781092 | 0.093334994 | 0.451907061 | 0.50590336 | 0.629824133 |
| Gdi2 | -0.000375391 | 0.995300499 | 0.99872501 | 1.011619751 | 1.010817297 |
| Gdpd1 | -0.087247777 | 0.098359957 | 0.461974211 | 0.181013602 | 0.261137654 |
| Gdpgp1 | 0.013382271 | 0.708633101 | 0.913767764 | 0.082934087 | 0.070129449 |
| Gem | -0.065576287 | 0.069770262 | 0.400867525 | 0.068344836 | 0.126030789 |
| Gemin2 | 0.062062848 | 0.141086927 | 0.528664246 | 0.12674566 | 0.077382793 |
| Gemin5 | 0.033482554 | 0.3144525 | 0.714975653 | 0.075270362 | 0.048844377 |
| Gemin6 | 0.003937909 | 0.941791595 | 0.986168593 | 0.207528326 | 0.205526335 |
| Gemin7 | 0.108425243 | 0.08464995 | 0.433706763 | 0.465024313 | 0.356852548 |
| Gemin8 | -0.016999288 | 0.568458279 | 0.860316006 | 0.052796576 | 0.069062393 |
| Get1 | -0.018961316 | 0.529494338 | 0.844323583 | 0.056080856 | 0.070830323 |
| Get3 | -0.079530696 | 0.183303543 | 0.590066231 | 0.315220667 | 0.394213835 |
| Get4 | -0.096542143 | 0.12245952 | 0.498631318 | 0.32157383 | 0.424458909 |
| Gfer | -0.011470059 | 0.844684433 | 0.955950886 | 0.276864029 | 0.287010964 |
| Gfm1 | -0.000274908 | 0.994324246 | 0.998695538 | 0.100473056 | 0.1025368 |
| Gfm2 | -0.022271211 | 0.624946023 | 0.88668351 | 0.137232736 | 0.149839017 |

| | | | | | |
|---|---|---|---|---|---|
| Gfod2 | 0.028300087 | 0.514945823 | 0.836801664 | 0.131937141 | 0.111440276 |
| Gfpt1 | -0.205972469 | 0.000909631 | 0.036086819 | 1.179915032 | 1.478980078 |
| Gfus | -0.156220655 | 0.00580966 | 0.113446612 | 0.215532005 | 0.368399671 |
| Gga1 | -0.117210874 | 0.025549291 | 0.254380218 | 0.194545951 | 0.290500779 |
| Gga2 | -0.023635932 | 0.713407974 | 0.915544794 | 0.39232184 | 0.394559979 |
| Gga3 | 0.012249816 | 0.739598407 | 0.922843014 | 0.092334911 | 0.079934679 |
| Ggact | 0.017036124 | 0.786421034 | 0.937336173 | 0.401433878 | 0.371415872 |
| Ggct | 0.104119348 | 0.043996635 | 0.329446347 | 0.19670475 | 0.106149118 |
| Ggcx | -0.02393811 | 0.576396652 | 0.863734428 | 0.122531242 | 0.143500963 |
| Ggh | -0.051373331 | 0.269271946 | 0.676978829 | 0.143169377 | 0.187177996 |
| Ggnbp1 | 0.0340995 | 0.323450284 | 0.720244873 | 0.079487942 | 0.040567936 |
| Ggnbp2 | 0.18408031 | 0.006011339 | 0.115144676 | 0.73225013 | 0.533977709 |
| Ggps1 | -0.108348152 | 0.05210768 | 0.356999852 | 0.242454199 | 0.34166401 |
| Ggt7 | 0.006259641 | 0.854354905 | 0.957593115 | 0.074745335 | 0.072546489 |
| Ghdc | 0.036983184 | 0.297804593 | 0.702656882 | 0.085738939 | 0.05185075 |
| Ghitm | -0.195323185 | 0.00197704 | 0.060683637 | 1.120574066 | 1.41031953 |
| Ghr | -0.136811192 | 0.047241168 | 0.34038737 | 0.680901994 | 0.832483532 |
| Gid4 | -0.013459144 | 0.794117433 | 0.939757086 | 0.189802668 | 0.201591865 |
| Gid8 | 0.002184407 | 0.96903066 | 0.993682135 | 0.24607481 | 0.253983286 |
| Gigyf1 | -0.05509076 | 0.334063503 | 0.729929513 | 0.24439051 | 0.293409124 |
| Gigyf2 | -0.122420924 | 0.056720811 | 0.368804883 | 0.376552228 | 0.495032322 |
| Gin1 | 0.018213258 | 0.65276098 | 0.896887331 | 0.111033447 | 0.097296762 |
| Ginm1 | -0.12431581 | 0.057330014 | 0.36957055 | 0.475421424 | 0.593870319 |
| Gins3 | -0.04817661 | 0.02459183 | 0.248318109 | 0.017711574 | 0.060939105 |
| Gins4 | 0.001208866 | 0.973438753 | 0.994079882 | 0.088716833 | 0.087739982 |
| Gipc1 | -0.040497866 | 0.296891756 | 0.702656882 | 0.101330914 | 0.141898968 |
| Gipc2 | 0.007806548 | 0.786953723 | 0.937461311 | 0.05290125 | 0.044086612 |
| Gipr | -0.081666131 | 0.097001694 | 0.45927202 | 0.16624041 | 0.23324227 |
| Git1 | 0.038374083 | 0.468758125 | 0.813719354 | 0.208013191 | 0.172639681 |
| Git2 | 0.044987261 | 0.412545592 | 0.781358105 | 0.21443162 | 0.175141813 |
| Gjd2 | 0.117260367 | 0.102498282 | 0.468903943 | 0.806019448 | 0.674497865 |
| Gk5 | 0.005493379 | 0.859829248 | 0.959220425 | 0.060508862 | 0.056405287 |
| Gkap1 | -0.01996171 | 0.755273479 | 0.926726954 | 0.37324564 | 0.381513533 |
| Gla | 0.054898123 | 0.262123521 | 0.668764912 | 0.150402139 | 0.099740206 |
| Glb1 | -0.098257642 | 0.004580314 | 0.099507822 | 0.06947064 | 0.160954191 |
| Glb1l2 | -0.017330174 | 0.730835122 | 0.920770724 | 0.176042292 | 0.180297096 |
| Glce | 0.011102552 | 0.852538839 | 0.957461155 | 0.280787998 | 0.249158127 |
| Gle1 | -0.050524741 | 0.295434645 | 0.702173225 | 0.149667109 | 0.196713241 |
| Glg1 | -0.053321954 | 0.434091325 | 0.796332921 | 0.852086761 | 0.914945757 |
| Glis2 | 0.028917293 | 0.344250466 | 0.737354232 | 0.060053259 | 0.034992292 |
| Glis3 | 0.041173941 | 0.421807247 | 0.787661271 | 0.187152734 | 0.146616719 |
| Glmn | -0.027796882 | 0.362124265 | 0.750073837 | 0.056314546 | 0.081621181 |
| Glmp | -0.010217132 | 0.866365942 | 0.962125879 | 0.343875646 | 0.358157541 |
| Glo1 | 0.00683966 | 0.918534796 | 0.980385833 | 0.451101865 | 0.432324067 |
| Glod4 | -0.028136089 | 0.636296432 | 0.89072626 | 0.289932256 | 0.313278753 |
| Glp1r | 0.184755965 | 0.002462222 | 0.067311494 | 1.595968097 | 1.298628314 |
| Glrb | -0.029365973 | 0.546196768 | 0.851446616 | 0.160165827 | 0.183094405 |
| Glrx | 0.013593914 | 0.823191639 | 0.950930859 | 0.317665885 | 0.319510112 |
| Glrx2 | -0.029053958 | 0.659551486 | 0.898227584 | 0.552785018 | 0.59495196 |
| Glrx3 | -0.077472512 | 0.215856363 | 0.629630897 | 0.362897168 | 0.452762969 |
| Glrx5 | -0.018051983 | 0.790220371 | 0.938632173 | 0.915239058 | 0.92359113 |
| Gls | -0.029217099 | 0.666564566 | 0.901197061 | 0.438662602 | 0.480560534 |
| Glt1d1 | -0.053920835 | 0.094015892 | 0.452959864 | 0.059230433 | 0.104861268 |
| Glt28d2 | -0.029549222 | 0.283433375 | 0.69129896 | 0.038634212 | 0.067389391 |
| Glt8d1 | 0.009227126 | 0.860007005 | 0.959312104 | 0.193132633 | 0.178263617 |
| Gltp | -0.074747447 | 0.080808635 | 0.426112224 | 0.114056639 | 0.186314968 |
| Glud1 | -0.07278847 | 0.2698593 | 0.677226912 | 0.447775885 | 0.5248895 |
| Glul | 0.11149216 | 0.150775932 | 0.54633142 | 1.017321701 | 0.856224667 |
| Glyr1 | -0.040708608 | 0.524312908 | 0.841062594 | 0.579635064 | 0.596750293 |
| Gm10033 | 0.027969641 | 0.581851059 | 0.866173066 | 0.18437774 | 0.15562306 |
| Gm10048 | -0.055793655 | 0.009252142 | 0.149508372 | 0.017883593 | 0.068274086 |
| Gm10076 | 0.099855598 | 0.003562163 | 0.083731698 | 2.934663758 | 2.60956805 |
| Gm10138 | -0.076108947 | 0.097644378 | 0.460898239 | 0.140939434 | 0.202619398 |
| Gm10177 | 0.037799656 | 0.325040597 | 0.720820254 | 0.101234484 | 0.072217566 |
| Gm10275 | -0.006744209 | 0.83031177 | 0.95325696 | 0.062530085 | 0.062430822 |

| | | | | | |
|---|---|---|---|---|---|
| Gm10399 | 0.000119741 | 0.996730809 | 0.999020084 | 0.054791627 | 0.05058692 |
| Gm10524 | 0.05865718 | 0.06318488 | 0.383440442 | 0.067834254 | 0.017972739 |
| Gm10657 | -0.04266898 | 0.035022872 | 0.294903156 | 0.017416748 | 0.054041274 |
| Gm10762 | -0.035025959 | 0.244109863 | 0.653891299 | 0.050865082 | 0.077772115 |
| Gm10941 | 0.001115979 | 0.984170969 | 0.99608406 | 0.225342304 | 0.217185991 |
| Gm11110 | -0.045813556 | 0.037436757 | 0.304994095 | 0.021782717 | 0.060030909 |
| Gm11423 | -0.020601221 | 0.549095649 | 0.852402509 | 0.073439764 | 0.089363503 |
| Gm11478 | -0.017990771 | 0.558874377 | 0.856615394 | 0.060602639 | 0.083952528 |
| Gm11789 | -0.030163263 | 0.466891998 | 0.813152001 | 0.106509604 | 0.136319641 |
| Gm11837 | -0.014886712 | 0.61660276 | 0.883226807 | 0.053589595 | 0.065140494 |
| Gm12216 | -0.018521815 | 0.499097717 | 0.830722138 | 0.045082491 | 0.058928761 |
| Gm12258 | -0.087872607 | 0.015031979 | 0.197813154 | 0.0763012 | 0.159803006 |
| Gm12346 | 0.046353846 | 0.234167966 | 0.644459522 | 0.102258296 | 0.062467712 |
| Gm12350 | 0.055845866 | 0.105192066 | 0.472399448 | 0.083814867 | 0.031455693 |
| Gm12940 | -0.006809761 | 0.81303601 | 0.948397417 | 0.051227521 | 0.051250766 |
| Gm12976 | 0.009121172 | 0.835517421 | 0.954911647 | 0.130235546 | 0.129324696 |
| Gm13436 | -0.020071819 | 0.38007725 | 0.762006446 | 0.027318224 | 0.047894451 |
| Gm13594 | -0.001666869 | 0.950747688 | 0.988114084 | 0.045982845 | 0.045782348 |
| Gm13611 | -0.02931362 | 0.340713491 | 0.734007579 | 0.057354273 | 0.089846578 |
| Gm13835 | 0.00762375 | 0.793192043 | 0.939604453 | 0.052641309 | 0.04446264 |
| Gm14167 | -0.089959848 | 0.00247695 | 0.067374388 | 0.047590653 | 0.117415816 |
| Gm14303 | 0.001642633 | 0.958124555 | 0.990272449 | 0.061373828 | 0.06111073 |
| Gm14305 | 0.002809427 | 0.918312897 | 0.980385833 | 0.044641712 | 0.042727104 |
| Gm14326 | -0.012021023 | 0.696916244 | 0.910970095 | 0.058109295 | 0.069243846 |
| Gm14418 | -0.043393684 | 0.079982715 | 0.424081623 | 0.031148854 | 0.068985901 |
| Gm15417 | 0.024111115 | 0.65928259 | 0.898224427 | 0.226200032 | 0.200445426 |
| Gm15427 | -0.045192733 | 0.159065479 | 0.556789205 | 0.06484637 | 0.097637071 |
| Gm15446 | 0.005707015 | 0.8484607 | 0.957111129 | 0.055789249 | 0.048150577 |
| Gm15500 | -0.217818595 | 1.22E-07 | 3.96E-05 | 0.090645768 | 0.292234984 |
| Gm15706 | -0.077915642 | 0.211211227 | 0.624058605 | 0.349918534 | 0.432181107 |
| Gm15860 | -0.023896188 | 0.337895738 | 0.732352032 | 0.034743326 | 0.055723337 |
| Gm15972 | 0.001256213 | 0.966701106 | 0.992504747 | 0.059675866 | 0.056618774 |
| Gm16062 | 0.133138971 | 0.022678889 | 0.238404929 | 0.235453707 | 0.131589768 |
| Gm16124 | -0.112619908 | 0.002420925 | 0.066938901 | 0.076512519 | 0.174841444 |
| Gm16174 | 0.105887122 | 0.094835968 | 0.454569536 | 0.396506247 | 0.267332448 |
| Gm16253 | -0.00284572 | 0.916436209 | 0.980132949 | 0.045510672 | 0.047445261 |
| Gm16286 | 0.004829237 | 0.942346309 | 0.986270063 | 0.541935987 | 0.564629176 |
| Gm16315 | 0.026822609 | 0.451024937 | 0.806495742 | 0.084305214 | 0.052671977 |
| Gm16536 | -0.011752202 | 0.658587901 | 0.898224427 | 0.042091175 | 0.05028572 |
| Gm1673 | -0.010307348 | 0.857634788 | 0.958476825 | 0.255398174 | 0.269898582 |
| Gm16740 | 0.052239194 | 0.100447454 | 0.465462186 | 0.067509448 | 0.019780887 |
| Gm17018 | -0.143332602 | 0.016304186 | 0.206355375 | 0.338020104 | 0.468576407 |
| Gm17066 | -0.012731507 | 0.64842923 | 0.895379629 | 0.046567171 | 0.058029406 |
| Gm17251 | -0.057673988 | 0.111087237 | 0.48205041 | 0.080582234 | 0.126578977 |
| Gm17491 | -0.050845273 | 0.103412548 | 0.469315384 | 0.056666017 | 0.100021663 |
| Gm17767 | 0.048858623 | 0.22969375 | 0.641131395 | 0.108250244 | 0.069310496 |
| Gm18811 | -0.125422902 | 4.86E-06 | 0.000762889 | 0.031775474 | 0.144355369 |
| Gm19412 | 0.01473547 | 0.683624221 | 0.909415018 | 0.0859727 | 0.07032087 |
| Gm19531 | -0.059394555 | 0.0182077 | 0.215000801 | 0.031944119 | 0.079515568 |
| Gm19705 | 0.048234384 | 0.063862136 | 0.38543852 | 0.047281462 | 0.009669102 |
| Gm19719 | -0.054463643 | 0.000730855 | 0.031755825 | 0.004809344 | 0.049685545 |
| Gm19938 | -0.018095452 | 0.489837166 | 0.826459641 | 0.037768631 | 0.050073377 |
| Gm2011 | 0.017568918 | 0.668200704 | 0.902353633 | 0.115192908 | 0.100559854 |
| Gm20208 | 0.035432935 | 0.535377548 | 0.846767168 | 0.248716979 | 0.214366443 |
| Gm20257 | -0.06377205 | 0.083923047 | 0.432188621 | 0.079534258 | 0.134016061 |
| Gm20300 | -0.001188339 | 0.976695574 | 0.994468977 | 0.108876803 | 0.107040907 |
| Gm20342 | -0.059462666 | 0.080419739 | 0.425115437 | 0.06459137 | 0.116038146 |
| Gm20400 | 0.004044101 | 0.89510606 | 0.972229731 | 0.059345847 | 0.060891502 |
| Gm20604 | 0.085544008 | 0.0625248 | 0.381496301 | 0.147273859 | 0.068853753 |
| Gm20712 | 0.013185683 | 0.718221872 | 0.91797181 | 0.087101984 | 0.072053628 |
| Gm2115 | -0.033929664 | 0.126246135 | 0.505736405 | 0.022316659 | 0.049588395 |
| Gm21926 | -0.020633649 | 0.554836163 | 0.853469818 | 0.082553657 | 0.104838104 |
| Gm21988 | -0.0423174 | 0.04607334 | 0.336808532 | 0.021297679 | 0.05914108 |
| Gm26532 | -0.0393794 | 0.103253875 | 0.469315384 | 0.027756018 | 0.062442097 |
| Gm26559 | 0.010098068 | 0.753798463 | 0.926726954 | 0.066033907 | 0.053863529 |

| | | | | | |
|---|---|---|---|---|---|
| Gm26670 | -0.00115488 | 0.964913982 | 0.991970325 | 0.041558722 | 0.04370257 |
| Gm26682 | -0.0150945 | 0.593160233 | 0.872442739 | 0.046210926 | 0.061707742 |
| Gm26782 | -0.139255498 | 0.001483535 | 0.050135475 | 0.117016477 | 0.237878282 |
| Gm26812 | -0.002836321 | 0.926503607 | 0.981818446 | 0.060516456 | 0.063381226 |
| Gm26881 | -0.011830366 | 0.736163294 | 0.922302218 | 0.076633598 | 0.086322716 |
| Gm26910 | -0.026549633 | 0.258797177 | 0.665948842 | 0.030103784 | 0.057671766 |
| Gm27033 | -0.006028531 | 0.921113072 | 0.980716231 | 0.259912442 | 0.2680864 |
| Gm27168 | 0.023496147 | 0.5001761 | 0.831130155 | 0.072277567 | 0.041165125 |
| Gm28438 | -0.593559995 | 2.48E-66 | 2.49E-62 | 0.00701287 | 0.575910503 |
| Gm29019 | 0.078108908 | 0.017648598 | 0.21172951 | 0.075199692 | 0.00682784 |
| Gm29054 | -0.01627112 | 0.557125109 | 0.855384435 | 0.042685193 | 0.06044765 |
| Gm2a | -0.093422981 | 0.03160827 | 0.282549988 | 0.111887762 | 0.19721308 |
| Gm30025 | -0.013720826 | 0.781108525 | 0.935482467 | 0.169474455 | 0.176898846 |
| Gm30173 | 0.097800757 | 0.012439727 | 0.178113462 | 0.111198158 | 0.029469724 |
| Gm30648 | 0.11637422 | 0.045752442 | 0.335685135 | 0.297171935 | 0.197392537 |
| Gm32031 | -0.074675203 | 0.2132855 | 0.627784916 | 0.293784157 | 0.351625449 |
| Gm32856 | -0.008575523 | 0.774555725 | 0.933391261 | 0.056234084 | 0.061202035 |
| Gm3294 | -0.04459693 | 0.263199139 | 0.669775695 | 0.099203561 | 0.130316851 |
| Gm33447 | -0.021570718 | 0.553386341 | 0.852854813 | 0.076119676 | 0.101948786 |
| Gm33989 | 0.037804098 | 0.452768866 | 0.807136288 | 0.17918834 | 0.149262553 |
| Gm34220 | -0.000198648 | 0.995588586 | 0.998772877 | 0.080456526 | 0.08638331 |
| Gm34552 | -0.01388941 | 0.594546353 | 0.873162794 | 0.038486172 | 0.048665373 |
| Gm34759 | -0.03480353 | 0.205475548 | 0.616549498 | 0.040583146 | 0.070028252 |
| Gm3625 | 0.019920948 | 0.60707398 | 0.878576426 | 0.10261629 | 0.088899434 |
| Gm36371 | 0.029561344 | 0.33539805 | 0.730736446 | 0.059871517 | 0.034638562 |
| Gm36738 | 0.022935507 | 0.713097016 | 0.915509496 | 0.357005916 | 0.324895071 |
| Gm36839 | 0.027077916 | 0.440835586 | 0.800354711 | 0.082953372 | 0.06292569 |
| Gm37033 | -0.030463329 | 0.380379912 | 0.762006446 | 0.074008971 | 0.096140893 |
| Gm37052 | -0.031827158 | 0.158634759 | 0.556789205 | 0.024901749 | 0.052382311 |
| Gm37060 | -0.023001389 | 0.358542748 | 0.747310348 | 0.034551503 | 0.051197723 |
| Gm37140 | -0.024162027 | 0.499214368 | 0.830722138 | 0.07777325 | 0.087562996 |
| Gm37254 | -0.002452868 | 0.944389087 | 0.987070339 | 0.078457278 | 0.079684445 |
| Gm37297 | -0.050364216 | 0.139547205 | 0.526950827 | 0.066707234 | 0.108945947 |
| Gm37376 | 0.061245075 | 0.315137527 | 0.714975653 | 0.2403899 | 0.189531293 |
| Gm37407 | -0.025522391 | 0.303677204 | 0.708405917 | 0.033442578 | 0.053962828 |
| Gm37452 | -0.027845376 | 0.47479258 | 0.817387411 | 0.096979245 | 0.116967073 |
| Gm37629 | 0.021740197 | 0.551596024 | 0.852505562 | 0.088544458 | 0.067129219 |
| Gm37728 | -0.0916912 | 0.001153416 | 0.041993113 | 0.037981632 | 0.122561972 |
| Gm37733 | -0.084462435 | 0.000604256 | 0.028340757 | 0.025069333 | 0.095391984 |
| Gm37834 | 0.005317993 | 0.875267012 | 0.965096393 | 0.075308941 | 0.069142855 |
| Gm37859 | -0.020319921 | 0.461261316 | 0.810249429 | 0.04530078 | 0.058966672 |
| Gm37899 | 0.020428549 | 0.66775328 | 0.901929709 | 0.149204159 | 0.12022859 |
| Gm379 | -0.030767465 | 0.558885699 | 0.856615394 | 0.183918747 | 0.213766737 |
| Gm38012 | -0.058263743 | 0.00236563 | 0.066061828 | 0.010837618 | 0.060120856 |
| Gm38020 | -0.067754487 | 0.007565682 | 0.133456507 | 0.03144659 | 0.088196542 |
| Gm38043 | -0.057895646 | 0.002122992 | 0.063221586 | 0.009919361 | 0.061459794 |
| Gm38070 | -0.143376775 | 0.065994677 | 0.390559302 | 0.427513898 | 0.532191864 |
| Gm38190 | -0.01592189 | 0.468906971 | 0.813836982 | 0.026026435 | 0.038645817 |
| Gm38285 | 0.016027613 | 0.617572435 | 0.883238152 | 0.064104673 | 0.048604607 |
| Gm38340 | -0.018267262 | 0.613708361 | 0.881481228 | 0.080249992 | 0.089545087 |
| Gm38365 | -0.01801758 | 0.451836415 | 0.806495742 | 0.030345903 | 0.046890666 |
| Gm38380 | -0.018495464 | 0.459615834 | 0.809611113 | 0.034199889 | 0.047171925 |
| Gm38394 | -0.011014263 | 0.815507836 | 0.949014742 | 0.144627247 | 0.159073744 |
| Gm3924 | -0.016842883 | 0.677358294 | 0.906975267 | 0.108329992 | 0.129634056 |
| Gm39469 | 0.034650591 | 0.243975843 | 0.653830568 | 0.058530381 | 0.035942839 |
| Gm4013 | -0.101275008 | 0.077804282 | 0.418285047 | 0.234055695 | 0.330051051 |
| Gm40309 | 0.016149606 | 0.586747944 | 0.869124722 | 0.056158789 | 0.041308279 |
| Gm40466 | -0.035451557 | 0.134242562 | 0.516942048 | 0.027046515 | 0.057419666 |
| Gm40477 | 0.018977424 | 0.543733547 | 0.850865857 | 0.061631311 | 0.040329016 |
| Gm41584 | -0.064223311 | 0.005210697 | 0.107592455 | 0.0237547 | 0.079413627 |
| Gm4204 | 0.072475184 | 0.054156964 | 0.362869235 | 0.099309289 | 0.037648895 |
| Gm42067 | 0.025688059 | 0.502442372 | 0.832222078 | 0.098266932 | 0.073921418 |
| Gm42440 | -0.04314746 | 0.037827817 | 0.306861584 | 0.018681807 | 0.053108097 |
| Gm42444 | 0.002342158 | 0.945504088 | 0.987084873 | 0.073327738 | 0.0696064 |
| Gm42446 | -0.01182572 | 0.757898529 | 0.927237784 | 0.092432185 | 0.09791652 |

| | | | | | |
|---|---|---|---|---|---|
| Gm42447 | 0.028400774 | 0.47954845 | 0.819939708 | 0.10996268 | 0.079582149 |
| Gm42448 | -0.004391372 | 0.903121874 | 0.974950771 | 0.085813826 | 0.08428247 |
| Gm42467 | -0.030243952 | 0.251685008 | 0.663127858 | 0.036954423 | 0.067225517 |
| Gm42685 | 0.007182802 | 0.840716834 | 0.954960614 | 0.081769095 | 0.067902476 |
| Gm42769 | -0.020852882 | 0.450659282 | 0.806495742 | 0.041646392 | 0.058886814 |
| Gm42770 | -0.040959332 | 0.201271687 | 0.611924411 | 0.061799037 | 0.096989343 |
| Gm42771 | -0.024710101 | 0.262069207 | 0.668764912 | 0.025093102 | 0.044903768 |
| Gm42826 | -0.021201437 | 0.5617333 | 0.857134531 | 0.085615791 | 0.100279663 |
| Gm42941 | -0.092158363 | 0.001071708 | 0.039987855 | 0.034638229 | 0.117531247 |
| Gm42970 | -0.036113919 | 0.283357189 | 0.69129896 | 0.06783042 | 0.10114583 |
| Gm42971 | 0.038158032 | 0.194225896 | 0.603543443 | 0.058551753 | 0.023744704 |
| Gm42986 | -0.054716663 | 0.015037505 | 0.197813154 | 0.021444327 | 0.068909989 |
| Gm42992 | 0.023659212 | 0.430139438 | 0.793295051 | 0.058097124 | 0.036955255 |
| Gm43048 | -0.051334255 | 0.034660672 | 0.293577356 | 0.029214861 | 0.073605276 |
| Gm43138 | -0.059638994 | 0.026551512 | 0.259238835 | 0.036415094 | 0.086063068 |
| Gm43201 | -0.036801304 | 0.21079577 | 0.623520777 | 0.048372277 | 0.079296915 |
| Gm43230 | -0.045396376 | 0.062282715 | 0.380250374 | 0.027646964 | 0.064308816 |
| Gm43305 | -0.043619226 | 0.199393161 | 0.610108047 | 0.069996033 | 0.107688757 |
| Gm43313 | -0.03541707 | 0.085539589 | 0.435299206 | 0.019047927 | 0.047574418 |
| Gm43316 | -0.04091964 | 0.064226098 | 0.386241667 | 0.022789092 | 0.055379305 |
| Gm4332 | -0.027401356 | 0.427234882 | 0.791172788 | 0.072843487 | 0.099994183 |
| Gm43336 | -0.030549618 | 0.275177724 | 0.68241738 | 0.045577921 | 0.069613245 |
| Gm43359 | -0.015752144 | 0.745570232 | 0.924566547 | 0.16025242 | 0.177083548 |
| Gm43360 | 0.005682079 | 0.863322298 | 0.960676137 | 0.068494516 | 0.057914937 |
| Gm43361 | -0.002957981 | 0.92180112 | 0.980859278 | 0.053053224 | 0.055089527 |
| Gm43362 | -0.12443368 | 0.016345967 | 0.206355375 | 0.186683037 | 0.303185158 |
| Gm43609 | 0.038841377 | 0.19234494 | 0.60309438 | 0.057472677 | 0.022461472 |
| Gm4366 | -0.047725167 | 0.036021736 | 0.298801785 | 0.024624047 | 0.061694147 |
| Gm43696 | -0.024475738 | 0.371715035 | 0.75760494 | 0.040373208 | 0.064573473 |
| Gm43775 | -0.035088512 | 0.156540838 | 0.554127149 | 0.033726966 | 0.062711928 |
| Gm43813 | -0.02738211 | 0.318887824 | 0.717175454 | 0.041700283 | 0.06684299 |
| Gm43843 | -0.028088746 | 0.305760164 | 0.709411642 | 0.041987424 | 0.06009891 |
| Gm43872 | -0.078026755 | 0.008889426 | 0.146996602 | 0.046702068 | 0.112989943 |
| Gm43874 | -0.054394014 | 0.055866911 | 0.367454907 | 0.045682612 | 0.087393936 |
| Gm43879 | 0.023782084 | 0.639722268 | 0.891345382 | 0.174827698 | 0.145281294 |
| Gm43880 | -0.048803012 | 0.132484895 | 0.513250014 | 0.061351099 | 0.105348941 |
| Gm43882 | -0.026878521 | 0.289358543 | 0.695472148 | 0.03409546 | 0.059295104 |
| Gm43885 | -0.051246207 | 0.163717152 | 0.563495619 | 0.081336084 | 0.13334608 |
| Gm43889 | -0.105165239 | 0.014136449 | 0.192781984 | 0.110985047 | 0.211169595 |
| Gm43890 | -0.037959149 | 0.106830503 | 0.474004524 | 0.026121154 | 0.057982068 |
| Gm44109 | 0.009631939 | 0.781944463 | 0.936001977 | 0.076322789 | 0.071091464 |
| Gm44164 | -0.035185659 | 0.169676581 | 0.57227323 | 0.034459962 | 0.062442193 |
| Gm4419 | 0.072201608 | 0.15660502 | 0.554127149 | 0.188154349 | 0.133944918 |
| Gm44250 | -0.090405723 | 0.006895079 | 0.126971675 | 0.061737465 | 0.139461782 |
| Gm44509 | -0.006296224 | 0.810441197 | 0.948175579 | 0.041468706 | 0.047295941 |
| Gm44797 | 0.026573595 | 0.409395554 | 0.779735046 | 0.067333501 | 0.042418026 |
| Gm44798 | -0.040869794 | 0.064975561 | 0.388652983 | 0.022718695 | 0.056209174 |
| Gm44799 | -0.015168316 | 0.55350896 | 0.852866048 | 0.03722711 | 0.050529638 |
| Gm44800 | -0.000248046 | 0.994516479 | 0.998695538 | 0.081353559 | 0.073283599 |
| Gm44829 | -0.022142072 | 0.413320812 | 0.781607961 | 0.04069162 | 0.061461826 |
| Gm44850 | 0.046310223 | 0.113815204 | 0.484874023 | 0.056981221 | 0.014593361 |
| Gm44862 | -0.013888201 | 0.595717406 | 0.873388199 | 0.038445332 | 0.051805794 |
| Gm44950 | 0.001119504 | 0.972281335 | 0.994070262 | 0.064959449 | 0.062680771 |
| Gm45051 | 0.044791611 | 0.161843144 | 0.560531277 | 0.072435975 | 0.041598603 |
| Gm45223 | -0.049423474 | 0.040153504 | 0.315774775 | 0.02705833 | 0.070238082 |
| Gm45486 | 0.040189976 | 0.156626355 | 0.554127149 | 0.054157253 | 0.019796922 |
| Gm45605 | -0.046440876 | 0.01228745 | 0.176947483 | 0.011526609 | 0.051141071 |
| Gm45606 | 0.040986519 | 0.197047704 | 0.607422546 | 0.067171404 | 0.030025428 |
| Gm45640 | -0.022766979 | 0.383348572 | 0.763886718 | 0.036242061 | 0.057232108 |
| Gm45716 | 0.02374609 | 0.673384521 | 0.905635862 | 0.220087751 | 0.207087747 |
| Gm45871 | 0.022242235 | 0.415648166 | 0.783472369 | 0.049620035 | 0.034471993 |
| Gm46123 | 0.004925948 | 0.871441903 | 0.964616742 | 0.058780573 | 0.05333502 |
| Gm46430 | -0.003371452 | 0.904227054 | 0.975255421 | 0.048703335 | 0.048762374 |
| Gm4673 | -0.053362318 | 0.031299323 | 0.281212368 | 0.027548671 | 0.082446202 |
| Gm47163 | 0.0760713 | 0.043483958 | 0.329086044 | 0.103106609 | 0.034350242 |

| | | | | | |
|---|---|---|---|---|---|
| Gm47283 | -0.04342425 | 0.146703565 | 0.536416643 | 0.051859553 | 0.092408864 |
| Gm47405 | -0.043853929 | 0.140541642 | 0.528664246 | 0.050605824 | 0.089887026 |
| Gm47585 | -0.02845714 | 0.198661265 | 0.609389286 | 0.022754707 | 0.04527556 |
| Gm47586 | -0.020261572 | 0.585509066 | 0.868186511 | 0.08630431 | 0.103716746 |
| Gm47615 | -0.046220911 | 0.029439745 | 0.273392198 | 0.018169836 | 0.05678741 |
| Gm47889 | -0.011435898 | 0.739567997 | 0.922843014 | 0.07602631 | 0.085334343 |
| Gm4791 | 0.047833895 | 0.280278272 | 0.687465228 | 0.129678461 | 0.083032138 |
| Gm48146 | -0.030454455 | 0.221739505 | 0.633858837 | 0.033878595 | 0.062449712 |
| Gm48225 | -0.0684051 | 0.017779753 | 0.212261786 | 0.042475949 | 0.098611297 |
| Gm48226 | 0.042287601 | 0.411367466 | 0.7808047 | 0.193581738 | 0.162868092 |
| Gm48266 | -0.030628154 | 0.342082252 | 0.735068249 | 0.062194316 | 0.086119122 |
| Gm48267 | 0.030047478 | 0.318129703 | 0.717175454 | 0.062440994 | 0.03410761 |
| Gm48435 | 0.058461697 | 0.04316723 | 0.328373709 | 0.056222863 | 0.008610274 |
| Gm48562 | -0.009966482 | 0.690631582 | 0.909796728 | 0.035382464 | 0.042194996 |
| Gm48623 | -0.027630218 | 0.280410745 | 0.687465228 | 0.037184439 | 0.062526145 |
| Gm49041 | -0.009398502 | 0.764688775 | 0.928757959 | 0.061480456 | 0.069465107 |
| Gm49066 | 0.028321979 | 0.466752622 | 0.813152001 | 0.100541956 | 0.074786758 |
| Gm4924 | -0.002047626 | 0.952174121 | 0.98838165 | 0.074146477 | 0.078541513 |
| Gm49336 | 0.00836687 | 0.859208864 | 0.959220425 | 0.149921984 | 0.136713661 |
| Gm49338 | -0.020453299 | 0.491762294 | 0.827270742 | 0.052724904 | 0.068478088 |
| Gm49359 | -0.054373159 | 0.060110508 | 0.378409279 | 0.049007325 | 0.09482685 |
| Gm49492 | -0.029343588 | 0.318620855 | 0.717175454 | 0.049720797 | 0.07897785 |
| Gm49494 | -0.020975928 | 0.438229045 | 0.799200693 | 0.041345011 | 0.058735521 |
| Gm49542 | -0.025416463 | 0.330205001 | 0.725699056 | 0.038593649 | 0.060693945 |
| Gm49599 | -0.057379677 | 0.386108197 | 0.766272928 | 0.413295716 | 0.475054102 |
| Gm49705 | -0.029040007 | 0.129760493 | 0.509514066 | 0.016786054 | 0.041756144 |
| Gm49741 | -0.003267081 | 0.893266631 | 0.9716581 | 0.035918272 | 0.041683927 |
| Gm49797 | -0.076449856 | 0.034848838 | 0.294178126 | 0.075679805 | 0.141665334 |
| Gm49839 | -0.024004819 | 0.369690489 | 0.756876906 | 0.037985919 | 0.056345881 |
| Gm49980 | -0.017979391 | 0.767003198 | 0.929423047 | 0.297280369 | 0.31377941 |
| Gm50069 | -0.044904115 | 0.013098914 | 0.183622622 | 0.0106218 | 0.047687236 |
| Gm50240 | -0.068528292 | 0.165675388 | 0.566763419 | 0.159966558 | 0.229613804 |
| Gm50268 | 0.01844343 | 0.538085003 | 0.848066852 | 0.056574154 | 0.041705287 |
| Gm50340 | -0.026922007 | 0.381792732 | 0.762597742 | 0.055041166 | 0.083916214 |
| Gm50353 | 0.02697617 | 0.35991969 | 0.74793249 | 0.055731799 | 0.033075474 |
| Gm50394 | -0.057710288 | 0.066800378 | 0.392751172 | 0.0542029 | 0.109274681 |
| Gm50431 | -0.048370146 | 0.029202125 | 0.272146451 | 0.022362419 | 0.062867801 |
| Gm5113 | -0.019247159 | 0.461953794 | 0.810513184 | 0.040339028 | 0.056409591 |
| Gm5141 | -0.034772197 | 0.099415204 | 0.463315751 | 0.021651153 | 0.051228399 |
| Gm52969 | -0.061527246 | 0.001551634 | 0.05174005 | 0.011756648 | 0.063372286 |
| Gm53012 | -0.0318512 | 0.212168618 | 0.625966024 | 0.034864269 | 0.060013849 |
| Gm5617 | -0.087969491 | 0.209501424 | 0.622488392 | 0.603376757 | 0.69651521 |
| Gm5860 | -0.071317547 | 0.172949778 | 0.575371867 | 0.178611532 | 0.23421886 |
| Gm5870 | -0.04659334 | 0.105653057 | 0.472566726 | 0.046267636 | 0.086095193 |
| Gm5914 | 0.102547607 | 0.029380642 | 0.273392198 | 0.159750709 | 0.069598114 |
| Gm6204 | -0.035937092 | 0.087247483 | 0.439557333 | 0.01913977 | 0.045826839 |
| Gm6563 | -0.029999345 | 0.229742824 | 0.641131395 | 0.034367353 | 0.060337193 |
| Gm6710 | -0.069010417 | 0.008108285 | 0.139651258 | 0.032546472 | 0.088116301 |
| Gm6863 | 0.027972321 | 0.333919606 | 0.729929513 | 0.054531651 | 0.029009467 |
| Gm7072 | -0.058736153 | 0.240287676 | 0.650744882 | 0.160366639 | 0.209124445 |
| Gm7160 | 0.017106798 | 0.643498184 | 0.892591387 | 0.089847928 | 0.075076763 |
| Gm7240 | -0.031122848 | 0.439759661 | 0.800048525 | 0.105649 | 0.131788507 |
| Gm7467 | 0.032508089 | 0.353152258 | 0.743880213 | 0.087403201 | 0.066373282 |
| Gm8186 | 0.027012504 | 0.401808189 | 0.775220138 | 0.069976724 | 0.047129928 |
| Gm8624 | -0.038329671 | 0.210421993 | 0.623103727 | 0.055928386 | 0.086651801 |
| Gm8730 | 0.000146287 | 0.99575476 | 0.998839749 | 0.047611407 | 0.047419008 |
| Gm9616 | 0.009156667 | 0.789401576 | 0.93843184 | 0.07773284 | 0.062845773 |
| Gm9794 | 0.009151546 | 0.862795216 | 0.960607386 | 0.206228154 | 0.191809821 |
| Gm9800 | 0.034688336 | 0.258076726 | 0.665948842 | 0.060212725 | 0.02992413 |
| Gm9828 | -0.005380199 | 0.84865541 | 0.957111129 | 0.049603872 | 0.052049975 |
| Gm9843 | -0.02174547 | 0.626624458 | 0.886996623 | 0.131543005 | 0.148942504 |
| Gm9856 | -0.131191327 | 0.001970704 | 0.0606747 | 0.107444486 | 0.22959387 |
| Gm9993 | -0.009311971 | 0.755282092 | 0.926726954 | 0.055745968 | 0.072831619 |
| Gmcl1 | 0.075790328 | 0.123324084 | 0.500122758 | 0.169499971 | 0.107471261 |
| Gmds | -0.049888875 | 0.414444074 | 0.782795478 | 0.34416517 | 0.377715374 |

| | | | | | |
|---|---|---|---|---|---|
| Gmeb1 | -0.001661531 | 0.964812347 | 0.991970325 | 0.09325432 | 0.101301442 |
| Gmeb2 | 0.005815678 | 0.886012023 | 0.969590273 | 0.109165937 | 0.10318885 |
| Gmfb | 0.080592102 | 0.179775845 | 0.586800052 | 0.329843693 | 0.269946214 |
| Gmnn | 0.005767795 | 0.855540887 | 0.957736324 | 0.061387382 | 0.055897992 |
| Gmppa | 0.057152791 | 0.318895492 | 0.717175454 | 0.262132195 | 0.205044 |
| Gmppb | -0.016573148 | 0.788533699 | 0.938071914 | 0.306854082 | 0.322260772 |
| Gmpr | -0.168962952 | 0.011320218 | 0.168327452 | 0.660112889 | 0.871674022 |
| Gmpr2 | -0.041372505 | 0.269182639 | 0.676978829 | 0.088990515 | 0.124459417 |
| Gmps | -0.007606858 | 0.907015134 | 0.977187883 | 0.420965594 | 0.431140457 |
| Gna11 | -0.003106434 | 0.964373824 | 0.991970325 | 0.883264044 | 0.877738017 |
| Gna12 | -0.018328771 | 0.594998846 | 0.873342705 | 0.07662842 | 0.090067418 |
| Gna13 | -0.108099078 | 0.089181811 | 0.443953048 | 0.413727298 | 0.540493665 |
| Gnai1 | -0.015623286 | 0.696393765 | 0.910970095 | 0.102095867 | 0.120343967 |
| Gnai2 | -0.216030288 | 4.60E-06 | 0.00074156 | 2.024287347 | 2.492959142 |
| Gnai3 | 0.141202392 | 0.042171477 | 0.324597482 | 0.808676991 | 0.639936961 |
| Gnao1 | 0.064122747 | 0.308207528 | 0.710654481 | 1.079331381 | 0.953260556 |
| Gnaq | -0.046708769 | 0.498668343 | 0.830722138 | 0.655420206 | 0.694462123 |
| Gnas | -0.072384375 | 0.0340822 | 0.291879728 | 2.730586787 | 2.925114584 |
| Gnaz | -0.019651127 | 0.771297972 | 0.932251553 | 0.51966965 | 0.553229555 |
| Gnb1 | 0.120985631 | 0.066345375 | 0.39159102 | 1.00639247 | 0.846298262 |
| Gnb2 | -0.009437774 | 0.864834658 | 0.961442912 | 1.42350278 | 1.437125496 |
| Gnb4 | -0.035233767 | 0.142998904 | 0.531979243 | 0.028227014 | 0.061788036 |
| Gnb5 | 0.031318139 | 0.533611768 | 0.84612966 | 0.188774214 | 0.162684982 |
| Gne | -0.024850375 | 0.607120853 | 0.878576426 | 0.163374138 | 0.186653502 |
| Gng12 | 0.146963233 | 0.004562208 | 0.099436985 | 2.087471148 | 1.783479659 |
| Gng2 | -0.032524274 | 0.18334879 | 0.590066231 | 0.029032059 | 0.056656057 |
| Gng4 | 0.059163793 | 0.418155903 | 0.784638399 | 0.727054016 | 0.644773684 |
| Gng5 | -0.214402525 | 2.83E-06 | 0.000490569 | 2.067467621 | 2.524279152 |
| Gng7 | -0.039389311 | 0.046154722 | 0.337043414 | 0.015754029 | 0.05001335 |
| Gnl1 | 0.110014598 | 0.066668775 | 0.392235932 | 0.333821738 | 0.242153984 |
| Gnl2 | -0.010095003 | 0.857114972 | 0.958302663 | 0.249088645 | 0.259437721 |
| Gnl3 | -0.095248065 | 0.166615704 | 0.568127031 | 0.60288178 | 0.699049021 |
| Gnl3l | -0.161897145 | 0.011371063 | 0.168833376 | 0.416032429 | 0.575413224 |
| Gnpat | 0.022776114 | 0.629752197 | 0.888005452 | 0.148169479 | 0.124076021 |
| Gnpda1 | 0.074115886 | 0.185243925 | 0.591560356 | 0.234661434 | 0.172937182 |
| Gnpda2 | -0.076603465 | 0.169642123 | 0.57227323 | 0.224026309 | 0.303357775 |
| Gnpnat1 | 0.067115954 | 0.262295927 | 0.668764912 | 0.312016677 | 0.261316393 |
| Gnptab | 0.002217734 | 0.941805982 | 0.986168593 | 0.05961557 | 0.057149325 |
| Gnptg | -0.087567575 | 0.196250506 | 0.606881632 | 0.572898321 | 0.658418616 |
| Gns | -0.131763828 | 0.041259876 | 0.320282582 | 0.803690342 | 0.947868012 |
| Golga1 | -0.014106205 | 0.783632619 | 0.936124803 | 0.197397638 | 0.214635322 |
| Golga2 | -0.01075555 | 0.853766638 | 0.957461155 | 0.299758741 | 0.312384164 |
| Golga3 | 0.004388444 | 0.94980095 | 0.98784658 | 0.523271597 | 0.513086801 |
| Golga4 | -0.141654498 | 0.012407376 | 0.177904051 | 1.502419916 | 1.72920389 |
| Golga5 | -0.181077495 | 0.004172054 | 0.092660198 | 0.317641099 | 0.509613101 |
| Golga7 | 0.01524011 | 0.799839994 | 0.941368905 | 0.314218699 | 0.313947737 |
| Golgb1 | -0.107320136 | 0.06225835 | 0.380250374 | 1.584522927 | 1.74820151 |
| Golim4 | -0.198991266 | 0.005457966 | 0.110446789 | 0.746241954 | 0.997535617 |
| Golm1 | 0.020391301 | 0.568938948 | 0.860524446 | 0.082914275 | 0.064214369 |
| Golm2 | -0.086565239 | 0.190283829 | 0.599459758 | 0.732543931 | 0.831578022 |
| Golph3 | 0.042172896 | 0.539943225 | 0.84917113 | 0.738287713 | 0.679046226 |
| Golph3l | -0.012119523 | 0.734759521 | 0.921732448 | 0.084952877 | 0.094134711 |
| Golt1b | 0.051154791 | 0.441024719 | 0.800354711 | 0.567377114 | 0.54546613 |
| Gon4l | -0.001507712 | 0.971391691 | 0.993892166 | 0.116506943 | 0.1178123 |
| Gon7 | 0.021642899 | 0.607768143 | 0.878733629 | 0.117929688 | 0.105341903 |
| Gopc | 0.040111379 | 0.449698216 | 0.806495742 | 0.212304883 | 0.184458437 |
| Gorab | -0.01578423 | 0.727296198 | 0.920455328 | 0.137616012 | 0.158034937 |
| Gorasp1 | -0.009233972 | 0.771473687 | 0.932251553 | 0.061827677 | 0.071092311 |
| Gorasp2 | 0.158590726 | 0.020507305 | 0.227971987 | 0.808513415 | 0.628070884 |
| Gosr1 | 0.006967055 | 0.895460476 | 0.972277888 | 0.211818565 | 0.214086594 |
| Gosr2 | 0.041035939 | 0.53119188 | 0.845476197 | 0.468773128 | 0.438072228 |
| Got1 | -0.093238763 | 0.11852831 | 0.491192747 | 0.267631317 | 0.347359146 |
| Got2 | -0.000289092 | 0.996428303 | 0.998992348 | 0.469611297 | 0.464104078 |
| Gpaa1 | -0.079507324 | 0.044056826 | 0.329446347 | 0.095870676 | 0.163851368 |
| Gpalpp1 | 0.014887 | 0.773815747 | 0.932949618 | 0.198686043 | 0.186784342 |

| | | | | | |
|---|---|---|---|---|---|
| Gpam | -0.006012985 | 0.851444887 | 0.957461155 | 0.064434256 | 0.072155809 |
| Gpank1 | 0.04023889 | 0.374371724 | 0.75956519 | 0.14382769 | 0.115150428 |
| Gpat4 | 0.087113443 | 0.122211602 | 0.497823804 | 0.252329111 | 0.166346988 |
| Gpatch1 | 0.054119629 | 0.283881576 | 0.691750274 | 0.190333213 | 0.145191935 |
| Gpatch11 | -0.001100215 | 0.984233934 | 0.99608406 | 0.245819232 | 0.24646653 |
| Gpatch2 | 0.091695568 | 0.135750396 | 0.51925561 | 0.356912663 | 0.26013388 |
| Gpatch2l | -0.013352945 | 0.709366225 | 0.914215305 | 0.086899631 | 0.103313575 |
| Gpatch4 | -0.018109529 | 0.675244382 | 0.906025597 | 0.117845214 | 0.133882813 |
| Gpatch8 | 0.056097798 | 0.399856523 | 0.774630365 | 0.492812295 | 0.431764837 |
| Gpbp1 | 0.070091687 | 0.281262423 | 0.68811697 | 0.477591772 | 0.435067308 |
| Gpbp1l1 | -0.153263994 | 0.007151503 | 0.128637347 | 0.232116714 | 0.376247654 |
| Gpc4 | -0.030028702 | 0.606211588 | 0.878341504 | 0.267334394 | 0.303065937 |
| Gpcpd1 | 0.017665088 | 0.730486427 | 0.920770724 | 0.19421276 | 0.186697488 |
| Gpd1 | -0.076039849 | 0.041008815 | 0.319321549 | 0.079944385 | 0.151642752 |
| Gpd1l | -0.062103456 | 0.034760313 | 0.29382176 | 0.047179024 | 0.101727804 |
| Gpd2 | 0.086217235 | 0.205223251 | 0.616549498 | 0.54679476 | 0.454585231 |
| Gphn | 0.098044768 | 0.088921671 | 0.443672569 | 0.274695872 | 0.178427983 |
| Gpi1 | -0.102683941 | 0.118534248 | 0.491192747 | 0.478161852 | 0.592578531 |
| Gpkow | -0.102610327 | 0.009644351 | 0.152441494 | 0.093606217 | 0.200623914 |
| Gpld1 | -0.01278961 | 0.618059146 | 0.883238152 | 0.038374082 | 0.050046213 |
| Gpn1 | 0.023567414 | 0.554725743 | 0.853469818 | 0.106627225 | 0.08713202 |
| Gpn2 | -0.008086445 | 0.821201399 | 0.950930859 | 0.083095552 | 0.090436692 |
| Gpn3 | 0.049385742 | 0.227916229 | 0.639451668 | 0.122885728 | 0.087175662 |
| Gpr107 | -0.024351613 | 0.688356737 | 0.909796728 | 0.31983757 | 0.342616432 |
| Gpr108 | -0.02924289 | 0.579552985 | 0.865234762 | 0.206658835 | 0.235661568 |
| Gpr137 | -0.025092923 | 0.54209314 | 0.850208191 | 0.111617607 | 0.136286071 |
| Gpr137b | -0.029587307 | 0.348382628 | 0.739576235 | 0.062558676 | 0.087168709 |
| Gpr137b-ps | -0.030785126 | 0.423551459 | 0.789048567 | 0.093033118 | 0.126176733 |
| Gpr137c | -0.023169906 | 0.346044766 | 0.738085711 | 0.031927156 | 0.052696453 |
| Gpr155 | -0.073448605 | 0.016360265 | 0.206355375 | 0.04861849 | 0.110843418 |
| Gpr158 | 0.036666894 | 0.560410913 | 0.856904336 | 0.362130641 | 0.320626165 |
| Gpr161 | 0.017644526 | 0.605848791 | 0.878178429 | 0.077666461 | 0.058494935 |
| Gpr180 | -0.032754953 | 0.536913835 | 0.847793898 | 0.200065551 | 0.233818158 |
| Gpr19 | 0.004083945 | 0.901609413 | 0.974269668 | 0.07068109 | 0.062939009 |
| Gpr27 | -0.019715323 | 0.745640809 | 0.924566547 | 0.30395836 | 0.333405238 |
| Gpr45 | -0.188345817 | 0.00416396 | 0.092660198 | 0.450736014 | 0.638923305 |
| Gpr62 | -0.044675049 | 0.056447288 | 0.367897032 | 0.026220894 | 0.064629908 |
| Gpr85 | -0.029012671 | 0.483592657 | 0.822768706 | 0.110642155 | 0.132257397 |
| Gpr89 | 0.054058372 | 0.261539788 | 0.668468258 | 0.17002838 | 0.121277071 |
| Gprasp1 | -0.100344424 | 0.144872179 | 0.533279057 | 0.446985658 | 0.536768478 |
| Gprasp2 | -0.033304568 | 0.56984197 | 0.86083487 | 0.262331933 | 0.295825424 |
| Gprc5c | 0.039475359 | 0.324994946 | 0.720820254 | 0.106681643 | 0.072932599 |
| Gprin1 | 0.066794574 | 0.039240581 | 0.312585484 | 0.070626699 | 0.009224659 |
| Gprin3 | -0.012162582 | 0.814611066 | 0.94873644 | 0.174704654 | 0.182796513 |
| Gps1 | 0.085195259 | 0.191319393 | 0.601966379 | 0.468531946 | 0.389243299 |
| Gps2 | 0.066992316 | 0.220765962 | 0.633858837 | 0.240866542 | 0.167756156 |
| Gpsm1 | 0.041277655 | 0.467858171 | 0.813188482 | 0.23775544 | 0.205206951 |
| Gpt2 | -0.056855534 | 0.233256893 | 0.643935869 | 0.146884881 | 0.195332328 |
| Gpx1 | 0.223279459 | 0.000802058 | 0.033128615 | 1.063031705 | 0.747206467 |
| Gpx2 | -0.260294639 | 0.000450404 | 0.023539853 | 0.607143542 | 0.939932613 |
| Gpx3 | 0.05427166 | 0.468276765 | 0.813213564 | 0.395699166 | 0.303903319 |
| Gpx4 | -0.215825148 | 0.000917548 | 0.03625761 | 1.084356746 | 1.436906015 |
| Gpx4-ps2 | -0.210000898 | 4.68E-05 | 0.004515775 | 0.160255995 | 0.359566096 |
| Gramd1a | 0.014548499 | 0.781137646 | 0.935482467 | 0.213348354 | 0.201044278 |
| Gramd1b | -0.019954214 | 0.630213247 | 0.888281191 | 0.114092731 | 0.120487802 |
| Gramd1c | -0.014598657 | 0.697862814 | 0.910970095 | 0.090785006 | 0.110978266 |
| Gramd3 | -0.110115949 | 0.098966502 | 0.463315751 | 0.491423149 | 0.637520087 |
| Grb10 | -9.47E-05 | 0.998506592 | 0.999402739 | 0.172831437 | 0.170631598 |
| Grb14 | 0.006967701 | 0.892073219 | 0.971118429 | 0.182931378 | 0.164423331 |
| Grb2 | -0.006789255 | 0.919085711 | 0.980436936 | 0.588265421 | 0.605762641 |
| Grb7 | -0.067126866 | 0.141368249 | 0.528664246 | 0.134666628 | 0.185801109 |
| Grcc10 | -0.098274887 | 0.11517274 | 0.487260227 | 0.327502313 | 0.416880119 |
| Grhpr | -0.005389231 | 0.921574625 | 0.98085171 | 0.221831502 | 0.212828051 |
| Gria2 | -0.114129115 | 0.040175788 | 0.315774775 | 0.203873269 | 0.299453166 |
| Grik5 | -0.074630262 | 0.0408663 | 0.319308264 | 0.081288741 | 0.144594333 |

| | | | | | |
|---|---|---|---|---|---|
| Grin1 | 0.091732648 | 0.039027575 | 0.311933003 | 0.142578396 | 0.058787369 |
| Grina | 0.125852583 | 0.069538117 | 0.400662503 | 0.69171677 | 0.54556859 |
| Grip1 | -0.052210207 | 0.283731443 | 0.691561408 | 0.156928305 | 0.20537393 |
| Gripap1 | 0.010320537 | 0.873536972 | 0.964940674 | 0.407020404 | 0.387543275 |
| Grk2 | 0.019949263 | 0.74561524 | 0.924566547 | 0.335415348 | 0.318182764 |
| Grk3 | -0.016550307 | 0.799622173 | 0.941260986 | 0.450448144 | 0.471914059 |
| Grk5 | 0.061587395 | 0.197231621 | 0.607747501 | 0.16353605 | 0.118651172 |
| Grk6 | -0.007246878 | 0.849514866 | 0.957111129 | 0.094359081 | 0.105447757 |
| Grn | 0.031293787 | 0.601654859 | 0.876557788 | 0.302319238 | 0.274367973 |
| Grpel1 | 0.009163951 | 0.89538526 | 0.972277888 | 0.931168875 | 0.914356409 |
| Grpel2 | 0.043893724 | 0.381767014 | 0.762597742 | 0.177045474 | 0.136154731 |
| Grsf1 | -0.033942001 | 0.601704888 | 0.876557788 | 0.484513007 | 0.509475076 |
| Grtp1 | -0.019560912 | 0.700644327 | 0.912109871 | 0.183616503 | 0.192295153 |
| Grwd1 | 0.022357823 | 0.474471036 | 0.817387411 | 0.061458066 | 0.051211729 |
| Gsap | -0.010786281 | 0.731580114 | 0.920770724 | 0.058177793 | 0.068942953 |
| Gsdma | -0.099185201 | 0.154141248 | 0.550967131 | 0.462453967 | 0.581500358 |
| Gsdme | -0.019396461 | 0.444301927 | 0.801932586 | 0.035106978 | 0.050671538 |
| Gse1 | 0.004151367 | 0.951484777 | 0.988311364 | 0.59994256 | 0.570691133 |
| Gsk3a | 0.021666966 | 0.703498796 | 0.912277105 | 0.266827291 | 0.255384835 |
| Gsk3b | 0.085641654 | 0.125683495 | 0.504406266 | 1.257115655 | 1.141994315 |
| Gskip | 0.02255697 | 0.642836694 | 0.892561795 | 0.169890373 | 0.150204701 |
| Gsn | 0.054422951 | 0.406429872 | 0.777612997 | 0.408756255 | 0.363735506 |
| Gspt1 | 0.087118835 | 0.176620181 | 0.581034664 | 1.157322364 | 1.021438315 |
| Gspt2 | -0.015198071 | 0.687410835 | 0.909796728 | 0.090413859 | 0.103733555 |
| Gsr | 0.03396565 | 0.615635679 | 0.882439418 | 0.467739738 | 0.442138841 |
| Gss | -0.050155529 | 0.317467967 | 0.716863138 | 0.178813389 | 0.227990287 |
| Gstm1 | -0.35354278 | 7.58E-07 | 0.000165426 | 0.553820865 | 1.04040394 |
| Gstm2 | -0.061849973 | 0.076894592 | 0.418199494 | 0.067730265 | 0.122204726 |
| Gstm4 | 0.013413265 | 0.628484626 | 0.887864718 | 0.049639222 | 0.040772275 |
| Gstm5 | -0.033193489 | 0.565573063 | 0.858087314 | 0.264695269 | 0.284433431 |
| Gsto1 | -0.089419162 | 0.014526817 | 0.193767197 | 0.077935734 | 0.154663839 |
| Gstp1 | -0.348738808 | 5.94E-07 | 0.000138661 | 0.369645323 | 0.798703358 |
| Gstp-ps | -0.073544988 | 0.003708447 | 0.085962309 | 0.028144839 | 0.096278568 |
| Gstt1 | -0.143038478 | 0.0169591 | 0.208345766 | 0.25683514 | 0.404345946 |
| Gstt2 | -0.060797706 | 0.112680951 | 0.483221516 | 0.090687291 | 0.142319849 |
| Gstz1 | -0.165835741 | 0.004863051 | 0.102275888 | 0.287749279 | 0.452322365 |
| Gt(ROSA)26Sor | 0.034849322 | 0.49512036 | 0.828944629 | 0.189829146 | 0.16119931 |
| Gtf2a1 | -0.021917421 | 0.706029011 | 0.912844672 | 0.257605389 | 0.270417693 |
| Gtf2a2 | -0.054573643 | 0.42087888 | 0.787234019 | 0.702674502 | 0.750840478 |
| Gtf2b | -0.066563749 | 0.308311395 | 0.710730702 | 0.440928127 | 0.520313509 |
| Gtf2e1 | 0.039861052 | 0.178322822 | 0.584719427 | 0.058644885 | 0.02911605 |
| Gtf2e2 | 0.048903999 | 0.400047688 | 0.774697097 | 0.291279305 | 0.264708222 |
| Gtf2f1 | -0.021078165 | 0.712205398 | 0.915109107 | 0.241155033 | 0.259504649 |
| Gtf2f2 | -0.022659991 | 0.691543622 | 0.909796728 | 0.244555833 | 0.27104042 |
| Gtf2h1 | 0.060967625 | 0.273074441 | 0.68112529 | 0.242144963 | 0.187343817 |
| Gtf2h2 | 0.024533682 | 0.426662996 | 0.79105114 | 0.061438559 | 0.037144627 |
| Gtf2h3 | -0.030914247 | 0.38997335 | 0.768386831 | 0.082300948 | 0.114075016 |
| Gtf2h4 | -0.00729126 | 0.844391789 | 0.95591703 | 0.089948466 | 0.097919997 |
| Gtf2h5 | -0.014328752 | 0.82903276 | 0.952883894 | 0.804364736 | 0.782421054 |
| Gtf2i | -0.13956545 | 0.039334686 | 0.312838547 | 0.496595795 | 0.639518125 |
| Gtf2ird1 | 0.075610607 | 0.081061494 | 0.426199169 | 0.134593295 | 0.069989056 |
| Gtf2ird2 | -0.095455663 | 0.000280528 | 0.016660718 | 0.030754815 | 0.11215973 |
| Gtf3a | -0.035028206 | 0.545610255 | 0.851259637 | 0.335915566 | 0.370892345 |
| Gtf3c1 | -0.012493549 | 0.830497488 | 0.95325696 | 0.28753861 | 0.301888257 |
| Gtf3c2 | 0.016544813 | 0.745464764 | 0.924566547 | 0.179881695 | 0.163303864 |
| Gtf3c3 | -0.013441335 | 0.78186735 | 0.936001977 | 0.166652363 | 0.175660025 |
| Gtf3c4 | 0.017715451 | 0.737116922 | 0.92276204 | 0.203111715 | 0.191511148 |
| Gtf3c6 | -0.026907572 | 0.570718203 | 0.861225331 | 0.157494854 | 0.182363854 |
| Gtpbp1 | 0.005014022 | 0.907979367 | 0.977726521 | 0.128092565 | 0.118186022 |
| Gtpbp10 | 0.028870593 | 0.3794984 | 0.762006446 | 0.070447783 | 0.047167617 |
| Gtpbp2 | -0.089151343 | 0.211975656 | 0.625764606 | 0.63493498 | 0.735818142 |
| Gtpbp3 | 0.00476263 | 0.877505095 | 0.966372465 | 0.060141118 | 0.057117616 |
| Gtpbp4 | 0.110268299 | 0.102715429 | 0.468979611 | 0.503330072 | 0.398296409 |
| Gtpbp6 | 0.019981639 | 0.685006264 | 0.909796728 | 0.168872265 | 0.155482401 |
| Gtpbp8 | 0.052173341 | 0.285154362 | 0.692407818 | 0.176942912 | 0.130938164 |

| | | | | | |
|---|---|---|---|---|---|
| Gucd1 | 0.118465539 | 0.01065686 | 0.163053211 | 0.163565606 | 0.063898049 |
| Gucy1b1 | 0.08402387 | 0.029715219 | 0.27412836 | 0.107608622 | 0.033372171 |
| Gucy2c | -0.000351554 | 0.993791169 | 0.998695538 | 0.136120143 | 0.127039421 |
| Guf1 | -0.049636999 | 0.295145481 | 0.701818335 | 0.153845422 | 0.193772105 |
| Guk1 | -0.072342657 | 0.232654496 | 0.643647512 | 1.205846743 | 1.297621412 |
| Gusb | -0.112800981 | 0.044881427 | 0.331476968 | 0.221806194 | 0.338524081 |
| Gxylt1 | 0.005243471 | 0.924460253 | 0.981375217 | 0.219629895 | 0.211325914 |
| Gyg | 0.092103225 | 0.093759571 | 0.45227176 | 0.227551298 | 0.144057943 |
| Gys1 | -0.042969749 | 0.207162327 | 0.618605371 | 0.065857672 | 0.103959651 |
| Gzf1 | 0.048052643 | 0.300488954 | 0.705994295 | 0.154737351 | 0.113228105 |
| H13 | 0.030525119 | 0.655051153 | 0.897454058 | 0.698545525 | 0.648706185 |
| H1f0 | -0.028712767 | 0.690518403 | 0.909796728 | 0.663571627 | 0.736911429 |
| H1f10 | -0.170694744 | 0.004880955 | 0.102275888 | 0.240812203 | 0.425111359 |
| H1f2 | -0.193312151 | 0.01833376 | 0.215223327 | 1.255731957 | 1.518951489 |
| H1f3 | 0.012391835 | 0.755574608 | 0.926726954 | 0.100536677 | 0.086264102 |
| H1f4 | -0.125454101 | 0.055442994 | 0.36562505 | 0.354208817 | 0.494095015 |
| H1f5 | -0.058628061 | 0.000797394 | 0.033072093 | 0.007312169 | 0.057829426 |
| H2ac18 | 0.082640461 | 0.22636052 | 0.638587289 | 0.643369709 | 0.533435821 |
| H2ac20 | -0.052982632 | 0.030941637 | 0.280493857 | 0.028404552 | 0.076125189 |
| H2ac24 | 0.285602351 | 4.85E-05 | 0.004588036 | 0.463963373 | 0.167022054 |
| H2ac6 | 0.043647304 | 0.16428227 | 0.563869641 | 0.065960359 | 0.031062931 |
| H2ac8 | 0.060885458 | 0.220143286 | 0.633234352 | 0.181884541 | 0.123356968 |
| H2aj | -0.0105005 | 0.838863017 | 0.954911647 | 1.672252928 | 1.667154988 |
| H2aw | 0.019692527 | 0.706992417 | 0.913266781 | 0.185540802 | 0.169272114 |
| H2ax | 0.044262186 | 0.441303354 | 0.800354711 | 0.258344407 | 0.223024042 |
| H2az1 | -0.143606757 | 0.001683905 | 0.05417102 | 2.031032073 | 2.328672496 |
| H2az2 | -0.085001232 | 0.233847601 | 0.644459522 | 0.72830054 | 0.797995919 |
| H2bc21 | -0.010663574 | 0.679835273 | 0.907783945 | 0.038339259 | 0.049875059 |
| H2bc4 | 0.153669858 | 0.015259847 | 0.198397782 | 1.895915117 | 1.614568767 |
| H2bc6 | 0.003708759 | 0.930088838 | 0.982596974 | 0.113175507 | 0.108658611 |
| H2bu2 | 0.022610428 | 0.628631925 | 0.887864718 | 0.134550886 | 0.109668842 |
| H2-D1 | -0.229243286 | 0.000266554 | 0.016131031 | 1.302379637 | 1.64027602 |
| H2-DMa | -0.030016282 | 0.207768216 | 0.619907724 | 0.02962314 | 0.05662115 |
| H2-K1 | -0.108602073 | 0.043750985 | 0.329086044 | 1.776805218 | 1.992446266 |
| H2-K2 | -0.016346189 | 0.607731065 | 0.878733629 | 0.063288896 | 0.078302974 |
| H2-Ke6 | 0.053820513 | 0.189599977 | 0.598067444 | 0.121457286 | 0.079163565 |
| H2-M3 | -0.037306451 | 0.409640011 | 0.780033541 | 0.137143086 | 0.157763828 |
| H2-Q2 | -0.149285521 | 0.022444242 | 0.237629595 | 0.419113652 | 0.579945627 |
| H2-Q4 | 0.084536143 | 0.111668729 | 0.482690499 | 0.206687315 | 0.122620581 |
| H2-T22 | 0.06764388 | 0.181161292 | 0.588832867 | 0.187386564 | 0.126635492 |
| H2-T23 | -0.197437599 | 0.000496123 | 0.025103854 | 0.220251674 | 0.418277998 |
| H3c15 | -0.034273332 | 0.126756277 | 0.506268505 | 0.023586136 | 0.050454394 |
| H3f3a | -0.143437393 | 0.002604863 | 0.069641377 | 1.855129737 | 2.13204838 |
| H3f3b | -0.213129384 | 3.16E-05 | 0.003337137 | 2.218967136 | 2.693875231 |
| H4c1 | 0.118818834 | 0.002302316 | 0.065344539 | 0.107768691 | 0 |
| H4c14 | -0.01854986 | 0.577372477 | 0.864551328 | 0.066556441 | 0.084442678 |
| H4c3 | -0.02733674 | 0.312443307 | 0.714349311 | 0.040998825 | 0.063465607 |
| H4c4 | -0.024151751 | 0.617264987 | 0.883238152 | 0.160024227 | 0.176894085 |
| H4c8 | 0.146114147 | 0.021807699 | 0.235078507 | 0.35268121 | 0.203092429 |
| H4c9 | 0.098993384 | 0.095262494 | 0.455812225 | 0.268345531 | 0.180128832 |
| H6pd | -0.010893252 | 0.728891407 | 0.920522999 | 0.063205137 | 0.067680434 |
| Habp4 | 0.041123891 | 0.426858113 | 0.79105114 | 0.202070905 | 0.169287435 |
| Hacd1 | 0.020221848 | 0.613096373 | 0.881106572 | 0.108191732 | 0.098475877 |
| Hacd2 | 0.031595659 | 0.453960193 | 0.807844867 | 0.121899377 | 0.095535758 |
| Hacd3 | 0.048923094 | 0.363011783 | 0.7514022 | 0.218319917 | 0.163092528 |
| Hace1 | -0.065164307 | 0.089820306 | 0.445385778 | 0.089374642 | 0.151234074 |
| Hacl1 | -0.060127575 | 0.000928723 | 0.036555283 | 0.009527385 | 0.06209793 |
| Hadh | -0.066517873 | 0.237210459 | 0.648388175 | 1.832239862 | 1.973902565 |
| Hadha | -0.087978086 | 0.171587868 | 0.574479873 | 0.452113396 | 0.533997522 |
| Hadhb | -0.082523478 | 0.200351251 | 0.610852219 | 0.424021788 | 0.523049073 |
| Hagh | -0.206639966 | 0.001998206 | 0.060960472 | 0.603141576 | 0.862406956 |
| Haghl | -0.073393044 | 0.133450457 | 0.514972026 | 0.155838926 | 0.214506945 |
| Hap1 | -0.264256595 | 0.000220148 | 0.014074076 | 0.800734208 | 1.1351188 |
| Hapln1 | -0.002627214 | 0.943778954 | 0.986918206 | 0.086542296 | 0.082864201 |
| Hapln4 | -0.051609911 | 0.288415585 | 0.694795498 | 0.164706031 | 0.204756473 |

| | | | | | |
|---|---|---|---|---|---|
| Harbi1 | 0.045317363 | 0.325538087 | 0.720833548 | 0.152136684 | 0.121676207 |
| Hars | -0.05899705 | 0.282012563 | 0.688868362 | 0.223598369 | 0.273378375 |
| Hars2 | -0.042452497 | 0.229500154 | 0.641012885 | 0.075841864 | 0.119282226 |
| Hat1 | -0.044719267 | 0.290816818 | 0.69667177 | 0.114298374 | 0.161411409 |
| Haus1 | 0.035241746 | 0.2543627 | 0.663146569 | 0.063659734 | 0.03435465 |
| Haus2 | -0.030744205 | 0.476317495 | 0.817948631 | 0.124077087 | 0.14435159 |
| Haus7 | -0.034906213 | 0.251604484 | 0.663127858 | 0.053496192 | 0.084177108 |
| Hax1 | -0.169182408 | 0.012780261 | 0.179909504 | 0.579369228 | 0.769535181 |
| Hbegf | 0.007682634 | 0.869332456 | 0.963610145 | 0.135117944 | 0.122476135 |
| Hbp1 | -0.087295056 | 0.125189288 | 0.503791124 | 0.238447492 | 0.335695 |
| Hbs1l | 0.057623132 | 0.379224918 | 0.762006446 | 0.42770869 | 0.380476994 |
| Hccs | -0.077337474 | 0.117000404 | 0.489713534 | 0.16317952 | 0.230323707 |
| Hcfc1 | 0.00867148 | 0.86680331 | 0.962125879 | 0.181105969 | 0.187714861 |
| Hcfc1r1 | 0.196422288 | 0.003975917 | 0.090217721 | 1.091557652 | 0.840318637 |
| Hcfc2 | -0.041564711 | 0.336750512 | 0.731477644 | 0.125032814 | 0.15753206 |
| Hcn2 | -0.072376112 | 0.078212029 | 0.41934516 | 0.105743813 | 0.166955053 |
| Hcn4 | 0.032777248 | 0.464363035 | 0.811633402 | 0.139356655 | 0.10860396 |
| Hdac1 | -0.038548837 | 0.446912648 | 0.804170357 | 0.176067278 | 0.205759274 |
| Hdac11 | 0.064017757 | 0.189204995 | 0.597561527 | 0.174254475 | 0.117412361 |
| Hdac2 | 0.015923119 | 0.790845706 | 0.938824685 | 0.335218554 | 0.317246224 |
| Hdac3 | 0.004454605 | 0.922975193 | 0.980859278 | 0.152094408 | 0.153810997 |
| Hdac4 | 0.064457167 | 0.196800872 | 0.60713211 | 0.18758089 | 0.131368711 |
| Hdac5 | 0.030663754 | 0.621167545 | 0.884242858 | 0.361087142 | 0.327279054 |
| Hdac6 | -0.056205203 | 0.399830016 | 0.774630365 | 0.453595028 | 0.498842522 |
| Hdac7 | -0.015053137 | 0.623887083 | 0.886265909 | 0.057712521 | 0.069581698 |
| Hdac8 | -0.017877962 | 0.466337433 | 0.813032623 | 0.033302534 | 0.045369828 |
| Hddc2 | -0.010647597 | 0.815935634 | 0.949036482 | 0.143154856 | 0.153290965 |
| Hddc3 | 0.040458358 | 0.163202844 | 0.562993622 | 0.058856099 | 0.023803934 |
| Hdgf | 0.040263663 | 0.554535319 | 0.853469818 | 0.793826246 | 0.736345297 |
| Hdgfl2 | -0.074207277 | 0.13440225 | 0.516977677 | 0.16927182 | 0.239469531 |
| Hdgfl3 | -0.041897419 | 0.523616321 | 0.841062594 | 0.404799569 | 0.468369593 |
| Hdhd5 | -0.081859357 | 0.003981912 | 0.090217721 | 0.041650727 | 0.110989761 |
| Hdlbp | -0.071304308 | 0.211443684 | 0.624561584 | 1.381807741 | 1.469610116 |
| Heatr1 | -0.078269991 | 0.013593237 | 0.189230674 | 0.052989648 | 0.11990508 |
| Heatr3 | 0.004564635 | 0.911722776 | 0.979026586 | 0.109020509 | 0.105685631 |
| Heatr5a | 0.036273675 | 0.333813333 | 0.729929513 | 0.097290478 | 0.061043527 |
| Heatr5b | -0.005539133 | 0.873450716 | 0.964940674 | 0.077500185 | 0.079988337 |
| Heatr6 | -0.068968229 | 0.188598226 | 0.596771878 | 0.192711159 | 0.267030262 |
| Heca | -0.099968761 | 0.092846609 | 0.451283978 | 0.288392866 | 0.375337772 |
| Hectd1 | -0.046189672 | 0.520158766 | 0.838864153 | 0.76725269 | 0.849008559 |
| Hectd3 | 0.046795145 | 0.17550647 | 0.579269463 | 0.082975686 | 0.04598707 |
| Hectd4 | 0.004831819 | 0.922612513 | 0.980859278 | 0.166833056 | 0.157978984 |
| Heg1 | 0.07974884 | 0.109116365 | 0.477972625 | 0.187454446 | 0.120551644 |
| Helb | -0.01311022 | 0.693051377 | 0.909848241 | 0.066553529 | 0.079599629 |
| Helq | -0.011741878 | 0.787783685 | 0.937804093 | 0.123671757 | 0.131023959 |
| Helz | -0.106044341 | 0.085063207 | 0.434272336 | 0.349054317 | 0.44576594 |
| Helz2 | -0.031209976 | 0.28348974 | 0.69129896 | 0.048888044 | 0.067554712 |
| Hemk1 | 0.01083185 | 0.705851206 | 0.912844672 | 0.051454189 | 0.041870977 |
| Hepacam2 | 0.129963829 | 0.049443174 | 0.34898814 | 0.684307501 | 0.518037923 |
| Herc1 | -0.001688583 | 0.980808982 | 0.995540893 | 0.624297784 | 0.609159238 |
| Herc2 | -0.055991638 | 0.391842062 | 0.769704269 | 0.44297363 | 0.49726713 |
| Herc3 | -0.026646031 | 0.50221629 | 0.832218079 | 0.097323307 | 0.121704884 |
| Herc4 | 0.029550709 | 0.566438308 | 0.858732162 | 0.198496344 | 0.159424761 |
| Herc6 | 0.003923272 | 0.930327098 | 0.982692866 | 0.133273882 | 0.120260176 |
| Herpud1 | -0.05298857 | 0.443527461 | 0.801522297 | 1.235420957 | 1.311327619 |
| Herpud2 | 0.01839738 | 0.744718748 | 0.924566547 | 0.261061787 | 0.246691075 |
| Hes1 | -0.005856982 | 0.872186318 | 0.964745713 | 0.084146446 | 0.087632764 |
| Hes6 | -0.04288461 | 0.489151896 | 0.826256745 | 0.355978598 | 0.411493004 |
| Hexa | -0.278019785 | 4.50E-05 | 0.004405397 | 0.661826452 | 1.019895196 |
| Hexb | -0.081717836 | 0.046567875 | 0.338417216 | 0.101681111 | 0.169482462 |
| Hexim1 | -0.010201959 | 0.842864474 | 0.955306089 | 0.181414635 | 0.186725768 |
| Hfe | -0.135939989 | 0.000843891 | 0.034153784 | 0.09381196 | 0.226119633 |
| Hgfac | 0.009092318 | 0.815394835 | 0.948993269 | 0.103523337 | 0.092064092 |
| Hgh1 | 0.051429741 | 0.085971814 | 0.436689826 | 0.059235348 | 0.01774999 |
| Hgs | 0.018909265 | 0.726966283 | 0.920455328 | 0.222882035 | 0.201403081 |

| | | | | | |
|---|---|---|---|---|---|
| Hgsnat | 0.010738154 | 0.799501886 | 0.941260986 | 0.12421414 | 0.107985557 |
| Hhat | -0.028945875 | 0.256485359 | 0.665229303 | 0.035234238 | 0.059550362 |
| Hibadh | 0.033507474 | 0.571058237 | 0.861225331 | 0.319890641 | 0.297855085 |
| Hibch | 0.001393953 | 0.973760794 | 0.994079882 | 0.122589456 | 0.117469219 |
| Hic2 | -0.01957356 | 0.504296465 | 0.832249514 | 0.051916104 | 0.064397703 |
| Hid1 | -0.155527559 | 0.00720244 | 0.128860765 | 0.26160763 | 0.408174976 |
| Hif1a | 0.07939123 | 0.235782497 | 0.646327697 | 0.473920815 | 0.401899311 |
| Hif1an | 0.007399703 | 0.872865647 | 0.964911254 | 0.149447502 | 0.145781416 |
| Higd2a | -0.017991602 | 0.79604763 | 0.939881197 | 0.655085474 | 0.669343648 |
| Hikeshi | 0.026795958 | 0.609412386 | 0.878855197 | 0.204986832 | 0.177407328 |
| Hilpda | 0.039837232 | 0.295810914 | 0.702426542 | 0.10106969 | 0.066844994 |
| Hinfp | 0.008450718 | 0.797428744 | 0.940690722 | 0.071527181 | 0.069521601 |
| Hint1 | -0.111028676 | 0.033979532 | 0.291747273 | 1.598377746 | 1.784372989 |
| Hint2 | -0.208857249 | 0.001448105 | 0.049437526 | 0.472679116 | 0.696064187 |
| Hint3 | -0.014172469 | 0.836293588 | 0.954911647 | 0.729021493 | 0.7404147 |
| Hip1 | -0.028670664 | 0.340820401 | 0.734080335 | 0.050688941 | 0.07526659 |
| Hip1r | -0.135395175 | 0.022175975 | 0.236913658 | 0.260956481 | 0.395644239 |
| Hipk1 | -0.020037897 | 0.729930624 | 0.920522999 | 0.262277158 | 0.278367025 |
| Hipk2 | 0.015564054 | 0.81005279 | 0.947942154 | 0.449540931 | 0.411605915 |
| Hipk3 | 0.070465151 | 0.28037567 | 0.687465228 | 0.420916655 | 0.344827837 |
| Hira | -0.043394328 | 0.264312245 | 0.67104216 | 0.099216792 | 0.136042869 |
| Hirip3 | 0.041307523 | 0.380701629 | 0.762006446 | 0.152299323 | 0.106196473 |
| Hivep1 | 0.023914367 | 0.684429178 | 0.909679945 | 0.265255862 | 0.237172426 |
| Hivep2 | 0.007968623 | 0.896318092 | 0.97233366 | 0.356031232 | 0.32454326 |
| Hivep3 | 0.113504067 | 0.078794402 | 0.421117898 | 0.41662001 | 0.300814648 |
| Hjurp | 0.047636041 | 0.284179887 | 0.691925959 | 0.138027089 | 0.09291451 |
| Hlcs | 0.089212451 | 0.045815845 | 0.335834191 | 0.148076258 | 0.070019101 |
| Hlf | 0.063298779 | 0.101189918 | 0.466962393 | 0.099032211 | 0.04290881 |
| Hltf | -0.044066943 | 0.419607526 | 0.786186437 | 0.216647226 | 0.244760646 |
| Hmbox1 | 0.042788674 | 0.366824929 | 0.753633555 | 0.153763251 | 0.117674432 |
| Hmbs | -0.08769556 | 0.117155439 | 0.489953809 | 0.215991572 | 0.302271351 |
| Hmces | 0.036733118 | 0.549268291 | 0.852402509 | 0.312454582 | 0.287256151 |
| Hmg20a | 0.020894265 | 0.67406767 | 0.905947671 | 0.168089935 | 0.143104165 |
| Hmg20b | -0.019044842 | 0.745938177 | 0.924566547 | 0.301844397 | 0.317586355 |
| Hmga1 | 0.067806329 | 0.053515799 | 0.360979891 | 0.085758291 | 0.023635074 |
| Hmgb1 | -0.172867825 | 0.004952151 | 0.10355155 | 1.171748913 | 1.460832289 |
| Hmgb2 | -0.025395161 | 0.679400181 | 0.907783945 | 0.31445371 | 0.359380201 |
| Hmgcl | 0.036554121 | 0.50097328 | 0.831764926 | 0.220915926 | 0.188328584 |
| Hmgcll1 | 0.050769991 | 0.175296499 | 0.578766765 | 0.100121454 | 0.059357444 |
| Hmgcr | 0.132935856 | 0.063154289 | 0.383440442 | 0.633937837 | 0.479681585 |
| Hmgcs1 | 0.070244227 | 0.223041318 | 0.634902356 | 0.270291715 | 0.206957963 |
| Hmgn1 | 0.134348646 | 0.042855746 | 0.327353972 | 1.042770511 | 0.87113493 |
| Hmgn2 | -0.060937495 | 0.346829383 | 0.738383053 | 0.414305161 | 0.482590761 |
| Hmgn3 | 0.034583808 | 0.570483349 | 0.86113346 | 1.299520242 | 1.256012491 |
| Hmgn5 | -0.079512727 | 0.133776725 | 0.515636323 | 0.19391951 | 0.263061057 |
| Hmgxb3 | 0.067041481 | 0.13012312 | 0.509514066 | 0.139175678 | 0.078322649 |
| Hmgxb4 | -0.08679524 | 0.144549813 | 0.533279057 | 0.281680121 | 0.359882984 |
| Hmox1 | -0.263709975 | 3.03E-07 | 8.23E-05 | 0.144155446 | 0.398649781 |
| Hmox2 | -0.088549043 | 0.177325743 | 0.582591976 | 0.493389934 | 0.572917394 |
| Hnf1a | 0.014681266 | 0.714561352 | 0.915970918 | 0.105011082 | 0.091900101 |
| Hnf1aos1 | -0.033037756 | 0.521373014 | 0.840205104 | 0.187353851 | 0.221671514 |
| Hnf1b | -0.078401357 | 0.025377661 | 0.2537682 | 0.071681949 | 0.143542725 |
| Hnf4a | -0.037924803 | 0.210827776 | 0.623520777 | 0.050953138 | 0.083738383 |
| Hnf4g | -0.022390984 | 0.560516485 | 0.856904336 | 0.096177083 | 0.110269416 |
| Hnrnpa0 | 0.00310038 | 0.962919219 | 0.991878382 | 0.560171485 | 0.56756095 |
| Hnrnpa1 | 0.015743732 | 0.821322613 | 0.950930859 | 0.487637091 | 0.493239431 |
| Hnrnpa2b1 | -0.038155027 | 0.556246196 | 0.855114576 | 1.265990244 | 1.279991348 |
| Hnrnpa3 | -0.022021164 | 0.729335668 | 0.920522999 | 1.293760169 | 1.328524186 |
| Hnrnpab | -0.060148965 | 0.366856331 | 0.753633555 | 0.936876009 | 1.002840561 |
| Hnrnpc | 0.153212327 | 0.014237831 | 0.193060412 | 1.026039171 | 0.822307895 |
| Hnrnpd | 0.012067421 | 0.862348827 | 0.960216905 | 0.916758609 | 0.893176246 |
| Hnrnpdl | -0.03745915 | 0.606388215 | 0.878341504 | 0.914134538 | 0.951479328 |
| Hnrnpf | -0.016574337 | 0.806819181 | 0.945532285 | 0.961742769 | 0.977357041 |
| Hnrnph1 | -0.119550278 | 0.066376616 | 0.39159102 | 0.679418892 | 0.805251787 |
| Hnrnph2 | -0.060127717 | 0.344760304 | 0.737445936 | 0.411375806 | 0.46577722 |

| | | | | | |
|---|---|---|---|---|---|
| Hnrnph3 | 0.111260274 | 0.099338333 | 0.463315751 | 0.572573157 | 0.484758794 |
| Hnrnpk | 0.026734629 | 0.633021526 | 0.889464068 | 1.411413603 | 1.407066774 |
| Hnrnpl | -0.015091486 | 0.813572443 | 0.948397417 | 0.489600685 | 0.484192304 |
| Hnrnpll | 0.103842582 | 0.061704688 | 0.380250374 | 0.233991612 | 0.14457225 |
| Hnrnpm | 0.006793593 | 0.917295885 | 0.980132949 | 0.463427516 | 0.451855989 |
| Hnrnpr | 0.081421442 | 0.220869234 | 0.633858837 | 0.626051138 | 0.549785315 |
| Hnrnpu | -0.191393373 | 0.00334989 | 0.081411238 | 1.206696951 | 1.457234777 |
| Hnrnpul1 | -0.010164646 | 0.871792986 | 0.964616742 | 0.413503922 | 0.415988302 |
| Hnrnpul2 | -0.017889576 | 0.798803343 | 0.941031591 | 0.74349102 | 0.741381416 |
| Homer1 | -0.023555377 | 0.456923837 | 0.808898849 | 0.060893192 | 0.081108098 |
| Homez | -0.041110761 | 0.172572145 | 0.575294174 | 0.053258032 | 0.084547487 |
| Hook1 | -0.052220618 | 0.441304891 | 0.800354711 | 0.692876254 | 0.72858408 |
| Hook2 | -0.069810545 | 0.168653876 | 0.57227323 | 0.168750336 | 0.228606219 |
| Hook3 | -0.252388903 | 0.000154247 | 0.010687095 | 0.399171793 | 0.649031792 |
| Hopx | 0.010996508 | 0.853859507 | 0.957461155 | 0.286556976 | 0.278018404 |
| Hp1bp3 | -0.209561948 | 0.002331194 | 0.06554117 | 0.623618661 | 0.82966031 |
| Hpcal1 | 0.053252109 | 0.35855457 | 0.747310348 | 0.301036867 | 0.24853026 |
| Hpf1 | 0.040891244 | 0.517455776 | 0.837694133 | 0.411730712 | 0.357306178 |
| Hpgds | 0.160464451 | 0.002129012 | 0.063221586 | 0.196873143 | 0.045552908 |
| Hprt | -0.106038485 | 0.114695948 | 0.486355399 | 0.536083857 | 0.638082826 |
| Hps1 | -0.028709328 | 0.404021444 | 0.77632362 | 0.075899855 | 0.103580449 |
| Hps3 | -0.017800363 | 0.654960545 | 0.897454058 | 0.104381321 | 0.116990955 |
| Hps4 | -0.047001845 | 0.360620623 | 0.749054824 | 0.183962422 | 0.225378137 |
| Hps5 | 0.041780533 | 0.192548251 | 0.60325934 | 0.070985456 | 0.036424108 |
| Hps6 | -0.010215072 | 0.68607002 | 0.909796728 | 0.038908597 | 0.04813701 |
| Hras | -0.028969021 | 0.655668998 | 0.897825804 | 0.516536885 | 0.545759035 |
| Hs1bp3 | -0.02692213 | 0.405604144 | 0.777214356 | 0.062287405 | 0.083099069 |
| Hs2st1 | -0.062350094 | 0.199151395 | 0.609601265 | 0.157128793 | 0.205915168 |
| Hs3st6 | 0.06394625 | 0.216702324 | 0.631259121 | 0.207696421 | 0.15938295 |
| Hs6st1 | -0.058241536 | 0.292036823 | 0.697566301 | 0.216982964 | 0.273467488 |
| Hsbp1 | 0.037190983 | 0.571290052 | 0.861225331 | 1.034502354 | 0.953547462 |
| Hsbp1l1 | -0.068408007 | 0.132458089 | 0.513250014 | 0.139613217 | 0.19983623 |
| Hscb | -0.086693231 | 0.056809734 | 0.368822317 | 0.132851446 | 0.220335969 |
| Hsd17b10 | -0.069857862 | 0.215913149 | 0.629630897 | 0.230696574 | 0.291296321 |
| Hsd17b11 | -0.032362544 | 0.53969909 | 0.84899357 | 0.199972469 | 0.220801658 |
| Hsd17b12 | 0.022034332 | 0.729683043 | 0.920522999 | 0.455965059 | 0.421960665 |
| Hsd17b4 | -0.085001329 | 0.054953533 | 0.364775849 | 0.126010167 | 0.193783863 |
| Hsd17b7 | 0.007832653 | 0.79523586 | 0.939757086 | 0.058667123 | 0.051584115 |
| Hsdl1 | -0.017996442 | 0.716588229 | 0.916579082 | 0.175378816 | 0.196778679 |
| Hsdl2 | -0.012302375 | 0.813339554 | 0.948397417 | 0.190409186 | 0.198986351 |
| Hsf1 | 0.019315715 | 0.684330101 | 0.909679945 | 0.151955633 | 0.133012046 |
| Hsf2 | 0.045094701 | 0.439421692 | 0.800048525 | 0.287733609 | 0.255770682 |
| Hsf4 | -0.02078103 | 0.43909507 | 0.799999494 | 0.040938172 | 0.062651034 |
| Hsp90aa1 | 0.26820582 | 1.04E-05 | 0.001327097 | 2.827450844 | 2.12240737 |
| Hsp90ab1 | 0.102196572 | 0.002245777 | 0.064959253 | 3.835684712 | 3.484072895 |
| Hsp90b1 | -0.056461855 | 0.144885661 | 0.533279057 | 2.811775691 | 2.899023565 |
| Hspa12a | 0.108771524 | 0.01081302 | 0.164475284 | 0.133484341 | 0.039239683 |
| Hspa13 | -0.141896518 | 0.021399355 | 0.232200356 | 0.324674319 | 0.462614824 |
| Hspa14 | 0.140122203 | 0.010570795 | 0.161983305 | 0.245623085 | 0.127014453 |
| Hspa1a | -0.114677435 | 0.149689429 | 0.544162548 | 0.323195961 | 0.405616672 |
| Hspa1b | -0.009598432 | 0.891144972 | 0.970742575 | 0.231247786 | 0.234786076 |
| Hspa2 | -0.122638497 | 0.010884266 | 0.165272884 | 0.133059843 | 0.258407565 |
| Hspa4 | -0.026015718 | 0.699699015 | 0.911507617 | 0.897372372 | 0.94685674 |
| Hspa4l | 0.288416155 | 8.16E-05 | 0.006856998 | 0.761971369 | 0.409943773 |
| Hspa5 | 0.14510333 | 0.000769844 | 0.032691088 | 3.131644592 | 2.747364936 |
| Hspa8 | -0.056526728 | 0.441162572 | 0.800354711 | 1.535235291 | 1.65337526 |
| Hspa9 | -0.141063352 | 0.018052443 | 0.213922515 | 1.622355737 | 1.88768074 |
| Hspb1 | 0.574447171 | 4.89E-07 | 0.00011975 | 1.068950966 | 0.227357905 |
| Hspb11 | -0.020300136 | 0.711105314 | 0.914546911 | 0.240553602 | 0.272853183 |
| Hspb6 | 0.018795806 | 0.723625368 | 0.919830015 | 0.206692904 | 0.189470183 |
| Hspb8 | 0.09072591 | 0.057114681 | 0.369250461 | 0.155327515 | 0.076015652 |
| Hspbp1 | -0.017999498 | 0.70848357 | 0.913767764 | 0.159215736 | 0.170429972 |
| Hspd1 | 0.387386457 | 1.30E-07 | 4.09E-05 | 1.96709896 | 1.243130697 |
| Hspe1 | 0.599956609 | 2.32E-18 | 5.83E-15 | 2.3158626 | 1.185149074 |
| Hspe1-rs1 | 0.131715936 | 0.002049296 | 0.062141346 | 0.139153346 | 0.028649837 |

| | | | | | |
|---|---|---|---|---|---|
| Hsph1 | 0.430549543 | 1.43E-05 | 0.001697747 | 1.49278031 | 0.734508483 |
| Htatip2 | -0.364956572 | 2.32E-09 | 1.45E-06 | 0.261783621 | 0.64775397 |
| Htatsf1 | -0.011451078 | 0.844732674 | 0.955950886 | 0.277661265 | 0.288314619 |
| Htra2 | -0.011258056 | 0.751189804 | 0.926186851 | 0.080825253 | 0.09051299 |
| Htt | 0.036870795 | 0.492487202 | 0.827850285 | 0.227487728 | 0.184790099 |
| Hus1 | -0.033863055 | 0.353714007 | 0.744256656 | 0.086026297 | 0.117335163 |
| Huwe1 | -0.214690736 | 0.00191405 | 0.059662472 | 0.588515756 | 0.8207319 |
| Hyal2 | -0.015784097 | 0.636427477 | 0.890755879 | 0.068817818 | 0.086460472 |
| Hyal3 | -0.04311681 | 0.032932621 | 0.287906455 | 0.017022135 | 0.05295843 |
| Hyi | -0.091611006 | 0.135951682 | 0.519431685 | 0.330505766 | 0.425787761 |
| Hyou1 | -0.050363439 | 0.466260073 | 0.813032623 | 0.725750136 | 0.763270305 |
| Hypk | 0.013122993 | 0.8106409 | 0.948188174 | 0.229849707 | 0.230043283 |
| Iah1 | 0.035862822 | 0.525936193 | 0.842051614 | 0.251278084 | 0.223596775 |
| Iapp | -0.056340503 | 1.08E-08 | 6.38E-06 | 8.951856438 | 9.311670536 |
| Iars | -0.046228426 | 0.463868654 | 0.811254677 | 0.391278339 | 0.416500912 |
| Iars2 | 0.060406608 | 0.275481508 | 0.682717999 | 0.237649461 | 0.183202071 |
| Ibtk | -0.095848117 | 0.084819891 | 0.433957325 | 0.226386059 | 0.327663699 |
| Ica1 | 0.005775107 | 0.929201105 | 0.982502239 | 0.525013916 | 0.512203481 |
| Ica1l | -0.103444581 | 0.042906272 | 0.327490687 | 0.166823485 | 0.260131446 |
| Ice1 | 0.075955294 | 0.168299423 | 0.571912906 | 0.239174185 | 0.156676104 |
| Ice2 | -0.057906002 | 0.048190495 | 0.344017069 | 0.046955824 | 0.095294497 |
| Icmt | -0.06625796 | 0.12392197 | 0.501565251 | 0.121100325 | 0.178598711 |
| Id2 | -0.235294305 | 0.003422123 | 0.082765892 | 0.6571008 | 0.929978723 |
| Id3 | -0.307000043 | 2.20E-08 | 9.59E-06 | 0.121648271 | 0.437333576 |
| Ide | 0.049894891 | 0.325579119 | 0.720833548 | 0.192590996 | 0.147152723 |
| Idh1 | 0.002597692 | 0.961596466 | 0.991441074 | 0.211523153 | 0.210932862 |
| Idh2 | 0.1855862 | 0.00763986 | 0.134528558 | 0.807802509 | 0.591722157 |
| Idh3a | -0.025283634 | 0.673759627 | 0.905654932 | 0.300672006 | 0.314121794 |
| Idh3b | 0.031345379 | 0.607266129 | 0.878640642 | 0.31280883 | 0.287640464 |
| Idh3g | -0.055515177 | 0.35582906 | 0.745326231 | 0.318928757 | 0.364232834 |
| Idi1 | 0.057810588 | 0.075159616 | 0.413795734 | 0.073299828 | 0.032166985 |
| Idnk | 0.170794235 | 0.010980346 | 0.165713947 | 0.557395003 | 0.381170834 |
| Ids | -0.065449943 | 0.335950282 | 0.730805382 | 0.76872141 | 0.828371784 |
| Idua | -0.055162906 | 0.107712949 | 0.47605234 | 0.071728917 | 0.12385659 |
| Ier2 | -0.388918471 | 9.63E-08 | 3.36E-05 | 0.454917669 | 0.892927397 |
| Ier3 | -0.09901693 | 0.009307529 | 0.149711006 | 0.082487915 | 0.177457166 |
| Ier3ip1 | 0.070563372 | 0.280740865 | 0.687895392 | 0.953231585 | 0.855604073 |
| Ier5 | -0.196136028 | 0.001229236 | 0.044063714 | 0.266826215 | 0.446045641 |
| Iffo1 | 0.029383474 | 0.511704457 | 0.835213712 | 0.142115339 | 0.120517604 |
| Ifi27 | 0.064240148 | 0.397919241 | 0.774311756 | 0.59635216 | 0.475204207 |
| Ifi30 | 0.005003421 | 0.929241684 | 0.982502239 | 0.250567113 | 0.243535841 |
| Ifi35 | 0.030961046 | 0.564096111 | 0.857926031 | 0.210393362 | 0.186896147 |
| Ifih1 | -0.008544681 | 0.845747543 | 0.956051742 | 0.122821804 | 0.1264701 |
| Ifit1 | 0.165929274 | 0.004204112 | 0.09314939 | 0.220329501 | 0.040323887 |
| Ifit3 | 0.079214457 | 0.037270749 | 0.304267772 | 0.088991182 | 0.008159573 |
| Ifitm2 | -0.187169946 | 0.000473002 | 0.024222066 | 0.17463378 | 0.339057942 |
| Ifnar1 | 0.031622093 | 0.423916281 | 0.78954309 | 0.104807859 | 0.084701538 |
| Ifnar2 | -0.057143719 | 0.153489384 | 0.549817979 | 0.100221983 | 0.162348637 |
| Ifngr1 | 0.035689323 | 0.306083529 | 0.709680069 | 0.082358601 | 0.058622562 |
| Ifngr2 | 0.036676444 | 0.390131604 | 0.76844731 | 0.126767558 | 0.09799755 |
| Ifrd1 | -0.157277128 | 0.025935406 | 0.256582976 | 0.524190511 | 0.685096527 |
| Ifrd2 | 0.018299105 | 0.496356453 | 0.830056214 | 0.046828411 | 0.038084157 |
| Ift122 | -0.083686014 | 0.046794639 | 0.339117542 | 0.109438791 | 0.178644247 |
| Ift172 | -0.084021331 | 0.016330939 | 0.206355375 | 0.070823567 | 0.139508029 |
| Ift20 | -0.178877076 | 0.010311685 | 0.15907731 | 0.875028966 | 1.100816207 |
| Ift22 | 0.078957742 | 0.154993461 | 0.552106544 | 0.2337024 | 0.157860938 |
| Ift27 | -0.038138515 | 0.394190426 | 0.770644587 | 0.131651369 | 0.157607151 |
| Ift43 | -0.047547832 | 0.135052837 | 0.518167175 | 0.057890543 | 0.089931919 |
| Ift46 | -0.09160847 | 0.046172631 | 0.337043414 | 0.134206944 | 0.215571151 |
| Ift52 | -0.001481153 | 0.976382495 | 0.994468977 | 0.172774032 | 0.166341667 |
| Ift57 | 0.069417057 | 0.338294794 | 0.732353334 | 0.792648107 | 0.682668158 |
| Ift74 | -0.04377449 | 0.477542019 | 0.818492016 | 0.327123386 | 0.352664332 |
| Ift80 | -0.076844999 | 0.033718443 | 0.290749155 | 0.075335795 | 0.141907328 |
| Ift81 | 0.024899103 | 0.695915564 | 0.910916082 | 0.400376217 | 0.360594076 |
| Ift88 | 0.017972375 | 0.713887308 | 0.915840939 | 0.168354923 | 0.159974696 |

| | | | | | |
|---|---|---|---|---|---|
| Iftap | -0.112340187 | 0.040120056 | 0.315774775 | 0.226463612 | 0.334351317 |
| Igbp1 | 0.007376579 | 0.888865978 | 0.970176435 | 0.201439119 | 0.19140819 |
| Igdcc4 | -0.014882872 | 0.557777428 | 0.855384435 | 0.034905214 | 0.048727728 |
| Igf1r | -0.144098587 | 0.059336194 | 0.375386569 | 1.485420485 | 1.709577754 |
| Igf2bp2 | 0.069060434 | 0.020931973 | 0.229383208 | 0.060082193 | 0 |
| Igf2r | 0.117229982 | 0.074078791 | 0.412383666 | 0.495168332 | 0.35914092 |
| Igfals | -0.046359551 | 0.298090458 | 0.703072054 | 0.134048734 | 0.181996414 |
| Igfbp7 | 0.006284883 | 0.929393076 | 0.982502239 | 0.559046711 | 0.571636309 |
| Ighmbp2 | 0.008262944 | 0.793175365 | 0.939604453 | 0.063522706 | 0.054864576 |
| Igip | 0.105148026 | 0.042309705 | 0.325136871 | 0.187086215 | 0.090213384 |
| Igsf1 | 0.033971493 | 0.272438245 | 0.680045428 | 0.06582639 | 0.043987436 |
| Igsf11 | -0.017168554 | 0.51962402 | 0.838531832 | 0.041367456 | 0.056609512 |
| Igsf8 | -0.005788574 | 0.906057189 | 0.976494793 | 0.169550239 | 0.170289807 |
| Igsf9b | -0.033070578 | 0.534496187 | 0.846347806 | 0.201927098 | 0.223175422 |
| Ik | 0.068871814 | 0.307972967 | 0.710440053 | 0.609808991 | 0.539777307 |
| Ikbip | -0.140674941 | 0.008435628 | 0.142539393 | 0.204645949 | 0.347112732 |
| Ikbkb | -0.061377183 | 0.178599097 | 0.584861057 | 0.140416396 | 0.183835712 |
| Ikbkg | -0.082398744 | 0.035126664 | 0.295034581 | 0.088313553 | 0.164209693 |
| Ikzf2 | 0.047725184 | 0.36442221 | 0.752769237 | 0.19715659 | 0.162057559 |
| Ikzf3 | 0.006339373 | 0.867703915 | 0.962549093 | 0.096571758 | 0.095405561 |
| Ikzf4 | -0.009636426 | 0.821458349 | 0.950930859 | 0.121018261 | 0.116504188 |
| Ikzf5 | -0.018050973 | 0.65499003 | 0.897454058 | 0.103283779 | 0.121208979 |
| Il10rb | 0.015452589 | 0.666535578 | 0.901197061 | 0.086774703 | 0.076390644 |
| Il11ra1 | -0.095046366 | 0.004830314 | 0.10205513 | 0.062705593 | 0.147058886 |
| Il13ra1 | 0.038385858 | 0.367249765 | 0.754003328 | 0.122865194 | 0.087025205 |
| Il15ra | 0.027984416 | 0.33785881 | 0.732352032 | 0.05520463 | 0.03230387 |
| Il17ra | 0.079054041 | 0.064773604 | 0.388281728 | 0.131313962 | 0.0624062 |
| Il17rc | -0.000349077 | 0.992167037 | 0.998695538 | 0.083641947 | 0.083069009 |
| Il17re | 0.051070473 | 0.106871798 | 0.474004524 | 0.066360655 | 0.021958019 |
| Il18bp | 0.010431134 | 0.764091705 | 0.928730554 | 0.081739559 | 0.06586567 |
| Il1r1 | -0.160397748 | 0.006962254 | 0.127751639 | 1.654926488 | 1.91624208 |
| Il3ra | -0.053825374 | 0.071775925 | 0.40586758 | 0.047802132 | 0.091204055 |
| Il6ra | -0.101388229 | 0.101928111 | 0.467829632 | 0.330018572 | 0.429255144 |
| Il6st | -0.033410286 | 0.603285195 | 0.876723966 | 0.394009697 | 0.438624593 |
| Ildr1 | -0.026105759 | 0.658371043 | 0.898224427 | 0.282606077 | 0.301113754 |
| Ilf2 | -0.009057127 | 0.880078934 | 0.967751911 | 0.32765872 | 0.344357645 |
| Ilf3 | -0.089491135 | 0.128114224 | 0.508260084 | 0.260796981 | 0.344923552 |
| Ilk | 0.076816699 | 0.18514202 | 0.591560356 | 0.287730596 | 0.209907072 |
| Ilkap | 0.126718657 | 0.058741768 | 0.372687182 | 0.482602058 | 0.350950115 |
| Ilrun | 0.114035745 | 0.091046337 | 0.445496137 | 0.586148773 | 0.472496203 |
| Ilvbl | -0.018592952 | 0.625664746 | 0.88683922 | 0.092849656 | 0.106826123 |
| Immp1l | -0.119450034 | 0.07488763 | 0.41344108 | 0.528220683 | 0.684384491 |
| Immp2l | -0.027472457 | 0.332263749 | 0.72806771 | 0.048164901 | 0.071353355 |
| Immt | -0.155392472 | 0.017132204 | 0.208919982 | 0.442257846 | 0.592053886 |
| Imp3 | 0.098828986 | 0.132557454 | 0.513250014 | 0.751120069 | 0.637471505 |
| Imp4 | 0.023587866 | 0.595517097 | 0.873342705 | 0.144330635 | 0.126691014 |
| Impa1 | -0.039951998 | 0.495906935 | 0.829846267 | 0.29223572 | 0.332592636 |
| Impact | -0.044030505 | 0.516896319 | 0.837583212 | 0.497379069 | 0.544830429 |
| Impdh1 | -0.149813404 | 0.00914126 | 0.1482335 | 0.254803821 | 0.399853977 |
| Impdh2 | -0.0871639 | 0.065653242 | 0.389687514 | 0.139600729 | 0.226571459 |
| Ina | 0.044891465 | 0.484744847 | 0.823246028 | 0.391912002 | 0.360586833 |
| Inafm1 | -0.05879703 | 0.289917504 | 0.69604181 | 0.22559924 | 0.276976677 |
| Inafm2 | 0.030103365 | 0.570542743 | 0.86113346 | 0.203881916 | 0.169782959 |
| Ing1 | 0.035792106 | 0.512343612 | 0.835753752 | 0.232193115 | 0.196729508 |
| Ing2 | 0.159475275 | 0.020816058 | 0.229342232 | 0.888242301 | 0.697125387 |
| Ing3 | -0.032678294 | 0.394893117 | 0.771549623 | 0.090016301 | 0.118996892 |
| Ing5 | -0.022631528 | 0.628255556 | 0.887864718 | 0.150119316 | 0.177335078 |
| Inha | 0.071855106 | 0.090723406 | 0.445385778 | 0.131095956 | 0.056658042 |
| Inip | 0.052179878 | 0.180019879 | 0.586833233 | 0.105468208 | 0.065380067 |
| Inka2 | 0.017543676 | 0.561049473 | 0.857134531 | 0.055305521 | 0.044138697 |
| Ino80 | -0.057786296 | 0.206346632 | 0.617685998 | 0.140442528 | 0.188394001 |
| Ino80b | -0.03066834 | 0.50717213 | 0.833462342 | 0.140764206 | 0.160150248 |
| Ino80c | -0.058245909 | 0.01572501 | 0.202348621 | 0.026658887 | 0.074717744 |
| Ino80d | -0.02055938 | 0.726528621 | 0.920263474 | 0.268185032 | 0.286312346 |
| Ino80e | 0.01811682 | 0.73723511 | 0.92276204 | 0.230817067 | 0.221671893 |

| | | | | | |
|---|---|---|---|---|---|
| Inpp1 | 0.035825809 | 0.401076281 | 0.774740795 | 0.128308305 | 0.098356809 |
| Inpp4a | 0.055060249 | 0.377960386 | 0.761937562 | 0.376082755 | 0.317393229 |
| Inpp5a | 0.090656031 | 0.114383118 | 0.485644398 | 0.279832664 | 0.191776162 |
| Inpp5b | -0.058150394 | 0.104870035 | 0.472399448 | 0.077110328 | 0.130363439 |
| Inpp5e | 0.059500704 | 0.042441195 | 0.325425723 | 0.059497801 | 0.00890831 |
| Inpp5f | 0.014669844 | 0.832636407 | 0.953797263 | 0.762339857 | 0.731661491 |
| Inpp5j | -0.004658503 | 0.890639664 | 0.970579007 | 0.073377616 | 0.074861321 |
| Inpp5k | -0.019546142 | 0.594470163 | 0.873162794 | 0.087975344 | 0.104105794 |
| Inppl1 | -0.038885948 | 0.399657611 | 0.774544012 | 0.142105508 | 0.172815322 |
| Ins1 | 0.011920901 | 0.423272456 | 0.788783074 | 10.42754545 | 10.25929699 |
| Ins2 | -0.01365544 | 0.155318814 | 0.552356396 | 11.75416259 | 11.82953309 |
| Insig1 | 0.158255023 | 0.008476483 | 0.142749092 | 0.297353743 | 0.165435893 |
| Insig2 | 0.072430681 | 0.274132113 | 0.681791282 | 0.488689334 | 0.434616677 |
| Insm1 | 0.083213513 | 0.199656715 | 0.610108047 | 1.332489329 | 1.210332222 |
| Insr | 0.019358852 | 0.690904987 | 0.909796728 | 0.164371885 | 0.148750005 |
| Insrr | -0.245794244 | 5.12E-05 | 0.004718243 | 0.286911772 | 0.525597825 |
| Ints1 | -0.024910115 | 0.496567975 | 0.830056214 | 0.08454363 | 0.101361518 |
| Ints10 | -0.052496291 | 0.195917372 | 0.606495975 | 0.103047588 | 0.15496932 |
| Ints11 | -0.005311394 | 0.897841302 | 0.972876254 | 0.114186881 | 0.121549292 |
| Ints12 | 0.084476637 | 0.01778541 | 0.212261786 | 0.091586195 | 0.023852891 |
| Ints13 | -0.021563091 | 0.620531322 | 0.883937014 | 0.124776616 | 0.13621231 |
| Ints14 | -0.048665473 | 0.122006192 | 0.49725002 | 0.059189849 | 0.098028831 |
| Ints2 | -0.012726794 | 0.756229699 | 0.926833377 | 0.105270271 | 0.116602439 |
| Ints3 | 0.016237361 | 0.763676371 | 0.928730554 | 0.208635861 | 0.182788197 |
| Ints4 | 0.018494077 | 0.631048105 | 0.888584432 | 0.101065294 | 0.083484888 |
| Ints5 | -0.003376669 | 0.926043306 | 0.981772637 | 0.083693448 | 0.091799428 |
| Ints6 | -0.11382543 | 0.047371916 | 0.340840086 | 0.232937842 | 0.323874195 |
| Ints6l | -0.166715579 | 0.000370914 | 0.020682602 | 0.137688389 | 0.280834639 |
| Ints7 | -0.021696827 | 0.480186164 | 0.819943608 | 0.058612805 | 0.073792633 |
| Ints8 | -0.020610897 | 0.6010361 | 0.876370606 | 0.099471729 | 0.115945067 |
| Ints9 | -0.008135728 | 0.835740276 | 0.954911647 | 0.102818473 | 0.112056764 |
| Intu | -0.048308656 | 0.219509809 | 0.63289494 | 0.09525392 | 0.132991638 |
| Invs | -0.065572513 | 0.062219059 | 0.380250374 | 0.072908426 | 0.134495791 |
| Ip6k1 | -0.039463611 | 0.534314361 | 0.846347806 | 0.407224135 | 0.441122127 |
| Ip6k2 | 0.008743256 | 0.893596404 | 0.971676636 | 0.54865327 | 0.556426079 |
| Ipmk | 0.134632663 | 0.041487913 | 0.321059507 | 0.421533025 | 0.279595957 |
| Ipo11 | 0.049524962 | 0.077058126 | 0.418199494 | 0.052363005 | 0.009476755 |
| Ipo13 | 0.045074895 | 0.215444959 | 0.629611185 | 0.091535979 | 0.052104422 |
| Ipo4 | -0.054423778 | 0.285426248 | 0.692407818 | 0.178590667 | 0.227532139 |
| Ipo5 | 0.142902015 | 0.019414079 | 0.221110882 | 0.355083919 | 0.214312881 |
| Ipo7 | -0.004267837 | 0.946062938 | 0.987084873 | 0.408273856 | 0.423521266 |
| Ipo8 | -0.034058714 | 0.45768024 | 0.808898849 | 0.135353704 | 0.170939541 |
| Ipo9 | -0.086264778 | 0.028985065 | 0.271130568 | 0.095837578 | 0.174691767 |
| Ipp | -0.006711317 | 0.857239064 | 0.958302663 | 0.089019088 | 0.096192861 |
| Ippk | 0.074781476 | 0.075796944 | 0.415113505 | 0.124172669 | 0.057248674 |
| Iqcb1 | 0.003470016 | 0.94631423 | 0.987084873 | 0.192673867 | 0.203820565 |
| Iqcc | -0.051463853 | 0.16664003 | 0.568127031 | 0.089391646 | 0.132466863 |
| Iqce | 0.01878061 | 0.588504577 | 0.86992264 | 0.082818664 | 0.061264579 |
| Iqgap1 | -0.023321165 | 0.702129034 | 0.912152193 | 1.149624457 | 1.202599335 |
| Iqgap2 | 0.012339579 | 0.695307842 | 0.910378884 | 0.060381324 | 0.042353714 |
| Irak1 | -0.064089806 | 0.339769394 | 0.733286616 | 0.571724374 | 0.647302446 |
| Irak1bp1 | -0.076541863 | 0.23575957 | 0.646327697 | 0.336297934 | 0.397327048 |
| Irak4 | -0.046333922 | 0.020427814 | 0.227815517 | 0.015360693 | 0.053799626 |
| Ireb2 | -0.115575707 | 0.082254253 | 0.428430691 | 0.41395941 | 0.534050633 |
| Irf1 | -0.132306749 | 0.011467108 | 0.169507156 | 0.165024345 | 0.297516864 |
| Irf2 | -0.009915784 | 0.879223259 | 0.967221631 | 0.439352929 | 0.448623731 |
| Irf2bp1 | 0.071631412 | 0.225687356 | 0.638019508 | 0.283906167 | 0.227866669 |
| Irf2bp2 | -0.017957884 | 0.782460222 | 0.936026395 | 0.952115355 | 1.019924045 |
| Irf2bpl | 0.042509646 | 0.540468321 | 0.849380522 | 0.532406554 | 0.475222674 |
| Irf3 | 0.045708963 | 0.310994436 | 0.713603232 | 0.143323585 | 0.104781609 |
| Irf6 | -0.000467954 | 0.993924986 | 0.998695538 | 0.360228016 | 0.370739814 |
| Irf7 | 0.008396961 | 0.806227009 | 0.945445148 | 0.074549778 | 0.059702323 |
| Irf9 | 0.003835272 | 0.926573006 | 0.981818446 | 0.119960173 | 0.111587669 |
| Irgm1 | 0.041948106 | 0.305468659 | 0.709411642 | 0.114794382 | 0.068972294 |
| Irgq | 0.024215947 | 0.626650698 | 0.886996623 | 0.168129563 | 0.142599297 |

| | | | | | |
|---|---|---|---|---|---|
| Irs2 | -0.096952409 | 0.16272893 | 0.562628407 | 0.478197762 | 0.567252762 |
| Irx2 | -0.039012827 | 0.132697398 | 0.513250014 | 0.028285486 | 0.062820407 |
| Isca1 | 0.0559956 | 0.260971462 | 0.66834369 | 0.178924384 | 0.130813819 |
| Isca2 | 0.074426629 | 0.217656552 | 0.632272789 | 0.330291379 | 0.253579797 |
| Iscu | -0.065940094 | 0.329444719 | 0.724538308 | 0.499969389 | 0.573577134 |
| Isg20 | 0.021005817 | 0.757080777 | 0.92705828 | 0.909796918 | 0.862192687 |
| Isg20l2 | -0.02853027 | 0.339227997 | 0.732701236 | 0.053801407 | 0.078889779 |
| Isl1 | 0.088985347 | 0.20714187 | 0.618605371 | 0.934726667 | 0.806834437 |
| Isoc1 | 0.098443329 | 0.136184559 | 0.519952162 | 0.553312798 | 0.456549301 |
| Isoc2b | -0.017453468 | 0.749332618 | 0.925326217 | 0.232970653 | 0.262659613 |
| Ist1 | 0.069670209 | 0.183613574 | 0.590066231 | 0.208516171 | 0.149129412 |
| Isy1 | 0.01789418 | 0.720463788 | 0.919412687 | 0.178488424 | 0.165000886 |
| Isyna1 | 0.010730221 | 0.738122188 | 0.922843014 | 0.065969157 | 0.051531945 |
| Itch | -0.064407201 | 0.279964856 | 0.687465228 | 0.298737355 | 0.364612124 |
| Itfg1 | -0.016953804 | 0.787610385 | 0.937804093 | 0.391664329 | 0.411193504 |
| Itfg2 | -0.034215233 | 0.408829786 | 0.779525942 | 0.115341728 | 0.144383732 |
| Itga2 | 0.003292807 | 0.915887737 | 0.980132949 | 0.056703173 | 0.053658389 |
| Itga3 | -0.000147182 | 0.996486592 | 0.998992348 | 0.07248944 | 0.067901425 |
| Itga6 | -0.068711223 | 0.249589374 | 0.661305641 | 0.255480563 | 0.297457027 |
| Itgav | 0.120414775 | 0.086478497 | 0.437271876 | 0.709701302 | 0.583557363 |
| Itgb1 | 0.218440267 | 0.001373213 | 0.047626161 | 0.878881601 | 0.613837766 |
| Itgb1bp1 | 0.013755926 | 0.80864785 | 0.946960503 | 0.250361922 | 0.23561367 |
| Itgb3bp | 0.017574183 | 0.749331371 | 0.925326217 | 0.240305045 | 0.230355307 |
| Itgb5 | 0.057182217 | 0.280633286 | 0.687842806 | 0.22515273 | 0.170937184 |
| Itgb8 | 0.051714357 | 0.299917435 | 0.70506682 | 0.174593733 | 0.11838211 |
| Itm2b | -0.109243498 | 0.009135907 | 0.1482335 | 2.335746427 | 2.592515222 |
| Itm2c | -0.024362304 | 0.722433503 | 0.919483582 | 0.891086464 | 0.906310287 |
| Itpa | 0.037718009 | 0.494677689 | 0.828842054 | 0.2382092 | 0.196616211 |
| Itpk1 | 0.072968541 | 0.04841698 | 0.344953921 | 0.093075312 | 0.026913983 |
| Itpkb | 0.142836082 | 0.020901809 | 0.229383208 | 0.381227134 | 0.24095592 |
| Itpkc | -0.002239869 | 0.928790342 | 0.982502239 | 0.037645636 | 0.040237449 |
| Itpr1 | -0.201360014 | 4.91E-05 | 0.004588036 | 0.134743819 | 0.312766431 |
| Itpr2 | -0.130458713 | 0.003336475 | 0.081282038 | 0.113554618 | 0.23008741 |
| Itpr3 | -0.061709 | 0.130639148 | 0.510509367 | 0.10315949 | 0.156123663 |
| Itprid2 | 0.114223924 | 0.070586802 | 0.402952703 | 0.413429013 | 0.298308617 |
| Itsn1 | -0.00065381 | 0.984285706 | 0.99608406 | 0.070923759 | 0.070618358 |
| Itsn2 | -0.066100252 | 0.268095014 | 0.676097903 | 0.299118796 | 0.360207927 |
| Ivd | -0.025115892 | 0.665569236 | 0.901006558 | 0.277529677 | 0.304165963 |
| Ivns1abp | 0.018054662 | 0.78696179 | 0.937461311 | 0.515008427 | 0.478342208 |
| Iws1 | 0.018131574 | 0.769653023 | 0.931397082 | 0.332678768 | 0.316384727 |
| Izumo4 | 0.004618387 | 0.922287643 | 0.980859278 | 0.152265883 | 0.150980393 |
| Jade1 | -0.00574798 | 0.934521195 | 0.983960188 | 0.910845164 | 0.905360054 |
| Jade2 | -0.029965831 | 0.501961241 | 0.832218079 | 0.133146888 | 0.171365764 |
| Jade3 | 0.023425848 | 0.654914484 | 0.897454058 | 0.194381876 | 0.173712595 |
| Jag2 | 0.050158237 | 0.188048243 | 0.595974902 | 0.099429139 | 0.054689462 |
| Jagn1 | -0.06506341 | 0.346110099 | 0.738085711 | 0.507157655 | 0.562951609 |
| Jak1 | 0.015599546 | 0.816860948 | 0.94911078 | 0.808657226 | 0.811466087 |
| Jak2 | 0.014957391 | 0.771969918 | 0.932402174 | 0.180812 | 0.15996048 |
| Jakmip1 | -0.121401971 | 0.009482559 | 0.151151979 | 0.128932169 | 0.241035022 |
| Jam2 | -0.004385163 | 0.910291666 | 0.978641543 | 0.096608929 | 0.099727007 |
| Jam3 | -0.002799198 | 0.927222402 | 0.981818446 | 0.059019142 | 0.06144563 |
| Jarid2 | -0.059739305 | 0.241358504 | 0.651739388 | 0.170759032 | 0.225846661 |
| Jazf1 | 0.027389536 | 0.384277617 | 0.764518225 | 0.063838131 | 0.040759988 |
| Jkamp | 0.094425644 | 0.112577198 | 0.483221516 | 0.364576012 | 0.28040716 |
| Jmjd1c | 0.00491892 | 0.943205252 | 0.986873983 | 0.538419658 | 0.537688735 |
| Jmjd6 | 0.053735877 | 0.340197274 | 0.733546237 | 0.25359674 | 0.201119834 |
| Jmjd8 | -0.005416773 | 0.905044073 | 0.975822039 | 0.14131064 | 0.141946306 |
| Jmy | -0.049022179 | 0.373390145 | 0.758519786 | 0.227015693 | 0.278017563 |
| Josd1 | 0.029917982 | 0.581562107 | 0.866173066 | 0.230619079 | 0.198501136 |
| Josd2 | -0.038365795 | 0.467656024 | 0.813188482 | 0.206506989 | 0.239335588 |
| Jph3 | -0.045781538 | 0.202451219 | 0.612972214 | 0.077024321 | 0.115632748 |
| Jph4 | -0.022190436 | 0.377937291 | 0.761937562 | 0.033808328 | 0.051684614 |
| Jpt1 | -0.014775319 | 0.770356387 | 0.931574344 | 0.177498373 | 0.193244033 |
| Jpt2 | 0.022482164 | 0.63582535 | 0.890315156 | 0.163427856 | 0.150755301 |
| Jpx | 0.05168396 | 0.055282044 | 0.365283656 | 0.052348369 | 0.008970781 |

| | | | | | |
|---|---|---|---|---|---|
| Jrkl | -0.000928993 | 0.978605963 | 0.995264774 | 0.077522534 | 0.083551935 |
| Jtb | -0.147717869 | 0.031134125 | 0.280770404 | 0.702420734 | 0.880686641 |
| Jun | -0.357325149 | 4.52E-05 | 0.004405397 | 0.876997253 | 1.383211248 |
| Junb | -0.351657206 | 9.54E-06 | 0.001260277 | 0.743545421 | 1.205946236 |
| Jund | -0.00117643 | 0.983360662 | 0.99608406 | 2.158346748 | 2.129469889 |
| Jup | 0.078478074 | 0.184758686 | 0.591385002 | 0.248075223 | 0.176435676 |
| Kalrn | -0.071924625 | 0.026803854 | 0.260688257 | 0.05789881 | 0.11814047 |
| Kank1 | -0.023124856 | 0.458968613 | 0.809370658 | 0.059636913 | 0.077422083 |
| Kansl1 | -0.014151795 | 0.813913128 | 0.948397417 | 0.32737197 | 0.334781221 |
| Kansl1l | 0.025914038 | 0.418806931 | 0.7851594 | 0.068115707 | 0.044586432 |
| Kansl2 | 0.033142295 | 0.521769295 | 0.840205104 | 0.193673299 | 0.167319241 |
| Kansl3 | -0.035525188 | 0.487434449 | 0.826112275 | 0.184556347 | 0.220888336 |
| Kantr | 0.055563329 | 0.245308911 | 0.655005464 | 0.15903857 | 0.101657783 |
| Kars | 0.054256111 | 0.347820593 | 0.739124045 | 0.299837123 | 0.231027835 |
| Kat14 | 0.004428829 | 0.929312521 | 0.982502239 | 0.186673339 | 0.181759873 |
| Kat2a | -0.060181952 | 0.065503877 | 0.389491953 | 0.062022158 | 0.114057424 |
| Kat2b | 0.077664678 | 0.153061899 | 0.549602359 | 0.225056653 | 0.154944268 |
| Kat5 | 0.014568826 | 0.751615909 | 0.926186851 | 0.144192171 | 0.134041399 |
| Kat6a | -0.096284378 | 0.119786688 | 0.493284734 | 0.314540728 | 0.404939793 |
| Kat6b | 0.005628756 | 0.919565622 | 0.980436936 | 0.250096356 | 0.249948613 |
| Kat7 | 0.072454838 | 0.193192182 | 0.60325934 | 0.243334505 | 0.18952063 |
| Kat8 | -0.013813134 | 0.698138316 | 0.911092742 | 0.081005604 | 0.103110706 |
| Katna1 | 0.099040489 | 0.088213421 | 0.441426313 | 0.303541609 | 0.217952712 |
| Katnal1 | 0.021211643 | 0.538625184 | 0.848357012 | 0.082878171 | 0.060155359 |
| Katnb1 | -0.050218055 | 0.055347559 | 0.365475957 | 0.034159493 | 0.073842298 |
| Katnbl1 | 0.00906689 | 0.824171411 | 0.951594208 | 0.110718129 | 0.097600105 |
| Katnip | -0.029062537 | 0.532823249 | 0.84612966 | 0.147553474 | 0.170441245 |
| Kazald1 | -0.075720211 | 0.003773503 | 0.086973441 | 0.031763815 | 0.096358443 |
| Kazn | 0.023983448 | 0.465695893 | 0.812791331 | 0.073106996 | 0.059794385 |
| Kbtbd2 | 0.03506289 | 0.519410415 | 0.838496482 | 0.233628948 | 0.198283113 |
| Kbtbd3 | -0.013175289 | 0.762280019 | 0.92872278 | 0.122744915 | 0.132365839 |
| Kbtbd4 | 0.057063451 | 0.131084122 | 0.510948091 | 0.104442096 | 0.063381675 |
| Kbtbd7 | -0.039146492 | 0.365261807 | 0.752802229 | 0.12015497 | 0.159660368 |
| Kcmf1 | 0.076658694 | 0.246598546 | 0.656353649 | 0.704163743 | 0.632301398 |
| Kcna2 | 0.043504502 | 0.205415374 | 0.616549498 | 0.077576975 | 0.035993936 |
| Kcnab3 | -0.005006748 | 0.896383716 | 0.97233366 | 0.087548214 | 0.085673761 |
| Kcnb1 | -0.07361335 | 0.249182086 | 0.66061449 | 0.363765867 | 0.445881953 |
| Kcnb2 | 0.083103689 | 0.102019562 | 0.467993757 | 0.191083784 | 0.115983776 |
| Kcnc3 | 0.027073723 | 0.534135498 | 0.846347806 | 0.125758899 | 0.096987054 |
| Kcnh1 | -0.027172583 | 0.566562753 | 0.858742125 | 0.153530867 | 0.163010574 |
| Kcnh2 | -0.09587994 | 0.054634705 | 0.363639613 | 0.167916856 | 0.243496595 |
| Kcnh6 | 0.009096625 | 0.750320968 | 0.92612569 | 0.053008431 | 0.044436998 |
| Kcnh8 | 0.067259506 | 0.02305732 | 0.240567904 | 0.059297317 | 0 |
| Kcnip1 | -0.131554925 | 0.047370799 | 0.340840086 | 0.55855592 | 0.724701597 |
| Kcnip4 | -0.038348277 | 0.145349519 | 0.533628684 | 0.03719194 | 0.07211321 |
| Kcnj11 | -0.052769019 | 0.362220573 | 0.750073837 | 0.252929397 | 0.309708993 |
| Kcnj12 | 0.052979115 | 0.110016956 | 0.479062989 | 0.073599126 | 0.023686103 |
| Kcnj6 | 0.017157729 | 0.54905107 | 0.852402509 | 0.051691898 | 0.038807859 |
| Kcnk1 | 0.026805271 | 0.672612267 | 0.905446529 | 0.367482902 | 0.338138603 |
| Kcnk12 | 0.062287818 | 0.226332459 | 0.638587289 | 0.190930621 | 0.143826731 |
| Kcnk16 | 0.151619864 | 0.018760968 | 0.217462874 | 0.399669807 | 0.229848598 |
| Kcnma1 | -0.091068924 | 0.169393216 | 0.57227323 | 0.3779947 | 0.461883399 |
| Kcnmb2 | -0.018751911 | 0.787180289 | 0.937573105 | 0.552285538 | 0.555250393 |
| Kcnn3 | -0.0270701 | 0.619881016 | 0.883866495 | 0.206152668 | 0.223961463 |
| Kcnq1ot1 | 0.002931399 | 0.97026209 | 0.993892166 | 1.021977433 | 0.976692467 |
| Kcnq4 | -0.035309794 | 0.143564744 | 0.532857888 | 0.030901674 | 0.062896203 |
| Kctd1 | -0.032580384 | 0.169861116 | 0.572496984 | 0.027950915 | 0.056057843 |
| Kctd10 | 0.024867511 | 0.457604485 | 0.808898849 | 0.076599021 | 0.058295222 |
| Kctd11 | 0.014392734 | 0.679840605 | 0.907783945 | 0.079810142 | 0.07098948 |
| Kctd12 | -0.021073293 | 0.640457194 | 0.891345382 | 0.131823078 | 0.146262382 |
| Kctd12b | -0.041291616 | 0.243745646 | 0.653788095 | 0.072534423 | 0.107173116 |
| Kctd13 | -0.041610724 | 0.461402829 | 0.810341242 | 0.245648045 | 0.301607164 |
| Kctd15 | -0.001255825 | 0.971810074 | 0.993892166 | 0.080775591 | 0.07841974 |
| Kctd18 | -0.047665373 | 0.204617973 | 0.616256046 | 0.089718563 | 0.134258893 |
| Kctd2 | -0.014380868 | 0.791527452 | 0.93896242 | 0.226023132 | 0.242925127 |

| | | | | | |
|---|---|---|---|---|---|
| Kctd20 | -0.126156695 | 0.051774756 | 0.355933715 | 0.366439364 | 0.502940931 |
| Kctd3 | -0.071443207 | 0.062597985 | 0.3817108 | 0.091069862 | 0.148004662 |
| Kctd5 | -0.045168608 | 0.2189553 | 0.632622127 | 0.088462371 | 0.126627884 |
| Kctd6 | 0.037423693 | 0.396846099 | 0.773576286 | 0.141739216 | 0.116342819 |
| Kctd8 | -0.011633761 | 0.74533432 | 0.924566547 | 0.08451299 | 0.092714745 |
| Kctd9 | -0.022563834 | 0.512586186 | 0.83587775 | 0.067825911 | 0.09035489 |
| Kdelr1 | 0.076329646 | 0.253696618 | 0.663146569 | 0.834497433 | 0.747480428 |
| Kdelr2 | -0.081164729 | 0.244218947 | 0.654008958 | 0.902265124 | 1.039184201 |
| Kdelr3 | 0.007336543 | 0.84773716 | 0.956938604 | 0.089767014 | 0.084198389 |
| Kdf1 | 0.018630093 | 0.621059235 | 0.884242858 | 0.100554059 | 0.087958446 |
| Kdm1a | 0.014075773 | 0.839945537 | 0.954911647 | 0.862862296 | 0.828509565 |
| Kdm1b | -0.03527905 | 0.291882244 | 0.697566301 | 0.065274181 | 0.097437257 |
| Kdm2a | -0.069768617 | 0.287957736 | 0.694589247 | 0.495550668 | 0.560135751 |
| Kdm2b | -0.002224064 | 0.974538311 | 0.994250968 | 0.492612279 | 0.506775629 |
| Kdm3a | -0.048360474 | 0.297468836 | 0.702656882 | 0.136878074 | 0.183821234 |
| Kdm3b | 0.046871579 | 0.289989685 | 0.69604181 | 0.131956634 | 0.078064872 |
| Kdm4a | 0.048856363 | 0.343869812 | 0.736696115 | 0.192730833 | 0.135951359 |
| Kdm4b | 0.00092381 | 0.979095299 | 0.995540893 | 0.082462503 | 0.090330655 |
| Kdm4c | -0.01022463 | 0.847171527 | 0.956905313 | 0.211656094 | 0.219585909 |
| Kdm5a | -0.120216193 | 0.062227488 | 0.380250374 | 0.413618724 | 0.539590745 |
| Kdm5b | -0.1188944 | 0.063463766 | 0.383957695 | 0.434548432 | 0.547004451 |
| Kdm5c | -0.006680377 | 0.911142862 | 0.978717991 | 0.283567512 | 0.293079405 |
| Kdm5d | 0.039176441 | 0.318483587 | 0.717175454 | 0.108039648 | 0.0686401 |
| Kdm6a | -0.00956542 | 0.853487919 | 0.957461155 | 0.183768354 | 0.196434054 |
| Kdm6b | -0.071930897 | 0.301516993 | 0.706960943 | 0.608695583 | 0.671343178 |
| Kdm7a | -0.044511303 | 0.475336898 | 0.817387411 | 0.322319949 | 0.369159968 |
| Kdsr | -0.01897099 | 0.612913101 | 0.881095502 | 0.08760969 | 0.107157018 |
| Keap1 | -0.0505669 | 0.456394283 | 0.808445095 | 0.54279413 | 0.593631154 |
| Khdc4 | -0.015937249 | 0.761285984 | 0.928506001 | 0.191790409 | 0.217111419 |
| Khdrbs1 | -0.068230938 | 0.303703363 | 0.708405917 | 0.490930688 | 0.564964673 |
| Khdrbs3 | 0.00086453 | 0.989864549 | 0.997717461 | 0.633006173 | 0.620540982 |
| Khk | -0.025065651 | 0.58892574 | 0.86992264 | 0.136888392 | 0.169636492 |
| Khnyn | -0.04154717 | 0.368665258 | 0.755778838 | 0.140885988 | 0.17871426 |
| Khsrp | -0.143934384 | 0.007875149 | 0.13698938 | 0.209979715 | 0.342972895 |
| Kidins220 | 0.025563147 | 0.708542528 | 0.913767764 | 0.652607379 | 0.638006198 |
| Kif12 | -0.073231171 | 0.217833945 | 0.632272789 | 0.28577927 | 0.343505781 |
| Kif13a | -0.034924316 | 0.295794508 | 0.702426542 | 0.066457602 | 0.093021343 |
| Kif13b | 0.109558738 | 0.064836124 | 0.388281728 | 0.31910061 | 0.224228614 |
| Kif16b | 0.006724518 | 0.87445125 | 0.964947297 | 0.116619559 | 0.113044929 |
| Kif19a | 0.024866184 | 0.450991121 | 0.806495742 | 0.071364871 | 0.047152387 |
| Kif1a | 0.000917346 | 0.989163196 | 0.997411191 | 0.726092033 | 0.714167323 |
| Kif1b | 0.127547532 | 0.057762937 | 0.370772606 | 1.139762524 | 0.960295971 |
| Kif1c | 0.072274423 | 0.06902467 | 0.399753362 | 0.112510611 | 0.044406587 |
| Kif21a | 0.082922748 | 0.202342153 | 0.612939912 | 1.257980969 | 1.15650987 |
| Kif2a | -0.01529665 | 0.803303245 | 0.943517638 | 0.352802933 | 0.378606368 |
| Kif3a | -0.032675098 | 0.491359981 | 0.827270742 | 0.148958682 | 0.176046696 |
| Kif3b | 0.064458384 | 0.234060221 | 0.644459522 | 0.217635639 | 0.158604149 |
| Kif5a | -0.047154282 | 0.008521791 | 0.143271724 | 0.010406956 | 0.050345393 |
| Kif5b | -0.138643423 | 0.001969863 | 0.0606747 | 2.07412851 | 2.348261815 |
| Kif5c | 0.061925152 | 0.356616855 | 0.745571198 | 0.48388889 | 0.438918251 |
| Kifap3 | -0.015362378 | 0.789891955 | 0.938440574 | 0.275672201 | 0.277097568 |
| Kifbp | 0.005418819 | 0.927175857 | 0.981818446 | 0.294159768 | 0.303241017 |
| Kifc3 | -0.088208424 | 0.023017336 | 0.240400628 | 0.091837654 | 0.159336116 |
| Kin | -0.066751777 | 0.305696241 | 0.709411642 | 0.390327499 | 0.438611982 |
| Kiz | -0.062215255 | 0.134411619 | 0.516977677 | 0.112105664 | 0.165488786 |
| Kl | 0.025783328 | 0.483556305 | 0.822768706 | 0.09214346 | 0.064333692 |
| Klc1 | 0.002525705 | 0.96649011 | 0.992504747 | 0.31016591 | 0.30876174 |
| Klc2 | 0.056162683 | 0.231921662 | 0.643345304 | 0.158287363 | 0.11243121 |
| Klc3 | -0.025824773 | 0.536270839 | 0.847520662 | 0.114142534 | 0.130759901 |
| Klc4 | -0.046858946 | 0.179700666 | 0.586745473 | 0.073001588 | 0.11196 |
| Klf10 | -0.018625237 | 0.705325799 | 0.912844672 | 0.161072239 | 0.173643234 |
| Klf11 | -0.062921585 | 0.071907273 | 0.40592424 | 0.072368854 | 0.127989125 |
| Klf13 | 0.109588783 | 0.09465782 | 0.454115053 | 0.51039261 | 0.407466926 |
| Klf16 | -0.026277573 | 0.349356885 | 0.740480699 | 0.043929746 | 0.070766701 |
| Klf3 | -0.044036278 | 0.425526433 | 0.790344696 | 0.229612939 | 0.269717813 |

| | | | | | |
|---|---|---|---|---|---|
| Klf4 | -0.020075177 | 0.615382718 | 0.882439418 | 0.098773097 | 0.109550082 |
| Klf5 | 0.029289714 | 0.612378598 | 0.880579368 | 0.239438852 | 0.198614829 |
| Klf6 | -0.030903761 | 0.535724912 | 0.847049612 | 0.147823076 | 0.197888959 |
| Klf7 | -0.008920236 | 0.88592335 | 0.969590273 | 0.313096729 | 0.315216291 |
| Klf9 | 0.002037 | 0.976733603 | 0.994468977 | 0.885109549 | 0.895099475 |
| Klhdc1 | -0.045695438 | 0.338915917 | 0.732653908 | 0.148798525 | 0.182107041 |
| Klhdc10 | -0.074117502 | 0.201922687 | 0.612724107 | 0.27814954 | 0.349573737 |
| Klhdc2 | -0.036544811 | 0.587332495 | 0.869235061 | 0.585512999 | 0.649113993 |
| Klhdc3 | -0.036685262 | 0.411935161 | 0.781143625 | 0.126440293 | 0.151754319 |
| Klhdc4 | -0.018557641 | 0.624550264 | 0.88668351 | 0.093406735 | 0.113022242 |
| Klhdc8a | -0.028524734 | 0.32640297 | 0.720866629 | 0.04776898 | 0.072433075 |
| Klhdc8b | -0.051950442 | 0.108930847 | 0.477972625 | 0.060219939 | 0.100124824 |
| Klhl11 | 0.024906171 | 0.51705346 | 0.837583212 | 0.100142764 | 0.081416539 |
| Klhl12 | 0.012206044 | 0.746004707 | 0.924566547 | 0.093852363 | 0.085637738 |
| Klhl15 | -0.029075907 | 0.410384138 | 0.780118483 | 0.081034064 | 0.108812369 |
| Klhl18 | -0.016216816 | 0.529176311 | 0.844323583 | 0.037921757 | 0.051954963 |
| Klhl2 | -0.027834165 | 0.499350931 | 0.830722138 | 0.11040961 | 0.137182575 |
| Klhl20 | 0.039345872 | 0.223501223 | 0.635670666 | 0.071969523 | 0.036540992 |
| Klhl21 | 0.009809575 | 0.821755425 | 0.950930859 | 0.131332331 | 0.132390345 |
| Klhl22 | -0.085016089 | 0.118598556 | 0.491192747 | 0.208356266 | 0.289071494 |
| Klhl24 | -0.003491319 | 0.949217469 | 0.987767085 | 0.238019891 | 0.247912202 |
| Klhl26 | 0.034330872 | 0.319854692 | 0.717256849 | 0.081689924 | 0.057018495 |
| Klhl28 | 0.013244313 | 0.762110108 | 0.92872278 | 0.129102148 | 0.11801972 |
| Klhl32 | 0.08665968 | 0.185894702 | 0.592701341 | 0.470877629 | 0.377194799 |
| Klhl36 | 0.009772624 | 0.791241188 | 0.938824685 | 0.092833025 | 0.08341277 |
| Klhl42 | -0.017222615 | 0.687869154 | 0.909796728 | 0.12173729 | 0.141110018 |
| Klhl7 | 0.037239823 | 0.546720351 | 0.851446616 | 0.330477396 | 0.308850226 |
| Klhl8 | 0.034844237 | 0.336768065 | 0.731477644 | 0.090097776 | 0.065215716 |
| Klhl9 | -0.020608943 | 0.746025694 | 0.924566547 | 0.409461126 | 0.417883523 |
| Kmt2a | -0.029746583 | 0.676792189 | 0.906695569 | 0.848489707 | 0.88509182 |
| Kmt2b | 0.086120984 | 0.101667234 | 0.467719786 | 0.222066835 | 0.150627918 |
| Kmt2c | -0.024466643 | 0.720910416 | 0.919412687 | 0.657206777 | 0.663035493 |
| Kmt2d | -0.019796886 | 0.722701484 | 0.919483582 | 0.241219361 | 0.266510971 |
| Kmt2e | -0.110296379 | 0.104649016 | 0.472076273 | 0.781942514 | 0.909477712 |
| Kmt5a | 0.191386409 | 0.004431532 | 0.097328864 | 0.525826514 | 0.336685525 |
| Kmt5b | -0.204894547 | 0.000487903 | 0.024858263 | 0.257003065 | 0.470748474 |
| Kmt5c | 0.037044607 | 0.228025061 | 0.639451668 | 0.062128665 | 0.029008334 |
| Knop1 | 0.061869294 | 0.324369088 | 0.720406991 | 0.379183441 | 0.310658173 |
| Kpna1 | -0.034356771 | 0.575068152 | 0.862723699 | 0.346995718 | 0.393258951 |
| Kpna2 | 0.058366312 | 0.144491544 | 0.533279057 | 0.111839365 | 0.059708397 |
| Kpna3 | 0.085588244 | 0.153746029 | 0.550142209 | 0.335937037 | 0.257534607 |
| Kpna4 | 0.152180232 | 0.015967794 | 0.203904258 | 0.38776813 | 0.242004698 |
| Kpna6 | 0.006856251 | 0.893818116 | 0.971749901 | 0.196570918 | 0.194616584 |
| Kpnb1 | 0.103027856 | 0.105894853 | 0.473025898 | 0.405067342 | 0.307751293 |
| Kras | -0.003809946 | 0.954683566 | 0.989066902 | 1.128451461 | 1.15931553 |
| Krba1 | 0.000713872 | 0.979674197 | 0.995540893 | 0.04859125 | 0.049071688 |
| Krcc1 | -0.08116954 | 0.246213805 | 0.655935044 | 0.575505563 | 0.64930005 |
| Kri1 | -0.046734394 | 0.198753764 | 0.609389286 | 0.077033146 | 0.114623108 |
| Krit1 | 0.012637313 | 0.842273401 | 0.955188334 | 0.392025919 | 0.376624675 |
| Krr1 | 0.066116251 | 0.244946245 | 0.654809396 | 0.2470309 | 0.190473667 |
| Krt10 | -0.029802824 | 0.618936274 | 0.883679002 | 0.305710853 | 0.340239568 |
| Krt18 | -0.073820138 | 0.284551743 | 0.691925959 | 1.379886741 | 1.507315147 |
| Krt222 | 0.025379293 | 0.401302918 | 0.774740795 | 0.060201911 | 0.034428034 |
| Krt7 | -0.24616926 | 9.83E-05 | 0.007650838 | 1.409843 | 1.859785483 |
| Krt8 | 0.244355434 | 0.003005806 | 0.076092349 | 1.278706125 | 0.883718627 |
| Krtcap2 | -0.115586059 | 0.014491579 | 0.193677739 | 1.887390756 | 2.095953091 |
| Krtcap3 | -0.029389055 | 0.557653609 | 0.855384435 | 0.183559057 | 0.206903592 |
| Ksr2 | -0.099200363 | 0.091702254 | 0.447237863 | 0.249082843 | 0.336824231 |
| Kti12 | 0.046693048 | 0.263532732 | 0.670148983 | 0.118787731 | 0.088505448 |
| Ktn1 | 0.104166629 | 0.13248969 | 0.513250014 | 1.080713091 | 0.933837324 |
| Kxd1 | -0.001332552 | 0.981338625 | 0.995719363 | 0.287865036 | 0.296782832 |
| Ky | -0.020995956 | 0.680477849 | 0.907783945 | 0.170772549 | 0.181482767 |
| Kyat1 | -0.001272881 | 0.968504864 | 0.993505636 | 0.06470152 | 0.063801279 |
| Kyat3 | -0.014625775 | 0.615035128 | 0.882439418 | 0.048527137 | 0.060159118 |
| L2hgdh | -0.052099899 | 0.131083214 | 0.510948091 | 0.070473116 | 0.119290884 |

| | | | | | |
|---|---|---|---|---|---|
| L3mbtl2 | 0.065159387 | 0.111763185 | 0.482690499 | 0.120937236 | 0.069541518 |
| Lacc1 | -0.019508547 | 0.501838258 | 0.832218079 | 0.049265946 | 0.061443075 |
| Lactb | 0.038385928 | 0.425881578 | 0.790724712 | 0.16339208 | 0.133752671 |
| Lactb2 | 0.190434268 | 0.002205055 | 0.064525187 | 0.367398895 | 0.187610315 |
| Lage3 | -0.000491375 | 0.993360582 | 0.998695538 | 0.297720943 | 0.310751645 |
| Lamc1 | 0.031335778 | 0.355556835 | 0.745326231 | 0.077334734 | 0.054836502 |
| Lamc2 | -0.076389127 | 0.039323258 | 0.312838547 | 0.077415806 | 0.139902629 |
| Lamp1 | -0.011006029 | 0.850524004 | 0.957244834 | 1.451125608 | 1.460230909 |
| Lamp2 | -0.003404758 | 0.960283721 | 0.99118854 | 0.883353995 | 0.866253079 |
| Lamtor1 | -0.05794529 | 0.396053449 | 0.772781583 | 0.676402343 | 0.736435018 |
| Lamtor2 | -0.092110341 | 0.145315883 | 0.533628684 | 1.016066811 | 1.126703536 |
| Lamtor3 | 0.10350592 | 0.119318428 | 0.492433827 | 0.559267128 | 0.453769862 |
| Lamtor4 | -0.036136609 | 0.590181176 | 0.87075199 | 0.84997776 | 0.902392232 |
| Lamtor5 | -0.005965975 | 0.929547067 | 0.982502239 | 0.536482294 | 0.558195287 |
| Lancl1 | -0.100438908 | 0.049040842 | 0.347612176 | 0.187123211 | 0.28363071 |
| Lancl2 | -0.026811373 | 0.410061468 | 0.780101937 | 0.062643194 | 0.08651502 |
| Lap3 | 0.092877199 | 0.087368425 | 0.439557333 | 0.234598561 | 0.15121169 |
| Laptm4a | -0.014997509 | 0.790694325 | 0.938824685 | 1.404597574 | 1.415667499 |
| Laptm4b | 0.141345111 | 0.01676326 | 0.207525869 | 0.300992388 | 0.174280197 |
| Larp1 | 0.087630738 | 0.194566375 | 0.603853652 | 0.772816243 | 0.669464089 |
| Larp1b | 0.057542102 | 0.375794921 | 0.760595784 | 0.476289418 | 0.421370201 |
| Larp4 | 0.104382621 | 0.118214584 | 0.491192747 | 0.665006224 | 0.545102391 |
| Larp4b | 0.025985813 | 0.705813178 | 0.912844672 | 0.733154401 | 0.685404053 |
| Larp7 | 0.016746129 | 0.685994488 | 0.909796728 | 0.114814672 | 0.099826062 |
| Lars | -0.187315687 | 0.006524269 | 0.122629384 | 0.837542465 | 1.054897919 |
| Lars2 | -0.082377436 | 0.215477875 | 0.629611185 | 0.433028419 | 0.514033202 |
| Las1l | -0.083098669 | 0.142248892 | 0.530762873 | 0.234160496 | 0.310908941 |
| Lasp1 | 0.009989739 | 0.870473906 | 0.964236463 | 0.317149786 | 0.313607123 |
| Lats1 | -0.08150074 | 0.18464901 | 0.59135996 | 0.314293487 | 0.391984914 |
| Lbh | -0.022972425 | 0.747385046 | 0.924930563 | 0.651716256 | 0.667339645 |
| Lbr | 0.091486967 | 0.07065824 | 0.402952703 | 0.195796443 | 0.114606395 |
| Lclat1 | 0.000684391 | 0.983480608 | 0.99608406 | 0.070493154 | 0.070260667 |
| Lcmt1 | -0.013210071 | 0.789953118 | 0.938440574 | 0.172916322 | 0.183894582 |
| Lcmt2 | -0.014484801 | 0.740641058 | 0.922901128 | 0.129729976 | 0.156111589 |
| Lcor | 0.034228425 | 0.505736753 | 0.832775992 | 0.199167383 | 0.178836829 |
| Lcorl | -0.066095055 | 0.335528496 | 0.730736446 | 0.506480585 | 0.57753516 |
| Ldah | -0.059023218 | 0.227781258 | 0.639451668 | 0.167440415 | 0.207256105 |
| Ldb1 | -0.056608059 | 0.314687619 | 0.714975653 | 0.23432134 | 0.284176256 |
| Ldhb | 0.069498847 | 0.118757435 | 0.49133074 | 0.142824008 | 0.084296256 |
| Ldhd | -0.002134022 | 0.935746661 | 0.984289378 | 0.043707628 | 0.043817996 |
| Ldlr | -0.039871012 | 0.494769591 | 0.828842054 | 0.273424198 | 0.302286948 |
| Ldlrad4 | -0.011507514 | 0.743944045 | 0.924566547 | 0.077542378 | 0.086512487 |
| Ldlrap1 | -0.053338905 | 0.069060442 | 0.399753362 | 0.049252645 | 0.093166091 |
| Lefty1 | 0.021245429 | 0.663660717 | 0.899599744 | 0.160372856 | 0.154289425 |
| Lekr1 | -0.032828926 | 0.240212542 | 0.650744882 | 0.041704139 | 0.070367793 |
| Lemd2 | 0.007190596 | 0.903685581 | 0.974985722 | 0.291517173 | 0.292937305 |
| Lemd3 | 0.008985474 | 0.822097606 | 0.950930859 | 0.10865245 | 0.103241171 |
| Leng1 | -0.042114988 | 0.313025407 | 0.714906079 | 0.110091165 | 0.159257589 |
| Leng8 | -0.019295358 | 0.686310903 | 0.909796728 | 0.160505424 | 0.171891061 |
| Leng9 | 0.043303875 | 0.267972232 | 0.676097903 | 0.102715739 | 0.069308532 |
| Leo1 | -0.073084155 | 0.198219524 | 0.609389286 | 0.231868679 | 0.310190445 |
| Leprot | -0.059197407 | 0.252226257 | 0.663127858 | 0.191835147 | 0.243589448 |
| Leprotl1 | -0.158078447 | 0.012975348 | 0.182144849 | 1.131825126 | 1.375305129 |
| Letm1 | 0.005742345 | 0.915933294 | 0.980132949 | 0.227062189 | 0.223208641 |
| Letm2 | 0.019991261 | 0.665633519 | 0.901006558 | 0.144747588 | 0.129685901 |
| Letmd1 | 0.054655526 | 0.141135535 | 0.528664246 | 0.096083886 | 0.050580834 |
| Lgals3bp | -0.236508499 | 0.000656944 | 0.029470643 | 0.721929965 | 1.017501884 |
| Lgals8 | -0.057310998 | 0.365653072 | 0.752886169 | 0.352282869 | 0.396791579 |
| Lgalsl | -0.050350397 | 0.369253127 | 0.756523935 | 0.230556745 | 0.278287782 |
| Lgi1 | 0.013003372 | 0.795643619 | 0.939757086 | 0.168941288 | 0.159181905 |
| Lgi2 | 0.100001564 | 0.080566404 | 0.425378745 | 0.266222641 | 0.17628647 |
| Lgmn | 0.003995949 | 0.891439281 | 0.970852438 | 0.056123979 | 0.052046825 |
| Lgr4 | -0.020096029 | 0.560923132 | 0.857134531 | 0.077910745 | 0.096666829 |
| Lhfpl4 | 0.023570089 | 0.652554577 | 0.896875696 | 0.201123572 | 0.181494977 |
| Lhfpl5 | -0.014963581 | 0.599083815 | 0.875637724 | 0.046613074 | 0.056673592 |

| | | | | | |
|---|---|---|---|---|---|
| Lhx1 | 0.006674932 | 0.88909765 | 0.97019712 | 0.150770228 | 0.149312289 |
| Lhx1os | -0.077098987 | 0.003785291 | 0.086973441 | 0.034177259 | 0.098062588 |
| Lias | -0.001016772 | 0.987314536 | 0.997268589 | 0.388751745 | 0.389793767 |
| Lifr | 0.119340229 | 0.052798395 | 0.359036238 | 0.357744134 | 0.236647919 |
| Lig1 | -0.005065664 | 0.859823553 | 0.959220425 | 0.05107989 | 0.054642081 |
| Lig3 | -0.08891828 | 0.040838635 | 0.319308264 | 0.111918474 | 0.197177929 |
| Lig4 | -0.006739306 | 0.844668401 | 0.955950886 | 0.078541726 | 0.085457509 |
| Lima1 | 0.045948214 | 0.400990637 | 0.774740795 | 0.215255046 | 0.159833927 |
| Limch1 | 0.054508735 | 0.408403084 | 0.7788603 | 0.369187457 | 0.3095083 |
| Limd1 | -0.054037578 | 0.388337319 | 0.767160379 | 0.35258277 | 0.388716435 |
| Limd2 | 0.135781324 | 0.031254379 | 0.281212368 | 0.375027104 | 0.251248089 |
| Limk2 | 0.059378087 | 0.104939358 | 0.472399448 | 0.091853442 | 0.038395927 |
| Lims1 | 0.033927871 | 0.511001626 | 0.834957869 | 0.186548567 | 0.16036416 |
| Lin37 | 0.016673805 | 0.738618942 | 0.922843014 | 0.166564601 | 0.168776696 |
| Lin52 | 0.0215482 | 0.457659714 | 0.808898849 | 0.054712644 | 0.03623728 |
| LIN54 | 0.137987236 | 0.002810991 | 0.073093048 | 0.162168499 | 0.047522582 |
| Lin7c | -0.019216274 | 0.767412523 | 0.929806796 | 0.441080471 | 0.459870507 |
| Lins1 | 0.02152966 | 0.430287831 | 0.793295051 | 0.049158623 | 0.041418412 |
| Lipa | 0.05375664 | 0.161262465 | 0.559461938 | 0.103485203 | 0.049148111 |
| Lipo3 | -0.121577525 | 0.000608771 | 0.02841969 | 0.069357957 | 0.169721582 |
| Litaf | 0.023827153 | 0.538791852 | 0.848423882 | 0.098330076 | 0.069064427 |
| Lix1l | -0.012322736 | 0.742001022 | 0.923430162 | 0.094827158 | 0.107137149 |
| Llgl1 | 0.008008876 | 0.785128722 | 0.936442895 | 0.052912257 | 0.044186019 |
| Llgl2 | -0.019803988 | 0.694189033 | 0.909986409 | 0.176506997 | 0.198233476 |
| Llph | 0.088170357 | 0.202069099 | 0.612724107 | 0.563301238 | 0.496604821 |
| Lman1 | -0.009955082 | 0.855371522 | 0.957736324 | 1.581183086 | 1.553064463 |
| Lman2 | 0.020334345 | 0.760131929 | 0.927859527 | 0.675470331 | 0.647672253 |
| Lman2l | -0.035297736 | 0.488200635 | 0.826256745 | 0.180171438 | 0.22648148 |
| Lmbr1 | 0.023481221 | 0.513958208 | 0.83659041 | 0.087351484 | 0.067847341 |
| Lmbrd1 | 0.028164552 | 0.662322935 | 0.898950007 | 0.435662773 | 0.411119897 |
| Lmbrd2 | -0.040938205 | 0.45006362 | 0.806495742 | 0.228401498 | 0.269625867 |
| Lmf1 | -0.029159603 | 0.422022364 | 0.787769847 | 0.084164134 | 0.106573213 |
| Lmf2 | -0.095186838 | 0.005793043 | 0.113342639 | 0.064644927 | 0.147500067 |
| Lmna | 0.078824541 | 0.260194848 | 0.667410092 | 0.620616505 | 0.531318483 |
| Lmnb1 | -0.01451514 | 0.670893336 | 0.904224039 | 0.071725162 | 0.087988018 |
| Lmnb2 | -0.00634372 | 0.823525769 | 0.951067558 | 0.050101498 | 0.057580184 |
| Lmo1 | -0.152847711 | 0.012171698 | 0.176287639 | 0.291530779 | 0.434807997 |
| Lmo4 | -0.040666703 | 0.163211997 | 0.562993622 | 0.046837589 | 0.077378912 |
| Lmo7 | -0.091809331 | 0.174765566 | 0.577992588 | 0.516847958 | 0.619936977 |
| Lmtk2 | -0.080397365 | 0.109033339 | 0.477972625 | 0.167249845 | 0.237751387 |
| Lncpint | -0.159298808 | 0.000150304 | 0.010687095 | 0.091541998 | 0.251457977 |
| Lnpep | 0.026403146 | 0.685573874 | 0.909796728 | 0.448489553 | 0.426585414 |
| Lnpk | 0.041878287 | 0.429944673 | 0.793263728 | 0.220099247 | 0.19511555 |
| Lnx1 | 0.102909377 | 0.002304669 | 0.065344539 | 0.085514276 | 0 |
| Lonp1 | -0.221261442 | 0.000786027 | 0.032786291 | 0.456114054 | 0.696539098 |
| Lonp2 | -0.025037898 | 0.694938413 | 0.910242368 | 0.394069449 | 0.431706047 |
| Lonrf1 | 0.015903065 | 0.691811512 | 0.909796728 | 0.103354518 | 0.090710417 |
| Lonrf2 | -0.025594184 | 0.454465214 | 0.807876964 | 0.072865619 | 0.098343577 |
| Loxl2 | -0.008930089 | 0.841228037 | 0.954960614 | 0.133020536 | 0.133061594 |
| Lpar6 | 0.017572452 | 0.572514585 | 0.861775874 | 0.060563347 | 0.041068737 |
| Lpcat1 | 0.060505438 | 0.131635416 | 0.512050022 | 0.113554203 | 0.059270332 |
| Lpcat3 | -0.053635478 | 0.348487572 | 0.73964258 | 0.261073318 | 0.31931291 |
| Lpgat1 | 0.137165917 | 0.04852087 | 0.345392885 | 0.77036998 | 0.595594794 |
| Lpin1 | 0.004061282 | 0.931814337 | 0.98340088 | 0.155239695 | 0.150171806 |
| Lpin2 | 0.092352721 | 0.081451267 | 0.427576549 | 0.205229899 | 0.120041486 |
| Lpl | -0.12518133 | 0.061994193 | 0.380250374 | 0.42451909 | 0.582335966 |
| Lpp | -0.11394898 | 0.105297307 | 0.472434351 | 0.655310219 | 0.79747919 |
| Lratd2 | 0.025083967 | 0.596610563 | 0.87393173 | 0.157828291 | 0.135030935 |
| Lrba | -0.152088533 | 0.009946993 | 0.155269009 | 0.255751709 | 0.394180262 |
| Lrch1 | -0.034143396 | 0.434537603 | 0.796628285 | 0.118323279 | 0.149576149 |
| Lrch2 | -0.027977551 | 0.355060862 | 0.745326231 | 0.051646567 | 0.070231055 |
| Lrch3 | 0.059988433 | 0.198925644 | 0.609389286 | 0.152743581 | 0.109425715 |
| Lrfn1 | -0.06357425 | 0.004605925 | 0.099848091 | 0.022341186 | 0.076910569 |
| Lrfn3 | -0.057312706 | 0.322254855 | 0.719780733 | 0.258954524 | 0.313175579 |
| Lrfn4 | 0.083547075 | 0.063255622 | 0.383440442 | 0.146485434 | 0.075867464 |

| | | | | | |
|---|---|---|---|---|---|
| Lrif1 | 0.084066593 | 0.184286303 | 0.590567014 | 0.353832963 | 0.270807385 |
| Lrig2 | 0.036683315 | 0.315029889 | 0.714975653 | 0.092743284 | 0.059923495 |
| Lrp10 | 0.022704831 | 0.71540848 | 0.916430034 | 0.379915142 | 0.364904706 |
| Lrp11 | 0.18285473 | 0.004761754 | 0.101438784 | 0.633310058 | 0.449628767 |
| Lrp12 | -0.040569436 | 0.358577925 | 0.747310348 | 0.128771211 | 0.179371634 |
| Lrp3 | -0.005979008 | 0.890399331 | 0.970579007 | 0.124390831 | 0.125987737 |
| Lrp6 | -0.002483017 | 0.966367732 | 0.992504747 | 0.283008897 | 0.273826442 |
| Lrp8 | -0.001035583 | 0.984150276 | 0.99608406 | 0.184643606 | 0.173774072 |
| Lrpap1 | -0.238923127 | 0.000199289 | 0.012906373 | 0.434398225 | 0.686477204 |
| Lrpprc | -0.040808282 | 0.503972988 | 0.832249514 | 0.296710468 | 0.337756965 |
| Lrrc1 | 0.041473458 | 0.372754902 | 0.75825687 | 0.150591304 | 0.116956529 |
| Lrrc10b | 0.129813591 | 0.073328244 | 0.410711821 | 0.837638728 | 0.68286669 |
| Lrrc14 | 0.088712944 | 0.051202078 | 0.355933715 | 0.145616086 | 0.070443565 |
| Lrrc20 | 0.046743707 | 0.254436329 | 0.663146569 | 0.117284805 | 0.070648011 |
| Lrrc24 | 0.021102963 | 0.508746244 | 0.834633222 | 0.068084379 | 0.048104291 |
| Lrrc27 | -0.03808374 | 0.055192549 | 0.365173116 | 0.016672723 | 0.049715226 |
| Lrrc28 | -0.020965813 | 0.404861329 | 0.776552851 | 0.034433674 | 0.051155608 |
| Lrrc4 | 0.000321933 | 0.99055951 | 0.997733826 | 0.04602016 | 0.048151826 |
| Lrrc40 | 0.102603715 | 0.014178145 | 0.193060412 | 0.130277833 | 0.044684418 |
| Lrrc41 | 0.133518061 | 0.040883859 | 0.319308264 | 0.423805645 | 0.284127988 |
| Lrrc42 | -0.001578995 | 0.975435111 | 0.994455356 | 0.182720777 | 0.185013309 |
| Lrrc45 | -0.06217685 | 0.036946714 | 0.303216817 | 0.049988927 | 0.099444363 |
| Lrrc47 | -0.006382353 | 0.909842585 | 0.978578013 | 0.259325069 | 0.266711287 |
| Lrrc49 | -0.020927385 | 0.490023295 | 0.82659234 | 0.052381775 | 0.072422885 |
| Lrrc51 | -0.023463296 | 0.395687528 | 0.772384513 | 0.045894658 | 0.069282226 |
| Lrrc57 | -0.001661819 | 0.965679446 | 0.992256509 | 0.100656289 | 0.100983424 |
| Lrrc58 | -0.111228778 | 0.1046968 | 0.472076273 | 0.707960084 | 0.826491032 |
| Lrrc59 | 0.133970178 | 0.033864118 | 0.291744277 | 0.403522606 | 0.277505154 |
| Lrrc61 | -0.023110874 | 0.418450393 | 0.784886311 | 0.046902226 | 0.066031238 |
| Lrrc73 | -0.027430794 | 0.653811506 | 0.897454058 | 0.317145944 | 0.350906765 |
| Lrrc8a | -0.039727875 | 0.174088051 | 0.577054745 | 0.047239122 | 0.086696604 |
| Lrrc8d | -0.021519308 | 0.723024984 | 0.919483582 | 0.339724702 | 0.346120216 |
| Lrrcc1 | 0.041148683 | 0.38155029 | 0.762568749 | 0.155480652 | 0.115572412 |
| Lrrfip1 | 0.180681898 | 0.000348173 | 0.019522993 | 0.192589631 | 0.036801257 |
| Lrrfip2 | 0.152407283 | 0.026637269 | 0.259571135 | 1.172442429 | 0.922346884 |
| Lrrn1 | -0.053853914 | 0.406382882 | 0.777612997 | 0.388268402 | 0.436170506 |
| Lrrn2 | -0.023904462 | 0.446903008 | 0.804170357 | 0.058953426 | 0.077107202 |
| Lrwd1 | -0.02469226 | 0.406961237 | 0.777736088 | 0.050413029 | 0.073648992 |
| Lsg1 | 0.078600722 | 0.168320287 | 0.571912906 | 0.228339781 | 0.159034041 |
| Lsm1 | 0.014997365 | 0.805234904 | 0.94472738 | 0.359238508 | 0.340541698 |
| Lsm10 | -0.020585286 | 0.70636582 | 0.912927342 | 0.235971263 | 0.247221382 |
| Lsm12 | -0.043902785 | 0.512227536 | 0.835753752 | 0.525106364 | 0.593464125 |
| Lsm14a | -0.040387576 | 0.506915441 | 0.833276466 | 0.346394542 | 0.385855165 |
| Lsm14b | -0.027684782 | 0.617913791 | 0.883238152 | 0.240728856 | 0.276068924 |
| Lsm2 | 0.017974658 | 0.7594726 | 0.927719223 | 0.291349612 | 0.267034925 |
| Lsm3 | 0.050438714 | 0.404791621 | 0.776552851 | 0.311647939 | 0.264943832 |
| Lsm4 | 0.047254173 | 0.496557392 | 0.830056214 | 0.71401996 | 0.656944214 |
| Lsm5 | 0.022694324 | 0.704677594 | 0.912507936 | 0.307291288 | 0.283221902 |
| Lsm6 | -0.067565358 | 0.304101581 | 0.708511506 | 0.857353399 | 0.926727758 |
| Lsm7 | -0.070253422 | 0.249645021 | 0.661305641 | 0.341969378 | 0.400573391 |
| Lsm8 | -0.059112167 | 0.329090539 | 0.724485236 | 0.326933447 | 0.383241219 |
| Lsr | -0.114266414 | 0.046912927 | 0.339316662 | 0.269024084 | 0.384514638 |
| Lss | -0.024003597 | 0.505163011 | 0.832249514 | 0.082198809 | 0.099799836 |
| Lta4h | -0.006540073 | 0.889085661 | 0.97019712 | 0.143493722 | 0.152608562 |
| Ltbp3 | -0.087623852 | 0.029536492 | 0.27358919 | 0.102609563 | 0.180886128 |
| Ltbr | -0.004082703 | 0.91953528 | 0.980436936 | 0.109834067 | 0.105481192 |
| Ltn1 | 0.022552436 | 0.691454883 | 0.909796728 | 0.255315067 | 0.238090425 |
| LTO1 | -0.015730987 | 0.691190176 | 0.909796728 | 0.102584328 | 0.115536782 |
| Ltv1 | -0.0548195 | 0.246380211 | 0.655946467 | 0.152409677 | 0.202086987 |
| Luc7l | -0.108312401 | 0.090296902 | 0.445385778 | 0.387873656 | 0.498137876 |
| Luc7l2 | -0.06902655 | 0.30307116 | 0.708026508 | 1.160625713 | 1.250375499 |
| Luc7l3 | -0.070121368 | 0.316435082 | 0.715693767 | 0.678097598 | 0.73279951 |
| Lurap1 | -0.027876858 | 0.254402354 | 0.663146569 | 0.029883587 | 0.054841483 |
| Lurap1l | 0.132154925 | 0.038960434 | 0.311933003 | 0.335987953 | 0.217946059 |
| Luzp1 | 0.206961496 | 0.000843818 | 0.034153784 | 0.361621122 | 0.17254083 |

| | | | | | |
|---|---|---|---|---|---|
| Lxn | -0.052120758 | 0.140952737 | 0.528664246 | 0.077561865 | 0.115312753 |
| Ly6e | -0.106221735 | 0.120573109 | 0.494691316 | 0.549699257 | 0.631288913 |
| Ly96 | -0.005720522 | 0.917238525 | 0.980132949 | 0.209180244 | 0.211111255 |
| Lyar | 0.052532331 | 0.393885933 | 0.770629121 | 0.328921986 | 0.274590036 |
| Lyn | -0.037889846 | 0.129996477 | 0.509514066 | 0.03216595 | 0.06142618 |
| Lypla1 | -0.019149813 | 0.715286044 | 0.916430034 | 0.194896412 | 0.210724643 |
| Lypla2 | 0.009070511 | 0.86694282 | 0.962125879 | 0.229628175 | 0.219820934 |
| Lyplal1 | -0.029862968 | 0.566470623 | 0.858732162 | 0.185573989 | 0.210741319 |
| Lyrm1 | -0.014476489 | 0.630858641 | 0.888561606 | 0.055636066 | 0.071880244 |
| Lyrm2 | -0.05773257 | 0.323733001 | 0.72034806 | 0.277308013 | 0.335231618 |
| Lyrm4 | -0.085634766 | 0.046275999 | 0.337552474 | 0.117328594 | 0.18772704 |
| Lyrm9 | 0.059407684 | 0.310597508 | 0.71348779 | 0.271727675 | 0.206026649 |
| Lysmd1 | -0.006287385 | 0.84229859 | 0.955188334 | 0.064308753 | 0.073696833 |
| Lysmd2 | 0.051525442 | 0.441462787 | 0.800354711 | 0.602184464 | 0.538537028 |
| Lysmd3 | -0.082409565 | 0.180228061 | 0.587198006 | 0.292926581 | 0.384112877 |
| Lysmd4 | 0.041532046 | 0.209913418 | 0.622606671 | 0.074004582 | 0.039807792 |
| Lyst | -0.014129558 | 0.795429523 | 0.939757086 | 0.219190783 | 0.234550183 |
| Lyve1 | 0.103921393 | 0.005603276 | 0.111587469 | 0.092639816 | 0.009224659 |
| Lzic | -0.001147063 | 0.975812806 | 0.994455356 | 0.090228663 | 0.100901544 |
| Lztfl1 | -0.108914214 | 0.07237713 | 0.407888408 | 0.337279955 | 0.428475286 |
| Lztr1 | -0.190073692 | 0.000154392 | 0.010687095 | 0.15198083 | 0.340281467 |
| Lzts2 | 0.136352163 | 0.01889384 | 0.218227362 | 0.277422053 | 0.167728151 |
| Lzts3 | -0.033434735 | 0.093815862 | 0.45227176 | 0.016467375 | 0.044178156 |
| M6pr | 0.026553826 | 0.682775026 | 0.909173483 | 0.45533509 | 0.441656231 |
| Macf1 | -0.083916869 | 0.20350183 | 0.614950615 | 1.059357799 | 1.16635182 |
| Macir | -0.02877699 | 0.608824085 | 0.878855197 | 0.227801705 | 0.271964817 |
| Maco1 | -0.020281122 | 0.765192629 | 0.928757959 | 0.595122173 | 0.616962339 |
| Macrod1 | -0.048138947 | 0.103018449 | 0.468979611 | 0.05184622 | 0.094527586 |
| Macrod2 | -0.005408637 | 0.915205024 | 0.980132949 | 0.18767336 | 0.187914713 |
| Macroh2a1 | 0.033455025 | 0.602372193 | 0.876614426 | 1.117850693 | 1.055686487 |
| Macroh2a2 | 0.080642043 | 0.162899763 | 0.562993622 | 0.287315554 | 0.225953333 |
| Mad1l1 | -0.050126049 | 0.03349514 | 0.290069649 | 0.02491548 | 0.072669655 |
| Mad2l1bp | -0.0350769 | 0.196752097 | 0.60713211 | 0.039664927 | 0.074997285 |
| Mad2l2 | -0.01692022 | 0.670499698 | 0.904104984 | 0.102249262 | 0.11361709 |
| Madd | -0.296126064 | 1.06E-05 | 0.001334578 | 0.749482278 | 1.111725742 |
| Maea | -0.00113089 | 0.98257136 | 0.995921574 | 0.198407264 | 0.200362833 |
| Maf1 | -0.123183225 | 0.022362711 | 0.237266944 | 0.191749783 | 0.30806393 |
| Mafa | 0.346370734 | 5.88E-06 | 0.000860632 | 2.237111163 | 1.629368477 |
| Mafb | -0.005040056 | 0.915080408 | 0.980132949 | 0.117655426 | 0.115038426 |
| Maff | -0.04547666 | 0.258774998 | 0.665948842 | 0.09808748 | 0.145873087 |
| Mafg | -0.009998028 | 0.880381166 | 0.967810606 | 0.50041957 | 0.524344299 |
| Mafk | -0.012569537 | 0.770330667 | 0.931574344 | 0.120709059 | 0.122597328 |
| Maged1 | -0.094840257 | 0.067693735 | 0.395254229 | 1.652010017 | 1.801216883 |
| Maged2 | -0.169671118 | 0.003905382 | 0.089087078 | 0.274621244 | 0.419459177 |
| Magee1 | -0.005193911 | 0.882761674 | 0.96861827 | 0.07944972 | 0.085181302 |
| Magef1 | 0.021695742 | 0.540584116 | 0.849380522 | 0.083008534 | 0.066769665 |
| Mageh1 | 0.032526182 | 0.629071148 | 0.887864718 | 0.562422981 | 0.518869996 |
| Magi1 | 0.030497444 | 0.400099085 | 0.774697097 | 0.085505632 | 0.06053288 |
| Magi2 | -0.003380311 | 0.949306328 | 0.987767085 | 0.207317342 | 0.214171863 |
| Magoh | 0.087575638 | 0.182679374 | 0.590066231 | 0.450955075 | 0.359573997 |
| Magohb | 0.014578166 | 0.748682926 | 0.925186829 | 0.145089896 | 0.134644215 |
| Magt1 | 0.025878585 | 0.697413137 | 0.910970095 | 0.817596967 | 0.785717163 |
| Maip1 | -0.007414301 | 0.890885804 | 0.970670952 | 0.225757724 | 0.237876242 |
| Mak16 | 0.001943621 | 0.975636168 | 0.994455356 | 0.389266601 | 0.38767056 |
| Mal2 | -0.044570992 | 0.167579707 | 0.570246024 | 0.062425825 | 0.099461016 |
| Malat1 | -0.021962213 | 0.51102383 | 0.834957869 | 6.695571151 | 6.609782971 |
| Malsu1 | 0.089730454 | 0.163372995 | 0.563109461 | 0.448612809 | 0.358675571 |
| Malt1 | 0.004504109 | 0.89061455 | 0.970579007 | 0.070081005 | 0.068557153 |
| Maml1 | -0.017281763 | 0.591564522 | 0.871500529 | 0.064006777 | 0.078839956 |
| Maml3 | 0.066974004 | 0.316525319 | 0.715693767 | 0.444093837 | 0.376718919 |
| Man1a | 0.001662904 | 0.979828868 | 0.995540893 | 0.583336631 | 0.577762951 |
| Man1a2 | 0.017534336 | 0.77835533 | 0.934523337 | 0.406789498 | 0.382828973 |
| Man1b1 | -0.030297946 | 0.449861551 | 0.806495742 | 0.102763328 | 0.131718534 |
| Man1c1 | 0.033265282 | 0.303103341 | 0.708026508 | 0.069527241 | 0.040653914 |
| Man2a1 | -0.092630142 | 0.031733173 | 0.283368203 | 0.110605674 | 0.188201498 |

| | | | | | |
|---|---|---|---|---|---|
| Man2a2 | -0.086772781 | 0.082824513 | 0.430310873 | 0.1605013 | 0.233105899 |
| Man2b1 | -0.046273726 | 0.303993239 | 0.708427238 | 0.127138342 | 0.167557462 |
| Man2b2 | -0.08567299 | 0.033636523 | 0.290545455 | 0.099109358 | 0.169410803 |
| Man2c1 | -0.123227962 | 0.002632476 | 0.069899887 | 0.094724779 | 0.207431008 |
| Manba | -0.11283735 | 0.000434913 | 0.023005843 | 0.055119501 | 0.150542506 |
| Manbal | 0.031816624 | 0.577765979 | 0.864673658 | 0.270422718 | 0.252181298 |
| Manea | 0.012412229 | 0.729671848 | 0.920522999 | 0.086985113 | 0.075131793 |
| Maneal | 0.03762729 | 0.271553381 | 0.67876599 | 0.080736932 | 0.047937671 |
| Manf | 0.089872676 | 0.137709231 | 0.522884425 | 1.870244923 | 1.621618127 |
| Maoa | 0.048157493 | 0.219417352 | 0.63289494 | 0.106538995 | 0.069425941 |
| Maob | 0.203277492 | 0.003232398 | 0.079642162 | 0.618840219 | 0.405960498 |
| Map10 | 0.035613138 | 0.229119804 | 0.640979764 | 0.059664054 | 0.027850895 |
| Map11 | 0.03279753 | 0.375340105 | 0.760147021 | 0.090892668 | 0.05873514 |
| Map1b | -0.135157793 | 0.023135353 | 0.240881268 | 1.475587827 | 1.705287615 |
| Map1lc3a | -0.058616399 | 0.392712081 | 0.769704269 | 0.797147848 | 0.902636694 |
| Map1lc3b | -0.01975351 | 0.765357476 | 0.928757959 | 1.052235549 | 1.070652007 |
| Map1s | 0.068040931 | 0.100502774 | 0.46546263 | 0.123110588 | 0.069627953 |
| Map2 | 0.171614593 | 0.015617906 | 0.201228402 | 0.729574404 | 0.540207391 |
| Map2k1 | 0.041717587 | 0.46169071 | 0.810421416 | 0.264337812 | 0.225112316 |
| Map2k2 | -0.028313743 | 0.682449609 | 0.909170768 | 0.661966959 | 0.697437365 |
| Map2k3 | -0.033145815 | 0.472081268 | 0.815861654 | 0.143301751 | 0.179635963 |
| Map2k3os | -0.049466871 | 0.017910319 | 0.212740677 | 0.018700348 | 0.062160426 |
| Map2k4 | 0.043906552 | 0.406955338 | 0.777736088 | 0.217691771 | 0.183155162 |
| Map2k5 | 0.01044924 | 0.803024455 | 0.943517638 | 0.116485058 | 0.112095475 |
| Map2k7 | 0.025939018 | 0.602622371 | 0.876632118 | 0.180459969 | 0.154306326 |
| Map3k1 | 0.093947424 | 0.116505295 | 0.488986751 | 0.280126455 | 0.171767346 |
| Map3k10 | 0.069521891 | 0.134839105 | 0.5179411 | 0.161173604 | 0.097465833 |
| Map3k11 | -0.010931138 | 0.780579961 | 0.935383906 | 0.103908496 | 0.114977267 |
| Map3k12 | 0.067482039 | 0.107528115 | 0.475444796 | 0.122369139 | 0.068172916 |
| Map3k15 | -0.051929634 | 0.388378144 | 0.767160379 | 0.290722592 | 0.335835073 |
| Map3k2 | -0.007094117 | 0.917446138 | 0.980132949 | 0.560394586 | 0.572989606 |
| Map3k20 | 0.076362397 | 0.031969686 | 0.283462667 | 0.088930743 | 0.023470235 |
| Map3k3 | 0.007245252 | 0.854461599 | 0.957596144 | 0.103604705 | 0.09500077 |
| Map3k4 | -0.002144941 | 0.973288356 | 0.994079882 | 0.385110924 | 0.371522521 |
| Map3k5 | 0.011993243 | 0.779739223 | 0.935139002 | 0.125265869 | 0.11966465 |
| Map3k7 | -0.017272355 | 0.75577039 | 0.926726954 | 0.22617561 | 0.241635842 |
| Map4 | -0.078794745 | 0.153491081 | 0.549817979 | 0.203681953 | 0.278387471 |
| Map4k2 | -0.015428622 | 0.64473033 | 0.893466123 | 0.07242718 | 0.082792377 |
| Map4k3 | 0.008190291 | 0.849778538 | 0.957111129 | 0.133800406 | 0.127866362 |
| Map4k4 | 0.047733042 | 0.458956211 | 0.809370658 | 0.405740325 | 0.349197298 |
| Map4k5 | 0.089589484 | 0.079267922 | 0.422322725 | 0.197206464 | 0.11801322 |
| Map6 | 0.003072751 | 0.930410511 | 0.982692866 | 0.080178326 | 0.077355714 |
| Map7 | -0.004670808 | 0.941444725 | 0.986168593 | 0.420127337 | 0.421326272 |
| Map7d1 | 0.072750016 | 0.269795152 | 0.677226912 | 0.421638117 | 0.34114758 |
| Map7d2 | -0.094580721 | 0.093162361 | 0.45188518 | 0.21904604 | 0.315014901 |
| Map9 | -0.033244041 | 0.568931436 | 0.860524446 | 0.262149744 | 0.287791189 |
| Mapk1 | 0.128593013 | 0.069940269 | 0.401596384 | 0.642641706 | 0.502315521 |
| Mapk10 | -0.059283211 | 0.233260527 | 0.643935869 | 0.161066426 | 0.214675952 |
| Mapk13 | -0.043681837 | 0.16421298 | 0.563869641 | 0.054204576 | 0.086288197 |
| Mapk14 | -0.011935601 | 0.853039197 | 0.957461155 | 0.474514477 | 0.489699668 |
| Mapk15 | -0.013849818 | 0.786993026 | 0.937461311 | 0.171104146 | 0.171908358 |
| Mapk1ip1 | 0.035580663 | 0.49387949 | 0.828842054 | 0.198363332 | 0.164492456 |
| Mapk1ip1l | -0.066171303 | 0.274045017 | 0.681791282 | 0.292216392 | 0.347897591 |
| Mapk3 | 0.146475637 | 0.027536887 | 0.264360019 | 0.581358608 | 0.42324194 |
| Mapk6 | -0.032162235 | 0.589463821 | 0.870321914 | 0.292621961 | 0.325524095 |
| Mapk8 | -0.020677533 | 0.668965184 | 0.903080505 | 0.15530354 | 0.178514043 |
| Mapk8ip1 | -0.047156559 | 0.434167119 | 0.796332921 | 0.354444486 | 0.381790045 |
| Mapk8ip2 | -0.028167076 | 0.66445642 | 0.900263106 | 0.405828225 | 0.414789148 |
| Mapk8ip3 | 0.189678052 | 0.002288786 | 0.06526292 | 0.399866782 | 0.212800736 |
| Mapk9 | -0.0512233 | 0.321739346 | 0.719227395 | 0.179920062 | 0.22387397 |
| Mapkap1 | -0.000386512 | 0.994222409 | 0.998695538 | 0.191379031 | 0.182908018 |
| Mapkapk2 | 0.121360066 | 0.06018412 | 0.378488731 | 0.411660544 | 0.285664956 |
| Mapkapk3 | 0.074396613 | 0.077895607 | 0.418318999 | 0.118519751 | 0.053148405 |
| Mapkbp1 | -0.047584191 | 0.11745809 | 0.490810511 | 0.051856613 | 0.096301547 |
| Mapre1 | 0.018391383 | 0.782570665 | 0.936026395 | 0.526822928 | 0.528622208 |

| | | | | | |
|---|---|---|---|---|---|
| Mapre2 | 0.10940846 | 0.105055579 | 0.472399448 | 0.667315963 | 0.556927876 |
| Mapre3 | 0.060531589 | 0.326837503 | 0.721457668 | 0.338323102 | 0.269778891 |
| Mapt | -0.009386403 | 0.873738979 | 0.964940674 | 0.275688751 | 0.288846995 |
| Marchf2 | -0.018611235 | 0.696900087 | 0.910970095 | 0.156418477 | 0.178374086 |
| Marchf4 | -0.008995883 | 0.811389159 | 0.948397417 | 0.089047888 | 0.091964708 |
| Marchf5 | -0.028729554 | 0.665745454 | 0.901036559 | 0.97747759 | 1.027542275 |
| Marchf6 | -0.254671396 | 0.000131105 | 0.00967571 | 0.551063997 | 0.818108118 |
| Marchf7 | 0.088341311 | 0.184342578 | 0.590567014 | 0.684211145 | 0.598615172 |
| Marchf8 | -0.027200428 | 0.694136819 | 0.909986409 | 0.606607649 | 0.646174212 |
| Marchf9 | -0.050360243 | 0.304668968 | 0.709410087 | 0.167898171 | 0.221435402 |
| Marcks | 0.126779538 | 0.079113236 | 0.421923248 | 0.731278508 | 0.57591921 |
| Marf1 | -0.025186755 | 0.662884636 | 0.899270729 | 0.284015801 | 0.325924465 |
| Mark1 | 0.026579741 | 0.519455129 | 0.838496482 | 0.116014022 | 0.097328753 |
| Mark2 | -0.046504912 | 0.488869622 | 0.826256745 | 0.480910979 | 0.523014359 |
| Mark3 | 0.010111835 | 0.88156486 | 0.968081674 | 0.515481858 | 0.510303073 |
| Mark4 | -0.001872816 | 0.962923479 | 0.991878382 | 0.107546161 | 0.11605218 |
| Mars1 | -0.118465335 | 0.057315181 | 0.36957055 | 0.375496435 | 0.488300789 |
| Marveld1 | 0.004374575 | 0.916118427 | 0.980132949 | 0.119932869 | 0.121829147 |
| Marveld2 | 0.003795533 | 0.915655063 | 0.980132949 | 0.085621994 | 0.083323044 |
| Marveld3 | 0.01514314 | 0.792293108 | 0.939347252 | 0.248276771 | 0.243287275 |
| Mast1 | 0.036048691 | 0.230702977 | 0.642320605 | 0.059113268 | 0.030053329 |
| Mast2 | -0.034773215 | 0.461269825 | 0.810249429 | 0.151778764 | 0.174147407 |
| Mast3 | 0.03341408 | 0.526094547 | 0.842170809 | 0.198001222 | 0.167359655 |
| Mast4 | -0.189803302 | 0.008928602 | 0.146996602 | 0.958341404 | 1.212345916 |
| Mat1a | -0.019214032 | 0.686912328 | 0.909796728 | 0.155460668 | 0.175070196 |
| Mat2a | -0.267025394 | 0.000144951 | 0.010542562 | 0.460743304 | 0.759284119 |
| Mat2b | -0.110841782 | 0.064039105 | 0.385820926 | 0.29129282 | 0.404290898 |
| Matr3 | 0.042419457 | 0.516248598 | 0.837427561 | 0.637926049 | 0.598081443 |
| Mau2 | -0.104400933 | 0.067182301 | 0.392969665 | 0.228277256 | 0.3241729 |
| Mavs | 0.047061247 | 0.105486275 | 0.472434351 | 0.056190666 | 0.018168094 |
| Max | -0.020131773 | 0.724367884 | 0.919830015 | 0.254612862 | 0.27754859 |
| Maz | -0.030897954 | 0.620109735 | 0.883937014 | 0.376148499 | 0.427483841 |
| Mbd1 | -0.035711402 | 0.47604742 | 0.817948631 | 0.174854552 | 0.213215519 |
| Mbd2 | -0.009321776 | 0.881356651 | 0.968081674 | 1.041972166 | 1.050713244 |
| Mbd3 | 0.009808719 | 0.874418881 | 0.964947297 | 0.391331787 | 0.379238177 |
| Mbd5 | 0.113765391 | 0.004643613 | 0.100256881 | 0.118751155 | 0.018323267 |
| Mbd6 | 0.007370274 | 0.884997958 | 0.969306472 | 0.192218219 | 0.177358823 |
| Mbip | -0.001880247 | 0.959169125 | 0.990654508 | 0.087936035 | 0.088631958 |
| Mblac1 | -0.021107874 | 0.373989403 | 0.759249926 | 0.029527497 | 0.050073637 |
| Mblac2 | -0.017345264 | 0.683198906 | 0.909173483 | 0.118544981 | 0.135709016 |
| Mbnl1 | -0.050976556 | 0.464402772 | 0.811633402 | 0.647197566 | 0.685517432 |
| Mbnl2 | -0.086724131 | 0.209861167 | 0.622606671 | 0.921761452 | 1.009310349 |
| Mboat2 | 0.092688204 | 0.160214835 | 0.558431893 | 0.489497704 | 0.408656543 |
| Mboat7 | 0.045339307 | 0.253309449 | 0.663146569 | 0.109716737 | 0.063999658 |
| Mbp | 0.092788499 | 0.159160241 | 0.556789205 | 0.470596636 | 0.382912899 |
| Mbtd1 | 0.019033674 | 0.669647264 | 0.903393762 | 0.137929097 | 0.112404113 |
| Mbtps1 | -0.052388447 | 0.34828863 | 0.739576235 | 0.231957064 | 0.281606091 |
| Mbtps2 | -0.079934427 | 0.030712229 | 0.279219789 | 0.079867899 | 0.149503066 |
| Mcat | -0.014037324 | 0.70238712 | 0.912152193 | 0.088238097 | 0.103590142 |
| Mcc | 0.017987699 | 0.693873928 | 0.909986409 | 0.144715328 | 0.127829579 |
| Mccc1 | -0.078587422 | 0.012025972 | 0.174689004 | 0.053245264 | 0.123585281 |
| Mcee | 0.038375211 | 0.52575658 | 0.842051614 | 0.315704312 | 0.285969706 |
| Mcf2l | -0.025444137 | 0.700918937 | 0.912152193 | 0.472331012 | 0.495532256 |
| Mcfd2 | -0.025452033 | 0.643210069 | 0.892561795 | 0.237693047 | 0.277686246 |
| Mcl1 | -0.094125371 | 0.152766898 | 0.549602359 | 0.447052864 | 0.568636274 |
| Mcm3ap | 0.039381027 | 0.32375113 | 0.72034806 | 0.110039497 | 0.078205774 |
| Mcm4 | 0.028085084 | 0.384090414 | 0.764448836 | 0.064283016 | 0.040699172 |
| Mcm7 | -0.047884391 | 0.087823897 | 0.440523963 | 0.04198471 | 0.085727547 |
| Mcm9 | -0.025164649 | 0.372387639 | 0.757837538 | 0.045678204 | 0.066543029 |
| Mcmbp | -0.022300219 | 0.658976345 | 0.898224427 | 0.181427337 | 0.202427314 |
| Mcoln1 | -0.015733755 | 0.580734561 | 0.866173066 | 0.04799459 | 0.05882567 |
| Mcph1 | 0.008376459 | 0.762239306 | 0.92872278 | 0.04968586 | 0.03979125 |
| Mcrip1 | 0.027105091 | 0.691867565 | 0.909796728 | 0.77948873 | 0.752284862 |
| Mcrip2 | 0.037256787 | 0.357854647 | 0.746823177 | 0.112116978 | 0.080108139 |
| Mcrs1 | -0.031252692 | 0.507655097 | 0.833674307 | 0.153806887 | 0.186131615 |

| | | | | | |
|---|---|---|---|---|---|
| Mcts1 | -0.086660239 | 0.150891029 | 0.546523037 | 0.295053921 | 0.381750215 |
| Mcts2 | 0.018759148 | 0.626992807 | 0.887105554 | 0.094481485 | 0.079640215 |
| Mcu | 0.011624916 | 0.725367192 | 0.92014727 | 0.073066869 | 0.063735028 |
| Mcub | 0.057074234 | 0.215351582 | 0.629611185 | 0.142054449 | 0.08871963 |
| Mcur1 | -0.103286537 | 0.068824976 | 0.399753362 | 0.235452605 | 0.330052696 |
| Mdc1 | -0.046911006 | 0.232912082 | 0.643935869 | 0.095009166 | 0.136928038 |
| Mdh1 | 0.070429088 | 0.23688915 | 0.647862778 | 1.430424729 | 1.297051157 |
| Mdh2 | -0.088519753 | 0.198831821 | 0.609389286 | 0.817272867 | 0.943486485 |
| Mdm1 | -0.197493093 | 0.00039034 | 0.02129261 | 0.211053176 | 0.386870592 |
| Mdm2 | -0.073562091 | 0.249186655 | 0.66061449 | 0.337843211 | 0.407784755 |
| Mdm4 | -0.283684137 | 4.43E-07 | 0.000111214 | 0.219671444 | 0.48395832 |
| Mdn1 | -0.034081456 | 0.427596074 | 0.791545709 | 0.120165741 | 0.148544994 |
| Mdp1 | -0.079593634 | 0.171594167 | 0.574479873 | 0.275856818 | 0.358133713 |
| Me1 | -0.05803263 | 0.275917275 | 0.683291806 | 0.184694579 | 0.247342441 |
| Me2 | -0.093074045 | 0.037194071 | 0.304267772 | 0.13316049 | 0.216673735 |
| Mea1 | -0.049418201 | 0.406722937 | 0.777736088 | 0.31156687 | 0.365486711 |
| Meaf6 | -0.015466134 | 0.766158007 | 0.92882091 | 0.186405843 | 0.198800287 |
| Mecp2 | 0.017107695 | 0.791071923 | 0.938824685 | 0.401646365 | 0.380209468 |
| Mecr | 0.026700831 | 0.340298177 | 0.733546237 | 0.051571071 | 0.029436847 |
| Med1 | 0.033538038 | 0.549737333 | 0.852455047 | 0.249204965 | 0.223974607 |
| Med10 | 0.014913578 | 0.811268782 | 0.948397417 | 0.388637942 | 0.38290846 |
| Med11 | 0.036569892 | 0.419251453 | 0.785665951 | 0.147006423 | 0.119026613 |
| Med12 | 0.000602533 | 0.983954489 | 0.99608406 | 0.055661009 | 0.05153039 |
| Med12l | 0.003114804 | 0.933229267 | 0.983806549 | 0.092949009 | 0.088444483 |
| Med13 | -0.111036394 | 0.105402554 | 0.472434351 | 0.4606919 | 0.578931881 |
| Med13l | 0.040725431 | 0.491923178 | 0.827270742 | 0.304805771 | 0.258982048 |
| Med14 | -0.00922988 | 0.835999244 | 0.954911647 | 0.130624756 | 0.14048368 |
| Med15 | 0.019833443 | 0.734215986 | 0.921626545 | 0.277711587 | 0.260302678 |
| Med16 | -0.010510876 | 0.802383835 | 0.9432568 | 0.111825763 | 0.120817943 |
| Med17 | 0.054693342 | 0.096954797 | 0.45927202 | 0.075586026 | 0.034323312 |
| Med18 | 0.001119231 | 0.968951855 | 0.993682135 | 0.052284708 | 0.050996286 |
| Med19 | 0.003616084 | 0.944672903 | 0.987070339 | 0.198108355 | 0.198541945 |
| Med20 | -0.002655714 | 0.943218304 | 0.986873983 | 0.0905028 | 0.095155119 |
| Med21 | -0.028008448 | 0.623562857 | 0.886173341 | 0.279184342 | 0.307494673 |
| Med22 | -0.016751788 | 0.731568761 | 0.920770724 | 0.166294368 | 0.179257125 |
| Med23 | 0.008876665 | 0.837037451 | 0.954911647 | 0.121058347 | 0.109771436 |
| Med24 | -0.044722221 | 0.153737333 | 0.550142209 | 0.056206202 | 0.090684908 |
| Med25 | 0.028402978 | 0.634210197 | 0.88965971 | 0.318319712 | 0.290275407 |
| Med26 | -0.001630629 | 0.959115894 | 0.990654508 | 0.063861184 | 0.068273362 |
| Med27 | 0.022404231 | 0.618835208 | 0.883660404 | 0.137086675 | 0.115040529 |
| Med28 | -0.102391222 | 0.111374356 | 0.482253844 | 0.86949015 | 1.01052603 |
| Med29 | -0.037393942 | 0.567171492 | 0.859145829 | 0.447352064 | 0.47159004 |
| Med30 | -0.060337307 | 0.299735887 | 0.70506682 | 0.26485527 | 0.326639124 |
| Med31 | 0.020882038 | 0.734176534 | 0.921626545 | 0.334867208 | 0.312562652 |
| Med4 | 0.007147188 | 0.887522103 | 0.970045467 | 0.184300997 | 0.180089389 |
| Med6 | 0.039695105 | 0.384167111 | 0.7644499 | 0.147893629 | 0.113247828 |
| Med7 | 0.005830327 | 0.918484093 | 0.980385833 | 0.275541569 | 0.276822539 |
| Med8 | -0.00739424 | 0.904210838 | 0.975255421 | 0.325416632 | 0.327101467 |
| Med9 | -0.042231858 | 0.450564162 | 0.806495742 | 0.240388924 | 0.294882107 |
| Med9os | 0.03885192 | 0.243303193 | 0.653166476 | 0.074917931 | 0.041371258 |
| Mef2a | 0.048876633 | 0.43114024 | 0.793555374 | 0.35942402 | 0.309970486 |
| Mef2c | -0.048932882 | 0.042045062 | 0.324121569 | 0.025481988 | 0.068481198 |
| Mef2d | 0.184033174 | 0.005141073 | 0.106393706 | 0.465569482 | 0.277399262 |
| Meg3 | -0.072644813 | 0.113360541 | 0.483758396 | 2.949586107 | 3.081595116 |
| Megf8 | -0.048435708 | 0.238840096 | 0.650445287 | 0.104251329 | 0.14264214 |
| Megf9 | 0.035850833 | 0.602254232 | 0.876569856 | 0.543390262 | 0.497963317 |
| Meig1 | -0.017133628 | 0.762553446 | 0.92872278 | 0.23421853 | 0.24170312 |
| Meis1 | -0.058160701 | 0.142650896 | 0.531472547 | 0.096043519 | 0.154937718 |
| Meis2 | -0.161863302 | 0.018271881 | 0.21500396 | 1.031843251 | 1.23104787 |
| Meis3 | 0.006988886 | 0.884830922 | 0.969283609 | 0.160718732 | 0.151610675 |
| Memo1 | -0.066904382 | 0.117848304 | 0.491192747 | 0.115972048 | 0.175359892 |
| Men1 | 0.018239675 | 0.680740601 | 0.907865189 | 0.140091274 | 0.127260314 |
| Mepce | -0.003434765 | 0.948586494 | 0.987651726 | 0.206959207 | 0.210775359 |
| Mesd | -0.080314424 | 0.135198628 | 0.518330263 | 0.2161878 | 0.282063512 |
| Met | 0.078876238 | 0.130338408 | 0.509823306 | 0.202393782 | 0.126960876 |

| | | | | | |
|---|---|---|---|---|---|
| Metap1 | 0.060502771 | 0.260007262 | 0.667410092 | 0.209994542 | 0.164880934 |
| Metap1d | -0.033111063 | 0.433444433 | 0.795625782 | 0.119148046 | 0.147563612 |
| Metap2 | 0.099426747 | 0.093111911 | 0.45188518 | 1.28366638 | 1.130952405 |
| Mettl1 | -0.258106857 | 0.000106328 | 0.008209368 | 0.39111848 | 0.65500378 |
| Mettl14 | 0.004911363 | 0.897576983 | 0.972811413 | 0.095385745 | 0.094001416 |
| Mettl16 | 0.02525712 | 0.64103311 | 0.891636547 | 0.207930557 | 0.19101538 |
| Mettl18 | 0.054398345 | 0.146273053 | 0.536014103 | 0.094507562 | 0.052379609 |
| Mettl21a | -0.050812699 | 0.092681798 | 0.450701165 | 0.053958903 | 0.099482655 |
| Mettl22 | -0.004619767 | 0.875715682 | 0.965151159 | 0.055504298 | 0.061190945 |
| Mettl23 | -0.004374621 | 0.941292844 | 0.986168593 | 0.307774209 | 0.332312495 |
| Mettl25 | -0.021933439 | 0.45754262 | 0.808898849 | 0.051833423 | 0.071781592 |
| Mettl26 | -0.072176562 | 0.303872425 | 0.708427238 | 0.75611409 | 0.867264183 |
| Mettl27 | -0.021965294 | 0.573859389 | 0.862168892 | 0.097916278 | 0.113242833 |
| Mettl3 | 0.065316667 | 0.058662615 | 0.372682944 | 0.086691103 | 0.033655708 |
| Mettl4 | -0.121303459 | 0.000839652 | 0.034153784 | 0.072356193 | 0.188623195 |
| Mettl5 | 0.034588158 | 0.431544215 | 0.793976916 | 0.13484307 | 0.101968965 |
| Mettl6 | 0.017710629 | 0.677155585 | 0.9069403 | 0.123209559 | 0.110574222 |
| Mettl7a1 | 0.138858671 | 0.037286974 | 0.304267772 | 0.418218481 | 0.282943696 |
| Mettl8 | -0.03499679 | 0.229367508 | 0.640979764 | 0.047996409 | 0.078676669 |
| Mettl9 | -0.102634415 | 0.128542199 | 0.508260084 | 0.608417097 | 0.727242599 |
| Mex3a | -0.046258743 | 0.010096579 | 0.156872081 | 0.009508361 | 0.049033895 |
| Mex3c | 0.133554039 | 0.004784319 | 0.101522651 | 0.164897749 | 0.051057989 |
| Mex3d | 0.01945781 | 0.576797252 | 0.863980874 | 0.080806729 | 0.068974804 |
| Mfap1a | 0.057787273 | 0.289338005 | 0.695472148 | 0.245302402 | 0.19802749 |
| Mfap1b | -0.009270246 | 0.877495736 | 0.966372465 | 0.305081302 | 0.318582737 |
| Mfap3 | -0.041201302 | 0.309752019 | 0.712742094 | 0.100860185 | 0.133593749 |
| Mff | -0.037865097 | 0.57104503 | 0.861225331 | 0.585964605 | 0.633208413 |
| Mfhas1 | 0.046557481 | 0.208702912 | 0.621219196 | 0.092136481 | 0.049205004 |
| Mfn1 | -0.035735945 | 0.387450141 | 0.767160379 | 0.114164311 | 0.140199784 |
| Mfn2 | 0.050996092 | 0.350470363 | 0.740908966 | 0.217727616 | 0.164410834 |
| Mfsd1 | -0.133464352 | 0.010807101 | 0.164475284 | 0.179830553 | 0.312181591 |
| Mfsd10 | 0.04230761 | 0.326271541 | 0.720866629 | 0.126703004 | 0.094698371 |
| Mfsd11 | 0.091413846 | 0.106587069 | 0.473998412 | 0.262092688 | 0.167824531 |
| Mfsd13a | 0.013121988 | 0.745134739 | 0.924566547 | 0.116086526 | 0.099871999 |
| Mfsd14a | -0.000355046 | 0.994823677 | 0.998704266 | 0.243955245 | 0.258751604 |
| Mfsd14b | -0.029854517 | 0.553187198 | 0.852763002 | 0.164740589 | 0.190991847 |
| Mfsd4b1 | -0.032008216 | 0.107013847 | 0.474424902 | 0.017344866 | 0.044324996 |
| Mfsd5 | 0.066719061 | 0.155677867 | 0.552356396 | 0.155660985 | 0.103443356 |
| Mfsd6 | 0.027936855 | 0.619137267 | 0.883797831 | 0.236511174 | 0.210506474 |
| Mfsd6l | 0.023577848 | 0.438291552 | 0.799200693 | 0.06179238 | 0.038632141 |
| Mfsd8 | -0.007583721 | 0.844938761 | 0.955950886 | 0.101565607 | 0.108303785 |
| Mga | 0.081644355 | 0.222943199 | 0.634822484 | 0.544179017 | 0.461595305 |
| Mgat1 | 0.03958967 | 0.440599869 | 0.80031914 | 0.205516259 | 0.178857321 |
| Mgat2 | 0.018572163 | 0.746542365 | 0.924631183 | 0.262298176 | 0.254031505 |
| Mgat3 | -0.018432037 | 0.68017903 | 0.907783945 | 0.134272248 | 0.15023956 |
| Mgat4a | -0.013392586 | 0.588924095 | 0.86992264 | 0.033409782 | 0.044573231 |
| Mgat4b | 0.085394298 | 0.109295775 | 0.477972625 | 0.227074088 | 0.154632786 |
| Mgat5 | -0.009326177 | 0.783122882 | 0.936124803 | 0.07204847 | 0.083053362 |
| Mgme1 | 0.092672198 | 0.027813738 | 0.26510449 | 0.121988585 | 0.046720148 |
| Mgmt | -0.040739267 | 0.228046094 | 0.639451668 | 0.063939125 | 0.107568659 |
| Mgrn1 | -0.03278225 | 0.509890374 | 0.834904701 | 0.176667732 | 0.202561914 |
| Mgst3 | 0.046863596 | 0.379856608 | 0.762006446 | 0.218790461 | 0.169697765 |
| Mia2 | -0.156073729 | 0.022886336 | 0.239281409 | 0.944246112 | 1.146527898 |
| Mia3 | -0.161661936 | 0.007864263 | 0.13698938 | 1.354257491 | 1.594345909 |
| Mib1 | -0.019698494 | 0.762690246 | 0.92872278 | 0.394233002 | 0.422856096 |
| Mib2 | -0.100658836 | 0.030035335 | 0.27505899 | 0.137358178 | 0.226301128 |
| Mical2 | 0.015180752 | 0.779293325 | 0.934723602 | 0.200690905 | 0.188442332 |
| Mical3 | 0.086340359 | 0.155403549 | 0.552356396 | 0.332803349 | 0.259934059 |
| Micall1 | -0.025913778 | 0.343277137 | 0.735897614 | 0.043004248 | 0.064466059 |
| Micos10 | 0.049380043 | 0.262990277 | 0.669775695 | 1.911242759 | 1.798587587 |
| Micos13 | -0.055241011 | 0.350000814 | 0.7406656 | 1.348174264 | 1.406494756 |
| Micu1 | -0.02897976 | 0.571752239 | 0.861404566 | 0.193950391 | 0.217142997 |
| Micu2 | 0.082481062 | 0.192398944 | 0.60309438 | 0.38700894 | 0.316186214 |
| Micu3 | -0.000272541 | 0.996575245 | 0.998992348 | 0.430420294 | 0.429929617 |
| Mid1 | 0.03099895 | 0.253843247 | 0.663146569 | 0.048641357 | 0.024793876 |

| | | | | | |
|---|---|---|---|---|---|
| Mid1ip1 | -0.094091815 | 0.172697155 | 0.575294174 | 0.504838819 | 0.616099032 |
| Mid2 | -0.064911228 | 0.170645106 | 0.572831081 | 0.140213869 | 0.196913478 |
| Mideas | -0.018668383 | 0.68200111 | 0.908884974 | 0.129682526 | 0.148250998 |
| Midn | 0.075005147 | 0.2098565 | 0.622606671 | 0.275148483 | 0.213781875 |
| Mief1 | -0.073382735 | 0.103429872 | 0.469315384 | 0.131771614 | 0.196810484 |
| Mien1 | -0.002554287 | 0.968520406 | 0.993505636 | 1.023375202 | 1.032651842 |
| Mier1 | -0.040669219 | 0.542945681 | 0.850428495 | 0.59414744 | 0.639621103 |
| Mier2 | 0.029072938 | 0.382449462 | 0.763150149 | 0.073626582 | 0.044868442 |
| Mier3 | 0.032700019 | 0.472469668 | 0.816209648 | 0.142852179 | 0.116310134 |
| Mif | 0.004779263 | 0.945581565 | 0.987084873 | 1.155497819 | 1.162724481 |
| Mif4gd | -0.041120929 | 0.318322892 | 0.717175454 | 0.110073768 | 0.150170493 |
| Miga1 | -0.006594925 | 0.895720946 | 0.97233366 | 0.178427868 | 0.18971316 |
| Miip | 0.002995383 | 0.917053408 | 0.980132949 | 0.050571685 | 0.045903924 |
| Mindy1 | -0.028323129 | 0.510664664 | 0.834957869 | 0.119694007 | 0.146178207 |
| Mindy2 | -0.071455288 | 0.280256826 | 0.687465228 | 0.588110131 | 0.632152176 |
| Mindy3 | -0.096461108 | 0.056219853 | 0.367897032 | 0.16345136 | 0.253102815 |
| Mink1 | -0.131242777 | 0.005715177 | 0.112697904 | 0.1345944 | 0.237587406 |
| Minpp1 | 0.012563689 | 0.757063969 | 0.92705828 | 0.109922078 | 0.097743333 |
| Mios | 0.030050125 | 0.460588903 | 0.810111566 | 0.113426629 | 0.089659023 |
| Mipep | -0.017888367 | 0.49347921 | 0.82875113 | 0.040368728 | 0.055221994 |
| Mipol1 | 0.090032274 | 0.058178849 | 0.371463807 | 0.167903763 | 0.094364626 |
| Mir22hg | -0.231498048 | 1.02E-05 | 0.001317415 | 0.165357287 | 0.373191447 |
| Mir6236 | -0.227618848 | 0.003534304 | 0.08334591 | 0.69902361 | 1.026204675 |
| Mir670hg | 0.155517215 | 0.028960222 | 0.271130568 | 0.727551959 | 0.543850793 |
| Mir703 | 0.052796701 | 0.100615498 | 0.465596015 | 0.073854993 | 0.034284295 |
| Mirg | 0.024152093 | 0.603609291 | 0.876762149 | 0.148442427 | 0.119958691 |
| Mis12 | 0.048449383 | 0.281362165 | 0.68811697 | 0.140574612 | 0.098847424 |
| Mitd1 | 0.074328981 | 0.154439744 | 0.551012197 | 0.203254252 | 0.131356074 |
| Mitf | -0.038472598 | 0.208361889 | 0.620388098 | 0.052228634 | 0.088686693 |
| Mkln1 | 0.05283091 | 0.428846833 | 0.792251095 | 0.478542647 | 0.428340334 |
| Mknk1 | 0.007117392 | 0.894099728 | 0.971851741 | 0.209714686 | 0.201626996 |
| Mknk2 | -0.012389536 | 0.759232454 | 0.927719223 | 0.106679896 | 0.117228358 |
| Mkrn1 | -0.11253616 | 0.096340005 | 0.457336038 | 0.529624163 | 0.657400774 |
| Mkrn2 | 0.105805985 | 0.057040992 | 0.369129876 | 0.252286245 | 0.167948263 |
| Mkrn2os | 0.105366782 | 0.077013991 | 0.418199494 | 0.291660973 | 0.197465087 |
| Mks1 | -0.019870382 | 0.484099729 | 0.822768706 | 0.046466663 | 0.063851456 |
| Mlec | -0.013980973 | 0.839233877 | 0.954911647 | 0.874157929 | 0.891858521 |
| Mlf2 | 0.116169662 | 0.090286224 | 0.445385778 | 0.729733333 | 0.596293871 |
| Mlh1 | -0.040010836 | 0.181434735 | 0.589059475 | 0.051298722 | 0.085445029 |
| Mlh3 | -0.031490458 | 0.537240011 | 0.847793898 | 0.174639788 | 0.214528175 |
| Mllt10 | -0.006469 | 0.917247505 | 0.980132949 | 0.328527694 | 0.327681012 |
| Mllt11 | 0.050300582 | 0.354646387 | 0.745150888 | 0.213672267 | 0.159107009 |
| Mllt3 | -0.017484648 | 0.75185511 | 0.926186851 | 0.22591616 | 0.245070104 |
| Mllt6 | -0.092871416 | 0.044132822 | 0.329446347 | 0.138826234 | 0.220388072 |
| Mlx | 0.123825688 | 0.037583063 | 0.30568979 | 0.328434062 | 0.206355262 |
| Mlxip | 0.030378822 | 0.518643077 | 0.838109801 | 0.154204985 | 0.127325944 |
| Mlxipl | -0.119759474 | 0.082000596 | 0.42777546 | 0.462339694 | 0.598030194 |
| Mlycd | 0.020201371 | 0.738794962 | 0.922843014 | 0.30290668 | 0.280751596 |
| Mmaa | 0.013444774 | 0.699818441 | 0.911507617 | 0.081649796 | 0.075626997 |
| Mmab | -0.041572067 | 0.165394426 | 0.566504119 | 0.052120673 | 0.091516278 |
| Mmachc | -0.046959221 | 0.081894338 | 0.42777546 | 0.039068419 | 0.077391189 |
| Mmadhc | 0.072310783 | 0.235161495 | 0.645247655 | 0.332481548 | 0.266534716 |
| Mmgt1 | -0.039177839 | 0.331651111 | 0.727636858 | 0.10134938 | 0.129968864 |
| Mmgt2 | 0.007882519 | 0.89287419 | 0.971493043 | 0.270040692 | 0.26953328 |
| Mms19 | -0.013490977 | 0.748884721 | 0.925186829 | 0.116292308 | 0.121801865 |
| Mmut | -0.071867095 | 0.161311432 | 0.559461938 | 0.177372709 | 0.248874019 |
| Mnat1 | 0.054779032 | 0.361650391 | 0.750013467 | 0.331103175 | 0.282557207 |
| Mns1 | 0.002978029 | 0.959710961 | 0.990908231 | 0.263565738 | 0.255979933 |
| Mnt | -0.077919304 | 0.091946487 | 0.448211214 | 0.128500589 | 0.200891293 |
| Mnx1 | 0.01925379 | 0.7103299 | 0.914486281 | 0.183512601 | 0.177087237 |
| Mob1a | -0.009076301 | 0.859748527 | 0.959220425 | 0.198428317 | 0.204686746 |
| Mob1b | -0.009231399 | 0.87851946 | 0.966959077 | 0.324321687 | 0.338806718 |
| Mob2 | 0.029293832 | 0.572602821 | 0.861775874 | 0.189333011 | 0.162445369 |
| Mob3c | -0.108410751 | 0.023912073 | 0.245655551 | 0.146141713 | 0.240630356 |
| Mob4 | 0.039230095 | 0.550261419 | 0.852505562 | 0.521813277 | 0.484934392 |

| | | | | | |
|---|---|---|---|---|---|
| Mocs2 | -0.041301842 | 0.537430785 | 0.847793898 | 0.703416573 | 0.750379536 |
| Mocs3 | 0.038185623 | 0.279745341 | 0.687465228 | 0.085314771 | 0.059531612 |
| Mogs | 0.029287706 | 0.592125313 | 0.872005806 | 0.223395257 | 0.190692024 |
| Mon1a | -0.027943791 | 0.407310441 | 0.777888922 | 0.070595757 | 0.093223015 |
| Mon1b | 0.00204546 | 0.954704276 | 0.989066902 | 0.083605812 | 0.080197431 |
| Mon2 | -0.062509338 | 0.327850765 | 0.722581934 | 0.391576704 | 0.476391366 |
| Morc2a | -0.092817187 | 0.033665873 | 0.290545455 | 0.1168529 | 0.196011501 |
| Morc3 | -0.055750968 | 0.271479859 | 0.67876599 | 0.174877694 | 0.235557097 |
| Morc4 | -0.023900657 | 0.581392068 | 0.866173066 | 0.111979619 | 0.125881595 |
| Morf4l1 | 0.007191277 | 0.878112373 | 0.966691068 | 1.900234439 | 1.872374638 |
| Morf4l2 | -0.0442642 | 0.43179622 | 0.79405252 | 1.530835798 | 1.608518935 |
| Morn1 | -0.03855608 | 0.053842633 | 0.361725909 | 0.015591375 | 0.045868607 |
| Morn2 | -0.179112499 | 0.001985444 | 0.060755792 | 0.266111851 | 0.427086164 |
| Morn4 | -0.059380025 | 0.063929244 | 0.385611669 | 0.058761917 | 0.109762106 |
| Mosmo | 0.01346207 | 0.798366757 | 0.940958114 | 0.209767176 | 0.194634039 |
| Mospd2 | 0.032154472 | 0.505471872 | 0.83252727 | 0.163114029 | 0.138435042 |
| Mospd3 | -0.074797886 | 0.023896292 | 0.245655551 | 0.06260505 | 0.122951012 |
| Mov10 | -0.012155386 | 0.680359877 | 0.907783945 | 0.052017154 | 0.062906175 |
| Mpc1 | 0.128249708 | 0.065279459 | 0.389310714 | 0.785049251 | 0.620392536 |
| Mpc2 | -0.076854898 | 0.234930181 | 0.64504573 | 0.938596095 | 1.021767303 |
| Mpdu1 | 0.030393972 | 0.525798676 | 0.842051614 | 0.163266462 | 0.131237246 |
| Mpdz | 0.083881773 | 0.090197877 | 0.445385778 | 0.188906746 | 0.111707768 |
| Mpg | -0.031699193 | 0.378369676 | 0.761977617 | 0.084143943 | 0.10779226 |
| Mphosph10 | 0.084134947 | 0.215662375 | 0.629611185 | 0.514203799 | 0.425077893 |
| Mphosph6 | 0.073358723 | 0.158175352 | 0.556468981 | 0.201563939 | 0.139248202 |
| Mphosph8 | 0.071414173 | 0.305052698 | 0.709410087 | 0.750045431 | 0.671826051 |
| Mphosph9 | -0.002615165 | 0.949856266 | 0.98784658 | 0.115057786 | 0.122525369 |
| Mplkip | 0.051198915 | 0.402961028 | 0.77525778 | 0.319578945 | 0.266258221 |
| Mpnd | -0.13065143 | 0.005278851 | 0.108299861 | 0.131811873 | 0.243112626 |
| Mpp2 | -0.021709813 | 0.611644041 | 0.880153582 | 0.121287042 | 0.132229049 |
| Mpp3 | 0.041828468 | 0.18193282 | 0.589454 | 0.06807494 | 0.030513502 |
| Mpp5 | 0.06945496 | 0.203274638 | 0.614800187 | 0.236592534 | 0.182386897 |
| Mpp6 | -0.006481023 | 0.915899398 | 0.980132949 | 0.314719751 | 0.321505538 |
| Mpp7 | 0.046623054 | 0.459331076 | 0.809393611 | 0.313563848 | 0.247397632 |
| Mppe1 | -0.099818746 | 0.001428109 | 0.048921251 | 0.049643932 | 0.140533236 |
| Mpped2 | -0.032845513 | 0.471043125 | 0.814586466 | 0.144241205 | 0.170911147 |
| Mprip | 0.055750141 | 0.382952021 | 0.763687454 | 0.407686246 | 0.352214701 |
| Mpst | 0.046977477 | 0.155544621 | 0.552356396 | 0.075557979 | 0.033463215 |
| Mpv17 | 0.028843612 | 0.62495772 | 0.88668351 | 0.284906231 | 0.252794401 |
| Mpv17l | 0.026628701 | 0.399465227 | 0.774544012 | 0.06711756 | 0.041735508 |
| Mpv17l2 | -0.076499432 | 0.171103302 | 0.573793465 | 0.220347524 | 0.295109914 |
| Mpz | -0.001361144 | 0.961052077 | 0.991375098 | 0.04848553 | 0.049312203 |
| Mpzl1 | 0.067052168 | 0.24547366 | 0.655159244 | 0.25963175 | 0.191849872 |
| Mr1 | -0.010766046 | 0.692663456 | 0.909796728 | 0.044867804 | 0.05429688 |
| Mras | 0.160391985 | 0.014536883 | 0.193767197 | 0.423397761 | 0.263005047 |
| Mre11a | 0.041358402 | 0.205810816 | 0.616737719 | 0.074618047 | 0.036881276 |
| Mreg | 0.037097599 | 0.41852262 | 0.784886311 | 0.137941674 | 0.103852486 |
| Mrfap1 | 0.075020138 | 0.06190888 | 0.380250374 | 2.206257324 | 2.083348009 |
| Mrgbp | -0.002602242 | 0.946966195 | 0.987501267 | 0.104273046 | 0.104176995 |
| Mrgpre | 0.0564515 | 0.112945557 | 0.483221516 | 0.090334314 | 0.041899223 |
| Mri1 | -0.011356948 | 0.775728466 | 0.933636662 | 0.10646796 | 0.121785188 |
| Mrln | 0.036308399 | 0.591382397 | 0.871477529 | 1.002191948 | 0.912265536 |
| Mrm1 | -0.034251449 | 0.163913707 | 0.563565555 | 0.030872952 | 0.061930183 |
| Mrm2 | 0.017321833 | 0.582998595 | 0.866383905 | 0.063372708 | 0.054526819 |
| Mrm3 | 0.041358173 | 0.181939672 | 0.589454 | 0.064698416 | 0.036054552 |
| Mroh1 | -0.061564328 | 0.192226503 | 0.60309438 | 0.145403382 | 0.197070295 |
| Mroh7 | -0.003626875 | 0.896229938 | 0.97233366 | 0.04922075 | 0.049650227 |
| Mrpl1 | 0.081047864 | 0.19469329 | 0.604060759 | 0.336452466 | 0.267463765 |
| Mrpl10 | -0.007175221 | 0.896753268 | 0.972419247 | 0.23318391 | 0.242819754 |
| Mrpl11 | -0.105936943 | 0.090497654 | 0.445385778 | 0.34183388 | 0.448231883 |
| Mrpl12 | 0.057727524 | 0.401155287 | 0.774740795 | 0.500864646 | 0.443144855 |
| Mrpl13 | -0.10412381 | 0.086280837 | 0.43696958 | 0.348822108 | 0.462244914 |
| Mrpl14 | 0.089548666 | 0.176470488 | 0.580732554 | 0.620764261 | 0.535588496 |
| Mrpl15 | 0.126544724 | 0.044213845 | 0.329453867 | 0.377220735 | 0.259903979 |
| Mrpl16 | 0.054733709 | 0.276084264 | 0.683536695 | 0.182032973 | 0.146130431 |

| | | | | | |
|---|---|---|---|---|---|
| Mrpl17 | 0.070878985 | 0.315563067 | 0.714975653 | 0.781813155 | 0.700524078 |
| Mrpl18 | 0.218576955 | 0.000620768 | 0.028602902 | 0.384747149 | 0.179576715 |
| Mrpl19 | 0.039533829 | 0.444568897 | 0.802110017 | 0.209341627 | 0.173541111 |
| Mrpl2 | -0.046256322 | 0.463731587 | 0.811254677 | 0.407063717 | 0.465709482 |
| Mrpl20 | 0.062800355 | 0.34221611 | 0.735068249 | 0.917200681 | 0.850135318 |
| Mrpl21 | -0.164112376 | 0.011119249 | 0.166572992 | 0.378166561 | 0.532059622 |
| Mrpl22 | 0.010367526 | 0.819210119 | 0.950128492 | 0.142541705 | 0.145042066 |
| Mrpl23 | -0.011457031 | 0.748803025 | 0.925186829 | 0.082382815 | 0.09233288 |
| Mrpl24 | -0.110596884 | 0.099555057 | 0.463642501 | 0.477910822 | 0.5838429 |
| Mrpl27 | -0.061288487 | 0.338106226 | 0.732352032 | 0.400203808 | 0.467451564 |
| Mrpl28 | 0.007597822 | 0.90318856 | 0.974950771 | 0.465325797 | 0.477489348 |
| Mrpl3 | -0.017393413 | 0.741200231 | 0.922901128 | 0.202048326 | 0.207634028 |
| Mrpl30 | -0.029887403 | 0.649100153 | 0.895557756 | 0.778917272 | 0.819297417 |
| Mrpl32 | 0.020414197 | 0.729768731 | 0.920522999 | 0.296899303 | 0.289715897 |
| Mrpl33 | 0.019375665 | 0.773910383 | 0.932949618 | 0.765046806 | 0.707101103 |
| Mrpl34 | -0.071744862 | 0.290225336 | 0.696221724 | 0.679180293 | 0.757706634 |
| Mrpl35 | 0.0087526 | 0.87222534 | 0.964745713 | 0.229532945 | 0.220508502 |
| Mrpl36 | -0.063783727 | 0.345590926 | 0.73792645 | 0.67868077 | 0.740393026 |
| Mrpl37 | 0.006411735 | 0.876993009 | 0.966020504 | 0.117280856 | 0.108244246 |
| Mrpl38 | -0.017729977 | 0.764904019 | 0.928757959 | 0.294963447 | 0.319920086 |
| Mrpl39 | -0.001589973 | 0.972851567 | 0.994079882 | 0.147543347 | 0.155820971 |
| Mrpl4 | -0.0517117 | 0.394698318 | 0.771487247 | 0.34027982 | 0.399812429 |
| Mrpl40 | -0.035912889 | 0.53644503 | 0.847520662 | 0.268769803 | 0.29933025 |
| Mrpl41 | 0.120446227 | 0.067183554 | 0.392969665 | 0.449586177 | 0.326016784 |
| Mrpl42 | -0.022797009 | 0.728656717 | 0.920519505 | 0.570217125 | 0.603853418 |
| Mrpl43 | -0.040448909 | 0.540348269 | 0.849380522 | 0.523363635 | 0.575401331 |
| Mrpl44 | -0.007997005 | 0.870866707 | 0.964504119 | 0.165978233 | 0.191046629 |
| Mrpl45 | 0.009139347 | 0.83842373 | 0.954911647 | 0.13512958 | 0.133262391 |
| Mrpl46 | 0.049011132 | 0.387967692 | 0.767160379 | 0.264950319 | 0.218218055 |
| Mrpl47 | -0.046405692 | 0.188275126 | 0.596501717 | 0.076177441 | 0.114566851 |
| Mrpl48 | -0.046421717 | 0.458835598 | 0.809370658 | 0.372141831 | 0.410325583 |
| Mrpl49 | 0.007307613 | 0.89048213 | 0.970579007 | 0.206112787 | 0.203340795 |
| Mrpl50 | 0.054686545 | 0.373225283 | 0.758519786 | 0.33273717 | 0.280631249 |
| Mrpl51 | 0.051766702 | 0.43766086 | 0.798546091 | 0.649512637 | 0.586667464 |
| Mrpl52 | -0.052107736 | 0.42868957 | 0.792251095 | 0.85271861 | 0.892348303 |
| Mrpl54 | 0.064068858 | 0.33464003 | 0.730169996 | 0.866795674 | 0.804319841 |
| Mrpl55 | -0.029824569 | 0.633546624 | 0.889501117 | 0.363674371 | 0.389485766 |
| Mrpl57 | -0.004297761 | 0.943685256 | 0.986918206 | 1.002453304 | 1.012092465 |
| Mrpl58 | -0.150723943 | 0.023847787 | 0.245655381 | 0.458879572 | 0.626274376 |
| Mrpl9 | -0.004973797 | 0.933398884 | 0.983869133 | 0.296155438 | 0.311031416 |
| Mrps10 | -0.053450331 | 0.314571422 | 0.714975653 | 0.193652052 | 0.257291244 |
| Mrps11 | -0.001318003 | 0.982013337 | 0.995803987 | 0.275000809 | 0.263282056 |
| Mrps12 | 0.030102699 | 0.658744974 | 0.898224427 | 1.016774022 | 0.967353674 |
| Mrps14 | -0.08739657 | 0.193533212 | 0.60325934 | 0.82408569 | 0.916144633 |
| Mrps15 | -0.001622279 | 0.978007255 | 0.995031396 | 0.281450311 | 0.28846788 |
| Mrps16 | 0.108074202 | 0.143144357 | 0.532282155 | 0.895203618 | 0.754104614 |
| Mrps17 | -0.04406203 | 0.49532467 | 0.829148384 | 0.378083797 | 0.43587211 |
| Mrps18a | -0.000912426 | 0.98904327 | 0.997411191 | 0.48110193 | 0.496288557 |
| Mrps18b | 0.037854034 | 0.533794262 | 0.84612966 | 0.301438341 | 0.269837062 |
| Mrps18c | 0.074924159 | 0.235000326 | 0.64504573 | 0.443843985 | 0.369335509 |
| Mrps2 | -0.069159369 | 0.051495006 | 0.355933715 | 0.071291174 | 0.132757226 |
| Mrps21 | 0.074407206 | 0.269131789 | 0.676978829 | 0.893581652 | 0.773306016 |
| Mrps22 | 0.01183328 | 0.760047912 | 0.927859527 | 0.099036929 | 0.087900805 |
| Mrps23 | 0.070685983 | 0.269171397 | 0.676978829 | 0.417454053 | 0.358004722 |
| Mrps24 | -0.013755446 | 0.83627755 | 0.954911647 | 0.697571611 | 0.711731061 |
| Mrps25 | 0.036400763 | 0.494758794 | 0.828842054 | 0.215473006 | 0.189596233 |
| Mrps26 | -0.023574296 | 0.718317286 | 0.91797181 | 0.55934366 | 0.58056949 |
| Mrps28 | -0.056799839 | 0.39132701 | 0.769704269 | 0.416492335 | 0.485011353 |
| Mrps30 | 0.035570838 | 0.476274702 | 0.817948631 | 0.177841545 | 0.154224164 |
| Mrps31 | 0.049667117 | 0.370317006 | 0.75730884 | 0.254518172 | 0.217668539 |
| Mrps33 | -0.095334499 | 0.134728514 | 0.517714432 | 1.014302765 | 1.159520629 |
| Mrps34 | 0.108076596 | 0.111181623 | 0.482251491 | 0.70045666 | 0.594957386 |
| Mrps35 | -0.0838444 | 0.104999807 | 0.472399448 | 0.185404822 | 0.262840431 |
| Mrps36 | 0.04428753 | 0.468150579 | 0.813213564 | 0.35063973 | 0.304174887 |
| Mrps5 | 0.109107643 | 0.084995857 | 0.434272336 | 0.382968786 | 0.265823848 |

| | | | | | |
|---|---|---|---|---|---|
| Mrps6 | -0.078368137 | 0.044881658 | 0.331476968 | 0.089678263 | 0.152462329 |
| Mrps7 | 0.04812483 | 0.451036819 | 0.806495742 | 0.408503835 | 0.372954164 |
| Mrps9 | 0.039646302 | 0.428001506 | 0.791658813 | 0.178114067 | 0.130511493 |
| Mrrf | 0.002264513 | 0.955629353 | 0.989278596 | 0.111474094 | 0.110692544 |
| Mrs2 | 0.024646717 | 0.661636882 | 0.898920864 | 0.251393912 | 0.235870616 |
| Mrtfb | 0.027831853 | 0.503789513 | 0.832249514 | 0.121547979 | 0.101701384 |
| Mrto4 | 0.016457565 | 0.742280525 | 0.923637085 | 0.177687286 | 0.175101652 |
| Msantd2 | 0.011121415 | 0.723550569 | 0.919830015 | 0.06306542 | 0.052941918 |
| Msantd3 | -0.034600767 | 0.288779745 | 0.695246414 | 0.066225538 | 0.093691924 |
| Msantd4 | -0.008161025 | 0.891127519 | 0.970742575 | 0.281975815 | 0.290158672 |
| Msh2 | 0.049353863 | 0.174887158 | 0.577992588 | 0.090078307 | 0.049309177 |
| Msh3 | 0.005524506 | 0.848668673 | 0.957111129 | 0.053834487 | 0.0438185 |
| Msh6 | 0.044806431 | 0.235023636 | 0.64504573 | 0.10087466 | 0.065574035 |
| Msi2 | -0.087239009 | 0.196449878 | 0.60713211 | 1.020774412 | 1.099247273 |
| Msl1 | -0.011880129 | 0.850214489 | 0.957111129 | 0.461188416 | 0.473043625 |
| Msl2 | 0.003691912 | 0.950059561 | 0.987860147 | 0.276145324 | 0.266026821 |
| Msl3 | 0.055652289 | 0.293216522 | 0.699053261 | 0.21022067 | 0.161540494 |
| Msl3l2 | 0.013854389 | 0.741393445 | 0.923017366 | 0.120250156 | 0.107764122 |
| Msmo1 | 0.036934937 | 0.582166335 | 0.866173066 | 0.46776939 | 0.415738791 |
| Msra | -0.037546418 | 0.167653663 | 0.570246024 | 0.039374593 | 0.07359064 |
| Msrb1 | -0.01827031 | 0.698603338 | 0.911319029 | 0.155817771 | 0.178196298 |
| Msrb2 | -0.046734479 | 0.108553599 | 0.477972625 | 0.047689185 | 0.088692168 |
| Msto1 | -0.05137231 | 0.039034358 | 0.311933003 | 0.031733039 | 0.074366536 |
| Mt1 | 0.908110272 | 5.16E-17 | 1.04E-13 | 4.199076966 | 1.958884449 |
| Mt2 | 0.946528856 | 9.14E-15 | 1.15E-11 | 2.285011638 | 0.643771747 |
| Mt3 | 0.096223544 | 0.036389676 | 0.30061167 | 0.140095701 | 0.046810552 |
| Mta1 | -0.057753048 | 0.288454004 | 0.694795498 | 0.198015787 | 0.242767754 |
| Mta2 | 0.009186011 | 0.847773046 | 0.956938604 | 0.162657796 | 0.146019304 |
| Mta3 | -0.052665494 | 0.313714507 | 0.714906079 | 0.190536049 | 0.242184299 |
| Mtag2 | -0.023878969 | 0.29748549 | 0.702656882 | 0.026875483 | 0.048135234 |
| Mtap | -0.147507852 | 0.001190316 | 0.043130693 | 0.12970431 | 0.264341496 |
| Mtarc2 | -0.010697738 | 0.871532092 | 0.964616742 | 0.472450643 | 0.480967602 |
| mt-Atp6 | -0.084820344 | 0.060888927 | 0.380250374 | 0.119322483 | 0.197819236 |
| mt-Atp8 | 0.004609517 | 0.945375486 | 0.987084873 | 0.458117427 | 0.405997221 |
| Mtch1 | -0.139379304 | 0.035929287 | 0.298528355 | 1.015878181 | 1.195696044 |
| Mtch2 | 0.024146567 | 0.71637843 | 0.916579082 | 0.613403905 | 0.576324218 |
| Mtcl1 | -0.025630183 | 0.674504368 | 0.906025597 | 0.291430467 | 0.31729923 |
| mt-Co1 | 0.064154127 | 0.37813473 | 0.761963116 | 2.131176106 | 1.939524423 |
| mt-Cytb | -0.177387305 | 0.002018339 | 0.061388095 | 3.970068375 | 4.404113837 |
| Mtdh | 0.069250795 | 0.245583343 | 0.655215315 | 1.27204548 | 1.147706031 |
| Mterf1a | 0.045028529 | 0.15006523 | 0.544542557 | 0.063755245 | 0.025431544 |
| Mterf3 | 0.003717844 | 0.943544858 | 0.986918206 | 0.203055458 | 0.211570654 |
| Mterf4 | 0.015271018 | 0.629398978 | 0.887881594 | 0.064752221 | 0.048398658 |
| Mtf1 | 0.081073848 | 0.164575809 | 0.564527879 | 0.281279361 | 0.21162093 |
| Mtf2 | -0.034294562 | 0.445370892 | 0.803123904 | 0.132604713 | 0.154747446 |
| Mtfmt | -0.006198976 | 0.840161751 | 0.954911647 | 0.058429183 | 0.068527393 |
| Mtfp1 | -0.059032068 | 0.090459461 | 0.445385778 | 0.072470928 | 0.12305774 |
| Mtfr1 | -0.037625838 | 0.500234858 | 0.831130155 | 0.22066901 | 0.265086767 |
| Mtfr1l | -0.002117459 | 0.971760189 | 0.993892166 | 0.333095627 | 0.330476008 |
| Mtg1 | -0.106323002 | 0.007177212 | 0.128762694 | 0.090970343 | 0.185609101 |
| Mtg2 | 0.038453831 | 0.398021413 | 0.774311756 | 0.146599588 | 0.111207224 |
| Mthfd1 | -0.012611682 | 0.817436801 | 0.949382903 | 0.223149634 | 0.237307138 |
| Mthfd1l | -0.050235106 | 0.285371577 | 0.692407818 | 0.139225506 | 0.181319563 |
| Mthfd2 | -0.077185634 | 0.255390324 | 0.664253092 | 1.201442842 | 1.256857889 |
| Mthfd2l | -0.202540166 | 4.94E-05 | 0.004588036 | 0.152655578 | 0.335928868 |
| Mthfr | -0.065398934 | 0.140719961 | 0.528664246 | 0.126501717 | 0.186727138 |
| Mthfs | -0.092527378 | 0.002266876 | 0.065256147 | 0.049601324 | 0.131068843 |
| Mthfsd | 0.02444219 | 0.498098927 | 0.830722138 | 0.087722318 | 0.060307855 |
| Mthfsl | -0.034170591 | 0.371222425 | 0.75760494 | 0.090760794 | 0.120389479 |
| Mtif2 | 0.076546429 | 0.183122037 | 0.590066231 | 0.260343821 | 0.187988963 |
| Mtif3 | 0.00082273 | 0.987584102 | 0.997268589 | 0.199879477 | 0.19672523 |
| Mtln | 0.049653636 | 0.463522136 | 0.811254677 | 0.530520281 | 0.479947848 |
| Mtm1 | -0.044646846 | 0.125513663 | 0.504113898 | 0.048709324 | 0.089046155 |
| Mtmr1 | 0.027048136 | 0.572685571 | 0.861775874 | 0.163357004 | 0.135820117 |
| Mtmr10 | 0.019533389 | 0.573595577 | 0.862168892 | 0.079555548 | 0.060366825 |

| | | | | | |
|---|---|---|---|---|---|
| Mtmr11 | -0.064911106 | 0.066644795 | 0.392235932 | 0.07330508 | 0.133483309 |
| Mtmr12 | 0.038415598 | 0.25218841 | 0.663127858 | 0.078676826 | 0.041912938 |
| Mtmr2 | 0.003434891 | 0.94620476 | 0.987084873 | 0.191897408 | 0.192617884 |
| Mtmr3 | 0.046830008 | 0.453650769 | 0.807844867 | 0.356044973 | 0.304634784 |
| Mtmr4 | -0.006351213 | 0.865488047 | 0.961791798 | 0.091231842 | 0.09295388 |
| Mtmr6 | -0.038927478 | 0.540394734 | 0.849380522 | 0.400324169 | 0.437213836 |
| Mtmr7 | 0.006520293 | 0.917079218 | 0.980132949 | 0.379159018 | 0.377151569 |
| Mtmr9 | 0.000553431 | 0.986776539 | 0.997268589 | 0.072394361 | 0.071344033 |
| mt-Nd1 | -0.134094037 | 0.056137697 | 0.367897032 | 2.816525887 | 3.006426309 |
| mt-Nd2 | -0.057481153 | 0.411114926 | 0.780671239 | 2.580505086 | 2.578635015 |
| mt-Nd3 | 0.814048336 | 7.07E-26 | 3.55E-22 | 0.936030693 | 0 |
| mt-Nd4 | -0.162566085 | 0.01156417 | 0.170690558 | 2.961230753 | 3.218623559 |
| mt-Nd4l | -0.036745651 | 0.121756215 | 0.496977281 | 0.027152165 | 0.059628648 |
| mt-Nd5 | -0.108140849 | 0.156182391 | 0.553727536 | 1.441238583 | 1.535355347 |
| mt-Nd6 | -0.059249054 | 0.224685388 | 0.637067124 | 0.139899845 | 0.205322601 |
| Mto1 | -0.042951411 | 0.158972981 | 0.556789205 | 0.057275843 | 0.097925123 |
| Mtor | -0.018793292 | 0.705583836 | 0.912844672 | 0.172885476 | 0.184189363 |
| Mtpap | -0.043057798 | 0.327452719 | 0.722182734 | 0.126909451 | 0.169404613 |
| Mtpn | 0.168047803 | 0.013644975 | 0.189413402 | 0.645384528 | 0.452911259 |
| Mtr | -0.020287811 | 0.484504918 | 0.823117106 | 0.051093433 | 0.069528009 |
| Mtres1 | 0.101398302 | 0.066287596 | 0.39159102 | 0.240065232 | 0.159455529 |
| Mtrex | 0.051533268 | 0.2897723 | 0.695966637 | 0.167776233 | 0.124838779 |
| Mtrf1 | -0.053218303 | 0.037209388 | 0.304267772 | 0.033613455 | 0.076031105 |
| Mtrf1l | 0.049747826 | 0.196504312 | 0.60713211 | 0.101200071 | 0.060236355 |
| mt-Rnr1 | 0.24731615 | 0.001791023 | 0.056529878 | 1.330671168 | 0.926442987 |
| mt-Rnr2 | 0.128249413 | 0.047706291 | 0.341775906 | 3.730931068 | 3.302072336 |
| Mtrr | -0.054138684 | 0.056717764 | 0.368804883 | 0.045384057 | 0.091431326 |
| Mtss1 | -0.08332221 | 0.189849005 | 0.598465598 | 0.389387022 | 0.482732747 |
| Mtss2 | -0.170591878 | 0.004468248 | 0.097920976 | 0.271859201 | 0.414884541 |
| Mttp | -0.120483314 | 0.004391024 | 0.096650684 | 0.107835091 | 0.219004704 |
| Mturn | -0.010536331 | 0.728187587 | 0.920519505 | 0.056589246 | 0.060630847 |
| Mtus1 | 0.035886417 | 0.533137219 | 0.84612966 | 0.25779506 | 0.224118 |
| Mtus2 | -0.086501594 | 0.003505806 | 0.08334591 | 0.042146826 | 0.1188239 |
| Mtx1 | 0.004071993 | 0.935563702 | 0.984289378 | 0.180526356 | 0.178634464 |
| Mtx2 | 0.010991733 | 0.858474253 | 0.958774461 | 0.358886981 | 0.347379063 |
| Muc4 | -0.056124815 | 0.280305323 | 0.687465228 | 0.182748078 | 0.205871238 |
| Mul1 | 0.078731817 | 0.133021271 | 0.513908582 | 0.204153152 | 0.135562855 |
| Mus81 | 0.083040533 | 0.028829357 | 0.270430142 | 0.10602058 | 0.038152421 |
| Mvb12a | -0.100893417 | 0.128733836 | 0.508260084 | 0.492112739 | 0.609021019 |
| Mvk | 0.029654349 | 0.278299065 | 0.685301206 | 0.049788941 | 0.029516592 |
| Mvp | -0.036351832 | 0.573757763 | 0.862168892 | 0.352996812 | 0.384986516 |
| Mxd1 | 0.035912429 | 0.407120261 | 0.777888922 | 0.128164002 | 0.092428797 |
| Mxd4 | -0.178487335 | 0.006360303 | 0.120222898 | 0.441719976 | 0.632388721 |
| Mxi1 | -0.010461699 | 0.833833268 | 0.954514657 | 0.168796437 | 0.176120059 |
| Mxra7 | -0.013479342 | 0.639627566 | 0.891345382 | 0.04961173 | 0.061809321 |
| Mxra8 | 0.043813839 | 0.172633817 | 0.575294174 | 0.069030824 | 0.035239822 |
| Myadm | 0.005489666 | 0.892572036 | 0.971348317 | 0.11201383 | 0.100660666 |
| Mybbp1a | 0.042706132 | 0.489017598 | 0.826256745 | 0.347248331 | 0.320641746 |
| Myc | -0.105766897 | 2.59E-06 | 0.000472189 | 0.017261942 | 0.105965253 |
| Mycbp | 0.082772991 | 0.106429687 | 0.473900883 | 0.210108752 | 0.124774305 |
| Mycbp2 | -0.011253215 | 0.868434727 | 0.96295459 | 1.005049206 | 0.997739383 |
| Myd88 | -0.002661079 | 0.925553275 | 0.981625089 | 0.049456257 | 0.04872492 |
| Mydgf | -0.073420685 | 0.271288624 | 0.67876599 | 0.735752622 | 0.810533866 |
| Myef2 | 0.017394101 | 0.777488259 | 0.934307164 | 0.334700285 | 0.31285253 |
| Myg1 | -0.080048122 | 0.126893844 | 0.506616353 | 0.185033396 | 0.254726691 |
| Myh10 | 0.019768115 | 0.682811916 | 0.909173483 | 0.162609219 | 0.137030219 |
| Myh14 | -0.010047255 | 0.82793642 | 0.952874813 | 0.140822938 | 0.149810631 |
| Myh9 | 0.088049074 | 0.193974445 | 0.60325934 | 0.491457023 | 0.391096673 |
| Myl12a | 0.038315101 | 0.532587848 | 0.84612966 | 0.345205131 | 0.316626523 |
| Myl12b | -0.063734906 | 0.201395135 | 0.611924411 | 0.172187255 | 0.249890538 |
| Myl6 | -0.053748297 | 0.27391564 | 0.681791282 | 1.745359427 | 1.824665256 |
| Mylip | -0.008642722 | 0.769866327 | 0.931407628 | 0.051947571 | 0.063759324 |
| Mynn | 0.046392366 | 0.354781104 | 0.745277928 | 0.174723727 | 0.141586031 |
| Myo10 | 0.12614619 | 0.063439954 | 0.383957695 | 0.47911736 | 0.343978375 |
| Myo18a | 0.037285843 | 0.534643814 | 0.846347806 | 0.3047875 | 0.268169165 |

| | | | | | |
|---|---|---|---|---|---|
| Myo1b | 0.205641178 | 4.37E-05 | 0.004389492 | 0.204769588 | 0.025956506 |
| Myo1c | 0.112287595 | 0.062718298 | 0.381980311 | 0.300253449 | 0.184140958 |
| Myo1d | -0.094909045 | 0.009022828 | 0.147945584 | 0.077943794 | 0.152530375 |
| Myo1e | 0.040617692 | 0.210473278 | 0.623103727 | 0.069424527 | 0.036246014 |
| Myo5a | -0.048391571 | 0.406266565 | 0.777612997 | 0.249710502 | 0.297485355 |
| Myo5b | -0.015785648 | 0.651652316 | 0.896468516 | 0.073225586 | 0.090007948 |
| Myo6 | -0.074469402 | 0.286082395 | 0.693090656 | 0.667557152 | 0.737070676 |
| Myo9a | -0.084908909 | 0.055153891 | 0.365158051 | 0.123777666 | 0.197263071 |
| Myo9b | 0.021438203 | 0.526781026 | 0.842463537 | 0.076850564 | 0.05354927 |
| Mypop | -0.014719641 | 0.732831577 | 0.921306823 | 0.125069357 | 0.139488838 |
| Myrip | -0.038580416 | 0.043268829 | 0.328509254 | 0.014305987 | 0.047591402 |
| Mysm1 | 0.023407099 | 0.722879632 | 0.919483582 | 0.46971519 | 0.428723173 |
| Myt1 | 0.025884526 | 0.570087614 | 0.86083487 | 0.143849232 | 0.114189479 |
| Myt1l | -0.111677498 | 0.058558393 | 0.372682944 | 0.256822825 | 0.35958668 |
| Mzt1 | -0.003280871 | 0.950604717 | 0.988114084 | 0.214348272 | 0.220679685 |
| Mzt2 | -0.053439567 | 0.255889134 | 0.664689244 | 0.13925567 | 0.188545366 |
| N4bp1 | 0.070576357 | 0.266174484 | 0.673899077 | 0.40421879 | 0.345812929 |
| N4bp2 | -0.175725646 | 2.11E-06 | 0.000400386 | 0.067504393 | 0.226897564 |
| N4bp2l1 | -0.160761202 | 0.002707322 | 0.071134541 | 0.186886533 | 0.326965464 |
| N4bp2l2 | 0.036032979 | 0.589019676 | 0.86992264 | 0.603288082 | 0.570644555 |
| N6amt1 | 0.000909315 | 0.986615091 | 0.997268589 | 0.207369202 | 0.202690042 |
| Naa10 | 0.067398445 | 0.139877317 | 0.527800236 | 0.156884028 | 0.092906944 |
| Naa15 | -0.023828607 | 0.726087311 | 0.92014727 | 0.531463785 | 0.558116055 |
| Naa16 | -0.001326996 | 0.978030878 | 0.995031396 | 0.156266326 | 0.161579782 |
| Naa20 | -0.012985588 | 0.847165902 | 0.956905313 | 0.703237681 | 0.710094828 |
| Naa25 | 0.052066585 | 0.255826543 | 0.664689244 | 0.149413915 | 0.106954769 |
| Naa30 | -0.031499691 | 0.559963645 | 0.856904336 | 0.208545403 | 0.243362946 |
| Naa35 | -0.080303593 | 0.214188443 | 0.628967058 | 0.397719085 | 0.486213341 |
| Naa38 | 0.081785727 | 0.215542894 | 0.629611185 | 0.602292335 | 0.500089684 |
| Naa40 | 0.046983272 | 0.288022362 | 0.694589247 | 0.133833841 | 0.085286279 |
| Naa50 | 0.010355726 | 0.880694994 | 0.967867694 | 0.585341747 | 0.573410961 |
| Naa60 | -0.021711348 | 0.672399138 | 0.905281039 | 0.195968956 | 0.213519218 |
| Naa80 | -0.013531062 | 0.697799199 | 0.910970095 | 0.079164615 | 0.08997134 |
| Naaa | -0.178242358 | 3.86E-07 | 0.000102034 | 0.055460443 | 0.219687745 |
| Nab1 | 0.123823207 | 0.056299852 | 0.367897032 | 0.373967795 | 0.252230411 |
| Nabp1 | 0.034837062 | 0.486981681 | 0.825787318 | 0.1666376 | 0.13470962 |
| Nabp2 | 0.003730584 | 0.952128446 | 0.98838165 | 0.384151662 | 0.380016854 |
| Naca | -0.107077599 | 0.012461195 | 0.17816668 | 2.032039895 | 2.271557773 |
| Nacc1 | 0.021261205 | 0.674476934 | 0.906025597 | 0.178708163 | 0.15959164 |
| Nacc2 | -0.020286147 | 0.499639348 | 0.830722138 | 0.055864136 | 0.068467967 |
| Nadk | 0.01033777 | 0.828774859 | 0.952874813 | 0.159347829 | 0.142152308 |
| Nadk2 | 0.002558697 | 0.946468548 | 0.987084873 | 0.095080354 | 0.09914102 |
| Nae1 | -0.010658947 | 0.864808309 | 0.961442912 | 0.406597273 | 0.416156584 |
| Naf1 | 0.006424762 | 0.880853563 | 0.967919035 | 0.122795752 | 0.119504586 |
| Naga | -0.138306999 | 0.036316255 | 0.300252265 | 0.435765055 | 0.573005419 |
| Nagk | -0.000738886 | 0.990204346 | 0.997733826 | 0.344069012 | 0.344044012 |
| Naglu | -0.01539578 | 0.673668226 | 0.905654932 | 0.083384594 | 0.093759238 |
| Nagpa | -0.066256149 | 0.029191258 | 0.272146451 | 0.05070146 | 0.108807987 |
| Naip2 | -0.056751526 | 0.198749847 | 0.609389286 | 0.125807591 | 0.173085874 |
| Nalcn | -0.068603522 | 0.091523418 | 0.446582669 | 0.096768218 | 0.148420472 |
| Nampt | 0.057917781 | 0.334329766 | 0.729969081 | 0.307573198 | 0.248469061 |
| Nanp | -0.025611858 | 0.46979582 | 0.814253264 | 0.076965584 | 0.099784241 |
| Nans | -0.004778439 | 0.936576615 | 0.98453816 | 0.347537916 | 0.339194681 |
| Nap1l1 | -0.074193994 | 0.242533836 | 0.653091138 | 0.851043113 | 0.949214767 |
| Nap1l2 | -0.091033751 | 0.042303442 | 0.325136871 | 0.122695526 | 0.203258544 |
| Nap1l3 | 0.06431535 | 0.079676841 | 0.423216727 | 0.096509256 | 0.038552257 |
| Nap1l4 | -0.010915941 | 0.864642101 | 0.961442912 | 0.475883866 | 0.485993427 |
| Nap1l5 | -0.064569128 | 0.346885787 | 0.738383053 | 0.359837885 | 0.423507174 |
| Napa | -0.041065048 | 0.528200273 | 0.843657883 | 0.918150796 | 0.987485452 |
| Napb | -0.008796783 | 0.85436349 | 0.957593115 | 0.158458051 | 0.171589359 |
| Napepld | 0.072913269 | 0.131723266 | 0.512050022 | 0.165014782 | 0.099770537 |
| Napg | -0.109702212 | 0.090241488 | 0.445385778 | 0.395506497 | 0.509164151 |
| Naprt | -0.029992669 | 0.222950011 | 0.634822484 | 0.032810466 | 0.061425893 |
| Narf | 0.044118526 | 0.115543936 | 0.487479818 | 0.052573612 | 0.017130354 |
| Nars | 0.095110626 | 0.120520059 | 0.494691316 | 1.301568711 | 1.154477968 |

| | | | | | |
|---|---|---|---|---|---|
| Nars2 | 0.00487498 | 0.890119733 | 0.970579007 | 0.080642017 | 0.080682189 |
| Nasp | 0.012325905 | 0.808747233 | 0.9469664 | 0.183748485 | 0.174819729 |
| Nat10 | -0.016953282 | 0.602582099 | 0.876632118 | 0.064396405 | 0.081449942 |
| Nat14 | -0.114033358 | 0.002180879 | 0.064192029 | 0.07575029 | 0.18079483 |
| Nat8f1 | 0.028781565 | 0.467789995 | 0.813188482 | 0.107829463 | 0.077338345 |
| Nat9 | -0.081892325 | 0.155540143 | 0.552356396 | 0.262671584 | 0.353206921 |
| Natd1 | -0.03563377 | 0.213172085 | 0.62773867 | 0.049420653 | 0.076625694 |
| Nav1 | -0.177723733 | 0.00677871 | 0.125996128 | 0.396551613 | 0.579495081 |
| Nav2 | -0.072497989 | 0.276409468 | 0.683667283 | 0.468642955 | 0.538104341 |
| Nav3 | 0.03415433 | 0.311184004 | 0.713746308 | 0.075825061 | 0.040003484 |
| Naxd | -0.032474565 | 0.598953054 | 0.875574105 | 0.386122304 | 0.413543513 |
| Naxe | -0.097170935 | 0.127187851 | 0.50738651 | 0.413037864 | 0.514425817 |
| Nbas | -0.195701882 | 0.00131426 | 0.04612319 | 0.304416595 | 0.492467919 |
| Nbdy | -0.005976186 | 0.921631212 | 0.98085171 | 0.324513778 | 0.328645674 |
| Nbea | 0.005614012 | 0.93464944 | 0.983960188 | 0.557929589 | 0.546024408 |
| Nbeal1 | -0.085673105 | 0.204495661 | 0.616256046 | 0.503931792 | 0.607549114 |
| Nbeal2 | 0.00829004 | 0.804836802 | 0.944702021 | 0.072430652 | 0.065377861 |
| Nbl1 | 0.011819841 | 0.814519813 | 0.94873644 | 0.175221366 | 0.182626801 |
| Nbn | 0.080833541 | 0.109248198 | 0.477972625 | 0.188316984 | 0.116183691 |
| Nbr1 | -0.148031746 | 0.020653252 | 0.227971987 | 0.429760279 | 0.580295616 |
| Ncald | -0.107739187 | 0.113744191 | 0.484874023 | 0.59859858 | 0.701556597 |
| Ncam1 | -0.201673143 | 0.003483321 | 0.083243072 | 0.444675365 | 0.659348269 |
| Ncapd3 | 0.021219201 | 0.571258089 | 0.861225331 | 0.092565837 | 0.072846415 |
| Ncaph2 | 0.036084766 | 0.451652107 | 0.806495742 | 0.159682937 | 0.137206915 |
| Ncbp1 | 0.059036406 | 0.263077264 | 0.669775695 | 0.221427064 | 0.167997313 |
| Ncbp2 | 0.101618883 | 0.093739729 | 0.45227176 | 0.340704395 | 0.254813782 |
| Ncbp3 | 0.021199861 | 0.696680158 | 0.910970095 | 0.219738346 | 0.217779415 |
| Ncdn | -0.046675564 | 0.24995972 | 0.661789952 | 0.10600549 | 0.14566072 |
| Nceh1 | -0.009514722 | 0.873069988 | 0.964940674 | 0.306798677 | 0.342148834 |
| Nck1 | 0.046491245 | 0.474322046 | 0.817387411 | 0.491086925 | 0.42467773 |
| Nck2 | 0.012546987 | 0.751956655 | 0.926186851 | 0.100646464 | 0.086835221 |
| Nckap1 | -0.05306768 | 0.432048809 | 0.794226501 | 0.959738153 | 1.042260977 |
| Nckap5 | -0.051594476 | 0.104145484 | 0.471172212 | 0.054449841 | 0.101003885 |
| Nckipsd | 0.02590978 | 0.44601256 | 0.803559158 | 0.077762876 | 0.052126371 |
| Ncl | -0.040969663 | 0.498045499 | 0.830722138 | 1.534410423 | 1.572804932 |
| Ncln | 0.023990037 | 0.568269932 | 0.860160655 | 0.122741658 | 0.099957103 |
| Ncoa1 | 0.044431088 | 0.501483484 | 0.832218079 | 0.477184005 | 0.433403279 |
| Ncoa2 | 0.032443981 | 0.58299225 | 0.866383905 | 0.276408785 | 0.25186402 |
| Ncoa3 | 0.089153368 | 0.103591673 | 0.469469748 | 0.250558643 | 0.16003213 |
| Ncoa4 | 0.127451761 | 0.022008971 | 0.23623533 | 0.244313897 | 0.127422198 |
| Ncoa5 | -0.023280719 | 0.536012196 | 0.84737036 | 0.089792928 | 0.108610241 |
| Ncoa6 | 0.000636048 | 0.990239356 | 0.997733826 | 0.194003695 | 0.199773156 |
| Ncoa7 | 0.115181997 | 0.097107801 | 0.459533711 | 0.621815819 | 0.468992686 |
| Ncor1 | -0.020541018 | 0.735157434 | 0.921955891 | 1.330763039 | 1.35638938 |
| Ncor2 | -0.035523081 | 0.5625494 | 0.857580244 | 0.350177501 | 0.365151552 |
| Ncstn | -0.112996943 | 0.038984503 | 0.311933003 | 0.219285959 | 0.310977219 |
| Ndel1 | 0.028601286 | 0.65475213 | 0.897454058 | 0.380398317 | 0.373847372 |
| Ndfip1 | -0.027155575 | 0.610896393 | 0.879838309 | 1.460034768 | 1.529908322 |
| Ndfip2 | 0.065642382 | 0.334313635 | 0.729969081 | 0.735289532 | 0.647134121 |
| Ndn | -0.135992529 | 0.011782938 | 0.172720733 | 0.215324399 | 0.322310085 |
| Ndrg1 | -0.073533007 | 0.2867602 | 0.693329451 | 0.479812417 | 0.567394167 |
| Ndrg3 | 0.077363958 | 0.235812696 | 0.646327697 | 0.430691478 | 0.348921243 |
| Ndrg4 | -0.103140675 | 0.004057912 | 0.091116924 | 0.071129563 | 0.162845151 |
| Ndst1 | 0.013592203 | 0.822317888 | 0.950930859 | 0.303934518 | 0.286709699 |
| Ndufa1 | 0.111765484 | 0.050754071 | 0.354829042 | 1.372882898 | 1.160401747 |
| Ndufa10 | -0.082804689 | 0.198791181 | 0.609389286 | 0.477023896 | 0.569968917 |
| Ndufa11 | -0.028776945 | 0.610898992 | 0.879838309 | 1.389042324 | 1.381971333 |
| Ndufa12 | 0.038865851 | 0.557385563 | 0.855384435 | 0.540186476 | 0.510083484 |
| Ndufa13 | -0.00252859 | 0.962535459 | 0.991757003 | 1.496874737 | 1.482074924 |
| Ndufa2 | -0.002294933 | 0.962005043 | 0.991441074 | 1.819866724 | 1.797662195 |
| Ndufa3 | -0.036357489 | 0.582565759 | 0.86638206 | 0.693332465 | 0.70976192 |
| Ndufa4 | 0.041368674 | 0.446158867 | 0.80367849 | 1.70570322 | 1.641132813 |
| Ndufa5 | 0.103966705 | 0.120703301 | 0.494691316 | 1.063506221 | 0.913315943 |
| Ndufa6 | -0.033041302 | 0.509993098 | 0.834904701 | 1.658612954 | 1.700646955 |
| Ndufa7 | 0.015515487 | 0.747201406 | 0.924930563 | 1.739254649 | 1.708980146 |

| | | | | | |
|---|---|---|---|---|---|
| Ndufa8 | 0.076460844 | 0.240556077 | 0.650744882 | 0.768237783 | 0.698145664 |
| Ndufa9 | 0.0730682 | 0.228500083 | 0.640093591 | 0.327183717 | 0.268367073 |
| Ndufab1 | -0.039440124 | 0.545766241 | 0.851259637 | 0.474984736 | 0.530588009 |
| Ndufaf1 | 0.012161197 | 0.691001593 | 0.909796728 | 0.060975584 | 0.054062898 |
| Ndufaf2 | 0.014746376 | 0.773436593 | 0.932714536 | 0.190768356 | 0.178140844 |
| Ndufaf3 | -0.004644062 | 0.941906338 | 0.986168593 | 0.446935188 | 0.469082622 |
| Ndufaf4 | 0.072471402 | 0.263036432 | 0.669775695 | 0.42072787 | 0.344994422 |
| Ndufaf5 | -0.022925652 | 0.448020997 | 0.804951652 | 0.055910593 | 0.08195774 |
| Ndufaf6 | 0.096468959 | 0.038119071 | 0.308052426 | 0.154357438 | 0.075814414 |
| Ndufaf7 | 0.051373943 | 0.335700484 | 0.730736446 | 0.223427846 | 0.179367181 |
| Ndufaf8 | 0.005378458 | 0.934062234 | 0.983960188 | 0.568258997 | 0.543596688 |
| Ndufb1 | 0.031077827 | 0.548568105 | 0.852186669 | 1.512743935 | 1.406000772 |
| Ndufb10 | 0.003073669 | 0.959928422 | 0.990928888 | 1.246461664 | 1.25658596 |
| Ndufb11 | -0.055440966 | 0.298802976 | 0.703845452 | 1.518911006 | 1.609021513 |
| Ndufb2 | -0.024037288 | 0.704566978 | 0.912507936 | 0.978865333 | 0.992819838 |
| Ndufb3 | 0.015605751 | 0.805698124 | 0.945049909 | 0.947051956 | 0.913338276 |
| Ndufb4 | -0.031560821 | 0.615463086 | 0.882439418 | 0.357830547 | 0.363527862 |
| Ndufb5 | -0.1098347 | 0.090542047 | 0.445385778 | 0.956722286 | 1.067142346 |
| Ndufb6 | -0.065587615 | 0.321593745 | 0.719227395 | 0.786768765 | 0.859115177 |
| Ndufb7 | -0.088128991 | 0.165355675 | 0.566504119 | 1.170356253 | 1.245911745 |
| Ndufb8 | -0.094740896 | 0.121148092 | 0.495300774 | 1.200020673 | 1.340210642 |
| Ndufb9 | 0.056892283 | 0.275046819 | 0.68241738 | 1.629466488 | 1.548743419 |
| Ndufc1 | 0.040971561 | 0.466285305 | 0.813032623 | 1.387222622 | 1.287602674 |
| Ndufc2 | -0.051217857 | 0.37858961 | 0.762006446 | 1.636269924 | 1.698190759 |
| Ndufs1 | -0.14884352 | 0.01425304 | 0.193060412 | 0.309842778 | 0.452420288 |
| Ndufs2 | 0.014168222 | 0.8386735 | 0.954911647 | 1.172696288 | 1.195178672 |
| Ndufs3 | -0.02305759 | 0.718911002 | 0.918613587 | 0.438783699 | 0.477135051 |
| Ndufs4 | 0.072391778 | 0.295342774 | 0.702121132 | 0.740613222 | 0.661598818 |
| Ndufs5 | 0.006316063 | 0.920557705 | 0.980436936 | 0.995900184 | 0.95827439 |
| Ndufs6 | -0.053004023 | 0.408126108 | 0.778479997 | 0.992348309 | 1.038341426 |
| Ndufs7 | -0.105975733 | 0.106798089 | 0.474004524 | 0.961235309 | 1.072762128 |
| Ndufs8 | -0.164327606 | 0.010317757 | 0.15907731 | 0.446939535 | 0.604387618 |
| Ndufv1 | -0.220382762 | 0.00063952 | 0.028919709 | 0.401764615 | 0.63354679 |
| Ndufv2 | 0.038681932 | 0.56795574 | 0.859814745 | 0.673939692 | 0.651169865 |
| Ndufv3 | 0.125791185 | 0.01766749 | 0.21172951 | 1.475907785 | 1.254866407 |
| Neat1 | -0.139003929 | 0.069362224 | 0.400472912 | 0.822256449 | 1.014193698 |
| Nebl | 0.037886498 | 0.241960256 | 0.652555608 | 0.07242905 | 0.038840449 |
| Necab2 | -0.034135414 | 0.443142173 | 0.801368502 | 0.12875327 | 0.153501219 |
| Necap1 | 0.040535501 | 0.490634398 | 0.826951175 | 0.287606003 | 0.253092272 |
| Necap2 | -0.042365004 | 0.263252478 | 0.669775695 | 0.089381871 | 0.125992314 |
| Nectin1 | 0.063390472 | 0.205133266 | 0.616549498 | 0.183790746 | 0.127137025 |
| Nectin2 | 0.026538433 | 0.604584141 | 0.877736706 | 0.18868254 | 0.157070027 |
| Nectin3 | 0.114626317 | 0.098222298 | 0.461543634 | 0.648665957 | 0.524146899 |
| Nedd4 | -0.11983809 | 0.060972088 | 0.380250374 | 1.154758515 | 1.322532417 |
| Nedd4l | 0.144405825 | 0.011314281 | 0.168327452 | 0.256230165 | 0.120714098 |
| Nedd8 | -0.102650928 | 0.087290514 | 0.439557333 | 1.02630611 | 1.169998436 |
| Nedd9 | -0.068528606 | 0.046939609 | 0.339316662 | 0.070235835 | 0.133953696 |
| Nefh | -0.012261513 | 0.651265193 | 0.896304503 | 0.04171724 | 0.056663759 |
| Nefm | 0.05568442 | 0.319340409 | 0.717256849 | 0.200094807 | 0.138431295 |
| Negr1 | 0.04264033 | 0.441594688 | 0.800354711 | 0.235629563 | 0.201186373 |
| Neil1 | 0.013373556 | 0.659134383 | 0.898224427 | 0.060277287 | 0.045649343 |
| Neil2 | -0.022502106 | 0.510897557 | 0.834957869 | 0.072601829 | 0.099231955 |
| Nek1 | -0.016717122 | 0.677637077 | 0.907222001 | 0.107918891 | 0.1156317 |
| Nek4 | -0.018603433 | 0.634344019 | 0.88965971 | 0.096924791 | 0.116779742 |
| Nek7 | -0.033158519 | 0.590124456 | 0.87075199 | 0.35468976 | 0.385650051 |
| Nek9 | 0.073152712 | 0.182290545 | 0.590019414 | 0.238393931 | 0.167202564 |
| Nelfa | -0.002695523 | 0.957730482 | 0.990137214 | 0.186459531 | 0.189435465 |
| Nelfb | -0.047208036 | 0.374361747 | 0.75956519 | 0.217131092 | 0.266311108 |
| Nelfcd | -0.100209167 | 0.005327976 | 0.10847241 | 0.073084031 | 0.163332017 |
| Nelfe | -0.126595821 | 0.033934565 | 0.291747273 | 0.282828983 | 0.396193202 |
| Nell1 | -0.078224532 | 0.218009988 | 0.632396893 | 0.406554969 | 0.47979789 |
| Nemf | -0.031951457 | 0.623926962 | 0.886265909 | 0.419911084 | 0.44561231 |
| Nemp1 | -0.008035021 | 0.793291129 | 0.939610935 | 0.057572981 | 0.066544278 |
| Nenf | -0.211374448 | 0.000597482 | 0.028287387 | 1.240358272 | 1.566711162 |
| Neo1 | 0.052983815 | 0.365003613 | 0.752802229 | 0.277716576 | 0.233193027 |

| | | | | | |
|---|---|---|---|---|---|
| Nepro | -0.062153128 | 0.014396525 | 0.193677739 | 0.030624047 | 0.082298217 |
| Net1 | 0.026353967 | 0.445345834 | 0.803123904 | 0.080589443 | 0.058658944 |
| Neu1 | 0.074249384 | 0.258608329 | 0.665948842 | 0.471143192 | 0.396183152 |
| Neurl1a | 0.071951995 | 0.113888627 | 0.48492417 | 0.149701231 | 0.087369366 |
| Neurl4 | -0.069344512 | 0.072513745 | 0.408428989 | 0.088510062 | 0.14981121 |
| Neurod1 | -0.141424802 | 0.019809284 | 0.224661898 | 1.329644107 | 1.546522762 |
| Nf1 | -0.09672459 | 0.079693097 | 0.423216727 | 0.217770644 | 0.302980186 |
| Nf2 | 0.015037231 | 0.759531837 | 0.927719223 | 0.172052162 | 0.153611239 |
| Nfasc | -0.096273749 | 0.098955358 | 0.463315751 | 0.272726938 | 0.330493901 |
| Nfat5 | -0.078539473 | 0.246100954 | 0.655935044 | 0.44113647 | 0.520272495 |
| Nfatc1 | -0.026931554 | 0.304889138 | 0.709410087 | 0.040071824 | 0.058362752 |
| Nfatc2 | -0.058937979 | 0.159205665 | 0.556789205 | 0.114383225 | 0.162609432 |
| Nfatc3 | 0.054818943 | 0.252038908 | 0.663127858 | 0.162571134 | 0.117998717 |
| Nfe2l1 | -0.043847772 | 0.488856329 | 0.826256745 | 1.094133756 | 1.152229498 |
| Nfe2l2 | 0.067808059 | 0.255009283 | 0.663605957 | 0.280817099 | 0.231299815 |
| Nfia | -0.033519348 | 0.466888163 | 0.813152001 | 0.134481832 | 0.171159626 |
| Nfic | 0.144168098 | 0.031915212 | 0.283462667 | 0.950671378 | 0.774855332 |
| Nfil3 | -0.028327933 | 0.615490777 | 0.882439418 | 0.226063906 | 0.243009951 |
| Nfix | -0.008359158 | 0.885950369 | 0.969590273 | 0.262202288 | 0.274237245 |
| Nfkb1 | 0.021352715 | 0.713184888 | 0.915509496 | 0.294762502 | 0.285788235 |
| Nfkb2 | -0.03754199 | 0.196678515 | 0.60713211 | 0.046540972 | 0.08499255 |
| Nfkbia | -0.090678619 | 0.232748742 | 0.643730814 | 0.5405056 | 0.684207211 |
| Nfkbib | 0.106190277 | 0.042338736 | 0.325136871 | 0.211362264 | 0.120291477 |
| Nfkbil1 | 0.054299148 | 0.359595307 | 0.747719327 | 0.313576986 | 0.278315075 |
| Nfkbiz | -0.082019067 | 0.123021378 | 0.499567826 | 0.185124799 | 0.263369615 |
| Nfrkb | 0.029728691 | 0.517968101 | 0.837813573 | 0.146551355 | 0.11636497 |
| Nfs1 | -0.101693242 | 0.032201003 | 0.284508332 | 0.142361737 | 0.225997361 |
| Nfu1 | -0.011602549 | 0.845784782 | 0.956051742 | 0.3343336 | 0.345346225 |
| Nfx1 | 0.081311594 | 0.184224719 | 0.590567014 | 0.327380493 | 0.242525082 |
| Nfxl1 | -0.040338479 | 0.453831192 | 0.807844867 | 0.20853675 | 0.255815405 |
| Nfyb | 0.032195829 | 0.604042877 | 0.877264992 | 0.326426885 | 0.293518417 |
| Nfyc | 0.012169962 | 0.762563466 | 0.92872278 | 0.110751874 | 0.104436993 |
| Ngdn | 0.089158099 | 0.159716841 | 0.557786338 | 0.374860563 | 0.295043171 |
| Ngly1 | -0.013634321 | 0.792791455 | 0.939573484 | 0.189445855 | 0.205221649 |
| Ngrn | -0.006627079 | 0.922367593 | 0.980859278 | 0.48627336 | 0.49480183 |
| Nhlrc1 | -0.014610304 | 0.578021239 | 0.864673658 | 0.04153564 | 0.057155316 |
| Nhlrc2 | 0.031635382 | 0.550705521 | 0.852505562 | 0.201679289 | 0.182039218 |
| Nhlrc3 | 0.087170475 | 0.052356905 | 0.357487248 | 0.138900983 | 0.064556169 |
| Nhp2 | -0.074573128 | 0.183451602 | 0.590066231 | 0.235549344 | 0.293647388 |
| Niban1 | -0.085887747 | 0.001564673 | 0.052002066 | 0.032682484 | 0.107115148 |
| Niban2 | 0.041161092 | 0.366176497 | 0.753107222 | 0.138369686 | 0.10777404 |
| Nicn1 | -0.047759762 | 0.274569669 | 0.682000399 | 0.130075127 | 0.177204095 |
| Nif3l1 | 0.006317053 | 0.873884096 | 0.964940674 | 0.10465567 | 0.098257474 |
| Nifk | -0.041489185 | 0.44320017 | 0.801368502 | 0.204308594 | 0.246244344 |
| Nim1k | 0.046376405 | 0.174242726 | 0.577186219 | 0.082046316 | 0.043022082 |
| Nin | -0.013773232 | 0.659101218 | 0.898224427 | 0.061124643 | 0.077530232 |
| Ninj1 | 0.042789815 | 0.509763344 | 0.834904701 | 0.894246167 | 0.866054463 |
| Nip7 | 0.039646713 | 0.502095852 | 0.832218079 | 0.280057098 | 0.247008392 |
| Nipa1 | 0.048922323 | 0.240515492 | 0.650744882 | 0.123516794 | 0.085242755 |
| Nipa2 | 0.024838767 | 0.671862013 | 0.90516497 | 0.296873172 | 0.27244941 |
| Nipal1 | -0.103350892 | 0.111329035 | 0.482253844 | 0.462174514 | 0.561326679 |
| Nipal2 | -0.112569006 | 0.070090046 | 0.402226298 | 0.355273695 | 0.457967597 |
| Nipal3 | -0.099109262 | 0.12573722 | 0.504406266 | 0.37803836 | 0.474052544 |
| Nipbl | 0.000188605 | 0.997814041 | 0.99938181 | 0.823172446 | 0.818084321 |
| Nipsnap1 | -0.004803566 | 0.923347009 | 0.980859278 | 0.18017193 | 0.184101599 |
| Nipsnap2 | -0.015762969 | 0.769384328 | 0.931397082 | 0.21263602 | 0.223033079 |
| Nipsnap3b | -0.043629552 | 0.378909084 | 0.762006446 | 0.162094891 | 0.205966642 |
| Nisch | -0.215464716 | 0.001267164 | 0.044967808 | 0.932579755 | 1.220588452 |
| Nit1 | -0.04590711 | 0.38848098 | 0.767160379 | 0.203575964 | 0.249392961 |
| Nit2 | -0.000444971 | 0.988311051 | 0.997411191 | 0.056419203 | 0.055533428 |
| Nkap | -0.025343902 | 0.653393596 | 0.897265224 | 0.249284175 | 0.269870527 |
| Nkapd1 | 0.025249876 | 0.633290518 | 0.889501117 | 0.205857474 | 0.183450216 |
| Nkiras1 | -0.049999734 | 0.427355805 | 0.791250732 | 0.34969998 | 0.388368444 |
| Nkiras2 | -0.031817329 | 0.476990958 | 0.818444859 | 0.132182668 | 0.164179482 |
| Nktr | -0.217713628 | 0.001204703 | 0.043494963 | 1.197110064 | 1.50419324 |

| | | | | | |
|---|---|---|---|---|---|
| Nkx2-2 | -0.044652097 | 0.545490808 | 0.851259637 | 0.682841347 | 0.739234698 |
| Nkx6-1 | 0.023723714 | 0.714332426 | 0.915840939 | 1.583928557 | 1.5326214 |
| Nlgn2 | -0.075467757 | 0.072643499 | 0.408930343 | 0.109241881 | 0.164115614 |
| Nlk | 0.105591564 | 0.087474646 | 0.439786592 | 0.363090699 | 0.254962987 |
| Nln | -0.040118547 | 0.170049802 | 0.572762895 | 0.047989353 | 0.084711776 |
| Nmd3 | 0.037103236 | 0.507732735 | 0.833674307 | 0.232105451 | 0.206223034 |
| Nme1 | 0.010675184 | 0.868489351 | 0.96295459 | 1.181282155 | 1.167298022 |
| Nme2 | 0.036394777 | 0.450258647 | 0.806495742 | 0.172058703 | 0.133791704 |
| Nme4 | -0.050187162 | 0.064804154 | 0.388281728 | 0.038382354 | 0.084436935 |
| Nme5 | -0.03502979 | 0.17603295 | 0.580243913 | 0.035455474 | 0.063376638 |
| Nme6 | 0.000237684 | 0.994219601 | 0.998695538 | 0.069681795 | 0.069819048 |
| Nme7 | 0.038597651 | 0.313826461 | 0.714906079 | 0.102107391 | 0.070976143 |
| Nmi | -0.091455032 | 0.059889724 | 0.378296512 | 0.142121662 | 0.237788747 |
| Nmnat1 | -0.030859699 | 0.315489429 | 0.714975653 | 0.056866717 | 0.086864817 |
| Nmnat2 | -0.07435899 | 0.043619161 | 0.329086044 | 0.080452283 | 0.146617916 |
| Nmnat3 | -0.066381647 | 0.123451228 | 0.500436178 | 0.111960087 | 0.168232559 |
| Nmral1 | -0.074409311 | 0.004644759 | 0.100256881 | 0.034601694 | 0.094945304 |
| Nmt1 | 0.174285312 | 0.005573546 | 0.111475656 | 0.381985991 | 0.219742033 |
| Nmt2 | 0.097450628 | 0.116534453 | 0.488986751 | 0.392835269 | 0.291823577 |
| Nnat | 0.156447376 | 0.008939613 | 0.146996602 | 0.214013555 | 0.057973693 |
| Noa1 | 0.026357598 | 0.539128193 | 0.848687213 | 0.130748256 | 0.113265208 |
| Nob1 | 0.020336784 | 0.692825784 | 0.909796728 | 0.192457582 | 0.172900393 |
| Noc2l | -0.003191777 | 0.953923196 | 0.989066902 | 0.222634116 | 0.227146353 |
| Noc3l | 0.014513511 | 0.728021535 | 0.920519505 | 0.10870265 | 0.095451946 |
| Noc4l | 0.046590624 | 0.100302074 | 0.465319642 | 0.056113834 | 0.019645343 |
| Noct | 0.057573249 | 0.199021429 | 0.609389286 | 0.131504906 | 0.079579953 |
| Nod1 | 0.015572135 | 0.647592123 | 0.895376633 | 0.075736478 | 0.057799098 |
| Nol10 | 0.0245343 | 0.392828408 | 0.769704269 | 0.053850003 | 0.034736415 |
| Nol11 | -0.072411908 | 0.193448661 | 0.60325934 | 0.229775612 | 0.293308846 |
| Nol12 | -0.12486191 | 0.034548883 | 0.29337321 | 0.282127245 | 0.415797951 |
| Nol3 | -0.073784343 | 0.090560972 | 0.445385778 | 0.119453493 | 0.189498869 |
| Nol4 | -0.137122129 | 0.017151546 | 0.208919982 | 0.246607849 | 0.388077519 |
| Nol4l | 0.050417386 | 0.108649716 | 0.477972625 | 0.069544503 | 0.026943217 |
| Nol6 | -0.048761596 | 0.158423271 | 0.556789205 | 0.073051993 | 0.118191801 |
| Nol7 | -0.068330702 | 0.315546548 | 0.714975653 | 0.750712359 | 0.836821238 |
| Nol8 | -0.021166317 | 0.710352603 | 0.914486281 | 0.221804996 | 0.250889789 |
| Nol9 | 0.015212353 | 0.669446799 | 0.903244727 | 0.08125556 | 0.068370298 |
| Nolc1 | -0.027502178 | 0.680216869 | 0.907783945 | 0.463182675 | 0.475329055 |
| Nom1 | 0.04550163 | 0.41105201 | 0.780671239 | 0.224902774 | 0.185931374 |
| Nomo1 | -0.172201778 | 0.000118271 | 0.008925455 | 0.11202908 | 0.271772216 |
| Nono | 0.010874737 | 0.868321309 | 0.96295459 | 0.557840215 | 0.550786075 |
| Nop10 | 0.014319605 | 0.828228437 | 0.952874813 | 1.103303778 | 1.101901776 |
| Nop14 | -0.037185294 | 0.557563458 | 0.855384435 | 0.354909091 | 0.395621012 |
| Nop16 | -0.089691823 | 0.040975296 | 0.319308264 | 0.118952664 | 0.208878155 |
| Nop2 | -0.036448882 | 0.260833123 | 0.66819348 | 0.062534992 | 0.103088046 |
| Nop53 | 0.015415541 | 0.818997815 | 0.950128492 | 0.511790309 | 0.499840989 |
| Nop56 | 0.031065692 | 0.578689944 | 0.865142348 | 0.234514794 | 0.206099065 |
| Nop58 | -0.038979125 | 0.586554994 | 0.86902362 | 0.551032433 | 0.577614943 |
| Nop9 | 0.022572365 | 0.474553748 | 0.817387411 | 0.064354178 | 0.045607413 |
| Norad | -0.144608418 | 0.03591398 | 0.298528355 | 0.78473781 | 0.983436931 |
| Nosip | -0.027236255 | 0.494719717 | 0.828842054 | 0.100539038 | 0.123242508 |
| Nostrin | -0.028228048 | 0.463479996 | 0.811254677 | 0.083832077 | 0.099522452 |
| Notch1 | 0.024898037 | 0.531614661 | 0.845612734 | 0.104215436 | 0.075458662 |
| Nova1 | -0.065089428 | 0.009081617 | 0.147974328 | 0.02924218 | 0.08259379 |
| Nova2 | 0.003077403 | 0.925665623 | 0.981625089 | 0.066586929 | 0.064628427 |
| Noxo1 | -0.017026954 | 0.470992761 | 0.814586466 | 0.031257345 | 0.043758086 |
| Npas2 | -0.063821103 | 0.002806662 | 0.073093048 | 0.017928927 | 0.07226533 |
| Npas4 | 0.008000414 | 0.823217917 | 0.950930859 | 0.076138395 | 0.068635279 |
| Npat | 0.007190454 | 0.892021088 | 0.971118429 | 0.199470557 | 0.200397193 |
| Npc1 | 0.009810104 | 0.820911226 | 0.950892784 | 0.129027507 | 0.117748091 |
| Npc2 | -0.096931209 | 0.135375103 | 0.518618255 | 0.98042009 | 1.119840523 |
| Npdc1 | -0.083829726 | 0.230972986 | 0.642359619 | 0.672959698 | 0.742772469 |
| Npepl1 | -0.022079106 | 0.748320869 | 0.924987261 | 0.487923963 | 0.532829218 |
| Npepps | 0.045172108 | 0.523384138 | 0.841062594 | 0.776144278 | 0.714664957 |
| Nphp1 | -0.079016024 | 0.054127805 | 0.362869235 | 0.105404409 | 0.167936854 |

| | | | | | |
|---|---|---|---|---|---|
| Nphp4 | -0.030496518 | 0.159071723 | 0.556789205 | 0.022237899 | 0.046310313 |
| Nploc4 | 0.004666732 | 0.93427271 | 0.983960188 | 0.244508681 | 0.225344829 |
| Npm1 | -0.012961998 | 0.826363858 | 0.952566917 | 1.547573462 | 1.5971789 |
| Npm3 | -0.051568833 | 0.350076478 | 0.7406656 | 0.233179225 | 0.28316856 |
| Npnt | -0.052918207 | 0.314306216 | 0.714975653 | 0.178539888 | 0.225151841 |
| Npr1 | -0.042416815 | 0.128841314 | 0.508260084 | 0.044493437 | 0.086563663 |
| Npr2 | -0.049853932 | 0.038958435 | 0.311933003 | 0.028708876 | 0.069039798 |
| Nprl2 | 0.001155342 | 0.975439018 | 0.994455356 | 0.093532009 | 0.096092365 |
| Nprl3 | 0.063251097 | 0.104923222 | 0.472399448 | 0.105797866 | 0.052193462 |
| Nptn | -0.045156607 | 0.45717617 | 0.808898849 | 0.329277437 | 0.378215333 |
| Nptxr | 0.123367943 | 0.055023477 | 0.364775849 | 0.394152815 | 0.263865107 |
| Npy | 0.182796427 | 0.009398407 | 0.150449465 | 0.250343643 | 0.035225664 |
| Nqo1 | -0.003811728 | 0.908969993 | 0.978154712 | 0.061142452 | 0.06861414 |
| Nqo2 | 0.052657967 | 0.157351175 | 0.554924229 | 0.09396609 | 0.046120913 |
| Nr1d1 | 0.094699657 | 0.06713797 | 0.392969665 | 0.176372152 | 0.098599494 |
| Nr1d2 | -0.053409123 | 0.402481638 | 0.77525778 | 0.340878409 | 0.394427939 |
| Nr1h2 | 0.076658357 | 0.126270343 | 0.505736405 | 0.174417572 | 0.104214272 |
| Nr1h3 | -0.060164462 | 0.079092325 | 0.421923248 | 0.063904999 | 0.117589713 |
| Nr1h4 | -0.034009372 | 0.283100103 | 0.691020364 | 0.06103258 | 0.087200847 |
| Nr2c1 | -0.001542857 | 0.958207236 | 0.990272449 | 0.053953811 | 0.052840341 |
| Nr2c2 | -0.080733474 | 0.13181786 | 0.512216749 | 0.20634155 | 0.262592351 |
| Nr2c2ap | 0.035454862 | 0.504352642 | 0.832249514 | 0.199569359 | 0.176703741 |
| Nr2f6 | 0.17562745 | 0.003652883 | 0.085463838 | 0.360899197 | 0.210319453 |
| Nr3c1 | 0.018695902 | 0.794996731 | 0.939757086 | 0.732318202 | 0.733038314 |
| Nr3c2 | 0.00949743 | 0.752590409 | 0.926272218 | 0.056003768 | 0.047409623 |
| Nr4a1 | -0.027024745 | 0.694947421 | 0.910242368 | 0.392297463 | 0.380698588 |
| Nr4a2 | -0.183902731 | 0.002691533 | 0.071134541 | 0.226488426 | 0.441092062 |
| Nr6a1 | 0.012313403 | 0.751377586 | 0.926186851 | 0.10062103 | 0.090579544 |
| Nras | 0.068052773 | 0.215644007 | 0.629611185 | 0.226996492 | 0.172234692 |
| Nrbf2 | -0.011699745 | 0.79103453 | 0.938824685 | 0.124647235 | 0.142156628 |
| Nrbp1 | -0.040318852 | 0.491079875 | 0.827146956 | 0.27215852 | 0.306982593 |
| Nrbp2 | -0.065940036 | 0.024027609 | 0.245835998 | 0.046511613 | 0.097605441 |
| Nrcam | 0.005556351 | 0.85391805 | 0.957461155 | 0.057611538 | 0.048134719 |
| Nrd1 | 0.014626096 | 0.832548975 | 0.953797263 | 0.788715409 | 0.74250365 |
| Nrde2 | -0.001355891 | 0.963640367 | 0.991970325 | 0.057866141 | 0.059840094 |
| Nrep | 0.022549044 | 0.51772543 | 0.837743618 | 0.080994901 | 0.063450056 |
| Nrf1 | -0.211028256 | 0.000305341 | 0.017715076 | 0.254322988 | 0.46110437 |
| Nrip1 | -0.068178443 | 0.279841707 | 0.687465228 | 0.351834108 | 0.442140083 |
| Nrip3 | 0.008472278 | 0.824285986 | 0.951617028 | 0.096318886 | 0.092314036 |
| Nrtn | -0.060887119 | 0.355300576 | 0.745326231 | 0.406121743 | 0.476426788 |
| Nrxn1 | -0.180564329 | 0.002884723 | 0.074240929 | 0.275325883 | 0.447322089 |
| Nsa2 | -0.141764586 | 0.033375728 | 0.289428903 | 0.663341303 | 0.823226275 |
| Nsd1 | -0.180789209 | 0.012671741 | 0.179596745 | 0.635425737 | 0.865131032 |
| Nsd2 | -0.104769088 | 0.08787688 | 0.440569553 | 0.344940173 | 0.454573773 |
| Nsd3 | 0.041378321 | 0.539745738 | 0.84899357 | 0.756826935 | 0.71604348 |
| Nsdhl | 0.036791544 | 0.189603528 | 0.598067444 | 0.05247125 | 0.018674818 |
| Nsf | 0.016756627 | 0.778852084 | 0.934641125 | 0.316126008 | 0.300940698 |
| Nsfl1c | -0.113755756 | 0.060116506 | 0.378409279 | 0.303680419 | 0.420237655 |
| Nsmaf | 0.009966935 | 0.847772387 | 0.956938604 | 0.196182212 | 0.192603364 |
| Nsmce1 | 0.056653463 | 0.30724401 | 0.709900582 | 0.252817921 | 0.208797589 |
| Nsmce2 | -0.090911685 | 0.084134299 | 0.432507481 | 0.193154382 | 0.26340822 |
| Nsmce3 | 0.110035234 | 0.100698234 | 0.465764134 | 0.492944113 | 0.381449821 |
| Nsmce4a | -0.038881816 | 0.552544546 | 0.852623515 | 0.447683387 | 0.498256069 |
| Nsmf | -0.019728963 | 0.54707375 | 0.851446616 | 0.066867592 | 0.082496301 |
| Nsrp1 | 0.234901222 | 0.000546845 | 0.026905298 | 0.488964704 | 0.248217612 |
| Nsun2 | -0.032698103 | 0.563556797 | 0.857926031 | 0.239072304 | 0.266914925 |
| Nsun3 | -0.016554201 | 0.697461556 | 0.910970095 | 0.120645406 | 0.149356957 |
| Nsun4 | -0.006302884 | 0.827491944 | 0.952851099 | 0.049923234 | 0.05053544 |
| Nsun5 | 0.027193947 | 0.394105736 | 0.770629121 | 0.065490362 | 0.043150649 |
| Nsun6 | -0.021282404 | 0.497243671 | 0.830145496 | 0.058247379 | 0.078773658 |
| Nt5c | -0.021235078 | 0.698564583 | 0.911319029 | 0.228432754 | 0.252110216 |
| Nt5c2 | -0.124537215 | 0.027607775 | 0.264407669 | 0.243632495 | 0.353925644 |
| Nt5c3 | 0.002643038 | 0.969382415 | 0.993839765 | 0.620361678 | 0.624350446 |
| Nt5c3b | -0.036891768 | 0.306113098 | 0.709680069 | 0.082079766 | 0.107519774 |
| Nt5dc1 | -0.030046484 | 0.349898871 | 0.7406656 | 0.060972361 | 0.089656081 |

| | | | | | |
|---|---|---|---|---|---|
| Nt5dc2 | -0.09989918 | 0.051380075 | 0.355933715 | 0.186463718 | 0.280638603 |
| Nt5dc3 | 0.098334766 | 0.043185543 | 0.328373709 | 0.177991442 | 0.09164462 |
| Nt5m | -0.025780076 | 0.634090412 | 0.88965971 | 0.226714745 | 0.260673654 |
| Ntan1 | -0.108645837 | 0.063232387 | 0.383440442 | 0.251668792 | 0.357894087 |
| Nthl1 | -0.03722822 | 0.095568398 | 0.455869783 | 0.023268673 | 0.054255792 |
| Ntmt1 | 0.051796076 | 0.296955483 | 0.702656882 | 0.177289233 | 0.13274546 |
| Ntpcr | 0.002951732 | 0.949381735 | 0.987767085 | 0.149117571 | 0.153161207 |
| Ntrk2 | -0.242626511 | 0.000444269 | 0.023346218 | 0.513767232 | 0.770779822 |
| Nuak1 | 0.021214997 | 0.639627191 | 0.891345382 | 0.136917095 | 0.118773751 |
| Nuak2 | 0.015403314 | 0.596785091 | 0.874059822 | 0.054082204 | 0.040480498 |
| Nub1 | 0.055436063 | 0.40195937 | 0.775220138 | 0.433510084 | 0.375119732 |
| Nubp1 | 0.013350226 | 0.753241925 | 0.926506029 | 0.118992758 | 0.107143733 |
| Nubp2 | 0.019606152 | 0.628250144 | 0.887864718 | 0.107793871 | 0.098073555 |
| Nubpl | -0.014180504 | 0.617382341 | 0.883238152 | 0.047405996 | 0.060063534 |
| Nucb1 | 0.037997651 | 0.577291438 | 0.864551328 | 0.59230633 | 0.558475672 |
| Nucb2 | -0.138290609 | 0.053749241 | 0.361340344 | 1.088076781 | 1.193018156 |
| Nucks1 | -0.067038815 | 0.303243728 | 0.708156655 | 0.959177453 | 1.026641356 |
| Nudc | 0.119113222 | 0.084446812 | 0.433265738 | 1.028508402 | 0.848413026 |
| Nudcd1 | -0.003482632 | 0.939277781 | 0.985603501 | 0.138245043 | 0.141890581 |
| Nudcd2 | -0.023416104 | 0.699372959 | 0.911319029 | 0.301337903 | 0.331231166 |
| Nudcd3 | 0.141362103 | 0.031762254 | 0.283375773 | 0.577273952 | 0.40716379 |
| Nudt11 | 0.001171841 | 0.964522425 | 0.991970325 | 0.041490709 | 0.039012853 |
| Nudt13 | 0.002933583 | 0.945746749 | 0.987084873 | 0.127103902 | 0.126787482 |
| Nudt14 | -0.043205614 | 0.377706223 | 0.761864421 | 0.163246823 | 0.204687602 |
| Nudt16 | 0.049816657 | 0.27476267 | 0.682115488 | 0.145168047 | 0.103602712 |
| Nudt16l1 | -0.052706428 | 0.277191781 | 0.684342381 | 0.167323756 | 0.213667148 |
| Nudt18 | 0.010248302 | 0.853905515 | 0.957461155 | 0.249287174 | 0.248047354 |
| Nudt19 | 0.000153829 | 0.997913457 | 0.99938181 | 0.284927361 | 0.28371317 |
| Nudt2 | 0.009695301 | 0.839044112 | 0.954911647 | 0.163711435 | 0.157413214 |
| Nudt21 | 0.015305317 | 0.772558996 | 0.932469974 | 0.209213733 | 0.195126442 |
| Nudt22 | 0.043661117 | 0.310087956 | 0.712953768 | 0.125607542 | 0.085564897 |
| Nudt3 | 0.092015465 | 0.114479846 | 0.485849564 | 0.288877755 | 0.201280406 |
| Nudt4 | 0.168445855 | 0.014014109 | 0.191923581 | 0.681560435 | 0.488489256 |
| Nudt5 | -0.066555205 | 0.251458455 | 0.663127858 | 0.281786134 | 0.347354993 |
| Nudt7 | -0.018414551 | 0.740621336 | 0.922901128 | 0.234948977 | 0.250334311 |
| Nudt9 | -0.036339631 | 0.506586969 | 0.833269979 | 0.219758399 | 0.248103535 |
| Nufip1 | 0.012647865 | 0.795581197 | 0.939757086 | 0.162058792 | 0.153810173 |
| Nufip2 | -0.002909932 | 0.964835031 | 0.991970325 | 0.475183234 | 0.477700794 |
| Numa1 | -0.001131623 | 0.986259677 | 0.997196869 | 0.452778782 | 0.435449483 |
| Numb | -0.002472866 | 0.947948146 | 0.987651726 | 0.09224275 | 0.098328451 |
| Numbl | 0.032118101 | 0.323353069 | 0.720244873 | 0.072614339 | 0.04727775 |
| Nup107 | -0.03520722 | 0.201731535 | 0.612641275 | 0.040695067 | 0.076745998 |
| Nup133 | -0.020975169 | 0.454025396 | 0.807844867 | 0.050233202 | 0.069420844 |
| Nup153 | 0.017747578 | 0.738291247 | 0.922843014 | 0.210199167 | 0.199856816 |
| Nup155 | -0.012759615 | 0.648003971 | 0.895376633 | 0.045067459 | 0.05548616 |
| Nup160 | -0.042481044 | 0.279426824 | 0.68716738 | 0.09820468 | 0.133812968 |
| Nup188 | -0.023157191 | 0.476081087 | 0.817948631 | 0.065853787 | 0.08317898 |
| Nup205 | -0.077686171 | 0.022360295 | 0.237266944 | 0.066905493 | 0.132898231 |
| Nup210 | 0.019708819 | 0.664256649 | 0.900113945 | 0.144444735 | 0.125152665 |
| Nup214 | 0.017152914 | 0.730824379 | 0.920770724 | 0.17146768 | 0.161710736 |
| Nup35 | 0.208435538 | 0.000180958 | 0.012028291 | 0.24305827 | 0.065135872 |
| Nup37 | -0.028186677 | 0.388302715 | 0.767160379 | 0.067600122 | 0.090331709 |
| Nup43 | 0.077410476 | 0.037984317 | 0.307458537 | 0.101101779 | 0.043029602 |
| Nup50 | 0.071461027 | 0.273454511 | 0.681426919 | 0.433769098 | 0.38145962 |
| Nup54 | 0.03027936 | 0.497104311 | 0.830056214 | 0.135641461 | 0.11006393 |
| Nup62 | -0.063570799 | 0.179041425 | 0.585380409 | 0.143147893 | 0.210621085 |
| Nup85 | -0.030061926 | 0.298755957 | 0.703845452 | 0.048882492 | 0.070484671 |
| Nup88 | 0.008061773 | 0.894444921 | 0.972016422 | 0.315064713 | 0.314551305 |
| Nup93 | 0.047919554 | 0.372090244 | 0.757676884 | 0.219558464 | 0.172736737 |
| Nup98 | -0.04116503 | 0.472236743 | 0.8159477 | 0.262134368 | 0.299401636 |
| Nupr1 | -0.962268525 | 1.96E-20 | 6.57E-17 | 1.172508832 | 3.032697456 |
| Nupr1l | -0.067449653 | 0.118115751 | 0.491192747 | 0.121546649 | 0.186723948 |
| Nus1 | 0.004884051 | 0.939393294 | 0.985603501 | 0.519802523 | 0.517530459 |
| Nvl | -0.095183689 | 0.061832872 | 0.380250374 | 0.179106369 | 0.264904145 |
| Nxf1 | 0.061728136 | 0.157991312 | 0.556211435 | 0.13225681 | 0.075906714 |

| | | | | | |
|---|---|---|---|---|---|
| Nxn | 0.076768118 | 0.046451356 | 0.338339813 | 0.109115409 | 0.043744247 |
| Nxph3 | 0.125389726 | 0.002781501 | 0.07270294 | 0.124713762 | 0.016551381 |
| Nxt1 | 0.079306422 | 0.128925991 | 0.508260084 | 0.21322844 | 0.144990989 |
| Nxt2 | 0.016818735 | 0.65795777 | 0.898224427 | 0.097534487 | 0.077158758 |
| Oard1 | 0.028045646 | 0.647036445 | 0.895025469 | 0.346846816 | 0.315922291 |
| Oasl2 | 0.099473475 | 0.027596245 | 0.264407669 | 0.13868601 | 0.044631264 |
| Oaz1 | 0.014414354 | 0.724188176 | 0.919830015 | 2.143821219 | 2.13903527 |
| Oaz2 | -0.134057945 | 0.033137749 | 0.287968471 | 0.32432558 | 0.451708435 |
| Obi1 | 0.00522936 | 0.870160276 | 0.96420829 | 0.065288582 | 0.058075276 |
| Obsl1 | -0.001754263 | 0.954726801 | 0.989066902 | 0.06109755 | 0.06103287 |
| Ocel1 | 0.025307801 | 0.505788826 | 0.832775992 | 0.099629682 | 0.076245409 |
| Ociad1 | -0.058326338 | 0.390166239 | 0.76844731 | 0.700702744 | 0.756397925 |
| Ociad2 | 0.163594226 | 0.016560886 | 0.206965324 | 0.99551492 | 0.773716265 |
| Ocln | 0.05602091 | 0.183569491 | 0.590066231 | 0.121503359 | 0.079023294 |
| Ocrl | -0.035074149 | 0.487956276 | 0.826183728 | 0.170108578 | 0.188036263 |
| Odad4 | -0.029182404 | 0.254434776 | 0.663146569 | 0.031446216 | 0.055154723 |
| Odc1 | 0.176223623 | 0.020383468 | 0.227815517 | 0.763999296 | 0.589481668 |
| Odf2 | -0.117902137 | 0.020668976 | 0.227971987 | 0.175603056 | 0.272589272 |
| Odf2l | -0.030683117 | 0.534608027 | 0.846347806 | 0.171374008 | 0.187128165 |
| Odr4 | 0.07705865 | 0.251419962 | 0.663127858 | 0.453826745 | 0.370430139 |
| Ofd1 | -0.059788076 | 0.253945186 | 0.663146569 | 0.184131847 | 0.246975648 |
| Oga | 0.042973561 | 0.524088853 | 0.841062594 | 0.486221599 | 0.421647237 |
| Ogdh | -0.037859313 | 0.584865616 | 0.867683611 | 0.625953951 | 0.661132803 |
| Ogdhl | -0.015329476 | 0.76264744 | 0.92872278 | 0.177854573 | 0.183526636 |
| Ogfod1 | -0.020253715 | 0.661576823 | 0.898920864 | 0.143079148 | 0.161030827 |
| Ogfod2 | 0.017028392 | 0.755996872 | 0.926726954 | 0.233621292 | 0.219625226 |
| Ogfod3 | -0.090774913 | 0.020271824 | 0.227608207 | 0.08614776 | 0.169743867 |
| Ogfr | -0.071856356 | 0.213768234 | 0.628127689 | 0.253458887 | 0.318968301 |
| Ogfrl1 | -0.026003707 | 0.428154926 | 0.791658813 | 0.064232865 | 0.085782276 |
| Ogt | -0.051592736 | 0.401407552 | 0.774793768 | 0.338792986 | 0.366798066 |
| Oip5os1 | 0.03936505 | 0.516493207 | 0.837427561 | 0.366839107 | 0.329048584 |
| Ola1 | -0.149269419 | 0.020372343 | 0.227815517 | 0.405972527 | 0.549710756 |
| Olfm1 | 0.027761974 | 0.458686003 | 0.809370658 | 0.096291544 | 0.069209912 |
| Olfm4 | -0.030652622 | 0.298230228 | 0.70313959 | 0.053248365 | 0.079702892 |
| Olfr1322 | -0.019117307 | 0.424168988 | 0.789771168 | 0.031408134 | 0.047395619 |
| Oma1 | -0.042128764 | 0.252885286 | 0.663146569 | 0.083067457 | 0.118293101 |
| Onecut2 | -0.038865238 | 0.1603791 | 0.558508561 | 0.041904759 | 0.077334678 |
| Opa1 | 0.118560553 | 0.048967954 | 0.347589359 | 0.334531057 | 0.215868305 |
| Opa3 | 0.064272866 | 0.211033396 | 0.623900499 | 0.200492016 | 0.149945581 |
| Ophn1 | 0.01457989 | 0.638101657 | 0.89125563 | 0.063644804 | 0.046658575 |
| Optn | -0.02123735 | 0.593398179 | 0.872481469 | 0.102016273 | 0.125907378 |
| Orai1 | 0.046887788 | 0.209308651 | 0.62245385 | 0.09758431 | 0.061778989 |
| Orai2 | -0.005155461 | 0.875401121 | 0.965096393 | 0.070404216 | 0.072979595 |
| Orai3 | -0.053615741 | 0.291116258 | 0.697026212 | 0.181797661 | 0.227331313 |
| Orc2 | -0.02043105 | 0.703834813 | 0.912277105 | 0.209729402 | 0.223130675 |
| Orc3 | 0.073006325 | 0.138980104 | 0.525848983 | 0.180852102 | 0.109090547 |
| Orc4 | 0.014870584 | 0.799348681 | 0.941260986 | 0.284296585 | 0.271482437 |
| Orc5 | 0.067396555 | 0.095584416 | 0.455869783 | 0.115000474 | 0.060170185 |
| Orc6 | 0.126642887 | 0.029579829 | 0.273633867 | 0.253836975 | 0.137403014 |
| Ormdl1 | 0.049672084 | 0.219112107 | 0.632622127 | 0.116077742 | 0.073478215 |
| Ormdl2 | 0.044039502 | 0.465875115 | 0.812791331 | 0.334661646 | 0.289837354 |
| Ormdl3 | -0.080174187 | 0.21995343 | 0.633217295 | 0.565470707 | 0.647347352 |
| Os9 | -0.104020411 | 0.137612097 | 0.522790545 | 0.678125916 | 0.779701236 |
| Osbp | 0.035321194 | 0.583994678 | 0.867093874 | 0.435030741 | 0.399549015 |
| Osbpl11 | 0.027832264 | 0.463042718 | 0.811092454 | 0.09733599 | 0.072924058 |
| Osbpl1a | 0.129170664 | 0.028267703 | 0.267663148 | 0.297070773 | 0.183848239 |
| Osbpl2 | 0.023413766 | 0.680307201 | 0.907783945 | 0.25619725 | 0.235263679 |
| Osbpl3 | 0.052955713 | 0.094612565 | 0.454115053 | 0.068489717 | 0.018571846 |
| Osbpl5 | 0.016836101 | 0.571230759 | 0.861225331 | 0.057234151 | 0.041945086 |
| Osbpl6 | 0.049240627 | 0.330467868 | 0.725898152 | 0.179327511 | 0.128178447 |
| Osbpl7 | 0.002637857 | 0.92868971 | 0.982502239 | 0.055738577 | 0.052395191 |
| Osbpl8 | 0.057448196 | 0.306595531 | 0.709680069 | 0.256488319 | 0.219815811 |
| Osbpl9 | 0.047623439 | 0.382983643 | 0.763687454 | 0.216200293 | 0.171295696 |
| Oscp1 | -0.040416489 | 0.127613651 | 0.508260084 | 0.038674743 | 0.073571884 |
| Oser1 | 0.045268479 | 0.435200771 | 0.796975049 | 0.255455204 | 0.220017805 |

| | | | | | |
|---|---|---|---|---|---|
| Osgep | 0.000413126 | 0.994024107 | 0.998695538 | 0.233734637 | 0.225298811 |
| Osgepl1 | 0.014015742 | 0.665175224 | 0.900993754 | 0.068181263 | 0.057566607 |
| Osgin1 | 0.001143213 | 0.980861203 | 0.995540893 | 0.151036199 | 0.15009712 |
| Osgin2 | 0.087857053 | 0.013779167 | 0.189915708 | 0.092453991 | 0.017721932 |
| Ost4 | 0.087919701 | 0.112553736 | 0.483221516 | 1.443290288 | 1.314760718 |
| Ostc | 0.026432448 | 0.649958918 | 0.895593446 | 1.303407677 | 1.26162911 |
| Ostf1 | -0.079665661 | 0.13387848 | 0.515830443 | 0.204573761 | 0.287176004 |
| Ostm1 | -0.054615826 | 0.31445259 | 0.714975653 | 0.217908294 | 0.260767582 |
| Otub1 | -0.098837896 | 0.108433211 | 0.477972625 | 0.366109551 | 0.453530762 |
| Otub2 | -0.040334846 | 0.312422019 | 0.714349311 | 0.097672826 | 0.134460776 |
| Otud3 | 0.041590625 | 0.411647976 | 0.781032788 | 0.175031186 | 0.14209441 |
| Otud4 | -0.030300197 | 0.626632333 | 0.886996623 | 0.33093574 | 0.363463157 |
| Otud5 | -0.002273902 | 0.968648235 | 0.993505636 | 0.266384545 | 0.280679511 |
| Otud6b | -0.01898447 | 0.712693801 | 0.915447617 | 0.187238866 | 0.202346208 |
| Otud7a | 0.01612924 | 0.621990149 | 0.884892293 | 0.072647537 | 0.062531269 |
| Otud7b | 0.07971251 | 0.162112288 | 0.560882811 | 0.264716657 | 0.190358496 |
| Otulin | 0.03268833 | 0.609228186 | 0.878855197 | 0.37622474 | 0.344681538 |
| Otulinl | 0.005211849 | 0.852737525 | 0.957461155 | 0.048942074 | 0.04596561 |
| Ovol2 | 0.114024898 | 0.04097155 | 0.319308264 | 0.223345133 | 0.127334422 |
| Oxa1l | 0.011851033 | 0.814846019 | 0.948794604 | 0.178576693 | 0.175170565 |
| Oxct1 | 0.024664689 | 0.720669831 | 0.919412687 | 0.627148489 | 0.580107096 |
| Oxld1 | -0.011979655 | 0.784492285 | 0.936423418 | 0.131612477 | 0.149530856 |
| Oxnad1 | -0.019929397 | 0.628288335 | 0.887864718 | 0.115535578 | 0.138619753 |
| Oxr1 | 0.039500869 | 0.504932069 | 0.832249514 | 0.284450356 | 0.246190263 |
| Oxsm | 0.030112513 | 0.32997211 | 0.725346052 | 0.064660136 | 0.037435124 |
| Oxsr1 | 0.042817191 | 0.398303437 | 0.774311756 | 0.191426847 | 0.153048817 |
| Oxtr | 0.103505895 | 0.008735036 | 0.146122588 | 0.105395584 | 0.007679646 |
| P2rx4 | -0.038967631 | 0.477149934 | 0.818444859 | 0.217208685 | 0.253479576 |
| P2ry1 | 0.118420865 | 0.023989551 | 0.245835998 | 0.213221795 | 0.104700002 |
| P3h1 | -0.058345127 | 0.05415974 | 0.362869235 | 0.048999316 | 0.0953404 |
| P3h3 | -0.038126655 | 0.069233552 | 0.400055935 | 0.019805243 | 0.049935654 |
| P3h4 | -0.119373797 | 0.016654327 | 0.207393899 | 0.157080712 | 0.265130758 |
| P4ha1 | -0.011845837 | 0.848779979 | 0.957111129 | 0.348986142 | 0.353788723 |
| P4ha2 | 0.034381664 | 0.501964187 | 0.832218079 | 0.193584919 | 0.159174874 |
| P4hb | -0.016749191 | 0.787541169 | 0.937804093 | 1.367616374 | 1.337616158 |
| P4htm | 0.039962839 | 0.430593262 | 0.793311823 | 0.183924328 | 0.142778116 |
| Pa2g4 | -0.07089388 | 0.293924666 | 0.699744278 | 0.46762021 | 0.544319053 |
| Pabpc1 | -0.10672068 | 0.062247767 | 0.380250374 | 1.257342902 | 1.42657787 |
| Pabpc1l | -0.061999896 | 0.005888907 | 0.113767137 | 0.022210275 | 0.080972732 |
| Pabpc4 | 0.022558939 | 0.592649029 | 0.872202098 | 0.119234391 | 0.103864985 |
| Pabpn1 | -0.102950749 | 0.046356249 | 0.33789228 | 0.173745118 | 0.272442649 |
| Pacc1 | 0.036764752 | 0.456480558 | 0.808445095 | 0.174944738 | 0.157259614 |
| Pacrg | 0.005195019 | 0.882256938 | 0.968472149 | 0.078176416 | 0.076419283 |
| Pacs1 | -0.008118013 | 0.845154687 | 0.955950886 | 0.117356933 | 0.119938501 |
| Pacs2 | -0.001712701 | 0.963452585 | 0.991970325 | 0.089694198 | 0.088423779 |
| Pacsin1 | 0.07146984 | 0.16637936 | 0.568127031 | 0.203470278 | 0.134511119 |
| Pacsin2 | -0.011788046 | 0.792892382 | 0.939582154 | 0.137548466 | 0.147641825 |
| Paf1 | -0.02918658 | 0.572144469 | 0.861494041 | 0.19554042 | 0.214934205 |
| Pafah1b1 | 0.075607903 | 0.192731073 | 0.60325934 | 1.459927601 | 1.345434448 |
| Pafah1b2 | 0.082612852 | 0.205328224 | 0.616549498 | 0.628394089 | 0.558820686 |
| Pafah1b3 | -0.002705429 | 0.96090712 | 0.991375098 | 0.232405032 | 0.232364227 |
| Pag1 | -0.048845083 | 0.279467694 | 0.68716738 | 0.13621759 | 0.179784907 |
| Paics | -0.151068706 | 0.01104272 | 0.166153611 | 0.313012005 | 0.45704719 |
| Paip1 | -0.02652822 | 0.650213155 | 0.895593446 | 0.321567061 | 0.367110066 |
| Paip2 | -0.13706386 | 0.034588547 | 0.293461747 | 1.029133841 | 1.205791805 |
| Paip2b | 0.06342214 | 0.293568947 | 0.69925611 | 0.329238685 | 0.266813211 |
| Pak1 | 0.048124038 | 0.1177107 | 0.491192747 | 0.061938396 | 0.021936373 |
| Pak1ip1 | 0.050278917 | 0.436344954 | 0.797595028 | 0.431042423 | 0.392841842 |
| Pak2 | 0.068014515 | 0.310662931 | 0.71348779 | 0.831062916 | 0.746465282 |
| Pak3 | -0.054437752 | 0.430919191 | 0.793421321 | 0.495740361 | 0.531537457 |
| Pak4 | 0.043093988 | 0.359742615 | 0.747719327 | 0.158306498 | 0.116387433 |
| Pam | -0.218681658 | 9.83E-05 | 0.007650838 | 1.866570144 | 2.253182795 |
| Pam16 | -0.165202778 | 0.012367032 | 0.177814713 | 0.628675118 | 0.818268777 |
| Pamr1 | 0.030955513 | 0.571915621 | 0.861494041 | 0.226059077 | 0.19404906 |
| Pan2 | -0.03189125 | 0.356618586 | 0.745571198 | 0.074495772 | 0.101753286 |

| | | | | | |
|---|---|---|---|---|---|
| Pan3 | -0.088210169 | 0.102951462 | 0.468979611 | 0.207495832 | 0.269285383 |
| Pank1 | 0.039361613 | 0.291411242 | 0.697147764 | 0.097472894 | 0.066349868 |
| Pank2 | 0.08533778 | 0.095936677 | 0.456574881 | 0.189664258 | 0.11264999 |
| Pank3 | 0.130315684 | 0.035189187 | 0.295084312 | 0.371302743 | 0.243581514 |
| Pank4 | 0.021249844 | 0.489320547 | 0.826402547 | 0.060780773 | 0.046912697 |
| Papola | 0.15560107 | 0.005997398 | 0.115097285 | 1.471463389 | 1.246084108 |
| Pappa2 | -0.175482199 | 0.012974729 | 0.182144849 | 0.351034704 | 0.522437833 |
| Papss1 | -0.019798472 | 0.710687871 | 0.914509508 | 0.210602857 | 0.226495143 |
| Papss2 | -0.06159314 | 0.369728751 | 0.756876906 | 0.881589729 | 0.979408091 |
| Paqr3 | -0.007529211 | 0.789041639 | 0.938231363 | 0.047794691 | 0.050823846 |
| Paqr9 | -0.035403545 | 0.240652794 | 0.650744882 | 0.052272397 | 0.084719832 |
| Pard3 | -0.026188174 | 0.644783647 | 0.893466123 | 0.240570551 | 0.260175212 |
| Pard3b | -0.053967035 | 0.122814349 | 0.499468242 | 0.072075564 | 0.114025106 |
| Pard6a | 0.090527176 | 0.116893332 | 0.489579867 | 0.289913053 | 0.213624333 |
| Pard6b | -0.040282579 | 0.199719976 | 0.610108047 | 0.055699731 | 0.091056921 |
| Pard6g | 0.092463108 | 0.020177185 | 0.227038569 | 0.108752073 | 0.03041699 |
| Parg | -0.053410223 | 0.346220956 | 0.738109544 | 0.23068586 | 0.285041767 |
| Park7 | 0.031637381 | 0.595088068 | 0.873342705 | 1.739010628 | 1.621511263 |
| Parl | -0.079310418 | 0.125164982 | 0.503791124 | 0.182667022 | 0.261823305 |
| Parm1 | 0.14516367 | 0.00798335 | 0.138153252 | 0.236150841 | 0.111652154 |
| Parn | 0.123670072 | 0.01462374 | 0.194386632 | 0.210583945 | 0.104303143 |
| Parp1 | 0.116735386 | 0.053397318 | 0.360908339 | 0.312580303 | 0.211440782 |
| Parp10 | 0.04217726 | 0.256826236 | 0.665256942 | 0.097035011 | 0.063421737 |
| Parp12 | 0.050967812 | 0.316303099 | 0.715674979 | 0.187340758 | 0.143882542 |
| Parp14 | 0.094308309 | 0.07942995 | 0.422526451 | 0.206999571 | 0.107742476 |
| Parp16 | -0.050455085 | 0.093442266 | 0.452015851 | 0.052085805 | 0.093066928 |
| Parp2 | -0.010708474 | 0.842645823 | 0.955306089 | 0.216789854 | 0.228974147 |
| Parp4 | -0.114156752 | 0.007123256 | 0.128637347 | 0.108488522 | 0.210804968 |
| Parp6 | 0.003610244 | 0.946155451 | 0.987084873 | 0.201185451 | 0.190166946 |
| Parp8 | -0.06113035 | 0.306907703 | 0.709680069 | 0.299012277 | 0.374361434 |
| Parp9 | 0.073022325 | 0.090103904 | 0.445385778 | 0.128793405 | 0.065223865 |
| Particl | -0.033738765 | 0.239999266 | 0.650744882 | 0.048654911 | 0.079906207 |
| Parva | 0.006369622 | 0.913488732 | 0.979979308 | 0.283648709 | 0.285007697 |
| Patj | 0.017403452 | 0.675366833 | 0.906025597 | 0.113615033 | 0.093697904 |
| Patl1 | 0.037696771 | 0.381119128 | 0.762430261 | 0.124959 | 0.091859263 |
| Patz1 | 0.012789493 | 0.736224506 | 0.922302218 | 0.097989958 | 0.088017868 |
| Paupar | -0.005753258 | 0.849939524 | 0.957111129 | 0.053864146 | 0.060667654 |
| Pawr | 0.071057779 | 0.188944715 | 0.596946223 | 0.229614766 | 0.172151474 |
| Pax6 | -0.052147963 | 0.400958512 | 0.774740795 | 1.419853187 | 1.4721426 |
| Pax6os1 | 0.034125285 | 0.35198419 | 0.742354553 | 0.091369725 | 0.056379998 |
| Paxbp1 | -0.097936446 | 0.116124572 | 0.488696992 | 0.299445494 | 0.393278903 |
| Paxip1 | 0.009274854 | 0.874110479 | 0.964947297 | 0.278861449 | 0.262808563 |
| Paxx | -0.051439842 | 0.276277095 | 0.683667283 | 0.144204974 | 0.200575177 |
| Pbdc1 | 0.104789667 | 0.089215233 | 0.443953048 | 0.369424078 | 0.267882661 |
| Pbld2 | -0.025158262 | 0.292912849 | 0.698685876 | 0.030157171 | 0.049121463 |
| Pbrm1 | 0.014277301 | 0.833513319 | 0.954366094 | 0.928245751 | 0.895386117 |
| Pbx1 | -0.091507568 | 0.146889787 | 0.536877806 | 0.390142445 | 0.488870846 |
| Pbx2 | 0.053333798 | 0.370044171 | 0.757234136 | 0.274601671 | 0.224552064 |
| Pbx3 | -0.089141325 | 0.128262181 | 0.508260084 | 0.280213205 | 0.378797856 |
| Pbx4 | -0.039413889 | 0.07032096 | 0.402558548 | 0.021375676 | 0.055228384 |
| Pbxip1 | -0.122100642 | 0.053016128 | 0.359299711 | 0.344966417 | 0.461741205 |
| Pcbd1 | -0.077212423 | 0.245917368 | 0.655932134 | 0.970000047 | 1.082021389 |
| Pcbd2 | -0.093308604 | 0.115900494 | 0.488163347 | 0.297962034 | 0.396497372 |
| Pcbp1 | 0.016152199 | 0.806643103 | 0.945532285 | 1.125861126 | 1.084278839 |
| Pcbp2 | 0.035881752 | 0.494297687 | 0.828842054 | 1.659912876 | 1.576350789 |
| Pcbp3 | 0.04945207 | 0.357266737 | 0.745983316 | 0.212212559 | 0.161221854 |
| Pcbp4 | 0.028338492 | 0.502949864 | 0.832249514 | 0.122650276 | 0.09789057 |
| Pcca | -0.036075991 | 0.445063444 | 0.802857978 | 0.148708221 | 0.181170099 |
| Pccb | -0.042637549 | 0.38013083 | 0.762006446 | 0.155120455 | 0.195522294 |
| Pcdh1 | -0.002629091 | 0.960377459 | 0.99118854 | 0.215461802 | 0.222433445 |
| Pcdh15 | -0.0032284 | 0.916450887 | 0.980132949 | 0.054427533 | 0.057100291 |
| Pcdhb22 | -0.060589743 | 0.000500226 | 0.025103854 | 0.006492615 | 0.059176375 |
| Pcdhga12 | -0.017923199 | 0.534282174 | 0.846347806 | 0.047352989 | 0.061050751 |
| Pced1a | -0.022737673 | 0.443684157 | 0.801522297 | 0.053201645 | 0.073521085 |
| Pcf11 | 0.007783035 | 0.905656408 | 0.976167673 | 0.401727768 | 0.398851859 |

| | | | | | |
|---|---|---|---|---|---|
| Pcgf1 | 0.007025949 | 0.837970313 | 0.954911647 | 0.075403992 | 0.072294252 |
| Pcgf2 | 0.013589005 | 0.731834453 | 0.920770724 | 0.108386616 | 0.096033726 |
| Pcgf3 | 0.024406509 | 0.60945155 | 0.878855197 | 0.154105607 | 0.131475382 |
| Pcgf5 | 5.25E-05 | 0.999360123 | 0.999858209 | 0.426648968 | 0.435603225 |
| Pcgf6 | -0.094019843 | 0.051535223 | 0.355933715 | 0.1501577 | 0.220421219 |
| Pcid2 | 0.03228125 | 0.452093742 | 0.806552593 | 0.125576918 | 0.095136441 |
| Pcif1 | 0.183376919 | 0.006573642 | 0.123096359 | 0.691272297 | 0.502097578 |
| Pck2 | -0.208017002 | 0.002467793 | 0.067311494 | 0.74213722 | 0.981242632 |
| Pclo | -0.100116074 | 0.12700528 | 0.506859642 | 1.077049836 | 1.203509151 |
| Pcm1 | 0.018938721 | 0.782784418 | 0.936114286 | 0.866681774 | 0.821444113 |
| Pcmt1 | -0.002279539 | 0.972517682 | 0.994079882 | 0.588272512 | 0.600393263 |
| Pcmtd1 | -0.062685335 | 0.342517476 | 0.735368536 | 0.87788392 | 0.951118773 |
| Pcmtd2 | -0.108648771 | 0.061740933 | 0.380250374 | 0.258577232 | 0.35631662 |
| Pcna | -0.020441212 | 0.733584029 | 0.921306823 | 0.299812854 | 0.311546107 |
| Pcnp | 0.063754634 | 0.306900898 | 0.709680069 | 0.816798582 | 0.757207572 |
| Pcnt | -0.023022814 | 0.53509923 | 0.846460359 | 0.086353242 | 0.103017696 |
| Pcnx | -0.000567921 | 0.992163824 | 0.998695538 | 0.265522462 | 0.262214527 |
| Pcnx3 | -0.000620847 | 0.988016905 | 0.997357506 | 0.118537607 | 0.113782076 |
| Pcnx4 | 0.021998357 | 0.569301106 | 0.860683115 | 0.098901582 | 0.079135917 |
| Pcp4 | 0.044409242 | 0.512304547 | 0.835753752 | 0.343590909 | 0.301106039 |
| Pcsk1 | -0.193671604 | 0.003994891 | 0.090307927 | 1.305451285 | 1.581559222 |
| Pcsk1n | -0.17099903 | 1.55E-05 | 0.00181139 | 3.17564187 | 3.643878921 |
| Pcsk2 | -0.145214069 | 4.95E-06 | 0.00076421 | 3.354380439 | 3.773245509 |
| Pcsk2os1 | -0.027480497 | 0.646976833 | 0.895025469 | 0.316994423 | 0.359982061 |
| Pcsk4 | -0.034051395 | 0.281620009 | 0.688579788 | 0.060580882 | 0.094039857 |
| Pcsk6 | -0.102169955 | 0.005580433 | 0.111475656 | 0.07327958 | 0.163604004 |
| Pcsk7 | -0.029448599 | 0.3762619 | 0.760785796 | 0.068472748 | 0.088938765 |
| Pcsk9 | 0.105070341 | 0.00933661 | 0.14984873 | 0.114112877 | 0.017442902 |
| Pctp | -0.031406135 | 0.27072225 | 0.678123091 | 0.044985966 | 0.074866549 |
| Pcx | -0.00934356 | 0.852435797 | 0.957461155 | 0.179539387 | 0.193238206 |
| Pcyox1 | -0.046690111 | 0.397898953 | 0.774311756 | 0.22605598 | 0.26958364 |
| Pcyox1l | -0.059639712 | 0.018642375 | 0.216996248 | 0.031893522 | 0.084961223 |
| Pcyt1a | 0.113346774 | 0.057548256 | 0.370501507 | 0.314812564 | 0.216527414 |
| Pcyt1b | -0.029310579 | 0.436800436 | 0.797968132 | 0.088611805 | 0.113985555 |
| Pcyt2 | 0.014460312 | 0.737945483 | 0.922843014 | 0.128206171 | 0.111775298 |
| Pdap1 | 0.208027439 | 0.000633871 | 0.028919709 | 1.252066477 | 0.974116596 |
| Pdcd10 | 0.151421605 | 0.025889445 | 0.256517626 | 0.621945634 | 0.457059532 |
| Pdcd11 | 0.000198485 | 0.997217188 | 0.999265301 | 0.251441191 | 0.247883001 |
| Pdcd2 | -0.035378188 | 0.497750108 | 0.830714638 | 0.195020615 | 0.232221176 |
| Pdcd2l | 0.048528586 | 0.29926112 | 0.704513802 | 0.155943963 | 0.114447461 |
| Pdcd4 | 0.141566678 | 0.035294514 | 0.295084312 | 1.284860124 | 1.062386452 |
| Pdcd5 | 0.01595553 | 0.803357749 | 0.943517638 | 0.681585115 | 0.651527707 |
| Pdcd5-ps | -0.025957243 | 0.200018697 | 0.61058019 | 0.020034931 | 0.040576049 |
| Pdcd6 | -0.075107361 | 0.274156582 | 0.681791282 | 0.737119251 | 0.829479084 |
| Pdcd6ip | -0.017484802 | 0.803014139 | 0.943517638 | 0.72986675 | 0.729960074 |
| Pdcd7 | 0.022910283 | 0.674325581 | 0.906025597 | 0.207528058 | 0.195829176 |
| Pdcl | -0.088560249 | 0.041478182 | 0.321059507 | 0.120803713 | 0.196161136 |
| Pdcl3 | -0.013547441 | 0.801673544 | 0.942642614 | 0.220120017 | 0.236786039 |
| Pde10a | -0.063269162 | 0.398771389 | 0.774321615 | 0.799617569 | 0.863298034 |
| Pde12 | 0.086346706 | 0.051255174 | 0.355933715 | 0.144091635 | 0.07891011 |
| Pde1c | 0.13846683 | 0.029422571 | 0.273392198 | 0.352598011 | 0.218197615 |
| Pde3b | -0.164704304 | 0.01398244 | 0.191923581 | 0.515200975 | 0.683439708 |
| Pde4a | -0.019559539 | 0.378369045 | 0.761977617 | 0.026438475 | 0.042738246 |
| Pde4b | -0.005184307 | 0.920830521 | 0.980519408 | 0.183676235 | 0.186484129 |
| Pde4d | -0.052768057 | 0.109479496 | 0.477972625 | 0.065946386 | 0.107902913 |
| Pde4dip | 0.09712432 | 0.137500334 | 0.522761688 | 0.544222679 | 0.465196206 |
| Pde5a | -0.150952767 | 0.028403905 | 0.268446318 | 0.616296933 | 0.775878579 |
| Pde6d | -0.048089914 | 0.392865893 | 0.769704269 | 0.242746405 | 0.297940719 |
| Pde7a | -0.035071423 | 0.243293372 | 0.653166476 | 0.055469412 | 0.086789217 |
| Pde7b | -0.053363678 | 0.011645332 | 0.171133529 | 0.017186666 | 0.062257538 |
| Pde8a | 0.074180027 | 0.236176211 | 0.646794168 | 0.369350226 | 0.295309085 |
| Pdgfa | 0.226387809 | 0.001080997 | 0.040036792 | 0.730102458 | 0.473586972 |
| Pdgfrl | -0.014864253 | 0.694205278 | 0.909986409 | 0.093072581 | 0.10662719 |
| Pdha1 | 0.035650179 | 0.563672368 | 0.857926031 | 0.363490144 | 0.326155467 |
| Pdhb | 0.071036778 | 0.262322154 | 0.668764912 | 0.448514608 | 0.383813404 |

| | | | | | |
|---|---|---|---|---|---|
| Pdhx | 0.034074568 | 0.311888562 | 0.71405691 | 0.077564982 | 0.044465813 |
| Pdia3 | 0.085806355 | 0.06126615 | 0.380250374 | 2.027804559 | 1.854374037 |
| Pdia4 | 0.049593911 | 0.467343157 | 0.813188482 | 0.570339398 | 0.50237302 |
| Pdia5 | -0.034857922 | 0.387215152 | 0.766866314 | 0.102542089 | 0.14825469 |
| Pdia6 | 0.228211972 | 0.000172386 | 0.011690799 | 1.766132706 | 1.360070743 |
| Pdik1l | 0.071273903 | 0.134269993 | 0.516942048 | 0.163085206 | 0.102120684 |
| Pdk1 | 0.061137646 | 0.124296394 | 0.502037386 | 0.115505234 | 0.064682286 |
| Pdk3 | 0.05687046 | 0.137313491 | 0.522249151 | 0.103215753 | 0.057670049 |
| Pdlim5 | 0.133366171 | 0.00710059 | 0.128637347 | 0.185901367 | 0.069316411 |
| Pdlim7 | 0.084671805 | 0.052938349 | 0.359257749 | 0.13293858 | 0.056116348 |
| Pdp2 | -0.014168816 | 0.658013987 | 0.898224427 | 0.063609725 | 0.078055739 |
| Pdpk1 | 0.031489441 | 0.638539328 | 0.89125563 | 0.657547575 | 0.626608276 |
| Pdpr | -0.025741784 | 0.278487037 | 0.685442974 | 0.030372403 | 0.05341496 |
| Pdrg1 | -0.089248081 | 0.14414592 | 0.533279057 | 0.335606895 | 0.418118328 |
| Pds5a | -0.053614081 | 0.378841891 | 0.762006446 | 0.34342607 | 0.381286132 |
| Pds5b | -0.068961835 | 0.234492447 | 0.644900587 | 0.25567742 | 0.307094838 |
| Pdss2 | -0.032216913 | 0.336740355 | 0.731477644 | 0.068429752 | 0.098100311 |
| Pdx1 | 0.034979507 | 0.612145165 | 0.880369826 | 1.01634834 | 0.991640556 |
| Pdxdc1 | -0.05356695 | 0.417619744 | 0.784482356 | 0.528756062 | 0.557100515 |
| Pdxk | 0.08693303 | 0.033124852 | 0.287968471 | 0.119246166 | 0.042158016 |
| Pdyn | 0.017851273 | 0.815022225 | 0.948889697 | 0.478713206 | 0.449927386 |
| Pdzd11 | -0.015030505 | 0.788274187 | 0.93787435 | 0.245660616 | 0.260910184 |
| Pdzd4 | 0.016672875 | 0.62899364 | 0.887864718 | 0.080256208 | 0.064344015 |
| Pdzd8 | -0.117099663 | 0.052309137 | 0.357487248 | 0.310226834 | 0.426952346 |
| Pdzk1ip1 | -0.025937206 | 0.694723778 | 0.910242368 | 0.474035703 | 0.502341989 |
| Pea15a | 0.25770405 | 0.000260024 | 0.015913799 | 0.635276562 | 0.349308948 |
| Pebp1 | -0.073270394 | 0.072738302 | 0.409234492 | 2.169419165 | 2.375651121 |
| Pecr | 0.0204693 | 0.63255059 | 0.889078599 | 0.131019216 | 0.122765405 |
| Pef1 | -0.071734499 | 0.141344828 | 0.528664246 | 0.159093841 | 0.220992415 |
| Peg13 | 0.129504555 | 0.01586486 | 0.203377154 | 0.220260606 | 0.100177735 |
| Peg3 | -0.156479018 | 0.001458684 | 0.049629872 | 2.302865485 | 2.627351946 |
| Peli1 | -0.02193594 | 0.678155354 | 0.907552705 | 0.194217378 | 0.214520897 |
| Peli3 | -0.003883039 | 0.900176237 | 0.973983939 | 0.062156121 | 0.068246722 |
| Pelp1 | -0.007375441 | 0.864432855 | 0.961442912 | 0.121050305 | 0.119729524 |
| Pemt | 0.002785273 | 0.918647299 | 0.980385833 | 0.047606242 | 0.042563138 |
| Pepd | 0.009437854 | 0.840086514 | 0.954911647 | 0.155087291 | 0.14033758 |
| Per1 | 0.086974698 | 0.067091416 | 0.392969665 | 0.155637732 | 0.070731696 |
| Per2 | 0.051667811 | 0.182678587 | 0.590066231 | 0.106476478 | 0.056106713 |
| Per3 | 0.080794003 | 0.098138475 | 0.461540022 | 0.171861778 | 0.096632286 |
| Perp | 0.105442068 | 0.014104027 | 0.192601517 | 0.139529523 | 0.052057295 |
| Pes1 | 0.0078161 | 0.870976516 | 0.964504119 | 0.164177397 | 0.165448963 |
| Pet100 | 0.104811122 | 0.080832686 | 0.426112224 | 0.344006567 | 0.240447199 |
| Pex1 | 0.008458597 | 0.825223721 | 0.952011047 | 0.096812291 | 0.088285049 |
| Pex10 | 0.028114519 | 0.342709572 | 0.735501771 | 0.060150337 | 0.042963228 |
| Pex11a | -0.043272582 | 0.404493069 | 0.776552851 | 0.197180054 | 0.228976518 |
| Pex11b | -0.019750938 | 0.691837863 | 0.909796728 | 0.169222738 | 0.18256644 |
| Pex11g | -0.008254064 | 0.837126641 | 0.954911647 | 0.107192754 | 0.11035623 |
| Pex12 | -0.009207819 | 0.82865732 | 0.952874813 | 0.119812686 | 0.13296427 |
| Pex13 | -0.101644081 | 0.051346082 | 0.355933715 | 0.18044805 | 0.285899868 |
| Pex14 | 0.099792978 | 0.032370939 | 0.285006239 | 0.163533363 | 0.076526558 |
| Pex16 | 0.012372163 | 0.812442818 | 0.948397417 | 0.198416212 | 0.195351366 |
| Pex19 | 0.045078333 | 0.326406297 | 0.720866629 | 0.152092469 | 0.112460017 |
| Pex2 | -0.046081635 | 0.417459852 | 0.784482356 | 0.24207197 | 0.291464812 |
| Pex26 | -0.017131015 | 0.648248604 | 0.895379629 | 0.088956656 | 0.102702268 |
| Pex3 | 0.025089353 | 0.652573798 | 0.896875696 | 0.233221804 | 0.207637779 |
| Pex5 | 0.057105219 | 0.182441131 | 0.590066231 | 0.130574744 | 0.083918116 |
| Pex5l | -0.072708782 | 0.228229238 | 0.639623601 | 0.317409953 | 0.391334816 |
| Pex6 | -0.073842623 | 0.106216572 | 0.473814268 | 0.137710413 | 0.196294794 |
| Pex7 | -0.017526943 | 0.777949086 | 0.934339474 | 0.357059468 | 0.375033517 |
| Pfas | -0.016756337 | 0.52962651 | 0.844323583 | 0.039312121 | 0.052397869 |
| Pfdn1 | -0.091255556 | 0.172855913 | 0.575371867 | 0.647834663 | 0.759560246 |
| Pfdn2 | -0.001842621 | 0.976944773 | 0.99458309 | 1.087324243 | 1.099504513 |
| Pfdn4 | 0.065812521 | 0.283735168 | 0.691561408 | 0.359683053 | 0.301085623 |
| Pfdn5 | -0.045609751 | 0.349579373 | 0.740550478 | 1.851656967 | 1.939167711 |
| Pfdn6 | -0.083847298 | 0.201954152 | 0.612724107 | 0.554713976 | 0.630956001 |

| | | | | | |
|---|---|---|---|---|---|
| Pfkfb2 | 0.052370764 | 0.256847785 | 0.665256942 | 0.150940256 | 0.107198968 |
| Pfkl | -0.070350826 | 0.104685509 | 0.472076273 | 0.122556192 | 0.191787968 |
| Pfkm | -0.011725944 | 0.766198855 | 0.92882091 | 0.10216563 | 0.111188039 |
| Pfkp | 0.06692866 | 0.209821439 | 0.622606671 | 0.214226635 | 0.151859976 |
| Pfn1 | 0.055366016 | 0.380677916 | 0.762006446 | 1.223751581 | 1.143638296 |
| Pfn2 | -0.042218637 | 0.395329546 | 0.771969388 | 0.170607454 | 0.208019122 |
| Pgam1 | -0.014267438 | 0.839128406 | 0.954911647 | 0.82671016 | 0.847326436 |
| Pgam5 | 0.014383203 | 0.7371431 | 0.92276204 | 0.1266362 | 0.113173919 |
| Pgap1 | 0.107504812 | 0.083570599 | 0.431740995 | 0.344136145 | 0.236002226 |
| Pgap2 | -0.104402644 | 0.019396972 | 0.221110882 | 0.127468546 | 0.221016534 |
| Pgap3 | -0.03099105 | 0.267714825 | 0.675818334 | 0.043455899 | 0.070678413 |
| Pgap4 | 0.028394226 | 0.470762012 | 0.814586466 | 0.106613782 | 0.088835588 |
| Pgap6 | -0.005472364 | 0.900234035 | 0.973983939 | 0.123262842 | 0.128381094 |
| Pgbd5 | 0.021575635 | 0.613477738 | 0.881402241 | 0.124919796 | 0.104496141 |
| Pgd | -0.058700778 | 0.159550079 | 0.557623868 | 0.1115934 | 0.182046656 |
| Pgf | 0.029473107 | 0.381951006 | 0.762762087 | 0.068566412 | 0.034482155 |
| Pgghg | -0.086597905 | 0.001003511 | 0.038152443 | 0.033331698 | 0.103746741 |
| Pggt1b | -0.004666791 | 0.935012086 | 0.984081583 | 0.252054125 | 0.265526061 |
| Pgk1 | -0.061516284 | 0.362850771 | 0.751223843 | 0.676310674 | 0.734039555 |
| Pgls | -0.184082669 | 0.005742882 | 0.11293531 | 0.842255287 | 1.041719071 |
| Pgm1 | -0.028355828 | 0.548367191 | 0.852076539 | 0.150399206 | 0.174719073 |
| Pgm2 | 0.025056425 | 0.542659356 | 0.85028523 | 0.113600531 | 0.095831247 |
| Pgm3 | -0.119485533 | 0.041888194 | 0.323160491 | 0.277006705 | 0.388821619 |
| Pgp | 0.160378833 | 0.019244383 | 0.220497571 | 0.65716928 | 0.485877805 |
| Pgpep1 | -0.010377659 | 0.816396989 | 0.949036482 | 0.136997269 | 0.144927468 |
| Pgrmc1 | -0.073257896 | 0.302370931 | 0.707681683 | 0.699572756 | 0.795015112 |
| Pgrmc2 | 0.001942112 | 0.978178219 | 0.995031396 | 0.759051966 | 0.757555241 |
| Pgs1 | 0.044134587 | 0.288135527 | 0.694614177 | 0.122918594 | 0.09115453 |
| Phactr1 | 0.044081488 | 0.545281708 | 0.851259637 | 0.965430913 | 0.91117283 |
| Phactr2 | -0.070249563 | 0.013576063 | 0.189230674 | 0.042207218 | 0.099716292 |
| Phactr4 | 0.061654774 | 0.2448342 | 0.654809396 | 0.207387016 | 0.140364161 |
| Phax | 0.040861594 | 0.533594494 | 0.84612966 | 0.427324414 | 0.391482298 |
| Phb | 0.033819334 | 0.43833813 | 0.799200693 | 0.136687421 | 0.110340365 |
| Phb2 | 0.108675444 | 0.125730163 | 0.504406266 | 0.642866579 | 0.537879528 |
| Phc1 | 0.0312217 | 0.433044781 | 0.795183034 | 0.111582155 | 0.083872032 |
| Phc2 | 0.101141882 | 0.11758345 | 0.491129875 | 0.39377421 | 0.304364854 |
| Phc3 | 0.009989546 | 0.856043076 | 0.958084785 | 0.226863214 | 0.215409302 |
| Phf1 | -0.059813639 | 0.28773385 | 0.694589247 | 0.237289758 | 0.298441842 |
| Phf10 | -0.037022017 | 0.593249098 | 0.872445596 | 0.535509943 | 0.558951785 |
| Phf12 | 0.052814883 | 0.36111411 | 0.749638535 | 0.265005102 | 0.209214369 |
| Phf13 | 0.084626101 | 0.038743459 | 0.31159303 | 0.124190967 | 0.054522349 |
| Phf14 | -0.060136427 | 0.361827052 | 0.750032543 | 0.478543868 | 0.537244564 |
| Phf2 | 0.049019701 | 0.393752856 | 0.770539562 | 0.261090208 | 0.220101786 |
| Phf20 | 0.033759769 | 0.594660828 | 0.873162794 | 0.3433193 | 0.308468334 |
| Phf20l1 | -0.011702114 | 0.86385025 | 0.96114233 | 0.928892692 | 0.918299548 |
| Phf21a | -0.120920911 | 0.044962225 | 0.331703493 | 0.275630187 | 0.388020565 |
| Phf21b | -0.002180278 | 0.95990321 | 0.990928888 | 0.126268009 | 0.128508893 |
| Phf23 | 0.084369578 | 0.113159963 | 0.483626277 | 0.220926639 | 0.145640788 |
| Phf3 | 0.024962385 | 0.705435425 | 0.912844672 | 0.604533713 | 0.593785132 |
| Phf5a | -0.066624473 | 0.3121469 | 0.714322488 | 0.477719896 | 0.546568321 |
| Phf6 | -0.082991028 | 0.051899763 | 0.356548882 | 0.111396343 | 0.192015496 |
| Phf7 | -0.065388992 | 0.076200064 | 0.416492204 | 0.088967614 | 0.148933466 |
| Phf8 | -0.018683265 | 0.678701938 | 0.907783945 | 0.132635557 | 0.14940808 |
| Phgdh | -0.169516105 | 0.016856404 | 0.208334711 | 0.504926953 | 0.696172518 |
| Phip | -0.058461139 | 0.417782521 | 0.784482356 | 0.70009989 | 0.752521684 |
| Phka1 | -0.06310587 | 0.057169993 | 0.369250461 | 0.065955494 | 0.117300369 |
| Phka2 | -0.022949214 | 0.414911392 | 0.783085706 | 0.046213604 | 0.062994896 |
| Phkb | -0.06845357 | 0.067805311 | 0.395649928 | 0.085457703 | 0.140948153 |
| Phkg2 | -0.077369809 | 0.098076713 | 0.461507723 | 0.14143594 | 0.210499951 |
| Phlda1 | 0.09353546 | 0.048774145 | 0.346704032 | 0.153100175 | 0.069378839 |
| Phlda3 | 0.01779476 | 0.676727417 | 0.906695569 | 0.10315798 | 0.08719892 |
| Phldb2 | 0.005185871 | 0.936833319 | 0.98453816 | 0.386905254 | 0.386290818 |
| Phlpp1 | 0.038030584 | 0.288184189 | 0.694614177 | 0.088993376 | 0.059063604 |
| Phlpp2 | -0.030196247 | 0.475285219 | 0.817387411 | 0.113685424 | 0.145071869 |
| Phospho2 | 0.056492032 | 0.335308943 | 0.730736446 | 0.278833518 | 0.225476333 |

| | | | | | |
|---|---|---|---|---|---|
| Phpt1 | -0.040026559 | 0.268458962 | 0.676166273 | 0.083008558 | 0.11729523 |
| Phrf1 | 0.001963879 | 0.970794553 | 0.993892166 | 0.206441251 | 0.203428658 |
| Phtf1 | 0.004856162 | 0.912433395 | 0.97958006 | 0.134419014 | 0.129577255 |
| Phyh | -0.11255347 | 0.096231575 | 0.457111365 | 0.768803277 | 0.941226704 |
| Phyhip | -0.015271566 | 0.611915195 | 0.880165207 | 0.05378499 | 0.071898856 |
| Phyhipl | 0.074928278 | 0.098946905 | 0.463315751 | 0.151829259 | 0.087780438 |
| Phykpl | -0.037252968 | 0.241848016 | 0.652555608 | 0.061002425 | 0.096586392 |
| Pi4k2a | 0.023175833 | 0.737974161 | 0.922843014 | 0.49728143 | 0.456177233 |
| Pi4k2b | -0.030853637 | 0.522477162 | 0.840670611 | 0.151531961 | 0.190544165 |
| Pi4ka | 0.139176313 | 0.025081352 | 0.251741532 | 0.385024964 | 0.26888747 |
| Pi4kb | 0.089208205 | 0.065874502 | 0.390314712 | 0.164300842 | 0.083126901 |
| Pias1 | 0.022573954 | 0.677362419 | 0.906975267 | 0.243667597 | 0.226206562 |
| Pias2 | -0.021246825 | 0.686398479 | 0.909796728 | 0.207793929 | 0.229257454 |
| Pias3 | 0.031776351 | 0.371109119 | 0.75760494 | 0.086838624 | 0.057309943 |
| Pias4 | 0.005094379 | 0.923393296 | 0.980859278 | 0.201315125 | 0.198722251 |
| Pibf1 | -0.07714878 | 0.127276787 | 0.507536958 | 0.173079771 | 0.244956454 |
| Picalm | 0.11332485 | 0.107348557 | 0.47507667 | 0.567169522 | 0.444357985 |
| Pick1 | -0.079136674 | 0.131479223 | 0.511891761 | 0.186865883 | 0.251770784 |
| Pid1 | 0.04016107 | 0.318330365 | 0.717175454 | 0.111761332 | 0.08081277 |
| Piga | -0.059140174 | 0.032653187 | 0.286486051 | 0.041516877 | 0.098721427 |
| Pigbos1 | -0.052537466 | 0.397813712 | 0.774311756 | 0.360964326 | 0.402275372 |
| Pigc | 0.044191538 | 0.20625705 | 0.617602033 | 0.084875696 | 0.053478459 |
| Pigf | 0.046321486 | 0.291444856 | 0.697147764 | 0.130965792 | 0.094253469 |
| Pigg | -0.01863814 | 0.56155258 | 0.857134531 | 0.064366961 | 0.080709385 |
| Pigh | -0.040446167 | 0.325006177 | 0.720820254 | 0.105560803 | 0.138852072 |
| Pigk | -0.189553048 | 0.000717833 | 0.031383916 | 0.2360867 | 0.411167255 |
| Pigl | -0.036729983 | 0.062131786 | 0.380250374 | 0.016598623 | 0.047970168 |
| Pigm | -0.006304501 | 0.835901629 | 0.954911647 | 0.055441079 | 0.064527136 |
| Pigo | 0.070110846 | 0.016174265 | 0.205235264 | 0.059812478 | 0 |
| Pigp | -0.176375764 | 0.009745881 | 0.153321961 | 0.641648566 | 0.829418211 |
| Pigq | -0.084804129 | 0.07411977 | 0.412383666 | 0.149782498 | 0.230061081 |
| Pigr | 0.024507616 | 0.494100998 | 0.828842054 | 0.076411692 | 0.054680838 |
| Pigs | -0.043250672 | 0.274572747 | 0.682000399 | 0.100385035 | 0.136458879 |
| Pigt | -0.129649048 | 0.010771189 | 0.164475284 | 0.175072296 | 0.287450206 |
| Pigu | 0.002510244 | 0.951942426 | 0.98838165 | 0.109875118 | 0.114255083 |
| Pigv | -0.040911583 | 0.253922415 | 0.663146569 | 0.078779268 | 0.109919116 |
| Pigx | -0.074744485 | 0.166188792 | 0.568127031 | 0.2015699 | 0.270090461 |
| Pigyl | 0.171442328 | 0.008900998 | 0.146996602 | 0.517111675 | 0.337015337 |
| Pih1d1 | -0.02913747 | 0.388893151 | 0.767614661 | 0.072603516 | 0.095554178 |
| Pik3c2a | -0.010759377 | 0.875362275 | 0.965096393 | 0.49882608 | 0.506481169 |
| Pik3c3 | 0.025464423 | 0.55308049 | 0.852729475 | 0.126701144 | 0.105836797 |
| Pik3ca | 0.042606924 | 0.466702076 | 0.813152001 | 0.276711041 | 0.243507608 |
| Pik3cb | -0.059397026 | 0.041671779 | 0.321985867 | 0.04589785 | 0.096365484 |
| Pik3ip1 | -0.011430687 | 0.694178019 | 0.909986409 | 0.049403595 | 0.054409203 |
| Pik3r1 | -0.205058964 | 0.002467932 | 0.067311494 | 0.516888572 | 0.716004801 |
| Pik3r2 | 0.026661862 | 0.510591176 | 0.834957869 | 0.110912045 | 0.086072191 |
| Pik3r3 | -0.031440624 | 0.494209698 | 0.828842054 | 0.145747982 | 0.180017801 |
| Pik3r4 | 0.015619106 | 0.712247917 | 0.915109107 | 0.12014663 | 0.106348356 |
| Pikfyve | 0.096349366 | 0.097833566 | 0.460975742 | 0.320143595 | 0.248846051 |
| Pim1 | 0.05913369 | 0.13048937 | 0.51021496 | 0.102830993 | 0.055281436 |
| Pim2 | 0.041436592 | 0.478139369 | 0.818590256 | 0.252556985 | 0.201492633 |
| Pim3 | -0.065757894 | 0.181400281 | 0.589059475 | 0.162634275 | 0.229667517 |
| Pin1 | 0.07795265 | 0.260102208 | 0.667410092 | 0.67590424 | 0.596502878 |
| Pin4 | -0.040054646 | 0.527272051 | 0.842966059 | 0.407039971 | 0.423503641 |
| Pink1 | -0.125134538 | 0.021052213 | 0.229924991 | 0.216857391 | 0.336602722 |
| Pinx1 | 0.026585576 | 0.454700603 | 0.807876964 | 0.085264779 | 0.065401206 |
| Pip4k2a | 0.001729563 | 0.971381861 | 0.993892166 | 0.161261753 | 0.160107693 |
| Pip4k2b | -0.011504441 | 0.706147059 | 0.912857825 | 0.058372023 | 0.0680771 |
| Pip4k2c | 0.003603657 | 0.952482034 | 0.98838165 | 0.316256441 | 0.298475907 |
| Pip4p1 | 0.023505053 | 0.691742977 | 0.909796728 | 0.312825745 | 0.302738552 |
| Pip4p2 | 0.010757853 | 0.83207233 | 0.953695326 | 0.188825568 | 0.198926846 |
| Pip5k1a | 0.029094545 | 0.61809951 | 0.883238152 | 0.282176071 | 0.250363379 |
| Pip5k1b | -0.028331645 | 0.662115833 | 0.898950007 | 0.444942818 | 0.479481485 |
| Pip5k1c | -0.021030228 | 0.653149191 | 0.897175096 | 0.150688945 | 0.170727987 |
| Pir | -0.098294367 | 6.38E-05 | 0.005670007 | 0.024357757 | 0.10794654 |

| | | | | | |
|---|---|---|---|---|---|
| Pisd | -0.064724901 | 0.121452191 | 0.49613986 | 0.109026526 | 0.165843638 |
| Pithd1 | -0.156762875 | 0.0205807 | 0.227971987 | 0.475562619 | 0.638739304 |
| Pitpna | -0.034995646 | 0.587845108 | 0.869466748 | 0.404207421 | 0.444213668 |
| Pitpnb | -0.06797517 | 0.310114706 | 0.712953768 | 0.495251329 | 0.574731305 |
| Pitpnc1 | -0.004938914 | 0.93305064 | 0.983777411 | 0.264134149 | 0.270710091 |
| Pitpnm1 | -0.061887498 | 0.152553368 | 0.549597327 | 0.116288147 | 0.168227816 |
| Pitpnm2 | -0.041259812 | 0.478170237 | 0.818590256 | 0.264322606 | 0.308611179 |
| Pitpnm3 | 0.091384526 | 0.06718501 | 0.392969665 | 0.18215976 | 0.096568629 |
| Pitrm1 | -0.061703033 | 0.173021507 | 0.57541977 | 0.134176215 | 0.189942885 |
| Pja1 | -0.035344873 | 0.550894261 | 0.852505562 | 0.286482811 | 0.320663598 |
| Pja2 | 0.02638321 | 0.707519016 | 0.913534231 | 0.72491608 | 0.721921884 |
| Pkd1 | 0.071510929 | 0.250323128 | 0.662228053 | 0.35921083 | 0.281185306 |
| Pkd2 | 0.03646334 | 0.521528854 | 0.840205104 | 0.256137684 | 0.226153184 |
| Pkdcc | -0.137902264 | 0.063278232 | 0.383440442 | 0.650288677 | 0.807848324 |
| Pkia | 0.048075201 | 0.30399483 | 0.708427238 | 0.150921162 | 0.108246144 |
| Pkib | 0.031090857 | 0.570403434 | 0.86113346 | 0.228215825 | 0.18971326 |
| Pkig | -0.049630136 | 0.407395942 | 0.777888922 | 0.339867895 | 0.405119114 |
| Pkm | -0.027820961 | 0.639215738 | 0.891345382 | 1.453170049 | 1.503512327 |
| Pkn1 | 0.071632868 | 0.215519001 | 0.629611185 | 0.282062593 | 0.220809867 |
| Pkn2 | 0.125619222 | 0.055097073 | 0.365022657 | 0.975817073 | 0.825004825 |
| Pknox1 | 0.036423022 | 0.47301716 | 0.81673374 | 0.17894928 | 0.139015015 |
| Pkp2 | -0.030845631 | 0.212766593 | 0.627177178 | 0.03249627 | 0.063544593 |
| Pkp3 | 0.017370646 | 0.571059165 | 0.861225331 | 0.061825724 | 0.044581177 |
| Pkp4 | 0.030141317 | 0.655790908 | 0.897854773 | 0.693146903 | 0.6388726 |
| Pla2g12a | -0.123911164 | 0.068062949 | 0.396257435 | 0.502803021 | 0.643384082 |
| Pla2g15 | 0.003028737 | 0.932241827 | 0.98343913 | 0.083725589 | 0.075892837 |
| Pla2g2d | -0.015950471 | 0.557715969 | 0.855384435 | 0.041285951 | 0.05542765 |
| Pla2g2f | 0.112253141 | 0.097006743 | 0.45927202 | 0.390515676 | 0.285464486 |
| Pla2g6 | -0.061567309 | 0.112410162 | 0.483221516 | 0.093017478 | 0.147190001 |
| Plaa | 0.042394161 | 0.477349547 | 0.818492016 | 0.301795692 | 0.265084571 |
| Plaat3 | -0.014785798 | 0.784908859 | 0.936423418 | 0.208166323 | 0.213660414 |
| Plag1 | 0.011564483 | 0.727305593 | 0.920455328 | 0.072440765 | 0.060102448 |
| Plagl1 | 0.012201369 | 0.682534795 | 0.909170768 | 0.055768491 | 0.050290257 |
| Plagl2 | 0.03075376 | 0.397150103 | 0.774018561 | 0.091753782 | 0.069508852 |
| Platr25 | -0.021172328 | 0.432778467 | 0.795079432 | 0.041541688 | 0.058016607 |
| Plbd2 | -0.019998411 | 0.537548342 | 0.847793898 | 0.065166665 | 0.077489258 |
| Plcb1 | 0.022342331 | 0.680494041 | 0.907783945 | 0.21792477 | 0.196291368 |
| Plcb4 | 0.00649614 | 0.902349348 | 0.974696557 | 0.203972327 | 0.194449259 |
| Plcg1 | 0.01250797 | 0.766228667 | 0.92882091 | 0.114843033 | 0.097000426 |
| Plch1 | 0.009495878 | 0.788019532 | 0.937804093 | 0.082068463 | 0.074397128 |
| Plcl2 | -0.026121081 | 0.59018888 | 0.87075199 | 0.161685249 | 0.185270449 |
| Plcxd2 | -0.004870708 | 0.917384143 | 0.980132949 | 0.144116793 | 0.159709834 |
| Plcxd3 | -0.071887112 | 0.136243318 | 0.519952162 | 0.158993391 | 0.225277043 |
| Pld3 | -0.093816229 | 0.113281397 | 0.483626277 | 0.314485233 | 0.391095486 |
| Plec | 0.013651253 | 0.788121231 | 0.937804093 | 0.189073726 | 0.172207662 |
| Plekha1 | -0.026555407 | 0.685915473 | 0.909796728 | 0.498775752 | 0.53500824 |
| Plekha2 | 0.006190691 | 0.884880513 | 0.969283609 | 0.11845775 | 0.113219162 |
| Plekha3 | -0.085586733 | 0.07898276 | 0.421675512 | 0.167986193 | 0.24969944 |
| Plekha5 | -0.173873443 | 0.001154737 | 0.041993113 | 0.188363517 | 0.352972323 |
| Plekha6 | -0.037610963 | 0.52999028 | 0.844635192 | 0.322013222 | 0.342737957 |
| Plekha8 | -0.017771497 | 0.647389302 | 0.895376633 | 0.099067078 | 0.113219983 |
| Plekhb1 | -0.048881706 | 0.022194133 | 0.236913658 | 0.019657813 | 0.05772442 |
| Plekhb2 | -0.010279496 | 0.856908787 | 0.958302663 | 0.24714198 | 0.25380679 |
| Plekhf2 | 0.018903782 | 0.630943366 | 0.888561606 | 0.104216441 | 0.093952943 |
| Plekhg3 | 0.057838876 | 0.23372149 | 0.644459522 | 0.166493878 | 0.116182186 |
| Plekhh1 | 0.031503648 | 0.402202475 | 0.77525778 | 0.093373543 | 0.067585563 |
| Plekhh2 | -0.016206025 | 0.618591577 | 0.883563919 | 0.065713571 | 0.075740284 |
| Plekhh3 | -0.058145195 | 0.129989774 | 0.509514066 | 0.094960472 | 0.151080849 |
| Plekhj1 | -0.073577873 | 0.256644818 | 0.665256942 | 0.444755131 | 0.51399971 |
| Plekhm1 | -0.033013485 | 0.260361024 | 0.667665713 | 0.049467504 | 0.078686208 |
| Plekhm2 | -0.003723697 | 0.922796003 | 0.980859278 | 0.09423342 | 0.099694383 |
| Plekhm3 | 0.023048673 | 0.499322576 | 0.830722138 | 0.080038864 | 0.06536816 |
| Plet1 | 0.0760243 | 0.049938541 | 0.351742555 | 0.095476986 | 0.027000433 |
| Plgrkt | -0.014788502 | 0.795304103 | 0.939757086 | 0.262379689 | 0.267889037 |
| Plin2 | 0.066041758 | 0.154087668 | 0.550967131 | 0.146648291 | 0.082728256 |

| | | | | | |
|---|---|---|---|---|---|
| Plk3 | 0.002130256 | 0.97294483 | 0.994079882 | 0.311163716 | 0.318075858 |
| Plod1 | 0.044997351 | 0.171543926 | 0.574479873 | 0.071832748 | 0.028107197 |
| Plod3 | -0.087995167 | 0.034500706 | 0.293212179 | 0.101208542 | 0.177457723 |
| Plp2 | 0.041749169 | 0.370123659 | 0.757234136 | 0.149709709 | 0.11596648 |
| Plpbp | -0.066925373 | 0.284720884 | 0.692115164 | 0.363891754 | 0.421136663 |
| Plpp2 | 0.033488562 | 0.421208614 | 0.787275765 | 0.115607684 | 0.092226106 |
| Plpp5 | -0.070643827 | 0.257422189 | 0.665256942 | 0.34975247 | 0.425704838 |
| Plpp6 | -0.066863913 | 0.129493434 | 0.509296863 | 0.124638474 | 0.186461896 |
| Plpp7 | -0.023165553 | 0.428029674 | 0.791658813 | 0.050084511 | 0.068527544 |
| Plppr1 | -0.016831149 | 0.492981208 | 0.828403211 | 0.034318693 | 0.047042846 |
| Plppr2 | -0.023254669 | 0.400122522 | 0.774697097 | 0.046797788 | 0.066746593 |
| Plppr4 | 0.050871346 | 0.259734468 | 0.667102369 | 0.137687859 | 0.09008042 |
| Plrg1 | -0.034851886 | 0.386228429 | 0.766272928 | 0.103771152 | 0.136224424 |
| Pls1 | -0.038659735 | 0.189336743 | 0.597789521 | 0.049408051 | 0.079989462 |
| Pls3 | -0.041939061 | 0.186216496 | 0.593124825 | 0.059626955 | 0.094842838 |
| Plscr1 | -0.038617332 | 0.443287526 | 0.801368502 | 0.167600299 | 0.200669038 |
| Plscr3 | -0.012035769 | 0.755989808 | 0.926726954 | 0.098767558 | 0.106526897 |
| Plut | 0.001285506 | 0.980753584 | 0.995540893 | 0.175483885 | 0.200089689 |
| Plxna2 | 0.117429043 | 0.051703489 | 0.355933715 | 0.302215509 | 0.176180336 |
| Plxnb1 | -0.020552127 | 0.551782614 | 0.852505562 | 0.074842358 | 0.087830191 |
| Plxnb2 | -0.066069918 | 0.328398387 | 0.723428541 | 0.5689012 | 0.609510942 |
| Pmepa1 | 0.128981501 | 0.041595038 | 0.321640523 | 0.327022503 | 0.214732667 |
| Pml | 0.032476403 | 0.379182674 | 0.762006446 | 0.090617784 | 0.053268652 |
| Pmm1 | -0.072796481 | 0.177402411 | 0.582653141 | 0.201028125 | 0.293070218 |
| Pmm2 | -0.052666875 | 0.246882572 | 0.656587274 | 0.13316626 | 0.171913712 |
| Pmp22 | 0.064313566 | 0.034994978 | 0.294903156 | 0.061239458 | 0.010952459 |
| Pmpca | -0.026852362 | 0.579457642 | 0.865234762 | 0.160531851 | 0.185064886 |
| Pmpcb | 0.012643488 | 0.836430267 | 0.954911647 | 0.318365233 | 0.31385477 |
| Pms2 | 0.040177605 | 0.186263769 | 0.593124825 | 0.058971499 | 0.026300682 |
| Pmvk | -0.136547428 | 0.060762744 | 0.380221737 | 0.557365083 | 0.705565364 |
| Pnck | 0.024739857 | 0.379411984 | 0.762006446 | 0.052126023 | 0.031057342 |
| Pnisr | -0.074297993 | 0.277984238 | 0.684996416 | 0.680890307 | 0.747246989 |
| Pnkd | -0.153661068 | 0.022541927 | 0.238161396 | 0.496390645 | 0.650631467 |
| Pnkp | -0.014645889 | 0.720830925 | 0.919412687 | 0.111601909 | 0.124144031 |
| Pnmal1 | -0.132976783 | 0.034365539 | 0.293212179 | 0.30901265 | 0.428629606 |
| Pnmal2 | -0.086675216 | 0.215471672 | 0.629611185 | 0.783781026 | 0.903351046 |
| Pnn | 0.107915998 | 0.127770142 | 0.508260084 | 0.917469203 | 0.777268526 |
| Pno1 | -0.129665042 | 0.02485849 | 0.250506687 | 0.237282712 | 0.370429185 |
| Pnp | -0.014573088 | 0.788121703 | 0.937804093 | 0.212762883 | 0.239027539 |
| Pnpla2 | -0.015185565 | 0.771891311 | 0.932402174 | 0.186760829 | 0.195956098 |
| Pnpla6 | -0.019686124 | 0.499483091 | 0.830722138 | 0.050228159 | 0.066743908 |
| Pnpla8 | -0.101405514 | 0.120055884 | 0.493583757 | 0.577440423 | 0.682989802 |
| Pnpo | -0.042204073 | 0.477188335 | 0.818444859 | 0.284532555 | 0.313666572 |
| Pnpt1 | -0.006228659 | 0.857170655 | 0.958302663 | 0.075476477 | 0.077083422 |
| Pnrc1 | -0.305801506 | 1.61E-05 | 0.001832428 | 0.66082037 | 1.046783479 |
| Pnrc2 | -0.145084324 | 0.026984752 | 0.261686913 | 0.392250381 | 0.527231378 |
| Poc1b | 0.000886476 | 0.981939833 | 0.995803987 | 0.103049461 | 0.103995319 |
| Poc5 | 0.021492075 | 0.58195953 | 0.866173066 | 0.103924023 | 0.083470151 |
| Podxl2 | -0.015187101 | 0.775064174 | 0.933555636 | 0.204250402 | 0.213501271 |
| Pofut1 | 0.003983445 | 0.894996727 | 0.972229731 | 0.055333542 | 0.048495305 |
| Pofut2 | 0.128136915 | 0.035990628 | 0.298790684 | 1.112856099 | 0.923349449 |
| Pogk | -0.043176749 | 0.428922233 | 0.792251095 | 0.218005794 | 0.255789615 |
| Poglut1 | -0.062856513 | 0.089889057 | 0.445385778 | 0.08632063 | 0.144145233 |
| Poglut3 | 0.07481172 | 0.051995269 | 0.35690059 | 0.106869351 | 0.040585867 |
| Pogz | -0.034453895 | 0.448126022 | 0.804951652 | 0.137450219 | 0.169870513 |
| Polb | -0.069746083 | 0.220221663 | 0.633234352 | 0.254948248 | 0.319870963 |
| Pold2 | -0.00626233 | 0.84045925 | 0.954911647 | 0.060260548 | 0.069124307 |
| Pold3 | 0.010400115 | 0.838870349 | 0.954911647 | 0.194284714 | 0.187799621 |
| Pold4 | -0.043179727 | 0.460968865 | 0.810249429 | 0.276186034 | 0.337412382 |
| Poldip2 | -0.027496294 | 0.607134213 | 0.878576426 | 0.212487334 | 0.241301017 |
| Poldip3 | -0.000444737 | 0.993918684 | 0.998695538 | 0.279902846 | 0.287347609 |
| Pole3 | -0.005973688 | 0.903435164 | 0.974950771 | 0.159492181 | 0.1670827 |
| Pole4 | -0.072172897 | 0.027847558 | 0.26510449 | 0.06111167 | 0.122362634 |
| Polg | -0.043377627 | 0.292923201 | 0.698685876 | 0.105723385 | 0.14101882 |
| Polk | 0.079862168 | 0.045931738 | 0.336263207 | 0.113143527 | 0.043812452 |

| | | | | | |
|---|---|---|---|---|---|
| Poll | 0.003015574 | 0.920502868 | 0.980436936 | 0.05645704 | 0.055574362 |
| Polr1a | -0.01285498 | 0.702085777 | 0.912152193 | 0.066924842 | 0.076765178 |
| Polr1b | -0.022983533 | 0.34982131 | 0.7406656 | 0.03194961 | 0.050496467 |
| Polr1c | 0.072638094 | 0.193909702 | 0.60325934 | 0.227651018 | 0.17138089 |
| Polr1d | -0.023655878 | 0.713004243 | 0.915509496 | 0.969605075 | 1.031447883 |
| Polr1f | -0.01331595 | 0.797437116 | 0.940690722 | 0.190975682 | 0.207710091 |
| Polr1h | -0.038296067 | 0.572918654 | 0.861868184 | 0.452602891 | 0.488321529 |
| Polr1has | 0.077166533 | 0.093447533 | 0.452015851 | 0.150871574 | 0.084602458 |
| Polr2a | 0.101020182 | 0.134714688 | 0.517714432 | 0.493105915 | 0.400824232 |
| Polr2b | -0.022184151 | 0.705968055 | 0.912844672 | 0.2946674 | 0.317632257 |
| Polr2c | 0.014894528 | 0.82451436 | 0.951625061 | 0.607044468 | 0.594939398 |
| Polr2d | 0.026499463 | 0.575540481 | 0.863006716 | 0.150252251 | 0.128403746 |
| Polr2e | 0.018335849 | 0.783551989 | 0.936124803 | 0.532738181 | 0.513229336 |
| Polr2f | -0.049634237 | 0.463851545 | 0.811254677 | 0.686361305 | 0.731050066 |
| Polr2g | -0.010399486 | 0.874875615 | 0.96500153 | 0.752076165 | 0.749018945 |
| Polr2h | 0.055113392 | 0.341771658 | 0.735000423 | 0.268197052 | 0.228518724 |
| Polr2i | 0.045544179 | 0.493518532 | 0.82875113 | 0.570585067 | 0.511527388 |
| Polr2j | 0.093154158 | 0.170609639 | 0.572831081 | 0.710252388 | 0.58011333 |
| Polr2k | -0.068765929 | 0.321659922 | 0.719227395 | 0.614430684 | 0.69691533 |
| Polr2l | 0.028841376 | 0.569668899 | 0.860834533 | 0.187727694 | 0.171649486 |
| Polr2m | -0.070076218 | 0.284949245 | 0.692407818 | 0.560565097 | 0.657565857 |
| Polr3a | 0.024932802 | 0.524200847 | 0.841062594 | 0.098136077 | 0.078486707 |
| Polr3c | -5.70E-05 | 0.998700057 | 0.999466149 | 0.079657967 | 0.085602046 |
| Polr3d | -0.077392935 | 0.200288898 | 0.610852219 | 0.298790801 | 0.380953612 |
| Polr3e | -0.052614079 | 0.14388052 | 0.533279057 | 0.07714142 | 0.128208223 |
| Polr3f | 0.002608908 | 0.955110706 | 0.989278596 | 0.142878569 | 0.136903022 |
| Polr3g | -0.024144144 | 0.356629204 | 0.745571198 | 0.039036838 | 0.063355829 |
| Polr3gl | -0.05515569 | 0.23988685 | 0.650744882 | 0.148277261 | 0.200681076 |
| Polr3h | -0.038732386 | 0.338120884 | 0.732352032 | 0.10363427 | 0.139510327 |
| Polr3k | -0.01267848 | 0.838248046 | 0.954911647 | 0.309229041 | 0.321577224 |
| Polrmt | -0.005111662 | 0.843773312 | 0.955524266 | 0.041043754 | 0.04525202 |
| Pom121 | 0.003243587 | 0.960912331 | 0.991375098 | 0.481881295 | 0.4767745 |
| Pomc | -0.04042893 | 0.43483093 | 0.796720977 | 0.191450379 | 0.222302561 |
| Pomgnt1 | -0.046162015 | 0.302153007 | 0.707585098 | 0.126682581 | 0.176453943 |
| Pomgnt2 | -0.00538343 | 0.883991227 | 0.968954079 | 0.087188464 | 0.086061586 |
| Pomp | -0.079022028 | 0.226158292 | 0.638587289 | 1.092983674 | 1.197536688 |
| Pomt1 | -0.005975602 | 0.81605847 | 0.949036482 | 0.040202792 | 0.046991868 |
| Pomt2 | -0.065284962 | 0.093663665 | 0.45227176 | 0.096159846 | 0.157514409 |
| Pon2 | -0.1262407 | 0.002615801 | 0.069641377 | 0.098553811 | 0.216100258 |
| Pon3 | -0.073158336 | 0.013783652 | 0.189915708 | 0.044365317 | 0.110528111 |
| Pop4 | 0.033300111 | 0.276723151 | 0.684050613 | 0.062572749 | 0.031930492 |
| Pop5 | 0.176838187 | 0.007045825 | 0.128637347 | 0.954819476 | 0.741976412 |
| Pop7 | -6.05E-06 | 0.999920618 | 0.99998562 | 0.333726577 | 0.345303262 |
| Por | 0.03784096 | 0.579803081 | 0.865479406 | 0.721627858 | 0.672599988 |
| Pot1a | -0.053644812 | 0.074721332 | 0.41344108 | 0.051547348 | 0.094796987 |
| Pou2f1 | -0.087349173 | 0.048269181 | 0.34433388 | 0.119881358 | 0.193694174 |
| Pou6f1 | 0.040002233 | 0.305537654 | 0.709411642 | 0.105804521 | 0.070870246 |
| Ppa1 | 0.020046732 | 0.750733299 | 0.926186851 | 0.434585918 | 0.412436804 |
| Ppa2 | 0.065741633 | 0.196832508 | 0.60713211 | 0.195966139 | 0.143081808 |
| Ppan | 0.056289085 | 0.240061633 | 0.650744882 | 0.160939971 | 0.11489411 |
| Ppard | 0.053461733 | 0.197286217 | 0.607747501 | 0.126740244 | 0.082453895 |
| Pparg | -0.016236811 | 0.708745845 | 0.913767764 | 0.12501742 | 0.153816244 |
| Ppat | 0.165463731 | 0.023949858 | 0.245792155 | 0.647930857 | 0.463309584 |
| Ppcs | 0.110295039 | 0.04537287 | 0.333876461 | 0.22782007 | 0.130412177 |
| Ppdpf | -0.039418191 | 0.503911068 | 0.832249514 | 0.314611173 | 0.351593154 |
| Ppfia1 | 0.066771563 | 0.31960652 | 0.717256849 | 0.492294095 | 0.425006706 |
| Ppfia3 | -0.0188815 | 0.472930031 | 0.816723798 | 0.037858464 | 0.052637419 |
| Ppfibp1 | -0.011387773 | 0.852405726 | 0.957461155 | 0.334345873 | 0.346029455 |
| Pphln1 | -0.030914354 | 0.51176291 | 0.835213712 | 0.15302349 | 0.189441617 |
| Ppia | -0.097496005 | 0.052608339 | 0.358472435 | 1.767205022 | 1.9891572 |
| Ppib | 0.051837153 | 0.453845402 | 0.807844867 | 0.867411434 | 0.793225933 |
| Ppid | 0.004041691 | 0.947455975 | 0.987651726 | 0.313039126 | 0.312587468 |
| Ppie | -0.054441744 | 0.186761413 | 0.593955736 | 0.106566818 | 0.161822791 |
| Ppif | 0.021499263 | 0.54353888 | 0.850865857 | 0.083008423 | 0.067913535 |
| Ppig | -0.008167781 | 0.900702181 | 0.974070444 | 1.083468006 | 1.053752898 |

| | | | | | |
|---|---|---|---|---|---|
| Ppih | -0.022270607 | 0.509930558 | 0.834904701 | 0.072469449 | 0.098880602 |
| Ppil1 | 0.063388269 | 0.089610385 | 0.445036831 | 0.096874677 | 0.0440354 |
| Ppil2 | -0.030934081 | 0.551059863 | 0.852505562 | 0.183754842 | 0.210400089 |
| Ppil3 | 0.059084692 | 0.101690026 | 0.467719786 | 0.091727119 | 0.040453174 |
| Ppil4 | 0.084965369 | 0.188373017 | 0.596623533 | 0.365631667 | 0.276457496 |
| Ppil6 | -0.028685359 | 0.262083199 | 0.668764912 | 0.034633158 | 0.056179474 |
| Ppip5k1 | 0.029947643 | 0.574321133 | 0.862168892 | 0.20943991 | 0.200276686 |
| Ppip5k2 | 0.054089998 | 0.280137438 | 0.687465228 | 0.179067358 | 0.125375942 |
| Ppl | 0.032634988 | 0.417717217 | 0.784482356 | 0.106415364 | 0.079350911 |
| Ppm1a | -0.075216027 | 0.256009405 | 0.664829597 | 0.538416347 | 0.621049778 |
| Ppm1b | -0.056476229 | 0.410839066 | 0.78053979 | 0.633949193 | 0.71959027 |
| Ppm1d | -0.00439633 | 0.923703998 | 0.980873574 | 0.137407729 | 0.146803711 |
| Ppm1e | 0.002579498 | 0.939077418 | 0.985520707 | 0.073274572 | 0.074741852 |
| Ppm1f | -0.081144161 | 0.00147201 | 0.049914078 | 0.029077813 | 0.099371184 |
| Ppm1g | 0.06444902 | 0.320107491 | 0.717489703 | 0.438738032 | 0.371865575 |
| Ppm1h | 0.027991209 | 0.565542617 | 0.858087314 | 0.166221211 | 0.152058138 |
| Ppm1k | -0.006303134 | 0.888552725 | 0.970176435 | 0.135827709 | 0.142382274 |
| Ppm1l | -0.00390214 | 0.953969863 | 0.989066902 | 0.598191604 | 0.598635101 |
| Ppme1 | -0.082122767 | 0.201434518 | 0.611924411 | 0.418391878 | 0.483544355 |
| Ppox | 0.053857648 | 0.135019223 | 0.518167175 | 0.091632521 | 0.047580922 |
| Ppp1ca | -0.054663718 | 0.392014995 | 0.769704269 | 0.909603078 | 0.982310679 |
| Ppp1cb | -0.076361645 | 0.22749347 | 0.639441223 | 1.062049774 | 1.160854657 |
| Ppp1cc | 0.024514102 | 0.671530249 | 0.904877555 | 0.263435364 | 0.261561055 |
| Ppp1r10 | -0.004552919 | 0.947964438 | 0.987651726 | 0.574051923 | 0.57024743 |
| Ppp1r11 | -0.034248811 | 0.607980114 | 0.87877121 | 0.700715857 | 0.738006204 |
| Ppp1r12a | -0.059823699 | 0.386960473 | 0.766664482 | 0.541399281 | 0.588144964 |
| Ppp1r12b | 0.019324576 | 0.633090695 | 0.889464068 | 0.112675857 | 0.09136177 |
| Ppp1r12c | 0.120947461 | 0.033607968 | 0.290545455 | 0.280234827 | 0.16918389 |
| Ppp1r13b | 0.036486019 | 0.533641489 | 0.84612966 | 0.295329136 | 0.265459605 |
| Ppp1r14b | -0.115842123 | 0.096393015 | 0.457336038 | 0.643988248 | 0.769469535 |
| Ppp1r15a | -0.339186828 | 9.73E-06 | 0.001268866 | 0.590306046 | 1.028534683 |
| Ppp1r15b | 0.043908054 | 0.470943532 | 0.814586466 | 0.346007778 | 0.316169668 |
| Ppp1r16a | -0.031095217 | 0.421393065 | 0.787357117 | 0.09954124 | 0.124077345 |
| Ppp1r1a | 0.156654821 | 0.02097188 | 0.229383208 | 1.265163518 | 1.071174041 |
| Ppp1r2 | 0.060302367 | 0.338613875 | 0.732629331 | 0.977139523 | 0.9372043 |
| Ppp1r21 | 0.0584592 | 0.212984028 | 0.627539904 | 0.151200119 | 0.100700111 |
| Ppp1r35 | 0.013817841 | 0.800502539 | 0.941927782 | 0.235066701 | 0.225094091 |
| Ppp1r37 | 0.00550395 | 0.931962655 | 0.98340088 | 0.46208444 | 0.460830961 |
| Ppp1r7 | -0.015709203 | 0.768961311 | 0.931268476 | 0.218892399 | 0.225110405 |
| Ppp1r8 | 0.009185649 | 0.842084487 | 0.955188334 | 0.144090354 | 0.137715232 |
| Ppp1r9a | -0.097662091 | 0.060674754 | 0.380221737 | 0.170351698 | 0.258107383 |
| Ppp1r9b | 0.084836202 | 0.090685217 | 0.445385778 | 0.191158215 | 0.115985346 |
| Ppp2ca | 0.002245561 | 0.973542126 | 0.994079882 | 0.828713131 | 0.813553048 |
| Ppp2cb | -0.128379893 | 0.025849915 | 0.256379045 | 0.253125935 | 0.3888544 |
| Ppp2r1a | -0.032532039 | 0.610307071 | 0.879490606 | 0.437700608 | 0.480033843 |
| Ppp2r1b | -0.087235851 | 0.02029584 | 0.227608207 | 0.084544527 | 0.165156351 |
| Ppp2r2a | 0.057697095 | 0.328258455 | 0.72332165 | 0.279667587 | 0.220696174 |
| Ppp2r2d | 0.160408736 | 0.022750068 | 0.238404929 | 0.744570027 | 0.513970769 |
| Ppp2r3a | 0.111383353 | 0.09563762 | 0.455869783 | 0.441773868 | 0.319909107 |
| Ppp2r3c | -0.069648779 | 0.152739545 | 0.549602359 | 0.15519026 | 0.212682152 |
| Ppp2r3d | -0.003221218 | 0.937370027 | 0.984964715 | 0.112523488 | 0.120380453 |
| Ppp2r5a | -0.071670716 | 0.288918527 | 0.69541373 | 0.700385036 | 0.795867129 |
| Ppp2r5b | 0.009428317 | 0.855528297 | 0.957736324 | 0.192439638 | 0.181224925 |
| Ppp2r5c | -0.047907715 | 0.463823977 | 0.811254677 | 0.585316617 | 0.622621561 |
| Ppp2r5d | -0.038182655 | 0.308942761 | 0.711182121 | 0.085076294 | 0.114904561 |
| Ppp2r5e | 0.008766626 | 0.889377037 | 0.970291013 | 0.372576184 | 0.368409255 |
| Ppp3ca | 0.040322302 | 0.546707529 | 0.851446616 | 0.424287678 | 0.3841059 |
| Ppp3cb | -0.036192023 | 0.579936756 | 0.865550219 | 0.43949818 | 0.473614243 |
| Ppp3cc | 0.038884503 | 0.190986686 | 0.601358592 | 0.059433993 | 0.026759151 |
| Ppp3r1 | 0.082224127 | 0.209681759 | 0.622606671 | 0.441068686 | 0.368219949 |
| Ppp4c | 0.019678351 | 0.744971081 | 0.924566547 | 0.3027231 | 0.293108054 |
| Ppp4r1 | 0.069136369 | 0.066834612 | 0.392751172 | 0.09778545 | 0.035112946 |
| Ppp4r1l-ps | -0.078764956 | 0.23742086 | 0.64861001 | 0.499346625 | 0.594543616 |
| Ppp4r2 | 0.106936941 | 0.1045341 | 0.471978751 | 0.493730328 | 0.373946899 |
| Ppp4r3a | 0.03948222 | 0.536430751 | 0.847520662 | 0.38995159 | 0.350158121 |

| | | | | | |
|---|---|---|---|---|---|
| Ppp4r3b | -0.05504258 | 0.41012203 | 0.780101937 | 0.466394363 | 0.517372135 |
| Ppp4r4 | 0.034617653 | 0.297775317 | 0.702656882 | 0.075445328 | 0.045187483 |
| Ppp5c | -0.048736926 | 0.357272273 | 0.745983316 | 0.201673078 | 0.252459011 |
| Ppp6c | 0.064548833 | 0.311845138 | 0.71405691 | 0.411240091 | 0.359329701 |
| Ppp6r1 | 0.03226916 | 0.551332153 | 0.852505562 | 0.21391658 | 0.179318935 |
| Ppp6r2 | -0.00835135 | 0.857088254 | 0.958302663 | 0.143988365 | 0.155447892 |
| Ppp6r3 | 0.023557741 | 0.70114923 | 0.912152193 | 0.340243192 | 0.321029105 |
| Pprc1 | 0.011883641 | 0.713854889 | 0.915840939 | 0.069492414 | 0.059829539 |
| Ppt1 | -0.064553468 | 0.200086063 | 0.610600127 | 0.170044639 | 0.225411471 |
| Ppt2 | -0.012612187 | 0.665050933 | 0.900946986 | 0.049008675 | 0.057796426 |
| Pptc7 | 0.059077737 | 0.246668743 | 0.656366431 | 0.184189531 | 0.128483095 |
| Ppwd1 | 0.026345837 | 0.646649445 | 0.894983519 | 0.256723934 | 0.239873337 |
| Ppy | 0.217944435 | 0.003374585 | 0.081813298 | 2.070059588 | 1.507232105 |
| Pqbp1 | -0.040521986 | 0.453635154 | 0.807844867 | 0.204727867 | 0.251268071 |
| Pradc1 | 0.029426939 | 0.435514288 | 0.796975049 | 0.094207384 | 0.068417 |
| Praf2 | 0.0420918 | 0.13085508 | 0.510610033 | 0.052181709 | 0.020890146 |
| Pramel12 | 0.050109473 | 0.138200463 | 0.523836119 | 0.082001929 | 0.044524632 |
| Prcc | 0.045132107 | 0.301745609 | 0.706960943 | 0.135341869 | 0.10048914 |
| Prdm10 | 0.029861646 | 0.338384768 | 0.732353334 | 0.065597715 | 0.037984892 |
| Prdm15 | 0.013685114 | 0.63372405 | 0.889606754 | 0.051224673 | 0.040221634 |
| Prdm2 | -0.018443471 | 0.763106143 | 0.928730554 | 0.351406081 | 0.363120333 |
| Prdm4 | 0.040619521 | 0.170396533 | 0.572762895 | 0.058308518 | 0.022135632 |
| Prdx1 | -0.168468386 | 0.00934595 | 0.14984873 | 1.3270712 | 1.612794007 |
| Prdx2 | -0.047632286 | 0.477909059 | 0.818590256 | 1.141516438 | 1.199130565 |
| Prdx3 | -0.072631815 | 0.225117378 | 0.637377467 | 0.276836734 | 0.357283279 |
| Prdx4 | -0.095793115 | 0.037556062 | 0.30568979 | 0.12831152 | 0.219288238 |
| Prdx5 | -0.064121795 | 0.314246902 | 0.714975653 | 0.943464772 | 1.009112685 |
| Prdx6 | 0.033598464 | 0.610404444 | 0.879504652 | 1.006006344 | 0.972153702 |
| Preb | -0.033248856 | 0.593447344 | 0.872481469 | 0.350580111 | 0.383352153 |
| Prelid1 | -0.037381607 | 0.549923032 | 0.852459771 | 0.406199604 | 0.436124581 |
| Prelid3b | 0.011358239 | 0.849439089 | 0.957111129 | 0.302109953 | 0.300569886 |
| Prep | 0.014423792 | 0.679166407 | 0.907783945 | 0.079431369 | 0.065578833 |
| Prepl | -0.011783534 | 0.85368625 | 0.957461155 | 0.431047114 | 0.425255285 |
| Prickle1 | -0.039384971 | 0.327600558 | 0.722347716 | 0.101378671 | 0.139389813 |
| Prim1 | -0.003262268 | 0.919839002 | 0.980436936 | 0.070378224 | 0.06984531 |
| Prkaa1 | 0.061912861 | 0.312314205 | 0.714349311 | 0.333414932 | 0.283864471 |
| Prkaa2 | 0.030870524 | 0.338917216 | 0.732653908 | 0.071010081 | 0.048793236 |
| Prkab1 | 0.036719691 | 0.387836886 | 0.767160379 | 0.122952786 | 0.095496932 |
| Prkab2 | 0.04692603 | 0.346505552 | 0.738383053 | 0.165766041 | 0.119630527 |
| Prkaca | -0.050457269 | 0.405454108 | 0.777075211 | 0.344613154 | 0.402558063 |
| Prkacb | -0.045126935 | 0.476410053 | 0.817948631 | 1.206372718 | 1.240779987 |
| Prkag1 | 0.028980662 | 0.595275919 | 0.873342705 | 0.220631916 | 0.197994743 |
| Prkag2 | 0.029616334 | 0.592300657 | 0.872005806 | 0.233337459 | 0.200674309 |
| Prkar1a | -0.085253379 | 0.109623691 | 0.477972625 | 1.761638426 | 1.917813836 |
| Prkar1b | -0.005116441 | 0.935389433 | 0.984268477 | 0.377926871 | 0.384205425 |
| Prkar2a | 0.082405776 | 0.205801597 | 0.616737719 | 0.446097722 | 0.365530945 |
| Prkar2b | 0.038305004 | 0.516150569 | 0.837427561 | 0.276227687 | 0.240370219 |
| Prkca | 0.091517161 | 0.169176361 | 0.57227323 | 0.498978629 | 0.406056022 |
| Prkcb | -0.013930572 | 0.827230426 | 0.952715064 | 0.374492668 | 0.376334383 |
| Prkcd | -0.008360406 | 0.771662552 | 0.9322553 | 0.050782462 | 0.061046365 |
| Prkce | 0.019396171 | 0.736944934 | 0.92276204 | 0.288644575 | 0.273592962 |
| Prkci | 0.004585254 | 0.910739021 | 0.978717991 | 0.112822668 | 0.110111668 |
| Prkcsh | -0.221367031 | 0.000766188 | 0.032691088 | 0.66032711 | 0.893354433 |
| Prkcz | -0.063258386 | 0.129044144 | 0.508493946 | 0.11280433 | 0.165261604 |
| Prkd1 | -0.017155397 | 0.71121211 | 0.914546911 | 0.148048912 | 0.15818185 |
| Prkd2 | -0.000674433 | 0.984373597 | 0.99608406 | 0.076684656 | 0.082509674 |
| Prkdc | 0.002775628 | 0.930534707 | 0.982720628 | 0.06106807 | 0.055280366 |
| Prkn | -0.04460577 | 0.151264421 | 0.546523037 | 0.055738476 | 0.092306362 |
| Prkra | -0.001721669 | 0.964989365 | 0.991970325 | 0.101422252 | 0.104675328 |
| Prkrip1 | 0.118476477 | 0.032127808 | 0.284111724 | 0.240474044 | 0.133181257 |
| Prkx | 0.024417825 | 0.595599364 | 0.873342705 | 0.141472073 | 0.118470161 |
| Prlr | -0.199863394 | 0.00232928 | 0.06554117 | 1.794508047 | 2.170386272 |
| Prmt1 | -0.121586821 | 0.054892656 | 0.364775849 | 0.354214562 | 0.468496335 |
| Prmt2 | -0.036201528 | 0.219592313 | 0.63289494 | 0.049891491 | 0.0853596 |
| Prmt3 | 0.017300356 | 0.649595254 | 0.895585658 | 0.098656679 | 0.083887304 |

| | | | | | |
|---|---|---|---|---|---|
| Prmt5 | 0.003783281 | 0.923256143 | 0.980859278 | 0.099057413 | 0.096096455 |
| Prmt7 | -0.089573681 | 0.01818959 | 0.215000801 | 0.084680811 | 0.162034041 |
| Prmt9 | 0.006904015 | 0.838346031 | 0.954911647 | 0.074418889 | 0.070956171 |
| Prnp | -0.130498612 | 0.009765444 | 0.153389301 | 1.728852339 | 1.981150728 |
| Prob1 | -0.004684651 | 0.866818886 | 0.962125879 | 0.04750801 | 0.050464906 |
| Prom1 | -0.058357775 | 0.369337042 | 0.756537937 | 0.377749568 | 0.432644462 |
| Prorp | 0.024067683 | 0.459076134 | 0.809370658 | 0.072755571 | 0.049030485 |
| Prorsd1 | -0.077543628 | 0.240488912 | 0.650744882 | 0.482710136 | 0.559648585 |
| Proscos | 0.075306262 | 0.060836567 | 0.380250374 | 0.107551323 | 0.043551322 |
| Proser1 | 0.000654799 | 0.988806526 | 0.997411191 | 0.144627784 | 0.14421782 |
| Prox1 | -0.005464495 | 0.932913986 | 0.983777411 | 0.364668316 | 0.373134222 |
| Prpf18 | 0.011839886 | 0.810339069 | 0.948166616 | 0.162197765 | 0.150170128 |
| Prpf19 | 0.026515381 | 0.686828553 | 0.909796728 | 0.465226385 | 0.448067088 |
| Prpf3 | 0.040990768 | 0.312408126 | 0.714349311 | 0.113096106 | 0.075751852 |
| Prpf31 | 0.084160913 | 0.061522037 | 0.380250374 | 0.144813432 | 0.070452474 |
| Prpf38a | 0.03297675 | 0.566359981 | 0.858732162 | 0.270833906 | 0.237409678 |
| Prpf38b | -0.021691227 | 0.756914435 | 0.92705828 | 0.502925109 | 0.50430158 |
| Prpf39 | -0.060692497 | 0.275616858 | 0.68288482 | 0.220539927 | 0.279802776 |
| Prpf4 | 0.090315467 | 0.089753195 | 0.445385778 | 0.220242666 | 0.137020555 |
| Prpf40a | -0.051690038 | 0.458430184 | 0.809370658 | 0.786089523 | 0.835079555 |
| Prpf40b | -0.068777365 | 0.044546198 | 0.330723271 | 0.07151504 | 0.125320458 |
| Prpf4b | -0.09919265 | 0.160972041 | 0.55925108 | 1.076055814 | 1.18022613 |
| Prpf6 | 0.074090232 | 0.248974277 | 0.660574894 | 0.433966371 | 0.368052531 |
| Prpf8 | -0.10645187 | 0.120610356 | 0.494691316 | 0.596608063 | 0.700800653 |
| Prps1 | -0.077509869 | 0.243844667 | 0.653830568 | 0.454104809 | 0.516288953 |
| Prps1l3 | -0.058277912 | 0.017033533 | 0.208804661 | 0.029496664 | 0.076709217 |
| Prps2 | -0.033550209 | 0.379035822 | 0.762006446 | 0.095794243 | 0.120725652 |
| Prpsap1 | -0.040086136 | 0.460311246 | 0.810111566 | 0.213368806 | 0.249542064 |
| Prpsap2 | -0.101349711 | 0.060437539 | 0.379132237 | 0.204983525 | 0.299742987 |
| Prr12 | 0.028094606 | 0.588600664 | 0.86992264 | 0.197134775 | 0.168272243 |
| Prr13 | -0.096394301 | 0.160398877 | 0.558508561 | 0.554012935 | 0.677224304 |
| Prr14 | -0.031699031 | 0.468304713 | 0.813213564 | 0.12504448 | 0.148404223 |
| Prr14l | -0.007614286 | 0.886440354 | 0.969612367 | 0.214333455 | 0.220242065 |
| Prr15l | 0.083851744 | 0.164628186 | 0.564527879 | 0.278016677 | 0.193725373 |
| Prr3 | 0.014195093 | 0.657931313 | 0.898224427 | 0.066138103 | 0.053990228 |
| Prr7 | 0.034588212 | 0.371695305 | 0.75760494 | 0.094849476 | 0.072417574 |
| Prrc1 | -0.173350423 | 0.010995864 | 0.165713947 | 0.631897745 | 0.8471645 |
| Prrc2a | -0.024468241 | 0.704615712 | 0.912507936 | 0.427070078 | 0.456924969 |
| Prrc2b | -0.213553279 | 0.001548388 | 0.05174005 | 0.801771425 | 1.061319343 |
| Prrc2c | -0.04498136 | 0.461241586 | 0.810249429 | 1.341400825 | 1.396022305 |
| Prrg2 | -0.011612087 | 0.843294727 | 0.955429414 | 0.304050993 | 0.313308735 |
| Prrt1 | -0.027798993 | 0.323163022 | 0.720244873 | 0.043090095 | 0.066668295 |
| Prss36 | 0.017161947 | 0.5337527 | 0.84612966 | 0.049653201 | 0.037776336 |
| Prss41 | -0.063055454 | 0.000161177 | 0.011080337 | 0.005562568 | 0.059345595 |
| Prss53 | -0.143922424 | 0.034335651 | 0.293212179 | 1.103331942 | 1.28254992 |
| Prune1 | 0.045817561 | 0.253750091 | 0.663146569 | 0.108598525 | 0.066480468 |
| Prune2 | -0.23176494 | 0.000621245 | 0.028602902 | 0.495566402 | 0.743859302 |
| Prxl2a | -0.083015545 | 0.156372399 | 0.554127149 | 0.285723516 | 0.355650918 |
| Prxl2b | -0.083037134 | 0.149965428 | 0.544377217 | 0.259013883 | 0.325605508 |
| Prxl2c | -0.085708448 | 0.143788505 | 0.533138243 | 0.291106799 | 0.368859712 |
| Psap | -0.148263327 | 0.005698801 | 0.112596197 | 1.642760369 | 1.898798174 |
| Psat1 | -0.078721774 | 0.182464674 | 0.590066231 | 0.262437209 | 0.326691274 |
| Psd | -0.012534296 | 0.713294635 | 0.9155164 | 0.070225887 | 0.080695219 |
| Psd3 | 0.033371335 | 0.511286235 | 0.835178052 | 0.181894834 | 0.152666073 |
| Psen1 | -0.069115996 | 0.144325896 | 0.533279057 | 0.154985845 | 0.215361736 |
| Psen2 | -0.114318498 | 0.01995935 | 0.225853434 | 0.1611078 | 0.260020281 |
| Psenen | 0.036804593 | 0.581400667 | 0.866173066 | 0.894552906 | 0.856628761 |
| Psip1 | -0.079942692 | 0.229150624 | 0.640979764 | 0.847929941 | 0.936082235 |
| Pskh1 | -0.018007646 | 0.640158856 | 0.891345382 | 0.099336493 | 0.114788198 |
| Psma1 | -0.017500402 | 0.795625282 | 0.939757086 | 0.768729841 | 0.783492643 |
| Psma2 | 0.020611391 | 0.733764737 | 0.921405813 | 1.152187206 | 1.122650072 |
| Psma3 | -0.07242475 | 0.272778455 | 0.680556141 | 1.130776691 | 1.274593116 |
| Psma4 | 0.07776456 | 0.25869352 | 0.665948842 | 0.790527818 | 0.717931892 |
| Psma5 | -0.16069409 | 0.012637976 | 0.179505076 | 0.476783082 | 0.665073994 |
| Psma6 | -0.021996178 | 0.7511036 | 0.926186851 | 0.744260102 | 0.805801739 |

| | | | | | |
|---|---|---|---|---|---|
| Psma7 | -0.140245403 | 0.014641456 | 0.194386632 | 1.453273614 | 1.712172753 |
| Psmb1 | 0.0203567 | 0.745730704 | 0.924566547 | 1.162188691 | 1.156156212 |
| Psmb10 | 0.131196655 | 0.056367604 | 0.367897032 | 0.447794916 | 0.303512137 |
| Psmb2 | -0.029334178 | 0.657718029 | 0.898224427 | 0.976224938 | 0.999824286 |
| Psmb3 | -0.088754157 | 0.203710824 | 0.614950615 | 0.734155636 | 0.834133583 |
| Psmb4 | -0.06100894 | 0.344519256 | 0.737445936 | 1.104655075 | 1.197433638 |
| Psmb5 | -0.143458819 | 0.03245673 | 0.285511129 | 0.648210337 | 0.823809997 |
| Psmb6 | -0.1320787 | 0.039751962 | 0.314071287 | 0.855287953 | 1.032197504 |
| Psmb7 | -0.029212397 | 0.656957609 | 0.898224427 | 0.579497681 | 0.614148209 |
| Psmb8 | 0.048677828 | 0.494939479 | 0.828918329 | 0.584942688 | 0.512984015 |
| Psmb9 | 0.116937693 | 0.035786528 | 0.298004804 | 0.244699315 | 0.138961213 |
| Psmc1 | 0.063605554 | 0.33227388 | 0.72806771 | 0.438148621 | 0.380457701 |
| Psmc2 | -0.063687807 | 0.365895813 | 0.75302167 | 0.694557398 | 0.778203976 |
| Psmc3 | 0.025312903 | 0.690640396 | 0.909796728 | 0.440541919 | 0.412268804 |
| Psmc4 | -0.06210081 | 0.34474175 | 0.737445936 | 0.43488765 | 0.499339504 |
| Psmc5 | -0.188113786 | 0.006356938 | 0.120222898 | 0.661356153 | 0.86824802 |
| Psmc6 | 0.125077048 | 0.064175506 | 0.386185094 | 0.760598215 | 0.635073648 |
| Psmd1 | -0.01901091 | 0.781113424 | 0.935482467 | 0.614153978 | 0.634410521 |
| Psmd10 | -0.119335762 | 0.014669113 | 0.194496547 | 0.14957882 | 0.271915162 |
| Psmd11 | -0.062806695 | 0.34360592 | 0.736445146 | 0.77699642 | 0.866814268 |
| Psmd12 | 0.071761111 | 0.310773244 | 0.71348779 | 0.952725469 | 0.863639243 |
| Psmd13 | 0.017645906 | 0.792238969 | 0.939347252 | 0.546180231 | 0.533177383 |
| Psmd14 | -0.051905281 | 0.393151081 | 0.769962419 | 0.360929965 | 0.404535238 |
| Psmd2 | 0.019127477 | 0.778773282 | 0.934641125 | 0.69177079 | 0.677027975 |
| Psmd3 | -0.071495933 | 0.296647619 | 0.702656882 | 0.532243655 | 0.601643887 |
| Psmd4 | -0.057448344 | 0.398604872 | 0.774321615 | 0.969916969 | 1.030161793 |
| Psmd5 | 0.000866676 | 0.987036261 | 0.997268589 | 0.206015103 | 0.207877313 |
| Psmd6 | -0.173770839 | 0.011287992 | 0.168327452 | 0.630439472 | 0.793803995 |
| Psmd7 | -0.012437917 | 0.848645568 | 0.957111129 | 1.049268504 | 1.056125318 |
| Psmd8 | -0.012499949 | 0.852777672 | 0.957461155 | 0.727201676 | 0.748085399 |
| Psmd9 | -0.046889703 | 0.402944879 | 0.77525778 | 0.239510212 | 0.284609958 |
| Psme1 | 0.030915879 | 0.656360395 | 0.89805697 | 0.673831998 | 0.603701965 |
| Psme2 | -0.101289442 | 0.143189516 | 0.532282155 | 0.899464388 | 1.019438945 |
| Psme3 | -0.01770928 | 0.747607003 | 0.924930563 | 0.231618073 | 0.240532697 |
| Psme3ip1 | -0.049692867 | 0.357972134 | 0.746823177 | 0.206424791 | 0.260547262 |
| Psme4 | -0.075323669 | 0.263466778 | 0.670148983 | 0.574629095 | 0.657370167 |
| Psmf1 | 0.023088013 | 0.675290645 | 0.906025597 | 0.24041212 | 0.218640198 |
| Psmg1 | -0.025960844 | 0.596073652 | 0.873655264 | 0.166579252 | 0.189957964 |
| Psmg2 | -0.01931578 | 0.739301041 | 0.922843014 | 0.26081533 | 0.277779771 |
| Psmg3 | -0.064122717 | 0.231215248 | 0.642677222 | 0.208673066 | 0.280371061 |
| Psmg4 | -0.026201735 | 0.687130662 | 0.909796728 | 0.475153409 | 0.491733723 |
| Pspc1 | -0.093612166 | 0.00251671 | 0.068270847 | 0.047979525 | 0.134591995 |
| Psph | -0.184106281 | 0.007255326 | 0.129345839 | 0.45488182 | 0.635179153 |
| Pstk | -0.076272951 | 0.030655291 | 0.279207941 | 0.075111327 | 0.142505466 |
| Ptar1 | -0.000843381 | 0.984876022 | 0.996391557 | 0.132639394 | 0.142687253 |
| Ptbp1 | -0.01586945 | 0.818656916 | 0.950128492 | 0.540815602 | 0.553815903 |
| Ptbp2 | 0.049317268 | 0.341906643 | 0.735000423 | 0.208376251 | 0.177127943 |
| Ptbp3 | -0.001283498 | 0.98337927 | 0.99608406 | 1.292604609 | 1.351703066 |
| Ptcd1 | -0.0098756 | 0.754213214 | 0.926726954 | 0.06095652 | 0.070609097 |
| Ptcd2 | 0.000666696 | 0.989302245 | 0.997451194 | 0.181557228 | 0.18162966 |
| Ptcd3 | -0.040564299 | 0.242883775 | 0.653166476 | 0.075805365 | 0.111902412 |
| Ptch1 | 0.01345663 | 0.74620904 | 0.924566547 | 0.118654815 | 0.110130586 |
| Ptdss1 | 0.068607478 | 0.109300787 | 0.477972625 | 0.12800917 | 0.068902747 |
| Ptdss2 | -0.017019301 | 0.634415785 | 0.88965971 | 0.082939513 | 0.09542865 |
| Pten | 0.049008363 | 0.453077066 | 0.807445758 | 0.81931241 | 0.785581242 |
| Ptges | -0.043202788 | 0.028543755 | 0.269514268 | 0.014470118 | 0.048937039 |
| Ptges2 | 0.106512969 | 0.022282205 | 0.236913658 | 0.165434717 | 0.078448692 |
| Ptges3 | -0.139980391 | 0.040312802 | 0.316388771 | 1.027841557 | 1.210097908 |
| Ptgr2 | 0.139114973 | 0.010466012 | 0.160622874 | 0.252878244 | 0.129946158 |
| Ptk2 | -0.008324852 | 0.845177373 | 0.955950886 | 0.121339876 | 0.135591187 |
| Ptk2b | 0.075022455 | 0.056436452 | 0.367897032 | 0.108172269 | 0.042904989 |
| Ptma | 0.06358902 | 0.054236226 | 0.362912667 | 2.885411978 | 2.729031833 |
| Ptms | -0.006959382 | 0.909311397 | 0.978321202 | 1.337982934 | 1.347992633 |
| Ptov1 | -0.105331914 | 0.117112409 | 0.489953809 | 0.5684546 | 0.686367583 |
| Ptp4a2 | 0.18265153 | 0.008414895 | 0.142539393 | 0.733981405 | 0.532847791 |

| | | | | | |
|---|---|---|---|---|---|
| Ptp4a3 | -0.049929295 | 0.35039961 | 0.740908966 | 0.192392835 | 0.238115139 |
| Ptpa | 0.07425471 | 0.222179925 | 0.633924544 | 0.406923209 | 0.341368528 |
| Ptpdc1 | 0.024737549 | 0.462360267 | 0.810535897 | 0.078144953 | 0.059712321 |
| Ptpmt1 | 0.003476824 | 0.953837133 | 0.989066902 | 0.293985756 | 0.289313133 |
| Ptpn1 | 0.071964597 | 0.299931452 | 0.70506682 | 0.667263834 | 0.567122019 |
| Ptpn11 | 0.054707534 | 0.426845052 | 0.79105114 | 0.876026358 | 0.834602065 |
| Ptpn12 | 0.081589952 | 0.103019455 | 0.468979611 | 0.176028117 | 0.090633339 |
| Ptpn13 | 0.026210871 | 0.569166027 | 0.860620982 | 0.140213619 | 0.116542286 |
| Ptpn2 | 0.017362673 | 0.783189898 | 0.936124803 | 0.31478219 | 0.298299659 |
| Ptpn21 | 0.068320755 | 0.19849956 | 0.609389286 | 0.207852354 | 0.154389394 |
| Ptpn23 | 0.013285052 | 0.780207269 | 0.935261001 | 0.160872425 | 0.142385686 |
| Ptpn3 | 0.011562586 | 0.714096287 | 0.915840939 | 0.065190046 | 0.056462773 |
| Ptpn4 | -0.064892882 | 0.201404218 | 0.611924411 | 0.17413783 | 0.225369902 |
| Ptpn6 | -0.028187812 | 0.561609765 | 0.857134531 | 0.158611495 | 0.18643166 |
| Ptpn9 | -0.037810536 | 0.48834552 | 0.826256745 | 0.23106881 | 0.256909579 |
| Ptpra | 0.032808749 | 0.586414725 | 0.86902362 | 0.339355713 | 0.305991966 |
| Ptprd | 0.009257523 | 0.812555195 | 0.948397417 | 0.101473589 | 0.095050445 |
| Ptprf | 0.095876088 | 0.15815555 | 0.556468981 | 1.181985939 | 1.026976784 |
| Ptprj | -0.021260407 | 0.662917617 | 0.899270729 | 0.166897706 | 0.18070319 |
| Ptprm | -0.004906235 | 0.915341954 | 0.980132949 | 0.145946396 | 0.144740462 |
| Ptprn | -0.348384751 | 1.84E-08 | 8.40E-06 | 1.338486048 | 1.881946588 |
| Ptprn2 | -0.119110464 | 0.059997505 | 0.378409279 | 0.400658957 | 0.522136033 |
| Ptprr | 0.012938902 | 0.698069002 | 0.911092742 | 0.074383332 | 0.05741052 |
| Ptprs | 0.045995793 | 0.276668996 | 0.684050613 | 0.120038683 | 0.082001309 |
| Ptprt | -0.0946632 | 0.047550445 | 0.341541257 | 0.138103151 | 0.22487759 |
| Ptpru | 0.008995899 | 0.795739546 | 0.939757086 | 0.078706492 | 0.073565776 |
| Ptrh2 | -0.032246029 | 0.533680849 | 0.84612966 | 0.191542892 | 0.214249755 |
| Ptrhd1 | 0.038407492 | 0.513033031 | 0.836298776 | 0.281594912 | 0.246089247 |
| Pts | -0.030571174 | 0.626117879 | 0.886938778 | 0.37327919 | 0.423428976 |
| Pttg1 | -0.114319774 | 0.016916687 | 0.208334711 | 0.139363299 | 0.243560301 |
| Pttg1ip | 0.046927516 | 0.448086012 | 0.804951652 | 0.364165676 | 0.327181276 |
| Puf60 | -0.186204909 | 0.004877431 | 0.102275888 | 0.414991462 | 0.611301674 |
| Pum1 | -0.082771925 | 0.197489579 | 0.607851242 | 0.634276196 | 0.720153081 |
| Pum2 | 0.046881367 | 0.480607203 | 0.820388721 | 0.717527623 | 0.650032342 |
| Pum3 | 0.101442241 | 0.083971378 | 0.432215755 | 0.309493441 | 0.223140698 |
| Pura | -0.046879496 | 0.442469965 | 0.801058989 | 1.231856759 | 1.301925135 |
| Purb | 0.046375205 | 0.481071499 | 0.820757204 | 1.07503776 | 1.012054559 |
| Purg | -0.034561586 | 0.185924974 | 0.592701341 | 0.033520369 | 0.067174116 |
| Pus1 | 0.068928582 | 0.109542527 | 0.477972625 | 0.132820152 | 0.07872268 |
| Pus10 | -0.008848688 | 0.771188976 | 0.932251553 | 0.056655757 | 0.059884171 |
| Pus3 | -0.037339044 | 0.311520176 | 0.71405691 | 0.086721112 | 0.121162707 |
| Pus7 | -0.010240942 | 0.806042286 | 0.945343121 | 0.114647527 | 0.124422373 |
| Pusl1 | 0.013121545 | 0.635548669 | 0.890176108 | 0.049623722 | 0.042573449 |
| Pvr | 0.061755332 | 0.044553025 | 0.330723271 | 0.066581708 | 0.012883206 |
| Pwp1 | -0.045776833 | 0.330584881 | 0.725898152 | 0.143518185 | 0.183507627 |
| Pwwp2a | -0.012115554 | 0.805385167 | 0.944793235 | 0.165096525 | 0.171976608 |
| Pwwp3a | 0.026968688 | 0.554683725 | 0.853469818 | 0.146412076 | 0.120000476 |
| Pwwp3b | 0.063268186 | 0.322070866 | 0.719780733 | 0.316981973 | 0.243571508 |
| Pxk | 0.089231963 | 0.169301604 | 0.57227323 | 0.445277375 | 0.352423561 |
| Pxmp2 | -0.048187032 | 0.006345101 | 0.120222898 | 0.009501698 | 0.049603218 |
| Pxmp4 | 0.007806853 | 0.851991353 | 0.957461155 | 0.114441003 | 0.108184385 |
| Pxylp1 | 0.049083331 | 0.135377088 | 0.518618255 | 0.074187484 | 0.031247111 |
| Pycard | 0.009825362 | 0.786817388 | 0.937461311 | 0.086327196 | 0.068620092 |
| Pycr1 | -0.11537476 | 0.038783714 | 0.311667042 | 0.217786488 | 0.324459562 |
| Pycr2 | 0.058092745 | 0.389131678 | 0.767934457 | 0.939991276 | 0.894023027 |
| Pycrl | -0.093524776 | 0.039753235 | 0.314071287 | 0.127371085 | 0.217997566 |
| Pygb | -0.026422638 | 0.221999708 | 0.633924544 | 0.023209895 | 0.044941945 |
| Pygo1 | -0.01650157 | 0.503288151 | 0.832249514 | 0.033149773 | 0.045541735 |
| Pygo2 | -0.050558466 | 0.159156074 | 0.556789205 | 0.071153425 | 0.116340317 |
| Pym1 | 0.010004745 | 0.77783154 | 0.934339474 | 0.084619395 | 0.077138252 |
| Pyroxd1 | 0.054529776 | 0.298925855 | 0.703969686 | 0.203732443 | 0.150706332 |
| Pyurf | -0.056107654 | 0.245105002 | 0.654809396 | 0.15087983 | 0.209352246 |
| Pyy | -0.024294799 | 0.81691422 | 0.94911078 | 1.603949886 | 1.538070366 |
| Qars | -0.054934891 | 0.266255716 | 0.673899077 | 0.163100204 | 0.215753354 |
| Qdpr | -0.098931166 | 0.147936655 | 0.539745621 | 0.699954693 | 0.827135035 |

| | | | | | |
|---|---|---|---|---|---|
| Qk | -0.010668934 | 0.819137676 | 0.950128492 | 0.14648326 | 0.16582473 |
| Qpctl | 3.62E-05 | 0.999303777 | 0.999858209 | 0.11519227 | 0.115864669 |
| Qrich1 | -0.075722743 | 0.130023175 | 0.509514066 | 0.167769052 | 0.236581746 |
| Qrsl1 | 0.002057879 | 0.947466899 | 0.987651726 | 0.058096751 | 0.059004732 |
| Qser1 | -0.029744893 | 0.605602939 | 0.87802905 | 0.235743319 | 0.261605481 |
| Qsox1 | -0.184575164 | 0.002591813 | 0.06955622 | 0.299803087 | 0.48179077 |
| Qsox2 | 0.015013205 | 0.798631173 | 0.940982298 | 0.277925334 | 0.254956531 |
| Qtrt1 | -0.018964725 | 0.668874978 | 0.903080193 | 0.134311053 | 0.157737819 |
| Qtrt2 | -0.031231392 | 0.383624326 | 0.763975666 | 0.076749243 | 0.110797099 |
| R3hcc1 | 0.023634115 | 0.61356617 | 0.881403127 | 0.151740235 | 0.1355557 |
| R3hcc1l | -0.013135954 | 0.697655407 | 0.910970095 | 0.071368286 | 0.085850149 |
| R3hdm1 | 0.022596972 | 0.751129256 | 0.926186851 | 0.68808011 | 0.655157105 |
| R3hdm2 | -0.117835348 | 0.071175969 | 0.404068555 | 0.711979692 | 0.859440187 |
| R3hdm4 | -0.060804928 | 0.290104538 | 0.696098314 | 0.264043907 | 0.304123486 |
| Rab10 | 0.034379943 | 0.605976101 | 0.878178429 | 0.817866492 | 0.771547481 |
| Rab10os | 0.013352859 | 0.775003647 | 0.933555636 | 0.153668488 | 0.136156777 |
| Rab11a | 0.010861224 | 0.875214069 | 0.965096393 | 0.714175005 | 0.70446965 |
| Rab11b | -0.041684923 | 0.50962499 | 0.834904701 | 0.380173009 | 0.415490299 |
| Rab11fip1 | -0.061858738 | 0.076783489 | 0.418199494 | 0.072334983 | 0.130096443 |
| Rab11fip2 | -0.082768565 | 0.1853599 | 0.591560356 | 0.339271038 | 0.426287652 |
| Rab11fip3 | -0.044233166 | 0.175941005 | 0.580131363 | 0.062155796 | 0.101165446 |
| Rab11fip5 | -0.024376948 | 0.309003211 | 0.711182121 | 0.030375709 | 0.050255814 |
| Rab12 | 0.109846983 | 0.074567734 | 0.41344108 | 0.374374519 | 0.272016466 |
| Rab13 | -0.085733097 | 0.00503335 | 0.104595715 | 0.04910075 | 0.128062977 |
| Rab14 | 0.110117997 | 0.051439815 | 0.355933715 | 1.479057832 | 1.305162217 |
| Rab18 | 0.046265041 | 0.492037399 | 0.827270742 | 0.709129969 | 0.662487513 |
| Rab19 | 0.038982396 | 0.194532868 | 0.603853652 | 0.061686031 | 0.026868867 |
| Rab1a | 0.071694605 | 0.280238276 | 0.687465228 | 1.065365747 | 0.987883302 |
| Rab1b | -0.041679055 | 0.48772045 | 0.826112275 | 0.333290217 | 0.364396716 |
| Rab20 | 0.04896372 | 0.099430321 | 0.463315751 | 0.059682288 | 0.018689498 |
| Rab21 | 0.111472079 | 0.107814224 | 0.476290212 | 0.827226924 | 0.675935893 |
| Rab22a | 0.023066836 | 0.739398141 | 0.922843014 | 0.692028337 | 0.679762753 |
| Rab23 | 0.053087927 | 0.175227942 | 0.578730783 | 0.108341287 | 0.05870313 |
| Rab24 | 0.083002148 | 0.208113298 | 0.620020309 | 0.443468027 | 0.355624684 |
| Rab25 | -0.079123687 | 0.221474621 | 0.633858837 | 0.450757817 | 0.529291799 |
| Rab26os | 0.031428648 | 0.340353176 | 0.733546237 | 0.072450534 | 0.042784601 |
| Rab27a | 0.045554665 | 0.515012724 | 0.836801664 | 0.616419157 | 0.53273114 |
| Rab28 | 0.025883849 | 0.693439508 | 0.909871028 | 0.460015574 | 0.444535598 |
| Rab2a | 0.018238095 | 0.725541295 | 0.92014727 | 1.717855381 | 1.679411572 |
| Rab2b | -0.012601348 | 0.689827778 | 0.909796728 | 0.061622464 | 0.069997245 |
| Rab33b | -0.042399709 | 0.429629569 | 0.79317385 | 0.215125922 | 0.268912462 |
| Rab34 | -0.067724268 | 0.216978932 | 0.631434486 | 0.210602707 | 0.269775903 |
| Rab35 | 0.050936322 | 0.277574526 | 0.684861238 | 0.154743226 | 0.108256358 |
| Rab36 | 0.02008521 | 0.627971095 | 0.887864718 | 0.115659011 | 0.095807776 |
| Rab37 | 0.051729837 | 0.384507544 | 0.764672523 | 0.292419679 | 0.247117962 |
| Rab39 | -0.026252887 | 0.241061513 | 0.651287862 | 0.025068787 | 0.048814399 |
| Rab39b | -0.056610082 | 0.367385606 | 0.754079617 | 0.379320012 | 0.449138146 |
| Rab3a | 0.02456463 | 0.712132669 | 0.915109107 | 0.657084722 | 0.642288491 |
| Rab3b | 0.099661595 | 0.104167693 | 0.471172212 | 0.30842521 | 0.211654207 |
| Rab3d | 0.040607973 | 0.542381493 | 0.850208191 | 0.523507881 | 0.482374219 |
| Rab3gap1 | -0.004342947 | 0.946269052 | 0.987084873 | 0.499889995 | 0.48152189 |
| Rab3gap2 | 0.06106409 | 0.32234337 | 0.719780733 | 0.335836157 | 0.287697392 |
| Rab3ip | -0.094815411 | 0.082481002 | 0.429166311 | 0.227991392 | 0.322627883 |
| Rab40c | 0.020080225 | 0.600253462 | 0.875944169 | 0.097043053 | 0.084348134 |
| Rab4a | 0.062464201 | 0.229391184 | 0.640979764 | 0.190778877 | 0.143480043 |
| Rab5a | 0.031799247 | 0.649512596 | 0.895585658 | 0.723795773 | 0.707721106 |
| Rab5b | -0.065180849 | 0.22433866 | 0.637067124 | 0.20208268 | 0.273053948 |
| Rab5c | 0.022753349 | 0.7148895 | 0.916116931 | 0.348425589 | 0.325115304 |
| Rab5if | 0.057860603 | 0.374589749 | 0.759807714 | 0.457347683 | 0.414937731 |
| Rab6a | 0.115659995 | 0.095322294 | 0.455812225 | 0.909561796 | 0.76609241 |
| Rab6b | 0.031337781 | 0.519644116 | 0.838531832 | 0.168093212 | 0.152468749 |
| Rab7 | 0.02545735 | 0.707563526 | 0.913534231 | 1.09190709 | 1.036971027 |
| Rab8a | 0.101979691 | 0.100292458 | 0.465319642 | 0.361230905 | 0.265650792 |
| Rab8b | 0.050535581 | 0.151239669 | 0.546523037 | 0.089597638 | 0.044538328 |
| Rab9 | 0.099481287 | 0.061939484 | 0.380250374 | 0.215729224 | 0.13160154 |

| | | | | | |
|---|---|---|---|---|---|
| Rabac1 | -0.079118247 | 0.135477989 | 0.518757369 | 1.573373933 | 1.714049542 |
| Rabep1 | 0.034982295 | 0.600154839 | 0.875927602 | 0.557704267 | 0.532471277 |
| Rabep2 | 0.031366664 | 0.33561896 | 0.730736446 | 0.072629977 | 0.047370731 |
| Rabepk | 0.030508615 | 0.430978683 | 0.793421321 | 0.105927543 | 0.07790194 |
| Rabgap1 | 0.00381754 | 0.950066155 | 0.987860147 | 0.346201375 | 0.341700913 |
| Rabgap1l | 0.017558326 | 0.733199409 | 0.921306823 | 0.192272954 | 0.171374554 |
| Rabgef1 | 0.08275578 | 0.071327467 | 0.404242678 | 0.155818573 | 0.089974608 |
| Rabggta | -0.019900292 | 0.549113478 | 0.852402509 | 0.069975343 | 0.08601717 |
| Rabggtb | 0.034615116 | 0.580317365 | 0.865989503 | 0.399349678 | 0.374297173 |
| Rabif | 0.001138333 | 0.978171108 | 0.995031396 | 0.11095888 | 0.117002406 |
| Rabl2 | -0.03788488 | 0.163765739 | 0.563495619 | 0.039030334 | 0.071359171 |
| Rabl3 | 0.034181702 | 0.345532133 | 0.73792645 | 0.090746002 | 0.068184647 |
| Rabl6 | 0.037813741 | 0.57606119 | 0.863450749 | 0.603231966 | 0.564770008 |
| Rac1 | -0.003051645 | 0.963506885 | 0.991970325 | 0.775959616 | 0.780005806 |
| Racgap1 | 0.045734525 | 0.129375395 | 0.5092317 | 0.062662725 | 0.026417564 |
| Rack1 | -0.132676754 | 0.002615686 | 0.069641377 | 1.978941065 | 2.260596437 |
| Rad1 | 0.085981064 | 0.03449365 | 0.293212179 | 0.118840435 | 0.047912969 |
| Rad17 | 0.113001153 | 0.06884129 | 0.399753362 | 0.347639648 | 0.235619891 |
| Rad18 | -0.00579546 | 0.847383683 | 0.956929571 | 0.056690664 | 0.061439719 |
| Rad21 | 0.015583532 | 0.819039363 | 0.950128492 | 0.677528989 | 0.656775346 |
| Rad23a | -0.077586439 | 0.238939162 | 0.650445287 | 0.511753727 | 0.601429304 |
| Rad23b | -0.008280833 | 0.900493816 | 0.974055009 | 0.657513798 | 0.688632651 |
| Rad50 | -0.00269158 | 0.957819845 | 0.990137214 | 0.186366299 | 0.176317143 |
| Rad51ap2 | 0.028512985 | 0.622290801 | 0.885194554 | 0.244951145 | 0.220906966 |
| Rad51d | 0.038800784 | 0.284370149 | 0.691925959 | 0.091676982 | 0.059715278 |
| Rad52 | -0.057296127 | 0.023304231 | 0.241886829 | 0.031122695 | 0.076107947 |
| Rad54l2 | 0.007151399 | 0.87491421 | 0.96500153 | 0.142442615 | 0.130560391 |
| Rad9b | 0.032400149 | 0.253402291 | 0.663146569 | 0.0541853 | 0.029730485 |
| Rae1 | 0.078031628 | 0.160742885 | 0.558841822 | 0.250547088 | 0.19203608 |
| Raf1 | 0.031255626 | 0.54728402 | 0.851449318 | 0.205635872 | 0.180871009 |
| Rai1 | -0.016294129 | 0.749825086 | 0.925706567 | 0.178562384 | 0.186152682 |
| Rai2 | -0.014740023 | 0.687593298 | 0.909796728 | 0.090201587 | 0.102901667 |
| Rala | 0.078347205 | 0.242389885 | 0.653091138 | 0.839893384 | 0.738020631 |
| Ralb | -0.047142663 | 0.33303283 | 0.728881491 | 0.152557338 | 0.192581787 |
| Ralbp1 | 0.098144018 | 0.115970824 | 0.488254681 | 1.010202094 | 0.848765516 |
| Ralgapa1 | -0.011802238 | 0.845939576 | 0.956051742 | 0.326930352 | 0.330446031 |
| Ralgapa2 | -0.06334641 | 0.245061929 | 0.654809396 | 0.213584313 | 0.272483453 |
| Ralgapb | -0.036574488 | 0.517857118 | 0.837803336 | 0.229904989 | 0.266791975 |
| Ralgds | 0.007002017 | 0.884226376 | 0.968954079 | 0.162361957 | 0.151837822 |
| Ralgps1 | 0.043605504 | 0.187106656 | 0.594488607 | 0.077438761 | 0.044425331 |
| Ralgps2 | -0.016149114 | 0.755287118 | 0.926726954 | 0.193331338 | 0.207750186 |
| Raly | 0.03879431 | 0.53322083 | 0.84612966 | 0.366834717 | 0.328289023 |
| Ramac | -0.049561772 | 0.414104912 | 0.782518584 | 0.324518444 | 0.379134087 |
| Ramp1 | 0.114612446 | 0.020599551 | 0.227971987 | 0.17020924 | 0.063242949 |
| Ramp2 | 0.042958927 | 0.365039531 | 0.752802229 | 0.150958127 | 0.120484149 |
| Ran | -0.026147705 | 0.684457801 | 0.909679945 | 1.049975503 | 1.07842381 |
| Ranbp1 | -0.124881808 | 0.050263222 | 0.353781178 | 0.446363192 | 0.557626122 |
| Ranbp10 | -0.033622863 | 0.429161286 | 0.792546794 | 0.120032413 | 0.142953302 |
| Ranbp2 | -0.095712305 | 0.170283417 | 0.572762895 | 0.729951817 | 0.847975851 |
| Ranbp3 | 0.081388619 | 0.11094557 | 0.481643894 | 0.185999097 | 0.118287671 |
| Ranbp6 | 0.006285588 | 0.863131229 | 0.960676137 | 0.088373429 | 0.082837548 |
| Ranbp9 | 0.160414673 | 0.005012806 | 0.104384928 | 0.281365393 | 0.139677091 |
| Rangap1 | 0.062905183 | 0.339231109 | 0.732701236 | 0.575958187 | 0.516691424 |
| Rap1a | 0.100605416 | 0.144896286 | 0.533279057 | 0.860557496 | 0.764054275 |
| Rap1b | -0.138336546 | 0.025492626 | 0.254305979 | 1.381250339 | 1.612465256 |
| Rap1gap | -0.041653945 | 0.38335134 | 0.763886718 | 0.141121493 | 0.185942813 |
| Rap1gap2 | 0.061989551 | 0.365687529 | 0.752886169 | 1.014328309 | 0.908097332 |
| Rap1gapos | 0.015549326 | 0.586481462 | 0.86902362 | 0.053217846 | 0.038174776 |
| Rap1gds1 | 0.030090279 | 0.48560504 | 0.824318763 | 0.12503435 | 0.101531724 |
| Rap2a | 0.067467411 | 0.174348191 | 0.577345033 | 0.170356833 | 0.114274956 |
| Rap2b | 0.024651904 | 0.504326676 | 0.832249514 | 0.088773678 | 0.073702963 |
| Rap2c | -0.020814702 | 0.538142651 | 0.848066852 | 0.074426861 | 0.09436023 |
| Rapgef1 | 0.039714372 | 0.441419428 | 0.800354711 | 0.189871032 | 0.146217359 |
| Rapgef2 | -0.038027506 | 0.353881308 | 0.744256656 | 0.106912854 | 0.140050036 |
| Rapgef4 | 0.00928197 | 0.883467944 | 0.968793593 | 0.40019895 | 0.38730264 |

| | | | | | |
|---|---|---|---|---|---|
| Rapgef6 | -0.095400383 | 0.072262178 | 0.407469371 | 0.201524453 | 0.281576845 |
| Raph1 | -0.002586875 | 0.955635299 | 0.989278596 | 0.147790798 | 0.146409046 |
| Rara | 0.0420224 | 0.248544351 | 0.659783035 | 0.089612469 | 0.048837071 |
| Rarg | 0.022751229 | 0.43060627 | 0.793311823 | 0.054503187 | 0.034233102 |
| Rars | -0.062989518 | 0.306189655 | 0.709680069 | 0.304545953 | 0.351109957 |
| Rars2 | -0.06541418 | 0.156792038 | 0.554322538 | 0.136180536 | 0.204207863 |
| Rasa1 | 0.079075964 | 0.204994482 | 0.616549498 | 0.39803329 | 0.329285139 |
| Rasa2 | 0.0872906 | 0.143432857 | 0.532606581 | 0.296897458 | 0.210955656 |
| Rasal2 | -0.001051682 | 0.983578769 | 0.99608406 | 0.184863206 | 0.182699009 |
| Rasd1 | -0.269688501 | 9.30E-05 | 0.007469648 | 0.371780592 | 0.626920398 |
| Rasd2 | -0.069430699 | 0.145644883 | 0.534102188 | 0.132752008 | 0.198475855 |
| Rasgef1b | 0.003040734 | 0.941505318 | 0.986168593 | 0.111457955 | 0.096930295 |
| Rasgrf1 | 0.03488598 | 0.625364299 | 0.88668351 | 0.643663189 | 0.577407473 |
| Rasgrf2 | -0.094049884 | 0.058351015 | 0.372056206 | 0.149911331 | 0.244451924 |
| Rassf1 | -0.090783907 | 0.086240795 | 0.43696958 | 0.179915437 | 0.255199083 |
| Rassf10 | 0.081729432 | 0.04551613 | 0.334195605 | 0.121795662 | 0.049591684 |
| Rassf3 | 0.002134479 | 0.971709943 | 0.993892166 | 0.338044235 | 0.341623652 |
| Rassf4 | -0.093576481 | 0.051077195 | 0.355933715 | 0.152111382 | 0.233584044 |
| Rassf6 | 0.03194699 | 0.250393838 | 0.662240566 | 0.05106424 | 0.023530904 |
| Rassf7 | 0.06111811 | 0.092435897 | 0.450107955 | 0.091678127 | 0.043470303 |
| Rassf8 | 0.100184839 | 0.008432921 | 0.142539393 | 0.108488291 | 0.03003178 |
| Rb1 | 0.081986038 | 0.105253026 | 0.472434351 | 0.18792422 | 0.108001895 |
| Rb1cc1 | -0.052189052 | 0.444219634 | 0.801932586 | 0.601496374 | 0.651244436 |
| Rbak | -0.052704753 | 0.066997663 | 0.392969665 | 0.046635003 | 0.09133159 |
| Rbbp4 | -0.032119263 | 0.628484208 | 0.887864718 | 0.502098228 | 0.546294059 |
| Rbbp5 | 0.0361298 | 0.305446967 | 0.709411642 | 0.086540856 | 0.06146721 |
| Rbbp6 | -0.025114315 | 0.707090895 | 0.913276453 | 0.819622802 | 0.864597705 |
| Rbbp7 | -0.055932835 | 0.426689683 | 0.79105114 | 0.739328013 | 0.800151192 |
| Rbbp8 | -0.010813903 | 0.754930665 | 0.926726954 | 0.07009468 | 0.082094601 |
| Rbbp9 | 0.01128323 | 0.721161975 | 0.919475778 | 0.065544567 | 0.057733596 |
| Rbck1 | 0.034587804 | 0.503248238 | 0.832249514 | 0.199477589 | 0.163558039 |
| Rbfa | 0.052920043 | 0.296185358 | 0.702600505 | 0.198462194 | 0.154385752 |
| Rbfox2 | 0.141124276 | 0.031162522 | 0.280770404 | 0.446936644 | 0.316060162 |
| Rbis | 0.052160494 | 0.417518774 | 0.784482356 | 0.465165566 | 0.407622079 |
| Rbks | -0.02883003 | 0.144045408 | 0.533279057 | 0.01710215 | 0.040183404 |
| Rbl2 | 0.067751765 | 0.174676623 | 0.577992588 | 0.180394956 | 0.118334832 |
| Rbm10 | 0.045666378 | 0.327174795 | 0.722043407 | 0.14997866 | 0.110872506 |
| Rbm11 | -0.055129438 | 0.002995138 | 0.076092349 | 0.009807291 | 0.057750825 |
| Rbm12 | 0.001853879 | 0.955894894 | 0.989308832 | 0.069755287 | 0.068940467 |
| Rbm12b2 | -0.006926026 | 0.827158562 | 0.952715064 | 0.060242137 | 0.07356485 |
| Rbm14 | 0.004375191 | 0.930123927 | 0.982596974 | 0.166907711 | 0.168060498 |
| Rbm15 | -0.051973383 | 0.220881185 | 0.633858837 | 0.111576125 | 0.16306924 |
| Rbm15b | 0.045114487 | 0.345148091 | 0.737702595 | 0.161183689 | 0.123787591 |
| Rbm17 | -0.119833001 | 0.078732435 | 0.421117898 | 0.479655969 | 0.595000588 |
| Rbm18 | -0.011810228 | 0.815942109 | 0.949036482 | 0.181867801 | 0.195383149 |
| Rbm19 | 0.036870632 | 0.242551895 | 0.653091138 | 0.068460547 | 0.032246802 |
| Rbm22 | -0.020111265 | 0.678940034 | 0.907783945 | 0.163029113 | 0.183613294 |
| Rbm25 | -0.085269151 | 0.220884094 | 0.633858837 | 0.915143945 | 0.999389486 |
| Rbm26 | -0.083412163 | 0.224568363 | 0.637067124 | 0.566614762 | 0.658567582 |
| Rbm27 | -0.006804615 | 0.917498877 | 0.980132949 | 0.471885575 | 0.504164377 |
| Rbm28 | -0.070869987 | 0.251373239 | 0.663127858 | 0.331040782 | 0.366367664 |
| Rbm3 | -0.116774726 | 0.083502852 | 0.431740995 | 0.688200418 | 0.803163634 |
| Rbm33 | 0.030758116 | 0.61797219 | 0.883238152 | 0.326862078 | 0.300009384 |
| Rbm34 | 0.00803518 | 0.872438802 | 0.964815806 | 0.174960712 | 0.170752449 |
| Rbm39 | -0.019773408 | 0.700749876 | 0.912128973 | 1.82824094 | 1.833917307 |
| Rbm4 | -0.024122188 | 0.409313506 | 0.779735046 | 0.049415037 | 0.066606898 |
| Rbm43 | 0.071921782 | 0.180838316 | 0.588545452 | 0.22750137 | 0.166416914 |
| Rbm47 | -0.008751851 | 0.895222394 | 0.972229731 | 0.47315018 | 0.495642534 |
| Rbm48 | -0.073059975 | 0.088943567 | 0.443672569 | 0.117382794 | 0.184063258 |
| Rbm4b | 0.016215727 | 0.763454914 | 0.928730554 | 0.216464958 | 0.199254433 |
| Rbm5 | -0.058766686 | 0.390878448 | 0.769527973 | 0.590865231 | 0.655845206 |
| Rbm6 | -0.106697106 | 0.084838042 | 0.433957325 | 0.311752457 | 0.400887678 |
| Rbm7 | 0.026213195 | 0.665342654 | 0.901006558 | 0.322722523 | 0.309349871 |
| Rbm8a | -0.12358054 | 0.077688487 | 0.418285047 | 0.634703634 | 0.754977518 |
| Rbms1 | -0.023128082 | 0.608727829 | 0.878855197 | 0.137073807 | 0.157756435 |

| | | | | | |
|---|---|---|---|---|---|
| Rbms2 | 0.027856794 | 0.365969882 | 0.75302167 | 0.065394706 | 0.045585743 |
| Rbmx | -0.083087829 | 0.075198109 | 0.413795734 | 0.1398016 | 0.21496826 |
| Rbmx2 | -0.081441913 | 0.096871505 | 0.45927202 | 0.153336442 | 0.227010274 |
| Rbmxl1 | -0.026434888 | 0.561848313 | 0.857134531 | 0.141960561 | 0.183401495 |
| Rbp2 | 0.042405471 | 0.176321183 | 0.580732554 | 0.061633126 | 0.024205761 |
| Rbp4 | 0.163903424 | 0.044647806 | 0.330723271 | 0.424478496 | 0.176157452 |
| Rbpj | 0.017114759 | 0.721668545 | 0.919475778 | 0.167891373 | 0.153562076 |
| Rbpms | 0.068554766 | 0.325230368 | 0.720833548 | 0.768971427 | 0.685553435 |
| Rbsn | 0.035091226 | 0.513337732 | 0.836423834 | 0.222695054 | 0.192684684 |
| Rbx1 | -0.012043008 | 0.831788722 | 0.953695326 | 1.3623669 | 1.377134646 |
| Rc3h1 | -0.046213831 | 0.425229382 | 0.790344696 | 0.264798042 | 0.311201808 |
| Rc3h2 | -0.033344452 | 0.616683617 | 0.883226807 | 0.446946813 | 0.478731951 |
| Rcan1 | -0.009641522 | 0.799945569 | 0.941382774 | 0.091982825 | 0.10141713 |
| Rcan2 | 0.045952972 | 0.457921855 | 0.809095362 | 0.355294741 | 0.304085893 |
| Rcan3 | -0.074042453 | 0.273529014 | 0.681426919 | 0.664633544 | 0.741253396 |
| Rcbtb1 | 0.056825886 | 0.386793711 | 0.766593284 | 0.386498676 | 0.326944848 |
| Rcbtb2 | 0.053853162 | 0.228873749 | 0.640603965 | 0.144980714 | 0.098081113 |
| Rcc1l | -0.012005714 | 0.745499509 | 0.924566547 | 0.087862544 | 0.100824904 |
| Rcc2 | -0.189809908 | 0.003523145 | 0.08334591 | 0.400190197 | 0.598030484 |
| Rce1 | 0.102599436 | 0.076965914 | 0.418199494 | 0.25417018 | 0.164797217 |
| Rchy1 | 0.131585301 | 0.044180012 | 0.329446347 | 0.441804904 | 0.324605612 |
| Rcl1 | 0.000254254 | 0.993548584 | 0.998695538 | 0.063411846 | 0.0684024 |
| Rcn1 | 0.00824503 | 0.82451826 | 0.951625061 | 0.093343711 | 0.089525107 |
| Rcn2 | -0.056915153 | 0.38117715 | 0.762430261 | 0.451242651 | 0.512706635 |
| Rcor1 | 0.041741427 | 0.483008104 | 0.822469365 | 0.296124429 | 0.257099671 |
| Rcor3 | 0.094540442 | 0.090091119 | 0.445385778 | 0.247481263 | 0.161431482 |
| Rdh11 | -0.001572516 | 0.961956327 | 0.991441074 | 0.07152559 | 0.073067945 |
| Rdh13 | 0.023548356 | 0.432763091 | 0.795079432 | 0.057329572 | 0.03768741 |
| Rdh14 | 0.025750165 | 0.649043494 | 0.895557756 | 0.274485239 | 0.25450431 |
| Rdm1 | -0.060618832 | 0.051555847 | 0.355933715 | 0.054964912 | 0.113104119 |
| Rdx | 0.029990178 | 0.640591374 | 0.891345382 | 0.969764608 | 0.921099346 |
| Rec8 | -0.012215552 | 0.777310796 | 0.934243618 | 0.120480956 | 0.130782551 |
| Recql | -0.004627576 | 0.893340734 | 0.9716581 | 0.07463129 | 0.08195091 |
| Recql5 | -0.025512631 | 0.325505551 | 0.720833548 | 0.038386128 | 0.060037627 |
| Reep2 | 0.040777095 | 0.210110445 | 0.622822957 | 0.075231102 | 0.042576373 |
| Reep3 | -0.115444879 | 0.06760905 | 0.394989542 | 0.37867204 | 0.492745321 |
| Reep4 | -0.026646598 | 0.30247628 | 0.707681683 | 0.036249262 | 0.060245913 |
| Reep5 | 0.011159266 | 0.834676151 | 0.954911647 | 1.419869997 | 1.427795195 |
| Reep6 | -0.213908519 | 0.001332512 | 0.046438985 | 0.492390689 | 0.703848919 |
| Rel | -0.057557113 | 0.169599497 | 0.57227323 | 0.105593887 | 0.159972055 |
| Rela | -0.004917196 | 0.91962273 | 0.980436936 | 0.161378256 | 0.164895637 |
| Relb | -0.015542902 | 0.672928342 | 0.905559579 | 0.081825044 | 0.102098507 |
| Relch | -0.052077514 | 0.345102847 | 0.737702595 | 0.230853754 | 0.282446505 |
| Rell1 | 0.11440639 | 0.028646048 | 0.269972194 | 0.220306171 | 0.122538078 |
| Relt | -0.085741288 | 0.021210696 | 0.23094225 | 0.07841084 | 0.150666682 |
| Rem2 | 0.061875506 | 0.060324631 | 0.378897573 | 0.079541377 | 0.025509499 |
| Repin1 | -0.011684644 | 0.748182848 | 0.924930563 | 0.08447901 | 0.097276894 |
| Reps1 | 0.048385975 | 0.407879752 | 0.77823964 | 0.329528055 | 0.293995805 |
| Reps2 | 0.040651569 | 0.436166541 | 0.797485381 | 0.202719167 | 0.156253878 |
| Rer1 | 0.000200307 | 0.997577411 | 0.999369645 | 0.800349377 | 0.776415641 |
| Rere | 0.154815229 | 0.018047321 | 0.213922515 | 0.694497905 | 0.531869047 |
| Resf1 | 0.056833539 | 0.382776976 | 0.763651859 | 0.345612976 | 0.291100294 |
| Resp18 | -0.032995618 | 0.623014651 | 0.885857477 | 2.690717814 | 2.763952527 |
| Ret | -0.035548566 | 0.525930677 | 0.842051614 | 0.231218257 | 0.2682424 |
| Retreg1 | 0.088625766 | 0.17867324 | 0.584913016 | 0.376872977 | 0.294117253 |
| Retreg2 | 0.077284268 | 0.258895325 | 0.665948842 | 0.579734285 | 0.508874821 |
| Retreg3 | 0.072851132 | 0.218128728 | 0.632396893 | 0.293692513 | 0.233868858 |
| Retsat | -0.054978639 | 0.020610995 | 0.227971987 | 0.02635395 | 0.072290772 |
| Rev1 | -0.010829159 | 0.81881154 | 0.950128492 | 0.152263874 | 0.163516963 |
| Rev3l | -0.046642418 | 0.491555156 | 0.827270742 | 0.444506226 | 0.472918418 |
| Rex1bd | -0.037814236 | 0.577702611 | 0.864673658 | 0.866565747 | 0.911678224 |
| Rexo1 | 0.112715064 | 0.053640581 | 0.361325633 | 0.295864984 | 0.19968687 |
| Rexo2 | -0.216155506 | 0.001789852 | 0.056529878 | 0.760950597 | 0.994506928 |
| Rexo4 | 0.104557501 | 0.024915104 | 0.25082538 | 0.164018096 | 0.072267499 |
| Rfc1 | -0.034029117 | 0.587132739 | 0.869181609 | 0.378903393 | 0.41045197 |

| | | | | | |
|---|---|---|---|---|---|
| Rfc2 | -0.024729673 | 0.642363429 | 0.892561795 | 0.208746947 | 0.238037162 |
| Rfc3 | -0.009232882 | 0.836389734 | 0.954911647 | 0.13671835 | 0.149973207 |
| Rfc4 | 0.072941009 | 0.022499984 | 0.237968744 | 0.0749333 | 0.01963691 |
| Rfc5 | 0.025704309 | 0.424262305 | 0.789771168 | 0.071429169 | 0.047667034 |
| Rfesd | 0.07564856 | 0.219366458 | 0.63289494 | 0.403125461 | 0.338225892 |
| Rffl | -0.020075809 | 0.53724386 | 0.847793898 | 0.064396918 | 0.083720091 |
| Rfk | -0.116572003 | 0.075174044 | 0.413795734 | 0.51686227 | 0.633038073 |
| Rflna | -0.0449416 | 0.514706703 | 0.836743564 | 0.490909679 | 0.52762366 |
| Rflnb | 0.131483551 | 0.027486367 | 0.264360019 | 0.252285925 | 0.143939611 |
| Rfng | 0.056161484 | 0.109555434 | 0.477972625 | 0.087111849 | 0.041321984 |
| Rfwd3 | -0.041122927 | 0.195569774 | 0.605843771 | 0.058892638 | 0.09621049 |
| Rfx3 | -0.060225486 | 0.217761787 | 0.632272789 | 0.160755552 | 0.211417407 |
| Rfx5 | 0.01650651 | 0.697261043 | 0.910970095 | 0.115814906 | 0.105286939 |
| Rfx6 | -0.101286635 | 0.081863475 | 0.42777546 | 0.24135086 | 0.337714845 |
| Rfx7 | 0.009829902 | 0.825161971 | 0.952011047 | 0.126837937 | 0.121149458 |
| Rfxank | -0.02600083 | 0.38346487 | 0.763961274 | 0.052998181 | 0.077548456 |
| Rfxap | -0.087920692 | 0.108183236 | 0.477710135 | 0.213984435 | 0.302656888 |
| Rgl1 | 0.031086845 | 0.505152896 | 0.832249514 | 0.148637501 | 0.122446205 |
| Rgl2 | -0.048487888 | 0.141170755 | 0.528664246 | 0.061207922 | 0.099784236 |
| Rgl3 | -0.014412307 | 0.658531402 | 0.898224427 | 0.066500825 | 0.075945932 |
| Rgmb | -0.002354049 | 0.953534175 | 0.989007183 | 0.107674959 | 0.109217146 |
| Rgp1 | -0.013271223 | 0.691662169 | 0.909796728 | 0.070661552 | 0.081570599 |
| Rgs11 | -0.078674607 | 0.174828444 | 0.577992588 | 0.250775356 | 0.312134194 |
| Rgs16 | -0.114667407 | 0.033278438 | 0.288940905 | 0.176342331 | 0.291979992 |
| Rgs17 | 0.069003744 | 0.301619848 | 0.706960943 | 0.388919032 | 0.312807984 |
| Rgs2 | 0.158192373 | 0.040936403 | 0.319308264 | 0.495830139 | 0.325474732 |
| Rgs3 | -0.011441972 | 0.757480636 | 0.927237784 | 0.082762952 | 0.100154342 |
| Rgs4 | 0.080958258 | 0.299944301 | 0.70506682 | 0.505218643 | 0.383887182 |
| Rgs7 | 0.09776839 | 0.08380341 | 0.431794059 | 0.260888995 | 0.179607474 |
| Rgs7bp | 0.027547329 | 0.598526405 | 0.875460438 | 0.188659263 | 0.157862779 |
| Rgs9 | -0.085451885 | 0.091033733 | 0.445496137 | 0.154787357 | 0.238142815 |
| Rhbdd1 | 0.019108263 | 0.705359214 | 0.912844672 | 0.18441033 | 0.1607125 |
| Rhbdd2 | 0.006467217 | 0.910124328 | 0.978641543 | 0.267381354 | 0.26311819 |
| Rhbdd3 | -0.022408318 | 0.581992189 | 0.866173066 | 0.10894908 | 0.122250505 |
| Rhbdl2 | 0.020131177 | 0.644929965 | 0.893466123 | 0.126200206 | 0.106754841 |
| Rhd | -0.025563995 | 0.315929993 | 0.714991959 | 0.033494384 | 0.053552018 |
| Rheb | 0.06515324 | 0.347875061 | 0.739124045 | 1.192560958 | 1.072410231 |
| Rhno1 | 0.039726893 | 0.186573188 | 0.593545195 | 0.061301177 | 0.028970965 |
| Rhoa | -0.034522299 | 0.556515589 | 0.85521955 | 1.271710346 | 1.322896041 |
| Rhob | -0.110723008 | 0.029625616 | 0.273726881 | 0.167231336 | 0.257627532 |
| Rhobtb1 | 0.019109062 | 0.758543242 | 0.927457489 | 0.400044731 | 0.385916369 |
| Rhobtb2 | 0.08322479 | 0.007145775 | 0.128637347 | 0.068248337 | 0 |
| Rhobtb3 | 0.113299361 | 0.05916598 | 0.3746681 | 0.31045273 | 0.201829676 |
| Rhoc | 0.032258821 | 0.436149546 | 0.797485381 | 0.116464411 | 0.086364445 |
| Rhog | 0.034330879 | 0.385847241 | 0.765970878 | 0.107593698 | 0.081923954 |
| Rhoq | -0.020253061 | 0.721864803 | 0.919475778 | 0.2731401 | 0.287111376 |
| Rhot1 | 0.045696115 | 0.371955955 | 0.75760494 | 0.194910461 | 0.14943874 |
| Rhot2 | -0.025339968 | 0.532367734 | 0.84612966 | 0.109555339 | 0.129757357 |
| Rhou | -0.091622216 | 0.10641712 | 0.473900883 | 0.220065536 | 0.315920206 |
| Rhpn2 | 0.108345954 | 0.061363333 | 0.380250374 | 0.282440608 | 0.185417198 |
| Rian | -0.150772355 | 0.03104118 | 0.280684975 | 1.058982775 | 1.233616112 |
| Ric1 | -0.002744499 | 0.949229919 | 0.987767085 | 0.120687687 | 0.130650718 |
| Ric3 | -0.016618675 | 0.739758383 | 0.922901128 | 0.180143914 | 0.195487904 |
| Ric8a | 0.080679363 | 0.129179927 | 0.508661798 | 0.211803813 | 0.147610446 |
| Ric8b | 0.037888363 | 0.496084784 | 0.830005497 | 0.231848213 | 0.202585445 |
| Rictor | -0.010086198 | 0.856734176 | 0.958302663 | 0.230864917 | 0.24964846 |
| Rida | -0.089522592 | 0.020869802 | 0.229383208 | 0.087914247 | 0.16244627 |
| Rif1 | -0.120699783 | 0.065854905 | 0.390314712 | 0.455506785 | 0.553281001 |
| Riiad1 | -0.024931732 | 0.560654655 | 0.856904336 | 0.117229018 | 0.140067648 |
| Rilpl1 | -0.040488647 | 0.242082809 | 0.652555608 | 0.073157512 | 0.109169814 |
| Rilpl2 | 0.029250057 | 0.596527279 | 0.87393173 | 0.235961814 | 0.221908401 |
| Rimbp2 | -0.029835767 | 0.640539078 | 0.891345382 | 0.459841936 | 0.477932751 |
| Rims2 | -0.166456411 | 0.008908261 | 0.146996602 | 0.346316976 | 0.47411491 |
| Rims3 | 0.13680333 | 0.024204616 | 0.24664135 | 0.350921541 | 0.228457578 |
| Ring1 | -0.042680853 | 0.380593986 | 0.762006446 | 0.161424737 | 0.206105886 |

| | | | | | |
|---|---|---|---|---|---|
| Rint1 | 0.082763422 | 0.147641273 | 0.539059825 | 0.281053064 | 0.198399602 |
| Riok1 | -0.047653879 | 0.425447477 | 0.790344696 | 0.278996531 | 0.330905267 |
| Riok2 | 0.048701325 | 0.290719303 | 0.69667177 | 0.149725521 | 0.103587914 |
| Riok3 | -0.092648956 | 0.154785025 | 0.551696483 | 0.802018713 | 0.914890089 |
| Riox1 | -0.010278441 | 0.745010567 | 0.924566547 | 0.06005785 | 0.07100036 |
| Riox2 | -0.031964481 | 0.437531859 | 0.798455866 | 0.109834664 | 0.142545578 |
| Ripk1 | -0.031674774 | 0.518942291 | 0.838342793 | 0.167574609 | 0.197799716 |
| Ripk2 | 0.01984442 | 0.566899237 | 0.858864944 | 0.079364086 | 0.061685106 |
| Ripk4 | -0.016682691 | 0.731671501 | 0.920770724 | 0.164594185 | 0.190470598 |
| Ripor1 | 0.018553328 | 0.708738951 | 0.913767764 | 0.174237976 | 0.157005257 |
| Ripply3 | 0.012904426 | 0.806696818 | 0.945532285 | 0.194488679 | 0.187312298 |
| Rit1 | -0.002372006 | 0.962608023 | 0.991757003 | 0.188538287 | 0.18960609 |
| Rit2 | -0.064523837 | 0.000424232 | 0.022649017 | 0.00864105 | 0.066661475 |
| Rita1 | -0.007695112 | 0.786341143 | 0.937336173 | 0.049191958 | 0.051590031 |
| Rlf | -0.080068655 | 0.172246156 | 0.575294174 | 0.259363201 | 0.349620905 |
| Rlim | -0.001792938 | 0.974741432 | 0.994357125 | 0.235247118 | 0.234052922 |
| Rmc1 | -0.010545062 | 0.795743495 | 0.939757086 | 0.109628378 | 0.128306645 |
| Rmdn1 | 0.017233776 | 0.727637104 | 0.920455328 | 0.172447226 | 0.155571849 |
| Rmdn3 | 0.024335337 | 0.657170813 | 0.898224427 | 0.245328328 | 0.21344631 |
| Rmi1 | 0.020461908 | 0.596361895 | 0.873822532 | 0.09784896 | 0.08350216 |
| Rmnd1 | -0.017465876 | 0.489578047 | 0.826459641 | 0.035411768 | 0.050699164 |
| Rmnd5a | -0.271339295 | 1.09E-05 | 0.001349633 | 0.329723689 | 0.589339956 |
| Rmnd5b | 0.019289094 | 0.636938821 | 0.890755879 | 0.115389783 | 0.107801163 |
| Rn18s | 0.062596308 | 0.035006817 | 0.294903156 | 4.58936215 | 4.334136195 |
| Rnase4 | -0.428330525 | 1.70E-08 | 8.40E-06 | 0.75947621 | 1.373648328 |
| Rnaseh1 | 0.039264972 | 0.361526381 | 0.750013467 | 0.130906537 | 0.100918337 |
| Rnaseh2a | -0.023445414 | 0.550973319 | 0.852505562 | 0.099858707 | 0.121551859 |
| Rnaseh2b | 0.017468755 | 0.614985013 | 0.882439418 | 0.081402003 | 0.068064358 |
| Rnaseh2c | 0.012298083 | 0.831025047 | 0.95325696 | 0.254417408 | 0.241356665 |
| Rnaset2a | -0.059017058 | 0.065885532 | 0.390314712 | 0.059638046 | 0.113580696 |
| Rnaset2b | 0.004015361 | 0.891405708 | 0.970852438 | 0.054423987 | 0.048841618 |
| Rnd3 | -0.005047428 | 0.869993949 | 0.964130426 | 0.051747896 | 0.054020956 |
| Rnf10 | 0.000800359 | 0.99035851 | 0.997733826 | 0.86128221 | 0.857176285 |
| Rnf103 | 0.014513692 | 0.806860071 | 0.945532285 | 0.305864798 | 0.289090861 |
| Rnf11 | -0.01415196 | 0.813209834 | 0.948397417 | 1.112368118 | 1.154699708 |
| Rnf111 | -0.008118395 | 0.886028793 | 0.969590273 | 0.245289975 | 0.244737258 |
| Rnf113a1 | 0.008309183 | 0.777000837 | 0.934230706 | 0.054543277 | 0.048168712 |
| Rnf113a2 | -0.025013392 | 0.617865471 | 0.883238152 | 0.172853723 | 0.197099531 |
| Rnf114 | 0.028612935 | 0.620267706 | 0.883937014 | 0.276620169 | 0.243657876 |
| Rnf115 | -0.009897109 | 0.884386601 | 0.968954079 | 0.590884333 | 0.607736897 |
| Rnf121 | 0.039428039 | 0.42513761 | 0.790344696 | 0.173689018 | 0.144908539 |
| Rnf123 | 0.031443891 | 0.265164902 | 0.672177989 | 0.052508446 | 0.021943285 |
| Rnf126 | -0.088725827 | 0.087501626 | 0.439786592 | 0.193006038 | 0.27608583 |
| Rnf128 | 0.131592244 | 0.043091862 | 0.328353923 | 0.414759503 | 0.2850189 |
| Rnf13 | -0.117127158 | 0.0193056 | 0.220946759 | 0.165044969 | 0.263293744 |
| Rnf130 | 0.116164035 | 0.070621922 | 0.402952703 | 1.169981654 | 1.013257629 |
| Rnf135 | 0.025040579 | 0.404211313 | 0.776339704 | 0.057273411 | 0.037898074 |
| Rnf138 | 0.072171588 | 0.240718305 | 0.650744882 | 0.311712467 | 0.267518128 |
| Rnf138rt1 | -0.048788587 | 0.33872606 | 0.732638051 | 0.1652378 | 0.206502987 |
| Rnf139 | 0.082247914 | 0.081903437 | 0.42777546 | 0.166632126 | 0.095566247 |
| Rnf14 | 0.091795267 | 0.184857053 | 0.591385002 | 0.618728731 | 0.525028512 |
| Rnf141 | -0.105393666 | 0.062017871 | 0.380250374 | 0.245528185 | 0.335012397 |
| Rnf145 | 0.099583458 | 0.048002295 | 0.343406298 | 0.190093716 | 0.095646989 |
| Rnf146 | -0.070686673 | 0.215226257 | 0.629611185 | 0.251307204 | 0.323154243 |
| Rnf149 | 0.097621478 | 0.018420545 | 0.2157189 | 0.126223821 | 0.040233886 |
| Rnf150 | 0.020505154 | 0.701705191 | 0.912152193 | 0.190518025 | 0.181031522 |
| Rnf157 | -0.059467259 | 0.054810397 | 0.364567232 | 0.054872279 | 0.104817767 |
| Rnf166 | -0.063633162 | 0.219634517 | 0.63289494 | 0.17809101 | 0.229511654 |
| Rnf167 | 0.070552369 | 0.248521147 | 0.659783035 | 0.334232244 | 0.267073665 |
| Rnf168 | 0.004031915 | 0.941575967 | 0.986168593 | 0.225268799 | 0.220876928 |
| Rnf169 | 0.019071139 | 0.666582344 | 0.901197061 | 0.133786981 | 0.12637169 |
| Rnf170 | 0.01283923 | 0.784869608 | 0.936423418 | 0.161587148 | 0.15351617 |
| Rnf181 | -0.050160026 | 0.353542764 | 0.744234211 | 0.214745689 | 0.263911596 |
| Rnf182 | -0.095303009 | 0.044622472 | 0.330723271 | 0.141791663 | 0.230861313 |
| Rnf185 | -0.059333004 | 0.219911495 | 0.633217295 | 0.158817122 | 0.206215051 |

| | | | | | |
|---|---|---|---|---|---|
| Rnf186 | 0.102075029 | 0.019062746 | 0.218929273 | 0.127170639 | 0.035842605 |
| Rnf187 | -0.024200048 | 0.713198042 | 0.915509496 | 0.523610491 | 0.57273858 |
| Rnf19a | -0.035106788 | 0.33992824 | 0.73341783 | 0.085758447 | 0.11790879 |
| Rnf19b | 0.045753686 | 0.398213962 | 0.774311756 | 0.223303382 | 0.18190125 |
| Rnf2 | 0.091955839 | 0.084095899 | 0.432507481 | 0.218241192 | 0.143814364 |
| Rnf20 | -0.004120218 | 0.948840418 | 0.987767085 | 0.461413376 | 0.465222455 |
| Rnf208 | -0.048596909 | 0.112916707 | 0.483221516 | 0.055222373 | 0.102813575 |
| Rnf213 | 0.054759904 | 0.170342755 | 0.572762895 | 0.109938032 | 0.058080275 |
| Rnf214 | -0.017500531 | 0.738188037 | 0.922843014 | 0.213957234 | 0.224816628 |
| Rnf215 | 0.013466342 | 0.728315513 | 0.920519505 | 0.103257825 | 0.095509426 |
| Rnf216 | 0.030170457 | 0.59620125 | 0.873714695 | 0.253668227 | 0.221633623 |
| Rnf217 | 0.026044122 | 0.641315919 | 0.891787413 | 0.228943684 | 0.202276648 |
| Rnf220 | 0.021604674 | 0.709055224 | 0.913931846 | 0.277960889 | 0.252399955 |
| Rnf227 | 0.101606213 | 0.128866919 | 0.508260084 | 0.446442935 | 0.333898788 |
| Rnf24 | 0.012496478 | 0.686682301 | 0.909796728 | 0.060087832 | 0.048362027 |
| Rnf25 | 0.060389817 | 0.176208495 | 0.580631865 | 0.145836374 | 0.091603615 |
| Rnf26 | 0.003458278 | 0.905518364 | 0.976167673 | 0.052723657 | 0.049780338 |
| Rnf31 | 0.08942683 | 0.034777349 | 0.29382176 | 0.128629945 | 0.050029957 |
| Rnf34 | 0.064055143 | 0.221504022 | 0.633858837 | 0.20267128 | 0.15702389 |
| Rnf38 | -0.041229333 | 0.297625707 | 0.702656882 | 0.099811687 | 0.136266539 |
| Rnf4 | 0.102157627 | 0.091078816 | 0.445496137 | 0.329662244 | 0.241653736 |
| Rnf40 | 0.076864819 | 0.047899533 | 0.342915559 | 0.108241203 | 0.04500794 |
| Rnf41 | 0.03855439 | 0.379379911 | 0.762006446 | 0.135947448 | 0.112607586 |
| Rnf44 | -0.05628072 | 0.222192542 | 0.633924544 | 0.136905747 | 0.184587198 |
| Rnf5 | -0.029254779 | 0.575021383 | 0.862723699 | 0.201233927 | 0.222520205 |
| Rnf6 | 0.080478286 | 0.216792885 | 0.631259121 | 0.525176821 | 0.441631786 |
| Rnf7 | 0.022758391 | 0.726023087 | 0.92014727 | 0.950643974 | 0.917229919 |
| Rnf8 | 0.102324935 | 0.041780925 | 0.322580882 | 0.198326866 | 0.113691379 |
| Rnft1 | 0.016596376 | 0.690442699 | 0.909796728 | 0.117857873 | 0.10716115 |
| Rngtt | -0.035229949 | 0.387627982 | 0.767160379 | 0.106333654 | 0.14195711 |
| Rnh1 | 0.069462495 | 0.30878128 | 0.711182121 | 0.784499649 | 0.706152412 |
| Rnmt | 0.047327587 | 0.404219517 | 0.776339704 | 0.248329325 | 0.204534612 |
| Rnpc3 | -0.034244257 | 0.454992625 | 0.807876964 | 0.137054325 | 0.168226703 |
| Rnpep | -0.016621089 | 0.733315387 | 0.921306823 | 0.168084686 | 0.178318721 |
| Rnpepl1 | -0.002223463 | 0.971642363 | 0.993892166 | 0.388396074 | 0.369935738 |
| Rnps1 | -0.074594633 | 0.26204924 | 0.668764912 | 0.4884248 | 0.567891187 |
| Ro60 | 0.073488748 | 0.268352123 | 0.676166273 | 0.610303636 | 0.553340671 |
| Robo1 | 0.028587155 | 0.572910434 | 0.861868184 | 0.182675449 | 0.164610816 |
| Robo2 | 0.023972322 | 0.72683101 | 0.920414187 | 0.493371488 | 0.468224696 |
| Rock1 | 0.083590957 | 0.157313829 | 0.554924229 | 1.263997628 | 1.145894477 |
| Rock2 | -0.032772092 | 0.643119773 | 0.892561795 | 0.632337558 | 0.642676507 |
| Rogdi | -0.163623671 | 0.009579197 | 0.151890044 | 0.329569019 | 0.486271088 |
| Rom1 | -0.031932752 | 0.364782794 | 0.752802229 | 0.079388627 | 0.102157109 |
| Romo1 | 0.214588915 | 3.28E-05 | 0.003427487 | 1.577381477 | 1.195440214 |
| Ropn1l | 0.004348884 | 0.878973399 | 0.967163177 | 0.052483769 | 0.051640846 |
| Ror2 | 0.079739159 | 0.056023134 | 0.367897032 | 0.122482791 | 0.054016955 |
| Rora | -0.094865679 | 0.161928279 | 0.560632679 | 0.578067425 | 0.680905371 |
| Rorc | -0.073498235 | 0.095407856 | 0.455842134 | 0.124101911 | 0.186615298 |
| Rp2 | -0.002287451 | 0.963803191 | 0.991970325 | 0.171623938 | 0.167825262 |
| Rp9 | 0.047009944 | 0.475296846 | 0.817387411 | 0.879935878 | 0.8110392 |
| Rpa1 | -0.109744282 | 0.044978425 | 0.331703493 | 0.194225583 | 0.329413472 |
| Rpa3 | 0.017932878 | 0.722445142 | 0.919483582 | 0.178853431 | 0.169490401 |
| Rpain | 0.034101491 | 0.264883352 | 0.671881274 | 0.060256534 | 0.030744295 |
| Rpap1 | -0.017657738 | 0.598730316 | 0.875574105 | 0.072217679 | 0.084735399 |
| Rpap2 | -0.010934555 | 0.780088042 | 0.935229775 | 0.100518047 | 0.106841126 |
| Rpap3 | -0.005220625 | 0.896617275 | 0.972419247 | 0.110777978 | 0.113974724 |
| Rpe | -0.030674323 | 0.546691459 | 0.851446616 | 0.185231295 | 0.217143779 |
| Rpf1 | -0.068426765 | 0.188918204 | 0.596946223 | 0.191021921 | 0.250378055 |
| Rpf2 | 0.02857507 | 0.490716469 | 0.826951175 | 0.117459099 | 0.094940599 |
| Rpgr | 0.012236596 | 0.754597056 | 0.926726954 | 0.102781559 | 0.090102536 |
| Rpgrip1 | 0.083478248 | 0.024019717 | 0.245835998 | 0.098057529 | 0.032728095 |
| Rpgrip1l | -0.001933819 | 0.94802848 | 0.987651726 | 0.052576601 | 0.055692162 |
| Rph3al | -0.043836356 | 0.504649519 | 0.832249514 | 0.385019839 | 0.421022372 |
| Rpia | 0.038583313 | 0.494278231 | 0.828842054 | 0.249624796 | 0.216895348 |
| Rpl10 | -0.106354887 | 0.001517108 | 0.05109802 | 2.902137451 | 3.213586654 |

| | | | | | |
|---|---|---|---|---|---|
| Rpl10a | -0.311643905 | 5.44E-06 | 0.000814735 | 0.945080742 | 1.398777185 |
| Rpl10a-ps1 | 0.015995217 | 0.605471688 | 0.87802905 | 0.063940758 | 0.049964753 |
| Rpl10-ps3 | -0.016217763 | 0.542684785 | 0.85028523 | 0.042536789 | 0.060544146 |
| Rpl11 | -0.135652595 | 8.20E-05 | 0.006856998 | 2.619435277 | 2.971566221 |
| Rpl12 | -0.181899929 | 9.83E-05 | 0.007650838 | 2.0342082 | 2.401730023 |
| Rpl13 | -0.135668972 | 4.09E-05 | 0.00414445 | 2.690212898 | 3.057269462 |
| Rpl13a | 0.01606333 | 0.813499206 | 0.948397417 | 0.745637341 | 0.738819953 |
| Rpl14 | -0.140492248 | 0.000293456 | 0.017224644 | 2.317566558 | 2.647772614 |
| Rpl15 | -0.027373505 | 0.650002587 | 0.895593446 | 1.241120379 | 1.304867524 |
| Rpl17 | -0.11534262 | 0.000946171 | 0.037078044 | 2.60981061 | 2.919934045 |
| Rpl18 | -0.127149697 | 0.001059605 | 0.039683797 | 2.196106988 | 2.490570259 |
| Rpl18a | -0.112180414 | 0.010002422 | 0.155789067 | 2.183409406 | 2.433469897 |
| Rpl19 | -0.103154211 | 0.000122298 | 0.009092661 | 3.206754767 | 3.534829532 |
| Rpl21 | -0.113430211 | 0.000805732 | 0.033143999 | 2.569357715 | 2.900437752 |
| Rpl22 | -0.097597559 | 0.040126379 | 0.315774775 | 1.786012982 | 1.937510064 |
| Rpl22l1 | -0.065244047 | 0.170907825 | 0.573329491 | 1.888228385 | 1.982149931 |
| Rpl23 | -0.100835066 | 0.000577557 | 0.027869893 | 2.929066261 | 3.194363901 |
| Rpl23a | -0.042908077 | 0.509960189 | 0.834904701 | 0.991609473 | 1.05325071 |
| Rpl24 | -0.063319008 | 0.339542329 | 0.733215653 | 0.597390761 | 0.693133412 |
| Rpl26 | -0.069742017 | 0.023782626 | 0.245330129 | 2.789855008 | 2.960286803 |
| Rpl27 | -0.003035696 | 0.956255211 | 0.989397512 | 1.482761271 | 1.460493896 |
| Rpl27a | 0.030515714 | 0.4231864 | 0.788769155 | 2.361324425 | 2.277197285 |
| Rpl28 | -0.034453266 | 0.245249926 | 0.655005464 | 2.819651882 | 2.914079205 |
| Rpl29 | -0.204055563 | 8.65E-05 | 0.007178374 | 1.64721769 | 2.055842463 |
| Rpl3 | -0.233306137 | 5.12E-06 | 0.000778614 | 1.910853019 | 2.388252526 |
| Rpl30 | 0.028367057 | 0.400593411 | 0.774740795 | 2.670655753 | 2.599860605 |
| Rpl31 | 0.027365575 | 0.582964421 | 0.866383905 | 1.698498225 | 1.641770919 |
| Rpl32 | -0.100441164 | 5.61E-05 | 0.00505496 | 3.30630162 | 3.594755565 |
| Rpl34 | -0.057791354 | 0.094996403 | 0.455121192 | 2.782823524 | 2.896599346 |
| Rpl35 | -0.04661961 | 0.280012591 | 0.687465228 | 2.095623666 | 2.150304111 |
| Rpl35a | 0.022582467 | 0.453917785 | 0.807844867 | 2.893264786 | 2.817847448 |
| Rpl36 | -0.038845571 | 0.207403594 | 0.619003827 | 2.867396053 | 2.938577371 |
| Rpl36a | -0.056541301 | 0.256907329 | 0.665256942 | 1.738458106 | 1.830099899 |
| Rpl36al | 0.019360022 | 0.747795928 | 0.924930563 | 1.340255334 | 1.28218522 |
| Rpl37 | 0.002916254 | 0.913292102 | 0.979979308 | 3.300030273 | 3.254146555 |
| Rpl37a | 0.054456715 | 0.026509674 | 0.259082375 | 3.439573715 | 3.23939182 |
| Rpl37rt | -0.01586521 | 0.695460898 | 0.910439681 | 0.110582286 | 0.130794189 |
| Rpl38 | 0.049847446 | 0.09092313 | 0.445385778 | 2.881872118 | 2.726973335 |
| Rpl38-ps2 | -0.034647049 | 0.12942754 | 0.509237248 | 0.025711132 | 0.055645441 |
| Rpl39 | -0.003087426 | 0.914767159 | 0.980132949 | 2.856387885 | 2.840470288 |
| Rpl3-ps1 | -0.075291085 | 0.036889197 | 0.303216817 | 0.074965809 | 0.142179355 |
| Rpl4 | -0.079136855 | 0.129086637 | 0.508493946 | 1.817723592 | 1.940020018 |
| Rpl41 | -0.011120093 | 0.615644801 | 0.882439418 | 3.885983504 | 3.918835534 |
| Rpl5 | -0.196540835 | 0.001635338 | 0.053297575 | 1.231057058 | 1.56038992 |
| Rpl6 | -0.089558988 | 0.018972651 | 0.218381306 | 2.377597739 | 2.587511902 |
| Rpl6l | -0.031486523 | 0.154449947 | 0.551012197 | 0.023127985 | 0.054530087 |
| Rpl7 | -0.076120463 | 0.124854555 | 0.503481386 | 1.603494019 | 1.755149884 |
| Rpl7a | -0.116185308 | 0.01889405 | 0.218227362 | 1.793880112 | 2.012746959 |
| Rpl7l1 | 0.030401244 | 0.620325658 | 0.883937014 | 0.33220113 | 0.310857356 |
| Rpl8 | -0.137241313 | 1.62E-05 | 0.001832428 | 2.978629949 | 3.350015137 |
| Rpl9 | -0.149867953 | 7.80E-07 | 0.000166207 | 2.961193753 | 3.405894221 |
| Rpl9-ps6 | -0.051790173 | 0.123237493 | 0.500122758 | 0.064369013 | 0.1139862 |
| Rplp0 | -0.086838453 | 0.0194026 | 0.221110882 | 2.473947149 | 2.673631488 |
| Rplp1 | 0.043286007 | 0.121858489 | 0.497192544 | 3.334434976 | 3.175603761 |
| Rplp2 | 0.005235414 | 0.874317564 | 0.964947297 | 2.808654439 | 2.773387737 |
| Rpn1 | 0.037127979 | 0.549242579 | 0.852402509 | 1.13227362 | 1.080617108 |
| Rpn2 | -0.030204587 | 0.658892212 | 0.898224427 | 1.001532836 | 1.017507343 |
| Rpp14 | 0.004041626 | 0.930809039 | 0.982803527 | 0.153035212 | 0.15773908 |
| Rpp21 | 0.137358486 | 0.043806531 | 0.329086044 | 0.520810167 | 0.380490656 |
| Rpp25l | 0.043860459 | 0.458033262 | 0.809095362 | 0.307998798 | 0.271165644 |
| Rpp30 | 0.045095802 | 0.285441334 | 0.692407818 | 0.122539608 | 0.084234324 |
| Rpp38 | 0.069501223 | 0.119077838 | 0.492069355 | 0.13388691 | 0.07414454 |
| Rpp40 | -0.019220897 | 0.469548772 | 0.814050152 | 0.039823528 | 0.053284945 |
| Rprd1a | -0.001312306 | 0.981897785 | 0.995803987 | 0.248461434 | 0.256860582 |
| Rprd1b | -0.070966644 | 0.292333208 | 0.698069506 | 0.497946258 | 0.564797968 |

| | | | | | |
|---|---|---|---|---|---|
| Rprd2 | -0.002031293 | 0.97314067 | 0.994079882 | 0.316677798 | 0.309591584 |
| Rprml | 0.015922154 | 0.695323755 | 0.910378884 | 0.102895088 | 0.105139477 |
| Rps10 | -0.027229286 | 0.680473101 | 0.907783945 | 0.929215722 | 0.98435125 |
| Rps10-ps1 | -0.061421408 | 0.044167992 | 0.329446347 | 0.05439352 | 0.113893729 |
| Rps10-ps2 | -0.072164151 | 0.080679289 | 0.425750801 | 0.105902257 | 0.171797865 |
| Rps11 | -0.096099074 | 0.004658137 | 0.100329878 | 2.479840733 | 2.72102068 |
| Rps11-ps1 | -0.094499449 | 0.005837974 | 0.113654038 | 0.071640913 | 0.147240135 |
| Rps11-ps2 | -0.017348355 | 0.604883154 | 0.877977183 | 0.069348466 | 0.08296938 |
| Rps12 | -0.069797928 | 0.053208282 | 0.360358652 | 2.723429513 | 2.861635945 |
| Rps13 | -0.108247629 | 0.001023462 | 0.038764098 | 2.595379496 | 2.867517445 |
| Rps14 | -0.074197639 | 0.001662968 | 0.053669478 | 3.463644366 | 3.672124137 |
| Rps15 | -0.02853775 | 0.453923555 | 0.807844867 | 2.588366617 | 2.609635549 |
| Rps15a | -0.035121333 | 0.346363241 | 0.738256073 | 2.330350968 | 2.416779314 |
| Rps16 | -0.079540792 | 0.02615962 | 0.257205918 | 2.562518354 | 2.76027219 |
| Rps17 | 0.015885682 | 0.649381461 | 0.895585658 | 2.408680245 | 2.35214166 |
| Rps18 | -0.150144133 | 6.93E-05 | 0.006092411 | 2.433079758 | 2.806014816 |
| Rps19 | -0.017730445 | 0.68479127 | 0.909796728 | 1.99096897 | 2.011883291 |
| Rps19bp1 | 0.073483842 | 0.254387381 | 0.663146569 | 0.400721701 | 0.319263754 |
| Rps2 | -0.134408406 | 0.000452644 | 0.023539853 | 2.369481509 | 2.740448658 |
| Rps20 | -0.094689105 | 0.005928015 | 0.113983697 | 2.854760291 | 3.064412695 |
| Rps21 | 0.034028816 | 0.291364542 | 0.697147764 | 3.430137443 | 3.246630753 |
| Rps23 | -0.051953594 | 0.118449705 | 0.491192747 | 2.679206507 | 2.77442082 |
| Rps23-ps1 | -0.026505696 | 0.466642339 | 0.813152001 | 0.081953049 | 0.107498859 |
| Rps24 | -0.121286404 | 2.00E-05 | 0.002221072 | 3.315212338 | 3.670357858 |
| Rps25 | -0.037038636 | 0.355910181 | 0.745326231 | 2.122532869 | 2.171971051 |
| Rps26 | -0.079027181 | 0.073629732 | 0.411252986 | 1.937724809 | 2.110761568 |
| Rps27 | -0.027443916 | 0.319446751 | 0.717256849 | 3.103465895 | 3.145541837 |
| Rps27a | -0.084815724 | 0.002089214 | 0.062782754 | 3.1658091 | 3.403189161 |
| Rps27a-ps3 | -0.010560354 | 0.744915197 | 0.924566547 | 0.067918063 | 0.07908721 |
| Rps27l | -0.002250591 | 0.964408818 | 0.991970325 | 1.840878908 | 1.825266896 |
| Rps27rt | 0.031746842 | 0.547313002 | 0.851449318 | 0.207596091 | 0.195236142 |
| Rps28 | 0.128378039 | 0.031442062 | 0.281771404 | 1.388165645 | 1.194168262 |
| Rps29 | 0.087280152 | 0.000855147 | 0.034332451 | 3.348322133 | 3.048707109 |
| Rps2-ps10 | -0.026754899 | 0.301235615 | 0.706666217 | 0.034995987 | 0.061310523 |
| Rps3 | -0.151774795 | 6.98E-05 | 0.006092411 | 2.4923733 | 2.856265683 |
| Rps3a1 | -0.189278118 | 1.37E-08 | 7.65E-06 | 2.871356457 | 3.415896585 |
| Rps4l | -0.023211908 | 0.75872876 | 0.927571324 | 0.782717399 | 0.849257824 |
| Rps4x | -0.216389211 | 8.05E-07 | 0.000166207 | 2.2673171 | 2.776091719 |
| Rps5 | -0.137430976 | 8.98E-05 | 0.007326725 | 2.538110517 | 2.893260098 |
| Rps6 | -0.07180526 | 0.153707947 | 0.550142209 | 1.593865331 | 1.722627866 |
| Rps6ka2 | -0.001640909 | 0.977290931 | 0.994733706 | 0.26435432 | 0.271668059 |
| Rps6ka3 | -0.068589088 | 0.226190734 | 0.638587289 | 0.249823045 | 0.299265805 |
| Rps6kb1 | 0.062970174 | 0.316770159 | 0.71592481 | 0.416793369 | 0.360670482 |
| Rps6kb2 | 0.060594353 | 0.169371107 | 0.57227323 | 0.140265018 | 0.083089654 |
| Rps6kc1 | -0.027679241 | 0.295821161 | 0.702426542 | 0.039354977 | 0.063644166 |
| Rps6kl1 | -0.024991879 | 0.340324191 | 0.733546237 | 0.038418821 | 0.06036493 |
| Rps6-ps4 | -0.04114086 | 0.236301848 | 0.646961714 | 0.074463775 | 0.113464167 |
| Rps7 | -0.162703804 | 0.000200264 | 0.012906373 | 1.81154504 | 2.162004797 |
| Rps8 | -0.099430169 | 0.000114025 | 0.008670195 | 3.452746964 | 3.728118467 |
| Rps9 | -0.159967914 | 0.0004355 | 0.023005843 | 2.157798283 | 2.471191486 |
| Rpsa | -0.112859761 | 0.004713727 | 0.100919217 | 2.325924906 | 2.603557685 |
| Rpsa-ps10 | -0.041139315 | 0.193433606 | 0.60325934 | 0.057729188 | 0.098215999 |
| Rptor | 0.101532152 | 0.016507428 | 0.206965324 | 0.128871117 | 0.042635358 |
| Rpusd1 | -0.036051307 | 0.382257986 | 0.76307148 | 0.10441495 | 0.137345832 |
| Rpusd3 | -0.036253803 | 0.182819698 | 0.590066231 | 0.040483552 | 0.072749935 |
| Rpusd4 | 0.058432979 | 0.080326711 | 0.425115437 | 0.078728287 | 0.031102229 |
| Rraga | 0.016520708 | 0.804681273 | 0.944629934 | 0.625188478 | 0.606152487 |
| Rragb | 0.062978901 | 0.101668904 | 0.467719786 | 0.103162141 | 0.044551767 |
| Rragc | 0.089050368 | 0.175669393 | 0.5796166 | 0.491955542 | 0.397192817 |
| Rragd | 0.009460817 | 0.888493504 | 0.970176435 | 0.569458719 | 0.580206805 |
| Rras | -0.102871575 | 0.01866002 | 0.216996248 | 0.114904518 | 0.212527313 |
| Rras2 | 0.07595695 | 0.086929703 | 0.438669394 | 0.134919417 | 0.067610307 |
| Rrbp1 | 0.065115116 | 0.181093881 | 0.588804431 | 1.913197898 | 1.754621266 |
| Rreb1 | 0.022509611 | 0.62536939 | 0.88668351 | 0.141447137 | 0.120951415 |
| Rrm1 | -0.00330119 | 0.922860984 | 0.980859278 | 0.075809297 | 0.081740096 |

| | | | | | |
|---|---|---|---|---|---|
| Rrm2b | 0.020060163 | 0.760880907 | 0.928280256 | 0.503762399 | 0.482430905 |
| Rrn3 | 0.080599121 | 0.087613191 | 0.43990675 | 0.161130874 | 0.091013902 |
| Rrnad1 | -0.001266187 | 0.970699183 | 0.993892166 | 0.077372617 | 0.077999606 |
| Rrp1 | -0.023875105 | 0.721665506 | 0.919475778 | 0.798334194 | 0.841072202 |
| Rrp12 | -0.022528553 | 0.504875923 | 0.832249514 | 0.067805089 | 0.084805493 |
| Rrp15 | -0.027274734 | 0.520378356 | 0.838907414 | 0.115343747 | 0.140623582 |
| Rrp1b | -0.004876708 | 0.913993694 | 0.980132949 | 0.141303598 | 0.151386462 |
| Rrp36 | 0.036305267 | 0.445562823 | 0.803149051 | 0.16072491 | 0.135560658 |
| Rrp7a | 0.046700344 | 0.428700118 | 0.792251095 | 0.289221713 | 0.250640797 |
| Rrp8 | 0.082471596 | 0.133667829 | 0.515414522 | 0.217701162 | 0.134261434 |
| Rrp9 | 0.007343729 | 0.821023808 | 0.95091345 | 0.068747237 | 0.065805359 |
| Rrs1 | 0.089991494 | 0.062150103 | 0.380250374 | 0.163692548 | 0.089054262 |
| Rsad1 | -0.020444036 | 0.40405645 | 0.77632362 | 0.034295253 | 0.053498201 |
| Rsad2 | 0.258623813 | 0.000524457 | 0.025930891 | 0.410209068 | 0.133879211 |
| Rsbn1 | 0.029768357 | 0.650043644 | 0.895593446 | 0.417046824 | 0.389377446 |
| Rsbn1l | -0.117513141 | 0.074515037 | 0.41344108 | 0.544284449 | 0.668144108 |
| Rsf1 | -0.115695473 | 0.097710396 | 0.460898239 | 0.649808705 | 0.790534876 |
| Rsl1d1 | 0.113343614 | 0.094315193 | 0.453645362 | 0.60974902 | 0.507096654 |
| Rsl24d1 | -0.077878463 | 0.11790123 | 0.491192747 | 0.167374035 | 0.248109913 |
| Rsph1 | -0.011064327 | 0.729926414 | 0.920522999 | 0.061163481 | 0.06600307 |
| Rsph3a | 0.044133116 | 0.146170151 | 0.535832654 | 0.0634873 | 0.027089404 |
| Rsph4a | -0.04718093 | 0.034286599 | 0.293212179 | 0.022258902 | 0.060158843 |
| Rsph9 | -0.04587123 | 0.337532075 | 0.732352032 | 0.15695082 | 0.195066206 |
| Rspo4 | 0.009719296 | 0.871874373 | 0.964616742 | 0.288925124 | 0.289182087 |
| Rspry1 | -0.070395205 | 0.192835829 | 0.60325934 | 0.200188484 | 0.26681579 |
| Rsrc1 | 0.023638355 | 0.691207137 | 0.909796728 | 0.317602683 | 0.297473696 |
| Rsrc2 | 0.145476116 | 0.029444751 | 0.273392198 | 1.134801786 | 0.927314765 |
| Rsrp1 | -0.231711084 | 0.001387945 | 0.047872176 | 0.874848557 | 1.172206122 |
| Rsu1 | -0.04384506 | 0.303395354 | 0.708181201 | 0.1165157 | 0.153905647 |
| Rtca | -0.086444288 | 0.059016556 | 0.373957814 | 0.133818063 | 0.225530693 |
| Rtcb | 0.030494461 | 0.655996924 | 0.89789619 | 0.67262875 | 0.628256137 |
| Rtf1 | 0.054485918 | 0.419753801 | 0.786284042 | 0.528615044 | 0.470522892 |
| Rtf2 | 0.032876541 | 0.593893383 | 0.872881519 | 0.329892569 | 0.29954253 |
| Rtl5 | 0.065343584 | 0.206433823 | 0.617762755 | 0.193797043 | 0.128438492 |
| Rtl6 | 0.012325395 | 0.754623275 | 0.926726954 | 0.105362715 | 0.098466907 |
| Rtl8a | 0.086258422 | 0.125159816 | 0.503791124 | 0.26628899 | 0.196477318 |
| Rtl8b | 0.019128076 | 0.691825218 | 0.909796728 | 0.1703662 | 0.163397468 |
| Rtn1 | -0.070484074 | 0.094646709 | 0.454115053 | 0.115594849 | 0.180588718 |
| Rtn2 | 0.072163341 | 0.166105656 | 0.568041726 | 0.19280767 | 0.128478234 |
| Rtn3 | 0.059020492 | 0.355642555 | 0.745326231 | 0.851698075 | 0.772588884 |
| Rtn4 | 0.026334057 | 0.682632131 | 0.909173483 | 1.427654736 | 1.370181049 |
| Rtn4ip1 | -0.041055887 | 0.155734226 | 0.552356396 | 0.048581369 | 0.081585219 |
| Rtn4rl1 | 0.03873516 | 0.478701974 | 0.819081437 | 0.234884286 | 0.203024644 |
| Rtp4 | 0.053548292 | 0.10984348 | 0.478722972 | 0.074548903 | 0.020970191 |
| Rtraf | -0.12347622 | 0.070875043 | 0.403272567 | 0.770193723 | 0.947635453 |
| Rubcn | 0.023528361 | 0.654459055 | 0.897454058 | 0.201845234 | 0.18259845 |
| Rufy1 | -0.029612088 | 0.562971063 | 0.857583554 | 0.182273128 | 0.211777512 |
| Rufy2 | 0.083307579 | 0.128716363 | 0.508260084 | 0.231120049 | 0.154081104 |
| Rufy3 | -0.124082365 | 0.065617576 | 0.389687514 | 0.518954269 | 0.614961472 |
| Rundc1 | 0.042898853 | 0.382047516 | 0.762803046 | 0.169887111 | 0.127150024 |
| Rundc3a | -0.185778499 | 0.003695027 | 0.085962309 | 0.717272979 | 0.9414143 |
| Rundc3b | -0.049761857 | 0.388110301 | 0.767160379 | 0.261579792 | 0.30281697 |
| Runx1t1 | -0.146456754 | 0.026117806 | 0.257205918 | 0.385617375 | 0.546635213 |
| Rusc1 | -0.052043231 | 0.323233397 | 0.720244873 | 0.200062225 | 0.244769986 |
| Ruvbl1 | -0.007921075 | 0.86069727 | 0.959655466 | 0.143380992 | 0.155613518 |
| Ruvbl2 | 0.009905559 | 0.822758018 | 0.950930859 | 0.130885262 | 0.118869539 |
| Rwdd1 | 0.040088196 | 0.484134581 | 0.822768706 | 1.327851698 | 1.259361044 |
| Rwdd2b | -0.033021879 | 0.153010162 | 0.549602359 | 0.026587274 | 0.054396178 |
| Rwdd3 | -0.063189928 | 0.011205204 | 0.167610475 | 0.030039129 | 0.084970778 |
| Rwdd4a | 0.084156429 | 0.215156218 | 0.629611185 | 0.668721541 | 0.589073726 |
| Rxra | 0.035245301 | 0.451409662 | 0.806495742 | 0.148646063 | 0.122133636 |
| Rxrb | -0.012478231 | 0.811492536 | 0.948397417 | 0.199230758 | 0.218973764 |
| Rxylt1 | 0.008765591 | 0.892898607 | 0.971493043 | 0.410389735 | 0.401044088 |
| Rybp | 0.111001815 | 0.094481276 | 0.454115053 | 0.474466843 | 0.361049449 |
| Ryk | 0.11869271 | 0.055584607 | 0.366077889 | 0.368486842 | 0.262135261 |

| | | | | | |
|---|---|---|---|---|---|
| S100a1 | -0.226996904 | 0.000953087 | 0.037078044 | 0.560983987 | 0.82496702 |
| S100a10 | 0.649852787 | 4.39E-15 | 6.30E-12 | 1.235564159 | 0.39810061 |
| S100a11 | -0.013945384 | 0.84039193 | 0.954911647 | 0.879064584 | 0.892433753 |
| S100a13 | -0.082735857 | 0.221529917 | 0.633858837 | 0.437938293 | 0.508778373 |
| S100a4 | -0.002052254 | 0.954265863 | 0.989066902 | 0.080013212 | 0.088550367 |
| S100a6 | -0.053150414 | 0.45232474 | 0.806821292 | 0.265660217 | 0.305562944 |
| S100pbp | -0.08174576 | 0.048147323 | 0.343953509 | 0.10623973 | 0.174925782 |
| S100z | 0.063001431 | 0.089507246 | 0.444861289 | 0.092678823 | 0.037438277 |
| Sac3d1 | 0.071418382 | 0.185194308 | 0.591560356 | 0.21200246 | 0.152347282 |
| Sacm1l | 0.002360714 | 0.971280211 | 0.993892166 | 0.450579147 | 0.442349595 |
| Sacs | -0.003413799 | 0.932292849 | 0.98343913 | 0.099207617 | 0.100355247 |
| Sae1 | -0.046860866 | 0.4009646 | 0.774740795 | 0.233981185 | 0.277727006 |
| Safb | -0.029855125 | 0.643156764 | 0.892561795 | 0.439186917 | 0.460448601 |
| Safb2 | 0.11499106 | 0.08428488 | 0.432888102 | 0.517314386 | 0.398223948 |
| Sall2 | 0.053805384 | 0.079887703 | 0.423801733 | 0.064049886 | 0.01833749 |
| Sall4 | 0.047473366 | 0.108702198 | 0.477972625 | 0.061240502 | 0.016423707 |
| Samd1 | -0.009120568 | 0.838676144 | 0.954911647 | 0.129337332 | 0.135898768 |
| Samd10 | -0.003598288 | 0.940373596 | 0.986159208 | 0.158175631 | 0.164913416 |
| Samd12 | 0.002276154 | 0.945794025 | 0.987084873 | 0.072429468 | 0.068979943 |
| Samd15 | -0.044914485 | 0.064383899 | 0.386943382 | 0.027769293 | 0.068526489 |
| Samd4 | 0.051700326 | 0.081893929 | 0.42777546 | 0.059222922 | 0.016610336 |
| Samd4b | 0.024752809 | 0.70395005 | 0.912277105 | 0.525003528 | 0.509348228 |
| Samd8 | 0.068929329 | 0.261736146 | 0.668764912 | 0.342903664 | 0.275114933 |
| Samd9l | 0.002794486 | 0.955668553 | 0.989278596 | 0.170245483 | 0.161156444 |
| Samhd1 | -0.010310329 | 0.795283306 | 0.939757086 | 0.103151686 | 0.109225286 |
| Samm50 | -0.002344423 | 0.969219282 | 0.993774026 | 0.336910123 | 0.335318236 |
| Sap130 | 0.062254672 | 0.089424602 | 0.444774394 | 0.096654471 | 0.045922541 |
| Sap18 | -0.044080981 | 0.504869261 | 0.832249514 | 0.465406042 | 0.507120408 |
| Sap30 | 0.075472682 | 0.081341286 | 0.427446328 | 0.134691978 | 0.071352203 |
| Sap30bp | 0.05467604 | 0.291957173 | 0.697566301 | 0.187052443 | 0.142676953 |
| Sap30l | -0.005143747 | 0.933778755 | 0.983869133 | 0.402389337 | 0.414190131 |
| Sar1a | -0.023181383 | 0.721895986 | 0.919475778 | 1.065880769 | 1.109902086 |
| Sar1b | -0.077582481 | 0.245046043 | 0.654809396 | 0.989195725 | 1.086858952 |
| Saraf | -0.118659959 | 0.037172764 | 0.304267772 | 1.547104625 | 1.735051513 |
| Sarnp | 0.01674352 | 0.798354432 | 0.940958114 | 0.456733218 | 0.442791366 |
| Sars | -0.135523249 | 0.024159446 | 0.246431256 | 1.370393402 | 1.572041505 |
| Sart1 | -0.042920369 | 0.402299874 | 0.77525778 | 0.187124155 | 0.230962878 |
| Sart3 | -0.05604211 | 0.210105826 | 0.622822957 | 0.133129226 | 0.184200317 |
| Sat1 | 0.029694712 | 0.646772003 | 0.895025469 | 0.341912461 | 0.326132023 |
| Sat2 | 0.03240136 | 0.268846793 | 0.676803426 | 0.058993649 | 0.02888242 |
| Satb1 | 0.0445611 | 0.417954019 | 0.784482356 | 0.224921692 | 0.186025025 |
| Sav1 | -0.024316707 | 0.547730131 | 0.851675805 | 0.107499871 | 0.129559927 |
| Saxo2 | -0.023711387 | 0.351688519 | 0.74208191 | 0.034398711 | 0.053838379 |
| Saysd1 | -0.01287357 | 0.836474977 | 0.954911647 | 0.371967866 | 0.378651133 |
| Sbds | -0.047869853 | 0.494042493 | 0.828842054 | 0.719533011 | 0.765552664 |
| Sbf1 | 0.046376576 | 0.232056843 | 0.643345304 | 0.106115878 | 0.061916661 |
| Sbf2 | -0.054174224 | 0.218531876 | 0.632622127 | 0.126012298 | 0.173411155 |
| Sbno1 | -0.105684647 | 0.12860708 | 0.508260084 | 0.725114765 | 0.840639771 |
| Sbno2 | 0.040859833 | 0.191006396 | 0.601358592 | 0.068119885 | 0.031575605 |
| Sc5d | -0.006885583 | 0.888069737 | 0.970045467 | 0.160833926 | 0.163181261 |
| Scaf1 | -0.050074173 | 0.364904249 | 0.752802229 | 0.222987954 | 0.266455623 |
| Scaf11 | -0.042949882 | 0.508555069 | 0.83445598 | 1.099593292 | 1.192605774 |
| Scaf4 | -0.077164271 | 0.118608043 | 0.491192747 | 0.168059699 | 0.241233064 |
| Scaf8 | -0.046758546 | 0.363520378 | 0.751765374 | 0.184578884 | 0.225632071 |
| Scai | -0.02400932 | 0.527806284 | 0.843341172 | 0.09219718 | 0.115530017 |
| Scamp1 | 0.01851552 | 0.752766912 | 0.926375857 | 0.287809257 | 0.251505568 |
| Scamp2 | 0.040482098 | 0.541854223 | 0.850208191 | 0.46898216 | 0.432677702 |
| Scamp3 | 0.064669973 | 0.30002395 | 0.70506682 | 0.370001363 | 0.319011237 |
| Scamp5 | 0.086898628 | 0.128005762 | 0.508260084 | 0.284392629 | 0.19856589 |
| Scand1 | 0.046587264 | 0.385295238 | 0.765415428 | 1.591511534 | 1.52982689 |
| Scap | -0.018751242 | 0.675277193 | 0.906025597 | 0.142405391 | 0.169309112 |
| Scaper | -0.084230447 | 0.136309534 | 0.520007144 | 0.233717633 | 0.310133991 |
| Scarb1 | 0.022530795 | 0.455032764 | 0.807876964 | 0.0589486 | 0.036059376 |
| Scarb2 | -0.013786984 | 0.83742378 | 0.954911647 | 0.684784434 | 0.706621326 |
| Scarf2 | 0.002323713 | 0.939512128 | 0.985603501 | 0.055951947 | 0.05390635 |

| | | | | | |
|---|---|---|---|---|---|
| Sccpdh | -0.020827015 | 0.611766155 | 0.880165207 | 0.110807521 | 0.134712493 |
| Scd2 | 0.109667599 | 0.05164949 | 0.355933715 | 0.224243563 | 0.125660038 |
| Scfd1 | 0.047794861 | 0.482214726 | 0.821803219 | 0.693614583 | 0.654467696 |
| Scfd2 | -0.056177205 | 0.06648113 | 0.39159102 | 0.051124696 | 0.099360306 |
| Scg2 | -0.081739936 | 0.000969575 | 0.037358974 | 4.252958281 | 4.467538653 |
| Scg3 | -0.065888193 | 0.031613394 | 0.282549988 | 3.337252698 | 3.541385476 |
| Scg5 | -0.164288896 | 7.97E-06 | 0.001126284 | 2.539133261 | 2.910824768 |
| Scgn | 0.106765116 | 0.142455495 | 0.531138857 | 0.80900341 | 0.689544684 |
| Sclt1 | -0.031879006 | 0.530609344 | 0.84508505 | 0.17684115 | 0.200936926 |
| Scly | -0.028724672 | 0.660876753 | 0.89868852 | 0.449136715 | 0.479221775 |
| Scmh1 | -0.031728616 | 0.506550058 | 0.833269979 | 0.153978812 | 0.183042934 |
| Scml4 | -0.009064811 | 0.76388025 | 0.928730554 | 0.05286466 | 0.06378134 |
| Scn1b | 0.046688283 | 0.258615494 | 0.665948842 | 0.112398482 | 0.066620591 |
| Scn3a | -0.094358517 | 0.035351911 | 0.295197277 | 0.117170217 | 0.210423462 |
| Scn8a | 0.006160138 | 0.90341344 | 0.974950771 | 0.188445988 | 0.183563006 |
| Scn9a | 0.033148713 | 0.562694749 | 0.857583554 | 0.263065394 | 0.232580323 |
| Scnm1 | 0.008382103 | 0.865592375 | 0.961801247 | 0.161474377 | 0.15631463 |
| Scnn1b | 0.041472984 | 0.544657672 | 0.851259637 | 0.537236328 | 0.485187638 |
| Sco1 | 0.049164693 | 0.090294681 | 0.445385778 | 0.05668455 | 0.018561198 |
| Sco2 | -0.000992477 | 0.979247464 | 0.995540893 | 0.096375925 | 0.101467184 |
| Scoc | -0.035933618 | 0.597881607 | 0.874644759 | 0.487794132 | 0.520127699 |
| Scp2 | -0.15432091 | 0.017100471 | 0.208804661 | 0.530794176 | 0.685304329 |
| Scpep1 | -0.079709511 | 0.194074366 | 0.60325934 | 0.310899489 | 0.387696511 |
| Scrib | 0.020760299 | 0.589020052 | 0.86992264 | 0.096965871 | 0.080348791 |
| Scrn1 | -0.022275742 | 0.713958898 | 0.915840939 | 0.328594268 | 0.358476108 |
| Scrn3 | 0.055802838 | 0.266483102 | 0.674047915 | 0.182918667 | 0.130539957 |
| Scrt1 | 0.034830966 | 0.413424811 | 0.781607961 | 0.122654713 | 0.097270794 |
| Scyl1 | -0.004799994 | 0.92885609 | 0.982502239 | 0.212066838 | 0.21456764 |
| Scyl2 | 0.058830998 | 0.25277027 | 0.663146569 | 0.186336807 | 0.13197455 |
| Scyl3 | -0.056625858 | 0.148640968 | 0.54192132 | 0.092716556 | 0.147801936 |
| Sdad1 | -0.077152921 | 0.129712185 | 0.509514066 | 0.175379819 | 0.242543282 |
| Sdc2 | -0.087169757 | 0.082664278 | 0.429897076 | 0.157698824 | 0.230906606 |
| Sdc3 | 0.136407525 | 0.002916582 | 0.074677892 | 0.155310881 | 0.038808654 |
| Sdc4 | 0.068094734 | 0.364127532 | 0.752624547 | 0.807596165 | 0.721605867 |
| Sdcbp | 0.013770995 | 0.837708771 | 0.954911647 | 0.641760795 | 0.618252196 |
| Sdcbp2 | 0.068909761 | 0.070596013 | 0.402952703 | 0.097020919 | 0.040298543 |
| Sdccag8 | 0.034005477 | 0.533328312 | 0.84612966 | 0.221853285 | 0.18725019 |
| Sde2 | 0.019453615 | 0.762039988 | 0.92872278 | 0.40215704 | 0.383807824 |
| Sdf2 | -0.042831089 | 0.513673 | 0.836550101 | 0.499218798 | 0.56185944 |
| Sdf2l1 | 0.381074081 | 5.92E-08 | 2.20E-05 | 1.138159145 | 0.632540976 |
| Sdf4 | -0.1542208 | 0.021214382 | 0.23094225 | 0.813263036 | 1.008335329 |
| Sdha | 0.01261931 | 0.85255378 | 0.957461155 | 0.65411547 | 0.635355228 |
| Sdhaf1 | 0.188555745 | 0.00402468 | 0.09071778 | 0.494294236 | 0.30977254 |
| Sdhaf2 | -0.016519077 | 0.747925044 | 0.924930563 | 0.182901507 | 0.209596725 |
| Sdhaf3 | 0.026478105 | 0.529341001 | 0.844323583 | 0.118098792 | 0.101877342 |
| Sdhaf4 | 0.027284797 | 0.647034778 | 0.895025469 | 0.283677347 | 0.258962933 |
| Sdhb | -0.058375056 | 0.387829662 | 0.767160379 | 0.643114244 | 0.70935552 |
| Sdhc | -0.044089495 | 0.510839048 | 0.834957869 | 0.582886756 | 0.609317066 |
| Sdhd | 0.022978905 | 0.740029436 | 0.922901128 | 0.543577129 | 0.512623056 |
| Sdk2 | 0.023713012 | 0.745540052 | 0.924566547 | 0.768256038 | 0.711878316 |
| Sdr39u1 | -0.087987944 | 0.082872464 | 0.430310873 | 0.178764927 | 0.258002529 |
| Sec11a | 0.095972642 | 0.166851145 | 0.568374737 | 0.690017177 | 0.602212165 |
| Sec11c | 0.018956844 | 0.729438137 | 0.920522999 | 1.658826758 | 1.575289528 |
| Sec13 | -0.014083762 | 0.829562465 | 0.953065609 | 0.536209138 | 0.546847551 |
| Sec14l1 | 0.11010919 | 0.085792692 | 0.436000632 | 0.388197049 | 0.269559417 |
| Sec16a | 0.09186761 | 0.158762426 | 0.556789205 | 0.41154607 | 0.313926671 |
| Sec16b | -0.027456888 | 0.506327035 | 0.833269979 | 0.114124337 | 0.129813255 |
| Sec22a | 0.006576146 | 0.886607607 | 0.969612367 | 0.145949799 | 0.144713932 |
| Sec22b | -0.060895594 | 0.384754893 | 0.764861331 | 0.758959191 | 0.84106476 |
| Sec22c | 0.040998768 | 0.257174141 | 0.665256942 | 0.089354317 | 0.050451271 |
| Sec23a | 0.056754908 | 0.215920051 | 0.629630897 | 0.14478763 | 0.107597929 |
| Sec23b | -0.011679832 | 0.854034096 | 0.957461155 | 0.375911631 | 0.376369863 |
| Sec23ip | 0.021532914 | 0.712081188 | 0.915109107 | 0.274433597 | 0.258638502 |
| Sec24a | -0.044809525 | 0.440013288 | 0.800219853 | 0.255044951 | 0.301068756 |
| Sec24b | 0.009973452 | 0.840409104 | 0.954911647 | 0.169235983 | 0.15103014 |

| | | | | | |
|---|---|---|---|---|---|
| Sec24c | -0.083230287 | 0.170095872 | 0.572762895 | 0.317184495 | 0.394224295 |
| Sec24d | -0.04300877 | 0.465789656 | 0.812791331 | 0.246349476 | 0.29032306 |
| Sec31a | -0.039549685 | 0.568681793 | 0.860399704 | 0.850638855 | 0.880530061 |
| Sec61a1 | -0.09605436 | 0.160480093 | 0.558508561 | 0.624003938 | 0.726725489 |
| Sec61a2 | 0.060747498 | 0.111883075 | 0.482998031 | 0.10519069 | 0.048779176 |
| Sec61b | -0.036337263 | 0.37197142 | 0.75760494 | 2.333441308 | 2.407918962 |
| Sec61g | 0.193774755 | 0.001600651 | 0.052831084 | 1.209331839 | 0.932282925 |
| Sec62 | 0.03092797 | 0.516344514 | 0.837427561 | 1.817592596 | 1.747955271 |
| Sec63 | -0.192459036 | 0.002900877 | 0.074465734 | 0.708739753 | 0.933212321 |
| Secisbp2 | 0.015001939 | 0.723070215 | 0.919483582 | 0.125003373 | 0.117587203 |
| Secisbp2l | -0.039137768 | 0.422580109 | 0.788224596 | 0.1620094 | 0.197461514 |
| Seh1l | 0.08406453 | 0.165654827 | 0.566763419 | 0.338303258 | 0.267393719 |
| Sel1l | 0.001072762 | 0.987470095 | 0.997268589 | 0.985455606 | 0.944503175 |
| Sel1l3 | 0.033924971 | 0.481303159 | 0.820999756 | 0.16122213 | 0.131938808 |
| Selenof | -0.024552574 | 0.674768907 | 0.906025597 | 1.311847733 | 1.334456556 |
| Selenoh | 0.020394751 | 0.734662868 | 0.921732448 | 0.3159738 | 0.300529275 |
| Selenoi | -0.027372359 | 0.583807718 | 0.867072813 | 0.172232957 | 0.197857289 |
| Selenok | 0.026263163 | 0.573104355 | 0.861889183 | 1.95055003 | 1.895857606 |
| Selenom | 0.082733373 | 0.117946846 | 0.491192747 | 1.78149684 | 1.614403976 |
| Selenon | 0.01819264 | 0.617587948 | 0.883238152 | 0.089302948 | 0.070478541 |
| Selenoo | -0.026261422 | 0.537837504 | 0.84785221 | 0.124799189 | 0.153162055 |
| Selenop | 0.054709124 | 0.413136283 | 0.781607961 | 1.190394689 | 1.086758839 |
| Selenos | -0.027220841 | 0.672220739 | 0.905281039 | 1.215314245 | 1.229381724 |
| Selenot | 0.057585423 | 0.402729996 | 0.77525778 | 0.957545898 | 0.881581239 |
| Selenow | 0.181191425 | 0.003920438 | 0.08922775 | 1.493501378 | 1.175395254 |
| Sem1 | 0.081959786 | 0.077162557 | 0.418199494 | 1.78352955 | 1.628586182 |
| Sema4b | 0.031439714 | 0.46989631 | 0.814258266 | 0.129068725 | 0.097707073 |
| Sema4d | -0.008095961 | 0.82235238 | 0.950930859 | 0.088265101 | 0.090197011 |
| Senp1 | 0.008095835 | 0.851094766 | 0.957461155 | 0.124148665 | 0.117239699 |
| Senp2 | 0.023729722 | 0.67632265 | 0.906611913 | 0.274462477 | 0.254805205 |
| Senp3 | -0.034840525 | 0.467098252 | 0.813188482 | 0.161710748 | 0.193114073 |
| Senp5 | 0.062585704 | 0.285125613 | 0.692407818 | 0.282057067 | 0.220327361 |
| Senp6 | 0.045500433 | 0.482970477 | 0.822469365 | 1.128179924 | 1.055963665 |
| Senp7 | 0.044021875 | 0.402940699 | 0.77525778 | 0.209510842 | 0.16742083 |
| Senp8 | -0.011531622 | 0.764210984 | 0.928730554 | 0.095950082 | 0.111487398 |
| Sephs1 | -0.031232099 | 0.38241583 | 0.763150149 | 0.080960951 | 0.116332214 |
| Sephs2 | 0.047298164 | 0.426232122 | 0.790724712 | 0.305961571 | 0.26942523 |
| Sepsecs | -0.053251877 | 0.319037012 | 0.717256849 | 0.206200968 | 0.256835964 |
| Septin10 | 0.032877723 | 0.377342592 | 0.761436991 | 0.096869034 | 0.070686871 |
| Septin11 | -0.098890052 | 0.112325635 | 0.483221516 | 0.326539351 | 0.425465244 |
| Septin2 | 0.024911032 | 0.448229031 | 0.804951652 | 0.070877164 | 0.04955568 |
| Septin3 | 0.011681657 | 0.673343269 | 0.905635862 | 0.049898155 | 0.036379288 |
| Septin7 | -0.063360429 | 0.348346597 | 0.739576235 | 0.811621064 | 0.882306115 |
| Septin8 | 0.001520168 | 0.960965374 | 0.991375098 | 0.06105212 | 0.05870538 |
| Septin9 | 0.061332975 | 0.323487564 | 0.720244873 | 0.338920025 | 0.281975215 |
| Serbp1 | 0.043768153 | 0.405857497 | 0.777402995 | 1.568099531 | 1.49642596 |
| Serf1 | -0.002544618 | 0.951473731 | 0.988311364 | 0.119267366 | 0.122619094 |
| Serf2 | 0.109545774 | 0.046957342 | 0.339316662 | 1.71575238 | 1.489755626 |
| Sergef | -0.032239928 | 0.410219988 | 0.780101937 | 0.101775346 | 0.129864438 |
| Serhl | -0.079009275 | 0.012604341 | 0.17944648 | 0.055948912 | 0.118508763 |
| Serinc1 | -0.147801273 | 0.016453703 | 0.206949651 | 1.153313354 | 1.397552556 |
| Serinc3 | 0.081632658 | 0.20451291 | 0.616256046 | 0.47080925 | 0.383174734 |
| Serinc5 | 0.002112595 | 0.956473454 | 0.989397512 | 0.100697525 | 0.092833241 |
| Serp1 | 0.034406755 | 0.611907544 | 0.880165207 | 1.290772105 | 1.244488727 |
| Serp2 | -0.032729751 | 0.370304409 | 0.75730884 | 0.084918983 | 0.112908645 |
| Serpina10 | 0.060829505 | 0.393675631 | 0.770538672 | 0.708235455 | 0.588088693 |
| Serpina7 | -0.157052812 | 0.004140519 | 0.092557659 | 0.146282312 | 0.318747037 |
| Serpinb6a | 0.180110603 | 0.007721933 | 0.135045705 | 0.818691369 | 0.617183714 |
| Serpinb9 | -0.05233486 | 0.10738687 | 0.47507667 | 0.061725918 | 0.103929228 |
| Serping1 | -0.086028676 | 0.083693986 | 0.431794059 | 0.154175436 | 0.22967271 |
| Serpinh1 | -0.021680191 | 0.447066474 | 0.804302957 | 0.043707283 | 0.058431354 |
| Serpini1 | 0.022257705 | 0.71264651 | 0.915447617 | 0.293635595 | 0.270188365 |
| Sertad1 | 0.007651815 | 0.882922044 | 0.96861827 | 0.185461007 | 0.181325838 |
| Sertad2 | 0.022227445 | 0.565143401 | 0.858087314 | 0.097401199 | 0.080875763 |
| Sertad3 | 0.010519875 | 0.73946189 | 0.922843014 | 0.064469281 | 0.059623133 |

| | | | | | |
|---|---|---|---|---|---|
| Sertad4 | -0.046659781 | 0.07932157 | 0.42236106 | 0.038524642 | 0.081307459 |
| Sesn1 | 0.043105458 | 0.292000492 | 0.697566301 | 0.112113544 | 0.07625086 |
| Sesn2 | -0.203853726 | 0.002324395 | 0.06554117 | 0.446887257 | 0.663560279 |
| Sesn3 | -0.111492091 | 0.111298629 | 0.482253844 | 0.604842892 | 0.727028728 |
| Sestd1 | 0.040791293 | 0.239418792 | 0.650744882 | 0.07860721 | 0.042236001 |
| Set | -0.082372587 | 0.19902033 | 0.609389286 | 1.121585988 | 1.248030408 |
| Setbp1 | 0.082240961 | 0.071088371 | 0.404027315 | 0.145690639 | 0.079273936 |
| Setd1a | 0.010897035 | 0.775646309 | 0.933636662 | 0.095908441 | 0.085638387 |
| Setd1b | -0.036925981 | 0.49954676 | 0.830722138 | 0.206784135 | 0.253926494 |
| Setd2 | 0.012539558 | 0.841301141 | 0.954960614 | 0.377059108 | 0.383144114 |
| Setd3 | -0.041604526 | 0.524608181 | 0.841229722 | 0.581619226 | 0.637621787 |
| Setd5 | 0.018266905 | 0.785708397 | 0.936931826 | 0.656902458 | 0.65379634 |
| Setd6 | 0.046713412 | 0.286889858 | 0.693357454 | 0.136715084 | 0.091030202 |
| Setd7 | 0.156015054 | 0.016708202 | 0.207483393 | 0.493613958 | 0.348016939 |
| Setdb1 | 0.04259775 | 0.33234841 | 0.72806771 | 0.133798143 | 0.104962581 |
| Setx | -0.035235209 | 0.530352062 | 0.845077576 | 0.222961187 | 0.243060305 |
| Sez6l | -0.18464661 | 0.008881675 | 0.146996602 | 0.483806516 | 0.681920479 |
| Sez6l2 | -0.255370179 | 0.000162946 | 0.011125798 | 0.910672134 | 1.257646677 |
| Sf1 | -0.020157948 | 0.757359811 | 0.927237784 | 0.461878108 | 0.493397526 |
| Sf3a1 | 0.099684818 | 0.07744833 | 0.418285047 | 0.267706438 | 0.175145215 |
| Sf3a2 | 0.014612328 | 0.76247314 | 0.92872278 | 0.169018121 | 0.160768225 |
| Sf3a3 | 0.012012741 | 0.823676215 | 0.951131865 | 0.234100375 | 0.230551576 |
| Sf3b1 | -0.000573226 | 0.993455659 | 0.998695538 | 0.873190091 | 0.883791568 |
| Sf3b2 | 0.006547284 | 0.922582773 | 0.980859278 | 1.120796084 | 1.095737824 |
| Sf3b3 | -0.041159179 | 0.511723677 | 0.835213712 | 0.366203639 | 0.407796833 |
| Sf3b4 | 0.041786772 | 0.342853807 | 0.735501771 | 0.133754241 | 0.098091149 |
| Sf3b5 | 0.209034173 | 0.002704262 | 0.071134541 | 0.846124386 | 0.626066759 |
| Sf3b6 | -0.02953663 | 0.66185366 | 0.898920864 | 1.029370491 | 1.079419154 |
| Sfi1 | -0.033858447 | 0.226716813 | 0.638720711 | 0.045689495 | 0.069101583 |
| Sfmbt1 | 0.02283788 | 0.702180483 | 0.912152193 | 0.295202739 | 0.270607784 |
| Sfn | 0.070270927 | 0.066461898 | 0.39159102 | 0.094981409 | 0.035759988 |
| Sfpq | -0.331938928 | 6.41E-07 | 0.000146286 | 0.467120302 | 0.802916623 |
| Sfr1 | -0.145264041 | 0.035445096 | 0.29572937 | 0.553551626 | 0.728702444 |
| Sfrp5 | -0.026234868 | 0.345693949 | 0.73792645 | 0.038697001 | 0.073873115 |
| Sfswap | 0.019144148 | 0.733212224 | 0.921306823 | 0.260399862 | 0.247091583 |
| Sft2d1 | -0.047950939 | 0.164256318 | 0.563869641 | 0.069948541 | 0.112467337 |
| Sft2d2 | -0.073621686 | 0.120824618 | 0.494986406 | 0.142971216 | 0.211125441 |
| Sft2d3 | 0.01750223 | 0.765848818 | 0.92882091 | 0.310615564 | 0.284596962 |
| Sfxn1 | 0.02608007 | 0.701076859 | 0.912152193 | 0.780985034 | 0.727596194 |
| Sfxn2 | -0.0438074 | 0.41116367 | 0.780671239 | 0.199775077 | 0.238732012 |
| Sfxn3 | -0.045295952 | 0.10157032 | 0.467719786 | 0.042279397 | 0.07911365 |
| Sgcb | -0.012965335 | 0.723063088 | 0.919483582 | 0.088885028 | 0.09737741 |
| Sgcz | -0.132756956 | 0.002284973 | 0.06526292 | 0.112114831 | 0.230953065 |
| Sgf29 | 0.018286771 | 0.641874303 | 0.892178303 | 0.106021239 | 0.096241601 |
| Sgip1 | -0.064661201 | 0.00297153 | 0.075891213 | 0.01834636 | 0.070627799 |
| Sgk2 | 0.055211303 | 0.12003966 | 0.493583757 | 0.083927146 | 0.03898255 |
| Sgk3 | 0.029010995 | 0.51743148 | 0.837694133 | 0.137586586 | 0.117144246 |
| Sgms1 | -0.034948252 | 0.443634596 | 0.801522297 | 0.136366734 | 0.164675424 |
| Sgms2 | 0.108566163 | 0.100975395 | 0.466401306 | 0.368291605 | 0.293393936 |
| Sgpl1 | 0.09043388 | 0.151482278 | 0.546576546 | 0.357568568 | 0.27331553 |
| Sgpp1 | -0.07720933 | 0.213370976 | 0.627852386 | 0.353322784 | 0.423799203 |
| Sgpp2 | 0.0197904 | 0.689022137 | 0.909796728 | 0.170718001 | 0.142423237 |
| Sgsm1 | 0.021761391 | 0.673683937 | 0.905654932 | 0.19843158 | 0.174633205 |
| Sgsm2 | -0.02145083 | 0.667690647 | 0.901929709 | 0.177304958 | 0.185957244 |
| Sgsm3 | -0.118187518 | 0.016937503 | 0.20833544 | 0.159078245 | 0.265458894 |
| Sgta | -0.082045101 | 0.180357606 | 0.587361872 | 0.347656935 | 0.414547751 |
| Sgtb | -0.04069216 | 0.318522268 | 0.717175454 | 0.100683606 | 0.146451027 |
| Sh2b1 | 0.018428752 | 0.661800176 | 0.898920864 | 0.121587303 | 0.101505767 |
| Sh2b3 | 0.01994733 | 0.502187109 | 0.832218079 | 0.056419136 | 0.038726785 |
| Sh2d3c | -0.065069876 | 0.038454776 | 0.310016536 | 0.055972641 | 0.107392799 |
| Sh2d5 | -0.037445505 | 0.123930916 | 0.501565251 | 0.029394341 | 0.059026131 |
| Sh3bgr | -0.000837003 | 0.977113326 | 0.994653798 | 0.050291395 | 0.047704879 |
| Sh3bgrl | -0.131662111 | 0.032264822 | 0.284812628 | 0.364430673 | 0.479802388 |
| Sh3bgrl3 | 0.15384676 | 0.023116738 | 0.240881268 | 0.489729471 | 0.323601666 |
| Sh3bp4 | 0.074218197 | 0.084157329 | 0.432507481 | 0.126912806 | 0.060614473 |

| | | | | | |
|---|---|---|---|---|---|
| Sh3bp5l | 0.013089554 | 0.670571827 | 0.904104984 | 0.060096031 | 0.048514626 |
| Sh3d19 | 0.044919286 | 0.435688163 | 0.796975049 | 0.247319387 | 0.203424727 |
| Sh3d21 | 0.035992799 | 0.216058167 | 0.629667778 | 0.056057906 | 0.027117923 |
| Sh3gl1 | 0.051581139 | 0.35090918 | 0.741476479 | 0.239512415 | 0.186489433 |
| Sh3gl2 | 0.056706851 | 0.17851772 | 0.584861057 | 0.122172002 | 0.066551822 |
| Sh3glb1 | 0.068752895 | 0.254231325 | 0.663146569 | 1.259400553 | 1.164269131 |
| Sh3glb2 | -0.101257509 | 0.022030115 | 0.23623533 | 0.128874987 | 0.222788041 |
| Sh3kbp1 | -0.076077615 | 0.14417566 | 0.533279057 | 0.191173834 | 0.260581037 |
| Sh3pxd2a | -0.095368148 | 0.172471532 | 0.575294174 | 0.604740887 | 0.712240609 |
| Sh3rf1 | -0.001766526 | 0.952537182 | 0.98838165 | 0.056091124 | 0.059292214 |
| Sh3yl1 | 0.007605719 | 0.880271077 | 0.967810606 | 0.188377049 | 0.18595548 |
| Sharpin | 0.017678414 | 0.693472222 | 0.909871028 | 0.139317558 | 0.120176912 |
| Shb | 0.089393815 | 0.114143984 | 0.485450496 | 0.258167699 | 0.181677712 |
| Shc1 | -0.103015553 | 0.086149855 | 0.436930819 | 0.28115971 | 0.369725513 |
| Shc2 | -0.108542433 | 0.000749814 | 0.032181025 | 0.054798829 | 0.154762589 |
| Shfl | 0.005357103 | 0.921453696 | 0.98085171 | 0.235594603 | 0.220146747 |
| Shisa2 | 0.088597522 | 0.039771307 | 0.314071287 | 0.131145378 | 0.0531905 |
| Shisa5 | 0.087253918 | 0.157281054 | 0.554924229 | 0.375284741 | 0.261061684 |
| Shisal1 | 0.075070137 | 0.04442204 | 0.330514466 | 0.096989727 | 0.026769491 |
| Shisal2b | 0.150151982 | 0.05247099 | 0.357919301 | 0.624597174 | 0.415676388 |
| Shkbp1 | -0.016289011 | 0.629137912 | 0.887864718 | 0.072425356 | 0.090992013 |
| Shld1 | 0.020367224 | 0.642560561 | 0.892561795 | 0.132610021 | 0.110284387 |
| Shld2 | -0.006291479 | 0.864886725 | 0.961442912 | 0.091578494 | 0.099456422 |
| Shld3 | -0.019697565 | 0.476882499 | 0.818444859 | 0.042485803 | 0.057642258 |
| Shmt2 | 0.000662637 | 0.990576624 | 0.997733826 | 0.230765268 | 0.210049947 |
| Shoc2 | -0.076905193 | 0.219687753 | 0.63289494 | 0.380299171 | 0.453156913 |
| Shprh | -0.034978126 | 0.514784448 | 0.836743564 | 0.199507518 | 0.226847629 |
| Shq1 | -0.016132245 | 0.541817117 | 0.850208191 | 0.041789574 | 0.054539491 |
| Shroom2 | -0.032096098 | 0.321652374 | 0.719227395 | 0.064851522 | 0.091873075 |
| Shroom3 | -0.022446772 | 0.606454057 | 0.878341504 | 0.128711376 | 0.142928716 |
| Shtn1 | -0.03404799 | 0.542522417 | 0.85028523 | 0.228129635 | 0.236275554 |
| Siae | -0.01719803 | 0.62189266 | 0.884879661 | 0.076458436 | 0.090948404 |
| Siah1a | 0.104872236 | 0.046920012 | 0.339316662 | 0.20361483 | 0.103303335 |
| Siah2 | -0.075851495 | 0.249098634 | 0.66061449 | 0.526712693 | 0.605572206 |
| Sidt2 | -0.018291474 | 0.69259713 | 0.909796728 | 0.145562224 | 0.154756988 |
| Sigirr | -0.005251991 | 0.867034828 | 0.962125879 | 0.060236016 | 0.062926913 |
| Sigmar1 | -0.044019055 | 0.287413143 | 0.694120722 | 0.112028755 | 0.157481996 |
| Sik1 | -0.097073558 | 0.008616951 | 0.144387881 | 0.07938346 | 0.156083354 |
| Sik2 | -0.059642373 | 0.109592474 | 0.477972625 | 0.082805843 | 0.133683796 |
| Sik3 | 0.088208588 | 0.163310866 | 0.563088685 | 0.413121877 | 0.334752494 |
| Sike1 | 0.065243341 | 0.274583861 | 0.682000399 | 0.323939995 | 0.256374033 |
| Sil1 | -0.08184542 | 0.185953624 | 0.592701341 | 0.351764784 | 0.440795027 |
| Sim1 | 0.003100273 | 0.931129444 | 0.982935026 | 0.080148847 | 0.089518218 |
| Simc1 | 0.040470759 | 0.183055964 | 0.590066231 | 0.062357475 | 0.030719072 |
| Sin3a | 0.005034933 | 0.927106904 | 0.981818446 | 0.221740984 | 0.205792973 |
| Sin3b | -0.069415631 | 0.308955717 | 0.711182121 | 0.711591293 | 0.784571893 |
| Sinhcaf | -0.059361615 | 0.21832509 | 0.632618221 | 0.150754299 | 0.213441526 |
| Sipa1l1 | -0.086289063 | 0.075194699 | 0.413795734 | 0.158125069 | 0.231170523 |
| Sipa1l3 | -0.026812602 | 0.626619074 | 0.886996623 | 0.214269165 | 0.235893165 |
| Sirpa | -0.064190925 | 0.311755954 | 0.71405691 | 0.398899761 | 0.474883376 |
| Sirt1 | -0.02737423 | 0.493842903 | 0.828842054 | 0.096863451 | 0.123599119 |
| Sirt2 | -0.095362598 | 0.082966403 | 0.430353379 | 0.222074982 | 0.301137669 |
| Sirt3 | -0.054669253 | 0.245068533 | 0.654809396 | 0.143292766 | 0.200475643 |
| Sirt4 | -0.045660533 | 0.081804601 | 0.42777546 | 0.038693289 | 0.080633818 |
| Sirt5 | -0.034014773 | 0.313338661 | 0.714906079 | 0.069280092 | 0.1001973 |
| Sirt6 | 0.063822522 | 0.05130504 | 0.355933715 | 0.074893505 | 0.021437311 |
| Sirt7 | -0.052424846 | 0.321742652 | 0.719227395 | 0.180341492 | 0.234488555 |
| Siva1 | -0.030889569 | 0.617775338 | 0.883238152 | 0.367767715 | 0.412248156 |
| Six4 | -8.70E-07 | 0.99998562 | 0.99998562 | 0.153354665 | 0.171723323 |
| Six5 | -0.009083739 | 0.750533662 | 0.92612569 | 0.048824153 | 0.061631645 |
| Skap1 | -0.10817258 | 0.094070978 | 0.452959864 | 0.358446683 | 0.452652582 |
| Skap2 | 0.073095339 | 0.112575683 | 0.483221516 | 0.158527419 | 0.092737075 |
| Ski | -0.054081264 | 0.306675216 | 0.709680069 | 0.195605959 | 0.250894867 |
| Skil | -0.058613859 | 0.35625914 | 0.745571198 | 0.351819698 | 0.405643148 |
| Skiv2l | 0.011292305 | 0.792499265 | 0.939347252 | 0.12772874 | 0.120692805 |

| | | | | | |
|---|---|---|---|---|---|
| Skp1 | -0.01762926 | 0.791317808 | 0.938824685 | 0.712566973 | 0.719065583 |
| Slain1 | -0.026633604 | 0.550425038 | 0.852505562 | 0.133052125 | 0.161307056 |
| Slain2 | 0.085094204 | 0.16181936 | 0.560531277 | 0.372041371 | 0.28323807 |
| Slbp | 0.043231621 | 0.402429916 | 0.77525778 | 0.196917994 | 0.153232832 |
| Slc10a3 | 0.011608796 | 0.68753723 | 0.909796728 | 0.055075682 | 0.044701466 |
| Slc10a7 | 0.079746411 | 0.017678854 | 0.21172951 | 0.081988539 | 0.014507486 |
| Slc11a2 | 0.053151458 | 0.239572886 | 0.650744882 | 0.149077064 | 0.110227953 |
| Slc12a2 | -0.045030849 | 0.30258249 | 0.707765194 | 0.118186594 | 0.164080894 |
| Slc12a6 | -0.109298321 | 0.011787756 | 0.172720733 | 0.111495737 | 0.20874782 |
| Slc12a7 | -0.010566572 | 0.843854447 | 0.955524266 | 0.203963006 | 0.195511965 |
| Slc15a4 | 0.038495145 | 0.313415167 | 0.714906079 | 0.101467083 | 0.072592737 |
| Slc16a10 | 0.043242997 | 0.521536828 | 0.840205104 | 0.545208899 | 0.496659984 |
| Slc16a11 | -0.007492302 | 0.822520291 | 0.950930859 | 0.07027775 | 0.085389203 |
| Slc16a12 | 0.011142476 | 0.856006673 | 0.958084785 | 0.359342531 | 0.330744695 |
| Slc16a6 | 0.01953045 | 0.724110024 | 0.919830015 | 0.22845527 | 0.21259894 |
| Slc16a9 | 0.031322674 | 0.518030391 | 0.837813573 | 0.1634718 | 0.131254531 |
| Slc17a5 | 0.010907097 | 0.813481744 | 0.948397417 | 0.146081426 | 0.137522418 |
| Slc18a1 | -0.041021789 | 0.1058971 | 0.473025898 | 0.032956657 | 0.063694757 |
| Slc19a1 | -0.020484271 | 0.421053796 | 0.787234019 | 0.035622472 | 0.050486139 |
| Slc19a2 | 0.022600644 | 0.452704859 | 0.807136288 | 0.056772361 | 0.039850474 |
| Slc1a4 | 0.033456402 | 0.508476199 | 0.83445598 | 0.177501862 | 0.146285484 |
| Slc20a1 | 0.046931233 | 0.235909122 | 0.646415467 | 0.107136004 | 0.073369503 |
| Slc20a2 | 0.007107082 | 0.810584671 | 0.948188174 | 0.057342125 | 0.053730371 |
| Slc22a15 | 0.029345112 | 0.424274797 | 0.789771168 | 0.09010234 | 0.069145589 |
| Slc22a17 | -0.124088606 | 0.017070282 | 0.208804661 | 0.177517157 | 0.287085688 |
| Slc22a23 | 0.058190483 | 0.352895409 | 0.743880213 | 0.357806258 | 0.308284005 |
| Slc23a2 | 0.05485569 | 0.25621697 | 0.664906869 | 0.165664677 | 0.106428076 |
| Slc24a2 | -0.038364131 | 0.561868715 | 0.857134531 | 0.495042091 | 0.508957549 |
| Slc24a5 | -0.028520452 | 0.475104506 | 0.817387411 | 0.105739746 | 0.139678243 |
| Slc25a1 | 0.278470362 | 8.06E-05 | 0.00685329 | 0.765132669 | 0.439627358 |
| Slc25a10 | -0.015862378 | 0.552382878 | 0.852623515 | 0.040861521 | 0.051596378 |
| Slc25a11 | -0.011734859 | 0.841069926 | 0.954960614 | 0.287794241 | 0.324639348 |
| Slc25a12 | 0.029260079 | 0.593146757 | 0.872442739 | 0.242856787 | 0.20958005 |
| Slc25a13 | -0.045284216 | 0.058657764 | 0.372682944 | 0.028100041 | 0.062615437 |
| Slc25a14 | -0.05366854 | 0.264495206 | 0.671066325 | 0.151479414 | 0.207997999 |
| Slc25a16 | 0.033183906 | 0.54231682 | 0.850208191 | 0.231805906 | 0.208068029 |
| Slc25a17 | -0.093972846 | 0.083274843 | 0.431284622 | 0.220466027 | 0.293683522 |
| Slc25a19 | 0.097579826 | 0.012725071 | 0.179750602 | 0.111914984 | 0.032458324 |
| Slc25a20 | -0.037551567 | 0.283473476 | 0.69129896 | 0.072406783 | 0.118447683 |
| Slc25a22 | -0.037756473 | 0.314670164 | 0.714975653 | 0.0890895 | 0.121245395 |
| Slc25a23 | -0.026615063 | 0.601634408 | 0.876557788 | 0.184240672 | 0.212022538 |
| Slc25a25 | 0.01716879 | 0.738111255 | 0.922843014 | 0.18373642 | 0.165665042 |
| Slc25a27 | 0.019986334 | 0.579547327 | 0.865234762 | 0.086369575 | 0.064982373 |
| Slc25a28 | 0.000452224 | 0.994465485 | 0.998695538 | 0.468031169 | 0.454585806 |
| Slc25a29 | -0.030678986 | 0.473283975 | 0.816838319 | 0.120456143 | 0.142222885 |
| Slc25a3 | -0.154971956 | 0.003012692 | 0.076092349 | 1.667272601 | 1.940648067 |
| Slc25a32 | -0.010109015 | 0.830084707 | 0.95325696 | 0.14931051 | 0.157289 |
| Slc25a35 | -0.061087516 | 0.023196026 | 0.241262711 | 0.035707066 | 0.091169618 |
| Slc25a36 | -0.029170938 | 0.652439492 | 0.896875696 | 0.404936861 | 0.43685058 |
| Slc25a37 | 0.004664368 | 0.88147904 | 0.968081674 | 0.059616725 | 0.059811262 |
| Slc25a38 | 0.030393959 | 0.359169023 | 0.747455833 | 0.070638008 | 0.047818919 |
| Slc25a39 | -0.154918052 | 0.01424246 | 0.193060412 | 0.888652151 | 1.076998212 |
| Slc25a4 | 0.122748856 | 0.023603375 | 0.244234094 | 1.593428125 | 1.409816155 |
| Slc25a40 | 0.062989981 | 0.14540943 | 0.533628684 | 0.135783745 | 0.073862265 |
| Slc25a42 | 0.066884246 | 0.090795653 | 0.445385778 | 0.111295878 | 0.052280156 |
| Slc25a44 | -0.015923341 | 0.784192053 | 0.936347328 | 0.269402336 | 0.283885273 |
| Slc25a46 | -0.019232558 | 0.749035386 | 0.925186829 | 0.303075201 | 0.32092818 |
| Slc25a5 | -0.145577463 | 0.030792754 | 0.279445638 | 0.941462294 | 1.135315728 |
| Slc25a51 | -0.047976237 | 0.484608547 | 0.823153832 | 0.560137122 | 0.591459176 |
| Slc25a53 | -0.036498134 | 0.273463055 | 0.681426919 | 0.065377736 | 0.100334941 |
| Slc26a2 | -0.007532766 | 0.838117675 | 0.954911647 | 0.090866167 | 0.091173863 |
| Slc27a1 | -0.012188121 | 0.701192032 | 0.912152193 | 0.062990657 | 0.07462225 |
| Slc27a4 | -0.019543806 | 0.628937216 | 0.887864718 | 0.104387207 | 0.123900669 |
| Slc29a1 | -0.030681199 | 0.607937882 | 0.87877121 | 0.278480591 | 0.319308065 |
| Slc29a3 | -0.018352425 | 0.542803596 | 0.850338644 | 0.056273052 | 0.070897339 |

| | | | | | |
|---|---|---|---|---|---|
| Slc29a4 | 0.026726499 | 0.420048373 | 0.786425204 | 0.072646097 | 0.047420598 |
| Slc2a2 | 0.408020002 | 8.11E-07 | 0.000166207 | 1.184084917 | 0.666127929 |
| Slc2a5 | 0.143407483 | 0.039540374 | 0.313480836 | 1.023058251 | 0.837368843 |
| Slc30a1 | 0.049923646 | 0.233251657 | 0.643935869 | 0.121706362 | 0.085224776 |
| Slc30a4 | 0.039123864 | 0.415816995 | 0.783472369 | 0.167309523 | 0.128898031 |
| Slc30a5 | 0.005205245 | 0.925966384 | 0.981772637 | 0.247831329 | 0.24517638 |
| Slc30a6 | 0.010418698 | 0.737052128 | 0.92276204 | 0.063406266 | 0.052957297 |
| Slc30a7 | 0.047475431 | 0.311716548 | 0.71405691 | 0.161856313 | 0.120402898 |
| Slc30a8 | -0.03779007 | 0.54217428 | 0.850208191 | 1.411620838 | 1.484282004 |
| Slc30a9 | -0.062547962 | 0.326490529 | 0.720866629 | 0.361327316 | 0.422995533 |
| Slc31a1 | 0.012871185 | 0.805029689 | 0.94470747 | 0.186450976 | 0.167583697 |
| Slc31a2 | 0.022818332 | 0.631555469 | 0.88887544 | 0.158578977 | 0.139100545 |
| Slc33a1 | -0.101541255 | 0.085748587 | 0.435997247 | 0.299179212 | 0.409002894 |
| Slc35a1 | -0.009318338 | 0.855490779 | 0.957736324 | 0.187714418 | 0.187012928 |
| Slc35a2 | 0.058751404 | 0.25649471 | 0.665229303 | 0.19947282 | 0.139093875 |
| Slc35a3 | 0.031312554 | 0.455969953 | 0.808152996 | 0.124655963 | 0.095179217 |
| Slc35a4 | 0.035815209 | 0.474261935 | 0.817387411 | 0.183435161 | 0.154452671 |
| Slc35a5 | 0.013309967 | 0.765436469 | 0.928757959 | 0.143430979 | 0.135664299 |
| Slc35b1 | 0.039643964 | 0.570069799 | 0.86083487 | 0.679309865 | 0.650137481 |
| Slc35b2 | 0.09500696 | 0.099256252 | 0.463315751 | 0.288403364 | 0.19911356 |
| Slc35b3 | 0.040937467 | 0.324688783 | 0.720676983 | 0.120549566 | 0.084266018 |
| Slc35b4 | -0.008908799 | 0.852084343 | 0.957461155 | 0.15696701 | 0.162939938 |
| Slc35c1 | -0.040799538 | 0.230491249 | 0.641909176 | 0.073141134 | 0.102683124 |
| Slc35c2 | -0.060111818 | 0.213777442 | 0.628127689 | 0.151489422 | 0.196045217 |
| Slc35d1 | -0.019638909 | 0.566900494 | 0.858864944 | 0.076864459 | 0.09450309 |
| Slc35d2 | -0.035425777 | 0.189469977 | 0.598022064 | 0.041155647 | 0.068114093 |
| Slc35d3 | -0.020537121 | 0.688798786 | 0.909796728 | 0.180334796 | 0.198277154 |
| Slc35e1 | 0.115365636 | 0.020322051 | 0.227647793 | 0.18498975 | 0.080267929 |
| Slc35e2 | 0.042757843 | 0.192988501 | 0.60325934 | 0.072996483 | 0.039735875 |
| Slc35e3 | -0.057705699 | 0.024774823 | 0.249914469 | 0.031458493 | 0.079603957 |
| Slc35e4 | -0.034358337 | 0.505088838 | 0.832249514 | 0.183621407 | 0.221644907 |
| Slc35f5 | -0.049597058 | 0.44991294 | 0.806495742 | 0.430645947 | 0.476816674 |
| Slc35g1 | -0.060201403 | 0.24050304 | 0.650744882 | 0.163512229 | 0.222906355 |
| Slc35g2 | 0.010859554 | 0.861391798 | 0.959896689 | 0.336187428 | 0.325229685 |
| Slc36a1 | 0.010379415 | 0.833671202 | 0.954437989 | 0.171846498 | 0.151981734 |
| Slc36a4 | 0.021173293 | 0.654610956 | 0.897454058 | 0.154982002 | 0.138158466 |
| Slc37a1 | 0.020913987 | 0.679257161 | 0.907783945 | 0.171480283 | 0.145860647 |
| Slc37a3 | -0.019371979 | 0.676458765 | 0.906611913 | 0.142923197 | 0.158990769 |
| Slc37a4 | 0.098996625 | 0.052731714 | 0.359036238 | 0.202648143 | 0.116388993 |
| Slc38a10 | -0.150731247 | 0.025819584 | 0.256331521 | 0.521355934 | 0.664181352 |
| Slc38a2 | -0.170630176 | 0.017428918 | 0.210563469 | 0.79728451 | 0.966232585 |
| Slc38a4 | 0.00642947 | 0.926807025 | 0.981818446 | 0.671358922 | 0.656715971 |
| Slc38a6 | -0.081717104 | 0.01616913 | 0.205235264 | 0.06370489 | 0.130293137 |
| Slc38a7 | 0.071753478 | 0.056383358 | 0.367897032 | 0.101647976 | 0.045400837 |
| Slc38a9 | -0.0463165 | 0.173460202 | 0.575630234 | 0.066207016 | 0.105264595 |
| Slc39a1 | -0.011608452 | 0.828806292 | 0.952874813 | 0.207040505 | 0.218892725 |
| Slc39a10 | -0.075851828 | 0.091188132 | 0.445813581 | 0.124167139 | 0.193337035 |
| Slc39a13 | -0.075895957 | 0.090716008 | 0.445385778 | 0.127447074 | 0.194810581 |
| Slc39a14 | 0.030817809 | 0.377045265 | 0.761412185 | 0.081221674 | 0.055632265 |
| Slc39a3 | 0.030381761 | 0.527833938 | 0.843341172 | 0.164038597 | 0.137501598 |
| Slc39a6 | -0.110967268 | 0.044498003 | 0.330723271 | 0.216507161 | 0.320169093 |
| Slc39a9 | 0.014570071 | 0.796595822 | 0.940196644 | 0.256920666 | 0.245125945 |
| Slc3a2 | -0.033869899 | 0.564828735 | 0.858087314 | 1.352552646 | 1.382190424 |
| Slc40a1 | -0.141798805 | 0.004726953 | 0.100945587 | 0.154609719 | 0.271661028 |
| Slc41a1 | -0.015799784 | 0.680709889 | 0.907865189 | 0.098528191 | 0.116152893 |
| Slc43a2 | -0.092457155 | 0.16109492 | 0.559461938 | 0.463097947 | 0.540109607 |
| Slc44a1 | -0.036442376 | 0.538253617 | 0.848108564 | 0.279406624 | 0.312299773 |
| Slc44a3 | -0.04044001 | 0.5036743 | 0.832249514 | 0.302747849 | 0.337563607 |
| Slc45a4 | 0.038821546 | 0.307060752 | 0.709680069 | 0.099764873 | 0.065443228 |
| Slc46a3 | -0.020095423 | 0.440517704 | 0.80031914 | 0.039306956 | 0.057066801 |
| Slc48a1 | -0.034158754 | 0.564070149 | 0.857926031 | 0.286970672 | 0.314796284 |
| Slc49a4 | 0.124193366 | 0.027046646 | 0.26190846 | 0.24078318 | 0.131736554 |
| Slc4a10 | 0.056413249 | 0.310846895 | 0.71348779 | 0.239520351 | 0.173082742 |
| Slc4a1ap | 0.048923747 | 0.321016819 | 0.718885724 | 0.182111823 | 0.140538181 |
| Slc4a2 | -0.075883141 | 0.101125798 | 0.466881156 | 0.140505866 | 0.204828764 |

| | | | | | |
|---|---|---|---|---|---|
| Slc4a3 | -0.050913074 | 0.229851645 | 0.641131395 | 0.11398268 | 0.149481328 |
| Slc4a4 | -0.051932466 | 0.215276297 | 0.629611185 | 0.104674102 | 0.159484413 |
| Slc4a7 | 0.030935351 | 0.666245134 | 0.901197061 | 0.964317563 | 0.918583731 |
| Slc4a8 | -0.017082927 | 0.683162408 | 0.909173483 | 0.113007096 | 0.131944861 |
| Slc50a1 | -0.038207132 | 0.569970992 | 0.86083487 | 0.598627733 | 0.642149201 |
| Slc5a3 | 0.05740053 | 0.083639465 | 0.431794059 | 0.074449074 | 0.025349736 |
| Slc5a6 | -0.016017876 | 0.626008966 | 0.886938778 | 0.067642764 | 0.082777085 |
| Slc66a1 | -0.015514101 | 0.551132874 | 0.852505562 | 0.039663232 | 0.055350721 |
| Slc66a2 | 0.034946063 | 0.489848402 | 0.826459641 | 0.187923024 | 0.158511518 |
| Slc6a17 | -0.195624306 | 0.000851706 | 0.034331622 | 0.26842077 | 0.449177623 |
| Slc6a19 | 0.056430939 | 0.126363527 | 0.505886017 | 0.092993542 | 0.040425987 |
| Slc6a6 | 0.038103486 | 0.561143369 | 0.857134531 | 1.156048009 | 1.101055688 |
| Slc6a8 | 0.116402058 | 0.014366527 | 0.193677739 | 0.165276461 | 0.060829685 |
| Slc6a9 | 0.005232106 | 0.852982146 | 0.957461155 | 0.050313883 | 0.043608379 |
| Slc7a1 | -0.042879916 | 0.498699127 | 0.830722138 | 0.384941979 | 0.422458065 |
| Slc7a11 | -0.102504894 | 0.002156376 | 0.063845264 | 0.056999323 | 0.137261891 |
| Slc7a14 | 0.056044401 | 0.421917893 | 0.787721334 | 1.084326352 | 0.97733104 |
| Slc7a2 | 0.035556757 | 0.584997011 | 0.867683611 | 0.463221264 | 0.436443912 |
| Slc7a4 | -0.048667706 | 0.160136451 | 0.558431893 | 0.074528351 | 0.115229796 |
| Slc7a5 | -0.005788633 | 0.928759555 | 0.982502239 | 0.395278109 | 0.39391299 |
| Slc7a6 | 0.009213546 | 0.757108371 | 0.92705828 | 0.057716986 | 0.047460637 |
| Slc7a6os | -0.074369196 | 0.08716554 | 0.439557333 | 0.118852084 | 0.181518241 |
| Slc7a8 | -0.07909899 | 0.186485231 | 0.593453477 | 0.294893429 | 0.36543479 |
| Slc8a1 | -0.203229836 | 0.000966629 | 0.037358974 | 0.305091144 | 0.495672163 |
| Slc9a1 | 0.054022681 | 0.079459126 | 0.422526451 | 0.065252922 | 0.017556798 |
| Slc9a3r1 | 0.073752158 | 0.097218654 | 0.459841484 | 0.138651516 | 0.07709115 |
| Slc9a3r2 | -0.040595771 | 0.233191908 | 0.643935869 | 0.06834118 | 0.098090068 |
| Slc9a6 | 0.047532987 | 0.238667943 | 0.650445287 | 0.115690568 | 0.069770931 |
| Slc9a7 | 0.074314943 | 0.048653789 | 0.346093609 | 0.10200846 | 0.038369294 |
| Slc9a8 | -0.051781201 | 0.226946748 | 0.638952177 | 0.121265908 | 0.163147992 |
| Slc9a9 | -0.058811319 | 0.034017451 | 0.291823211 | 0.038479854 | 0.089095864 |
| Slco1a6 | -0.011728084 | 0.735953552 | 0.922302218 | 0.072146348 | 0.081008326 |
| Slco2a1 | 0.033981526 | 0.409934345 | 0.780101937 | 0.11454534 | 0.083024624 |
| Slco3a1 | 0.127309312 | 0.018679362 | 0.216996248 | 0.23352291 | 0.128014021 |
| Slf1 | -0.037639256 | 0.352981604 | 0.743880213 | 0.110788047 | 0.143526449 |
| Slf2 | 0.051141698 | 0.332311113 | 0.72806771 | 0.201202728 | 0.157721277 |
| Slirp | -0.041480013 | 0.537535817 | 0.847793898 | 0.548354107 | 0.551663595 |
| Slitrk6 | 0.073802147 | 0.179858531 | 0.586833233 | 0.225416092 | 0.149990945 |
| Slk | -0.097508835 | 0.128861752 | 0.508260084 | 0.384033023 | 0.491175902 |
| Slmap | 0.006677719 | 0.902063109 | 0.974597139 | 0.210210953 | 0.202205186 |
| Sltm | 0.089495402 | 0.198269025 | 0.609389286 | 0.777456176 | 0.67109975 |
| Slu7 | 0.035104276 | 0.545594251 | 0.851259637 | 0.289771271 | 0.272095215 |
| Slx1b | -0.056972392 | 0.095488302 | 0.455869783 | 0.073882557 | 0.119814955 |
| Slx4 | -0.008354234 | 0.751664557 | 0.926186851 | 0.04104405 | 0.044319754 |
| Slx4ip | 0.01260665 | 0.731496696 | 0.920770724 | 0.090100006 | 0.076005871 |
| Smad1 | -2.48E-06 | 0.999970331 | 0.99998562 | 0.454376744 | 0.445757639 |
| Smad2 | -0.02152685 | 0.679138894 | 0.907783945 | 0.181240818 | 0.207921068 |
| Smad4 | -0.044978405 | 0.487353666 | 0.826112275 | 0.494109842 | 0.545147475 |
| Smad5 | 0.043765854 | 0.448314559 | 0.804961222 | 0.270533256 | 0.228832689 |
| Smad7 | -0.174964441 | 0.002227448 | 0.064615296 | 0.21936959 | 0.387637563 |
| Smap1 | 0.068484587 | 0.317705937 | 0.717070945 | 0.660851726 | 0.595467695 |
| Smap2 | 0.169851657 | 0.006409653 | 0.120927987 | 1.307117138 | 1.045445656 |
| Smarca1 | -0.048647937 | 0.154380994 | 0.551012197 | 0.066769659 | 0.102246534 |
| Smarca2 | -0.116479094 | 0.077206701 | 0.418199494 | 0.483488226 | 0.58855289 |
| Smarca4 | -0.100361761 | 0.149453726 | 0.543502554 | 0.765172195 | 0.871480256 |
| Smarca5 | 0.018044632 | 0.786000723 | 0.937083312 | 1.029083723 | 0.993446558 |
| Smarcad1 | 0.021959591 | 0.706221083 | 0.912857825 | 0.274733499 | 0.248195545 |
| Smarcal1 | -0.067895275 | 0.044251834 | 0.329492329 | 0.067488738 | 0.124494861 |
| Smarcb1 | 0.023746576 | 0.690666079 | 0.909796728 | 0.279281845 | 0.260553136 |
| Smarcc1 | -0.044629144 | 0.45727772 | 0.808898849 | 0.294248469 | 0.335052209 |
| Smarcc2 | -0.013719037 | 0.839493633 | 0.954911647 | 0.568404405 | 0.560051161 |
| Smarcd1 | 0.003098878 | 0.957148463 | 0.98987052 | 0.245455687 | 0.242695655 |
| Smarcd2 | 0.070177463 | 0.219988912 | 0.633217295 | 0.278182711 | 0.213046319 |
| Smarce1 | 0.127882793 | 0.03526963 | 0.295084312 | 0.335387491 | 0.218871681 |
| Smc1a | -0.006146174 | 0.92451954 | 0.981375217 | 0.421574053 | 0.418200159 |

| | | | | | |
|---|---|---|---|---|---|
| Smc3 | -0.029750308 | 0.648348686 | 0.895379629 | 0.612916733 | 0.654256648 |
| Smc4 | 0.023908492 | 0.657736767 | 0.898224427 | 0.200727965 | 0.185397242 |
| Smc5 | -0.043127389 | 0.394044511 | 0.770629121 | 0.177695762 | 0.219201835 |
| Smc6 | -0.117653872 | 0.085201967 | 0.434759607 | 0.696238694 | 0.84563868 |
| Smchd1 | -0.160438084 | 0.01109101 | 0.166398305 | 0.369009066 | 0.529149639 |
| Smco4 | 0.042283273 | 0.132202121 | 0.513250014 | 0.05312482 | 0.022876297 |
| Smcr8 | 0.07770352 | 0.043974278 | 0.329446347 | 0.110337031 | 0.043488583 |
| Smdt1 | 0.078692682 | 0.105168481 | 0.472399448 | 1.840750119 | 1.677385579 |
| Smg1 | 0.031264802 | 0.620525684 | 0.883937014 | 0.402528236 | 0.376847204 |
| Smg5 | 0.019820545 | 0.670624849 | 0.904104984 | 0.150702782 | 0.128901682 |
| Smg6 | -0.097672031 | 0.07719764 | 0.418199494 | 0.22208658 | 0.314690199 |
| Smg7 | 0.059694739 | 0.333562056 | 0.729562509 | 0.365326546 | 0.309528548 |
| Smg8 | 0.007251073 | 0.794387837 | 0.939757086 | 0.049113463 | 0.044941779 |
| Smg9 | -0.043895621 | 0.261365408 | 0.668468258 | 0.096682668 | 0.13326654 |
| Smim1 | -0.102827533 | 0.121965324 | 0.49725002 | 0.620850477 | 0.715919056 |
| Smim10l1 | 0.114662673 | 0.088085251 | 0.441393741 | 0.959817309 | 0.838833925 |
| Smim11 | 0.024630526 | 0.708343974 | 0.913767764 | 0.53016245 | 0.516965455 |
| Smim12 | 0.054375105 | 0.233336132 | 0.643935869 | 0.148611022 | 0.103922925 |
| Smim13 | -0.055077362 | 0.329147422 | 0.724485236 | 0.257799702 | 0.307573019 |
| Smim14 | -0.218984388 | 0.00026868 | 0.016148166 | 1.03419702 | 1.330369059 |
| Smim15 | 0.057400228 | 0.277296051 | 0.684342381 | 0.222399081 | 0.16736945 |
| Smim19 | 0.051831477 | 0.412442237 | 0.781358105 | 0.406582456 | 0.351198921 |
| Smim20 | 0.148971083 | 0.034450108 | 0.293212179 | 0.561782053 | 0.404939717 |
| Smim22 | 0.011834502 | 0.857287996 | 0.958302663 | 0.808368015 | 0.737345543 |
| Smim26 | 0.06896114 | 0.30652653 | 0.709680069 | 0.671302526 | 0.581976955 |
| Smim27 | -0.060497966 | 0.331666696 | 0.727636858 | 0.326538039 | 0.368890323 |
| Smim4 | 0.095969886 | 0.149214345 | 0.543025517 | 0.60276022 | 0.471740031 |
| Smim7 | -0.08004503 | 0.239160566 | 0.650706044 | 0.580436576 | 0.648726203 |
| Smim8 | 0.01950515 | 0.736098131 | 0.922302218 | 0.271022038 | 0.247426185 |
| Smn1 | 0.021400723 | 0.688188588 | 0.909796728 | 0.201833594 | 0.183643809 |
| Smndc1 | 0.009870563 | 0.880107546 | 0.967751911 | 0.392078387 | 0.388375777 |
| Smoc1 | -0.07218538 | 0.019430088 | 0.221110882 | 0.055436118 | 0.115268893 |
| Smpd1 | 0.088212874 | 0.136652613 | 0.520249727 | 0.332907723 | 0.251666377 |
| Smpd2 | -0.006149762 | 0.899056517 | 0.973823054 | 0.162586342 | 0.167603291 |
| Smpd3 | 0.056199752 | 0.198008988 | 0.609076376 | 0.132439679 | 0.090515512 |
| Smpd4 | 0.040185228 | 0.195465163 | 0.605706651 | 0.064160493 | 0.024270742 |
| Smpdl3a | -0.037027307 | 0.449750913 | 0.806495742 | 0.165858426 | 0.203704221 |
| Sms | -0.024019353 | 0.68745128 | 0.909796728 | 0.301869597 | 0.311799485 |
| Smu1 | 0.004105988 | 0.94566189 | 0.987084873 | 0.295023248 | 0.305588502 |
| Smug1 | -0.009724865 | 0.81738701 | 0.949382903 | 0.115520642 | 0.126131384 |
| Smurf1 | -0.015478793 | 0.578945687 | 0.865228985 | 0.044645283 | 0.059725659 |
| Smurf2 | 0.039441954 | 0.430682088 | 0.793311823 | 0.183372616 | 0.14804501 |
| Smyd2 | -0.052012017 | 0.366943029 | 0.753633555 | 0.266936619 | 0.309853907 |
| Smyd3 | -0.053411293 | 0.170384238 | 0.572762895 | 0.095221883 | 0.145716139 |
| Smyd4 | -0.039935794 | 0.323928693 | 0.720406991 | 0.105187715 | 0.148631047 |
| Smyd5 | -0.019144095 | 0.534690788 | 0.846347806 | 0.059030585 | 0.080886667 |
| Snai2 | -0.052424301 | 0.009790019 | 0.153535036 | 0.016328341 | 0.058747331 |
| Snap23 | -0.073825574 | 0.128383024 | 0.508260084 | 0.153782894 | 0.222330925 |
| Snap25 | -0.313805963 | 5.92E-06 | 0.000860632 | 0.724201231 | 1.101391847 |
| Snap29 | -0.0514845 | 0.284455397 | 0.691925959 | 0.151572473 | 0.201414446 |
| Snap47 | 0.02188068 | 0.726328925 | 0.92014727 | 0.379900004 | 0.38084087 |
| Snap91 | 0.013507633 | 0.750510702 | 0.92612569 | 0.120809325 | 0.102992286 |
| Snapc1 | 0.071353062 | 0.240225054 | 0.650744882 | 0.297235888 | 0.230704859 |
| Snapc2 | -0.009381031 | 0.829883116 | 0.953254388 | 0.125481147 | 0.13041736 |
| Snapc3 | -0.084593787 | 0.01837276 | 0.215429197 | 0.07530854 | 0.143397023 |
| Snapc4 | 0.033929753 | 0.242105008 | 0.652555608 | 0.056145236 | 0.02491172 |
| Snapc5 | -0.003611841 | 0.947120693 | 0.987559775 | 0.233520561 | 0.241497771 |
| Snapin | 0.01717477 | 0.700616054 | 0.912109871 | 0.133515875 | 0.11613445 |
| Sncb | -0.048502744 | 0.406707198 | 0.777736088 | 0.268737294 | 0.310321477 |
| Snd1 | -0.231860787 | 0.000638593 | 0.028919709 | 0.528685346 | 0.786574596 |
| Snf8 | -0.110505891 | 0.086376609 | 0.436976828 | 0.496654084 | 0.619829254 |
| Snhg1 | 0.054524552 | 0.455089803 | 0.807876964 | 0.699491604 | 0.656826774 |
| Snhg10 | 0.038428671 | 0.251577268 | 0.663127858 | 0.074206661 | 0.043884591 |
| Snhg11 | -0.287657746 | 6.99E-06 | 0.001002924 | 0.227391028 | 0.549416903 |
| Snhg12 | -0.117567059 | 0.08139479 | 0.427503666 | 0.528448268 | 0.66883628 |

| | | | | | |
|---|---|---|---|---|---|
| Snhg14 | -0.01963766 | 0.45002641 | 0.806495742 | 0.038534482 | 0.052661514 |
| Snhg15 | -0.097025127 | 0.052868503 | 0.359257749 | 0.146865739 | 0.251486227 |
| Snhg16 | -0.065439937 | 0.028748839 | 0.270181023 | 0.043718806 | 0.11100015 |
| Snhg17 | 0.013922551 | 0.679471663 | 0.907783945 | 0.075112939 | 0.062338375 |
| Snhg18 | -0.000522519 | 0.992508936 | 0.998695538 | 0.208049165 | 0.198670417 |
| Snhg20 | -0.035345601 | 0.547661712 | 0.851675805 | 0.296871911 | 0.325813254 |
| Snhg3 | -0.0598521 | 0.357080045 | 0.745983316 | 0.404922697 | 0.457731467 |
| Snhg4 | -0.144807916 | 0.007088848 | 0.128637347 | 0.196579972 | 0.32389755 |
| Snhg5 | -0.054291185 | 0.287976316 | 0.694589247 | 0.182652585 | 0.239312616 |
| Snhg6 | 0.023103563 | 0.726667396 | 0.920323111 | 0.448777423 | 0.404961014 |
| Snhg8 | -0.08166287 | 0.233879428 | 0.644459522 | 0.758115781 | 0.826474022 |
| Snhg9 | 0.019842478 | 0.723863227 | 0.919830015 | 0.248314723 | 0.221779156 |
| Snip1 | -0.05296874 | 0.211139584 | 0.624030627 | 0.115577431 | 0.163999092 |
| Snph | 0.096392112 | 0.047571453 | 0.341541257 | 0.173113372 | 0.090810317 |
| Snrk | -0.022320704 | 0.654895853 | 0.897454058 | 0.177713258 | 0.192738509 |
| Snrnp200 | 0.077519906 | 0.23143313 | 0.643104741 | 0.438614355 | 0.354707089 |
| Snrnp25 | 0.021555793 | 0.501712239 | 0.832218079 | 0.064470051 | 0.047365853 |
| Snrnp27 | 0.086394169 | 0.193492467 | 0.60325934 | 0.587911111 | 0.495646552 |
| Snrnp35 | -0.033940794 | 0.392190311 | 0.769704269 | 0.097616649 | 0.125553077 |
| Snrnp40 | 0.014383395 | 0.775056283 | 0.933555636 | 0.196034044 | 0.197191861 |
| Snrnp48 | 0.14222852 | 0.029547742 | 0.27358919 | 0.471419936 | 0.340263316 |
| Snrnp70 | -0.17819519 | 0.009548362 | 0.151640678 | 0.758111104 | 0.94292828 |
| Snrpa | -0.193163416 | 0.000153754 | 0.010687095 | 0.168245132 | 0.358391323 |
| Snrpa1 | -0.053720713 | 0.355815301 | 0.745326231 | 0.27211113 | 0.317957581 |
| Snrpb | -0.104064313 | 0.115386347 | 0.487313382 | 0.598794667 | 0.713094039 |
| Snrpb2 | -0.045891878 | 0.435010944 | 0.796898128 | 0.304468911 | 0.349374582 |
| Snrpc | -0.005197278 | 0.935058888 | 0.984081583 | 0.36916285 | 0.383494557 |
| Snrpd1 | 0.134054068 | 0.056946633 | 0.368959821 | 0.696149731 | 0.557503458 |
| Snrpd2 | -0.015046448 | 0.82457738 | 0.951625061 | 0.797547942 | 0.808798054 |
| Snrpd3 | 0.097311234 | 0.149149441 | 0.542986195 | 0.526092091 | 0.4177505 |
| Snrpe | -0.018905243 | 0.780589379 | 0.935383906 | 0.791529972 | 0.808076114 |
| Snrpf | 0.085378048 | 0.206930522 | 0.618511511 | 0.776073415 | 0.672483627 |
| Snrpg | 0.131539284 | 0.035308983 | 0.295084312 | 0.909866215 | 0.743989888 |
| Snrpn | -0.072217605 | 0.159559206 | 0.557623868 | 0.176199026 | 0.25849254 |
| Sntb2 | 0.046969693 | 0.385414922 | 0.765415428 | 0.209759235 | 0.166466493 |
| Sntg1 | 0.053289883 | 0.219583935 | 0.63289494 | 0.129227615 | 0.083379123 |
| Snu13 | 0.10881427 | 0.110320124 | 0.479758704 | 0.6013755 | 0.486527895 |
| Snupn | 0.047250445 | 0.191512374 | 0.602043906 | 0.088595677 | 0.051603581 |
| Snw1 | -0.022230537 | 0.739242733 | 0.922843014 | 0.534759849 | 0.565531787 |
| Snx1 | 0.067603597 | 0.205614604 | 0.616737719 | 0.228572438 | 0.170760532 |
| Snx10 | 0.079048948 | 0.263897358 | 0.670396806 | 0.656367355 | 0.554443196 |
| Snx11 | -0.06221548 | 0.032848637 | 0.287697882 | 0.050033045 | 0.10437545 |
| Snx12 | -0.010963735 | 0.822384397 | 0.950930859 | 0.161172272 | 0.17038992 |
| Snx13 | 0.135556393 | 0.012287775 | 0.176947483 | 0.250491741 | 0.138335042 |
| Snx14 | -0.071355707 | 0.149826031 | 0.544321624 | 0.161328024 | 0.231472966 |
| Snx15 | -0.02789892 | 0.579440825 | 0.865234762 | 0.186255233 | 0.21635826 |
| Snx16 | -0.006175959 | 0.897853218 | 0.972876254 | 0.162109242 | 0.167646866 |
| Snx17 | -0.079367067 | 0.193815544 | 0.60325934 | 0.336116485 | 0.418262292 |
| Snx18 | -0.005822306 | 0.896194314 | 0.97233366 | 0.134481497 | 0.150417504 |
| Snx19 | 0.021047461 | 0.625260111 | 0.88668351 | 0.123920248 | 0.10425438 |
| Snx2 | -0.024710084 | 0.696849651 | 0.910970095 | 0.3951446 | 0.414748222 |
| Snx21 | -0.043947948 | 0.353996693 | 0.744256656 | 0.151885396 | 0.194389433 |
| Snx24 | 0.035659537 | 0.470297741 | 0.814364464 | 0.172967454 | 0.146050084 |
| Snx25 | -0.000543306 | 0.987351355 | 0.997268589 | 0.07336261 | 0.077981478 |
| Snx27 | -0.021823487 | 0.71081973 | 0.914531044 | 0.267508281 | 0.281409617 |
| Snx3 | 0.12111371 | 0.052338774 | 0.357487248 | 0.970904376 | 0.837940035 |
| Snx30 | 0.055288243 | 0.355816114 | 0.745326231 | 0.308171411 | 0.268021713 |
| Snx32 | -0.004321732 | 0.908288124 | 0.977744305 | 0.094043388 | 0.091554238 |
| Snx33 | -0.02232896 | 0.391837667 | 0.769704269 | 0.040173395 | 0.059433375 |
| Snx4 | 0.000803017 | 0.988944935 | 0.997411191 | 0.262944746 | 0.276977098 |
| Snx5 | -0.003388643 | 0.959667122 | 0.990908231 | 0.515960059 | 0.523563051 |
| Snx6 | 0.05058842 | 0.446649095 | 0.804157614 | 0.587112811 | 0.541362541 |
| Snx7 | -0.05552191 | 0.062115702 | 0.380250374 | 0.047651955 | 0.097355636 |
| Snx8 | -0.056181807 | 0.165671668 | 0.566763419 | 0.10463652 | 0.162731743 |
| Snx9 | 0.008517493 | 0.878993972 | 0.967163177 | 0.231195756 | 0.226497048 |

| | | | | | |
|---|---|---|---|---|---|
| Soat1 | -0.072044518 | 0.172830221 | 0.575371867 | 0.184256278 | 0.240864777 |
| Sobp | 0.090244891 | 0.172095412 | 0.575294174 | 0.446859396 | 0.36555953 |
| Socs2 | 0.028633594 | 0.624181143 | 0.886501504 | 0.269607863 | 0.237471829 |
| Socs3 | -0.108063084 | 0.014476871 | 0.193677739 | 0.117551841 | 0.209294732 |
| Socs4 | 0.011027954 | 0.841548932 | 0.954960614 | 0.241124559 | 0.229788535 |
| Socs5 | 0.026286806 | 0.510968424 | 0.834957869 | 0.112371308 | 0.081615111 |
| Socs6 | -0.017550074 | 0.719974621 | 0.918919795 | 0.156976636 | 0.171008239 |
| Socs7 | -0.00795411 | 0.886488604 | 0.969612367 | 0.232726206 | 0.236020492 |
| Sod1 | -0.245493027 | 0.000266788 | 0.016131031 | 0.893093035 | 1.202356146 |
| Sod2 | 0.026620046 | 0.699360895 | 0.911319029 | 0.787633546 | 0.784574614 |
| Soga1 | 0.030861571 | 0.3418861 | 0.735000423 | 0.073062507 | 0.052215236 |
| Soga3 | 0.033479424 | 0.583478322 | 0.866840131 | 0.363023903 | 0.335072552 |
| Son | -0.034631408 | 0.513335904 | 0.836423834 | 1.702810713 | 1.737253114 |
| Sorbs2 | 0.008490559 | 0.900410756 | 0.974055009 | 0.74122777 | 0.708906064 |
| Sord | -0.028204273 | 0.348290172 | 0.739576235 | 0.055714822 | 0.079596064 |
| Sorl1 | 0.050104631 | 0.434625535 | 0.796628285 | 0.386780701 | 0.334902192 |
| Sort1 | -0.035945274 | 0.563231512 | 0.857617435 | 0.324415547 | 0.344831336 |
| Sos1 | -0.096028127 | 0.097463 | 0.460576232 | 0.271508197 | 0.358708541 |
| Sos2 | 0.069891873 | 0.223107122 | 0.634909606 | 0.265454344 | 0.196536331 |
| Sowaha | 0.034726759 | 0.289345648 | 0.695472148 | 0.070673051 | 0.041902685 |
| Sowahc | -0.045298777 | 0.510439987 | 0.834957869 | 0.496537231 | 0.547171064 |
| Sox12 | -0.044050936 | 0.319860681 | 0.717256849 | 0.124591716 | 0.157015343 |
| Sox4 | 0.144910805 | 0.044711107 | 0.330947919 | 0.524799418 | 0.366980649 |
| Sp1 | -0.008098719 | 0.876924051 | 0.966020504 | 0.192895968 | 0.202191791 |
| Sp2 | 0.017543803 | 0.599841046 | 0.875897759 | 0.072413723 | 0.058748262 |
| Sp3 | 0.072061079 | 0.238383182 | 0.650177173 | 0.346463774 | 0.277713597 |
| Sp3os | -0.055330602 | 0.269971556 | 0.677226912 | 0.171409591 | 0.225575064 |
| Sp4 | -0.039706964 | 0.438320326 | 0.799200693 | 0.180549794 | 0.210061559 |
| Spa17 | 0.037088423 | 0.465481356 | 0.812791331 | 0.191025355 | 0.156515043 |
| Spaca6 | 0.028533981 | 0.494512649 | 0.828842054 | 0.121476239 | 0.099131323 |
| Spag7 | 0.028517914 | 0.647943914 | 0.895376633 | 0.335623497 | 0.316682855 |
| Spag9 | -0.019845453 | 0.777113895 | 0.934230706 | 0.656514715 | 0.684808975 |
| Spast | -0.072252097 | 0.221430522 | 0.633858837 | 0.271887186 | 0.341838664 |
| Spata13 | -0.027407572 | 0.378838869 | 0.762006446 | 0.059313415 | 0.081945584 |
| Spata2 | -0.055570974 | 0.15119685 | 0.546523037 | 0.091486009 | 0.137307481 |
| Spata24 | -0.094353112 | 0.000778437 | 0.032691088 | 0.037177044 | 0.117682132 |
| Spata5 | -0.015350508 | 0.732969995 | 0.921306823 | 0.130810541 | 0.150272892 |
| SPATA5L1 | -0.03111493 | 0.212482287 | 0.626707233 | 0.033014415 | 0.057598697 |
| Spata6 | 0.003715079 | 0.903474326 | 0.974950771 | 0.059699503 | 0.055929489 |
| Spata7 | 0.072361629 | 0.103017264 | 0.468979611 | 0.139287698 | 0.077066688 |
| Spats2 | -0.035267217 | 0.318839582 | 0.717175454 | 0.080340739 | 0.108503324 |
| Spc25 | 0.146915625 | 0.046493026 | 0.33839775 | 0.919277942 | 0.703717559 |
| Spcs1 | 0.134191146 | 0.033045906 | 0.287968471 | 1.515855217 | 1.287213463 |
| Spcs2 | -0.135470691 | 0.027212076 | 0.262622702 | 1.391422348 | 1.607146317 |
| Spcs3 | 0.130057394 | 0.052093228 | 0.356999852 | 0.82457339 | 0.674625397 |
| Specc1 | 0.054747603 | 0.096415568 | 0.457336038 | 0.076279829 | 0.030013561 |
| Specc1l | -0.026508817 | 0.695164963 | 0.910378884 | 0.438550128 | 0.467832851 |
| Spef1 | 0.044846564 | 0.115297702 | 0.487260227 | 0.056053675 | 0.016618799 |
| Spen | -0.090857216 | 0.183414437 | 0.590066231 | 0.438696108 | 0.523940311 |
| Spg11 | -0.069888025 | 0.020094132 | 0.226612137 | 0.052336276 | 0.115543995 |
| Spg20 | 0.006627273 | 0.890132197 | 0.970579007 | 0.161393561 | 0.157519903 |
| Spg21 | -0.047966285 | 0.414751711 | 0.783085706 | 0.287464203 | 0.32788814 |
| Spg7 | -0.027019554 | 0.579182485 | 0.865234762 | 0.161879195 | 0.182291559 |
| Sphk2 | 0.043912787 | 0.364694496 | 0.752802229 | 0.166113822 | 0.131886482 |
| Sphkap | -0.030547982 | 0.605375996 | 0.87802905 | 1.473742304 | 1.517967933 |
| Spice1 | 0.009804167 | 0.774850944 | 0.933555636 | 0.073240551 | 0.056260473 |
| Spidr | -0.05036661 | 0.070249514 | 0.402558548 | 0.037799227 | 0.083315582 |
| Spin1 | 0.049444334 | 0.307077866 | 0.709680069 | 0.177354574 | 0.141329048 |
| Spin2c | -0.052537705 | 0.002282981 | 0.06526292 | 0.00776051 | 0.051935815 |
| Spindoc | 0.013258522 | 0.690582729 | 0.909796728 | 0.071123229 | 0.06047068 |
| Spint1 | -0.068788959 | 0.304956829 | 0.709410087 | 0.71070187 | 0.760071925 |
| Spint2 | -0.141010718 | 0.027719683 | 0.264936073 | 1.177398962 | 1.41527488 |
| Spire1 | 0.00302409 | 0.958364244 | 0.990297691 | 0.250984435 | 0.250380475 |
| Spns1 | -0.017376008 | 0.691694833 | 0.909796728 | 0.125401683 | 0.145208487 |
| Spock1 | -0.071516523 | 0.163507725 | 0.563380378 | 0.162863372 | 0.254206312 |

| | | | | | |
|---|---|---|---|---|---|
| Spock2 | -0.069373038 | 0.287816283 | 0.694589247 | 1.120237969 | 1.160039221 |
| Spon2 | -0.130769528 | 0.000151769 | 0.010687095 | 0.061375978 | 0.177021885 |
| Spop | -0.120130955 | 0.070297435 | 0.402558548 | 0.472674693 | 0.621150406 |
| Spopl | 0.104997038 | 0.043776699 | 0.329086044 | 0.200649022 | 0.107114877 |
| Spout1 | 0.038445732 | 0.34309203 | 0.735815108 | 0.113333867 | 0.080591793 |
| Spp1 | -0.305510814 | 0.001077115 | 0.040036792 | 0.521449013 | 1.000955613 |
| Sppl2a | -0.026530389 | 0.672387989 | 0.905281039 | 0.343409787 | 0.364957754 |
| Sppl3 | -0.058678885 | 0.286809194 | 0.693329451 | 0.224999237 | 0.287260935 |
| Spr | -0.007625255 | 0.895563412 | 0.972284474 | 0.266436201 | 0.273776604 |
| Spred2 | 0.049171247 | 0.238316108 | 0.65017091 | 0.120416678 | 0.079916446 |
| Spred3 | -0.007179232 | 0.797406342 | 0.940690722 | 0.047090583 | 0.047677819 |
| Spring1 | 0.000416984 | 0.992371414 | 0.998695538 | 0.129808518 | 0.132712634 |
| Sprtn | 0.007308159 | 0.828190667 | 0.952874813 | 0.072671957 | 0.064331508 |
| Spry2 | 0.062317848 | 0.077166703 | 0.418199494 | 0.085514715 | 0.029489089 |
| Spryd3 | -0.00750425 | 0.894856752 | 0.972229731 | 0.243494221 | 0.247598606 |
| Spryd4 | -0.013569609 | 0.667483908 | 0.901808587 | 0.060541989 | 0.072650641 |
| Spryd7 | 0.104002473 | 0.070855869 | 0.403272567 | 0.286380665 | 0.195460498 |
| Spsb1 | -0.041252575 | 0.132480842 | 0.513250014 | 0.040693169 | 0.072647067 |
| Spsb2 | -0.032609873 | 0.595810827 | 0.873397585 | 0.301663561 | 0.318348079 |
| Spsb3 | 0.001244919 | 0.98009993 | 0.995540893 | 0.183401253 | 0.185826855 |
| Sptan1 | 0.110149456 | 0.088982054 | 0.443672569 | 0.413095074 | 0.311363035 |
| Sptbn1 | -0.093339464 | 0.17195806 | 0.575294174 | 0.769858992 | 0.889427259 |
| Sptbn2 | -0.030492528 | 0.571061475 | 0.861225331 | 0.201726291 | 0.216798331 |
| Sptbn4 | -0.002310392 | 0.965760325 | 0.992256509 | 0.225156578 | 0.223381116 |
| Sptlc1 | 0.033495578 | 0.495765843 | 0.829748503 | 0.179209361 | 0.14474591 |
| Sptlc2 | 0.031273964 | 0.533371082 | 0.84612966 | 0.181790019 | 0.150246891 |
| Sptssa | -0.016870354 | 0.796297892 | 0.939955539 | 0.466875442 | 0.48913124 |
| Spty2d1 | 0.031559922 | 0.637636454 | 0.891210491 | 0.424203801 | 0.402087141 |
| Sqle | 0.033617331 | 0.625744282 | 0.88683922 | 0.659487588 | 0.604105906 |
| Sqor | -0.018914235 | 0.749759814 | 0.925706567 | 0.309260974 | 0.338151583 |
| Sqstm1 | -0.262002155 | 1.20E-05 | 0.001459181 | 1.66323754 | 2.196154095 |
| Sra1 | 0.013298068 | 0.838591753 | 0.954911647 | 1.046685176 | 1.054552362 |
| Srbd1 | 0.019539801 | 0.586461379 | 0.86902362 | 0.082380611 | 0.062889118 |
| Src | 0.010422627 | 0.797483048 | 0.940690722 | 0.111258315 | 0.091441969 |
| Srcap | 0.009357109 | 0.790899783 | 0.938824685 | 0.082743715 | 0.073287058 |
| Srd5a3 | 0.021180814 | 0.628028751 | 0.887864718 | 0.12675115 | 0.106925376 |
| Srebf1 | 0.010623923 | 0.827086941 | 0.952715064 | 0.171665256 | 0.162304109 |
| Srebf2 | 0.057979118 | 0.358677853 | 0.747363423 | 0.341122179 | 0.275272453 |
| Srek1 | 0.053605442 | 0.410356408 | 0.780118483 | 0.540197991 | 0.456128157 |
| Srek1ip1 | 0.046363153 | 0.467315433 | 0.813188482 | 0.416031768 | 0.3606944 |
| Srf | 0.016123221 | 0.587053628 | 0.869181609 | 0.056160738 | 0.045179805 |
| Srfbp1 | 0.047434603 | 0.253582902 | 0.663146569 | 0.118899277 | 0.086993249 |
| Srgap1 | 0.124675167 | 0.017184685 | 0.209069918 | 0.216179084 | 0.099892815 |
| Sri | 0.207736009 | 0.002465025 | 0.067311494 | 0.650136599 | 0.414220265 |
| Srm | 0.024017821 | 0.698973094 | 0.911319029 | 0.309736733 | 0.291789253 |
| Srp14 | -0.029834656 | 0.608564003 | 0.878855197 | 1.262401325 | 1.304975679 |
| Srp19 | -0.030555832 | 0.647918368 | 0.895376633 | 0.539101003 | 0.57642328 |
| Srp54a | 0.076397797 | 0.205845508 | 0.616737719 | 0.318995029 | 0.24238241 |
| Srp68 | 0.056194427 | 0.398708779 | 0.774321615 | 0.485285802 | 0.440423106 |
| Srp72 | -0.132079791 | 0.046567888 | 0.338417216 | 0.733112101 | 0.881314773 |
| Srp9 | -0.033925102 | 0.55290308 | 0.852623515 | 1.400709514 | 1.446903009 |
| Srpk1 | -0.027344691 | 0.66701087 | 0.901533545 | 0.41731311 | 0.438957593 |
| Srpk2 | -0.006584521 | 0.923380276 | 0.980859278 | 0.950675777 | 0.998782318 |
| Srpr | -0.067940974 | 0.251508273 | 0.663127858 | 1.348102271 | 1.445510201 |
| Srprb | 0.003654172 | 0.949386178 | 0.987767085 | 0.248011941 | 0.253660549 |
| Srr | -0.037194778 | 0.313440348 | 0.714906079 | 0.084474933 | 0.115673258 |
| Srrd | -0.004254922 | 0.920523261 | 0.980436936 | 0.124111698 | 0.131704588 |
| Srrm1 | 0.060890337 | 0.365754006 | 0.752886169 | 1.104039665 | 1.020303988 |
| Srrm2 | -0.135719247 | 0.013737096 | 0.189915708 | 1.684639308 | 1.919445055 |
| Srrm3 | 0.001476085 | 0.976159851 | 0.994468977 | 0.171331155 | 0.1685224 |
| Srrm4 | -0.033105101 | 0.254642088 | 0.66334523 | 0.047180688 | 0.072207433 |
| Srrt | -0.059175037 | 0.21862051 | 0.632622127 | 0.146526082 | 0.196969948 |
| Srsf1 | -0.010752172 | 0.859797812 | 0.959220425 | 0.319804937 | 0.334856903 |
| Srsf10 | -0.099217424 | 0.155010087 | 0.552106544 | 0.56277323 | 0.667152531 |
| Srsf11 | -0.005633696 | 0.934490856 | 0.983960188 | 0.935784233 | 0.95630906 |

| | | | | | |
|---|---|---|---|---|---|
| Srsf2 | -0.090142467 | 0.191523981 | 0.602043906 | 0.522312522 | 0.627827858 |
| Srsf3 | -0.38509184 | 4.47E-08 | 1.72E-05 | 0.476360506 | 0.886605571 |
| Srsf4 | 0.065865698 | 0.268083388 | 0.676097903 | 0.302011737 | 0.234479986 |
| Srsf5 | -0.210365615 | 0.001962305 | 0.0606747 | 0.869927924 | 1.120238799 |
| Srsf6 | -0.317437541 | 7.45E-07 | 0.000165426 | 0.342893749 | 0.652175401 |
| Srsf7 | -0.108936289 | 0.122803822 | 0.499468242 | 0.748088517 | 0.867288309 |
| Srsf9 | 0.127763187 | 0.051764956 | 0.355933715 | 0.677162626 | 0.554070708 |
| Srxn1 | -0.257603059 | 0.00019237 | 0.012702738 | 0.426318693 | 0.733586535 |
| Ss18 | -0.046117076 | 0.263040508 | 0.669775695 | 0.109033039 | 0.154370111 |
| Ss18l2 | 0.053231814 | 0.425527821 | 0.790344696 | 1.249040398 | 1.152951429 |
| Ssb | -0.032580158 | 0.600902608 | 0.876370606 | 1.091984761 | 1.132185043 |
| Ssbp1 | -0.083446348 | 0.201891363 | 0.612724107 | 0.533363618 | 0.60271098 |
| Ssbp2 | -0.05080819 | 0.299314781 | 0.704513802 | 0.162389174 | 0.199762436 |
| Ssbp3 | 0.002984574 | 0.964159476 | 0.991970325 | 0.504280111 | 0.497021554 |
| Ssbp4 | -0.154308913 | 0.017462016 | 0.210563469 | 0.392445096 | 0.5514179 |
| Ssh1 | 0.026203352 | 0.631585595 | 0.88887544 | 0.225230948 | 0.194789977 |
| Ssh2 | -0.038745982 | 0.383248802 | 0.763886718 | 0.133300966 | 0.166453109 |
| Ssna1 | -0.055472439 | 0.371564533 | 0.75760494 | 0.378260722 | 0.441759349 |
| Sspn | -0.000841994 | 0.976461101 | 0.994468977 | 0.050586383 | 0.055709659 |
| Ssr1 | 0.043966944 | 0.509264704 | 0.834904701 | 0.73183662 | 0.690931658 |
| Ssr2 | -0.063485245 | 0.334577677 | 0.730169996 | 1.011003335 | 1.08258548 |
| Ssr3 | -0.032555543 | 0.632788421 | 0.889288348 | 0.802094866 | 0.823006004 |
| Ssr4 | 0.034159907 | 0.482544214 | 0.822151804 | 2.521479544 | 2.304942491 |
| Ssrp1 | -0.15006267 | 0.025693368 | 0.255331022 | 0.454833961 | 0.622548699 |
| Sst | 0.236605006 | 0.009057479 | 0.147945584 | 0.730614732 | 0.321935167 |
| Sstr3 | 0.071097375 | 0.246279381 | 0.655935044 | 0.332332281 | 0.264124218 |
| Ssu72 | 0.134434618 | 0.049103527 | 0.347612176 | 0.690178948 | 0.558226354 |
| Ssx2ip | 0.05500645 | 0.376751384 | 0.761269164 | 0.375137497 | 0.309585441 |
| St13 | 0.185987531 | 0.004257678 | 0.094128449 | 1.398542523 | 1.093219284 |
| St14 | -0.052155021 | 0.304986928 | 0.709410087 | 0.16645278 | 0.209716114 |
| St18 | 0.045283383 | 0.508911333 | 0.8346818 | 0.699963532 | 0.645642504 |
| St3gal1 | 0.050852538 | 0.193105268 | 0.60325934 | 0.100586792 | 0.055078853 |
| St3gal3 | 0.021394095 | 0.623596922 | 0.886173341 | 0.132043331 | 0.11510022 |
| St3gal5 | -0.042193294 | 0.522424006 | 0.840670611 | 0.420471452 | 0.455374767 |
| St3gal6 | -0.003662208 | 0.920211309 | 0.980436936 | 0.083567644 | 0.084633251 |
| St6gal1 | -0.083687914 | 0.020176228 | 0.227038569 | 0.076443172 | 0.142788391 |
| St6galnac4 | 0.017363081 | 0.637982829 | 0.89125563 | 0.091999696 | 0.078067276 |
| St6galnac6 | 0.042910243 | 0.224371842 | 0.637067124 | 0.086261949 | 0.049065624 |
| St7 | -0.017031039 | 0.549704099 | 0.852455047 | 0.045991581 | 0.06223237 |
| St7l | 0.077006322 | 0.137883598 | 0.52303011 | 0.196080966 | 0.12563653 |
| St8sia1 | 0.026810571 | 0.44276592 | 0.801368502 | 0.079375471 | 0.050131107 |
| St8sia3 | 0.04074825 | 0.361190754 | 0.749642598 | 0.136360452 | 0.1002309 |
| St8sia5 | 0.044537679 | 0.425525666 | 0.790344696 | 0.227973614 | 0.185382146 |
| Stag1 | 0.049901886 | 0.437186721 | 0.798116245 | 0.349097781 | 0.279287121 |
| Stag2 | -0.024345851 | 0.707805559 | 0.913729182 | 0.444891249 | 0.464476269 |
| Stam | 0.002091251 | 0.963764912 | 0.991970325 | 0.146459678 | 0.149756165 |
| Stam2 | 0.111169156 | 0.084317649 | 0.432888102 | 0.44163031 | 0.336356894 |
| Stambp | 0.094514958 | 0.114944705 | 0.486793251 | 0.287125179 | 0.188993159 |
| Stambpl1 | -0.019821833 | 0.504152782 | 0.832249514 | 0.051713172 | 0.067770171 |
| Star | -0.061839209 | 0.081007089 | 0.426199169 | 0.062110146 | 0.115652403 |
| Stard10 | 0.154480932 | 0.022877122 | 0.239281409 | 0.860229584 | 0.678908872 |
| Stard13 | -0.051371591 | 0.215136009 | 0.629611185 | 0.108909213 | 0.149712134 |
| Stard3 | -0.00974483 | 0.775903225 | 0.933654329 | 0.076474496 | 0.085717971 |
| Stard3nl | 0.061403897 | 0.334959108 | 0.73038987 | 0.396679118 | 0.346125562 |
| Stard4 | 0.066573996 | 0.056971395 | 0.368959821 | 0.088114197 | 0.032746929 |
| Stard5 | -0.102807865 | 0.041451635 | 0.321059507 | 0.163425631 | 0.254415845 |
| Stard7 | 0.014398652 | 0.820175976 | 0.950589638 | 0.404655762 | 0.398047556 |
| Stat1 | 0.095260622 | 0.058382836 | 0.372056206 | 0.182637273 | 0.094177927 |
| Stat2 | -0.071110955 | 0.063845866 | 0.38543852 | 0.081131947 | 0.139498295 |
| Stat3 | 0.085704189 | 0.202125089 | 0.612724107 | 0.963899185 | 0.835378172 |
| Stat5a | 0.037268018 | 0.268307147 | 0.676166273 | 0.074758715 | 0.042488518 |
| Stat5b | 0.100138194 | 0.077399604 | 0.418285047 | 0.266958178 | 0.176905772 |
| Stat6 | 0.024282541 | 0.477843756 | 0.818590256 | 0.078036166 | 0.060283173 |
| Stau1 | 0.042639734 | 0.435468059 | 0.796975049 | 0.249319802 | 0.194184517 |
| Stau2 | -0.101030492 | 0.078009732 | 0.418637956 | 0.247532572 | 0.352789074 |

| | | | | | |
|---|---|---|---|---|---|
| Stbd1 | -0.019826344 | 0.626363824 | 0.886996623 | 0.106040781 | 0.11807993 |
| Steap1 | -0.217033069 | 1.81E-08 | 8.40E-06 | 0.064604992 | 0.261532742 |
| Steap2 | -0.02106349 | 0.701401393 | 0.912152193 | 0.232136738 | 0.232048488 |
| Stim1 | 0.015516416 | 0.822927274 | 0.950930859 | 0.545071855 | 0.510720017 |
| Stim2 | 0.010358402 | 0.853825992 | 0.957461155 | 0.246567877 | 0.234542737 |
| Stimate | -0.022411172 | 0.484135895 | 0.822768706 | 0.064533739 | 0.083855041 |
| Stip1 | 0.192143954 | 0.005864111 | 0.113654038 | 0.494090016 | 0.293950491 |
| Stk11 | 0.087728361 | 0.188509249 | 0.59667844 | 0.557180199 | 0.471750328 |
| Stk11ip | 0.002615045 | 0.9278403 | 0.982252198 | 0.052234061 | 0.047194366 |
| Stk16 | -0.03451089 | 0.569618694 | 0.860834533 | 0.362263432 | 0.392858479 |
| Stk19 | -0.111072714 | 0.031958968 | 0.283462667 | 0.185045924 | 0.281842583 |
| Stk24 | 0.057268773 | 0.252150905 | 0.663127858 | 0.180473651 | 0.14404484 |
| Stk25 | 0.008692636 | 0.881476499 | 0.968081674 | 0.290670942 | 0.289797188 |
| Stk26 | -0.045148812 | 0.462898701 | 0.811092454 | 0.325621699 | 0.363138441 |
| Stk3 | 0.028959597 | 0.425030965 | 0.790344696 | 0.08779699 | 0.058325932 |
| Stk32a | -0.208179786 | 0.002409403 | 0.066938901 | 0.593759246 | 0.841012123 |
| Stk35 | 0.02370454 | 0.705351922 | 0.912844672 | 0.287108146 | 0.265509023 |
| Stk38 | -0.025565167 | 0.683261491 | 0.909173483 | 0.355556234 | 0.370386991 |
| Stk38l | 0.100052643 | 0.051087254 | 0.355933715 | 0.193652969 | 0.100356228 |
| Stk39 | -0.013454172 | 0.775534444 | 0.933636662 | 0.159304461 | 0.162780609 |
| Stk4 | -0.006249187 | 0.906457558 | 0.976716534 | 0.217659532 | 0.220317976 |
| Stk40 | -0.054588665 | 0.398507589 | 0.774321615 | 0.40933379 | 0.43564766 |
| Stmn3 | -0.105713221 | 0.075580411 | 0.414321131 | 0.256796603 | 0.360199424 |
| Stoml1 | -0.077451598 | 0.009613802 | 0.152198314 | 0.047624314 | 0.117682364 |
| Stoml2 | 0.049653945 | 0.386629275 | 0.766593284 | 0.270616973 | 0.225893685 |
| Stox1 | 0.054591402 | 0.258883808 | 0.665948842 | 0.17619833 | 0.133546349 |
| Stox2 | 0.010346054 | 0.843617534 | 0.955471586 | 0.201013217 | 0.192190287 |
| Strada | 0.017988816 | 0.577653333 | 0.864673658 | 0.067069125 | 0.051827398 |
| Stradb | 0.090894657 | 0.009048031 | 0.147945584 | 0.089162859 | 0.008502086 |
| Strap | -0.039902444 | 0.561399575 | 0.857134531 | 0.668646038 | 0.711295044 |
| Strbp | -0.006801259 | 0.920519479 | 0.980436936 | 0.580120443 | 0.596173182 |
| Strip1 | -0.033841284 | 0.397978793 | 0.774311756 | 0.10289438 | 0.13691555 |
| Strn | -0.010855275 | 0.811364546 | 0.948397417 | 0.139960331 | 0.149639463 |
| Strn3 | 0.005326684 | 0.936476 | 0.98453816 | 0.568423449 | 0.552885848 |
| Strn4 | 0.038337809 | 0.44732086 | 0.804616393 | 0.186674562 | 0.160260601 |
| Stt3a | -0.0487807 | 0.482157354 | 0.821803219 | 0.599980866 | 0.66798706 |
| Stt3b | -0.130006463 | 0.026020424 | 0.256801368 | 0.296312242 | 0.418579395 |
| Stub1 | -0.117305593 | 0.073915584 | 0.412240835 | 0.868391027 | 1.026428615 |
| Stx12 | 0.107436481 | 0.092665604 | 0.450701165 | 0.428411125 | 0.319302325 |
| Stx16 | -0.096445555 | 0.110243722 | 0.479634259 | 0.294290646 | 0.38074598 |
| Stx17 | 0.007452236 | 0.876194956 | 0.965459301 | 0.158983462 | 0.154995416 |
| Stx18 | -0.074731365 | 0.188950697 | 0.596946223 | 0.252521263 | 0.332954999 |
| Stx2 | 0.032027243 | 0.286961486 | 0.69336361 | 0.058760469 | 0.027050611 |
| Stx3 | 0.015100609 | 0.813711564 | 0.948397417 | 0.400730428 | 0.39037097 |
| Stx4a | 0.025897761 | 0.658007722 | 0.898224427 | 0.271820698 | 0.239733399 |
| Stx5a | 0.005893164 | 0.923093505 | 0.980859278 | 0.33935223 | 0.32735886 |
| Stx6 | -0.01703985 | 0.767847778 | 0.930221865 | 0.262016071 | 0.288716622 |
| Stx7 | 0.07595111 | 0.257136266 | 0.665256942 | 0.797798009 | 0.714316452 |
| Stx8 | -0.030241539 | 0.609948595 | 0.879352778 | 0.301605652 | 0.317116667 |
| Stxbp1 | 0.02715827 | 0.688056929 | 0.909796728 | 0.748880961 | 0.709675144 |
| Stxbp2 | 0.008296116 | 0.866579015 | 0.962125879 | 0.169215642 | 0.160671477 |
| Stxbp3 | 0.001498102 | 0.97945899 | 0.995540893 | 0.259518773 | 0.252663305 |
| Stxbp4 | -0.013437247 | 0.655512256 | 0.897825804 | 0.057193083 | 0.063682578 |
| Stxbp5 | 0.00230273 | 0.966794751 | 0.992504747 | 0.244594929 | 0.251009017 |
| Stxbp5l | -0.037998514 | 0.247297278 | 0.657515967 | 0.055591992 | 0.081603536 |
| Stxbp6 | 0.039595715 | 0.45528869 | 0.808087105 | 0.216847589 | 0.189836795 |
| Styx | -0.029612644 | 0.256681436 | 0.665256942 | 0.038673805 | 0.063820836 |
| Sub1 | -0.001367316 | 0.98278489 | 0.995982628 | 1.2489753 | 1.247012883 |
| Sucla2 | 0.108927059 | 0.08378063 | 0.431794059 | 0.378330571 | 0.272717087 |
| Suclg1 | 0.023055213 | 0.740670125 | 0.922901128 | 0.698499157 | 0.675413069 |
| Suclg2 | -0.040349215 | 0.455550019 | 0.808152996 | 0.205525537 | 0.233985685 |
| Suco | 0.036717679 | 0.584457265 | 0.867267529 | 0.669097895 | 0.647589247 |
| Suds3 | -0.080666511 | 0.214507772 | 0.629611185 | 0.483514733 | 0.54926093 |
| Sugp1 | -0.225198682 | 3.39E-05 | 0.003504394 | 0.192025594 | 0.399654261 |
| Sugp2 | -0.058657447 | 0.040919323 | 0.319308264 | 0.045070966 | 0.092922216 |

| | | | | | |
|---|---|---|---|---|---|
| Sugt1 | 0.001566452 | 0.980663486 | 0.995540893 | 0.4206772 | 0.416295666 |
| Sult1c2 | -0.060059241 | 0.037253064 | 0.304267772 | 0.045559715 | 0.097138126 |
| Sult1d1 | -0.063343167 | 0.254713653 | 0.66334523 | 0.197636466 | 0.244412182 |
| Sult2b1 | -0.003945519 | 0.910535301 | 0.978693812 | 0.080091451 | 0.085018229 |
| Sult4a1 | -0.011260562 | 0.692884945 | 0.909796728 | 0.048310702 | 0.057361675 |
| Sumf1 | -0.003661007 | 0.936865992 | 0.98453816 | 0.144541374 | 0.145284989 |
| Sumf2 | 0.040348739 | 0.166396961 | 0.568127031 | 0.058318181 | 0.025230736 |
| Sumo1 | 0.119061189 | 0.067986837 | 0.396044041 | 1.103963748 | 0.97002424 |
| Sumo2 | 0.085433177 | 0.123016784 | 0.499567826 | 1.507081432 | 1.392338673 |
| Sumo3 | 0.015553845 | 0.819835201 | 0.950575887 | 0.716339341 | 0.683124375 |
| Sun1 | 0.030033569 | 0.611206909 | 0.87989943 | 0.287828752 | 0.273670596 |
| Sun2 | -0.060093928 | 0.074713414 | 0.41344108 | 0.068736411 | 0.121426089 |
| Suox | -0.039356582 | 0.464263013 | 0.811633402 | 0.210752572 | 0.241552309 |
| Supt16 | 0.062249194 | 0.363636633 | 0.751765374 | 0.613265043 | 0.543041595 |
| Supt20 | 0.027568455 | 0.578715326 | 0.865142348 | 0.178550231 | 0.153675373 |
| Supt3 | -0.00811618 | 0.797737759 | 0.940696966 | 0.062406623 | 0.071486264 |
| Supt4a | 0.015418477 | 0.811725984 | 0.948397417 | 0.374878655 | 0.364985116 |
| Supt5 | 0.081273566 | 0.218384046 | 0.632618221 | 0.491901861 | 0.412212965 |
| Supt6 | -0.050874644 | 0.462401967 | 0.810535897 | 0.591247681 | 0.640621109 |
| Supt7l | -0.014974799 | 0.587546912 | 0.869281892 | 0.044906977 | 0.057421595 |
| Supv3l1 | 0.019719399 | 0.571682881 | 0.861404566 | 0.079790615 | 0.063896356 |
| Surf1 | -0.004139416 | 0.946353044 | 0.987084873 | 0.328012976 | 0.337534294 |
| Surf2 | 0.066918568 | 0.219151852 | 0.632622127 | 0.236959613 | 0.183795055 |
| Surf4 | -0.066740364 | 0.321464364 | 0.719227395 | 0.906047293 | 0.98969842 |
| Surf6 | 0.026376163 | 0.458499446 | 0.809370658 | 0.083568875 | 0.062834106 |
| Susd4 | 0.026864844 | 0.341809783 | 0.735000423 | 0.052616105 | 0.025972995 |
| Susd6 | 0.076829936 | 0.226472959 | 0.638587289 | 0.355643769 | 0.27437103 |
| Suz12 | -0.00847969 | 0.898504193 | 0.973371461 | 0.51020562 | 0.524484681 |
| Sv2a | 0.10809102 | 0.100262144 | 0.465319642 | 0.422734219 | 0.300610968 |
| Sv2b | 0.043593236 | 0.143699452 | 0.533004951 | 0.061143528 | 0.020616715 |
| Svbp | -0.040957945 | 0.297688102 | 0.702656882 | 0.099363878 | 0.131962285 |
| Svil | 0.080414701 | 0.115190431 | 0.487260227 | 0.191601532 | 0.113606381 |
| Svip | 0.077767655 | 0.256198037 | 0.664906869 | 0.586622944 | 0.480930167 |
| Swi5 | -0.168977048 | 0.007687356 | 0.134995868 | 1.198383826 | 1.419364593 |
| Swt1 | 0.105789706 | 0.099233765 | 0.463315751 | 0.375903828 | 0.263670545 |
| Syap1 | -0.078140363 | 0.201173653 | 0.611914909 | 0.327866656 | 0.393782772 |
| Sybu | 0.044019383 | 0.240048663 | 0.650744882 | 0.094505718 | 0.062438348 |
| Syce2 | -0.035835175 | 0.402896008 | 0.77525778 | 0.11975937 | 0.162031472 |
| Syde2 | -0.049085396 | 0.016031553 | 0.203940048 | 0.014618491 | 0.058977598 |
| Syf2 | -0.100220469 | 0.142007957 | 0.53012127 | 0.608067903 | 0.714127233 |
| Sympk | 0.052331387 | 0.346943428 | 0.738383053 | 0.240790889 | 0.199728143 |
| Syn1 | 0.114402587 | 0.051299212 | 0.355933715 | 0.30455116 | 0.191479023 |
| Syn2 | 0.139456931 | 0.020286616 | 0.227608207 | 0.338765994 | 0.209567406 |
| Syncrip | -0.058107239 | 0.375274047 | 0.760147021 | 0.438642032 | 0.487327676 |
| Syndig1l | 0.179540435 | 0.008395001 | 0.142539393 | 0.608652408 | 0.419031868 |
| Syne1 | -0.169279431 | 0.001376067 | 0.047626161 | 0.178942043 | 0.349176196 |
| Syne2 | 0.096748094 | 0.077577052 | 0.418285047 | 0.227350168 | 0.146750369 |
| Syne4 | 0.067258051 | 0.28645496 | 0.693329451 | 0.37340837 | 0.297790629 |
| Syngr1 | 0.012722755 | 0.802121924 | 0.94305936 | 0.187807963 | 0.168480961 |
| Syngr2 | -0.018918236 | 0.740550405 | 0.922901128 | 0.275500119 | 0.297828307 |
| Synj1 | 0.118806513 | 0.050552661 | 0.354499194 | 0.355223327 | 0.236756106 |
| Synj2bp | -0.002319064 | 0.969934975 | 0.993892166 | 0.343901549 | 0.348176142 |
| Synpo | 0.002621838 | 0.942329434 | 0.986270063 | 0.081997549 | 0.077531283 |
| Synpr | 0.01057403 | 0.798410671 | 0.940958114 | 0.117258072 | 0.112736171 |
| Synrg | 0.10886954 | 0.108627735 | 0.477972625 | 0.81694137 | 0.694061306 |
| Syp | -0.139865038 | 0.039577167 | 0.313524882 | 0.708293451 | 0.879548917 |
| Sypl | 0.074365553 | 0.234005366 | 0.644459522 | 0.346497211 | 0.281457151 |
| Sys1 | 0.129675548 | 0.029926155 | 0.274935828 | 1.173666803 | 1.008610439 |
| Syt11 | -0.050038747 | 0.444322375 | 0.801932586 | 0.463222064 | 0.518232373 |
| Syt13 | -0.198636029 | 0.002057951 | 0.062198787 | 1.305445954 | 1.589208273 |
| Syt14 | 0.052226448 | 0.407249457 | 0.777888922 | 0.366965863 | 0.320177184 |
| Syt4 | -0.052261572 | 0.421487838 | 0.787357117 | 1.539723317 | 1.619003547 |
| Syt5 | -0.042146498 | 0.448187187 | 0.804951652 | 0.225863412 | 0.256246472 |
| Syt7 | 0.025003781 | 0.718295184 | 0.91797181 | 1.121741519 | 1.055979088 |
| Syt9 | -0.016016116 | 0.609445235 | 0.878855197 | 0.060010998 | 0.071295054 |

| | | | | | |
|---|---|---|---|---|---|
| Sytl1 | -0.05446514 | 0.155717664 | 0.552356396 | 0.088695766 | 0.133982051 |
| Sytl4 | 0.341004306 | 3.17E-06 | 0.000530034 | 1.064696156 | 0.643888559 |
| Sytl5 | -0.020415012 | 0.585422907 | 0.868186511 | 0.084145377 | 0.105965348 |
| Syvn1 | -0.07802506 | 0.089152725 | 0.443953048 | 0.132520359 | 0.198932533 |
| Szrd1 | 0.005324883 | 0.924623204 | 0.981375217 | 0.24996389 | 0.232721072 |
| T2 | 0.010963663 | 0.76647886 | 0.929011994 | 0.089869727 | 0.077553989 |
| Tab1 | 0.071963794 | 0.054588651 | 0.363639613 | 0.0981677 | 0.042583633 |
| Tab2 | 0.117404536 | 0.080463663 | 0.425115437 | 0.502258035 | 0.367583296 |
| Tab3 | -0.002602016 | 0.93457283 | 0.983960188 | 0.064018678 | 0.066726728 |
| Tacc1 | -0.134649699 | 0.052491503 | 0.357919301 | 0.631176854 | 0.769134351 |
| Tacc2 | 0.056175108 | 0.377114403 | 0.761412185 | 0.344206035 | 0.296773438 |
| Taco1 | -0.00167587 | 0.955390278 | 0.989278596 | 0.056111725 | 0.064470643 |
| Taco1os | -0.039030182 | 0.063201651 | 0.383440442 | 0.019452942 | 0.05208755 |
| Tada1 | -0.006977109 | 0.878638447 | 0.966984001 | 0.137587915 | 0.145381788 |
| Tada2a | -0.03623236 | 0.346926104 | 0.738383053 | 0.094074714 | 0.129955118 |
| Tada2b | 0.013417019 | 0.782181282 | 0.936026395 | 0.163092392 | 0.143636176 |
| Tada3 | 0.056964196 | 0.252110728 | 0.663127858 | 0.179385214 | 0.126516942 |
| Taf1 | -0.005071412 | 0.937796402 | 0.985021197 | 0.433128772 | 0.436543545 |
| Taf10 | -0.019242294 | 0.771313011 | 0.932251553 | 0.912170134 | 0.927599637 |
| Taf11 | 0.030166147 | 0.623115406 | 0.885857477 | 0.355270578 | 0.332955162 |
| Taf12 | -0.015301637 | 0.760380675 | 0.927895542 | 0.181168299 | 0.205039032 |
| Taf13 | 0.052341839 | 0.20363143 | 0.614950615 | 0.117696056 | 0.074336076 |
| Taf15 | -0.120340615 | 0.091375257 | 0.4462832 | 0.608568319 | 0.747551289 |
| Taf1a | 0.021811579 | 0.515885613 | 0.837427561 | 0.073476487 | 0.061521777 |
| Taf1b | -0.000556017 | 0.987924902 | 0.997357506 | 0.08711827 | 0.089748909 |
| Taf1d | 0.020370651 | 0.721832232 | 0.919475778 | 0.273621183 | 0.267775271 |
| Taf2 | -0.012906715 | 0.765025212 | 0.928757959 | 0.125971827 | 0.138862536 |
| Taf3 | 0.086678086 | 0.149895882 | 0.544321624 | 0.325197601 | 0.237848636 |
| Taf4 | -0.006303679 | 0.845469365 | 0.955950886 | 0.066303621 | 0.070138539 |
| Taf5 | 0.052575761 | 0.132563423 | 0.513250014 | 0.080280221 | 0.033484169 |
| Taf5l | 0.023587226 | 0.592341894 | 0.872005806 | 0.129995481 | 0.111026127 |
| Taf6 | 0.082101152 | 0.054381916 | 0.36337543 | 0.133475951 | 0.061274795 |
| Taf7 | -0.058563143 | 0.35432343 | 0.744784139 | 0.300171055 | 0.370542027 |
| Taf8 | 0.030044768 | 0.298124406 | 0.703072054 | 0.05556838 | 0.028451647 |
| Taf9b | -0.060869708 | 0.137751596 | 0.522884425 | 0.107611173 | 0.161920755 |
| Tafa5 | 0.117584564 | 0.035061679 | 0.294914843 | 0.259048994 | 0.155705642 |
| Tagln2 | -0.106555947 | 0.108997809 | 0.477972625 | 1.150679363 | 1.305960386 |
| Taldo1 | -0.170531864 | 0.00926514 | 0.149508372 | 0.531599338 | 0.733280719 |
| Tamalin | 0.052362844 | 0.11259486 | 0.483221516 | 0.074934652 | 0.036511622 |
| Tanc1 | -0.004416875 | 0.920269172 | 0.980436936 | 0.123292521 | 0.12877844 |
| Tanc2 | -0.053370983 | 0.349080396 | 0.740431094 | 0.250672395 | 0.30644583 |
| Tango2 | 0.023045231 | 0.560426118 | 0.856904336 | 0.101945396 | 0.080850865 |
| Tank | -0.024212159 | 0.565203073 | 0.858087314 | 0.113432035 | 0.129442416 |
| Taok1 | -0.046303573 | 0.492060011 | 0.827270742 | 0.549801105 | 0.587063218 |
| Taok2 | 0.054395472 | 0.363264075 | 0.751765374 | 0.329654869 | 0.262934658 |
| Taok3 | -0.073458784 | 0.277876591 | 0.684996416 | 0.589504217 | 0.671717449 |
| Tap1 | 0.048052445 | 0.216789406 | 0.631259121 | 0.107254639 | 0.063537202 |
| Tap2 | 0.034943352 | 0.516714084 | 0.837438925 | 0.20705698 | 0.183043872 |
| Tapbp | -0.233724937 | 0.00074671 | 0.032181025 | 0.698833524 | 0.94672187 |
| Tapbpl | 0.037758717 | 0.527704396 | 0.843341172 | 0.284821844 | 0.256040089 |
| Tapt1 | 0.078385079 | 0.094689214 | 0.454115053 | 0.157319604 | 0.089098034 |
| Tarbp2 | -0.024505726 | 0.460594007 | 0.810111566 | 0.071456282 | 0.095651179 |
| Tardbp | -0.04988278 | 0.43422637 | 0.796332921 | 0.455119399 | 0.519464208 |
| Tars | -0.008966399 | 0.895963785 | 0.97233366 | 0.562267489 | 0.577010706 |
| Tars2 | 0.037058563 | 0.253460428 | 0.663146569 | 0.068577668 | 0.034886752 |
| Tarsl2 | 0.038374655 | 0.349514522 | 0.740550478 | 0.114994957 | 0.081980541 |
| Tasor | 0.001764688 | 0.979055844 | 0.995540893 | 0.476113598 | 0.493655745 |
| Tasor2 | -0.055014121 | 0.378232487 | 0.761977617 | 0.357792062 | 0.403015753 |
| Tasp1 | -0.026200996 | 0.491023559 | 0.827146956 | 0.089927378 | 0.118257872 |
| Tat | 0.032460092 | 0.443953783 | 0.801865057 | 0.12059923 | 0.091562782 |
| Tatdn1 | 0.010832569 | 0.822726067 | 0.950930859 | 0.162938056 | 0.158653745 |
| Tatdn2 | 0.05850146 | 0.182062205 | 0.589470435 | 0.135295653 | 0.091728725 |
| Tatdn3 | -0.045128106 | 0.10912397 | 0.477972625 | 0.044719057 | 0.0783653 |
| Tax1bp1 | -0.037840728 | 0.546859162 | 0.851446616 | 1.185236059 | 1.249483995 |
| Tax1bp3 | 0.017790964 | 0.599818136 | 0.875897759 | 0.075570326 | 0.059953766 |

| | | | | | |
|---|---|---|---|---|---|
| Taz | -0.14270879 | 0.008213024 | 0.140469359 | 0.204025255 | 0.323904477 |
| Tbc1d1 | 0.15382401 | 0.021985648 | 0.23623533 | 0.481619817 | 0.321492241 |
| Tbc1d10a | 0.00122554 | 0.975671147 | 0.994455356 | 0.108548508 | 0.110130543 |
| Tbc1d10b | 0.01848956 | 0.724374863 | 0.919830015 | 0.205065153 | 0.195174727 |
| Tbc1d12 | 0.072433933 | 0.115813338 | 0.48800104 | 0.151230016 | 0.092400203 |
| Tbc1d13 | -0.048835636 | 0.122021202 | 0.49725002 | 0.058401443 | 0.100911809 |
| Tbc1d14 | 0.026587954 | 0.565299744 | 0.858087314 | 0.151419616 | 0.12188984 |
| Tbc1d15 | -0.128124259 | 0.061697422 | 0.380250374 | 0.552415413 | 0.712385033 |
| Tbc1d16 | -0.001691928 | 0.978086896 | 0.995031396 | 0.324594226 | 0.319311549 |
| Tbc1d17 | -0.003383318 | 0.948499388 | 0.987651726 | 0.194017677 | 0.212032613 |
| Tbc1d19 | -0.046975299 | 0.306044049 | 0.709680069 | 0.137769075 | 0.171778263 |
| Tbc1d20 | 0.178331109 | 0.005776301 | 0.113235802 | 0.394035824 | 0.215954926 |
| Tbc1d22a | 0.010496108 | 0.75798543 | 0.927237784 | 0.076388357 | 0.06704262 |
| Tbc1d22b | 0.066003774 | 0.178222498 | 0.584581441 | 0.17626021 | 0.113042645 |
| Tbc1d23 | -0.022436779 | 0.680070538 | 0.907783945 | 0.21875434 | 0.249902982 |
| Tbc1d24 | -0.002987406 | 0.942889036 | 0.986735195 | 0.116136881 | 0.119856548 |
| Tbc1d30 | -0.080837846 | 0.032040095 | 0.283836215 | 0.085107485 | 0.151191468 |
| Tbc1d31 | -0.046650335 | 0.198520258 | 0.609389286 | 0.076504222 | 0.128123071 |
| Tbc1d32 | 0.008160009 | 0.814174624 | 0.948563053 | 0.076496769 | 0.073248107 |
| Tbc1d4 | 0.062883995 | 0.028079146 | 0.266632345 | 0.055827926 | 0 |
| Tbc1d5 | -0.04183019 | 0.30284305 | 0.708026508 | 0.104773351 | 0.142729051 |
| Tbc1d7 | 0.033261007 | 0.355782058 | 0.745326231 | 0.088810867 | 0.063742946 |
| Tbc1d8 | -0.014562955 | 0.737686651 | 0.922843014 | 0.126655058 | 0.131672653 |
| Tbc1d8b | -0.024006135 | 0.620399047 | 0.883937014 | 0.161556807 | 0.181211393 |
| Tbc1d9 | 0.04223762 | 0.389911171 | 0.768386831 | 0.168738804 | 0.134005878 |
| Tbc1d9b | 0.02429514 | 0.661014331 | 0.89875384 | 0.236710212 | 0.219254817 |
| Tbca | -0.026414064 | 0.67940195 | 0.907783945 | 1.049194632 | 1.082006972 |
| Tbcb | -0.132343768 | 0.050374005 | 0.354312467 | 0.575946362 | 0.714070177 |
| Tbcc | 0.111265685 | 0.062120018 | 0.380250374 | 0.292463293 | 0.191410946 |
| Tbccd1 | -0.010314187 | 0.790877638 | 0.938824685 | 0.096439432 | 0.10107885 |
| Tbcd | 0.003097623 | 0.950968304 | 0.988114084 | 0.17631898 | 0.173730959 |
| Tbce | -0.037663827 | 0.473314475 | 0.816838319 | 0.195782992 | 0.226345759 |
| Tbcel | 0.021501965 | 0.517279931 | 0.837694133 | 0.073128819 | 0.054780374 |
| Tbck | 0.05810512 | 0.233918978 | 0.644459522 | 0.173295674 | 0.116508551 |
| Tbk1 | -0.011102808 | 0.828510427 | 0.952874813 | 0.179989464 | 0.188512682 |
| Tbl1x | 0.038050937 | 0.594003093 | 0.872914941 | 0.807258994 | 0.755754076 |
| Tbl1xr1 | -0.039600201 | 0.266903681 | 0.674279448 | 0.075157877 | 0.116439425 |
| Tbl2 | -0.053809896 | 0.226472674 | 0.638587289 | 0.135669421 | 0.184093961 |
| Tbl3 | -0.093603837 | 0.007105919 | 0.128637347 | 0.068761247 | 0.153336153 |
| Tbp | 0.015224288 | 0.691415806 | 0.909796728 | 0.095365187 | 0.084769738 |
| Tbpl1 | -0.028762314 | 0.611139594 | 0.87989943 | 0.239787725 | 0.266279176 |
| Tbrg1 | 0.033964904 | 0.597062951 | 0.874211647 | 1.257935462 | 1.184758459 |
| Tbrg4 | 0.010934071 | 0.702619129 | 0.912196119 | 0.052562891 | 0.049307226 |
| Tcaf1 | -0.03754436 | 0.551567583 | 0.852505562 | 0.437881992 | 0.454309602 |
| Tcaim | -0.006716155 | 0.872877878 | 0.964911254 | 0.117606468 | 0.12767799 |
| Tcea1 | -0.027615143 | 0.69690653 | 0.910970095 | 0.643164651 | 0.701521505 |
| Tceal1 | -0.08167061 | 0.126458904 | 0.505886017 | 0.211515918 | 0.28205997 |
| Tceal3 | -0.224462113 | 0.001033471 | 0.038850011 | 0.522821275 | 0.752765654 |
| Tceal8 | 0.042013261 | 0.536355056 | 0.847520662 | 0.49660984 | 0.456776857 |
| Tceal9 | -0.031211843 | 0.471038185 | 0.814586466 | 2.22271232 | 2.268041499 |
| Tceanc | 0.009715998 | 0.79520752 | 0.939757086 | 0.090172391 | 0.080401185 |
| Tceanc2 | 0.005028286 | 0.898463875 | 0.973371461 | 0.099977247 | 0.090310972 |
| Tcerg1 | 0.04698748 | 0.436946583 | 0.797968132 | 0.325634935 | 0.277899112 |
| Tcf12 | -0.064585403 | 0.264362898 | 0.67104216 | 0.266680013 | 0.325576944 |
| Tcf20 | 0.012697478 | 0.825905528 | 0.952251932 | 0.270919633 | 0.254623453 |
| Tcf24 | -0.03519433 | 0.470592769 | 0.814509333 | 0.153714854 | 0.184825924 |
| Tcf25 | -0.022434823 | 0.694354458 | 0.910059506 | 1.468335518 | 1.477134483 |
| Tcf3 | 0.051837795 | 0.206186113 | 0.617573862 | 0.118668123 | 0.070549995 |
| Tcf4 | 0.049467879 | 0.460789177 | 0.810249429 | 0.501208274 | 0.467741589 |
| Tcf7l1 | -0.046893221 | 0.060259912 | 0.378728077 | 0.029197519 | 0.068810736 |
| Tcf7l2 | -0.115639316 | 0.003299355 | 0.080573298 | 0.088301958 | 0.19401011 |
| Tchp | 0.038837236 | 0.368587538 | 0.755773876 | 0.128812776 | 0.09354228 |
| Tcim | -0.071566497 | 0.044020335 | 0.329446347 | 0.063594294 | 0.137131122 |
| Tcn2 | -0.097614676 | 0.085567932 | 0.435299206 | 0.248658572 | 0.347674613 |
| Tcof1 | -0.033671902 | 0.524241494 | 0.841062594 | 0.195534586 | 0.225864965 |

| | | | | | |
|---|---|---|---|---|---|
| Tcp1 | -0.012555958 | 0.856622497 | 0.958302663 | 0.757386091 | 0.77113549 |
| Tcp11l1 | 0.039112827 | 0.477950863 | 0.818590256 | 0.240238749 | 0.19282719 |
| Tcp11l2 | -0.06209468 | 0.057594662 | 0.370562578 | 0.062010392 | 0.115112962 |
| Tcta | -0.016565676 | 0.795056317 | 0.939757086 | 0.400252713 | 0.400733842 |
| Tctn1 | -0.032192598 | 0.154483244 | 0.551012197 | 0.025108016 | 0.051768059 |
| Tctn2 | -0.031453234 | 0.276342094 | 0.683667283 | 0.049531354 | 0.072901141 |
| Tctn3 | -0.050619989 | 0.077682282 | 0.418285047 | 0.046067931 | 0.089988691 |
| Tdg | 0.059925835 | 0.239365785 | 0.650744882 | 0.185631879 | 0.135399981 |
| Tdp1 | -0.028032172 | 0.325128642 | 0.720833548 | 0.047708494 | 0.070284322 |
| Tdp2 | -0.00825897 | 0.865176723 | 0.961658778 | 0.158625354 | 0.164774801 |
| Tdrd3 | 0.020237624 | 0.55168009 | 0.852505562 | 0.076036251 | 0.060952373 |
| Tdrd7 | 0.032036203 | 0.455952783 | 0.808152996 | 0.130926557 | 0.104333427 |
| Tdrkh | -0.018718478 | 0.634966821 | 0.889733629 | 0.102601965 | 0.12720286 |
| Tdrp | -0.07306058 | 0.006721182 | 0.125391263 | 0.033803848 | 0.101519853 |
| Tead1 | 0.060606429 | 0.35565672 | 0.745326231 | 0.427112404 | 0.37102073 |
| Tec | 0.006999463 | 0.853325804 | 0.957461155 | 0.093068102 | 0.083957714 |
| Tecpr1 | -0.133526451 | 0.002523985 | 0.06828366 | 0.120586985 | 0.238074247 |
| Tecpr2 | -0.001888016 | 0.957226589 | 0.98987052 | 0.080240437 | 0.077598245 |
| Tecr | -0.122806274 | 0.07699361 | 0.418199494 | 0.680967491 | 0.803629429 |
| Tef | -0.106915335 | 0.046701202 | 0.338929835 | 0.187683498 | 0.274486236 |
| Tefm | 0.062119542 | 0.056058545 | 0.367897032 | 0.075727654 | 0.02539718 |
| Telo2 | -0.025855703 | 0.327333782 | 0.722182734 | 0.039258578 | 0.063193389 |
| Ten1 | -0.013037767 | 0.837715164 | 0.954911647 | 0.426737949 | 0.443115037 |
| Tenm2 | -0.031345177 | 0.518714564 | 0.838109801 | 0.156442419 | 0.180970038 |
| Tenm3 | -0.058747045 | 0.153065225 | 0.549602359 | 0.109440962 | 0.165121616 |
| Tenm4 | -0.174310411 | 1.46E-06 | 0.000288034 | 0.059208046 | 0.214316389 |
| Tent2 | 0.034265042 | 0.422489909 | 0.788202829 | 0.119242 | 0.09045049 |
| Tent4a | 0.053142066 | 0.114379689 | 0.485644398 | 0.077652518 | 0.033487625 |
| Tent4b | 0.016249388 | 0.736628151 | 0.922692718 | 0.154785901 | 0.142124275 |
| Tent5a | -0.015041393 | 0.855267772 | 0.957736324 | 1.438374116 | 1.39389559 |
| Tent5c | 0.033514431 | 0.582261074 | 0.866185623 | 0.322927111 | 0.279461856 |
| Tep1 | -0.007556696 | 0.830943898 | 0.95325696 | 0.08046626 | 0.086377727 |
| Terf1 | -0.074171182 | 0.113923602 | 0.48492417 | 0.138471501 | 0.209659889 |
| Terf2 | -0.01179611 | 0.835927706 | 0.954911647 | 0.245195756 | 0.265603545 |
| Terf2ip | -0.006059298 | 0.90719367 | 0.977187883 | 0.197613037 | 0.216621482 |
| Tert | -0.0538457 | 0.012737197 | 0.179750602 | 0.019313445 | 0.062831738 |
| Tes | 0.047654355 | 0.459060761 | 0.809370658 | 0.456882187 | 0.405303286 |
| Tesc | -0.020798545 | 0.347722201 | 0.739124045 | 0.025259861 | 0.043434921 |
| Tesk1 | -0.079402184 | 0.146381371 | 0.536193937 | 0.21281001 | 0.281315743 |
| Tet1 | 0.019397312 | 0.511503907 | 0.835198424 | 0.056048957 | 0.043134727 |
| Tet2 | 0.056861763 | 0.319355885 | 0.717256849 | 0.253295937 | 0.193356211 |
| Tet3 | -0.019282867 | 0.731004631 | 0.920770724 | 0.247905363 | 0.264861793 |
| Tex10 | 0.011775586 | 0.764002277 | 0.928730554 | 0.101398351 | 0.099473131 |
| Tex2 | 0.039440326 | 0.270975515 | 0.678418869 | 0.085912664 | 0.050142784 |
| Tex261 | -0.049582935 | 0.438459702 | 0.799277158 | 0.50495222 | 0.550545402 |
| Tex264 | 0.013948693 | 0.812946168 | 0.948397417 | 0.311734178 | 0.307935041 |
| Tex30 | 0.049314492 | 0.327453783 | 0.722182734 | 0.182558212 | 0.141674218 |
| Tex9 | -0.046216272 | 0.238999922 | 0.650445287 | 0.094032066 | 0.133544407 |
| Tfam | 0.036217389 | 0.543949042 | 0.851058306 | 0.310217088 | 0.273467578 |
| Tfb1m | -0.023099394 | 0.346077306 | 0.738085711 | 0.033365501 | 0.056898201 |
| Tfb2m | 0.00960021 | 0.835520975 | 0.954911647 | 0.14879638 | 0.143008019 |
| Tfcp2 | 0.036336694 | 0.348707009 | 0.73995185 | 0.104969335 | 0.074752504 |
| Tfdp1 | -0.056350062 | 0.253056196 | 0.663146569 | 0.168100826 | 0.212518929 |
| Tfdp2 | -0.120393759 | 0.014591881 | 0.194242323 | 0.157439625 | 0.272941608 |
| Tfe3 | -0.04795861 | 0.351360751 | 0.74181907 | 0.179734876 | 0.221171938 |
| Tfg | -0.036560308 | 0.555711076 | 0.854553711 | 0.40406698 | 0.446935601 |
| Tfip11 | -0.04404397 | 0.271319342 | 0.67876599 | 0.098972161 | 0.136343677 |
| Tfpt | -0.032505149 | 0.391234569 | 0.769704269 | 0.094601879 | 0.128292153 |
| Tfrc | 0.192807698 | 0.00822915 | 0.140469359 | 0.630813812 | 0.430023899 |
| Tgds | 0.025810239 | 0.518260929 | 0.837916389 | 0.113410822 | 0.101403162 |
| Tgfbr1 | 0.077758937 | 0.102169682 | 0.468127803 | 0.158007779 | 0.083459075 |
| Tgfbr3 | 0.014489943 | 0.679623724 | 0.907783945 | 0.080699717 | 0.062009936 |
| Tgfbrap1 | 0.038464372 | 0.478062069 | 0.818590256 | 0.230341839 | 0.194695835 |
| Tgoln1 | -0.066673604 | 0.225660419 | 0.638019508 | 1.452949957 | 1.553491873 |
| Tgs1 | -0.042579842 | 0.416007411 | 0.78353657 | 0.18415133 | 0.212863528 |

| | | | | | |
|---|---|---|---|---|---|
| Tha1 | -0.034717824 | 0.221790598 | 0.633858837 | 0.046814414 | 0.077047599 |
| Thada | -0.062966016 | 0.047197306 | 0.340345488 | 0.058304122 | 0.116297479 |
| Thap1 | -0.086758419 | 0.049774126 | 0.350830687 | 0.120578258 | 0.195786908 |
| Thap11 | 0.030721878 | 0.58752622 | 0.869281892 | 0.248503865 | 0.219279881 |
| Thap12 | 0.13503424 | 0.010011353 | 0.155789067 | 0.218582176 | 0.105861386 |
| Thap2 | 0.022709113 | 0.648452179 | 0.895379629 | 0.180640819 | 0.15802383 |
| Thap3 | 0.055676502 | 0.323489478 | 0.720244873 | 0.2680696 | 0.220408267 |
| Thap4 | 0.032827007 | 0.417698045 | 0.784482356 | 0.113370588 | 0.091552165 |
| Thap6 | -0.055854921 | 0.06609964 | 0.390719721 | 0.052448075 | 0.0999507 |
| Thap7 | 0.140739514 | 0.011843634 | 0.173034289 | 0.265503403 | 0.152869354 |
| Thbs3 | -0.016844101 | 0.513694435 | 0.836550101 | 0.035619711 | 0.049847439 |
| Them4 | -0.009373253 | 0.854054969 | 0.957461155 | 0.176649988 | 0.184982194 |
| Them6 | -0.011757531 | 0.78163073 | 0.935961303 | 0.118738365 | 0.135188609 |
| Thnsl1 | 0.075905507 | 0.092176002 | 0.44911191 | 0.14293464 | 0.082040434 |
| Thoc1 | -0.025343505 | 0.650314521 | 0.89561016 | 0.229818175 | 0.24781822 |
| Thoc2 | -0.001710534 | 0.979943311 | 0.995540893 | 0.496306508 | 0.491415253 |
| Thoc2l | 0.077219197 | 0.249435542 | 0.661099693 | 0.707379539 | 0.611408617 |
| Thoc3 | 0.032239361 | 0.565160292 | 0.858087314 | 0.249449661 | 0.225427639 |
| Thoc5 | -0.006037166 | 0.866897526 | 0.962125879 | 0.085398995 | 0.088335387 |
| Thoc6 | -0.001025916 | 0.974042511 | 0.994149348 | 0.062530458 | 0.062335403 |
| Thoc7 | 0.01965555 | 0.772070621 | 0.932411602 | 0.718852138 | 0.691101042 |
| Thop1 | -0.014964999 | 0.65201894 | 0.896672002 | 0.070530125 | 0.090461173 |
| Thra | 0.046554843 | 0.402843906 | 0.77525778 | 0.244347712 | 0.208969739 |
| Thrap3 | -0.024390359 | 0.722059189 | 0.919475778 | 0.582050854 | 0.614112312 |
| Thrb | 0.02783315 | 0.540830602 | 0.849634802 | 0.140659341 | 0.112915095 |
| Thumpd1 | -0.026605535 | 0.652693593 | 0.896887331 | 0.264706927 | 0.275462831 |
| Thumpd2 | -0.034195216 | 0.193449367 | 0.60325934 | 0.037232991 | 0.066741053 |
| Thumpd3 | 0.058942507 | 0.248463622 | 0.659783035 | 0.188864319 | 0.144713876 |
| Thyn1 | 0.003821597 | 0.937816335 | 0.985021197 | 0.168968132 | 0.163494088 |
| Tia1 | -0.011087906 | 0.866050285 | 0.962125879 | 0.485195828 | 0.49317666 |
| Tial1 | -0.059577814 | 0.357264226 | 0.745983316 | 0.387166028 | 0.459300615 |
| Ticam1 | -0.001105535 | 0.9707294 | 0.993892166 | 0.056991793 | 0.057062127 |
| Tigar | -0.024289103 | 0.584164878 | 0.8671012 | 0.134095928 | 0.151294713 |
| Tigd2 | 0.04267777 | 0.488623671 | 0.826256745 | 0.363870191 | 0.323753523 |
| Timm10 | 0.077491184 | 0.144668006 | 0.533279057 | 0.222934062 | 0.161429181 |
| Timm10b | -0.01622293 | 0.803012599 | 0.943517638 | 0.512923269 | 0.540762721 |
| Timm13 | 0.100958474 | 0.08532432 | 0.434941698 | 1.312935696 | 1.161056468 |
| Timm17a | 0.142387888 | 0.036467889 | 0.301010036 | 0.603105803 | 0.455086587 |
| Timm17b | -0.070105237 | 0.279945763 | 0.687465228 | 0.436731237 | 0.518358346 |
| Timm21 | -0.029912094 | 0.333423069 | 0.729464185 | 0.054094014 | 0.079533056 |
| Timm22 | 0.008120961 | 0.887736726 | 0.970045467 | 0.307315397 | 0.291537195 |
| Timm23 | 0.052581014 | 0.435224318 | 0.796975049 | 0.700826192 | 0.64182619 |
| Timm29 | 0.028474912 | 0.5948722 | 0.8732971 | 0.213621264 | 0.192643571 |
| Timm44 | 0.051706407 | 0.356614067 | 0.745571198 | 0.231742947 | 0.1718725 |
| Timm50 | 0.035928532 | 0.517049908 | 0.837583212 | 0.236952993 | 0.204336698 |
| Timm8a1 | 0.053932174 | 0.183251501 | 0.590066231 | 0.112643057 | 0.078484894 |
| Timm8b | 0.294303859 | 3.26E-06 | 0.000535636 | 1.382880614 | 0.961921444 |
| Timm9 | 0.02799747 | 0.642591626 | 0.892561795 | 0.332399724 | 0.308708855 |
| Timmdc1 | -0.051465033 | 0.201187526 | 0.611914909 | 0.102617353 | 0.146036243 |
| Timp2 | -0.099197256 | 0.008101165 | 0.139651258 | 0.078349363 | 0.161843502 |
| Tinf2 | 0.008540177 | 0.795148682 | 0.939757086 | 0.068410871 | 0.060779215 |
| Tiparp | 0.018421338 | 0.746391534 | 0.924631183 | 0.222090335 | 0.20745364 |
| Tipin | -0.024266478 | 0.650141411 | 0.895593446 | 0.210941841 | 0.235510251 |
| Tiprl | -0.050785247 | 0.450997478 | 0.806495742 | 0.481865267 | 0.541958331 |
| Tjap1 | -0.032180332 | 0.372320859 | 0.757837538 | 0.080082628 | 0.108985343 |
| Tjp1 | -0.047049891 | 0.476354593 | 0.817948631 | 0.461158829 | 0.495116256 |
| Tjp2 | 0.071954988 | 0.223753155 | 0.636206917 | 0.269988745 | 0.201206024 |
| Tjp3 | 0.053779454 | 0.077491449 | 0.418285047 | 0.062949177 | 0.018338136 |
| Tk2 | -0.000188249 | 0.996585159 | 0.998992348 | 0.126916987 | 0.138603112 |
| Tkfc | -0.035469061 | 0.254191445 | 0.663146569 | 0.057887774 | 0.090986898 |
| Tkt | -0.171272533 | 0.005220437 | 0.107592455 | 0.338845998 | 0.511602671 |
| Tlcd1 | -0.035955 | 0.281021938 | 0.68791143 | 0.067001572 | 0.106607043 |
| Tlcd3a | -0.027882902 | 0.269970624 | 0.677226912 | 0.035066136 | 0.058971384 |
| Tlcd3b | -0.080073218 | 0.028594773 | 0.26974223 | 0.076587086 | 0.146919273 |
| Tlcd4 | -0.021028612 | 0.738565043 | 0.922843014 | 0.366538271 | 0.387052477 |

| | | | | | |
|---|---|---|---|---|---|
| Tle1 | -0.048396187 | 0.369239685 | 0.756523935 | 0.203407093 | 0.249163337 |
| Tle2 | -0.023103659 | 0.460335499 | 0.810111566 | 0.059254387 | 0.083208991 |
| Tle3 | 0.043970147 | 0.513831959 | 0.836550101 | 0.421273868 | 0.388745807 |
| Tle4 | 0.025422467 | 0.473690978 | 0.817334768 | 0.083176487 | 0.062678285 |
| Tle5 | -0.051282842 | 0.230850742 | 0.642359619 | 1.993526324 | 2.109565732 |
| Tle6 | -0.043445751 | 0.076227576 | 0.416492204 | 0.029362238 | 0.067751394 |
| Tlk1 | -0.079197854 | 0.246742655 | 0.656389087 | 0.725789165 | 0.812778454 |
| Tlk2 | 0.109105829 | 0.083122859 | 0.430942219 | 0.342802106 | 0.235898158 |
| Tln1 | 0.080005724 | 0.221708318 | 0.633858837 | 0.460747296 | 0.394401581 |
| Tln2 | 0.021340065 | 0.467692278 | 0.813188482 | 0.057174521 | 0.040163331 |
| Tlnrd1 | 0.034309653 | 0.392149164 | 0.769704269 | 0.112727924 | 0.090561461 |
| Tlr2 | -0.044724568 | 0.257654375 | 0.665656876 | 0.093745029 | 0.129132716 |
| Tm2d1 | -0.173190475 | 0.008473802 | 0.142749092 | 0.584402737 | 0.777914911 |
| Tm2d2 | -0.017151534 | 0.797101991 | 0.940683443 | 0.638547322 | 0.660295863 |
| Tm2d3 | 0.013735598 | 0.832063997 | 0.953695326 | 0.554770084 | 0.544770278 |
| Tm4sf4 | 0.194032512 | 0.000569407 | 0.027743399 | 1.821403645 | 1.497386581 |
| Tm7sf3 | -0.047522623 | 0.368247587 | 0.755518937 | 0.197543516 | 0.241512044 |
| Tm9sf2 | -0.006007903 | 0.928737869 | 0.982502239 | 0.827431739 | 0.835123431 |
| Tm9sf3 | -0.100051985 | 0.128702977 | 0.508260084 | 0.993602894 | 1.109001407 |
| Tm9sf4 | -0.082078344 | 0.197335512 | 0.607747501 | 0.397786881 | 0.466471182 |
| Tma16 | -0.060322558 | 0.163645968 | 0.563469839 | 0.112509176 | 0.167209723 |
| Tma7 | 0.038841508 | 0.451193489 | 0.806495742 | 1.504075278 | 1.408922525 |
| Tmbim1 | 0.003376163 | 0.944617404 | 0.987070339 | 0.167721092 | 0.157582398 |
| Tmbim4 | -0.028348727 | 0.675591102 | 0.906025597 | 0.884686084 | 0.919369339 |
| Tmbim6 | -0.292240429 | 5.16E-07 | 0.000123208 | 1.288878229 | 1.771877361 |
| Tmc4 | -0.042501868 | 0.470427933 | 0.814364464 | 0.281017259 | 0.321323143 |
| Tmc6 | -0.010639185 | 0.676724251 | 0.906695569 | 0.037537097 | 0.045694151 |
| Tmc7 | -0.033170503 | 0.249794028 | 0.661525768 | 0.047353503 | 0.078176757 |
| Tmcc1 | -0.024598631 | 0.602193185 | 0.876569856 | 0.15344045 | 0.178420154 |
| Tmcc2 | 0.059798774 | 0.102941077 | 0.468979611 | 0.093547831 | 0.041222001 |
| Tmcc3 | -0.040133607 | 0.49962909 | 0.830722138 | 0.288296851 | 0.322791676 |
| Tmco1 | -0.073004404 | 0.261089382 | 0.66834369 | 0.650094985 | 0.712585482 |
| Tmco3 | -0.059568788 | 0.287991104 | 0.694589247 | 0.223370378 | 0.282616346 |
| Tmco6 | 0.005882697 | 0.84314451 | 0.955367063 | 0.05532039 | 0.049830614 |
| Tmed1 | 0.048902982 | 0.114916082 | 0.486793251 | 0.067601264 | 0.027691716 |
| Tmed10 | -0.178987629 | 0.000243359 | 0.015171416 | 1.627126489 | 1.951403165 |
| Tmed2 | 0.059777453 | 0.298293153 | 0.70313959 | 1.257084458 | 1.181047836 |
| Tmed3 | -0.073600515 | 0.225725543 | 0.638019508 | 1.407026984 | 1.505362835 |
| Tmed4 | -0.024550162 | 0.719841385 | 0.918866588 | 0.618686724 | 0.649717187 |
| Tmed5 | -0.133497996 | 0.040316951 | 0.316388771 | 0.459863814 | 0.597874731 |
| Tmed6 | -0.047364731 | 0.117831576 | 0.491192747 | 0.05538077 | 0.091775687 |
| Tmed7 | -0.082978879 | 0.20535081 | 0.616549498 | 0.547438373 | 0.633234473 |
| Tmed8 | 0.174670543 | 0.005492547 | 0.110700196 | 0.396031291 | 0.225120289 |
| Tmed9 | 0.004529336 | 0.940591964 | 0.986168593 | 1.338067432 | 1.354255369 |
| Tmem101 | -0.019727796 | 0.702403537 | 0.912152193 | 0.187140032 | 0.206458802 |
| Tmem102 | 0.03778314 | 0.306916404 | 0.709680069 | 0.091878158 | 0.059142066 |
| Tmem104 | 0.053315211 | 0.123309011 | 0.500122758 | 0.082685285 | 0.036329642 |
| Tmem106a | 0.026947837 | 0.447992525 | 0.804951652 | 0.085388345 | 0.063389719 |
| Tmem106b | -0.146077339 | 0.017899626 | 0.212740677 | 0.307215315 | 0.454014749 |
| Tmem106c | 0.012396798 | 0.828579574 | 0.952874813 | 0.252292285 | 0.236724204 |
| Tmem107 | 0.091970136 | 0.157985483 | 0.556211435 | 0.489953101 | 0.397380473 |
| Tmem108 | -0.020910894 | 0.732042314 | 0.920855835 | 0.304533084 | 0.322852816 |
| Tmem109 | -0.078691289 | 0.152248735 | 0.548949837 | 0.22392267 | 0.297839675 |
| Tmem11 | -0.086680922 | 0.157923163 | 0.556211435 | 0.319978286 | 0.41466651 |
| Tmem115 | 0.097560571 | 0.064419886 | 0.386943382 | 0.223275596 | 0.149399714 |
| Tmem117 | 0.047608211 | 0.307337385 | 0.709952896 | 0.154094924 | 0.104671957 |
| Tmem120a | 0.033756115 | 0.323327003 | 0.720244873 | 0.077541627 | 0.048751314 |
| Tmem121 | -0.051634334 | 0.10151316 | 0.467719786 | 0.057420045 | 0.103933271 |
| Tmem123 | 0.041470632 | 0.38058477 | 0.762006446 | 0.16310494 | 0.133374259 |
| Tmem126a | -0.057711333 | 0.386504602 | 0.766593284 | 0.69957998 | 0.753086238 |
| Tmem126b | 0.022318145 | 0.675202602 | 0.906025597 | 0.198948663 | 0.18947555 |
| Tmem127 | -0.073384255 | 0.209728927 | 0.622606671 | 0.271832129 | 0.332930901 |
| Tmem128 | 0.035133961 | 0.574159902 | 0.862168892 | 0.369208644 | 0.332162462 |
| Tmem129 | 0.055958602 | 0.175072545 | 0.578407878 | 0.119181932 | 0.079237871 |
| Tmem130 | -0.104482956 | 0.03181501 | 0.283462667 | 0.148166569 | 0.240551068 |

| | | | | | |
|---|---|---|---|---|---|
| Tmem131 | -0.051937923 | 0.35939155 | 0.747719327 | 0.241169271 | 0.291318768 |
| Tmem131l | -0.010398474 | 0.828888114 | 0.952874813 | 0.1537586 | 0.165354661 |
| Tmem132a | 0.086801207 | 0.01220575 | 0.176526096 | 0.086989105 | 0.008502086 |
| Tmem132b | 0.0714536 | 0.172043096 | 0.575294174 | 0.202622951 | 0.128567365 |
| Tmem134 | 0.00570959 | 0.928799216 | 0.982502239 | 0.632887011 | 0.616654825 |
| Tmem135 | -0.019409052 | 0.691660801 | 0.909796728 | 0.169873198 | 0.187560499 |
| Tmem138 | 0.01592731 | 0.68087098 | 0.90791843 | 0.101536503 | 0.083566197 |
| Tmem140 | 0.021373067 | 0.597591424 | 0.874521003 | 0.105916407 | 0.082013932 |
| Tmem141 | -0.078089667 | 0.10184838 | 0.467829632 | 0.147984635 | 0.218654437 |
| Tmem143 | -0.015737529 | 0.630937665 | 0.888561606 | 0.06447007 | 0.079869799 |
| Tmem144 | 0.018527181 | 0.542067529 | 0.850208191 | 0.058832682 | 0.048438759 |
| Tmem147 | 0.013720434 | 0.839738378 | 0.954911647 | 0.692473468 | 0.670472597 |
| Tmem14c | -0.072994692 | 0.264418825 | 0.67104216 | 0.637733844 | 0.728835491 |
| Tmem150a | -0.039758668 | 0.191821371 | 0.602601283 | 0.05574609 | 0.088806934 |
| Tmem150c | -0.02044255 | 0.654985286 | 0.897454058 | 0.137001189 | 0.156295893 |
| Tmem151a | -0.00632958 | 0.850962269 | 0.957461155 | 0.069072702 | 0.07744431 |
| Tmem158 | 0.065443674 | 0.019992379 | 0.225972421 | 0.055878611 | 0.008860735 |
| Tmem159 | 0.021978632 | 0.617694217 | 0.883238152 | 0.136328766 | 0.121970105 |
| Tmem160 | -0.170355821 | 0.01223155 | 0.176644707 | 0.541688129 | 0.720244865 |
| Tmem161a | 0.03523304 | 0.450643809 | 0.806495742 | 0.158873608 | 0.128778659 |
| Tmem161b | 0.007149661 | 0.828729102 | 0.952874813 | 0.070784408 | 0.06240703 |
| Tmem163 | 0.060979936 | 0.358772822 | 0.747406146 | 0.640508639 | 0.594858985 |
| Tmem164 | -0.022423998 | 0.535035545 | 0.846460359 | 0.083052806 | 0.10750325 |
| Tmem165 | 0.028991914 | 0.616609912 | 0.883226807 | 0.272979892 | 0.241173738 |
| Tmem167 | 0.023840832 | 0.726204089 | 0.92014727 | 1.037584518 | 0.991819301 |
| Tmem167b | -0.01992592 | 0.730709748 | 0.920770724 | 0.289984166 | 0.319464742 |
| Tmem168 | 0.069980861 | 0.057823428 | 0.370772606 | 0.097751456 | 0.036595422 |
| Tmem17 | -0.035572047 | 0.140301813 | 0.528607094 | 0.028606816 | 0.064754488 |
| Tmem170 | -0.046611689 | 0.184000103 | 0.590326848 | 0.076592708 | 0.121260101 |
| Tmem170b | 0.018873179 | 0.658985741 | 0.898224427 | 0.12171135 | 0.107364162 |
| Tmem175 | 0.029528203 | 0.37424553 | 0.75956519 | 0.071764377 | 0.041758489 |
| Tmem176a | -0.016084952 | 0.812474807 | 0.948397417 | 0.835795423 | 0.827194436 |
| Tmem176b | -0.115344817 | 0.077751372 | 0.418285047 | 0.988203673 | 1.093118916 |
| Tmem178b | 0.127268588 | 0.011819755 | 0.17293714 | 0.188728351 | 0.076106042 |
| Tmem179 | -0.025126673 | 0.36020201 | 0.748364226 | 0.040411611 | 0.064420011 |
| Tmem179b | -0.026268631 | 0.675899423 | 0.906225288 | 0.363495056 | 0.414912201 |
| Tmem18 | 0.028811354 | 0.440434293 | 0.80031914 | 0.096337887 | 0.072927765 |
| Tmem181a | -0.085523704 | 0.051047416 | 0.355933715 | 0.121828092 | 0.198466206 |
| Tmem181b-ps | -0.01220382 | 0.78216069 | 0.936026395 | 0.131621577 | 0.163145971 |
| Tmem183a | 0.013142243 | 0.816219887 | 0.949036482 | 0.253950473 | 0.245354692 |
| Tmem184a | -0.111674282 | 0.07322808 | 0.410379808 | 0.33766588 | 0.437878619 |
| Tmem184b | 0.112868906 | 0.051603306 | 0.355933715 | 0.289150451 | 0.19185547 |
| Tmem184c | -0.037785796 | 0.480618132 | 0.820388721 | 0.210867208 | 0.242749812 |
| Tmem185a | -0.014969009 | 0.764626132 | 0.928757959 | 0.177600218 | 0.179356736 |
| Tmem185b | 0.027353452 | 0.354434606 | 0.744861838 | 0.056739165 | 0.036300672 |
| Tmem186 | 0.013337334 | 0.644786883 | 0.893466123 | 0.0542041 | 0.043728617 |
| Tmem189 | 0.05155967 | 0.18750755 | 0.594820883 | 0.106532059 | 0.064222023 |
| Tmem19 | 0.063730012 | 0.204640968 | 0.616256046 | 0.18989469 | 0.136233971 |
| Tmem191c | 0.038234514 | 0.306883392 | 0.709680069 | 0.094326629 | 0.064617045 |
| Tmem192 | -0.068422635 | 0.221004934 | 0.633858837 | 0.232472357 | 0.298427138 |
| Tmem198 | 0.020386767 | 0.619943363 | 0.883866495 | 0.110214662 | 0.094942123 |
| Tmem199 | -0.063183515 | 0.278901096 | 0.686110367 | 0.294072643 | 0.35897851 |
| Tmem201 | -0.033407464 | 0.20397884 | 0.615554905 | 0.036722798 | 0.06701921 |
| Tmem203 | 0.087585814 | 0.114288582 | 0.485644398 | 0.254264607 | 0.181591577 |
| Tmem205 | -0.074186313 | 0.222446401 | 0.634095002 | 0.3308943 | 0.411045459 |
| Tmem208 | -0.038542959 | 0.565617781 | 0.858087314 | 0.483780166 | 0.534422572 |
| Tmem209 | -0.020935397 | 0.542304862 | 0.850208191 | 0.071622367 | 0.091604752 |
| Tmem212 | 0.009444285 | 0.850067153 | 0.957111129 | 0.18039261 | 0.156122581 |
| Tmem214 | -0.192381293 | 0.00060056 | 0.028299632 | 0.221190083 | 0.388011174 |
| Tmem215 | -0.04232145 | 0.560071896 | 0.856904336 | 0.671184679 | 0.703022325 |
| Tmem216 | -0.089008793 | 0.031911216 | 0.283462667 | 0.106703484 | 0.186005662 |
| Tmem218 | -0.000929921 | 0.988362542 | 0.997411191 | 0.400468369 | 0.406133971 |
| Tmem219 | -0.102216456 | 0.111614525 | 0.482690499 | 0.447615008 | 0.547235762 |
| Tmem222 | -0.024559833 | 0.663410244 | 0.899593114 | 0.253086716 | 0.279047755 |
| Tmem223 | -0.005875653 | 0.928418412 | 0.982502239 | 0.43103726 | 0.429594223 |

| | | | | | |
|---|---|---|---|---|---|
| Tmem229b | 0.059973566 | 0.2582931 | 0.665948842 | 0.208364568 | 0.154581365 |
| Tmem230 | -0.112557743 | 0.037849421 | 0.306861584 | 0.224101979 | 0.344765617 |
| Tmem231 | -0.001285452 | 0.970707204 | 0.993892166 | 0.08023491 | 0.080754578 |
| Tmem234 | 0.083533168 | 0.200546935 | 0.610932587 | 1.060728713 | 0.952057402 |
| Tmem237 | -0.053128868 | 0.124230666 | 0.501973913 | 0.069739539 | 0.113637141 |
| Tmem238 | -0.111674541 | 0.081971622 | 0.42777546 | 0.570838873 | 0.66891388 |
| Tmem240 | -0.001700738 | 0.952449088 | 0.98838165 | 0.050687886 | 0.052628483 |
| Tmem242 | 0.007754242 | 0.886914195 | 0.969712176 | 0.228770803 | 0.220722181 |
| Tmem243 | 0.141851952 | 0.03046725 | 0.277999808 | 0.446079871 | 0.29560572 |
| Tmem245 | 0.162479926 | 0.016012216 | 0.203940048 | 0.489618983 | 0.344071507 |
| Tmem248 | 0.033433936 | 0.623113 | 0.885857477 | 0.658227794 | 0.599039053 |
| Tmem25 | -0.052081142 | 0.024570362 | 0.248318109 | 0.024530226 | 0.067414957 |
| Tmem250-ps | 0.099970162 | 0.065267468 | 0.389310714 | 0.231978691 | 0.143380645 |
| Tmem251 | 0.031534858 | 0.595474785 | 0.873342705 | 0.302541257 | 0.277040455 |
| Tmem254a | 0.090776858 | 0.075494832 | 0.414292853 | 0.20028371 | 0.113576387 |
| Tmem255a | 0.065531422 | 0.061132866 | 0.380250374 | 0.082979874 | 0.023439202 |
| Tmem256 | -0.045769883 | 0.487640042 | 0.826112275 | 1.200986741 | 1.235785456 |
| Tmem258 | -0.052198082 | 0.409347443 | 0.779735046 | 1.025237925 | 1.068688189 |
| Tmem259 | -0.052990596 | 0.349020986 | 0.740431094 | 0.232976725 | 0.281787107 |
| Tmem260 | 0.064253203 | 0.096013222 | 0.456656576 | 0.108369389 | 0.04985 |
| Tmem263 | 0.017515212 | 0.777300083 | 0.934243618 | 0.345902825 | 0.338259207 |
| Tmem267 | -0.005803911 | 0.841436685 | 0.954960614 | 0.049147202 | 0.058757406 |
| Tmem268 | -0.012482926 | 0.681240231 | 0.908154905 | 0.055760368 | 0.068487717 |
| Tmem28 | -0.019632834 | 0.552632721 | 0.852623515 | 0.068426936 | 0.084161627 |
| Tmem29 | 0.027768761 | 0.441151742 | 0.800354711 | 0.086570807 | 0.066365282 |
| Tmem30a | -0.070918903 | 0.273647522 | 0.681538507 | 0.962642967 | 1.058904394 |
| Tmem30b | -0.045348627 | 0.263025192 | 0.669775695 | 0.106826549 | 0.155354175 |
| Tmem33 | 0.025246953 | 0.698453989 | 0.911319029 | 0.523163615 | 0.486840754 |
| Tmem37 | -0.050956971 | 0.029917226 | 0.274935828 | 0.02528596 | 0.065102218 |
| Tmem38a | 0.078986072 | 0.043764141 | 0.329086044 | 0.114890086 | 0.04921037 |
| Tmem38b | 0.046695053 | 0.252453129 | 0.663146569 | 0.11534774 | 0.080122032 |
| Tmem39a | -0.017778796 | 0.750936947 | 0.926186851 | 0.220156698 | 0.239178936 |
| Tmem39b | -0.068064034 | 0.019401029 | 0.221110882 | 0.046922982 | 0.105495196 |
| Tmem41b | -0.012148415 | 0.829320098 | 0.953043946 | 0.248569154 | 0.262703494 |
| Tmem42 | -0.036847532 | 0.447453374 | 0.804698299 | 0.158944957 | 0.197872393 |
| Tmem43 | 0.024728639 | 0.575087895 | 0.862723699 | 0.133236319 | 0.109616832 |
| Tmem50a | -0.071013694 | 0.277285728 | 0.684342381 | 0.694355892 | 0.775352734 |
| Tmem50b | 0.035725586 | 0.516540204 | 0.837427561 | 0.252233231 | 0.213241736 |
| Tmem51 | 0.073287666 | 0.111763746 | 0.482690499 | 0.148508557 | 0.079528489 |
| Tmem53 | -0.070274169 | 0.021103431 | 0.230233845 | 0.050594586 | 0.110956263 |
| Tmem59 | -0.105056644 | 0.074062082 | 0.412383666 | 1.361565721 | 1.52134136 |
| Tmem59l | -0.075834526 | 0.248965834 | 0.660574894 | 0.320103486 | 0.39432966 |
| Tmem60 | 0.160680159 | 0.011935362 | 0.17412097 | 0.393984045 | 0.242674801 |
| Tmem62 | -0.104260943 | 0.006355099 | 0.120222898 | 0.088125769 | 0.182475334 |
| Tmem63a | -0.02816798 | 0.591155592 | 0.871281744 | 0.192002754 | 0.212171253 |
| Tmem63b | -0.043516867 | 0.460892834 | 0.810249429 | 0.276633617 | 0.31024807 |
| Tmem64 | -0.117702385 | 0.007559037 | 0.133456507 | 0.120255668 | 0.218991341 |
| Tmem65 | 0.124658392 | 0.034441079 | 0.293212179 | 1.457866123 | 1.279871857 |
| Tmem67 | 0.024272121 | 0.423572151 | 0.789048567 | 0.061543843 | 0.033845055 |
| Tmem68 | 0.051314859 | 0.227138529 | 0.639312792 | 0.125436709 | 0.086591445 |
| Tmem70 | -0.011403006 | 0.845271718 | 0.955950886 | 0.27807872 | 0.293229047 |
| Tmem71 | 0.067223701 | 0.149140093 | 0.542986195 | 0.159694591 | 0.093328794 |
| Tmem80 | -0.170034537 | 0.000419975 | 0.022541679 | 0.14649738 | 0.301447123 |
| Tmem86a | -0.03590437 | 0.297385218 | 0.702656882 | 0.069767598 | 0.103638793 |
| Tmem87a | 0.013165857 | 0.719415704 | 0.918741515 | 0.09038103 | 0.076068671 |
| Tmem87b | 0.000838554 | 0.987730103 | 0.997268589 | 0.220331105 | 0.223090382 |
| Tmem8b | -0.00979077 | 0.752424939 | 0.926186851 | 0.060962024 | 0.069057234 |
| Tmem9 | -0.035630818 | 0.494759532 | 0.828842054 | 0.200641061 | 0.228661184 |
| Tmem94 | 0.061981523 | 0.158946763 | 0.556789205 | 0.134943352 | 0.077780369 |
| Tmem97 | -0.022053551 | 0.660276934 | 0.898481506 | 0.167430459 | 0.187928761 |
| Tmem9b | 0.098782398 | 0.098470708 | 0.462278063 | 0.338380162 | 0.244067027 |
| Tmf1 | 0.180942291 | 0.006555703 | 0.122989887 | 0.676537314 | 0.500098466 |
| Tmigd3 | 0.035364508 | 0.440400335 | 0.80031914 | 0.139816999 | 0.116851761 |
| Tmod1 | 0.060511166 | 0.345681496 | 0.73792645 | 0.388039406 | 0.320192969 |
| Tmod2 | -0.113067805 | 0.097465768 | 0.460576232 | 0.788184996 | 0.930458344 |

| | | | | | |
|---|---|---|---|---|---|
| Tmod3 | -0.04981536 | 0.320726222 | 0.718701556 | 0.171405423 | 0.21912593 |
| Tmpo | 0.006098379 | 0.927234935 | 0.981818446 | 0.529559951 | 0.52027268 |
| Tmprss2 | -0.014205186 | 0.687470771 | 0.909796728 | 0.078497352 | 0.088160242 |
| Tmprss4 | 0.012801364 | 0.800775899 | 0.942028562 | 0.174935062 | 0.156530721 |
| Tmsb10 | 0.089282114 | 0.213540908 | 0.628127689 | 0.727389139 | 0.584232562 |
| Tmsb15b1 | -0.011447748 | 0.772840817 | 0.932469974 | 0.102010155 | 0.107076573 |
| Tmsb15b2 | 0.274968002 | 8.93E-05 | 0.007326725 | 0.478594597 | 0.192397962 |
| Tmsb4x | 0.110624629 | 0.061960467 | 0.380250374 | 1.992274593 | 1.80361875 |
| Tmtc1 | 0.005778058 | 0.901549459 | 0.974269668 | 0.146123362 | 0.125526346 |
| Tmtc2 | -0.073469341 | 0.067072463 | 0.392969665 | 0.093485215 | 0.157324426 |
| Tmtc3 | 0.034529691 | 0.506556372 | 0.833269979 | 0.197535785 | 0.164741982 |
| Tmtc4 | 0.045752483 | 0.161438485 | 0.559709178 | 0.074426744 | 0.038931453 |
| Tmub1 | -0.003954592 | 0.938376528 | 0.985197198 | 0.187789014 | 0.204893326 |
| Tmub2 | 0.019573693 | 0.70228284 | 0.912152193 | 0.185698682 | 0.168441829 |
| Tmx1 | -0.073426011 | 0.271684985 | 0.67876599 | 0.490819581 | 0.562939661 |
| Tmx2 | 0.08374013 | 0.128887793 | 0.508260084 | 0.243737753 | 0.175310657 |
| Tmx3 | -0.058624022 | 0.325592039 | 0.720833548 | 0.328973735 | 0.382632728 |
| Tmx4 | 0.145592673 | 0.024389291 | 0.247768538 | 0.726321882 | 0.570948064 |
| Tnfaip1 | 0.101752177 | 0.059834065 | 0.37818294 | 0.221109169 | 0.131864056 |
| Tnfaip3 | -0.062324633 | 0.001610672 | 0.052831084 | 0.011195686 | 0.065002006 |
| Tnfaip8 | 0.145333698 | 0.010139383 | 0.157293649 | 0.263143457 | 0.131012867 |
| Tnfrsf11a | 0.050394819 | 0.14494224 | 0.533279057 | 0.083586258 | 0.040627117 |
| Tnfrsf12a | 0.334115042 | 1.44E-05 | 0.001697747 | 0.538671226 | 0.205145074 |
| Tnfrsf1a | 0.079792688 | 0.162025984 | 0.560777516 | 0.281930705 | 0.208502002 |
| Tnfrsf21 | 0.033977543 | 0.490904891 | 0.827129829 | 0.173325639 | 0.142873981 |
| Tnfrsf23 | 0.013303792 | 0.752238352 | 0.926186851 | 0.116922537 | 0.112817548 |
| Tnfrsf9 | 0.083971965 | 0.051444155 | 0.355933715 | 0.130554622 | 0.060422091 |
| Tnfsf12 | -0.028412433 | 0.528860432 | 0.844309234 | 0.137172487 | 0.157819235 |
| Tnik | 0.028442285 | 0.56463841 | 0.858087314 | 0.175288885 | 0.149499191 |
| Tnip1 | 0.005265153 | 0.873897652 | 0.964940674 | 0.068446151 | 0.067993249 |
| Tnip2 | 0.021080965 | 0.554080675 | 0.853092152 | 0.086634723 | 0.064191102 |
| Tnk1 | 0.008458593 | 0.813763036 | 0.948397417 | 0.088430859 | 0.079707387 |
| Tnk2 | 0.018751815 | 0.673147357 | 0.905559579 | 0.133958209 | 0.119533013 |
| Tnks | -0.018670802 | 0.774268797 | 0.933157531 | 0.430477383 | 0.46769734 |
| Tnks1bp1 | -0.007127883 | 0.84587147 | 0.956051742 | 0.085454797 | 0.085092862 |
| Tnks2 | 0.080090059 | 0.243048961 | 0.653166476 | 0.711229434 | 0.622411833 |
| Tnnc1 | 0.011865282 | 0.759589178 | 0.927719223 | 0.094447447 | 0.082676638 |
| Tnpo1 | 0.057511168 | 0.381280536 | 0.762430537 | 0.595979215 | 0.525468133 |
| Tnpo2 | -0.057795907 | 0.169765645 | 0.572367409 | 0.120957317 | 0.170252703 |
| Tnpo3 | -0.049985976 | 0.420436193 | 0.786922065 | 0.372280599 | 0.406833029 |
| Tnr | -0.019575251 | 0.688790301 | 0.909796728 | 0.15695071 | 0.177069976 |
| Tnrc18 | -0.030437021 | 0.599852617 | 0.875897759 | 0.268924186 | 0.295502572 |
| Tnrc6a | 0.024410967 | 0.727688854 | 0.920455328 | 0.70531017 | 0.670794677 |
| Tnrc6b | -0.054453341 | 0.41259842 | 0.781358105 | 0.486185929 | 0.535047234 |
| Tnrc6c | -0.128938933 | 0.056235762 | 0.367897032 | 0.615235606 | 0.741716124 |
| Tns2 | 0.069066001 | 0.050536684 | 0.354499194 | 0.089933559 | 0.032062643 |
| Tns3 | 0.037940676 | 0.50350855 | 0.832249514 | 0.271061269 | 0.242288199 |
| Tob1 | -0.133975976 | 0.04254846 | 0.325502204 | 0.47198616 | 0.611237719 |
| Tob2 | -0.151892113 | 0.011058146 | 0.166153611 | 0.258922703 | 0.390866917 |
| Toe1 | 0.019883506 | 0.565237153 | 0.858087314 | 0.079152232 | 0.065065171 |
| Togaram1 | 0.014588228 | 0.817526794 | 0.949382903 | 0.332349199 | 0.33720839 |
| Tollip | 0.107932174 | 0.073887109 | 0.412240835 | 0.308922411 | 0.198432407 |
| Tom1 | 0.04366876 | 0.266640387 | 0.674088763 | 0.107814032 | 0.074821857 |
| Tom1l1 | 0.034517468 | 0.417315142 | 0.784482356 | 0.123915053 | 0.096698307 |
| Tom1l2 | -0.036573522 | 0.466627217 | 0.813152001 | 0.164580479 | 0.187666863 |
| Tomm20 | 0.153368837 | 0.005310544 | 0.108344012 | 1.414546812 | 1.198145692 |
| Tomm22 | 0.013244805 | 0.840343528 | 0.954911647 | 0.587004998 | 0.568242612 |
| Tomm34 | -0.075159146 | 0.116408256 | 0.488986751 | 0.154658227 | 0.225166601 |
| Tomm40 | -0.16983552 | 0.005373342 | 0.109174563 | 0.328956446 | 0.504188912 |
| Tomm40l | 0.023256448 | 0.552187308 | 0.852623515 | 0.104536819 | 0.089246293 |
| Tomm5 | 0.123542917 | 0.066943157 | 0.392969665 | 0.572256325 | 0.443767988 |
| Tomm6 | -0.021790012 | 0.499127076 | 0.830722138 | 0.064956872 | 0.08443408 |
| Tomm7 | 0.115563399 | 0.032958505 | 0.287906455 | 1.517300145 | 1.318708216 |
| Tomm70a | 0.062523595 | 0.353227241 | 0.743882043 | 0.496528225 | 0.443231164 |
| Top1 | 0.065838634 | 0.285479142 | 0.692407818 | 1.201467395 | 1.089426572 |

| | | | | | |
|---|---|---|---|---|---|
| Top2b | -0.020567689 | 0.745379591 | 0.924566547 | 0.466912217 | 0.484801006 |
| Top3b | 0.005868534 | 0.890704716 | 0.970579007 | 0.128883861 | 0.123057534 |
| Topbp1 | 0.076622935 | 0.049289196 | 0.348262299 | 0.109234805 | 0.048504421 |
| Topors | 0.110046755 | 0.080940243 | 0.426199169 | 0.384216015 | 0.281748539 |
| Tor1a | -0.031528822 | 0.600741901 | 0.876274737 | 0.306837174 | 0.347554986 |
| Tor1aip1 | -0.006268887 | 0.923173225 | 0.980859278 | 0.404681697 | 0.411277711 |
| Tor1aip2 | -0.047551316 | 0.468052728 | 0.813213564 | 0.459400148 | 0.488431909 |
| Tor1b | -0.063937254 | 0.351182655 | 0.741599055 | 0.631935783 | 0.702116094 |
| Tor2a | 0.038046944 | 0.521765825 | 0.840205104 | 0.294919446 | 0.256324316 |
| Tor3a | 0.087842353 | 0.181623095 | 0.589059475 | 0.507103223 | 0.403358666 |
| Tor4a | -0.027293576 | 0.213207246 | 0.62773867 | 0.023886956 | 0.043952113 |
| Tox | 0.051717306 | 0.29079824 | 0.69667177 | 0.171094716 | 0.112786803 |
| Tox3 | -0.128017105 | 0.077019002 | 0.418199494 | 0.755451502 | 0.913228036 |
| Tox4 | -0.053803671 | 0.402784497 | 0.77525778 | 0.449188312 | 0.506014267 |
| Tpcn1 | 0.054068268 | 0.217721593 | 0.632272789 | 0.136811966 | 0.085522639 |
| Tpd52 | -0.115286749 | 0.081968383 | 0.42777546 | 0.968375425 | 1.118323657 |
| Tpd52l1 | -0.025977959 | 0.564143848 | 0.857926031 | 0.133414837 | 0.158780631 |
| Tpd52l2 | 0.079705929 | 0.230944548 | 0.642359619 | 0.504534393 | 0.420848879 |
| Tpgs1 | -0.063485904 | 0.286762966 | 0.693329451 | 0.315374359 | 0.386384004 |
| Tpgs2 | -0.011243087 | 0.753203702 | 0.926506029 | 0.081902658 | 0.096226766 |
| Tpi1 | 0.115624484 | 0.089859125 | 0.445385778 | 0.859732587 | 0.715436402 |
| Tpm1 | 0.18264332 | 0.008787717 | 0.146759253 | 0.590941421 | 0.38247245 |
| Tpm3 | 0.128561164 | 0.055527612 | 0.365942644 | 0.553767691 | 0.419599315 |
| Tpm4 | 0.087448605 | 0.230115448 | 0.641131395 | 0.742556145 | 0.622594516 |
| Tpp1 | 0.05789329 | 0.342294237 | 0.735068249 | 0.316946949 | 0.256940097 |
| Tpp2 | 0.013259424 | 0.825332411 | 0.952011047 | 0.324142315 | 0.308680381 |
| Tppp3 | 0.140992023 | 0.004567147 | 0.099436985 | 0.173675225 | 0.036780864 |
| Tpr | -0.023046012 | 0.733547307 | 0.921306823 | 1.323333727 | 1.336049289 |
| Tpra1 | 0.006788859 | 0.847337227 | 0.956929571 | 0.079955778 | 0.077108808 |
| Tprgl | -0.071872523 | 0.238701808 | 0.650445287 | 0.346866614 | 0.425550848 |
| Tprkb | -0.086127011 | 0.119802329 | 0.493284734 | 0.21577866 | 0.291362643 |
| Tpst1 | -0.040272212 | 0.335862604 | 0.730805382 | 0.109619043 | 0.14536329 |
| Tpst2 | -0.082288198 | 0.18967198 | 0.5980954 | 0.340447986 | 0.435475863 |
| Tpt1 | -0.032319869 | 0.140012074 | 0.528110178 | 3.812330409 | 3.945530579 |
| Tpt1-ps3 | 0.028860919 | 0.616196723 | 0.883033482 | 0.286696404 | 0.247320057 |
| Tra2a | -0.105186816 | 0.129828519 | 0.509514066 | 0.536671731 | 0.652290285 |
| Tra2b | -0.132177889 | 0.050647788 | 0.354499194 | 0.803826799 | 0.952101188 |
| Trabd | 0.030370348 | 0.662295426 | 0.898950007 | 0.641757263 | 0.610651184 |
| Tradd | -0.080960761 | 0.018293652 | 0.21500396 | 0.066737298 | 0.13694104 |
| Traf2 | 0.006674478 | 0.82572754 | 0.952186044 | 0.055278814 | 0.048167743 |
| Traf3 | -0.027400028 | 0.609515894 | 0.878855197 | 0.193539738 | 0.214728941 |
| Traf3ip1 | -0.014282878 | 0.692358488 | 0.909796728 | 0.086563204 | 0.097431435 |
| Traf3ip2 | -0.082100446 | 0.024522454 | 0.248116801 | 0.077265259 | 0.14753564 |
| Traf4 | 0.002605573 | 0.929773017 | 0.982502239 | 0.054010874 | 0.056043661 |
| Traf5 | 0.016414706 | 0.549708422 | 0.852455047 | 0.048515456 | 0.035187849 |
| Traf6 | 0.012938018 | 0.765336829 | 0.928757959 | 0.125829371 | 0.116066453 |
| Traf7 | -0.048327574 | 0.292549479 | 0.698292299 | 0.136413972 | 0.171744242 |
| Trafd1 | 0.012292029 | 0.846695811 | 0.956583279 | 0.347022526 | 0.324447887 |
| Trak1 | 0.132212587 | 0.027366624 | 0.263607301 | 0.31854389 | 0.179739306 |
| Trak2 | 0.026873772 | 0.557391099 | 0.855384435 | 0.143342953 | 0.117786066 |
| Tram1 | -0.017230358 | 0.77912687 | 0.934664917 | 1.194355055 | 1.203745343 |
| Trap1 | -0.005011791 | 0.886130256 | 0.969590273 | 0.080497636 | 0.085335584 |
| Trappc1 | -0.012621703 | 0.848752726 | 0.957111129 | 0.546321241 | 0.551164761 |
| Trappc10 | -0.077803606 | 0.214038378 | 0.628710329 | 0.414802483 | 0.484584252 |
| Trappc11 | 0.046403243 | 0.434054986 | 0.796332921 | 0.30439386 | 0.256837349 |
| Trappc12 | -0.010794138 | 0.820433327 | 0.950668356 | 0.149565551 | 0.161057442 |
| Trappc13 | -0.026260058 | 0.648717633 | 0.895430849 | 0.262175952 | 0.285552553 |
| Trappc2 | -0.0170703 | 0.737195545 | 0.92276204 | 0.177488769 | 0.1883349 |
| Trappc2l | -0.07970065 | 0.230917927 | 0.642359619 | 0.867156707 | 0.938871122 |
| Trappc3 | 0.008517026 | 0.892264038 | 0.971118429 | 0.392497555 | 0.375152463 |
| Trappc4 | -0.000818446 | 0.990415115 | 0.997733826 | 0.621355566 | 0.644073429 |
| Trappc5 | -0.075098214 | 0.206865764 | 0.618502137 | 0.299587592 | 0.370209571 |
| Trappc6a | -0.069306506 | 0.257417999 | 0.665256942 | 0.331893122 | 0.399831551 |
| Trappc6b | -0.055333927 | 0.403974757 | 0.77632362 | 0.70040944 | 0.751711666 |
| Trappc8 | 0.074749473 | 0.144873718 | 0.533279057 | 0.197511183 | 0.13578326 |

| | | | | | |
|---|---|---|---|---|---|
| Trappc9 | -0.05411206 | 0.263215947 | 0.669775695 | 0.160321616 | 0.201930373 |
| Trerf1 | 0.077336804 | 0.078760685 | 0.421117898 | 0.141634495 | 0.076667002 |
| Triap1 | 0.005961605 | 0.926115705 | 0.981772637 | 0.386610834 | 0.384600654 |
| Trib1 | -0.07610909 | 0.267971122 | 0.676097903 | 0.451116445 | 0.516611265 |
| Trib2 | -0.019263301 | 0.463362227 | 0.811254677 | 0.037977548 | 0.05285882 |
| Trib3 | -0.180445048 | 0.008111655 | 0.139651258 | 0.469064396 | 0.63568855 |
| Trim11 | -0.001116534 | 0.974261481 | 0.994170647 | 0.077757321 | 0.081980177 |
| Trim12c | 0.067516688 | 0.038921698 | 0.311933003 | 0.077186859 | 0.016344088 |
| Trim13 | -0.012220349 | 0.631831822 | 0.88887544 | 0.036722788 | 0.048594701 |
| Trim2 | 0.085239025 | 0.130847044 | 0.510610033 | 0.218617441 | 0.12857132 |
| Trim21 | 0.029572243 | 0.311753924 | 0.71405691 | 0.054473072 | 0.028511411 |
| Trim23 | 0.030465192 | 0.4542715 | 0.807876964 | 0.111989637 | 0.087668741 |
| Trim24 | 0.006153755 | 0.920542426 | 0.980436936 | 0.322110655 | 0.3191575 |
| Trim25 | 0.000776468 | 0.988528628 | 0.997411191 | 0.223656315 | 0.217856852 |
| Trim26 | 0.008007012 | 0.888808536 | 0.970176435 | 0.259860223 | 0.257102254 |
| Trim27 | -0.09623235 | 0.096114363 | 0.456770768 | 0.258236489 | 0.360523593 |
| Trim28 | 0.097902446 | 0.086847143 | 0.438473226 | 0.278808582 | 0.19620364 |
| Trim3 | 0.055614671 | 0.146511247 | 0.536299559 | 0.101353503 | 0.050696203 |
| Trim32 | -0.009449123 | 0.755117666 | 0.926726954 | 0.05510968 | 0.061862661 |
| Trim33 | 0.049080866 | 0.419168366 | 0.785656935 | 0.344300604 | 0.303276547 |
| Trim34a | 0.024236372 | 0.416906246 | 0.784482356 | 0.058114444 | 0.034031051 |
| Trim35 | -0.093337009 | 0.183776115 | 0.590259477 | 0.575806736 | 0.670101962 |
| Trim37 | -0.020477438 | 0.729531933 | 0.920522999 | 0.270990042 | 0.28717877 |
| Trim39 | -0.027705986 | 0.530464047 | 0.84508505 | 0.128652945 | 0.157176826 |
| Trim41 | -0.019699117 | 0.731287192 | 0.920770724 | 0.244640862 | 0.250582766 |
| Trim44 | -0.103710624 | 0.130157823 | 0.509514066 | 0.629333899 | 0.736202744 |
| Trim46 | -0.025316312 | 0.409842253 | 0.780101937 | 0.057228257 | 0.079369013 |
| Trim56 | -0.05581778 | 0.38960194 | 0.768258286 | 0.366156447 | 0.420437841 |
| Trim66 | -0.085118003 | 0.000971475 | 0.037358974 | 0.028654228 | 0.102587918 |
| Trim7 | 0.016668936 | 0.553416698 | 0.852854813 | 0.050377169 | 0.038045707 |
| Trim8 | 0.097970544 | 0.070728827 | 0.403126197 | 0.230407917 | 0.145623712 |
| Trim9 | 0.041388543 | 0.256237897 | 0.664906869 | 0.090332697 | 0.052769503 |
| Trio | 0.119025361 | 0.060695413 | 0.380221737 | 0.394122014 | 0.269302467 |
| Triobp | 0.001985077 | 0.959390104 | 0.990780788 | 0.098540031 | 0.100040456 |
| Trip10 | -0.069551908 | 0.008224825 | 0.140469359 | 0.033191202 | 0.093563251 |
| Trip11 | -0.068768576 | 0.322853143 | 0.720244873 | 0.993800326 | 1.08546085 |
| Trip12 | 0.014840792 | 0.82013221 | 0.950589638 | 0.541339837 | 0.537066074 |
| Trip4 | -0.005305469 | 0.918849211 | 0.980385833 | 0.195352591 | 0.193228175 |
| Triqk | -0.159315091 | 2.19E-05 | 0.002386658 | 0.073026823 | 0.216178314 |
| Trir | -0.142428974 | 0.026578147 | 0.259246704 | 0.700596774 | 0.847406181 |
| Trit1 | 0.03235853 | 0.351160168 | 0.741599055 | 0.077785453 | 0.04628235 |
| Trmo | -0.053029813 | 0.018888418 | 0.218227362 | 0.022585114 | 0.068402497 |
| Trmt1 | -0.033420871 | 0.432859326 | 0.795079432 | 0.123298117 | 0.15460754 |
| Trmt10a | 0.026297227 | 0.544421303 | 0.851259637 | 0.129407293 | 0.107837484 |
| Trmt10b | -0.004964892 | 0.896746585 | 0.972419247 | 0.095934184 | 0.109269135 |
| Trmt10c | 0.088254128 | 0.106270368 | 0.473814268 | 0.228543562 | 0.153468009 |
| Trmt11 | 0.02353892 | 0.578852654 | 0.865218777 | 0.126319113 | 0.110584815 |
| Trmt112 | 0.025922783 | 0.697465644 | 0.910970095 | 0.703952921 | 0.690970097 |
| Trmt12 | -0.028852937 | 0.433551042 | 0.795675956 | 0.080641622 | 0.112759199 |
| Trmt13 | -0.019522173 | 0.608751464 | 0.878855197 | 0.090154129 | 0.105580843 |
| Trmt1l | 0.00626456 | 0.893097069 | 0.97160365 | 0.152500704 | 0.143662186 |
| Trmt2a | 0.020729988 | 0.662210151 | 0.898950007 | 0.156084642 | 0.143402908 |
| Trmt2b | 0.025921875 | 0.451556689 | 0.806495742 | 0.079276417 | 0.057505963 |
| Trmt6 | -0.022494992 | 0.737698532 | 0.922843014 | 0.454059805 | 0.461265667 |
| Trmt61a | 0.032579388 | 0.37662782 | 0.761269164 | 0.090330489 | 0.061266583 |
| Trmt61b | -0.031779314 | 0.452690693 | 0.807136288 | 0.114486168 | 0.150759898 |
| Trmt9b | -0.040648879 | 0.186983815 | 0.594286432 | 0.053947787 | 0.08421094 |
| Trmu | -0.019236148 | 0.597304022 | 0.874309534 | 0.082468481 | 0.100937545 |
| Trnau1ap | 0.012708692 | 0.81656661 | 0.949036482 | 0.22377075 | 0.226283629 |
| Trnt1 | 0.048454527 | 0.404574738 | 0.776552851 | 0.270677693 | 0.229336988 |
| Trp53 | -0.02526859 | 0.645939738 | 0.894371244 | 0.215146843 | 0.236803878 |
| Trp53bp1 | -0.064919122 | 0.25703708 | 0.665256942 | 0.244490849 | 0.305130953 |
| Trp53bp2 | 0.031600495 | 0.529418804 | 0.844323583 | 0.183386219 | 0.163203204 |
| Trp53cor1 | -0.031656117 | 0.501031259 | 0.831764926 | 0.14694752 | 0.168547507 |
| Trp53i11 | 0.022522222 | 0.603495654 | 0.876723966 | 0.128931778 | 0.11253908 |

| | | | | | |
|---|---|---|---|---|---|
| Trp53i13 | -0.173881035 | 0.003170764 | 0.078774653 | 0.250555291 | 0.425990832 |
| Trp53inp1 | -0.065372439 | 0.332961166 | 0.728881491 | 0.381515907 | 0.45398118 |
| Trp53inp2 | 0.066360693 | 0.318212635 | 0.717175454 | 0.715161573 | 0.643018184 |
| Trp53rka | 0.006909144 | 0.8734044 | 0.964940674 | 0.124619205 | 0.118849923 |
| Trp53rkb | -0.044237736 | 0.308967402 | 0.711182121 | 0.128538731 | 0.164695951 |
| Trpc1 | 0.028137827 | 0.657222398 | 0.898224427 | 0.386996839 | 0.356330884 |
| Trpc4 | -0.007376204 | 0.794179201 | 0.939757086 | 0.047540496 | 0.048044401 |
| Trpc4ap | -0.002274357 | 0.972571742 | 0.994079882 | 0.404651301 | 0.42484556 |
| Trpm3 | -0.011130204 | 0.808420326 | 0.946804528 | 0.139789887 | 0.14515063 |
| Trpm4 | 0.03407742 | 0.544782882 | 0.851259637 | 0.245176561 | 0.211625438 |
| Trpm5 | 0.135834618 | 0.034329791 | 0.293212179 | 0.341414688 | 0.232412803 |
| Trpm7 | -0.060248289 | 0.25587121 | 0.664689244 | 0.202794901 | 0.255394939 |
| Trps1 | 0.050895249 | 0.294501569 | 0.700619162 | 0.16791345 | 0.126556824 |
| Trpt1 | -0.083428612 | 0.006987434 | 0.127979693 | 0.055040551 | 0.129396026 |
| Trrap | -0.072848127 | 0.167298579 | 0.569598316 | 0.200849358 | 0.267720158 |
| Trub1 | 0.023078987 | 0.605505338 | 0.87802905 | 0.136522924 | 0.113724294 |
| Trub2 | 0.035064098 | 0.375803742 | 0.760595784 | 0.108706513 | 0.071505464 |
| Tsc1 | -0.062704562 | 0.236068476 | 0.646675571 | 0.189492492 | 0.243156393 |
| Tsc2 | -0.118100105 | 0.002994466 | 0.076092349 | 0.095016312 | 0.18757098 |
| Tsc22d1 | -0.024667773 | 0.682048583 | 0.908884974 | 1.460449384 | 1.4892871 |
| Tsc22d2 | -0.038124158 | 0.515741259 | 0.837349565 | 0.264345314 | 0.29371779 |
| Tsc22d3 | -0.006459834 | 0.90355605 | 0.974950771 | 0.201007264 | 0.215075554 |
| Tsc22d4 | 0.008899081 | 0.889723855 | 0.970458414 | 0.499560354 | 0.488822436 |
| Tsen15 | -0.006096596 | 0.893909219 | 0.971749901 | 0.152614504 | 0.157334514 |
| Tsen2 | -0.02032191 | 0.5514047 | 0.852505562 | 0.072077551 | 0.087348672 |
| Tsen34 | -0.054717705 | 0.323421209 | 0.720244873 | 0.220308815 | 0.28413532 |
| Tsen54 | 0.013184633 | 0.699047574 | 0.911319029 | 0.076506113 | 0.070527142 |
| Tsfm | -0.064095353 | 0.223287517 | 0.63524286 | 0.199865521 | 0.249143618 |
| Tsg101 | -0.084650028 | 0.179487646 | 0.586240644 | 0.379105388 | 0.468516247 |
| Tsga10 | 0.040217019 | 0.266537065 | 0.674047915 | 0.084875126 | 0.043707066 |
| Tshz1 | 0.129870284 | 0.067958692 | 0.396044041 | 0.955155398 | 0.776883194 |
| Tshz2 | -0.106147028 | 0.114366113 | 0.485644398 | 0.643126661 | 0.765388982 |
| Tshz3 | 0.038646872 | 0.491656508 | 0.827270742 | 0.242274292 | 0.206823362 |
| Tsn | 0.043937919 | 0.501909256 | 0.832218079 | 0.831310532 | 0.786252128 |
| Tsnax | -0.101640416 | 0.080474179 | 0.425115437 | 0.27433487 | 0.372978489 |
| Tspan13 | 0.087074214 | 0.14237388 | 0.531031821 | 1.31459861 | 1.16854351 |
| Tspan14 | -0.029312753 | 0.179969802 | 0.586833233 | 0.023136169 | 0.051084457 |
| Tspan15 | -0.10384262 | 0.118412967 | 0.491192747 | 0.470965872 | 0.574714183 |
| Tspan17 | 0.035826843 | 0.455649817 | 0.808152996 | 0.16277241 | 0.124413812 |
| Tspan2 | 0.003472436 | 0.95424938 | 0.989066902 | 0.31859055 | 0.333732235 |
| Tspan3 | -0.045288661 | 0.502770133 | 0.832249514 | 0.778546288 | 0.837141865 |
| Tspan31 | 0.100754126 | 0.11764695 | 0.491190698 | 0.438869314 | 0.340266335 |
| Tspan33 | -0.022863058 | 0.671558524 | 0.904877555 | 0.206978358 | 0.231207134 |
| Tspan5 | 0.01049605 | 0.882402889 | 0.968472149 | 0.778951929 | 0.787959815 |
| Tspan7 | -0.002720335 | 0.967231777 | 0.992750317 | 0.811722366 | 0.811664365 |
| Tspan8 | -0.006040908 | 0.9147131 | 0.980132949 | 0.179477901 | 0.17789921 |
| Tspo | -0.014967146 | 0.799654521 | 0.941260986 | 0.277678053 | 0.293518488 |
| Tspyl1 | -0.044354661 | 0.48630892 | 0.825197169 | 0.406588047 | 0.457219596 |
| Tspyl2 | 0.042344014 | 0.4985594 | 0.830722138 | 0.325505144 | 0.309593336 |
| Tspyl4 | 0.001246544 | 0.983742287 | 0.99608406 | 1.092472428 | 1.07693375 |
| Tspyl5 | 0.01633118 | 0.584277594 | 0.867129116 | 0.057175129 | 0.043007831 |
| Tsr1 | -0.040435364 | 0.46258514 | 0.810715393 | 0.206029084 | 0.250748719 |
| Tsr2 | -0.030765471 | 0.544026112 | 0.851058306 | 0.186612419 | 0.214237906 |
| Tsr3 | -0.04620121 | 0.403760329 | 0.77632362 | 0.235318084 | 0.29821713 |
| Tssc4 | -0.021442567 | 0.764629812 | 0.928757959 | 0.67949815 | 0.725754005 |
| Tstd2 | -0.019700211 | 0.61938277 | 0.8838136 | 0.097552136 | 0.119323493 |
| Tstd3 | 0.089338959 | 0.032074563 | 0.283890993 | 0.127772052 | 0.049709627 |
| Ttbk2 | 0.008290698 | 0.860449629 | 0.959592547 | 0.146002997 | 0.137348757 |
| Ttc1 | -0.024187988 | 0.691980846 | 0.909796728 | 0.32543122 | 0.352682814 |
| Ttc13 | 0.002192987 | 0.942172293 | 0.986270063 | 0.057879948 | 0.05448874 |
| Ttc14 | -0.079814591 | 0.258328919 | 0.665948842 | 0.66795038 | 0.742187951 |
| Ttc17 | -0.010790142 | 0.728827063 | 0.920522999 | 0.061392875 | 0.070488585 |
| Ttc19 | 0.084173564 | 0.172296085 | 0.575294174 | 0.345388669 | 0.266123808 |
| Ttc21b | -0.001246718 | 0.971483154 | 0.993892166 | 0.073002356 | 0.072763655 |
| Ttc26 | -0.014390468 | 0.683080867 | 0.909173483 | 0.07909792 | 0.087794172 |

| | | | | | |
|---|---|---|---|---|---|
| Ttc28 | 0.092469293 | 0.214674377 | 0.629611185 | 0.683246619 | 0.567475773 |
| Ttc3 | 0.023971575 | 0.727529124 | 0.920455328 | 0.896166102 | 0.867678516 |
| Ttc30b | -0.032598177 | 0.215735576 | 0.629630897 | 0.038232246 | 0.06286147 |
| Ttc32 | -0.011699718 | 0.804946867 | 0.94470747 | 0.147036368 | 0.152577316 |
| Ttc33 | -0.14192931 | 0.014462588 | 0.193677739 | 0.263030314 | 0.406498604 |
| Ttc37 | 0.000470208 | 0.993366451 | 0.998695538 | 0.23530278 | 0.24747872 |
| Ttc38 | -0.028721807 | 0.267564052 | 0.675673686 | 0.037261946 | 0.062428537 |
| Ttc39a | 0.042884065 | 0.456234836 | 0.808445095 | 0.269034087 | 0.224917199 |
| Ttc39b | -0.045111168 | 0.497046249 | 0.830056214 | 0.481504886 | 0.518217917 |
| Ttc39c | 0.031840381 | 0.324296645 | 0.720406991 | 0.069632621 | 0.039069471 |
| Ttc4 | -0.091972312 | 0.089101391 | 0.443953048 | 0.195793032 | 0.271888594 |
| Ttc5 | 0.07164832 | 0.118675143 | 0.491192747 | 0.156158463 | 0.088743503 |
| Ttc7 | -0.010745581 | 0.78871934 | 0.938142932 | 0.103869094 | 0.114853961 |
| Ttc7b | 0.0121189 | 0.745113751 | 0.924566547 | 0.087748892 | 0.075046598 |
| Ttc8 | -0.079672617 | 0.088700077 | 0.443367864 | 0.137599376 | 0.213342482 |
| Ttc9 | -0.024823594 | 0.551599786 | 0.852505562 | 0.116139389 | 0.132649541 |
| Ttc9c | -0.109172527 | 0.022279936 | 0.236913658 | 0.149967608 | 0.24551884 |
| Ttf1 | 0.045449559 | 0.392815311 | 0.769704269 | 0.208795578 | 0.165890768 |
| Tti1 | 0.056669912 | 0.105117351 | 0.472399448 | 0.088560176 | 0.042145646 |
| Tti2 | 0.013811507 | 0.715467346 | 0.916430034 | 0.095758979 | 0.089142434 |
| Ttl | 0.009292066 | 0.794860341 | 0.939757086 | 0.084479484 | 0.085116922 |
| Ttll1 | -0.02625359 | 0.483626848 | 0.822768706 | 0.088431122 | 0.114710615 |
| Ttll13 | -0.057524828 | 0.010426634 | 0.160509392 | 0.023031504 | 0.073524141 |
| Ttll5 | 0.029930651 | 0.301237344 | 0.706666217 | 0.055972951 | 0.027621797 |
| Ttll7 | -0.067816395 | 0.326241156 | 0.720866629 | 0.655480392 | 0.71937391 |
| Ttpal | -0.049501495 | 0.090364041 | 0.445385778 | 0.045541187 | 0.088047279 |
| Ttr | 0.158692292 | 0.002867313 | 0.074173238 | 2.740542726 | 2.338398576 |
| Ttyh1 | -0.056398273 | 0.34693905 | 0.738383053 | 0.22608471 | 0.289410683 |
| Ttyh2 | 0.080420518 | 0.17995916 | 0.586833233 | 0.295156723 | 0.211801384 |
| Tuba1a | 0.096146274 | 0.159209427 | 0.556789205 | 0.494293885 | 0.394634483 |
| Tuba1b | 0.037381513 | 0.497903794 | 0.830722138 | 0.219499952 | 0.19551491 |
| Tuba1c | -0.037916583 | 0.155183347 | 0.552356396 | 0.037013734 | 0.071357526 |
| Tuba4a | 0.01796018 | 0.78949073 | 0.93843184 | 0.469617158 | 0.456223817 |
| Tubb2a | -0.004510616 | 0.941671764 | 0.986168593 | 0.306575377 | 0.320653611 |
| Tubb3 | -0.101315811 | 0.070435116 | 0.40282465 | 0.201758702 | 0.292378173 |
| Tubb4b | -0.13516072 | 0.052022075 | 0.35690059 | 0.578063864 | 0.737119324 |
| Tubb5 | 0.067657855 | 0.302848592 | 0.708026508 | 0.423443572 | 0.362174135 |
| Tubb6 | 0.046400998 | 0.178179268 | 0.584581441 | 0.077335562 | 0.037174845 |
| Tubg1 | 0.077125373 | 0.167659262 | 0.570246024 | 0.250817943 | 0.191932646 |
| Tubg2 | 0.022457883 | 0.424503144 | 0.789923742 | 0.050709868 | 0.033678219 |
| Tubgcp2 | -0.00187808 | 0.954427374 | 0.989066902 | 0.069752336 | 0.067900801 |
| Tubgcp3 | 0.020619931 | 0.634617944 | 0.88965971 | 0.128413601 | 0.118923236 |
| Tubgcp4 | 0.016844937 | 0.625325908 | 0.88668351 | 0.074302484 | 0.064673048 |
| Tubgcp5 | -0.040868709 | 0.35951948 | 0.747719327 | 0.134738981 | 0.169117915 |
| Tubgcp6 | -0.013850735 | 0.621373108 | 0.884390511 | 0.0469657 | 0.053776581 |
| Tufm | 0.055747678 | 0.169026007 | 0.57227323 | 0.117054113 | 0.072331433 |
| Tuft1 | 0.068737521 | 0.093274869 | 0.45188518 | 0.118863991 | 0.06454215 |
| Tug1 | 0.029304247 | 0.6195124 | 0.883866495 | 0.284232882 | 0.257792923 |
| Tulp3 | 0.02943273 | 0.386218021 | 0.766272928 | 0.076565223 | 0.053189501 |
| Tulp4 | 0.001465802 | 0.982253685 | 0.995846488 | 0.910776481 | 0.903536295 |
| Tusc1 | -0.053891909 | 0.131723937 | 0.512050022 | 0.080355813 | 0.125755257 |
| Tusc2 | 0.157957143 | 0.00686158 | 0.126831819 | 0.314864921 | 0.17617528 |
| TUSC3 | 0.000136703 | 0.998363499 | 0.99938181 | 0.758249181 | 0.756006208 |
| Tut1 | 0.013439441 | 0.725482688 | 0.92014727 | 0.093996689 | 0.08476855 |
| Tut4 | -0.059659422 | 0.387826939 | 0.767160379 | 0.649492524 | 0.702638877 |
| Tut7 | 0.067660367 | 0.301697233 | 0.706960943 | 0.490384513 | 0.416253153 |
| Tvp23b | 0.032306503 | 0.62902887 | 0.887864718 | 0.500144183 | 0.471014002 |
| Twf1 | -0.076356905 | 0.264029314 | 0.670562304 | 0.620810878 | 0.704116241 |
| Twf2 | 0.009156163 | 0.87782771 | 0.966621692 | 0.275392009 | 0.26342704 |
| Twnk | -0.005398719 | 0.842991905 | 0.955306089 | 0.045885035 | 0.055076226 |
| Twsg1 | -0.023156345 | 0.740220665 | 0.922901128 | 1.023625831 | 1.020861839 |
| Txlna | -0.080732915 | 0.187836156 | 0.595675039 | 0.301463782 | 0.384135519 |
| Txlng | -0.074895616 | 0.113072614 | 0.483557659 | 0.152533743 | 0.216323735 |
| Txn1 | 0.076710043 | 0.276768909 | 0.684050613 | 0.942409114 | 0.834948076 |
| Txn2 | -0.076081083 | 0.244640462 | 0.654809396 | 0.509623645 | 0.607970315 |

| | | | | | |
|---|---|---|---|---|---|
| Txndc11 | 0.050058242 | 0.312145148 | 0.714322488 | 0.176978546 | 0.140030396 |
| Txndc12 | -0.063774548 | 0.302458252 | 0.707681683 | 0.325277934 | 0.402232834 |
| Txndc15 | -0.041953364 | 0.497107492 | 0.830056214 | 0.315983537 | 0.348096832 |
| Txndc16 | -0.015879708 | 0.667238192 | 0.901719367 | 0.084546962 | 0.099169862 |
| Txndc17 | 0.154662496 | 0.020491866 | 0.227971987 | 0.981940435 | 0.80293595 |
| Txndc5 | -0.127598519 | 0.026336615 | 0.257893268 | 0.245983928 | 0.36797745 |
| Txndc9 | 0.118077149 | 0.06605412 | 0.390680729 | 0.471470056 | 0.377305468 |
| Txnip | -0.391846853 | 1.60E-05 | 0.001832428 | 0.965103299 | 1.468377014 |
| Txnl1 | -0.031866683 | 0.616462567 | 0.883226807 | 1.120486696 | 1.172830113 |
| Txnl4a | 0.261091574 | 0.000176965 | 0.011841314 | 0.711115425 | 0.4481374 |
| Txnrd1 | -0.219201218 | 0.000576065 | 0.027869893 | 1.163967258 | 1.500555974 |
| Txnrd2 | 0.008868604 | 0.813855514 | 0.948397417 | 0.092918626 | 0.080838211 |
| Tyk2 | -0.01534609 | 0.590773935 | 0.871005673 | 0.048399356 | 0.063634379 |
| Tyms | -0.068300481 | 0.001639906 | 0.053297575 | 0.017852366 | 0.076783362 |
| Tysnd1 | -0.029898358 | 0.478569038 | 0.819081437 | 0.113492973 | 0.150980234 |
| Tyw1 | -0.071726551 | 0.065345161 | 0.389471129 | 0.093236578 | 0.162049723 |
| Tyw5 | 0.003711259 | 0.914114619 | 0.980132949 | 0.07155929 | 0.064785346 |
| U2af1 | 0.031320278 | 0.645415505 | 0.893892014 | 0.586811378 | 0.566508726 |
| U2af2 | 0.063758642 | 0.308119894 | 0.710615665 | 0.373957762 | 0.312845723 |
| U2surp | 0.138252475 | 0.056770304 | 0.368804883 | 0.978563837 | 0.813340488 |
| Uaca | 0.08486796 | 0.203299972 | 0.614800187 | 0.477123356 | 0.393912509 |
| Uap1 | 0.081144839 | 0.105944507 | 0.473027145 | 0.18832509 | 0.122294262 |
| Uap1l1 | 0.063922499 | 0.319510226 | 0.717256849 | 0.367303207 | 0.286215684 |
| Uba1 | -0.042401514 | 0.524668533 | 0.841229722 | 0.445991595 | 0.505869342 |
| Uba2 | -0.027889794 | 0.631179791 | 0.888645191 | 0.284949544 | 0.320664322 |
| Uba3 | 0.067307905 | 0.221744469 | 0.633858837 | 0.245555308 | 0.192798269 |
| Uba5 | 0.127527339 | 0.056295541 | 0.367897032 | 0.452667904 | 0.338027419 |
| Uba52 | 0.21067616 | 0.000232232 | 0.014633967 | 1.581861115 | 1.204385491 |
| Uba6 | -0.062989028 | 0.268721249 | 0.676657093 | 0.230888759 | 0.29660433 |
| Ubac1 | 0.009533002 | 0.846153626 | 0.956185988 | 0.1615992 | 0.154395081 |
| Ubac2 | 0.062716655 | 0.046596851 | 0.338417216 | 0.072195761 | 0.019843358 |
| Ubald1 | 0.057336103 | 0.259365582 | 0.666816687 | 0.192658293 | 0.145789614 |
| Ubald2 | -0.069960127 | 0.141447888 | 0.528664246 | 0.150861239 | 0.222680917 |
| Ubap1 | -0.00153476 | 0.975300877 | 0.994455356 | 0.163911983 | 0.166883059 |
| Ubap2 | 0.035507804 | 0.545164715 | 0.851259637 | 0.279025139 | 0.235998899 |
| Ubap2l | 0.071386241 | 0.304835942 | 0.709410087 | 0.874409308 | 0.795653105 |
| Ubb | -0.087727765 | 0.074184484 | 0.41251505 | 2.461848092 | 2.647197111 |
| Ubb-ps | 0.061195673 | 0.136198189 | 0.519952162 | 0.119579332 | 0.066497619 |
| Ubc | -0.097635437 | 0.031881609 | 0.283462667 | 2.410448934 | 2.656907037 |
| Ube2a | -0.013475124 | 0.83333191 | 0.954303562 | 0.399331256 | 0.405814105 |
| Ube2b | -0.03795759 | 0.567764193 | 0.859654428 | 1.175305954 | 1.237337097 |
| Ube2d1 | -0.032527647 | 0.491951558 | 0.827270742 | 0.153003221 | 0.178762911 |
| Ube2d2a | 0.164388561 | 0.012026512 | 0.174689004 | 0.753635292 | 0.590474912 |
| Ube2d3 | 0.065132612 | 0.281925043 | 0.688868362 | 1.32973864 | 1.277857037 |
| Ube2d-ps | 0.050993255 | 0.261285143 | 0.668468258 | 0.147233291 | 0.104183065 |
| Ube2e1 | -0.181037749 | 0.006218492 | 0.118659697 | 0.467802666 | 0.654323922 |
| Ube2e2 | 0.092171681 | 0.049529009 | 0.349348321 | 0.162382522 | 0.089325904 |
| Ube2e3 | 0.030714583 | 0.643091018 | 0.892561795 | 0.624041007 | 0.608479069 |
| Ube2f | 0.062891385 | 0.300776437 | 0.706227193 | 0.332292229 | 0.270760945 |
| Ube2g1 | -0.056330014 | 0.269583117 | 0.677215248 | 0.180702882 | 0.232117389 |
| Ube2g2 | 0.001287353 | 0.982590475 | 0.995921574 | 0.284074106 | 0.301068197 |
| Ube2h | 0.003617003 | 0.9563634 | 0.989397512 | 0.533891167 | 0.519327994 |
| Ube2i | 0.159183966 | 0.020626719 | 0.227971987 | 0.696300816 | 0.523807491 |
| Ube2j1 | 0.039707243 | 0.528714698 | 0.844309234 | 0.383541324 | 0.34005196 |
| Ube2j2 | 0.013765676 | 0.830416741 | 0.95325696 | 0.421391022 | 0.41973904 |
| Ube2k | 0.015576698 | 0.816749597 | 0.94911078 | 0.838401714 | 0.837333955 |
| Ube2l3 | -0.076247735 | 0.262111393 | 0.668764912 | 0.663776015 | 0.767890615 |
| Ube2l6 | 0.026109176 | 0.425066052 | 0.790344696 | 0.068653159 | 0.042993673 |
| Ube2m | -0.031979934 | 0.648136818 | 0.895379629 | 0.62929837 | 0.669183697 |
| Ube2n | 0.01290947 | 0.84349765 | 0.95544362 | 0.508697777 | 0.519826063 |
| Ube2o | 0.107072014 | 0.10935704 | 0.477972625 | 0.471144112 | 0.372096047 |
| Ube2q1 | -0.10767953 | 0.088953378 | 0.443672569 | 0.403301111 | 0.507223733 |
| Ube2q2 | -0.057477969 | 0.354927246 | 0.745282523 | 0.366180203 | 0.413618673 |
| Ube2r2 | -0.026735013 | 0.694787654 | 0.910242368 | 0.718854164 | 0.776728779 |
| Ube2s | 0.028634467 | 0.334190356 | 0.729969081 | 0.059588124 | 0.035766474 |

| | | | | | |
|---|---|---|---|---|---|
| Ube2v1 | 0.059622775 | 0.04541276 | 0.333925179 | 0.061050568 | 0.006913716 |
| Ube2v2 | -0.010030238 | 0.868549159 | 0.96295459 | 0.322821776 | 0.334113897 |
| Ube2w | -0.07016098 | 0.28608893 | 0.693090656 | 0.414879339 | 0.490788064 |
| Ube2z | 0.08876986 | 0.164322104 | 0.563869641 | 0.388895621 | 0.295296878 |
| Ube3a | 0.084030582 | 0.170589019 | 0.572831081 | 1.085557011 | 0.985269486 |
| Ube3b | -0.006230512 | 0.874794119 | 0.96500153 | 0.101028372 | 0.107664149 |
| Ube3c | 0.02516991 | 0.692519045 | 0.909796728 | 0.38212887 | 0.345944094 |
| Ube4a | 0.092308056 | 0.077671542 | 0.418285047 | 0.201820103 | 0.123950819 |
| Ube4b | 0.058749017 | 0.371721159 | 0.75760494 | 0.508675096 | 0.436196638 |
| Ubfd1 | -0.134027431 | 0.015330976 | 0.198807499 | 0.215354182 | 0.338341861 |
| Ubl3 | -0.026680548 | 0.635275856 | 0.889918181 | 1.437698919 | 1.477272512 |
| Ubl4a | -0.038082722 | 0.537522445 | 0.847793898 | 0.376845735 | 0.42318349 |
| Ubl5 | 0.05370571 | 0.266211728 | 0.673899077 | 1.579235258 | 1.438520481 |
| Ubl7 | -0.063112701 | 0.112645398 | 0.483221516 | 0.095837209 | 0.153868887 |
| Ublcp1 | -0.033629439 | 0.438692623 | 0.799411376 | 0.127037041 | 0.156166336 |
| Ubn1 | 0.085143987 | 0.22735873 | 0.639441223 | 0.67767835 | 0.592314775 |
| Ubn2 | -0.223824689 | 0.001291954 | 0.045499435 | 0.740723197 | 1.001364115 |
| Ubp1 | 0.012820589 | 0.816138058 | 0.949036482 | 0.236938359 | 0.217518312 |
| Ubqln1 | 0.046422467 | 0.483164868 | 0.822511157 | 0.502458367 | 0.443992496 |
| Ubqln2 | -0.085041863 | 0.200908954 | 0.611438197 | 0.469741168 | 0.586027265 |
| Ubqln4 | -0.0390309 | 0.356950324 | 0.745931793 | 0.114422696 | 0.147501397 |
| Ubr1 | 0.024174527 | 0.66636208 | 0.901197061 | 0.248395267 | 0.229359737 |
| Ubr2 | 0.002300107 | 0.973570642 | 0.994079882 | 0.681952387 | 0.68042898 |
| Ubr3 | -0.017584653 | 0.789965568 | 0.938440574 | 0.533580443 | 0.557147475 |
| Ubr4 | -0.183204123 | 0.007148153 | 0.128637347 | 0.936028001 | 1.151754572 |
| Ubr5 | -0.007696329 | 0.909763827 | 0.978578013 | 0.615094492 | 0.622033736 |
| Ubr7 | 0.023888116 | 0.656000076 | 0.89789619 | 0.216308592 | 0.196171394 |
| Ubtd1 | 0.044722475 | 0.184833241 | 0.591385002 | 0.078054662 | 0.046354292 |
| Ubtd2 | 0.046897837 | 0.146936668 | 0.536877806 | 0.071989126 | 0.033808564 |
| Ubtf | -0.014469552 | 0.836787043 | 0.954911647 | 0.63016378 | 0.679673383 |
| Ubxn1 | -0.025192298 | 0.703854078 | 0.912277105 | 0.911240402 | 0.92196702 |
| Ubxn10 | -0.006209011 | 0.880504386 | 0.967810606 | 0.112672014 | 0.114775674 |
| Ubxn2a | 0.129261261 | 0.071605598 | 0.40513269 | 0.712314147 | 0.584047068 |
| Ubxn2b | -0.037297484 | 0.295974458 | 0.702458178 | 0.080859326 | 0.114462949 |
| Ubxn4 | -0.028525099 | 0.629004107 | 0.887864718 | 1.354770545 | 1.379342545 |
| Ubxn6 | -0.256391778 | 2.57E-05 | 0.00273918 | 0.30711368 | 0.579988772 |
| Ubxn7 | 0.001236467 | 0.980725233 | 0.995540893 | 0.180592948 | 0.186517842 |
| Ubxn8 | -0.03272365 | 0.31302384 | 0.714906079 | 0.060881445 | 0.089659554 |
| Uchl1 | -0.086882189 | 0.000395953 | 0.021482072 | 0.023473572 | 0.098605992 |
| Uchl3 | -0.09006034 | 0.115296846 | 0.487260227 | 0.214946426 | 0.31011638 |
| Uchl5 | 0.078956747 | 0.181509122 | 0.589059475 | 0.275050592 | 0.208371609 |
| Uck1 | -0.030656309 | 0.512443062 | 0.835780144 | 0.146207835 | 0.179416008 |
| Uck2 | 0.038167898 | 0.48562288 | 0.824318763 | 0.238480815 | 0.207502868 |
| Uckl1 | -0.059816533 | 0.230148094 | 0.641131395 | 0.178143106 | 0.230829057 |
| Ucn3 | 0.18093734 | 0.021363078 | 0.232117183 | 0.974152133 | 0.751796273 |
| Ucp2 | -0.187799103 | 0.007561293 | 0.133456507 | 0.631369991 | 0.829450724 |
| Ufc1 | -0.063422474 | 0.315718405 | 0.714975653 | 0.991971692 | 1.067562756 |
| Ufd1 | -0.00167966 | 0.978169721 | 0.995031396 | 0.348279465 | 0.341984283 |
| Ufl1 | -0.128488597 | 0.061934236 | 0.380250374 | 0.515801757 | 0.645180122 |
| Ufm1 | 0.032216494 | 0.576819352 | 0.863980874 | 1.175259843 | 1.092150766 |
| Ufsp1 | 0.01028781 | 0.764146408 | 0.928730554 | 0.075008864 | 0.071865142 |
| Ufsp2 | -0.15640386 | 0.019722826 | 0.223934391 | 0.561163222 | 0.710115211 |
| Ugcg | 0.034645022 | 0.482143596 | 0.821803219 | 0.176140266 | 0.145192188 |
| Ugdh | -0.144828176 | 0.001959484 | 0.0606747 | 0.128659993 | 0.27072261 |
| Uggt1 | 0.046184592 | 0.484042564 | 0.822768706 | 0.516234613 | 0.468537768 |
| Uggt2 | -0.021265455 | 0.527347204 | 0.842966059 | 0.070105198 | 0.090761134 |
| Ugp2 | -0.067053203 | 0.217175648 | 0.631640679 | 0.212507777 | 0.282533991 |
| Ugt2b34 | -0.214563548 | 0.000587047 | 0.028192291 | 0.263213693 | 0.498736122 |
| Uhmk1 | 0.16683531 | 0.004172802 | 0.092660198 | 0.311506985 | 0.161100661 |
| Uhrf1bp1 | 0.020370712 | 0.608543859 | 0.878855197 | 0.106064804 | 0.085903241 |
| Uhrf1bp1l | 0.097157807 | 0.103650987 | 0.469469748 | 0.295115494 | 0.195959092 |
| Uhrf2 | 0.009910426 | 0.871658409 | 0.964616742 | 0.342872379 | 0.343651175 |
| Uimc1 | -0.020532374 | 0.703245557 | 0.912277105 | 0.207621104 | 0.219537729 |
| Ulk1 | -0.048512581 | 0.259729378 | 0.667102369 | 0.112392125 | 0.159193408 |
| Ulk2 | -0.084585236 | 0.10089067 | 0.466401306 | 0.187439917 | 0.261980285 |

| | | | | | |
|---|---|---|---|---|---|
| Ulk3 | -0.02669405 | 0.286563311 | 0.693329451 | 0.034568263 | 0.056795822 |
| Umad1 | 0.000136839 | 0.997481187 | 0.999369645 | 0.123148636 | 0.133446931 |
| Umps | 0.060680912 | 0.28134642 | 0.68811697 | 0.265345765 | 0.220931912 |
| Unc119 | 0.042834591 | 0.355246313 | 0.745326231 | 0.159035046 | 0.117189455 |
| Unc119b | -0.027220523 | 0.603465505 | 0.876723966 | 0.196497244 | 0.203641491 |
| Unc13a | 0.038101517 | 0.478880047 | 0.819246469 | 0.221929982 | 0.1683454 |
| Unc13b | -0.057718289 | 0.111719477 | 0.482690499 | 0.079698354 | 0.12678627 |
| Unc45a | -0.001338068 | 0.981433861 | 0.995719363 | 0.241367836 | 0.231705746 |
| Unc50 | 0.014499984 | 0.82306123 | 0.950930859 | 0.438623317 | 0.418094361 |
| Unc5a | -0.023117825 | 0.559668437 | 0.856904336 | 0.099050892 | 0.12326455 |
| Unc5b | -0.027986048 | 0.233112339 | 0.643935869 | 0.028481602 | 0.054632527 |
| Unc5c | 0.03648282 | 0.254776911 | 0.66334523 | 0.065376224 | 0.029123647 |
| Unc79 | -0.027155414 | 0.519448575 | 0.838496482 | 0.115628471 | 0.14606682 |
| Unc80 | -0.188680776 | 0.006436811 | 0.121212521 | 0.616638037 | 0.833346174 |
| Unc93b1 | -0.024745225 | 0.564057969 | 0.857926031 | 0.122389636 | 0.155643007 |
| Unk | -0.017089969 | 0.578211207 | 0.864775129 | 0.058386486 | 0.072704503 |
| Upb1 | -0.029575776 | 0.665495874 | 0.901006558 | 0.576928893 | 0.598333509 |
| Upf1 | -0.06639544 | 0.289014084 | 0.695472148 | 0.313393234 | 0.376064687 |
| Upf2 | -0.026169307 | 0.666352998 | 0.901197061 | 0.300558534 | 0.314793107 |
| Upf3a | -0.036051218 | 0.469353385 | 0.814050152 | 0.167278372 | 0.197043561 |
| Upf3b | -0.004484512 | 0.948220254 | 0.987651726 | 0.71602653 | 0.682643745 |
| Upp1 | 0.054432483 | 0.330562447 | 0.725898152 | 0.239530683 | 0.190374759 |
| Upp2 | 0.00716963 | 0.871099914 | 0.964504119 | 0.137287212 | 0.124215541 |
| Uqcc1 | -0.029116529 | 0.54679958 | 0.851446616 | 0.152911017 | 0.183410898 |
| Uqcc2 | 0.134112868 | 0.002217399 | 0.064615296 | 2.184381336 | 1.911057189 |
| Uqcc3 | 0.05197372 | 0.436585571 | 0.797879526 | 0.517866711 | 0.460843656 |
| Uqcr10 | -0.011134936 | 0.826437453 | 0.952566917 | 1.717820726 | 1.695944787 |
| Uqcr11 | -0.027270179 | 0.615694256 | 0.882439418 | 1.542991115 | 1.564008605 |
| Uqcrb | -0.02548928 | 0.624828827 | 0.88668351 | 1.572879459 | 1.593948327 |
| Uqcrc1 | -0.104475787 | 0.106790209 | 0.474004524 | 0.419192465 | 0.510198747 |
| Uqcrc2 | -0.097407807 | 0.128734497 | 0.508260084 | 0.709923047 | 0.818639639 |
| Uqcrfs1 | -0.056308611 | 0.387847324 | 0.767160379 | 0.793439494 | 0.862439224 |
| Uqcrh | 0.040740851 | 0.399598425 | 0.774544012 | 1.716432508 | 1.636271924 |
| Uqcrq | -0.094619305 | 0.007693298 | 0.134995868 | 2.51869067 | 2.724815231 |
| Urb1 | 0.040921112 | 0.196199321 | 0.606881632 | 0.066146411 | 0.029111181 |
| Urb2 | -0.03301483 | 0.179107625 | 0.585380409 | 0.032701154 | 0.060370697 |
| Urgcp | 0.008660686 | 0.858626653 | 0.958837975 | 0.166956786 | 0.158944807 |
| Uri1 | -0.18333883 | 0.007308872 | 0.130069404 | 0.616704837 | 0.813259776 |
| Urm1 | -0.103211348 | 0.043402524 | 0.329026536 | 0.17410265 | 0.265786066 |
| Urod | -0.09196896 | 0.040703799 | 0.319175023 | 0.134839194 | 0.219988269 |
| Uros | -0.006506423 | 0.838299141 | 0.954911647 | 0.065046519 | 0.074641024 |
| Use1 | -0.211710961 | 0.00188743 | 0.059335578 | 0.632680461 | 0.867255991 |
| Usf1 | 0.00030197 | 0.995441367 | 0.99872501 | 0.205081415 | 0.210400257 |
| Usf2 | 0.00624654 | 0.924694319 | 0.981375217 | 0.451405383 | 0.44765773 |
| Usf3 | -0.035956373 | 0.441852655 | 0.800518971 | 0.146085099 | 0.179969613 |
| Ush1c | -0.035683031 | 0.523797332 | 0.841062594 | 0.238910724 | 0.272275241 |
| Uso1 | 0.037625779 | 0.57155187 | 0.861404566 | 0.84675959 | 0.80978922 |
| Usp1 | -0.107305819 | 0.053321053 | 0.360635718 | 0.209699157 | 0.31266192 |
| Usp10 | 0.063245427 | 0.273839399 | 0.681791282 | 0.278587253 | 0.219255628 |
| Usp11 | -0.000173412 | 0.99521187 | 0.99872501 | 0.05155904 | 0.051741619 |
| Usp12 | 0.012255805 | 0.822141138 | 0.950930859 | 0.223639659 | 0.212392786 |
| Usp14 | 0.111600869 | 0.095301378 | 0.455812225 | 0.510392619 | 0.392770955 |
| Usp15 | 0.018815954 | 0.776250324 | 0.933751738 | 0.460919893 | 0.453810107 |
| Usp16 | 0.078406607 | 0.251930046 | 0.663127858 | 0.742198827 | 0.650842774 |
| Usp18 | 0.01003171 | 0.793609268 | 0.939757086 | 0.09436959 | 0.083586227 |
| Usp19 | -0.039977601 | 0.529269145 | 0.844323583 | 0.384153401 | 0.406659546 |
| Usp2 | 0.038475556 | 0.193447649 | 0.60325934 | 0.057846077 | 0.021523517 |
| Usp20 | -0.028427526 | 0.568685029 | 0.860399704 | 0.177271829 | 0.200915088 |
| Usp21 | -0.003034643 | 0.918756466 | 0.980385833 | 0.055424202 | 0.055001103 |
| Usp22 | -0.023200284 | 0.709662182 | 0.914361916 | 0.355229275 | 0.399974163 |
| Usp24 | 0.079178155 | 0.136576453 | 0.520249727 | 0.217276103 | 0.144881885 |
| Usp25 | -0.017971921 | 0.654201596 | 0.897454058 | 0.105640446 | 0.118963481 |
| Usp29 | 0.057824099 | 0.331229489 | 0.727045202 | 0.303839168 | 0.240870062 |
| Usp3 | 0.108533988 | 0.090089989 | 0.445385778 | 0.402454662 | 0.299125546 |
| Usp30 | -0.018865478 | 0.60189661 | 0.876557788 | 0.085120553 | 0.098405353 |

| | | | | | |
|---|---|---|---|---|---|
| Usp31 | -0.028120109 | 0.587342053 | 0.869235061 | 0.185085588 | 0.207862789 |
| Usp32 | 0.045047935 | 0.293580278 | 0.69925611 | 0.12927187 | 0.090382314 |
| Usp33 | 0.057616854 | 0.396690729 | 0.773573897 | 0.521187258 | 0.459341706 |
| Usp34 | -0.071211722 | 0.277830985 | 0.684996416 | 0.830561792 | 0.927718743 |
| Usp35 | 0.047477491 | 0.421560094 | 0.787357117 | 0.289835005 | 0.233559348 |
| Usp36 | -0.039128926 | 0.510843605 | 0.834957869 | 0.26570039 | 0.302286365 |
| Usp37 | 0.03882706 | 0.457128319 | 0.808898849 | 0.202388911 | 0.173331919 |
| Usp38 | -0.105461785 | 0.029500009 | 0.27358919 | 0.148092759 | 0.242781024 |
| Usp39 | 0.06579168 | 0.141196356 | 0.528664246 | 0.143487861 | 0.092185273 |
| Usp4 | -0.008935158 | 0.872636957 | 0.964911254 | 0.232522994 | 0.243916441 |
| Usp40 | -0.036981182 | 0.374869218 | 0.760042036 | 0.11043731 | 0.145628706 |
| Usp42 | 0.040687903 | 0.371352807 | 0.75760494 | 0.13954766 | 0.106006238 |
| Usp45 | 0.132520412 | 0.023603036 | 0.244234094 | 0.287204832 | 0.159086419 |
| Usp46 | 0.052117689 | 0.378864307 | 0.762006446 | 0.3149293 | 0.255634265 |
| Usp47 | -0.053577002 | 0.421565921 | 0.787357117 | 0.519725501 | 0.561122557 |
| Usp48 | -0.025766498 | 0.673008546 | 0.905559579 | 0.346657861 | 0.355852462 |
| Usp5 | -0.045407833 | 0.376039412 | 0.760642399 | 0.177541675 | 0.218944528 |
| Usp50 | 0.044556957 | 0.474878238 | 0.817387411 | 0.332097388 | 0.282781203 |
| Usp51 | 0.0285937 | 0.3502056 | 0.740782636 | 0.059770232 | 0.032961161 |
| Usp53 | 0.02473326 | 0.606360747 | 0.878341504 | 0.151800295 | 0.130089709 |
| Usp54 | -0.041670064 | 0.057302596 | 0.36957055 | 0.02166464 | 0.057209138 |
| Usp6nl | -0.048585347 | 0.258751074 | 0.665948842 | 0.114698069 | 0.160734892 |
| Usp7 | 0.040891585 | 0.543725564 | 0.850865857 | 0.503083777 | 0.464709196 |
| Usp8 | 0.007785577 | 0.904789289 | 0.975652137 | 0.393080374 | 0.384154771 |
| Usp9x | -0.084345648 | 0.218394155 | 0.632618221 | 0.96054313 | 1.05843402 |
| Uspl1 | 0.070466084 | 0.228626296 | 0.640268452 | 0.26523561 | 0.194985765 |
| Ust | 0.032708047 | 0.342744248 | 0.735501771 | 0.08333897 | 0.060235877 |
| Utp11 | 0.067153283 | 0.243171956 | 0.653166476 | 0.249918944 | 0.202624539 |
| Utp14a | -0.06534534 | 0.160117919 | 0.558431893 | 0.134633872 | 0.202642416 |
| Utp14b | -0.037559411 | 0.123820636 | 0.50152854 | 0.029101934 | 0.066514937 |
| Utp15 | 0.013718469 | 0.745744604 | 0.924566547 | 0.11992236 | 0.103486781 |
| Utp18 | 0.080848437 | 0.173137082 | 0.575613413 | 0.303367756 | 0.22518573 |
| Utp20 | -0.018451832 | 0.566741433 | 0.858864944 | 0.061366766 | 0.076042877 |
| Utp23 | -0.03212862 | 0.467885173 | 0.813188482 | 0.129850961 | 0.15820656 |
| Utp25 | -0.080341249 | 0.016599291 | 0.206965324 | 0.060029913 | 0.13585962 |
| Utp3 | 0.095854194 | 0.150556279 | 0.545732529 | 0.519609813 | 0.413398929 |
| Utp4 | -0.013845925 | 0.731423546 | 0.920770724 | 0.105493573 | 0.128903976 |
| Utp6 | -0.023269477 | 0.580686517 | 0.866173066 | 0.113412506 | 0.132153042 |
| Utrn | 0.088274492 | 0.193294298 | 0.60325934 | 0.528911758 | 0.454331933 |
| Uty | -0.051727642 | 0.286380711 | 0.693329451 | 0.164502959 | 0.20502945 |
| Uvrag | 0.045551291 | 0.399268454 | 0.774544012 | 0.231623265 | 0.193565208 |
| Uvssa | -0.019732409 | 0.735513814 | 0.922102442 | 0.260240932 | 0.270052871 |
| Uxs1 | 0.030990908 | 0.451170635 | 0.806495742 | 0.118739249 | 0.095940257 |
| Uxt | -0.077947291 | 0.166853898 | 0.568374737 | 0.243767383 | 0.31439881 |
| Vac14 | 0.018264948 | 0.639020956 | 0.891345382 | 0.094617398 | 0.081082956 |
| Vamp1 | -0.028048665 | 0.237518735 | 0.648678718 | 0.029270239 | 0.058058154 |
| Vamp2 | -0.141888966 | 0.038018745 | 0.307489238 | 0.675082009 | 0.863517657 |
| Vamp3 | 0.014828411 | 0.803620065 | 0.943715291 | 0.341995147 | 0.345848786 |
| Vamp4 | 0.0222928 | 0.721314323 | 0.919475778 | 0.427389422 | 0.410717361 |
| Vamp5 | -0.012859392 | 0.632215089 | 0.888980506 | 0.04026869 | 0.049113544 |
| Vamp8 | 0.031852758 | 0.603305712 | 0.876723966 | 1.080296169 | 1.048106188 |
| Vapa | -0.003337218 | 0.957879083 | 0.990137214 | 0.948122483 | 1.003939768 |
| Vapb | -0.058199153 | 0.371756312 | 0.75760494 | 1.086545094 | 1.18509339 |
| Vars | -0.196088327 | 0.000320045 | 0.018285574 | 0.201254495 | 0.38149114 |
| Vars2 | -0.040890859 | 0.168971316 | 0.57227323 | 0.050811334 | 0.086494313 |
| Vasp | -0.006465063 | 0.899347078 | 0.97386413 | 0.185897218 | 0.187702141 |
| Vat1 | 0.152348054 | 0.016379983 | 0.206355375 | 0.392318819 | 0.251467415 |
| Vat1l | -0.014757152 | 0.759067509 | 0.927719223 | 0.156941102 | 0.159107443 |
| Vbp1 | 0.023874242 | 0.697791753 | 0.910970095 | 0.384325513 | 0.360426335 |
| Vcl | -0.018450103 | 0.714949081 | 0.916116931 | 0.175116384 | 0.192088314 |
| Vcp | -0.10384856 | 0.085311767 | 0.434941698 | 1.131930039 | 1.312402304 |
| Vcpip1 | -0.004159751 | 0.945734101 | 0.987084873 | 0.327035229 | 0.347010988 |
| Vdac1 | 0.036019652 | 0.605540227 | 0.87802905 | 0.96705092 | 0.907154933 |
| Vdac2 | -0.039552197 | 0.551916025 | 0.852505562 | 0.844851462 | 0.893350368 |
| Vdac3 | -0.253935593 | 0.000200597 | 0.012906373 | 0.753635552 | 1.0674157 |

| | | | | | |
|---|---|---|---|---|---|
| Vdr | 0.076919211 | 0.270052499 | 0.677226912 | 0.529236283 | 0.445267262 |
| Vegfa | 0.029230919 | 0.63038445 | 0.888397743 | 1.362672096 | 1.345145061 |
| Vegfb | 0.011759642 | 0.840064368 | 0.954911647 | 0.303249172 | 0.289538826 |
| Vezf1 | 0.004746717 | 0.94452578 | 0.987070339 | 0.631653201 | 0.619715442 |
| Vezt | 0.03301274 | 0.451837901 | 0.806495742 | 0.132896176 | 0.109979528 |
| Vgf | -0.249759248 | 0.000149693 | 0.010687095 | 0.289764825 | 0.538930254 |
| Vgll4 | 0.027925487 | 0.523885484 | 0.841062594 | 0.13216458 | 0.107260574 |
| Vhl | 0.041426644 | 0.332264994 | 0.72806771 | 0.128642202 | 0.092160946 |
| Vil1 | 0.031050702 | 0.599517145 | 0.875897759 | 0.276776349 | 0.242446438 |
| Vipas39 | 0.036973439 | 0.427090379 | 0.79105114 | 0.153082568 | 0.118159799 |
| Vipr1 | 0.029833588 | 0.555011512 | 0.853608727 | 0.174122262 | 0.150571334 |
| Virma | 0.013944341 | 0.804351802 | 0.944353613 | 0.258566473 | 0.255438499 |
| Vkorc1 | -0.115441852 | 0.065491552 | 0.389491953 | 0.368234471 | 0.500056413 |
| Vkorc1l1 | 0.041409198 | 0.468568466 | 0.813530823 | 0.254579762 | 0.215341581 |
| Vldlr | 0.015454532 | 0.823080092 | 0.950930859 | 0.653988794 | 0.625379109 |
| Vma21 | 0.043150168 | 0.490380424 | 0.826674417 | 0.348149812 | 0.313527451 |
| Vmac | -0.038299436 | 0.443287532 | 0.801368502 | 0.174752118 | 0.210990865 |
| Vmp1 | -0.015905153 | 0.810016444 | 0.947942154 | 0.418799022 | 0.44981402 |
| Vopp1 | 0.014948907 | 0.59978085 | 0.875897759 | 0.053250317 | 0.038179303 |
| Vps11 | 0.084922132 | 0.112851649 | 0.483221516 | 0.221279782 | 0.147595029 |
| Vps13a | -0.065617289 | 0.304822403 | 0.709410087 | 0.390486034 | 0.453320238 |
| Vps13b | -0.182967789 | 0.000379529 | 0.020930399 | 0.165063956 | 0.326535734 |
| Vps13c | -0.132086313 | 0.039452977 | 0.313282855 | 0.371718023 | 0.495602275 |
| Vps13d | 0.009965625 | 0.819174726 | 0.950128492 | 0.128943804 | 0.115357158 |
| Vps16 | -0.053963144 | 0.073970882 | 0.412240835 | 0.053605703 | 0.10361795 |
| Vps18 | 0.016887764 | 0.709631691 | 0.914361916 | 0.139597852 | 0.132247012 |
| Vps25 | -0.041611223 | 0.053264086 | 0.360493347 | 0.020516486 | 0.055882737 |
| Vps26a | -0.005109041 | 0.918300358 | 0.980385833 | 0.170677363 | 0.174150554 |
| Vps26b | -0.065455674 | 0.203623598 | 0.614950615 | 0.1908458 | 0.252581902 |
| Vps26c | 0.070299004 | 0.106706778 | 0.474004524 | 0.138467463 | 0.078543578 |
| Vps28 | -0.0609826 | 0.354931798 | 0.745282523 | 0.546706603 | 0.62634447 |
| Vps29 | -0.050855067 | 0.441027602 | 0.800354711 | 0.669517171 | 0.709777857 |
| Vps33a | 0.012096738 | 0.742703918 | 0.923846726 | 0.090618787 | 0.080462417 |
| Vps33b | -0.009146992 | 0.769783176 | 0.931407628 | 0.062917012 | 0.070445624 |
| Vps35 | 0.055933932 | 0.380072399 | 0.762006446 | 0.918809471 | 0.841250642 |
| Vps35l | -0.111086201 | 0.013543595 | 0.189064067 | 0.126276762 | 0.220882336 |
| Vps36 | -0.006032972 | 0.923602522 | 0.980869592 | 0.380184604 | 0.387320883 |
| Vps37a | 0.164453429 | 0.015586297 | 0.201079258 | 0.972447464 | 0.79846881 |
| Vps37b | -0.005007707 | 0.920679156 | 0.980462248 | 0.169094704 | 0.183095526 |
| Vps37c | 0.027947501 | 0.582118865 | 0.866173066 | 0.176884467 | 0.154782206 |
| Vps37d | 0.016771862 | 0.628615608 | 0.887864718 | 0.07584178 | 0.060031018 |
| Vps39 | -0.022890523 | 0.617777175 | 0.883238152 | 0.140038841 | 0.158465981 |
| Vps41 | -0.03514658 | 0.552012402 | 0.852523231 | 0.287317337 | 0.321893856 |
| Vps45 | -0.071803458 | 0.120659368 | 0.494691316 | 0.134166651 | 0.19395656 |
| Vps4a | -0.032229744 | 0.586872412 | 0.869180817 | 0.286970393 | 0.315088211 |
| Vps4b | -0.076160391 | 0.252090084 | 0.663127858 | 0.603141885 | 0.658824766 |
| Vps50 | 0.095003388 | 0.069057755 | 0.399753362 | 0.203207675 | 0.122849221 |
| Vps51 | -0.07335403 | 0.153323577 | 0.549817979 | 0.180760602 | 0.228409116 |
| Vps52 | 0.026764103 | 0.602679269 | 0.876632118 | 0.197968128 | 0.170135862 |
| Vps53 | 0.043368679 | 0.477529537 | 0.818492016 | 0.351209708 | 0.311908834 |
| Vps54 | -0.051582633 | 0.243383742 | 0.653166476 | 0.130072503 | 0.169306702 |
| Vps72 | -0.05201256 | 0.301267783 | 0.706666217 | 0.17152295 | 0.225873286 |
| Vps8 | 0.151961056 | 0.010282468 | 0.159021773 | 0.279381705 | 0.141645307 |
| Vps9d1 | 0.067652723 | 0.076687712 | 0.418199494 | 0.104603005 | 0.04909938 |
| Vrk1 | 0.046082118 | 0.441199022 | 0.800354711 | 0.314630854 | 0.266507121 |
| Vrk2 | -0.028070818 | 0.188447644 | 0.596671608 | 0.02215998 | 0.044905019 |
| Vrk3 | 0.006052464 | 0.899115098 | 0.973823054 | 0.150505695 | 0.146100847 |
| Vsig10 | 0.008626648 | 0.84299552 | 0.955306089 | 0.125343896 | 0.112568869 |
| Vsnl1 | -0.010968324 | 0.654220034 | 0.897454058 | 0.033816219 | 0.043694378 |
| Vstm2l | 0.073048088 | 0.265201239 | 0.672177989 | 0.523951299 | 0.464799742 |
| Vta1 | -0.06049387 | 0.319850367 | 0.717256849 | 0.331651152 | 0.381478367 |
| Vti1a | -0.114086724 | 0.004552661 | 0.099436985 | 0.095635433 | 0.193551351 |
| Vti1b | -0.102576796 | 0.131163621 | 0.511059496 | 0.570990968 | 0.694585736 |
| Vwa5a | 0.058901965 | 0.191739043 | 0.602531237 | 0.140424249 | 0.081266495 |
| Vwa5b2 | -0.001820638 | 0.973739724 | 0.994079882 | 0.223918057 | 0.23044975 |

| | | | | | |
|---|---|---|---|---|---|
| Vwa8 | -0.088440682 | 0.061848736 | 0.380250374 | 0.13993111 | 0.222009905 |
| Wac | 0.12189142 | 0.074297952 | 0.412917246 | 0.663977505 | 0.532166023 |
| Wapl | 0.038067951 | 0.58102246 | 0.866173066 | 0.739982427 | 0.68963915 |
| Wars | 0.105169461 | 0.099413633 | 0.463315751 | 0.410576295 | 0.29580252 |
| Wasf2 | 0.049749733 | 0.434552398 | 0.796628285 | 0.412807315 | 0.36238414 |
| Washc1 | 0.030959476 | 0.51132501 | 0.835178052 | 0.153715015 | 0.140684971 |
| Washc2 | -0.010878748 | 0.855409308 | 0.957736324 | 0.293017673 | 0.310139616 |
| Washc3 | -0.127693829 | 0.031154047 | 0.280770404 | 0.323946019 | 0.451726394 |
| Washc4 | 0.140286464 | 0.012751064 | 0.179750602 | 0.261067009 | 0.137454202 |
| Washc5 | -0.017518626 | 0.746837101 | 0.92465005 | 0.22876083 | 0.230792004 |
| Wasl | 0.005979799 | 0.929688049 | 0.982502239 | 0.724089432 | 0.723908172 |
| Wbp1 | 0.116550111 | 0.028690463 | 0.27013713 | 0.228381808 | 0.125986953 |
| Wbp11 | 0.092204488 | 0.194462613 | 0.603853652 | 0.660550313 | 0.56068083 |
| Wbp1l | 0.006947735 | 0.871695417 | 0.964616742 | 0.121717529 | 0.114655022 |
| Wbp2 | 0.007442813 | 0.910286636 | 0.978641543 | 0.581288665 | 0.56030559 |
| Wbp4 | -0.038657705 | 0.560252205 | 0.856904336 | 0.675922829 | 0.721459099 |
| Wdcp | 0.018634456 | 0.669339353 | 0.903227201 | 0.132603731 | 0.117875762 |
| Wdfy1 | -0.075765572 | 0.219106653 | 0.632622127 | 0.361672441 | 0.438270785 |
| Wdfy2 | -0.062477976 | 0.167728541 | 0.570288402 | 0.128234351 | 0.193422863 |
| Wdfy3 | -0.031848229 | 0.546841463 | 0.851446616 | 0.199395364 | 0.219320992 |
| Wdr1 | 0.026961669 | 0.664141336 | 0.900079204 | 0.36342828 | 0.32359286 |
| Wdr11 | -0.060929647 | 0.199632031 | 0.610108047 | 0.149751921 | 0.204178737 |
| Wdr12 | 0.039430684 | 0.475350236 | 0.817387411 | 0.236929029 | 0.21177595 |
| Wdr13 | -0.039912872 | 0.365413217 | 0.752802229 | 0.125251082 | 0.154919509 |
| Wdr18 | -0.105542893 | 0.029949813 | 0.274935828 | 0.157626551 | 0.2529881 |
| Wdr20 | -0.030544973 | 0.455032026 | 0.807876964 | 0.102902774 | 0.132313677 |
| Wdr24 | -0.004879016 | 0.891892504 | 0.971118429 | 0.086234062 | 0.08887528 |
| Wdr25 | -0.017600483 | 0.544396333 | 0.851259637 | 0.047244215 | 0.057585074 |
| Wdr26 | -0.019234564 | 0.777109553 | 0.934230706 | 0.559782188 | 0.572452088 |
| Wdr27 | 0.031122285 | 0.383569795 | 0.763975666 | 0.085801579 | 0.058288064 |
| Wdr3 | -0.028006301 | 0.4753129 | 0.817387411 | 0.098735158 | 0.126244443 |
| Wdr33 | 0.126918904 | 0.060082964 | 0.378409279 | 0.65217784 | 0.511426169 |
| Wdr35 | -0.080973891 | 0.100323972 | 0.465319642 | 0.158428002 | 0.224001953 |
| Wdr36 | 0.059529202 | 0.21539025 | 0.629611185 | 0.161930238 | 0.109432793 |
| Wdr37 | 0.078611766 | 0.11971937 | 0.493284734 | 0.189809491 | 0.131380511 |
| Wdr4 | -0.010064485 | 0.749018146 | 0.925186829 | 0.061475068 | 0.068893001 |
| Wdr41 | 0.051603758 | 0.18639301 | 0.593348124 | 0.107451277 | 0.058264416 |
| Wdr43 | 0.058517015 | 0.371740003 | 0.75760494 | 0.399526999 | 0.349111476 |
| Wdr44 | -0.047090502 | 0.202045706 | 0.612724107 | 0.083251622 | 0.124264927 |
| Wdr45 | -0.048667544 | 0.1746117 | 0.577992588 | 0.07998676 | 0.12114336 |
| Wdr45b | -0.060460263 | 0.303500465 | 0.708261839 | 0.302608377 | 0.352909417 |
| Wdr46 | -0.012851462 | 0.725776774 | 0.92014727 | 0.086315011 | 0.093622906 |
| Wdr47 | 0.007105093 | 0.879701956 | 0.96752517 | 0.14940788 | 0.141341128 |
| Wdr48 | -0.082692093 | 0.048951721 | 0.347589359 | 0.106066801 | 0.193854455 |
| Wdr5 | 0.069146359 | 0.069764242 | 0.400867525 | 0.103141277 | 0.044521634 |
| Wdr53 | 0.02822545 | 0.451810878 | 0.806495742 | 0.091962512 | 0.077090045 |
| Wdr54 | -0.009645383 | 0.756345914 | 0.926833377 | 0.059989103 | 0.065887688 |
| Wdr55 | 0.035348793 | 0.270263514 | 0.677481242 | 0.06864939 | 0.037544351 |
| Wdr59 | 0.015132507 | 0.611396932 | 0.879924148 | 0.057598219 | 0.044910848 |
| Wdr5b | -0.04778851 | 0.043117512 | 0.328353923 | 0.02729831 | 0.068807994 |
| Wdr6 | -0.181307037 | 0.003057625 | 0.076723447 | 0.333802954 | 0.491127443 |
| Wdr61 | -0.02401351 | 0.703875311 | 0.912277105 | 0.4240964 | 0.460649335 |
| Wdr7 | -0.011501198 | 0.822181461 | 0.950930859 | 0.17425345 | 0.182640994 |
| Wdr70 | 0.028832607 | 0.530931148 | 0.845463419 | 0.141152216 | 0.118678727 |
| Wdr73 | 0.030254237 | 0.351983921 | 0.742354553 | 0.072313496 | 0.047723391 |
| Wdr74 | 0.043408406 | 0.435469438 | 0.796975049 | 0.22161649 | 0.186092899 |
| Wdr75 | 0.003029999 | 0.959099241 | 0.990654508 | 0.276532878 | 0.272567239 |
| Wdr77 | -0.079618445 | 0.026253513 | 0.257330576 | 0.071450365 | 0.135396628 |
| Wdr81 | -0.013661413 | 0.650453771 | 0.895679037 | 0.05384308 | 0.068684822 |
| Wdr82 | 0.001263024 | 0.978587434 | 0.995264774 | 0.148900595 | 0.143831886 |
| Wdr83 | -0.015025036 | 0.684071572 | 0.909648432 | 0.088457425 | 0.106466101 |
| Wdr83os | -0.162026405 | 0.014766608 | 0.195426645 | 0.597198202 | 0.760604958 |
| Wdr89 | 0.060573166 | 0.250991251 | 0.663121659 | 0.219556191 | 0.152710007 |
| Wdr90 | 0.011384927 | 0.725465911 | 0.92014727 | 0.067507992 | 0.05203945 |
| Wdr92 | -0.020724363 | 0.556619412 | 0.85521955 | 0.077172386 | 0.096313123 |

| | | | | | |
|---|---|---|---|---|---|
| Wdsub1 | 0.079622811 | 0.156601627 | 0.554127149 | 0.244066913 | 0.185390953 |
| Wdtc1 | -0.019263155 | 0.655400325 | 0.89780989 | 0.120893533 | 0.133825348 |
| Wdyhv1 | -0.033260971 | 0.419894637 | 0.786284042 | 0.110577544 | 0.135597018 |
| Wee1 | 0.033639786 | 0.371242935 | 0.75760494 | 0.100779304 | 0.07321766 |
| Wfs1 | -0.089794293 | 0.168967723 | 0.57227323 | 0.403696027 | 0.481210164 |
| Whamm | 0.084056214 | 0.128856128 | 0.508260084 | 0.228819009 | 0.15314044 |
| Whrn | -0.008847265 | 0.841175954 | 0.954960614 | 0.129302257 | 0.141775708 |
| Wipf2 | -0.02203496 | 0.691121192 | 0.909796728 | 0.224655468 | 0.2364771 |
| Wipi1 | -0.221489151 | 0.001752893 | 0.055676541 | 0.418377543 | 0.63638453 |
| Wipi2 | -0.128247698 | 0.03299769 | 0.287968471 | 0.29635895 | 0.41900953 |
| Wiz | 0.030905707 | 0.523606213 | 0.841062594 | 0.160128406 | 0.132320075 |
| Wls | -0.199512465 | 0.002369459 | 0.066061828 | 1.077226974 | 1.364600358 |
| Wnk1 | -0.013962724 | 0.820160585 | 0.950589638 | 0.950864102 | 0.939330851 |
| Wnk2 | 0.039436534 | 0.18498705 | 0.591498892 | 0.060562557 | 0.026613455 |
| Wnk3 | -0.021868865 | 0.537476382 | 0.847793898 | 0.06932294 | 0.08561944 |
| Wnk4 | -0.003959829 | 0.93879001 | 0.985324082 | 0.181005508 | 0.175568129 |
| Wnt4 | -0.118990839 | 0.058068739 | 0.37129897 | 1.715488933 | 1.990081426 |
| Wrap53 | -0.024235502 | 0.331758735 | 0.727679726 | 0.036704865 | 0.062032204 |
| Wrap73 | 0.000843484 | 0.981073672 | 0.995588708 | 0.0830209 | 0.081571791 |
| Wrn | 0.054785434 | 0.157209894 | 0.554924229 | 0.107079764 | 0.059452262 |
| Wrnip1 | 0.060349459 | 0.239801607 | 0.650744882 | 0.20255995 | 0.155927334 |
| Wsb1 | -0.017184227 | 0.582151658 | 0.866173066 | 0.058873583 | 0.071037991 |
| Wsb2 | 0.042830727 | 0.52569482 | 0.842051614 | 0.804245961 | 0.765806721 |
| Wtap | 0.104432831 | 0.086641289 | 0.43765406 | 0.333282855 | 0.255096285 |
| Wtip | -0.121159693 | 0.000175452 | 0.011818903 | 0.057635786 | 0.161923183 |
| Wwox | -0.022212134 | 0.469960958 | 0.814258266 | 0.057294261 | 0.073302925 |
| Wwp1 | 0.015297541 | 0.728531925 | 0.920519505 | 0.129760737 | 0.11750189 |
| Wwp2 | 0.000693102 | 0.989071123 | 0.997411191 | 0.175173722 | 0.170931872 |
| Xab2 | 0.037192221 | 0.39916339 | 0.774544012 | 0.139671941 | 0.110164114 |
| Xaf1 | 0.117136758 | 0.007723048 | 0.135045705 | 0.13967742 | 0.033928643 |
| Xbp1 | -0.11670856 | 0.088223691 | 0.441426313 | 1.109618855 | 1.250071282 |
| Xiap | -0.033429781 | 0.552860473 | 0.852623515 | 0.252118743 | 0.270576732 |
| Xk | -0.00027368 | 0.994272534 | 0.998695538 | 0.098652273 | 0.103237605 |
| Xkr6 | -0.05117073 | 0.289611278 | 0.695912952 | 0.158867265 | 0.206471487 |
| Xpa | -0.123773785 | 0.017698621 | 0.21172951 | 0.175835139 | 0.289505255 |
| Xpc | 0.025089725 | 0.607606562 | 0.878733629 | 0.176519379 | 0.162523211 |
| Xpnpep1 | -0.047889478 | 0.266543407 | 0.674047915 | 0.122676017 | 0.161848251 |
| Xpnpep3 | 0.06714777 | 0.14516956 | 0.533628684 | 0.14876349 | 0.095500917 |
| Xpo1 | 0.11805887 | 0.045171916 | 0.332885843 | 0.308409917 | 0.196042356 |
| Xpo4 | -0.051512971 | 0.272777412 | 0.680556141 | 0.140495313 | 0.181413498 |
| Xpo5 | 0.051857835 | 0.272369506 | 0.680042969 | 0.157960557 | 0.113124816 |
| Xpo6 | -0.116967823 | 0.015235809 | 0.198397782 | 0.150749112 | 0.245442271 |
| Xpo7 | -0.016155185 | 0.791255614 | 0.938824685 | 0.319614825 | 0.329539649 |
| Xpot | 0.02075383 | 0.756006805 | 0.926726954 | 0.475284773 | 0.444986164 |
| Xpr1 | 0.026626333 | 0.647755458 | 0.895376633 | 0.286692631 | 0.257732053 |
| Xrcc1 | -0.032885054 | 0.46051691 | 0.810111566 | 0.131017606 | 0.158962733 |
| Xrcc3 | -0.00026965 | 0.991814655 | 0.998579967 | 0.042794674 | 0.041303824 |
| Xrcc4 | 0.077250707 | 0.05769271 | 0.370772606 | 0.118596624 | 0.060699465 |
| Xrcc5 | -0.056132683 | 0.290898813 | 0.69667177 | 0.19640694 | 0.242392436 |
| Xrcc6 | -0.041088624 | 0.407921125 | 0.77823964 | 0.168646258 | 0.19976439 |
| Xrn1 | 0.02573717 | 0.69223787 | 0.909796728 | 0.418477403 | 0.386575137 |
| Xrn2 | 0.036156272 | 0.572154954 | 0.861494041 | 0.36654554 | 0.338858068 |
| Yae1d1 | -0.073781888 | 0.193350808 | 0.60325934 | 0.238405079 | 0.312244935 |
| Yaf2 | 0.119106987 | 0.073214034 | 0.410379808 | 0.480043061 | 0.371032521 |
| Yars | -0.013764325 | 0.832452444 | 0.953797263 | 0.433812532 | 0.451180961 |
| Yars2 | 0.03942614 | 0.27686267 | 0.68411389 | 0.089987814 | 0.062616907 |
| Ybey | 7.42E-05 | 0.998181194 | 0.99938181 | 0.064770821 | 0.067960276 |
| Ybx1 | -0.106281197 | 0.05541848 | 0.36562505 | 1.575739048 | 1.746989049 |
| Ybx3 | 0.111690811 | 0.10172684 | 0.467719786 | 0.782753861 | 0.682791906 |
| Ydjc | -0.030398875 | 0.364329215 | 0.752732058 | 0.070707773 | 0.093091753 |
| Yeats2 | 0.031180775 | 0.391918896 | 0.769704269 | 0.093365426 | 0.063344536 |
| Yeats4 | -0.051214594 | 0.401747387 | 0.775220138 | 0.326558998 | 0.361268826 |
| Yes1 | -0.060401885 | 0.185325336 | 0.591560356 | 0.134404604 | 0.187045975 |
| Yif1a | 0.012194916 | 0.851904121 | 0.957461155 | 0.498863648 | 0.492973696 |
| Yif1b | 0.024602814 | 0.666104315 | 0.901197061 | 0.275357515 | 0.25745879 |

| | | | | | |
|---|---|---|---|---|---|
| Yipf1 | -0.096302389 | 0.105467028 | 0.472434351 | 0.329645778 | 0.408060627 |
| Yipf2 | -0.059491446 | 0.232292094 | 0.643543798 | 0.166649832 | 0.218642267 |
| Yipf3 | 0.01285235 | 0.844964137 | 0.955950886 | 0.50119002 | 0.477631245 |
| Yipf4 | -0.146498508 | 0.023635701 | 0.244316717 | 0.474800965 | 0.633430435 |
| Yipf5 | -0.013573313 | 0.839729706 | 0.954911647 | 0.551307877 | 0.560186776 |
| Yipf6 | -0.001109419 | 0.98651196 | 0.997268589 | 0.42699555 | 0.418889159 |
| Yju2 | 0.065214873 | 0.115230746 | 0.487260227 | 0.117267654 | 0.059146562 |
| Ykt6 | 0.050257939 | 0.375864809 | 0.760595784 | 0.255695688 | 0.208197536 |
| Ylpm1 | -0.007517057 | 0.909746404 | 0.978578013 | 0.430052438 | 0.42777304 |
| Yme1l1 | 0.03030516 | 0.661422238 | 0.898920864 | 0.592254457 | 0.562458306 |
| Yod1 | 0.02741963 | 0.590392373 | 0.870796216 | 0.180849469 | 0.161405349 |
| Ypel2 | 0.01029697 | 0.868357855 | 0.96295459 | 0.344416848 | 0.337023807 |
| Ypel3 | -0.167613573 | 0.013524269 | 0.189057229 | 0.714584437 | 0.909499955 |
| Ypel4 | -0.01241268 | 0.772674555 | 0.932469974 | 0.121906497 | 0.130667195 |
| Ypel5 | -0.074708322 | 0.226422574 | 0.638587289 | 0.357138895 | 0.427821636 |
| Yrdc | 0.071764919 | 0.268257703 | 0.676166273 | 0.41726762 | 0.351595377 |
| Ythdc1 | 0.141556035 | 0.039443067 | 0.313282855 | 0.758656636 | 0.615486446 |
| Ythdc2 | -0.109648602 | 0.015255007 | 0.198397782 | 0.12418142 | 0.22388666 |
| Ythdf1 | 0.032498248 | 0.574569474 | 0.862412713 | 0.276459882 | 0.254462245 |
| Ythdf2 | -0.037171746 | 0.5079368 | 0.833850451 | 0.233696315 | 0.273036574 |
| Ythdf3 | 0.047757696 | 0.437469439 | 0.798455866 | 0.399708705 | 0.363259327 |
| Ywhab | -0.177232051 | 0.009941519 | 0.155269009 | 0.554819241 | 0.708829338 |
| Ywhae | -0.045399594 | 0.378057936 | 0.761961347 | 1.562257488 | 1.643982006 |
| Ywhag | 0.078687905 | 0.281039114 | 0.68791143 | 0.792723628 | 0.685311806 |
| Ywhah | 0.154116584 | 0.029077471 | 0.271741694 | 0.935527337 | 0.736718158 |
| Ywhaq | -0.008736328 | 0.853092323 | 0.957461155 | 1.891435684 | 1.94749641 |
| Ywhaz | 0.02326996 | 0.70595209 | 0.912844672 | 1.213980056 | 1.15934168 |
| Yy1 | -0.033400317 | 0.627748329 | 0.887864718 | 0.792956729 | 0.863008951 |
| Zadh2 | 0.12293541 | 0.008593338 | 0.144232999 | 0.169241876 | 0.061548551 |
| Zbed3 | -0.030112889 | 0.467503167 | 0.813188482 | 0.111705129 | 0.137526323 |
| Zbed5 | -0.033782678 | 0.157404531 | 0.554924229 | 0.025930878 | 0.05731 |
| Zbtb1 | 0.023306784 | 0.621893108 | 0.884879661 | 0.150865611 | 0.133451561 |
| Zbtb10 | 0.016591893 | 0.660161829 | 0.898481506 | 0.09249685 | 0.087634168 |
| Zbtb11 | 0.028644656 | 0.634063224 | 0.88965971 | 0.294935112 | 0.276269191 |
| Zbtb11os1 | -0.096221707 | 0.009065113 | 0.147945584 | 0.079245492 | 0.171462093 |
| Zbtb12 | -0.074378904 | 0.08037449 | 0.425115437 | 0.115341018 | 0.175839649 |
| Zbtb14 | 0.092425623 | 0.077646913 | 0.418285047 | 0.215016824 | 0.12940547 |
| Zbtb17 | -0.020923085 | 0.461981131 | 0.810513184 | 0.048233666 | 0.066539956 |
| Zbtb18 | -0.014773964 | 0.773436223 | 0.932714536 | 0.185733773 | 0.191487362 |
| Zbtb2 | 0.040173153 | 0.196750779 | 0.60713211 | 0.064770903 | 0.032259962 |
| Zbtb20 | -0.180582508 | 0.006893547 | 0.126971675 | 1.31525828 | 1.5542911 |
| Zbtb21 | -0.031791185 | 0.529166101 | 0.844323583 | 0.183889672 | 0.213731685 |
| Zbtb22 | 0.070028346 | 0.067840343 | 0.395649928 | 0.105607315 | 0.053721306 |
| Zbtb24 | -0.046218565 | 0.285600116 | 0.692407818 | 0.118948704 | 0.162800392 |
| Zbtb25 | 0.017158719 | 0.560205964 | 0.856904336 | 0.055914383 | 0.03864067 |
| Zbtb26 | 0.010627565 | 0.716421426 | 0.916579082 | 0.055019119 | 0.048801294 |
| Zbtb33 | 0.052758598 | 0.2542435 | 0.663146569 | 0.152166088 | 0.108204023 |
| Zbtb37 | -0.065410249 | 0.032292196 | 0.284812628 | 0.052369314 | 0.109122756 |
| Zbtb38 | 0.083704611 | 0.183374939 | 0.590066231 | 0.40117603 | 0.326037931 |
| Zbtb39 | -0.000317191 | 0.991458045 | 0.998321067 | 0.050824257 | 0.051542172 |
| Zbtb4 | -0.018258379 | 0.773026704 | 0.932556373 | 0.35741728 | 0.347371527 |
| Zbtb40 | 0.024925568 | 0.509796857 | 0.834904701 | 0.09762158 | 0.077554799 |
| Zbtb41 | -0.008528561 | 0.860157227 | 0.959373052 | 0.164737058 | 0.174696642 |
| Zbtb43 | 0.009194701 | 0.834358344 | 0.954794181 | 0.126205209 | 0.120085774 |
| Zbtb44 | -0.033284099 | 0.564636037 | 0.858087314 | 0.244453225 | 0.276915421 |
| Zbtb45 | 0.026719311 | 0.371270307 | 0.75760494 | 0.060557499 | 0.040248003 |
| Zbtb5 | -0.053953294 | 0.118322011 | 0.491192747 | 0.070704368 | 0.122149798 |
| Zbtb6 | 0.029209075 | 0.543338267 | 0.850777876 | 0.154841903 | 0.130797631 |
| Zbtb7a | 0.072910976 | 0.29555494 | 0.702292835 | 0.860911691 | 0.750104845 |
| Zbtb7b | -0.021578833 | 0.660588131 | 0.89868852 | 0.16287309 | 0.174805083 |
| Zbtb7c | -0.052784108 | 0.051456868 | 0.355933715 | 0.037673177 | 0.078675352 |
| Zbtb8os | -0.041592144 | 0.452822999 | 0.807136288 | 0.238397687 | 0.278346663 |
| Zc2hc1a | -0.026045885 | 0.51933107 | 0.838496482 | 0.108676437 | 0.133904352 |
| Zc2hc1c | 0.080758771 | 0.054570638 | 0.363639613 | 0.122150893 | 0.054694874 |
| Zc3h10 | 0.024779224 | 0.63185074 | 0.88887544 | 0.188873406 | 0.158465074 |

| | | | | | |
|---|---|---|---|---|---|
| Zc3h12b | -0.039713888 | 0.156743481 | 0.554322538 | 0.045048222 | 0.077364969 |
| Zc3h12c | -0.074729935 | 0.120350435 | 0.494254219 | 0.150654526 | 0.214642721 |
| Zc3h13 | 0.126505226 | 0.061093747 | 0.380250374 | 0.476057919 | 0.35574126 |
| Zc3h14 | 0.04908515 | 0.45925136 | 0.809393611 | 0.554170633 | 0.515388069 |
| Zc3h15 | 0.125805009 | 0.043746716 | 0.329086044 | 1.142530398 | 0.964766993 |
| Zc3h18 | -0.029549354 | 0.619789089 | 0.883866495 | 0.305214218 | 0.348568547 |
| Zc3h3 | 0.089917591 | 0.130895648 | 0.510610033 | 0.280160611 | 0.198448604 |
| Zc3h4 | 0.00664439 | 0.867443052 | 0.962366078 | 0.107517181 | 0.102245741 |
| Zc3h6 | -0.036954985 | 0.504875249 | 0.832249514 | 0.202296718 | 0.235930418 |
| Zc3h7a | 0.003503598 | 0.954774257 | 0.989066902 | 0.362519365 | 0.355136208 |
| Zc3h7b | 0.04652076 | 0.367816411 | 0.754809512 | 0.19932268 | 0.15065152 |
| Zc3h8 | 0.046239784 | 0.213592569 | 0.628127689 | 0.088551588 | 0.047001642 |
| Zc3hav1 | -0.046754103 | 0.462335697 | 0.810535897 | 0.385811871 | 0.441469563 |
| Zc3hc1 | -0.00953051 | 0.719113215 | 0.918638072 | 0.039650812 | 0.048601098 |
| Zcchc10 | 0.033918553 | 0.449824718 | 0.806495742 | 0.141267734 | 0.109016777 |
| Zcchc12 | -0.060946961 | 0.218879294 | 0.632622127 | 0.153988235 | 0.202588291 |
| Zcchc14 | -0.035449214 | 0.515680998 | 0.837349565 | 0.223818016 | 0.25548215 |
| Zcchc17 | -0.075200912 | 0.243947074 | 0.653830568 | 0.404852005 | 0.499532849 |
| Zcchc18 | -0.032210843 | 0.643450563 | 0.892591387 | 0.718759669 | 0.760014047 |
| Zcchc2 | 0.037802729 | 0.476137924 | 0.817948631 | 0.209999825 | 0.181714157 |
| Zcchc3 | 0.033069809 | 0.43972619 | 0.800048525 | 0.127544617 | 0.102352345 |
| Zcchc4 | -0.037008107 | 0.180248785 | 0.587198006 | 0.040188853 | 0.069557107 |
| Zcchc7 | -0.12702894 | 0.05582419 | 0.367414687 | 0.46225328 | 0.571361295 |
| Zcchc8 | -0.00685628 | 0.889211948 | 0.970216363 | 0.172738471 | 0.180368419 |
| Zcchc9 | 0.023610655 | 0.697216908 | 0.910970095 | 0.334638214 | 0.307832114 |
| Zcrb1 | 0.02091656 | 0.722394936 | 0.919483582 | 1.328630595 | 1.254353668 |
| Zdbf2 | -0.159093515 | 0.018259354 | 0.21500396 | 0.454061309 | 0.619007228 |
| Zdhhc1 | -0.020357386 | 0.586593108 | 0.86902362 | 0.086947086 | 0.106056166 |
| Zdhhc12 | 0.020465858 | 0.61839922 | 0.883414883 | 0.113036514 | 0.096245412 |
| Zdhhc13 | -0.011091575 | 0.721340751 | 0.919475778 | 0.062292087 | 0.075912892 |
| Zdhhc14 | -0.043562519 | 0.239942585 | 0.650744882 | 0.08873478 | 0.12904047 |
| Zdhhc16 | 0.000521714 | 0.987177753 | 0.997268589 | 0.069081682 | 0.071728055 |
| Zdhhc17 | 0.028354396 | 0.561316204 | 0.857134531 | 0.163613812 | 0.138601234 |
| Zdhhc18 | 0.027889966 | 0.385343572 | 0.765415428 | 0.070939814 | 0.048231112 |
| Zdhhc2 | 0.0445076 | 0.50022456 | 0.831130155 | 1.1425757 | 1.071588947 |
| Zdhhc20 | 0.067502451 | 0.273213488 | 0.681302802 | 0.33874905 | 0.279230854 |
| Zdhhc21 | -0.05030776 | 0.221303849 | 0.633858837 | 0.101809055 | 0.149827702 |
| Zdhhc22 | 0.024686947 | 0.507284769 | 0.833462342 | 0.08999836 | 0.071717269 |
| Zdhhc23 | -0.075978906 | 0.087193623 | 0.439557333 | 0.121027253 | 0.177670671 |
| Zdhhc24 | -0.007358396 | 0.813106762 | 0.948397417 | 0.06016718 | 0.06291682 |
| Zdhhc3 | -0.055555997 | 0.383023079 | 0.763687454 | 0.384406884 | 0.445184571 |
| Zdhhc4 | -0.020537271 | 0.584948217 | 0.867683611 | 0.088507414 | 0.104986619 |
| Zdhhc5 | -0.022951345 | 0.69338156 | 0.909871028 | 0.254810679 | 0.279929924 |
| Zdhhc6 | -0.027589094 | 0.546919585 | 0.851446616 | 0.140491722 | 0.170417947 |
| Zdhhc7 | 0.069526169 | 0.094093984 | 0.452959864 | 0.123884877 | 0.069550169 |
| Zdhhc8 | -0.000115283 | 0.998386112 | 0.99938181 | 0.247410011 | 0.248966053 |
| Zeb1 | -0.0398165 | 0.442839919 | 0.801368502 | 0.175549672 | 0.214553727 |
| Zer1 | -0.021396789 | 0.62240307 | 0.885228796 | 0.123951918 | 0.142476918 |
| Zfand1 | -0.019151553 | 0.629298556 | 0.887864718 | 0.10538032 | 0.125521464 |
| Zfand2a | -0.002712954 | 0.934121606 | 0.983960188 | 0.069267022 | 0.071334844 |
| Zfand2b | 0.085833776 | 0.144395035 | 0.533279057 | 0.272166725 | 0.184439667 |
| Zfand3 | 0.037983084 | 0.528173049 | 0.843657883 | 0.343379806 | 0.31773284 |
| Zfand5 | -0.113569574 | 0.097871805 | 0.460975742 | 0.649505967 | 0.789844282 |
| Zfand6 | 0.021138728 | 0.754535884 | 0.926726954 | 0.640767164 | 0.632059071 |
| Zfas1 | 0.015537561 | 0.822219067 | 0.950930859 | 0.570600285 | 0.549685111 |
| Zfc3h1 | -0.106059878 | 0.04600114 | 0.336525833 | 0.181523261 | 0.276768602 |
| Zfhx2 | -0.081817607 | 0.231121658 | 0.642595035 | 0.802736777 | 0.889391288 |
| Zfhx3 | 0.060777171 | 0.366925562 | 0.753633555 | 0.459212547 | 0.380448444 |
| Zfp1 | 0.027451089 | 0.534424898 | 0.846347806 | 0.133778984 | 0.111688073 |
| Zfp106 | -0.008163969 | 0.903503749 | 0.974950771 | 0.655910322 | 0.647972194 |
| Zfp110 | -0.019188983 | 0.689721774 | 0.909796728 | 0.160034987 | 0.188238312 |
| Zfp111 | -0.018275862 | 0.47284167 | 0.816711726 | 0.036451698 | 0.048817266 |
| Zfp113 | -0.010096891 | 0.812508889 | 0.948397417 | 0.122429541 | 0.143021997 |
| Zfp119b | 0.04640319 | 0.094695906 | 0.454115053 | 0.051860351 | 0.015725051 |
| Zfp12 | -0.065355135 | 0.083570024 | 0.431740995 | 0.088859396 | 0.153752442 |

| | | | | | |
|---|---|---|---|---|---|
| Zfp120 | -0.023709563 | 0.634520002 | 0.88965971 | 0.163744597 | 0.200199897 |
| Zfp125 | -0.048025166 | 0.08609136 | 0.436930819 | 0.042589795 | 0.082397475 |
| Zfp131 | 0.027511327 | 0.634903727 | 0.889733629 | 0.277736112 | 0.247819131 |
| Zfp142 | 0.001392365 | 0.971996082 | 0.993981119 | 0.104539225 | 0.103952792 |
| Zfp143 | 0.010320631 | 0.783545201 | 0.936124803 | 0.090475505 | 0.080996514 |
| Zfp146 | -0.019467689 | 0.623697606 | 0.88619095 | 0.103682067 | 0.121868563 |
| Zfp148 | -0.049546474 | 0.464500173 | 0.811662297 | 0.635101486 | 0.675686457 |
| Zfp157 | 0.043889184 | 0.314045069 | 0.714975653 | 0.135459738 | 0.10419813 |
| Zfp160 | -0.017185735 | 0.68131976 | 0.908154905 | 0.109939927 | 0.12716802 |
| Zfp169 | -0.047133419 | 0.02729459 | 0.263165994 | 0.019193493 | 0.059021326 |
| Zfp174 | -0.037827373 | 0.251181982 | 0.663127858 | 0.067409362 | 0.101541561 |
| Zfp180 | 0.002063091 | 0.95761426 | 0.990137214 | 0.096255658 | 0.099049541 |
| Zfp182 | -0.063945224 | 0.041131893 | 0.319536229 | 0.0532804 | 0.111502981 |
| Zfp2 | -0.048570108 | 0.166564395 | 0.568127031 | 0.076428319 | 0.113770862 |
| Zfp207 | 0.043935036 | 0.483058375 | 0.822469365 | 0.347152613 | 0.311281598 |
| Zfp212 | 0.006704472 | 0.838129217 | 0.954911647 | 0.070317281 | 0.065290191 |
| Zfp213 | 0.005912603 | 0.827633345 | 0.952851099 | 0.045403227 | 0.042296945 |
| Zfp217 | -0.017833575 | 0.523201693 | 0.841062594 | 0.047114102 | 0.057647302 |
| Zfp219 | -0.03268427 | 0.402697493 | 0.77525778 | 0.095015048 | 0.124678675 |
| Zfp229 | -0.026243704 | 0.25563474 | 0.664689244 | 0.026431072 | 0.048749606 |
| Zfp235 | 0.009395384 | 0.786833308 | 0.937461311 | 0.080282947 | 0.072518981 |
| Zfp236 | 0.064739062 | 0.158630395 | 0.556789205 | 0.149215006 | 0.095139263 |
| Zfp24 | 0.09386512 | 0.128847804 | 0.508260084 | 0.347938703 | 0.252576057 |
| Zfp251 | -0.015827253 | 0.744868655 | 0.924566547 | 0.165511316 | 0.177131962 |
| Zfp26 | -0.021027736 | 0.574124589 | 0.862168892 | 0.08573867 | 0.104970766 |
| Zfp260 | -0.065494278 | 0.240116731 | 0.650744882 | 0.231338411 | 0.280502991 |
| Zfp263 | 0.017921989 | 0.710273995 | 0.914486281 | 0.161681685 | 0.142195187 |
| Zfp266 | -0.019464182 | 0.746558843 | 0.924631183 | 0.312186077 | 0.331415835 |
| Zfp267 | 0.004105479 | 0.913454842 | 0.979979308 | 0.089209672 | 0.08566958 |
| Zfp27 | -0.021595599 | 0.391725942 | 0.769704269 | 0.035627421 | 0.05049911 |
| Zfp275 | -0.00828122 | 0.823460401 | 0.951067558 | 0.088052587 | 0.096527452 |
| Zfp277 | 0.019320704 | 0.592624474 | 0.872202098 | 0.089439863 | 0.082807962 |
| Zfp28 | 0.054723308 | 0.047689452 | 0.341775906 | 0.052418957 | 0.008162079 |
| Zfp280c | -0.020348188 | 0.526448068 | 0.842199436 | 0.064532568 | 0.084035671 |
| Zfp280d | -0.050612139 | 0.413408197 | 0.781607961 | 0.339851435 | 0.390230044 |
| Zfp281 | -0.001620882 | 0.970288716 | 0.993892166 | 0.132348539 | 0.135297218 |
| Zfp282 | 0.045408525 | 0.257335679 | 0.665256942 | 0.10999627 | 0.071600967 |
| Zfp287 | -0.040680088 | 0.158254621 | 0.556552777 | 0.04901765 | 0.084126461 |
| Zfp292 | 0.024504753 | 0.731352857 | 0.920770724 | 0.598332263 | 0.568641941 |
| Zfp30 | 0.11700701 | 0.012021779 | 0.174689004 | 0.161220499 | 0.055954658 |
| Zfp316 | -0.031014584 | 0.258663636 | 0.665948842 | 0.043947366 | 0.070365784 |
| Zfp317 | -0.035254887 | 0.339723512 | 0.733286616 | 0.085688787 | 0.113909805 |
| Zfp318 | -0.002741821 | 0.959061574 | 0.990654508 | 0.205628949 | 0.212213892 |
| Zfp319 | -0.021855426 | 0.531342962 | 0.845517106 | 0.078236874 | 0.101581004 |
| Zfp322a | 0.006372602 | 0.887711303 | 0.970045467 | 0.140158411 | 0.137043471 |
| Zfp326 | -0.072384616 | 0.209096639 | 0.62220663 | 0.263835647 | 0.330079708 |
| Zfp329 | -0.013456789 | 0.819667293 | 0.950548887 | 0.30955385 | 0.314746642 |
| Zfp330 | 0.046645786 | 0.475969318 | 0.817948631 | 0.406930931 | 0.369512377 |
| Zfp334 | -0.025642592 | 0.37464266 | 0.759807714 | 0.048421711 | 0.067610709 |
| Zfp335 | -0.017478805 | 0.632513665 | 0.889078599 | 0.085779911 | 0.099946834 |
| Zfp346 | -0.016517996 | 0.691240214 | 0.909796728 | 0.108021225 | 0.124710793 |
| Zfp35 | 0.011199486 | 0.736119474 | 0.922302218 | 0.071678032 | 0.063071481 |
| Zfp358 | 0.031063456 | 0.499387791 | 0.830722138 | 0.149274114 | 0.12161021 |
| Zfp36 | -0.102459944 | 0.018762862 | 0.217462874 | 0.116223304 | 0.196517402 |
| Zfp362 | -0.009867909 | 0.763275651 | 0.928730554 | 0.06732759 | 0.077522586 |
| Zfp367 | 0.038718884 | 0.27517638 | 0.68241738 | 0.083505139 | 0.053976256 |
| Zfp369 | -0.01100648 | 0.692037883 | 0.909796728 | 0.046392945 | 0.054504748 |
| Zfp36l2 | 0.040304796 | 0.538664864 | 0.848357012 | 0.371428676 | 0.318096087 |
| Zfp37 | -0.044112208 | 0.229530767 | 0.641012885 | 0.081977814 | 0.126263213 |
| Zfp384 | -0.03344694 | 0.351502218 | 0.741961674 | 0.081114018 | 0.110718938 |
| Zfp386 | 0.015448598 | 0.752054896 | 0.926186851 | 0.165758229 | 0.150691079 |
| Zfp395 | 0.050692411 | 0.307903747 | 0.710440053 | 0.175811239 | 0.125415664 |
| Zfp397 | 0.078621783 | 0.171366632 | 0.574292783 | 0.282614634 | 0.223983002 |
| Zfp398 | 0.003864685 | 0.946128622 | 0.987084873 | 0.256936287 | 0.254775015 |
| Zfp40 | 0.034816921 | 0.252248895 | 0.663127858 | 0.064926693 | 0.034809319 |

| | | | | | |
|---|---|---|---|---|---|
| Zfp407 | -0.12963042 | 0.00070534 | 0.031325216 | 0.080851525 | 0.19110587 |
| Zfp410 | -0.021323339 | 0.638464652 | 0.89125563 | 0.137759114 | 0.161920844 |
| Zfp414 | 0.011358306 | 0.693542922 | 0.909871028 | 0.053686857 | 0.047125979 |
| Zfp420 | -0.02698814 | 0.35092856 | 0.741476479 | 0.046947001 | 0.067685005 |
| Zfp422 | 0.022591771 | 0.663513781 | 0.899593114 | 0.212241661 | 0.191301475 |
| Zfp426 | 0.004097994 | 0.932110739 | 0.98343913 | 0.16576614 | 0.161681185 |
| Zfp428 | 0.016619244 | 0.603399891 | 0.876723966 | 0.070438465 | 0.059976186 |
| Zfp438 | 0.016494583 | 0.649594107 | 0.895585658 | 0.088696371 | 0.072879213 |
| Zfp442 | 0.024139561 | 0.393265107 | 0.770035482 | 0.054169919 | 0.034525372 |
| Zfp444 | -0.010828307 | 0.783572309 | 0.936124803 | 0.099270277 | 0.107241611 |
| Zfp445 | -0.025884043 | 0.680498795 | 0.907783945 | 0.391252516 | 0.403854216 |
| Zfp449 | -0.007972893 | 0.764948954 | 0.928757959 | 0.041228262 | 0.048849539 |
| Zfp451 | -0.004647087 | 0.93879189 | 0.985324082 | 0.311817601 | 0.328668376 |
| Zfp458 | -0.00954267 | 0.734627832 | 0.921732448 | 0.047179223 | 0.055612684 |
| Zfp462 | 0.043096601 | 0.420470563 | 0.786922065 | 0.214985237 | 0.177668444 |
| Zfp467 | 0.025517627 | 0.583754855 | 0.867072813 | 0.142604895 | 0.111540751 |
| Zfp507 | -0.001949253 | 0.962530207 | 0.991757003 | 0.114910126 | 0.112788495 |
| Zfp51 | 0.024916316 | 0.445539342 | 0.803149051 | 0.068777837 | 0.046409286 |
| Zfp511 | 0.063290055 | 0.220247198 | 0.633234352 | 0.195591262 | 0.135510098 |
| Zfp512 | -0.035080142 | 0.417545326 | 0.784482356 | 0.119829054 | 0.156359457 |
| Zfp512b | -0.104185656 | 0.018289191 | 0.21500396 | 0.113076032 | 0.207784803 |
| Zfp513 | -0.015310508 | 0.687227239 | 0.909796728 | 0.092084697 | 0.105670781 |
| Zfp516 | -0.109869465 | 0.116654147 | 0.489152833 | 0.600467702 | 0.719431636 |
| Zfp518a | 0.008309542 | 0.836876746 | 0.954911647 | 0.109778649 | 0.100123444 |
| Zfp518b | -0.042459925 | 0.118651125 | 0.491192747 | 0.041418814 | 0.0771994 |
| Zfp52 | -0.053862407 | 0.116878596 | 0.489579867 | 0.072333031 | 0.117885512 |
| Zfp523 | 0.029007619 | 0.475308979 | 0.817387411 | 0.115015727 | 0.08858939 |
| Zfp524 | 0.014354155 | 0.741073228 | 0.922901128 | 0.126692677 | 0.113344789 |
| Zfp53 | 0.004815945 | 0.870331545 | 0.964236463 | 0.055395237 | 0.059696914 |
| Zfp553 | 0.072481182 | 0.134433769 | 0.516977677 | 0.167898252 | 0.115196864 |
| Zfp560 | -0.145083031 | 0.020037423 | 0.226226793 | 0.354897473 | 0.486186343 |
| Zfp563 | 0.03347381 | 0.322224314 | 0.719780733 | 0.075235205 | 0.046104374 |
| Zfp566 | 0.042687726 | 0.176467085 | 0.580732554 | 0.066113666 | 0.028033162 |
| Zfp568 | -0.121201849 | 0.014952509 | 0.197256514 | 0.148766037 | 0.265245542 |
| Zfp574 | 0.088387394 | 0.015109415 | 0.197980673 | 0.094071532 | 0.02483254 |
| Zfp579 | -0.012280317 | 0.745970135 | 0.924566547 | 0.093151073 | 0.105481342 |
| Zfp580 | -0.061692762 | 0.018919789 | 0.218273469 | 0.03210094 | 0.090835341 |
| Zfp592 | 0.018360128 | 0.684995061 | 0.909796728 | 0.144129893 | 0.13219506 |
| Zfp593 | -0.020735109 | 0.675458781 | 0.906025597 | 0.17025894 | 0.182870598 |
| Zfp597 | -0.006021175 | 0.882826081 | 0.96861827 | 0.105429309 | 0.113349827 |
| Zfp598 | 0.042743433 | 0.396358942 | 0.773119057 | 0.189147869 | 0.148129849 |
| Zfp60 | -0.039659856 | 0.393621059 | 0.770538672 | 0.140466427 | 0.175815837 |
| Zfp605 | -0.021323079 | 0.593134854 | 0.872442739 | 0.108327928 | 0.124450219 |
| Zfp606 | -0.092445689 | 0.00660766 | 0.123502952 | 0.063560969 | 0.147213999 |
| Zfp608 | -0.083209446 | 0.163227203 | 0.562993622 | 0.280448612 | 0.375761789 |
| Zfp609 | -0.018149194 | 0.727438211 | 0.920455328 | 0.18899737 | 0.203065632 |
| Zfp61 | 0.047389951 | 0.163849209 | 0.563565555 | 0.077185887 | 0.036498943 |
| Zfp612 | 0.018829 | 0.765865655 | 0.92882091 | 0.401133759 | 0.385625887 |
| Zfp617 | 0.006339634 | 0.871068305 | 0.964504119 | 0.099378224 | 0.096314173 |
| Zfp618 | -0.019729796 | 0.53310329 | 0.84612966 | 0.060435668 | 0.078561683 |
| Zfp62 | -0.019739992 | 0.702392088 | 0.912152193 | 0.18801544 | 0.211235224 |
| Zfp622 | 0.140613901 | 0.042546419 | 0.325502204 | 0.521872704 | 0.394534466 |
| Zfp626 | 0.051717576 | 0.054070687 | 0.362869235 | 0.050156579 | 0.007385677 |
| Zfp629 | 0.016563056 | 0.567471266 | 0.859340539 | 0.053818963 | 0.038006908 |
| Zfp637 | 0.091810908 | 0.093285453 | 0.45188518 | 0.245815513 | 0.172897744 |
| Zfp638 | -0.045930559 | 0.491770651 | 0.827270742 | 0.479760898 | 0.5200628 |
| Zfp639 | 0.109011831 | 0.069069754 | 0.399753362 | 0.335882163 | 0.223813203 |
| Zfp64 | -0.00282026 | 0.939817523 | 0.985679047 | 0.092972297 | 0.099504041 |
| Zfp641 | -0.036767536 | 0.224671167 | 0.637067124 | 0.054956178 | 0.079854568 |
| Zfp644 | -0.059468417 | 0.364202194 | 0.752624547 | 0.435678823 | 0.495889213 |
| Zfp646 | -0.006702799 | 0.89968428 | 0.973983939 | 0.208741193 | 0.211539819 |
| Zfp65 | 0.041332768 | 0.228075732 | 0.639451668 | 0.077945181 | 0.038553598 |
| Zfp651 | -0.019473745 | 0.651119193 | 0.89622646 | 0.114282013 | 0.124700754 |
| Zfp652 | -0.090811247 | 0.166365596 | 0.568127031 | 0.419920401 | 0.519922762 |
| Zfp654 | 0.069828242 | 0.180931722 | 0.588627104 | 0.190507935 | 0.128358589 |

| | | | | | |
|---|---|---|---|---|---|
| Zfp655 | -0.025380699 | 0.601383048 | 0.876557788 | 0.156924747 | 0.18337579 |
| Zfp664 | -0.04503534 | 0.490387514 | 0.826674417 | 0.512094777 | 0.545544868 |
| Zfp667 | -0.011129062 | 0.857991118 | 0.9586857 | 0.329219781 | 0.328311014 |
| Zfp668 | 0.055184716 | 0.052325185 | 0.357487248 | 0.056972841 | 0.010952459 |
| Zfp672 | 0.132885239 | 0.016897019 | 0.208334711 | 0.249102496 | 0.1283372 |
| Zfp68 | -0.01111834 | 0.840370861 | 0.954911647 | 0.234095772 | 0.25006088 |
| Zfp687 | -0.016651783 | 0.668246942 | 0.902353633 | 0.096723022 | 0.116512154 |
| Zfp688 | 0.056797231 | 0.270927338 | 0.678418869 | 0.200375009 | 0.155987449 |
| Zfp689 | -0.020809751 | 0.349722958 | 0.7406656 | 0.025528958 | 0.045280042 |
| Zfp692 | -0.019556139 | 0.400764452 | 0.774740795 | 0.029778107 | 0.04657505 |
| Zfp703 | 0.006171467 | 0.850010402 | 0.957111129 | 0.064969571 | 0.060267667 |
| Zfp704 | 0.060265612 | 0.325512323 | 0.720833548 | 0.342307492 | 0.279837987 |
| Zfp706 | -0.023970791 | 0.710126018 | 0.914486281 | 0.936256185 | 0.970921622 |
| Zfp707 | 0.0247382 | 0.421107846 | 0.787234019 | 0.061639368 | 0.040473298 |
| Zfp710 | 0.01443974 | 0.719003852 | 0.918615283 | 0.109613364 | 0.090937289 |
| Zfp719 | 0.047280958 | 0.237887149 | 0.649177085 | 0.110241161 | 0.06955741 |
| Zfp729a | -0.057662526 | 0.184032351 | 0.590326848 | 0.118921196 | 0.183420146 |
| Zfp729b | 0.042860175 | 0.254217635 | 0.663146569 | 0.098594375 | 0.070883032 |
| Zfp738 | -0.014920147 | 0.754738632 | 0.926726954 | 0.149718266 | 0.162786888 |
| Zfp74 | -0.010773263 | 0.780536742 | 0.935383906 | 0.100004579 | 0.109444533 |
| Zfp740 | -0.035023041 | 0.443267559 | 0.801368502 | 0.134740995 | 0.167858669 |
| Zfp746 | -0.052828045 | 0.297455277 | 0.702656882 | 0.183438698 | 0.231079898 |
| Zfp747 | -0.037045683 | 0.163606311 | 0.563469839 | 0.040370744 | 0.070705463 |
| Zfp748 | 0.049040646 | 0.15110362 | 0.546523037 | 0.080675201 | 0.042672257 |
| Zfp758 | -0.038460637 | 0.337621793 | 0.732352032 | 0.101750064 | 0.138638274 |
| Zfp768 | 0.065590212 | 0.076441743 | 0.417435132 | 0.096263486 | 0.040116535 |
| Zfp770 | 0.030234044 | 0.353469825 | 0.744234211 | 0.067958456 | 0.045665902 |
| Zfp771 | 0.097211648 | 0.095799214 | 0.456136959 | 0.280345486 | 0.207278363 |
| Zfp775 | 0.049205788 | 0.129928214 | 0.509514066 | 0.071952968 | 0.034493968 |
| Zfp777 | 0.013251951 | 0.712982952 | 0.915509496 | 0.084454456 | 0.076664446 |
| Zfp780b | -0.009866634 | 0.806317671 | 0.945445148 | 0.108901584 | 0.118272001 |
| Zfp787 | 0.097721688 | 0.104330352 | 0.47169538 | 0.316152309 | 0.226735226 |
| Zfp788 | -0.008450631 | 0.853765172 | 0.957461155 | 0.139492211 | 0.146675158 |
| Zfp790 | -0.016040804 | 0.487678344 | 0.826112275 | 0.029665894 | 0.044189176 |
| Zfp799 | 0.037628005 | 0.253910611 | 0.663146569 | 0.07565445 | 0.038818596 |
| Zfp800 | 0.0363818 | 0.543616115 | 0.850865857 | 0.320658529 | 0.276378593 |
| Zfp809 | 0.050164204 | 0.269498498 | 0.677215248 | 0.14250195 | 0.099485909 |
| Zfp81 | -0.040340385 | 0.27057685 | 0.678097339 | 0.081528338 | 0.114116486 |
| Zfp810 | -0.019958489 | 0.640342245 | 0.891345382 | 0.119559137 | 0.127262974 |
| Zfp820 | -0.073923213 | 0.003187347 | 0.078991107 | 0.026740933 | 0.093559138 |
| Zfp821 | 0.014281656 | 0.742413707 | 0.923637085 | 0.131782389 | 0.115822675 |
| Zfp827 | -0.044725822 | 0.310128141 | 0.712953768 | 0.126512645 | 0.158889397 |
| Zfp830 | -0.048484493 | 0.340427942 | 0.733549861 | 0.171019012 | 0.212994783 |
| Zfp839 | 0.003170953 | 0.91604844 | 0.980132949 | 0.058954505 | 0.054572948 |
| Zfp84 | -0.071606414 | 0.088891724 | 0.443672569 | 0.110641707 | 0.174943758 |
| Zfp846 | 0.008499144 | 0.782617665 | 0.936026395 | 0.060485636 | 0.056767648 |
| Zfp862-ps | -0.027581 | 0.457239427 | 0.808898849 | 0.086391413 | 0.115196818 |
| Zfp865 | 0.029965223 | 0.415065852 | 0.783085706 | 0.093294334 | 0.061898062 |
| Zfp866 | 0.003146757 | 0.916678668 | 0.980132949 | 0.05533677 | 0.056868058 |
| Zfp868 | 0.022997174 | 0.499658219 | 0.830722138 | 0.07725352 | 0.061197125 |
| Zfp869 | 0.000422577 | 0.994684796 | 0.998704266 | 0.401277401 | 0.410983262 |
| Zfp87 | 0.011927875 | 0.724536823 | 0.919830015 | 0.077013643 | 0.070588324 |
| Zfp871 | 0.028379051 | 0.666896922 | 0.901500931 | 0.509061553 | 0.4736172 |
| Zfp874a | -0.010048361 | 0.714329575 | 0.915840939 | 0.044120623 | 0.055930597 |
| Zfp882 | -0.050146715 | 0.022071131 | 0.236422565 | 0.022204507 | 0.063759243 |
| Zfp9 | -0.002264814 | 0.968260459 | 0.993505636 | 0.236156821 | 0.242595887 |
| Zfp90 | 0.026863405 | 0.401954469 | 0.775220138 | 0.06803942 | 0.042647746 |
| Zfp91 | -0.028768526 | 0.639460542 | 0.891345382 | 0.410190976 | 0.431820481 |
| Zfp92 | -0.078831866 | 0.058043402 | 0.37129897 | 0.10730545 | 0.17036651 |
| Zfp930 | -0.007332265 | 0.797741361 | 0.940696966 | 0.050408244 | 0.057892595 |
| Zfp931 | 0.057520029 | 0.138080841 | 0.523580431 | 0.104934813 | 0.051331658 |
| Zfp932 | 0.043202482 | 0.178891278 | 0.585380409 | 0.071475208 | 0.036534443 |
| Zfp937 | -0.044908231 | 0.074968892 | 0.41344108 | 0.033919167 | 0.072070337 |
| Zfp938 | 0.041713133 | 0.477418832 | 0.818492016 | 0.304488121 | 0.270113526 |
| Zfp94 | -0.042723238 | 0.064040815 | 0.385820926 | 0.024665304 | 0.062107431 |

| | | | | | |
|---|---|---|---|---|---|
| Zfp941 | -0.019312791 | 0.581949529 | 0.866173066 | 0.077091592 | 0.095217109 |
| Zfp942 | 0.006977493 | 0.86063778 | 0.959655466 | 0.098700913 | 0.09437449 |
| Zfp943 | -0.047570842 | 0.20454524 | 0.616256046 | 0.088800079 | 0.129101906 |
| Zfp944 | -0.049479797 | 0.17343688 | 0.575630234 | 0.087156087 | 0.13033779 |
| Zfp945 | -0.054817234 | 0.247489829 | 0.657853659 | 0.146340683 | 0.188070659 |
| Zfp946 | 0.008761001 | 0.811753377 | 0.948397417 | 0.086530736 | 0.07614845 |
| Zfp948 | 0.030343373 | 0.39896989 | 0.774544012 | 0.086396129 | 0.067575895 |
| Zfp949 | 0.018060529 | 0.644682299 | 0.893466123 | 0.10348656 | 0.085040889 |
| Zfp950 | -0.028616701 | 0.650989503 | 0.896170847 | 0.379245725 | 0.393044515 |
| Zfp952 | 0.030732842 | 0.411219372 | 0.780671239 | 0.098444212 | 0.078236483 |
| Zfp954 | 0.008986389 | 0.759726511 | 0.927720653 | 0.05781225 | 0.053734626 |
| Zfp955b | 0.007453742 | 0.831794326 | 0.953695326 | 0.080455573 | 0.071058127 |
| Zfp956 | -0.000382484 | 0.988744808 | 0.997411191 | 0.043645203 | 0.041400088 |
| Zfp958 | 0.002987335 | 0.932946531 | 0.983777411 | 0.079904797 | 0.081077509 |
| Zfp959 | 0.042397297 | 0.222523658 | 0.634095002 | 0.080023771 | 0.046646891 |
| Zfp961 | -0.006145491 | 0.8617364 | 0.960067516 | 0.080199187 | 0.08520315 |
| Zfp974 | -0.020997618 | 0.390935851 | 0.769527973 | 0.032688948 | 0.05392779 |
| Zfp975 | 0.034136525 | 0.252725294 | 0.663146569 | 0.058881585 | 0.032123066 |
| Zfp976 | -0.014346114 | 0.640473817 | 0.891345382 | 0.057476501 | 0.068359505 |
| Zfp983 | 0.048208728 | 0.177768431 | 0.583473427 | 0.087674809 | 0.042797472 |
| Zfp994 | 0.007613781 | 0.874457628 | 0.964947297 | 0.153564043 | 0.154479302 |
| Zfp995 | 0.019409819 | 0.477876742 | 0.818590256 | 0.04852369 | 0.029986101 |
| Zfpl1 | 0.059082447 | 0.307585564 | 0.710199288 | 0.269211474 | 0.217982277 |
| Zfr | -0.040702423 | 0.539605002 | 0.84899357 | 0.969373025 | 1.027389049 |
| Zfr2 | -0.033051857 | 0.338076455 | 0.732352032 | 0.073421649 | 0.105890916 |
| Zfx | 0.045370218 | 0.376241025 | 0.760785796 | 0.19849931 | 0.157796354 |
| Zfyve1 | -0.009284119 | 0.728560929 | 0.920519505 | 0.042789487 | 0.048933618 |
| Zfyve16 | -0.042882965 | 0.414141548 | 0.782518584 | 0.19766994 | 0.244317757 |
| Zfyve19 | 0.013130298 | 0.722048485 | 0.919475778 | 0.091750313 | 0.080585189 |
| Zfyve21 | 0.031281667 | 0.604501021 | 0.877736706 | 0.33264891 | 0.314858299 |
| Zfyve26 | 0.066260531 | 0.040124677 | 0.315774775 | 0.06875752 | 0.010161188 |
| Zfyve27 | -0.016367137 | 0.701425759 | 0.912152193 | 0.122233798 | 0.136838748 |
| Zfyve9 | -0.054229953 | 0.366479726 | 0.75345289 | 0.302264996 | 0.337399835 |
| Zgpat | -0.05083498 | 0.238842339 | 0.650445287 | 0.118336315 | 0.167939758 |
| Zhx1 | -0.099982122 | 0.098175545 | 0.461540022 | 0.319798014 | 0.416803333 |
| Zhx2 | 0.01032304 | 0.79776951 | 0.940696966 | 0.107719432 | 0.095629575 |
| Zhx3 | 0.052510737 | 0.209365766 | 0.62245385 | 0.129317697 | 0.082339399 |
| Zim1 | 0.029740777 | 0.438620941 | 0.799411376 | 0.097577173 | 0.072169536 |
| Zkscan1 | -0.116291746 | 0.058450422 | 0.372250559 | 0.316242675 | 0.427004298 |
| Zkscan14 | 0.003235472 | 0.919149066 | 0.980436936 | 0.062024016 | 0.066316941 |
| Zkscan17 | 0.044694579 | 0.233726053 | 0.644459522 | 0.099454295 | 0.06134736 |
| Zkscan3 | -0.00858274 | 0.888764955 | 0.970176435 | 0.341470861 | 0.353368272 |
| Zkscan6 | 0.01977947 | 0.624470905 | 0.88668351 | 0.108622113 | 0.09793213 |
| Zkscan7 | -0.029368191 | 0.356283096 | 0.745571198 | 0.061403772 | 0.087214206 |
| Zkscan8 | 0.005547181 | 0.883842737 | 0.968954079 | 0.097019703 | 0.088150565 |
| Zmat1 | -0.018420962 | 0.574275101 | 0.862168892 | 0.064320626 | 0.082244967 |
| Zmat2 | 0.058681288 | 0.38516487 | 0.765373153 | 0.611626325 | 0.563021378 |
| Zmat3 | -0.024960249 | 0.644555457 | 0.893466123 | 0.224950721 | 0.245773495 |
| Zmat5 | -0.035443652 | 0.486867859 | 0.825733812 | 0.191024081 | 0.224862155 |
| Zmiz1 | 0.037963146 | 0.562818682 | 0.857583554 | 0.441543898 | 0.379631982 |
| Zmiz2 | 0.060743129 | 0.280906527 | 0.68791143 | 0.262584303 | 0.200661141 |
| Zmpste24 | -0.089923519 | 0.147066807 | 0.537157767 | 0.332219323 | 0.41458989 |
| Zmym1 | 0.015973155 | 0.661281749 | 0.898920864 | 0.087236411 | 0.074932827 |
| Zmym2 | 0.033509336 | 0.560153822 | 0.856904336 | 0.270802054 | 0.248502504 |
| Zmym3 | -0.010727628 | 0.752403866 | 0.926186851 | 0.073436444 | 0.081705528 |
| Zmym4 | -0.018485373 | 0.761542635 | 0.928598328 | 0.308556596 | 0.323049093 |
| Zmym5 | -0.007567749 | 0.907776442 | 0.977612891 | 0.382048038 | 0.388741272 |
| Zmym6 | -0.025693659 | 0.407922561 | 0.77823964 | 0.059024821 | 0.086403286 |
| Zmynd11 | -0.076895175 | 0.25054419 | 0.662289186 | 0.586209975 | 0.639467798 |
| Zmynd19 | 0.066339987 | 0.145403882 | 0.533628684 | 0.141909427 | 0.083744062 |
| Zmynd8 | 0.086418023 | 0.225487717 | 0.638019508 | 0.755978815 | 0.672265225 |
| Znfx1 | 0.050240322 | 0.398761172 | 0.774321615 | 0.296315401 | 0.257342421 |
| Znhit1 | 0.048911536 | 0.439458812 | 0.800048525 | 0.432129928 | 0.382764522 |
| Znhit2 | -0.004192235 | 0.941954598 | 0.986168593 | 0.247197775 | 0.253706914 |
| Znhit3 | -0.098858492 | 0.057145631 | 0.369250461 | 0.175285857 | 0.259264222 |

| | | | | | |
|---|---|---|---|---|---|
| Znhit6 | 0.04884563 | 0.335640565 | 0.730736446 | 0.177659237 | 0.134082766 |
| Znrf1 | 0.012182648 | 0.794254501 | 0.939757086 | 0.148241357 | 0.131649039 |
| Znrf2 | -0.009567252 | 0.882101111 | 0.968458636 | 0.468769581 | 0.48636198 |
| Znrf3 | -0.009289144 | 0.751809727 | 0.926186851 | 0.050626101 | 0.062558248 |
| Zpr1 | -0.053120722 | 0.33629439 | 0.731236306 | 0.211338819 | 0.282440313 |
| Zranb1 | 0.002047044 | 0.976714392 | 0.994468977 | 0.74032894 | 0.747158405 |
| Zranb2 | -0.160439356 | 0.018947983 | 0.218347772 | 0.749434947 | 0.909513472 |
| Zrsr1 | 0.008077721 | 0.881392927 | 0.968081674 | 0.223793402 | 0.213808043 |
| Zrsr2 | 0.053598446 | 0.39904789 | 0.774544012 | 0.439243744 | 0.391118503 |
| Zscan12 | 0.010569084 | 0.758275388 | 0.927355924 | 0.078029706 | 0.070555837 |
| Zscan18 | -0.027234001 | 0.240565692 | 0.650744882 | 0.027064414 | 0.04895386 |
| Zscan21 | 0.014395107 | 0.649683465 | 0.895585658 | 0.067006027 | 0.052452813 |
| Zscan22 | 0.003477469 | 0.927189611 | 0.981818446 | 0.094904313 | 0.0917225 |
| Zscan26 | -0.038880382 | 0.506922995 | 0.833276466 | 0.257368944 | 0.309986671 |
| Zscan29 | 0.066610323 | 0.074798531 | 0.41344108 | 0.098808091 | 0.041606409 |
| Zswim1 | 0.053220641 | 0.176469585 | 0.580732554 | 0.106450862 | 0.061918257 |
| Zswim5 | 0.053189275 | 0.103347084 | 0.469315384 | 0.075005003 | 0.030477854 |
| Zswim7 | -0.054792029 | 0.353039727 | 0.743880213 | 0.277168631 | 0.338861357 |
| Zswim8 | -0.017905569 | 0.766892288 | 0.929400857 | 0.310976548 | 0.319549673 |
| Zup1 | -0.066394046 | 0.222093151 | 0.633924544 | 0.206163452 | 0.259213026 |
| Zw10 | 0.033897026 | 0.373986948 | 0.759249926 | 0.10105261 | 0.073744702 |
| Zwint | -0.090965433 | 0.197955042 | 0.609076376 | 0.676215511 | 0.769717493 |
| Zxdb | -0.032559171 | 0.460299663 | 0.810111566 | 0.130863312 | 0.16540185 |
| Zxdc | -0.048128579 | 0.358411963 | 0.747310348 | 0.189315719 | 0.237922406 |
| Zyg11b | -0.15579867 | 0.015450601 | 0.199744575 | 0.440203796 | 0.602748158 |
| Zyx | 0.004889382 | 0.86488244 | 0.961442912 | 0.051390559 | 0.043265725 |
| Zzef1 | -0.029112043 | 0.600628851 | 0.876237177 | 0.236810442 | 0.264841532 |
| Zzz3 | -0.021361518 | 0.738197272 | 0.922843014 | 0.396117426 | 0.400221414 |
