## Supplementary data file 3 for "Sorting receptor SORCS2 facilitates a protective stress response in pancreatic islets"

| GeneID | Log2FC | p-value | FDR | Average (Sorcs2-/-) | Average (Sorcs2+/+) |
|---|---|---|---|---|---|
| 0610009B22Rik | 0.0701661 | 0.050547746 | 0.4231507 | 0.602008529 | 0.475790572 |
| 0610009E02Rik | 0.010642308 | 0.561193434 | 0.8926240 | 0.078473313 | 0.064646662 |
| 0610009L18Rik | -0.056718103 | 0.015444662 | 0.2428067 | 0.089960997 | 0.130135154 |
| 0610010F05Rik | -0.037549902 | 0.111090633 | 0.5671739 | 0.106379027 | 0.13409742 |
| 0610010K14Rik | 0.02221618 | 0.408035802 | 0.8270770 | 0.223179793 | 0.167257029 |
| 0610012G03Rik | 0.06113849 | 0.082114616 | 0.5148480 | 0.656324354 | 0.587456218 |
| 0610030E20Rik | -0.009283843 | 0.73903754 | 0.9474350 | 0.180639342 | 0.192191333 |
| 0610040B10Rik | 0.012359196 | 0.504515354 | 0.8695566 | 0.072721658 | 0.072344159 |
| 0610040J01Rik | 0.029454325 | 0.403570815 | 0.8264703 | 0.457962182 | 0.40658122 |
| 1010001B22Rik | -0.005801252 | 0.72084688 | 0.9445652 | 0.05304201 | 0.056359409 |
| 1110002L01Rik | -0.015445943 | 0.397996682 | 0.8238469 | 0.064677023 | 0.074816189 |
| 1110004F10Rik | 0.021632975 | 0.5375551 | 0.8814126 | 0.875242562 | 0.80125819 |
| 1110008P14Rik | 0.055250103 | 0.130501102 | 0.5943851 | 0.795787667 | 0.734414907 |
| 1110018N20Rik | -0.053249067 | 0.027623367 | 0.3181899 | 0.104163385 | 0.14346521 |
| 1110019D14Rik | -0.038117811 | 0.009445857 | 0.1947602 | 0.023774294 | 0.052514175 |
| 1110020A21Rik | -0.042008078 | 0.078063343 | 0.5048164 | 0.107887326 | 0.132991918 |
| 1110032A03Rik | -0.026002565 | 0.425486258 | 0.8325096 | 0.314325937 | 0.332472498 |
| 1110035H17Rik | -0.00734953 | 0.639154355 | 0.9160042 | 0.048989 | 0.051935886 |
| 1110038B12Rik | -0.0602803 | 0.045424159 | 0.4035299 | 0.187095911 | 0.242606761 |
| 1110038F14Rik | 0.020593142 | 0.530931094 | 0.8776448 | 0.396226631 | 0.349658076 |
| 1110046J04Rik | 0.006199814 | 0.669816728 | 0.9239496 | 0.044351988 | 0.042636519 |
| 1110051M20Rik | 0.00024943 | 0.991040306 | 0.9982478 | 0.119931216 | 0.115023978 |
| 1110059E24Rik | -0.000197498 | 0.994016868 | 0.9985888 | 0.181768291 | 0.162680818 |
| 1110059G10Rik | 0.025810467 | 0.360547698 | 0.8012480 | 0.230097736 | 0.183569915 |
| 1110065P20Rik | 0.012778359 | 0.715412804 | 0.9431411 | 0.772159903 | 0.724549569 |
| 1190007I07Rik | -0.033775029 | 0.240271238 | 0.7120404 | 0.19958748 | 0.227582006 |
| 1300002E11Rik | 0.003557721 | 0.911705064 | 0.9840185 | 0.337018263 | 0.311015156 |
| 1500004A13Rik | 0.009911864 | 0.650361178 | 0.9177795 | 0.121722999 | 0.098298647 |
| 1500009L16Rik | 0.016267439 | 0.540518987 | 0.8834978 | 0.156673084 | 0.178694568 |
| 1500011B03Rik | -0.039553541 | 0.262710855 | 0.7275735 | 0.927280975 | 0.916318781 |
| 1500015A07Rik | -0.012251295 | 0.655067882 | 0.9206439 | 0.180322295 | 0.181876205 |
| 1500026H17Rik | 0.003218894 | 0.882794297 | 0.9799360 | 0.098276357 | 0.117175621 |
| 1600012H06Rik | 0.00018431 | 0.993428011 | 0.9985888 | 0.122840501 | 0.111598119 |
| 1600014C10Rik | -0.013359652 | 0.338883952 | 0.7834245 | 0.036476055 | 0.041116036 |
| 1600020E01Rik | -0.001146429 | 0.962047797 | 0.9963994 | 0.128672898 | 0.132426164 |
| 1700007L15Rik | 0.005597837 | 0.791800043 | 0.9624054 | 0.101382274 | 0.095670509 |
| 1700008J07Rik | -0.018695281 | 0.235119656 | 0.7069981 | 0.047585237 | 0.055850866 |
| 1700017B05Rik | 0.000706086 | 0.960415292 | 0.9958628 | 0.043401346 | 0.041873789 |
| 1700019D03Rik | 0.053630751 | 0.089680166 | 0.5302360 | 0.334534612 | 0.279397641 |
| 1700025G04Rik | -0.003437564 | 0.851100204 | 0.9749822 | 0.073890664 | 0.067565329 |
| 1700029J07Rik | -0.027370317 | 0.093799654 | 0.5333348 | 0.04386179 | 0.063543017 |
| 1700037C18Rik | -0.017125955 | 0.29249205 | 0.7526152 | 0.044662401 | 0.058117035 |
| 1700037H04Rik | -0.008731482 | 0.683514124 | 0.9309982 | 0.123086905 | 0.099844214 |
| 1700052K11Rik | -0.016551885 | 0.382611505 | 0.8129162 | 0.074799457 | 0.07802314 |
| 1700056E22Rik | 0.051623322 | 0.020407097 | 0.2774595 | 0.152237355 | 0.098928787 |
| 1700084E18Rik | 0.002430406 | 0.877136183 | 0.9785583 | 0.052802961 | 0.049472858 |
| 1700086L19Rik | -0.157879715 | 0.00021766 | 0.0172735 | 1.64365784 | 1.678825388 |
| 1700086O06Rik | 0.010250555 | 0.535266921 | 0.8795949 | 0.062607495 | 0.055504033 |
| 1700086P04Rik | -0.009272289 | 0.54907471 | 0.8871105 | 0.04583872 | 0.04952408 |
| 1700088E04Rik | 0.010211859 | 0.63634571 | 0.9153541 | 0.101597263 | 0.107649853 |
| 1700096K18Rik | -0.01779286 | 0.519160459 | 0.8741795 | 0.190844333 | 0.18891988 |
| 1700110K17Rik | -0.000393224 | 0.983554535 | 0.9976072 | 0.063348072 | 0.083908483 |
| 1700113A16Rik | 0.052957708 | 0.00497537 | 0.1410161 | 0.109101387 | 0.063972228 |
| 1700123O20Rik | 0.004056012 | 0.893032327 | 0.9822826 | 0.279575403 | 0.259108267 |
| 1810008I18Rik | -0.040849792 | 0.223269655 | 0.6955958 | 0.245789302 | 0.352904929 |
| 1810010H24Rik | 0.018174831 | 0.240071862 | 0.7118791 | 0.061636481 | 0.045532295 |
| 1810013L24Rik | 0.035687959 | 0.191757779 | 0.6700281 | 0.212453733 | 0.180024281 |
| 1810014B01Rik | 0.000888568 | 0.953459904 | 0.9954690 | 0.045784048 | 0.048253671 |
| 1810021B22Rik | -0.000487313 | 0.976069683 | 0.9976072 | 0.05866329 | 0.05339729 |
| 1810024B03Rik | 0.061928606 | 0.000401241 | 0.0261862 | 0.097833847 | 0.053226387 |
| 1810030O07Rik | -0.014888717 | 0.578684608 | 0.9003073 | 0.176485478 | 0.180794511 |
| 1810037I17Rik | 9.71E-05 | 0.996929819 | 0.9994073 | 1.647311346 | 1.621809893 |
| 1810044D09Rik | 0.008946444 | 0.69069828 | 0.9346237 | 0.109726591 | 0.111692655 |
| 1810058I24Rik | 0.027013643 | 0.443243187 | 0.8431541 | 0.634131307 | 0.57712072 |

| | | | | | |
|---|---|---|---|---|---|
| 1810059H22Rik | -0.005659773 | 0.773547297 | 0.9557311 | 0.078299818 | 0.090518252 |
| 1810062G17Rik | -0.007489171 | 0.66612827 | 0.9230765 | 0.054653968 | 0.06755166 |
| 2010001A14Rik | -0.022322052 | 0.154072768 | 0.6304934 | 0.037435003 | 0.056183302 |
| 2010204K13Rik | 0.049646982 | 0.071532513 | 0.4888240 | 0.214560166 | 0.163578024 |
| 2010315B03Rik | -0.030735471 | 0.161204028 | 0.6386357 | 0.090161546 | 0.119052115 |
| 2010320M18Rik | -0.003658525 | 0.904921406 | 0.9831707 | 0.264758004 | 0.253744821 |
| 2210016F16Rik | -0.000692083 | 0.983205179 | 0.9976072 | 0.362271737 | 0.36670632 |
| 2210016L21Rik | 0.041462044 | 0.255196037 | 0.7226790 | 0.579024723 | 0.504838534 |
| 2210408F21Rik | 0.00050694 | 0.986247848 | 0.9976072 | 0.214214817 | 0.228967929 |
| 2210408I21Rik | 0.016279383 | 0.296285181 | 0.7545964 | 0.060382575 | 0.044911221 |
| 2300009A05Rik | 0.009390068 | 0.792290919 | 0.9624460 | 0.922761156 | 0.856829296 |
| 2310001H17Rik | -0.064049716 | 0.000589976 | 0.0344268 | 0.049057773 | 0.080278256 |
| 2310009B15Rik | 0.015147827 | 0.63929966 | 0.9160042 | 0.298498497 | 0.284466168 |
| 2310010J17Rik | 0.033947969 | 0.278896106 | 0.7395480 | 0.329104329 | 0.277607394 |
| 2310011J03Rik | -0.002449868 | 0.934084309 | 0.9904988 | 0.251395925 | 0.244667288 |
| 2310015A10Rik | -0.03934144 | 0.052459552 | 0.4319240 | 0.063364468 | 0.097438722 |
| 2310022A10Rik | 0.05534039 | 0.002579469 | 0.0927113 | 0.111658577 | 0.061888467 |
| 2310022B05Rik | -0.00389719 | 0.90445621 | 0.9831707 | 0.327433156 | 0.312670437 |
| 2310030G06Rik | 0.000713897 | 0.98258876 | 0.9976072 | 0.366346805 | 0.345820428 |
| 2310033P09Rik | -0.028575002 | 0.402145517 | 0.8254259 | 0.375524449 | 0.357133936 |
| 2310034G01Rik | -0.022458324 | 0.350558233 | 0.7927303 | 0.135426399 | 0.140740576 |
| 2310039H08Rik | 0.084444023 | 0.018381345 | 0.2642651 | 0.933457535 | 0.812354978 |
| 2310040G24Rik | -0.014507215 | 0.413864099 | 0.8300990 | 0.051019355 | 0.071919564 |
| 2310057M21Rik | 0.031511704 | 0.175006159 | 0.6534142 | 0.151303989 | 0.112933517 |
| 2310061I04Rik | 0.049660235 | 0.083858454 | 0.5184165 | 0.252941528 | 0.202776473 |
| 2310068J16Rik | 0.000731885 | 0.963747337 | 0.9966421 | 0.054041152 | 0.053937814 |
| 2410002F23Rik | -0.024726476 | 0.374287705 | 0.8087297 | 0.184135139 | 0.187147386 |
| 2410004B18Rik | -0.01165274 | 0.659948242 | 0.9211780 | 0.194608828 | 0.174848866 |
| 2410006H16Rik | 0.013841293 | 0.692939405 | 0.9359725 | 1.29945774 | 1.321444631 |
| 2410021H03Rik | -0.005890881 | 0.751201016 | 0.9506053 | 0.051487878 | 0.072742369 |
| 2410022M11Rik | 0.015546286 | 0.458924604 | 0.8522649 | 0.112711027 | 0.098422894 |
| 2510002D24Rik | -0.049698172 | 0.125217697 | 0.5870522 | 0.280092535 | 0.319457328 |
| 2510009E07Rik | -0.042601829 | 0.096866062 | 0.5379319 | 0.130151267 | 0.170907777 |
| 2510039O18Rik | 0.019311036 | 0.497003578 | 0.8684629 | 0.228482626 | 0.221302592 |
| 2510046G10Rik | -0.019470973 | 0.463143321 | 0.8524425 | 0.177914117 | 0.172690778 |
| 2610001J05Rik | 0.014852462 | 0.671593078 | 0.9251775 | 0.447209739 | 0.420616499 |
| 2610002M06Rik | -0.000832611 | 0.977873492 | 0.9976072 | 0.268691752 | 0.226151295 |
| 2610005L07Rik | -0.013723296 | 0.469663097 | 0.8567594 | 0.076789381 | 0.080512916 |
| 2610008E11Rik | -0.011088456 | 0.645441442 | 0.9169695 | 0.135379848 | 0.129838425 |
| 2610035D17Rik | -0.014717984 | 0.520573965 | 0.8741795 | 0.121743458 | 0.108329295 |
| 2610037D02Rik | -0.038491014 | 0.074690195 | 0.4966285 | 0.095028226 | 0.103427791 |
| 2610044O15Rik8 | 0.028723665 | 0.092607186 | 0.5325584 | 0.079176611 | 0.051606112 |
| 2610301B20Rik | 0.034087185 | 0.172398564 | 0.6522284 | 0.183049301 | 0.137454864 |
| 2610306M01Rik | -0.005507611 | 0.826684854 | 0.9716906 | 0.168379273 | 0.152816457 |
| 2610507B11Rik | -0.056401358 | 0.06216503 | 0.4629707 | 0.227890532 | 0.277404119 |
| 2700033N17Rik | 0.038904963 | 0.018870085 | 0.2677986 | 0.072931849 | 0.049284454 |
| 2700046G09Rik | -0.014179545 | 0.538514761 | 0.8816335 | 0.117950311 | 0.127911875 |
| 2700049A03Rik | 0.006266764 | 0.733342631 | 0.9464134 | 0.081335951 | 0.068111139 |
| 2700062C07Rik | -0.034392368 | 0.206169281 | 0.6797565 | 0.174009387 | 0.189614956 |
| 2700081O15Rik | -0.023722935 | 0.13029045 | 0.5939711 | 0.041154835 | 0.055961325 |
| 2700097O09Rik | 0.023957791 | 0.245060802 | 0.7141607 | 0.118584891 | 0.083890516 |
| 2810001G20Rik | -0.019521605 | 0.401815901 | 0.8254259 | 0.119782881 | 0.130605617 |
| 2810004N23Rik | 0.017964444 | 0.568910013 | 0.8956532 | 0.280138253 | 0.28012278 |
| 2810006K23Rik | -0.01749697 | 0.459987991 | 0.8524343 | 0.121878251 | 0.133293434 |
| 2810013P06Rik | -0.104521371 | 0.00111585 | 0.0519682 | 0.233632966 | 0.314353311 |
| 2810025M15Rik | -0.06055602 | 0.036753779 | 0.3626279 | 0.182774407 | 0.234656819 |
| 2810402E24Rik | 0.011865178 | 0.627625528 | 0.9128095 | 0.147566271 | 0.136966082 |
| 2810403D21Rik | -0.026086992 | 0.125065016 | 0.5870522 | 0.04674598 | 0.06338456 |
| 2810454H06Rik | -0.028126119 | 0.094734224 | 0.5350704 | 0.042505338 | 0.06069685 |
| 2900005J15Rik | -0.002442151 | 0.891927653 | 0.9819597 | 0.06303698 | 0.072990909 |
| 2900026A02Rik | -0.023189589 | 0.492980044 | 0.8669301 | 0.322026095 | 0.340398303 |
| 2900027M19Rik | -0.006237457 | 0.829297566 | 0.9722601 | 0.179158272 | 0.187129845 |
| 2900052L18Rik | -0.006951605 | 0.745884171 | 0.9485717 | 0.100066597 | 0.099445497 |
| 2900076A07Rik | 0.018252042 | 0.411173744 | 0.8290007 | 0.124148977 | 0.105632394 |
| 2900092N22Rik | 0.006793624 | 0.672904532 | 0.9260839 | 0.051888521 | 0.051050017 |
| 2900093K20Rik | -0.001828076 | 0.951273948 | 0.9950060 | 0.257283634 | 0.250266884 |

| | | | | | |
|---|---|---|---|---|---|
| 3010003L21Rik | 0.012961611 | 0.522765538 | 0.8750611 | 0.087636249 | 0.080791408 |
| 3110009E18Rik | -0.025131523 | 0.170426595 | 0.6507045 | 0.059813464 | 0.078545305 |
| 3110040N11Rik | 0.046702043 | 0.13623409 | 0.6038615 | 0.326314001 | 0.255533904 |
| 3110056K07Rik | -0.03286382 | 0.19436705 | 0.6717191 | 0.14099117 | 0.156220788 |
| 3110080O07Rik | 0.014520938 | 0.347890933 | 0.7917953 | 0.059402135 | 0.042503283 |
| 3110082I17Rik | -0.001710147 | 0.944260495 | 0.9936932 | 0.155998562 | 0.130300923 |
| 3222401L13Rik | -0.038345462 | 0.142730268 | 0.6158696 | 0.111652472 | 0.166712675 |
| 3830406C13Rik | -0.007363882 | 0.812037722 | 0.9673015 | 0.283977829 | 0.259175519 |
| 3930402G23Rik | -0.038618261 | 0.038997147 | 0.3786326 | 0.038753756 | 0.076273048 |
| 4632404H12Rik | 0.008165298 | 0.582168295 | 0.9018999 | 0.053656735 | 0.038316202 |
| 4632415L05Rik | -0.020481089 | 0.369206905 | 0.8056719 | 0.105911386 | 0.11894955 |
| 4632427E13Rik | 0.025738185 | 0.45751795 | 0.8518352 | 0.377113323 | 0.321559444 |
| 4632428C04Rik | -0.014484488 | 0.364978198 | 0.8050191 | 0.051866999 | 0.054032393 |
| 4732440D04Rik | -0.02804654 | 0.087872683 | 0.5278913 | 0.044065392 | 0.061875069 |
| 4732491K20Rik | -0.010492407 | 0.509082606 | 0.8704067 | 0.053856243 | 0.052134101 |
| 4833420G17Rik | -0.024416544 | 0.409110453 | 0.8276447 | 0.231357707 | 0.220578455 |
| 4833439L19Rik | 0.160137914 | 1.06E-05 | 0.0016229 | 0.659539354 | 0.520499866 |
| 4921524J17Rik | 0.014281934 | 0.627584653 | 0.9128095 | 0.248212184 | 0.229309679 |
| 4930402H24Rik | -0.024909075 | 0.390485332 | 0.8191191 | 0.189037898 | 0.221505631 |
| 4930430F08Rik | -0.017046348 | 0.44267333 | 0.8431541 | 0.107659354 | 0.111044868 |
| 4930431P19Rik | -0.00290457 | 0.898794557 | 0.9825441 | 0.124255967 | 0.116819958 |
| 4930453N24Rik | 0.013416188 | 0.634232176 | 0.9144743 | 0.216390258 | 0.206026609 |
| 4930481A15Rik | -0.028472673 | 0.096297801 | 0.5369725 | 0.050549866 | 0.069079591 |
| 4930503L19Rik | -0.013938347 | 0.489423678 | 0.8651997 | 0.082872661 | 0.091298668 |
| 4930522L14Rik | 0.007515508 | 0.699618952 | 0.9372491 | 0.096891671 | 0.080060995 |
| 4930563E22Rik | 0.014951473 | 0.276569484 | 0.7367264 | 0.048009026 | 0.033553512 |
| 4930570G19Rik | -0.009573554 | 0.55210817 | 0.8894073 | 0.053127611 | 0.05365925 |
| 4930578M01Rik | -0.03710603 | 0.055261452 | 0.4435223 | 0.054610669 | 0.084634201 |
| 4930579K19Rik | 0.011226824 | 0.674911799 | 0.9265887 | 0.193162105 | 0.165939429 |
| 4930581F22Rik | -0.041282418 | 0.023461111 | 0.2972960 | 0.042833016 | 0.076616807 |
| 4930589O11Rik | -0.009232336 | 0.57409875 | 0.8974946 | 0.049645316 | 0.049232228 |
| 4931406C07Rik | 0.004269507 | 0.889920511 | 0.9819597 | 0.22479118 | 0.236702556 |
| 4931440P22Rik | -0.034136104 | 0.046236912 | 0.4073656 | 0.049715372 | 0.065297813 |
| 4932422M17Rik | 0.020813278 | 0.1544347 | 0.6304934 | 0.055817922 | 0.037360766 |
| 4932438A13Rik | -0.014572865 | 0.68233097 | 0.9304086 | 0.444492353 | 0.478658693 |
| 4933412E12Rik | -0.010944904 | 0.448459439 | 0.8463296 | 0.043298936 | 0.042745839 |
| 4933421O10Rik | -0.003776728 | 0.901843125 | 0.9825441 | 0.263514329 | 0.263729294 |
| 4933434E20Rik | 0.054372261 | 0.093179427 | 0.5330587 | 0.349620933 | 0.267750215 |
| 5031425E22Rik | -0.028586299 | 0.364041969 | 0.8049171 | 0.276374007 | 0.277819611 |
| 5031439G07Rik | -0.031207234 | 0.106067474 | 0.5617171 | 0.059773704 | 0.085644998 |
| 5033406O09Rik | -0.028537564 | 0.047443025 | 0.4106761 | 0.02414894 | 0.048992618 |
| 5033430I15Rik | 0.002289306 | 0.909580984 | 0.9837373 | 0.08464619 | 0.089124619 |
| 5330406M23Rik | -0.038738633 | 0.034666199 | 0.3548902 | 0.044640301 | 0.077845518 |
| 5330431K02Rik | -0.045253722 | 0.022793341 | 0.2926894 | 0.057301598 | 0.093336315 |
| 5330434G04Rik | -0.083550526 | 0.004786485 | 0.1372310 | 0.151698697 | 0.218712119 |
| 5330438D12Rik | -0.0795196 | 0.003380603 | 0.1111729 | 0.146461072 | 0.18498352 |
| 5330439M10Rik | -0.007904081 | 0.802387385 | 0.9647143 | 0.205531023 | 0.218052427 |
| 5430403G16Rik | -0.024332423 | 0.278852538 | 0.7395480 | 0.115464613 | 0.114372651 |
| 5430405H02Rik | -0.017360807 | 0.453749058 | 0.8490482 | 0.115042011 | 0.118833861 |
| 5430416N02Rik | 0.019037016 | 0.519365468 | 0.8741795 | 0.216608077 | 0.21576329 |
| 5530601H04Rik | -0.002377028 | 0.90635382 | 0.9835423 | 0.105192811 | 0.086348865 |
| 5730409E04Rik | -0.004686958 | 0.781418361 | 0.9590847 | 0.062473826 | 0.06164486 |
| 5730455P16Rik | 0.030727087 | 0.261984363 | 0.7275735 | 0.214034336 | 0.173627151 |
| 5730480H06Rik | 0.018874373 | 0.343444283 | 0.7872955 | 0.104678101 | 0.079035897 |
| 5830444B04Rik | 0.019863923 | 0.218219104 | 0.6897120 | 0.0656995 | 0.046785087 |
| 5830448L01Rik | -0.018497485 | 0.227728747 | 0.7015153 | 0.045607161 | 0.05397909 |
| 5830454E08Rik | 0.004917741 | 0.811459418 | 0.9673015 | 0.101684378 | 0.090624249 |
| 5930430L01Rik | -0.015603328 | 0.437947131 | 0.8399914 | 0.078644596 | 0.087808772 |
| 6030458C11Rik | -0.024035829 | 0.254464047 | 0.7223071 | 0.077447038 | 0.101450763 |
| 6330418K02Rik | -0.000364841 | 0.985361434 | 0.9976072 | 0.087010406 | 0.08386245 |
| 6430503K07Rik | 0.009985945 | 0.582763242 | 0.9019790 | 0.079000024 | 0.063845477 |
| 6430548M08Rik | 0.034584334 | 0.196545709 | 0.6730962 | 0.180350229 | 0.164600366 |
| 6430590A07Rik | -0.020550759 | 0.354518496 | 0.7969131 | 0.103968388 | 0.108275745 |
| 6720427I07Rik | -0.022938039 | 0.165054706 | 0.6420707 | 0.046602968 | 0.062078975 |
| 6820431F20Rik | -0.016235268 | 0.667592803 | 0.9230765 | 0.594044032 | 0.561330334 |
| 7SK | -0.007522938 | 0.644309435 | 0.9168391 | 0.048228332 | 0.053089532 |

| | | | | | |
|---|---|---|---|---|---|
| 8030462N17Rik | 0.051521578 | 0.027041026 | 0.31558468 | 0.171255344 | 0.115845651 |
| 8430429K09Rik | -0.00326386 | 0.857467412 | 0.97631891 | 0.067113585 | 0.069091462 |
| 9130016M20Rik | -0.005661546 | 0.759262302 | 0.95215339 | 0.080262748 | 0.073045406 |
| 9130023H24Rik | -0.009355468 | 0.572543077 | 0.89697209 | 0.052923301 | 0.061989976 |
| 9130401M01Rik | -0.004782673 | 0.870455584 | 0.97801781 | 0.24486038 | 0.234915017 |
| 9230114K14Rik | 0.003892621 | 0.861190681 | 0.97696750 | 0.111535793 | 0.110506955 |
| 9330020H09Rik | -0.014763044 | 0.497641518 | 0.86846291 | 0.096827355 | 0.108791769 |
| 9330151L19Rik | -0.027827873 | 0.067305178 | 0.48137517 | 0.038317744 | 0.052855815 |
| 9330160F10Rik | -0.050876968 | 0.014509827 | 0.23719126 | 0.064986402 | 0.103220786 |
| 9330162012Rik | -0.007321364 | 0.658401757 | 0.92117801 | 0.053113364 | 0.057821664 |
| 9330162G02Rik | -0.002324992 | 0.910858938 | 0.98374748 | 0.090598361 | 0.091394816 |
| 9430038I01Rik | -0.040547465 | 0.027475102 | 0.31818990 | 0.057055776 | 0.08159102 |
| 9430065F17Rik | 0.004696386 | 0.781213949 | 0.95908471 | 0.064511683 | 0.058523042 |
| 9430081H08Rik | -0.023745443 | 0.271428834 | 0.73420300 | 0.086226597 | 0.103819951 |
| 9530068E07Rik | -0.030592749 | 0.385246901 | 0.81398280 | 0.483487842 | 0.490051952 |
| 9530077C05Rik | -0.021030779 | 0.2319521 | 0.70301400 | 0.051261047 | 0.072985774 |
| 9930017N22Rik | 0.000995252 | 0.955044039 | 0.99546908 | 0.064708317 | 0.065215958 |
| 9930021J03Rik | 0.027655904 | 0.416260403 | 0.83009907 | 0.374938295 | 0.332958317 |
| A030001D20Rik | 0.014453604 | 0.528037541 | 0.87711527 | 0.131323705 | 0.113472625 |
| A130010J15Rik | -0.042199537 | 0.017593719 | 0.25824014 | 0.045659069 | 0.076021888 |
| A130014A01Rik | -0.035945707 | 0.090556184 | 0.53210319 | 0.08122859 | 0.103273191 |
| A1cf | -0.030774937 | 0.23033504 | 0.70197349 | 0.150701121 | 0.14715941 |
| A230051N06Rik | -0.085244771 | 0.005958354 | 0.15454384 | 0.22597878 | 0.273276279 |
| A230057D06Rik | -0.035724532 | 0.310813742 | 0.76793831 | 0.269388378 | 0.348111514 |
| A2ml1 | 0.015966244 | 0.610558582 | 0.90908029 | 0.283372065 | 0.214049303 |
| A330023F24Rik | -0.060659191 | 0.004109672 | 0.12660851 | 0.055572905 | 0.118067997 |
| A330076H08Rik | -0.004299768 | 0.917610384 | 0.98481302 | 0.632767254 | 0.720675858 |
| A430005L14Rik | -0.027704216 | 0.425169211 | 0.83244607 | 0.390538829 | 0.403920084 |
| A430018G15Rik | -0.016694662 | 0.391989929 | 0.82016249 | 0.078730118 | 0.085019375 |
| A430033K04Rik | -0.013696861 | 0.446605782 | 0.84435474 | 0.059134924 | 0.072432902 |
| A530017D24Rik | -0.024365778 | 0.105166024 | 0.55938174 | 0.032134363 | 0.05314033 |
| A530058N18Rik | 0.016537966 | 0.573809771 | 0.89749469 | 0.219942906 | 0.244338374 |
| A730020E08Rik | -0.005551482 | 0.756920968 | 0.95215339 | 0.072148214 | 0.067978975 |
| A730063M14Rik | -0.037879465 | 0.019251182 | 0.26955860 | 0.029080792 | 0.064504309 |
| A930005H10Rik | -0.013620928 | 0.563244562 | 0.89298161 | 0.129348865 | 0.134827551 |
| A930029G22Rik | -0.028499873 | 0.054984045 | 0.44294400 | 0.035632387 | 0.050496661 |
| AA414768 | -0.007510134 | 0.637935014 | 0.91600428 | 0.053523374 | 0.056277908 |
| Aaas | 0.021925348 | 0.192746688 | 0.67002811 | 0.081867429 | 0.049704838 |
| Aacs | 0.017831658 | 0.415400018 | 0.83009907 | 0.120727448 | 0.101210763 |
| Aagab | 0.008434252 | 0.73879578 | 0.94743504 | 0.167566814 | 0.153310673 |
| Aak1 | -0.007019457 | 0.832255027 | 0.97234942 | 0.321836467 | 0.35485045 |
| Aamdc | -0.014309943 | 0.523007824 | 0.87506119 | 0.108576252 | 0.114739435 |
| Aamp | -0.080090876 | 0.024996646 | 0.30415400 | 0.69764616 | 0.731952544 |
| Aar2 | 0.031889174 | 0.148034629 | 0.62556790 | 0.133058055 | 0.097385449 |
| Aard | 0.038339879 | 0.150448573 | 0.62803209 | 0.19865826 | 0.105305458 |
| Aars | 0.100118771 | 0.006212442 | 0.15761490 | 0.610623584 | 0.539007231 |
| Aars2 | 0.033356388 | 0.01479208 | 0.23898609 | 0.060617341 | 0.028994249 |
| Aasdh | -0.026151763 | 0.126472904 | 0.58893500 | 0.051189191 | 0.066269863 |
| Aasdhppt | 0.018238964 | 0.522556843 | 0.87506119 | 0.230128103 | 0.198136558 |
| Aatf | 0.049347427 | 0.031933701 | 0.33852287 | 0.151745744 | 0.110780478 |
| Aatk | 0.008492817 | 0.740707449 | 0.94790800 | 0.175531137 | 0.146422463 |
| Abat | -0.024049062 | 0.339036466 | 0.78342458 | 0.12730803 | 0.161968992 |
| Abca1 | -7.62E-05 | 0.998039491 | 0.99940731 | 0.23546401 | 0.27863005 |
| Abca2 | -0.00987565 | 0.665276788 | 0.92307657 | 0.111363578 | 0.127283364 |
| Abca3 | -0.031541312 | 0.266490912 | 0.73073994 | 0.190255043 | 0.226195027 |
| Abca5 | -0.079894152 | 0.010465774 | 0.20174069 | 0.204812844 | 0.287077277 |
| Abca7 | -0.018593099 | 0.418908371 | 0.83009907 | 0.115674366 | 0.120749345 |
| Abca8b | -0.000363888 | 0.983634033 | 0.99760728 | 0.06932154 | 0.06484704 |
| Abcb10 | 0.032042118 | 0.263598877 | 0.72838464 | 0.225910782 | 0.200922242 |
| Abcb4 | 0.021057752 | 0.219597556 | 0.69101621 | 0.061872252 | 0.063073615 |
| Abcb7 | 0.01700473 | 0.448514738 | 0.84632969 | 0.122953061 | 0.111513678 |
| Abcb8 | -0.054033421 | 0.010524171 | 0.20174069 | 0.079332651 | 0.115832045 |
| Abcb9 | -0.006356258 | 0.800410299 | 0.96470534 | 0.147592409 | 0.14640547 |
| Abcc1 | -0.028675102 | 0.103700125 | 0.55665868 | 0.054283438 | 0.073110092 |
| Abcc4 | -0.024358033 | 0.123838343 | 0.58525434 | 0.030252139 | 0.066727361 |
| Abcc5 | -0.018328353 | 0.490870997 | 0.86581549 | 0.168834762 | 0.166791317 |

| | | | | | |
|---|---|---|---|---|---|
| Abcc8 | -0.077043744 | 0.015147871 | 0.2415866 | 1.428491763 | 1.550810043 |
| Abcd1 | 0.027213551 | 0.133075477 | 0.5988251 | 0.084273534 | 0.063464801 |
| Abcd3 | 0.036147194 | 0.330535703 | 0.7779360 | 0.661952102 | 0.626685422 |
| Abce1 | 0.103901602 | 0.001839171 | 0.0745064 | 0.410767871 | 0.322291234 |
| Abcf1 | 0.009729917 | 0.785594674 | 0.9602516 | 0.513296171 | 0.488887926 |
| Abcf2 | 0.011079814 | 0.687962252 | 0.9329576 | 0.200546473 | 0.176935679 |
| Abcf3 | -0.028961881 | 0.310315837 | 0.7677033 | 0.208415967 | 0.22728962 |
| Abcg1 | -0.036451433 | 0.190041844 | 0.6699489 | 0.17550828 | 0.207853315 |
| Abcg2 | -0.051602644 | 0.017289592 | 0.2570999 | 0.083753071 | 0.108185206 |
| Abhd10 | -0.014380081 | 0.53291834 | 0.8788944 | 0.116429714 | 0.126631683 |
| Abhd11 | -0.070183056 | 0.00881845 | 0.1869209 | 0.151672894 | 0.198605442 |
| Abhd12 | -0.031028199 | 0.269001171 | 0.7337525 | 0.197008154 | 0.206341587 |
| Abhd13 | 0.001323219 | 0.954424799 | 0.9954690 | 0.126764595 | 0.117866368 |
| Abhd14a | -0.004319763 | 0.886468184 | 0.9810089 | 0.264387551 | 0.236820925 |
| Abhd14b | -0.033441521 | 0.212256316 | 0.6822059 | 0.160844815 | 0.188680571 |
| Abhd16a | 0.006758332 | 0.837002235 | 0.9726619 | 0.338042944 | 0.324699318 |
| Abhd17a | -0.011628568 | 0.735583754 | 0.9468380 | 0.533334026 | 0.491618788 |
| Abhd17b | 0.003353143 | 0.894448367 | 0.9824361 | 0.147687719 | 0.144799477 |
| Abhd17c | 0.039652428 | 0.205290772 | 0.6795076 | 0.26102164 | 0.257712144 |
| Abhd18 | -0.010675418 | 0.58342068 | 0.9020469 | 0.072226821 | 0.08397741 |
| Abhd2 | 0.045222342 | 0.141050467 | 0.6138718 | 0.27255494 | 0.227582893 |
| Abhd4 | 0.003090509 | 0.86542378 | 0.9770148 | 0.0742312 | 0.067161497 |
| Abhd5 | -0.007063971 | 0.780776555 | 0.9590847 | 0.151989941 | 0.149455743 |
| Abhd6 | -0.0116566 | 0.571177836 | 0.8965907 | 0.091571125 | 0.104111834 |
| Abhd8 | -0.004632629 | 0.881178441 | 0.9795705 | 0.309244311 | 0.280869446 |
| Abi1 | -0.015606705 | 0.608903614 | 0.9088672 | 0.250350058 | 0.268822041 |
| Abi2 | 0.026674011 | 0.355626196 | 0.7969131 | 0.236122173 | 0.223780768 |
| Abitram | 0.001416683 | 0.941877902 | 0.9930310 | 0.08082328 | 0.084300342 |
| Abl1 | 0.032078386 | 0.085206996 | 0.5211664 | 0.092079766 | 0.06794144 |
| Abl2 | -0.0445332 | 0.063316553 | 0.4680329 | 0.103355437 | 0.140339974 |
| Ablim1 | -0.016342604 | 0.440412105 | 0.8417896 | 0.084166981 | 0.112912329 |
| Abr | 0.01418823 | 0.638473873 | 0.9160042 | 0.272289453 | 0.239482554 |
| Abracl | 0.012707182 | 0.737143046 | 0.9469752 | 0.719760373 | 0.76302046 |
| Abraxas1 | 0.003176008 | 0.830280505 | 0.9722601 | 0.047040812 | 0.045394475 |
| Abraxas2 | 0.034023073 | 0.279396012 | 0.7404778 | 0.304956744 | 0.297274938 |
| Abt1 | -0.048510323 | 0.061506604 | 0.4613744 | 0.146864608 | 0.169574361 |
| Abtb1 | -0.03859803 | 0.030866673 | 0.3353750 | 0.046705296 | 0.077343554 |
| Acaa1a | 0.006488984 | 0.849729058 | 0.9748841 | 0.525762085 | 0.466484993 |
| Acaa2 | 0.021614117 | 0.261209627 | 0.7275735 | 0.095372683 | 0.074600577 |
| Acaca | -0.029321622 | 0.161422519 | 0.6392460 | 0.083505505 | 0.108815226 |
| Acad11 | -0.015297885 | 0.4190405 | 0.8300990 | 0.072130558 | 0.079689633 |
| Acad8 | 0.001806677 | 0.931449005 | 0.9893956 | 0.107629794 | 0.097250501 |
| Acad9 | -0.037922268 | 0.026182719 | 0.3097265 | 0.038951523 | 0.071733392 |
| Acadl | 0.018506427 | 0.596046915 | 0.9068020 | 0.468324556 | 0.402673582 |
| Acadm | -0.038401621 | 0.113292435 | 0.5696203 | 0.112536716 | 0.146719923 |
| Acads | -0.019833924 | 0.434080593 | 0.8377586 | 0.156144601 | 0.157999824 |
| Acadsb | 0.060402322 | 0.066369863 | 0.4776087 | 0.359721128 | 0.303768513 |
| Acadvl | -0.033089979 | 0.307396294 | 0.765611 | 0.338233898 | 0.336410657 |
| Acap2 | 0.033611266 | 0.214015235 | 0.6835257 | 0.197652746 | 0.170180742 |
| Acap3 | -0.004952959 | 0.757383526 | 0.9521533 | 0.045961386 | 0.055840173 |
| Acat1 | -0.072484072 | 0.044334023 | 0.4009056 | 0.668690088 | 0.70930085 |
| Acbd3 | 0.069692752 | 0.057263072 | 0.4491407 | 0.90808427 | 0.845305042 |
| Acbd4 | 0.000748929 | 0.965071211 | 0.9966421 | 0.060135529 | 0.062487723 |
| Acbd5 | 0.011191528 | 0.759673595 | 0.9521533 | 0.498364986 | 0.494855781 |
| Acbd6 | 0.03129572 | 0.378920519 | 0.8104552 | 0.568868665 | 0.498760804 |
| Acd | -0.0230912 | 0.438836212 | 0.8403314 | 0.235286033 | 0.239905912 |
| Ace2 | -0.040624272 | 0.318934746 | 0.7710043 | 0.557201039 | 0.493362241 |
| Acer2 | -0.032935207 | 0.112880317 | 0.5696203 | 0.079793066 | 0.097386426 |
| Acer3 | -0.002284174 | 0.897372691 | 0.9825441 | 0.071964211 | 0.065720705 |
| Ache | 0.001247576 | 0.953200423 | 0.9954690 | 0.088769657 | 0.087839376 |
| Acin1 | -0.013723451 | 0.718349373 | 0.9438444 | 0.735908525 | 0.753044345 |
| Acly | -0.068587266 | 0.031403128 | 0.3385228 | 1.616323618 | 1.931463677 |
| Aco1 | 0.011257005 | 0.601241888 | 0.9079493 | 0.117586063 | 0.098323768 |
| Aco2 | -0.043185866 | 0.237528473 | 0.7098511 | 0.703355319 | 0.697814243 |
| Acot13 | 0.0742371 | 0.044963971 | 0.4025655 | 0.62192716 | 0.547420835 |
| Acot2 | 0.008367773 | 0.615715311 | 0.9112316 | 0.068420485 | 0.055017076 |

| | | | | | |
|---|---|---|---|---|---|
| Acot8 | -0.015670869 | 0.560061653 | 0.8925646 | 0.179377771 | 0.17545799 |
| Acox1 | 0.027020082 | 0.38985314 | 0.8181390 | 0.320717697 | 0.297890471 |
| Acp1 | -0.034975591 | 0.313956397 | 0.7705213 | 0.834215607 | 0.810389793 |
| Acp2 | -0.01442782 | 0.554410895 | 0.8902995 | 0.134256535 | 0.13835351 |
| Acp6 | -0.013719438 | 0.449564744 | 0.8476871 | 0.070207269 | 0.070566937 |
| Acsf3 | 0.016543675 | 0.438873691 | 0.8403314 | 0.107027402 | 0.101104383 |
| Acsl1 | -0.005890949 | 0.767452521 | 0.9544221 | 0.087076872 | 0.08691915 |
| Acsl3 | -0.000754148 | 0.962165287 | 0.9964171 | 0.048503471 | 0.053794828 |
| Acsl4 | 0.024684128 | 0.441283393 | 0.8426431 | 0.278703789 | 0.252024415 |
| Acsl5 | -0.011529336 | 0.664535942 | 0.9230765 | 0.164624457 | 0.187345692 |
| Acsl6 | 0.004660662 | 0.759573452 | 0.9521533 | 0.045334379 | 0.049194099 |
| Acss2 | -0.004169367 | 0.818646061 | 0.9698995 | 0.065709967 | 0.072473198 |
| Actb | -0.018358909 | 0.477300507 | 0.8607489 | 2.412301018 | 2.357604299 |
| Actg1 | 0.006563041 | 0.792747725 | 0.9624460 | 2.69470186 | 2.595970349 |
| Actl6a | 0.024496715 | 0.435672918 | 0.8389295 | 0.333105122 | 0.278355542 |
| Actl6b | -0.020458675 | 0.45891559 | 0.8522649 | 0.183296793 | 0.17986964 |
| Actn1 | 0.0014565 | 0.959185157 | 0.9957785 | 0.207390661 | 0.215258076 |
| Actn4 | 0.011603242 | 0.724195568 | 0.9445652 | 0.292718386 | 0.298534552 |
| Actr10 | -0.016973442 | 0.620078795 | 0.9121505 | 0.674695874 | 0.653445432 |
| Actr1a | 0.019987553 | 0.471657667 | 0.8585034 | 0.224981931 | 0.193593148 |
| Actr1b | -0.061465948 | 0.070232835 | 0.4855119 | 0.33775799 | 0.378811324 |
| Actr2 | -0.001898396 | 0.955846923 | 0.9954690 | 0.436963933 | 0.435611378 |
| Actr3 | -0.018326722 | 0.613891103 | 0.9108386 | 0.708016586 | 0.668074635 |
| Actr5 | -0.002903059 | 0.86922522 | 0.9776398 | 0.063735934 | 0.064076728 |
| Actr6 | -0.050532742 | 0.01361939 | 0.2295523 | 0.062854092 | 0.103538182 |
| Actr8 | -0.002154684 | 0.934622263 | 0.9906456 | 0.172448841 | 0.166982677 |
| Acvr1 | -0.031754412 | 0.165486119 | 0.6425136 | 0.099380196 | 0.125589485 |
| Acvr1b | 0.019305149 | 0.256773295 | 0.7248694 | 0.074001561 | 0.057168926 |
| Acvr1c | 0.011488484 | 0.599972687 | 0.9078234 | 0.114575932 | 0.09363552 |
| Acvr2a | -0.011667456 | 0.620423012 | 0.9121505 | 0.12652847 | 0.127971892 |
| Acvr2b | 0.028312458 | 0.204600332 | 0.6781273 | 0.144214576 | 0.104052671 |
| Acyp1 | 0.021875027 | 0.527792696 | 0.8768553 | 0.481279553 | 0.441455065 |
| Acyp2 | -0.017183485 | 0.456314018 | 0.8505514 | 0.09728454 | 0.136516155 |
| Adal | -0.006600282 | 0.665829579 | 0.9230765 | 0.0390676 | 0.050291089 |
| Adam10 | 0.026201324 | 0.313201382 | 0.7697557 | 0.178939883 | 0.152087609 |
| Adam15 | 0.000152585 | 0.9935374 | 0.9985888 | 0.071190038 | 0.078507722 |
| Adam17 | 0.044846656 | 0.013502741 | 0.2292511 | 0.098204057 | 0.056585926 |
| Adam22 | 0.073466254 | 0.003467182 | 0.1131396 | 0.174785513 | 0.141526014 |
| Adam23 | 0.002523239 | 0.858761961 | 0.9769675 | 0.043847928 | 0.039292247 |
| Adam32 | 0.013342404 | 0.435464733 | 0.8389295 | 0.06559691 | 0.053180416 |
| Adam9 | 0.048642417 | 0.0297089 | 0.3298864 | 0.126326025 | 0.107255668 |
| Adamts18 | -0.07557393 | 0.007170519 | 0.1680001 | 0.133274645 | 0.176118109 |
| Adar | -0.01239225 | 0.610818569 | 0.9091253 | 0.12738501 | 0.13843906 |
| Adarb1 | -0.014785768 | 0.47429023 | 0.8599815 | 0.077029585 | 0.102408162 |
| Adat1 | 0.005489723 | 0.728929799 | 0.9459020 | 0.060331395 | 0.049933395 |
| Adck1 | -0.024890235 | 0.132102157 | 0.5978345 | 0.046697444 | 0.062912013 |
| Adck2 | 0.017870138 | 0.245880953 | 0.7152900 | 0.066918111 | 0.04472144 |
| Adck5 | 0.004202611 | 0.866214254 | 0.9770148 | 0.155687109 | 0.143833672 |
| Adcy6 | -0.078308035 | 0.002552426 | 0.0927113 | 0.123025403 | 0.173531772 |
| Adcy9 | -0.017144611 | 0.531192025 | 0.8776448 | 0.18138264 | 0.197534711 |
| Add1 | -0.058287033 | 0.055545321 | 0.4449332 | 0.243615952 | 0.272605125 |
| Add3 | -0.053902379 | 0.068129706 | 0.4830926 | 0.196484147 | 0.225109912 |
| Adgb | 0.011810726 | 0.405755904 | 0.8270770 | 0.048475902 | 0.038231583 |
| Adgrb3 | 0.001139012 | 0.952830195 | 0.9953744 | 0.071984051 | 0.076671552 |
| Adgrg1 | 0.019297836 | 0.563392035 | 0.8930727 | 0.383093581 | 0.359230781 |
| Adgrl1 | -0.044890624 | 0.150566671 | 0.6280320 | 0.241446237 | 0.285411786 |
| Adh1 | 0.004810664 | 0.819556403 | 0.9699062 | 0.102094967 | 0.09616989 |
| Adh5 | -0.086039381 | 0.010171941 | 0.1999013 | 0.407242553 | 0.43049327 |
| Adi1 | 0.026316658 | 0.420881811 | 0.8300990 | 0.316722891 | 0.328321278 |
| Adipor1 | 0.00844582 | 0.807533253 | 0.9653807 | 0.449227403 | 0.429976533 |
| Adipor2 | -0.027163298 | 0.3740468 | 0.8087297 | 0.258821265 | 0.264402475 |
| Adk | -0.074701509 | 0.038191885 | 0.3732645 | 0.508391554 | 0.517120469 |
| Adm2 | -0.036026874 | 0.1541269 | 0.6304934 | 0.102088927 | 0.15310669 |
| Adnp | -0.018696565 | 0.335474445 | 0.7810153 | 0.069975082 | 0.090484093 |
| Adnp2 | 0.036229112 | 0.04278307 | 0.3959774 | 0.094689172 | 0.059896849 |
| Ado | 0.021208453 | 0.369060833 | 0.8056719 | 0.141791719 | 0.122316281 |

| | | | | | |
|---|---|---|---|---|---|
| Adora1 | 0.035440414 | 0.022036811 | 0.2895010 | 0.062817365 | 0.043101674 |
| Adora2a | 0.014360206 | 0.530258493 | 0.8776448 | 0.122176707 | 0.103958227 |
| Adora2b | 0.053900696 | 0.000461042 | 0.0291307 | 0.069493307 | 0.036775685 |
| Adpgk | 0.00919875 | 0.584170989 | 0.9021406 | 0.064250917 | 0.058650789 |
| Adprh | 0.01177611 | 0.712065416 | 0.9419834 | 0.316754286 | 0.3054943 |
| Adprm | 0.013735306 | 0.546613947 | 0.8858459 | 0.13248091 | 0.117500718 |
| Adprs | 0.017285637 | 0.535970666 | 0.8799251 | 0.226887943 | 0.187612843 |
| Adra2a | -0.012668064 | 0.665723033 | 0.9230765 | 0.165443769 | 0.241636283 |
| Adrb1 | -0.053727392 | 0.016274818 | 0.2514444 | 0.091899205 | 0.10835195 |
| Adrm1 | -0.020413641 | 0.543034302 | 0.8840682 | 0.390952344 | 0.390201107 |
| Adsl | 0.03671388 | 0.107505056 | 0.5633723 | 0.14156326 | 0.110552142 |
| Adss | 0.01927824 | 0.491403922 | 0.8663100 | 0.219344463 | 0.173423897 |
| Adssl1 | 0.025792207 | 0.093506617 | 0.5330951 | 0.057965584 | 0.042077258 |
| Aebp2 | -0.014061422 | 0.624645996 | 0.9121505 | 0.187048582 | 0.212897814 |
| Aen | 0.002804394 | 0.930034236 | 0.9885288 | 0.290062567 | 0.263973901 |
| Afdn | -0.061180406 | 0.080398903 | 0.5098712 | 0.409860357 | 0.468399428 |
| Aff1 | -0.009799083 | 0.714917083 | 0.9430821 | 0.157623849 | 0.18116434 |
| Aff2 | 0.009431078 | 0.647178888 | 0.9169695 | 0.092190409 | 0.086377902 |
| Aff4 | -0.01210764 | 0.743058908 | 0.9479080 | 0.70923351 | 0.744407156 |
| Afg3l1 | 0.006460734 | 0.835342602 | 0.9723494 | 0.255587309 | 0.243820563 |
| Afg3l2 | -0.014200047 | 0.624678861 | 0.9121505 | 0.222912497 | 0.236533239 |
| Aftph | -0.01823572 | 0.600846167 | 0.9078437 | 0.386185366 | 0.40353489 |
| Aga | -0.079085897 | 0.032234403 | 0.3393267 | 0.731024438 | 0.793690791 |
| Agap1 | -0.060858128 | 0.096291901 | 0.5369725 | 0.434187447 | 0.559577735 |
| Agap3 | -0.021465392 | 0.458235612 | 0.8521621 | 0.228249395 | 0.224538698 |
| Agbl3 | -0.029516828 | 0.209728626 | 0.6807660 | 0.125641842 | 0.135100085 |
| Agbl5 | 0.021337994 | 0.138389894 | 0.6079839 | 0.055225803 | 0.038225482 |
| Agfg1 | -0.00683212 | 0.850436094 | 0.9748841 | 0.46762488 | 0.457841327 |
| Agfg2 | -0.001180009 | 0.95510577 | 0.9954690 | 0.092200516 | 0.091876104 |
| Aggf1 | -0.003772094 | 0.902588021 | 0.9825441 | 0.281075722 | 0.251878466 |
| Agk | 0.006291123 | 0.74432323 | 0.9484504 | 0.087925381 | 0.078757623 |
| Agl | 0.011587746 | 0.665733132 | 0.9230765 | 0.181342577 | 0.176092766 |
| Ago1 | -0.030676986 | 0.176569475 | 0.6536659 | 0.113290876 | 0.12884044 |
| Ago2 | 0.005774167 | 0.869181018 | 0.9776398 | 0.408635523 | 0.424218604 |
| Ago3 | -0.038225142 | 0.188321501 | 0.6668854 | 0.185276575 | 0.214678481 |
| Ago4 | -0.026270538 | 0.175993596 | 0.6536659 | 0.073194 | 0.083823939 |
| Agpat1 | -0.060794245 | 0.022163999 | 0.2898221 | 0.13778283 | 0.178322899 |
| Agpat2 | 0.027289614 | 0.254492916 | 0.7223071 | 0.152973461 | 0.127150828 |
| Agpat3 | 0.026770715 | 0.444560056 | 0.8431541 | 0.47000822 | 0.433372921 |
| Agpat4 | -0.008588946 | 0.763334733 | 0.9532128 | 0.217035503 | 0.195427666 |
| Agpat5 | 0.020378566 | 0.461598144 | 0.8524425 | 0.196230176 | 0.192901436 |
| Agps | 0.018069537 | 0.417888303 | 0.8300990 | 0.125791619 | 0.107038082 |
| Agt | -0.038045227 | 0.106637928 | 0.5623243 | 0.11235919 | 0.122408021 |
| Agtpbp1 | 0.021495356 | 0.505637069 | 0.8695566 | 0.311795907 | 0.29227288 |
| Agtrap | 0.001093792 | 0.963787828 | 0.9966421 | 0.141687689 | 0.141909927 |
| Ahctf1 | -0.014443601 | 0.617921913 | 0.9115017 | 0.207224133 | 0.217107225 |
| Ahcyl1 | 0.032424645 | 0.3734741 | 0.8085689 | 0.562477626 | 0.533291925 |
| Ahcyl2 | -0.001161215 | 0.972291754 | 0.9976072 | 0.367309997 | 0.377228217 |
| Ahdc1 | -0.043293513 | 0.077300645 | 0.5022647 | 0.122002268 | 0.145000358 |
| Ahi1 | -0.076544119 | 0.04100663 | 0.3901716 | 0.816348751 | 0.946457251 |
| Ahsa1 | -0.019931383 | 0.563221614 | 0.8929816 | 0.447066271 | 0.423247255 |
| Ahsa2 | 0.011790057 | 0.664257818 | 0.9230765 | 0.197045989 | 0.174102195 |
| AI413582 | -0.053582063 | 0.105788747 | 0.5617171 | 0.280614619 | 0.352646054 |
| AI467606 | -0.00129682 | 0.940652325 | 0.9929301 | 0.067779845 | 0.057987043 |
| AI480526 | -0.039427659 | 0.063471818 | 0.4688312 | 0.07621833 | 0.105381035 |
| AI506816 | -0.034051156 | 0.198718466 | 0.6730962 | 0.174213948 | 0.16996644 |
| AI597479 | 0.016556278 | 0.339256837 | 0.7837512 | 0.074791684 | 0.059889215 |
| AI661453 | -0.015833862 | 0.478494538 | 0.8607489 | 0.108097374 | 0.115546034 |
| AI837181 | -0.009833837 | 0.708248458 | 0.9412948 | 0.179745812 | 0.172812389 |
| AI987944 | 0.024884905 | 0.300468579 | 0.7595584 | 0.158958541 | 0.127889811 |
| Aida | 0.019430828 | 0.520193099 | 0.8741795 | 0.229257544 | 0.240836292 |
| Aifm1 | -0.015006577 | 0.497200854 | 0.8684629 | 0.096396637 | 0.114141547 |
| Aifm2 | 0.011043942 | 0.528810697 | 0.8776448 | 0.06820539 | 0.058708172 |
| Aig1 | -0.053078689 | 0.138147815 | 0.6079839 | 0.495130669 | 0.547872484 |
| Aimp1 | -0.002212763 | 0.950927296 | 0.9948532 | 0.813263259 | 0.752573312 |
| Aimp2 | -0.000622965 | 0.977729075 | 0.9976072 | 0.126700735 | 0.114164026 |

| | | | | | |
|---|---|---|---|---|---|
| Aip | -0.037161779 | 0.25032808 | 0.7207210 | 0.344728516 | 0.340536631 |
| Airn | -0.056153528 | 0.046321421 | 0.4073656 | 0.164544969 | 0.203774108 |
| Ak1 | -0.026403874 | 0.232519255 | 0.7039417 | 0.096732375 | 0.114972569 |
| Ak2 | 0.040343312 | 0.227947407 | 0.7015153 | 0.409244015 | 0.354925052 |
| Ak3 | -0.003049206 | 0.87838645 | 0.9785583 | 0.085129766 | 0.085565438 |
| Ak6 | 0.016916318 | 0.592257959 | 0.9060200 | 0.319682803 | 0.268940684 |
| Akain1 | -0.025823731 | 0.309485582 | 0.7673813 | 0.112380196 | 0.177069849 |
| Akap1 | -0.000784102 | 0.962784345 | 0.9965380 | 0.055460616 | 0.063966468 |
| Akap10 | 0.001374607 | 0.934384396 | 0.9906052 | 0.06537675 | 0.056265394 |
| Akap11 | -0.024064363 | 0.508074807 | 0.8704067 | 0.427859053 | 0.47004105 |
| Akap12 | -0.014172575 | 0.411961005 | 0.8297979 | 0.058296343 | 0.057768966 |
| Akap13 | -1.36E-05 | 0.999708697 | 0.9997086 | 0.428080391 | 0.481900959 |
| Akap17b | -0.0887998 | 0.006420089 | 0.1603078 | 0.228466168 | 0.280490906 |
| Akap7 | 0.012272706 | 0.567230515 | 0.8952062 | 0.117055271 | 0.099559241 |
| Akap8 | -0.013469734 | 0.636466278 | 0.9153541 | 0.18901966 | 0.201688623 |
| Akap8l | -0.070879043 | 0.057283468 | 0.4491407 | 0.540476038 | 0.578079035 |
| Akap9 | -0.049259487 | 0.175712967 | 0.6536659 | 1.148188754 | 1.200028468 |
| Akip1 | 0.006665406 | 0.724268446 | 0.9445652 | 0.082825631 | 0.077954829 |
| Akirin1 | -0.025638842 | 0.479433409 | 0.8610807 | 0.510653454 | 0.553174178 |
| Akirin2 | 0.018227287 | 0.598489911 | 0.9075513 | 0.448682184 | 0.392462155 |
| Akr1a1 | 0.055568736 | 0.096058804 | 0.5368469 | 1.203223666 | 1.045488301 |
| Akr1b10 | -0.004697754 | 0.824935191 | 0.9715054 | 0.093660081 | 0.096060533 |
| Akr1b3 | -0.017875436 | 0.620282182 | 0.9121505 | 0.446672823 | 0.450606401 |
| Akr1c12 | 0.002962841 | 0.909708726 | 0.9837373 | 0.147946166 | 0.161387909 |
| Akr1c13 | 0.026702498 | 0.334331269 | 0.7804836 | 0.197842033 | 0.186786251 |
| Akr1c14 | -0.027727213 | 0.124368401 | 0.5852543 | 0.045706847 | 0.081367483 |
| Akr1c19 | -0.00941889 | 0.795749003 | 0.9638376 | 0.438124549 | 0.394668247 |
| Akr1e1 | 0.028597036 | 0.302566385 | 0.7614050 | 0.215705053 | 0.178995893 |
| Akr7a5 | -0.022274886 | 0.475481949 | 0.8603832 | 0.24334013 | 0.251997747 |
| Akt1 | 0.02623265 | 0.407135544 | 0.8270770 | 0.307726295 | 0.282508655 |
| Akt1s1 | 0.023937511 | 0.466614939 | 0.8553370 | 0.379675308 | 0.318766293 |
| Akt2 | -0.001638292 | 0.957374294 | 0.9956630 | 0.248644736 | 0.247087781 |
| Akt3 | 0.01876297 | 0.476530745 | 0.8607489 | 0.177857412 | 0.166846998 |
| Aktip | -0.037479222 | 0.23304168 | 0.7051391 | 0.287878475 | 0.290667282 |
| Alad | 0.010140435 | 0.70728937 | 0.9408354 | 0.193970086 | 0.178395948 |
| Alas1 | -0.020115568 | 0.479378425 | 0.8610807 | 0.195237541 | 0.185233592 |
| Alcam | -0.046549481 | 0.224661109 | 0.6968297 | 0.99902981 | 0.983367827 |
| Aldh18a1 | -0.018278519 | 0.580941328 | 0.9015399 | 0.333853409 | 0.346636214 |
| Aldh1l2 | 0.052066248 | 0.113860429 | 0.5696977 | 0.272290847 | 0.276345925 |
| Aldh2 | -0.004144375 | 0.86458119 | 0.9769675 | 0.148064325 | 0.141143703 |
| Aldh3a2 | 0.020332297 | 0.370268723 | 0.8065581 | 0.126430871 | 0.113114195 |
| Aldh4a1 | -0.016765067 | 0.313155724 | 0.7697557 | 0.048662891 | 0.059376568 |
| Aldh5a1 | -0.004930092 | 0.862165668 | 0.9769675 | 0.182519657 | 0.194169413 |
| Aldh6a1 | -0.017007258 | 0.411241222 | 0.8290007 | 0.098313563 | 0.096179404 |
| Aldh7a1 | 0.041653259 | 0.08985149 | 0.5302360 | 0.183700473 | 0.121997309 |
| Aldh9a1 | -0.004871989 | 0.838951573 | 0.9727216 | 0.131686601 | 0.134945618 |
| Aldoa | 0.014529256 | 0.599595536 | 0.9078234 | 1.852128591 | 1.784198158 |
| Aldoc | 0.020219474 | 0.317551009 | 0.7709420 | 0.101365913 | 0.080313477 |
| Alg1 | -0.010026336 | 0.6842185 | 0.9311703 | 0.140577183 | 0.138248174 |
| Alg10b | 0.037893699 | 0.151591879 | 0.6281105 | 0.185267083 | 0.163305429 |
| Alg11 | -0.002234384 | 0.919887084 | 0.9854513 | 0.120049407 | 0.112950315 |
| Alg12 | -0.013478353 | 0.519315499 | 0.8741795 | 0.089559293 | 0.09237743 |
| Alg14 | 0.008641752 | 0.657662955 | 0.9211780 | 0.086770681 | 0.079360394 |
| Alg2 | -0.023794692 | 0.402328195 | 0.8254592 | 0.200550387 | 0.211918776 |
| Alg3 | 0.006174999 | 0.787195727 | 0.9608311 | 0.124669917 | 0.114283232 |
| Alg5 | 0.040110753 | 0.191196345 | 0.6700281 | 0.303851098 | 0.255563534 |
| Alg6 | -0.003688715 | 0.83405781 | 0.9723494 | 0.064694635 | 0.06753282 |
| Alg9 | -0.02230488 | 0.156515726 | 0.6316663 | 0.036536664 | 0.058423322 |
| Alkbh1 | -0.023323125 | 0.508358551 | 0.8704067 | 0.356971799 | 0.42944803 |
| Alkbh2 | -0.018604998 | 0.412303422 | 0.8297979 | 0.111042778 | 0.115001795 |
| Alkbh3 | -0.036610022 | 0.149599831 | 0.6264374 | 0.134915769 | 0.160074611 |
| Alkbh5 | 0.062068788 | 0.093817475 | 0.5333348 | 0.69786965 | 0.650868612 |
| Alkbh6 | -0.013679621 | 0.676648701 | 0.9278898 | 0.363589702 | 0.347895192 |
| Alkbh7 | -0.051131451 | 0.064298641 | 0.4700387 | 0.191461464 | 0.209570903 |
| Alkbh8 | -0.021095048 | 0.309499361 | 0.7673813 | 0.08892847 | 0.099590255 |
| Alms1 | -0.007980632 | 0.614749176 | 0.9108386 | 0.042796877 | 0.053914749 |

| | | | | | |
|---|---|---|---|---|---|
| Alox5ap | -0.006895456 | 0.728269065 | 0.9458355 | 0.06801968 | 0.088945073 |
| Als2 | -0.022127325 | 0.167479324 | 0.6446304 | 0.03994011 | 0.056293301 |
| Alyref | 0.050845446 | 0.146505554 | 0.6225768 | 0.6177054 | 0.50872156 |
| Alyref2 | -0.042958557 | 0.047879037 | 0.4133681 | 0.09643006 | 0.117463079 |
| Amacr | 0.007046004 | 0.700302915 | 0.9375165 | 0.081473525 | 0.071893205 |
| Ambp | -0.004494728 | 0.858860355 | 0.9769675 | 0.13649911 | 0.136165053 |
| Ambra1 | -0.007743263 | 0.707745882 | 0.9408790 | 0.09324013 | 0.09606843 |
| Amd1 | 0.003065163 | 0.891088509 | 0.9819597 | 0.128558388 | 0.108925341 |
| Amd2 | -0.013308587 | 0.471531474 | 0.8585034 | 0.065073151 | 0.073880095 |
| Amdhd2 | 0.042132962 | 0.041437429 | 0.3917471 | 0.126134405 | 0.081922201 |
| Amer1 | -0.017163596 | 0.33722892 | 0.7832617 | 0.057416074 | 0.068687495 |
| Amer2 | 0.025986211 | 0.38224713 | 0.8126882 | 0.258664492 | 0.222210722 |
| Amfr | 0.025403146 | 0.438305384 | 0.8403201 | 0.374991746 | 0.360311574 |
| Amigo1 | 0.026752912 | 0.29103083 | 0.7514382 | 0.164012888 | 0.146602312 |
| Amigo2 | 0.011970182 | 0.594786646 | 0.9067325 | 0.112892882 | 0.112849359 |
| Ammecr1l | 0.007511813 | 0.735408607 | 0.9468380 | 0.119552296 | 0.105060485 |
| Amn1 | -0.017291596 | 0.551746066 | 0.8893234 | 0.22147573 | 0.217705719 |
| Amotl1 | -0.068233085 | 0.041185219 | 0.3905902 | 0.260362297 | 0.30446179 |
| Amotl2 | -0.013185202 | 0.501970343 | 0.8695566 | 0.081103426 | 0.084375749 |
| Ampd2 | -0.01064434 | 0.722991741 | 0.9445652 | 0.223133404 | 0.251158684 |
| Amph | -0.002579163 | 0.899714222 | 0.9825441 | 0.09038859 | 0.094371352 |
| Amt | 0.020271416 | 0.256184683 | 0.7238259 | 0.089760272 | 0.061420353 |
| Amz2 | -0.037177813 | 0.252656762 | 0.7223071 | 0.292375626 | 0.318909629 |
| Anapc1 | 0.013219058 | 0.642018243 | 0.9166712 | 0.215572291 | 0.192931054 |
| Anapc10 | -0.040246218 | 0.085867097 | 0.5232196 | 0.119710682 | 0.136127725 |
| Anapc11 | -0.017048382 | 0.625893996 | 0.9128095 | 0.607739524 | 0.570943764 |
| Anapc13 | 0.146814709 | 1.70E-05 | 0.0023782 | 1.087809354 | 0.922787364 |
| Anapc15 | 0.020299018 | 0.30536726 | 0.7646271 | 0.101684276 | 0.08102101 |
| Anapc16 | 0.023916389 | 0.496626964 | 0.8684629 | 0.503707158 | 0.493289412 |
| Anapc2 | -0.023477882 | 0.340953015 | 0.7851366 | 0.140611837 | 0.153612777 |
| Anapc4 | 0.018303178 | 0.528599347 | 0.8774607 | 0.251210788 | 0.222617356 |
| Anapc5 | -0.001186273 | 0.974642058 | 0.9976072 | 0.580203658 | 0.566772575 |
| Anapc7 | -0.030824861 | 0.253213549 | 0.7223071 | 0.168324799 | 0.192398529 |
| Ang | 0.008819206 | 0.824503336 | 0.971271 | 0.662330467 | 0.61961729 |
| Angel2 | -0.04311034 | 0.169072469 | 0.6471109 | 0.247264199 | 0.287628117 |
| Angptl6 | -0.032925496 | 0.2913797 | 0.7516725 | 0.21423816 | 0.286388472 |
| Ank | -0.026976229 | 0.509848876 | 0.8711607 | 0.939806045 | 0.895544958 |
| Ank2 | 0.057515664 | 0.103157915 | 0.5553141 | 0.405230461 | 0.384654606 |
| Ank3 | -0.010148788 | 0.74490132 | 0.9485717 | 0.253195413 | 0.256043341 |
| Ankfy1 | -0.011978856 | 0.630171836 | 0.9130564 | 0.136856625 | 0.142728027 |
| Ankhd1 | 0.016325081 | 0.627859656 | 0.9128095 | 0.389898194 | 0.357475701 |
| Ankib1 | -0.066459854 | 0.012389314 | 0.2195296 | 0.132259359 | 0.186527283 |
| Ankle2 | -0.006014981 | 0.808045896 | 0.9656445 | 0.145277755 | 0.146646421 |
| Ankmy2 | -0.039714759 | 0.107405039 | 0.5633723 | 0.127507526 | 0.151319453 |
| Ankra2 | 0.020676948 | 0.514865511 | 0.8733714 | 0.294917173 | 0.306347512 |
| Ankrd10 | 0.001571017 | 0.964447314 | 0.9966421 | 0.421515637 | 0.489233132 |
| Ankrd11 | -0.015394562 | 0.684682344 | 0.9313273 | 0.921932441 | 0.943646269 |
| Ankrd12 | 0.000901406 | 0.98142386 | 0.9976072 | 0.827935284 | 0.842583952 |
| Ankrd13a | -0.003792793 | 0.865854634 | 0.9770148 | 0.125583806 | 0.108906354 |
| Ankrd13b | -0.012508325 | 0.381250238 | 0.8119369 | 0.03573923 | 0.046331489 |
| Ankrd13c | -0.006598879 | 0.833639816 | 0.9723494 | 0.287623454 | 0.273984327 |
| Ankrd16 | -0.00403398 | 0.814838365 | 0.9685114 | 0.05839775 | 0.064588807 |
| Ankrd17 | 0.003649338 | 0.924557663 | 0.9878036 | 0.5754344 | 0.562006141 |
| Ankrd24 | -0.040168092 | 0.159657171 | 0.6368311 | 0.180825791 | 0.20966788 |
| Ankrd26 | 0.004630523 | 0.843739247 | 0.9734889 | 0.120516454 | 0.118387773 |
| Ankrd27 | 0.009008291 | 0.591236963 | 0.9060200 | 0.066006198 | 0.057442897 |
| Ankrd28 | -0.016304414 | 0.365907796 | 0.8050191 | 0.062527312 | 0.073189141 |
| Ankrd33b | -0.048393586 | 0.078897755 | 0.5062520 | 0.153528252 | 0.196095088 |
| Ankrd37 | 0.00954322 | 0.6042309 | 0.9087139 | 0.075541734 | 0.070910331 |
| Ankrd39 | 0.00230816 | 0.926777102 | 0.9879210 | 0.152983811 | 0.145420984 |
| Ankrd40 | -0.009838755 | 0.757409259 | 0.9521533 | 0.295323097 | 0.292557199 |
| Ankrd44 | 0.055827079 | 0.032157672 | 0.3392328 | 0.190108893 | 0.144652609 |
| Ankrd46 | 0.053123595 | 0.017896363 | 0.2587928 | 0.15131679 | 0.100246976 |
| Ankrd49 | 0.034570418 | 0.144522712 | 0.6193215 | 0.149219895 | 0.114522563 |
| Ankrd50 | -0.026482662 | 0.173333801 | 0.6525507 | 0.062744322 | 0.087631979 |
| Ankrd54 | -0.055174801 | 0.089832426 | 0.5302360 | 0.259058812 | 0.312950957 |

| | | | | | |
|---|---|---|---|---|---|
| Anks1 | 0.00198131 | 0.92099367 | 0.9861175 | 0.08783338 | 0.092103508 |
| Anks3 | -0.031242271 | 0.17967294 | 0.6550070 | 0.119678648 | 0.125916113 |
| Anks4b | 0.0236569 | 0.29121372 | 0.7516725 | 0.108978422 | 0.109547543 |
| Ankzf1 | -0.014077302 | 0.420259464 | 0.8300990 | 0.061350424 | 0.067175018 |
| Ano10 | -0.003786005 | 0.830245749 | 0.9722601 | 0.069583723 | 0.067268137 |
| Ano6 | 0.018280703 | 0.459659785 | 0.8524343 | 0.149297069 | 0.141638883 |
| Ano8 | -0.012167422 | 0.47459053 | 0.8600971 | 0.053271628 | 0.065397603 |
| Anp32a | -0.018860095 | 0.591410435 | 0.9060200 | 1.00648565 | 0.950528519 |
| Anp32b | 0.057310074 | 0.097379049 | 0.5384203 | 1.17972493 | 1.055461308 |
| Anp32e | -0.088598826 | 0.016798847 | 0.2546713 | 0.624248144 | 0.677780896 |
| Anpep | -0.002330277 | 0.912336641 | 0.9840393 | 0.099476599 | 0.091935 |
| Antkmt | 0.027363609 | 0.185999652 | 0.6640243 | 0.120429817 | 0.090137357 |
| Anxa11 | 0.019159453 | 0.499266484 | 0.8690742 | 0.210091702 | 0.204077765 |
| Anxa4 | 0.022423556 | 0.547075085 | 0.8858508 | 0.545862362 | 0.533669237 |
| Anxa5 | 0.007775285 | 0.798562335 | 0.9644698 | 1.554493431 | 1.480235853 |
| Anxa6 | -0.02323368 | 0.502576281 | 0.8695566 | 0.392723249 | 0.389524936 |
| Anxa7 | -0.05507485 | 0.114054406 | 0.5696977 | 0.378198814 | 0.404984305 |
| Aopep | -0.015404006 | 0.665058149 | 0.9230765 | 0.346617931 | 0.417126968 |
| Aox1 | -0.015433385 | 0.317266669 | 0.7708266 | 0.039704906 | 0.052316416 |
| Ap1ar | 0.02170898 | 0.38033218 | 0.8117350 | 0.158952906 | 0.135860076 |
| Ap1b1 | 0.02474923 | 0.400091757 | 0.8244516 | 0.24850962 | 0.219531117 |
| Ap1g1 | -0.00667259 | 0.838105239 | 0.9727216 | 0.297544117 | 0.273764784 |
| Ap1m1 | 0.007004614 | 0.79049575 | 0.9621438 | 0.16754781 | 0.153276711 |
| Ap1m2 | 0.005185282 | 0.857526898 | 0.9763189 | 0.229491655 | 0.212347991 |
| Ap1s1 | 0.039726772 | 0.27267423 | 0.7345096 | 0.551579912 | 0.465454014 |
| Ap1s2 | 0.065055527 | 0.087506982 | 0.5270608 | 0.76317323 | 0.66049855 |
| Ap2a1 | 0.007682184 | 0.808500885 | 0.9658392 | 0.340056379 | 0.310147081 |
| Ap2a2 | 0.055524365 | 0.094251447 | 0.5334417 | 0.392127067 | 0.323027152 |
| Ap2b1 | -0.010409445 | 0.748979442 | 0.9501120 | 0.336480838 | 0.343317923 |
| Ap2m1 | -0.036641153 | 0.314067714 | 0.7705987 | 0.67006869 | 0.669388941 |
| Ap2s1 | -0.018320639 | 0.59180608 | 0.9060200 | 0.759490842 | 0.716763917 |
| Ap3b1 | -0.00627102 | 0.845132119 | 0.9734889 | 0.28584425 | 0.283128775 |
| Ap3b2 | 0.016679242 | 0.630724927 | 0.9132784 | 0.42691978 | 0.411358161 |
| Ap3d1 | -0.032676847 | 0.327057303 | 0.7778857 | 0.312044232 | 0.354265574 |
| Ap3m1 | -0.01249513 | 0.679296996 | 0.9288316 | 0.261200188 | 0.245493886 |
| Ap3m2 | 0.015852965 | 0.396368531 | 0.8236229 | 0.082016694 | 0.069878608 |
| Ap3s1 | 0.059377696 | 0.079123391 | 0.5073717 | 0.630697984 | 0.521054829 |
| Ap3s2 | 0.03147032 | 0.257508328 | 0.7250597 | 0.214610978 | 0.181860606 |
| Ap4b1 | 0.011298783 | 0.438451112 | 0.8403201 | 0.053381907 | 0.039114271 |
| Ap4m1 | -0.036368064 | 0.044427555 | 0.4009562 | 0.045628264 | 0.078981627 |
| Ap4s1 | 0.012872477 | 0.63175382 | 0.9133517 | 0.18026847 | 0.172933543 |
| Ap5m1 | -0.024835885 | 0.280099778 | 0.7408618 | 0.095869625 | 0.123309632 |
| Ap5s1 | 0.03878537 | 0.050204361 | 0.4218341 | 0.111634919 | 0.073741084 |
| Apaf1 | 0.017001851 | 0.261785699 | 0.7275735 | 0.060570259 | 0.039223333 |
| Apba1 | -0.019334963 | 0.25210357 | 0.7221678 | 0.039185139 | 0.067439105 |
| Apba3 | 0.006510335 | 0.723922332 | 0.9445652 | 0.089276793 | 0.06906599 |
| Apbb1 | -0.023989152 | 0.410662839 | 0.8285083 | 0.243468568 | 0.233533441 |
| Apbb2 | 0.003048991 | 0.902244255 | 0.9825441 | 0.141415043 | 0.137909565 |
| Apbb3 | -0.016487399 | 0.379985275 | 0.8116825 | 0.071510519 | 0.079067213 |
| Apc | -0.089604874 | 0.007565995 | 0.1721437 | 0.281001419 | 0.359290001 |
| Apeh | -0.007142576 | 0.772273162 | 0.9552333 | 0.137900152 | 0.146429267 |
| Apex1 | 0.021357098 | 0.415907081 | 0.8300990 | 0.190684928 | 0.152820049 |
| Aph1a | -0.015355069 | 0.617182365 | 0.9115017 | 0.246793283 | 0.264214101 |
| Aph1b | 0.016517442 | 0.366688857 | 0.8053794 | 0.084107506 | 0.06621747 |
| Api5 | 0.046430043 | 0.15628305 | 0.6315293 | 0.366274692 | 0.324324985 |
| Apip | -0.039015071 | 0.174357254 | 0.6527657 | 0.167585498 | 0.216789399 |
| Aplp1 | -0.122424385 | 2.45E-06 | 0.0004494 | 2.131895094 | 2.070611207 |
| Aplp2 | -0.049719545 | 0.172154377 | 0.6520594 | 0.447797432 | 0.453069001 |
| Apmap | -0.032737155 | 0.199554033 | 0.6730962 | 0.143403612 | 0.157124227 |
| Apoa2 | -0.036671838 | 0.140636984 | 0.6130953 | 0.14325202 | 0.146196544 |
| Apobec1 | 0.050279219 | 0.076698046 | 0.5015054 | 0.234473928 | 0.177684665 |
| Apobec3 | -0.052919983 | 0.021786966 | 0.2874025 | 0.095001505 | 0.130696174 |
| Apod | -0.032219001 | 0.048713621 | 0.4158684 | 0.045020182 | 0.05806125 |
| Apold1 | 0.000596956 | 0.970191456 | 0.9976072 | 0.061199178 | 0.047027841 |
| Apoo | -0.005593587 | 0.848014525 | 0.9744357 | 0.25154729 | 0.232347278 |
| Apool | 0.011953713 | 0.647887066 | 0.9169695 | 0.172546814 | 0.145946229 |

| | | | | | |
|---|---|---|---|---|---|
| App | -0.125439383 | 0.001640244 | 0.0704381 | 1.033236357 | 1.128986753 |
| Appbp2 | -0.040926316 | 0.251046954 | 0.7207210 | 0.466023166 | 0.461126529 |
| Appl1 | -0.014596894 | 0.684048067 | 0.9311703 | 0.453641158 | 0.441920589 |
| Appl2 | -0.041221912 | 0.056763679 | 0.4473248 | 0.082050596 | 0.118880172 |
| Aprt | -0.012812397 | 0.72443212 | 0.9445652 | 0.597568534 | 0.608829256 |
| Aptx | 0.022772741 | 0.330079213 | 0.7778857 | 0.149154542 | 0.121765694 |
| Aqp11 | 0.009553175 | 0.576220768 | 0.8994610 | 0.062535025 | 0.063567084 |
| Aqr | 0.000428998 | 0.988176494 | 0.9976143 | 0.213335237 | 0.21910648 |
| Araf | 0.004492142 | 0.898583219 | 0.9825441 | 0.485560779 | 0.502092022 |
| Arap1 | 0.003370964 | 0.873703818 | 0.9784331 | 0.106674735 | 0.102032392 |
| Arc | -0.049935635 | 0.058435468 | 0.4517548 | 0.099503287 | 0.151082436 |
| Arcn1 | -0.046650922 | 0.206792234 | 0.6797565 | 0.76277171 | 0.792596406 |
| Arel1 | 0.002419584 | 0.913940156 | 0.9845236 | 0.116293103 | 0.109629256 |
| Arf1 | 0.010614222 | 0.731495914 | 0.9459973 | 1.459016339 | 1.39466513 |
| Arf2 | -0.007901333 | 0.761398196 | 0.9528631 | 0.167503299 | 0.165935865 |
| Arf3 | 0.063165001 | 0.076973241 | 0.5015240 | 0.649416184 | 0.597244966 |
| Arf4 | -0.029838325 | 0.397905192 | 0.8238469 | 1.3025752 | 1.228513974 |
| Arf5 | 0.017349855 | 0.478842672 | 0.8608564 | 1.863901631 | 1.735476623 |
| Arf6 | -0.043867004 | 0.128648141 | 0.5914729 | 0.201608131 | 0.216495278 |
| Arfgap1 | -0.005731628 | 0.861829037 | 0.9769675 | 0.326852559 | 0.325097052 |
| Arfgap2 | -0.059404705 | 0.065687964 | 0.4756383 | 0.302754924 | 0.331329001 |
| Arfgap3 | -0.062809256 | 0.068431315 | 0.4833869 | 0.360687306 | 0.40222586 |
| Arfgef1 | -0.042387133 | 0.240100212 | 0.7118791 | 0.724353978 | 0.760512668 |
| Arfgef2 | -0.012739743 | 0.627856686 | 0.9128095 | 0.160184728 | 0.173227216 |
| Arfgef3 | 0.018441015 | 0.627580129 | 0.9128095 | 0.792218009 | 0.836780335 |
| Arfip1 | 0.023661119 | 0.46334379 | 0.8524425 | 0.34157228 | 0.300370505 |
| Arfip2 | -0.060883869 | 0.075694062 | 0.4969457 | 0.361593353 | 0.392626855 |
| Arfrp1 | 0.042592573 | 0.195243736 | 0.6719932 | 0.416551634 | 0.350268105 |
| Arg1 | -0.084231307 | 0.02595818 | 0.3089760 | 0.258339266 | 0.327334946 |
| Arglu1 | -0.097273907 | 0.015373481 | 0.2425875 | 0.672169549 | 0.76538208 |
| Arhgap1 | -0.005691688 | 0.838449844 | 0.9727216 | 0.197812247 | 0.19238979 |
| Arhgap12 | -0.006067946 | 0.824996169 | 0.9715054 | 0.176316627 | 0.182873129 |
| Arhgap17 | -0.067492719 | 0.006648578 | 0.1617967 | 0.099823669 | 0.156268656 |
| Arhgap18 | -0.010134453 | 0.57180867 | 0.8966165 | 0.058729023 | 0.074289812 |
| Arhgap21 | -0.004973345 | 0.863511252 | 0.9769675 | 0.191232378 | 0.202299615 |
| Arhgap23 | -0.026769553 | 0.090355034 | 0.5319418 | 0.044591472 | 0.057676438 |
| Arhgap24 | 0.00921274 | 0.761055808 | 0.9526414 | 0.234134836 | 0.221784715 |
| Arhgap26 | -0.006732716 | 0.659834216 | 0.9211780 | 0.041823187 | 0.05150751 |
| Arhgap29 | -0.04460983 | 0.06545503 | 0.4746446 | 0.111628134 | 0.128405864 |
| Arhgap32 | -0.026518265 | 0.354132588 | 0.7967782 | 0.197151859 | 0.213211631 |
| Arhgap35 | 0.045082454 | 0.075389147 | 0.4966285 | 0.17081149 | 0.134559945 |
| Arhgap44 | 0.00125868 | 0.950476381 | 0.9948462 | 0.084671946 | 0.087930418 |
| Arhgap5 | -0.006387195 | 0.856531155 | 0.9763189 | 0.991197948 | 1.012760517 |
| Arhgap6 | 0.008650628 | 0.781657447 | 0.9590847 | 0.243247002 | 0.208550248 |
| Arhgdia | -0.007513424 | 0.841472835 | 0.9734589 | 0.80251069 | 0.74955404 |
| Arhgdig | 0.014366603 | 0.632474115 | 0.9139320 | 0.229479786 | 0.245232664 |
| Arhgef1 | -0.000673774 | 0.981310813 | 0.9976072 | 0.199430434 | 0.194690483 |
| Arhgef11 | -0.031227032 | 0.167655913 | 0.6446304 | 0.089665894 | 0.131962771 |
| Arhgef12 | 0.026153128 | 0.472941665 | 0.8595788 | 0.598689911 | 0.61718403 |
| Arhgef16 | 0.003358489 | 0.909381806 | 0.9837373 | 0.237724518 | 0.232329493 |
| Arhgef17 | 0.054809166 | 0.051560987 | 0.4287384 | 0.232031016 | 0.182999445 |
| Arhgef18 | -0.005718278 | 0.771460178 | 0.9552333 | 0.081790828 | 0.087658323 |
| Arhgef2 | -0.018696716 | 0.567369768 | 0.8952062 | 0.26847065 | 0.2860988 |
| Arhgef25 | -0.040381377 | 0.01444375 | 0.2371912 | 0.044563032 | 0.066296158 |
| Arhgef28 | -0.046485112 | 0.088767364 | 0.5288722 | 0.152703481 | 0.180671598 |
| Arhgef5 | -0.008209373 | 0.781671055 | 0.9590847 | 0.209680006 | 0.224771001 |
| Arhgef7 | 0.026886023 | 0.192734708 | 0.6700281 | 0.106612707 | 0.082331343 |
| Arhgef9 | -0.038021069 | 0.092192312 | 0.5325584 | 0.111350864 | 0.118314017 |
| Arid1a | -0.036138024 | 0.316182877 | 0.7705987 | 0.502512125 | 0.524497493 |
| Arid1b | -0.056303389 | 0.052062432 | 0.4313858 | 0.182436005 | 0.232626422 |
| Arid2 | -0.002955563 | 0.919532353 | 0.9853630 | 0.244253627 | 0.24370282 |
| Arid3a | -0.010013706 | 0.666654536 | 0.9230765 | 0.110348083 | 0.123759065 |
| Arid3b | 0.001545151 | 0.917966138 | 0.9849890 | 0.046007522 | 0.047652236 |
| Arid4a | 0.079123018 | 0.017220082 | 0.2570999 | 0.389536474 | 0.289349207 |
| Arid4b | -0.087304839 | 0.017181778 | 0.2570787 | 0.645408238 | 0.68044264 |
| Arid5a | -0.016066915 | 0.334006045 | 0.7804836 | 0.039034651 | 0.066331797 |

| | | | | | |
|---|---|---|---|---|---|
| Arid5b | 0.031405696 | 0.352567833 | 0.7952929 | 0.308699489 | 0.297568052 |
| Arih1 | 0.038561898 | 0.234555297 | 0.7064894 | 0.348473096 | 0.311036668 |
| Arih2 | -0.041401712 | 0.150793415 | 0.6280320 | 0.19671132 | 0.225679589 |
| Arl1 | -0.075149467 | 0.025872947 | 0.3088084 | 1.224683803 | 1.214859798 |
| Arl13b | -0.021278742 | 0.412781761 | 0.8299138 | 0.14948289 | 0.163175759 |
| Arl14ep | 0.068911819 | 0.006436103 | 0.1603078 | 0.188937946 | 0.132823316 |
| Arl15 | 0.009611788 | 0.507592067 | 0.8704067 | 0.053608134 | 0.039233013 |
| Arl16 | -0.035326676 | 0.144808917 | 0.6193638 | 0.132792373 | 0.14558188 |
| Arl2 | 0.043561775 | 0.203590814 | 0.6772740 | 0.519471475 | 0.455094817 |
| Arl2bp | -0.032406366 | 0.329762413 | 0.7778857 | 0.389595009 | 0.401605508 |
| Arl3 | -0.002770979 | 0.933668388 | 0.9903410 | 1.037596437 | 0.994009216 |
| Arl4a | -0.081429613 | 0.031672845 | 0.3385228 | 0.6701173 | 0.772507306 |
| Arl4d | -0.065711632 | 0.00712351 | 0.1680001 | 0.09531337 | 0.145277282 |
| Arl5a | -0.024076267 | 0.477490276 | 0.8607489 | 0.344834478 | 0.340877991 |
| Arl5b | -0.010020329 | 0.614345418 | 0.9108386 | 0.07961305 | 0.090769451 |
| Arl6 | -0.003951092 | 0.894728782 | 0.9824361 | 0.264124956 | 0.235520679 |
| Arl6ip1 | -0.007935541 | 0.828151263 | 0.9718751 | 0.868297023 | 0.84218936 |
| Arl6ip4 | 0.044739638 | 0.209455909 | 0.6807660 | 0.551654598 | 0.478141125 |
| Arl6ip5 | 0.048737833 | 0.150895885 | 0.6280320 | 0.488255433 | 0.417369378 |
| Arl6ip6 | 0.037104849 | 0.152468782 | 0.6281105 | 0.204309504 | 0.15175046 |
| Arl8a | -0.045446518 | 0.149321525 | 0.6264374 | 0.254922465 | 0.302867686 |
| Arl8b | 0.020377737 | 0.554387651 | 0.8902995 | 0.424070922 | 0.452119622 |
| Armc1 | 0.002235561 | 0.944928403 | 0.9936932 | 0.330690078 | 0.313422064 |
| Armc10 | -0.011169309 | 0.751885847 | 0.9507594 | 0.390255549 | 0.407533231 |
| Armc5 | 0.002503645 | 0.915622456 | 0.9845236 | 0.13629035 | 0.131050271 |
| Armc6 | -0.021387203 | 0.206025393 | 0.6797565 | 0.052723631 | 0.059976774 |
| Armc8 | -0.030283752 | 0.254845449 | 0.7223665 | 0.156162511 | 0.187372301 |
| Armc9 | -0.015684053 | 0.426964062 | 0.8337565 | 0.083127277 | 0.090285289 |
| Armcx1 | -0.025888119 | 0.368128445 | 0.8056719 | 0.213479493 | 0.214078365 |
| Armcx2 | -0.000817753 | 0.97930354 | 0.9976072 | 0.302726374 | 0.26609581 |
| Armcx3 | 0.043455218 | 0.221795938 | 0.6933883 | 0.584705476 | 0.519509573 |
| Armcx4 | -0.011880239 | 0.644495367 | 0.9168391 | 0.140467756 | 0.155403166 |
| Armcx5 | 0.017138582 | 0.489640253 | 0.8651997 | 0.144720042 | 0.137764906 |
| Armcx6 | -0.001015143 | 0.95964141 | 0.9957785 | 0.087140372 | 0.088280721 |
| Armh3 | -0.024114999 | 0.173244662 | 0.6525507 | 0.046854458 | 0.068982482 |
| Armt1 | -0.012077209 | 0.561614989 | 0.8926240 | 0.087190049 | 0.099908787 |
| Arnt | -0.021669612 | 0.500233795 | 0.8693704 | 0.255607652 | 0.280824746 |
| Arnt2 | -0.029359669 | 0.168837162 | 0.6466658 | 0.099419088 | 0.104544263 |
| Arntl | 0.011618533 | 0.563820223 | 0.8931805 | 0.096694931 | 0.081075308 |
| Arpc1a | -0.044384163 | 0.194985503 | 0.6719932 | 1.290712257 | 1.187441957 |
| Arpc1b | -0.051424525 | 0.132506875 | 0.5988251 | 0.343053409 | 0.360570502 |
| Arpc2 | 0.005292618 | 0.851339138 | 0.9749822 | 1.517131286 | 1.432319469 |
| Arpc3 | -0.031312541 | 0.316733533 | 0.7705987 | 1.219302673 | 1.150711299 |
| Arpc4 | 0.008847321 | 0.775525046 | 0.9558538 | 0.29238626 | 0.250645739 |
| Arpc5 | 0.027532253 | 0.433012568 | 0.8374897 | 0.98633205 | 0.89558309 |
| Arpc5l | 0.022475969 | 0.504897317 | 0.8695566 | 1.207753499 | 1.233680526 |
| Arpin | -0.014067958 | 0.543185098 | 0.8840682 | 0.115780524 | 0.125463671 |
| Arpp19 | 0.072044134 | 0.027656294 | 0.3181899 | 1.07304791 | 1.00543667 |
| Arrb2 | 0.008197324 | 0.709646363 | 0.9418251 | 0.129883446 | 0.107066412 |
| Arrdc1 | -0.046023403 | 0.039261854 | 0.3799003 | 0.096622043 | 0.122397218 |
| Arrdc3 | 0.000774636 | 0.973780852 | 0.9976072 | 0.1412833 | 0.126663 |
| Arrdc4 | 0.066038506 | 0.028947958 | 0.3264172 | 0.295490809 | 0.202591748 |
| Arsa | 0.007256741 | 0.692533087 | 0.9358575 | 0.077853737 | 0.076143385 |
| Arsb | -0.008514182 | 0.778850206 | 0.9581793 | 0.257243977 | 0.240330502 |
| ARVCF | 0.001211921 | 0.944718335 | 0.9936932 | 0.06769904 | 0.058355447 |
| Arx | -0.003067016 | 0.937618403 | 0.9918079 | 0.568644696 | 0.461773234 |
| Arxes1 | 0.025178254 | 0.082871718 | 0.5173615 | 0.057957018 | 0.03970533 |
| Arxes2 | 0.004929688 | 0.839859743 | 0.9728408 | 0.137831215 | 0.143747663 |
| As3mt | 0.012088397 | 0.712144048 | 0.9419834 | 0.345428807 | 0.31041914 |
| Asah1 | 0.062054744 | 0.09345448 | 0.5330951 | 0.618174614 | 0.566577754 |
| Asah2 | 0.047214263 | 0.057969013 | 0.4508199 | 0.155160722 | 0.141358201 |
| Asap1 | -0.009017672 | 0.757824554 | 0.9521533 | 0.204290593 | 0.208734939 |
| Asap2 | 0.010700191 | 0.560803943 | 0.8926240 | 0.081069147 | 0.066828064 |
| Asb1 | 0.00126271 | 0.945857753 | 0.9942681 | 0.078608572 | 0.075435277 |
| Asb13 | 0.007955 | 0.753895646 | 0.9518448 | 0.160346344 | 0.141637564 |
| Asb3 | 7.63E-05 | 0.996558372 | 0.9994073 | 0.073220356 | 0.067331036 |

| | | | | | |
|---|---|---|---|---|---|
| Asb4 | -0.00897682 | 0.621333066 | 0.9121505 | 0.058878136 | 0.060860778 |
| Asb6 | -0.015036819 | 0.435345335 | 0.8389295 | 0.080627384 | 0.081158258 |
| Asb7 | 0.022825 | 0.30065582 | 0.7596805 | 0.120863064 | 0.096474427 |
| Asb8 | 0.001070418 | 0.966915683 | 0.9976072 | 0.16992251 | 0.159610072 |
| Ascc1 | -0.011713751 | 0.694596933 | 0.9361958 | 0.232426882 | 0.224268211 |
| Ascc2 | -0.031696179 | 0.212012311 | 0.6822059 | 0.124562289 | 0.155401597 |
| Ascc3 | 0.047830828 | 0.205534154 | 0.6797565 | 0.606653964 | 0.564463593 |
| Ascl1 | 0.012427306 | 0.522951074 | 0.8750611 | 0.082279025 | 0.057178498 |
| Asf1a | -0.01924428 | 0.506336594 | 0.8695566 | 0.216327291 | 0.217472045 |
| Ash1l | -0.090204849 | 0.019580464 | 0.2709040 | 0.811710562 | 0.909681703 |
| Ash2l | 0.010769283 | 0.716512421 | 0.9431411 | 0.2434613 | 0.219428327 |
| Asic1 | 0.017687294 | 0.309706236 | 0.7676876 | 0.070176348 | 0.062631062 |
| Asl | 0.006429031 | 0.762404902 | 0.9530061 | 0.110938721 | 0.100417119 |
| Asns | 0.111618234 | 0.005437464 | 0.1488982 | 1.261348517 | 1.178167953 |
| Asnsd1 | 0.027916161 | 0.214916179 | 0.6844200 | 0.137264787 | 0.105089041 |
| Asph | -0.040662921 | 0.178903833 | 0.6549987 | 0.220205023 | 0.249001829 |
| Asphd1 | -0.001737139 | 0.936234218 | 0.9912468 | 0.115436258 | 0.105918346 |
| Aspscr1 | -0.016663968 | 0.524423201 | 0.8758385 | 0.164901618 | 0.171341865 |
| Asrgl1 | -0.039007241 | 0.206956334 | 0.6797565 | 0.267668829 | 0.271977562 |
| Aste1 | 0.015491908 | 0.296895038 | 0.7545964 | 0.052031303 | 0.04257765 |
| Asxl1 | 0.062076469 | 0.011748111 | 0.2135511 | 0.191567159 | 0.117019374 |
| Asxl2 | -0.054754047 | 0.115322247 | 0.5724670 | 0.376533342 | 0.406440292 |
| Atad1 | 0.014187385 | 0.659826801 | 0.9211780 | 0.324022715 | 0.314855653 |
| Atad2 | 0.005234838 | 0.775597202 | 0.9558538 | 0.073075036 | 0.065588434 |
| Atad2b | -0.006487034 | 0.824029915 | 0.9710592 | 0.217160851 | 0.208838888 |
| Atad3a | -0.019384033 | 0.42791992 | 0.8340365 | 0.119027627 | 0.138955318 |
| Atat1 | -0.023380026 | 0.305146643 | 0.7646008 | 0.112451117 | 0.120426741 |
| Ate1 | -0.055133514 | 0.125424349 | 0.5871682 | 0.402309302 | 0.460098131 |
| Atf1 | 0.021721685 | 0.493698183 | 0.8674510 | 0.338755782 | 0.298180966 |
| Atf2 | 0.001958659 | 0.954459031 | 0.9954690 | 0.377623096 | 0.374882703 |
| Atf3 | -0.117784886 | 0.02972969 | 0.3298864 | 0.564196784 | 0.85978399 |
| Atf4 | -0.08395287 | 0.008839699 | 0.1869718 | 1.329400488 | 1.459649449 |
| Atf5 | 0.051331255 | 0.235456316 | 0.7075815 | 0.886518358 | 1.021970414 |
| Atf6 | 0.039770556 | 0.244599845 | 0.7137971 | 0.355867669 | 0.362769602 |
| Atf6b | -0.047352464 | 0.052930073 | 0.4342980 | 0.118357795 | 0.148600502 |
| Atf7 | -0.014107975 | 0.566782626 | 0.8949756 | 0.135051432 | 0.134517375 |
| Atf7ip | -0.007209455 | 0.810586478 | 0.9668615 | 0.239491212 | 0.243624378 |
| Atg10 | -0.001369508 | 0.959950018 | 0.9958628 | 0.191671278 | 0.185355419 |
| Atg101 | 0.018660179 | 0.491926798 | 0.8664619 | 0.191729385 | 0.173745603 |
| Atg12 | 0.066402546 | 0.028178537 | 0.3216698 | 0.287568786 | 0.218287799 |
| Atg13 | -0.052011387 | 0.059860486 | 0.4573692 | 0.159254133 | 0.207683723 |
| Atg14 | 0.014484606 | 0.493259309 | 0.8669617 | 0.111242436 | 0.095438384 |
| Atg16l1 | -0.011542396 | 0.626918458 | 0.9128095 | 0.116243332 | 0.131402129 |
| Atg16l2 | 0.046802069 | 0.031492667 | 0.3385228 | 0.136156051 | 0.090939288 |
| Atg2a | 0.006866747 | 0.741051092 | 0.9479080 | 0.10554943 | 0.092189743 |
| Atg2b | 0.002138043 | 0.928712294 | 0.9881953 | 0.12433962 | 0.126697131 |
| Atg3 | -0.004467638 | 0.8950351 | 0.9824361 | 0.479665183 | 0.450466202 |
| Atg4b | -0.034146259 | 0.166262791 | 0.6429285 | 0.135851381 | 0.148202154 |
| Atg4c | 0.013102539 | 0.443732413 | 0.8431541 | 0.073518064 | 0.057890177 |
| Atg4d | -0.014826452 | 0.605635481 | 0.9087139 | 0.197840701 | 0.208658558 |
| Atg5 | -0.007754053 | 0.800168072 | 0.9647053 | 0.260991743 | 0.270868798 |
| Atg7 | -0.00395349 | 0.827512831 | 0.9717006 | 0.071454193 | 0.069109688 |
| Atic | -0.034203821 | 0.242875115 | 0.7132719 | 0.211658069 | 0.23015582 |
| Atl1 | 0.011168724 | 0.502828939 | 0.8695566 | 0.069391706 | 0.05350804 |
| Atl2 | 0.001912207 | 0.942145127 | 0.9931828 | 0.166960656 | 0.168961173 |
| Atl3 | 0.015368883 | 0.598788432 | 0.9075513 | 0.238690395 | 0.208647778 |
| Atm | -0.00485522 | 0.850163233 | 0.9748841 | 0.152508827 | 0.154223283 |
| Atmin | -0.014241458 | 0.488802122 | 0.8646428 | 0.09015635 | 0.095539184 |
| Atn1 | -0.042361996 | 0.145983504 | 0.6216370 | 0.202765588 | 0.228818514 |
| Atoh8 | -0.027954741 | 0.146033815 | 0.6216370 | 0.077742708 | 0.074747999 |
| Atox1 | 0.083205672 | 0.017297709 | 0.2570999 | 1.012463705 | 0.975902374 |
| Atp11a | -0.051231677 | 0.04367222 | 0.3986436 | 0.118855972 | 0.165685757 |
| Atp11b | 0.004221124 | 0.905158359 | 0.9832544 | 0.425973341 | 0.414604167 |
| Atp13a1 | -0.031299916 | 0.205630774 | 0.6797565 | 0.12100771 | 0.153440084 |
| Atp13a2 | 0.036560576 | 0.122884982 | 0.5833237 | 0.159843974 | 0.116754686 |
| Atp13a3 | -0.028642847 | 0.369037601 | 0.8056719 | 0.2407805 | 0.26961417 |

| | | | | | |
|---|---|---|---|---|---|
| Atp1a1 | 0.098198575 | 0.002609645 | 0.0934573 | 1.657948641 | 1.559017608 |
| Atp1b1 | 0.175484019 | 5.61E-07 | 0.0001589 | 1.346658175 | 1.168465628 |
| Atp1b2 | 0.016055004 | 0.209024095 | 0.6807660 | 0.040681392 | 0.031621234 |
| Atp1b3 | -0.005043377 | 0.87515678 | 0.9784331 | 0.32239785 | 0.339813708 |
| Atp23 | 0.005202791 | 0.73745782 | 0.9470008 | 0.059803885 | 0.049936915 |
| Atp2a2 | 0.102439223 | 0.002199089 | 0.0836783 | 1.527739825 | 1.523302859 |
| Atp2a3 | 0.074590044 | 0.036721464 | 0.3626279 | 0.455423012 | 0.446281997 |
| Atp2b1 | 0.033610892 | 0.353968307 | 0.7967782 | 1.034704805 | 1.000106124 |
| Atp2b2 | -0.06065274 | 0.019026701 | 0.2683814 | 0.114289945 | 0.170953927 |
| Atp2b4 | -0.046235178 | 0.093753265 | 0.5333348 | 0.178659789 | 0.18883238 |
| Atp2c1 | -0.024929386 | 0.436799995 | 0.8392515 | 0.281798761 | 0.346110247 |
| Atp5a1 | 0.009207095 | 0.71557373 | 0.9431411 | 1.669369422 | 1.644789812 |
| Atp5b | 0.018411709 | 0.503776148 | 0.8695566 | 1.796008693 | 1.678455923 |
| Atp5c1 | 0.037798857 | 0.136362273 | 0.6041597 | 1.924766004 | 1.773536193 |
| Atp5d | -0.006221265 | 0.792748514 | 0.9624460 | 1.931463583 | 1.83131277 |
| Atp5e | 0.053234643 | 0.00076049 | 0.0405594 | 3.209790474 | 2.997480245 |
| Atp5g1 | 0.058332187 | 0.023991994 | 0.3005057 | 2.170345938 | 2.027090462 |
| Atp5g2 | 0.059832643 | 0.071651401 | 0.4888240 | 0.387072093 | 0.301852615 |
| Atp5g3 | 0.046448501 | 0.135520763 | 0.6028321 | 1.335426371 | 1.214200216 |
| Atp5h | -0.044670666 | 0.03254049 | 0.3412279 | 2.137025577 | 2.099838939 |
| Atp5j | 0.029417214 | 0.201608157 | 0.6752150 | 2.079670544 | 1.918785521 |
| Atp5j2 | 0.091078716 | 5.37E-05 | 0.0058575 | 2.122346706 | 1.925904464 |
| Atp5k | 0.154245166 | 3.82E-06 | 0.0006414 | 1.255237857 | 1.062913089 |
| Atp5l | 0.15463428 | 1.04E-06 | 0.0002286 | 1.551000283 | 1.360690069 |
| Atp5md | 0.163593628 | 6.75E-11 | 6.08E-08 | 2.015040116 | 1.711475375 |
| Atp5mpl | 0.135959909 | 1.76E-05 | 0.0024169 | 1.29470029 | 1.049099078 |
| Atp5o | 0.04313575 | 0.218316115 | 0.6897120 | 0.935245668 | 0.823069957 |
| Atp5pb | 0.099600624 | 0.000646256 | 0.0359849 | 1.615301094 | 1.413915965 |
| Atp6ap1 | -0.047691522 | 0.171924452 | 0.6516968 | 0.773810383 | 0.791535971 |
| Atp6ap2 | -0.050044024 | 0.102407305 | 0.5529567 | 1.324354567 | 1.372857143 |
| Atp6v0a1 | -0.040509581 | 0.215734484 | 0.6854856 | 0.295052668 | 0.346432176 |
| Atp6v0a2 | 0.018850651 | 0.468610545 | 0.8563037 | 0.175532549 | 0.157597965 |
| Atp6v0b | 0.000625012 | 0.981503931 | 0.9976072 | 1.704204107 | 1.697516974 |
| Atp6v0c | -0.006418837 | 0.7672782 | 0.9544221 | 2.107375288 | 2.064278763 |
| Atp6v0d1 | -0.063414473 | 0.073006862 | 0.4903372 | 0.663412551 | 0.711958541 |
| Atp6v0e | 0.058879935 | 0.071300397 | 0.4881297 | 1.32203911 | 1.267756727 |
| Atp6v0e2 | 0.026412758 | 0.447670961 | 0.8455660 | 0.451900492 | 0.389184305 |
| Atp6v1a | 0.09156394 | 0.011569908 | 0.2120789 | 0.490471663 | 0.467948938 |
| Atp6v1b2 | -0.003442399 | 0.926519491 | 0.9879210 | 0.719149952 | 0.744989776 |
| Atp6v1c1 | 0.026144016 | 0.465816545 | 0.8549201 | 0.557157492 | 0.516128575 |
| Atp6v1d | 0.006311119 | 0.85986808 | 0.9769675 | 0.794588169 | 0.792510115 |
| Atp6v1e1 | 0.051876922 | 0.133501355 | 0.5988251 | 1.026069516 | 0.99124484 |
| Atp6v1f | -0.034742278 | 0.163468212 | 0.6409504 | 1.713339047 | 1.689557795 |
| Atp6v1g1 | 0.00215613 | 0.938258432 | 0.9923791 | 1.518764215 | 1.475659106 |
| Atp6v1h | -4.80E-05 | 0.998889746 | 0.9996989 | 0.445190247 | 0.433750271 |
| Atp7a | 0.019118888 | 0.452277667 | 0.8484983 | 0.154365139 | 0.15171483 |
| Atp8a1 | 0.010718089 | 0.62232174 | 0.9121505 | 0.106762725 | 0.109078387 |
| Atp8a2 | -0.014710538 | 0.358117051 | 0.7992173 | 0.041568176 | 0.05661547 |
| Atp8b1 | 0.085204116 | 0.023585235 | 0.2980452 | 0.643193238 | 0.586910051 |
| Atp8b2 | -0.012786394 | 0.568999332 | 0.8956532 | 0.106605045 | 0.108384263 |
| Atp9a | 0.031307347 | 0.329845648 | 0.7778857 | 0.323792019 | 0.309257882 |
| Atp9b | 0.003329282 | 0.881218546 | 0.9795705 | 0.116031918 | 0.109876358 |
| Atpaf1 | 0.008014289 | 0.707519937 | 0.9408790 | 0.107491582 | 0.103279891 |
| Atpaf2 | -0.006537207 | 0.747341189 | 0.9496370 | 0.092922432 | 0.09239031 |
| Atpif1 | 0.030972623 | 0.113689609 | 0.5696977 | 2.335789205 | 2.190010911 |
| Atpsckmt | -0.010718524 | 0.655059378 | 0.9206439 | 0.125016612 | 0.137277424 |
| Atr | -0.013192135 | 0.564392518 | 0.8933738 | 0.116821216 | 0.108454227 |
| Atraid | -0.143152717 | 5.75E-05 | 0.0061312 | 0.790854532 | 0.963741125 |
| Atrip | 0.022337624 | 0.281029416 | 0.7408618 | 0.11438638 | 0.084812741 |
| Atrn | -0.006352482 | 0.773641266 | 0.9557311 | 0.105077341 | 0.105083449 |
| Atrnl1 | 0.015897742 | 0.289150903 | 0.7494616 | 0.059773921 | 0.040220865 |
| Atrx | -0.031260732 | 0.383654772 | 0.8129787 | 1.062933072 | 1.085293318 |
| Atxn1 | 0.008829427 | 0.749109929 | 0.9501560 | 0.1892572 | 0.173801146 |
| Atxn10 | -0.032255047 | 0.348086648 | 0.7917953 | 1.021336464 | 0.925346612 |
| Atxn1l | -0.000500402 | 0.979122955 | 0.9976072 | 0.075839863 | 0.076230712 |
| Atxn2 | -0.033873264 | 0.30828088 | 0.7662606 | 0.345567536 | 0.354950182 |

| | | | | | |
|---|---|---|---|---|---|
| Atxn2l | 0.042346319 | 0.228704047 | 0.70196775 | 0.489505376 | 0.460987745 |
| Atxn3 | 0.007774391 | 0.670972747 | 0.92470826 | 0.077423887 | 0.069316867 |
| Atxn7 | 0.003599555 | 0.901008213 | 0.98254414 | 0.214884964 | 0.199135018 |
| Atxn7l1 | -0.014413761 | 0.377428524 | 0.81017979 | 0.055115861 | 0.058176126 |
| Atxn7l2 | 0.046201107 | 0.002440055 | 0.08998271 | 0.07047876 | 0.03546257 |
| Atxn7l3 | -0.058053517 | 0.022836861 | 0.29268948 | 0.127522907 | 0.169845835 |
| Atxn7l3b | 0.000480299 | 0.989016994 | 0.99808365 | 1.015378659 | 1.041375835 |
| AU022252 | 0.001309645 | 0.933629239 | 0.99034101 | 0.053365236 | 0.05189557 |
| AU040320 | -0.008955421 | 0.724958315 | 0.94487071 | 0.140005023 | 0.165379173 |
| AU041133 | -0.019632891 | 0.388874528 | 0.81702876 | 0.10324471 | 0.122149379 |
| Auh | 0.071007323 | 0.044467058 | 0.40095622 | 0.524803677 | 0.460908528 |
| Aup1 | 0.005090096 | 0.864503016 | 0.97696756 | 0.221337628 | 0.245885514 |
| Aurkaip1 | 0.051693907 | 0.152353983 | 0.62811059 | 0.739181809 | 0.708068838 |
| Auts2 | -0.048215131 | 0.244619305 | 0.71379715 | 0.7514583 | 0.807461452 |
| AV356131 | -0.004959314 | 0.856916147 | 0.97631893 | 0.210867879 | 0.196143641 |
| Aven | -0.028726272 | 0.116657595 | 0.57274726 | 0.052186666 | 0.077818524 |
| Avl9 | -0.03863116 | 0.25175361 | 0.72191687 | 0.333518854 | 0.366773605 |
| Avpi1 | -0.026554149 | 0.405855115 | 0.82707706 | 0.197710274 | 0.280440301 |
| Avpr1b | -0.013893104 | 0.646866612 | 0.91696956 | 0.230090954 | 0.195841924 |
| AW146154 | 0.004590125 | 0.761767 | 0.95292921 | 0.054511404 | 0.045883235 |
| AW209491 | 0.00429104 | 0.857327333 | 0.97631893 | 0.143007917 | 0.129411055 |
| AW549877 | -0.006166731 | 0.819374619 | 0.96990621 | 0.16598815 | 0.182573771 |
| AW551984 | 0.010838672 | 0.429146568 | 0.83403655 | 0.049186281 | 0.034620902 |
| AW554918 | -0.015067774 | 0.347790422 | 0.79179535 | 0.044576236 | 0.055074701 |
| Axin1 | -0.027658921 | 0.314364195 | 0.77059875 | 0.196857197 | 0.202240035 |
| Axin2 | 0.014776402 | 0.277397289 | 0.73794076 | 0.048663914 | 0.031681405 |
| Azi2 | -0.013777536 | 0.695560695 | 0.93634989 | 0.405420993 | 0.401186547 |
| Azin1 | -0.022458881 | 0.515130663 | 0.87337141 | 0.399019672 | 0.415461658 |
| B230206L02Rik | -0.011434407 | 0.472765097 | 0.85957296 | 0.054714512 | 0.051646333 |
| B230216N24Rik | -0.086553385 | 0.001088297 | 0.05116544 | 0.065101768 | 0.154397434 |
| B230217C12Rik | -0.033065628 | 0.201036684 | 0.67465626 | 0.136214521 | 0.15792348 |
| B230219D22Rik | 0.090422941 | 0.016216318 | 0.25135291 | 0.804643241 | 0.68260684 |
| B230307C23Rik | -0.023194683 | 0.21202237 | 0.68220598 | 0.057326131 | 0.081656478 |
| B230322F03Rik | -0.004355489 | 0.812561564 | 0.96730151 | 0.085357224 | 0.071519566 |
| B230354K17Rik | -0.060982825 | 0.005753743 | 0.15302179 | 0.075118817 | 0.120307417 |
| B230369F24Rik | 0.011732968 | 0.415268977 | 0.83009907 | 0.048239225 | 0.039040808 |
| B2m | -0.018040133 | 0.648447723 | 0.91723961 | 0.429296906 | 0.471270531 |
| B330016D10Rik | -0.011468611 | 0.621033344 | 0.91215057 | 0.117824127 | 0.117737201 |
| B3galnt1 | -0.034801746 | 0.172217303 | 0.65205946 | 0.128007008 | 0.15868989 |
| B3galnt2 | -0.023460895 | 0.187037763 | 0.66551566 | 0.053963004 | 0.072495002 |
| B3galt2 | -0.014236188 | 0.427504247 | 0.83403655 | 0.053107569 | 0.073043627 |
| B3galt4 | 0.009822834 | 0.459674326 | 0.85243431 | 0.045449786 | 0.032871988 |
| B3galt5 | -0.003823118 | 0.794248894 | 0.96256135 | 0.042201388 | 0.046100053 |
| B3galt6 | -0.006834974 | 0.735300997 | 0.94683808 | 0.091586486 | 0.092963086 |
| B3gat3 | -0.011069216 | 0.741856436 | 0.94790806 | 0.398059789 | 0.36984703 |
| B3gnt9 | 0.015197487 | 0.393534334 | 0.82089394 | 0.086746232 | 0.058419441 |
| B4galt1 | -0.040036084 | 0.047823956 | 0.41325229 | 0.065846808 | 0.092491315 |
| B4galt2 | 0.011394781 | 0.463164081 | 0.85244258 | 0.065699683 | 0.044389697 |
| B4galt3 | 0.036336225 | 0.275002686 | 0.73531715 | 0.42103508 | 0.355161731 |
| B4galt5 | -0.004271497 | 0.822114427 | 0.97069202 | 0.072582387 | 0.079689802 |
| B4galt6 | -0.016116324 | 0.651569494 | 0.91874716 | 0.461104963 | 0.467125844 |
| B4galt7 | -0.007727798 | 0.69351678 | 0.93597259 | 0.088392375 | 0.085291256 |
| B630019K06Rik | -0.014965268 | 0.510138374 | 0.87116073 | 0.119223425 | 0.114700533 |
| B830012L14Rik | -0.033259417 | 0.206010913 | 0.67975655 | 0.13671723 | 0.158384559 |
| B830017H08Rik | 0.052674334 | 0.224773572 | 0.69682971 | 0.603573227 | 0.782002523 |
| B930095G15Rik | -0.018874118 | 0.387360912 | 0.81655597 | 0.096202884 | 0.108524457 |
| B9d1 | -0.003835812 | 0.898261339 | 0.98254414 | 0.244690133 | 0.246912496 |
| B9d2 | 0.002728242 | 0.923681389 | 0.98759639 | 0.21420574 | 0.203795253 |
| Babam1 | 0.012960457 | 0.705761223 | 0.94013813 | 0.419302335 | 0.392002895 |
| Babam2 | 0.032504546 | 0.319282862 | 0.77100435 | 0.382178436 | 0.334663982 |
| Bace1 | -0.073074685 | 0.009278005 | 0.19303689 | 0.155450946 | 0.202436608 |
| Bace2 | 0.048875813 | 0.270619253 | 0.73420306 | 0.707896595 | 0.910420217 |
| Bach1 | -0.003730928 | 0.898341673 | 0.98254414 | 0.196941627 | 0.214578391 |
| Bad | -0.007133315 | 0.833313218 | 0.97234941 | 0.382013162 | 0.377856126 |
| Bag1 | -0.001062826 | 0.974123363 | 0.99760728 | 1.252616034 | 1.152783511 |
| Bag2 | -0.046948296 | 0.042921476 | 0.39681364 | 0.08584881 | 0.137982587 |

| | | | | | |
|---|---|---|---|---|---|
| Bag3 | -0.018464489 | 0.471126411 | 0.8580121 | 0.13051483 | 0.162431931 |
| Bag4 | 0.029649592 | 0.319865227 | 0.7712841 | 0.242147179 | 0.216204268 |
| Bag5 | -0.023528119 | 0.474614073 | 0.8600971 | 0.348783686 | 0.351082003 |
| Bag6 | 0.004506784 | 0.886691606 | 0.9811467 | 0.340678966 | 0.302311055 |
| Bahd1 | 0.037043329 | 0.076155734 | 0.4986566 | 0.125645704 | 0.087153358 |
| Baiap2l1 | 0.022999052 | 0.463278049 | 0.8524425 | 0.282330134 | 0.250158518 |
| Baiap3 | -0.021928807 | 0.38813455 | 0.8170287 | 0.142111743 | 0.147462478 |
| Bak1 | -0.003524884 | 0.826267994 | 0.9716906 | 0.053154115 | 0.053360655 |
| Bambi | -0.033183041 | 0.406411355 | 0.8270770 | 0.590073768 | 0.56587768 |
| Banf1 | -0.000646627 | 0.985143482 | 0.9976072 | 0.501587434 | 0.465858545 |
| Banp | 0.057455693 | 0.053002902 | 0.4345361 | 0.257082282 | 0.218425387 |
| Bap1 | 0.024974558 | 0.20285279 | 0.6770860 | 0.097552081 | 0.080390702 |
| Basp1 | -0.054124806 | 0.196104754 | 0.6730962 | 0.472742237 | 0.469619358 |
| Bax | -0.077915726 | 0.025109481 | 0.3045061 | 1.189885041 | 1.194260282 |
| Baz1a | -0.008583948 | 0.691890586 | 0.9355149 | 0.098189789 | 0.105316411 |
| Baz1b | -0.077214567 | 0.041051727 | 0.3901716 | 0.482127962 | 0.550463547 |
| Baz2a | -0.051421353 | 0.144834715 | 0.6193638 | 0.451049509 | 0.467574897 |
| Baz2b | -0.014262405 | 0.702201543 | 0.9387923 | 0.763063596 | 0.760157359 |
| BB218582 | -0.017102403 | 0.433374389 | 0.8375363 | 0.106346127 | 0.108811073 |
| Bbc3 | 0.046331838 | 0.069219458 | 0.4835495 | 0.174356384 | 0.137001487 |
| Bbip1 | -0.027806926 | 0.453261292 | 0.8490009 | 0.659936217 | 0.637269661 |
| Bbs1 | 0.012318325 | 0.348253792 | 0.7919939 | 0.044224403 | 0.03217841 |
| Bbs2 | -0.024314943 | 0.237980629 | 0.7104326 | 0.081316452 | 0.092356385 |
| Bbs4 | -0.033451621 | 0.151863086 | 0.6281105 | 0.124085821 | 0.132837811 |
| Bbs5 | 0.002661761 | 0.873689444 | 0.9784331 | 0.065263105 | 0.056540538 |
| Bbs7 | -0.011997223 | 0.553693451 | 0.8902995 | 0.097174687 | 0.088796502 |
| Bbx | -0.063197998 | 0.08890245 | 0.5292814 | 0.817420875 | 0.838998554 |
| BC002059 | 0.011954478 | 0.450685363 | 0.8478942 | 0.058628849 | 0.046801327 |
| BC003965 | 0.005671671 | 0.824700885 | 0.9713883 | 0.17582819 | 0.161130583 |
| BC004004 | 0.033584811 | 0.340220505 | 0.7844658 | 0.512852411 | 0.449272944 |
| BC005537 | -0.067057738 | 0.07473081 | 0.4966285 | 0.617365354 | 0.692068847 |
| BC005624 | -0.013119472 | 0.716715483 | 0.9431474 | 0.578887354 | 0.553083007 |
| BC024063 | 0.005475503 | 0.668634212 | 0.9230765 | 0.042266507 | 0.029673771 |
| BC029722 | -0.018468962 | 0.481766058 | 0.8620306 | 0.16012237 | 0.17722197 |
| BC031181 | 0.071777777 | 0.039970105 | 0.3834656 | 0.903452465 | 0.824399614 |
| BC051142 | -0.030812581 | 0.111036559 | 0.5671739 | 0.060967837 | 0.088353478 |
| BC051226 | 0.031779845 | 0.151436537 | 0.6281105 | 0.132009593 | 0.102413482 |
| Bcam | -0.032076707 | 0.075656174 | 0.4969457 | 0.050166699 | 0.070230216 |
| Bcap29 | 0.011437322 | 0.643105303 | 0.9166712 | 0.14754683 | 0.139280017 |
| Bcap31 | 0.029801617 | 0.387370201 | 0.8165559 | 0.596276986 | 0.525484603 |
| Bcar1 | -0.019719359 | 0.463124847 | 0.8524425 | 0.166618788 | 0.185820408 |
| Bcas2 | 0.003462205 | 0.919016275 | 0.9851211 | 0.928362816 | 0.895775817 |
| Bcas3 | -0.003325491 | 0.835449898 | 0.9723494 | 0.050513573 | 0.053699408 |
| Bcat2 | -0.025305299 | 0.086735187 | 0.5250566 | 0.025801748 | 0.054103273 |
| Bccip | 0.049589686 | 0.168792052 | 0.6466658 | 0.737594668 | 0.638923081 |
| Bcdin3d | 0.024316139 | 0.276249102 | 0.7364662 | 0.135478176 | 0.10524892 |
| Bckdha | -0.028113638 | 0.295430739 | 0.7543559 | 0.168002741 | 0.185293512 |
| Bckdhb | 0.004682969 | 0.770678821 | 0.9552333 | 0.055473646 | 0.053992677 |
| Bckdk | 0.003540692 | 0.90875331 | 0.9837373 | 0.270226389 | 0.272787738 |
| Bcl10 | -0.031310519 | 0.273339174 | 0.7351955 | 0.196734548 | 0.205025303 |
| Bcl11a | -0.023696899 | 0.296676932 | 0.7545964 | 0.112547692 | 0.115107435 |
| Bcl2 | -0.02872742 | 0.131870113 | 0.5978345 | 0.072859612 | 0.08262472 |
| Bcl2l1 | -0.01785017 | 0.523505035 | 0.8753025 | 0.189160072 | 0.197737797 |
| Bcl2l11 | -0.027523564 | 0.272313245 | 0.7342806 | 0.142203626 | 0.146755742 |
| Bcl2l13 | 0.009897392 | 0.686105547 | 0.9318810 | 0.150293423 | 0.137616221 |
| Bcl2l2 | -0.024107501 | 0.349091362 | 0.7927303 | 0.136386307 | 0.170222304 |
| Bcl3 | -0.057476155 | 0.049206373 | 0.4182752 | 0.1603567 | 0.219238235 |
| Bcl6b | 0.013099824 | 0.372557272 | 0.8085689 | 0.052087784 | 0.040684779 |
| Bcl7a | -0.049917888 | 0.083689429 | 0.5183040 | 0.17872471 | 0.215353424 |
| Bcl7b | 0.006380688 | 0.85497287 | 0.9755431 | 0.596956185 | 0.566702018 |
| Bcl7c | -0.044832033 | 0.167244833 | 0.6445488 | 0.303893224 | 0.301358974 |
| Bcl9 | -0.048541232 | 0.101910519 | 0.5515288 | 0.197478152 | 0.253201671 |
| Bcl9l | -0.019151753 | 0.387764985 | 0.8170287 | 0.093550718 | 0.11091201 |
| Bclaf1 | -0.066303844 | 0.08035895 | 0.5098712 | 0.801828066 | 0.882522626 |
| Bclaf3 | -0.000177813 | 0.992807462 | 0.9985888 | 0.092061854 | 0.086238633 |
| Bcor | -0.029984552 | 0.368899229 | 0.8056719 | 0.308174561 | 0.353351796 |

| | | | | | |
|---|---|---|---|---|---|
| Bcorl1 | 0.008804397 | 0.663638933 | 0.9230765 | 0.098241275 | 0.083964101 |
| Bcr | -0.019857613 | 0.497378298 | 0.8684629 | 0.227695683 | 0.240904908 |
| Bcs1l | 0.012403482 | 0.55665778 | 0.8915944 | 0.112907419 | 0.094656289 |
| Bdh1 | -0.02946131 | 0.084188354 | 0.5186471 | 0.039282424 | 0.073283639 |
| Bdnf | -0.020622909 | 0.237571156 | 0.7098511 | 0.050530659 | 0.063327552 |
| Bdp1 | -0.071868335 | 0.032366153 | 0.3397589 | 0.296222874 | 0.347742059 |
| Becn1 | 0.033048516 | 0.36079328 | 0.8014037 | 0.519829615 | 0.472760389 |
| Bend5 | -0.007490008 | 0.714823178 | 0.9430821 | 0.093637116 | 0.084786216 |
| Bend7 | -0.08441181 | 0.026421465 | 0.3109145 | 0.482167246 | 0.494854941 |
| Bet1 | 0.039221351 | 0.287341211 | 0.7471316 | 0.695138554 | 0.617068067 |
| Bet1l | 0.009741105 | 0.782117873 | 0.9590847 | 0.468550416 | 0.405749406 |
| Bex1 | -0.037925264 | 0.304338356 | 0.7634877 | 0.998128239 | 1.021923384 |
| Bex2 | -0.013415536 | 0.668613591 | 0.9230765 | 1.608360442 | 1.655205009 |
| Bex3 | -0.053964748 | 0.006106343 | 0.1568368 | 2.454337152 | 2.53306732 |
| Bex4 | -0.033729089 | 0.371273243 | 0.8074114 | 0.826920964 | 0.846233977 |
| Bfar | -0.034555033 | 0.262403873 | 0.7275735 | 0.229107905 | 0.2596455 |
| Bhlha15 | -0.004680801 | 0.805385194 | 0.9647143 | 0.075427221 | 0.07739392 |
| Bhlhb9 | -0.003930938 | 0.901321953 | 0.9825441 | 0.305712488 | 0.289412167 |
| Bhlhe40 | -0.032645291 | 0.159189256 | 0.6357726 | 0.085864104 | 0.130514879 |
| Bhlhe41 | 0.06707315 | 0.033530828 | 0.3464852 | 0.326465223 | 0.266085036 |
| Bicc1 | -0.018303695 | 0.520244429 | 0.8741795 | 0.189227694 | 0.191170781 |
| Bicd1 | 0.029808565 | 0.38711434 | 0.8163635 | 0.395478322 | 0.363271113 |
| Bicd2 | 0.009056554 | 0.60890355 | 0.9088672 | 0.071582075 | 0.063784621 |
| Bicdl1 | -0.014171016 | 0.456088779 | 0.8502914 | 0.07202998 | 0.078581541 |
| Bicral | 0.041835995 | 0.081517494 | 0.5135726 | 0.161613634 | 0.12375457 |
| Bik | -0.070735655 | 0.004285871 | 0.1302787 | 0.115442889 | 0.142609674 |
| Bin1 | -0.038757555 | 0.094214958 | 0.5334417 | 0.098683247 | 0.137028448 |
| Bin3 | -0.013780202 | 0.520210445 | 0.8741795 | 0.101407931 | 0.10280416 |
| Birc2 | -0.062409607 | 0.051940257 | 0.4313858 | 0.260813103 | 0.294957674 |
| Birc3 | -0.008764981 | 0.58488443 | 0.9023908 | 0.043146677 | 0.054962177 |
| Birc6 | 0.040301531 | 0.265486316 | 0.7306783 | 0.669808062 | 0.620148121 |
| Bivm | 0.030757544 | 0.04743629 | 0.4106761 | 0.072673438 | 0.044040729 |
| Blcap | -0.002650514 | 0.921695232 | 0.9862621 | 0.173563202 | 0.184661087 |
| Blmh | -0.002928775 | 0.929587452 | 0.9882657 | 0.34368562 | 0.329754151 |
| Bloc1s1 | 0.004062252 | 0.770029429 | 0.9552333 | 0.041899655 | 0.037922902 |
| Bloc1s2 | -0.00062966 | 0.984534239 | 0.9976072 | 0.343682849 | 0.305003241 |
| Bloc1s4 | -0.034594389 | 0.115758844 | 0.5724670 | 0.104373091 | 0.117902657 |
| Bloc1s5 | 0.010716668 | 0.614549361 | 0.9108386 | 0.115506157 | 0.101444607 |
| Bloc1s6 | -0.016123236 | 0.523238765 | 0.8750824 | 0.146794664 | 0.15048923 |
| Blvra | 0.002321689 | 0.907019075 | 0.9836812 | 0.09549504 | 0.085469283 |
| Blvrb | -0.056720155 | 0.047093434 | 0.4106761 | 0.193184474 | 0.226063318 |
| Blzf1 | 0.018642879 | 0.453783054 | 0.8490482 | 0.155952773 | 0.139051027 |
| Bmi1 | 0.007021188 | 0.809732772 | 0.9661473 | 0.214377706 | 0.218748749 |
| Bmp1 | -0.019416053 | 0.450100762 | 0.8478268 | 0.130729413 | 0.16635933 |
| Bmpr1a | -0.030513422 | 0.370618968 | 0.8066125 | 0.354752529 | 0.370495646 |
| Bmpr2 | -0.023566733 | 0.400075959 | 0.8244516 | 0.168068103 | 0.193449067 |
| Bms1 | 0.008845485 | 0.704767389 | 0.9400796 | 0.128502553 | 0.123131458 |
| Bmt2 | -0.044017691 | 0.134058067 | 0.6006009 | 0.195494563 | 0.213005504 |
| Bmyc | 0.058928838 | 0.093339022 | 0.5330587 | 0.475619711 | 0.393918095 |
| Bnip1 | 0.007586928 | 0.737362711 | 0.9470008 | 0.123256923 | 0.11280542 |
| Bnip2 | -0.009850964 | 0.669734172 | 0.9239496 | 0.116965189 | 0.119980627 |
| Bnip3 | -0.029699466 | 0.394424589 | 0.8214190 | 0.453191578 | 0.50396982 |
| Bnip3l | 0.056288897 | 0.091651038 | 0.5325584 | 0.527654449 | 0.404643209 |
| Bod1 | -0.006978086 | 0.844364206 | 0.9734889 | 0.617714189 | 0.559648544 |
| Bod1l | -0.039732461 | 0.275258523 | 0.7356525 | 0.617918777 | 0.656803941 |
| Bola1 | 0.045209386 | 0.221400564 | 0.6929153 | 0.56937225 | 0.511087275 |
| Bola2 | 0.120737146 | 0.000706179 | 0.0384906 | 0.890314485 | 0.7792868 |
| Bola3 | 0.021800393 | 0.537086335 | 0.8807896 | 0.784177736 | 0.718824996 |
| Bop1 | -0.026888462 | 0.205508534 | 0.6797565 | 0.086721926 | 0.104629981 |
| Borcs5 | -0.035873524 | 0.05666822 | 0.4473248 | 0.063346892 | 0.083833667 |
| Borcs6 | -0.057352296 | 0.041143447 | 0.3905674 | 0.175083693 | 0.202698947 |
| Borcs7 | -0.000175225 | 0.995636128 | 0.9994073 | 0.312245411 | 0.312546903 |
| Borcs8 | 0.022559816 | 0.442420079 | 0.8431541 | 0.244936338 | 0.221612214 |
| Bpgm | -0.01903822 | 0.42965308 | 0.8340365 | 0.127901521 | 0.141495837 |
| Bphl | 0.019104674 | 0.553179722 | 0.8900377 | 0.332954143 | 0.310546955 |
| Bpnt1 | 0.012381204 | 0.69320894 | 0.9359725 | 0.272516521 | 0.263767872 |

| | | | | | |
|---|---|---|---|---|---|
| Bpnt2 | -0.007563122 | 0.830186446 | 0.9722601 | 0.50890416 | 0.512772237 |
| Bptf | -0.000718468 | 0.984962904 | 0.9976072 | 0.729183743 | 0.757169442 |
| Braf | 0.009797698 | 0.786010951 | 0.9602516 | 0.516220808 | 0.521694335 |
| Brap | -0.015191849 | 0.574287117 | 0.8974946 | 0.160909751 | 0.177342834 |
| Brcc3 | 0.015667035 | 0.555960494 | 0.8911667 | 0.190392543 | 0.172375848 |
| Brd1 | -0.051517647 | 0.157137589 | 0.6321187 | 0.509859779 | 0.55698238 |
| Brd2 | -0.151301446 | 2.72E-05 | 0.0035037 | 0.846247031 | 1.06824982 |
| Brd3 | -0.067101217 | 0.053817145 | 0.4390346 | 0.356928557 | 0.412292339 |
| Brd3os | -0.016243344 | 0.401473595 | 0.8254259 | 0.069677972 | 0.085361744 |
| Brd4 | -0.049425335 | 0.180741567 | 0.6569883 | 0.505694552 | 0.537876114 |
| Brd7 | -0.011610545 | 0.692917069 | 0.9359725 | 0.221816159 | 0.226183823 |
| Brd8 | 0.009615909 | 0.794556033 | 0.9627453 | 0.607792773 | 0.558165578 |
| Brd9 | -0.019859644 | 0.42807106 | 0.8340365 | 0.124992693 | 0.150378157 |
| Brdt | -0.002116627 | 0.887664613 | 0.9812414 | 0.05287331 | 0.043651376 |
| Brf1 | -0.025980209 | 0.243408999 | 0.7132719 | 0.093346368 | 0.113727941 |
| Brf2 | -0.020038938 | 0.257422529 | 0.7250597 | 0.064684383 | 0.071370903 |
| Bri3 | 0.008747613 | 0.805762278 | 0.9647143 | 1.009074557 | 0.959729228 |
| Bri3bp | 0.006639016 | 0.781870802 | 0.9590847 | 0.135651165 | 0.126040555 |
| Brip1os | -0.030085894 | 0.184882559 | 0.6635437 | 0.108406121 | 0.113994444 |
| Brix1 | 0.0271527 | 0.422591976 | 0.8314897 | 0.412488619 | 0.360144676 |
| Brk1 | -0.014723548 | 0.647584593 | 0.9169695 | 0.914456387 | 0.928168273 |
| Brms1l | -0.009565252 | 0.605756612 | 0.9087139 | 0.073457944 | 0.074555873 |
| Brox | 0.008223501 | 0.75257802 | 0.9507898 | 0.165829044 | 0.15961465 |
| Brpf1 | -0.002039806 | 0.926612164 | 0.9879210 | 0.104528199 | 0.108039417 |
| Brpf3 | -0.023313527 | 0.252645863 | 0.7223071 | 0.074539233 | 0.093853555 |
| Brsk1 | -0.011292138 | 0.518593214 | 0.8741622 | 0.057668732 | 0.065522936 |
| Brsk2 | -0.017888143 | 0.222499305 | 0.6949600 | 0.027726396 | 0.051972797 |
| Brwd1 | -0.051304504 | 0.128138361 | 0.5912244 | 0.328366001 | 0.384095318 |
| Brwd3 | -0.00713378 | 0.772241853 | 0.9552333 | 0.135886626 | 0.13184654 |
| Bscl2 | 0.005858131 | 0.843476548 | 0.9734889 | 0.221271091 | 0.22453947 |
| Bsdc1 | -0.044585023 | 0.141454546 | 0.6143551 | 0.220572503 | 0.259296807 |
| Bsg | -0.070128998 | 0.000496276 | 0.0305779 | 2.552426656 | 2.527972721 |
| Bsn | -0.008287685 | 0.663212865 | 0.9230058 | 0.073331885 | 0.081498738 |
| Bspry | -0.007010505 | 0.662403647 | 0.9227576 | 0.054183816 | 0.05862997 |
| Bst2 | 0.06750037 | 0.01500328 | 0.2400524 | 0.194259917 | 0.142701656 |
| Btaf1 | -1.61E-05 | 0.99950113 | 0.9996989 | 0.159997592 | 0.147221702 |
| Btbd1 | -0.022528773 | 0.517793088 | 0.8738804 | 0.495475016 | 0.438013558 |
| Btbd10 | -0.010740454 | 0.678969544 | 0.9287614 | 0.163030356 | 0.153628318 |
| Btbd17 | -0.004712838 | 0.857292952 | 0.9763189 | 0.157413332 | 0.168545223 |
| Btbd2 | -0.036603533 | 0.14464707 | 0.6193638 | 0.139051213 | 0.156270792 |
| Btbd3 | 0.025215421 | 0.274366752 | 0.7351955 | 0.137035438 | 0.118295493 |
| Btbd6 | 0.02033227 | 0.334642299 | 0.7804836 | 0.111327748 | 0.091091737 |
| Btbd7 | 0.003055206 | 0.928756276 | 0.9881953 | 0.378396206 | 0.364157079 |
| Btbd8 | -0.016516012 | 0.533883798 | 0.8793171 | 0.132536863 | 0.160455939 |
| Btbd9 | -0.044473103 | 0.1429297 | 0.6161940 | 0.225285079 | 0.261417628 |
| Btd | 0.002763789 | 0.841903898 | 0.9734889 | 0.041955403 | 0.039570594 |
| Btf3 | -0.017012481 | 0.459924666 | 0.8524343 | 2.072869868 | 2.018638156 |
| Btf3l4 | -0.026252516 | 0.3460057 | 0.7894150 | 0.183193711 | 0.192428405 |
| Btf3-ps18 | -0.019670325 | 0.274354932 | 0.7351955 | 0.057964351 | 0.073489381 |
| Btg1 | -0.035990897 | 0.35782578 | 0.7989722 | 0.552057379 | 0.592331728 |
| Btg2 | -0.185587595 | 2.45E-07 | 8.37E-05 | 1.869887932 | 2.28450903 |
| Btg3 | 0.036116771 | 0.340241283 | 0.7844658 | 1.23812304 | 1.05762808 |
| Btrc | 0.024155812 | 0.449864029 | 0.8478268 | 0.326434375 | 0.274416464 |
| Bub3 | -0.037440161 | 0.269650895 | 0.7340360 | 0.432470212 | 0.423097049 |
| Bud13 | 0.027332125 | 0.113091269 | 0.5696203 | 0.084177367 | 0.054733705 |
| Bud23 | -0.018198219 | 0.531764484 | 0.8778671 | 0.232913703 | 0.220245085 |
| Bud31 | 0.006860426 | 0.843210216 | 0.9734889 | 0.519079747 | 0.480924654 |
| Bysl | -0.020419054 | 0.322448683 | 0.7735649 | 0.08718578 | 0.096775065 |
| Bzw1 | -0.056445068 | 0.121602685 | 0.5819916 | 0.528836571 | 0.555201231 |
| Bzw2 | -0.010278819 | 0.705328234 | 0.9401381 | 0.178202893 | 0.180643385 |
| C030006K11Rik | -0.046500707 | 0.0520968 | 0.4313858 | 0.107790238 | 0.13668872 |
| C030014I23Rik | 0.021156746 | 0.25762978 | 0.7250597 | 0.081628866 | 0.072026943 |
| C030034I22Rik | -0.003826447 | 0.87210118 | 0.9784261 | 0.133664825 | 0.122753554 |
| C130036L24Rik | 0.008156271 | 0.640294711 | 0.9165200 | 0.076185607 | 0.065024391 |
| C130051F05Rik | 0.00042748 | 0.979129143 | 0.9976072 | 0.052704045 | 0.051002722 |
| C130089K02Rik | -0.106628166 | 0.00021731 | 0.0172735 | 0.128182418 | 0.209497061 |

| | | | | | |
|---|---|---|---|---|---|
| C1d | 0.055648012 | 0.110591183 | 0.5655888 | 0.54117035 | 0.448357757 |
| C1galt1 | 0.038924808 | 0.2849289 | 0.7450791 | 0.556399924 | 0.521487473 |
| C1galt1c1 | 0.00435735 | 0.884897738 | 0.9803788 | 0.285030589 | 0.230604393 |
| C1qa | -0.01078754 | 0.706023223 | 0.9402269 | 0.210096351 | 0.172726186 |
| C1qbp | -0.029752799 | 0.365789082 | 0.8050191 | 0.3239044 | 0.363720389 |
| C1qtnf12 | 0.000138819 | 0.993439135 | 0.9985888 | 0.05346874 | 0.060413027 |
| C1qtnf4 | 0.038333651 | 0.280712483 | 0.7408618 | 1.181973778 | 1.075414881 |
| C1qtnf6 | -0.029268035 | 0.066890133 | 0.4795761 | 0.040074323 | 0.059878878 |
| C230096K16Rik | 0.001135198 | 0.938816763 | 0.9925463 | 0.047138828 | 0.043518151 |
| C2cd2l | -0.002109387 | 0.928593501 | 0.9881953 | 0.128828071 | 0.130594195 |
| C2cd3 | 0.001159705 | 0.961950994 | 0.9963994 | 0.140426626 | 0.136006737 |
| C2cd4a | 0.00026 | 0.989746237 | 0.9980836 | 0.061625162 | 0.078908563 |
| C2cd4b | -0.073029759 | 0.134852635 | 0.6013051 | 1.059399731 | 1.237347665 |
| C2cd4c | 0.004004357 | 0.771078877 | 0.9552333 | 0.039414246 | 0.037977173 |
| C2cd4d | 0.021790158 | 0.078441943 | 0.5056175 | 0.045008997 | 0.025126661 |
| C2cd5 | -0.060512115 | 0.014874474 | 0.2391487 | 0.10454444 | 0.159565366 |
| C330007P06Rik | 0.019686875 | 0.407775871 | 0.8270770 | 0.151877604 | 0.132908465 |
| C330013E15Rik | -0.036350833 | 0.031385003 | 0.3385228 | 0.039582587 | 0.069024728 |
| C330018D20Rik | -0.020728316 | 0.194417553 | 0.6717191 | 0.041600622 | 0.058681967 |
| C730034F03Rik | -0.026355049 | 0.112831167 | 0.5696203 | 0.048217018 | 0.059952324 |
| C77080 | -0.0136909 | 0.522567698 | 0.8750611 | 0.092597629 | 0.104216427 |
| C87436 | -0.023740803 | 0.373220248 | 0.8085689 | 0.17400179 | 0.179020348 |
| C8b | -0.032864005 | 0.33872675 | 0.7834245 | 0.401224532 | 0.335291831 |
| C920006O11Rik | -0.000519594 | 0.979685004 | 0.9976072 | 0.091575072 | 0.088949508 |
| C920021L13Rik | -0.011325274 | 0.473451778 | 0.8597092 | 0.043933082 | 0.055514528 |
| C9orf72 | 0.007963316 | 0.728334091 | 0.9458355 | 0.126028814 | 0.116638238 |
| Caap1 | -0.012283067 | 0.630097749 | 0.9130564 | 0.151713554 | 0.160742811 |
| Cab39 | -0.006684815 | 0.839063122 | 0.9727216 | 0.323773764 | 0.339607466 |
| Cab39l | 0.009574946 | 0.730079596 | 0.9459020 | 0.20069479 | 0.180183039 |
| Cabcoco1 | -0.00408182 | 0.778907093 | 0.9581793 | 0.046125984 | 0.044827247 |
| Cabin1 | -0.102179994 | 0.000632899 | 0.0356725 | 0.163429311 | 0.255626212 |
| Cables2 | 0.002398971 | 0.890824183 | 0.9819597 | 0.072898803 | 0.065608224 |
| Cabp1 | -0.027299571 | 0.317123213 | 0.7706668 | 0.127411034 | 0.206523238 |
| Cabp7 | 0.002409081 | 0.908690681 | 0.9837373 | 0.091559327 | 0.101944915 |
| Cacfd1 | -0.014699569 | 0.648854572 | 0.9172569 | 0.282575965 | 0.298065684 |
| Cacna1a | -0.018433518 | 0.566234877 | 0.8947196 | 0.255847285 | 0.281541237 |
| Cacna1c | -0.004225859 | 0.866854362 | 0.9772002 | 0.140705357 | 0.151315559 |
| Cacna1d | -0.052020667 | 0.143441516 | 0.6165078 | 0.399179803 | 0.453547282 |
| Cacna2d1 | 0.026368593 | 0.473261796 | 0.8597092 | 0.66175954 | 0.663070372 |
| Cacna2d4 | -0.017992924 | 0.306882881 | 0.7652785 | 0.063932232 | 0.068443634 |
| Cacnb1 | -0.000545446 | 0.969484383 | 0.9976072 | 0.045850062 | 0.040644661 |
| Cacnb2 | -0.030485215 | 0.263415768 | 0.7283846 | 0.148594991 | 0.184895753 |
| Cacnb3 | -0.062732617 | 0.035216923 | 0.3577036 | 0.220258951 | 0.251803961 |
| Cacng4 | -0.003498389 | 0.84314404 | 0.9734889 | 0.071632247 | 0.059255426 |
| Cactin | 0.019542518 | 0.310524046 | 0.7677462 | 0.100858315 | 0.075210279 |
| Cacul1 | -0.008301398 | 0.742454381 | 0.9479080 | 0.163776302 | 0.156090639 |
| Cacybp | -0.001219966 | 0.972171037 | 0.9976072 | 0.958081835 | 0.899811656 |
| Cad | 0.020667143 | 0.327640053 | 0.7778857 | 0.111546481 | 0.095874778 |
| Cadm1 | 0.060801338 | 0.047177936 | 0.4106761 | 0.295938644 | 0.195246527 |
| Cadps | -0.053446243 | 0.150404896 | 0.6280320 | 0.442244858 | 0.46924829 |
| Cadps2 | -0.016224628 | 0.358842702 | 0.7999369 | 0.052312997 | 0.070465814 |
| Calb1 | 0.001819852 | 0.965095199 | 0.9966421 | 0.6091655 | 0.471728851 |
| Calca | -0.02465495 | 0.211404533 | 0.6822059 | 0.074247331 | 0.086374602 |
| Calcoco1 | 0.001615964 | 0.942864571 | 0.9934906 | 0.124162588 | 0.107583321 |
| Cald1 | -0.102262777 | 0.002714425 | 0.0964810 | 0.306339871 | 0.336151646 |
| Calm1 | 0.010325811 | 0.482450274 | 0.8620306 | 3.19512331 | 3.150528932 |
| Calm2 | -0.027676788 | 0.280175041 | 0.7408618 | 2.040898358 | 1.950832894 |
| Calm3 | -0.053793478 | 0.109480215 | 0.5655888 | 1.224054292 | 1.23961401 |
| Calr | 0.057474414 | 0.07007517 | 0.4850436 | 2.060006477 | 1.874510236 |
| Calu | -0.002906851 | 0.904482928 | 0.9831707 | 0.13080229 | 0.132827986 |
| Caly | -0.024960924 | 0.182863009 | 0.6607594 | 0.056533655 | 0.076391092 |
| Camk1 | -0.05529924 | 0.054649092 | 0.4421851 | 0.19540265 | 0.234148593 |
| Camk2b | -0.089950903 | 0.014924102 | 0.2393294 | 0.617508152 | 0.6779948 |
| Camk2d | -0.050409362 | 0.131494684 | 0.5972652 | 0.312152374 | 0.333402722 |
| Camk2g | -0.012907552 | 0.549586522 | 0.8874817 | 0.101982626 | 0.106311836 |
| Camk2n1 | 0.043006798 | 0.098416547 | 0.5417825 | 2.208390638 | 2.195100412 |

| | | | | | |
|---|---|---|---|---|---|
| Camk2n2 | -0.045218647 | 0.223621515 | 0.6956178 | 0.943822649 | 0.941042136 |
| Camkk2 | -0.00703962 | 0.712408526 | 0.9421288 | 0.070754463 | 0.076590884 |
| Camkmt | 0.01249251 | 0.564571071 | 0.8935138 | 0.116414419 | 0.100486117 |
| Caml | 0.004981603 | 0.871172238 | 0.9782269 | 0.268597234 | 0.264453155 |
| Camsap1 | 0.000392318 | 0.989215832 | 0.9980836 | 0.206849742 | 0.215098586 |
| Camsap2 | -0.013954123 | 0.578031769 | 0.8997914 | 0.139709886 | 0.148851031 |
| Camsap3 | -0.022403202 | 0.132632414 | 0.5988251 | 0.034247279 | 0.053340551 |
| Camta1 | -0.035231821 | 0.284030691 | 0.7434491 | 0.310844251 | 0.35191511 |
| Camta2 | 0.027400474 | 0.257627774 | 0.7250597 | 0.157979885 | 0.131533356 |
| Cand1 | -0.057569588 | 0.092024834 | 0.5325584 | 0.333475594 | 0.366968657 |
| Cand2 | -0.016119097 | 0.369947105 | 0.8065581 | 0.069366212 | 0.069932463 |
| Cant1 | 0.014643551 | 0.565375061 | 0.8943433 | 0.155939753 | 0.156007795 |
| Canx | -0.024070884 | 0.449982698 | 0.8478268 | 1.382180116 | 1.394141884 |
| Cap1 | 0.073085185 | 0.018856967 | 0.2677986 | 0.342248847 | 0.25132839 |
| Capn1 | 0.011203197 | 0.559418254 | 0.8923733 | 0.087353569 | 0.078059514 |
| Capn10 | -0.012301481 | 0.520899364 | 0.8741795 | 0.083379368 | 0.084648697 |
| Capn15 | -0.029153942 | 0.268293052 | 0.7323870 | 0.151572786 | 0.17122635 |
| Capn2 | -0.06306601 | 0.022551799 | 0.2914468 | 0.181929593 | 0.204008205 |
| Capn5 | -0.002524467 | 0.900969161 | 0.9825441 | 0.084117499 | 0.090607512 |
| Capn7 | 0.009624462 | 0.730934722 | 0.9459020 | 0.204674015 | 0.196336279 |
| Capns1 | -0.008889112 | 0.796314837 | 0.9642793 | 1.093961465 | 1.022714562 |
| Caprin1 | 0.049091637 | 0.197639793 | 0.6730962 | 0.670140917 | 0.615882271 |
| Capsl | 0.022124162 | 0.489777834 | 0.8652887 | 0.307726622 | 0.275632 |
| Capza1 | -0.020881232 | 0.212844046 | 0.6827572 | 0.048149359 | 0.061263649 |
| Capza2 | 0.00418258 | 0.91028982 | 0.9837373 | 0.698413038 | 0.66139947 |
| Capzb | -0.016611143 | 0.642154951 | 0.9166712 | 0.715653124 | 0.674020323 |
| Car10 | 0.041623565 | 0.072792829 | 0.4903372 | 0.120728938 | 0.121738777 |
| Car8 | -0.043764167 | 0.143066017 | 0.6162812 | 0.154494361 | 0.21074889 |
| Card19 | -0.053706198 | 0.124295157 | 0.5852543 | 0.37309186 | 0.404878517 |
| Card6 | -0.00401828 | 0.800425599 | 0.9647053 | 0.051116944 | 0.051057978 |
| Carf | -0.018080036 | 0.374649818 | 0.8091718 | 0.08166609 | 0.096053981 |
| Carhsp1 | 0.012972556 | 0.593587758 | 0.9061761 | 0.150605421 | 0.132379498 |
| Carm1 | -0.008905462 | 0.692492293 | 0.9358575 | 0.11552695 | 0.11455493 |
| Carmil1 | -0.048384082 | 0.029612506 | 0.3296925 | 0.082880015 | 0.118588893 |
| Carmil2 | -0.029289319 | 0.106445843 | 0.5621561 | 0.051078963 | 0.074621332 |
| Carmil3 | 0.013382068 | 0.401858453 | 0.8254259 | 0.053097719 | 0.052194802 |
| Carnmt1 | 0.00022833 | 0.993840451 | 0.9985888 | 0.261639038 | 0.23467785 |
| Cars | 0.086756923 | 0.015067755 | 0.2406958 | 0.470260038 | 0.388823012 |
| Cars2 | 0.001255266 | 0.955429899 | 0.9954690 | 0.119492598 | 0.111952983 |
| Casc3 | -0.03799293 | 0.211716797 | 0.6822059 | 0.218152143 | 0.271788785 |
| Casd1 | -0.041853163 | 0.115523034 | 0.5724670 | 0.155173384 | 0.183696263 |
| Cask | -0.004230738 | 0.887376932 | 0.9812414 | 0.249487786 | 0.242367263 |
| Casp2 | 0.02958165 | 0.055929556 | 0.4449332 | 0.070143904 | 0.04334677 |
| Casp3 | 0.018062682 | 0.343648143 | 0.7872955 | 0.084899297 | 0.074439721 |
| Casp6 | 0.001227087 | 0.947786075 | 0.9945018 | 0.079887385 | 0.070997555 |
| Casp8 | -0.022349817 | 0.178088951 | 0.6538489 | 0.047344577 | 0.059739808 |
| Casp8ap2 | 0.002326543 | 0.936554303 | 0.9912468 | 0.208329167 | 0.201918667 |
| Casr | -0.017558486 | 0.227859576 | 0.7015153 | 0.035105295 | 0.04896315 |
| Cast | 0.003847283 | 0.875879184 | 0.9785583 | 0.136020745 | 0.146092733 |
| Castor2 | -0.103039535 | 0.010537361 | 0.2017406 | 1.047170938 | 1.07208631 |
| Casz1 | -0.084018769 | 0.006456648 | 0.1603078 | 0.193062729 | 0.284781905 |
| Cbarp | 0.020390748 | 0.551882554 | 0.8893234 | 0.378610881 | 0.357703857 |
| Cbfa2t2 | -0.04351772 | 0.074138839 | 0.4942589 | 0.116159087 | 0.136773343 |
| Cbfa2t3 | -0.061403705 | 0.076784483 | 0.5015054 | 0.353533533 | 0.418086326 |
| Cbfb | -0.033564318 | 0.174211292 | 0.6527657 | 0.133432222 | 0.148416814 |
| Cbl | -0.076473144 | 0.011418295 | 0.2105633 | 0.19062312 | 0.2574875 |
| Cblb | -0.030145387 | 0.148741096 | 0.6264374 | 0.07304949 | 0.098735359 |
| Cblc | -0.01263619 | 0.598793877 | 0.9075513 | 0.133540671 | 0.134171108 |
| Cbll1 | -0.005567579 | 0.848593366 | 0.9747621 | 0.217928879 | 0.22344453 |
| Cbr1 | 0.017504261 | 0.598515361 | 0.9075513 | 0.362011829 | 0.333367396 |
| Cbr4 | -0.00038215 | 0.985223508 | 0.9976072 | 0.103187379 | 0.092286254 |
| Cbs | -0.057765608 | 0.030133847 | 0.3311437 | 0.130237696 | 0.184265691 |
| Cbwd1 | 0.010774992 | 0.683808965 | 0.9311703 | 0.194671542 | 0.160523933 |
| Cbx1 | 0.049147167 | 0.15698339 | 0.6320110 | 0.658331266 | 0.592333036 |
| Cbx3 | 0.054065355 | 0.135130156 | 0.6021972 | 0.720318521 | 0.572211418 |
| Cbx4 | 0.01572905 | 0.6032971 | 0.9087139 | 0.226854238 | 0.229816023 |

| | | | | | |
|---|---|---|---|---|---|
| Cbx5 | 0.006681752 | 0.854492393 | 0.9755431 | 0.570882523 | 0.52056303 |
| Cbx6 | 0.04575845 | 0.184238669 | 0.6628862 | 0.455634093 | 0.395013382 |
| Cbx7 | -0.031240004 | 0.215923974 | 0.6858680 | 0.136854582 | 0.160138936 |
| Cbx8 | -0.012218813 | 0.499037914 | 0.8690742 | 0.066265089 | 0.070860109 |
| Cby1 | 0.007852944 | 0.785825113 | 0.9602516 | 0.233093801 | 0.205779989 |
| Cc2d1a | -0.022709841 | 0.243366672 | 0.7132719 | 0.07610318 | 0.09096514 |
| Cc2d1b | -0.000687976 | 0.974964949 | 0.9976072 | 0.099500435 | 0.106327496 |
| Cc2d2a | 0.021972667 | 0.362962709 | 0.8038825 | 0.146083743 | 0.125364053 |
| Ccar1 | 0.01123459 | 0.75697887 | 0.9521533 | 0.461215241 | 0.449768985 |
| Ccar2 | 0.034708575 | 0.093338236 | 0.5330587 | 0.135655913 | 0.086850838 |
| Ccdc107 | 0.001982884 | 0.956504165 | 0.9955934 | 0.457565105 | 0.444421112 |
| Ccdc112 | -0.036597803 | 0.136729593 | 0.6052465 | 0.114677373 | 0.151679963 |
| Ccdc113 | -0.014285407 | 0.450767595 | 0.8478942 | 0.068622541 | 0.081995915 |
| Ccdc115 | -0.033343675 | 0.259902653 | 0.7263475 | 0.228748681 | 0.242505553 |
| Ccdc117 | 0.039907933 | 0.064037512 | 0.4700387 | 0.116302247 | 0.093063603 |
| Ccdc12 | -0.024087607 | 0.490975247 | 0.8658154 | 0.688042846 | 0.684286103 |
| Ccdc122 | -0.034382606 | 0.008099659 | 0.1773700 | 0.010218652 | 0.046082745 |
| Ccdc124 | 0.079034602 | 0.024852843 | 0.3041540 | 0.638241458 | 0.520287307 |
| Ccdc126 | 0.028504485 | 0.088996113 | 0.5292814 | 0.075677015 | 0.05084606 |
| Ccdc127 | -0.043327463 | 0.196295266 | 0.6730962 | 0.349883855 | 0.393034916 |
| Ccdc130 | -0.003853852 | 0.86876821 | 0.9776398 | 0.122821716 | 0.119586561 |
| Ccdc134 | -0.018532426 | 0.313256436 | 0.7697557 | 0.060460379 | 0.071769525 |
| Ccdc136 | -0.025868062 | 0.203500587 | 0.6772740 | 0.076814182 | 0.092561068 |
| Ccdc137 | 0.003772313 | 0.846486972 | 0.9739657 | 0.087273647 | 0.081776528 |
| Ccdc148 | 0.005628751 | 0.7356752 | 0.9468380 | 0.059531759 | 0.055489417 |
| Ccdc149 | -0.019944351 | 0.229878125 | 0.7019677 | 0.050245107 | 0.064552839 |
| Ccdc157 | -0.0052124 | 0.746266753 | 0.9486117 | 0.049188766 | 0.055286872 |
| Ccdc160 | -0.016116576 | 0.281491793 | 0.7414759 | 0.036230109 | 0.051281246 |
| Ccdc162 | 0.01536309 | 0.427539208 | 0.8340365 | 0.07235237 | 0.08149145 |
| Ccdc163 | -0.012639603 | 0.478261488 | 0.8607489 | 0.0590016 | 0.067416043 |
| Ccdc166 | 0.046891168 | 0.021938001 | 0.2890105 | 0.12326331 | 0.078633534 |
| Ccdc167 | -0.031126682 | 0.277663352 | 0.7382158 | 0.174546538 | 0.208302083 |
| Ccdc173 | 0.002180761 | 0.898933656 | 0.9825441 | 0.060028066 | 0.061703933 |
| Ccdc174 | -0.029292147 | 0.376962333 | 0.8101797 | 0.355309209 | 0.366551968 |
| Ccdc181 | 0.029924781 | 0.213522962 | 0.6830745 | 0.164597283 | 0.126218031 |
| Ccdc184 | 0.016570504 | 0.531753408 | 0.8778671 | 0.179789903 | 0.157144796 |
| Ccdc186 | -0.032602975 | 0.362752642 | 0.8035967 | 0.452314683 | 0.45421945 |
| Ccdc189 | -0.023129822 | 0.152253126 | 0.6281105 | 0.039335077 | 0.061236941 |
| Ccdc191 | 0.007027084 | 0.799533007 | 0.9647053 | 0.217907707 | 0.188902238 |
| Ccdc22 | 0.010227124 | 0.418992978 | 0.8300990 | 0.040607364 | 0.029823035 |
| Ccdc25 | -0.003098231 | 0.927894948 | 0.9880547 | 0.340496564 | 0.355542399 |
| Ccdc28a | -0.024181681 | 0.225515369 | 0.6979011 | 0.078084636 | 0.094773486 |
| Ccdc28b | -0.025827593 | 0.350735254 | 0.7927303 | 0.184720412 | 0.200330449 |
| Ccdc30 | -0.019707422 | 0.354087828 | 0.7967782 | 0.088357312 | 0.108044411 |
| Ccdc32 | -0.032257248 | 0.186318072 | 0.6644983 | 0.132127878 | 0.151359849 |
| Ccdc34 | 0.027845934 | 0.379020055 | 0.8104934 | 0.283155551 | 0.264679544 |
| Ccdc43 | 0.023440135 | 0.326933098 | 0.7778857 | 0.151132098 | 0.136948945 |
| Ccdc47 | 0.016948142 | 0.657292422 | 0.9211780 | 0.628243399 | 0.652897261 |
| Ccdc50 | 0.055483788 | 0.062359748 | 0.4637246 | 0.280111211 | 0.222774457 |
| Ccdc58 | 0.002483538 | 0.925864688 | 0.9879210 | 0.200803421 | 0.17453931 |
| Ccdc59 | 0.02265128 | 0.534649079 | 0.8795949 | 0.797292767 | 0.774050294 |
| Ccdc6 | 0.033284022 | 0.20771748 | 0.6807334 | 0.17800152 | 0.165214155 |
| Ccdc66 | -0.003664689 | 0.872633521 | 0.9784331 | 0.116830111 | 0.114475008 |
| Ccdc71 | 0.00433444 | 0.828742486 | 0.9722601 | 0.102119363 | 0.087974572 |
| Ccdc71l | -0.048948021 | 0.004102043 | 0.1266085 | 0.036516113 | 0.06945873 |
| Ccdc77 | -0.00645456 | 0.719031845 | 0.9441159 | 0.069025137 | 0.071118843 |
| Ccdc8 | 0.023337988 | 0.168135027 | 0.6452222 | 0.078567332 | 0.053387717 |
| Ccdc82 | 0.009043204 | 0.698636196 | 0.9369692 | 0.13278737 | 0.126251379 |
| Ccdc84 | -0.036604884 | 0.072573319 | 0.4902939 | 0.078406374 | 0.093797963 |
| Ccdc85b | 0.008268111 | 0.805905998 | 0.9647143 | 0.396490317 | 0.380565611 |
| Ccdc86 | -0.022497384 | 0.496246915 | 0.8684629 | 0.325592814 | 0.300087447 |
| Ccdc88a | -0.070676311 | 0.068826628 | 0.4835495 | 0.568667124 | 0.630642426 |
| Ccdc88c | 0.022388974 | 0.113717711 | 0.5696977 | 0.054304053 | 0.033095381 |
| Ccdc9 | 0.000958527 | 0.969582477 | 0.9976072 | 0.154141012 | 0.137068306 |
| Ccdc90b | 0.028380492 | 0.282160709 | 0.7418590 | 0.206226649 | 0.158891634 |
| Ccdc91 | -0.005789383 | 0.827717484 | 0.9717109 | 0.203178709 | 0.177473446 |

| | | | | | |
|---|---|---|---|---|---|
| Ccdc92 | 0.047776669 | 0.094818514 | 0.5350704 | 0.240749394 | 0.188526431 |
| Ccdc92b | -0.022717651 | 0.192483672 | 0.6700281 | 0.057177944 | 0.070440492 |
| Ccdc93 | -0.009934287 | 0.542732592 | 0.8840682 | 0.048961771 | 0.057619068 |
| Ccdc97 | -0.013671134 | 0.555991007 | 0.8911667 | 0.107728145 | 0.124538216 |
| Ccl25 | -0.042323533 | 0.03523663 | 0.3577036 | 0.064503506 | 0.093778274 |
| Ccl27a | 0.043716945 | 0.151544218 | 0.6281105 | 0.280358149 | 0.241419263 |
| Ccm2 | 0.034329575 | 0.177815032 | 0.6538187 | 0.180795279 | 0.152224697 |
| Ccnc | 0.001486624 | 0.954155952 | 0.9954690 | 0.175825741 | 0.163836007 |
| Ccnd1 | -0.017422694 | 0.666581716 | 0.9230765 | 0.811326191 | 0.843863796 |
| Ccnd2 | 0.083094002 | 0.000962919 | 0.0471160 | 2.642260113 | 2.433485055 |
| Ccnd3 | 0.0162615 | 0.599493849 | 0.9078234 | 0.282686019 | 0.259140024 |
| Ccndbp1 | -0.080642358 | 0.00433693 | 0.1311656 | 0.166202909 | 0.227250413 |
| Ccng1 | 0.032976866 | 0.359116882 | 0.8000088 | 0.451214161 | 0.391390988 |
| Ccng2 | -0.004532944 | 0.781611218 | 0.9590847 | 0.053416406 | 0.056943311 |
| Ccnh | 0.013268617 | 0.672015414 | 0.9255022 | 0.308754344 | 0.283024545 |
| Ccni | -0.005266182 | 0.885278471 | 0.9804580 | 0.660289101 | 0.642026457 |
| Ccnk | 0.006514669 | 0.792737906 | 0.9624460 | 0.147707307 | 0.139376124 |
| Ccnl1 | -0.044175645 | 0.253294582 | 0.7223071 | 0.581759894 | 0.634475202 |
| Ccnl2 | -0.030913624 | 0.397740752 | 0.8238469 | 0.668445985 | 0.623951099 |
| Ccnq | 0.004415341 | 0.85324106 | 0.9755431 | 0.13422638 | 0.133485829 |
| Ccnt1 | 0.04643395 | 0.18887662 | 0.6674941 | 0.439383196 | 0.404786635 |
| Ccnt2 | -0.011676419 | 0.724507548 | 0.9445652 | 0.331969389 | 0.340469806 |
| Ccny | -0.025700907 | 0.441011575 | 0.8424484 | 0.345387615 | 0.369003044 |
| Ccp110 | -0.036280577 | 0.059372786 | 0.4548580 | 0.057973442 | 0.085855723 |
| Ccpg1 | -0.030816558 | 0.328078507 | 0.7778857 | 0.271208647 | 0.294417426 |
| Ccpg1os | -0.007121792 | 0.798962308 | 0.9646672 | 0.212230222 | 0.202776571 |
| Ccs | 0.004790009 | 0.871207567 | 0.9782269 | 0.247342548 | 0.234298825 |
| Ccser1 | 0.011508136 | 0.564303615 | 0.8933738 | 0.093621195 | 0.083558877 |
| Ccser2 | -0.015844566 | 0.520185498 | 0.8741795 | 0.132834664 | 0.145289481 |
| Cct2 | -0.062192296 | 0.07239486 | 0.4902095 | 0.895692671 | 0.85942949 |
| Cct3 | -0.097942878 | 0.00542202 | 0.1488982 | 0.635713845 | 0.654778386 |
| Cct4 | -0.025365003 | 0.479241838 | 0.8610807 | 0.704556043 | 0.677327596 |
| Cct5 | -0.034963216 | 0.306854844 | 0.7652785 | 0.878223826 | 0.821367853 |
| Cct6a | -0.011711283 | 0.748951907 | 0.9501120 | 0.864830928 | 0.782833323 |
| Cct7 | -0.034357544 | 0.293171605 | 0.7533684 | 1.153984065 | 1.116886204 |
| Cct8 | -0.017178827 | 0.623177298 | 0.9121505 | 0.826781125 | 0.746787834 |
| Ccz1 | -0.03718522 | 0.219611794 | 0.6910162 | 0.252824771 | 0.262736118 |
| Cd151 | 0.072484416 | 0.010537158 | 0.2017406 | 0.262673239 | 0.168451008 |
| Cd164 | 0.019758639 | 0.45749599 | 0.8518352 | 1.789286377 | 1.902653935 |
| Cd164l2 | 0.012559003 | 0.510738048 | 0.8716066 | 0.079911452 | 0.071831385 |
| Cd200 | 0.087432898 | 0.016953799 | 0.2555884 | 0.578651495 | 0.46191616 |
| Cd24a | 0.007136348 | 0.84334453 | 0.9734889 | 0.188757267 | 0.232195556 |
| Cd274 | -0.023989893 | 0.210420751 | 0.6817825 | 0.061001859 | 0.089170306 |
| Cd276 | 0.016415451 | 0.322147475 | 0.7733024 | 0.063256605 | 0.054150534 |
| Cd2ap | 0.031302622 | 0.40699727 | 0.8270770 | 0.725353004 | 0.669852949 |
| Cd2bp2 | -0.016423775 | 0.532442833 | 0.8784022 | 0.161990756 | 0.167813396 |
| Cd320 | -0.038049671 | 0.120940071 | 0.5811445 | 0.151387963 | 0.159422256 |
| Cd3eap | -0.047945642 | 0.082223815 | 0.5148480 | 0.161922544 | 0.192957644 |
| Cd44 | 0.048383663 | 0.010446197 | 0.2017406 | 0.090467225 | 0.06939621 |
| Cd47 | 0.0206439 | 0.537060976 | 0.8807896 | 1.089911845 | 1.0806615 |
| Cd59a | -0.051059829 | 0.143325025 | 0.6165078 | 0.323610699 | 0.338831337 |
| Cd59b | -0.023990755 | 0.113690317 | 0.5696977 | 0.033340555 | 0.053014105 |
| Cd63 | 0.022978095 | 0.282747742 | 0.7424186 | 2.506102855 | 2.469726795 |
| Cd81 | -0.061808443 | 0.092042918 | 0.5325584 | 1.375080565 | 1.24107746 |
| Cd82 | -0.017445677 | 0.593308407 | 0.9060585 | 0.289692959 | 0.315299286 |
| Cd9 | 0.048412634 | 0.083154284 | 0.5178207 | 0.207075723 | 0.137962007 |
| Cd99l2 | -0.020198923 | 0.374344927 | 0.8087297 | 0.103904098 | 0.118293988 |
| Cda | 0.036513306 | 0.247841468 | 0.7177068 | 0.304080071 | 0.283348345 |
| Cdadc1 | -0.019985138 | 0.518827189 | 0.8741795 | 0.238601586 | 0.261650899 |
| Cdc123 | 0.017792821 | 0.598866237 | 0.9075513 | 0.454640179 | 0.388879416 |
| Cdc14b | -0.041974265 | 0.115706846 | 0.5724670 | 0.140691475 | 0.181522264 |
| Cdc16 | 0.076716011 | 0.013316142 | 0.2285400 | 0.344825138 | 0.258187379 |
| Cdc23 | 0.041921685 | 0.110228047 | 0.5655888 | 0.201081546 | 0.146591136 |
| Cdc25a | 0.010038428 | 0.711826795 | 0.9419834 | 0.185693701 | 0.17050602 |
| Cdc26 | 0.013374456 | 0.703215844 | 0.9391887 | 0.645746619 | 0.556349649 |
| Cdc27 | -0.001171003 | 0.961901163 | 0.9963994 | 0.152894424 | 0.140108764 |

| | | | | | |
|---|---|---|---|---|---|
| Cdc34 | 0.024654556 | 0.481226629 | 0.86201507 | 0.403504245 | 0.381774905 |
| Cdc37 | -0.091598464 | 0.011867391 | 0.21457162 | 0.575203201 | 0.624855386 |
| Cdc37l1 | -0.000849788 | 0.980893857 | 0.99760728 | 0.442005597 | 0.46778621 |
| Cdc40 | -0.046176324 | 0.139618399 | 0.61218711 | 0.252951453 | 0.298070129 |
| Cdc42 | -0.018896197 | 0.563818958 | 0.89318055 | 1.186278844 | 1.055910091 |
| Cdc42bpa | -0.022450913 | 0.52714716 | 0.87685539 | 0.367146156 | 0.383070067 |
| Cdc42bpb | 0.011072322 | 0.684293573 | 0.93117031 | 0.196103252 | 0.182497024 |
| Cdc42bpg | -0.019223945 | 0.290905062 | 0.75130908 | 0.061610563 | 0.078194885 |
| Cdc42ep3 | -0.062309293 | 0.031268013 | 0.33825375 | 0.186222656 | 0.191227545 |
| Cdc42ep4 | -1.14E-05 | 0.999418199 | 0.99969890 | 0.058642835 | 0.049998424 |
| Cdc42ep5 | -0.030942389 | 0.057221901 | 0.44914073 | 0.041589263 | 0.063532257 |
| Cdc42se1 | -0.040491135 | 0.162626835 | 0.64007035 | 0.208694017 | 0.234289691 |
| Cdc42se2 | -0.032301466 | 0.323816558 | 0.77553362 | 0.290396296 | 0.335692781 |
| Cdc5l | 0.070156542 | 0.04741776 | 0.41067610 | 0.64552759 | 0.519298322 |
| Cdc73 | 0.078158165 | 0.020145793 | 0.27527033 | 0.434406326 | 0.356691742 |
| Cdca4 | 0.009066305 | 0.526903141 | 0.87685539 | 0.047526052 | 0.03912206 |
| Cdcp1 | 0.010151768 | 0.689496025 | 0.93401619 | 0.160659926 | 0.149789452 |
| Cdh1 | 0.120866664 | 0.000440456 | 0.02837224 | 0.485016069 | 0.407832129 |
| Cdh2 | -0.020779411 | 0.569930801 | 0.89625342 | 0.494620892 | 0.516067506 |
| Cdh22 | -0.000897152 | 0.950163403 | 0.99484627 | 0.038465846 | 0.045509712 |
| Cdh4 | -0.014494962 | 0.317931083 | 0.77100438 | 0.027491037 | 0.048997021 |
| Cdh8 | -6.45E-05 | 0.997634485 | 0.99940731 | 0.103521961 | 0.111989552 |
| Cdhr1 | 0.021844489 | 0.219253142 | 0.69069265 | 0.065985469 | 0.065822364 |
| Cdin1 | 0.008107869 | 0.6052827 | 0.90871392 | 0.063325894 | 0.047972666 |
| Cdip1 | 0.042616756 | 0.170531123 | 0.65070459 | 0.287003385 | 0.233360932 |
| Cdipt | -0.001363791 | 0.967211061 | 0.99760728 | 0.342312296 | 0.35608841 |
| Cdk10 | -0.024607305 | 0.238537967 | 0.71097411 | 0.097447774 | 0.102496219 |
| Cdk11b | 0.089404719 | 0.016747715 | 0.25467130 | 0.594102583 | 0.507011492 |
| Cdk12 | -0.058717266 | 0.106702737 | 0.56232438 | 0.477596721 | 0.510313166 |
| Cdk13 | -0.030589303 | 0.377296683 | 0.81017979 | 0.372141419 | 0.379135308 |
| Cdk14 | -0.046460577 | 0.084808699 | 0.52022650 | 0.143343674 | 0.189334853 |
| Cdk16 | -0.024558207 | 0.440374636 | 0.84178961 | 0.261917896 | 0.282790536 |
| Cdk17 | 0.027349176 | 0.190017911 | 0.66994891 | 0.115784775 | 0.090917505 |
| Cdk18 | 0.014427539 | 0.297351767 | 0.75509628 | 0.048018392 | 0.036317754 |
| Cdk2ap1 | 0.025697576 | 0.392522852 | 0.82044388 | 0.333690596 | 0.250068389 |
| Cdk2ap2 | -0.123741526 | 0.001555136 | 0.06793680 | 0.648916219 | 0.706147302 |
| Cdk4 | 0.03558108 | 0.320725329 | 0.77218975 | 0.629761322 | 0.545891147 |
| Cdk5 | -0.002498506 | 0.92620526 | 0.98792100 | 0.178777532 | 0.173481165 |
| Cdk5r2 | -0.005868109 | 0.825589244 | 0.97169062 | 0.178148106 | 0.168554237 |
| Cdk5rap2 | -0.010903826 | 0.562677125 | 0.89262402 | 0.064972062 | 0.078958621 |
| Cdk5rap3 | -0.094825637 | 0.007937439 | 0.17584088 | 0.53390666 | 0.583968402 |
| Cdk7 | 0.017469692 | 0.526499009 | 0.87685539 | 0.223734308 | 0.184941935 |
| Cdk8 | -0.002697675 | 0.939822108 | 0.99282169 | 0.440046214 | 0.417461333 |
| Cdk9 | -0.025662416 | 0.413785356 | 0.83009907 | 0.293399018 | 0.286417838 |
| Cdkal1 | 0.002292468 | 0.909796724 | 0.98373739 | 0.100342686 | 0.087898972 |
| Cdkl1 | 0.030982911 | 0.138367518 | 0.60798394 | 0.111681488 | 0.088340606 |
| Cdkl2 | -0.03297776 | 0.198265304 | 0.67309622 | 0.132360848 | 0.168115324 |
| Cdkl3 | 0.034293832 | 0.024537885 | 0.30395880 | 0.074127417 | 0.038991232 |
| Cdkl5 | 0.020256647 | 0.289858536 | 0.74977699 | 0.090547603 | 0.073093965 |
| Cdkn1a | 0.116061144 | 0.018217656 | 0.26267318 | 0.9366917 | 0.891760284 |
| Cdkn1b | -0.048373489 | 0.191318357 | 0.67002811 | 0.539947882 | 0.529391284 |
| Cdkn2aip | -0.024943208 | 0.41787396 | 0.83009907 | 0.253315547 | 0.248398796 |
| Cdkn2aipnl | -0.005301767 | 0.821677059 | 0.97069202 | 0.131703101 | 0.127591661 |
| Cdkn2b | 0.011273974 | 0.633686742 | 0.9144743 | 0.133423401 | 0.120285858 |
| Cdkn2c | -0.047836282 | 0.035979112 | 0.35996902 | 0.098383121 | 0.123157345 |
| Cdkn2d | -0.010401736 | 0.701589414 | 0.93818771 | 0.196908074 | 0.185056093 |
| Cdo1 | -0.007862034 | 0.720614779 | 0.94452957 | 0.138048332 | 0.104692113 |
| Cdon | -0.003696006 | 0.813842461 | 0.96779150 | 0.046031493 | 0.05434194 |
| Cdpf1 | -0.019284749 | 0.477209404 | 0.86074894 | 0.183709344 | 0.186130496 |
| Cdr2 | -0.027925099 | 0.246250789 | 0.71608931 | 0.125678911 | 0.144934228 |
| Cdr2l | -0.000620798 | 0.968311214 | 0.99760728 | 0.051852308 | 0.047895659 |
| Cds1 | 0.010570025 | 0.631400137 | 0.91335177 | 0.104796086 | 0.1053316 |
| Cds2 | 0.005695411 | 0.864657416 | 0.97696750 | 0.36592766 | 0.358577802 |
| Cdv3 | 0.036419702 | 0.32276337 | 0.77375848 | 0.565546503 | 0.521058671 |
| Cdyl | -0.009127644 | 0.693865164 | 0.93597259 | 0.113516681 | 0.12399813 |
| Cdyl2 | -0.001952216 | 0.899398061 | 0.98254414 | 0.055703219 | 0.049279785 |

| | | | | | |
|---|---|---|---|---|---|
| Cebpb | -0.01641344 | 0.668672204 | 0.9230765 | 1.115274641 | 1.199198132 |
| Cebpd | -0.069386248 | 0.106739598 | 0.5623243 | 0.442552394 | 0.574717127 |
| Cebpg | 0.110694907 | 0.003640931 | 0.1157629 | 0.696960527 | 0.606639307 |
| Cebpz | 0.038724307 | 0.292851396 | 0.7531983 | 0.560337179 | 0.524898736 |
| Cebpzos | 0.040580141 | 0.24569192 | 0.7152900 | 0.557438334 | 0.498811228 |
| Celf1 | 0.036183605 | 0.323981956 | 0.7755552 | 0.616747976 | 0.590258025 |
| Celf2 | 0.029195289 | 0.31686213 | 0.7705987 | 0.229823974 | 0.204579024 |
| Celf3 | -0.048803653 | 0.178617611 | 0.6545573 | 0.500431863 | 0.551508874 |
| Celf4 | -0.060330586 | 0.106084006 | 0.5617171 | 0.521428207 | 0.593469308 |
| Celf6 | -0.015583142 | 0.57063877 | 0.8962534 | 0.183501864 | 0.190608752 |
| Celsr2 | 0.032930926 | 0.091232334 | 0.5321282 | 0.093634592 | 0.07711926 |
| Celsr3 | 0.020748451 | 0.343639177 | 0.7872955 | 0.118597931 | 0.104643815 |
| Cemip2 | -0.008805976 | 0.54222109 | 0.8836953 | 0.029287359 | 0.048409104 |
| Cenpb | -0.033988133 | 0.337992316 | 0.7834245 | 0.447151017 | 0.457602095 |
| Cenpc1 | 0.029715264 | 0.241381326 | 0.7128617 | 0.170798836 | 0.142783778 |
| Cenpj | 0.001449015 | 0.93511582 | 0.9908021 | 0.070524082 | 0.067388276 |
| Cenpq | -0.000745624 | 0.969810784 | 0.9976072 | 0.087784179 | 0.084350283 |
| Cenps | -0.001357922 | 0.956035816 | 0.9954690 | 0.148834948 | 0.140193154 |
| Cenpv | -0.001851699 | 0.958803774 | 0.9957785 | 0.734082783 | 0.678604719 |
| Cenpw | 0.019434358 | 0.29404549 | 0.7533922 | 0.088249704 | 0.065727106 |
| Cenpx | -0.011717354 | 0.73485556 | 0.9468380 | 0.575932401 | 0.541170687 |
| Cep104 | 0.009648865 | 0.712185105 | 0.9419834 | 0.165686431 | 0.158027842 |
| Cep112 | -0.010493406 | 0.522256313 | 0.8750611 | 0.051143982 | 0.055092896 |
| Cep120 | 0.006015189 | 0.798407575 | 0.9644698 | 0.129522232 | 0.125398392 |
| Cep126 | -0.004689767 | 0.856389442 | 0.9763189 | 0.150548212 | 0.155750166 |
| Cep131 | 0.027365509 | 0.058711738 | 0.4520948 | 0.062448422 | 0.036803161 |
| Cep162 | -0.033118966 | 0.161101472 | 0.6384844 | 0.096972209 | 0.132486906 |
| Cep164 | 0.020531201 | 0.479936961 | 0.8612472 | 0.230681825 | 0.217384229 |
| Cep170 | 0.025546741 | 0.417020293 | 0.8300990 | 0.279057346 | 0.251036589 |
| Cep170b | -0.04356834 | 0.174072013 | 0.6527657 | 0.267745689 | 0.302849906 |
| Cep19 | 0.016225506 | 0.506161365 | 0.8695566 | 0.148158171 | 0.142907088 |
| Cep192 | -0.013190239 | 0.512638707 | 0.8722771 | 0.080364547 | 0.092087919 |
| Cep20 | -0.034843364 | 0.288221936 | 0.7480799 | 0.324211171 | 0.333595104 |
| Cep250 | 0.006347564 | 0.643983151 | 0.9168391 | 0.039292223 | 0.037491885 |
| Cep290 | 0.020156692 | 0.534904838 | 0.8795949 | 0.271459914 | 0.269492939 |
| Cep295 | -0.021937396 | 0.275498923 | 0.7356525 | 0.080411673 | 0.091299965 |
| Cep350 | 0.017873377 | 0.597138751 | 0.9075513 | 0.357839745 | 0.333911403 |
| Cep41 | 0.01257523 | 0.466276106 | 0.8552993 | 0.076964792 | 0.058291342 |
| Cep43 | -0.008388476 | 0.682501739 | 0.9304590 | 0.088685622 | 0.087630019 |
| Cep44 | 0.001522674 | 0.950031919 | 0.9948462 | 0.14623375 | 0.131807091 |
| Cep57 | 0.041494312 | 0.192155158 | 0.6700281 | 0.35699974 | 0.289614833 |
| Cep57l1 | -0.006886742 | 0.648595018 | 0.9172569 | 0.045652527 | 0.045948814 |
| Cep63 | -0.051333293 | 0.023094349 | 0.2949903 | 0.091746157 | 0.124078314 |
| Cep68 | -0.026615065 | 0.142422996 | 0.6157288 | 0.05824444 | 0.076319933 |
| Cep70 | -0.019544361 | 0.495122946 | 0.8682869 | 0.214524776 | 0.207540343 |
| Cep78 | -0.01866369 | 0.427915463 | 0.8340365 | 0.112028866 | 0.123876462 |
| Cep83 | 0.01497556 | 0.605464176 | 0.9087139 | 0.224056718 | 0.213866058 |
| Cep83os | -0.014479907 | 0.397123435 | 0.8238469 | 0.050444599 | 0.067577035 |
| Cep85 | -0.027858956 | 0.22135708 | 0.6929153 | 0.106159197 | 0.12672423 |
| Cep89 | -0.011076904 | 0.615009203 | 0.9108526 | 0.095752509 | 0.103809621 |
| Cep97 | -0.01908127 | 0.310716839 | 0.7678901 | 0.067081321 | 0.080037533 |
| Cept1 | 0.004958433 | 0.810516384 | 0.9668615 | 0.095367101 | 0.088396656 |
| Cerk | 0.030451476 | 0.322733288 | 0.7737584 | 0.28093304 | 0.260268357 |
| Cerkl | -0.010141091 | 0.715103939 | 0.9431411 | 0.177093402 | 0.194702247 |
| Cers2 | 0.003947568 | 0.900814618 | 0.9825441 | 0.334319582 | 0.295875807 |
| Cers4 | -0.061497742 | 0.020829076 | 0.2799949 | 0.135997519 | 0.174752005 |
| Cers5 | -0.050924543 | 0.053758244 | 0.4389150 | 0.146903081 | 0.187196654 |
| Cers6 | 0.033544419 | 0.175332799 | 0.6534142 | 0.164499379 | 0.135316631 |
| Cert1 | 0.021427183 | 0.500703989 | 0.8693704 | 0.314551147 | 0.303472096 |
| Cetn2 | 0.029530138 | 0.394009651 | 0.8211293 | 0.453638674 | 0.384909717 |
| Cetn3 | 0.052924431 | 0.134290041 | 0.6006119 | 0.629645803 | 0.495716194 |
| Cetn4 | -0.029033107 | 0.239738484 | 0.7118791 | 0.120510067 | 0.147146363 |
| Cfap126 | 0.01809706 | 0.485239778 | 0.8629715 | 0.174491402 | 0.150262232 |
| Cfap20 | -0.089396869 | 0.009970486 | 0.1987218 | 0.510840394 | 0.590339028 |
| Cfap298 | -0.027332135 | 0.416957769 | 0.8300990 | 0.371851261 | 0.385490424 |
| Cfap36 | 0.012052474 | 0.730994601 | 0.9459020 | 0.945527818 | 0.915941439 |

| | | | | | |
|---|---|---|---|---|---|
| Cfap410 | 0.005697832 | 0.84534853 | 0.9734889 | 0.239434461 | 0.220349813 |
| Cfap97 | 0.027707371 | 0.374097967 | 0.8087297 | 0.308099281 | 0.26824149 |
| Cfdp1 | 0.007978089 | 0.822842703 | 0.9710592 | 0.771409937 | 0.670717401 |
| Cfi | -0.055210853 | 0.00662239 | 0.1617967 | 0.06228992 | 0.089889007 |
| Cfl1 | -0.022517354 | 0.42107445 | 0.8300990 | 1.61881433 | 1.532260987 |
| Cfl2 | -0.009326232 | 0.801232971 | 0.9647143 | 0.563063771 | 0.526498459 |
| Cflar | -0.018586362 | 0.588365943 | 0.9044624 | 0.354790006 | 0.355929411 |
| Cggbp1 | 0.079351783 | 0.027863574 | 0.3199151 | 0.580732214 | 0.50887678 |
| Cgn | -0.023821433 | 0.36754989 | 0.8056719 | 0.166862056 | 0.168264533 |
| Cgref1 | -0.017061316 | 0.302301999 | 0.7611258 | 0.038042611 | 0.060978359 |
| Cgrrf1 | 0.017955871 | 0.488624977 | 0.8644836 | 0.168534736 | 0.167228603 |
| Chac1 | 0.116611143 | 0.003082777 | 0.1032885 | 0.459957236 | 0.413188137 |
| Chac2 | 0.04259223 | 0.030019142 | 0.3308280 | 0.111549985 | 0.075080592 |
| Champ1 | 0.006339576 | 0.77165827 | 0.9552333 | 0.117350395 | 0.105089987 |
| Chaserr | -0.002709664 | 0.940286795 | 0.9928458 | 0.560303337 | 0.531915717 |
| Chchd1 | 0.030854886 | 0.351600864 | 0.7941440 | 1.054298274 | 0.977689748 |
| Chchd10 | 0.046895033 | 0.24087154 | 0.7120404 | 1.073568598 | 1.093724346 |
| Chchd2 | 0.034756324 | 0.057058414 | 0.4485283 | 2.973066699 | 2.685568123 |
| Chchd3 | 0.018231206 | 0.561835582 | 0.8926240 | 0.36051974 | 0.306261637 |
| Chchd4 | -0.051636349 | 0.099145703 | 0.5433551 | 0.261453577 | 0.296521836 |
| Chchd5 | -0.005053963 | 0.860948352 | 0.9769675 | 0.20511181 | 0.209519362 |
| Chchd6 | -0.064157071 | 0.064531113 | 0.4711545 | 0.441069235 | 0.459911014 |
| Chchd7 | -0.008681321 | 0.802834424 | 0.9647143 | 0.862175633 | 0.803886855 |
| Chd1 | 0.008123875 | 0.798604349 | 0.9644698 | 0.30845141 | 0.285139603 |
| Chd1l | 0.004276834 | 0.784690025 | 0.9601202 | 0.056641108 | 0.051022965 |
| Chd2 | -0.007585828 | 0.832334768 | 0.9723494 | 0.413480913 | 0.394355926 |
| Chd3 | -0.01407251 | 0.711199251 | 0.9419834 | 0.690586358 | 0.672330211 |
| Chd4 | 0.024687849 | 0.502277821 | 0.8695566 | 1.116884965 | 1.085236218 |
| Chd5 | -0.0278552 | 0.236845279 | 0.7095050 | 0.096578471 | 0.142148525 |
| Chd6 | -0.022062695 | 0.499619464 | 0.8692081 | 0.311572802 | 0.328174091 |
| Chd7 | -0.0455224 | 0.159459382 | 0.6362980 | 0.243412806 | 0.326710118 |
| Chd8 | -0.004101268 | 0.902692281 | 0.9825441 | 0.366296187 | 0.357223508 |
| Chd9 | -0.076932051 | 0.036462738 | 0.3620803 | 0.508184807 | 0.582412749 |
| Cherp | 0.000598507 | 0.982556917 | 0.9976072 | 0.194956714 | 0.189648639 |
| Chfr | -0.010170396 | 0.667592203 | 0.9230765 | 0.127326536 | 0.129484308 |
| Chga | -0.031524338 | 0.005559242 | 0.1502661 | 5.285302641 | 5.382559226 |
| Chgb | -0.037192399 | 0.102508816 | 0.5529567 | 3.049984066 | 3.340101556 |
| Chic1 | 0.144685837 | 2.86E-05 | 0.0035979 | 1.61177972 | 1.405893554 |
| Chic2 | -0.004882215 | 0.88510816 | 0.9803788 | 0.420211474 | 0.40335154 |
| Chid1 | -0.026565721 | 0.443321139 | 0.8431541 | 0.444555621 | 0.423589837 |
| Chka | -0.121090237 | 0.002461295 | 0.0904297 | 0.630518714 | 0.811023888 |
| Chkb | -0.005763449 | 0.811163926 | 0.9671569 | 0.129057936 | 0.131279439 |
| Chm | -0.006093194 | 0.812815897 | 0.9673015 | 0.14272838 | 0.155351415 |
| Chmp1a | -0.00297245 | 0.926961455 | 0.9879210 | 0.34401214 | 0.314526344 |
| Chmp1b | -0.010605331 | 0.67430187 | 0.9264646 | 0.141244369 | 0.147771998 |
| Chmp2a | -0.011080001 | 0.729226153 | 0.9459020 | 1.329792952 | 1.281157555 |
| Chmp2b | -0.045973425 | 0.153200066 | 0.6290333 | 0.308420948 | 0.303616992 |
| Chmp3 | 2.54E-05 | 0.999420451 | 0.9996989 | 0.568715679 | 0.547029642 |
| Chmp4b | 0.069544924 | 0.038397542 | 0.3747897 | 1.040041609 | 0.916739282 |
| Chmp4c | 0.052539654 | 0.033971251 | 0.3488333 | 0.165850443 | 0.132957715 |
| Chmp5 | -0.0040238 | 0.905474852 | 0.9832544 | 0.946813607 | 0.902426127 |
| Chmp6 | 0.014349068 | 0.518980492 | 0.8741795 | 0.120332089 | 0.111991694 |
| Chmp7 | -0.01408107 | 0.601744745 | 0.9081899 | 0.191723781 | 0.184508413 |
| Chn1 | 0.006867823 | 0.708962443 | 0.9414811 | 0.075420638 | 0.069628755 |
| Chordc1 | -0.014895112 | 0.674190977 | 0.9264405 | 0.441511767 | 0.452304625 |
| Chp1 | -0.082403477 | 0.010960451 | 0.2047427 | 0.270885036 | 0.322207426 |
| Chpf | -0.002106741 | 0.948488418 | 0.9945042 | 0.332537828 | 0.338197966 |
| Chpf2 | -0.007817381 | 0.752478699 | 0.9507898 | 0.138133733 | 0.137640419 |
| Chpt1 | 0.010354043 | 0.65997364 | 0.9211780 | 0.146737654 | 0.128415321 |
| Chrac1 | 0.028927041 | 0.405314257 | 0.8270770 | 0.582584118 | 0.550631223 |
| Chrm3 | 0.013769651 | 0.407613438 | 0.8270770 | 0.057186712 | 0.06097303 |
| Chrna1os | 0.045535058 | 0.035827751 | 0.3593642 | 0.131956797 | 0.089509028 |
| Chrnb2 | -0.005406514 | 0.748923762 | 0.9501120 | 0.061919442 | 0.062975865 |
| Chst11 | 0.02756816 | 0.226877236 | 0.7003768 | 0.111844189 | 0.1257883 |
| Chst12 | -0.004840043 | 0.776277191 | 0.9561298 | 0.056808388 | 0.065098914 |
| Chtop | 0.026014591 | 0.448608534 | 0.8463296 | 0.440703169 | 0.390701684 |

| | | | | | |
|---|---|---|---|---|---|
| Chuk | 0.070955788 | 0.021031507 | 0.2811759 | 0.299990376 | 0.239611285 |
| Churc1 | -0.017529208 | 0.618215949 | 0.9115017 | 0.877685255 | 0.824075476 |
| Ciao1 | 0.004211527 | 0.853914878 | 0.9755431 | 0.133913707 | 0.124070097 |
| Ciao2a | 0.009267909 | 0.76330143 | 0.9532128 | 0.286592086 | 0.263406055 |
| Ciao2b | -0.046853051 | 0.146606814 | 0.6225768 | 0.350322688 | 0.351057949 |
| Ciao3 | 0.026864064 | 0.198479802 | 0.6730962 | 0.118871702 | 0.092770404 |
| Ciapin1 | -0.011048323 | 0.721962862 | 0.9445652 | 0.247685353 | 0.250814285 |
| Cib1 | 0.011714195 | 0.721686194 | 0.9445652 | 0.423271773 | 0.402020748 |
| Cib2 | 0.020291758 | 0.532360402 | 0.8784022 | 0.323416704 | 0.331903842 |
| Cib3 | 0.000152209 | 0.994576207 | 0.9988050 | 0.08973264 | 0.116275362 |
| Cibar1 | -0.023199033 | 0.523285185 | 0.8750824 | 0.565054507 | 0.542931215 |
| Cibar2 | 0.015137076 | 0.368678224 | 0.8056719 | 0.067082893 | 0.054114441 |
| Cic | 0.002821759 | 0.91377111 | 0.9845236 | 0.161955586 | 0.165730251 |
| Cinp | 0.022287918 | 0.337597243 | 0.7834245 | 0.135951181 | 0.117982412 |
| Cipc | 0.026014954 | 0.191231135 | 0.6700281 | 0.103293708 | 0.077707564 |
| Cir1 | 0.046005923 | 0.068981991 | 0.4835495 | 0.190459817 | 0.151019314 |
| Cirbp | 0.00761049 | 0.813468595 | 0.9674680 | 1.184919645 | 1.17910104 |
| Cisd1 | 0.037839712 | 0.224528625 | 0.6966918 | 1.256810485 | 1.145052402 |
| Cisd2 | -0.007745366 | 0.829758575 | 0.9722601 | 0.575026534 | 0.573154442 |
| Cisd3 | -0.029006913 | 0.314623658 | 0.7705987 | 0.202590569 | 0.217429177 |
| Cish | -0.009337952 | 0.714699395 | 0.9430747 | 0.148853064 | 0.146151954 |
| Cit | -0.044972488 | 0.098928549 | 0.5433551 | 0.158269949 | 0.155556101 |
| Cited2 | 0.059641162 | 0.129801712 | 0.5936051 | 0.509002143 | 0.458255119 |
| Cited4 | -0.025225516 | 0.196989737 | 0.6730962 | 0.067131015 | 0.088119527 |
| Ciz1 | -0.017306751 | 0.469400301 | 0.8567527 | 0.125904203 | 0.131946011 |
| Ckap4 | 0.026789007 | 0.461094305 | 0.8524425 | 0.422772377 | 0.379990811 |
| Ckap5 | -0.010448708 | 0.743286597 | 0.9479080 | 0.296753131 | 0.282166026 |
| Ckb | 0.005741917 | 0.897242392 | 0.9825441 | 0.500721359 | 0.600914922 |
| Ckmt1 | -0.012200132 | 0.602380304 | 0.9086609 | 0.11955899 | 0.126548423 |
| Cks1b | 0.023750563 | 0.377727319 | 0.8101797 | 0.204679936 | 0.16880636 |
| Clasp1 | 0.019573428 | 0.460713265 | 0.8524425 | 0.170231977 | 0.162023668 |
| Clasp2 | -0.035352398 | 0.256226468 | 0.7238259 | 0.233902634 | 0.260498011 |
| Clasrp | -0.021420155 | 0.241364578 | 0.7128617 | 0.053359184 | 0.07687181 |
| Clba1 | -0.024449024 | 0.235832053 | 0.7079390 | 0.092219941 | 0.101682099 |
| Clcc1 | -0.023521437 | 0.344218652 | 0.7876929 | 0.137073361 | 0.149679388 |
| Clcn3 | -0.026868983 | 0.458196529 | 0.8521621 | 0.773548962 | 0.805483236 |
| Clcn4 | -0.045046277 | 0.163879287 | 0.6410026 | 0.299241478 | 0.323817574 |
| Clcn5 | -0.044150204 | 0.133588706 | 0.5988251 | 0.181294717 | 0.219711395 |
| Clcn6 | -0.018337755 | 0.246300077 | 0.7160893 | 0.037253099 | 0.05549573 |
| Clcn7 | -0.011331136 | 0.465227087 | 0.8543229 | 0.040291424 | 0.052758883 |
| Cldn12 | 0.009718979 | 0.637403685 | 0.9159850 | 0.097055636 | 0.093177044 |
| Cldn3 | -0.00120637 | 0.972797301 | 0.9976072 | 0.429867953 | 0.412337041 |
| Cldn4 | 0.02379411 | 0.531326732 | 0.8776448 | 0.505367787 | 0.462766033 |
| Cldn6 | -0.043043382 | 0.096376624 | 0.5371101 | 0.176564056 | 0.163133382 |
| Cldn7 | 0.121602556 | 0.002260404 | 0.0855847 | 1.161520223 | 1.093149566 |
| Cldnd1 | -0.038822124 | 0.257762564 | 0.7250597 | 0.405630413 | 0.408637189 |
| Clec11a | 0.019432866 | 0.349580595 | 0.7927303 | 0.115344344 | 0.08646293 |
| Clec2d | -0.046304883 | 0.029717991 | 0.3298864 | 0.078811918 | 0.091863836 |
| Clic1 | -0.051577578 | 0.134820181 | 0.6013051 | 1.081451104 | 1.081941662 |
| Clic4 | 0.029940255 | 0.420597439 | 0.8300990 | 0.578772366 | 0.584665638 |
| Clint1 | 0.035334371 | 0.338997006 | 0.7834245 | 0.578287982 | 0.555677842 |
| Clip1 | -0.005324142 | 0.881871739 | 0.9797222 | 0.460140889 | 0.4868672 |
| Clip2 | 0.017456962 | 0.254174966 | 0.7223071 | 0.064906344 | 0.043609815 |
| Clip3 | -0.022572529 | 0.355602395 | 0.7969131 | 0.140714403 | 0.148857788 |
| Clk1 | 0.003256283 | 0.933951392 | 0.9904637 | 0.611343434 | 0.594842477 |
| Clk2 | -0.041171991 | 0.088050307 | 0.5278913 | 0.119864498 | 0.138130335 |
| Clk3 | -0.074537938 | 0.010032142 | 0.1990377 | 0.186246122 | 0.230938952 |
| Clk4 | -0.002846297 | 0.933025991 | 0.9902233 | 0.345042067 | 0.324140756 |
| Cln3 | 0.030043851 | 0.167810755 | 0.6449137 | 0.114769593 | 0.0988504 |
| Cln5 | -0.002885123 | 0.902854131 | 0.9825441 | 0.120350132 | 0.127448292 |
| Cln6 | 0.02675348 | 0.409415338 | 0.8278434 | 0.293387142 | 0.300224804 |
| Cln8 | 0.010538134 | 0.542774559 | 0.8840682 | 0.069394481 | 0.061879596 |
| Clns1a | 0.067130453 | 0.046298165 | 0.4073656 | 0.433646779 | 0.36120107 |
| Clock | -0.02605103 | 0.45057871 | 0.8478942 | 0.409411897 | 0.44469748 |
| Clp1 | 0.009821543 | 0.646922256 | 0.9169695 | 0.114045912 | 0.09661986 |
| Clpb | -0.03211894 | 0.253791155 | 0.7223071 | 0.17547052 | 0.200549517 |

| | | | | | |
|---|---|---|---|---|---|
| Clpp | -0.005959918 | 0.868325117 | 0.97756927 | 0.617296153 | 0.611800899 |
| Clptm1 | -0.035220885 | 0.257103231 | 0.72505979 | 0.255124303 | 0.292335668 |
| Clptm1l | -0.023086858 | 0.53173212 | 0.87786714 | 0.477448714 | 0.481459885 |
| Clpx | 0.019914325 | 0.554370049 | 0.89029959 | 0.390911851 | 0.366000352 |
| Clstn1 | -0.003000551 | 0.913848003 | 0.98452363 | 0.170935352 | 0.200745166 |
| Clstn3 | -0.059481727 | 0.0086593 | 0.18461360 | 0.082397154 | 0.129983899 |
| Clta | 0.074480279 | 0.007958927 | 0.17584088 | 1.507647105 | 1.377856787 |
| Cltb | 0.011719262 | 0.698600695 | 0.93696920 | 1.322719875 | 1.244359507 |
| Cltc | 0.010021623 | 0.777214054 | 0.95645247 | 1.166214948 | 1.156731919 |
| Cltrn | -0.196675639 | 1.06E-11 | 1.31E-08 | 2.830037485 | 3.039754421 |
| Clu | 0.06787951 | 0.14741809 | 0.62411421 | 0.673384644 | 0.658116973 |
| Cluap1 | 0.012901226 | 0.595242761 | 0.9067325 | 0.155994278 | 0.127961296 |
| Cluh | -0.022392897 | 0.405476984 | 0.82707700 | 0.150929533 | 0.182321178 |
| Clybl | 0.008917227 | 0.586956315 | 0.90357024 | 0.062439369 | 0.055537143 |
| Cmas | -0.041667931 | 0.254891371 | 0.72236653 | 0.64962678 | 0.680352803 |
| Cmbl | -0.01122688 | 0.655439979 | 0.92080099 | 0.141238875 | 0.150575245 |
| Cmc1 | 0.007794695 | 0.809013928 | 0.96587052 | 0.353648853 | 0.334093604 |
| Cmc2 | -0.006933999 | 0.779586294 | 0.95865824 | 0.152569338 | 0.146523707 |
| Cmc4 | -0.024872778 | 0.314495956 | 0.77059879 | 0.147252779 | 0.152754985 |
| Cmip | -0.022693706 | 0.516030593 | 0.87357053 | 0.46904395 | 0.507668695 |
| Cmpk1 | -0.018402253 | 0.613915264 | 0.91083863 | 0.764373473 | 0.740013938 |
| Cmpk2 | 0.038601605 | 0.047362727 | 0.41067610 | 0.099119927 | 0.072509837 |
| Cmss1 | -0.000154366 | 0.994941092 | 0.99907033 | 0.139164372 | 0.133512937 |
| Cmtm4 | 0.018299472 | 0.601767415 | 0.90818998 | 0.423323333 | 0.427279314 |
| Cmtm6 | -0.008920323 | 0.722723925 | 0.94456521 | 0.164143328 | 0.158810798 |
| Cmtm7 | 0.012439419 | 0.483838743 | 0.86248860 | 0.082997349 | 0.061626393 |
| Cmtm8 | 0.005904707 | 0.770177973 | 0.95523334 | 0.101727696 | 0.083986933 |
| Cmtr1 | 0.003955193 | 0.80112225 | 0.96471431 | 0.047522953 | 0.051554644 |
| Cmtr2 | 0.018201378 | 0.293239854 | 0.75336844 | 0.080131205 | 0.060310708 |
| Cnbp | -0.008616773 | 0.793422852 | 0.96245081 | 1.223780302 | 1.170518016 |
| Cndp2 | -0.005550095 | 0.837744165 | 0.97272164 | 0.192035582 | 0.187238543 |
| Cnep1r1 | 0.035931153 | 0.179073928 | 0.65500709 | 0.195103703 | 0.162709963 |
| Cnih1 | 0.045716308 | 0.15177751 | 0.62811059 | 0.356926467 | 0.288608711 |
| Cnih4 | 0.073249061 | 0.004747134 | 0.13717421 | 0.211513331 | 0.146528652 |
| Cnksr1 | -0.021086509 | 0.249151134 | 0.71927281 | 0.054364928 | 0.080304475 |
| Cnksr3 | 0.026497161 | 0.087570223 | 0.52712171 | 0.062381537 | 0.044297723 |
| Cnn3 | 0.026833472 | 0.408683532 | 0.82764474 | 0.284475063 | 0.30983415 |
| Cnnm2 | -0.001943857 | 0.926997089 | 0.98792100 | 0.10068885 | 0.104183069 |
| Cnnm3 | -0.029262415 | 0.136001947 | 0.60329191 | 0.070635033 | 0.088586327 |
| Cnot1 | -0.032475683 | 0.330928784 | 0.77810228 | 0.333342837 | 0.377082077 |
| Cnot10 | -0.036730973 | 0.220158305 | 0.69145345 | 0.227215773 | 0.244098261 |
| Cnot11 | -0.022866387 | 0.376383167 | 0.81017979 | 0.143090805 | 0.16612137 |
| Cnot2 | -0.016715574 | 0.580238259 | 0.90105878 | 0.248861596 | 0.244990813 |
| Cnot3 | -0.015346473 | 0.634129504 | 0.9144743 | 0.29435388 | 0.29211108 |
| Cnot4 | -0.018970175 | 0.579656441 | 0.90087519 | 0.404583035 | 0.406175071 |
| Cnot6 | -0.027553558 | 0.362371434 | 0.80304783 | 0.222575695 | 0.241706025 |
| Cnot6l | -0.002832454 | 0.935617585 | 0.99106529 | 0.944763934 | 0.87616941 |
| Cnot7 | 0.01754014 | 0.607567861 | 0.90871392 | 0.505434651 | 0.426261178 |
| Cnot8 | 0.001562169 | 0.944010583 | 0.99358991 | 0.120715616 | 0.107093131 |
| Cnot9 | 0.010315174 | 0.584944169 | 0.90239083 | 0.086915845 | 0.078257199 |
| Cnppd1 | -0.027403255 | 0.238905711 | 0.71126790 | 0.110162347 | 0.130833182 |
| Cnpy2 | -0.03488767 | 0.204339312 | 0.67805984 | 1.497530206 | 1.506838082 |
| Cnpy3 | -0.028645351 | 0.319387709 | 0.77106980 | 0.233749801 | 0.221430421 |
| Cnpy4 | -0.067173124 | 0.000267508 | 0.02010364 | 0.037375299 | 0.083209681 |
| Cnrip1 | -0.029060955 | 0.305961234 | 0.76509590 | 0.175686276 | 0.196616768 |
| Cnst | -0.029743562 | 0.148309269 | 0.62556941 | 0.076478888 | 0.100267877 |
| Cntfr | 0.028199238 | 0.11118244 | 0.56719141 | 0.069719795 | 0.061731476 |
| Cntln | -0.022089172 | 0.423794897 | 0.83204620 | 0.168649122 | 0.19828479 |
| Cntn1 | -0.010808784 | 0.709691868 | 0.94182519 | 0.149744604 | 0.224806227 |
| Cntnap2 | -0.000423993 | 0.983485491 | 0.99760728 | 0.070682025 | 0.100915699 |
| Cntrl | -0.000626171 | 0.977089316 | 0.99760728 | 0.105295138 | 0.110021084 |
| Coa3 | 0.077668888 | 0.006511455 | 0.16108138 | 1.636602604 | 1.434766187 |
| Coa4 | -0.037050814 | 0.103487476 | 0.55611904 | 0.110918717 | 0.128668669 |
| Coa5 | 0.019162349 | 0.522069594 | 0.87496711 | 0.256090019 | 0.226642992 |
| Coa6 | -0.045151765 | 0.120113216 | 0.57973049 | 0.219525454 | 0.22477388 |
| Coa7 | 0.025068768 | 0.24762515 | 0.71770680 | 0.123471626 | 0.097425571 |

| | | | | | |
|---|---|---|---|---|---|
| Coa8 | 0.026850689 | 0.431928368 | 0.8362079 | 0.464620558 | 0.425952169 |
| Coasy | 0.005126777 | 0.868479642 | 0.9775692 | 0.274342047 | 0.266676722 |
| Cobl | -0.018048844 | 0.317780432 | 0.7709420 | 0.064483119 | 0.069632186 |
| Cobll1 | 0.056516771 | 0.047397315 | 0.4106761 | 0.233354784 | 0.201707667 |
| Cog1 | 0.012828018 | 0.628626985 | 0.9130564 | 0.189378574 | 0.162318678 |
| Cog2 | 0.01482308 | 0.580493804 | 0.9013145 | 0.187858541 | 0.170773653 |
| Cog3 | 0.013754662 | 0.508576337 | 0.8704067 | 0.104690355 | 0.093291691 |
| Cog4 | -0.05426259 | 0.034004219 | 0.3488333 | 0.125065312 | 0.166251297 |
| Cog5 | -0.008417055 | 0.743469657 | 0.9479080 | 0.145609149 | 0.1587372 |
| Cog6 | -0.003497912 | 0.916033564 | 0.9845236 | 0.356614298 | 0.350741592 |
| Cog7 | 0.001587477 | 0.957623145 | 0.9956630 | 0.249119395 | 0.244333341 |
| Cog8 | 0.011219463 | 0.65620708 | 0.9211780 | 0.177651478 | 0.143594613 |
| Coil | 0.007507682 | 0.687180847 | 0.9326630 | 0.082575275 | 0.072539802 |
| Col1a2 | -0.003778792 | 0.811620102 | 0.9673015 | 0.056337025 | 0.043575718 |
| Col27a1 | -0.066127568 | 0.013433602 | 0.2292511 | 0.107171967 | 0.178545059 |
| Col6a6 | 0.006137047 | 0.697648317 | 0.9368717 | 0.047842709 | 0.055882511 |
| Col9a2 | 0.005441934 | 0.735565873 | 0.9468380 | 0.060891693 | 0.044205391 |
| Colgalt1 | -0.005908338 | 0.832754011 | 0.9723494 | 0.208854463 | 0.198839456 |
| Commd1 | 0.029970606 | 0.377504117 | 0.8101797 | 0.464659138 | 0.39723341 |
| Commd10 | 0.026083669 | 0.104310217 | 0.5580045 | 0.070637745 | 0.046891149 |
| Commd2 | 0.010593151 | 0.710770514 | 0.9419834 | 0.231342617 | 0.20541294 |
| Commd3 | 0.041723828 | 0.252019451 | 0.7221354 | 0.728476881 | 0.608239026 |
| Commd4 | -0.06198039 | 0.07252713 | 0.4902939 | 0.394956602 | 0.404637063 |
| Commd6 | -0.048600477 | 0.154890389 | 0.6304934 | 0.618706431 | 0.649135164 |
| Commd7 | -0.068179047 | 0.061700847 | 0.4613744 | 0.546825113 | 0.606691066 |
| Commd8 | -0.025746328 | 0.315338876 | 0.7705987 | 0.153465863 | 0.165139032 |
| Commd9 | -0.06550846 | 0.050350271 | 0.4222102 | 0.375088031 | 0.394847253 |
| Comt | -0.001162063 | 0.97352404 | 0.9976072 | 0.501302212 | 0.526211018 |
| Comtd1 | -0.009973686 | 0.623306466 | 0.9121505 | 0.087135248 | 0.095197367 |
| Cop1 | -0.035948503 | 0.224854044 | 0.6968297 | 0.220184877 | 0.228350737 |
| Copa | 0.009111176 | 0.800220324 | 0.9647053 | 0.958064295 | 0.959559121 |
| Copb1 | 0.004148047 | 0.902910526 | 0.9825441 | 0.969678164 | 0.939905617 |
| Copb2 | -0.052784702 | 0.129816702 | 0.5936051 | 1.25575279 | 1.207976396 |
| Cope | -0.046614426 | 0.111883568 | 0.5679582 | 1.450026937 | 1.477745455 |
| Copg1 | -0.012802602 | 0.725491383 | 0.9450332 | 0.636536423 | 0.627013358 |
| Copg2 | -0.024006317 | 0.361831658 | 0.8029910 | 0.162831855 | 0.17259528 |
| Coprs | 0.008001126 | 0.785852469 | 0.9602516 | 0.255346702 | 0.243485382 |
| Cops2 | -0.007253512 | 0.823646491 | 0.9710592 | 0.320465838 | 0.304962259 |
| Cops3 | -0.011387271 | 0.751198697 | 0.9506053 | 0.569328394 | 0.549850756 |
| Cops4 | -0.017264403 | 0.624723134 | 0.9121505 | 0.574634056 | 0.531065088 |
| Cops5 | -0.038674222 | 0.280976939 | 0.7408618 | 0.434943257 | 0.452720784 |
| Cops6 | -0.046849583 | 0.166272178 | 0.6429285 | 0.480593142 | 0.46740187 |
| Cops7a | -0.007884709 | 0.81506551 | 0.9685493 | 0.449697353 | 0.41892265 |
| Cops7b | 0.017461381 | 0.225692219 | 0.6979011 | 0.055641045 | 0.038417721 |
| Cops8 | -0.013175313 | 0.698064505 | 0.9369692 | 0.36931804 | 0.360135664 |
| Cops9 | 0.086664378 | 0.006134343 | 0.1568368 | 1.260248727 | 1.159181746 |
| Copz1 | 0.006315846 | 0.863043357 | 0.9769675 | 0.630779266 | 0.575780218 |
| Copz2 | -0.018651245 | 0.593466059 | 0.9061387 | 0.673777302 | 0.631826116 |
| Coq10a | -0.00800949 | 0.803598069 | 0.9647143 | 0.305605395 | 0.303958897 |
| Coq10b | -0.106624592 | 0.003166573 | 0.1050582 | 0.337023612 | 0.415115942 |
| Coq2 | 0.054891808 | 0.115294989 | 0.5724670 | 0.491890908 | 0.408583984 |
| Coq4 | 0.000365178 | 0.984443358 | 0.9976072 | 0.07890622 | 0.075532359 |
| Coq5 | 0.033832221 | 0.270355746 | 0.7340360 | 0.281663161 | 0.244166845 |
| Coq6 | -0.01167884 | 0.532617218 | 0.8785438 | 0.081511956 | 0.078082767 |
| Coq7 | 0.017585216 | 0.581629088 | 0.9018069 | 0.338109222 | 0.280106352 |
| Coq8a | 0.014270923 | 0.416188011 | 0.8300990 | 0.078133279 | 0.061507994 |
| Coq8b | 0.009198403 | 0.713954266 | 0.9428150 | 0.151687824 | 0.140279789 |
| Coq9 | -0.015605817 | 0.609391506 | 0.9088672 | 0.243633677 | 0.257216271 |
| Coro1c | 0.001288886 | 0.960525802 | 0.9958628 | 0.164631977 | 0.162483711 |
| Coro2b | -0.008236457 | 0.753750192 | 0.9517823 | 0.161854378 | 0.16488694 |
| Coro7 | 0.016337449 | 0.36174772 | 0.8029844 | 0.073466665 | 0.067114317 |
| Cotl1 | 0.009180335 | 0.764977101 | 0.9540090 | 0.287915573 | 0.25378845 |
| Cox10 | 0.021964476 | 0.253993308 | 0.7223071 | 0.088991482 | 0.078383684 |
| Cox11 | 0.019267529 | 0.309406068 | 0.7673813 | 0.089831749 | 0.0693478 |
| Cox14 | 0.02054325 | 0.531250366 | 0.8776448 | 1.160827421 | 1.081456006 |
| Cox15 | 0.012452669 | 0.460905328 | 0.8524425 | 0.07073804 | 0.055867877 |

| | | | | | |
|---|---|---|---|---|---|
| Cox16 | 0.071349448 | 0.015514494 | 0.2435186 | 0.261718544 | 0.192852449 |
| Cox17 | 0.103155105 | 0.00061377 | 0.0350454 | 1.445470668 | 1.284656731 |
| Cox18 | -0.040578261 | 0.171720445 | 0.6515402 | 0.217786737 | 0.239447436 |
| Cox19 | -0.017939678 | 0.571559047 | 0.8966165 | 0.297921066 | 0.295595195 |
| Cox20 | 0.005487633 | 0.877276498 | 0.9785583 | 0.466959968 | 0.463845572 |
| Cox4i1 | 0.028387306 | 0.115709807 | 0.5724670 | 2.50566097 | 2.361673299 |
| Cox5a | 0.053791412 | 0.106922858 | 0.5626921 | 1.010908108 | 0.949506666 |
| Cox5b | 0.059077961 | 0.020765875 | 0.2798878 | 1.844417525 | 1.678887905 |
| Cox6a1 | 0.108293472 | 0.000164871 | 0.0138603 | 1.723471764 | 1.480964311 |
| Cox6a2 | 0.050333756 | 0.198311974 | 0.6730962 | 0.267996097 | 0.319714256 |
| Cox6b1 | 0.057825779 | 0.005284346 | 0.1464265 | 2.102418662 | 1.962169999 |
| Cox6c | 0.09099317 | 2.16E-05 | 0.0028168 | 2.244703129 | 2.069720001 |
| Cox7a2 | 0.085797334 | 2.87E-05 | 0.0035979 | 2.326790548 | 2.102393774 |
| Cox7a2l | -0.027771102 | 0.322004462 | 0.7733024 | 1.714969265 | 1.709849584 |
| Cox7b | 0.072510483 | 0.013474198 | 0.2292511 | 1.403614678 | 1.306459562 |
| Cox7c | 0.145846855 | 4.70E-13 | 1.17E-09 | 2.682095686 | 2.326388117 |
| Cox8a | 0.081009148 | 1.12E-07 | 4.28E-05 | 3.035311995 | 2.779368964 |
| Cpa2 | 0.002258224 | 0.88986866 | 0.9819597 | 0.052190036 | 0.049052078 |
| Cpd | 0.146426144 | 1.61E-05 | 0.0023189 | 0.454542395 | 0.331120611 |
| Cpe | -0.012764369 | 0.29613598 | 0.7545964 | 3.51654539 | 3.423343943 |
| Cpeb1 | -0.027282993 | 0.395384044 | 0.8225618 | 0.262645217 | 0.295458991 |
| Cpeb2 | -0.006635793 | 0.701445638 | 0.9381877 | 0.053630617 | 0.064396164 |
| Cpeb3 | -0.016850897 | 0.514725558 | 0.8733714 | 0.140652145 | 0.157583524 |
| Cpeb4 | 9.65E-05 | 0.997623546 | 0.9994073 | 0.312569809 | 0.330170381 |
| Cplane1 | -0.066903722 | 0.017853214 | 0.2587928 | 0.16144544 | 0.207022459 |
| Cplx2 | 0.012458203 | 0.740787044 | 0.9479080 | 1.24556758 | 1.191599989 |
| Cpm | -0.030507636 | 0.11040285 | 0.5655888 | 0.054566732 | 0.087306226 |
| Cpn1 | -0.027751769 | 0.413562317 | 0.8300990 | 0.463361324 | 0.422563166 |
| Cpne3 | 0.047917527 | 0.143074157 | 0.6162812 | 0.338386464 | 0.302288207 |
| Cpne4 | 0.015863154 | 0.328477318 | 0.7778857 | 0.052012359 | 0.050149191 |
| Cpne8 | 0.034014883 | 0.164704672 | 0.6420707 | 0.174559046 | 0.12252006 |
| Cpox | -0.017380252 | 0.535786663 | 0.8798218 | 0.192963451 | 0.207539842 |
| Cpped1 | -0.04197605 | 0.017786462 | 0.2587928 | 0.050726874 | 0.07743444 |
| Cpq | -0.129519522 | 0.000324907 | 0.0226977 | 1.259229787 | 1.234863653 |
| Cpsf1 | 0.004300179 | 0.826279568 | 0.9716906 | 0.090455653 | 0.080590156 |
| Cpsf2 | 0.023299121 | 0.374843802 | 0.8092384 | 0.176326087 | 0.164516727 |
| Cpsf3 | -0.006535597 | 0.815793361 | 0.9686999 | 0.211710857 | 0.196744431 |
| Cpsf4 | -0.004227684 | 0.790505057 | 0.9621438 | 0.050886188 | 0.053545545 |
| Cpsf6 | -0.047189361 | 0.150930294 | 0.6280320 | 0.290683181 | 0.339412716 |
| Cpsf7 | -0.027240257 | 0.38011481 | 0.8117350 | 0.247181874 | 0.262207713 |
| Cpt1a | -0.00094186 | 0.963509828 | 0.9966421 | 0.085685249 | 0.095076471 |
| Cpt1c | -0.019357413 | 0.297880972 | 0.7557491 | 0.066924485 | 0.073300868 |
| Cpt2 | 0.023985189 | 0.383863911 | 0.8129787 | 0.223469676 | 0.185460593 |
| Cptp | 0.012186781 | 0.45469737 | 0.8499020 | 0.066395648 | 0.051334738 |
| Cr1l | -0.01872393 | 0.559926506 | 0.8925646 | 0.335257087 | 0.319422296 |
| Cracdl | 0.021834541 | 0.133536727 | 0.5988251 | 0.046119155 | 0.039527251 |
| Cradd | -0.006101577 | 0.759475133 | 0.9521533 | 0.096007728 | 0.08529073 |
| Cramp1l | 0.013949364 | 0.506502089 | 0.8695566 | 0.102247672 | 0.096301744 |
| Crat | 0.008286503 | 0.566719902 | 0.8949756 | 0.051091319 | 0.041237804 |
| Crb3 | 0.02665145 | 0.451679893 | 0.8481913 | 0.442041988 | 0.430778477 |
| Crbn | -0.052605883 | 0.033907696 | 0.3485640 | 0.122552956 | 0.158399696 |
| Crcp | -0.019653507 | 0.463103377 | 0.8524425 | 0.185558293 | 0.171507082 |
| Creb1 | 0.030380861 | 0.352059194 | 0.7949982 | 0.340707799 | 0.302163807 |
| Creb3 | 0.036170204 | 0.284819818 | 0.7450791 | 0.420047357 | 0.3660229 |
| Creb3l1 | 0.063376476 | 0.019303219 | 0.2695586 | 0.193565568 | 0.181523037 |
| Creb3l2 | 0.025768871 | 0.480599715 | 0.8617579 | 0.584896822 | 0.559542251 |
| Crebbp | -0.032689437 | 0.364852323 | 0.8050191 | 0.451091627 | 0.471585568 |
| Crebl2 | 0.005127226 | 0.819616833 | 0.9699062 | 0.114170854 | 0.112617094 |
| Crebrf | -0.014335298 | 0.651114539 | 0.9182621 | 0.260030368 | 0.274878523 |
| Crebzf | -0.04223987 | 0.17514698 | 0.6534142 | 0.240595116 | 0.276121371 |
| Creg1 | -0.01614718 | 0.663822365 | 0.9230765 | 1.01369285 | 1.186815118 |
| Creld1 | -0.015196848 | 0.608454395 | 0.9088672 | 0.258256286 | 0.236444238 |
| Creld2 | 0.08688086 | 0.027562738 | 0.3181899 | 0.618170916 | 0.502553441 |
| Crem | 0.025140691 | 0.505321056 | 0.8695566 | 0.491691702 | 0.491983825 |
| Crip1 | 0.006006773 | 0.875658247 | 0.9784331 | 0.450014135 | 0.502313762 |
| Cript | -0.000747374 | 0.982903562 | 0.9976072 | 0.533234824 | 0.496800965 |

| | | | | | |
|---|---|---|---|---|---|
| Crk | 0.023245532 | 0.418529541 | 0.83009907 | 0.225840789 | 0.208568864 |
| Crkl | 0.011957518 | 0.613443814 | 0.91083865 | 0.129694514 | 0.126709321 |
| Crlf1 | -0.003901773 | 0.858667247 | 0.97696750 | 0.092998478 | 0.115502726 |
| Crlf2 | -0.020483298 | 0.450957266 | 0.84789420 | 0.198516781 | 0.19813777 |
| Crlf3 | -0.016197708 | 0.310319116 | 0.76770334 | 0.04374431 | 0.054523198 |
| Crls1 | 0.016937001 | 0.583791329 | 0.90214067 | 0.277091544 | 0.252982137 |
| Crmp1 | -0.022372968 | 0.272199536 | 0.73428063 | 0.083339753 | 0.097649543 |
| Crnkl1 | 0.020675282 | 0.422370595 | 0.83148973 | 0.190369499 | 0.152043497 |
| Crocc | 0.027567206 | 0.073346339 | 0.49161870 | 0.064754374 | 0.045661155 |
| Crot | 0.032150375 | 0.155083413 | 0.63076150 | 0.124552134 | 0.111020832 |
| Crtap | -0.023937097 | 0.18343868 | 0.66159407 | 0.063696867 | 0.07500975 |
| Crtc1 | 0.029820064 | 0.142829249 | 0.61602875 | 0.111103169 | 0.085739407 |
| Crtc2 | 0.020339915 | 0.315159548 | 0.77059875 | 0.102169617 | 0.088023173 |
| Crtc3 | 0.012634351 | 0.679782736 | 0.92892004 | 0.243618634 | 0.252020842 |
| Cry1 | -0.042940914 | 0.083981394 | 0.51841657 | 0.12257726 | 0.158523135 |
| Cry2 | 0.001286709 | 0.967465537 | 0.99760728 | 0.259529803 | 0.271743289 |
| Cryba2 | 0.001808933 | 0.969058014 | 0.99760728 | 1.234383243 | 1.036185651 |
| Crybb3 | -0.089086291 | 0.011485731 | 0.21125160 | 0.078577594 | 0.244930947 |
| Crybg3 | -0.006633441 | 0.763633328 | 0.95322041 | 0.095188811 | 0.107813159 |
| Cryl1 | 0.043529838 | 0.029076804 | 0.32641720 | 0.102838281 | 0.077607798 |
| Cryz | 0.018303206 | 0.316146219 | 0.77059875 | 0.082668378 | 0.065051848 |
| Cryzl1 | -0.033524144 | 0.28250524 | 0.74228118 | 0.249159723 | 0.273421396 |
| Cryzl2 | -0.00615003 | 0.802145451 | 0.96471431 | 0.1340049 | 0.143037575 |
| Cs | -0.010237969 | 0.718056672 | 0.94382369 | 0.204945902 | 0.207716706 |
| Csad | -0.013586853 | 0.622410783 | 0.91215057 | 0.18780315 | 0.192562678 |
| Csde1 | 0.04851329 | 0.174263392 | 0.65276574 | 1.086703155 | 1.049334187 |
| Cse1l | 0.038790058 | 0.10571197 | 0.56171712 | 0.164391692 | 0.129411414 |
| Csf2ra | -0.023917878 | 0.403176383 | 0.82647037 | 0.191792354 | 0.20596935 |
| Csgalnact2 | 0.025096346 | 0.233641691 | 0.70513913 | 0.111333739 | 0.084407878 |
| Csk | 0.000903174 | 0.971951206 | 0.99760728 | 0.172947991 | 0.148921146 |
| Csmd2 | -0.023064209 | 0.191410205 | 0.67002813 | 0.050876533 | 0.06765211 |
| Csnk1a1 | 0.049056108 | 0.111480816 | 0.56757614 | 1.439719494 | 1.374805612 |
| Csnk1d | -0.039473306 | 0.23141577 | 0.70261419 | 0.29864822 | 0.323479501 |
| Csnk1e | 0.022019117 | 0.481233819 | 0.86201507 | 0.283460743 | 0.258358947 |
| Csnk1g1 | -0.029456273 | 0.312526275 | 0.76891385 | 0.196287043 | 0.213987933 |
| Csnk1g2 | -0.002074204 | 0.951636228 | 0.99515668 | 0.406002742 | 0.391451096 |
| Csnk1g3 | -0.00698133 | 0.835479265 | 0.97234942 | 0.378539655 | 0.35944387 |
| Csnk2a1 | 0.060630901 | 0.085339639 | 0.52128640 | 0.541574742 | 0.471017497 |
| Csnk2a2 | 0.09703955 | 0.005463604 | 0.14889821 | 0.521082619 | 0.441977781 |
| Csnk2b | -0.022800412 | 0.520430781 | 0.87417954 | 0.600024538 | 0.589114763 |
| Cspp1 | 0.04427956 | 0.239249897 | 0.71158843 | 0.512789761 | 0.476223483 |
| Csrnp1 | -0.034842023 | 0.250322772 | 0.72072102 | 0.171346951 | 0.213292009 |
| Csrp1 | 0.02552533 | 0.177287374 | 0.65366590 | 0.101863616 | 0.067162218 |
| Csrp2 | -0.016222031 | 0.43768541 | 0.83999140 | 0.095187448 | 0.088632814 |
| Cst3 | -0.065766619 | 1.78E-05 | 0.00241691 | 3.964057342 | 3.890959625 |
| Cst6 | -0.010030258 | 0.617551066 | 0.91150170 | 0.079289696 | 0.096000199 |
| Cstb | 0.060855179 | 0.081962686 | 0.51460110 | 1.003295615 | 0.890089791 |
| Cstf1 | 0.022172551 | 0.204904227 | 0.67868111 | 0.080162332 | 0.062194063 |
| Cstf2 | -0.014509773 | 0.464912325 | 0.85390303 | 0.067498932 | 0.09022972 |
| Cstf2t | 0.004964735 | 0.8289995 | 0.97226010 | 0.124098963 | 0.113458158 |
| Cstf3 | -0.039404775 | 0.206803487 | 0.67975655 | 0.257351098 | 0.274348335 |
| Ctbp1 | -0.000118553 | 0.997264397 | 0.99940731 | 0.608386121 | 0.568467083 |
| Ctbp2 | -0.023117186 | 0.340300121 | 0.78446580 | 0.139469261 | 0.134773371 |
| Ctbs | -0.024390812 | 0.253559692 | 0.72230717 | 0.08909057 | 0.108047968 |
| Ctc1 | 0.006688563 | 0.716263201 | 0.94314111 | 0.080595113 | 0.068577359 |
| Ctcf | -0.003847533 | 0.916857977 | 0.98454547 | 0.605689257 | 0.589048037 |
| Ctdnep1 | -0.083902697 | 0.013842154 | 0.23116862 | 0.33800256 | 0.402354326 |
| Ctdp1 | 0.009142332 | 0.589393004 | 0.90465390 | 0.062891198 | 0.057999438 |
| Ctdsp1 | -0.033553123 | 0.164263396 | 0.64155213 | 0.120272647 | 0.142812736 |
| Ctdsp2 | 0.011405777 | 0.706974188 | 0.94083543 | 0.279257805 | 0.262992268 |
| Ctdspl | 0.002631346 | 0.940383335 | 0.99284589 | 0.473266381 | 0.430863469 |
| Ctdspl2 | 0.019029967 | 0.49923145 | 0.86907428 | 0.202245178 | 0.192002916 |
| Ctf1 | -0.031254465 | 0.13604476 | 0.60329191 | 0.074885906 | 0.10158453 |
| Ctif | -0.006051698 | 0.705591417 | 0.94013812 | 0.047185129 | 0.05855143 |
| Ctnna1 | -0.061772928 | 0.096027664 | 0.53684695 | 0.45878661 | 0.514060714 |
| Ctnnal1 | 0.025560658 | 0.360569168 | 0.80124805 | 0.178140977 | 0.182242359 |

| | | | | | |
|---|---|---|---|---|---|
| Ctnnb1 | -0.046149456 | 0.182908601 | 0.6607594 | 0.796952325 | 0.852776676 |
| Ctnnbip1 | 0.027436713 | 0.381880762 | 0.8126135 | 0.289619769 | 0.251318938 |
| Ctnnbl1 | 0.061411292 | 0.016948361 | 0.2555884 | 0.202216189 | 0.148521905 |
| Ctnnd1 | 0.011605866 | 0.689096453 | 0.9337299 | 0.240091054 | 0.238544172 |
| Ctnnd2 | -0.028884723 | 0.281403036 | 0.7414390 | 0.164117485 | 0.182779351 |
| Ctps | 0.013484522 | 0.500343397 | 0.8693704 | 0.099431694 | 0.080429123 |
| Ctps2 | -0.009231765 | 0.718653428 | 0.9440039 | 0.163637954 | 0.159684813 |
| Ctr9 | -0.01745956 | 0.61791557 | 0.9115017 | 0.383835116 | 0.369084411 |
| Ctsa | -0.023658696 | 0.42995963 | 0.8340944 | 0.245075498 | 0.256241051 |
| Ctsb | -0.059915513 | 0.048531497 | 0.4157447 | 1.353943022 | 1.347576378 |
| Ctsd | 0.010358954 | 0.774586149 | 0.9558538 | 0.862907349 | 0.823020199 |
| Ctsf | -0.065224185 | 0.046654038 | 0.4094577 | 0.262573223 | 0.329840411 |
| Ctsl | -0.011142747 | 0.71617833 | 0.9431411 | 1.794690188 | 1.736231697 |
| Ctso | 0.003023538 | 0.927012065 | 0.9879210 | 0.376988024 | 0.348538203 |
| Ctsz | 0.038867686 | 0.380828096 | 0.8117350 | 1.625505731 | 1.367027058 |
| Cttn | 0.025908114 | 0.419933586 | 0.8300990 | 0.34541746 | 0.291475452 |
| Cttnbp2 | -0.006793281 | 0.796702012 | 0.9644244 | 0.119638751 | 0.174707381 |
| Ctu1 | 0.010821138 | 0.412201604 | 0.8297979 | 0.042124154 | 0.033007536 |
| Ctu2 | -0.033028863 | 0.124415335 | 0.5852543 | 0.091066699 | 0.109784948 |
| Ctxn1 | 0.033139852 | 0.039634355 | 0.3820921 | 0.068361645 | 0.04479611 |
| Ctxn2 | -0.028113059 | 0.367858364 | 0.8056719 | 0.330262388 | 0.258371831 |
| Cuedc1 | -0.00010577 | 0.99740873 | 0.9994073 | 0.288366321 | 0.286790411 |
| Cuedc2 | -0.054228437 | 0.104697425 | 0.5580045 | 0.35603711 | 0.372643848 |
| Cul1 | -0.016071423 | 0.652305463 | 0.9190271 | 0.515946925 | 0.50732158 |
| Cul2 | -0.002552696 | 0.926791997 | 0.9879210 | 0.185144632 | 0.184410703 |
| Cul3 | 0.005655708 | 0.875638871 | 0.9784331 | 0.724260807 | 0.678501615 |
| Cul4a | -0.028966412 | 0.360509081 | 0.8012480 | 0.282662505 | 0.297054961 |
| Cul4b | -0.022294731 | 0.413487463 | 0.8300990 | 0.182452885 | 0.182209056 |
| Cul5 | -0.041948197 | 0.230623525 | 0.7022964 | 0.487481782 | 0.508390544 |
| Cul7 | 0.005557554 | 0.81983219 | 0.9700304 | 0.155596445 | 0.132889229 |
| Cul9 | -0.049983673 | 0.035738105 | 0.3591914 | 0.097355885 | 0.145578715 |
| Cuta | 0.011933271 | 0.739328761 | 0.9475634 | 0.863546072 | 0.815411031 |
| Cutc | 0.008785694 | 0.659449839 | 0.9211780 | 0.098280613 | 0.08847534 |
| Cux1 | -0.029468816 | 0.34022397 | 0.7844658 | 0.260327621 | 0.277784786 |
| Cux2 | -0.003000895 | 0.883756223 | 0.9801988 | 0.094938319 | 0.093190453 |
| Cwc15 | -0.00076972 | 0.982533161 | 0.9976072 | 0.620214109 | 0.623758739 |
| Cwc22 | 0.035540314 | 0.079916887 | 0.5081894 | 0.110256542 | 0.07819613 |
| Cwc25 | -0.030291284 | 0.296758123 | 0.7545964 | 0.192838656 | 0.216840572 |
| Cwc27 | 0.015348045 | 0.536115711 | 0.8799251 | 0.148079311 | 0.143076582 |
| Cwf19l1 | -0.000945438 | 0.949316895 | 0.9948462 | 0.048587335 | 0.044120666 |
| Cwf19l2 | 0.010790352 | 0.740189444 | 0.9479080 | 0.289788752 | 0.28289834 |
| Cxadr | -0.052879978 | 0.049798013 | 0.4200648 | 0.136764263 | 0.192043222 |
| Cxxc1 | -0.001965686 | 0.93695362 | 0.9913161 | 0.147968187 | 0.141108673 |
| Cxxc4 | 0.071047094 | 0.043682017 | 0.3986436 | 0.499845173 | 0.443914385 |
| Cxxc5 | 0.035574527 | 0.344636601 | 0.7879223 | 0.72292349 | 0.664144374 |
| Cyb561 | -0.01215555 | 0.712659273 | 0.9421288 | 0.332609147 | 0.342751331 |
| Cyb561d2 | 0.026305052 | 0.252776089 | 0.7223071 | 0.137527554 | 0.11751274 |
| Cyb5a | -0.068706884 | 0.041034261 | 0.3901716 | 0.294561146 | 0.394010951 |
| Cyb5b | 0.027809824 | 0.410158361 | 0.8283287 | 0.388258009 | 0.382740498 |
| Cyb5d2 | -0.023078288 | 0.199016822 | 0.6730962 | 0.059757599 | 0.074665555 |
| Cyb5r1 | -0.057452047 | 0.091114455 | 0.5321282 | 0.200160849 | 0.315522396 |
| Cyb5r3 | -0.061441158 | 0.064084908 | 0.4700387 | 1.193940394 | 1.199811748 |
| Cyb5r4 | -0.003503716 | 0.894834698 | 0.9824361 | 0.156658136 | 0.166232051 |
| Cyb5rl | -0.034264979 | 0.037802848 | 0.3701917 | 0.044915092 | 0.062840514 |
| Cybc1 | -0.019915125 | 0.230187572 | 0.7019677 | 0.04533276 | 0.059829811 |
| Cyc1 | 0.000572072 | 0.98762995 | 0.9976072 | 0.619019161 | 0.590433504 |
| Cycs | 0.088582569 | 0.018669666 | 0.2672483 | 1.035950286 | 0.824254819 |
| Cyfip1 | 0.003578549 | 0.902854803 | 0.9825441 | 0.259853997 | 0.229031569 |
| Cyfip2 | 0.057416919 | 0.128375231 | 0.5914729 | 0.664206641 | 0.688436536 |
| Cyhr1 | -0.047579873 | 0.172726217 | 0.6524920 | 0.534761143 | 0.558131622 |
| Cyld | 0.03325092 | 0.291890001 | 0.7520906 | 0.312482087 | 0.278151518 |
| Cyp20a1 | 0.008634793 | 0.754811969 | 0.9519143 | 0.183382665 | 0.173049423 |
| Cyp2j6 | -0.018306232 | 0.243452554 | 0.7132719 | 0.042095083 | 0.055226666 |
| Cyp2u1 | -0.019346502 | 0.256257745 | 0.7238259 | 0.04699535 | 0.065903661 |
| Cyp4f13 | -0.056840201 | 0.008672376 | 0.1846136 | 0.076851912 | 0.121412547 |
| Cyp4f16 | -0.023623631 | 0.138891477 | 0.6093779 | 0.043987012 | 0.05990649 |

| | | | | | |
|---|---|---|---|---|---|
| Cyp4f17 | -0.016776751 | 0.273658374 | 0.7351955 | 0.045178322 | 0.051156267 |
| Cyp4v3 | -0.04797757 | 0.067616044 | 0.4814530 | 0.136487458 | 0.164253005 |
| Cyp51 | 0.018214536 | 0.506120134 | 0.8695566 | 0.205986199 | 0.177017421 |
| Cyp7b1 | 0.009450066 | 0.517462964 | 0.8738804 | 0.053997214 | 0.040918033 |
| Cyren | 0.052487449 | 0.023340069 | 0.2964577 | 0.157331152 | 0.106427776 |
| Cyrib | 0.029732489 | 0.379759736 | 0.8115177 | 0.421524343 | 0.355903945 |
| Cystm1 | -0.02361282 | 0.468204157 | 0.8561686 | 1.551381575 | 1.414166383 |
| Cyth1 | -0.003148995 | 0.891236028 | 0.9819597 | 0.121805338 | 0.125104663 |
| Cyth2 | -0.057050669 | 0.069981507 | 0.4848331 | 0.281300139 | 0.30839699 |
| Cyth3 | -0.031337948 | 0.069035001 | 0.4835495 | 0.045979706 | 0.073131134 |
| Czib | -0.030289385 | 0.359577876 | 0.8003169 | 0.325597014 | 0.328234547 |
| D030056L22Rik | 0.015543984 | 0.503811685 | 0.8695566 | 0.132010012 | 0.111784316 |
| D10Wsu102e | 0.003048145 | 0.917411878 | 0.9848120 | 0.227160314 | 0.200642999 |
| D11Wsu47e | -0.023699763 | 0.127310969 | 0.5898761 | 0.037699428 | 0.057484269 |
| D130020L05Rik | 0.022304993 | 0.366960226 | 0.8053794 | 0.160210795 | 0.137176749 |
| D16Ertd472e | 0.013035151 | 0.408851307 | 0.8276447 | 0.063108483 | 0.042997959 |
| D17H6S53E | 0.015356524 | 0.384376524 | 0.8135348 | 0.089875991 | 0.05738357 |
| D230025D16Rik | 0.007670898 | 0.742426178 | 0.9479080 | 0.146955009 | 0.121551208 |
| D330023K18Rik | 0.010308301 | 0.713770478 | 0.9428150 | 0.214541049 | 0.197730831 |
| D330050I16Rik | -0.026809884 | 0.137489675 | 0.6070876 | 0.050142203 | 0.073927469 |
| D3Ertd751e | -0.040890619 | 0.106227956 | 0.5617171 | 0.125416368 | 0.161883841 |
| D5Ertd579e | 0.031956305 | 0.333387749 | 0.7804836 | 0.340248932 | 0.331248255 |
| D630024D03Rik | -0.000415774 | 0.985404156 | 0.9976072 | 0.102827481 | 0.126574411 |
| D630039A03Rik | -0.007541671 | 0.643229409 | 0.9166712 | 0.056144245 | 0.057904807 |
| D630045J12Rik | 0.014842327 | 0.329890528 | 0.7778857 | 0.052658986 | 0.044366633 |
| D6Wsu163e | 8.91E-05 | 0.995896188 | 0.9994073 | 0.064889295 | 0.063913222 |
| D730003I15Rik | 0.00530317 | 0.883170919 | 0.9800934 | 0.431069842 | 0.473123631 |
| D830050J10Rik | -0.026883173 | 0.094239057 | 0.5334417 | 0.038712819 | 0.06036192 |
| D8Ertd738e | 0.016822351 | 0.581069849 | 0.9015399 | 1.34961467 | 1.301108879 |
| Daam1 | -0.016082408 | 0.66386113 | 0.9230765 | 0.486560314 | 0.537314111 |
| Dab2ip | -0.017622413 | 0.534982292 | 0.8795949 | 0.181371907 | 0.200604468 |
| Dach1 | -0.061817828 | 0.055843377 | 0.4449332 | 0.250662427 | 0.310190297 |
| Dact1 | -0.009548525 | 0.610106951 | 0.9090802 | 0.073931894 | 0.073013564 |
| Dact2 | 0.001749007 | 0.936845142 | 0.9913070 | 0.131538918 | 0.100433273 |
| Dact3 | 0.006594336 | 0.648176618 | 0.9170676 | 0.051434586 | 0.039008109 |
| Dad1 | 0.016257089 | 0.446975434 | 0.8448925 | 2.364680916 | 2.28480705 |
| Dag1 | -0.030986751 | 0.334463 | 0.7804836 | 0.268113757 | 0.305904076 |
| Dagla | -0.006166523 | 0.698712485 | 0.9369692 | 0.048329899 | 0.056148562 |
| Daglb | -0.004791603 | 0.765626967 | 0.9541481 | 0.0518761 | 0.055822137 |
| Dalrd3 | -0.030692418 | 0.290442379 | 0.7508961 | 0.212468419 | 0.222360484 |
| Dancr | -0.019529649 | 0.473654281 | 0.8597092 | 0.174004977 | 0.17708331 |
| Dand5 | -0.026424137 | 0.128104076 | 0.5912244 | 0.050379147 | 0.071692641 |
| Dap | 0.040273388 | 0.233411717 | 0.7051391 | 1.278687388 | 1.168973998 |
| Dap3 | 0.031083554 | 0.372040604 | 0.8085689 | 0.523773504 | 0.446134735 |
| Dapk1 | 0.054531282 | 0.070730508 | 0.4879008 | 0.273582238 | 0.220020925 |
| Dapk3 | -0.020842501 | 0.534884994 | 0.8795949 | 0.332748719 | 0.32880918 |
| Dapl1 | 0.119101382 | 0.019238129 | 0.2695586 | 0.595410149 | 0.475128676 |
| Dapp1 | -0.002976262 | 0.88509686 | 0.9803788 | 0.08818716 | 0.094461723 |
| Dars | 0.012688036 | 0.680048261 | 0.9290839 | 0.298749238 | 0.268554281 |
| Dars2 | 0.011047327 | 0.44396292 | 0.8431541 | 0.051230839 | 0.03899623 |
| Daxx | 0.00494246 | 0.841427212 | 0.9734589 | 0.149268004 | 0.1349873 |
| Dazap1 | 0.054456939 | 0.113592257 | 0.5696977 | 0.473153503 | 0.401681125 |
| Dazap2 | -0.040675344 | 0.228342293 | 0.7015153 | 0.364147296 | 0.389762652 |
| Dbi | 0.091737225 | 0.010196583 | 0.1999013 | 0.500827175 | 0.418609371 |
| Dbnl | -0.0613962 | 0.036253779 | 0.3610818 | 0.216834007 | 0.247531372 |
| Dbp | 0.035680661 | 0.209749211 | 0.6807660 | 0.218159451 | 0.161431462 |
| Dbpht2 | -0.178842901 | 4.98E-05 | 0.0055597 | 0.737781239 | 0.776359519 |
| Dbr1 | -0.012550012 | 0.604602599 | 0.9087139 | 0.139638574 | 0.137587105 |
| Dbt | -0.015583846 | 0.396512653 | 0.8237498 | 0.056351371 | 0.078665617 |
| Dcaf1 | -0.001976013 | 0.943737051 | 0.9935293 | 0.199509752 | 0.192260414 |
| Dcaf10 | 0.006911822 | 0.797011777 | 0.9644244 | 0.173108907 | 0.170299352 |
| Dcaf11 | -0.033042923 | 0.334198621 | 0.7804836 | 0.402752351 | 0.39277267 |
| Dcaf12 | 0.013658773 | 0.549666141 | 0.8874817 | 0.124206922 | 0.117461014 |
| Dcaf12l1 | 0.042148904 | 0.092123832 | 0.5325584 | 0.147829704 | 0.13256714 |
| Dcaf13 | 0.028848113 | 0.343261506 | 0.7872955 | 0.283388821 | 0.233616788 |
| Dcaf15 | 0.005428711 | 0.772178714 | 0.9552333 | 0.0821566 | 0.073449034 |

| | | | | | |
|---|---|---|---|---|---|
| Dcaf17 | -0.034463743 | 0.183779135 | 0.6617383 | 0.134811456 | 0.164786086 |
| Dcaf5 | 0.004483261 | 0.868048842 | 0.9775280 | 0.184883848 | 0.177246069 |
| Dcaf6 | 0.014240488 | 0.635031872 | 0.9150953 | 0.253743799 | 0.232941968 |
| Dcaf7 | 0.029613678 | 0.374976348 | 0.8093484 | 0.372726606 | 0.356116585 |
| Dcaf8 | -0.011525833 | 0.745931293 | 0.9485717 | 0.557452699 | 0.545847935 |
| Dcakd | 0.017178528 | 0.32620374 | 0.7765637 | 0.081659389 | 0.060716505 |
| Dcbld1 | 0.010683285 | 0.531166263 | 0.8776448 | 0.065245429 | 0.058778367 |
| Dcbld2 | 0.017051008 | 0.31834273 | 0.7710043 | 0.065454546 | 0.05446907 |
| Dcdc2a | -0.021372264 | 0.306543014 | 0.7652785 | 0.059055931 | 0.097658327 |
| Dclk2 | -0.054021771 | 0.006175171 | 0.1574747 | 0.055392527 | 0.091974774 |
| Dclre1a | -0.007806837 | 0.620055891 | 0.9121505 | 0.050014187 | 0.053059643 |
| Dclre1c | -0.033422487 | 0.032063802 | 0.3392328 | 0.035585728 | 0.057003714 |
| Dcp1a | 0.012634592 | 0.496543843 | 0.8684629 | 0.081464414 | 0.069812286 |
| Dcp1b | 0.026574592 | 0.116387978 | 0.5724670 | 0.078051053 | 0.051945114 |
| Dcp2 | -0.009750894 | 0.698966443 | 0.9369928 | 0.154371893 | 0.155420703 |
| Dcps | -0.013044064 | 0.641779435 | 0.9166712 | 0.197924297 | 0.191154463 |
| Dctn1 | -0.042429353 | 0.1799622 | 0.6550362 | 0.278697733 | 0.314038269 |
| Dctn2 | -0.023608942 | 0.505752358 | 0.8695566 | 0.62158835 | 0.611151998 |
| Dctn3 | 0.033283011 | 0.302235818 | 0.7611258 | 1.167663233 | 1.05948054 |
| Dctn4 | 0.018546795 | 0.575711234 | 0.8989541 | 0.343953491 | 0.308497339 |
| Dctn5 | 0.007560066 | 0.75154327 | 0.9506053 | 0.142254439 | 0.121945907 |
| Dctn6 | 0.022714994 | 0.500596295 | 0.8693704 | 0.40875138 | 0.359260393 |
| Dctpp1 | 0.002690552 | 0.921417776 | 0.9861327 | 0.194373451 | 0.188852348 |
| Dcun1d1 | 0.001395588 | 0.955712476 | 0.9954690 | 0.164273058 | 0.149581801 |
| Dcun1d2 | -0.018562805 | 0.358227703 | 0.7992844 | 0.078112821 | 0.091539458 |
| Dcun1d3 | 0.016350147 | 0.490067585 | 0.8655242 | 0.129025997 | 0.126147898 |
| Dcun1d4 | 0.025176965 | 0.206285095 | 0.6797565 | 0.10140308 | 0.080586778 |
| Dcun1d5 | 0.019667554 | 0.589106533 | 0.9046539 | 0.554631336 | 0.48939287 |
| Dcx | 0.00798714 | 0.646491006 | 0.9169695 | 0.059694634 | 0.06385491 |
| Dcxr | 0.010952347 | 0.607127842 | 0.9087139 | 0.10642649 | 0.096182409 |
| Dda1 | 0.046850337 | 0.17327268 | 0.6525507 | 0.438411095 | 0.365675085 |
| Ddb1 | -0.055671783 | 0.129042139 | 0.5918160 | 0.723163699 | 0.747755232 |
| Ddc | 0.114518623 | 0.006940329 | 0.1662996 | 0.728655901 | 0.617599833 |
| Ddhd1 | -0.083441908 | 0.002303883 | 0.0868993 | 0.125052488 | 0.188630053 |
| Ddhd2 | -0.079222256 | 0.015382292 | 0.2425875 | 0.283628497 | 0.345629627 |
| Ddi2 | 0.022649448 | 0.44395742 | 0.8431541 | 0.246253768 | 0.22295649 |
| Ddit3 | 0.088987973 | 0.036047287 | 0.3601098 | 0.7817826 | 0.793974729 |
| Ddit4 | -0.066785891 | 0.047955686 | 0.4136699 | 0.264047696 | 0.325596749 |
| Ddo | -0.002631255 | 0.852857554 | 0.9755431 | 0.046491873 | 0.042124881 |
| Ddost | -0.070384022 | 0.025976418 | 0.3089760 | 1.500991169 | 1.483993733 |
| Ddr1 | -0.069936524 | 0.041062422 | 0.3901716 | 0.367208383 | 0.403298131 |
| Ddrgk1 | 0.007864107 | 0.82933178 | 0.9722601 | 0.811799634 | 0.786067693 |
| Ddt | -0.038109741 | 0.296405205 | 0.7545964 | 0.595472553 | 0.607443565 |
| Ddx1 | 0.015583693 | 0.656551318 | 0.9211780 | 0.542170746 | 0.486577171 |
| Ddx10 | 0.050790404 | 0.036774566 | 0.3626279 | 0.166286935 | 0.117694535 |
| Ddx17 | -0.126622831 | 0.000610527 | 0.0350454 | 0.52068549 | 0.666881865 |
| Ddx18 | 0.049352857 | 0.119446997 | 0.5783300 | 0.310720395 | 0.27496134 |
| Ddx19a | 0.018884692 | 0.43357173 | 0.8377544 | 0.153728167 | 0.129346365 |
| Ddx19b | 0.008051516 | 0.685978899 | 0.9318810 | 0.085072691 | 0.090188629 |
| Ddx20 | 0.006237115 | 0.690676031 | 0.9346237 | 0.053769299 | 0.049692213 |
| Ddx21 | -0.023589182 | 0.476018432 | 0.8607231 | 0.301966889 | 0.31765138 |
| Ddx23 | 0.052066208 | 0.090586625 | 0.5321031 | 0.279133491 | 0.226252758 |
| Ddx24 | 0.016702214 | 0.626981746 | 0.9128095 | 1.029868197 | 1.052580454 |
| Ddx25 | 0.010471194 | 0.64728818 | 0.9169695 | 0.136656048 | 0.116314935 |
| Ddx27 | -0.00855578 | 0.762072854 | 0.9529513 | 0.20844142 | 0.203412055 |
| Ddx28 | -0.003074123 | 0.845802492 | 0.9736268 | 0.051610499 | 0.05257354 |
| Ddx31 | 0.004078225 | 0.829756205 | 0.9722601 | 0.082470397 | 0.077088296 |
| Ddx39a | -0.018955805 | 0.461115602 | 0.8524425 | 0.157057985 | 0.155050561 |
| Ddx39b | -0.131139026 | 0.000357747 | 0.0241418 | 0.688321717 | 0.747553634 |
| Ddx3x | 0.026067347 | 0.501576088 | 0.8695566 | 0.732486617 | 0.73661458 |
| Ddx3y | -0.024829694 | 0.500280252 | 0.8693704 | 0.450081437 | 0.464085074 |
| Ddx41 | -0.070334155 | 0.050404101 | 0.4223046 | 0.467040092 | 0.527732634 |
| Ddx42 | 0.025935933 | 0.47967061 | 0.8610807 | 0.575007336 | 0.556673685 |
| Ddx46 | -0.026047065 | 0.461979103 | 0.8524425 | 0.389935305 | 0.415324954 |
| Ddx47 | -0.059524187 | 0.045073076 | 0.4027410 | 0.195146487 | 0.237811168 |
| Ddx49 | -0.008483312 | 0.772467931 | 0.9552333 | 0.215406903 | 0.211492072 |

| | | | | | |
|---|---|---|---|---|---|
| Ddx5 | -0.064956274 | 0.002658007 | 0.0948468 | 2.548458253 | 2.641786088 |
| Ddx50 | -0.084204891 | 0.023494201 | 0.2972960 | 1.138830854 | 1.180469857 |
| Ddx52 | -0.018717392 | 0.495639724 | 0.8684629 | 0.188204289 | 0.191573566 |
| Ddx54 | 0.016995862 | 0.529542504 | 0.8776448 | 0.207689201 | 0.181760175 |
| Ddx55 | -0.017229622 | 0.514992216 | 0.8733714 | 0.176584325 | 0.170161174 |
| Ddx56 | 0.031569328 | 0.220682456 | 0.6925561 | 0.193488864 | 0.147198411 |
| Ddx58 | -0.019770165 | 0.391547723 | 0.8200150 | 0.103149009 | 0.121655903 |
| Ddx59 | 0.035345955 | 0.020730814 | 0.2798878 | 0.072584382 | 0.039392004 |
| Ddx6 | -0.060953871 | 0.100733988 | 0.5480085 | 1.043311072 | 1.054389053 |
| Ddx60 | 0.014335208 | 0.36997597 | 0.8065581 | 0.05356679 | 0.047059559 |
| Deaf1 | 0.019140779 | 0.341094832 | 0.7851366 | 0.101232233 | 0.08268618 |
| Decr1 | 0.060390721 | 0.005966763 | 0.1545438 | 0.154235011 | 0.101109238 |
| Decr2 | -0.033362527 | 0.101212405 | 0.5492489 | 0.079567342 | 0.097478829 |
| Dedd | -0.031180614 | 0.205835338 | 0.6797565 | 0.139463385 | 0.151949393 |
| Dedd2 | -0.020377467 | 0.329518353 | 0.7778857 | 0.084835746 | 0.100767647 |
| Def8 | -0.009290754 | 0.628872494 | 0.9130564 | 0.073605561 | 0.081556473 |
| Defb1 | 0.040189225 | 0.383802105 | 0.8129787 | 0.690553709 | 0.735489106 |
| Degs1 | 0.023299511 | 0.527203274 | 0.8768553 | 0.609660233 | 0.592012495 |
| Dek | -0.00503913 | 0.892261593 | 0.9820058 | 0.615050672 | 0.559859962 |
| Dele1 | 0.00293827 | 0.909451422 | 0.9837373 | 0.181096841 | 0.159834212 |
| Denn2b | -0.012057962 | 0.608772703 | 0.9088672 | 0.104787766 | 0.128508752 |
| Dennd10 | 0.010431298 | 0.678363664 | 0.9281886 | 0.171635205 | 0.144395387 |
| Dennd11 | -0.030518143 | 0.334360592 | 0.7804836 | 0.255805941 | 0.284091184 |
| Dennd1a | 0.002505261 | 0.924572304 | 0.9878036 | 0.165456387 | 0.162099405 |
| Dennd1b | -0.056373509 | 0.03526554 | 0.3577036 | 0.139033796 | 0.180097128 |
| Dennd2a | -0.018269335 | 0.261509861 | 0.7275735 | 0.05213722 | 0.051859776 |
| Dennd2d | -0.050385604 | 0.006129068 | 0.1568368 | 0.040522294 | 0.083464951 |
| Dennd4a | -0.034532675 | 0.231327017 | 0.7026141 | 0.179565561 | 0.224167584 |
| Dennd4c | 0.047475717 | 0.149395717 | 0.6264374 | 0.361234231 | 0.333971705 |
| Dennd5a | -0.005369676 | 0.847756196 | 0.9744069 | 0.196816917 | 0.192838527 |
| Dennd5b | -0.028171704 | 0.367655809 | 0.8056719 | 0.246710322 | 0.257723113 |
| Dennd6a | -0.007837394 | 0.792311631 | 0.9624460 | 0.231956327 | 0.228427251 |
| Denr | 0.065528293 | 0.069880517 | 0.4847655 | 1.02597001 | 0.947526516 |
| Depdc5 | -0.018268493 | 0.270718636 | 0.7342030 | 0.050351547 | 0.062347145 |
| Depp1 | -0.091903976 | 0.00797914 | 0.1758957 | 0.138224279 | 0.253232876 |
| Deptor | 0.008309044 | 0.818456114 | 0.9697903 | 0.504875192 | 0.484670326 |
| Dera | 0.026078569 | 0.259191296 | 0.7263213 | 0.147819359 | 0.113008049 |
| Derl1 | -0.010518528 | 0.780921998 | 0.9590847 | 0.606125751 | 0.664892601 |
| Derl2 | 0.032876935 | 0.345103405 | 0.7880440 | 0.448995133 | 0.404086563 |
| Desi1 | 0.035320865 | 0.321626527 | 0.7732349 | 0.514322953 | 0.45750659 |
| Desi2 | 0.096624477 | 0.007529431 | 0.1721437 | 0.562920648 | 0.449944098 |
| Dexi | 0.006043895 | 0.810716948 | 0.9668615 | 0.173493721 | 0.157678409 |
| Dffa | -0.000670436 | 0.974534154 | 0.9976072 | 0.093373905 | 0.095568189 |
| Dffb | 0.018018083 | 0.301760025 | 0.7609383 | 0.068690553 | 0.060264749 |
| Dgat1 | 0.021171835 | 0.237035113 | 0.7095317 | 0.089957189 | 0.064409035 |
| Dgcr2 | -0.047068079 | 0.084390185 | 0.5191841 | 0.149032766 | 0.195642377 |
| Dgcr6 | 0.028056272 | 0.398485984 | 0.8239850 | 0.393747976 | 0.363810313 |
| Dgcr8 | 0.005072602 | 0.791341014 | 0.9623764 | 0.081192642 | 0.074891171 |
| Dgkd | 0.028990844 | 0.306737208 | 0.7652785 | 0.217319541 | 0.197413422 |
| Dgke | 0.0090359 | 0.655112143 | 0.9206439 | 0.089549808 | 0.092869675 |
| Dgkg | 0.007071622 | 0.733762663 | 0.9464134 | 0.077500037 | 0.098553125 |
| Dgkh | 0.035944756 | 0.115976465 | 0.5724670 | 0.124336595 | 0.112941587 |
| Dgkq | -0.013609463 | 0.4232651 | 0.8317437 | 0.049928214 | 0.063440185 |
| Dgkz | 0.022553229 | 0.392434259 | 0.8204315 | 0.177681873 | 0.161812505 |
| Dguok | -0.051017884 | 0.130703913 | 0.5950357 | 0.427689545 | 0.451612447 |
| Dhcr7 | 0.015749979 | 0.321986474 | 0.7733024 | 0.069322932 | 0.046576256 |
| Dhdds | -0.029136467 | 0.315876594 | 0.7705987 | 0.21161367 | 0.22918713 |
| Dhdh | 0.019713606 | 0.337732929 | 0.7834245 | 0.109657445 | 0.08743487 |
| Dhfr | -0.033946941 | 0.0678247 | 0.4817235 | 0.057158648 | 0.076387247 |
| Dhps | -0.025217002 | 0.43845737 | 0.8403201 | 0.298398192 | 0.300648032 |
| Dhrs1 | -0.007953666 | 0.738413585 | 0.9474220 | 0.129271187 | 0.130901834 |
| Dhrs11 | 0.011809629 | 0.373051249 | 0.8085689 | 0.045319784 | 0.031813507 |
| Dhrs13 | 0.034287274 | 0.026284052 | 0.3100330 | 0.0728299 | 0.039973234 |
| Dhrs4 | 0.022130263 | 0.377294029 | 0.8101797 | 0.159479979 | 0.148917505 |
| Dhrs7 | 0.018597884 | 0.617151562 | 0.9115017 | 0.579815267 | 0.582372742 |
| Dhrs7b | -0.011158639 | 0.619962346 | 0.9121505 | 0.10866911 | 0.112063588 |

| | | | | | |
|---|---|---|---|---|---|
| Dhx15 | 0.037446173 | 0.286746557 | 0.74685807 | 0.41887665 | 0.393814942 |
| Dhx16 | -0.000984692 | 0.961663928 | 0.99635575 | 0.089964482 | 0.093185418 |
| Dhx29 | 0.024877554 | 0.350461987 | 0.79273039 | 0.188307222 | 0.166206356 |
| Dhx30 | -0.008879632 | 0.773508918 | 0.95573117 | 0.292205371 | 0.275079099 |
| Dhx32 | 0.016781436 | 0.631141035 | 0.91335177 | 0.40574114 | 0.383500009 |
| Dhx33 | -0.016007187 | 0.311990876 | 0.76843798 | 0.050793333 | 0.053957707 |
| Dhx34 | -0.03243652 | 0.068271049 | 0.48338697 | 0.054538699 | 0.07493972 |
| Dhx36 | 0.075666799 | 0.041370838 | 0.39160182 | 0.678586675 | 0.5848961 |
| Dhx38 | -0.02706296 | 0.207207382 | 0.67995277 | 0.088058594 | 0.107415271 |
| Dhx40 | 0.002123004 | 0.950170811 | 0.99484627 | 0.386837155 | 0.348621402 |
| Dhx57 | -0.012447378 | 0.603828877 | 0.90871392 | 0.126060556 | 0.130451999 |
| Dhx8 | -0.031717197 | 0.204800119 | 0.67856285 | 0.127196454 | 0.152043767 |
| Dhx9 | -0.057451367 | 0.100955693 | 0.54873570 | 0.350711036 | 0.396879574 |
| Diablo | -0.034431227 | 0.258064513 | 0.72511021 | 0.263125591 | 0.264490342 |
| Diaph1 | -0.043941867 | 0.191590812 | 0.67002813 | 0.370789714 | 0.374490162 |
| Diaph2 | 0.013767105 | 0.580612868 | 0.90135831 | 0.141699296 | 0.143734345 |
| Dicer1 | 0.018093296 | 0.604845741 | 0.90871392 | 0.522957286 | 0.515618465 |
| Dido1 | -0.008227486 | 0.785072551 | 0.96025161 | 0.246115625 | 0.241456147 |
| Dimt1 | 0.023903197 | 0.333847309 | 0.78048367 | 0.163118355 | 0.139098792 |
| Dio1 | 0.066439564 | 0.012802463 | 0.22335359 | 0.185056071 | 0.158697179 |
| Dip2b | 0.006577364 | 0.807083576 | 0.96529303 | 0.162419188 | 0.175172878 |
| Dip2c | 0.031729061 | 0.295895593 | 0.75459642 | 0.275655362 | 0.243626407 |
| Dipk1b | 0.016430841 | 0.421059393 | 0.83009907 | 0.09663906 | 0.096982876 |
| Dipk2a | 0.028784369 | 0.203341288 | 0.67727403 | 0.133994776 | 0.108433449 |
| Dis3 | -0.019862365 | 0.349197886 | 0.79273039 | 0.088852094 | 0.102438707 |
| Dis3l | 0.003175085 | 0.834945109 | 0.97234942 | 0.049056756 | 0.04944125 |
| Dis3l2 | 0.005227863 | 0.786366032 | 0.96044700 | 0.085840076 | 0.074923032 |
| Disp2 | -0.070184309 | 0.049301329 | 0.41872361 | 0.437402569 | 0.537573566 |
| Dixdc1 | -0.04471337 | 0.086178166 | 0.52379073 | 0.154354336 | 0.157899015 |
| Dkc1 | 0.006421351 | 0.792730058 | 0.96244609 | 0.142079 | 0.143430794 |
| Dlat | 0.078720445 | 0.014671942 | 0.23765848 | 0.37174744 | 0.283477183 |
| Dld | -0.017802074 | 0.59318273 | 0.90605850 | 0.363424482 | 0.347040855 |
| Dleu2 | -0.01205265 | 0.664476342 | 0.92307657 | 0.168420428 | 0.173513027 |
| Dlg1 | -0.02375857 | 0.401850084 | 0.82542593 | 0.179945062 | 0.205346223 |
| Dlg3 | 0.030028322 | 0.157793104 | 0.63244751 | 0.118738888 | 0.095372955 |
| Dlg4 | -0.001629087 | 0.95433959 | 0.99546908 | 0.192381744 | 0.194200411 |
| Dlg5 | -0.009666603 | 0.61763815 | 0.91150170 | 0.06710695 | 0.085324465 |
| Dlgap1 | 0.013232195 | 0.439347152 | 0.84056388 | 0.059451322 | 0.062838872 |
| Dlgap4 | 0.008119004 | 0.780220906 | 0.95908199 | 0.228796517 | 0.222900181 |
| Dlk1 | 0.024208841 | 0.373253191 | 0.80856898 | 0.124892975 | 0.090855444 |
| Dlst | -0.047988105 | 0.122846731 | 0.58332371 | 0.25477969 | 0.282703946 |
| Dmac1 | 0.009287007 | 0.768035385 | 0.95442217 | 0.302176791 | 0.277881561 |
| Dmac2 | 0.027529634 | 0.19076573 | 0.67002813 | 0.125857357 | 0.087673869 |
| Dmac2l | 0.021100532 | 0.384634 | 0.81360648 | 0.174265915 | 0.125236221 |
| Dmap1 | -0.028426531 | 0.166502029 | 0.64292850 | 0.074518338 | 0.102052181 |
| Dmd | -0.054500908 | 0.006615082 | 0.16179671 | 0.065905185 | 0.092307264 |
| Dmpk | 0.014711706 | 0.556929169 | 0.89159442 | 0.133934014 | 0.150142984 |
| Dmrtc1a | -0.029238827 | 0.221022491 | 0.69262049 | 0.147918564 | 0.134494918 |
| Dmtf1 | 0.104072736 | 0.000465967 | 0.02925567 | 0.298985598 | 0.189910321 |
| Dmwd | 0.012899356 | 0.643113692 | 0.91667127 | 0.218459296 | 0.18453323 |
| Dmxl1 | 0.0075157 | 0.834287141 | 0.97234942 | 0.404751739 | 0.397187486 |
| Dmxl2 | 0.088906716 | 0.003060968 | 0.10328859 | 0.282631778 | 0.23533337 |
| Dnah11 | -0.009481158 | 0.618056413 | 0.91150170 | 0.071640351 | 0.066764859 |
| Dnaja1 | -0.065650378 | 0.07120719 | 0.4881297 | 0.786725182 | 0.815603139 |
| Dnaja2 | -0.001405064 | 0.968477136 | 0.99760728 | 0.724093201 | 0.67397262 |
| Dnaja3 | 0.00881439 | 0.76159452 | 0.95286310 | 0.241954305 | 0.22192608 |
| Dnaja4 | -0.042090317 | 0.209788481 | 0.68076602 | 0.308335494 | 0.371230872 |
| Dnajb1 | -0.158261156 | 9.65E-05 | 0.00903437 | 0.294368524 | 0.533343545 |
| Dnajb11 | 0.006363567 | 0.864392112 | 0.97696750 | 0.46391155 | 0.467402799 |
| Dnajb12 | -0.013119974 | 0.652939995 | 0.91920173 | 0.221270144 | 0.21346732 |
| Dnajb14 | -0.009209116 | 0.802620125 | 0.96471431 | 0.550365016 | 0.557681186 |
| Dnajb2 | 0.017692605 | 0.591893771 | 0.90602008 | 0.375809785 | 0.340991262 |
| Dnajb3 | -0.000390892 | 0.981614676 | 0.99760728 | 0.067996746 | 0.058221412 |
| Dnajb4 | -0.059296346 | 0.061601956 | 0.46137448 | 0.226973554 | 0.293992035 |
| Dnajb6 | 0.014939848 | 0.660521535 | 0.92117801 | 0.92166048 | 0.8656187 |
| Dnajb9 | -0.070746349 | 0.072042422 | 0.48882409 | 1.062075274 | 1.141789345 |

| | | | | | |
|---|---|---|---|---|---|
| Dnajc1 | 0.02487296 | 0.500401948 | 0.8693704 | 0.638605882 | 0.624095694 |
| Dnajc10 | -0.010782262 | 0.752262663 | 0.9507898 | 0.356492146 | 0.399844287 |
| Dnajc11 | -0.029761843 | 0.171319991 | 0.6511472 | 0.087122505 | 0.106870254 |
| Dnajc12 | -0.023010474 | 0.535557415 | 0.8795949 | 0.826793771 | 0.759942872 |
| Dnajc13 | 0.012581997 | 0.622655568 | 0.9121505 | 0.164467348 | 0.159535249 |
| Dnajc14 | 0.028932682 | 0.267860002 | 0.7323841 | 0.211453934 | 0.167722933 |
| Dnajc15 | 0.064459656 | 0.055754098 | 0.4449332 | 0.470980092 | 0.403715916 |
| Dnajc17 | 0.007475252 | 0.745661726 | 0.9485717 | 0.126190793 | 0.117540345 |
| Dnajc18 | 0.032270199 | 0.201993582 | 0.6752150 | 0.186464993 | 0.14783827 |
| Dnajc19 | 0.038277438 | 0.272313492 | 0.7342806 | 0.697990746 | 0.65697314 |
| Dnajc2 | -0.018999307 | 0.571751688 | 0.8966165 | 0.351099442 | 0.35813 |
| Dnajc21 | 0.013726756 | 0.7064273 | 0.9403863 | 0.823753703 | 0.806164994 |
| Dnajc24 | 0.072878899 | 0.020743887 | 0.2798878 | 0.320728294 | 0.274846426 |
| Dnajc25 | 0.001822861 | 0.950051815 | 0.9948462 | 0.238327527 | 0.209214437 |
| Dnajc27 | 0.005794216 | 0.748671958 | 0.9501120 | 0.073856923 | 0.066144215 |
| Dnajc3 | -0.025618849 | 0.397985564 | 0.8238469 | 1.633004791 | 1.658409132 |
| Dnajc30 | 0.014589261 | 0.597249251 | 0.9075513 | 0.219029734 | 0.184523316 |
| Dnajc4 | -0.036538277 | 0.254968637 | 0.7223665 | 0.296516994 | 0.322234476 |
| Dnajc5 | -0.016149902 | 0.639540896 | 0.9160042 | 0.421350452 | 0.426715313 |
| Dnajc6 | -0.026527865 | 0.223296216 | 0.6955958 | 0.087494017 | 0.114837067 |
| Dnajc7 | -0.093327998 | 0.01120786 | 0.2082059 | 0.414928709 | 0.488149599 |
| Dnajc8 | 0.030215806 | 0.400616097 | 0.8246755 | 0.626693931 | 0.573995859 |
| Dnajc9 | -0.003086086 | 0.923474694 | 0.9874818 | 0.295562669 | 0.293517436 |
| Dnal1 | 0.005361362 | 0.797470657 | 0.9644244 | 0.097922572 | 0.090030347 |
| Dnal4 | 0.032107279 | 0.216720796 | 0.6872330 | 0.185044927 | 0.164764905 |
| Dnase2a | 0.041020223 | 0.211185785 | 0.6822059 | 0.303703823 | 0.255742515 |
| Dner | 0.009107424 | 0.622454594 | 0.9121505 | 0.078183802 | 0.074276915 |
| Dnlz | -0.002687142 | 0.939442277 | 0.9927844 | 0.506106764 | 0.484470909 |
| Dnm1l | 0.003861324 | 0.903749598 | 0.9828103 | 0.272511744 | 0.262988565 |
| Dnm2 | 0.015677171 | 0.634720736 | 0.9147798 | 0.403939672 | 0.367751828 |
| Dnm3 | -0.092227715 | 0.004620273 | 0.1360032 | 0.221869106 | 0.311846694 |
| Dnmt1 | -0.001183847 | 0.963154407 | 0.9966421 | 0.148548977 | 0.153062965 |
| Dnmt3a | -0.038959968 | 0.285274927 | 0.7451828 | 0.450793711 | 0.479802889 |
| Dnpep | -0.076421398 | 0.038423502 | 0.3747897 | 0.512692844 | 0.521952898 |
| Dnttip1 | 0.063749601 | 0.053226544 | 0.4352904 | 0.374903903 | 0.306424377 |
| Dnttip2 | 0.010036391 | 0.771577715 | 0.9552333 | 0.429550236 | 0.409936194 |
| Dock11 | -0.000636594 | 0.984410602 | 0.9976072 | 0.309698119 | 0.247656991 |
| Dock5 | 0.004723882 | 0.790052666 | 0.9621068 | 0.060148291 | 0.064282903 |
| Dock6 | -0.001757988 | 0.90855046 | 0.9837373 | 0.048224994 | 0.047686721 |
| Dock7 | 0.024789836 | 0.365431688 | 0.8050191 | 0.195467826 | 0.1715848 |
| Dock8 | -0.003846826 | 0.854485021 | 0.9755431 | 0.096058671 | 0.090667848 |
| Dock9 | 0.011115582 | 0.590125826 | 0.9049386 | 0.098368872 | 0.093312337 |
| Dohh | -0.020683841 | 0.546664921 | 0.8858459 | 0.463289831 | 0.437268284 |
| Dolk | -0.025633832 | 0.187650686 | 0.6660088 | 0.067225612 | 0.085845648 |
| Dop1a | -0.045275817 | 0.122649392 | 0.5833237 | 0.186770339 | 0.22077321 |
| Dop1b | -0.058154186 | 0.009282117 | 0.1930368 | 0.078774956 | 0.126159526 |
| Dot1l | 0.037571227 | 0.118702283 | 0.5777984 | 0.152827568 | 0.12069709 |
| Dpagt1 | -0.009669011 | 0.675071624 | 0.9265887 | 0.116207755 | 0.123338182 |
| Dpf2 | 0.007681299 | 0.812331619 | 0.9673015 | 0.323815462 | 0.31196021 |
| Dpf3 | -0.034203104 | 0.04255754 | 0.3952910 | 0.045883224 | 0.065419201 |
| Dph3 | 0.052803157 | 0.143417893 | 0.6165078 | 0.761828181 | 0.666557363 |
| Dph5 | 0.00292179 | 0.888103286 | 0.9812414 | 0.101472342 | 0.092512376 |
| Dph6 | -0.039981119 | 0.155651896 | 0.6312943 | 0.165908098 | 0.216011517 |
| Dph7 | -0.015498136 | 0.395030061 | 0.8222195 | 0.060813322 | 0.075409565 |
| Dpm1 | -0.085962592 | 0.015914598 | 0.2474560 | 0.428018636 | 0.451447815 |
| Dpm2 | 0.009027846 | 0.795117069 | 0.9631898 | 0.475088665 | 0.439633206 |
| Dpm3 | 0.172434273 | 2.17E-09 | 1.66E-06 | 1.724264348 | 1.524034756 |
| Dpp10 | -0.042983996 | 0.06610956 | 0.4772671 | 0.118363655 | 0.12342624 |
| Dpp3 | -0.008874293 | 0.731101111 | 0.9459020 | 0.170005021 | 0.162419816 |
| Dpp4 | 0.029744583 | 0.275468562 | 0.7356525 | 0.177496225 | 0.162155847 |
| Dpp7 | -0.043362609 | 0.099249954 | 0.5433551 | 0.137578362 | 0.173925139 |
| Dpp8 | -0.035503741 | 0.237394743 | 0.7098511 | 0.20307972 | 0.245993763 |
| Dpp9 | 0.002671556 | 0.933454926 | 0.9903410 | 0.340244814 | 0.303172835 |
| Dpy19l4 | 0.000701222 | 0.971565652 | 0.9976072 | 0.081657317 | 0.086253368 |
| Dpy30 | 0.016274665 | 0.641712897 | 0.9166712 | 0.621905675 | 0.558661699 |
| Dpysl2 | 0.079166527 | 0.020506317 | 0.2778998 | 0.47716562 | 0.363002977 |

| | | | | | |
|---|---|---|---|---|---|
| Dr1 | -0.00610484 | 0.804344761 | 0.9647143 | 0.140086827 | 0.139626183 |
| Dram2 | 0.013946388 | 0.589530623 | 0.9046775 | 0.171054932 | 0.154936864 |
| Drap1 | -0.037308243 | 0.242566187 | 0.7132719 | 1.190414281 | 1.165042066 |
| Drc1 | -0.024208175 | 0.32260467 | 0.7737520 | 0.157892866 | 0.129589294 |
| Drg1 | 0.007167372 | 0.827313345 | 0.9717006 | 0.380430745 | 0.340140217 |
| Drg2 | -0.017083723 | 0.510625021 | 0.8716066 | 0.164595645 | 0.169311924 |
| Drosha | -0.066361786 | 0.004401628 | 0.1311882 | 0.082253238 | 0.134873653 |
| Dsg2 | 0.009044337 | 0.726615828 | 0.9451913 | 0.152865244 | 0.162759468 |
| Dsp | 0.002313977 | 0.943842937 | 0.9935293 | 0.265940086 | 0.307190546 |
| Dst | -0.082708894 | 0.012271445 | 0.2185506 | 1.463837245 | 1.625507164 |
| Dstn | 0.078254091 | 0.011589885 | 0.2120789 | 1.518706913 | 1.34362042 |
| Dstyk | -0.00386391 | 0.874947703 | 0.9784331 | 0.149226614 | 0.139587339 |
| Dtd1 | -0.017680703 | 0.594544543 | 0.9065829 | 0.402049822 | 0.389764228 |
| Dtd2 | 0.028884469 | 0.296134925 | 0.7545964 | 0.215293102 | 0.185474676 |
| Dtnb | -0.08713916 | 0.001031635 | 0.0494387 | 0.123900251 | 0.190401925 |
| Dtnbos | 0.010287746 | 0.498246426 | 0.8684629 | 0.047060346 | 0.047368187 |
| Dtnbp1 | -0.057183602 | 0.056383558 | 0.4471022 | 0.215873513 | 0.255587512 |
| Dtwd1 | -0.038857745 | 0.032179242 | 0.3392328 | 0.055374343 | 0.077942582 |
| Dtx2 | -0.026778891 | 0.073447515 | 0.4917333 | 0.030549087 | 0.052276083 |
| Dtx3 | -0.036810339 | 0.24272649 | 0.7132719 | 0.278957587 | 0.300707639 |
| Dtx3l | 0.010381504 | 0.614771922 | 0.9108386 | 0.102469456 | 0.091743186 |
| Dtymk | 0.040152182 | 0.190811597 | 0.6700281 | 0.295412315 | 0.258920847 |
| Dus1l | -0.012379253 | 0.585402802 | 0.9024935 | 0.118150041 | 0.122992273 |
| Dus2 | 0.013672305 | 0.542693969 | 0.8840682 | 0.119662041 | 0.105925153 |
| Dus3l | 0.00285883 | 0.873894098 | 0.9784331 | 0.079970429 | 0.069106384 |
| Dusp1 | -0.114703829 | 0.013565564 | 0.2292511 | 1.134571613 | 1.354466019 |
| Dusp10 | -0.01125844 | 0.750596992 | 0.9505837 | 0.324159336 | 0.312888682 |
| Dusp11 | 0.066667166 | 0.066908608 | 0.4795761 | 0.624041247 | 0.555597029 |
| Dusp12 | 0.04986691 | 0.041357798 | 0.3916018 | 0.185948504 | 0.129562298 |
| Dusp16 | -0.023036993 | 0.373407312 | 0.8085689 | 0.15295324 | 0.154117464 |
| Dusp18 | -0.063395552 | 0.019828625 | 0.2724376 | 0.115668942 | 0.183498863 |
| Dusp19 | 0.011159817 | 0.658780141 | 0.9211780 | 0.175171674 | 0.14502343 |
| Dusp22 | -0.017013351 | 0.497241402 | 0.8684629 | 0.14614836 | 0.160027213 |
| Dusp23 | -0.009687424 | 0.664808799 | 0.9230765 | 0.097298737 | 0.11826696 |
| Dusp26 | -0.042135728 | 0.155547612 | 0.6312943 | 0.218784114 | 0.216814216 |
| Dusp28 | -0.00517601 | 0.849458841 | 0.9748841 | 0.185717616 | 0.178743803 |
| Dusp3 | 0.02725212 | 0.368467933 | 0.8056719 | 0.263314935 | 0.238193473 |
| Dusp4 | 0.032046487 | 0.101711053 | 0.5510917 | 0.096844721 | 0.075881601 |
| Dusp5 | -0.140456409 | 0.00032899 | 0.0228222 | 0.336604517 | 0.53901761 |
| Dusp7 | 0.021808047 | 0.26348696 | 0.7283846 | 0.09269893 | 0.08026046 |
| Dusp8 | -0.015586508 | 0.655123566 | 0.9206439 | 0.364814127 | 0.374828729 |
| Dut | -0.036554871 | 0.325650231 | 0.7763099 | 0.599314982 | 0.610601921 |
| Dvl1 | -0.02138688 | 0.367692497 | 0.8056719 | 0.121507809 | 0.138458424 |
| Dvl2 | -0.016930631 | 0.340356962 | 0.7844658 | 0.061208897 | 0.067322901 |
| Dvl3 | -0.034217926 | 0.192768174 | 0.6700281 | 0.145685635 | 0.173621861 |
| Dxo | 0.021557611 | 0.39869203 | 0.8239850 | 0.191851795 | 0.15465593 |
| Dym | -0.01037125 | 0.652528779 | 0.9191465 | 0.115172827 | 0.125820627 |
| Dync1h1 | -0.047600056 | 0.193402669 | 0.6710847 | 0.579740714 | 0.635325184 |
| Dync1i1 | 0.014789033 | 0.25800749 | 0.7251102 | 0.045975448 | 0.029682494 |
| Dync1i2 | -0.024436664 | 0.497957129 | 0.8684629 | 0.787908811 | 0.77217875 |
| Dync1li1 | 0.008676622 | 0.775313838 | 0.9558538 | 0.243662973 | 0.237782685 |
| Dync1li2 | -0.001549102 | 0.959388892 | 0.9957785 | 0.257301033 | 0.251858886 |
| Dync2h1 | -0.048227581 | 0.173641362 | 0.6527657 | 0.374006821 | 0.38878816 |
| Dync2i1 | 0.025552868 | 0.182173821 | 0.6595419 | 0.091504302 | 0.072648084 |
| Dync2i2 | -0.042803928 | 0.070742943 | 0.4879008 | 0.122967002 | 0.137243276 |
| Dync2li1 | 0.000703312 | 0.983019183 | 0.9976072 | 0.383005042 | 0.341750711 |
| Dynll1 | -0.085624312 | 0.017829762 | 0.2587928 | 0.949072554 | 0.980977738 |
| Dynll2 | 0.01531695 | 0.631796359 | 0.9133517 | 1.144994155 | 1.134540598 |
| Dynlrb1 | -0.01570041 | 0.637195115 | 0.9158312 | 1.193950863 | 1.094857293 |
| Dynlt1a | -0.013452736 | 0.607794043 | 0.9087139 | 0.184865977 | 0.161968008 |
| Dynlt1b | 0.008429654 | 0.682074401 | 0.9303225 | 0.101036819 | 0.091897845 |
| Dynlt1f | 0.046819717 | 0.125234759 | 0.5870522 | 0.394797699 | 0.260962753 |
| Dynlt2a2 | -0.003371336 | 0.86337081 | 0.9769675 | 0.080711071 | 0.082103014 |
| Dynlt2b | -0.053663398 | 0.091336827 | 0.5321282 | 0.338031179 | 0.342229705 |
| Dynlt3 | 0.04841132 | 0.154374372 | 0.6304934 | 0.391774073 | 0.363417589 |
| Dyrk1a | -0.001446281 | 0.958462058 | 0.9957785 | 0.193522617 | 0.185573675 |

| | | | | | |
|---|---|---|---|---|---|
| Dyrk1b | -0.011384003 | 0.472910582 | 0.85957884 | 0.047524013 | 0.055577331 |
| Dyrk2 | 0.060269554 | 0.076154089 | 0.49865668 | 0.382241631 | 0.319501258 |
| Dyrk3 | -9.78E-05 | 0.996429907 | 0.99940731 | 0.092615569 | 0.107789739 |
| Dzip1l | 0.010185898 | 0.486375473 | 0.86373583 | 0.053660033 | 0.043676385 |
| Dzip3 | 0.004466004 | 0.862578673 | 0.97696750 | 0.162233925 | 0.149772486 |
| E030022I16Rik | -0.028089838 | 0.085930068 | 0.52324897 | 0.038296608 | 0.063893578 |
| E030030I06Rik | -0.012991608 | 0.547569424 | 0.88626018 | 0.099434425 | 0.100132758 |
| E130102H24Rik | -0.018403324 | 0.315218299 | 0.77059875 | 0.06026046 | 0.078242173 |
| E130307A14Rik | -0.009236845 | 0.531243628 | 0.87764481 | 0.045362628 | 0.044852122 |
| E130308A19Rik | 0.019093644 | 0.27032726 | 0.73403607 | 0.079728636 | 0.057954378 |
| E130309D02Rik | 0.029176514 | 0.165159037 | 0.64207072 | 0.117892455 | 0.088601946 |
| E130311K13Rik | 0.003667662 | 0.855075373 | 0.97554315 | 0.088690977 | 0.087997224 |
| E130317F20Rik | -0.01907583 | 0.26668836 | 0.73087655 | 0.053301269 | 0.068902777 |
| E230001N04Rik | 0.006807505 | 0.683862319 | 0.93117032 | 0.066458123 | 0.054603133 |
| E230020A03Rik | -0.042915181 | 0.0750921 | 0.49662859 | 0.098715162 | 0.133295419 |
| E2f3 | -0.001493834 | 0.929421654 | 0.98819537 | 0.062211958 | 0.058861065 |
| E2f4 | -0.029813277 | 0.296328976 | 0.75459642 | 0.21874269 | 0.21859937 |
| E2f5 | 0.017581616 | 0.485341393 | 0.86297153 | 0.152468173 | 0.144814837 |
| E2f6 | 0.020617451 | 0.312327124 | 0.76861450 | 0.102786814 | 0.08991063 |
| E4f1 | -0.004113219 | 0.823617713 | 0.97105924 | 0.083191272 | 0.075654698 |
| Eaf1 | 0.003301785 | 0.895185519 | 0.98243614 | 0.137050703 | 0.158225496 |
| Eapp | 0.040428007 | 0.224841384 | 0.69682977 | 0.338860926 | 0.304380718 |
| Ears2 | -0.009215041 | 0.66910878 | 0.92355073 | 0.117252106 | 0.101834718 |
| Ebag9 | -0.003265855 | 0.910006567 | 0.98373739 | 0.224155938 | 0.202328805 |
| Ebna1bp2 | -0.005591703 | 0.871825045 | 0.97822694 | 0.514785791 | 0.471285521 |
| Ebp | 0.021999698 | 0.49777024 | 0.86846292 | 0.331001013 | 0.301654196 |
| Ebpl | 0.015268254 | 0.626757395 | 0.91280959 | 0.296337575 | 0.275206164 |
| Ecd | -0.018356386 | 0.581093407 | 0.90153997 | 0.311231103 | 0.331453751 |
| Ece1 | 0.039260643 | 0.296699667 | 0.75459642 | 0.844455221 | 0.821568454 |
| Ech1 | -0.011994107 | 0.703368262 | 0.93918879 | 0.290576241 | 0.301422846 |
| Echdc1 | 0.036380266 | 0.120990599 | 0.58114450 | 0.157476058 | 0.119847886 |
| Echdc2 | -0.003466645 | 0.890586052 | 0.98195975 | 0.139058369 | 0.144039621 |
| Echs1 | -0.043704434 | 0.202321878 | 0.67576869 | 0.424236702 | 0.433376934 |
| Eci1 | -0.047137548 | 0.155223263 | 0.63081309 | 0.331661125 | 0.342199413 |
| Eci2 | -0.047843721 | 0.192570502 | 0.67002811 | 0.688866111 | 0.681564125 |
| Ecpas | -0.019412367 | 0.588721153 | 0.90446241 | 0.456860713 | 0.469517278 |
| Ecsit | -0.017944012 | 0.527664807 | 0.87685539 | 0.202108936 | 0.200425611 |
| Edc3 | -0.007572293 | 0.67523285 | 0.92658872 | 0.073232093 | 0.070884468 |
| Edc4 | -0.02242828 | 0.353313599 | 0.79619965 | 0.135030837 | 0.141006649 |
| Edem1 | -0.000817272 | 0.975847396 | 0.99760728 | 0.145139853 | 0.196061598 |
| Edem2 | -0.033794888 | 0.334459376 | 0.78048367 | 0.448658463 | 0.448066446 |
| Edem3 | -0.015692024 | 0.530726194 | 0.87764481 | 0.132085425 | 0.153511764 |
| Edf1 | 0.063722866 | 0.014243922 | 0.23475005 | 1.746901102 | 1.597579074 |
| Edn3 | -0.01762205 | 0.610479619 | 0.90908029 | 0.338942257 | 0.271653229 |
| Edrf1 | 0.020486169 | 0.353557398 | 0.79656810 | 0.122232229 | 0.101311914 |
| Eea1 | 0.02156517 | 0.558388169 | 0.89198239 | 0.522059755 | 0.522595015 |
| Eed | 0.064637322 | 0.015313022 | 0.24258752 | 0.205524874 | 0.152085706 |
| Eef1a1 | -0.041838116 | 0.000565898 | 0.03361503 | 4.344088725 | 4.285613905 |
| Eef1a2 | -0.12174801 | 0.002201617 | 0.08367830 | 0.622809479 | 0.609725052 |
| Eef1akmt1 | -0.01478025 | 0.629152368 | 0.91305644 | 0.250445705 | 0.258807638 |
| Eef1akmt2 | -0.006032553 | 0.813030117 | 0.96730151 | 0.163899064 | 0.158819654 |
| Eef1akmt4 | -0.026601554 | 0.070803296 | 0.48790080 | 0.034799751 | 0.050227174 |
| Eef1aknmt | -0.0307051 | 0.062641894 | 0.46512543 | 0.053949216 | 0.062227512 |
| Eef1b2 | -0.075797967 | 9.24E-05 | 0.00889470 | 2.551787272 | 2.539917761 |
| Eef1d | -0.00564642 | 0.852524357 | 0.97554315 | 1.371597503 | 1.26455857 |
| Eef1e1 | 0.027157468 | 0.306625437 | 0.76527857 | 0.185779175 | 0.162425519 |
| Eef1g | -0.046266618 | 0.134794915 | 0.60130514 | 1.560585064 | 1.463410984 |
| Eef2 | -0.046014001 | 0.022517188 | 0.29144683 | 2.524608231 | 2.477275412 |
| Eef2kmt | 0.002768556 | 0.935287913 | 0.99082188 | 0.419638759 | 0.363292514 |
| Eefsec | -0.028979941 | 0.150197455 | 0.62803209 | 0.079149336 | 0.094103784 |
| Efcab1 | -0.002473748 | 0.944996429 | 0.99369321 | 0.604916229 | 0.587882013 |
| Efcab10 | 0.014351577 | 0.441907927 | 0.84287090 | 0.079983381 | 0.073604816 |
| Efcab14 | 0.005868789 | 0.868319953 | 0.97756927 | 0.477999764 | 0.469687116 |
| Efcab15 | 0.080017917 | 0.025398054 | 0.30576297 | 0.529824939 | 0.429982274 |
| Efcab2 | -0.005906166 | 0.755429173 | 0.95213280 | 0.080946753 | 0.076857021 |
| Efcc1 | -0.010219785 | 0.570587017 | 0.89625342 | 0.068311312 | 0.061959365 |

| | | | | | |
|---|---|---|---|---|---|
| Efemp2 | -0.041267725 | 0.075343233 | 0.49662859 | 0.093958894 | 0.130539577 |
| Efhd2 | 0.008547494 | 0.77077764 | 0.95523334 | 0.230876547 | 0.215331179 |
| Efl1 | -0.012954697 | 0.565624806 | 0.89438911 | 0.102080375 | 0.116610703 |
| Efna1 | -0.006562242 | 0.756169483 | 0.95215335 | 0.09714101 | 0.093514924 |
| Efna5 | 0.042768077 | 0.067502146 | 0.48145306 | 0.135579305 | 0.11401241 |
| Efnb1 | -0.004061446 | 0.874463615 | 0.97843318 | 0.145253171 | 0.146513617 |
| Efr3a | -0.001175391 | 0.973823629 | 0.99760728 | 0.48771167 | 0.44913093 |
| Eftud2 | 0.018704874 | 0.444712414 | 0.84315416 | 0.151759315 | 0.133636786 |
| Egf | 4.68E-05 | 0.997494975 | 0.99940731 | 0.044793526 | 0.046724202 |
| Egflam | 0.006119609 | 0.756273619 | 0.95215335 | 0.109117709 | 0.070153125 |
| Egln1 | -0.050286151 | 0.132634178 | 0.59882516 | 0.340436459 | 0.356865313 |
| Egln2 | 0.009540824 | 0.770537479 | 0.95523334 | 0.447084195 | 0.409480732 |
| Egr1 | -0.07193524 | 0.065278415 | 0.47405701 | 1.745339477 | 1.918642288 |
| Egr2 | -0.017272561 | 0.360642397 | 0.80124805 | 0.069655656 | 0.072561772 |
| Egr4 | -0.026256325 | 0.186698209 | 0.66476891 | 0.048052038 | 0.071551114 |
| Ehbp1 | -0.048453293 | 0.019261066 | 0.26955866 | 0.07646509 | 0.105335877 |
| Ehd1 | 0.010715649 | 0.645870078 | 0.91696956 | 0.137705705 | 0.120996992 |
| Ehd3 | 0.015268999 | 0.301626127 | 0.76093836 | 0.050252058 | 0.041632614 |
| Ehd4 | -0.011437762 | 0.582980196 | 0.90197908 | 0.094797546 | 0.095915323 |
| Ehf | -0.049169385 | 0.110318532 | 0.56558881 | 0.178973244 | 0.214692393 |
| Ehmt1 | 0.011847207 | 0.699771319 | 0.93731184 | 0.245591847 | 0.244988353 |
| Ehmt2 | -0.043172502 | 0.19908647 | 0.67309621 | 0.390272527 | 0.405408196 |
| Ei24 | -0.025428818 | 0.455843825 | 0.85029141 | 0.575179653 | 0.525527802 |
| Eid1 | 0.061492817 | 0.080968181 | 0.51255576 | 1.256873144 | 1.156349772 |
| Eid2 | -0.048220948 | 0.112623355 | 0.56962035 | 0.270491517 | 0.279607923 |
| Eid2b | 0.006596071 | 0.804423149 | 0.96471431 | 0.197160279 | 0.180528878 |
| Eif1 | 0.018720192 | 0.101062525 | 0.54873576 | 3.983700421 | 3.89488047 |
| Eif1a | 0.043390167 | 0.237786578 | 0.71028089 | 0.626312442 | 0.565743084 |
| Eif1ad | 0.000845389 | 0.977172326 | 0.99760728 | 0.227002419 | 0.236044387 |
| Eif1ax | -0.012296017 | 0.717754972 | 0.94356338 | 0.427913096 | 0.413341975 |
| Eif1b | -0.026535942 | 0.415932082 | 0.83009907 | 1.215782126 | 1.149509211 |
| Eif2a | -0.046199617 | 0.156820177 | 0.63176836 | 0.352009161 | 0.357209895 |
| Eif2ak1 | -0.03909197 | 0.09242333 | 0.53255841 | 0.094241713 | 0.131128585 |
| Eif2ak2 | 0.04102552 | 0.131390076 | 0.59726521 | 0.212492592 | 0.161253136 |
| Eif2ak3 | -0.011201447 | 0.665564907 | 0.92307657 | 0.149299663 | 0.162804103 |
| Eif2ak4 | -0.012830733 | 0.495151831 | 0.86828696 | 0.068026414 | 0.076686166 |
| Eif2b1 | 0.028832088 | 0.267373673 | 0.73168145 | 0.194975052 | 0.157983253 |
| Eif2b2 | 0.015643039 | 0.617246559 | 0.91150176 | 0.290187182 | 0.267695035 |
| Eif2b3 | 0.006628787 | 0.805564646 | 0.96471431 | 0.195276486 | 0.179897932 |
| Eif2b4 | 0.03883302 | 0.120163457 | 0.57973049 | 0.174329116 | 0.127876893 |
| Eif2b5 | 0.040956135 | 0.164333061 | 0.64155211 | 0.253331109 | 0.211714808 |
| Eif2d | -0.012287214 | 0.634091962 | 0.9144743 | 0.160594943 | 0.163144675 |
| Eif2s1 | -0.063932986 | 0.065183359 | 0.47405701 | 0.483141856 | 0.491559664 |
| Eif2s2 | 0.105666521 | 0.000229242 | 0.01790612 | 1.872743448 | 1.696170217 |
| Eif2s3x | 0.02254659 | 0.444408325 | 0.84315416 | 0.232107666 | 0.217284239 |
| Eif2s3y | -0.016049975 | 0.640455326 | 0.91652002 | 0.416993799 | 0.389104287 |
| Eif3a | 0.029414953 | 0.378129236 | 0.81019153 | 1.367283555 | 1.311268257 |
| Eif3b | 0.000242022 | 0.994435663 | 0.99876498 | 0.52430469 | 0.491608275 |
| Eif3c | 0.056364183 | 0.067626606 | 0.48145306 | 1.619973907 | 1.644431289 |
| Eif3d | 0.002615682 | 0.940277883 | 0.99284589 | 0.546473418 | 0.530873578 |
| Eif3e | 0.004335783 | 0.902496066 | 0.98254414 | 0.999626332 | 0.912989641 |
| Eif3f | -0.038492087 | 0.176516054 | 0.65366596 | 1.572419002 | 1.442498657 |
| Eif3g | -0.054973009 | 0.12399254 | 0.58525434 | 0.662583709 | 0.68564495 |
| Eif3h | -0.077328818 | 0.007708871 | 0.17340589 | 1.368027582 | 1.355409225 |
| Eif3i | -0.042773892 | 0.191892507 | 0.67002811 | 1.023571413 | 0.969259234 |
| Eif3j1 | 0.056253563 | 0.109450403 | 0.56558881 | 0.560964159 | 0.509890754 |
| Eif3k | -0.023080781 | 0.478043806 | 0.86074894 | 1.339053453 | 1.253760465 |
| Eif3l | -0.023886184 | 0.475244474 | 0.86029656 | 0.423893366 | 0.418506223 |
| Eif3m | 0.050639725 | 0.162882174 | 0.64007035 | 0.914612145 | 0.796088285 |
| Eif4a1 | -0.001430052 | 0.964239629 | 0.99664211 | 1.472606984 | 1.344525258 |
| Eif4a2 | -0.033197707 | 0.295940858 | 0.75459641 | 1.300229185 | 1.309635908 |
| Eif4a3 | 0.006818511 | 0.846820003 | 0.97396571 | 0.48002971 | 0.43566195 |
| Eif4b | -0.018086822 | 0.629969151 | 0.91305644 | 0.803700628 | 0.742754242 |
| Eif4e | -0.013342327 | 0.701654702 | 0.93818771 | 0.821941425 | 0.769536587 |
| Eif4e2 | -0.005183456 | 0.859630013 | 0.97696756 | 0.252382351 | 0.239841786 |
| Eif4e3 | -0.032358989 | 0.172610763 | 0.65230429 | 0.11370943 | 0.137821168 |

| | | | | | |
|---|---|---|---|---|---|
| Eif4ebp1 | 0.175425397 | 0.000106505 | 0.00978269 | 1.042306103 | 0.898470687 |
| Eif4ebp2 | 0.053826104 | 0.086842218 | 0.5250566 | 0.344518261 | 0.278507751 |
| Eif4enif1 | 0.000898577 | 0.973495805 | 0.9976072 | 0.17446571 | 0.177895912 |
| Eif4g1 | -0.03188582 | 0.374821337 | 0.8092384 | 0.819070212 | 0.827780087 |
| Eif4g2 | -0.036868632 | 0.247729141 | 0.7177068 | 1.29180846 | 1.379479612 |
| Eif4g3 | -0.03676206 | 0.318649515 | 0.7710043 | 0.581792272 | 0.663487778 |
| Eif4h | -0.025821108 | 0.467614149 | 0.8558546 | 0.723685733 | 0.688191217 |
| Eif5 | 0.011529707 | 0.708541055 | 0.9414057 | 1.476265474 | 1.455767461 |
| Eif5a | -0.054275377 | 0.031534373 | 0.3385228 | 2.025087865 | 1.990674572 |
| Eif5b | 0.064307639 | 0.078481846 | 0.5056175 | 0.960759129 | 0.885629163 |
| Eif6 | -0.019248697 | 0.602468066 | 0.9086609 | 0.619436709 | 0.617821319 |
| Eipr1 | 0.000350858 | 0.987269784 | 0.9976072 | 0.117877434 | 0.110988037 |
| Elac1 | -0.004481419 | 0.790567996 | 0.9621438 | 0.055049533 | 0.060975307 |
| Elac2 | -0.016301315 | 0.389723133 | 0.8180392 | 0.070903754 | 0.080858823 |
| Elapor1 | -0.113057982 | 0.001173417 | 0.0541535 | 1.139936614 | 1.18352137 |
| Elapor2 | 0.149691175 | 8.33E-07 | 0.0002071 | 0.321095187 | 0.198415639 |
| Elavl1 | 0.032627295 | 0.339471881 | 0.7840654 | 0.414536347 | 0.367655075 |
| Elavl4 | 0.0652726 | 0.044501291 | 0.4009562 | 0.311775674 | 0.289110116 |
| Elf1 | -0.000968134 | 0.976412409 | 0.9976072 | 0.346149179 | 0.328286614 |
| Elf2 | -0.005360399 | 0.878569865 | 0.9785583 | 0.449942497 | 0.444772027 |
| Elf3 | 0.011588645 | 0.652635469 | 0.9191465 | 0.15876299 | 0.142913864 |
| Elk3 | 0.055398561 | 0.015246528 | 0.2423807 | 0.141039403 | 0.105335215 |
| Elk4 | 0.016161683 | 0.481427372 | 0.8620306 | 0.13603785 | 0.110986822 |
| Ell | 0.012165454 | 0.588633908 | 0.9044624 | 0.144942295 | 0.112356805 |
| Ell2 | 0.011632667 | 0.725650406 | 0.9450332 | 0.307479944 | 0.315190899 |
| Elmo1 | 0.004023479 | 0.830274944 | 0.9722601 | 0.064602057 | 0.078980584 |
| Elmo2 | -0.00829101 | 0.629309913 | 0.9130564 | 0.058107103 | 0.065735087 |
| Elmo3 | -0.009417122 | 0.668532454 | 0.9230765 | 0.106850227 | 0.108065233 |
| Elmod1 | 0.01879674 | 0.372691253 | 0.8085689 | 0.113773467 | 0.09069111 |
| Elmod2 | 0.034741673 | 0.0966085 | 0.5373016 | 0.116009262 | 0.08559851 |
| Elmod3 | -0.024710295 | 0.100762864 | 0.5480085 | 0.030386144 | 0.055058976 |
| Eloa | -0.021516227 | 0.515468697 | 0.8734806 | 0.36114281 | 0.342981938 |
| Elob | 0.122706983 | 2.83E-06 | 0.0004924 | 1.88344384 | 1.673970158 |
| Eloc | 0.016524576 | 0.61678572 | 0.9115017 | 1.119990013 | 1.047436451 |
| Elof1 | -0.000279636 | 0.993728167 | 0.9985888 | 0.731951867 | 0.703716057 |
| Elovl1 | -0.022024652 | 0.362387888 | 0.8030478 | 0.136376546 | 0.142230695 |
| Elovl2 | 0.016475488 | 0.371311595 | 0.8074114 | 0.076358456 | 0.070228904 |
| Elovl5 | 0.040279219 | 0.1263444 | 0.5889350 | 0.181340685 | 0.160526623 |
| Elovl6 | 0.006026385 | 0.798277179 | 0.9644698 | 0.123738175 | 0.124529936 |
| Elovl7 | -0.004153082 | 0.826720656 | 0.9716906 | 0.076033137 | 0.077987765 |
| Elp1 | 0.041754736 | 0.128848079 | 0.5914729 | 0.216884679 | 0.172944373 |
| Elp2 | -0.027670822 | 0.270379821 | 0.7340360 | 0.143997688 | 0.151225251 |
| Elp3 | -0.04059786 | 0.184615809 | 0.6630661 | 0.225228438 | 0.258285584 |
| Elp4 | 0.012016462 | 0.650070815 | 0.9175779 | 0.17762055 | 0.164100953 |
| Elp5 | -0.019249537 | 0.499213713 | 0.8690742 | 0.190893296 | 0.201322561 |
| Elp6 | -0.009725854 | 0.56656466 | 0.8949556 | 0.05493729 | 0.060998409 |
| Emb | 0.087392471 | 0.019046424 | 0.2683814 | 0.991298829 | 0.914728335 |
| Emc1 | -0.029287749 | 0.282060584 | 0.7418354 | 0.169659855 | 0.180409024 |
| Emc10 | -0.0798039 | 0.026854806 | 0.3138777 | 0.861517733 | 0.895846513 |
| Emc2 | -0.015437149 | 0.639207274 | 0.9160042 | 0.363067095 | 0.346040257 |
| Emc3 | -0.036434117 | 0.23688928 | 0.7095050 | 0.260075391 | 0.27482325 |
| Emc4 | 0.035654208 | 0.264108707 | 0.7291840 | 0.316564164 | 0.296458514 |
| Emc6 | 0.012588598 | 0.709160346 | 0.9414976 | 0.507566917 | 0.488739715 |
| Emc7 | -0.01113669 | 0.747944942 | 0.9496497 | 0.959606715 | 0.945654639 |
| Emc8 | -0.036052851 | 0.215052148 | 0.6845467 | 0.218369279 | 0.225487144 |
| Emc9 | 0.043866893 | 0.145579021 | 0.6208701 | 0.296208963 | 0.239090409 |
| Emd | -0.000160629 | 0.996357498 | 0.9994073 | 0.479836743 | 0.492240426 |
| Emg1 | 0.001453449 | 0.967692348 | 0.9976072 | 0.74210426 | 0.69047252 |
| Emilin1 | 0.031928934 | 0.106178979 | 0.5617171 | 0.091666769 | 0.080477499 |
| Eml1 | -0.002450466 | 0.894098284 | 0.9824361 | 0.072879222 | 0.075428695 |
| Eml2 | -0.008784285 | 0.734813674 | 0.9468380 | 0.158078605 | 0.167545875 |
| Eml3 | -0.024984015 | 0.242801664 | 0.7132719 | 0.0858682 | 0.102198129 |
| Eml4 | 0.01415596 | 0.636166862 | 0.9153541 | 0.235001546 | 0.216827489 |
| Eml5 | 0.004016488 | 0.91000215 | 0.9837373 | 0.383604785 | 0.401291076 |
| Eml6 | 0.00929185 | 0.626926166 | 0.9128095 | 0.087730327 | 0.073788699 |
| Emsy | -0.041435241 | 0.155469107 | 0.6312943 | 0.188544025 | 0.2176682 |

| | | | | | |
|---|---|---|---|---|---|
| Enah | 0.01091119 | 0.766136264 | 0.9544221 | 0.467401681 | 0.462617291 |
| Enc1 | 0.011560187 | 0.460833617 | 0.8524425 | 0.055074768 | 0.046884841 |
| Endod1 | -0.009598758 | 0.587979715 | 0.9044624 | 0.059472285 | 0.07161156 |
| Endog | 0.005678045 | 0.854723041 | 0.9755431 | 0.277772633 | 0.27944041 |
| Endov | 0.015371082 | 0.363572729 | 0.8046911 | 0.080513495 | 0.055358409 |
| Enho | 0.1029116 | 0.01878875 | 0.2677936 | 0.723344528 | 0.84490899 |
| Eno1 | 0.032939593 | 0.322073383 | 0.7733024 | 0.385795298 | 0.340472421 |
| Eno2 | 0.012423508 | 0.624116407 | 0.9121505 | 0.149002773 | 0.146487848 |
| Eno3 | 0.013243718 | 0.338306115 | 0.7834245 | 0.049928106 | 0.035562141 |
| Enoph1 | -0.02063206 | 0.315181013 | 0.7705987 | 0.08470098 | 0.09312037 |
| Enox2 | 0.02023565 | 0.235864624 | 0.7079390 | 0.075419583 | 0.057690045 |
| Enpep | 0.011168149 | 0.541554563 | 0.8835017 | 0.066230205 | 0.071073688 |
| Enpp2 | -0.004890686 | 0.891797544 | 0.9819597 | 0.37675884 | 0.482949475 |
| Enpp4 | 0.007486021 | 0.77948195 | 0.9586487 | 0.178458134 | 0.170904749 |
| Enpp5 | -0.037497084 | 0.262786415 | 0.7275735 | 0.326030539 | 0.360611266 |
| Ensa | -0.016680124 | 0.602030751 | 0.9084492 | 0.273122353 | 0.27896099 |
| ENSMUSG00000095041 | -0.189010117 | 8.99E-07 | 0.0002072 | 0.547580216 | 0.732371899 |
| ENSMUSG00000115801 | -0.020538408 | 0.554424329 | 0.8902995 | 0.261256088 | 0.296607473 |
| ENSMUSG00000117901 | -0.013760419 | 0.377773355 | 0.8101797 | 0.033377872 | 0.056237461 |
| Entpd2 | 0.037982 | 0.005934328 | 0.1545438 | 0.057056322 | 0.033445162 |
| Entpd3 | -0.001288017 | 0.972019959 | 0.9976072 | 0.366448352 | 0.472743159 |
| Entpd5 | -0.016212956 | 0.487006524 | 0.8640311 | 0.121287629 | 0.125681947 |
| Entpd6 | -0.006790345 | 0.669956634 | 0.9239496 | 0.04272378 | 0.055721037 |
| Entpd7 | 0.030277976 | 0.300022878 | 0.7590275 | 0.234069949 | 0.21866369 |
| Entr1 | -0.024343175 | 0.407818333 | 0.8270770 | 0.220253768 | 0.244201147 |
| Eny2 | 0.020520399 | 0.557625835 | 0.8919136 | 0.958829481 | 0.888070925 |
| Eola1 | -0.000108651 | 0.996671433 | 0.9994073 | 0.163810712 | 0.154317441 |
| Ep300 | -0.025670903 | 0.461913485 | 0.8524425 | 0.377428581 | 0.395941685 |
| Ep400 | 0.014588917 | 0.644652497 | 0.9168391 | 0.321349797 | 0.30520443 |
| Epb41 | 0.013872215 | 0.631627377 | 0.9133517 | 0.20588813 | 0.21747651 |
| Epb41l1 | 0.038840572 | 0.038981995 | 0.3786326 | 0.097904029 | 0.07084739 |
| Epb41l2 | -0.010439641 | 0.595777006 | 0.9068020 | 0.091893663 | 0.076491105 |
| Epb41l3 | 0.039164471 | 0.296235972 | 0.7545964 | 0.576035832 | 0.576326928 |
| Epb41l4aos | -0.018848817 | 0.500939675 | 0.8693704 | 0.176828448 | 0.201879684 |
| Epb41l4b | 0.028295274 | 0.350235687 | 0.7927303 | 0.269962485 | 0.238261254 |
| Epb41l5 | 0.001744368 | 0.950519457 | 0.9948462 | 0.199129733 | 0.189749779 |
| Epc1 | 0.018242168 | 0.573594284 | 0.8973494 | 0.286726084 | 0.278527575 |
| Epc2 | 0.004002417 | 0.898272872 | 0.9825441 | 0.278766499 | 0.263497893 |
| Epcam | -0.010596377 | 0.696288269 | 0.9363862 | 2.419678259 | 2.303554443 |
| Epg5 | -0.011955105 | 0.442753748 | 0.8431541 | 0.041891816 | 0.050378546 |
| Ephx1 | -0.011194851 | 0.534453436 | 0.8795949 | 0.055495295 | 0.073763514 |
| Ephx4 | -0.012024172 | 0.49588121 | 0.8684629 | 0.067049701 | 0.06830201 |
| Epm2aip1 | -0.085389947 | 0.01424592 | 0.2347500 | 0.416092682 | 0.468285794 |
| Epn1 | -0.011846037 | 0.738127962 | 0.9474220 | 0.487578915 | 0.488101439 |
| Epn2 | -0.028249751 | 0.240115684 | 0.7118791 | 0.115946336 | 0.133173014 |
| EPPK1 | -0.0139354 | 0.376258779 | 0.8101773 | 0.041947105 | 0.051469339 |
| Eprs | 0.003526087 | 0.905293256 | 0.9832544 | 1.694826421 | 1.747800857 |
| Eps15 | -0.040566695 | 0.069581392 | 0.4837345 | 0.083156551 | 0.12095275 |
| Eps15l1 | 0.021324392 | 0.305093397 | 0.7646008 | 0.122952234 | 0.088640745 |
| Eps8 | -0.000337186 | 0.990182783 | 0.9980836 | 0.176186367 | 0.191700337 |
| Eps8l2 | 0.014208458 | 0.629963077 | 0.9130564 | 0.239787087 | 0.224695733 |
| Eral1 | 0.022592296 | 0.133327278 | 0.5988251 | 0.063534061 | 0.040949495 |
| Erap1 | 0.018092869 | 0.529529319 | 0.8776448 | 0.233758173 | 0.210632436 |
| Erbin | -0.018367702 | 0.567690688 | 0.8954510 | 0.291953514 | 0.307961546 |
| Erc1 | -0.024162863 | 0.324555414 | 0.7757552 | 0.11708639 | 0.141320792 |
| Ercc1 | -0.006558684 | 0.759532473 | 0.9521533 | 0.10692838 | 0.101940862 |
| Ercc2 | -0.008149904 | 0.645028528 | 0.9169695 | 0.065264208 | 0.067984923 |
| Ercc3 | 0.024713231 | 0.17642971 | 0.6536659 | 0.084440432 | 0.067264387 |
| Ercc4 | -0.005198074 | 0.724872309 | 0.9448707 | 0.046584563 | 0.044259742 |
| Ercc5 | -0.028555902 | 0.280319012 | 0.7408618 | 0.165162918 | 0.172505394 |
| Ercc6 | 0.008695844 | 0.65164361 | 0.9187471 | 0.081758181 | 0.075779001 |
| Ercc6l2 | -0.006170819 | 0.815542246 | 0.9686999 | 0.167553722 | 0.168528817 |
| Erf | -0.016809755 | 0.410239815 | 0.8283287 | 0.085773876 | 0.09536848 |
| Erg28 | -0.012323815 | 0.695938752 | 0.9363862 | 0.319749303 | 0.278096007 |
| Ergic1 | 0.009071243 | 0.787687618 | 0.9608781 | 0.368984905 | 0.382491577 |
| Ergic2 | 0.060900308 | 0.078844333 | 0.5062367 | 0.518575896 | 0.465232756 |

| | | | | | |
|---|---|---|---|---|---|
| Ergic3 | -0.087609113 | 0.018460964 | 0.2650257 | 0.599955699 | 0.653591257 |
| Erh | 0.085500601 | 0.012127484 | 0.2175490 | 0.49979701 | 0.377697135 |
| Eri1 | -0.015892187 | 0.518254591 | 0.8740369 | 0.147170029 | 0.141537972 |
| Eri3 | -0.027441119 | 0.405789522 | 0.8270770 | 0.33657677 | 0.325222777 |
| Erich1 | 0.039212058 | 0.15209479 | 0.6281105 | 0.200040334 | 0.161527966 |
| Erich3 | -0.010806973 | 0.65952217 | 0.9211780 | 0.112332702 | 0.138475949 |
| Erlec1 | -0.055697314 | 0.126238247 | 0.5889350 | 0.725041527 | 0.813001993 |
| Erlin1 | -0.008046778 | 0.674465338 | 0.9265608 | 0.065692555 | 0.081757799 |
| Erlin2 | 0.036990668 | 0.146198801 | 0.6219091 | 0.177524172 | 0.150608618 |
| Ermard | -0.007863782 | 0.639236137 | 0.9160042 | 0.055832116 | 0.059908084 |
| Ermp1 | -0.006242851 | 0.716358188 | 0.9431411 | 0.059426855 | 0.064962074 |
| Ern1 | 0.020103392 | 0.562839043 | 0.8926240 | 0.37305283 | 0.390162271 |
| Ero1a | -0.01269395 | 0.710450986 | 0.9419370 | 0.355996123 | 0.385198659 |
| Ero1b | 0.166319707 | 7.24E-05 | 0.0073280 | 1.119504061 | 1.066719783 |
| Erp29 | 0.031311804 | 0.397647471 | 0.8238469 | 0.765631533 | 0.738737372 |
| Erp44 | -0.045041286 | 0.230075086 | 0.7019677 | 0.657783932 | 0.663224175 |
| Errfi1 | -0.028860704 | 0.463721199 | 0.8526625 | 0.382858465 | 0.437110305 |
| Esam | -0.000563707 | 0.973456646 | 0.9976072 | 0.06380499 | 0.056714579 |
| Esco1 | 0.007526792 | 0.798488675 | 0.9644698 | 0.247686118 | 0.240037352 |
| Esd | -0.01660697 | 0.622571187 | 0.9121505 | 1.227037276 | 1.191705517 |
| Esf1 | 0.027239777 | 0.429065124 | 0.8340365 | 0.400276103 | 0.376881983 |
| Esrp1 | -0.000263112 | 0.993900008 | 0.9985888 | 0.339574695 | 0.359854885 |
| Esrra | 0.001968526 | 0.918892563 | 0.9851211 | 0.085729813 | 0.07830607 |
| Ess2 | -0.014216901 | 0.489587667 | 0.8651997 | 0.083823205 | 0.088754219 |
| Esyt1 | 0.103559732 | 0.000813336 | 0.0426893 | 0.329505045 | 0.250833664 |
| Esyt2 | -0.039640706 | 0.178936142 | 0.6549987 | 0.208431302 | 0.242482723 |
| Etaa1 | -0.013964856 | 0.478286004 | 0.8607489 | 0.07450898 | 0.087382186 |
| Etf1 | 0.008984704 | 0.803471961 | 0.9647143 | 0.567508452 | 0.544151402 |
| Etfa | -0.015859039 | 0.633942862 | 0.9144743 | 0.380138885 | 0.371711769 |
| Etfb | -0.08801696 | 0.012400833 | 0.2195296 | 0.500033677 | 0.527409987 |
| Etfdh | -0.023269216 | 0.307578672 | 0.7656663 | 0.107935319 | 0.1280871 |
| Etfrf1 | 0.008727872 | 0.765034452 | 0.9540090 | 0.233405867 | 0.215453411 |
| Ethe1 | 0.012533785 | 0.56033238 | 0.8925646 | 0.113681702 | 0.09830393 |
| Etl4 | -0.047482983 | 0.115517737 | 0.5724670 | 0.218699724 | 0.24956327 |
| Etnk1 | -0.01113197 | 0.758049888 | 0.9521533 | 0.449038272 | 0.487618016 |
| Etohd2 | -0.013108463 | 0.405939444 | 0.8270770 | 0.050624262 | 0.052436885 |
| Ets2 | -0.016452186 | 0.644386449 | 0.9168391 | 0.380605383 | 0.399405547 |
| Etv1 | -0.074490691 | 0.092784259 | 0.5326503 | 1.479399766 | 1.428445278 |
| Etv3 | -0.027378139 | 0.257973312 | 0.7251102 | 0.117976083 | 0.141099122 |
| Etv5 | 0.01118883 | 0.517334267 | 0.8738218 | 0.056737318 | 0.065288944 |
| Etv6 | -0.011802647 | 0.549462318 | 0.8874415 | 0.085217398 | 0.08406973 |
| Eva1b | 0.002681761 | 0.941087208 | 0.9929301 | 0.519008311 | 0.499355998 |
| Eva1c | -0.009160361 | 0.553858053 | 0.8902995 | 0.045076254 | 0.052861745 |
| Evi5 | 0.005349102 | 0.863123652 | 0.9769675 | 0.279524839 | 0.260822171 |
| Evi5l | -0.017507119 | 0.283367259 | 0.7427848 | 0.051254208 | 0.056683159 |
| Evl | -0.00546047 | 0.843686776 | 0.9734889 | 0.20414533 | 0.183734619 |
| Evpl | -0.000233306 | 0.98770013 | 0.9976072 | 0.046990648 | 0.048256663 |
| Ewsr1 | -0.02858887 | 0.426918396 | 0.8337565 | 0.562297555 | 0.585918052 |
| Exd2 | -0.02259291 | 0.250823332 | 0.7207210 | 0.072228528 | 0.087116292 |
| Exoc1 | 0.006998368 | 0.761944897 | 0.9529513 | 0.134134254 | 0.117675447 |
| Exoc2 | -0.027428101 | 0.253255874 | 0.7223071 | 0.122054216 | 0.145455635 |
| Exoc3 | 0.007515925 | 0.827408839 | 0.9717006 | 0.422172145 | 0.386939961 |
| Exoc4 | -0.032860593 | 0.337025006 | 0.7829714 | 0.335743845 | 0.346004072 |
| Exoc5 | -0.018332284 | 0.53968637 | 0.8829112 | 0.233277205 | 0.251936031 |
| Exoc6 | 0.022378693 | 0.310072991 | 0.7677033 | 0.128683925 | 0.09916163 |
| Exoc6b | -0.027874503 | 0.350671475 | 0.7927303 | 0.215464625 | 0.246487747 |
| Exoc7 | -0.011611432 | 0.70539769 | 0.9401381 | 0.263365892 | 0.273426287 |
| Exoc8 | -0.011829876 | 0.510876679 | 0.8716066 | 0.065867415 | 0.070419206 |
| Exosc1 | -0.007677603 | 0.802211513 | 0.9647143 | 0.251808801 | 0.245565739 |
| Exosc10 | -0.007855047 | 0.738323841 | 0.9474220 | 0.123893836 | 0.126768047 |
| Exosc2 | -0.020546829 | 0.270048289 | 0.7340360 | 0.057561309 | 0.077132317 |
| Exosc3 | 0.001356311 | 0.96384665 | 0.9966421 | 0.280752166 | 0.272328218 |
| Exosc4 | -9.11E-05 | 0.997874577 | 0.9994073 | 0.449981468 | 0.43564547 |
| Exosc5 | -0.074276971 | 0.019919252 | 0.2733042 | 0.291399486 | 0.329491575 |
| Exosc7 | 0.017511939 | 0.570452101 | 0.8962534 | 0.330447226 | 0.308138868 |
| Exosc8 | 0.001899418 | 0.940343155 | 0.9928458 | 0.182357443 | 0.146404306 |

| | | | | | |
|---|---|---|---|---|---|
| Exosc9 | 0.002743189 | 0.915753841 | 0.98452361 | 0.169786305 | 0.148415248 |
| Exph5 | -0.054586872 | 0.036543649 | 0.36215084 | 0.123739137 | 0.17204528 |
| Ext1 | -0.005408201 | 0.803563694 | 0.96471431 | 0.093199758 | 0.105728472 |
| Ext2 | -0.010157757 | 0.692017016 | 0.93551490 | 0.150138161 | 0.163560376 |
| Extl2 | -0.000992357 | 0.964622462 | 0.99664211 | 0.10940805 | 0.117877324 |
| Extl3 | -0.021594475 | 0.293638909 | 0.75336844 | 0.074303412 | 0.098727809 |
| Eya1 | 0.012489568 | 0.380682234 | 0.81173500 | 0.047240728 | 0.037613976 |
| Eya3 | 0.033625845 | 0.111688527 | 0.56757614 | 0.11409737 | 0.09404242 |
| Ezh1 | 0.01824369 | 0.481800006 | 0.86203068 | 0.172883631 | 0.156707059 |
| Ezh2 | -0.02958109 | 0.045105372 | 0.40274101 | 0.021623858 | 0.05364483 |
| Ezr | 0.006950673 | 0.781006701 | 0.95908472 | 0.139575697 | 0.135473929 |
| F11r | 0.0194618 | 0.534211751 | 0.87956524 | 0.319447243 | 0.264761536 |
| F3 | 0.010983136 | 0.682677771 | 0.93049780 | 0.176860557 | 0.156214804 |
| F8a | 0.022397824 | 0.412449952 | 0.82979790 | 0.19124724 | 0.19346062 |
| Faah | -0.033810467 | 0.163216501 | 0.64043239 | 0.113556281 | 0.139370679 |
| Faap100 | -0.006297708 | 0.782800696 | 0.95908628 | 0.124390713 | 0.119957081 |
| Faap20 | -0.015579816 | 0.63403394 | 0.9144743 | 0.351764817 | 0.340120763 |
| Faap24 | 0.00303667 | 0.856716918 | 0.97631892 | 0.061688497 | 0.059603682 |
| Fabp3 | 0.005987844 | 0.812161357 | 0.96730151 | 0.136168054 | 0.140220824 |
| Fabp5 | -0.048001723 | 0.035626066 | 0.35919149 | 0.055024751 | 0.109583267 |
| Fadd | -0.016212961 | 0.266477511 | 0.73073994 | 0.04276483 | 0.04673398 |
| Fads1 | 0.013635037 | 0.453883774 | 0.84904829 | 0.079799411 | 0.068414712 |
| Fads2 | 0.042183744 | 0.013719858 | 0.23053228 | 0.095422754 | 0.045325517 |
| Fads6 | -0.027131374 | 0.239097064 | 0.71141058 | 0.103711326 | 0.127204791 |
| Faf1 | 0.006536975 | 0.838390452 | 0.97272164 | 0.354910071 | 0.314428086 |
| Faf2 | 0.018050307 | 0.599520253 | 0.90782341 | 0.437289668 | 0.407811192 |
| Fah | 0.015391493 | 0.319665487 | 0.77128410 | 0.055009522 | 0.046551004 |
| Fahd1 | -0.033561266 | 0.24267991 | 0.71327199 | 0.217452303 | 0.214031868 |
| Fahd2a | 0.003545673 | 0.896447132 | 0.98254414 | 0.201868496 | 0.184943617 |
| Faim | 0.001731594 | 0.947026296 | 0.99450181 | 0.174718373 | 0.174190813 |
| Fam102a | -0.059551518 | 0.048433757 | 0.41562531 | 0.21297147 | 0.274135178 |
| Fam104a | 0.035801337 | 0.306578006 | 0.76527857 | 0.925222192 | 0.851708343 |
| Fam110a | -0.047271352 | 0.039752082 | 0.38219230 | 0.088895541 | 0.133939137 |
| Fam110b | -0.034212966 | 0.108533041 | 0.56368177 | 0.082769213 | 0.111994594 |
| Fam114a1 | 0.042409653 | 0.205729477 | 0.67975655 | 0.381244627 | 0.329750401 |
| Fam114a2 | -0.036552706 | 0.310331695 | 0.76770334 | 0.444919605 | 0.469060558 |
| Fam117a | 0.014334993 | 0.60987553 | 0.90908029 | 0.211546859 | 0.191557334 |
| Fam117b | 0.011868951 | 0.640822888 | 0.91667127 | 0.164460611 | 0.148571824 |
| Fam118a | -0.102957214 | 0.000443484 | 0.02838290 | 0.161127723 | 0.230758925 |
| Fam118b | -0.029912704 | 0.081535562 | 0.51357269 | 0.050299913 | 0.068429941 |
| Fam120a | -0.076109329 | 0.035491858 | 0.35919149 | 0.553566116 | 0.612359303 |
| Fam120b | -0.03083041 | 0.309161702 | 0.76738130 | 0.236035597 | 0.269360371 |
| Fam120c | -0.009330353 | 0.633112505 | 0.91411960 | 0.074830964 | 0.084748589 |
| Fam122a | -0.034441477 | 0.338711464 | 0.78342458 | 0.501222378 | 0.562699525 |
| Fam126a | -0.031030904 | 0.069988016 | 0.48483318 | 0.046556754 | 0.064967262 |
| Fam126b | 0.03451018 | 0.266313181 | 0.73073994 | 0.293214299 | 0.265742745 |
| Fam131a | 0.017359012 | 0.507907201 | 0.87040677 | 0.184688722 | 0.160444411 |
| Fam133b | 0.04255255 | 0.185217268 | 0.66376783 | 0.312825001 | 0.268627056 |
| Fam135a | 0.030081825 | 0.421053878 | 0.83009907 | 0.593097005 | 0.604025692 |
| Fam135b | -0.034518162 | 0.065238838 | 0.47405701 | 0.040412934 | 0.087659956 |
| Fam136a | 0.023330602 | 0.379377962 | 0.81108392 | 0.197108043 | 0.16167394 |
| Fam13b | -0.049002233 | 0.065189663 | 0.47405701 | 0.151549343 | 0.187843545 |
| Fam149a | 0.01348654 | 0.578560091 | 0.90028487 | 0.154358222 | 0.128175617 |
| Fam149b | -0.002253804 | 0.930591168 | 0.98901482 | 0.165856568 | 0.159241613 |
| Fam151a | 0.114911481 | 0.000614709 | 0.03504548 | 0.303193093 | 0.282011259 |
| Fam155a | -0.017702049 | 0.501310863 | 0.86955665 | 0.153278375 | 0.167606792 |
| Fam160a2 | -0.018542374 | 0.427947131 | 0.83403655 | 0.117233935 | 0.130644249 |
| Fam160b1 | -0.040448745 | 0.194339609 | 0.67171917 | 0.272097912 | 0.294152331 |
| Fam160b2 | -0.012130663 | 0.55045355 | 0.88830567 | 0.089861631 | 0.096468693 |
| Fam161a | -0.014242472 | 0.370532681 | 0.80661258 | 0.043417176 | 0.053145932 |
| Fam162a | 0.009139426 | 0.767661204 | 0.95442217 | 0.274350743 | 0.259262418 |
| Fam166c | -0.011433486 | 0.416800708 | 0.83009907 | 0.043646507 | 0.039043841 |
| Fam168a | -0.011113892 | 0.703679102 | 0.93918879 | 0.222700829 | 0.237995822 |
| Fam168b | 0.021102385 | 0.37621277 | 0.81017735 | 0.154133194 | 0.131450978 |
| Fam169a | -0.016343685 | 0.434834284 | 0.83856067 | 0.081977122 | 0.103027745 |
| Fam171a1 | 0.011202305 | 0.559109415 | 0.89212890 | 0.085254356 | 0.077425303 |

| | | | | | |
|---|---|---|---|---|---|
| Fam171b | -0.021655769 | 0.43705279 | 0.8394017 | 0.16020935 | 0.200378933 |
| Fam172a | -0.004942882 | 0.878127907 | 0.9785583 | 0.336075551 | 0.298567208 |
| Fam174a | -0.006665214 | 0.85209107 | 0.9753915 | 0.772729696 | 0.718160571 |
| Fam174b | 0.009530653 | 0.793708583 | 0.9624508 | 0.575754099 | 0.542808715 |
| Fam174c | -0.022716594 | 0.415547997 | 0.8300990 | 0.18320144 | 0.205213532 |
| Fam183b | 0.010807944 | 0.767153547 | 0.9544221 | 1.020144582 | 0.928281285 |
| Fam184a | -0.012268655 | 0.623498919 | 0.9121505 | 0.131898423 | 0.149858568 |
| Fam185a | 0.005338694 | 0.745820284 | 0.9485717 | 0.060891595 | 0.053840477 |
| Fam187b | -0.01315252 | 0.562699921 | 0.8926240 | 0.110645518 | 0.123830566 |
| Fam189b | 0.000772781 | 0.960376464 | 0.9958628 | 0.048408586 | 0.051274423 |
| Fam193a | -0.02177399 | 0.5037357 | 0.8695566 | 0.297143316 | 0.295937699 |
| Fam193b | -0.084080797 | 0.000883135 | 0.0451582 | 0.103486882 | 0.166027312 |
| Fam199x | 0.015309961 | 0.48795845 | 0.8642517 | 0.118715317 | 0.106539293 |
| Fam204a | 0.001433313 | 0.967704874 | 0.9976072 | 0.449270794 | 0.447909964 |
| Fam207a | 0.016780363 | 0.502010393 | 0.8695566 | 0.159370905 | 0.151026239 |
| Fam20a | 0.000264628 | 0.992366163 | 0.9985888 | 0.186959654 | 0.17239387 |
| Fam20b | -0.004479267 | 0.888929568 | 0.9816521 | 0.302867103 | 0.304249772 |
| Fam20c | -0.042454139 | 0.203580247 | 0.6772740 | 0.373305197 | 0.378454412 |
| Fam210a | -0.026802331 | 0.22019168 | 0.6914534 | 0.102601265 | 0.110754338 |
| Fam210b | -0.096308956 | 0.014091976 | 0.2345510 | 0.412461734 | 0.550829001 |
| Fam214a | 0.00964463 | 0.713887509 | 0.9428150 | 0.171785469 | 0.169741156 |
| Fam214b | -0.015396328 | 0.538602214 | 0.8816335 | 0.135919196 | 0.152508164 |
| Fam216a | -0.095805202 | 0.00471475 | 0.1367553 | 0.363154264 | 0.426621147 |
| Fam219a | 0.037509891 | 0.112045788 | 0.5679582 | 0.149597225 | 0.121477602 |
| Fam219b | 0.018957584 | 0.420164727 | 0.8300990 | 0.145134762 | 0.119527366 |
| Fam220a | -0.039080455 | 0.033143078 | 0.3453564 | 0.046182283 | 0.076046957 |
| Fam222a | -0.02874394 | 0.269200359 | 0.7338465 | 0.149117193 | 0.162411988 |
| Fam222b | 0.021860012 | 0.438994152 | 0.8403747 | 0.230401453 | 0.196602473 |
| Fam229b | 0.000274194 | 0.989758011 | 0.9980836 | 0.10696676 | 0.099524381 |
| Fam234a | 0.029565939 | 0.357717553 | 0.7989722 | 0.295578638 | 0.270480116 |
| Fam234b | -0.007934543 | 0.727207979 | 0.9452179 | 0.10828317 | 0.110713732 |
| Fam241a | -0.007821197 | 0.628664717 | 0.9130564 | 0.047929263 | 0.05664532 |
| Fam241b | 0.033504834 | 0.251064979 | 0.7207210 | 0.2623885 | 0.219232905 |
| Fam32a | 0.01317578 | 0.693272006 | 0.9359725 | 0.419702244 | 0.368755627 |
| Fam3a | -0.022843449 | 0.430885532 | 0.8351669 | 0.21802741 | 0.219235899 |
| Fam3c | -0.037327621 | 0.290792755 | 0.7513090 | 0.479791717 | 0.423658311 |
| Fam43a | -0.022630729 | 0.259932844 | 0.7263475 | 0.087652978 | 0.082737554 |
| Fam50a | 0.016099707 | 0.635999468 | 0.9153541 | 0.370386998 | 0.345655201 |
| Fam53a | 0.01699133 | 0.5464377 | 0.8858459 | 0.226376443 | 0.191263255 |
| Fam53b | 0.036633705 | 0.006061105 | 0.1565785 | 0.056116938 | 0.028686819 |
| Fam53c | -0.00891054 | 0.726370654 | 0.9451202 | 0.141803811 | 0.144984355 |
| Fam71e1 | -0.001439504 | 0.945010274 | 0.9936932 | 0.103191854 | 0.098558178 |
| Fam76a | 0.00579712 | 0.866852037 | 0.9772002 | 0.443702748 | 0.425654916 |
| Fam76b | 0.035412173 | 0.195019211 | 0.6719932 | 0.203315478 | 0.173100407 |
| Fam81a | 0.003253193 | 0.89198023 | 0.9819597 | 0.143853576 | 0.120969592 |
| Fam83g | -0.042537725 | 0.049042048 | 0.4175940 | 0.072077933 | 0.11159414 |
| Fam83h | 0.011326946 | 0.524564733 | 0.8758385 | 0.073957917 | 0.063289487 |
| Fam8a1 | -0.022146974 | 0.427722666 | 0.8340365 | 0.208514613 | 0.200818692 |
| Fam91a1 | 0.002542947 | 0.934252787 | 0.9905715 | 0.270073657 | 0.266130476 |
| Fam98a | 0.04995302 | 0.10434711 | 0.5580045 | 0.290687716 | 0.235319954 |
| Fam98b | 0.032344494 | 0.305619625 | 0.7646271 | 0.303064086 | 0.276457091 |
| Fam98c | -0.018555272 | 0.503206319 | 0.8695566 | 0.195123627 | 0.196584279 |
| Fance | -0.004843246 | 0.827477459 | 0.9717006 | 0.126366838 | 0.110867617 |
| Fancf | -0.001572269 | 0.913927293 | 0.9845236 | 0.047078021 | 0.043858866 |
| Fancl | -0.007557524 | 0.677059388 | 0.9280496 | 0.075800403 | 0.071856884 |
| Fancm | 0.002483076 | 0.897150476 | 0.9825441 | 0.080742798 | 0.078353259 |
| Far1 | -0.015835772 | 0.631448809 | 0.9133517 | 0.352891113 | 0.345359629 |
| Farp1 | 0.040625472 | 0.116363244 | 0.5724670 | 0.180370966 | 0.150521255 |
| Farp2 | -0.009943141 | 0.490085658 | 0.8655242 | 0.040626654 | 0.044097684 |
| Fars2 | 0.002518753 | 0.929176266 | 0.9881953 | 0.221269629 | 0.200618856 |
| Farsa | -0.055169711 | 0.039008325 | 0.3786326 | 0.148906049 | 0.191819787 |
| Farsb | -0.007725801 | 0.803816887 | 0.9647143 | 0.299347683 | 0.285553578 |
| Fasn | -0.04227439 | 0.108311599 | 0.5636817 | 0.149908332 | 0.168803543 |
| Fastk | 0.017451049 | 0.583122151 | 0.9020071 | 0.297681462 | 0.271223642 |
| Fastkd1 | -0.019783072 | 0.239908894 | 0.7118791 | 0.049895217 | 0.065464639 |
| Fastkd2 | -0.021118907 | 0.265750195 | 0.7306783 | 0.072369066 | 0.085050294 |

| | | | | | |
|---|---|---|---|---|---|
| Fat1 | -0.004744149 | 0.843786062 | 0.9734889 | 0.130271159 | 0.1276268 |
| Fau | 0.024991972 | 0.039760335 | 0.3821923 | 3.938999101 | 3.755499058 |
| Fbf1 | -0.027453201 | 0.118770849 | 0.5777984 | 0.05365248 | 0.07033235 |
| Fbh1 | -0.028282139 | 0.194473536 | 0.6717191 | 0.103525876 | 0.114320634 |
| Fbl | 0.019051841 | 0.556158791 | 0.8912586 | 0.375963687 | 0.311436997 |
| Fbp1 | -0.103355997 | 0.018971197 | 0.2683814 | 0.444932714 | 0.466959723 |
| Fbrs | 0.044135357 | 0.073462579 | 0.4917333 | 0.168706983 | 0.125422043 |
| Fbrsl1 | -0.00668038 | 0.796803771 | 0.9644244 | 0.155002183 | 0.174215548 |
| Fbxl12 | -0.011243379 | 0.648795418 | 0.9172569 | 0.146941825 | 0.144188529 |
| Fbxl14 | 0.023961535 | 0.325800338 | 0.7763291 | 0.171125508 | 0.13858704 |
| Fbxl15 | -0.015754556 | 0.603025189 | 0.9087139 | 0.227281263 | 0.239236535 |
| Fbxl16 | -0.04789855 | 0.139853499 | 0.6125151 | 0.30731081 | 0.371009408 |
| Fbxl17 | -0.017840285 | 0.482937833 | 0.8620306 | 0.150391157 | 0.16070757 |
| Fbxl18 | 0.011441864 | 0.511495125 | 0.8720873 | 0.071299078 | 0.061504632 |
| Fbxl19 | -0.021437694 | 0.145806411 | 0.6213461 | 0.033866247 | 0.050928863 |
| Fbxl2 | 0.008818119 | 0.584262439 | 0.9021406 | 0.057706372 | 0.053987269 |
| Fbxl20 | 0.010266817 | 0.619622809 | 0.9121505 | 0.107484582 | 0.090566644 |
| Fbxl3 | 0.004153839 | 0.881107098 | 0.9795705 | 0.192712072 | 0.183470405 |
| Fbxl4 | -0.040262704 | 0.031887401 | 0.3385228 | 0.058614074 | 0.087217747 |
| Fbxl5 | -0.007261596 | 0.823927638 | 0.9710592 | 0.31822256 | 0.327085111 |
| Fbxl6 | -0.024579668 | 0.261276226 | 0.7275735 | 0.101243901 | 0.113632663 |
| Fbxo11 | 0.037781406 | 0.208674153 | 0.6807660 | 0.280691351 | 0.231765911 |
| Fbxo16 | 0.008191481 | 0.624568532 | 0.9121505 | 0.069694662 | 0.057074586 |
| Fbxo17 | -0.000276338 | 0.987606829 | 0.9976072 | 0.065577395 | 0.072238854 |
| Fbxo2 | 0.039983778 | 0.209936718 | 0.6808016 | 0.280571796 | 0.276630451 |
| Fbxo21 | 0.023941553 | 0.405374778 | 0.8270770 | 0.241083337 | 0.220649088 |
| Fbxo22 | -0.028055865 | 0.324492739 | 0.7757552 | 0.216997951 | 0.226156829 |
| Fbxo25 | -0.028735053 | 0.135253612 | 0.6024767 | 0.062321589 | 0.093164427 |
| Fbxo28 | 0.001804879 | 0.94493689 | 0.9936932 | 0.173122936 | 0.160501269 |
| Fbxo3 | 0.000234784 | 0.994372277 | 0.9987649 | 0.361638176 | 0.350291769 |
| Fbxo30 | 0.011400068 | 0.541590138 | 0.8835017 | 0.086202424 | 0.071400924 |
| Fbxo31 | 0.03034185 | 0.165693428 | 0.6425780 | 0.133746345 | 0.105386695 |
| Fbxo32 | 0.014570464 | 0.4648676 | 0.8539030 | 0.089638749 | 0.08765238 |
| Fbxo33 | -0.034048099 | 0.283715717 | 0.7429936 | 0.238853622 | 0.271386351 |
| Fbxo34 | -0.005927389 | 0.844903287 | 0.9734889 | 0.26680273 | 0.254441463 |
| Fbxo36 | -0.001646227 | 0.955189071 | 0.9954690 | 0.245509342 | 0.226405075 |
| Fbxo38 | -0.005441057 | 0.833309016 | 0.9723494 | 0.159538021 | 0.157816697 |
| Fbxo4 | -0.010990907 | 0.642634324 | 0.9166712 | 0.139096983 | 0.129134093 |
| Fbxo42 | 0.017505892 | 0.435675047 | 0.8389295 | 0.140086633 | 0.112811 |
| Fbxo44 | -0.003392452 | 0.898407511 | 0.9825441 | 0.166252039 | 0.167676955 |
| Fbxo45 | 0.014800163 | 0.488049459 | 0.8642517 | 0.113674024 | 0.094084621 |
| Fbxo46 | -0.026998983 | 0.136717954 | 0.6052465 | 0.057961455 | 0.069793597 |
| Fbxo6 | 0.050950234 | 0.039281474 | 0.3799003 | 0.176031333 | 0.137439626 |
| Fbxo7 | 0.001132401 | 0.959050081 | 0.9957785 | 0.112034953 | 0.113828262 |
| Fbxo8 | 0.010960872 | 0.667138413 | 0.9230765 | 0.170281086 | 0.154097574 |
| Fbxo9 | -0.031509827 | 0.346635878 | 0.7899443 | 0.332660871 | 0.369192377 |
| Fbxw11 | 0.054909698 | 0.05111043 | 0.4267806 | 0.242272497 | 0.199715641 |
| Fbxw2 | -0.005987719 | 0.860068602 | 0.9769675 | 0.393999395 | 0.371557092 |
| Fbxw4 | -0.016158685 | 0.527303117 | 0.8768553 | 0.152471403 | 0.151348998 |
| Fbxw5 | 0.031086639 | 0.2795649 | 0.7405297 | 0.251437443 | 0.212455264 |
| Fbxw7 | 0.047935058 | 0.131270227 | 0.5972652 | 0.298948377 | 0.279455991 |
| Fbxw8 | -0.011709273 | 0.638407063 | 0.9160042 | 0.135452174 | 0.141645811 |
| Fbxw9 | 0.004525482 | 0.848244062 | 0.9745866 | 0.147996232 | 0.120370578 |
| Fcf1 | -0.003503674 | 0.905743854 | 0.9832544 | 0.256803669 | 0.246338496 |
| Fcho2 | 0.031025158 | 0.354974646 | 0.7969131 | 0.341586031 | 0.312406145 |
| Fchsd2 | 0.016599584 | 0.443147136 | 0.8431541 | 0.127796955 | 0.095923948 |
| Fdft1 | 0.043420206 | 0.154353208 | 0.6304934 | 0.276547921 | 0.218733699 |
| Fdps | -0.008638598 | 0.802666557 | 0.9647143 | 0.366637719 | 0.322138644 |
| Fdx1 | -0.032934385 | 0.343024276 | 0.7872955 | 0.488280576 | 0.509274659 |
| Fdxacb1 | -0.001785723 | 0.908366558 | 0.9837373 | 0.05553454 | 0.049424634 |
| Fdxr | 0.001625229 | 0.931977131 | 0.9894277 | 0.084367095 | 0.082318259 |
| Fech | -0.037762518 | 0.199447557 | 0.6730962 | 0.221611777 | 0.241556432 |
| Fem1a | -0.023983427 | 0.339007294 | 0.7834245 | 0.141987301 | 0.154740389 |
| Fem1b | -0.012464797 | 0.68341346 | 0.9309982 | 0.255077443 | 0.261826351 |
| Fem1c | -0.020212319 | 0.419042684 | 0.8300990 | 0.137296578 | 0.147403069 |
| Fen1 | 0.005062456 | 0.734783092 | 0.9468380 | 0.056198296 | 0.043926157 |

| | | | | | |
|---|---|---|---|---|---|
| Fer | 0.015017435 | 0.518718105 | 0.87417954 | 0.133277553 | 0.114288156 |
| Fermt2 | -0.00556861 | 0.879001615 | 0.97864152 | 0.479588479 | 0.512833212 |
| Fev | -0.033998935 | 0.188026906 | 0.66686690 | 0.155656013 | 0.153002654 |
| Fez1 | 0.002943136 | 0.91075043 | 0.98373739 | 0.164147552 | 0.160489737 |
| Fez2 | 0.02453925 | 0.185614111 | 0.66376783 | 0.087882852 | 0.068352699 |
| Ffar1 | 0.080350341 | 0.006608625 | 0.16179671 | 0.239507086 | 0.206281085 |
| Ffar2 | -0.092609492 | 0.00085802 | 0.04456311 | 0.117208555 | 0.196824449 |
| Ffar4 | -0.027661987 | 0.344772879 | 0.78799761 | 0.228568572 | 0.204427911 |
| Fgb | -0.006933241 | 0.81646929 | 0.96882480 | 0.107798126 | 0.128929712 |
| Fgd1 | -0.027455256 | 0.083080058 | 0.51775319 | 0.037360361 | 0.059433577 |
| Fgd2 | -0.061155792 | 0.030792528 | 0.33530392 | 0.180882471 | 0.218562696 |
| Fgd4 | -0.033754492 | 0.160271248 | 0.63703375 | 0.111275617 | 0.130318243 |
| Fgd6 | 0.030160338 | 0.124009904 | 0.58525434 | 0.090473926 | 0.084052523 |
| Fgf12 | -0.025972798 | 0.450456282 | 0.84789420 | 0.359926423 | 0.351983068 |
| Fgf9 | -0.015739772 | 0.503906228 | 0.86955665 | 0.106583636 | 0.131400378 |
| Fgfbp3 | -0.004635495 | 0.766394673 | 0.95442217 | 0.043470653 | 0.052134551 |
| Fgfr1 | 0.013777893 | 0.610244558 | 0.90908029 | 0.190547686 | 0.153169068 |
| Fgfr1op2 | -0.001127512 | 0.975434675 | 0.99760728 | 0.560375866 | 0.574586325 |
| Fggy | 0.014288613 | 0.438729031 | 0.84033149 | 0.086230693 | 0.071216524 |
| Fgl2 | -0.046567712 | 0.265735844 | 0.73067832 | 0.515176617 | 0.541771931 |
| Fh1 | 0.051932601 | 0.195031099 | 0.67199328 | 0.633582992 | 0.643466139 |
| Fhad1 | -0.021770036 | 0.225683255 | 0.69790112 | 0.061558198 | 0.074534147 |
| Fhdc1 | -0.013842532 | 0.497886169 | 0.86846292 | 0.065356124 | 0.099815752 |
| Fhit | -0.009116645 | 0.524534362 | 0.87583859 | 0.036251848 | 0.04298918 |
| Fhl1 | 0.011914016 | 0.638897297 | 0.91600428 | 0.151061308 | 0.149646418 |
| Fhl2 | -0.023922038 | 0.171817471 | 0.65154025 | 0.047883967 | 0.064569802 |
| Fhod3 | 0.030139065 | 0.2152673 | 0.68465900 | 0.139468948 | 0.137129381 |
| Fibp | -0.075760844 | 0.026161865 | 0.30972651 | 0.396404173 | 0.424989858 |
| Ficd | 0.002181995 | 0.929298485 | 0.98819537 | 0.145439575 | 0.145880106 |
| Fig4 | 0.010180856 | 0.530421596 | 0.87764481 | 0.058560067 | 0.053425469 |
| Filip1 | 0.033147186 | 0.209587879 | 0.68076602 | 0.148935355 | 0.163140232 |
| Fip1l1 | -0.014244229 | 0.70372886 | 0.93918879 | 0.762234087 | 0.757339808 |
| Firre | -0.013682835 | 0.524912641 | 0.87583859 | 0.083647943 | 0.106978557 |
| Fis1 | -0.01437301 | 0.662752033 | 0.92286639 | 1.052228159 | 0.974510903 |
| Fitm2 | -0.003944946 | 0.88425901 | 0.98028002 | 0.168297469 | 0.204705557 |
| Fiz1 | 0.007025822 | 0.774067889 | 0.95582697 | 0.149398755 | 0.129461991 |
| Fkbp11 | 0.015752802 | 0.713109174 | 0.94245177 | 0.61695161 | 0.608397057 |
| Fkbp14 | -0.003579558 | 0.835654926 | 0.97234942 | 0.062400029 | 0.065524872 |
| Fkbp15 | -0.038195122 | 0.15503396 | 0.63076150 | 0.154129147 | 0.175101376 |
| Fkbp1a | 0.001603063 | 0.956765373 | 0.99560605 | 1.430269082 | 1.348723361 |
| Fkbp1b | 0.030884845 | 0.378388849 | 0.81019153 | 0.299322328 | 0.36424209 |
| Fkbp2 | 0.092306865 | 0.002723255 | 0.09648105 | 1.730197117 | 1.602377283 |
| Fkbp3 | 0.004838715 | 0.895146461 | 0.98243614 | 0.809026329 | 0.728544351 |
| Fkbp4 | -0.029001183 | 0.433237539 | 0.83753632 | 0.864468529 | 0.81784386 |
| Fkbp5 | 0.007874749 | 0.803114032 | 0.96471431 | 0.287050857 | 0.278462794 |
| Fkbp7 | 0.005241055 | 0.768155914 | 0.95442217 | 0.0762575 | 0.066071376 |
| Fkbp8 | -0.057961409 | 0.104004871 | 0.55799259 | 0.491872136 | 0.522005637 |
| Fkbp9 | -0.003336268 | 0.927371006 | 0.98792100 | 0.642439441 | 0.613568119 |
| Fkbpl | 0.006312226 | 0.747839168 | 0.94963703 | 0.093335277 | 0.080391258 |
| Fkrp | -0.01590993 | 0.428552756 | 0.83403655 | 0.074974164 | 0.090879035 |
| Fktn | -0.004995114 | 0.844962667 | 0.97348890 | 0.137972447 | 0.152894883 |
| Flad1 | 0.002142004 | 0.921126687 | 0.98611750 | 0.10344717 | 0.106020737 |
| Flcn | 0.007470566 | 0.738858253 | 0.94743504 | 0.116570577 | 0.118591507 |
| Flii | -0.028459347 | 0.325699115 | 0.77630994 | 0.225408858 | 0.231918954 |
| Flnb | 0.043738962 | 0.198664806 | 0.67309622 | 0.372928419 | 0.374055278 |
| Flot1 | -0.051675742 | 0.113601116 | 0.56969774 | 0.267458906 | 0.32137736 |
| Flot2 | -0.026128091 | 0.352367768 | 0.79525680 | 0.17522193 | 0.200065537 |
| Flrt1 | -0.016899905 | 0.528331915 | 0.87731044 | 0.141352186 | 0.189357253 |
| Flt3l | -0.025854024 | 0.095394728 | 0.53517678 | 0.028331072 | 0.059664609 |
| Flvcr1 | -0.010524665 | 0.562453455 | 0.89262402 | 0.071825774 | 0.073826345 |
| Flywch1 | -0.03694082 | 0.283201338 | 0.74278487 | 0.360623293 | 0.410254545 |
| Flywch2 | 0.024749365 | 0.40638755 | 0.82707770 | 0.280061801 | 0.233033835 |
| Fmc1 | -0.00690233 | 0.842717615 | 0.97348890 | 0.528673637 | 0.543812891 |
| Fmn2 | -0.052137188 | 0.137150467 | 0.60683882 | 0.323470828 | 0.361227374 |
| Fmnl2 | 0.020909649 | 0.485421486 | 0.86297153 | 0.214090793 | 0.229417157 |
| Fmo5 | -0.044496546 | 0.009463147 | 0.19476020 | 0.047627766 | 0.065029459 |

| | | | | | |
|---|---|---|---|---|---|
| Fmr1 | -0.011079512 | 0.730862478 | 0.9459020 | 0.273503363 | 0.285863178 |
| Fn3k | -0.037384558 | 0.03495991 | 0.3567924 | 0.042787925 | 0.076988371 |
| Fnbp1 | -0.038390875 | 0.047386626 | 0.4106761 | 0.064396883 | 0.087972348 |
| Fnbp1l | -0.048972746 | 0.06307898 | 0.4666245 | 0.136125766 | 0.169600128 |
| Fnbp4 | -0.040827102 | 0.199398393 | 0.6730962 | 0.246057148 | 0.270064648 |
| Fndc10 | -0.026302104 | 0.088976487 | 0.5292814 | 0.039177359 | 0.057760333 |
| Fndc3a | -0.019071611 | 0.551631904 | 0.8893234 | 1.257119726 | 1.262367671 |
| Fndc3b | 0.015392447 | 0.667927324 | 0.9230765 | 0.494306032 | 0.511220792 |
| Fndc5 | 0.023379802 | 0.203519159 | 0.6772740 | 0.091971455 | 0.063928562 |
| Fnip1 | -0.009889178 | 0.76903332 | 0.9552333 | 0.306492987 | 0.326473731 |
| Fnip2 | 0.057929947 | 0.022509213 | 0.2914468 | 0.17538801 | 0.14533996 |
| Fnta | -0.032698425 | 0.311698614 | 0.7684379 | 0.346423024 | 0.340485056 |
| Fntb | 0.004408475 | 0.812182524 | 0.9673015 | 0.079178044 | 0.07541388 |
| Focad | -0.01952721 | 0.291678731 | 0.7519368 | 0.061701449 | 0.076099493 |
| Folr1 | 0.012878863 | 0.740796702 | 0.9479080 | 0.387769001 | 0.353848984 |
| Fos | -0.184669487 | 6.73E-08 | 2.90E-05 | 2.370612762 | 2.792975272 |
| Fosb | -0.222646778 | 3.31E-06 | 0.0005666 | 0.556028579 | 0.934198813 |
| Fosl2 | -0.053960074 | 0.049516964 | 0.4192321 | 0.089485948 | 0.173108283 |
| Foxa2 | -0.003389047 | 0.925631125 | 0.9879210 | 0.58022187 | 0.574995774 |
| Foxa3 | 0.011071345 | 0.696950948 | 0.9364422 | 0.223683724 | 0.190508517 |
| Foxj2 | 0.007095291 | 0.726044691 | 0.9450680 | 0.088888729 | 0.096256898 |
| Foxj3 | 0.03081458 | 0.258410382 | 0.7251572 | 0.209194805 | 0.186754522 |
| Foxk1 | 0.026794946 | 0.140665832 | 0.6130953 | 0.087376362 | 0.064672743 |
| Foxk2 | 0.034099202 | 0.193524432 | 0.6710847 | 0.190504428 | 0.162843156 |
| Foxn2 | 0.020313707 | 0.356040509 | 0.7972735 | 0.125461033 | 0.103915966 |
| Foxn3 | -0.05392828 | 0.072605014 | 0.4902939 | 0.219442787 | 0.259604332 |
| Foxo1 | 0.023468687 | 0.468123846 | 0.8561686 | 0.29252607 | 0.287650889 |
| Foxo3 | 0.031824713 | 0.19717859 | 0.6730962 | 0.164477477 | 0.1421705 |
| Foxo6 | 0.0201666 | 0.199227692 | 0.6730962 | 0.05974039 | 0.047757835 |
| Foxp1 | -0.073791807 | 0.056019138 | 0.4452803 | 0.767016792 | 0.80585488 |
| Foxp2 | -0.026063477 | 0.145496906 | 0.6207868 | 0.049612542 | 0.06610215 |
| Foxp4 | -0.030248907 | 0.325249907 | 0.7763099 | 0.236166287 | 0.258262038 |
| Foxred1 | 0.001090865 | 0.954673217 | 0.9954690 | 0.085075344 | 0.078601464 |
| Fpgs | -0.015231121 | 0.357201904 | 0.7986123 | 0.047681321 | 0.066253247 |
| Fpgt | 0.003552432 | 0.880179394 | 0.9792571 | 0.12780605 | 0.122528721 |
| Fra10ac1 | 0.018596072 | 0.584275567 | 0.9021406 | 0.403493681 | 0.352032853 |
| Frg1 | 0.0296486 | 0.339995329 | 0.7844658 | 0.348221078 | 0.279921154 |
| Frk | -0.017083981 | 0.46093773 | 0.8524425 | 0.101706881 | 0.127829434 |
| Frmd4a | -0.022062915 | 0.301434654 | 0.7608403 | 0.078619624 | 0.104507452 |
| Frmd4b | -0.034142162 | 0.20393685 | 0.6775129 | 0.15679895 | 0.168351548 |
| Frmd5 | -0.009925327 | 0.527351734 | 0.8768553 | 0.03842423 | 0.059406057 |
| Frrs1 | 0.000124064 | 0.993893452 | 0.9985888 | 0.047375584 | 0.058712046 |
| Frrs1l | 0.011120572 | 0.591821782 | 0.9060200 | 0.111685095 | 0.084921473 |
| Frs2 | -0.005033401 | 0.838773239 | 0.9727216 | 0.139218095 | 0.143059177 |
| Fry | -0.030728415 | 0.250666541 | 0.7207210 | 0.157482061 | 0.186941359 |
| Fryl | 0.022336011 | 0.517809423 | 0.8738804 | 0.395547267 | 0.35596432 |
| Frzb | 0.037861707 | 0.209762407 | 0.6807660 | 0.216489355 | 0.245612697 |
| Fsd1l | 0.006177943 | 0.73367592 | 0.9464134 | 0.078488626 | 0.069440669 |
| Fth1 | -0.069517964 | 5.56E-07 | 0.0001589 | 3.307767762 | 3.472861516 |
| Ftl1 | -0.023923031 | 0.157551393 | 0.6324475 | 3.493304659 | 3.532843709 |
| Ftl1-ps1 | -0.019107699 | 0.294519173 | 0.7536588 | 0.057526435 | 0.079470454 |
| Fto | -0.018941574 | 0.519764195 | 0.8741795 | 0.205731011 | 0.214531545 |
| Ftsj1 | 0.020505367 | 0.234613525 | 0.7064894 | 0.077571519 | 0.057208912 |
| Ftsj3 | -0.00129174 | 0.962548355 | 0.9964418 | 0.196495725 | 0.178013914 |
| Ftx | -0.023930245 | 0.322183352 | 0.7733024 | 0.129921747 | 0.137702791 |
| Fubp1 | -0.091377989 | 0.016556035 | 0.2530598 | 0.765740189 | 0.779439851 |
| Fubp3 | -0.014455153 | 0.568634477 | 0.8956532 | 0.152224866 | 0.150158061 |
| Fuca1 | 0.015584886 | 0.651915719 | 0.9189792 | 0.441904676 | 0.42108112 |
| Fuca2 | -0.037858221 | 0.136021997 | 0.6032919 | 0.139078009 | 0.157181215 |
| Fundc1 | -0.009340668 | 0.756567297 | 0.9521533 | 0.25301057 | 0.24247358 |
| Fundc2 | -0.070776871 | 0.049174126 | 0.4182752 | 0.634993837 | 0.668081563 |
| Fuom | 0.067388728 | 0.058744991 | 0.4520948 | 0.624214684 | 0.580704485 |
| Furin | -0.044161554 | 0.101718651 | 0.5510917 | 0.166037381 | 0.188261307 |
| Fus | -0.092549614 | 0.021102011 | 0.2817388 | 0.893836043 | 1.011356804 |
| Fut11 | -0.011392526 | 0.561436584 | 0.8926240 | 0.078316852 | 0.087015554 |
| Fut8 | -0.02207312 | 0.266711001 | 0.7308765 | 0.072796055 | 0.090821649 |

| | | | | | |
|---|---|---|---|---|---|
| Fuz | 0.006623443 | 0.745410224 | 0.9485717 | 0.102808582 | 0.091205336 |
| Fxn | 0.000533307 | 0.983318939 | 0.9976072 | 0.155182004 | 0.157493882 |
| Fxr1 | 0.01806068 | 0.610211846 | 0.9090802 | 0.705553126 | 0.62109158 |
| Fxr2 | 0.036325073 | 0.283263196 | 0.7427848 | 0.428787371 | 0.35999873 |
| Fxyd3 | -0.021450771 | 0.570492514 | 0.8962534 | 0.289853256 | 0.324449563 |
| Fxyd5 | -0.029211966 | 0.063900835 | 0.4700387 | 0.029832325 | 0.056791692 |
| Fxyd6 | 0.049442487 | 0.219063022 | 0.6903129 | 0.676122151 | 0.614734599 |
| Fyco1 | 0.02136196 | 0.420593258 | 0.8300990 | 0.18743657 | 0.174590698 |
| Fyn | -0.027177592 | 0.150859807 | 0.6280320 | 0.062688713 | 0.079635096 |
| Fyttd1 | -0.011531531 | 0.7499142 | 0.9504537 | 0.475445509 | 0.491649572 |
| Fzd3 | 0.055495178 | 0.063525119 | 0.4688758 | 0.27644465 | 0.24572355 |
| Fzd6 | -0.006023537 | 0.706987402 | 0.9408354 | 0.043363333 | 0.054096654 |
| Fzr1 | -0.003549226 | 0.890320887 | 0.9819597 | 0.169417194 | 0.154505524 |
| G2e3 | -0.043552981 | 0.075440471 | 0.4966285 | 0.117620244 | 0.142393502 |
| G3bp1 | -0.02994103 | 0.415868127 | 0.8300990 | 0.642304093 | 0.636563195 |
| G3bp2 | 0.007549872 | 0.833441468 | 0.9723494 | 0.837217514 | 0.81953631 |
| G6pc2 | 0.237593188 | 8.95E-07 | 0.0002072 | 1.005393883 | 0.954742018 |
| G6pc3 | -0.04678425 | 0.124783891 | 0.5863837 | 0.246147276 | 0.264227622 |
| G6pdx | 0.024886745 | 0.203410089 | 0.6772740 | 0.088934185 | 0.083474887 |
| G730003C15Rik | -0.014818826 | 0.289207565 | 0.7494616 | 0.036836387 | 0.040923416 |
| Gaa | -0.096759695 | 0.00560155 | 0.1509982 | 0.403570715 | 0.504297221 |
| Gab1 | -0.038902966 | 0.011753926 | 0.2135511 | 0.024443346 | 0.054992235 |
| Gab2 | -0.013550797 | 0.486302239 | 0.8637358 | 0.077684056 | 0.084278418 |
| Gabarap | -0.004888776 | 0.791339557 | 0.9623764 | 2.436286904 | 2.347984728 |
| Gabarapl1 | -0.036206168 | 0.315802875 | 0.7705987 | 0.69259271 | 0.72207589 |
| Gabarapl2 | -0.004411203 | 0.86614329 | 0.9770148 | 1.731093061 | 1.597572297 |
| Gabbr1 | -0.037996889 | 0.212493947 | 0.6826230 | 0.232443797 | 0.245494116 |
| Gabbr2 | 0.018303173 | 0.519670608 | 0.8741795 | 0.161661891 | 0.206824212 |
| Gabpa | -0.04278132 | 0.037222778 | 0.3659563 | 0.067493784 | 0.096933244 |
| Gabpb1 | 0.072134633 | 0.000881527 | 0.0451582 | 0.149504108 | 0.082885254 |
| Gabpb2 | -0.054529304 | 0.044803692 | 0.4018558 | 0.132081378 | 0.184898623 |
| Gabrb3 | -0.010370647 | 0.733852032 | 0.9464134 | 0.25309326 | 0.256026949 |
| Gad1 | 0.050151281 | 0.043794689 | 0.3989378 | 0.135596448 | 0.130458869 |
| Gadd45a | -0.074567798 | 0.050852631 | 0.4253440 | 0.411323568 | 0.460302951 |
| Gadd45b | 0.043757638 | 0.292352446 | 0.7526152 | 0.457820131 | 0.455789241 |
| Gadd45g | -0.055257719 | 0.218251928 | 0.6897120 | 1.389228775 | 1.490869312 |
| Gadd45gip1 | 0.000283338 | 0.994058998 | 0.9985888 | 0.646458467 | 0.620069332 |
| Gak | 0.011626633 | 0.674979956 | 0.9265887 | 0.174625304 | 0.182377242 |
| Gal3st1 | 0.012528136 | 0.508689027 | 0.8704067 | 0.09176207 | 0.074978545 |
| Galc | -0.008175063 | 0.710581427 | 0.9419370 | 0.099460852 | 0.106693711 |
| Gale | 0.035976712 | 0.209631732 | 0.6807660 | 0.259902251 | 0.187496603 |
| Galk2 | 0.000601794 | 0.98137684 | 0.9976072 | 0.167692437 | 0.156175158 |
| Galns | -0.040844919 | 0.127642236 | 0.5905834 | 0.152204656 | 0.172669263 |
| Galnt1 | 0.015186726 | 0.653822865 | 0.9200648 | 0.350307923 | 0.342847411 |
| Galnt10 | 0.017140566 | 0.350226045 | 0.7927303 | 0.079709713 | 0.07096475 |
| Galnt11 | 0.013674248 | 0.646209089 | 0.9169695 | 0.264884804 | 0.230409318 |
| Galnt12 | 0.0011354 | 0.954654881 | 0.9954690 | 0.101346722 | 0.08799638 |
| Galnt13 | -0.031416765 | 0.146964769 | 0.6232627 | 0.104341613 | 0.103165123 |
| Galnt14 | -0.012808314 | 0.554821784 | 0.8902995 | 0.082919773 | 0.116748867 |
| Galnt18 | 0.001805806 | 0.915747518 | 0.9845236 | 0.060104978 | 0.064874301 |
| Galnt4 | 0.018128908 | 0.21465304 | 0.6842380 | 0.055105838 | 0.040669453 |
| Galnt7 | 0.006763948 | 0.754570473 | 0.9519143 | 0.117000097 | 0.096980909 |
| Galt | -0.021703831 | 0.280622462 | 0.7408618 | 0.080573921 | 0.090303851 |
| Gan | 0.000739781 | 0.967914897 | 0.9976072 | 0.073440011 | 0.067662745 |
| Ganab | -0.036096818 | 0.304222263 | 0.7634877 | 0.399205406 | 0.425178256 |
| Ganc | -0.03436564 | 0.160370747 | 0.6370337 | 0.107736118 | 0.14926828 |
| Gap43 | 0.007889386 | 0.730381587 | 0.9459020 | 0.082233421 | 0.087413002 |
| Gapdh | -0.042011205 | 0.107969709 | 0.5636817 | 2.077921025 | 2.019964107 |
| Gapvd1 | 0.019264858 | 0.574324199 | 0.8974946 | 0.348728352 | 0.352361953 |
| Gar1 | -0.023210517 | 0.453155268 | 0.8489707 | 0.279418793 | 0.267947848 |
| Garre1 | -0.029186895 | 0.199669905 | 0.6732581 | 0.108081407 | 0.127401745 |
| Gars | 0.059334846 | 0.094830766 | 0.5350704 | 1.17546197 | 1.065784676 |
| Gart | -0.011609113 | 0.622479748 | 0.9121505 | 0.119620072 | 0.12890807 |
| Gas2 | -0.062434448 | 0.01774558 | 0.2587928 | 0.139178155 | 0.169977979 |
| Gas2l1 | -0.053772072 | 0.005119624 | 0.1434651 | 0.060965551 | 0.094434956 |
| Gas5 | 0.043253962 | 0.079733817 | 0.5076761 | 2.235367094 | 2.104035368 |

| | | | | | |
|---|---|---|---|---|---|
| Gas8 | -0.024815265 | 0.247935262 | 0.7177068 | 0.090476398 | 0.110686709 |
| Gata6 | 0.020794159 | 0.554668452 | 0.8902995 | 0.430459138 | 0.360285239 |
| Gatad1 | -0.086007988 | 0.017563764 | 0.2582401 | 0.46370276 | 0.48589855 |
| Gatad2a | -0.048968944 | 0.113138645 | 0.5696203 | 0.237382751 | 0.283812635 |
| Gatad2b | -0.000763866 | 0.983249928 | 0.9976072 | 0.481847284 | 0.467198474 |
| Gatb | -0.013254833 | 0.388824719 | 0.8170287 | 0.047805394 | 0.049939413 |
| Gatc | 0.003022135 | 0.875006169 | 0.9784331 | 0.078713872 | 0.081243842 |
| Gatd1 | 0.004446864 | 0.891801462 | 0.9819597 | 0.377194013 | 0.348960804 |
| Gatd3a | 0.013528995 | 0.58230997 | 0.9018999 | 0.16602693 | 0.133513407 |
| Gatm | -0.038447018 | 0.23828374 | 0.7109096 | 0.256616607 | 0.265226923 |
| Gba | -0.036744531 | 0.126854208 | 0.5893234 | 0.122958648 | 0.138405842 |
| Gbe1 | -0.008031584 | 0.656261256 | 0.9211780 | 0.064416392 | 0.071368054 |
| Gbf1 | -0.017356694 | 0.639494446 | 0.9160042 | 0.608828451 | 0.616915721 |
| Gbp7 | 0.032924227 | 0.149140642 | 0.6264374 | 0.133136226 | 0.103245039 |
| Gc | -0.170850729 | 6.69E-05 | 0.0070648 | 0.843329501 | 0.968914078 |
| Gca | -0.025083802 | 0.35107658 | 0.7931404 | 0.166238323 | 0.160846336 |
| Gcat | 0.009163458 | 0.663986834 | 0.9230765 | 0.105425836 | 0.09218444 |
| Gcc1 | -0.028733262 | 0.19068827 | 0.6700281 | 0.09410575 | 0.1136484 |
| Gcc2 | 0.020629089 | 0.575292892 | 0.8986689 | 0.716570525 | 0.664448401 |
| Gcdh | 0.005734769 | 0.810721213 | 0.9668615 | 0.149414302 | 0.129987809 |
| Gcfc2 | -0.01156104 | 0.478531296 | 0.8607489 | 0.051888898 | 0.057708565 |
| Gcg | -0.051703336 | 0.125293552 | 0.5870522 | 10.08279846 | 9.847983946 |
| Gcgr | 0.006112003 | 0.745625932 | 0.9485717 | 0.067875763 | 0.076035282 |
| Gch1 | -0.047377336 | 0.163028426 | 0.6400703 | 1.345182279 | 1.502309388 |
| Gck | -0.00551734 | 0.838969744 | 0.9727216 | 0.169602546 | 0.179057414 |
| Gclc | -0.081277415 | 0.010376362 | 0.2017406 | 0.210819151 | 0.292919199 |
| Gclm | -0.132003314 | 5.34E-05 | 0.0058575 | 1.298710964 | 1.45355063 |
| Gcn1 | 0.037860249 | 0.24322442 | 0.7132719 | 0.368009308 | 0.306620133 |
| Gcnt2 | 0.054957171 | 0.075128654 | 0.4966285 | 0.268883519 | 0.240285448 |
| Gcsh | -0.073835913 | 0.036669206 | 0.3626279 | 0.435548231 | 0.450698256 |
| Gdap1 | -0.015162943 | 0.663318987 | 0.9230058 | 0.405411728 | 0.452575351 |
| Gdap1l1 | -0.037736828 | 0.240804001 | 0.7120404 | 0.278021628 | 0.29484 |
| Gdap2 | -0.012182376 | 0.626516773 | 0.9128095 | 0.133683229 | 0.157724142 |
| Gde1 | -0.08979057 | 0.016409546 | 0.2525257 | 0.653079312 | 0.780682102 |
| Gdf15 | -0.032038894 | 0.435786688 | 0.8389295 | 0.297826211 | 0.355454486 |
| Gdi1 | -0.008059432 | 0.825197451 | 0.9716271 | 0.615364468 | 0.59135858 |
| Gdi2 | 0.002331249 | 0.945519509 | 0.9940179 | 1.080848893 | 0.975189146 |
| Gdpd1 | 0.00034594 | 0.989390493 | 0.9980836 | 0.154857358 | 0.166347582 |
| Gem | -0.072574714 | 0.014605059 | 0.2375117 | 0.095806107 | 0.212722824 |
| Gemin2 | 0.006153443 | 0.760178016 | 0.9521533 | 0.090854063 | 0.089047195 |
| Gemin5 | 0.023210161 | 0.212973032 | 0.6827572 | 0.095444986 | 0.065786919 |
| Gemin6 | -0.027442144 | 0.359045726 | 0.8000088 | 0.251701923 | 0.241140253 |
| Gemin7 | -0.029695165 | 0.372411867 | 0.8085689 | 0.407266344 | 0.396498504 |
| Gemin8 | 0.011003306 | 0.501938209 | 0.8695566 | 0.062262714 | 0.055051804 |
| Get1 | -0.008844479 | 0.582156946 | 0.9018999 | 0.057925101 | 0.056088658 |
| Get3 | -0.025723011 | 0.446536297 | 0.8443547 | 0.337666449 | 0.355494952 |
| Get4 | 0.016976523 | 0.591386771 | 0.9060200 | 0.299100082 | 0.281249395 |
| Gfer | 0.003322583 | 0.914842409 | 0.9845236 | 0.301847593 | 0.270394743 |
| Gfm1 | -0.015785608 | 0.456053325 | 0.8502914 | 0.102216955 | 0.099180425 |
| Gfm2 | 0.028319996 | 0.209713668 | 0.6807660 | 0.127881109 | 0.108825103 |
| Gfod1 | 0.013442557 | 0.426341168 | 0.8333604 | 0.074056656 | 0.051783468 |
| Gfod2 | -0.036171512 | 0.112439148 | 0.5693702 | 0.102869433 | 0.12136856 |
| Gfpt1 | -0.026229716 | 0.432557014 | 0.8370982 | 1.155010855 | 1.234870585 |
| Gfra1 | 0.019170542 | 0.35937712 | 0.8002291 | 0.124833449 | 0.083930653 |
| Gfra3 | -0.03988128 | 0.079700623 | 0.5076761 | 0.117823073 | 0.111320244 |
| Gfus | 0.00344189 | 0.907128612 | 0.9836812 | 0.240981622 | 0.238071379 |
| Gga1 | -0.017263686 | 0.541267135 | 0.8835017 | 0.2029713 | 0.208847274 |
| Gga2 | 0.013788623 | 0.640938936 | 0.9166712 | 0.232836086 | 0.239240898 |
| Gga3 | -0.007006908 | 0.705767 | 0.9401381 | 0.069398866 | 0.074078277 |
| Ggact | 0.108772161 | 0.000258832 | 0.0196314 | 0.308516561 | 0.212515638 |
| Ggct | 0.003906668 | 0.884209156 | 0.9802800 | 0.194463821 | 0.164855506 |
| Ggcx | -0.060206283 | 0.00835092 | 0.1793097 | 0.082585817 | 0.13366192 |
| Ggh | -0.041185456 | 0.187447721 | 0.6659251 | 0.264388328 | 0.285880227 |
| Ggnbp1 | 0.03379005 | 0.034704263 | 0.3549137 | 0.075407837 | 0.04372724 |
| Ggnbp2 | -0.056234304 | 0.110445431 | 0.5655888 | 0.551497037 | 0.591958436 |
| Ggps1 | -0.020841227 | 0.501953312 | 0.8695566 | 0.281346601 | 0.28686831 |

| | | | | | |
|---|---|---|---|---|---|
| Ggt7 | 0.010178684 | 0.470420467 | 0.8573527 | 0.041628463 | 0.040680114 |
| Ghdc | -0.016530178 | 0.352874092 | 0.7957515 | 0.063199398 | 0.07524253 |
| Ghitm | 0.110925077 | 0.004888827 | 0.1389603 | 0.898875454 | 0.891719881 |
| Ghr | -0.059797037 | 0.094320238 | 0.5334417 | 0.368860187 | 0.466888633 |
| Gid4 | 0.009631297 | 0.725964773 | 0.9450680 | 0.202145622 | 0.179233185 |
| Gid8 | -0.016226232 | 0.565821101 | 0.8944932 | 0.201851148 | 0.208603039 |
| Gigyf1 | -0.050902058 | 0.102485263 | 0.5529567 | 0.241506276 | 0.267943008 |
| Gigyf2 | -0.018580306 | 0.583904887 | 0.9021406 | 0.337663181 | 0.342374724 |
| Gin1 | 0.012872899 | 0.495238638 | 0.8682869 | 0.088379572 | 0.071506956 |
| Ginm1 | -0.015775847 | 0.659218033 | 0.9211780 | 0.571201759 | 0.566962103 |
| Gins4 | -0.025140384 | 0.262087595 | 0.7275735 | 0.122336843 | 0.120860527 |
| Gipc1 | -0.016529169 | 0.53866743 | 0.8816335 | 0.177294257 | 0.168731216 |
| Gipc2 | -0.002928559 | 0.875522458 | 0.9784331 | 0.083185597 | 0.072988277 |
| Gipr | -0.014271442 | 0.515989946 | 0.8735705 | 0.097995821 | 0.11313102 |
| Git1 | -0.002170535 | 0.935846204 | 0.9912016 | 0.185743363 | 0.186391597 |
| Git2 | -0.037638558 | 0.194756012 | 0.6719932 | 0.166763092 | 0.205387327 |
| Gjd2 | 0.142788911 | 1.87E-05 | 0.0024893 | 0.364956133 | 0.285372347 |
| Gkap1 | -0.045349889 | 0.18848041 | 0.6668854 | 0.339594781 | 0.368335764 |
| Gla | -0.006058373 | 0.784461816 | 0.9601202 | 0.105513914 | 0.111975432 |
| Glb1 | -0.043300488 | 0.086030149 | 0.5232489 | 0.122728256 | 0.162606276 |
| Glb1l2 | -0.013883304 | 0.5938856 | 0.9062213 | 0.143943668 | 0.162911361 |
| Glce | 0.004184591 | 0.86095926 | 0.9769675 | 0.119257294 | 0.13815311 |
| Gle1 | -0.03351137 | 0.261412283 | 0.7275735 | 0.211718019 | 0.227934077 |
| Glg1 | -0.0041045 | 0.912149386 | 0.9840393 | 0.682122821 | 0.716583919 |
| Glis2 | -0.02353642 | 0.124405693 | 0.5852543 | 0.030678827 | 0.053933544 |
| Glis3 | 0.031602981 | 0.151778765 | 0.6281105 | 0.118375312 | 0.111320985 |
| Glmn | -0.025894158 | 0.168053052 | 0.6451572 | 0.069554974 | 0.079264291 |
| Glmp | -0.008608284 | 0.797852679 | 0.9644698 | 0.402587797 | 0.387434792 |
| Glo1 | -0.052236164 | 0.145480853 | 0.6207868 | 0.776292418 | 0.733249226 |
| Glod4 | 0.036878773 | 0.267095988 | 0.7313133 | 0.502314807 | 0.419470032 |
| Glp1r | 0.063967476 | 0.137512693 | 0.6070876 | 0.55235757 | 0.682161454 |
| Glrb | 0.00701701 | 0.797478071 | 0.9644244 | 0.196655602 | 0.185204669 |
| Glrx | -0.010723016 | 0.685324631 | 0.9316044 | 0.152417549 | 0.178404365 |
| Glrx2 | 0.002941567 | 0.934907311 | 0.9907594 | 0.559509978 | 0.517497772 |
| Glrx3 | -0.053491521 | 0.137403674 | 0.6070876 | 0.598081747 | 0.580605045 |
| Glrx5 | -0.029903159 | 0.407880482 | 0.8270770 | 0.691984822 | 0.71334952 |
| Gls | -0.114348363 | 0.007560659 | 0.1721437 | 0.904044192 | 0.894070307 |
| Glt1d1 | -0.012261516 | 0.452821622 | 0.8486662 | 0.044803366 | 0.06066779 |
| Glt8d1 | -0.031681642 | 0.294380967 | 0.7536152 | 0.247981163 | 0.260102323 |
| Gltp | 0.000432571 | 0.986025398 | 0.9976072 | 0.148090242 | 0.135929632 |
| Glud1 | -0.032306123 | 0.393730427 | 0.8210059 | 0.797059873 | 0.722535582 |
| Glul | 0.050609994 | 0.183676525 | 0.6617383 | 0.577510447 | 0.544554515 |
| Glyr1 | 0.033528526 | 0.337418057 | 0.7834245 | 0.494717794 | 0.451590418 |
| Gm10033 | -0.004744811 | 0.855536088 | 0.9757403 | 0.163400819 | 0.150400482 |
| Gm10039 | 0.02296994 | 0.18916678 | 0.6680435 | 0.073941968 | 0.061377283 |
| Gm10076 | 0.114232383 | 4.40E-12 | 7.28E-09 | 3.177377099 | 2.963433245 |
| Gm10130 | -0.006398099 | 0.720527821 | 0.9445295 | 0.063553411 | 0.061262559 |
| Gm10138 | -0.002418934 | 0.919605947 | 0.9853630 | 0.122263758 | 0.131386265 |
| Gm10177 | 0.043317098 | 0.091123952 | 0.5321282 | 0.188223933 | 0.134086124 |
| Gm10275 | -0.006654978 | 0.719559666 | 0.9444523 | 0.061832233 | 0.07841664 |
| Gm10399 | -0.012618334 | 0.56921507 | 0.8956532 | 0.108867734 | 0.113079423 |
| Gm10524 | -0.002805512 | 0.866088491 | 0.9770148 | 0.058142256 | 0.059447272 |
| Gm10762 | 0.001667967 | 0.931833359 | 0.9893956 | 0.090393943 | 0.079389751 |
| Gm10941 | 0.020646334 | 0.399332445 | 0.8239850 | 0.118535001 | 0.13845874 |
| Gm11423 | -0.027128226 | 0.204374892 | 0.6780598 | 0.09090333 | 0.103105453 |
| Gm11478 | 0.04172418 | 0.084499753 | 0.5191841 | 0.197236784 | 0.124411683 |
| Gm11789 | 0.029993758 | 0.045082659 | 0.4027410 | 0.055461117 | 0.041973832 |
| Gm11808 | 0.018197911 | 0.191212196 | 0.6700281 | 0.061485718 | 0.032625563 |
| Gm11837 | -0.056307716 | 0.06958561 | 0.4837345 | 0.236827279 | 0.254581578 |
| Gm12254 | -0.007085393 | 0.658289612 | 0.9211780 | 0.057452112 | 0.051965064 |
| Gm12258 | -0.017654229 | 0.32023993 | 0.7719458 | 0.053838782 | 0.072813072 |
| Gm12350 | -0.004031119 | 0.833401451 | 0.9723494 | 0.075171676 | 0.083714536 |
| Gm12940 | -0.003281154 | 0.839256901 | 0.9727382 | 0.052299527 | 0.054040851 |
| Gm12976 | -0.031223771 | 0.137305774 | 0.6070876 | 0.072006744 | 0.105063368 |
| Gm13205 | 0.026709312 | 0.038780785 | 0.3777809 | 0.054054037 | 0.026588321 |
| Gm13268 | 0.020708743 | 0.142220021 | 0.6152737 | 0.054453791 | 0.034319839 |

| | | | | | |
|---|---|---|---|---|---|
| Gm13421 | -0.009739434 | 0.516573984 | 0.8736890 | 0.046857617 | 0.048020957 |
| Gm13436 | -0.018789588 | 0.344003464 | 0.7875638 | 0.081820354 | 0.090741527 |
| Gm13594 | 0.008917724 | 0.520038069 | 0.8741795 | 0.049709619 | 0.033837303 |
| Gm13611 | 0.025815593 | 0.266244172 | 0.7307399 | 0.152317804 | 0.119664947 |
| Gm13835 | -0.00067713 | 0.963752843 | 0.9966421 | 0.044921512 | 0.047654382 |
| Gm14004 | 0.01222586 | 0.49444212 | 0.8678106 | 0.081968954 | 0.062407344 |
| Gm14167 | -0.015560769 | 0.329097565 | 0.7778857 | 0.039821474 | 0.060657301 |
| Gm14295 | 0.026455417 | 0.083949553 | 0.5184165 | 0.064766534 | 0.045992635 |
| Gm14296 | 0.006683775 | 0.668017557 | 0.9230765 | 0.055809539 | 0.047920119 |
| Gm14303 | 0.009320167 | 0.608955034 | 0.9088672 | 0.081597668 | 0.069074218 |
| Gm14326 | 0.038588612 | 0.007490913 | 0.1721437 | 0.064795916 | 0.03038597 |
| Gm14403 | 0.011609913 | 0.433280309 | 0.8375363 | 0.052301276 | 0.042615895 |
| Gm14586 | 0.004290027 | 0.801578866 | 0.9647143 | 0.06732582 | 0.061084083 |
| Gm14966 | 0.003933599 | 0.773132543 | 0.9556977 | 0.04374821 | 0.036124705 |
| Gm15417 | 0.012783262 | 0.66793643 | 0.9230765 | 0.231789388 | 0.246360228 |
| Gm15427 | -0.042843618 | 0.048642257 | 0.4158684 | 0.088443685 | 0.119007746 |
| Gm15441 | 0.007755401 | 0.63151696 | 0.9133517 | 0.06522946 | 0.052715069 |
| Gm15446 | -0.017068192 | 0.319194741 | 0.7710043 | 0.055309318 | 0.06123687 |
| Gm15492 | 0.046640384 | 6.96E-05 | 0.0072372 | 0.053653917 | 0.016934314 |
| Gm15500 | -0.054034327 | 0.068756488 | 0.4835495 | 0.217911387 | 0.247793672 |
| Gm15706 | 0.035803342 | 0.282042388 | 0.7418354 | 0.342409873 | 0.312868607 |
| Gm15972 | 0.021292699 | 0.198495909 | 0.6730962 | 0.068189789 | 0.054048695 |
| Gm16062 | 0.038008395 | 0.025025328 | 0.3041540 | 0.071526088 | 0.050701061 |
| Gm16124 | -0.034311244 | 0.171084275 | 0.6507063 | 0.140136617 | 0.150028359 |
| Gm16174 | 0.06474546 | 0.028272163 | 0.3219975 | 0.22280385 | 0.206705889 |
| Gm16253 | 0.003738452 | 0.800941616 | 0.9647143 | 0.043970693 | 0.046007109 |
| Gm16286 | 0.008182619 | 0.811679845 | 0.9673015 | 0.690608667 | 0.649345723 |
| Gm16315 | -0.012839651 | 0.436001271 | 0.8390644 | 0.044887655 | 0.064981638 |
| Gm1673 | 0.075089779 | 0.044225196 | 0.4007863 | 0.609375934 | 0.486878432 |
| Gm16740 | 0.000344847 | 0.985119754 | 0.9976072 | 0.075347999 | 0.072046603 |
| Gm16973 | -0.00101491 | 0.947389071 | 0.9945018 | 0.047626498 | 0.045502257 |
| Gm17018 | 0.051107132 | 0.125227725 | 0.5870522 | 0.395817381 | 0.352955555 |
| Gm17251 | 0.029129138 | 0.160560281 | 0.6371283 | 0.114924164 | 0.086864508 |
| Gm17491 | -0.053524965 | 0.001093585 | 0.0511715 | 0.033193463 | 0.067016473 |
| Gm17733 | 0.018313497 | 0.146736081 | 0.6228591 | 0.043502496 | 0.027610076 |
| Gm18811 | -0.027748553 | 0.057623781 | 0.4501007 | 0.023690214 | 0.050885909 |
| Gm19196 | 0.013316658 | 0.325948696 | 0.7763291 | 0.047167622 | 0.033103195 |
| Gm19412 | -0.014721793 | 0.376807689 | 0.8101797 | 0.044873463 | 0.062701507 |
| Gm19938 | -0.022614012 | 0.233648421 | 0.7051391 | 0.064358985 | 0.077997936 |
| Gm2011 | -0.031931794 | 0.163050181 | 0.6400703 | 0.101858618 | 0.125597829 |
| Gm20208 | 0.013881501 | 0.654667972 | 0.9206439 | 0.265326833 | 0.251630331 |
| Gm20257 | -0.020853533 | 0.327642675 | 0.7778857 | 0.092552154 | 0.105433551 |
| Gm20300 | -0.057629301 | 0.014933967 | 0.2393294 | 0.100753011 | 0.139302636 |
| Gm20342 | -0.043101878 | 0.02326348 | 0.2958637 | 0.054906519 | 0.077790014 |
| Gm20554 | -0.012746528 | 0.591671928 | 0.9060200 | 0.143919037 | 0.116958847 |
| Gm20604 | -0.051276017 | 0.027661497 | 0.3181899 | 0.107325992 | 0.141790472 |
| Gm20721 | 0.005761668 | 0.741606763 | 0.9479080 | 0.071067194 | 0.056812168 |
| Gm20939 | 0.017641032 | 0.181528751 | 0.6583478 | 0.048477257 | 0.028936496 |
| Gm21596 | 0.032643461 | 0.010354028 | 0.2017406 | 0.05432686 | 0.022574692 |
| Gm21926 | -0.021534099 | 0.285388359 | 0.7451828 | 0.086148882 | 0.097701749 |
| Gm26532 | -0.050049405 | 0.023165067 | 0.2949903 | 0.060028947 | 0.115744535 |
| Gm26559 | -0.002937862 | 0.850659709 | 0.9748841 | 0.052943787 | 0.052092377 |
| Gm26670 | 0.005997282 | 0.666869169 | 0.9230765 | 0.046063808 | 0.038080839 |
| Gm26782 | -0.039986038 | 0.12476898 | 0.5863837 | 0.146921791 | 0.173114866 |
| Gm26881 | -0.039028329 | 0.061800387 | 0.4614965 | 0.079712619 | 0.105815069 |
| Gm26910 | 1.89E-05 | 0.99888893 | 0.9996989 | 0.041320147 | 0.035809046 |
| Gm27033 | 0.039549232 | 0.064934188 | 0.4732896 | 0.107238819 | 0.098459435 |
| Gm28438 | -0.558896788 | 2.12E-53 | 1.05E-49 | 0.008573137 | 0.602517552 |
| Gm28791 | 0.01890958 | 0.190468478 | 0.6700281 | 0.052698069 | 0.037557523 |
| Gm2a | -0.020719517 | 0.475067039 | 0.8602394 | 0.239334076 | 0.221070624 |
| Gm30025 | -0.019504794 | 0.416488821 | 0.8300990 | 0.128012032 | 0.134111853 |
| Gm30551 | 0.006498524 | 0.658275487 | 0.9211780 | 0.056665518 | 0.037147898 |
| Gm30648 | -0.025778497 | 0.418194695 | 0.8300990 | 0.247333292 | 0.267035623 |
| Gm32031 | -0.016288333 | 0.583684851 | 0.9021406 | 0.214447692 | 0.246210791 |
| Gm32856 | -0.011892239 | 0.429289139 | 0.8340365 | 0.036856629 | 0.050128416 |
| Gm3294 | 0.021679863 | 0.265785092 | 0.7306783 | 0.092322817 | 0.077089087 |

| | | | | | |
|---|---|---|---|---|---|
| Gm33989 | -0.036067092 | 0.181004555 | 0.65735074 | 0.144734288 | 0.169228026 |
| Gm34220 | 0.017772005 | 0.408727147 | 0.82764474 | 0.119065743 | 0.097038837 |
| Gm34961 | -0.025102507 | 0.088441383 | 0.52847714 | 0.028373624 | 0.050427705 |
| Gm3625 | 0.019820149 | 0.365991553 | 0.80501911 | 0.123256621 | 0.09436695 |
| Gm36371 | 0.012953652 | 0.424412123 | 0.8324361 | 0.061961291 | 0.051147291 |
| Gm36608 | 0.003556204 | 0.782014733 | 0.95908472 | 0.038099264 | 0.032583427 |
| Gm36738 | 0.076304056 | 0.029081062 | 0.32641720 | 0.428702149 | 0.372105272 |
| Gm36839 | -0.016521369 | 0.407846114 | 0.82707770 | 0.074958736 | 0.087290383 |
| Gm37140 | -0.013611111 | 0.485268762 | 0.86297153 | 0.066230294 | 0.083036694 |
| Gm37254 | -0.018779059 | 0.257790918 | 0.72505979 | 0.045353305 | 0.061037691 |
| Gm37297 | -0.030361626 | 0.036994539 | 0.36407324 | 0.022544089 | 0.051591527 |
| Gm37376 | -0.010846722 | 0.701602167 | 0.93818771 | 0.152687999 | 0.148797246 |
| Gm37452 | 0.016724081 | 0.516852619 | 0.87375221 | 0.157998087 | 0.143388267 |
| Gm37465 | -0.018340554 | 0.287447265 | 0.74713163 | 0.049896775 | 0.065238624 |
| Gm37517 | -0.026298496 | 0.110361001 | 0.56558881 | 0.046558568 | 0.056905821 |
| Gm37728 | -0.043239014 | 0.014685953 | 0.23765848 | 0.035199677 | 0.076781261 |
| Gm37733 | 0.001447963 | 0.926011764 | 0.98792100 | 0.046342352 | 0.047781113 |
| Gm37834 | 0.010396716 | 0.515635349 | 0.87348064 | 0.060025136 | 0.049807507 |
| Gm37859 | -0.049948454 | 0.046265054 | 0.40736569 | 0.119704493 | 0.145239994 |
| Gm37899 | -0.026756688 | 0.354683819 | 0.79691318 | 0.16804463 | 0.183878038 |
| Gm379 | -0.086167239 | 0.013463691 | 0.22925110 | 0.257585881 | 0.326418878 |
| Gm38070 | -0.039096138 | 0.355218409 | 0.79691318 | 0.298069051 | 0.451483867 |
| Gm38157 | -0.017171628 | 0.463243825 | 0.85244258 | 0.10835543 | 0.113466792 |
| Gm38285 | 0.026957762 | 0.092989931 | 0.53282111 | 0.07121836 | 0.044741967 |
| Gm38340 | -0.031119187 | 0.146550364 | 0.62257688 | 0.078131467 | 0.098315815 |
| Gm38394 | -0.107484098 | 0.000244964 | 0.01883750 | 0.151344474 | 0.225033872 |
| Gm3924 | 0.007162176 | 0.720001205 | 0.94445237 | 0.080413879 | 0.094753645 |
| Gm39469 | -0.00017454 | 0.990638273 | 0.99808369 | 0.050418857 | 0.040687479 |
| Gm4013 | -0.053765457 | 0.078398434 | 0.50561759 | 0.200329905 | 0.270869029 |
| Gm40309 | -0.03758385 | 0.035113034 | 0.35770363 | 0.047699559 | 0.069476081 |
| Gm4045 | 0.019372656 | 0.147019631 | 0.62326271 | 0.054280429 | 0.02920088 |
| Gm4204 | 0.000432597 | 0.980575458 | 0.99760728 | 0.063351904 | 0.067781433 |
| Gm42067 | -0.00888408 | 0.66024814 | 0.92117801 | 0.076542726 | 0.098018259 |
| Gm42372 | -0.027623229 | 0.138210615 | 0.60798394 | 0.073129171 | 0.073496579 |
| Gm42447 | 0.017185592 | 0.310581216 | 0.76774624 | 0.061160172 | 0.060834838 |
| Gm42448 | 0.032516555 | 0.017729411 | 0.25879289 | 0.04905244 | 0.033412826 |
| Gm42467 | -0.028796732 | 0.067164133 | 0.48071298 | 0.035235088 | 0.058970842 |
| Gm42685 | 0.046053087 | 0.005942981 | 0.15454384 | 0.078904307 | 0.047797985 |
| Gm42770 | 0.012769279 | 0.375888275 | 0.81017735 | 0.047619746 | 0.038746938 |
| Gm42826 | -0.005308325 | 0.789744077 | 0.96184914 | 0.073730954 | 0.084889604 |
| Gm42841 | 0.001908437 | 0.908200696 | 0.98373739 | 0.054543602 | 0.05375564 |
| Gm4285 | 0.007744054 | 0.552113934 | 0.88940731 | 0.043693241 | 0.032300028 |
| Gm42937 | -0.014407245 | 0.41816262 | 0.83009907 | 0.053360236 | 0.068622507 |
| Gm42941 | -0.031295681 | 0.08874519 | 0.52887221 | 0.052463833 | 0.07436662 |
| Gm42992 | -0.015050571 | 0.492889905 | 0.86693011 | 0.101127163 | 0.100227944 |
| Gm42993 | -0.057414891 | 0.001870692 | 0.07452714 | 0.041400494 | 0.07868269 |
| Gm43305 | -0.008787887 | 0.667756523 | 0.92307657 | 0.094194942 | 0.093965001 |
| Gm4332 | -0.012923228 | 0.599424835 | 0.90782341 | 0.133262278 | 0.139646896 |
| Gm43336 | -0.031622883 | 0.116713046 | 0.57274720 | 0.068396542 | 0.090119025 |
| Gm43359 | -0.04782723 | 0.042325786 | 0.39529100 | 0.088962348 | 0.127233399 |
| Gm43362 | -0.048311625 | 0.165283733 | 0.64207071 | 0.345910042 | 0.386777153 |
| Gm43737 | 0.030520733 | 0.01537036 | 0.24258752 | 0.053988917 | 0.02221431 |
| Gm43813 | -0.009448109 | 0.62950792 | 0.91305644 | 0.070720274 | 0.075752643 |
| Gm43879 | 0.016767296 | 0.329636067 | 0.77788578 | 0.058109481 | 0.063767258 |
| Gm43889 | 0.001620324 | 0.914445943 | 0.98452363 | 0.038238953 | 0.053006904 |
| Gm43909 | 0.010084591 | 0.456979147 | 0.85163124 | 0.046995673 | 0.034215914 |
| Gm44109 | 0.009040042 | 0.630732953 | 0.91327848 | 0.087513828 | 0.074184359 |
| Gm44164 | -0.029849662 | 0.087487079 | 0.52706087 | 0.045145341 | 0.070235575 |
| Gm4419 | 0.003259971 | 0.865808987 | 0.97701481 | 0.082868409 | 0.084207499 |
| Gm44250 | -0.009932633 | 0.606352251 | 0.90871393 | 0.066912516 | 0.081490475 |
| Gm44518 | -0.023421184 | 0.108588086 | 0.56368171 | 0.030209331 | 0.048611853 |
| Gm44639 | -0.038619638 | 0.01456144 | 0.23719120 | 0.026284044 | 0.05858056 |
| Gm44797 | -0.009226325 | 0.636101095 | 0.91535418 | 0.072749379 | 0.080283766 |
| Gm44799 | 0.002793183 | 0.877551494 | 0.97855831 | 0.069766735 | 0.067453888 |
| Gm44800 | -0.021233629 | 0.2874434 | 0.74713163 | 0.076441381 | 0.090042412 |
| Gm44801 | -0.025244433 | 0.173169 | 0.65255079 | 0.05832386 | 0.075761335 |

| | | | | | |
|---|---|---|---|---|---|
| Gm44829 | -0.003062512 | 0.863639083 | 0.9769675 | 0.065371303 | 0.060362431 |
| Gm44863 | 0.014904302 | 0.333989914 | 0.7804836 | 0.055711229 | 0.043678169 |
| Gm44950 | -0.006732249 | 0.711583534 | 0.9419834 | 0.070316946 | 0.072589486 |
| Gm45051 | 0.020098207 | 0.316286735 | 0.7705987 | 0.100328368 | 0.077007445 |
| Gm45159 | -0.015623945 | 0.403438584 | 0.8264703 | 0.056940554 | 0.068931676 |
| Gm45223 | -0.01594311 | 0.380660641 | 0.8117350 | 0.064395318 | 0.068923714 |
| Gm45486 | 0.002323482 | 0.882279392 | 0.9798677 | 0.050227868 | 0.048451453 |
| Gm45716 | -0.006498294 | 0.693739081 | 0.9359725 | 0.049630285 | 0.064242368 |
| Gm45847 | 0.003508267 | 0.815038627 | 0.9685493 | 0.048548699 | 0.044854906 |
| Gm45871 | -0.029380425 | 0.108843987 | 0.5644183 | 0.062648567 | 0.075145911 |
| Gm46404 | -0.011022205 | 0.472685379 | 0.8595729 | 0.041737092 | 0.052615359 |
| Gm46430 | -0.018160052 | 0.233403773 | 0.7051391 | 0.042510327 | 0.05166555 |
| Gm46565 | 0.015857332 | 0.27422034 | 0.7351955 | 0.050148098 | 0.041193011 |
| Gm47163 | -0.002249937 | 0.917264968 | 0.9848120 | 0.107013267 | 0.108141677 |
| Gm47283 | -0.028348193 | 0.277943311 | 0.7382158 | 0.153255782 | 0.154171062 |
| Gm47586 | -0.038030747 | 0.060949078 | 0.4605600 | 0.06480426 | 0.098333041 |
| Gm4791 | 0.03150438 | 0.042510506 | 0.3952910 | 0.062779421 | 0.045293476 |
| Gm48226 | -0.038707317 | 0.193139593 | 0.6710839 | 0.222618352 | 0.235948293 |
| Gm48266 | 0.00487938 | 0.793827976 | 0.9624508 | 0.081551119 | 0.069557669 |
| Gm48267 | -0.035259851 | 0.064226313 | 0.4700387 | 0.064152207 | 0.079728195 |
| Gm48562 | 0.00068529 | 0.966964379 | 0.9976072 | 0.056353202 | 0.055407657 |
| Gm48582 | -0.019440511 | 0.287477895 | 0.7471316 | 0.053902983 | 0.070935811 |
| Gm4890 | 0.010455868 | 0.434021971 | 0.8377586 | 0.045276203 | 0.033726199 |
| Gm48933 | 0.023303807 | 0.054044007 | 0.4398596 | 0.043135451 | 0.023628134 |
| Gm49041 | -0.024333009 | 0.301251597 | 0.7606291 | 0.117146532 | 0.124226242 |
| Gm49066 | -0.006154191 | 0.779774302 | 0.9587705 | 0.109402561 | 0.108842201 |
| Gm4924 | -0.001316731 | 0.952362441 | 0.9951566 | 0.114022727 | 0.10895947 |
| Gm49336 | 0.01849458 | 0.418364254 | 0.8300990 | 0.120704539 | 0.108377547 |
| Gm49338 | -0.008270546 | 0.609314361 | 0.9088672 | 0.051060674 | 0.05882673 |
| Gm49359 | 0.003040862 | 0.824502782 | 0.971271 | 0.046771236 | 0.034439441 |
| Gm49494 | -0.012282549 | 0.443154767 | 0.8431541 | 0.037038904 | 0.061233641 |
| Gm49599 | -0.070473565 | 0.061327934 | 0.4611772 | 0.431931015 | 0.475107536 |
| Gm49602 | 0.004036035 | 0.788658668 | 0.9615897 | 0.049788974 | 0.044907257 |
| Gm49797 | 0.000700747 | 0.970528644 | 0.9976072 | 0.078579589 | 0.076377463 |
| Gm49980 | -0.036234426 | 0.239040286 | 0.7114105 | 0.225437682 | 0.253643475 |
| Gm50240 | -0.03875076 | 0.174575677 | 0.6530130 | 0.20147629 | 0.215394229 |
| Gm50340 | 0.008349802 | 0.622783167 | 0.9121505 | 0.06424845 | 0.059851902 |
| Gm50394 | 0.009924429 | 0.584717007 | 0.9023908 | 0.069025286 | 0.068404417 |
| Gm50455 | 0.010535903 | 0.434626714 | 0.8384863 | 0.046073654 | 0.035017657 |
| Gm5617 | 0.073052733 | 0.044442225 | 0.4009562 | 0.819215469 | 0.721742892 |
| Gm5860 | 0.022831359 | 0.316384036 | 0.7705987 | 0.098989166 | 0.112601798 |
| Gm5870 | 0.008679551 | 0.643011877 | 0.9166712 | 0.071322115 | 0.079113133 |
| Gm5914 | 0.003341016 | 0.887506587 | 0.9812414 | 0.128466358 | 0.131053785 |
| Gm609 | -0.021722371 | 0.258497048 | 0.7251953 | 0.065448459 | 0.080718922 |
| Gm6563 | 0.015846675 | 0.34482839 | 0.7879976 | 0.066957326 | 0.055778785 |
| Gm6658 | -0.023731496 | 0.142134539 | 0.6152737 | 0.044329359 | 0.058936969 |
| Gm6863 | 0.000542279 | 0.976595463 | 0.9976072 | 0.065999769 | 0.073956346 |
| Gm7072 | -0.063958625 | 0.019025517 | 0.2683814 | 0.146931291 | 0.194202014 |
| Gm7160 | 0.007877229 | 0.626693346 | 0.9128095 | 0.057322861 | 0.053791157 |
| Gm7240 | 0.00134066 | 0.954369546 | 0.9954690 | 0.12202298 | 0.122149915 |
| Gm7467 | -0.005240524 | 0.787599986 | 0.9608781 | 0.081695123 | 0.080687953 |
| Gm8186 | 0.01883975 | 0.250534753 | 0.7207210 | 0.070612303 | 0.052143021 |
| Gm8624 | -0.024054378 | 0.25303446 | 0.7223071 | 0.089226027 | 0.101535239 |
| Gm8730 | -0.057039004 | 0.012737658 | 0.2232465 | 0.089754387 | 0.126886951 |
| Gm8797 | -0.032091486 | 0.047245342 | 0.4106761 | 0.035182081 | 0.062228448 |
| Gm9294 | -0.035891872 | 0.02651275 | 0.3112502 | 0.035993135 | 0.062598283 |
| Gm9385 | 0.014636726 | 0.482003216 | 0.8620306 | 0.110287985 | 0.093394434 |
| Gm9616 | -0.018054411 | 0.497283813 | 0.8684629 | 0.166421381 | 0.169595006 |
| Gm9794 | -0.068333078 | 0.043645592 | 0.3986436 | 0.31800741 | 0.364317175 |
| Gm9828 | 0.002754996 | 0.841992799 | 0.9734889 | 0.045596379 | 0.036121533 |
| Gm9843 | -0.013980842 | 0.607429103 | 0.9087139 | 0.178938082 | 0.180622681 |
| Gm9856 | 0.000843774 | 0.974803853 | 0.9976072 | 0.186430426 | 0.165744364 |
| Gm9920 | 0.007124483 | 0.621348863 | 0.9121505 | 0.047719688 | 0.039602775 |
| Gm9993 | 0.004476 | 0.803051788 | 0.9647143 | 0.072992466 | 0.062605468 |
| Gmcl1 | -0.012250754 | 0.520242085 | 0.8741795 | 0.076133596 | 0.081120152 |
| Gmds | -0.047217585 | 0.183764742 | 0.6617383 | 0.423239463 | 0.44895686 |

| | | | | | |
|---|---|---|---|---|---|
| Gmeb1 | -0.02044733 | 0.231291199 | 0.70261419 | 0.05472773 | 0.066671208 |
| Gmeb2 | -0.045888982 | 0.023669282 | 0.29872682 | 0.062692476 | 0.101028588 |
| Gmfb | -0.062782836 | 0.047284865 | 0.41067610 | 0.283878754 | 0.306148977 |
| Gmnn | 0.000613123 | 0.965240048 | 0.99668790 | 0.045559168 | 0.040414465 |
| Gmppa | -0.092660698 | 0.00578788 | 0.15327813 | 0.355372271 | 0.392560955 |
| Gmppb | -0.025209397 | 0.473887502 | 0.85972275 | 0.378659748 | 0.374086643 |
| Gmpr | 0.02399279 | 0.505904502 | 0.86955665 | 0.707741516 | 0.682713537 |
| Gmpr2 | 0.015515939 | 0.504198953 | 0.86955665 | 0.138547955 | 0.11927243 |
| Gmps | -0.041788597 | 0.233441577 | 0.70513913 | 0.466189705 | 0.464383775 |
| Gna11 | -0.067032251 | 0.052168838 | 0.43146469 | 0.841237191 | 0.895672672 |
| Gna12 | -0.022475717 | 0.304008069 | 0.76348775 | 0.104107226 | 0.114492374 |
| Gna13 | 0.016249107 | 0.616116595 | 0.91140420 | 0.296005126 | 0.30114344 |
| Gnai1 | -0.006422322 | 0.854779688 | 0.97554315 | 0.473441793 | 0.394923745 |
| Gnai2 | -0.050323368 | 0.025491376 | 0.30614340 | 1.902665083 | 2.06218869 |
| Gnai3 | 0.044799963 | 0.179843029 | 0.65500705 | 0.420484273 | 0.387209292 |
| Gnao1 | -0.008439149 | 0.824022719 | 0.97105924 | 0.888784051 | 0.865374379 |
| Gnaq | 0.014997347 | 0.68564963 | 0.93160448 | 0.534661989 | 0.518013386 |
| Gnas | -0.103240881 | 2.42E-08 | 1.27E-05 | 4.668677436 | 4.639408922 |
| Gnat2 | -0.009472103 | 0.577114726 | 0.89946101 | 0.053745501 | 0.063957786 |
| Gnaz | -0.021751534 | 0.547098402 | 0.88585084 | 0.399171289 | 0.430813282 |
| Gnb1 | 0.093436905 | 0.007119446 | 0.16800019 | 1.066089354 | 0.872310799 |
| Gnb2 | -0.048900316 | 0.055610273 | 0.44493323 | 1.75672152 | 1.704274238 |
| Gnb4 | -0.032066069 | 0.094017589 | 0.53344171 | 0.072182099 | 0.083538616 |
| Gnb5 | -0.029671988 | 0.286647354 | 0.74685807 | 0.212296473 | 0.207041068 |
| Gne | 0.011795557 | 0.621800133 | 0.91215057 | 0.141869205 | 0.127693144 |
| Gng12 | 0.14286564 | 0.000649326 | 0.03598497 | 1.001722199 | 1.014156503 |
| Gng2 | -0.073592789 | 0.007015119 | 0.16768669 | 0.144499598 | 0.182670712 |
| Gng4 | 0.047367613 | 0.17557749 | 0.65366590 | 0.485070722 | 0.476965069 |
| Gng5 | 0.037039505 | 0.077759506 | 0.50383690 | 2.470232573 | 2.422387048 |
| Gnl1 | -0.037881136 | 0.235888805 | 0.70793905 | 0.347150607 | 0.334981966 |
| Gnl2 | -0.003659883 | 0.904775702 | 0.98317077 | 0.272324539 | 0.245766355 |
| Gnl3 | -0.06218896 | 0.103034475 | 0.55531410 | 0.508304839 | 0.590001707 |
| Gnl3l | 0.030482719 | 0.382186189 | 0.81268823 | 0.390637427 | 0.372213478 |
| Gnpat | -0.077159439 | 0.001790104 | 0.07386815 | 0.101885201 | 0.156085578 |
| Gnpda1 | -0.021317826 | 0.358616432 | 0.79979204 | 0.106571433 | 0.130148167 |
| Gnpda2 | 0.012422867 | 0.698200331 | 0.93696920 | 0.325319297 | 0.305974587 |
| Gnpnat1 | 0.003599461 | 0.915803613 | 0.98452361 | 0.339818836 | 0.33205443 |
| Gnptab | -0.025757698 | 0.081940458 | 0.51460110 | 0.030901304 | 0.051579561 |
| Gnptg | -0.103664253 | 0.003807077 | 0.11951329 | 0.758502132 | 0.826509936 |
| Gns | -0.056887788 | 0.116296223 | 0.57246708 | 0.544492178 | 0.673371875 |
| Golga1 | 0.045418203 | 0.107643769 | 0.56337238 | 0.23644123 | 0.188762112 |
| Golga2 | 0.00630781 | 0.843273566 | 0.97348890 | 0.304736591 | 0.275111333 |
| Golga3 | 0.051093982 | 0.138838722 | 0.60937791 | 0.42251316 | 0.387141245 |
| Golga4 | -0.063064719 | 0.087026818 | 0.52540553 | 0.991054998 | 1.106034928 |
| Golga5 | 0.001023031 | 0.974261865 | 0.99760728 | 0.285373882 | 0.280699493 |
| Golga7 | -0.0837571 | 0.00783139 | 0.17497159 | 0.264595543 | 0.312721469 |
| Golga7b | 0.025190533 | 0.152910563 | 0.62862147 | 0.084557589 | 0.055972459 |
| Golgb1 | -0.072729529 | 0.02228765 | 0.29053020 | 1.516922889 | 1.624478456 |
| Golim4 | -0.008708514 | 0.819036018 | 0.96990621 | 0.586230657 | 0.651770566 |
| Golm1 | -0.020503588 | 0.223007165 | 0.69559587 | 0.042994674 | 0.067163264 |
| Golm2 | 0.007709298 | 0.833126829 | 0.97234942 | 0.583018017 | 0.620826987 |
| Golph3 | 0.038668813 | 0.294680686 | 0.75365882 | 0.615538112 | 0.5917487 |
| Golph3l | 0.001482583 | 0.950834192 | 0.99485327 | 0.152156597 | 0.129495516 |
| Golt1b | 0.020739648 | 0.561032585 | 0.89262402 | 0.593348327 | 0.505149245 |
| Gon4l | 0.016888541 | 0.465442437 | 0.85440210 | 0.136896481 | 0.114128112 |
| Gon7 | -0.012077714 | 0.607355343 | 0.90871392 | 0.142873808 | 0.130358486 |
| Gopc | -0.030992684 | 0.291841983 | 0.75209060 | 0.210828049 | 0.217251595 |
| Gorab | -0.022237594 | 0.391821397 | 0.82001503 | 0.155789711 | 0.168517324 |
| Gorasp1 | -0.000141005 | 0.993859138 | 0.99858888 | 0.072525798 | 0.069280699 |
| Gorasp2 | -0.081104302 | 0.022204115 | 0.28982211 | 0.635093767 | 0.720205246 |
| Gosr1 | 0.038030475 | 0.112697795 | 0.56962035 | 0.153933389 | 0.122755286 |
| Gosr2 | -0.043890932 | 0.206903676 | 0.67975655 | 0.387842525 | 0.437028058 |
| Got1 | -0.031222794 | 0.329570206 | 0.77788578 | 0.233222652 | 0.276704762 |
| Got2 | 0.053799032 | 0.11571696 | 0.57246708 | 0.405375331 | 0.34642833 |
| Gpaa1 | 0.024694174 | 0.281631117 | 0.74164605 | 0.149801058 | 0.11566423 |
| Gpalpp1 | 0.022943072 | 0.325141044 | 0.77630994 | 0.132080835 | 0.11451092 |

| | | | | | |
|---|---|---|---|---|---|
| Gpank1 | -0.001207195 | 0.956110519 | 0.9954690 | 0.112585913 | 0.107357738 |
| Gpat4 | -0.008607248 | 0.760330499 | 0.9522129 | 0.20120478 | 0.207136739 |
| Gpatch1 | -0.000401371 | 0.987577004 | 0.9976072 | 0.16253137 | 0.152997359 |
| Gpatch11 | 0.051565009 | 0.106718007 | 0.5623243 | 0.334457593 | 0.270944247 |
| Gpatch2 | -0.039631115 | 0.173176188 | 0.6525507 | 0.181791721 | 0.231001381 |
| Gpatch2l | 0.012680967 | 0.454961026 | 0.8499460 | 0.074709023 | 0.055067713 |
| Gpatch4 | 0.005278183 | 0.797165111 | 0.9644244 | 0.093825941 | 0.089396794 |
| Gpatch8 | 0.002186006 | 0.951135536 | 0.9949662 | 0.441403048 | 0.433044711 |
| Gpbp1 | -0.024539829 | 0.479582736 | 0.8610807 | 0.47515902 | 0.466836307 |
| Gpbp1l1 | -0.016037078 | 0.615481007 | 0.9111433 | 0.306150332 | 0.302446533 |
| Gpc4 | -0.046174356 | 0.169647561 | 0.6482680 | 0.374181176 | 0.392642899 |
| Gpcpd1 | -0.024940148 | 0.338528496 | 0.7834245 | 0.136295607 | 0.161372346 |
| Gpd1 | -0.017787438 | 0.316254935 | 0.7705987 | 0.059207829 | 0.067894708 |
| Gpd2 | 0.079717026 | 0.02742093 | 0.3181899 | 0.532591173 | 0.417830492 |
| Gphn | 0.048763642 | 0.084851471 | 0.5202265 | 0.222857395 | 0.178958182 |
| Gpi1 | -0.056401543 | 0.120212261 | 0.5797304 | 0.710401776 | 0.700818045 |
| Gpkow | 0.000367242 | 0.984868921 | 0.9976072 | 0.079160691 | 0.077749449 |
| Gpld1 | -0.089180553 | 7.43E-05 | 0.0073749 | 0.066319708 | 0.129920821 |
| Gpn1 | 0.024246365 | 0.294694514 | 0.7536588 | 0.138319082 | 0.116926047 |
| Gpn2 | -0.006353225 | 0.731667685 | 0.9460562 | 0.084676726 | 0.075861986 |
| Gpn3 | -0.00261701 | 0.91644522 | 0.9845285 | 0.148859159 | 0.140408961 |
| Gpr107 | -0.044255569 | 0.15232183 | 0.6281105 | 0.232602893 | 0.259665463 |
| Gpr108 | 0.002580639 | 0.934502611 | 0.9906246 | 0.303304354 | 0.281751474 |
| Gpr119 | -0.042557213 | 0.208913239 | 0.6807660 | 0.319370024 | 0.265258695 |
| Gpr135 | 0.023394204 | 0.063964164 | 0.4700387 | 0.051916314 | 0.024583497 |
| Gpr137 | -0.017715851 | 0.480837661 | 0.8617579 | 0.151005883 | 0.146859561 |
| Gpr137b | -0.013488352 | 0.495131034 | 0.8682869 | 0.088530335 | 0.088887035 |
| Gpr137b-ps | -0.015909041 | 0.459919858 | 0.8524343 | 0.098979974 | 0.104555101 |
| Gpr137c | 0.003017078 | 0.837899681 | 0.9727216 | 0.052186857 | 0.040440637 |
| Gpr146 | -0.024059354 | 0.164443389 | 0.6417303 | 0.041880171 | 0.064138772 |
| Gpr158 | 0.020961405 | 0.346581642 | 0.7899443 | 0.097621224 | 0.120699072 |
| Gpr180 | -0.008770116 | 0.782156797 | 0.9590847 | 0.301743808 | 0.275513287 |
| Gpr19 | 0.004184574 | 0.844888637 | 0.9734889 | 0.106197615 | 0.099745875 |
| Gpr27 | -0.043584112 | 0.162999209 | 0.6400703 | 0.264907055 | 0.293914658 |
| Gpr45 | 0.030664614 | 0.287336147 | 0.7471316 | 0.190440879 | 0.21208522 |
| Gpr85 | 0.000660467 | 0.977674047 | 0.9976072 | 0.120281001 | 0.125532309 |
| Gpr89 | 0.006934699 | 0.791584511 | 0.9624054 | 0.179204115 | 0.16617737 |
| Gprasp1 | -0.043142635 | 0.259639142 | 0.7263475 | 0.562732536 | 0.582083722 |
| Gprasp2 | -0.048617716 | 0.128487049 | 0.5914729 | 0.249230612 | 0.290357626 |
| Gprc5c | 0.00565349 | 0.797747032 | 0.9644698 | 0.119309138 | 0.101068792 |
| Gprin1 | 0.011707626 | 0.552719043 | 0.8900045 | 0.098922163 | 0.074136995 |
| Gprin3 | -0.005247529 | 0.801541646 | 0.9647143 | 0.087927067 | 0.098438137 |
| Gps1 | -0.011566848 | 0.733656726 | 0.9464134 | 0.588332519 | 0.543265131 |
| Gps2 | 0.042495894 | 0.16711458 | 0.6442971 | 0.277270258 | 0.238595292 |
| Gpsm1 | -0.041581229 | 0.115775502 | 0.5724670 | 0.141426477 | 0.176482536 |
| Gpt2 | 0.027280383 | 0.196848042 | 0.6730962 | 0.098465197 | 0.099271911 |
| Gpx1 | -0.059732464 | 0.128088631 | 0.5912244 | 0.807453087 | 0.897130571 |
| Gpx2 | -0.068478568 | 0.079424507 | 0.5076205 | 0.483443213 | 0.640838332 |
| Gpx3 | -0.122515867 | 0.000658003 | 0.0360629 | 2.649867469 | 2.559151652 |
| Gpx4 | -0.078911744 | 0.004770677 | 0.1371742 | 1.545570786 | 1.611303266 |
| Gpx4-ps2 | -0.056245412 | 0.058002689 | 0.4508199 | 0.201506503 | 0.259711175 |
| Gramd1a | -0.050694294 | 0.095341415 | 0.5351767 | 0.220629157 | 0.244487319 |
| Gramd1b | 0.007765668 | 0.642114631 | 0.9166712 | 0.059371927 | 0.060702422 |
| Gramd1c | 0.032402495 | 0.040022196 | 0.3835943 | 0.072674776 | 0.043716176 |
| Gramd3 | 0.001466045 | 0.966831457 | 0.9976072 | 0.404654934 | 0.420777492 |
| Grb10 | 0.012952431 | 0.725638825 | 0.9450332 | 0.442943671 | 0.383236357 |
| Grb14 | -0.054365796 | 0.069315265 | 0.4835495 | 0.207399899 | 0.237308137 |
| Grb2 | -0.028049705 | 0.418810996 | 0.8300990 | 0.458356907 | 0.470468702 |
| Grb7 | -0.006298727 | 0.812312311 | 0.9673015 | 0.175704035 | 0.17126284 |
| Grcc10 | 0.054985802 | 0.112015391 | 0.5679582 | 0.470040711 | 0.4006534 |
| Grhpr | -0.020690597 | 0.486186671 | 0.8637127 | 0.242282237 | 0.237483189 |
| Gria2 | -0.123049531 | 0.01079126 | 0.2035157 | 0.709146252 | 0.790185584 |
| Gria3 | -0.018662025 | 0.559064501 | 0.8921289 | 0.274761424 | 0.247822338 |
| Grik5 | -0.019863873 | 0.405077431 | 0.8270770 | 0.128692236 | 0.140014361 |
| Grin1 | 0.002755291 | 0.881137274 | 0.9795705 | 0.059437496 | 0.075062071 |
| Grina | -0.020582597 | 0.568348243 | 0.8955308 | 0.492544999 | 0.508038339 |

| | | | | | |
|---|---|---|---|---|---|
| Grip1 | -0.006125421 | 0.699400631 | 0.93720988 | 0.042575381 | 0.0580907 |
| Gripap1 | -0.059588128 | 0.089549494 | 0.53023604 | 0.406761976 | 0.464343265 |
| Grk2 | 0.046054695 | 0.119321852 | 0.57827370 | 0.27133745 | 0.238557955 |
| Grk3 | -0.010372205 | 0.733060968 | 0.94641343 | 0.223407083 | 0.244665821 |
| Grk5 | -0.009940885 | 0.589864349 | 0.90467754 | 0.06714644 | 0.077239523 |
| Grk6 | -0.010855223 | 0.589382222 | 0.90465390 | 0.088848111 | 0.08855141 |
| Grn | 0.006783839 | 0.785383253 | 0.96025161 | 0.147456355 | 0.148124282 |
| Grpel1 | 0.019135611 | 0.591139504 | 0.90602008 | 0.784598502 | 0.734411734 |
| Grpel2 | -0.033704928 | 0.16642537 | 0.64292850 | 0.120472487 | 0.145099212 |
| Grsf1 | -0.037752191 | 0.266866004 | 0.73109935 | 0.402075695 | 0.418025094 |
| Grtp1 | -0.0243721 | 0.347939399 | 0.79179535 | 0.145195481 | 0.166144292 |
| Grwd1 | -0.002063928 | 0.890883441 | 0.98195975 | 0.054065349 | 0.046805611 |
| Gsap | 0.004355206 | 0.792756727 | 0.96244605 | 0.063648241 | 0.056075251 |
| Gsdma | -0.100691596 | 0.003577212 | 0.11494803 | 0.220520512 | 0.402791341 |
| Gse1 | -0.054238799 | 0.14014599 | 0.61301372 | 0.432664546 | 0.495342943 |
| Gsk3a | 0.005392294 | 0.857754746 | 0.97635422 | 0.268440712 | 0.258814381 |
| Gsk3b | 0.10576233 | 0.00278795 | 0.09798352 | 1.024898326 | 0.918401673 |
| Gskip | 0.064829769 | 0.009976157 | 0.19872185 | 0.180861841 | 0.127864548 |
| Gsn | 0.023200589 | 0.504431099 | 0.86955665 | 0.39456927 | 0.360429573 |
| Gspt1 | 0.036870241 | 0.331094256 | 0.77830687 | 0.822047787 | 0.760432466 |
| Gspt2 | -0.024382057 | 0.388782911 | 0.81702870 | 0.206612533 | 0.198187916 |
| Gsr | -0.032431918 | 0.336179314 | 0.78216775 | 0.347444958 | 0.36892412 |
| Gss | -0.01054358 | 0.633418862 | 0.91436483 | 0.109393564 | 0.112738663 |
| Gstm1 | -0.073032673 | 0.080119329 | 0.50882442 | 0.438824279 | 0.575106372 |
| Gstm2 | -0.014076558 | 0.404621228 | 0.82707700 | 0.041443897 | 0.064310154 |
| Gstm4 | 0.027715537 | 0.101832273 | 0.55140619 | 0.077438771 | 0.051412396 |
| Gstm5 | 0.054821616 | 0.129158801 | 0.59207731 | 0.596431805 | 0.500348724 |
| Gsto1 | 0.003277768 | 0.877654654 | 0.97855831 | 0.084970373 | 0.099578825 |
| Gsto2 | 0.011167998 | 0.484426268 | 0.86275947 | 0.064416616 | 0.050346465 |
| Gstp1 | 0.036388504 | 0.395881196 | 0.82306793 | 0.88892831 | 0.871852077 |
| Gstp-ps | -0.000951891 | 0.965043334 | 0.99664211 | 0.113473687 | 0.105173772 |
| Gstt1 | -0.044633428 | 0.21984585 | 0.69102371 | 0.498173703 | 0.493317756 |
| Gstt2 | -0.091935029 | 0.001016691 | 0.04895909 | 0.147459319 | 0.20519707 |
| Gstz1 | -0.028696672 | 0.406710064 | 0.82707700 | 0.533270583 | 0.482766166 |
| Gt(ROSA)26Sor | 0.001028984 | 0.974671918 | 0.99760728 | 0.309669601 | 0.283517788 |
| Gtf2a1 | -0.0367866 | 0.180103744 | 0.65516280 | 0.171537139 | 0.195412718 |
| Gtf2a2 | -0.01613765 | 0.663267719 | 0.92300582 | 0.705669352 | 0.677837404 |
| Gtf2b | -0.044543639 | 0.210764767 | 0.68220598 | 0.433361259 | 0.49823589 |
| Gtf2e1 | 0.01283653 | 0.443047304 | 0.84315410 | 0.074681081 | 0.05506467 |
| Gtf2e2 | -0.004689854 | 0.87971509 | 0.97910621 | 0.315351974 | 0.280288311 |
| Gtf2f1 | -0.099008485 | 0.000930132 | 0.04619851 | 0.195558756 | 0.252477298 |
| Gtf2f2 | 0.005356548 | 0.838869122 | 0.97272164 | 0.172559635 | 0.169181208 |
| Gtf2h1 | -0.032388289 | 0.240051374 | 0.71187913 | 0.157606672 | 0.188344045 |
| Gtf2h2 | 0.008382025 | 0.626702035 | 0.91280959 | 0.076531612 | 0.061165537 |
| Gtf2h3 | -0.013257222 | 0.550007029 | 0.88788767 | 0.106038172 | 0.108118015 |
| Gtf2h4 | -0.06386991 | 0.00197746 | 0.07815301 | 0.073338637 | 0.107845177 |
| Gtf2h5 | 0.010141555 | 0.775773535 | 0.95585380 | 0.918816323 | 0.900568891 |
| Gtf2i | 0.000571772 | 0.987352107 | 0.99760728 | 0.542018204 | 0.524114879 |
| Gtf2ird1 | -0.021265643 | 0.328034704 | 0.77788578 | 0.09807884 | 0.107942506 |
| Gtf3a | 0.014451593 | 0.65430446 | 0.92040559 | 0.342252864 | 0.315310656 |
| Gtf3c1 | -0.020159892 | 0.503179593 | 0.86955665 | 0.238035036 | 0.241759995 |
| Gtf3c2 | 0.022181023 | 0.417536891 | 0.83009907 | 0.202955956 | 0.17907348 |
| Gtf3c3 | 0.025483466 | 0.28893542 | 0.74934362 | 0.159299201 | 0.126274626 |
| Gtf3c4 | -0.023984326 | 0.302455793 | 0.76131983 | 0.098876963 | 0.132068739 |
| Gtf3c5 | -0.005977306 | 0.733782129 | 0.94641343 | 0.058452723 | 0.06841775 |
| Gtf3c6 | 0.015724296 | 0.598874114 | 0.90755130 | 0.267196439 | 0.230434861 |
| Gtpbp1 | 0.009445799 | 0.622229796 | 0.91215057 | 0.092834778 | 0.078339514 |
| Gtpbp10 | -0.000993748 | 0.955542973 | 0.99546908 | 0.067766829 | 0.06707369 |
| Gtpbp2 | 0.053546316 | 0.132810964 | 0.59882510 | 0.403022393 | 0.399265735 |
| Gtpbp3 | 0.042114637 | 0.011499583 | 0.21125160 | 0.086148896 | 0.048125213 |
| Gtpbp4 | -0.038233569 | 0.268026645 | 0.73238419 | 0.441827405 | 0.443678906 |
| Gtpbp6 | 0.059727622 | 0.046724409 | 0.40945771 | 0.302976932 | 0.221493619 |
| Gtpbp8 | 0.044174848 | 0.062889299 | 0.46589983 | 0.15681976 | 0.12067229 |
| Gucd1 | 0.005439581 | 0.81268629 | 0.96730151 | 0.118086002 | 0.117183615 |
| Gucy1b1 | 0.003435398 | 0.870181622 | 0.97801782 | 0.109157322 | 0.089372791 |
| Gucy2c | 0.012773221 | 0.433318677 | 0.83753631 | 0.051676322 | 0.056768726 |

| | | | | | |
|---|---|---|---|---|---|
| Guf1 | 0.015277099 | 0.506287345 | 0.86955669 | 0.133490764 | 0.115034583 |
| Guk1 | -0.027145877 | 0.369212157 | 0.80567191 | 1.544184717 | 1.478601675 |
| Gusb | -0.028585012 | 0.35393566 | 0.79677824 | 0.253016735 | 0.282266155 |
| Gxylt1 | -0.014257588 | 0.576792197 | 0.89946101 | 0.140771298 | 0.149667678 |
| Gyg | 0.043248946 | 0.092546386 | 0.53255842 | 0.189889701 | 0.145207459 |
| Gys1 | -0.004391716 | 0.805443164 | 0.96471431 | 0.062996754 | 0.068666152 |
| Gzf1 | 0.004843732 | 0.847103488 | 0.97406590 | 0.153425373 | 0.150948069 |
| H13 | -0.123498953 | 0.000758825 | 0.04055940 | 0.622556064 | 0.739753941 |
| H1f0 | 0.023326048 | 0.560588274 | 0.89262401 | 0.657613784 | 0.601483309 |
| H1f10 | -0.037813445 | 0.324392075 | 0.77575529 | 0.501549095 | 0.476704811 |
| H1f2 | -0.057355547 | 0.192428975 | 0.67002813 | 0.718446164 | 0.871938714 |
| H1f3 | 0.003269982 | 0.84414904 | 0.97348890 | 0.046164404 | 0.060741046 |
| H1f4 | -0.033010072 | 0.378770025 | 0.81030809 | 0.38197831 | 0.430041447 |
| H2ac18 | 0.080950457 | 0.018846346 | 0.26779862 | 0.374795418 | 0.372987143 |
| H2ac24 | 0.09273264 | 0.000844613 | 0.04409760 | 0.224739798 | 0.177714609 |
| H2ac8 | -0.002245902 | 0.91617123 | 0.98452361 | 0.090679558 | 0.109074663 |
| H2aj | 0.050446887 | 0.070884733 | 0.48790080 | 1.647538654 | 1.626235555 |
| H2aw | 0.005400335 | 0.812877914 | 0.96730151 | 0.133523349 | 0.110587388 |
| H2ax | -0.031514851 | 0.383581422 | 0.81297870 | 0.49708782 | 0.477693425 |
| H2az1 | 0.029864078 | 0.165214839 | 0.64207071 | 2.480848171 | 2.356791146 |
| H2az2 | 0.001773792 | 0.961704266 | 0.99635579 | 0.920797615 | 0.857343253 |
| H2bc4 | 0.093238531 | 0.022883943 | 0.29291447 | 1.091429255 | 1.061396766 |
| H2bc6 | 0.021468847 | 0.206908045 | 0.67975659 | 0.065604992 | 0.059819569 |
| H2bu2 | -0.009302885 | 0.822250514 | 0.97069202 | 0.815056975 | 0.680485356 |
| H2-D1 | -0.069574377 | 0.016470179 | 0.25252577 | 1.854890149 | 1.850124349 |
| H2-K1 | -0.010779372 | 0.660033933 | 0.92117802 | 2.18705122 | 2.139758793 |
| H2-K2 | -0.040355138 | 0.055254987 | 0.44352231 | 0.08245001 | 0.10319157 |
| H2-Ke6 | 0.024744402 | 0.321755238 | 0.77323490 | 0.191387849 | 0.137987298 |
| H2-M3 | 0.021894922 | 0.45360654 | 0.84904829 | 0.237118706 | 0.207549927 |
| H2-Q2 | -0.06225151 | 0.091347323 | 0.53212820 | 0.578784799 | 0.604175137 |
| H2-Q4 | 0.054474134 | 0.01756527 | 0.25824014 | 0.158928432 | 0.100393006 |
| H2-T22 | -0.000821573 | 0.97442992 | 0.99760728 | 0.163318702 | 0.152188695 |
| H2-T23 | -0.101520918 | 0.007707641 | 0.17340589 | 0.593382122 | 0.606358921 |
| H3f3a | -0.027666919 | 0.15312921 | 0.62900279 | 2.566857538 | 2.444298388 |
| H3f3b | -0.023788244 | 0.221299852 | 0.69291531 | 2.822687608 | 2.814868416 |
| H4c4 | 0.003886489 | 0.867524729 | 0.97738161 | 0.114034301 | 0.115326713 |
| H4c8 | 0.056103694 | 0.024107207 | 0.30156809 | 0.153087743 | 0.128393736 |
| H4c9 | 0.003228454 | 0.902423178 | 0.98254414 | 0.145523878 | 0.158152351 |
| H6pd | 0.013265999 | 0.323958952 | 0.77555520 | 0.040378665 | 0.037199408 |
| Habp4 | -0.005439939 | 0.837982438 | 0.97272164 | 0.171539295 | 0.170965067 |
| Hacd1 | 0.0091109 | 0.605514084 | 0.90871391 | 0.066672023 | 0.061573516 |
| Hacd2 | 0.070405607 | 0.00279694 | 0.09798351 | 0.180783723 | 0.111080253 |
| Hacd3 | -0.008176078 | 0.762938923 | 0.95320007 | 0.191909602 | 0.179103298 |
| Hace1 | -0.028873566 | 0.114428857 | 0.57070601 | 0.057987552 | 0.076699658 |
| Hadh | 0.155760149 | 0.000536808 | 0.03227350 | 0.81630806 | 0.871964021 |
| Hadha | 0.007832847 | 0.827333685 | 0.97170067 | 0.495899435 | 0.473849019 |
| Hadhb | -0.03349189 | 0.316036923 | 0.77059879 | 0.582955322 | 0.584080221 |
| Hagh | -0.045701888 | 0.176925433 | 0.65366590 | 0.93695347 | 0.91320365 |
| Haghl | 0.000572124 | 0.98379886 | 0.99760728 | 0.187352807 | 0.196652735 |
| Hap1 | -0.084316682 | 0.035576227 | 0.35919149 | 0.72851545 | 0.923435439 |
| Hapln1 | -0.009942772 | 0.765437422 | 0.95403181 | 0.31622335 | 0.248412357 |
| Hapln4 | 0.013915925 | 0.515342525 | 0.87348064 | 0.08832009 | 0.105836792 |
| Harbi1 | -0.011267421 | 0.604946869 | 0.90871391 | 0.10125348 | 0.107519493 |
| Hars | -0.036340144 | 0.177863941 | 0.65381879 | 0.154205465 | 0.184334223 |
| Hars2 | 0.014540492 | 0.450881989 | 0.84789420 | 0.099243432 | 0.076058051 |
| Hat1 | 0.028132905 | 0.277742739 | 0.73821581 | 0.185094533 | 0.148399634 |
| Haus1 | -0.021817903 | 0.168266436 | 0.64547681 | 0.048803558 | 0.057549789 |
| Haus2 | -0.027248668 | 0.262852853 | 0.72757354 | 0.116036195 | 0.135753576 |
| Haus7 | -0.002044911 | 0.912819878 | 0.98420329 | 0.087147039 | 0.074805951 |
| Hax1 | -0.000301596 | 0.993504405 | 0.99858888 | 0.576856946 | 0.575226001 |
| Hbegf | -0.018784432 | 0.47308225 | 0.85967684 | 0.114001201 | 0.147823003 |
| Hbp1 | -0.039247736 | 0.216774394 | 0.68723301 | 0.272547873 | 0.294276736 |
| Hbs1l | 0.039413498 | 0.209402686 | 0.68076601 | 0.307644488 | 0.271206178 |
| Hccs | 0.005865486 | 0.835022224 | 0.97234941 | 0.1921185 | 0.185943966 |
| Hcfc1 | -0.004995951 | 0.861937786 | 0.97696750 | 0.211610879 | 0.211415108 |
| Hcfc1r1 | 0.038419861 | 0.271369058 | 0.73420300 | 1.292221152 | 1.11741822 |

| | | | | | |
|---|---|---|---|---|---|
| Hcfc2 | -0.028074117 | 0.23724935 | 0.7098511 | 0.120634951 | 0.145658132 |
| Hcn2 | 0.043978717 | 0.056908807 | 0.4480439 | 0.163264549 | 0.112477975 |
| Hcn3 | -0.004581169 | 0.759865073 | 0.9521533 | 0.049714505 | 0.042255635 |
| Hcn4 | 0.023236795 | 0.110187881 | 0.5655888 | 0.04899022 | 0.043763379 |
| Hdac1 | -0.069633691 | 0.024528504 | 0.3039588 | 0.255268087 | 0.308756491 |
| Hdac11 | -0.022398572 | 0.397962279 | 0.8238469 | 0.160133145 | 0.173878549 |
| Hdac2 | -0.020842654 | 0.514323297 | 0.8733459 | 0.329884389 | 0.311162049 |
| Hdac3 | -0.026499074 | 0.33371736 | 0.7804836 | 0.21448823 | 0.198808568 |
| Hdac4 | 0.012916287 | 0.597735222 | 0.9075513 | 0.143362699 | 0.129674473 |
| Hdac5 | -0.000591299 | 0.984805579 | 0.9976072 | 0.276472458 | 0.278971973 |
| Hdac6 | -0.034970074 | 0.342647141 | 0.7872431 | 0.466287879 | 0.478283761 |
| Hdac7 | -0.012923116 | 0.411058081 | 0.8289685 | 0.044917419 | 0.053667927 |
| Hdac8 | -0.01734831 | 0.320863523 | 0.7721897 | 0.053836394 | 0.068509457 |
| Hddc2 | 0.057718544 | 0.005841762 | 0.1537142 | 0.119480449 | 0.088025343 |
| Hddc3 | 0.005778189 | 0.666135262 | 0.9230765 | 0.046698844 | 0.032959132 |
| Hdgf | 0.012329405 | 0.726947255 | 0.9452179 | 0.865594977 | 0.820331045 |
| Hdgfl2 | 0.046754417 | 0.099977016 | 0.5455291 | 0.256621252 | 0.188889258 |
| Hdgfl3 | -0.059837116 | 0.071230572 | 0.4881297 | 0.278926566 | 0.339012336 |
| Hdhd5 | -0.000154377 | 0.993028268 | 0.9985888 | 0.06214845 | 0.069628043 |
| Hdlbp | 0.008633475 | 0.817230853 | 0.9693806 | 1.204258744 | 1.164317672 |
| Heatr1 | -0.000997929 | 0.949948495 | 0.9948462 | 0.058458849 | 0.057577942 |
| Heatr3 | -0.003368973 | 0.876603165 | 0.9785583 | 0.103867241 | 0.105689022 |
| Heatr5a | -0.008640333 | 0.675879292 | 0.9269628 | 0.081049129 | 0.091784339 |
| Heatr5b | 0.008497885 | 0.677804686 | 0.9281886 | 0.093882142 | 0.088374029 |
| Heatr6 | -0.020901426 | 0.451989304 | 0.8484352 | 0.192502015 | 0.189631437 |
| Hebp1 | 0.025905481 | 0.066000031 | 0.4768538 | 0.061727114 | 0.032337856 |
| Heca | 0.004333915 | 0.882419176 | 0.9798677 | 0.221441962 | 0.228414106 |
| Hectd1 | 0.014596086 | 0.687086471 | 0.9326625 | 0.714606536 | 0.691228002 |
| Hectd2 | 0.008273258 | 0.67352626 | 0.9261686 | 0.086403756 | 0.077089313 |
| Hectd3 | -0.019318082 | 0.238663698 | 0.7109741 | 0.046360817 | 0.05839653 |
| Hectd4 | -0.019244569 | 0.436304094 | 0.8390644 | 0.136819323 | 0.147576197 |
| Heg1 | 0.009245642 | 0.728341423 | 0.9458355 | 0.182830415 | 0.163105817 |
| Helb | -0.011144154 | 0.468520252 | 0.8563037 | 0.045692082 | 0.050423443 |
| Helq | 0.050682738 | 0.010776426 | 0.2035157 | 0.109560821 | 0.07705529 |
| Helz | -0.028142069 | 0.415253522 | 0.8300990 | 0.426560093 | 0.429120332 |
| Helz2 | 0.011234131 | 0.475126226 | 0.8602394 | 0.057325191 | 0.049029227 |
| Hepacam2 | 0.060762831 | 0.090345795 | 0.5319418 | 0.466800676 | 0.417090745 |
| Herc1 | 0.018448081 | 0.613140416 | 0.9108386 | 0.567515927 | 0.561146357 |
| Herc2 | -0.031707758 | 0.374363591 | 0.8087297 | 0.433269175 | 0.463869834 |
| Herc3 | -0.087205525 | 0.000637987 | 0.0357561 | 0.107810447 | 0.180828279 |
| Herc4 | -0.007760506 | 0.756909743 | 0.9521533 | 0.139768115 | 0.148603003 |
| Herc6 | -0.018734408 | 0.273758179 | 0.7351955 | 0.049824049 | 0.063983413 |
| Herpud1 | 0.035825503 | 0.33894011 | 0.7834245 | 1.131942114 | 1.136140091 |
| Herpud2 | 0.029103346 | 0.343582562 | 0.7872955 | 0.276087453 | 0.235709283 |
| Hes1 | -0.029204634 | 0.425895909 | 0.8328183 | 0.316713598 | 0.315458955 |
| Hes6 | -0.022135153 | 0.503032584 | 0.8695566 | 0.307898426 | 0.32005853 |
| Hexa | -0.047577479 | 0.179670083 | 0.6550070 | 0.91800887 | 0.946554732 |
| Hexb | -0.016603536 | 0.547960611 | 0.8863001 | 0.194456578 | 0.188376236 |
| Hexim1 | -0.026874447 | 0.393044323 | 0.8206236 | 0.229290852 | 0.257982235 |
| Hfe | -0.045020172 | 0.114836792 | 0.5715910 | 0.168102229 | 0.218065178 |
| Hgfac | -0.001219078 | 0.966068744 | 0.9972322 | 0.205482621 | 0.193167114 |
| Hgh1 | -0.009805851 | 0.532188934 | 0.8782755 | 0.050411484 | 0.050328191 |
| Hgs | -0.013188566 | 0.626110275 | 0.9128095 | 0.173952617 | 0.185697449 |
| Hgsnat | 0.04156983 | 0.054048779 | 0.4398596 | 0.13169351 | 0.096451222 |
| Hhatl | 0.002306736 | 0.874610378 | 0.9784331 | 0.049942768 | 0.039069038 |
| Hhex | -0.030393589 | 0.135819559 | 0.6032919 | 0.03916007 | 0.084030901 |
| Hibadh | -0.019711324 | 0.567445229 | 0.8952062 | 0.503232397 | 0.466602168 |
| Hibch | 0.016474377 | 0.482818659 | 0.8620306 | 0.140865838 | 0.119179747 |
| Hid1 | -0.046996025 | 0.179862823 | 0.6550070 | 0.393433149 | 0.421709132 |
| Hif1a | -0.034041678 | 0.327492318 | 0.7778857 | 0.390063209 | 0.430174173 |
| Hif1an | -0.005641618 | 0.802359712 | 0.9647143 | 0.113638081 | 0.113274961 |
| Higd1a | -0.105800367 | 0.013206569 | 0.2274733 | 1.282865211 | 1.198386401 |
| Higd2a | 0.048122885 | 0.181267193 | 0.6577068 | 0.721710106 | 0.664594196 |
| Hikeshi | -0.006627618 | 0.82987674 | 0.9722601 | 0.279061733 | 0.25731769 |
| Hilpda | -0.034725951 | 0.156353315 | 0.6315293 | 0.122546898 | 0.137904515 |
| Hinfp | 0.001811545 | 0.930736651 | 0.9890634 | 0.097785758 | 0.092920834 |

| | | | | | |
|---|---|---|---|---|---|
| Hint1 | 0.009689696 | 0.657975473 | 0.9211780 | 2.227108439 | 2.097282907 |
| Hint2 | -0.053131734 | 0.15346921 | 0.6298548 | 0.722664094 | 0.757084162 |
| Hint3 | 0.041846037 | 0.249352758 | 0.7192728 | 0.612959495 | 0.576781214 |
| Hip1 | 0.005602681 | 0.712075506 | 0.9419834 | 0.049008606 | 0.045082463 |
| Hip1r | -0.053613228 | 0.086224825 | 0.5237907 | 0.237499124 | 0.288616412 |
| Hipk1 | 0.022427661 | 0.471128223 | 0.8580121 | 0.286696884 | 0.256004372 |
| Hipk2 | -0.004812526 | 0.888161962 | 0.9812414 | 0.337972315 | 0.357511208 |
| Hipk3 | -0.009093597 | 0.789249412 | 0.9618371 | 0.341092908 | 0.352986839 |
| Hira | -0.000104804 | 0.995866378 | 0.9994073 | 0.085976256 | 0.083250212 |
| Hirip3 | 0.008175252 | 0.743108298 | 0.9479080 | 0.147958777 | 0.142122355 |
| Hivep1 | 0.041495454 | 0.122961883 | 0.5833237 | 0.196163865 | 0.160023655 |
| Hivep2 | 0.051509847 | 0.069275402 | 0.4835495 | 0.222965969 | 0.188802804 |
| Hivep3 | -0.046916028 | 0.128811534 | 0.5914729 | 0.252327566 | 0.291805105 |
| Hjurp | 0.002844847 | 0.911625507 | 0.9840185 | 0.15758501 | 0.15644114 |
| Hk1 | 0.003484088 | 0.846754649 | 0.9739657 | 0.080069666 | 0.065395692 |
| Hlcs | 0.001176975 | 0.958656553 | 0.9957785 | 0.112730842 | 0.116453924 |
| Hlf | -0.012878412 | 0.468981657 | 0.8563037 | 0.059940467 | 0.067995105 |
| Hltf | 0.040822414 | 0.187378172 | 0.6659251 | 0.264826625 | 0.231745268 |
| Hmbox1 | 0.024004978 | 0.281101103 | 0.7408618 | 0.126221316 | 0.104979844 |
| Hmbs | 0.015714712 | 0.568210305 | 0.8955308 | 0.2284363 | 0.196314595 |
| Hmces | -0.040298908 | 0.212080193 | 0.6822059 | 0.308148251 | 0.32511915 |
| Hmg20a | -0.017965524 | 0.487322639 | 0.8640311 | 0.147284746 | 0.1640162 |
| Hmg20b | 0.005327571 | 0.872368297 | 0.9784331 | 0.354740281 | 0.331531772 |
| Hmgb1 | -0.023946747 | 0.429781414 | 0.8340365 | 1.397880483 | 1.315230872 |
| Hmgb2 | -0.019754377 | 0.57334496 | 0.8973494 | 0.420831096 | 0.401692932 |
| Hmgb3 | -0.020762029 | 0.390976761 | 0.8196300 | 0.137192393 | 0.126095658 |
| Hmgcl | -0.024277272 | 0.405258059 | 0.8270770 | 0.255913603 | 0.24968399 |
| Hmgcll1 | 0.026836052 | 0.141172231 | 0.6139045 | 0.088802013 | 0.068777619 |
| Hmgcr | -0.047321455 | 0.193662755 | 0.6710847 | 0.459273634 | 0.48287325 |
| Hmgcs1 | 0.01751478 | 0.581380194 | 0.9017028 | 0.287830932 | 0.248929859 |
| Hmgn1 | 0.093491077 | 0.00524556 | 0.1461684 | 1.416297985 | 1.150859325 |
| Hmgn2 | 0.092182 | 0.010773021 | 0.2035157 | 0.622242838 | 0.471801884 |
| Hmgn3 | 0.087130089 | 0.007197589 | 0.1680001 | 1.398913856 | 1.244746997 |
| Hmgn5 | 0.005186522 | 0.877126282 | 0.9785583 | 0.351334126 | 0.298449086 |
| Hmgxb3 | -0.024071404 | 0.261905706 | 0.7275735 | 0.090558326 | 0.109091471 |
| Hmgxb4 | 0.015877515 | 0.607902661 | 0.9087393 | 0.27954395 | 0.247484254 |
| Hmox1 | -0.076942317 | 0.01688137 | 0.2552792 | 0.150473065 | 0.265611328 |
| Hmox2 | 0.008538806 | 0.804663673 | 0.9647143 | 0.476200551 | 0.445446122 |
| Hnf1a | 0.025484277 | 0.170720456 | 0.6507063 | 0.087520494 | 0.070343542 |
| Hnf1aos1 | -0.008799546 | 0.762708382 | 0.9532000 | 0.222605429 | 0.21461022 |
| Hnf1b | 0.006693722 | 0.716412824 | 0.9431411 | 0.076025392 | 0.068653506 |
| Hnf4a | -0.013411485 | 0.409650193 | 0.8279808 | 0.044352226 | 0.060480017 |
| Hnf4g | 0.011896813 | 0.44977599 | 0.8478268 | 0.054121036 | 0.051609064 |
| Hnrnpa0 | -0.149796688 | 5.46E-05 | 0.0058902 | 0.632696833 | 0.737173725 |
| Hnrnpa1 | -0.042746811 | 0.270063172 | 0.7340360 | 0.766895918 | 0.716757487 |
| Hnrnpa2b1 | -0.057784836 | 0.069564177 | 0.4837345 | 1.537527096 | 1.569750667 |
| Hnrnpa3 | -0.004860534 | 0.881815044 | 0.9797222 | 1.502695043 | 1.385927074 |
| Hnrnpab | -0.005344923 | 0.881947523 | 0.9797222 | 1.02918683 | 0.981966257 |
| Hnrnpc | -0.070511279 | 0.051431105 | 0.4282799 | 0.881722234 | 0.898776977 |
| Hnrnpd | -0.037760518 | 0.305554803 | 0.7646271 | 0.906839808 | 0.925833352 |
| Hnrnpdl | -0.088854455 | 0.010778389 | 0.2035157 | 0.996601555 | 1.067794064 |
| Hnrnpf | -0.031626456 | 0.363919267 | 0.8049171 | 1.005913293 | 0.943403278 |
| Hnrnph1 | -0.000107499 | 0.997720753 | 0.9994073 | 0.69792049 | 0.683828251 |
| Hnrnph2 | -0.031763914 | 0.366958391 | 0.8053794 | 0.415961631 | 0.425346924 |
| Hnrnph3 | -0.02492819 | 0.477746667 | 0.8607489 | 0.635634454 | 0.613596076 |
| Hnrnpk | -0.026465758 | 0.350065552 | 0.7927303 | 1.490004468 | 1.411393819 |
| Hnrnpl | 0.002537494 | 0.942349262 | 0.9932112 | 0.478796135 | 0.454221206 |
| Hnrnpll | -0.00596369 | 0.84195655 | 0.9734889 | 0.247741903 | 0.233398466 |
| Hnrnpm | -0.052426926 | 0.141091524 | 0.6138718 | 0.391328783 | 0.438740098 |
| Hnrnpr | -0.012707118 | 0.723269226 | 0.9445652 | 0.585514518 | 0.552283307 |
| Hnrnpu | 0.004934452 | 0.892318609 | 0.9820058 | 1.232290633 | 1.154151537 |
| Hnrnpul1 | -0.010857515 | 0.747480887 | 0.9496370 | 0.384516918 | 0.38243777 |
| Hnrnpul2 | -0.008440785 | 0.827319544 | 0.9717006 | 0.578564915 | 0.575500984 |
| Homer1 | -0.032240225 | 0.108124274 | 0.5636817 | 0.06723817 | 0.094633268 |
| Homez | 0.009624009 | 0.639380944 | 0.9160042 | 0.115925533 | 0.088272872 |
| Hook1 | -0.016142184 | 0.662734566 | 0.9228663 | 0.592314824 | 0.607316422 |

| | | | | | |
|---|---|---|---|---|---|
| Hook2 | -0.011678437 | 0.658207146 | 0.9211780 | 0.158808589 | 0.165466202 |
| Hook3 | -0.038435299 | 0.294312755 | 0.7536152 | 0.472223512 | 0.484615901 |
| Hopx | -0.006948861 | 0.815973483 | 0.9686999 | 0.2085207 | 0.231217736 |
| Hp1bp3 | -0.002441348 | 0.947867745 | 0.9945018 | 0.633380617 | 0.627984841 |
| Hpcal1 | -0.021211921 | 0.416858715 | 0.8300990 | 0.14025417 | 0.176387575 |
| Hpf1 | -0.0003468 | 0.991956714 | 0.9985765 | 0.490385533 | 0.451409117 |
| Hpgds | -0.000590898 | 0.972882831 | 0.9976072 | 0.047591279 | 0.066324347 |
| Hprt | 0.002915588 | 0.936348116 | 0.9912468 | 0.860462453 | 0.780519829 |
| Hps1 | 0.00392593 | 0.874873061 | 0.9784331 | 0.151067265 | 0.135100985 |
| Hps3 | -0.003517004 | 0.827451956 | 0.9717006 | 0.043398615 | 0.056687419 |
| Hps4 | 0.016049928 | 0.49732149 | 0.8684629 | 0.135234801 | 0.127732505 |
| Hps5 | -0.022706866 | 0.131384191 | 0.5972652 | 0.033643337 | 0.05355033 |
| Hras | -0.018762164 | 0.597607556 | 0.9075513 | 0.581788572 | 0.576833476 |
| Hs2st1 | 0.001962452 | 0.943820514 | 0.9935293 | 0.202758133 | 0.191121283 |
| Hs3st6 | 0.028948657 | 0.196627008 | 0.6730962 | 0.123007641 | 0.112481761 |
| Hs6st1 | -0.008943212 | 0.782559141 | 0.9590862 | 0.292013644 | 0.306880728 |
| Hsbp1 | -0.019510021 | 0.525725891 | 0.8765043 | 1.215311769 | 1.186862424 |
| Hsbp1l1 | 0.000744498 | 0.980313609 | 0.9976072 | 0.250439009 | 0.23194239 |
| Hscb | 0.02660011 | 0.271110121 | 0.7342030 | 0.144086043 | 0.129918302 |
| Hsd17b10 | -0.047061339 | 0.140834642 | 0.6133099 | 0.274940917 | 0.318036452 |
| Hsd17b11 | -0.013648014 | 0.533044311 | 0.8789560 | 0.098014514 | 0.11380092 |
| Hsd17b12 | -0.038531144 | 0.259105165 | 0.7262851 | 0.367088634 | 0.391678208 |
| Hsd17b4 | 0.029277515 | 0.272394428 | 0.7342806 | 0.202642958 | 0.163649692 |
| Hsd17b7 | 0.006594931 | 0.711644755 | 0.9419834 | 0.06856393 | 0.063417504 |
| Hsdl1 | 0.004935349 | 0.836098069 | 0.9723494 | 0.132717773 | 0.126681517 |
| Hsdl2 | -0.068431603 | 0.014232749 | 0.2347500 | 0.187371822 | 0.225889741 |
| Hsf1 | -0.022761164 | 0.350990497 | 0.7931265 | 0.130452154 | 0.140887774 |
| Hsf2 | 0.000674274 | 0.977680692 | 0.9976072 | 0.125635153 | 0.131170941 |
| Hsp90aa1 | -0.026444012 | 0.397428519 | 0.8238469 | 1.7662796 | 1.794728676 |
| Hsp90ab1 | -0.023495709 | 0.103605693 | 0.5564528 | 3.362763939 | 3.404608602 |
| Hsp90b1 | -0.005079832 | 0.799687859 | 0.9647053 | 3.03312971 | 2.970036479 |
| Hspa13 | -0.077048127 | 0.033688569 | 0.3468530 | 0.387240156 | 0.46167915 |
| Hspa14 | 0.027467173 | 0.338509019 | 0.7834245 | 0.240460032 | 0.200093489 |
| Hspa1a | -0.164065175 | 3.82E-05 | 0.0044635 | 0.12800508 | 0.36810197 |
| Hspa1b | -0.069293319 | 0.011896612 | 0.2145716 | 0.069912684 | 0.160001454 |
| Hspa2 | -0.076235907 | 0.024931387 | 0.3041540 | 0.343228368 | 0.364273664 |
| Hspa4 | -0.00794833 | 0.837302137 | 0.9727177 | 0.826294361 | 0.783884201 |
| Hspa4l | -0.022540591 | 0.517294009 | 0.8738218 | 0.396168045 | 0.398770264 |
| Hspa5 | -0.07467097 | 0.003804448 | 0.1195132 | 2.745052232 | 2.848227357 |
| Hspa8 | -0.058758273 | 0.045295465 | 0.4034756 | 1.898721431 | 1.790186915 |
| Hspa9 | 0.098314114 | 0.009423321 | 0.1947486 | 1.195715111 | 1.179460524 |
| Hspb1 | 0.053402471 | 0.061412822 | 0.4611772 | 0.14887892 | 0.132558083 |
| Hspb11 | 0.047068415 | 0.1128758 | 0.5696203 | 0.271203134 | 0.220155672 |
| Hspb6 | 0.004250882 | 0.84741758 | 0.9742716 | 0.103991784 | 0.114691504 |
| Hspb8 | -0.034491121 | 0.093010954 | 0.5328211 | 0.074891038 | 0.093440395 |
| Hspbap1 | -0.018504949 | 0.235368668 | 0.7075324 | 0.04188141 | 0.056420354 |
| Hspbp1 | 0.009211623 | 0.734370413 | 0.9468380 | 0.19317721 | 0.181718686 |
| Hspd1 | 0.055864787 | 0.108588294 | 0.5636817 | 1.150896876 | 1.010216207 |
| Hspe1 | 0.059841825 | 0.068631567 | 0.4835495 | 1.498539319 | 1.320979093 |
| Hsph1 | 0.039664957 | 0.26286528 | 0.7275735 | 0.392120919 | 0.37768026 |
| Htatip2 | -0.075581855 | 0.02422645 | 0.3022973 | 0.262680626 | 0.350317016 |
| Htatsf1 | -0.059840375 | 0.072017274 | 0.4888240 | 0.317091368 | 0.349835862 |
| Htra2 | -0.017310378 | 0.388157326 | 0.8170287 | 0.079720985 | 0.090623173 |
| Htt | 0.024017395 | 0.392140193 | 0.8201624 | 0.200556167 | 0.183693967 |
| Hus1 | 0.000929908 | 0.959603679 | 0.9957785 | 0.082288111 | 0.06979323 |
| Huwe1 | -0.091287446 | 0.012069849 | 0.2169074 | 0.598897729 | 0.683859976 |
| Hyal1 | -0.015711538 | 0.377746665 | 0.8101797 | 0.060810187 | 0.068404613 |
| Hyal2 | 0.016106586 | 0.414361542 | 0.8300990 | 0.104404012 | 0.080722454 |
| Hyi | -0.060428921 | 0.087931309 | 0.5278913 | 0.364073718 | 0.418913758 |
| Hyou1 | 0.036338393 | 0.365701685 | 0.8050191 | 0.74559772 | 0.724835657 |
| Hypk | 0.058953235 | 0.055055447 | 0.4429440 | 0.321630111 | 0.246063405 |
| Iah1 | 0.01162887 | 0.731221166 | 0.9459020 | 0.323611313 | 0.323211626 |
| Iapp | 0.065229772 | 0.026185407 | 0.3097265 | 7.174385403 | 7.650590343 |
| Iars | 0.030173675 | 0.34984322 | 0.7927303 | 0.313327761 | 0.291667819 |
| Iars2 | 0.001487945 | 0.957036887 | 0.9956630 | 0.181344533 | 0.191349084 |
| Ibtk | -0.006268068 | 0.843709395 | 0.9734889 | 0.272778045 | 0.277787521 |

| | | | | | |
|---|---|---|---|---|---|
| Ica1 | -0.080443622 | 0.024312403 | 0.3027057 | 0.464291802 | 0.528411561 |
| Ica1l | 0.011024023 | 0.543988306 | 0.8842645 | 0.066469216 | 0.077410481 |
| Ice1 | -0.041186317 | 0.116743433 | 0.5727472 | 0.138274937 | 0.176631863 |
| Ice2 | 0.02565054 | 0.11544174 | 0.5724670 | 0.073242381 | 0.049628534 |
| Icmt | 0.017672275 | 0.407322507 | 0.8270770 | 0.130825279 | 0.09962085 |
| Id1 | -0.149014128 | 0.008268863 | 0.1790985 | 1.161643573 | 1.24506219 |
| Id2 | -0.024596102 | 0.611850311 | 0.9100887 | 0.99483642 | 1.07049969 |
| Id3 | -0.132269895 | 0.010029599 | 0.1990377 | 1.501548139 | 1.540568312 |
| Id4 | -0.021109505 | 0.299689724 | 0.7585920 | 0.072463894 | 0.083543312 |
| Ide | -0.025809105 | 0.32043183 | 0.7720873 | 0.151347782 | 0.165824189 |
| Idh1 | -0.021029333 | 0.397854859 | 0.8238469 | 0.135325307 | 0.143002596 |
| Idh2 | 0.027750377 | 0.443735915 | 0.8431541 | 0.574065973 | 0.505903694 |
| Idh3a | 0.016858687 | 0.598550068 | 0.9075513 | 0.344117355 | 0.309475793 |
| Idh3b | -0.031852032 | 0.349547714 | 0.7927303 | 0.346251809 | 0.372271855 |
| Idh3g | 0.029321145 | 0.395514612 | 0.8225618 | 0.448896714 | 0.398006256 |
| Idi1 | 0.039068127 | 0.094083039 | 0.5334417 | 0.148078581 | 0.096268339 |
| Idnk | -0.026078114 | 0.450432405 | 0.8478942 | 0.392144793 | 0.403988191 |
| Ids | -0.014514777 | 0.698446001 | 0.9369692 | 0.55095435 | 0.610711794 |
| Idua | 0.019342178 | 0.174377947 | 0.6527657 | 0.051692835 | 0.038029529 |
| Ier2 | -0.201128268 | 1.47E-06 | 0.0002912 | 1.358059467 | 1.583092648 |
| Ier3 | -0.048591133 | 0.067490367 | 0.4814530 | 0.111809323 | 0.156782895 |
| Ier3ip1 | 0.091455953 | 0.006279767 | 0.1585121 | 1.079503939 | 0.942656032 |
| Ier5 | -0.035058656 | 0.275750468 | 0.7357967 | 0.231526324 | 0.278654299 |
| Ier5l | 0.022660087 | 0.294766145 | 0.7536588 | 0.117739758 | 0.085112003 |
| Iffo1 | -0.010828465 | 0.653934256 | 0.9200648 | 0.127033123 | 0.134366779 |
| Iffo2 | -0.037303137 | 0.010277903 | 0.2010982 | 0.029412659 | 0.05107884 |
| Ifi27 | 0.13572318 | 0.000904582 | 0.0458299 | 0.513828431 | 0.421781009 |
| Ifi30 | -0.001471343 | 0.95752883 | 0.9956630 | 0.181182512 | 0.187965674 |
| Ifi35 | 0.006164624 | 0.836989537 | 0.9726619 | 0.230039761 | 0.219588597 |
| Ifih1 | -0.040581363 | 0.292548836 | 0.7526152 | 0.336986247 | 0.32609618 |
| Ifit1 | 0.006505159 | 0.747677132 | 0.9496370 | 0.088256464 | 0.078203263 |
| Ifitm2 | -0.078849792 | 0.040990467 | 0.3901716 | 0.722544759 | 0.655296518 |
| Ifnar1 | -0.002980421 | 0.871821867 | 0.9782269 | 0.073621242 | 0.072412954 |
| Ifnar2 | 0.001127478 | 0.969094428 | 0.9976072 | 0.265983982 | 0.230061142 |
| Ifngr1 | 0.011943005 | 0.522944014 | 0.8750611 | 0.083778917 | 0.069855187 |
| Ifngr2 | -0.023623931 | 0.174214148 | 0.6527657 | 0.050017435 | 0.068213483 |
| Ifrd1 | -0.000658026 | 0.98599632 | 0.9976072 | 0.434390686 | 0.484531376 |
| Ifrd2 | 0.008931558 | 0.641545986 | 0.9166712 | 0.088504902 | 0.072562132 |
| Ift122 | -0.035955192 | 0.104643915 | 0.5580045 | 0.098528001 | 0.11588529 |
| Ift172 | -0.00720808 | 0.690052423 | 0.9343871 | 0.063750901 | 0.072203722 |
| Ift20 | -0.022316662 | 0.477927519 | 0.8607489 | 1.192363557 | 1.224859279 |
| Ift22 | 0.065909633 | 0.030752791 | 0.3353039 | 0.317124708 | 0.243784398 |
| Ift27 | -0.01870742 | 0.511920241 | 0.8720873 | 0.219604105 | 0.209533665 |
| Ift43 | 0.007145101 | 0.72355559 | 0.9445652 | 0.106280486 | 0.08719625 |
| Ift46 | -0.04938104 | 0.061942678 | 0.4617923 | 0.161125356 | 0.17551219 |
| Ift52 | 0.01058449 | 0.726235228 | 0.9450680 | 0.278100124 | 0.238735799 |
| Ift57 | 0.098409543 | 0.009964503 | 0.1987218 | 0.681053556 | 0.577384784 |
| Ift74 | -0.013183994 | 0.684297544 | 0.9311703 | 0.301354302 | 0.315279197 |
| Ift80 | -0.001013318 | 0.962300291 | 0.9964199 | 0.10259747 | 0.102226128 |
| Ift81 | 0.06563577 | 0.043199638 | 0.3976662 | 0.338137996 | 0.2991664 |
| Ift88 | -0.027060102 | 0.23972974 | 0.7118791 | 0.107635841 | 0.133710774 |
| Iftap | 0.088534879 | 0.006245191 | 0.1580415 | 0.398155692 | 0.28932804 |
| Igbp1 | 0.020833617 | 0.450509035 | 0.8478942 | 0.214379568 | 0.191019754 |
| Igf1r | 0.080930151 | 0.076525536 | 0.5007475 | 0.78466971 | 0.958886483 |
| Igf2bp2 | -0.013721252 | 0.419359276 | 0.8300990 | 0.058485692 | 0.059889359 |
| Igf2r | 0.030922854 | 0.335423823 | 0.7810153 | 0.337476386 | 0.326517328 |
| Igfals | 0.001784145 | 0.931643732 | 0.9893956 | 0.09717923 | 0.09753985 |
| Igfbp7 | -0.046436818 | 0.27805548 | 0.7383057 | 1.433971507 | 1.299111465 |
| Ighmbp2 | 0.01019948 | 0.51807372 | 0.8740291 | 0.053746677 | 0.049064424 |
| Igip | 0.029343982 | 0.274630764 | 0.7351955 | 0.191867439 | 0.160561903 |
| Igsf21 | 0.00849864 | 0.678298482 | 0.9281886 | 0.096766257 | 0.080890362 |
| Igsf8 | -0.003490387 | 0.878209554 | 0.9785583 | 0.111925602 | 0.124461248 |
| Igsf9b | 0.009348276 | 0.67798131 | 0.9281886 | 0.109171608 | 0.114232969 |
| Igtp | 0.005066583 | 0.715382331 | 0.9431411 | 0.048059901 | 0.038802004 |
| Ik | 0.038757431 | 0.271852423 | 0.7342806 | 0.588553483 | 0.509095619 |
| Ikbip | -0.013640255 | 0.694462642 | 0.9361958 | 0.43668927 | 0.389101905 |

| | | | | | |
|---|---|---|---|---|---|
| Ikbkb | -0.005788557 | 0.805484621 | 0.9647143 | 0.115520369 | 0.126058688 |
| Ikbkg | -0.021789946 | 0.272368682 | 0.7342806 | 0.070642724 | 0.089986186 |
| Ikzf2 | 0.006934039 | 0.759850468 | 0.9521533 | 0.107191999 | 0.108425839 |
| Ikzf3 | -0.006280301 | 0.645569445 | 0.9169695 | 0.027952189 | 0.04510396 |
| Ikzf4 | -0.003198189 | 0.902453302 | 0.9825441 | 0.143742117 | 0.158878391 |
| Ikzf5 | 0.029265121 | 0.197367245 | 0.6730962 | 0.127978153 | 0.108138693 |
| Il10rb | 0.015107647 | 0.430392972 | 0.8345383 | 0.08750162 | 0.075815401 |
| Il11ra1 | 0.006674817 | 0.75703577 | 0.9521533 | 0.114190948 | 0.104628443 |
| Il13ra1 | 0.012681773 | 0.531130148 | 0.8776448 | 0.096160865 | 0.088581112 |
| Il15ra | 0.01441902 | 0.374021764 | 0.8087297 | 0.065341374 | 0.051419717 |
| Il17ra | 0.004319022 | 0.827642083 | 0.9717109 | 0.085910314 | 0.087719038 |
| Il17rc | 0.038302052 | 0.035553167 | 0.3591914 | 0.096544903 | 0.062822525 |
| Il18 | -0.002211425 | 0.905703638 | 0.9832544 | 0.074895735 | 0.072409799 |
| Il18bp | -0.018973928 | 0.383259284 | 0.8129787 | 0.077792337 | 0.112701069 |
| Il1r1 | 0.074023405 | 0.067592453 | 0.4814530 | 1.434926708 | 1.422184826 |
| Il3ra | -0.018062701 | 0.399149802 | 0.8239850 | 0.101417516 | 0.104761953 |
| Il6ra | -0.01251669 | 0.695082659 | 0.9362759 | 0.277607978 | 0.266476246 |
| Il6st | 0.046205191 | 0.167315226 | 0.6445697 | 0.402458031 | 0.35042153 |
| Ildr1 | -0.026991563 | 0.333942922 | 0.7804836 | 0.183860565 | 0.201365209 |
| Ilf2 | 0.003952308 | 0.908746722 | 0.9837373 | 0.440396103 | 0.404431534 |
| Ilf3 | -0.083311587 | 0.010540599 | 0.2017406 | 0.254548425 | 0.320533056 |
| Ilk | 0.038872124 | 0.208693179 | 0.6807660 | 0.297675425 | 0.24630318 |
| Ilkap | -0.01644118 | 0.643962554 | 0.9168391 | 0.510170614 | 0.488537239 |
| Ilrun | -0.047773328 | 0.146567584 | 0.6225768 | 0.309500636 | 0.369073357 |
| Ilvbl | -0.001577314 | 0.936145039 | 0.9912468 | 0.0889994 | 0.089532424 |
| Immp1l | 0.069422404 | 0.050142217 | 0.4218341 | 0.816361603 | 0.683620675 |
| Immp2l | -0.035074465 | 0.077398915 | 0.5022647 | 0.073497398 | 0.092556541 |
| Immt | 0.003514416 | 0.921220989 | 0.9861175 | 0.490153567 | 0.464435618 |
| Imp3 | -0.054225506 | 0.119792195 | 0.5793947 | 0.823839466 | 0.825194254 |
| Imp4 | -0.008115738 | 0.75119043 | 0.9506053 | 0.149778291 | 0.152822878 |
| Impa1 | -0.036709873 | 0.245159345 | 0.7142380 | 0.32145217 | 0.334645051 |
| Impact | -0.020245119 | 0.574134498 | 0.8974946 | 0.491585163 | 0.51966527 |
| Impdh1 | -0.021046669 | 0.479310868 | 0.8610807 | 0.246933754 | 0.246836237 |
| Impdh2 | 0.027769918 | 0.314886507 | 0.7705987 | 0.229801487 | 0.176728005 |
| Ina | -0.033882926 | 0.343355722 | 0.7872955 | 0.329604789 | 0.358056554 |
| Inafm1 | 0.03648523 | 0.2691673 | 0.7338465 | 0.427741468 | 0.354857048 |
| Inafm2 | 0.004220857 | 0.877519363 | 0.9785583 | 0.187499509 | 0.183399218 |
| Incenp | 0.021028152 | 0.112797015 | 0.5696203 | 0.052759307 | 0.027765919 |
| Ing1 | -0.023927631 | 0.48754457 | 0.8640311 | 0.34825924 | 0.362095227 |
| Ing2 | 0.000957473 | 0.979504109 | 0.9976072 | 0.588983181 | 0.61095616 |
| Ing3 | 0.032485147 | 0.226295554 | 0.6993308 | 0.206153294 | 0.165242213 |
| Ing5 | 0.037847382 | 0.139767064 | 0.6124069 | 0.171554986 | 0.140400583 |
| Inha | -0.001056345 | 0.960500939 | 0.9958628 | 0.106119859 | 0.105846666 |
| Inip | 0.012034859 | 0.493028809 | 0.8669301 | 0.071071262 | 0.061081051 |
| Ino80 | 0.01412986 | 0.595644018 | 0.9068020 | 0.196408481 | 0.160906838 |
| Ino80b | 0.055967332 | 0.052248836 | 0.4314646 | 0.263684073 | 0.199120338 |
| Ino80c | 0.005132108 | 0.733724081 | 0.9464134 | 0.055954227 | 0.045273636 |
| Ino80d | -0.036229953 | 0.261195051 | 0.7275735 | 0.26907994 | 0.292405693 |
| Ino80e | 0.004747875 | 0.867845056 | 0.9775204 | 0.238762724 | 0.207734418 |
| Inpp1 | 0.01215684 | 0.558111671 | 0.8919622 | 0.113620361 | 0.095150518 |
| Inpp4a | -0.051920743 | 0.09250623 | 0.5325584 | 0.230309932 | 0.273201924 |
| Inpp5a | 0.010849205 | 0.668132825 | 0.9230765 | 0.145678205 | 0.157023354 |
| Inpp5b | -0.004755745 | 0.797945813 | 0.9644698 | 0.072048601 | 0.077581722 |
| Inpp5e | -0.003342749 | 0.835817574 | 0.9723494 | 0.053063973 | 0.054364882 |
| Inpp5f | -0.084398274 | 0.02231876 | 0.2905539 | 0.418286552 | 0.58808356 |
| Inpp5k | -0.006425649 | 0.755122817 | 0.9519403 | 0.092875695 | 0.092249132 |
| Inppl1 | -0.00985929 | 0.688198877 | 0.9329669 | 0.128782492 | 0.146329098 |
| Ins1 | 0.109879297 | 0.002805174 | 0.0979835 | 7.038309206 | 7.852239951 |
| Ins2 | 0.094203042 | 0.001556489 | 0.0679368 | 8.452024791 | 9.293722201 |
| Insig1 | 0.044974349 | 0.135514967 | 0.6028321 | 0.270181224 | 0.196690557 |
| Insig2 | 0.056395314 | 0.072007953 | 0.4888240 | 0.316369279 | 0.27574384 |
| Insm1 | 0.023418724 | 0.551325024 | 0.8890026 | 0.754778111 | 0.77650014 |
| Insr | -0.011627063 | 0.637123927 | 0.9158312 | 0.129457788 | 0.144342754 |
| Insrr | -0.047279715 | 0.110284214 | 0.5655888 | 0.162933482 | 0.24269705 |
| Ints1 | -0.011719138 | 0.606854951 | 0.9087139 | 0.117462381 | 0.120561086 |
| Ints10 | -0.010374534 | 0.622066696 | 0.9121505 | 0.09297599 | 0.097785854 |

| | | | | | |
|---|---|---|---|---|---|
| Ints11 | 0.037969822 | 0.093086279 | 0.5328424 | 0.141580588 | 0.110819675 |
| Ints12 | -0.01845277 | 0.365327465 | 0.8050191 | 0.07558532 | 0.094697797 |
| Ints13 | 0.016643225 | 0.507899812 | 0.8704067 | 0.160157923 | 0.13513332 |
| Ints14 | 0.006297846 | 0.702417578 | 0.9388187 | 0.064644295 | 0.055756017 |
| Ints2 | 0.038515767 | 0.06293405 | 0.4658998 | 0.120548635 | 0.082918257 |
| Ints3 | 0.03093904 | 0.239899128 | 0.7118791 | 0.193649295 | 0.172078329 |
| Ints4 | -0.015601416 | 0.335285266 | 0.7809414 | 0.04982455 | 0.060364451 |
| Ints5 | 0.035177295 | 0.056184936 | 0.4462406 | 0.096156886 | 0.065326338 |
| Ints6 | -0.014971451 | 0.629695567 | 0.9130564 | 0.272950005 | 0.257362348 |
| Ints6l | -0.054467709 | 0.030143427 | 0.3311437 | 0.114550286 | 0.156201127 |
| Ints7 | 0.003825301 | 0.810948789 | 0.9670167 | 0.052964822 | 0.053516048 |
| Ints8 | -0.021178767 | 0.349184364 | 0.7927303 | 0.117049402 | 0.117055469 |
| Ints9 | 0.016910125 | 0.40974109 | 0.8279958 | 0.102202722 | 0.085929254 |
| Intu | -0.006233448 | 0.717530168 | 0.9435179 | 0.057891003 | 0.066903102 |
| Invs | 0.007213672 | 0.712769848 | 0.9421288 | 0.095151361 | 0.083597714 |
| Ip6k1 | -0.004482982 | 0.894418105 | 0.9824361 | 0.442198544 | 0.420572893 |
| Ip6k2 | 0.040833533 | 0.243295926 | 0.7132719 | 0.502290029 | 0.447670512 |
| Ipmk | -0.012979799 | 0.65551377 | 0.9208009 | 0.230321754 | 0.222646238 |
| Ipo11 | 0.004675177 | 0.757499406 | 0.9521533 | 0.055853921 | 0.045470404 |
| Ipo13 | 0.062211852 | 0.000270186 | 0.0201522 | 0.09587625 | 0.051312108 |
| Ipo4 | 0.024776808 | 0.382854702 | 0.8129787 | 0.208018554 | 0.195068563 |
| Ipo5 | -0.041360209 | 0.179617196 | 0.6550070 | 0.270909618 | 0.286191751 |
| Ipo7 | 0.004883039 | 0.876650787 | 0.9785583 | 0.269163722 | 0.27447435 |
| Ipo8 | -0.002011005 | 0.939338466 | 0.9927804 | 0.172928848 | 0.167462877 |
| Ipo9 | -0.016178836 | 0.474381689 | 0.8599901 | 0.104500602 | 0.118968561 |
| Ipp | -0.014502328 | 0.538411484 | 0.8816335 | 0.127510686 | 0.123703668 |
| Ippk | 0.029559945 | 0.128743384 | 0.5914729 | 0.105907813 | 0.0717254 |
| Iqcb1 | -0.005737509 | 0.843863342 | 0.9734889 | 0.251645864 | 0.232035232 |
| Iqcc | 0.004721929 | 0.84143434 | 0.9734589 | 0.141767156 | 0.122404825 |
| Iqce | 0.023911388 | 0.173049059 | 0.6525507 | 0.078830393 | 0.062715753 |
| Iqgap1 | -0.01650336 | 0.648922327 | 0.9172569 | 0.54217326 | 0.622342198 |
| Iqsec1 | -0.001944453 | 0.909292761 | 0.9837373 | 0.061594912 | 0.062766168 |
| Irak1 | -0.017769843 | 0.6221456 | 0.9121505 | 0.641594173 | 0.652679894 |
| Irak1bp1 | -0.020834219 | 0.513511045 | 0.8730127 | 0.289337742 | 0.311251345 |
| Ireb2 | -0.025324565 | 0.445096527 | 0.8432629 | 0.289078191 | 0.351926079 |
| Irf1 | -0.053188091 | 0.088594505 | 0.5284771 | 0.21285075 | 0.256290477 |
| Irf2 | 0.040880773 | 0.217052984 | 0.6872330 | 0.400782372 | 0.342754615 |
| Irf2bp1 | -0.113709064 | 9.38E-05 | 0.0089506 | 0.19539501 | 0.26467499 |
| Irf2bp2 | -0.026984765 | 0.473420072 | 0.8597092 | 0.855399374 | 0.885966646 |
| Irf2bpl | 0.010087182 | 0.773931999 | 0.9558269 | 0.420073327 | 0.411904603 |
| Irf3 | -0.026342185 | 0.30661549 | 0.7652785 | 0.139111113 | 0.160836135 |
| Irf6 | 0.088238848 | 0.005931978 | 0.1545438 | 0.352251769 | 0.268270689 |
| Irf7 | -0.00792206 | 0.690492107 | 0.9346237 | 0.057107838 | 0.076794742 |
| Irf9 | -0.033639023 | 0.076793026 | 0.5015054 | 0.052397583 | 0.083971884 |
| Irgm1 | -0.003350565 | 0.857505309 | 0.9763189 | 0.071061828 | 0.076069403 |
| Irgq | -0.013914475 | 0.583217614 | 0.9020133 | 0.148587776 | 0.157201483 |
| Irs2 | -0.109816762 | 0.006464027 | 0.1603078 | 0.408399453 | 0.602113878 |
| Irx1 | -0.02274633 | 0.579775566 | 0.9008751 | 0.661659591 | 0.525142662 |
| Irx2 | -0.056733271 | 0.163825523 | 0.6410026 | 0.649640693 | 0.55626789 |
| Isca1 | -0.022230025 | 0.423623616 | 0.8319830 | 0.187639898 | 0.194669235 |
| Isca2 | -2.08E-05 | 0.999485079 | 0.9996989 | 0.341841036 | 0.317912403 |
| Iscu | 0.024862979 | 0.484023369 | 0.8625857 | 0.756076002 | 0.675645316 |
| Isg20 | 0.028472148 | 0.45829484 | 0.8521621 | 0.877121947 | 0.839644019 |
| Isg20l2 | -0.015105633 | 0.403253308 | 0.8264703 | 0.067883045 | 0.070961593 |
| Isl1 | 0.026923332 | 0.477973686 | 0.8607489 | 1.329282583 | 1.234830018 |
| Isoc1 | 0.056021753 | 0.096223215 | 0.5369725 | 0.370873926 | 0.346209026 |
| Isoc2b | 0.016004836 | 0.583308044 | 0.9020133 | 0.237100267 | 0.223200576 |
| Ist1 | -0.025520017 | 0.383054401 | 0.8129787 | 0.208978738 | 0.227100109 |
| Isy1 | -0.009101886 | 0.723869895 | 0.9445652 | 0.161871925 | 0.152581529 |
| Isyna1 | -0.021147514 | 0.426896201 | 0.8337565 | 0.174468226 | 0.168157552 |
| Itch | 0.010855017 | 0.738961955 | 0.9474350 | 0.308364849 | 0.286794624 |
| Itfg1 | -0.03758198 | 0.293992593 | 0.7533922 | 0.537228085 | 0.521769386 |
| Itfg2 | -0.014416805 | 0.608154117 | 0.9088672 | 0.187622956 | 0.189992592 |
| Itga3 | 0.014557708 | 0.428412514 | 0.8340365 | 0.078051942 | 0.065414921 |
| Itga6 | -0.033603147 | 0.191310866 | 0.6700281 | 0.10741242 | 0.173581723 |
| Itgav | 0.060039343 | 0.075377518 | 0.4966285 | 0.35544005 | 0.364816055 |

| | | | | | |
|---|---|---|---|---|---|
| Itgb1 | 0.047643228 | 0.207428755 | 0.6804541 | 0.913512095 | 0.815006111 |
| Itgb1bp1 | -0.034863119 | 0.223495727 | 0.6955958 | 0.197669792 | 0.214927345 |
| Itgb3bp | -0.010022526 | 0.752148945 | 0.9507898 | 0.301465068 | 0.299575712 |
| Itgb5 | -0.029232084 | 0.233526304 | 0.7051391 | 0.120001263 | 0.154775423 |
| Itgb8 | 0.047582198 | 0.004054088 | 0.1260707 | 0.074455546 | 0.051911096 |
| Itm2b | -0.095276048 | 4.89E-08 | 2.42E-05 | 3.53478984 | 3.600218134 |
| Itm2c | -0.08103812 | 0.017623849 | 0.2582401 | 1.447607082 | 1.401558445 |
| Itpa | -0.006099842 | 0.839447034 | 0.9727382 | 0.247230112 | 0.239263815 |
| Itpk1 | -0.023369341 | 0.210445383 | 0.6817825 | 0.053707397 | 0.075178072 |
| Itpkb | -0.013001753 | 0.585438088 | 0.9024935 | 0.088871439 | 0.134984489 |
| Itpkc | 0.001245295 | 0.938929164 | 0.9925593 | 0.054026511 | 0.051871423 |
| Itpr1 | -0.125669748 | 0.001724635 | 0.0726586 | 0.616352389 | 0.618194675 |
| Itpr3 | 0.001477092 | 0.948631684 | 0.9945493 | 0.113435508 | 0.124557365 |
| Itprid2 | 0.070582097 | 0.005367729 | 0.1483227 | 0.180506041 | 0.143926719 |
| Itsn1 | 0.014451487 | 0.354921218 | 0.7969131 | 0.055639445 | 0.048919296 |
| Itsn2 | -0.047910751 | 0.142174306 | 0.6152737 | 0.26394762 | 0.290371167 |
| Ivd | 0.008851356 | 0.782468459 | 0.9590862 | 0.316430947 | 0.288629284 |
| Ivns1abp | -0.007043261 | 0.851953265 | 0.9753463 | 0.545995429 | 0.548389496 |
| Iws1 | 0.01238128 | 0.684909547 | 0.9313643 | 0.274505633 | 0.247580909 |
| Izumo4 | -0.035421605 | 0.159939574 | 0.6369331 | 0.125381567 | 0.159972456 |
| Jade1 | -0.020849011 | 0.578152941 | 0.8997924 | 0.436520093 | 0.53148218 |
| Jade2 | 0.011650055 | 0.561022039 | 0.8926240 | 0.090533516 | 0.093438928 |
| Jade3 | 0.067148405 | 0.009942346 | 0.1987218 | 0.200895848 | 0.140829625 |
| Jagn1 | -0.008113828 | 0.808868297 | 0.9658428 | 0.429166297 | 0.42233808 |
| Jak1 | 0.043432896 | 0.218760968 | 0.6901690 | 0.615904842 | 0.599703766 |
| Jak2 | -0.037782477 | 0.090727929 | 0.5321031 | 0.080312179 | 0.121527926 |
| Jakmip1 | -0.007530551 | 0.694759705 | 0.9362759 | 0.06458506 | 0.089156767 |
| Jam2 | 0.004442764 | 0.815538883 | 0.9686999 | 0.073919793 | 0.08071012 |
| Jam3 | -7.16E-05 | 0.997073203 | 0.9994073 | 0.093608098 | 0.082239895 |
| Jarid2 | 0.015962469 | 0.574029265 | 0.8974946 | 0.199502262 | 0.178025334 |
| Jazf1 | 0.004428265 | 0.780651594 | 0.9590847 | 0.060724794 | 0.048965419 |
| Jkamp | -0.002367529 | 0.939877873 | 0.9928216 | 0.306990284 | 0.275988108 |
| Jmjd1c | -0.065096941 | 0.075188429 | 0.4966285 | 0.376935397 | 0.466552639 |
| Jmjd6 | -0.02884956 | 0.37732518 | 0.8101797 | 0.306876487 | 0.313317553 |
| Jmjd8 | 0.020007028 | 0.424997533 | 0.8324460 | 0.165640335 | 0.141540441 |
| Jmy | -0.025749322 | 0.373298816 | 0.8085689 | 0.214055476 | 0.221314784 |
| Josd1 | -0.068956643 | 0.010767056 | 0.2035157 | 0.134971659 | 0.197189372 |
| Josd2 | 0.042746072 | 0.182304839 | 0.6595419 | 0.384626771 | 0.317601757 |
| Jph3 | -0.000829092 | 0.961516433 | 0.9963557 | 0.06318112 | 0.060729443 |
| Jpt1 | -0.049408806 | 0.10465523 | 0.5580045 | 0.239284156 | 0.264640124 |
| Jpt2 | 0.035560739 | 0.197914807 | 0.6730962 | 0.215964422 | 0.171743871 |
| Jpx | 0.002643843 | 0.854467083 | 0.9755431 | 0.049550829 | 0.039249955 |
| Jrkl | -0.008618336 | 0.679755922 | 0.9289200 | 0.099492766 | 0.090755872 |
| Jtb | -0.008708368 | 0.805666781 | 0.9647143 | 0.832571576 | 0.837729782 |
| Jun | -0.278123494 | 6.26E-12 | 8.87E-09 | 1.753476784 | 2.307413656 |
| Junb | -0.094340539 | 0.014554173 | 0.2371912 | 2.175761857 | 2.25751756 |
| Jund | -0.096072056 | 3.13E-05 | 0.0038515 | 2.422567262 | 2.584127184 |
| Jup | -0.081993807 | 0.007748419 | 0.1737612 | 0.204867752 | 0.255878543 |
| Kalrn | -0.015275827 | 0.417135173 | 0.8300990 | 0.067703158 | 0.075996602 |
| Kank1 | 0.01204369 | 0.494383885 | 0.8678106 | 0.077761896 | 0.061438411 |
| Kansl1 | -0.053004474 | 0.108134725 | 0.5636817 | 0.288949304 | 0.333069109 |
| Kansl1l | -0.02795759 | 0.156094189 | 0.6312943 | 0.067602969 | 0.086174214 |
| Kansl2 | -0.022663501 | 0.396198071 | 0.8234412 | 0.17222598 | 0.18554368 |
| Kansl3 | -0.014493716 | 0.582717355 | 0.9019790 | 0.15573317 | 0.168815303 |
| Kantr | 0.017830652 | 0.467455868 | 0.8558546 | 0.151782434 | 0.135293506 |
| Kars | 0.012802283 | 0.671119046 | 0.9247813 | 0.250596004 | 0.244846034 |
| Kat14 | -0.013012112 | 0.541341763 | 0.8835017 | 0.094252971 | 0.105842107 |
| Kat2a | -0.008811213 | 0.61543081 | 0.9111433 | 0.064282816 | 0.066237185 |
| Kat2b | 0.002385573 | 0.926120579 | 0.9879210 | 0.164507229 | 0.155582761 |
| Kat5 | -0.020656051 | 0.410512144 | 0.8283974 | 0.14231862 | 0.154059673 |
| Kat6a | -0.032963785 | 0.268150559 | 0.7323870 | 0.213808618 | 0.24693427 |
| Kat6b | -0.024579461 | 0.362350298 | 0.8030478 | 0.146417797 | 0.190786598 |
| Kat7 | 0.004961941 | 0.85642243 | 0.9763189 | 0.194665921 | 0.181282215 |
| Kat8 | -0.017378207 | 0.447938079 | 0.8457452 | 0.117977091 | 0.120711814 |
| Katna1 | -0.03806732 | 0.201799954 | 0.6752150 | 0.231143794 | 0.262046382 |
| Katnal1 | -0.021717835 | 0.326449189 | 0.7769616 | 0.104250898 | 0.106464407 |

| | | | | | |
|---|---|---|---|---|---|
| Katnb1 | -0.008570833 | 0.616030659 | 0.9114042 | 0.060393898 | 0.060243635 |
| Katnbl1 | 0.002336176 | 0.914646971 | 0.9845236 | 0.112969123 | 0.10410885 |
| Katnip | -0.056389603 | 0.057401978 | 0.4494298 | 0.187926819 | 0.231931886 |
| Kazald1 | -0.042988049 | 0.015175424 | 0.2416375 | 0.048399812 | 0.07284516 |
| Kazn | 0.018865311 | 0.309963557 | 0.7677033 | 0.088042761 | 0.066915922 |
| Kbtbd2 | 0.01483109 | 0.600937167 | 0.9078437 | 0.204081076 | 0.183575982 |
| Kbtbd3 | 0.002910457 | 0.910696171 | 0.9837373 | 0.16592782 | 0.157608873 |
| Kbtbd4 | -0.009282709 | 0.61838775 | 0.9115017 | 0.066983326 | 0.074576625 |
| Kbtbd7 | 0.027467222 | 0.162904254 | 0.6400703 | 0.105902577 | 0.081342131 |
| Kcmf1 | -0.031079911 | 0.383952141 | 0.8129787 | 0.538865815 | 0.54239946 |
| Kcnab3 | 0.017458343 | 0.175744725 | 0.6536659 | 0.037919484 | 0.03026668 |
| Kcnb1 | -0.023108108 | 0.478018028 | 0.8607489 | 0.294747796 | 0.313009967 |
| Kcnb2 | -0.028842904 | 0.155908792 | 0.6312943 | 0.063159611 | 0.095275239 |
| Kcnc3 | -0.04976246 | 0.074230513 | 0.4945377 | 0.155511125 | 0.190011097 |
| Kcnh1 | 0.005077759 | 0.772432572 | 0.9552333 | 0.052868498 | 0.065430686 |
| Kcnh2 | -0.074732196 | 0.008519204 | 0.1821346 | 0.169823949 | 0.240570272 |
| Kcnh6 | -0.006197526 | 0.709515496 | 0.9418251 | 0.051344048 | 0.05918094 |
| Kcnip1 | 0.051991986 | 0.090757924 | 0.5321031 | 0.232225398 | 0.25064002 |
| Kcnip3 | -0.006220911 | 0.772825327 | 0.9555561 | 0.120287598 | 0.091503878 |
| Kcnip4 | -0.013302021 | 0.4219412 | 0.8311470 | 0.051619209 | 0.0574388 |
| Kcnj11 | 0.033601173 | 0.320780213 | 0.7721897 | 0.36880187 | 0.317627523 |
| Kcnj3 | -0.001899665 | 0.907475816 | 0.9837373 | 0.061275302 | 0.049587343 |
| Kcnj6 | -0.029149067 | 0.112389094 | 0.5693702 | 0.061603842 | 0.079046789 |
| Kcnk1 | -0.058768978 | 0.088355541 | 0.5284771 | 0.360560107 | 0.394059012 |
| Kcnk12 | 0.001747969 | 0.946782329 | 0.9945018 | 0.175060742 | 0.160483926 |
| Kcnk16 | 0.132869161 | 0.000119382 | 0.0107661 | 0.497103616 | 0.343420438 |
| Kcnk3 | 0.052886106 | 0.123550145 | 0.5850202 | 0.454579952 | 0.304779167 |
| Kcnma1 | -0.037854409 | 0.24460933 | 0.7137971 | 0.282339387 | 0.317426259 |
| Kcnmb2 | 0.009273378 | 0.791950846 | 0.9624099 | 0.348150961 | 0.368628486 |
| Kcnn3 | 0.053020047 | 0.007874212 | 0.1751394 | 0.112041178 | 0.070261576 |
| Kcnq1ot1 | -0.038512583 | 0.444442717 | 0.8431541 | 0.981044603 | 1.050619392 |
| Kcnu1 | 0.022244988 | 0.069833171 | 0.4847655 | 0.046522556 | 0.024674101 |
| Kctd10 | -0.01478861 | 0.416035999 | 0.8300990 | 0.061351147 | 0.069976556 |
| Kctd11 | 0.019432217 | 0.307039833 | 0.7654775 | 0.093106079 | 0.069986326 |
| Kctd12 | -0.033480187 | 0.27035132 | 0.7340360 | 0.20289964 | 0.223261659 |
| Kctd12b | -0.051244284 | 0.121964497 | 0.5819916 | 0.211512448 | 0.230385471 |
| Kctd13 | -0.017245765 | 0.599724886 | 0.9078234 | 0.379688189 | 0.357558021 |
| Kctd15 | 0.000665487 | 0.970464623 | 0.9976072 | 0.059694833 | 0.071888478 |
| Kctd18 | -0.027757676 | 0.12433776 | 0.5852543 | 0.053195128 | 0.079410204 |
| Kctd2 | -0.050250243 | 0.099034888 | 0.5433551 | 0.246875403 | 0.263945199 |
| Kctd20 | -0.017460614 | 0.601544882 | 0.9081304 | 0.334456775 | 0.323092182 |
| Kctd3 | -0.011669248 | 0.578828277 | 0.9003073 | 0.086952324 | 0.099454776 |
| Kctd5 | -0.002499451 | 0.898933478 | 0.9825441 | 0.083015797 | 0.085260693 |
| Kctd6 | -0.011699742 | 0.571864178 | 0.8966165 | 0.090045242 | 0.098418648 |
| Kctd8 | 0.017030581 | 0.334343592 | 0.7804836 | 0.067087519 | 0.058734949 |
| Kctd9 | 0.018015683 | 0.336698247 | 0.7826493 | 0.085096544 | 0.071691899 |
| Kdelr1 | -0.012314215 | 0.731155845 | 0.9459020 | 0.989751039 | 0.909976439 |
| Kdelr2 | 0.001213756 | 0.973593356 | 0.9976072 | 1.242178335 | 1.104575049 |
| Kdelr3 | -0.014889942 | 0.602537505 | 0.9086609 | 0.212635973 | 0.183903326 |
| Kdf1 | -0.010582685 | 0.636240631 | 0.9153541 | 0.114561823 | 0.11437528 |
| Kdm1a | -0.006612346 | 0.858992639 | 0.9769675 | 0.718984765 | 0.704251856 |
| Kdm2a | -0.032827262 | 0.359060397 | 0.8000088 | 0.49509602 | 0.493879653 |
| Kdm2b | -0.066565697 | 0.025162883 | 0.3047811 | 0.1537584 | 0.262035084 |
| Kdm3a | -0.015281449 | 0.399839136 | 0.8244516 | 0.06511024 | 0.076055447 |
| Kdm3b | 0.003194792 | 0.892078765 | 0.9819597 | 0.121876035 | 0.118599305 |
| Kdm4a | -0.015882818 | 0.533139852 | 0.8789674 | 0.141706777 | 0.162658867 |
| Kdm4b | -0.016012503 | 0.400415577 | 0.8246612 | 0.063620196 | 0.080753194 |
| Kdm4c | -0.024103518 | 0.393005014 | 0.8206236 | 0.191221512 | 0.192932293 |
| Kdm5a | -0.071343734 | 0.044680507 | 0.4015733 | 0.393469602 | 0.436691905 |
| Kdm5b | 0.042769761 | 0.230317121 | 0.7019734 | 0.579570966 | 0.499957917 |
| Kdm5c | 0.004999142 | 0.861454941 | 0.9769675 | 0.217405816 | 0.200656822 |
| Kdm5d | 0.002351334 | 0.908988737 | 0.9837373 | 0.099968908 | 0.092285677 |
| Kdm6a | -0.013213116 | 0.608994967 | 0.9088672 | 0.15889282 | 0.162860066 |
| Kdm6b | 0.02222137 | 0.54130737 | 0.8835017 | 0.570716297 | 0.575730033 |
| Kdm7a | -0.07157012 | 0.04157367 | 0.3921210 | 0.400713447 | 0.439836125 |
| Kdsr | -0.037310303 | 0.06131892 | 0.4611772 | 0.070036352 | 0.096476568 |

| | | | | | |
|---|---|---|---|---|---|
| Keap1 | -0.025608545 | 0.467274188 | 0.85585468 | 0.498530661 | 0.515271685 |
| Khdc4 | -0.041512092 | 0.153629077 | 0.63001258 | 0.193891988 | 0.22378071 |
| Khdrbs1 | -0.043397164 | 0.239563459 | 0.71187911 | 0.489307205 | 0.48952669 |
| Khdrbs3 | 0.000852113 | 0.981743765 | 0.99760728 | 0.848778909 | 0.751752013 |
| Khk | 0.057450294 | 0.009906133 | 0.19872185 | 0.153333066 | 0.099095051 |
| Khnyn | -0.001821471 | 0.938741263 | 0.99254631 | 0.136778132 | 0.126060398 |
| Khsrp | -0.031752829 | 0.209102754 | 0.68076601 | 0.14291265 | 0.163692867 |
| Kidins220 | 0.013905963 | 0.689895296 | 0.93430188 | 0.39466217 | 0.410789292 |
| Kif12 | -0.058336073 | 0.071292554 | 0.4881297 | 0.260385477 | 0.318559317 |
| Kif13a | -0.016324534 | 0.354744427 | 0.79691318 | 0.06413681 | 0.065876886 |
| Kif13b | 0.000305179 | 0.991501175 | 0.99834460 | 0.208842572 | 0.21082105 |
| Kif16b | -0.014231454 | 0.515630759 | 0.87348064 | 0.091434072 | 0.107597708 |
| Kif1a | 0.026171765 | 0.46306736 | 0.85244258 | 0.570492471 | 0.554285881 |
| Kif1b | -0.040458732 | 0.283539493 | 0.74292439 | 0.737807457 | 0.775205554 |
| Kif1c | 0.029237052 | 0.118136661 | 0.57587994 | 0.089035264 | 0.069556591 |
| Kif21a | -0.055995883 | 0.119513774 | 0.57833000 | 1.177053971 | 1.154440968 |
| Kif2a | 0.004949076 | 0.880344267 | 0.97925711 | 0.407832484 | 0.376682697 |
| Kif3a | -0.036121473 | 0.19711948 | 0.67309622 | 0.184608157 | 0.197736984 |
| Kif3b | -0.020407161 | 0.419537479 | 0.83009907 | 0.131445076 | 0.155641141 |
| Kif3c | 0.033037368 | 0.011608758 | 0.21207897 | 0.053560255 | 0.028483415 |
| Kif5b | -0.008925551 | 0.750286875 | 0.95045378 | 1.736474126 | 1.833350088 |
| Kif5c | -0.00050322 | 0.988228679 | 0.99761431 | 0.332329873 | 0.351640696 |
| Kifap3 | 0.026272469 | 0.397598979 | 0.82384691 | 0.287601999 | 0.262649891 |
| Kifbp | -0.041253082 | 0.160280418 | 0.63703375 | 0.210844033 | 0.233469242 |
| Kifc3 | -0.043450926 | 0.072199865 | 0.48927244 | 0.116458579 | 0.138956082 |
| Kin | 0.004652187 | 0.900593141 | 0.98254414 | 0.603265271 | 0.573007037 |
| Kiz | -0.029818648 | 0.249301386 | 0.71927285 | 0.149460429 | 0.163254838 |
| Kl | -0.032299812 | 0.192102063 | 0.67002811 | 0.134153479 | 0.144163887 |
| Klb | 0.013946215 | 0.529709798 | 0.87764481 | 0.12917681 | 0.100875939 |
| Klc1 | 0.010889082 | 0.711071979 | 0.94198349 | 0.232820992 | 0.215940654 |
| Klc2 | -0.015297744 | 0.495625013 | 0.86846291 | 0.112736666 | 0.11850043 |
| Klc3 | -0.063708481 | 0.021717337 | 0.28737460 | 0.185695297 | 0.205633546 |
| Klc4 | 0.020795095 | 0.258303259 | 0.72511021 | 0.08695938 | 0.064793812 |
| Klf10 | -0.021160677 | 0.503398701 | 0.86955665 | 0.238883639 | 0.260015282 |
| Klf11 | -0.012472485 | 0.564317946 | 0.89337381 | 0.100316066 | 0.107137003 |
| Klf12 | -0.007240036 | 0.658556775 | 0.92117801 | 0.053745937 | 0.05628367 |
| Klf13 | 0.026485703 | 0.455432288 | 0.85029141 | 0.430821911 | 0.401465585 |
| Klf16 | -0.009321181 | 0.568516035 | 0.89561361 | 0.054140365 | 0.054978411 |
| Klf2 | -0.058395238 | 0.010488855 | 0.20174069 | 0.026657837 | 0.111014986 |
| Klf3 | -0.006542598 | 0.818946986 | 0.96990621 | 0.189050402 | 0.198977105 |
| Klf4 | -0.236070049 | 1.31E-06 | 0.00026581 | 0.660818872 | 1.016903103 |
| Klf5 | 0.012091675 | 0.61788361 | 0.91150170 | 0.119012397 | 0.132445555 |
| Klf6 | -0.122134954 | 0.000354012 | 0.02405344 | 0.171848798 | 0.317785814 |
| Klf7 | -0.0062707 | 0.854081312 | 0.97554315 | 0.356518589 | 0.355593602 |
| Klf9 | 0.023214537 | 0.541852098 | 0.88367200 | 0.786237195 | 0.792638176 |
| Klhdc1 | -0.014906312 | 0.536601167 | 0.88043061 | 0.124059654 | 0.141246537 |
| Klhdc10 | 0.004005292 | 0.897657477 | 0.98254414 | 0.260970832 | 0.262134049 |
| Klhdc2 | 0.015738245 | 0.665920773 | 0.92307657 | 0.645094114 | 0.577033265 |
| Klhdc3 | -0.008275762 | 0.716960788 | 0.94314747 | 0.114592202 | 0.12402258 |
| Klhdc4 | -0.000247835 | 0.990514269 | 0.99808365 | 0.100465989 | 0.093381355 |
| Klhdc8b | 0.05275532 | 0.062222287 | 0.46304957 | 0.251542904 | 0.171113041 |
| Klhl11 | -0.030651441 | 0.099176036 | 0.54335515 | 0.054808168 | 0.077128042 |
| Klhl12 | 0.000349713 | 0.987035224 | 0.99760728 | 0.097047081 | 0.099623211 |
| Klhl13 | -0.046965823 | 0.080043421 | 0.50866799 | 0.174705582 | 0.164943762 |
| Klhl15 | 0.005309759 | 0.782766787 | 0.95908628 | 0.086085741 | 0.077797062 |
| Klhl18 | 0.004927595 | 0.737211942 | 0.94697528 | 0.046431949 | 0.043235353 |
| Klhl2 | 0.018849399 | 0.381192564 | 0.81193690 | 0.114868335 | 0.096297708 |
| Klhl21 | 0.020041757 | 0.365261143 | 0.80501911 | 0.119807045 | 0.104778247 |
| Klhl22 | -0.025930792 | 0.293345934 | 0.75336844 | 0.122080459 | 0.146488912 |
| Klhl24 | -0.074945864 | 0.01436494 | 0.23631874 | 0.226588398 | 0.275208984 |
| Klhl26 | -0.02596561 | 0.184087741 | 0.66260899 | 0.081300899 | 0.086737081 |
| Klhl28 | 0.037316406 | 0.085718631 | 0.52263601 | 0.119275862 | 0.093721541 |
| Klhl32 | 0.024001011 | 0.401484922 | 0.82542591 | 0.199524953 | 0.205409439 |
| Klhl36 | -0.030009109 | 0.057824145 | 0.45081999 | 0.036609817 | 0.057026628 |
| Klhl42 | -0.005040378 | 0.806465023 | 0.96480137 | 0.103944559 | 0.089706261 |
| Klhl7 | -0.107266543 | 0.00233316 | 0.08767020 | 0.427333852 | 0.449103051 |

| | | | | | |
|---|---|---|---|---|---|
| Klhl8 | 0.002687154 | 0.861157877 | 0.9769675 | 0.047343045 | 0.052464922 |
| Klhl9 | 0.034836756 | 0.321765517 | 0.7732349 | 0.475118893 | 0.393717227 |
| Kmt2a | -0.07806485 | 0.046004396 | 0.4073656 | 0.842457408 | 0.882363127 |
| Kmt2b | -0.026707481 | 0.365763003 | 0.8050191 | 0.211247204 | 0.230451172 |
| Kmt2c | 0.014486205 | 0.700403114 | 0.9375165 | 0.586797931 | 0.559274485 |
| Kmt2d | 0.001799561 | 0.954032146 | 0.9954690 | 0.307703376 | 0.272721742 |
| Kmt2e | -0.044526532 | 0.221797419 | 0.6933883 | 0.954459928 | 0.962738189 |
| Kmt5a | 0.024100849 | 0.475810259 | 0.8606925 | 0.37834488 | 0.344114706 |
| Kmt5b | -0.072015355 | 0.024999086 | 0.3041540 | 0.252916106 | 0.318573275 |
| Kmt5c | 0.002782399 | 0.854996118 | 0.9755431 | 0.062713381 | 0.045016487 |
| Knop1 | 0.018646054 | 0.559933728 | 0.8925646 | 0.301249116 | 0.297054574 |
| Kpna1 | -0.029110335 | 0.336703986 | 0.7826493 | 0.239331151 | 0.255354003 |
| Kpna2 | -0.030243883 | 0.042540878 | 0.3952910 | 0.024719767 | 0.055313264 |
| Kpna3 | 0.062831916 | 0.029951393 | 0.3308280 | 0.271854338 | 0.205483615 |
| Kpna4 | -0.031210693 | 0.355530969 | 0.7969131 | 0.351192682 | 0.349530323 |
| Kpna6 | -0.001801097 | 0.942222195 | 0.9931828 | 0.14818593 | 0.148526787 |
| Kpnb1 | 0.031132929 | 0.331950957 | 0.7793175 | 0.38549604 | 0.345960228 |
| Kptn | 0.015757502 | 0.378342259 | 0.8101915 | 0.078698204 | 0.062702639 |
| Kras | 0.068303153 | 0.0607498 | 0.4600290 | 1.022052037 | 0.937734615 |
| Krba1 | 0.011539934 | 0.514026128 | 0.8733342 | 0.082807559 | 0.063824582 |
| Krcc1 | -0.035112813 | 0.337876599 | 0.7834245 | 0.647283458 | 0.653668868 |
| Kremen1 | 0.000187434 | 0.990357934 | 0.9980836 | 0.056836613 | 0.048496508 |
| Kri1 | -0.032261298 | 0.091820287 | 0.5325584 | 0.052837124 | 0.088108767 |
| Krit1 | -0.056592675 | 0.10735727 | 0.5633723 | 0.381707783 | 0.407409969 |
| Krr1 | -0.028732192 | 0.272775543 | 0.7345096 | 0.168432235 | 0.174241224 |
| Krt1 | -0.067844033 | 0.002903764 | 0.1000185 | 0.113947734 | 0.12333031 |
| Krt10 | 0.004655246 | 0.885771577 | 0.9807851 | 0.325407157 | 0.30751473 |
| Krt18 | -0.070707045 | 0.031440031 | 0.3385228 | 1.380678335 | 1.506739616 |
| Krt222 | 0.026482421 | 0.1112084 | 0.5671914 | 0.063085795 | 0.055487716 |
| Krt7 | -0.043262965 | 0.190276551 | 0.6700281 | 1.36326001 | 1.476256579 |
| Krt8 | -0.071464497 | 0.058803147 | 0.4521916 | 0.952292248 | 1.051721509 |
| Krtcap2 | 0.056072732 | 0.031931937 | 0.3385228 | 2.10717058 | 2.110840846 |
| Krtcap3 | 0.050122902 | 0.120768433 | 0.5810004 | 0.335710133 | 0.270122752 |
| Ksr2 | -0.013550864 | 0.673526276 | 0.9261686 | 0.293769118 | 0.285360272 |
| Kti12 | -0.018008182 | 0.500445468 | 0.8693704 | 0.166291158 | 0.16773743 |
| Ktn1 | 0.054092509 | 0.138239143 | 0.6079839 | 0.782674642 | 0.75327448 |
| Kxd1 | -0.041336278 | 0.193965137 | 0.6716564 | 0.313847714 | 0.309133684 |
| Ky | 0.04653542 | 0.013356883 | 0.2288433 | 0.089209338 | 0.068437543 |
| L2hgdh | -0.025969074 | 0.152088323 | 0.6281105 | 0.059198004 | 0.075952911 |
| L3mbtl2 | -0.004146972 | 0.834601173 | 0.9723494 | 0.090694957 | 0.083083877 |
| L3mbtl3 | 0.009068133 | 0.546328341 | 0.8858413 | 0.058564776 | 0.041848058 |
| Lactb | -0.014471596 | 0.601080957 | 0.9078445 | 0.187767097 | 0.191997044 |
| Lactb2 | 0.006952168 | 0.798765732 | 0.9645472 | 0.177268275 | 0.180635559 |
| Lage3 | -0.005508811 | 0.865865256 | 0.9770148 | 0.377210583 | 0.351601619 |
| Lamc1 | 0.009859982 | 0.570445723 | 0.8962534 | 0.069984063 | 0.06137681 |
| Lamp1 | 0.018796527 | 0.506546833 | 0.8695566 | 1.563789907 | 1.4463173 |
| Lamp2 | -0.12843114 | 0.000322934 | 0.0226977 | 0.916772884 | 1.070872058 |
| Lamtor1 | 0.001855575 | 0.959336973 | 0.9957785 | 0.738282441 | 0.702927835 |
| Lamtor2 | 0.037836019 | 0.213155154 | 0.6827572 | 1.381866794 | 1.281711128 |
| Lamtor3 | 0.004086174 | 0.90556406 | 0.9832544 | 0.447938229 | 0.409015475 |
| Lamtor4 | 0.007794956 | 0.821116088 | 0.9705476 | 0.943338321 | 0.920264447 |
| Lamtor5 | -0.070471552 | 0.042791111 | 0.3959774 | 0.53368306 | 0.562198479 |
| Lancl1 | -0.042499875 | 0.106463458 | 0.5621561 | 0.147752687 | 0.185995568 |
| Lancl2 | 0.000973586 | 0.955578102 | 0.9954690 | 0.066391681 | 0.064523373 |
| Lap3 | -0.056534206 | 0.08301817 | 0.5177531 | 0.334566842 | 0.32570631 |
| Laptm4a | 0.034378038 | 0.24827403 | 0.7184592 | 1.448886842 | 1.33245643 |
| Laptm4b | 0.025230345 | 0.422988284 | 0.8315789 | 0.286028336 | 0.256147802 |
| Large2 | -0.015968079 | 0.376853325 | 0.8101797 | 0.057159607 | 0.074366694 |
| Larp1 | 0.02335794 | 0.508305412 | 0.8704067 | 0.501772665 | 0.485775505 |
| Larp1b | 0.01404608 | 0.678737447 | 0.9285719 | 0.383979704 | 0.408066431 |
| Larp4 | -0.025239371 | 0.486519664 | 0.8637358 | 0.525423696 | 0.525019242 |
| Larp4b | -0.014289435 | 0.689697828 | 0.9341619 | 0.517615618 | 0.536960177 |
| Larp7 | 0.04910993 | 0.052280299 | 0.4314646 | 0.191012805 | 0.130327034 |
| Lars | 0.025582783 | 0.502270443 | 0.8695566 | 0.811726374 | 0.837513705 |
| Lars2 | -0.07140843 | 0.045600826 | 0.4046155 | 0.327651398 | 0.391519338 |
| Las1l | -0.038348479 | 0.220983914 | 0.6926204 | 0.2270919 | 0.27748664 |

| | | | | | |
|---|---|---|---|---|---|
| Lasp1 | -0.036425181 | 0.268295802 | 0.7323870 | 0.356498174 | 0.369729595 |
| Lats1 | -0.067699946 | 0.02535118 | 0.3055695 | 0.206640254 | 0.269789101 |
| Lbh | 0.11531154 | 0.00180357 | 0.0739314 | 0.502650831 | 0.419149036 |
| Lbr | 0.019916982 | 0.353256526 | 0.7961996 | 0.115825778 | 0.097717564 |
| Lca5 | -0.006794175 | 0.679834627 | 0.9289200 | 0.053888793 | 0.055095138 |
| Lclat1 | 0.015153518 | 0.408218463 | 0.8272782 | 0.078894688 | 0.064580991 |
| Lcmt1 | -0.044403463 | 0.097425868 | 0.5384203 | 0.172563601 | 0.188149465 |
| Lcmt2 | 0.020553474 | 0.350071901 | 0.7927303 | 0.11991131 | 0.107801592 |
| Lcor | -0.015511618 | 0.561561929 | 0.8926240 | 0.160307891 | 0.178621577 |
| Lcorl | 0.029125545 | 0.409125019 | 0.8276447 | 0.426124368 | 0.405879561 |
| Ldah | -0.004311269 | 0.870450459 | 0.9780178 | 0.16245855 | 0.162712885 |
| Ldb1 | -0.015342686 | 0.578061574 | 0.8997914 | 0.177645763 | 0.195157745 |
| Ldhb | 0.037162667 | 0.075495556 | 0.4966285 | 0.119599099 | 0.085671633 |
| Ldhd | -0.020037319 | 0.206366821 | 0.6797565 | 0.042549592 | 0.054842278 |
| Ldlr | -0.033749565 | 0.317001664 | 0.7706462 | 0.280969214 | 0.308249099 |
| Ldlrad4 | 0.006311746 | 0.710071542 | 0.9419370 | 0.060516692 | 0.060502779 |
| Ldlrap1 | -0.012098697 | 0.624747089 | 0.9121505 | 0.142463275 | 0.139338955 |
| Lefty1 | 0.024932255 | 0.406788194 | 0.8270770 | 0.291081904 | 0.222729811 |
| Lekr1 | 0.000334488 | 0.983175421 | 0.9976072 | 0.053257682 | 0.050133373 |
| Lemd2 | 0.016000437 | 0.587221095 | 0.9038375 | 0.253782284 | 0.225307963 |
| Lemd3 | -0.029274388 | 0.184498896 | 0.6628862 | 0.095549139 | 0.11643048 |
| Leng1 | 0.026790846 | 0.227995609 | 0.7015153 | 0.141876056 | 0.106899805 |
| Leng8 | -0.025252164 | 0.344550845 | 0.7879223 | 0.148038155 | 0.175976318 |
| Leng9 | 0.024437822 | 0.266153316 | 0.7307399 | 0.12028869 | 0.098311389 |
| Leo1 | -0.039578132 | 0.223459574 | 0.6955958 | 0.283809784 | 0.301628446 |
| Leprot | -0.025236696 | 0.364078964 | 0.8049171 | 0.21288455 | 0.200859776 |
| Leprotl1 | 0.003174859 | 0.933140961 | 0.9902394 | 0.744765606 | 0.846013129 |
| Letm1 | 0.06795439 | 0.011976133 | 0.2156138 | 0.209688974 | 0.169150259 |
| Letm2 | -0.01091482 | 0.603345169 | 0.9087139 | 0.089538471 | 0.102568973 |
| Letmd1 | -0.022199744 | 0.275294169 | 0.7356525 | 0.081118182 | 0.095750804 |
| Lgals3bp | -0.08847788 | 0.016719561 | 0.2546713 | 1.191338978 | 1.294859596 |
| Lgals8 | 0.012301847 | 0.692553469 | 0.9358575 | 0.298962538 | 0.271114017 |
| Lgalsl | -0.032471943 | 0.286655841 | 0.7468580 | 0.216036903 | 0.25854249 |
| Lgi1 | -0.013299073 | 0.508963067 | 0.8704067 | 0.071578238 | 0.091146322 |
| Lgi2 | 0.074950356 | 0.006360644 | 0.1597407 | 0.221971952 | 0.146974298 |
| Lgmn | -0.034519803 | 0.177595646 | 0.6538187 | 0.13525728 | 0.158720158 |
| Lgr4 | 0.020807749 | 0.178783575 | 0.6549235 | 0.067612333 | 0.041664 |
| Lhfpl4 | -0.009911874 | 0.696152678 | 0.9363862 | 0.145338566 | 0.155582255 |
| Lhfpl5 | -0.011575577 | 0.425191936 | 0.8324460 | 0.037673903 | 0.047751534 |
| Lhx1 | -0.033684397 | 0.114734926 | 0.5713707 | 0.058190609 | 0.100298249 |
| Lias | 0.000918782 | 0.977300707 | 0.9976072 | 0.37405884 | 0.357885564 |
| Lifr | 0.070095391 | 0.008080926 | 0.1773512 | 0.217865951 | 0.14597986 |
| Lig1 | 0.007177024 | 0.72334045 | 0.9445652 | 0.093141776 | 0.080489822 |
| Lig3 | -0.019518126 | 0.380825925 | 0.8117350 | 0.097616269 | 0.113470177 |
| Lig4 | 0.01365454 | 0.450151596 | 0.8478268 | 0.075258565 | 0.066190153 |
| Lima1 | 0.006470359 | 0.790493444 | 0.9621438 | 0.13325039 | 0.138017313 |
| Limch1 | 0.01672345 | 0.389077003 | 0.8170287 | 0.073363287 | 0.086376351 |
| Limd1 | -0.044452749 | 0.193654283 | 0.6710847 | 0.317129467 | 0.360396894 |
| Limd2 | 0.062293736 | 0.052716904 | 0.4332656 | 0.32784013 | 0.266284382 |
| Limk2 | 0.012945461 | 0.579014976 | 0.9003073 | 0.134880529 | 0.115514544 |
| Lims1 | 0.006548072 | 0.805393441 | 0.9647143 | 0.173659498 | 0.159912234 |
| Lin37 | 0.0137066 | 0.593854023 | 0.9062213 | 0.171723281 | 0.158282612 |
| Lin52 | 0.014228239 | 0.399366956 | 0.8239850 | 0.069414839 | 0.055437477 |
| LIN54 | 0.005711627 | 0.787397849 | 0.9608311 | 0.097937909 | 0.096186482 |
| Lin7c | -0.077853276 | 0.023749126 | 0.2989737 | 0.383942535 | 0.445597883 |
| Lins1 | -0.007096322 | 0.642906571 | 0.9166712 | 0.042565715 | 0.050331267 |
| Lipa | 1.39E-05 | 0.999321992 | 0.9996989 | 0.058550523 | 0.055131026 |
| Lipo3 | 0.004832716 | 0.800452586 | 0.9647053 | 0.087782977 | 0.078712094 |
| Lipt2 | -0.019146007 | 0.218134122 | 0.6897120 | 0.044447819 | 0.049287943 |
| Litaf | 0.013048259 | 0.491036084 | 0.8658154 | 0.081819738 | 0.071554783 |
| Lix1l | -0.021782987 | 0.365930381 | 0.8050191 | 0.143346408 | 0.133977957 |
| Llgl1 | -0.036010504 | 0.016815494 | 0.2546713 | 0.02884468 | 0.052951484 |
| Llgl2 | 0.006390552 | 0.808893417 | 0.9658428 | 0.184814553 | 0.157233418 |
| Llph | -0.001741108 | 0.960359821 | 0.9958628 | 0.450625847 | 0.437102661 |
| Lman1 | -0.043225438 | 0.177331542 | 0.6536659 | 1.597338056 | 1.670170938 |
| Lman2 | -0.044721881 | 0.229609236 | 0.7019677 | 0.638413551 | 0.668649559 |

| | | | | | |
|---|---|---|---|---|---|
| Lman2l | 0.020327249 | 0.463548383 | 0.8525027 | 0.217218281 | 0.189428051 |
| Lmbr1 | 0.039697738 | 0.032112274 | 0.3392328 | 0.100766796 | 0.065082405 |
| Lmbr1l | -0.002894936 | 0.864066652 | 0.9769675 | 0.067369774 | 0.059843493 |
| Lmbrd1 | 0.072091246 | 0.014181655 | 0.2347500 | 0.266400626 | 0.218109575 |
| Lmbrd2 | 0.01525325 | 0.59525171 | 0.9067325 | 0.212959586 | 0.210529454 |
| Lmf1 | -0.006311139 | 0.782736919 | 0.9590862 | 0.113396215 | 0.121296421 |
| Lmf2 | -0.014402474 | 0.415689211 | 0.8300990 | 0.057595971 | 0.076245732 |
| Lmna | -0.088677172 | 0.01760339 | 0.2582401 | 0.619936045 | 0.673254763 |
| Lmnb1 | -0.009651739 | 0.614911518 | 0.9108439 | 0.081293696 | 0.078429709 |
| Lmnb2 | 0.026766415 | 0.088543604 | 0.5284771 | 0.06675194 | 0.04592741 |
| Lmo1 | 0.036846286 | 0.155963242 | 0.6312943 | 0.144785261 | 0.160282666 |
| Lmo4 | 0.019399631 | 0.369678427 | 0.8063346 | 0.109314746 | 0.090857652 |
| Lmo7 | 0.011610792 | 0.72383753 | 0.9445652 | 0.296513676 | 0.322000629 |
| Lmtk2 | -0.009902 | 0.660654319 | 0.9211780 | 0.102208476 | 0.118902862 |
| Lncpint | -0.031634505 | 0.214369069 | 0.6837977 | 0.113241061 | 0.149090957 |
| Lnpep | 0.014919635 | 0.666103831 | 0.9230765 | 0.373489351 | 0.354089633 |
| Lnpk | -0.009796143 | 0.687747938 | 0.9327945 | 0.133731152 | 0.135364296 |
| Lnx1 | 0.002728046 | 0.878302321 | 0.9785583 | 0.06734908 | 0.066902613 |
| Lonp1 | -0.047834865 | 0.18596789 | 0.6640243 | 0.3459592 | 0.423449816 |
| Lonp2 | 0.01758853 | 0.592292362 | 0.9060200 | 0.320667676 | 0.302086916 |
| Lonrf1 | 0.030906018 | 0.080647 | 0.5108673 | 0.080419738 | 0.057440777 |
| Lonrf2 | 0.001940692 | 0.926507576 | 0.9879210 | 0.121145148 | 0.092001653 |
| Loxl2 | 0.020387091 | 0.161037908 | 0.6384844 | 0.050699874 | 0.041960649 |
| Lpar6 | -0.006605881 | 0.70733374 | 0.9408354 | 0.07129611 | 0.062304241 |
| Lpcat1 | -0.031118236 | 0.200560458 | 0.6739701 | 0.132770314 | 0.14122644 |
| Lpcat3 | 0.000480498 | 0.986719617 | 0.9976072 | 0.219865164 | 0.209271809 |
| Lpgat1 | 0.107711633 | 0.002399592 | 0.0890983 | 0.46498298 | 0.386029066 |
| Lpin1 | 0.066021607 | 0.009917243 | 0.1987218 | 0.195908939 | 0.142569407 |
| Lpin2 | -0.043087096 | 0.165560036 | 0.6425491 | 0.227735569 | 0.253893064 |
| Lpl | -0.106618701 | 0.011419668 | 0.2105633 | 0.757657827 | 0.767367809 |
| Lpp | -0.018821564 | 0.618365234 | 0.9115017 | 0.770833764 | 0.758694475 |
| Lratd1 | -0.025724613 | 0.21599617 | 0.6858777 | 0.084837909 | 0.087687118 |
| Lratd2 | 0.029541347 | 0.181642952 | 0.6583478 | 0.11418864 | 0.103103478 |
| Lrba | -0.028877856 | 0.390757739 | 0.8193912 | 0.320579903 | 0.334185159 |
| Lrch1 | -0.012696684 | 0.545528896 | 0.8852685 | 0.098230759 | 0.100911562 |
| Lrch2 | 0.027686467 | 0.104741355 | 0.5580045 | 0.077837189 | 0.056379605 |
| Lrch3 | 0.045394358 | 0.058220375 | 0.4508199 | 0.164506656 | 0.114437378 |
| Lrfn3 | -0.036216455 | 0.234721882 | 0.7064894 | 0.238421695 | 0.272727173 |
| Lrfn4 | 0.001236892 | 0.948246357 | 0.9945018 | 0.079290385 | 0.079550537 |
| Lrif1 | -0.008207878 | 0.800709607 | 0.9647143 | 0.315764566 | 0.298738024 |
| Lrig1 | -0.03101594 | 0.068988634 | 0.4835495 | 0.043019579 | 0.065737305 |
| Lrig2 | 0.015331948 | 0.431598159 | 0.8357317 | 0.098128701 | 0.084460957 |
| Lrp1 | -0.002956049 | 0.881007989 | 0.9795705 | 0.082393786 | 0.080257684 |
| Lrp10 | 0.081818961 | 0.010758912 | 0.2035157 | 0.360202696 | 0.297954591 |
| Lrp11 | -0.03710658 | 0.302652075 | 0.7614274 | 0.579895911 | 0.574859311 |
| Lrp12 | 0.001868886 | 0.941589796 | 0.9929301 | 0.142348055 | 0.149372449 |
| Lrp3 | -0.011361658 | 0.673299499 | 0.9261686 | 0.187636859 | 0.1816568 |
| Lrp6 | 0.034474235 | 0.283107395 | 0.7427848 | 0.307524066 | 0.267765245 |
| Lrp8 | -0.032121474 | 0.284434972 | 0.7442877 | 0.206140348 | 0.224448101 |
| Lrpap1 | 0.031466424 | 0.374141285 | 0.8087297 | 0.820846271 | 0.713489714 |
| Lrpprc | -0.013577838 | 0.705172127 | 0.9401381 | 0.35702147 | 0.357593094 |
| Lrrc1 | -0.035219959 | 0.13007474 | 0.5938064 | 0.10221475 | 0.126646728 |
| Lrrc10b | -0.03630968 | 0.332074137 | 0.7793175 | 0.51678377 | 0.577718758 |
| Lrrc14 | 0.002118941 | 0.924262496 | 0.9878036 | 0.125500508 | 0.112959378 |
| Lrrc20 | -0.00310879 | 0.876579916 | 0.9785583 | 0.078925574 | 0.090234962 |
| Lrrc24 | 0.012478762 | 0.516741585 | 0.8737134 | 0.084837614 | 0.072127064 |
| Lrrc40 | 0.013619053 | 0.497817319 | 0.8684629 | 0.102381617 | 0.08545386 |
| Lrrc41 | 0.029508294 | 0.358501695 | 0.7997159 | 0.372741389 | 0.323094594 |
| Lrrc42 | 0.010481324 | 0.660856269 | 0.9212611 | 0.153428547 | 0.130221615 |
| Lrrc45 | 0.001065936 | 0.950339121 | 0.9948462 | 0.066814055 | 0.058363556 |
| Lrrc47 | -0.003909044 | 0.8906824 | 0.9819597 | 0.208602984 | 0.209621534 |
| Lrrc49 | 0.003166328 | 0.861581323 | 0.9769675 | 0.078054012 | 0.067293399 |
| Lrrc51 | -0.034126908 | 0.023154605 | 0.2949903 | 0.03091743 | 0.05546223 |
| Lrrc57 | -0.032806399 | 0.084047447 | 0.5185016 | 0.061661455 | 0.084244894 |
| Lrrc58 | -0.006222125 | 0.858667087 | 0.9769675 | 0.536974985 | 0.558000042 |
| Lrrc59 | -0.01748422 | 0.614218225 | 0.9108386 | 0.410740641 | 0.416969759 |

| | | | | | |
|---|---|---|---|---|---|
| Lrrc61 | 0.002264649 | 0.872368516 | 0.9784331 | 0.041220737 | 0.040931722 |
| Lrrc73 | 0.0295872 | 0.359509981 | 0.8003169 | 0.350048949 | 0.29276627 |
| Lrrc8a | 0.013676781 | 0.442542604 | 0.8431541 | 0.079650489 | 0.062875929 |
| Lrrc8b | 0.000383778 | 0.981118925 | 0.9976072 | 0.055203583 | 0.052471663 |
| Lrrc8d | -0.02747639 | 0.373346498 | 0.8085689 | 0.238949089 | 0.280234894 |
| Lrrcc1 | 0.004718482 | 0.851211465 | 0.9749822 | 0.169261372 | 0.145115444 |
| Lrrfip1 | 0.037105793 | 0.003092411 | 0.1032885 | 0.049456063 | 0.024848504 |
| Lrrfip2 | 0.045089665 | 0.223544319 | 0.6955958 | 0.523759804 | 0.534491555 |
| Lrrn1 | -0.045228624 | 0.174353385 | 0.6527657 | 0.296139715 | 0.319083747 |
| Lrwd1 | 0.022369807 | 0.171754036 | 0.6515402 | 0.075506296 | 0.052078115 |
| Lsg1 | -0.021116668 | 0.453013941 | 0.8488663 | 0.216504976 | 0.214153946 |
| Lsm1 | 0.056099792 | 0.06985419 | 0.4847655 | 0.330729107 | 0.268733663 |
| Lsm10 | 0.046221478 | 0.095419448 | 0.5351767 | 0.234001465 | 0.186600375 |
| Lsm12 | 0.060752206 | 0.073672716 | 0.4920747 | 0.451827049 | 0.383015961 |
| Lsm14a | -0.017796982 | 0.579028332 | 0.9003073 | 0.28367944 | 0.294246808 |
| Lsm14b | -0.040184447 | 0.159747136 | 0.6369225 | 0.171764409 | 0.19655673 |
| Lsm2 | 0.005399813 | 0.869597324 | 0.9777384 | 0.3701917 | 0.329320531 |
| Lsm3 | 0.019398884 | 0.548040818 | 0.8863001 | 0.351243804 | 0.306973044 |
| Lsm4 | -0.058110889 | 0.093028849 | 0.5328211 | 1.009152038 | 0.991602031 |
| Lsm5 | -0.025979908 | 0.433985721 | 0.8377586 | 0.392062568 | 0.390601746 |
| Lsm6 | -0.002079634 | 0.948223158 | 0.9945018 | 1.224546508 | 1.180600649 |
| Lsm7 | 0.054425058 | 0.10749021 | 0.5633723 | 0.436949474 | 0.374845255 |
| Lsm8 | -0.017187087 | 0.598985204 | 0.9075810 | 0.377099363 | 0.341089484 |
| Lsr | -0.111093706 | 0.001008509 | 0.0488020 | 0.314560055 | 0.366030686 |
| Lst1 | 0.027293777 | 0.107499194 | 0.5633723 | 0.078112011 | 0.051664347 |
| Lta4h | 0.008678416 | 0.735861244 | 0.9468380 | 0.164758273 | 0.14530401 |
| Ltbp3 | -0.083005314 | 0.00813199 | 0.1775389 | 0.24573606 | 0.294682253 |
| Ltbr | -0.03493634 | 0.109729753 | 0.5655888 | 0.085233841 | 0.111366933 |
| Ltn1 | -0.034500044 | 0.224150776 | 0.6962539 | 0.18391924 | 0.212698476 |
| LTO1 | -0.006313535 | 0.759751788 | 0.9521533 | 0.099103316 | 0.097237306 |
| Ltv1 | -0.023155141 | 0.333417073 | 0.7804836 | 0.119185204 | 0.142736589 |
| Luc7l | -0.032789751 | 0.383574199 | 0.8129787 | 0.520471209 | 0.517531623 |
| Luc7l2 | -0.063789257 | 0.0972026 | 0.5383862 | 1.143085513 | 1.227452904 |
| Luc7l3 | -0.046943577 | 0.213015445 | 0.6827572 | 0.585675661 | 0.644898755 |
| Lurap1l | 0.01473224 | 0.560817594 | 0.8926240 | 0.158135209 | 0.14401157 |
| Luzp1 | -9.68E-05 | 0.997459063 | 0.9994073 | 0.254381159 | 0.240639758 |
| Lxn | -0.039001599 | 0.042991738 | 0.3970931 | 0.057194004 | 0.091121319 |
| Ly6e | -0.041464581 | 0.273513066 | 0.7351955 | 0.631696435 | 0.676695676 |
| Ly6h | 0.003953744 | 0.871463393 | 0.9782269 | 0.083557395 | 0.098502593 |
| Ly96 | -0.033079747 | 0.239300305 | 0.7115884 | 0.180316435 | 0.203568112 |
| Lyar | -0.014167195 | 0.657591872 | 0.9211780 | 0.283950216 | 0.277943414 |
| Lypla1 | -0.048543711 | 0.046967573 | 0.4105007 | 0.122760751 | 0.15193062 |
| Lypla2 | 0.00443931 | 0.880330622 | 0.9792571 | 0.241722035 | 0.24366077 |
| Lyplal1 | -0.031985524 | 0.291972436 | 0.7521076 | 0.270219764 | 0.256185449 |
| Lyrm1 | 0.01381063 | 0.385144891 | 0.8139828 | 0.059998361 | 0.050415032 |
| Lyrm2 | -0.041989326 | 0.199441509 | 0.6730962 | 0.299955299 | 0.351219222 |
| Lyrm4 | 0.026714349 | 0.269960588 | 0.7340360 | 0.176794554 | 0.134940791 |
| Lyrm7 | -0.012394586 | 0.397465302 | 0.8238469 | 0.041621979 | 0.045379136 |
| Lyrm9 | 0.071993759 | 0.017336666 | 0.2570999 | 0.28772214 | 0.236337299 |
| Lysmd2 | 0.001524904 | 0.964911365 | 0.9966421 | 0.535191053 | 0.486403858 |
| Lysmd3 | -0.052595246 | 0.11616455 | 0.5724670 | 0.288249619 | 0.334742619 |
| Lyst | -0.018040604 | 0.524921869 | 0.8758385 | 0.175734011 | 0.192363319 |
| Lzic | -0.009664537 | 0.586913457 | 0.9035702 | 0.068632725 | 0.070053071 |
| Lztfl1 | -0.032972548 | 0.341202016 | 0.7851366 | 0.407759744 | 0.410944746 |
| Lztr1 | 0.004961463 | 0.86389424 | 0.9769675 | 0.202163321 | 0.200132116 |
| Lzts2 | -0.029543588 | 0.287479986 | 0.7471316 | 0.178083364 | 0.192662238 |
| M6pr | 0.02921647 | 0.415300629 | 0.8300990 | 0.518368489 | 0.449402465 |
| Macf1 | -0.037157536 | 0.324263156 | 0.7757552 | 0.612541127 | 0.718579497 |
| Macir | 0.022771678 | 0.318876975 | 0.7710043 | 0.121523343 | 0.115195329 |
| Maco1 | 0.017992444 | 0.625283593 | 0.9121784 | 0.661371625 | 0.618944941 |
| Macrod1 | -0.030499071 | 0.088494808 | 0.5284771 | 0.056869323 | 0.072802829 |
| Macrod2 | 0.027820876 | 0.294929795 | 0.7536588 | 0.190411968 | 0.156561408 |
| Macroh2a1 | 0.053654239 | 0.071901665 | 0.4888240 | 1.328417488 | 1.189235261 |
| Macroh2a2 | -0.003890801 | 0.89975487 | 0.9825441 | 0.282515233 | 0.258083127 |
| Mad2l1bp | -0.00737926 | 0.668558539 | 0.9230765 | 0.06277433 | 0.060510738 |
| Mad2l2 | -0.007599609 | 0.738454337 | 0.9474220 | 0.111544112 | 0.111878288 |

| | | | | | |
|---|---|---|---|---|---|
| Madd | 0.003941809 | 0.916074865 | 0.9845236 | 0.540716285 | 0.62786404 |
| Maea | 0.040155338 | 0.14127406 | 0.6139045 | 0.219065645 | 0.175083401 |
| Maf1 | -0.036568812 | 0.228023985 | 0.7015153 | 0.258132971 | 0.267484378 |
| Mafa | 0.165779412 | 0.001735898 | 0.0726586 | 0.815046995 | 0.801946516 |
| Mafb | -0.099567669 | 0.046948145 | 0.4105007 | 1.969876657 | 1.787739069 |
| Maff | -0.050729805 | 0.087073255 | 0.5254055 | 0.126270121 | 0.215233367 |
| Mafg | -0.054242209 | 0.149920284 | 0.6272497 | 0.577946642 | 0.616662077 |
| Mafk | -0.021631193 | 0.31665315 | 0.7705987 | 0.084689394 | 0.107886695 |
| Maged1 | -0.058149451 | 0.024201384 | 0.3022973 | 1.839213883 | 1.858345866 |
| Maged2 | -0.004100512 | 0.911268293 | 0.9839753 | 0.615509717 | 0.57675559 |
| Magee1 | -0.008148043 | 0.706210332 | 0.9403498 | 0.114401146 | 0.098860754 |
| Magef1 | -0.021978653 | 0.328396416 | 0.7778857 | 0.111793272 | 0.120611438 |
| Mageh1 | 0.025041465 | 0.521829185 | 0.8748598 | 1.025546862 | 0.882379951 |
| Magi2 | -0.034058093 | 0.275775435 | 0.7357967 | 0.240629678 | 0.259397686 |
| Magi3 | -0.002654438 | 0.884411082 | 0.9802800 | 0.071436822 | 0.067785989 |
| Magoh | 0.034454831 | 0.335816938 | 0.7816292 | 0.665870071 | 0.56801594 |
| Magohb | 0.040643956 | 0.138514898 | 0.6082637 | 0.248233523 | 0.187823878 |
| Magt1 | -0.029707725 | 0.421411664 | 0.8304337 | 0.725266247 | 0.783462751 |
| Maip1 | -0.017884964 | 0.501226041 | 0.8695566 | 0.176321883 | 0.183182278 |
| Mak16 | 0.065872048 | 0.043969772 | 0.3997984 | 0.366210487 | 0.286607264 |
| Mal2 | -0.013156032 | 0.462334139 | 0.8524425 | 0.057703605 | 0.069304129 |
| Malat1 | -0.037268131 | 0.202429329 | 0.6759000 | 5.974094728 | 6.258361994 |
| Malsu1 | 0.04884908 | 0.132065539 | 0.5978345 | 0.378862998 | 0.322592525 |
| Malt1 | 0.003422819 | 0.854368302 | 0.9755431 | 0.071234963 | 0.069829693 |
| Maml1 | -0.009762543 | 0.57002481 | 0.8962534 | 0.05716898 | 0.061668341 |
| Maml3 | -0.011313614 | 0.752381781 | 0.9507898 | 0.39584603 | 0.398212479 |
| Mamld1 | -0.022550272 | 0.36865142 | 0.8056719 | 0.148642598 | 0.136861796 |
| Man1a | 0.003279437 | 0.928270777 | 0.9880613 | 0.50294847 | 0.504362489 |
| Man1a2 | -0.062887293 | 0.061318238 | 0.4611772 | 0.338947597 | 0.416434347 |
| Man1b1 | -0.001573949 | 0.942656851 | 0.9934299 | 0.114618254 | 0.109617203 |
| Man1c1 | 0.019839489 | 0.444865783 | 0.8431541 | 0.206308854 | 0.155598713 |
| Man2a1 | -0.005833199 | 0.864812371 | 0.9769675 | 0.345125874 | 0.326214458 |
| Man2a2 | -0.065952974 | 0.010450477 | 0.2017406 | 0.107273985 | 0.176459205 |
| Man2b1 | -0.000814696 | 0.974375258 | 0.9976072 | 0.161794507 | 0.154098223 |
| Man2b2 | -0.016463901 | 0.499093873 | 0.8690742 | 0.137303868 | 0.137983656 |
| Man2c1 | -0.001987763 | 0.934929402 | 0.9907594 | 0.153929476 | 0.133358104 |
| Manba | 0.013556413 | 0.394656492 | 0.8216143 | 0.056345675 | 0.052221294 |
| Manbal | -0.030092699 | 0.318140406 | 0.7710043 | 0.23727489 | 0.25061599 |
| Manea | -0.001321251 | 0.943486657 | 0.9935293 | 0.071081862 | 0.076289237 |
| Maneal | -0.013269337 | 0.4773855 | 0.8607489 | 0.070352515 | 0.07737736 |
| Manf | -0.08408929 | 0.010179686 | 0.1999013 | 1.967322975 | 2.013743126 |
| Mansc1 | 0.02308798 | 0.073619697 | 0.4920747 | 0.049926442 | 0.028067237 |
| Maoa | -0.014866089 | 0.506489798 | 0.8695566 | 0.112799632 | 0.110430119 |
| Maob | 0.103136114 | 0.007341012 | 0.1701468 | 0.629073063 | 0.485326799 |
| Map10 | 0.003579014 | 0.819452485 | 0.9699062 | 0.053155936 | 0.047427293 |
| Map11 | -0.007016738 | 0.732908384 | 0.9464134 | 0.092481565 | 0.092737005 |
| Map1b | -0.076651688 | 0.018307262 | 0.2635820 | 1.298342597 | 1.374388214 |
| Map1lc3a | -0.003096933 | 0.927660479 | 0.9879470 | 1.06763853 | 1.04242952 |
| Map1lc3b | -0.024571249 | 0.474976617 | 0.8602394 | 0.999946904 | 0.956135791 |
| Map1s | -0.029011803 | 0.218544851 | 0.6899952 | 0.103701426 | 0.131837484 |
| Map2 | -0.052445865 | 0.162448426 | 0.6400703 | 0.518343487 | 0.552863455 |
| Map2k1 | -0.031768993 | 0.30149834 | 0.7608403 | 0.24406811 | 0.249829797 |
| Map2k2 | 0.024079185 | 0.482470142 | 0.8620306 | 0.779455691 | 0.735102174 |
| Map2k3 | -0.04478127 | 0.042678871 | 0.3958356 | 0.084259774 | 0.117984004 |
| Map2k4 | 0.034560737 | 0.17444478 | 0.6527696 | 0.178766963 | 0.144152732 |
| Map2k5 | 0.003041846 | 0.892776403 | 0.9821882 | 0.127099786 | 0.112261394 |
| Map2k7 | -0.025910532 | 0.350580678 | 0.7927303 | 0.175155472 | 0.189277529 |
| Map3k1 | 0.010802115 | 0.665604783 | 0.9230765 | 0.15400986 | 0.144710993 |
| Map3k10 | 0.003923452 | 0.876876156 | 0.9785583 | 0.162070907 | 0.154579613 |
| Map3k11 | -0.015818102 | 0.417325948 | 0.8300990 | 0.073790561 | 0.088664734 |
| Map3k12 | 0.016307581 | 0.372753972 | 0.8085689 | 0.085709813 | 0.067671609 |
| Map3k15 | -0.084310514 | 0.0323583 | 0.3397589 | 0.60492273 | 0.610671062 |
| Map3k2 | -0.024026399 | 0.520841686 | 0.8741795 | 0.494813027 | 0.518640331 |
| Map3k20 | -0.016649687 | 0.292978845 | 0.7533307 | 0.04098746 | 0.052346759 |
| Map3k21 | -0.010087375 | 0.543546979 | 0.8842220 | 0.056300483 | 0.06104854 |
| Map3k3 | -0.010907624 | 0.638605746 | 0.9160042 | 0.12779167 | 0.119519997 |

| | | | | | |
|---|---|---|---|---|---|
| Map3k4 | 0.021089369 | 0.522047351 | 0.8749671 | 0.322680323 | 0.303694103 |
| Map3k5 | -0.016975115 | 0.517948509 | 0.8739665 | 0.153292379 | 0.152766448 |
| Map3k7 | -0.015963825 | 0.564320672 | 0.8933738 | 0.170496252 | 0.184352186 |
| Map4 | -0.007533138 | 0.782834844 | 0.9590862 | 0.176225127 | 0.185050148 |
| Map4k2 | 0.016051746 | 0.408010579 | 0.8270770 | 0.093261877 | 0.076363173 |
| Map4k3 | -0.014611776 | 0.497484116 | 0.8684629 | 0.091905169 | 0.106679223 |
| Map4k4 | -0.065597599 | 0.046768511 | 0.4094824 | 0.289108632 | 0.348402797 |
| Map4k5 | 0.009742609 | 0.698221277 | 0.9369692 | 0.139677213 | 0.135501992 |
| Map6 | -0.021535317 | 0.177374795 | 0.6536659 | 0.042757 | 0.057328453 |
| Map7 | -0.012044708 | 0.712606927 | 0.9421288 | 0.326995253 | 0.328660769 |
| Map7d1 | -0.018900613 | 0.525398886 | 0.8762536 | 0.212511605 | 0.233015144 |
| Map7d2 | -0.06350709 | 0.043296634 | 0.3976875 | 0.208301988 | 0.26806004 |
| Map9 | 0.041744063 | 0.188440935 | 0.6668854 | 0.309378956 | 0.251841727 |
| Mapk1 | 0.020901602 | 0.544461633 | 0.8842656 | 0.481048103 | 0.453417848 |
| Mapk10 | -0.038231198 | 0.110146753 | 0.5655888 | 0.097316369 | 0.144785236 |
| Mapk13 | 0.022295797 | 0.125340382 | 0.5870522 | 0.051432354 | 0.040004324 |
| Mapk14 | -0.047149724 | 0.148888264 | 0.6264374 | 0.27738653 | 0.353500206 |
| Mapk15 | -0.074892122 | 0.029295197 | 0.3280004 | 0.281198275 | 0.335024628 |
| Mapk1ip1 | -0.027428567 | 0.317682089 | 0.7709420 | 0.187885502 | 0.202055667 |
| Mapk1ip1l | -0.033941367 | 0.262963574 | 0.7276284 | 0.245033074 | 0.258892876 |
| Mapk3 | -0.011263967 | 0.751268059 | 0.9506053 | 0.637505645 | 0.611874341 |
| Mapk4 | 0.00958732 | 0.677055974 | 0.9280496 | 0.12620777 | 0.106336362 |
| Mapk6 | -0.019074496 | 0.484432487 | 0.8627594 | 0.170949754 | 0.176768055 |
| Mapk7 | 0.001132185 | 0.944919468 | 0.9936932 | 0.059424629 | 0.055168766 |
| Mapk8 | -0.044286522 | 0.093407503 | 0.5330951 | 0.139404227 | 0.16832363 |
| Mapk8ip1 | 0.015285669 | 0.631693295 | 0.9133517 | 0.373169527 | 0.323815377 |
| Mapk8ip2 | -0.045260529 | 0.162883496 | 0.6400703 | 0.289838365 | 0.349346099 |
| Mapk8ip3 | -0.046729841 | 0.144851222 | 0.6193638 | 0.247512274 | 0.278763658 |
| Mapk9 | -0.000882789 | 0.972770801 | 0.9976072 | 0.158315035 | 0.154702397 |
| Mapkap1 | 0.014680536 | 0.561154684 | 0.8926240 | 0.16357323 | 0.149826688 |
| Mapkapk2 | 0.004807366 | 0.874874504 | 0.9784331 | 0.246846743 | 0.253538578 |
| Mapkbp1 | 0.005511192 | 0.721710466 | 0.9445652 | 0.052051442 | 0.049776455 |
| Mapre1 | 0.022882502 | 0.515922608 | 0.8735705 | 0.490853332 | 0.438914587 |
| Mapre2 | 0.102308481 | 0.002195928 | 0.0836783 | 0.478292518 | 0.400304277 |
| Mapre3 | -0.040224594 | 0.233338466 | 0.7051391 | 0.383084937 | 0.371293141 |
| Mapt | -0.024248765 | 0.275410777 | 0.7356525 | 0.083131553 | 0.125799651 |
| Marchf2 | -0.025182978 | 0.369149671 | 0.8056719 | 0.201838061 | 0.198795983 |
| Marchf4 | 0.009327292 | 0.58299157 | 0.9019790 | 0.060821312 | 0.063432374 |
| Marchf5 | -0.026285633 | 0.468844385 | 0.8563037 | 0.709809402 | 0.729156564 |
| Marchf6 | -0.018883866 | 0.596420336 | 0.9069085 | 0.446875949 | 0.485831289 |
| Marchf7 | 0.001005842 | 0.978294949 | 0.9976072 | 0.576451985 | 0.551089099 |
| Marchf8 | -0.006551695 | 0.857160437 | 0.9763189 | 0.510062176 | 0.517543547 |
| Marchf9 | 0.000442527 | 0.987751893 | 0.9976072 | 0.22952189 | 0.213393527 |
| Marcks | 0.031932851 | 0.403828308 | 0.8266563 | 0.749279259 | 0.691673139 |
| Marcksl1 | 0.042823256 | 0.048232995 | 0.4148910 | 0.135637227 | 0.077699088 |
| Marf1 | 0.027994892 | 0.377941782 | 0.8101915 | 0.325559713 | 0.276283718 |
| Mark1 | -0.022139349 | 0.31350716 | 0.7699903 | 0.099409692 | 0.114373864 |
| Mark2 | -0.014210505 | 0.667407052 | 0.9230765 | 0.343029441 | 0.371640135 |
| Mark3 | -0.1181956 | 0.001042222 | 0.0497059 | 0.358722626 | 0.460822054 |
| Mark4 | -0.029634581 | 0.167630975 | 0.6446304 | 0.090597585 | 0.104674948 |
| Mars1 | 0.012728969 | 0.724034182 | 0.9445652 | 0.453758846 | 0.436650817 |
| Marveld1 | 0.005109959 | 0.828121256 | 0.9718751 | 0.13536885 | 0.124760656 |
| Marveld2 | 0.003517959 | 0.829397993 | 0.9722601 | 0.060342118 | 0.053774987 |
| Marveld3 | -0.024612674 | 0.362423907 | 0.8030478 | 0.16812294 | 0.181462794 |
| Mast2 | -0.08643858 | 0.002563974 | 0.0927113 | 0.162968106 | 0.229652369 |
| Mast3 | 0.01108204 | 0.650074377 | 0.9175779 | 0.147066129 | 0.14674001 |
| Mast4 | 0.009982165 | 0.792990998 | 0.9624508 | 0.821773122 | 0.844831665 |
| Mat1a | 0.035946551 | 0.219634496 | 0.6910162 | 0.250511666 | 0.193469801 |
| Mat2a | -0.127788818 | 0.000931422 | 0.0461985 | 0.334798725 | 0.544032627 |
| Mat2b | -0.018021079 | 0.593215867 | 0.9060585 | 0.364311971 | 0.363060411 |
| Matr3 | 0.038458611 | 0.298009566 | 0.7558821 | 0.695078048 | 0.640025878 |
| Mau2 | -0.037376402 | 0.237331029 | 0.7098511 | 0.259258772 | 0.269679189 |
| Mavs | 0.022175194 | 0.15192001 | 0.6281105 | 0.066788933 | 0.044076913 |
| Max | -0.03614635 | 0.23695465 | 0.7095050 | 0.239714674 | 0.261195591 |
| Maz | 0.012793088 | 0.700273276 | 0.9375165 | 0.393088543 | 0.345166855 |
| Mb21d2 | -0.021182898 | 0.186421254 | 0.6644983 | 0.040054342 | 0.056250029 |

| | | | | | |
|---|---|---|---|---|---|
| Mbd1 | -0.037984207 | 0.157621 | 0.6324475 | 0.149194208 | 0.185823406 |
| Mbd2 | 0.050152002 | 0.15665551 | 0.6317683 | 0.848045063 | 0.778417537 |
| Mbd3 | 0.043282071 | 0.228963495 | 0.7019677 | 0.633834314 | 0.527760591 |
| Mbd5 | -0.01331438 | 0.596094346 | 0.9068020 | 0.147365825 | 0.141891016 |
| Mbd6 | 0.006103424 | 0.825863856 | 0.9716906 | 0.189924175 | 0.191157317 |
| Mbip | -0.012068253 | 0.577941517 | 0.8997914 | 0.104335237 | 0.107936713 |
| Mblac1 | 0.012359376 | 0.390780222 | 0.8193912 | 0.049973865 | 0.038951674 |
| Mblac2 | 0.011038768 | 0.624655552 | 0.9121505 | 0.143471374 | 0.107791808 |
| Mbnl1 | 0.031744064 | 0.364354544 | 0.8049882 | 0.455122013 | 0.468165068 |
| Mbnl2 | -0.038903917 | 0.274155323 | 0.7351955 | 1.236211669 | 1.213756578 |
| Mboat2 | 0.022811856 | 0.448219363 | 0.8461153 | 0.24246662 | 0.249538279 |
| Mboat7 | -0.025198826 | 0.28928814 | 0.7494746 | 0.124378456 | 0.13573626 |
| Mbp | 0.015235621 | 0.644386929 | 0.9168391 | 0.332336651 | 0.331218107 |
| Mbtd1 | -0.020572593 | 0.46664157 | 0.8553370 | 0.187097506 | 0.195406464 |
| Mbtps1 | -0.03847443 | 0.206605792 | 0.6797565 | 0.260844744 | 0.259852854 |
| Mbtps2 | -0.013257872 | 0.506110126 | 0.8695566 | 0.088638606 | 0.089476141 |
| Mcat | 0.01619592 | 0.435731753 | 0.8389295 | 0.136032599 | 0.089486979 |
| Mcc | -0.017762291 | 0.300167859 | 0.7590275 | 0.045496329 | 0.068751236 |
| Mccc1 | 0.002695051 | 0.891631344 | 0.9819597 | 0.091939904 | 0.083195723 |
| Mccc2 | 0.028870602 | 0.097605333 | 0.5391118 | 0.086591202 | 0.058867483 |
| Mcee | -0.072361539 | 0.027148547 | 0.3164672 | 0.354073191 | 0.392234874 |
| Mcf2l | 0.009554937 | 0.767299487 | 0.9544221 | 0.32482805 | 0.312839426 |
| Mcfd2 | -0.008875363 | 0.75954527 | 0.9521533 | 0.211818291 | 0.213792107 |
| Mcl1 | -0.009626035 | 0.787154437 | 0.9608311 | 0.529216152 | 0.518629446 |
| Mcm3ap | 0.01081756 | 0.606493092 | 0.9087139 | 0.102324379 | 0.09138714 |
| Mcm4 | -0.001030843 | 0.951829308 | 0.9951566 | 0.061726008 | 0.060358921 |
| Mcm7 | -0.017650878 | 0.346103449 | 0.7894564 | 0.062943213 | 0.07687413 |
| Mcm9 | -0.036102875 | 0.033257639 | 0.3454615 | 0.041222088 | 0.0664531 |
| Mcmbp | 0.001511445 | 0.949313405 | 0.9948462 | 0.142087349 | 0.127361926 |
| Mcph1 | 0.001701183 | 0.916336044 | 0.9845236 | 0.055930767 | 0.054467008 |
| Mcrip1 | 0.011198482 | 0.754326626 | 0.9519043 | 0.855582597 | 0.840649243 |
| Mcrip2 | -0.007290333 | 0.71979996 | 0.9444523 | 0.094295914 | 0.091453858 |
| Mcrs1 | 0.005130565 | 0.853906604 | 0.9755431 | 0.213427797 | 0.190492424 |
| Mctp2 | -0.021027361 | 0.400733644 | 0.8247464 | 0.141635968 | 0.132858098 |
| Mcts1 | 0.013329392 | 0.695802067 | 0.9363862 | 0.48447678 | 0.420169961 |
| Mcts2 | 0.010031566 | 0.572430152 | 0.8969368 | 0.083866403 | 0.062821589 |
| Mcu | -0.00615153 | 0.750986787 | 0.9506053 | 0.073675017 | 0.083140502 |
| Mcub | -0.007888429 | 0.771616407 | 0.9552333 | 0.198394877 | 0.179254791 |
| Mcur1 | -0.013156192 | 0.674805971 | 0.9265887 | 0.264378139 | 0.274715424 |
| Mdc1 | 0.010303576 | 0.618689176 | 0.9116750 | 0.095425907 | 0.091952636 |
| Mdh1 | 0.040810478 | 0.19405308 | 0.6716564 | 1.30361708 | 1.268726304 |
| Mdh2 | -0.043735998 | 0.200993796 | 0.6746562 | 1.055781063 | 1.016923133 |
| Mdm1 | 0.075518443 | 0.006670852 | 0.1617967 | 0.21693836 | 0.169960883 |
| Mdm2 | -0.031017067 | 0.330004905 | 0.7778857 | 0.267022206 | 0.287161327 |
| Mdm4 | -0.005295992 | 0.867750334 | 0.9775204 | 0.279805089 | 0.290261666 |
| Mdn1 | -0.019189906 | 0.380517394 | 0.8117350 | 0.088420741 | 0.109574291 |
| Mdp1 | -0.041019548 | 0.170613171 | 0.6507045 | 0.209946688 | 0.25968695 |
| Me1 | -0.063260912 | 0.053140193 | 0.4352904 | 0.274130736 | 0.294292726 |
| Me2 | 0.032594584 | 0.166961218 | 0.6442066 | 0.161505986 | 0.116714533 |
| Mea1 | -0.018144997 | 0.605837679 | 0.9087139 | 0.46160365 | 0.446006194 |
| Meaf6 | -0.079806041 | 0.00672648 | 0.1627480 | 0.217733091 | 0.256922896 |
| Mecp2 | 0.011592866 | 0.728098287 | 0.9458355 | 0.346315579 | 0.33288762 |
| Mecr | 0.029009666 | 0.137647495 | 0.6074124 | 0.106421295 | 0.074964578 |
| Med1 | 0.042769234 | 0.140576186 | 0.6130953 | 0.242948191 | 0.200650432 |
| Med10 | 0.047330092 | 0.144198572 | 0.6184391 | 0.387278892 | 0.323760357 |
| Med11 | 0.010300315 | 0.63117584 | 0.9133517 | 0.122449671 | 0.092767103 |
| Med12l | -0.018695101 | 0.376783007 | 0.8101797 | 0.092631337 | 0.100237532 |
| Med13 | 0.035357549 | 0.311819293 | 0.7684379 | 0.424188131 | 0.39090166 |
| Med13l | 0.009392324 | 0.757622243 | 0.9521533 | 0.249849092 | 0.233977002 |
| Med14 | -0.00225104 | 0.927694217 | 0.9879470 | 0.135915619 | 0.145577308 |
| Med15 | 0.000248471 | 0.993036816 | 0.9985888 | 0.209977713 | 0.210741335 |
| Med16 | -0.00850675 | 0.699086413 | 0.9370270 | 0.102262259 | 0.109219562 |
| Med17 | -0.02447442 | 0.177125127 | 0.6536659 | 0.065535778 | 0.07082018 |
| Med18 | -0.007778539 | 0.572168251 | 0.8968097 | 0.04221506 | 0.039623427 |
| Med19 | -0.025496794 | 0.414431785 | 0.8300990 | 0.268574822 | 0.265522703 |
| Med20 | -0.020199344 | 0.354655882 | 0.7969131 | 0.102521567 | 0.110639455 |

| | | | | | |
|---|---|---|---|---|---|
| Med21 | 0.010666642 | 0.75623241 | 0.9521533 | 0.46511651 | 0.41330639 |
| Med22 | 0.03434141 | 0.123841697 | 0.5852543 | 0.126400934 | 0.106068819 |
| Med23 | 0.008529173 | 0.681528389 | 0.9302093 | 0.099964702 | 0.096704861 |
| Med24 | -0.009058314 | 0.635794018 | 0.9153541 | 0.076817911 | 0.07978003 |
| Med25 | -0.049412129 | 0.114706685 | 0.5713707 | 0.269689674 | 0.300442171 |
| Med26 | -0.027038707 | 0.133577491 | 0.5988251 | 0.058057022 | 0.077720855 |
| Med27 | 0.029020713 | 0.208414303 | 0.6807660 | 0.141705091 | 0.118121877 |
| Med28 | 0.031348648 | 0.318331162 | 0.7710043 | 1.32865549 | 1.180371907 |
| Med29 | 0.030361462 | 0.378577697 | 0.8102958 | 0.520046909 | 0.47704505 |
| Med30 | -0.01271632 | 0.695214989 | 0.9362759 | 0.352789227 | 0.345853465 |
| Med31 | -0.008507256 | 0.79430718 | 0.9625613 | 0.396310765 | 0.378714318 |
| Med4 | -0.059567966 | 0.024543453 | 0.3039588 | 0.143478032 | 0.179721794 |
| Med6 | -0.007463239 | 0.770904255 | 0.9552333 | 0.160153353 | 0.161726162 |
| Med7 | 0.014380728 | 0.612671095 | 0.9105164 | 0.224968692 | 0.200684911 |
| Med8 | 0.003521384 | 0.906979032 | 0.9836812 | 0.265363934 | 0.251517869 |
| Med9 | 0.014582631 | 0.62767523 | 0.9128095 | 0.294171066 | 0.238270976 |
| Med9os | 0.009505746 | 0.62986914 | 0.9130564 | 0.088672557 | 0.081671459 |
| Mef2a | 0.030100084 | 0.371246485 | 0.8074114 | 0.344679251 | 0.322401553 |
| Mef2d | -0.016860329 | 0.600473725 | 0.9078437 | 0.297304794 | 0.327094111 |
| Meg3 | -0.087384436 | 0.011878488 | 0.2145716 | 2.890890312 | 3.138427629 |
| Megf8 | -0.016110222 | 0.487213723 | 0.8640311 | 0.117999171 | 0.127865763 |
| Megf9 | -0.012145829 | 0.720049732 | 0.9444523 | 0.334243734 | 0.354949079 |
| Meig1 | 0.062733511 | 0.026863128 | 0.3138777 | 0.247639565 | 0.180072218 |
| Meis1 | -0.00034073 | 0.987469818 | 0.9976072 | 0.095238238 | 0.104134133 |
| Meis2 | 0.062962418 | 0.083902856 | 0.5184165 | 1.170115652 | 1.11954262 |
| Meis3 | -0.039836768 | 0.157521802 | 0.6324475 | 0.180070485 | 0.204805334 |
| Memo1 | 0.001314853 | 0.958646385 | 0.9957785 | 0.150643614 | 0.143566971 |
| Men1 | 0.000586908 | 0.978018414 | 0.9976072 | 0.103441493 | 0.101137785 |
| Meox1 | 0.00589944 | 0.698759898 | 0.9369692 | 0.054234346 | 0.044935579 |
| Mepce | -0.031084115 | 0.286772419 | 0.7468580 | 0.199905134 | 0.228776069 |
| Mesd | -0.009408148 | 0.761056793 | 0.9526414 | 0.288193688 | 0.272069805 |
| Met | 0.00496194 | 0.799969866 | 0.9647053 | 0.080925867 | 0.081055007 |
| Metap1 | -0.014278635 | 0.606210741 | 0.9087139 | 0.192254368 | 0.19461713 |
| Metap1d | -0.032401498 | 0.165771156 | 0.6426142 | 0.113099937 | 0.131153122 |
| Metap2 | 0.073526021 | 0.02754812 | 0.3181899 | 1.091783459 | 1.032501475 |
| Mettl1 | -0.03235795 | 0.378600947 | 0.8102958 | 0.60884108 | 0.614335864 |
| Mettl14 | 0.018339633 | 0.319805812 | 0.7712841 | 0.075606658 | 0.067517469 |
| Mettl15 | 0.005158367 | 0.731891694 | 0.9461673 | 0.054412285 | 0.043958171 |
| Mettl16 | 0.016382151 | 0.592874309 | 0.9060585 | 0.304693502 | 0.251289667 |
| Mettl18 | 0.002992338 | 0.853751382 | 0.9755431 | 0.061155884 | 0.054828557 |
| Mettl2 | -0.029732715 | 0.059564964 | 0.4559293 | 0.035122363 | 0.058854736 |
| Mettl21a | -0.019041632 | 0.325686908 | 0.7763099 | 0.074296594 | 0.084068381 |
| Mettl22 | 0.009655491 | 0.624418324 | 0.9121505 | 0.096568383 | 0.082929423 |
| Mettl23 | 0.043912485 | 0.178467351 | 0.6542483 | 0.366442397 | 0.311972675 |
| Mettl25 | -0.010198994 | 0.584950098 | 0.9023908 | 0.067164734 | 0.078012966 |
| Mettl26 | -0.017827283 | 0.614606273 | 0.9108386 | 0.949012705 | 0.932578812 |
| Mettl27 | -0.038080374 | 0.151114835 | 0.6280470 | 0.155851945 | 0.175073008 |
| Mettl3 | 0.00702723 | 0.736423678 | 0.9469752 | 0.102089813 | 0.093369523 |
| Mettl4 | 0.013373665 | 0.553170664 | 0.8900377 | 0.123238503 | 0.106016261 |
| Mettl5 | -0.022745941 | 0.366337189 | 0.8052437 | 0.148521163 | 0.149320658 |
| Mettl6 | -0.002434189 | 0.912335983 | 0.9840393 | 0.107832308 | 0.107495986 |
| Mettl7a1 | 0.01240227 | 0.649305309 | 0.9174061 | 0.161821053 | 0.181149566 |
| Mettl8 | -0.025898648 | 0.129031282 | 0.5918160 | 0.05023201 | 0.063730551 |
| Mettl9 | -0.041028411 | 0.27143068 | 0.7342030 | 0.715866457 | 0.72395363 |
| Mex3b | -0.02436694 | 0.121981939 | 0.5819916 | 0.037430422 | 0.053985648 |
| Mex3c | 0.013452493 | 0.6000663 | 0.9078234 | 0.170993416 | 0.152652733 |
| Mex3d | 0.00636052 | 0.704596768 | 0.9400796 | 0.064679585 | 0.054577525 |
| Mfap1a | -0.043817528 | 0.089167489 | 0.5299829 | 0.128544905 | 0.175262547 |
| Mfap1b | 0.024898544 | 0.373648832 | 0.8087297 | 0.222844234 | 0.201176129 |
| Mfap2 | -0.012661472 | 0.627747472 | 0.9128095 | 0.196999881 | 0.149784192 |
| Mfap3 | -0.063618842 | 0.004197626 | 0.1286848 | 0.08377855 | 0.122996281 |
| Mff | -0.008268634 | 0.815319909 | 0.9686999 | 0.595114853 | 0.585064251 |
| Mfge8 | -0.010296326 | 0.604616683 | 0.9087139 | 0.068636996 | 0.079535338 |
| Mfhas1 | 0.024462918 | 0.16587785 | 0.6427766 | 0.077224231 | 0.061917244 |
| Mfn1 | -0.019928888 | 0.389482927 | 0.8177080 | 0.120272752 | 0.122325935 |
| Mfn2 | 0.02375476 | 0.343279974 | 0.7872955 | 0.161012023 | 0.144246498 |

| | | | | | |
|---|---|---|---|---|---|
| Mfsd1 | 0.020652309 | 0.416943478 | 0.83009907 | 0.173542208 | 0.152622336 |
| Mfsd10 | -0.0193046 | 0.417870231 | 0.83009907 | 0.128466299 | 0.135120243 |
| Mfsd11 | 0.047510341 | 0.074958058 | 0.49662859 | 0.206627545 | 0.161000018 |
| Mfsd13a | 0.034884243 | 0.02895903 | 0.32641726 | 0.075436877 | 0.048867302 |
| Mfsd14a | 0.02713478 | 0.341021233 | 0.78513662 | 0.234666725 | 0.195931066 |
| Mfsd14b | 0.002557306 | 0.91770278 | 0.98481302 | 0.151333903 | 0.140907201 |
| Mfsd5 | -0.00363672 | 0.86184591 | 0.97696756 | 0.097662584 | 0.098356398 |
| Mfsd6 | -0.002608287 | 0.92732856 | 0.98792100 | 0.206384051 | 0.207586627 |
| Mfsd8 | 0.003093493 | 0.863833036 | 0.97696756 | 0.074268197 | 0.06889255 |
| Mga | -0.001432397 | 0.967604281 | 0.99760728 | 0.399590908 | 0.420391325 |
| Mgat1 | -0.002941125 | 0.904962062 | 0.98317077 | 0.133733298 | 0.139609541 |
| Mgat2 | -0.020293849 | 0.564132188 | 0.89337382 | 0.40466079 | 0.367047717 |
| Mgat3 | -0.031954513 | 0.153527131 | 0.62985489 | 0.106004183 | 0.1193773 |
| Mgat4b | 0.006664765 | 0.805674624 | 0.96471431 | 0.189530268 | 0.165963317 |
| Mgat5 | 0.004969325 | 0.838861952 | 0.97272164 | 0.15047482 | 0.129198271 |
| Mgme1 | 0.001825157 | 0.903260696 | 0.98277125 | 0.048237959 | 0.046312635 |
| Mgmt | -0.02368796 | 0.357669642 | 0.79897222 | 0.143190759 | 0.150633123 |
| Mgrn1 | -0.038744349 | 0.114136854 | 0.56982264 | 0.115927895 | 0.152002752 |
| Mgst3 | 0.010071502 | 0.727180808 | 0.94521791 | 0.210252318 | 0.20317929 |
| Mia2 | -0.051270957 | 0.17845154 | 0.65424838 | 0.672826832 | 0.771251135 |
| Mia3 | -0.050797135 | 0.164785051 | 0.64207072 | 0.986903816 | 1.144441753 |
| Mib1 | -0.041128915 | 0.203769233 | 0.67740978 | 0.293262429 | 0.344601682 |
| Mib2 | -0.014882779 | 0.567263064 | 0.89520621 | 0.152172309 | 0.159728112 |
| Mical2 | -0.029566906 | 0.240551034 | 0.71204045 | 0.120179444 | 0.133894133 |
| Mical3 | -0.029231352 | 0.367625025 | 0.80567193 | 0.28768335 | 0.29241259 |
| Micall1 | 0.001282455 | 0.938598045 | 0.99252660 | 0.061850271 | 0.054499352 |
| Micos10 | 0.099358377 | 9.65E-05 | 0.00903437 | 1.845042666 | 1.678541649 |
| Micos13 | 0.009836988 | 0.730868639 | 0.94590205 | 1.512351278 | 1.502587832 |
| Micu1 | -0.032437352 | 0.199437679 | 0.67309621 | 0.134724775 | 0.16252287 |
| Micu2 | 0.019690865 | 0.539715111 | 0.88291120 | 0.345504631 | 0.297669317 |
| Micu3 | 0.030031442 | 0.388800881 | 0.81702870 | 0.467615826 | 0.429963449 |
| Mid1 | -0.011367718 | 0.451361473 | 0.84801240 | 0.039372095 | 0.051742688 |
| Mid1ip1 | -0.051134487 | 0.183472309 | 0.66159407 | 0.784561591 | 0.748273006 |
| Mid2 | -0.020435983 | 0.31751455 | 0.77094201 | 0.076856256 | 0.1032945 |
| Mideas | -0.017670883 | 0.409578485 | 0.82798083 | 0.084374202 | 0.109938609 |
| Midn | 0.004524534 | 0.895071946 | 0.98243614 | 0.339700691 | 0.372786333 |
| Mief1 | -0.050126263 | 0.019441834 | 0.27081127 | 0.078942564 | 0.110757998 |
| Mien1 | -0.052802213 | 0.090937907 | 0.53212820 | 1.197279636 | 1.185108139 |
| Mier1 | 0.012610329 | 0.73102659 | 0.94590205 | 0.528658773 | 0.502776912 |
| Mier2 | -0.026205856 | 0.127303063 | 0.58987613 | 0.050121856 | 0.065920866 |
| Mier3 | -0.011139828 | 0.613808109 | 0.91083865 | 0.102622347 | 0.107486397 |
| Mif | 0.090902743 | 0.015406264 | 0.24258752 | 1.283029586 | 1.120464869 |
| Mif4gd | -0.015718632 | 0.448986446 | 0.84675770 | 0.08455256 | 0.104210243 |
| Miga1 | -0.024252197 | 0.323340801 | 0.77495548 | 0.123105176 | 0.147241059 |
| Miga2 | -0.00076023 | 0.960731743 | 0.99597229 | 0.049459233 | 0.047323278 |
| Miip | -0.021882518 | 0.203896595 | 0.67751291 | 0.052311279 | 0.067094675 |
| Mindy1 | -0.033786122 | 0.122878257 | 0.58332371 | 0.083378303 | 0.118195687 |
| Mindy2 | 0.050109917 | 0.156086965 | 0.63129431 | 0.469835903 | 0.412774542 |
| Mindy3 | -0.005284201 | 0.835075532 | 0.97234942 | 0.15265552 | 0.142496356 |
| Mink1 | 0.008200204 | 0.727122727 | 0.94521791 | 0.139104238 | 0.125642909 |
| Minpp1 | -0.008616764 | 0.649844729 | 0.91757794 | 0.075306406 | 0.078181096 |
| Mios | 0.004891046 | 0.814486269 | 0.96832499 | 0.104001708 | 0.094008587 |
| Mipep | -0.014528302 | 0.414954334 | 0.83009907 | 0.067061763 | 0.070246871 |
| Mipol1 | 0.006214062 | 0.770382643 | 0.95523334 | 0.105156638 | 0.097450953 |
| Mir22hg | -0.05993984 | 0.072826872 | 0.49033721 | 0.290665369 | 0.324692132 |
| Mir6236 | -0.28179796 | 9.24E-10 | 7.64E-07 | 0.727699715 | 0.902896202 |
| Mir670hg | 0.221426303 | 2.92E-11 | 3.22E-08 | 0.399179375 | 0.265078345 |
| Mir703 | 0.015424433 | 0.533472283 | 0.87904420 | 0.165845987 | 0.136015415 |
| Mirg | -0.051005684 | 0.032102671 | 0.33923281 | 0.100053231 | 0.138924505 |
| Mis12 | 0.034630554 | 0.12792975 | 0.59122443 | 0.137939952 | 0.101466153 |
| Mitd1 | 0.055539983 | 0.023724231 | 0.29897377 | 0.172313986 | 0.132739271 |
| Mkln1 | -0.127209975 | 0.000798058 | 0.04211027 | 0.457059654 | 0.544748419 |
| Mknk1 | 0.004611056 | 0.86416097 | 0.97696756 | 0.166910921 | 0.175875934 |
| Mknk2 | -0.010132559 | 0.632092368 | 0.91355493 | 0.091831849 | 0.097455768 |
| Mkrn1 | -0.001092167 | 0.975071676 | 0.99760728 | 0.586093309 | 0.583074355 |
| Mkrn2 | 0.013652228 | 0.595173081 | 0.9067325 | 0.170600884 | 0.152026639 |

| | | | | | |
|---|---|---|---|---|---|
| Mkrn2os | 0.026138136 | 0.395935504 | 0.8230679 | 0.289806714 | 0.236932065 |
| Mks1 | -0.029642803 | 0.060491488 | 0.4593891 | 0.0426838 | 0.056165055 |
| Mlec | -0.005297841 | 0.883336016 | 0.9801670 | 0.830949037 | 0.87009498 |
| Mlf2 | 0.045855994 | 0.193380125 | 0.6710847 | 0.755418788 | 0.685392074 |
| Mlh1 | -0.001282599 | 0.941649059 | 0.9929301 | 0.07261132 | 0.065603068 |
| Mlh3 | -0.027829736 | 0.268867349 | 0.7337452 | 0.135422719 | 0.152999848 |
| Mllt10 | -0.030411202 | 0.383936135 | 0.8129787 | 0.432884898 | 0.435176197 |
| Mllt11 | 0.007842291 | 0.786109214 | 0.9602516 | 0.24336543 | 0.214722254 |
| Mllt3 | -0.008648049 | 0.760014156 | 0.9521533 | 0.207574636 | 0.20642403 |
| Mllt6 | 0.075179361 | 0.004203013 | 0.1286848 | 0.21405923 | 0.153099393 |
| Mlx | 0.047768287 | 0.12594022 | 0.5882766 | 0.326595237 | 0.268241662 |
| Mlxip | -0.06161876 | 0.009688499 | 0.1970097 | 0.090840928 | 0.150763527 |
| Mlxipl | -0.012539021 | 0.703411607 | 0.9391887 | 0.222910895 | 0.289588552 |
| Mlycd | -0.015614127 | 0.608331322 | 0.9088672 | 0.275874536 | 0.260918136 |
| Mmaa | 0.000692139 | 0.966478431 | 0.9974475 | 0.064789721 | 0.059569094 |
| Mmadhc | 0.017324086 | 0.58156265 | 0.9018069 | 0.318132892 | 0.278295347 |
| Mmgt1 | -0.016065 | 0.368803207 | 0.8056719 | 0.057122706 | 0.075568905 |
| Mmgt2 | 0.00101739 | 0.973577372 | 0.9976072 | 0.267259237 | 0.258032519 |
| Mmp24 | 0.019220487 | 0.197759431 | 0.6730962 | 0.055514657 | 0.04060911 |
| Mms19 | -0.001360099 | 0.949508971 | 0.9948462 | 0.101542769 | 0.104923139 |
| Mmut | 0.002839768 | 0.915000698 | 0.9845236 | 0.171925649 | 0.163214435 |
| Mnat1 | 0.032425934 | 0.274165617 | 0.7351955 | 0.24404582 | 0.227777723 |
| Mns1 | -0.028436676 | 0.25957372 | 0.7263475 | 0.131900449 | 0.159565574 |
| Mnt | -0.017467032 | 0.458785999 | 0.8522649 | 0.130827814 | 0.130708552 |
| Mnx1 | 0.009627159 | 0.584944232 | 0.9023908 | 0.057811908 | 0.069056422 |
| Mob1a | -0.018477538 | 0.393867675 | 0.8210059 | 0.093325119 | 0.109220099 |
| Mob1b | -0.016330807 | 0.579938409 | 0.9008751 | 0.206974544 | 0.228817106 |
| Mob2 | -0.017388795 | 0.537886408 | 0.8816335 | 0.213628701 | 0.203183717 |
| Mob3c | 0.025062797 | 0.289933927 | 0.7497769 | 0.142885297 | 0.122324392 |
| Mob4 | 0.028750537 | 0.418008395 | 0.8300990 | 0.615612778 | 0.513788132 |
| Mocs2 | 0.008613994 | 0.80474449 | 0.9647143 | 0.532651139 | 0.537741752 |
| Mocs3 | -0.023497683 | 0.222944407 | 0.6955958 | 0.070044264 | 0.086800556 |
| Mogs | 0.054588238 | 0.077415603 | 0.5022647 | 0.322365844 | 0.249535637 |
| Mon1a | -0.021133457 | 0.25475032 | 0.7223665 | 0.057494708 | 0.081384307 |
| Mon1b | 0.009647208 | 0.527736439 | 0.8768553 | 0.054286109 | 0.046492746 |
| Mon2 | -0.023785835 | 0.475461702 | 0.8603832 | 0.332281108 | 0.348855193 |
| Morc2a | -0.034329018 | 0.171006289 | 0.6507063 | 0.125094247 | 0.15259445 |
| Morc3 | -0.025805246 | 0.33073189 | 0.7779915 | 0.1561832 | 0.166075564 |
| Morc4 | -0.000540257 | 0.971171297 | 0.9976072 | 0.044344733 | 0.048411624 |
| Morf4l1 | 0.011792935 | 0.657668778 | 0.9211780 | 1.741321672 | 1.711462206 |
| Morf4l2 | 0.009582929 | 0.742976172 | 0.9479080 | 1.384775488 | 1.311291611 |
| Morn2 | -0.014311761 | 0.679301807 | 0.9288316 | 0.486913838 | 0.445538854 |
| Morn4 | 0.013099504 | 0.417594017 | 0.8300990 | 0.060644143 | 0.05090684 |
| Mosmo | -0.018858137 | 0.46908584 | 0.8563363 | 0.160443624 | 0.164718568 |
| Mospd2 | -0.024009325 | 0.265387222 | 0.7306783 | 0.088137735 | 0.107688313 |
| Mospd3 | -0.017532526 | 0.442123016 | 0.8429483 | 0.110678185 | 0.117791352 |
| Mov10 | 0.008417373 | 0.628731565 | 0.9130564 | 0.073648776 | 0.060060705 |
| Mpc1 | 0.096624135 | 0.007409382 | 0.1713311 | 0.763333043 | 0.633854219 |
| Mpc2 | 0.048506764 | 0.113155776 | 0.5696203 | 1.375110188 | 1.203848868 |
| Mpdu1 | -0.06745253 | 0.013302334 | 0.2285400 | 0.149521132 | 0.197847738 |
| Mpdz | -0.026290145 | 0.260464594 | 0.7270142 | 0.099487777 | 0.135267711 |
| Mpg | 0.00811072 | 0.741420168 | 0.9479080 | 0.157046747 | 0.140973503 |
| Mphosph10 | -0.025396851 | 0.45559012 | 0.8502914 | 0.331286024 | 0.356716725 |
| Mphosph6 | -0.003793147 | 0.902260137 | 0.9825441 | 0.277203782 | 0.25055506 |
| Mphosph8 | 0.041487052 | 0.247500119 | 0.7177068 | 0.57809451 | 0.533969816 |
| Mphosph9 | -0.025389075 | 0.273384991 | 0.7351955 | 0.112252237 | 0.124937412 |
| Mpi | -0.013193679 | 0.433841883 | 0.8377586 | 0.050309294 | 0.064199664 |
| Mplkip | -0.053068952 | 0.090656631 | 0.5321031 | 0.279160077 | 0.313182004 |
| Mpnd | -0.035118268 | 0.200520727 | 0.6739701 | 0.174245475 | 0.197926098 |
| Mpp2 | -0.065833631 | 0.039427667 | 0.3804693 | 0.239428315 | 0.267927434 |
| Mpp5 | -0.008906651 | 0.738327135 | 0.9474220 | 0.15683899 | 0.16506508 |
| Mpp6 | 0.018891717 | 0.50346911 | 0.8695566 | 0.190380568 | 0.193612924 |
| Mpp7 | -0.069430161 | 0.04250926 | 0.3952910 | 0.289405828 | 0.333281923 |
| Mppe1 | 0.01340096 | 0.425757031 | 0.8327109 | 0.071161781 | 0.059091514 |
| Mpped2 | -0.027989444 | 0.325888443 | 0.7763291 | 0.184238378 | 0.202167855 |
| Mprip | -0.01964629 | 0.570184851 | 0.8962534 | 0.430516766 | 0.436776206 |

| | | | | | |
|---|---|---|---|---|---|
| Mpst | 0.000526996 | 0.98041442 | 0.99760728 | 0.111793507 | 0.09853709 |
| Mpv17 | -0.069661299 | 0.029855369 | 0.33082809 | 0.279566943 | 0.320587523 |
| Mpv17l | -0.013281043 | 0.571154119 | 0.89659073 | 0.12466794 | 0.122904353 |
| Mpv17l2 | -0.068164515 | 0.022126465 | 0.28982211 | 0.231909698 | 0.267554218 |
| Mpz | -0.018070908 | 0.341492885 | 0.78525948 | 0.067898218 | 0.081250397 |
| Mpzl1 | 0.00246713 | 0.929995289 | 0.98852883 | 0.189794492 | 0.183888391 |
| Mr1 | -0.012249282 | 0.409233617 | 0.82764474 | 0.03844649 | 0.049784246 |
| Mras | -0.00393319 | 0.882305637 | 0.97986777 | 0.152949936 | 0.169687965 |
| Mre11a | -0.00964724 | 0.589800574 | 0.90467754 | 0.06623241 | 0.068592906 |
| Mrfap1 | -0.014689834 | 0.47033488 | 0.85735279 | 2.274622075 | 2.203116018 |
| Mrgbp | -0.004624956 | 0.847025032 | 0.97406590 | 0.141357597 | 0.128898017 |
| Mrgpre | -0.014159431 | 0.311989422 | 0.76843798 | 0.034481951 | 0.0461397 |
| Mri1 | -0.003681762 | 0.895335881 | 0.98249247 | 0.203721032 | 0.189758336 |
| Mrln | 0.025380052 | 0.519265988 | 0.87417954 | 0.764016464 | 0.77532909 |
| Mrm1 | 0.005570242 | 0.744943382 | 0.94857174 | 0.069362623 | 0.060309507 |
| Mrm2 | -0.010465918 | 0.505407825 | 0.86955663 | 0.054427645 | 0.051936883 |
| Mrm3 | 0.008773119 | 0.60757079 | 0.90871392 | 0.071673036 | 0.058863013 |
| Mroh1 | -0.033975815 | 0.100600815 | 0.54800859 | 0.073178642 | 0.109591493 |
| Mrpl1 | 0.01632384 | 0.606701358 | 0.90871392 | 0.328984059 | 0.2869046 |
| Mrpl10 | -0.01821489 | 0.539992778 | 0.88307424 | 0.227672734 | 0.23951182 |
| Mrpl11 | 0.024444256 | 0.467360567 | 0.85585468 | 0.449385364 | 0.40607426 |
| Mrpl12 | 0.076192398 | 0.020693527 | 0.27988788 | 0.40474529 | 0.3230519 |
| Mrpl13 | -0.017399678 | 0.619341789 | 0.91215057 | 0.458991345 | 0.455100402 |
| Mrpl14 | 0.051318943 | 0.142176742 | 0.61527370 | 0.734166117 | 0.643079046 |
| Mrpl15 | -0.06516624 | 0.043280921 | 0.39768759 | 0.293609401 | 0.321422129 |
| Mrpl16 | -0.002635101 | 0.926272146 | 0.98792100 | 0.233461982 | 0.213854863 |
| Mrpl17 | -0.088913609 | 0.009675035 | 0.19700979 | 0.863188618 | 0.872787202 |
| Mrpl18 | 0.056716836 | 0.058169664 | 0.45081999 | 0.292098586 | 0.229149535 |
| Mrpl19 | -7.42E-05 | 0.997940863 | 0.99940731 | 0.202226936 | 0.207540315 |
| Mrpl2 | -0.059967095 | 0.085670816 | 0.52263603 | 0.443365073 | 0.478598695 |
| Mrpl20 | -0.04032815 | 0.244762629 | 0.71379719 | 0.873796722 | 0.886180947 |
| Mrpl21 | 0.027185952 | 0.445332376 | 0.84339388 | 0.582630707 | 0.534588965 |
| Mrpl22 | 0.037047431 | 0.163933854 | 0.64100269 | 0.222169936 | 0.166044766 |
| Mrpl23 | 0.019459615 | 0.364809725 | 0.80501911 | 0.123834369 | 0.097769264 |
| Mrpl23-ps1 | 0.004758645 | 0.736738343 | 0.94697528 | 0.046740043 | 0.039228386 |
| Mrpl24 | -0.036453716 | 0.311414456 | 0.76843798 | 0.563713396 | 0.576458032 |
| Mrpl27 | 0.00636616 | 0.854138793 | 0.97554315 | 0.464758209 | 0.4473504 |
| Mrpl28 | 0.062228277 | 0.075072458 | 0.49662859 | 0.637698212 | 0.545397388 |
| Mrpl3 | 0.009591701 | 0.71619073 | 0.94314111 | 0.183684238 | 0.16679977 |
| Mrpl30 | -0.063447421 | 0.060526369 | 0.45938911 | 0.802753714 | 0.835876487 |
| Mrpl32 | -0.054988627 | 0.093938057 | 0.53344171 | 0.328689636 | 0.355345545 |
| Mrpl33 | 0.237151269 | 4.32E-11 | 4.28E-08 | 1.120370996 | 0.810265069 |
| Mrpl34 | 0.05462126 | 0.107579878 | 0.56337238 | 0.698577231 | 0.613183214 |
| Mrpl35 | 0.045925359 | 0.140326601 | 0.61309530 | 0.336608184 | 0.271514851 |
| Mrpl36 | -0.017170871 | 0.638249454 | 0.91600428 | 0.873807873 | 0.828156471 |
| Mrpl37 | -0.017606731 | 0.402096552 | 0.82542591 | 0.09579166 | 0.09851371 |
| Mrpl38 | -0.018091127 | 0.602125533 | 0.90845401 | 0.418950547 | 0.399195854 |
| Mrpl39 | -0.013967429 | 0.565318651 | 0.89434330 | 0.132320245 | 0.146187012 |
| Mrpl4 | 0.063398525 | 0.058102081 | 0.45081999 | 0.462834386 | 0.395377828 |
| Mrpl40 | -0.012016239 | 0.699411772 | 0.93720988 | 0.274738518 | 0.270087068 |
| Mrpl41 | -0.019004759 | 0.586656167 | 0.90352882 | 0.624589947 | 0.572675882 |
| Mrpl42 | -0.03993682 | 0.26285866 | 0.72757354 | 0.569901658 | 0.610676307 |
| Mrpl43 | -0.046175242 | 0.185277957 | 0.66376783 | 0.520082157 | 0.517082513 |
| Mrpl44 | 0.033458575 | 0.207079065 | 0.67975655 | 0.214845576 | 0.165326762 |
| Mrpl45 | -0.017149194 | 0.469497185 | 0.85675949 | 0.121337537 | 0.128217628 |
| Mrpl46 | 0.001026086 | 0.970972576 | 0.99760728 | 0.213514869 | 0.205361126 |
| Mrpl47 | -0.019078346 | 0.37624184 | 0.81017731 | 0.090504976 | 0.110510617 |
| Mrpl48 | 0.053888625 | 0.117757226 | 0.57463969 | 0.516387425 | 0.448761727 |
| Mrpl49 | 0.019420584 | 0.48834533 | 0.86429717 | 0.241448575 | 0.193598507 |
| Mrpl50 | 0.026350102 | 0.386310762 | 0.81546943 | 0.315364239 | 0.256348566 |
| Mrpl51 | -0.077500852 | 0.029931004 | 0.33082809 | 0.562627101 | 0.609008649 |
| Mrpl52 | 0.078177107 | 0.017086652 | 0.2562701 | 1.178065741 | 1.081268013 |
| Mrpl54 | 0.054687288 | 0.124425546 | 0.58525434 | 0.965169118 | 0.885727894 |
| Mrpl55 | 0.014401844 | 0.657766854 | 0.92117801 | 0.417272075 | 0.39186148 |
| Mrpl57 | -0.030524811 | 0.356714279 | 0.79824174 | 1.000522307 | 1.010243424 |
| Mrpl58 | 0.034305776 | 0.320535667 | 0.77215001 | 0.560918515 | 0.513768525 |

| | | | | | |
|---|---|---|---|---|---|
| Mrpl9 | 0.022364794 | 0.463956159 | 0.8529364 | 0.313360855 | 0.264597527 |
| Mrps10 | 0.009445883 | 0.773626719 | 0.9557311 | 0.359670741 | 0.298261143 |
| Mrps11 | -0.039492775 | 0.220835794 | 0.6926204 | 0.323985484 | 0.349292858 |
| Mrps12 | 0.092972759 | 0.003534439 | 0.1149480 | 1.059546824 | 0.912977288 |
| Mrps14 | 0.049621317 | 0.165275369 | 0.6420707 | 0.796245369 | 0.733958648 |
| Mrps15 | -0.005055356 | 0.878775118 | 0.9785583 | 0.401356344 | 0.367431262 |
| Mrps16 | 0.019563406 | 0.583527431 | 0.9020713 | 1.031452567 | 0.9501479 |
| Mrps17 | -0.021453154 | 0.54315423 | 0.8840682 | 0.503621809 | 0.494508635 |
| Mrps18a | -0.021377709 | 0.540835258 | 0.8835017 | 0.568392367 | 0.555773435 |
| Mrps18b | -0.009505628 | 0.767280641 | 0.9544221 | 0.317974405 | 0.299167101 |
| Mrps18c | 0.070662798 | 0.041709436 | 0.3925594 | 0.49366564 | 0.410738401 |
| Mrps2 | -0.01030277 | 0.560223459 | 0.8925646 | 0.057438667 | 0.069130594 |
| Mrps21 | 0.147899485 | 1.09E-05 | 0.0016229 | 1.119979048 | 0.956830723 |
| Mrps22 | 0.013676107 | 0.496145394 | 0.8684629 | 0.094919034 | 0.085905351 |
| Mrps23 | 0.019996545 | 0.547876339 | 0.8863001 | 0.409269145 | 0.360167311 |
| Mrps24 | -0.000832982 | 0.981771352 | 0.9976072 | 0.735753452 | 0.71007004 |
| Mrps25 | 0.025055272 | 0.354486866 | 0.7969131 | 0.207883394 | 0.1855624 |
| Mrps26 | -0.0180671 | 0.60542577 | 0.9087139 | 0.529563138 | 0.530764998 |
| Mrps27 | -0.004592445 | 0.779474458 | 0.9586487 | 0.055107495 | 0.056399194 |
| Mrps28 | 0.013302257 | 0.702505187 | 0.9388187 | 0.604023782 | 0.532131675 |
| Mrps30 | -0.018194118 | 0.462771794 | 0.8524425 | 0.157919461 | 0.148444906 |
| Mrps31 | -0.014479886 | 0.634546363 | 0.9147798 | 0.274648694 | 0.273571095 |
| Mrps33 | 0.006255191 | 0.843001009 | 0.9734889 | 1.285089376 | 1.181606575 |
| Mrps34 | -0.022481288 | 0.541302738 | 0.8835017 | 0.599721946 | 0.601350844 |
| Mrps35 | -0.014537392 | 0.587638292 | 0.9043394 | 0.192777679 | 0.180760872 |
| Mrps36 | 0.076375887 | 0.033630296 | 0.3468530 | 0.539689106 | 0.411908122 |
| Mrps5 | 0.049847796 | 0.106801491 | 0.5623517 | 0.301624452 | 0.270861624 |
| Mrps6 | 0.012287598 | 0.646918453 | 0.9169695 | 0.192802849 | 0.171152811 |
| Mrps7 | -0.056689673 | 0.089636676 | 0.5302360 | 0.396370258 | 0.409321492 |
| Mrps9 | -0.02036887 | 0.482758412 | 0.8620306 | 0.217282552 | 0.226886483 |
| Mrrf | -0.028590657 | 0.199414748 | 0.6730962 | 0.101868765 | 0.118372118 |
| Mrs2 | 0.003642288 | 0.89670055 | 0.9825441 | 0.183096196 | 0.202672325 |
| Mrtfa | 0.018682086 | 0.211486469 | 0.6822059 | 0.060330382 | 0.041726237 |
| Mrtfb | 0.012477797 | 0.570126487 | 0.8962534 | 0.120858759 | 0.107805909 |
| Mrto4 | -0.003865901 | 0.878358911 | 0.9785583 | 0.155493398 | 0.150507485 |
| Msantd2 | -0.014828151 | 0.425113465 | 0.8324460 | 0.06685611 | 0.074256889 |
| Msantd3 | -0.014661425 | 0.454851807 | 0.8499020 | 0.075223819 | 0.087053632 |
| Msantd4 | -0.037718626 | 0.2031851 | 0.6772740 | 0.240266422 | 0.242416318 |
| Msh2 | 0.031749335 | 0.14269709 | 0.6158696 | 0.136088158 | 0.097652339 |
| Msh3 | -0.036205537 | 0.065873861 | 0.4762891 | 0.062383544 | 0.087414758 |
| Msh6 | -0.014553165 | 0.406130225 | 0.8270770 | 0.063893962 | 0.065403005 |
| Msi1 | 0.013189984 | 0.496111993 | 0.8684629 | 0.08727893 | 0.075405832 |
| Msi2 | 0.01562634 | 0.68216411 | 0.9303225 | 0.976885232 | 0.97735674 |
| Msl1 | -0.012200782 | 0.728478839 | 0.9458355 | 0.444858639 | 0.431525155 |
| Msl2 | 0.025194319 | 0.399115837 | 0.8239850 | 0.24179573 | 0.208395564 |
| Msl3 | -0.025153253 | 0.366266656 | 0.8052437 | 0.177902757 | 0.190617474 |
| Msl3l2 | -0.01190442 | 0.651025365 | 0.9182621 | 0.178320824 | 0.167793108 |
| Msmo1 | 0.015665164 | 0.64504578 | 0.9169695 | 0.341444929 | 0.350514498 |
| Msrb1 | 0.021447867 | 0.424684505 | 0.8324460 | 0.196072788 | 0.16638485 |
| Msrb2 | 0.013263782 | 0.50851598 | 0.8704067 | 0.103572285 | 0.08674645 |
| Msto1 | 0.034980555 | 0.02114828 | 0.2819770 | 0.071871176 | 0.04099943 |
| Mt1 | 0.089138924 | 0.09060933 | 0.5321031 | 1.183320958 | 1.490005552 |
| Mt2 | 0.060306434 | 0.147131383 | 0.6234700 | 0.359729682 | 0.406437497 |
| Mta1 | 0.000747224 | 0.979528927 | 0.9976072 | 0.230386651 | 0.211923456 |
| Mta2 | 0.011937073 | 0.645955549 | 0.9169695 | 0.171290139 | 0.150528241 |
| Mta3 | -0.033647619 | 0.254595271 | 0.7223071 | 0.215531977 | 0.236993487 |
| Mtap | 0.008248242 | 0.726572351 | 0.9451913 | 0.139234686 | 0.122114664 |
| Mtarc2 | 0.003930808 | 0.908233775 | 0.9837373 | 0.440130306 | 0.429574599 |
| mt-Atp6 | 0.002297702 | 0.9276686 | 0.9879470 | 0.127915892 | 0.150615216 |
| mt-Atp8 | 0.025007665 | 0.441534937 | 0.8426454 | 0.270925902 | 0.273616662 |
| Mtch1 | -0.03936147 | 0.196363382 | 0.6730962 | 1.303406824 | 1.274775888 |
| Mtch2 | 0.013376283 | 0.700949176 | 0.9377499 | 0.588326409 | 0.542706115 |
| Mtcl1 | 0.008461655 | 0.738013575 | 0.9474220 | 0.148566843 | 0.158061057 |
| mt-Co1 | 0.007844563 | 0.859021936 | 0.9769675 | 1.654675522 | 1.790235981 |
| mt-Cytb | -0.084349487 | 0.021572811 | 0.2863286 | 3.921259202 | 4.313161713 |
| Mtdh | 0.0766491 | 0.028230092 | 0.3218879 | 1.120198521 | 1.06441232 |

| | | | | | |
|---|---|---|---|---|---|
| Mterf1a | -0.024213852 | 0.168676842 | 0.64655110 | 0.050458411 | 0.073494234 |
| Mterf3 | -0.000471829 | 0.986494127 | 0.99760728 | 0.20572806 | 0.202086641 |
| Mterf4 | -0.018698094 | 0.301261286 | 0.76062915 | 0.060295698 | 0.074873916 |
| Mtf1 | -0.012947081 | 0.664988136 | 0.92307657 | 0.234887251 | 0.246608765 |
| Mtf2 | -0.039652181 | 0.158530144 | 0.63448003 | 0.162996731 | 0.193010385 |
| Mtfmt | -0.018550254 | 0.298988664 | 0.75739723 | 0.063637404 | 0.069936004 |
| Mtfp1 | -0.008178123 | 0.736947559 | 0.94697528 | 0.141650674 | 0.140775757 |
| Mtfr1 | 0.002634635 | 0.918500059 | 0.98512115 | 0.178867964 | 0.163093767 |
| Mtfr1l | -0.036711637 | 0.269902975 | 0.73403607 | 0.428187702 | 0.417008654 |
| Mtg1 | 0.03439864 | 0.149139798 | 0.62643745 | 0.168379242 | 0.12301403 |
| Mtg2 | -0.033427565 | 0.187042088 | 0.66551560 | 0.135207615 | 0.15780312 |
| Mthfd1 | -0.002255671 | 0.931625809 | 0.98939560 | 0.176059028 | 0.167163738 |
| Mthfd1l | -0.013555564 | 0.516639087 | 0.87368901 | 0.080686629 | 0.104910575 |
| Mthfd2 | 0.159784328 | 0.000198787 | 0.01603224 | 1.074589942 | 0.978495252 |
| Mthfd2l | -0.048705553 | 0.104453519 | 0.55800455 | 0.218772267 | 0.23837617 |
| Mthfr | -0.007720436 | 0.689414871 | 0.93401619 | 0.071061905 | 0.081842029 |
| Mthfs | -0.025598709 | 0.282284 | 0.74198650 | 0.143973525 | 0.129466312 |
| Mthfsd | -0.02418478 | 0.293752938 | 0.75336844 | 0.116199123 | 0.123202813 |
| Mthfsl | -0.008153102 | 0.744284794 | 0.94845041 | 0.164775348 | 0.144613008 |
| Mtif2 | 0.040849435 | 0.143440205 | 0.61650780 | 0.224746575 | 0.172976984 |
| Mtif3 | 0.048525582 | 0.071156821 | 0.4881297 | 0.231226171 | 0.163781162 |
| Mtln | 0.063061036 | 0.079675765 | 0.50767610 | 0.587890029 | 0.536567932 |
| Mtm1 | 0.004225329 | 0.806447988 | 0.96480137 | 0.066024259 | 0.060958228 |
| Mtmr1 | -0.018459543 | 0.405538545 | 0.82707770 | 0.10027007 | 0.118139142 |
| Mtmr10 | 0.008491881 | 0.594222038 | 0.90652070 | 0.051247587 | 0.053265807 |
| Mtmr11 | -0.017127154 | 0.419842153 | 0.83009907 | 0.10687525 | 0.10550263 |
| Mtmr12 | -0.021716499 | 0.203472771 | 0.67727401 | 0.048744426 | 0.065609708 |
| Mtmr14 | -0.011585265 | 0.516960792 | 0.87378617 | 0.06042929 | 0.06657194 |
| Mtmr2 | -0.009445145 | 0.693400527 | 0.93597259 | 0.132497841 | 0.134169689 |
| Mtmr3 | -0.040494435 | 0.16706096 | 0.64429717 | 0.20096763 | 0.239971195 |
| Mtmr4 | -0.045962371 | 0.02251436 | 0.29144683 | 0.062474755 | 0.09875457 |
| Mtmr6 | 0.019909173 | 0.502961816 | 0.86955665 | 0.248405895 | 0.23551443 |
| Mtmr7 | -0.03852062 | 0.251623271 | 0.72183425 | 0.342403343 | 0.333832971 |
| Mtmr9 | -0.000860506 | 0.959965132 | 0.99586287 | 0.062445794 | 0.06158148 |
| mt-Nd1 | -0.066116769 | 0.119502342 | 0.57833000 | 2.738920698 | 3.087749617 |
| mt-Nd2 | -0.0525849 | 0.245796261 | 0.71529004 | 2.2020059 | 2.513511169 |
| mt-Nd3 | 0.79390696 | 1.13E-132 | 1.12E-128 | 0.82253181 | 0.008437543 |
| mt-Nd4 | -0.052093596 | 0.192568747 | 0.67002813 | 2.814759262 | 3.101375473 |
| mt-Nd5 | 0.005668406 | 0.899676178 | 0.98254414 | 1.259967298 | 1.352196362 |
| mt-Nd6 | 0.027895855 | 0.213323561 | 0.68300605 | 0.11352653 | 0.093964367 |
| Mto1 | 0.003797685 | 0.844954688 | 0.97348890 | 0.087044895 | 0.07971751 |
| Mtor | 0.010221814 | 0.662084757 | 0.92245510 | 0.129126731 | 0.126018376 |
| Mtpap | 0.01602266 | 0.487672392 | 0.8640311 | 0.144999173 | 0.113098605 |
| Mtpn | 0.043154135 | 0.211976871 | 0.68220598 | 0.476030029 | 0.432031823 |
| Mtr | -0.014785142 | 0.386259412 | 0.81546943 | 0.055560744 | 0.060805102 |
| Mtres1 | -0.017518763 | 0.535079823 | 0.87959492 | 0.218538828 | 0.210994136 |
| Mtrex | -0.025504478 | 0.32533308 | 0.77630994 | 0.147722068 | 0.154299392 |
| Mtrf1 | -0.013146316 | 0.379663671 | 0.81151774 | 0.044837659 | 0.048346153 |
| Mtrf1l | 0.014528066 | 0.510122891 | 0.87116072 | 0.124724154 | 0.106271906 |
| mt-Rnr1 | 0.010439626 | 0.818106098 | 0.96972308 | 0.836839338 | 0.85936099 |
| mt-Rnr2 | 0.023911658 | 0.584154763 | 0.90214067 | 2.918448634 | 2.991589287 |
| Mtrr | 0.023075257 | 0.117611957 | 0.57445131 | 0.060088335 | 0.037318233 |
| Mtss1 | -0.061889433 | 0.06845559 | 0.48338697 | 0.280402464 | 0.345219281 |
| Mtss2 | 0.007653006 | 0.775619927 | 0.95585380 | 0.156510437 | 0.168256756 |
| Mttp | -0.035487878 | 0.100761983 | 0.54800855 | 0.090295442 | 0.11449995 |
| Mturn | -0.037498523 | 0.024655712 | 0.30415400 | 0.040594403 | 0.067776729 |
| Mtus1 | 0.006898054 | 0.776636491 | 0.95633490 | 0.129843289 | 0.136629597 |
| Mtus2 | -0.009044422 | 0.666706635 | 0.92307657 | 0.095077665 | 0.090706368 |
| Mtx1 | -0.000829469 | 0.977267908 | 0.99760728 | 0.258773934 | 0.229732483 |
| Mtx2 | -0.0146607 | 0.659702091 | 0.92117802 | 0.402343615 | 0.385841631 |
| Muc4 | 0.031847503 | 0.085316736 | 0.52128640 | 0.077035763 | 0.070245237 |
| Mul1 | -0.003375095 | 0.8977774 | 0.98254414 | 0.168436968 | 0.16973924 |
| Mus81 | -0.025480013 | 0.17295743 | 0.65255078 | 0.059875269 | 0.082303471 |
| Mvb12a | 0.041003627 | 0.26039533 | 0.72701428 | 0.718990846 | 0.643041558 |
| Mvd | 0.005421056 | 0.732545137 | 0.94620413 | 0.054064467 | 0.047806615 |
| Mvk | 0.007743319 | 0.622871401 | 0.91215057 | 0.064680992 | 0.049186772 |

| | | | | | |
|---|---|---|---|---|---|
| Mvp | 0.012180714 | 0.70358932 | 0.93918879 | 0.296479012 | 0.294148724 |
| Mxd1 | 0.006182134 | 0.803386287 | 0.96471431 | 0.141222824 | 0.130604422 |
| Mxd4 | -0.041184975 | 0.267165282 | 0.73131339 | 0.6569292 | 0.659856006 |
| Mxi1 | -0.013985909 | 0.61357104 | 0.91083869 | 0.20092857 | 0.181727068 |
| Myadm | -0.019375384 | 0.313027719 | 0.76975571 | 0.076568289 | 0.085899038 |
| Mybbp1a | 0.031577194 | 0.341391746 | 0.78524569 | 0.342964125 | 0.313075999 |
| Mycbp | 0.021780204 | 0.512871226 | 0.87244634 | 0.397314143 | 0.332915504 |
| Mycbp2 | -0.043869854 | 0.247293302 | 0.71764088 | 0.903562658 | 0.935012701 |
| Myd88 | -0.010002611 | 0.556228075 | 0.89125868 | 0.056305372 | 0.062737161 |
| Mydgf | -0.072078062 | 0.0599566 | 0.45751497 | 0.926783934 | 0.950785567 |
| Myef2 | 0.044304016 | 0.208089405 | 0.68076601 | 0.460162351 | 0.387715939 |
| Myg1 | -0.010867127 | 0.728636648 | 0.94583558 | 0.297612487 | 0.292397605 |
| Myh10 | 0.014294775 | 0.598316158 | 0.90755130 | 0.19761183 | 0.16605253 |
| Myh14 | 0.007206649 | 0.743275484 | 0.94790800 | 0.116587942 | 0.106240812 |
| Myh9 | -0.036868245 | 0.316753425 | 0.77059879 | 0.434976444 | 0.444311112 |
| Myl12a | 0.038917535 | 0.291424661 | 0.75167259 | 0.694826096 | 0.567969885 |
| Myl12b | -0.045868332 | 0.169569266 | 0.64821854 | 0.384121008 | 0.344741307 |
| Myl6 | 0.044495785 | 0.044731713 | 0.40157339 | 2.287764835 | 2.089367885 |
| Mylip | -0.005434603 | 0.76004244 | 0.95215339 | 0.059870157 | 0.06667384 |
| Mynn | -0.003460005 | 0.89161128 | 0.98195979 | 0.160679118 | 0.151524587 |
| Myo10 | 0.019336383 | 0.489069369 | 0.86496131 | 0.194696316 | 0.197295894 |
| Myo18a | -0.018589879 | 0.509894795 | 0.87116071 | 0.182589145 | 0.188239693 |
| Myo1b | -0.049359739 | 0.12632802 | 0.58893500 | 0.301858949 | 0.291603902 |
| Myo1c | 0.016304117 | 0.569166836 | 0.89565324 | 0.206035775 | 0.194809645 |
| Myo1d | -0.040550034 | 0.081362231 | 0.51354914 | 0.096377473 | 0.128956106 |
| Myo1e | 0.0144701 | 0.423991316 | 0.83204620 | 0.077776094 | 0.058944543 |
| Myo5a | 0.018522071 | 0.510096379 | 0.87116071 | 0.190820184 | 0.181310139 |
| Myo5b | -0.020405411 | 0.142573829 | 0.61586961 | 0.027333387 | 0.046229642 |
| Myo6 | -0.029379293 | 0.419461689 | 0.83009907 | 0.4178818 | 0.467217872 |
| Myo9a | -0.043579452 | 0.091016171 | 0.53212820 | 0.126869251 | 0.154797728 |
| Myo9b | -0.005288554 | 0.784260487 | 0.96012021 | 0.077503353 | 0.07867869 |
| Mypop | 0.009411776 | 0.672336943 | 0.92581659 | 0.11711113 | 0.10754555 |
| Mysm1 | 0.024652945 | 0.484970816 | 0.86297151 | 0.431619859 | 0.402613803 |
| Myt1 | 0.003327645 | 0.897521198 | 0.98254414 | 0.164599007 | 0.157392077 |
| Myt1l | -0.08299566 | 0.016463508 | 0.25252577 | 0.29430127 | 0.359801155 |
| Mzt1 | 0.07710067 | 0.009184641 | 0.19221864 | 0.304989545 | 0.221873504 |
| Mzt2 | -0.015317955 | 0.567307089 | 0.89520621 | 0.171570907 | 0.180621156 |
| N4bp1 | -0.047827591 | 0.162629686 | 0.64007039 | 0.376272462 | 0.402097708 |
| N4bp2 | 0.004368968 | 0.85766993 | 0.97635421 | 0.138354662 | 0.135004863 |
| N4bp2l1 | -0.022545992 | 0.442633871 | 0.84315410 | 0.214007868 | 0.229199872 |
| N4bp2l2 | 0.008864203 | 0.804785989 | 0.96471431 | 0.586874405 | 0.601505719 |
| N4bp2os | 0.003712375 | 0.850223079 | 0.97488410 | 0.097299086 | 0.081082927 |
| N6amt1 | -0.029945499 | 0.3048493 | 0.76424180 | 0.188236692 | 0.23813382 |
| Naa10 | 0.027765103 | 0.317518085 | 0.77094201 | 0.223019041 | 0.17982728 |
| Naa15 | -0.035030683 | 0.329864761 | 0.77788578 | 0.398407552 | 0.414716631 |
| Naa16 | 0.005931912 | 0.821247687 | 0.97054760 | 0.184869111 | 0.167882912 |
| Naa20 | 0.027417809 | 0.423926937 | 0.83204620 | 0.526482422 | 0.521155948 |
| Naa25 | 0.01706949 | 0.497591422 | 0.86846292 | 0.164742509 | 0.137867171 |
| Naa30 | 0.025100434 | 0.305392182 | 0.76462719 | 0.15869275 | 0.132630593 |
| Naa35 | 0.01748544 | 0.598688042 | 0.90755130 | 0.334748154 | 0.315229589 |
| Naa38 | 0.052085617 | 0.158756042 | 0.63502410 | 0.693216624 | 0.622657495 |
| Naa40 | -0.025898436 | 0.220566221 | 0.69241041 | 0.080566938 | 0.104634819 |
| Naa50 | 0.061408224 | 0.07011619 | 0.48504363 | 0.405874894 | 0.346898488 |
| Naa60 | 0.010272584 | 0.660570704 | 0.92117802 | 0.132208181 | 0.125792657 |
| Naa80 | 0.027493444 | 0.164074436 | 0.64129960 | 0.102560309 | 0.078791101 |
| Naaa | -0.017896712 | 0.317037034 | 0.77064625 | 0.051511667 | 0.072867 |
| Nab1 | -0.004786984 | 0.8603796 | 0.97696750 | 0.177135005 | 0.187551144 |
| Nab2 | -0.015300173 | 0.332693552 | 0.78003309 | 0.043736575 | 0.055177525 |
| Nabp1 | -0.019843924 | 0.30219577 | 0.76112584 | 0.056229048 | 0.088188878 |
| Nabp2 | -0.018249108 | 0.582325502 | 0.90189991 | 0.393191237 | 0.393844765 |
| Naca | -0.017919098 | 0.373308427 | 0.80856898 | 2.616987092 | 2.504063354 |
| Nacc1 | -0.005417901 | 0.819055696 | 0.96990621 | 0.134505739 | 0.126587279 |
| Nacc2 | -0.000996132 | 0.951834109 | 0.99515668 | 0.057969057 | 0.058587116 |
| Nadk | 0.005471456 | 0.815916533 | 0.96869997 | 0.123282351 | 0.127539814 |
| Nadk2 | -0.02935065 | 0.118971479 | 0.57803248 | 0.06779202 | 0.0771059 |
| Nae1 | -0.062221381 | 0.066393387 | 0.4776087 | 0.379067321 | 0.399910698 |

| | | | | | |
|---|---|---|---|---|---|
| Naf1 | -0.030372118 | 0.15082718 | 0.62803209 | 0.085423341 | 0.10154019 |
| Naga | -0.027940089 | 0.430973125 | 0.83517354 | 0.437337371 | 0.463844671 |
| Nagk | -0.017245725 | 0.622842676 | 0.91215057 | 0.456734629 | 0.440906283 |
| Naglu | 0.010280651 | 0.585831287 | 0.9029041 | 0.086937217 | 0.074635918 |
| Nagpa | -0.014773016 | 0.369176572 | 0.80567193 | 0.061348931 | 0.057784269 |
| Naip2 | 0.025522293 | 0.191856234 | 0.67002813 | 0.090688074 | 0.077651531 |
| Nalcn | -0.00413311 | 0.873123409 | 0.97843318 | 0.157305973 | 0.154511476 |
| Nampt | -0.033151505 | 0.200245637 | 0.67383058 | 0.137727828 | 0.166752449 |
| Nanp | 0.051381752 | 0.00185284 | 0.07450649 | 0.092663242 | 0.043050154 |
| Nans | 0.025442938 | 0.464493615 | 0.85360812 | 0.449220376 | 0.393856386 |
| Nap1l1 | 0.048601455 | 0.175241693 | 0.65341424 | 0.767337028 | 0.711997285 |
| Nap1l2 | -0.018359617 | 0.482640013 | 0.86203068 | 0.158586608 | 0.171431915 |
| Nap1l3 | -0.013080629 | 0.596438622 | 0.90690851 | 0.139321883 | 0.133554915 |
| Nap1l4 | -0.041337286 | 0.22976271 | 0.70196775 | 0.45649507 | 0.430574714 |
| Nap1l5 | -0.088901972 | 0.035676038 | 0.35919149 | 0.788686332 | 0.810825718 |
| Napa | 0.011614042 | 0.738985273 | 0.94743504 | 0.949585371 | 0.892100614 |
| Napb | 0.031673303 | 0.187427507 | 0.66592513 | 0.141218076 | 0.116438866 |
| Napepld | 0.031873487 | 0.187524982 | 0.66592513 | 0.155644751 | 0.122012971 |
| Napg | -0.003753507 | 0.909747634 | 0.98373739 | 0.388052175 | 0.37127135 |
| Nars | 0.080904132 | 0.037475969 | 0.36808080 | 0.989993729 | 0.962042303 |
| Nars2 | -0.013477846 | 0.41526215 | 0.83009907 | 0.047654046 | 0.061171891 |
| Nasp | -0.017259444 | 0.541516033 | 0.88350175 | 0.177297087 | 0.194770285 |
| Nat10 | 0.03359542 | 0.096627641 | 0.53730168 | 0.114254075 | 0.082955946 |
| Nat14 | 0.012290049 | 0.596960464 | 0.90742381 | 0.145905894 | 0.112244263 |
| Nat8f1 | 0.007816462 | 0.696772105 | 0.93644228 | 0.094788215 | 0.082679658 |
| Nat9 | 0.031804995 | 0.319114595 | 0.77100438 | 0.374870525 | 0.30340824 |
| Nav1 | -0.043552841 | 0.209436453 | 0.68076601 | 0.36445526 | 0.400422694 |
| Nav2 | -0.049287773 | 0.201584745 | 0.67521501 | 0.486875921 | 0.526560326 |
| Nav3 | 0.014180302 | 0.412543062 | 0.82979790 | 0.062666819 | 0.057190613 |
| Naxd | -0.018911716 | 0.595533755 | 0.90680201 | 0.484542753 | 0.474977248 |
| Naxe | -0.056096083 | 0.118528261 | 0.57750508 | 0.700315183 | 0.693892731 |
| Nbas | 0.010832898 | 0.730263326 | 0.94590205 | 0.273712821 | 0.290613059 |
| Nbdy | 0.012197779 | 0.7436616 | 0.94790800 | 0.834462218 | 0.710611998 |
| Nbea | 0.002814185 | 0.93983437 | 0.99282169 | 0.533119523 | 0.518406473 |
| Nbeal1 | -0.040326393 | 0.278255252 | 0.73863850 | 0.45272582 | 0.479432212 |
| Nbeal2 | 0.003225178 | 0.872469614 | 0.97843318 | 0.098361462 | 0.088325974 |
| Nbl1 | 0.003693433 | 0.874714205 | 0.97843318 | 0.109536188 | 0.121350871 |
| Nbn | 0.006947336 | 0.731526747 | 0.94599730 | 0.10000564 | 0.089621156 |
| Nbr1 | -0.024838899 | 0.473706749 | 0.85970928 | 0.400541701 | 0.423767803 |
| Ncald | -0.00060272 | 0.987328927 | 0.99760728 | 0.490337522 | 0.526906061 |
| Ncam1 | -0.026520891 | 0.480569954 | 0.86175794 | 0.81549 | 0.762517905 |
| Ncam2 | 0.006813711 | 0.762368234 | 0.95300611 | 0.085580802 | 0.093515869 |
| Ncapd3 | -0.000442606 | 0.984546426 | 0.99760728 | 0.123393791 | 0.122455393 |
| Ncaph2 | 0.030358411 | 0.244690401 | 0.71379715 | 0.195399466 | 0.155781296 |
| Ncbp1 | 0.010101936 | 0.704057314 | 0.93950074 | 0.190810155 | 0.17366264 |
| Ncbp2 | -0.023974409 | 0.464328613 | 0.85346300 | 0.3508735 | 0.344636318 |
| Ncbp3 | -0.004140657 | 0.877393068 | 0.97855833 | 0.166201025 | 0.177713226 |
| Ncdn | 0.033481895 | 0.092157477 | 0.53255842 | 0.108451526 | 0.078793395 |
| Nceh1 | 0.033477396 | 0.237488631 | 0.70985110 | 0.216758912 | 0.21646114 |
| Nck1 | -0.003457549 | 0.91879156 | 0.98512115 | 0.383486922 | 0.384503386 |
| Nck2 | 0.006859833 | 0.61860813 | 0.91167507 | 0.041194703 | 0.037061681 |
| Nckap1 | -0.022932826 | 0.524615874 | 0.87583859 | 0.725419024 | 0.753175901 |
| Nckipsd | 0.023817031 | 0.211313781 | 0.68220598 | 0.09803205 | 0.066415903 |
| Ncl | 0.055506167 | 0.104713525 | 0.55800455 | 1.428018705 | 1.344224176 |
| Ncln | -0.022997415 | 0.358725524 | 0.79985551 | 0.130323016 | 0.151295334 |
| Ncoa1 | -0.015591132 | 0.647703575 | 0.91696950 | 0.339657859 | 0.370514808 |
| Ncoa2 | -0.021444495 | 0.451713208 | 0.84819137 | 0.187984685 | 0.216262128 |
| Ncoa3 | 0.021715593 | 0.35621507 | 0.79748442 | 0.141328164 | 0.130594971 |
| Ncoa4 | 0.042433934 | 0.094313705 | 0.53344171 | 0.177924162 | 0.155405449 |
| Ncoa5 | 0.002816459 | 0.890969602 | 0.98195975 | 0.101623959 | 0.093413924 |
| Ncoa6 | -0.012709842 | 0.626496615 | 0.91280959 | 0.146345284 | 0.16444146 |
| Ncoa7 | 0.055760628 | 0.140630652 | 0.61309530 | 0.635688628 | 0.573180009 |
| Ncor1 | -0.037856219 | 0.24088788 | 0.71204045 | 1.335746306 | 1.348455147 |
| Ncor2 | 0.021531914 | 0.500899655 | 0.86937040 | 0.314938635 | 0.282161992 |
| Ncs1 | 0.020699742 | 0.194954188 | 0.67199328 | 0.074376168 | 0.043586606 |
| Ncstn | -0.015888124 | 0.624040483 | 0.91215057 | 0.328550004 | 0.333464891 |

| | | | | | |
|---|---|---|---|---|---|
| Ndel1 | -0.041467229 | 0.212253756 | 0.6822059 | 0.295791906 | 0.32709135 |
| Ndfip1 | -0.059317401 | 0.017732856 | 0.2587928 | 1.744344977 | 1.735596823 |
| Ndfip2 | -0.01816432 | 0.612132374 | 0.9101248 | 0.532459666 | 0.532732483 |
| Ndn | -0.041954246 | 0.247340139 | 0.7176408 | 0.574374364 | 0.546396875 |
| Ndrg1 | -0.041322159 | 0.254512827 | 0.7223071 | 0.57355796 | 0.582398138 |
| Ndrg3 | -0.010281098 | 0.766003114 | 0.9543771 | 0.444183456 | 0.417870405 |
| Ndrg4 | -0.033218968 | 0.052058906 | 0.4313858 | 0.040554914 | 0.071803647 |
| Ndst1 | -0.033631103 | 0.252446836 | 0.7223071 | 0.196605048 | 0.23440331 |
| Ndst2 | 0.015969983 | 0.230823287 | 0.7024275 | 0.044972437 | 0.031245596 |
| Ndst4 | -0.005117455 | 0.856891641 | 0.9763189 | 0.212630442 | 0.169629877 |
| Ndufa1 | 0.088000377 | 0.001516019 | 0.0671379 | 1.586820977 | 1.471626767 |
| Ndufa10 | -0.021692436 | 0.543903953 | 0.8842645 | 0.525618575 | 0.531332646 |
| Ndufa11 | 0.061686509 | 0.027551209 | 0.3181899 | 1.611105215 | 1.522341846 |
| Ndufa12 | 0.031998165 | 0.382422246 | 0.8126882 | 0.515337394 | 0.487476737 |
| Ndufa13 | 0.023361582 | 0.318898717 | 0.7710043 | 1.940809623 | 1.885901655 |
| Ndufa2 | 0.113316572 | 2.53E-06 | 0.0004554 | 2.008630728 | 1.852197808 |
| Ndufa3 | 0.134193398 | 0.000225616 | 0.0177627 | 0.770836684 | 0.613273672 |
| Ndufa4 | 0.108051253 | 8.85E-05 | 0.0086926 | 1.856672609 | 1.610808279 |
| Ndufa5 | 0.180571863 | 1.88E-08 | 1.10E-05 | 1.445603362 | 1.18967322 |
| Ndufa6 | 0.062046408 | 0.007513654 | 0.1721437 | 2.087403705 | 1.860443011 |
| Ndufa7 | 0.06267104 | 0.005494015 | 0.1489751 | 2.085173829 | 1.92522366 |
| Ndufa8 | -0.043797101 | 0.189792007 | 0.6697747 | 0.886151008 | 0.86083633 |
| Ndufa9 | 0.07479692 | 0.022006472 | 0.2895010 | 0.421656489 | 0.316761717 |
| Ndufab1 | 0.031475119 | 0.366283254 | 0.8052437 | 0.717421075 | 0.584667081 |
| Ndufaf1 | -0.004529855 | 0.790942681 | 0.9622456 | 0.063937707 | 0.06376135 |
| Ndufaf2 | 0.020410324 | 0.49056604 | 0.8658154 | 0.261985209 | 0.239119219 |
| Ndufaf3 | -0.018252733 | 0.600761096 | 0.9078437 | 0.491918925 | 0.480839618 |
| Ndufaf4 | 0.01557316 | 0.598756063 | 0.9075513 | 0.240117414 | 0.230067191 |
| Ndufaf5 | 0.012327202 | 0.426760004 | 0.8337565 | 0.061458025 | 0.047572869 |
| Ndufaf6 | -0.0100549 | 0.652267077 | 0.9190271 | 0.096353904 | 0.114535487 |
| Ndufaf7 | 0.008788338 | 0.729607024 | 0.9459020 | 0.163745505 | 0.150710283 |
| Ndufaf8 | 0.101386205 | 0.003625563 | 0.1157629 | 0.690034083 | 0.600357736 |
| Ndufb1 | 0.116625596 | 3.20E-05 | 0.0038728 | 1.834529412 | 1.640959937 |
| Ndufb10 | 0.005406265 | 0.842381201 | 0.9734889 | 1.522490289 | 1.423991965 |
| Ndufb11 | 0.014877629 | 0.483845708 | 0.8624886 | 2.140437055 | 2.01419774 |
| Ndufb2 | 0.117256945 | 0.000138781 | 0.0120763 | 1.379339468 | 1.213872046 |
| Ndufb3 | 0.045987172 | 0.169453163 | 0.6480244 | 1.201648568 | 1.091364283 |
| Ndufb4 | 0.050638466 | 0.158239858 | 0.6339819 | 0.559338887 | 0.50843956 |
| Ndufb5 | 0.038629876 | 0.230653409 | 0.7022964 | 1.345787833 | 1.219144624 |
| Ndufb6 | 0.025325977 | 0.455472372 | 0.8502914 | 1.094149923 | 0.98755149 |
| Ndufb7 | 0.045417872 | 0.126859318 | 0.5893234 | 1.377377481 | 1.308360838 |
| Ndufb8 | 0.083172484 | 0.007618085 | 0.1729322 | 1.41254298 | 1.243593743 |
| Ndufb9 | 0.029464608 | 0.24361547 | 0.7132719 | 1.701805696 | 1.595493894 |
| Ndufc1 | 0.132681764 | 8.07E-06 | 0.0012505 | 1.502990686 | 1.342032816 |
| Ndufc2 | 0.082660616 | 0.003132666 | 0.1042820 | 1.772074208 | 1.670978247 |
| Ndufs1 | 0.06121459 | 0.058232134 | 0.4508199 | 0.391165257 | 0.309980305 |
| Ndufs2 | 0.024534938 | 0.488439515 | 0.8643096 | 1.178241478 | 1.076765322 |
| Ndufs3 | 0.042367316 | 0.229523432 | 0.7019677 | 0.676490205 | 0.565617882 |
| Ndufs4 | 0.006177396 | 0.861372961 | 0.9769675 | 0.790305367 | 0.746091005 |
| Ndufs5 | 0.126230477 | 7.01E-05 | 0.0072372 | 1.254874737 | 1.036004006 |
| Ndufs5-ps | 0.005066467 | 0.702388952 | 0.9388187 | 0.045079662 | 0.032142671 |
| Ndufs6 | -0.04613346 | 0.177282993 | 0.6536659 | 1.097822377 | 1.125033954 |
| Ndufs7 | 0.041710752 | 0.184346581 | 0.6628862 | 1.154214751 | 1.076663458 |
| Ndufs8 | -0.051511769 | 0.152973009 | 0.6286214 | 0.603009762 | 0.631610705 |
| Ndufv1 | 0.016200462 | 0.649291822 | 0.9174061 | 0.653765868 | 0.62625642 |
| Ndufv2 | -0.029794239 | 0.403723393 | 0.8266121 | 0.870088008 | 0.812222301 |
| Ndufv3 | 0.120970084 | 3.14E-05 | 0.0038515 | 1.519917936 | 1.361418372 |
| Neat1 | -0.158992965 | 0.000123257 | 0.0110153 | 0.394779393 | 0.684837556 |
| Neb | -0.014056154 | 0.452208996 | 0.8484983 | 0.056852254 | 0.075034285 |
| Nebl | 0.005843023 | 0.687721435 | 0.9327945 | 0.038993904 | 0.046089009 |
| Necab2 | -0.086494918 | 0.001852319 | 0.0745064 | 0.140208317 | 0.20327858 |
| Necap1 | 0.017869744 | 0.524864017 | 0.8758385 | 0.223860485 | 0.204768676 |
| Necap2 | -0.023064035 | 0.306064395 | 0.7651609 | 0.094441842 | 0.115047245 |
| Nectin1 | 0.009976468 | 0.520469963 | 0.8741795 | 0.050462246 | 0.050481424 |
| Nectin2 | -0.007213918 | 0.749618493 | 0.9504425 | 0.108703746 | 0.115424498 |
| Nectin3 | 0.024478006 | 0.466860925 | 0.8554230 | 0.3228912 | 0.333157918 |

| | | | | | |
|---|---|---|---|---|---|
| Nedd4 | 0.006040696 | 0.860136295 | 0.9769675 | 1.258538782 | 1.186828901 |
| Nedd4l | -0.030260269 | 0.272113564 | 0.7342806 | 0.181952766 | 0.186768925 |
| Nedd8 | 0.033515207 | 0.264996422 | 0.7306700 | 1.237281576 | 1.131017622 |
| Nedd9 | -0.009754266 | 0.725035923 | 0.9448707 | 0.16972502 | 0.171213454 |
| Nefm | -0.054156944 | 0.171654921 | 0.6515402 | 0.402457958 | 0.390238643 |
| Negr1 | 0.017325987 | 0.504501296 | 0.8695566 | 0.167716476 | 0.151155278 |
| Neil1 | -0.016806966 | 0.30216662 | 0.7611258 | 0.05069246 | 0.05902036 |
| Neil2 | 0.006631239 | 0.638139011 | 0.9160042 | 0.044055393 | 0.040923603 |
| Nek1 | 0.011901465 | 0.589790371 | 0.9046775 | 0.112708904 | 0.102316569 |
| Nek4 | -0.015777454 | 0.417973314 | 0.8300990 | 0.068501109 | 0.086557457 |
| Nek7 | -0.034281286 | 0.265142558 | 0.7306700 | 0.276655135 | 0.306822094 |
| Nek9 | -0.065242603 | 0.01279924 | 0.2233535 | 0.111850744 | 0.171480274 |
| Nelfa | -0.008336871 | 0.745948408 | 0.9485717 | 0.156614729 | 0.164354344 |
| Nelfb | 0.003684651 | 0.901956728 | 0.9825441 | 0.255284753 | 0.241311932 |
| Nelfcd | -0.043669346 | 0.08812379 | 0.5278913 | 0.14545912 | 0.156345847 |
| Nelfe | 0.039766158 | 0.236679743 | 0.7095050 | 0.445544495 | 0.365294196 |
| Nell1 | 0.01859251 | 0.581359443 | 0.9017028 | 0.368079608 | 0.346719592 |
| Nemf | 0.015637634 | 0.67139314 | 0.9250305 | 0.430158851 | 0.435466451 |
| Nemp1 | 0.002586241 | 0.898209449 | 0.9825441 | 0.100862282 | 0.08814089 |
| Nenf | -0.068024851 | 0.00765804 | 0.1734058 | 1.888886672 | 1.919319518 |
| Neo1 | -0.006742655 | 0.830831597 | 0.9722601 | 0.289168301 | 0.287985678 |
| Net1 | 0.012350449 | 0.422947915 | 0.8315789 | 0.051866376 | 0.04574369 |
| Neu1 | -0.072961376 | 0.033360627 | 0.3456484 | 0.370170976 | 0.431418338 |
| Neurl1a | -0.015531173 | 0.524899642 | 0.8758385 | 0.136676755 | 0.141923333 |
| Neurl4 | -0.009986037 | 0.66830288 | 0.9230765 | 0.113292561 | 0.122517604 |
| Neurod1 | 0.084661546 | 0.02041922 | 0.2774595 | 1.421359145 | 1.293403119 |
| Nf1 | -0.023328872 | 0.447818062 | 0.8456796 | 0.23362617 | 0.251230456 |
| Nf2 | -0.036269469 | 0.148130962 | 0.6255679 | 0.128776804 | 0.158878614 |
| Nfasc | -0.062362034 | 0.059891402 | 0.4573692 | 0.308413059 | 0.383633509 |
| Nfat5 | -0.126532552 | 0.000387531 | 0.0256287 | 0.30869093 | 0.424918969 |
| Nfatc1 | -0.018597545 | 0.248483864 | 0.7185162 | 0.039398648 | 0.060125748 |
| Nfatc2 | 0.004520695 | 0.822994952 | 0.9710592 | 0.096073014 | 0.090781378 |
| Nfatc3 | -0.010484653 | 0.665225215 | 0.9230765 | 0.135682244 | 0.138421685 |
| Nfe2l1 | 0.084201826 | 0.020084798 | 0.2748154 | 0.779088419 | 0.777443413 |
| Nfe2l2 | -0.000667207 | 0.983064142 | 0.9976072 | 0.268515653 | 0.252811608 |
| Nfia | 0.046543212 | 0.073004244 | 0.4903372 | 0.201087576 | 0.137902257 |
| Nfic | -0.000175413 | 0.996206578 | 0.9994073 | 0.615880919 | 0.637461681 |
| Nfil3 | 0.012828276 | 0.69357439 | 0.9359725 | 0.325170493 | 0.312378077 |
| Nfix | 0.000849884 | 0.964168322 | 0.9966421 | 0.063589918 | 0.07578584 |
| Nfkb1 | 0.065260852 | 0.023495978 | 0.2972960 | 0.241816759 | 0.193501422 |
| Nfkbia | -0.064826204 | 0.151985717 | 0.6281105 | 0.510622987 | 0.614369184 |
| Nfkbib | -0.040042856 | 0.117130214 | 0.5735102 | 0.136059905 | 0.170726982 |
| Nfkbil1 | 0.032539513 | 0.254463996 | 0.7223071 | 0.24396272 | 0.219564769 |
| Nfkbiz | -0.021812288 | 0.443954304 | 0.8431541 | 0.158841714 | 0.204989281 |
| Nfrkb | -0.010946023 | 0.667526055 | 0.9230765 | 0.142993268 | 0.151826382 |
| Nfs1 | 0.028757006 | 0.264495662 | 0.7300492 | 0.175496469 | 0.150653917 |
| Nfu1 | -0.01666204 | 0.63375764 | 0.9144743 | 0.410182577 | 0.410598677 |
| Nfx1 | -0.02718883 | 0.399309766 | 0.8239850 | 0.283370589 | 0.302738335 |
| Nfxl1 | -0.037715943 | 0.186020532 | 0.6640243 | 0.174723211 | 0.225956224 |
| Nfya | -0.005120425 | 0.759548195 | 0.9521533 | 0.057854465 | 0.055516756 |
| Nfyb | 0.037748223 | 0.251082176 | 0.7207210 | 0.342789894 | 0.305661671 |
| Nfyc | 0.034768697 | 0.092457571 | 0.5325584 | 0.123022181 | 0.083384882 |
| Ngdn | -0.011925866 | 0.708654138 | 0.9414057 | 0.333609913 | 0.304699278 |
| Ngly1 | 0.023050063 | 0.397702564 | 0.8238469 | 0.230595084 | 0.188434763 |
| Ngrn | 0.0239629 | 0.490417699 | 0.8658154 | 0.507921446 | 0.450374723 |
| Nhlrc2 | -0.018933289 | 0.502416833 | 0.8695566 | 0.18875441 | 0.198812959 |
| Nhlrc3 | -0.011856227 | 0.518430363 | 0.8740648 | 0.063327735 | 0.074407958 |
| Nhp2 | -0.037025615 | 0.253868476 | 0.7223071 | 0.299668578 | 0.323819067 |
| Nhsl1 | 0.018716039 | 0.419245689 | 0.8300990 | 0.131402394 | 0.105348362 |
| Niban2 | 0.006600546 | 0.739836525 | 0.9479080 | 0.087117837 | 0.08847456 |
| Nicn1 | -0.001728325 | 0.943852912 | 0.9935293 | 0.15090955 | 0.144507306 |
| Nif3l1 | 0.03143354 | 0.052272622 | 0.4314646 | 0.075690791 | 0.048781915 |
| Nifk | 0.023129016 | 0.407610392 | 0.8270770 | 0.215588301 | 0.183583243 |
| Nim1k | 0.00140987 | 0.940855551 | 0.9929301 | 0.080702591 | 0.072153664 |
| Nin | -0.004650107 | 0.764438198 | 0.9537450 | 0.043394161 | 0.05133314 |
| Ninj1 | 0.09788736 | 0.003639386 | 0.1157629 | 1.012068348 | 0.885827911 |

| | | | | | |
|---|---|---|---|---|---|
| Nip7 | -0.014853617 | 0.645819948 | 0.9169695 | 0.292443269 | 0.290558924 |
| Nipa1 | -0.014183155 | 0.455949629 | 0.8502914 | 0.075720562 | 0.079885396 |
| Nipa2 | 0.013608327 | 0.65799207 | 0.9211780 | 0.263301099 | 0.253164805 |
| Nipal1 | -0.022829227 | 0.520982812 | 0.8741795 | 0.398042743 | 0.424443788 |
| Nipal2 | -0.056099079 | 0.079615227 | 0.5076761 | 0.252850364 | 0.303872915 |
| Nipal3 | 0.027988084 | 0.372797968 | 0.8085689 | 0.292723794 | 0.26964728 |
| Nipbl | 0.010298251 | 0.784840214 | 0.9601202 | 0.791970398 | 0.757549401 |
| Nipsnap1 | 0.00710973 | 0.796004394 | 0.9640292 | 0.22078315 | 0.197844161 |
| Nipsnap2 | 0.036789386 | 0.210301848 | 0.6817628 | 0.236911297 | 0.215630609 |
| Nipsnap3b | 0.006762209 | 0.818309022 | 0.9697318 | 0.286597561 | 0.245212119 |
| Nisch | -0.024152762 | 0.503679739 | 0.8695566 | 0.934124763 | 0.959915461 |
| Nit1 | 0.031797703 | 0.255710656 | 0.7233104 | 0.243525924 | 0.192282237 |
| Nit2 | -0.004513409 | 0.807770268 | 0.9655477 | 0.080237185 | 0.071411161 |
| Nkap | 0.040317589 | 0.15855605 | 0.6344800 | 0.242131601 | 0.200967987 |
| Nkapd1 | -0.019570812 | 0.49840317 | 0.8684629 | 0.212427888 | 0.21675108 |
| Nkapl | -0.010147142 | 0.589147815 | 0.9046539 | 0.069065023 | 0.072268878 |
| Nkiras1 | -0.05842222 | 0.075429563 | 0.4966285 | 0.316145833 | 0.354397932 |
| Nkiras2 | 0.050841474 | 0.031849475 | 0.3385228 | 0.16231294 | 0.112482774 |
| Nkrf | 0.02475181 | 0.058620212 | 0.4520948 | 0.05041252 | 0.029450979 |
| Nktr | -0.115895514 | 0.001788463 | 0.0738681 | 1.229987695 | 1.401178616 |
| Nkx2-2 | 0.023120244 | 0.536046047 | 0.8799251 | 0.559167366 | 0.549622636 |
| Nkx6-1 | 0.052803792 | 0.205833052 | 0.6797565 | 0.64685207 | 0.710225925 |
| Nle1 | -0.009622748 | 0.519711533 | 0.8741795 | 0.043210188 | 0.047393157 |
| Nlgn2 | -0.061579826 | 0.033243155 | 0.3454615 | 0.174465981 | 0.214313253 |
| Nlk | 0.019743495 | 0.530364536 | 0.8776448 | 0.283741006 | 0.2733487 |
| Nmd3 | -0.028677168 | 0.349238898 | 0.7927303 | 0.226367608 | 0.236525278 |
| Nme1 | 0.006310978 | 0.835761347 | 0.9723494 | 1.65069144 | 1.547830974 |
| Nme2 | 0.033738051 | 0.217713435 | 0.6886853 | 0.198290974 | 0.175694107 |
| Nme4 | -0.012068404 | 0.443787201 | 0.8431541 | 0.051364879 | 0.055055851 |
| Nme5 | -0.031258985 | 0.042229436 | 0.3952037 | 0.038408811 | 0.055943674 |
| Nme6 | -0.006605231 | 0.71529845 | 0.9431411 | 0.077180903 | 0.0702495 |
| Nme7 | 0.008900101 | 0.612637534 | 0.9105164 | 0.071611665 | 0.062112441 |
| Nmi | 0.002377444 | 0.923791067 | 0.9876072 | 0.160320485 | 0.150508864 |
| Nmnat1 | -0.011944595 | 0.419164687 | 0.8300990 | 0.037139998 | 0.049026042 |
| Nmnat3 | -0.000255817 | 0.990004934 | 0.9980836 | 0.090306176 | 0.091039013 |
| Nmral1 | 0.008457844 | 0.610505663 | 0.9090802 | 0.062206151 | 0.056207083 |
| Nmt1 | -0.001951888 | 0.952147321 | 0.9951566 | 0.328685916 | 0.318743286 |
| Nmt2 | 0.041200707 | 0.185314283 | 0.6637678 | 0.29579374 | 0.243869046 |
| Nnat | 0.076514221 | 9.19E-07 | 0.0002072 | 0.083494362 | 0.024916482 |
| Noa1 | 0.013003376 | 0.538454494 | 0.8816335 | 0.121882436 | 0.097308259 |
| Nob1 | 0.01506086 | 0.558112374 | 0.8919622 | 0.164473654 | 0.160153913 |
| Noc2l | -0.040817345 | 0.120059775 | 0.5797304 | 0.135604914 | 0.170882317 |
| Noc3l | 0.045403988 | 0.03083855 | 0.3353750 | 0.126650145 | 0.083894923 |
| Noc4l | -0.022863198 | 0.247217054 | 0.7176408 | 0.076438348 | 0.091820553 |
| Noct | 0.014596613 | 0.545080138 | 0.8848298 | 0.1543692 | 0.123245102 |
| Nod1 | -0.012694412 | 0.420684065 | 0.8300990 | 0.043190179 | 0.055842452 |
| Nol10 | -0.004998011 | 0.741051506 | 0.9479080 | 0.050630577 | 0.04498549 |
| Nol11 | 0.059390122 | 0.048691684 | 0.4158684 | 0.293115116 | 0.23416653 |
| Nol12 | 0.001926038 | 0.952757116 | 0.9953744 | 0.406179068 | 0.360973619 |
| Nol3 | -0.015425609 | 0.415855736 | 0.8300990 | 0.054231361 | 0.088517857 |
| Nol4 | 0.011649872 | 0.712013775 | 0.9419834 | 0.277414677 | 0.267190455 |
| Nol4l | 0.010196076 | 0.557346402 | 0.8916648 | 0.066045155 | 0.063100974 |
| Nol6 | 0.00076194 | 0.970425846 | 0.9976072 | 0.090003529 | 0.087162652 |
| Nol7 | 0.093590913 | 0.010122281 | 0.1996282 | 1.010431882 | 0.849503063 |
| Nol8 | 0.001705862 | 0.955135103 | 0.9954690 | 0.214104177 | 0.218693105 |
| Nol9 | -0.020325751 | 0.244164027 | 0.7132719 | 0.053651698 | 0.069052525 |
| Nolc1 | -0.008577023 | 0.804425186 | 0.9647143 | 0.363232299 | 0.378316977 |
| Nom1 | -0.01086813 | 0.693436062 | 0.9359725 | 0.191721623 | 0.183450752 |
| Nomo1 | -0.019903568 | 0.430759369 | 0.8350855 | 0.142701195 | 0.152820582 |
| Nono | -0.034353746 | 0.334775676 | 0.7804875 | 0.557090635 | 0.565678433 |
| Nop10 | 0.067799676 | 0.055161197 | 0.4434352 | 1.24700349 | 1.136542441 |
| Nop14 | -0.007152214 | 0.830800745 | 0.9722601 | 0.314169523 | 0.315597446 |
| Nop16 | -0.003249003 | 0.901121329 | 0.9825441 | 0.178980948 | 0.165235634 |
| Nop2 | -0.007986554 | 0.670965934 | 0.9247082 | 0.077216729 | 0.078381545 |
| Nop53 | -0.056010533 | 0.126432034 | 0.5889350 | 0.519376741 | 0.540024485 |
| Nop56 | -0.060228181 | 0.048516944 | 0.4157447 | 0.226748889 | 0.254127613 |

| | | | | | |
|---|---|---|---|---|---|
| Nop58 | -0.031716671 | 0.392831731 | 0.82062368 | 0.55020595 | 0.567454415 |
| Nop9 | 0.000341982 | 0.98493279 | 0.99760728 | 0.074527883 | 0.067973931 |
| Norad | -0.001147976 | 0.975830159 | 0.99760728 | 0.828100585 | 0.812609556 |
| Nosip | -0.008365898 | 0.739234684 | 0.94756347 | 0.156139776 | 0.148663618 |
| Nostrin | -0.0608884 | 0.105924433 | 0.56171711 | 0.371351559 | 0.367292007 |
| Notch1 | -0.008093156 | 0.733085265 | 0.94641343 | 0.117461313 | 0.122406581 |
| Nova2 | -0.038703574 | 0.089517126 | 0.53023604 | 0.100908483 | 0.12166118 |
| Npas4 | -0.034852873 | 0.126514164 | 0.58893500 | 0.072571074 | 0.114476356 |
| Npat | -0.012255068 | 0.634226233 | 0.9144743 | 0.145893696 | 0.152129234 |
| Npc1 | -0.01221584 | 0.559829594 | 0.89256468 | 0.095845055 | 0.101195114 |
| Npc2 | -0.037241823 | 0.188637164 | 0.66688548 | 1.725724644 | 1.621039214 |
| Npdc1 | -0.021780257 | 0.519388972 | 0.87417954 | 0.956203995 | 0.91997177 |
| Npepl1 | -0.064942271 | 0.062805449 | 0.46571225 | 0.327913441 | 0.399788623 |
| Npepps | 0.006797635 | 0.850459099 | 0.97488410 | 0.601276677 | 0.584355771 |
| Nphp1 | 0.02095423 | 0.384460806 | 0.81353481 | 0.153644846 | 0.132510994 |
| Nploc4 | -0.001766387 | 0.945277464 | 0.99386883 | 0.152016535 | 0.160621249 |
| Npm1 | -0.023056927 | 0.397088841 | 0.82384692 | 2.044968948 | 1.936550505 |
| Npm3 | 0.027531737 | 0.325367878 | 0.77630994 | 0.223140709 | 0.184885337 |
| Npnt | 0.016616434 | 0.388750069 | 0.81702870 | 0.078618618 | 0.084126213 |
| Npr2 | -0.000733504 | 0.964250073 | 0.99664213 | 0.053113007 | 0.056247572 |
| Nprl2 | -0.038169556 | 0.135338594 | 0.60258470 | 0.142313105 | 0.161629989 |
| Nprl3 | -0.020394415 | 0.273683688 | 0.73519553 | 0.058100222 | 0.080957061 |
| Nptn | 0.047112796 | 0.156859424 | 0.63176830 | 0.440006434 | 0.366981647 |
| Nptx1 | -0.005169999 | 0.783027675 | 0.95913695 | 0.080485174 | 0.07184475 |
| Nptxr | 0.055255176 | 0.041465177 | 0.39174719 | 0.206427987 | 0.164529299 |
| Nr1d1 | 0.036321693 | 0.071124935 | 0.4881297 | 0.106527081 | 0.073997071 |
| Nr1d2 | 0.012349381 | 0.641020298 | 0.91667127 | 0.181622852 | 0.165849891 |
| Nr1h2 | -0.012730679 | 0.630489759 | 0.91319293 | 0.176329029 | 0.174286623 |
| Nr1h4 | -0.004765164 | 0.768127863 | 0.95442217 | 0.049978121 | 0.058095296 |
| Nr2c2 | 0.046989907 | 0.083579909 | 0.51830408 | 0.22994244 | 0.173683775 |
| Nr2c2ap | -0.049307234 | 0.083572178 | 0.51830408 | 0.205737836 | 0.216121745 |
| Nr2f6 | -0.022242647 | 0.480850736 | 0.86175794 | 0.274621191 | 0.280619547 |
| Nr3c1 | -0.023080842 | 0.530248885 | 0.87764481 | 0.539284044 | 0.543099193 |
| Nr3c2 | 0.009273274 | 0.568263853 | 0.89553080 | 0.061319982 | 0.055346497 |
| Nr4a1 | -0.003976268 | 0.920929332 | 0.98611750 | 0.403283468 | 0.515609215 |
| Nr4a2 | -0.086082134 | 0.075255994 | 0.49662859 | 0.732690802 | 0.815451268 |
| Nr6a1 | -0.057108757 | 0.00972442 | 0.19727248 | 0.074714183 | 0.1178686 |
| Nras | 0.017708911 | 0.508789934 | 0.87040677 | 0.183807397 | 0.16533331 |
| Nrbf2 | 0.000679896 | 0.977042859 | 0.99760728 | 0.127120225 | 0.127759803 |
| Nrbp1 | 0.021640228 | 0.479622813 | 0.86108079 | 0.280227445 | 0.240655119 |
| Nrbp2 | -0.021851254 | 0.185611576 | 0.66376783 | 0.042314612 | 0.060245691 |
| Nrcam | -0.002518163 | 0.890555174 | 0.98195975 | 0.053739379 | 0.068023819 |
| Nrd1 | -0.01336095 | 0.724411925 | 0.94456522 | 0.643663934 | 0.643925759 |
| Nrde2 | -0.018296345 | 0.217016517 | 0.68723302 | 0.034406347 | 0.049057896 |
| Nrep | 0.00627391 | 0.696626031 | 0.93638620 | 0.054475173 | 0.055175555 |
| Nrf1 | -0.012861793 | 0.631664351 | 0.91335177 | 0.139917729 | 0.175136917 |
| Nrg2 | 0.007833729 | 0.603074608 | 0.90871392 | 0.051017083 | 0.040739058 |
| Nrip1 | 0.004477366 | 0.899422201 | 0.98254414 | 0.357295096 | 0.358923379 |
| Nrsn1 | -0.00861256 | 0.642241193 | 0.91667127 | 0.06997401 | 0.073118739 |
| Nrtn | 0.019622215 | 0.558343504 | 0.89198239 | 0.292645989 | 0.326628152 |
| Nrxn1 | -0.034961035 | 0.28052519 | 0.74086184 | 0.227553936 | 0.306484865 |
| Nrxn2 | 0.025798458 | 0.089744601 | 0.53023604 | 0.060019777 | 0.037733318 |
| Nsa2 | 0.0547811 | 0.130026528 | 0.59380645 | 0.986743839 | 0.821733891 |
| Nsd1 | -0.060506762 | 0.120529286 | 0.58083458 | 0.5876514 | 0.626263978 |
| Nsd2 | -0.011979469 | 0.716261511 | 0.94314111 | 0.30389845 | 0.300092777 |
| Nsd3 | -0.023057627 | 0.546157041 | 0.88584131 | 0.676415433 | 0.674690451 |
| Nsdhl | 0.017001383 | 0.283667947 | 0.74299364 | 0.072279625 | 0.045990122 |
| Nsf | 0.021119635 | 0.506888156 | 0.86995337 | 0.306876147 | 0.301555573 |
| Nsfl1c | 0.047122859 | 0.152115498 | 0.62811059 | 0.417314194 | 0.328313139 |
| Nsmaf | -0.005658249 | 0.83010082 | 0.97226010 | 0.159308632 | 0.164236088 |
| Nsmce1 | 0.036428195 | 0.252994994 | 0.72230717 | 0.349481846 | 0.28341763 |
| Nsmce2 | 0.01610716 | 0.597979893 | 0.90755130 | 0.269009585 | 0.24179413 |
| Nsmce3 | -0.016806074 | 0.617929325 | 0.91150170 | 0.427425327 | 0.402036909 |
| Nsmce4a | 0.005499085 | 0.878513071 | 0.97855832 | 0.568022676 | 0.519053519 |
| Nsmf | 0.024648328 | 0.171138381 | 0.6507063 | 0.08981754 | 0.064491557 |
| Nsrp1 | 0.048289822 | 0.154798379 | 0.63049349 | 0.426731398 | 0.354325683 |

| | | | | | |
|---|---|---|---|---|---|
| Nsun2 | -0.01486987 | 0.611539087 | 0.9097882 | 0.246442807 | 0.220198655 |
| Nsun3 | -0.005518478 | 0.763575976 | 0.9532204 | 0.073781469 | 0.072272108 |
| Nsun4 | 0.019155489 | 0.258320515 | 0.7251102 | 0.072896921 | 0.056554759 |
| Nsun5 | -0.033275019 | 0.039201628 | 0.3799003 | 0.034670684 | 0.059868665 |
| Nsun6 | -0.023923158 | 0.185682865 | 0.6637744 | 0.058327888 | 0.069420192 |
| Nt5c | 0.011330147 | 0.716287866 | 0.9431411 | 0.282138888 | 0.259394568 |
| Nt5c2 | 0.023317001 | 0.432788237 | 0.8373823 | 0.245595302 | 0.230416852 |
| Nt5c3 | -0.023429925 | 0.495759895 | 0.8684629 | 0.400469109 | 0.449562109 |
| Nt5c3b | -0.009660382 | 0.666637753 | 0.9230765 | 0.115500449 | 0.116313741 |
| Nt5dc1 | 0.016233679 | 0.296594224 | 0.7545964 | 0.062606063 | 0.045806578 |
| Nt5dc2 | 0.002393197 | 0.933735841 | 0.9903410 | 0.24307321 | 0.203029876 |
| Nt5dc3 | -0.008718818 | 0.667836175 | 0.9230765 | 0.083167493 | 0.092618289 |
| Nt5m | -0.045176896 | 0.150405588 | 0.6280320 | 0.271633472 | 0.290759735 |
| Ntan1 | -0.006187427 | 0.850527422 | 0.9748841 | 0.343102092 | 0.317975373 |
| Ntmt1 | -0.008018894 | 0.772440702 | 0.9552333 | 0.208792749 | 0.18280057 |
| Ntn4 | -0.022115051 | 0.182398751 | 0.6596411 | 0.056965799 | 0.057477748 |
| Ntpcr | 0.010169026 | 0.68124085 | 0.9299864 | 0.155065108 | 0.140080137 |
| Ntrk2 | 0.032436847 | 0.341619926 | 0.7853157 | 0.298762583 | 0.344094471 |
| Nuak1 | 0.015679703 | 0.493122101 | 0.8669301 | 0.133006842 | 0.112301224 |
| Nuak2 | -0.011602104 | 0.435112789 | 0.8389295 | 0.04151949 | 0.049089886 |
| Nub1 | 0.004189864 | 0.899071339 | 0.9825441 | 0.33049791 | 0.301061852 |
| Nubp1 | 0.029016932 | 0.251083445 | 0.7207210 | 0.170086205 | 0.141839851 |
| Nubp2 | -0.003758617 | 0.872899401 | 0.9784331 | 0.145727794 | 0.125883391 |
| Nubpl | 0.003271776 | 0.859990772 | 0.9769675 | 0.080818744 | 0.074677116 |
| Nucb1 | 0.005522951 | 0.875267673 | 0.9784331 | 0.926126784 | 0.845578595 |
| Nucb2 | -0.052280384 | 0.176390126 | 0.6536659 | 0.729664176 | 0.913503095 |
| Nucks1 | 0.001215354 | 0.972813435 | 0.9976072 | 0.851679766 | 0.84831813 |
| Nudc | -0.020803525 | 0.556714189 | 0.8915944 | 0.83421864 | 0.801363218 |
| Nudcd1 | -0.044188692 | 0.078012816 | 0.5048164 | 0.126748293 | 0.15690237 |
| Nudcd2 | -0.003326235 | 0.921858717 | 0.9862854 | 0.430692488 | 0.392435093 |
| Nudcd3 | 0.012355345 | 0.724515803 | 0.9445652 | 0.4553305 | 0.464010788 |
| Nudt10 | 0.008705709 | 0.754998495 | 0.9519143 | 0.233293162 | 0.177846014 |
| Nudt11 | -0.094020343 | 0.015915019 | 0.2474560 | 0.458147831 | 0.431933319 |
| Nudt13 | -0.044896631 | 0.108228178 | 0.5636817 | 0.174214514 | 0.199959253 |
| Nudt14 | -0.004963019 | 0.865126687 | 0.9770148 | 0.23043387 | 0.225138879 |
| Nudt16 | 0.02358546 | 0.3345484 | 0.7804836 | 0.163693422 | 0.132499975 |
| Nudt16l1 | -0.012764928 | 0.635623392 | 0.9153541 | 0.182693105 | 0.188339336 |
| Nudt18 | 0.067777348 | 0.020919918 | 0.2808194 | 0.30426812 | 0.21350479 |
| Nudt19 | -0.032712961 | 0.231303227 | 0.7026141 | 0.173325625 | 0.198100503 |
| Nudt2 | -0.001443218 | 0.95664429 | 0.9955934 | 0.182745451 | 0.168543705 |
| Nudt21 | 0.086528318 | 0.002891577 | 0.1000185 | 0.3184551 | 0.209190324 |
| Nudt22 | -0.030737557 | 0.163029866 | 0.6400703 | 0.101658321 | 0.118927681 |
| Nudt3 | -0.024185651 | 0.440734486 | 0.8422435 | 0.281253507 | 0.29559791 |
| Nudt4 | 0.053771667 | 0.102060602 | 0.5520398 | 0.39111153 | 0.359374575 |
| Nudt5 | 0.01462956 | 0.652824781 | 0.9192017 | 0.335084812 | 0.308973503 |
| Nudt6 | -0.01010978 | 0.527727835 | 0.8768553 | 0.049775376 | 0.055325378 |
| Nudt7 | -0.001155084 | 0.964639554 | 0.9966421 | 0.155685886 | 0.15723168 |
| Nudt9 | -0.023949296 | 0.335282829 | 0.7809414 | 0.138529567 | 0.157344099 |
| Nufip1 | 0.002245259 | 0.92253952 | 0.9868009 | 0.131392888 | 0.122810552 |
| Nufip2 | -0.021118518 | 0.576078304 | 0.8993857 | 0.505349699 | 0.507071513 |
| Numa1 | -0.057091514 | 0.110666117 | 0.5655888 | 0.37908322 | 0.41287649 |
| Numb | -0.008785355 | 0.636605047 | 0.9153677 | 0.072565789 | 0.075491608 |
| Numbl | 0.040222032 | 0.028022912 | 0.3208303 | 0.106266051 | 0.062025377 |
| Nup107 | 0.012582977 | 0.459840133 | 0.8524343 | 0.080491872 | 0.053752408 |
| Nup133 | -0.028186342 | 0.079351108 | 0.5076205 | 0.034247873 | 0.060833199 |
| Nup153 | -0.012327724 | 0.642877633 | 0.9166712 | 0.169146521 | 0.16949474 |
| Nup160 | 0.025522418 | 0.212295352 | 0.6822059 | 0.110392655 | 0.085540982 |
| Nup188 | -0.009930767 | 0.568346211 | 0.8955308 | 0.061651113 | 0.069935432 |
| Nup205 | 0.038561265 | 0.046684631 | 0.4094577 | 0.102447174 | 0.068798065 |
| Nup210 | -0.003129371 | 0.9019028 | 0.9825441 | 0.149863333 | 0.153792289 |
| Nup214 | -0.020887219 | 0.451144769 | 0.8479265 | 0.187860246 | 0.191679138 |
| Nup35 | -0.021914501 | 0.388460561 | 0.8170287 | 0.151359421 | 0.159138444 |
| Nup37 | -0.025169653 | 0.14330127 | 0.6165078 | 0.054212827 | 0.069156137 |
| Nup43 | -0.022224057 | 0.234958315 | 0.7068170 | 0.069236018 | 0.07747583 |
| Nup50 | 0.010562871 | 0.741227274 | 0.9479080 | 0.320216807 | 0.29877704 |
| Nup54 | 0.034792982 | 0.156850307 | 0.6317683 | 0.165504929 | 0.130625234 |

| | | | | | |
|---|---|---|---|---|---|
| Nup62 | 0.012851017 | 0.618187324 | 0.9115017 | 0.157986386 | 0.144474541 |
| Nup85 | 0.031627676 | 0.082128578 | 0.5148480 | 0.094438124 | 0.06024207 |
| Nup88 | 0.013011864 | 0.657139678 | 0.9211780 | 0.246486731 | 0.219314068 |
| Nup93 | 0.011221166 | 0.623468805 | 0.9121505 | 0.10532735 | 0.125432519 |
| Nup98 | -0.029867657 | 0.329337976 | 0.7778857 | 0.279823161 | 0.27325554 |
| Nupl2 | -0.008342437 | 0.597994702 | 0.9075513 | 0.043254999 | 0.049312444 |
| Nupr1 | -0.039522288 | 0.511888259 | 0.8720873 | 0.865849126 | 1.479237307 |
| Nupr1l | -0.014690905 | 0.469838784 | 0.8568606 | 0.083138393 | 0.100439962 |
| Nus1 | 0.032592524 | 0.315461278 | 0.7705987 | 0.325858287 | 0.311817461 |
| Nvl | -0.050047485 | 0.06823063 | 0.4833869 | 0.17758715 | 0.200569365 |
| Nxf1 | 0.00092923 | 0.968994548 | 0.9976072 | 0.144167783 | 0.131183741 |
| Nxn | 0.026570042 | 0.188208082 | 0.6668854 | 0.113006649 | 0.082731163 |
| Nxph1 | 0.014708377 | 0.562766937 | 0.8926240 | 0.185740943 | 0.127283463 |
| Nxt1 | -0.001252213 | 0.961130512 | 0.9961577 | 0.169746313 | 0.164006343 |
| Nxt2 | -0.006093017 | 0.737028913 | 0.9469752 | 0.069615944 | 0.07204273 |
| Nyap1 | -0.012671962 | 0.461146585 | 0.8524425 | 0.061862025 | 0.061245624 |
| Nynrin | -0.001963689 | 0.91664659 | 0.9845316 | 0.086653392 | 0.071595614 |
| Oaf | -0.000487142 | 0.977373682 | 0.9976072 | 0.072522372 | 0.063087083 |
| Oard1 | 0.00348542 | 0.919082285 | 0.9851211 | 0.421776842 | 0.39843217 |
| Oat | -0.005133443 | 0.877268568 | 0.9785583 | 0.299793751 | 0.251973655 |
| Oaz1 | 0.022116032 | 0.215572044 | 0.6852119 | 2.780083341 | 2.55703877 |
| Oaz2 | -0.031196015 | 0.340469076 | 0.7845419 | 0.359678691 | 0.345314744 |
| Obi1 | 0.026568405 | 0.179615938 | 0.6550070 | 0.11062178 | 0.076419589 |
| Obsl1 | 0.026553903 | 0.154356342 | 0.6304934 | 0.097059231 | 0.065085506 |
| Ocel1 | -0.019998692 | 0.330219953 | 0.7778857 | 0.082720819 | 0.099086238 |
| Ociad1 | 0.052262893 | 0.156466671 | 0.6316663 | 0.741821948 | 0.64883215 |
| Ociad2 | -0.037479765 | 0.327896028 | 0.7778857 | 0.667654603 | 0.741473242 |
| Ocrl | -0.068170567 | 0.068707402 | 0.4835495 | 0.51043067 | 0.521268869 |
| Odad1 | 0.017588719 | 0.162997495 | 0.6400703 | 0.043212026 | 0.026983109 |
| Odc1 | -0.026675431 | 0.502524796 | 0.8695566 | 0.473367988 | 0.575434575 |
| Odf2 | -0.010171735 | 0.723518 | 0.9445652 | 0.209100211 | 0.20872041 |
| Odf2l | 0.012358621 | 0.620551146 | 0.9121505 | 0.142365038 | 0.131372782 |
| Odr4 | -0.002394333 | 0.940837393 | 0.9929301 | 0.317381472 | 0.334199627 |
| Ofd1 | -0.001228431 | 0.964658694 | 0.9966421 | 0.190196301 | 0.189115963 |
| Oga | 0.060295757 | 0.064234223 | 0.4700387 | 0.336990251 | 0.28695366 |
| Ogdh | -0.010438672 | 0.774324446 | 0.9558538 | 0.584928517 | 0.624230956 |
| Ogdhl | 0.022832311 | 0.330027692 | 0.7778857 | 0.124418696 | 0.122977607 |
| Ogfod1 | -0.003979817 | 0.862165755 | 0.9769675 | 0.11788336 | 0.120391134 |
| Ogfod2 | 0.045244703 | 0.151123827 | 0.6280470 | 0.328870922 | 0.27978534 |
| Ogfod3 | -0.036696954 | 0.20194878 | 0.6752150 | 0.196697763 | 0.215852587 |
| Ogfr | 0.050178123 | 0.102213728 | 0.5522658 | 0.318237504 | 0.243105094 |
| Ogfrl1 | -0.012247985 | 0.434751449 | 0.8385606 | 0.038229889 | 0.052127585 |
| Ogt | -0.069094319 | 0.037545257 | 0.3683965 | 0.277995831 | 0.352302744 |
| Oip5os1 | 0.023092726 | 0.452304344 | 0.8484983 | 0.272319738 | 0.240683976 |
| Ola1 | -0.002362675 | 0.947154131 | 0.9945018 | 0.521460563 | 0.494678768 |
| Olfm1 | 0.012413277 | 0.682555523 | 0.9304590 | 0.265757474 | 0.207547975 |
| Olfm3 | 0.007778933 | 0.72250444 | 0.9445652 | 0.115917591 | 0.088159512 |
| Oma1 | 0.041791259 | 0.04432059 | 0.4009056 | 0.12219392 | 0.084533478 |
| Onecut2 | -0.024272264 | 0.211026431 | 0.6822059 | 0.071814588 | 0.078285136 |
| Opa1 | 0.011941122 | 0.684593828 | 0.9313273 | 0.245695875 | 0.223286991 |
| Opa3 | 0.008273753 | 0.76801787 | 0.9544221 | 0.206621838 | 0.189045071 |
| Ophn1 | 0.000265232 | 0.987110991 | 0.9976072 | 0.060594573 | 0.054879758 |
| Optn | -0.026356487 | 0.151498737 | 0.6281105 | 0.059949503 | 0.082681195 |
| Orai1 | 0.015645779 | 0.451599456 | 0.8481913 | 0.104591093 | 0.085855666 |
| Orai2 | 0.010014738 | 0.512543919 | 0.8722771 | 0.050209775 | 0.04862346 |
| Orai3 | 0.026605402 | 0.311446242 | 0.7684379 | 0.200402756 | 0.153057646 |
| Orc2 | -8.38E-05 | 0.997684449 | 0.9994073 | 0.211196855 | 0.210791569 |
| Orc3 | -0.046698217 | 0.055930619 | 0.4449332 | 0.115914886 | 0.150688355 |
| Orc4 | 0.00746167 | 0.80675093 | 0.9650270 | 0.270825598 | 0.242481742 |
| Orc5 | 0.028896205 | 0.111956712 | 0.5679582 | 0.090355467 | 0.065296369 |
| Orc6 | 0.010052651 | 0.705674827 | 0.9401381 | 0.184066659 | 0.161783469 |
| Ormdl1 | -0.010414021 | 0.624181645 | 0.9121505 | 0.10437965 | 0.101230298 |
| Ormdl2 | 0.004880014 | 0.877883444 | 0.9785583 | 0.288704365 | 0.282959744 |
| Ormdl3 | -0.06213063 | 0.078611853 | 0.5057194 | 0.431869284 | 0.493835411 |
| Os9 | -0.070905281 | 0.058518653 | 0.4517548 | 0.708324043 | 0.755921363 |
| Osbp | -0.010785051 | 0.73562273 | 0.9468380 | 0.292469191 | 0.304712187 |

| | | | | | |
|---|---|---|---|---|---|
| Osbpl11 | 0.001709758 | 0.917503365 | 0.9848120 | 0.059703969 | 0.056326274 |
| Osbpl1a | -0.009429655 | 0.749838596 | 0.9504537 | 0.218098109 | 0.227748816 |
| Osbpl2 | 0.022926332 | 0.428615172 | 0.8340365 | 0.228547375 | 0.209339762 |
| Osbpl6 | 0.026576359 | 0.299997561 | 0.7590275 | 0.173985556 | 0.138943678 |
| Osbpl8 | 0.013536193 | 0.604414002 | 0.9087139 | 0.17000245 | 0.15282739 |
| Osbpl9 | 0.014306492 | 0.576910471 | 0.8994610 | 0.176757847 | 0.151908269 |
| Oscp1 | 0.008361791 | 0.645700053 | 0.9169695 | 0.080244078 | 0.069312105 |
| Oser1 | -0.008356905 | 0.791850098 | 0.9624054 | 0.324819154 | 0.307286827 |
| Osgep | -0.043879132 | 0.15053529 | 0.6280320 | 0.291285983 | 0.298468165 |
| Osgepl1 | -0.000500168 | 0.973203241 | 0.9976072 | 0.046713717 | 0.04564216 |
| Osgin1 | -0.018707665 | 0.436290588 | 0.8390644 | 0.113405754 | 0.133524685 |
| Osgin2 | -0.00775399 | 0.665841164 | 0.9230765 | 0.073020583 | 0.066637438 |
| Ost4 | 0.070937624 | 0.009366573 | 0.1943120 | 1.600249313 | 1.437904496 |
| Ostc | -0.023514494 | 0.410524365 | 0.8283974 | 1.872279456 | 1.802178429 |
| Ostf1 | 0.086880971 | 0.005028774 | 0.1417199 | 0.382011938 | 0.266089151 |
| Ostm1 | 0.039122425 | 0.082189685 | 0.5148480 | 0.120421399 | 0.107239808 |
| Otub1 | 0.062482201 | 0.078431626 | 0.5056175 | 0.490220387 | 0.410707261 |
| Otub2 | 0.051522323 | 0.007190466 | 0.1680001 | 0.104196849 | 0.073148064 |
| Otud3 | 0.005680494 | 0.74540437 | 0.9485717 | 0.068898307 | 0.064114993 |
| Otud4 | -0.015847832 | 0.577104632 | 0.8994610 | 0.180073569 | 0.212545675 |
| Otud5 | 0.04103704 | 0.167859603 | 0.6449137 | 0.264170086 | 0.224171914 |
| Otud6b | -0.018980384 | 0.475552537 | 0.8603832 | 0.17012503 | 0.165443667 |
| Otud7b | 0.004226202 | 0.875516069 | 0.9784331 | 0.164956229 | 0.16905302 |
| Otulin | -0.000386559 | 0.990073719 | 0.9980836 | 0.284135227 | 0.285812729 |
| Otulinl | 0.008670272 | 0.552430423 | 0.8897726 | 0.049061906 | 0.043755161 |
| Ovol2 | 0.050092832 | 0.001173691 | 0.0541535 | 0.066266202 | 0.045549768 |
| Oxa1l | -0.035268788 | 0.198231104 | 0.6730962 | 0.170953823 | 0.187487153 |
| Oxct1 | 0.087807035 | 0.01227057 | 0.2185506 | 0.500034798 | 0.413020611 |
| Oxld1 | 0.018868672 | 0.385840821 | 0.8148905 | 0.125760829 | 0.106009571 |
| Oxnad1 | -0.008062011 | 0.694571836 | 0.9361958 | 0.094329629 | 0.09393465 |
| Oxr1 | 0.00633173 | 0.830013018 | 0.9722601 | 0.215906443 | 0.210080063 |
| Oxsm | 0.014939128 | 0.331430288 | 0.7785433 | 0.054349697 | 0.046528168 |
| Oxsr1 | -0.037064053 | 0.12137273 | 0.5815270 | 0.112968679 | 0.139459144 |
| Oxtr | -0.009980384 | 0.800091622 | 0.9647053 | 0.494572445 | 0.418631423 |
| P2rx4 | -0.017587488 | 0.583013093 | 0.9019790 | 0.291063921 | 0.273546821 |
| P2ry1 | 0.040923368 | 0.01055478 | 0.2017406 | 0.065318926 | 0.049395358 |
| P3h1 | -0.019039208 | 0.318136998 | 0.7710043 | 0.072147038 | 0.079926535 |
| P3h3 | -0.006081976 | 0.698960133 | 0.9369928 | 0.046988395 | 0.048357589 |
| P3h4 | -0.016218996 | 0.635632108 | 0.9153541 | 0.442444358 | 0.374170768 |
| P4ha1 | 0.02364596 | 0.4616892 | 0.8524425 | 0.319814063 | 0.290999052 |
| P4ha2 | 0.035544839 | 0.159879048 | 0.6369331 | 0.178777473 | 0.145264115 |
| P4hb | 0.041254232 | 0.244867478 | 0.7138070 | 1.212964599 | 1.195181624 |
| P4htm | 0.019559766 | 0.454106183 | 0.8491486 | 0.170597866 | 0.165299229 |
| Pa2g4 | -0.018650587 | 0.604787787 | 0.9087139 | 0.574830838 | 0.553988812 |
| Pabpc1 | 0.028088555 | 0.407322896 | 0.8270770 | 1.282533334 | 1.204599402 |
| Pabpc1l | -0.026995818 | 0.132096956 | 0.5978345 | 0.052617765 | 0.072780274 |
| Pabpc4 | -0.023899653 | 0.309989337 | 0.7677033 | 0.118125572 | 0.127470123 |
| Pabpn1 | -0.069577546 | 0.025660276 | 0.3074274 | 0.237252577 | 0.27306998 |
| Pacc1 | -0.00698865 | 0.751360286 | 0.9506053 | 0.095343127 | 0.109227663 |
| Pacrg | 0.008078152 | 0.649110829 | 0.9173927 | 0.078799492 | 0.067802298 |
| Pacs1 | -0.025011616 | 0.216614524 | 0.6872330 | 0.073535435 | 0.098736683 |
| Pacs2 | -0.03815592 | 0.048117172 | 0.4147022 | 0.062903849 | 0.088646578 |
| Pacsin1 | 0.031202364 | 0.23411753 | 0.7056961 | 0.180704155 | 0.163826968 |
| Pacsin2 | -0.002578538 | 0.915217213 | 0.9845236 | 0.137668791 | 0.13997592 |
| Paf1 | -0.032000098 | 0.271150753 | 0.7342030 | 0.198078267 | 0.236863637 |
| Pafah1b1 | 0.004690734 | 0.894490865 | 0.9824361 | 1.142085756 | 1.138111395 |
| Pafah1b2 | 0.056724681 | 0.117278368 | 0.5739523 | 0.638377177 | 0.559189494 |
| Pafah1b3 | -0.003805395 | 0.910590252 | 0.9837373 | 0.410748103 | 0.388788256 |
| Pag1 | 0.003562019 | 0.812383361 | 0.9673015 | 0.041812389 | 0.04780863 |
| Paics | -0.023661582 | 0.455719612 | 0.8502914 | 0.320509967 | 0.31840852 |
| Paip1 | 0.041891015 | 0.201589826 | 0.6752150 | 0.394381462 | 0.315473815 |
| Paip2 | -0.021874553 | 0.479757901 | 0.8610816 | 1.296295586 | 1.220354963 |
| Paip2b | 0.021962245 | 0.441502754 | 0.8426454 | 0.211415883 | 0.20745735 |
| Pak1 | 0.051171802 | 0.00015482 | 0.0131265 | 0.063032221 | 0.025795566 |
| Pak1ip1 | -0.028215318 | 0.418649483 | 0.8300990 | 0.458983177 | 0.472505511 |
| Pak2 | -0.079809011 | 0.029352515 | 0.3282716 | 0.637008278 | 0.702733465 |

| | | | | | |
|---|---|---|---|---|---|
| Pak3 | -0.043703053 | 0.307402078 | 0.765611 | 1.092315546 | 1.023572946 |
| Pak4 | -0.053335412 | 0.022062779 | 0.2895010 | 0.096425084 | 0.136674609 |
| Pak5 | 0.023019799 | 0.160113848 | 0.6370337 | 0.075768095 | 0.043951167 |
| Pak6 | -0.01133762 | 0.542243292 | 0.8836953 | 0.072325325 | 0.071821181 |
| Palld | 0.003694208 | 0.787816173 | 0.9609168 | 0.044206608 | 0.035168301 |
| Pam | -0.108778789 | 6.16E-07 | 0.0001651 | 2.707836753 | 2.789293165 |
| Pam16 | 0.163091802 | 4.81E-06 | 0.0007953 | 0.915499417 | 0.697372936 |
| Pamr1 | 0.016057759 | 0.564676485 | 0.8935381 | 0.178030544 | 0.190765339 |
| Pan2 | -0.01942931 | 0.242830385 | 0.7132719 | 0.042326805 | 0.065246798 |
| Pan3 | -0.045543348 | 0.115806755 | 0.5724670 | 0.199677706 | 0.219232252 |
| Pank1 | 0.016711322 | 0.43374118 | 0.8377586 | 0.116645757 | 0.091893492 |
| Pank2 | -0.015616635 | 0.527756606 | 0.8768553 | 0.130016011 | 0.145673375 |
| Pank3 | -0.053389152 | 0.073528917 | 0.4918454 | 0.208627327 | 0.254433354 |
| Pank4 | -0.005695473 | 0.718661104 | 0.9440039 | 0.049084034 | 0.052593262 |
| Panx2 | -0.010868972 | 0.492812929 | 0.8669301 | 0.043916985 | 0.053205276 |
| Papola | 0.044932918 | 0.190864931 | 0.6700281 | 1.117708099 | 1.047660548 |
| Papolg | -0.016852945 | 0.309090954 | 0.7673813 | 0.050234792 | 0.064126366 |
| Pappa2 | -0.127476388 | 0.004681332 | 0.1365847 | 1.044590615 | 1.081051595 |
| Papss1 | 0.018012855 | 0.540678223 | 0.8835017 | 0.247024663 | 0.220835315 |
| Papss2 | 0.066364954 | 0.076106777 | 0.4986566 | 0.431284631 | 0.463293617 |
| Paqr9 | -0.010399403 | 0.525049115 | 0.8758385 | 0.050630321 | 0.057500267 |
| Pard3 | 0.01608691 | 0.470844095 | 0.8579671 | 0.110700403 | 0.113311163 |
| Pard3b | -0.028585776 | 0.107632792 | 0.5633723 | 0.045268407 | 0.078503738 |
| Pard6a | 0.010201601 | 0.741729672 | 0.9479080 | 0.282008036 | 0.272861314 |
| Parg | 0.030356225 | 0.261618238 | 0.7275735 | 0.204383698 | 0.168820668 |
| Park7 | 0.034928455 | 0.189161773 | 0.6680435 | 1.769792264 | 1.654956027 |
| Parl | 0.039869706 | 0.197709438 | 0.6730962 | 0.329025889 | 0.274322987 |
| Parm1 | 0.01539239 | 0.439317581 | 0.8405638 | 0.087921799 | 0.085714087 |
| Parn | 0.042593786 | 0.111500601 | 0.5675761 | 0.205266466 | 0.155824214 |
| Parp1 | -0.020770815 | 0.493712572 | 0.8674510 | 0.236312446 | 0.247375964 |
| Parp10 | 0.002137676 | 0.903429964 | 0.9827712 | 0.071403457 | 0.068058647 |
| Parp11 | 0.010669403 | 0.539866504 | 0.8830133 | 0.074350273 | 0.056193912 |
| Parp12 | -0.025487464 | 0.228945687 | 0.7019677 | 0.079301656 | 0.106322265 |
| Parp14 | 0.011253065 | 0.582048505 | 0.9018999 | 0.097876271 | 0.088657325 |
| Parp16 | -0.000402225 | 0.982657208 | 0.9976072 | 0.066733114 | 0.070476594 |
| Parp2 | 0.003558389 | 0.903614579 | 0.9827712 | 0.259395231 | 0.237165615 |
| Parp4 | -0.008183865 | 0.717679732 | 0.9435633 | 0.110961255 | 0.117213921 |
| Parp6 | -0.041791436 | 0.148862228 | 0.6264374 | 0.208763703 | 0.221398396 |
| Parp8 | 0.018523342 | 0.543546962 | 0.8842220 | 0.277926476 | 0.250836038 |
| Parp9 | -0.037214066 | 0.042510744 | 0.3952910 | 0.043906423 | 0.076787571 |
| Particl | -0.033246148 | 0.052466586 | 0.4319240 | 0.047001852 | 0.069716178 |
| Parva | 0.081012376 | 0.006873524 | 0.1654321 | 0.298029428 | 0.228719233 |
| Patj | 0.015788041 | 0.484756061 | 0.8629715 | 0.124062418 | 0.108853901 |
| Patl1 | -0.018822096 | 0.354903983 | 0.7969131 | 0.080852953 | 0.093799159 |
| Patz1 | 0.011325645 | 0.617173028 | 0.9115017 | 0.124984907 | 0.112655542 |
| Pawr | 0.004043419 | 0.899535313 | 0.9825441 | 0.297910409 | 0.263059118 |
| Pax6 | -0.007449646 | 0.823558168 | 0.9710592 | 1.195937609 | 1.24298727 |
| Pax6os1 | -0.024492012 | 0.133257095 | 0.5988251 | 0.035216668 | 0.06249943 |
| Paxbp1 | 0.031384688 | 0.311024912 | 0.7680774 | 0.271145042 | 0.237515639 |
| Paxip1 | -0.018126326 | 0.454807589 | 0.8499020 | 0.116710741 | 0.149063139 |
| Paxx | -0.019606962 | 0.52079545 | 0.8741795 | 0.277702991 | 0.258986316 |
| Pbdc1 | 0.019316349 | 0.562637145 | 0.8926240 | 0.402548089 | 0.358930742 |
| Pbrm1 | 0.011469868 | 0.763339023 | 0.9532128 | 0.83555513 | 0.826312 |
| Pbx1 | -0.093194402 | 0.01310858 | 0.2265455 | 0.578716262 | 0.598676107 |
| Pbx2 | 0.004402551 | 0.88507604 | 0.9803788 | 0.277367824 | 0.257478838 |
| Pbx3 | 0.002566043 | 0.925513896 | 0.9879210 | 0.165219956 | 0.188293974 |
| Pbxip1 | -0.122727146 | 0.001457763 | 0.0648475 | 0.453143498 | 0.571212062 |
| Pcbd1 | -0.046780822 | 0.114636548 | 0.5713707 | 1.448321674 | 1.370588305 |
| Pcbd2 | -0.05997585 | 0.092674804 | 0.5326385 | 0.472809937 | 0.502192959 |
| Pcbp1 | -0.000667223 | 0.983192729 | 0.9976072 | 1.290341782 | 1.210153466 |
| Pcbp2 | 0.026380504 | 0.364515789 | 0.8050031 | 1.508602402 | 1.495730322 |
| Pcbp3 | -0.008527764 | 0.713685212 | 0.9428150 | 0.114478916 | 0.136936037 |
| Pcbp4 | -0.044571419 | 0.092050652 | 0.5325584 | 0.176883647 | 0.173664357 |
| Pcca | -0.011207062 | 0.622835875 | 0.9121505 | 0.11610268 | 0.130317148 |
| Pccb | 0.01324163 | 0.625978167 | 0.9128095 | 0.190635067 | 0.167874524 |
| Pcdh1 | -0.040785813 | 0.113796693 | 0.5696977 | 0.132111501 | 0.165192026 |

| | | | | | |
|---|---|---|---|---|---|
| Pcdhga12 | -0.021187338 | 0.218839599 | 0.6901690 | 0.040548331 | 0.065828733 |
| Pced1a | -0.004369578 | 0.78965484 | 0.9618491 | 0.05591754 | 0.056503267 |
| Pcf11 | -0.01272918 | 0.710502529 | 0.9419370 | 0.3381498 | 0.328219045 |
| Pcgf1 | -0.014439649 | 0.406094365 | 0.8270770 | 0.05990281 | 0.068437759 |
| Pcgf2 | -0.060829833 | 0.010112369 | 0.1996282 | 0.088723885 | 0.143681663 |
| Pcgf3 | -0.012069806 | 0.606744595 | 0.9087139 | 0.124055904 | 0.128548998 |
| Pcgf5 | -0.057684047 | 0.084151213 | 0.5186471 | 0.325146111 | 0.354897079 |
| Pcgf6 | -0.005771801 | 0.841369276 | 0.9734589 | 0.212037569 | 0.208012844 |
| Pcid2 | -0.042036567 | 0.035996903 | 0.3599690 | 0.065358671 | 0.10161184 |
| Pcif1 | 0.01494096 | 0.680683595 | 0.9296958 | 0.593941056 | 0.570388271 |
| Pck2 | 0.043226179 | 0.260046134 | 0.7264594 | 0.658103016 | 0.679700627 |
| Pclo | -0.051360426 | 0.19253219 | 0.6700281 | 0.469158223 | 0.637444733 |
| Pcm1 | 0.051439639 | 0.177838729 | 0.6538187 | 0.794736182 | 0.732938544 |
| Pcmt1 | -0.111102202 | 0.00169577 | 0.0718890 | 0.644313124 | 0.694498513 |
| Pcmtd1 | -0.002012378 | 0.956511659 | 0.9955934 | 0.635487727 | 0.656633884 |
| Pcmtd2 | -0.020122237 | 0.521810935 | 0.8748598 | 0.295371841 | 0.288377738 |
| Pcna | 0.006319332 | 0.841084138 | 0.9734589 | 0.295998149 | 0.288069856 |
| Pcnp | -0.000534627 | 0.988261682 | 0.9976143 | 0.763089279 | 0.71544791 |
| Pcnt | -0.014761395 | 0.515074744 | 0.8733714 | 0.108543874 | 0.112745015 |
| Pcnx | -0.034225342 | 0.204571571 | 0.6781273 | 0.145975994 | 0.187767761 |
| Pcnx3 | -0.038175961 | 0.140152735 | 0.6130137 | 0.137550096 | 0.17431268 |
| Pcnx4 | 0.015281379 | 0.374097151 | 0.8087297 | 0.07256026 | 0.060670251 |
| Pcp4 | 0.086361499 | 0.006887449 | 0.1654321 | 0.242403439 | 0.201217921 |
| Pcp4l1 | -0.036470641 | 0.048264545 | 0.4148910 | 0.045237165 | 0.073552896 |
| Pcsk1 | 0.103910849 | 0.010629647 | 0.2027809 | 0.58710591 | 0.702877019 |
| Pcsk1n | -0.06246841 | 8.35E-07 | 0.0002071 | 4.873885543 | 4.836425518 |
| Pcsk2 | -0.061127719 | 6.62E-08 | 2.90E-05 | 4.362407678 | 4.346932911 |
| Pcsk2os1 | -0.050957835 | 0.137853226 | 0.6074257 | 0.379682708 | 0.390834986 |
| Pcsk4 | -0.00260698 | 0.866819418 | 0.9772002 | 0.048734252 | 0.050734025 |
| Pcsk6 | -0.038172467 | 0.043548535 | 0.3986436 | 0.052239706 | 0.093000233 |
| Pcsk7 | -0.007265596 | 0.724335377 | 0.9445652 | 0.090032211 | 0.092879597 |
| Pcx | -0.026516702 | 0.286124878 | 0.7459550 | 0.112231746 | 0.157753805 |
| Pcyox1 | 0.033556685 | 0.190505042 | 0.6700281 | 0.165098589 | 0.145702385 |
| Pcyox1l | 0.0313205 | 0.08845126 | 0.5284771 | 0.098749789 | 0.064700582 |
| Pcyt1a | -0.025879033 | 0.357070131 | 0.7985624 | 0.18017724 | 0.205896738 |
| Pcyt1b | 0.018563231 | 0.297345003 | 0.7550962 | 0.072501928 | 0.065476777 |
| Pcyt2 | 0.015458537 | 0.536307917 | 0.8800950 | 0.161874878 | 0.137074773 |
| Pdap1 | 0.045245562 | 0.1964458 | 0.6730962 | 1.179632478 | 1.098987697 |
| Pdcd10 | 0.024958371 | 0.478144827 | 0.8607489 | 0.670090185 | 0.595103819 |
| Pdcd11 | 0.004522937 | 0.869010355 | 0.9776398 | 0.178614519 | 0.190887514 |
| Pdcd2 | 0.018798658 | 0.494060425 | 0.8678106 | 0.210169628 | 0.18749871 |
| Pdcd2l | -0.002474105 | 0.91506216 | 0.9845236 | 0.135596968 | 0.127371757 |
| Pdcd4 | 0.06744514 | 0.064816933 | 0.4727823 | 1.118449347 | 0.988945392 |
| Pdcd5 | 0.104975809 | 0.003048108 | 0.1032885 | 0.855522197 | 0.67130347 |
| Pdcd6 | -0.070357846 | 0.043762757 | 0.3989378 | 0.704878562 | 0.744113608 |
| Pdcd6ip | 0.040065175 | 0.271102104 | 0.7342030 | 0.603888891 | 0.569486909 |
| Pdcd7 | 0.016131125 | 0.547779292 | 0.8863001 | 0.20155742 | 0.166793635 |
| Pdcl | 0.008470401 | 0.755514131 | 0.9521328 | 0.179793733 | 0.16956407 |
| Pdcl3 | 0.01072976 | 0.741682556 | 0.9479080 | 0.359205897 | 0.303295771 |
| Pde10a | -0.079696975 | 0.086896396 | 0.5250566 | 0.878844281 | 1.056591534 |
| Pde12 | -0.023969931 | 0.318412827 | 0.7710043 | 0.117697432 | 0.13223914 |
| Pde1c | 0.043186109 | 0.173508793 | 0.6527657 | 0.254937981 | 0.213510534 |
| Pde3b | -0.014510698 | 0.67089261 | 0.9247082 | 0.456581698 | 0.450923252 |
| Pde4b | 0.004411499 | 0.861306249 | 0.9769675 | 0.136070493 | 0.148780348 |
| Pde4d | -0.019825133 | 0.504964146 | 0.8695566 | 0.195696154 | 0.194790572 |
| Pde4dip | -0.001455133 | 0.966331154 | 0.9973993 | 0.365861506 | 0.39626793 |
| Pde5a | 0.038278278 | 0.229524824 | 0.7019677 | 0.28075341 | 0.325524913 |
| Pde6d | -0.046967406 | 0.115832441 | 0.5724670 | 0.227533791 | 0.261202925 |
| Pde7a | -0.029945144 | 0.143899701 | 0.6175769 | 0.083992798 | 0.094483058 |
| Pde8a | 0.007337097 | 0.817974816 | 0.9696833 | 0.292692277 | 0.288736472 |
| Pde8b | 0.007718448 | 0.650831603 | 0.9182548 | 0.068357777 | 0.058092097 |
| Pdgfa | -0.047854917 | 0.172133625 | 0.6520594 | 0.362550364 | 0.414610459 |
| Pdgfrl | -0.046858699 | 0.086838604 | 0.5250566 | 0.184593888 | 0.187246683 |
| Pdha1 | 0.020410692 | 0.541957358 | 0.8836720 | 0.352804407 | 0.327168466 |
| Pdhb | -0.035658223 | 0.297653932 | 0.7555596 | 0.420889782 | 0.426726184 |
| Pdhx | -0.003718388 | 0.834576469 | 0.9723494 | 0.064114299 | 0.069952848 |

| | | | | | |
|---|---|---|---|---|---|
| Pdia3 | 0.021078258 | 0.381895623 | 0.8126135 | 2.450390901 | 2.256497462 |
| Pdia4 | -0.062194128 | 0.125937102 | 0.5882766 | 0.723962335 | 0.734818368 |
| Pdia5 | -0.018910183 | 0.475946612 | 0.8607231 | 0.189423809 | 0.170147075 |
| Pdia6 | 0.034447401 | 0.293744272 | 0.7533684 | 1.904127726 | 1.722165961 |
| Pdik1l | 0.04323954 | 0.054585247 | 0.4421851 | 0.138893374 | 0.108818929 |
| Pdk1 | -0.02444008 | 0.259926289 | 0.7263475 | 0.091477378 | 0.106713846 |
| Pdk2 | 0.014090606 | 0.333519314 | 0.7804836 | 0.055572451 | 0.039169032 |
| Pdk3 | -0.01936422 | 0.328865337 | 0.7778857 | 0.075366199 | 0.09181345 |
| Pdk4 | -0.108273552 | 0.002574376 | 0.0927113 | 0.316905598 | 0.302470002 |
| Pdlim5 | 0.025170079 | 0.263808892 | 0.7285590 | 0.125152849 | 0.108641164 |
| Pdlim7 | 0.00070715 | 0.972556589 | 0.9976072 | 0.094394514 | 0.090143983 |
| Pdp2 | 0.014842246 | 0.386967679 | 0.8163635 | 0.076435184 | 0.057036864 |
| Pdpk1 | -0.000996351 | 0.97675811 | 0.9976072 | 0.406414238 | 0.422551908 |
| Pdrg1 | 0.024558279 | 0.49765037 | 0.8684629 | 0.481437962 | 0.452580066 |
| Pds5a | 0.007599993 | 0.813090799 | 0.9673015 | 0.310779269 | 0.305612996 |
| Pds5b | 0.003590794 | 0.911404628 | 0.9840154 | 0.304647582 | 0.296435432 |
| Pdss2 | -0.027578202 | 0.141595716 | 0.6144486 | 0.058665487 | 0.078819333 |
| Pdx1 | 0.037311394 | 0.316687314 | 0.7705987 | 0.402741234 | 0.474414714 |
| Pdxdc1 | 0.029447157 | 0.403492127 | 0.8264703 | 0.481510056 | 0.450763488 |
| Pdxk | 0.008863431 | 0.641505849 | 0.9166712 | 0.075357555 | 0.081449423 |
| Pdyn | -0.003161139 | 0.929178961 | 0.9881953 | 0.174288441 | 0.269408961 |
| Pdzd11 | -0.036957545 | 0.237962712 | 0.7104326 | 0.312792738 | 0.307993558 |
| Pdzd4 | -0.029417944 | 0.114332096 | 0.5705102 | 0.057356057 | 0.083503621 |
| Pdzd8 | -0.005158459 | 0.862639234 | 0.9769675 | 0.234334019 | 0.24537984 |
| Pdzk1ip1 | -0.000401001 | 0.990444878 | 0.9980836 | 0.38075389 | 0.381568553 |
| Pea15a | 0.007152183 | 0.813137837 | 0.9673015 | 0.236943808 | 0.261810159 |
| Pebp1 | -0.00162508 | 0.928246129 | 0.9880613 | 2.664593037 | 2.495440487 |
| Pecr | -0.004742764 | 0.84997073 | 0.9748841 | 0.153593188 | 0.147419102 |
| Pef1 | 0.00726979 | 0.796405305 | 0.9642793 | 0.230374304 | 0.20444746 |
| Peg10 | 0.013524288 | 0.629691841 | 0.9130564 | 0.159583017 | 0.107852943 |
| Peg13 | -0.031950994 | 0.161002278 | 0.6384844 | 0.082496036 | 0.12637781 |
| Peg3 | -0.068691848 | 0.015869575 | 0.2474560 | 2.551342823 | 2.666517598 |
| Peli1 | -0.000968378 | 0.968662462 | 0.9976072 | 0.134062558 | 0.143728619 |
| Pelp1 | -0.024719548 | 0.288002776 | 0.7477694 | 0.113793487 | 0.12891872 |
| Pemt | 0.022369205 | 0.137730009 | 0.6074257 | 0.059133426 | 0.044493138 |
| Pepd | -0.012026836 | 0.677233035 | 0.9280496 | 0.213351283 | 0.202936506 |
| Per1 | -0.021587017 | 0.396940062 | 0.8238469 | 0.143370655 | 0.156241528 |
| Per2 | 0.025148472 | 0.205163426 | 0.6793128 | 0.096120393 | 0.076677134 |
| Per3 | 0.003132357 | 0.899576294 | 0.9825441 | 0.147331764 | 0.128656644 |
| Perp | 0.011233023 | 0.536788693 | 0.8805926 | 0.074359875 | 0.06843095 |
| Pes1 | 0.028871443 | 0.228301227 | 0.7015153 | 0.153891756 | 0.121104163 |
| Pet100 | 0.000749638 | 0.982293708 | 0.9976072 | 0.359966457 | 0.36250466 |
| Pex1 | 0.008001686 | 0.644423359 | 0.9168391 | 0.073009818 | 0.060868908 |
| Pex10 | 0.010088889 | 0.572430004 | 0.8969368 | 0.081016485 | 0.063401146 |
| Pex11a | -0.007077676 | 0.775264515 | 0.9558538 | 0.138768309 | 0.149490432 |
| Pex11b | 0.007389086 | 0.78139387 | 0.9590847 | 0.206195617 | 0.183536494 |
| Pex11g | -0.035777825 | 0.089652142 | 0.5302360 | 0.070626773 | 0.110361102 |
| Pex12 | 0.030333765 | 0.143658363 | 0.6169224 | 0.114284613 | 0.083580893 |
| Pex13 | 0.033614698 | 0.188224882 | 0.6668854 | 0.175283375 | 0.146964611 |
| Pex14 | 0.002033949 | 0.936080267 | 0.9912468 | 0.174568152 | 0.155323059 |
| Pex16 | -0.014741836 | 0.598285598 | 0.9075513 | 0.20341893 | 0.202933906 |
| Pex19 | -0.00958104 | 0.710110617 | 0.9419370 | 0.169704161 | 0.161807499 |
| Pex2 | -0.041170738 | 0.188604816 | 0.6668854 | 0.258370346 | 0.28914511 |
| Pex26 | -0.007407748 | 0.700858695 | 0.9377499 | 0.079680655 | 0.082675412 |
| Pex3 | -0.024206402 | 0.400488359 | 0.8246612 | 0.200593756 | 0.219037883 |
| Pex5 | -0.01523546 | 0.525061707 | 0.8758385 | 0.129629554 | 0.134432663 |
| Pex5l | 0.038894341 | 0.201826092 | 0.6752150 | 0.247332887 | 0.213396966 |
| Pex6 | -0.022517834 | 0.41977911 | 0.8300990 | 0.209635447 | 0.203125975 |
| Pex7 | 0.00096024 | 0.978184942 | 0.9976072 | 0.40565318 | 0.391164598 |
| Pfas | 0.033124329 | 0.018926504 | 0.2682156 | 0.065800501 | 0.03232225 |
| Pfdn1 | 0.021373399 | 0.538337018 | 0.8816335 | 0.866092736 | 0.804424069 |
| Pfdn2 | 0.024291252 | 0.453966952 | 0.8490482 | 1.146689181 | 1.063182678 |
| Pfdn4 | 0.012346535 | 0.696870966 | 0.9364422 | 0.330944722 | 0.296148607 |
| Pfdn5 | 0.000175114 | 0.992302091 | 0.9985888 | 2.580141647 | 2.470190567 |
| Pfdn6 | 0.011484538 | 0.742093799 | 0.9479080 | 0.790237101 | 0.73297453 |
| Pfkfb2 | 0.018445599 | 0.350241848 | 0.7927303 | 0.094317601 | 0.081998374 |

| | | | | | |
|---|---|---|---|---|---|
| Pfkl | -0.020694364 | 0.481826176 | 0.86203068 | 0.22607914 | 0.210519896 |
| Pfkm | 0.04304178 | 0.043212701 | 0.39766629 | 0.132093154 | 0.090270292 |
| Pfkp | 0.021738352 | 0.406189929 | 0.82707700 | 0.167992317 | 0.144965701 |
| Pfn1 | 0.010891289 | 0.735859493 | 0.94683808 | 1.542581794 | 1.418667492 |
| Pfn2 | -0.049197036 | 0.094945052 | 0.53509361 | 0.227582301 | 0.236820805 |
| Pgam1 | 0.028585887 | 0.413981912 | 0.83009907 | 1.011145089 | 0.932032966 |
| Pgam5 | 0.006833297 | 0.770589666 | 0.95523334 | 0.12789618 | 0.120713797 |
| Pgap1 | -0.010207149 | 0.748115705 | 0.94974504 | 0.245028764 | 0.255618989 |
| Pgap2 | -0.030910521 | 0.21123951 | 0.68220598 | 0.134304044 | 0.146131625 |
| Pgap4 | -0.00485796 | 0.835981044 | 0.97234941 | 0.141106511 | 0.127245934 |
| Pgap6 | -0.027747507 | 0.191873848 | 0.67002811 | 0.078511176 | 0.107845967 |
| Pgbd5 | 0.004724697 | 0.834330653 | 0.97234941 | 0.11975868 | 0.115548043 |
| Pgd | 0.003993476 | 0.878828239 | 0.97855831 | 0.160718744 | 0.169002975 |
| Pgf | 0.018453053 | 0.353101961 | 0.79608441 | 0.089369636 | 0.077770426 |
| Pgghg | -0.026441542 | 0.21264067 | 0.68265224 | 0.090514628 | 0.105649429 |
| Pggt1b | -0.036959911 | 0.215579283 | 0.68521194 | 0.216085829 | 0.239204293 |
| Pgk1 | 0.120061086 | 0.000977436 | 0.04753019 | 0.664139832 | 0.541192056 |
| Pgls | -0.075337156 | 0.022198128 | 0.28982211 | 1.075439958 | 1.16470566 |
| Pgm1 | -0.031696165 | 0.260156428 | 0.72656299 | 0.17651397 | 0.193296615 |
| Pgm2 | 0.00368214 | 0.872986223 | 0.97843318 | 0.132254798 | 0.115343688 |
| Pgm3 | -0.064041907 | 0.054837781 | 0.44226891 | 0.331580858 | 0.368705358 |
| Pgp | 0.006966647 | 0.847792947 | 0.97440690 | 0.685488432 | 0.631762535 |
| Pgpep1 | -0.042647126 | 0.104794606 | 0.55800455 | 0.146231157 | 0.171590836 |
| Pgrmc1 | -0.115462292 | 0.001661486 | 0.07104281 | 1.040364467 | 1.088439493 |
| Pgrmc2 | 0.009497252 | 0.799222819 | 0.96470534 | 0.600130798 | 0.659207352 |
| Pgs1 | 0.007637705 | 0.740610281 | 0.94790800 | 0.131446619 | 0.11732641 |
| Phactr1 | 0.053158492 | 0.157237397 | 0.63226387 | 0.503809449 | 0.52207778 |
| Phactr2 | -0.019688267 | 0.154602356 | 0.63049349 | 0.024419289 | 0.046667621 |
| Phactr4 | -0.079081232 | 0.002553998 | 0.09271130 | 0.10932752 | 0.184916171 |
| Phax | 0.026696736 | 0.425585365 | 0.83253930 | 0.414248762 | 0.375772529 |
| Phb | -0.051933449 | 0.025817021 | 0.30880844 | 0.112740966 | 0.13556315 |
| Phb2 | -0.013641568 | 0.706348045 | 0.94038631 | 0.647833214 | 0.590680659 |
| Phc1 | 0.01493303 | 0.513901199 | 0.87333427 | 0.125989018 | 0.108985566 |
| Phc2 | -0.005436191 | 0.872783621 | 0.97843318 | 0.375830577 | 0.349465716 |
| Phc3 | -0.031542414 | 0.251108637 | 0.72072101 | 0.165942425 | 0.189307084 |
| Pheta1 | -0.014168905 | 0.344939016 | 0.78804400 | 0.043720449 | 0.049212491 |
| Phf1 | -0.005540032 | 0.851327413 | 0.97498221 | 0.235496987 | 0.231976355 |
| Phf10 | 0.051881877 | 0.175337399 | 0.65341424 | 0.506108318 | 0.477833635 |
| Phf12 | -0.061547895 | 0.05281006 | 0.43367201 | 0.233950176 | 0.282948092 |
| Phf13 | -0.003945058 | 0.874712603 | 0.97843318 | 0.138303921 | 0.137448835 |
| Phf14 | -0.05142903 | 0.140519668 | 0.61309530 | 0.385961198 | 0.422855016 |
| Phf2 | 0.022896705 | 0.421364348 | 0.83043379 | 0.216272114 | 0.206462996 |
| Phf20 | 0.006353654 | 0.837024823 | 0.97266190 | 0.289431296 | 0.255886937 |
| Phf20l1 | -0.005546176 | 0.881529158 | 0.97969630 | 0.52766208 | 0.562002571 |
| Phf21a | 0.002983872 | 0.923070692 | 0.98722250 | 0.292395578 | 0.277508795 |
| Phf21b | -0.008988788 | 0.684788461 | 0.93132731 | 0.096025155 | 0.112017091 |
| Phf23 | -0.017904457 | 0.525666131 | 0.87650434 | 0.201508202 | 0.199642562 |
| Phf3 | -0.127712487 | 0.000612969 | 0.03504548 | 0.498211404 | 0.585663427 |
| Phf5a | -0.040927087 | 0.244110905 | 0.71327199 | 0.558507046 | 0.586682932 |
| Phf6 | 0.025554128 | 0.276371597 | 0.73659490 | 0.154428735 | 0.122479577 |
| Phf7 | -0.004440216 | 0.829437374 | 0.97226010 | 0.09146095 | 0.092522684 |
| Phf8 | -0.016463989 | 0.535020348 | 0.87959491 | 0.161773297 | 0.166970739 |
| Phgdh | 0.089446849 | 0.027681239 | 0.31818990 | 0.467969154 | 0.441997647 |
| Phip | -0.097219729 | 0.013056055 | 0.22626311 | 0.749497039 | 0.833693465 |
| Phka1 | 0.003825097 | 0.83254523 | 0.97234941 | 0.074349051 | 0.066862663 |
| Phka2 | -0.006748984 | 0.696450663 | 0.93638620 | 0.060066678 | 0.064564588 |
| Phkb | 0.004176489 | 0.855543275 | 0.97574031 | 0.121674149 | 0.122540044 |
| Phkg2 | -0.0146407 | 0.558195744 | 0.89196227 | 0.15290994 | 0.15002935 |
| Phlda1 | -0.030820793 | 0.248731607 | 0.71874169 | 0.118218846 | 0.166321107 |
| Phlda3 | -0.019471096 | 0.42965405 | 0.83403655 | 0.102280297 | 0.114412054 |
| Phldb2 | -0.015131898 | 0.619706167 | 0.91215057 | 0.240145447 | 0.250119396 |
| Phlpp1 | -0.01386828 | 0.441161322 | 0.84257229 | 0.064035972 | 0.072390297 |
| Phlpp2 | -0.010284764 | 0.632121073 | 0.91355491 | 0.09693866 | 0.102703505 |
| Phospho2 | 0.034071246 | 0.241864321 | 0.71327199 | 0.251730924 | 0.204718807 |
| Phpt1 | 0.011976337 | 0.59457949 | 0.90658291 | 0.124559152 | 0.108829805 |
| Phrf1 | -0.020495826 | 0.437920956 | 0.83999140 | 0.165970549 | 0.171929009 |

| | | | | | |
|---|---|---|---|---|---|
| Phtf1 | -0.008753078 | 0.725259422 | 0.9450275 | 0.163699794 | 0.145289579 |
| Phyh | 0.005859984 | 0.869811137 | 0.9777883 | 0.771504789 | 0.728520759 |
| Phyhipl | 0.002894055 | 0.920348526 | 0.9858392 | 0.217027063 | 0.201928357 |
| Phykpl | -0.021325815 | 0.247941656 | 0.7177068 | 0.058327747 | 0.078358734 |
| Pi4k2a | 0.027349161 | 0.398542658 | 0.8239850 | 0.337539633 | 0.317772747 |
| Pi4k2b | -0.017910192 | 0.42999586 | 0.8340944 | 0.108515392 | 0.124392825 |
| Pi4ka | -0.015793693 | 0.623133869 | 0.9121505 | 0.317603322 | 0.312881795 |
| Pi4kb | -0.028157121 | 0.195922715 | 0.6727425 | 0.088730291 | 0.115222946 |
| Pias1 | 0.038773573 | 0.213147886 | 0.6827572 | 0.305594972 | 0.243567996 |
| Pias2 | -0.019525635 | 0.428385537 | 0.8340365 | 0.133968604 | 0.150434254 |
| Pias3 | 0.012233905 | 0.546757085 | 0.8858459 | 0.107002274 | 0.082700196 |
| Pias4 | -0.063506163 | 0.0113047 | 0.2096123 | 0.105766349 | 0.160061353 |
| Pibf1 | -0.029581577 | 0.200135864 | 0.6738305 | 0.11024834 | 0.129380341 |
| Picalm | 0.016626641 | 0.629335689 | 0.9130564 | 0.423576607 | 0.393829723 |
| Pick1 | -0.014229567 | 0.61981073 | 0.9121505 | 0.239101807 | 0.218923754 |
| Pid1 | -0.024872098 | 0.198618745 | 0.6730962 | 0.066796145 | 0.086459705 |
| Pigb | -0.00111231 | 0.941578181 | 0.9929301 | 0.046841739 | 0.04869964 |
| Pigbos1 | 0.027180391 | 0.436658373 | 0.8391598 | 0.448007445 | 0.415736731 |
| Pigc | 0.01186306 | 0.517757049 | 0.8738804 | 0.086310925 | 0.069194894 |
| Pigf | 0.003787435 | 0.879954942 | 0.9792337 | 0.168919299 | 0.149877517 |
| Pigg | -0.024386284 | 0.127067107 | 0.5893647 | 0.038820924 | 0.059621023 |
| Pigh | 0.007033071 | 0.711704075 | 0.9419834 | 0.079432336 | 0.076446397 |
| Pigk | -0.032206716 | 0.320295288 | 0.7719458 | 0.353219251 | 0.35259018 |
| Pigm | -0.006513218 | 0.728557847 | 0.9458355 | 0.074141539 | 0.074727761 |
| Pign | -0.004633438 | 0.77505084 | 0.9558538 | 0.05130246 | 0.053517174 |
| Pigo | 0.021158596 | 0.13303358 | 0.5988251 | 0.055665379 | 0.035147538 |
| Pigp | -0.001307175 | 0.97072109 | 0.9976072 | 0.831944768 | 0.80585347 |
| Pigq | -0.013997925 | 0.609240928 | 0.9088672 | 0.174783692 | 0.193442176 |
| Pigs | 0.000500225 | 0.985047172 | 0.9976072 | 0.166018832 | 0.165246568 |
| Pigt | -0.059702953 | 0.042023821 | 0.3943957 | 0.215686692 | 0.244095697 |
| Pigu | 0.022888458 | 0.345504061 | 0.7886332 | 0.157253891 | 0.13370365 |
| Pigv | 0.013946 | 0.413213166 | 0.8300990 | 0.069820088 | 0.058024358 |
| Pigx | -0.048020816 | 0.122666106 | 0.5833237 | 0.274818389 | 0.290109651 |
| Pigyl | 0.037710794 | 0.274681052 | 0.7351955 | 0.527762702 | 0.45273239 |
| Pih1d1 | -0.002313423 | 0.916341381 | 0.9845236 | 0.115328268 | 0.106745609 |
| Pik3c2a | -0.016466411 | 0.615816199 | 0.9112316 | 0.280748519 | 0.306133387 |
| Pik3c2g | -0.0577042 | 0.005431643 | 0.1488982 | 0.052486477 | 0.095390719 |
| Pik3c3 | -0.00108933 | 0.959060243 | 0.9957785 | 0.101918743 | 0.102815246 |
| Pik3ca | -0.029219946 | 0.299444163 | 0.7583574 | 0.188622066 | 0.202647401 |
| Pik3ip1 | -0.011965745 | 0.481932792 | 0.8620306 | 0.062326207 | 0.059216691 |
| Pik3r1 | -0.03994266 | 0.220940447 | 0.6926204 | 0.245747136 | 0.325695512 |
| Pik3r2 | -0.008306528 | 0.696065044 | 0.9363862 | 0.096044854 | 0.103218385 |
| Pik3r3 | 0.008114597 | 0.705428533 | 0.9401381 | 0.103851175 | 0.103043113 |
| Pik3r4 | -0.020957984 | 0.332822828 | 0.7801518 | 0.087980521 | 0.113774656 |
| Pikfyve | 0.003276394 | 0.90497302 | 0.9831707 | 0.185827336 | 0.192267631 |
| Pim1 | -0.031452 | 0.143935264 | 0.6175769 | 0.08700077 | 0.106022958 |
| Pim2 | 0.041291948 | 0.163822722 | 0.6410026 | 0.225714415 | 0.189680027 |
| Pim3 | -0.054021722 | 0.091443551 | 0.5323474 | 0.243736931 | 0.285181958 |
| Pin1 | -0.028319913 | 0.414302312 | 0.8300990 | 0.674139422 | 0.662985075 |
| Pin4 | -0.006452705 | 0.849056901 | 0.9748841 | 0.403958143 | 0.403666492 |
| Pink1 | -0.00126505 | 0.969278167 | 0.9976072 | 0.384298022 | 0.345086918 |
| Pinx1 | -0.013746379 | 0.438163422 | 0.8402437 | 0.061439306 | 0.06573602 |
| Pip4k2a | -0.002168317 | 0.902846781 | 0.9825441 | 0.066110324 | 0.069242892 |
| Pip4k2b | 0.007176601 | 0.647390258 | 0.9169695 | 0.053764507 | 0.052778836 |
| Pip4k2c | 0.008516509 | 0.785197583 | 0.9602516 | 0.313109014 | 0.293997172 |
| Pip4p1 | 0.003200506 | 0.921737552 | 0.9862621 | 0.358944012 | 0.327994646 |
| Pip4p2 | 0.017455244 | 0.520640511 | 0.8741795 | 0.194518147 | 0.171978102 |
| Pip5k1a | 0.033617011 | 0.289645201 | 0.7496864 | 0.323959582 | 0.289114895 |
| Pip5k1b | -0.048428156 | 0.115719361 | 0.5724670 | 0.195233807 | 0.289483676 |
| Pip5k1c | 0.008084364 | 0.741579885 | 0.9479080 | 0.140128523 | 0.137970804 |
| Pir | -0.018024084 | 0.228072643 | 0.7015153 | 0.031541892 | 0.052802702 |
| Pisd | 0.001842798 | 0.93516938 | 0.9908021 | 0.114470702 | 0.11891524 |
| Pithd1 | 0.022167404 | 0.546297035 | 0.8858413 | 0.677362775 | 0.607264223 |
| Pitpna | -0.026049025 | 0.427414457 | 0.8340365 | 0.351558601 | 0.355422629 |
| Pitpnb | 0.050413534 | 0.152416759 | 0.6281105 | 0.556795763 | 0.486236611 |
| Pitpnc1 | -0.077125359 | 0.028326915 | 0.3222511 | 0.340465028 | 0.392185696 |

| | | | | | |
|---|---|---|---|---|---|
| Pitpnm1 | 0.007554978 | 0.776986056 | 0.9564524 | 0.166091588 | 0.161510656 |
| Pitpnm2 | -0.025953785 | 0.411356062 | 0.8290638 | 0.285206674 | 0.276922027 |
| Pitpnm3 | 0.010556323 | 0.588601119 | 0.9044624 | 0.080997775 | 0.079580989 |
| Pitrm1 | -0.024195698 | 0.305819612 | 0.7649345 | 0.115005635 | 0.1356162 |
| Pja1 | 0.002105581 | 0.948923174 | 0.9947498 | 0.386351839 | 0.351461198 |
| Pja2 | -0.002599321 | 0.943749002 | 0.9935293 | 0.839347292 | 0.798843116 |
| Pkd1 | -0.021993423 | 0.486547968 | 0.8637358 | 0.270404578 | 0.297385528 |
| Pkd2 | 0.003970769 | 0.888894348 | 0.9816521 | 0.198725776 | 0.204885907 |
| Pkdcc | -0.002097111 | 0.954630228 | 0.9954690 | 0.598339389 | 0.579608918 |
| Pkia | -0.035940832 | 0.100423949 | 0.5473657 | 0.080528013 | 0.112042884 |
| Pkib | 0.023961638 | 0.328771231 | 0.7778857 | 0.141893128 | 0.139709071 |
| Pkig | 0.032383933 | 0.340218994 | 0.7844658 | 0.470156001 | 0.413738136 |
| Pkm | 0.032941554 | 0.271500565 | 0.7342030 | 1.513708951 | 1.436917985 |
| Pkn1 | -0.004881789 | 0.874098979 | 0.9784331 | 0.25140328 | 0.25838114 |
| Pkn2 | -0.000769123 | 0.983565297 | 0.9976072 | 0.633790372 | 0.649576495 |
| Pknox1 | -0.028427488 | 0.269865769 | 0.7340360 | 0.121189969 | 0.151576717 |
| Pkp2 | -0.002284108 | 0.891883585 | 0.9819597 | 0.058969791 | 0.057644741 |
| Pkp3 | 0.002280721 | 0.876988062 | 0.9785583 | 0.045328661 | 0.047172206 |
| Pkp4 | 0.010786917 | 0.75290374 | 0.9510766 | 0.384511177 | 0.392900655 |
| Pla2g12a | -0.067622464 | 0.072026704 | 0.4888240 | 0.942389623 | 0.959861668 |
| Pla2g15 | 0.00175835 | 0.924319934 | 0.9878036 | 0.068471678 | 0.073950523 |
| Pla2g2d | -0.028712211 | 0.085980518 | 0.5232489 | 0.047565847 | 0.062132855 |
| Pla2g2f | 0.00431061 | 0.894405809 | 0.9824361 | 0.306096166 | 0.281205203 |
| Pla2g6 | -0.034347399 | 0.202200414 | 0.6755904 | 0.177763254 | 0.188679266 |
| Plaa | 0.016501867 | 0.566487453 | 0.8949556 | 0.221194958 | 0.211798447 |
| Plaat3 | -0.077878869 | 0.026347706 | 0.3104147 | 0.463460282 | 0.449339836 |
| Plag1 | -0.030471243 | 0.097199975 | 0.5383862 | 0.060083498 | 0.077357434 |
| Plagl2 | 0.013517778 | 0.483181956 | 0.8620306 | 0.086924258 | 0.07845609 |
| Plbd2 | 0.024579811 | 0.130272149 | 0.5939711 | 0.065393798 | 0.05006957 |
| Plcb1 | 0.008973926 | 0.716026705 | 0.9431411 | 0.138634332 | 0.133607637 |
| Plcb4 | -0.008446037 | 0.770124223 | 0.9552333 | 0.1936558 | 0.198118118 |
| Plcg1 | -0.0229153 | 0.23017976 | 0.7019677 | 0.063240409 | 0.085335431 |
| Plcl2 | 0.031702798 | 0.179416882 | 0.6550070 | 0.144330237 | 0.117120467 |
| Plcxd2 | -0.017372466 | 0.512914021 | 0.8724463 | 0.150975787 | 0.167878484 |
| Plcxd3 | -0.024516121 | 0.316805016 | 0.7705987 | 0.121590792 | 0.139026999 |
| Pld1 | -0.021387088 | 0.190111512 | 0.6699489 | 0.042393846 | 0.059550334 |
| Pld3 | -0.005377697 | 0.887331816 | 0.9812414 | 0.513954648 | 0.496395783 |
| Plec | 0.006701082 | 0.804539032 | 0.9647143 | 0.17792814 | 0.169815945 |
| Plekha1 | -0.057246394 | 0.121891056 | 0.5819916 | 0.44002309 | 0.468066129 |
| Plekha2 | 0.001456661 | 0.943165402 | 0.9935293 | 0.085666932 | 0.089485155 |
| Plekha3 | -0.04388994 | 0.108135058 | 0.5636817 | 0.155720049 | 0.200445861 |
| Plekha5 | -0.00844166 | 0.759585888 | 0.9521533 | 0.191731121 | 0.186465792 |
| Plekha6 | 0.017568388 | 0.562385446 | 0.8926240 | 0.265149158 | 0.256824112 |
| Plekha8 | 0.005336492 | 0.801941754 | 0.9647143 | 0.107775956 | 0.098394764 |
| Plekhb1 | -0.07363318 | 0.000717472 | 0.0388924 | 0.074341091 | 0.122172029 |
| Plekhb2 | 0.016670921 | 0.582881598 | 0.9019790 | 0.262258341 | 0.240075982 |
| Plekhf2 | 0.00237586 | 0.913797814 | 0.9845236 | 0.112117675 | 0.103552565 |
| Plekhg3 | -0.03238275 | 0.158349869 | 0.6341666 | 0.10402093 | 0.125679195 |
| Plekhh2 | -0.013227935 | 0.54042019 | 0.8834819 | 0.095484604 | 0.100915209 |
| Plekhh3 | -0.040127778 | 0.083605682 | 0.5183040 | 0.111753542 | 0.132422904 |
| Plekhj1 | -0.036641423 | 0.296446546 | 0.7545964 | 0.571448615 | 0.549363023 |
| Plekhm1 | -0.016130203 | 0.310323966 | 0.7677033 | 0.04459741 | 0.057676559 |
| Plekhm2 | -0.034386785 | 0.116548403 | 0.5727304 | 0.090240196 | 0.115052441 |
| Plekhm3 | -0.018935459 | 0.199793653 | 0.6734464 | 0.036357167 | 0.051120978 |
| Plekho1 | 0.002343785 | 0.873631842 | 0.9784331 | 0.048395984 | 0.041158876 |
| Plgrkt | -0.033565445 | 0.311919741 | 0.7684379 | 0.288868955 | 0.323205201 |
| Plin2 | -0.035010691 | 0.104598653 | 0.5580045 | 0.091133263 | 0.113497317 |
| Plin3 | 0.023786076 | 0.328541296 | 0.7778857 | 0.166703647 | 0.12188706 |
| Plk2 | -0.020955739 | 0.452014152 | 0.8484352 | 0.142796934 | 0.169961048 |
| Plk3 | -0.026864845 | 0.270759471 | 0.7342030 | 0.092371892 | 0.153908169 |
| Plod1 | 0.002028051 | 0.901406075 | 0.9825441 | 0.059526238 | 0.05713688 |
| Plod3 | -0.023876675 | 0.388183677 | 0.8170287 | 0.19244798 | 0.18657451 |
| Plp2 | 0.044775317 | 0.067759159 | 0.4817235 | 0.18548821 | 0.126447905 |
| Plpbp | -0.051775671 | 0.1421866 | 0.6152737 | 0.389429147 | 0.429636089 |
| Plpp2 | 0.023910996 | 0.164294951 | 0.6415521 | 0.072319253 | 0.058259304 |
| Plpp5 | -0.064509238 | 0.052386562 | 0.4319240 | 0.28683577 | 0.383498153 |

| | | | | | |
|---|---|---|---|---|---|
| Plpp6 | -0.010545562 | 0.667917217 | 0.9230765 | 0.137941493 | 0.145739962 |
| Plppr2 | -0.034889779 | 0.039950048 | 0.3834656 | 0.033422115 | 0.070675975 |
| Plppr4 | -0.00733018 | 0.630375463 | 0.9131607 | 0.033998641 | 0.055779312 |
| Plrg1 | -0.001351907 | 0.954415854 | 0.9954690 | 0.142299586 | 0.12468241 |
| Pls1 | -6.83E-05 | 0.997230868 | 0.9994073 | 0.090099446 | 0.081033291 |
| Pls3 | -0.044049934 | 0.073228413 | 0.4912860 | 0.116324045 | 0.145066502 |
| Plscr1 | 0.004634603 | 0.845424133 | 0.9734889 | 0.143682751 | 0.127535634 |
| Plscr3 | 0.015288914 | 0.48825744 | 0.8642971 | 0.125241627 | 0.104434253 |
| Plut | -0.001995354 | 0.914486673 | 0.9845236 | 0.048757187 | 0.069652869 |
| Plxna2 | -0.017094072 | 0.534199424 | 0.8795652 | 0.170687836 | 0.188100475 |
| Plxnb1 | 0.035864069 | 0.081380973 | 0.5135491 | 0.118114777 | 0.080385657 |
| Plxnb2 | 0.028213283 | 0.428547899 | 0.8340365 | 0.421317859 | 0.43905785 |
| Pmepa1 | 0.019089127 | 0.42973091 | 0.8340365 | 0.117753534 | 0.127441317 |
| Pml | -0.008382412 | 0.600986542 | 0.9078437 | 0.048081548 | 0.054231857 |
| Pmm1 | -0.049003793 | 0.070922677 | 0.4879008 | 0.138556017 | 0.191496703 |
| Pmm2 | -0.028810828 | 0.232193977 | 0.7033173 | 0.118896194 | 0.143902138 |
| Pmpca | 0.019004751 | 0.491778066 | 0.8664619 | 0.21057365 | 0.178156885 |
| Pmpcb | -0.021743589 | 0.506265878 | 0.8695566 | 0.391456379 | 0.380599554 |
| Pms2 | -0.003921053 | 0.831517334 | 0.9723494 | 0.072582894 | 0.06937881 |
| Pmvk | -0.033424494 | 0.395210874 | 0.8224233 | 0.500458966 | 0.582606842 |
| Pnck | -0.0194281 | 0.308796118 | 0.7669648 | 0.077131361 | 0.078399552 |
| Pnisr | -0.074986885 | 0.059713104 | 0.4567108 | 0.620983026 | 0.682000349 |
| Pnkd | 0.051555283 | 0.134493651 | 0.6009806 | 0.56034759 | 0.491084942 |
| Pnkp | -0.027733547 | 0.196611921 | 0.6730962 | 0.085908928 | 0.111411204 |
| Pnmal1 | -0.104054477 | 0.001625497 | 0.0701084 | 0.281329206 | 0.368302038 |
| Pnmal2 | -0.099534817 | 0.007540685 | 0.1721437 | 0.889885454 | 0.94270242 |
| Pnn | -0.089151991 | 0.023976596 | 0.3005057 | 0.72625783 | 0.790616558 |
| Pno1 | 0.010383486 | 0.722635721 | 0.9445652 | 0.247027344 | 0.216702533 |
| Pnoc | 0.027860357 | 0.219834945 | 0.6910237 | 0.1102823 | 0.070428598 |
| Pnp | -0.037658493 | 0.212612478 | 0.6826522 | 0.242587218 | 0.253067336 |
| Pnpla2 | 0.044742168 | 0.125527573 | 0.5873743 | 0.269742423 | 0.220935387 |
| Pnpla6 | -0.009249564 | 0.607294736 | 0.9087139 | 0.065165565 | 0.071684129 |
| Pnpla8 | 0.008841811 | 0.799967443 | 0.9647053 | 0.570598833 | 0.542602265 |
| Pnpo | -0.051567758 | 0.089982287 | 0.5306922 | 0.207682286 | 0.2492043 |
| Pnpt1 | 0.019468153 | 0.293543146 | 0.7533684 | 0.093855458 | 0.067301296 |
| Pnrc1 | -0.041748016 | 0.245823495 | 0.7152900 | 1.146465307 | 1.114585412 |
| Pnrc2 | -0.019545853 | 0.548422626 | 0.8867730 | 0.32291126 | 0.348140573 |
| Poc1b | 0.008059927 | 0.726180875 | 0.9450680 | 0.123634223 | 0.116143208 |
| Poc5 | 0.018785105 | 0.254273583 | 0.7223071 | 0.068952552 | 0.053993476 |
| Podxl2 | 0.013192514 | 0.626238185 | 0.9128095 | 0.176142313 | 0.173754902 |
| Pofut1 | -0.028172026 | 0.140009584 | 0.6129281 | 0.063645267 | 0.084027302 |
| Pofut2 | 0.103151667 | 0.002839567 | 0.0988368 | 0.865110126 | 0.823489919 |
| Pogk | 0.010189434 | 0.638701003 | 0.9160042 | 0.106118608 | 0.101248908 |
| Poglut1 | -0.017006281 | 0.489473102 | 0.8651997 | 0.13059413 | 0.142418176 |
| Poglut3 | -0.010650323 | 0.490886661 | 0.8658154 | 0.045228939 | 0.05137723 |
| Pogz | -0.030261297 | 0.205928844 | 0.6797565 | 0.108871998 | 0.136605489 |
| Pola1 | -0.024361547 | 0.095886855 | 0.5368118 | 0.024995527 | 0.050613214 |
| Polb | 0.012610817 | 0.68829229 | 0.9329669 | 0.302526879 | 0.267819559 |
| Pold2 | -0.017053555 | 0.412585032 | 0.8297979 | 0.092167585 | 0.095837472 |
| Pold3 | -0.006796149 | 0.794933941 | 0.9630855 | 0.166244405 | 0.166060223 |
| Pold4 | -0.006998347 | 0.844553199 | 0.9734889 | 0.473721351 | 0.453894429 |
| Poldip2 | 0.04609475 | 0.116375977 | 0.5724670 | 0.257057849 | 0.215285957 |
| Poldip3 | -0.028219356 | 0.362010755 | 0.8030292 | 0.251521682 | 0.263916905 |
| Pole3 | -0.032415102 | 0.195394908 | 0.6719932 | 0.138816548 | 0.15303505 |
| Pole4 | 0.034201582 | 0.134381323 | 0.6007493 | 0.139934846 | 0.104420212 |
| Polg | 0.049227948 | 0.023123634 | 0.2949903 | 0.136016933 | 0.088405639 |
| Polg2 | -0.00426115 | 0.770127977 | 0.9552333 | 0.043262779 | 0.043705461 |
| Polh | 0.013450383 | 0.265828433 | 0.7306783 | 0.043969306 | 0.02382314 |
| Polk | 0.011349035 | 0.52613281 | 0.8768553 | 0.078363707 | 0.065307681 |
| Poll | 0.021364934 | 0.119299803 | 0.5782737 | 0.0487572 | 0.034961593 |
| Polr1a | 0.030578582 | 0.148092256 | 0.6255679 | 0.116730469 | 0.090717225 |
| Polr1c | -0.034988853 | 0.206247017 | 0.6797565 | 0.175450464 | 0.194239801 |
| Polr1d | 0.05882034 | 0.089619693 | 0.5302360 | 1.022990716 | 0.892565338 |
| Polr1f | -0.024439331 | 0.413384377 | 0.8300990 | 0.235193135 | 0.240049675 |
| Polr1h | -0.002362735 | 0.946406292 | 0.9945018 | 0.499128194 | 0.499646834 |
| Polr1has | -0.001445537 | 0.949551781 | 0.9948462 | 0.12412827 | 0.118851601 |

| | | | | | |
|---|---|---|---|---|---|
| Polr2a | -0.029176816 | 0.367823148 | 0.8056719 | 0.270169232 | 0.308481121 |
| Polr2b | -0.056138246 | 0.060140456 | 0.4581503 | 0.226441835 | 0.254524011 |
| Polr2c | -0.017714611 | 0.624264218 | 0.9121505 | 0.60518829 | 0.594186456 |
| Polr2d | -0.023405543 | 0.388588908 | 0.8170287 | 0.196553172 | 0.185847843 |
| Polr2e | -0.040685941 | 0.253636012 | 0.7223071 | 0.711744531 | 0.691980939 |
| Polr2f | 0.019908568 | 0.574439845 | 0.8975339 | 0.843791805 | 0.800985326 |
| Polr2g | -0.06351854 | 0.077311309 | 0.5022647 | 0.757559832 | 0.785021046 |
| Polr2h | -0.041870856 | 0.183743508 | 0.6617383 | 0.286862336 | 0.285092756 |
| Polr2i | 0.093315496 | 0.008013887 | 0.1762699 | 0.643377903 | 0.555515003 |
| Polr2j | 0.011483887 | 0.745935066 | 0.9485717 | 0.587624705 | 0.565276917 |
| Polr2k | 0.018319621 | 0.614089717 | 0.9108386 | 0.797394103 | 0.745545454 |
| Polr2l | 0.050683749 | 0.049697473 | 0.4195735 | 0.197016473 | 0.150135358 |
| Polr2m | -0.00751506 | 0.833807163 | 0.9723494 | 0.576030086 | 0.55865644 |
| Polr3a | -0.021930506 | 0.262831874 | 0.7275735 | 0.065767644 | 0.086820055 |
| Polr3c | 0.056620522 | 0.007258668 | 0.1690281 | 0.136012622 | 0.083102204 |
| Polr3d | 0.080093224 | 0.017808413 | 0.2587928 | 0.43390131 | 0.334806363 |
| Polr3e | -0.020334183 | 0.239417165 | 0.7117225 | 0.050683432 | 0.067742819 |
| Polr3f | -0.04379937 | 0.057319775 | 0.4491407 | 0.099733859 | 0.123950574 |
| Polr3g | -0.013596917 | 0.340707144 | 0.7849082 | 0.03300873 | 0.047363055 |
| Polr3gl | -0.009300117 | 0.759741473 | 0.9521533 | 0.282273095 | 0.259547215 |
| Polr3h | -0.028535552 | 0.211765883 | 0.6822059 | 0.105158887 | 0.126077847 |
| Polr3k | 0.003033654 | 0.92813653 | 0.9880613 | 0.373695051 | 0.347305503 |
| Pom121 | -0.034991582 | 0.272899092 | 0.7346428 | 0.289433251 | 0.324813704 |
| Pomc | 0.00849093 | 0.755798013 | 0.9521533 | 0.175443366 | 0.190049985 |
| Pomgnt1 | -0.010215729 | 0.696524378 | 0.9363862 | 0.17031825 | 0.166709564 |
| Pomgnt2 | -0.001201996 | 0.962368858 | 0.9964199 | 0.171938074 | 0.148648384 |
| Pomp | 0.094277183 | 0.003420373 | 0.1119805 | 1.476866448 | 1.238485539 |
| Pomt2 | -0.030537078 | 0.155685079 | 0.6312943 | 0.09113351 | 0.110294351 |
| Pon2 | -0.003742891 | 0.884300529 | 0.9802800 | 0.144939946 | 0.152239441 |
| Pon3 | -0.050831166 | 0.026851807 | 0.3138777 | 0.091490892 | 0.136302937 |
| Pop4 | -0.021581921 | 0.195848963 | 0.6727425 | 0.047964704 | 0.062537861 |
| Pop5 | 0.056592495 | 0.11628538 | 0.5724670 | 0.834827775 | 0.734578193 |
| Pop7 | -0.012906574 | 0.691459007 | 0.9351429 | 0.374171359 | 0.348508155 |
| Por | -0.016805535 | 0.632675171 | 0.9139562 | 0.42370576 | 0.496966687 |
| Pot1a | -0.030423326 | 0.071684747 | 0.4888240 | 0.04125723 | 0.065472808 |
| Pot1b | -0.021037102 | 0.174775181 | 0.6534142 | 0.045859933 | 0.051202357 |
| Pou2f1 | -0.02082935 | 0.404990662 | 0.8270770 | 0.135280795 | 0.153706285 |
| Pou3f4 | -0.005494085 | 0.772269464 | 0.9552333 | 0.083473192 | 0.064459984 |
| Pou6f2 | -0.023443487 | 0.382910242 | 0.8129787 | 0.179778946 | 0.1542634 |
| Ppa1 | -0.022621507 | 0.554122248 | 0.8902995 | 0.698201968 | 0.640777192 |
| Ppa2 | -0.048460757 | 0.0997136 | 0.5446910 | 0.241243654 | 0.257636601 |
| Ppan | -0.037484414 | 0.18447838 | 0.6628862 | 0.190455549 | 0.198315875 |
| Ppard | -0.006287387 | 0.790335945 | 0.9621438 | 0.125741426 | 0.121219283 |
| Pparg | -0.000202476 | 0.989728577 | 0.9980836 | 0.043652318 | 0.059071552 |
| Ppat | -0.007321623 | 0.838324742 | 0.9727216 | 0.360008072 | 0.384675737 |
| Ppcdc | -0.006293439 | 0.750398595 | 0.9504537 | 0.088789528 | 0.085698014 |
| Ppcs | -0.012361123 | 0.635844457 | 0.9153541 | 0.166114815 | 0.176227116 |
| Ppdpf | -0.057628936 | 0.109228686 | 0.5649366 | 0.535663737 | 0.510047629 |
| Ppfia1 | -0.009306589 | 0.782092978 | 0.9590847 | 0.338594099 | 0.372111418 |
| Ppfia3 | 0.004288347 | 0.80140738 | 0.9647143 | 0.06555014 | 0.058338887 |
| Ppfibp1 | -0.057738576 | 0.085025166 | 0.5209234 | 0.297146808 | 0.332301019 |
| Pphln1 | 0.018156051 | 0.456013247 | 0.8502914 | 0.151006108 | 0.124954235 |
| Ppia | 0.023171734 | 0.288026919 | 0.7477694 | 2.293122304 | 2.158522382 |
| Ppib | 0.005136702 | 0.889255973 | 0.9819033 | 1.168224724 | 1.070117552 |
| Ppid | -0.043060243 | 0.182291285 | 0.6595419 | 0.30977413 | 0.319325995 |
| Ppie | -0.006857478 | 0.812203606 | 0.9673015 | 0.224498973 | 0.220010944 |
| Ppif | 0.0005542 | 0.977692368 | 0.9976072 | 0.097742875 | 0.08324517 |
| Ppig | 0.014525223 | 0.679504799 | 0.9289200 | 1.02756944 | 0.979060156 |
| Ppih | -0.021814113 | 0.276146395 | 0.7363903 | 0.08679455 | 0.092016232 |
| Ppil1 | -0.062567847 | 0.004805228 | 0.1373713 | 0.090126487 | 0.124318958 |
| Ppil2 | -0.042331031 | 0.154876782 | 0.6304934 | 0.234424152 | 0.246669484 |
| Ppil3 | -0.014139855 | 0.534602212 | 0.8795949 | 0.131925515 | 0.119645659 |
| Ppil4 | 0.014122812 | 0.661719299 | 0.9220754 | 0.326837081 | 0.295101881 |
| Ppip5k1 | -0.042394327 | 0.08452444 | 0.5191841 | 0.117004402 | 0.140812744 |
| Ppip5k2 | -0.026925815 | 0.293561345 | 0.7533684 | 0.132368802 | 0.163295471 |
| Ppl | -0.015862798 | 0.396802261 | 0.8238469 | 0.066331917 | 0.070204386 |

| | | | | | |
|---|---|---|---|---|---|
| Ppm1a | 0.020533083 | 0.553582095 | 0.8902995 | 0.513774143 | 0.464696929 |
| Ppm1b | 0.00566204 | 0.88002706 | 0.9792337 | 0.61597692 | 0.610736504 |
| Ppm1d | -0.002535194 | 0.912517932 | 0.9840393 | 0.125998569 | 0.11823715 |
| Ppm1g | -0.027057322 | 0.447529103 | 0.8455660 | 0.461575785 | 0.463482577 |
| Ppm1h | -0.058452403 | 0.009382606 | 0.1943120 | 0.074774197 | 0.127314263 |
| Ppm1k | -0.000883696 | 0.966058067 | 0.9972322 | 0.085562888 | 0.097450509 |
| Ppm1l | 0.056032713 | 0.128684025 | 0.5914729 | 0.554536249 | 0.512202661 |
| Ppme1 | 0.05242077 | 0.121405493 | 0.5815270 | 0.420152247 | 0.352178982 |
| Ppox | -0.001444964 | 0.946259582 | 0.9944798 | 0.114325972 | 0.104358106 |
| Ppp1ca | -0.065067195 | 0.050220373 | 0.4218341 | 1.024388719 | 1.031181872 |
| Ppp1cb | 0.054279642 | 0.117780006 | 0.5746396 | 1.075061855 | 0.934830139 |
| Ppp1cc | -0.007911415 | 0.798107361 | 0.9644698 | 0.275138266 | 0.256559078 |
| Ppp1r10 | 0.073633547 | 0.042427393 | 0.3952910 | 0.519228312 | 0.435856423 |
| Ppp1r11 | 0.020537018 | 0.576686604 | 0.8994610 | 0.79843787 | 0.736430334 |
| Ppp1r12a | -0.069674938 | 0.066567814 | 0.4782962 | 0.487318219 | 0.520530401 |
| Ppp1r12b | -0.031586381 | 0.150250398 | 0.6280320 | 0.094645225 | 0.11254864 |
| Ppp1r12c | -0.021607969 | 0.459739737 | 0.8524343 | 0.225578168 | 0.219813451 |
| Ppp1r13b | -0.025916849 | 0.366967249 | 0.8053794 | 0.206717863 | 0.22356778 |
| Ppp1r14b | -0.095361546 | 0.007196859 | 0.1680001 | 1.045217161 | 1.076989267 |
| Ppp1r15a | -0.151710419 | 0.000366072 | 0.0245367 | 0.939635901 | 1.099963873 |
| Ppp1r15b | 0.006936724 | 0.825761494 | 0.9716906 | 0.309988956 | 0.291381873 |
| Ppp1r16a | -0.028559229 | 0.216316862 | 0.6866762 | 0.12264408 | 0.128199672 |
| Ppp1r1a | 0.193034519 | 2.60E-06 | 0.0004613 | 0.588181708 | 0.503409981 |
| Ppp1r2 | -0.004270746 | 0.902308919 | 0.9825441 | 0.866061967 | 0.80878961 |
| Ppp1r21 | -0.011586375 | 0.562430152 | 0.8926240 | 0.073994674 | 0.094553389 |
| Ppp1r35 | 0.008639469 | 0.787455369 | 0.9608311 | 0.362919151 | 0.326775172 |
| Ppp1r37 | 0.020861764 | 0.543882941 | 0.8842645 | 0.530449985 | 0.447896015 |
| Ppp1r3e | -0.008214666 | 0.637833948 | 0.9160042 | 0.060318028 | 0.066785156 |
| Ppp1r7 | -0.011029656 | 0.712692951 | 0.9421288 | 0.242103127 | 0.241773632 |
| Ppp1r8 | 0.03091607 | 0.141936755 | 0.6152737 | 0.105453516 | 0.095423617 |
| Ppp1r9a | 0.003014003 | 0.926535064 | 0.9879210 | 0.296084331 | 0.27427009 |
| Ppp1r9b | -0.036946648 | 0.181594235 | 0.6583478 | 0.173224808 | 0.189442232 |
| Ppp2ca | 0.018037022 | 0.612906452 | 0.9107297 | 0.76477059 | 0.717119824 |
| Ppp2cb | -0.000968238 | 0.972188829 | 0.9976072 | 0.202681831 | 0.198439229 |
| Ppp2r1a | -0.039478162 | 0.255832509 | 0.7234488 | 0.479415058 | 0.487945581 |
| Ppp2r1b | 0.005927496 | 0.769331256 | 0.9552333 | 0.099565029 | 0.087789521 |
| Ppp2r2a | 0.033114911 | 0.329048755 | 0.7778857 | 0.376843185 | 0.322365213 |
| Ppp2r2b | 0.026006805 | 0.30407834 | 0.7634877 | 0.188366861 | 0.125011526 |
| Ppp2r2d | 0.101243039 | 0.004423829 | 0.1313903 | 0.543036391 | 0.468254114 |
| Ppp2r3a | 0.025008973 | 0.382354854 | 0.8126882 | 0.220603827 | 0.201460861 |
| Ppp2r3c | 0.005380599 | 0.819061652 | 0.9699062 | 0.127423496 | 0.122413611 |
| Ppp2r3d | -0.025024091 | 0.32122447 | 0.7726834 | 0.143507123 | 0.148885794 |
| Ppp2r5a | -0.02031146 | 0.586769191 | 0.9035626 | 0.620621694 | 0.628540491 |
| Ppp2r5b | -0.033064067 | 0.257046004 | 0.7250597 | 0.236312877 | 0.235299783 |
| Ppp2r5c | -0.01159464 | 0.749623269 | 0.9504425 | 0.542403009 | 0.526662625 |
| Ppp2r5d | -0.025833775 | 0.111732633 | 0.5675761 | 0.043960464 | 0.059582507 |
| Ppp2r5e | -0.019948143 | 0.541059829 | 0.8835017 | 0.285257841 | 0.303425103 |
| Ppp3ca | -0.023101685 | 0.496756062 | 0.8684629 | 0.352032932 | 0.382132107 |
| Ppp3cb | 0.039240881 | 0.259815247 | 0.7263475 | 0.511436309 | 0.43641324 |
| Ppp3cc | -0.048771622 | 0.009671097 | 0.1970097 | 0.05751189 | 0.084417876 |
| Ppp3r1 | 0.029896723 | 0.316405718 | 0.7705987 | 0.282521374 | 0.257848743 |
| Ppp4c | -0.058596395 | 0.082261512 | 0.5148480 | 0.370680104 | 0.396315024 |
| Ppp4r1 | 0.002617215 | 0.885099238 | 0.9803788 | 0.06490189 | 0.06947013 |
| Ppp4r1l-ps | -0.033391531 | 0.264846636 | 0.7306700 | 0.190373158 | 0.26175033 |
| Ppp4r2 | -0.001011751 | 0.975482843 | 0.9976072 | 0.373800361 | 0.347286419 |
| Ppp4r3a | 0.009500129 | 0.775926293 | 0.9558538 | 0.333648925 | 0.317734065 |
| Ppp4r3b | 0.024518872 | 0.491819736 | 0.8664619 | 0.462966401 | 0.411815686 |
| Ppp4r4 | 0.004021855 | 0.831605532 | 0.9723494 | 0.084317307 | 0.071196836 |
| Ppp5c | -0.053354931 | 0.066821548 | 0.4795761 | 0.213946042 | 0.23769845 |
| Ppp6c | 0.009244464 | 0.787301095 | 0.9608311 | 0.400024027 | 0.372733319 |
| Ppp6r1 | 0.007199719 | 0.766662637 | 0.9544221 | 0.138336027 | 0.141605415 |
| Ppp6r2 | -0.006085683 | 0.812964292 | 0.9673015 | 0.163528694 | 0.159496941 |
| Ppp6r3 | -0.041104452 | 0.188514956 | 0.6668854 | 0.264193908 | 0.274622633 |
| Pprc1 | -0.007915907 | 0.652985348 | 0.9192017 | 0.066967786 | 0.064452357 |
| Ppt1 | 0.0317371 | 0.282563196 | 0.7422811 | 0.277617873 | 0.215338214 |
| Ppt2 | -0.000612645 | 0.973411979 | 0.9976072 | 0.085480721 | 0.070219571 |

| | | | | | |
|---|---|---|---|---|---|
| Pptc7 | 0.005752376 | 0.801759555 | 0.9647143 | 0.123433956 | 0.117337583 |
| Ppwd1 | 0.004021219 | 0.885433003 | 0.9805196 | 0.196540046 | 0.196056727 |
| Ppy | -0.014161903 | 0.839070477 | 0.9727216 | 4.661498179 | 4.760705671 |
| Pqbp1 | -0.029298412 | 0.310568749 | 0.7677462 | 0.183631966 | 0.217709637 |
| Pradc1 | 0.000230507 | 0.989583357 | 0.9980836 | 0.070947103 | 0.067674282 |
| Praf2 | 0.003807796 | 0.836350259 | 0.9725231 | 0.080466517 | 0.071889847 |
| Pramel12 | 0.001211471 | 0.94838581 | 0.9945018 | 0.079810941 | 0.073401963 |
| Prcc | -0.010862021 | 0.639842972 | 0.9162786 | 0.124014156 | 0.121993673 |
| Prdm2 | -0.007526321 | 0.819629416 | 0.9699062 | 0.289692019 | 0.302752173 |
| Prdm4 | 0.001918759 | 0.905700346 | 0.9832544 | 0.057636424 | 0.049428164 |
| Prdx1 | -0.011995121 | 0.668401747 | 0.9230765 | 1.772054845 | 1.707516938 |
| Prdx2 | 0.014258904 | 0.609454137 | 0.9088672 | 1.803693423 | 1.640690286 |
| Prdx3 | -0.027583945 | 0.39692268 | 0.8238469 | 0.325703046 | 0.320790969 |
| Prdx4 | -0.097178104 | 0.009239919 | 0.1929684 | 0.50475985 | 0.512334504 |
| Prdx5 | -0.115446422 | 0.000183028 | 0.0150053 | 1.282998124 | 1.366753101 |
| Prdx6 | -0.000949791 | 0.978169057 | 0.9976072 | 1.018856193 | 1.009865764 |
| Preb | -0.009840888 | 0.775041263 | 0.9558538 | 0.409124544 | 0.400465252 |
| Prelid1 | -0.000694904 | 0.983862286 | 0.9976072 | 0.458862916 | 0.423970348 |
| Prelid3b | -0.03688419 | 0.285037829 | 0.7450791 | 0.474351703 | 0.452096035 |
| Prep | -0.003674113 | 0.806434298 | 0.9648013 | 0.042769443 | 0.049482277 |
| Prepl | 0.02592283 | 0.425337584 | 0.8325096 | 0.330217404 | 0.330085225 |
| Prex1 | -0.026606863 | 0.170580173 | 0.6507045 | 0.064093413 | 0.082019249 |
| Prickle1 | 0.018475978 | 0.370232033 | 0.8065581 | 0.107341955 | 0.090919591 |
| Prim1 | -0.021936781 | 0.199998085 | 0.6737970 | 0.049020301 | 0.070313399 |
| Primpol | 0.002930115 | 0.839529137 | 0.9727382 | 0.044430156 | 0.040094564 |
| Prkaa1 | -0.022969511 | 0.439129322 | 0.8404713 | 0.214603962 | 0.229479008 |
| Prkaa2 | -0.001836627 | 0.924833658 | 0.9879210 | 0.077393254 | 0.082661201 |
| Prkab1 | -0.010008405 | 0.611924604 | 0.9100887 | 0.079446386 | 0.085071483 |
| Prkab2 | 0.000665767 | 0.977968334 | 0.9976072 | 0.133784359 | 0.132444581 |
| Prkaca | 0.030577272 | 0.355630164 | 0.7969131 | 0.429927188 | 0.340629755 |
| Prkacb | 0.06138666 | 0.097101894 | 0.5383862 | 0.919746578 | 0.880771095 |
| Prkag1 | -0.039530962 | 0.19365871 | 0.6710847 | 0.27727813 | 0.272561609 |
| Prkag2 | -0.019409375 | 0.383568184 | 0.8129787 | 0.103655632 | 0.115555997 |
| Prkar1a | -0.079286465 | 0.003725305 | 0.1176911 | 1.780256581 | 1.969971258 |
| Prkar1b | 0.01871248 | 0.57353459 | 0.8973494 | 0.32799925 | 0.320183075 |
| Prkar2a | 0.013705119 | 0.658269278 | 0.9211780 | 0.291948038 | 0.291705258 |
| Prkar2b | 0.027352375 | 0.274454712 | 0.7351955 | 0.141956231 | 0.148841968 |
| Prkca | 0.076376646 | 0.014501666 | 0.2371912 | 0.331748137 | 0.268864759 |
| Prkcb | 0.039450869 | 0.13597109 | 0.6032919 | 0.180594971 | 0.173676553 |
| Prkcd | -0.023136012 | 0.186612731 | 0.6647031 | 0.052406789 | 0.072576434 |
| Prkce | -0.013992556 | 0.642559307 | 0.9166712 | 0.229751148 | 0.249789576 |
| Prkci | -0.011541888 | 0.556987538 | 0.8915944 | 0.071125893 | 0.085662308 |
| Prkcsh | 0.009282459 | 0.797579503 | 0.9644244 | 0.661486914 | 0.662635761 |
| Prkcz | -0.046142716 | 0.059379156 | 0.4548580 | 0.120867658 | 0.15261094 |
| Prkd1 | -0.003864173 | 0.884426032 | 0.9802800 | 0.164240512 | 0.165171953 |
| Prkd2 | 0.010375744 | 0.576362731 | 0.8994610 | 0.087000666 | 0.068051759 |
| Prkdc | -0.012902494 | 0.480888122 | 0.8617579 | 0.065853726 | 0.073669248 |
| Prkn | -0.009271977 | 0.613661254 | 0.9108386 | 0.063242503 | 0.073753203 |
| Prkra | -0.075917804 | 0.006304235 | 0.1587259 | 0.161125535 | 0.201441133 |
| Prkrip1 | 0.044099342 | 0.110251827 | 0.5655888 | 0.221622009 | 0.186967004 |
| Prkx | 0.001659732 | 0.932652119 | 0.9899584 | 0.083971264 | 0.085976334 |
| Prlr | 0.140995437 | 0.003308298 | 0.1093943 | 0.861472403 | 0.943233456 |
| Prmt1 | -0.049203564 | 0.157594108 | 0.6324475 | 0.678132254 | 0.645323553 |
| Prmt2 | -0.058070572 | 0.010997472 | 0.2047427 | 0.094247689 | 0.126941337 |
| Prmt3 | 0.009715622 | 0.557380435 | 0.8916648 | 0.064121009 | 0.057049911 |
| Prmt5 | -0.010427612 | 0.659367716 | 0.9211780 | 0.137066895 | 0.130939784 |
| Prmt6 | -0.015506111 | 0.375796876 | 0.8101773 | 0.062741996 | 0.070591308 |
| Prmt7 | -0.023253097 | 0.296214041 | 0.7545964 | 0.108679581 | 0.121566864 |
| Prmt9 | 0.004782974 | 0.807167899 | 0.9652930 | 0.086586501 | 0.079076542 |
| Prnp | -0.078439847 | 0.004761412 | 0.1371742 | 1.503585033 | 1.683891981 |
| Prom1 | -0.043917186 | 0.161626551 | 0.6394225 | 0.205814903 | 0.278708947 |
| Prorp | -0.015309447 | 0.370133087 | 0.8065581 | 0.052556114 | 0.064843926 |
| Prorsd1 | 0.026579227 | 0.458019464 | 0.8521621 | 0.521438795 | 0.452493329 |
| Proser1 | 0.010837803 | 0.656249047 | 0.9211780 | 0.140236016 | 0.130652935 |
| Prox1 | 0.003227274 | 0.932676536 | 0.9899584 | 0.572291734 | 0.535321366 |
| Prox1os | -0.018727582 | 0.391570808 | 0.8200150 | 0.105892622 | 0.106455521 |

| | | | | | |
|---|---|---|---|---|---|
| Prpf18 | -0.000749642 | 0.976075101 | 0.9976072 | 0.167889495 | 0.155947519 |
| Prpf19 | 0.045183568 | 0.169083829 | 0.6471109 | 0.399859025 | 0.348422146 |
| Prpf3 | 0.019813659 | 0.381217964 | 0.8119369 | 0.132509954 | 0.106263023 |
| Prpf31 | -0.026481641 | 0.307323403 | 0.765611 | 0.161967101 | 0.164377037 |
| Prpf38a | 0.00523262 | 0.85335839 | 0.9755431 | 0.229388384 | 0.197563334 |
| Prpf38b | -0.048962523 | 0.177377969 | 0.6536659 | 0.431992032 | 0.474640678 |
| Prpf39 | -0.072878931 | 0.017101896 | 0.2562701 | 0.190415725 | 0.268108936 |
| Prpf4 | 0.002699865 | 0.921215265 | 0.9861175 | 0.187044322 | 0.184190085 |
| Prpf40a | -0.022674499 | 0.542881342 | 0.8840682 | 0.713336248 | 0.714700894 |
| Prpf40b | -0.013340126 | 0.592181903 | 0.9060200 | 0.148989803 | 0.144547645 |
| Prpf4b | -0.065590742 | 0.09511687 | 0.5351767 | 0.849066621 | 0.95476988 |
| Prpf6 | 0.056543699 | 0.068321652 | 0.4833869 | 0.300249889 | 0.253206515 |
| Prpf8 | 0.003278167 | 0.926954506 | 0.9879210 | 0.498330676 | 0.483489273 |
| Prps1 | -0.000623832 | 0.986068333 | 0.9976072 | 0.52870354 | 0.508291497 |
| Prps1l3 | -0.009710092 | 0.52290545 | 0.8750611 | 0.044662935 | 0.049771564 |
| Prps2 | 0.022839233 | 0.247732731 | 0.7177068 | 0.094650933 | 0.084737721 |
| Prpsap1 | -0.003249038 | 0.90163431 | 0.9825441 | 0.172855911 | 0.174639885 |
| Prpsap2 | -0.022693134 | 0.30176729 | 0.7609383 | 0.097303636 | 0.113988322 |
| Prr12 | -0.035017037 | 0.187408771 | 0.6659251 | 0.144375891 | 0.174705276 |
| Prr13 | -0.087861161 | 0.011672477 | 0.2128510 | 0.675339427 | 0.708854379 |
| Prr14 | -0.024951449 | 0.328078796 | 0.7778857 | 0.147290358 | 0.152021329 |
| Prr14l | 0.056038572 | 0.029940787 | 0.3308280 | 0.207641515 | 0.134729491 |
| Prr15 | 0.012577252 | 0.485632858 | 0.8630379 | 0.087636904 | 0.059593336 |
| Prr15l | 0.034093113 | 0.095592672 | 0.5357510 | 0.102147963 | 0.084145147 |
| Prr16 | 0.046326538 | 0.000108121 | 0.0098399 | 0.061387796 | 0.016321653 |
| Prr3 | -0.019706212 | 0.404871957 | 0.8270770 | 0.125489016 | 0.128857002 |
| Prr7 | -0.009779509 | 0.62266826 | 0.9121505 | 0.083620345 | 0.087488159 |
| Prrc1 | -0.006389454 | 0.864247217 | 0.9769675 | 0.688295342 | 0.70297363 |
| Prrc2a | -0.071696993 | 0.043867436 | 0.3992339 | 0.438439476 | 0.504633926 |
| Prrc2b | -0.002456438 | 0.947950058 | 0.9945018 | 0.689105916 | 0.708358546 |
| Prrc2c | -0.016635815 | 0.642381778 | 0.9166712 | 1.076876787 | 1.123906463 |
| Prrg2 | -0.002121041 | 0.947404255 | 0.9945018 | 0.322006856 | 0.317002408 |
| Prrt1 | -0.031669866 | 0.081058972 | 0.5125557 | 0.053808485 | 0.075545904 |
| Prss36 | -0.026552164 | 0.160202586 | 0.6370337 | 0.067748438 | 0.077200022 |
| Prss53 | 0.062406243 | 0.127499076 | 0.5901963 | 0.503056366 | 0.583748699 |
| Prune1 | -0.003459666 | 0.871688186 | 0.9782269 | 0.104850018 | 0.103273434 |
| Prune2 | 0.042857604 | 0.227732241 | 0.7015153 | 0.508125761 | 0.448510742 |
| Prxl2a | 0.094360163 | 0.009084108 | 0.1909202 | 0.540362653 | 0.409993721 |
| Prxl2b | -0.031526645 | 0.344119523 | 0.7876478 | 0.352219313 | 0.368720949 |
| Prxl2c | -0.058343865 | 0.067050491 | 0.4802461 | 0.285264066 | 0.316544042 |
| Psap | -0.0195993 | 0.494409134 | 0.8678106 | 1.476995023 | 1.537987132 |
| Psat1 | 0.051166683 | 0.118986122 | 0.5780324 | 0.307699678 | 0.300285194 |
| Psd | 0.004517971 | 0.813010176 | 0.9673015 | 0.080388285 | 0.083125801 |
| Psd3 | 0.008747273 | 0.722620754 | 0.9445652 | 0.157839039 | 0.136062755 |
| Psen1 | 0.000597162 | 0.979534436 | 0.9976072 | 0.130348209 | 0.119372864 |
| Psen2 | 0.015048494 | 0.611244876 | 0.9094869 | 0.270748749 | 0.235962816 |
| Psenen | 0.010516893 | 0.758195459 | 0.9521533 | 1.088995928 | 0.999096683 |
| Psip1 | -0.060805484 | 0.092734289 | 0.5326503 | 0.885589705 | 0.897695548 |
| Pskh1 | -0.024853602 | 0.228416798 | 0.7015153 | 0.089313746 | 0.097867876 |
| Psma1 | -0.042593188 | 0.224192503 | 0.6962539 | 0.76985399 | 0.739525807 |
| Psma2 | 0.012448644 | 0.698481488 | 0.9369692 | 1.39115475 | 1.302687155 |
| Psma3 | -0.021905103 | 0.467993582 | 0.8561686 | 1.436909088 | 1.352424811 |
| Psma4 | 0.028208151 | 0.428140955 | 0.8340365 | 0.842023941 | 0.730991514 |
| Psma5 | -0.050856186 | 0.159005947 | 0.6355208 | 0.637274715 | 0.620741593 |
| Psma6 | 0.055306149 | 0.116183835 | 0.5724670 | 0.842990104 | 0.766659265 |
| Psma7 | -0.011739325 | 0.656417995 | 0.9211780 | 1.758751471 | 1.731234982 |
| Psmb1 | -0.01351037 | 0.615750387 | 0.9112316 | 1.601126423 | 1.492788775 |
| Psmb10 | -0.024535842 | 0.462746492 | 0.8524425 | 0.357796853 | 0.353450066 |
| Psmb2 | 0.037929887 | 0.254627843 | 0.7223071 | 1.237032268 | 1.127435862 |
| Psmb3 | 0.004696786 | 0.887618484 | 0.9812414 | 1.062624839 | 0.955386341 |
| Psmb4 | -0.066906906 | 0.022741495 | 0.2926013 | 1.367952254 | 1.379503365 |
| Psmb5 | 0.015597727 | 0.653646462 | 0.9200018 | 1.031074727 | 0.922403606 |
| Psmb6 | -0.068334168 | 0.024794856 | 0.3041540 | 1.284192989 | 1.28886831 |
| Psmb7 | 0.021458958 | 0.549041208 | 0.8871105 | 0.610474731 | 0.551932365 |
| Psmb8 | -0.005988398 | 0.867994218 | 0.9775280 | 0.471724028 | 0.478027487 |
| Psmb9 | -0.04395262 | 0.113454268 | 0.5696977 | 0.166017949 | 0.191495732 |

| | | | | | |
|---|---|---|---|---|---|
| Psmc1 | 0.026417476 | 0.444737509 | 0.84315410 | 0.499087976 | 0.433600579 |
| Psmc2 | -0.110579686 | 0.001869764 | 0.07452714 | 0.600174325 | 0.693192376 |
| Psmc3 | -0.023126792 | 0.49082623 | 0.86581549 | 0.383111616 | 0.38542461 |
| Psmc4 | -0.022778246 | 0.513831019 | 0.87333427 | 0.484504322 | 0.483140404 |
| Psmc5 | -0.02121502 | 0.554740837 | 0.89029959 | 0.827468387 | 0.819046471 |
| Psmc6 | 0.109337389 | 0.001794579 | 0.07386815 | 0.693488295 | 0.574881716 |
| Psmd1 | 0.003077862 | 0.931490978 | 0.98939560 | 0.685050232 | 0.642073414 |
| Psmd10 | 0.006578351 | 0.819941464 | 0.97004404 | 0.221455421 | 0.210916196 |
| Psmd11 | -0.013691878 | 0.703711773 | 0.93918879 | 0.779289012 | 0.762929551 |
| Psmd12 | -0.030552232 | 0.389010643 | 0.81702870 | 0.787942697 | 0.796668368 |
| Psmd13 | 0.04297195 | 0.229901115 | 0.70196775 | 0.660499477 | 0.563458736 |
| Psmd14 | -0.005107372 | 0.875211865 | 0.97843318 | 0.341320828 | 0.320162128 |
| Psmd2 | 0.029187232 | 0.413017474 | 0.83005133 | 0.725516539 | 0.633941621 |
| Psmd3 | -0.048005493 | 0.166561867 | 0.64292850 | 0.469086313 | 0.498295154 |
| Psmd4 | 0.052863187 | 0.139655186 | 0.61218711 | 0.902095684 | 0.847995757 |
| Psmd5 | -0.017011134 | 0.507437814 | 0.87040677 | 0.153050497 | 0.1652326 |
| Psmd6 | -0.015576753 | 0.671747484 | 0.92526173 | 0.693660433 | 0.666596508 |
| Psmd7 | 0.030321231 | 0.368219443 | 0.80567193 | 0.969678079 | 0.94719429 |
| Psmd8 | 0.044688857 | 0.208591847 | 0.68076602 | 0.945362357 | 0.825416955 |
| Psmd9 | 0.006929426 | 0.821882798 | 0.97069202 | 0.291844188 | 0.272748578 |
| Psme1 | 0.010419024 | 0.776519514 | 0.95630957 | 0.925253867 | 0.85113237 |
| Psme2 | 0.035872045 | 0.29522744 | 0.75410785 | 1.171611926 | 1.05867889 |
| Psme3 | -0.0224643 | 0.42332924 | 0.83174372 | 0.178214549 | 0.196886526 |
| Psme3ip1 | -0.024157869 | 0.408008124 | 0.82707700 | 0.209388156 | 0.22809875 |
| Psme4 | -0.031584583 | 0.364285556 | 0.80498822 | 0.404198109 | 0.446941846 |
| Psmf1 | 0.03197971 | 0.249341303 | 0.71927285 | 0.233291579 | 0.182402781 |
| Psmg1 | -0.014137163 | 0.641734646 | 0.91667127 | 0.248989056 | 0.243640813 |
| Psmg2 | -0.047708473 | 0.109559875 | 0.56558881 | 0.226878491 | 0.26132904 |
| Psmg3 | 0.003191204 | 0.911150555 | 0.98395531 | 0.240443447 | 0.210851591 |
| Psmg4 | -0.010356411 | 0.774104223 | 0.95582697 | 0.547606266 | 0.542555381 |
| Pspc1 | -0.008814049 | 0.678323927 | 0.92818862 | 0.090440929 | 0.097004348 |
| Psph | 0.079406061 | 0.057951596 | 0.45081999 | 0.551505017 | 0.544251661 |
| Pstk | -0.003428613 | 0.870624749 | 0.97809711 | 0.099091965 | 0.099210088 |
| Ptar1 | 0.003043362 | 0.891934926 | 0.98195975 | 0.119707888 | 0.110305459 |
| Ptbp1 | 0.016719752 | 0.642368784 | 0.91667127 | 0.557481631 | 0.54938787 |
| Ptbp2 | -0.026611913 | 0.307698742 | 0.76577308 | 0.136932645 | 0.162316856 |
| Ptbp3 | -0.028425593 | 0.457365196 | 0.85183522 | 0.8988947 | 1.001330167 |
| Ptcd1 | 0.007561326 | 0.627447346 | 0.91280959 | 0.058421398 | 0.046253436 |
| Ptcd2 | 0.013820793 | 0.600988897 | 0.90784374 | 0.189105676 | 0.165928061 |
| Ptcd3 | 0.026058109 | 0.173123469 | 0.65255078 | 0.096174962 | 0.072730718 |
| Ptch1 | -0.017734056 | 0.488334499 | 0.86429717 | 0.14973272 | 0.144664582 |
| Ptdss1 | 0.030939335 | 0.162398033 | 0.64007035 | 0.128771813 | 0.102507154 |
| Ptdss2 | -0.021925786 | 0.234736815 | 0.70648944 | 0.063603516 | 0.075874181 |
| Pten | -0.025892544 | 0.476261002 | 0.86072310 | 0.499273564 | 0.522116728 |
| Ptges2 | -0.013872776 | 0.521605859 | 0.87478108 | 0.102272901 | 0.105291691 |
| Ptges3 | 0.003332834 | 0.918363575 | 0.98512115 | 1.237291367 | 1.137957588 |
| Ptgr2 | -0.003205886 | 0.912454567 | 0.98403934 | 0.240819715 | 0.226359729 |
| Ptk2 | 0.011726574 | 0.622660841 | 0.91215057 | 0.135554836 | 0.135417255 |
| Ptk2b | -0.028215063 | 0.200732629 | 0.67432024 | 0.093131661 | 0.108417717 |
| Ptma | -0.018480963 | 0.305383655 | 0.76462715 | 2.794010849 | 2.818009852 |
| Ptms | -0.019949508 | 0.51199968 | 0.87208730 | 1.429479397 | 1.403815162 |
| Ptov1 | -0.080355376 | 0.022563626 | 0.29144683 | 1.010992548 | 0.981297193 |
| Ptp4a2 | 0.018848839 | 0.613991217 | 0.91083865 | 0.653907108 | 0.627894204 |
| Ptp4a3 | -0.036787745 | 0.25829537 | 0.72511022 | 0.311135372 | 0.310487865 |
| Ptpa | 0.010199229 | 0.768137433 | 0.95442217 | 0.402112608 | 0.365864347 |
| Ptpdc1 | -0.013039936 | 0.492636284 | 0.86693013 | 0.075211159 | 0.078484196 |
| Ptpmt1 | -0.019895541 | 0.528314769 | 0.87731044 | 0.307737413 | 0.306341961 |
| Ptpn1 | 0.00807787 | 0.823726494 | 0.97105924 | 0.488022071 | 0.499852483 |
| Ptpn11 | -0.022124797 | 0.549006299 | 0.88711058 | 0.603837398 | 0.641535647 |
| Ptpn12 | 0.016254784 | 0.445853663 | 0.84373680 | 0.112460031 | 0.098442272 |
| Ptpn13 | 0.024317026 | 0.271061817 | 0.73420300 | 0.128434449 | 0.093518777 |
| Ptpn2 | 0.019747814 | 0.526536244 | 0.87685539 | 0.274206118 | 0.250159544 |
| Ptpn21 | -0.00295509 | 0.897394072 | 0.98254414 | 0.120230019 | 0.120281357 |
| Ptpn23 | 0.017932391 | 0.425202014 | 0.83244601 | 0.129901647 | 0.1084593 |
| Ptpn3 | 0.025464092 | 0.130454249 | 0.59438518 | 0.081372739 | 0.051505691 |
| Ptpn4 | -0.051525305 | 0.055830169 | 0.44493323 | 0.157926148 | 0.189417158 |

| | | | | | |
|---|---|---|---|---|---|
| Ptpn6 | 0.005084263 | 0.823862111 | 0.9710592 | 0.124633373 | 0.11905942 |
| Ptpn9 | 0.002289203 | 0.938424659 | 0.9924491 | 0.215613507 | 0.215302282 |
| Ptpra | -0.038107205 | 0.229640169 | 0.7019677 | 0.276603167 | 0.296524108 |
| Ptprd | -0.077725612 | 0.025593106 | 0.3069934 | 0.331040373 | 0.33737274 |
| Ptprf | 0.006882761 | 0.84904835 | 0.9748841 | 0.865634782 | 0.897510448 |
| Ptprj | -0.024442167 | 0.337783256 | 0.7834245 | 0.136491039 | 0.163040237 |
| Ptprk | -0.03219429 | 0.128687077 | 0.5914729 | 0.097047482 | 0.098024482 |
| Ptprm | 0.014614875 | 0.418687172 | 0.8300990 | 0.06296219 | 0.074786814 |
| Ptprn | -0.103960525 | 0.000350395 | 0.0239718 | 1.585120136 | 1.821320578 |
| Ptprn2 | -0.038832644 | 0.259709269 | 0.7263475 | 0.339929272 | 0.369030984 |
| Ptprr | -1.39E-05 | 0.999360674 | 0.9996989 | 0.058786013 | 0.062518742 |
| Ptprs | -0.00035987 | 0.981969549 | 0.9976072 | 0.047465839 | 0.056047635 |
| Ptprt | -0.010634671 | 0.723492133 | 0.9445652 | 0.215283393 | 0.229474184 |
| Ptpru | 0.001058056 | 0.95784114 | 0.9957785 | 0.08793537 | 0.081273871 |
| Ptrh2 | 0.046405662 | 0.079425221 | 0.5076205 | 0.202571034 | 0.159394916 |
| Ptrhd1 | -0.011400257 | 0.690962665 | 0.9348540 | 0.204565121 | 0.211744845 |
| Pts | -0.037237433 | 0.311997259 | 0.7684379 | 0.50101547 | 0.537633316 |
| Pttg1 | -0.005822502 | 0.840289544 | 0.9729978 | 0.226701046 | 0.20377424 |
| Pttg1ip | 0.041473215 | 0.187782222 | 0.6662373 | 0.339458152 | 0.299785864 |
| Puf60 | -0.01844765 | 0.592769545 | 0.9060585 | 0.5180909 | 0.509875243 |
| Pum1 | -0.01703464 | 0.644582833 | 0.9168391 | 0.534566846 | 0.543402547 |
| Pum2 | 0.02326004 | 0.52905619 | 0.8776448 | 0.576819998 | 0.567318953 |
| Pum3 | -0.003399496 | 0.907826074 | 0.9837373 | 0.235574703 | 0.225591584 |
| Pura | -0.057380474 | 0.077543614 | 0.5027664 | 1.351275131 | 1.365296115 |
| Purb | -0.005593353 | 0.881723612 | 0.9797222 | 0.890157534 | 0.874772882 |
| Pus1 | -0.027922787 | 0.19171665 | 0.6700281 | 0.090831837 | 0.107581727 |
| Pus10 | -0.017655636 | 0.287669417 | 0.7472324 | 0.044638273 | 0.058953541 |
| Pus3 | -0.019493612 | 0.332228316 | 0.7793535 | 0.080705826 | 0.090576689 |
| Pus7 | 0.019838774 | 0.386819056 | 0.8162614 | 0.133508247 | 0.113308462 |
| Pusl1 | -0.040527409 | 0.025276001 | 0.3050747 | 0.057077717 | 0.080271652 |
| Pwp1 | 0.011215544 | 0.651993572 | 0.9189792 | 0.16159643 | 0.143671335 |
| Pwp2 | 0.000670841 | 0.964763265 | 0.9966421 | 0.045010638 | 0.047956374 |
| Pwwp2a | -0.021686143 | 0.437124267 | 0.8394017 | 0.18373251 | 0.185903747 |
| Pwwp3a | -0.022976167 | 0.281820305 | 0.7417588 | 0.087089036 | 0.106595684 |
| Pwwp3b | -0.011173446 | 0.753107966 | 0.9512135 | 0.319921266 | 0.309549474 |
| Pxk | 0.008803754 | 0.775388547 | 0.9558538 | 0.272222857 | 0.26719066 |
| Pxmp4 | -0.020341866 | 0.313893052 | 0.7705213 | 0.08173256 | 0.095854984 |
| Pxn | -0.03654807 | 0.015675689 | 0.2456600 | 0.029074994 | 0.055002703 |
| Pxylp1 | 0.02281413 | 0.186420327 | 0.6644983 | 0.07460069 | 0.058116468 |
| Pycard | -0.003166023 | 0.880610961 | 0.9794439 | 0.093168041 | 0.096355458 |
| Pycr1 | 0.00678528 | 0.83410653 | 0.9723494 | 0.301025383 | 0.298199544 |
| Pycr2 | -0.004669451 | 0.893315953 | 0.9823405 | 0.918916302 | 0.871338145 |
| Pycrl | -0.040155508 | 0.118821456 | 0.5777984 | 0.149298915 | 0.169466112 |
| Pygb | -0.035520579 | 0.046015639 | 0.4073656 | 0.054682292 | 0.072324809 |
| Pygl | 0.029122285 | 0.02908799 | 0.3264172 | 0.058648539 | 0.027477954 |
| Pygo1 | 0.017043634 | 0.251162557 | 0.7207210 | 0.052666559 | 0.041955862 |
| Pygo2 | -0.014185429 | 0.491717738 | 0.8664619 | 0.082609427 | 0.095895036 |
| Pym1 | 0.021278012 | 0.26061383 | 0.7271077 | 0.098299766 | 0.069516417 |
| Pyroxd1 | 0.038583688 | 0.149527825 | 0.6264374 | 0.20166559 | 0.160133392 |
| Pyurf | 0.028812873 | 0.281823508 | 0.7417588 | 0.200707455 | 0.165422321 |
| Pyy | -0.110777466 | 0.005714594 | 0.1523891 | 6.888701894 | 7.283345264 |
| Qars | -0.053441246 | 0.068463579 | 0.4833869 | 0.209576093 | 0.229337851 |
| Qdpr | -0.037521562 | 0.307549385 | 0.7656663 | 0.702678239 | 0.705946964 |
| Qk | -0.008475138 | 0.730811885 | 0.9459020 | 0.136605761 | 0.13904246 |
| Qpct | -0.018864706 | 0.238610977 | 0.7109741 | 0.038853288 | 0.053368313 |
| Qpctl | -0.032225704 | 0.197502753 | 0.6730962 | 0.139142463 | 0.152201492 |
| Qrich1 | -0.033368395 | 0.274469879 | 0.7351955 | 0.219564002 | 0.245288658 |
| Qser1 | 0.005513159 | 0.837610157 | 0.9727216 | 0.186267679 | 0.17432126 |
| Qsox1 | -0.074019938 | 0.030478939 | 0.3340895 | 0.383764609 | 0.431982807 |
| Qsox2 | -0.045502187 | 0.109168289 | 0.5649188 | 0.17548405 | 0.224847609 |
| Qtrt1 | -0.004187501 | 0.871644768 | 0.9782269 | 0.16492011 | 0.160375003 |
| Qtrt2 | -0.026442876 | 0.097984813 | 0.5409067 | 0.033183346 | 0.059263276 |
| R3hcc1 | 0.034174554 | 0.094877911 | 0.5350704 | 0.104117774 | 0.087719624 |
| R3hcc1l | -0.01966865 | 0.302844584 | 0.7616612 | 0.068525478 | 0.082287335 |
| R3hdm1 | -0.016133271 | 0.657165577 | 0.9211780 | 0.558479185 | 0.573037034 |
| R3hdm2 | 0.05082797 | 0.163628786 | 0.6410026 | 0.646346504 | 0.591126958 |

| | | | | | |
|---|---|---|---|---|---|
| R3hdm4 | -0.013465185 | 0.677401933 | 0.9281529 | 0.327959872 | 0.332303411 |
| R74862 | -0.013171312 | 0.401067311 | 0.8252619 | 0.052613913 | 0.04954703 |
| Rab10 | -0.00621058 | 0.863410426 | 0.9769675 | 0.621736424 | 0.612546697 |
| Rab10os | 0.019502214 | 0.399854968 | 0.8244516 | 0.131523209 | 0.12103385 |
| Rab11a | -0.038256771 | 0.285572263 | 0.7452977 | 0.715735049 | 0.699558151 |
| Rab11b | 0.00927278 | 0.788910941 | 0.9616609 | 0.472943316 | 0.429049471 |
| Rab11fip1 | -0.008417118 | 0.61542842 | 0.9111433 | 0.055457233 | 0.063899482 |
| Rab11fip2 | -0.023753209 | 0.496356155 | 0.8684629 | 0.384393798 | 0.379933739 |
| Rab11fip3 | -0.003558473 | 0.840286161 | 0.9729978 | 0.066545842 | 0.071229535 |
| Rab12 | -0.035184671 | 0.332298473 | 0.7793535 | 0.519959652 | 0.503535086 |
| Rab13 | -0.014645131 | 0.468716949 | 0.8563037 | 0.092439385 | 0.090625276 |
| Rab14 | -0.012919894 | 0.663151387 | 0.9230058 | 1.354137116 | 1.351065972 |
| Rab18 | 0.013129119 | 0.714593563 | 0.9430747 | 0.718656552 | 0.667798731 |
| Rab19 | -0.024083978 | 0.132599289 | 0.5988251 | 0.034917014 | 0.062802792 |
| Rab1a | -0.013421964 | 0.703638991 | 0.9391887 | 0.955038596 | 0.91767898 |
| Rab1b | 0.024715474 | 0.461757534 | 0.8524425 | 0.375261813 | 0.354006273 |
| Rab20 | -0.022067575 | 0.277947205 | 0.7382158 | 0.083984108 | 0.088612944 |
| Rab21 | -0.081751551 | 0.025433832 | 0.3058225 | 0.598218547 | 0.64118713 |
| Rab22a | -0.010346801 | 0.75729366 | 0.9521533 | 0.34203584 | 0.378933362 |
| Rab23 | -0.014929621 | 0.424441713 | 0.8324361 | 0.07255703 | 0.079228972 |
| Rab24 | 0.049315349 | 0.157075405 | 0.6321187 | 0.488631558 | 0.44235505 |
| Rab25 | -0.051115386 | 0.157775983 | 0.6324475 | 0.590417236 | 0.602248461 |
| Rab26os | -0.039969969 | 0.072601855 | 0.4902939 | 0.085208243 | 0.11576933 |
| Rab27a | 0.007414062 | 0.832072454 | 0.9723494 | 0.442474451 | 0.442440868 |
| Rab27b | 0.010986181 | 0.47211732 | 0.8590766 | 0.053670281 | 0.040709897 |
| Rab28 | 0.003624418 | 0.91447712 | 0.9845236 | 0.464667498 | 0.428148085 |
| Rab2a | 0.028123354 | 0.282076928 | 0.7418354 | 1.727977931 | 1.629769185 |
| Rab2b | -0.011134066 | 0.527394498 | 0.8768553 | 0.057306545 | 0.068726766 |
| Rab33b | -0.021414098 | 0.461884382 | 0.8524425 | 0.220006141 | 0.235518151 |
| Rab34 | -0.04167492 | 0.190831428 | 0.6700281 | 0.293674305 | 0.282573905 |
| Rab35 | -0.004495057 | 0.846111714 | 0.9738285 | 0.120758667 | 0.122917137 |
| Rab36 | -0.010794186 | 0.601424661 | 0.9080871 | 0.08480867 | 0.097877797 |
| Rab37 | -0.018283103 | 0.565455802 | 0.8943433 | 0.286346147 | 0.277285505 |
| Rab39b | 0.01267918 | 0.688345295 | 0.9329669 | 0.298613233 | 0.294066088 |
| Rab3a | 0.025816644 | 0.469584265 | 0.8567594 | 0.566421148 | 0.525545925 |
| Rab3b | -0.017757454 | 0.544846062 | 0.8845945 | 0.181612512 | 0.221414087 |
| Rab3c | -0.006850042 | 0.813472923 | 0.9674680 | 0.237246146 | 0.18828823 |
| Rab3d | 0.007566009 | 0.80735167 | 0.9653373 | 0.259550594 | 0.276733147 |
| Rab3gap1 | -0.001777694 | 0.959409627 | 0.9957785 | 0.398386515 | 0.403293531 |
| Rab3gap2 | 0.014064196 | 0.64000953 | 0.9162786 | 0.25222629 | 0.239560787 |
| Rab3ip | -0.017321802 | 0.562615649 | 0.8926240 | 0.283407974 | 0.247351098 |
| Rab40b | -0.049661094 | 0.020445863 | 0.2774595 | 0.093066412 | 0.114710105 |
| Rab40c | 0.002851663 | 0.899881816 | 0.9825441 | 0.125887566 | 0.116460356 |
| Rab4a | 0.008180516 | 0.75989929 | 0.9521533 | 0.178498628 | 0.169530336 |
| Rab5a | -0.056508691 | 0.123484536 | 0.5849888 | 0.639696834 | 0.685969263 |
| Rab5b | 0.006426522 | 0.826396867 | 0.9716906 | 0.256645919 | 0.231217964 |
| Rab5c | 0.024761242 | 0.42949975 | 0.8340365 | 0.33299342 | 0.313105017 |
| Rab5if | -0.011282707 | 0.737246986 | 0.9469752 | 0.430769198 | 0.414636045 |
| Rab6a | 0.050852051 | 0.171021304 | 0.6507063 | 0.694442299 | 0.6161577 |
| Rab6b | 0.027009724 | 0.238184578 | 0.7108276 | 0.126870169 | 0.117803956 |
| Rab7 | 0.064517909 | 0.073712005 | 0.4920747 | 0.813078975 | 0.74793254 |
| Rab8a | 0.019592807 | 0.521354081 | 0.8745066 | 0.283459425 | 0.268434665 |
| Rab8b | 0.018916561 | 0.236464308 | 0.7091069 | 0.062983798 | 0.048143597 |
| Rab9 | -0.031630331 | 0.243524875 | 0.7132719 | 0.185127327 | 0.190831686 |
| Rabac1 | 0.017617497 | 0.470996502 | 0.8580121 | 1.911963883 | 1.822002819 |
| Rabep1 | -0.002916788 | 0.931819673 | 0.9893956 | 0.395043689 | 0.405188513 |
| Rabep2 | -0.019485193 | 0.360612633 | 0.8012480 | 0.099816678 | 0.097896066 |
| Rabepk | -0.021126892 | 0.331200062 | 0.7783711 | 0.093706342 | 0.110505109 |
| Rabgap1 | 0.021659574 | 0.504466981 | 0.8695566 | 0.349843211 | 0.306142531 |
| Rabgap1l | -0.054346401 | 0.057060767 | 0.4485283 | 0.172652991 | 0.200735679 |
| Rabgef1 | 0.033754512 | 0.095178722 | 0.5351767 | 0.109118211 | 0.082996673 |
| Rabggta | 0.014175313 | 0.398920084 | 0.8239850 | 0.071116577 | 0.055564231 |
| Rabggtb | 0.003779008 | 0.915701164 | 0.9845236 | 0.478304109 | 0.435638893 |
| Rabif | 0.007023 | 0.76055992 | 0.9522598 | 0.135028178 | 0.115472378 |
| Rabl2 | 0.004804016 | 0.751213883 | 0.9506053 | 0.056202447 | 0.045934142 |
| Rabl3 | 0.019829839 | 0.312112124 | 0.7684379 | 0.094923428 | 0.074807153 |

| | | | | | |
|---|---|---|---|---|---|
| Rabl6 | -0.051152425 | 0.174219923 | 0.65276574 | 0.574838234 | 0.595142537 |
| Rac1 | 0.013123021 | 0.718238417 | 0.94382369 | 0.743284772 | 0.6816502 |
| Rack1 | -0.113078553 | 1.13E-08 | 7.35E-06 | 2.701688486 | 2.758685626 |
| Rad1 | 0.023942122 | 0.263563705 | 0.72838464 | 0.127528528 | 0.092921274 |
| Rad17 | -0.013718362 | 0.669919679 | 0.92394964 | 0.310258525 | 0.304553261 |
| Rad18 | -0.056119799 | 0.014828 | 0.23901395 | 0.098841577 | 0.122070918 |
| Rad21 | 0.034938448 | 0.345717941 | 0.78893995 | 0.623993988 | 0.58767857 |
| Rad23a | -0.039727797 | 0.258754865 | 0.72571339 | 0.621227116 | 0.599760627 |
| Rad23b | -0.027406574 | 0.441655693 | 0.84270521 | 0.511838684 | 0.525198415 |
| Rad50 | 0.047655468 | 0.067839494 | 0.48172351 | 0.192435565 | 0.14894877 |
| Rad51ap2 | -0.015399685 | 0.5099955 | 0.87116071 | 0.085030771 | 0.121448748 |
| Rad51c | 0.027699761 | 0.017372986 | 0.25722391 | 0.04214825 | 0.021895795 |
| Rad51d | -0.018823394 | 0.369579158 | 0.80629541 | 0.10096261 | 0.101581462 |
| Rad54l2 | -0.048611468 | 0.017242415 | 0.25709990 | 0.066262795 | 0.102561193 |
| Rad9b | 0.008445606 | 0.559444666 | 0.89237335 | 0.048952896 | 0.040222677 |
| Rae1 | 0.02499602 | 0.390218146 | 0.81873181 | 0.236689341 | 0.210575541 |
| Raf1 | -0.033551658 | 0.240784436 | 0.71204045 | 0.18340285 | 0.206139294 |
| Rai1 | 0.016723062 | 0.514621411 | 0.87337141 | 0.158745021 | 0.14649821 |
| Rai2 | 0.051563081 | 0.043214139 | 0.39766629 | 0.205305248 | 0.13098299 |
| Rala | 0.026000444 | 0.45435281 | 0.84944963 | 0.837305087 | 0.758329104 |
| Ralb | -0.028396705 | 0.207929239 | 0.68074521 | 0.112824001 | 0.121493859 |
| Ralbp1 | 0.081561235 | 0.019979415 | 0.27375110 | 1.050627618 | 0.933427528 |
| Ralgapa1 | 0.047515705 | 0.143499649 | 0.61650780 | 0.356274912 | 0.303255806 |
| Ralgapa2 | -0.019287933 | 0.544722775 | 0.88453919 | 0.282614152 | 0.272489101 |
| Ralgapb | 0.003514601 | 0.895042456 | 0.98243614 | 0.188948011 | 0.173369644 |
| Ralgds | -0.03975015 | 0.110438796 | 0.56558881 | 0.117687484 | 0.153601232 |
| Ralgps1 | -0.023356359 | 0.200517107 | 0.67397010 | 0.060268958 | 0.074257911 |
| Ralgps2 | -0.022402236 | 0.407926661 | 0.82707700 | 0.155517384 | 0.171429452 |
| Raly | 0.032314896 | 0.327574469 | 0.77788578 | 0.42228696 | 0.34584401 |
| Ramac | 0.059182371 | 0.0685953 | 0.48354951 | 0.43178424 | 0.327049318 |
| Ramp1 | 0.052204795 | 0.019715458 | 0.27201299 | 0.133517832 | 0.092139229 |
| Ramp2 | 0.002420203 | 0.918590632 | 0.98512119 | 0.11868655 | 0.13434596 |
| Ran | 0.061564771 | 0.066585394 | 0.47829624 | 1.300469632 | 1.16152013 |
| Ranbp1 | 0.023763372 | 0.526790496 | 0.87685539 | 0.819016708 | 0.712253513 |
| Ranbp10 | 0.019090591 | 0.412640458 | 0.82979790 | 0.142115576 | 0.120844664 |
| Ranbp2 | -0.061892042 | 0.110613027 | 0.56558881 | 0.57421345 | 0.629169698 |
| Ranbp3 | -0.003945524 | 0.887958228 | 0.98124141 | 0.21298088 | 0.19540774 |
| Ranbp6 | 0.045214781 | 0.033469047 | 0.34620745 | 0.145836723 | 0.088488066 |
| Ranbp9 | -0.005838116 | 0.835100973 | 0.97234941 | 0.197305204 | 0.19802152 |
| Rangap1 | 0.00588241 | 0.87334894 | 0.97843318 | 0.489104859 | 0.486808038 |
| Rangrf | -0.009150447 | 0.554612269 | 0.89029959 | 0.044888358 | 0.05045653 |
| Rap1a | -0.009042251 | 0.793906359 | 0.96245081 | 0.761493142 | 0.727948943 |
| Rap1b | 0.046216549 | 0.141512245 | 0.61435511 | 1.454150209 | 1.358455355 |
| Rap1gap | -0.070068723 | 0.030756126 | 0.33530391 | 0.244755361 | 0.295255279 |
| Rap1gap2 | 0.019277187 | 0.607771197 | 0.90871391 | 0.719163991 | 0.733598723 |
| Rap1gds1 | -0.006381449 | 0.735140521 | 0.94683808 | 0.067613764 | 0.07303551 |
| Rap2a | -0.038057568 | 0.137332427 | 0.60708763 | 0.141769316 | 0.158873428 |
| Rap2b | 0.028383215 | 0.117525781 | 0.57431317 | 0.087341606 | 0.064069958 |
| Rap2c | 0.003442704 | 0.839372433 | 0.97273829 | 0.062133223 | 0.059278761 |
| Rapgef1 | 0.038354947 | 0.086635645 | 0.52505662 | 0.138559297 | 0.108254771 |
| Rapgef2 | -0.002320676 | 0.915576199 | 0.98452363 | 0.113577317 | 0.105571427 |
| Rapgef4 | -0.069347991 | 0.026858387 | 0.31387771 | 0.221038707 | 0.291522323 |
| Rapgef6 | -0.000338259 | 0.990259375 | 0.99808365 | 0.198682757 | 0.193248041 |
| Raph1 | -0.000942069 | 0.959641345 | 0.99577853 | 0.065164582 | 0.074274617 |
| Rara | 0.024969158 | 0.084583403 | 0.51922488 | 0.055843366 | 0.037775049 |
| Rarg | -0.007903771 | 0.643333207 | 0.91667127 | 0.061627806 | 0.063275971 |
| Rars | -0.038784061 | 0.234988156 | 0.70681701 | 0.301712515 | 0.320373039 |
| Rars2 | 0.03334917 | 0.152886485 | 0.62862147 | 0.152370991 | 0.116000019 |
| Rasa1 | 0.01326708 | 0.681680822 | 0.93028941 | 0.330291163 | 0.307163123 |
| Rasa2 | -0.04936746 | 0.083743103 | 0.51830408 | 0.179527365 | 0.215518725 |
| Rasal2 | 0.028450287 | 0.244180614 | 0.71327199 | 0.156567655 | 0.135572312 |
| Rasd1 | -0.137079375 | 0.001883972 | 0.07475600 | 0.511245545 | 0.667513485 |
| Rasd2 | -0.021374436 | 0.206803482 | 0.67975659 | 0.030302696 | 0.066884952 |
| Rasgrf1 | 0.02987152 | 0.349076916 | 0.79273039 | 0.254846106 | 0.321433721 |
| Rasgrf2 | -0.083273926 | 0.007692616 | 0.17340589 | 0.204403952 | 0.277286819 |
| Rassf1 | -0.018173371 | 0.579727983 | 0.90087519 | 0.30993881 | 0.301922158 |

| | | | | | |
|---|---|---|---|---|---|
| Rassf10 | 0.002975481 | 0.875303934 | 0.9784331 | 0.075733701 | 0.074869581 |
| Rassf3 | 0.029459568 | 0.3557973 | 0.7970888 | 0.298267113 | 0.269437738 |
| Rassf4 | 0.018387528 | 0.41049786 | 0.8283974 | 0.117943104 | 0.112200906 |
| Rassf7 | -0.01417354 | 0.52290986 | 0.8750611 | 0.115476212 | 0.114527552 |
| Rb1 | 0.053731814 | 0.03058257 | 0.3348555 | 0.178169546 | 0.129547 |
| Rb1cc1 | -0.031384344 | 0.38522228 | 0.8139828 | 0.489137777 | 0.520302922 |
| Rbak | -0.040127906 | 0.010812684 | 0.2035210 | 0.032065056 | 0.057646846 |
| Rbbp4 | -0.043024706 | 0.231836659 | 0.7030140 | 0.645252429 | 0.60174642 |
| Rbbp5 | -0.011766686 | 0.530463805 | 0.8776448 | 0.068206642 | 0.077046474 |
| Rbbp6 | 0.042018469 | 0.262141184 | 0.7275735 | 0.871588528 | 0.811052923 |
| Rbbp7 | -0.12317554 | 0.000546577 | 0.0326629 | 1.115525906 | 1.149509508 |
| Rbbp8 | -0.013957941 | 0.457515638 | 0.8518352 | 0.063473431 | 0.07332345 |
| Rbbp9 | -0.000574978 | 0.973307654 | 0.9976072 | 0.061806257 | 0.062695826 |
| Rbck1 | -0.029629675 | 0.336543818 | 0.7826493 | 0.255882449 | 0.261032727 |
| Rbfa | -0.037458258 | 0.155172279 | 0.6308130 | 0.156120787 | 0.187936159 |
| Rbfox2 | 0.053101009 | 0.086411098 | 0.5246010 | 0.309919008 | 0.257376568 |
| Rbis | 0.063142977 | 0.058945796 | 0.4529374 | 0.422413341 | 0.381702167 |
| Rbl1 | 0.018434646 | 0.187559963 | 0.6659251 | 0.053118593 | 0.03489424 |
| Rbl2 | 0.022434635 | 0.37052515 | 0.8066125 | 0.158172905 | 0.135356663 |
| Rbm10 | 0.065931777 | 0.026460562 | 0.3110056 | 0.29814735 | 0.205590385 |
| Rbm11 | -0.026497127 | 0.145052748 | 0.6196913 | 0.059498092 | 0.074403781 |
| Rbm12 | 0.030548789 | 0.081068548 | 0.5125557 | 0.078912735 | 0.059734948 |
| Rbm12b2 | -0.006908036 | 0.673982915 | 0.9264090 | 0.055468321 | 0.059088314 |
| Rbm14 | 0.01941162 | 0.330689831 | 0.7779915 | 0.102032159 | 0.078474129 |
| Rbm15 | 0.038008549 | 0.122021515 | 0.5819916 | 0.170519282 | 0.125620833 |
| Rbm15b | 0.017090362 | 0.50366105 | 0.8695566 | 0.166617441 | 0.150865086 |
| Rbm17 | 0.02959661 | 0.364523601 | 0.8050031 | 0.368648645 | 0.350581555 |
| Rbm18 | 0.004944253 | 0.85468447 | 0.9755431 | 0.186863695 | 0.178545042 |
| Rbm19 | 0.009554039 | 0.553223031 | 0.8900377 | 0.061866859 | 0.050086609 |
| Rbm22 | 0.013555394 | 0.643524604 | 0.9167845 | 0.232471154 | 0.212854839 |
| Rbm24 | -0.013073933 | 0.45278626 | 0.8486662 | 0.059261022 | 0.061357138 |
| Rbm25 | 0.028726073 | 0.446302314 | 0.8442637 | 1.025105801 | 0.945530633 |
| Rbm26 | -0.030327052 | 0.394283823 | 0.8213556 | 0.447441847 | 0.474848313 |
| Rbm27 | -0.009839256 | 0.771481925 | 0.9552333 | 0.349936121 | 0.364857441 |
| Rbm28 | 0.02621471 | 0.449962668 | 0.8478268 | 0.413042449 | 0.361535334 |
| Rbm3 | 0.004717031 | 0.901513242 | 0.9825441 | 0.771944129 | 0.820909727 |
| Rbm33 | 0.024064007 | 0.472145782 | 0.8590766 | 0.338527123 | 0.296771654 |
| Rbm34 | -0.030238398 | 0.250460555 | 0.7207210 | 0.157779238 | 0.168471925 |
| Rbm39 | 0.036870923 | 0.185599018 | 0.6637678 | 1.922839771 | 1.883134664 |
| Rbm43 | -0.023991527 | 0.419668475 | 0.8300990 | 0.231808752 | 0.228725091 |
| Rbm47 | 0.003410906 | 0.915973809 | 0.9845236 | 0.304579361 | 0.296655959 |
| Rbm48 | 0.052516426 | 0.033218959 | 0.3454615 | 0.170615153 | 0.133906447 |
| Rbm4b | -0.05456397 | 0.074688911 | 0.4966285 | 0.251482007 | 0.260450485 |
| Rbm5 | -0.065236738 | 0.088078685 | 0.5278913 | 0.535984657 | 0.574779396 |
| Rbm6 | -0.002488105 | 0.941385586 | 0.9929301 | 0.340661934 | 0.332922751 |
| Rbm7 | 0.007461112 | 0.818252439 | 0.9697318 | 0.357688358 | 0.314358987 |
| Rbm8a | -0.041660053 | 0.2309739 | 0.7024275 | 0.72924204 | 0.743923108 |
| Rbms1 | 0.011049777 | 0.646238726 | 0.9169695 | 0.13590935 | 0.123812896 |
| Rbms2 | -0.02434308 | 0.172522432 | 0.6522284 | 0.058317301 | 0.068745306 |
| Rbms3 | -0.01827184 | 0.466592181 | 0.8553370 | 0.136511199 | 0.13777221 |
| Rbmx | -0.029079906 | 0.273092551 | 0.7349642 | 0.168074728 | 0.179570509 |
| Rbmx2 | 0.026515177 | 0.408511598 | 0.8276447 | 0.334711913 | 0.259895126 |
| Rbmxl1 | 0.003016274 | 0.911933367 | 0.9840393 | 0.214782208 | 0.178468072 |
| Rbp4 | 0.026026772 | 0.571843722 | 0.8966165 | 3.311976995 | 2.951246943 |
| Rbpj | -0.025419084 | 0.334695316 | 0.7804836 | 0.141505331 | 0.169917195 |
| Rbpms | -0.016054734 | 0.679244079 | 0.9288316 | 0.72085132 | 0.706082006 |
| Rbsn | -0.037276983 | 0.155908469 | 0.6312943 | 0.141453557 | 0.18142845 |
| Rbx1 | 0.050923065 | 0.064546269 | 0.4711545 | 1.546203785 | 1.448892301 |
| Rc3h1 | -0.005098281 | 0.867296534 | 0.9773465 | 0.256865218 | 0.243300278 |
| Rc3h2 | 0.003648653 | 0.914897434 | 0.9845236 | 0.3607452 | 0.356482518 |
| Rcan1 | -0.000758735 | 0.968236321 | 0.9976072 | 0.082240832 | 0.079082053 |
| Rcan2 | 0.028696951 | 0.334055606 | 0.7804836 | 0.255598196 | 0.222661716 |
| Rcan3 | -0.004310832 | 0.902565363 | 0.9825441 | 0.463361554 | 0.485735911 |
| Rcbtb1 | 0.0138378 | 0.639929687 | 0.9162786 | 0.237338052 | 0.228961041 |
| Rcbtb2 | 0.01755066 | 0.459209276 | 0.8522649 | 0.15225008 | 0.121992856 |
| Rcc1l | -0.025407195 | 0.204354877 | 0.6780598 | 0.077992158 | 0.092063619 |

| | | | | | |
|---|---|---|---|---|---|
| Rcc2 | 0.008814398 | 0.80110507 | 0.9647143 | 0.40312966 | 0.439715698 |
| Rccd1 | -0.038758915 | 0.016696086 | 0.2546713 | 0.037776967 | 0.05831815 |
| Rce1 | -0.026399383 | 0.363361023 | 0.8044168 | 0.220923605 | 0.227759883 |
| Rchy1 | -0.048665699 | 0.126596448 | 0.5890416 | 0.279659796 | 0.325573435 |
| Rcl1 | -0.017364092 | 0.385182121 | 0.8139828 | 0.083633209 | 0.089651179 |
| Rcn1 | 0.005697622 | 0.863682852 | 0.9769675 | 0.415901711 | 0.312015802 |
| Rcn2 | -0.013442477 | 0.716994293 | 0.9431474 | 0.586349951 | 0.555728671 |
| Rcn3 | 0.027067371 | 0.492367937 | 0.8669301 | 0.580267494 | 0.434295792 |
| Rcor1 | -0.01800193 | 0.5627057 | 0.8926240 | 0.249183964 | 0.250570604 |
| Rcor3 | -0.030038995 | 0.308383845 | 0.7663245 | 0.198280156 | 0.230215938 |
| Rdh11 | 0.007991345 | 0.625269485 | 0.9121784 | 0.059058908 | 0.05260971 |
| Rdh13 | -0.016375277 | 0.285453117 | 0.7451828 | 0.036595431 | 0.052467782 |
| Rdh14 | -0.023835301 | 0.431446248 | 0.8356213 | 0.243940354 | 0.261519445 |
| Rdm1 | -0.030116154 | 0.185408022 | 0.6637678 | 0.10358095 | 0.119922582 |
| Rdx | 0.018851853 | 0.589251203 | 0.9046539 | 0.896899921 | 0.844206815 |
| Rec8 | -0.021990309 | 0.229523478 | 0.7019677 | 0.052414482 | 0.080050298 |
| Recql | -0.015900517 | 0.439593254 | 0.8407104 | 0.094997106 | 0.090986031 |
| Reep2 | 0.015026819 | 0.439679404 | 0.8407131 | 0.098515857 | 0.07810327 |
| Reep3 | 0.028660913 | 0.382303971 | 0.8126882 | 0.338943396 | 0.318944538 |
| Reep5 | 0.04262466 | 0.175098275 | 0.6534142 | 1.444007647 | 1.387272262 |
| Reep6 | -0.05394622 | 0.121674383 | 0.5819916 | 0.350871621 | 0.427582681 |
| Rel | -0.024512374 | 0.20003351 | 0.6737970 | 0.057096437 | 0.082014965 |
| Rela | -0.029693857 | 0.210742888 | 0.6822059 | 0.111927101 | 0.139336207 |
| Relb | -0.016197919 | 0.333740652 | 0.7804836 | 0.040104459 | 0.057173344 |
| Relch | 0.030452046 | 0.307396504 | 0.765611 | 0.24076299 | 0.212492775 |
| Rell1 | -0.012582647 | 0.529364498 | 0.8776448 | 0.075505341 | 0.090577602 |
| Rell2 | -0.03437148 | 0.056523415 | 0.4471389 | 0.055656168 | 0.072515085 |
| Relt | 0.025074301 | 0.229350609 | 0.7019677 | 0.116286305 | 0.091998447 |
| Rem2 | 0.006187525 | 0.773888179 | 0.9558269 | 0.113209171 | 0.094265635 |
| Repin1 | 0.009905538 | 0.571702559 | 0.8966165 | 0.078575138 | 0.060961734 |
| Reps1 | -0.042205942 | 0.116261683 | 0.5724670 | 0.136968609 | 0.176678473 |
| Reps2 | 0.026673603 | 0.242873116 | 0.7132719 | 0.119613843 | 0.117299019 |
| Rer1 | 0.029975823 | 0.39552622 | 0.8225618 | 0.806336739 | 0.764338359 |
| Rere | 0.043829868 | 0.218947772 | 0.6901690 | 0.47488229 | 0.433742788 |
| Resf1 | 0.02870371 | 0.39313614 | 0.8206236 | 0.315128411 | 0.306490146 |
| Resp18 | -0.037734996 | 0.021726915 | 0.2873746 | 4.707510326 | 4.579680453 |
| Ret | -0.005537264 | 0.815683789 | 0.9686999 | 0.110708467 | 0.139892407 |
| Retreg1 | 0.051579823 | 0.069489682 | 0.4837345 | 0.204973697 | 0.175939326 |
| Retreg2 | -0.076578527 | 0.029014719 | 0.3264172 | 0.501978087 | 0.537032203 |
| Retreg3 | 0.009111929 | 0.734740933 | 0.9468380 | 0.188328082 | 0.171779746 |
| Rev1 | 0.003155338 | 0.906227315 | 0.9835423 | 0.17614557 | 0.166057576 |
| Rev3l | -0.015221989 | 0.657815542 | 0.9211780 | 0.354518611 | 0.363703058 |
| Rex1bd | 0.02792962 | 0.43435423 | 0.8381237 | 0.808431696 | 0.789930325 |
| Rexo1 | -0.012999683 | 0.668591774 | 0.9230765 | 0.253235022 | 0.258837147 |
| Rexo2 | -0.004155793 | 0.909472722 | 0.9837373 | 0.883746597 | 0.915400122 |
| Rexo4 | 0.009745812 | 0.687464647 | 0.9327945 | 0.15612812 | 0.12944311 |
| Rfc1 | -0.009657758 | 0.769445071 | 0.9552333 | 0.337951235 | 0.343780742 |
| Rfc2 | 0.063535591 | 0.030764165 | 0.3353039 | 0.283475755 | 0.20588534 |
| Rfc3 | -0.027124102 | 0.228287252 | 0.7015153 | 0.102469296 | 0.114994828 |
| Rfc4 | 0.011446062 | 0.559053255 | 0.8921289 | 0.098119646 | 0.075214981 |
| Rfc5 | -0.011127744 | 0.509082669 | 0.8704067 | 0.058640906 | 0.06137341 |
| Rfesd | -0.03501787 | 0.243279708 | 0.7132719 | 0.232719785 | 0.249977193 |
| Rffl | -0.027299025 | 0.060842071 | 0.4603763 | 0.024871726 | 0.051988922 |
| Rfk | -0.005660115 | 0.869343254 | 0.9776539 | 0.439869648 | 0.45351011 |
| Rflna | 0.062033356 | 0.126954059 | 0.5893234 | 0.614988823 | 0.544467268 |
| Rflnb | 0.030225415 | 0.178101409 | 0.6538489 | 0.123976876 | 0.102825747 |
| Rfng | 0.004113393 | 0.820911853 | 0.9704976 | 0.078719712 | 0.065907696 |
| Rft1 | -0.004619659 | 0.76587873 | 0.9543420 | 0.053030321 | 0.050232262 |
| Rfwd3 | 0.019460805 | 0.251797619 | 0.7219168 | 0.073225555 | 0.053053016 |
| Rfx2 | -0.002886361 | 0.862982691 | 0.9769675 | 0.062198404 | 0.059247277 |
| Rfx3 | -0.03968859 | 0.116566811 | 0.5727304 | 0.119413844 | 0.153020289 |
| Rfx5 | -0.00399956 | 0.840414809 | 0.9730292 | 0.077776417 | 0.088067295 |
| Rfx6 | -0.003721326 | 0.899826735 | 0.9825441 | 0.221675648 | 0.223424175 |
| Rfx7 | -0.054923403 | 0.051462676 | 0.4282799 | 0.163888303 | 0.20171559 |
| Rfxank | -7.25E-05 | 0.996349805 | 0.9994073 | 0.056139596 | 0.053228424 |
| Rfxap | 0.015143202 | 0.647226153 | 0.9169695 | 0.381211386 | 0.322171615 |

| | | | | | |
|---|---|---|---|---|---|
| Rgl1 | -0.00661559 | 0.730483918 | 0.9459020 | 0.073949198 | 0.086064991 |
| Rgl2 | -0.003476294 | 0.86483386 | 0.9769675 | 0.09121283 | 0.089881641 |
| Rgl3 | 0.008806883 | 0.65040227 | 0.9177795 | 0.08843652 | 0.075673429 |
| Rgmb | -0.013238726 | 0.497967475 | 0.8684629 | 0.072334478 | 0.082725551 |
| Rgp1 | -0.001424122 | 0.92935925 | 0.9881953 | 0.050128557 | 0.059135977 |
| Rgs11 | -0.04316551 | 0.097392829 | 0.5384203 | 0.117807559 | 0.181465077 |
| Rgs16 | 0.011109755 | 0.645292538 | 0.9169695 | 0.130261338 | 0.12729965 |
| Rgs17 | -0.007135126 | 0.86160205 | 0.9769675 | 0.586991165 | 0.583757501 |
| Rgs2 | 0.012588816 | 0.719695887 | 0.9444523 | 0.284643693 | 0.315599543 |
| Rgs3 | -0.027798429 | 0.079831069 | 0.5079693 | 0.03896459 | 0.058391388 |
| Rgs4 | 0.004363338 | 0.926378089 | 0.9879210 | 0.586878595 | 0.577313651 |
| Rgs7 | 0.042768254 | 0.110101364 | 0.5655888 | 0.196932607 | 0.167997845 |
| Rgs7bp | -0.012496253 | 0.614789516 | 0.9108386 | 0.134011973 | 0.135576174 |
| Rgs9 | -0.057476912 | 0.047560338 | 0.4113326 | 0.168960459 | 0.232557765 |
| Rhbdd1 | 0.000341627 | 0.987564933 | 0.9976072 | 0.108522192 | 0.111260952 |
| Rhbdd2 | -0.018654741 | 0.546169015 | 0.8858413 | 0.289761016 | 0.287399283 |
| Rhbdd3 | 0.009929674 | 0.612058422 | 0.9101248 | 0.091594733 | 0.084932632 |
| Rhbdl2 | -0.003654313 | 0.819569556 | 0.9699062 | 0.047301268 | 0.058522024 |
| Rhd | -0.01670683 | 0.263031802 | 0.7276284 | 0.039014865 | 0.049044666 |
| Rheb | 0.049422215 | 0.134222933 | 0.6006009 | 1.069815594 | 0.965325122 |
| Rhno1 | 0.0434594 | 0.029564799 | 0.3295312 | 0.125774102 | 0.076512971 |
| Rhoa | 0.019571425 | 0.494302355 | 0.8678106 | 1.616431895 | 1.450768977 |
| Rhob | -0.134491089 | 0.000423678 | 0.0274698 | 0.33993239 | 0.458088615 |
| Rhobtb1 | 0.038845143 | 0.247880897 | 0.7177068 | 0.356860936 | 0.333382765 |
| Rhobtb2 | -0.002944815 | 0.84012062 | 0.9729978 | 0.041487029 | 0.046817676 |
| Rhobtb3 | -0.079075515 | 0.009147439 | 0.1918448 | 0.216218603 | 0.263314725 |
| Rhoc | -0.00566405 | 0.841388996 | 0.9734589 | 0.21206595 | 0.18261918 |
| Rhog | 0.002120056 | 0.915399299 | 0.9845236 | 0.095247402 | 0.084159442 |
| Rhoq | -0.003074866 | 0.910645263 | 0.9837373 | 0.163288287 | 0.191570717 |
| Rhot1 | -0.047025735 | 0.06105206 | 0.4605600 | 0.126350022 | 0.157365698 |
| Rhot2 | -0.004302852 | 0.836697959 | 0.9726619 | 0.094028017 | 0.099443732 |
| Rhou | -0.049426554 | 0.199245012 | 0.6730962 | 0.389917478 | 0.438716527 |
| Rhpn2 | 0.043650247 | 0.075364495 | 0.4966285 | 0.16272619 | 0.131706385 |
| Rian | 0.011777515 | 0.758629295 | 0.9521533 | 1.053389013 | 1.067555002 |
| Ribc1 | 0.009747891 | 0.482169639 | 0.8620306 | 0.048845384 | 0.034698913 |
| Ric1 | 0.007556763 | 0.752378495 | 0.9507898 | 0.131753664 | 0.123998767 |
| Ric3 | 0.015013682 | 0.561380767 | 0.8926240 | 0.159703388 | 0.156359893 |
| Ric8a | -0.003672295 | 0.893799691 | 0.9824361 | 0.187337116 | 0.17942919 |
| Ric8b | 0.037067362 | 0.192644229 | 0.6700281 | 0.216625853 | 0.190558706 |
| Rictor | 0.022661544 | 0.460275344 | 0.8524425 | 0.262532427 | 0.246882796 |
| Rida | -0.011380952 | 0.659181061 | 0.9211780 | 0.161081112 | 0.16755678 |
| Rif1 | 0.004034691 | 0.907731731 | 0.9837373 | 0.402046843 | 0.399162342 |
| Riiad1 | 0.008210677 | 0.769933097 | 0.9552333 | 0.20423895 | 0.188208112 |
| Rilpl1 | 0.01231677 | 0.563656718 | 0.8931805 | 0.114181572 | 0.098010468 |
| Rilpl2 | 0.026976283 | 0.356663957 | 0.7982417 | 0.269012107 | 0.208500252 |
| Rimbp2 | -0.070938749 | 0.035210555 | 0.3577036 | 0.305822084 | 0.366366886 |
| Rims2 | -0.0548815 | 0.123853247 | 0.5852543 | 0.372056532 | 0.413713111 |
| Rims3 | 0.017766687 | 0.414745511 | 0.8300990 | 0.098498393 | 0.103717507 |
| Ring1 | -0.016889278 | 0.577123924 | 0.8994610 | 0.261328114 | 0.248906273 |
| Rint1 | 0.012016291 | 0.718971757 | 0.9441159 | 0.379769993 | 0.344288517 |
| Riok1 | -0.003250564 | 0.914171601 | 0.9845236 | 0.255518912 | 0.252848627 |
| Riok2 | 0.034383113 | 0.127081783 | 0.5893647 | 0.137129959 | 0.106054814 |
| Riok3 | 0.041740048 | 0.241638945 | 0.7131979 | 0.646286343 | 0.605363603 |
| Riox1 | -0.010349281 | 0.55649092 | 0.8915358 | 0.066858691 | 0.066873161 |
| Riox2 | -0.001060233 | 0.960186649 | 0.9958628 | 0.107639031 | 0.102407024 |
| Ripk1 | 0.034164016 | 0.18539004 | 0.6637678 | 0.181668975 | 0.147514262 |
| Ripk2 | 0.021176589 | 0.230097015 | 0.7019677 | 0.077696489 | 0.062112433 |
| Ripk4 | -0.024928304 | 0.332069336 | 0.7793175 | 0.14312974 | 0.157303695 |
| Ripor1 | -0.027985388 | 0.254231584 | 0.7223071 | 0.126258108 | 0.144713144 |
| Ripply3 | 0.005019717 | 0.845838353 | 0.9736268 | 0.179383395 | 0.149207418 |
| Rit1 | 0.012731187 | 0.661486572 | 0.9218806 | 0.238010963 | 0.22423632 |
| Rita1 | 0.007868086 | 0.595317417 | 0.9067325 | 0.050034597 | 0.044759662 |
| Rlf | -0.025090485 | 0.407272711 | 0.8270770 | 0.217776904 | 0.245089343 |
| Rlim | -0.029947601 | 0.360070585 | 0.8010608 | 0.245425948 | 0.280562681 |
| Rmc1 | 0.010721456 | 0.568059893 | 0.8955308 | 0.081465531 | 0.072006739 |
| Rmdn1 | 0.006319036 | 0.804207834 | 0.9647143 | 0.168333797 | 0.150568278 |

| | | | | | |
|---|---|---|---|---|---|
| Rmdn3 | -0.016526582 | 0.51612872 | 0.8735705 | 0.149173587 | 0.162100774 |
| Rmi1 | 0.003292289 | 0.863215125 | 0.9769675 | 0.079303191 | 0.07840633 |
| Rmnd1 | 0.014698741 | 0.381639159 | 0.8124164 | 0.075053458 | 0.056028573 |
| Rmnd5a | -0.011019874 | 0.745629965 | 0.9485717 | 0.379866253 | 0.388652007 |
| Rmnd5b | -0.030594782 | 0.177268333 | 0.6536659 | 0.104957876 | 0.124028364 |
| Rn18s | -0.151420983 | 2.30E-12 | 4.56E-09 | 4.120040052 | 4.502080273 |
| Rnase4 | -0.100813618 | 0.014150214 | 0.2347500 | 0.832110827 | 0.990383069 |
| Rnaseh1 | 0.028993392 | 0.211729984 | 0.6822059 | 0.140245841 | 0.119092786 |
| Rnaseh2a | 0.017661821 | 0.485132025 | 0.8629715 | 0.171410237 | 0.140665605 |
| Rnaseh2b | 0.05133575 | 0.009528415 | 0.1956974 | 0.11966141 | 0.069786876 |
| Rnaseh2c | -0.021611161 | 0.555576757 | 0.8910341 | 0.580675275 | 0.541237277 |
| Rnaset2a | 0.013432492 | 0.511014573 | 0.8716066 | 0.102404042 | 0.08926023 |
| Rnaset2b | 0.007127908 | 0.699943625 | 0.9374160 | 0.084695437 | 0.068667583 |
| Rnd2 | -0.01725976 | 0.316391328 | 0.7705987 | 0.059467601 | 0.065135692 |
| Rnd3 | -0.031983297 | 0.144921095 | 0.6193956 | 0.078664428 | 0.101567637 |
| Rnf10 | 0.045237463 | 0.192592955 | 0.6700281 | 0.863319548 | 0.768034111 |
| Rnf103 | 0.025663612 | 0.39992775 | 0.8244516 | 0.279065635 | 0.268834825 |
| Rnf11 | 0.022680621 | 0.487125529 | 0.8640311 | 1.181109886 | 1.089439429 |
| Rnf111 | -0.038462731 | 0.190363792 | 0.6700281 | 0.209684448 | 0.237630203 |
| Rnf113a1 | 0.003171736 | 0.848365676 | 0.9746134 | 0.067632631 | 0.055394723 |
| Rnf113a2 | -0.066313738 | 0.028915041 | 0.3264172 | 0.226878328 | 0.267662873 |
| Rnf114 | -0.02450906 | 0.412554274 | 0.8297979 | 0.214770557 | 0.23323452 |
| Rnf115 | -0.025465009 | 0.438886845 | 0.8403314 | 0.392423892 | 0.404751986 |
| Rnf121 | 0.043044842 | 0.117824308 | 0.5746396 | 0.21342082 | 0.173136357 |
| Rnf123 | 0.015230342 | 0.349685605 | 0.7927303 | 0.065056197 | 0.055826195 |
| Rnf126 | 0.048652469 | 0.101053735 | 0.5487357 | 0.271124445 | 0.2185756 |
| Rnf128 | 0.000293626 | 0.992750171 | 0.9985888 | 0.324071001 | 0.305261438 |
| Rnf13 | 0.021800084 | 0.437802388 | 0.8399914 | 0.218202904 | 0.19010312 |
| Rnf130 | 0.050002752 | 0.12744132 | 0.5901963 | 1.523252237 | 1.288912224 |
| Rnf135 | -0.010132087 | 0.482129761 | 0.8620306 | 0.040382573 | 0.045421013 |
| Rnf138 | -0.011814353 | 0.712021995 | 0.9419834 | 0.336033181 | 0.310720185 |
| Rnf138rt1 | -0.075471587 | 0.060178416 | 0.4581503 | 0.537052711 | 0.53487807 |
| Rnf139 | -0.012351389 | 0.629566194 | 0.9130564 | 0.149893736 | 0.158470929 |
| Rnf14 | 0.033753233 | 0.323537068 | 0.7751516 | 0.402532167 | 0.390737237 |
| Rnf141 | -0.018585918 | 0.510787158 | 0.8716066 | 0.190334489 | 0.209496614 |
| Rnf145 | 0.023627795 | 0.375123368 | 0.8094896 | 0.195732618 | 0.157359836 |
| Rnf146 | 0.003956155 | 0.895719335 | 0.9825441 | 0.2581213 | 0.245034938 |
| Rnf149 | 0.01508044 | 0.466616519 | 0.8553370 | 0.112092501 | 0.090532761 |
| Rnf150 | -0.002750579 | 0.883618181 | 0.9801988 | 0.061695798 | 0.080839487 |
| Rnf157 | -0.04368275 | 0.006099644 | 0.1568368 | 0.026732718 | 0.063957468 |
| Rnf166 | -0.020889978 | 0.443885526 | 0.8431541 | 0.17589404 | 0.186132387 |
| Rnf167 | 0.018537321 | 0.577975666 | 0.8997914 | 0.366433556 | 0.332252019 |
| Rnf168 | 0.014924075 | 0.580985169 | 0.9015399 | 0.199372524 | 0.173993536 |
| Rnf169 | -0.007834217 | 0.71060066 | 0.9419370 | 0.093051065 | 0.09360135 |
| Rnf170 | 0.015949969 | 0.52935393 | 0.8776448 | 0.159131927 | 0.13694336 |
| Rnf181 | -0.008624345 | 0.78424735 | 0.9601202 | 0.316694232 | 0.294951626 |
| Rnf182 | 0.017228822 | 0.330803256 | 0.7779915 | 0.063020262 | 0.067011241 |
| Rnf185 | -0.008481575 | 0.708879113 | 0.9414811 | 0.116738554 | 0.121186939 |
| Rnf186 | 0.028034873 | 0.145815912 | 0.6213461 | 0.094582614 | 0.074627468 |
| Rnf187 | -0.031055167 | 0.378160154 | 0.8101915 | 0.684082486 | 0.626050692 |
| Rnf19a | -0.018102558 | 0.41205575 | 0.8297979 | 0.109203222 | 0.109767762 |
| Rnf19b | -0.016133773 | 0.469891345 | 0.8568606 | 0.104521565 | 0.122922892 |
| Rnf2 | 0.001381469 | 0.955850774 | 0.9954690 | 0.163584368 | 0.151718824 |
| Rnf20 | 0.017783217 | 0.600243325 | 0.9078234 | 0.362174804 | 0.342203868 |
| Rnf208 | -0.003802784 | 0.862991529 | 0.9769675 | 0.109582778 | 0.105730105 |
| Rnf212b | -0.009749125 | 0.553028977 | 0.8900377 | 0.060661418 | 0.049964991 |
| Rnf213 | -0.039993696 | 0.028832878 | 0.3264172 | 0.047629525 | 0.079765108 |
| Rnf214 | -0.027393307 | 0.324349469 | 0.7757552 | 0.187552658 | 0.208969585 |
| Rnf215 | 0.024541564 | 0.303018278 | 0.7617692 | 0.159656959 | 0.124252316 |
| Rnf216 | 0.010241875 | 0.726948218 | 0.9452179 | 0.248251982 | 0.220313421 |
| Rnf217 | -0.026151973 | 0.319544179 | 0.7712599 | 0.139559142 | 0.171944509 |
| Rnf220 | 0.016429313 | 0.58597748 | 0.9029041 | 0.253168648 | 0.236744321 |
| Rnf227 | 0.020919733 | 0.515091683 | 0.8733714 | 0.300504301 | 0.278925353 |
| Rnf24 | 0.016506594 | 0.286063935 | 0.7459550 | 0.053677319 | 0.047490642 |
| Rnf25 | 0.022206917 | 0.305598406 | 0.7646271 | 0.126379058 | 0.105340631 |
| Rnf26 | -0.000180796 | 0.989357576 | 0.9980836 | 0.040170517 | 0.03799516 |

| | | | | | |
|---|---|---|---|---|---|
| Rnf31 | 0.001316285 | 0.947506831 | 0.9945018 | 0.092747035 | 0.084333128 |
| Rnf32 | 0.006792926 | 0.707676481 | 0.9408790 | 0.07568729 | 0.064123312 |
| Rnf34 | 0.015606623 | 0.547137914 | 0.8858508 | 0.155559578 | 0.150467286 |
| Rnf38 | 0.011074555 | 0.62448641 | 0.9121505 | 0.125765973 | 0.111331655 |
| Rnf4 | 0.004473084 | 0.890602815 | 0.9819597 | 0.305129504 | 0.294049238 |
| Rnf40 | -0.00506269 | 0.793902179 | 0.9624508 | 0.081141611 | 0.083507819 |
| Rnf41 | 0.00673397 | 0.777163342 | 0.9564524 | 0.137703805 | 0.128770023 |
| Rnf44 | 0.028540932 | 0.214246175 | 0.6837977 | 0.13944626 | 0.112476057 |
| Rnf5 | -0.017219122 | 0.622231194 | 0.9121505 | 0.49867182 | 0.454286956 |
| Rnf6 | 0.005277633 | 0.883745789 | 0.9801988 | 0.452339381 | 0.444833162 |
| Rnf7 | 0.038789553 | 0.267784216 | 0.7323841 | 1.001460656 | 0.890531272 |
| Rnf8 | 0.011135861 | 0.700576097 | 0.9375165 | 0.24394868 | 0.215827085 |
| Rnft1 | -0.001750079 | 0.926817866 | 0.9879210 | 0.078152499 | 0.080310547 |
| Rngtt | -0.007014005 | 0.765403025 | 0.9540318 | 0.131471256 | 0.124480824 |
| Rnh1 | -0.026799155 | 0.453841473 | 0.8490482 | 0.844492433 | 0.803781101 |
| Rnmt | -0.030302694 | 0.328830704 | 0.7778857 | 0.244397919 | 0.25564795 |
| Rnpc3 | -0.012110447 | 0.605521622 | 0.9087139 | 0.120714177 | 0.12562911 |
| Rnpep | -0.017677802 | 0.499367278 | 0.8690742 | 0.156676933 | 0.169768854 |
| Rnpepl1 | 0.010904299 | 0.736958864 | 0.9469752 | 0.318528801 | 0.29895455 |
| Rnps1 | 0.027406991 | 0.441955257 | 0.8428709 | 0.516012826 | 0.485303838 |
| Ro60 | -0.028488981 | 0.425044714 | 0.8324460 | 0.461563858 | 0.47435898 |
| Robo1 | 0.011819671 | 0.467712594 | 0.8558769 | 0.057142536 | 0.058365455 |
| Robo2 | 0.050124079 | 0.174267976 | 0.6527657 | 0.435876003 | 0.389528665 |
| Rock1 | 0.008728652 | 0.809685512 | 0.9661473 | 1.092100911 | 1.067326881 |
| Rock2 | -0.024313279 | 0.506569343 | 0.8695566 | 0.465996877 | 0.47902971 |
| Rogdi | -0.04671455 | 0.164895956 | 0.6420707 | 0.33443122 | 0.384476036 |
| Rom1 | 0.001340574 | 0.951951136 | 0.9951566 | 0.119302268 | 0.11264418 |
| Romo1 | 0.167579347 | 2.31E-08 | 1.27E-05 | 1.552672734 | 1.339274955 |
| Ropn1l | 0.004378787 | 0.799637131 | 0.9647053 | 0.067939512 | 0.060804803 |
| Ror2 | -0.006912829 | 0.754501231 | 0.9519143 | 0.101627108 | 0.105330261 |
| Rora | -0.047826171 | 0.180693691 | 0.6569883 | 0.458058861 | 0.501644014 |
| Rorc | -0.005670633 | 0.837843559 | 0.9727216 | 0.189912302 | 0.183418689 |
| Rp2 | -0.028379242 | 0.271770339 | 0.7342806 | 0.16323647 | 0.17253681 |
| Rp9 | 0.043337945 | 0.208535189 | 0.6807660 | 0.93188229 | 0.862394695 |
| Rpa1 | 0.030604058 | 0.290821204 | 0.7513090 | 0.243788175 | 0.197112485 |
| Rpa2 | -0.005143461 | 0.786858109 | 0.9608311 | 0.085414355 | 0.075733457 |
| Rpa3 | -0.004139401 | 0.886156943 | 0.9809263 | 0.236419553 | 0.211776533 |
| Rpain | -0.012096924 | 0.45880654 | 0.8522649 | 0.051416416 | 0.056022671 |
| Rpap1 | -0.032051081 | 0.093764362 | 0.5333348 | 0.062987865 | 0.085411967 |
| Rpap2 | 0.035760029 | 0.096050523 | 0.5368469 | 0.132473796 | 0.093164272 |
| Rpap3 | 0.035258746 | 0.107040199 | 0.5630110 | 0.132075918 | 0.097975588 |
| Rpe | 0.01710707 | 0.514299018 | 0.8733459 | 0.188832178 | 0.15985751 |
| Rpf1 | 0.013471668 | 0.6251608 | 0.9121784 | 0.205267252 | 0.179771703 |
| Rpf2 | -0.01220573 | 0.63021144 | 0.9130564 | 0.169544305 | 0.15211061 |
| Rpgr | -0.017128853 | 0.393168835 | 0.8206236 | 0.082844766 | 0.091771675 |
| Rpgrip1 | 0.038040842 | 0.054642828 | 0.4421851 | 0.117857292 | 0.065519752 |
| Rpgrip1l | 0.015291094 | 0.365252783 | 0.8050191 | 0.073049412 | 0.05285464 |
| Rph3al | 0.008510121 | 0.802051623 | 0.9647143 | 0.370290691 | 0.360761632 |
| Rpia | -0.071839583 | 0.018706815 | 0.2673942 | 0.219687579 | 0.253221427 |
| Rpl10 | -0.032882711 | 0.011403097 | 0.2105633 | 3.777668811 | 3.627680051 |
| Rpl10a | -0.078873296 | 0.002049058 | 0.0803441 | 2.046466475 | 1.89657608 |
| Rpl10a-ps1 | -0.01239396 | 0.642657937 | 0.9166712 | 0.190929522 | 0.166335307 |
| Rpl10-ps3 | 0.00451336 | 0.822120748 | 0.9706920 | 0.108977119 | 0.082406826 |
| Rpl11 | -0.009718578 | 0.471617591 | 0.8585034 | 3.694925586 | 3.510485956 |
| Rpl12 | -0.073131362 | 0.000145967 | 0.0125912 | 2.733077316 | 2.79210596 |
| Rpl13 | -0.040480743 | 0.001558978 | 0.0679368 | 4.00049387 | 3.848167596 |
| Rpl13a | -0.055737168 | 0.103420963 | 0.5560628 | 1.192620686 | 1.075716137 |
| Rpl14 | -0.051731629 | 0.000579868 | 0.0340372 | 3.332313808 | 3.254954616 |
| Rpl14-ps1 | -0.01869856 | 0.29886614 | 0.7572802 | 0.052668333 | 0.074397527 |
| Rpl15 | 0.010837215 | 0.685231913 | 0.9316044 | 2.057366198 | 1.816851944 |
| Rpl17 | -0.03749344 | 0.013069433 | 0.2262631 | 3.577942475 | 3.436003265 |
| Rpl18 | -0.003439393 | 0.821152797 | 0.9705476 | 3.190437634 | 3.004524112 |
| Rpl18a | -0.036105913 | 0.025198384 | 0.3048389 | 2.986202404 | 2.9154321 |
| Rpl19 | -0.030033186 | 0.004643711 | 0.1362887 | 4.303705727 | 4.188317409 |
| Rpl19-ps11 | -0.030435965 | 0.095436263 | 0.5351767 | 0.058951945 | 0.076364964 |
| Rpl21 | -0.026663371 | 0.046258817 | 0.4073656 | 3.563779205 | 3.442043914 |

| | | | | | |
|---|---|---|---|---|---|
| Rpl22 | -0.003639926 | 0.851496326 | 0.9750483 | 2.455105888 | 2.393685254 |
| Rpl22l1 | 0.03696923 | 0.051980499 | 0.4313858 | 2.753126191 | 2.574083275 |
| Rpl23 | -0.01728995 | 0.161703328 | 0.6394225 | 3.989375274 | 3.904654451 |
| Rpl23a | -0.013907132 | 0.673958519 | 0.9264090 | 1.661791084 | 1.526496085 |
| Rpl24 | 0.070359913 | 0.041825489 | 0.3932785 | 1.262594182 | 0.950186159 |
| Rpl26 | -0.025450081 | 0.042070231 | 0.3944581 | 3.645444679 | 3.581351747 |
| Rpl27 | 0.133269359 | 9.00E-08 | 3.72E-05 | 2.092210011 | 1.789335793 |
| Rpl27a | 0.051810022 | 0.002407093 | 0.0890983 | 3.153789935 | 2.908561446 |
| Rpl28 | 0.022775015 | 0.097180238 | 0.5383862 | 3.539372317 | 3.353690118 |
| Rpl29 | -0.038553466 | 0.04424003 | 0.4007863 | 2.694548047 | 2.510087819 |
| Rpl3 | -0.06177271 | 0.000198628 | 0.0160322 | 2.973772966 | 2.940356358 |
| Rpl30 | 0.043983662 | 0.001417731 | 0.0633508 | 3.560266056 | 3.347403847 |
| Rpl31 | 0.057613178 | 0.017456188 | 0.2580346 | 2.152678356 | 2.005401039 |
| Rpl32 | -0.040995094 | 0.00032398 | 0.0226977 | 4.070718166 | 4.102047194 |
| Rpl34 | 0.024403205 | 0.060732772 | 0.4600290 | 3.698420846 | 3.55412709 |
| Rpl35 | 0.09646454 | 5.28E-07 | 0.0001588 | 2.822695528 | 2.554141764 |
| Rpl35a | 0.046536598 | 0.000501256 | 0.0306941 | 3.809778532 | 3.569782113 |
| Rpl36 | 0.057411979 | 3.33E-05 | 0.0039788 | 3.526477285 | 3.360362224 |
| Rpl36a | 0.037824089 | 0.060306809 | 0.4587757 | 2.527957742 | 2.399133398 |
| Rpl36al | 0.043231706 | 0.156105338 | 0.6312943 | 1.692450426 | 1.578288575 |
| Rpl36a-ps2 | 0.006236514 | 0.694057375 | 0.9361045 | 0.059832421 | 0.050384113 |
| Rpl37 | 0.067638901 | 5.66E-08 | 2.67E-05 | 3.963309292 | 3.742231679 |
| Rpl37a | 0.061394658 | 1.77E-07 | 6.26E-05 | 4.061919967 | 3.833515791 |
| Rpl37rt | 0.000535775 | 0.984237435 | 0.9976072 | 0.188777784 | 0.171967773 |
| Rpl38 | 0.056094436 | 4.99E-05 | 0.0055597 | 3.553563979 | 3.332380327 |
| Rpl39 | 0.082125803 | 9.80E-08 | 3.89E-05 | 3.338472659 | 3.120118004 |
| Rpl3-ps1 | -0.073684271 | 0.002959321 | 0.1015794 | 0.115793376 | 0.162910255 |
| Rpl4 | -0.08452706 | 3.43E-05 | 0.0040504 | 2.343393993 | 2.352610277 |
| Rpl41 | 0.077352156 | 8.83E-14 | 2.92E-10 | 4.537754979 | 4.187775365 |
| Rpl5 | -0.129413253 | 1.07E-06 | 0.0002318 | 2.056582891 | 2.021230415 |
| Rpl6 | 0.003600932 | 0.806357764 | 0.9648013 | 3.306064234 | 3.101387988 |
| Rpl6l | -0.033141194 | 0.109066663 | 0.5646875 | 0.09508769 | 0.100669003 |
| Rpl7 | -0.065727754 | 0.000917957 | 0.0461985 | 2.528373831 | 2.410960242 |
| Rpl7a | -0.063698283 | 0.000525108 | 0.0319574 | 2.761259764 | 2.628412002 |
| Rpl7l1 | -0.006318977 | 0.848925514 | 0.9748841 | 0.378666846 | 0.351397986 |
| Rpl7-ps9 | -0.002132235 | 0.91015475 | 0.9837373 | 0.083632748 | 0.075498165 |
| Rpl8 | -0.06686701 | 2.35E-09 | 1.67E-06 | 4.079511542 | 4.104288805 |
| Rpl9 | -0.030212552 | 0.010844832 | 0.2035210 | 4.28223139 | 4.134170616 |
| Rpl9-ps6 | -0.033268424 | 0.164748075 | 0.6420707 | 0.111743372 | 0.140989138 |
| Rplp0 | -0.057200202 | 0.000105847 | 0.0097826 | 3.385832047 | 3.32026599 |
| Rplp1 | -0.019056847 | 0.126351458 | 0.5889350 | 4.120560419 | 4.038414132 |
| Rplp2 | -0.002076207 | 0.882750833 | 0.9799360 | 3.472155113 | 3.400979884 |
| Rpn1 | 0.120081872 | 0.000937805 | 0.0462837 | 1.254060201 | 1.065428896 |
| Rpn2 | -0.014240625 | 0.681888235 | 0.9303225 | 1.252236086 | 1.247239594 |
| Rpp14 | 0.003758629 | 0.86353738 | 0.9769675 | 0.1186346 | 0.105534402 |
| Rpp21 | -0.005001316 | 0.887467161 | 0.9812414 | 0.491856958 | 0.500564551 |
| Rpp25l | -0.014939793 | 0.616643966 | 0.9115017 | 0.246308052 | 0.244739982 |
| Rpp30 | -0.012035835 | 0.617341536 | 0.9115017 | 0.139371368 | 0.132693679 |
| Rpp38 | 0.001662067 | 0.944712543 | 0.9936932 | 0.151373578 | 0.130061886 |
| Rprd1a | 0.035941891 | 0.208477132 | 0.6807660 | 0.227104145 | 0.185511623 |
| Rprd1b | -0.011731589 | 0.725732192 | 0.9450332 | 0.343505634 | 0.364448055 |
| Rprd2 | 0.018712075 | 0.574311981 | 0.8974946 | 0.338052406 | 0.309230451 |
| Rps10 | -0.026375881 | 0.380375163 | 0.8117350 | 1.622923307 | 1.443500545 |
| Rps10-ps1 | -0.00733134 | 0.75962751 | 0.9521533 | 0.173795166 | 0.131125951 |
| Rps10-ps2 | -0.027325449 | 0.375426883 | 0.8099683 | 0.299873035 | 0.250812481 |
| Rps11 | 0.005917122 | 0.707230109 | 0.9408354 | 3.096149463 | 3.004718644 |
| Rps11-ps1 | 0.004523209 | 0.854964913 | 0.9755431 | 0.147669289 | 0.135907416 |
| Rps11-ps2 | 0.01307501 | 0.544098855 | 0.8842645 | 0.102384821 | 0.101338603 |
| Rps12 | -0.031028341 | 0.022479603 | 0.2914468 | 3.557621384 | 3.532952414 |
| Rps13 | -0.015527885 | 0.243479305 | 0.7132719 | 3.635729971 | 3.514036351 |
| Rps14 | -0.029497145 | 0.015737831 | 0.2462449 | 4.032871418 | 4.058573808 |
| Rps15 | -0.054429145 | 0.002897468 | 0.1000185 | 3.087978863 | 3.174419875 |
| Rps15a | 0.006751893 | 0.64823369 | 0.9170676 | 3.376844502 | 3.185648806 |
| Rps15a-ps5 | 0.028457408 | 0.055827877 | 0.4449332 | 0.069561886 | 0.036468736 |
| Rps16 | -0.019876054 | 0.165302806 | 0.6420707 | 3.615500181 | 3.472481623 |
| Rps17 | 0.004133651 | 0.820138863 | 0.9700462 | 2.905433933 | 2.879108853 |

| | | | | | |
|---|---|---|---|---|---|
| Rps18 | -0.045991588 | 0.0020491 | 0.0803441 | 3.382265653 | 3.362927659 |
| Rps19 | 0.053405091 | 0.008314174 | 0.1793097 | 2.621494964 | 2.400952566 |
| Rps19bp1 | 0.016656189 | 0.613450871 | 0.9108386 | 0.33106537 | 0.313147046 |
| Rps2 | -0.05699339 | 0.000232774 | 0.0180400 | 3.612333011 | 3.437931644 |
| Rps20 | -0.021618803 | 0.083754177 | 0.5183040 | 3.87531648 | 3.807899318 |
| Rps21 | 0.056092805 | 1.16E-06 | 0.0002449 | 4.193758997 | 3.954297556 |
| Rps23 | 0.017869243 | 0.197495257 | 0.6730962 | 3.56319479 | 3.383208732 |
| Rps23-ps1 | -0.051784571 | 0.048872732 | 0.4165098 | 0.166320773 | 0.187101085 |
| Rps24 | -0.018104198 | 0.076917117 | 0.5015240 | 4.586649637 | 4.459791751 |
| Rps25 | -0.030203331 | 0.111680007 | 0.5675761 | 2.624748853 | 2.629408384 |
| Rps26 | 0.015866278 | 0.452606008 | 0.8486662 | 2.832266196 | 2.582029743 |
| Rps26-ps1 | 0.002518804 | 0.899704206 | 0.9825441 | 0.09158793 | 0.087407922 |
| Rps27 | 0.066913301 | 4.03E-07 | 0.0001290 | 3.66453302 | 3.478467041 |
| Rps27a | -0.008491035 | 0.467030326 | 0.8555754 | 4.266966886 | 4.095763067 |
| Rps27a-ps3 | 0.00571636 | 0.832710223 | 0.9723494 | 0.206228333 | 0.175174986 |
| Rps27l | 0.091519278 | 0.000282234 | 0.0207389 | 2.094916788 | 1.901878574 |
| Rps27rt | 0.097511855 | 0.001223648 | 0.0559382 | 0.316486037 | 0.229423563 |
| Rps28 | 0.075507221 | 0.007093097 | 0.1680001 | 1.733451351 | 1.575830876 |
| Rps29 | 0.045278804 | 7.37E-05 | 0.0073749 | 3.971540168 | 3.804887537 |
| Rps2-ps10 | -0.01689111 | 0.377811668 | 0.8101797 | 0.071771093 | 0.08405698 |
| Rps3 | -0.065143367 | 9.16E-07 | 0.0002072 | 3.474219265 | 3.502732195 |
| Rps3a1 | -0.056311731 | 5.31E-06 | 0.0008490 | 4.276416882 | 4.185134151 |
| Rps4l | 0.019947585 | 0.621803934 | 0.9121505 | 0.732548056 | 0.689470849 |
| Rps4x | -0.038630782 | 0.008161065 | 0.1775389 | 3.515618644 | 3.395022326 |
| Rps5 | -0.050583139 | 0.000486608 | 0.0301697 | 3.607506966 | 3.54041624 |
| Rps6 | -0.051404829 | 0.016468143 | 0.2525257 | 2.203494458 | 2.084056648 |
| Rps6ka2 | 0.021973562 | 0.429797869 | 0.8340365 | 0.203357788 | 0.194060738 |
| Rps6ka3 | 0.04966793 | 0.07834948 | 0.5056175 | 0.229703744 | 0.188357224 |
| Rps6kb1 | -0.023964626 | 0.452625892 | 0.8486662 | 0.300541003 | 0.322972813 |
| Rps6kb2 | 0.007414329 | 0.74370645 | 0.9479080 | 0.121292125 | 0.116074114 |
| Rps6kc1 | 0.01648555 | 0.252504763 | 0.7223071 | 0.054774855 | 0.039994001 |
| Rps6kl1 | 0.003429023 | 0.830249568 | 0.9722601 | 0.058978505 | 0.049284478 |
| Rps6-ps4 | -0.024502415 | 0.328312738 | 0.7778857 | 0.153502592 | 0.145970498 |
| Rps7 | -0.006421618 | 0.718810987 | 0.9440758 | 2.89772549 | 2.65965318 |
| Rps7-ps3 | -0.011846492 | 0.401861403 | 0.8254259 | 0.045248825 | 0.04270676 |
| Rps8 | 0.002324087 | 0.847478826 | 0.9742716 | 4.016416579 | 3.938028643 |
| Rps9 | -0.015742921 | 0.304199515 | 0.7634877 | 3.308699362 | 3.169045899 |
| Rpsa | -0.066779501 | 5.29E-06 | 0.0008490 | 3.494719696 | 3.376732504 |
| Rpsa-ps10 | -0.047494491 | 0.070396016 | 0.4863011 | 0.149426302 | 0.166892429 |
| Rptor | -0.011015405 | 0.638104588 | 0.9160042 | 0.128457971 | 0.129129958 |
| Rpusd1 | -0.00413359 | 0.826684489 | 0.9716906 | 0.068341357 | 0.078693509 |
| Rpusd4 | -0.000492095 | 0.97762911 | 0.9976072 | 0.068666149 | 0.066814951 |
| Rraga | -0.085520344 | 0.016252199 | 0.2514444 | 0.702841248 | 0.710226747 |
| Rragb | -0.01728524 | 0.549016622 | 0.8871105 | 0.207855302 | 0.189106518 |
| Rragc | 0.035083892 | 0.294915571 | 0.7536588 | 0.36346816 | 0.327586354 |
| Rragd | -0.014299467 | 0.649602727 | 0.9174778 | 0.233053157 | 0.292761712 |
| Rras | -0.013513477 | 0.577525264 | 0.8997914 | 0.13550018 | 0.141652707 |
| Rras2 | -0.014808219 | 0.461726884 | 0.8524425 | 0.084355344 | 0.089064613 |
| Rrbp1 | -0.020174377 | 0.484074097 | 0.8625857 | 1.795718935 | 1.841967896 |
| Rreb1 | -0.0257945 | 0.262538722 | 0.7275735 | 0.096012357 | 0.128857397 |
| Rrm1 | -0.017771935 | 0.34770488 | 0.7917953 | 0.073986537 | 0.082373821 |
| Rrm2b | -0.021496314 | 0.533629454 | 0.8790442 | 0.411211724 | 0.419915385 |
| Rrn3 | 0.009236406 | 0.698089436 | 0.9369692 | 0.143617451 | 0.129458538 |
| Rrnad1 | 0.007348078 | 0.681907642 | 0.9303225 | 0.073371891 | 0.065453647 |
| Rrp1 | -0.00117001 | 0.974147796 | 0.9976072 | 0.796818867 | 0.786865945 |
| Rrp15 | -0.006560202 | 0.774846963 | 0.9558538 | 0.128406586 | 0.116019519 |
| Rrp1b | 0.003377377 | 0.888751445 | 0.9816521 | 0.145553808 | 0.13248511 |
| Rrp36 | -0.041605909 | 0.095175866 | 0.5351767 | 0.138528818 | 0.157557834 |
| Rrp7a | -0.067986736 | 0.028040316 | 0.3208303 | 0.274231828 | 0.29753771 |
| Rrp8 | 0.003348539 | 0.902403154 | 0.9825441 | 0.172465386 | 0.179372101 |
| Rrp9 | 0.022364164 | 0.279518779 | 0.7405297 | 0.113386815 | 0.087183873 |
| Rrs1 | -0.032109971 | 0.233964493 | 0.7056961 | 0.17199031 | 0.180915857 |
| Rsad2 | 0.082964949 | 1.30E-05 | 0.0018933 | 0.104131675 | 0.047619992 |
| Rsbn1 | -0.002161513 | 0.950462959 | 0.9948462 | 0.370919107 | 0.382935218 |
| Rsbn1l | -0.078451096 | 0.032866677 | 0.3439213 | 0.465697784 | 0.561051341 |
| Rsf1 | -0.035772743 | 0.349026511 | 0.7927303 | 0.562701586 | 0.582924737 |

| | | | | | |
|---|---|---|---|---|---|
| Rsl1d1 | 0.050452799 | 0.159334728 | 0.6360565 | 0.542867745 | 0.487062905 |
| Rsl24d1 | 0.006818427 | 0.78038299 | 0.9590847 | 0.156350569 | 0.133209943 |
| Rsph1 | -0.029871162 | 0.149862988 | 0.6272497 | 0.082211806 | 0.096147833 |
| Rsph3a | -0.000882544 | 0.958174952 | 0.9957785 | 0.060962249 | 0.057798932 |
| Rsph3b | 0.018385732 | 0.222015345 | 0.6936668 | 0.0608307 | 0.042910847 |
| Rsph9 | 0.00560254 | 0.839685301 | 0.9727523 | 0.186387935 | 0.179340797 |
| Rspo4 | 0.039501704 | 0.134714308 | 0.6013051 | 0.170312908 | 0.158873314 |
| Rspry1 | -0.049688649 | 0.092476427 | 0.5325584 | 0.198762054 | 0.235944493 |
| Rsrc1 | -0.046048746 | 0.165220767 | 0.6420707 | 0.348469457 | 0.371839191 |
| Rsrc2 | -0.00029069 | 0.993406985 | 0.9985888 | 0.940837866 | 0.946845331 |
| Rsrp1 | -0.116721942 | 0.001563282 | 0.0679368 | 1.003597457 | 1.11139711 |
| Rsu1 | -0.071266269 | 0.007158559 | 0.1680001 | 0.141675431 | 0.179824448 |
| Rtca | 0.008452832 | 0.730784149 | 0.9459020 | 0.163648366 | 0.137770332 |
| Rtcb | 0.051772105 | 0.131937869 | 0.5978345 | 0.564152522 | 0.484803117 |
| Rtf1 | -0.02881097 | 0.412549687 | 0.8297979 | 0.435572297 | 0.452859017 |
| Rtf2 | -0.016788101 | 0.604790259 | 0.9087139 | 0.334557379 | 0.328773829 |
| Rtkn | 0.012837265 | 0.473817695 | 0.8597227 | 0.077015725 | 0.058183336 |
| Rtl5 | -0.006841941 | 0.759594414 | 0.9521533 | 0.102537794 | 0.116658003 |
| Rtl6 | -0.027896317 | 0.149274435 | 0.6264374 | 0.065234271 | 0.088822418 |
| Rtl8a | -0.069839837 | 0.025049781 | 0.3041540 | 0.263598291 | 0.286087156 |
| Rtl8b | 0.012976052 | 0.647208024 | 0.9169695 | 0.23272478 | 0.188595772 |
| Rtn1 | 0.009017109 | 0.755006262 | 0.9519143 | 0.237976359 | 0.208326888 |
| Rtn2 | -0.031920859 | 0.203591847 | 0.6772740 | 0.156923446 | 0.159957379 |
| Rtn3 | 0.026844076 | 0.451348914 | 0.8480124 | 0.929555265 | 0.845615233 |
| Rtn4 | 0.024717545 | 0.487553264 | 0.8640311 | 0.843407799 | 0.891416754 |
| Rtn4rl1 | -0.016099863 | 0.504998202 | 0.8695566 | 0.116918374 | 0.139289201 |
| Rtraf | 0.022787219 | 0.49972293 | 0.8692357 | 1.076296023 | 0.930678262 |
| Rubcn | 0.012785616 | 0.637881615 | 0.9160042 | 0.189635588 | 0.178179062 |
| Rufy1 | -0.033636108 | 0.221425566 | 0.6929153 | 0.174606781 | 0.197150375 |
| Rufy2 | -0.028994264 | 0.309455768 | 0.7673813 | 0.186155622 | 0.20435091 |
| Rufy3 | -0.034442335 | 0.316370856 | 0.7705987 | 0.359557369 | 0.405432044 |
| Rundc1 | -0.018590832 | 0.478666408 | 0.8608358 | 0.153142775 | 0.167474696 |
| Rundc3a | -0.033352266 | 0.337839376 | 0.7834245 | 0.776479128 | 0.798476792 |
| Rundc3b | 0.040176278 | 0.11547865 | 0.5724670 | 0.183005929 | 0.157073661 |
| Runx1t1 | 0.025794495 | 0.478440447 | 0.8607489 | 0.399146713 | 0.379409431 |
| Rusc1 | -0.037339066 | 0.193989452 | 0.6716564 | 0.190224323 | 0.212153954 |
| Ruvbl1 | -0.026738864 | 0.29240799 | 0.7526152 | 0.162140895 | 0.160231364 |
| Ruvbl2 | 0.01614296 | 0.50157493 | 0.8695566 | 0.152790216 | 0.133353545 |
| Rwdd1 | 0.074819731 | 0.037657342 | 0.3691312 | 0.923575953 | 0.877146588 |
| Rwdd2a | 0.008152769 | 0.605849566 | 0.9087139 | 0.057879435 | 0.048415868 |
| Rwdd3 | -0.029860304 | 0.093274506 | 0.5330587 | 0.053106502 | 0.073260592 |
| Rwdd4a | -0.001378805 | 0.97030687 | 0.9976072 | 0.551342893 | 0.573631267 |
| Rxra | -0.045517713 | 0.077277438 | 0.5022647 | 0.134803725 | 0.169612981 |
| Rxrb | -0.068901617 | 0.024592431 | 0.3041540 | 0.238486707 | 0.275601992 |
| Rxylt1 | -0.007130547 | 0.824356687 | 0.971271 | 0.313991733 | 0.316479587 |
| Rybp | 0.039002012 | 0.243992949 | 0.7132719 | 0.376152323 | 0.361634355 |
| Ryk | 0.015405965 | 0.614773397 | 0.9108386 | 0.277144068 | 0.266633752 |
| S100a1 | -0.047672971 | 0.197748267 | 0.6730962 | 0.383000433 | 0.516624935 |
| S100a10 | 0.175626376 | 3.00E-07 | 9.92E-05 | 0.400750911 | 0.303406473 |
| S100a11 | -0.021473863 | 0.507270999 | 0.8704067 | 1.611860267 | 1.459989382 |
| S100a13 | 0.001787816 | 0.957345179 | 0.9956630 | 0.357175612 | 0.391006786 |
| S100a16 | -0.005551452 | 0.738088452 | 0.9474220 | 0.049427046 | 0.054304538 |
| S100a6 | 0.00820132 | 0.713874748 | 0.9428150 | 0.072233548 | 0.087241234 |
| S100pbp | -0.011627976 | 0.641475976 | 0.9166712 | 0.149122508 | 0.144447189 |
| Saal1 | 0.008728049 | 0.549078529 | 0.8871105 | 0.054871985 | 0.041850518 |
| Sac3d1 | 0.00750075 | 0.785403375 | 0.9602516 | 0.216645906 | 0.188844088 |
| Sacm1l | 0.004982613 | 0.879346274 | 0.9788055 | 0.363741873 | 0.342511588 |
| Sacs | 0.000502924 | 0.976957926 | 0.9976072 | 0.060143227 | 0.062853318 |
| Sae1 | -0.025743401 | 0.40032807 | 0.8246612 | 0.26240432 | 0.264461182 |
| Safb | -0.055392 | 0.108364672 | 0.5636817 | 0.395684794 | 0.410753696 |
| Safb2 | -0.076457139 | 0.024622462 | 0.3041540 | 0.304396134 | 0.387880461 |
| Sall2 | -0.008571024 | 0.586600716 | 0.9035288 | 0.05461723 | 0.054334722 |
| Sall4 | -0.030565446 | 0.060036329 | 0.4577712 | 0.040586751 | 0.060777735 |
| Samd1 | 0.005029272 | 0.835050757 | 0.9723494 | 0.152786109 | 0.1240709 |
| Samd10 | 0.006426121 | 0.801614633 | 0.9647143 | 0.180726416 | 0.154880008 |
| Samd12 | 0.037027437 | 0.048166226 | 0.4147647 | 0.09213154 | 0.068141162 |

| | | | | | |
|---|---|---|---|---|---|
| Samd14 | -0.000954537 | 0.958896223 | 0.9957785 | 0.087354084 | 0.07104031 |
| Samd4 | 0.003656648 | 0.844337333 | 0.9734889 | 0.071749492 | 0.072208267 |
| Samd4b | 0.043346822 | 0.20754234 | 0.6806016 | 0.411994771 | 0.410098207 |
| Samd5 | 0.011971194 | 0.616302597 | 0.9114903 | 0.140331226 | 0.108701744 |
| Samd8 | 0.00710301 | 0.805771743 | 0.9647143 | 0.212504634 | 0.206268252 |
| Samd9l | 0.01583343 | 0.488058282 | 0.8642517 | 0.131966683 | 0.102755739 |
| Samhd1 | -0.015534509 | 0.539191392 | 0.8823455 | 0.150084662 | 0.147125434 |
| Samm50 | -0.024893752 | 0.458823986 | 0.8522649 | 0.389703902 | 0.384471418 |
| Sap130 | 0.002344142 | 0.910688392 | 0.9837373 | 0.100191335 | 0.094154991 |
| Sap18 | 0.022571695 | 0.544197061 | 0.8842645 | 0.662012001 | 0.572169128 |
| Sap30 | -0.040934428 | 0.090983079 | 0.5321282 | 0.125036305 | 0.142925164 |
| Sap30bp | -0.000589592 | 0.981619746 | 0.9976072 | 0.167110321 | 0.162515696 |
| Sap30l | -0.043384701 | 0.173792847 | 0.6527657 | 0.301870194 | 0.321714273 |
| Sar1a | 0.017583479 | 0.617305186 | 0.9115017 | 0.958972317 | 0.920529409 |
| Sar1b | -0.011772409 | 0.730566522 | 0.9459020 | 0.860525936 | 0.859974716 |
| Saraf | 0.017470613 | 0.577778566 | 0.8997914 | 1.327907351 | 1.35099211 |
| Sarnp | 0.061591307 | 0.092213305 | 0.5325584 | 0.709910044 | 0.584233725 |
| Sars | 0.016120698 | 0.623637304 | 0.9121505 | 1.274844806 | 1.302447814 |
| Sart1 | -0.065057802 | 0.021360906 | 0.2840485 | 0.170992909 | 0.218209473 |
| Sart3 | -0.004405005 | 0.860589351 | 0.9769675 | 0.1569803 | 0.144160499 |
| Sash1 | -0.031950039 | 0.149088421 | 0.6264374 | 0.09769187 | 0.112585855 |
| Sat1 | 0.014281546 | 0.688768146 | 0.9334125 | 0.327219905 | 0.341924689 |
| Sat2 | -0.004574316 | 0.833504357 | 0.9723494 | 0.111949984 | 0.107034853 |
| Satb1 | 0.003879389 | 0.892781414 | 0.9821882 | 0.212317615 | 0.199012022 |
| Sav1 | -0.014003191 | 0.540225154 | 0.8833086 | 0.11805312 | 0.125378279 |
| Saxo2 | 0.005470829 | 0.685590807 | 0.9316044 | 0.043648298 | 0.036849113 |
| Saysd1 | -0.012398924 | 0.695402874 | 0.9362759 | 0.307555246 | 0.295430421 |
| Sbds | -0.034372013 | 0.347713737 | 0.7917953 | 0.64562403 | 0.658656361 |
| Sbf1 | -0.010695061 | 0.57900264 | 0.9003073 | 0.078089232 | 0.086000984 |
| Sbf2 | -0.037453948 | 0.113909024 | 0.5696977 | 0.107747836 | 0.140440629 |
| Sbno1 | 0.006593213 | 0.862294213 | 0.9769675 | 0.695239457 | 0.67723152 |
| Sbno2 | -0.023026551 | 0.256624351 | 0.7246551 | 0.080374861 | 0.089336299 |
| Sc5d | 0.022342404 | 0.383335759 | 0.8129787 | 0.179486774 | 0.145311161 |
| Scaf1 | -0.041378975 | 0.165307322 | 0.6420707 | 0.22920872 | 0.24748741 |
| Scaf11 | 0.031769923 | 0.394108771 | 0.8211634 | 1.012866624 | 0.959171217 |
| Scaf4 | -0.002885399 | 0.903519932 | 0.9827712 | 0.120776053 | 0.134154172 |
| Scaf8 | 0.016800226 | 0.568877234 | 0.8956532 | 0.224236494 | 0.200699438 |
| Scai | -0.010485755 | 0.633156833 | 0.9141196 | 0.09860322 | 0.104376818 |
| Scamp1 | 0.046169437 | 0.113756738 | 0.5696977 | 0.247961386 | 0.22594252 |
| Scamp2 | 0.005522248 | 0.871204906 | 0.9782269 | 0.454688773 | 0.428608498 |
| Scamp3 | 0.021545202 | 0.481613036 | 0.8620306 | 0.297584108 | 0.265462137 |
| Scamp5 | 0.012638338 | 0.607208858 | 0.9087139 | 0.142450727 | 0.144674628 |
| Scand1 | 0.055171141 | 0.031902255 | 0.3385228 | 1.532972552 | 1.45608013 |
| Scap | -0.0049389 | 0.849763679 | 0.9748841 | 0.15686301 | 0.169652939 |
| Scaper | 0.034050234 | 0.269909353 | 0.7340360 | 0.263562722 | 0.245563307 |
| Scarb1 | 0.01275412 | 0.388509386 | 0.8170287 | 0.055719525 | 0.0409052 |
| Scarb2 | 0.023198662 | 0.506300147 | 0.8695566 | 0.45083907 | 0.47208332 |
| Sccpdh | -0.033994099 | 0.145193655 | 0.6200262 | 0.109840881 | 0.129759884 |
| Scd2 | 0.045357972 | 0.123015445 | 0.5833237 | 0.228522803 | 0.177914792 |
| Scfd1 | 0.000472979 | 0.989817119 | 0.9980836 | 0.83972945 | 0.783183179 |
| Scfd2 | -0.006425132 | 0.736272579 | 0.9469498 | 0.07907426 | 0.082111636 |
| Scg2 | -0.007203211 | 0.747824764 | 0.9496370 | 3.48091828 | 3.834712875 |
| Scg3 | -0.006858577 | 0.719973521 | 0.9444523 | 2.826239284 | 2.960486058 |
| Scg5 | -0.067088976 | 4.23E-07 | 0.0001311 | 3.569542128 | 3.58717269 |
| Scgn | 0.031156858 | 0.413827082 | 0.8300990 | 0.846683884 | 0.737590366 |
| Scin | 0.00867492 | 0.565958194 | 0.8945674 | 0.063387461 | 0.038417319 |
| Sclt1 | -0.005597296 | 0.835610692 | 0.9723494 | 0.164357276 | 0.170923496 |
| Scly | -0.051612933 | 0.129307853 | 0.5922132 | 0.339607094 | 0.391034839 |
| Scmh1 | -0.030812024 | 0.250067248 | 0.7207210 | 0.163618591 | 0.177799836 |
| Scn1b | 0.00440088 | 0.767807758 | 0.9544221 | 0.046022208 | 0.045974638 |
| Scn3a | 0.029233792 | 0.37059288 | 0.8066125 | 0.288953267 | 0.248191297 |
| Scn8a | -0.030972705 | 0.208850564 | 0.6807660 | 0.127794204 | 0.14552707 |
| Scn9a | -0.035019761 | 0.253880212 | 0.7223071 | 0.196602565 | 0.243286983 |
| Scnm1 | -0.035636889 | 0.192064299 | 0.6700281 | 0.187653933 | 0.189850256 |
| Scnn1b | -0.01464747 | 0.685541849 | 0.9316044 | 0.442784027 | 0.467420564 |
| Sco2 | 0.012101409 | 0.512463033 | 0.8722771 | 0.077638173 | 0.069987574 |

| | | | | | |
|---|---|---|---|---|---|
| Scoc | -0.000403753 | 0.991203805 | 0.99824789 | 0.519719439 | 0.48618317 |
| Scp2 | 0.037807993 | 0.28765221 | 0.74723242 | 0.781016256 | 0.645326056 |
| Scpep1 | -0.075104352 | 0.01238026 | 0.21952968 | 0.191563234 | 0.263972984 |
| Scrib | -0.008234034 | 0.695410212 | 0.93627599 | 0.094779833 | 0.099650269 |
| Scrn1 | -0.01396521 | 0.647588721 | 0.91696950 | 0.235151799 | 0.253509606 |
| Scrn2 | -0.019513173 | 0.266176372 | 0.73073994 | 0.057198208 | 0.069980135 |
| Scrn3 | 0.005329962 | 0.84510403 | 0.97348890 | 0.207040904 | 0.188167266 |
| Scrt1 | -0.02473902 | 0.221608218 | 0.69326821 | 0.073773576 | 0.093879669 |
| Scyl1 | -0.012985746 | 0.660703796 | 0.92117801 | 0.247534287 | 0.245758924 |
| Scyl2 | -0.064958371 | 0.006663904 | 0.16179671 | 0.097434155 | 0.14976104 |
| Scyl3 | 0.007351348 | 0.719646307 | 0.94445237 | 0.090623219 | 0.088620303 |
| Sdad1 | -0.026678408 | 0.300121036 | 0.75902757 | 0.151512838 | 0.155311393 |
| Sdc2 | -0.048165892 | 0.173709211 | 0.65276574 | 0.482487558 | 0.455684417 |
| Sdc4 | 0.003659864 | 0.923956539 | 0.98767767 | 0.840366913 | 0.819893291 |
| Sdcbp | -0.024417527 | 0.494880528 | 0.86828690 | 0.636966049 | 0.628245302 |
| Sdccag8 | 0.059728484 | 0.024948822 | 0.30415400 | 0.213950016 | 0.161367165 |
| Sde2 | -0.024732361 | 0.462233228 | 0.85244258 | 0.353629908 | 0.37186517 |
| Sdf2 | -0.00621685 | 0.859661354 | 0.97696750 | 0.605827605 | 0.574864086 |
| Sdf2l1 | 0.137941962 | 0.001745761 | 0.07276447 | 0.996641365 | 0.797503551 |
| Sdf4 | -0.087829243 | 0.003384498 | 0.11117293 | 1.415123359 | 1.423937067 |
| Sdha | -0.014019775 | 0.690553415 | 0.93462378 | 0.62459971 | 0.601643783 |
| Sdhaf1 | -0.016126331 | 0.648807552 | 0.91725697 | 0.438565883 | 0.414486979 |
| Sdhaf2 | 0.028268252 | 0.261924268 | 0.72757354 | 0.181801521 | 0.143469724 |
| Sdhaf3 | 0.039785544 | 0.058008577 | 0.45081999 | 0.125481506 | 0.086554444 |
| Sdhaf4 | -0.003094922 | 0.928299574 | 0.98806134 | 0.443170477 | 0.421279751 |
| Sdhb | 0.004009715 | 0.906405686 | 0.98354237 | 0.88072378 | 0.797242675 |
| Sdhc | 0.080887805 | 0.023967848 | 0.30050578 | 0.624904972 | 0.530077295 |
| Sdhd | 0.009480738 | 0.793273574 | 0.96245081 | 0.678932262 | 0.591028462 |
| Sdk1 | -0.008492432 | 0.622648659 | 0.91215057 | 0.061365019 | 0.062568307 |
| Sdk2 | 0.00845619 | 0.803522605 | 0.96471431 | 0.300761603 | 0.379180741 |
| Sdr39u1 | -0.041094409 | 0.120558308 | 0.58083458 | 0.170893732 | 0.184754387 |
| Sdsl | -0.045604645 | 0.019473914 | 0.27081127 | 0.063047152 | 0.091464919 |
| Sec11a | -0.011608351 | 0.74353665 | 0.94790800 | 0.829041935 | 0.80386176 |
| Sec11c | 0.034384638 | 0.308186041 | 0.76626060 | 1.477802309 | 1.500244578 |
| Sec13 | 0.05844119 | 0.096199342 | 0.53697250 | 0.615611421 | 0.525773575 |
| Sec14l1 | 0.001607129 | 0.954150755 | 0.99546908 | 0.182778072 | 0.198944036 |
| Sec16a | -0.045397201 | 0.182830418 | 0.66075941 | 0.346570614 | 0.383779147 |
| Sec16b | -0.014408789 | 0.426200449 | 0.83324959 | 0.050722395 | 0.077847897 |
| Sec22a | -0.006727654 | 0.790923335 | 0.96224569 | 0.172177692 | 0.1605404 |
| Sec22b | 0.019112977 | 0.594419248 | 0.90658293 | 0.944139767 | 0.850325315 |
| Sec22c | -0.002060405 | 0.908788361 | 0.98373739 | 0.069882235 | 0.071770375 |
| Sec23a | 0.011493436 | 0.645833053 | 0.91696950 | 0.152119364 | 0.13942176 |
| Sec23b | -0.082320589 | 0.021763705 | 0.28740251 | 0.440069349 | 0.490926263 |
| Sec23ip | 0.021658623 | 0.473676882 | 0.85970928 | 0.256896515 | 0.239538467 |
| Sec24a | -0.00081964 | 0.979551136 | 0.99760728 | 0.268047426 | 0.26603297 |
| Sec24b | 0.009641854 | 0.72111414 | 0.94456522 | 0.182313683 | 0.170547022 |
| Sec24c | 0.020461582 | 0.496833787 | 0.86846292 | 0.267426874 | 0.248968423 |
| Sec24d | 0.018184409 | 0.570275868 | 0.89625342 | 0.31989905 | 0.294429205 |
| Sec31a | 0.006766737 | 0.849286795 | 0.97488410 | 0.789983129 | 0.772313004 |
| Sec61a1 | -0.124628263 | 0.000905511 | 0.04582992 | 0.629139788 | 0.755097557 |
| Sec61a2 | -0.017341443 | 0.417470535 | 0.83009907 | 0.091661332 | 0.10726383 |
| Sec61b | 0.038441906 | 0.043442693 | 0.39842159 | 2.844188665 | 2.721698297 |
| Sec61g | 0.130726804 | 0.000334632 | 0.02305244 | 1.151943153 | 1.014819051 |
| Sec62 | 0.016875851 | 0.547798468 | 0.88630011 | 1.552672418 | 1.569305323 |
| Sec63 | -0.011451496 | 0.76325115 | 0.95321287 | 0.632074612 | 0.647879439 |
| Secisbp2 | -0.051849923 | 0.045328867 | 0.40347568 | 0.135870862 | 0.172085237 |
| Secisbp2l | -0.037949482 | 0.114016567 | 0.56969774 | 0.110751802 | 0.14349112 |
| Seh1l | 0.031871483 | 0.269771248 | 0.73403607 | 0.2469805 | 0.207334031 |
| Sel1l | 0.089562331 | 0.012967404 | 0.22528310 | 0.484943874 | 0.47561432 |
| Sel1l3 | -0.013683749 | 0.595136367 | 0.9067325 | 0.161423926 | 0.15819635 |
| Selenof | 0.023499424 | 0.405877644 | 0.82707700 | 1.581260041 | 1.47444516 |
| Selenoh | -0.040806636 | 0.235746161 | 0.70793905 | 0.420478952 | 0.42719494 |
| Selenoi | -0.004335548 | 0.887792849 | 0.98124141 | 0.275024936 | 0.236320454 |
| Selenok | -0.007777095 | 0.730011656 | 0.94590205 | 2.195230563 | 2.152753685 |
| Selenom | 0.072645301 | 0.01281131 | 0.22335359 | 1.713638726 | 1.650393041 |
| Selenon | 0.025070183 | 0.341675689 | 0.78531570 | 0.187417554 | 0.154296132 |

| | | | | | |
|---|---|---|---|---|---|
| Selenoo | 0.011299282 | 0.627631145 | 0.91280959 | 0.140685769 | 0.119752226 |
| Selenop | 0.077993767 | 0.066210682 | 0.47728867 | 0.962083518 | 0.877216236 |
| Selenos | 0.02300909 | 0.511774935 | 0.87208730 | 1.178525328 | 1.186890803 |
| Selenot | 0.070449052 | 0.058048103 | 0.45081999 | 0.813226524 | 0.759506104 |
| Selenow | 0.185328901 | 1.59E-06 | 0.00030991 | 1.265717512 | 1.061352751 |
| Sem1 | 0.096272173 | 4.21E-05 | 0.00486194 | 2.033757488 | 1.794957402 |
| Sema3e | -0.02980371 | 0.218860236 | 0.69016901 | 0.133923891 | 0.133051586 |
| Sema4a | -0.020935613 | 0.19248853 | 0.67002811 | 0.044694179 | 0.057985605 |
| Sema4b | 0.00632658 | 0.7457148 | 0.94857174 | 0.089023969 | 0.081275244 |
| Sema4d | -0.002570535 | 0.890158493 | 0.98195975 | 0.070199533 | 0.075057829 |
| Sema6d | -0.040985798 | 0.021027531 | 0.28117593 | 0.038992041 | 0.075594792 |
| Senp1 | 0.016871048 | 0.424669973 | 0.83244601 | 0.108701503 | 0.092708252 |
| Senp2 | 0.005699188 | 0.849634187 | 0.97488410 | 0.245083879 | 0.230777046 |
| Senp3 | 0.017076332 | 0.492847951 | 0.86693011 | 0.158403685 | 0.146647916 |
| Senp5 | -0.017112678 | 0.531364389 | 0.87764481 | 0.183664221 | 0.191523082 |
| Senp6 | -0.013303541 | 0.7105374 | 0.94193701 | 1.000996277 | 1.002901634 |
| Senp7 | -0.028874934 | 0.320844133 | 0.77218975 | 0.218986986 | 0.224541307 |
| Senp8 | -0.01955988 | 0.339989874 | 0.78446580 | 0.101424092 | 0.094774396 |
| Sephs1 | -0.019396886 | 0.325405658 | 0.77630994 | 0.086512375 | 0.089631756 |
| Sephs2 | -0.035697974 | 0.246447344 | 0.71614119 | 0.257566801 | 0.283373152 |
| Sepsecs | -0.003512561 | 0.896406642 | 0.98254414 | 0.185006137 | 0.175734832 |
| Septin10 | 0.010934688 | 0.636324848 | 0.91535418 | 0.140930437 | 0.113873071 |
| Septin11 | 0.000649719 | 0.982144013 | 0.99760728 | 0.197854057 | 0.213091315 |
| Septin3 | 0.014708913 | 0.352590547 | 0.79529291 | 0.054440173 | 0.047946108 |
| Septin7 | -0.049519003 | 0.170897596 | 0.6507063 | 0.895044881 | 0.895449797 |
| Septin8 | -0.03348377 | 0.11045127 | 0.56558881 | 0.082489895 | 0.102672272 |
| Septin9 | 0.014030918 | 0.653969484 | 0.92006485 | 0.257260303 | 0.262079564 |
| Serbp1 | 0.0381471 | 0.223832035 | 0.69605447 | 1.438636252 | 1.362033666 |
| Serf1 | 0.023204869 | 0.367883689 | 0.80567193 | 0.187226923 | 0.152984638 |
| Serf2 | 0.141699905 | 5.77E-07 | 0.00015894 | 1.937806951 | 1.604263939 |
| Sergef | -0.013876404 | 0.53351031 | 0.87904420 | 0.103273827 | 0.116658659 |
| Serhl | -0.025295645 | 0.11305718 | 0.56962035 | 0.045644807 | 0.057192726 |
| Serinc1 | -0.076420814 | 0.032967951 | 0.34461750 | 0.912461779 | 1.054157153 |
| Serinc3 | -0.011867219 | 0.722378482 | 0.94456522 | 0.357057041 | 0.382894948 |
| Serinc5 | -0.051804084 | 0.020185938 | 0.27543948 | 0.094172308 | 0.123387372 |
| Serp1 | -0.015189128 | 0.678067601 | 0.92818862 | 1.145686907 | 1.14599516 |
| Serp2 | -0.034779553 | 0.154583448 | 0.63049349 | 0.131359764 | 0.144534769 |
| Serpina10 | -0.072341817 | 0.072207143 | 0.48927244 | 0.534047303 | 0.648214993 |
| Serpina7 | -0.106796415 | 0.020288436 | 0.27645781 | 0.334586443 | 0.448641862 |
| Serpinb6a | 0.019653977 | 0.561789505 | 0.89262402 | 0.390178219 | 0.411295101 |
| Serpinb9 | -0.018659815 | 0.350025396 | 0.79273039 | 0.076967546 | 0.087199125 |
| Serpine1 | -0.036140057 | 0.100373723 | 0.54736570 | 0.046029513 | 0.085602541 |
| Serpine2 | -0.061912088 | 0.009775832 | 0.19791071 | 0.078356432 | 0.119780458 |
| Serping1 | -0.126540592 | 0.000569904 | 0.03365149 | 0.373315308 | 0.460264603 |
| Serpinh1 | -0.020518348 | 0.304355632 | 0.76348775 | 0.083958545 | 0.087121842 |
| Serpini1 | -0.00084606 | 0.975698226 | 0.99760728 | 0.167607968 | 0.189750091 |
| Sertad1 | -0.048569093 | 0.171555961 | 0.65154025 | 0.358446241 | 0.412571043 |
| Sertad2 | -0.026078354 | 0.243820009 | 0.71327199 | 0.090206518 | 0.117697389 |
| Sertad3 | -0.012651518 | 0.609083865 | 0.90886720 | 0.162302275 | 0.14156333 |
| Sertm1 | -0.002978653 | 0.852427137 | 0.97554315 | 0.068349358 | 0.047745053 |
| Sesn1 | -0.030428626 | 0.162719722 | 0.64007035 | 0.106886113 | 0.113142916 |
| Sesn2 | -0.00486799 | 0.893671069 | 0.98243614 | 0.40317527 | 0.414637811 |
| Sesn3 | -0.010755441 | 0.77714218 | 0.95645247 | 0.806243471 | 0.750872274 |
| Sestd1 | 0.01494552 | 0.377166525 | 0.81017979 | 0.078068706 | 0.054384992 |
| Set | -0.018695722 | 0.597719567 | 0.90755130 | 1.085062116 | 1.096173716 |
| Setbp1 | -0.008083087 | 0.772319544 | 0.95523334 | 0.182094798 | 0.188033039 |
| Setd1a | -0.00058521 | 0.977109104 | 0.99760728 | 0.081346929 | 0.089051337 |
| Setd1b | 0.024690171 | 0.45919865 | 0.85226491 | 0.362299395 | 0.319487482 |
| Setd2 | -0.041156466 | 0.242224914 | 0.71327199 | 0.288436247 | 0.32453776 |
| Setd3 | 0.027289437 | 0.44482038 | 0.84315410 | 0.566981374 | 0.536315042 |
| Setd4 | 0.001711479 | 0.908564762 | 0.98373739 | 0.047075355 | 0.044780725 |
| Setd5 | 0.029493068 | 0.432508597 | 0.83709824 | 0.638636249 | 0.574323695 |
| Setd6 | -0.000349285 | 0.986855552 | 0.99760728 | 0.097207381 | 0.103261478 |
| Setd7 | -0.003745625 | 0.906580233 | 0.98362418 | 0.266605002 | 0.274310858 |
| Setdb1 | -0.020122053 | 0.335217649 | 0.78094149 | 0.092915824 | 0.096740384 |
| Setx | 0.01562854 | 0.591222398 | 0.90602008 | 0.207618133 | 0.195282152 |

| | | | | | |
|---|---|---|---|---|---|
| Sez6l | -0.060592394 | 0.083282744 | 0.5179716 | 0.246065413 | 0.401418397 |
| Sez6l2 | -0.05330612 | 0.103197173 | 0.5553141 | 1.287454168 | 1.356334355 |
| Sf1 | 0.044301227 | 0.210836529 | 0.6822059 | 0.492679106 | 0.417896553 |
| Sf3a1 | 0.016500229 | 0.56118494 | 0.8926240 | 0.230772374 | 0.203464102 |
| Sf3a2 | -0.017108222 | 0.510108088 | 0.8711607 | 0.167531667 | 0.160962289 |
| Sf3a3 | -0.047745682 | 0.104607533 | 0.5580045 | 0.211461356 | 0.24255904 |
| Sf3b1 | -0.057347306 | 0.121114834 | 0.5811445 | 0.942478725 | 0.957240114 |
| Sf3b2 | -0.040277497 | 0.190744415 | 0.6700281 | 1.270629629 | 1.241339601 |
| Sf3b3 | -0.043530254 | 0.221856298 | 0.6933883 | 0.443061481 | 0.474385726 |
| Sf3b4 | -0.026176278 | 0.350583136 | 0.7927303 | 0.21580664 | 0.211257876 |
| Sf3b5 | 0.015168226 | 0.664038724 | 0.9230765 | 0.908138069 | 0.82743431 |
| Sf3b6 | 0.004628471 | 0.885936805 | 0.9808586 | 1.110341304 | 1.092546554 |
| Sfi1 | 0.022468741 | 0.137895451 | 0.6074257 | 0.061678716 | 0.04068059 |
| Sfmbt1 | -0.019377129 | 0.491857846 | 0.8664619 | 0.182594754 | 0.208535912 |
| Sfpq | -0.011763845 | 0.757974002 | 0.9521533 | 0.587569254 | 0.579963789 |
| Sfr1 | -0.095744694 | 0.008328428 | 0.1793097 | 0.89294041 | 0.92314227 |
| Sfrp5 | -0.12986848 | 0.00096417 | 0.0471160 | 0.787402073 | 0.700419851 |
| Sfswap | -0.003233837 | 0.910531401 | 0.9837373 | 0.204619569 | 0.206938359 |
| Sft2d1 | 0.024135993 | 0.184787687 | 0.6634433 | 0.093166889 | 0.063980593 |
| Sft2d2 | 0.024269749 | 0.262437764 | 0.7275735 | 0.113085326 | 0.098511585 |
| Sft2d3 | -0.012098206 | 0.710283586 | 0.9419370 | 0.293636311 | 0.294843685 |
| Sfxn1 | 0.014894019 | 0.677550387 | 0.9281886 | 0.610471164 | 0.622210424 |
| Sfxn2 | -0.043685575 | 0.124392052 | 0.5852543 | 0.164134001 | 0.206237312 |
| Sfxn3 | -0.01272734 | 0.439550805 | 0.8407104 | 0.046235071 | 0.059980612 |
| Sgcb | 0.035272776 | 0.133436076 | 0.5988251 | 0.152506488 | 0.116542343 |
| Sgce | 0.065257237 | 0.017338697 | 0.2570999 | 0.252990458 | 0.155387664 |
| Sgf29 | -0.00820853 | 0.686390463 | 0.9319728 | 0.088151871 | 0.09134524 |
| Sgk2 | -0.017795626 | 0.328748446 | 0.7778857 | 0.06000898 | 0.076746981 |
| Sgk3 | 0.014588474 | 0.512621151 | 0.8722771 | 0.116896609 | 0.10450494 |
| Sgms1 | -0.033710116 | 0.229968178 | 0.7019677 | 0.165135577 | 0.193051722 |
| Sgms2 | 0.113555838 | 0.000299379 | 0.0218370 | 0.342688428 | 0.227365951 |
| Sgpl1 | -0.009746874 | 0.735785845 | 0.9468380 | 0.183850601 | 0.212939317 |
| Sgpp1 | 0.050033957 | 0.094416395 | 0.5336812 | 0.262189671 | 0.23264918 |
| Sgpp2 | -0.052263125 | 0.024772058 | 0.3041540 | 0.084087676 | 0.13113524 |
| Sgsm1 | -0.033345535 | 0.230155055 | 0.7019677 | 0.185514157 | 0.201980392 |
| Sgsm2 | 0.077600122 | 0.002510847 | 0.0919099 | 0.201835995 | 0.152447559 |
| Sgsm3 | 0.0093224 | 0.699629998 | 0.9372491 | 0.144531908 | 0.133548135 |
| Sgta | -0.009657482 | 0.780056536 | 0.9589987 | 0.450954915 | 0.423051645 |
| Sgtb | -0.017627208 | 0.409171726 | 0.8276447 | 0.079731976 | 0.104536887 |
| Sh2b1 | 0.024354828 | 0.255347588 | 0.7226955 | 0.122012688 | 0.095486146 |
| Sh2b3 | 0.020180599 | 0.184403532 | 0.6628862 | 0.058856766 | 0.041878102 |
| Sh2d3c | -0.007037167 | 0.693838406 | 0.9359725 | 0.072955843 | 0.071786278 |
| Sh3bgr | 0.01519708 | 0.493153906 | 0.8669301 | 0.133339219 | 0.092355556 |
| Sh3bgrl | -0.046477213 | 0.179408724 | 0.6550070 | 0.411779865 | 0.43940043 |
| Sh3bgrl3 | -0.024678164 | 0.50398617 | 0.8695566 | 0.479399958 | 0.472026255 |
| Sh3bp4 | -0.046092287 | 0.070908859 | 0.4879008 | 0.134755167 | 0.165710204 |
| Sh3bp5l | -0.002006221 | 0.912868363 | 0.9842032 | 0.073907771 | 0.073745384 |
| Sh3d19 | 0.005560287 | 0.826936089 | 0.9717006 | 0.141684064 | 0.147859861 |
| Sh3d21 | 0.002350521 | 0.878251587 | 0.9785583 | 0.055025993 | 0.044920436 |
| Sh3gl1 | -0.059301175 | 0.06792039 | 0.4819529 | 0.303457068 | 0.32700553 |
| Sh3gl2 | 0.022925894 | 0.23193661 | 0.7030140 | 0.098149878 | 0.078464448 |
| Sh3glb1 | 0.010180282 | 0.759076501 | 0.9521533 | 1.152513553 | 1.100805244 |
| Sh3glb2 | -0.008769372 | 0.741795146 | 0.9479080 | 0.170265064 | 0.178790598 |
| Sh3kbp1 | 0.000123236 | 0.996461679 | 0.9994073 | 0.173823118 | 0.184342447 |
| Sh3pxd2a | 0.038522495 | 0.209901433 | 0.6808016 | 0.227873061 | 0.261084098 |
| Sh3yl1 | -0.009399661 | 0.742267424 | 0.9479080 | 0.212308431 | 0.204647074 |
| Sharpin | -0.01859089 | 0.447267413 | 0.8452834 | 0.128707962 | 0.143394093 |
| Shb | -0.057028548 | 0.031671661 | 0.3385228 | 0.138046995 | 0.185416062 |
| Shc1 | -0.12517077 | 0.000278373 | 0.0206079 | 0.239559248 | 0.386742411 |
| Shc2 | -0.02260179 | 0.21485606 | 0.6844200 | 0.059403147 | 0.074106247 |
| Shfl | 0.017937226 | 0.550984425 | 0.8887423 | 0.246676995 | 0.242011746 |
| Shisa5 | 0.068465824 | 0.04022801 | 0.3851948 | 0.405017245 | 0.323775611 |
| Shisal1 | 0.017286341 | 0.419688315 | 0.8300990 | 0.115426102 | 0.09226053 |
| Shisal2b | 0.100070403 | 0.019561077 | 0.2709040 | 0.401715132 | 0.388999563 |
| Shkbp1 | -0.012403411 | 0.555869808 | 0.8911667 | 0.098038526 | 0.100787447 |
| Shld1 | -0.022639304 | 0.316160002 | 0.7705987 | 0.11062177 | 0.125464468 |

| | | | | | |
|---|---|---|---|---|---|
| Shld2 | -0.015909454 | 0.504371798 | 0.86955665 | 0.134745277 | 0.12771212 |
| Shld3 | -0.047036077 | 0.00372109 | 0.11769117 | 0.031617664 | 0.064641492 |
| Shmt2 | 0.076415219 | 0.016156804 | 0.25082237 | 0.309527825 | 0.249087771 |
| Shoc2 | -0.014154932 | 0.663161189 | 0.92300582 | 0.302910477 | 0.308719265 |
| Shpk | 0.005523817 | 0.714146164 | 0.9428174 | 0.047672144 | 0.044972125 |
| Shprh | -0.024903497 | 0.298705822 | 0.75706738 | 0.116480398 | 0.140027344 |
| Shq1 | -0.013474127 | 0.447672665 | 0.84556603 | 0.061764125 | 0.065286304 |
| Shroom2 | 0.007790144 | 0.682790256 | 0.93052333 | 0.086369839 | 0.07562368 |
| Shroom3 | 0.019843911 | 0.425418427 | 0.83250960 | 0.15188679 | 0.130583733 |
| Shtn1 | 0.01715596 | 0.513258298 | 0.87273265 | 0.178049065 | 0.162435735 |
| Siae | 0.014464285 | 0.484821654 | 0.86297153 | 0.109516813 | 0.092837514 |
| Siah1a | -0.024667579 | 0.368763273 | 0.80567193 | 0.181317796 | 0.194476358 |
| Siah1b | 0.003025722 | 0.84302496 | 0.97348890 | 0.054147547 | 0.046091363 |
| Siah2 | 0.056321904 | 0.09860961 | 0.54254427 | 0.387315646 | 0.372270023 |
| Sidt2 | -0.01517106 | 0.530582415 | 0.87764481 | 0.124986775 | 0.136767752 |
| Sigirr | 0.017504208 | 0.253017504 | 0.72230717 | 0.060275578 | 0.047542061 |
| Sigmar1 | 0.001744249 | 0.943014935 | 0.99349067 | 0.148156148 | 0.131100359 |
| Sik1 | -0.06794641 | 0.031228873 | 0.33819915 | 0.172153119 | 0.251809663 |
| Sik2 | -0.015164471 | 0.511674451 | 0.87208730 | 0.108183182 | 0.114634122 |
| Sik3 | 0.024710994 | 0.43028696 | 0.83449592 | 0.282090581 | 0.252398645 |
| Sike1 | -0.047293134 | 0.092889786 | 0.53282112 | 0.194439303 | 0.223303586 |
| Sil1 | -0.007247484 | 0.837156036 | 0.97266190 | 0.507582943 | 0.496278866 |
| Sim1 | -0.04983821 | 0.058714486 | 0.45209489 | 0.139659088 | 0.158877134 |
| Simc1 | 0.003980809 | 0.822130079 | 0.97069202 | 0.069078123 | 0.066933758 |
| Sin3a | 0.021808449 | 0.462377753 | 0.85244258 | 0.238296935 | 0.220802157 |
| Sin3b | -0.002272094 | 0.950069288 | 0.99484627 | 0.654063404 | 0.624792103 |
| Sinhcaf | -0.039393862 | 0.152722061 | 0.62862147 | 0.156638992 | 0.180181885 |
| Sipa1l1 | 0.002912059 | 0.916548631 | 0.98453163 | 0.194412311 | 0.186119278 |
| Sipa1l3 | -0.008876729 | 0.730429761 | 0.94590205 | 0.153587502 | 0.156846862 |
| Sirpa | 0.042305978 | 0.134182489 | 0.60060093 | 0.207700861 | 0.194016116 |
| Sirt1 | 0.010196094 | 0.68114776 | 0.92998645 | 0.142815205 | 0.134536667 |
| Sirt2 | -0.055271871 | 0.069399445 | 0.48373458 | 0.265294866 | 0.283371266 |
| Sirt3 | -0.018249147 | 0.492869409 | 0.86693011 | 0.182810104 | 0.180836316 |
| Sirt4 | -0.021503763 | 0.188389472 | 0.66688548 | 0.044159413 | 0.060333289 |
| Sirt5 | -0.009338241 | 0.603193026 | 0.90871392 | 0.069312027 | 0.068257902 |
| Sirt6 | 0.012231898 | 0.490980018 | 0.86581549 | 0.078485242 | 0.061035404 |
| Sirt7 | -0.000455087 | 0.986545692 | 0.99760728 | 0.173144318 | 0.16801843 |
| Siva1 | -0.010021063 | 0.761618102 | 0.95286310 | 0.405949251 | 0.377089338 |
| Six4 | 0.008439833 | 0.687624744 | 0.93279457 | 0.10170126 | 0.09156556 |
| Six5 | -0.004262914 | 0.797594552 | 0.96442442 | 0.061509338 | 0.058274339 |
| Skap1 | 0.03023113 | 0.396901065 | 0.82384693 | 0.522736906 | 0.455335566 |
| Skap2 | 0.055912546 | 0.048683346 | 0.41586843 | 0.243150358 | 0.199720947 |
| Ski | 0.019764425 | 0.461339186 | 0.85244258 | 0.181034937 | 0.158965635 |
| Skil | -0.122684883 | 0.002351402 | 0.08788808 | 0.417237505 | 0.541373988 |
| Skiv2l | -0.012292952 | 0.638477003 | 0.91600428 | 0.155530901 | 0.158235066 |
| Skp1 | 0.058670128 | 0.104405304 | 0.55800455 | 0.850895875 | 0.737795497 |
| Slain1 | 0.000542586 | 0.980432386 | 0.99760728 | 0.11761646 | 0.106854525 |
| Slain2 | 0.022112225 | 0.510963277 | 0.87160669 | 0.364205325 | 0.339351336 |
| Slbp | -0.019626646 | 0.516455742 | 0.87368901 | 0.261604023 | 0.253906964 |
| Slc10a3 | -0.017152066 | 0.319768859 | 0.77128410 | 0.055812259 | 0.063240245 |
| Slc10a7 | 0.035164587 | 0.071078289 | 0.4881297 | 0.103091147 | 0.070115239 |
| Slc11a2 | 0.034962694 | 0.135687582 | 0.60305593 | 0.154174289 | 0.122559511 |
| Slc12a2 | -0.037845951 | 0.069062139 | 0.48354952 | 0.077245756 | 0.099987299 |
| Slc12a6 | -0.018005929 | 0.470151966 | 0.85717830 | 0.138752382 | 0.149792567 |
| Slc12a7 | 0.006414753 | 0.743006238 | 0.94790800 | 0.074556175 | 0.092105092 |
| Slc15a4 | -0.023426098 | 0.197140721 | 0.67309622 | 0.055199339 | 0.076068562 |
| Slc16a10 | -0.071187506 | 0.064063583 | 0.47003870 | 0.673546922 | 0.696306582 |
| Slc16a11 | -0.024915114 | 0.394479885 | 0.82141907 | 0.222718199 | 0.202123905 |
| Slc16a12 | -0.092349536 | 0.009944527 | 0.19872185 | 0.333203308 | 0.399049039 |
| Slc16a6 | -0.004810229 | 0.808834249 | 0.96584288 | 0.066050378 | 0.093016365 |
| Slc16a9 | -0.006133967 | 0.754906558 | 0.95191435 | 0.072284392 | 0.085273115 |
| Slc17a5 | 0.026079042 | 0.279071729 | 0.73981602 | 0.149756981 | 0.12914481 |
| Slc19a1 | 0.019020831 | 0.147356964 | 0.62411422 | 0.044885675 | 0.03007526 |
| Slc1a4 | 0.042761544 | 0.044943376 | 0.40256551 | 0.11741179 | 0.095781221 |
| Slc20a1 | -0.005990839 | 0.754274955 | 0.95190435 | 0.068132907 | 0.078230767 |
| Slc20a2 | -0.005277241 | 0.736710119 | 0.94697528 | 0.04589858 | 0.053669129 |

| | | | | | |
|---|---|---|---|---|---|
| Slc22a15 | -0.002876804 | 0.882930279 | 0.9799360 | 0.084059755 | 0.081895628 |
| Slc22a17 | 0.012413026 | 0.669766414 | 0.9239496 | 0.231628908 | 0.205435538 |
| Slc22a23 | 0.06246737 | 0.019320178 | 0.2695586 | 0.188189505 | 0.168652784 |
| Slc23a2 | 0.020543485 | 0.335154999 | 0.7809414 | 0.100633882 | 0.096928862 |
| Slc24a2 | 0.008694904 | 0.787980608 | 0.9609992 | 0.28806139 | 0.31536075 |
| Slc24a5 | 0.015360426 | 0.558953936 | 0.8921289 | 0.212253167 | 0.147304189 |
| Slc25a1 | 0.086592626 | 0.018038917 | 0.2604746 | 0.536572459 | 0.490495064 |
| Slc25a11 | -0.06984819 | 0.03880633 | 0.3777809 | 0.357537573 | 0.368618466 |
| Slc25a12 | 0.022083777 | 0.420569793 | 0.8300990 | 0.203453572 | 0.191645828 |
| Slc25a14 | -0.009813587 | 0.719952423 | 0.9444523 | 0.198637921 | 0.185336111 |
| Slc25a16 | 0.023568528 | 0.369071676 | 0.8056719 | 0.172002151 | 0.162677288 |
| Slc25a17 | -0.002257086 | 0.944462707 | 0.9936932 | 0.314304315 | 0.303927579 |
| Slc25a19 | -0.008248646 | 0.638842808 | 0.9160042 | 0.062926944 | 0.070043257 |
| Slc25a20 | 1.00E-05 | 0.999598187 | 0.9996989 | 0.091425923 | 0.082657736 |
| Slc25a22 | 0.002710664 | 0.901857886 | 0.9825441 | 0.114180947 | 0.103050372 |
| Slc25a23 | 0.001536525 | 0.956046301 | 0.9954690 | 0.206177927 | 0.20283394 |
| Slc25a25 | -0.068007345 | 0.004403801 | 0.1311882 | 0.083705078 | 0.159570021 |
| Slc25a27 | -0.050201827 | 0.054518421 | 0.4421851 | 0.143100694 | 0.160920989 |
| Slc25a28 | -0.040434383 | 0.223379779 | 0.6955958 | 0.338438453 | 0.385028988 |
| Slc25a29 | -0.033116572 | 0.086464477 | 0.5246040 | 0.052762487 | 0.093670826 |
| Slc25a3 | -0.036656204 | 0.075231797 | 0.4966285 | 2.201248808 | 2.156496107 |
| Slc25a30 | -0.021528739 | 0.207921697 | 0.6807452 | 0.048905974 | 0.066039576 |
| Slc25a32 | -0.009476952 | 0.673096919 | 0.9260917 | 0.115708598 | 0.111501371 |
| Slc25a33 | -0.017391603 | 0.48256597 | 0.8620306 | 0.157496747 | 0.129543735 |
| Slc25a36 | -0.011232784 | 0.730301999 | 0.9459020 | 0.302463738 | 0.314917741 |
| Slc25a37 | -0.028669614 | 0.049614778 | 0.4192321 | 0.02507091 | 0.052229483 |
| Slc25a38 | -0.013383031 | 0.436667665 | 0.8391598 | 0.061967702 | 0.067718219 |
| Slc25a39 | -0.006160657 | 0.852911536 | 0.9755431 | 1.237468188 | 1.143447266 |
| Slc25a4 | 0.044332599 | 0.104950262 | 0.5585335 | 1.661548973 | 1.512757097 |
| Slc25a40 | 0.050761527 | 0.004703087 | 0.1367553 | 0.093719886 | 0.065516923 |
| Slc25a42 | -0.019667829 | 0.265227159 | 0.7306700 | 0.051148262 | 0.066902427 |
| Slc25a44 | 0.009412195 | 0.739962253 | 0.9479080 | 0.200338427 | 0.202767043 |
| Slc25a46 | 0.006370563 | 0.822864277 | 0.9710592 | 0.221745811 | 0.213423954 |
| Slc25a5 | -0.041468849 | 0.214502228 | 0.6839801 | 1.517379675 | 1.472364315 |
| Slc25a51 | -0.037345757 | 0.298378314 | 0.7564305 | 0.489804991 | 0.506500002 |
| Slc25a53 | -0.008538729 | 0.697425088 | 0.9366987 | 0.112731826 | 0.102676072 |
| Slc26a2 | -0.008513087 | 0.676841016 | 0.9280252 | 0.076497617 | 0.097915624 |
| Slc27a1 | -0.03736012 | 0.04958115 | 0.4192321 | 0.060923182 | 0.085199485 |
| Slc27a4 | -0.008595518 | 0.640396474 | 0.9165200 | 0.077484375 | 0.07733842 |
| Slc29a1 | 0.040050168 | 0.195321914 | 0.6719932 | 0.29680437 | 0.258129766 |
| Slc29a4 | -0.036918314 | 0.088420481 | 0.5284771 | 0.086564464 | 0.113499095 |
| Slc2a2 | 0.198641167 | 2.27E-06 | 0.0004245 | 0.464909922 | 0.354644192 |
| Slc2a3 | 0.012815874 | 0.401255055 | 0.8254259 | 0.066158134 | 0.036931396 |
| Slc2a5 | -0.045189763 | 0.240631368 | 0.7120404 | 0.764679419 | 0.810539933 |
| Slc2a8 | -0.013254796 | 0.476712229 | 0.8607489 | 0.065165669 | 0.081137671 |
| Slc30a1 | -0.028939526 | 0.11707918 | 0.5735102 | 0.049052109 | 0.082777007 |
| Slc30a4 | 0.000794077 | 0.974265656 | 0.9976072 | 0.141986872 | 0.138033007 |
| Slc30a5 | 0.011449354 | 0.695031147 | 0.9362759 | 0.25589519 | 0.227653328 |
| Slc30a6 | -0.029569128 | 0.081540019 | 0.5135726 | 0.045360323 | 0.066621612 |
| Slc30a7 | 0.005244616 | 0.830759577 | 0.9722601 | 0.141586724 | 0.13838434 |
| Slc30a8 | -0.020093638 | 0.559028177 | 0.8921289 | 1.998729762 | 1.82180351 |
| Slc30a9 | -0.037781148 | 0.263712433 | 0.7284954 | 0.370968248 | 0.385125207 |
| Slc31a1 | -0.006243287 | 0.791341164 | 0.9623764 | 0.125364849 | 0.133210261 |
| Slc31a2 | 0.015805472 | 0.566848008 | 0.8949756 | 0.23237854 | 0.170389904 |
| Slc33a1 | -0.070951567 | 0.027277004 | 0.3172190 | 0.253678187 | 0.328761447 |
| Slc35a1 | 0.009485361 | 0.733468971 | 0.9464134 | 0.195611828 | 0.189371713 |
| Slc35a2 | -0.004751168 | 0.864892002 | 0.9769675 | 0.194118988 | 0.188714267 |
| Slc35a3 | 0.003257123 | 0.868383562 | 0.9775692 | 0.084832727 | 0.086364717 |
| Slc35a4 | 0.007410438 | 0.766829567 | 0.9544221 | 0.165794762 | 0.150575477 |
| Slc35a5 | -0.036109653 | 0.129558313 | 0.5930865 | 0.119268647 | 0.140926299 |
| Slc35b1 | 0.078122874 | 0.040738855 | 0.3893347 | 0.699418635 | 0.604150718 |
| Slc35b2 | -0.028632179 | 0.288383985 | 0.7482199 | 0.182132061 | 0.190811301 |
| Slc35b3 | -0.024673753 | 0.328491753 | 0.7778857 | 0.134728178 | 0.150597974 |
| Slc35b4 | 0.055543001 | 0.029033177 | 0.3264172 | 0.18649775 | 0.137792275 |
| Slc35c1 | -0.034087351 | 0.098341317 | 0.5417261 | 0.073129944 | 0.103222272 |
| Slc35c2 | -0.015034549 | 0.55476894 | 0.8902995 | 0.150255939 | 0.156490307 |

| | | | | | |
|---|---|---|---|---|---|
| Slc35d1 | -0.01569854 | 0.38441994 | 0.8135348 | 0.066120134 | 0.070967192 |
| Slc35d3 | 0.023728659 | 0.253623639 | 0.7223071 | 0.109798455 | 0.088164062 |
| Slc35e1 | -0.012524664 | 0.576985861 | 0.8994610 | 0.105554579 | 0.119640495 |
| Slc35e2 | -0.015245639 | 0.354770415 | 0.7969131 | 0.049034347 | 0.060038493 |
| Slc35e4 | 0.010250163 | 0.661439055 | 0.9218806 | 0.134780352 | 0.132126849 |
| Slc35f5 | -0.003691027 | 0.90016836 | 0.9825441 | 0.216529373 | 0.233167253 |
| Slc35f6 | -0.014073826 | 0.365378599 | 0.8050191 | 0.04360899 | 0.052919163 |
| Slc35g1 | -0.003066736 | 0.893164249 | 0.9822826 | 0.119165783 | 0.117640304 |
| Slc35g2 | 0.052589305 | 0.13486935 | 0.6013051 | 0.475625025 | 0.393731566 |
| Slc36a1 | 0.004634902 | 0.817381506 | 0.9694433 | 0.09215608 | 0.095356792 |
| Slc36a4 | 0.040605818 | 0.092595224 | 0.5325584 | 0.150978943 | 0.121015691 |
| Slc37a1 | -0.003115888 | 0.908683098 | 0.9837373 | 0.173398557 | 0.180573501 |
| Slc37a3 | 0.036160892 | 0.161905864 | 0.6398829 | 0.189122997 | 0.154733916 |
| Slc37a4 | 0.006369331 | 0.781500175 | 0.9590847 | 0.123457527 | 0.1147791 |
| Slc38a10 | -0.058762323 | 0.097347521 | 0.5384203 | 0.590022664 | 0.612581917 |
| Slc38a2 | 0.007230758 | 0.854882421 | 0.9755431 | 0.751144697 | 0.800440403 |
| Slc38a4 | 0.102722455 | 0.008512904 | 0.1821346 | 0.944768737 | 0.768894119 |
| Slc38a5 | -0.034506417 | 0.36007364 | 0.8010608 | 0.387501383 | 0.282421584 |
| Slc38a6 | -0.011313825 | 0.504274129 | 0.8695566 | 0.050125598 | 0.061638479 |
| Slc38a7 | 0.018045216 | 0.282620566 | 0.7422811 | 0.066030072 | 0.056823389 |
| Slc38a9 | -0.021377246 | 0.269017939 | 0.7337525 | 0.06967182 | 0.082429102 |
| Slc39a1 | 0.05058596 | 0.053716449 | 0.4389150 | 0.210896181 | 0.143626221 |
| Slc39a10 | -0.009386924 | 0.710630706 | 0.9419370 | 0.15166796 | 0.143210157 |
| Slc39a11 | -0.011390183 | 0.629690707 | 0.9130564 | 0.147412218 | 0.125340239 |
| Slc39a13 | -0.052982956 | 0.035591183 | 0.3591914 | 0.129275568 | 0.154828378 |
| Slc39a14 | -0.010074367 | 0.610974274 | 0.9092206 | 0.074879564 | 0.086023825 |
| Slc39a3 | 0.036443529 | 0.127764194 | 0.5908721 | 0.156015425 | 0.119096189 |
| Slc39a6 | -0.030300093 | 0.29352254 | 0.7533684 | 0.202139324 | 0.218060012 |
| Slc39a9 | -0.051186483 | 0.065256549 | 0.4740570 | 0.156436506 | 0.209302956 |
| Slc3a2 | -0.06780048 | 0.046055654 | 0.4073656 | 1.281804088 | 1.34880462 |
| Slc40a1 | -0.132704995 | 0.000470124 | 0.0293309 | 0.383299266 | 0.466155036 |
| Slc41a1 | -0.006171187 | 0.7462249 | 0.9486117 | 0.076886256 | 0.079400506 |
| Slc41a2 | 0.034714775 | 0.106032947 | 0.5617171 | 0.136073766 | 0.083693793 |
| Slc43a2 | -0.090754911 | 0.012687705 | 0.2227646 | 0.427482905 | 0.504583849 |
| Slc44a1 | -0.046306751 | 0.083487824 | 0.5183040 | 0.145834052 | 0.191525722 |
| Slc44a3 | -0.038008038 | 0.198917605 | 0.6730962 | 0.19955441 | 0.235772903 |
| Slc45a4 | 0.004603245 | 0.831511861 | 0.9723494 | 0.101808129 | 0.099999455 |
| Slc46a3 | -0.008748879 | 0.569105667 | 0.8956532 | 0.047151764 | 0.051136141 |
| Slc48a1 | -0.053106658 | 0.111642671 | 0.5675761 | 0.373991779 | 0.396013785 |
| Slc49a4 | 0.059187522 | 0.009691611 | 0.1970097 | 0.159181031 | 0.101878264 |
| Slc4a10 | 0.021420212 | 0.272693255 | 0.7345096 | 0.081481819 | 0.080433778 |
| Slc4a1ap | -0.040773648 | 0.085214915 | 0.5211664 | 0.107115642 | 0.145571803 |
| Slc4a2 | -0.024700476 | 0.357099083 | 0.7985624 | 0.161398877 | 0.173220524 |
| Slc4a3 | -0.072979288 | 0.010104275 | 0.1996282 | 0.171033107 | 0.221554576 |
| Slc4a4 | -0.031960927 | 0.209561145 | 0.6807660 | 0.11293294 | 0.149076979 |
| Slc4a7 | -0.013616472 | 0.720425557 | 0.9445295 | 0.502411525 | 0.608925994 |
| Slc4a8 | 0.008215387 | 0.705020874 | 0.9401381 | 0.10506789 | 0.100298244 |
| Slc50a1 | -0.041187891 | 0.265309844 | 0.7306700 | 0.666477706 | 0.686191122 |
| Slc52a2 | -0.001286976 | 0.936795759 | 0.9913070 | 0.053570584 | 0.056165126 |
| Slc5a6 | -0.020120429 | 0.216674131 | 0.6872330 | 0.035646759 | 0.06406452 |
| Slc66a2 | -0.009749737 | 0.738606413 | 0.9474350 | 0.213134852 | 0.214966592 |
| Slc6a17 | -0.033641343 | 0.28524055 | 0.7451828 | 0.264933887 | 0.283875973 |
| Slc6a6 | 0.047864958 | 0.217686928 | 0.6886853 | 0.721855117 | 0.696938828 |
| Slc6a8 | -0.010442539 | 0.657265643 | 0.9211780 | 0.107718202 | 0.129082667 |
| Slc6a9 | 0.028181191 | 0.090560772 | 0.5321031 | 0.071439485 | 0.051560424 |
| Slc7a1 | 0.098018392 | 0.002114395 | 0.0819328 | 0.318313038 | 0.251921 |
| Slc7a11 | 0.008201703 | 0.610771988 | 0.9091253 | 0.049149527 | 0.05392591 |
| Slc7a14 | 0.060900328 | 0.09814141 | 0.5411688 | 0.638487518 | 0.626472766 |
| Slc7a2 | -0.09287988 | 0.025900063 | 0.3088084 | 0.784623149 | 0.765407583 |
| Slc7a4 | 0.009623744 | 0.621012152 | 0.9121505 | 0.085669247 | 0.078523586 |
| Slc7a5 | 0.157394004 | 1.19E-06 | 0.0002449 | 0.380328839 | 0.274409276 |
| Slc7a6 | 0.00652873 | 0.721519767 | 0.9445652 | 0.083856271 | 0.069171798 |
| Slc7a6os | -0.031589295 | 0.194823808 | 0.6719932 | 0.121938897 | 0.143898292 |
| Slc7a8 | -0.106524585 | 0.006884515 | 0.1654321 | 0.677113527 | 0.67190969 |
| Slc8a1 | 0.007140212 | 0.82860851 | 0.9722601 | 0.29769364 | 0.307505146 |
| Slc9a1 | -0.025482995 | 0.122461803 | 0.5833237 | 0.040567059 | 0.066539302 |

| | | | | | |
|---|---|---|---|---|---|
| Slc9a3r1 | -0.025863903 | 0.302248576 | 0.7611258 | 0.140174248 | 0.145213872 |
| Slc9a3r2 | -0.012910999 | 0.445131997 | 0.8432629 | 0.055036529 | 0.065679761 |
| Slc9a6 | 0.034752428 | 0.090197891 | 0.5316477 | 0.112787642 | 0.085166311 |
| Slc9a8 | -0.038713581 | 0.089825388 | 0.5302360 | 0.09569489 | 0.12551914 |
| Slco1a6 | -0.011078142 | 0.455969763 | 0.8502914 | 0.031148898 | 0.050540268 |
| Slco2a1 | 0.03975744 | 0.011802896 | 0.2140488 | 0.077108213 | 0.041306702 |
| Slco3a1 | 0.002580105 | 0.915626642 | 0.9845236 | 0.136371998 | 0.141330872 |
| Slf1 | -0.011565894 | 0.646650294 | 0.9169695 | 0.150432729 | 0.141708546 |
| Slf2 | 0.021144259 | 0.433961587 | 0.8377586 | 0.185154807 | 0.172774727 |
| Slirp | 0.112697717 | 0.002062584 | 0.0805544 | 0.641592631 | 0.56792679 |
| Slitrk6 | 0.034016529 | 0.113258784 | 0.5696203 | 0.099938776 | 0.09362002 |
| Slk | 0.017804285 | 0.607471163 | 0.9087139 | 0.398549963 | 0.362121756 |
| Slmap | 0.018597684 | 0.487326094 | 0.8640311 | 0.181166808 | 0.162154432 |
| Sltm | 0.086180917 | 0.022738808 | 0.2926013 | 0.830724948 | 0.714943368 |
| Slu7 | -0.007494932 | 0.814240177 | 0.9681484 | 0.310414458 | 0.302694908 |
| Slx1b | 0.035922588 | 0.024914208 | 0.3041540 | 0.070407049 | 0.046773589 |
| Slx4ip | -0.02741753 | 0.20160246 | 0.6752150 | 0.087641726 | 0.107113512 |
| Smad1 | -0.01528304 | 0.659422511 | 0.9211780 | 0.390581797 | 0.401007232 |
| Smad2 | -0.026682336 | 0.28064762 | 0.7408618 | 0.140637661 | 0.147379461 |
| Smad4 | -0.042497849 | 0.226794113 | 0.7003768 | 0.423913553 | 0.440451753 |
| Smad5 | -0.01911062 | 0.465355184 | 0.8544000 | 0.14308731 | 0.169180241 |
| Smad7 | -0.108103339 | 0.002756156 | 0.0972991 | 0.328059863 | 0.429062535 |
| Smap1 | 0.045167204 | 0.211558944 | 0.6822059 | 0.637183919 | 0.571374 |
| Smap2 | 0.12542124 | 0.000629237 | 0.0356687 | 0.937222564 | 0.789075279 |
| Smarca1 | -0.050619707 | 0.231924754 | 0.7030140 | 0.664336208 | 0.575199094 |
| Smarca2 | -0.04326085 | 0.272727653 | 0.7345096 | 0.486510575 | 0.540768614 |
| Smarca4 | -0.017344045 | 0.628823477 | 0.9130564 | 0.727944421 | 0.738719887 |
| Smarca5 | -0.023541008 | 0.522360678 | 0.8750611 | 0.855424766 | 0.859504141 |
| Smarcad1 | 0.050240702 | 0.099364894 | 0.5436843 | 0.267865116 | 0.221959281 |
| Smarcal1 | -0.01287177 | 0.521075924 | 0.8741879 | 0.07805975 | 0.092516475 |
| Smarcb1 | -0.018946344 | 0.560372262 | 0.8925646 | 0.363689598 | 0.356613908 |
| Smarcc1 | -0.073750361 | 0.028490779 | 0.3233736 | 0.298341587 | 0.342023536 |
| Smarcc2 | -0.034470507 | 0.329935317 | 0.7778857 | 0.432509126 | 0.483932171 |
| Smarcd1 | -0.015433189 | 0.621744313 | 0.9121505 | 0.250927816 | 0.262683756 |
| Smarcd2 | -0.003818558 | 0.879295159 | 0.9788055 | 0.135841541 | 0.155943316 |
| Smarce1 | -0.009215363 | 0.767272796 | 0.9544221 | 0.271226377 | 0.267150684 |
| Smc1a | -0.045480929 | 0.206080362 | 0.6797565 | 0.55229622 | 0.556059268 |
| Smc2 | -0.004498223 | 0.787299604 | 0.9608311 | 0.047896692 | 0.057816018 |
| Smc3 | 0.042434933 | 0.257726358 | 0.7250597 | 0.658838894 | 0.577297934 |
| Smc4 | 0.009710828 | 0.742028975 | 0.9479080 | 0.242572826 | 0.204831979 |
| Smc5 | -0.035367824 | 0.213789463 | 0.6831238 | 0.176177954 | 0.199579239 |
| Smc6 | -0.05759544 | 0.117522916 | 0.5743131 | 0.734851311 | 0.747463589 |
| Smchd1 | -0.037634491 | 0.294141944 | 0.7533922 | 0.419397555 | 0.431084855 |
| Smco4 | -0.047377581 | 0.019523049 | 0.2708652 | 0.068192241 | 0.092295381 |
| Smcr8 | 0.005791382 | 0.743364518 | 0.9479080 | 0.06858469 | 0.064299416 |
| Smdt1 | 0.040706038 | 0.085561951 | 0.5223227 | 2.001657171 | 1.851235639 |
| Smg1 | -0.041921943 | 0.199154416 | 0.6730962 | 0.283514945 | 0.301649997 |
| Smg5 | -0.017501472 | 0.483265224 | 0.8620306 | 0.123795533 | 0.140038874 |
| Smg6 | 0.001947916 | 0.950914548 | 0.9948532 | 0.272904767 | 0.256291102 |
| Smg7 | 0.044538999 | 0.128286964 | 0.5914729 | 0.262153895 | 0.215663132 |
| Smg8 | 0.021460591 | 0.104136838 | 0.5580045 | 0.048784227 | 0.030506852 |
| Smg9 | -5.27E-05 | 0.99785643 | 0.9994073 | 0.08745026 | 0.081678622 |
| Smim1 | 0.092899842 | 0.00658607 | 0.1617967 | 0.394472043 | 0.36589973 |
| Smim10l1 | -0.00087095 | 0.980860596 | 0.9976072 | 0.772329091 | 0.786368715 |
| Smim10l2a | 0.013097763 | 0.35792667 | 0.7989722 | 0.046476632 | 0.040103006 |
| Smim11 | 0.006640648 | 0.85162544 | 0.9750836 | 0.644089082 | 0.603403134 |
| Smim12 | 0.041286289 | 0.061688387 | 0.4613744 | 0.144164182 | 0.104055818 |
| Smim13 | 0.004187181 | 0.888090067 | 0.9812414 | 0.225655002 | 0.227736529 |
| Smim14 | -0.051351338 | 0.106222611 | 0.5617171 | 1.325383972 | 1.357184511 |
| Smim15 | -0.027663752 | 0.316582473 | 0.7705987 | 0.183827267 | 0.199522494 |
| Smim19 | 0.011318416 | 0.740268511 | 0.9479080 | 0.366090587 | 0.359930335 |
| Smim20 | 0.013053248 | 0.69640997 | 0.9363862 | 0.394968516 | 0.366233042 |
| Smim22 | 0.100613489 | 0.005076936 | 0.1426719 | 0.96303285 | 0.834477021 |
| Smim26 | 0.115828704 | 0.001368268 | 0.0616964 | 0.655371745 | 0.571915815 |
| Smim27 | 0.017240664 | 0.592589942 | 0.9060561 | 0.307818471 | 0.298566529 |
| Smim4 | 0.094791333 | 0.008750346 | 0.1858745 | 0.554612833 | 0.503069877 |

| | | | | | |
|---|---|---|---|---|---|
| Smim6 | -0.00585955 | 0.747478877 | 0.9496370 | 0.07651993 | 0.066975068 |
| Smim7 | -0.021490763 | 0.554949314 | 0.8903602 | 0.816519904 | 0.770046741 |
| Smim8 | 0.022623697 | 0.463488104 | 0.8525027 | 0.310472101 | 0.265183408 |
| Smn1 | -0.044587584 | 0.12130076 | 0.5815270 | 0.216214682 | 0.230669845 |
| Smndc1 | 0.001933518 | 0.953417527 | 0.9954690 | 0.343012723 | 0.345834374 |
| Smox | 0.018597016 | 0.270266575 | 0.7340360 | 0.078007531 | 0.052792153 |
| Smpd1 | 0.026411744 | 0.414689889 | 0.8300990 | 0.355085537 | 0.302222049 |
| Smpd2 | -0.013623028 | 0.535559813 | 0.8795949 | 0.103254657 | 0.112453731 |
| Smpd3 | 0.020418753 | 0.373441902 | 0.8085689 | 0.121640738 | 0.111777603 |
| Smpd4 | 0.012303246 | 0.408994548 | 0.8276447 | 0.04996553 | 0.044562429 |
| Smpdl3a | -0.007035998 | 0.781935125 | 0.9590847 | 0.156456952 | 0.154241069 |
| Sms | 0.003846158 | 0.903548697 | 0.9827712 | 0.278154753 | 0.289121991 |
| Smu1 | -0.019597169 | 0.543801554 | 0.8842645 | 0.334073831 | 0.31928881 |
| Smug1 | -0.021025694 | 0.241287756 | 0.7128617 | 0.054150434 | 0.072910646 |
| Smurf1 | -0.017120716 | 0.254098821 | 0.7223071 | 0.037715372 | 0.049563828 |
| Smurf2 | -0.014533899 | 0.599661612 | 0.9078234 | 0.172330911 | 0.173509235 |
| Smyd2 | 0.019121771 | 0.555497996 | 0.8910341 | 0.341581264 | 0.317710548 |
| Smyd3 | -0.011104336 | 0.665209791 | 0.9230765 | 0.146401949 | 0.153263952 |
| Smyd4 | -0.011068876 | 0.638655089 | 0.9160042 | 0.122462715 | 0.127284906 |
| Smyd5 | -0.002189025 | 0.904067581 | 0.9829946 | 0.079953993 | 0.069694626 |
| Snai2 | -0.040478235 | 0.004850509 | 0.1382673 | 0.016454483 | 0.051212678 |
| Snap23 | 0.020499621 | 0.437670223 | 0.8399914 | 0.186531185 | 0.159606768 |
| Snap25 | -0.074534869 | 0.044162267 | 0.4007863 | 0.620874915 | 0.759178771 |
| Snap29 | -0.026228964 | 0.284039559 | 0.7434491 | 0.128457802 | 0.146749596 |
| Snap47 | 0.004140985 | 0.902568225 | 0.9825441 | 0.44012457 | 0.402298296 |
| Snap91 | -0.001859958 | 0.927237129 | 0.9879210 | 0.079241691 | 0.087409977 |
| Snapc1 | -0.025811406 | 0.380446689 | 0.8117350 | 0.216495881 | 0.231705708 |
| Snapc2 | -0.005642513 | 0.780790154 | 0.9590847 | 0.098993376 | 0.090532024 |
| Snapc3 | -0.026197062 | 0.19462935 | 0.6719932 | 0.084485845 | 0.093400053 |
| Snapc5 | -0.016146275 | 0.595094689 | 0.9067325 | 0.251270925 | 0.247291296 |
| Snapin | 0.025868408 | 0.231466047 | 0.7026141 | 0.124600227 | 0.101224442 |
| Sncb | 0.064100239 | 0.02628402 | 0.3100330 | 0.2490563 | 0.192308362 |
| Snd1 | -0.048743874 | 0.176753251 | 0.6536659 | 0.544773557 | 0.580514201 |
| Snf8 | 0.024645702 | 0.503693414 | 0.8695566 | 0.714542431 | 0.631406565 |
| Snhg1 | 0.064243055 | 0.096596711 | 0.5373016 | 0.812917427 | 0.736265239 |
| Snhg10 | -0.018727164 | 0.321562015 | 0.7732349 | 0.065021812 | 0.081661022 |
| Snhg11 | -0.195036317 | 7.08E-05 | 0.0072372 | 0.298984313 | 0.553746557 |
| Snhg12 | -0.037920665 | 0.315596648 | 0.7705987 | 0.550401179 | 0.626557822 |
| Snhg14 | -0.029304259 | 0.078712793 | 0.5057194 | 0.036402457 | 0.062356249 |
| Snhg15 | -0.015356075 | 0.621949672 | 0.9121505 | 0.213987507 | 0.243294362 |
| Snhg16 | -0.03456563 | 0.089764251 | 0.5302360 | 0.071058716 | 0.098519659 |
| Snhg17 | 0.008237785 | 0.675152795 | 0.9265887 | 0.092993047 | 0.081405422 |
| Snhg18 | 0.024027491 | 0.462371672 | 0.8524425 | 0.307758247 | 0.286429093 |
| Snhg20 | -0.11556255 | 0.00079656 | 0.0421102 | 0.29445876 | 0.379134793 |
| Snhg3 | 0.027640928 | 0.43878859 | 0.8403314 | 0.517572002 | 0.479586353 |
| Snhg4 | -0.098512977 | 0.003042681 | 0.1032885 | 0.260503168 | 0.341182964 |
| Snhg5 | -0.019547463 | 0.429210258 | 0.8340365 | 0.112510357 | 0.151417263 |
| Snhg6 | 0.07726686 | 0.020830273 | 0.2799949 | 0.389631377 | 0.342859127 |
| Snhg8 | -0.002984357 | 0.928974064 | 0.9881953 | 1.329448256 | 1.28566878 |
| Snhg9 | -0.07174137 | 0.01362967 | 0.2295523 | 0.153744508 | 0.249831871 |
| Snip1 | 0.004602935 | 0.849889459 | 0.9748841 | 0.150521412 | 0.129428814 |
| Snph | -0.035557967 | 0.071704905 | 0.4888240 | 0.05871141 | 0.094466377 |
| Snrk | -0.026599896 | 0.306770299 | 0.7652785 | 0.150584706 | 0.165116629 |
| Snrnp200 | 0.055463154 | 0.099673869 | 0.5446910 | 0.381310812 | 0.31471701 |
| Snrnp25 | 0.028034227 | 0.179510436 | 0.6550070 | 0.127493412 | 0.086761784 |
| Snrnp27 | -0.021005878 | 0.557953609 | 0.8919622 | 0.575426428 | 0.55022274 |
| Snrnp35 | -0.004557004 | 0.833053642 | 0.9723494 | 0.112330769 | 0.103529823 |
| Snrnp40 | -0.027343823 | 0.326109773 | 0.7765263 | 0.209058379 | 0.202228494 |
| Snrnp48 | 0.007748533 | 0.820075981 | 0.9700462 | 0.393583272 | 0.360631916 |
| Snrnp70 | -0.109199838 | 0.004228656 | 0.1290716 | 0.821622064 | 0.91980757 |
| Snrpa | -0.02150184 | 0.52629888 | 0.8768553 | 0.425046555 | 0.380095394 |
| Snrpa1 | 0.021946881 | 0.515496967 | 0.8734806 | 0.400673044 | 0.342788961 |
| Snrpb | 0.032857994 | 0.342759838 | 0.7872603 | 0.948845577 | 0.858276588 |
| Snrpb2 | 0.010871347 | 0.7048702 | 0.9400796 | 0.241129982 | 0.214264259 |
| Snrpc | -0.060499849 | 0.082568818 | 0.5164455 | 0.429912221 | 0.422429913 |
| Snrpd1 | -0.003966547 | 0.910564329 | 0.9837373 | 0.78893511 | 0.727958698 |

| | | | | | |
|---|---|---|---|---|---|
| Snrpd2 | 0.095539586 | 0.006209548 | 0.1576149 | 1.02309297 | 0.867348674 |
| Snrpd3 | 0.010771215 | 0.763915878 | 0.9533331 | 0.693665351 | 0.616600428 |
| Snrpe | 0.078775117 | 0.016538522 | 0.2530598 | 1.077721613 | 0.937520557 |
| Snrpf | 0.031908028 | 0.350076951 | 0.7927303 | 0.987012326 | 0.900721687 |
| Snrpg | 0.122953712 | 0.000398927 | 0.0261862 | 1.103655664 | 0.928219606 |
| Snrpn | -0.018812695 | 0.559532485 | 0.8923733 | 0.334563783 | 0.300040561 |
| Snta1 | -0.011404112 | 0.476703826 | 0.8607489 | 0.054915626 | 0.052689974 |
| Sntb2 | -0.030169327 | 0.216969784 | 0.6872330 | 0.118746861 | 0.139342058 |
| Snu13 | 0.028746389 | 0.428709084 | 0.8340365 | 0.840152052 | 0.726469395 |
| Snupn | 0.049136447 | 0.01949186 | 0.2708112 | 0.126740116 | 0.086060865 |
| Snw1 | -0.013176178 | 0.720679872 | 0.9445295 | 0.557066847 | 0.542055566 |
| Snx1 | -0.000774502 | 0.979630899 | 0.9976072 | 0.247804198 | 0.240304459 |
| Snx10 | 0.045184909 | 0.211547061 | 0.6822059 | 0.486495901 | 0.474151845 |
| Snx11 | -0.009663053 | 0.592948312 | 0.9060585 | 0.062634667 | 0.074872153 |
| Snx12 | 0.008483305 | 0.745513993 | 0.9485717 | 0.175487109 | 0.159627737 |
| Snx13 | -0.030529611 | 0.225240634 | 0.6973742 | 0.137608939 | 0.164537081 |
| Snx14 | -0.023516724 | 0.391789846 | 0.8200150 | 0.183891019 | 0.190626503 |
| Snx15 | -0.016997079 | 0.58241954 | 0.9019047 | 0.288702768 | 0.276146147 |
| Snx16 | 0.032755085 | 0.194369176 | 0.6717191 | 0.195362586 | 0.145766224 |
| Snx17 | -0.062736548 | 0.06196025 | 0.4617923 | 0.361471805 | 0.402898557 |
| Snx18 | 0.047471838 | 0.132966963 | 0.5988251 | 0.293766796 | 0.224623134 |
| Snx19 | 0.005224433 | 0.793687804 | 0.9624508 | 0.088257583 | 0.086655862 |
| Snx2 | 0.043015009 | 0.193681011 | 0.6710847 | 0.400493845 | 0.351426352 |
| Snx21 | -0.014767819 | 0.619483374 | 0.9121505 | 0.25765593 | 0.2325165 |
| Snx24 | -0.042102144 | 0.177345708 | 0.6536659 | 0.277436966 | 0.27614535 |
| Snx25 | -0.02037439 | 0.265250648 | 0.7306700 | 0.062786835 | 0.076718936 |
| Snx27 | 0.001492214 | 0.957967611 | 0.9957785 | 0.195398537 | 0.198952802 |
| Snx3 | 0.00448861 | 0.887577595 | 0.9812414 | 1.146114513 | 1.057134982 |
| Snx30 | -0.051673332 | 0.068950852 | 0.4835495 | 0.169259194 | 0.216085813 |
| Snx32 | -0.011721939 | 0.546909609 | 0.8858508 | 0.069205758 | 0.085525833 |
| Snx4 | -0.031057122 | 0.303428585 | 0.7626074 | 0.251646994 | 0.261533996 |
| Snx5 | 0.056016114 | 0.098140274 | 0.5411688 | 0.410974473 | 0.355663698 |
| Snx6 | -0.000809638 | 0.981341717 | 0.9976072 | 0.532197329 | 0.512829154 |
| Snx7 | 0.015151523 | 0.341407714 | 0.7852456 | 0.066511157 | 0.04957564 |
| Snx8 | 0.008843185 | 0.598541726 | 0.9075513 | 0.065072256 | 0.057239698 |
| Snx9 | 0.009744444 | 0.748921005 | 0.9501120 | 0.274521072 | 0.243881692 |
| Soat1 | 0.028319956 | 0.231096573 | 0.7025675 | 0.119163697 | 0.126436448 |
| Sobp | -0.001204331 | 0.97012482 | 0.9976072 | 0.290139511 | 0.298269878 |
| Socs1 | -0.028771139 | 0.064162818 | 0.4700387 | 0.028636992 | 0.057402113 |
| Socs2 | -0.00057276 | 0.985681394 | 0.9976072 | 0.262478484 | 0.283773671 |
| Socs3 | 0.040769114 | 0.274600621 | 0.7351955 | 0.401994865 | 0.334710784 |
| Socs4 | 0.025546624 | 0.364055065 | 0.8049171 | 0.199562878 | 0.172623983 |
| Socs5 | -0.012881286 | 0.538395455 | 0.8816335 | 0.087780347 | 0.101791776 |
| Socs6 | -0.015237248 | 0.545332524 | 0.8850946 | 0.140305234 | 0.155632109 |
| Socs7 | -0.051689668 | 0.063915038 | 0.4700387 | 0.158560552 | 0.206289147 |
| Sod1 | -0.039764121 | 0.255515751 | 0.7229652 | 1.021962869 | 1.121175758 |
| Sod2 | -0.018510811 | 0.608506978 | 0.9088672 | 0.884723335 | 0.887849836 |
| Soga3 | -0.01536516 | 0.647563218 | 0.9169695 | 0.34314249 | 0.35936533 |
| Son | 0.007675174 | 0.797579678 | 0.9644244 | 1.669006753 | 1.69452391 |
| Sorbs1 | -0.013743885 | 0.482713261 | 0.8620306 | 0.07116076 | 0.084573334 |
| Sorbs2 | 0.08972713 | 0.013452652 | 0.2292511 | 0.479129103 | 0.445308938 |
| Sorcs2 | -0.070703881 | 1.88E-05 | 0.0024893 | 0.020949429 | 0.069336527 |
| Sord | -0.030855095 | 0.054750699 | 0.4422689 | 0.042288369 | 0.061702895 |
| Sorl1 | -0.013770554 | 0.596783209 | 0.9072933 | 0.110160996 | 0.170274191 |
| Sort1 | -0.033108173 | 0.30822198 | 0.7662606 | 0.298117457 | 0.33599913 |
| Sos1 | -0.026346374 | 0.329491547 | 0.7778857 | 0.169950206 | 0.185227615 |
| Sos2 | -0.008456961 | 0.7670751 | 0.9544221 | 0.197935603 | 0.212207218 |
| Sowahc | -0.0253287 | 0.466094941 | 0.8551251 | 0.284594341 | 0.358415109 |
| Sox12 | -0.027390981 | 0.236935313 | 0.7095050 | 0.106406075 | 0.121367681 |
| Sox4 | -0.023623924 | 0.511216914 | 0.8718019 | 0.331110958 | 0.340036661 |
| Sox5 | 0.018475464 | 0.192254569 | 0.6700281 | 0.054334951 | 0.03281813 |
| Sp1 | -0.010739217 | 0.705950339 | 0.9402269 | 0.208648717 | 0.200648212 |
| Sp2 | -0.000745535 | 0.963220897 | 0.9966421 | 0.054303333 | 0.056947839 |
| Sp3 | -0.037052941 | 0.241731609 | 0.7132592 | 0.278171017 | 0.294557319 |
| Sp3os | -0.054072457 | 0.058261212 | 0.4508199 | 0.15837031 | 0.220460666 |
| Sp4 | -0.0353087 | 0.181036515 | 0.6573507 | 0.156816931 | 0.171690471 |

| | | | | | |
|---|---|---|---|---|---|
| Spa17 | 0.019303718 | 0.524216143 | 0.87583859 | 0.259133986 | 0.230910326 |
| Spaca6 | 0.008807671 | 0.687503815 | 0.93279457 | 0.106536752 | 0.108264811 |
| Spag7 | 0.045228563 | 0.191693768 | 0.67002813 | 0.489889278 | 0.390935251 |
| Spag9 | -0.018592181 | 0.617350115 | 0.91150170 | 0.565254983 | 0.574396907 |
| Spast | -0.05667883 | 0.050007706 | 0.42111753 | 0.182327978 | 0.230630348 |
| Spata13 | -0.014771073 | 0.33830228 | 0.78342458 | 0.040519553 | 0.051711349 |
| Spata2 | -0.008609824 | 0.644084081 | 0.91683910 | 0.068398788 | 0.077338896 |
| Spata24 | 0.014267851 | 0.443696192 | 0.84315410 | 0.083111364 | 0.06955982 |
| Spata5 | -0.005200045 | 0.803947273 | 0.96471431 | 0.096147734 | 0.096323508 |
| SPATA5L1 | 0.021399054 | 0.231779749 | 0.70301400 | 0.087210586 | 0.059818397 |
| Spata6 | -0.022213361 | 0.242377796 | 0.71327199 | 0.077866992 | 0.079758372 |
| Spata7 | -0.032643791 | 0.215162988 | 0.68454677 | 0.135113547 | 0.171633689 |
| Spats2 | -0.008959905 | 0.691352936 | 0.93514293 | 0.113526635 | 0.112270761 |
| Spc24 | -0.021189729 | 0.207719772 | 0.68073344 | 0.051674756 | 0.06431717 |
| Spc25 | 0.049341897 | 0.269560437 | 0.73403607 | 0.958822082 | 0.862720433 |
| Spcs1 | 0.052820494 | 0.05589141 | 0.44493321 | 1.695080717 | 1.549226322 |
| Spcs2 | -0.083729796 | 0.007086492 | 0.16800019 | 1.67363528 | 1.743645578 |
| Spcs3 | 0.012410355 | 0.742823208 | 0.94790800 | 0.676401483 | 0.683413639 |
| Specc1 | 0.001850256 | 0.912483524 | 0.98403934 | 0.063985398 | 0.052872099 |
| Specc1l | -0.003665018 | 0.91046057 | 0.98373739 | 0.299731862 | 0.295953904 |
| Spen | 0.066457447 | 0.048411735 | 0.41562531 | 0.385623654 | 0.336449791 |
| Spg11 | 0.006375306 | 0.68613497 | 0.93188101 | 0.060706128 | 0.048186636 |
| Spg20 | 0.040818687 | 0.144794407 | 0.61936384 | 0.220760591 | 0.194194122 |
| Spg21 | 0.013268645 | 0.675801549 | 0.92696288 | 0.332003216 | 0.302558781 |
| Spg7 | -0.014360887 | 0.632972487 | 0.91411960 | 0.279226658 | 0.266291888 |
| Sphk2 | 0.045885165 | 0.065439194 | 0.47464465 | 0.174219 | 0.132703753 |
| Sphkap | 0.051148846 | 0.160566617 | 0.63712831 | 1.618275105 | 1.4757862 |
| Spice1 | -0.009078397 | 0.579426946 | 0.90078597 | 0.056389579 | 0.055608212 |
| Spidr | 0.016158097 | 0.257492902 | 0.72505979 | 0.056004134 | 0.036060487 |
| Spin1 | 0.051475077 | 0.041670052 | 0.39255940 | 0.178112987 | 0.137148074 |
| Spin2c | -0.032076257 | 0.031617319 | 0.33852287 | 0.032438306 | 0.052528264 |
| Spindoc | 0.003525411 | 0.854580448 | 0.97554319 | 0.086548163 | 0.075519238 |
| Spint1 | -0.0142906 | 0.697050011 | 0.93644859 | 0.584789307 | 0.579785062 |
| Spint2 | -0.138827774 | 1.19E-08 | 7.35E-06 | 2.231345938 | 2.264071259 |
| Spire1 | 0.014884629 | 0.592403423 | 0.90605029 | 0.21462687 | 0.198529845 |
| Spire2 | -0.036387256 | 0.032703302 | 0.34257311 | 0.046593101 | 0.069267992 |
| Spns1 | 0.022205131 | 0.346247605 | 0.78960373 | 0.144454406 | 0.124412655 |
| Spock1 | -0.060268521 | 0.005278608 | 0.14642658 | 0.051831102 | 0.129603259 |
| Spock2 | 0.086559731 | 0.026195619 | 0.30972651 | 0.530134674 | 0.581754957 |
| Spon2 | -0.057692552 | 0.036198634 | 0.36089491 | 0.165595716 | 0.190067414 |
| Spop | 0.019102446 | 0.585891512 | 0.9029041 | 0.537496955 | 0.48910322 |
| Spopl | 0.007716468 | 0.764638031 | 0.95387440 | 0.160852896 | 0.149475971 |
| Spout1 | -0.028577576 | 0.230091407 | 0.70196779 | 0.129319298 | 0.134097422 |
| Spp1 | -0.260233841 | 9.16E-05 | 0.00889470 | 0.643074999 | 1.188001125 |
| Sppl2a | 0.09537457 | 0.003580539 | 0.11494801 | 0.363459854 | 0.301778326 |
| Sppl3 | 0.002791776 | 0.919838756 | 0.98545139 | 0.186899071 | 0.189819336 |
| Spr | -0.010991117 | 0.684030964 | 0.93117031 | 0.174245977 | 0.179565726 |
| Spred2 | 0.012619239 | 0.543516796 | 0.88422204 | 0.087638005 | 0.09326499 |
| Spred3 | -0.012102925 | 0.480484808 | 0.86175794 | 0.057430567 | 0.063890676 |
| Spring1 | 0.017244147 | 0.422618067 | 0.83148971 | 0.11003332 | 0.104032701 |
| Sprtn | 0.036207837 | 0.049480388 | 0.41923219 | 0.096168738 | 0.065218272 |
| Spry2 | 0.017398322 | 0.372362552 | 0.80856898 | 0.098097627 | 0.073632894 |
| Spryd3 | -0.00622975 | 0.849793412 | 0.97488410 | 0.356237405 | 0.320070375 |
| Spryd4 | -0.001112397 | 0.953477741 | 0.99546908 | 0.083539039 | 0.077822657 |
| Spryd7 | -0.009427005 | 0.742704636 | 0.94790800 | 0.213713135 | 0.218753394 |
| Spsb1 | -0.049197403 | 0.044697299 | 0.40157339 | 0.107359469 | 0.141749832 |
| Spsb2 | -0.010828348 | 0.751668182 | 0.95060539 | 0.38104342 | 0.353493109 |
| Spsb3 | -0.042034387 | 0.109788607 | 0.56558881 | 0.148494059 | 0.185929141 |
| Sptan1 | -0.064074097 | 0.069285745 | 0.48354951 | 0.342899723 | 0.382935139 |
| Sptbn1 | 0.004990547 | 0.898991376 | 0.98254414 | 0.84412519 | 0.781570148 |
| Sptbn2 | -0.015651722 | 0.563778574 | 0.89318059 | 0.158172059 | 0.176124055 |
| Sptbn4 | -0.090182888 | 0.000873685 | 0.04514038 | 0.124827078 | 0.197621691 |
| Sptlc1 | 0.052698805 | 0.033104028 | 0.34531229 | 0.165927092 | 0.128181239 |
| Sptlc2 | -0.024500033 | 0.276511492 | 0.73672644 | 0.100909871 | 0.119900764 |
| Sptssa | -0.023834122 | 0.505043467 | 0.86955669 | 0.523652053 | 0.502751962 |
| Spty2d1 | -0.043398902 | 0.232024263 | 0.70301791 | 0.419789232 | 0.454597467 |

| | | | | | |
|---|---|---|---|---|---|
| Sqle | -0.01881094 | 0.592209672 | 0.90602008 | 0.348373172 | 0.420223717 |
| Sqor | -0.08318513 | 0.007314203 | 0.16992247 | 0.22800447 | 0.294776995 |
| Sqstm1 | -0.051708917 | 0.151780578 | 0.62811059 | 1.238555154 | 1.456348196 |
| Sra1 | 0.040087163 | 0.218924328 | 0.69016901 | 1.310069466 | 1.155861277 |
| Srbd1 | 0.031197392 | 0.107791026 | 0.56343910 | 0.10457121 | 0.074071533 |
| Src | -0.025282454 | 0.177888792 | 0.65381875 | 0.060808482 | 0.085350123 |
| Srcap | -0.002650435 | 0.893952267 | 0.98243614 | 0.09150433 | 0.081403347 |
| Srd5a3 | -0.005295493 | 0.800410695 | 0.96470534 | 0.098767184 | 0.09624389 |
| Srebf1 | 0.008096333 | 0.716578086 | 0.94314111 | 0.114919313 | 0.103305969 |
| Srebf2 | 0.014608439 | 0.659195556 | 0.92117802 | 0.31639084 | 0.283746869 |
| Srek1 | -0.002826106 | 0.939051567 | 0.99258301 | 0.530014561 | 0.511051293 |
| Srek1ip1 | 0.109830045 | 0.000323358 | 0.02269773 | 0.330147179 | 0.255390457 |
| Srf | -0.014839848 | 0.378745761 | 0.81030809 | 0.052358824 | 0.063582909 |
| Srfbp1 | -0.011902679 | 0.587891154 | 0.90446241 | 0.11273631 | 0.114949162 |
| Srgap1 | -0.006208176 | 0.806262385 | 0.96480137 | 0.13810225 | 0.150441513 |
| Srgap2 | -0.019315909 | 0.330126388 | 0.77788578 | 0.075245539 | 0.087320696 |
| Sri | 0.123187558 | 0.000731369 | 0.03943032 | 0.771327405 | 0.57373873 |
| Srm | -0.028750945 | 0.420738435 | 0.83009907 | 0.415254263 | 0.403260931 |
| Srp14 | 0.017581934 | 0.5078243 | 0.87040677 | 1.613613302 | 1.543948682 |
| Srp19 | 0.017744565 | 0.629577837 | 0.91305644 | 0.610838383 | 0.553340583 |
| Srp54a | 0.03812568 | 0.21101526 | 0.68220598 | 0.261105274 | 0.233654123 |
| Srp68 | -0.017119999 | 0.623461808 | 0.91215057 | 0.442734345 | 0.419958683 |
| Srp72 | -0.104998742 | 0.004566415 | 0.13522041 | 0.870232188 | 0.950202309 |
| Srp9 | 0.021583741 | 0.445760043 | 0.84373680 | 1.55744268 | 1.480886578 |
| Srpk1 | -0.01527215 | 0.65523021 | 0.92066341 | 0.414592972 | 0.435016198 |
| Srpk2 | -0.010568463 | 0.769378135 | 0.95523334 | 0.739059693 | 0.762909645 |
| Srpr | -0.005432772 | 0.867068193 | 0.97720023 | 1.298238389 | 1.31262 |
| Srprb | -0.013573268 | 0.666462741 | 0.92307657 | 0.294008744 | 0.285690647 |
| Srr | -0.017012459 | 0.363366132 | 0.80441687 | 0.064781725 | 0.080224464 |
| Srrd | -0.011182536 | 0.580031786 | 0.90087917 | 0.091381894 | 0.09405332 |
| Srrm1 | -0.007630962 | 0.822079383 | 0.97069202 | 1.218984173 | 1.178061267 |
| Srrm2 | -0.051596002 | 0.05658604 | 0.4472777 | 1.75611093 | 1.807290931 |
| Srrm3 | 0.028986438 | 0.274538131 | 0.73519555 | 0.189842242 | 0.165281013 |
| Srrt | -0.026383775 | 0.383530323 | 0.81297870 | 0.241581207 | 0.226332832 |
| Srsf1 | -0.047714269 | 0.176951558 | 0.65366590 | 0.390328491 | 0.421126016 |
| Srsf10 | -0.016962183 | 0.645280187 | 0.91696950 | 0.604535078 | 0.598428923 |
| Srsf11 | -0.103535715 | 0.005959625 | 0.15454384 | 0.885475821 | 0.96027952 |
| Srsf2 | -0.096548403 | 0.013734333 | 0.23053228 | 0.809991767 | 0.840911363 |
| Srsf3 | -0.070844487 | 0.045350341 | 0.40347568 | 1.06463183 | 1.079141121 |
| Srsf4 | 0.028731645 | 0.356810661 | 0.79827734 | 0.293971416 | 0.256184462 |
| Srsf5 | -0.128570346 | 0.000166558 | 0.01388450 | 1.149843122 | 1.299048421 |
| Srsf6 | -0.11789781 | 0.001062042 | 0.05016880 | 0.490543264 | 0.591649437 |
| Srsf7 | -0.052904639 | 0.176902473 | 0.65366590 | 1.156245999 | 1.132184392 |
| Srsf9 | -0.059838434 | 0.092330276 | 0.53255842 | 0.75872838 | 0.765451975 |
| Srxn1 | -0.076603419 | 0.040324699 | 0.38574832 | 0.379919356 | 0.474518594 |
| Ss18 | -0.023432232 | 0.309505327 | 0.76738130 | 0.118032545 | 0.125128197 |
| Ss18l2 | 0.093809143 | 0.010412046 | 0.20174069 | 1.084091794 | 1.040047411 |
| Ssb | 0.007385368 | 0.823469631 | 0.97105924 | 1.188889129 | 1.121950511 |
| Ssbp1 | 0.036197851 | 0.313718405 | 0.77031840 | 0.717988227 | 0.655622291 |
| Ssbp2 | 0.005193193 | 0.86057536 | 0.97696750 | 0.241267924 | 0.21111378 |
| Ssbp3 | 0.001976731 | 0.953352238 | 0.99546908 | 0.401121228 | 0.390631006 |
| Ssbp4 | -0.052696466 | 0.142690454 | 0.61586962 | 0.495266974 | 0.550006846 |
| Ssh1 | 0.022865144 | 0.328353015 | 0.77788578 | 0.13413802 | 0.118868416 |
| Ssh2 | 0.011151217 | 0.593322181 | 0.90605850 | 0.100014811 | 0.094955281 |
| Ssh3 | 0.006223781 | 0.683085888 | 0.93079835 | 0.059872144 | 0.043013871 |
| Ssna1 | -0.070795392 | 0.035703499 | 0.35919149 | 0.410412842 | 0.427473068 |
| Ssr1 | 0.021227994 | 0.575464122 | 0.89870971 | 0.940469533 | 0.824573748 |
| Ssr2 | -0.036151931 | 0.240319764 | 0.71204045 | 1.790336867 | 1.654563541 |
| Ssr3 | -0.032973239 | 0.36649829 | 0.80537944 | 0.78554503 | 0.771344338 |
| Ssr4 | -0.025199409 | 0.149558962 | 0.62643745 | 2.823353906 | 2.841706898 |
| Ssrp1 | -0.048461687 | 0.173152605 | 0.65255078 | 0.489124228 | 0.509041151 |
| Sst | -0.015441181 | 0.869857074 | 0.97778834 | 1.600162301 | 1.992564742 |
| Sstr2 | 0.005235615 | 0.797127024 | 0.96442442 | 0.096689383 | 0.074002002 |
| Sstr3 | -0.008568159 | 0.755563502 | 0.95213280 | 0.18061677 | 0.189649653 |
| Ssu72 | 0.014295456 | 0.695278132 | 0.93627595 | 0.751901997 | 0.678487544 |
| Ssx2ip | 0.035110171 | 0.178226577 | 0.65384898 | 0.171457651 | 0.151420847 |

| | | | | | |
|---|---|---|---|---|---|
| St13 | 0.058522842 | 0.099920384 | 0.5455202 | 1.061467581 | 0.953465117 |
| St14 | 0.009477867 | 0.697319009 | 0.9366830 | 0.153229623 | 0.134069604 |
| St18 | -0.069682665 | 0.055531548 | 0.4449332 | 0.422753242 | 0.526300667 |
| St3gal3 | -0.011035222 | 0.552843557 | 0.8900045 | 0.067438486 | 0.076275024 |
| St3gal5 | 0.013129247 | 0.665172195 | 0.9230765 | 0.237764111 | 0.245228634 |
| St3gal6 | -0.006214375 | 0.8163312 | 0.9688248 | 0.196914715 | 0.165869088 |
| St6gal1 | -0.088534883 | 0.00065721 | 0.0360629 | 0.102014234 | 0.1645858 |
| St6galnac4 | 0.020832461 | 0.291281444 | 0.7516725 | 0.100396798 | 0.081492564 |
| St6galnac5 | -0.008413782 | 0.678088726 | 0.9281886 | 0.090841466 | 0.080671977 |
| St6galnac6 | 0.019509448 | 0.40201083 | 0.8254259 | 0.155937847 | 0.116317251 |
| St7l | 0.011456077 | 0.649633493 | 0.9174778 | 0.162723319 | 0.145497672 |
| St8sia3 | -0.034592731 | 0.225014569 | 0.6968918 | 0.195314618 | 0.201484588 |
| St8sia5 | -0.007387659 | 0.721593209 | 0.9445652 | 0.066787898 | 0.104742165 |
| Stag1 | 0.043057551 | 0.186488819 | 0.6645003 | 0.332314783 | 0.300288786 |
| Stag2 | 0.041638341 | 0.238836096 | 0.7112679 | 0.419976031 | 0.374781874 |
| Stam | 0.016765515 | 0.413476694 | 0.8300990 | 0.102796026 | 0.086089273 |
| Stam2 | 0.028679876 | 0.352336618 | 0.7952568 | 0.276588411 | 0.25853916 |
| Stambp | 0.013425415 | 0.65951529 | 0.9211780 | 0.283082072 | 0.257960433 |
| Stambpl1 | -0.025814802 | 0.087743311 | 0.5276030 | 0.030977778 | 0.051746822 |
| Star | -0.03550822 | 0.05677238 | 0.4473248 | 0.022274317 | 0.076509438 |
| Stard10 | -0.005700835 | 0.875629663 | 0.9784331 | 0.856179596 | 0.793799733 |
| Stard13 | 0.002538618 | 0.894748435 | 0.9824361 | 0.078257823 | 0.079588947 |
| Stard3 | 0.000307322 | 0.987248377 | 0.9976072 | 0.079467134 | 0.078788987 |
| Stard3nl | -0.040693368 | 0.219577786 | 0.6910162 | 0.331853075 | 0.369496837 |
| Stard4 | -0.026173154 | 0.131441983 | 0.5972652 | 0.051934018 | 0.067783651 |
| Stard5 | 0.019045934 | 0.482905634 | 0.8620306 | 0.1725323 | 0.17484345 |
| Stard7 | 0.029240783 | 0.344516716 | 0.7879223 | 0.290046774 | 0.260841293 |
| Stat1 | -0.006109231 | 0.740059535 | 0.9479080 | 0.072758613 | 0.073666169 |
| Stat2 | 0.01039481 | 0.572862261 | 0.8973304 | 0.076480996 | 0.068714595 |
| Stat3 | -0.005141177 | 0.888861998 | 0.9816521 | 0.640765104 | 0.705483168 |
| Stat5a | 0.005074885 | 0.741915837 | 0.9479080 | 0.054806422 | 0.046337978 |
| Stat5b | 0.025359069 | 0.339027066 | 0.7834245 | 0.180368655 | 0.170030135 |
| Stat6 | -0.008237075 | 0.672708884 | 0.9259431 | 0.079649356 | 0.083910145 |
| Stau1 | 0.018016063 | 0.510886699 | 0.8716066 | 0.208691261 | 0.180436353 |
| Stau2 | 0.003170782 | 0.92130414 | 0.9861175 | 0.299675516 | 0.298790831 |
| Steap1 | -0.013111449 | 0.547336295 | 0.8860274 | 0.064736828 | 0.103746135 |
| Steap2 | -0.01028666 | 0.721372971 | 0.9445652 | 0.210801655 | 0.216696117 |
| Stim1 | -0.006442872 | 0.836102879 | 0.9723494 | 0.262717166 | 0.318006381 |
| Stim2 | 0.075011892 | 0.005459194 | 0.1488982 | 0.216511013 | 0.149777068 |
| Stip1 | 0.008225491 | 0.809477142 | 0.9661473 | 0.426766215 | 0.376339101 |
| Stk11 | 0.016828083 | 0.643312375 | 0.9166712 | 0.561218568 | 0.507716652 |
| Stk11ip | -0.004848546 | 0.763894736 | 0.9533331 | 0.047404086 | 0.054706784 |
| Stk16 | -0.04514984 | 0.17787731 | 0.6538187 | 0.419927406 | 0.405972989 |
| Stk19 | 0.019247303 | 0.528596076 | 0.8774607 | 0.260621475 | 0.240315675 |
| Stk24 | -0.061561645 | 0.024902886 | 0.3041540 | 0.156105993 | 0.200113381 |
| Stk25 | 0.038361193 | 0.223975726 | 0.6962539 | 0.343938251 | 0.278742722 |
| Stk26 | 0.015140563 | 0.45778271 | 0.8520083 | 0.085303456 | 0.099607706 |
| Stk3 | 0.013594937 | 0.466801838 | 0.8554230 | 0.088084065 | 0.068618192 |
| Stk32a | -0.037418252 | 0.314457989 | 0.7705987 | 0.479550388 | 0.582875782 |
| Stk32b | -0.014812438 | 0.318075733 | 0.7710043 | 0.044425319 | 0.047578585 |
| Stk35 | 0.011778017 | 0.627924663 | 0.9128095 | 0.126642382 | 0.137718844 |
| Stk38 | 0.037544526 | 0.266513822 | 0.7307399 | 0.38068446 | 0.325665561 |
| Stk38l | 0.011758447 | 0.634666292 | 0.9147798 | 0.148335524 | 0.138152175 |
| Stk39 | 0.004192116 | 0.86604316 | 0.9770148 | 0.14165501 | 0.141817587 |
| Stk4 | -0.005019618 | 0.842617857 | 0.9734889 | 0.150157418 | 0.153778673 |
| Stk40 | 0.017207728 | 0.607128418 | 0.9087139 | 0.328318159 | 0.327613357 |
| Stmn3 | -0.025014391 | 0.499340321 | 0.8690742 | 0.457044766 | 0.491059231 |
| Stn1 | 0.020941721 | 0.103225739 | 0.5553141 | 0.044488158 | 0.027849131 |
| Stoml1 | -0.009104332 | 0.586103648 | 0.9029582 | 0.059315456 | 0.060394874 |
| Stoml2 | -0.035506905 | 0.267108288 | 0.7313133 | 0.312515807 | 0.317214417 |
| Ston2 | -0.009901722 | 0.588710077 | 0.9044624 | 0.074451892 | 0.075759971 |
| Stox1 | 0.008882232 | 0.669880134 | 0.9239496 | 0.09528312 | 0.098281465 |
| Stox2 | -0.02653676 | 0.376707716 | 0.8101797 | 0.214519755 | 0.229341026 |
| Strada | -0.013438772 | 0.437706644 | 0.8399914 | 0.058655396 | 0.062601763 |
| Stradb | -0.000197355 | 0.991140457 | 0.9982478 | 0.071944882 | 0.068565967 |
| Strap | 0.039564065 | 0.252586616 | 0.7223071 | 0.548476446 | 0.475763968 |

| | | | | | |
|---|---|---|---|---|---|
| Strbp | 0.034027323 | 0.345116139 | 0.7880440 | 0.548333855 | 0.488993342 |
| Strip1 | 0.014495838 | 0.508159555 | 0.8704067 | 0.120379668 | 0.102492919 |
| Strn | 0.004276244 | 0.80025221 | 0.9647053 | 0.06227659 | 0.063986099 |
| Strn3 | -0.022192032 | 0.527298774 | 0.8768553 | 0.45701489 | 0.462553873 |
| Strn4 | -0.038332861 | 0.153727807 | 0.6301569 | 0.160585026 | 0.18968468 |
| Stt3a | -0.031459417 | 0.3917995 | 0.8200150 | 0.623154485 | 0.638732658 |
| Stt3b | -0.067879021 | 0.031755193 | 0.3385228 | 0.275988682 | 0.305588435 |
| Stub1 | -0.063972211 | 0.054184897 | 0.4403011 | 1.219923955 | 1.185124676 |
| Stx12 | -0.005026025 | 0.876121853 | 0.9785583 | 0.321587807 | 0.324170885 |
| Stx16 | -0.068538571 | 0.063690762 | 0.4697489 | 0.397980512 | 0.430816112 |
| Stx17 | -0.052222479 | 0.041876588 | 0.3933861 | 0.135664984 | 0.163704009 |
| Stx18 | 0.021602987 | 0.472233355 | 0.8590784 | 0.246791492 | 0.235373561 |
| Stx1a | -0.004936261 | 0.783069575 | 0.9591369 | 0.068928353 | 0.066901937 |
| Stx3 | -0.014167687 | 0.646759853 | 0.9169695 | 0.260843256 | 0.28244505 |
| Stx4a | -0.063872814 | 0.054051296 | 0.4398596 | 0.291381359 | 0.351219929 |
| Stx5a | -0.021989343 | 0.53025952 | 0.8776448 | 0.431612472 | 0.418871924 |
| Stx6 | 0.038735133 | 0.186268994 | 0.6644983 | 0.273681039 | 0.220861162 |
| Stx7 | 0.010825907 | 0.762196434 | 0.9529857 | 0.555917432 | 0.559612728 |
| Stx8 | 0.001419296 | 0.965467626 | 0.9968191 | 0.407534191 | 0.36452027 |
| Stxbp1 | 0.048701888 | 0.172493082 | 0.6522284 | 0.530371178 | 0.505627048 |
| Stxbp2 | -0.058225028 | 0.028107648 | 0.3212302 | 0.13308226 | 0.183941031 |
| Stxbp3 | 0.006641582 | 0.83178297 | 0.9723494 | 0.294000453 | 0.26866532 |
| Stxbp4 | 0.014023093 | 0.461745377 | 0.8524425 | 0.092656124 | 0.073081729 |
| Stxbp5 | -0.007215775 | 0.793988732 | 0.9624508 | 0.175673869 | 0.196594991 |
| Stxbp5l | -0.051015591 | 0.094116523 | 0.5334417 | 0.18653215 | 0.2167877 |
| Stxbp6 | -0.018370249 | 0.460738188 | 0.8524425 | 0.134207271 | 0.1500844 |
| Sub1 | 0.052194447 | 0.057528666 | 0.4500665 | 1.499454007 | 1.362262698 |
| Sucla2 | 0.031462045 | 0.343849064 | 0.7873921 | 0.401442468 | 0.344794383 |
| Suclg1 | -0.035521145 | 0.308722602 | 0.7669648 | 0.77341926 | 0.743017656 |
| Suclg2 | 0.032175803 | 0.289413949 | 0.7496047 | 0.262117308 | 0.237913137 |
| Suco | -0.023663757 | 0.51487711 | 0.8733714 | 0.529888311 | 0.548795938 |
| Suds3 | 0.024472653 | 0.477068008 | 0.8607489 | 0.482753835 | 0.440944354 |
| Sugp1 | 0.025396667 | 0.33463764 | 0.7804836 | 0.191359761 | 0.156431505 |
| Sugp2 | -0.011903 | 0.523156793 | 0.8750824 | 0.073522325 | 0.077326586 |
| Sugt1 | -0.065591418 | 0.069187992 | 0.4835495 | 0.497744048 | 0.532169717 |
| Sult1d1 | 0.072234938 | 0.019828335 | 0.2724376 | 0.263610083 | 0.201716963 |
| Sult2b1 | -0.002530475 | 0.869027495 | 0.9776398 | 0.042830662 | 0.052751485 |
| Sult4a1 | -0.002867932 | 0.86066086 | 0.9769675 | 0.057839807 | 0.05597977 |
| Sumf1 | 0.060245727 | 0.031579999 | 0.3385228 | 0.23055051 | 0.168833122 |
| Sumf2 | 0.004841836 | 0.775956774 | 0.9558538 | 0.070084901 | 0.060195565 |
| Sumo1 | 0.02251108 | 0.484234897 | 0.8627173 | 1.094338896 | 1.017030751 |
| Sumo2 | 0.038995111 | 0.123459027 | 0.5849888 | 1.933363384 | 1.715259884 |
| Sumo3 | 0.017737018 | 0.625006394 | 0.9121764 | 0.687050591 | 0.647206387 |
| Sun1 | -0.009347236 | 0.758394669 | 0.9521533 | 0.240752611 | 0.238369999 |
| Suox | -0.071547167 | 0.014547861 | 0.2371912 | 0.19753666 | 0.23984049 |
| Supt16 | 0.036574011 | 0.322401228 | 0.7735649 | 0.560840194 | 0.495153672 |
| Supt20 | 0.004715554 | 0.858576674 | 0.9769675 | 0.173011376 | 0.163677171 |
| Supt3 | -0.021759511 | 0.32357891 | 0.7751516 | 0.108656701 | 0.110505187 |
| Supt4a | 0.013457226 | 0.684768391 | 0.9313273 | 0.332463769 | 0.326900985 |
| Supt5 | -0.065287561 | 0.069268605 | 0.4835495 | 0.389736032 | 0.459718588 |
| Supt6 | -0.023678785 | 0.529400797 | 0.8776448 | 0.455804697 | 0.483069593 |
| Supt7l | 0.02289956 | 0.176867032 | 0.6536659 | 0.07856627 | 0.055490101 |
| Supv3l1 | 0.044236104 | 0.009593476 | 0.1966266 | 0.090410889 | 0.051647315 |
| Surf1 | 0.010480978 | 0.744913631 | 0.9485717 | 0.388865852 | 0.352481098 |
| Surf2 | 0.008681158 | 0.750294933 | 0.9504537 | 0.197236218 | 0.180291627 |
| Surf4 | 0.060902787 | 0.096747008 | 0.5376640 | 0.958205079 | 0.902325626 |
| Surf6 | 0.021006699 | 0.240699132 | 0.7120404 | 0.082266367 | 0.061779563 |
| Susd6 | -0.02340209 | 0.392081324 | 0.8201624 | 0.160298527 | 0.201788623 |
| Suv39h2 | 0.025087663 | 0.057616477 | 0.4501007 | 0.050876582 | 0.028665636 |
| Suz12 | -0.021668294 | 0.544138124 | 0.8842645 | 0.402040933 | 0.414795166 |
| Sv2a | 0.016779983 | 0.595961274 | 0.9068020 | 0.25965228 | 0.292700279 |
| Sv2b | 0.025963115 | 0.183088531 | 0.6609309 | 0.102179113 | 0.069868964 |
| Svbp | 0.008270846 | 0.685407251 | 0.9316044 | 0.106239696 | 0.088990049 |
| Svil | 0.021595163 | 0.352414207 | 0.7952568 | 0.12022411 | 0.115982619 |
| Svip | 0.147880582 | 7.86E-06 | 0.0012370 | 0.394237575 | 0.299452723 |
| Svop | 0.030356256 | 0.072736805 | 0.4903372 | 0.080884506 | 0.050323216 |

| | | | | | |
|---|---|---|---|---|---|
| Swi5 | -0.028844611 | 0.270072612 | 0.73403607 | 1.787319026 | 1.710322794 |
| Swsap1 | -0.015790477 | 0.350471176 | 0.79273039 | 0.052116 | 0.060492385 |
| Swt1 | -0.036091163 | 0.25958305 | 0.72634759 | 0.266417394 | 0.287280876 |
| Syap1 | -0.040212091 | 0.192029843 | 0.67002813 | 0.260296131 | 0.297326979 |
| Sybu | -0.019041457 | 0.401711 | 0.82542593 | 0.108956957 | 0.115666479 |
| Syce2 | 0.001840102 | 0.936470743 | 0.99124689 | 0.119774389 | 0.119959182 |
| Syf2 | -0.056331542 | 0.108818243 | 0.56441837 | 0.55106808 | 0.577002705 |
| Sympk | 0.047452847 | 0.108362111 | 0.56368177 | 0.27109482 | 0.230032604 |
| Syn1 | -0.006922788 | 0.807900135 | 0.96558660 | 0.205536253 | 0.214578468 |
| Syn2 | 0.032437087 | 0.329758683 | 0.77788578 | 0.332167728 | 0.299546776 |
| Syncrip | -0.015724658 | 0.660166092 | 0.92117803 | 0.434154689 | 0.441941693 |
| Syndig1l | 0.010930658 | 0.727534842 | 0.94551888 | 0.236580297 | 0.278438458 |
| Syne1 | -0.086607027 | 0.004678808 | 0.13658474 | 0.190201303 | 0.271992294 |
| Syne2 | 0.007155439 | 0.751578669 | 0.95060533 | 0.102308626 | 0.109179274 |
| Syne4 | -0.054723625 | 0.092016264 | 0.53255842 | 0.28859495 | 0.335055307 |
| Syngr1 | 0.02528152 | 0.265033744 | 0.73067008 | 0.123658647 | 0.112871963 |
| Syngr2 | 0.020898881 | 0.494431907 | 0.86781063 | 0.271365562 | 0.249231817 |
| Synj1 | 0.007697275 | 0.789658398 | 0.96184914 | 0.230952787 | 0.214971843 |
| Synj2bp | -0.00942222 | 0.775460989 | 0.95585380 | 0.353783124 | 0.340435661 |
| Synpo | -0.040174733 | 0.013542952 | 0.22925110 | 0.032497832 | 0.064168853 |
| Synpr | -0.014130775 | 0.452675955 | 0.84866625 | 0.064460542 | 0.082131899 |
| Synrg | 0.029777704 | 0.417260088 | 0.83009907 | 0.562983402 | 0.572329568 |
| Syp | 0.046034521 | 0.197744341 | 0.67309621 | 0.551294756 | 0.548100349 |
| Sypl | -0.027245555 | 0.398634289 | 0.82398500 | 0.322692295 | 0.335735006 |
| Sys1 | 0.062549609 | 0.062814818 | 0.46571225 | 1.093142869 | 0.981967506 |
| Syt1 | 0.004139079 | 0.804616538 | 0.96471431 | 0.05601119 | 0.053942785 |
| Syt11 | -0.056404929 | 0.120948748 | 0.58114450 | 0.411724806 | 0.462614077 |
| Syt13 | -0.070450408 | 0.043456871 | 0.39842159 | 1.105607522 | 1.28606212 |
| Syt14 | -0.025367838 | 0.415980041 | 0.83009907 | 0.250915037 | 0.279359974 |
| Syt4 | 0.017649318 | 0.660144183 | 0.92117803 | 0.906659094 | 1.070719762 |
| Syt5 | -0.028636051 | 0.403492253 | 0.82647037 | 0.312748058 | 0.32139304 |
| Syt7 | 0.022332301 | 0.527081975 | 0.87685539 | 0.99837539 | 0.972747713 |
| Syt9 | -0.030607713 | 0.140652224 | 0.61309530 | 0.080620277 | 0.095207621 |
| Sytl1 | -0.057295328 | 0.005496462 | 0.14897511 | 0.052246064 | 0.11285046 |
| Sytl2 | -0.027267962 | 0.285701031 | 0.74543773 | 0.14259109 | 0.1328713 |
| Sytl4 | 0.099683845 | 0.001855152 | 0.07450649 | 0.301196922 | 0.298030022 |
| Sytl5 | 0.020000565 | 0.477471444 | 0.86074894 | 0.236630549 | 0.183959928 |
| Syvn1 | -0.009206648 | 0.717492294 | 0.94351793 | 0.155799027 | 0.160186971 |
| Szrd1 | -0.014142375 | 0.637204347 | 0.91583123 | 0.244117082 | 0.249754349 |
| Szt2 | -0.013297515 | 0.404757823 | 0.82707700 | 0.050920669 | 0.0563786 |
| Tab1 | -0.003600979 | 0.84321421 | 0.97348890 | 0.076628813 | 0.068992734 |
| Tab2 | -0.041184251 | 0.242166365 | 0.71327199 | 0.369142598 | 0.407692759 |
| Tab3 | -0.031948422 | 0.030216063 | 0.33157449 | 0.025198598 | 0.053059749 |
| Tacc1 | 0.034891779 | 0.346551296 | 0.78994438 | 0.695023477 | 0.679107814 |
| Tacc2 | 0.052272449 | 0.056434449 | 0.44713897 | 0.185342072 | 0.172891381 |
| Taco1 | -0.003525879 | 0.793658459 | 0.96245087 | 0.044951757 | 0.035948111 |
| Tada1 | -0.028482126 | 0.211239921 | 0.68220598 | 0.118282151 | 0.126164039 |
| Tada2a | -0.014280219 | 0.477717359 | 0.86074894 | 0.089407518 | 0.08946969 |
| Tada2b | 0.017451922 | 0.416993372 | 0.83009907 | 0.104227716 | 0.104675842 |
| Tada3 | -0.047597244 | 0.106193409 | 0.56171713 | 0.214627688 | 0.230371314 |
| Taf1 | 0.010871048 | 0.759679348 | 0.95215335 | 0.413189223 | 0.40804103 |
| Taf10 | 0.009969328 | 0.762050908 | 0.95295130 | 1.215616105 | 1.127513541 |
| Taf11 | -0.05622397 | 0.083091036 | 0.51775319 | 0.324844871 | 0.356143992 |
| Taf12 | 0.009958896 | 0.722738276 | 0.94456523 | 0.214781662 | 0.191532699 |
| Taf13 | -0.01267727 | 0.562818202 | 0.89262403 | 0.109769577 | 0.111560056 |
| Taf15 | 0.041171121 | 0.274808974 | 0.73519553 | 0.509816632 | 0.496753373 |
| Taf1a | -0.030749412 | 0.079520401 | 0.50762057 | 0.050260727 | 0.06805813 |
| Taf1b | -0.0294109 | 0.096431995 | 0.53711700 | 0.053494216 | 0.072585483 |
| Taf1d | -0.022047171 | 0.512504435 | 0.87227718 | 0.31795617 | 0.339465698 |
| Taf2 | 0.017553054 | 0.424808367 | 0.83244603 | 0.119975403 | 0.103771425 |
| Taf3 | -0.016227675 | 0.606105718 | 0.90871393 | 0.251892163 | 0.265679613 |
| Taf4 | -0.05202686 | 0.002190086 | 0.08367830 | 0.031666741 | 0.073570226 |
| Taf5 | 0.009072135 | 0.680325761 | 0.92933510 | 0.116928702 | 0.10140867 |
| Taf5l | -0.009344414 | 0.647854517 | 0.91696950 | 0.090037926 | 0.095812247 |
| Taf6 | -0.013391134 | 0.598308099 | 0.90755130 | 0.154650673 | 0.16046855 |
| Taf6l | 0.034491511 | 0.016800375 | 0.25467130 | 0.066202185 | 0.03379481 |

| | | | | | |
|---|---|---|---|---|---|
| Taf7 | -0.02016983 | 0.544465209 | 0.8842656 | 0.310411862 | 0.307538349 |
| Taf8 | -0.004984732 | 0.796902554 | 0.9644244 | 0.095350733 | 0.082427285 |
| Taf9b | 0.000919315 | 0.969090621 | 0.9976072 | 0.148098743 | 0.133522705 |
| Tafa5 | 0.043066941 | 0.079383511 | 0.5076205 | 0.155540236 | 0.130815862 |
| Tagap1 | -0.005577143 | 0.769725903 | 0.9552333 | 0.090802303 | 0.080138062 |
| Tagln2 | -0.133751296 | 0.000259246 | 0.0196314 | 1.086615358 | 1.289923946 |
| Taldo1 | 0.078304033 | 0.033329196 | 0.3456484 | 0.810057897 | 0.708797414 |
| Tanc1 | -0.036867702 | 0.120716854 | 0.5810004 | 0.119690636 | 0.137644852 |
| Tanc2 | -0.008275675 | 0.768049386 | 0.9544221 | 0.199226208 | 0.191943343 |
| Tango2 | -0.005883571 | 0.748517603 | 0.9501120 | 0.077454998 | 0.074695733 |
| Tank | -0.028935409 | 0.175407473 | 0.6534142 | 0.083601362 | 0.109133744 |
| Taok1 | -0.035588262 | 0.334411661 | 0.7804836 | 0.519196269 | 0.55284397 |
| Taok2 | -0.070822409 | 0.036500031 | 0.3620803 | 0.300004443 | 0.361995804 |
| Taok3 | -0.044064013 | 0.224954953 | 0.6968918 | 0.573855315 | 0.585904155 |
| Tap1 | 0.016919054 | 0.289745971 | 0.7496864 | 0.064038955 | 0.049713998 |
| Tap2 | -0.020310634 | 0.421517055 | 0.8304765 | 0.14289496 | 0.155950429 |
| Tapbp | -0.002371268 | 0.946218349 | 0.9944798 | 0.672581332 | 0.697172924 |
| Tapbpl | -0.000806546 | 0.978384778 | 0.9976072 | 0.211373158 | 0.230067373 |
| Tapt1 | -0.01038373 | 0.643099674 | 0.9166712 | 0.118962834 | 0.116904694 |
| Tarbp2 | -0.02463861 | 0.230979702 | 0.7024275 | 0.089905065 | 0.098332622 |
| Tardbp | 0.030117441 | 0.402710614 | 0.8259022 | 0.479249492 | 0.442665833 |
| Tars | 0.055109707 | 0.128509298 | 0.5914729 | 0.504205134 | 0.49436748 |
| Tars2 | 0.020833335 | 0.279770637 | 0.7407935 | 0.095850511 | 0.073826453 |
| Tarsl2 | -0.008566949 | 0.632667284 | 0.9139562 | 0.060031497 | 0.070611843 |
| Tasor | -0.029421648 | 0.420333594 | 0.8300990 | 0.398638437 | 0.428273404 |
| Tasor2 | -0.044360981 | 0.142449357 | 0.6157288 | 0.227260963 | 0.246704382 |
| Tasp1 | -0.009490552 | 0.612242531 | 0.9101522 | 0.070829078 | 0.075918133 |
| Tat | -0.003615168 | 0.83304152 | 0.9723494 | 0.054785951 | 0.061062156 |
| Tatdn1 | -0.039344383 | 0.149055337 | 0.6264374 | 0.182358022 | 0.195322028 |
| Tatdn2 | -0.002771246 | 0.897156387 | 0.9825441 | 0.109133019 | 0.097817942 |
| Tatdn3 | -0.029024736 | 0.088070572 | 0.5278913 | 0.041375997 | 0.064979959 |
| Tax1bp1 | -0.045715966 | 0.170833859 | 0.6507063 | 1.179067566 | 1.200852533 |
| Tax1bp3 | -0.04177477 | 0.079488236 | 0.5076205 | 0.120456599 | 0.129237758 |
| Taz | -0.094199109 | 0.001828684 | 0.0745064 | 0.169962371 | 0.26011632 |
| Tbc1d1 | -0.0220821 | 0.485556725 | 0.8630379 | 0.265104224 | 0.279192589 |
| Tbc1d10a | -0.03607299 | 0.098351689 | 0.5417261 | 0.096508328 | 0.115833931 |
| Tbc1d10b | -0.026777265 | 0.277142668 | 0.7376464 | 0.141763883 | 0.152893898 |
| Tbc1d12 | 0.00804919 | 0.730292087 | 0.9459020 | 0.139543881 | 0.119971243 |
| Tbc1d14 | 0.031542657 | 0.117075483 | 0.5735102 | 0.10885375 | 0.086325501 |
| Tbc1d15 | 0.044446806 | 0.228378115 | 0.7015153 | 0.528633207 | 0.502365307 |
| Tbc1d16 | 0.076374942 | 0.01747977 | 0.2580346 | 0.356800033 | 0.274081531 |
| Tbc1d17 | -0.027969958 | 0.348051976 | 0.7917953 | 0.231686366 | 0.229137264 |
| Tbc1d19 | -0.004368887 | 0.84673782 | 0.9739657 | 0.115012555 | 0.114235045 |
| Tbc1d20 | 0.032798454 | 0.273652605 | 0.7351955 | 0.271567697 | 0.237775897 |
| Tbc1d22a | 0.019865058 | 0.279813857 | 0.7407935 | 0.092672023 | 0.063463758 |
| Tbc1d22b | 0.034951984 | 0.091332372 | 0.5321282 | 0.116187142 | 0.092608064 |
| Tbc1d23 | -0.019657016 | 0.515113172 | 0.8733714 | 0.223276585 | 0.230684302 |
| Tbc1d24 | 0.012937835 | 0.483320743 | 0.8620306 | 0.072041294 | 0.074110157 |
| Tbc1d30 | -0.011421102 | 0.483328091 | 0.8620306 | 0.050763884 | 0.058026556 |
| Tbc1d31 | 0.019921241 | 0.268067865 | 0.7323841 | 0.076543669 | 0.06307129 |
| Tbc1d32 | -0.020354331 | 0.312117 | 0.7684379 | 0.073651327 | 0.089855492 |
| Tbc1d5 | 0.008091013 | 0.702677611 | 0.9389226 | 0.10047405 | 0.100490681 |
| Tbc1d7 | 0.00322013 | 0.876175523 | 0.9785583 | 0.109541958 | 0.093191902 |
| Tbc1d8 | -0.010318314 | 0.58520622 | 0.9024598 | 0.067021224 | 0.079439168 |
| Tbc1d8b | 0.051618509 | 0.040986481 | 0.3901716 | 0.1857532 | 0.137553438 |
| Tbc1d9 | -1.76E-05 | 0.999327343 | 0.9996989 | 0.099886214 | 0.09397682 |
| Tbc1d9b | 0.039684919 | 0.214173771 | 0.6837977 | 0.314788375 | 0.264672636 |
| Tbca | 0.053472014 | 0.065520897 | 0.4747752 | 1.501497588 | 1.340740831 |
| Tbcb | -0.027053894 | 0.455668248 | 0.8502914 | 0.860304337 | 0.839288927 |
| Tbcc | -0.005315586 | 0.864214051 | 0.9769675 | 0.274207474 | 0.262237995 |
| Tbccd1 | 0.025023813 | 0.20302861 | 0.6772740 | 0.100564335 | 0.074453141 |
| Tbcd | 0.021558847 | 0.384658712 | 0.8136064 | 0.154450437 | 0.14339143 |
| Tbce | -0.038221571 | 0.162171901 | 0.6400703 | 0.165346442 | 0.193891845 |
| Tbcel | -0.064362334 | 0.012426044 | 0.2195296 | 0.133464067 | 0.166022671 |
| Tbck | -0.034770343 | 0.214376119 | 0.6837977 | 0.18989386 | 0.195099919 |
| Tbk1 | -0.004743647 | 0.835811207 | 0.9723494 | 0.123768041 | 0.117326476 |

| | | | | | |
|---|---|---|---|---|---|
| Tbl1x | 0.029610677 | 0.441539431 | 0.8426454 | 0.807609173 | 0.735691867 |
| Tbl1xr1 | 0.005476652 | 0.802554709 | 0.9647143 | 0.112073931 | 0.102849227 |
| Tbl2 | 0.004845198 | 0.826342278 | 0.9716906 | 0.106319305 | 0.108680859 |
| Tbl3 | -0.017546258 | 0.387806537 | 0.8170287 | 0.081865336 | 0.096516121 |
| Tbp | -0.015742817 | 0.393137947 | 0.8206236 | 0.062970555 | 0.072371558 |
| Tbpl1 | 0.003091212 | 0.897242622 | 0.9825441 | 0.134045542 | 0.138916636 |
| Tbrg1 | 0.028787195 | 0.386361529 | 0.8154694 | 1.228069191 | 1.136819045 |
| Tbrg4 | -0.004097454 | 0.789455606 | 0.9618491 | 0.050781148 | 0.047172546 |
| Tcaf1 | 0.033269685 | 0.357891047 | 0.7989722 | 0.510602614 | 0.452957343 |
| Tcaim | -0.011871264 | 0.508999997 | 0.8704067 | 0.064926169 | 0.071637999 |
| Tcea1 | 0.072163654 | 0.048765263 | 0.4159513 | 0.575965083 | 0.522173146 |
| Tcea2 | 0.014338813 | 0.28714285 | 0.7471316 | 0.047675274 | 0.033494652 |
| Tceal1 | -0.025799962 | 0.42390819 | 0.8320462 | 0.321240337 | 0.315189773 |
| Tceal3 | -0.013569337 | 0.722043826 | 0.9445652 | 0.592618951 | 0.5944363 |
| Tceal8 | 0.020016067 | 0.596231513 | 0.9068716 | 0.723513225 | 0.650750232 |
| Tceal9 | 0.051844461 | 0.007860581 | 0.1751394 | 2.558819998 | 2.420422971 |
| Tceanc | -0.003230538 | 0.845744326 | 0.9736268 | 0.058645515 | 0.057554846 |
| Tceanc2 | 0.025243872 | 0.185381168 | 0.6637678 | 0.096786666 | 0.072187886 |
| Tcerg1 | -0.000969758 | 0.976846344 | 0.9976072 | 0.329872478 | 0.318801647 |
| Tcf12 | -0.037158794 | 0.248741496 | 0.7187416 | 0.260964586 | 0.277744448 |
| Tcf20 | 0.018983778 | 0.487301429 | 0.8640311 | 0.188889854 | 0.178740628 |
| Tcf24 | 0.024859011 | 0.154552883 | 0.6304934 | 0.073786035 | 0.068692688 |
| Tcf25 | 0.022701259 | 0.424126558 | 0.8321470 | 1.477720379 | 1.447713805 |
| Tcf3 | -0.031661442 | 0.176267574 | 0.6536659 | 0.103678872 | 0.131950012 |
| Tcf4 | 0.010849324 | 0.758572889 | 0.9521533 | 0.419612714 | 0.41451031 |
| Tcf7l2 | -0.016012724 | 0.47948182 | 0.8610807 | 0.116924063 | 0.11777988 |
| Tchp | -0.011582361 | 0.605295709 | 0.9087139 | 0.103534459 | 0.119533968 |
| Tcim | 0.03340725 | 0.061033495 | 0.4605600 | 0.070234069 | 0.053433466 |
| Tcn2 | -0.01664412 | 0.588237565 | 0.9044624 | 0.243932769 | 0.277288601 |
| Tcof1 | -0.021700512 | 0.446204458 | 0.8442396 | 0.178488328 | 0.209727295 |
| Tcp1 | -0.015800622 | 0.658025066 | 0.9211780 | 0.883509738 | 0.833005703 |
| Tcp11l1 | 0.039383582 | 0.125783591 | 0.5882766 | 0.184504729 | 0.151785743 |
| Tcp11l2 | -0.036879419 | 0.105835055 | 0.5617171 | 0.107829107 | 0.120545062 |
| Tcta | 8.42E-05 | 0.997953977 | 0.9994073 | 0.3686115 | 0.349167625 |
| Tctn1 | 0.004483479 | 0.773385181 | 0.9557311 | 0.061645983 | 0.045671845 |
| Tctn2 | 0.013609221 | 0.41162812 | 0.8294436 | 0.065181358 | 0.058923735 |
| Tctn3 | 0.01580139 | 0.314515717 | 0.7705987 | 0.065481505 | 0.048672554 |
| Tdg | 0.048718386 | 0.03814046 | 0.3731295 | 0.166130847 | 0.111346234 |
| Tdp1 | 0.001257692 | 0.949039835 | 0.9947670 | 0.09086542 | 0.082310686 |
| Tdp2 | -0.014753581 | 0.588453686 | 0.9044624 | 0.181315555 | 0.17734157 |
| Tdrd3 | -0.022628466 | 0.278623016 | 0.7393789 | 0.091514643 | 0.10003984 |
| Tdrd7 | 0.029625292 | 0.154066706 | 0.6304934 | 0.118310986 | 0.088040568 |
| Tdrkh | -0.029370771 | 0.276712747 | 0.7369101 | 0.192382324 | 0.185383573 |
| Tdrp | 0.027262917 | 0.081613149 | 0.5137071 | 0.074319072 | 0.042202567 |
| Tead1 | -0.004408236 | 0.893686535 | 0.9824361 | 0.314770828 | 0.311241147 |
| Tec | 0.031587368 | 0.044601233 | 0.4014920 | 0.063846108 | 0.046852344 |
| Tecpr1 | -0.010822462 | 0.682174017 | 0.9303225 | 0.16661613 | 0.16724191 |
| Tecpr2 | -0.011892623 | 0.496561159 | 0.8684629 | 0.054826699 | 0.070629679 |
| Tecr | 0.026335733 | 0.457233508 | 0.8518352 | 1.042450655 | 0.939733974 |
| Tef | 0.048000388 | 0.034848948 | 0.3560263 | 0.143907814 | 0.096589486 |
| Tefm | -0.014296088 | 0.45368444 | 0.8490482 | 0.076206554 | 0.080496195 |
| Telo2 | 0.017495928 | 0.218531749 | 0.6899952 | 0.055642496 | 0.036752666 |
| Ten1 | -0.018165814 | 0.606384642 | 0.9087139 | 0.462944578 | 0.466637818 |
| Tenm2 | -0.023750642 | 0.391517333 | 0.8200150 | 0.158100345 | 0.183851758 |
| Tenm3 | 0.033707059 | 0.071843029 | 0.4888240 | 0.084891751 | 0.071874379 |
| Tenm4 | -0.034812207 | 0.124128402 | 0.5852543 | 0.078254273 | 0.123892576 |
| Tent2 | -0.023070427 | 0.253431441 | 0.7223071 | 0.077908005 | 0.092844885 |
| Tent4a | -0.004001783 | 0.838220623 | 0.9727216 | 0.09171979 | 0.084303014 |
| Tent4b | -0.022021667 | 0.297858795 | 0.7557491 | 0.089405292 | 0.103249881 |
| Tent5a | -0.026231035 | 0.557183389 | 0.8916372 | 0.702624101 | 0.92286849 |
| Tent5c | -0.029920083 | 0.375786834 | 0.8101773 | 0.317876501 | 0.337169033 |
| Tent5d | 0.009236056 | 0.694214789 | 0.9361896 | 0.157496794 | 0.098801622 |
| Tep1 | -0.021918091 | 0.275701451 | 0.7357967 | 0.077540227 | 0.093491966 |
| Terf1 | -0.027752476 | 0.213771212 | 0.6831238 | 0.104540955 | 0.119448984 |
| Terf2 | 0.023227208 | 0.398613593 | 0.8239850 | 0.197870653 | 0.183126208 |
| Terf2ip | 0.033069613 | 0.242351295 | 0.7132719 | 0.214537229 | 0.189725873 |

| | | | | | |
|---|---|---|---|---|---|
| Tes | 0.049238896 | 0.099210421 | 0.54335515 | 0.24078552 | 0.228479481 |
| Tesc | 0.015002755 | 0.47693529 | 0.86074894 | 0.110965922 | 0.082344557 |
| Tesk1 | -0.043321825 | 0.14607217 | 0.62163703 | 0.21059561 | 0.247163563 |
| Tet1 | -0.01936952 | 0.260693905 | 0.72710774 | 0.050014427 | 0.065702335 |
| Tet2 | -0.011951938 | 0.643759251 | 0.91683910 | 0.146763883 | 0.163308379 |
| Tet3 | 0.040422581 | 0.092496603 | 0.53255842 | 0.136152163 | 0.126669739 |
| Tex10 | -0.037088017 | 0.132901103 | 0.59882510 | 0.117613489 | 0.14725085 |
| Tex2 | -0.021742602 | 0.232542051 | 0.70394170 | 0.054683697 | 0.078595795 |
| Tex261 | -0.001653388 | 0.962918579 | 0.99657301 | 0.504771972 | 0.483403882 |
| Tex264 | -0.004681067 | 0.871668324 | 0.97822694 | 0.212061501 | 0.224048351 |
| Tex30 | 0.016305664 | 0.537988414 | 0.88163354 | 0.191779847 | 0.163769536 |
| Tex9 | -0.020093856 | 0.377383705 | 0.81017979 | 0.09912718 | 0.114464265 |
| Tfam | -0.020538683 | 0.494987937 | 0.86828690 | 0.240465601 | 0.249224794 |
| Tfb2m | 0.002493262 | 0.910722639 | 0.98373739 | 0.106413461 | 0.10588951 |
| Tfcp2 | -0.001977816 | 0.918716142 | 0.98512115 | 0.087653046 | 0.084212463 |
| Tfdp1 | -0.005996857 | 0.817788187 | 0.96961898 | 0.157376046 | 0.160071186 |
| Tfdp2 | -0.049485931 | 0.099048991 | 0.54335515 | 0.203233059 | 0.24225921 |
| Tfe3 | 0.025461167 | 0.402561984 | 0.82576817 | 0.247457494 | 0.220463535 |
| Tfg | -0.038406467 | 0.272385759 | 0.73428062 | 0.498341176 | 0.48735135 |
| Tfip11 | 0.000244009 | 0.990611446 | 0.99808363 | 0.096432216 | 0.093760429 |
| Tfpi | -0.010658547 | 0.605088381 | 0.90871392 | 0.095383403 | 0.076732311 |
| Tfpt | 0.015681321 | 0.474242337 | 0.85998155 | 0.121855639 | 0.105675853 |
| Tfrc | 0.026179554 | 0.468217238 | 0.85616860 | 0.390837502 | 0.375715955 |
| Tgds | 0.035255202 | 0.110045998 | 0.56558881 | 0.139658702 | 0.09621479 |
| Tgfb2 | 0.022757783 | 0.108483784 | 0.56368171 | 0.056358333 | 0.030424636 |
| Tgfbr1 | 0.059224951 | 0.008180196 | 0.17756573 | 0.149968822 | 0.097259138 |
| Tgfbr3 | -0.015145041 | 0.336872492 | 0.78280044 | 0.044971016 | 0.052937286 |
| Tgfbrap1 | 0.010138036 | 0.673435438 | 0.92616863 | 0.141088474 | 0.131444974 |
| Tgoln1 | -0.075420991 | 0.016984109 | 0.25558849 | 1.268357725 | 1.379261722 |
| Tgs1 | -0.006293795 | 0.831649751 | 0.97234941 | 0.236497137 | 0.222252938 |
| Tha1 | -0.042246651 | 0.046688199 | 0.40945771 | 0.086951626 | 0.107798311 |
| Thada | -0.009854179 | 0.570904913 | 0.89652948 | 0.061808504 | 0.064246925 |
| Thap1 | 0.013041991 | 0.592542473 | 0.90605611 | 0.149407081 | 0.131095741 |
| Thap11 | 0.010945201 | 0.709052957 | 0.94148110 | 0.254059798 | 0.21932845 |
| Thap12 | 0.074370737 | 0.003063636 | 0.10328859 | 0.193339148 | 0.131513799 |
| Thap2 | -0.016568512 | 0.546777708 | 0.88584590 | 0.194513074 | 0.184824732 |
| Thap3 | 0.020073901 | 0.501611836 | 0.86955663 | 0.271886832 | 0.236584788 |
| Thap4 | -0.040484344 | 0.064260531 | 0.47003870 | 0.100784395 | 0.11917324 |
| Thap6 | 0.007786493 | 0.672559781 | 0.92586630 | 0.073734907 | 0.070161302 |
| Thap7 | 0.033119229 | 0.230191643 | 0.70196775 | 0.235543814 | 0.18610054 |
| Thbs3 | -0.042681775 | 0.039297114 | 0.37990034 | 0.081403753 | 0.10018024 |
| Them4 | -0.020635028 | 0.349781459 | 0.79273039 | 0.092666832 | 0.116355393 |
| Them6 | -0.011420109 | 0.525037713 | 0.87583859 | 0.06152032 | 0.073537796 |
| Thnsl1 | -0.011542544 | 0.584299784 | 0.90214067 | 0.095458027 | 0.099094472 |
| Thnsl2 | -0.004256087 | 0.77587557 | 0.95585380 | 0.046670116 | 0.045331911 |
| Thoc1 | -0.01851959 | 0.477832363 | 0.86074894 | 0.153096968 | 0.166440015 |
| Thoc2 | 0.044568307 | 0.224405176 | 0.69652670 | 0.547649039 | 0.487895458 |
| Thoc2l | 0.033085725 | 0.393187939 | 0.82062368 | 0.769524914 | 0.723801256 |
| Thoc3 | 0.009463837 | 0.747789669 | 0.94963702 | 0.256689195 | 0.245931361 |
| Thoc5 | -0.004698294 | 0.820286889 | 0.97010561 | 0.098867234 | 0.096135845 |
| Thoc6 | 0.024199292 | 0.213681768 | 0.68312382 | 0.105258233 | 0.070625479 |
| Thoc7 | 0.041492137 | 0.240452854 | 0.71204049 | 0.861103192 | 0.750833168 |
| Thop1 | -0.033422168 | 0.101690413 | 0.55109170 | 0.076740533 | 0.099537699 |
| Thra | -0.009637353 | 0.772166585 | 0.95523334 | 0.375607418 | 0.330024728 |
| Thrap3 | 0.034552857 | 0.347892805 | 0.79179535 | 0.589406311 | 0.521265852 |
| Thrb | -0.011716917 | 0.581789746 | 0.90189991 | 0.090857311 | 0.100396833 |
| Thsd7a | 0.002738862 | 0.889630508 | 0.98195979 | 0.093541759 | 0.074799522 |
| Thumpd1 | -0.00919036 | 0.767824541 | 0.95442217 | 0.279006236 | 0.274825509 |
| Thumpd3 | -0.008835383 | 0.720369182 | 0.94452957 | 0.152464215 | 0.144203879 |
| Thyn1 | 0.047637801 | 0.148667352 | 0.62643749 | 0.394651512 | 0.300080143 |
| Tia1 | -0.065763771 | 0.072993411 | 0.49033721 | 0.44739504 | 0.503691815 |
| Tial1 | -0.026490497 | 0.436274911 | 0.83906441 | 0.417130769 | 0.43922786 |
| Ticam1 | -0.051160028 | 0.012473205 | 0.21977651 | 0.064701906 | 0.100641659 |
| Tigar | 0.006442324 | 0.725346728 | 0.94502751 | 0.070690549 | 0.071192313 |
| Tigd2 | -0.02771294 | 0.400526012 | 0.82466127 | 0.31813015 | 0.325095412 |
| Timm10 | -0.014914153 | 0.624678383 | 0.91215057 | 0.249743965 | 0.253082496 |

| | | | | | |
|---|---|---|---|---|---|
| Timm10b | 0.03480962 | 0.318921211 | 0.77100438 | 0.611247202 | 0.534173989 |
| Timm13 | 0.047806252 | 0.104753331 | 0.55800455 | 1.368059618 | 1.304035725 |
| Timm17a | 0.035843619 | 0.312177929 | 0.76843798 | 0.551516845 | 0.512922856 |
| Timm17b | 0.002084807 | 0.9523899 | 0.99515668 | 0.725456289 | 0.64932187 |
| Timm21 | -0.015040364 | 0.378234897 | 0.81019153 | 0.06272463 | 0.062506904 |
| Timm22 | -0.016227838 | 0.57207269 | 0.89680168 | 0.205553314 | 0.229119783 |
| Timm23 | 0.109947075 | 0.003548223 | 0.11494803 | 0.794937688 | 0.609898587 |
| Timm29 | -0.041875264 | 0.119327246 | 0.57827370 | 0.152747444 | 0.188969615 |
| Timm44 | -0.005393752 | 0.850763326 | 0.97488410 | 0.200352207 | 0.201722165 |
| Timm50 | -0.015739424 | 0.600552051 | 0.90784374 | 0.255522619 | 0.260818424 |
| Timm8a1 | 0.004110298 | 0.823114842 | 0.97105924 | 0.070857538 | 0.073684536 |
| Timm8b | 0.122084587 | 0.000451858 | 0.02873352 | 0.847643678 | 0.737612778 |
| Timm9 | -0.021540491 | 0.507378065 | 0.87040677 | 0.330971672 | 0.332671258 |
| Timmdc1 | 0.003276034 | 0.893092838 | 0.98228263 | 0.158646827 | 0.132792071 |
| Timp2 | -0.078298131 | 0.016298262 | 0.25144441 | 0.26834937 | 0.308917077 |
| Tinf2 | 0.024347822 | 0.133418195 | 0.59882510 | 0.068856103 | 0.05185964 |
| Tiparp | -0.047000845 | 0.175204968 | 0.65341424 | 0.271640085 | 0.337320213 |
| Tipin | -0.008230212 | 0.76972208 | 0.95523334 | 0.189922591 | 0.20279364 |
| Tiprl | -0.030830654 | 0.357662784 | 0.79897222 | 0.460503622 | 0.429371315 |
| Tjap1 | -0.010440146 | 0.550599739 | 0.88830567 | 0.063563412 | 0.06657097 |
| Tjp1 | 0.071306458 | 0.03648465 | 0.36208030 | 0.387490908 | 0.326486909 |
| Tjp2 | 0.016372559 | 0.444724219 | 0.84315410 | 0.115004681 | 0.104336388 |
| Tk2 | 0.007861914 | 0.722223578 | 0.94456522 | 0.110856247 | 0.109053772 |
| Tkfc | -0.0035795 | 0.850724524 | 0.97488410 | 0.083426795 | 0.072691814 |
| Tkt | -0.006437173 | 0.854238506 | 0.97554319 | 0.446213926 | 0.435592198 |
| Tlcd1 | 0.007793236 | 0.691383866 | 0.93514293 | 0.092195149 | 0.081314509 |
| Tlcd3b | -0.038790993 | 0.107324772 | 0.56337238 | 0.126237078 | 0.14083456 |
| Tlcd4 | 0.036061556 | 0.204453901 | 0.67809518 | 0.231159099 | 0.20588624 |
| Tle1 | -0.013401672 | 0.646168913 | 0.91696950 | 0.207284577 | 0.220034884 |
| Tle2 | 0.003635741 | 0.827882111 | 0.97178920 | 0.062486873 | 0.057912391 |
| Tle3 | 0.001915588 | 0.942961418 | 0.99349067 | 0.143929158 | 0.170971726 |
| Tle4 | -0.031124243 | 0.102154758 | 0.55224800 | 0.05523958 | 0.080861751 |
| Tle5 | -0.024857116 | 0.195154361 | 0.67199328 | 2.410589167 | 2.325899663 |
| Tle6 | -0.008370224 | 0.647847247 | 0.91696950 | 0.07865519 | 0.070861543 |
| Tlk1 | 0.016043757 | 0.66063232 | 0.92117803 | 0.534878807 | 0.530258653 |
| Tlk2 | 0.054669031 | 0.073247184 | 0.49128604 | 0.296569774 | 0.222888657 |
| Tln1 | -0.021710717 | 0.535363 | 0.87959493 | 0.425759944 | 0.435562858 |
| Tlnrd1 | -0.013118185 | 0.56926348 | 0.89565324 | 0.123083242 | 0.119735768 |
| Tlr2 | -9.06E-05 | 0.995314718 | 0.99934433 | 0.047352706 | 0.051834403 |
| Tm2d1 | 0.008995707 | 0.802259863 | 0.96471433 | 0.726395971 | 0.685804476 |
| Tm2d2 | -0.031867359 | 0.364198813 | 0.80498823 | 0.664599324 | 0.658133132 |
| Tm2d3 | 0.033113702 | 0.34119137 | 0.78513662 | 0.607956847 | 0.535068499 |
| Tm4sf4 | 0.048907892 | 0.074131494 | 0.49425893 | 2.449642798 | 2.198324108 |
| Tm7sf3 | -0.008044125 | 0.788879622 | 0.96166093 | 0.256787849 | 0.23586378 |
| Tm9sf1 | -0.011384876 | 0.502055048 | 0.86955665 | 0.061223883 | 0.05922221 |
| Tm9sf2 | 0.029297546 | 0.419512399 | 0.83009907 | 0.727060775 | 0.708481967 |
| Tm9sf3 | 0.012141576 | 0.736902685 | 0.94697528 | 0.908699052 | 0.885139912 |
| Tm9sf4 | -0.10642422 | 0.002109906 | 0.08193280 | 0.36433134 | 0.444532703 |
| Tma16 | 0.035424654 | 0.11066328 | 0.56558883 | 0.140762651 | 0.101246147 |
| Tma7 | 0.069890626 | 0.006420694 | 0.16030787 | 1.755465989 | 1.610153726 |
| Tmbim1 | -0.020245258 | 0.474005942 | 0.85978038 | 0.189226785 | 0.203358686 |
| Tmbim4 | 0.041430307 | 0.250989423 | 0.72072102 | 0.964311046 | 0.902817254 |
| Tmbim6 | -0.10160332 | 0.000309144 | 0.02238473 | 1.372343153 | 1.518160508 |
| Tmc4 | -0.067456181 | 0.042695983 | 0.39583565 | 0.309332527 | 0.362293204 |
| Tmc6 | -0.003392887 | 0.844220722 | 0.97348890 | 0.071630842 | 0.062752738 |
| Tmcc1 | -0.088696699 | 0.001243753 | 0.05642600 | 0.143039352 | 0.205998352 |
| Tmcc2 | 0.003758059 | 0.871424767 | 0.97822694 | 0.133823212 | 0.120332447 |
| Tmcc3 | 0.01280253 | 0.600204287 | 0.90782343 | 0.125530256 | 0.141240092 |
| Tmco1 | 0.017473479 | 0.624804756 | 0.91215057 | 0.629966735 | 0.616161396 |
| Tmco3 | 0.016166944 | 0.573470219 | 0.89734948 | 0.220977932 | 0.208664761 |
| Tmco6 | 0.006290061 | 0.750358569 | 0.95045378 | 0.099843972 | 0.079815162 |
| Tmed1 | -0.004886319 | 0.832902915 | 0.97234943 | 0.131336766 | 0.116122991 |
| Tmed10 | -0.025062795 | 0.302075672 | 0.76112584 | 1.962952599 | 1.930879015 |
| Tmed2 | 0.056675548 | 0.075557979 | 0.49670984 | 1.473860356 | 1.29096382 |
| Tmed3 | -0.076837435 | 0.002128327 | 0.08215177 | 2.202159007 | 2.174216405 |
| Tmed4 | -0.00611354 | 0.866737412 | 0.97720023 | 0.778137044 | 0.728760327 |

| | | | | | |
|---|---|---|---|---|---|
| Tmed5 | 0.032149802 | 0.36106996 | 0.8018388 | 0.438578592 | 0.428320174 |
| Tmed6 | 0.009537322 | 0.621391 | 0.9121505 | 0.075848398 | 0.07738998 |
| Tmed7 | 0.002429221 | 0.947196701 | 0.9945018 | 0.59841315 | 0.572905706 |
| Tmed8 | -0.00308795 | 0.927240473 | 0.9879210 | 0.341276868 | 0.325086334 |
| Tmed9 | -0.03897047 | 0.203729048 | 0.6774097 | 1.464828144 | 1.480491948 |
| Tmeff1 | -0.0226334 | 0.234030701 | 0.7056961 | 0.073996678 | 0.079206739 |
| Tmem101 | 0.003454882 | 0.892429081 | 0.9820184 | 0.167896462 | 0.162196584 |
| Tmem102 | 0.022857717 | 0.207035889 | 0.6797565 | 0.078792875 | 0.064079727 |
| Tmem104 | -0.000862354 | 0.957182715 | 0.9956630 | 0.0531966 | 0.053277674 |
| Tmem106b | 0.018410809 | 0.571215063 | 0.8965907 | 0.350053574 | 0.336899464 |
| Tmem106c | -0.018159457 | 0.58500761 | 0.9023908 | 0.354023663 | 0.336255387 |
| Tmem107 | -0.032417449 | 0.376238237 | 0.8101773 | 0.668318235 | 0.615172595 |
| Tmem108 | -0.10511857 | 0.005010043 | 0.1415943 | 0.360043942 | 0.45927791 |
| Tmem109 | 0.030840317 | 0.272369731 | 0.7342806 | 0.227435587 | 0.190227298 |
| Tmem11 | -0.056581878 | 0.07157382 | 0.4888240 | 0.280733075 | 0.311571814 |
| Tmem115 | 0.043603794 | 0.126818041 | 0.5893234 | 0.231645383 | 0.194975771 |
| Tmem117 | -0.009694279 | 0.64766534 | 0.9169695 | 0.100476129 | 0.097088657 |
| Tmem120a | 0.017616684 | 0.348459392 | 0.7922798 | 0.088435735 | 0.072370676 |
| Tmem121 | 0.024623326 | 0.154800533 | 0.6304934 | 0.071510562 | 0.062016973 |
| Tmem123 | 0.025536675 | 0.296880276 | 0.7545964 | 0.160927753 | 0.137942873 |
| Tmem126a | 0.004795693 | 0.898763025 | 0.9825441 | 0.654239312 | 0.65953437 |
| Tmem126b | -0.019691152 | 0.460607715 | 0.8524425 | 0.182084664 | 0.178449287 |
| Tmem127 | 0.007770581 | 0.784342037 | 0.9601202 | 0.201539725 | 0.20818172 |
| Tmem128 | 0.095989518 | 0.004091976 | 0.1266085 | 0.418184644 | 0.338789614 |
| Tmem129 | -0.036238002 | 0.054722347 | 0.4422689 | 0.059734718 | 0.083953589 |
| Tmem130 | -0.038026495 | 0.176724392 | 0.6536659 | 0.163479107 | 0.198431122 |
| Tmem131 | -0.048836122 | 0.098986941 | 0.5433551 | 0.196860907 | 0.232291507 |
| Tmem131l | 0.028124299 | 0.274885305 | 0.7352014 | 0.188546285 | 0.144004342 |
| Tmem132a | 0.020514367 | 0.152061851 | 0.6281105 | 0.053012963 | 0.038053553 |
| Tmem132b | 0.055522231 | 0.00436857 | 0.1311882 | 0.104157282 | 0.074845467 |
| Tmem134 | -0.051523418 | 0.149539011 | 0.6264374 | 0.913896425 | 0.887656582 |
| Tmem135 | 0.048380441 | 0.047366492 | 0.4106761 | 0.156405228 | 0.129879865 |
| Tmem138 | 0.00502844 | 0.805380056 | 0.9647143 | 0.094234387 | 0.085605898 |
| Tmem140 | -0.023202277 | 0.162322971 | 0.6400703 | 0.042859101 | 0.063017752 |
| Tmem141 | 0.002922849 | 0.918530506 | 0.9851211 | 0.213968267 | 0.201347226 |
| Tmem143 | -0.009860089 | 0.609587349 | 0.9089292 | 0.082042268 | 0.080613164 |
| Tmem147 | 0.112911342 | 0.0016872 | 0.0718327 | 0.735616326 | 0.622000343 |
| Tmem14c | -0.040312321 | 0.257503397 | 0.7250597 | 0.692946419 | 0.70628325 |
| Tmem150a | -0.011572746 | 0.407448306 | 0.8270770 | 0.03693847 | 0.043424693 |
| Tmem150c | 0.024934709 | 0.189776457 | 0.6697747 | 0.086363252 | 0.081519072 |
| Tmem151a | 0.012493317 | 0.444413813 | 0.8431541 | 0.066098275 | 0.054594378 |
| Tmem159 | -0.018419559 | 0.448674178 | 0.8463296 | 0.131680584 | 0.142104107 |
| Tmem160 | 0.030911707 | 0.40986356 | 0.8280746 | 0.614675008 | 0.6142528 |
| Tmem161a | -0.007618999 | 0.774556682 | 0.9558538 | 0.164869642 | 0.176689058 |
| Tmem161b | -0.008207162 | 0.609356444 | 0.9088672 | 0.060947286 | 0.05270211 |
| Tmem163 | 0.037500428 | 0.261258544 | 0.7275735 | 0.357145854 | 0.356586889 |
| Tmem164 | 0.014598719 | 0.304763719 | 0.7642204 | 0.046482179 | 0.040031317 |
| Tmem165 | 0.022037309 | 0.477749142 | 0.8607489 | 0.286582667 | 0.256010782 |
| Tmem167 | 0.044275243 | 0.21151624 | 0.6822059 | 1.023230142 | 0.970733089 |
| Tmem167b | 0.032818586 | 0.275925556 | 0.7359993 | 0.267654044 | 0.230109756 |
| Tmem168 | -0.031800769 | 0.160369593 | 0.6370337 | 0.101086559 | 0.116126672 |
| Tmem17 | -0.016193803 | 0.288426717 | 0.7482199 | 0.046154651 | 0.04898735 |
| Tmem170b | -0.02261158 | 0.213820514 | 0.6831238 | 0.056713386 | 0.074069512 |
| Tmem175 | 0.015150553 | 0.46294568 | 0.8524425 | 0.107213923 | 0.089993524 |
| Tmem176a | -0.094605491 | 0.004356498 | 0.1311882 | 1.271742401 | 1.329181287 |
| Tmem176b | -0.006960799 | 0.830066058 | 0.9722601 | 1.305293927 | 1.269292115 |
| Tmem178b | -0.016820659 | 0.516049938 | 0.8735705 | 0.140494966 | 0.154470943 |
| Tmem179 | -0.032262005 | 0.093700107 | 0.5333348 | 0.04826612 | 0.083588116 |
| Tmem179b | 0.043056016 | 0.199495674 | 0.6730962 | 0.422153994 | 0.364785983 |
| Tmem18 | 0.022868077 | 0.148445293 | 0.6258297 | 0.067480472 | 0.046049156 |
| Tmem181a | 0.015019089 | 0.498175658 | 0.8684629 | 0.126906416 | 0.104068924 |
| Tmem181b-ps | -0.013609959 | 0.516613276 | 0.8736890 | 0.092131605 | 0.101299269 |
| Tmem183a | 0.049513821 | 0.12292437 | 0.5833237 | 0.375341958 | 0.287537101 |
| Tmem184a | 0.031310487 | 0.277839301 | 0.7382158 | 0.232922545 | 0.232369095 |
| Tmem184b | 0.021858391 | 0.453342542 | 0.8490009 | 0.253532099 | 0.213027455 |
| Tmem184c | -0.01820707 | 0.517155362 | 0.8738172 | 0.191100872 | 0.200334059 |

| | | | | | |
|---|---|---|---|---|---|
| Tmem185a | 0.006263667 | 0.799370599 | 0.96470534 | 0.144442061 | 0.134828191 |
| Tmem185b | 0.005840584 | 0.736149194 | 0.94691390 | 0.080081988 | 0.061560034 |
| Tmem186 | -0.000489093 | 0.976619041 | 0.99760728 | 0.061398969 | 0.056122018 |
| Tmem189 | -0.007297386 | 0.678289522 | 0.92818862 | 0.058196126 | 0.069502672 |
| Tmem19 | -0.018503813 | 0.418681754 | 0.83009907 | 0.103786438 | 0.125236784 |
| Tmem191c | 0.00044929 | 0.978196559 | 0.99760728 | 0.052208279 | 0.058352551 |
| Tmem192 | -0.039520823 | 0.149460815 | 0.62643745 | 0.154435189 | 0.198092492 |
| Tmem198 | 0.029469575 | 0.206637073 | 0.67975655 | 0.143661859 | 0.12070134 |
| Tmem199 | 0.063933411 | 0.041583806 | 0.39212105 | 0.350122445 | 0.278303738 |
| Tmem201 | -0.034964603 | 0.025898056 | 0.30880844 | 0.032884453 | 0.056262621 |
| Tmem203 | 0.015505947 | 0.588189299 | 0.90446241 | 0.234632518 | 0.207561229 |
| Tmem205 | -0.013921047 | 0.679775266 | 0.92892004 | 0.410352592 | 0.413227667 |
| Tmem208 | -0.036057395 | 0.31896517 | 0.77100438 | 0.616400359 | 0.624529265 |
| Tmem209 | -0.014158215 | 0.413010264 | 0.83005132 | 0.059587535 | 0.066057762 |
| Tmem212 | -0.014757749 | 0.57779807 | 0.89979143 | 0.141530515 | 0.169931696 |
| Tmem214 | -0.042959728 | 0.166106524 | 0.64292850 | 0.268141702 | 0.281241428 |
| Tmem215 | 0.094592698 | 0.004403084 | 0.13118829 | 0.286445513 | 0.276520906 |
| Tmem216 | -0.00020172 | 0.993275605 | 0.99858888 | 0.142787465 | 0.125063806 |
| Tmem218 | 0.018631478 | 0.575347406 | 0.89866891 | 0.349229783 | 0.322149288 |
| Tmem219 | 0.019135596 | 0.571352647 | 0.89661650 | 0.493497662 | 0.4675095 |
| Tmem222 | -0.095333451 | 0.002981165 | 0.10197640 | 0.252371757 | 0.325459018 |
| Tmem223 | -0.003686745 | 0.917470386 | 0.98481209 | 0.530330617 | 0.501974638 |
| Tmem229b | 0.045636796 | 0.013803277 | 0.23090811 | 0.097489422 | 0.064893385 |
| Tmem230 | -0.046025006 | 0.141284685 | 0.61390454 | 0.279142185 | 0.286381641 |
| Tmem231 | 0.002694697 | 0.867027531 | 0.97720021 | 0.056362644 | 0.057066812 |
| Tmem234 | 0.01236698 | 0.731262387 | 0.94590209 | 0.960814529 | 0.916214677 |
| Tmem237 | 0.005685982 | 0.817596706 | 0.96958270 | 0.150720938 | 0.133911616 |
| Tmem238 | 0.034904856 | 0.327621837 | 0.77788578 | 0.506419641 | 0.523992966 |
| Tmem240 | -0.006073015 | 0.711164957 | 0.94198349 | 0.065489481 | 0.056436678 |
| Tmem242 | 0.05031756 | 0.078038089 | 0.5048164 | 0.286985242 | 0.201367981 |
| Tmem243 | 0.089901145 | 0.003080922 | 0.10328859 | 0.330842444 | 0.237119432 |
| Tmem245 | 0.011210947 | 0.727847596 | 0.94580141 | 0.308888126 | 0.287489753 |
| Tmem248 | 0.08554575 | 0.015281764 | 0.24255210 | 0.619590388 | 0.555992045 |
| Tmem250-ps | 0.059414861 | 0.054524952 | 0.44218514 | 0.315182292 | 0.239186666 |
| Tmem251 | -0.004354574 | 0.886195731 | 0.98092631 | 0.26273663 | 0.252321798 |
| Tmem254a | 0.095162096 | 1.10E-05 | 0.00162295 | 0.175912169 | 0.086144672 |
| Tmem256 | 0.112699637 | 0.000925862 | 0.04619851 | 1.22594177 | 1.138842415 |
| Tmem258 | 0.033474203 | 0.289746619 | 0.74968641 | 1.491015769 | 1.393007774 |
| Tmem259 | -0.044406395 | 0.132580035 | 0.59882510 | 0.212694331 | 0.248620948 |
| Tmem260 | -0.000585491 | 0.979869748 | 0.99760728 | 0.118242148 | 0.117813689 |
| Tmem263 | 0.040069739 | 0.261757843 | 0.72757354 | 0.52872141 | 0.437753268 |
| Tmem267 | -0.033631643 | 0.042117886 | 0.39453201 | 0.036165104 | 0.063630565 |
| Tmem268 | -0.02773847 | 0.056747167 | 0.44732480 | 0.026348805 | 0.052799258 |
| Tmem28 | -0.01060749 | 0.568202427 | 0.89553080 | 0.072252146 | 0.073748918 |
| Tmem29 | 0.022842301 | 0.254099246 | 0.72230717 | 0.108389763 | 0.082255553 |
| Tmem30a | 0.005671102 | 0.874679996 | 0.97843318 | 0.875912066 | 0.844104734 |
| Tmem30b | 0.030358747 | 0.154306118 | 0.63049349 | 0.116912231 | 0.09769801 |
| Tmem33 | -0.041570319 | 0.216889919 | 0.68723302 | 0.392220296 | 0.426882531 |
| Tmem35b | -0.001312901 | 0.931571712 | 0.98939560 | 0.049659025 | 0.046601145 |
| Tmem37 | -0.045632074 | 0.019317173 | 0.26955860 | 0.069497678 | 0.090459387 |
| Tmem38a | 0.006357331 | 0.808466782 | 0.96583920 | 0.18335124 | 0.160340874 |
| Tmem38b | 0.018534215 | 0.315934779 | 0.77059875 | 0.087859711 | 0.067611061 |
| Tmem39a | 0.034116892 | 0.274759574 | 0.73519553 | 0.314453551 | 0.260958894 |
| Tmem39b | -0.020361173 | 0.163271524 | 0.64043239 | 0.03346696 | 0.050041878 |
| Tmem41b | 0.006188121 | 0.845075823 | 0.97348890 | 0.24796511 | 0.254915284 |
| Tmem42 | 0.014766946 | 0.605420366 | 0.90871392 | 0.223613289 | 0.210583206 |
| Tmem43 | -0.029200746 | 0.148073356 | 0.62556790 | 0.076179549 | 0.099086122 |
| Tmem50a | 0.059619295 | 0.078679207 | 0.50571949 | 0.820087635 | 0.757991632 |
| Tmem50b | 0.045763217 | 0.113199784 | 0.56962039 | 0.237743542 | 0.194907178 |
| Tmem51 | 0.059466948 | 0.031941271 | 0.33852287 | 0.232410772 | 0.175802558 |
| Tmem53 | -0.007106278 | 0.654511901 | 0.92056689 | 0.05408675 | 0.049857298 |
| Tmem59 | -0.003972714 | 0.896338832 | 0.98254414 | 1.523061467 | 1.492706582 |
| Tmem59l | -0.052694524 | 0.195701142 | 0.67268029 | 0.804473299 | 0.765529674 |
| Tmem60 | -0.019748227 | 0.533490281 | 0.87904420 | 0.29172409 | 0.287126711 |
| Tmem62 | -0.018350812 | 0.324663665 | 0.77575525 | 0.06389374 | 0.079987508 |
| Tmem63a | -0.007251011 | 0.762896381 | 0.95320007 | 0.132048513 | 0.134331233 |

| | | | | | |
|---|---|---|---|---|---|
| Tmem63b | -0.025241082 | 0.398137312 | 0.82384691 | 0.212519202 | 0.247218709 |
| Tmem64 | -0.015886888 | 0.518235197 | 0.87403698 | 0.132093892 | 0.135587591 |
| Tmem65 | 0.093125007 | 0.013779471 | 0.23089924 | 0.806026784 | 0.768167676 |
| Tmem67 | 0.010097945 | 0.537780719 | 0.88163354 | 0.061918152 | 0.055663861 |
| Tmem68 | -0.006573514 | 0.732293892 | 0.94620411 | 0.079266841 | 0.081403952 |
| Tmem70 | 0.019367164 | 0.514052301 | 0.87333427 | 0.278674591 | 0.254585682 |
| Tmem71 | 0.015118777 | 0.379826298 | 0.81151774 | 0.065926378 | 0.058579769 |
| Tmem80 | -0.062129813 | 0.022808581 | 0.29268948 | 0.172415292 | 0.204710558 |
| Tmem86a | -0.008320206 | 0.690664015 | 0.93462378 | 0.106116238 | 0.093454118 |
| Tmem87a | 0.038026962 | 0.058499386 | 0.45175489 | 0.108749276 | 0.079263278 |
| Tmem87b | 0.006150243 | 0.833583431 | 0.97234941 | 0.237085018 | 0.216370017 |
| Tmem8b | 0.002998182 | 0.869232168 | 0.97763980 | 0.08199351 | 0.065588109 |
| Tmem9 | 0.03130848 | 0.262840592 | 0.72757354 | 0.239209001 | 0.195954945 |
| Tmem94 | -0.025715369 | 0.159008339 | 0.63552084 | 0.053376506 | 0.079501736 |
| Tmem97 | -0.047505517 | 0.07547239 | 0.49662859 | 0.137991447 | 0.182383124 |
| Tmem98 | 0.017231068 | 0.244055559 | 0.71327199 | 0.053763199 | 0.043672901 |
| Tmem9b | -0.017915001 | 0.516466516 | 0.87368901 | 0.182669152 | 0.1945827 |
| Tmf1 | 0.001027369 | 0.976951548 | 0.99760728 | 0.483642234 | 0.478520461 |
| Tmigd3 | -0.019915942 | 0.313052865 | 0.76975571 | 0.059256869 | 0.086263558 |
| Tmod1 | 0.022908086 | 0.343598303 | 0.78729551 | 0.137420133 | 0.133509892 |
| Tmod2 | 0.029503466 | 0.422452714 | 0.83148971 | 0.613727438 | 0.599370336 |
| Tmod3 | -0.027255292 | 0.306680523 | 0.76527857 | 0.157769636 | 0.171747467 |
| Tmpo | 0.003224231 | 0.92713624 | 0.98792100 | 0.465118776 | 0.434551344 |
| Tmprss4 | 0.018271177 | 0.260717968 | 0.72710774 | 0.055506874 | 0.057898786 |
| Tmsb10 | 0.152472298 | 0.000135862 | 0.01199599 | 1.401733965 | 1.10955065 |
| Tmsb15b1 | -0.020053078 | 0.604144777 | 0.90871391 | 0.600363503 | 0.478456128 |
| Tmsb15b2 | 0.107026412 | 0.031740338 | 0.33852287 | 1.931095001 | 1.519896395 |
| Tmsb15l | -0.056485113 | 0.076998093 | 0.50152401 | 0.301718475 | 0.25993277 |
| Tmsb4x | -0.015080055 | 0.58850999 | 0.90446241 | 2.953541434 | 2.72628749 |
| Tmtc1 | -0.008847411 | 0.732513769 | 0.94620411 | 0.137015039 | 0.155145185 |
| Tmtc2 | -0.02580044 | 0.197008068 | 0.67309621 | 0.058944016 | 0.095899138 |
| Tmtc3 | 0.025345073 | 0.28076133 | 0.74086184 | 0.138215981 | 0.129041894 |
| Tmtc4 | 0.026746791 | 0.121149882 | 0.58114450 | 0.078687617 | 0.054285075 |
| Tmub1 | -0.025602137 | 0.345166479 | 0.78804400 | 0.16504891 | 0.188605431 |
| Tmub2 | 0.040201221 | 0.130228748 | 0.59397110 | 0.198197017 | 0.159968247 |
| Tmx1 | -0.083981798 | 0.012437065 | 0.21952968 | 0.38235456 | 0.459063681 |
| Tmx2 | 0.029206249 | 0.331326195 | 0.77848314 | 0.289271975 | 0.232638963 |
| Tmx3 | 0.056592745 | 0.075228215 | 0.49662859 | 0.316873401 | 0.278416023 |
| Tmx4 | -0.013000773 | 0.740896197 | 0.94790800 | 0.805587273 | 0.774897805 |
| Tnfaip1 | 0.012702462 | 0.670604821 | 0.92458649 | 0.265194903 | 0.22471168 |
| Tnfaip8 | -0.010005947 | 0.643597525 | 0.91678451 | 0.094347325 | 0.097969013 |
| Tnfrsf12a | 0.027704284 | 0.508889191 | 0.87040677 | 0.757336061 | 0.623464114 |
| Tnfrsf1a | -0.044731823 | 0.137833992 | 0.60742578 | 0.188532747 | 0.245802998 |
| Tnfrsf21 | -0.019415645 | 0.329079998 | 0.77788578 | 0.059537521 | 0.094834869 |
| Tnfrsf23 | -0.018369543 | 0.404768762 | 0.82707700 | 0.086431089 | 0.105134136 |
| Tnfsf12 | -0.004086355 | 0.878632244 | 0.97855831 | 0.182467901 | 0.168425727 |
| Tnik | -0.037638312 | 0.091352261 | 0.53212820 | 0.082838779 | 0.124382438 |
| Tnip1 | 0.027186239 | 0.197470312 | 0.67309621 | 0.111587111 | 0.084384543 |
| Tnip2 | -0.022864586 | 0.243595057 | 0.71327199 | 0.068912972 | 0.089331728 |
| Tnk1 | 0.001505403 | 0.941458303 | 0.99293011 | 0.094867873 | 0.090385352 |
| Tnk2 | 0.042613615 | 0.084227872 | 0.51864710 | 0.173202552 | 0.129984655 |
| Tnks | 0.002380908 | 0.940401209 | 0.99284589 | 0.316887883 | 0.319787431 |
| Tnks1bp1 | -0.05193376 | 0.012841615 | 0.22348910 | 0.06821233 | 0.10780328 |
| Tnks2 | 0.002834931 | 0.939690186 | 0.99282169 | 0.650931601 | 0.634336599 |
| Tnpo1 | -0.004631292 | 0.890347714 | 0.98195979 | 0.385051319 | 0.40420135 |
| Tnpo2 | -0.007461186 | 0.761576106 | 0.95286310 | 0.145605463 | 0.143140146 |
| Tnpo3 | -0.056211978 | 0.081759919 | 0.51430461 | 0.296001367 | 0.326136039 |
| Tnr | -0.06549341 | 0.055626919 | 0.44493321 | 0.293119672 | 0.32552147 |
| Tnrc18 | -0.022083292 | 0.48376569 | 0.86248860 | 0.253366083 | 0.278200216 |
| Tnrc6a | -0.001643179 | 0.965005617 | 0.99664211 | 0.634943047 | 0.633730448 |
| Tnrc6b | -0.04061228 | 0.302898569 | 0.76166129 | 0.671085965 | 0.68706194 |
| Tnrc6c | -0.030883835 | 0.420573947 | 0.83009907 | 0.635189789 | 0.674472936 |
| Tns2 | -0.013962565 | 0.399340259 | 0.82398500 | 0.056706472 | 0.058680826 |
| Tns3 | -0.005078419 | 0.869180772 | 0.97763980 | 0.248204655 | 0.232048543 |
| Tob1 | -0.088943534 | 0.022692552 | 0.29260131 | 0.655415665 | 0.717141235 |
| Tob2 | -0.014856602 | 0.677639698 | 0.92818861 | 0.354931898 | 0.364109662 |

| | | | | | |
|---|---|---|---|---|---|
| Toe1 | 0.01023644 | 0.610605038 | 0.90908029 | 0.094018142 | 0.086376785 |
| Togaram1 | -0.030305446 | 0.306638774 | 0.76527857 | 0.199009926 | 0.229623561 |
| Tollip | 0.018244046 | 0.476240647 | 0.86072310 | 0.161604884 | 0.148102416 |
| Tom1 | 0.01669952 | 0.370223982 | 0.80655813 | 0.085669091 | 0.07375531 |
| Tom1l1 | -0.006965785 | 0.7181743 | 0.94382369 | 0.08083307 | 0.07994654 |
| Tom1l2 | 0.020993559 | 0.405634312 | 0.82707700 | 0.174031172 | 0.144982042 |
| Tomm20 | 0.073346166 | 0.015792394 | 0.24670952 | 1.329588328 | 1.211304584 |
| Tomm22 | 0.016954981 | 0.639399749 | 0.91600428 | 0.735131651 | 0.647667971 |
| Tomm34 | -0.036296986 | 0.201857051 | 0.67521501 | 0.200654967 | 0.216852803 |
| Tomm40 | 0.043552787 | 0.197153996 | 0.67309622 | 0.478206883 | 0.403863689 |
| Tomm40l | -0.012749331 | 0.569666096 | 0.89614457 | 0.116138338 | 0.120727181 |
| Tomm5 | -0.027077561 | 0.437132021 | 0.83940179 | 0.597957506 | 0.568724266 |
| Tomm6 | 0.067648339 | 0.005794284 | 0.15327813 | 0.195970832 | 0.120892945 |
| Tomm7 | 0.118210608 | 6.38E-07 | 0.00016650 | 1.947211111 | 1.715859032 |
| Tomm70a | -0.003479714 | 0.916767586 | 0.98454547 | 0.364464255 | 0.352808437 |
| Top1 | -0.034656054 | 0.309951225 | 0.76770334 | 1.225674833 | 1.228598787 |
| Top2b | -0.023928553 | 0.535399415 | 0.87959492 | 0.592045573 | 0.559821585 |
| Top3b | -0.049762922 | 0.049961426 | 0.42108525 | 0.12625102 | 0.157141879 |
| Topbp1 | -0.011262882 | 0.557066764 | 0.89159443 | 0.076422985 | 0.080029307 |
| Topors | 0.008073618 | 0.815877266 | 0.96869997 | 0.396281502 | 0.380435216 |
| Tor1a | 0.003136218 | 0.92313266 | 0.98722250 | 0.327469122 | 0.291211823 |
| Tor1aip1 | -0.039156356 | 0.214721075 | 0.68423805 | 0.235327092 | 0.28087184 |
| Tor1aip2 | -0.063558298 | 0.062635311 | 0.46512543 | 0.330919918 | 0.395050799 |
| Tor1b | -0.062242425 | 0.083206275 | 0.51782073 | 0.507298799 | 0.578471666 |
| Tor2a | 0.016431474 | 0.613214922 | 0.91083865 | 0.358206825 | 0.32549348 |
| Tor3a | 0.026615692 | 0.353934765 | 0.79677824 | 0.205145457 | 0.210920331 |
| Tox | 0.069174145 | 0.024809833 | 0.30415400 | 0.294598808 | 0.229452124 |
| Tox3 | -0.062673538 | 0.107803179 | 0.56343910 | 0.754987925 | 0.797875613 |
| Tox4 | 0.02355456 | 0.51844731 | 0.87406480 | 0.504825009 | 0.472958486 |
| Tpcn1 | 0.012437287 | 0.550624151 | 0.88830567 | 0.107682171 | 0.090676184 |
| Tpd52 | -0.005938323 | 0.862567047 | 0.97696750 | 0.90156054 | 0.894498801 |
| Tpd52l1 | 0.006275933 | 0.771276747 | 0.95523334 | 0.096517817 | 0.107505895 |
| Tpd52l2 | -0.009870143 | 0.764354757 | 0.95374505 | 0.374605006 | 0.368466129 |
| Tpgs1 | -0.025939422 | 0.408749049 | 0.82764474 | 0.273719449 | 0.300814431 |
| Tpgs2 | -0.008799482 | 0.619042868 | 0.91206078 | 0.059736502 | 0.066928365 |
| Tpi1 | -0.016846337 | 0.657296668 | 0.92117802 | 0.818866797 | 0.772914789 |
| Tpm1 | -0.024627233 | 0.520598059 | 0.87417954 | 0.55016695 | 0.562926465 |
| Tpm3 | -0.014092387 | 0.68127133 | 0.92998645 | 0.48316885 | 0.447674976 |
| Tpm4 | 0.017168861 | 0.64163452 | 0.91667127 | 0.503767533 | 0.494168929 |
| Tpp1 | 0.027820188 | 0.373467233 | 0.80856898 | 0.310885846 | 0.283568971 |
| Tpp2 | 0.01202351 | 0.72436306 | 0.94456522 | 0.377552623 | 0.330348053 |
| Tpr | -0.004995657 | 0.886420254 | 0.98100890 | 1.173206291 | 1.16872816 |
| Tpra1 | 0.033338283 | 0.056298325 | 0.44678350 | 0.082160102 | 0.059234052 |
| Tprgl | -0.115704291 | 0.00105524 | 0.05008603 | 0.498297076 | 0.559660653 |
| Tprkb | 0.027242529 | 0.330287795 | 0.77788578 | 0.229090688 | 0.19009823 |
| Tpst1 | -0.02734965 | 0.317737604 | 0.77094201 | 0.195802003 | 0.187407461 |
| Tpst2 | 0.031014229 | 0.377161086 | 0.81017979 | 0.479763792 | 0.420247461 |
| Tpt1 | -0.040142937 | 0.000150516 | 0.01287168 | 4.562522901 | 4.525610843 |
| Tpt1-ps3 | -0.059195206 | 0.082692695 | 0.51656897 | 0.34148214 | 0.375322838 |
| Tra2a | -0.074946875 | 0.060580999 | 0.45945222 | 0.610785119 | 0.693463339 |
| Tra2b | -0.002538422 | 0.947069677 | 0.99450182 | 0.840429832 | 0.862791284 |
| Trabd | -0.033129288 | 0.369113617 | 0.80567193 | 0.542955201 | 0.551866118 |
| Tradd | 0.001234705 | 0.952421129 | 0.99515668 | 0.1095775 | 0.086564108 |
| Traf2 | -0.010363118 | 0.558022781 | 0.89196227 | 0.058291851 | 0.066510905 |
| Traf3 | -0.004950864 | 0.836581457 | 0.97266190 | 0.127615539 | 0.130969595 |
| Traf3ip1 | -0.007112229 | 0.691967223 | 0.93551490 | 0.059474363 | 0.070518039 |
| Traf3ip2 | 0.030403185 | 0.116397792 | 0.57246708 | 0.092663696 | 0.07602831 |
| Traf4 | 0.036011484 | 0.061827514 | 0.46149650 | 0.112846005 | 0.069467086 |
| Traf6 | -0.018960523 | 0.343802425 | 0.78739211 | 0.066768892 | 0.094382843 |
| Traf7 | 0.0267586 | 0.278683248 | 0.73937891 | 0.167044656 | 0.136554536 |
| Trafd1 | 0.073982633 | 0.012201901 | 0.21841537 | 0.254999576 | 0.198391491 |
| Trak1 | -0.024809522 | 0.399963605 | 0.82445164 | 0.19949043 | 0.233970126 |
| Trak2 | -0.023746986 | 0.259538161 | 0.72634755 | 0.085402403 | 0.108050588 |
| Tram1 | -0.088421406 | 0.008348573 | 0.17930979 | 1.011778083 | 1.120769823 |
| Trap1 | -0.035727723 | 0.125119735 | 0.58705221 | 0.102246497 | 0.13675983 |
| Trappc1 | 0.028845612 | 0.418280252 | 0.83009907 | 0.59103162 | 0.535580854 |

| | | | | | |
|---|---|---|---|---|---|
| Trappc10 | -0.038084645 | 0.228025239 | 0.7015153 | 0.261696798 | 0.296264031 |
| Trappc11 | -0.015129851 | 0.627760822 | 0.9128095 | 0.256261129 | 0.271074329 |
| Trappc12 | 0.004144329 | 0.866076668 | 0.9770148 | 0.146786511 | 0.146224801 |
| Trappc13 | 0.0318117 | 0.286857714 | 0.7468841 | 0.283038256 | 0.236823064 |
| Trappc2 | 6.18E-05 | 0.998198349 | 0.9994073 | 0.207583708 | 0.195149448 |
| Trappc2l | 0.01620764 | 0.60739456 | 0.9087139 | 1.160375896 | 1.092804592 |
| Trappc3 | -0.016782901 | 0.623727402 | 0.9121505 | 0.477241207 | 0.435669044 |
| Trappc4 | -0.042038686 | 0.233316888 | 0.7051391 | 0.641895066 | 0.617103038 |
| Trappc5 | -0.010205781 | 0.741847898 | 0.9479080 | 0.278637243 | 0.289347099 |
| Trappc6a | -0.049149654 | 0.166214408 | 0.6429285 | 0.510405626 | 0.5411649 |
| Trappc6b | 0.03617891 | 0.296970206 | 0.7545964 | 0.776116594 | 0.709762228 |
| Trappc8 | -0.017830149 | 0.47329369 | 0.8597092 | 0.13575 | 0.147301184 |
| Trappc9 | -0.014414469 | 0.585234322 | 0.9024598 | 0.167877409 | 0.175661699 |
| Trerf1 | -0.020478815 | 0.428396044 | 0.8340365 | 0.154682214 | 0.155797472 |
| Triap1 | -0.000802687 | 0.980173626 | 0.9976072 | 0.333777507 | 0.323039845 |
| Trib1 | 0.007889205 | 0.833361254 | 0.9723494 | 0.468012343 | 0.453244214 |
| Trib3 | 0.026988025 | 0.422846151 | 0.8315789 | 0.273716335 | 0.311419506 |
| Trim11 | -0.009815011 | 0.608931365 | 0.9088672 | 0.074598079 | 0.081179084 |
| Trim12a | 0.012474463 | 0.431345836 | 0.8356213 | 0.070797412 | 0.044286021 |
| Trim12c | 0.002841076 | 0.87028255 | 0.9780178 | 0.06890716 | 0.061790118 |
| Trim2 | -0.01651789 | 0.565665066 | 0.8943891 | 0.203750149 | 0.202076097 |
| Trim23 | 0.003082346 | 0.86706649 | 0.9772002 | 0.081247472 | 0.070359644 |
| Trim24 | -0.051619534 | 0.122514227 | 0.5833237 | 0.266190763 | 0.301914117 |
| Trim25 | 0.006930469 | 0.789059062 | 0.9617232 | 0.162958837 | 0.151234285 |
| Trim26 | 0.009672524 | 0.736028718 | 0.9468817 | 0.224274098 | 0.201799577 |
| Trim27 | -0.009579606 | 0.773097565 | 0.9556977 | 0.340037074 | 0.322754058 |
| Trim28 | -0.014256508 | 0.652668156 | 0.9191465 | 0.280600542 | 0.278812809 |
| Trim3 | 0.001611495 | 0.926416897 | 0.9879210 | 0.06601921 | 0.064388041 |
| Trim32 | -0.014125544 | 0.391371207 | 0.8200150 | 0.053720574 | 0.059040627 |
| Trim33 | 0.028483658 | 0.3551493 | 0.7969131 | 0.283283127 | 0.255863946 |
| Trim35 | -0.000391379 | 0.991869185 | 0.9985765 | 0.949655029 | 0.867938849 |
| Trim37 | 0.030697684 | 0.296882575 | 0.7545964 | 0.237942408 | 0.232138858 |
| Trim39 | 0.004378957 | 0.865557362 | 0.9770148 | 0.154551408 | 0.150793428 |
| Trim41 | 0.017676704 | 0.524323533 | 0.8758385 | 0.218277249 | 0.199059633 |
| Trim44 | -0.047310103 | 0.212090303 | 0.6822059 | 0.510023947 | 0.571147724 |
| Trim46 | -0.028605223 | 0.177252096 | 0.6536659 | 0.08700188 | 0.105307997 |
| Trim56 | -0.0343619 | 0.291596901 | 0.7519213 | 0.268533972 | 0.304865198 |
| Trim62 | -0.024193201 | 0.120015796 | 0.5797304 | 0.039368548 | 0.055797561 |
| Trim8 | -0.031353248 | 0.250730138 | 0.7207210 | 0.175981906 | 0.192295496 |
| Trim9 | -0.041126793 | 0.035183029 | 0.3577036 | 0.062510466 | 0.090464743 |
| Trio | -0.03403458 | 0.248924425 | 0.7188734 | 0.202537291 | 0.244386839 |
| Triobp | 0.004350888 | 0.86961559 | 0.9777384 | 0.185920079 | 0.163202129 |
| Trip10 | -0.010809219 | 0.464580844 | 0.8536102 | 0.037029017 | 0.04895164 |
| Trip11 | -0.007839147 | 0.8412105 | 0.9734589 | 0.773909516 | 0.804634773 |
| Trip12 | -0.063165786 | 0.071833496 | 0.4888240 | 0.45218099 | 0.509246685 |
| Trip4 | -0.01282592 | 0.62493846 | 0.9121764 | 0.169607669 | 0.175078482 |
| Triqk | 0.031569575 | 0.289747759 | 0.7496864 | 0.298321362 | 0.230487001 |
| Trir | 0.013783606 | 0.681261494 | 0.9299864 | 1.020420653 | 0.931599578 |
| Trit1 | -0.013917756 | 0.483145739 | 0.8620306 | 0.087192324 | 0.087715714 |
| Trmt1 | -0.001925657 | 0.941682106 | 0.9929301 | 0.173184577 | 0.161487375 |
| Trmt10a | -0.015575785 | 0.506254719 | 0.8695566 | 0.112638914 | 0.126287848 |
| Trmt10b | 0.025215436 | 0.258835916 | 0.7257355 | 0.132207232 | 0.109694052 |
| Trmt10c | -0.026188469 | 0.355638575 | 0.7969131 | 0.218725195 | 0.211400003 |
| Trmt11 | -0.00316507 | 0.887691229 | 0.9812414 | 0.116388768 | 0.113203483 |
| Trmt112 | 0.004710644 | 0.895928369 | 0.9825441 | 0.822875261 | 0.785275203 |
| Trmt12 | -0.001258771 | 0.94705623 | 0.9945018 | 0.080601041 | 0.077328569 |
| Trmt13 | 0.010447239 | 0.656830056 | 0.9211780 | 0.139394103 | 0.118220347 |
| Trmt1l | 0.024264427 | 0.314233586 | 0.7705987 | 0.149312143 | 0.128911874 |
| Trmt2a | 0.01930314 | 0.402263994 | 0.8254592 | 0.140543607 | 0.113252505 |
| Trmt2b | -0.023956974 | 0.22611424 | 0.6989882 | 0.06746342 | 0.085860401 |
| Trmt5 | 0.008050368 | 0.595579223 | 0.9068020 | 0.050758206 | 0.041580475 |
| Trmt6 | 0.040236616 | 0.248424213 | 0.7185162 | 0.428164822 | 0.407223246 |
| Trmt61a | 0.033008702 | 0.061034797 | 0.4605600 | 0.084117994 | 0.058311487 |
| Trmt61b | -0.005559186 | 0.834588996 | 0.9723494 | 0.155437503 | 0.165251198 |
| Trmt9b | 0.017100242 | 0.417516258 | 0.8300990 | 0.123138146 | 0.091182259 |
| Trmu | -0.00959213 | 0.589122325 | 0.9046539 | 0.065641543 | 0.069798145 |

| | | | | | |
|---|---|---|---|---|---|
| Trnau1ap | 0.011148293 | 0.720605479 | 0.9445295 | 0.298760804 | 0.261138242 |
| Trnt1 | 0.039211169 | 0.179806058 | 0.6550070 | 0.268894742 | 0.213441011 |
| Trp53 | 0.023755035 | 0.462226914 | 0.8524425 | 0.313288228 | 0.275354345 |
| Trp53bp1 | -5.65E-05 | 0.998524993 | 0.9996334 | 0.245447721 | 0.241588212 |
| Trp53bp2 | -0.014340922 | 0.53198223 | 0.8780804 | 0.108477337 | 0.121421594 |
| Trp53cor1 | -0.022276238 | 0.416859834 | 0.8300990 | 0.145115748 | 0.182482619 |
| Trp53i11 | -0.007455249 | 0.75416267 | 0.9519043 | 0.123434903 | 0.129569224 |
| Trp53i13 | -0.003404179 | 0.922384101 | 0.9867411 | 0.445188652 | 0.389155883 |
| Trp53inp1 | -0.063549987 | 0.081924286 | 0.5146011 | 0.299693109 | 0.380552136 |
| Trp53inp2 | -0.023010504 | 0.504543516 | 0.8695566 | 0.493699725 | 0.546527962 |
| Trp53rka | -0.027074461 | 0.233294905 | 0.7051391 | 0.117818822 | 0.124667542 |
| Trp53rkb | 0.002028991 | 0.937477522 | 0.9917646 | 0.164339928 | 0.153621635 |
| Trpc1 | -0.042695192 | 0.178185729 | 0.6538489 | 0.242932352 | 0.30426484 |
| Trpc4ap | -0.046711418 | 0.166565161 | 0.6429285 | 0.366869947 | 0.376346247 |
| Trpm3 | 0.02982804 | 0.075999177 | 0.4986189 | 0.063216315 | 0.056704734 |
| Trpm4 | -0.016112354 | 0.500682068 | 0.8693704 | 0.109345504 | 0.136056469 |
| Trpm5 | 0.076259424 | 0.000535075 | 0.0322735 | 0.127172766 | 0.087255354 |
| Trpm7 | 0.043575374 | 0.12839983 | 0.5914729 | 0.228013604 | 0.18227122 |
| Trps1 | 0.026418368 | 0.258204848 | 0.7251102 | 0.126775903 | 0.123307949 |
| Trpt1 | -0.009893063 | 0.533622159 | 0.8790442 | 0.051452657 | 0.054693759 |
| Trrap | -0.011739317 | 0.707648941 | 0.9408790 | 0.272122585 | 0.277743821 |
| Trub1 | -0.016727964 | 0.504596303 | 0.8695566 | 0.14085853 | 0.153878316 |
| Trub2 | -0.027365478 | 0.167964981 | 0.6450687 | 0.076622219 | 0.092238494 |
| Tsc1 | -0.015267909 | 0.550438535 | 0.8883056 | 0.142691913 | 0.1583032 |
| Tsc2 | -0.018784187 | 0.427165556 | 0.8339858 | 0.127839676 | 0.139550578 |
| Tsc22d1 | -0.026632377 | 0.460219779 | 0.8524425 | 1.39998228 | 1.448155593 |
| Tsc22d2 | 0.000587073 | 0.985468656 | 0.9976072 | 0.271638661 | 0.268085124 |
| Tsc22d3 | 0.004327349 | 0.863427152 | 0.9769675 | 0.142634306 | 0.140252184 |
| Tsc22d4 | -0.050334456 | 0.169382609 | 0.6480044 | 0.47806517 | 0.521130823 |
| Tsen15 | -0.009651697 | 0.76055866 | 0.9522598 | 0.316834483 | 0.290998744 |
| Tsen2 | -0.004181651 | 0.835063455 | 0.9723494 | 0.096724072 | 0.089740656 |
| Tsen34 | -0.031264332 | 0.3305444 | 0.7779360 | 0.371317191 | 0.349688496 |
| Tsen54 | 0.009381472 | 0.617588184 | 0.9115017 | 0.093005959 | 0.070502348 |
| Tsfm | 0.014302574 | 0.606063524 | 0.9087139 | 0.217804707 | 0.197818385 |
| Tsg101 | 0.020493293 | 0.55799341 | 0.8919622 | 0.529701453 | 0.448458427 |
| Tshz1 | 0.020947219 | 0.554027958 | 0.8902995 | 0.516585476 | 0.536460268 |
| Tshz2 | -0.008435073 | 0.830672463 | 0.9722601 | 0.769813857 | 0.808510545 |
| Tshz3 | -0.070092306 | 0.021261543 | 0.2831067 | 0.206400511 | 0.254623608 |
| Tsn | 0.009529762 | 0.784753805 | 0.9601202 | 0.841648343 | 0.787613705 |
| Tsnax | -0.015981973 | 0.626907434 | 0.9128095 | 0.420613923 | 0.389401305 |
| Tspan12 | -0.045323927 | 0.036988013 | 0.3640732 | 0.10807649 | 0.108520105 |
| Tspan13 | 0.075129107 | 0.030047992 | 0.3308280 | 1.121415164 | 1.031455222 |
| Tspan15 | -0.020508473 | 0.549384079 | 0.8874415 | 0.360179558 | 0.379836668 |
| Tspan17 | -0.009511521 | 0.725991609 | 0.9450680 | 0.184960529 | 0.177323357 |
| Tspan2 | -0.030264597 | 0.338461087 | 0.7834245 | 0.239647753 | 0.271514147 |
| Tspan3 | 0.045556142 | 0.204225413 | 0.6780598 | 1.002852711 | 0.878148113 |
| Tspan31 | 0.018903845 | 0.576523151 | 0.8994610 | 0.452059185 | 0.4036326 |
| Tspan33 | -0.019659459 | 0.482189867 | 0.8620306 | 0.184196838 | 0.201981314 |
| Tspan5 | 0.000349068 | 0.992134087 | 0.9985765 | 0.453981198 | 0.487270141 |
| Tspan7 | -0.050671142 | 0.163844252 | 0.6410026 | 1.061611099 | 1.033139638 |
| Tspan8 | -0.078677763 | 0.087309796 | 0.5265125 | 0.41486041 | 0.504370391 |
| Tspan9 | 0.031135748 | 0.034315457 | 0.3516625 | 0.062420747 | 0.033472788 |
| Tspo | -0.033592556 | 0.368787811 | 0.8056719 | 0.516830169 | 0.516160506 |
| Tspyl1 | 0.000952846 | 0.978389175 | 0.9976072 | 0.473154462 | 0.43622531 |
| Tspyl2 | -0.02128235 | 0.526751562 | 0.8768553 | 0.332112194 | 0.328532621 |
| Tspyl4 | 0.075367773 | 0.046183335 | 0.4073656 | 0.699210201 | 0.705134687 |
| Tspyl5 | -0.010919155 | 0.505029798 | 0.8695566 | 0.045388388 | 0.058247281 |
| Tsr1 | 0.033425559 | 0.244013272 | 0.7132719 | 0.227649178 | 0.193772273 |
| Tsr2 | 0.000412915 | 0.988018966 | 0.9976143 | 0.206728799 | 0.178900012 |
| Tsr3 | 0.026469089 | 0.372188856 | 0.8085689 | 0.293568121 | 0.242686631 |
| Tssc4 | 0.011750569 | 0.757845222 | 0.9521533 | 0.632424672 | 0.612793549 |
| Tstd2 | -0.024737866 | 0.199460262 | 0.6730962 | 0.058496495 | 0.08451428 |
| Tstd3 | 0.00359252 | 0.853166266 | 0.9755431 | 0.087232294 | 0.079109536 |
| Ttbk2 | 0.024897045 | 0.293378515 | 0.7533684 | 0.134000737 | 0.113919577 |
| Ttc1 | 0.038291212 | 0.238385769 | 0.7109741 | 0.365540083 | 0.335216033 |
| Ttc13 | -0.025604561 | 0.122030509 | 0.5819916 | 0.047965652 | 0.06506521 |

| | | | | | |
|---|---|---|---|---|---|
| Ttc14 | -0.019807236 | 0.616358598 | 0.9114903 | 0.607931689 | 0.645831683 |
| Ttc17 | 0.021019572 | 0.185454079 | 0.6637678 | 0.061822565 | 0.050637348 |
| Ttc19 | -0.019622937 | 0.555638839 | 0.8910341 | 0.334583907 | 0.351921096 |
| Ttc21b | 0.008049768 | 0.675046076 | 0.9265887 | 0.083658217 | 0.073743347 |
| Ttc22 | -0.014229815 | 0.405611283 | 0.8270770 | 0.059625901 | 0.063784984 |
| Ttc26 | -0.016075462 | 0.304394564 | 0.7634877 | 0.035083379 | 0.056690306 |
| Ttc27 | -0.013773054 | 0.332325145 | 0.7793535 | 0.033542592 | 0.046260699 |
| Ttc28 | 0.04696605 | 0.141994414 | 0.6152737 | 0.272239685 | 0.278526451 |
| Ttc3 | -0.043232065 | 0.266083027 | 0.7307399 | 0.883199038 | 0.88447476 |
| Ttc32 | -0.072098675 | 0.009950156 | 0.1987218 | 0.168737537 | 0.220159284 |
| Ttc33 | -0.051021524 | 0.118763785 | 0.5777984 | 0.306885791 | 0.332830603 |
| Ttc37 | 0.043704916 | 0.156354456 | 0.6315293 | 0.270877877 | 0.227242896 |
| Ttc39a | -0.038524603 | 0.200247235 | 0.6738305 | 0.228860649 | 0.250235989 |
| Ttc39b | -0.017833131 | 0.609746049 | 0.9090292 | 0.466151177 | 0.466555805 |
| Ttc4 | 0.017140115 | 0.534553976 | 0.8795949 | 0.205096682 | 0.189217483 |
| Ttc5 | 0.004448063 | 0.853874903 | 0.9755431 | 0.148833857 | 0.139797481 |
| Ttc7 | 0.00708309 | 0.68349607 | 0.9309982 | 0.06666748 | 0.061520928 |
| Ttc7b | 0.018692069 | 0.362113876 | 0.8030478 | 0.108269298 | 0.086817506 |
| Ttc8 | 0.013837455 | 0.60998512 | 0.9090802 | 0.187179864 | 0.180128466 |
| Ttc9 | -0.026400375 | 0.163810233 | 0.6410026 | 0.053150171 | 0.084573219 |
| Ttc9c | -0.029234972 | 0.255330949 | 0.7226955 | 0.152524319 | 0.1635658 |
| Ttf1 | -0.034569493 | 0.246744996 | 0.7165428 | 0.21606769 | 0.227739167 |
| Tti1 | -0.022780068 | 0.223315959 | 0.6955958 | 0.062283113 | 0.085844271 |
| Tti2 | 0.004782555 | 0.798547458 | 0.9644698 | 0.084853505 | 0.073102391 |
| Ttl | 0.022740665 | 0.155688935 | 0.6312943 | 0.064486489 | 0.046061915 |
| Ttll1 | -0.00471654 | 0.774388472 | 0.9558538 | 0.05753136 | 0.055073165 |
| Ttll12 | -0.007793081 | 0.605701772 | 0.9087139 | 0.041252913 | 0.049231816 |
| Ttll5 | -0.029421562 | 0.083668858 | 0.5183040 | 0.05016207 | 0.063866048 |
| Ttll7 | -0.030711181 | 0.404763573 | 0.8270770 | 0.597191981 | 0.668648143 |
| Ttpal | -0.022581204 | 0.168601766 | 0.6465131 | 0.040779333 | 0.063649667 |
| Ttr | -0.01458807 | 0.652115156 | 0.9190200 | 6.401501819 | 6.032157939 |
| Ttyh1 | -0.058326441 | 0.046114871 | 0.4073656 | 0.102696873 | 0.191278883 |
| Ttyh2 | -0.003124072 | 0.882896428 | 0.9799360 | 0.086949205 | 0.102596434 |
| Tuba1a | -0.049962957 | 0.193491747 | 0.6710847 | 0.967902771 | 0.877198352 |
| Tuba1b | 0.010231061 | 0.749464887 | 0.9504425 | 0.323020125 | 0.269800858 |
| Tuba1c | -0.016954207 | 0.4173143 | 0.8300990 | 0.076893054 | 0.09380939 |
| Tuba4a | -0.080045243 | 0.035872597 | 0.3594506 | 0.585955595 | 0.669535469 |
| Tubb2a | -0.011169738 | 0.760186956 | 0.9521533 | 0.486006166 | 0.433428737 |
| Tubb2b | 0.014655758 | 0.296271726 | 0.7545964 | 0.049975338 | 0.035392571 |
| Tubb3 | 0.016668423 | 0.40203972 | 0.8254259 | 0.067211294 | 0.083187271 |
| Tubb4a | -0.027618161 | 0.157699669 | 0.6324475 | 0.072379965 | 0.084405805 |
| Tubb4b | -0.077899596 | 0.053215194 | 0.4352904 | 0.805408403 | 0.864452909 |
| Tubb5 | 0.059008704 | 0.134227044 | 0.6006009 | 0.723459055 | 0.590950326 |
| Tubb6 | -0.0458369 | 0.138379431 | 0.6079839 | 0.220027708 | 0.232651994 |
| Tubg1 | -0.044600542 | 0.13344049 | 0.5988251 | 0.234841603 | 0.248005906 |
| Tubg2 | -0.014249378 | 0.393565685 | 0.8208939 | 0.059965904 | 0.062453595 |
| Tubgcp2 | -0.015947991 | 0.477580914 | 0.8607489 | 0.117398937 | 0.114445814 |
| Tubgcp3 | -0.008621628 | 0.675622835 | 0.9269628 | 0.084871012 | 0.099852004 |
| Tubgcp4 | -0.00581166 | 0.778304577 | 0.9576756 | 0.090123755 | 0.091181288 |
| Tubgcp5 | -0.040555425 | 0.044239304 | 0.4007863 | 0.07142009 | 0.098873984 |
| Tufm | -0.022528727 | 0.342995584 | 0.7872955 | 0.138821202 | 0.134449542 |
| Tuft1 | -0.053291383 | 0.021590104 | 0.2863286 | 0.102909786 | 0.130596055 |
| Tug1 | 0.037983547 | 0.212828756 | 0.6827572 | 0.275351366 | 0.242145275 |
| Tulp3 | 0.000536127 | 0.978629988 | 0.9976072 | 0.090093466 | 0.089466334 |
| Tulp4 | 0.041356682 | 0.273884581 | 0.7351955 | 0.649062948 | 0.616194944 |
| Tusc1 | -0.018585392 | 0.271503459 | 0.7342030 | 0.052120209 | 0.065750714 |
| Tusc2 | 0.015292223 | 0.607141444 | 0.9087139 | 0.281200584 | 0.241845365 |
| TUSC3 | -0.025331744 | 0.459358874 | 0.8523830 | 0.846311352 | 0.821961804 |
| Tut1 | -0.018098357 | 0.307828098 | 0.7658523 | 0.054443717 | 0.067557011 |
| Tut4 | 0.013211664 | 0.716792063 | 0.9431474 | 0.707981321 | 0.682883819 |
| Tut7 | -0.11745788 | 0.003564891 | 0.1149480 | 0.496869629 | 0.596743376 |
| Tvp23b | -0.018729952 | 0.607527538 | 0.9087139 | 0.725969826 | 0.682100876 |
| Twf1 | -0.008202771 | 0.820782927 | 0.9704608 | 0.510264318 | 0.518832994 |
| Twf2 | 0.033024915 | 0.254251721 | 0.7223071 | 0.249490495 | 0.207233006 |
| Twnk | -0.025845193 | 0.111741553 | 0.5675761 | 0.042529501 | 0.061194646 |
| Twsg1 | 0.10260603 | 0.006536908 | 0.1613087 | 0.889232656 | 0.758933273 |

| | | | | | |
|---|---|---|---|---|---|
| Txlna | 0.017108379 | 0.59991831 | 0.9078234 | 0.341736878 | 0.314905068 |
| Txlng | -0.020750701 | 0.328498976 | 0.7778857 | 0.087146931 | 0.106060778 |
| Txn1 | -0.001871138 | 0.956132004 | 0.9954690 | 1.212137695 | 1.1695291 |
| Txn2 | -0.001205869 | 0.972804787 | 0.9976072 | 0.492115887 | 0.454892076 |
| Txndc11 | -0.013681401 | 0.658343769 | 0.9211780 | 0.265507044 | 0.262318147 |
| Txndc12 | -0.062665388 | 0.05796419 | 0.4508199 | 0.417337715 | 0.424724664 |
| Txndc15 | -0.008409489 | 0.805813042 | 0.9647143 | 0.391647196 | 0.396341817 |
| Txndc16 | -0.011799816 | 0.493143774 | 0.8669301 | 0.058022705 | 0.066453842 |
| Txndc17 | 0.042918561 | 0.209094352 | 0.6807660 | 1.120242186 | 1.005277528 |
| Txndc5 | -0.05104355 | 0.15063546 | 0.6280320 | 0.491290771 | 0.485227637 |
| Txndc9 | -0.021128309 | 0.530369159 | 0.8776448 | 0.45977871 | 0.445341826 |
| Txnip | 0.084247463 | 0.050301861 | 0.4221611 | 1.507495948 | 1.277884113 |
| Txnl1 | -0.071793603 | 0.03370628 | 0.3468530 | 1.097865466 | 1.14527121 |
| Txnl4a | -0.026795574 | 0.445178252 | 0.8432629 | 0.384347411 | 0.410110327 |
| Txnrd1 | -0.115565236 | 0.001245696 | 0.0564260 | 1.13937264 | 1.334233562 |
| Txnrd2 | 0.011199971 | 0.636300939 | 0.9153541 | 0.143254423 | 0.129319651 |
| Tyk2 | -0.008680507 | 0.593673282 | 0.9061761 | 0.055581755 | 0.056435501 |
| Tyms | -0.014164551 | 0.359312463 | 0.8002291 | 0.045789918 | 0.052079692 |
| Tysnd1 | 0.00264191 | 0.913328697 | 0.9845236 | 0.150620837 | 0.13502686 |
| Tyw1 | -0.003951589 | 0.860431941 | 0.9769675 | 0.114548577 | 0.12047574 |
| Tyw5 | 0.028566621 | 0.162966816 | 0.6400703 | 0.113342517 | 0.085351299 |
| U2af1 | -0.058049404 | 0.1228962 | 0.5833237 | 0.688267802 | 0.683069322 |
| U2af2 | 0.010071381 | 0.755823559 | 0.9521533 | 0.37427166 | 0.337553746 |
| U2surp | 0.058614383 | 0.11918248 | 0.5782737 | 0.75010848 | 0.695535223 |
| Uaca | -0.00969784 | 0.722187272 | 0.9445652 | 0.176208474 | 0.189348327 |
| Uap1 | 0.019047144 | 0.47477795 | 0.8602369 | 0.197644686 | 0.160171901 |
| Uap1l1 | -0.01416061 | 0.62332055 | 0.9121505 | 0.201230145 | 0.23194379 |
| Uba1 | -0.020159608 | 0.573353427 | 0.8973494 | 0.482961137 | 0.500891974 |
| Uba2 | 0.004814666 | 0.878359326 | 0.9785583 | 0.315056625 | 0.27906307 |
| Uba3 | -0.011357594 | 0.675789643 | 0.9269628 | 0.177417342 | 0.183155031 |
| Uba5 | -0.036842312 | 0.311009858 | 0.7680774 | 0.460947135 | 0.470495954 |
| Uba52 | 0.103725192 | 0.000318364 | 0.0226977 | 1.989463902 | 1.828694622 |
| Uba6 | 0.000329197 | 0.991156991 | 0.9982478 | 0.242501305 | 0.226934258 |
| Ubac1 | -0.013360305 | 0.554313628 | 0.8902995 | 0.126102189 | 0.119464438 |
| Ubac2 | -0.016804281 | 0.387073343 | 0.8163635 | 0.078562346 | 0.083814911 |
| Ubald1 | -0.016858595 | 0.49982637 | 0.8692632 | 0.14107203 | 0.144716428 |
| Ubald2 | -0.006830147 | 0.831079233 | 0.9723494 | 0.327931754 | 0.313155151 |
| Ubap1 | 0.000849743 | 0.969765702 | 0.9976072 | 0.122668629 | 0.112154793 |
| Ubap2 | 0.03002601 | 0.295257552 | 0.7541078 | 0.22212866 | 0.192326453 |
| Ubap2l | -0.001953441 | 0.957274198 | 0.9956630 | 0.914154197 | 0.888246004 |
| Ubb | 0.026841572 | 0.086909575 | 0.5250566 | 3.135588312 | 2.96245174 |
| Ubb-ps | 0.035514173 | 0.133553127 | 0.5988251 | 0.162956951 | 0.117361165 |
| Ubc | -0.052034839 | 0.010870468 | 0.2035210 | 2.591898032 | 2.691265906 |
| Ube2a | 0.023765556 | 0.489319792 | 0.8651997 | 0.441645875 | 0.387822865 |
| Ube2b | -0.038657971 | 0.194116829 | 0.6716564 | 1.323381394 | 1.288244092 |
| Ube2d1 | -0.053254805 | 0.035631182 | 0.3591914 | 0.126288164 | 0.163395237 |
| Ube2d2a | -0.068842631 | 0.061307015 | 0.4611772 | 0.744433948 | 0.748871881 |
| Ube2d3 | -0.024114803 | 0.445577023 | 0.8436961 | 1.394465014 | 1.325768216 |
| Ube2d-ps | 0.02130582 | 0.36668955 | 0.8053794 | 0.147836215 | 0.119896641 |
| Ube2e1 | -0.070432739 | 0.036691562 | 0.3626279 | 0.484607733 | 0.514394813 |
| Ube2e2 | 0.002381505 | 0.920751656 | 0.9861175 | 0.137952519 | 0.135180675 |
| Ube2e3 | 0.049149102 | 0.177385966 | 0.6536659 | 0.668942268 | 0.562115876 |
| Ube2f | 0.006231839 | 0.844793114 | 0.9734889 | 0.34982846 | 0.296141894 |
| Ube2g1 | -0.036435354 | 0.19543353 | 0.6719932 | 0.199650498 | 0.209675802 |
| Ube2g2 | 0.013074518 | 0.686255062 | 0.9319165 | 0.33698049 | 0.29013227 |
| Ube2h | 0.002350286 | 0.946904308 | 0.9945018 | 0.377784782 | 0.369350304 |
| Ube2i | -0.006316873 | 0.86082955 | 0.9769675 | 0.844102163 | 0.765705924 |
| Ube2j1 | 0.0331612 | 0.330276005 | 0.7778857 | 0.397841483 | 0.343179902 |
| Ube2j2 | -0.031211208 | 0.331777661 | 0.7791748 | 0.335645272 | 0.334958499 |
| Ube2k | 0.035358511 | 0.327627995 | 0.7778857 | 0.786245955 | 0.751502048 |
| Ube2l3 | 0.046760777 | 0.182074138 | 0.6594287 | 0.872139247 | 0.732789688 |
| Ube2l6 | -0.013065397 | 0.476132338 | 0.8607231 | 0.062202374 | 0.07070767 |
| Ube2m | 0.028789719 | 0.416460833 | 0.8300990 | 0.580410076 | 0.564881067 |
| Ube2n | 0.031633326 | 0.361967805 | 0.8030292 | 0.660485584 | 0.580977408 |
| Ube2o | -0.00640019 | 0.843595759 | 0.9734889 | 0.302877542 | 0.319611903 |
| Ube2q1 | -0.08349851 | 0.012219812 | 0.2184153 | 0.389399697 | 0.44201235 |

| | | | | | |
|---|---|---|---|---|---|
| Ube2q2 | -0.071852104 | 0.031066678 | 0.3368103 | 0.311888023 | 0.364251238 |
| Ube2r2 | -0.025751506 | 0.480701303 | 0.8617579 | 0.656572029 | 0.629854194 |
| Ube2s | 0.033598451 | 0.087756556 | 0.5276030 | 0.10335059 | 0.070006943 |
| Ube2v2 | -0.018002667 | 0.607720168 | 0.9087139 | 0.484615095 | 0.449798165 |
| Ube2w | -0.027590533 | 0.405660616 | 0.8270770 | 0.391242651 | 0.39215349 |
| Ube2z | -0.007176869 | 0.825435512 | 0.9716906 | 0.316423566 | 0.315711913 |
| Ube3a | 0.031675256 | 0.37322404 | 0.8085689 | 0.909252423 | 0.845587685 |
| Ube3b | -0.016037829 | 0.406223425 | 0.8270770 | 0.07528822 | 0.082834741 |
| Ube3c | 0.045953127 | 0.109024289 | 0.5646875 | 0.226435713 | 0.205050677 |
| Ube4a | -0.027596944 | 0.277212306 | 0.7376464 | 0.140045089 | 0.16683998 |
| Ube4b | 0.009016989 | 0.794021921 | 0.9624508 | 0.429876613 | 0.411801924 |
| Ubfd1 | -0.004484516 | 0.890332641 | 0.9819597 | 0.340418997 | 0.322142071 |
| Ubl3 | -0.00101943 | 0.973146928 | 0.9976072 | 1.504497232 | 1.463607394 |
| Ubl4a | 0.036655098 | 0.261536948 | 0.7275735 | 0.347429147 | 0.315583105 |
| Ubl5 | 0.070550962 | 0.005644609 | 0.1517466 | 1.819445722 | 1.69220701 |
| Ubl7 | -0.030785113 | 0.199003166 | 0.6730962 | 0.130718039 | 0.13387021 |
| Ublcp1 | 0.024937428 | 0.280326505 | 0.7408618 | 0.136701371 | 0.114329238 |
| Ubn1 | -0.006653321 | 0.857287079 | 0.9763189 | 0.705379978 | 0.682683274 |
| Ubn2 | -0.053549568 | 0.152186676 | 0.6281105 | 0.81775391 | 0.833549209 |
| Ubp1 | -0.006642546 | 0.805877443 | 0.9647143 | 0.178821223 | 0.181769757 |
| Ubqln1 | -0.038638582 | 0.234572975 | 0.7064894 | 0.308958316 | 0.34208596 |
| Ubqln2 | -0.090106187 | 0.014841996 | 0.2390139 | 0.660740667 | 0.710846275 |
| Ubqln4 | -0.008737592 | 0.633055095 | 0.9141196 | 0.064479857 | 0.073661085 |
| Ubr1 | -0.058908271 | 0.051192102 | 0.4270537 | 0.208416311 | 0.246038294 |
| Ubr2 | -0.063244522 | 0.09223051 | 0.5325584 | 0.649739698 | 0.734734807 |
| Ubr3 | 0.06393584 | 0.06736387 | 0.4814478 | 0.471560386 | 0.408586668 |
| Ubr4 | -0.005377991 | 0.8905619 | 0.9819597 | 0.794166606 | 0.903949816 |
| Ubr5 | -0.053560431 | 0.148025161 | 0.6255679 | 0.531151679 | 0.568131412 |
| Ubr7 | -0.005767695 | 0.816761625 | 0.9690557 | 0.149610845 | 0.148113551 |
| Ubtd1 | -0.001652899 | 0.918568934 | 0.9851211 | 0.058214955 | 0.050731511 |
| Ubtd2 | 0.024715226 | 0.071484724 | 0.4888240 | 0.051883002 | 0.032159147 |
| Ubtf | 0.018207632 | 0.617131673 | 0.9115017 | 0.503094714 | 0.472579974 |
| Ubxn1 | -0.067278339 | 0.036097793 | 0.3602516 | 1.206140816 | 1.196932747 |
| Ubxn10 | -0.016376902 | 0.378022411 | 0.8101915 | 0.075058652 | 0.079049166 |
| Ubxn2a | -0.027725114 | 0.443808894 | 0.8431541 | 0.487386607 | 0.511263308 |
| Ubxn2b | -0.020362895 | 0.303694384 | 0.7630821 | 0.075060863 | 0.092483542 |
| Ubxn4 | -0.051605362 | 0.125957615 | 0.5882766 | 1.121300546 | 1.159798492 |
| Ubxn6 | -0.063937307 | 0.0768492 | 0.5015240 | 0.485377869 | 0.528768042 |
| Ubxn7 | 0.005832545 | 0.826656157 | 0.9716906 | 0.175172946 | 0.16199916 |
| Ubxn8 | 0.004520403 | 0.808448293 | 0.9658392 | 0.083914226 | 0.072168666 |
| Uchl1 | -0.041350489 | 0.135576461 | 0.6028321 | 0.142450341 | 0.15325959 |
| Uchl3 | 0.013917318 | 0.651096601 | 0.9182621 | 0.281090683 | 0.248684726 |
| Uchl5 | -0.000797197 | 0.980167749 | 0.9976072 | 0.296975103 | 0.27069665 |
| Uck1 | 0.020768846 | 0.446422116 | 0.8443293 | 0.230052903 | 0.17910287 |
| Uck2 | 0.000427106 | 0.987382791 | 0.9976072 | 0.177812207 | 0.181563465 |
| Uckl1 | -0.004996992 | 0.862540049 | 0.9769675 | 0.231765656 | 0.226880273 |
| Ucn3 | 0.164719566 | 1.69E-05 | 0.0023782 | 0.396454591 | 0.328930612 |
| Ucp2 | -0.1207003 | 0.001201132 | 0.0551630 | 0.932222284 | 1.012102993 |
| Uevld | 0.003307358 | 0.816381344 | 0.9688248 | 0.048211568 | 0.041002156 |
| Ufc1 | 0.019432122 | 0.541189966 | 0.8835017 | 1.292528427 | 1.214899914 |
| Ufd1 | 0.027722171 | 0.364827447 | 0.8050191 | 0.280588292 | 0.247508125 |
| Ufl1 | 0.086661468 | 0.018594691 | 0.2665597 | 0.588126797 | 0.51719055 |
| Ufm1 | 0.055079319 | 0.111683682 | 0.5675761 | 1.023040795 | 0.923324328 |
| Ufsp1 | 0.008107086 | 0.670353368 | 0.9243682 | 0.084278955 | 0.076527572 |
| Ufsp2 | 0.019826677 | 0.592191237 | 0.9060200 | 0.682475082 | 0.666144781 |
| Ugcg | 0.000443142 | 0.986058922 | 0.9976072 | 0.153623567 | 0.15548496 |
| Ugdh | -0.06608497 | 0.052617303 | 0.4328056 | 0.298559468 | 0.341196685 |
| Uggt1 | -0.064013371 | 0.054801541 | 0.4422689 | 0.318109145 | 0.393427641 |
| Uggt2 | 0.009182719 | 0.662487873 | 0.9227576 | 0.104694493 | 0.09232886 |
| Ugp2 | -0.005439781 | 0.849771712 | 0.9748841 | 0.2267806 | 0.213636475 |
| Ugt2b34 | -0.187471003 | 4.60E-05 | 0.0052477 | 0.895183057 | 0.990712674 |
| Ugt2b35 | -0.02028884 | 0.294057473 | 0.7533922 | 0.062853046 | 0.076038916 |
| Ugt8a | -0.005488085 | 0.745510739 | 0.9485717 | 0.067104544 | 0.054684286 |
| Uhmk1 | 0.009570245 | 0.733102546 | 0.9464134 | 0.206826062 | 0.192472263 |
| Uhrf1bp1 | -0.000879251 | 0.962590921 | 0.9964418 | 0.073749101 | 0.077939095 |
| Uhrf1bp1l | 0.041392102 | 0.174358298 | 0.6527657 | 0.261937689 | 0.226866392 |

| | | | | | |
|---|---|---|---|---|---|
| Uhrf2 | 0.001335823 | 0.96876335 | 0.99760728 | 0.375522558 | 0.364752638 |
| Uimc1 | -0.007225251 | 0.808771646 | 0.96584288 | 0.244063097 | 0.237965472 |
| Ulk1 | -0.010405506 | 0.673954892 | 0.92640901 | 0.152395186 | 0.146662727 |
| Ulk2 | -0.029214095 | 0.29739528 | 0.75509628 | 0.152782352 | 0.198128647 |
| Ulk3 | 0.027862591 | 0.039330409 | 0.37990034 | 0.054256497 | 0.028787443 |
| Ulk4 | -0.003309452 | 0.831578482 | 0.97234941 | 0.046526918 | 0.050653926 |
| Umad1 | 0.048972902 | 0.043676833 | 0.39864361 | 0.168378374 | 0.126331028 |
| Umps | 0.062178167 | 0.028905528 | 0.32641726 | 0.261634278 | 0.186847171 |
| Unc119 | -0.021327983 | 0.458052454 | 0.85216216 | 0.20168417 | 0.212645335 |
| Unc119b | 0.040783869 | 0.094989907 | 0.53509361 | 0.154456843 | 0.132633874 |
| Unc13a | 0.039367446 | 0.148270749 | 0.62556941 | 0.212155017 | 0.174798911 |
| Unc13b | -0.01232355 | 0.593288785 | 0.90605856 | 0.105509157 | 0.117232076 |
| Unc45a | 0.004708047 | 0.878387619 | 0.97855831 | 0.267986265 | 0.243417549 |
| Unc50 | -0.012434443 | 0.720919924 | 0.94456521 | 0.534943084 | 0.523275661 |
| Unc5a | -0.012164637 | 0.591477755 | 0.90602008 | 0.115372512 | 0.1188319 |
| Unc5c | -0.057621803 | 0.005712055 | 0.15238917 | 0.066707181 | 0.100414183 |
| Unc79 | -0.038051281 | 0.156770806 | 0.63176836 | 0.167480133 | 0.175548757 |
| Unc80 | -0.022349389 | 0.562732935 | 0.89262401 | 0.670062436 | 0.69179599 |
| Unc93b1 | -0.006447596 | 0.793315532 | 0.96245081 | 0.143715315 | 0.142045582 |
| Unk | -0.021291047 | 0.177553384 | 0.65381879 | 0.040569918 | 0.056074072 |
| Upb1 | -0.088101632 | 0.025279377 | 0.30507471 | 0.640813755 | 0.6913247 |
| Upf1 | -0.001435123 | 0.961211926 | 0.99615778 | 0.233812944 | 0.231891904 |
| Upf2 | -0.011978741 | 0.708104218 | 0.94122924 | 0.288214453 | 0.272838958 |
| Upf3a | -0.011087544 | 0.692847095 | 0.93597259 | 0.203602764 | 0.186596473 |
| Upf3b | 0.010982978 | 0.772305762 | 0.95523334 | 0.557484853 | 0.566628017 |
| Upk3a | 0.021513283 | 0.484715255 | 0.86297151 | 0.183297306 | 0.127003845 |
| Upp1 | -0.033361629 | 0.382258058 | 0.81268821 | 0.594708611 | 0.531374962 |
| Upp2 | -0.06158382 | 0.019658223 | 0.27160107 | 0.133929384 | 0.18501585 |
| Uqcc1 | -0.015999298 | 0.506287228 | 0.86955669 | 0.122335997 | 0.141390443 |
| Uqcc2 | 0.054047819 | 0.028450045 | 0.32328114 | 2.004560212 | 1.923418896 |
| Uqcc3 | -0.015208654 | 0.665110733 | 0.92307657 | 0.436634981 | 0.45345844 |
| Uqcr10 | 0.132096057 | 1.83E-06 | 0.00034860 | 1.760572264 | 1.600540178 |
| Uqcr11 | 0.074295598 | 0.003972296 | 0.12391569 | 1.919424951 | 1.77459673 |
| Uqcrb | 0.094772634 | 0.000178349 | 0.01474359 | 2.026708903 | 1.811544172 |
| Uqcrc1 | -0.012246809 | 0.729950846 | 0.94590209 | 0.539533111 | 0.538931999 |
| Uqcrc2 | 0.000530115 | 0.987634907 | 0.99760728 | 0.866884281 | 0.810899478 |
| Uqcrfs1 | -0.002805961 | 0.936588418 | 0.99124689 | 1.107404935 | 0.991006892 |
| Uqcrh | 0.052455649 | 0.023818301 | 0.29946456 | 2.128270293 | 1.956542491 |
| Uqcrq | 0.139115705 | 1.47E-07 | 5.40E-05 | 2.388124985 | 2.279910833 |
| Urb1 | 0.007789646 | 0.645279713 | 0.91696956 | 0.065137598 | 0.056734898 |
| Urgcp | 0.018219592 | 0.496686247 | 0.86846291 | 0.194623063 | 0.169407831 |
| Uri1 | -0.015092634 | 0.675085986 | 0.92658871 | 0.527023862 | 0.562344863 |
| Urm1 | -0.068794765 | 0.013208128 | 0.22747331 | 0.165829181 | 0.216349583 |
| Urod | 0.02464531 | 0.423334079 | 0.83174371 | 0.310128665 | 0.259481336 |
| Uros | -0.004276629 | 0.817822788 | 0.96961898 | 0.081505622 | 0.075546548 |
| Usb1 | -0.009912368 | 0.538426271 | 0.88163354 | 0.043045133 | 0.05781532 |
| Use1 | -0.091675147 | 0.011000796 | 0.20474270 | 0.742197685 | 0.813943508 |
| Usf1 | 0.002493091 | 0.93184709 | 0.98939560 | 0.23673662 | 0.214457728 |
| Usf2 | -0.041862051 | 0.224247098 | 0.69625390 | 0.404125488 | 0.425601301 |
| Usf3 | -0.02160955 | 0.355959109 | 0.79727124 | 0.103182254 | 0.12582012 |
| Ush1c | -0.009635098 | 0.754322447 | 0.95190435 | 0.254696613 | 0.253928245 |
| Uso1 | 0.052472343 | 0.154692476 | 0.63049349 | 0.77324248 | 0.725382597 |
| Usp1 | -0.042263117 | 0.126938288 | 0.58932347 | 0.163169435 | 0.198990345 |
| Usp10 | 0.008116795 | 0.788313448 | 0.96128699 | 0.251053282 | 0.233510597 |
| Usp11 | 0.019184596 | 0.393851308 | 0.82100599 | 0.139578525 | 0.108073794 |
| Usp12 | -0.008783511 | 0.740238777 | 0.94790806 | 0.170903585 | 0.170747499 |
| Usp14 | -0.011376436 | 0.739570458 | 0.9477508 | 0.404823739 | 0.391472354 |
| Usp15 | -0.024523675 | 0.485942347 | 0.86343329 | 0.397216573 | 0.394997094 |
| Usp16 | -0.059597111 | 0.113106314 | 0.56962039 | 0.444706201 | 0.513437137 |
| Usp18 | -0.017467539 | 0.368834127 | 0.80567191 | 0.085240105 | 0.084647685 |
| Usp19 | -0.064489307 | 0.049393458 | 0.41914721 | 0.26617398 | 0.333996595 |
| Usp2 | 0.009769402 | 0.55108998 | 0.88876810 | 0.048330942 | 0.055010413 |
| Usp20 | 0.00749919 | 0.782743929 | 0.95908628 | 0.192961011 | 0.181997149 |
| Usp21 | -0.008580673 | 0.622831412 | 0.91215057 | 0.058635592 | 0.067438651 |
| Usp22 | -0.007216281 | 0.82259238 | 0.97098009 | 0.325614188 | 0.314915905 |
| Usp24 | -0.033358977 | 0.217115755 | 0.68723301 | 0.159472592 | 0.181301655 |

| | | | | | |
|---|---|---|---|---|---|
| Usp25 | -0.007287993 | 0.734523075 | 0.94683808 | 0.100997862 | 0.104071915 |
| Usp29 | -0.109994195 | 0.001568301 | 0.06793680 | 0.307736392 | 0.384453336 |
| Usp3 | 0.043040234 | 0.175814114 | 0.65366590 | 0.324496996 | 0.267129932 |
| Usp30 | -0.009812779 | 0.60759241 | 0.90871391 | 0.080917034 | 0.075072262 |
| Usp31 | 0.029644524 | 0.196441213 | 0.67309621 | 0.133781295 | 0.117362663 |
| Usp32 | 0.00704052 | 0.752580845 | 0.95078980 | 0.107517218 | 0.110778562 |
| Usp33 | -0.0425291 | 0.198508914 | 0.67309621 | 0.29385124 | 0.346319229 |
| Usp34 | -0.089285247 | 0.014663397 | 0.23765848 | 0.796335468 | 0.839586797 |
| Usp35 | 0.012732772 | 0.596077204 | 0.90680201 | 0.134256712 | 0.135047611 |
| Usp36 | 0.017297023 | 0.589743503 | 0.90467754 | 0.281071416 | 0.263015361 |
| Usp37 | -0.022861505 | 0.361207489 | 0.80196470 | 0.140975577 | 0.150010776 |
| Usp38 | -0.056241931 | 0.009036331 | 0.19031931 | 0.072834212 | 0.115308302 |
| Usp39 | 0.006409147 | 0.763486562 | 0.95322041 | 0.102333835 | 0.094914571 |
| Usp4 | -0.013295298 | 0.64315315 | 0.91667127 | 0.204549601 | 0.208241346 |
| Usp40 | -0.028307869 | 0.220107392 | 0.69145345 | 0.110242293 | 0.119658283 |
| Usp42 | -0.006347591 | 0.784500162 | 0.96012021 | 0.115871008 | 0.125288354 |
| Usp45 | -0.047845488 | 0.069023742 | 0.48354951 | 0.138327984 | 0.179846463 |
| Usp46 | 0.044315288 | 0.128115612 | 0.59122443 | 0.254055469 | 0.208952707 |
| Usp47 | 0.013267373 | 0.700152707 | 0.93751653 | 0.45069959 | 0.418700888 |
| Usp48 | 0.037784497 | 0.250748737 | 0.72072101 | 0.364129975 | 0.318914393 |
| Usp5 | -0.074086026 | 0.00795764 | 0.17584088 | 0.156386456 | 0.211464698 |
| Usp50 | 0.049691444 | 0.119883077 | 0.57955171 | 0.279484513 | 0.27505523 |
| Usp51 | 0.021891192 | 0.280284242 | 0.74086184 | 0.103937364 | 0.077414013 |
| Usp53 | 0.004535865 | 0.870834982 | 0.97822251 | 0.210597424 | 0.183182242 |
| Usp6nl | -0.000213719 | 0.991347561 | 0.99829131 | 0.074107107 | 0.088387134 |
| Usp7 | -0.025811036 | 0.478851378 | 0.86085641 | 0.504308526 | 0.504648723 |
| Usp8 | 0.054311364 | 0.108944405 | 0.56464394 | 0.404788258 | 0.350535836 |
| Usp9x | -0.062228102 | 0.099098295 | 0.54335515 | 0.985400825 | 1.007088136 |
| Uspl1 | -0.000934788 | 0.974329091 | 0.99760728 | 0.205818266 | 0.206502811 |
| Utp11 | 0.033100187 | 0.298193971 | 0.75615649 | 0.335176224 | 0.281584752 |
| Utp14a | 0.005914298 | 0.804205206 | 0.96471431 | 0.124028336 | 0.12924946 |
| Utp14b | -0.029612346 | 0.050950489 | 0.42580357 | 0.028070396 | 0.052249586 |
| Utp15 | -0.002310954 | 0.907117938 | 0.98368121 | 0.094781375 | 0.083361051 |
| Utp18 | 0.085870745 | 0.00521858 | 0.14582621 | 0.322680687 | 0.222971597 |
| Utp20 | 0.006900312 | 0.707222232 | 0.94083541 | 0.075271623 | 0.065410277 |
| Utp23 | -0.025280137 | 0.246577669 | 0.71626661 | 0.099175743 | 0.112561084 |
| Utp25 | 0.027356114 | 0.131597377 | 0.59745811 | 0.091125627 | 0.061707097 |
| Utp3 | 0.023939021 | 0.500721503 | 0.86937040 | 0.535286827 | 0.48285715 |
| Utp4 | -0.022287162 | 0.311235457 | 0.76840610 | 0.094366021 | 0.110503553 |
| Utp6 | 0.007065497 | 0.757012034 | 0.95215335 | 0.120970402 | 0.12043554 |
| Utrn | 0.005740552 | 0.874713641 | 0.97843318 | 0.468130692 | 0.43863574 |
| Uty | 0.032675917 | 0.209057153 | 0.68076601 | 0.17923419 | 0.144584799 |
| Uvrag | -0.010622019 | 0.700585694 | 0.93751651 | 0.177426827 | 0.186633863 |
| Uvssa | 0.013555288 | 0.642504598 | 0.91667127 | 0.2092681 | 0.193767886 |
| Uxs1 | -0.011780564 | 0.573312242 | 0.89734948 | 0.094377272 | 0.097659932 |
| Uxt | 0.101203064 | 0.001389744 | 0.06238128 | 0.371286092 | 0.271630339 |
| Vac14 | -0.004677915 | 0.790762749 | 0.96224569 | 0.060125642 | 0.069016469 |
| Vamp2 | -0.010181091 | 0.77502299 | 0.95585380 | 0.678369022 | 0.683898384 |
| Vamp3 | 0.004848028 | 0.881447239 | 0.97969630 | 0.359569475 | 0.297982194 |
| Vamp4 | 0.027631044 | 0.422617552 | 0.83148971 | 0.533505058 | 0.487005664 |
| Vamp8 | 0.061084792 | 0.072906458 | 0.49033721 | 1.097413678 | 0.98563448 |
| Vapa | 0.046196433 | 0.165697034 | 0.64257801 | 1.151111214 | 1.007698519 |
| Vapb | 0.074103407 | 0.033688561 | 0.34685300 | 0.776545131 | 0.737039313 |
| Vars | -0.04383942 | 0.177115117 | 0.65366590 | 0.29753254 | 0.319911379 |
| Vars2 | -0.00576223 | 0.735594409 | 0.94683808 | 0.064445497 | 0.058937457 |
| Vasp | 0.016259887 | 0.577776709 | 0.89979141 | 0.246719718 | 0.229565498 |
| Vat1 | 0.005552987 | 0.864135278 | 0.97696750 | 0.326971013 | 0.29360869 |
| Vat1l | -0.005613706 | 0.83214683 | 0.97234941 | 0.176380087 | 0.176237815 |
| Vbp1 | 0.036769489 | 0.27429998 | 0.73519551 | 0.418721759 | 0.362462422 |
| Vcl | -0.002736663 | 0.913712033 | 0.98452361 | 0.148602986 | 0.144872903 |
| Vcp | 0.021860037 | 0.477430159 | 0.86074894 | 1.329305448 | 1.240375195 |
| Vcpip1 | 0.003363208 | 0.91158392 | 0.98401851 | 0.271684689 | 0.247090843 |
| Vcpkmt | -0.015405497 | 0.344616017 | 0.78792235 | 0.053237675 | 0.057520571 |
| Vdac1 | -0.048797251 | 0.170896106 | 0.6507063 | 0.711528261 | 0.779316754 |
| Vdac2 | -0.06272569 | 0.061405193 | 0.46117728 | 1.037420722 | 1.036217834 |
| Vdac3 | 0.031064276 | 0.399067324 | 0.82398500 | 0.953978824 | 0.911380635 |

| | | | | | |
|---|---|---|---|---|---|
| Vdr | 0.042123369 | 0.22417583 | 0.6962539 | 0.391208814 | 0.358058533 |
| Vegfa | 0.07740736 | 0.033380163 | 0.3456484 | 1.053562237 | 1.038187383 |
| Vegfb | 0.00071144 | 0.982124248 | 0.9976072 | 0.315163053 | 0.311769892 |
| Vezf1 | 0.052373614 | 0.147472149 | 0.6241142 | 0.542412705 | 0.47686714 |
| Vezt | -0.00410362 | 0.833546234 | 0.9723494 | 0.079664293 | 0.081813635 |
| Vgf | -0.137555326 | 0.004294472 | 0.1302787 | 0.649837276 | 0.749487929 |
| Vgll4 | -0.0160772 | 0.552759623 | 0.8900045 | 0.184462089 | 0.180448976 |
| Vhl | -0.013760662 | 0.519430463 | 0.8741795 | 0.10115994 | 0.106380319 |
| Vil1 | 0.045930066 | 0.03003029 | 0.3308280 | 0.111521809 | 0.094816938 |
| Vipas39 | -0.004303691 | 0.855958026 | 0.9761011 | 0.138100402 | 0.131385549 |
| Vipr1 | 0.003504135 | 0.898073065 | 0.9825441 | 0.190427665 | 0.179528862 |
| Virma | -0.011077298 | 0.711822371 | 0.9419834 | 0.254258015 | 0.232150377 |
| Vkorc1 | 0.100899289 | 0.004590407 | 0.1355263 | 0.798565317 | 0.555718536 |
| Vkorc1l1 | -0.013714973 | 0.628731111 | 0.9130564 | 0.204483984 | 0.218094967 |
| Vldlr | 0.054864215 | 0.107676819 | 0.5633723 | 0.350588782 | 0.370834015 |
| Vma21 | 0.001706153 | 0.960952879 | 0.9960974 | 0.440146477 | 0.392147813 |
| Vmac | -0.010160703 | 0.704856792 | 0.9400796 | 0.185265685 | 0.183889644 |
| Vmp1 | -0.025800864 | 0.465897026 | 0.8549201 | 0.379642522 | 0.417086432 |
| Vopp1 | -0.021090941 | 0.259584594 | 0.7263475 | 0.062110479 | 0.078355658 |
| Vps11 | -0.028842084 | 0.318515255 | 0.7710043 | 0.229243753 | 0.2334973 |
| Vps13a | -0.002224153 | 0.948251358 | 0.9945018 | 0.415807083 | 0.407192679 |
| Vps13b | -0.006229112 | 0.839575129 | 0.9727382 | 0.241360383 | 0.242302292 |
| Vps13c | 0.022651011 | 0.45904105 | 0.8522649 | 0.247987228 | 0.268968765 |
| Vps13d | -0.008300004 | 0.715858212 | 0.9431411 | 0.110958428 | 0.118481411 |
| Vps16 | -0.018080877 | 0.381083112 | 0.8119369 | 0.092196924 | 0.097027265 |
| Vps18 | -0.022762587 | 0.256940816 | 0.7250597 | 0.071675042 | 0.094519772 |
| Vps26a | -0.004432813 | 0.851340323 | 0.9749822 | 0.127820208 | 0.12860438 |
| Vps26b | -0.009905877 | 0.659215608 | 0.9211780 | 0.114110628 | 0.119455444 |
| Vps26c | -0.020404965 | 0.373843486 | 0.8087297 | 0.126229602 | 0.125452745 |
| Vps28 | -0.019830448 | 0.573477469 | 0.8973494 | 0.914896176 | 0.850434392 |
| Vps29 | -0.002069434 | 0.954254061 | 0.9954690 | 0.674169644 | 0.639798047 |
| Vps33a | -0.015523849 | 0.418614344 | 0.8300990 | 0.070389559 | 0.08667075 |
| Vps33b | 0.017336948 | 0.281109274 | 0.7408618 | 0.063490896 | 0.051854434 |
| Vps35 | 0.035682026 | 0.316168602 | 0.7705987 | 0.603463501 | 0.564091097 |
| Vps35l | 0.04706326 | 0.080438352 | 0.5098712 | 0.199743268 | 0.162616207 |
| Vps36 | -0.014885217 | 0.656220812 | 0.9211780 | 0.337160271 | 0.355800309 |
| Vps37a | 0.011769334 | 0.756927911 | 0.9521533 | 0.830385812 | 0.784017529 |
| Vps37b | -0.007413292 | 0.791818881 | 0.9624054 | 0.193460966 | 0.197717499 |
| Vps37c | 0.03994639 | 0.121767938 | 0.5819916 | 0.193248899 | 0.147308573 |
| Vps37d | -0.011723813 | 0.631420502 | 0.9133517 | 0.143999936 | 0.137466591 |
| Vps39 | -0.011588109 | 0.558943118 | 0.8921289 | 0.076704091 | 0.087677035 |
| Vps41 | -0.080620656 | 0.017004877 | 0.2555884 | 0.352313217 | 0.38999278 |
| Vps45 | 0.050202527 | 0.066252672 | 0.4772886 | 0.213294249 | 0.172946602 |
| Vps4a | 0.004114487 | 0.884212357 | 0.9802800 | 0.21086575 | 0.201152117 |
| Vps4b | 0.028297585 | 0.431456927 | 0.8356213 | 0.455830421 | 0.425191743 |
| Vps50 | 0.013404655 | 0.605116834 | 0.9087139 | 0.176250106 | 0.15573844 |
| Vps51 | -0.023566043 | 0.388337798 | 0.8170287 | 0.179855374 | 0.19275 |
| Vps52 | 0.05819676 | 0.017887622 | 0.2587928 | 0.181624948 | 0.123428596 |
| Vps53 | 0.026706451 | 0.393547853 | 0.8208939 | 0.325064857 | 0.278875072 |
| Vps54 | -0.013868429 | 0.562488342 | 0.8926240 | 0.125588614 | 0.139551391 |
| Vps72 | -0.007061941 | 0.807399601 | 0.9653373 | 0.222421253 | 0.22232987 |
| Vps8 | 0.044965921 | 0.024320213 | 0.3027057 | 0.109351128 | 0.075004798 |
| Vps9d1 | -0.01009424 | 0.628854016 | 0.9130564 | 0.096661256 | 0.101697536 |
| Vrk1 | 0.06003515 | 0.029527129 | 0.3294815 | 0.214877147 | 0.171298417 |
| Vrk3 | 0.031038733 | 0.252307094 | 0.7223071 | 0.217891451 | 0.179502737 |
| Vsig10 | -0.00640297 | 0.791627878 | 0.9624054 | 0.13441294 | 0.136149328 |
| Vstm2l | 0.060271386 | 0.045146762 | 0.4027480 | 0.221070557 | 0.213120808 |
| Vta1 | 0.001135087 | 0.972884077 | 0.9976072 | 0.412533721 | 0.37112288 |
| Vti1a | -0.033271665 | 0.1170097 | 0.5735102 | 0.084145569 | 0.108855794 |
| Vti1b | 0.042651846 | 0.228172012 | 0.7015153 | 0.741006584 | 0.643911222 |
| Vwa5a | 0.049024377 | 0.03136557 | 0.3385228 | 0.139257322 | 0.102869077 |
| Vwa5b2 | -0.025406001 | 0.416965263 | 0.8300990 | 0.26039776 | 0.264030585 |
| Vwa8 | 0.001441477 | 0.948288158 | 0.9945018 | 0.107665063 | 0.115661804 |
| Vwde | -0.073272524 | 0.009033817 | 0.1903193 | 0.153891274 | 0.184309717 |
| Wac | 0.007197774 | 0.844519561 | 0.9734889 | 0.498815762 | 0.485426743 |
| Wapl | -0.042376839 | 0.24880635 | 0.7187416 | 0.57013964 | 0.590827357 |

| | | | | | |
|---|---|---|---|---|---|
| Wars | -0.026170845 | 0.42092544 | 0.8300990 | 0.208998811 | 0.291700769 |
| Wasf2 | 0.009440345 | 0.77714494 | 0.9564524 | 0.394561823 | 0.377697288 |
| Washc1 | 0.023107294 | 0.299613491 | 0.7585920 | 0.136033666 | 0.108019297 |
| Washc2 | -0.027069676 | 0.383217518 | 0.8129787 | 0.231217741 | 0.253076537 |
| Washc3 | 0.03842323 | 0.235932109 | 0.7079390 | 0.392754724 | 0.346483667 |
| Washc4 | 0.007233543 | 0.770773231 | 0.9552333 | 0.139244706 | 0.143365316 |
| Washc5 | -0.006318035 | 0.786034831 | 0.9602516 | 0.110916838 | 0.134194348 |
| Wasl | 0.012195957 | 0.728378114 | 0.9458355 | 0.673643916 | 0.626644978 |
| Wbp1 | 0.000470876 | 0.98823274 | 0.9976143 | 0.306094958 | 0.279424441 |
| Wbp11 | -0.000906519 | 0.980037966 | 0.9976072 | 0.55089176 | 0.538549085 |
| Wbp1l | -0.032013664 | 0.157641675 | 0.6324475 | 0.094965289 | 0.11604508 |
| Wbp2 | -0.054625808 | 0.122624613 | 0.5833237 | 0.530141384 | 0.551030826 |
| Wbp4 | -0.034636597 | 0.337669829 | 0.7834245 | 0.648517963 | 0.639837777 |
| Wdcp | -0.001171436 | 0.956652946 | 0.9955934 | 0.102383475 | 0.105663239 |
| Wdfy1 | -0.120830112 | 0.000136648 | 0.0119959 | 0.227139278 | 0.341529983 |
| Wdfy2 | -0.017867712 | 0.375981039 | 0.8101773 | 0.081149113 | 0.097175928 |
| Wdfy3 | -0.013777714 | 0.60263424 | 0.9086687 | 0.162748329 | 0.169700025 |
| Wdpcp | 0.003848339 | 0.830617318 | 0.9722601 | 0.073686314 | 0.06722906 |
| Wdr1 | -0.025411944 | 0.405370203 | 0.8270770 | 0.213139145 | 0.254067304 |
| Wdr11 | -0.003091687 | 0.896970963 | 0.9825441 | 0.130758015 | 0.13659769 |
| Wdr12 | -0.017253707 | 0.524141303 | 0.8758385 | 0.171822914 | 0.188450171 |
| Wdr13 | -0.004162695 | 0.86243149 | 0.9769675 | 0.132508815 | 0.132147302 |
| Wdr18 | -0.037916619 | 0.208439266 | 0.6807660 | 0.216970185 | 0.244045246 |
| Wdr20 | -0.025106819 | 0.285804858 | 0.7455125 | 0.117938819 | 0.129214549 |
| Wdr24 | -0.01930633 | 0.29408968 | 0.7533922 | 0.058078539 | 0.077295141 |
| Wdr26 | 0.048169318 | 0.175358969 | 0.6534142 | 0.552046115 | 0.494712798 |
| Wdr27 | 0.003468639 | 0.86289388 | 0.9769675 | 0.096941946 | 0.084713968 |
| Wdr3 | 0.0175592 | 0.403374639 | 0.8264703 | 0.115633495 | 0.090234752 |
| Wdr33 | 0.001838613 | 0.959103899 | 0.9957785 | 0.530804258 | 0.512497089 |
| Wdr35 | -0.012195102 | 0.594264531 | 0.9065207 | 0.112243618 | 0.119279133 |
| Wdr36 | -0.019944036 | 0.429599273 | 0.8340365 | 0.133635646 | 0.155518655 |
| Wdr37 | 0.004167812 | 0.865985078 | 0.9770148 | 0.14189296 | 0.143426571 |
| Wdr4 | 0.01891348 | 0.27777955 | 0.7382158 | 0.074292369 | 0.060991505 |
| Wdr41 | 0.000496899 | 0.980488953 | 0.9976072 | 0.097583993 | 0.089056993 |
| Wdr43 | 0.020657507 | 0.529556965 | 0.8776448 | 0.332410343 | 0.308530789 |
| Wdr44 | -0.000216921 | 0.992113651 | 0.9985765 | 0.113251486 | 0.106731089 |
| Wdr45 | -0.023695827 | 0.223355965 | 0.6955958 | 0.071767846 | 0.085165591 |
| Wdr45b | 0.020362436 | 0.527434125 | 0.8768553 | 0.343629858 | 0.289998664 |
| Wdr46 | -0.021035683 | 0.338992376 | 0.7834245 | 0.115781746 | 0.112424995 |
| Wdr47 | -0.008166698 | 0.724053114 | 0.9445652 | 0.117970818 | 0.124957163 |
| Wdr48 | 0.010892415 | 0.654964132 | 0.9206439 | 0.145425797 | 0.131083887 |
| Wdr5 | 0.007550074 | 0.696590225 | 0.9363862 | 0.085617345 | 0.080260316 |
| Wdr53 | -0.028057415 | 0.198134275 | 0.6730962 | 0.098442196 | 0.112611724 |
| Wdr54 | 0.000365132 | 0.983381782 | 0.9976072 | 0.067636175 | 0.063442539 |
| Wdr55 | -0.00691029 | 0.698461682 | 0.9369692 | 0.076777331 | 0.068197528 |
| Wdr59 | 0.035584083 | 0.054194328 | 0.4403011 | 0.098447289 | 0.060476819 |
| Wdr5b | 0.002045197 | 0.88247366 | 0.9798677 | 0.045269286 | 0.037147093 |
| Wdr6 | -0.026620399 | 0.444073546 | 0.8431541 | 0.402602456 | 0.410625261 |
| Wdr61 | -0.034325091 | 0.313454813 | 0.7699903 | 0.441963731 | 0.430782395 |
| Wdr7 | 0.055710189 | 0.007173015 | 0.1680001 | 0.123532766 | 0.087924889 |
| Wdr70 | -5.40E-05 | 0.998162097 | 0.9994073 | 0.132555422 | 0.1282729 |
| Wdr73 | 0.007953537 | 0.677186217 | 0.9280496 | 0.08545493 | 0.077048309 |
| Wdr74 | -0.025081141 | 0.405792866 | 0.8270770 | 0.247770164 | 0.255971619 |
| Wdr75 | -0.046077692 | 0.124408693 | 0.5852543 | 0.224593092 | 0.246680417 |
| Wdr76 | 0.017027871 | 0.152589218 | 0.6283458 | 0.039532761 | 0.025640974 |
| Wdr77 | 0.031841696 | 0.131778446 | 0.5978345 | 0.126293241 | 0.086760225 |
| Wdr82 | -0.004540004 | 0.857493994 | 0.9763189 | 0.155824868 | 0.153181442 |
| Wdr83 | -0.011582045 | 0.636503343 | 0.9153541 | 0.145738597 | 0.14078305 |
| Wdr83os | -0.005976751 | 0.865432246 | 0.9770148 | 0.979127403 | 0.905229086 |
| Wdr89 | -0.018644949 | 0.535218673 | 0.8795949 | 0.226917209 | 0.245451315 |
| Wdr90 | -0.023344138 | 0.267447272 | 0.7316814 | 0.077459498 | 0.110055029 |
| Wdr92 | 0.005747766 | 0.750305204 | 0.9504537 | 0.075785321 | 0.06817206 |
| Wdsub1 | -0.033121845 | 0.208070583 | 0.6807660 | 0.144834192 | 0.17514318 |
| Wdtc1 | 0.009178297 | 0.717058288 | 0.9431474 | 0.165354739 | 0.149455227 |
| Wdyhv1 | -0.017895982 | 0.459048321 | 0.8522649 | 0.134733631 | 0.135192952 |
| Wee1 | -0.009237961 | 0.695798854 | 0.9363862 | 0.112760473 | 0.119964468 |

| | | | | | |
|---|---|---|---|---|---|
| Wfs1 | 0.020796522 | 0.496063676 | 0.8684629 | 0.244756942 | 0.272200259 |
| Whamm | -0.071262947 | 0.019259491 | 0.2695586 | 0.238445651 | 0.271557917 |
| Whrn | -0.017116266 | 0.307884988 | 0.7658523 | 0.044820658 | 0.068376602 |
| Wipf1 | 0.024099666 | 0.030775178 | 0.3353039 | 0.042104028 | 0.017730509 |
| Wipf2 | -0.012642356 | 0.658286867 | 0.9211780 | 0.196530011 | 0.205194386 |
| Wipi1 | -0.0139659 | 0.714702054 | 0.9430747 | 0.358878585 | 0.467409582 |
| Wipi2 | -0.019771942 | 0.542052226 | 0.8836742 | 0.312479386 | 0.317966062 |
| Wiz | 0.013393322 | 0.546314828 | 0.8858413 | 0.121439961 | 0.110915582 |
| Wls | 0.007546166 | 0.846392088 | 0.9739657 | 0.706439553 | 0.77623091 |
| Wnk1 | -0.064208819 | 0.07272115 | 0.4903372 | 0.923615538 | 0.964181847 |
| Wnk2 | -0.024963343 | 0.243993524 | 0.7132719 | 0.087509619 | 0.097767773 |
| Wnk3 | -0.095476484 | 0.024374307 | 0.3029989 | 0.503328997 | 0.489876807 |
| Wnk4 | -0.022796027 | 0.459159643 | 0.8522649 | 0.271666007 | 0.253129059 |
| Wnt4 | -0.097315161 | 0.003896211 | 0.1219255 | 1.872451137 | 1.937664684 |
| Wrap53 | -0.015856545 | 0.336728557 | 0.7826493 | 0.049795683 | 0.05878144 |
| Wrap73 | 0.001104695 | 0.95742237 | 0.9956630 | 0.10613711 | 0.094487305 |
| Wrn | -0.029654457 | 0.207824512 | 0.6807452 | 0.120475698 | 0.130203853 |
| Wrnip1 | -0.011324158 | 0.673058913 | 0.9260917 | 0.176738707 | 0.175838949 |
| Wsb1 | 0.007611819 | 0.686785236 | 0.9323812 | 0.084450771 | 0.071826665 |
| Wsb2 | -0.00939349 | 0.789597111 | 0.9618491 | 0.588294114 | 0.57600576 |
| Wtap | 0.010585751 | 0.735899361 | 0.9468380 | 0.286204749 | 0.258174829 |
| Wtip | -0.042304618 | 0.059162132 | 0.4538966 | 0.11633266 | 0.119696766 |
| Wwc1 | -0.030640735 | 0.176616274 | 0.6536659 | 0.113272746 | 0.111342156 |
| Wwox | -0.014043995 | 0.526061222 | 0.8768553 | 0.102479213 | 0.111824034 |
| Wwp1 | 0.016169196 | 0.440993261 | 0.8424484 | 0.115736402 | 0.094701495 |
| Wwp2 | -0.022184284 | 0.324178328 | 0.7757552 | 0.102604273 | 0.122414226 |
| Xab2 | 0.000961565 | 0.967601067 | 0.9976072 | 0.126632117 | 0.120286476 |
| Xbp1 | -0.07471128 | 0.024725886 | 0.3041540 | 1.177263278 | 1.23529135 |
| Xiap | -0.006523815 | 0.822191291 | 0.9706920 | 0.221899966 | 0.21290933 |
| Xk | -0.001540157 | 0.919459964 | 0.9853630 | 0.045019319 | 0.051536899 |
| Xkr4 | 0.000329991 | 0.982368977 | 0.9976072 | 0.049941116 | 0.042215254 |
| Xkr6 | 0.003158 | 0.906081805 | 0.9835136 | 0.181100403 | 0.174839604 |
| Xlr3a | -0.004257071 | 0.784601654 | 0.9601202 | 0.048961038 | 0.043446679 |
| Xpa | -0.021791041 | 0.48091653 | 0.8617579 | 0.257200988 | 0.255735393 |
| Xpc | -0.041245912 | 0.085070153 | 0.5209234 | 0.093970348 | 0.14121189 |
| Xpnpep1 | -0.023773329 | 0.271010491 | 0.7342030 | 0.094578247 | 0.112334005 |
| Xpnpep3 | -0.015829826 | 0.444707026 | 0.8431541 | 0.087001879 | 0.09533864 |
| Xpo1 | -0.02255088 | 0.426777234 | 0.8337565 | 0.206331793 | 0.210756036 |
| Xpo4 | 0.062178078 | 0.019822382 | 0.2724376 | 0.204642833 | 0.153294352 |
| Xpo5 | 0.001643526 | 0.943287914 | 0.9935293 | 0.110405188 | 0.118620799 |
| Xpo6 | -0.018425902 | 0.517133663 | 0.8738172 | 0.19049417 | 0.204631107 |
| Xpo7 | -0.009425435 | 0.768255117 | 0.9544258 | 0.303032206 | 0.2924033 |
| Xpot | 0.045463945 | 0.171391973 | 0.6511713 | 0.337888684 | 0.333514006 |
| Xpr1 | 0.031778294 | 0.246462301 | 0.7161411 | 0.199884223 | 0.178469525 |
| Xrcc1 | 0.03205308 | 0.174867682 | 0.6534142 | 0.151350652 | 0.122391247 |
| Xrcc4 | -0.001308549 | 0.952259981 | 0.9951566 | 0.120497156 | 0.108289512 |
| Xrcc5 | -0.024075757 | 0.388923186 | 0.8170287 | 0.185252466 | 0.197827123 |
| Xrcc6 | -0.030506718 | 0.265136199 | 0.7306700 | 0.183096333 | 0.187907117 |
| Xrn1 | 0.003290368 | 0.925092412 | 0.9879210 | 0.387054887 | 0.356570684 |
| Xrn2 | -0.02045443 | 0.554396054 | 0.8902995 | 0.350696502 | 0.366168759 |
| Yae1d1 | 0.013037645 | 0.663410434 | 0.9230058 | 0.252260653 | 0.232035165 |
| Yaf2 | -0.000753481 | 0.982981943 | 0.9976072 | 0.483787064 | 0.452688758 |
| Yars | 0.07338408 | 0.0279741 | 0.3208127 | 0.39113411 | 0.313871215 |
| Yars2 | 0.02373321 | 0.289130584 | 0.7494616 | 0.134148272 | 0.105990854 |
| Ybey | -0.004586566 | 0.787388752 | 0.9608311 | 0.057588784 | 0.065078096 |
| Ybx1 | -0.029726238 | 0.222857472 | 0.6955958 | 2.153280954 | 2.063574154 |
| Ybx3 | -0.010968795 | 0.767565063 | 0.9544221 | 0.551351321 | 0.557176188 |
| Ydjc | -0.035484219 | 0.078538933 | 0.5056175 | 0.065300363 | 0.10187034 |
| Yeats2 | -0.007791693 | 0.732279198 | 0.9462041 | 0.115683652 | 0.110344413 |
| Yeats4 | 0.052797914 | 0.121693824 | 0.5819916 | 0.436636979 | 0.384101818 |
| Yes1 | -0.008369365 | 0.723949015 | 0.9445652 | 0.118984594 | 0.131775048 |
| Yif1a | 0.00472406 | 0.896393875 | 0.9825441 | 0.646718049 | 0.586942505 |
| Yif1b | 0.031202898 | 0.343319153 | 0.7872955 | 0.358380335 | 0.309124781 |
| Yipf1 | 0.039180561 | 0.226916446 | 0.7003768 | 0.373791709 | 0.348866016 |
| Yipf2 | -0.000352744 | 0.990293612 | 0.9980836 | 0.229236599 | 0.2262587 |
| Yipf3 | -0.047027744 | 0.185759363 | 0.6638086 | 0.517900289 | 0.540139492 |

| | | | | | |
|---|---|---|---|---|---|
| Yipf4 | -0.031843859 | 0.373270795 | 0.80856898 | 0.634361144 | 0.6055388 |
| Yipf5 | 0.083611995 | 0.020987354 | 0.28117593 | 0.649948638 | 0.537706582 |
| Yipf6 | 0.039766692 | 0.226459829 | 0.69962052 | 0.332472872 | 0.295425084 |
| Yju2 | 0.02342058 | 0.26149963 | 0.72757354 | 0.117137551 | 0.087678465 |
| Ykt6 | -0.002055617 | 0.941461074 | 0.99293011 | 0.195631833 | 0.193221549 |
| Ylpm1 | -0.03941114 | 0.221052064 | 0.69262049 | 0.297895304 | 0.326991935 |
| Yme1l1 | 0.023706195 | 0.498210488 | 0.86846292 | 0.464194428 | 0.412103933 |
| Yod1 | 0.028939381 | 0.251925508 | 0.72207484 | 0.181721115 | 0.14256214 |
| Ypel2 | -0.099013878 | 0.005817891 | 0.15349329 | 0.309761228 | 0.394694138 |
| Ypel3 | -0.045082841 | 0.213370542 | 0.68300605 | 0.978241678 | 0.928964597 |
| Ypel4 | 0.00881655 | 0.732465829 | 0.94620413 | 0.169681926 | 0.149579267 |
| Ypel5 | 0.044321057 | 0.179764571 | 0.65500705 | 0.372848607 | 0.318556454 |
| Yrdc | -0.039688823 | 0.248511023 | 0.71851627 | 0.384257084 | 0.403431807 |
| Ythdc1 | 0.079416731 | 0.03095645 | 0.33598247 | 0.751294039 | 0.663968928 |
| Ythdc2 | -0.050556228 | 0.043190394 | 0.39766629 | 0.119023369 | 0.148975438 |
| Ythdf1 | 0.023591683 | 0.45104211 | 0.84789420 | 0.299753184 | 0.253992703 |
| Ythdf2 | 0.035511114 | 0.25501286 | 0.72236652 | 0.28680604 | 0.250019343 |
| Ythdf3 | -0.025376226 | 0.409977234 | 0.82813564 | 0.255103614 | 0.271535525 |
| Ywhab | 0.040485013 | 0.265793417 | 0.73067832 | 0.785563386 | 0.72196404 |
| Ywhae | 0.015969467 | 0.542920317 | 0.88406828 | 1.808143138 | 1.685953888 |
| Ywhag | 0.048198307 | 0.180002902 | 0.65503624 | 0.536579477 | 0.522946686 |
| Ywhah | -0.0160821 | 0.664573035 | 0.92307657 | 0.800818162 | 0.77782747 |
| Ywhaq | 0.018748614 | 0.414150437 | 0.83009907 | 1.882384376 | 1.802148858 |
| Ywhaz | 0.052728731 | 0.150994924 | 0.62803758 | 0.783199576 | 0.824393678 |
| Yy1 | -0.030811742 | 0.406977491 | 0.82707700 | 0.712298006 | 0.718853116 |
| Zadh2 | 0.013384072 | 0.541961744 | 0.88367200 | 0.114935838 | 0.106037645 |
| Zbed3 | -0.027720665 | 0.244792129 | 0.71379715 | 0.126561387 | 0.137682055 |
| Zbtb1 | -0.048864082 | 0.05122923 | 0.42705374 | 0.116179005 | 0.153250914 |
| Zbtb10 | -0.000400774 | 0.985518269 | 0.99760728 | 0.103255148 | 0.107060956 |
| Zbtb11 | 0.058708366 | 0.061586906 | 0.46137448 | 0.311767576 | 0.241100773 |
| Zbtb11os1 | -0.051332344 | 0.014186582 | 0.23475005 | 0.067503597 | 0.109614175 |
| Zbtb12 | -0.035264961 | 0.089318246 | 0.53023604 | 0.077471066 | 0.098043356 |
| Zbtb14 | -0.022614151 | 0.36062222 | 0.80124805 | 0.131960602 | 0.145302263 |
| Zbtb17 | -0.017745941 | 0.285327174 | 0.74518287 | 0.049986038 | 0.0652042 |
| Zbtb18 | 0.004688502 | 0.809286297 | 0.96607943 | 0.07689764 | 0.08587831 |
| Zbtb2 | -0.030297947 | 0.092519292 | 0.53255842 | 0.047787506 | 0.075418396 |
| Zbtb20 | -0.05269665 | 0.129914611 | 0.59362180 | 1.502897268 | 1.58316228 |
| Zbtb21 | -0.006236874 | 0.803937988 | 0.96471431 | 0.142675161 | 0.142783137 |
| Zbtb22 | -0.025749958 | 0.201710645 | 0.67521501 | 0.079781185 | 0.091120543 |
| Zbtb24 | 0.023054646 | 0.31926781 | 0.77100438 | 0.147696218 | 0.118527469 |
| Zbtb25 | -0.017388522 | 0.300530922 | 0.75955840 | 0.052495885 | 0.063980962 |
| Zbtb33 | -0.007500116 | 0.724295166 | 0.94456522 | 0.103938314 | 0.101693166 |
| Zbtb37 | -0.010756743 | 0.568373178 | 0.89553080 | 0.065121502 | 0.074311096 |
| Zbtb38 | -0.041529005 | 0.234107861 | 0.70569610 | 0.354235281 | 0.379963732 |
| Zbtb39 | 0.011981799 | 0.387896866 | 0.81702870 | 0.046958572 | 0.038093064 |
| Zbtb4 | -0.029876296 | 0.398115308 | 0.82384692 | 0.417123318 | 0.42880076 |
| Zbtb40 | 0.00621562 | 0.694426973 | 0.93619580 | 0.057700443 | 0.050783486 |
| Zbtb41 | -0.00872954 | 0.73108557 | 0.94590205 | 0.143841978 | 0.159272326 |
| Zbtb43 | 0.016034186 | 0.457757648 | 0.85200834 | 0.115662137 | 0.097009716 |
| Zbtb44 | -0.005631858 | 0.847951045 | 0.97443578 | 0.220290189 | 0.216461137 |
| Zbtb45 | -0.005284115 | 0.729011116 | 0.94590205 | 0.049755184 | 0.050523878 |
| Zbtb48 | 0.01832719 | 0.242336864 | 0.71327199 | 0.063176985 | 0.045066988 |
| Zbtb5 | 0.00211424 | 0.912268697 | 0.98403934 | 0.07557324 | 0.076269027 |
| Zbtb6 | -0.006242033 | 0.803358187 | 0.96471431 | 0.150447661 | 0.141506467 |
| Zbtb7a | -0.082198772 | 0.027202637 | 0.31672554 | 0.717935159 | 0.797350467 |
| Zbtb7b | -0.01850917 | 0.445842654 | 0.84373680 | 0.116526382 | 0.151323191 |
| Zbtb7c | -0.017352036 | 0.370950736 | 0.80715755 | 0.075840288 | 0.081905259 |
| Zbtb8a | 0.005078413 | 0.729163658 | 0.94590205 | 0.054970301 | 0.043690268 |
| Zbtb8os | -0.001627884 | 0.959270174 | 0.99577853 | 0.318756321 | 0.313229713 |
| Zc2hc1a | 0.002563417 | 0.914802097 | 0.98452361 | 0.148413123 | 0.131064464 |
| Zc2hc1c | 0.009627684 | 0.605743136 | 0.90871392 | 0.083599731 | 0.069539738 |
| Zc3h10 | -0.008888735 | 0.746106418 | 0.94861170 | 0.169713399 | 0.180099486 |
| Zc3h12b | -0.019984356 | 0.183459339 | 0.66159407 | 0.039720946 | 0.051572085 |
| Zc3h12c | -0.034763304 | 0.106480995 | 0.56215618 | 0.08070037 | 0.111883678 |
| Zc3h13 | 0.005078081 | 0.887623006 | 0.98124141 | 0.415383282 | 0.394452435 |
| Zc3h14 | -0.07202173 | 0.045437798 | 0.40352995 | 0.491618252 | 0.531666977 |

| | | | | | |
|---|---|---|---|---|---|
| Zc3h15 | 0.058462084 | 0.091289945 | 0.5321282 | 0.887357858 | 0.81830097 |
| Zc3h18 | 0.040925124 | 0.195897669 | 0.6727425 | 0.303209593 | 0.246838878 |
| Zc3h3 | 0.027446448 | 0.268073289 | 0.7323841 | 0.130932468 | 0.123286005 |
| Zc3h4 | -0.001640042 | 0.941335593 | 0.9929301 | 0.107278207 | 0.108880769 |
| Zc3h6 | -0.023588597 | 0.422998731 | 0.8315789 | 0.214824977 | 0.22306708 |
| Zc3h7a | -0.041085266 | 0.213072896 | 0.6827572 | 0.290923328 | 0.321695493 |
| Zc3h7b | 0.009732773 | 0.73194436 | 0.9461673 | 0.221308444 | 0.209718817 |
| Zc3h8 | -0.028525841 | 0.146967269 | 0.6232627 | 0.065221454 | 0.088266213 |
| Zc3hav1 | -0.025407008 | 0.432988515 | 0.8374897 | 0.305802802 | 0.316690023 |
| Zcchc10 | -0.031764518 | 0.144529173 | 0.6193215 | 0.087992346 | 0.106287574 |
| Zcchc12 | -0.019683063 | 0.50197115 | 0.8695566 | 0.150995205 | 0.190890592 |
| Zcchc14 | 0.034437364 | 0.2070421 | 0.6797565 | 0.205302548 | 0.172988046 |
| Zcchc17 | -0.010521566 | 0.762944418 | 0.9532000 | 0.461009927 | 0.451118976 |
| Zcchc18 | 0.032442292 | 0.399172954 | 0.8239850 | 1.210072388 | 1.018748925 |
| Zcchc2 | -0.038662521 | 0.113949034 | 0.5696977 | 0.111945363 | 0.147424534 |
| Zcchc3 | 0.05249389 | 0.066153458 | 0.4772671 | 0.253216219 | 0.193932314 |
| Zcchc7 | -0.057058945 | 0.131963588 | 0.5978345 | 0.458922005 | 0.514159413 |
| Zcchc8 | 0.005314671 | 0.834719121 | 0.9723494 | 0.159086024 | 0.152179111 |
| Zcchc9 | 0.008408217 | 0.785826422 | 0.9602516 | 0.268585194 | 0.273246616 |
| Zcrb1 | 0.02612718 | 0.367592963 | 0.8056719 | 1.551376712 | 1.424403435 |
| Zdbf2 | -0.044497914 | 0.281079619 | 0.7408618 | 0.725753023 | 0.730623536 |
| Zdhhc1 | 0.028335128 | 0.209577523 | 0.6807660 | 0.140737002 | 0.106559399 |
| Zdhhc12 | -0.004461871 | 0.846625513 | 0.9739657 | 0.126414987 | 0.124698568 |
| Zdhhc13 | -0.006960866 | 0.667623687 | 0.9230765 | 0.053394666 | 0.054095914 |
| Zdhhc14 | 0.001755783 | 0.929021411 | 0.9881953 | 0.084056456 | 0.082378792 |
| Zdhhc16 | 0.032215812 | 0.108409415 | 0.5636817 | 0.117417523 | 0.078885632 |
| Zdhhc17 | -0.058948943 | 0.018785199 | 0.2677936 | 0.118024858 | 0.159206901 |
| Zdhhc18 | -0.012110334 | 0.467583621 | 0.8558546 | 0.057373235 | 0.060063697 |
| Zdhhc2 | 0.054600717 | 0.14832049 | 0.6255694 | 0.577555771 | 0.616881925 |
| Zdhhc20 | -0.002662206 | 0.933267912 | 0.9902682 | 0.270192164 | 0.268958241 |
| Zdhhc21 | -0.026591819 | 0.243419348 | 0.7132719 | 0.098605351 | 0.119280683 |
| Zdhhc23 | -0.004917415 | 0.784232357 | 0.9601202 | 0.057864545 | 0.07197786 |
| Zdhhc24 | 0.002220888 | 0.874873799 | 0.9784331 | 0.044696099 | 0.042644025 |
| Zdhhc3 | -0.005191096 | 0.867396217 | 0.9773478 | 0.258399179 | 0.276673689 |
| Zdhhc4 | 0.003118864 | 0.896169892 | 0.9825441 | 0.144796734 | 0.123328611 |
| Zdhhc5 | 0.015580305 | 0.566233286 | 0.8947196 | 0.194712455 | 0.185173935 |
| Zdhhc6 | -0.005564705 | 0.785941745 | 0.9602516 | 0.092580963 | 0.093496573 |
| Zdhhc7 | 0.004037252 | 0.826300729 | 0.9716906 | 0.07461241 | 0.072216014 |
| Zdhhc8 | -0.017378236 | 0.514238205 | 0.8733459 | 0.171287261 | 0.193155293 |
| Zeb1 | 0.002613653 | 0.924395613 | 0.9878036 | 0.17979565 | 0.175758323 |
| Zer1 | 0.019774014 | 0.442324303 | 0.8431541 | 0.167476146 | 0.148975199 |
| Zfand1 | -0.00786531 | 0.743362845 | 0.9479080 | 0.148573952 | 0.13590347 |
| Zfand2a | -0.005666507 | 0.726787726 | 0.9452179 | 0.051753852 | 0.056285701 |
| Zfand2b | -0.051876477 | 0.082662606 | 0.5165689 | 0.211415278 | 0.260219699 |
| Zfand3 | 0.019810076 | 0.531160425 | 0.8776448 | 0.304976584 | 0.281253668 |
| Zfand5 | -0.103589666 | 0.005704351 | 0.1523891 | 0.702802746 | 0.78174632 |
| Zfand6 | -0.009441759 | 0.793797037 | 0.9624508 | 0.693563028 | 0.650088763 |
| Zfas1 | -0.074595677 | 0.056521921 | 0.4471389 | 0.513170496 | 0.61686452 |
| Zfc3h1 | -0.028571026 | 0.283201363 | 0.7427848 | 0.146594989 | 0.163423232 |
| Zfhx2 | 0.049239244 | 0.182974412 | 0.6607594 | 0.741159068 | 0.776693717 |
| Zfhx3 | -0.020093022 | 0.530752044 | 0.8776448 | 0.258542877 | 0.29337107 |
| Zfp1 | -0.015316851 | 0.573598092 | 0.8973494 | 0.191571207 | 0.180038856 |
| Zfp101 | 0.015028483 | 0.255951486 | 0.7235790 | 0.04528359 | 0.030060278 |
| Zfp106 | 0.027333223 | 0.451033771 | 0.8478942 | 0.539752768 | 0.532662195 |
| Zfp110 | -0.020841483 | 0.468964179 | 0.8563037 | 0.211598334 | 0.214908877 |
| Zfp111 | -0.011584603 | 0.436132722 | 0.8390644 | 0.041640845 | 0.049430431 |
| Zfp113 | 0.031667268 | 0.166371518 | 0.6429285 | 0.142728717 | 0.106235895 |
| Zfp119a | -0.014727328 | 0.391821698 | 0.8200150 | 0.068136539 | 0.063663609 |
| Zfp119b | -0.008047432 | 0.599912536 | 0.9078234 | 0.051492977 | 0.047500314 |
| Zfp12 | -0.002197818 | 0.898328518 | 0.9825441 | 0.067639864 | 0.06278889 |
| Zfp120 | -0.034627564 | 0.176518355 | 0.6536659 | 0.160741502 | 0.163017935 |
| Zfp125 | -0.007275222 | 0.637558008 | 0.9160042 | 0.050942878 | 0.048163658 |
| Zfp131 | 0.021375645 | 0.511754905 | 0.8720873 | 0.321139319 | 0.307732334 |
| Zfp142 | 0.024062851 | 0.251108956 | 0.7207210 | 0.106755922 | 0.092674105 |
| Zfp143 | -0.018120533 | 0.330459869 | 0.7779360 | 0.071208233 | 0.077329064 |
| Zfp146 | -0.057439771 | 0.01355101 | 0.2292511 | 0.094897043 | 0.129588055 |

| | | | | | |
|---|---|---|---|---|---|
| Zfp148 | -0.032449932 | 0.381618935 | 0.8124164 | 0.599311817 | 0.613163602 |
| Zfp157 | -0.014639964 | 0.556938462 | 0.8915944 | 0.148696029 | 0.142708593 |
| Zfp160 | -0.012275154 | 0.541451147 | 0.8835017 | 0.079003487 | 0.08997149 |
| Zfp180 | -0.021714777 | 0.228358655 | 0.7015153 | 0.055284741 | 0.077544431 |
| Zfp182 | 0.005631474 | 0.756128978 | 0.9521533 | 0.08275496 | 0.069962574 |
| Zfp2 | -0.00211608 | 0.898365165 | 0.9825441 | 0.055173299 | 0.05719152 |
| Zfp207 | -0.01373014 | 0.688343523 | 0.9329669 | 0.373117348 | 0.380616395 |
| Zfp212 | -0.008381263 | 0.674074623 | 0.9264090 | 0.08099544 | 0.086836415 |
| Zfp213 | -0.002811075 | 0.852356197 | 0.9755431 | 0.044851322 | 0.048993224 |
| Zfp217 | -0.004064629 | 0.826389745 | 0.9716906 | 0.078197581 | 0.072372431 |
| Zfp219 | -0.047688855 | 0.039679746 | 0.3821583 | 0.10224853 | 0.129211938 |
| Zfp235 | -0.00096095 | 0.954761276 | 0.9954690 | 0.06075925 | 0.059393186 |
| Zfp236 | -0.029298503 | 0.190045972 | 0.6699489 | 0.096642137 | 0.113583358 |
| Zfp24 | 0.027893962 | 0.384879533 | 0.8139000 | 0.322946781 | 0.279903483 |
| Zfp251 | -0.02445461 | 0.32500936 | 0.7763099 | 0.12889205 | 0.152645722 |
| Zfp26 | -0.016925904 | 0.436448829 | 0.8390644 | 0.099266703 | 0.111671878 |
| Zfp260 | -0.012128187 | 0.713920769 | 0.9428150 | 0.33897744 | 0.315406878 |
| Zfp263 | -0.003516374 | 0.883759912 | 0.9801988 | 0.136905979 | 0.12903629 |
| Zfp266 | -0.087207401 | 0.012668735 | 0.2227646 | 0.35321971 | 0.39688347 |
| Zfp267 | -0.032348445 | 0.095890175 | 0.5368118 | 0.06499995 | 0.08853842 |
| Zfp275 | -0.020725361 | 0.33620598 | 0.7821677 | 0.093931275 | 0.11110186 |
| Zfp277 | 0.004649462 | 0.830529882 | 0.9722601 | 0.117088513 | 0.105031183 |
| Zfp280b | 0.003169127 | 0.842493196 | 0.9734889 | 0.055728224 | 0.051588407 |
| Zfp280c | 0.0036906 | 0.822150983 | 0.9706920 | 0.059779105 | 0.053230069 |
| Zfp280d | -0.046281956 | 0.179711768 | 0.6550070 | 0.34361102 | 0.367724137 |
| Zfp281 | -0.025800843 | 0.266052756 | 0.7307399 | 0.117810413 | 0.127387263 |
| Zfp282 | 0.019254421 | 0.350217386 | 0.7927303 | 0.097101923 | 0.089311452 |
| Zfp287 | -0.016040724 | 0.342672993 | 0.7872431 | 0.052404767 | 0.059970003 |
| Zfp292 | -0.071321098 | 0.055044226 | 0.4429440 | 0.566925883 | 0.600347475 |
| Zfp30 | -0.016274493 | 0.498352813 | 0.8684629 | 0.124554699 | 0.13317589 |
| Zfp316 | -0.000548481 | 0.972525656 | 0.9976072 | 0.055597032 | 0.0513465 |
| Zfp317 | 0.006741991 | 0.71111361 | 0.9419834 | 0.080896158 | 0.067700719 |
| Zfp318 | 0.022745215 | 0.422562407 | 0.8314897 | 0.220924952 | 0.200159119 |
| Zfp319 | -0.005424778 | 0.751393639 | 0.9506053 | 0.059322036 | 0.064803786 |
| Zfp322a | -0.004161855 | 0.884723086 | 0.9803788 | 0.212052616 | 0.194592802 |
| Zfp324 | -0.030797563 | 0.06171814 | 0.4613744 | 0.044329674 | 0.060465701 |
| Zfp326 | -0.046150774 | 0.174136676 | 0.6527657 | 0.360377562 | 0.367788253 |
| Zfp329 | -0.068368035 | 0.042353276 | 0.3952910 | 0.29985029 | 0.338062657 |
| Zfp330 | -0.040107577 | 0.225631844 | 0.6979011 | 0.337578061 | 0.374463063 |
| Zfp334 | -0.02095814 | 0.16763847 | 0.6446304 | 0.035203947 | 0.051584272 |
| Zfp335 | 0.013856112 | 0.487659828 | 0.8640311 | 0.09787658 | 0.081354119 |
| Zfp346 | 0.01196585 | 0.46886695 | 0.8563037 | 0.063785033 | 0.05527593 |
| Zfp35 | 0.005362447 | 0.750032173 | 0.9504537 | 0.060294393 | 0.054048456 |
| Zfp354c | 0.030409368 | 0.02606851 | 0.3097001 | 0.062058509 | 0.028938637 |
| Zfp358 | -0.041845098 | 0.086697069 | 0.5250566 | 0.141713354 | 0.155696934 |
| Zfp36 | -0.04189856 | 0.319158725 | 0.7710043 | 0.558577787 | 0.591022047 |
| Zfp362 | 0.024214006 | 0.152828682 | 0.6286214 | 0.071939086 | 0.060016536 |
| Zfp365 | 0.01634035 | 0.280076868 | 0.7408618 | 0.059082319 | 0.038202091 |
| Zfp367 | 0.011508052 | 0.460609314 | 0.8524425 | 0.061780239 | 0.048446696 |
| Zfp369 | 0.004709667 | 0.765430007 | 0.9540318 | 0.057226728 | 0.050737485 |
| Zfp36l2 | -0.092387446 | 0.008148418 | 0.1775389 | 0.287336435 | 0.390607927 |
| Zfp37 | 0.012659838 | 0.551871851 | 0.8893234 | 0.11402071 | 0.094515415 |
| Zfp384 | -0.003293893 | 0.855425536 | 0.9757403 | 0.069684025 | 0.069979478 |
| Zfp386 | 0.03548441 | 0.181948908 | 0.6592159 | 0.206926269 | 0.16022859 |
| Zfp39 | -0.005832391 | 0.714123303 | 0.9428174 | 0.050734377 | 0.052584572 |
| Zfp395 | -0.066900266 | 0.031531879 | 0.3385228 | 0.246906898 | 0.274343277 |
| Zfp397 | -0.031250158 | 0.311924089 | 0.7684379 | 0.261041966 | 0.262889229 |
| Zfp398 | -0.01168561 | 0.672520048 | 0.9258663 | 0.183206309 | 0.187028573 |
| Zfp40 | 0.004817211 | 0.76543134 | 0.9540318 | 0.055945945 | 0.05439887 |
| Zfp407 | -0.027175701 | 0.198824141 | 0.6730962 | 0.080770633 | 0.101086365 |
| Zfp408 | 0.011065241 | 0.4364065 | 0.8390644 | 0.050263971 | 0.040244127 |
| Zfp410 | -0.035680177 | 0.162445744 | 0.6400703 | 0.13907599 | 0.157782702 |
| Zfp414 | -0.007506509 | 0.668295521 | 0.9230765 | 0.069143512 | 0.065062866 |
| Zfp420 | 0.010596819 | 0.46331226 | 0.8524425 | 0.049280493 | 0.038433203 |
| Zfp422 | -0.003605183 | 0.895145389 | 0.9824361 | 0.199872075 | 0.189680216 |
| Zfp426 | -0.0191701 | 0.435697618 | 0.8389295 | 0.128613097 | 0.139729488 |

| | | | | | |
|---|---|---|---|---|---|
| Zfp428 | -0.021859505 | 0.230600172 | 0.7022964 | 0.066901415 | 0.072123665 |
| Zfp442 | -0.015832701 | 0.407042617 | 0.8270770 | 0.082068421 | 0.078211747 |
| Zfp444 | -0.000599529 | 0.980221439 | 0.9976072 | 0.148313622 | 0.13142607 |
| Zfp445 | -0.006348898 | 0.850578813 | 0.9748841 | 0.332014307 | 0.345283266 |
| Zfp449 | -0.04242122 | 0.007759703 | 0.1737612 | 0.03495893 | 0.060546988 |
| Zfp451 | -0.012943088 | 0.693656465 | 0.9359725 | 0.301248321 | 0.301354576 |
| Zfp458 | 0.005696966 | 0.722708634 | 0.9445652 | 0.062112616 | 0.048769073 |
| Zfp462 | 0.006106806 | 0.82350551 | 0.9710592 | 0.170042941 | 0.168512298 |
| Zfp467 | -0.000476243 | 0.987010409 | 0.9976072 | 0.213583995 | 0.202269158 |
| Zfp507 | -0.028472577 | 0.22881257 | 0.7019677 | 0.109422995 | 0.128705489 |
| Zfp51 | -0.014663744 | 0.354093333 | 0.7967782 | 0.055749783 | 0.053747729 |
| Zfp511 | -0.035679186 | 0.227730034 | 0.7015153 | 0.232327321 | 0.240047975 |
| Zfp512 | 0.022101481 | 0.365488872 | 0.8050191 | 0.165158692 | 0.131271297 |
| Zfp512b | -0.050783763 | 0.03225656 | 0.3393267 | 0.102309411 | 0.140109878 |
| Zfp513 | -0.009873015 | 0.649962303 | 0.9175779 | 0.106040647 | 0.103913266 |
| Zfp516 | -0.029443489 | 0.429694009 | 0.8340365 | 0.637666842 | 0.642405464 |
| Zfp518a | -0.022369394 | 0.294853632 | 0.7536588 | 0.088740458 | 0.102211168 |
| Zfp52 | -0.005236288 | 0.767792362 | 0.9544221 | 0.068381704 | 0.067204019 |
| Zfp523 | -0.017921852 | 0.500834954 | 0.8693704 | 0.183613478 | 0.167994528 |
| Zfp524 | -0.005222054 | 0.802975368 | 0.9647143 | 0.100646751 | 0.100348744 |
| Zfp53 | 0.015074896 | 0.324691112 | 0.7757552 | 0.067037855 | 0.042741063 |
| Zfp532 | 0.009011838 | 0.582185214 | 0.9018999 | 0.062963467 | 0.053820011 |
| Zfp553 | 0.039215454 | 0.1030889 | 0.5553141 | 0.167925165 | 0.125956939 |
| Zfp560 | -0.030111209 | 0.363652275 | 0.8046911 | 0.396930897 | 0.431042392 |
| Zfp563 | -0.018301256 | 0.241502578 | 0.7130076 | 0.039366909 | 0.055462279 |
| Zfp568 | 0.047047397 | 0.155891541 | 0.6312943 | 0.343136516 | 0.272791793 |
| Zfp574 | -0.009414539 | 0.600905935 | 0.9078437 | 0.061618263 | 0.073341575 |
| Zfp579 | 0.016610864 | 0.427372145 | 0.8340365 | 0.111090581 | 0.090618905 |
| Zfp580 | 0.014567584 | 0.417644866 | 0.8300990 | 0.088675941 | 0.061688282 |
| Zfp592 | -0.020676547 | 0.333578649 | 0.7804836 | 0.091262969 | 0.108865265 |
| Zfp593 | -0.006622593 | 0.814781952 | 0.9685114 | 0.193845672 | 0.197017205 |
| Zfp597 | 0.005983905 | 0.771571196 | 0.9552333 | 0.108951296 | 0.093996399 |
| Zfp598 | 0.00311743 | 0.904117226 | 0.9829946 | 0.156909535 | 0.161216122 |
| Zfp60 | -0.026067255 | 0.245550792 | 0.7151684 | 0.105157612 | 0.123659209 |
| Zfp605 | -0.027545865 | 0.213529655 | 0.6830745 | 0.104839926 | 0.115154612 |
| Zfp606 | 0.041944785 | 0.029199291 | 0.3272960 | 0.110050398 | 0.072055309 |
| Zfp608 | -0.007042772 | 0.820413118 | 0.9701392 | 0.239971134 | 0.25047783 |
| Zfp609 | -0.020568995 | 0.355525932 | 0.7969131 | 0.09822548 | 0.114450362 |
| Zfp61 | 0.024868361 | 0.177999083 | 0.6538489 | 0.102273523 | 0.066238069 |
| Zfp612 | -0.024295911 | 0.5079602 | 0.8704067 | 0.437619768 | 0.443463582 |
| Zfp617 | -0.013792689 | 0.487288076 | 0.8640311 | 0.084399356 | 0.092901267 |
| Zfp618 | -0.016918605 | 0.512151133 | 0.8721955 | 0.154333557 | 0.153384042 |
| Zfp62 | 0.015397794 | 0.560332139 | 0.8925646 | 0.173786526 | 0.147558529 |
| Zfp622 | -0.062311912 | 0.060480969 | 0.4593891 | 0.303169272 | 0.369559815 |
| Zfp626 | -0.009724546 | 0.54436686 | 0.8842656 | 0.050867856 | 0.05506209 |
| Zfp628 | -0.021855448 | 0.20151782 | 0.6752150 | 0.046600279 | 0.062909494 |
| Zfp629 | -0.017435389 | 0.252780589 | 0.7223071 | 0.034212218 | 0.052812779 |
| Zfp637 | 0.005622311 | 0.842777967 | 0.9734889 | 0.224347494 | 0.199511757 |
| Zfp638 | 0.008697402 | 0.817154046 | 0.9693806 | 0.539602431 | 0.516516632 |
| Zfp639 | 0.004938053 | 0.872684872 | 0.9784331 | 0.25289273 | 0.266350087 |
| Zfp64 | -0.034389755 | 0.078544023 | 0.5056175 | 0.063868896 | 0.089353573 |
| Zfp641 | 0.018161672 | 0.282963125 | 0.7427848 | 0.071852477 | 0.052983833 |
| Zfp644 | -0.040994517 | 0.280913552 | 0.7408618 | 0.53596996 | 0.560078152 |
| Zfp646 | 0.035138374 | 0.172524934 | 0.6522284 | 0.176790325 | 0.141084808 |
| Zfp65 | 0.001068825 | 0.95077646 | 0.9948532 | 0.062317979 | 0.065022919 |
| Zfp651 | 0.025309479 | 0.316545257 | 0.7705987 | 0.168590981 | 0.133709017 |
| Zfp652 | -0.030755611 | 0.377773894 | 0.8101797 | 0.382821069 | 0.396567084 |
| Zfp653 | -0.018207655 | 0.219818102 | 0.6910237 | 0.039636936 | 0.048278035 |
| Zfp654 | -0.016999059 | 0.461394531 | 0.8524425 | 0.10014381 | 0.122574525 |
| Zfp655 | 0.000604777 | 0.979478884 | 0.9976072 | 0.128371133 | 0.126602192 |
| Zfp664 | -0.011208546 | 0.753699912 | 0.9517823 | 0.46463403 | 0.473057844 |
| Zfp667 | -0.003753229 | 0.900263984 | 0.9825441 | 0.224090276 | 0.23213735 |
| Zfp668 | 0.017622246 | 0.19528591 | 0.6719932 | 0.048573865 | 0.032256911 |
| Zfp672 | 0.032754302 | 0.229209226 | 0.7019677 | 0.201831954 | 0.174428873 |
| Zfp68 | -0.0033069 | 0.904676798 | 0.9831707 | 0.198077575 | 0.189445119 |
| Zfp687 | -0.019545448 | 0.416055114 | 0.8300990 | 0.120003983 | 0.136614397 |

| | | | | | |
|---|---|---|---|---|---|
| Zfp688 | -0.030377838 | 0.271841253 | 0.7342806 | 0.191320641 | 0.188989466 |
| Zfp692 | -0.00176491 | 0.908077969 | 0.9837373 | 0.049707618 | 0.047115969 |
| Zfp704 | -0.050061503 | 0.181212538 | 0.6577068 | 0.46948648 | 0.49755444 |
| Zfp706 | 0.013113316 | 0.71472139 | 0.9430747 | 0.882277052 | 0.827226769 |
| Zfp707 | 0.007853172 | 0.695411418 | 0.9362759 | 0.097539355 | 0.084824093 |
| Zfp709 | -0.046943894 | 0.010873605 | 0.2035210 | 0.04582669 | 0.08183797 |
| Zfp710 | 0.014170788 | 0.513110917 | 0.8726316 | 0.120961649 | 0.101210684 |
| Zfp715 | 0.002042679 | 0.891395197 | 0.9819597 | 0.048835987 | 0.045985382 |
| Zfp719 | -0.014207616 | 0.474952299 | 0.8602394 | 0.08359952 | 0.090457376 |
| Zfp729a | -0.035708067 | 0.067655796 | 0.4814530 | 0.064055111 | 0.093052936 |
| Zfp729b | -0.049393392 | 0.001729963 | 0.0726586 | 0.025507293 | 0.060205369 |
| Zfp738 | -0.02575679 | 0.270941216 | 0.7342030 | 0.108322362 | 0.129394891 |
| Zfp74 | 0.021355036 | 0.301147557 | 0.7606291 | 0.108351198 | 0.091893482 |
| Zfp740 | 0.023972809 | 0.357377637 | 0.7988251 | 0.179501275 | 0.152942992 |
| Zfp746 | -0.022848449 | 0.365209209 | 0.8050191 | 0.14221135 | 0.147283238 |
| Zfp747 | -0.029760481 | 0.064071247 | 0.4700387 | 0.039777747 | 0.059835992 |
| Zfp748 | 0.036455734 | 0.036430296 | 0.3620803 | 0.083107719 | 0.05513076 |
| Zfp758 | -0.01830036 | 0.321140534 | 0.7726689 | 0.062820735 | 0.074506556 |
| Zfp763 | 0.018026884 | 0.18080424 | 0.6569883 | 0.047701579 | 0.032719893 |
| Zfp768 | 0.006740942 | 0.708581287 | 0.9414057 | 0.080042878 | 0.069413982 |
| Zfp770 | -0.026007566 | 0.159199518 | 0.6357726 | 0.062021376 | 0.075743558 |
| Zfp771 | 0.007322087 | 0.809573815 | 0.9661473 | 0.278876957 | 0.250080514 |
| Zfp775 | -0.003985697 | 0.770896357 | 0.9552333 | 0.041272912 | 0.039075017 |
| Zfp777 | -0.014068751 | 0.429473214 | 0.8340365 | 0.063815996 | 0.069880609 |
| Zfp780b | 0.047254928 | 0.029503066 | 0.3294815 | 0.147054548 | 0.091449679 |
| Zfp783 | -0.01140517 | 0.490435692 | 0.8658154 | 0.05527562 | 0.056217497 |
| Zfp787 | 0.001423492 | 0.963542036 | 0.9966421 | 0.289807456 | 0.265739571 |
| Zfp788 | 0.019877139 | 0.423523946 | 0.8319519 | 0.167303041 | 0.139677791 |
| Zfp799 | -0.011875162 | 0.49951711 | 0.8691825 | 0.066760768 | 0.06485509 |
| Zfp800 | -0.1172583 | 0.000373461 | 0.0248639 | 0.248545162 | 0.347720845 |
| Zfp809 | 0.011196324 | 0.590464834 | 0.9053185 | 0.099145314 | 0.090863429 |
| Zfp81 | 0.028667121 | 0.14083871 | 0.6133099 | 0.103502911 | 0.073420253 |
| Zfp810 | -0.035030395 | 0.12985112 | 0.5936051 | 0.107363824 | 0.121760903 |
| Zfp820 | -0.008360029 | 0.628953573 | 0.9130564 | 0.061985166 | 0.06027225 |
| Zfp821 | -0.037447857 | 0.099685858 | 0.5446910 | 0.109304692 | 0.12277585 |
| Zfp827 | 0.004462156 | 0.84322697 | 0.9734889 | 0.119661221 | 0.113345481 |
| Zfp830 | -0.079214907 | 0.007485626 | 0.1721437 | 0.189732465 | 0.250230156 |
| Zfp84 | 0.002732686 | 0.897679353 | 0.9825441 | 0.099992429 | 0.097175882 |
| Zfp846 | -0.017040149 | 0.355199046 | 0.7969131 | 0.058867155 | 0.077898016 |
| Zfp862-ps | -0.015846452 | 0.42930369 | 0.8340365 | 0.099832523 | 0.088423205 |
| Zfp865 | -0.007022223 | 0.716194435 | 0.9431411 | 0.075965505 | 0.083155259 |
| Zfp866 | -0.025512421 | 0.161724927 | 0.6394225 | 0.05970966 | 0.077098984 |
| Zfp868 | -0.011647322 | 0.508885065 | 0.8704067 | 0.064461188 | 0.065677503 |
| Zfp869 | 0.007200125 | 0.826302441 | 0.9716906 | 0.327024931 | 0.315639208 |
| Zfp87 | -0.025746206 | 0.160414346 | 0.6370337 | 0.059260014 | 0.079388522 |
| Zfp871 | -0.08750315 | 0.013965897 | 0.2328431 | 0.423335254 | 0.498250493 |
| Zfp874b | 0.023837145 | 0.077412485 | 0.5022647 | 0.0493537 | 0.028792727 |
| Zfp882 | -0.006053491 | 0.667868217 | 0.9230765 | 0.040710665 | 0.043176682 |
| Zfp9 | -0.006167142 | 0.845357765 | 0.9734889 | 0.265931673 | 0.258959996 |
| Zfp90 | -0.037719342 | 0.049561753 | 0.4192321 | 0.06079284 | 0.085129595 |
| Zfp91 | 0.066781306 | 0.035813457 | 0.3593642 | 0.344824081 | 0.289127437 |
| Zfp930 | 0.000634572 | 0.967890128 | 0.9976072 | 0.063363937 | 0.047695415 |
| Zfp931 | -0.018449474 | 0.375546152 | 0.8100495 | 0.089422337 | 0.093756683 |
| Zfp932 | 0.00385094 | 0.83840837 | 0.9727216 | 0.097070611 | 0.074595822 |
| Zfp933 | -0.002037564 | 0.889432415 | 0.9819597 | 0.046707397 | 0.043562129 |
| Zfp937 | -0.026565498 | 0.1174175 | 0.5743131 | 0.050691538 | 0.06490189 |
| Zfp938 | 0.056816365 | 0.084466679 | 0.5191841 | 0.354409738 | 0.293990071 |
| Zfp941 | -0.025344686 | 0.249871408 | 0.7205594 | 0.101407109 | 0.112006357 |
| Zfp942 | -0.026329165 | 0.170214273 | 0.6501831 | 0.068807405 | 0.083314205 |
| Zfp943 | -0.033487193 | 0.12111213 | 0.5811445 | 0.085602141 | 0.106651168 |
| Zfp944 | -0.036057886 | 0.065871073 | 0.4762891 | 0.06718 | 0.092396794 |
| Zfp945 | -0.043166846 | 0.140287241 | 0.6130953 | 0.231066176 | 0.251780825 |
| Zfp946 | -0.003017129 | 0.874110252 | 0.9784331 | 0.076903066 | 0.077490163 |
| Zfp948 | 0.029482711 | 0.129280854 | 0.5922132 | 0.100748085 | 0.07353147 |
| Zfp949 | 0.016545817 | 0.441997424 | 0.8428709 | 0.126785328 | 0.098841463 |
| Zfp950 | -0.016755781 | 0.636020317 | 0.9153541 | 0.41379076 | 0.420677159 |

| | | | | | |
|---|---|---|---|---|---|
| Zfp952 | -0.021797735 | 0.283411365 | 0.7427848 | 0.078270478 | 0.092439746 |
| Zfp954 | 0.024109223 | 0.202019975 | 0.6752150 | 0.101323548 | 0.069265899 |
| Zfp955b | -0.006550676 | 0.756842777 | 0.9521533 | 0.104837938 | 0.099065946 |
| Zfp956 | 0.004680606 | 0.781710823 | 0.9590847 | 0.066800715 | 0.058288858 |
| Zfp958 | 0.024349466 | 0.271551496 | 0.7342030 | 0.129911082 | 0.105502822 |
| Zfp959 | 0.020612065 | 0.243080499 | 0.7132719 | 0.081652005 | 0.055934738 |
| Zfp961 | -0.013063227 | 0.366813671 | 0.8053794 | 0.039521687 | 0.046099738 |
| Zfp974 | 0.019046664 | 0.197244893 | 0.6730962 | 0.057341147 | 0.041305 |
| Zfp975 | 0.01681941 | 0.262460213 | 0.7275735 | 0.05360315 | 0.041065381 |
| Zfp976 | -0.025501979 | 0.15420403 | 0.6304934 | 0.057374401 | 0.070083775 |
| Zfp979 | -0.019076847 | 0.257602535 | 0.7250597 | 0.054532604 | 0.063152788 |
| Zfp983 | 0.015911779 | 0.385578859 | 0.8145107 | 0.089211864 | 0.06661279 |
| Zfp994 | 0.009645312 | 0.703498373 | 0.9391887 | 0.159057285 | 0.146851898 |
| Zfp995 | 0.001173102 | 0.948202188 | 0.9945018 | 0.079609113 | 0.066849719 |
| Zfpl1 | 0.045271441 | 0.159808482 | 0.6369225 | 0.354934855 | 0.301463513 |
| Zfr | 0.050217031 | 0.179150053 | 0.6550070 | 0.831303452 | 0.782651934 |
| Zfr2 | -0.031541159 | 0.133135424 | 0.5988251 | 0.078993251 | 0.103271176 |
| Zfx | -0.040162718 | 0.152021981 | 0.6281105 | 0.175183131 | 0.20157422 |
| Zfyve1 | 0.013955609 | 0.435494133 | 0.8389295 | 0.073308318 | 0.064567719 |
| Zfyve16 | 0.048082142 | 0.08121015 | 0.5131240 | 0.218264008 | 0.170961749 |
| Zfyve19 | -0.025852179 | 0.292444272 | 0.7526152 | 0.135053075 | 0.143797041 |
| Zfyve21 | -0.026177168 | 0.392291372 | 0.8203057 | 0.252614648 | 0.272062339 |
| Zfyve26 | -0.009325342 | 0.605308837 | 0.9087139 | 0.06519627 | 0.068877565 |
| Zfyve27 | -0.019533536 | 0.410756626 | 0.8285290 | 0.128538399 | 0.136045132 |
| Zfyve9 | -0.006753213 | 0.816229191 | 0.9688248 | 0.207947703 | 0.204707379 |
| Zgpat | 0.035924492 | 0.059110989 | 0.4538552 | 0.106576981 | 0.071538211 |
| Zhx1 | 0.004736163 | 0.876820623 | 0.9785583 | 0.245513056 | 0.261398801 |
| Zhx2 | -0.004150668 | 0.837085256 | 0.9726619 | 0.079438644 | 0.089980916 |
| Zhx3 | -0.018945855 | 0.380410847 | 0.8117350 | 0.084026638 | 0.103832199 |
| Zim1 | 0.014334977 | 0.648132738 | 0.9170676 | 0.276549672 | 0.231159074 |
| Zkscan1 | -0.015119186 | 0.622892461 | 0.9121505 | 0.281117147 | 0.279141105 |
| Zkscan14 | -0.065937713 | 0.002356676 | 0.0878880 | 0.081578922 | 0.117623913 |
| Zkscan17 | -0.024320026 | 0.215138389 | 0.6845467 | 0.071610165 | 0.089748662 |
| Zkscan2 | -0.00271545 | 0.85430633 | 0.9755431 | 0.048789242 | 0.041781891 |
| Zkscan3 | -0.069674864 | 0.033035841 | 0.3449637 | 0.356693721 | 0.371762504 |
| Zkscan6 | 0.020753983 | 0.290837224 | 0.7513090 | 0.106374401 | 0.078129521 |
| Zkscan7 | -0.002181728 | 0.906853021 | 0.9836812 | 0.069308156 | 0.074299116 |
| Zkscan8 | 0.00247096 | 0.89850632 | 0.9825441 | 0.082920666 | 0.077164232 |
| Zmat1 | -0.02340803 | 0.238586455 | 0.7109741 | 0.086312382 | 0.082576418 |
| Zmat2 | 0.013265636 | 0.708711508 | 0.9414057 | 0.692980406 | 0.646213824 |
| Zmat3 | 0.015937337 | 0.579869942 | 0.9008751 | 0.218723667 | 0.196481027 |
| Zmat5 | -0.037714353 | 0.200346853 | 0.6739371 | 0.207215412 | 0.23860653 |
| Zmiz1 | 0.001119596 | 0.972160416 | 0.9976072 | 0.298289396 | 0.302060318 |
| Zmiz2 | -0.045029734 | 0.109668049 | 0.5655888 | 0.17655902 | 0.211422324 |
| Zmpste24 | 0.027380113 | 0.365752448 | 0.8050191 | 0.264974734 | 0.249483684 |
| Zmym1 | -0.008972427 | 0.657640733 | 0.9211780 | 0.095063149 | 0.088268846 |
| Zmym2 | -0.016837407 | 0.559059818 | 0.8921289 | 0.195209103 | 0.206443286 |
| Zmym3 | 0.003692168 | 0.862055996 | 0.9769675 | 0.107368344 | 0.100721108 |
| Zmym4 | 0.057370376 | 0.095291664 | 0.5351767 | 0.37744148 | 0.328912491 |
| Zmym5 | -0.037859253 | 0.276947404 | 0.7373371 | 0.379645382 | 0.402175927 |
| Zmym6 | -0.01676653 | 0.340311967 | 0.7844658 | 0.053440559 | 0.066098846 |
| Zmynd11 | -0.043484111 | 0.222736644 | 0.6954823 | 0.474291621 | 0.514651931 |
| Zmynd19 | -0.007635455 | 0.732146142 | 0.9462041 | 0.123403543 | 0.111707716 |
| Zmynd8 | 0.056213287 | 0.119307099 | 0.5782737 | 0.560635527 | 0.514680415 |
| Znfx1 | 0.018959161 | 0.4725944 | 0.8595729 | 0.176379115 | 0.165046848 |
| Znhit1 | 0.011288378 | 0.754642482 | 0.9519143 | 0.623001806 | 0.56046105 |
| Znhit2 | 0.040148458 | 0.166116435 | 0.6429285 | 0.254530586 | 0.2009182 |
| Znhit3 | 0.001773632 | 0.947389662 | 0.9945018 | 0.1881826 | 0.166477232 |
| Znhit6 | -0.036511477 | 0.161699698 | 0.6394225 | 0.149985094 | 0.169038276 |
| Znrf1 | -0.046675965 | 0.096903664 | 0.5379319 | 0.187118258 | 0.206047949 |
| Znrf2 | 0.002043326 | 0.951778984 | 0.9951566 | 0.358376847 | 0.370835472 |
| Znrf3 | 0.005304748 | 0.747153585 | 0.9496173 | 0.059958461 | 0.054887964 |
| Zpr1 | -0.063825018 | 0.027653522 | 0.3181899 | 0.207771042 | 0.23063864 |
| Zranb1 | 0.044109813 | 0.23091658 | 0.7024275 | 0.557595048 | 0.519448224 |
| Zranb2 | -0.016961847 | 0.649540259 | 0.9174778 | 0.809573184 | 0.818536636 |
| Zrsr1 | 0.018513514 | 0.468964705 | 0.8563037 | 0.158445035 | 0.155335431 |

| | | | | | |
|---|---|---|---|---|---|
| Zrsr2 | 0.0192177 | 0.576435136 | 0.89946101 | 0.379321627 | 0.360503636 |
| Zscan12 | -0.000484435 | 0.976394911 | 0.99760728 | 0.061281928 | 0.054520137 |
| Zscan18 | -0.019767764 | 0.229185995 | 0.70196775 | 0.039222895 | 0.064010017 |
| Zscan2 | 0.015483705 | 0.382751118 | 0.81297870 | 0.083944729 | 0.061177077 |
| Zscan21 | -0.020458491 | 0.200875821 | 0.67457283 | 0.039335842 | 0.058098697 |
| Zscan22 | 0.01046094 | 0.517752305 | 0.87388048 | 0.061731395 | 0.052387262 |
| Zscan26 | -0.003720805 | 0.90137905 | 0.98254414 | 0.256889737 | 0.246378923 |
| Zscan29 | 0.004867893 | 0.792680921 | 0.96244605 | 0.080378519 | 0.069601585 |
| Zswim1 | 0.020008934 | 0.284991908 | 0.74507912 | 0.090971438 | 0.071271344 |
| Zswim5 | 0.004272905 | 0.765000972 | 0.95400902 | 0.044683296 | 0.040848959 |
| Zswim7 | -0.028881054 | 0.325705848 | 0.77630994 | 0.206432503 | 0.229987104 |
| Zswim8 | -0.058022797 | 0.07934695 | 0.50762057 | 0.289836225 | 0.353280177 |
| Zswim9 | 0.010673545 | 0.485155805 | 0.86297153 | 0.057424759 | 0.044043815 |
| Zup1 | -0.014503958 | 0.614816089 | 0.91083865 | 0.206740128 | 0.207119045 |
| Zw10 | 0.026452525 | 0.167392837 | 0.64461837 | 0.097613163 | 0.077154509 |
| Zwint | -0.022805684 | 0.530301231 | 0.87764481 | 1.143201299 | 1.057515198 |
| Zxdb | 0.040421774 | 0.120653754 | 0.58100041 | 0.205310865 | 0.156237213 |
| Zxdc | -0.021331483 | 0.419350871 | 0.83009907 | 0.16117908 | 0.184109988 |
| Zyg11b | 0.020092407 | 0.531229307 | 0.87764481 | 0.276597844 | 0.287339644 |
| Zyx | -0.01721936 | 0.314903304 | 0.77059875 | 0.049609719 | 0.063630485 |
| Zzef1 | -0.036706816 | 0.236394495 | 0.70910699 | 0.235265045 | 0.25247138 |
| Zzz3 | 0.037869061 | 0.242733735 | 0.71327199 | 0.322197936 | 0.296643802 |
